# Epha1 is a cell surface marker for neuromesodermal progenitors and their early mesoderm derivatives

**DOI:** 10.1101/584524

**Authors:** Luisa de Lemos, André Dias, Ana Nóvoa, Moisés Mallo

**Author notes:** correspondence to Moisés Mallo.

## Abstract

The vertebrate body is built during embryonic development by the sequential addition of new tissue as the embryo grows at its caudal end. During this process, the neuro-mesodermal progenitors (NMPs) generate the postcranial neural tube and paraxial mesoderm. Recently, several approaches have been designed to determine their molecular fingerprint but a simple method to isolate NMPs from embryos without the need for transgenic markers is still missing. We isolated NMPs using a genetic strategy that exploits their self-renew properties, and searched their transcriptome for cell surface markers. We found a distinct Epha1 expression profile in progenitor-containing areas of the mouse embryo, consisting of two cell subpopulations with different Epha1 expression levels. We show that Sox2^+^/T^+^ cells are preferentially associated with the Epha1 compartment, indicating that NMPs might be contained within this cell pool. Transcriptional profiling showed enrichment of high Epha1-expressing cells in known NMP and early mesoderm markers. Also, tail bud cells with lower Epha1 levels contained a molecular signature suggesting the presence of notochord progenitors. Our results thus indicate that Epha1 could represent a valuable cell surface marker for different subsets of axial progenitors, most particularly for NMPs taking mesodermal fates.

## INTRODUCTION

Vertebrate body axis extension occurs in a head to tail sequence and relies on populations of cells with self-renewing properties, collectively known as axial progenitors. (Aires et al., 2018; Dias and Aires, 2020; Stern et al., 2006; Steventon and Martinez Arias, 2017). These progenitors, initially located in the caudal epiblast and later in the tailbud, include several cell pools classified according to their potential. The neuro-mesodermal progenitors (NMPs) are one of those cell populations, consisting of bipotent cells that generate both neural and mesodermal derivatives (Cambray and Wilson, 2002; Cambray and Wilson, 2007; Henrique et al., 2015; Tzouanacou et al., 2009; Wilson et al., 2009). The lateral and paraxial mesoderm progenitors are an additional cell population within the epiblast, whose potential is restricted to the production of mesodermal derivatives (Wymeersch et al., 2016). Recently, transcriptome data analyses led to the identification of notochord progenitors, which maintain a stable molecular signature during axial elongation (including genes like *Noto*, *Shh* and *Foxa2*), possibly acting as a stable progenitor niche particularly for NMPs (Wymeersch et al., 2019). Since the initial identification of axial progenitors, increasing efforts have been conducted to determine the precise molecular characteristics of each different cell population, with particular focus on NMPs. Combined mapping and expression studies indicate that the early neural marker *Sox2* and the mesodermal transcription factor *T* (Brachyury) are mainly co-expressed in regions known to contain NMPs (Cambray and Wilson, 2007; Martin and Kimelman, 2012; Olivera-Martinez et al., 2012; Tsakiridis et al., 2014; Wymeersch et al., 2016). More recently, the introduction of improved high throughput techniques allowing *in vivo* and *in vitro* transcriptomic analyses, and most particularly single-cell RNA sequencing (scRNA-seq), has shed light on the role of these key transcription factors and provided a deeper understanding of other molecular players (e.g. *Wnt3a*, *Tbx6* and *Cdx2*) and gene regulatory networks involved in the control of NMP maintenance and differentiation both in the embryo and in *in vitro* models (Dias et al., 2020; Edri et al., 2019; Gouti et al., 2017; Koch et al., 2017; Wymeersch et al., 2019). Interestingly, some of these studies have shown that the NMP transcriptome changes extensively over time, including the activation of an incomplete epithelial to mesenchymal transition (EMT) when entering the tail bud (Dias et al., 2020; Wymeersch et al., 2019). It has also been reported expression of *Tbx6* in a subpopulation of the tail NMPs and that this expression is involved in NMP cell fate decisions (Javali et al., 2017). In addition, gene expression and lineage tracing experiments indicated that *Nkx1-2* (formerly *Sax1*) is present in NMPs and early neural and mesodermal progenitors throughout axial extension (Albors et al., 2018).

Despite all these studies, it is still not possible to isolate NMPs in a physiologically active form without relying on previous modifications to introduce reporter genes, mostly because the molecules typically used to identify those progenitors are transcription factors. In the present work, we searched for cell surface markers specifically enriched in the axial progenitor population that could facilitate isolation of NMPs using conventional cell sorting approaches. For this, we exploited the intrinsic self-renewal properties of these progenitors to obtain from the tail bud a cell population highly enriched in NMPs. In particular, we recovered the cells containing the fluorescent marker that had been genetically introduced into the progenitors when they were still engaged in axial elongation at much earlier stages of development. Analysis of the transcript content of these cells provided several genes encoding cytoplasmic membrane proteins that might allow sorting active NMPs. From these molecules, Epha1 stood out as the best candidate, as its expression is mostly limited to the progenitor-containing areas of the mouse embryo throughout axial extension and due to the existence of fluorescent-activated cell sorting (FACS)-tested antibodies that could facilitate the development of protocols for NMP isolation from embryonic tissues. Using one of these Epha1 antibodies, we found that tail bud Epha1-positive cells can be divided in two subpopulations, Epha1^High^ and Epha1^Low^, on the basis of their Epha1 expression levels according to FACS plots. While cells containing low Epha1 expression levels were also obtained from tissues containing differentiated NMP derivatives, Epha1^High^ cells were exclusively found in the tail bud. Variations in Epha1 immunoreactivity were also observed by immunofluorescence, with stronger signals being detected in regions of the caudal epiblast and the tail bud known to contain NMPs and their early mesoderm derivatives. Epha1 positive cells from the tail bud, and most prominently those in the Epha1^High^ compartment, were highly enriched in Sox2^+^/T^+^ cells, the most frequently used molecular signature of NMPs (Cambray and Wilson, 2007; Koch et al., 2017; Wymeersch et al., 2016), suggesting that Epha1 could be a marker for these progenitors. Analysis of the transcriptomes obtained from tail bud Epha1^High^ and Epha1^Low^ cells revealed enrichment of both cell pools in NMP markers, further indicating a connection between Epha1 and axial progenitors. These transcriptome analyses also showed that Epha1^High^ and Epha1^Low^ cells represent different cell populations. In particular, Epha1^High^ cells expressed markers associated with NMPs entering mesodermal routes. Conversely, Epha1^Low^ cells included a molecular signature congruent with the existence of notochord progenitors within this cell compartment. Together, our results indicate that Epha1 could be a valuable cell surface marker for the isolation of different subsets of mouse embryonic axial progenitors and that high Epha1 values might be a molecular signature of NMPs entering the mesodermal progenitor compartment.

## RESULTS

### Labelling and isolation of long-term NMPs and their immediate descendants from developing mouse embryos

Most NMPs transcriptome analyses using embryonic tissue have been performed on microdissected regions of the embryo containing NMPs but that also include other cell types (e.g. some that already entered in mesoderm differentiation routes), or on cells isolated on the basis of the expression of a gene enriched in these progenitors (Dias et al., 2020; Gouti et al., 2017; Koch et al., 2017; Wymeersch et al., 2019). Here we used an unbiased approach to isolate NMPs exploiting their self-renewing properties. In particular, as the NMPs in the tail bud derive from those located in the node streak border at earlier developmental stages (Cambray and Wilson, 2007), we designed a genetic strategy to label these progenitors when at the caudal epiblast and isolated them from the tail bud at later developmental stages. This system combines the *Cdx2P-Cre^ERT^* transgene (Jurberg et al., 2013) with the *ROSA26*-*YFP-Cre* reporter (Srinivas et al., 2001). By administering a single low tamoxifen dose at embryonic day (E) 7.5 we induced a short pulse of permanent YFP label into a subset of axial progenitors that allowed following their fate at later developmental stages. We have previously used this system with the *ROSA26-βgal-Cre* reporter (Soriano, 1999), which proved the efficiency of the method, as estimated by the presence of labelled cells in the embryonic tissues posterior to the position of Cre-mediated recombination, all the way down to the tail tip (Aires et al., 2019) (Figure 1A–C).

**Figure 1.**
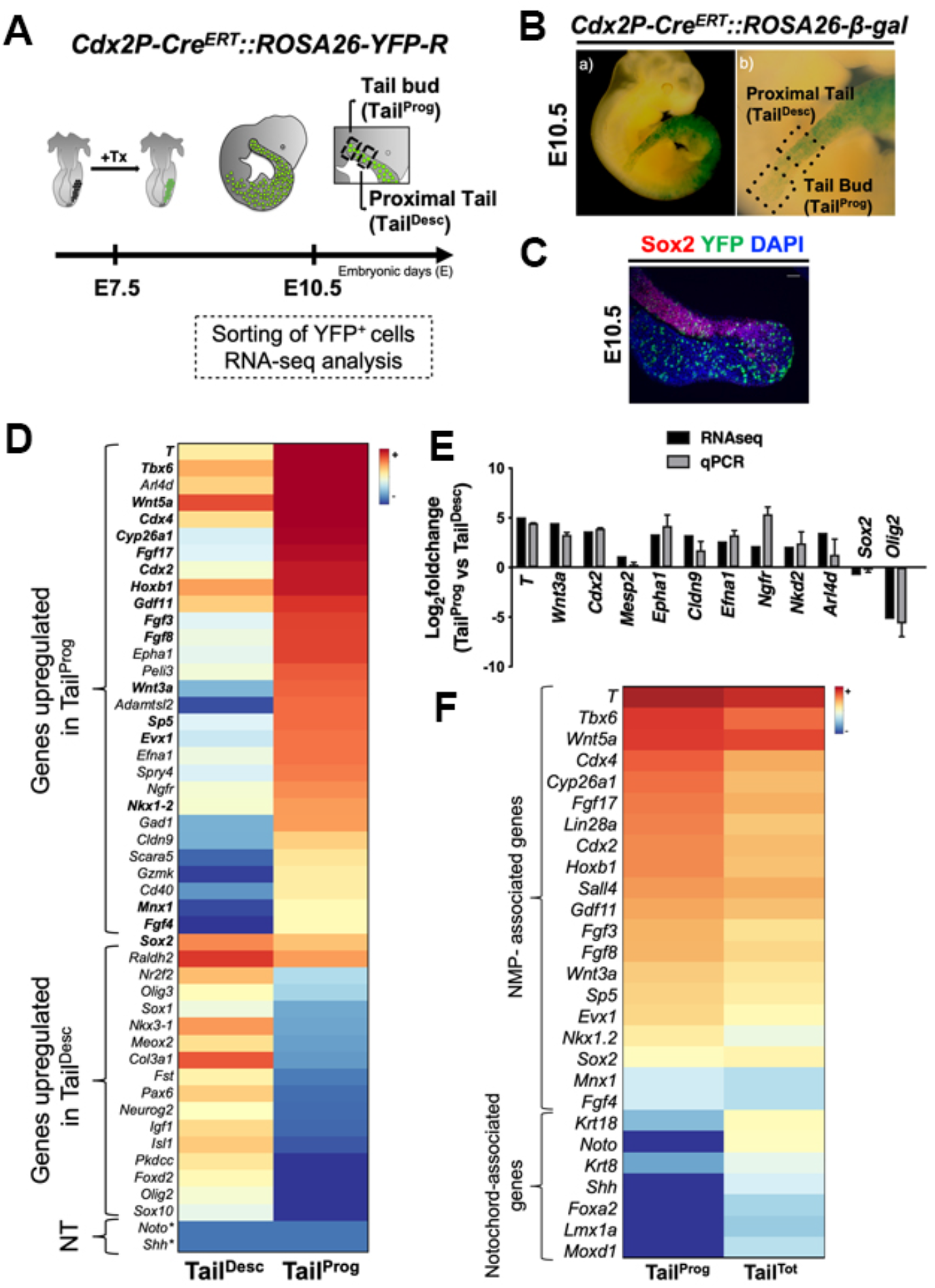
Labeling, isolation and transcriptomic analysis of long term axial progenitors and their immediate descendants. **A.** Schematic representation of the genetic labelling strategy used to induce a permanent YFP label in axial progenitors in *Cdx2P-Cre^ERT^∷ROSA26-YFP* reporter embryos. Tx: tamoxifen. **B.** β-Gal staining of E10.5 embryos labeled with the same genetic scheme but with ROSA26-reporter to show contribution of labeled cells until the tail bud. b) shows a magnification of the tail region. Dotted squares, in (b), delimited the regions from where Tail^Prog^ and Tail^Desc^ populations were obtained. **C.** Immunofluorescence for Sox2 (magenta) and YFP (green) in sagittal sections of *Cdx2P-Cre^ERT^∷ROSA26-YFP* tails. DAPI is shown in blue. Scale bar: 100 μm. **D.** Heat map displaying several differentially expressed genes between Tail^Prog^ and Tail^Desc^ cell populations. Genes associated with NMPs are highlighted in bold. NT represent notochord genes. The color key bar represents the average of normalized counts on logarithmic scale. Note: values labeled with (*) are equal to zero, therefore logarithmic scale could not be applicable. **E**. Validation of RNA-seq data by RT-qPCR comparing the data obtained by RNA-seq (black bars) and by RT-qPCR (grey bars). The error bars represent the standard deviation error of three independent replicates. **F**. Heat maps comparing transcriptome data from Tail^Prog^ YFP cells and from the entire tailbud (Tail^Tot^) (same region) (obtained from Aires et al., 2019). Tail^Prog^ is highly enriched in NMPs but seem to lack notochord progenitors. The color key bar represents the average of normalized counts on logarithmic scale.

Descendants of progenitors labelled with YFP at E7.5 were then recovered from the tail region at E10.5 by FACS. Two sets of YFP-positive cells were recovered: a first set was isolated from the tail bud (Tail^Prog^), which is expected to be highly enriched in progenitors; a second set was recovered from more anterior tail regions, where labelled cells are already part of the tissues derived from the progenitors (Tail^Desc^). Comparison of the transcriptome of these two cell pools, obtained by RNA sequencing (RNA-seq), identified 2465 genes showing differential expression (*p*-value <0,05) between the two cell groups (Table S1). Of these, 847 genes were highly expressed in the Tail^Prog^, whereas 611 genes were up-regulated in the Tail^Desc^ sample. A selection of 12 differentially expressed genes was then used to validate the RNA-seq data by reverse transcription-quantitative polymerase chain reaction (RT-qPCR) (Figure 1E).

Initial analysis of the differentially expressed genes revealed high enrichment of the Tail^Prog^ cells in factors that have been linked to neuro-mesodermal identity (Figure 1D). For instance, *Cdx2*, *Cdx4* and *T*, known to be highly expressed in NMPs and proven to be essential for their activity (Amin et al., 2016; Chawengsaksophak et al., 2004; Herrmann et al., 1990; Savory et al., 2011; van Rooijen et al., 2012), were among the most strongly up-regulated genes in the Tail^Prog^ compartment (Figure 1D). Similarly, other genes whose expression has been shown to be enriched in tail bud NMPs, including *Mnx1*, *Nkx1-2*, *Fgf8*, *Fgf4*, *Fgf3*, *Evx1*, *Cyp26a*, *Hoxb1*, *Sp5*, *Gdf11*, *Wnt5a* or *Wnt3a* (Abu-Abed et al., 2001; Aires et al., 2019; Albors et al., 2018; Boulet and Capecchi, 2012; Cambray and Wilson, 2007; Dush and Martin, 1992; Greco et al., 1996; Harrison et al., 2000; Javali et al., 2017; McPherron et al., 1999; Murphy and Hill, 1991; Naiche et al., 2011; Robinton et al., 2019; Sakai et al., 2001; Takada et al., 1994; Wymeersch et al., 2019; Yamaguchi et al., 1999), also showed significant differential expression in the Tail^Prog^ cell pool (Figure 1D). In addition, we also found other genes previously not linked to NMPs like *Epha1*, *Arl4d*, *Efna1*, *Cldn9*, *Gad1* and *Scara5*, to be highly expressed in the progenitor compartment (Figure 1D).

Conversely, we found enrichment of the Tail^Desc^ cell pool in markers for neural and mesodermal NMP derivatives, including *Ngn2*, *Sox1*, *Olig2*, *Olig3*, *Sox10*, *Meox2*, *Raldh2*, *Pax6*, *Fst* and *Nr2f2* (Albano et al., 1994; Aubert et al., 2003; Candia et al., 1992; Gradwohl et al., 1996; Jonk et al., 1994; Kuhlbrodt et al., 1998; Niederreither et al., 1997; Takeichi et al., 2002; Walther and Gruss, 1991) (Figure 1D), indicating that the Tail^Prog^ cells have the differentiation potential expected from NMPs. *Sox2*, one of the components of the typical NMP signature (Aires et al., 2018; Koch et al., 2017; Wymeersch et al., 2016), was expressed at slightly higher levels in Tail^Desc^ than in Tail^Prog^ cells (Figure 1D). This is not surprising, as *Sox2* is highly expressed in the neural tube, which is one of the tissues from which the Tail^Desc^ cells were recovered. Interestingly, both Tail^Prog^ and Tail^Desc^ datasets do not contain notochord markers such as *Shh*, *Noto* or *Foxa2*, which were readily found in a dataset obtained from similar stage unsorted tail buds (Aires et al., 2019) (Tail^Tot^) (Figure 1F; Table S2), indicating that the Tail^Prog^ cell population is highly enriched in NMPs and not in other axial progenitors. Together, these data indicate that the genetic lineage tracing strategy here described labels tail NMPs and their neural and mesodermal descendants, and that the isolated Tail^Prog^ population is highly enriched in NMPs.

### *Epha1* is expressed in axial progenitors and early mesodermal-fated cells

From the genes differentially expressed (*p*-value <0,05) between the Tail^Prog^ and the Tail^Desc^ cell populations, we concentrated on those coding for membrane proteins that could be used to isolate physiologically active NMPs without previous genomic modifications (e.g. transgenic reporters). Gene Ontology (GO) categorization (Ashburner et al., 2000) of genes differentially up-regulated in Tail^Prog^ cells with a log_2_ Fold change >2 and q value <0,05, identified 61 genes assigned to the category “membrane” (GO:0005886). From these, we further selected the 16 genes that had >10 reads in the Tail^Prog^ RNA-seq dataset (Figure 2A), as, for these datasets, this was the read number empirically determined to be the lower threshold level allowing mRNA or protein detection in the embryo using conventional methods. Expression analyses at E10.5 by *in situ* hybridization (ISH) revealed that the staining patterns for some of those genes included a strong signal in the tail region, although these patterns differed among the various genes (Figure 2B). In addition, for most of them, tail bud expression represented a subset of more complex expression patterns that included other embryonic regions.

**Figure 2.**
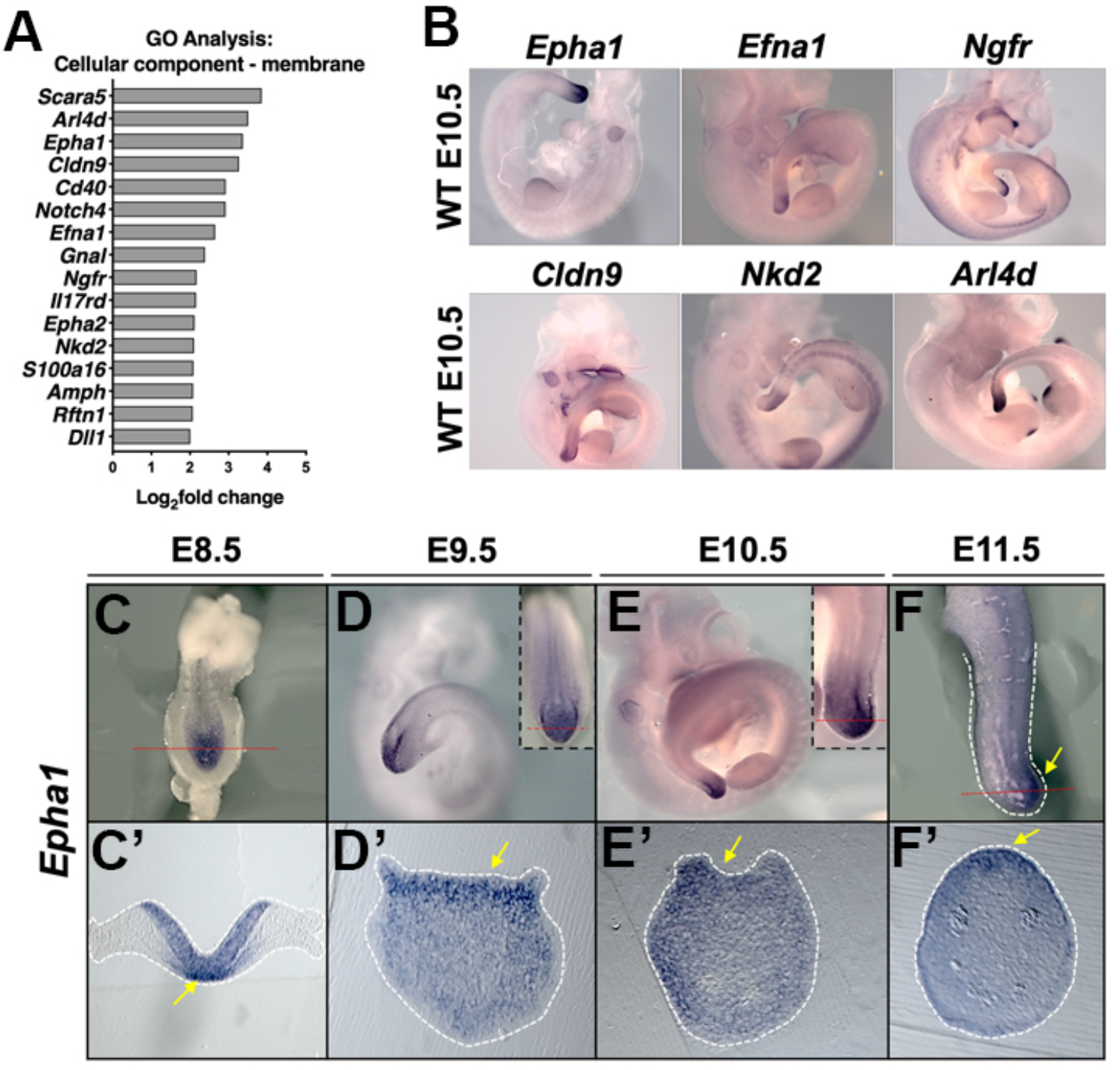
Identification of potential cell-surface markers for axial progenitors. **A**. Differentially expressed genes in Tail^Prog^ cells encoding cell membrane proteins according to gene ontology classification. **B**. Whole-mount *in situ* hybridization in E10.5 embryos using probes for *Epha1*, *Efna1*, *Ngfr*, *Cldn9*, *Nkd2* and *Arl4d*. **C-F**. Whole-mount *in situ* hybridization for *Epha1* in wild type embryos from E8.5 to E11.5 showing high *Epha1* expression in the caudal epiblast and in the tailbud progenitor regions. Inlets show magnifications of the posterior regions. **C’-F’**. Transversal sections of these regions (dotted red lines) made for each timepoint indicating that *Epha1* is still expressed in early mesoderm cells. Yellow arrows indicate the regions where stronger *Epha1* expression is observed (e.g. primitive streak in the E8.5 embryo).

From these genes, we focused on *Epha1* based not only on its expression pattern, apparently restricted to the progenitor zone at different developmental stages, but also on the availability of FACS-validated antibodies, able to provide reliable data with cells obtained from solid embryonic tissues. By ISH we detected *Epha1* expression already at E8.5, when it is mostly observed in the caudal lateral epiblast, the area containing NMPs (Cambray and Wilson, 2007; Wymeersch et al., 2016), as well as in cells entering in the mesodermal compartment (Figure 2C, C’). *Epha1* expression was also observed in the caudal epiblast of E9.5 embryos, fading anteriorly when entering the regions corresponding to the presomitic mesoderm and the caudal neural tube (Figure 2D, D’), and at E10.5 and E11.5, *Epha1* expression was essentially restricted to the tailbud (Figure 2E, E’). Analysis of Epha1 by immunostaining confirmed the expression patterns obtained by ISH, although slightly broader, possibly derived from higher stability of the protein than of the mRNA. In addition, although this technique is not purely quantitative, it showed that the Epha1 protein levels were not uniform in the positive domain. In particular, in E8.5 embryos Epha1 expression was particularly strong in the region of the primitive streak (PS) (Figure 3A - white arrow), known to contain progenitor cells undergoing an EMT to generate the mesodermal layer (Acloque et al., 2009; Hay, 1968; Wilson et al., 2009) and at E10.5 in the region abutting the posterior end of the neural tube, positive for both T and Sox2, as well as mesenchyme caudal to this region thought to contain mesoderm progenitors (Dias et al., 2020; McGrew et al., 2008; Wymeersch et al., 2016; Wymeersch et al., 2019) (Figure 3B; white arrows). These data suggest that, at various developmental stages, *Epha1* is strongly expressed in regions containing NMPs and their early mesoderm derivatives. Consistent with this observation, using a gastrulation/early organogenesis stage (Theiler stage 12) mouse scRNA-seq dataset (Pijuan-Sala et al., 2019), we also found *Epha1* to be highly expressed in the NMP and caudal mesoderm clusters (Figure 4), overlapping with the region co-expressing the main NMP-associated genes (*Sox2*, *T and Nkx1-2)* (Albors et al., 2018; Cambray and Wilson, 2007; Koch et al., 2017; Martin and Kimelman, 2012; Olivera-Martinez et al., 2012; Tsakiridis et al., 2014; Wymeersch et al., 2016), further suggesting an association between Epha1 expression and NMPs and their early mesoderm derivatives.

**Figure 3.**
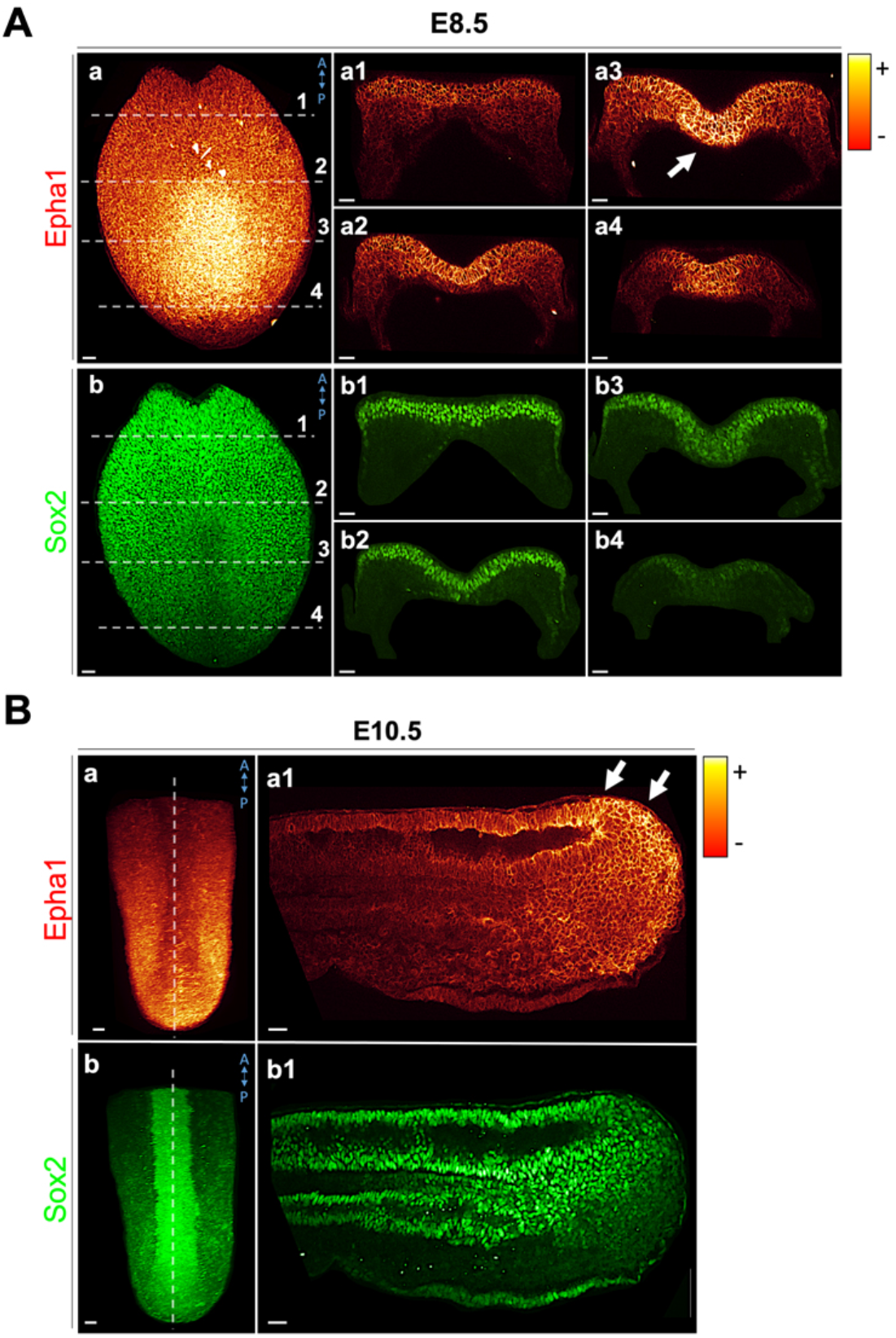
High Epha1 expression labels NMPs and early mesoderm cells. Whole-mount two-photon immunofluorescence staining for Epha1 (red/yellow gradient) and Sox2 (green) in E8.5 (**A**) and E10.5 (**B**) embryos. Transversal and sagittal sections (dotted white lines) are also shown. High expression of Epha1 (white arrows) was observed in regions known to contain NMPs and early mesoderm cells (e.g. primitive streak at E8.5). Scale bars: 50μm.

**Figure 4.**
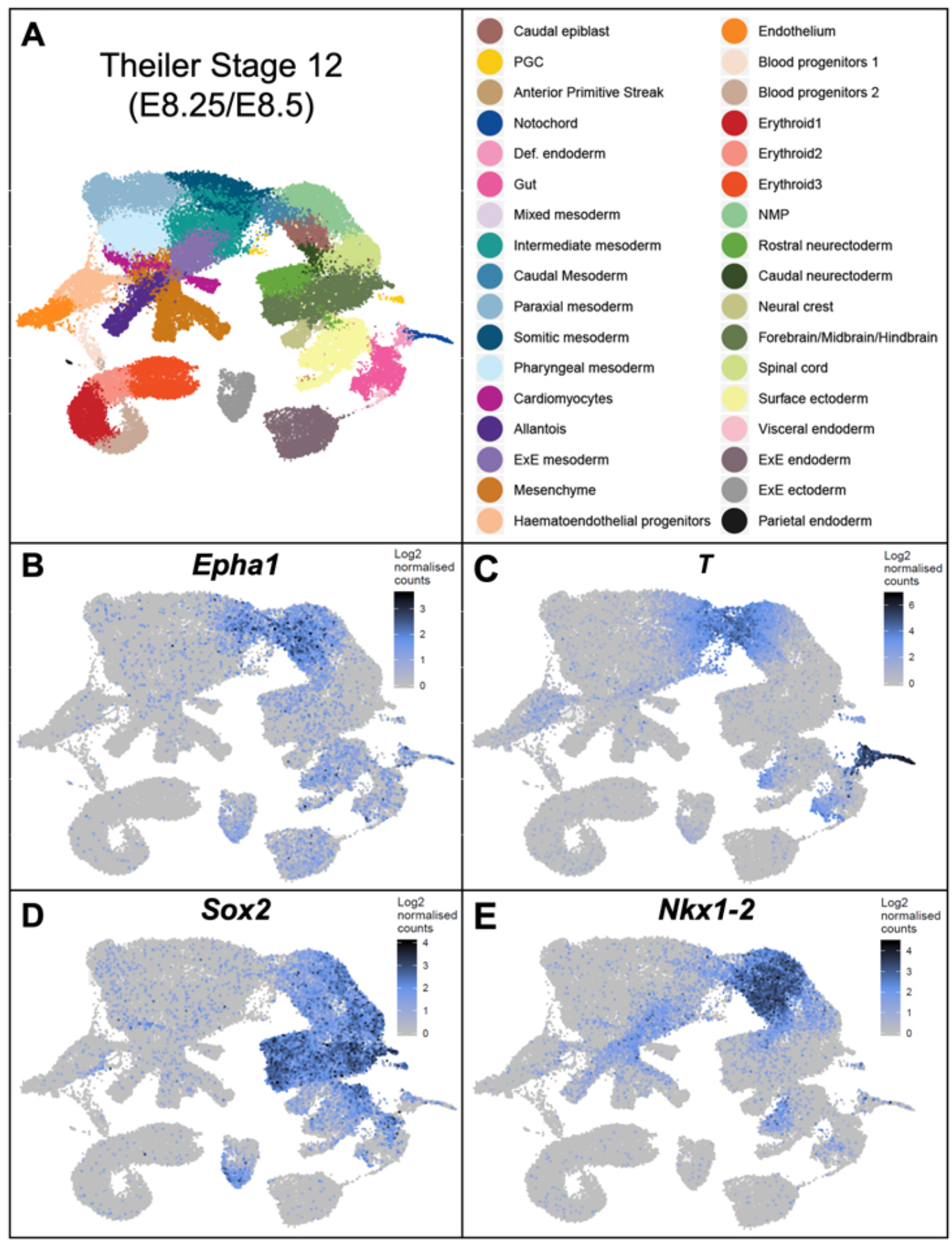
Distribution of *Epha1* expression in the single-cell UMAP of Theiler Stage 12 embryos. Analysis of the scRNA-seq dataset of Theiler stage 12 (around E8.5) mouse embryos. **A**. UMAP (Uniform Manifold Approximation and Projection). *Epha1* expression (**B**) overlaps with the regions co-expressing *T* (**C**), *Sox2* (**D**) and *Nkx1-2* (**E**), which are known to contain NMPs and their early mesoderm derivatives [clusters termed “NMP” (light green), “caudal epiblast” (brown) and “caudal mesoderm” (medium sky blue)].

### High Epha1 expressing cells are enriched in T and Sox2 double-positive cells

To test this hypothesis, we performed FACS analysis of cells obtained from the tail bud and adjacent anterior tail region of E10.5 embryos using an antibody against Epha1. Both areas contained a high proportion of Epha1 positive cells (Figure 5A–C), which fits with the immunofluorescence data. Interestingly, however, the staining patterns in the two embryonic regions were different, as cells from the tail bud included a population with higher staining intensity that was never observed in the FACS plots obtained from the anterior tail region (Figure 5B). These results indicate the existence of cells with different amounts of Epha1 in the tail bud, consistent with the not uniform staining intensities observed by immunofluorescence. Based on their Epha1 content, we grouped Epha1-positive tail bud cells in two subpopulations, which will be referred to as Epha1^High^ and Epha1^Low^ (Supplementary Figure 3). We also obtained similar Epha1^high^ and Epha1^low^ cell compartments in the progenitor-containing area of E8.5 embryos, although the proportion of Epha1^high^ cells was lower in this tissue than in the tail bud from E10.5 embryos (Figure 6A).

**Figure 5.**
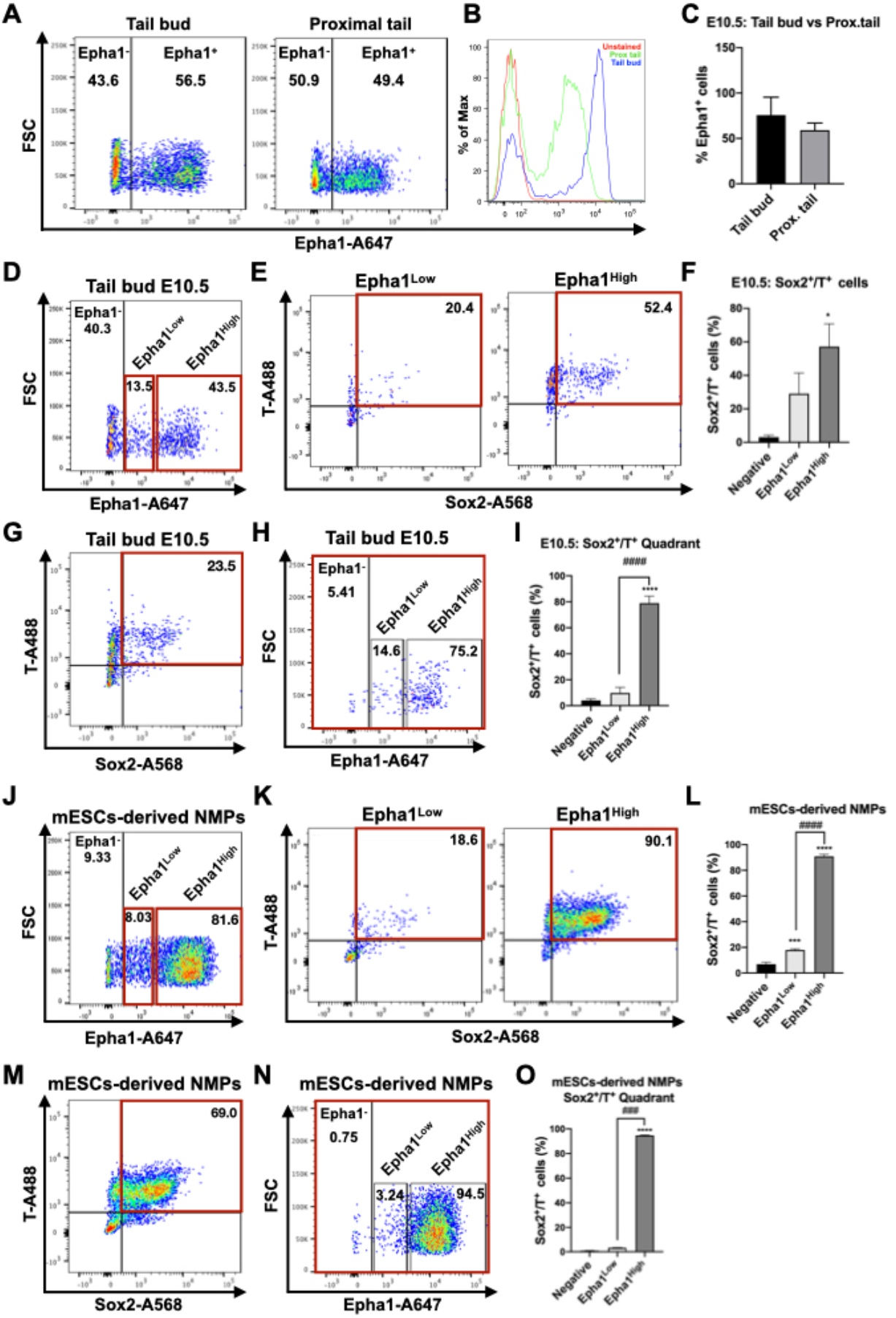
Flow cytometry analysis of Epha1, Sox2 and T expression in E10.5 embryos and mESCs-derived NMPS. **A.** FACS analysis showing fluorescence spread into Epha1-channel from the tail bud and proximal tail of E10.5 embryos. **B.** Single parameter histogram showing the different fluorescent intensity of Epha1-positive cells from tail bud (blue), proximal tail (green) and unstained control (red). **C.** Percentage of Epha1-positive cells found in the tail bud and proximal tail of E10.5 embryos. **D.** FACS dot-plots displaying Epha1 subpopulations of E10.5 tail bud. **E.** FACS profiles of Sox2 and T expression in cells from the Epha1^Low^ and Epha1^High^ compartments indicated in D. **F.** Quantification of Sox2^+^/T^+^ cells within the different Epha1 subpopulations of E10.5 tail bud. **G.** FACS dot-plot displaying Sox2 and T expression in cells from E10.5 tail buds. **H.** FACS profile showing distribution of Sox2^+^/T^+^ cells from the red window in G among the Epha1 compartment. **I.** Quantification of Sox2^+^/T^+^ cells from E10.5 tail buds between the different Epha1 subpopulations. **J.** FACS dot-plot displaying Epha1 subpopulations in mESCs-derived NMPs. **K.** FACS profiles showing Sox2 and T expression in cells from the Epha1^Low^ and Epha1^High^ windows defined in J. **L.** Quantification of Sox2^+^/T^+^ cells in the different Epha1 subpopulations of mESCs-derived NMPs. **M.** FACS dot-plot displaying Sox2 and T expression in mESCs-derived NMPs. **N.** FACS profiles showing the distribution of Sox2^+^/T^+^ cells from the gate indicated in M among the Epha1 compartments. **O.** Quantification of the distribution of Sox2^+^/T^+^ cells from mESCs-derived NMPs in the different Epha1 subpopulations. Quantification in C, F, I, L and O derives from at least 3 independent experiments, using one-way analysis of variance ANOVA to determine statistical significance. (*p<0,05, ***p<0,001 and ****p<0,0001 vs Negative; ###p<0,001 and ####p<0,0001 vs Epha1^Low^). Error bars indicate the standard deviation (SD). The FACS plots show the results from a representative experiment.

**Figure 6.**
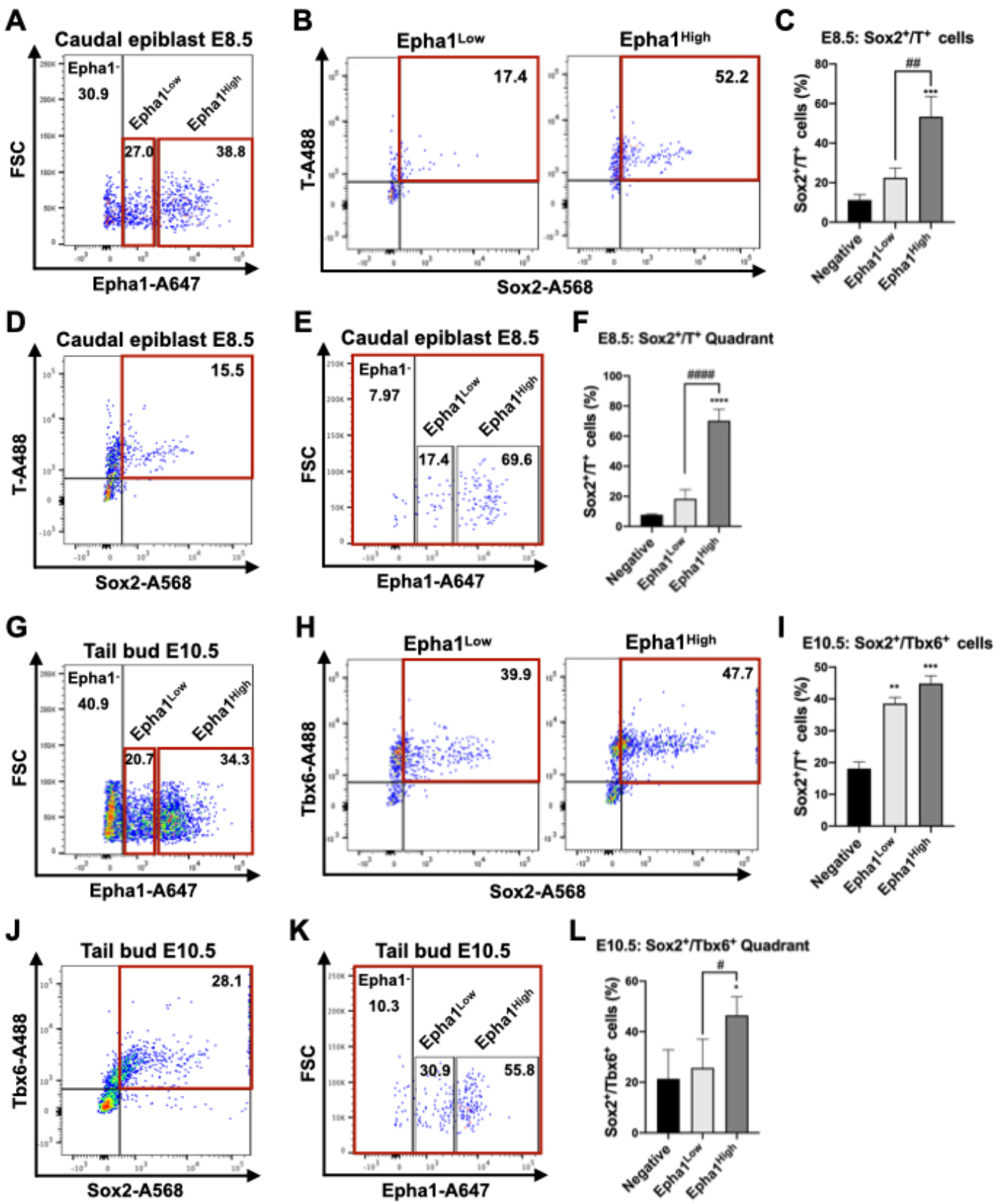
Flow cytometry analysis of Epha1 subpopulations that co-express Sox2 and T in E8.5 embryos or Sox2 and Tbx6 in E10.5 tail buds. **A.** FACS dot-plot displaying Epha1 subpopulations of E8.5 posterior ends. **B.** FACS profiles of Sox2 and T expression in cells from the Epha1^Low^ and Epha1^High^ compartments indicated in A. **C.** Quantification of Sox2^+^/T^+^ cells within the different Epha1 subpopulations from posterior ends of E8.5. **D.** FACS dot-plot displaying Sox2 and T expression in cells from the posterior regions of E8.5 embryos. **E.** FACS profile showing distribution of Sox2^+^/T^+^ cells from the red window in D among the Epha1 compartments. **F.** Quantification of the distribution of Sox2^+^/T^+^ cells from the posterior regions of E8.5 embryos between the different Epha1 subpopulations. **G.** FACS dot-plot displaying Epha1 subpopulations in the tail buds of E10.5 embryos **H.** FACS profiles showing Sox2 and Tbx6 expression in cells from the Epha1^Low^ and Epha1^High^ windows defined in G. **I.** Quantification of Sox2+/Tbx6+ cells from E10.5 tail buds among the different Epha1 subpopulations. **J.** FACS dot-plot displaying Sox2 and Tbx6 expression in cells from the tail buds of E10.5 embryos. **K.** FACS profiles showing distribution of Sox2+/Tbx6+ cells from the red window in J among the various Epha1 compartments. **L.** Quantification of the distribution of Sox2+/Tbx6+ cells from the posterior regions of E8.5 embryos between the different Epha1 subpopulations. Quantification in C, F, I, and L derives from at least 3 independent experiments, using one-way analysis of variance ANOVA to determine statistical significance. (*p<0,05, ***p<0,001 and ****p<0,0001 vs Negative; ###p<0,001 and ####p<0,0001 vs Epha1^Low^). Error bars indicate the standard deviation (SD). The FACS plots show the results from a representative experiment.

These observations prompted us to assess the distribution of Sox2^+^/T^+^ cells, a common criterion used to identify the NMPs (Cambray and Wilson, 2007; Koch et al., 2017; Wymeersch et al., 2016), among the different Epha1 compartments obtained from the tail bud of E10.5 embryos. We observed that the proportion of cells co-expressing Sox2 and T differed significantly among the three cell compartments, being about 57% for Epha1^High^ cells, 29% for Epha1^Low^ cells and was almost absent from the Epha1-negative pool (3%), consistent with NMPs being enriched in the Epha1^High^ compartment (Figure 5D–F; Table S3). We then performed a complementary analysis by first isolating Sox2^+^/T^+^ cells from the E10.5 tail bud and analyzed their distribution among the different Epha1 compartments. We observed that about 79% of the cells co-expressing Sox2 and T were Epha1^High^, whereas only 9.8% and 4% of these cells were Epha1^Low^ or Epha1-negative, respectively (Figure 5G–I; Table S4). These results are consistent with the Epha1^High^ subpopulation being indeed enriched in NMPs.

A similar distribution of Sox2^+^/T^+^ cells among the various Epha1 compartments and of Epha1^High^ and Epha1^Low^ from the Sox2^+^/T^+^ gated cells was also observed at E8.5 (Figure 6). In particular, 53,4% of Epha1^High^ and 22,5% of Epha1^Low^ were found within the Sox2^+^/T^+^ cells (Figure 6A–C; Table S3), and 70% and 18,5% of the Sox2^+^/T^+^ cells showed Epha1^High^ and Epha1^Low^ profiles, respectively (Figure 6D–F; Table S4). In addition, NMPs obtained from embryonic stem (ES) cell cultures according to standard inducing conditions (Gouti et al., 2014; Turner et al., 2014) were preferentially found in the Epha1^High^ compartment, from which 91% were positive for both T and Sox2 (Figure 5J–L; Table S3), further suggesting NMP enrichment also in this Epha1-positive compartment of *in vitro* differentiated ES cells. Consistent with this, 94% of Sox2^+^/T^+^ cells obtained from ES cell-derived NMPs mapped to the Epha1^High^ compartment (Figure 5M–O; Table S4). Together, taking the simultaneous expression of T and Sox2 as the NMP defining criterion, these results suggest that these cells are preferentially Epha1^High^, although some NMPs can also have an Epha1^Low^ phenotype, at least in embryonic tissues. This might reflect heterogeneity in the NMP progenitors when defined as Sox2^+^/T^+^ cells. Indeed, the cell distribution within the Sox2^+^/T^+^ gate from both Epha1^High^ and Epha1^Low^ cells shows different levels of Sox2 and T expression (Figures 5 and 6).

It has recently been reported that a subset of tail bud NMPs co-express Tbx6 and Sox2 (Javali et al., 2017). Analysis of Tbx6^+^/Sox2^+^ cells from E10.5 tail buds revealed that a high proportion of these cells also expressed Epha1, 46,5% of them entering the Epha1^High^ and 27% the Epha1^Low^ compartments (Figure 6J–L; Tables S3 and S4). This observation gives additional support to the connection between Epha1 and NMPs in the tail bud.

### High Epha1 labels NMPs entering mesodermal fates

To further characterize the Epha1-positive cell populations in the tail bud we isolated the Epha1^High^ and Epha1^Low^ compartments from this region of E10.5 embryos and analyzed their transcript content by RNA-seq. Both cell compartments showed enrichment in genes that have been associated with NMPs (e.g. *T*, *Wnt3a*, *Tbx6* and *Nkx1-2*) to levels similar to those observed in Tail^Prog^ cells, an effect particularly clear in Epha1^High^ cells (Figure 7A). However, Epha1^High^ and Epha1^Low^ cells seemed to represent two different cell compartments, as 875 genes showed differential expression (*p*-value <0,05) between these two Epha1 cell populations (Table S5). These include several early mesoderm related genes, like *Tbx6*, *Dll1*, *Msgn1*, *Cited1, Lfng, Snai1 or Wnt3a* (Carver et al., 2001; Dias et al., 2020; Gouti et al., 2017; Koch et al., 2017; Serth et al., 2003; Wymeersch et al., 2019) upregulated in the Epha1^High^ compartment (Figure 7C), thus suggesting that this cell population might contain NMPs already entering mesodermal fates. Interestingly, we also found that these cells display a more mesenchymal state than that observed in the axial progenitor pool (Tail^Prog^) (Figure 7D), thus suggesting that tail bud NMPs might still undergo EMT during mesoderm formation.

**Figure 7.**
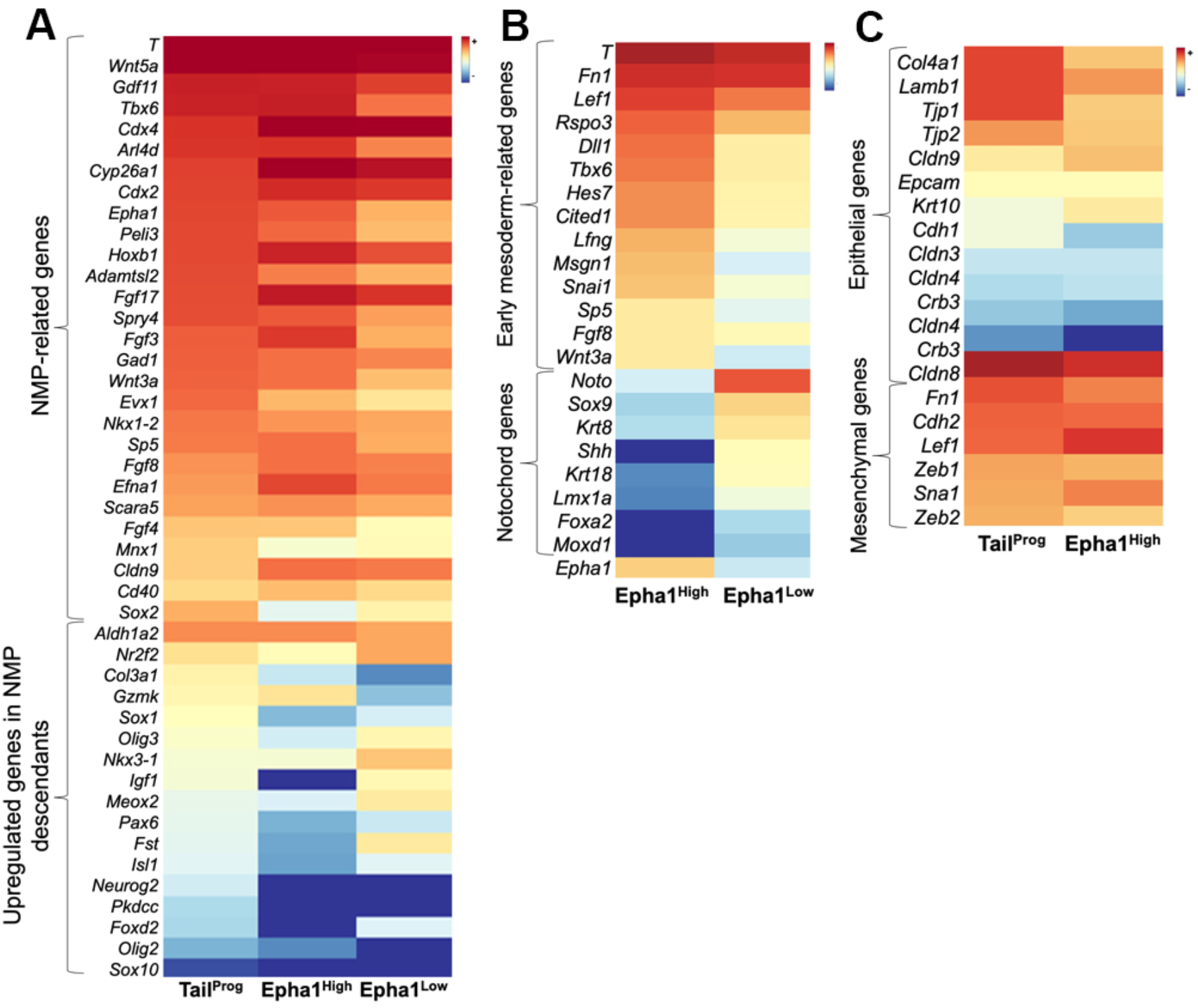
Isolation of two different axial progenitor cell populations inside the Epha1 positive domain. **A.** Heatmap showing expression of genes associated with NMPs and their descendants in Epha1^High^, Epha1^Low^ cells taking as reference their levels in Tail^Prog^ cells. The Epha1 cell compartments are enriched in NMP markers and the levels of differentiated markers are even lower than in Tail^Prog^ cells, more evidently in Epha1^High^ cells. The color key bar represents the average of normalized counts on logarithmic scale. **B.** Heatmap highlighting some differentially expressed genes between the two Epha1 cell populations. Epha1^High^ cells show high expression of several genes associated with mesoderm progenitors. The Epha1^Low^ compartment is enriched in notochord markers (e.g. *Noto*, *Shh* and *Foxa2*). **C**. Heatmap comparing epithelial and mesenchymal markers between the Epha1^High^ and Tail^Prog^ cell populations. Epha1^High^ seem to have a more complete mesenchymal phenotype, once several epithelial-associated genes are downregulated in the cell population. The color key bar represents the average of normalized counts on logarithmic scale.

The transcriptome profile of Epha1^Low^ cells also showed substantial enrichment in some NMP markers, which is consistent with the presence of Sox2+/T+ cells in this cell compartment (Figure 5). Interestingly, these cells also contained high abundance of transcripts for *Shh*, *Noto*, *Foxa2*, *Krt8* and *Krt18*, not present in either the Epha1^High^ or the Tail^Prog^ cell populations (Figures 7A and 1D, F). These genes are commonly expressed in the notochord (Abdelkhalek et al., 2004; Ang et al., 1993; Echelard et al., 1993; Rodrigues-Pinto et al., 2016; Wymeersch et al., 2019), thus suggesting that the tail bud Epha1^Low^ cell population might also be enriched in notochord progenitors.

Together, our results indicate that *Epha1* is a valuable cell surface marker that can be used to label and isolate axial progenitors, and that higher Epha1 expression might be a hallmark of NMPs entering the mesodermal progenitor compartment.

## DISCUSSION

In this study, we have used a genetic strategy to label and isolate physiologically active NMPs from the tail bud of mouse embryos and analyzed their mRNA content using high throughput methods. Although the Cre driver used to label the cells is not expected to be exclusive for the progenitors at early stages, it is expected that from the cells labeled by a pulse of Cre activity only bona fide progenitors will carry the reporter to the tail bud. Indeed, labeled cells can be observed along the whole axis caudal to the region where the reporter was activated (Aires et al., 2019), Figure 1B. Both the enrichment of tail bud cells isolated with this strategy (Tail^Prog^) in known markers for NMPs, as well as the high concordance between our dataset and previously published data for NMPs obtained from embryos or from *in vitro* differentiated human or mouse embryonic stem cells (Dias et al., 2020; Gouti et al., 2017; Koch et al., 2017; Verrier et al., 2018; Wymeersch et al., 2019), provide support to our lineage tracing strategy as a non-biased, reliable tail bud axial progenitor labelling methodology. However, it should be noted that the labeling conditions used to restrict the time frame of effective cell labelling also reduced the number of effectively labelled progenitors. As a consequence, the tail bud cells isolated and analyzed with this approach represent just a fraction of the actual progenitors and, therefore, it is possible that the tail bud NMP molecular fingerprint obtained in these experiments is not fully comprehensive.

Our initial goal was to identify cell surface markers that could facilitate isolation of physiologically active NMPs. From the list of new genes that we found to be differentially expressed between NMPs and their derivatives, only Epha1 showed expression restricted to the progenitor containing areas. Whether any of the other candidate surface molecules, alone or in combination, could help developing protocols for NMP isolation will require additional work.

An important finding from our experiments was that the levels of Epha1 are not uniform in the caudal epiblast or the tail bud cells expressing this molecule. This was clearer from the FACS profiles showing the existence of at least two populations of Epha1 positive cells in this embryonic region. The observation that the cells with higher Epha1 content are exclusive of the tail bud (not observed in cells from more anterior areas of the tail, containing derivatives of the progenitors) could indicate that high Epha1 content could be a molecular fingerprint of NMPs. Indeed, there is high correlation between high Epha1 and the Sox2^+^/T^+^ phenotype. Unfortunately, the association between Epha1 expression levels and NMP identity might be more complex. In particular, a significant number of Sox2^+^/T^+^ cells were also found in cells with lower Epha1 expression levels, indicating that some NMPs could be within the Epha1^Low^ compartment. Consistent with this, the transcriptome profile of tail bud Epha1^Low^ cells show a considerable enrichment in genes known to be expressed in NMPs. However, the FACS profiles of these cells also indicate that, if NMPs are present within this cell population, they should represent just a small subset of the cells in the Epha1^Low^ compartment. Interestingly, the transcriptomic profile of these cells also indicate that they might contain precursors for the notochord, which are clearly not present in the Epha1^high^ compartment. In addition, it should be noted that the Epha1 protein can be detected both by immunofluorescence or FACS analysis in areas where its mRNA is already below detection levels, indicating higher protein than transcript stability. Therefore, tail bud cells with low Epha1 levels might also include cells in already more advanced differentiation stages that are in the process of losing their Epha1 molecules. Indeed, more anterior regions of the embryonic tail also contain a significant number of cells with low Epha1 expression levels, despite not containing detectable amounts of its mRNA.

The question is then to understand the identity of cells containing high Epha1 expression levels. The high proportion of tail bud Sox2^+^/T^+^ and Sox2^+^/Tbx6^+^ cells within the Epha1^High^ compartment indicates that this cell population is enriched in NMPs and possibly also in their immediate derivatives. The enrichment of these cells in NMP markers observed in the RNA-seq dataset is consistent with this hypothesis. Interestingly, the transcriptome profile of these cells also showed significant enrichment in mesoderm-associated genes, thus indicating that at least part of these cells could represent NMPs entering mesodermal fates. Indeed, scRNA-seq analysis of Theiler stage 12 (around E8.5) (Pijuan-Sala et al., 2019) indicates that *Epha1* is strongly expressed in cells that were classified as belonging to NMP and caudal mesodermal clusters, and that colocalize with the Sox2^+^/T^+^/Nkx1-2^+^ domain. The distribution of higher Epha1 immunoreactivity in the caudal tissue of E8.5 embryos and in the tail bud is also consistent with this hypothesis, as it is associated with areas representing early steps of mesodermal formation (e.g. primitive streak) (McGrew et al., 2008; Wymeersch et al., 2016; Wymeersch et al., 2019). Similarly, the general patterns observed within the Sox2^+^/T^+^ gate in the FACS plots from Epha1^High^ cells might indicate the presence of cells at early stages in the progression from NMPs to mesodermal progenitors, as the cell distribution in the Sox2 expression axis include a significant proportion of cells with relatively low Sox2 contents but comparatively high T expression. Interestingly, this pattern is not observed in the Sox2^+^/T^+^ cells from the Epha1^Low^ gate, further highlighting the differences between the Epha1^High^ and Epha1^Low^ compartments. Therefore, we suggest that Epha1 is associated with axial progenitors, including both NMPs and those for the notochord, and that a transient increase in Epha1 expression is a hallmark of NMPs entering the mesodermal progenitor compartment that will generate paraxial mesoderm during axial extension. Interestingly, tail bud Epha1^High^ cells seem to contain a more complete mesenchymal profile than tail bud NMPs, which have been shown to contain an intermediate epithelial/mesenchymal signature derived from an incomplete EMT (Dias et al., 2020). This finding might indicate that in the tail bud, NMP mesodermal differentiation includes completion of the partial EMT characteristic of the neuro-mesodermal progenitors. Additional work, particularly at single-cell resolution, will be required to further explore this hypothesis.

In conclusion, our results indicate that, while low Epha1 expression levels might label a heterogeneous cell population, including various progenitors and differentiating tissues, the Epha1^High^ phenotype seems to identify a specific subset of NMPs already taking mesodermal routes and might thus provide a useful criterion to isolate and study NMP-derived mesodermal progenitors.

## EXPERIMENTAL PROCEDURES

### Mice and embryos

In this work embryo staging was defined according to the standard timed mating approach, considering E0.5 the morning on which a mating plug was found. To isolate axial progenitors from developing embryos, matings were set up between *Cdx2P-Cre^ERT^* transgenics (Jurberg et al., 2013) and *ROSA26*-*YFP-Cre* reporter (Srinivas et al., 2001) mice. Pregnant females were treated at E7.5 with a single intraperitoneal injection of 200 μl of a 1 mg/ml solution of tamoxifen (Sigma #T5648) in corn oil. Embryos were then collected at E10.5 by caesarean section, dissected in ice-cold phosphate-buffered saline (PBS) (Biowest #L0615) and processed for cell sorting (see below). Wild type embryos were collected at E8.5, E9.5 and E10.5, dissected in ice-cold PBS, fixed in 4% paraformaldehyde (PFA) (Sigma #P6148) in PBS and processed for *in situ* hybridization, immunofluorescence or flow cytometry studies. Mice were genotyped as described in Aires et al, 2019, using primers listed in Table S6.

All animal procedures were performed in accordance with Portuguese (Portaria 1005/92) and European (directive 2010/63/EU) legislations and guidance on animal use in bioscience research. The project was reviewed and approved by the Ethics Committee of “Instituto Gulbenkian de Ciência” and by the Portuguese National Entity “Direcção Geral de Alimentação Veterinária” (license reference: 014308).

### Expression analysis on embryos

Whole mount *in situ* hybridization was performed according to the protocol described in (Aires et al., 2019). Probes for *Efna1*, *Epha1*, *Ngfr*, *Cldn9*, *Nkd2*, *Arl4d* were prepared by amplifying cDNA fragments and cloning them into appropriate vectors for in vitro transcription. The sequences of all primers used to amplify these cDNAs are listed in Table S6. Whole mount-stained embryos were embedded in gelatin and sectioned in a vibratome following the protocol described in (Dias et al., 2020).

Whole-mount immunostaining was done according to the protocol described in (Osorno et al., 2012). RapiClear 1.52 (SunJin laboratory) was used for embryo clearing. Primary antibodies (used at 1:200 dilution): rabbit anti-Sox2 (Abcam, AB92494) and goat anti-Epha1 antibody (R&D systems #AF3034). Secondary antibodies (all used at 1:1000 dilution): donkey anti-goat 488 (Molecular Probes, A-11055) and donkey anti-rabbit 568 (Molecular Probes, A10042). Images were acquired with a Prairie two-photon system, and the image dataset pre-processing was performed as described in (Dias et al., 2020). However, no deconvolution was performed in order to reduce possible pre-processing interferences that could misrepresent the real Epha1 expression levels in the tail bud.

Immunofluorescence staining of mouse sections was performed as described in (Aires et al., 2019). Primary antibodies (used at 1:200) chicken anti-GFP (Abcam, AB13970) and rabbit anti-Sox2 (Abcam, AB92494). Secondary antibodies (1:1000): goat anti-chicken 488 (Thermo Fisher, A-11039) and donkey anti-rabbit 568 (Molecular Probes, A10042). Samples were analyzed using a Leica Sp5 live confocal.

The β-galactosidase staining was performed as described in (Aires et al., 2019).

### Mouse ES cell culture and differentiation

CJ7 mouse embryonic stem (ES) cells (Swiatek and Gridley, 1993) were maintained in ES cell medium [DMEM High Glucose (Biowest #S17532L0102), 15% Defined fetal bovine serum (Hyclone^TM^, GE Healthcare #SH30070.03), 1% MEM non-essential amino acid solution (Sigma #M-7145), 2 mM L-glutamine, 1% EmbryoMax Nucleosides (Millipore #ES-008-D), 100 U/ml Penicillin and 100 μg/ml Streptomycin (Sigma #P7539), 0.1 mM β-mercaptoethanol and 1000 U/ml LIF (Millipore #ESG1107)] on mitomycin C-inactivated primary mouse embryo fibroblasts. To start differentiation, ES cells were derived into NMP-like cells using a protocol adapted from (Gouti et al., 2014). Briefly, ES cells were removed from feeders by dissociation using 0.05 % trypsin-EDTA solution (Sigma #59417C) and seeded at a density of 5.000 cells/cm^2^ on CellBIND^®^Surface dishes (Corning #3294) in N2B27 medium [Dulbecco’s Modified Eagle Medium/F12 (Gibco #21331-020) and Neurobasal medium (Gibco #21103-049) (1:1), 40 μg/ml BSA, 0.1 mM β-mercaptoethanol and supplemented with 1x N-2 (Gibco #LS17502048) and 1x B-27 minus vitamin A (Gibco #LS12587001)]. Cells were grown in N2B27 medium with 10 ng/ml bFgf (Peprotech #100-18B) for 3 days (D1-D3). Neuromesodermal identity was induced by the addition of 5 μM CHIR99021 (Abcam #ab120890) from D2 to D3. After differentiation cells were detached with Accutase solution and processed for FACS analysis as described above.

### Cell Sorting for RNA-sequencing by Fluorescence-Activated Cell sorting (FACS)

Two regions were collected from E10.5 *Cdx2P-Cre^ERT^*∷*ROSA26*-*YFP* embryos: the tail buds and a more proximal region of the tail tip (up to the third somite). To obtain a single-cell suspension, tissue was incubated on ice for 5 minutes in Accutase solution (Sigma #A6964). Digestion was terminated by adding two volumes of PBS/10% donkey serum (DS) (Biowest S2170) and washed twice with PBS/10% DS. Cells were then resuspended in PBS/2% DS and filtered through a 100 μm cell strainer. Cells were sorted according to their YFP-positive fluorescence in a MoFlo sorter (Beckman Coulter) using a 488 nm excitation laser with detector filter of 520/40. The YFP parameters were set using cells from *Cdx2P-Cre^ERT^* tails dissected in parallel to serve as YFP negative control. YFP-positive cells were collected directly in TRI Reagent^®^ (Sigma #T9424) and stored at −80°C. The collected YFP-positive cells from the tail tip and the proximal tail were designated as Tail^Prog^ and Tail^Desc^, respectively.

To isolate Epha1 cell populations, single-cell suspensions were obtained from tail buds and proximal tail regions of E10.5 or from the caudal epiblast of E8.5 wild type embryos as mentioned above. Cells were incubated with 100 μl of blocking solution (10% DS with 1:100 dilution of 2.4G2 anti-mouse Fc block in PBS) on ice for 30 minutes and then stained on ice for 1 hour with a 1:100 dilution of goat anti-Epha1 antibody (R&D systems #AF3034). After two washes with PBS/10% DS, cells were incubated with a 1:1500 dilution of donkey anti-goat A647 antibody (Thermo Fisher #A-21447) for an additional hour on ice and subsequently washed twice with PBS/10% DS. Stained cells were resuspended in PBS/2% DS, filtered through a cell strainer and sorted on a FACSAria IIu (BD Biosciences). Epha1-positive cells were identified using a 633 nm excitation laser with filter detection of 660/20. Gating conditions of Epha1^High^ and Epha1^Low^ cell populations (illustrated in supplementary Figure 1) were based on an apparent separation in total Epha1-positive cells histogram. For RNA-seq analyses, sorted cells were collected directly in TRI Reagent^®^ and kept at −80°C until further use. For all RNA-seq experiments, 15.000-20.000 purified-sorted cells per sample were collected, to obtain a sufficient concentration of high-quality RNA.

### RNA-sequencing analysis

Total RNA was isolated from the TRI Reagent^®^ suspension following the manufacturer’s protocol, with the addition of 10 μg RNase-free glycogen (Roche #10901393001) in the isopropanol step. RNA samples were then resuspended in RNase-free water. RNA concentration and purity were determined on an AATI Fragment Analyzer (Agilent). For the Tail^Desc^ and Tail^Prog^ samples, RNA-seq libraries were prepared from two biological replicates using TruSeq® Stranded mRNA sample Prep Kit (Illumina #20020594) and sequenced using Illumina HiSeq 2500 system at the CRG Genomics Unit (Barcelona, Spain). At least 25 million single end 50 bases reads were generated for each library. Read alignments were performed by TopHat2 v2.0.9 (Kim et al., 2013) with Bowtie2 v2.1.0.0. (Langmead and Salzberg, 2012) Differential expression analysis between Tail^Desc^, Tail^Prog^ and Tail^Tot^ (Aires et al., 2019) was performed using CuffDiff v2.1.1 (Trapnell et al., 2013).

RNA-seq from Epha1^High^ and Epha1^Low^ cells was performed using two separate biological replicates. Libraries were prepared from total RNA using the SMART-Seq2 protocol (Picelli et al., 2014). Sequencing was performed on Illumina NextSeq500 at the IGC Genomics Facility, generating 20-25 million single-end 75 bases reads per sample. Read alignments were performed as above. Read count normalization and differential expression between Epha1^High^ and Epha1^Low^ samples were analyzed using the DESeq2 R package (Love et al., 2014). To faithfully compare all samples (Tail^Contr^, Tail^Prog^, Epha1^High^ and Epha1^Low^) from the above-mentioned RNA-seq independent experiments, normalization and differential expression were performed using the DESeq2 R package.

The sequencing data of the RNA-seq experiments was deposited in the NCBI trace and Short-read Archive (SRA), accession numbers PRJNA527654 and PRJNA527619.

### Quantitative RT-qPCR

Quantitative RT-qPCR was done as described in (Aires et al., 2019). The sequences of the primers used are given in Table S6.

### Protein expression profile analysis by FACS

Cells obtained from embryos and stained with Epha1 antibodies as described above were then washed twice with PBS/10% DS and processed for staining using True-Nuclear^TM^ transcription factor Buffer Set (BioLegend #424401) according to the manufacturer’s instructions and following the protocol described in (Aires et al., 2019). The following primary antibodies were used: rabbit anti-Sox2 (Abcam #ab92494) (1:200) and mouse anti-T/Bras (Santa Cruz #sc-166962) (1:100), or rabbit anti-Sox2 (1:200) and mouse anti-Tbx6 (Santa Cruz #517027) (1:100). Donkey anti-rabbit A568 (Thermo Fisher Scientific #A10042) (1:1500) and donkey anti-mouse Alexa Fluor^®^ 488 (Abcam #ab150105) (1:1500) were used as secondary antibodies. For multicolor FACS analysis, single stain compensation controls were used to correct the spectral overlap between different fluorophores. Gating conditions of Epha1^High^ and Epha1^Low^ subsets were based on an apparent separation in total Epha1-positive cells histogram (A647 channel). Quadrant gates were established according to fluorescence levels detected by the control samples processed without primary antibodies. Flow cytometry data was analyzed using FlowJo^TM^ 10 (BD, Biosciences) software. Quadrant averages were calculated using at least 3 independent experiments and one-way analysis of variance ANOVA was used to determine statistical significance.

### Single-cell analysis

The public single-cell molecular map of mouse gastrulation and early organogenesis (Pijuan-Sala et al., 2019) was used to assess the gene expression of *Epha1*, *T*, *Sox2* and *Nkx1-2* in Theiler stage 12 wild type mouse embryos.

## Supporting information

Table S1 containing datasets

Table S2 containing datasets

## AUTHOR CONTRIBUTION

Conceptualization: L. dL. and M.M.; Methodology: L.dL., A.D. and M.M.; Investigation: L.dL., A.D. and A.N.; Writing-Original Draft: L. dL., A.D. and M.M.; Funding Acquisition: M.M.; Supervision: M.M.

## ACKNOWLEDGMENTS

We would like to thank the SunJin laboratory for the RapiClear test sample, the IGC animal house, Flow Cytometry, Genomics and Imaging Facilities for their expert services, advice and assistance, and Daniel Sobral from the IGC Bioinformatics Unit for assistance with RNA-seq analysis. We would also acknowledge all members of the Mallo laboratory for helpful discussions and comments throughout the course of the project and Rita Aires and Ana Casaca for reading the manuscript. This work has been supported by grants PTDC/BEX-BID/0899/2014 (FCT, Portugal), LISBOA-01-0145-FEDER-030254 (FCT, Portugal) and SCML-MC-60-2014 (from Santa Casa da Misericórdia de Lisboa, Portugal) to M.M., PhD fellowship PD/BD/128426/2017 to A.D. and the research infrastructure Congento, project LISBOA-01-0145-FEDER-022170.

## REFERENCES

Abdelkhalek, H. B., Beckers, A., Schuster-gossler, K., Pavlova, M. N., Burkhardt, H., Lickert, H., Rossant, J., Reinhardt, R., Schalkwyk, L. C., Herrmann, B. G., et al. (2004). The mouse homeobox gene Not is required for caudal notochord development and affected by the truncate mutation Hanaa. Genes Dev. 18, 1725–1736.

Abu-Abed, S., Dollé, P., Metzger, D., Beckett, B., Chambon, P. and Petkovich, M. (2001). The retinoic acid-metabolizing enzyme, CYP26A1, is essential for normal hindbrain patterning, vertebral identity, and development of posterior structures. Genes Dev. 15, 226–240.

Acloque, H., Adams, M. S., Fishwick, K., Bronner-Fraser, M. and Nieto, M. A. (2009). Epithelial-mesenchymal transitions : the importance of changing cell state in development and disease. J. Clin. Invest. 119, 1438–1449.

Aires, R., Dias, A. and Mallo, M. (2018). Deconstructing the molecular mechanisms shaping the vertebrate body plan. Curr. Opin. Cell Biol. 55, 81–86.

Aires, R., de Lemos, L., Nóvoa, A., Jurberg, A. D., Mascrez, B., Duboule, D. and Mallo, M. (2019). Tail Bud Progenitor Activity Relies on a Network Comprising Gdf11, Lin28, and Hox13 Genes. Dev. Cell 48, 383–395.

Albano, R. M., Arkell, R., Beddington, R. S. and Smith, J. C. (1994). Expression of inhibin subunits and follistatin during postimplantation mouse development: decidual expression of activin and expression of follistatin in primitive streak, somites and hindbrain. Development 120, 803–13.

Albors, A. R., Halley, P. A. and Storey, K. G. (2018). Lineage tracing of axial progenitors using Nkx1-2CreER T2 mice defines their trunk and tail contributions. Development 145, dev164319.

Amin, S., Neijts, R., Simmini, S., van Rooijen, C., Tan, S. C., Kester, L., van Oudenaarden, A., Creyghton, M. P. and Deschamps, J. (2016). Cdx and T Brachyury Co-activate Growth Signaling in the Embryonic Axial Progenitor Niche. Cell Rep. 17, 3165–3177.

Ang, S.-L., Wierda, A., Wong, D., Stevens, K. A., Cascio, S., Rossant, J. and Zaret, K. S. (1993). The formation and maintenance of the definitive endoderm lineage in the mouse: involvement of HNF3/forkhead proteins. Development 119, 1301–1315.

Ashburner, M., Ball, C. A., Blake, J. A., Botstein, D., Butler, H., Cherry, J. M., Davis, A. P., Dolinski, K., Dwight, S. S., Eppig, J. T., et al. (2000). Gene Ontology: tool for the unification of biology. Nat. Genet. 25, 25–29.

Aubert, J., Stavridis, M. P., Tweedie, S., Reilly, M. O., Vierlinger, K., Li, M., Ghazal, P., Pratt, T., Mason, J. O., Roy, D., et al. (2003). Screening for mammalian neural genes via fluorescence-activated cell sorter purification of neural precursors from Sox1–gfp knock-in mice. Proc Natl Acad Sci U S A 100, 11836–11841.

Boulet, A. M. and Capecchi, M. R. (2012). Signaling by FGF4 and FGF8 is required for axial elongation of the mouse embryo. Dev. Biol. 371, 235–245.

Cambray, N. and Wilson, V. (2002). Axial progenitors with extensive potency are localised to the mouse chordoneural hinge. Development 129, 4855–4866.

Cambray, N. and Wilson, V. (2007). Two distinct sources for a population of maturing axial progenitors. Development 134, 2829–2840.

Candia, A. F., Hu, J., Crosby, J., Lalley, P. A., Noden, D., Nadeau, J. H. and Wright, C. V. E. (1992). Mox-1 and Mox-2 define a novel homeobox gene subfamily and are differentially expressed during early mesodermal patterning in mouse embryos. Development 116, 1123–1136.

Carver, E. A., Jiang, R., Lan, Y., Oram, K. F. and Gridley, T. (2001). The mouse Snail gene encodes a key regulator of the epithelial-mesenchymal transition. Mol Cell Biol 21, 8184–8188.

Chawengsaksophak, K., de Graaff, W., Rossant, J., Deschamps, J. and Beck, F. (2004). Cdx2 is essential for axial elongation in mouse development. Proc. Natl. Acad. Sci. U. S. A. 101, 7641–7645.

Dias, A. and Aires, R. (2020). Axial Stem Cells and the Formation of the Vertebrate Body. In Concepts and Applications of Stem Cell Biology. (ed. Rodrigues, G.) and Roelen, B. A. J.), pp. 131–158. Springer, Cham.

Dias, A., Lozovska, A., Wymeersch, F. J., Nóvoa, A., Binagui-Casas, A., Sobral, D., Martins, G. G., Wilson, V. and Mallo, M. (2020). A TgfbRI/Snai1-dependent developmental module at the core of vertebrate axial elongation. Elife 9, e56615.

Dush, M. K. and Martin, G. R. (1992). Analysis of mouse Evx genes: Evx-1 displays graded expression in the primitive streak. Dev. Biol. 151, 273–287.

Echelard, Y., Epstein, D. J., St-Jacques, B., Shen, L., Mohler, J., McMahon, J. A. and McMahon, A. P. (1993). Sonic hedgehog, a member of a family of putative signaling molecules, is implicated in the regulation of CNS polarity. Cell 75, 1417–1430.

Edri, S., Hayward, P., Jawaid, W. and Martinez Arias, A. (2019). Neuro-mesodermal progenitors (NMPs): a comparative study between pluripotent stem cells and embryo-derived populations. Development 146, dev180190.

Gouti, M., Tsakiridis, A., Wymeersch, F. J., Huang, Y., Kleinjung, J., Wilson, V. and Briscoe, J. (2014). In vitro generation of neuromesodermal progenitors reveals distinct roles for wnt signalling in the specification of spinal cord and paraxial mesoderm identity. PLoS Biol. 12, e1001937.

Gouti, M., Delile, J., Stamataki, D., Wymeersch, F. J., Huang, Y., Kleinjung, J., Wilson, V. and Briscoe, J. (2017). A gene regulatory network balances neural and mesoderm specification during vertebrate trunk development. Dev. Cell 41, 1–19.

Gradwohl, G., Fode, C. and Guillemot, F. (1996). Restricted expression of a novel murine atonal-related bHLH protein in undifferentiated neural precursors. Dev. Biol. 180, 227–241.

Greco, T. L., Takada, S., Newhouse, M. M., McMahon, J. A., McMahon, A. P. and Camper, S. A. (1996). Analysis of the vestigial tail mutation demonstrates that Wnt-3a gene dosage regulates mouse axial development. Genes Dev. 10, 313–324.

Harrison, S. M., Houzelstein, D., Dunwoodie, S. L. and Beddington, R. S. P. (2000). Sp5, a new member of the Sp1 family, is dynamically expressed during development and genetically interacts with Brachyury. Dev. Biol. 227, 358–372.

Hay, B. (1968). Organization and fine structure of epithelium and mesenchyme in the developing chick embryo. In Epithelial-Mesenchymal Interactions (ed. Fleischmajer, R.) and Billingham, R. E.), pp. 31–55. Philadelphia: Williams & Wilkins Co.

Henrique, D., Abranches, E., Verrier, L. and Storey, K. G. (2015). Neuromesodermal progenitors and the making of the spinal cord. Development 142, 2864–2875.

Herrmann, B. G., Labeit, S., Poustka, A., King, T. R. and Lehrach, H. (1990). Cloning of the T gene required in mesoderm formation in the mouse. Nature 343, 617–622.

Javali, A., Misra, A., Leonavicius, K., Acharya, D., Vyas, B. and Sambasivan, R. (2017). Co-expression of Tbx6 and Sox2 identifies a novel transient neuromesoderm progenitor cell state. Development 144, 4522–4529.

Jonk, L. J. C., de Jonge, M. E. J., Pals, C. E. G. M., Wissink, S., Vervaart, J. M. A., Schoorlemmer, J. and Kruijer, W. (1994). Cloning and expression during development of three murine members of the COUP family of nuclear orphan receptors. Mech. Dev. 47, 81–97.

Jurberg, A. D., Aires, R., Varela-Lasheras, I., Nóvoa, A. and Mallo, M. (2013). Switching axial progenitors from producing trunk to tail tissues in vertebrate embryos. Dev. Cell 25, 451–462.

Koch, F., Scholze, M., Wittler, L., Schifferl, D., Sudheer, S., Grote, P., Timmermann, B., Macura, K. and Herrmann, B. G. (2017). Antagonistic Activities of Sox2 and Brachyury Control the Fate Choice of Neuro-Mesodermal Progenitors. Dev. Cell 42, 514–526.

Kuhlbrodt, K., Herbarth, B., Sock, E., Hermans-Borgmeyer, I. and Wegner, M. (1998). Sox10, a novel transcriptional modulator in glial cells. J. Neurosci. 18, 237–250.

Langmead, B. and Salzberg, S. L. (2012). Fast gapped-read alignment with Bowtie 2. Nat. Methods 9, 357–359.

Love, M. I., Huber, W. and Anders, S. (2014). Moderated estimation of fold change and dispersion for RNA-seq data with DESeq2. Genome Biol. 15, 550.

Martin, B. L. and Kimelman, D. (2012). Canonical Wnt Signaling Dynamically Controls Multiple Stem Cell Fate Decisions during Vertebrate Body Formation. Dev. Cell 22, 223–232.

McGrew, M. J., Sherman, A., Lillico, S. G., Ellard, F. M., Radcliffe, P. A., Gilhooley, H. J., Mitrophanous, K. A., Cambray, N., Wilson, V. and Sang, H. (2008). Localised axial progenitor cell populations in the avian tail bud are not committed to a posterior Hox identity. Development 135, 2289–99.

McPherron, A. C., Lawle, A. M. and Lee, S.-J. (1999). Regulation of anterior/posterior patterning of the axial skeleton by growth/differentiation factor 11. Nat. Genet. 22, 260–264.

Murphy, P. and Hill, R. E. (1991). Expression of the mouse labial-like homeobox-containing genes, Hox 2.9 and Hox 1.6, during segmentation of the hindbrain. Development 111, 61–74.

Naiche, L. A., Holder, N. and Lewandoski, M. (2011). FGF4 and FGF8 comprise the wavefront activity that controls somitogenesis. Proc. Natl. Acad. Sci. U. S. A. 108, 4018–4023.

Niederreither, K., McCaffery, P., Dräger, U. C., Chambon, P. and Dollé, P. (1997). Restricted expression and retinoic acid-induced downregulation of the retinaldehyde dehydrogenase type 2 (RALDH-2) gene during mouse development. Mech. Dev. 62, 67–78.

Olivera-Martinez, I., Harada, H., Halley, P. A. and Storey, K. G. (2012). Loss of FGF-Dependent Mesoderm Identity and Rise of Endogenous Retinoid Signalling Determine Cessation of Body Axis Elongation. PLoS Biol. 10,.

Osorno, R., Tsakiridis, A., Wong, F., Cambray, N., Economou, C., Wilkie, R., Blin, G., Scotting, P. J., Chambers, I. and Wilson, V. (2012). The developmental dismantling of pluripotency is reversed by ectopic Oct4 expression. Development 139, 2288–2298.

Picelli, S., Faridani, O. R., Björklund, Å. K., Winberg, G., Sagasser, S. and Sandberg, R. (2014). Full-length RNA-seq from single cells using Smart-seq2. Nat. Protoc. 9, 171–181.

Pijuan-Sala, B., Griffiths, J. A., Guibentif, C., Hiscock, T. W., Jawaid, W., Calero-Nieto, F. J., Mulas, C., Ibarra-Soria, X., Tyser, R. C. V., Ho, D. L. L., et al. (2019). A single-cell molecular map of mouse gastrulation and early organogenesis. Nature 566, 490–495.

Robinton, D. A., Chal, J., Lummertz da Rocha, E., Han, A., Yermalovich, A. V., Oginuma, M., Schlaeger, T. M., Sousa, P., Rodriguez, A., Urbach, A., et al. (2019). The Lin28/let-7 Pathway Regulates the Mammalian Caudal Body Axis Elongation Program. Dev. Cell 48, 396–405.

Rodrigues-Pinto, R., Berry, A., Piper-Hanley, K., Hanley, N., Richardson, S. M. and Hoyland, J. A. (2016). Spatiotemporal analysis of putative notochordal cell markers reveals CD24 and keratins 8, 18, and 19 as notochord-specific markers during early human intervertebral disc development. J. Orthop. Res. 34, 1327–1340.

Sakai, Y., Meno, C., Fujii, H., Nishino, J., Shiratori, H., Saijoh, Y., Rossant, J. and Hamada, H. (2001). The retinoic acid-inactivating enzyme CYP26 is essential for establishing an uneven distribution of retinoic acid along the anterio-posterior axis within the mouse embryo. Genes Dev. 15, 213–225.

Savory, J. G. A., Mansfield, M., Rijli, F. M. and Lohnes, D. (2011). Cdx mediates neural tube closure through transcriptional regulation of the planar cell polarity gene *Ptk7*. Development 138, 1361–1370.

Serth, K., Schuster-Gossler, K., Cordes, R. and Gossler, A. (2003). Transcriptional oscillation of Lunatic fringe is essential for somitogenesis. Genes Dev. 17, 912–925.

Soriano, P. (1999). Generalized lacZ expression with the ROSA26 Cre reporter strain. Nat. Genet. 21, 70–71.

Srinivas, S., Watanabe, T., Lin, C.-S., William, C. M., Tanabe, Y., Jessell, T. M. and Costantini, F. (2001). Cre reporter strains produced by targeted insertion of EYFP and ECFP into the ROSA26 locus. BMC Dev Biol 1, 4.

Stern, C. D., Charité, J., Deschamps, J., Duboule, D., Durston, A. J., Kmita, M., Nicolas, J. F., Palmeirim, I., Smith, J. C. and Wolpert, L. (2006). Head-tail patterning of the vertebrate embryo: One, two or many unresolved problems? Int. J. Dev. Biol. 50, 3–15.

Steventon, B. and Martinez Arias, A. (2017). Evo-engineering and the cellular and molecular origins of the vertebrate spinal cord. Dev. Biol. 432, 3–13.

Swiatek, P. J. and Gridley, T. (1993). Perinatal lethality and defects in hindbrain development in mice homozygous for a targeted mutation of the zinc finger gene Krox20. Genes Dev. 7, 2071–2084.

Takada, S., Stark, K. L., Shea, M. J., Vassileva, G., McMahon, J. A. and McMahon, A. P. (1994). Wnt-3a regulates somite and tailbud formation in the mouse embryo. Genes Dev. 8, 174–189.

Takeichi, M., Okubo, K., Uchida, T., Ohtsuki, T., Chisaka, O., Takebayashi, H., Ikenaka, K., Kawamoto, S. and Nabeshima, Y. (2002). Non-overlapping expression of Olig3 and Olig2 in the embryonic neural tube. Mech. Dev. 113, 169–174.

Trapnell, C., Hendrickson, D. G., Sauvageau, M., Goff, L., Rinn, J. L. and Pachter, L. (2013). Differential analysis of gene regulation at transcript resolution with RNA-seq. Nat. Biotechnol. 31, 46–53.

Tsakiridis, A., Huang, Y., Blin, G., Skylaki, S., Wymeersch, F., Osorno, R. R., Economou, C., Karagianni, E., Zhao, S., Lowell, S., et al. (2014). Distinct Wnt-driven primitive streak-like populations reflect in vivo lineage precursors. Development 141, 1209–1221.

Turner, D. a, Hayward, P. C., Baillie-Johnson, P., Rué, P., Broome, R., Faunes, F. and Martinez Arias, A. (2014). Wnt/β-catenin and FGF signalling direct the specification and maintenance of a neuromesodermal axial progenitor in ensembles of mouse embryonic stem cells. Development 141, 4243–4353.

Tzouanacou, E., Wegener, A., Wymeersch, F. J., Wilson, V. and Nicolas, J. F. (2009). Redefining the Progression of Lineage Segregations during Mammalian Embryogenesis by Clonal Analysis. Dev. Cell 17, 365–376.

van Rooijen, C., Simmini, S., Bialecka, M., Neijts, R., van de Ven, C., Beck, F. and Deschamps, J. (2012). Evolutionarily conserved requirement of Cdx for post-occipital tissue emergence. Development 139, 2576–2583.

Verrier, L., Davidson, L., Gierliński, M., Dady, A. and Storey, K. G. (2018). Neural differentiation, selection and transcriptomic profiling of human neuromesodermal progenitor-like cells in vitro. Development 145, dev166215.

Walther, C. and Gruss, P. (1991). Pax-6, a murine paired box gene, is expressed in the developing CNS. Development 113, 1435–1449.

Wilson, V., Olivera-Martinez, I. and Storey, K. G. (2009). Stem cells, signals and vertebrate body axis extension. Development 136, 2133–2133.

Wymeersch, F. J., Huang, Y., Blin, G., Cambray, N., Wilkie, R., Wong, F. C. K. and Wilson, V. (2016). Position-dependent plasticity of distinct progenitor types in the primitive streak. Elife 5, e10042.

Wymeersch, F. J., Skylaki, S., Huang, Y., Watson, J. A., Economou, C., Marek-Johnston, C., Tomlinson, S. R. and Wilson, V. (2019). Transcriptionally dynamic progenitor populations organised around a stable niche drive axial patterning. Development 146, dev168161.

Yamaguchi, T. P., Bradley, A., McMahon, A. P. and Jones, S. (1999). A Wnt5a pathway underlies outgrowth of multiple structures in the vertebrate embryo. Development 126, 1211–1223.

