## Supplemental figure and tables for "Epha1 is a cell surface marker for neuromesodermal progenitors and their early mesoderm derivatives"

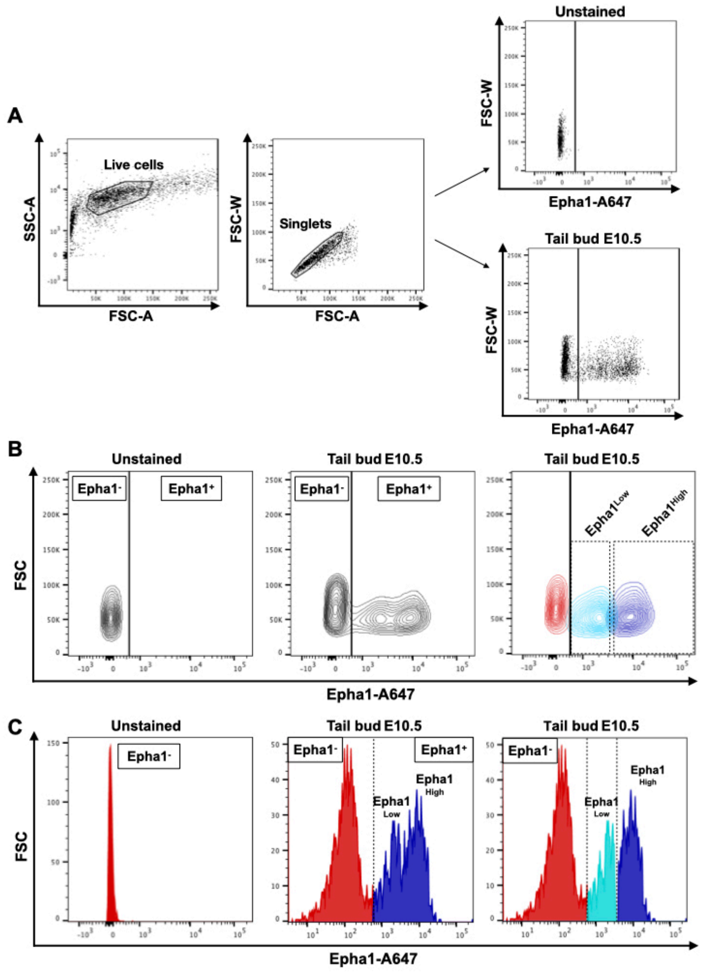


**Figure S1. Gating strategy for FACS analysis used to define Epha1 positive cells from the tail buds of E10.5 embryos. A.** Representative example of exclusion of debris and doublets cells (using forward scatter and side scatter gates) and definition of Epha1 gating. **B.** Representative gating scheme illustrating two subpopulations for Epha1-positive cells: Epha1$^{Low}$ (light blue) and Epha1$^{High}$ (dark blue). **C.** Representative histogram showing two peaks with different fluorescent intensity in the Epha1 channel, corresponding to the defined Epha1 subsets: Epha1$^{Low}$ and Epha1$^{High}$. All gates were firstly set using unstained control.

**Table S3.** FACS data of T/Sox2 and Tbx6/Sox2 subpopulations within the different Epha1 domains in E10.5 and E8.5 embryos and in mESC-derived NMPs (mean ± SD).

| Sample | Epha1 channel | Distribution in Sox2, T and Tbx6 channels | | | |
|---|---|---|---|---|---|
| | | $Sox2^-/T^-$ | $Sox2^-/T^+$ | $Sox2^+/T^+$ | $Sox2^+/T^-$ |
| **E10.5 embryo** | Epha1$^{Neg}$ | 58±2,55 | 33,60±1,41 | **3,07±1,26** | 5,34±0,15 |
| | Epha1$^{Low}$ | 51,87±13,83 | 15,92±6,62 | **29,10±12,30** | 9,22±6,09 |
| | Epha1$^{High}$ | 11,78±6,51 | 36,90±5,37 | **57,17±13,59** | 2,42±1,45 |
| **E8.5 embryo** | Epha1$^{Neg}$ | 55,07±9,98 | 25,67±1,55 | **11,17±2,75** | 8,12±8,78 |
| | Epha1$^{Low}$ | 46,40±13,48 | 21,87±12,35 | **22,50±4,87** | 9,22±6,09 |
| | Epha1$^{High}$ | 15,98±11,81 | 25,47±13,07 | **53,37±10,20** | 5,17±7,49 |
| **mESCs-derived NMPs** | Epha1$^{Neg}$ | 58,17±0,8 | 0,57±0,37 | **6,85±1,65** | 34,4±1,25 |
| | Epha1$^{Low}$ | 59,83±3,52 | 4,05±2,09 | **17,97±0,93** | 18,13±2,28 |
| | Epha1$^{High}$ | 1,61±0,6 | 5,72±0,57 | **90,83±1,63** | 1,86±0,67 |
| | | $Sox2^-/Tbx6^-$ | $Sox2^-/Tbx6^+$ | $Sox2^+/Tbx6^+$ | $Sox2^+/Tbx6^-$ |
| **E10.5 embryo** | Epha1$^{Neg}$ | 17,80±3,39 | 68,15±10,11 | **18,10±2,12** | 3,02±0,78 |
| | Epha1$^{Low}$ | 51,25±4,31 | 35,2±14,71 | **38,55±1,91** | 11,34±9,14 |
| | Epha1$^{High}$ | 23,55±4,31 | 27,20±3,54 | **44,87±2,41** | 2,37±0,01 |

**Table S4.** FACS data of Epha1 subpopulations within the different T/Sox2 and Tbx6/Sox2 quadrants from embryos at E10.5, E8.5 and mESC-derived NMPs (mean ± SD).

| Sample | Quadrant (Q) | Epha1 Channel | | |
|---|---|---|---|---|
| | | **Epha1$^{Neg}$** | **Epha1$^{Low}$** | **Epha1$^{High}$** |
| **E10.5 embryos** | Q1: $T^+/Sox2^-$ | 40,30±1,75 | 8,75±2,79 | 46,70±2,51 |
| | **Q2: $T^+/Sox2^+$** | **4,04±1,25** | **9,79±4,30** | **78,97±5,44** |
| | Q3: $T^-/Sox2^-$ | 67,03±5,33 | 21,47±3,74 | 6,33±2,45 |
| | Q4: $T^-/Sox2^+$ | 48,73±8,90 | 32,27±2,29 | 10,10±2,59 |
| **E8.5 Embryos** | Q1: $T^+/Sox2^-$ | 20,47±2,36 | 29,67±3,67 | 46,00±6,24 |
| | **Q2: $T^+/Sox2^+$** | **7,76±0,50** | **18,40±6,06** | **70,30±7,47** |
| | Q3: $T^-/Sox2^-$ | 42,20±18,35 | 39,17±7,31 | 12,85±10,97 |
| | Q4: $T^-/Sox2^+$ | 36,93±20,69 | 42,17±2,36 | 20,27±17,56 |
| **mESCs-derived NMPs** | Q1: $T^+/Sox2^-$ | 0,76±0,4 | 9,11±1,11 | 88,37±1,72 |
| | **Q2: $T^+/Sox2^+$** | **0,79±0,15** | **3,33±0,24** | **94,73±0,40** |
| | Q3: $T^-/Sox2^-$ | 35,60±0,92 | 49,77±3,76 | 9,45±1,97 |
| | Q4: $T^-/Sox2^+$ | 48,37±5,73 | 31,17±3,72 | 17,00±3,15 |
| **E10.5 embryos** | Q1: $Tbx6^+/Sox2^-$ | 52,03±22,37 | 18,44±11,52 | 25,33±15,30 |
| | **Q2: $Tbx6^+/Sox2^+$** | **21,40±13,98** | **27,17±13,40** | **46,47±9,06** |
| | Q3: $Tbx6^-/Sox2^-$ | 53,77±12,79 | 18,60±3,27 | 24,87±11,38 |
| | Q4: $Tbx6^-/Sox2^+$ | 40,23±4,98 | 33,27±7,77 | 29,07±8,03 |

**Table S6.** List of primers used in this work.

| | | |
|---|---|---|
| **Primers for genotyping (sequence 5’ to 3’)** | | |
| ***cre*** | Forward | CGAGTGATGAGGTTCGCAAG |
| | Reverse | CACCAGCTTGCATGATCT |
| ***YFP wild type Allele*** | Forward | CTGGCTTCTGAGGACCG |
| | Reverse | CAGGACAACGCCCACACA |
| ***YFP mutant Allele*** | Forward | AGGGCGAGGAGCTGTTCA |
| | Reverse | TGAAGTCGATGCCCTTCAG |
| **Primers for RT-qPCR (sequence 5’ to 3’)** | | |
| ***β-Actin*** | Forward | ATGAAGATCCTGACCGAGCG |
| | Reverse | TACTTGCGCTCAGGAGGAGC |
| ***Arl4d*** | Forward | GCCTCGAGGGCTGAAGACACCCCAGCTT |
| | Reverse | CTGAATTCGCCTTGCTGATCCGGTGTAA |
| ***Cdx2*** | Forward | GCGAAACCTGTGCGAGTGGATG |
| | Reverse | TTTCCTCTCCTTGGCTCTGCG |
| ***Cldn9*** | Forward | GCCTCGAGGGCTGGCTAGGAACTTTGGT |
| | Reverse | CTGAATTCGGACACGTACAGCAGAGGAG |
| ***Efna1*** | Forward | GCCTCGAGCTCTCTTGGGTCTGTGCTGC |
| | Reverse | CTGAATTCGTACTTCCGGGTCATCTGCTT |
| ***Epha1*** | Forward | GCCTCGAGCAAGATTGCAAGACTGTGGC |
| | Reverse | CTGAATTCCCTCCCACATTACAATCCCA |
| ***Mesp2*** | Forward | GCCATGAGTAGTGGGGTGTC |
| | Reverse | GTCAGCGGCTCTTTCTAGGG |
| ***Ngfr*** | Forward | GCCTCGAGTGCCTGGACAGTGTTACGTT |
| | Reverse | CTGAATTCAGGAATGAGGTTGTCAGCGG |
| ***Nkd2*** | Forward | GCCTCGAGGGAGAGAGAGTCCCGAAGGG |
| | Reverse | CTGAATTCACATGTCCTCTCTGGTGACTT |
| ***Olig2*** | Forward | TTACAGACCGAGCCAACACC |
| | Reverse | TCAACCTTCCGAATGTGAATTAGA |
| ***Sox2*** | Forward | TTTGTCCGAGACCGAGAAGC |
| | Reverse | CTCCGGGAAGCGTGTACTTA |
| ***T*** | Forward | ACCCAGCTCTAAGGAACCAC |
| | Reverse | GCTGGCGTTATGACTCACAG |
| ***Wnt3a*** | Forward | ATTGAATTTGGAGGAATGGT |
| | Reverse | CTTGAAGTACGTGTAACGTG |
| **Primers for in situ hybridization probes (sequence 5’ to 3’)** | | |
| ***Arl4d*** | Forward | GCCTCGAGGGCTGAAGACACCCCAGCTT |
| | Reverse | CTGAATTCGCCTTGCTGATCCGGTGTAA |
| ***Cldn9*** | Forward | GCCTCGAGGGCTGGCTAGGAACTTTGGT |
| | Reverse | CTGAATTCGGACACGTACAGCAGAGGAG |
| ***Efna1*** | Forward | GCCTCGAGCTCTCTTGGGTCTGTGCTGC |
| | Reverse | CTGAATTCGTACTTCCGGGTCATCTGCTT |
| ***Epha1*** | Forward | GCCTCGAGCAAGATTGCAAGACTGTGGC |
| | Reverse | CTGAATTCCCTCCCACATTACAATCCCA |
| ***Ngfr*** | Forward | GCCTCGAGTGCCTGGACAGTGTTACGTT |
| | Reverse | CTGAATTCAGGAATGAGGTTGTCAGCGG |
| ***Nkd2*** | Forward | GCCTCGAGGGAGAGAGAGTCCCGAAGGG |
| | Reverse | CTGAATTCACATGTCCTCTCTGGTGACTT |
