## Supplementary material for "Epha1 is a cell surface marker for neuromesodermal progenitors and their early mesoderm derivatives": Table S1 containing datasets

| gene | $Tail^{Desc}$ | $Tail^{Prog}$ | log2(fold_change) | p_value | q_value |
|---|---|---|---|---|---|
| AI646519 | 2,38425 | 0 | -inf | 5,00E-005 | 0,000828682 |
| Itih2 | 1,58598 | 0 | -inf | 5,00E-005 | 0,000828682 |
| LOC102636514 | 1,14393 | 0 | -inf | 5,00E-005 | 0,000828682 |
| Sst | 1,06651 | 0 | -inf | 5,00E-005 | 0,000828682 |
| Spp1 | 0,984743 | 0 | -inf | 5,00E-005 | 0,000828682 |
| 1110017D15Rik | 0,702737 | 0 | -inf | 5,00E-005 | 0,000828682 |
| Adamtsl2 | 0,720218 | 41,5535 | 5,85039 | 5,00E-005 | 0,000828682 |
| T | 11,6298 | 387,129 | 5,05692 | 5,00E-005 | 0,000828682 |
| Tg | 0,0978718 | 2,52635 | 4,69002 | 5,00E-005 | 0,000828682 |
| Cyp26a1 | 4,47831 | 114,255 | 4,67316 | 5,00E-005 | 0,000828682 |
| Wnt3a | 1,91882 | 43,3302 | 4,49708 | 5,00E-005 | 0,000828682 |
| Msgn1 | 17,1417 | 370,957 | 4,43567 | 5,00E-005 | 0,000828682 |
| Fgf17 | 4,8635 | 100,671 | 4,37151 | 5,00E-005 | 0,000828682 |
| Fgf4 | 0,423431 | 8,68437 | 4,35822 | 5,00E-005 | 0,000828682 |
| Mixl1 | 0,19945 | 4,05406 | 4,34527 | 5,00E-005 | 0,000828682 |
| Avpr1b | 0,234972 | 4,53124 | 4,26935 | 5,00E-005 | 0,000828682 |
| Kcnh5 | 0,12496 | 2,08628 | 4,0614 | 5,00E-005 | 0,000828682 |
| Lypd6b | 0,573637 | 9,07675 | 3,98397 | 5,00E-005 | 0,000828682 |
| Gad1 | 1,76913 | 27,0266 | 3,93327 | 5,00E-005 | 0,000828682 |
| Scara5 | 0,874159 | 12,6114 | 3,85069 | 5,00E-005 | 0,000828682 |
| Oasl2 | 0,273807 | 3,84163 | 3,81049 | 5,00E-005 | 0,000828682 |
| Mnx1 | 0,690573 | 9,17673 | 3,73211 | 5,00E-005 | 0,000828682 |
| Ptgs1 | 0,588638 | 7,45215 | 3,6622 | 5,00E-005 | 0,000828682 |
| Cdx2 | 7,10774 | 89,681 | 3,65734 | 5,00E-005 | 0,000828682 |
| Fgf3 | 5,11287 | 60,2707 | 3,55925 | 5,00E-005 | 0,000828682 |
| 5730457N03Rik | 3,70173 | 42,2513 | 3,51273 | 5,00E-005 | 0,000828682 |
| Arl4d | 16,9365 | 192,416 | 3,50601 | 5,00E-005 | 0,000828682 |
| Tmprss2 | 0,41677 | 4,33724 | 3,37945 | 5,00E-005 | 0,000828682 |
| Epha1 | 5,44836 | 56,1742 | 3,36601 | 5,00E-005 | 0,000828682 |
| Evx1 | 3,94065 | 38,8081 | 3,29985 | 5,00E-005 | 0,000828682 |
| Fgf8 | 6,16785 | 59,368 | 3,26685 | 5,00E-005 | 0,000828682 |
| Cldn9 | 1,74932 | 16,8368 | 3,26675 | 5,00E-005 | 0,000828682 |
| Cdx4 | 14,6594 | 135,285 | 3,2061 | 5,00E-005 | 0,000828682 |
| Hbb-y | 9,17446 | 84,3205 | 3,20019 | 5,00E-005 | 0,000828682 |
| Gabrb2 | 0,131336 | 1,17016 | 3,15538 | 5,00E-005 | 0,000828682 |
| Hbb-bh1 | 6,11632 | 54,3906 | 3,15262 | 5,00E-005 | 0,000828682 |
| Lzts1 | 0,502106 | 4,29288 | 3,09588 | 5,00E-005 | 0,000828682 |
| Egfr | 0,173531 | 1,46901 | 3,08158 | 5,00E-005 | 0,000828682 |
| Lmo2 | 19,2711 | 162,931 | 3,07975 | 5,00E-005 | 0,000828682 |
| Adamts17 | 0,72806 | 6,05739 | 3,05657 | 5,00E-005 | 0,000828682 |
| Ripk4 | 0,241981 | 2,01056 | 3,05463 | 5,00E-005 | 0,000828682 |
| Fstl4 | 0,386719 | 3,18131 | 3,04026 | 5,00E-005 | 0,000828682 |
| Tbx6 | 24,2716 | 198,974 | 3,03524 | 5,00E-005 | 0,000828682 |
| Igsf21 | 0,849528 | 6,8777 | 3,01719 | 5,00E-005 | 0,000828682 |
| Sp5 | 4,93199 | 39,8336 | 3,01375 | 5,00E-005 | 0,000828682 |
| Cpa6 | 0,352995 | 2,83613 | 3,0062 | 5,00E-005 | 0,000828682 |
| Exph5 | 0,0807415 | 0,635671 | 2,9769 | 5,00E-005 | 0,000828682 |
| Uap1l1 | 2,91883 | 22,8054 | 2,96591 | 5,00E-005 | 0,000828682 |
| Dmtn | 1,20202 | 9,19104 | 2,93476 | 5,00E-005 | 0,000828682 |

| | | | | | |
|---|---|---|---|---|---|
| Cd40 | 1,34419 | 10,1952 | 2,92308 | 5,00E-005 | 0,000828682 |
| Notch4 | 2,51295 | 19,0307 | 2,92087 | 5,00E-005 | 0,000828682 |
| Spry4 | 4,65284 | 34,8557 | 2,90521 | 5,00E-005 | 0,000828682 |
| Nod2 | 0,328254 | 2,34375 | 2,83593 | 5,00E-005 | 0,000828682 |
| Peli3 | 6,61883 | 47,1273 | 2,83191 | 5,00E-005 | 0,000828682 |
| Tll2 | 1,09256 | 7,68301 | 2,81396 | 5,00E-005 | 0,000828682 |
| Elf3 | 1,04455 | 7,26625 | 2,79832 | 5,00E-005 | 0,000828682 |
| Hes7 | 16,8414 | 116,91 | 2,79531 | 5,00E-005 | 0,000828682 |
| Hba-x | 7,34465 | 50,2881 | 2,77545 | 5,00E-005 | 0,000828682 |
| Pcp4l1 | 0,548399 | 3,71544 | 2,76024 | 5,00E-005 | 0,000828682 |
| Otx1 | 0,29417 | 1,94417 | 2,72443 | 5,00E-005 | 0,000828682 |
| Sv2b | 1,7862 | 11,7118 | 2,71299 | 5,00E-005 | 0,000828682 |
| Epor | 0,749054 | 4,8363 | 2,69076 | 5,00E-005 | 0,000828682 |
| Rspo3 | 17,7535 | 112,499 | 2,66373 | 5,00E-005 | 0,000828682 |
| S1pr5 | 1,43289 | 9,04735 | 2,65857 | 5,00E-005 | 0,000828682 |
| Chst7 | 4,92552 | 31,0857 | 2,6579 | 5,00E-005 | 0,000828682 |
| Efna1 | 6,17267 | 38,6717 | 2,64731 | 5,00E-005 | 0,000828682 |
| Wdr72 | 0,20241 | 1,23293 | 2,60674 | 5,00E-005 | 0,000828682 |
| Pde1c | 0,95793 | 5,79715 | 2,59735 | 5,00E-005 | 0,000828682 |
| Fgf9 | 0,96793 | 5,81839 | 2,58765 | 5,00E-005 | 0,000828682 |
| Irgm1 | 0,747965 | 4,46933 | 2,57902 | 5,00E-005 | 0,000828682 |
| Prr18 | 0,390048 | 2,32241 | 2,5739 | 5,00E-005 | 0,000828682 |
| Pla2g7 | 3,72687 | 22,1815 | 2,57332 | 5,00E-005 | 0,000828682 |
| Cdh20 | 0,745197 | 4,34955 | 2,54517 | 5,00E-005 | 0,000828682 |
| Cmklr1 | 0,647498 | 3,7507 | 2,53421 | 5,00E-005 | 0,000828682 |
| Cited1 | 28,6118 | 164,542 | 2,52378 | 5,00E-005 | 0,000828682 |
| Eogt | 12,3372 | 67,4187 | 2,45014 | 5,00E-005 | 0,000828682 |
| Trim9 | 5,98826 | 32,0372 | 2,41954 | 5,00E-005 | 0,000828682 |
| Esr1 | 0,740432 | 3,93278 | 2,40911 | 5,00E-005 | 0,000828682 |
| Cdhr1 | 0,997451 | 5,27297 | 2,4023 | 5,00E-005 | 0,000828682 |
| Klhl32 | 0,705512 | 3,72397 | 2,4001 | 5,00E-005 | 0,000828682 |
| Lamc2 | 0,285106 | 1,50162 | 2,39695 | 5,00E-005 | 0,000828682 |
| Gnal | 2,69161 | 14,0906 | 2,38819 | 5,00E-005 | 0,000828682 |
| Nptx2 | 4,08403 | 20,7099 | 2,34226 | 5,00E-005 | 0,000828682 |
| 6330419J24Rik | 1,35727 | 6,81655 | 2,32834 | 5,00E-005 | 0,000828682 |
| Rnf43 | 0,416102 | 2,06406 | 2,31047 | 5,00E-005 | 0,000828682 |
| Inpp4b | 0,336311 | 1,66537 | 2,30797 | 5,00E-005 | 0,000828682 |
| Endod1 | 1,63872 | 8,09187 | 2,3039 | 5,00E-005 | 0,000828682 |
| Urad | 6,18122 | 30,2 | 2,28859 | 5,00E-005 | 0,000828682 |
| Nfatc1 | 0,829619 | 4,0465 | 2,28616 | 5,00E-005 | 0,000828682 |
| B230323A14Rik | 2,23241 | 10,8424 | 2,28001 | 5,00E-005 | 0,000828682 |
| Dgkg | 0,577698 | 2,77734 | 2,26532 | 5,00E-005 | 0,000828682 |
| Hottip | 1,19476 | 5,73393 | 2,2628 | 5,00E-005 | 0,000828682 |
| Unc5d | 0,156085 | 0,744257 | 2,25347 | 5,00E-005 | 0,000828682 |
| Dusp4 | 27,6122 | 131,178 | 2,24814 | 5,00E-005 | 0,000828682 |
| Myom2 | 0,264146 | 1,24848 | 2,24077 | 5,00E-005 | 0,000828682 |
| Sct | 5,50081 | 25,8949 | 2,23495 | 5,00E-005 | 0,000828682 |
| Btg2 | 16,7174 | 78,0724 | 2,22346 | 5,00E-005 | 0,000828682 |
| Itgb8 | 1,62685 | 7,5763 | 2,21941 | 5,00E-005 | 0,000828682 |
| Dll3 | 36,3577 | 169,044 | 2,21707 | 5,00E-005 | 0,000828682 |

| | | | | | |
|---|---|---|---|---|---|
| Irf1 | 2,33335 | 10,7682 | 2,2063 | 5,00E-005 | 0,000828682 |
| Fermt1 | 0,416873 | 1,91162 | 2,19712 | 5,00E-005 | 0,000828682 |
| Cntn2 | 0,323624 | 1,45398 | 2,16761 | 5,00E-005 | 0,000828682 |
| Ngfr | 6,92856 | 31,1091 | 2,16671 | 5,00E-005 | 0,000828682 |
| Etv4 | 18,6811 | 83,761 | 2,1647 | 5,00E-005 | 0,000828682 |
| Il17rd | 11,7495 | 52,2845 | 2,15378 | 5,00E-005 | 0,000828682 |
| 2610528A11Rik | 32,3547 | 142,237 | 2,13625 | 5,00E-005 | 0,000828682 |
| Car3 | 6,4267 | 28,133 | 2,13011 | 5,00E-005 | 0,000828682 |
| Dync1i1 | 0,644696 | 2,81424 | 2,12605 | 5,00E-005 | 0,000828682 |
| Epha2 | 5,06553 | 21,9777 | 2,11726 | 5,00E-005 | 0,000828682 |
| Slc15a2 | 0,779888 | 3,36176 | 2,10788 | 5,00E-005 | 0,000828682 |
| Nkd2 | 23,0523 | 99,217 | 2,10568 | 5,00E-005 | 0,000828682 |
| Pmaip1 | 11,1425 | 47,9245 | 2,10469 | 5,00E-005 | 0,000828682 |
| Tpm1 | 389,815 | 1675,5 | 2,10373 | 5,00E-005 | 0,000828682 |
| Ptprb | 0,258603 | 1,1071 | 2,09797 | 5,00E-005 | 0,000828682 |
| S100a16 | 2,85161 | 12,1518 | 2,09132 | 5,00E-005 | 0,000828682 |
| Gna14 | 0,649291 | 2,75501 | 2,08512 | 5,00E-005 | 0,000828682 |
| Amph | 3,10355 | 13,1335 | 2,08126 | 5,00E-005 | 0,000828682 |
| Rftn1 | 11,5459 | 48,5749 | 2,07283 | 5,00E-005 | 0,000828682 |
| Greb1 | 26,4431 | 107,943 | 2,02931 | 5,00E-005 | 0,000828682 |
| Loxl4 | 0,307674 | 1,2538 | 2,02683 | 5,00E-005 | 0,000828682 |
| Rimbp2 | 0,641978 | 2,5893 | 2,01197 | 5,00E-005 | 0,000828682 |
| Dll1 | 44,1993 | 177,656 | 2,00699 | 5,00E-005 | 0,000828682 |
| Pax2 | 0,605754 | 2,42284 | 1,99989 | 5,00E-005 | 0,000828682 |
| Gucy1a2 | 1,39371 | 5,56758 | 1,99812 | 5,00E-005 | 0,000828682 |
| Ets2 | 37,8342 | 150,989 | 1,99668 | 5,00E-005 | 0,000828682 |
| Etv5 | 8,47795 | 33,6875 | 1,99043 | 5,00E-005 | 0,000828682 |
| Rgs9 | 1,53736 | 6,06244 | 1,97944 | 5,00E-005 | 0,000828682 |
| Ndrg2 | 2,55499 | 10,0502 | 1,97584 | 5,00E-005 | 0,000828682 |
| Nkx1-2 | 7,21487 | 28,0127 | 1,95703 | 5,00E-005 | 0,000828682 |
| Robo4 | 0,732044 | 2,82798 | 1,94977 | 5,00E-005 | 0,000828682 |
| Cdkn1a | 3,58707 | 13,8038 | 1,94418 | 5,00E-005 | 0,000828682 |
| Vwa2 | 9,57728 | 36,654 | 1,93628 | 5,00E-005 | 0,000828682 |
| Hoxc13 | 17,6186 | 67,3001 | 1,93351 | 5,00E-005 | 0,000828682 |
| Ankrd34b | 0,390696 | 1,48333 | 1,92472 | 5,00E-005 | 0,000828682 |
| Gdf11 | 17,4457 | 66,2024 | 1,92401 | 5,00E-005 | 0,000828682 |
| Mapkapk3 | 1,31433 | 4,94475 | 1,91157 | 5,00E-005 | 0,000828682 |
| Dlgap2 | 0,556546 | 2,08741 | 1,90714 | 5,00E-005 | 0,000828682 |
| Fgf15 | 14,0127 | 52,2721 | 1,8993 | 5,00E-005 | 0,000828682 |
| Krt19 | 2,38331 | 8,85722 | 1,89389 | 5,00E-005 | 0,000828682 |
| Aim1 | 0,175841 | 0,652798 | 1,89237 | 5,00E-005 | 0,000828682 |
| Dennd2c | 0,602166 | 2,23291 | 1,89069 | 5,00E-005 | 0,000828682 |
| Erap1 | 2,35804 | 8,70878 | 1,88488 | 5,00E-005 | 0,000828682 |
| Trim36 | 1,46976 | 5,41595 | 1,88163 | 5,00E-005 | 0,000828682 |
| Kcnab3 | 4,82249 | 17,6495 | 1,87178 | 5,00E-005 | 0,000828682 |
| Nfam1 | 1,16865 | 4,27133 | 1,86985 | 5,00E-005 | 0,000828682 |
| Wnt8a | 1,30259 | 4,74103 | 1,86381 | 5,00E-005 | 0,000828682 |
| Kitl | 8,13178 | 29,5804 | 1,863 | 5,00E-005 | 0,000828682 |
| Gbp7 | 0,340606 | 1,22944 | 1,85183 | 5,00E-005 | 0,000828682 |
| Mira | 26,3392 | 95,0117 | 1,85089 | 5,00E-005 | 0,000828682 |

| | | | | | |
|---|---|---|---|---|---|
| Sema5b | 3,30647 | 11,8696 | 1,84391 | 5,00E-005 | 0,000828682 |
| Zfpm1 | 1,90006 | 6,80073 | 1,83964 | 5,00E-005 | 0,000828682 |
| Mpzl2 | 3,56139 | 12,5492 | 1,81708 | 5,00E-005 | 0,000828682 |
| Notum | 1,78738 | 6,24468 | 1,80478 | 5,00E-005 | 0,000828682 |
| Egln3 | 7,3742 | 25,6312 | 1,79734 | 5,00E-005 | 0,000828682 |
| Foxb1 | 13,7941 | 47,9388 | 1,79715 | 5,00E-005 | 0,000828682 |
| Rassf9 | 1,23152 | 4,26845 | 1,79327 | 5,00E-005 | 0,000828682 |
| Col12a1 | 0,217628 | 0,747328 | 1,77988 | 5,00E-005 | 0,000828682 |
| Dysf | 0,877939 | 2,9876 | 1,76679 | 5,00E-005 | 0,000828682 |
| Rbm20 | 0,805066 | 2,7367 | 1,76526 | 5,00E-005 | 0,000828682 |
| Wnt5a | 55,1647 | 186,793 | 1,75963 | 5,00E-005 | 0,000828682 |
| Vstm2b | 9,55348 | 32,2099 | 1,7534 | 5,00E-005 | 0,000828682 |
| Bhlhe40 | 1,29268 | 4,34093 | 1,74764 | 5,00E-005 | 0,000828682 |
| Nim1k | 1,07296 | 3,593 | 1,7436 | 5,00E-005 | 0,000828682 |
| C77370 | 0,222893 | 0,744821 | 1,74054 | 5,00E-005 | 0,000828682 |
| Pik3c2b | 5,07426 | 16,9272 | 1,73807 | 5,00E-005 | 0,000828682 |
| Spock2 | 18,0048 | 59,9568 | 1,73554 | 5,00E-005 | 0,000828682 |
| Grhl3 | 1,36311 | 4,51562 | 1,72803 | 5,00E-005 | 0,000828682 |
| Atf7ip2 | 1,37615 | 4,55742 | 1,72758 | 5,00E-005 | 0,000828682 |
| 1300002K09Rik | 4,59984 | 15,083 | 1,71327 | 5,00E-005 | 0,000828682 |
| Smc6 | 31,8076 | 103,749 | 1,70566 | 5,00E-005 | 0,000828682 |
| Agpat9 | 2,85162 | 9,27189 | 1,70108 | 5,00E-005 | 0,000828682 |
| Aplnr | 4,77943 | 15,5193 | 1,69916 | 5,00E-005 | 0,000828682 |
| Ptk7 | 106,529 | 344,417 | 1,69291 | 5,00E-005 | 0,000828682 |
| Mmp17 | 4,01126 | 12,9237 | 1,68789 | 5,00E-005 | 0,000828682 |
| Hoxb1 | 27,3047 | 87,6444 | 1,68251 | 5,00E-005 | 0,000828682 |
| Tcea3 | 5,50226 | 17,6137 | 1,6786 | 5,00E-005 | 0,000828682 |
| Lrrc55 | 0,782497 | 2,49743 | 1,67429 | 5,00E-005 | 0,000828682 |
| Dock8 | 0,333464 | 1,06379 | 1,6736 | 5,00E-005 | 0,000828682 |
| Ccdc171 | 0,79158 | 2,47595 | 1,64517 | 5,00E-005 | 0,000828682 |
| Rab8b | 12,7804 | 39,966 | 1,64484 | 5,00E-005 | 0,000828682 |
| B3gnt7 | 3,23583 | 10,0931 | 1,64115 | 5,00E-005 | 0,000828682 |
| Itga1 | 2,67899 | 8,33184 | 1,63694 | 5,00E-005 | 0,000828682 |
| Nrarp | 30,8363 | 94,8818 | 1,6215 | 5,00E-005 | 0,000828682 |
| Hoxc8 | 8,17342 | 25,0545 | 1,61606 | 5,00E-005 | 0,000828682 |
| Fam132b | 1,62952 | 4,98569 | 1,61335 | 5,00E-005 | 0,000828682 |
| Parp12 | 1,69053 | 5,16933 | 1,6125 | 5,00E-005 | 0,000828682 |
| Slc2a3 | 70,449 | 214,956 | 1,60939 | 5,00E-005 | 0,000828682 |
| Fam222a | 2,79167 | 8,4748 | 1,60205 | 5,00E-005 | 0,000828682 |
| Rell1 | 12,7401 | 38,6495 | 1,60107 | 5,00E-005 | 0,000828682 |
| Csdc2 | 1,33109 | 4,02602 | 1,59674 | 5,00E-005 | 0,000828682 |
| Hoxc6 | 18,4851 | 55,5966 | 1,58864 | 5,00E-005 | 0,000828682 |
| Maff | 2,53384 | 7,60801 | 1,58619 | 5,00E-005 | 0,000828682 |
| Hoxa13 | 6,81253 | 20,454 | 1,58612 | 5,00E-005 | 0,000828682 |
| Rhbg | 6,0613 | 18,1348 | 1,58106 | 5,00E-005 | 0,000828682 |
| Hoxc5 | 6,76402 | 20,1496 | 1,5748 | 5,00E-005 | 0,000828682 |
| Fabp7 | 62,8222 | 186,8 | 1,57215 | 5,00E-005 | 0,000828682 |
| Glis3 | 1,25905 | 3,72377 | 1,56443 | 5,00E-005 | 0,000828682 |
| Ephb1 | 0,913097 | 2,69914 | 1,56366 | 5,00E-005 | 0,000828682 |
| Cdx1 | 2,54452 | 7,52069 | 1,56347 | 5,00E-005 | 0,000828682 |

| | | | | | |
|---|---|---|---|---|---|
| Snph | 1,67927 | 4,90193 | 1,54551 | 5,00E-005 | 0,000828682 |
| AW551984 | 4,8376 | 14,0618 | 1,53942 | 5,00E-005 | 0,000828682 |
| Afap1l2 | 0,746151 | 2,15939 | 1,53308 | 5,00E-005 | 0,000828682 |
| Rnf128 | 1,73598 | 5,00031 | 1,52627 | 5,00E-005 | 0,000828682 |
| Cacna1a | 2,21229 | 6,34444 | 1,51995 | 5,00E-005 | 0,000828682 |
| Pcmtd1 | 6,35127 | 18,2071 | 1,51939 | 5,00E-005 | 0,000828682 |
| Spry2 | 15,2472 | 43,4501 | 1,51082 | 5,00E-005 | 0,000828682 |
| Sema4d | 3,93245 | 11,1409 | 1,50237 | 5,00E-005 | 0,000828682 |
| Socs2 | 6,62848 | 18,5203 | 1,48236 | 5,00E-005 | 0,000828682 |
| Efna3 | 9,30526 | 25,8972 | 1,47668 | 5,00E-005 | 0,000828682 |
| Hopx | 2,95198 | 8,21037 | 1,47577 | 5,00E-005 | 0,000828682 |
| Wnt5b | 13,0466 | 36,2569 | 1,47459 | 5,00E-005 | 0,000828682 |
| Ddx58 | 0,821103 | 2,27801 | 1,47214 | 5,00E-005 | 0,000828682 |
| Adcy5 | 0,675725 | 1,87046 | 1,46889 | 5,00E-005 | 0,000828682 |
| Smtnl2 | 4,72732 | 13,0746 | 1,46768 | 5,00E-005 | 0,000828682 |
| Emp1 | 7,90677 | 21,7472 | 1,45967 | 5,00E-005 | 0,000828682 |
| Rbm24 | 2,62433 | 7,20083 | 1,45621 | 5,00E-005 | 0,000828682 |
| D430041D05Rik | 0,882111 | 2,41806 | 1,45482 | 5,00E-005 | 0,000828682 |
| Pde4b | 4,32797 | 11,8544 | 1,45366 | 5,00E-005 | 0,000828682 |
| Cdh3 | 4,8547 | 13,2699 | 1,4507 | 5,00E-005 | 0,000828682 |
| Col13a1 | 8,91589 | 24,038 | 1,43086 | 5,00E-005 | 0,000828682 |
| Arhgef5 | 1,75324 | 4,72117 | 1,42912 | 5,00E-005 | 0,000828682 |
| Sall3 | 8,65324 | 23,2955 | 1,42874 | 5,00E-005 | 0,000828682 |
| Csf1 | 3,48656 | 9,35418 | 1,42381 | 5,00E-005 | 0,000828682 |
| Wnt11 | 1,43665 | 3,851 | 1,42253 | 5,00E-005 | 0,000828682 |
| Arhgap31 | 3,92149 | 10,4923 | 1,41985 | 5,00E-005 | 0,000828682 |
| Hoxa7 | 9,77536 | 26,0451 | 1,41379 | 5,00E-005 | 0,000828682 |
| Ifitm1 | 184,717 | 490,816 | 1,40986 | 5,00E-005 | 0,000828682 |
| Ypel3 | 18,4832 | 49,1093 | 1,40978 | 5,00E-005 | 0,000828682 |
| Tln2 | 8,08266 | 21,4435 | 1,40764 | 5,00E-005 | 0,000828682 |
| Mmp2 | 28,6987 | 75,9681 | 1,40441 | 5,00E-005 | 0,000828682 |
| Clstn2 | 2,06995 | 5,46808 | 1,40143 | 5,00E-005 | 0,000828682 |
| Trim25 | 1,0812 | 2,8502 | 1,39842 | 5,00E-005 | 0,000828682 |
| Rhpn2 | 5,02971 | 13,2272 | 1,39496 | 5,00E-005 | 0,000828682 |
| Fam131b | 5,90668 | 15,5198 | 1,39369 | 5,00E-005 | 0,000828682 |
| Lama1 | 24,543 | 64,2085 | 1,38745 | 5,00E-005 | 0,000828682 |
| E330009J07Rik | 5,66766 | 14,8185 | 1,38657 | 5,00E-005 | 0,000828682 |
| Lef1 | 48,7965 | 127,378 | 1,38427 | 5,00E-005 | 0,000828682 |
| Alpl | 7,40302 | 19,2904 | 1,3817 | 5,00E-005 | 0,000828682 |
| Lfng | 23,9363 | 61,8702 | 1,37004 | 5,00E-005 | 0,000828682 |
| Ptprm | 1,84059 | 4,75495 | 1,36926 | 5,00E-005 | 0,000828682 |
| Doc2g | 2,94401 | 7,6051 | 1,36918 | 5,00E-005 | 0,000828682 |
| Hoxc4 | 14,4414 | 37,299 | 1,36892 | 5,00E-005 | 0,000828682 |
| Mylip | 18,2085 | 46,955 | 1,36667 | 5,00E-005 | 0,000828682 |
| Atxn7 | 4,41762 | 11,3851 | 1,36581 | 5,00E-005 | 0,000828682 |
| Dgkk | 1,20993 | 3,10727 | 1,36073 | 5,00E-005 | 0,000828682 |
| Ccnjl | 16,8485 | 43,2634 | 1,36052 | 5,00E-005 | 0,000828682 |
| Sept8 | 32,8576 | 84,1913 | 1,35744 | 5,00E-005 | 0,000828682 |
| Optn | 1,60009 | 4,08701 | 1,3529 | 5,00E-005 | 0,000828682 |
| Adamts3 | 1,26659 | 3,23468 | 1,35268 | 5,00E-005 | 0,000828682 |

| | | | | | |
|---|---|---|---|---|---|
| Ripk3 | 7,7114 | 19,5574 | 1,34265 | 5,00E-005 | 0,000828682 |
| Car13 | 9,64521 | 24,3068 | 1,33348 | 5,00E-005 | 0,000828682 |
| Hoxc12 | 40,347 | 101,179 | 1,32638 | 5,00E-005 | 0,000828682 |
| Hspb8 | 4,00485 | 10,0106 | 1,32171 | 5,00E-005 | 0,000828682 |
| Schip1 | 11,0453 | 27,5477 | 1,3185 | 5,00E-005 | 0,000828682 |
| Dusp6 | 54,9793 | 136,976 | 1,31696 | 5,00E-005 | 0,000828682 |
| Ddit4 | 40,0624 | 99,6086 | 1,31402 | 5,00E-005 | 0,000828682 |
| Chic1 | 2,71256 | 6,73973 | 1,31303 | 5,00E-005 | 0,000828682 |
| Hoxb13 | 20,9009 | 51,6943 | 1,30644 | 5,00E-005 | 0,000828682 |
| Kdm7a | 8,07553 | 19,9099 | 1,30185 | 5,00E-005 | 0,000828682 |
| Sp8 | 6,15936 | 15,1431 | 1,29781 | 5,00E-005 | 0,000828682 |
| Gm11538 | 2,37904 | 5,849 | 1,29781 | 5,00E-005 | 0,000828682 |
| Pgbd5 | 2,12587 | 5,2229 | 1,2968 | 5,00E-005 | 0,000828682 |
| Gm53 | 33,8388 | 82,7297 | 1,28973 | 5,00E-005 | 0,000828682 |
| Hoxa11os | 79,4114 | 193,193 | 1,28262 | 5,00E-005 | 0,000828682 |
| Pdzd2 | 1,2854 | 3,12056 | 1,27959 | 5,00E-005 | 0,000828682 |
| Itga5 | 23,2234 | 56,2414 | 1,27605 | 5,00E-005 | 0,000828682 |
| Adam19 | 12,5488 | 30,3695 | 1,27508 | 5,00E-005 | 0,000828682 |
| Engase | 2,90691 | 6,99577 | 1,267 | 5,00E-005 | 0,000828682 |
| Ptpn3 | 0,793103 | 1,90565 | 1,2647 | 5,00E-005 | 0,000828682 |
| Slit3 | 8,71609 | 20,8653 | 1,25935 | 5,00E-005 | 0,000828682 |
| Aspg | 2,89926 | 6,93899 | 1,25904 | 5,00E-005 | 0,000828682 |
| Nbeal1 | 1,23451 | 2,95223 | 1,25787 | 5,00E-005 | 0,000828682 |
| Cdk18 | 2,22436 | 5,29452 | 1,25111 | 5,00E-005 | 0,000828682 |
| Lmo7 | 1,41238 | 3,35765 | 1,24932 | 5,00E-005 | 0,000828682 |
| Trps1 | 6,1806 | 14,5916 | 1,23932 | 5,00E-005 | 0,000828682 |
| Map6 | 2,97177 | 6,98706 | 1,23336 | 5,00E-005 | 0,000828682 |
| Il17ra | 3,23298 | 7,57445 | 1,22828 | 5,00E-005 | 0,000828682 |
| Zbtb16 | 5,13367 | 12,0173 | 1,22705 | 5,00E-005 | 0,000828682 |
| Ptpn6 | 2,50452 | 5,86122 | 1,22667 | 5,00E-005 | 0,000828682 |
| Zyx | 44,4737 | 103,552 | 1,21933 | 5,00E-005 | 0,000828682 |
| Ddx3y | 3,70542 | 8,59853 | 1,21445 | 5,00E-005 | 0,000828682 |
| Lrrk2 | 1,23674 | 2,86892 | 1,21397 | 5,00E-005 | 0,000828682 |
| Dock6 | 9,11431 | 21,0735 | 1,20923 | 5,00E-005 | 0,000828682 |
| Gprc5a | 4,35897 | 9,99586 | 1,19734 | 5,00E-005 | 0,000828682 |
| Rarg | 31,7032 | 72,6008 | 1,19536 | 5,00E-005 | 0,000828682 |
| Mogat2 | 27,9426 | 63,7712 | 1,19044 | 5,00E-005 | 0,000828682 |
| Pcdh19 | 13,0896 | 29,7469 | 1,18432 | 5,00E-005 | 0,000828682 |
| Arap3 | 1,75897 | 3,99594 | 1,1838 | 5,00E-005 | 0,000828682 |
| Gbx2 | 15,7325 | 35,7312 | 1,18344 | 5,00E-005 | 0,000828682 |
| Hoxc10 | 88,0394 | 199,797 | 1,18232 | 5,00E-005 | 0,000828682 |
| Naaa | 22,5554 | 51,1552 | 1,18141 | 5,00E-005 | 0,000828682 |
| Qsox1 | 11,9536 | 26,8478 | 1,16736 | 5,00E-005 | 0,000828682 |
| Jade2 | 1,23617 | 2,77556 | 1,1669 | 5,00E-005 | 0,000828682 |
| Pmepa1 | 2,00197 | 4,48353 | 1,16322 | 5,00E-005 | 0,000828682 |
| Epb4.1l2 | 50,5839 | 113,064 | 1,16039 | 5,00E-005 | 0,000828682 |
| Fbln2 | 10,1887 | 22,7726 | 1,16033 | 5,00E-005 | 0,000828682 |
| Mesp2 | 5,21535 | 11,5717 | 1,14976 | 5,00E-005 | 0,000828682 |
| Ppfibp1 | 28,2249 | 62,5359 | 1,14771 | 5,00E-005 | 0,000828682 |
| Kif26b | 19,9107 | 44,0514 | 1,14564 | 5,00E-005 | 0,000828682 |

| | | | | | |
|---|---|---|---|---|---|
| Atp1b2 | 2,45355 | 5,40343 | 1,139 | 5,00E-005 | 0,000828682 |
| Cxcr4 | 18,3063 | 40,2203 | 1,13558 | 5,00E-005 | 0,000828682 |
| Rps6ka5 | 2,54837 | 5,59278 | 1,13399 | 5,00E-005 | 0,000828682 |
| Ezr | 28,2249 | 61,4129 | 1,12157 | 5,00E-005 | 0,000828682 |
| Lypd6 | 4,64154 | 10,0883 | 1,12001 | 5,00E-005 | 0,000828682 |
| Smug1 | 10,4752 | 22,7242 | 1,11726 | 5,00E-005 | 0,000828682 |
| Blnk | 14,6264 | 31,5804 | 1,11045 | 5,00E-005 | 0,000828682 |
| Lcor | 1,65783 | 3,57643 | 1,10923 | 5,00E-005 | 0,000828682 |
| Hotair | 11,0979 | 23,9216 | 1,10802 | 5,00E-005 | 0,000828682 |
| Spred3 | 2,81366 | 6,04683 | 1,10373 | 5,00E-005 | 0,000828682 |
| Parp8 | 5,39207 | 11,588 | 1,10371 | 5,00E-005 | 0,000828682 |
| Heg1 | 12,5243 | 26,8392 | 1,09961 | 5,00E-005 | 0,000828682 |
| Antxr1 | 16,3959 | 35,1035 | 1,09828 | 5,00E-005 | 0,000828682 |
| Lrp8 | 7,66029 | 16,3135 | 1,0906 | 5,00E-005 | 0,000828682 |
| Epha4 | 24,4467 | 51,9969 | 1,08879 | 5,00E-005 | 0,000828682 |
| Afap1l1 | 2,37145 | 5,04245 | 1,08835 | 5,00E-005 | 0,000828682 |
| Prickle1 | 24,0596 | 51,1316 | 1,0876 | 5,00E-005 | 0,000828682 |
| Fn1 | 179,162 | 380,422 | 1,08633 | 5,00E-005 | 0,000828682 |
| Slc16a2 | 3,92632 | 8,3333 | 1,08571 | 5,00E-005 | 0,000828682 |
| Fat1 | 29,5866 | 62,6906 | 1,08331 | 5,00E-005 | 0,000828682 |
| Adamts9 | 8,77078 | 18,5356 | 1,07952 | 5,00E-005 | 0,000828682 |
| Pcolce2 | 12,0926 | 25,5262 | 1,07785 | 5,00E-005 | 0,000828682 |
| Pea15a | 58,7902 | 123,368 | 1,06932 | 5,00E-005 | 0,000828682 |
| Stc2 | 6,35895 | 13,3063 | 1,06525 | 5,00E-005 | 0,000828682 |
| Dlc1 | 17,7497 | 37,1385 | 1,06512 | 5,00E-005 | 0,000828682 |
| Susd4 | 10,3535 | 21,6387 | 1,0635 | 5,00E-005 | 0,000828682 |
| St6galnac4 | 8,13362 | 16,9875 | 1,0625 | 5,00E-005 | 0,000828682 |
| Fgd4 | 3,24476 | 6,77261 | 1,0616 | 5,00E-005 | 0,000828682 |
| Irx2 | 5,31216 | 11,0701 | 1,0593 | 5,00E-005 | 0,000828682 |
| Gcnt2 | 6,09092 | 12,6903 | 1,05899 | 5,00E-005 | 0,000828682 |
| Magi1 | 15,9764 | 33,2773 | 1,05859 | 5,00E-005 | 0,000828682 |
| Dhrs3 | 9,37066 | 19,4427 | 1,053 | 5,00E-005 | 0,000828682 |
| Sall1 | 77,4493 | 160,508 | 1,05132 | 5,00E-005 | 0,000828682 |
| Arid3b | 55,9807 | 115,864 | 1,04943 | 5,00E-005 | 0,000828682 |
| Smoc1 | 45,2855 | 93,4952 | 1,04584 | 5,00E-005 | 0,000828682 |
| Adamts20 | 4,28591 | 8,80414 | 1,03858 | 5,00E-005 | 0,000828682 |
| Atp1a2 | 11,3901 | 23,3003 | 1,03256 | 5,00E-005 | 0,000828682 |
| Unc5b | 13,1935 | 26,9567 | 1,03081 | 5,00E-005 | 0,000828682 |
| Zic3 | 19,6497 | 40,002 | 1,02556 | 5,00E-005 | 0,000828682 |
| Jag1 | 6,85643 | 13,9411 | 1,02382 | 5,00E-005 | 0,000828682 |
| Hoxd4 | 59,027 | 119,933 | 1,02278 | 5,00E-005 | 0,000828682 |
| Mtss1 | 10,4191 | 21,1563 | 1,02185 | 5,00E-005 | 0,000828682 |
| Spred2 | 20,2946 | 41,1209 | 1,01878 | 5,00E-005 | 0,000828682 |
| Smim3 | 7,75859 | 15,6953 | 1,01647 | 5,00E-005 | 0,000828682 |
| Tcirg1 | 5,10641 | 10,3219 | 1,01533 | 5,00E-005 | 0,000828682 |
| Tgfb1i1 | 3,55476 | 7,17791 | 1,01381 | 5,00E-005 | 0,000828682 |
| Ptprj | 3,04492 | 6,14745 | 1,01358 | 5,00E-005 | 0,000828682 |
| Atp10a | 2,11153 | 4,25172 | 1,00976 | 5,00E-005 | 0,000828682 |
| 2610018G03Rik | 9,84353 | 19,8048 | 1,0086 | 5,00E-005 | 0,000828682 |
| B3gnt2 | 4,36734 | 8,77068 | 1,00594 | 5,00E-005 | 0,000828682 |

| | | | | | |
|---|---|---|---|---|---|
| Hoxb9 | 49,7365 | 99,5908 | 1,00171 | 5,00E-005 | 0,000828682 |
| Sh3bp1 | 14,2558 | 28,5388 | 1,00138 | 5,00E-005 | 0,000828682 |
| Hoxd12 | 46,0816 | 92,2098 | 1,00073 | 5,00E-005 | 0,000828682 |
| Crb2 | 11,9634 | 23,866 | 0,996332 | 5,00E-005 | 0,000828682 |
| Sulf1 | 39,0631 | 77,7153 | 0,992393 | 5,00E-005 | 0,000828682 |
| Hoxd11 | 59,61 | 118,577 | 0,992202 | 5,00E-005 | 0,000828682 |
| Cachd1 | 28,0958 | 55,8656 | 0,991607 | 5,00E-005 | 0,000828682 |
| Dhtkd1 | 3,12392 | 6,17247 | 0,982492 | 5,00E-005 | 0,000828682 |
| Prkag2 | 20,8528 | 41,0847 | 0,978356 | 5,00E-005 | 0,000828682 |
| Hoxb4 | 14,0977 | 27,7593 | 0,977515 | 5,00E-005 | 0,000828682 |
| Mllt3 | 30,8894 | 60,7381 | 0,975489 | 5,00E-005 | 0,000828682 |
| Hoxc9 | 18,0251 | 35,4141 | 0,974314 | 5,00E-005 | 0,000828682 |
| Gng2 | 20,0878 | 39,2851 | 0,967665 | 5,00E-005 | 0,000828682 |
| Lhpp | 38,5113 | 75,1155 | 0,963831 | 5,00E-005 | 0,000828682 |
| Slc1a4 | 13,9942 | 27,2495 | 0,961402 | 5,00E-005 | 0,000828682 |
| Rab11fip1 | 4,1226 | 8,02103 | 0,960234 | 5,00E-005 | 0,000828682 |
| Slit2 | 5,85225 | 11,3415 | 0,954551 | 5,00E-005 | 0,000828682 |
| Zfp36l2 | 15,0325 | 28,8487 | 0,940422 | 5,00E-005 | 0,000828682 |
| Sema6a | 11,128 | 21,3312 | 0,938776 | 5,00E-005 | 0,000828682 |
| Nkd1 | 18,957 | 36,2033 | 0,933387 | 5,00E-005 | 0,000828682 |
| Furin | 8,61551 | 16,436 | 0,931853 | 5,00E-005 | 0,000828682 |
| Smoc2 | 9,84182 | 18,6769 | 0,924261 | 5,00E-005 | 0,000828682 |
| Pde9a | 14,5356 | 27,5095 | 0,920343 | 5,00E-005 | 0,000828682 |
| Per3 | 2,52031 | 4,75914 | 0,9171 | 5,00E-005 | 0,000828682 |
| BC005561 | 5,05337 | 9,53521 | 0,916018 | 5,00E-005 | 0,000828682 |
| Zdhhc9 | 6,97721 | 13,1482 | 0,914143 | 5,00E-005 | 0,000828682 |
| Ccdc88a | 8,91813 | 16,8039 | 0,913979 | 5,00E-005 | 0,000828682 |
| Ptpru | 3,02559 | 5,69777 | 0,913182 | 5,00E-005 | 0,000828682 |
| F11r | 9,36265 | 17,6159 | 0,911891 | 5,00E-005 | 0,000828682 |
| Foxo4 | 37,4194 | 70,3559 | 0,910886 | 5,00E-005 | 0,000828682 |
| Hoxb3 | 32,9767 | 61,9234 | 0,909035 | 5,00E-005 | 0,000828682 |
| Gja1 | 180,959 | 338,564 | 0,903763 | 5,00E-005 | 0,000828682 |
| Sema4g | 11,8941 | 22,2271 | 0,902072 | 5,00E-005 | 0,000828682 |
| Sp4 | 5,21271 | 9,72537 | 0,89972 | 5,00E-005 | 0,000828682 |
| Pik3c2a | 9,1275 | 16,9714 | 0,894816 | 5,00E-005 | 0,000828682 |
| A630007B06Rik | 11,961 | 22,1212 | 0,887092 | 5,00E-005 | 0,000828682 |
| Rin2 | 6,91346 | 12,7703 | 0,885306 | 5,00E-005 | 0,000828682 |
| Cenpf | 16,8169 | 30,9431 | 0,87971 | 5,00E-005 | 0,000828682 |
| Pdia5 | 22,589 | 41,5113 | 0,877886 | 5,00E-005 | 0,000828682 |
| Kat6b | 5,51848 | 10,1304 | 0,876344 | 5,00E-005 | 0,000828682 |
| Hoxd9 | 18,0985 | 33,2068 | 0,875603 | 5,00E-005 | 0,000828682 |
| Bmpr2 | 6,91215 | 12,6558 | 0,872587 | 5,00E-005 | 0,000828682 |
| Creg1 | 9,78295 | 17,8641 | 0,868725 | 5,00E-005 | 0,000828682 |
| Sulf2 | 125,161 | 228,224 | 0,866667 | 5,00E-005 | 0,000828682 |
| Casc5 | 6,88306 | 12,5106 | 0,862032 | 5,00E-005 | 0,000828682 |
| Dst | 5,57896 | 10,0779 | 0,85313 | 5,00E-005 | 0,000828682 |
| Prnp | 13,7248 | 24,7789 | 0,85233 | 5,00E-005 | 0,000828682 |
| Dpysl5 | 26,1902 | 47,1865 | 0,849345 | 5,00E-005 | 0,000828682 |
| Myh9 | 43,263 | 77,938 | 0,849194 | 5,00E-005 | 0,000828682 |
| Pdlim5 | 13,3522 | 23,9993 | 0,845914 | 5,00E-005 | 0,000828682 |

| | | | | | |
|---|---|---|---|---|---|
| Igf1r | 17,2755 | 31,0246 | 0,844691 | 5,00E-005 | 0,000828682 |
| Ephb4 | 41,1724 | 73,7858 | 0,841665 | 5,00E-005 | 0,000828682 |
| Cyld | 3,55354 | 6,35751 | 0,839205 | 5,00E-005 | 0,000828682 |
| Scai | 3,70107 | 6,60978 | 0,836659 | 5,00E-005 | 0,000828682 |
| Akap9 | 6,53977 | 11,6452 | 0,832426 | 5,00E-005 | 0,000828682 |
| Bod1l | 7,96959 | 14,1176 | 0,824921 | 5,00E-005 | 0,000828682 |
| Krba1 | 8,06365 | 14,2836 | 0,824857 | 5,00E-005 | 0,000828682 |
| Shf | 33,6797 | 59,5982 | 0,82339 | 5,00E-005 | 0,000828682 |
| Rif1 | 21,6021 | 38,1834 | 0,821778 | 5,00E-005 | 0,000828682 |
| Serinc5 | 17,3559 | 30,6599 | 0,820923 | 5,00E-005 | 0,000828682 |
| Has2 | 21,1858 | 37,0646 | 0,806944 | 5,00E-005 | 0,000828682 |
| Tenm4 | 21,352 | 37,3173 | 0,805473 | 5,00E-005 | 0,000828682 |
| Laptm4b | 57,5877 | 100,351 | 0,801218 | 5,00E-005 | 0,000828682 |
| Stra6 | 15,7633 | 27,4459 | 0,800022 | 5,00E-005 | 0,000828682 |
| Slc37a3 | 23,2355 | 40,4533 | 0,799924 | 5,00E-005 | 0,000828682 |
| Wls | 74,6049 | 129,615 | 0,796889 | 5,00E-005 | 0,000828682 |
| Cdc14b | 12,8982 | 22,333 | 0,792009 | 5,00E-005 | 0,000828682 |
| Cgnl1 | 16,233 | 28,0276 | 0,787922 | 5,00E-005 | 0,000828682 |
| Smarca1 | 7,99444 | 13,7964 | 0,787227 | 5,00E-005 | 0,000828682 |
| Axin2 | 18,8731 | 32,4397 | 0,781429 | 5,00E-005 | 0,000828682 |
| Naa60 | 34,5317 | 59,2076 | 0,77786 | 5,00E-005 | 0,000828682 |
| Dcbld2 | 29,4437 | 50,2297 | 0,770582 | 5,00E-005 | 0,000828682 |
| Lrrfip1 | 18,0466 | 30,6006 | 0,761836 | 5,00E-005 | 0,000828682 |
| Rfx3 | 6,72076 | 11,3952 | 0,761729 | 5,00E-005 | 0,000828682 |
| Ccnl1 | 51,5393 | 87,2057 | 0,758749 | 5,00E-005 | 0,000828682 |
| Dnmt3a | 35,1051 | 59,1639 | 0,753039 | 5,00E-005 | 0,000828682 |
| Lamc1 | 24,84 | 41,8066 | 0,751069 | 5,00E-005 | 0,000828682 |
| Hoxd10 | 71,8879 | 120,538 | 0,745666 | 5,00E-005 | 0,000828682 |
| Tcf7 | 21,7685 | 36,4004 | 0,74171 | 5,00E-005 | 0,000828682 |
| Limch1 | 27,2529 | 45,2186 | 0,730505 | 5,00E-005 | 0,000828682 |
| Snai1 | 39,4214 | 65,0314 | 0,722159 | 5,00E-005 | 0,000828682 |
| Mical1 | 22,925 | 37,1674 | 0,697114 | 5,00E-005 | 0,000828682 |
| Amot | 20,2629 | 32,5429 | 0,683503 | 5,00E-005 | 0,000828682 |
| D17Wsu92e | 38,2373 | 60,3545 | 0,658481 | 5,00E-005 | 0,000828682 |
| Ttc28 | 8,09431 | 4,88892 | -0,727391 | 5,00E-005 | 0,000828682 |
| Fhl1 | 64,9297 | 38,963 | -0,736773 | 5,00E-005 | 0,000828682 |
| Arrb1 | 12,7205 | 7,56918 | -0,748941 | 5,00E-005 | 0,000828682 |
| Tmem119 | 36,9591 | 21,898 | -0,755135 | 5,00E-005 | 0,000828682 |
| Foxp1 | 16,492 | 9,68557 | -0,767857 | 5,00E-005 | 0,000828682 |
| Ebf2 | 21,7362 | 12,7521 | -0,768198 | 5,00E-005 | 0,000828682 |
| Net1 | 30,0797 | 17,604 | -0,772888 | 5,00E-005 | 0,000828682 |
| Tcf15 | 116,223 | 67,5435 | -0,783004 | 5,00E-005 | 0,000828682 |
| Hdgfrp3 | 17,484 | 10,1127 | -0,789873 | 5,00E-005 | 0,000828682 |
| Mrc2 | 34,3234 | 19,8136 | -0,792702 | 5,00E-005 | 0,000828682 |
| Cer1 | 39,0247 | 22,4471 | -0,797859 | 5,00E-005 | 0,000828682 |
| Ptn | 82,827 | 47,5997 | -0,799149 | 5,00E-005 | 0,000828682 |
| Frem1 | 17,2125 | 9,88772 | -0,799745 | 5,00E-005 | 0,000828682 |
| Plce1 | 5,27545 | 3,02655 | -0,801621 | 5,00E-005 | 0,000828682 |
| Fstl1 | 151,101 | 86,5437 | -0,80401 | 5,00E-005 | 0,000828682 |
| Efnb3 | 22,8561 | 13,08 | -0,805218 | 5,00E-005 | 0,000828682 |

| | | | | | |
|---|---|---|---|---|---|
| Eno3 | 67,5424 | 37,5434 | -0,847235 | 5,00E-005 | 0,000828682 |
| Adamts1 | 7,26935 | 4,03245 | -0,850168 | 5,00E-005 | 0,000828682 |
| Pik3r1 | 9,07751 | 5,02036 | -0,854506 | 5,00E-005 | 0,000828682 |
| Rcn3 | 76,2256 | 41,9415 | -0,861895 | 5,00E-005 | 0,000828682 |
| Col1a2 | 27,3675 | 15,0427 | -0,863404 | 5,00E-005 | 0,000828682 |
| Marcks | 127,647 | 70,1051 | -0,864573 | 5,00E-005 | 0,000828682 |
| Stxbp6 | 11,4137 | 6,22746 | -0,874057 | 5,00E-005 | 0,000828682 |
| Lsp1 | 14,276 | 7,78751 | -0,874355 | 5,00E-005 | 0,000828682 |
| Pvrl1 | 6,46685 | 3,52012 | -0,877439 | 5,00E-005 | 0,000828682 |
| Colec12 | 38,6011 | 20,8893 | -0,885874 | 5,00E-005 | 0,000828682 |
| Pvrl2 | 46,611 | 25,1518 | -0,89001 | 5,00E-005 | 0,000828682 |
| Slc1a3 | 7,86643 | 4,23687 | -0,892712 | 5,00E-005 | 0,000828682 |
| Arl4c | 12,4284 | 6,60848 | -0,911255 | 5,00E-005 | 0,000828682 |
| Flrt2 | 4,85827 | 2,57476 | -0,916005 | 5,00E-005 | 0,000828682 |
| Zfp36l1 | 52,1987 | 27,652 | -0,916628 | 5,00E-005 | 0,000828682 |
| Plk2 | 22,2326 | 11,7413 | -0,921082 | 5,00E-005 | 0,000828682 |
| Uaca | 10,0618 | 5,28838 | -0,927994 | 5,00E-005 | 0,000828682 |
| Dock5 | 6,95869 | 3,65714 | -0,9281 | 5,00E-005 | 0,000828682 |
| Celsr2 | 4,55814 | 2,39213 | -0,930154 | 5,00E-005 | 0,000828682 |
| Zfhx4 | 4,29585 | 2,22898 | -0,946564 | 5,00E-005 | 0,000828682 |
| Itga8 | 8,13696 | 4,20968 | -0,95078 | 5,00E-005 | 0,000828682 |
| Zadh2 | 23,39 | 11,951 | -0,968756 | 5,00E-005 | 0,000828682 |
| Sfrp1 | 23,4083 | 11,9311 | -0,972302 | 5,00E-005 | 0,000828682 |
| Klhl13 | 12,8179 | 6,52991 | -0,973027 | 5,00E-005 | 0,000828682 |
| Pbx3 | 12,0694 | 6,1024 | -0,98391 | 5,00E-005 | 0,000828682 |
| Zfhx3 | 2,06008 | 1,03972 | -0,986511 | 5,00E-005 | 0,000828682 |
| Igfbp4 | 245,838 | 124,058 | -0,986687 | 5,00E-005 | 0,000828682 |
| Meox1 | 217,344 | 108,476 | -1,00261 | 5,00E-005 | 0,000828682 |
| Efna5 | 6,02014 | 3,00211 | -1,00382 | 5,00E-005 | 0,000828682 |
| Dusp9 | 67,2929 | 33,488 | -1,00681 | 5,00E-005 | 0,000828682 |
| Bmi1 | 18,1511 | 9,02989 | -1,00727 | 5,00E-005 | 0,000828682 |
| Utrn | 16,292 | 8,10506 | -1,00727 | 5,00E-005 | 0,000828682 |
| Parvb | 13,6528 | 6,7877 | -1,0082 | 5,00E-005 | 0,000828682 |
| Lrrn3 | 6,71897 | 3,33892 | -1,00886 | 5,00E-005 | 0,000828682 |
| Nuak2 | 8,60098 | 4,24997 | -1,01705 | 5,00E-005 | 0,000828682 |
| Tcf7l2 | 5,09735 | 2,50215 | -1,02658 | 5,00E-005 | 0,000828682 |
| Spsb4 | 51,1514 | 25,0259 | -1,03135 | 5,00E-005 | 0,000828682 |
| Dab2 | 13,7576 | 6,72769 | -1,03205 | 5,00E-005 | 0,000828682 |
| Ednra | 12,2406 | 5,96864 | -1,03619 | 5,00E-005 | 0,000828682 |
| Ptprd | 6,51339 | 3,16594 | -1,04078 | 5,00E-005 | 0,000828682 |
| Slc6a17 | 3,51564 | 1,70859 | -1,04098 | 5,00E-005 | 0,000828682 |
| Rgmb | 14,1042 | 6,8531 | -1,04129 | 5,00E-005 | 0,000828682 |
| Ier5l | 20,535 | 9,91009 | -1,05112 | 5,00E-005 | 0,000828682 |
| Robo1 | 16,3735 | 7,89336 | -1,05265 | 5,00E-005 | 0,000828682 |
| Gypc | 19,1001 | 9,19426 | -1,05477 | 5,00E-005 | 0,000828682 |
| Nrip1 | 12,7203 | 6,08977 | -1,06268 | 5,00E-005 | 0,000828682 |
| Msx1 | 23,0149 | 11,0132 | -1,06334 | 5,00E-005 | 0,000828682 |
| Wscd1 | 6,07407 | 2,9065 | -1,06338 | 5,00E-005 | 0,000828682 |
| Itga6 | 11,357 | 5,43287 | -1,0638 | 5,00E-005 | 0,000828682 |
| Alx3 | 11,4459 | 5,44505 | -1,07182 | 5,00E-005 | 0,000828682 |

| | | | | | |
|---|---|---|---|---|---|
| Tbx2 | 7,87576 | 3,7195 | -1,08231 | 5,00E-005 | 0,000828682 |
| Six1 | 36,3957 | 17,184 | -1,08271 | 5,00E-005 | 0,000828682 |
| Dtx4 | 14,4831 | 6,81862 | -1,08681 | 5,00E-005 | 0,000828682 |
| Nid1 | 10,6322 | 4,99774 | -1,08909 | 5,00E-005 | 0,000828682 |
| Met | 15,8599 | 7,40795 | -1,09823 | 5,00E-005 | 0,000828682 |
| Thsd4 | 6,70457 | 3,12002 | -1,10359 | 5,00E-005 | 0,000828682 |
| Pappa | 3,63361 | 1,6852 | -1,10848 | 5,00E-005 | 0,000828682 |
| Tubb3 | 12,2247 | 5,65671 | -1,11176 | 5,00E-005 | 0,000828682 |
| Hsd11b2 | 17,5754 | 8,12542 | -1,11305 | 5,00E-005 | 0,000828682 |
| Mpped2 | 6,94697 | 3,1547 | -1,13888 | 5,00E-005 | 0,000828682 |
| Fkbp14 | 7,97572 | 3,58824 | -1,15234 | 5,00E-005 | 0,000828682 |
| Ppap2b | 32,8378 | 14,7688 | -1,1528 | 5,00E-005 | 0,000828682 |
| Aldh1a2 | 65,9811 | 29,6524 | -1,1539 | 5,00E-005 | 0,000828682 |
| Prkce | 3,59287 | 1,61341 | -1,15503 | 5,00E-005 | 0,000828682 |
| Fbn2 | 106,47 | 47,621 | -1,16077 | 5,00E-005 | 0,000828682 |
| Dact3 | 14,7347 | 6,57962 | -1,16314 | 5,00E-005 | 0,000828682 |
| Sema3a | 8,31936 | 3,7091 | -1,1654 | 5,00E-005 | 0,000828682 |
| Alx4 | 12,4052 | 5,50374 | -1,17246 | 5,00E-005 | 0,000828682 |
| Peg10 | 126,557 | 56,0649 | -1,17462 | 5,00E-005 | 0,000828682 |
| Dusp23 | 8,92368 | 3,95288 | -1,17473 | 5,00E-005 | 0,000828682 |
| Fmn2 | 2,178 | 0,962801 | -1,17769 | 5,00E-005 | 0,000828682 |
| Syt7 | 2,04867 | 0,888778 | -1,20479 | 5,00E-005 | 0,000828682 |
| Slc38a4 | 40,956 | 17,7408 | -1,207 | 5,00E-005 | 0,000828682 |
| Tshz1 | 15,6355 | 6,74551 | -1,21283 | 5,00E-005 | 0,000828682 |
| Sec24d | 3,77382 | 1,61983 | -1,22018 | 5,00E-005 | 0,000828682 |
| Cdkn1c | 82,4703 | 35,2913 | -1,22456 | 5,00E-005 | 0,000828682 |
| Map1b | 18,9849 | 8,08249 | -1,23198 | 5,00E-005 | 0,000828682 |
| C630043F03Rik | 7,29921 | 3,08504 | -1,24245 | 5,00E-005 | 0,000828682 |
| Dclk2 | 7,5397 | 3,18603 | -1,24275 | 5,00E-005 | 0,000828682 |
| Eef2k | 8,48229 | 3,57622 | -1,24602 | 5,00E-005 | 0,000828682 |
| Cpa2 | 19,4884 | 8,19979 | -1,24896 | 5,00E-005 | 0,000828682 |
| Grrp1 | 23,076 | 9,61597 | -1,26289 | 5,00E-005 | 0,000828682 |
| Adamts12 | 3,94037 | 1,64148 | -1,26334 | 5,00E-005 | 0,000828682 |
| Gli1 | 31,522 | 13,1312 | -1,26336 | 5,00E-005 | 0,000828682 |
| Cpm | 7,58143 | 3,15386 | -1,26535 | 5,00E-005 | 0,000828682 |
| Arhgef26 | 4,77485 | 1,97901 | -1,27068 | 5,00E-005 | 0,000828682 |
| Mafb | 3,25749 | 1,34049 | -1,281 | 5,00E-005 | 0,000828682 |
| Ldb2 | 10,586 | 4,34644 | -1,28425 | 5,00E-005 | 0,000828682 |
| Ckb | 55,8409 | 22,8844 | -1,28696 | 5,00E-005 | 0,000828682 |
| Tbx3 | 13,6948 | 5,59253 | -1,29205 | 5,00E-005 | 0,000828682 |
| Col18a1 | 48,1515 | 19,5064 | -1,30363 | 5,00E-005 | 0,000828682 |
| Usp29 | 1,52484 | 0,617521 | -1,3041 | 5,00E-005 | 0,000828682 |
| Sobp | 1,92467 | 0,778917 | -1,30507 | 5,00E-005 | 0,000828682 |
| Heyl | 5,43878 | 2,19567 | -1,30862 | 5,00E-005 | 0,000828682 |
| Igdcc3 | 109,245 | 44,0549 | -1,31019 | 5,00E-005 | 0,000828682 |
| Fgfr3 | 6,98099 | 2,80494 | -1,31546 | 5,00E-005 | 0,000828682 |
| Peg3 | 182,597 | 73,0119 | -1,32246 | 5,00E-005 | 0,000828682 |
| Fzd2 | 48,5704 | 18,8783 | -1,36335 | 5,00E-005 | 0,000828682 |
| Pcdh18 | 21,8385 | 8,47998 | -1,36474 | 5,00E-005 | 0,000828682 |
| Hs6st2 | 9,43995 | 3,6496 | -1,37104 | 5,00E-005 | 0,000828682 |

| | | | | | |
|---|---|---|---|---|---|
| Ssc5d | 7,19925 | 2,77801 | -1,3738 | 5,00E-005 | 0,000828682 |
| Kctd12 | 14,129 | 5,45179 | -1,37386 | 5,00E-005 | 0,000828682 |
| Mdfi | 24,8264 | 9,55954 | -1,37686 | 5,00E-005 | 0,000828682 |
| Ntn1 | 35,8939 | 13,794 | -1,3797 | 5,00E-005 | 0,000828682 |
| Psd2 | 2,83176 | 1,08354 | -1,38594 | 5,00E-005 | 0,000828682 |
| Tns3 | 19,9083 | 7,60463 | -1,38842 | 5,00E-005 | 0,000828682 |
| Sh3bp5 | 32,3012 | 12,2705 | -1,39639 | 5,00E-005 | 0,000828682 |
| Wnt16 | 15,9623 | 6,03847 | -1,40241 | 5,00E-005 | 0,000828682 |
| Aim2 | 3,33332 | 1,24118 | -1,42525 | 5,00E-005 | 0,000828682 |
| Dmrt2 | 20,8678 | 7,76653 | -1,42593 | 5,00E-005 | 0,000828682 |
| Tns1 | 7,83866 | 2,91454 | -1,42734 | 5,00E-005 | 0,000828682 |
| Csrnp3 | 2,09232 | 0,77654 | -1,42997 | 5,00E-005 | 0,000828682 |
| Rdh10 | 17,7646 | 6,58304 | -1,43218 | 5,00E-005 | 0,000828682 |
| Nedd9 | 12,7276 | 4,71222 | -1,43348 | 5,00E-005 | 0,000828682 |
| Ildr2 | 6,18697 | 2,28613 | -1,43633 | 5,00E-005 | 0,000828682 |
| Srgap3 | 4,27537 | 1,57824 | -1,43773 | 5,00E-005 | 0,000828682 |
| Nr6a1 | 4,46975 | 1,64591 | -1,44131 | 5,00E-005 | 0,000828682 |
| Fzd4 | 6,23456 | 2,28482 | -1,44821 | 5,00E-005 | 0,000828682 |
| Pdgfrb | 29,5146 | 10,7928 | -1,45135 | 5,00E-005 | 0,000828682 |
| Id3 | 315,598 | 114,837 | -1,4585 | 5,00E-005 | 0,000828682 |
| Gli2 | 12,2548 | 4,41888 | -1,4716 | 5,00E-005 | 0,000828682 |
| Klf3 | 9,66277 | 3,47784 | -1,47425 | 5,00E-005 | 0,000828682 |
| Scube3 | 9,21529 | 3,29316 | -1,48456 | 5,00E-005 | 0,000828682 |
| Twist2 | 21,1594 | 7,55389 | -1,48601 | 5,00E-005 | 0,000828682 |
| Galnt16 | 10,565 | 3,76533 | -1,48845 | 5,00E-005 | 0,000828682 |
| Adamts15 | 5,48175 | 1,93961 | -1,49887 | 5,00E-005 | 0,000828682 |
| Plch1 | 18,418 | 6,51083 | -1,5002 | 5,00E-005 | 0,000828682 |
| Postn | 3,11195 | 1,09622 | -1,50528 | 5,00E-005 | 0,000828682 |
| Fam195a | 11,4793 | 4,04355 | -1,50534 | 5,00E-005 | 0,000828682 |
| Slc35f1 | 2,50176 | 0,879583 | -1,50805 | 5,00E-005 | 0,000828682 |
| Tgfbi | 29,4077 | 10,3329 | -1,50895 | 5,00E-005 | 0,000828682 |
| Dnm1 | 8,87086 | 3,08633 | -1,52318 | 5,00E-005 | 0,000828682 |
| Metrn | 105,764 | 36,5781 | -1,53179 | 5,00E-005 | 0,000828682 |
| Ets1 | 27,0631 | 9,31106 | -1,53931 | 5,00E-005 | 0,000828682 |
| Angpt1 | 6,35549 | 2,17138 | -1,54939 | 5,00E-005 | 0,000828682 |
| Npr3 | 1,04735 | 0,357666 | -1,55006 | 5,00E-005 | 0,000828682 |
| Meis2 | 63,9354 | 21,7426 | -1,55609 | 5,00E-005 | 0,000828682 |
| Apln | 3,45786 | 1,1759 | -1,55611 | 5,00E-005 | 0,000828682 |
| Pcolce | 56,7755 | 19,297 | -1,55689 | 5,00E-005 | 0,000828682 |
| Gpm6b | 3,38585 | 1,14658 | -1,56218 | 5,00E-005 | 0,000828682 |
| Ncald | 2,71634 | 0,915598 | -1,56888 | 5,00E-005 | 0,000828682 |
| Elavl4 | 3,52474 | 1,18742 | -1,56969 | 5,00E-005 | 0,000828682 |
| Fam181b | 38,9405 | 13,0955 | -1,5722 | 5,00E-005 | 0,000828682 |
| Col23a1 | 21,6816 | 7,27989 | -1,57448 | 5,00E-005 | 0,000828682 |
| Col14a1 | 1,80134 | 0,601974 | -1,5813 | 5,00E-005 | 0,000828682 |
| Tgfb2 | 6,45166 | 2,14165 | -1,59095 | 5,00E-005 | 0,000828682 |
| Fam198b | 1,52199 | 0,50412 | -1,59412 | 5,00E-005 | 0,000828682 |
| Gas1 | 65,1685 | 21,5044 | -1,59954 | 5,00E-005 | 0,000828682 |
| Col2a1 | 70,2208 | 22,9593 | -1,61282 | 5,00E-005 | 0,000828682 |
| Epha7 | 9,24421 | 3,00855 | -1,61948 | 5,00E-005 | 0,000828682 |

| | | | | | |
|---|---|---|---|---|---|
| Gdf10 | 10,5222 | 3,42398 | -1,61969 | 5,00E-005 | 0,000828682 |
| Serpinf1 | 40,6943 | 13,21 | -1,62319 | 5,00E-005 | 0,000828682 |
| 1700011H14Rik | 7,33095 | 2,37477 | -1,62621 | 5,00E-005 | 0,000828682 |
| Pgm5 | 8,6369 | 2,78191 | -1,63444 | 5,00E-005 | 0,000828682 |
| Zfp9 | 4,52228 | 1,45299 | -1,63803 | 5,00E-005 | 0,000828682 |
| Bmp4 | 24,0962 | 7,73437 | -1,63945 | 5,00E-005 | 0,000828682 |
| Arid5b | 1,61987 | 0,519077 | -1,64186 | 5,00E-005 | 0,000828682 |
| Tppp3 | 16,9038 | 5,41291 | -1,64287 | 5,00E-005 | 0,000828682 |
| Tox2 | 7,07119 | 2,24714 | -1,65386 | 5,00E-005 | 0,000828682 |
| Fgfbp3 | 7,63029 | 2,4124 | -1,66127 | 5,00E-005 | 0,000828682 |
| Kdr | 4,46238 | 1,40443 | -1,66783 | 5,00E-005 | 0,000828682 |
| Bgn | 8,23849 | 2,58961 | -1,66965 | 5,00E-005 | 0,000828682 |
| Tgfbr3 | 2,53095 | 0,790366 | -1,67909 | 5,00E-005 | 0,000828682 |
| Cdo1 | 9,23328 | 2,88209 | -1,67973 | 5,00E-005 | 0,000828682 |
| Cdc42ep1 | 14,9771 | 4,66603 | -1,68249 | 5,00E-005 | 0,000828682 |
| Fmod | 6,92001 | 2,12701 | -1,70195 | 5,00E-005 | 0,000828682 |
| Olig3 | 8,88869 | 2,71476 | -1,71114 | 5,00E-005 | 0,000828682 |
| Sox21 | 2,84585 | 0,868236 | -1,7127 | 5,00E-005 | 0,000828682 |
| Cd248 | 13,2183 | 3,99986 | -1,72451 | 5,00E-005 | 0,000828682 |
| Meis1 | 37,0892 | 11,2033 | -1,72708 | 5,00E-005 | 0,000828682 |
| Slit1 | 3,41791 | 1,02767 | -1,73373 | 5,00E-005 | 0,000828682 |
| Scube1 | 17,4373 | 5,2397 | -1,73462 | 5,00E-005 | 0,000828682 |
| Wfikkn1 | 11,1797 | 3,30534 | -1,75801 | 5,00E-005 | 0,000828682 |
| Slc7a11 | 0,977182 | 0,288417 | -1,76047 | 5,00E-005 | 0,000828682 |
| A730017C20Rik | 3,20779 | 0,938063 | -1,77382 | 5,00E-005 | 0,000828682 |
| Id2 | 201,822 | 58,2761 | -1,79211 | 5,00E-005 | 0,000828682 |
| Sepp1 | 54,4947 | 15,719 | -1,79361 | 5,00E-005 | 0,000828682 |
| Lmcd1 | 3,75781 | 1,08356 | -1,79412 | 5,00E-005 | 0,000828682 |
| Col1a1 | 12,0853 | 3,48372 | -1,79455 | 5,00E-005 | 0,000828682 |
| Fzd8 | 2,55415 | 0,734345 | -1,79832 | 5,00E-005 | 0,000828682 |
| Igfbp5 | 55,9054 | 15,9144 | -1,81266 | 5,00E-005 | 0,000828682 |
| Ahnak | 0,938837 | 0,265293 | -1,82329 | 5,00E-005 | 0,000828682 |
| Nrtn | 6,78602 | 1,91234 | -1,82723 | 5,00E-005 | 0,000828682 |
| Sox6 | 3,45152 | 0,971099 | -1,82954 | 5,00E-005 | 0,000828682 |
| Synpo | 1,55031 | 0,433216 | -1,8394 | 5,00E-005 | 0,000828682 |
| Itga9 | 10,5489 | 2,89821 | -1,86385 | 5,00E-005 | 0,000828682 |
| Col26a1 | 63,7254 | 17,0726 | -1,90018 | 5,00E-005 | 0,000828682 |
| Leprel1 | 7,83542 | 2,09333 | -1,90421 | 5,00E-005 | 0,000828682 |
| Cthrc1 | 11,4173 | 3,03269 | -1,91255 | 5,00E-005 | 0,000828682 |
| Acss1 | 2,69791 | 0,715019 | -1,91579 | 5,00E-005 | 0,000828682 |
| Pdgfc | 9,33817 | 2,46161 | -1,92354 | 5,00E-005 | 0,000828682 |
| Npnt | 4,04907 | 1,06728 | -1,92365 | 5,00E-005 | 0,000828682 |
| Sox5 | 4,74074 | 1,24779 | -1,92574 | 5,00E-005 | 0,000828682 |
| Magel2 | 5,17743 | 1,35846 | -1,93026 | 5,00E-005 | 0,000828682 |
| Miat | 20,042 | 5,25202 | -1,93208 | 5,00E-005 | 0,000828682 |
| Acvrl1 | 3,99054 | 1,03567 | -1,94602 | 5,00E-005 | 0,000828682 |
| Mecom | 4,70026 | 1,2087 | -1,95928 | 5,00E-005 | 0,000828682 |
| Sox1 | 6,33335 | 1,62821 | -1,95968 | 5,00E-005 | 0,000828682 |
| Nova2 | 1,18546 | 0,303545 | -1,96547 | 5,00E-005 | 0,000828682 |
| Cyp1b1 | 1,588 | 0,403937 | -1,97501 | 5,00E-005 | 0,000828682 |

| | | | | | |
|---|---|---|---|---|---|
| Figf | 3,78373 | 0,951708 | -1,99122 | 5,00E-005 | 0,000828682 |
| Camk2a | 2,15872 | 0,542274 | -1,99308 | 5,00E-005 | 0,000828682 |
| Enpp2 | 3,27937 | 0,815477 | -2,0077 | 5,00E-005 | 0,000828682 |
| Cped1 | 9,46784 | 2,34221 | -2,01516 | 5,00E-005 | 0,000828682 |
| Bmp5 | 6,42536 | 1,55886 | -2,04328 | 5,00E-005 | 0,000828682 |
| Dmd | 1,37003 | 0,330046 | -2,05346 | 5,00E-005 | 0,000828682 |
| Slc38a3 | 8,93255 | 2,15025 | -2,05457 | 5,00E-005 | 0,000828682 |
| Clmp | 18,3447 | 4,39989 | -2,05983 | 5,00E-005 | 0,000828682 |
| Ebf1 | 4,12401 | 0,986672 | -2,0634 | 5,00E-005 | 0,000828682 |
| Sertad4 | 6,14249 | 1,4581 | -2,07474 | 5,00E-005 | 0,000828682 |
| Fgfr2 | 31,2229 | 7,41039 | -2,07498 | 5,00E-005 | 0,000828682 |
| Cxcl12 | 80,7962 | 19,1041 | -2,08041 | 5,00E-005 | 0,000828682 |
| Ppfibp2 | 9,87476 | 2,32436 | -2,08691 | 5,00E-005 | 0,000828682 |
| Bmp3 | 2,66302 | 0,62321 | -2,09527 | 5,00E-005 | 0,000828682 |
| Brsk2 | 1,95675 | 0,457673 | -2,09607 | 5,00E-005 | 0,000828682 |
| Des | 8,17886 | 1,89159 | -2,1123 | 5,00E-005 | 0,000828682 |
| Chl1 | 2,6243 | 0,597472 | -2,13499 | 5,00E-005 | 0,000828682 |
| Twist1 | 99,2084 | 22,4955 | -2,14082 | 5,00E-005 | 0,000828682 |
| Mest | 1013,06 | 224,529 | -2,17375 | 5,00E-005 | 0,000828682 |
| Arg1 | 73,1445 | 15,9985 | -2,19281 | 5,00E-005 | 0,000828682 |
| Col9a1 | 28,8113 | 6,27988 | -2,19782 | 5,00E-005 | 0,000828682 |
| Ccnd2 | 130,836 | 27,8032 | -2,23443 | 5,00E-005 | 0,000828682 |
| Tmem200b | 3,69015 | 0,779131 | -2,24374 | 5,00E-005 | 0,000828682 |
| Nrp1 | 20,6141 | 4,09791 | -2,33067 | 5,00E-005 | 0,000828682 |
| Mir6236 | 9956,69 | 1944,74 | -2,35608 | 5,00E-005 | 0,000828682 |
| Prss23 | 3,07226 | 0,59852 | -2,35983 | 5,00E-005 | 0,000828682 |
| L1cam | 0,931176 | 0,177427 | -2,39183 | 5,00E-005 | 0,000828682 |
| Crnde | 10,5538 | 1,99593 | -2,40262 | 5,00E-005 | 0,000828682 |
| Adcy4 | 1,70077 | 0,319223 | -2,41356 | 5,00E-005 | 0,000828682 |
| Lrrc32 | 2,06807 | 0,385988 | -2,42166 | 5,00E-005 | 0,000828682 |
| Foxp4 | 40,3115 | 7,47299 | -2,43143 | 5,00E-005 | 0,000828682 |
| Ptgfr | 2,53442 | 0,463862 | -2,44989 | 5,00E-005 | 0,000828682 |
| Eya2 | 14,4 | 2,60723 | -2,46548 | 5,00E-005 | 0,000828682 |
| Snai2 | 25,5791 | 4,62927 | -2,46611 | 5,00E-005 | 0,000828682 |
| Zim1 | 9,08646 | 1,60077 | -2,50496 | 5,00E-005 | 0,000828682 |
| Plxnc1 | 1,58874 | 0,278721 | -2,51099 | 5,00E-005 | 0,000828682 |
| Fli1 | 12,4298 | 2,17826 | -2,51256 | 5,00E-005 | 0,000828682 |
| Ccdc160 | 6,78928 | 1,18765 | -2,51515 | 5,00E-005 | 0,000828682 |
| Prmt8 | 2,72339 | 0,473274 | -2,52466 | 5,00E-005 | 0,000828682 |
| Tnfaip6 | 6,1788 | 1,06352 | -2,53848 | 5,00E-005 | 0,000828682 |
| Sorcs2 | 2,47861 | 0,422238 | -2,5534 | 5,00E-005 | 0,000828682 |
| Plat | 13,4423 | 2,24202 | -2,58391 | 5,00E-005 | 0,000828682 |
| Foxp2 | 2,7237 | 0,451319 | -2,59335 | 5,00E-005 | 0,000828682 |
| Foxf2 | 6,55912 | 1,07879 | -2,60409 | 5,00E-005 | 0,000828682 |
| Hhip | 2,39567 | 0,387292 | -2,62894 | 5,00E-005 | 0,000828682 |
| Ptchd2 | 0,630625 | 0,101504 | -2,63525 | 5,00E-005 | 0,000828682 |
| Rab3il1 | 13,4471 | 2,11304 | -2,66991 | 5,00E-005 | 0,000828682 |
| Wnt4 | 2,9998 | 0,468852 | -2,67766 | 5,00E-005 | 0,000828682 |
| Skida1 | 3,43007 | 0,531642 | -2,68971 | 5,00E-005 | 0,000828682 |
| Nell2 | 9,12849 | 1,38674 | -2,71868 | 5,00E-005 | 0,000828682 |

| | | | | | |
|---|---|---|---|---|---|
| Thbs1 | 3,0964 | 0,464878 | -2,73567 | 5,00E-005 | 0,000828682 |
| Hdac9 | 1,06849 | 0,160407 | -2,73576 | 5,00E-005 | 0,000828682 |
| Scube2 | 9,35714 | 1,40148 | -2,73911 | 5,00E-005 | 0,000828682 |
| Rubie | 4,55814 | 0,680756 | -2,74324 | 5,00E-005 | 0,000828682 |
| Nr2f2 | 20,763 | 3,05607 | -2,76427 | 5,00E-005 | 0,000828682 |
| Msx3 | 8,76466 | 1,27424 | -2,78206 | 5,00E-005 | 0,000828682 |
| Cd36 | 13,428 | 1,94582 | -2,78679 | 5,00E-005 | 0,000828682 |
| Col6a3 | 0,760419 | 0,109511 | -2,79572 | 5,00E-005 | 0,000828682 |
| Srrm4 | 0,8431 | 0,118846 | -2,82661 | 5,00E-005 | 0,000828682 |
| Prrx2 | 65,6949 | 9,25207 | -2,82793 | 5,00E-005 | 0,000828682 |
| Nrcam | 5,63661 | 0,791396 | -2,83236 | 5,00E-005 | 0,000828682 |
| Eya1 | 21,3335 | 2,95876 | -2,85005 | 5,00E-005 | 0,000828682 |
| Tmem30b | 9,04254 | 1,23894 | -2,86763 | 5,00E-005 | 0,000828682 |
| Sox9 | 16,2868 | 2,23048 | -2,86828 | 5,00E-005 | 0,000828682 |
| Irx3 | 20,1442 | 2,75369 | -2,87093 | 5,00E-005 | 0,000828682 |
| Aff3 | 9,88962 | 1,33544 | -2,8886 | 5,00E-005 | 0,000828682 |
| Prrx1 | 51,5126 | 6,89605 | -2,90108 | 5,00E-005 | 0,000828682 |
| Fam181a | 11,5438 | 1,53004 | -2,91547 | 5,00E-005 | 0,000828682 |
| Slc40a1 | 1,75839 | 0,231876 | -2,92283 | 5,00E-005 | 0,000828682 |
| Runx1t1 | 1,67381 | 0,218975 | -2,9343 | 5,00E-005 | 0,000828682 |
| Rfx4 | 6,54186 | 0,848429 | -2,94684 | 5,00E-005 | 0,000828682 |
| Parm1 | 13,985 | 1,76665 | -2,98479 | 5,00E-005 | 0,000828682 |
| Asb4 | 15,5281 | 1,95388 | -2,99047 | 5,00E-005 | 0,000828682 |
| Irx5 | 6,67245 | 0,834228 | -2,9997 | 5,00E-005 | 0,000828682 |
| Pag1 | 3,35877 | 0,411957 | -3,02737 | 5,00E-005 | 0,000828682 |
| Spon1 | 5,9806 | 0,721875 | -3,05047 | 5,00E-005 | 0,000828682 |
| Uncx | 17,0554 | 2,04962 | -3,0568 | 5,00E-005 | 0,000828682 |
| Pgf | 10,6569 | 1,27241 | -3,06615 | 5,00E-005 | 0,000828682 |
| Pou3f2 | 4,5608 | 0,539567 | -3,07941 | 5,00E-005 | 0,000828682 |
| Crym | 5,40509 | 0,63577 | -3,08774 | 5,00E-005 | 0,000828682 |
| Gpr124 | 15,575 | 1,82637 | -3,09218 | 5,00E-005 | 0,000828682 |
| Dpep1 | 3,53244 | 0,411316 | -3,10235 | 5,00E-005 | 0,000828682 |
| Abca9 | 2,18935 | 0,24395 | -3,16584 | 5,00E-005 | 0,000828682 |
| Grem1 | 4,05486 | 0,450633 | -3,16963 | 5,00E-005 | 0,000828682 |
| Ptx3 | 15,3748 | 1,70651 | -3,17145 | 5,00E-005 | 0,000828682 |
| Sncaip | 10,2787 | 1,13688 | -3,1765 | 5,00E-005 | 0,000828682 |
| Gdnf | 2,14903 | 0,237651 | -3,17677 | 5,00E-005 | 0,000828682 |
| Neurog2 | 8,28381 | 0,911309 | -3,18428 | 5,00E-005 | 0,000828682 |
| Epha3 | 2,0137 | 0,221101 | -3,18707 | 5,00E-005 | 0,000828682 |
| Ptges | 3,20565 | 0,349045 | -3,19913 | 5,00E-005 | 0,000828682 |
| Ndnf | 7,35709 | 0,797726 | -3,20517 | 5,00E-005 | 0,000828682 |
| Unc5c | 2,35894 | 0,251217 | -3,23113 | 5,00E-005 | 0,000828682 |
| Meox2 | 14,3184 | 1,47921 | -3,27497 | 5,00E-005 | 0,000828682 |
| Fez1 | 21,7195 | 2,23797 | -3,27873 | 5,00E-005 | 0,000828682 |
| Fst | 10,4971 | 1,05958 | -3,30843 | 5,00E-005 | 0,000828682 |
| Tnc | 2,53607 | 0,249943 | -3,34292 | 5,00E-005 | 0,000828682 |
| Dnm3os | 1,05404 | 0,103726 | -3,34509 | 5,00E-005 | 0,000828682 |
| Tbx4 | 2,01543 | 0,188227 | -3,42054 | 5,00E-005 | 0,000828682 |
| Rhbdl3 | 2,10158 | 0,192134 | -3,45129 | 5,00E-005 | 0,000828682 |
| Foxd3 | 4,46499 | 0,362118 | -3,62412 | 5,00E-005 | 0,000828682 |

| | | | | | |
|---|---|---|---|---|---|
| Ebf3 | 5,82196 | 0,389886 | -3,90038 | 5,00E-005 | 0,000828682 |
| Rarb | 4,91129 | 0,327096 | -3,90832 | 5,00E-005 | 0,000828682 |
| Tbx18 | 3,87715 | 0,258 | -3,90956 | 5,00E-005 | 0,000828682 |
| Nkx6-1 | 9,39715 | 0,611695 | -3,94134 | 5,00E-005 | 0,000828682 |
| Cldn1 | 4,26326 | 0,277232 | -3,94279 | 5,00E-005 | 0,000828682 |
| Osr1 | 8,6757 | 0,547286 | -3,98661 | 5,00E-005 | 0,000828682 |
| Robo2 | 8,21496 | 0,51582 | -3,99331 | 5,00E-005 | 0,000828682 |
| Moxd1 | 5,92445 | 0,360523 | -4,03852 | 5,00E-005 | 0,000828682 |
| Igf1 | 15,156 | 0,884529 | -4,09884 | 5,00E-005 | 0,000828682 |
| Ednrb | 2,83377 | 0,159683 | -4,14944 | 5,00E-005 | 0,000828682 |
| Nr2f1 | 19,1311 | 1,04764 | -4,19071 | 5,00E-005 | 0,000828682 |
| Nkx3-1 | 28,7892 | 1,5725 | -4,1944 | 5,00E-005 | 0,000828682 |
| Pax6 | 17,2448 | 0,93305 | -4,20806 | 5,00E-005 | 0,000828682 |
| Ddr2 | 4,09697 | 0,211303 | -4,27717 | 5,00E-005 | 0,000828682 |
| Itih5 | 6,89553 | 0,337756 | -4,35161 | 5,00E-005 | 0,000828682 |
| Gfra1 | 2,37772 | 0,10128 | -4,55316 | 5,00E-005 | 0,000828682 |
| Isl1 | 18,1031 | 0,770939 | -4,55348 | 5,00E-005 | 0,000828682 |
| Foxd2 | 9,31207 | 0,368923 | -4,65771 | 5,00E-005 | 0,000828682 |
| Foxd2os | 7,28562 | 0,279762 | -4,70278 | 5,00E-005 | 0,000828682 |
| Pkdcc | 12,5852 | 0,4478 | -4,81273 | 5,00E-005 | 0,000828682 |
| Mybpc1 | 4,8215 | 0,155228 | -4,95702 | 5,00E-005 | 0,000828682 |
| Col3a1 | 50,0665 | 1,39277 | -5,16781 | 5,00E-005 | 0,000828682 |
| Olig2 | 7,02792 | 0,191407 | -5,19838 | 5,00E-005 | 0,000828682 |
| Zic1 | 7,3623 | 0,170876 | -5,42913 | 5,00E-005 | 0,000828682 |
| Pax1 | 26,8099 | 0,452744 | -5,88793 | 5,00E-005 | 0,000828682 |
| Dmgdh | 0,201392 | 2,63974 | 3,71232 | 0,0001 | 0,00156822 |
| Dnajc22 | 1,70538 | 4,87339 | 1,51483 | 0,0001 | 0,00156822 |
| Chodl | 1,19217 | 3,09045 | 1,37423 | 0,0001 | 0,00156822 |
| Art5 | 4,0018 | 9,01854 | 1,17225 | 0,0001 | 0,00156822 |
| Cyp27a1 | 4,5437 | 9,67289 | 1,09008 | 0,0001 | 0,00156822 |
| Piezo1 | 6,50555 | 13,3365 | 1,03564 | 0,0001 | 0,00156822 |
| 2010111I01Rik | 35,6056 | 70,1288 | 0,977902 | 0,0001 | 0,00156822 |
| Ptar1 | 7,74267 | 15,1289 | 0,966403 | 0,0001 | 0,00156822 |
| Dsc2 | 3,02164 | 5,71707 | 0,919945 | 0,0001 | 0,00156822 |
| Rab15 | 7,58888 | 13,8802 | 0,871071 | 0,0001 | 0,00156822 |
| Prickle2 | 1,83885 | 3,35108 | 0,865825 | 0,0001 | 0,00156822 |
| Sorbs1 | 3,64542 | 6,48988 | 0,832105 | 0,0001 | 0,00156822 |
| Kcnq1ot1 | 0,454556 | 0,805938 | 0,826211 | 0,0001 | 0,00156822 |
| Etl4 | 6,23225 | 10,9718 | 0,815981 | 0,0001 | 0,00156822 |
| Zfp518a | 5,31475 | 9,31844 | 0,810088 | 0,0001 | 0,00156822 |
| Dync2h1 | 4,53082 | 7,82913 | 0,78908 | 0,0001 | 0,00156822 |
| Skil | 9,67077 | 16,6684 | 0,78541 | 0,0001 | 0,00156822 |
| Ddx26b | 15,5313 | 26,5478 | 0,773411 | 0,0001 | 0,00156822 |
| Lrp4 | 8,09533 | 13,8327 | 0,772919 | 0,0001 | 0,00156822 |
| Tmem50b | 15,1325 | 25,6993 | 0,764081 | 0,0001 | 0,00156822 |
| Lpp | 5,678 | 9,63671 | 0,763157 | 0,0001 | 0,00156822 |
| Arhgap5 | 19,6424 | 33,1672 | 0,75579 | 0,0001 | 0,00156822 |
| Mllt6 | 17,0872 | 28,1096 | 0,718149 | 0,0001 | 0,00156822 |
| Irs1 | 6,9071 | 11,2486 | 0,703598 | 0,0001 | 0,00156822 |
| Dlg1 | 37,2246 | 58,9274 | 0,662685 | 0,0001 | 0,00156822 |

| | | | | | |
|---|---|---|---|---|---|
| Ubn2 | 8,25234 | 12,8683 | 0,640951 | 0,0001 | 0,00156822 |
| Zfp516 | 27,2573 | 17,5851 | -0,683094 | 0,0001 | 0,00156822 |
| Cadm1 | 61,8823 | 38,0614 | -0,701197 | 0,0001 | 0,00156822 |
| Pbx1 | 35,0699 | 21,379 | -0,714037 | 0,0001 | 0,00156822 |
| Cdh11 | 75,1851 | 45,5644 | -0,722539 | 0,0001 | 0,00156822 |
| Timp3 | 22,0619 | 13,0185 | -0,760995 | 0,0001 | 0,00156822 |
| Sfrp2 | 59,1349 | 33,3439 | -0,826585 | 0,0001 | 0,00156822 |
| Klhl29 | 3,63437 | 2,02386 | -0,844599 | 0,0001 | 0,00156822 |
| Lgals1 | 66,318 | 36,6056 | -0,857336 | 0,0001 | 0,00156822 |
| Tle1 | 22,1321 | 11,2971 | -0,970193 | 0,0001 | 0,00156822 |
| Ramp2 | 27,4628 | 13,5986 | -1,01402 | 0,0001 | 0,00156822 |
| Pard3b | 1,92647 | 0,844109 | -1,19046 | 0,0001 | 0,00156822 |
| Dact2 | 4,43764 | 1,75826 | -1,33564 | 0,0001 | 0,00156822 |
| Plcd1 | 3,99852 | 1,54165 | -1,37499 | 0,0001 | 0,00156822 |
| Col9a2 | 3,25797 | 1,20976 | -1,42925 | 0,0001 | 0,00156822 |
| Ptprn2 | 1,2117 | 0,379843 | -1,67356 | 0,0001 | 0,00156822 |
| Scrn1 | 3,77192 | 1,07208 | -1,81488 | 0,0001 | 0,00156822 |
| Six2 | 3,08869 | 0,836682 | -1,88424 | 0,0001 | 0,00156822 |
| Cldn11 | 3,09514 | 0,375462 | -3,04326 | 0,0001 | 0,00156822 |
| Gzmk | 0,634708 | 10,6106 | 4,06327 | 0,00015 | 0,0022611 |
| Svopl | 0,387869 | 1,92349 | 2,31009 | 0,00015 | 0,0022611 |
| Oxtr | 0,185864 | 0,919659 | 2,30686 | 0,00015 | 0,0022611 |
| Ifitm5 | 2,55195 | 11,5011 | 2,1721 | 0,00015 | 0,0022611 |
| Zscan10 | 0,632693 | 2,04873 | 1,69515 | 0,00015 | 0,0022611 |
| Gxylt2 | 1,12733 | 3,63773 | 1,69013 | 0,00015 | 0,0022611 |
| Gatsl3 | 1,96698 | 5,5164 | 1,48774 | 0,00015 | 0,0022611 |
| Cldn6 | 3,91847 | 9,1295 | 1,22025 | 0,00015 | 0,0022611 |
| Pde10a | 1,83006 | 3,23301 | 0,820987 | 0,00015 | 0,0022611 |
| Adamts10 | 11,5408 | 20,0345 | 0,795745 | 0,00015 | 0,0022611 |
| Phip | 14,5577 | 24,8179 | 0,769601 | 0,00015 | 0,0022611 |
| Rock2 | 16,1232 | 27,194 | 0,754153 | 0,00015 | 0,0022611 |
| Apcdd1 | 13,9758 | 23,3661 | 0,741482 | 0,00015 | 0,0022611 |
| Plekhg3 | 7,96501 | 13,2905 | 0,73865 | 0,00015 | 0,0022611 |
| Rap2c | 31,5628 | 51,4399 | 0,704661 | 0,00015 | 0,0022611 |
| Ankrd50 | 8,74842 | 14,1779 | 0,696553 | 0,00015 | 0,0022611 |
| Maml3 | 10,7802 | 17,425 | 0,692775 | 0,00015 | 0,0022611 |
| Mfap2 | 115,373 | 71,249 | -0,695368 | 0,00015 | 0,0022611 |
| Prtg | 11,639 | 6,8793 | -0,758633 | 0,00015 | 0,0022611 |
| Sox2 | 33,1559 | 19,3327 | -0,778223 | 0,00015 | 0,0022611 |
| Deptor | 3,96112 | 2,27098 | -0,802595 | 0,00015 | 0,0022611 |
| Rassf2 | 5,65198 | 3,14499 | -0,845701 | 0,00015 | 0,0022611 |
| Plekhg1 | 3,20876 | 1,76519 | -0,862193 | 0,00015 | 0,0022611 |
| Crip2 | 14,1044 | 7,08061 | -0,9942 | 0,00015 | 0,0022611 |
| Kcne1l | 12,1683 | 5,99678 | -1,02087 | 0,00015 | 0,0022611 |
| Gadd45a | 20,2586 | 9,61273 | -1,07552 | 0,00015 | 0,0022611 |
| Hes5 | 8,39297 | 3,26616 | -1,36159 | 0,00015 | 0,0022611 |
| Plk3 | 3,04223 | 1,15055 | -1,4028 | 0,00015 | 0,0022611 |
| Pou3f3 | 2,22578 | 0,57677 | -1,94824 | 0,00015 | 0,0022611 |
| Gata3 | 1,39278 | 0,309082 | -2,1719 | 0,00015 | 0,0022611 |
| Tmem100 | 2,70513 | 0,572264 | -2,24094 | 0,00015 | 0,0022611 |

| | | | | | |
|---|---|---|---|---|---|
| 6430573F11Rik | 2,89997 | 0,567278 | -2,35391 | 0,00015 | 0,0022611 |
| Prdm8 | 2,86536 | 0,126012 | -4,50708 | 0,00015 | 0,0022611 |
| Gpr83 | 0,263564 | 2,1701 | 3,04154 | 0,0002 | 0,00294897 |
| Rab19 | 0,75688 | 3,72777 | 2,30018 | 0,0002 | 0,00294897 |
| H2-Q10 | 0,753056 | 3,22659 | 2,09918 | 0,0002 | 0,00294897 |
| Arc | 0,441513 | 1,8882 | 2,09649 | 0,0002 | 0,00294897 |
| Alcam | 0,423914 | 1,31739 | 1,63584 | 0,0002 | 0,00294897 |
| Rora | 1,89094 | 5,087 | 1,42771 | 0,0002 | 0,00294897 |
| Stambpl1 | 1,81221 | 4,31337 | 1,25106 | 0,0002 | 0,00294897 |
| Ptpn4 | 3,19893 | 5,84177 | 0,868817 | 0,0002 | 0,00294897 |
| Egflam | 3,6745 | 6,5645 | 0,837139 | 0,0002 | 0,00294897 |
| Akna | 3,45377 | 6,05103 | 0,809011 | 0,0002 | 0,00294897 |
| Man1c1 | 4,08411 | 7,12394 | 0,802653 | 0,0002 | 0,00294897 |
| Vcan | 60,0973 | 102,41 | 0,76899 | 0,0002 | 0,00294897 |
| Tubb6 | 36,6133 | 60,1491 | 0,716175 | 0,0002 | 0,00294897 |
| Cacna2d1 | 7,22261 | 11,7104 | 0,697201 | 0,0002 | 0,00294897 |
| Setx | 10,0021 | 16,2119 | 0,696743 | 0,0002 | 0,00294897 |
| Lars2 | 97,0621 | 54,6573 | -0,828495 | 0,0002 | 0,00294897 |
| Sh2d3c | 3,55092 | 1,43807 | -1,30406 | 0,0002 | 0,00294897 |
| Elavl3 | 1,5446 | 0,57221 | -1,43262 | 0,0002 | 0,00294897 |
| Tmem132e | 1,50585 | 0,101496 | -3,89109 | 0,0002 | 0,00294897 |
| Pik3cd | 0,916568 | 2,04963 | 1,16105 | 0,00025 | 0,00361964 |
| Epcam | 2,68571 | 5,88907 | 1,13274 | 0,00025 | 0,00361964 |
| Shb | 6,83324 | 11,7248 | 0,778927 | 0,00025 | 0,00361964 |
| Myadm | 11,6319 | 19,6671 | 0,757704 | 0,00025 | 0,00361964 |
| Nhsl1 | 6,07895 | 10,1685 | 0,742215 | 0,00025 | 0,00361964 |
| Nipbl | 13,5482 | 22,2519 | 0,71583 | 0,00025 | 0,00361964 |
| Hoxd8 | 25,5331 | 41,8087 | 0,711435 | 0,00025 | 0,00361964 |
| Trip10 | 22,6638 | 36,9978 | 0,707051 | 0,00025 | 0,00361964 |
| Arfgef1 | 15,6811 | 25,2696 | 0,68837 | 0,00025 | 0,00361964 |
| Insr | 8,6881 | 13,9415 | 0,682278 | 0,00025 | 0,00361964 |
| S1pr2 | 23,1481 | 13,9538 | -0,730234 | 0,00025 | 0,00361964 |
| Itm2a | 10,5399 | 5,21119 | -1,01618 | 0,00025 | 0,00361964 |
| Fbxl16 | 2,2507 | 0,952078 | -1,24122 | 0,00025 | 0,00361964 |
| Kctd12b | 1,19928 | 0,428795 | -1,48381 | 0,00025 | 0,00361964 |
| Phactr3 | 0,7764 | 0,214219 | -1,85771 | 0,00025 | 0,00361964 |
| Btbd17 | 2,83292 | 0,265465 | -3,4157 | 0,00025 | 0,00361964 |
| Fam163b | 0,12254 | 2,70664 | 4,46518 | 0,0003 | 0,00423833 |
| Arhgap22 | 0,219134 | 1,85766 | 3,0836 | 0,0003 | 0,00423833 |
| Wdr65 | 0,203311 | 0,972998 | 2,25875 | 0,0003 | 0,00423833 |
| Bzrap1 | 0,259715 | 0,760407 | 1,54985 | 0,0003 | 0,00423833 |
| Evpl | 0,398395 | 1,1002 | 1,46549 | 0,0003 | 0,00423833 |
| Tmem63a | 0,826641 | 2,18006 | 1,39903 | 0,0003 | 0,00423833 |
| Mgst1 | 10,4155 | 20,3509 | 0,966367 | 0,0003 | 0,00423833 |
| Nyap1 | 2,99682 | 5,55897 | 0,891387 | 0,0003 | 0,00423833 |
| Dsp | 4,50985 | 7,55437 | 0,744231 | 0,0003 | 0,00423833 |
| Hoxd1 | 15,3095 | 25,5155 | 0,736949 | 0,0003 | 0,00423833 |
| Atrx | 23,548 | 38,6042 | 0,713153 | 0,0003 | 0,00423833 |
| Rian | 59,3679 | 95,4434 | 0,684963 | 0,0003 | 0,00423833 |
| Brca2 | 6,07795 | 9,70727 | 0,675479 | 0,0003 | 0,00423833 |

| | | | | | |
|---|---|---|---|---|---|
| Plxna2 | 10,773 | 16,9062 | 0,650138 | 0,0003 | 0,00423833 |
| Tgif1 | 50,8806 | 79,0583 | 0,635803 | 0,0003 | 0,00423833 |
| Cmtm3 | 32,7649 | 19,5031 | -0,748446 | 0,0003 | 0,00423833 |
| Phlda1 | 20,8209 | 11,9897 | -0,796241 | 0,0003 | 0,00423833 |
| Myocd | 2,15274 | 1,03215 | -1,06052 | 0,0003 | 0,00423833 |
| Nrg1 | 2,60368 | 0,685951 | -1,92438 | 0,0003 | 0,00423833 |
| Esrrg | 0,728973 | 0,120549 | -2,59625 | 0,0003 | 0,00423833 |
| Gm16551 | 1,71731 | 0,217344 | -2,98209 | 0,0003 | 0,00423833 |
| Nkx2-9 | 6,09702 | 0,380783 | -4,00106 | 0,0003 | 0,00423833 |
| Gpr133 | 0,168812 | 0,765823 | 2,1816 | 0,00035 | 0,00483294 |
| Sost | 1,18169 | 3,23244 | 1,45177 | 0,00035 | 0,00483294 |
| 9330159F19Rik | 0,673521 | 1,45657 | 1,11278 | 0,00035 | 0,00483294 |
| Tnfrsf1b | 1,06257 | 2,27877 | 1,10069 | 0,00035 | 0,00483294 |
| Meg3 | 49,6179 | 88,3556 | 0,832461 | 0,00035 | 0,00483294 |
| Ddhd1 | 3,52614 | 6,01878 | 0,771381 | 0,00035 | 0,00483294 |
| Plag1 | 4,59264 | 7,80612 | 0,765282 | 0,00035 | 0,00483294 |
| Ifngr2 | 12,373 | 20,9392 | 0,759012 | 0,00035 | 0,00483294 |
| Sh3bp4 | 19,1446 | 30,5425 | 0,673884 | 0,00035 | 0,00483294 |
| Wwp1 | 9,54639 | 15,2077 | 0,671779 | 0,00035 | 0,00483294 |
| Gpr125 | 76,7685 | 122,087 | 0,669323 | 0,00035 | 0,00483294 |
| Notch1 | 28,2603 | 44,158 | 0,643897 | 0,00035 | 0,00483294 |
| Cd24a | 315,48 | 200,894 | -0,651114 | 0,00035 | 0,00483294 |
| Myc | 38,2514 | 24,0974 | -0,666633 | 0,00035 | 0,00483294 |
| Tshz2 | 6,63176 | 3,88655 | -0,770902 | 0,00035 | 0,00483294 |
| Cntfr | 24,0665 | 13,5891 | -0,824574 | 0,00035 | 0,00483294 |
| Megf9 | 9,09267 | 5,1102 | -0,831323 | 0,00035 | 0,00483294 |
| Emid1 | 8,42879 | 4,41749 | -0,932098 | 0,00035 | 0,00483294 |
| Dpp4 | 1,1218 | 0,41523 | -1,43383 | 0,00035 | 0,00483294 |
| Ampd3 | 0,921243 | 0,201005 | -2,19635 | 0,00035 | 0,00483294 |
| Myf5 | 1,78073 | 0,377654 | -2,23733 | 0,00035 | 0,00483294 |
| A230077H06Rik | 0,252139 | 1,40681 | 2,48014 | 0,0004 | 0,00544136 |
| Ppef2 | 1,33228 | 3,31576 | 1,31544 | 0,0004 | 0,00544136 |
| Pik3ap1 | 1,46556 | 3,42846 | 1,22611 | 0,0004 | 0,00544136 |
| Nipal3 | 1,39949 | 2,83048 | 1,01615 | 0,0004 | 0,00544136 |
| Acta2 | 8,5468 | 14,9466 | 0,806362 | 0,0004 | 0,00544136 |
| Per1 | 4,99852 | 8,45913 | 0,759008 | 0,0004 | 0,00544136 |
| Cep170 | 14,8699 | 23,559 | 0,663884 | 0,0004 | 0,00544136 |
| Jund | 36,4741 | 22,9653 | -0,667416 | 0,0004 | 0,00544136 |
| Wdr86 | 10,7983 | 5,47056 | -0,98105 | 0,0004 | 0,00544136 |
| Astn1 | 1,58692 | 0,768941 | -1,04529 | 0,0004 | 0,00544136 |
| Lama4 | 1,61882 | 0,734315 | -1,14047 | 0,0004 | 0,00544136 |
| Ano1 | 2,1413 | 0,945727 | -1,17899 | 0,0004 | 0,00544136 |
| Dcdc2a | 0,90793 | 0,369556 | -1,29679 | 0,0004 | 0,00544136 |
| D930028M14Rik | 5,11595 | 1,35958 | -1,91184 | 0,0004 | 0,00544136 |
| Ptprn | 0,186019 | 1,04036 | 2,48356 | 0,00045 | 0,00602568 |
| Ddx43 | 0,345441 | 1,76033 | 2,34933 | 0,00045 | 0,00602568 |
| 2700086A05Rik | 26,8794 | 115,888 | 2,10815 | 0,00045 | 0,00602568 |
| Mical2 | 0,497972 | 1,23471 | 1,31003 | 0,00045 | 0,00602568 |
| Ranbp2 | 25,366 | 40,7764 | 0,684836 | 0,00045 | 0,00602568 |
| Shc1 | 18,2667 | 28,7857 | 0,656134 | 0,00045 | 0,00602568 |

| | | | | | |
|---|---|---|---|---|---|
| Igsf3 | 12,6089 | 19,724 | 0,64551 | 0,00045 | 0,00602568 |
| Vasp | 42,1401 | 65,8094 | 0,643101 | 0,00045 | 0,00602568 |
| Rsf1 | 8,25183 | 12,7786 | 0,630945 | 0,00045 | 0,00602568 |
| Zcchc24 | 18,7967 | 11,7968 | -0,67208 | 0,00045 | 0,00602568 |
| Axl | 5,81752 | 3,33133 | -0,804308 | 0,00045 | 0,00602568 |
| Slc7a2 | 1,83001 | 0,962366 | -0,927195 | 0,00045 | 0,00602568 |
| Plxdc2 | 2,15217 | 1,08563 | -0,987267 | 0,00045 | 0,00602568 |
| Dpf3 | 3,9727 | 1,45386 | -1,45023 | 0,00045 | 0,00602568 |
| Asic4 | 1,45334 | 0,246974 | -2,55694 | 0,00045 | 0,00602568 |
| Krt82 | 0,251243 | 1,96386 | 2,96653 | 0,0005 | 0,00656499 |
| Samd3 | 0,415596 | 1,93611 | 2,2199 | 0,0005 | 0,00656499 |
| Kcnh1 | 0,332896 | 0,859428 | 1,36831 | 0,0005 | 0,00656499 |
| Slc22a21 | 1,11773 | 2,83245 | 1,34148 | 0,0005 | 0,00656499 |
| Ybx2 | 2,10272 | 4,95578 | 1,23686 | 0,0005 | 0,00656499 |
| Atg16l2 | 3,33263 | 6,53088 | 0,970619 | 0,0005 | 0,00656499 |
| Clstn3 | 2,29624 | 4,39847 | 0,937726 | 0,0005 | 0,00656499 |
| Palm3 | 4,5564 | 8,40747 | 0,883778 | 0,0005 | 0,00656499 |
| Hebp1 | 13,3908 | 24,0248 | 0,843282 | 0,0005 | 0,00656499 |
| Tsku | 9,7302 | 16,3166 | 0,745798 | 0,0005 | 0,00656499 |
| Kcp | 4,5186 | 7,56958 | 0,74434 | 0,0005 | 0,00656499 |
| Snx11 | 10,2605 | 16,8923 | 0,719262 | 0,0005 | 0,00656499 |
| Nsg2 | 14,2669 | 23,2137 | 0,702301 | 0,0005 | 0,00656499 |
| Oaf | 37,8076 | 24,1331 | -0,647661 | 0,0005 | 0,00656499 |
| Dusp7 | 17,2283 | 10,5 | -0,714382 | 0,0005 | 0,00656499 |
| Slc25a12 | 12,5112 | 7,38608 | -0,760335 | 0,0005 | 0,00656499 |
| Tet2 | 2,78267 | 1,58096 | -0,815672 | 0,0005 | 0,00656499 |
| Gap43 | 13,1761 | 6,71379 | -0,97272 | 0,0005 | 0,00656499 |
| Smad9 | 1,40047 | 0,610579 | -1,19767 | 0,0005 | 0,00656499 |
| Rrad | 0,766337 | 3,06898 | 2,00171 | 0,00055 | 0,0071123 |
| Ier3 | 1,01784 | 3,94882 | 1,95591 | 0,00055 | 0,0071123 |
| Nell1 | 0,740746 | 1,91162 | 1,36775 | 0,00055 | 0,0071123 |
| F5 | 0,455413 | 1,04663 | 1,20051 | 0,00055 | 0,0071123 |
| Lix1 | 4,34572 | 7,59798 | 0,806022 | 0,00055 | 0,0071123 |
| Steap2 | 1,86771 | 3,20887 | 0,7808 | 0,00055 | 0,0071123 |
| Ccdc136 | 6,94783 | 11,3258 | 0,704983 | 0,00055 | 0,0071123 |
| Pde5a | 4,70225 | 7,60908 | 0,694371 | 0,00055 | 0,0071123 |
| Acadl | 21,2278 | 34,0263 | 0,680691 | 0,00055 | 0,0071123 |
| Ifitm2 | 224,23 | 343,947 | 0,617203 | 0,00055 | 0,0071123 |
| Lin28b | 24,1394 | 15,7746 | -0,613784 | 0,00055 | 0,0071123 |
| Ank3 | 7,01398 | 4,26564 | -0,717469 | 0,00055 | 0,0071123 |
| Fam101a | 3,43559 | 1,22141 | -1,49201 | 0,00055 | 0,0071123 |
| Prdm6 | 1,7744 | 0,590143 | -1,58819 | 0,00055 | 0,0071123 |
| Sema3d | 0,768533 | 0,0576953 | -3,73558 | 0,00055 | 0,0071123 |
| Cdc42ep5 | 6,49839 | 14,1645 | 1,12412 | 0,0006 | 0,00765851 |
| Actn3 | 1,63144 | 3,44536 | 1,07851 | 0,0006 | 0,00765851 |
| Acsl6 | 4,25297 | 7,75033 | 0,865789 | 0,0006 | 0,00765851 |
| Hoxb7 | 28,3362 | 46,761 | 0,72266 | 0,0006 | 0,00765851 |
| Tpr | 42,4629 | 69,4589 | 0,709955 | 0,0006 | 0,00765851 |
| Zfp217 | 9,78181 | 15,5151 | 0,665499 | 0,0006 | 0,00765851 |
| Mllt4 | 38,0129 | 58,1734 | 0,613872 | 0,0006 | 0,00765851 |

| | | | | | |
|---|---|---|---|---|---|
| Lpgat1 | 21,5191 | 32,4771 | 0,593802 | 0,0006 | 0,00765851 |
| Tcf12 | 137,432 | 88,022 | -0,642782 | 0,0006 | 0,00765851 |
| Plekha8 | 10,3625 | 6,57312 | -0,65672 | 0,0006 | 0,00765851 |
| P4ha3 | 1,94184 | 0,624465 | -1,63673 | 0,0006 | 0,00765851 |
| Rgs8 | 0,76544 | 0,21294 | -1,84584 | 0,0006 | 0,00765851 |
| Nhlh2 | 1,13163 | 0,176278 | -2,68248 | 0,0006 | 0,00765851 |
| Tmem35 | 0,927904 | 2,88449 | 1,63627 | 0,00065 | 0,00825564 |
| Dkk2 | 0,653981 | 1,69782 | 1,37636 | 0,00065 | 0,00825564 |
| 4932438A13Rik | 3,12481 | 5,0003 | 0,678245 | 0,00065 | 0,00825564 |
| 6330403A02Rik | 1,75867 | 0,860023 | -1,03204 | 0,00065 | 0,00825564 |
| Adra1a | 0,716891 | 0,0666163 | -3,42781 | 0,00065 | 0,00825564 |
| Synj2 | 0,47706 | 2,63324 | 2,4646 | 0,0007 | 0,00881217 |
| 2700054A10Rik | 1,1361 | 2,3301 | 1,03631 | 0,0007 | 0,00881217 |
| Phc2 | 63,3857 | 42,074 | -0,591228 | 0,0007 | 0,00881217 |
| Tspan9 | 23,1102 | 14,9832 | -0,625177 | 0,0007 | 0,00881217 |
| Emilin1 | 29,636 | 18,9589 | -0,644478 | 0,0007 | 0,00881217 |
| Hspa12a | 3,33183 | 1,82113 | -0,871476 | 0,0007 | 0,00881217 |
| Foxl1 | 4,28119 | 2,09292 | -1,0325 | 0,0007 | 0,00881217 |
| Ror1 | 2,04885 | 0,894917 | -1,19499 | 0,0007 | 0,00881217 |
| Fcgrt | 2,2594 | 0,542794 | -2,05746 | 0,0007 | 0,00881217 |
| Prlr | 0,0286933 | 0,59914 | 4,38411 | 0,00075 | 0,00933172 |
| Tceal6 | 2,76147 | 6,82819 | 1,30607 | 0,00075 | 0,00933172 |
| Spint1 | 1,44677 | 3,46538 | 1,26018 | 0,00075 | 0,00933172 |
| Zbtb38 | 1,93244 | 3,70384 | 0,938595 | 0,00075 | 0,00933172 |
| Cap2 | 3,67796 | 6,39068 | 0,797065 | 0,00075 | 0,00933172 |
| Adrbk2 | 2,3262 | 3,94064 | 0,760455 | 0,00075 | 0,00933172 |
| Peak1 | 3,50954 | 5,62058 | 0,679435 | 0,00075 | 0,00933172 |
| Zdhhc18 | 12,1836 | 19,463 | 0,675798 | 0,00075 | 0,00933172 |
| Sall4 | 48,5157 | 74,3303 | 0,615501 | 0,00075 | 0,00933172 |
| Sdk1 | 0,619632 | 0,223795 | -1,46923 | 0,00075 | 0,00933172 |
| Wt1 | 1,10406 | 0,162734 | -2,76223 | 0,00075 | 0,00933172 |
| Wnt1 | 2,85105 | 0,15713 | -4,18146 | 0,00075 | 0,00933172 |
| Lefty2 | 0,219629 | 1,52892 | 2,79937 | 0,0008 | 0,00982989 |
| Fbxo25 | 8,5084 | 14,7968 | 0,798321 | 0,0008 | 0,00982989 |
| Ptpn14 | 10,1519 | 15,7798 | 0,636331 | 0,0008 | 0,00982989 |
| Gabarapl1 | 21,3521 | 33,0723 | 0,631246 | 0,0008 | 0,00982989 |
| Jarid2 | 20,7235 | 31,5242 | 0,605193 | 0,0008 | 0,00982989 |
| Pax3 | 30,2582 | 19,5171 | -0,632589 | 0,0008 | 0,00982989 |
| Fbxl7 | 6,61759 | 3,98809 | -0,730607 | 0,0008 | 0,00982989 |
| Draxin | 5,85338 | 3,5 | -0,741914 | 0,0008 | 0,00982989 |
| Apba2 | 3,01075 | 1,41602 | -1,08828 | 0,0008 | 0,00982989 |
| Glrx | 6,34763 | 2,87024 | -1,14505 | 0,0008 | 0,00982989 |
| Foxa2 | 2,44548 | 0,930692 | -1,39374 | 0,0008 | 0,00982989 |
| Vwa1 | 1,34688 | 0,394822 | -1,77035 | 0,0008 | 0,00982989 |
| Tbx22 | 1,92636 | 0,186634 | -3,36759 | 0,0008 | 0,00982989 |
| Rgs4 | 0,358663 | 1,24759 | 1,79844 | 0,00085 | 0,0102866 |
| Eps8l1 | 0,525802 | 1,37661 | 1,38853 | 0,00085 | 0,0102866 |
| Mfsd2a | 6,11734 | 10,5776 | 0,790037 | 0,00085 | 0,0102866 |
| Hey2 | 8,21791 | 13,8941 | 0,757634 | 0,00085 | 0,0102866 |
| Ncoa7 | 5,15028 | 8,64147 | 0,746625 | 0,00085 | 0,0102866 |

| | | | | | |
|---|---|---|---|---|---|
| Dxo | 33,0835 | 51,4845 | 0,638026 | 0,00085 | 0,0102866 |
| Ttbk2 | 5,83616 | 8,98715 | 0,622842 | 0,00085 | 0,0102866 |
| Apc | 10,4532 | 15,7741 | 0,593621 | 0,00085 | 0,0102866 |
| Klhdc10 | 35,9564 | 23,7743 | -0,596848 | 0,00085 | 0,0102866 |
| Lhfpl2 | 25,3376 | 16,4144 | -0,626319 | 0,00085 | 0,0102866 |
| Asxl3 | 4,78149 | 2,9862 | -0,679151 | 0,00085 | 0,0102866 |
| Commd3 | 69,8565 | 43,4588 | -0,684747 | 0,00085 | 0,0102866 |
| 1190002N15Rik | 10,5468 | 6,49988 | -0,698316 | 0,00085 | 0,0102866 |
| Chchd6 | 20,9351 | 11,7315 | -0,835535 | 0,00085 | 0,0102866 |
| Depdc7 | 4,87977 | 2,20824 | -1,14392 | 0,00085 | 0,0102866 |
| Tgm2 | 1,16911 | 0,36451 | -1,68139 | 0,00085 | 0,0102866 |
| Myrfl | 0,400287 | 1,27801 | 1,6748 | 0,0009 | 0,0107899 |
| Tgfa | 0,674938 | 1,59053 | 1,23668 | 0,0009 | 0,0107899 |
| Adcy7 | 0,859107 | 1,71157 | 0,994414 | 0,0009 | 0,0107899 |
| Wdr47 | 2,9412 | 5,19565 | 0,8209 | 0,0009 | 0,0107899 |
| Med12 | 15,8872 | 24,1595 | 0,60473 | 0,0009 | 0,0107899 |
| Csrp2 | 273,359 | 415,633 | 0,604512 | 0,0009 | 0,0107899 |
| Sox12 | 30,7997 | 20,2012 | -0,608473 | 0,0009 | 0,0107899 |
| Gfra2 | 1,76913 | 0,743108 | -1,25139 | 0,0009 | 0,0107899 |
| Tfap2c | 1,58491 | 0,500644 | -1,66255 | 0,0009 | 0,0107899 |
| Osr2 | 2,1522 | 0,332967 | -2,69236 | 0,0009 | 0,0107899 |
| Sema7a | 0,631304 | 1,58696 | 1,32986 | 0,00095 | 0,0113048 |
| Dnaaf3 | 1,77429 | 3,86045 | 1,12153 | 0,00095 | 0,0113048 |
| Junb | 2,79216 | 5,78823 | 1,05174 | 0,00095 | 0,0113048 |
| Fbxl14 | 16,8984 | 28,1499 | 0,73624 | 0,00095 | 0,0113048 |
| Agtrap | 9,81277 | 15,5047 | 0,659973 | 0,00095 | 0,0113048 |
| Fermt2 | 53,0857 | 80,7342 | 0,604856 | 0,00095 | 0,0113048 |
| Cttnbp2 | 1,75325 | 0,871031 | -1,00924 | 0,00095 | 0,0113048 |
| Btbd11 | 1,08634 | 0,431355 | -1,33252 | 0,00095 | 0,0113048 |
| BC089491 | 0,66327 | 2,91437 | 2,13552 | 0,001 | 0,0117688 |
| Lpcat2 | 0,629072 | 1,77938 | 1,50008 | 0,001 | 0,0117688 |
| Rab27a | 0,734081 | 1,83991 | 1,32562 | 0,001 | 0,0117688 |
| Capg | 2,73245 | 6,22311 | 1,18744 | 0,001 | 0,0117688 |
| Slc26a11 | 4,41371 | 7,83761 | 0,828423 | 0,001 | 0,0117688 |
| Adamts6 | 2,82518 | 4,93741 | 0,80541 | 0,001 | 0,0117688 |
| Cdk6 | 6,31582 | 10,6193 | 0,749652 | 0,001 | 0,0117688 |
| Kbtbd8 | 3,77507 | 6,31805 | 0,742977 | 0,001 | 0,0117688 |
| Bicc1 | 5,90111 | 9,76997 | 0,727368 | 0,001 | 0,0117688 |
| Zfml | 29,0879 | 45,6104 | 0,648944 | 0,001 | 0,0117688 |
| Atoh8 | 11,3264 | 6,6608 | -0,765925 | 0,001 | 0,0117688 |
| Adam11 | 1,80343 | 0,85621 | -1,07471 | 0,001 | 0,0117688 |
| Osbpl10 | 4,0854 | 7,66344 | 0,907514 | 0,00105 | 0,0122449 |
| Rnf19b | 11,8515 | 18,9907 | 0,680224 | 0,00105 | 0,0122449 |
| Gcc2 | 8,32146 | 13,0432 | 0,648384 | 0,00105 | 0,0122449 |
| Ktn1 | 40,9812 | 62,1838 | 0,601577 | 0,00105 | 0,0122449 |
| Fxyd6 | 75,8339 | 49,8729 | -0,604586 | 0,00105 | 0,0122449 |
| Arhgap29 | 8,14045 | 5,07118 | -0,682789 | 0,00105 | 0,0122449 |
| Tenc1 | 4,22024 | 2,45657 | -0,78068 | 0,00105 | 0,0122449 |
| Dhrs7 | 3,01914 | 0,811572 | -1,89535 | 0,00105 | 0,0122449 |
| C130021I20Rik | 0,807645 | 0,155507 | -2,37674 | 0,00105 | 0,0122449 |

| | | | | | |
|---|---|---|---|---|---|
| Foxf1 | 1,47467 | 0,178269 | -3,04826 | 0,00105 | 0,0122449 |
| Usp53 | 1,74041 | 3,34264 | 0,941557 | 0,0011 | 0,0127815 |
| H2-D1 | 7,22232 | 12,5379 | 0,795766 | 0,0011 | 0,0127815 |
| Fndc3c1 | 56,7157 | 37,3347 | -0,603231 | 0,0011 | 0,0127815 |
| Dnajc15 | 78,5173 | 48,0325 | -0,709 | 0,0011 | 0,0127815 |
| Pacsin1 | 0,496796 | 1,23942 | 1,31894 | 0,00115 | 0,0132903 |
| Smpdl3a | 3,7637 | 7,30363 | 0,956461 | 0,00115 | 0,0132903 |
| Cyp39a1 | 6,60276 | 10,6875 | 0,694781 | 0,00115 | 0,0132903 |
| Tm9sf4 | 32,765 | 49,3543 | 0,59102 | 0,00115 | 0,0132903 |
| Lin28a | 62,5818 | 94,1685 | 0,5895 | 0,00115 | 0,0132903 |
| Igsf9 | 28,2676 | 42,2626 | 0,58023 | 0,00115 | 0,0132903 |
| Grik4 | 0,888503 | 1,96858 | 1,14771 | 0,0012 | 0,0137689 |
| D8Ertd82e | 9,24732 | 14,3273 | 0,631655 | 0,0012 | 0,0137689 |
| Ankrd26 | 11,9599 | 18,2801 | 0,612071 | 0,0012 | 0,0137689 |
| Ppp1r12a | 26,7276 | 40,2069 | 0,589117 | 0,0012 | 0,0137689 |
| Fignl2 | 65,0008 | 43,5595 | -0,57747 | 0,0012 | 0,0137689 |
| Sox4 | 48,4852 | 32,457 | -0,579015 | 0,0012 | 0,0137689 |
| Emilin2 | 20,2549 | 13,1762 | -0,620345 | 0,0012 | 0,0137689 |
| Usp25 | 15,261 | 9,91356 | -0,622372 | 0,0012 | 0,0137689 |
| Lhfpl3 | 0,141779 | 1,20603 | 3,08855 | 0,00125 | 0,0142787 |
| Rasgrp1 | 0,264133 | 0,802617 | 1,60345 | 0,00125 | 0,0142787 |
| Emp2 | 1,20504 | 2,45692 | 1,02777 | 0,00125 | 0,0142787 |
| Apela | 39,7237 | 61,8659 | 0,639144 | 0,00125 | 0,0142787 |
| Grb14 | 1,76956 | 0,233905 | -2,91939 | 0,00125 | 0,0142787 |
| Kcns3 | 0,311959 | 1,09324 | 1,80919 | 0,0013 | 0,0147448 |
| Slc1a2 | 0,367521 | 0,845987 | 1,20281 | 0,0013 | 0,0147448 |
| Ankrd12 | 1,17961 | 2,10453 | 0,835192 | 0,0013 | 0,0147448 |
| Braf | 7,7064 | 11,5796 | 0,587456 | 0,0013 | 0,0147448 |
| Nnat | 341,178 | 222,699 | -0,615431 | 0,0013 | 0,0147448 |
| Dkk3 | 3,33477 | 1,76111 | -0,921099 | 0,0013 | 0,0147448 |
| Llgl2 | 3,38547 | 1,78479 | -0,923602 | 0,0013 | 0,0147448 |
| Eda | 2,37929 | 1,25273 | -0,925457 | 0,0013 | 0,0147448 |
| Kirrel3 | 1,64209 | 3,11292 | 0,92274 | 0,00135 | 0,0152714 |
| Podxl | 9,52043 | 6,05355 | -0,653246 | 0,00135 | 0,0152714 |
| Tshz3 | 2,71444 | 1,46466 | -0,890082 | 0,00135 | 0,0152714 |
| Esco1 | 3,88888 | 6,35949 | 0,709556 | 0,0014 | 0,0157537 |
| Scara3 | 6,74728 | 10,9195 | 0,69453 | 0,0014 | 0,0157537 |
| D430019H16Rik | 4,66796 | 7,34955 | 0,654864 | 0,0014 | 0,0157537 |
| Mbd2 | 16,1828 | 9,79821 | -0,723869 | 0,0014 | 0,0157537 |
| Asic2 | 1,05276 | 0,237878 | -2,14589 | 0,0014 | 0,0157537 |
| Adh1 | 2,27617 | 0,293095 | -2,95717 | 0,0014 | 0,0157537 |
| Lgals3bp | 0,452178 | 1,56106 | 1,78756 | 0,00145 | 0,0162451 |
| Col17a1 | 0,188184 | 0,606446 | 1,68823 | 0,00145 | 0,0162451 |
| Firre | 10,0417 | 15,1786 | 0,596027 | 0,00145 | 0,0162451 |
| Basp1 | 125,38 | 84,8003 | -0,564162 | 0,00145 | 0,0162451 |
| Gm7694 | 1,5803 | 0,706169 | -1,16211 | 0,00145 | 0,0162451 |
| Tshr | 0,0834645 | 0,69679 | 3,06149 | 0,0015 | 0,0167031 |
| Dopey2 | 1,1189 | 2,80256 | 1,32466 | 0,0015 | 0,0167031 |
| Sp6 | 0,572314 | 1,40653 | 1,29726 | 0,0015 | 0,0167031 |
| Med12l | 1,5475 | 3,17483 | 1,03673 | 0,0015 | 0,0167031 |

| | | | | | |
|---|---|---|---|---|---|
| Tmem173 | 6,45279 | 10,7989 | 0,742893 | 0,0015 | 0,0167031 |
| Casd1 | 34,1732 | 22,4853 | -0,603886 | 0,0015 | 0,0167031 |
| 2610034B18Rik | 11,4756 | 6,84497 | -0,745459 | 0,0015 | 0,0167031 |
| Helz2 | 1,67197 | 2,74337 | 0,714399 | 0,00155 | 0,0172151 |
| Col4a1 | 66,7336 | 44,9884 | -0,56886 | 0,00155 | 0,0172151 |
| Pax9 | 15,7965 | 10,0219 | -0,656444 | 0,00155 | 0,0172151 |
| Dsg2 | 0,288584 | 0,743948 | 1,36621 | 0,0016 | 0,0176482 |
| Abca4 | 0,383512 | 0,827397 | 1,10931 | 0,0016 | 0,0176482 |
| Abcb1b | 0,669111 | 1,44155 | 1,10731 | 0,0016 | 0,0176482 |
| Fgd5 | 1,6468 | 2,91554 | 0,824099 | 0,0016 | 0,0176482 |
| Nfib | 1,84175 | 2,99216 | 0,700114 | 0,0016 | 0,0176482 |
| Lrrcc1 | 11,4162 | 18,4784 | 0,694755 | 0,0016 | 0,0176482 |
| Cbl | 11,8897 | 18,3701 | 0,627647 | 0,0016 | 0,0176482 |
| Chrna6 | 1,11079 | 0,148256 | -2,90543 | 0,0016 | 0,0176482 |
| BC064078 | 0,376639 | 5,81987 | 3,94973 | 0,00165 | 0,0181373 |
| Myl2 | 1,15881 | 5,55495 | 2,26113 | 0,00165 | 0,0181373 |
| Rras | 11,2245 | 19,8989 | 0,826035 | 0,00165 | 0,0181373 |
| Enpp3 | 1,38266 | 0,468761 | -1,56052 | 0,00165 | 0,0181373 |
| Mmp25 | 0,179452 | 0,891854 | 2,31321 | 0,0017 | 0,0185913 |
| Il17d | 1,94491 | 4,59201 | 1,23942 | 0,0017 | 0,0185913 |
| Upf2 | 13,3593 | 20,4789 | 0,616296 | 0,0017 | 0,0185913 |
| Rcn2 | 119,226 | 175,829 | 0,560473 | 0,0017 | 0,0185913 |
| Tspan13 | 19,0479 | 12,014 | -0,664923 | 0,0017 | 0,0185913 |
| Pkia | 5,4503 | 3,16933 | -0,782156 | 0,0017 | 0,0185913 |
| Pm20d1 | 1,62814 | 3,08831 | 0,923593 | 0,00175 | 0,0190085 |
| 2610307P16Rik | 3,2594 | 5,8903 | 0,853737 | 0,00175 | 0,0190085 |
| Lrch1 | 3,08543 | 5,12447 | 0,73193 | 0,00175 | 0,0190085 |
| Arhgap24 | 8,74627 | 13,7408 | 0,651727 | 0,00175 | 0,0190085 |
| Rock1 | 9,84871 | 15,1151 | 0,61798 | 0,00175 | 0,0190085 |
| Atr | 9,32114 | 13,9234 | 0,578931 | 0,00175 | 0,0190085 |
| Pitrm1 | 82,6506 | 123,05 | 0,574144 | 0,00175 | 0,0190085 |
| Pitx1 | 1,64259 | 0,592445 | -1,47122 | 0,00175 | 0,0190085 |
| Egf | 0,396457 | 0,979011 | 1,30416 | 0,0018 | 0,0194691 |
| Taok1 | 8,73754 | 12,9886 | 0,571942 | 0,0018 | 0,0194691 |
| Rgma | 25,4688 | 17,3271 | -0,555704 | 0,0018 | 0,0194691 |
| Clip2 | 12,866 | 8,39435 | -0,616071 | 0,0018 | 0,0194691 |
| Tor3a | 3,38165 | 1,78261 | -0,923732 | 0,0018 | 0,0194691 |
| Fam172a | 8,72771 | 5,49399 | -0,667749 | 0,00185 | 0,0199595 |
| Ddit4l | 3,31127 | 1,60104 | -1,04837 | 0,00185 | 0,0199595 |
| Fndc5 | 2,06597 | 0,885147 | -1,22283 | 0,00185 | 0,0199595 |
| Lhx5 | 0,35131 | 1,15168 | 1,71292 | 0,0019 | 0,0203789 |
| Itpr3 | 0,813196 | 1,47732 | 0,861311 | 0,0019 | 0,0203789 |
| A830080D01Rik | 5,93401 | 9,7319 | 0,713713 | 0,0019 | 0,0203789 |
| St3gal1 | 7,00623 | 10,6748 | 0,607504 | 0,0019 | 0,0203789 |
| Cd63 | 46,6646 | 70,9676 | 0,604833 | 0,0019 | 0,0203789 |
| Epb4.1l3 | 61,202 | 89,7844 | 0,552887 | 0,0019 | 0,0203789 |
| Map1a | 1,96857 | 1,01056 | -0,962002 | 0,0019 | 0,0203789 |
| Sacs | 0,829421 | 1,85875 | 1,16415 | 0,00195 | 0,0208281 |
| H2-Q4 | 6,61432 | 11,3012 | 0,772815 | 0,00195 | 0,0208281 |
| Trip11 | 4,71286 | 7,34175 | 0,639521 | 0,00195 | 0,0208281 |

| | | | | | |
|---|---|---|---|---|---|
| Atl2 | 25,7355 | 38,0298 | 0,563371 | 0,00195 | 0,0208281 |
| Lmx1a | 0,841803 | 0,163541 | -2,36383 | 0,00195 | 0,0208281 |
| S100a6 | 13,6212 | 24,6188 | 0,853909 | 0,002 | 0,0212913 |
| Bdp1 | 7,85429 | 11,7146 | 0,57675 | 0,002 | 0,0212913 |
| Smurf1 | 10,8876 | 16,2286 | 0,575858 | 0,002 | 0,0212913 |
| Cyth3 | 14,1073 | 9,23122 | -0,611846 | 0,002 | 0,0212913 |
| Camkk1 | 1,3761 | 2,74094 | 0,994087 | 0,00205 | 0,0217514 |
| Cenpe | 17,3268 | 26,0448 | 0,587991 | 0,00205 | 0,0217514 |
| Stox2 | 16,2102 | 10,9659 | -0,563876 | 0,00205 | 0,0217514 |
| Phgdh | 18,0482 | 11,3994 | -0,662901 | 0,00205 | 0,0217514 |
| Skap1 | 0,242378 | 1,18639 | 2,29124 | 0,0021 | 0,0221901 |
| Plcxd1 | 1,21219 | 3,95607 | 1,70645 | 0,0021 | 0,0221901 |
| Anxa4 | 3,66541 | 6,86524 | 0,905336 | 0,0021 | 0,0221901 |
| Gmpr | 5,98628 | 10,571 | 0,820378 | 0,0021 | 0,0221901 |
| Myl9 | 24,7851 | 38,8172 | 0,647225 | 0,0021 | 0,0221901 |
| Ppp4r4 | 1,86965 | 3,47407 | 0,893863 | 0,00215 | 0,0226438 |
| Hoxb8 | 10,2072 | 15,816 | 0,631803 | 0,00215 | 0,0226438 |
| Trabd2b | 1,05618 | 0,523088 | -1,01373 | 0,00215 | 0,0226438 |
| Celf3 | 0,736377 | 0,121184 | -2,60324 | 0,00215 | 0,0226438 |
| St14 | 0,199361 | 0,717395 | 1,84738 | 0,0022 | 0,023038 |
| Mgam | 0,294314 | 0,729905 | 1,31035 | 0,0022 | 0,023038 |
| Usp9x | 12,6908 | 18,6348 | 0,554217 | 0,0022 | 0,023038 |
| Nr2f6 | 42,6545 | 28,5483 | -0,579293 | 0,0022 | 0,023038 |
| Zcchc3 | 28,1461 | 18,7657 | -0,584841 | 0,0022 | 0,023038 |
| Efs | 15,8452 | 10,4689 | -0,597935 | 0,0022 | 0,023038 |
| Nup210 | 9,46675 | 6,15769 | -0,62048 | 0,0022 | 0,023038 |
| Smagp | 0,507289 | 2,27209 | 2,16314 | 0,00225 | 0,0234659 |
| Sipa1l3 | 6,62705 | 9,99254 | 0,592484 | 0,00225 | 0,0234659 |
| Gulp1 | 11,4019 | 7,21631 | -0,659942 | 0,00225 | 0,0234659 |
| Erbb3 | 1,14488 | 0,459643 | -1,31661 | 0,00225 | 0,0234659 |
| Bai3 | 0,628643 | 0,226316 | -1,4739 | 0,00225 | 0,0234659 |
| Timp4 | 1,34293 | 3,52795 | 1,39345 | 0,0023 | 0,0239096 |
| Jdp2 | 5,48155 | 9,76582 | 0,833158 | 0,0023 | 0,0239096 |
| Whrn | 5,47659 | 8,60895 | 0,65256 | 0,0023 | 0,0239096 |
| Bre | 24,9668 | 15,511 | -0,686717 | 0,0023 | 0,0239096 |
| Mgll | 0,788371 | 1,6356 | 1,05287 | 0,00235 | 0,0243308 |
| Aldoc | 2,53874 | 5,02331 | 0,984526 | 0,00235 | 0,0243308 |
| Thra | 9,71262 | 15,2882 | 0,654488 | 0,00235 | 0,0243308 |
| Rasa3 | 11,5815 | 17,4049 | 0,587676 | 0,00235 | 0,0243308 |
| Mga | 12,5086 | 18,6299 | 0,574694 | 0,00235 | 0,0243308 |
| Lrch2 | 7,07584 | 11,041 | 0,6419 | 0,0024 | 0,0247685 |
| Ucp2 | 5,83595 | 9,05839 | 0,634286 | 0,0024 | 0,0247685 |
| Ankrd6 | 5,75898 | 3,61025 | -0,673715 | 0,0024 | 0,0247685 |
| Gm266 | 5,42447 | 2,42298 | -1,1627 | 0,0024 | 0,0247685 |
| Gas2l3 | 2,75173 | 4,38981 | 0,673819 | 0,00245 | 0,0251429 |
| Esyt1 | 4,90458 | 7,80056 | 0,669447 | 0,00245 | 0,0251429 |
| Srf | 21,2887 | 31,385 | 0,559984 | 0,00245 | 0,0251429 |
| Tmsb4x | 665,189 | 457,078 | -0,541325 | 0,00245 | 0,0251429 |
| Tox3 | 4,81054 | 2,78068 | -0,79076 | 0,00245 | 0,0251429 |
| Pou2f2 | 2,12382 | 1,05969 | -1,00302 | 0,00245 | 0,0251429 |

| | | | | | |
|---|---|---|---|---|---|
| Gm15055 | 9,75987 | 3,41398 | -1,5154 | 0,00245 | 0,0251429 |
| Itga7 | 0,1086 | 0,596393 | 2,45725 | 0,0025 | 0,0255538 |
| Sema3g | 0,347189 | 1,00962 | 1,54002 | 0,0025 | 0,0255538 |
| Vamp8 | 16,6077 | 8,88745 | -0,902014 | 0,0025 | 0,0255538 |
| Cntnap2 | 2,27792 | 1,13745 | -1,00192 | 0,0025 | 0,0255538 |
| Wnt7a | 1,05314 | 0,0880929 | -3,57952 | 0,0025 | 0,0255538 |
| Egr2 | 0,231883 | 0,922081 | 1,9915 | 0,00255 | 0,0259408 |
| Tmem229a | 1,34523 | 2,46365 | 0,872948 | 0,00255 | 0,0259408 |
| Kif20b | 16,2796 | 24,3705 | 0,582067 | 0,00255 | 0,0259408 |
| Sfxn1 | 61,7285 | 89,3471 | 0,533484 | 0,00255 | 0,0259408 |
| Mageh1 | 31,0097 | 20,1925 | -0,618901 | 0,00255 | 0,0259408 |
| Fkbp7 | 10,3496 | 5,19073 | -0,995566 | 0,00255 | 0,0259408 |
| Traf5 | 0,567153 | 1,60518 | 1,50093 | 0,0026 | 0,0263867 |
| Myo9a | 4,8466 | 7,32769 | 0,596386 | 0,0026 | 0,0263867 |
| Sorl1 | 5,44355 | 3,50927 | -0,633376 | 0,0026 | 0,0263867 |
| Rp2h | 9,71934 | 16,9206 | 0,799851 | 0,00265 | 0,0268093 |
| Atm | 8,65502 | 12,5837 | 0,539951 | 0,00265 | 0,0268093 |
| Loxl1 | 1,06499 | 0,373985 | -1,50979 | 0,00265 | 0,0268093 |
| Vtn | 1,55749 | 0,329581 | -2,24052 | 0,00265 | 0,0268093 |
| Lingo1 | 2,95759 | 5,09874 | 0,785719 | 0,0027 | 0,0271865 |
| Pcdh7 | 1,59751 | 2,74457 | 0,780756 | 0,0027 | 0,0271865 |
| Sowahc | 4,89341 | 7,5727 | 0,629969 | 0,0027 | 0,0271865 |
| Rbm38 | 30,1477 | 19,8509 | -0,602844 | 0,0027 | 0,0271865 |
| Abcd2 | 2,16706 | 1,20237 | -0,849859 | 0,0027 | 0,0271865 |
| Ddc | 2,6565 | 1,16606 | -1,18788 | 0,0027 | 0,0271865 |
| Mgat4a | 1,04877 | 2,02719 | 0,950786 | 0,00275 | 0,0275817 |
| Bcorl1 | 6,69204 | 10,1848 | 0,605898 | 0,00275 | 0,0275817 |
| Dixdc1 | 13,4744 | 19,7128 | 0,548909 | 0,00275 | 0,0275817 |
| Lbx2 | 6,54047 | 2,75277 | -1,24851 | 0,00275 | 0,0275817 |
| Dbx1 | 2,95556 | 0,127943 | -4,52986 | 0,00275 | 0,0275817 |
| Nmi | 1,59289 | 3,6632 | 1,20146 | 0,0028 | 0,0279956 |
| Spry1 | 11,915 | 18,3489 | 0,622917 | 0,0028 | 0,0279956 |
| Tmf1 | 6,59155 | 9,84791 | 0,579199 | 0,0028 | 0,0279956 |
| 6330403K07Rik | 51,4237 | 75,7693 | 0,55918 | 0,0028 | 0,0279956 |
| Htra1 | 0,393195 | 1,44041 | 1,87316 | 0,00285 | 0,0284291 |
| Arid4b | 12,6771 | 18,7751 | 0,56659 | 0,00285 | 0,0284291 |
| Rc3h2 | 6,6163 | 9,78119 | 0,563987 | 0,00285 | 0,0284291 |
| Edar | 0,0815947 | 0,843954 | 3,37062 | 0,0029 | 0,0288158 |
| Crlf1 | 0,681758 | 2,02222 | 1,56861 | 0,0029 | 0,0288158 |
| Dennd4a | 2,24092 | 3,54007 | 0,659689 | 0,0029 | 0,0288158 |
| Clmn | 1,96394 | 3,01791 | 0,619797 | 0,0029 | 0,0288158 |
| Thoc2 | 13,7491 | 20,8155 | 0,598317 | 0,0029 | 0,0288158 |
| Cers4 | 2,85589 | 4,85164 | 0,76453 | 0,00295 | 0,0291545 |
| Cxadr | 21,1 | 32,0027 | 0,600956 | 0,00295 | 0,0291545 |
| Sparc | 53,8979 | 37,1306 | -0,537618 | 0,00295 | 0,0291545 |
| Tspyl4 | 6,88737 | 4,34155 | -0,665742 | 0,00295 | 0,0291545 |
| Coro2b | 2,26062 | 1,15743 | -0,965791 | 0,00295 | 0,0291545 |
| Mpped1 | 1,28402 | 0,48908 | -1,39253 | 0,00295 | 0,0291545 |
| Rcan2 | 0,8732 | 0,316061 | -1,46611 | 0,00295 | 0,0291545 |
| S100a1 | 3,09868 | 8,40713 | 1,43996 | 0,003 | 0,0296031 |

| | | | | | |
|---|---|---|---|---|---|
| Acer2 | 4,53472 | 8,61297 | 0,925499 | 0,003 | 0,0296031 |
| Dgkh | 0,738123 | 1,50952 | 1,03216 | 0,00305 | 0,0300502 |
| Olig1 | 1,84671 | 0,121829 | -3,92203 | 0,00305 | 0,0300502 |
| Itgb2l | 0,099232 | 1,15242 | 3,53771 | 0,0031 | 0,030426 |
| Plek | 0,269891 | 0,769356 | 1,51128 | 0,0031 | 0,030426 |
| Gnpda1 | 6,73404 | 10,7339 | 0,672631 | 0,0031 | 0,030426 |
| Dnmt3b | 15,6889 | 23,0165 | 0,552927 | 0,0031 | 0,030426 |
| Cav1 | 1,1326 | 0,384462 | -1,55872 | 0,0031 | 0,030426 |
| Pygl | 11,5849 | 17,455 | 0,591394 | 0,00315 | 0,0308695 |
| Mgp | 6,37884 | 1,72127 | -1,88982 | 0,00315 | 0,0308695 |
| Phf5a | 86,7229 | 59,4866 | -0,543847 | 0,0032 | 0,0313117 |
| St6galnac3 | 6,11009 | 3,82046 | -0,677446 | 0,0032 | 0,0313117 |
| Mir219-1 | 0 | 71,1615 | inf | 0,00325 | 0,031608 |
| Tnni1 | 5,63448 | 10,5446 | 0,904147 | 0,00325 | 0,031608 |
| Tmem151b | 3,35449 | 5,32833 | 0,667593 | 0,00325 | 0,031608 |
| Tex2 | 4,46791 | 6,97141 | 0,641852 | 0,00325 | 0,031608 |
| Sapcd2 | 8,99177 | 13,607 | 0,597669 | 0,00325 | 0,031608 |
| Zfp292 | 10,7826 | 15,9315 | 0,563184 | 0,00325 | 0,031608 |
| Tec | 0,765806 | 0,170433 | -2,16777 | 0,00325 | 0,031608 |
| Nhlh1 | 1,27443 | 0,116716 | -3,44878 | 0,00325 | 0,031608 |
| Otop1 | 0,0915812 | 0,846046 | 3,20761 | 0,0033 | 0,0319973 |
| Atp2c1 | 33,2328 | 49,7179 | 0,58116 | 0,0033 | 0,0319973 |
| Boc | 16,4048 | 11,0452 | -0,570699 | 0,0033 | 0,0319973 |
| Gyltl1b | 3,65041 | 1,94855 | -0,905658 | 0,0033 | 0,0319973 |
| Wnt2b | 0,254266 | 0,828507 | 1,70417 | 0,00335 | 0,0324331 |
| Dach1 | 10,4196 | 7,01133 | -0,571544 | 0,00335 | 0,0324331 |
| Brwd3 | 4,69115 | 7,11006 | 0,59992 | 0,0034 | 0,0328675 |
| Med13 | 10,1095 | 14,6611 | 0,536278 | 0,0034 | 0,0328675 |
| Mirg | 9,12123 | 15,1081 | 0,728026 | 0,00345 | 0,0332008 |
| Dpy19l4 | 7,58398 | 11,6351 | 0,617458 | 0,00345 | 0,0332008 |
| Casp8ap2 | 15,5233 | 22,8323 | 0,55664 | 0,00345 | 0,0332008 |
| Celsr1 | 14,188 | 20,6189 | 0,539298 | 0,00345 | 0,0332008 |
| Nrep | 182,881 | 126,169 | -0,535552 | 0,00345 | 0,0332008 |
| Plcl2 | 7,1816 | 4,56358 | -0,654139 | 0,00345 | 0,0332008 |
| Stc1 | 0,294532 | 0,840652 | 1,51308 | 0,0035 | 0,0336315 |
| Sema5a | 6,44451 | 9,32395 | 0,532871 | 0,0035 | 0,0336315 |
| Mki67 | 49,4044 | 72,4683 | 0,55271 | 0,00355 | 0,0340609 |
| Pknox2 | 34,6147 | 23,6712 | -0,548255 | 0,00355 | 0,0340609 |
| Entpd1 | 0,390346 | 0,932103 | 1,25574 | 0,0036 | 0,0344633 |
| Epha5 | 0,297868 | 0,646641 | 1,11829 | 0,0036 | 0,0344633 |
| Tmem198b | 10,5092 | 16,0223 | 0,608421 | 0,0036 | 0,0344633 |
| Nuak1 | 8,86916 | 13,1156 | 0,56441 | 0,00365 | 0,0348639 |
| Gm1673 | 73,429 | 44,3826 | -0,726356 | 0,00365 | 0,0348639 |
| Tfap2a | 0,774143 | 0,19987 | -1,95354 | 0,00365 | 0,0348639 |
| Lipg | 2,56217 | 4,29929 | 0,746734 | 0,0037 | 0,0351322 |
| Cecr2 | 16,9175 | 24,5409 | 0,536671 | 0,0037 | 0,0351322 |
| Col5a1 | 17,2177 | 11,9095 | -0,531771 | 0,0037 | 0,0351322 |
| Efnb2 | 24,3372 | 16,7331 | -0,540462 | 0,0037 | 0,0351322 |
| Pygo1 | 7,05248 | 3,9754 | -0,827033 | 0,0037 | 0,0351322 |
| Lrrtm4 | 1,63582 | 0,79955 | -1,03275 | 0,0037 | 0,0351322 |

| | | | | | |
|---|---|---|---|---|---|
| Chrna4 | 1,05454 | 0,470553 | -1,16419 | 0,0037 | 0,0351322 |
| Nkx3-2 | 2,29262 | 0,181589 | -3,65825 | 0,0037 | 0,0351322 |
| Alms1 | 4,98483 | 7,26083 | 0,542592 | 0,00375 | 0,0355543 |
| Ooep | 2,61406 | 0,796992 | -1,71366 | 0,00375 | 0,0355543 |
| Letmd1 | 12,696 | 8,23944 | -0,623757 | 0,0038 | 0,0359752 |
| Emcn | 3,98451 | 1,87287 | -1,08915 | 0,0038 | 0,0359752 |
| 1700017B05Rik | 11,013 | 7,42859 | -0,568047 | 0,00385 | 0,0363948 |
| Cyp7b1 | 1,28206 | 0,403604 | -1,66745 | 0,00385 | 0,0363948 |
| AU015836 | 0,104409 | 0,793109 | 2,92527 | 0,0039 | 0,0367322 |
| Tmc8 | 0,158156 | 0,641624 | 2,02038 | 0,0039 | 0,0367322 |
| Stat3 | 3,5215 | 5,54631 | 0,65534 | 0,0039 | 0,0367322 |
| Tspan18 | 5,33302 | 3,2906 | -0,696602 | 0,0039 | 0,0367322 |
| Prdm12 | 2,6373 | 0,0918228 | -4,84407 | 0,0039 | 0,0367322 |
| Cpne5 | 0,459564 | 1,04739 | 1,18846 | 0,00395 | 0,0370399 |
| Slc2a6 | 1,21229 | 2,5573 | 1,07688 | 0,00395 | 0,0370399 |
| Wrnip1 | 24,8999 | 36,0376 | 0,533365 | 0,00395 | 0,0370399 |
| Notch2 | 19,3136 | 13,4189 | -0,525349 | 0,00395 | 0,0370399 |
| Sox13 | 22,2237 | 15,1368 | -0,55404 | 0,00395 | 0,0370399 |
| Myb | 1,09477 | 0,465862 | -1,23266 | 0,00395 | 0,0370399 |
| Cygb | 1,35281 | 0,470161 | -1,52473 | 0,004 | 0,0374814 |
| Cyp4f15 | 0,396139 | 1,26898 | 1,67959 | 0,00405 | 0,0377843 |
| Dock10 | 0,277316 | 0,61556 | 1,15037 | 0,00405 | 0,0377843 |
| Wdr81 | 2,0138 | 3,23128 | 0,682188 | 0,00405 | 0,0377843 |
| Fam212a | 26,921 | 40,8879 | 0,602943 | 0,00405 | 0,0377843 |
| Smc5 | 24,5524 | 35,6323 | 0,537319 | 0,00405 | 0,0377843 |
| Jph2 | 1,11393 | 0,500825 | -1,15328 | 0,00405 | 0,0377843 |
| Cck | 0,763382 | 3,88293 | 2,34667 | 0,0041 | 0,0381675 |
| T2 | 0,620417 | 2,19231 | 1,82114 | 0,0041 | 0,0381675 |
| St6galnac5 | 1,21432 | 0,309247 | -1,97332 | 0,0041 | 0,0381675 |
| Pygm | 0,690994 | 1,56357 | 1,1781 | 0,00415 | 0,038549 |
| Nab1 | 2,0149 | 3,33442 | 0,726731 | 0,00415 | 0,038549 |
| Oat | 94,8206 | 135,984 | 0,520167 | 0,00415 | 0,038549 |
| Mst1r | 0,140982 | 0,670721 | 2,2502 | 0,0042 | 0,0389289 |
| Tmbim1 | 2,64392 | 6,68964 | 1,33925 | 0,0042 | 0,0389289 |
| Tnik | 0,857725 | 0,389128 | -1,14027 | 0,0042 | 0,0389289 |
| Tmed5 | 7,23993 | 10,9451 | 0,596236 | 0,00425 | 0,0393355 |
| Rhou | 22,0484 | 15,1474 | -0,541602 | 0,00425 | 0,0393355 |
| Plod2 | 36,2522 | 25,073 | -0,531932 | 0,0043 | 0,0396837 |
| Snhg3 | 117,356 | 75,7263 | -0,632027 | 0,0043 | 0,0396837 |
| Pdzd4 | 4,72637 | 2,88871 | -0,710305 | 0,0043 | 0,0396837 |
| Ankrd44 | 2,20903 | 1,3101 | -0,753734 | 0,0043 | 0,0396837 |
| Tmem180 | 2,12792 | 3,92072 | 0,881675 | 0,00435 | 0,0400013 |
| Wnk3 | 3,01499 | 4,5192 | 0,583913 | 0,00435 | 0,0400013 |
| Hoxa9 | 42,2999 | 60,3617 | 0,512982 | 0,00435 | 0,0400013 |
| Edn3 | 5,61228 | 3,37965 | -0,731712 | 0,00435 | 0,0400013 |
| Sim1 | 0,929916 | 0,472606 | -0,976461 | 0,00435 | 0,0400013 |
| Fryl | 7,71584 | 11,0974 | 0,524326 | 0,00445 | 0,0408915 |
| Rap1gap | 0,515734 | 1,20597 | 1,22549 | 0,0045 | 0,0411447 |
| Ppfia4 | 0,641381 | 1,19434 | 0,896962 | 0,0045 | 0,0411447 |
| Cd2ap | 8,95751 | 13,1993 | 0,55929 | 0,0045 | 0,0411447 |

| | | | | | |
|---|---|---|---|---|---|
| Hsd17b7 | 9,50006 | 13,9126 | 0,550383 | 0,0045 | 0,0411447 |
| Tmtc3 | 8,40933 | 12,274 | 0,545546 | 0,0045 | 0,0411447 |
| Mtap | 91,1536 | 63,7866 | -0,515046 | 0,0045 | 0,0411447 |
| Tsen54 | 15,7228 | 10,1757 | -0,627736 | 0,0045 | 0,0411447 |
| Gpr98 | 0,670853 | 0,397148 | -0,756318 | 0,00455 | 0,0415722 |
| AI504432 | 1,28907 | 2,24601 | 0,801035 | 0,0046 | 0,0419097 |
| Rhbdf1 | 8,89517 | 13,474 | 0,599086 | 0,0046 | 0,0419097 |
| Prkdc | 3,79714 | 5,53878 | 0,544653 | 0,0046 | 0,0419097 |
| Epb4.1l5 | 13,509 | 19,3567 | 0,518908 | 0,0046 | 0,0419097 |
| Fgfr1 | 63,7224 | 91,1757 | 0,516849 | 0,00465 | 0,0422452 |
| Mbnl2 | 4,58166 | 2,84643 | -0,686717 | 0,00465 | 0,0422452 |
| Eya4 | 1,33113 | 0,629201 | -1,08106 | 0,00465 | 0,0422452 |
| Alx1 | 0,882489 | 0,191568 | -2,20372 | 0,00465 | 0,0422452 |
| Lnpep | 5,36287 | 8,06868 | 0,589327 | 0,0047 | 0,042639 |
| Lmo3 | 1,10542 | 0,185339 | -2,57636 | 0,0047 | 0,042639 |
| Adam2 | 0,270473 | 0,925941 | 1,77544 | 0,00475 | 0,0430318 |
| Rragd | 32,1062 | 22,3657 | -0,521565 | 0,00475 | 0,0430318 |
| Snhg4 | 23,812 | 15,0966 | -0,657461 | 0,0048 | 0,0434234 |
| Glrb | 1,16955 | 0,471722 | -1,30995 | 0,0048 | 0,0434234 |
| Col27a1 | 2,59597 | 3,98114 | 0,616909 | 0,00485 | 0,0438448 |
| Dmxl1 | 4,25765 | 6,23046 | 0,549281 | 0,0049 | 0,0442656 |
| Exoc6 | 3,08383 | 5,17506 | 0,746853 | 0,00495 | 0,0446544 |
| Cntrob | 14,9831 | 21,7929 | 0,540521 | 0,00495 | 0,0446544 |
| Mical3 | 5,42962 | 8,31937 | 0,615624 | 0,005 | 0,0450421 |
| Mir17hg | 22,499 | 15,3332 | -0,553201 | 0,005 | 0,0450421 |
| Eml2 | 6,3283 | 9,9562 | 0,653778 | 0,00505 | 0,0453969 |
| Klf10 | 6,22228 | 9,54438 | 0,617207 | 0,00505 | 0,0453969 |
| S100a10 | 74,1442 | 48,9005 | -0,600483 | 0,00505 | 0,0453969 |
| Mpp5 | 9,75992 | 14,3039 | 0,551464 | 0,0051 | 0,0457502 |
| Sned1 | 1,57043 | 0,826573 | -0,925944 | 0,0051 | 0,0457502 |
| Sfmbt2 | 0,910273 | 0,467471 | -0,961421 | 0,0051 | 0,0457502 |
| Rps6ka3 | 29,1787 | 20,4341 | -0,513936 | 0,00515 | 0,0461665 |
| Igsf8 | 18,9593 | 28,0041 | 0,562735 | 0,0052 | 0,0465172 |
| Fign | 4,2175 | 2,84251 | -0,569225 | 0,0052 | 0,0465172 |
| Tmeff2 | 0,66314 | 0,168509 | -1,97649 | 0,0052 | 0,0465172 |
| Stx3 | 4,56091 | 7,17767 | 0,654192 | 0,00525 | 0,0468338 |
| Shroom4 | 5,188 | 7,82126 | 0,592221 | 0,00525 | 0,0468338 |
| Bzw2 | 180,767 | 127,852 | -0,499657 | 0,00525 | 0,0468338 |
| Mgat5 | 7,96955 | 5,05975 | -0,655432 | 0,00525 | 0,0468338 |
| Zbtb41 | 4,44933 | 6,60208 | 0,569334 | 0,0053 | 0,0472142 |
| Arhgap19 | 15,8574 | 11,0683 | -0,518719 | 0,0053 | 0,0472142 |
| Uty | 2,54652 | 4,03472 | 0,66394 | 0,00535 | 0,0475605 |
| Eif5a2 | 7,00094 | 10,2415 | 0,548804 | 0,00535 | 0,0475605 |
| Mysm1 | 5,17134 | 7,55375 | 0,546655 | 0,00535 | 0,0475605 |
| Mmp16 | 2,79629 | 4,4593 | 0,673306 | 0,0054 | 0,0479385 |
| Celf2 | 1,27239 | 0,71778 | -0,825922 | 0,0054 | 0,0479385 |
| Frrs1 | 2,12846 | 3,76679 | 0,823524 | 0,00545 | 0,0482822 |
| Adam23 | 2,4829 | 4,12965 | 0,733992 | 0,00545 | 0,0482822 |
| Arid4a | 6,7288 | 9,87826 | 0,553907 | 0,00545 | 0,0482822 |
| 3930402G23Rik | 0,21571 | 0,784755 | 1,86315 | 0,0055 | 0,0485575 |

| | | | | | |
|---|---|---|---|---|---|
| Hdac11 | 8,54857 | 13,0456 | 0,609809 | 0,0055 | 0,0485575 |
| Herc2 | 10,5141 | 14,9968 | 0,512323 | 0,0055 | 0,0485575 |
| Tek | 0,854865 | 0,381933 | -1,16238 | 0,0055 | 0,0485575 |
| Thbd | 0,629657 | 0,192982 | -1,7061 | 0,0055 | 0,0485575 |
| Sytl2 | 0,31885 | 0,749764 | 1,23356 | 0,00555 | 0,0488979 |
| Rapgef6 | 8,71676 | 12,4661 | 0,516142 | 0,00555 | 0,0488979 |
| Suox | 4,86516 | 2,78829 | -0,803104 | 0,00555 | 0,0488979 |
| Syk | 0,912313 | 1,64826 | 0,853348 | 0,0056 | 0,0493046 |
| Tgm3 | 0,0499766 | 3,10354 | 5,95652 | 0,00565 | 0,0495408 |
| Fam126b | 1,4437 | 2,31037 | 0,678358 | 0,00565 | 0,0495408 |
| Irx1 | 13,1018 | 19,4691 | 0,571422 | 0,00565 | 0,0495408 |
| Wrn | 4,34612 | 6,43845 | 0,566986 | 0,00565 | 0,0495408 |
| Tnfrsf19 | 46,155 | 65,1878 | 0,498114 | 0,00565 | 0,0495408 |
| Vangl1 | 5,26938 | 3,36488 | -0,647078 | 0,00565 | 0,0495408 |
| Piwil4 | 0,299967 | 0,902326 | 1,58884 | 0,0057 | 0,049911 |
| Chd3 | 37,9797 | 53,655 | 0,498483 | 0,0057 | 0,049911 |
| Ablim2 | 0,0157033 | 1,0683 | 6,0881 | 0,00575 | 0,0502459 |
| Ikzf3 | 0,232699 | 0,639557 | 1,45861 | 0,00575 | 0,0502459 |
| Plaur | 1,6101 | 0,394271 | -2,02989 | 0,00575 | 0,0502459 |
| Slc26a10 | 0,510063 | 1,38424 | 1,44035 | 0,00585 | 0,0510502 |
| Grik1 | 0,696426 | 1,4175 | 1,02531 | 0,00585 | 0,0510502 |
| Cdkl5 | 1,57787 | 2,80067 | 0,827792 | 0,0059 | 0,0514166 |
| Fbln7 | 1,95623 | 3,44749 | 0,817472 | 0,0059 | 0,0514166 |
| Olfml2a | 1,2833 | 0,482889 | -1,4101 | 0,00595 | 0,0518171 |
| Fcho2 | 8,92245 | 13,0139 | 0,544545 | 0,006 | 0,0521817 |
| 4930412O13Rik | 1,07603 | 0,491874 | -1,12936 | 0,006 | 0,0521817 |
| Rapgefl1 | 1,08025 | 1,9515 | 0,853215 | 0,00605 | 0,0524742 |
| Ptprk | 22,0763 | 31,4306 | 0,509671 | 0,00605 | 0,0524742 |
| Nipal1 | 0,693435 | 0,264518 | -1,3904 | 0,00605 | 0,0524742 |
| Tcf21 | 2,28258 | 0,288436 | -2,98434 | 0,00605 | 0,0524742 |
| Zfp148 | 8,85215 | 12,6913 | 0,519745 | 0,0061 | 0,0527297 |
| Add3 | 52,8952 | 37,4062 | -0,499858 | 0,0061 | 0,0527297 |
| Sema3f | 15,8667 | 10,8943 | -0,542425 | 0,0061 | 0,0527297 |
| D630003M21Rik | 1,85449 | 1,01238 | -0,873278 | 0,0061 | 0,0527297 |
| Pitx2 | 0,795641 | 0,179942 | -2,14459 | 0,0061 | 0,0527296 |
| Gng4 | 1,05265 | 0,417063 | -1,33569 | 0,00615 | 0,0530903 |
| Dleu7 | 1,68956 | 0,406812 | -2,05421 | 0,00615 | 0,0530903 |
| B230216G23Rik | 1,12052 | 2,56004 | 1,192 | 0,0062 | 0,0534141 |
| Myo1e | 3,88373 | 5,95267 | 0,616093 | 0,0062 | 0,0534141 |
| Mrps33 | 6,56769 | 3,43871 | -0,933518 | 0,0062 | 0,0534141 |
| Pitpnc1 | 10,3745 | 7,14788 | -0,537453 | 0,00625 | 0,0537726 |
| Tns4 | 0,937662 | 0,438066 | -1,09792 | 0,00625 | 0,0537726 |
| Aldh9a1 | 16,5212 | 11,3391 | -0,543018 | 0,0063 | 0,0541302 |
| Arxes2 | 17,1833 | 10,9832 | -0,64571 | 0,0063 | 0,0541302 |
| Fkbp1b | 2,26929 | 5,00359 | 1,14073 | 0,00635 | 0,054414 |
| Nanos1 | 1,75964 | 2,98658 | 0,763215 | 0,00635 | 0,054414 |
| Gas7 | 1,28191 | 2,11278 | 0,720849 | 0,00635 | 0,054414 |
| Fam115c | 2,43381 | 3,9722 | 0,706722 | 0,00635 | 0,054414 |
| Cyp46a1 | 1,01651 | 0,295813 | -1,78086 | 0,0064 | 0,0547693 |
| Neurog1 | 3,4726 | 0,131617 | -4,7216 | 0,0064 | 0,0547693 |

| | | | | | |
|---|---|---|---|---|---|
| C030039L03Rik | 0,757513 | 1,42699 | 0,913631 | 0,00645 | 0,0551237 |
| Tbcel | 7,94511 | 11,5083 | 0,534534 | 0,00645 | 0,0551236 |
| Cep290 | 3,03726 | 6,06994 | 0,998909 | 0,0065 | 0,0554402 |
| Rabgap1l | 2,23044 | 3,79699 | 0,767528 | 0,0065 | 0,0554402 |
| Lima1 | 30,9647 | 21,9119 | -0,498911 | 0,0065 | 0,0554402 |
| Xiap | 15,5469 | 22,1159 | 0,508463 | 0,00655 | 0,0557924 |
| Vkorc1 | 21,3721 | 12,7134 | -0,749384 | 0,00655 | 0,0557924 |
| Vps13b | 3,5567 | 5,11144 | 0,523189 | 0,0066 | 0,0561438 |
| Bcor | 22,2441 | 31,2435 | 0,490134 | 0,0066 | 0,0561438 |
| Lor | 0,77902 | 1,90369 | 1,28906 | 0,00665 | 0,0563449 |
| Sema6c | 4,1966 | 6,43421 | 0,616543 | 0,00665 | 0,0563449 |
| Phf20l1 | 8,70439 | 12,4695 | 0,518586 | 0,00665 | 0,0563449 |
| Stag2 | 29,4243 | 41,6469 | 0,501199 | 0,00665 | 0,0563449 |
| Cotl1 | 98,7596 | 70,0613 | -0,495304 | 0,00665 | 0,0563449 |
| Tmem41a | 21,2044 | 13,7143 | -0,628682 | 0,00665 | 0,0563449 |
| 2310030G06Rik | 0,837101 | 2,32791 | 1,47556 | 0,0067 | 0,0566563 |
| Col24a1 | 0,677054 | 1,21952 | 0,848968 | 0,0067 | 0,0566563 |
| Ern1 | 2,31914 | 3,81211 | 0,717003 | 0,0067 | 0,0566563 |
| Il17rc | 1,02456 | 2,08999 | 1,02849 | 0,00675 | 0,0570039 |
| H3f3a | 32,7674 | 21,6755 | -0,596194 | 0,00675 | 0,0570039 |
| Pcp4 | 53,1481 | 34,6085 | -0,618889 | 0,0068 | 0,0573884 |
| Dusp26 | 1,51262 | 3,04282 | 1,00836 | 0,0069 | 0,0581177 |
| Mapkbp1 | 4,52288 | 6,64312 | 0,554619 | 0,0069 | 0,0581177 |
| Lrrc8d | 10,4556 | 7,15767 | -0,546712 | 0,0069 | 0,0581177 |
| Tdrd7 | 4,62501 | 2,85908 | -0,693908 | 0,007 | 0,0589213 |
| Rnf182 | 2,14703 | 3,6155 | 0,751849 | 0,00705 | 0,0591868 |
| Kcnip3 | 3,10482 | 5,03146 | 0,696467 | 0,00705 | 0,0591868 |
| A230046K03Rik | 10,3539 | 14,893 | 0,524462 | 0,00705 | 0,0591868 |
| Rab11fip4 | 8,64091 | 5,68752 | -0,603384 | 0,00705 | 0,0591868 |
| Rln1 | 0,547432 | 3,71733 | 2,76352 | 0,00715 | 0,0599087 |
| Col4a2 | 35,261 | 25,086 | -0,491191 | 0,00715 | 0,0599087 |
| Ralgds | 12,0089 | 8,13742 | -0,561463 | 0,00715 | 0,0599087 |
| Asap3 | 2,61717 | 4,02453 | 0,620809 | 0,0072 | 0,0601704 |
| Vps13a | 2,05846 | 3,10257 | 0,5919 | 0,0072 | 0,0601704 |
| H2-Ke6 | 27,5017 | 17,7965 | -0,627927 | 0,0072 | 0,0601704 |
| Tmem51 | 3,06887 | 1,55659 | -0,979316 | 0,0072 | 0,0601704 |
| Spint2 | 6,65774 | 11,0131 | 0,726123 | 0,00725 | 0,0603916 |
| Cpn1 | 7,35524 | 11,587 | 0,655669 | 0,00725 | 0,0603916 |
| 2810474O19Rik | 8,32346 | 11,9213 | 0,518287 | 0,00725 | 0,0603916 |
| Sox3 | 11,2163 | 7,28266 | -0,623064 | 0,00725 | 0,0603916 |
| Gm715 | 2,16742 | 0,765863 | -1,50082 | 0,00725 | 0,0603916 |
| Cep112 | 2,93348 | 4,67975 | 0,673815 | 0,0073 | 0,0607292 |
| Hs3st3a1 | 3,49493 | 1,95661 | -0,836907 | 0,0073 | 0,0607292 |
| Ptpn5 | 0,477155 | 1,08705 | 1,18789 | 0,00735 | 0,0609081 |
| C1ql1 | 2,90979 | 5,48785 | 0,915326 | 0,00735 | 0,0609081 |
| BC018507 | 23,0283 | 32,2722 | 0,486883 | 0,00735 | 0,0609081 |
| Arhgap33 | 7,91124 | 5,34357 | -0,566099 | 0,00735 | 0,0609081 |
| Lrrtm3 | 1,19581 | 0,586306 | -1,02826 | 0,00735 | 0,0609081 |
| Cldn10 | 1,81865 | 0,504896 | -1,84881 | 0,00735 | 0,0609081 |
| 2610020H08Rik | 9,30071 | 14,084 | 0,598641 | 0,0074 | 0,0612434 |

| | | | | | |
|---|---|---|---|---|---|
| E2f2 | 4,97731 | 3,25861 | -0,61111 | 0,0074 | 0,0612434 |
| Cited4 | 0,276969 | 1,65632 | 2,58018 | 0,00745 | 0,0615381 |
| Suco | 11,6543 | 16,3993 | 0,492778 | 0,00745 | 0,0615381 |
| Atf3 | 2,58211 | 1,26495 | -1,02947 | 0,00745 | 0,0615381 |
| Cep68 | 22,0497 | 30,8403 | 0,484061 | 0,0075 | 0,0618714 |
| Rerg | 1,28925 | 0,51685 | -1,31872 | 0,0075 | 0,0618714 |
| A930007I19Rik | 0,234596 | 0,870382 | 1,89147 | 0,00755 | 0,0621241 |
| Stxbp1 | 6,55332 | 9,62482 | 0,554534 | 0,00755 | 0,0621241 |
| Zfp329 | 5,78361 | 8,29945 | 0,521046 | 0,00755 | 0,0621241 |
| Rps6ka1 | 24,5003 | 34,2796 | 0,48455 | 0,00755 | 0,0621241 |
| Klk8 | 4,89175 | 8,41887 | 0,783277 | 0,00765 | 0,0629065 |
| Ttyh1 | 0,290583 | 0,757623 | 1,38253 | 0,0077 | 0,0631962 |
| Zfp119b | 1,17995 | 2,44155 | 1,04908 | 0,0077 | 0,0631962 |
| Sstr2 | 1,34718 | 0,32458 | -2,0533 | 0,0077 | 0,0631962 |
| Ash1l | 9,24969 | 12,9275 | 0,48297 | 0,00775 | 0,0635659 |
| Nrk | 1,34024 | 0,764921 | -0,809104 | 0,0078 | 0,0639351 |
| Itprip | 11,3001 | 16,2893 | 0,527591 | 0,00785 | 0,0643038 |
| Sec14l2 | 3,37248 | 5,49381 | 0,703996 | 0,0079 | 0,0646721 |
| Lad1 | 0,686964 | 1,42746 | 1,05514 | 0,008 | 0,065449 |
| Kif21a | 4,35846 | 2,93854 | -0,568716 | 0,00805 | 0,0658161 |
| Zfp7 | 13,029 | 18,7015 | 0,521431 | 0,00815 | 0,0665065 |
| Auts2 | 15,9431 | 22,2624 | 0,481676 | 0,00815 | 0,0665065 |
| Dpysl4 | 14,0564 | 9,4963 | -0,565792 | 0,00815 | 0,0665065 |
| Kcnk1 | 2,7321 | 4,66702 | 0,772488 | 0,0082 | 0,0668295 |
| Ndp | 0,975135 | 0,241487 | -2,01366 | 0,0082 | 0,0668295 |
| Lsm14b | 70,3703 | 50,7334 | -0,472032 | 0,00825 | 0,0671516 |
| Id4 | 2,67328 | 1,25871 | -1,08667 | 0,00825 | 0,0671517 |
| Nr1d1 | 2,08343 | 3,57346 | 0,77836 | 0,0083 | 0,0675158 |
| Tnip1 | 5,97159 | 9,09415 | 0,606824 | 0,00835 | 0,0678365 |
| Uba6 | 9,09697 | 12,9595 | 0,510548 | 0,00835 | 0,0678365 |
| Dse | 16,3451 | 11,496 | -0,507729 | 0,0084 | 0,0681564 |
| Tfpi | 20,7124 | 13,7686 | -0,589113 | 0,0084 | 0,0681564 |
| Ankrd45 | 1,80056 | 3,00363 | 0,738262 | 0,00845 | 0,0684322 |
| Atp6ap2 | 20,8033 | 29,5518 | 0,506429 | 0,00845 | 0,0684322 |
| Smc2 | 43,2446 | 60,7106 | 0,489427 | 0,00845 | 0,0684322 |
| Zfp654 | 3,32401 | 5,13982 | 0,628795 | 0,0085 | 0,0685774 |
| Foxc2 | 55,5569 | 39,6861 | -0,485334 | 0,0085 | 0,0685774 |
| Rpl35a | 232,325 | 164,646 | -0,496779 | 0,0085 | 0,0685774 |
| Cdc42ep3 | 3,206 | 1,71388 | -0,903509 | 0,0085 | 0,0685774 |
| Prdm1 | 0,882883 | 0,433053 | -1,02768 | 0,0085 | 0,0685774 |
| 5930403L14Rik | 1,0099 | 0,392826 | -1,36224 | 0,0085 | 0,0685774 |
| Sgtb | 3,95349 | 6,20518 | 0,650345 | 0,00855 | 0,0688077 |
| Rgs2 | 5,74428 | 3,62372 | -0,664654 | 0,00855 | 0,0688077 |
| Fam81a | 1,18829 | 0,512136 | -1,21428 | 0,00855 | 0,0688077 |
| Lrmp | 0,922007 | 0,255808 | -1,84972 | 0,00855 | 0,0688077 |
| Adamts14 | 1,56706 | 2,56173 | 0,709061 | 0,0086 | 0,0691666 |
| Klhl28 | 2,80754 | 4,69029 | 0,74037 | 0,00865 | 0,0694817 |
| Ackr3 | 9,8905 | 6,3498 | -0,639332 | 0,00865 | 0,0694817 |
| Sh3glb1 | 22,2483 | 31,5038 | 0,50183 | 0,0087 | 0,0698395 |
| Tmem106a | 0,368363 | 1,00853 | 1,45305 | 0,00875 | 0,0700218 |

| | | | | | |
|---|---|---|---|---|---|
| Apobec3 | 2,0577 | 3,5952 | 0,805041 | 0,00875 | 0,0700218 |
| Ap1s1 | 65,2015 | 46,1217 | -0,499461 | 0,00875 | 0,0700218 |
| Crispld1 | 12,3709 | 8,52531 | -0,537123 | 0,00875 | 0,0700218 |
| Mfng | 5,04467 | 2,91618 | -0,790679 | 0,00875 | 0,0700218 |
| Fzd7 | 48,4676 | 34,8735 | -0,474892 | 0,0088 | 0,0702904 |
| Stk17b | 3,15603 | 1,86804 | -0,75659 | 0,0088 | 0,0702904 |
| Lgr5 | 0,680109 | 0,299895 | -1,18131 | 0,0088 | 0,0702904 |
| Pnliprp2 | 1,21529 | 2,63975 | 1,1191 | 0,00885 | 0,0706458 |
| Lefty1 | 0,151484 | 1,40026 | 3,20845 | 0,0089 | 0,0709125 |
| Bmpr1b | 1,57219 | 2,55487 | 0,700476 | 0,0089 | 0,0709125 |
| Lhx6 | 1,1685 | 0,554954 | -1,07422 | 0,0089 | 0,0709125 |
| Nefl | 3,77702 | 2,09942 | -0,847258 | 0,00895 | 0,0712667 |
| Cnr1 | 1,12872 | 1,89635 | 0,748537 | 0,009 | 0,0715316 |
| Glipr2 | 16,6522 | 24,0818 | 0,532235 | 0,009 | 0,0715316 |
| Ckap5 | 52,7375 | 73,0376 | 0,46981 | 0,009 | 0,0715316 |
| Cmbl | 2,28173 | 0,665327 | -1,77799 | 0,00905 | 0,0718845 |
| Zbtb37 | 2,40949 | 3,86298 | 0,680986 | 0,00915 | 0,072544 |
| Gli3 | 20,8503 | 15,0519 | -0,470122 | 0,00915 | 0,072544 |
| Fgf10 | 0,737579 | 0,335068 | -1,13835 | 0,00915 | 0,072544 |
| Bik | 0,482545 | 2,23696 | 2,2128 | 0,0092 | 0,0728954 |
| Cyp4f17 | 1,85577 | 3,49666 | 0,913962 | 0,00925 | 0,0732011 |
| Lhfp | 5,0417 | 7,97093 | 0,660838 | 0,00925 | 0,0732011 |
| Fam199x | 4,55482 | 6,58417 | 0,531608 | 0,0093 | 0,0735061 |
| Ripply1 | 4,92194 | 1,92232 | -1,35638 | 0,0093 | 0,0735061 |
| Gpc1 | 41,7362 | 57,667 | 0,466445 | 0,00935 | 0,0738558 |
| Susd5 | 0,861412 | 1,58231 | 0,877259 | 0,0094 | 0,0740683 |
| Egr1 | 4,22563 | 6,77286 | 0,680598 | 0,0094 | 0,0740683 |
| Rbp1 | 26,8247 | 37,4367 | 0,480891 | 0,0094 | 0,0740683 |
| Actn1 | 63,2692 | 87,9215 | 0,474712 | 0,0094 | 0,0740683 |
| Cux2 | 2,69138 | 1,66972 | -0,68874 | 0,00945 | 0,0744166 |
| Megf10 | 1,10347 | 0,63519 | -0,796787 | 0,0095 | 0,0747644 |
| Plagl2 | 30,7866 | 42,7706 | 0,474315 | 0,00955 | 0,0750198 |
| Ctnnd2 | 1,46401 | 0,852621 | -0,779951 | 0,00955 | 0,0750198 |
| Chst1 | 1,48864 | 0,712737 | -1,06256 | 0,00955 | 0,0750198 |
| Esf1 | 25,4309 | 35,8522 | 0,495481 | 0,0096 | 0,0752743 |
| Hyou1 | 73,9761 | 102,108 | 0,46497 | 0,0096 | 0,0752743 |
| Dab1 | 2,37703 | 0,829498 | -1,51885 | 0,0096 | 0,0752743 |
| Fut8 | 19,4201 | 13,6392 | -0,509792 | 0,00965 | 0,075574 |
| Vamp5 | 1,00961 | 0,240261 | -2,07113 | 0,00965 | 0,075574 |
| Erich2 | 0,405497 | 2,77358 | 2,77399 | 0,0097 | 0,0758729 |
| Pigv | 6,89942 | 10,639 | 0,624814 | 0,0097 | 0,0758729 |
| Hpse | 0,415781 | 0,98141 | 1,23903 | 0,00975 | 0,0761248 |
| Gpr157 | 0,395146 | 0,820798 | 1,05464 | 0,00975 | 0,0761248 |
| Tle4 | 24,2624 | 33,7313 | 0,475363 | 0,00975 | 0,0761248 |
| Rnf122 | 18,6035 | 13,1132 | -0,504553 | 0,0098 | 0,0764686 |
| Klc4 | 12,8872 | 18,6468 | 0,532989 | 0,00985 | 0,076812 |
| B3gnt8 | 0,183806 | 0,822226 | 2,16135 | 0,00995 | 0,0774506 |
| Hook3 | 6,36784 | 8,88614 | 0,480754 | 0,00995 | 0,0774506 |
| Bub1 | 54,4215 | 75,4175 | 0,470723 | 0,00995 | 0,0774506 |
| 2010010A06Rik | 0,365378 | 7,06718 | 4,27367 | 0,01 | 0,0776984 |

| | | | | | |
|---|---|---|---|---|---|
| Ppp2r3a | 5,2716 | 7,49856 | 0,508371 | 0,01 | 0,0776984 |
| Scg3 | 0,920471 | 0,0990856 | -3,21563 | 0,01 | 0,0776984 |
| Cdh2 | 47,3548 | 65,5905 | 0,469975 | 0,01005 | 0,0779924 |
| Adcyap1r1 | 0,867973 | 0,464355 | -0,902421 | 0,01005 | 0,0779924 |
| Slc38a6 | 4,53182 | 6,9442 | 0,615717 | 0,0101 | 0,0782857 |
| Gen1 | 10,424 | 14,8602 | 0,511547 | 0,0101 | 0,0782857 |
| Glb1l2 | 1,28593 | 2,23742 | 0,799026 | 0,01015 | 0,078389 |
| Adcy1 | 1,10644 | 1,68986 | 0,610973 | 0,01015 | 0,078389 |
| Ccdc18 | 3,83508 | 5,82404 | 0,602766 | 0,01015 | 0,078389 |
| 03/07/16 | 31,5049 | 44,1112 | 0,48557 | 0,01015 | 0,078389 |
| Lrrn1 | 15,6748 | 11,0066 | -0,510073 | 0,01015 | 0,078389 |
| C130026L21Rik | 0,983499 | 0,199845 | -2,29904 | 0,01015 | 0,078389 |
| Selenbp1 | 1,83236 | 3,46074 | 0,917372 | 0,0102 | 0,0785859 |
| Pcdh8 | 42,2527 | 58,312 | 0,464749 | 0,0102 | 0,0785859 |
| Fabp5 | 75,5099 | 52,9914 | -0,510908 | 0,0102 | 0,0785859 |
| Rhpn1 | 5,39797 | 3,34106 | -0,69211 | 0,0102 | 0,0785859 |
| Kcnc3 | 1,49526 | 2,42503 | 0,697612 | 0,01025 | 0,0788291 |
| Mycbp2 | 7,4797 | 10,343 | 0,467599 | 0,01025 | 0,0788291 |
| Ccdc40 | 3,83614 | 2,49019 | -0,623399 | 0,01025 | 0,0788291 |
| Wnt10a | 0,0995269 | 1,31484 | 3,72365 | 0,0103 | 0,0789769 |
| Serac1 | 2,44705 | 3,91609 | 0,678371 | 0,0103 | 0,0789769 |
| Higd1a | 12,0361 | 7,95693 | -0,597079 | 0,0103 | 0,0789769 |
| Anpep | 4,2091 | 2,65534 | -0,664616 | 0,0103 | 0,0789769 |
| Irf5 | 1,70262 | 0,807708 | -1,07585 | 0,0103 | 0,0789769 |
| Gck | 0,245256 | 0,761916 | 1,63534 | 0,01035 | 0,0792655 |
| Otud1 | 0,410406 | 1,02423 | 1,31942 | 0,01035 | 0,0792656 |
| Actn2 | 0,348693 | 0,872941 | 1,32393 | 0,0104 | 0,0795535 |
| Nufip2 | 10,4213 | 14,4799 | 0,474515 | 0,0104 | 0,0795535 |
| Spin4 | 1,9949 | 3,18209 | 0,673658 | 0,0105 | 0,0802227 |
| Fbxl17 | 2,36069 | 3,3382 | 0,499862 | 0,0105 | 0,0802228 |
| Osbpl2 | 7,71898 | 11,4082 | 0,563586 | 0,01055 | 0,0805568 |
| Ndufs4 | 66,4064 | 47,6049 | -0,480212 | 0,0106 | 0,0808904 |
| Dtx3l | 0,966709 | 1,79398 | 0,892009 | 0,01065 | 0,0810791 |
| Zfp503 | 32,1986 | 23,2922 | -0,467151 | 0,01065 | 0,0810791 |
| Cdon | 10,5128 | 7,51148 | -0,484982 | 0,01065 | 0,0810791 |
| Acot1 | 7,53843 | 4,54636 | -0,729553 | 0,01065 | 0,0810791 |
| Hspa1a | 2,54604 | 4,43484 | 0,800627 | 0,0107 | 0,0814114 |
| 5430421N21Rik | 0,193324 | 2,2494 | 3,54045 | 0,01075 | 0,08155 |
| Fam19a4 | 0,741408 | 1,60986 | 1,1186 | 0,01075 | 0,08155 |
| Timm50 | 112,166 | 155,538 | 0,471634 | 0,01075 | 0,08155 |
| Fam114a1 | 4,43919 | 2,76556 | -0,682725 | 0,01075 | 0,08155 |
| Gca | 0,672346 | 0,254681 | -1,40051 | 0,01075 | 0,08155 |
| Myo5c | 0,577928 | 1,02693 | 0,829373 | 0,0108 | 0,0818325 |
| Plekha1 | 24,852 | 34,4514 | 0,471199 | 0,0108 | 0,0818325 |
| Rnd1 | 0,909295 | 1,84326 | 1,01944 | 0,01085 | 0,0819693 |
| Chst11 | 6,35875 | 9,02136 | 0,504601 | 0,01085 | 0,0819693 |
| Acvr1b | 14,6646 | 20,5734 | 0,488437 | 0,01085 | 0,0819693 |
| Far1 | 27,3277 | 37,9796 | 0,474862 | 0,01085 | 0,0819693 |
| Ostf1 | 5,2515 | 3,02666 | -0,795004 | 0,01085 | 0,0819693 |
| Creld1 | 9,20146 | 13,7193 | 0,576275 | 0,0109 | 0,0822985 |

| | | | | | |
|---|---|---|---|---|---|
| Dusp2 | 0,489821 | 1,39297 | 1,50784 | 0,01095 | 0,0825788 |
| Pcm1 | 24,8978 | 34,7912 | 0,482702 | 0,01095 | 0,0825788 |
| Stk38l | 6,80184 | 9,72072 | 0,515138 | 0,011 | 0,0829072 |
| Cbr3 | 1,14242 | 2,70235 | 1,24212 | 0,01105 | 0,0832351 |
| Rnf39 | 0,233887 | 1,28495 | 2,45782 | 0,0111 | 0,0835627 |
| Slc45a4 | 5,8129 | 8,48249 | 0,545232 | 0,01115 | 0,0838407 |
| Dctpp1 | 276,191 | 200,153 | -0,464563 | 0,01115 | 0,0838407 |
| Agap1 | 7,19921 | 5,16566 | -0,478885 | 0,0112 | 0,084118 |
| Ift27 | 40,5445 | 28,0907 | -0,529415 | 0,0112 | 0,084118 |
| Tdrd1 | 0,642414 | 1,1825 | 0,880267 | 0,01125 | 0,0844441 |
| Tbc1d9 | 3,20472 | 2,10405 | -0,607026 | 0,0113 | 0,0847203 |
| Nr1h5 | 4,24449 | 2,57052 | -0,723533 | 0,0113 | 0,0847203 |
| Sec31b | 0,446888 | 0,93461 | 1,06445 | 0,0114 | 0,0854201 |
| Bcl11b | 1,42266 | 2,21098 | 0,636099 | 0,01145 | 0,0857447 |
| Tyro3 | 17,6333 | 24,3849 | 0,467682 | 0,0115 | 0,0860688 |
| Ocln | 1,0498 | 1,88613 | 0,845319 | 0,0116 | 0,0866656 |
| Setd1b | 8,73439 | 12,1224 | 0,472892 | 0,0116 | 0,0866656 |
| Plxnb1 | 20,0572 | 27,4696 | 0,453718 | 0,0116 | 0,0866656 |
| Sgpl1 | 30,9877 | 42,6084 | 0,459442 | 0,01165 | 0,0869885 |
| Fam161a | 1,27113 | 2,47436 | 0,960945 | 0,01175 | 0,0876331 |
| 1810011O10Rik | 14,6299 | 9,42464 | -0,63441 | 0,01175 | 0,0876331 |
| Cbln2 | 0,838392 | 0,232117 | -1,85277 | 0,01185 | 0,0883276 |
| Tmx4 | 5,52469 | 7,92679 | 0,520842 | 0,0119 | 0,0886488 |
| Bai2 | 0,651625 | 1,16582 | 0,839226 | 0,01195 | 0,0887635 |
| Mum1l1 | 12,4523 | 17,4547 | 0,487207 | 0,01195 | 0,0887635 |
| Fam217b | 1,53728 | 0,696583 | -1,14202 | 0,01195 | 0,0887635 |
| Hgf | 1,47407 | 0,652977 | -1,1747 | 0,01195 | 0,0887635 |
| Dok4 | 4,90417 | 1,83092 | -1,42144 | 0,01195 | 0,0887635 |
| H2-Bl | 0,827594 | 2,17206 | 1,39207 | 0,012 | 0,0889803 |
| 1700030J22Rik | 0,835023 | 1,60052 | 0,93865 | 0,012 | 0,0889803 |
| Usp34 | 14,694 | 22,189 | 0,594619 | 0,012 | 0,0889803 |
| Ccdc64 | 0,671329 | 1,5608 | 1,2172 | 0,0121 | 0,0895665 |
| Ltbp1 | 14,9086 | 10,8292 | -0,461221 | 0,0121 | 0,0895665 |
| Pigc | 7,82867 | 5,26733 | -0,571694 | 0,0121 | 0,0895665 |
| Gramd4 | 11,3434 | 15,8266 | 0,480497 | 0,01215 | 0,0898848 |
| Stbd1 | 4,95412 | 7,81563 | 0,657735 | 0,0122 | 0,0901507 |
| Tcp11l2 | 2,22784 | 1,19727 | -0,895899 | 0,0122 | 0,0901507 |
| Fam102a | 6,29544 | 4,26518 | -0,561699 | 0,01225 | 0,090416 |
| 2810405F15Rik | 1,36271 | 0,442226 | -1,62363 | 0,01225 | 0,090416 |
| Crhbp | 0,203271 | 0,968386 | 2,25218 | 0,0124 | 0,0914705 |
| Arhgap10 | 20,0818 | 27,6532 | 0,461556 | 0,01245 | 0,0917866 |
| Ccdc177 | 0,772033 | 0,36696 | -1,07304 | 0,0125 | 0,0920494 |
| Lurap1l | 1,3594 | 0,522262 | -1,38013 | 0,0125 | 0,0920494 |
| Fam13c | 2,20406 | 4,41884 | 1,0035 | 0,01255 | 0,0923645 |
| Prr5 | 8,90947 | 13,2685 | 0,574593 | 0,0126 | 0,0925732 |
| Mtss1l | 17,3749 | 24,083 | 0,471011 | 0,0126 | 0,0925732 |
| Chn2 | 2,06782 | 1,08418 | -0,931502 | 0,0126 | 0,0925732 |
| Rbpms2 | 13,2103 | 19,0135 | 0,525356 | 0,0127 | 0,0931479 |
| Strbp | 7,70235 | 10,5 | 0,447024 | 0,0127 | 0,0931479 |
| Frat2 | 2,00926 | 1,01513 | -0,985005 | 0,0127 | 0,0931479 |

| | | | | | |
|---|---|---|---|---|---|
| Snap25 | 0,724684 | 0,142506 | -2,34633 | 0,01275 | 0,0934611 |
| Car2 | 5,00348 | 7,91712 | 0,662043 | 0,0128 | 0,0937205 |
| Gatad2b | 16,7048 | 23,8789 | 0,515474 | 0,0128 | 0,0937205 |
| Ticam2 | 0,570058 | 1,16545 | 1,03171 | 0,01285 | 0,0938189 |
| Tspo | 15,8388 | 24,2491 | 0,614473 | 0,01285 | 0,0938189 |
| Rbm41 | 4,23769 | 6,15767 | 0,539105 | 0,01285 | 0,0938189 |
| Chchd3 | 89,7698 | 65,6014 | -0,452503 | 0,01285 | 0,0938189 |
| Bcl9l | 7,16071 | 5,02075 | -0,5122 | 0,01285 | 0,0938189 |
| Prkar2b | 18,6356 | 13,3677 | -0,479309 | 0,0129 | 0,0941304 |
| Slc50a1 | 15,1635 | 9,50124 | -0,674415 | 0,013 | 0,0948061 |
| Ism2 | 0,415132 | 1,03711 | 1,32093 | 0,01305 | 0,0950627 |
| Aprt | 139,842 | 101,645 | -0,460261 | 0,01305 | 0,0950627 |
| Zmym2 | 24,3225 | 33,5121 | 0,462394 | 0,01315 | 0,0957368 |
| 6720416L17Rik | 11,2941 | 16,3091 | 0,530109 | 0,01325 | 0,0964101 |
| 2410004P03Rik | 0,203602 | 0,806691 | 1,98627 | 0,0133 | 0,0965002 |
| Parp4 | 1,1023 | 1,77522 | 0,687481 | 0,0133 | 0,0965002 |
| Nav2 | 3,04633 | 4,30444 | 0,498755 | 0,0133 | 0,0965002 |
| Usp2 | 1,11975 | 0,55832 | -1,00402 | 0,0133 | 0,0965002 |
| Pipox | 1,06021 | 0,340579 | -1,63828 | 0,0133 | 0,0965002 |
| Nckap5l | 10,1994 | 14,2019 | 0,477592 | 0,0134 | 0,0971708 |
| Tro | 22,3357 | 30,7075 | 0,459242 | 0,01345 | 0,0974233 |
| Fzd9 | 3,32332 | 1,89829 | -0,807923 | 0,01345 | 0,0974233 |
| Eid1 | 76,675 | 56,2637 | -0,446551 | 0,0135 | 0,0977302 |
| Jph4 | 0,525718 | 1,00264 | 0,931437 | 0,01355 | 0,0979264 |
| Kcnj15 | 0,598884 | 1,04697 | 0,805873 | 0,01355 | 0,0979264 |
| A830010M20Rik | 1,0812 | 1,72813 | 0,676569 | 0,01355 | 0,0979264 |
| Hoxa1 | 12,3983 | 17,5438 | 0,500816 | 0,01365 | 0,0985381 |
| Hpse2 | 0,712521 | 0,152067 | -2,22823 | 0,01365 | 0,0985381 |
| Pnmal1 | 1,43081 | 0,576413 | -1,31166 | 0,01375 | 0,0992042 |
| Tapbp | 3,40787 | 5,32691 | 0,644429 | 0,0138 | 0,0993414 |
| Nol8 | 15,6249 | 21,4832 | 0,459357 | 0,0138 | 0,0993414 |
| Sh3kbp1 | 5,66211 | 3,91896 | -0,530869 | 0,0138 | 0,0993414 |
| Scarf2 | 9,33026 | 6,39991 | -0,543866 | 0,0138 | 0,0993414 |
| Adrb2 | 0,34538 | 0,933224 | 1,43404 | 0,01385 | 0,0995338 |
| Suz12 | 37,3259 | 50,4869 | 0,435732 | 0,01385 | 0,0995338 |
| Bex1 | 74,9177 | 53,3726 | -0,489209 | 0,01385 | 0,0995338 |
| Plekhg2 | 28,1909 | 38,4869 | 0,449136 | 0,0139 | 0,0998372 |
| Cox5b | 74,5897 | 51,4299 | -0,536371 | 0,01395 | 0,10014 |
| Sema3e | 1,87152 | 2,82287 | 0,592952 | 0,014 | 0,100275 |
| D930016D06Rik | 7,19007 | 10,4452 | 0,538758 | 0,014 | 0,100275 |
| Fam135a | 12,2434 | 16,8985 | 0,464893 | 0,014 | 0,100275 |
| Pip4k2a | 8,32825 | 5,70274 | -0,546358 | 0,014 | 0,100275 |
| Nomo1 | 34,8806 | 47,5316 | 0,446462 | 0,0141 | 0,100878 |
| S1pr1 | 2,89796 | 1,73465 | -0,740394 | 0,0141 | 0,100878 |
| Arl3 | 130,403 | 94,7251 | -0,461162 | 0,01415 | 0,10118 |
| Parp9 | 0,875749 | 1,87972 | 1,10193 | 0,0142 | 0,101481 |
| D330045A20Rik | 0,884348 | 1,59979 | 0,855195 | 0,01425 | 0,101668 |
| Grb7 | 3,38577 | 1,99285 | -0,764649 | 0,01425 | 0,101668 |
| Bdh2 | 3,79806 | 1,91942 | -0,984592 | 0,01425 | 0,101668 |
| Slc7a3 | 0,668416 | 1,41053 | 1,07742 | 0,0143 | 0,101742 |

| | | | | | |
|---|---|---|---|---|---|
| Sgms2 | 0,359878 | 0,696255 | 0,952105 | 0,0143 | 0,101742 |
| Synm | 3,28422 | 4,67981 | 0,510896 | 0,0143 | 0,101742 |
| Dnm1l | 32,8337 | 44,7694 | 0,447335 | 0,0143 | 0,101742 |
| Irs4 | 0,607826 | 0,304289 | -0,998215 | 0,0143 | 0,101742 |
| Tdrp | 0,872931 | 1,71186 | 0,971625 | 0,0144 | 0,102226 |
| Acot9 | 4,45952 | 7,24272 | 0,699644 | 0,0144 | 0,102226 |
| Hoxb2 | 34,5639 | 47,9547 | 0,472406 | 0,0144 | 0,102226 |
| Zmynd8 | 34,5282 | 47,2275 | 0,451852 | 0,0144 | 0,102226 |
| Arhgap26 | 0,474924 | 0,826656 | 0,79959 | 0,01445 | 0,102525 |
| Usp32 | 4,93962 | 6,91549 | 0,485432 | 0,0145 | 0,102823 |
| Erbb2 | 14,1387 | 19,9419 | 0,496152 | 0,01455 | 0,10312 |
| Pon3 | 4,15425 | 6,70419 | 0,690473 | 0,0146 | 0,10336 |
| Rmst | 0,643294 | 0,200799 | -1,67973 | 0,0146 | 0,10336 |
| Thbs2 | 0,507387 | 0,90962 | 0,842177 | 0,01465 | 0,103486 |
| Vwa5a | 2,68957 | 4,06855 | 0,59714 | 0,01465 | 0,103486 |
| Abhd2 | 4,71159 | 7,01923 | 0,575097 | 0,01465 | 0,103486 |
| 8430419L09Rik | 5,61759 | 8,01398 | 0,512569 | 0,01465 | 0,103486 |
| Per2 | 0,941321 | 1,54744 | 0,717124 | 0,01475 | 0,104078 |
| Twf1 | 26,569 | 36,3165 | 0,450886 | 0,01475 | 0,104078 |
| Csnk1g1 | 7,99442 | 11,1049 | 0,474127 | 0,0148 | 0,104373 |
| Hbegf | 1,75675 | 3,01619 | 0,779823 | 0,01485 | 0,104611 |
| Prdm13 | 1,39932 | 0,0573235 | -4,60946 | 0,01485 | 0,104611 |
| Rbks | 2,52907 | 1,02202 | -1,30719 | 0,0149 | 0,104905 |
| Cdk20 | 2,81827 | 4,70784 | 0,740255 | 0,01495 | 0,105142 |
| Epn2 | 13,5047 | 18,5037 | 0,454346 | 0,01495 | 0,105142 |
| Nfat5 | 8,22811 | 11,4382 | 0,475223 | 0,01505 | 0,105729 |
| Smim11 | 90,0639 | 61,8932 | -0,541168 | 0,01505 | 0,105729 |
| Zfp395 | 16,1523 | 22,1699 | 0,456859 | 0,0151 | 0,105964 |
| Qdpr | 62,5643 | 45,4017 | -0,462592 | 0,0151 | 0,105964 |
| Ptges2 | 11,866 | 8,03091 | -0,563202 | 0,0152 | 0,106608 |
| Vash1 | 14,4488 | 19,6874 | 0,446325 | 0,01525 | 0,106725 |
| Ldhb | 67,5166 | 48,6472 | -0,472884 | 0,01525 | 0,106725 |
| Rgl1 | 10,4979 | 7,46139 | -0,492588 | 0,01525 | 0,106725 |
| Gng8 | 2,9131 | 1,18738 | -1,29478 | 0,01525 | 0,106725 |
| Pamr1 | 0,880749 | 0,380882 | -1,20939 | 0,0153 | 0,107017 |
| Trim56 | 2,99635 | 4,58059 | 0,612327 | 0,01535 | 0,107249 |
| Traf3ip2 | 2,85825 | 1,24854 | -1,19489 | 0,01535 | 0,107249 |
| Zfp62 | 21,5914 | 29,5632 | 0,453347 | 0,0154 | 0,107482 |
| Adk | 69,4286 | 51,1381 | -0,441132 | 0,0154 | 0,107482 |
| Skap2 | 56,4227 | 76,7343 | 0,443594 | 0,0155 | 0,108062 |
| Lonrf2 | 2,59813 | 1,67619 | -0,632294 | 0,0155 | 0,108062 |
| Ncoa1 | 1,95731 | 2,89437 | 0,564381 | 0,01565 | 0,108989 |
| Rhob | 28,7989 | 39,4759 | 0,454957 | 0,01565 | 0,108989 |
| Cnksr2 | 2,84905 | 4,19164 | 0,557034 | 0,0157 | 0,109219 |
| Lasp1 | 35,9479 | 48,5564 | 0,433754 | 0,0157 | 0,109219 |
| Fnbp1 | 11,2813 | 15,8061 | 0,486555 | 0,0158 | 0,109795 |
| Tox | 0,86519 | 0,375226 | -1,20526 | 0,0158 | 0,109795 |
| Slc7a7 | 1,63293 | 6,92928 | 2,08524 | 0,01585 | 0,110083 |
| Cyp26c1 | 0,119095 | 2,03893 | 4,09763 | 0,0159 | 0,110371 |
| Ret | 1,15191 | 1,79841 | 0,642696 | 0,01595 | 0,110658 |

| | | | | | |
|---|---|---|---|---|---|
| Jph1 | 1,20455 | 1,9526 | 0,696901 | 0,01605 | 0,111231 |
| Erc1 | 11,9695 | 16,2903 | 0,444643 | 0,01605 | 0,111231 |
| Serpinb1a | 0,276349 | 0,866828 | 1,64925 | 0,0161 | 0,111337 |
| Alas2 | 0,339781 | 0,956943 | 1,49383 | 0,0161 | 0,111337 |
| Gulo | 0,42896 | 1,04228 | 1,28083 | 0,0161 | 0,111337 |
| Serping1 | 6,23956 | 3,88541 | -0,683379 | 0,0161 | 0,111337 |
| Mink1 | 10,991 | 15,0997 | 0,458203 | 0,0162 | 0,111969 |
| Micu2 | 39,4672 | 53,4565 | 0,437712 | 0,01625 | 0,112254 |
| Zbtb8b | 0,338452 | 0,773671 | 1,19277 | 0,01635 | 0,112884 |
| Fn3k | 0,423195 | 1,69147 | 1,99888 | 0,0164 | 0,113107 |
| Sgk1 | 5,17968 | 7,76019 | 0,58323 | 0,0164 | 0,113107 |
| Pitpnm1 | 2,59076 | 4,96089 | 0,937225 | 0,0165 | 0,113553 |
| Anxa3 | 6,86835 | 10,4855 | 0,610366 | 0,0165 | 0,113553 |
| Bcar3 | 4,72373 | 6,94159 | 0,555339 | 0,0165 | 0,113553 |
| Slc12a7 | 23,4849 | 31,8898 | 0,441366 | 0,0165 | 0,113553 |
| Xrn1 | 7,6688 | 10,6259 | 0,470511 | 0,0166 | 0,11418 |
| Josd2 | 18,9537 | 12,4617 | -0,604976 | 0,01665 | 0,114462 |
| Ninl | 14,4018 | 19,5945 | 0,444195 | 0,0167 | 0,114622 |
| Smg1 | 14,664 | 19,8156 | 0,434358 | 0,0167 | 0,114622 |
| Ralgps2 | 19,9778 | 14,7836 | -0,434401 | 0,0167 | 0,114622 |
| Cltb | 4,51541 | 7,30997 | 0,695011 | 0,01675 | 0,114903 |
| Fam189b | 10,5498 | 14,8458 | 0,492828 | 0,01695 | 0,116151 |
| Gm608 | 5,7231 | 7,7919 | 0,445179 | 0,01695 | 0,116151 |
| Fam174a | 5,87004 | 3,7336 | -0,652803 | 0,017 | 0,116431 |
| Dlg4 | 11,0345 | 15,3227 | 0,473648 | 0,01705 | 0,116712 |
| Dlk1 | 35,2962 | 26,1585 | -0,432232 | 0,0171 | 0,116991 |
| Crtac1 | 0,306022 | 0,779393 | 1,34872 | 0,0172 | 0,117425 |
| Dennd4b | 6,25365 | 8,66768 | 0,470946 | 0,0172 | 0,117425 |
| Sik1 | 8,38013 | 11,5779 | 0,466325 | 0,0172 | 0,117425 |
| Ptgfrn | 39,5111 | 53,2047 | 0,429294 | 0,0172 | 0,117425 |
| Isg20 | 0,422783 | 1,32526 | 1,64829 | 0,01725 | 0,117579 |
| Parp3 | 0,731822 | 1,42404 | 0,960423 | 0,01725 | 0,117579 |
| Tef | 8,28127 | 11,5039 | 0,474196 | 0,01725 | 0,117579 |
| Fxyd5 | 0,273124 | 1,42511 | 2,38345 | 0,0173 | 0,117794 |
| Fgfr4 | 1,09373 | 0,541484 | -1,01426 | 0,0173 | 0,117794 |
| Atp13a3 | 18,2932 | 24,6761 | 0,431806 | 0,01735 | 0,118072 |
| Sfn | 0,80234 | 1,76878 | 1,14047 | 0,0174 | 0,118224 |
| Vsig10 | 0,754315 | 1,34093 | 0,829997 | 0,0174 | 0,118224 |
| Slain1 | 4,15764 | 6,33887 | 0,60846 | 0,0174 | 0,118224 |
| Mrpl30 | 116,139 | 84,5911 | -0,457281 | 0,0175 | 0,118841 |
| Rasgrp2 | 0,260302 | 0,762805 | 1,55113 | 0,01755 | 0,119117 |
| Atp9a | 8,22437 | 11,544 | 0,489164 | 0,0176 | 0,119267 |
| Psme2b | 30,4523 | 42,5631 | 0,483053 | 0,0176 | 0,119267 |
| Ubl3 | 14,091 | 10,0346 | -0,489791 | 0,0176 | 0,119267 |
| Dock4 | 2,46995 | 3,53873 | 0,51875 | 0,01765 | 0,119543 |
| Slc41a1 | 8,2817 | 11,4324 | 0,465134 | 0,0177 | 0,119818 |
| Cx3cl1 | 2,3645 | 3,72417 | 0,655385 | 0,01775 | 0,120093 |
| Tbc1d8b | 0,834305 | 1,38307 | 0,729226 | 0,0178 | 0,120305 |
| Ccdc15 | 2,23015 | 3,4875 | 0,645055 | 0,0178 | 0,120305 |
| Zdhhc20 | 14,6083 | 19,9317 | 0,44828 | 0,01785 | 0,120516 |

| | | | | | |
|---|---|---|---|---|---|
| Arhgdib | 1,5057 | 0,459453 | -1,71245 | 0,01785 | 0,120516 |
| Gm16386 | 1,62548 | 2,62387 | 0,690831 | 0,0179 | 0,120726 |
| Setbp1 | 10,3194 | 13,8938 | 0,429082 | 0,0179 | 0,120726 |
| Pon2 | 11,6476 | 7,93058 | -0,554538 | 0,01795 | 0,121 |
| Rictor | 6,56537 | 8,9755 | 0,451116 | 0,018 | 0,121273 |
| Klf9 | 1,86767 | 3,01918 | 0,692919 | 0,01805 | 0,121546 |
| Setd2 | 19,8874 | 26,8653 | 0,433894 | 0,01815 | 0,122155 |
| Acsl1 | 4,05699 | 5,87352 | 0,533816 | 0,0182 | 0,122428 |
| Igsf1 | 0,823653 | 1,65485 | 1,0066 | 0,01825 | 0,122571 |
| Tmem87b | 9,88285 | 13,5695 | 0,457369 | 0,01825 | 0,122571 |
| St8sia1 | 1,92737 | 1,28914 | -0,580231 | 0,01825 | 0,122571 |
| Palm2 | 0,758682 | 0,442486 | -0,777864 | 0,0183 | 0,122843 |
| Wdr59 | 5,36655 | 7,53394 | 0,489409 | 0,0184 | 0,12332 |
| Dab2ip | 22,4054 | 30,1746 | 0,429488 | 0,0184 | 0,12332 |
| Slc25a23 | 10,9332 | 7,79925 | -0,487315 | 0,0184 | 0,12332 |
| Mesp1 | 3,90095 | 6,77018 | 0,795369 | 0,01845 | 0,123591 |
| Orai1 | 3,25861 | 5,25449 | 0,689293 | 0,0185 | 0,123861 |
| Rab11fip5 | 0,947301 | 1,55839 | 0,718167 | 0,0186 | 0,124466 |
| Gm16702 | 0,140886 | 4,85729 | 5,10756 | 0,0187 | 0,124939 |
| Zfp81 | 8,99753 | 12,9762 | 0,528264 | 0,0187 | 0,124939 |
| Tiam1 | 9,40062 | 12,945 | 0,461572 | 0,0187 | 0,124939 |
| Gatsl2 | 3,4674 | 5,03113 | 0,53703 | 0,0188 | 0,125542 |
| Upp1 | 1,52842 | 0,116994 | -3,70753 | 0,01885 | 0,125811 |
| Prkca | 0,90806 | 1,40899 | 0,633802 | 0,0189 | 0,126013 |
| Atp11c | 9,46669 | 12,9637 | 0,453543 | 0,0189 | 0,126013 |
| Zdbf2 | 5,33885 | 7,25686 | 0,442817 | 0,01895 | 0,126215 |
| Krt5 | 0,729333 | 0,148576 | -2,29538 | 0,01895 | 0,126215 |
| Ppp1r3c | 0,637241 | 1,26341 | 0,987407 | 0,019 | 0,126351 |
| Soga3 | 5,49973 | 7,86131 | 0,51541 | 0,019 | 0,126351 |
| Tmub1 | 9,53616 | 6,06715 | -0,65239 | 0,019 | 0,126351 |
| Smpd3 | 1,13162 | 1,82654 | 0,690726 | 0,01905 | 0,126553 |
| Fktn | 9,9071 | 13,6925 | 0,466853 | 0,01905 | 0,126553 |
| Mab21l1 | 4,49884 | 0,637241 | -2,81964 | 0,01915 | 0,127151 |
| Slc6a13 | 1,31336 | 0,07701 | -4,09207 | 0,0192 | 0,127417 |
| Ly6h | 1,84546 | 0,624918 | -1,56224 | 0,01925 | 0,127683 |
| Ttc38 | 1,70657 | 2,69425 | 0,658785 | 0,01945 | 0,12881 |
| Pgm1 | 15,7088 | 21,791 | 0,472155 | 0,01945 | 0,12881 |
| Npat | 6,74019 | 9,23306 | 0,454021 | 0,01945 | 0,12881 |
| Gde1 | 31,5769 | 22,7113 | -0,475457 | 0,01955 | 0,129405 |
| Ppm1k | 2,00976 | 2,98635 | 0,571359 | 0,0196 | 0,129669 |
| Zbtb44 | 4,70163 | 6,42509 | 0,450557 | 0,01975 | 0,130594 |
| Znf512b | 11,5922 | 8,05439 | -0,525301 | 0,0198 | 0,130857 |
| Nrn1 | 0,318493 | 0,962263 | 1,59517 | 0,01985 | 0,131053 |
| Socs5 | 4,88441 | 3,34704 | -0,545298 | 0,01985 | 0,131053 |
| Mid1ip1 | 19,4167 | 26,5766 | 0,452862 | 0,0199 | 0,131248 |
| Ptch1 | 44,2705 | 33,0175 | -0,423114 | 0,0199 | 0,131248 |
| Prkcdbp | 4,65508 | 7,78374 | 0,741656 | 0,01995 | 0,13151 |
| Uxs1 | 17,6665 | 25,2256 | 0,513877 | 0,02005 | 0,131831 |
| Abcc1 | 7,96062 | 10,9099 | 0,454683 | 0,02005 | 0,131831 |
| Mis18bp1 | 20,1272 | 27,4355 | 0,4469 | 0,02005 | 0,131831 |

| | | | | | |
|---|---|---|---|---|---|
| Zfp703 | 16,2795 | 11,8419 | -0,45916 | 0,02005 | 0,131831 |
| Dtnb | 14,4619 | 10,2669 | -0,494254 | 0,02005 | 0,131831 |
| Txlnb | 0,614198 | 1,09764 | 0,837633 | 0,0201 | 0,132092 |
| Hif1a | 49,4803 | 66,0258 | 0,416175 | 0,0202 | 0,132681 |
| Prex2 | 4,35009 | 5,92753 | 0,446385 | 0,02025 | 0,132941 |
| Parva | 24,4524 | 18,1068 | -0,433441 | 0,0204 | 0,133857 |
| Ttyh2 | 1,41597 | 0,711484 | -0,992887 | 0,02045 | 0,134048 |
| C1qtnf7 | 1,02809 | 0,229972 | -2,16043 | 0,02045 | 0,134048 |
| Abca8b | 0,676218 | 0,366708 | -0,882856 | 0,0205 | 0,134307 |
| Tulp3 | 24,9983 | 33,5845 | 0,425967 | 0,02055 | 0,134566 |
| Elf4 | 2,25578 | 1,49221 | -0,596173 | 0,0206 | 0,134756 |
| Itga3 | 1,25038 | 0,732723 | -0,771025 | 0,0206 | 0,134756 |
| A630089N07Rik | 4,13638 | 6,41924 | 0,634034 | 0,02065 | 0,134946 |
| Calcrl | 2,41513 | 1,55244 | -0,63756 | 0,02065 | 0,134946 |
| 2310014L17Rik | 0,791649 | 1,42595 | 0,848994 | 0,02075 | 0,135461 |
| Hdac10 | 5,29622 | 7,78747 | 0,556193 | 0,02075 | 0,135461 |
| Greb1l | 57,5558 | 77,6198 | 0,431465 | 0,02085 | 0,136045 |
| Sept5 | 10,1781 | 14,3968 | 0,500281 | 0,0209 | 0,136094 |
| Ly75 | 7,60605 | 10,4008 | 0,451474 | 0,0209 | 0,136094 |
| Wdr1 | 59,11 | 79,1794 | 0,421723 | 0,0209 | 0,136094 |
| Cacna1g | 2,64917 | 1,83159 | -0,532443 | 0,0209 | 0,136094 |
| Abtb2 | 2,5676 | 1,6426 | -0,644438 | 0,02105 | 0,137001 |
| B4galnt3 | 0,286205 | 0,692888 | 1,27557 | 0,02115 | 0,137443 |
| Zfp738 | 3,02824 | 4,43482 | 0,550396 | 0,02115 | 0,137443 |
| Lrrc17 | 0,825855 | 0,290964 | -1,50505 | 0,02115 | 0,137443 |
| Dnajb5 | 7,8113 | 11,1927 | 0,518919 | 0,02125 | 0,138023 |
| Iqsec2 | 0,500026 | 0,884301 | 0,822533 | 0,02135 | 0,138532 |
| Dapk1 | 14,2655 | 19,0978 | 0,420874 | 0,02135 | 0,138532 |
| G2e3 | 10,7837 | 14,514 | 0,42859 | 0,02145 | 0,139111 |
| Rreb1 | 5,8727 | 7,98532 | 0,443326 | 0,02155 | 0,139618 |
| Kcnk2 | 0,937424 | 0,47978 | -0,966328 | 0,02155 | 0,139618 |
| Rassf8 | 9,75689 | 14,0016 | 0,521096 | 0,0216 | 0,139871 |
| D230030E09Rik | 0,204552 | 4,93994 | 4,59396 | 0,02165 | 0,140054 |
| Szt2 | 6,40419 | 8,64045 | 0,43209 | 0,02165 | 0,140054 |
| Hs3st1 | 0,698203 | 1,53354 | 1,13515 | 0,02175 | 0,14063 |
| Usp47 | 57,0114 | 75,6654 | 0,408385 | 0,0219 | 0,141528 |
| Tex15 | 0,465857 | 0,781558 | 0,746464 | 0,02205 | 0,142354 |
| C230035I16Rik | 3,22364 | 1,03487 | -1,63925 | 0,02205 | 0,142354 |
| Tspan2 | 5,05875 | 7,15249 | 0,499664 | 0,02215 | 0,142856 |
| Top1mt | 5,16887 | 3,25791 | -0,665902 | 0,02215 | 0,142856 |
| Pnma2 | 0,438188 | 0,825601 | 0,913895 | 0,0223 | 0,143607 |
| 5830418K08Rik | 12,0652 | 18,4432 | 0,612237 | 0,0223 | 0,143607 |
| Fam149a | 5,92273 | 4,24638 | -0,480029 | 0,0223 | 0,143607 |
| Frmd4a | 23,5459 | 31,3583 | 0,413369 | 0,02245 | 0,1445 |
| Bicd1 | 0,82047 | 1,26319 | 0,622548 | 0,0225 | 0,144677 |
| Lama5 | 5,5979 | 7,50419 | 0,422809 | 0,0225 | 0,144677 |
| Gstt1 | 0,464519 | 1,56376 | 1,75121 | 0,02255 | 0,144853 |
| 03/03/16 | 4,97215 | 7,58111 | 0,608539 | 0,02255 | 0,144853 |
| Smad4 | 45,5137 | 34,0267 | -0,419632 | 0,0226 | 0,145102 |
| S100a11 | 34,7489 | 51,4544 | 0,566327 | 0,0227 | 0,145671 |

| | | | | | |
|---|---|---|---|---|---|
| Zfp229 | 2,74875 | 4,0574 | 0,561781 | 0,02275 | 0,145846 |
| Zfp946 | 7,08549 | 10,1327 | 0,516074 | 0,02275 | 0,145846 |
| Nme4 | 60,647 | 44,0336 | -0,461831 | 0,023 | 0,147301 |
| Zfp185 | 1,86314 | 1,1215 | -0,732309 | 0,023 | 0,147301 |
| Klf15 | 2,34618 | 1,35229 | -0,794914 | 0,02305 | 0,147548 |
| Phtf2 | 7,19782 | 9,85858 | 0,45382 | 0,0231 | 0,147573 |
| Rpsa | 13,6788 | 9,59286 | -0,511904 | 0,0231 | 0,147573 |
| Ggct | 5,86658 | 3,54699 | -0,725926 | 0,0231 | 0,147573 |
| Cbx8 | 4,51879 | 2,67962 | -0,753907 | 0,0231 | 0,147573 |
| Tssc1 | 16,273 | 22,5661 | 0,471674 | 0,02315 | 0,147745 |
| Atcay | 0,639618 | 0,283535 | -1,17368 | 0,02315 | 0,147745 |
| Slc39a11 | 4,31597 | 6,38368 | 0,564706 | 0,0232 | 0,147844 |
| Hey1 | 29,0559 | 39,0846 | 0,427772 | 0,0232 | 0,147844 |
| Fbxo21 | 38,0768 | 50,7926 | 0,415705 | 0,0232 | 0,147844 |
| Nog | 2,22647 | 3,6897 | 0,728746 | 0,02325 | 0,148015 |
| Tmem17 | 6,01438 | 9,31646 | 0,631365 | 0,02325 | 0,148015 |
| Fam83g | 0,558665 | 0,981723 | 0,813332 | 0,0233 | 0,14826 |
| Arntl | 2,30732 | 3,59289 | 0,638926 | 0,0234 | 0,148675 |
| Dzip3 | 14,0582 | 18,9295 | 0,429224 | 0,0234 | 0,148675 |
| Tjp1 | 29,1409 | 38,5403 | 0,403324 | 0,0234 | 0,148675 |
| Zfp715 | 4,85564 | 6,83848 | 0,494013 | 0,0235 | 0,149163 |
| Rgs3 | 1,0324 | 0,570107 | -0,856692 | 0,0235 | 0,149163 |
| Scd2 | 373,426 | 520,035 | 0,477784 | 0,0236 | 0,149649 |
| Galnt5 | 1,05801 | 0,555129 | -0,930453 | 0,0236 | 0,149649 |
| Tbxa2r | 0,200747 | 0,671042 | 1,74103 | 0,0237 | 0,150209 |
| Pdlim4 | 45,4067 | 61,1226 | 0,4288 | 0,02375 | 0,150303 |
| Chchd1 | 220,843 | 164,891 | -0,421513 | 0,02375 | 0,150303 |
| Cd93 | 0,768555 | 0,442889 | -0,795205 | 0,02375 | 0,150303 |
| Zfp677 | 0,864301 | 1,56261 | 0,854354 | 0,0238 | 0,150323 |
| Ehbp1 | 6,12208 | 8,43785 | 0,462855 | 0,0238 | 0,150323 |
| Ston2 | 3,35171 | 2,40684 | -0,477757 | 0,0238 | 0,150323 |
| Dctn3 | 40,9218 | 29,2903 | -0,482445 | 0,0238 | 0,150323 |
| Caprin2 | 9,18676 | 12,6237 | 0,458503 | 0,02385 | 0,150491 |
| Zswim6 | 6,86739 | 4,98093 | -0,463347 | 0,02385 | 0,150491 |
| Frmd5 | 2,57232 | 1,65756 | -0,634004 | 0,0239 | 0,150732 |
| Atpif1 | 449,556 | 337,6 | -0,413184 | 0,02395 | 0,150973 |
| Serinc2 | 17,1895 | 23,6115 | 0,457965 | 0,024 | 0,151214 |
| Rb1cc1 | 5,45153 | 7,47913 | 0,456208 | 0,02405 | 0,15138 |
| Ski | 36,4084 | 48,0585 | 0,400522 | 0,02405 | 0,15138 |
| Rab28 | 42,2527 | 31,2498 | -0,4352 | 0,0241 | 0,151546 |
| Rn45s | 567,913 | 389,302 | -0,544781 | 0,0241 | 0,151546 |
| Dbt | 12,7629 | 17,3355 | 0,441778 | 0,02415 | 0,151712 |
| Pctp | 16,3875 | 11,6983 | -0,486292 | 0,02415 | 0,151712 |
| Gm11627 | 2,63445 | 0,803783 | -1,71262 | 0,02425 | 0,152266 |
| 2810025M15Rik | 52,2896 | 34,1702 | -0,613785 | 0,02435 | 0,152819 |
| Eln | 0,866531 | 1,46876 | 0,761279 | 0,0244 | 0,153058 |
| Klhdc7a | 0,417506 | 0,742553 | 0,830698 | 0,02465 | 0,154475 |
| Ly6e | 34,346 | 45,8975 | 0,418272 | 0,02465 | 0,154475 |
| Jrkl | 4,94964 | 7,15698 | 0,532027 | 0,0247 | 0,154562 |
| Lphn1 | 14,3936 | 19,1059 | 0,408583 | 0,0247 | 0,154562 |

| | | | | | |
|---|---|---|---|---|---|
| Rpl7a | 120,603 | 89,8391 | -0,424852 | 0,0247 | 0,154562 |
| 2610044O15Rik8 | 4,0226 | 6,2571 | 0,637367 | 0,0248 | 0,155036 |
| Fam92a | 43,7558 | 32,1815 | -0,443242 | 0,0248 | 0,155036 |
| Rgs9bp | 0,448071 | 0,762067 | 0,76619 | 0,0249 | 0,155586 |
| Slc25a21 | 1,34183 | 0,679467 | -0,981728 | 0,025 | 0,156134 |
| Odc1 | 51,1698 | 38,499 | -0,410473 | 0,02505 | 0,156371 |
| Rtn2 | 9,22988 | 6,0069 | -0,619691 | 0,0251 | 0,15653 |
| Chsy3 | 0,813079 | 0,420292 | -0,952003 | 0,0251 | 0,15653 |
| Pid1 | 1,6375 | 2,68606 | 0,713994 | 0,02515 | 0,156614 |
| Krtcap2 | 217,86 | 161,769 | -0,429465 | 0,02515 | 0,156614 |
| Tmem216 | 26,8571 | 18,8253 | -0,512633 | 0,02515 | 0,156614 |
| Foxn1 | 0,0405185 | 0,682708 | 4,07462 | 0,0252 | 0,156773 |
| Ajuba | 35,9564 | 27,0614 | -0,410012 | 0,0252 | 0,156773 |
| Slc5a3 | 3,0736 | 4,67112 | 0,603841 | 0,0254 | 0,157864 |
| Fbn1 | 4,53594 | 3,31606 | -0,451932 | 0,0254 | 0,157864 |
| Bnc2 | 0,69149 | 0,394986 | -0,807909 | 0,0255 | 0,158332 |
| A830082K12Rik | 1,74612 | 0,091587 | -4,25287 | 0,0255 | 0,158332 |
| Rcan3 | 2,23078 | 3,28277 | 0,557364 | 0,02555 | 0,158336 |
| Fam208a | 5,77878 | 8,47384 | 0,552251 | 0,02555 | 0,158336 |
| Fzd10 | 34,8703 | 46,4863 | 0,414804 | 0,02555 | 0,158336 |
| Caly | 1,33588 | 0,44593 | -1,58291 | 0,02555 | 0,158336 |
| Heph | 18,3877 | 24,4052 | 0,408447 | 0,02565 | 0,158802 |
| Hmgn2 | 58,2378 | 43,1435 | -0,432815 | 0,02565 | 0,158802 |
| Cd83 | 12,6363 | 17,4859 | 0,468618 | 0,0257 | 0,159035 |
| Rasgrp4 | 0,674025 | 1,39837 | 1,05288 | 0,02575 | 0,159191 |
| App | 48,0404 | 36,1992 | -0,408292 | 0,02575 | 0,159191 |
| Zfp964 | 1,38664 | 2,51157 | 0,856998 | 0,0258 | 0,159346 |
| Sh3rf1 | 9,84896 | 7,28651 | -0,434743 | 0,0258 | 0,159346 |
| Mir196a-1 | 508,77 | 1100,68 | 1,11331 | 0,02585 | 0,159578 |
| Cox19 | 58,1436 | 42,0023 | -0,469152 | 0,02595 | 0,160042 |
| Vat1l | 1,70536 | 1,02833 | -0,729771 | 0,02595 | 0,160042 |
| Esco2 | 14,3788 | 19,6607 | 0,451372 | 0,026 | 0,160273 |
| Hspbp1 | 43,484 | 32,2129 | -0,432844 | 0,02605 | 0,160504 |
| Dicer1 | 33,8991 | 44,8231 | 0,402997 | 0,0261 | 0,160657 |
| Coro1a | 2,12811 | 1,10047 | -0,951454 | 0,0261 | 0,160657 |
| Fam189a2 | 0,469351 | 0,980378 | 1,06267 | 0,02615 | 0,160657 |
| Arhgef10 | 5,64092 | 7,72192 | 0,453028 | 0,02615 | 0,160657 |
| Slc43a3 | 18,4606 | 25,0024 | 0,437613 | 0,02615 | 0,160657 |
| Rpl18a | 61,7732 | 46,2028 | -0,419 | 0,02615 | 0,160657 |
| Rad50 | 13,4225 | 17,8893 | 0,414441 | 0,02635 | 0,161808 |
| Rnf19a | 10,8222 | 14,5761 | 0,429607 | 0,0265 | 0,162574 |
| Rpl3 | 62,7206 | 46,8785 | -0,420013 | 0,0265 | 0,162574 |
| Chm | 4,51976 | 6,33469 | 0,487028 | 0,02655 | 0,162803 |
| Prss22 | 0,156132 | 0,988684 | 2,66274 | 0,02665 | 0,16326 |
| Ints12 | 15,7447 | 11,5194 | -0,450802 | 0,02665 | 0,16326 |
| Gm4349 | 0,167027 | 1,27899 | 2,93685 | 0,0267 | 0,163488 |
| Sgsh | 2,99361 | 4,33628 | 0,534575 | 0,0268 | 0,164022 |
| Fzd6 | 1,88849 | 1,17089 | -0,689627 | 0,02685 | 0,16425 |
| Bcl2l11 | 21,197 | 28,5371 | 0,428983 | 0,0269 | 0,164477 |
| Arhgap42 | 10,104 | 7,50249 | -0,429483 | 0,02695 | 0,164704 |

| | | | | | |
|---|---|---|---|---|---|
| Zscan26 | 18,9844 | 25,1753 | 0,407195 | 0,027 | 0,164931 |
| Mov10 | 8,06992 | 11,0592 | 0,454616 | 0,02715 | 0,16569 |
| Asphd2 | 9,65238 | 6,49926 | -0,570609 | 0,02715 | 0,16569 |
| Tesk2 | 6,63911 | 11,137 | 0,746295 | 0,0272 | 0,165916 |
| Casp6 | 27,4018 | 37,146 | 0,438938 | 0,02735 | 0,166752 |
| Acsl5 | 13,5001 | 9,86794 | -0,452153 | 0,0274 | 0,166977 |
| Dgka | 1,29714 | 2,24797 | 0,793292 | 0,02745 | 0,167123 |
| Nudt9 | 52,8024 | 39,4111 | -0,422001 | 0,02745 | 0,167123 |
| Nap1l2 | 1,0994 | 1,90805 | 0,795383 | 0,02755 | 0,167573 |
| Zdhhc1 | 4,20609 | 2,68887 | -0,645477 | 0,02755 | 0,167573 |
| H2-Eb1 | 0,154138 | 0,718017 | 2,2198 | 0,0276 | 0,167718 |
| Snap91 | 3,08582 | 2,0048 | -0,622196 | 0,0276 | 0,167718 |
| Msantd3 | 43,204 | 32,1704 | -0,425431 | 0,02765 | 0,167942 |
| Fam83h | 1,22795 | 1,92384 | 0,647733 | 0,02785 | 0,168997 |
| Gpr56 | 5,68221 | 3,95579 | -0,522488 | 0,02785 | 0,168997 |
| Ppm1f | 17,9344 | 23,7417 | 0,404695 | 0,0279 | 0,16922 |
| Fam120b | 10,0369 | 13,4431 | 0,421557 | 0,02795 | 0,169444 |
| Nfkbie | 1,63587 | 2,67984 | 0,712082 | 0,0281 | 0,170272 |
| Psd3 | 4,5257 | 6,06045 | 0,421284 | 0,0284 | 0,171766 |
| Btbd7 | 15,9107 | 21,1339 | 0,409562 | 0,0284 | 0,171766 |
| Morc3 | 17,529 | 23,2563 | 0,407876 | 0,0284 | 0,171766 |
| Sh3pxd2a | 11,8818 | 15,6721 | 0,399438 | 0,0284 | 0,171766 |
| Tmem106c | 13,4585 | 9,32049 | -0,530037 | 0,02845 | 0,171987 |
| Slc29a1 | 11,355 | 7,99195 | -0,506704 | 0,02855 | 0,172429 |
| Podxl2 | 11,2383 | 7,83239 | -0,520895 | 0,02855 | 0,172429 |
| Dcun1d1 | 5,22016 | 3,68135 | -0,50386 | 0,0287 | 0,173253 |
| Triobp | 11,5943 | 16,1006 | 0,473703 | 0,02875 | 0,173474 |
| Pigg | 4,56423 | 6,47147 | 0,503722 | 0,0288 | 0,173694 |
| Rgs7 | 0,419624 | 0,901618 | 1,10342 | 0,02885 | 0,173913 |
| Athl1 | 7,80635 | 10,76 | 0,462953 | 0,0289 | 0,17397 |
| Nhp2 | 148,518 | 110,283 | -0,429427 | 0,0289 | 0,17397 |
| Haus4 | 21,9778 | 15,8554 | -0,47107 | 0,0289 | 0,17397 |
| A930004D18Rik | 1,3088 | 0,594882 | -1,13757 | 0,02895 | 0,174189 |
| Tapbpl | 0,582825 | 1,28408 | 1,1396 | 0,029 | 0,174244 |
| Pkd1 | 6,24577 | 9,68842 | 0,633383 | 0,029 | 0,174244 |
| Cd109 | 2,01629 | 1,3468 | -0,582172 | 0,029 | 0,174244 |
| Zfp951 | 1,05084 | 1,78648 | 0,765567 | 0,02915 | 0,174982 |
| Rhbdd2 | 11,7057 | 15,7228 | 0,425643 | 0,02915 | 0,174982 |
| Sntb2 | 5,40578 | 7,44531 | 0,461829 | 0,0293 | 0,1758 |
| Cep170b | 2,89045 | 4,0047 | 0,470399 | 0,0294 | 0,176318 |
| Hoxc11 | 45,4322 | 61,2377 | 0,430704 | 0,02945 | 0,176535 |
| Tph1 | 1,98128 | 0,600134 | -1,72308 | 0,0295 | 0,176752 |
| Klc2 | 16,5443 | 22,1132 | 0,418574 | 0,02955 | 0,176886 |
| Islr | 12,8747 | 9,20245 | -0,484451 | 0,02955 | 0,176886 |
| Apbb1ip | 0,314747 | 0,720386 | 1,19458 | 0,0296 | 0,176938 |
| Tmem30a | 23,059 | 30,4985 | 0,403409 | 0,0296 | 0,176938 |
| C030046E11Rik | 13,2222 | 17,4272 | 0,398372 | 0,0296 | 0,176938 |
| BC052040 | 10,5282 | 7,55191 | -0,479345 | 0,02965 | 0,177072 |
| 1810014B01Rik | 2,64211 | 1,48699 | -0,829298 | 0,02965 | 0,177072 |
| Clybl | 13,9387 | 9,38238 | -0,571072 | 0,0297 | 0,177205 |

| | | | | | |
|---|---|---|---|---|---|
| Akr1c13 | 2,35335 | 1,15997 | -1,02063 | 0,0297 | 0,177205 |
| Nme6 | 33,7498 | 24,3585 | -0,47045 | 0,0298 | 0,177719 |
| Fras1 | 2,24758 | 3,03565 | 0,433632 | 0,02985 | 0,177935 |
| 4732456N10Rik | 0,557298 | 1,12043 | 1,00753 | 0,02995 | 0,178365 |
| Tubb4a | 8,10742 | 11,3557 | 0,486105 | 0,02995 | 0,178365 |
| Fes | 1,04041 | 1,78666 | 0,780115 | 0,03 | 0,17858 |
| Stxbp5 | 2,502 | 3,45946 | 0,467463 | 0,03015 | 0,17939 |
| Col7a1 | 0,706053 | 1,07977 | 0,612869 | 0,03025 | 0,179901 |
| Tob1 | 7,97099 | 11,1177 | 0,480032 | 0,03035 | 0,180245 |
| Slc9a1 | 7,80899 | 10,4898 | 0,425774 | 0,03035 | 0,180245 |
| Onecut1 | 0,676174 | 0,0954369 | -2,82478 | 0,03035 | 0,180245 |
| Rps11 | 1303,31 | 990,374 | -0,396133 | 0,03045 | 0,180718 |
| Igdcc4 | 5,56633 | 4,07344 | -0,450479 | 0,0305 | 0,180718 |
| Scn2b | 2,56184 | 1,67049 | -0,616914 | 0,0305 | 0,180718 |
| Naalad2 | 0,951953 | 0,47075 | -1,01593 | 0,0305 | 0,180718 |
| Pou3f4 | 1,26453 | 0,319953 | -1,98268 | 0,0305 | 0,180718 |
| Tom1l2 | 6,37497 | 4,58344 | -0,475987 | 0,0307 | 0,181819 |
| Neat1 | 0,394115 | 0,803757 | 1,02814 | 0,03075 | 0,181947 |
| Akap11 | 17,287 | 22,7584 | 0,396713 | 0,03075 | 0,181947 |
| Penk | 0,657692 | 1,47803 | 1,16819 | 0,0308 | 0,182159 |
| Gm15446 | 0,458936 | 1,1415 | 1,31457 | 0,0309 | 0,182498 |
| Yes1 | 14,5464 | 19,5422 | 0,425939 | 0,0309 | 0,182498 |
| Sec31a | 52,0062 | 39,7473 | -0,387826 | 0,0309 | 0,182498 |
| Zfp423 | 17,5302 | 23,0965 | 0,397832 | 0,03095 | 0,182541 |
| Plin2 | 36,5389 | 27,4595 | -0,412129 | 0,03095 | 0,182541 |
| Mamdc2 | 0,773674 | 0,383459 | -1,01265 | 0,03095 | 0,182541 |
| Rlf | 12,6138 | 16,564 | 0,393055 | 0,03105 | 0,183047 |
| Zfp462 | 22,3981 | 17,0773 | -0,391296 | 0,0311 | 0,183173 |
| Slc35g1 | 6,67901 | 4,76692 | -0,486578 | 0,0311 | 0,183173 |
| Rims2 | 0,783954 | 1,24991 | 0,672977 | 0,03125 | 0,183972 |
| Prim1 | 87,5039 | 66,5849 | -0,394152 | 0,03135 | 0,184391 |
| Zmiz2 | 17,6911 | 13,325 | -0,408884 | 0,03135 | 0,184391 |
| Tgfb1 | 1,21868 | 2,12689 | 0,803418 | 0,03145 | 0,184895 |
| Ncoa2 | 11,51 | 15,1017 | 0,391819 | 0,0316 | 0,185692 |
| Cenpw | 20,2587 | 27,5148 | 0,441665 | 0,03165 | 0,1859 |
| Farp1 | 36,2691 | 27,7747 | -0,38497 | 0,0317 | 0,186109 |
| Igtp | 0,699367 | 1,38387 | 0,984582 | 0,032 | 0,187784 |
| Diap2 | 0,720106 | 1,10765 | 0,621227 | 0,0322 | 0,188785 |
| Efcab1 | 1,89251 | 0,839964 | -1,1719 | 0,0322 | 0,188785 |
| Amotl2 | 32,2785 | 42,1765 | 0,385864 | 0,03225 | 0,188819 |
| Ctdspl | 6,40143 | 4,65428 | -0,459836 | 0,03225 | 0,188819 |
| Aard | 5,17786 | 3,20397 | -0,692495 | 0,03225 | 0,188819 |
| D930007P13Rik | 1,68908 | 3,02322 | 0,839852 | 0,03235 | 0,189318 |
| Dock9 | 1,89875 | 2,68258 | 0,498567 | 0,0324 | 0,189524 |
| Gpr50 | 0,0775987 | 1,21768 | 3,97195 | 0,03245 | 0,189558 |
| Zfp592 | 10,0791 | 13,2268 | 0,392108 | 0,03245 | 0,189558 |
| Olfml1 | 2,40731 | 0,513703 | -2,22842 | 0,03245 | 0,189558 |
| Epb4.1l4b | 2,78483 | 4,14036 | 0,57217 | 0,0327 | 0,190844 |
| Dio3 | 1,07479 | 0,490931 | -1,13046 | 0,0327 | 0,190844 |
| Snord37 | 1564,88 | 1452,25 | -0,107761 | 0,0328 | 0,191341 |

| | | | | | |
|---|---|---|---|---|---|
| Pnp2 | 0,14998 | 1,01508 | 2,75875 | 0,03285 | 0,191545 |
| Arhgef15 | 0,957942 | 1,54938 | 0,693678 | 0,0329 | 0,19175 |
| Klk15 | 0,233137 | 2,13067 | 3,19205 | 0,03295 | 0,191954 |
| Appl2 | 12,0105 | 16,1422 | 0,426547 | 0,033 | 0,192158 |
| Stard13 | 2,10375 | 3,02123 | 0,522171 | 0,03305 | 0,192217 |
| Cldn7 | 1,35542 | 2,63813 | 0,960778 | 0,0331 | 0,192217 |
| Dlg5 | 18,4087 | 24,0963 | 0,388422 | 0,0331 | 0,192217 |
| Plod1 | 18,7835 | 14,0953 | -0,41425 | 0,0331 | 0,192217 |
| Ttl | 10,2067 | 7,60441 | -0,424606 | 0,0331 | 0,192217 |
| Akr1e1 | 24,3636 | 17,9852 | -0,437919 | 0,0331 | 0,192217 |
| Col8a1 | 1,3144 | 1,98926 | 0,597826 | 0,03315 | 0,19242 |
| Atp5o | 242,141 | 185,174 | -0,386964 | 0,03325 | 0,192913 |
| Ptpdc1 | 3,52098 | 4,9625 | 0,495088 | 0,03335 | 0,193318 |
| Ago1 | 26,2607 | 34,3257 | 0,386385 | 0,03335 | 0,193318 |
| Ric3 | 5,02759 | 7,33419 | 0,544768 | 0,0334 | 0,193433 |
| Snx30 | 5,60517 | 4,14122 | -0,436703 | 0,0334 | 0,193433 |
| Tle6 | 1,90905 | 3,1161 | 0,706885 | 0,0335 | 0,19375 |
| Uso1 | 36,2282 | 47,2129 | 0,382067 | 0,0335 | 0,19375 |
| Tmem37 | 2,22023 | 0,987638 | -1,16865 | 0,0335 | 0,19375 |
| Kdm8 | 7,91581 | 10,9858 | 0,472829 | 0,03355 | 0,193865 |
| B3gat1 | 0,832938 | 0,436149 | -0,933387 | 0,03355 | 0,193865 |
| Mdk | 1447,89 | 1097,62 | -0,399579 | 0,03375 | 0,194932 |
| Gyg | 20,0276 | 14,6636 | -0,449756 | 0,0339 | 0,195711 |
| St3gal6 | 7,62396 | 11,0196 | 0,531465 | 0,03395 | 0,195823 |
| Rpl13 | 91,207 | 68,5362 | -0,412278 | 0,03395 | 0,195823 |
| Ttc39a | 0,77876 | 1,42976 | 0,876522 | 0,03415 | 0,196888 |
| Ifnlr1 | 0,348299 | 0,675595 | 0,955834 | 0,03425 | 0,197376 |
| Sap30l | 16,4024 | 22,389 | 0,448888 | 0,0343 | 0,197575 |
| Chchd10 | 2,51886 | 1,19125 | -1,0803 | 0,03435 | 0,197775 |
| 4632428N05Rik | 0,443021 | 0,790666 | 0,835692 | 0,0344 | 0,197885 |
| Mmp14 | 39,3095 | 29,7877 | -0,400162 | 0,0344 | 0,197885 |
| Nfkbid | 0,438327 | 0,971202 | 1,14776 | 0,03445 | 0,197995 |
| Ndufb4 | 20,9456 | 12,8067 | -0,709749 | 0,03445 | 0,197995 |
| Zfp597 | 2,74265 | 3,83237 | 0,482665 | 0,03455 | 0,198392 |
| Kif13a | 9,21822 | 6,95118 | -0,407231 | 0,03455 | 0,198392 |
| Fam185a | 6,53091 | 9,01164 | 0,464506 | 0,0346 | 0,198501 |
| Efr3b | 3,42298 | 4,68216 | 0,451923 | 0,0346 | 0,198501 |
| Prr14l | 5,55021 | 7,3177 | 0,398849 | 0,03485 | 0,199846 |
| Gabarap | 154,036 | 118,221 | -0,381778 | 0,0349 | 0,199954 |
| Pdp1 | 3,9273 | 2,74733 | -0,515505 | 0,0349 | 0,199954 |
| Trim21 | 1,86322 | 2,88365 | 0,630098 | 0,035 | 0,200348 |
| Kifap3 | 22,6113 | 17,2203 | -0,392936 | 0,035 | 0,200348 |
| Hist1h2bb | 1,44895 | 4,01421 | 1,47011 | 0,03505 | 0,200455 |
| Krt8 | 1,825 | 3,03133 | 0,732056 | 0,03505 | 0,200455 |
| Mospd2 | 6,6702 | 9,15529 | 0,456875 | 0,0351 | 0,200562 |
| 2410089E03Rik | 6,85621 | 9,05149 | 0,400744 | 0,0351 | 0,200562 |
| Asah2 | 0,610479 | 1,02579 | 0,748728 | 0,03525 | 0,20124 |
| Sema6d | 2,96923 | 4,13091 | 0,476367 | 0,03525 | 0,20124 |
| Edil3 | 0,594111 | 0,974423 | 0,713815 | 0,0353 | 0,201347 |
| Agrn | 33,9037 | 44,095 | 0,379174 | 0,0353 | 0,201347 |

| | | | | | |
|---|---|---|---|---|---|
| Ocrl | 10,1979 | 13,4444 | 0,398732 | 0,03545 | 0,202022 |
| Nos1ap | 0,945696 | 0,416851 | -1,18184 | 0,03545 | 0,202022 |
| Ndufb9 | 180,354 | 137,122 | -0,395371 | 0,0355 | 0,202218 |
| Odf2 | 58,6097 | 76,7141 | 0,388353 | 0,03555 | 0,202413 |
| Morf4l2 | 186,366 | 142,667 | -0,385492 | 0,03565 | 0,202892 |
| Mbd5 | 3,78732 | 5,11223 | 0,432777 | 0,0358 | 0,203565 |
| Fcho1 | 0,685059 | 0,337129 | -1,02293 | 0,0358 | 0,203565 |
| Rac3 | 11,2233 | 16,2372 | 0,532807 | 0,03585 | 0,203669 |
| Rit1 | 9,07188 | 6,48634 | -0,483997 | 0,03585 | 0,203669 |
| Msx2 | 6,35413 | 9,00138 | 0,502452 | 0,0359 | 0,203772 |
| Sirt5 | 6,53833 | 4,19746 | -0,639404 | 0,0359 | 0,203772 |
| Hook1 | 3,63676 | 5,08446 | 0,483441 | 0,03595 | 0,203876 |
| Cwf19l2 | 8,2596 | 11,1011 | 0,426563 | 0,03595 | 0,203876 |
| Gpc3 | 389,345 | 510,615 | 0,391185 | 0,0361 | 0,204636 |
| Hp | 0,120647 | 0,963283 | 2,99717 | 0,0362 | 0,205112 |
| Npc1 | 9,05431 | 12,2885 | 0,440633 | 0,0363 | 0,205497 |
| Asrgl1 | 16,7758 | 12,387 | -0,437555 | 0,0363 | 0,205497 |
| B4galt5 | 8,86373 | 11,8348 | 0,417045 | 0,03635 | 0,205599 |
| Fxyd1 | 2,62298 | 0,931785 | -1,49314 | 0,03635 | 0,205599 |
| Ldoc1l | 7,27459 | 5,37331 | -0,437055 | 0,0364 | 0,205791 |
| Tes | 17,5173 | 13,0041 | -0,429807 | 0,0365 | 0,206265 |
| Anxa5 | 37,8092 | 49,6319 | 0,392528 | 0,03655 | 0,206457 |
| Tulp1 | 0,283986 | 0,726631 | 1,3554 | 0,0366 | 0,206557 |
| Zgrf1 | 7,76492 | 10,2296 | 0,397705 | 0,0366 | 0,206557 |
| Rnaseh2c | 94,407 | 70,635 | -0,41851 | 0,03665 | 0,206749 |
| Zswim4 | 15,9217 | 20,8437 | 0,388619 | 0,0368 | 0,207503 |
| Anks6 | 1,96675 | 2,94648 | 0,583181 | 0,03685 | 0,207603 |
| 3110009E18Rik | 10,2578 | 5,5996 | -0,873322 | 0,03685 | 0,207603 |
| Rpl14 | 310,873 | 238,645 | -0,381462 | 0,0369 | 0,207793 |
| Atf2 | 22,0412 | 28,8221 | 0,386976 | 0,03695 | 0,207984 |
| Ago3 | 2,05783 | 2,86428 | 0,477046 | 0,037 | 0,208174 |
| Irf9 | 5,36856 | 7,5984 | 0,501162 | 0,03705 | 0,208181 |
| Fnbp1l | 41,4629 | 54,0096 | 0,381394 | 0,03705 | 0,208181 |
| Mpv17l | 0,89211 | 0,483972 | -0,882299 | 0,03705 | 0,208181 |
| Zfp65 | 4,03977 | 5,6523 | 0,484567 | 0,0372 | 0,208932 |
| Nphp4 | 2,18721 | 3,13642 | 0,520026 | 0,03735 | 0,209591 |
| Slc39a3 | 4,04838 | 2,79579 | -0,534086 | 0,03735 | 0,209591 |
| Agtpbp1 | 6,04144 | 4,3638 | -0,469308 | 0,0375 | 0,210341 |
| AI987944 | 2,38079 | 3,57887 | 0,58806 | 0,0378 | 0,211931 |
| AW822252 | 0,240181 | 0,974703 | 2,02084 | 0,03785 | 0,211934 |
| Smc3 | 51,2127 | 66,7201 | 0,38162 | 0,03785 | 0,211934 |
| Dock7 | 37,3274 | 48,433 | 0,375755 | 0,03785 | 0,211934 |
| Vav2 | 12,6235 | 16,7963 | 0,412028 | 0,038 | 0,212681 |
| Zfp607 | 2,47612 | 3,61355 | 0,545339 | 0,03805 | 0,212775 |
| Adcy2 | 1,07289 | 0,641159 | -0,74275 | 0,03805 | 0,212775 |
| A930024E05Rik | 2,38032 | 3,77286 | 0,664502 | 0,03815 | 0,213148 |
| Zfp945 | 2,62131 | 3,63386 | 0,471217 | 0,03815 | 0,213148 |
| Slc1a5 | 8,25543 | 11,3906 | 0,464429 | 0,0383 | 0,213707 |
| Sox11 | 44,4462 | 34,3475 | -0,371856 | 0,0383 | 0,213707 |
| Rnf130 | 32,1566 | 23,0222 | -0,482087 | 0,0383 | 0,213707 |

| | | | | | |
|---|---|---|---|---|---|
| Pcmtd2 | 13,2592 | 17,5065 | 0,400904 | 0,0384 | 0,214172 |
| Polr2f | 269,472 | 206,956 | -0,380811 | 0,03855 | 0,214915 |
| Mtap7d3 | 1,61436 | 2,54301 | 0,655577 | 0,03885 | 0,216399 |
| Man2a1 | 14,3161 | 18,6457 | 0,381211 | 0,03885 | 0,216399 |
| Frmd8 | 14,7127 | 19,4242 | 0,400789 | 0,03925 | 0,218533 |
| Chrd | 1,03077 | 1,70902 | 0,729458 | 0,03945 | 0,219551 |
| Gad1os | 0,683546 | 4,67832 | 2,77488 | 0,0395 | 0,219734 |
| Fsd1l | 3,60342 | 4,98126 | 0,467146 | 0,03955 | 0,219813 |
| Tmem62 | 0,933651 | 0,493505 | -0,919818 | 0,03955 | 0,219813 |
| Zfp800 | 5,0755 | 6,96303 | 0,456166 | 0,0396 | 0,219813 |
| Foxn3 | 8,20623 | 11,2168 | 0,450867 | 0,0396 | 0,219813 |
| Neurog3 | 1,29861 | 0,0955744 | -3,76421 | 0,0396 | 0,219813 |
| 1110001J03Rik | 49,6616 | 34,0445 | -0,544706 | 0,03965 | 0,219996 |
| AK129341 | 3,71243 | 5,21687 | 0,490822 | 0,0397 | 0,220178 |
| 5930412G12Rik | 1,37495 | 2,23992 | 0,70407 | 0,03975 | 0,22036 |
| Larp4 | 11,3367 | 14,7307 | 0,377825 | 0,03985 | 0,220819 |
| Hes3 | 10,7411 | 7,62983 | -0,493419 | 0,0399 | 0,221001 |
| Tecr | 168,905 | 130,938 | -0,367337 | 0,03995 | 0,221182 |
| Pml | 6,23912 | 8,2949 | 0,410881 | 0,0401 | 0,221917 |
| Jade3 | 7,90687 | 10,4523 | 0,402641 | 0,04015 | 0,222002 |
| Tcf19 | 20,863 | 15,4106 | -0,437023 | 0,04015 | 0,222002 |
| Jak1 | 23,0093 | 30,2809 | 0,396193 | 0,0402 | 0,222183 |
| Cyb561d2 | 3,23312 | 2,00614 | -0,688501 | 0,04035 | 0,22282 |
| Ap3m2 | 1,26121 | 0,750856 | -0,748201 | 0,04035 | 0,22282 |
| Ciart | 4,63041 | 7,31545 | 0,659807 | 0,04045 | 0,223084 |
| Cxx1c | 22,4424 | 30,7628 | 0,45496 | 0,04045 | 0,223084 |
| Msi2 | 30,5425 | 39,5451 | 0,372679 | 0,04045 | 0,223084 |
| Uqcrq | 221,303 | 168,553 | -0,392823 | 0,0405 | 0,223263 |
| Sumo3 | 98,7498 | 76,3144 | -0,371824 | 0,04055 | 0,223443 |
| Pkn3 | 5,33535 | 8,00727 | 0,585726 | 0,0406 | 0,223527 |
| Golgb1 | 10,8105 | 14,0345 | 0,376554 | 0,0406 | 0,223527 |
| Jade1 | 13,1209 | 17,0019 | 0,373827 | 0,04065 | 0,22361 |
| Gadd45g | 22,3828 | 16,103 | -0,475061 | 0,04065 | 0,22361 |
| Sparcl1 | 92,7625 | 120,049 | 0,37201 | 0,04075 | 0,223776 |
| Rcbtb2 | 90,2998 | 70,124 | -0,364813 | 0,04075 | 0,223776 |
| Sept11 | 134,806 | 103,534 | -0,380781 | 0,04075 | 0,223776 |
| Pafah1b3 | 113,017 | 86,2055 | -0,390693 | 0,04075 | 0,223776 |
| Fhod1 | 10,0742 | 7,50701 | -0,424362 | 0,0409 | 0,224504 |
| Hfm1 | 1,41433 | 0,737888 | -0,938644 | 0,04095 | 0,224682 |
| Pclo | 0,522113 | 0,753052 | 0,528387 | 0,041 | 0,22475 |
| Gpr108 | 11,6713 | 15,8499 | 0,441504 | 0,04105 | 0,22475 |
| Mfap4 | 51,151 | 66,532 | 0,379286 | 0,04105 | 0,22475 |
| Pld1 | 0,967904 | 0,587987 | -0,71908 | 0,04105 | 0,22475 |
| Sox2ot | 2,62353 | 0,30103 | -3,12353 | 0,04105 | 0,22475 |
| C130083M11Rik | 0,351814 | 0,800774 | 1,18658 | 0,0411 | 0,224832 |
| Erc2 | 1,46352 | 2,11431 | 0,530739 | 0,0411 | 0,224832 |
| Vsig8 | 0,19262 | 0,589358 | 1,61339 | 0,04115 | 0,224995 |
| Gm8615 | 1,09437 | 1,89749 | 0,793993 | 0,0412 | 0,224995 |
| Zfp827 | 5,75918 | 7,58338 | 0,396978 | 0,0412 | 0,224995 |
| Pkn2 | 29,1019 | 37,619 | 0,370347 | 0,0412 | 0,224995 |

| | | | | | |
|---|---|---|---|---|---|
| Kremen1 | 6,67816 | 8,85795 | 0,407523 | 0,0413 | 0,225445 |
| Gm527 | 2,22637 | 3,68148 | 0,725594 | 0,04135 | 0,225526 |
| E2f8 | 13,4876 | 17,6781 | 0,390327 | 0,04135 | 0,225526 |
| Car15 | 1,27597 | 2,48896 | 0,963943 | 0,0414 | 0,225607 |
| Rsrc1 | 18,4175 | 13,9696 | -0,398789 | 0,0414 | 0,225607 |
| Rbm15 | 13,8646 | 18,2976 | 0,400245 | 0,0416 | 0,226601 |
| Gstm6 | 1,49016 | 2,76171 | 0,89009 | 0,04175 | 0,227225 |
| Sgcd | 0,821469 | 0,263983 | -1,63776 | 0,04175 | 0,227225 |
| Mphosph9 | 8,3248 | 10,8129 | 0,37726 | 0,0418 | 0,227401 |
| Kxd1 | 27,8723 | 20,5752 | -0,437922 | 0,04185 | 0,227576 |
| Ubl4b | 0,789491 | 0,118408 | -2,73716 | 0,0419 | 0,227751 |
| Hapln3 | 4,402 | 2,73939 | -0,684306 | 0,042 | 0,228198 |
| Shisa3 | 12,4493 | 9,40281 | -0,404901 | 0,0422 | 0,229188 |
| Mir6910 | 156,867 | 290,378 | 0,888393 | 0,04225 | 0,229265 |
| Gkap1 | 9,24833 | 6,38788 | -0,533854 | 0,04225 | 0,229265 |
| Edem1 | 14,5262 | 18,8507 | 0,375964 | 0,0424 | 0,229982 |
| Ccdc102a | 10,8696 | 7,9407 | -0,452964 | 0,0425 | 0,230427 |
| Zfp593 | 16,8227 | 12,481 | -0,430673 | 0,0427 | 0,231413 |
| Gltscr1l | 3,07891 | 4,18149 | 0,441597 | 0,043 | 0,232941 |
| Mvb12a | 23,1768 | 16,7697 | -0,466823 | 0,0431 | 0,233359 |
| Nefm | 2,56602 | 3,75551 | 0,549475 | 0,04315 | 0,233359 |
| Map4k2 | 4,09835 | 5,59353 | 0,448716 | 0,04315 | 0,233359 |
| Rasgrp3 | 4,66224 | 6,30656 | 0,435829 | 0,04315 | 0,233359 |
| 4933439K11Rik | 0,751025 | 2,0824 | 1,47131 | 0,04335 | 0,234243 |
| Gm17296 | 7,42111 | 9,84175 | 0,40728 | 0,04335 | 0,234243 |
| H2-K1 | 10,1266 | 13,9547 | 0,462596 | 0,0434 | 0,234316 |
| Pex7 | 8,99125 | 12,3841 | 0,461898 | 0,0434 | 0,234316 |
| Ogfrl1 | 3,25529 | 4,48595 | 0,462626 | 0,04355 | 0,234928 |
| Ak3 | 36,9671 | 28,6088 | -0,36978 | 0,04355 | 0,234928 |
| Psrc1 | 46,0071 | 35,4591 | -0,375699 | 0,04365 | 0,235342 |
| Kbtbd11 | 4,55527 | 6,03554 | 0,405945 | 0,0437 | 0,235342 |
| Mcoln3 | 6,27171 | 4,21467 | -0,573441 | 0,0437 | 0,235342 |
| 2610016A17Rik | 4,91079 | 2,02139 | -1,28061 | 0,0437 | 0,235342 |
| Thap7 | 13,7349 | 9,56937 | -0,52135 | 0,0438 | 0,235781 |
| Zfp850 | 4,37988 | 5,87349 | 0,423324 | 0,0439 | 0,23622 |
| Purb | 25,5667 | 32,8476 | 0,361525 | 0,04395 | 0,23639 |
| Mlf1 | 9,59141 | 13,752 | 0,519825 | 0,044 | 0,236461 |
| Crispld2 | 1,06018 | 0,640284 | -0,72752 | 0,044 | 0,236461 |
| Spata33 | 1,11378 | 3,2052 | 1,52496 | 0,0441 | 0,236573 |
| Emilin3 | 2,85605 | 4,09942 | 0,521398 | 0,0441 | 0,236573 |
| Mtmr11 | 6,93926 | 9,40945 | 0,439329 | 0,0441 | 0,236573 |
| Fam57a | 14,9947 | 21,1439 | 0,495792 | 0,04415 | 0,236573 |
| Socs7 | 9,00782 | 11,7013 | 0,377414 | 0,04415 | 0,236573 |
| Lmbrd1 | 5,24729 | 3,89056 | -0,431595 | 0,04415 | 0,236573 |
| Fhod3 | 5,05771 | 3,10762 | -0,702671 | 0,04415 | 0,236573 |
| Ccdc55 | 10,7778 | 14,2588 | 0,403788 | 0,0442 | 0,236742 |
| Plac1 | 1,47691 | 2,79357 | 0,919529 | 0,0444 | 0,237615 |
| Fat4 | 5,46809 | 4,21548 | -0,375341 | 0,0444 | 0,237615 |
| Gpr153 | 5,06537 | 6,89648 | 0,445191 | 0,04445 | 0,237783 |
| 1700123M08Rik | 1,00468 | 0,23609 | -2,08933 | 0,04455 | 0,238219 |

| | | | | | |
|---|---|---|---|---|---|
| Fam174b | 3,99154 | 5,65515 | 0,50262 | 0,0446 | 0,238288 |
| Psma7 | 369,207 | 287,937 | -0,358673 | 0,0446 | 0,238288 |
| Gcat | 23,068 | 17,4266 | -0,404601 | 0,04475 | 0,23889 |
| Tspan4 | 12,4512 | 6,91047 | -0,849425 | 0,04475 | 0,23889 |
| Pdpn | 19,6776 | 26,0175 | 0,402926 | 0,0449 | 0,239591 |
| AI848285 | 0,251324 | 0,678223 | 1,43221 | 0,04495 | 0,239758 |
| 2810408I11Rik | 20,253 | 13,8496 | -0,548284 | 0,045 | 0,239925 |
| Fndc3b | 6,5788 | 8,58912 | 0,384685 | 0,0451 | 0,240259 |
| Ctnnbip1 | 22,0706 | 16,8685 | -0,3878 | 0,0451 | 0,240259 |
| Pcbd1 | 21,8863 | 15,2972 | -0,516756 | 0,04515 | 0,240425 |
| C1galt1 | 7,25958 | 9,9985 | 0,461825 | 0,0453 | 0,241124 |
| Eno4 | 0,336224 | 0,702771 | 1,06363 | 0,04535 | 0,241156 |
| Depdc1a | 1,9594 | 2,96749 | 0,598835 | 0,04535 | 0,241156 |
| D6Ertd527e | 0,0504018 | 0,807648 | 4,00218 | 0,0454 | 0,241156 |
| Ifi35 | 0,195338 | 0,724304 | 1,89062 | 0,0454 | 0,241156 |
| Got1l1 | 0,757727 | 0,217642 | -1,79972 | 0,0454 | 0,241156 |
| Usp35 | 1,26867 | 1,93034 | 0,605545 | 0,04545 | 0,241222 |
| Hspa1b | 4,90557 | 7,02378 | 0,517826 | 0,04545 | 0,241222 |
| Ankrd52 | 21,8947 | 28,0839 | 0,359159 | 0,04555 | 0,241652 |
| Reck | 11,1939 | 8,51577 | -0,394508 | 0,04565 | 0,242083 |
| Ifrd2 | 31,3377 | 23,8551 | -0,393601 | 0,04585 | 0,243043 |
| Acat1 | 7,98419 | 5,88159 | -0,440941 | 0,04595 | 0,243372 |
| Nab2 | 7,25098 | 4,6267 | -0,648191 | 0,04595 | 0,243372 |
| Myof | 0,610539 | 0,941253 | 0,624499 | 0,04625 | 0,244759 |
| D7Ertd715e | 7,3386 | 10,2497 | 0,482004 | 0,04625 | 0,244759 |
| Vcam1 | 26,3149 | 34,0511 | 0,371819 | 0,0463 | 0,244922 |
| Gbp9 | 0,341285 | 0,652646 | 0,935325 | 0,0464 | 0,24535 |
| Adamts7 | 2,53585 | 1,76824 | -0,520161 | 0,04645 | 0,245513 |
| Ntm | 2,88411 | 1,79725 | -0,682334 | 0,0465 | 0,245676 |
| Prokr2 | 0,603008 | 0,983026 | 0,705052 | 0,0466 | 0,245901 |
| Ndrg3 | 21,3955 | 27,798 | 0,377672 | 0,0466 | 0,245901 |
| D030045P18Rik | 1,7438 | 0,066321 | -4,71663 | 0,0466 | 0,245901 |
| Gxylt1 | 14,4727 | 18,6581 | 0,366468 | 0,04665 | 0,246063 |
| S1pr3 | 46,8609 | 60,2297 | 0,362089 | 0,04685 | 0,247017 |
| Minos1 | 39,2741 | 30,4677 | -0,366295 | 0,0469 | 0,247179 |
| Neu3 | 0,900662 | 1,45507 | 0,692028 | 0,04705 | 0,247766 |
| Mgat3 | 9,52669 | 7,22975 | -0,398029 | 0,04705 | 0,247766 |
| Tcea2 | 5,99252 | 3,83174 | -0,645165 | 0,0472 | 0,248454 |
| Csgalnact2 | 4,07976 | 5,61052 | 0,459651 | 0,04725 | 0,248513 |
| Gsto1 | 96,2404 | 123,905 | 0,364517 | 0,04725 | 0,248513 |
| Zkscan6 | 4,76025 | 6,7331 | 0,500231 | 0,04735 | 0,248835 |
| Apopt1 | 34,6453 | 25,7216 | -0,429677 | 0,04735 | 0,248835 |
| H2afy2 | 187,698 | 146,699 | -0,355558 | 0,04745 | 0,249156 |
| Jakmip2 | 3,21236 | 2,2123 | -0,538085 | 0,04745 | 0,249156 |
| Zbed6 | 26,5551 | 39,5698 | 0,575414 | 0,0475 | 0,249317 |
| Gp1ba | 0,32395 | 0,675952 | 1,06115 | 0,04755 | 0,249477 |
| Thbs4 | 0,422043 | 0,788468 | 0,901661 | 0,04765 | 0,2499 |
| Zfand5 | 13,4076 | 10,3648 | -0,371351 | 0,04775 | 0,250322 |
| Pold2 | 51,808 | 39,6301 | -0,386577 | 0,0479 | 0,250903 |
| Gjb2 | 0,685974 | 0,309055 | -1,15029 | 0,0479 | 0,250903 |

| | | | | | |
|---|---|---|---|---|---|
| Ptrf | 0,627187 | 1,08665 | 0,792921 | 0,04795 | 0,251062 |
| Ptgis | 16,8574 | 22,4178 | 0,41126 | 0,04805 | 0,251381 |
| Map4k4 | 86,7854 | 111,661 | 0,363601 | 0,04805 | 0,251381 |
| Svep1 | 3,25013 | 4,27644 | 0,395913 | 0,04815 | 0,251596 |
| Wapal | 30,2586 | 38,8475 | 0,360478 | 0,04815 | 0,251596 |
| Pds5a | 55,9861 | 71,8474 | 0,359868 | 0,04815 | 0,251596 |
| Nudt8 | 11,8172 | 17,3275 | 0,552178 | 0,04845 | 0,252958 |
| Dmpk | 8,73561 | 11,7227 | 0,42433 | 0,04845 | 0,252958 |
| Dock11 | 8,78077 | 11,4766 | 0,38628 | 0,0485 | 0,253116 |
| Prcp | 8,80815 | 11,6868 | 0,407964 | 0,04855 | 0,25317 |
| B9d2 | 12,6856 | 8,68986 | -0,545784 | 0,04855 | 0,25317 |
| BC048502 | 0,618163 | 1,80208 | 1,5436 | 0,04865 | 0,253589 |
| Kcna3 | 1,16523 | 1,99291 | 0,774266 | 0,04875 | 0,254007 |
| Chchd2 | 118,548 | 92,1435 | -0,363523 | 0,04885 | 0,254425 |
| Ubxn7 | 12,6851 | 16,2664 | 0,358757 | 0,0494 | 0,257185 |
| Ccdc28a | 2,21998 | 3,76104 | 0,760585 | 0,0495 | 0,257496 |
| Ubxn2a | 5,99943 | 8,25765 | 0,460906 | 0,0495 | 0,257496 |
| 0610010K14Rik | 100,307 | 76,7959 | -0,385319 | 0,04965 | 0,258172 |
| Il15ra | 0,25567 | 0,593652 | 1,21534 | 0,04975 | 0,258482 |
| Pfkfb1 | 3,48328 | 2,21542 | -0,652861 | 0,04975 | 0,258482 |
| Jak2 | 6,19916 | 8,14811 | 0,394395 | 0,0498 | 0,258533 |
| Ppt2 | 24,3603 | 31,8948 | 0,388792 | 0,0498 | 0,258533 |
| Zdhhc13 | 17,2266 | 22,5804 | 0,390431 | 0,0499 | 0,258947 |
| Slco5a1 | 1,57634 | 2,17434 | 0,464 | 0,0502 | 0,260293 |
| Hes1 | 29,0043 | 37,9733 | 0,388718 | 0,0502 | 0,260293 |
| Hist2h2be | 2,68357 | 1,74685 | -0,619397 | 0,05025 | 0,260342 |
| Acsbg1 | 1,48679 | 0,89243 | -0,736388 | 0,05025 | 0,260342 |
| Fam132a | 10,6711 | 7,42098 | -0,524026 | 0,05045 | 0,261167 |
| Tmem26 | 1,63934 | 1,09632 | -0,580455 | 0,05045 | 0,261167 |
| Igf2bp3 | 80,8932 | 63,2749 | -0,354385 | 0,0505 | 0,261321 |
| Dclk1 | 3,11415 | 4,19878 | 0,431133 | 0,0507 | 0,26225 |
| Ppp1cb | 92,6907 | 118,595 | 0,355551 | 0,05075 | 0,262297 |
| Lyrm7 | 1,37537 | 0,772531 | -0,832153 | 0,05075 | 0,262297 |
| Akr1b8 | 8,65048 | 12,2243 | 0,498902 | 0,0508 | 0,262344 |
| Tgfb3 | 3,45613 | 2,40225 | -0,524771 | 0,0508 | 0,262344 |
| Gprin1 | 2,1297 | 1,42658 | -0,578091 | 0,05085 | 0,262496 |
| Atp1b1 | 6,33487 | 10,268 | 0,696762 | 0,0509 | 0,262649 |
| Cyb561 | 1,50615 | 2,35193 | 0,642976 | 0,05095 | 0,262801 |
| Abcc10 | 1,90714 | 2,71181 | 0,507851 | 0,05105 | 0,263211 |
| 5031439G07Rik | 15,8718 | 12,3074 | -0,36693 | 0,0511 | 0,263363 |
| Zfp663 | 0,388168 | 0,737552 | 0,926064 | 0,0513 | 0,264182 |
| Tnks1bp1 | 10,8433 | 14,0121 | 0,369877 | 0,0513 | 0,264182 |
| Ndufa12 | 105,447 | 79,8283 | -0,401552 | 0,05135 | 0,264333 |
| Cryz | 15,0642 | 10,7732 | -0,48367 | 0,0514 | 0,264484 |
| Smc4 | 82,4508 | 105,616 | 0,357225 | 0,05145 | 0,264529 |
| Pdlim1 | 7,79582 | 5,36525 | -0,539054 | 0,05145 | 0,264529 |
| Lhx9 | 0,313935 | 0,677428 | 1,1096 | 0,0515 | 0,26468 |
| Ints6 | 10,8091 | 13,971 | 0,370197 | 0,0516 | 0,265088 |
| Sorbs2 | 0,637232 | 0,386461 | -0,721495 | 0,0518 | 0,266009 |
| Ablim1 | 5,80906 | 7,62013 | 0,39151 | 0,0519 | 0,266416 |

| | | | | | |
|---|---|---|---|---|---|
| Rpl36al | 81,3797 | 60,344 | -0,431457 | 0,05195 | 0,266566 |
| Tubd1 | 6,58455 | 4,61143 | -0,51387 | 0,0521 | 0,267229 |
| Eaf1 | 31,1492 | 39,8133 | 0,354054 | 0,0522 | 0,267635 |
| Rnf123 | 8,9387 | 12,2091 | 0,449822 | 0,0524 | 0,268553 |
| Slc4a4 | 2,97212 | 2,19017 | -0,440447 | 0,05255 | 0,269214 |
| Cth | 18,3767 | 24,1006 | 0,391194 | 0,05265 | 0,269618 |
| Ndfip1 | 113,297 | 145,46 | 0,360512 | 0,0527 | 0,269659 |
| Lyrm4 | 7,9505 | 5,48719 | -0,534977 | 0,0527 | 0,269659 |
| Slc35a1 | 30,326 | 39,0852 | 0,366068 | 0,0528 | 0,270063 |
| Lemd3 | 7,31445 | 9,54119 | 0,383419 | 0,05285 | 0,270211 |
| Yod1 | 1,72767 | 2,93155 | 0,762835 | 0,05295 | 0,270399 |
| Mbp | 3,92841 | 2,82236 | -0,477041 | 0,05295 | 0,270399 |
| 1600027J07Rik | 3,45438 | 1,25617 | -1,45939 | 0,05295 | 0,270399 |
| Cdh6 | 7,4837 | 10,0995 | 0,432462 | 0,053 | 0,270547 |
| Zyg11b | 9,9939 | 12,7776 | 0,354494 | 0,05305 | 0,270587 |
| Kif5b | 41,1682 | 52,6012 | 0,353564 | 0,05305 | 0,270587 |
| Rpl27a | 83,5034 | 65,0632 | -0,359993 | 0,05315 | 0,270989 |
| Hoxd3 | 32,222 | 48,6817 | 0,595335 | 0,05335 | 0,271901 |
| 9230114K14Rik | 0,623688 | 1,15952 | 0,894627 | 0,05345 | 0,272302 |
| Loxl3 | 2,69901 | 1,16198 | -1,21585 | 0,0535 | 0,272449 |
| S100a13 | 4,45363 | 7,30857 | 0,714607 | 0,05365 | 0,272887 |
| Col16a1 | 1,05212 | 1,55974 | 0,568007 | 0,05365 | 0,272887 |
| Foxo3 | 7,66639 | 10,2383 | 0,417359 | 0,05365 | 0,272887 |
| Sema4b | 3,61702 | 4,94808 | 0,452066 | 0,05375 | 0,273179 |
| Yeats4 | 116,767 | 91,7023 | -0,348598 | 0,05375 | 0,273179 |
| Slc35c1 | 4,41645 | 3,10138 | -0,509977 | 0,0538 | 0,273325 |
| Plcg2 | 2,09023 | 1,42071 | -0,557052 | 0,05385 | 0,273471 |
| Ccdc28b | 13,6371 | 9,28286 | -0,554899 | 0,05395 | 0,27387 |
| Msrb2 | 15,3367 | 11,0423 | -0,47395 | 0,054 | 0,274016 |
| Snord82 | 288,707 | 302,996 | 0,0696921 | 0,05405 | 0,274161 |
| Lphn3 | 1,16952 | 1,7046 | 0,543517 | 0,0543 | 0,27532 |
| Cnot6l | 6,7384 | 8,67067 | 0,363738 | 0,05435 | 0,275356 |
| Prex1 | 13,0876 | 16,704 | 0,351988 | 0,05435 | 0,275356 |
| Nnt | 17,439 | 13,5653 | -0,362397 | 0,0545 | 0,275789 |
| A430005L14Rik | 43,461 | 33,1539 | -0,390545 | 0,0545 | 0,275789 |
| Ptgr1 | 10,0776 | 7,3922 | -0,447075 | 0,0545 | 0,275789 |
| Prdm11 | 1,88282 | 2,96309 | 0,654205 | 0,05455 | 0,275934 |
| Zfp61 | 15,0646 | 11,3831 | -0,404269 | 0,0546 | 0,275969 |
| Alox5ap | 1,63307 | 0,657547 | -1,31242 | 0,0546 | 0,275969 |
| Pcx | 7,97099 | 12,1413 | 0,607094 | 0,0547 | 0,276365 |
| Mir150 | 0 | 486,198 | inf | 0,0549 | 0,277157 |
| Agpat2 | 7,08986 | 9,94465 | 0,488164 | 0,0549 | 0,277157 |
| Hspg2 | 13,4817 | 17,1194 | 0,344622 | 0,055 | 0,277553 |
| Snrk | 5,38309 | 7,10729 | 0,400864 | 0,05505 | 0,277696 |
| Nemf | 21,7209 | 27,934 | 0,36294 | 0,05515 | 0,277982 |
| Slc2a1 | 141,596 | 110,813 | -0,353653 | 0,05515 | 0,277982 |
| C430049B03Rik | 4,51958 | 3,21323 | -0,492165 | 0,0553 | 0,278629 |
| Nudt4 | 66,1322 | 51,8898 | -0,349901 | 0,0554 | 0,279023 |
| 1190005I06Rik | 2,69131 | 4,82098 | 0,841019 | 0,0555 | 0,279417 |
| Zcchc18 | 4,80125 | 6,78288 | 0,498486 | 0,05555 | 0,279449 |

| | | | | | |
|---|---|---|---|---|---|
| Crtc2 | 16,4275 | 21,302 | 0,374882 | 0,05555 | 0,279449 |
| Plvap | 4,45054 | 2,96595 | -0,585487 | 0,0556 | 0,279591 |
| Ankrd34a | 0,691957 | 1,20093 | 0,795401 | 0,05565 | 0,279733 |
| Cox6c | 324,587 | 253,867 | -0,354535 | 0,0559 | 0,280879 |
| Diap3 | 5,96041 | 7,96748 | 0,418712 | 0,05605 | 0,281523 |
| Rpl37 | 110,198 | 86,2158 | -0,354078 | 0,05615 | 0,281804 |
| Rpl31 | 23,9824 | 17,3549 | -0,466633 | 0,05615 | 0,281804 |
| Ndn | 82,3785 | 64,7893 | -0,346512 | 0,05645 | 0,283199 |
| Lgr4 | 18,4844 | 14,4703 | -0,353213 | 0,0565 | 0,283339 |
| Tmem42 | 8,09477 | 5,09744 | -0,667218 | 0,0566 | 0,28373 |
| Taf1 | 16,6373 | 21,0783 | 0,341338 | 0,0567 | 0,28412 |
| Prkcd | 17,8184 | 23,005 | 0,368582 | 0,0568 | 0,28451 |
| Nrxn2 | 1,94747 | 2,67058 | 0,455551 | 0,0569 | 0,2849 |
| Akr1b10 | 6,43127 | 9,21404 | 0,51873 | 0,057 | 0,285289 |
| Cecr5 | 14,5319 | 19,0978 | 0,39418 | 0,05705 | 0,285317 |
| Ube2l6 | 3,88969 | 2,4586 | -0,661818 | 0,05705 | 0,285317 |
| Kank4 | 1,49488 | 0,97847 | -0,611431 | 0,0572 | 0,285955 |
| Rps19 | 134,419 | 104,046 | -0,369515 | 0,0573 | 0,286344 |
| Msmo1 | 90,4565 | 71,0734 | -0,347914 | 0,05755 | 0,287481 |
| Rps26 | 1029,1 | 812,73 | -0,340538 | 0,05765 | 0,287756 |
| Rad51d | 4,15438 | 2,6961 | -0,623756 | 0,05765 | 0,287757 |
| Zfp369 | 4,81023 | 6,57865 | 0,451685 | 0,0577 | 0,287894 |
| Gab1 | 9,36154 | 7,20367 | -0,378013 | 0,05775 | 0,28792 |
| Rpl37a | 54,8501 | 40,7812 | -0,42759 | 0,05775 | 0,28792 |
| Ube3a | 18,831 | 24,1296 | 0,357688 | 0,0578 | 0,288057 |
| Pld2 | 0,980433 | 1,51181 | 0,624784 | 0,05795 | 0,288692 |
| Cspp1 | 12,1955 | 15,7134 | 0,365643 | 0,058 | 0,288829 |
| Hmgxb3 | 16,5333 | 20,9676 | 0,342792 | 0,05805 | 0,288854 |
| Rps21 | 645,836 | 507,305 | -0,348312 | 0,05805 | 0,288854 |
| Fgd3 | 0,869052 | 0,495022 | -0,811951 | 0,0582 | 0,289488 |
| Gatm | 1,78048 | 1,09659 | -0,699247 | 0,0583 | 0,289873 |
| Tmem139 | 0,491958 | 1,18412 | 1,26721 | 0,0585 | 0,290642 |
| Nkiras2 | 32,5771 | 25,3881 | -0,359705 | 0,0585 | 0,290642 |
| Arl5a | 22,0858 | 28,0898 | 0,34693 | 0,05855 | 0,290778 |
| Cep55 | 28,3467 | 36,2321 | 0,354087 | 0,0586 | 0,290801 |
| Sltm | 51,8873 | 65,7454 | 0,341508 | 0,0586 | 0,290801 |
| Mmp23 | 1,05335 | 1,91191 | 0,860026 | 0,05865 | 0,290937 |
| Zfp748 | 3,75556 | 5,04951 | 0,427113 | 0,0587 | 0,29096 |
| Hmbs | 31,6957 | 24,5681 | -0,367497 | 0,0587 | 0,29096 |
| Layn | 0,32636 | 0,759644 | 1,21886 | 0,0588 | 0,291343 |
| Pglyrp3 | 1,3298 | 2,4574 | 0,88592 | 0,05895 | 0,291973 |
| Mrpl27 | 43,3538 | 32,3362 | -0,423007 | 0,05905 | 0,292243 |
| Tmem160 | 28,7065 | 21,0294 | -0,448968 | 0,05905 | 0,292243 |
| Slc24a3 | 1,65033 | 2,42175 | 0,553298 | 0,0591 | 0,292377 |
| Ssna1 | 76,8314 | 59,4585 | -0,369814 | 0,05915 | 0,292512 |
| Ccpg1os | 9,90043 | 14,6293 | 0,563294 | 0,0592 | 0,292646 |
| Lrrc51 | 5,69366 | 3,53069 | -0,689405 | 0,0595 | 0,294016 |
| Lpin1 | 1,70977 | 2,39795 | 0,487997 | 0,05955 | 0,294037 |
| Pcca | 18,273 | 14,0923 | -0,374807 | 0,05955 | 0,294037 |
| Zfp941 | 0,723276 | 1,14936 | 0,668215 | 0,05965 | 0,294304 |

| | | | | | |
|---|---|---|---|---|---|
| Hist1h1e | 1,63895 | 0,611362 | -1,42267 | 0,05965 | 0,294304 |
| Snord66 | 408,824 | 1447,84 | 1,82435 | 0,05975 | 0,294344 |
| Atp2a2 | 108,938 | 138,275 | 0,344034 | 0,05975 | 0,294344 |
| Ccdc92 | 3,19357 | 2,12892 | -0,585049 | 0,05975 | 0,294344 |
| Amer2 | 2,71282 | 1,78461 | -0,604189 | 0,05975 | 0,294344 |
| Etv6 | 18,1771 | 23,0811 | 0,344591 | 0,05985 | 0,294611 |
| Tnfaip8l1 | 1,36412 | 0,800571 | -0,768867 | 0,05985 | 0,294611 |
| Ppapdc2 | 2,22635 | 1,46886 | -0,599979 | 0,05995 | 0,29499 |
| Ero1l | 8,27527 | 10,721 | 0,373563 | 0,06005 | 0,295369 |
| Pir | 8,01115 | 5,65735 | -0,501882 | 0,0601 | 0,295501 |
| Cox6a1 | 314,557 | 246,911 | -0,349332 | 0,0602 | 0,29588 |
| Gltscr2 | 71,624 | 56,0387 | -0,35402 | 0,06025 | 0,296012 |
| Bpgm | 22,9617 | 29,5844 | 0,365604 | 0,0606 | 0,297276 |
| Mtmr6 | 10,3956 | 13,3935 | 0,365559 | 0,0606 | 0,297276 |
| Lmna | 1,84063 | 1,09487 | -0,749439 | 0,0606 | 0,297276 |
| Pklr | 1,07817 | 0,622018 | -0,793552 | 0,0606 | 0,297276 |
| Slc25a5 | 211,965 | 167,351 | -0,340948 | 0,0607 | 0,297653 |
| Lrrc20 | 6,32356 | 8,51386 | 0,429077 | 0,06075 | 0,29767 |
| Cyb5r1 | 17,5507 | 23,1187 | 0,39753 | 0,06075 | 0,29767 |
| Tmem65 | 9,87792 | 12,8046 | 0,374382 | 0,0608 | 0,297688 |
| Onecut2 | 0,772284 | 0,537601 | -0,522598 | 0,0608 | 0,297688 |
| Srxn1 | 7,78215 | 5,75339 | -0,435758 | 0,0609 | 0,298064 |
| Mir219b | 174,635 | 0 | -inf | 0,06095 | 0,298195 |
| Rpl41 | 497,566 | 393,255 | -0,339423 | 0,061 | 0,298326 |
| Gpc4 | 13,9468 | 18,1513 | 0,38014 | 0,06105 | 0,298456 |
| Fjx1 | 11,8024 | 15,4387 | 0,38747 | 0,0611 | 0,298587 |
| Dusp16 | 6,71317 | 5,10245 | -0,395802 | 0,06125 | 0,299206 |
| Smtn | 9,85962 | 12,8119 | 0,377881 | 0,0615 | 0,300313 |
| Trpm7 | 20,1025 | 25,502 | 0,343238 | 0,06155 | 0,300328 |
| Morc4 | 3,34963 | 2,41457 | -0,472235 | 0,06155 | 0,300328 |
| Ppip5k1 | 1,94567 | 3,1518 | 0,695906 | 0,06165 | 0,300588 |
| Mtmr12 | 6,90741 | 8,98496 | 0,379367 | 0,06165 | 0,300588 |
| 2610035F20Rik | 5,96538 | 8,69028 | 0,542787 | 0,0617 | 0,300717 |
| Ypel1 | 15,9296 | 10,5303 | -0,597154 | 0,06175 | 0,300847 |
| Dennd1b | 1,96972 | 2,6429 | 0,424126 | 0,06185 | 0,301105 |
| Tulp4 | 3,6632 | 4,71255 | 0,363404 | 0,06185 | 0,301105 |
| Sall2 | 31,1546 | 24,6643 | -0,337017 | 0,0619 | 0,301234 |
| Cacnb3 | 13,417 | 17,3146 | 0,367927 | 0,06195 | 0,301363 |
| Fam32a | 46,0171 | 36,0822 | -0,350886 | 0,06205 | 0,301621 |
| Mxra7 | 12,397 | 9,17513 | -0,434186 | 0,06205 | 0,301621 |
| Gm684 | 1,36127 | 0,680542 | -1,00019 | 0,06225 | 0,302478 |
| Tyk2 | 3,56513 | 4,789 | 0,425771 | 0,0625 | 0,303578 |
| Mcur1 | 3,74694 | 5,0163 | 0,420911 | 0,06255 | 0,303706 |
| Strn | 6,8909 | 8,9854 | 0,38289 | 0,0627 | 0,304319 |
| Pros1 | 13,1011 | 10,1055 | -0,374542 | 0,06275 | 0,304447 |
| Kcnip4 | 0,699127 | 1,42438 | 1,02671 | 0,0628 | 0,304459 |
| Ndufb5 | 137,315 | 108,315 | -0,342246 | 0,0628 | 0,304459 |
| Fam89b | 52,6296 | 40,9732 | -0,361194 | 0,0629 | 0,304829 |
| Fmnl2 | 14,332 | 18,1761 | 0,342797 | 0,06295 | 0,304956 |
| Slc1a6 | 0,715886 | 1,27754 | 0,83557 | 0,06305 | 0,304991 |

| | | | | | |
|---|---|---|---|---|---|
| Rxra | 16,1746 | 12,6997 | -0,348929 | 0,06305 | 0,304991 |
| Aadat | 2,92431 | 1,86844 | -0,646262 | 0,06305 | 0,304991 |
| Slc16a13 | 3,98406 | 5,58384 | 0,487018 | 0,0631 | 0,304991 |
| Lrch3 | 16,2631 | 20,6403 | 0,343857 | 0,0631 | 0,304991 |
| Akap2 | 18,9172 | 14,9897 | -0,335727 | 0,0631 | 0,304991 |
| Elmsan1 | 7,28131 | 9,30579 | 0,353931 | 0,06315 | 0,305118 |
| Dcaf12l1 | 1,09732 | 0,673119 | -0,70505 | 0,0633 | 0,305728 |
| Creb3l2 | 12,2686 | 9,46589 | -0,374166 | 0,06335 | 0,305854 |
| Fam13b | 18,4164 | 23,5383 | 0,354021 | 0,0635 | 0,306463 |
| Abhd14b | 3,9879 | 5,88642 | 0,56176 | 0,06365 | 0,307071 |
| Pgpep1 | 7,37575 | 5,65256 | -0,383885 | 0,06375 | 0,307438 |
| Lmtk2 | 7,44062 | 9,50958 | 0,353959 | 0,0638 | 0,307448 |
| Rev3l | 8,26939 | 10,5543 | 0,351982 | 0,0638 | 0,307448 |
| D630033O11Rik | 0,788971 | 2,2078 | 1,48457 | 0,06385 | 0,307573 |
| Zfp26 | 7,21482 | 9,14628 | 0,342223 | 0,064 | 0,30818 |
| Adipor2 | 34,4518 | 43,3643 | 0,331929 | 0,06415 | 0,308787 |
| Jmjd1c | 10,6341 | 13,4433 | 0,338193 | 0,0642 | 0,308911 |
| Dock1 | 18,0587 | 22,8525 | 0,339661 | 0,06445 | 0,309882 |
| D1Ertd622e | 5,90787 | 4,29016 | -0,461605 | 0,06445 | 0,309882 |
| Map6d1 | 0,737421 | 1,19603 | 0,697691 | 0,0645 | 0,310006 |
| Slc25a27 | 5,72494 | 7,70043 | 0,427679 | 0,06455 | 0,31013 |
| Gcnt1 | 2,71601 | 3,69939 | 0,4458 | 0,0646 | 0,310138 |
| Zfp871 | 13,3588 | 16,8586 | 0,335691 | 0,0646 | 0,310138 |
| Arhgap21 | 14,8662 | 18,724 | 0,332849 | 0,0647 | 0,310502 |
| Hprt | 80,3731 | 63,4281 | -0,341592 | 0,06495 | 0,311585 |
| Adcy3 | 3,81758 | 6,09803 | 0,675685 | 0,065 | 0,311592 |
| Icam2 | 1,50417 | 0,692453 | -1,11918 | 0,065 | 0,311592 |
| Ap2s1 | 154,016 | 121,658 | -0,340249 | 0,06515 | 0,312194 |
| Phf21b | 4,9877 | 3,6628 | -0,445431 | 0,0652 | 0,312317 |
| Ugcg | 23,1667 | 29,3336 | 0,340504 | 0,0653 | 0,31268 |
| Raph1 | 9,39002 | 11,8908 | 0,340647 | 0,0655 | 0,31352 |
| Crtc3 | 2,97818 | 2,16467 | -0,460283 | 0,06575 | 0,314487 |
| Slc16a7 | 0,397838 | 0,768943 | 0,950695 | 0,0658 | 0,314487 |
| Ofd1 | 4,85525 | 6,38911 | 0,396069 | 0,0658 | 0,314487 |
| Megf6 | 5,32433 | 4,08546 | -0,382101 | 0,0658 | 0,314487 |
| Katnal1 | 4,33949 | 3,28636 | -0,401034 | 0,0659 | 0,314496 |
| Cdkn3 | 14,6594 | 10,0817 | -0,540081 | 0,0659 | 0,314496 |
| Epb4.1l4a | 2,20019 | 1,50977 | -0,543301 | 0,0659 | 0,314496 |
| Ngly1 | 4,52785 | 3,09646 | -0,548205 | 0,0659 | 0,314496 |
| Slc25a25 | 8,83002 | 11,6015 | 0,39382 | 0,06595 | 0,314618 |
| Gnb5 | 7,12305 | 5,18819 | -0,457264 | 0,06605 | 0,314743 |
| 2010204K13Rik | 24,7991 | 17,1908 | -0,528655 | 0,06605 | 0,314743 |
| Msx1os | 3,98205 | 2,04781 | -0,959432 | 0,06605 | 0,314743 |
| Dennd6b | 6,00555 | 7,8036 | 0,377844 | 0,0661 | 0,314865 |
| Arid1a | 48,1154 | 60,5628 | 0,331932 | 0,0664 | 0,316176 |
| Ythdf3 | 32,7024 | 41,1919 | 0,332963 | 0,0665 | 0,316535 |
| Fbxo10 | 7,67464 | 10,9081 | 0,507224 | 0,06665 | 0,317131 |
| Lrfn1 | 6,47255 | 8,72164 | 0,430265 | 0,06675 | 0,317489 |
| Galc | 8,95714 | 11,5687 | 0,369115 | 0,0669 | 0,317967 |
| Xpo4 | 19,6482 | 24,9879 | 0,346834 | 0,0669 | 0,317967 |

| | | | | | |
|---|---|---|---|---|---|
| Gm13152 | 2,10098 | 3,4233 | 0,704329 | 0,06695 | 0,318087 |
| Dirc2 | 5,21611 | 3,93146 | -0,407908 | 0,06705 | 0,318444 |
| Arhgef25 | 43,3199 | 54,5689 | 0,333049 | 0,0671 | 0,318564 |
| Hdx | 2,17466 | 3,04285 | 0,484629 | 0,0672 | 0,318802 |
| Dhcr7 | 27,9068 | 21,9454 | -0,346698 | 0,0672 | 0,318802 |
| Slc6a8 | 9,1631 | 11,7863 | 0,363201 | 0,06725 | 0,318922 |
| Pwwp2a | 24,7715 | 31,3197 | 0,33839 | 0,06735 | 0,319278 |
| Tnfrsf12a | 2,1546 | 10,7537 | 2,31935 | 0,0674 | 0,319397 |
| Srek1 | 33,7476 | 42,5156 | 0,333208 | 0,0675 | 0,319635 |
| Adam22 | 1,12816 | 0,720857 | -0,646181 | 0,0675 | 0,319635 |
| Arvcf | 14,378 | 10,4569 | -0,459407 | 0,06755 | 0,319753 |
| Slc20a2 | 21,0865 | 27,0602 | 0,359855 | 0,0676 | 0,319872 |
| Maged2 | 116,614 | 146,102 | 0,325235 | 0,06765 | 0,31999 |
| Palld | 20,1416 | 25,5999 | 0,345957 | 0,06785 | 0,320818 |
| Mad2l2 | 24,089 | 18,3113 | -0,395635 | 0,0679 | 0,320818 |
| Ttc14 | 10,7952 | 13,7375 | 0,347726 | 0,06795 | 0,320818 |
| Fbxl5 | 30,4383 | 38,3918 | 0,33491 | 0,06795 | 0,320818 |
| Ndufaf2 | 88,4589 | 68,6006 | -0,366786 | 0,06795 | 0,320818 |
| Atp5g1 | 4,74201 | 2,84206 | -0,738562 | 0,068 | 0,320936 |
| Dzip1l | 12,8321 | 16,3997 | 0,353914 | 0,0683 | 0,322233 |
| Fgf2 | 0,297273 | 1,317 | 2,1474 | 0,0685 | 0,322821 |
| Atp5h | 107,16 | 82,8854 | -0,370579 | 0,0685 | 0,322821 |
| Actc1 | 17,3769 | 13,0095 | -0,417607 | 0,0685 | 0,322821 |
| Efhd1 | 0,447273 | 0,908348 | 1,02209 | 0,06855 | 0,322936 |
| Fbxl18 | 5,57362 | 7,47996 | 0,424414 | 0,0686 | 0,322936 |
| Gpr85 | 1,2616 | 0,777311 | -0,698685 | 0,0686 | 0,322936 |
| Lsr | 24,4673 | 31,1951 | 0,350465 | 0,0687 | 0,323288 |
| Tbc1d12 | 3,3413 | 4,50625 | 0,431519 | 0,06885 | 0,323757 |
| Rfx7 | 16,0428 | 20,1889 | 0,331636 | 0,06885 | 0,323757 |
| Slc7a6 | 51,5043 | 64,5976 | 0,326788 | 0,069 | 0,324343 |
| Cul7 | 31,0319 | 38,9009 | 0,32605 | 0,06905 | 0,324456 |
| Malat1 | 60,495 | 76,2341 | 0,333622 | 0,0691 | 0,324456 |
| Coa3 | 110,98 | 87,0296 | -0,350726 | 0,0691 | 0,324456 |
| Homer1 | 12,9285 | 16,7627 | 0,374703 | 0,06925 | 0,324923 |
| Rnf14 | 25,8656 | 32,684 | 0,337545 | 0,06925 | 0,324923 |
| Mapk8ip1 | 21,1376 | 16,6427 | -0,344923 | 0,0693 | 0,325039 |
| Tnks | 18,0195 | 22,6701 | 0,331233 | 0,0695 | 0,325857 |
| Pmel | 0,526401 | 1,21212 | 1,2033 | 0,0696 | 0,326207 |
| Frmd4b | 12,6578 | 16,0205 | 0,3399 | 0,06965 | 0,326322 |
| Enoph1 | 28,4962 | 22,2437 | -0,357371 | 0,06985 | 0,327135 |
| Zfc3h1 | 11,4804 | 14,4695 | 0,333839 | 0,0699 | 0,327135 |
| Akap1 | 19,052 | 15,0034 | -0,344651 | 0,0699 | 0,327135 |
| Tlr5 | 0,405936 | 0,727158 | 0,841014 | 0,07 | 0,327364 |
| Acad10 | 0,523399 | 0,862293 | 0,720266 | 0,07 | 0,327364 |
| Nudt18 | 0,594412 | 0,959144 | 0,690284 | 0,07025 | 0,328294 |
| Prdm10 | 4,44283 | 5,95241 | 0,421993 | 0,07025 | 0,328294 |
| Vps28 | 53,0065 | 40,9751 | -0,371422 | 0,0703 | 0,328408 |
| Hus1 | 17,1559 | 13,5859 | -0,336596 | 0,07035 | 0,328522 |
| Hunk | 17,8751 | 22,5185 | 0,333154 | 0,07045 | 0,32863 |
| Atp5g3 | 795,657 | 632,693 | -0,330642 | 0,07045 | 0,32863 |

| | | | | | |
|---|---|---|---|---|---|
| Ywhaq | 51,3758 | 40,8096 | -0,332183 | 0,07045 | 0,32863 |
| Rhof | 0,650486 | 1,14844 | 0,82009 | 0,07055 | 0,328977 |
| Car8 | 0,633756 | 0,389622 | -0,701852 | 0,07065 | 0,329323 |
| Gm14420 | 0,760606 | 1,15496 | 0,602623 | 0,0707 | 0,329437 |
| Ube2l3 | 73,8643 | 58,7375 | -0,330596 | 0,07085 | 0,329896 |
| Prpsap2 | 9,31569 | 6,81452 | -0,45105 | 0,07085 | 0,329896 |
| Tnnc1 | 0,452521 | 1,37186 | 1,60008 | 0,07095 | 0,330228 |
| Smox | 11,2144 | 27,2013 | 1,27833 | 0,071 | 0,330228 |
| St6galnac2 | 0,755181 | 1,32115 | 0,806902 | 0,07105 | 0,330228 |
| Camk4 | 0,661141 | 0,921702 | 0,479343 | 0,07105 | 0,330228 |
| Mxd4 | 11,3417 | 8,12502 | -0,481197 | 0,07105 | 0,330228 |
| Ier5 | 11,9457 | 15,2654 | 0,353766 | 0,0711 | 0,330334 |
| Pik3cb | 2,24247 | 3,06586 | 0,451203 | 0,07115 | 0,330334 |
| Gale | 12,9748 | 17,1626 | 0,403557 | 0,07115 | 0,330334 |
| Nr1h3 | 1,74389 | 2,69082 | 0,625735 | 0,0716 | 0,332062 |
| Eml3 | 19,7237 | 15,4904 | -0,348556 | 0,0716 | 0,332062 |
| Hlx | 1,06865 | 0,591814 | -0,852575 | 0,0716 | 0,332062 |
| C1qtnf1 | 0,925208 | 1,39149 | 0,588778 | 0,0717 | 0,332406 |
| Tpst2 | 12,4987 | 9,4397 | -0,40496 | 0,07175 | 0,332518 |
| Mboat2 | 14,9277 | 19,057 | 0,35233 | 0,0718 | 0,332612 |
| Pdlim3 | 2,09757 | 1,26849 | -0,725609 | 0,07185 | 0,332612 |
| Stmn3 | 0,845089 | 0,259035 | -1,70596 | 0,07185 | 0,332612 |
| Psmc1 | 140,713 | 112,394 | -0,324195 | 0,0719 | 0,332612 |
| Tuba1b | 136,154 | 108,585 | -0,32641 | 0,0719 | 0,332612 |
| Siah2 | 5,85246 | 7,83734 | 0,421322 | 0,072 | 0,332715 |
| Adh5 | 203,611 | 163,023 | -0,320736 | 0,072 | 0,332715 |
| Gnptab | 6,72681 | 5,19806 | -0,37195 | 0,072 | 0,332715 |
| Ccne1 | 59,3258 | 47,1886 | -0,33022 | 0,07205 | 0,332826 |
| Rpl12 | 10,4084 | 15,5697 | 0,580988 | 0,0722 | 0,33338 |
| Arl6 | 25,2305 | 19,4717 | -0,373793 | 0,07225 | 0,33338 |
| Ahcyl2 | 6,96322 | 8,91483 | 0,356453 | 0,0723 | 0,33338 |
| Zfyve16 | 7,77267 | 9,92239 | 0,352277 | 0,0723 | 0,33338 |
| Rps15a | 39,6077 | 31,5803 | -0,326758 | 0,0723 | 0,33338 |
| Glis2 | 18,1133 | 14,2272 | -0,348398 | 0,07245 | 0,333943 |
| Loh12cr1 | 5,60331 | 3,90709 | -0,520184 | 0,0725 | 0,333943 |
| 4930500L23Rik | 1,22413 | 0,236018 | -2,37479 | 0,0725 | 0,333943 |
| Irs2 | 7,86741 | 10,1252 | 0,363993 | 0,07255 | 0,334053 |
| Phf13 | 16,6745 | 21,2212 | 0,347865 | 0,0727 | 0,334623 |
| Immp1l | 12,7608 | 8,99251 | -0,504921 | 0,07285 | 0,335194 |
| Naa15 | 46,4867 | 58,3938 | 0,328996 | 0,0729 | 0,335303 |
| Nktr | 32,8428 | 41,187 | 0,326612 | 0,0731 | 0,336103 |
| Dusp3 | 2,39312 | 1,68154 | -0,509107 | 0,0732 | 0,336442 |
| Map3k2 | 3,06659 | 3,94215 | 0,362346 | 0,07325 | 0,336551 |
| Eif3a | 138,844 | 175,433 | 0,337462 | 0,07345 | 0,337349 |
| Mccc2 | 21,1477 | 16,5005 | -0,35799 | 0,0735 | 0,337458 |
| 2700046A07Rik | 0,543915 | 1,00087 | 0,8798 | 0,0736 | 0,33776 |
| Pdpr | 9,33661 | 12,2228 | 0,388604 | 0,07365 | 0,33776 |
| D330023K18Rik | 5,1988 | 2,70331 | -0,943452 | 0,07365 | 0,33776 |
| Celf4 | 1,33702 | 1,9329 | 0,531742 | 0,0737 | 0,33776 |
| Shprh | 8,03684 | 10,164 | 0,338763 | 0,0737 | 0,33776 |

| | | | | | |
|---|---|---|---|---|---|
| Stxbp3b | 2,59611 | 4,16812 | 0,683049 | 0,07375 | 0,33776 |
| Churc1 | 35,8933 | 26,5283 | -0,43618 | 0,07375 | 0,33776 |
| Fbxo30 | 13,6369 | 17,1599 | 0,331528 | 0,0738 | 0,337868 |
| Adamts4 | 3,03965 | 4,14117 | 0,446133 | 0,0739 | 0,338205 |
| G6pc3 | 13,8243 | 18,1482 | 0,392617 | 0,074 | 0,338421 |
| Fam160b2 | 7,55797 | 9,73696 | 0,365472 | 0,074 | 0,338421 |
| Ssbp2 | 9,09403 | 7,15596 | -0,345774 | 0,07405 | 0,338529 |
| Irf2 | 4,72131 | 6,45573 | 0,451393 | 0,07415 | 0,338866 |
| Mtmr1 | 2,6693 | 1,93649 | -0,46302 | 0,0742 | 0,338974 |
| Entpd6 | 11,9943 | 9,17267 | -0,386935 | 0,07425 | 0,339081 |
| Crabp2 | 534,166 | 426,675 | -0,324151 | 0,07445 | 0,339874 |
| Myo18a | 8,06832 | 6,36232 | -0,342717 | 0,07455 | 0,340209 |
| Dnpep | 37,387 | 29,3999 | -0,346723 | 0,0747 | 0,340772 |
| A530054K11Rik | 1,72265 | 2,39404 | 0,474815 | 0,0748 | 0,340972 |
| Zfp280d | 19,9562 | 25,2825 | 0,3413 | 0,0748 | 0,340972 |
| Zfp422 | 52,5906 | 41,9391 | -0,326509 | 0,07485 | 0,340972 |
| Tspan7 | 34,5939 | 27,183 | -0,347811 | 0,07485 | 0,340972 |
| Map2k6 | 59,8432 | 47,6923 | -0,327432 | 0,07495 | 0,341306 |
| Hoxa10 | 86,1725 | 107,645 | 0,320979 | 0,07505 | 0,34164 |
| Zfp420 | 3,0638 | 4,19236 | 0,45244 | 0,0752 | 0,34208 |
| Speg | 18,6024 | 13,5204 | -0,460357 | 0,0752 | 0,34208 |
| Zkscan7 | 4,32864 | 5,94195 | 0,457023 | 0,07525 | 0,342186 |
| Clock | 4,29788 | 7,01566 | 0,706954 | 0,07535 | 0,342277 |
| P2rx3 | 1,02562 | 1,51421 | 0,562072 | 0,07535 | 0,342277 |
| Zfp160 | 8,6987 | 11,1842 | 0,36259 | 0,07535 | 0,342277 |
| Gm13826 | 294,514 | 236,163 | -0,318555 | 0,0755 | 0,342715 |
| Lrrc8b | 3,18209 | 2,22494 | -0,516208 | 0,0755 | 0,342716 |
| U2surp | 31,6173 | 39,5592 | 0,323298 | 0,0757 | 0,343502 |
| Mcam | 1,73689 | 1,13953 | -0,608069 | 0,0758 | 0,343818 |
| Cables2 | 9,33227 | 12,0595 | 0,369875 | 0,07585 | 0,343818 |
| Rasa2 | 5,61036 | 7,18056 | 0,356003 | 0,07585 | 0,343818 |
| Vrk3 | 21,5547 | 16,7592 | -0,363048 | 0,0759 | 0,343923 |
| Ssh1 | 5,00242 | 6,65381 | 0,411553 | 0,07595 | 0,344028 |
| Itgav | 10,6882 | 13,4441 | 0,330943 | 0,076 | 0,344133 |
| Dera | 43,5017 | 34,3694 | -0,339948 | 0,07605 | 0,344238 |
| Fam179b | 9,80306 | 12,3422 | 0,332292 | 0,0761 | 0,344342 |
| Ccdc157 | 4,98486 | 6,77564 | 0,442804 | 0,07625 | 0,3449 |
| Ptms | 224,543 | 280,422 | 0,320608 | 0,07635 | 0,345108 |
| Sdhb | 158,836 | 126,877 | -0,32411 | 0,07635 | 0,345108 |
| Tmem88 | 4,15842 | 7,68816 | 0,886604 | 0,07655 | 0,345891 |
| Fosl2 | 0,529214 | 0,808875 | 0,612065 | 0,0766 | 0,345995 |
| Fut10 | 19,9877 | 25,1695 | 0,332562 | 0,07665 | 0,346045 |
| Gm5577 | 4,60383 | 6,56472 | 0,511899 | 0,0767 | 0,346045 |
| N4bp2 | 23,2834 | 29,0539 | 0,319427 | 0,0767 | 0,346045 |
| Smarcd3 | 26,3969 | 33,5284 | 0,345016 | 0,07675 | 0,346045 |
| Cope | 101,029 | 80,2304 | -0,332542 | 0,07675 | 0,346045 |
| Tmx3 | 16,2522 | 20,4494 | 0,331419 | 0,0768 | 0,346045 |
| Ylpm1 | 25,9765 | 32,3987 | 0,31873 | 0,0768 | 0,346045 |
| Antxr2 | 13,7672 | 10,8183 | -0,347765 | 0,07685 | 0,346149 |
| Zkscan8 | 7,28618 | 9,37398 | 0,363499 | 0,07695 | 0,346356 |

| | | | | | |
|---|---|---|---|---|---|
| Ptpn12 | 29,0365 | 36,3985 | 0,326007 | 0,07695 | 0,346356 |
| Slc4a7 | 12,2562 | 15,3497 | 0,324693 | 0,07705 | 0,346684 |
| 2210018M11Rik | 19,6469 | 24,6938 | 0,329843 | 0,07715 | 0,347013 |
| Igfbp3 | 20,4345 | 25,9268 | 0,343438 | 0,0772 | 0,347116 |
| Birc3 | 1,63618 | 1,06957 | -0,613302 | 0,07725 | 0,347219 |
| Ccdc23 | 58,9243 | 45,5652 | -0,37093 | 0,07745 | 0,347977 |
| Npbwr1 | 0,425938 | 0,722203 | 0,761762 | 0,0775 | 0,347977 |
| Ndufa9 | 104,427 | 83,5162 | -0,322366 | 0,0775 | 0,347977 |
| Lxn | 17,7968 | 31,163 | 0,80822 | 0,07755 | 0,34808 |
| Ephb3 | 10,98 | 8,59472 | -0,353354 | 0,0776 | 0,348182 |
| Mgea5 | 59,5148 | 74,0965 | 0,316158 | 0,0777 | 0,348509 |
| Ism1 | 17,5398 | 22,2445 | 0,34282 | 0,0779 | 0,349284 |
| BC048507 | 51,1254 | 38,6405 | -0,403925 | 0,07795 | 0,349346 |
| Lca5 | 9,12204 | 11,5448 | 0,339819 | 0,078 | 0,349346 |
| Med14 | 16,2197 | 20,3009 | 0,323797 | 0,078 | 0,349346 |
| Wipf2 | 6,09142 | 7,72059 | 0,341933 | 0,07805 | 0,349346 |
| Ppp1r14a | 26,8899 | 18,9906 | -0,501774 | 0,07805 | 0,349346 |
| Papd7 | 15,9312 | 20,0509 | 0,331808 | 0,07815 | 0,349672 |
| Leprel4 | 10,5346 | 13,7898 | 0,388462 | 0,0782 | 0,349773 |
| Aagab | 26,4776 | 33,299 | 0,330702 | 0,0784 | 0,350423 |
| Aff2 | 3,47211 | 2,57032 | -0,433862 | 0,0784 | 0,350423 |
| Cplx2 | 5,96556 | 4,58494 | -0,379755 | 0,0785 | 0,350748 |
| Tnnt1 | 1,2636 | 2,27182 | 0,846306 | 0,07865 | 0,351296 |
| Enc1 | 24,5013 | 30,5808 | 0,319766 | 0,0789 | 0,35229 |
| Efr3a | 12,9359 | 16,2148 | 0,325926 | 0,07895 | 0,352391 |
| Fxr2 | 64,0052 | 79,6625 | 0,315711 | 0,079 | 0,352491 |
| Gas2 | 11,2764 | 8,56011 | -0,397604 | 0,0791 | 0,352815 |
| Nsmce1 | 44,4282 | 34,7278 | -0,355385 | 0,07935 | 0,353807 |
| 2810428I15Rik | 48,1377 | 36,6572 | -0,393072 | 0,0794 | 0,353907 |
| Plekhm1 | 3,15667 | 4,17143 | 0,402141 | 0,07955 | 0,354329 |
| Cox7a2l | 54,5343 | 42,8023 | -0,349477 | 0,07955 | 0,354329 |
| Ranbp6 | 8,28334 | 10,5629 | 0,35072 | 0,0796 | 0,354429 |
| Phc3 | 3,63476 | 4,61073 | 0,343133 | 0,07965 | 0,354528 |
| Saa3 | 0,290572 | 1,82765 | 2,65302 | 0,07975 | 0,35485 |
| Hif3a | 1,52747 | 2,0708 | 0,439042 | 0,08005 | 0,355938 |
| Tnk2 | 16,1398 | 20,2308 | 0,325934 | 0,08005 | 0,355938 |
| Fbxl3 | 11,5695 | 14,6481 | 0,340388 | 0,0801 | 0,356037 |
| Eri3 | 58,5215 | 46,0333 | -0,346291 | 0,08015 | 0,356136 |
| Chrdl1 | 1,78812 | 2,4907 | 0,478111 | 0,08045 | 0,35732 |
| Galnt12 | 0,701415 | 1,19459 | 0,768178 | 0,0805 | 0,35732 |
| C1d | 24,9364 | 19,8307 | -0,330519 | 0,0805 | 0,35732 |
| Kmt2a | 12,2035 | 15,2102 | 0,317741 | 0,0806 | 0,35764 |
| Pgap1 | 1,14447 | 1,53978 | 0,428045 | 0,08065 | 0,357738 |
| 2810454H06Rik | 0,57317 | 0,966835 | 0,754307 | 0,08075 | 0,357934 |
| Dnm2 | 24,7256 | 32,6144 | 0,399506 | 0,08075 | 0,357934 |
| D1Pas1 | 0,931852 | 1,42948 | 0,617313 | 0,0808 | 0,358032 |
| Bmp2 | 2,99327 | 2,15845 | -0,471726 | 0,08085 | 0,35813 |
| Dhodh | 16,8474 | 13,1592 | -0,356456 | 0,0809 | 0,358228 |
| Pabpc4 | 25,3854 | 20,2051 | -0,329274 | 0,08115 | 0,359211 |
| Mypop | 2,64348 | 1,77708 | -0,572927 | 0,0812 | 0,359308 |

| | | | | | |
|---|---|---|---|---|---|
| Nop10 | 156,99 | 125,277 | -0,325551 | 0,08145 | 0,36029 |
| Ndufv2 | 95,0935 | 76,1753 | -0,320023 | 0,08155 | 0,360608 |
| Dok5 | 0,949087 | 0,499116 | -0,927167 | 0,08175 | 0,36134 |
| Fuca2 | 4,0875 | 5,38446 | 0,397581 | 0,0818 | 0,36134 |
| Hgs | 24,1024 | 19,1779 | -0,329736 | 0,0818 | 0,36134 |
| 2010107G23Rik | 2,8071 | 1,84056 | -0,608934 | 0,08205 | 0,362319 |
| Kctd14 | 0,337489 | 0,648913 | 0,943185 | 0,08215 | 0,362511 |
| Tnrc18 | 24,5858 | 19,6767 | -0,321337 | 0,08215 | 0,362511 |
| Trim61 | 0,768864 | 0,34978 | -1,13628 | 0,08225 | 0,362828 |
| Phlda2 | 1,39501 | 2,71096 | 0,95853 | 0,08235 | 0,363086 |
| Hoxa5 | 47,2106 | 66,047 | 0,484381 | 0,08245 | 0,363086 |
| Tmem261 | 81,4017 | 63,9002 | -0,349239 | 0,08245 | 0,363086 |
| Alg14 | 21,5155 | 15,9337 | -0,433298 | 0,08245 | 0,363086 |
| Cd44 | 1,61107 | 1,12209 | -0,521835 | 0,08245 | 0,363086 |
| Rpl13a | 736,878 | 590,776 | -0,318816 | 0,0826 | 0,363621 |
| Zfp949 | 2,91497 | 3,97055 | 0,445859 | 0,0827 | 0,363812 |
| Atg16l1 | 15,0474 | 18,9602 | 0,33346 | 0,0827 | 0,363812 |
| Arf4 | 177,768 | 143,409 | -0,309863 | 0,08275 | 0,363907 |
| Csrnp1 | 2,59904 | 1,80072 | -0,529407 | 0,08285 | 0,364222 |
| Fnip1 | 4,18314 | 5,38055 | 0,363169 | 0,0829 | 0,364317 |
| Capn6 | 17,5749 | 13,936 | -0,334703 | 0,0831 | 0,365071 |
| Fggy | 2,64343 | 3,83319 | 0,536133 | 0,0833 | 0,365824 |
| Srsf12 | 1,40505 | 2,07486 | 0,562393 | 0,0834 | 0,365888 |
| Mmp15 | 7,76897 | 6,04207 | -0,36268 | 0,0834 | 0,365888 |
| Ccdc181 | 4,5393 | 3,18432 | -0,511483 | 0,0834 | 0,365888 |
| Mxra8 | 9,64002 | 7,30516 | -0,400121 | 0,08345 | 0,365982 |
| Osbpl5 | 2,65845 | 1,91725 | -0,471545 | 0,0835 | 0,366076 |
| Tatdn3 | 4,29992 | 6,23308 | 0,535634 | 0,0836 | 0,366233 |
| Spc24 | 69,5837 | 55,6829 | -0,321517 | 0,0836 | 0,366233 |
| Phlpp2 | 5,13471 | 6,49736 | 0,339571 | 0,08365 | 0,366233 |
| Dnajb11 | 64,3184 | 80,1317 | 0,317142 | 0,08365 | 0,366233 |
| Tet1 | 3,50172 | 4,393 | 0,327144 | 0,0837 | 0,366327 |
| Chst3 | 2,09178 | 2,79596 | 0,418611 | 0,0838 | 0,366639 |
| Hmgb1 | 15,1287 | 11,8286 | -0,355 | 0,084 | 0,367389 |
| Ldlrap1 | 4,80013 | 6,43161 | 0,422108 | 0,08405 | 0,367451 |
| Qsox2 | 12,9722 | 16,5533 | 0,351704 | 0,0841 | 0,367451 |
| Habp4 | 7,60304 | 5,73962 | -0,405622 | 0,0841 | 0,367451 |
| Rab37 | 1,00251 | 0,571655 | -0,810397 | 0,08415 | 0,367544 |
| Snrpe | 263,295 | 210,173 | -0,325101 | 0,0842 | 0,367605 |
| Myo7a | 4,29358 | 5,45939 | 0,346559 | 0,08425 | 0,367605 |
| Cep192 | 17,6248 | 21,9045 | 0,313617 | 0,08425 | 0,367605 |
| Senp7 | 10,6258 | 14,39 | 0,437499 | 0,0843 | 0,367698 |
| Syne2 | 8,69559 | 6,99062 | -0,314864 | 0,08435 | 0,367791 |
| Fam69c | 1,17491 | 0,688721 | -0,770561 | 0,0844 | 0,367884 |
| Paox | 2,30587 | 1,46706 | -0,652387 | 0,08445 | 0,367977 |
| Tubb2a-ps2 | 99,4415 | 78,5283 | -0,340635 | 0,0845 | 0,36807 |
| Crybg3 | 3,7063 | 2,49024 | -0,573695 | 0,0846 | 0,36838 |
| Ntpcr | 8,51496 | 6,0265 | -0,498679 | 0,08465 | 0,368473 |
| Aatf | 33,7601 | 26,7707 | -0,334665 | 0,0847 | 0,368566 |
| Bbs4 | 8,48918 | 10,9971 | 0,373425 | 0,08475 | 0,368658 |

| | | | | | |
|---|---|---|---|---|---|
| Mfsd4 | 1,67935 | 2,45076 | 0,545321 | 0,08485 | 0,368968 |
| Rps18 | 362,172 | 291,166 | -0,314836 | 0,08505 | 0,369712 |
| 2310009B15Rik | 8,73786 | 5,50517 | -0,666494 | 0,0851 | 0,369804 |
| B3galnt1 | 7,30483 | 5,44405 | -0,42417 | 0,08525 | 0,370331 |
| B3galt1 | 0,948546 | 0,633296 | -0,582838 | 0,08565 | 0,371943 |
| Rep15 | 2,17334 | 3,53984 | 0,703768 | 0,0859 | 0,372902 |
| Plekhg4 | 1,26735 | 1,81533 | 0,518416 | 0,086 | 0,37321 |
| Nutf2 | 19,6201 | 15,335 | -0,35551 | 0,08605 | 0,373265 |
| Itpripl1 | 3,59854 | 4,80943 | 0,418457 | 0,0861 | 0,373265 |
| G0s2 | 3,3551 | 1,90977 | -0,812955 | 0,0861 | 0,373265 |
| Kctd11 | 4,93021 | 6,55419 | 0,410769 | 0,08625 | 0,373536 |
| Ccndbp1 | 15,6996 | 20,2784 | 0,369218 | 0,08625 | 0,373536 |
| Fam43a | 6,62906 | 5,04596 | -0,393675 | 0,08625 | 0,373536 |
| E130309D02Rik | 30,1321 | 24,0143 | -0,327406 | 0,08645 | 0,37424 |
| Sord | 4,74034 | 6,41467 | 0,436385 | 0,0865 | 0,37424 |
| Nphp3 | 4,27822 | 5,5371 | 0,372119 | 0,0865 | 0,37424 |
| Ggh | 5,20723 | 3,55369 | -0,551199 | 0,0866 | 0,374546 |
| Snord95 | 126,221 | 234,273 | 0,892239 | 0,08675 | 0,374942 |
| Bnip2 | 21,252 | 26,2903 | 0,306926 | 0,08675 | 0,374942 |
| Tbc1d16 | 21,9077 | 17,6611 | -0,310862 | 0,0868 | 0,375032 |
| Fbxo48 | 2,39463 | 3,43943 | 0,522368 | 0,0869 | 0,375338 |
| Zc3hc1 | 49,0299 | 39,3806 | -0,316175 | 0,08715 | 0,376291 |
| Wdr44 | 0,881076 | 1,30913 | 0,571269 | 0,0873 | 0,376305 |
| Bcl11a | 3,36097 | 4,93928 | 0,555423 | 0,0873 | 0,376305 |
| Cyp26b1 | 1,03149 | 1,48094 | 0,521778 | 0,0873 | 0,376305 |
| Maml2 | 7,46851 | 9,45986 | 0,340998 | 0,0873 | 0,376305 |
| Sdc2 | 12,6384 | 15,947 | 0,335464 | 0,0873 | 0,376305 |
| Polr3g | 12,4295 | 9,7637 | -0,348274 | 0,0875 | 0,376914 |
| Rbpms | 18,7924 | 13,7996 | -0,445525 | 0,0875 | 0,376914 |
| Nfe2l1 | 47,4249 | 58,6235 | 0,305833 | 0,08755 | 0,377003 |
| Plscr3 | 18,8086 | 14,9019 | -0,335896 | 0,08765 | 0,37718 |
| Nicn1 | 13,3559 | 10,3269 | -0,371067 | 0,08765 | 0,37718 |
| Pecam1 | 1,53953 | 1,00742 | -0,61183 | 0,0877 | 0,377269 |
| Mir1934 | 0 | 72,1509 | inf | 0,0879 | 0,377711 |
| Pdzd8 | 14,3504 | 17,8217 | 0,312541 | 0,0879 | 0,377711 |
| Zfp955b | 4,70029 | 6,1924 | 0,39775 | 0,08795 | 0,377711 |
| Syngr2 | 15,2552 | 19,725 | 0,370722 | 0,08795 | 0,377711 |
| Sun1 | 22,6926 | 28,2619 | 0,316638 | 0,08795 | 0,377711 |
| Dnajc13 | 18,5785 | 22,9825 | 0,306903 | 0,08815 | 0,378443 |
| Numbl | 6,8994 | 5,23361 | -0,398664 | 0,0883 | 0,37896 |
| Ifitm3 | 8,20878 | 12,0347 | 0,551962 | 0,08845 | 0,379477 |
| Zfp90 | 11,4966 | 14,6707 | 0,351728 | 0,0885 | 0,379525 |
| Atp11a | 11,136 | 13,8262 | 0,312172 | 0,08855 | 0,379525 |
| Rpp14 | 24,2966 | 19,3145 | -0,331076 | 0,08855 | 0,379525 |
| Pdzrn3 | 25,6696 | 31,8979 | 0,313402 | 0,0886 | 0,379613 |
| B4galt6 | 9,89716 | 7,88592 | -0,327736 | 0,08865 | 0,3797 |
| Casp9 | 8,52121 | 10,8143 | 0,343814 | 0,0887 | 0,379788 |
| Slc43a2 | 1,00544 | 1,40073 | 0,478347 | 0,08885 | 0,379923 |
| Hmgn1 | 385,679 | 311,769 | -0,306923 | 0,08885 | 0,379923 |
| Klhl24 | 8,64466 | 6,86182 | -0,333219 | 0,08885 | 0,379923 |

| | | | | | |
|---|---|---|---|---|---|
| Radil | 1,71234 | 1,10702 | -0,629294 | 0,08885 | 0,379923 |
| Glrx3 | 39,3768 | 31,4145 | -0,325916 | 0,0889 | 0,38001 |
| Hspb11 | 30,1936 | 21,9701 | -0,458703 | 0,08895 | 0,380097 |
| Eif2b2 | 63,6001 | 51,2044 | -0,312761 | 0,08915 | 0,380825 |
| Hectd2 | 1,31861 | 1,85002 | 0,488517 | 0,08925 | 0,381125 |
| Rundc3b | 1,56648 | 1,06373 | -0,558394 | 0,0894 | 0,381638 |
| Mtus1 | 1,71115 | 2,37258 | 0,471485 | 0,08945 | 0,381725 |
| Art1 | 0,59034 | 1,15231 | 0,964906 | 0,0895 | 0,381811 |
| Ccdc124 | 100,773 | 80,6631 | -0,321128 | 0,08955 | 0,381897 |
| Mapk13 | 0,738463 | 0,315676 | -1,22608 | 0,0897 | 0,38241 |
| Eif2s3y | 15,0445 | 19,2452 | 0,355268 | 0,08975 | 0,382496 |
| Zbtb9 | 11,957 | 15,1922 | 0,345474 | 0,08985 | 0,382795 |
| Dpy19l3 | 6,14044 | 7,71278 | 0,328908 | 0,08995 | 0,38284 |
| Qser1 | 18,0557 | 22,3377 | 0,307028 | 0,08995 | 0,38284 |
| Emb | 18,1413 | 14,3646 | -0,33676 | 0,08995 | 0,38284 |
| Tlcd2 | 4,34215 | 2,9544 | -0,555546 | 0,09 | 0,382925 |
| Atp6v0e2 | 9,27959 | 6,95714 | -0,415567 | 0,09005 | 0,383011 |
| Diap1 | 10,5424 | 13,1909 | 0,323346 | 0,0901 | 0,383055 |
| Tm7sf3 | 10,8433 | 13,6968 | 0,337034 | 0,09015 | 0,383055 |
| Sfxn2 | 11,3935 | 8,95 | -0,348253 | 0,09015 | 0,383055 |
| Atg4d | 10,5251 | 15,1135 | 0,521999 | 0,09025 | 0,383353 |
| Kctd7 | 0,919974 | 1,35336 | 0,556883 | 0,0903 | 0,383439 |
| Pard6g | 34,0918 | 42,2636 | 0,309989 | 0,09035 | 0,383524 |
| Lrrn2 | 0,895418 | 0,554145 | -0,692298 | 0,09045 | 0,383822 |
| H2-Ke2 | 151,694 | 119,688 | -0,341888 | 0,0906 | 0,384289 |
| Lmtk3 | 5,38844 | 6,90373 | 0,357509 | 0,09065 | 0,384289 |
| Atox1 | 165,018 | 129,99 | -0,344219 | 0,09065 | 0,384289 |
| Ece2 | 31,9753 | 24,4029 | -0,389908 | 0,0907 | 0,384374 |
| Sipa1 | 7,80257 | 6,08316 | -0,35913 | 0,09075 | 0,384459 |
| Pigx | 57,5731 | 45,7166 | -0,332677 | 0,0908 | 0,384544 |
| St6gal1 | 11,1369 | 13,9926 | 0,329306 | 0,09105 | 0,385475 |
| Slc7a5 | 23,3264 | 28,9902 | 0,313603 | 0,09115 | 0,385771 |
| Clcn3 | 8,41283 | 10,5311 | 0,323988 | 0,0912 | 0,385855 |
| Irf2bp2 | 8,54252 | 10,7627 | 0,333305 | 0,09125 | 0,38594 |
| Dnajc3 | 21,603 | 26,8124 | 0,311669 | 0,0913 | 0,385981 |
| 1110004F10Rik | 97,0717 | 78,6338 | -0,3039 | 0,09135 | 0,385981 |
| A930006K02Rik | 1,595 | 0,697874 | -1,19252 | 0,09135 | 0,385981 |
| Dok2 | 1,84016 | 2,77977 | 0,595135 | 0,09145 | 0,386149 |
| Nfu1 | 23,4106 | 17,6561 | -0,406996 | 0,09145 | 0,386149 |
| Spock3 | 1,964 | 1,34313 | -0,548189 | 0,09155 | 0,386444 |
| Prdx6 | 184,639 | 149,364 | -0,305877 | 0,0916 | 0,386485 |
| Cacna1d | 0,368097 | 1,00806 | 1,45343 | 0,09165 | 0,386485 |
| Rgag4 | 5,29736 | 7,92737 | 0,581568 | 0,09165 | 0,386485 |
| Hoxb5 | 8,60799 | 11,299 | 0,39245 | 0,09175 | 0,386522 |
| Aurkaip1 | 64,84 | 51,9381 | -0,320091 | 0,09175 | 0,386522 |
| Ap1s3 | 0,631738 | 1,01277 | 0,680902 | 0,0918 | 0,386522 |
| Ghrl | 2,16517 | 0,876481 | -1,30469 | 0,0918 | 0,386522 |
| Rbm27 | 16,8424 | 20,818 | 0,305737 | 0,09185 | 0,386522 |
| Pebp1 | 108,825 | 87,9549 | -0,307172 | 0,09185 | 0,386522 |
| Emc1 | 8,2441 | 10,3214 | 0,324207 | 0,0919 | 0,386522 |

| | | | | | |
|---|---|---|---|---|---|
| Dmtf1 | 35,9616 | 44,8294 | 0,317986 | 0,0919 | 0,386522 |
| Stxbp2 | 7,41972 | 9,57298 | 0,367604 | 0,0921 | 0,38711 |
| Cd34 | 3,93029 | 2,78916 | -0,494804 | 0,0921 | 0,38711 |
| Tmcc3 | 1,28655 | 0,905861 | -0,506144 | 0,0922 | 0,387403 |
| Tomm5 | 46,7005 | 35,7434 | -0,385762 | 0,0925 | 0,388448 |
| Ptprg | 11,4225 | 14,1458 | 0,308495 | 0,09255 | 0,388448 |
| Mphosph8 | 41,4974 | 52,3263 | 0,334516 | 0,0926 | 0,388448 |
| Slc16a1 | 38,2073 | 47,2327 | 0,305936 | 0,0926 | 0,388448 |
| Rhobtb2 | 6,17896 | 4,83623 | -0,353481 | 0,0926 | 0,388448 |
| Gtf3c6 | 95,156 | 76,5517 | -0,31386 | 0,09265 | 0,38853 |
| Tor4a | 0,45185 | 0,748395 | 0,727956 | 0,0929 | 0,389451 |
| Sssca1 | 89,5909 | 71,2389 | -0,330686 | 0,09295 | 0,389534 |
| Arhgap20 | 0,759966 | 0,486738 | -0,642789 | 0,09305 | 0,389825 |
| Zfp385b | 1,02072 | 1,52822 | 0,582261 | 0,09315 | 0,390117 |
| Acot7 | 35,5822 | 44,531 | 0,323655 | 0,0932 | 0,390199 |
| l7Rn6 | 52,0008 | 40,6528 | -0,355181 | 0,09335 | 0,390699 |
| Eva1b | 19,8243 | 15,2956 | -0,37415 | 0,0936 | 0,391618 |
| Clcn1 | 0,518863 | 0,829348 | 0,676623 | 0,0937 | 0,391909 |
| Zic5 | 2,79768 | 3,91069 | 0,483193 | 0,09385 | 0,39228 |
| Ajap1 | 3,57349 | 4,77884 | 0,419329 | 0,09385 | 0,39228 |
| Thbs3 | 15,4923 | 12,2951 | -0,333462 | 0,09415 | 0,393278 |
| Tmem9b | 9,6048 | 7,18301 | -0,419167 | 0,09415 | 0,393278 |
| Taf6l | 16,011 | 20,9202 | 0,385836 | 0,0944 | 0,394194 |
| Tjap1 | 28,751 | 35,6775 | 0,311403 | 0,09455 | 0,394644 |
| Cyb5rl | 4,60044 | 6,39993 | 0,476285 | 0,0946 | 0,394644 |
| Sqstm1 | 36,2835 | 44,9094 | 0,307706 | 0,0946 | 0,394644 |
| C330027C09Rik | 26,9781 | 33,5327 | 0,313778 | 0,09465 | 0,394724 |
| Tbk1 | 14,2536 | 17,8858 | 0,327482 | 0,0948 | 0,395221 |
| Spag7 | 104,572 | 84,6856 | -0,304304 | 0,09485 | 0,395301 |
| Ppp1r11 | 39,707 | 31,7308 | -0,323508 | 0,09505 | 0,396006 |
| Pou3f3os | 6,92468 | 0,248896 | -4,79813 | 0,09525 | 0,396453 |
| Hand2 | 2,67908 | 0,041913 | -5,9982 | 0,09525 | 0,396453 |
| Sox10 | 5,62716 | 0,033928 | -7,37379 | 0,09525 | 0,396453 |
| Tmem181a | 2,33299 | 3,19923 | 0,455545 | 0,0954 | 0,396948 |
| Cebpa | 1,30933 | 0,843458 | -0,634443 | 0,09545 | 0,397027 |
| Capn15 | 12,9182 | 16,2244 | 0,32876 | 0,09555 | 0,397186 |
| Por | 36,1198 | 44,5793 | 0,303582 | 0,09555 | 0,397186 |
| Tmtc4 | 8,28905 | 6,45765 | -0,360198 | 0,0956 | 0,397265 |
| Uvssa | 2,56437 | 3,30604 | 0,366502 | 0,09565 | 0,397344 |
| Cnn3 | 202,98 | 164,733 | -0,301213 | 0,0958 | 0,397838 |
| Mgat2 | 22,693 | 28,2507 | 0,316038 | 0,09585 | 0,397917 |
| 2810468N07Rik | 3,3899 | 2,22189 | -0,609453 | 0,0959 | 0,397996 |
| Pcna | 166,831 | 135,281 | -0,302432 | 0,09595 | 0,398075 |
| Col11a1 | 8,29388 | 6,63043 | -0,322944 | 0,09615 | 0,398775 |
| 0610010F05Rik | 9,09536 | 11,426 | 0,329118 | 0,0964 | 0,399683 |
| Armcx3 | 1,49632 | 1,01644 | -0,557889 | 0,09675 | 0,401005 |
| Wnk1 | 38,351 | 47,2183 | 0,300082 | 0,09695 | 0,401522 |
| Rere | 24,3951 | 29,9947 | 0,298114 | 0,09695 | 0,401522 |
| Rps5 | 1845,52 | 1483,48 | -0,315045 | 0,097 | 0,401522 |
| Gng11 | 10,299 | 7,23742 | -0,508951 | 0,097 | 0,401522 |

| | | | | | |
|---|---|---|---|---|---|
| Dennd2a | 29,5118 | 36,3084 | 0,299014 | 0,0971 | 0,401806 |
| Nup50 | 41,0197 | 33,2779 | -0,30175 | 0,09715 | 0,401883 |
| Trim44 | 46,6769 | 57,3245 | 0,296443 | 0,0972 | 0,401961 |
| Tmem242 | 99,8753 | 80,5277 | -0,310644 | 0,0973 | 0,402244 |
| Pgm2l1 | 5,07591 | 4,04778 | -0,326536 | 0,0974 | 0,402398 |
| Dusp10 | 0,86522 | 0,509403 | -0,764259 | 0,0974 | 0,402398 |
| A2m | 1,93075 | 1,39023 | -0,473835 | 0,0976 | 0,403095 |
| Ahi1 | 8,71646 | 10,9068 | 0,323409 | 0,09765 | 0,403172 |
| Noxo1 | 0,865067 | 1,37862 | 0,672337 | 0,09775 | 0,403455 |
| Letm2 | 1,76821 | 2,58119 | 0,545749 | 0,0979 | 0,403501 |
| Mprip | 37,7406 | 46,3482 | 0,296397 | 0,0979 | 0,403501 |
| Scand1 | 39,7404 | 30,604 | -0,376888 | 0,0979 | 0,403501 |
| Ripply3 | 2,45873 | 3,60202 | 0,550892 | 0,09795 | 0,403501 |
| Bbs12 | 8,50926 | 10,961 | 0,365273 | 0,09795 | 0,403501 |
| Epc2 | 14,0257 | 17,4324 | 0,313694 | 0,09795 | 0,403501 |
| Osbpl8 | 3,97143 | 5,03005 | 0,340915 | 0,09805 | 0,403783 |
| Zfp953 | 0,750296 | 1,10299 | 0,555887 | 0,0981 | 0,40386 |
| Bhlhe41 | 0,380787 | 0,587993 | 0,626817 | 0,09815 | 0,403936 |
| Rasl10b | 2,10595 | 1,48273 | -0,506212 | 0,0982 | 0,403959 |
| Cdk2 | 63,5554 | 78,1149 | 0,297583 | 0,09825 | 0,403959 |
| Eef1d | 208,196 | 169,432 | -0,297235 | 0,09825 | 0,403959 |
| Tmsb10 | 354,513 | 436,688 | 0,300762 | 0,09845 | 0,404651 |
| Carf | 1,44791 | 2,57559 | 0,830933 | 0,09855 | 0,404933 |
| Wbp5 | 179,57 | 145,818 | -0,300371 | 0,09865 | 0,405084 |
| Plekhj1 | 55,4606 | 44,4631 | -0,318852 | 0,09865 | 0,405084 |
| Mir677 | 41,3978 | 159,903 | 1,94957 | 0,09885 | 0,405775 |
| Wdr5b | 4,64042 | 3,29613 | -0,493481 | 0,09895 | 0,406056 |
| Lenep | 5,03863 | 3,03238 | -0,732579 | 0,09905 | 0,406336 |
| Mir703 | 3018,67 | 2164,71 | -0,479737 | 0,09915 | 0,406616 |
| Map3k5 | 0,53363 | 0,796946 | 0,578641 | 0,0993 | 0,406971 |
| Ssu72 | 56,1626 | 45,1506 | -0,314864 | 0,0993 | 0,406971 |
| Dctn6 | 53,4076 | 42,4484 | -0,331334 | 0,09945 | 0,407403 |
| Ndufa4 | 581,811 | 473,234 | -0,297998 | 0,0995 | 0,407403 |
| Sfswap | 32,5256 | 40,0241 | 0,299292 | 0,0996 | 0,407403 |
| Lta4h | 69,1412 | 56,1887 | -0,299266 | 0,0996 | 0,407403 |
| Ppp2r3d | 4,30452 | 5,75406 | 0,418728 | 0,09965 | 0,407403 |
| Peg13 | 4,46452 | 3,11743 | -0,518147 | 0,09965 | 0,407403 |
| Cnih2 | 7,06855 | 3,75772 | -0,911558 | 0,09965 | 0,407403 |
| Tmem82 | 0,406847 | 0,87853 | 1,11061 | 0,0997 | 0,407403 |
| Marveld2 | 0,344304 | 0,611687 | 0,829111 | 0,0997 | 0,407403 |
| Psme2 | 8,97486 | 12,5209 | 0,480378 | 0,09975 | 0,407403 |
| Pnmal2 | 0,958635 | 1,38595 | 0,531821 | 0,0998 | 0,407403 |
| Cdyl | 9,19085 | 11,5747 | 0,332708 | 0,0998 | 0,407403 |
| Mir717 | 0 | 31,9666 | inf | 0,09985 | 0,407403 |
| Cd38 | 1,30035 | 1,89034 | 0,53974 | 0,09985 | 0,407403 |
| Gemin7 | 78,2808 | 62,4907 | -0,325017 | 0,1 | 0,407886 |
| Psme1 | 45,3022 | 56,7582 | 0,325248 | 0,1002 | 0,408442 |
| Ids | 4,17322 | 3,21023 | -0,378484 | 0,1002 | 0,408442 |
| Serpini1 | 1,01064 | 0,646217 | -0,645175 | 0,1003 | 0,408663 |
| Cops5 | 105,554 | 85,7389 | -0,299955 | 0,10035 | 0,408663 |

| | | | | | |
|---|---|---|---|---|---|
| Enox2 | 4,05863 | 3,07208 | -0,401776 | 0,10035 | 0,408663 |
| Stk3 | 18,3473 | 22,8964 | 0,31955 | 0,10045 | 0,408941 |
| Mdga1 | 1,86469 | 2,43877 | 0,387217 | 0,1009 | 0,410643 |
| Rpl6 | 75,5228 | 60,9047 | -0,310357 | 0,10105 | 0,411123 |
| Znrf3 | 4,36413 | 5,48733 | 0,330412 | 0,10135 | 0,412212 |
| Hoxb6 | 25,0567 | 31,6091 | 0,335148 | 0,1019 | 0,414318 |
| Xrcc6bp1 | 11,6093 | 8,51797 | -0,446693 | 0,10205 | 0,414796 |
| Sema4c | 29,5822 | 36,3946 | 0,298994 | 0,1021 | 0,414868 |
| Abi3bp | 1,02907 | 0,686258 | -0,584523 | 0,1023 | 0,415549 |
| Cdk12 | 17,5855 | 21,6786 | 0,301884 | 0,1025 | 0,416098 |
| Nhs | 3,60638 | 2,75863 | -0,3866 | 0,1025 | 0,416098 |
| Pcyt1a | 16,6501 | 13,4865 | -0,304009 | 0,10265 | 0,416575 |
| Rwdd2b | 7,35853 | 5,54063 | -0,409368 | 0,1027 | 0,416646 |
| Hdac8 | 8,87202 | 6,6608 | -0,413567 | 0,10275 | 0,416717 |
| Actr5 | 10,1298 | 7,88216 | -0,361946 | 0,1028 | 0,416788 |
| Mkl2 | 6,36743 | 8,20427 | 0,365664 | 0,10285 | 0,416859 |
| Pak4 | 36,7436 | 29,8631 | -0,29913 | 0,10295 | 0,417133 |
| Myo6 | 1,71005 | 1,27979 | -0,418127 | 0,1031 | 0,417609 |
| 9830147E19Rik | 2,20491 | 2,95107 | 0,420521 | 0,10315 | 0,417679 |
| Cage1 | 0,413987 | 0,714928 | 0,788215 | 0,1032 | 0,41775 |
| Hist1h1b | 2,29636 | 3,80091 | 0,726994 | 0,1033 | 0,417891 |
| Ago2 | 21,0614 | 25,9181 | 0,299359 | 0,1033 | 0,417891 |
| Abca2 | 3,43737 | 4,34441 | 0,337856 | 0,1035 | 0,418436 |
| Tcf7l1 | 24,5733 | 19,8814 | -0,305669 | 0,1035 | 0,418436 |
| Zfp821 | 29,1222 | 36,1162 | 0,310531 | 0,10355 | 0,418506 |
| Psmg3 | 51,7184 | 40,9181 | -0,337938 | 0,1039 | 0,419789 |
| Fanca | 13,2711 | 17,1157 | 0,367026 | 0,1041 | 0,420332 |
| Tmem120a | 15,8473 | 12,111 | -0,387924 | 0,1041 | 0,420332 |
| Aplp1 | 3,5524 | 4,81264 | 0,438037 | 0,1042 | 0,420408 |
| Wwc1 | 4,60419 | 3,47745 | -0,404919 | 0,1042 | 0,420408 |
| Agps | 26,4022 | 21,5277 | -0,294459 | 0,10425 | 0,420408 |
| Trim27 | 38,8095 | 31,5667 | -0,298004 | 0,10425 | 0,420408 |
| Fam101b | 14,8247 | 18,4204 | 0,313302 | 0,10435 | 0,420547 |
| 2810032G03Rik | 1,89424 | 1,10863 | -0,772846 | 0,10435 | 0,420547 |
| Lat | 0,613513 | 0,223626 | -1,45601 | 0,10445 | 0,420818 |
| Pcbp3 | 14,4368 | 11,3528 | -0,346705 | 0,1045 | 0,420824 |
| Mir23a | 103,582 | 0 | -inf | 0,10455 | 0,420824 |
| Kif1a | 2,21102 | 1,69577 | -0,382773 | 0,10455 | 0,420824 |
| Ceacam1 | 0,522312 | 1,07588 | 1,04253 | 0,1047 | 0,421032 |
| Micall1 | 5,79661 | 7,23332 | 0,319449 | 0,1047 | 0,421032 |
| Kat2b | 0,920683 | 0,604989 | -0,605795 | 0,1047 | 0,421032 |
| Dgat2 | 7,51687 | 9,71447 | 0,370004 | 0,10475 | 0,421038 |
| Coq7 | 49,1968 | 38,9728 | -0,336097 | 0,1048 | 0,421038 |
| Atg10 | 3,39365 | 2,31121 | -0,554191 | 0,1048 | 0,421038 |
| BC039771 | 0,770455 | 1,2543 | 0,703102 | 0,10505 | 0,42191 |
| Focad | 9,31722 | 11,554 | 0,310417 | 0,10515 | 0,42218 |
| Sf3b6 | 133,531 | 108,781 | -0,295752 | 0,1055 | 0,423452 |
| Fbxo3 | 35,546 | 45,3742 | 0,352182 | 0,10555 | 0,423521 |
| Commd9 | 20,2658 | 15,6175 | -0,375887 | 0,10565 | 0,423789 |
| Cldn26 | 0,0607168 | 0,792691 | 3,70659 | 0,10575 | 0,424058 |

| | | | | | |
|---|---|---|---|---|---|
| Adpgk | 33,8689 | 41,6457 | 0,298207 | 0,1059 | 0,424527 |
| Wwc2 | 11,3048 | 13,9787 | 0,306303 | 0,10595 | 0,424594 |
| Sgsm1 | 4,76554 | 6,10476 | 0,357294 | 0,106 | 0,424662 |
| Mbnl3 | 1,4989 | 1,9628 | 0,389007 | 0,10605 | 0,42473 |
| Nek8 | 5,14299 | 6,69234 | 0,379903 | 0,1061 | 0,424797 |
| Fam193b | 26,1216 | 31,9849 | 0,29215 | 0,10625 | 0,425265 |
| Rasa1 | 18,6967 | 23,0501 | 0,301992 | 0,1064 | 0,425535 |
| 3010026O09Rik | 13,7266 | 10,2927 | -0,415342 | 0,1064 | 0,425535 |
| Cdc27 | 34,8921 | 42,6944 | 0,291146 | 0,10645 | 0,425535 |
| Myd88 | 1,95073 | 1,27485 | -0,61368 | 0,10645 | 0,425535 |
| 2700094K13Rik | 408,375 | 334,572 | -0,287579 | 0,10655 | 0,425802 |
| Ppic | 76,5598 | 62,1827 | -0,300075 | 0,1067 | 0,426268 |
| Zfp780b | 2,39255 | 3,12836 | 0,386855 | 0,10675 | 0,426335 |
| Atg3 | 18,2539 | 14,4939 | -0,33276 | 0,1068 | 0,426402 |
| Zfp428 | 41,1605 | 32,8546 | -0,325166 | 0,1069 | 0,426669 |
| Ndufa13 | 73,5722 | 59,4583 | -0,307282 | 0,10695 | 0,426735 |
| Ccdc42 | 0,856859 | 1,50568 | 0,813283 | 0,1071 | 0,42687 |
| Jmy | 8,6445 | 10,616 | 0,296382 | 0,1071 | 0,42687 |
| Sptlc2 | 14,3869 | 17,8691 | 0,312719 | 0,10715 | 0,42687 |
| Myl12b | 85,228 | 105,109 | 0,302485 | 0,10715 | 0,42687 |
| Emc2 | 42,5201 | 33,9204 | -0,325993 | 0,10715 | 0,42687 |
| Sirt2 | 27,4493 | 22,0107 | -0,318563 | 0,1072 | 0,426936 |
| Six4 | 8,14461 | 10,1035 | 0,310938 | 0,1074 | 0,427467 |
| 2810408M09Rik | 16,2589 | 12,7715 | -0,348303 | 0,1074 | 0,427467 |
| Pitpnm2 | 4,73053 | 3,73186 | -0,342106 | 0,1077 | 0,428462 |
| Slc44a1 | 8,51196 | 11,035 | 0,374527 | 0,10775 | 0,428462 |
| Decr2 | 9,43685 | 7,29139 | -0,372111 | 0,10775 | 0,428462 |
| BC020402 | 0,882845 | 1,38722 | 0,651961 | 0,1078 | 0,428528 |
| 2700062C07Rik | 21,0238 | 16,1393 | -0,381441 | 0,10805 | 0,429388 |
| Cpeb3 | 0,983245 | 0,687199 | -0,516823 | 0,10825 | 0,43005 |
| Usp46 | 22,8688 | 18,5055 | -0,305426 | 0,1083 | 0,430115 |
| Zfp532 | 20,7676 | 26,0476 | 0,326821 | 0,10845 | 0,430444 |
| Rab4a | 8,21236 | 6,0723 | -0,435553 | 0,10845 | 0,430444 |
| Fam214a | 2,06057 | 1,50507 | -0,45321 | 0,1085 | 0,43051 |
| Cend1 | 1,65555 | 1,01933 | -0,69969 | 0,1086 | 0,430773 |
| Abhd8 | 23,5391 | 18,8316 | -0,321905 | 0,1088 | 0,431167 |
| Epb4.1l1 | 3,76989 | 2,94014 | -0,358637 | 0,1088 | 0,431167 |
| Tal2 | 0,998951 | 0,0856102 | -3,54456 | 0,1088 | 0,431167 |
| Elk4 | 4,73789 | 6,11727 | 0,368644 | 0,10895 | 0,431624 |
| Pdp2 | 6,0403 | 10,5699 | 0,807265 | 0,10905 | 0,431624 |
| Ascc3 | 5,97576 | 7,41704 | 0,311722 | 0,10905 | 0,431624 |
| Fundc2 | 17,1258 | 13,7872 | -0,312838 | 0,10905 | 0,431624 |
| Gm7244 | 4,3102 | 6,54434 | 0,602492 | 0,1091 | 0,431689 |
| Tmem245 | 6,20246 | 7,70805 | 0,313525 | 0,10915 | 0,431753 |
| Fth1 | 324,462 | 265,769 | -0,287879 | 0,1092 | 0,431818 |
| Srpx | 0,94528 | 0,56731 | -0,736604 | 0,10925 | 0,431883 |
| C030037D09Rik | 1,99771 | 3,15978 | 0,661479 | 0,10935 | 0,432012 |
| Col22a1 | 2,20079 | 2,86218 | 0,379097 | 0,10935 | 0,432012 |
| Pls3 | 31,5667 | 38,5772 | 0,289345 | 0,10955 | 0,432669 |
| Reep2 | 3,87823 | 5,27798 | 0,444588 | 0,10975 | 0,433058 |

| | | | | | |
|---|---|---|---|---|---|
| Flywch2 | 18,122 | 24,2861 | 0,422385 | 0,10975 | 0,433058 |
| Rnf6 | 10,6637 | 13,3404 | 0,323094 | 0,10975 | 0,433058 |
| Edf1 | 152,235 | 123,491 | -0,301896 | 0,10985 | 0,43332 |
| Ripply2 | 54,7972 | 41,7925 | -0,390858 | 0,1099 | 0,433384 |
| Sf3b5 | 154,971 | 126,015 | -0,298402 | 0,11 | 0,433645 |
| Ctf1 | 3,23047 | 4,61578 | 0,514831 | 0,11005 | 0,433709 |
| BC005537 | 61,9285 | 75,6888 | 0,289476 | 0,11015 | 0,433969 |
| Med22 | 47,5037 | 37,7237 | -0,332569 | 0,11025 | 0,43423 |
| Atxn2 | 22,2156 | 27,1648 | 0,290166 | 0,1103 | 0,434294 |
| Vps26a | 24,9075 | 20,1979 | -0,302378 | 0,1105 | 0,434948 |
| Smim22 | 0,639086 | 0 | -inf | 0,11055 | 0,435011 |
| Grhl1 | 0,355905 | 0,602658 | 0,75985 | 0,1107 | 0,435334 |
| Rsbn1l | 7,45871 | 9,39719 | 0,333304 | 0,1107 | 0,435334 |
| Capn1 | 2,00378 | 2,77368 | 0,469079 | 0,11075 | 0,43539 |
| Frmd7 | 1,00527 | 1,4157 | 0,493922 | 0,1108 | 0,43539 |
| Gls2 | 1,38914 | 2,66504 | 0,939963 | 0,11085 | 0,43539 |
| Aldh3a2 | 8,62096 | 10,8667 | 0,333999 | 0,11085 | 0,43539 |
| Tmem98 | 63,4074 | 51,5031 | -0,299993 | 0,1109 | 0,435453 |
| 2700049A03Rik | 9,53846 | 11,7863 | 0,305283 | 0,11095 | 0,435516 |
| Fgf18 | 31,2509 | 25,0368 | -0,319851 | 0,11105 | 0,435775 |
| Acacb | 0,439137 | 0,62986 | 0,52036 | 0,11115 | 0,435901 |
| Lamtor4 | 19,4981 | 14,7428 | -0,403322 | 0,11115 | 0,435901 |
| Zfp41 | 41,1262 | 33,7018 | -0,287229 | 0,1116 | 0,437452 |
| Trappc2 | 2,28952 | 3,80419 | 0,732543 | 0,11165 | 0,437452 |
| Fmr1 | 25,7181 | 31,4718 | 0,291275 | 0,1117 | 0,437452 |
| Ap1m1 | 64,5131 | 52,682 | -0,292282 | 0,11175 | 0,437452 |
| Esd | 48,7635 | 39,7807 | -0,293732 | 0,11175 | 0,437452 |
| Rasl11a | 15,6075 | 11,875 | -0,394303 | 0,11175 | 0,437452 |
| Matr3 | 112,336 | 138,485 | 0,301907 | 0,112 | 0,438225 |
| Ptov1 | 163,422 | 134,257 | -0,283606 | 0,11205 | 0,438225 |
| Lrrc34 | 1,32551 | 0,80409 | -0,721114 | 0,11205 | 0,438225 |
| Dnajc16 | 6,43051 | 8,11371 | 0,335429 | 0,11215 | 0,438349 |
| Inip | 23,0712 | 18,7366 | -0,300235 | 0,11215 | 0,438349 |
| Tekt2 | 0,350095 | 0,733628 | 1,0673 | 0,11235 | 0,438863 |
| Polr2j | 178,472 | 144,795 | -0,301692 | 0,11235 | 0,438863 |
| Fuom | 1,6135 | 1,07441 | -0,586646 | 0,11245 | 0,43912 |
| Ankrd9 | 1,63429 | 1,10278 | -0,567519 | 0,1125 | 0,439181 |
| Hipk2 | 18,5243 | 22,8359 | 0,301886 | 0,11265 | 0,439499 |
| Shd | 9,60234 | 7,24565 | -0,406273 | 0,11265 | 0,439499 |
| Cox4i2 | 7,40549 | 14,5594 | 0,975284 | 0,11285 | 0,440146 |
| Nhsl2 | 4,06963 | 3,11298 | -0,386603 | 0,1129 | 0,440207 |
| Rab5b | 9,3132 | 11,6717 | 0,325667 | 0,11295 | 0,440268 |
| Arfgef2 | 4,70531 | 5,83989 | 0,31165 | 0,113 | 0,440329 |
| Ddah2 | 173,467 | 142,421 | -0,284499 | 0,1131 | 0,440585 |
| Tsc22d3 | 31,2115 | 25,2993 | -0,302978 | 0,1132 | 0,440841 |
| Hoxd13 | 62,9583 | 76,7073 | 0,284967 | 0,11325 | 0,440902 |
| Hsd17b4 | 38,2483 | 31,1657 | -0,295436 | 0,1133 | 0,440962 |
| Fancm | 10,5915 | 12,9521 | 0,29028 | 0,1134 | 0,441218 |
| Acer3 | 4,99224 | 6,36432 | 0,35032 | 0,11355 | 0,441667 |
| Zfp101 | 13,9123 | 17,3923 | 0,322088 | 0,1136 | 0,441715 |

| | | | | | |
|---|---|---|---|---|---|
| Ing2 | 10,8232 | 13,5598 | 0,325208 | 0,11365 | 0,441715 |
| Hist3h2a | 7,73559 | 11,6096 | 0,585737 | 0,1137 | 0,441715 |
| Mcm4 | 144,274 | 118,252 | -0,286941 | 0,1137 | 0,441715 |
| Casc4 | 14,4432 | 17,7594 | 0,298197 | 0,11375 | 0,441763 |
| Rrm1 | 170,194 | 139,285 | -0,289143 | 0,1138 | 0,441763 |
| Ncor1 | 32,0768 | 39,9206 | 0,315605 | 0,11385 | 0,441763 |
| Lrp5 | 11,3727 | 9,19064 | -0,307331 | 0,11385 | 0,441763 |
| Nipsnap1 | 28,8882 | 23,3303 | -0,308277 | 0,11395 | 0,442017 |
| Usp42 | 13,8651 | 16,9895 | 0,293181 | 0,1142 | 0,442853 |
| Pald1 | 16,4436 | 13,3623 | -0,299351 | 0,11445 | 0,443688 |
| Socs3 | 3,88084 | 2,8778 | -0,431406 | 0,1146 | 0,444136 |
| Lrrc28 | 6,42508 | 8,24039 | 0,358998 | 0,1147 | 0,444255 |
| Snrpd2 | 276,44 | 225,275 | -0,295276 | 0,1147 | 0,444255 |
| Ears2 | 12,8991 | 16,2855 | 0,33632 | 0,11475 | 0,444314 |
| Srp19 | 94,1298 | 76,3091 | -0,302797 | 0,11505 | 0,445341 |
| Fam98a | 31,1903 | 25,4784 | -0,291826 | 0,11515 | 0,445594 |
| Hells | 79,526 | 97,3369 | 0,29156 | 0,11525 | 0,445637 |
| Prdm9 | 0,871521 | 1,27919 | 0,553622 | 0,1153 | 0,445637 |
| Kif3a | 13,6859 | 16,7286 | 0,289629 | 0,1153 | 0,445637 |
| Add2 | 1,7207 | 1,21709 | -0,499555 | 0,1153 | 0,445637 |
| F930015N05Rik | 2,17719 | 1,36168 | -0,677078 | 0,11545 | 0,446082 |
| Aldoart2 | 14,0204 | 17,7724 | 0,342113 | 0,1155 | 0,446141 |
| Ubr3 | 16,4306 | 19,934 | 0,278847 | 0,11565 | 0,446359 |
| Mob3b | 2,92898 | 3,7463 | 0,355069 | 0,1157 | 0,446359 |
| Card10 | 0,760296 | 1,09016 | 0,519904 | 0,11575 | 0,446359 |
| B130024G19Rik | 1,20726 | 0,189291 | -2,67306 | 0,11575 | 0,446359 |
| Fam159a | 1,45817 | 0,0998367 | -3,86845 | 0,1158 | 0,446359 |
| AW549542 | 1,16472 | 0,0443369 | -4,71533 | 0,1158 | 0,446359 |
| Dbx2 | 1,08984 | 0,0335691 | -5,02084 | 0,1158 | 0,446359 |
| Ap3s2 | 20,4101 | 16,7334 | -0,286551 | 0,1159 | 0,44661 |
| Osbpl7 | 2,97088 | 3,92446 | 0,401604 | 0,116 | 0,446786 |
| Bola2 | 266,023 | 212,898 | -0,321392 | 0,11605 | 0,446786 |
| Agtr1a | 1,08303 | 0,67028 | -0,692232 | 0,11605 | 0,446786 |
| Ppp1r7 | 31,5991 | 25,9369 | -0,284876 | 0,11625 | 0,447421 |
| Mapk9 | 13,04 | 15,9758 | 0,292944 | 0,11655 | 0,448365 |
| S100a3 | 0,187044 | 0,832189 | 2,15353 | 0,1166 | 0,448365 |
| Gdpd3 | 1,60325 | 0,883396 | -0,859864 | 0,1166 | 0,448365 |
| Slc16a3 | 65,538 | 79,9127 | 0,286094 | 0,11675 | 0,448673 |
| Zbtb10 | 6,12987 | 4,93136 | -0,31387 | 0,11675 | 0,448673 |
| Clcf1 | 1,45754 | 2,55371 | 0,809058 | 0,1172 | 0,450206 |
| Il18 | 0,692576 | 1,38904 | 1,00405 | 0,11725 | 0,450206 |
| Rpl18 | 302,952 | 248,827 | -0,283943 | 0,1173 | 0,450206 |
| Cbwd1 | 15,1062 | 18,9022 | 0,323417 | 0,11735 | 0,450206 |
| Psmb7 | 343,734 | 282,993 | -0,280527 | 0,11735 | 0,450206 |
| Hspe1 | 176,395 | 144,651 | -0,286234 | 0,1174 | 0,450206 |
| Arf2 | 25,9998 | 21,165 | -0,296821 | 0,1174 | 0,450206 |
| Slc39a6 | 47,5673 | 57,8998 | 0,283587 | 0,11745 | 0,450206 |
| Acbd6 | 45,7951 | 36,8405 | -0,313901 | 0,1175 | 0,450206 |
| Zcchc2 | 3,86717 | 3,048 | -0,343411 | 0,1175 | 0,450206 |
| Nmral1 | 116,375 | 95,6817 | -0,282469 | 0,11775 | 0,451029 |

| | | | | | |
|---|---|---|---|---|---|
| Mrps15 | 73,4398 | 59,4684 | -0,30444 | 0,1179 | 0,451469 |
| Ufc1 | 83,4876 | 67,8052 | -0,300165 | 0,11805 | 0,451752 |
| Prim2 | 41,4226 | 33,6967 | -0,297812 | 0,1181 | 0,451752 |
| Hps6 | 1,87491 | 2,61949 | 0,482463 | 0,11815 | 0,451752 |
| Cdc34 | 98,2508 | 80,4845 | -0,287758 | 0,11815 | 0,451752 |
| Oxr1 | 11,8211 | 9,51281 | -0,313419 | 0,11815 | 0,451752 |
| Pycrl | 59,4839 | 48,4646 | -0,295569 | 0,11825 | 0,451787 |
| Cars2 | 10,2516 | 7,91122 | -0,373874 | 0,11825 | 0,451787 |
| Xylt2 | 13,5372 | 16,7163 | 0,304325 | 0,1183 | 0,451787 |
| Tmem132c | 64,1401 | 77,9042 | 0,280474 | 0,1183 | 0,451787 |
| Pik3ca | 10,4324 | 12,7356 | 0,287804 | 0,1184 | 0,452034 |
| Mrpl18 | 106,951 | 87,3383 | -0,292261 | 0,11845 | 0,452091 |
| D3Ertd751e | 5,37705 | 7,36294 | 0,453466 | 0,11865 | 0,452585 |
| D3Ertd254e | 4,92938 | 6,13588 | 0,315865 | 0,11865 | 0,452585 |
| Slc25a4 | 462,663 | 380,506 | -0,282043 | 0,1187 | 0,452641 |
| Rps14 | 873,925 | 719,331 | -0,280854 | 0,1188 | 0,452675 |
| Caskin2 | 10,0501 | 8,11283 | -0,308938 | 0,1188 | 0,452675 |
| Fam195b | 87,0651 | 71,144 | -0,291352 | 0,11885 | 0,452675 |
| Fam162b | 1,38445 | 0,242515 | -2,51317 | 0,11885 | 0,452675 |
| Rbm26 | 28,7877 | 36,7155 | 0,350934 | 0,1189 | 0,452731 |
| Crebl2 | 0,64824 | 1,03266 | 0,671769 | 0,119 | 0,452977 |
| Snrpg | 223,815 | 180,589 | -0,309601 | 0,11925 | 0,453794 |
| Ssh2 | 2,2717 | 2,88045 | 0,34252 | 0,1193 | 0,453849 |
| Ide | 24,6596 | 29,9928 | 0,282466 | 0,11935 | 0,453905 |
| Rapgef5 | 1,28523 | 0,902158 | -0,51057 | 0,11975 | 0,455156 |
| Atg9b | 0,647994 | 0,368857 | -0,812921 | 0,11975 | 0,455156 |
| Abca5 | 2,29124 | 1,78364 | -0,361304 | 0,1198 | 0,455212 |
| Spag4 | 3,03153 | 4,30355 | 0,505481 | 0,11985 | 0,455267 |
| Slc8a2 | 1,12621 | 0,780108 | -0,529726 | 0,11995 | 0,455512 |
| Tsc22d2 | 5,08081 | 6,25548 | 0,300061 | 0,12005 | 0,455757 |
| Purg | 16,2976 | 20,565 | 0,335532 | 0,12015 | 0,456001 |
| Secisbp2l | 9,74547 | 11,9069 | 0,288997 | 0,12025 | 0,456246 |
| Gm4890 | 1,16012 | 1,69167 | 0,544167 | 0,1204 | 0,45668 |
| Stam | 18,1134 | 22,1368 | 0,289389 | 0,12045 | 0,456683 |
| Rhoq | 20,1437 | 16,0657 | -0,326346 | 0,12055 | 0,456683 |
| BC049715 | 0,309598 | 0,838238 | 1,43696 | 0,1206 | 0,456683 |
| Gata2 | 0,85708 | 0,543385 | -0,657455 | 0,1206 | 0,456683 |
| 4930447C04Rik | 0,79652 | 1,29061 | 0,696276 | 0,12065 | 0,456683 |
| Otud3 | 5,68734 | 7,56952 | 0,412449 | 0,12065 | 0,456683 |
| Etfb | 94,8328 | 77,5722 | -0,289845 | 0,12065 | 0,456683 |
| Pop4 | 73,6283 | 60,0975 | -0,292956 | 0,1209 | 0,457494 |
| Tram2 | 3,47981 | 4,91894 | 0,49934 | 0,121 | 0,457738 |
| Ociad2 | 3,25385 | 4,42883 | 0,444778 | 0,12105 | 0,457792 |
| Uspl1 | 36,7548 | 30,201 | -0,283333 | 0,1212 | 0,458224 |
| Ostm1 | 2,58808 | 1,87632 | -0,463977 | 0,12125 | 0,458278 |
| Ppp1r26 | 4,52029 | 5,67273 | 0,327625 | 0,12145 | 0,458899 |
| Eif3j1,Eif3j2 | 1,30252 | 4,67823 | 1,84466 | 0,12175 | 0,459761 |
| Plekha7 | 4,06822 | 3,14787 | -0,370022 | 0,12175 | 0,459761 |
| Prob1 | 0,973214 | 1,36014 | 0,482925 | 0,1218 | 0,459815 |
| C1qtnf6 | 5,45914 | 4,14329 | -0,397898 | 0,12185 | 0,459868 |

| | | | | | |
|---|---|---|---|---|---|
| Slc39a10 | 34,5223 | 28,3746 | -0,282931 | 0,1219 | 0,459922 |
| Traf3ip1 | 10,0158 | 12,4274 | 0,311253 | 0,12195 | 0,459975 |
| Eme2 | 6,60348 | 8,73848 | 0,404156 | 0,1221 | 0,460405 |
| Klhl20 | 7,55726 | 9,41599 | 0,317249 | 0,12215 | 0,460458 |
| D030056L22Rik | 28,2575 | 34,9266 | 0,30569 | 0,12245 | 0,461454 |
| Pcyt1b | 1,1964 | 0,847064 | -0,49816 | 0,12255 | 0,461695 |
| Map9 | 3,44589 | 4,34351 | 0,333987 | 0,1226 | 0,461748 |
| Pqlc3 | 1,05676 | 1,62606 | 0,62173 | 0,123 | 0,462763 |
| Acsl4 | 11,3342 | 13,875 | 0,291799 | 0,123 | 0,462763 |
| Gatc | 29,142 | 23,6143 | -0,303437 | 0,123 | 0,462763 |
| Cyp4f16 | 4,76774 | 6,28311 | 0,398173 | 0,12305 | 0,462763 |
| Mpzl1 | 61,2139 | 74,3296 | 0,280079 | 0,12305 | 0,462763 |
| Ampd2 | 27,0187 | 22,2424 | -0,280642 | 0,1231 | 0,462816 |
| Tomm40l | 11,9047 | 9,5524 | -0,317599 | 0,12335 | 0,46362 |
| Col6a1 | 1,31484 | 0,927356 | -0,503693 | 0,12345 | 0,463776 |
| Slc46a1 | 6,71974 | 8,70361 | 0,373207 | 0,1235 | 0,463776 |
| Syngr1 | 1,17747 | 0,488334 | -1,26975 | 0,1235 | 0,463776 |
| Cep97 | 9,42023 | 13,4757 | 0,51653 | 0,12375 | 0,464579 |
| 2810442N19Rik | 1,4654 | 0 | -inf | 0,12385 | 0,464682 |
| Slbp | 80,172 | 66,0294 | -0,279991 | 0,12385 | 0,464682 |
| Lrrc16b | 4,32508 | 3,38099 | -0,355279 | 0,12395 | 0,464907 |
| Timm9 | 61,8328 | 49,7203 | -0,314539 | 0,124 | 0,464907 |
| Aven | 33,3229 | 41,4609 | 0,315239 | 0,12415 | 0,464907 |
| Sqle | 76,2302 | 62,7885 | -0,279864 | 0,12415 | 0,464907 |
| Utp14b | 8,83015 | 13,8197 | 0,646213 | 0,1242 | 0,464907 |
| Ophn1 | 3,46645 | 4,47151 | 0,367302 | 0,1242 | 0,464907 |
| Lrfn4 | 25,7221 | 32,5644 | 0,340289 | 0,1242 | 0,464907 |
| Nkx6-2 | 1,92677 | 0,121709 | -3,98468 | 0,1242 | 0,464907 |
| Syt14 | 3,07189 | 4,28261 | 0,479361 | 0,1243 | 0,465146 |
| Snora15 | 31,362 | 20,0096 | -0,64832 | 0,12455 | 0,465946 |
| St3gal3 | 14,6719 | 18,2242 | 0,312793 | 0,1246 | 0,465997 |
| Slc25a14 | 6,36249 | 4,69685 | -0,437899 | 0,125 | 0,467322 |
| Arxes1 | 12,8634 | 10,0419 | -0,357237 | 0,12505 | 0,467322 |
| Mastl | 7,96734 | 9,83931 | 0,304458 | 0,1251 | 0,467322 |
| Nt5dc3 | 0,610343 | 0,409005 | -0,577502 | 0,1251 | 0,467322 |
| Mef2a | 8,68105 | 10,6544 | 0,295503 | 0,12515 | 0,467372 |
| BC024139 | 1,70642 | 0,0873977 | -4,28723 | 0,1252 | 0,467423 |
| Rest | 14,2144 | 17,3465 | 0,287292 | 0,12545 | 0,46822 |
| Bcas2 | 104,593 | 86,1277 | -0,280234 | 0,12575 | 0,469067 |
| Prdx3 | 71,2741 | 58,5037 | -0,284851 | 0,12575 | 0,469067 |
| Ormdl1 | 21,9814 | 17,7099 | -0,31173 | 0,1259 | 0,46949 |
| Mrps7 | 75,7667 | 62,1892 | -0,2849 | 0,126 | 0,46964 |
| Eif3k | 378,904 | 313,539 | -0,273187 | 0,12605 | 0,46964 |
| Rpl22l1 | 307,463 | 252,617 | -0,283461 | 0,12605 | 0,46964 |
| Atp11b | 15,6379 | 19,0608 | 0,285559 | 0,1262 | 0,470062 |
| Haus3 | 29,0889 | 35,782 | 0,298765 | 0,12635 | 0,470484 |
| Mrs2 | 14,952 | 18,5942 | 0,314516 | 0,12645 | 0,470583 |
| Ntn4 | 1,19799 | 0,769623 | -0,638389 | 0,12645 | 0,470583 |
| Pfn2 | 31,741 | 25,8126 | -0,298272 | 0,1266 | 0,471005 |
| Psma4 | 118,126 | 97,6088 | -0,27524 | 0,1268 | 0,471612 |

| | | | | | |
|---|---|---|---|---|---|
| Fbxl4 | 3,40049 | 4,5496 | 0,419996 | 0,12695 | 0,471809 |
| Ptplad2 | 0,685217 | 0,373037 | -0,877242 | 0,12695 | 0,471809 |
| Dennd1c | 0,7042 | 1,09399 | 0,635544 | 0,127 | 0,471809 |
| Atp5e | 664,727 | 549,472 | -0,274715 | 0,127 | 0,471809 |
| Ifitm7 | 0,687657 | 2,28869 | 1,73476 | 0,12715 | 0,472229 |
| Gm2027 | 2,16127 | 1,5482 | -0,481291 | 0,12725 | 0,472464 |
| Sept7 | 88,0779 | 106,694 | 0,276632 | 0,12735 | 0,472611 |
| Pwp1 | 56,448 | 46,5576 | -0,277905 | 0,1274 | 0,472611 |
| Ethe1 | 8,57414 | 6,45276 | -0,410076 | 0,1274 | 0,472611 |
| Fcrl6 | 0,917187 | 0,533989 | -0,780405 | 0,12755 | 0,472942 |
| Gstm1 | 7,38677 | 9,85416 | 0,415788 | 0,1276 | 0,472942 |
| Tpbg | 6,18822 | 7,80701 | 0,335245 | 0,1276 | 0,472942 |
| 1700049G17Rik | 2,01225 | 2,63608 | 0,389587 | 0,12775 | 0,473361 |
| Cdc42bpb | 22,3314 | 27,0543 | 0,276784 | 0,1278 | 0,47341 |
| Cntrl | 14,7992 | 18,0401 | 0,285687 | 0,12785 | 0,473458 |
| Ppm1h | 1,02706 | 0,745126 | -0,462961 | 0,128 | 0,473877 |
| Gm13251 | 0,596018 | 0,906685 | 0,605244 | 0,12815 | 0,474255 |
| Gna13 | 16,2255 | 19,6941 | 0,279497 | 0,1282 | 0,474255 |
| Ttc30a1 | 4,89806 | 3,74648 | -0,386674 | 0,12825 | 0,474255 |
| Mme | 1,58697 | 1,17939 | -0,428237 | 0,12825 | 0,474255 |
| Ppp1r9a | 5,60717 | 4,56799 | -0,295712 | 0,1283 | 0,474303 |
| Ptbp3 | 16,2803 | 19,7269 | 0,277031 | 0,12835 | 0,474351 |
| Slc9a5 | 7,07764 | 8,74305 | 0,304868 | 0,1285 | 0,474769 |
| Mylk | 2,06072 | 1,58127 | -0,382066 | 0,12865 | 0,475186 |
| Art3 | 0,766993 | 0,0592156 | -3,69516 | 0,12895 | 0,476068 |
| Drd1a | 0,0155277 | 2,1413 | 7,10749 | 0,129 | 0,476068 |
| 6530402F18Rik | 0,617645 | 0,923285 | 0,579999 | 0,129 | 0,476068 |
| Camk1g | 4,35132 | 3,25955 | -0,41678 | 0,1291 | 0,4763 |
| Alkbh7 | 20,3226 | 15,6422 | -0,377641 | 0,12915 | 0,476347 |
| Uhmk1 | 9,32204 | 11,8494 | 0,346091 | 0,12955 | 0,477685 |
| Ppp1r9b | 23,3866 | 19,2276 | -0,282506 | 0,1297 | 0,477924 |
| Rps3 | 815,361 | 664,83 | -0,294453 | 0,1297 | 0,477924 |
| Galnt4 | 3,80907 | 5,63204 | 0,564218 | 0,1298 | 0,477924 |
| Ncoa4 | 1,23203 | 1,69461 | 0,459921 | 0,12985 | 0,477924 |
| Zfp236 | 7,64208 | 9,30014 | 0,283287 | 0,12985 | 0,477924 |
| Romo1 | 136,522 | 103,933 | -0,393485 | 0,12985 | 0,477924 |
| H2-M3 | 0,806333 | 1,36534 | 0,759808 | 0,1299 | 0,477924 |
| Lyn | 1,21678 | 1,71042 | 0,491282 | 0,12995 | 0,477924 |
| Lzts2 | 47,1697 | 38,9276 | -0,277066 | 0,12995 | 0,477924 |
| Tssc4 | 44,0495 | 35,9766 | -0,292069 | 0,13 | 0,477971 |
| Lrp3 | 23,861 | 19,6809 | -0,277854 | 0,13005 | 0,478018 |
| Wbp1l | 21,8484 | 17,9606 | -0,282687 | 0,1301 | 0,478064 |
| Ruvbl2 | 142,984 | 118,451 | -0,27156 | 0,13035 | 0,478709 |
| Frk | 0,87056 | 0,599462 | -0,538274 | 0,13035 | 0,478709 |
| Tmem136 | 5,3431 | 6,75032 | 0,337279 | 0,13055 | 0,479306 |
| Gnl3l | 16,9999 | 20,6894 | 0,283367 | 0,13075 | 0,479723 |
| Slitrk5 | 4,00311 | 3,13897 | -0,350828 | 0,1308 | 0,479723 |
| Pspn | 2,61597 | 1,16833 | -1,16291 | 0,1308 | 0,479723 |
| Zbtb7a | 2,58508 | 3,30467 | 0,354298 | 0,13095 | 0,479723 |
| Tmem201 | 23,4286 | 28,3135 | 0,273217 | 0,13095 | 0,479723 |

| | | | | | |
|---|---|---|---|---|---|
| Nabp2 | 90,5289 | 74,512 | -0,280906 | 0,13095 | 0,479723 |
| Dagla | 0,670703 | 0,457788 | -0,550996 | 0,13095 | 0,479723 |
| Idh2 | 196,558 | 163,055 | -0,269592 | 0,131 | 0,479723 |
| H2afv | 130,126 | 107,877 | -0,270518 | 0,131 | 0,479723 |
| Pola2 | 57,4587 | 47,596 | -0,271686 | 0,13125 | 0,480364 |
| Stx17 | 4,05731 | 3,21792 | -0,334397 | 0,13125 | 0,480364 |
| Man2a2 | 13,6605 | 16,5537 | 0,277143 | 0,13135 | 0,480456 |
| Tmem263 | 11,4174 | 9,26682 | -0,301086 | 0,13135 | 0,480456 |
| Snrpb2 | 135,834 | 112,464 | -0,27239 | 0,1315 | 0,480867 |
| Ebf4 | 1,20492 | 1,73105 | 0,522706 | 0,13155 | 0,480913 |
| Lrrc18 | 0,243701 | 0,596907 | 1,2924 | 0,1316 | 0,480959 |
| Zc2hc1c | 1,39723 | 1,00386 | -0,47701 | 0,13165 | 0,481004 |
| Prkx | 4,58724 | 5,85717 | 0,352578 | 0,13175 | 0,481096 |
| Rbx1 | 67,4546 | 55,6452 | -0,27766 | 0,13175 | 0,481096 |
| Flt4 | 0,495446 | 0,725682 | 0,550611 | 0,1318 | 0,481141 |
| 3110045C21Rik | 5,90386 | 9,19166 | 0,638667 | 0,13205 | 0,481916 |
| Tpmt | 5,98756 | 4,63855 | -0,368292 | 0,1321 | 0,481916 |
| Trim6 | 2,33495 | 3,0879 | 0,403235 | 0,13215 | 0,481916 |
| Mettl7a1 | 7,0672 | 5,38289 | -0,392759 | 0,1322 | 0,481916 |
| Gm4285 | 1,19394 | 0,669654 | -0,834239 | 0,1322 | 0,481916 |
| Mybl2 | 14,1843 | 11,579 | -0,292783 | 0,13235 | 0,482326 |
| Gm16907 | 2,31653 | 5,01961 | 1,11561 | 0,1324 | 0,482371 |
| 4930451G09Rik | 0,971732 | 1,48204 | 0,608953 | 0,13245 | 0,482416 |
| Tpd52 | 9,17115 | 7,19479 | -0,350151 | 0,1325 | 0,482461 |
| Zfx | 8,18353 | 9,99017 | 0,287787 | 0,13275 | 0,483234 |
| Rims4 | 0,302608 | 0,808723 | 1,4182 | 0,1328 | 0,483279 |
| Rufy3 | 11,4633 | 14,0368 | 0,292192 | 0,133 | 0,483732 |
| Plp1 | 2,83561 | 2,12645 | -0,415213 | 0,133 | 0,483732 |
| Pold1 | 39,1265 | 32,4591 | -0,269524 | 0,13305 | 0,483777 |
| Porcn | 3,09617 | 4,22188 | 0,447402 | 0,13345 | 0,485094 |
| Rpl39 | 448,685 | 371,04 | -0,27413 | 0,13375 | 0,486047 |
| Mplkip | 10,9554 | 8,78544 | -0,318456 | 0,134 | 0,486817 |
| Ubash3b | 2,19168 | 1,59237 | -0,46086 | 0,1341 | 0,486949 |
| Samd4 | 2,02956 | 1,56557 | -0,374474 | 0,13415 | 0,486949 |
| Lrp11 | 1,28634 | 0,895008 | -0,523295 | 0,13415 | 0,486949 |
| Fbl | 43,745 | 35,5439 | -0,299515 | 0,13425 | 0,487174 |
| Banp | 8,87874 | 10,8218 | 0,285507 | 0,1343 | 0,487217 |
| Snhg11 | 4,0679 | 5,06946 | 0,317548 | 0,1344 | 0,487442 |
| Top3b | 21,2446 | 25,793 | 0,279887 | 0,13445 | 0,487486 |
| Atp8a2 | 0,66063 | 0,968711 | 0,552223 | 0,1345 | 0,487529 |
| Mdh1 | 176,086 | 146,407 | -0,266295 | 0,13475 | 0,488108 |
| Extl3 | 25,7659 | 31,1484 | 0,273696 | 0,1348 | 0,488108 |
| Zdhhc2 | 2,82694 | 3,80133 | 0,427265 | 0,13485 | 0,488108 |
| Sec13 | 162,493 | 135,129 | -0,266045 | 0,13485 | 0,488108 |
| Marcksl1 | 568,79 | 468,208 | -0,280748 | 0,13485 | 0,488108 |
| Dancr | 17,0195 | 13,1374 | -0,373508 | 0,1352 | 0,489142 |
| Rsad1 | 3,61144 | 2,79537 | -0,369533 | 0,13525 | 0,489142 |
| Sv2a | 1,19483 | 0,838443 | -0,511024 | 0,13525 | 0,489142 |
| Prpf40b | 22,4718 | 27,2849 | 0,279986 | 0,1353 | 0,489185 |
| Arhgef40 | 23,0306 | 29,7285 | 0,368294 | 0,1354 | 0,489271 |

| | | | | | |
|---|---|---|---|---|---|
| Rnf149 | 22,1366 | 27,0153 | 0,287345 | 0,1354 | 0,489271 |
| Gm5127 | 0,720577 | 0,143511 | -2,32799 | 0,13555 | 0,48958 |
| Ndufc2 | 166,112 | 136,932 | -0,278692 | 0,1356 | 0,48958 |
| Arhgef28 | 0,93448 | 0,668317 | -0,483632 | 0,1356 | 0,48958 |
| Phldb1 | 17,6963 | 21,4221 | 0,275653 | 0,13565 | 0,489622 |
| Elf2 | 21,2801 | 17,5507 | -0,277971 | 0,1358 | 0,490026 |
| Unkl | 1,76624 | 2,62879 | 0,573721 | 0,1359 | 0,490249 |
| 4632434I11Rik | 10,2695 | 12,6628 | 0,302239 | 0,1362 | 0,491193 |
| Sept10 | 22,4362 | 27,5839 | 0,297998 | 0,13685 | 0,493398 |
| Rps6 | 15,7025 | 12,3758 | -0,343472 | 0,1373 | 0,494882 |
| Bbs1 | 4,77715 | 5,93074 | 0,312061 | 0,13745 | 0,495144 |
| Oser1 | 3,10856 | 2,17865 | -0,512814 | 0,13745 | 0,495144 |
| 03/08/16 | 13,596 | 16,5419 | 0,282947 | 0,13765 | 0,495725 |
| Col6a2 | 1,60101 | 2,17624 | 0,442855 | 0,1377 | 0,495766 |
| Eif4ebp1 | 63,7353 | 52,0673 | -0,291717 | 0,13785 | 0,496167 |
| Ibtk | 18,6286 | 22,5512 | 0,27569 | 0,1379 | 0,496208 |
| Rcn1 | 115,086 | 95,713 | -0,265925 | 0,138 | 0,49633 |
| BC051226 | 3,34708 | 2,0931 | -0,677258 | 0,13805 | 0,49633 |
| Fam46c | 0,917407 | 0,55306 | -0,730126 | 0,13805 | 0,49633 |
| Il15 | 0,760997 | 0,37831 | -1,00832 | 0,13815 | 0,49655 |
| Prps2 | 17,2391 | 14,1484 | -0,285049 | 0,1382 | 0,496591 |
| Rpl29 | 75,1625 | 61,1541 | -0,297564 | 0,1383 | 0,496811 |
| Tex40 | 0,365056 | 0,926692 | 1,34397 | 0,1384 | 0,496892 |
| Mast2 | 22,6422 | 27,2469 | 0,26708 | 0,1384 | 0,496892 |
| Cpt1c | 6,11594 | 7,74522 | 0,340731 | 0,1385 | 0,496973 |
| Khdrbs3 | 53,2722 | 44,0918 | -0,272873 | 0,1385 | 0,496973 |
| Inca1 | 0,500884 | 0,944165 | 0,914564 | 0,13855 | 0,497013 |
| Mrps26 | 45,049 | 36,4608 | -0,305148 | 0,1387 | 0,497273 |
| Irak1bp1 | 10,3356 | 8,12086 | -0,34792 | 0,1387 | 0,497273 |
| Chml | 6,47431 | 8,0935 | 0,322037 | 0,1389 | 0,497851 |
| Mamdc4 | 3,13309 | 4,01389 | 0,357413 | 0,1393 | 0,499145 |
| Tvp23b | 11,0936 | 8,70952 | -0,34906 | 0,1395 | 0,499722 |
| Zfp942 | 7,3672 | 9,2255 | 0,324511 | 0,13955 | 0,499762 |
| Nf1 | 7,4912 | 9,04851 | 0,272485 | 0,1397 | 0,50016 |
| Bcat1 | 20,8997 | 17,3639 | -0,267386 | 0,1398 | 0,500238 |
| Rab2a | 51,1837 | 42,3814 | -0,272253 | 0,1398 | 0,500238 |
| Rab34 | 63,1985 | 52,2419 | -0,274682 | 0,1399 | 0,500457 |
| Acd | 25,8416 | 20,744 | -0,317003 | 0,14005 | 0,500592 |
| Abcd4 | 4,56035 | 5,88772 | 0,368562 | 0,1401 | 0,500592 |
| Fam117b | 11,0573 | 13,4419 | 0,28173 | 0,1401 | 0,500592 |
| Zfp389 | 0,846824 | 1,55118 | 0,873228 | 0,14015 | 0,500592 |
| Uchl1 | 42,0915 | 51,499 | 0,291017 | 0,1402 | 0,500592 |
| Camta1 | 7,66724 | 9,88386 | 0,366368 | 0,14025 | 0,500592 |
| Ankrd13b | 19,245 | 15,8021 | -0,284366 | 0,14025 | 0,500592 |
| Lrrc59 | 137,775 | 112,82 | -0,288292 | 0,14025 | 0,500592 |
| Mis18a | 46,0313 | 37,7215 | -0,287227 | 0,1403 | 0,500631 |
| Mblac1 | 4,02497 | 2,84155 | -0,5023 | 0,14045 | 0,501027 |
| Pds5b | 29,1089 | 35,0142 | 0,26648 | 0,1409 | 0,502151 |
| Arhgap28 | 30,2225 | 25,1061 | -0,267584 | 0,1409 | 0,502151 |
| Mir6904 | 0 | 206,081 | inf | 0,14095 | 0,502151 |

| | | | | | |
|---|---|---|---|---|---|
| Mif | 335,63 | 278,445 | -0,26948 | 0,14095 | 0,502151 |
| Abhd17c | 34,213 | 28,2493 | -0,276329 | 0,141 | 0,502151 |
| Oscp1 | 7,23951 | 5,60883 | -0,368194 | 0,141 | 0,502151 |
| Tab3 | 4,33666 | 5,36583 | 0,307218 | 0,14105 | 0,50219 |
| Hibch | 11,7012 | 9,21727 | -0,344249 | 0,14115 | 0,502406 |
| Ikzf2 | 0,675526 | 0,492128 | -0,456977 | 0,1412 | 0,502445 |
| Fpgt | 3,76509 | 4,81044 | 0,353484 | 0,1413 | 0,502661 |
| D130040H23Rik | 1,23601 | 1,82453 | 0,561834 | 0,1414 | 0,502878 |
| Tsc22d1 | 141,611 | 116,029 | -0,287455 | 0,1415 | 0,503094 |
| Trak2 | 7,08038 | 8,63996 | 0,287198 | 0,14155 | 0,503132 |
| D8Ertd738e | 224,198 | 186,097 | -0,268722 | 0,1416 | 0,50317 |
| Olfm4 | 4,52001 | 6,05191 | 0,421065 | 0,1417 | 0,503386 |
| Rybp | 6,25065 | 7,73049 | 0,306553 | 0,14185 | 0,50364 |
| Oxnad1 | 15,4268 | 12,3789 | -0,317549 | 0,14185 | 0,50364 |
| 2610035D17Rik | 4,64239 | 3,44112 | -0,431991 | 0,14205 | 0,504211 |
| Apex2 | 15,6165 | 19,2735 | 0,303547 | 0,1423 | 0,504958 |
| Trove2 | 6,08966 | 7,42354 | 0,285744 | 0,1424 | 0,505173 |
| Tmem39b | 21,8382 | 17,7579 | -0,29839 | 0,1426 | 0,505743 |
| Msl3 | 17,3708 | 14,2591 | -0,284781 | 0,14275 | 0,506033 |
| Morn4 | 21,2955 | 26,0676 | 0,29171 | 0,1428 | 0,506033 |
| Ralgapa1 | 10,2203 | 12,3201 | 0,269565 | 0,1428 | 0,506033 |
| Atad2 | 44,1597 | 52,9234 | 0,261177 | 0,1429 | 0,506247 |
| Kpna6 | 21,1967 | 25,4808 | 0,265576 | 0,14295 | 0,506285 |
| Setd4 | 2,83397 | 3,88887 | 0,456528 | 0,1431 | 0,506508 |
| Mbd3 | 151,638 | 126,003 | -0,267172 | 0,1431 | 0,506508 |
| Stk36 | 2,86622 | 2,22657 | -0,364325 | 0,14315 | 0,506508 |
| Ndufaf7 | 13,1195 | 10,5299 | -0,317221 | 0,1432 | 0,506508 |
| Spink2 | 0,96136 | 1,95372 | 1,02307 | 0,14325 | 0,506508 |
| Sepw1 | 421,79 | 351,545 | -0,262816 | 0,14325 | 0,506508 |
| Uqcrb | 103,498 | 83,8942 | -0,302965 | 0,14335 | 0,506722 |
| Taf11 | 64,1031 | 52,8297 | -0,279045 | 0,14355 | 0,507289 |
| Zfat | 3,32864 | 2,58329 | -0,365721 | 0,14365 | 0,507334 |
| Bhmt2 | 0,0350557 | 0,904026 | 4,68864 | 0,1438 | 0,507334 |
| Cort | 0,59477 | 1,59849 | 1,42631 | 0,1438 | 0,507334 |
| Susd1 | 1,61165 | 2,19471 | 0,445493 | 0,1438 | 0,507334 |
| Atp2b4 | 1,608 | 2,05277 | 0,352306 | 0,1438 | 0,507334 |
| Jun | 38,5192 | 31,7493 | -0,278854 | 0,1438 | 0,507334 |
| Nudt10 | 5,78151 | 4,35303 | -0,409428 | 0,1439 | 0,507547 |
| Zmym5 | 11,6147 | 14,0382 | 0,273409 | 0,14395 | 0,507584 |
| Kctd3 | 21,2339 | 25,6009 | 0,269825 | 0,144 | 0,507621 |
| Kcnd1 | 0,598865 | 0,984806 | 0,717608 | 0,14415 | 0,507907 |
| Uggt1 | 22,7145 | 27,243 | 0,262273 | 0,1442 | 0,507907 |
| Sowaha | 0,847046 | 0,564157 | -0,586344 | 0,1442 | 0,507907 |
| Rpp40 | 7,88253 | 6,26764 | -0,330736 | 0,14435 | 0,508296 |
| Slc35e3 | 5,64252 | 4,46206 | -0,338629 | 0,1445 | 0,508545 |
| Prkar1b | 1,34401 | 0,914874 | -0,5549 | 0,1445 | 0,508545 |
| Armc10 | 33,6217 | 27,6715 | -0,280994 | 0,14455 | 0,508581 |
| Zc3h7a | 14,6335 | 17,7322 | 0,277102 | 0,14485 | 0,509357 |
| Nbl1 | 4,81327 | 3,57941 | -0,427295 | 0,14485 | 0,509357 |
| Arhgef10l | 9,61028 | 7,86495 | -0,289142 | 0,14505 | 0,509921 |

| | | | | | |
|---|---|---|---|---|---|
| Prrt1 | 0,336097 | 0,805113 | 1,26032 | 0,14515 | 0,509993 |
| Ank2 | 6,46505 | 5,26111 | -0,297293 | 0,14515 | 0,509993 |
| Kmt2d | 9,13826 | 10,9881 | 0,265948 | 0,1454 | 0,510592 |
| Trp53inp1 | 2,7782 | 2,16199 | -0,36179 | 0,1454 | 0,510592 |
| Nhlrc2 | 11,8595 | 14,4426 | 0,28428 | 0,1457 | 0,510877 |
| Pef1 | 30,4148 | 24,8677 | -0,290501 | 0,1457 | 0,510877 |
| Batf3 | 2,35833 | 1,31306 | -0,844828 | 0,1457 | 0,510877 |
| Arl2bp | 107,765 | 89,9076 | -0,261373 | 0,14575 | 0,510877 |
| Aldh5a1 | 5,95758 | 4,83156 | -0,302239 | 0,14575 | 0,510877 |
| Svil | 9,13024 | 11,0064 | 0,26962 | 0,1458 | 0,510877 |
| Ndufa8 | 102,272 | 84,0842 | -0,282498 | 0,1458 | 0,510877 |
| Mcrs1 | 12,0605 | 9,55979 | -0,335242 | 0,1458 | 0,510877 |
| 4930523C07Rik | 0,391114 | 0,607336 | 0,634906 | 0,1459 | 0,510949 |
| Kars | 203,758 | 169,93 | -0,261913 | 0,1459 | 0,510949 |
| AU041133 | 2,23327 | 3,03472 | 0,44241 | 0,14615 | 0,511476 |
| Gpx3 | 63,7776 | 76,6471 | 0,265181 | 0,1462 | 0,511476 |
| Fam210a | 10,5248 | 12,6437 | 0,264622 | 0,1462 | 0,511476 |
| Ubr2 | 14,5268 | 17,3848 | 0,259104 | 0,14625 | 0,511476 |
| Perp | 1,27052 | 0,803482 | -0,661077 | 0,14625 | 0,511476 |
| Nr2c2ap | 19,7571 | 13,5204 | -0,547229 | 0,1464 | 0,511861 |
| Zkscan4 | 1,48997 | 1,02408 | -0,540954 | 0,1466 | 0,512421 |
| Mrpl2 | 58,3998 | 47,9286 | -0,285077 | 0,14685 | 0,513015 |
| Mapk8ip2 | 1,86911 | 1,36728 | -0,451042 | 0,14685 | 0,513015 |
| Lap3 | 95,6741 | 79,8551 | -0,260745 | 0,1469 | 0,51305 |
| Stoml1 | 5,39185 | 6,94817 | 0,365852 | 0,14695 | 0,513085 |
| Ogdhl | 11,4893 | 14,0008 | 0,28522 | 0,147 | 0,51312 |
| Gbe1 | 6,83145 | 8,5493 | 0,323616 | 0,14715 | 0,513364 |
| Plcb3 | 13,8409 | 16,737 | 0,274111 | 0,14715 | 0,513364 |
| Zfp445 | 42,5955 | 51,0899 | 0,262336 | 0,1473 | 0,513748 |
| Dyrk1b | 6,72107 | 5,24653 | -0,357328 | 0,14765 | 0,514829 |
| Zfp326 | 43,714 | 52,4043 | 0,261591 | 0,14785 | 0,515383 |
| Vcpkmt | 18,2168 | 14,2444 | -0,354878 | 0,14795 | 0,515383 |
| Bag2 | 27,6538 | 22,7181 | -0,283637 | 0,148 | 0,515383 |
| Trmt2b | 3,91343 | 2,56727 | -0,608201 | 0,148 | 0,515383 |
| Aldh1l1 | 0,503936 | 0,777676 | 0,625929 | 0,14805 | 0,515383 |
| Naaladl1 | 1,65728 | 1,14211 | -0,537118 | 0,14805 | 0,515383 |
| Atp10d | 1,4294 | 1,86013 | 0,379992 | 0,14845 | 0,516635 |
| Gse1 | 19,8266 | 24,8222 | 0,324195 | 0,1485 | 0,516669 |
| Rbm12b1 | 2,04875 | 2,7279 | 0,413052 | 0,14885 | 0,517746 |
| 2410004B18Rik | 33,8911 | 27,3313 | -0,310357 | 0,1489 | 0,51778 |
| Ptpn23 | 15,7234 | 18,8995 | 0,265432 | 0,149 | 0,517987 |
| Elmo3 | 0,596854 | 0,896295 | 0,586596 | 0,14915 | 0,518368 |
| Osbpl6 | 3,31787 | 4,22171 | 0,347569 | 0,14925 | 0,518575 |
| Clta | 211,074 | 176,715 | -0,256325 | 0,1495 | 0,519187 |
| Nup210l | 0,672733 | 0,928879 | 0,465458 | 0,1496 | 0,519187 |
| Zfp449 | 4,18943 | 5,27878 | 0,33345 | 0,14965 | 0,519187 |
| Snf8 | 38,8235 | 31,3891 | -0,306669 | 0,14965 | 0,519187 |
| Dnajb4 | 10,3943 | 8,36594 | -0,313188 | 0,14965 | 0,519187 |
| Ica1 | 3,22111 | 2,28986 | -0,492299 | 0,1497 | 0,519187 |
| D11Wsu47e | 12,7069 | 15,6461 | 0,300186 | 0,14975 | 0,519187 |

| | | | | | |
|---|---|---|---|---|---|
| Grb10 | 402,635 | 328,842 | -0,292079 | 0,14975 | 0,519187 |
| Pdgfd | 0,683555 | 1,10537 | 0,693397 | 0,14985 | 0,519394 |
| Prodh | 6,87545 | 5,36121 | -0,358896 | 0,14995 | 0,5196 |
| Prps1 | 48,3731 | 40,2522 | -0,265135 | 0,1501 | 0,519979 |
| Snapc1 | 13,3873 | 16,467 | 0,298708 | 0,15015 | 0,520012 |
| Btf3 | 171,188 | 142,966 | -0,25991 | 0,1503 | 0,520035 |
| Rnf187 | 111,509 | 93,1112 | -0,260139 | 0,1503 | 0,520035 |
| Rnpep | 33,3073 | 40,0347 | 0,265411 | 0,15035 | 0,520035 |
| Dcaf6 | 4,81797 | 6,05566 | 0,329858 | 0,1504 | 0,520035 |
| Hat1 | 152,733 | 127,548 | -0,25997 | 0,1504 | 0,520035 |
| Mrpl42 | 110,517 | 90,9164 | -0,281655 | 0,1504 | 0,520035 |
| Maob | 2,47068 | 1,79238 | -0,463036 | 0,15045 | 0,520068 |
| Fam58b | 16,7154 | 13,2083 | -0,33973 | 0,15065 | 0,520619 |
| Cdc123 | 119,159 | 99,3096 | -0,26288 | 0,1508 | 0,520717 |
| Mrps11 | 49,5608 | 40,5907 | -0,288051 | 0,1508 | 0,520717 |
| Synrg | 3,95431 | 3,19835 | -0,306098 | 0,1508 | 0,520717 |
| Npm3 | 33,0353 | 40,889 | 0,307704 | 0,15085 | 0,520749 |
| Eif5a | 820,746 | 681,201 | -0,268855 | 0,1509 | 0,520782 |
| Rrp1b | 32,0345 | 38,3829 | 0,260837 | 0,151 | 0,520987 |
| Igip | 0,888804 | 1,46417 | 0,720148 | 0,1511 | 0,521191 |
| Ly6a | 0,280049 | 0,597789 | 1,09396 | 0,1512 | 0,521396 |
| Nrbp2 | 1,20749 | 0,808495 | -0,578697 | 0,1513 | 0,521601 |
| Samhd1 | 23,651 | 28,3384 | 0,260856 | 0,15135 | 0,521633 |
| Has3 | 0,779439 | 1,06191 | 0,446155 | 0,15145 | 0,521697 |
| Lactb | 10,3197 | 12,8795 | 0,319679 | 0,15145 | 0,521697 |
| Psmb6 | 320,495 | 268,288 | -0,256518 | 0,1516 | 0,522074 |
| Jph3 | 1,04307 | 1,43985 | 0,46509 | 0,1517 | 0,522278 |
| Zfp930 | 10,8277 | 13,3071 | 0,29746 | 0,152 | 0,52317 |
| Birc6 | 13,597 | 16,2654 | 0,25852 | 0,15215 | 0,523546 |
| Pim3 | 14,6396 | 17,8691 | 0,287596 | 0,1525 | 0,52461 |
| Agpat4 | 16,3516 | 13,2609 | -0,302253 | 0,15275 | 0,525329 |
| Il13ra1 | 0,875794 | 1,22784 | 0,487464 | 0,153 | 0,525907 |
| Rapgef2 | 4,00314 | 4,93597 | 0,302202 | 0,153 | 0,525907 |
| Mpc2 | 19,4983 | 15,1236 | -0,36655 | 0,15315 | 0,526281 |
| Trim63 | 1,16811 | 1,72538 | 0,562746 | 0,15325 | 0,526374 |
| Tbc1d10a | 9,51662 | 11,9191 | 0,324758 | 0,1533 | 0,526374 |
| Gpr180 | 10,4063 | 8,44324 | -0,301588 | 0,1533 | 0,526374 |
| Tmem161a | 16,0411 | 19,4723 | 0,279652 | 0,15345 | 0,526748 |
| Vav3 | 0,607887 | 0,909469 | 0,581222 | 0,1535 | 0,526778 |
| Faxc | 8,59418 | 7,14498 | -0,26643 | 0,1537 | 0,527213 |
| BC051142 | 0,393134 | 0,705838 | 0,844315 | 0,15375 | 0,527213 |
| AI462493 | 36,2959 | 29,0978 | -0,3189 | 0,15375 | 0,527213 |
| Sbf1 | 21,7074 | 25,9389 | 0,256935 | 0,1539 | 0,527446 |
| 1500009L16Rik | 15,3865 | 12,1071 | -0,345813 | 0,1539 | 0,527446 |
| 4921531C22Rik | 1,64486 | 1,10622 | -0,572325 | 0,15395 | 0,527476 |
| Fis1 | 134,357 | 112,058 | -0,261823 | 0,15405 | 0,527487 |
| Mrps27 | 48,4228 | 40,3661 | -0,262543 | 0,15405 | 0,527487 |
| Rps2 | 283,418 | 237,208 | -0,25678 | 0,1541 | 0,527487 |
| Fam78a | 0,425273 | 0,644214 | 0,59915 | 0,1542 | 0,527487 |
| Plagl1 | 47,7945 | 57,2513 | 0,260463 | 0,1542 | 0,527487 |

| | | | | | |
|---|---|---|---|---|---|
| Igbp1 | 61,0718 | 50,7326 | -0,267594 | 0,1542 | 0,527487 |
| Cpt1b | 0,95133 | 1,81626 | 0,932951 | 0,15435 | 0,52786 |
| Ankdd1b | 1,12374 | 0,78328 | -0,520705 | 0,1544 | 0,52789 |
| Haus6 | 25,9093 | 31,0423 | 0,260767 | 0,15445 | 0,52792 |
| Zcchc17 | 40,2354 | 33,1511 | -0,27941 | 0,15455 | 0,528011 |
| Rarres2 | 0,693445 | 0 | -inf | 0,1546 | 0,528011 |
| Cnnm3 | 8,23099 | 9,99398 | 0,279994 | 0,1546 | 0,528011 |
| Rplp0 | 786,558 | 652,502 | -0,269569 | 0,15465 | 0,528041 |
| Sh3bgrl2 | 6,47828 | 8,03286 | 0,310301 | 0,1547 | 0,528071 |
| Gm14403 | 1,08238 | 1,49449 | 0,465456 | 0,15475 | 0,528081 |
| Srrm2 | 135,043 | 164,702 | 0,286435 | 0,15485 | 0,528081 |
| Ddah1 | 6,14412 | 7,56876 | 0,30085 | 0,1549 | 0,528081 |
| Trak1 | 9,23985 | 7,58368 | -0,284971 | 0,1549 | 0,528081 |
| Hist1h1d | 1,94984 | 3,12641 | 0,681147 | 0,15495 | 0,528081 |
| Sarnp | 16,7636 | 13,1468 | -0,350621 | 0,15495 | 0,528081 |
| Camsap3 | 2,9633 | 3,76626 | 0,345929 | 0,155 | 0,528112 |
| Cadm4 | 5,69133 | 4,39099 | -0,374218 | 0,1551 | 0,528312 |
| A330021E22Rik | 0,860193 | 1,23119 | 0,517325 | 0,1552 | 0,528372 |
| Kdm6a | 15,7491 | 18,8733 | 0,261084 | 0,1552 | 0,528372 |
| Fkbpl | 8,86316 | 11,5164 | 0,377796 | 0,15525 | 0,528402 |
| Vprbp | 20,1746 | 24,1242 | 0,257941 | 0,1554 | 0,528492 |
| Rbm25 | 74,2203 | 88,7384 | 0,257745 | 0,1554 | 0,528492 |
| Ipo7 | 110,489 | 131,833 | 0,254811 | 0,1554 | 0,528492 |
| Perm1 | 0,748623 | 1,06248 | 0,505129 | 0,1556 | 0,529032 |
| Med8 | 51,816 | 43,2667 | -0,260141 | 0,15575 | 0,529321 |
| Scarna10 | 94,8965 | 0 | -inf | 0,15585 | 0,529321 |
| S100pbp | 27,5246 | 33,4382 | 0,280778 | 0,15585 | 0,529321 |
| Ccnd1 | 114,244 | 136,448 | 0,256233 | 0,15585 | 0,529321 |
| Ufm1 | 8,99029 | 7,39849 | -0,281137 | 0,1559 | 0,52935 |
| Snrpc | 62,3276 | 50,9144 | -0,291795 | 0,15595 | 0,52938 |
| Gfpt1 | 17,1983 | 14,3382 | -0,262406 | 0,15605 | 0,529579 |
| Nit2 | 36,0693 | 29,6127 | -0,284554 | 0,1564 | 0,530486 |
| Arl6ip5 | 25,3124 | 20,5696 | -0,29933 | 0,1564 | 0,530486 |
| Whsc1l1 | 18,328 | 22,5116 | 0,296618 | 0,15665 | 0,530913 |
| Crmp1 | 14,6281 | 17,7215 | 0,276753 | 0,15665 | 0,530913 |
| Zfp937 | 1,73649 | 1,33726 | -0,376896 | 0,15665 | 0,530913 |
| Rtbdn | 0,471971 | 0,805169 | 0,770594 | 0,1568 | 0,531281 |
| Aplp2 | 81,2162 | 96,828 | 0,253657 | 0,15685 | 0,53131 |
| Asph | 27,5382 | 22,5839 | -0,286141 | 0,15715 | 0,532186 |
| Ei24 | 76,4841 | 64,2241 | -0,252046 | 0,1572 | 0,532215 |
| Sapcd1 | 1,17975 | 2,38664 | 1,0165 | 0,15745 | 0,532639 |
| Pnrc2 | 92,9822 | 77,8184 | -0,256844 | 0,15745 | 0,532639 |
| Cdk4 | 916,566 | 762,222 | -0,266027 | 0,15745 | 0,532639 |
| Ttpa | 2,35259 | 1,74377 | -0,43204 | 0,15755 | 0,532837 |
| Ttc17 | 5,7551 | 7,06073 | 0,294975 | 0,15765 | 0,533035 |
| Itgb1bp2 | 0,777661 | 1,30914 | 0,751404 | 0,15775 | 0,533092 |
| Stub1 | 110,347 | 89,7016 | -0,298839 | 0,15775 | 0,533092 |
| Ltbp4 | 2,65966 | 3,34673 | 0,331508 | 0,15805 | 0,533965 |
| Adarb1 | 2,37843 | 2,97424 | 0,322514 | 0,1583 | 0,534444 |
| Ccdc66 | 11,3391 | 15,1178 | 0,414943 | 0,15835 | 0,534444 |

| | | | | | |
|---|---|---|---|---|---|
| Pfn1 | 721,645 | 604,127 | -0,256438 | 0,15835 | 0,534444 |
| Erbb2ip | 26,6747 | 31,9876 | 0,262043 | 0,1584 | 0,534444 |
| Cops7a | 59,5159 | 49,4473 | -0,267384 | 0,1584 | 0,534444 |
| Gnb2l1 | 1442,81 | 1192,4 | -0,275013 | 0,15845 | 0,534472 |
| Scaf8 | 28,3173 | 33,7268 | 0,25221 | 0,15855 | 0,534528 |
| Serpinb9 | 0,629589 | 0,406506 | -0,631133 | 0,15855 | 0,534528 |
| Cdk5r1 | 2,08709 | 1,57792 | -0,403464 | 0,15865 | 0,534725 |
| Foxj3 | 14,0915 | 16,8935 | 0,26165 | 0,1589 | 0,535286 |
| Smco4 | 19,9307 | 15,712 | -0,343125 | 0,1589 | 0,535286 |
| Capns2 | 0,171511 | 0,929657 | 2,43839 | 0,15905 | 0,53551 |
| Cit | 5,62262 | 6,8253 | 0,279649 | 0,15905 | 0,53551 |
| Pla2g6 | 6,91621 | 5,54404 | -0,319043 | 0,1591 | 0,535538 |
| Necab2 | 9,57923 | 7,62839 | -0,328531 | 0,1592 | 0,535649 |
| Rasa4 | 0,749843 | 1,10524 | 0,559698 | 0,1593 | 0,535649 |
| Adprh | 106,992 | 89,5851 | -0,256171 | 0,1593 | 0,535649 |
| 2810417H13Rik | 116,694 | 95,7851 | -0,284852 | 0,1593 | 0,535649 |
| Slc15a4 | 10,9443 | 13,3858 | 0,290524 | 0,1595 | 0,536068 |
| Pisd-ps2 | 2,15285 | 3,0456 | 0,500477 | 0,15955 | 0,536068 |
| Slc39a7 | 103,74 | 87,0797 | -0,252566 | 0,15955 | 0,536068 |
| Pank3 | 14,5491 | 17,3574 | 0,254616 | 0,1596 | 0,536096 |
| Ehhadh | 1,12369 | 1,56081 | 0,474051 | 0,15965 | 0,536123 |
| Cldn12 | 16,7138 | 20,0456 | 0,262241 | 0,1598 | 0,536486 |
| Spast | 19,7212 | 16,4787 | -0,259138 | 0,15995 | 0,536849 |
| Ell | 7,41257 | 9,13708 | 0,301759 | 0,16 | 0,536877 |
| Hist1h2ai | 0,389409 | 1,42181 | 1,86837 | 0,1602 | 0,537407 |
| Ntmt1 | 51,1047 | 42,2718 | -0,27376 | 0,16025 | 0,537435 |
| B630005N14Rik | 8,52046 | 10,3431 | 0,279665 | 0,1606 | 0,538468 |
| Ralb | 24,304 | 29,2941 | 0,269417 | 0,16065 | 0,538494 |
| AF357399 | 101,365 | 0 | -inf | 0,1608 | 0,538716 |
| Mob1a | 44,6705 | 53,5863 | 0,26254 | 0,1608 | 0,538716 |
| Slc27a6 | 1,64047 | 2,244 | 0,451963 | 0,16085 | 0,538743 |
| Rps24 | 175,041 | 147,066 | -0,251229 | 0,1609 | 0,538769 |
| Lancl2 | 17,2643 | 14,3058 | -0,271184 | 0,16095 | 0,538796 |
| Ctgf | 9,85133 | 12,1442 | 0,30188 | 0,16145 | 0,540214 |
| Bbs5 | 9,93052 | 7,73763 | -0,359977 | 0,1615 | 0,540214 |
| Slc39a8 | 2,09349 | 1,56478 | -0,419955 | 0,1615 | 0,540214 |
| Taf4b | 3,2385 | 4,04211 | 0,319784 | 0,16165 | 0,540575 |
| Atf6b | 50,2783 | 59,9971 | 0,254956 | 0,16175 | 0,540654 |
| Psmd14 | 144,529 | 121,318 | -0,252565 | 0,1618 | 0,540654 |
| Hmgcl | 25,9422 | 21,1998 | -0,29125 | 0,1618 | 0,540654 |
| Cmss1 | 34,5731 | 27,347 | -0,338265 | 0,16185 | 0,54068 |
| Tmod2 | 3,64209 | 2,98346 | -0,287777 | 0,16205 | 0,541066 |
| Copz2 | 11,1952 | 8,47289 | -0,401956 | 0,16205 | 0,541066 |
| Asap1 | 19,4072 | 23,1702 | 0,25568 | 0,16215 | 0,541119 |
| Shkbp1 | 9,98353 | 8,04691 | -0,311115 | 0,16215 | 0,541119 |
| Cep131 | 18,086 | 21,6823 | 0,261648 | 0,16235 | 0,541416 |
| Arrb2 | 20,4024 | 16,5473 | -0,302147 | 0,16235 | 0,541416 |
| Enho | 9,72068 | 7,38955 | -0,39557 | 0,1624 | 0,541416 |
| Otud4 | 30,0064 | 35,6168 | 0,24729 | 0,16245 | 0,541416 |
| Lsm3 | 339,878 | 284,809 | -0,255022 | 0,16245 | 0,541416 |

| | | | | | |
|---|---|---|---|---|---|
| 3110082I17Rik | 27,7574 | 22,6404 | -0,293972 | 0,16255 | 0,541467 |
| Ttc7 | 1,18038 | 0,858046 | -0,460121 | 0,16255 | 0,541468 |
| Nedd4l | 1,76954 | 3,0555 | 0,788036 | 0,1629 | 0,542493 |
| 6030458C11Rik | 20,0943 | 24,2568 | 0,271603 | 0,16295 | 0,542518 |
| Cep152 | 7,7183 | 9,33844 | 0,274899 | 0,163 | 0,542544 |
| Fam171a2 | 11,9073 | 9,74868 | -0,288566 | 0,1631 | 0,542736 |
| Zfp808 | 1,97515 | 2,63518 | 0,415943 | 0,1632 | 0,542928 |
| Cda | 0,735126 | 1,39753 | 0,926818 | 0,16325 | 0,542953 |
| Lrrk1 | 2,51394 | 3,12212 | 0,312575 | 0,1634 | 0,543221 |
| Gng7 | 1,41704 | 1,91506 | 0,434507 | 0,1635 | 0,543221 |
| Narg2 | 12,1736 | 14,6917 | 0,271244 | 0,1635 | 0,543221 |
| Nat8l | 1,96867 | 1,53783 | -0,356327 | 0,1635 | 0,543221 |
| Psap | 89,618 | 106,499 | 0,248981 | 0,1637 | 0,543604 |
| Lipt2 | 12,571 | 9,87117 | -0,348808 | 0,1637 | 0,543604 |
| Rab3b | 1,44998 | 1,95534 | 0,431389 | 0,1639 | 0,544128 |
| Rtn1 | 5,56427 | 7,29399 | 0,390515 | 0,1646 | 0,546292 |
| Arl8a | 13,7718 | 16,9452 | 0,299157 | 0,16465 | 0,546292 |
| Zfp706 | 45,872 | 38,5147 | -0,252203 | 0,16475 | 0,546292 |
| Tuft1 | 1,68143 | 2,29164 | 0,446695 | 0,1648 | 0,546292 |
| Prpf4b | 60,5449 | 72,1388 | 0,252769 | 0,1648 | 0,546292 |
| Lypd2 | 0,789034 | 1,87271 | 1,24697 | 0,16485 | 0,546292 |
| Adam15 | 3,54152 | 2,7157 | -0,383042 | 0,16485 | 0,546292 |
| Rpp25l | 58,9694 | 48,5445 | -0,280659 | 0,165 | 0,546648 |
| Gimap6 | 0,997196 | 0,588065 | -0,761902 | 0,16515 | 0,547003 |
| Abcb8 | 17,0892 | 14,1465 | -0,272641 | 0,1653 | 0,547315 |
| Ppp1r15a | 9,21362 | 11,364 | 0,302631 | 0,16535 | 0,547315 |
| Snrpa1 | 153,94 | 129,37 | -0,250868 | 0,1654 | 0,547315 |
| Necap2 | 18,8447 | 15,4145 | -0,289865 | 0,16545 | 0,547315 |
| 4930558J18Rik | 1,30307 | 1,85685 | 0,510939 | 0,1655 | 0,547315 |
| Rft1 | 8,26679 | 10,2355 | 0,308178 | 0,1655 | 0,547315 |
| Nradd | 20,6865 | 25,5219 | 0,303046 | 0,16565 | 0,54767 |
| Ccdc159 | 1,23469 | 2,07267 | 0,747335 | 0,1658 | 0,548024 |
| Ier2 | 49,8577 | 41,4799 | -0,265403 | 0,1659 | 0,548214 |
| Uck2 | 115,438 | 96,8269 | -0,253635 | 0,1662 | 0,549064 |
| 2010107E04Rik | 681,033 | 570,883 | -0,254529 | 0,16625 | 0,549087 |
| Tub | 3,03569 | 2,18683 | -0,473183 | 0,1663 | 0,549111 |
| Cramp1l | 12,0446 | 14,384 | 0,256076 | 0,1664 | 0,5493 |
| Rps4x | 360,976 | 303,93 | -0,248164 | 0,1665 | 0,549347 |
| 2900097C17Rik | 12,008 | 9,97302 | -0,267897 | 0,1665 | 0,549347 |
| Helb | 4,30621 | 5,76259 | 0,4203 | 0,1666 | 0,549536 |
| Zfp451 | 23,1017 | 27,9281 | 0,273718 | 0,16665 | 0,549559 |
| Pbrm1 | 33,6698 | 40,0048 | 0,248717 | 0,1667 | 0,549583 |
| Snora75 | 58,7635 | 161,933 | 1,46241 | 0,16675 | 0,549607 |
| Cdk5rap2 | 22,1209 | 26,3766 | 0,253848 | 0,1668 | 0,54963 |
| Rbm33 | 14,8085 | 17,5846 | 0,247894 | 0,16695 | 0,549983 |
| Gtf2a1 | 7,22591 | 8,72712 | 0,272326 | 0,16705 | 0,550006 |
| Eapp | 42,0292 | 34,5735 | -0,281725 | 0,16715 | 0,550006 |
| Gm4961 | 1,18902 | 0,795259 | -0,580282 | 0,16715 | 0,550006 |
| Rsrc2 | 80,8898 | 96,4561 | 0,253916 | 0,16725 | 0,550006 |
| Gm5088 | 16,329 | 13,5401 | -0,270198 | 0,16725 | 0,550006 |

| | | | | | |
|---|---|---|---|---|---|
| Prorsd1 | 6,48585 | 5,06482 | -0,356786 | 0,16725 | 0,550006 |
| Ndufs8 | 98,4535 | 81,8081 | -0,267199 | 0,1673 | 0,550006 |
| Wnk2 | 1,62373 | 1,25809 | -0,368073 | 0,1673 | 0,550006 |
| Dgke | 3,37632 | 4,18737 | 0,310592 | 0,1674 | 0,550053 |
| Itgb3bp | 7,91682 | 6,37335 | -0,312869 | 0,1674 | 0,550053 |
| Cnih4 | 14,3322 | 11,8642 | -0,27265 | 0,1675 | 0,550241 |
| 09/15/16 | 167,383 | 140,87 | -0,248789 | 0,16765 | 0,550592 |
| Rbm43 | 5,09065 | 3,91113 | -0,380263 | 0,1678 | 0,550849 |
| Cyba | 30,6467 | 38,0048 | 0,310446 | 0,1679 | 0,550849 |
| Pdxdc1 | 34,1944 | 28,0347 | -0,286544 | 0,1679 | 0,550849 |
| Galm | 5,03802 | 3,89227 | -0,372246 | 0,1679 | 0,550849 |
| Prtn3 | 0,958284 | 0,461741 | -1,05337 | 0,16795 | 0,550873 |
| Thap11 | 46,6921 | 39,0987 | -0,25606 | 0,16805 | 0,55106 |
| Aamdc | 13,179 | 10,0204 | -0,395308 | 0,1682 | 0,551293 |
| Alg3 | 22,9332 | 18,7506 | -0,290499 | 0,16825 | 0,551293 |
| Wwox | 2,90186 | 2,14809 | -0,433923 | 0,16825 | 0,551293 |
| Stradb | 7,84619 | 6,26145 | -0,325496 | 0,16845 | 0,551807 |
| Rps3a1 | 184,248 | 155,129 | -0,24818 | 0,16855 | 0,551853 |
| Yif1b | 33,0298 | 26,9877 | -0,291467 | 0,16855 | 0,551853 |
| Lamtor3 | 15,4912 | 12,349 | -0,327048 | 0,1686 | 0,551876 |
| Ccdc34 | 57,7816 | 48,5485 | -0,251184 | 0,16875 | 0,552226 |
| Mss51 | 1,60641 | 2,2556 | 0,489677 | 0,16885 | 0,552412 |
| 1110008F13Rik | 111,792 | 94,051 | -0,249296 | 0,16895 | 0,552598 |
| Polr3k | 40,7841 | 34,2219 | -0,253087 | 0,1691 | 0,552807 |
| Zbed3 | 18,2217 | 14,9724 | -0,283345 | 0,1691 | 0,552807 |
| Zfp51 | 3,78858 | 4,88662 | 0,367181 | 0,1692 | 0,552993 |
| Dstyk | 6,26203 | 7,56119 | 0,271982 | 0,16935 | 0,553342 |
| Zbtb33 | 12,0627 | 14,4526 | 0,260781 | 0,1695 | 0,553454 |
| Rps8 | 207,306 | 174,363 | -0,249664 | 0,1695 | 0,553454 |
| Plxna3 | 10,2155 | 12,1883 | 0,254738 | 0,16955 | 0,553454 |
| Sar1b | 41,443 | 34,3061 | -0,272666 | 0,1696 | 0,553454 |
| Mrps21 | 186,7 | 154,111 | -0,276755 | 0,1696 | 0,553454 |
| Ppargc1a | 0,782992 | 0,573677 | -0,448758 | 0,1699 | 0,554292 |
| Clic4 | 107,51 | 127,683 | 0,248098 | 0,17005 | 0,5545 |
| Slitrk2 | 0,830883 | 0,620991 | -0,420073 | 0,17005 | 0,5545 |
| Zfp64 | 9,8217 | 11,9975 | 0,288687 | 0,1702 | 0,554707 |
| Rnf146 | 21,6404 | 26,1495 | 0,27306 | 0,1702 | 0,554707 |
| Mrpl43 | 50,131 | 41,5929 | -0,269367 | 0,17025 | 0,554729 |
| Hmga2 | 202,248 | 168,957 | -0,259472 | 0,1703 | 0,554751 |
| Tusc2 | 19,9914 | 16,3348 | -0,291435 | 0,17045 | 0,555098 |
| Nme1 | 296,598 | 250,015 | -0,246494 | 0,1705 | 0,55512 |
| Sdf4 | 31,5919 | 26,6675 | -0,244477 | 0,17065 | 0,555468 |
| Plekhf2 | 8,11172 | 9,95093 | 0,294824 | 0,1707 | 0,555489 |
| Clasp2 | 14,7319 | 17,6517 | 0,260866 | 0,17075 | 0,555511 |
| Mcm7 | 136,27 | 114,78 | -0,247595 | 0,1709 | 0,555858 |
| Zfp560 | 2,33132 | 2,96055 | 0,344722 | 0,17095 | 0,55588 |
| Tbca | 246,733 | 206,269 | -0,258425 | 0,171 | 0,555902 |
| Tax1bp1 | 52,2914 | 62,0438 | 0,246714 | 0,17115 | 0,556248 |
| Clns1a | 63,6237 | 53,5819 | -0,247819 | 0,17125 | 0,556432 |
| Rab20 | 0,86907 | 1,64676 | 0,922086 | 0,17145 | 0,5568 |

| | | | | | |
|---|---|---|---|---|---|
| Fopnl | 41,8365 | 34,7986 | -0,265734 | 0,17145 | 0,5568 |
| Ntn3 | 2,33288 | 1,58536 | -0,557302 | 0,17165 | 0,557309 |
| Ilf2 | 257,52 | 216,196 | -0,252343 | 0,17185 | 0,557676 |
| Fam110b | 5,25084 | 4,04776 | -0,375425 | 0,17185 | 0,557676 |
| Snord67 | 0 | 16,1249 | inf | 0,17205 | 0,558042 |
| Cox6b1 | 547,111 | 460,671 | -0,248096 | 0,17205 | 0,558042 |
| Msn | 64,0156 | 75,7208 | 0,242265 | 0,1722 | 0,558388 |
| Dqx1 | 1,83782 | 2,43538 | 0,406152 | 0,17225 | 0,558409 |
| Cacng7 | 21,0721 | 26,4154 | 0,326043 | 0,1723 | 0,55843 |
| Esrp1 | 0,687968 | 0,974013 | 0,501599 | 0,1725 | 0,558937 |
| Rps20 | 813,26 | 685,956 | -0,2456 | 0,1727 | 0,559443 |
| Pdcd6 | 51,207 | 42,8306 | -0,257697 | 0,1728 | 0,559485 |
| Psmc3ip | 49,4736 | 38,3849 | -0,366122 | 0,1728 | 0,559485 |
| Me1 | 2,26401 | 2,94146 | 0,377652 | 0,17305 | 0,560153 |
| Gm10033 | 2,32204 | 3,26058 | 0,489738 | 0,17315 | 0,560194 |
| Mccc1 | 14,1144 | 16,963 | 0,265229 | 0,17315 | 0,560194 |
| Rnaseh2b | 38,1165 | 31,7003 | -0,265919 | 0,1733 | 0,560538 |
| Txn2 | 137,808 | 115,852 | -0,250378 | 0,17335 | 0,560558 |
| Acbd3 | 19,3789 | 23,1206 | 0,254688 | 0,1737 | 0,561467 |
| Coasy | 16,2955 | 13,4597 | -0,275832 | 0,17375 | 0,561467 |
| Pnp | 19,7065 | 23,6333 | 0,262151 | 0,17385 | 0,561467 |
| Crebbp | 9,02215 | 10,7859 | 0,257599 | 0,17385 | 0,561467 |
| Smdt1 | 167,939 | 139,933 | -0,263201 | 0,17385 | 0,561467 |
| Dnajc27 | 8,55382 | 10,2651 | 0,263108 | 0,1739 | 0,561487 |
| Ndufs6 | 95,0878 | 78,0008 | -0,285772 | 0,17395 | 0,561507 |
| Cog1 | 13,6333 | 17,1481 | 0,330921 | 0,17435 | 0,562657 |
| Mrpl46 | 53,054 | 44,0142 | -0,269492 | 0,17445 | 0,562838 |
| Scfd2 | 4,09053 | 3,24319 | -0,334876 | 0,1746 | 0,56318 |
| Mrpl28 | 111,106 | 93,4589 | -0,249535 | 0,17475 | 0,563523 |
| Selk | 51,8096 | 42,8576 | -0,27367 | 0,1748 | 0,563542 |
| Zfp760 | 6,33101 | 7,74186 | 0,290244 | 0,1749 | 0,563723 |
| Pcf11 | 14,0987 | 16,7903 | 0,252068 | 0,1752 | 0,564548 |
| Ubl4 | 23,4805 | 19,0634 | -0,30066 | 0,17525 | 0,564567 |
| Adra2a | 1,29272 | 0,943936 | -0,453653 | 0,1753 | 0,564587 |
| Ccnt2 | 31,9412 | 37,8884 | 0,246339 | 0,1755 | 0,564947 |
| Arpc1a | 79,0673 | 66,5105 | -0,249499 | 0,1755 | 0,564947 |
| Cdc42bpa | 7,76641 | 6,50768 | -0,255104 | 0,1757 | 0,565449 |
| Rabggta | 9,51084 | 11,6497 | 0,292652 | 0,1759 | 0,565951 |
| Aff4 | 15,5742 | 18,4203 | 0,242132 | 0,17625 | 0,566935 |
| Lrig3 | 28,9462 | 34,3157 | 0,245493 | 0,17645 | 0,567293 |
| Akap12 | 6,51177 | 5,39354 | -0,271817 | 0,17645 | 0,567293 |
| Ube3c | 27,3397 | 32,3306 | 0,241904 | 0,1766 | 0,567633 |
| Bmf | 12,3841 | 10,3327 | -0,261275 | 0,17665 | 0,567639 |
| Ryr1 | 0,470204 | 0,60814 | 0,371117 | 0,1767 | 0,567639 |
| Eif3e | 106,312 | 89,6811 | -0,245427 | 0,17675 | 0,567639 |
| Fam98c | 5,51787 | 7,6532 | 0,47195 | 0,1769 | 0,567639 |
| D630045M09Rik | 0,703566 | 0,353446 | -0,993196 | 0,1769 | 0,567639 |
| Lrrtm1 | 0,746453 | 0,138929 | -2,42571 | 0,1769 | 0,567639 |
| Rtn4rl2 | 1,09993 | 0,640351 | -0,780478 | 0,17695 | 0,567639 |
| Mib1 | 22,5518 | 26,7466 | 0,24611 | 0,177 | 0,567639 |

| | | | | | |
|---|---|---|---|---|---|
| Pnisr | 45,6119 | 54,0082 | 0,243769 | 0,177 | 0,567639 |
| Nxt2 | 3,15096 | 4,08514 | 0,374591 | 0,17725 | 0,568299 |
| Slc8a1 | 0,747677 | 0,93405 | 0,321084 | 0,1774 | 0,568637 |
| Pdia4 | 135,844 | 160,844 | 0,243713 | 0,1775 | 0,568816 |
| Scaf11 | 46,7097 | 55,249 | 0,242227 | 0,1776 | 0,568994 |
| A930013F10Rik | 3,17218 | 4,96918 | 0,647535 | 0,17775 | 0,569333 |
| Psmc3 | 210,717 | 177,98 | -0,243589 | 0,178 | 0,569991 |
| Fgd2 | 0,391539 | 0,622546 | 0,669024 | 0,17815 | 0,570329 |
| AA388235 | 3,63203 | 2,87013 | -0,339663 | 0,1783 | 0,570667 |
| Cbx4 | 1,014 | 0,749289 | -0,436466 | 0,1784 | 0,57072 |
| Inadl | 4,35692 | 5,46492 | 0,326894 | 0,17845 | 0,57072 |
| Cdh15 | 0,770108 | 0,0203295 | -5,24341 | 0,17845 | 0,57072 |
| Rpl22 | 203,934 | 172,555 | -0,241047 | 0,1785 | 0,570737 |
| Dkk1 | 19,1969 | 23,0843 | 0,266038 | 0,17865 | 0,570932 |
| Timm13 | 73,7215 | 62,0573 | -0,248485 | 0,17865 | 0,570932 |
| Birc5 | 196,924 | 166,114 | -0,245466 | 0,1788 | 0,571269 |
| Zfp644 | 11,8975 | 14,1911 | 0,254331 | 0,17895 | 0,571606 |
| Slc8b1 | 2,08872 | 2,7473 | 0,395394 | 0,17915 | 0,572103 |
| Ctrl | 1,05483 | 1,77314 | 0,749297 | 0,17925 | 0,572137 |
| Nmt2 | 15,8296 | 18,8064 | 0,248594 | 0,17925 | 0,572137 |
| Mid2 | 0,843672 | 1,12147 | 0,410638 | 0,1795 | 0,572793 |
| Hoxa6 | 36,3871 | 61,021 | 0,74588 | 0,17965 | 0,573129 |
| Zbtb21 | 5,46641 | 6,62237 | 0,276753 | 0,1798 | 0,573356 |
| Rcan1 | 24,675 | 20,6558 | -0,2565 | 0,1799 | 0,573356 |
| Psmg1 | 77,4031 | 64,719 | -0,258201 | 0,1799 | 0,573356 |
| Il11ra1 | 7,48237 | 5,75594 | -0,378442 | 0,1799 | 0,573356 |
| Zfp54 | 1,08686 | 0,739873 | -0,554817 | 0,18025 | 0,574329 |
| Bud31 | 201,937 | 166,993 | -0,274112 | 0,18035 | 0,574505 |
| Nckap1 | 61,0218 | 71,9871 | 0,238413 | 0,18045 | 0,57468 |
| Myo9b | 14,0326 | 16,6096 | 0,243235 | 0,1806 | 0,574873 |
| 5930430L01Rik | 0,825364 | 0,569207 | -0,536078 | 0,1806 | 0,574873 |
| Ctps2 | 11,6912 | 17,36 | 0,570343 | 0,18075 | 0,575114 |
| Fyttd1 | 34,7736 | 41,0649 | 0,239913 | 0,1808 | 0,575114 |
| 1700084E18Rik | 1,32264 | 2,76321 | 1,06293 | 0,1809 | 0,575114 |
| Mars2 | 4,34691 | 5,4478 | 0,325685 | 0,1809 | 0,575114 |
| Abl2 | 10,3276 | 12,2375 | 0,244793 | 0,1809 | 0,575114 |
| Mrpl14 | 54,2369 | 44,1326 | -0,29743 | 0,18125 | 0,575957 |
| Vapb | 15,3732 | 12,9782 | -0,244318 | 0,1813 | 0,575958 |
| Cdc7 | 31,5173 | 26,5728 | -0,246194 | 0,1813 | 0,575957 |
| Camkk2 | 7,03312 | 8,4632 | 0,26704 | 0,18155 | 0,576562 |
| 4930506C21Rik | 0,292879 | 0,86748 | 1,56652 | 0,1816 | 0,576562 |
| Dscc1 | 19,9946 | 16,2787 | -0,296623 | 0,18165 | 0,576562 |
| Rnaseh2a | 25,569 | 21,3231 | -0,261976 | 0,18175 | 0,576562 |
| Ranbp1 | 679,956 | 575,277 | -0,241184 | 0,1818 | 0,576562 |
| Mettl16 | 37,2547 | 31,4394 | -0,244852 | 0,1818 | 0,576562 |
| Kansl1 | 30,0581 | 35,4623 | 0,238533 | 0,18185 | 0,576562 |
| Thsd1 | 0,837914 | 0,593352 | -0,497914 | 0,18185 | 0,576562 |
| Gls | 22,7488 | 26,9241 | 0,243111 | 0,18195 | 0,576737 |
| Helz | 10,8623 | 12,9102 | 0,249174 | 0,18225 | 0,577545 |
| Zmynd19 | 52,4278 | 43,8366 | -0,258194 | 0,1823 | 0,577561 |

| | | | | | |
|---|---|---|---|---|---|
| Cep350 | 5,52017 | 6,54666 | 0,246046 | 0,1824 | 0,577735 |
| 2810459M11Rik | 0,501346 | 0,743828 | 0,569162 | 0,18255 | 0,577925 |
| Dctn5 | 62,8674 | 52,9457 | -0,247797 | 0,18255 | 0,577925 |
| Fert2 | 9,23794 | 11,1724 | 0,27429 | 0,1828 | 0,578574 |
| Cspg4 | 1,15025 | 1,46637 | 0,350295 | 0,18285 | 0,578589 |
| Suv420h1 | 15,3522 | 18,3555 | 0,257765 | 0,18295 | 0,578763 |
| Kif16b | 8,17121 | 6,80865 | -0,263182 | 0,18305 | 0,578936 |
| Mdm4 | 24,3 | 28,787 | 0,244461 | 0,1831 | 0,578952 |
| R3hcc1 | 40,3943 | 33,8385 | -0,255485 | 0,18335 | 0,579599 |
| Ccl25 | 4,61627 | 6,20407 | 0,426486 | 0,18355 | 0,580089 |
| Ckap2l | 39,5742 | 33,4424 | -0,242882 | 0,1838 | 0,580736 |
| Pear1 | 0,735132 | 1,01333 | 0,463028 | 0,18405 | 0,581239 |
| Nid2 | 61,8052 | 74,5831 | 0,27112 | 0,18405 | 0,581239 |
| Plekho1 | 85,202 | 71,9363 | -0,244168 | 0,1841 | 0,581254 |
| C030034I22Rik | 0,693322 | 1,00884 | 0,541099 | 0,1842 | 0,581427 |
| Mrpl47 | 50,2415 | 41,5396 | -0,274393 | 0,1845 | 0,58223 |
| Mtg2 | 14,4171 | 11,7011 | -0,301145 | 0,1846 | 0,582403 |
| Zfp637 | 36,2475 | 29,9242 | -0,276568 | 0,1847 | 0,582575 |
| Tada1 | 30,586 | 22,0186 | -0,474151 | 0,1848 | 0,582747 |
| Xrcc4 | 5,12517 | 6,65981 | 0,377882 | 0,18495 | 0,582933 |
| Npdc1 | 4,94828 | 3,72389 | -0,410118 | 0,18495 | 0,582933 |
| Slc2a8 | 4,86996 | 6,19987 | 0,348327 | 0,18505 | 0,583105 |
| Lrrc8c | 4,10134 | 3,2096 | -0,353704 | 0,1851 | 0,58312 |
| Armc5 | 6,12179 | 7,63926 | 0,31948 | 0,18515 | 0,583134 |
| Evi5 | 10,9497 | 13,0232 | 0,250189 | 0,1852 | 0,583148 |
| Eef1a1 | 1647,3 | 1355,27 | -0,281528 | 0,1853 | 0,58332 |
| D930015E06Rik | 28,4198 | 24,0325 | -0,241908 | 0,1854 | 0,583491 |
| Ank | 5,32248 | 4,28592 | -0,312492 | 0,1855 | 0,583663 |
| Zer1 | 9,72074 | 11,6324 | 0,259011 | 0,18575 | 0,584306 |
| Actr2 | 59,2688 | 69,9588 | 0,239231 | 0,1859 | 0,584635 |
| Mir3535 | 27,4387 | 57,0723 | 1,05658 | 0,18595 | 0,584649 |
| Kcna6 | 1,48425 | 1,89707 | 0,354041 | 0,1861 | 0,58469 |
| Naa30 | 10,6805 | 12,7633 | 0,257018 | 0,1861 | 0,58469 |
| Eif4e3 | 10,1785 | 8,32453 | -0,290088 | 0,1861 | 0,58469 |
| Slc20a1 | 52,0763 | 43,9981 | -0,243185 | 0,18625 | 0,584889 |
| Lias | 31,5159 | 26,3336 | -0,259176 | 0,1863 | 0,584889 |
| Asb13 | 9,9865 | 8,11042 | -0,300202 | 0,1863 | 0,584889 |
| Myo1c | 4,40133 | 3,59042 | -0,293787 | 0,18635 | 0,584903 |
| Mllt10 | 20,1182 | 16,4017 | -0,294657 | 0,1865 | 0,58523 |
| Echs1 | 104,864 | 88,7209 | -0,241169 | 0,18675 | 0,585872 |
| Slc25a28 | 12,1015 | 14,869 | 0,29713 | 0,1868 | 0,585885 |
| Ndufb8 | 320,729 | 271,51 | -0,240345 | 0,187 | 0,586021 |
| Fem1c | 19,7011 | 23,4649 | 0,252232 | 0,18705 | 0,586021 |
| Dld | 96,0529 | 81,4166 | -0,238507 | 0,18705 | 0,586021 |
| Rplp1 | 1795,28 | 1519,33 | -0,240775 | 0,18705 | 0,586021 |
| Ndufb2 | 115,609 | 95,4455 | -0,276504 | 0,1872 | 0,586021 |
| Rtfdc1 | 33,0356 | 27,8046 | -0,248698 | 0,18725 | 0,586021 |
| Hist1h4b | 1,53913 | 0,564099 | -1,44809 | 0,18725 | 0,586021 |
| Zcchc9 | 21,321 | 25,7072 | 0,269896 | 0,1873 | 0,586021 |
| Cul5 | 13,4764 | 15,947 | 0,242851 | 0,1873 | 0,586021 |

| | | | | | |
|---|---|---|---|---|---|
| Sertad3 | 6,55378 | 4,95396 | -0,403747 | 0,1873 | 0,586021 |
| Eif3g | 246,434 | 208,817 | -0,238964 | 0,1874 | 0,58619 |
| Cab39l | 19,245 | 16,0979 | -0,257608 | 0,18745 | 0,586204 |
| Krt7 | 0,870824 | 1,34086 | 0,622703 | 0,18765 | 0,586543 |
| Stxbp3a | 7,88429 | 9,66366 | 0,293589 | 0,18765 | 0,586543 |
| Fam214b | 4,24628 | 3,36231 | -0,336749 | 0,1877 | 0,586557 |
| Tceal5 | 2,70972 | 1,81034 | -0,581887 | 0,18785 | 0,586883 |
| Psmd10 | 22,38 | 18,4449 | -0,278991 | 0,188 | 0,587208 |
| B3gnt1 | 14,0068 | 11,4693 | -0,288347 | 0,18815 | 0,587534 |
| Wdyhv1 | 19,3015 | 15,7356 | -0,294675 | 0,1882 | 0,587547 |
| Ankib1 | 6,8892 | 8,22879 | 0,256343 | 0,18825 | 0,58756 |
| Calm2 | 206,087 | 174,701 | -0,238367 | 0,1883 | 0,587573 |
| Scarna13 | 1,47653 | 0 | -inf | 0,1884 | 0,587742 |
| Sidt2 | 14,1824 | 16,8089 | 0,24512 | 0,18845 | 0,587755 |
| Fech | 9,04629 | 7,41963 | -0,285979 | 0,1885 | 0,587768 |
| Snord42b | 0 | 2645,8 | inf | 0,1887 | 0,588249 |
| Arhgap17 | 15,061 | 17,9685 | 0,254652 | 0,1888 | 0,588417 |
| Mrpl13 | 195,999 | 165,635 | -0,242838 | 0,18895 | 0,588742 |
| Zbtb8a | 3,46201 | 4,47869 | 0,371467 | 0,189 | 0,588755 |
| Kdm5d | 7,04419 | 8,44115 | 0,261005 | 0,18915 | 0,588936 |
| 1110038B12Rik | 299,648 | 252,959 | -0,244364 | 0,18915 | 0,588936 |
| Tex26 | 0,587825 | 1,03843 | 0,820944 | 0,18945 | 0,589727 |
| Slc39a13 | 6,66443 | 5,30005 | -0,330475 | 0,18955 | 0,589734 |
| Satb1 | 9,94718 | 8,36661 | -0,249644 | 0,18965 | 0,589734 |
| Coa5 | 36,7117 | 31,1292 | -0,237972 | 0,1897 | 0,589734 |
| Hist1h2be | 0,248515 | 0,592316 | 1,25303 | 0,18975 | 0,589734 |
| Frmd6 | 12,2665 | 14,593 | 0,250554 | 0,18975 | 0,589734 |
| Raet1d | 5,44624 | 3,79505 | -0,52114 | 0,1898 | 0,589734 |
| Chkb | 16,6873 | 20,2807 | 0,28136 | 0,18985 | 0,589734 |
| Snord15a | 11,7966 | 18,4672 | 0,646594 | 0,1899 | 0,589734 |
| Fancd2os | 1,35064 | 0,860998 | -0,649563 | 0,1899 | 0,589734 |
| Krit1 | 10,1632 | 12,0503 | 0,245711 | 0,18995 | 0,589734 |
| Galt | 5,94917 | 7,47534 | 0,32945 | 0,19 | 0,589734 |
| Isy1 | 66,0013 | 55,7199 | -0,244302 | 0,19005 | 0,589734 |
| Ndufb7 | 184,277 | 154,737 | -0,252056 | 0,19005 | 0,589734 |
| Stamos | 0,914763 | 1,61427 | 0,819415 | 0,1901 | 0,589747 |
| Ccdc109b | 0,41157 | 0,751611 | 0,868849 | 0,19015 | 0,589759 |
| Asap2 | 12,4369 | 15,0003 | 0,270368 | 0,19025 | 0,589927 |
| Arhgef3 | 1,97773 | 2,55366 | 0,36872 | 0,1905 | 0,590417 |
| Ubr7 | 46,0176 | 39,0215 | -0,237914 | 0,1905 | 0,590417 |
| Rab6b | 1,94909 | 2,47888 | 0,346884 | 0,1906 | 0,590584 |
| Phf21a | 13,2149 | 15,6463 | 0,243659 | 0,1907 | 0,590608 |
| C2cd2l | 3,07386 | 2,44435 | -0,3306 | 0,1907 | 0,590608 |
| Cacna1c | 2,33999 | 1,88608 | -0,311114 | 0,19075 | 0,590621 |
| Mina | 21,5942 | 17,5038 | -0,302982 | 0,19085 | 0,590788 |
| Arhgef18 | 7,31158 | 8,74349 | 0,258025 | 0,191 | 0,591109 |
| Ntng2 | 0,412674 | 0,821014 | 0,992403 | 0,1911 | 0,591194 |
| Ecscr | 2,8004 | 3,9268 | 0,487719 | 0,19115 | 0,591194 |
| 1700052N19Rik | 22,9199 | 19,1744 | -0,257419 | 0,1912 | 0,591194 |
| Asna1 | 94,1466 | 79,6622 | -0,241013 | 0,19125 | 0,591194 |

| | | | | | |
|---|---|---|---|---|---|
| Ube2e1 | 66,9748 | 56,1464 | -0,254425 | 0,1913 | 0,591194 |
| Vps72 | 98,2666 | 83,2427 | -0,239378 | 0,19135 | 0,591194 |
| Ndufa1 | 145,767 | 121,881 | -0,258184 | 0,19135 | 0,591194 |
| Polq | 7,54245 | 8,93732 | 0,244808 | 0,1914 | 0,591206 |
| Nudc | 168,282 | 142,831 | -0,236572 | 0,1917 | 0,591848 |
| Tmem159 | 7,44926 | 5,78842 | -0,363927 | 0,1917 | 0,591848 |
| Chp1 | 28,9694 | 24,3915 | -0,248149 | 0,1919 | 0,592073 |
| Klf8 | 0,605373 | 0,842071 | 0,476119 | 0,19195 | 0,592073 |
| Sppl2a | 10,8788 | 12,861 | 0,241483 | 0,19195 | 0,592073 |
| 2810004N23Rik | 86,7616 | 73,3822 | -0,241627 | 0,192 | 0,592073 |
| Sirpa | 8,80446 | 10,5706 | 0,263755 | 0,19205 | 0,592073 |
| Serp2 | 5,4375 | 3,87389 | -0,48916 | 0,19205 | 0,592073 |
| Ppip5k2 | 22,8139 | 26,8872 | 0,237006 | 0,1921 | 0,592085 |
| Zfp317 | 15,4348 | 18,3221 | 0,247395 | 0,19225 | 0,592405 |
| Spire1 | 15,2355 | 18,0084 | 0,241233 | 0,19235 | 0,592571 |
| Cep57 | 34,9224 | 41,1429 | 0,236488 | 0,1925 | 0,592891 |
| Zfp473 | 1,98726 | 2,55249 | 0,361122 | 0,19255 | 0,592902 |
| Ecm1 | 1,99311 | 2,69169 | 0,433493 | 0,19285 | 0,593667 |
| Tmtc2 | 2,11971 | 1,6757 | -0,339101 | 0,1929 | 0,593667 |
| Vps45 | 8,48937 | 6,91749 | -0,295407 | 0,193 | 0,593667 |
| Fkbp9 | 70,0589 | 59,618 | -0,232822 | 0,19305 | 0,593667 |
| Zfhx2 | 1,34308 | 1,05482 | -0,348552 | 0,1931 | 0,593667 |
| Mtrf1l | 7,0312 | 8,62084 | 0,294058 | 0,19325 | 0,593667 |
| Bptf | 19,6985 | 23,2212 | 0,237363 | 0,1933 | 0,593667 |
| Knstrn | 36,3176 | 30,7916 | -0,238132 | 0,1933 | 0,593667 |
| Tubb2a | 7,05164 | 8,87755 | 0,332203 | 0,19335 | 0,593667 |
| Slc4a3 | 4,90473 | 5,98124 | 0,286271 | 0,19335 | 0,593667 |
| Glrx5 | 100,571 | 84,8733 | -0,244837 | 0,19335 | 0,593667 |
| Baz2a | 20,4728 | 24,1138 | 0,236154 | 0,1934 | 0,593667 |
| Cdt1 | 54,4557 | 46,1909 | -0,237473 | 0,1934 | 0,593667 |
| 2610306M01Rik | 6,88771 | 5,1818 | -0,410571 | 0,1935 | 0,593832 |
| Sdhc | 135,428 | 114,953 | -0,236482 | 0,19355 | 0,593843 |
| Wdr62 | 6,32841 | 7,63298 | 0,270403 | 0,1936 | 0,593855 |
| Rnd2 | 10,0281 | 7,44527 | -0,429658 | 0,19365 | 0,593866 |
| Slc9b2 | 2,0597 | 2,73819 | 0,410788 | 0,1937 | 0,593878 |
| Rbl2 | 2,71486 | 3,63425 | 0,420779 | 0,19395 | 0,594502 |
| Commd4 | 36,7665 | 30,1099 | -0,288154 | 0,1943 | 0,595432 |
| Brms1l | 18,3208 | 15,3159 | -0,258453 | 0,1944 | 0,595487 |
| Fam96a | 90,6998 | 76,7907 | -0,240167 | 0,19445 | 0,595487 |
| Atp5j2 | 123,239 | 102,29 | -0,268796 | 0,1945 | 0,595487 |
| Nup153 | 47,6888 | 56,0271 | 0,232474 | 0,19455 | 0,595487 |
| Piezo2 | 0,847828 | 0,660361 | -0,360517 | 0,19455 | 0,595487 |
| Zfp719 | 4,51547 | 5,47674 | 0,278441 | 0,1946 | 0,595498 |
| Efcab14 | 10,9704 | 13,0625 | 0,251816 | 0,19465 | 0,595509 |
| Snord53 | 0 | 1122,95 | inf | 0,1947 | 0,59552 |
| Brd2 | 96,058 | 112,748 | 0,23113 | 0,19495 | 0,596022 |
| Eif4g3 | 50,2343 | 58,9482 | 0,230776 | 0,195 | 0,596022 |
| Tpd52l2 | 35,6794 | 30,2974 | -0,235898 | 0,19505 | 0,596022 |
| Mtx1 | 41,1241 | 34,6137 | -0,248639 | 0,19505 | 0,596022 |
| Uba3 | 44,9339 | 52,9475 | 0,23676 | 0,1952 | 0,596197 |

| | | | | | |
|---|---|---|---|---|---|
| Tpgs1 | 29,4998 | 24,3251 | -0,278264 | 0,1952 | 0,596197 |
| Urah | 0,309895 | 0,883897 | 1,5121 | 0,1953 | 0,59636 |
| Cldn5 | 2,7906 | 1,98309 | -0,492824 | 0,19535 | 0,596371 |
| Ppp1r13l | 3,3196 | 4,1771 | 0,331493 | 0,1954 | 0,596381 |
| Nck2 | 13,6635 | 16,3335 | 0,257508 | 0,19565 | 0,596825 |
| Crabp1 | 300,587 | 255,892 | -0,232247 | 0,1957 | 0,596825 |
| Hoxa3 | 13,7294 | 20,1771 | 0,555445 | 0,1959 | 0,596825 |
| Sergef | 6,2353 | 4,82819 | -0,368976 | 0,1959 | 0,596825 |
| Bclaf1 | 79,1427 | 93,2345 | 0,236407 | 0,19595 | 0,596825 |
| Snx3 | 79,5925 | 67,5587 | -0,236491 | 0,19595 | 0,596825 |
| Zfp935 | 4,97671 | 6,21363 | 0,320245 | 0,196 | 0,596825 |
| Zfp862-ps | 6,46172 | 7,69884 | 0,252724 | 0,196 | 0,596825 |
| Ptprf | 41,6005 | 48,9167 | 0,233727 | 0,19605 | 0,596825 |
| Pim1 | 30,4252 | 36,2625 | 0,253213 | 0,1961 | 0,596825 |
| Nudt21 | 10,6752 | 8,36585 | -0,351677 | 0,1961 | 0,596825 |
| Mboat7 | 12,5698 | 15,0022 | 0,25522 | 0,19615 | 0,596825 |
| Fndc3a | 18,5646 | 21,8445 | 0,234718 | 0,19615 | 0,596825 |
| Gpr63 | 2,52744 | 3,30288 | 0,386049 | 0,1963 | 0,597008 |
| Fam192a | 50,8441 | 43,0645 | -0,239582 | 0,19635 | 0,597008 |
| 1190002F15Rik | 14,2849 | 10,7536 | -0,409678 | 0,19635 | 0,597008 |
| Casq1 | 0,643857 | 0,995466 | 0,628633 | 0,1964 | 0,597019 |
| Unc93b1 | 0,919756 | 1,33224 | 0,534528 | 0,19645 | 0,597029 |
| Icmt | 17,3006 | 20,3898 | 0,237029 | 0,1965 | 0,59704 |
| Slc35b4 | 13,4902 | 16,0589 | 0,251463 | 0,19655 | 0,59705 |
| Zfp52 | 0,633018 | 0,912656 | 0,527826 | 0,1966 | 0,597061 |
| Sar1a | 98,8161 | 84,23 | -0,230412 | 0,19665 | 0,597071 |
| Gdpgp1 | 2,21781 | 1,70658 | -0,378031 | 0,1967 | 0,597082 |
| Rnf11 | 23,1429 | 27,5492 | 0,251441 | 0,19695 | 0,597699 |
| Dctd | 54,6704 | 46,2611 | -0,24096 | 0,197 | 0,59771 |
| Rpl4 | 1352,12 | 1132,64 | -0,255536 | 0,19705 | 0,59772 |
| Adamts19 | 1,82983 | 1,41752 | -0,368343 | 0,1971 | 0,59773 |
| 1110019D14Rik | 1,264 | 1,8367 | 0,539118 | 0,19715 | 0,597741 |
| Rbfox2 | 36,564 | 31,0522 | -0,23573 | 0,1973 | 0,598054 |
| Vps37a | 6,24156 | 8,86516 | 0,50624 | 0,1975 | 0,598377 |
| Zfp442 | 2,25683 | 1,69466 | -0,413303 | 0,1975 | 0,598377 |
| Rbm12b2 | 5,7002 | 6,93931 | 0,28378 | 0,1976 | 0,598539 |
| Ube2e3 | 117,792 | 100,07 | -0,235231 | 0,19775 | 0,598721 |
| Cd1d1 | 3,24354 | 2,43056 | -0,416283 | 0,1978 | 0,598721 |
| Fbxo32 | 0,694422 | 0,515887 | -0,428758 | 0,1978 | 0,598721 |
| AA543186 | 0,976379 | 1,43857 | 0,559126 | 0,19785 | 0,598731 |
| Lig1 | 101,395 | 86,026 | -0,237147 | 0,19815 | 0,599234 |
| Mir1839 | 281,387 | 0 | -inf | 0,1982 | 0,599234 |
| Erg | 0,844488 | 1,23119 | 0,543903 | 0,19825 | 0,599234 |
| Ftl1 | 352,137 | 300,177 | -0,230325 | 0,19825 | 0,599234 |
| Aar2 | 19,1379 | 16,1517 | -0,244742 | 0,19825 | 0,599234 |
| Ppp1r21 | 9,47715 | 11,3936 | 0,265698 | 0,19845 | 0,599698 |
| Alkbh8 | 14,2922 | 17,1335 | 0,261592 | 0,1985 | 0,599707 |
| Lypla2 | 67,0167 | 56,8896 | -0,236355 | 0,19855 | 0,599717 |
| Gba | 13,7345 | 11,2419 | -0,28892 | 0,1986 | 0,599727 |
| Rgp1 | 19,3033 | 23,7133 | 0,296853 | 0,19865 | 0,599737 |

| | | | | | |
|---|---|---|---|---|---|
| Rabgap1 | 21,3233 | 25,0302 | 0,231241 | 0,1988 | 0,600049 |
| Snx15 | 10,1455 | 8,1201 | -0,321269 | 0,1989 | 0,600209 |
| Shc2 | 12,7931 | 10,7295 | -0,253776 | 0,19905 | 0,600521 |
| Ikzf4 | 1,52995 | 2,00266 | 0,388433 | 0,1991 | 0,60053 |
| Tmem101 | 16,3643 | 19,7691 | 0,272694 | 0,1992 | 0,600549 |
| Chaf1b | 57,4258 | 48,784 | -0,23529 | 0,1992 | 0,600549 |
| 2410004I01Rik | 1,36231 | 0,805454 | -0,758184 | 0,19935 | 0,60086 |
| Ndufb6 | 137,744 | 115,602 | -0,252821 | 0,1994 | 0,60087 |
| Ppapdc1a | 0,380092 | 0,759835 | 0,999337 | 0,19945 | 0,600879 |
| Mpc1 | 1,28483 | 2,30166 | 0,841089 | 0,19955 | 0,600908 |
| Klf16 | 10,9914 | 13,1916 | 0,26324 | 0,1996 | 0,600908 |
| Rpa2 | 107,07 | 91,0415 | -0,233959 | 0,1996 | 0,600908 |
| Arap1 | 2,60251 | 3,27311 | 0,330757 | 0,19965 | 0,600917 |
| Emp3 | 4,46458 | 6,1835 | 0,469898 | 0,1997 | 0,600927 |
| Vcl | 25,2305 | 29,6745 | 0,234052 | 0,2 | 0,601689 |
| Mrps22 | 60,0746 | 50,7137 | -0,24438 | 0,20005 | 0,601698 |
| Tti1 | 17,8637 | 21,1221 | 0,241726 | 0,2001 | 0,601707 |
| Casp7 | 5,70988 | 7,07029 | 0,308309 | 0,2003 | 0,602167 |
| Pnck | 0,607193 | 0,955331 | 0,653845 | 0,20035 | 0,602177 |
| Pip4k2b | 28,0364 | 23,8981 | -0,230405 | 0,20045 | 0,602195 |
| Tmem165 | 69,2077 | 56,1041 | -0,302827 | 0,20045 | 0,602195 |
| Gpm6a | 0,600283 | 0,862827 | 0,523429 | 0,2008 | 0,602964 |
| Jmjd7 | 11,6893 | 15,7443 | 0,429638 | 0,2008 | 0,602964 |
| Disp1 | 4,9417 | 5,97291 | 0,273427 | 0,2009 | 0,602982 |
| Psma2 | 233,43 | 198,722 | -0,232236 | 0,2009 | 0,602982 |
| Zfp787 | 15,0972 | 12,4727 | -0,275504 | 0,201 | 0,603009 |
| Zmym3 | 15,277 | 18,6707 | 0,289415 | 0,20105 | 0,603009 |
| Sgpp2 | 2,498 | 1,0818 | -1,20733 | 0,20105 | 0,603009 |
| Tmem74b | 0,720257 | 1,14912 | 0,673946 | 0,2011 | 0,603018 |
| Akr1c18 | 1,07943 | 0,635516 | -0,764273 | 0,20115 | 0,603027 |
| Fbxw7 | 8,68552 | 10,3786 | 0,256928 | 0,20125 | 0,603045 |
| Tmem208 | 46,4259 | 38,5021 | -0,269995 | 0,20125 | 0,603045 |
| Cpe | 28,2886 | 23,8338 | -0,247211 | 0,2013 | 0,603054 |
| Rrp7a | 28,4274 | 23,686 | -0,263247 | 0,20145 | 0,603362 |
| Sort1 | 14,3977 | 16,9241 | 0,233244 | 0,2015 | 0,603371 |
| Chd6 | 14,2551 | 16,7572 | 0,233306 | 0,2016 | 0,60353 |
| Ly96 | 1,78726 | 0,92782 | -0,94583 | 0,2017 | 0,603688 |
| Gpam | 10,6105 | 8,87457 | -0,25775 | 0,2018 | 0,603715 |
| Avl9 | 6,42601 | 7,63634 | 0,248958 | 0,20185 | 0,603715 |
| Mamld1 | 0,689662 | 0,468197 | -0,558774 | 0,20185 | 0,603715 |
| Uckl1 | 9,05849 | 12,8735 | 0,507067 | 0,20195 | 0,603873 |
| Fiz1 | 31,9724 | 27,1295 | -0,236961 | 0,2021 | 0,604066 |
| Impa2 | 11,3806 | 9,12771 | -0,318255 | 0,20215 | 0,604066 |
| Tstd3 | 14,4191 | 10,5593 | -0,449477 | 0,2022 | 0,604066 |
| Dusp28 | 4,76845 | 6,13366 | 0,363226 | 0,20225 | 0,604066 |
| Rfx2 | 1,5935 | 1,19121 | -0,419774 | 0,20225 | 0,604066 |
| Mex3c | 30,7742 | 26,1853 | -0,232967 | 0,20265 | 0,604996 |
| Mlec | 111,702 | 131,915 | 0,239959 | 0,2027 | 0,604996 |
| Coq10a | 3,49879 | 4,43239 | 0,34123 | 0,20275 | 0,604996 |
| Zfp318 | 8,04708 | 9,76019 | 0,278445 | 0,20275 | 0,604996 |

| | | | | | |
|---|---|---|---|---|---|
| Slc9a3r2 | 4,02541 | 3,0938 | -0,379757 | 0,2028 | 0,605004 |
| Mturn | 1,4044 | 1,07345 | -0,387691 | 0,203 | 0,60546 |
| Prdx6b | 4,05799 | 3,20741 | -0,339358 | 0,2031 | 0,605618 |
| Mkx | 1,89874 | 2,46644 | 0,377388 | 0,20315 | 0,605626 |
| Rps19bp1 | 59,0082 | 49,3921 | -0,256635 | 0,20325 | 0,605783 |
| Zfp866 | 10,3628 | 12,2568 | 0,242175 | 0,2034 | 0,606089 |
| Gramd1a | 32,4262 | 38,1886 | 0,235981 | 0,2035 | 0,606247 |
| Mpp3 | 6,37353 | 5,18367 | -0,298121 | 0,20365 | 0,606412 |
| Scrn3 | 3,79066 | 3,00861 | -0,33335 | 0,20365 | 0,606412 |
| Peli2 | 11,3303 | 9,59449 | -0,239907 | 0,2038 | 0,606718 |
| Itga10 | 1,08496 | 1,41412 | 0,382266 | 0,20415 | 0,607204 |
| Mrpl40 | 64,129 | 54,1181 | -0,244867 | 0,20415 | 0,607204 |
| Tmem53 | 0,935225 | 1,50117 | 0,682705 | 0,2042 | 0,607204 |
| Spred1 | 24,9678 | 29,625 | 0,246751 | 0,2042 | 0,607204 |
| Pbx2 | 100,888 | 86,1974 | -0,227044 | 0,2042 | 0,607204 |
| She | 0,666922 | 0,888069 | 0,413154 | 0,20505 | 0,60959 |
| Heatr5b | 6,35886 | 7,57517 | 0,252511 | 0,20515 | 0,609605 |
| Imp4 | 49,0788 | 41,7374 | -0,233758 | 0,20515 | 0,609605 |
| Ndufa5 | 126,423 | 106,459 | -0,247967 | 0,2052 | 0,609612 |
| Gpr107 | 15,194 | 17,9525 | 0,240685 | 0,20535 | 0,609917 |
| Aldh2 | 57,3632 | 48,9406 | -0,229094 | 0,20545 | 0,610072 |
| Zufsp | 6,3944 | 7,88638 | 0,302553 | 0,2056 | 0,610242 |
| Ctsl | 117,651 | 137,722 | 0,227248 | 0,20565 | 0,610242 |
| Setd1a | 24,8844 | 29,1256 | 0,227044 | 0,20565 | 0,610242 |
| Dbndd1 | 4,43085 | 5,70997 | 0,365899 | 0,2057 | 0,61025 |
| Ctxn1 | 37,5848 | 31,541 | -0,252923 | 0,2059 | 0,610702 |
| Dmxl2 | 3,15541 | 3,76469 | 0,2547 | 0,20615 | 0,611019 |
| Zfp362 | 54,8804 | 64,3262 | 0,229115 | 0,20615 | 0,611019 |
| Psmd7 | 192,791 | 164,799 | -0,226329 | 0,20615 | 0,611019 |
| Tcfl5 | 1,66588 | 2,36107 | 0,50316 | 0,2063 | 0,611048 |
| Hmgn5 | 19,7944 | 23,607 | 0,254123 | 0,2063 | 0,611048 |
| Cyb5d2 | 1,18737 | 1,63008 | 0,457182 | 0,20635 | 0,611048 |
| Smn1 | 76,7755 | 65,3787 | -0,231824 | 0,20635 | 0,611048 |
| Pth1r | 41,9916 | 35,7741 | -0,231187 | 0,2064 | 0,611055 |
| Pbx4 | 2,97891 | 2,16195 | -0,462451 | 0,20645 | 0,611062 |
| Mcm5 | 141,808 | 120,998 | -0,228955 | 0,2067 | 0,61166 |
| Slc35a4 | 39,143 | 32,9807 | -0,247131 | 0,20675 | 0,611667 |
| Atp5a1 | 598,038 | 503,96 | -0,246926 | 0,2069 | 0,61197 |
| Kdm3a | 30,1468 | 35,2998 | 0,227652 | 0,20715 | 0,612568 |
| Rcc1 | 99,453 | 85,2135 | -0,222932 | 0,2073 | 0,61287 |
| Mrps34 | 50,2397 | 41,9959 | -0,25858 | 0,20745 | 0,613172 |
| Pet117 | 0 | 3,0559 | inf | 0,2077 | 0,613635 |
| Rem2 | 1,22644 | 1,72606 | 0,493014 | 0,20775 | 0,613635 |
| Cenpi | 14,8582 | 17,5961 | 0,243995 | 0,20775 | 0,613635 |
| Lats1 | 10,9268 | 12,8425 | 0,233058 | 0,20785 | 0,613789 |
| Tecpr1 | 5,50297 | 6,63924 | 0,270808 | 0,2081 | 0,614386 |
| Bcl2l1 | 17,9155 | 21,4328 | 0,25861 | 0,2083 | 0,614693 |
| 1300002E11Rik | 13,2868 | 9,749 | -0,446666 | 0,2083 | 0,614693 |
| Abca3 | 1,24978 | 0,951676 | -0,393129 | 0,20835 | 0,6147 |
| Cfl1 | 716,936 | 609,526 | -0,234156 | 0,2084 | 0,614706 |

| | | | | | |
|---|---|---|---|---|---|
| Gk5 | 2,30054 | 2,96939 | 0,368195 | 0,20855 | 0,615007 |
| Dcun1d5 | 105,047 | 89,4891 | -0,231246 | 0,20865 | 0,615025 |
| Ankrd11 | 37,9629 | 44,3794 | 0,2253 | 0,2087 | 0,615025 |
| Nat2 | 2,55969 | 1,82807 | -0,485643 | 0,2087 | 0,615025 |
| Stat6 | 5,18016 | 4,03743 | -0,359558 | 0,20875 | 0,615031 |
| Lmbrd2 | 4,35315 | 7,64491 | 0,812439 | 0,2089 | 0,615332 |
| 3110043O21Rik | 7,16107 | 8,62868 | 0,268965 | 0,20895 | 0,615338 |
| Klhl11 | 7,04158 | 8,58462 | 0,285854 | 0,20905 | 0,615491 |
| 1700071M16Rik | 0,226273 | 0,871084 | 1,94475 | 0,20925 | 0,615938 |
| Cenpp | 30,0753 | 24,8769 | -0,27377 | 0,2093 | 0,615944 |
| Reep3 | 8,84488 | 10,4581 | 0,241711 | 0,20935 | 0,61595 |
| Eef1b2 | 444,376 | 377,732 | -0,234417 | 0,20945 | 0,616103 |
| Itpr2 | 1,28477 | 1,0354 | -0,311319 | 0,2095 | 0,616109 |
| Mir215 | 33,0087 | 0 | -inf | 0,20955 | 0,616114 |
| Dnase1l1 | 1,44406 | 2,04083 | 0,499028 | 0,2096 | 0,61612 |
| Cdk8 | 27,4822 | 23,2654 | -0,240311 | 0,2098 | 0,616567 |
| Hoxa4 | 7,87111 | 14,6389 | 0,895168 | 0,20995 | 0,616866 |
| 2310039H08Rik | 14,3069 | 11,1629 | -0,358 | 0,21 | 0,616872 |
| 1110051M20Rik | 21,3999 | 17,9812 | -0,251115 | 0,21015 | 0,617052 |
| Zc3h6 | 1,15859 | 1,50369 | 0,376135 | 0,2102 | 0,617052 |
| Plb1 | 0,502331 | 1,00105 | 0,9948 | 0,2103 | 0,617052 |
| Ccnb1 | 32,1251 | 27,2233 | -0,23886 | 0,2103 | 0,617052 |
| Rc3h1 | 7,68472 | 9,01341 | 0,230081 | 0,21035 | 0,617052 |
| Plec | 3,07096 | 2,58472 | -0,248678 | 0,21035 | 0,617052 |
| Metap1 | 46,2532 | 39,4958 | -0,227854 | 0,2104 | 0,617058 |
| Iqcb1 | 32,2059 | 37,8157 | 0,231659 | 0,2105 | 0,61721 |
| Thumpd3 | 43,3023 | 50,96 | 0,234922 | 0,2108 | 0,617948 |
| Slc25a33 | 9,66737 | 11,7651 | 0,283321 | 0,2109 | 0,6181 |
| Dhx32 | 9,35025 | 6,01218 | -0,637116 | 0,2111 | 0,61843 |
| Polr2k | 3,0864 | 4,54445 | 0,558182 | 0,21125 | 0,61843 |
| Mnd1 | 6,15924 | 4,56471 | -0,432229 | 0,21125 | 0,61843 |
| Hspb6 | 2,15555 | 2,96437 | 0,45967 | 0,21135 | 0,61843 |
| Acy3 | 2,23428 | 3,02322 | 0,436274 | 0,21135 | 0,61843 |
| Akap10 | 11,9492 | 14,1381 | 0,242682 | 0,21135 | 0,61843 |
| Ndufa10 | 133,006 | 113,497 | -0,228848 | 0,21135 | 0,61843 |
| Ptprz1 | 3,72421 | 4,45038 | 0,256993 | 0,2114 | 0,618435 |
| Plbd2 | 13,5773 | 16,0028 | 0,237127 | 0,21155 | 0,618733 |
| Ankhd1 | 19,8501 | 23,2444 | 0,22774 | 0,21165 | 0,618743 |
| Wrb | 19,9614 | 16,8062 | -0,248214 | 0,21165 | 0,618743 |
| Rpl31-ps12 | 0,496755 | 1,27861 | 1,36396 | 0,21175 | 0,618895 |
| Trim8 | 13,6331 | 16,1374 | 0,243294 | 0,2122 | 0,619927 |
| Rps9 | 1019,16 | 870,915 | -0,226777 | 0,2122 | 0,619927 |
| Gpbp1l1 | 11,7819 | 14,016 | 0,2505 | 0,21235 | 0,620224 |
| Mus81 | 9,41438 | 11,4331 | 0,28028 | 0,21245 | 0,620375 |
| Naca | 149,766 | 127,207 | -0,235531 | 0,2126 | 0,620672 |
| Cyc1 | 178,845 | 153,008 | -0,225105 | 0,21275 | 0,620969 |
| Sema4f | 0,484675 | 0,686634 | 0,502523 | 0,2129 | 0,621265 |
| Mrps18b | 26,1358 | 21,5478 | -0,278487 | 0,213 | 0,621416 |
| Nono | 143,439 | 128,796 | -0,22321 | 0,21325 | 0,621862 |
| Scyl1 | 31,1335 | 26,3776 | -0,239153 | 0,21325 | 0,621862 |

| | | | | | |
|---|---|---|---|---|---|
| Dnajc9 | 126,481 | 108,232 | -0,22479 | 0,21335 | 0,621893 |
| Rai1 | 20,766 | 25,2825 | 0,283917 | 0,2134 | 0,621893 |
| Ndufaf3 | 10,5572 | 8,33234 | -0,341429 | 0,21345 | 0,621893 |
| D630045J12Rik | 7,65095 | 8,95787 | 0,227516 | 0,21355 | 0,621893 |
| Rragb | 12,0079 | 9,97556 | -0,267517 | 0,21355 | 0,621893 |
| Six5 | 9,98042 | 8,29209 | -0,267364 | 0,2136 | 0,621893 |
| Efcc1 | 4,05861 | 3,22293 | -0,332612 | 0,2136 | 0,621893 |
| Fam110a | 9,97856 | 12,1971 | 0,289633 | 0,2139 | 0,622497 |
| Hsbp1 | 96,5677 | 82,4178 | -0,228584 | 0,214 | 0,622497 |
| Entpd5 | 2,50905 | 3,09744 | 0,303938 | 0,21405 | 0,622497 |
| Cnot6 | 63,3965 | 73,8934 | 0,221043 | 0,21405 | 0,622497 |
| Rpl14-ps1 | 1106,56 | 944,367 | -0,228657 | 0,21405 | 0,622497 |
| Mtmr4 | 28,9733 | 33,8559 | 0,224682 | 0,21415 | 0,622646 |
| Fbxo36 | 2,44133 | 1,62027 | -0,591432 | 0,2142 | 0,622651 |
| Spire2 | 1,73657 | 2,29955 | 0,405106 | 0,2146 | 0,623531 |
| H2afy3 | 29,9071 | 24,8348 | -0,268124 | 0,2146 | 0,623531 |
| Ubxn6 | 31,0314 | 26,2266 | -0,242699 | 0,2147 | 0,62368 |
| Best1 | 0,767619 | 0,498028 | -0,624165 | 0,21475 | 0,623684 |
| Smarcad1 | 25,9517 | 30,2983 | 0,223409 | 0,21485 | 0,623833 |
| Tsr2 | 20,4557 | 16,4323 | -0,315968 | 0,21495 | 0,623982 |
| Lsm4 | 153,003 | 130,673 | -0,227598 | 0,215 | 0,623986 |
| Ppp1r3g | 0,763119 | 1,09884 | 0,526001 | 0,2151 | 0,623994 |
| Socs1 | 1,8688 | 2,61932 | 0,487082 | 0,2151 | 0,623994 |
| Kif3c | 5,85578 | 4,77333 | -0,294865 | 0,2152 | 0,624143 |
| Mgst3 | 6,93575 | 4,96653 | -0,481815 | 0,21525 | 0,624147 |
| Trim23 | 5,1151 | 6,15507 | 0,267012 | 0,2153 | 0,624151 |
| Polr2i | 73,137 | 61,4093 | -0,252145 | 0,21545 | 0,624163 |
| Fcer1g | 0,927912 | 0,374993 | -1,30713 | 0,21545 | 0,624163 |
| Slc25a48 | 0,713946 | 0,0328922 | -4,44 | 0,21545 | 0,624163 |
| Adc | 1,3777 | 1,89565 | 0,460426 | 0,21555 | 0,624171 |
| St3gal2 | 25,0771 | 21,4584 | -0,224833 | 0,21555 | 0,624171 |
| Plekha3 | 10,3832 | 12,3947 | 0,255469 | 0,2156 | 0,624174 |
| Rarres1 | 0,409496 | 0,715924 | 0,805959 | 0,21575 | 0,624468 |
| Nbea | 4,78065 | 5,68881 | 0,250918 | 0,21585 | 0,624475 |
| 1110058L19Rik | 27,4474 | 22,0744 | -0,314295 | 0,21585 | 0,624475 |
| Inpp5k | 6,6214 | 5,38951 | -0,296982 | 0,2159 | 0,624479 |
| Cbln1 | 0,974373 | 1,35363 | 0,474284 | 0,21605 | 0,624491 |
| H2-Ab1 | 4,23801 | 5,58989 | 0,399431 | 0,21605 | 0,624491 |
| Nedd8 | 401,65 | 344,167 | -0,222829 | 0,21605 | 0,624491 |
| Kif18a | 9,06461 | 10,8432 | 0,258469 | 0,2161 | 0,624494 |
| Ercc6l2 | 13,986 | 16,4342 | 0,232718 | 0,2162 | 0,624643 |
| Ccdc12 | 31,0005 | 25,4878 | -0,282485 | 0,21635 | 0,624806 |
| Rara | 11,5627 | 9,71316 | -0,25147 | 0,21645 | 0,624806 |
| Bag4 | 11,7945 | 13,9078 | 0,237787 | 0,2165 | 0,624806 |
| Dgcr6 | 44,9114 | 38,0478 | -0,239269 | 0,2165 | 0,624806 |
| Stra13 | 47,7414 | 39,3175 | -0,280069 | 0,2165 | 0,624806 |
| Slc19a3 | 0,697424 | 0,980548 | 0,491551 | 0,21655 | 0,62481 |
| Ulk3 | 3,11569 | 3,91223 | 0,32844 | 0,21665 | 0,624958 |
| Spag9 | 21,35 | 24,9759 | 0,226299 | 0,2168 | 0,62525 |
| Hars | 78,2722 | 66,9841 | -0,224682 | 0,21685 | 0,625253 |

| | | | | | |
|---|---|---|---|---|---|
| LOC100503496 | 0,190945 | 1,28041 | 2,74538 | 0,2169 | 0,625257 |
| 4930599N23Rik | 0 | 0,762124 | inf | 0,217 | 0,625405 |
| Slc35f6 | 3,87425 | 4,79892 | 0,308792 | 0,2171 | 0,625552 |
| Mir6921 | 0 | 201,135 | inf | 0,2173 | 0,625847 |
| Scarb2 | 13,2896 | 11,0593 | -0,265035 | 0,2173 | 0,625847 |
| Acot11 | 0,688743 | 0,909551 | 0,40119 | 0,2175 | 0,626142 |
| Itm2b | 41,3633 | 35,1621 | -0,234332 | 0,2175 | 0,626142 |
| Smad7 | 5,53919 | 6,63757 | 0,260981 | 0,2176 | 0,626166 |
| Mir7033 | 0 | 171,198 | inf | 0,21765 | 0,626166 |
| Ppdpf | 38,6292 | 32,0435 | -0,269663 | 0,21775 | 0,626166 |
| Atp5k | 682,319 | 580,456 | -0,233259 | 0,2178 | 0,626166 |
| Vcpip1 | 9,78531 | 11,5039 | 0,233437 | 0,21785 | 0,626166 |
| Akr1a1 | 301,234 | 258,178 | -0,222518 | 0,21785 | 0,626166 |
| P4htm | 2,21707 | 1,63021 | -0,443598 | 0,21785 | 0,626166 |
| Agl | 4,11934 | 3,4673 | -0,248601 | 0,21805 | 0,626601 |
| Gm15800 | 4,75922 | 5,57675 | 0,228698 | 0,2183 | 0,627179 |
| Zfr | 43,0182 | 50,0828 | 0,219366 | 0,21835 | 0,627182 |
| Rab13 | 33,504 | 28,012 | -0,258287 | 0,2191 | 0,629054 |
| Wdpcp | 4,16084 | 3,28896 | -0,339242 | 0,2191 | 0,629054 |
| Cipc | 14,8005 | 12,5555 | -0,237321 | 0,2193 | 0,629068 |
| Glt28d2 | 0,636585 | 0,420359 | -0,59873 | 0,2193 | 0,629068 |
| Mir6352 | 0 | 7,73561 | inf | 0,21935 | 0,629068 |
| Arl6ip4 | 59,5155 | 50,5041 | -0,236867 | 0,21935 | 0,629068 |
| Mtm1 | 0,628192 | 0,41515 | -0,597571 | 0,21935 | 0,629068 |
| Ppwd1 | 16,9341 | 21,0132 | 0,311359 | 0,2194 | 0,629071 |
| Zfp467 | 3,88962 | 5,14567 | 0,403731 | 0,21945 | 0,629074 |
| Fam208b | 15,8924 | 23,2489 | 0,548828 | 0,2195 | 0,629076 |
| Dhrs4 | 32,9197 | 27,6331 | -0,252557 | 0,2196 | 0,629085 |
| Idh1 | 100,225 | 86,0673 | -0,219709 | 0,21965 | 0,629085 |
| Tesk1 | 10,3819 | 8,74144 | -0,248129 | 0,21965 | 0,629085 |
| 1700120K04Rik | 2,03879 | 1,08352 | -0,911982 | 0,2197 | 0,629087 |
| Med30 | 65,9696 | 55,7576 | -0,242632 | 0,21975 | 0,62909 |
| Gtf3c1 | 31,2631 | 36,5001 | 0,223438 | 0,2198 | 0,629093 |
| Fzd1 | 24,868 | 21,2897 | -0,224137 | 0,22005 | 0,629668 |
| Vps8 | 5,55912 | 6,64663 | 0,257765 | 0,22025 | 0,6301 |
| Ssx2ip | 12,3771 | 14,5967 | 0,237961 | 0,2204 | 0,630388 |
| Trpc1 | 1,1135 | 1,49448 | 0,424539 | 0,2205 | 0,630393 |
| Hist1h2bc | 5,74528 | 4,09404 | -0,488851 | 0,2205 | 0,630393 |
| Hk2 | 32,2194 | 37,5865 | 0,222283 | 0,2208 | 0,63111 |
| H6pd | 4,14916 | 5,00418 | 0,270316 | 0,22125 | 0,631978 |
| Pspc1 | 57,8631 | 67,4116 | 0,220354 | 0,22125 | 0,631978 |
| Msi1 | 56,1238 | 47,8921 | -0,228824 | 0,2213 | 0,631978 |
| Dynll1 | 123,801 | 106,317 | -0,219651 | 0,22135 | 0,631978 |
| 4930503E24Rik | 22,297 | 18,5279 | -0,267152 | 0,22135 | 0,631978 |
| Dbndd2 | 0,995231 | 1,31821 | 0,40548 | 0,2216 | 0,632551 |
| Nin | 14,1479 | 12,0346 | -0,233404 | 0,22165 | 0,632553 |
| Lrp12 | 14,4849 | 17,0303 | 0,233561 | 0,2217 | 0,632555 |
| Coq9 | 18,2267 | 15,2079 | -0,261238 | 0,2218 | 0,6327 |
| Ndufs2 | 163,857 | 140,774 | -0,219057 | 0,2219 | 0,632844 |
| Ndufaf4 | 17,3862 | 20,4327 | 0,232942 | 0,222 | 0,632989 |

| | | | | | |
|---|---|---|---|---|---|
| Fhdc1 | 3,09506 | 2,53334 | -0,288929 | 0,22205 | 0,632991 |
| Dip2c | 4,4681 | 5,29092 | 0,243855 | 0,22215 | 0,633135 |
| Ptcd2 | 30,7536 | 26,0288 | -0,240644 | 0,2224 | 0,633707 |
| Snord85 | 0 | 162,066 | inf | 0,22255 | 0,633993 |
| Mid1 | 2,23847 | 1,76736 | -0,340919 | 0,2226 | 0,633995 |
| Cpped1 | 7,90449 | 6,50213 | -0,281759 | 0,2228 | 0,634424 |
| Fpgs | 16,8744 | 20,3022 | 0,266798 | 0,2229 | 0,634434 |
| Atad5 | 12,6358 | 14,7554 | 0,223729 | 0,22305 | 0,634434 |
| Gpsm1 | 31,0306 | 25,63 | -0,275859 | 0,22305 | 0,634434 |
| Vps39 | 11,0944 | 13,0407 | 0,233179 | 0,2231 | 0,634434 |
| Drap1 | 128,635 | 110,191 | -0,223279 | 0,2231 | 0,634434 |
| Prrg1 | 1,3136 | 1,7034 | 0,374889 | 0,22315 | 0,634434 |
| Slc9a3r1 | 44,6956 | 52,2523 | 0,225362 | 0,22315 | 0,634434 |
| Gpr27 | 1,0916 | 1,64961 | 0,595677 | 0,2232 | 0,634436 |
| AW549877 | 18,9904 | 22,1881 | 0,224517 | 0,22325 | 0,634438 |
| Rhod | 15,6352 | 18,9797 | 0,279659 | 0,2234 | 0,634583 |
| Vps13d | 3,15708 | 3,71319 | 0,234072 | 0,2234 | 0,634583 |
| Dlx5 | 1,75359 | 2,44514 | 0,479608 | 0,2235 | 0,634586 |
| 3010001F23Rik | 1,64317 | 1,02211 | -0,684929 | 0,2235 | 0,634586 |
| Oxct1 | 63,2912 | 54,3958 | -0,218509 | 0,2236 | 0,634729 |
| Elk1 | 12,9276 | 15,2463 | 0,238005 | 0,2237 | 0,634732 |
| Ythdc1 | 47,2967 | 55,2401 | 0,223977 | 0,2237 | 0,634732 |
| Kdsr | 2,78661 | 3,38507 | 0,280675 | 0,22395 | 0,635026 |
| Yy2 | 5,60052 | 7,48522 | 0,418485 | 0,224 | 0,635026 |
| Rnf34 | 41,9524 | 49,0213 | 0,224653 | 0,224 | 0,635026 |
| Slc35f2 | 2,51092 | 3,20462 | 0,35194 | 0,22405 | 0,635026 |
| Tceal3 | 2,02662 | 2,8617 | 0,4978 | 0,2241 | 0,635026 |
| Anapc7 | 43,3511 | 37,1487 | -0,222757 | 0,22415 | 0,635026 |
| Shmt1 | 61,9329 | 53,0593 | -0,223102 | 0,22415 | 0,635026 |
| Gng12 | 23,9449 | 27,9196 | 0,221564 | 0,2247 | 0,636443 |
| Apaf1 | 11,6205 | 9,93184 | -0,226536 | 0,22495 | 0,63687 |
| St3gal5 | 12,0509 | 10,0278 | -0,265138 | 0,22495 | 0,63687 |
| Uqcrfs1 | 153,739 | 132,095 | -0,218909 | 0,22505 | 0,637013 |
| Cdca8 | 133,765 | 114,82 | -0,220329 | 0,2252 | 0,637158 |
| Mtch2 | 48,6643 | 41,7798 | -0,220057 | 0,22525 | 0,637158 |
| Tob2 | 8,36995 | 9,89823 | 0,241951 | 0,2253 | 0,637158 |
| Wdr46 | 48,9617 | 41,99 | -0,221607 | 0,2253 | 0,637158 |
| 3110002H16Rik | 6,0421 | 8,59895 | 0,509111 | 0,22565 | 0,637866 |
| Ndufv3 | 127,561 | 105,768 | -0,270293 | 0,22565 | 0,637867 |
| Zeb1 | 15,9348 | 18,5823 | 0,221744 | 0,22575 | 0,637868 |
| Gm6548 | 33,2924 | 28,3649 | -0,231088 | 0,22575 | 0,637868 |
| Uckl1os | 0,693815 | 2,43825 | 1,81322 | 0,2259 | 0,638151 |
| Sec23a | 29,2892 | 25,1399 | -0,220393 | 0,22595 | 0,638152 |
| Acss3 | 0,587674 | 0,369365 | -0,66997 | 0,226 | 0,638153 |
| Mrpl20 | 203,216 | 173,945 | -0,224377 | 0,22605 | 0,638153 |
| 1810044D09Rik | 7,0604 | 4,82578 | -0,548989 | 0,22615 | 0,638295 |
| Tank | 14,3093 | 17,0029 | 0,248832 | 0,2262 | 0,638296 |
| Puf60 | 148,924 | 128,15 | -0,216735 | 0,2263 | 0,638438 |
| Mir6963 | 0 | 88,0666 | inf | 0,2265 | 0,638583 |
| Zfp605 | 3,10605 | 3,87622 | 0,319567 | 0,22655 | 0,638583 |

| | | | | | |
|---|---|---|---|---|---|
| Hip1 | 27,0798 | 31,4899 | 0,21767 | 0,22655 | 0,638583 |
| Pla2g4c | 1,52752 | 1,97257 | 0,368886 | 0,22665 | 0,638583 |
| 1810043H04Rik | 39,0848 | 31,4179 | -0,31502 | 0,22665 | 0,638583 |
| Gngt2 | 1,00848 | 0,443619 | -1,18479 | 0,22665 | 0,638583 |
| Zcchc8 | 28,5249 | 33,2001 | 0,218963 | 0,2267 | 0,638583 |
| Sos2 | 8,19395 | 9,63487 | 0,233705 | 0,22685 | 0,638725 |
| Commd8 | 18,3574 | 15,6424 | -0,230894 | 0,22685 | 0,638725 |
| Mark2 | 18,5072 | 21,5782 | 0,221489 | 0,22715 | 0,639289 |
| Rpl15 | 51,1706 | 43,8454 | -0,222888 | 0,22715 | 0,639289 |
| Tctn1 | 6,54957 | 8,12414 | 0,310814 | 0,2272 | 0,63929 |
| Reep5 | 16,2803 | 19,1601 | 0,234971 | 0,2275 | 0,639853 |
| Ube2d3 | 61,5673 | 52,8551 | -0,22012 | 0,22755 | 0,639853 |
| Mgat5b | 1,66055 | 1,2854 | -0,369445 | 0,22755 | 0,639853 |
| Zswim3 | 2,94287 | 3,7157 | 0,336407 | 0,22765 | 0,639994 |
| Smap1 | 39,8236 | 33,9041 | -0,232167 | 0,22775 | 0,639995 |
| Plin3 | 23,4357 | 19,889 | -0,236738 | 0,22775 | 0,639995 |
| Snrnp35 | 24,5702 | 20,4601 | -0,264096 | 0,2278 | 0,639995 |
| Tgfbr2 | 2,22245 | 1,77968 | -0,320531 | 0,22795 | 0,640136 |
| Avpi1 | 8,05802 | 6,24705 | -0,367251 | 0,22795 | 0,640136 |
| Mir6972 | 0 | 365,166 | inf | 0,2283 | 0,640979 |
| Adrbk1 | 28,3672 | 33,0101 | 0,218686 | 0,2285 | 0,6414 |
| Ino80d | 4,92594 | 5,74086 | 0,220868 | 0,22855 | 0,6414 |
| Mafg | 15,4051 | 18,386 | 0,255202 | 0,2287 | 0,64154 |
| Ptpn21 | 3,71913 | 4,46496 | 0,263681 | 0,22875 | 0,64154 |
| Crim1 | 1,90259 | 1,37496 | -0,468573 | 0,22875 | 0,64154 |
| Mir7115 | 0 | 579,406 | inf | 0,22885 | 0,64154 |
| Gstt3 | 0,432303 | 0,685115 | 0,664305 | 0,2289 | 0,64154 |
| Cox7b | 137,398 | 118,133 | -0,217946 | 0,2289 | 0,64154 |
| Mir7006 | 315,832 | 0 | -inf | 0,229 | 0,64154 |
| Hnrnpc | 232,537 | 199,164 | -0,223502 | 0,22905 | 0,64154 |
| Psma6 | 322,362 | 272,157 | -0,244243 | 0,22905 | 0,64154 |
| Prkra | 60,8272 | 52,1383 | -0,222373 | 0,2292 | 0,641819 |
| Adprm | 11,5486 | 14,012 | 0,278935 | 0,2293 | 0,641819 |
| Sult5a1 | 1,16214 | 0,707475 | -0,716033 | 0,2294 | 0,641819 |
| Snora78 | 17,0344 | 40,2549 | 1,24072 | 0,22945 | 0,641819 |
| Agap3 | 28,9059 | 33,7268 | 0,222532 | 0,22945 | 0,641819 |
| Tspan17 | 1,36049 | 0,920353 | -0,563872 | 0,22945 | 0,641819 |
| Vps13c | 2,64954 | 3,14043 | 0,24522 | 0,2295 | 0,641819 |
| Mir6973b | 47,2529 | 0 | -inf | 0,22985 | 0,642518 |
| Mrps18c | 95,001 | 79,6725 | -0,253861 | 0,22985 | 0,642518 |
| Ppp2r5a | 13,6445 | 11,2651 | -0,27646 | 0,22995 | 0,642658 |
| Ggt7 | 1,98212 | 2,56058 | 0,369424 | 0,23015 | 0,642796 |
| Cnksr3 | 17,081 | 20,0087 | 0,228233 | 0,23015 | 0,642796 |
| Hadh | 73,0065 | 84,8516 | 0,216917 | 0,23015 | 0,642796 |
| Mir6977 | 196,743 | 0 | -inf | 0,2302 | 0,642796 |
| Mir6406 | 0 | 6,71844 | inf | 0,2305 | 0,643354 |
| Polr2h | 96,4964 | 82,0361 | -0,234216 | 0,2305 | 0,643354 |
| Sh3bgrl | 49,0927 | 57,3665 | 0,224701 | 0,2309 | 0,644189 |
| Stard5 | 4,44069 | 3,54558 | -0,32476 | 0,2309 | 0,644189 |
| Ube2o | 16,4336 | 19,1575 | 0,221262 | 0,23095 | 0,644189 |

| | | | | | |
|---|---|---|---|---|---|
| Eif3i | 281,994 | 243,148 | -0,213834 | 0,231 | 0,644189 |
| 5430417L22Rik | 4,12023 | 3,32358 | -0,309986 | 0,23115 | 0,644467 |
| Mlxip | 15,6574 | 18,2069 | 0,217642 | 0,23125 | 0,644603 |
| Coq4 | 14,4021 | 12,0319 | -0,259425 | 0,2313 | 0,644603 |
| Mrps12 | 70,3939 | 59,4056 | -0,244851 | 0,23135 | 0,644603 |
| 9430038I01Rik | 10,4964 | 8,38742 | -0,323601 | 0,2314 | 0,644604 |
| Uqcrc2 | 163,16 | 140,565 | -0,215045 | 0,2316 | 0,644881 |
| Tomm22 | 126,846 | 108,926 | -0,219723 | 0,2316 | 0,644881 |
| Hba-a1,Hba-a2 | 0,31879 | 3,63248 | 3,51027 | 0,23175 | 0,645018 |
| Setd7 | 14,4678 | 16,7847 | 0,214308 | 0,23175 | 0,645018 |
| Tor1b | 24,7896 | 28,8896 | 0,220811 | 0,23185 | 0,645157 |
| Ppil3 | 54,8787 | 46,3771 | -0,242835 | 0,23215 | 0,64585 |
| 2510009E07Rik | 7,74646 | 9,13949 | 0,238576 | 0,2322 | 0,64585 |
| Dap3 | 22,378 | 19,2326 | -0,218524 | 0,2323 | 0,645983 |
| Sbno1 | 17,0646 | 19,8065 | 0,214969 | 0,23235 | 0,645983 |
| Sdhd | 169,676 | 146,164 | -0,215193 | 0,23245 | 0,645983 |
| Fchsd2 | 12,2948 | 14,4004 | 0,228058 | 0,2325 | 0,645983 |
| Gpank1 | 12,0208 | 9,79451 | -0,295487 | 0,2325 | 0,645983 |
| Rilpl2 | 13,0847 | 10,6493 | -0,297121 | 0,23255 | 0,645983 |
| Lamp2 | 25,341 | 29,8018 | 0,233927 | 0,2327 | 0,64626 |
| Arhgef4 | 1,41368 | 1,04749 | -0,432511 | 0,2328 | 0,646397 |
| 2700089E24Rik | 25,0029 | 21,1923 | -0,23856 | 0,23285 | 0,646397 |
| Prdx5 | 69,4964 | 58,8267 | -0,240466 | 0,233 | 0,646672 |
| Maneal | 2,79531 | 1,77912 | -0,651848 | 0,23305 | 0,646672 |
| Eif1a | 49,2314 | 57,1592 | 0,215406 | 0,23315 | 0,64681 |
| Nrros | 0,881123 | 1,17246 | 0,412129 | 0,2333 | 0,647086 |
| Hint2 | 20,8744 | 26,0959 | 0,322088 | 0,2334 | 0,647223 |
| Zscan2 | 10,0023 | 8,31153 | -0,26714 | 0,23355 | 0,647358 |
| Glrx2 | 6,01117 | 4,98714 | -0,269432 | 0,23355 | 0,647358 |
| Kif5c | 9,35703 | 10,9305 | 0,224241 | 0,2336 | 0,647358 |
| Xpo1 | 110,156 | 128,573 | 0,223035 | 0,23375 | 0,647489 |
| Zfp407 | 6,2352 | 7,30465 | 0,228379 | 0,2338 | 0,647489 |
| Matn2 | 2,11343 | 1,65343 | -0,354127 | 0,23385 | 0,647489 |
| Cish | 1,40643 | 1,01872 | -0,465286 | 0,2339 | 0,647489 |
| Atg2b | 6,49278 | 7,5783 | 0,223038 | 0,23395 | 0,647489 |
| Pacsin2 | 21,9746 | 25,5713 | 0,218688 | 0,23395 | 0,647489 |
| 9130024F11Rik | 0,579344 | 0,822996 | 0,506466 | 0,23405 | 0,647615 |
| Spg21 | 23,9208 | 20,3804 | -0,231086 | 0,23425 | 0,647615 |
| Gramd1b | 1,61699 | 1,22072 | -0,40557 | 0,23425 | 0,647615 |
| Sec11c | 10,4896 | 13,3522 | 0,348116 | 0,23435 | 0,647615 |
| Zmat5 | 33,8232 | 28,0045 | -0,272353 | 0,23435 | 0,647615 |
| Spata5 | 18,5783 | 21,7234 | 0,225633 | 0,2344 | 0,647615 |
| Ftsj1 | 11,4895 | 9,72382 | -0,240722 | 0,2344 | 0,647615 |
| Rusc1 | 4,17973 | 5,2521 | 0,329485 | 0,23445 | 0,647615 |
| Zcchc7 | 11,4584 | 13,6101 | 0,248282 | 0,23445 | 0,647615 |
| Smurf2 | 19,0379 | 22,1199 | 0,216475 | 0,2346 | 0,647887 |
| Asxl2 | 6,12501 | 7,16619 | 0,226495 | 0,23465 | 0,647887 |
| Retsat | 4,7642 | 5,94946 | 0,320525 | 0,2347 | 0,647887 |
| Mir8118 | 5,39034 | 0 | -inf | 0,2349 | 0,6483 |
| D430020J02Rik | 4,66154 | 5,67823 | 0,284635 | 0,2351 | 0,648573 |

| | | | | | |
|---|---|---|---|---|---|
| Traf4 | 42,7706 | 36,8675 | -0,21427 | 0,2351 | 0,648573 |
| Slc29a3 | 1,49649 | 1,87839 | 0,327917 | 0,2352 | 0,648709 |
| Clpp | 53,763 | 45,8498 | -0,229697 | 0,23535 | 0,648975 |
| Sc5d | 23,215 | 19,7407 | -0,233885 | 0,2354 | 0,648975 |
| Tsga10 | 1,80094 | 2,38226 | 0,403584 | 0,2355 | 0,648975 |
| Amica1 | 0,799436 | 0,0188832 | -5,40381 | 0,2355 | 0,648975 |
| Uggt2 | 5,63651 | 6,64741 | 0,23799 | 0,2356 | 0,648975 |
| Mertk | 3,43186 | 2,76688 | -0,310733 | 0,2356 | 0,648975 |
| Snord88a | 219,42 | 0 | -inf | 0,2358 | 0,649369 |
| Pet100 | 16,4527 | 13,5371 | -0,281406 | 0,23595 | 0,649369 |
| Mir1b | 0 | 22,5111 | inf | 0,2361 | 0,649369 |
| Mark1 | 7,43638 | 8,76928 | 0,237859 | 0,2361 | 0,649369 |
| Uqcc2 | 162,011 | 137,849 | -0,233007 | 0,23615 | 0,649369 |
| Gorab | 11,9944 | 10,0561 | -0,254295 | 0,23615 | 0,649369 |
| Nol7 | 50,7107 | 40,5912 | -0,321124 | 0,23615 | 0,649369 |
| Blm | 28,705 | 33,2726 | 0,213031 | 0,2362 | 0,649369 |
| Dyx1c1 | 1,13871 | 1,5718 | 0,465009 | 0,23625 | 0,649369 |
| Cdc37 | 156,644 | 134,915 | -0,215438 | 0,23625 | 0,649369 |
| Tma7 | 71,2718 | 60,0014 | -0,248335 | 0,2363 | 0,649369 |
| Git2 | 7,06545 | 8,33301 | 0,238058 | 0,2365 | 0,649495 |
| Cdk14 | 6,48683 | 5,47656 | -0,244243 | 0,2365 | 0,649495 |
| Lsm2 | 129,344 | 111,114 | -0,219181 | 0,23655 | 0,649495 |
| Gclc | 16,092 | 13,7152 | -0,230567 | 0,23655 | 0,649495 |
| Ticam1 | 2,47696 | 3,12311 | 0,334416 | 0,23665 | 0,649495 |
| Rab10 | 42,2415 | 36,2495 | -0,2207 | 0,2367 | 0,649495 |
| Slc37a2 | 1,42573 | 1,10572 | -0,366709 | 0,2367 | 0,649495 |
| Mrpl12 | 162,13 | 139,696 | -0,214854 | 0,2368 | 0,649625 |
| Mir6407 | 15,1634 | 0 | -inf | 0,2369 | 0,649625 |
| Arhgap4 | 1,21178 | 1,59663 | 0,397898 | 0,2369 | 0,649625 |
| Fam3c | 10,6135 | 12,5763 | 0,244808 | 0,23695 | 0,649625 |
| Mpdu1 | 36,8845 | 31,3684 | -0,233701 | 0,2371 | 0,649757 |
| Lysmd2 | 5,94837 | 4,61085 | -0,367462 | 0,2371 | 0,649757 |
| Mir1957b | 0 | 19,9412 | inf | 0,23715 | 0,649757 |
| Cox5a | 334,883 | 289,07 | -0,212236 | 0,2373 | 0,650027 |
| Muc1 | 1,38018 | 0,998623 | -0,466845 | 0,23735 | 0,650027 |
| Cycs | 1,71109 | 1,30652 | -0,389188 | 0,23745 | 0,650162 |
| Ythdc2 | 5,96841 | 7,02601 | 0,235359 | 0,23755 | 0,650293 |
| Abcc4 | 1,80649 | 2,22647 | 0,301567 | 0,2376 | 0,650293 |
| Errfi1 | 18,1126 | 15,4269 | -0,231542 | 0,23765 | 0,650293 |
| Cep85 | 16,0979 | 18,7956 | 0,223524 | 0,23775 | 0,650416 |
| Stard8 | 0,947551 | 0,722173 | -0,391859 | 0,2378 | 0,650416 |
| Clasrp | 27,9416 | 32,6813 | 0,226055 | 0,23785 | 0,650416 |
| Mir6538 | 42,2685 | 0 | -inf | 0,23795 | 0,650416 |
| Ash2l | 62,7075 | 54,2239 | -0,209707 | 0,238 | 0,650416 |
| Tigd5 | 1,17184 | 0,902999 | -0,375978 | 0,238 | 0,650416 |
| Kdm5b | 34,0542 | 39,444 | 0,211972 | 0,23805 | 0,650416 |
| 1500011K16Rik | 12,3065 | 9,69454 | -0,34418 | 0,2383 | 0,650959 |
| Atp5s | 5,13635 | 4,0097 | -0,357249 | 0,23835 | 0,650959 |
| Narf | 12,4887 | 10,6585 | -0,228616 | 0,2385 | 0,65123 |
| Hddc2 | 73,6509 | 62,5778 | -0,235051 | 0,2388 | 0,651908 |

| | | | | | |
|---|---|---|---|---|---|
| C530008M17Rik | 14,2921 | 16,6024 | 0,216176 | 0,23885 | 0,651908 |
| Zfp799 | 10,5532 | 12,3857 | 0,231003 | 0,23895 | 0,652042 |
| Mir6920 | 0 | 139,735 | inf | 0,23915 | 0,65231 |
| Pnpla8 | 11,9735 | 14,0344 | 0,229117 | 0,23915 | 0,65231 |
| Gtl3 | 60,0534 | 51,4488 | -0,223108 | 0,2393 | 0,65244 |
| Kbtbd3 | 2,02196 | 2,61451 | 0,370782 | 0,23935 | 0,65244 |
| H3f3b | 239,872 | 207,221 | -0,211098 | 0,23935 | 0,65244 |
| Dpm3 | 38,5244 | 31,0804 | -0,309768 | 0,2395 | 0,65271 |
| Cyp4f13 | 0,478774 | 0,755677 | 0,658425 | 0,2396 | 0,652844 |
| Soga1 | 15,9501 | 18,4981 | 0,213809 | 0,23975 | 0,653114 |
| Lgals3 | 0,0245889 | 0,613336 | 4,6406 | 0,2399 | 0,653384 |
| Phf10 | 86,4712 | 101,67 | 0,233599 | 0,24005 | 0,653651 |
| Urb1 | 9,12025 | 10,6236 | 0,220126 | 0,2401 | 0,653651 |
| Huwe1 | 61,3978 | 71,6537 | 0,222854 | 0,24035 | 0,654051 |
| Echdc3 | 0,6808 | 1,02806 | 0,594626 | 0,2404 | 0,654051 |
| Akap7 | 6,84677 | 5,70331 | -0,263623 | 0,2404 | 0,654051 |
| Cenpm | 22,5711 | 18,8052 | -0,263343 | 0,2407 | 0,65428 |
| Mir7035 | 41,4193 | 0 | -inf | 0,24075 | 0,65428 |
| Elp5 | 80,5306 | 65,1914 | -0,304855 | 0,24075 | 0,65428 |
| Nes | 52,9501 | 61,3006 | 0,211267 | 0,2408 | 0,65428 |
| Zfp113 | 6,59042 | 7,72598 | 0,229348 | 0,24085 | 0,65428 |
| Adipor1 | 54,4606 | 63,1811 | 0,21428 | 0,2409 | 0,65428 |
| Fscn1 | 286,316 | 246,542 | -0,215774 | 0,2409 | 0,65428 |
| Mtmr3 | 16,8961 | 19,6227 | 0,215833 | 0,241 | 0,65428 |
| Fen1 | 92,8169 | 80,2619 | -0,209672 | 0,241 | 0,65428 |
| Mir6387 | 0 | 10,4817 | inf | 0,2411 | 0,65428 |
| Ppp1r36 | 0,995388 | 1,45328 | 0,545986 | 0,2411 | 0,65428 |
| Prrc1 | 16,2504 | 18,8992 | 0,217851 | 0,24115 | 0,65428 |
| Bscl2 | 12,5593 | 10,363 | -0,27731 | 0,24115 | 0,65428 |
| Mir6915 | 0 | 168,351 | inf | 0,2413 | 0,65428 |
| Aplf | 5,53296 | 4,55461 | -0,280726 | 0,2413 | 0,65428 |
| Polr3gl | 9,40771 | 7,6041 | -0,307065 | 0,2413 | 0,65428 |
| Mir7067 | 0 | 535,577 | inf | 0,2414 | 0,654407 |
| Pltp | 11,1963 | 7,7597 | -0,528945 | 0,24145 | 0,654407 |
| Tmem170b | 2,21127 | 2,66305 | 0,268204 | 0,2415 | 0,654407 |
| Angptl1 | 1,99417 | 0,211865 | -3,23457 | 0,2419 | 0,65505 |
| Snord92 | 77,7465 | 0 | -inf | 0,24195 | 0,65505 |
| Nudt2 | 37,2631 | 31,2092 | -0,255775 | 0,242 | 0,65505 |
| Lyplal1 | 7,79806 | 6,29809 | -0,308201 | 0,242 | 0,65505 |
| Vat1 | 6,12366 | 5,04582 | -0,279307 | 0,2421 | 0,65505 |
| AA415398 | 0,810902 | 1,16757 | 0,525915 | 0,24225 | 0,65505 |
| Pdia6 | 346,411 | 403,435 | 0,219852 | 0,24225 | 0,65505 |
| Polr1c | 102,916 | 88,6838 | -0,214727 | 0,24225 | 0,65505 |
| Polk | 3,00424 | 3,66658 | 0,287435 | 0,24235 | 0,65505 |
| Dap | 21,2762 | 25,141 | 0,240802 | 0,24235 | 0,65505 |
| Klhl22 | 32,904 | 28,3127 | -0,216815 | 0,24235 | 0,65505 |
| Aarsd1 | 74,5174 | 64,1195 | -0,216816 | 0,24235 | 0,65505 |
| Casp3 | 67,3931 | 77,8449 | 0,208002 | 0,2425 | 0,655165 |
| Mir8093 | 8,47996 | 0 | -inf | 0,24255 | 0,655165 |
| Lmo1 | 2,14696 | 2,97062 | 0,468469 | 0,2426 | 0,655165 |

| | | | | | |
|---|---|---|---|---|---|
| Mir7078 | 196,743 | 0 | -inf | 0,24265 | 0,655165 |
| Trmt112 | 25,0493 | 20,1449 | -0,31436 | 0,24265 | 0,655165 |
| Armc8 | 11,6619 | 13,6812 | 0,230391 | 0,2427 | 0,655165 |
| Mrpl51 | 26,1645 | 22,1995 | -0,237083 | 0,24275 | 0,655165 |
| Clpb | 9,78565 | 8,31243 | -0,235398 | 0,24285 | 0,655297 |
| Ndufs3 | 27,9146 | 23,2524 | -0,263642 | 0,243 | 0,655516 |
| 2310067B10Rik | 12,7376 | 10,9179 | -0,222403 | 0,24305 | 0,655516 |
| Nfx1 | 41,8734 | 48,4523 | 0,210532 | 0,2432 | 0,655516 |
| Rps28 | 1,63354 | 3,4263 | 1,06865 | 0,24325 | 0,655516 |
| Grpel1 | 33,7137 | 28,6341 | -0,235601 | 0,24325 | 0,655516 |
| Mir3100 | 266,297 | 0 | -inf | 0,2433 | 0,655516 |
| Dtnbp1 | 45,8116 | 53,5193 | 0,224347 | 0,24345 | 0,655516 |
| Pdap1 | 156,63 | 135,579 | -0,208235 | 0,24345 | 0,655516 |
| Cfl2 | 52,0428 | 44,9328 | -0,211928 | 0,24345 | 0,655516 |
| Slc2a5 | 1,33314 | 1,73417 | 0,379418 | 0,2435 | 0,655516 |
| Mir1188 | 0 | 26,8737 | inf | 0,24375 | 0,655516 |
| Pank4 | 24,2216 | 28,2749 | 0,223231 | 0,24375 | 0,655516 |
| Tk1 | 40,422 | 34,3406 | -0,235226 | 0,24375 | 0,655516 |
| Nagpa | 5,24198 | 4,21774 | -0,31364 | 0,24375 | 0,655516 |
| Spice1 | 11,211 | 13,1286 | 0,227796 | 0,2438 | 0,655516 |
| Rnf44 | 59,2343 | 68,4784 | 0,209214 | 0,2438 | 0,655516 |
| Metap2 | 43,0259 | 37,1084 | -0,213461 | 0,2438 | 0,655516 |
| Cfb | 1,09111 | 1,45769 | 0,417895 | 0,24395 | 0,655782 |
| Mir6945 | 0 | 420,429 | inf | 0,24405 | 0,655913 |
| Bach2 | 4,24727 | 3,60278 | -0,237423 | 0,2442 | 0,656179 |
| Ncor2 | 19,7638 | 22,9246 | 0,214037 | 0,24435 | 0,656301 |
| Cct5 | 543,565 | 467,851 | -0,216405 | 0,24435 | 0,656301 |
| Clgn | 1,05289 | 0,741374 | -0,506089 | 0,2444 | 0,656301 |
| Dhcr24 | 82,7225 | 95,8481 | 0,21247 | 0,24445 | 0,656301 |
| Ptp4a1 | 3,82762 | 3,13909 | -0,286103 | 0,24465 | 0,656691 |
| Xab2 | 40,0474 | 34,5738 | -0,212031 | 0,2447 | 0,656691 |
| Mir6976 | 446,217 | 0 | -inf | 0,24475 | 0,656691 |
| Cbs | 1,59006 | 1,19416 | -0,413082 | 0,2448 | 0,656691 |
| Mir6955 | 112,141 | 0 | -inf | 0,24505 | 0,65708 |
| Nrip2 | 0,898715 | 1,27615 | 0,505858 | 0,24505 | 0,65708 |
| Rbmxl1 | 33,4414 | 28,661 | -0,222548 | 0,2451 | 0,65708 |
| Dnase1l2 | 1,48369 | 2,10128 | 0,502073 | 0,24515 | 0,65708 |
| Uqcr11 | 211,544 | 181,107 | -0,224115 | 0,24525 | 0,657211 |
| Ccdc47 | 20,767 | 24,1667 | 0,218727 | 0,2455 | 0,657743 |
| Mir7005 | 261,511 | 0 | -inf | 0,2457 | 0,657812 |
| Rfk | 27,7462 | 23,8173 | -0,220282 | 0,2457 | 0,657812 |
| Mir6943 | 0 | 229,596 | inf | 0,2458 | 0,657812 |
| Pfkfb2 | 3,07155 | 3,81825 | 0,313946 | 0,24585 | 0,657812 |
| H2-Q7,H2-Q9 | 7,25223 | 9,06644 | 0,322112 | 0,2459 | 0,657812 |
| Mab21l2 | 3,15958 | 0,0170319 | -7,53535 | 0,2459 | 0,657812 |
| AF357355 | 189,793 | 0 | -inf | 0,24595 | 0,657812 |
| Rap2b | 9,58897 | 8,16633 | -0,231687 | 0,24595 | 0,657812 |
| Cox4i1 | 623,421 | 539,917 | -0,207468 | 0,246 | 0,657812 |
| Coil | 23,3505 | 19,9844 | -0,224582 | 0,24605 | 0,657812 |
| Mir1938 | 55,8906 | 0 | -inf | 0,24615 | 0,657812 |

| | | | | | |
|---|---|---|---|---|---|
| Pepd | 27,3051 | 23,3142 | -0,227961 | 0,2462 | 0,657812 |
| Cetn3 | 138,507 | 119,683 | -0,210735 | 0,24625 | 0,657812 |
| Ccdc6 | 14,715 | 17,0551 | 0,212915 | 0,2463 | 0,657812 |
| Zfand1 | 3,85864 | 2,99412 | -0,365958 | 0,2463 | 0,657812 |
| Mpdz | 27,0258 | 31,2187 | 0,208073 | 0,24635 | 0,657812 |
| Pdpk1 | 11,5249 | 13,351 | 0,212186 | 0,2464 | 0,657812 |
| Kntc1 | 17,5823 | 20,3334 | 0,209729 | 0,2465 | 0,657812 |
| Clec2l | 1,20627 | 0,798249 | -0,595641 | 0,2465 | 0,657812 |
| Mir681 | 0 | 30,7392 | inf | 0,24695 | 0,658876 |
| Pla2g4b | 2,93612 | 4,39056 | 0,580494 | 0,24705 | 0,659005 |
| Crlf3 | 22,8177 | 26,5974 | 0,221133 | 0,24735 | 0,659246 |
| Htatip2 | 0,79501 | 0,499865 | -0,669435 | 0,24735 | 0,659246 |
| Ccdc114 | 2,28845 | 2,91366 | 0,348456 | 0,2474 | 0,659246 |
| Map2k2 | 78,7306 | 68,151 | -0,20819 | 0,2474 | 0,659246 |
| Nmnat3 | 0,756032 | 1,07318 | 0,505374 | 0,24745 | 0,659246 |
| Mir199b | 0 | 9,68609 | inf | 0,2475 | 0,659246 |
| Oxa1l | 32,5618 | 27,8026 | -0,227963 | 0,2475 | 0,659246 |
| Mir7013 | 0 | 241,945 | inf | 0,24765 | 0,659509 |
| Kif23 | 38,6452 | 44,6935 | 0,209778 | 0,24805 | 0,6603 |
| Slc25a32 | 4,44042 | 3,61581 | -0,296377 | 0,24805 | 0,6603 |
| Zc3h7b | 50,7217 | 58,5094 | 0,206065 | 0,2482 | 0,660562 |
| Ccz1 | 69,0066 | 59,7628 | -0,207486 | 0,24835 | 0,660824 |
| Krt39 | 0,0232169 | 1,69352 | 6,18871 | 0,24855 | 0,661082 |
| Myo19 | 12,0136 | 14,8286 | 0,303708 | 0,24855 | 0,661082 |
| Sat2 | 6,99034 | 8,96212 | 0,358479 | 0,24865 | 0,661207 |
| Rangap1 | 93,6956 | 81,1824 | -0,206813 | 0,2487 | 0,661207 |
| Gdpd5 | 1,58474 | 1,22744 | -0,368595 | 0,249 | 0,661867 |
| Slco2b1 | 0,0409135 | 0,778415 | 4,24989 | 0,2491 | 0,661911 |
| Abhd14a | 22,2388 | 26,406 | 0,247792 | 0,2492 | 0,661911 |
| Map2 | 3,79701 | 3,16655 | -0,261954 | 0,2492 | 0,661911 |
| Mir6385 | 0 | 11,8485 | inf | 0,24935 | 0,661911 |
| Zfp60 | 11,7978 | 13,7659 | 0,222574 | 0,24935 | 0,661911 |
| Bfar | 17,5635 | 14,8251 | -0,244533 | 0,24935 | 0,661911 |
| 1700088E04Rik | 1,80537 | 0,207297 | -3,12252 | 0,24945 | 0,661911 |
| Apool | 45,567 | 38,6659 | -0,236927 | 0,2495 | 0,661911 |
| Mir6395 | 56,1882 | 0 | -inf | 0,24955 | 0,661911 |
| Tmem167 | 11,8339 | 10,2109 | -0,212816 | 0,2496 | 0,661911 |
| Cep57l1 | 9,94358 | 8,25736 | -0,268085 | 0,24965 | 0,661911 |
| Gm14305 | 0,847285 | 1,23328 | 0,541576 | 0,24975 | 0,661911 |
| Ift80 | 15,9415 | 18,5082 | 0,215376 | 0,24975 | 0,661911 |
| Pyurf | 5,45747 | 4,6145 | -0,242057 | 0,24975 | 0,661911 |
| Nifk | 90,6554 | 78,4795 | -0,208078 | 0,24985 | 0,661911 |
| Dapk3 | 22,4888 | 19,1333 | -0,233122 | 0,24995 | 0,661911 |
| Psmg4 | 57,2716 | 47,5685 | -0,267812 | 0,25 | 0,661911 |
| Bckdhb | 28,4702 | 24,181 | -0,235579 | 0,25005 | 0,661911 |
| Acsf3 | 8,54905 | 7,09297 | -0,269374 | 0,25005 | 0,661911 |
| Mroh1 | 3,0596 | 4,6201 | 0,59458 | 0,2501 | 0,661911 |
| Cuta | 85,6162 | 73,0125 | -0,229741 | 0,2501 | 0,661911 |
| Plcb4 | 4,18076 | 5,03872 | 0,269292 | 0,2502 | 0,662039 |
| Nudcd2 | 109,522 | 94,7888 | -0,208428 | 0,2503 | 0,662167 |

| | | | | | |
|---|---|---|---|---|---|
| Tnni3 | 1,55239 | 2,32768 | 0,584397 | 0,2504 | 0,662295 |
| Nol12 | 20,1554 | 17,1658 | -0,23163 | 0,2506 | 0,662613 |
| Lamb1 | 45,2918 | 52,3027 | 0,207637 | 0,25085 | 0,662613 |
| Fxn | 14,2791 | 11,6696 | -0,291148 | 0,25085 | 0,662613 |
| Tfap2b | 1,91951 | 0,051765 | -5,21262 | 0,25085 | 0,662613 |
| Mir1956 | 399,445 | 0 | -inf | 0,2509 | 0,662613 |
| Fam69b | 10,9273 | 9,0463 | -0,272541 | 0,2509 | 0,662613 |
| Lnp | 6,41613 | 7,4664 | 0,218709 | 0,25095 | 0,662613 |
| Tie1 | 1,07243 | 0,805068 | -0,413699 | 0,25115 | 0,662613 |
| Leprel2 | 11,9421 | 10,122 | -0,238557 | 0,2512 | 0,662613 |
| Trmt5 | 22,4717 | 26,3505 | 0,229719 | 0,25125 | 0,662613 |
| Pdcd2l | 23,936 | 20,1943 | -0,245236 | 0,25125 | 0,662613 |
| 4632415L05Rik | 5,99509 | 5,04539 | -0,248815 | 0,25125 | 0,662613 |
| Prrc2b | 48,8581 | 56,3981 | 0,207049 | 0,2514 | 0,662613 |
| Hmgb1-rs17 | 0,560415 | 0,885673 | 0,660277 | 0,25145 | 0,662613 |
| Ctsh | 2,51103 | 1,87831 | -0,418847 | 0,25145 | 0,662613 |
| Saa1 | 1,5581 | 0,0665609 | -4,54897 | 0,25145 | 0,662613 |
| Fibin | 0,735977 | 0,0143832 | -5,6772 | 0,25145 | 0,662613 |
| Neurod4 | 1,82872 | 0,00854021 | -7,74235 | 0,25145 | 0,662613 |
| Tmcc2 | 8,62962 | 10,1716 | 0,237185 | 0,2516 | 0,662736 |
| Siae | 1,94793 | 1,52792 | -0,350373 | 0,2516 | 0,662736 |
| Sec61g | 0,0399875 | 0,594111 | 3,89311 | 0,2517 | 0,662863 |
| Ptplb | 6,42997 | 7,62865 | 0,246617 | 0,25195 | 0,663249 |
| Apip | 44,6651 | 37,8143 | -0,240217 | 0,25195 | 0,663249 |
| Prrc2a | 118,783 | 138,184 | 0,218259 | 0,2521 | 0,663359 |
| Capn2 | 41,2505 | 47,6431 | 0,207856 | 0,25215 | 0,663359 |
| Slc22a17 | 21,1623 | 18,1019 | -0,225357 | 0,25215 | 0,663359 |
| Fam105a | 1,11006 | 1,45681 | 0,392174 | 0,25225 | 0,663359 |
| Olfm2 | 2,43281 | 1,81729 | -0,420834 | 0,25225 | 0,663359 |
| Fcf1 | 42,204 | 35,3216 | -0,256828 | 0,2524 | 0,663617 |
| Tubb2b | 76,8314 | 66,5479 | -0,207304 | 0,2526 | 0,664007 |
| Fam109b | 0,90332 | 1,21714 | 0,430182 | 0,2527 | 0,664133 |
| 9930021J03Rik | 7,33965 | 8,54182 | 0,218833 | 0,2529 | 0,664518 |
| Atp6v1c1 | 24,7051 | 21,0993 | -0,227618 | 0,25295 | 0,664518 |
| Map3k6 | 0,485902 | 0,665007 | 0,452704 | 0,2532 | 0,664898 |
| Zfp120 | 2,16565 | 2,69043 | 0,31304 | 0,25325 | 0,664898 |
| Tonsl | 10,7986 | 12,609 | 0,223607 | 0,25325 | 0,664898 |
| Gpx7 | 30,8118 | 26,1286 | -0,237855 | 0,2534 | 0,665156 |
| Ngfrap1 | 338,643 | 294,046 | -0,203725 | 0,2535 | 0,665282 |
| Dync1i2 | 83,1196 | 72,1591 | -0,204007 | 0,2536 | 0,665408 |
| Slco3a1 | 1,41305 | 1,09751 | -0,36458 | 0,2537 | 0,665534 |
| Tmem231 | 7,9918 | 9,4789 | 0,246199 | 0,25385 | 0,665792 |
| Mir7014 | 223,109 | 0 | -inf | 0,25405 | 0,665903 |
| Psmb4 | 296,394 | 256,889 | -0,20637 | 0,25405 | 0,665903 |
| Hdgf | 244,628 | 211,671 | -0,208761 | 0,25405 | 0,665903 |
| Setd5 | 50,7134 | 58,4439 | 0,204684 | 0,2541 | 0,665903 |
| Mfsd8 | 2,42822 | 1,91685 | -0,341164 | 0,25425 | 0,66616 |
| Mir6970 | 279,907 | 0 | -inf | 0,25445 | 0,666548 |
| Grhpr | 34,9887 | 40,9109 | 0,225593 | 0,25465 | 0,666904 |
| Zfp790 | 5,09922 | 6,03385 | 0,242803 | 0,2548 | 0,666904 |

| | | | | | |
|---|---|---|---|---|---|
| Golga4 | 16,8332 | 19,4517 | 0,208588 | 0,2548 | 0,666904 |
| Def8 | 17,6697 | 15,2406 | -0,21336 | 0,25485 | 0,666904 |
| Fam86 | 7,02717 | 5,42597 | -0,373063 | 0,25485 | 0,666904 |
| Nol11 | 46,9103 | 40,6445 | -0,206845 | 0,25495 | 0,666904 |
| Pawr | 19,8774 | 16,8704 | -0,236635 | 0,25495 | 0,666904 |
| Dus1l | 42,3177 | 36,5429 | -0,211667 | 0,2551 | 0,667161 |
| Pdcd5 | 34,9665 | 42,0199 | 0,2651 | 0,2552 | 0,667286 |
| Thoc5 | 42,4465 | 36,7399 | -0,208296 | 0,25545 | 0,667646 |
| Cep120 | 11,5662 | 9,91068 | -0,222858 | 0,25555 | 0,667646 |
| Rgl3 | 3,22568 | 4,00309 | 0,311514 | 0,2556 | 0,667646 |
| Pign | 7,28592 | 8,48743 | 0,220218 | 0,2556 | 0,667646 |
| 2610318N02Rik | 25,4865 | 21,6047 | -0,23839 | 0,25565 | 0,667646 |
| Ogfod2 | 4,69098 | 3,67858 | -0,350741 | 0,25565 | 0,667646 |
| Mir8091 | 8,00883 | 30,374 | 1,92317 | 0,25585 | 0,66787 |
| Wdr76 | 14,362 | 12,366 | -0,215874 | 0,25585 | 0,66787 |
| Hnrnph3 | 87,3088 | 73,7843 | -0,242813 | 0,2559 | 0,66787 |
| Tpd52l1 | 18,6274 | 14,6948 | -0,342118 | 0,25595 | 0,66787 |
| Lsm1 | 17,9852 | 15,3825 | -0,225527 | 0,25605 | 0,66787 |
| Uhrf1 | 122,572 | 141,284 | 0,204975 | 0,2561 | 0,66787 |
| Gmpr2 | 41,7367 | 35,9189 | -0,216574 | 0,2561 | 0,66787 |
| Rpl8 | 2031,27 | 1742,53 | -0,221198 | 0,25625 | 0,667973 |
| Gzmm | 0,708895 | 1,10903 | 0,645651 | 0,2563 | 0,667973 |
| Zfp598 | 30,125 | 34,861 | 0,210651 | 0,25635 | 0,667973 |
| Slc26a2 | 4,7576 | 5,67771 | 0,255074 | 0,2564 | 0,667973 |
| Cs | 152,664 | 132,438 | -0,205043 | 0,2564 | 0,667973 |
| Kbtbd7 | 4,37238 | 5,20222 | 0,250708 | 0,25665 | 0,668353 |
| Banf1 | 398,225 | 334,463 | -0,251736 | 0,25665 | 0,668353 |
| Hoxa2 | 16,3915 | 19,3185 | 0,237038 | 0,25685 | 0,668597 |
| Pgp | 40,4215 | 34,4294 | -0,231482 | 0,25685 | 0,668597 |
| Zbtbd6 | 1,59255 | 1,22248 | -0,381535 | 0,2569 | 0,668597 |
| Copz1 | 148,301 | 128,917 | -0,20209 | 0,257 | 0,668716 |
| Slc25a30 | 6,04415 | 4,86086 | -0,314327 | 0,25705 | 0,668716 |
| Fam117a | 11,4653 | 13,5426 | 0,24023 | 0,2572 | 0,668835 |
| Mrps14 | 27,9963 | 24,0217 | -0,220899 | 0,2572 | 0,668835 |
| 1700108F19Rik | 1,09911 | 0,63399 | -0,793803 | 0,2573 | 0,668959 |
| Psmd13 | 146,01 | 126,909 | -0,202273 | 0,2574 | 0,669084 |
| Sh2b3 | 5,2558 | 6,38132 | 0,279945 | 0,2575 | 0,669208 |
| Fip1l1 | 45,4603 | 52,6419 | 0,211603 | 0,25765 | 0,669321 |
| Fgfr1op | 23,2605 | 20,1215 | -0,209143 | 0,25765 | 0,669321 |
| Txndc5 | 132,43 | 115,039 | -0,203118 | 0,2577 | 0,669321 |
| Riiad1 | 1,00035 | 0,479967 | -1,05949 | 0,25785 | 0,669575 |
| Mtcp1 | 10,1626 | 8,18831 | -0,311628 | 0,258 | 0,669829 |
| Tmem260 | 6,91537 | 5,88589 | -0,232546 | 0,2582 | 0,670212 |
| Msrb3 | 7,47017 | 6,28796 | -0,248549 | 0,2583 | 0,670336 |
| Plekhh2 | 4,84862 | 5,67263 | 0,226444 | 0,2584 | 0,670448 |
| Fxyd2 | 0,698143 | 0,315335 | -1,14664 | 0,25845 | 0,670448 |
| Swap70 | 16,5515 | 19,1748 | 0,212252 | 0,2585 | 0,670448 |
| Ptma | 244,665 | 212,499 | -0,203352 | 0,25875 | 0,670961 |
| G6pd2 | 5,54041 | 4,4187 | -0,326371 | 0,2589 | 0,671214 |
| Hmgn3 | 48,1679 | 40,9112 | -0,235578 | 0,259 | 0,671284 |

| | | | | | |
|---|---|---|---|---|---|
| Ubr1 | 8,97521 | 10,368 | 0,208123 | 0,25905 | 0,671284 |
| Mapk8ip3 | 17,1335 | 19,9024 | 0,216121 | 0,25925 | 0,671284 |
| Fignl1 | 51,9148 | 45,0914 | -0,203293 | 0,25925 | 0,671284 |
| Wbp1 | 53,1053 | 45,8163 | -0,212995 | 0,2593 | 0,671284 |
| Thoc6 | 47,1434 | 39,8969 | -0,24078 | 0,25935 | 0,671284 |
| Wdsub1 | 5,82856 | 7,09541 | 0,283745 | 0,2594 | 0,671284 |
| Cct3 | 426,909 | 369,073 | -0,210024 | 0,2594 | 0,671284 |
| Snrpd1 | 259,01 | 224,773 | -0,20454 | 0,25945 | 0,671284 |
| Rabac1 | 21,4725 | 17,7136 | -0,277636 | 0,25945 | 0,671284 |
| Glb1l | 4,5256 | 6,02608 | 0,413109 | 0,2596 | 0,67153 |
| Tmem132d | 0,663175 | 0,486634 | -0,446552 | 0,25965 | 0,67153 |
| Eea1 | 3,05738 | 3,6242 | 0,245365 | 0,2598 | 0,671647 |
| Aspscr1 | 35,1207 | 40,7396 | 0,21411 | 0,2598 | 0,671647 |
| Hint1 | 660,014 | 573,407 | -0,202936 | 0,2599 | 0,671771 |
| Sumf2 | 15,2344 | 17,8703 | 0,230233 | 0,2603 | 0,671918 |
| Carhsp1 | 38,1786 | 43,9884 | 0,204357 | 0,2604 | 0,671918 |
| Caap1 | 13,8783 | 11,729 | -0,242754 | 0,2604 | 0,671918 |
| Copb2 | 109,426 | 95,144 | -0,201773 | 0,26045 | 0,671918 |
| Zfp931 | 1,32959 | 1,79854 | 0,435846 | 0,2605 | 0,671918 |
| Espl1 | 24,6882 | 28,4486 | 0,204533 | 0,2605 | 0,671918 |
| 1700021F05Rik | 35,1903 | 29,7024 | -0,244599 | 0,26055 | 0,671918 |
| Mfsd9 | 1,02103 | 0,758527 | -0,428748 | 0,26055 | 0,671918 |
| 2900009J06Rik | 4,36607 | 2,81137 | -0,635062 | 0,26055 | 0,671918 |
| Ppp1r12b | 2,20719 | 2,74993 | 0,317181 | 0,2607 | 0,671918 |
| Timm21 | 34,9549 | 40,7593 | 0,221633 | 0,2607 | 0,671918 |
| Ost4 | 72,3509 | 58,2625 | -0,312443 | 0,2607 | 0,671918 |
| Mrpl22 | 27,3261 | 22,5788 | -0,275308 | 0,26075 | 0,671918 |
| Car9 | 0,681012 | 0,98679 | 0,535063 | 0,26085 | 0,671918 |
| Ppp1r14b | 83,2004 | 71,735 | -0,213913 | 0,26085 | 0,671918 |
| Eci1 | 36,5422 | 31,1703 | -0,22939 | 0,26085 | 0,671918 |
| Cenpv | 28,9689 | 24,4456 | -0,244929 | 0,26085 | 0,671918 |
| 1110038F14Rik | 38,6229 | 32,9082 | -0,231008 | 0,2609 | 0,671918 |
| Fam63b | 1,23092 | 1,50309 | 0,288204 | 0,2611 | 0,672298 |
| Tbc1d32 | 2,02993 | 2,53143 | 0,318526 | 0,2613 | 0,672672 |
| AI846148 | 26,0031 | 30,4424 | 0,227399 | 0,26135 | 0,672672 |
| Ddx3x | 198,643 | 230,398 | 0,213945 | 0,2615 | 0,672788 |
| Zfand2b | 9,54508 | 7,76568 | -0,297644 | 0,2615 | 0,672788 |
| Atxn2l | 71,5185 | 82,1321 | 0,199629 | 0,2618 | 0,673425 |
| Cby1 | 30,5109 | 25,862 | -0,238491 | 0,26195 | 0,673509 |
| Tacc3 | 94,7012 | 82,3271 | -0,202014 | 0,262 | 0,673509 |
| Med31 | 16,3117 | 13,3881 | -0,284962 | 0,262 | 0,673509 |
| Gipc1 | 57,7025 | 50,0052 | -0,206555 | 0,26205 | 0,673509 |
| Ccnj | 19,5741 | 22,665 | 0,211521 | 0,2621 | 0,673509 |
| Lyrm2 | 34,3673 | 29,4422 | -0,223151 | 0,26215 | 0,673509 |
| Trdmt1 | 2,67226 | 2,10998 | -0,34083 | 0,2622 | 0,673509 |
| Cacybp | 129,37 | 112,488 | -0,201729 | 0,2623 | 0,673631 |
| Angel1 | 4,34124 | 5,18902 | 0,257355 | 0,26245 | 0,673874 |
| Ncaph2 | 55,786 | 48,3696 | -0,205803 | 0,2625 | 0,673874 |
| Zfp143 | 16,7215 | 19,4215 | 0,215944 | 0,26285 | 0,674625 |
| Aldh18a1 | 47,3173 | 41,1307 | -0,202153 | 0,26295 | 0,674625 |

| | | | | | |
|---|---|---|---|---|---|
| Tyms-ps | 13,4356 | 11,1467 | -0,269444 | 0,26295 | 0,674625 |
| Rab31 | 18,6345 | 16,0553 | -0,214922 | 0,26305 | 0,674746 |
| Atg4c | 5,79454 | 6,93092 | 0,258353 | 0,2633 | 0,675097 |
| Mir7-1 | 20,0456 | 81,3387 | 2,02066 | 0,2634 | 0,675097 |
| Cttnbp2nl | 17,6983 | 20,4101 | 0,20567 | 0,2634 | 0,675097 |
| Tfg | 70,0162 | 60,8612 | -0,202166 | 0,2634 | 0,675097 |
| Ankrd13c | 13,4379 | 15,6915 | 0,223677 | 0,26345 | 0,675097 |
| Tspan11 | 0,957152 | 0,729713 | -0,39142 | 0,2636 | 0,675347 |
| Ube2z | 39,972 | 45,9311 | 0,200482 | 0,2637 | 0,675454 |
| Tbc1d2b | 5,60193 | 4,7744 | -0,230605 | 0,26375 | 0,675454 |
| Icosl | 1,28756 | 1,67136 | 0,376381 | 0,2638 | 0,675454 |
| 1810058I24Rik | 72,4033 | 62,5023 | -0,212146 | 0,2639 | 0,675576 |
| Pnpo | 7,8549 | 6,51335 | -0,270194 | 0,26405 | 0,675818 |
| Wee1 | 25,7422 | 29,6745 | 0,205088 | 0,2641 | 0,675818 |
| Rsbn1 | 8,17553 | 10,2878 | 0,331547 | 0,2642 | 0,675925 |
| 2310040G24Rik | 1,43299 | 2,26156 | 0,658289 | 0,2643 | 0,675925 |
| Zfp59 | 2,62697 | 3,21094 | 0,289595 | 0,2643 | 0,675925 |
| Rhebl1 | 10,3176 | 12,553 | 0,282922 | 0,2646 | 0,676423 |
| Zfp472 | 7,62699 | 9,1142 | 0,257001 | 0,2646 | 0,676423 |
| Hivep1 | 7,48494 | 6,47011 | -0,210201 | 0,265 | 0,677175 |
| Tmem9 | 41,1544 | 35,4394 | -0,215693 | 0,265 | 0,677175 |
| Psmb2 | 296,925 | 258,639 | -0,199156 | 0,2652 | 0,677551 |
| Prrc2c | 46,4077 | 53,4749 | 0,204497 | 0,26535 | 0,677665 |
| Mpv17l2 | 8,27995 | 6,40078 | -0,371374 | 0,26535 | 0,677665 |
| Pced1b | 1,115 | 1,6482 | 0,563857 | 0,26545 | 0,677785 |
| Nsa2 | 12,1835 | 9,67703 | -0,332291 | 0,26555 | 0,677906 |
| Slc16a4 | 0,743001 | 1,04035 | 0,485635 | 0,26595 | 0,678649 |
| Gadd45b | 1,30163 | 1,84759 | 0,505324 | 0,266 | 0,678649 |
| Zfp85 | 1,79449 | 2,36976 | 0,401163 | 0,266 | 0,678649 |
| Ddx19b | 7,67368 | 8,88288 | 0,21346 | 0,2662 | 0,679024 |
| Hspa14 | 78,3746 | 67,9167 | -0,206621 | 0,2667 | 0,680165 |
| Klhl36 | 1,38479 | 1,02298 | -0,436882 | 0,2668 | 0,680246 |
| Brd4 | 36,3241 | 41,981 | 0,208809 | 0,26695 | 0,680246 |
| Gng5 | 117,436 | 100,658 | -0,222415 | 0,26695 | 0,680246 |
| 9630033F20Rik | 6,58281 | 5,54866 | -0,246563 | 0,26695 | 0,680246 |
| Gm5129 | 0,0552326 | 0,976164 | 4,14353 | 0,267 | 0,680246 |
| Rtkn2 | 2,61574 | 2,12595 | -0,299112 | 0,26705 | 0,680246 |
| Lrrc9 | 0,445207 | 0,6519 | 0,550174 | 0,26715 | 0,680365 |
| B930003M22Rik | 1,13778 | 0,739896 | -0,620827 | 0,26735 | 0,680724 |
| Baiap2l1 | 7,34778 | 8,92252 | 0,280142 | 0,2674 | 0,680724 |
| Coa7 | 12,9903 | 11,1003 | -0,22684 | 0,26745 | 0,680724 |
| Prrg2 | 5,08715 | 6,40177 | 0,331614 | 0,26775 | 0,681027 |
| Cant1 | 18,6023 | 21,5862 | 0,214627 | 0,26785 | 0,681027 |
| Neurl4 | 27,0988 | 31,1615 | 0,201536 | 0,26785 | 0,681027 |
| Itm2c | 62,3947 | 54,2829 | -0,200926 | 0,26785 | 0,681027 |
| Snhg1 | 591,861 | 514,564 | -0,201907 | 0,268 | 0,681027 |
| Ptk2 | 50,618 | 58,0311 | 0,197176 | 0,26805 | 0,681027 |
| Cops8 | 90,2415 | 78,6122 | -0,199037 | 0,26805 | 0,681027 |
| Dars2 | 18,549 | 16,0274 | -0,210797 | 0,26805 | 0,681027 |
| Zfp948 | 5,95632 | 7,14242 | 0,261991 | 0,2681 | 0,681027 |

| | | | | | |
|---|---|---|---|---|---|
| P4ha2 | 13,9314 | 11,8212 | -0,236967 | 0,2681 | 0,681027 |
| Rufy2 | 7,01321 | 9,00925 | 0,361332 | 0,26835 | 0,681392 |
| Nqo1 | 2,33906 | 1,76312 | -0,407801 | 0,26835 | 0,681392 |
| Rab33a | 1,32987 | 0,883747 | -0,589575 | 0,26845 | 0,681511 |
| Srebf1 | 18,8084 | 23,8167 | 0,340595 | 0,26865 | 0,681876 |
| Psmf1 | 13,9607 | 16,1676 | 0,211731 | 0,2687 | 0,681876 |
| Zc3h11a | 49,1111 | 58,9848 | 0,264295 | 0,2688 | 0,681987 |
| Spty2d1 | 5,83938 | 6,8075 | 0,22131 | 0,26885 | 0,681987 |
| Ccdc84 | 8,08708 | 10,4313 | 0,367226 | 0,269 | 0,682176 |
| Tpt1 | 37,853 | 44,515 | 0,233884 | 0,2691 | 0,682176 |
| Mrpl45 | 34,0486 | 29,5572 | -0,204087 | 0,2691 | 0,682176 |
| Ak6 | 59,688 | 48,0411 | -0,313171 | 0,2692 | 0,682176 |
| Lrfn3 | 5,81993 | 4,83489 | -0,267518 | 0,26925 | 0,682176 |
| Kifc2 | 0,518162 | 0,724948 | 0,484474 | 0,2693 | 0,682176 |
| Nt5c | 33,8724 | 27,9243 | -0,278588 | 0,2693 | 0,682176 |
| Fancd2 | 13,1483 | 15,1771 | 0,20702 | 0,26935 | 0,682176 |
| Slc35e4 | 1,52012 | 1,17397 | -0,372788 | 0,26945 | 0,682295 |
| Ptpmt1 | 27,826 | 23,7237 | -0,230105 | 0,2696 | 0,68254 |
| Cep89 | 17,376 | 20,1783 | 0,215708 | 0,26975 | 0,682785 |
| Mrgpre | 0,651094 | 0,868175 | 0,41512 | 0,26985 | 0,682903 |
| Pnpla2 | 3,35306 | 4,12413 | 0,298611 | 0,27 | 0,683148 |
| Gcfc2 | 8,04571 | 10,7805 | 0,422128 | 0,27015 | 0,683393 |
| Dpy30 | 172,131 | 149,764 | -0,200816 | 0,27025 | 0,683511 |
| Galnt11 | 14,4001 | 16,756 | 0,218602 | 0,2706 | 0,684126 |
| Ppp2r1a | 161,953 | 141,075 | -0,199109 | 0,2706 | 0,684126 |
| Arhgef12 | 12,9428 | 14,8575 | 0,199041 | 0,2708 | 0,684362 |
| Cald1 | 47,6387 | 41,4835 | -0,199595 | 0,2708 | 0,684362 |
| Bbc3 | 3,55895 | 4,44175 | 0,319678 | 0,27115 | 0,685112 |
| Coro1b | 53,5653 | 46,5859 | -0,201405 | 0,27125 | 0,685186 |
| Arpc1b | 17,2447 | 20,3001 | 0,235335 | 0,2714 | 0,685186 |
| Aaed1 | 10,1831 | 12,4963 | 0,295322 | 0,27145 | 0,685186 |
| Hp1bp3 | 54,4585 | 47,523 | -0,196531 | 0,27145 | 0,685186 |
| 1500011B03Rik | 10,0846 | 8,08172 | -0,319416 | 0,2715 | 0,685186 |
| Slc10a3 | 7,74692 | 5,55722 | -0,47926 | 0,2715 | 0,685186 |
| Zc2hc1a | 7,77527 | 9,13858 | 0,233078 | 0,2717 | 0,685556 |
| Ppp3ca | 21,2111 | 18,2622 | -0,215963 | 0,2718 | 0,685665 |
| Itpk1 | 10,8024 | 12,6442 | 0,227115 | 0,27185 | 0,685665 |
| Sptan1 | 42,3505 | 48,5001 | 0,195608 | 0,27195 | 0,685782 |
| BC025920 | 2,30643 | 2,95042 | 0,355263 | 0,27245 | 0,686764 |
| Ppih | 26,4596 | 21,8615 | -0,275401 | 0,27245 | 0,686764 |
| Arsi | 1,28452 | 0,90924 | -0,498499 | 0,2725 | 0,686764 |
| Zfp609 | 18,0405 | 20,7108 | 0,199142 | 0,27265 | 0,686963 |
| H2-Q8 | 2,46107 | 3,99545 | 0,699071 | 0,27275 | 0,686963 |
| Ankle2 | 19,2968 | 22,1247 | 0,197295 | 0,2728 | 0,686963 |
| Rab8a | 65,0402 | 56,6602 | -0,198998 | 0,2728 | 0,686963 |
| Cnbp | 424,674 | 367,974 | -0,206751 | 0,27285 | 0,686963 |
| Eif4b | 182,171 | 158,191 | -0,203625 | 0,2729 | 0,686963 |
| Kdm6b | 18,0099 | 20,7174 | 0,202054 | 0,27305 | 0,687035 |
| Cables1 | 2,36743 | 2,92408 | 0,30466 | 0,2731 | 0,687035 |
| Josd1 | 19,7914 | 17,1203 | -0,209172 | 0,27315 | 0,687035 |

| | | | | | |
|---|---|---|---|---|---|
| Sec61a2 | 37,3031 | 42,9342 | 0,202831 | 0,2732 | 0,687035 |
| Srcrb4d | 1,08879 | 0,807305 | -0,431545 | 0,2732 | 0,687035 |
| Aftph | 9,96387 | 11,5732 | 0,216015 | 0,27325 | 0,687035 |
| 4930539J05Rik | 3,01032 | 4,4749 | 0,571938 | 0,2734 | 0,687125 |
| Nsmce2 | 65,2595 | 56,5825 | -0,205832 | 0,2734 | 0,687125 |
| Ppp1r3b | 1,11224 | 0,862232 | -0,36732 | 0,27345 | 0,687125 |
| Haus7 | 54,7748 | 63,0856 | 0,203799 | 0,2735 | 0,687125 |
| Ccdc106 | 1,93907 | 2,58788 | 0,416409 | 0,2738 | 0,687744 |
| Slx4ip | 6,31668 | 8,49272 | 0,427061 | 0,274 | 0,687977 |
| Ssbp4 | 16,07 | 13,5446 | -0,246653 | 0,274 | 0,687977 |
| Kmt2b | 17,7991 | 20,4116 | 0,197584 | 0,2742 | 0,688209 |
| Fbxl19 | 50,8773 | 58,291 | 0,196251 | 0,2742 | 0,688209 |
| Rsl1 | 1,94898 | 2,50405 | 0,361541 | 0,2744 | 0,688554 |
| Stk39 | 26,5323 | 23,0383 | -0,203716 | 0,2746 | 0,688554 |
| Rab22a | 17,0902 | 19,907 | 0,220108 | 0,2747 | 0,688554 |
| Trim41 | 20,8725 | 24,0164 | 0,202418 | 0,2747 | 0,688554 |
| 2700069I18Rik | 1,46019 | 0,0736366 | -4,30959 | 0,2747 | 0,688554 |
| Cenpb | 44,7964 | 33,0616 | -0,438228 | 0,27475 | 0,688554 |
| Gnb2 | 171,831 | 150,12 | -0,194877 | 0,2748 | 0,688554 |
| C330021F23Rik | 1,4819 | 1,98218 | 0,419634 | 0,2749 | 0,688554 |
| Foxj2 | 2,40049 | 2,89563 | 0,270548 | 0,2749 | 0,688554 |
| Elac1 | 3,47644 | 2,90561 | -0,258772 | 0,27495 | 0,688554 |
| Bhlhe22 | 0,602691 | 0,819782 | 0,443822 | 0,2751 | 0,688554 |
| Ccdc174 | 16,7052 | 19,5606 | 0,227655 | 0,2751 | 0,688554 |
| Ubr4 | 17,829 | 20,4985 | 0,201292 | 0,2751 | 0,688554 |
| Gpd2 | 10,9773 | 9,52285 | -0,205059 | 0,27515 | 0,688554 |
| Bcl2l12 | 30,6613 | 25,7188 | -0,253595 | 0,2752 | 0,688554 |
| Trmt13 | 4,69815 | 6,3532 | 0,435391 | 0,27525 | 0,688554 |
| Ralbp1 | 24,8914 | 28,5748 | 0,199098 | 0,27525 | 0,688554 |
| Tmem186 | 12,7418 | 10,9254 | -0,221886 | 0,27545 | 0,688768 |
| Zfpl1 | 21,9879 | 18,6294 | -0,239125 | 0,2755 | 0,688768 |
| Lin7c | 36,6667 | 41,9977 | 0,19584 | 0,27555 | 0,688768 |
| Igf2r | 50,6341 | 44,0106 | -0,202257 | 0,27555 | 0,688768 |
| Snx18 | 19,6387 | 17,0806 | -0,201339 | 0,2757 | 0,689008 |
| Acvr1 | 10,8198 | 12,6244 | 0,222537 | 0,27595 | 0,689489 |
| Srgap1 | 2,34514 | 2,78458 | 0,247782 | 0,276 | 0,689489 |
| B4galt1 | 13,9248 | 16,0361 | 0,203669 | 0,27615 | 0,68973 |
| Slc43a1 | 16,7816 | 19,4609 | 0,213694 | 0,27635 | 0,689789 |
| Dcakd | 182,723 | 159,694 | -0,194349 | 0,27635 | 0,689789 |
| Pmm1 | 19,4851 | 16,4236 | -0,246599 | 0,2764 | 0,689789 |
| Cmpk1 | 33,8615 | 29,2746 | -0,209999 | 0,27645 | 0,689789 |
| Golt1b | 35,9777 | 31,3492 | -0,198671 | 0,2765 | 0,689789 |
| Eif1ax | 268,398 | 233,868 | -0,198679 | 0,2765 | 0,689789 |
| Upf3a | 33,8787 | 29,1274 | -0,218001 | 0,27655 | 0,689789 |
| Gm12191 | 321,214 | 265,899 | -0,272654 | 0,2767 | 0,689944 |
| Hace1 | 10,7686 | 12,5132 | 0,216612 | 0,2768 | 0,689944 |
| Pcsk9 | 17,9027 | 15,4857 | -0,209241 | 0,27685 | 0,689944 |
| Upf3b | 19,777 | 16,9872 | -0,219376 | 0,2769 | 0,689944 |
| Arl4a | 9,59633 | 8,21792 | -0,22371 | 0,2769 | 0,689944 |
| Hist1h4k | 0,671341 | 1,56114 | 1,21748 | 0,277 | 0,689944 |

| | | | | | |
|---|---|---|---|---|---|
| Mir6948 | 704,096 | 1460,24 | 1,05236 | 0,2771 | 0,689944 |
| Nfrkb | 10,6185 | 12,2524 | 0,206481 | 0,27715 | 0,689944 |
| Arf6 | 34,6229 | 30,1929 | -0,197516 | 0,27715 | 0,689944 |
| Serf2 | 23,708 | 20,5567 | -0,205765 | 0,27715 | 0,689944 |
| Lbr | 58,3761 | 66,8543 | 0,195642 | 0,27735 | 0,690174 |
| Bckdha | 26,3708 | 22,6888 | -0,21696 | 0,27735 | 0,690174 |
| Stk19 | 10,1705 | 12,4036 | 0,286374 | 0,2775 | 0,690279 |
| Mrpl4 | 39,7244 | 34,3192 | -0,211007 | 0,2775 | 0,690279 |
| Olfm1 | 14,8195 | 17,518 | 0,241336 | 0,27775 | 0,690586 |
| Mtcl1 | 5,25381 | 6,099 | 0,215207 | 0,27775 | 0,690586 |
| Donson | 22,0221 | 25,4978 | 0,211421 | 0,27785 | 0,690586 |
| Ttc3 | 81,7481 | 94,1199 | 0,203315 | 0,27785 | 0,690586 |
| Herc3 | 1,88659 | 2,41965 | 0,359016 | 0,27795 | 0,690586 |
| Dcun1d4 | 17,1965 | 14,9249 | -0,204393 | 0,27795 | 0,690586 |
| Zbtb34 | 4,40067 | 5,14196 | 0,224597 | 0,278 | 0,690586 |
| Acsf2 | 5,23174 | 4,35767 | -0,263733 | 0,27815 | 0,690824 |
| Supv3l1 | 26,1161 | 22,511 | -0,214309 | 0,27835 | 0,69091 |
| Blmh | 126,717 | 110,652 | -0,195582 | 0,2784 | 0,69091 |
| Rhoc | 63,2336 | 54,6867 | -0,209502 | 0,2784 | 0,69091 |
| Rbbp7 | 124,769 | 104,606 | -0,254293 | 0,2784 | 0,69091 |
| Gnaz | 0,751746 | 1,02875 | 0,452573 | 0,2785 | 0,691025 |
| Igf2 | 223,157 | 193,411 | -0,20639 | 0,2786 | 0,691139 |
| Pycr2 | 50,6151 | 58,2202 | 0,201952 | 0,27905 | 0,692112 |
| Cnrip1 | 8,53584 | 6,99845 | -0,286498 | 0,2791 | 0,692112 |
| Atp5c1 | 254,047 | 222,379 | -0,192073 | 0,27945 | 0,692836 |
| Leng9 | 0,638467 | 0,94134 | 0,560103 | 0,2795 | 0,692836 |
| Actl6a | 126,939 | 110,988 | -0,193736 | 0,2798 | 0,693436 |
| Wipi1 | 7,80181 | 6,43975 | -0,276804 | 0,27985 | 0,693436 |
| Ccdc71l | 6,04965 | 5,14295 | -0,234256 | 0,28015 | 0,694045 |
| Tanc2 | 4,03797 | 4,66693 | 0,208844 | 0,28025 | 0,694159 |
| Tpgs2 | 12,4591 | 10,431 | -0,256325 | 0,2804 | 0,694396 |
| Txnrd3 | 8,32838 | 7,05263 | -0,239874 | 0,28055 | 0,694633 |
| Usp12 | 10,8902 | 12,5924 | 0,209523 | 0,2807 | 0,694829 |
| H13 | 26,2095 | 31,5525 | 0,267663 | 0,2808 | 0,694829 |
| Trim62 | 7,30274 | 6,19263 | -0,237885 | 0,28085 | 0,694829 |
| 2610507I01Rik | 3,41786 | 4,15746 | 0,282609 | 0,2809 | 0,694829 |
| Rprd2 | 9,07044 | 10,415 | 0,199416 | 0,2809 | 0,694829 |
| Mrrf | 10,9997 | 8,01385 | -0,456894 | 0,281 | 0,694943 |
| Set | 87,4028 | 76,4274 | -0,19359 | 0,2812 | 0,695262 |
| Mknk2 | 50,1175 | 43,8011 | -0,194346 | 0,28125 | 0,695262 |
| Atp13a1 | 30,0629 | 34,3934 | 0,194148 | 0,2813 | 0,695262 |
| Mier2 | 21,1349 | 24,3703 | 0,205499 | 0,2814 | 0,695262 |
| Stard7 | 60,0108 | 52,4043 | -0,195535 | 0,2814 | 0,695262 |
| Cnnm4 | 3,32448 | 3,96236 | 0,253232 | 0,2816 | 0,69558 |
| Fam196b | 0,951606 | 0,730494 | -0,381493 | 0,28165 | 0,69558 |
| Il3ra | 3,18307 | 4,06596 | 0,353178 | 0,2817 | 0,69558 |
| Brd1 | 21,077 | 24,1293 | 0,195113 | 0,28175 | 0,69558 |
| L2hgdh | 9,11012 | 7,7881 | -0,226198 | 0,2818 | 0,69558 |
| Elovl1 | 15,1445 | 21,6876 | 0,518074 | 0,28215 | 0,69631 |
| Gne | 7,45045 | 6,2721 | -0,248378 | 0,2823 | 0,69636 |

| | | | | | |
|---|---|---|---|---|---|
| Notch3 | 30,6945 | 35,071 | 0,192301 | 0,28235 | 0,69636 |
| Vti1b | 42,5361 | 36,825 | -0,208002 | 0,28235 | 0,69636 |
| Zfp239 | 9,15336 | 7,74571 | -0,240905 | 0,2824 | 0,69636 |
| Lrrc42 | 48,6324 | 55,7801 | 0,197832 | 0,2825 | 0,69636 |
| Pccb | 18,8975 | 21,8901 | 0,212085 | 0,28255 | 0,69636 |
| Glt8d2 | 1,24276 | 0,69441 | -0,839686 | 0,28255 | 0,69636 |
| Zmat2 | 80,7841 | 70,6266 | -0,19386 | 0,28265 | 0,696377 |
| Mpzl3 | 0,931274 | 1,41387 | 0,602369 | 0,2829 | 0,696377 |
| Ddx1 | 167,313 | 146,454 | -0,192098 | 0,28295 | 0,696377 |
| Pycr1 | 4,10465 | 4,94042 | 0,267374 | 0,283 | 0,696377 |
| Nr4a2 | 1,51765 | 1,91105 | 0,332522 | 0,28305 | 0,696377 |
| Pde4a | 1,60863 | 1,98163 | 0,300856 | 0,28305 | 0,696377 |
| Sec14l1 | 26,0385 | 30,1259 | 0,210354 | 0,28305 | 0,696377 |
| Pnn | 106,365 | 121,73 | 0,194661 | 0,28305 | 0,696377 |
| Strip1 | 33,4224 | 38,2919 | 0,196224 | 0,2831 | 0,696377 |
| Mir3064 | 1226,33 | 801,803 | -0,613031 | 0,2831 | 0,696377 |
| Gpd1 | 0,965612 | 1,26355 | 0,387962 | 0,2832 | 0,69649 |
| Ndufa2 | 163,986 | 140,623 | -0,22174 | 0,28335 | 0,696725 |
| Akap17b | 2,052 | 2,45377 | 0,257971 | 0,28355 | 0,696949 |
| Apoc1 | 0,95391 | 0,235289 | -2,01942 | 0,28355 | 0,696949 |
| Sec61b | 132,064 | 114,051 | -0,211565 | 0,2837 | 0,696998 |
| Ccser2 | 6,05975 | 7,0064 | 0,209414 | 0,2838 | 0,696998 |
| Mkln1 | 30,445 | 34,8602 | 0,195375 | 0,2838 | 0,696998 |
| Ppm1l | 3,94306 | 3,28949 | -0,261452 | 0,2838 | 0,696998 |
| Ttc7b | 12,3869 | 10,7017 | -0,21098 | 0,28385 | 0,696998 |
| Pemt | 4,22082 | 5,41938 | 0,360605 | 0,28395 | 0,696998 |
| Zdhhc7 | 13,7268 | 11,8006 | -0,218137 | 0,28395 | 0,696998 |
| Fem1b | 37,5898 | 42,9749 | 0,193152 | 0,2842 | 0,697201 |
| Mif4gd | 6,60351 | 8,06958 | 0,289261 | 0,28425 | 0,697201 |
| Sccpdh | 31,2289 | 35,9612 | 0,20356 | 0,28425 | 0,697201 |
| Tmem60 | 25,4924 | 21,4382 | -0,249886 | 0,28425 | 0,697201 |
| Irf2bpl | 5,90503 | 6,91292 | 0,227349 | 0,2845 | 0,69735 |
| Serpinb6a | 16,9597 | 20,8498 | 0,297921 | 0,28455 | 0,69735 |
| Nanos3 | 0,787846 | 1,31913 | 0,743598 | 0,2846 | 0,69735 |
| Dlgap4 | 25,1169 | 21,8606 | -0,200321 | 0,2846 | 0,69735 |
| Atn1 | 23,004 | 26,3796 | 0,197541 | 0,2847 | 0,69735 |
| Armcx4 | 29,779 | 26,0359 | -0,193793 | 0,2847 | 0,69735 |
| Mrps16 | 69,3804 | 59,109 | -0,231151 | 0,2847 | 0,69735 |
| Flna | 128,994 | 148,921 | 0,207245 | 0,28475 | 0,69735 |
| Trappc10 | 10,8674 | 12,5041 | 0,202395 | 0,2848 | 0,69735 |
| B230118H07Rik | 10,8212 | 9,20562 | -0,233268 | 0,28495 | 0,697562 |
| Trnau1ap | 39,8042 | 34,2781 | -0,215635 | 0,285 | 0,697562 |
| Hnrnph2 | 52,6996 | 46,0288 | -0,195255 | 0,28505 | 0,697562 |
| Acsm3 | 0,55113 | 1,19292 | 1,11404 | 0,28525 | 0,697631 |
| Clu | 4,03306 | 4,98502 | 0,305726 | 0,28525 | 0,697631 |
| Klf7 | 2,47544 | 1,85436 | -0,416768 | 0,28525 | 0,697631 |
| H2-M5 | 1,10035 | 1,59172 | 0,532619 | 0,2853 | 0,697631 |
| Tbccd1 | 13,3195 | 15,4432 | 0,21343 | 0,28535 | 0,697631 |
| Rimklb | 12,8437 | 14,81 | 0,20551 | 0,28575 | 0,69821 |
| Mrpl34 | 107,35 | 92,7484 | -0,210926 | 0,28575 | 0,69821 |

| | | | | | |
|---|---|---|---|---|---|
| Acaa2 | 23,241 | 19,91 | -0,223178 | 0,28575 | 0,69821 |
| Fbxl20 | 5,76655 | 6,65616 | 0,206982 | 0,2859 | 0,698443 |
| Ptges3 | 44,897 | 39,1231 | -0,198599 | 0,286 | 0,698544 |
| Ddhd2 | 8,48939 | 9,86328 | 0,216407 | 0,28605 | 0,698544 |
| Pdk1 | 15,9966 | 18,3625 | 0,198994 | 0,28635 | 0,699013 |
| Mrps18a | 68,1745 | 58,9973 | -0,208583 | 0,2866 | 0,699013 |
| Neto2 | 6,59718 | 5,6678 | -0,21906 | 0,2866 | 0,699013 |
| Mcu | 4,82068 | 5,74465 | 0,252982 | 0,28665 | 0,699013 |
| Uqcr10 | 341,766 | 298,163 | -0,196909 | 0,28665 | 0,699013 |
| Elp4 | 13,5493 | 11,7355 | -0,207337 | 0,28665 | 0,699013 |
| Evc2 | 14,6542 | 16,849 | 0,201351 | 0,2867 | 0,699013 |
| Homer3 | 38,6484 | 32,4723 | -0,251202 | 0,28675 | 0,699013 |
| Tead4 | 1,47432 | 1,86617 | 0,340036 | 0,28695 | 0,699013 |
| AW554918 | 5,92636 | 6,88797 | 0,216932 | 0,28695 | 0,699013 |
| Arl8b | 33,6568 | 29,372 | -0,196458 | 0,28705 | 0,699013 |
| Gigyf1 | 15,7107 | 17,9882 | 0,195303 | 0,2871 | 0,699013 |
| Slc12a9 | 4,79155 | 5,68554 | 0,246803 | 0,28715 | 0,699013 |
| Ppm1d | 16,8314 | 14,5431 | -0,210827 | 0,28725 | 0,699013 |
| Tmed3 | 16,422 | 13,871 | -0,243557 | 0,28725 | 0,699013 |
| Cdc26 | 57,0182 | 49,0674 | -0,216658 | 0,2873 | 0,699013 |
| Scyl2 | 16,8328 | 19,3681 | 0,20241 | 0,28735 | 0,699013 |
| Fgfr1op2 | 24,6092 | 28,231 | 0,198081 | 0,28735 | 0,699013 |
| Usp48 | 27,9281 | 31,882 | 0,191026 | 0,28735 | 0,699013 |
| Cuedc2 | 36,506 | 31,7008 | -0,203615 | 0,28735 | 0,699013 |
| Kpna3 | 43,6899 | 49,9557 | 0,193352 | 0,28745 | 0,699013 |
| Ngdn | 47,9223 | 41,4254 | -0,210179 | 0,28745 | 0,699013 |
| Ptpn11 | 53,8234 | 61,3582 | 0,189023 | 0,2875 | 0,699013 |
| Chrna3 | 0,695251 | 0,501395 | -0,471586 | 0,28755 | 0,699013 |
| Hrsp12 | 29,8488 | 25,4303 | -0,231128 | 0,2877 | 0,699113 |
| Prss35 | 1,19997 | 0,905194 | -0,4067 | 0,2877 | 0,699113 |
| Rasd1 | 1,09166 | 2,20804 | 1,01624 | 0,28785 | 0,699345 |
| Mrps17 | 57,0999 | 49,5224 | -0,205407 | 0,288 | 0,699555 |
| Jmjd7-pla2g4b | 2,75115 | 3,81339 | 0,47104 | 0,2881 | 0,699555 |
| Zfp672 | 12,0408 | 8,80978 | -0,450758 | 0,2881 | 0,699555 |
| Rpap1 | 18,7293 | 21,4238 | 0,193921 | 0,28825 | 0,699654 |
| Pop1 | 25,5341 | 22,2455 | -0,198913 | 0,28825 | 0,699654 |
| Pou2f1 | 12,1842 | 13,9382 | 0,194026 | 0,2884 | 0,699753 |
| Ssr2 | 148,313 | 129,749 | -0,192929 | 0,2884 | 0,699753 |
| Rad54l2 | 20,1981 | 23,0866 | 0,192838 | 0,28855 | 0,699941 |
| Pmf1 | 103,536 | 90,2948 | -0,197413 | 0,28865 | 0,699941 |
| Ivd | 40,2288 | 35,0801 | -0,197577 | 0,28865 | 0,699941 |
| Trip6 | 43,9765 | 38,2787 | -0,20019 | 0,2887 | 0,699941 |
| Zbtb12 | 71,209 | 62,2718 | -0,19348 | 0,28875 | 0,69994 |
| Ppil1 | 110,289 | 96,3562 | -0,194836 | 0,28885 | 0,700051 |
| Wdr36 | 47,2104 | 41,3172 | -0,192364 | 0,289 | 0,700271 |
| Iqsec1 | 7,16873 | 8,26996 | 0,206163 | 0,28905 | 0,700271 |
| Zfp618 | 7,63457 | 8,97415 | 0,233229 | 0,2892 | 0,70037 |
| Iqce | 6,09524 | 7,1209 | 0,224377 | 0,2892 | 0,70037 |
| Ypel4 | 1,06803 | 1,678 | 0,651782 | 0,28945 | 0,700677 |
| Rab39b | 3,35825 | 2,75743 | -0,284383 | 0,2895 | 0,700677 |

| | | | | | |
|---|---|---|---|---|---|
| AW146154 | 1,19049 | 1,5151 | 0,347856 | 0,28955 | 0,700677 |
| Brsk1 | 11,5018 | 13,4605 | 0,226876 | 0,2896 | 0,700677 |
| Fam84b | 4,18954 | 3,49142 | -0,262977 | 0,2896 | 0,700677 |
| Dnajc12 | 9,48469 | 11,4603 | 0,27297 | 0,2897 | 0,700775 |
| Atad2b | 11,3622 | 12,993 | 0,193488 | 0,28975 | 0,700775 |
| Cd3eap | 25,0959 | 21,0143 | -0,256076 | 0,2899 | 0,700995 |
| Cdk13 | 15,0717 | 17,217 | 0,191995 | 0,28995 | 0,700995 |
| Ict1 | 75,4193 | 65,2169 | -0,209689 | 0,2902 | 0,701335 |
| Taf12 | 36,516 | 31,5761 | -0,209694 | 0,2902 | 0,701335 |
| Sugt1 | 65,2876 | 57,0672 | -0,194147 | 0,2904 | 0,701554 |
| Lpl | 18,069 | 15,736 | -0,199449 | 0,2904 | 0,701554 |
| Rec8 | 1,43037 | 1,07261 | -0,415262 | 0,29055 | 0,701773 |
| Timm8b | 213,521 | 177,708 | -0,264867 | 0,2906 | 0,701773 |
| Egln2 | 34,2386 | 29,749 | -0,202782 | 0,2907 | 0,701882 |
| Fbf1 | 16,8339 | 14,6891 | -0,196617 | 0,29115 | 0,702446 |
| Btbd3 | 9,3308 | 8,0794 | -0,207752 | 0,29115 | 0,702446 |
| Atp5f1 | 121,83 | 97,6933 | -0,318536 | 0,2913 | 0,702446 |
| Uqcrh | 458,678 | 401,968 | -0,190401 | 0,29135 | 0,702446 |
| Tmod3 | 34,3082 | 30,0298 | -0,192159 | 0,2914 | 0,702446 |
| Aimp1 | 107,109 | 93,6314 | -0,194015 | 0,2914 | 0,702446 |
| Il1rap | 1,32952 | 1,05113 | -0,338965 | 0,29145 | 0,702446 |
| Abat | 7,44978 | 8,63391 | 0,212816 | 0,29155 | 0,702446 |
| Nudt6 | 2,90233 | 2,15859 | -0,427118 | 0,2916 | 0,702446 |
| Apex1 | 274,972 | 238,92 | -0,202758 | 0,29165 | 0,702446 |
| 6030419C18Rik | 1,75197 | 1,34342 | -0,383071 | 0,29165 | 0,702446 |
| Ina | 0,777273 | 0,117194 | -2,72952 | 0,29165 | 0,702446 |
| Samd1 | 37,23 | 32,4746 | -0,197151 | 0,2917 | 0,702446 |
| Mlycd | 3,31457 | 2,64854 | -0,323623 | 0,2917 | 0,702446 |
| Zgpat | 26,7141 | 32,5723 | 0,286042 | 0,29185 | 0,702675 |
| Swi5 | 496,137 | 433,946 | -0,193223 | 0,29195 | 0,702784 |
| Snx27 | 22,9395 | 26,1929 | 0,191341 | 0,2922 | 0,703254 |
| Abi1 | 19,007 | 22,0668 | 0,215349 | 0,2923 | 0,703362 |
| Cds2 | 11,2674 | 13,326 | 0,24209 | 0,29245 | 0,703591 |
| Sgms1 | 4,91185 | 4,12276 | -0,252654 | 0,2927 | 0,704037 |
| Rdh5 | 0,91523 | 1,32905 | 0,53819 | 0,29275 | 0,704037 |
| Abl1 | 50,5183 | 44,3218 | -0,18879 | 0,2928 | 0,704037 |
| Trp53i11 | 28,8656 | 25,1266 | -0,200137 | 0,29305 | 0,704471 |
| Tmem199 | 37,4425 | 43,1879 | 0,20595 | 0,2931 | 0,704471 |
| Gnao1 | 4,67581 | 3,92731 | -0,251676 | 0,29315 | 0,704471 |
| 2310069G16Rik | 0,613762 | 0,194054 | -1,66122 | 0,2932 | 0,704471 |
| Rexo2 | 48,559 | 42,0581 | -0,207356 | 0,2936 | 0,705288 |
| Uxt | 7,12079 | 5,23016 | -0,445182 | 0,29365 | 0,705288 |
| Gm20939 | 0,705546 | 0,966337 | 0,453787 | 0,29375 | 0,705384 |
| Rnf121 | 22,0896 | 25,6053 | 0,213081 | 0,2938 | 0,705384 |
| Wdr83os | 53,0189 | 45,3368 | -0,225825 | 0,29395 | 0,705588 |
| Soat1 | 11,3948 | 9,87586 | -0,206394 | 0,294 | 0,705588 |
| Ppp1r1a | 35,6478 | 30,8011 | -0,210833 | 0,29405 | 0,705588 |
| Rnu11 | 31,2006 | 90,003 | 1,5284 | 0,2942 | 0,705816 |
| Dym | 18,8051 | 21,6823 | 0,205394 | 0,29445 | 0,706284 |
| Arntl2 | 1,33015 | 1,70646 | 0,359422 | 0,2946 | 0,706379 |

| | | | | | |
|---|---|---|---|---|---|
| Ndor1 | 11,0413 | 12,6874 | 0,200487 | 0,2946 | 0,706379 |
| Syngr4 | 0,459765 | 0,798231 | 0,795909 | 0,29475 | 0,706607 |
| Syt9 | 2,27313 | 2,81153 | 0,306673 | 0,295 | 0,707062 |
| Ino80b | 22,0902 | 18,7044 | -0,240026 | 0,29505 | 0,707062 |
| Lpar6 | 30,4398 | 35,4467 | 0,219692 | 0,2952 | 0,707289 |
| Galnt18 | 0,900733 | 0,658612 | -0,451671 | 0,29555 | 0,707995 |
| Ppia | 0,918846 | 1,47635 | 0,684144 | 0,2958 | 0,708449 |
| Hmgcll1 | 2,01582 | 1,62329 | -0,312442 | 0,29585 | 0,708449 |
| Syce2 | 31,2147 | 24,1563 | -0,369828 | 0,29605 | 0,708784 |
| Gm6194 | 0,474857 | 0,718416 | 0,597327 | 0,2961 | 0,708784 |
| Tmem120b | 3,51266 | 2,79933 | -0,327481 | 0,2963 | 0,70913 |
| Cdk5 | 16,8884 | 14,5302 | -0,216974 | 0,2964 | 0,709224 |
| Mtch1 | 126,582 | 111,163 | -0,187386 | 0,29645 | 0,709224 |
| Rnf111 | 9,23141 | 10,6253 | 0,202882 | 0,2968 | 0,709929 |
| Taz | 18,4773 | 21,4021 | 0,211997 | 0,2971 | 0,710292 |
| Txnip | 12,8749 | 14,9074 | 0,211464 | 0,2972 | 0,710292 |
| Wdr24 | 11,3451 | 13,1245 | 0,210187 | 0,29725 | 0,710292 |
| Ppid | 90,4751 | 79,4229 | -0,187967 | 0,2973 | 0,710292 |
| Ccdc112 | 4,75674 | 5,6702 | 0,253425 | 0,2974 | 0,710292 |
| Srgap2 | 12,8173 | 14,6014 | 0,188018 | 0,29745 | 0,710292 |
| Tmeff1 | 68,1292 | 59,9101 | -0,185474 | 0,29745 | 0,710292 |
| Tmed4 | 44,3068 | 38,6195 | -0,198198 | 0,29745 | 0,710292 |
| Serf1 | 120,978 | 105,237 | -0,201102 | 0,29745 | 0,710292 |
| Pm20d2 | 0,735254 | 0,928576 | 0,336777 | 0,2977 | 0,710612 |
| Hdac4 | 4,76448 | 5,59723 | 0,232396 | 0,29775 | 0,710612 |
| Phf11c | 1,36284 | 0,973905 | -0,484762 | 0,29775 | 0,710612 |
| 3000002C10Rik | 4,1768 | 5,23808 | 0,326642 | 0,29795 | 0,710811 |
| Scmh1 | 11,54 | 13,315 | 0,206408 | 0,298 | 0,710811 |
| Lphn2 | 41,1143 | 46,7628 | 0,185723 | 0,298 | 0,710811 |
| Dopey1 | 2,42892 | 3,10159 | 0,352694 | 0,29815 | 0,710905 |
| Prpf38b | 63,2175 | 72,0156 | 0,187986 | 0,29815 | 0,710905 |
| Tle3 | 34,2131 | 39,0246 | 0,189834 | 0,2984 | 0,711369 |
| Map2k4 | 36,4877 | 41,5868 | 0,188719 | 0,29875 | 0,712071 |
| Mtmr7 | 2,37243 | 4,27361 | 0,849091 | 0,2989 | 0,712296 |
| Zfp783 | 3,24098 | 3,96097 | 0,289422 | 0,299 | 0,712402 |
| C77080 | 3,41884 | 4,92006 | 0,525168 | 0,2994 | 0,713037 |
| Nupl1 | 42,2469 | 48,15 | 0,18869 | 0,2994 | 0,713037 |
| Rab3gap1 | 8,79279 | 10,1723 | 0,21026 | 0,29945 | 0,713037 |
| Fkbp4 | 416,274 | 364,055 | -0,193377 | 0,29955 | 0,713037 |
| Usp31 | 5,97227 | 6,84103 | 0,195936 | 0,2996 | 0,713037 |
| Stt3b | 61,1783 | 69,5028 | 0,184051 | 0,2996 | 0,713037 |
| Fbxo9 | 11,7068 | 13,7058 | 0,22744 | 0,2997 | 0,713143 |
| Sipa1l1 | 17,1477 | 19,5694 | 0,19058 | 0,2999 | 0,713419 |
| Ubald2 | 8,75172 | 10,4811 | 0,260158 | 0,29995 | 0,713419 |
| Zfp788 | 6,30157 | 7,36132 | 0,224253 | 0,3 | 0,713419 |
| Phf8 | 9,20625 | 10,5617 | 0,198155 | 0,3001 | 0,713419 |
| Fgfrl1 | 25,0773 | 28,677 | 0,193512 | 0,3001 | 0,713419 |
| Sgsm2 | 3,38349 | 2,84509 | -0,250039 | 0,30015 | 0,713419 |
| Gm14378 | 2,08027 | 1,49553 | -0,476107 | 0,30035 | 0,713762 |
| Mocs3 | 3,4658 | 4,29827 | 0,310569 | 0,3006 | 0,714092 |

| | | | | | |
|---|---|---|---|---|---|
| Slx4 | 8,95627 | 10,2963 | 0,201148 | 0,3006 | 0,714092 |
| Srsf3 | 115,008 | 101,067 | -0,186418 | 0,30075 | 0,714184 |
| Tet3 | 11,3536 | 9,95692 | -0,189374 | 0,30075 | 0,714184 |
| Filip1l | 3,88695 | 2,79904 | -0,473704 | 0,301 | 0,714632 |
| Tomm6 | 151,552 | 132,85 | -0,190007 | 0,30105 | 0,714632 |
| 1810021B22Rik | 0,547571 | 1,01973 | 0,897062 | 0,3015 | 0,715408 |
| Gtpbp3 | 13,4533 | 11,6393 | -0,208948 | 0,3015 | 0,715408 |
| Npm2 | 1,12673 | 1,6485 | 0,549011 | 0,30155 | 0,715408 |
| Gpr162 | 2,62632 | 3,21379 | 0,29123 | 0,3016 | 0,715408 |
| N4bp2l2 | 10,3158 | 11,7484 | 0,187611 | 0,3019 | 0,715987 |
| Pvr | 2,38023 | 1,91786 | -0,311604 | 0,30215 | 0,716447 |
| Wdr34 | 10,8104 | 9,13302 | -0,243255 | 0,3024 | 0,716803 |
| Mettl21a | 6,83418 | 5,78179 | -0,241253 | 0,3025 | 0,716803 |
| Fbxw17 | 10,1564 | 8,55502 | -0,247544 | 0,30255 | 0,716803 |
| Cuedc1 | 20,1114 | 24,2138 | 0,267818 | 0,30265 | 0,716803 |
| Nfya | 46,8759 | 41,2421 | -0,184727 | 0,30265 | 0,716803 |
| Rsph1 | 0,743695 | 0,452392 | -0,717139 | 0,30265 | 0,716803 |
| Ctdsp1 | 24,1804 | 21,0876 | -0,197442 | 0,30275 | 0,716803 |
| Pomgnt2 | 3,74846 | 3,07282 | -0,286732 | 0,30275 | 0,716803 |
| Chst2 | 5,88977 | 5,09661 | -0,208672 | 0,30285 | 0,716803 |
| Gcnt4 | 1,23142 | 0,987149 | -0,31898 | 0,3029 | 0,716803 |
| Nxn | 64,4216 | 73,24 | 0,185087 | 0,3031 | 0,716803 |
| Mrpl55 | 42,5906 | 36,4814 | -0,223375 | 0,3031 | 0,716803 |
| Ppp3cc | 2,99066 | 3,71058 | 0,311177 | 0,30315 | 0,716803 |
| Heatr3 | 44,5845 | 39,1377 | -0,187981 | 0,30315 | 0,716803 |
| 1700052K11Rik | 3,34917 | 4,07665 | 0,283582 | 0,3032 | 0,716803 |
| Jam2 | 2,95627 | 3,49821 | 0,242839 | 0,30325 | 0,716803 |
| Pcnp | 23,0924 | 20,0397 | -0,204557 | 0,30325 | 0,716803 |
| Fsip1 | 1,7733 | 1,33887 | -0,405421 | 0,30345 | 0,717144 |
| Zfp429 | 1,90711 | 1,47625 | -0,369448 | 0,30355 | 0,717248 |
| Nat14 | 2,13391 | 1,63414 | -0,384971 | 0,30365 | 0,71731 |
| Ddrgk1 | 28,0635 | 24,0551 | -0,222348 | 0,3037 | 0,71731 |
| Arfgap1 | 28,3458 | 24,7598 | -0,195135 | 0,30375 | 0,71731 |
| Inpp1 | 5,07522 | 4,14778 | -0,291131 | 0,3038 | 0,71731 |
| Acss2 | 2,89685 | 2,3635 | -0,293565 | 0,30395 | 0,717344 |
| Atp7b | 0,82248 | 1,03928 | 0,337526 | 0,304 | 0,717344 |
| Kansl3 | 28,6566 | 32,6054 | 0,186242 | 0,30415 | 0,717344 |
| Alkbh5 | 24,0359 | 27,3372 | 0,185673 | 0,30415 | 0,717344 |
| Pogz | 18,2592 | 20,7532 | 0,184711 | 0,30415 | 0,717344 |
| Thoc3 | 55,3407 | 48,5634 | -0,188473 | 0,30415 | 0,717344 |
| Cops4 | 114,446 | 100,608 | -0,18592 | 0,30455 | 0,718142 |
| Uqcrc1 | 141,463 | 124,666 | -0,182354 | 0,3046 | 0,718142 |
| Rad51 | 63,9615 | 56,1793 | -0,187166 | 0,3047 | 0,718231 |
| Clcn2 | 7,13691 | 8,32905 | 0,222852 | 0,30475 | 0,718231 |
| Bok | 21,9494 | 25,428 | 0,212237 | 0,3049 | 0,718453 |
| Arl10 | 17,8811 | 21,0949 | 0,238463 | 0,305 | 0,718556 |
| Chrnb4 | 1,18143 | 0,922282 | -0,35726 | 0,30525 | 0,719013 |
| Slc12a6 | 4,37671 | 5,32523 | 0,282996 | 0,30545 | 0,719238 |
| Tmem176a | 10,3686 | 12,5232 | 0,272385 | 0,30555 | 0,719238 |
| Trappc13 | 28,6594 | 24,9363 | -0,20076 | 0,30555 | 0,719238 |

| | | | | | |
|---|---|---|---|---|---|
| Ttll11 | 0,426859 | 0,592478 | 0,473003 | 0,30575 | 0,719238 |
| 03/09/16 | 4,75808 | 3,88966 | -0,290736 | 0,30575 | 0,719238 |
| Eif4h | 271,876 | 238,553 | -0,188638 | 0,3058 | 0,719238 |
| Cmtr1 | 31,6431 | 36,6712 | 0,212755 | 0,30585 | 0,719238 |
| Exosc8 | 101,272 | 88,9435 | -0,18728 | 0,30585 | 0,719238 |
| Gm12504 | 0,998523 | 0,726627 | -0,458581 | 0,30585 | 0,719238 |
| Rims3 | 1,2503 | 0,822208 | -0,604694 | 0,30595 | 0,719341 |
| Ndrg4 | 2,38578 | 1,92808 | -0,307295 | 0,30615 | 0,719665 |
| 2010308F09Rik | 0,462385 | 0,976868 | 1,07907 | 0,3062 | 0,719665 |
| Tcf20 | 17,7221 | 20,2037 | 0,189068 | 0,3064 | 0,719786 |
| Hps1 | 5,1588 | 4,31864 | -0,256458 | 0,3064 | 0,719786 |
| Hdhd2 | 22,7601 | 19,8054 | -0,200611 | 0,30645 | 0,719786 |
| Crebrf | 0,779677 | 0,963269 | 0,305063 | 0,30655 | 0,719786 |
| B4galnt4 | 25,7226 | 29,3487 | 0,190258 | 0,3066 | 0,719786 |
| Rars | 118,414 | 104,29 | -0,183242 | 0,3066 | 0,719786 |
| Gm14057 | 0,891478 | 0,444862 | -1,00284 | 0,30665 | 0,719786 |
| Thtpa | 10,4775 | 8,91624 | -0,232786 | 0,3067 | 0,719786 |
| Pxdn | 73,1241 | 83,2464 | 0,187042 | 0,30685 | 0,719977 |
| Pramef8 | 5,98307 | 6,96362 | 0,218949 | 0,30695 | 0,719977 |
| Mtf2 | 36,5967 | 41,6299 | 0,185906 | 0,30695 | 0,719977 |
| Rras2 | 29,2979 | 33,4977 | 0,193262 | 0,3071 | 0,720197 |
| Psmd6 | 170,092 | 150,044 | -0,180931 | 0,3072 | 0,7203 |
| Heca | 6,5703 | 7,63784 | 0,217205 | 0,3074 | 0,720637 |
| Arl2 | 59,0616 | 51,2725 | -0,204036 | 0,3075 | 0,720725 |
| Mettl15 | 2,13845 | 1,66494 | -0,361096 | 0,30755 | 0,720726 |
| Arsa | 3,60387 | 4,29807 | 0,254142 | 0,3077 | 0,720931 |
| Atp6v0a1 | 14,672 | 16,7493 | 0,19104 | 0,30775 | 0,720931 |
| Tgm4 | 3,44145 | 2,83063 | -0,281891 | 0,30795 | 0,721253 |
| Tmem28 | 1,06004 | 1,46784 | 0,469583 | 0,308 | 0,721253 |
| Sesn2 | 5,82868 | 4,90678 | -0,248391 | 0,3081 | 0,721356 |
| Tspyl3 | 1,93197 | 2,38382 | 0,303207 | 0,3083 | 0,721429 |
| Fam184a | 1,61987 | 1,98713 | 0,294803 | 0,3083 | 0,721429 |
| Rpl32 | 1091,72 | 922,742 | -0,242604 | 0,3083 | 0,721429 |
| Zfp959 | 4,3257 | 5,23837 | 0,276185 | 0,30875 | 0,72235 |
| Psma1 | 221,305 | 194,887 | -0,183399 | 0,30895 | 0,722686 |
| Praf2 | 23,7771 | 20,399 | -0,221076 | 0,30915 | 0,722744 |
| Evi5l | 4,17814 | 3,3473 | -0,319864 | 0,30915 | 0,722744 |
| Tmem179b | 20,3442 | 27,129 | 0,415218 | 0,3092 | 0,722744 |
| Acot10 | 3,91575 | 4,84283 | 0,306561 | 0,3092 | 0,722744 |
| A430033K04Rik | 3,30251 | 2,73351 | -0,272807 | 0,3093 | 0,722846 |
| Bcr | 10,5584 | 9,25915 | -0,189443 | 0,30945 | 0,722933 |
| Ccdc94 | 11,8002 | 9,87334 | -0,257197 | 0,30945 | 0,722933 |
| 2610001J05Rik | 27,1434 | 23,6064 | -0,201426 | 0,30975 | 0,723502 |
| Acp6 | 12,0846 | 14,0812 | 0,2206 | 0,30985 | 0,723604 |
| Nkain4 | 1,016 | 0,629825 | -0,689872 | 0,3101 | 0,724026 |
| Epm2aip1 | 18,1839 | 20,6511 | 0,183554 | 0,3102 | 0,724026 |
| Lrp6 | 18,4073 | 16,2044 | -0,183894 | 0,3102 | 0,724026 |
| Ep300 | 16,9088 | 19,1954 | 0,182979 | 0,31055 | 0,724711 |
| AA474331 | 2,09513 | 2,92761 | 0,48268 | 0,3107 | 0,724737 |
| Anp32b | 410,623 | 361,596 | -0,183436 | 0,31075 | 0,724737 |

| | | | | | |
|---|---|---|---|---|---|
| Mark4 | 13,9548 | 16,1397 | 0,209856 | 0,31085 | 0,724737 |
| Txn1 | 230,202 | 202,61 | -0,184194 | 0,31085 | 0,724737 |
| Zpr1 | 28,5616 | 25,1373 | -0,184247 | 0,31085 | 0,724737 |
| Ddi2 | 27,1601 | 31,3031 | 0,204821 | 0,3109 | 0,724737 |
| Acp1 | 14,2262 | 12,3982 | -0,198416 | 0,31115 | 0,725188 |
| Armc7 | 3,4957 | 2,47778 | -0,496531 | 0,3113 | 0,725405 |
| Gpr137b | 2,80101 | 2,28561 | -0,293367 | 0,31155 | 0,725724 |
| Mafk | 2,25533 | 1,82095 | -0,308649 | 0,31155 | 0,725724 |
| Tmem8b | 2,11301 | 2,53576 | 0,263118 | 0,3118 | 0,726175 |
| Gpatch8 | 19,7472 | 22,4141 | 0,182763 | 0,3121 | 0,726726 |
| Znrd1 | 92,7142 | 80,7745 | -0,19889 | 0,31215 | 0,726726 |
| 3110040N11Rik | 13,9317 | 11,4824 | -0,278943 | 0,31235 | 0,72706 |
| Glg1 | 44,5721 | 50,6509 | 0,184446 | 0,3126 | 0,72751 |
| Pla2g3 | 0,720654 | 0,964179 | 0,419994 | 0,3128 | 0,727843 |
| Mapre3 | 4,03587 | 4,91276 | 0,283653 | 0,3129 | 0,727944 |
| Fam227a | 0,745738 | 0,584269 | -0,352036 | 0,3131 | 0,728277 |
| Fam76b | 16,5357 | 18,8952 | 0,192438 | 0,31325 | 0,728346 |
| Cdpf1 | 16,0934 | 13,3977 | -0,264489 | 0,3133 | 0,728346 |
| Amigo1 | 1,44144 | 1,18155 | -0,28683 | 0,3133 | 0,728346 |
| Prr12 | 16,3409 | 18,5713 | 0,184588 | 0,31345 | 0,728499 |
| Gnrh1 | 1,55423 | 0,872767 | -0,832535 | 0,3135 | 0,728499 |
| Rad23b | 107,86 | 95,1004 | -0,18164 | 0,31355 | 0,728499 |
| Nsun4 | 5,3853 | 6,28713 | 0,223374 | 0,3136 | 0,728499 |
| Car14 | 101,765 | 115,953 | 0,1883 | 0,31365 | 0,728499 |
| Slc25a35 | 0,579461 | 0,910364 | 0,651732 | 0,31375 | 0,728552 |
| Nphp1 | 27,6928 | 31,491 | 0,185428 | 0,31385 | 0,728552 |
| Myh10 | 92,7089 | 81,174 | -0,191691 | 0,3139 | 0,728552 |
| Mrpl36 | 48,3507 | 41,932 | -0,205488 | 0,3139 | 0,728552 |
| Ccrn4l | 7,63662 | 8,86203 | 0,214702 | 0,314 | 0,728653 |
| Eml1 | 23,8605 | 19,3882 | -0,29944 | 0,31435 | 0,729317 |
| Znrf2 | 6,55697 | 5,55959 | -0,238052 | 0,3144 | 0,729317 |
| Zfp558 | 2,93405 | 3,4952 | 0,252483 | 0,3146 | 0,729649 |
| Kctd21 | 1,37995 | 1,08618 | -0,345363 | 0,315 | 0,730445 |
| Nprl3 | 10,3834 | 12,4668 | 0,263815 | 0,3153 | 0,730927 |
| Pax8 | 0,637508 | 0,86687 | 0,443372 | 0,31535 | 0,730927 |
| Slc19a2 | 5,99082 | 6,98658 | 0,221833 | 0,31545 | 0,730927 |
| Pam16 | 1,14721 | 1,88489 | 0,716345 | 0,31555 | 0,730927 |
| Rae1 | 71,6486 | 63,0733 | -0,18391 | 0,31555 | 0,730927 |
| Eif6 | 81,8792 | 71,9748 | -0,186005 | 0,31555 | 0,730927 |
| Arhgef1 | 23,4749 | 20,6381 | -0,185807 | 0,31575 | 0,731259 |
| Dip2a | 5,18338 | 5,96022 | 0,201472 | 0,3159 | 0,731474 |
| Cul4b | 37,7657 | 42,8519 | 0,182282 | 0,3161 | 0,731772 |
| Hadhb | 7,07872 | 5,93834 | -0,253428 | 0,3162 | 0,731772 |
| Ikbke | 1,23269 | 0,965546 | -0,352397 | 0,3162 | 0,731772 |
| Pisd | 22,1531 | 19,3507 | -0,195119 | 0,31635 | 0,731987 |
| Git1 | 27,5868 | 24,2506 | -0,18596 | 0,31655 | 0,732302 |
| Rita1 | 7,25355 | 8,66664 | 0,256787 | 0,3166 | 0,732302 |
| Spopl | 6,2381 | 7,35518 | 0,237653 | 0,3169 | 0,73255 |
| Snrnp48 | 30,1317 | 34,5649 | 0,198028 | 0,31695 | 0,73255 |
| Ctr9 | 34,4612 | 30,3682 | -0,18241 | 0,317 | 0,73255 |

| | | | | | |
|---|---|---|---|---|---|
| Thada | 6,52646 | 7,45727 | 0,192346 | 0,31705 | 0,73255 |
| Src | 19,7102 | 22,3673 | 0,182448 | 0,31705 | 0,73255 |
| Mrpl54 | 132,799 | 115,993 | -0,195215 | 0,31705 | 0,73255 |
| Cox14 | 40,7486 | 34,91 | -0,223109 | 0,31715 | 0,732616 |
| Prkd2 | 5,59659 | 6,52985 | 0,222501 | 0,3172 | 0,732616 |
| Mtfr2 | 8,7978 | 7,49841 | -0,230557 | 0,31725 | 0,732616 |
| Tmem219 | 7,87021 | 6,3471 | -0,310305 | 0,3174 | 0,732831 |
| Rabl2 | 28,1234 | 24,4849 | -0,199878 | 0,3176 | 0,73316 |
| Fntb | 17,5752 | 15,2939 | -0,200582 | 0,3178 | 0,733474 |
| Ext2 | 28,2104 | 32,0466 | 0,183945 | 0,31785 | 0,733474 |
| Serbp1 | 169,929 | 148,843 | -0,19114 | 0,31805 | 0,733539 |
| Tubgcp5 | 11,3185 | 12,9551 | 0,194842 | 0,3181 | 0,733539 |
| Ppt1 | 23,5026 | 20,5303 | -0,195064 | 0,3181 | 0,733539 |
| Sdhaf1 | 28,3018 | 24,2737 | -0,221499 | 0,31815 | 0,733539 |
| Ppib | 465,665 | 527,646 | 0,180277 | 0,3183 | 0,733539 |
| 1700124L16Rik | 2,14869 | 1,04854 | -1,03507 | 0,3183 | 0,733539 |
| Terf2ip | 11,9558 | 10,4209 | -0,19823 | 0,31835 | 0,733539 |
| Cxcl17 | 0,329447 | 1,1714 | 1,83012 | 0,3184 | 0,733539 |
| Usp51 | 3,46426 | 2,82387 | -0,294875 | 0,3184 | 0,733539 |
| Eda2r | 2,71054 | 2,20936 | -0,294954 | 0,31845 | 0,733539 |
| 2410004N09Rik | 77,6142 | 66,1005 | -0,231659 | 0,31885 | 0,734278 |
| Gm9958 | 1,92963 | 2,48311 | 0,363822 | 0,31895 | 0,734278 |
| Mgat4b | 18,0367 | 15,7087 | -0,199372 | 0,31895 | 0,734278 |
| Tfb1m | 17,2923 | 14,4052 | -0,263539 | 0,319 | 0,734278 |
| Map2k7 | 25,5937 | 29,0254 | 0,181529 | 0,31915 | 0,734492 |
| Artn | 0,722271 | 0,513743 | -0,491495 | 0,3193 | 0,734672 |
| 2810008D09Rik | 21,6965 | 17,8783 | -0,279253 | 0,31935 | 0,734672 |
| Emg1 | 189,944 | 165,539 | -0,198402 | 0,3194 | 0,734672 |
| Alkbh4 | 8,42046 | 7,07249 | -0,251682 | 0,31955 | 0,734868 |
| 4930427A07Rik | 11,5869 | 10,077 | -0,201435 | 0,3196 | 0,734868 |
| Slc35a3 | 5,30479 | 6,16361 | 0,216481 | 0,3199 | 0,735426 |
| Rps6ka6 | 28,3206 | 32,125 | 0,181843 | 0,32005 | 0,735639 |
| Acat2 | 76,0618 | 67,1094 | -0,180659 | 0,32015 | 0,735737 |
| Zfp382 | 3,16577 | 3,86743 | 0,288818 | 0,3203 | 0,735831 |
| Actr1a | 59,0211 | 52,0264 | -0,181988 | 0,3204 | 0,735831 |
| Vps51 | 19,1048 | 16,4151 | -0,218915 | 0,32045 | 0,735831 |
| Gal3st4 | 2,67659 | 1,82779 | -0,550293 | 0,3205 | 0,735831 |
| Mir5133 | 89,017 | 343,836 | 1,94957 | 0,3206 | 0,735831 |
| Tuba4a | 0,652833 | 1,57111 | 1,267 | 0,3206 | 0,735831 |
| R3hcc1l | 12,2125 | 13,9683 | 0,193805 | 0,3206 | 0,735831 |
| Lrrfip2 | 15,9031 | 18,1298 | 0,189056 | 0,32065 | 0,735832 |
| Rps27,Rps27rt | 4,93541 | 6,51299 | 0,400151 | 0,3209 | 0,736142 |
| Stk25 | 32,245 | 36,7255 | 0,187704 | 0,3209 | 0,736142 |
| 0610011F06Rik | 26,7422 | 22,7183 | -0,235269 | 0,3211 | 0,736469 |
| Ak2 | 123,255 | 108,776 | -0,180287 | 0,3213 | 0,736779 |
| Atp5j | 279,218 | 237,133 | -0,235697 | 0,32135 | 0,736779 |
| Tdrkh | 8,7743 | 10,1803 | 0,21443 | 0,32145 | 0,736859 |
| Rpa1 | 91,3464 | 80,8653 | -0,175827 | 0,3215 | 0,736859 |
| Parp1 | 166,241 | 146,55 | -0,181884 | 0,32165 | 0,736889 |
| Pfkp | 3,27488 | 6,068 | 0,889778 | 0,3217 | 0,736889 |

| | | | | | |
|---|---|---|---|---|---|
| N4bp1 | 18,0438 | 20,4284 | 0,17907 | 0,32175 | 0,736889 |
| Slc25a16 | 6,10816 | 5,21412 | -0,228314 | 0,32175 | 0,736889 |
| Zfp414 | 22,1615 | 18,9745 | -0,223993 | 0,3218 | 0,736889 |
| A830018L16Rik | 0,873912 | 1,13578 | 0,378118 | 0,3219 | 0,736986 |
| Brpf1 | 17,9893 | 20,4214 | 0,182942 | 0,32205 | 0,737066 |
| Rp9 | 38,858 | 33,8257 | -0,200093 | 0,32205 | 0,737066 |
| Angptl4 | 10,9494 | 9,36078 | -0,226147 | 0,3223 | 0,737456 |
| Spg20 | 34,847 | 30,7394 | -0,180947 | 0,3224 | 0,737456 |
| Ube2r2 | 60,4294 | 53,4193 | -0,177889 | 0,32245 | 0,737456 |
| Pick1 | 12,6037 | 10,8419 | -0,217218 | 0,32245 | 0,737456 |
| Dek | 162,311 | 184,562 | 0,18534 | 0,32265 | 0,737599 |
| Eva1a | 2,42374 | 1,90885 | -0,344536 | 0,32265 | 0,737599 |
| Scnm1 | 42,4001 | 36,6441 | -0,210485 | 0,3227 | 0,737599 |
| Klf5 | 0,468907 | 0,695813 | 0,569397 | 0,32275 | 0,737599 |
| Tapt1 | 16,0359 | 18,2913 | 0,189858 | 0,3228 | 0,737599 |
| Eif3f | 188,375 | 166,614 | -0,177101 | 0,3231 | 0,738153 |
| Ubn1 | 21,2449 | 18,7617 | -0,179327 | 0,32345 | 0,738689 |
| Wwtr1 | 12,7504 | 11,2115 | -0,185567 | 0,32345 | 0,738689 |
| Kdm3b | 24,1691 | 27,3484 | 0,178293 | 0,32365 | 0,738883 |
| Tmem258 | 327,83 | 288,276 | -0,1855 | 0,32365 | 0,738883 |
| Tmem206 | 4,32426 | 3,37997 | -0,355443 | 0,32385 | 0,739079 |
| Zfp612 | 3,45544 | 4,04519 | 0,227341 | 0,32405 | 0,739079 |
| Galnt7 | 24,2199 | 27,4115 | 0,178591 | 0,3241 | 0,739079 |
| Taf1d | 85,0226 | 102,499 | 0,269685 | 0,32425 | 0,739079 |
| Tmem194 | 10,7182 | 12,3671 | 0,206445 | 0,3243 | 0,739079 |
| Hist1h1a | 1,27756 | 1,91523 | 0,584132 | 0,32435 | 0,739079 |
| Hspa5 | 551,935 | 630,951 | 0,193029 | 0,32435 | 0,739079 |
| Hnrnpdl | 188,417 | 166,596 | -0,177576 | 0,32435 | 0,739079 |
| Cdc37l1 | 15,8508 | 18,155 | 0,195811 | 0,3244 | 0,739079 |
| Myzap | 0,985376 | 0,729784 | -0,433206 | 0,3244 | 0,739079 |
| Alas1 | 12,6702 | 14,5413 | 0,198719 | 0,32455 | 0,739079 |
| Tceb1 | 134,375 | 118,644 | -0,17963 | 0,32455 | 0,739079 |
| Alyref2 | 3,90906 | 3,09384 | -0,33742 | 0,32455 | 0,739079 |
| Il10rb | 1,45047 | 1,87329 | 0,369053 | 0,3246 | 0,739079 |
| Arhgap11a | 25,4253 | 28,732 | 0,176395 | 0,3246 | 0,739079 |
| Cdca4 | 71,1965 | 62,8929 | -0,178908 | 0,3247 | 0,739175 |
| Ubxn1 | 184,068 | 162,328 | -0,181324 | 0,3248 | 0,739272 |
| Xrcc5 | 31,9703 | 28,1951 | -0,181288 | 0,3249 | 0,739316 |
| Ndufa6 | 154,693 | 135,507 | -0,191042 | 0,32495 | 0,739316 |
| Ube2v2 | 2,60312 | 2,20356 | -0,240405 | 0,325 | 0,739316 |
| Med10 | 76,3973 | 66,9675 | -0,190061 | 0,32505 | 0,739316 |
| Tmem150a | 6,74653 | 9,45622 | 0,487119 | 0,3252 | 0,739395 |
| Galk1 | 79,5314 | 70,2896 | -0,178213 | 0,3252 | 0,739395 |
| Scaf4 | 22,1828 | 25,1274 | 0,17982 | 0,3253 | 0,739491 |
| Cdc16 | 110,099 | 97,2898 | -0,178441 | 0,3255 | 0,739815 |
| Pik3r3 | 16,8713 | 14,8785 | -0,181341 | 0,3258 | 0,740365 |
| Mphosph10 | 51,0914 | 57,8324 | 0,178797 | 0,32605 | 0,740785 |
| Glud1 | 44,183 | 39,0746 | -0,177263 | 0,3261 | 0,740785 |
| Plekhm3 | 2,18163 | 2,54039 | 0,219645 | 0,3263 | 0,741108 |
| Ryk | 42,9444 | 48,4823 | 0,174988 | 0,32665 | 0,741576 |

| | | | | | |
|---|---|---|---|---|---|
| Rfc5 | 41,3293 | 36,3694 | -0,18444 | 0,3268 | 0,741576 |
| Zfp882 | 2,77579 | 3,2477 | 0,226519 | 0,3269 | 0,741576 |
| Ppif | 38,4243 | 33,5847 | -0,194218 | 0,3269 | 0,741576 |
| Cript | 39,1976 | 34,1952 | -0,196974 | 0,3269 | 0,741576 |
| Shroom3 | 16,1925 | 14,2604 | -0,183308 | 0,327 | 0,741576 |
| Ficd | 0,757175 | 0,980833 | 0,37338 | 0,32705 | 0,741576 |
| Abcd1 | 4,33842 | 5,09046 | 0,230627 | 0,3271 | 0,741576 |
| Smad5 | 39,4319 | 44,4706 | 0,173489 | 0,3271 | 0,741576 |
| Pbdc1 | 41,1471 | 36,1372 | -0,187305 | 0,32715 | 0,741576 |
| Tmem189 | 16,4712 | 19,0022 | 0,206215 | 0,3272 | 0,741576 |
| Ldlr | 30,1556 | 35,0522 | 0,178051 | 0,3272 | 0,741576 |
| Trim3 | 6,72304 | 7,83279 | 0,220412 | 0,32745 | 0,741976 |
| Fam219a | 1,32533 | 0,995792 | -0,41244 | 0,3275 | 0,741976 |
| Jam3 | 61,8829 | 54,6921 | -0,17821 | 0,32755 | 0,741976 |
| Sypl | 15,2354 | 17,2477 | 0,178978 | 0,3277 | 0,742185 |
| L3mbtl3 | 16,4734 | 18,7509 | 0,186825 | 0,32785 | 0,742376 |
| Akt2 | 13,4983 | 11,7875 | -0,195522 | 0,3279 | 0,742376 |
| Trmt1 | 39,4393 | 34,7119 | -0,184204 | 0,328 | 0,742436 |
| Cwc22 | 12,1807 | 13,9175 | 0,192304 | 0,3281 | 0,742435 |
| Creb1 | 16,6641 | 18,8665 | 0,179087 | 0,3281 | 0,742436 |
| Zfp786 | 2,20316 | 1,78016 | -0,307569 | 0,32845 | 0,743096 |
| Dram2 | 6,75036 | 7,86583 | 0,220636 | 0,3286 | 0,743287 |
| Limd1 | 25,3816 | 22,4303 | -0,178335 | 0,32865 | 0,743287 |
| Tprn | 2,88082 | 3,46994 | 0,268433 | 0,32875 | 0,743328 |
| Hras | 24,4025 | 21,3995 | -0,189451 | 0,32885 | 0,743328 |
| Cfp | 5,60947 | 6,78101 | 0,273636 | 0,3289 | 0,743328 |
| Ncapd3 | 18,874 | 21,3531 | 0,178049 | 0,3289 | 0,743328 |
| Ppa2 | 41,1882 | 36,0356 | -0,19281 | 0,3291 | 0,74351 |
| Plekhg5 | 2,7611 | 3,27579 | 0,246603 | 0,3292 | 0,74351 |
| Fam115a | 30,782 | 27,192 | -0,178902 | 0,32935 | 0,74351 |
| Zfp408 | 8,74171 | 10,026 | 0,19776 | 0,32945 | 0,74351 |
| Eml4 | 47,4438 | 42,0244 | -0,174992 | 0,32945 | 0,74351 |
| Ccne2 | 23,9787 | 29,3897 | 0,293557 | 0,32955 | 0,74351 |
| Usp39 | 71,9201 | 63,7253 | -0,17453 | 0,3296 | 0,74351 |
| Kctd17 | 22,1169 | 19,2017 | -0,203913 | 0,3296 | 0,74351 |
| Trpt1 | 5,15568 | 4,00875 | -0,36301 | 0,3296 | 0,74351 |
| Pigo | 7,70845 | 8,83364 | 0,196568 | 0,32965 | 0,74351 |
| Pole2 | 30,5934 | 26,8275 | -0,189507 | 0,32965 | 0,74351 |
| Stard10 | 5,38888 | 6,59591 | 0,291586 | 0,32975 | 0,74351 |
| Hist1h2bk | 3,03307 | 2,00334 | -0,59837 | 0,32975 | 0,74351 |
| Tubgcp2 | 37,1272 | 32,5269 | -0,190844 | 0,32985 | 0,74351 |
| Fam151b | 0,757848 | 0,502975 | -0,59142 | 0,32985 | 0,74351 |
| B230217C12Rik | 8,42792 | 10,1003 | 0,261145 | 0,33025 | 0,744281 |
| Sde2 | 24,497 | 27,7537 | 0,180077 | 0,33035 | 0,744376 |
| Psenen | 1,0774 | 1,68645 | 0,646441 | 0,3305 | 0,744452 |
| Tm6sf1 | 3,23244 | 5,01294 | 0,633031 | 0,3305 | 0,744452 |
| Tsnax | 51,6893 | 45,7119 | -0,177296 | 0,3308 | 0,744735 |
| Atg9a | 9,32562 | 8,12955 | -0,198023 | 0,3308 | 0,744735 |
| Pcsk4 | 0,606335 | 0,437082 | -0,472211 | 0,3308 | 0,744735 |
| 1110057K04Rik | 30,7985 | 27,0534 | -0,187052 | 0,3309 | 0,74483 |

| | | | | | |
|---|---|---|---|---|---|
| Acrbp | 5,136 | 6,24835 | 0,282829 | 0,3313 | 0,745432 |
| Tmem107 | 15,4328 | 18,1433 | 0,233436 | 0,33135 | 0,745432 |
| Chd1l | 15,5541 | 13,6238 | -0,191159 | 0,33135 | 0,745432 |
| Ano6 | 15,799 | 13,9948 | -0,17494 | 0,3314 | 0,745432 |
| Mmp28 | 0,663596 | 0,475778 | -0,480018 | 0,33165 | 0,745864 |
| Cyp51 | 68,5302 | 77,3103 | 0,17392 | 0,3321 | 0,746428 |
| Cdc25b | 8,7348 | 4,94327 | -0,821308 | 0,33215 | 0,746428 |
| Ankmy2 | 8,07527 | 6,91212 | -0,224383 | 0,3322 | 0,746428 |
| 2310047M10Rik | 4,69258 | 5,63939 | 0,265156 | 0,33225 | 0,746428 |
| Prelid2 | 21,5385 | 25,5114 | 0,244224 | 0,33225 | 0,746428 |
| Zfp27 | 5,97239 | 6,92819 | 0,21417 | 0,33225 | 0,746428 |
| Ece1 | 13,5379 | 15,3488 | 0,18113 | 0,33235 | 0,746522 |
| Stk38 | 37,4992 | 42,3484 | 0,175448 | 0,3326 | 0,746934 |
| 3110021N24Rik | 2,54007 | 3,88008 | 0,611217 | 0,33265 | 0,746934 |
| Lamtor2 | 94,8524 | 82,7169 | -0,197503 | 0,33305 | 0,747701 |
| Arl1 | 57,435 | 50,1511 | -0,19565 | 0,33325 | 0,74787 |
| Ing3 | 10,8265 | 9,45849 | -0,194885 | 0,3333 | 0,74787 |
| Pcsk6 | 1,46728 | 1,19568 | -0,295308 | 0,3333 | 0,74787 |
| Meis3 | 15,9369 | 18,3323 | 0,202015 | 0,3334 | 0,747963 |
| Tmem220 | 1,0997 | 1,46243 | 0,411266 | 0,3335 | 0,748057 |
| Zfp367 | 7,78 | 6,74397 | -0,206171 | 0,33365 | 0,748244 |
| Bahd1 | 5,4449 | 6,27247 | 0,204129 | 0,3337 | 0,748244 |
| Pofut1 | 14,2335 | 12,5693 | -0,179387 | 0,3338 | 0,748337 |
| Nfic | 1,32572 | 1,59026 | 0,262484 | 0,33395 | 0,748543 |
| Gosr2 | 37,275 | 32,9708 | -0,177021 | 0,33405 | 0,748636 |
| Trip12 | 43,0334 | 48,5401 | 0,17372 | 0,3342 | 0,748842 |
| Hykk | 1,84737 | 1,53887 | -0,263601 | 0,3343 | 0,748935 |
| Cmpk2 | 0,769813 | 0,583291 | -0,400292 | 0,3344 | 0,749028 |
| Spaca6 | 5,40852 | 6,66536 | 0,301448 | 0,3346 | 0,749345 |
| Lifr | 5,00009 | 5,81724 | 0,218382 | 0,33475 | 0,74955 |
| Tsc1 | 11,1675 | 12,6143 | 0,175748 | 0,33485 | 0,749643 |
| Eif2b1 | 57,8807 | 51,2312 | -0,176062 | 0,33495 | 0,749661 |
| Strap | 144,52 | 128,135 | -0,173603 | 0,335 | 0,749661 |
| Snrpb | 462,853 | 410,431 | -0,173414 | 0,33505 | 0,749661 |
| 1700029J07Rik | 1,33237 | 1,67226 | 0,327808 | 0,33515 | 0,749661 |
| Sema3c | 5,34641 | 4,6356 | -0,205816 | 0,33515 | 0,749661 |
| Dennd4c | 3,90089 | 4,47047 | 0,196624 | 0,3354 | 0,75007 |
| Akt1s1 | 48,5085 | 42,6988 | -0,184043 | 0,33545 | 0,75007 |
| Cenpj | 10,8915 | 12,3942 | 0,186466 | 0,33565 | 0,750237 |
| Fbrs | 16,1942 | 18,4462 | 0,18784 | 0,3357 | 0,750237 |
| Atg14 | 7,72561 | 6,68243 | -0,209275 | 0,3357 | 0,750237 |
| Cxxc5 | 33,903 | 38,3241 | 0,176837 | 0,3359 | 0,750532 |
| Cyr61 | 39,7792 | 45,2696 | 0,18653 | 0,33605 | 0,750532 |
| Ift43 | 39,2293 | 33,8743 | -0,211741 | 0,33605 | 0,750532 |
| Eif5b | 34,5406 | 39,4197 | 0,190623 | 0,33615 | 0,750532 |
| Chrac1 | 39,816 | 34,5356 | -0,205264 | 0,33615 | 0,750532 |
| Ndufa11 | 18,6296 | 16,3643 | -0,187045 | 0,3362 | 0,750532 |
| Hdac2 | 205,444 | 182,091 | -0,174081 | 0,33625 | 0,750532 |
| Mier3 | 9,52768 | 10,8352 | 0,185532 | 0,3363 | 0,750532 |
| Wdr6 | 93,8116 | 106,173 | 0,178585 | 0,33665 | 0,751182 |

| | | | | | |
|---|---|---|---|---|---|
| Zswim1 | 6,51148 | 7,5819 | 0,219575 | 0,33685 | 0,751459 |
| Hacl1 | 2,13038 | 2,60679 | 0,29116 | 0,3369 | 0,751459 |
| C920021L13Rik | 0,385479 | 0,729773 | 0,920795 | 0,33695 | 0,751459 |
| Zfp82 | 5,34886 | 6,33293 | 0,243641 | 0,33715 | 0,751775 |
| Ogfr | 77,7315 | 68,954 | -0,172865 | 0,33725 | 0,751867 |
| Fem1a | 8,72648 | 7,66822 | -0,186509 | 0,33735 | 0,751959 |
| Tubg2 | 0,580414 | 0,828112 | 0,512743 | 0,3375 | 0,752163 |
| Ntrk3 | 1,25089 | 1,54799 | 0,307436 | 0,3378 | 0,752701 |
| Phb2 | 283,87 | 251,665 | -0,173728 | 0,33795 | 0,752774 |
| Pigl | 10,589 | 9,0209 | -0,231228 | 0,33795 | 0,752774 |
| Zscan20 | 3,58308 | 4,14411 | 0,209864 | 0,33805 | 0,752866 |
| Klhl34 | 1,92943 | 2,37164 | 0,297715 | 0,3382 | 0,752938 |
| Rfc3 | 83,2885 | 73,6699 | -0,177042 | 0,3382 | 0,752938 |
| Lsm6 | 23,912 | 21,163 | -0,176188 | 0,33855 | 0,753587 |
| Tex261 | 29,9006 | 26,2114 | -0,189983 | 0,3387 | 0,75379 |
| Tcf4 | 24,7331 | 27,8471 | 0,171087 | 0,33885 | 0,753993 |
| Elovl5 | 55,9185 | 49,6318 | -0,172058 | 0,3392 | 0,754562 |
| Fam49a | 5,94517 | 6,81693 | 0,197403 | 0,3394 | 0,754562 |
| Ndnl2 | 30,3268 | 34,6606 | 0,192704 | 0,3394 | 0,754562 |
| Slc35b1 | 63,8966 | 72,3771 | 0,179793 | 0,3394 | 0,754562 |
| Bach1 | 15,0862 | 13,335 | -0,178013 | 0,3394 | 0,754562 |
| Tagln3 | 23,9345 | 20,7493 | -0,206025 | 0,3396 | 0,754569 |
| Aldh1l2 | 0,690107 | 0,856657 | 0,311897 | 0,33975 | 0,754569 |
| Rhoa | 66,3642 | 58,794 | -0,174736 | 0,33975 | 0,754569 |
| Nup37 | 62,2242 | 54,7976 | -0,183363 | 0,33985 | 0,754569 |
| Son | 83,4249 | 94,2919 | 0,176656 | 0,3399 | 0,754569 |
| Herc4 | 23,3555 | 26,3055 | 0,171599 | 0,3399 | 0,754569 |
| Atp6v0e | 43,7546 | 37,9501 | -0,20533 | 0,3399 | 0,754569 |
| Zmym4 | 25,0759 | 28,5759 | 0,188494 | 0,34 | 0,754569 |
| Hsf2 | 25,3191 | 28,6683 | 0,179231 | 0,34 | 0,754569 |
| Rsph3a | 2,06452 | 2,57108 | 0,316574 | 0,34005 | 0,754569 |
| Kidins220 | 8,77134 | 7,72558 | -0,183154 | 0,34005 | 0,754569 |
| Gabarapl2 | 8,00764 | 6,55823 | -0,288071 | 0,3402 | 0,754771 |
| Ybx1 | 243,394 | 215,871 | -0,173127 | 0,34045 | 0,755057 |
| Rbpj | 8,1987 | 7,20458 | -0,186481 | 0,34045 | 0,755057 |
| Mtx2 | 143,485 | 127,17 | -0,174142 | 0,34055 | 0,755057 |
| Aggf1 | 32,963 | 29,1557 | -0,177067 | 0,34065 | 0,755057 |
| 2610305D13Rik | 1,06185 | 0,808961 | -0,392435 | 0,34065 | 0,755057 |
| Birc2 | 18,143 | 20,5876 | 0,182366 | 0,3407 | 0,755057 |
| Lgmn | 21,768 | 24,8102 | 0,188726 | 0,34075 | 0,755057 |
| Tasp1 | 8,08268 | 9,41442 | 0,220038 | 0,3408 | 0,755057 |
| Armc1 | 37,0031 | 32,8203 | -0,173056 | 0,3409 | 0,755149 |
| Mrpl16 | 46,7874 | 41,0975 | -0,187071 | 0,341 | 0,75524 |
| Ttc13 | 20,371 | 23,0884 | 0,180654 | 0,34115 | 0,755441 |
| Aimp2 | 83,9709 | 70,2221 | -0,257966 | 0,3414 | 0,755845 |
| Lysmd4 | 4,71601 | 5,50311 | 0,222681 | 0,34145 | 0,755845 |
| 2200002D01Rik | 3,2945 | 6,44839 | 0,968879 | 0,3416 | 0,755896 |
| 5730507C01Rik | 0,888899 | 1,15333 | 0,375715 | 0,34165 | 0,755896 |
| Cyb561a3 | 5,13425 | 5,9785 | 0,21963 | 0,34165 | 0,755896 |
| R3hdm1 | 21,4445 | 24,1995 | 0,17437 | 0,34185 | 0,756019 |

| | | | | | |
|---|---|---|---|---|---|
| Trappc4 | 63,4455 | 56,0382 | -0,179109 | 0,34185 | 0,756019 |
| Sdc4 | 3,87819 | 3,22724 | -0,265084 | 0,3419 | 0,756019 |
| Ybx3 | 162,508 | 144,398 | -0,170463 | 0,34195 | 0,756019 |
| Emc6 | 56,0498 | 49,5051 | -0,179131 | 0,342 | 0,756019 |
| Ddx11 | 16,8613 | 19,0787 | 0,17825 | 0,3422 | 0,756141 |
| Sirt1 | 18,8045 | 21,2467 | 0,176162 | 0,34225 | 0,756141 |
| Luc7l3 | 73,5215 | 82,7308 | 0,170258 | 0,34225 | 0,756141 |
| Gm13498 | 10,4349 | 8,99825 | -0,213697 | 0,3423 | 0,756141 |
| Kbtbd4 | 19,2018 | 16,8528 | -0,188254 | 0,34235 | 0,756141 |
| Ttk | 29,0065 | 32,7237 | 0,173958 | 0,3425 | 0,756172 |
| Arf5 | 93,6225 | 82,748 | -0,178131 | 0,3426 | 0,756172 |
| Kti12 | 43,4239 | 36,3755 | -0,255524 | 0,3426 | 0,756172 |
| Dpp6 | 0,699393 | 0,547381 | -0,353558 | 0,3426 | 0,756172 |
| Tmem41b | 4,25797 | 3,63145 | -0,229622 | 0,3428 | 0,756464 |
| Micall2 | 3,04621 | 2,54305 | -0,260452 | 0,34285 | 0,756464 |
| Chrna7 | 2,9013 | 2,35714 | -0,29966 | 0,34325 | 0,757216 |
| Mad2l1bp | 15,6812 | 18,1829 | 0,213546 | 0,34335 | 0,757306 |
| Ap4e1 | 6,37297 | 7,25607 | 0,187223 | 0,34355 | 0,757387 |
| Cebpb | 0,576319 | 0,832919 | 0,531309 | 0,3436 | 0,757387 |
| Nif3l1 | 41,405 | 36,6698 | -0,175212 | 0,34365 | 0,757387 |
| 4930556M19Rik | 0,913171 | 1,21882 | 0,416533 | 0,3437 | 0,757387 |
| Mob1b | 6,75661 | 7,78863 | 0,205069 | 0,3437 | 0,757387 |
| Tbl1xr1 | 8,74967 | 7,7371 | -0,177436 | 0,34375 | 0,757387 |
| Eri2 | 13,7335 | 15,6375 | 0,18731 | 0,3438 | 0,757387 |
| Psmb3 | 78,6219 | 68,9428 | -0,189532 | 0,344 | 0,757568 |
| Hfe | 0,783759 | 0,544276 | -0,526072 | 0,344 | 0,757568 |
| Psmc4 | 229,113 | 203,729 | -0,169411 | 0,34415 | 0,757708 |
| 2310036O22Rik | 200,529 | 178,019 | -0,171778 | 0,34425 | 0,757708 |
| Fam118b | 24,9119 | 19,6052 | -0,345601 | 0,34425 | 0,757708 |
| Lnx2 | 4,67214 | 5,39757 | 0,208227 | 0,3443 | 0,757708 |
| Cgn | 1,14085 | 1,38909 | 0,284033 | 0,3445 | 0,757998 |
| Arhgef17 | 17,0065 | 15,0973 | -0,171801 | 0,34455 | 0,757998 |
| Abcc5 | 20,3006 | 23,82 | 0,230649 | 0,3447 | 0,758068 |
| Slmap | 17,2793 | 19,4973 | 0,174229 | 0,3447 | 0,758068 |
| Nhp2l1 | 11,5238 | 9,76698 | -0,238631 | 0,34485 | 0,758268 |
| Zfp825 | 5,34287 | 6,44675 | 0,270958 | 0,345 | 0,758468 |
| Grtp1 | 4,77459 | 5,81253 | 0,283789 | 0,34515 | 0,758668 |
| Rusc2 | 1,10995 | 0,899895 | -0,302661 | 0,3453 | 0,758867 |
| Uba5 | 29,419 | 26,0389 | -0,176083 | 0,3456 | 0,759136 |
| Cln5 | 14,2221 | 12,4182 | -0,195684 | 0,3456 | 0,759136 |
| 5730409E04Rik | 2,7958 | 2,30957 | -0,275634 | 0,3456 | 0,759136 |
| Rpl21 | 4,79682 | 3,98888 | -0,266095 | 0,3458 | 0,759426 |
| Rmi2 | 4,61114 | 3,95387 | -0,22186 | 0,34585 | 0,759426 |
| Nip7 | 38,7918 | 34,4168 | -0,172637 | 0,346 | 0,759625 |
| Adcy6 | 18,2493 | 20,5778 | 0,173246 | 0,3461 | 0,759714 |
| Stmn1 | 45,7727 | 40,1894 | -0,187673 | 0,34625 | 0,759914 |
| Rcbtb1 | 48,1714 | 42,8574 | -0,168631 | 0,34635 | 0,760003 |
| Tor1a | 30,5812 | 26,8687 | -0,186719 | 0,3465 | 0,760202 |
| Timm17b | 25,8492 | 22,7051 | -0,1871 | 0,34675 | 0,760588 |
| Slc30a4 | 6,3352 | 5,53522 | -0,19475 | 0,3469 | 0,760588 |

| | | | | | |
|---|---|---|---|---|---|
| Ldha | 986,175 | 1122,43 | 0,186709 | 0,347 | 0,760588 |
| Trrap | 23,3621 | 26,306 | 0,171223 | 0,347 | 0,760588 |
| Col9a3 | 1,15577 | 1,52252 | 0,397607 | 0,34705 | 0,760588 |
| Satb2 | 2,18678 | 2,57483 | 0,235665 | 0,34715 | 0,760588 |
| Ddx41 | 42,3631 | 37,6005 | -0,172056 | 0,34715 | 0,760588 |
| Rpl17 | 8,65709 | 7,01853 | -0,302713 | 0,34715 | 0,760588 |
| Tomm70a | 90,5309 | 80,5526 | -0,168479 | 0,3473 | 0,760766 |
| Ndufaf5 | 7,70277 | 6,64131 | -0,21391 | 0,34735 | 0,760766 |
| Chchd4 | 58,5114 | 51,7118 | -0,178224 | 0,34775 | 0,761471 |
| Pura | 5,19445 | 6,21625 | 0,259075 | 0,3478 | 0,761471 |
| Gpr19 | 5,38725 | 4,47672 | -0,267106 | 0,34785 | 0,761471 |
| Prkch | 1,89404 | 1,54307 | -0,295659 | 0,34805 | 0,761758 |
| Wdr55 | 44,4201 | 39,2259 | -0,179404 | 0,3481 | 0,761758 |
| Cnpy4 | 36,4005 | 32,1779 | -0,177887 | 0,34835 | 0,762045 |
| Dync2li1 | 19,6302 | 16,9768 | -0,20951 | 0,34835 | 0,762045 |
| G3bp1 | 149,833 | 133,292 | -0,168764 | 0,3486 | 0,762297 |
| Ramp3 | 1,23596 | 0,877805 | -0,493657 | 0,34865 | 0,762297 |
| Peo1 | 29,3424 | 26,0335 | -0,172621 | 0,34875 | 0,762297 |
| Hmgcr | 81,9568 | 92,2128 | 0,170104 | 0,3488 | 0,762297 |
| Sptlc1 | 18,6119 | 21,1114 | 0,181798 | 0,3489 | 0,762297 |
| Srpk2 | 10,7631 | 13,0257 | 0,275267 | 0,349 | 0,762297 |
| Ikzf5 | 5,91608 | 6,78406 | 0,197508 | 0,349 | 0,762297 |
| Abt1 | 7,71306 | 6,75278 | -0,191821 | 0,349 | 0,762297 |
| Stx18 | 13,9866 | 11,544 | -0,276901 | 0,349 | 0,762297 |
| Bahcc1 | 14,8538 | 13,1929 | -0,171071 | 0,34915 | 0,762474 |
| Ttll5 | 6,62729 | 7,56701 | 0,191305 | 0,3492 | 0,762474 |
| Mettl9 | 130,787 | 116,2 | -0,170605 | 0,3494 | 0,762487 |
| Camk2n2 | 3,61097 | 2,39947 | -0,589671 | 0,3494 | 0,762487 |
| Rcsd1 | 6,34881 | 7,40493 | 0,222001 | 0,3495 | 0,762487 |
| Manf | 82,7991 | 93,0472 | 0,168347 | 0,3495 | 0,762487 |
| Polr2a | 67,9292 | 76,441 | 0,170314 | 0,3497 | 0,762487 |
| Gapvd1 | 25,3278 | 28,4866 | 0,169562 | 0,34975 | 0,762487 |
| Pygb | 23,7297 | 21,0246 | -0,174613 | 0,34975 | 0,762487 |
| Gab2 | 2,30641 | 1,95941 | -0,235228 | 0,34975 | 0,762487 |
| Tnpo2 | 36,8093 | 41,5141 | 0,173531 | 0,3498 | 0,762487 |
| Zfp454 | 8,89925 | 7,64289 | -0,219566 | 0,3498 | 0,762487 |
| Scp2 | 12,1784 | 10,6439 | -0,1943 | 0,3499 | 0,762575 |
| Canx | 172,818 | 197,135 | 0,172666 | 0,35015 | 0,762799 |
| Zfp507 | 16,2726 | 18,323 | 0,171217 | 0,3502 | 0,762799 |
| Shfm1 | 695,273 | 619,334 | -0,166862 | 0,35025 | 0,762799 |
| Ccl27a | 8,24471 | 12,1496 | 0,559372 | 0,3503 | 0,762799 |
| Etv2 | 0,815507 | 1,19588 | 0,552307 | 0,3503 | 0,762799 |
| Fam136a | 69,7839 | 61,9455 | -0,171896 | 0,3504 | 0,762887 |
| Ccdc51 | 5,17863 | 6,07543 | 0,230415 | 0,3505 | 0,762955 |
| Vps54 | 19,0856 | 22,6171 | 0,244929 | 0,35055 | 0,762955 |
| Ap5m1 | 11,3893 | 14,1816 | 0,316337 | 0,35065 | 0,763043 |
| Arhgap44 | 2,05318 | 2,93577 | 0,515875 | 0,35095 | 0,763086 |
| Smim14 | 5,18294 | 6,12465 | 0,240856 | 0,3511 | 0,763086 |
| Scamp1 | 13,616 | 15,4129 | 0,178828 | 0,35115 | 0,763086 |
| Psmd11 | 35,5072 | 31,3409 | -0,180063 | 0,35115 | 0,763086 |

| | | | | | |
|---|---|---|---|---|---|
| Klhl15 | 2,51807 | 3,0516 | 0,277245 | 0,35125 | 0,763086 |
| Stag1 | 30,271 | 34,0299 | 0,168866 | 0,35125 | 0,763086 |
| Upf1 | 34,1785 | 38,5315 | 0,17295 | 0,3513 | 0,763086 |
| Efcab2 | 2,97261 | 3,56731 | 0,26311 | 0,35135 | 0,763086 |
| Pcnt | 13,3637 | 15,0091 | 0,167524 | 0,3514 | 0,763086 |
| Wasf3 | 14,4663 | 12,6517 | -0,193371 | 0,3514 | 0,763086 |
| Isoc1 | 52,1181 | 46,4078 | -0,167417 | 0,35145 | 0,763086 |
| Abhd11 | 16,6567 | 14,1902 | -0,231207 | 0,3515 | 0,763086 |
| Ppp1r10 | 23,3839 | 26,3094 | 0,170065 | 0,35155 | 0,763086 |
| AI314180 | 29,438 | 26,2118 | -0,167463 | 0,35165 | 0,763086 |
| Asb1 | 3,30288 | 3,80578 | 0,204465 | 0,3517 | 0,763086 |
| Mrpl35 | 12,5763 | 11,0542 | -0,186116 | 0,3517 | 0,763086 |
| 1600020E01Rik | 26,0218 | 30,6791 | 0,237532 | 0,35175 | 0,763086 |
| Tmem97 | 29,0561 | 33,0831 | 0,187258 | 0,35175 | 0,763086 |
| Zfpm2 | 0,55968 | 0,706188 | 0,335451 | 0,3518 | 0,763086 |
| Ip6k1 | 27,6634 | 31,1074 | 0,169279 | 0,3519 | 0,763153 |
| Wfs1 | 2,5719 | 3,05603 | 0,248827 | 0,35195 | 0,763153 |
| Ulk2 | 9,0579 | 10,2509 | 0,1785 | 0,3522 | 0,763396 |
| Mrpl39 | 52,0156 | 46,1906 | -0,171347 | 0,3522 | 0,763396 |
| Mia3 | 20,9281 | 23,5231 | 0,168639 | 0,35225 | 0,763396 |
| Ttc1 | 39,8617 | 35,1624 | -0,180969 | 0,3523 | 0,763396 |
| Eral1 | 17,4032 | 15,2494 | -0,190601 | 0,3525 | 0,763638 |
| Tspyl2 | 17,403 | 19,7498 | 0,1825 | 0,35255 | 0,763638 |
| Tars | 90,3947 | 80,462 | -0,167931 | 0,3526 | 0,763638 |
| Intu | 4,50502 | 5,1441 | 0,191383 | 0,35265 | 0,763638 |
| Gjc1 | 74,7036 | 83,8831 | 0,167203 | 0,35275 | 0,763705 |
| Nme2 | 335,167 | 298,174 | -0,168726 | 0,3528 | 0,763705 |
| Necap1 | 24,8312 | 28,1083 | 0,178841 | 0,35295 | 0,763901 |
| Fam126a | 11,7573 | 9,02961 | -0,380817 | 0,3531 | 0,764097 |
| Dtx3 | 51,8234 | 59,6674 | 0,203337 | 0,3532 | 0,764184 |
| Tsen34 | 45,7145 | 40,4091 | -0,177973 | 0,3533 | 0,764251 |
| Nudt11 | 7,01812 | 6,03465 | -0,217815 | 0,35335 | 0,764251 |
| Dyrk3 | 7,23521 | 6,15537 | -0,233189 | 0,3535 | 0,764447 |
| Il4ra | 0,555547 | 0,698292 | 0,329922 | 0,3536 | 0,764534 |
| Mbip | 20,8038 | 18,1172 | -0,199489 | 0,3537 | 0,764601 |
| Ggta1 | 6,80086 | 7,81158 | 0,199897 | 0,35375 | 0,764601 |
| Elovl6 | 29,4469 | 33,0826 | 0,167961 | 0,35385 | 0,764688 |
| Eif2ak1 | 39,4978 | 33,7213 | -0,228114 | 0,35395 | 0,764754 |
| Rbms2 | 13,3789 | 11,8405 | -0,176221 | 0,354 | 0,764754 |
| Gnai1 | 9,57317 | 10,9279 | 0,190951 | 0,35415 | 0,764929 |
| Tmem80 | 16,8416 | 12,4963 | -0,430532 | 0,3542 | 0,764929 |
| Psmb10 | 6,77752 | 8,10314 | 0,257723 | 0,3546 | 0,765664 |
| Myg1 | 44,8125 | 39,6079 | -0,178111 | 0,35485 | 0,766075 |
| Rrnad1 | 10,344 | 12,7472 | 0,301379 | 0,355 | 0,76627 |
| Rpa3 | 76,6114 | 67,2215 | -0,188637 | 0,35515 | 0,766422 |
| Ran | 161,521 | 144,17 | -0,163949 | 0,3552 | 0,766422 |
| Cited2 | 64,8189 | 72,807 | 0,167663 | 0,35525 | 0,766422 |
| Apoo | 1,50045 | 1,0835 | -0,469695 | 0,35545 | 0,766641 |
| Rpl36 | 39,1692 | 46,5223 | 0,248203 | 0,3555 | 0,766641 |
| Tifa | 0,569145 | 0,780092 | 0,454849 | 0,3556 | 0,766641 |

| | | | | | |
|---|---|---|---|---|---|
| Ankrd24 | 2,43195 | 2,89437 | 0,251138 | 0,3556 | 0,766641 |
| Sap30bp | 36,5797 | 32,4556 | -0,172573 | 0,35565 | 0,766641 |
| Pcbp4 | 195,842 | 174,381 | -0,167447 | 0,35575 | 0,766664 |
| 4933407K13Rik | 2,36964 | 2,82322 | 0,252674 | 0,35585 | 0,766664 |
| Dmwd | 36,0355 | 40,5186 | 0,169165 | 0,3559 | 0,766664 |
| Fkbp11 | 18,4225 | 15,4849 | -0,250609 | 0,3559 | 0,766664 |
| Oaz1 | 24,8042 | 21,5189 | -0,204982 | 0,3562 | 0,767182 |
| Cyfip2 | 3,14706 | 3,93966 | 0,324064 | 0,3566 | 0,767914 |
| Gm10560 | 1,97833 | 1,40713 | -0,491524 | 0,3567 | 0,768001 |
| 03/02/16 | 6,18026 | 5,12642 | -0,269718 | 0,3568 | 0,768087 |
| 9930014A18Rik | 1,57606 | 2,01657 | 0,355584 | 0,357 | 0,768367 |
| Bard1 | 7,82384 | 8,88678 | 0,183783 | 0,35705 | 0,768367 |
| Bmyc | 33,9577 | 29,4557 | -0,205194 | 0,3572 | 0,768561 |
| Syvn1 | 22,828 | 27,1639 | 0,250883 | 0,3573 | 0,768647 |
| Gins4 | 64,8262 | 57,6025 | -0,170446 | 0,3574 | 0,768734 |
| 2900056M20Rik | 1,29742 | 1,57405 | 0,278832 | 0,3575 | 0,768798 |
| Syde1 | 21,7912 | 24,5672 | 0,172992 | 0,35755 | 0,768798 |
| C030006K11Rik | 5,59716 | 4,67614 | -0,259377 | 0,35765 | 0,768821 |
| Mief2 | 4,78776 | 5,61388 | 0,229647 | 0,3578 | 0,768821 |
| Zfp397 | 18,3133 | 20,5811 | 0,168429 | 0,3578 | 0,768821 |
| Sdcbp | 59,8558 | 67,1067 | 0,164966 | 0,3578 | 0,768821 |
| Lhfpl4 | 1,81148 | 1,50972 | -0,262891 | 0,35795 | 0,768993 |
| Paip2b | 42,0267 | 47,202 | 0,167543 | 0,358 | 0,768993 |
| Vcp | 61,1211 | 54,5588 | -0,163857 | 0,3581 | 0,769079 |
| Zfp963 | 3,54824 | 4,2303 | 0,253656 | 0,3582 | 0,769122 |
| 2010315B03Rik | 1,42656 | 1,13845 | -0,325465 | 0,35825 | 0,769122 |
| H19 | 741,751 | 652,973 | -0,183911 | 0,3583 | 0,769122 |
| Trim26 | 15,3752 | 17,3862 | 0,177337 | 0,3586 | 0,769637 |
| Higd2a | 99,1084 | 112,269 | 0,179886 | 0,35875 | 0,76983 |
| Otud7b | 9,90067 | 11,1223 | 0,167854 | 0,3589 | 0,770023 |
| Smg7 | 30,462 | 34,1804 | 0,166159 | 0,35905 | 0,770195 |
| Timm23 | 2,5223 | 1,94587 | -0,374326 | 0,3591 | 0,770195 |
| Gars | 131,436 | 117,249 | -0,164783 | 0,3592 | 0,77028 |
| Ppig | 18,3571 | 20,6503 | 0,16983 | 0,3593 | 0,77028 |
| Zfp521 | 22,42 | 19,9605 | -0,167639 | 0,35935 | 0,77028 |
| Sh3tc2 | 1,52211 | 1,82653 | 0,263033 | 0,3594 | 0,77028 |
| Xylb | 4,4487 | 5,14734 | 0,210443 | 0,3595 | 0,77028 |
| Recql | 20,4284 | 17,6684 | -0,20941 | 0,3595 | 0,77028 |
| Cdc25a | 57,4511 | 51,3681 | -0,161461 | 0,35975 | 0,770644 |
| Slfn9 | 4,05861 | 4,70569 | 0,213421 | 0,3598 | 0,770644 |
| Lztfl1 | 8,67714 | 9,90149 | 0,190426 | 0,35985 | 0,770644 |
| Tbc1d10b | 16,2734 | 14,3921 | -0,177244 | 0,35995 | 0,770729 |
| Sgcb | 6,90768 | 6,0177 | -0,198988 | 0,36015 | 0,770985 |
| Fam213a | 17,1301 | 14,9108 | -0,200174 | 0,3602 | 0,770985 |
| Eef1g | 124,07 | 110,645 | -0,165217 | 0,36025 | 0,770985 |
| Btbd9 | 5,33426 | 4,68383 | -0,187598 | 0,3605 | 0,771392 |
| Hibadh | 42,2215 | 47,5529 | 0,171554 | 0,3607 | 0,771562 |
| Ispd | 3,27797 | 2,74283 | -0,257138 | 0,3607 | 0,771562 |
| Zbtb14 | 8,83856 | 7,73214 | -0,192944 | 0,3608 | 0,771648 |
| Rasgef1b | 2,37744 | 1,96044 | -0,278233 | 0,3609 | 0,771733 |

| | | | | | |
|---|---|---|---|---|---|
| Aco1 | 32,0496 | 35,9533 | 0,165819 | 0,36115 | 0,772095 |
| Zfp655 | 10,6562 | 12,1727 | 0,191953 | 0,3612 | 0,772095 |
| Bphl | 13,9662 | 11,9659 | -0,223012 | 0,36125 | 0,772095 |
| Det1 | 10,2647 | 8,91073 | -0,204071 | 0,36135 | 0,772115 |
| Stt3a | 90,7365 | 81,0755 | -0,162416 | 0,3614 | 0,772115 |
| Larp1b | 1,95403 | 2,43884 | 0,319743 | 0,3615 | 0,772115 |
| Pias3 | 11,3366 | 13,0082 | 0,198441 | 0,3615 | 0,772115 |
| Exosc2 | 80,6043 | 71,9086 | -0,164693 | 0,36165 | 0,772307 |
| Blvra | 7,46606 | 8,9264 | 0,257732 | 0,36185 | 0,772605 |
| 4833439L19Rik | 52,707 | 60,653 | 0,202586 | 0,3621 | 0,772882 |
| Mrps9 | 51,1244 | 45,3207 | -0,173844 | 0,3621 | 0,772882 |
| Sfmbt1 | 6,86791 | 6,07532 | -0,176911 | 0,36245 | 0,7735 |
| Fah | 1,32904 | 1,71762 | 0,370032 | 0,36255 | 0,773563 |
| Rrm2 | 257,601 | 229,737 | -0,165157 | 0,3626 | 0,773563 |
| Lage3 | 41,3044 | 35,9956 | -0,198474 | 0,36275 | 0,773754 |
| Bag1 | 123,626 | 110,334 | -0,164106 | 0,36295 | 0,77403 |
| Gm5141 | 2,64641 | 2,2059 | -0,262666 | 0,363 | 0,77403 |
| Rassf3 | 10,3315 | 9,07663 | -0,186817 | 0,3631 | 0,774092 |
| Ppox | 18,1191 | 20,6236 | 0,186788 | 0,36315 | 0,774092 |
| Ndfip2 | 21,1074 | 23,8414 | 0,175723 | 0,3633 | 0,774151 |
| Tm9sf3 | 46,0722 | 51,6383 | 0,164546 | 0,36345 | 0,774151 |
| Zfp563 | 1,67019 | 1,41284 | -0,241414 | 0,36345 | 0,774151 |
| Btrc | 8,71788 | 9,83231 | 0,173552 | 0,3635 | 0,774151 |
| Cd302 | 9,59929 | 8,15343 | -0,235521 | 0,36355 | 0,774151 |
| 1600016N20Rik | 3,01432 | 3,88329 | 0,365447 | 0,3636 | 0,774151 |
| Hn1 | 119,362 | 106,52 | -0,164211 | 0,3636 | 0,774151 |
| Hsp90b1 | 563,344 | 639,163 | 0,182168 | 0,3637 | 0,774236 |
| Sptbn2 | 4,54257 | 3,9962 | -0,184878 | 0,3641 | 0,774914 |
| Zfp579 | 7,97932 | 6,87122 | -0,215698 | 0,36415 | 0,774914 |
| Acbd5 | 6,40368 | 7,33156 | 0,19522 | 0,3642 | 0,774914 |
| Wsb1 | 97,1103 | 86,6484 | -0,164451 | 0,3644 | 0,775189 |
| Mir1895 | 528,237 | 86,6223 | -2,60838 | 0,36445 | 0,775189 |
| Copa | 87,4088 | 77,0024 | -0,182874 | 0,3647 | 0,775441 |
| Cdk19 | 4,91195 | 5,59407 | 0,187602 | 0,36475 | 0,775441 |
| Ywhag | 108,064 | 121,136 | 0,164744 | 0,36475 | 0,775441 |
| Glt1d1 | 3,9249 | 3,20129 | -0,294002 | 0,365 | 0,775795 |
| Ppfia1 | 21,2994 | 23,8226 | 0,161515 | 0,3652 | 0,775795 |
| 1700001G17Rik | 0,904553 | 1,31682 | 0,541785 | 0,3653 | 0,775795 |
| Ammecr1l | 22,7279 | 25,4782 | 0,164803 | 0,3653 | 0,775795 |
| Top3a | 14,2086 | 16,0221 | 0,173301 | 0,36535 | 0,775795 |
| Gm13034 | 0,769636 | 1,10259 | 0,51865 | 0,3654 | 0,775795 |
| Zswim7 | 10,9371 | 13,1375 | 0,264465 | 0,3654 | 0,775795 |
| Dvl2 | 33,2319 | 37,2416 | 0,164345 | 0,3654 | 0,775795 |
| E030030I06Rik | 2,56497 | 3,30473 | 0,365587 | 0,36575 | 0,775957 |
| Clcn6 | 8,31284 | 9,50752 | 0,193726 | 0,36575 | 0,775957 |
| Mipep | 17,3128 | 15,2615 | -0,181937 | 0,36575 | 0,775957 |
| Yipf5 | 22,7328 | 20,1136 | -0,176602 | 0,3658 | 0,775957 |
| Yipf6 | 6,12607 | 5,37197 | -0,189511 | 0,36585 | 0,775957 |
| Spns1 | 24,1381 | 21,4379 | -0,171148 | 0,3659 | 0,775957 |
| Esrra | 6,91307 | 5,66703 | -0,286735 | 0,3659 | 0,775957 |

| | | | | | |
|---|---|---|---|---|---|
| Siva1 | 93,6723 | 82,7776 | -0,178382 | 0,3661 | 0,776253 |
| Crtap | 17,5847 | 15,3945 | -0,191901 | 0,3663 | 0,776365 |
| Gclm | 6,66258 | 5,68456 | -0,229034 | 0,3663 | 0,776365 |
| Wdr25 | 3,21297 | 2,68083 | -0,261227 | 0,36635 | 0,776365 |
| Wdfy2 | 7,3108 | 8,487 | 0,215225 | 0,3665 | 0,776365 |
| 0610012G03Rik | 21,2736 | 18,6084 | -0,193106 | 0,3666 | 0,776365 |
| Tmsb15l | 9,77076 | 12,3112 | 0,333431 | 0,36665 | 0,776365 |
| Tomm7 | 114,356 | 93,4362 | -0,291473 | 0,3667 | 0,776365 |
| Ube2g1 | 30,1273 | 26,8826 | -0,164396 | 0,36675 | 0,776365 |
| Pms1 | 10,012 | 11,4034 | 0,187737 | 0,36685 | 0,776365 |
| Phox2a | 0,0472577 | 0,67946 | 3,84577 | 0,3669 | 0,776365 |
| Usp22 | 58,0754 | 51,8986 | -0,162231 | 0,3669 | 0,776365 |
| Atp6v1e1 | 75,05 | 66,8759 | -0,166365 | 0,3669 | 0,776365 |
| BC029214 | 2,41802 | 1,81661 | -0,412577 | 0,36695 | 0,776365 |
| Foxk2 | 38,3659 | 42,9188 | 0,161783 | 0,367 | 0,776365 |
| 3110057O12Rik | 2,20254 | 1,80005 | -0,291132 | 0,36715 | 0,776504 |
| Cnot8 | 38,1068 | 33,8628 | -0,170346 | 0,36725 | 0,776504 |
| 2610002M06Rik | 2,84644 | 2,45374 | -0,214176 | 0,36735 | 0,776504 |
| Agbl3 | 1,36946 | 0,981566 | -0,480454 | 0,36735 | 0,776504 |
| Rnf220 | 50,4878 | 56,5148 | 0,162693 | 0,36745 | 0,776504 |
| Zbtb3 | 2,58926 | 3,14655 | 0,281227 | 0,3675 | 0,776504 |
| Gpbp1 | 49,3307 | 55,1828 | 0,161735 | 0,3675 | 0,776504 |
| Trim16 | 0,838371 | 1,04064 | 0,311816 | 0,36755 | 0,776504 |
| Mknk1 | 7,80944 | 6,76591 | -0,206935 | 0,36795 | 0,777221 |
| Ttc23 | 26,3273 | 23,3112 | -0,175536 | 0,3681 | 0,77732 |
| Mcmdc2 | 0,49235 | 0,666 | 0,435839 | 0,3682 | 0,77732 |
| Nufip1 | 17,2934 | 15,3705 | -0,170058 | 0,36825 | 0,77732 |
| Ssb | 251,118 | 282,549 | 0,170133 | 0,3683 | 0,77732 |
| R3hdm2 | 18,2745 | 20,4842 | 0,164683 | 0,3683 | 0,77732 |
| Pagr1a | 52,0132 | 46,2634 | -0,169007 | 0,3685 | 0,777614 |
| Etv1 | 0,799515 | 0,630313 | -0,343058 | 0,3686 | 0,777675 |
| Ttc39c | 4,99639 | 4,24182 | -0,236205 | 0,36865 | 0,777675 |
| Flcn | 11,8032 | 10,4145 | -0,180579 | 0,3688 | 0,777841 |
| Scfd1 | 30,6584 | 27,201 | -0,172625 | 0,36885 | 0,777841 |
| DQ267102 | 239,602 | 492,001 | 1,03802 | 0,36895 | 0,777924 |
| Pank1 | 5,15702 | 5,84068 | 0,1796 | 0,3692 | 0,778323 |
| Ireb2 | 35,6632 | 39,9721 | 0,164558 | 0,3694 | 0,778617 |
| Tyms | 31,1725 | 27,8605 | -0,162052 | 0,36965 | 0,778993 |
| Prdx2 | 630,709 | 564,187 | -0,160802 | 0,3697 | 0,778993 |
| Mir682 | 11075 | 9581,1 | -0,209048 | 0,3698 | 0,779054 |
| Zfp839 | 4,12987 | 4,79107 | 0,214251 | 0,36985 | 0,779053 |
| D330041H03Rik | 1,12727 | 1,70581 | 0,597629 | 0,37005 | 0,779174 |
| Ttc12 | 1,20487 | 1,47118 | 0,288097 | 0,37005 | 0,779174 |
| Agfg1 | 27,6471 | 31,0249 | 0,166302 | 0,3701 | 0,779174 |
| Vil1 | 0,138045 | 1,20365 | 3,12419 | 0,37015 | 0,779174 |
| Abhd12 | 20,2931 | 22,9458 | 0,177243 | 0,3703 | 0,779211 |
| Pgd | 114,099 | 102,003 | -0,161678 | 0,3703 | 0,779211 |
| Metap1d | 15,329 | 13,241 | -0,211254 | 0,37035 | 0,779211 |
| Sf3b2 | 191,636 | 170,763 | -0,166375 | 0,37065 | 0,779511 |
| Btaf1 | 17,2535 | 19,276 | 0,159919 | 0,3707 | 0,779511 |

| | | | | | |
|---|---|---|---|---|---|
| Zcwpw1 | 4,48308 | 3,78793 | -0,24308 | 0,3707 | 0,779511 |
| 4931406P16Rik | 12,153 | 13,6379 | 0,166308 | 0,37075 | 0,779511 |
| Hipk3 | 7,98708 | 7,10265 | -0,169311 | 0,3708 | 0,779511 |
| 5730455P16Rik | 9,40365 | 10,5969 | 0,172353 | 0,37095 | 0,779511 |
| Gpi1 | 82,4666 | 73,7975 | -0,160237 | 0,371 | 0,779511 |
| Wrap53 | 25,8775 | 22,8866 | -0,177195 | 0,371 | 0,779511 |
| Gosr1 | 11,438 | 12,8826 | 0,171595 | 0,3711 | 0,779511 |
| Morf4l1 | 24,8966 | 22,0554 | -0,174818 | 0,3711 | 0,779511 |
| Kpnb1 | 243,967 | 215,962 | -0,175913 | 0,3712 | 0,779525 |
| Hist1h2bp | 1,0664 | 1,76098 | 0,723631 | 0,3713 | 0,779525 |
| Snta1 | 1,40302 | 1,09752 | -0,354288 | 0,3713 | 0,779525 |
| Slc35a2 | 9,46303 | 8,17961 | -0,210271 | 0,37135 | 0,779525 |
| Ccdc14 | 9,14444 | 10,3382 | 0,177014 | 0,37145 | 0,77954 |
| Rad23a | 34,1019 | 29,2515 | -0,221345 | 0,3715 | 0,77954 |
| Pcnx | 7,80389 | 8,73429 | 0,162497 | 0,3716 | 0,77954 |
| BC003965 | 15,1311 | 13,3963 | -0,175682 | 0,3716 | 0,77954 |
| Sap30 | 61,8693 | 55,0144 | -0,169413 | 0,37225 | 0,780753 |
| Kmt2e | 15,5911 | 17,9788 | 0,205577 | 0,3723 | 0,780753 |
| Nkiras1 | 1,74067 | 2,47505 | 0,50781 | 0,3726 | 0,781186 |
| BC037034 | 15,4597 | 17,7176 | 0,196672 | 0,3727 | 0,781186 |
| Ano8 | 8,86011 | 7,21191 | -0,296944 | 0,3727 | 0,781186 |
| Lmf1 | 9,85559 | 8,49683 | -0,214017 | 0,37275 | 0,781186 |
| Ldlrad3 | 23,6505 | 26,4974 | 0,163979 | 0,3729 | 0,781372 |
| Snhg8 | 41,2354 | 36,0523 | -0,193791 | 0,3732 | 0,781873 |
| Man2b2 | 2,82418 | 3,32132 | 0,233924 | 0,3735 | 0,782374 |
| Xkr8 | 1,79951 | 1,50125 | -0,261447 | 0,3737 | 0,782665 |
| Nsmce4a | 133,753 | 119,66 | -0,160622 | 0,37385 | 0,782828 |
| Ppap2a | 7,3746 | 11,6388 | 0,658304 | 0,3739 | 0,782828 |
| Naa38 | 88,5404 | 76,5479 | -0,209974 | 0,3741 | 0,783119 |
| Rpl10a | 2,94059 | 3,77602 | 0,360763 | 0,37455 | 0,78391 |
| Slc38a7 | 6,49646 | 5,65597 | -0,19988 | 0,3746 | 0,78391 |
| Rbbp8 | 22,9355 | 25,6837 | 0,163276 | 0,37475 | 0,784096 |
| Xpnpep1 | 51,3631 | 45,9685 | -0,160085 | 0,3749 | 0,784108 |
| Sgk3 | 1,6634 | 1,96061 | 0,237164 | 0,37495 | 0,784108 |
| Asxl1 | 32,0996 | 35,8621 | 0,159906 | 0,37495 | 0,784108 |
| Cbx2 | 53,5685 | 47,9445 | -0,16002 | 0,375 | 0,784108 |
| Chmp2b | 20,912 | 18,4353 | -0,181855 | 0,37515 | 0,784142 |
| Zfp758 | 2,61564 | 3,11338 | 0,251319 | 0,3752 | 0,784142 |
| Med28 | 19,8302 | 17,7115 | -0,163015 | 0,37525 | 0,784142 |
| Alpk1 | 0,624391 | 0,493691 | -0,338841 | 0,3754 | 0,784142 |
| Cux1 | 33,3089 | 38,0956 | 0,193715 | 0,3755 | 0,784142 |
| Rpp30 | 44,3208 | 39,2945 | -0,173655 | 0,3755 | 0,784142 |
| Zfp819 | 0,631125 | 0,83504 | 0,403921 | 0,37555 | 0,784142 |
| Pvt1 | 3,53761 | 4,66986 | 0,400603 | 0,37565 | 0,784142 |
| Cyb561d1 | 1,85908 | 1,55538 | -0,25732 | 0,37565 | 0,784142 |
| Elof1 | 86,2459 | 67,9164 | -0,344697 | 0,37565 | 0,784142 |
| Daam2 | 14,6899 | 13,106 | -0,164591 | 0,3757 | 0,784142 |
| H2-T9 | 1,41545 | 2,13642 | 0,593929 | 0,37575 | 0,784142 |
| Dnajc10 | 51,9957 | 58,0082 | 0,157862 | 0,37595 | 0,784432 |
| Gucd1 | 15,713 | 13,8973 | -0,177154 | 0,37615 | 0,784675 |

| | | | | | |
|---|---|---|---|---|---|
| LOC102634401 | 4,46661 | 6,61285 | 0,566091 | 0,3762 | 0,784675 |
| Zfp764 | 2,80734 | 2,33719 | -0,264428 | 0,37625 | 0,784675 |
| Tmem5 | 30,2069 | 26,6387 | -0,181356 | 0,37635 | 0,784733 |
| Eif1ad | 54,9665 | 45,6697 | -0,267314 | 0,3764 | 0,784733 |
| 4930430F08Rik | 5,06607 | 6,82591 | 0,430154 | 0,37655 | 0,784918 |
| Dhdh | 3,81504 | 2,46428 | -0,63053 | 0,3768 | 0,785311 |
| Tmem147 | 63,0107 | 55,7957 | -0,175442 | 0,37695 | 0,785449 |
| Tefm | 14,6861 | 16,9027 | 0,202806 | 0,37705 | 0,785449 |
| Fkbp2 | 65,4512 | 57,5567 | -0,185435 | 0,37705 | 0,785449 |
| 4921524J17Rik | 8,68594 | 10,0872 | 0,21577 | 0,37715 | 0,78553 |
| Uchl5 | 37,4946 | 33,3573 | -0,168679 | 0,3773 | 0,785562 |
| Slc35c2 | 23,0308 | 20,41 | -0,174289 | 0,37735 | 0,785562 |
| Apoa1bp | 58,5427 | 51,8192 | -0,176004 | 0,3774 | 0,785562 |
| Atxn10 | 137,713 | 123,338 | -0,159043 | 0,3775 | 0,785562 |
| Skp1a | 302,784 | 270,92 | -0,160425 | 0,3775 | 0,785562 |
| Fars2 | 8,41305 | 7,23074 | -0,218485 | 0,3776 | 0,785562 |
| Repin1 | 16,6252 | 18,7221 | 0,171374 | 0,37765 | 0,785562 |
| Crkl | 35,6526 | 39,7866 | 0,158273 | 0,37775 | 0,785562 |
| Suds3 | 61,2112 | 54,8437 | -0,158471 | 0,37775 | 0,785562 |
| Dhx57 | 13,8615 | 15,5808 | 0,168685 | 0,3778 | 0,785562 |
| Golga1 | 8,07959 | 9,10106 | 0,171751 | 0,3779 | 0,785562 |
| Whsc1 | 68,8343 | 61,6334 | -0,159416 | 0,3779 | 0,785562 |
| Rhbdd1 | 4,51405 | 5,20943 | 0,206703 | 0,3783 | 0,786138 |
| Sbf2 | 16,0737 | 17,9619 | 0,160237 | 0,3783 | 0,786138 |
| Cd82 | 1,8162 | 2,25823 | 0,314271 | 0,37845 | 0,786149 |
| Ogt | 56,0338 | 62,4568 | 0,156561 | 0,37845 | 0,786149 |
| Tarsl2 | 4,37257 | 3,75682 | -0,218972 | 0,3785 | 0,786149 |
| Triap1 | 41,6932 | 36,8224 | -0,179229 | 0,37855 | 0,786149 |
| 4930453N24Rik | 33,7341 | 29,8937 | -0,174368 | 0,37875 | 0,786437 |
| Brat1 | 6,366 | 7,28506 | 0,194554 | 0,379 | 0,786574 |
| Orai2 | 1,51667 | 1,24923 | -0,279865 | 0,379 | 0,786574 |
| Calr3 | 1,05859 | 0,790719 | -0,420909 | 0,379 | 0,786574 |
| Rnf26 | 13,7883 | 12,1229 | -0,185707 | 0,3791 | 0,786654 |
| Tcf3 | 136,03 | 121,818 | -0,159203 | 0,3792 | 0,786734 |
| Mrps23 | 61,0969 | 51,793 | -0,238344 | 0,3795 | 0,787229 |
| Npm3-ps1 | 40,7525 | 46,0904 | 0,177579 | 0,3797 | 0,787342 |
| Nlrp5-ps | 0,978709 | 0,756734 | -0,371094 | 0,3797 | 0,787342 |
| Nmnat2 | 1,08014 | 1,30769 | 0,275812 | 0,3798 | 0,787342 |
| Wars | 63,333 | 56,4865 | -0,165052 | 0,3798 | 0,787342 |
| Gtf2a2 | 89,8725 | 80,1773 | -0,164685 | 0,3801 | 0,787799 |
| Parg | 22,6178 | 25,27 | 0,159965 | 0,38015 | 0,787799 |
| H2-T22,H2-T9 | 1,27851 | 9,67E-005 | -13,6903 | 0,3803 | 0,787799 |
| Syn1 | 0,591938 | 1,13968 | 0,945111 | 0,3804 | 0,787799 |
| Drg2 | 63,791 | 57,0556 | -0,160984 | 0,3804 | 0,787799 |
| Rbmx | 104,925 | 94,1439 | -0,156415 | 0,38045 | 0,787799 |
| Mcmbp | 45,3079 | 40,6004 | -0,158271 | 0,38045 | 0,787799 |
| 4930519F09Rik | 4,22751 | 3,4921 | -0,275715 | 0,38085 | 0,7885 |
| E4f1 | 14,1017 | 15,9293 | 0,175813 | 0,38105 | 0,788786 |
| Uhrf1bp1l | 12,2354 | 13,6884 | 0,161898 | 0,38115 | 0,788866 |
| Mrpl38 | 48,3684 | 43,0952 | -0,166539 | 0,3815 | 0,789463 |

| | | | | | |
|---|---|---|---|---|---|
| Ahsa1 | 218,347 | 196,303 | -0,153542 | 0,3818 | 0,789957 |
| Adnp2 | 14,7875 | 16,5602 | 0,163336 | 0,38195 | 0,790012 |
| Ptp4a3 | 5,72394 | 4,98569 | -0,199216 | 0,38195 | 0,790012 |
| Telo2 | 15,3578 | 17,246 | 0,167295 | 0,38215 | 0,790083 |
| 9530077C05Rik | 2,57037 | 2,11505 | -0,281286 | 0,38215 | 0,790083 |
| Tecpr2 | 5,07922 | 5,71369 | 0,169816 | 0,3824 | 0,790083 |
| Rdx | 99,0113 | 110,399 | 0,157064 | 0,3824 | 0,790083 |
| Tmem126b | 25,1627 | 21,832 | -0,204841 | 0,3824 | 0,790083 |
| Kin | 32,0857 | 28,3852 | -0,176796 | 0,38255 | 0,790083 |
| Gstm2 | 2,3765 | 1,82607 | -0,380093 | 0,38255 | 0,790083 |
| Akt3 | 22,7361 | 25,4175 | 0,160835 | 0,3826 | 0,790083 |
| Btg1 | 14,6301 | 13,0533 | -0,164527 | 0,3826 | 0,790083 |
| Dnaaf2 | 19,4691 | 17,1814 | -0,18034 | 0,3826 | 0,790083 |
| Mepce | 21,403 | 23,9941 | 0,164865 | 0,3827 | 0,790138 |
| Tmem135 | 5,73494 | 5,00386 | -0,196737 | 0,38275 | 0,790138 |
| Krt18 | 3,35924 | 4,0925 | 0,284846 | 0,38285 | 0,790193 |
| Nr2c2 | 7,42902 | 8,77728 | 0,240601 | 0,3829 | 0,790193 |
| Snord96a | 984,341 | 1089,29 | 0,146155 | 0,38305 | 0,790225 |
| Pex10 | 8,74527 | 7,51518 | -0,218695 | 0,38305 | 0,790225 |
| Mrpl9 | 56,2631 | 50,2823 | -0,162138 | 0,3831 | 0,790225 |
| Mast4 | 4,31345 | 4,85295 | 0,170021 | 0,3833 | 0,790445 |
| Fam20b | 25,0761 | 22,4249 | -0,161213 | 0,38335 | 0,790445 |
| As3mt | 22,2028 | 19,6124 | -0,178973 | 0,3834 | 0,790445 |
| Sh3yl1 | 4,24739 | 5,02887 | 0,24366 | 0,38355 | 0,790445 |
| Spata1 | 1,19209 | 1,68716 | 0,501099 | 0,3836 | 0,790445 |
| Dnajb14 | 4,28778 | 3,59381 | -0,254716 | 0,38365 | 0,790445 |
| Atf6 | 5,75259 | 6,47936 | 0,171638 | 0,3837 | 0,790445 |
| Traf3 | 7,35635 | 8,25682 | 0,166595 | 0,3837 | 0,790445 |
| Abcg2 | 35,3939 | 39,5258 | 0,159295 | 0,3839 | 0,790706 |
| Samm50 | 83,4124 | 74,7425 | -0,158332 | 0,38395 | 0,790706 |
| Stk10 | 1,42872 | 1,69218 | 0,244156 | 0,38405 | 0,790785 |
| Pex2 | 30,1448 | 26,7662 | -0,171495 | 0,3843 | 0,791077 |
| Qpct | 1,02067 | 0,772933 | -0,401102 | 0,3844 | 0,791077 |
| AI414108 | 1,88962 | 1,59786 | -0,241953 | 0,38445 | 0,791077 |
| Plk1 | 66,2797 | 59,4318 | -0,157332 | 0,3845 | 0,791077 |
| Trpc3 | 1,10077 | 0,894372 | -0,29957 | 0,3845 | 0,791077 |
| Nt5c3 | 22,3365 | 19,7285 | -0,179119 | 0,385 | 0,791931 |
| Arv1 | 12,8939 | 14,9024 | 0,208861 | 0,3851 | 0,791931 |
| Pidd1 | 11,9115 | 10,5321 | -0,177569 | 0,3851 | 0,791931 |
| Sh3gl3 | 5,82538 | 6,86222 | 0,236324 | 0,3852 | 0,792009 |
| 1700003E16Rik | 0,548993 | 0,7442 | 0,438903 | 0,38535 | 0,792167 |
| Stard3nl | 35,3908 | 39,867 | 0,171818 | 0,3854 | 0,792167 |
| Bet1l | 9,91828 | 8,54443 | -0,215106 | 0,38565 | 0,792529 |
| Nosip | 23,9921 | 21,3572 | -0,167838 | 0,3857 | 0,792529 |
| Ube2m | 140,008 | 125,406 | -0,158912 | 0,38595 | 0,792916 |
| Osgin2 | 13,2914 | 15,012 | 0,175629 | 0,38615 | 0,7932 |
| H2-Q5 | 0,934479 | 1,28495 | 0,459482 | 0,38625 | 0,793278 |
| Rpf2 | 38,3346 | 34,1472 | -0,166878 | 0,38635 | 0,793356 |
| Pex11g | 2,88251 | 2,26973 | -0,344803 | 0,38645 | 0,793435 |
| Memo1 | 56,3067 | 63,0308 | 0,16275 | 0,38655 | 0,793513 |

| | | | | | |
|---|---|---|---|---|---|
| Rpl23 | 163,145 | 146,313 | -0,157094 | 0,38665 | 0,793543 |
| Spop | 46,4341 | 41,5855 | -0,159104 | 0,3867 | 0,793543 |
| Smg6 | 16,6413 | 18,5692 | 0,158149 | 0,38675 | 0,793543 |
| Klhl42 | 2,98601 | 2,60311 | -0,197986 | 0,38695 | 0,793802 |
| Fam171a1 | 20,4395 | 18,2915 | -0,160191 | 0,387 | 0,793802 |
| Coq6 | 24,1475 | 21,3727 | -0,176108 | 0,3871 | 0,793856 |
| Stx1a | 4,4084 | 5,19851 | 0,237845 | 0,38715 | 0,793856 |
| Nolc1 | 118,856 | 106,92 | -0,152674 | 0,38745 | 0,794095 |
| Ak4 | 14,7042 | 16,4364 | 0,16066 | 0,38755 | 0,794095 |
| Psmc5 | 276,955 | 248,736 | -0,155034 | 0,38755 | 0,794095 |
| Klhl23 | 14,1642 | 12,6483 | -0,163306 | 0,38755 | 0,794095 |
| Tmem143 | 5,40111 | 6,28437 | 0,218512 | 0,3876 | 0,794095 |
| Max | 46,3238 | 41,4903 | -0,15898 | 0,38765 | 0,794095 |
| Cpsf6 | 43,5992 | 48,5533 | 0,155265 | 0,3877 | 0,794095 |
| Zfp658 | 3,7845 | 4,37925 | 0,210583 | 0,388 | 0,79446 |
| Grcc10 | 218,723 | 195,844 | -0,1594 | 0,3881 | 0,79446 |
| Mettl1 | 48,1109 | 42,5759 | -0,176328 | 0,3881 | 0,79446 |
| Sec22a | 5,07366 | 5,7865 | 0,189665 | 0,3882 | 0,79446 |
| Dhx8 | 35,3435 | 39,3535 | 0,155049 | 0,38825 | 0,79446 |
| Rmnd1 | 4,46358 | 3,76258 | -0,24648 | 0,38825 | 0,79446 |
| Ccdc141 | 0,659782 | 0,802927 | 0,283279 | 0,38835 | 0,794465 |
| Ikbkb | 17,2768 | 19,3191 | 0,161197 | 0,3884 | 0,794465 |
| Nfkb2 | 3,10633 | 3,65476 | 0,234563 | 0,38845 | 0,794465 |
| Zfp446 | 4,18401 | 4,77717 | 0,191271 | 0,3885 | 0,794465 |
| Smarca5-ps | 4,34956 | 4,99484 | 0,199566 | 0,38865 | 0,794518 |
| Supt3 | 15,0136 | 13,0683 | -0,20019 | 0,38865 | 0,794518 |
| Abr | 17,9198 | 20,029 | 0,160537 | 0,3889 | 0,794878 |
| Cfdp1 | 210,676 | 189,142 | -0,155556 | 0,38895 | 0,794878 |
| Bub1b | 91,3272 | 81,9821 | -0,155735 | 0,38935 | 0,795116 |
| Mir1901 | 2,95364 | 225,599 | 6,25512 | 0,3894 | 0,795116 |
| Igfbp2 | 120,566 | 134,542 | 0,15823 | 0,3894 | 0,795116 |
| Rap2a | 11,1581 | 9,92328 | -0,169197 | 0,3894 | 0,795116 |
| Gdi1 | 43,6084 | 48,7055 | 0,159479 | 0,38945 | 0,795116 |
| Sept6 | 39,3506 | 35,2648 | -0,158156 | 0,3895 | 0,795116 |
| Car12 | 2,75666 | 2,3452 | -0,233207 | 0,3895 | 0,795116 |
| Mapk6 | 73,1423 | 65,8294 | -0,151974 | 0,3903 | 0,796622 |
| Dscr3 | 28,334 | 25,2688 | -0,165176 | 0,39045 | 0,796628 |
| Ube2i | 15,5602 | 13,838 | -0,169224 | 0,3906 | 0,796628 |
| Ccdc30 | 1,04849 | 1,30276 | 0,313266 | 0,3907 | 0,796628 |
| Pex11a | 2,73352 | 2,08875 | -0,388122 | 0,3907 | 0,796628 |
| Cbx1 | 168,163 | 151,06 | -0,154741 | 0,39075 | 0,796628 |
| Slc19a1 | 14,4015 | 12,7169 | -0,179469 | 0,3908 | 0,796628 |
| Dpm2 | 44,1676 | 38,9987 | -0,179562 | 0,3908 | 0,796628 |
| A930005H10Rik | 5,97312 | 4,80586 | -0,313692 | 0,3908 | 0,796628 |
| Cdc42 | 188,394 | 169,148 | -0,155466 | 0,3909 | 0,796706 |
| Gpkow | 15,1177 | 16,9324 | 0,163555 | 0,39105 | 0,79686 |
| Nudt1 | 41,2913 | 36,5024 | -0,177843 | 0,3911 | 0,79686 |
| Lipo1 | 2,24173 | 2,67802 | 0,256555 | 0,39125 | 0,796863 |
| Raly | 323,628 | 290,717 | -0,154721 | 0,39125 | 0,796863 |
| Vps37c | 9,11273 | 10,3699 | 0,186447 | 0,3913 | 0,796863 |

| | | | | | |
|---|---|---|---|---|---|
| Angel2 | 29,2349 | 32,6182 | 0,157989 | 0,39135 | 0,796863 |
| 4930503L19Rik | 24,5353 | 27,45 | 0,161944 | 0,39145 | 0,796916 |
| Bola1 | 24,8369 | 21,4255 | -0,213158 | 0,3915 | 0,796916 |
| Tagln2 | 115,096 | 103,255 | -0,156631 | 0,39165 | 0,79707 |
| Chd2 | 11,1166 | 12,633 | 0,184478 | 0,3917 | 0,79707 |
| Tomm40 | 106,482 | 95,6471 | -0,154815 | 0,3918 | 0,797147 |
| 4930526I15Rik | 5,45856 | 6,53095 | 0,258772 | 0,3919 | 0,797224 |
| 2310034G01Rik | 0,648351 | 1,00439 | 0,631477 | 0,3922 | 0,797507 |
| Pcdh1 | 1,24131 | 1,49828 | 0,271442 | 0,3922 | 0,797507 |
| Fancl | 20,6395 | 17,9599 | -0,200631 | 0,39225 | 0,797507 |
| Zfp574 | 9,84984 | 11,07 | 0,168482 | 0,3923 | 0,797507 |
| Foxo1 | 4,09915 | 3,59781 | -0,188206 | 0,39235 | 0,797507 |
| BC018242 | 3,76892 | 4,40505 | 0,225006 | 0,3925 | 0,797586 |
| Pfdn5 | 210,517 | 188,86 | -0,156621 | 0,3926 | 0,797586 |
| Tnrc6a | 26,1865 | 29,1279 | 0,153583 | 0,39265 | 0,797586 |
| Dhx58 | 0,803263 | 1,02781 | 0,355624 | 0,3927 | 0,797586 |
| Zkscan1 | 9,59547 | 10,6964 | 0,156698 | 0,3927 | 0,797586 |
| Ptbp2 | 69,2844 | 62,2208 | -0,155133 | 0,3928 | 0,797663 |
| Pnkd | 20,0051 | 16,6667 | -0,263397 | 0,3931 | 0,798146 |
| Slc35e2 | 7,74706 | 8,69802 | 0,167039 | 0,3932 | 0,798223 |
| Abcf2 | 93,7208 | 84,4627 | -0,150054 | 0,39355 | 0,798531 |
| Nsun3 | 3,37465 | 4,09808 | 0,280211 | 0,39365 | 0,798531 |
| Jtb | 36,5296 | 32,3226 | -0,176524 | 0,39365 | 0,798531 |
| Cbfb | 61,9513 | 55,6267 | -0,155356 | 0,39375 | 0,798531 |
| Slc35a5 | 2,44087 | 2,84881 | 0,222963 | 0,3938 | 0,798531 |
| Gstcd | 23,5625 | 26,3354 | 0,160506 | 0,3938 | 0,798531 |
| Gtf2f1 | 67,824 | 60,9329 | -0,154576 | 0,39385 | 0,798531 |
| Hs3st3b1 | 5,28128 | 4,66947 | -0,177628 | 0,39385 | 0,798531 |
| Mir8102 | 8,00883 | 21,4057 | 1,41833 | 0,39415 | 0,798963 |
| Zfp623 | 16,5755 | 14,7244 | -0,170844 | 0,3942 | 0,798963 |
| Gal3st3 | 1,60972 | 1,30843 | -0,298972 | 0,39425 | 0,798964 |
| Zfp652 | 12,6427 | 14,2488 | 0,172541 | 0,39435 | 0,79904 |
| Spin2c | 10,7416 | 9,24117 | -0,217056 | 0,3945 | 0,799218 |
| Creb3l4 | 1,71677 | 2,13962 | 0,317659 | 0,3946 | 0,799269 |
| Ppp1r37 | 25,0579 | 27,947 | 0,157427 | 0,39465 | 0,799269 |
| Hsdl2 | 8,66414 | 9,84116 | 0,183773 | 0,39475 | 0,799345 |
| Phf3 | 19,3 | 21,4669 | 0,153512 | 0,3949 | 0,799523 |
| Mir3960 | 79,829 | 296,334 | 1,89224 | 0,395 | 0,799599 |
| Cltc | 74,9104 | 88,6575 | 0,243078 | 0,39525 | 0,799979 |
| Gm12070 | 1,63961 | 1,28829 | -0,347895 | 0,39535 | 0,80003 |
| Pcgf3 | 23,1798 | 25,8854 | 0,159271 | 0,3954 | 0,80003 |
| 2810002D19Rik | 7,39192 | 6,30279 | -0,229959 | 0,39555 | 0,800207 |
| Shisa5 | 9,32618 | 8,08162 | -0,206642 | 0,3959 | 0,800539 |
| Manea | 10,9426 | 12,276 | 0,165883 | 0,39605 | 0,800539 |
| Cul3 | 70,3143 | 78,1618 | 0,152644 | 0,39605 | 0,800539 |
| Sumo1 | 54,9827 | 48,9603 | -0,167364 | 0,39615 | 0,800539 |
| Eif2s3x | 5,67827 | 4,96399 | -0,193951 | 0,39615 | 0,800539 |
| Tsacc | 5,66703 | 7,32734 | 0,370699 | 0,3963 | 0,800539 |
| Ccdc22 | 18,7623 | 16,5133 | -0,184205 | 0,3963 | 0,800539 |
| Chst15 | 13,5063 | 12,0532 | -0,164215 | 0,39635 | 0,800539 |

| | | | | | |
|---|---|---|---|---|---|
| Ssfa2 | 7,02757 | 6,24488 | -0,170353 | 0,39635 | 0,800539 |
| Atat1 | 25,0272 | 28,2292 | 0,173695 | 0,39645 | 0,800539 |
| Ruvbl1 | 24,6283 | 21,872 | -0,171227 | 0,39645 | 0,800539 |
| Rpl24 | 0,840494 | 1,29134 | 0,619559 | 0,3966 | 0,800539 |
| Ccdc50 | 30,9361 | 34,3772 | 0,152159 | 0,3966 | 0,800539 |
| Copg1 | 70,1274 | 62,3735 | -0,169045 | 0,3966 | 0,800539 |
| Fastkd1 | 11,2536 | 9,95223 | -0,177292 | 0,39665 | 0,800539 |
| C1galt1c1 | 7,44889 | 8,67262 | 0,219441 | 0,39675 | 0,800615 |
| Bex2 | 108,38 | 97,2045 | -0,157 | 0,39685 | 0,800643 |
| Psma5 | 37,725 | 33,5773 | -0,168033 | 0,3969 | 0,800643 |
| Cry1 | 13,333 | 14,9772 | 0,167768 | 0,397 | 0,800643 |
| Nme5 | 1,08388 | 1,53193 | 0,49914 | 0,3971 | 0,800643 |
| Cdyl2 | 1,39213 | 1,60935 | 0,209183 | 0,39715 | 0,800643 |
| Ccp110 | 27,7473 | 30,8662 | 0,153679 | 0,39715 | 0,800643 |
| Lpar1 | 28,8502 | 32,0869 | 0,153402 | 0,3973 | 0,800643 |
| Ipo4 | 53,7679 | 48,3864 | -0,152145 | 0,39735 | 0,800643 |
| Rpap2 | 10,7745 | 8,89709 | -0,276217 | 0,3974 | 0,800643 |
| Slc6a9 | 4,38305 | 3,71318 | -0,23928 | 0,39745 | 0,800643 |
| Endov | 9,47853 | 10,6083 | 0,162461 | 0,3976 | 0,800643 |
| Rfesd | 5,72504 | 4,94411 | -0,211575 | 0,3976 | 0,800643 |
| Prdx4 | 154,151 | 138,708 | -0,152299 | 0,39765 | 0,800643 |
| Rnf166 | 13,1386 | 11,5397 | -0,187196 | 0,3977 | 0,800643 |
| Ctla2a | 0,634096 | 0,423672 | -0,581753 | 0,3977 | 0,800643 |
| Denr | 77,3465 | 69,688 | -0,150425 | 0,3979 | 0,800719 |
| Spats2l | 2,3368 | 2,73639 | 0,22774 | 0,39795 | 0,800719 |
| Ifngr1 | 9,90571 | 11,2743 | 0,186706 | 0,398 | 0,800719 |
| Aldh1b1 | 7,39093 | 8,46311 | 0,195432 | 0,39805 | 0,800719 |
| Srsf1 | 209,207 | 187,244 | -0,160011 | 0,39805 | 0,800719 |
| Wdfy1 | 2,6149 | 3,03322 | 0,214095 | 0,39835 | 0,800921 |
| Senp5 | 19,0268 | 21,1501 | 0,152634 | 0,39835 | 0,800921 |
| Rcc2 | 149,809 | 134,499 | -0,155529 | 0,3984 | 0,800921 |
| Ppp1r35 | 29,6597 | 26,0709 | -0,186066 | 0,3984 | 0,800921 |
| Snord35b | 220,847 | 203,384 | -0,11884 | 0,39865 | 0,801273 |
| Bet1 | 28,2884 | 25,1354 | -0,170486 | 0,3987 | 0,801273 |
| Nt5e | 0,604023 | 0,76428 | 0,339497 | 0,3989 | 0,801549 |
| Ppie | 70,9004 | 63,5665 | -0,157527 | 0,39905 | 0,801725 |
| C2cd5 | 12,7629 | 11,4222 | -0,160113 | 0,39925 | 0,802001 |
| Hax1 | 19,3606 | 22,2402 | 0,200049 | 0,39945 | 0,802152 |
| Rlim | 12,8184 | 14,2553 | 0,153279 | 0,39945 | 0,802152 |
| Idh3g | 84,4085 | 75,8239 | -0,154735 | 0,39965 | 0,802327 |
| Mterfd2 | 10,0808 | 8,94328 | -0,172731 | 0,39975 | 0,802327 |
| Slc38a2 | 90,9528 | 81,9127 | -0,151031 | 0,3998 | 0,802327 |
| Tiam2 | 1,69626 | 2,14976 | 0,341821 | 0,39985 | 0,802327 |
| Zfp553 | 32,9533 | 29,6034 | -0,154662 | 0,39985 | 0,802327 |
| Timm44 | 59,3713 | 53,3155 | -0,155209 | 0,39995 | 0,802402 |
| Erp44 | 28,9738 | 32,2676 | 0,155337 | 0,40025 | 0,802753 |
| Edem3 | 11,6048 | 12,919 | 0,154768 | 0,40025 | 0,802753 |
| Stoml2 | 100,909 | 90,766 | -0,152837 | 0,40065 | 0,803153 |
| Phb | 23,1424 | 20,5879 | -0,168744 | 0,40065 | 0,803153 |
| Mir378b | 64,4071 | 120,881 | 0,908291 | 0,4007 | 0,803153 |

| | | | | | |
|---|---|---|---|---|---|
| Kdm4b | 17,1574 | 15,3894 | -0,156903 | 0,4007 | 0,803153 |
| Zzz3 | 18,2522 | 20,297 | 0,153189 | 0,4009 | 0,803403 |
| Tsc22d4 | 18,8605 | 21,1223 | 0,163399 | 0,40095 | 0,803403 |
| Prrg3 | 3,79024 | 3,33421 | -0,184942 | 0,40115 | 0,803476 |
| Kdm4c | 4,76293 | 4,17575 | -0,189811 | 0,40115 | 0,803476 |
| Llph | 19,9807 | 17,4904 | -0,192044 | 0,4012 | 0,803476 |
| Gpatch4 | 46,5602 | 41,807 | -0,155355 | 0,4013 | 0,803476 |
| Ddx46 | 46,6278 | 51,7189 | 0,149501 | 0,4014 | 0,803476 |
| Ak1 | 5,90243 | 6,81722 | 0,207876 | 0,4015 | 0,803476 |
| Pdf | 38,3562 | 34,2279 | -0,164287 | 0,4015 | 0,803476 |
| Bccip | 131,832 | 117,637 | -0,164349 | 0,4015 | 0,803476 |
| Nop9 | 24,5697 | 22,0746 | -0,154493 | 0,40155 | 0,803476 |
| Prr24 | 4,07488 | 4,84521 | 0,249803 | 0,4017 | 0,803625 |
| Hadha | 44,4928 | 40,042 | -0,15206 | 0,40175 | 0,803625 |
| Dnajc24 | 23,7097 | 20,8654 | -0,184366 | 0,40195 | 0,803875 |
| Klhl21 | 6,63917 | 7,50462 | 0,176775 | 0,402 | 0,803875 |
| St8sia2 | 3,21759 | 2,81098 | -0,194906 | 0,4021 | 0,803949 |
| Myrf | 3,33015 | 3,78639 | 0,185233 | 0,4022 | 0,804024 |
| Gldc | 11,2967 | 10,0568 | -0,167726 | 0,40245 | 0,804297 |
| Spg11 | 6,70218 | 7,62219 | 0,185575 | 0,4026 | 0,804297 |
| Zbtb40 | 4,87126 | 5,46534 | 0,166017 | 0,4026 | 0,804297 |
| Slc44a2 | 16,7226 | 18,6504 | 0,157407 | 0,4026 | 0,804297 |
| Gemin8 | 5,21208 | 4,42502 | -0,236176 | 0,40265 | 0,804297 |
| Clspn | 33,5277 | 37,2001 | 0,149952 | 0,40295 | 0,804771 |
| Steap3 | 0,816642 | 1,02378 | 0,326137 | 0,4031 | 0,80482 |
| Slc35e1 | 15,0837 | 16,7605 | 0,152079 | 0,4031 | 0,80482 |
| Snrpd3 | 728,806 | 657,105 | -0,149412 | 0,40335 | 0,805043 |
| Atp9b | 12,2164 | 13,6026 | 0,155061 | 0,4034 | 0,805043 |
| Dbi | 416,244 | 374,868 | -0,151047 | 0,4034 | 0,805043 |
| Snora30 | 11,7589 | 39,14 | 1,73489 | 0,40365 | 0,805416 |
| Psip1 | 171,352 | 191,197 | 0,158098 | 0,40375 | 0,805491 |
| Axin1 | 30,6694 | 33,9748 | 0,147663 | 0,40385 | 0,805565 |
| Erlec1 | 7,89088 | 8,9023 | 0,173992 | 0,404 | 0,805588 |
| Ankrd46 | 29,7757 | 26,6853 | -0,15809 | 0,404 | 0,805588 |
| Rab36 | 3,09289 | 2,61426 | -0,242554 | 0,40405 | 0,805588 |
| Mapk1ip1l | 45,3232 | 50,3229 | 0,150966 | 0,4043 | 0,805676 |
| Parp11 | 9,12089 | 8,09201 | -0,172676 | 0,40435 | 0,805676 |
| Wdr45 | 8,47471 | 7,29816 | -0,215632 | 0,4045 | 0,805676 |
| Zdhhc21 | 4,2729 | 4,7941 | 0,166045 | 0,4046 | 0,805676 |
| Exosc1 | 33,242 | 29,7172 | -0,161707 | 0,4046 | 0,805676 |
| Ust | 3,61141 | 4,13299 | 0,194623 | 0,4048 | 0,805676 |
| Exosc10 | 50,9139 | 45,8809 | -0,150167 | 0,4048 | 0,805676 |
| Smarcd2 | 27,8414 | 31,0103 | 0,155516 | 0,4049 | 0,805676 |
| Sft2d1 | 0,975585 | 1,49718 | 0,617906 | 0,40495 | 0,805676 |
| Pelp1 | 34,7049 | 38,5344 | 0,151009 | 0,405 | 0,805676 |
| Pa2g4 | 238,069 | 214,226 | -0,152251 | 0,405 | 0,805676 |
| Rgs10 | 4,75279 | 5,78655 | 0,283929 | 0,4051 | 0,805676 |
| Gga2 | 43,0309 | 38,8704 | -0,1467 | 0,4051 | 0,805676 |
| Dhfr | 32,8404 | 28,9985 | -0,179495 | 0,4051 | 0,805676 |
| Lyrm1 | 2,83393 | 2,2151 | -0,35543 | 0,40515 | 0,805676 |

| | | | | | |
|---|---|---|---|---|---|
| Tmem150c | 0,67629 | 0,525381 | -0,364277 | 0,4052 | 0,805676 |
| Tollip | 22,6879 | 25,2424 | 0,153922 | 0,40525 | 0,805676 |
| Pdcd6ip | 21,6365 | 24,0032 | 0,149756 | 0,40525 | 0,805676 |
| Fasn | 74,8553 | 83,4356 | 0,156558 | 0,40535 | 0,805676 |
| Pstk | 28,4045 | 25,0921 | -0,178889 | 0,40535 | 0,805676 |
| H2-DMa | 0,741483 | 0,984932 | 0,409611 | 0,4055 | 0,805849 |
| Grwd1 | 68,1091 | 61,2788 | -0,152459 | 0,40575 | 0,805946 |
| Slc35d2 | 0,869243 | 0,668379 | -0,379093 | 0,40575 | 0,805946 |
| 1700001O22Rik | 1,06106 | 0,768686 | -0,465039 | 0,40575 | 0,805946 |
| Tango2 | 5,85028 | 4,97372 | -0,234179 | 0,4058 | 0,805946 |
| Cyb5r4 | 16,1629 | 14,4068 | -0,165932 | 0,40605 | 0,806317 |
| Cep41 | 7,99707 | 7,07204 | -0,177345 | 0,4062 | 0,80649 |
| Dnaic1 | 0,809801 | 0,526255 | -0,621806 | 0,4063 | 0,806564 |
| Ranbp9 | 23,6553 | 27,6823 | 0,226801 | 0,4064 | 0,806612 |
| Tmem203 | 7,98411 | 6,66611 | -0,260286 | 0,40645 | 0,806612 |
| Zfp36 | 2,24998 | 2,72912 | 0,278521 | 0,4066 | 0,806734 |
| Zfp182 | 1,585 | 1,8488 | 0,222102 | 0,40665 | 0,806734 |
| Dot1l | 17,6891 | 19,6472 | 0,151465 | 0,4067 | 0,806734 |
| Osbpl1a | 15,4015 | 17,2399 | 0,162678 | 0,40685 | 0,806782 |
| Hps3 | 4,69895 | 4,0724 | -0,206461 | 0,40685 | 0,806782 |
| Jup | 1,66545 | 1,95121 | 0,228457 | 0,40715 | 0,807226 |
| Rpl10 | 5,6793 | 7,10661 | 0,323449 | 0,4072 | 0,807226 |
| Rad51ap1 | 42,2547 | 37,886 | -0,157445 | 0,4073 | 0,8073 |
| AI506816 | 16,3307 | 22,9561 | 0,491287 | 0,40745 | 0,807327 |
| Col4a6 | 2,00032 | 2,30092 | 0,201979 | 0,4075 | 0,807327 |
| Fads3 | 2,03778 | 1,71143 | -0,251791 | 0,40755 | 0,807327 |
| Thop1 | 60,4246 | 54,4736 | -0,149578 | 0,4077 | 0,807327 |
| Dtymk | 191,45 | 171,815 | -0,156113 | 0,4077 | 0,807327 |
| Pacs2 | 19,161 | 21,2551 | 0,149634 | 0,40795 | 0,807327 |
| Txndc16 | 5,54015 | 6,28294 | 0,181513 | 0,40815 | 0,807327 |
| Fer1l5 | 1,01304 | 1,1972 | 0,240967 | 0,40825 | 0,807327 |
| Hspa8 | 342,336 | 307,916 | -0,152876 | 0,40825 | 0,807327 |
| Slc4a2 | 14,247 | 12,7741 | -0,157436 | 0,40825 | 0,807327 |
| Fam229b | 4,1384 | 5,15683 | 0,317413 | 0,40835 | 0,807327 |
| 5031425E22Rik | 5,39972 | 6,197 | 0,198685 | 0,4084 | 0,807327 |
| Efemp2 | 49,6372 | 55,133 | 0,151495 | 0,40845 | 0,807327 |
| Peg12 | 27,459 | 24,6222 | -0,157324 | 0,40845 | 0,807327 |
| Snx7 | 5,04238 | 5,85086 | 0,214544 | 0,4085 | 0,807327 |
| Senp6 | 41,1345 | 45,5718 | 0,147793 | 0,4085 | 0,807327 |
| Mcm6 | 211,699 | 190,378 | -0,153148 | 0,40855 | 0,807327 |
| Zfp777 | 14,4506 | 16,1679 | 0,161998 | 0,4086 | 0,807327 |
| Anxa2 | 9,75665 | 11,2603 | 0,206789 | 0,40865 | 0,807327 |
| Fam118a | 14,3079 | 16,0415 | 0,165004 | 0,40875 | 0,807327 |
| Ccbe1 | 1,06636 | 0,894234 | -0,253965 | 0,40875 | 0,807327 |
| Rab6a | 49,7809 | 55,3553 | 0,153129 | 0,4088 | 0,807327 |
| Abhd17b | 16,3363 | 14,5681 | -0,165266 | 0,4088 | 0,807327 |
| Fancb | 7,40549 | 8,40732 | 0,183051 | 0,40895 | 0,807327 |
| Scamp2 | 13,3307 | 11,8172 | -0,173867 | 0,40895 | 0,807327 |
| Ift20 | 38,6099 | 34,0836 | -0,179894 | 0,40895 | 0,807327 |
| 1700021K19Rik | 6,05514 | 5,38445 | -0,169361 | 0,40905 | 0,807375 |

| | | | | | |
|---|---|---|---|---|---|
| Selt | 27,5134 | 30,5587 | 0,15145 | 0,4091 | 0,807375 |
| Nup43 | 79,9266 | 71,8938 | -0,152809 | 0,40945 | 0,807912 |
| Appl1 | 10,7711 | 11,9794 | 0,153387 | 0,4097 | 0,807912 |
| Osgep | 26,608 | 23,746 | -0,164176 | 0,4097 | 0,807912 |
| Msrb1 | 24,211 | 21,1665 | -0,193876 | 0,4097 | 0,807912 |
| 1700024F13Rik | 2,65573 | 1,95366 | -0,442927 | 0,4097 | 0,807912 |
| Fam149b | 23,7265 | 21,1485 | -0,165946 | 0,40975 | 0,807912 |
| Bcl2l2 | 9,42983 | 10,6048 | 0,169416 | 0,41005 | 0,808379 |
| Zkscan2 | 2,24202 | 2,59677 | 0,211918 | 0,4103 | 0,808747 |
| Fam120a | 40,0686 | 44,4185 | 0,148688 | 0,41045 | 0,808841 |
| Ddx39 | 122,619 | 110,679 | -0,147803 | 0,41055 | 0,808841 |
| Pex16 | 7,56788 | 6,42078 | -0,237141 | 0,41075 | 0,808841 |
| Acadm | 14,9914 | 13,3014 | -0,172554 | 0,4108 | 0,808841 |
| Lpcat4 | 6,82024 | 7,85263 | 0,203354 | 0,4109 | 0,808841 |
| Dedd | 18,1254 | 20,7106 | 0,19236 | 0,4109 | 0,808841 |
| Nek6 | 19,007 | 17,0893 | -0,153431 | 0,41095 | 0,808841 |
| Kcnab2 | 0,997593 | 0,814374 | -0,292761 | 0,41095 | 0,808841 |
| Serp1 | 103,396 | 90,989 | -0,184421 | 0,4111 | 0,808841 |
| Spc25 | 81,7742 | 73,5051 | -0,153802 | 0,41115 | 0,808841 |
| Mkrn1 | 30,1675 | 33,503 | 0,151293 | 0,4112 | 0,808841 |
| Vps25 | 101,043 | 90,8405 | -0,153564 | 0,4112 | 0,808841 |
| Hnrnpab | 648,264 | 578,62 | -0,163967 | 0,4112 | 0,808841 |
| Zfp933 | 3,33178 | 3,80881 | 0,193048 | 0,41125 | 0,808841 |
| Oma1 | 10,649 | 9,34708 | -0,188128 | 0,4114 | 0,808841 |
| Ctbp1 | 135,784 | 122,56 | -0,147824 | 0,41145 | 0,808841 |
| Ehd3 | 5,87443 | 6,63931 | 0,176585 | 0,41165 | 0,808841 |
| Dhx36 | 31,2356 | 34,6711 | 0,150541 | 0,41165 | 0,808841 |
| 1810022K09Rik | 76,2213 | 56,0488 | -0,443511 | 0,4117 | 0,808841 |
| Rcor1 | 18,1113 | 20,1941 | 0,157044 | 0,41185 | 0,808841 |
| Acot13 | 38,7428 | 34,0199 | -0,18755 | 0,41185 | 0,808841 |
| Tbc1d8 | 0,964836 | 1,28376 | 0,412016 | 0,4119 | 0,808841 |
| Nol10 | 33,4556 | 37,094 | 0,148939 | 0,4119 | 0,808841 |
| Eif3d | 201,596 | 181,933 | -0,148059 | 0,4119 | 0,808841 |
| Ptrhd1 | 5,38763 | 6,56764 | 0,285723 | 0,41195 | 0,808841 |
| Ehmt2 | 66,976 | 60,5344 | -0,14589 | 0,412 | 0,808841 |
| Nr1h2 | 21,6921 | 24,2481 | 0,160699 | 0,41205 | 0,808841 |
| Parl | 50,1185 | 44,8526 | -0,160151 | 0,4122 | 0,808986 |
| Ipmk | 13,9155 | 15,4726 | 0,153026 | 0,41225 | 0,808986 |
| Usp36 | 14,4718 | 16,071 | 0,151212 | 0,4125 | 0,809229 |
| Rbm6 | 38,4936 | 42,5956 | 0,146085 | 0,4125 | 0,809229 |
| Pask | 16,2748 | 18,1007 | 0,153412 | 0,4126 | 0,809302 |
| Cdk2ap2 | 24,1976 | 27,5025 | 0,1847 | 0,4131 | 0,810035 |
| Tmem209 | 21,28 | 19,1064 | -0,155443 | 0,4131 | 0,810035 |
| Cisd3 | 8,96784 | 7,463 | -0,265005 | 0,41345 | 0,810571 |
| Sacm1l | 16,6842 | 18,5953 | 0,156451 | 0,4135 | 0,810571 |
| Rnase4 | 1,57389 | 1,97713 | 0,329079 | 0,4137 | 0,810839 |
| Tspan31 | 67,8366 | 61,1254 | -0,15029 | 0,414 | 0,811303 |
| Lypla1 | 33,0367 | 29,7064 | -0,153298 | 0,4143 | 0,811513 |
| Cklf | 12,0688 | 14,4754 | 0,262314 | 0,4144 | 0,811513 |
| Gpatch2l | 6,66932 | 7,51592 | 0,172411 | 0,4144 | 0,811513 |

| | | | | | |
|---|---|---|---|---|---|
| Syngr3 | 0,557076 | 0,749568 | 0,428185 | 0,41445 | 0,811513 |
| Sos1 | 7,78848 | 8,64132 | 0,149911 | 0,41455 | 0,811513 |
| Rtcb | 148,359 | 134,135 | -0,145409 | 0,41455 | 0,811513 |
| Ccbl2 | 4,30534 | 3,68368 | -0,224982 | 0,41455 | 0,811513 |
| Steap1 | 5,04528 | 5,99864 | 0,249702 | 0,41485 | 0,811853 |
| Rpusd1 | 9,24891 | 8,06071 | -0,198377 | 0,41485 | 0,811853 |
| Map3k4 | 19,2021 | 21,2594 | 0,146834 | 0,41495 | 0,811899 |
| Rnf180 | 0,591743 | 0,737307 | 0,317294 | 0,415 | 0,811899 |
| Gtse1 | 33,5025 | 37,1918 | 0,150713 | 0,4154 | 0,812531 |
| Trp53 | 115,869 | 104,65 | -0,146916 | 0,41545 | 0,812531 |
| Smpd1 | 12,1148 | 10,74 | -0,173774 | 0,41555 | 0,812603 |
| Rnf144a | 25,7434 | 23,2471 | -0,147149 | 0,41575 | 0,81287 |
| Pacrgl | 12,1789 | 10,6493 | -0,193624 | 0,41585 | 0,812942 |
| Hspd1 | 198,859 | 179,684 | -0,146281 | 0,41605 | 0,813183 |
| Carkd | 33,7276 | 30,1629 | -0,161153 | 0,4161 | 0,813183 |
| 2210016F16Rik | 11,2273 | 9,97846 | -0,170128 | 0,4167 | 0,813957 |
| Ano10 | 9,83444 | 11,0855 | 0,172756 | 0,41675 | 0,813957 |
| Kirrel | 24,7571 | 22,4031 | -0,144144 | 0,41675 | 0,813957 |
| Lonp2 | 22,8172 | 20,4088 | -0,160936 | 0,41675 | 0,813957 |
| Bag5 | 26,4653 | 23,7217 | -0,157898 | 0,41715 | 0,814614 |
| Mapk7 | 35,2182 | 39,0647 | 0,149546 | 0,4173 | 0,814704 |
| Isca2 | 38,2478 | 33,9466 | -0,172109 | 0,41735 | 0,814704 |
| Mutyh | 12,2407 | 13,9037 | 0,183783 | 0,4174 | 0,814704 |
| E330033B04Rik | 0,688509 | 0,825305 | 0,26145 | 0,41745 | 0,814704 |
| Plekhh1 | 3,28469 | 3,83582 | 0,22378 | 0,4176 | 0,814873 |
| Cep19 | 19,7845 | 17,6243 | -0,16681 | 0,41785 | 0,815086 |
| Elovl2 | 14,194 | 12,7156 | -0,158674 | 0,4179 | 0,815086 |
| Hcfc2 | 6,87051 | 6,00443 | -0,194389 | 0,4179 | 0,815086 |
| Inf2 | 1,45646 | 1,70805 | 0,229885 | 0,4181 | 0,815228 |
| Usp38 | 12,517 | 11,2397 | -0,155296 | 0,4181 | 0,815228 |
| Cpq | 4,24019 | 3,60037 | -0,235986 | 0,41835 | 0,815346 |
| Arl6ip1 | 150,153 | 135,682 | -0,146204 | 0,41855 | 0,815346 |
| Swt1 | 6,75714 | 7,61426 | 0,172291 | 0,41865 | 0,815346 |
| Rad1 | 30,598 | 27,352 | -0,16179 | 0,4187 | 0,815346 |
| Actr3b | 1,35769 | 1,06445 | -0,351047 | 0,4187 | 0,815346 |
| Fat3 | 4,15604 | 4,61788 | 0,15202 | 0,41875 | 0,815346 |
| Smyd2 | 78,9476 | 87,5062 | 0,148489 | 0,41875 | 0,815346 |
| Cetn2 | 59,6806 | 53,624 | -0,154383 | 0,41875 | 0,815346 |
| Rassf1 | 20,9944 | 23,4928 | 0,162209 | 0,4189 | 0,815346 |
| Fam89a | 3,49134 | 2,88449 | -0,275461 | 0,41895 | 0,815346 |
| Rab1b | 82,4569 | 74,5747 | -0,144954 | 0,419 | 0,815346 |
| 1700008O03Rik | 0,496707 | 0,765402 | 0,623823 | 0,41905 | 0,815346 |
| Adck2 | 3,91344 | 4,47701 | 0,194098 | 0,41905 | 0,815346 |
| Fastkd2 | 16,6371 | 14,9084 | -0,15828 | 0,41905 | 0,815346 |
| Xist | 39,3421 | 43,8923 | 0,157893 | 0,41935 | 0,8154 |
| Trim39 | 14,2931 | 15,9408 | 0,157406 | 0,41945 | 0,8154 |
| Smek2 | 20,5518 | 22,7708 | 0,147923 | 0,41945 | 0,8154 |
| Zfp260 | 41,2182 | 37,2061 | -0,147743 | 0,41945 | 0,8154 |
| Stx8 | 4,1381 | 3,62527 | -0,19088 | 0,41955 | 0,8154 |
| Rab11a | 38,6086 | 42,8126 | 0,149114 | 0,4196 | 0,8154 |

| | | | | | |
|---|---|---|---|---|---|
| Mylpf | 3,22607 | 2,50167 | -0,366884 | 0,4196 | 0,8154 |
| Pias4 | 25,5063 | 28,5375 | 0,162005 | 0,41965 | 0,8154 |
| Dkc1 | 119,477 | 107,821 | -0,148091 | 0,41965 | 0,8154 |
| Ykt6 | 68,2552 | 61,732 | -0,144921 | 0,42005 | 0,815824 |
| Evc | 10,3013 | 9,22692 | -0,158906 | 0,42015 | 0,815824 |
| Ctsb | 21,3887 | 18,6617 | -0,196766 | 0,4202 | 0,815824 |
| Fosb | 0,765716 | 0,945834 | 0,304779 | 0,42025 | 0,815824 |
| Uros | 8,92277 | 10,1989 | 0,192852 | 0,42025 | 0,815824 |
| Klhl12 | 19,9661 | 22,1915 | 0,152451 | 0,42025 | 0,815824 |
| Rasl11b | 24,0672 | 26,8844 | 0,159698 | 0,4205 | 0,81601 |
| Coprs | 50,9339 | 57,0385 | 0,163308 | 0,42055 | 0,81601 |
| Fam187b | 0,733282 | 0,99655 | 0,442575 | 0,4206 | 0,81601 |
| Plekha5 | 14,6765 | 16,2726 | 0,148941 | 0,4206 | 0,81601 |
| 8430427H17Rik | 11,9897 | 13,2991 | 0,149532 | 0,42095 | 0,816565 |
| Sptbn1 | 29,457 | 32,5804 | 0,145392 | 0,42105 | 0,816609 |
| Tamm41 | 37,2446 | 33,2991 | -0,161549 | 0,4211 | 0,816609 |
| Snrnp40 | 139,743 | 126,471 | -0,143977 | 0,4213 | 0,81675 |
| Zbtb18 | 23,468 | 21,1806 | -0,147947 | 0,4213 | 0,81675 |
| Sema4a | 0,722333 | 0,895915 | 0,310697 | 0,42145 | 0,816908 |
| Papss2 | 0,522401 | 0,65864 | 0,334332 | 0,42165 | 0,816908 |
| Arhgap12 | 11,3147 | 12,5815 | 0,153109 | 0,42165 | 0,816908 |
| Pik3r2 | 29,4028 | 26,5264 | -0,148525 | 0,42165 | 0,816908 |
| Mkrn3 | 1,96395 | 1,63583 | -0,263733 | 0,4217 | 0,816908 |
| 9130011E15Rik | 8,4908 | 9,57131 | 0,172815 | 0,4219 | 0,817172 |
| Katna1 | 36,0601 | 40,0362 | 0,150904 | 0,42205 | 0,817276 |
| Zfp619 | 1,46892 | 1,70849 | 0,217964 | 0,42215 | 0,817276 |
| Armcx1 | 19,8115 | 17,7174 | -0,16117 | 0,42215 | 0,817276 |
| Mtmr9 | 20,3435 | 18,2415 | -0,157346 | 0,4223 | 0,817276 |
| Lman2l | 20,4637 | 18,3348 | -0,158487 | 0,4223 | 0,817276 |
| Rpl7l1 | 80,9551 | 73,2091 | -0,145099 | 0,42245 | 0,817276 |
| Atxn7l3b | 101,059 | 91,33 | -0,146035 | 0,42265 | 0,817276 |
| Mphosph6 | 42,5425 | 47,4846 | 0,158555 | 0,4227 | 0,817276 |
| Fbxo8 | 5,32722 | 4,6783 | -0,187398 | 0,4227 | 0,817276 |
| Fam20c | 2,72052 | 3,17194 | 0,221484 | 0,42275 | 0,817276 |
| Cttn | 66,5056 | 73,5855 | 0,145948 | 0,42285 | 0,817276 |
| Atrnl1 | 7,6698 | 8,52257 | 0,152099 | 0,4229 | 0,817276 |
| Ptprs | 48,8366 | 44,1762 | -0,144691 | 0,423 | 0,817276 |
| Angptl6 | 1,78571 | 1,40326 | -0,347715 | 0,423 | 0,817276 |
| Rprm | 7,52483 | 8,69815 | 0,209049 | 0,42305 | 0,817276 |
| Odf2l | 12,7541 | 14,332 | 0,168287 | 0,42305 | 0,817276 |
| Ncoa6 | 18,2262 | 20,196 | 0,148054 | 0,4231 | 0,817276 |
| 4933426M11Rik | 7,30227 | 6,51814 | -0,163886 | 0,4231 | 0,817276 |
| Prosc | 21,658 | 19,3212 | -0,164717 | 0,4232 | 0,817335 |
| Tex30 | 39,6196 | 35,4691 | -0,159653 | 0,42335 | 0,817335 |
| Hist1h4d | 2,12738 | 3,25252 | 0,612483 | 0,4235 | 0,817335 |
| Slc25a36 | 15,6451 | 17,4496 | 0,157484 | 0,4235 | 0,817335 |
| Skp2 | 65,0529 | 58,6745 | -0,148881 | 0,4237 | 0,817335 |
| Fam103a1 | 4,67287 | 3,93578 | -0,24766 | 0,4239 | 0,817335 |
| Ect2 | 35,6009 | 39,399 | 0,146248 | 0,42395 | 0,817335 |
| Pja2 | 28,5643 | 31,5768 | 0,144653 | 0,42395 | 0,817335 |

| | | | | | |
|---|---|---|---|---|---|
| Pdcd2 | 30,7552 | 34,4754 | 0,164741 | 0,42405 | 0,817335 |
| Ormdl3 | 25,2862 | 28,1663 | 0,155622 | 0,42405 | 0,817335 |
| Gprasp1 | 28,5364 | 31,6906 | 0,151252 | 0,42405 | 0,817335 |
| Capns1 | 93,0495 | 84,1933 | -0,144294 | 0,42405 | 0,817335 |
| Fxyd3 | 0,137735 | 0,607063 | 2,13995 | 0,4241 | 0,817335 |
| Bcar1 | 25,3554 | 28,1075 | 0,148663 | 0,4241 | 0,817335 |
| Gpc2 | 35,9737 | 32,4402 | -0,14916 | 0,4241 | 0,817335 |
| Fam83d | 16,4843 | 18,421 | 0,160264 | 0,42415 | 0,817335 |
| 2510039O18Rik | 15,8743 | 17,6886 | 0,156126 | 0,42425 | 0,817404 |
| Orc6 | 94,0808 | 85,0499 | -0,145591 | 0,42455 | 0,81786 |
| Zfp93 | 1,98125 | 2,32603 | 0,231457 | 0,4248 | 0,818218 |
| Ctps | 96,6894 | 106,736 | 0,14262 | 0,4249 | 0,818288 |
| Creld2 | 46,7782 | 51,9591 | 0,15154 | 0,42505 | 0,818305 |
| Nadk2 | 20,7561 | 18,5348 | -0,163303 | 0,42505 | 0,818305 |
| Rnf181 | 51,7036 | 46,1066 | -0,165292 | 0,4251 | 0,818305 |
| Cast | 5,20894 | 5,93913 | 0,189263 | 0,4252 | 0,818348 |
| Atrn | 6,35897 | 7,06881 | 0,152676 | 0,42525 | 0,818348 |
| Khnyn | 20,3016 | 22,4906 | 0,147729 | 0,42545 | 0,81861 |
| Frzb | 41,8891 | 46,412 | 0,147922 | 0,4256 | 0,818749 |
| Zc3hav1l | 18,8496 | 17,0428 | -0,145373 | 0,42565 | 0,818749 |
| Prkrip1 | 21,958 | 19,603 | -0,163674 | 0,42575 | 0,818819 |
| Sod2 | 16,3961 | 14,749 | -0,152736 | 0,42605 | 0,819123 |
| Plk4 | 37,6309 | 41,5579 | 0,143206 | 0,4261 | 0,819123 |
| Rhbdf2 | 1,65515 | 1,39604 | -0,24562 | 0,4261 | 0,819123 |
| Sap130 | 26,7533 | 29,5429 | 0,143096 | 0,4262 | 0,819193 |
| Wdr35 | 12,5377 | 11,2665 | -0,154232 | 0,4264 | 0,819401 |
| Mlh3 | 4,33255 | 4,8713 | 0,169091 | 0,4265 | 0,819401 |
| Orc1 | 17,3301 | 15,5629 | -0,155167 | 0,4265 | 0,819401 |
| 6720489N17Rik | 6,64601 | 7,56082 | 0,186054 | 0,4266 | 0,819471 |
| Msc | 1,56193 | 1,24982 | -0,321613 | 0,4267 | 0,819487 |
| Fam213b | 17,8825 | 15,4416 | -0,211734 | 0,42675 | 0,819487 |
| Snx13 | 3,87691 | 4,36394 | 0,170724 | 0,4268 | 0,819487 |
| Ensa | 75,7353 | 68,3405 | -0,148225 | 0,4271 | 0,819902 |
| Arcn1 | 82,0076 | 74,2707 | -0,142964 | 0,4273 | 0,819902 |
| Atad1 | 53,7808 | 48,6476 | -0,144722 | 0,4273 | 0,819902 |
| Mesdc1 | 15,7681 | 14,1685 | -0,154314 | 0,4273 | 0,819902 |
| Mrpl33 | 189,099 | 169,767 | -0,155588 | 0,42735 | 0,819902 |
| Prpf31 | 56,4162 | 51,1719 | -0,140758 | 0,4274 | 0,819902 |
| Usp8 | 24,4482 | 27,0086 | 0,143691 | 0,4275 | 0,819918 |
| Brpf3 | 13,1722 | 14,5938 | 0,147863 | 0,42755 | 0,819918 |
| Nkain3 | 0,721406 | 0,878955 | 0,284978 | 0,4276 | 0,819918 |
| Ankrd13a | 31,2562 | 28,2418 | -0,146307 | 0,4278 | 0,820179 |
| Slc3a2 | 58,4931 | 64,4872 | 0,140745 | 0,42795 | 0,820222 |
| Rhobtb3 | 45,0751 | 40,81 | -0,14341 | 0,42795 | 0,820222 |
| Tra2b | 67,912 | 61,4749 | -0,143671 | 0,42805 | 0,820291 |
| Sod1 | 278,19 | 251,878 | -0,143344 | 0,42845 | 0,820881 |
| Nub1 | 22,0035 | 24,4068 | 0,149549 | 0,4285 | 0,820881 |
| Ccna2 | 132,657 | 118,718 | -0,160162 | 0,42855 | 0,820881 |
| H2afj | 10,7977 | 12,2188 | 0,178375 | 0,4287 | 0,820923 |
| Pin1 | 28,9548 | 26,1861 | -0,145001 | 0,4287 | 0,820923 |

| | | | | | |
|---|---|---|---|---|---|
| Zfp410 | 37,8388 | 41,8047 | 0,143799 | 0,4289 | 0,821183 |
| Mvb12b | 14,5321 | 16,0603 | 0,144248 | 0,42905 | 0,821199 |
| Chtop | 80,6076 | 73,1461 | -0,140134 | 0,4291 | 0,821199 |
| Plekho2 | 3,06242 | 2,6824 | -0,191151 | 0,4291 | 0,821199 |
| Dcps | 73,8433 | 65,9987 | -0,16203 | 0,42925 | 0,821336 |
| Map4 | 34,9042 | 38,5635 | 0,143833 | 0,4293 | 0,821336 |
| Rpl5 | 14,1607 | 11,892 | -0,251909 | 0,42945 | 0,821501 |
| Marveld1 | 4,21501 | 3,66581 | -0,201406 | 0,42975 | 0,821818 |
| Coro7 | 7,1832 | 8,34593 | 0,216445 | 0,42985 | 0,821818 |
| Hs2st1 | 22,748 | 25,1397 | 0,14423 | 0,42985 | 0,821818 |
| Rbm3 | 27,3838 | 24,2527 | -0,175178 | 0,4299 | 0,821818 |
| Cnot7 | 58,0793 | 65,1841 | 0,166496 | 0,43 | 0,821818 |
| Alg9 | 15,1457 | 16,8865 | 0,156966 | 0,43 | 0,821818 |
| Ndufv1 | 89,6399 | 81,2335 | -0,142067 | 0,4301 | 0,821886 |
| 4933427D14Rik | 10,2595 | 11,4206 | 0,154672 | 0,43025 | 0,82205 |
| Phrf1 | 32,7362 | 36,1332 | 0,142436 | 0,43035 | 0,822119 |
| Vim | 278,555 | 252,108 | -0,14392 | 0,4305 | 0,822161 |
| Csrp1 | 33,3295 | 29,9726 | -0,153156 | 0,4305 | 0,822161 |
| Pdk2 | 2,19342 | 1,83228 | -0,259542 | 0,431 | 0,822871 |
| Mir7082 | 94,4213 | 34,3317 | -1,45957 | 0,431 | 0,822871 |
| Mycl | 27,4306 | 30,3252 | 0,14473 | 0,43125 | 0,823159 |
| Arl15 | 9,51065 | 10,6315 | 0,160722 | 0,43135 | 0,823159 |
| Wars2 | 3,65161 | 3,19772 | -0,191488 | 0,4314 | 0,823159 |
| 1500004A13Rik | 1,61936 | 2,12929 | 0,39495 | 0,4315 | 0,823159 |
| Tceal8 | 74,6182 | 67,5323 | -0,143949 | 0,4315 | 0,823159 |
| Prdm2 | 21,204 | 23,4577 | 0,145724 | 0,43155 | 0,823159 |
| Zfp184 | 13,3361 | 14,8784 | 0,157891 | 0,4316 | 0,823159 |
| Peg3os | 36,0305 | 8,17592 | -2,13977 | 0,4317 | 0,823227 |
| Stk32c | 0,837594 | 0,591591 | -0,501651 | 0,4318 | 0,823295 |
| Tmem110 | 6,77537 | 7,6031 | 0,166289 | 0,4319 | 0,823364 |
| Wfdc2 | 7,6796 | 9,18842 | 0,258786 | 0,432 | 0,823432 |
| Ralgapb | 13,0608 | 14,351 | 0,141706 | 0,4322 | 0,823568 |
| Klhdc2 | 105,836 | 95,9251 | -0,141856 | 0,4322 | 0,823568 |
| Chd7 | 28,1881 | 25,5552 | -0,141467 | 0,4325 | 0,823895 |
| Pdhb | 55,7857 | 50,3458 | -0,148024 | 0,4325 | 0,823895 |
| Sclt1 | 12,4642 | 13,9099 | 0,158321 | 0,4327 | 0,824031 |
| Moap1 | 1,00853 | 0,471487 | -1,09697 | 0,4327 | 0,824031 |
| Dnajc14 | 19,3141 | 21,5622 | 0,158855 | 0,433 | 0,82448 |
| Pkp2 | 2,16807 | 2,54746 | 0,232645 | 0,43315 | 0,824644 |
| Larp4b | 19,6434 | 21,7087 | 0,14423 | 0,43335 | 0,824752 |
| Naa50 | 61,9137 | 68,3496 | 0,142674 | 0,43335 | 0,824752 |
| Gm11110 | 3,42355 | 2,55585 | -0,421692 | 0,4334 | 0,824752 |
| Cenpn | 35,6612 | 32,1105 | -0,151308 | 0,4336 | 0,824983 |
| Mfsd10 | 24,0776 | 21,5928 | -0,157138 | 0,43365 | 0,824983 |
| E430018J23Rik | 3,248 | 3,80104 | 0,226844 | 0,4338 | 0,825132 |
| Shisa4 | 15,55 | 13,6398 | -0,189088 | 0,43395 | 0,825132 |
| Mdfic | 3,49437 | 3,03458 | -0,203536 | 0,43395 | 0,825132 |
| Wdr61 | 100,079 | 90,6797 | -0,142287 | 0,43405 | 0,825132 |
| Nmrk1 | 0,790758 | 0,591678 | -0,418423 | 0,43405 | 0,825132 |
| 2810029C07Rik | 1,5571 | 1,8946 | 0,283038 | 0,4342 | 0,825268 |

| | | | | | |
|---|---|---|---|---|---|
| Gm4532 | 5,93537 | 7,16654 | 0,271939 | 0,43425 | 0,825268 |
| Exoc4 | 17,8405 | 16,1048 | -0,147665 | 0,43445 | 0,825499 |
| Asl | 12,5279 | 11,0239 | -0,184504 | 0,4345 | 0,825499 |
| Dbn1 | 130,034 | 118,032 | -0,139711 | 0,43465 | 0,825634 |
| Otub2 | 5,0938 | 5,80081 | 0,187511 | 0,4347 | 0,825634 |
| Ldb1 | 116,983 | 106,092 | -0,140979 | 0,43485 | 0,825727 |
| Nfe2l2 | 19,2859 | 17,3262 | -0,154592 | 0,43495 | 0,825727 |
| Nfxl1 | 21,7539 | 24,0471 | 0,144584 | 0,43505 | 0,825727 |
| Mex3d | 17,808 | 16,0343 | -0,151357 | 0,43515 | 0,825727 |
| 4933412E12Rik | 1,9785 | 1,59413 | -0,311638 | 0,43515 | 0,825727 |
| Prpf6 | 52,3089 | 47,4025 | -0,142093 | 0,4352 | 0,825727 |
| Mapre2 | 29,8428 | 27,0867 | -0,139797 | 0,43525 | 0,825727 |
| Mybl1 | 3,09012 | 3,51777 | 0,186999 | 0,43535 | 0,825727 |
| Fuk | 6,96525 | 7,80458 | 0,164146 | 0,43535 | 0,825727 |
| Smek1 | 40,0093 | 44,1923 | 0,14346 | 0,4355 | 0,825727 |
| Dcun1d3 | 3,78779 | 4,261 | 0,169833 | 0,4356 | 0,825727 |
| Fdxr | 8,43532 | 7,40273 | -0,188386 | 0,4356 | 0,825727 |
| Prkag1 | 44,5113 | 40,1422 | -0,149051 | 0,43565 | 0,825727 |
| Xntrpc | 0,904474 | 0,25011 | -1,85452 | 0,43565 | 0,825727 |
| Fance | 13,0652 | 11,5688 | -0,175492 | 0,43585 | 0,825984 |
| Pgrmc2 | 34,5607 | 31,3138 | -0,142335 | 0,43595 | 0,826051 |
| Il18bp | 5,19028 | 6,06358 | 0,224358 | 0,43605 | 0,826119 |
| Prpf40a | 74,337 | 81,8452 | 0,138816 | 0,43625 | 0,826332 |
| Kcnn1 | 0,694259 | 0,560452 | -0,308883 | 0,4363 | 0,826332 |
| Eif2b5 | 79,6718 | 72,4026 | -0,138027 | 0,43655 | 0,826332 |
| Impad1 | 33,2565 | 36,6368 | 0,139657 | 0,43665 | 0,826332 |
| Dcxr | 4,23887 | 3,46993 | -0,288776 | 0,43665 | 0,826332 |
| Car4 | 7,17717 | 6,15502 | -0,221653 | 0,4367 | 0,826332 |
| Nudt12 | 2,71794 | 2,29899 | -0,241514 | 0,4367 | 0,826332 |
| Tpm2 | 17,464 | 15,6263 | -0,160408 | 0,43675 | 0,826332 |
| BC028528 | 6,06768 | 8,77278 | 0,531888 | 0,4368 | 0,826332 |
| St6gal2 | 1,0097 | 1,2508 | 0,308927 | 0,4369 | 0,826332 |
| Nexn | 1,11606 | 1,35792 | 0,28299 | 0,4369 | 0,826332 |
| Adsl | 33,1944 | 30,1122 | -0,140591 | 0,437 | 0,826332 |
| Tmem43 | 27,1301 | 24,5511 | -0,144105 | 0,437 | 0,826332 |
| Slc30a9 | 34,3839 | 37,9084 | 0,140784 | 0,43715 | 0,826439 |
| Zfp280c | 16,6416 | 18,4122 | 0,145868 | 0,43725 | 0,826439 |
| Ttc32 | 10,188 | 8,75734 | -0,218311 | 0,43725 | 0,826439 |
| Tbc1d1 | 20,5463 | 18,6655 | -0,138505 | 0,4375 | 0,826763 |
| Gm16576 | 0,509499 | 0,619659 | 0,282394 | 0,43755 | 0,826763 |
| Pes1 | 40,7868 | 36,9706 | -0,141724 | 0,43775 | 0,82687 |
| Zfp277 | 17,118 | 15,3088 | -0,161159 | 0,43775 | 0,82687 |
| Idh3b | 95,008 | 80,1979 | -0,244484 | 0,4378 | 0,82687 |
| Egln1 | 27,7385 | 25,1206 | -0,143021 | 0,43825 | 0,827598 |
| Lcorl | 4,48338 | 6,14646 | 0,455168 | 0,4385 | 0,827826 |
| Gmfb | 38,9732 | 42,9065 | 0,138714 | 0,4385 | 0,827826 |
| Sept4 | 0,452062 | 0,624983 | 0,467298 | 0,43865 | 0,827987 |
| Lepre1 | 15,0742 | 13,5664 | -0,152035 | 0,43875 | 0,828054 |
| Shroom1 | 0,5277 | 0,662616 | 0,328456 | 0,43895 | 0,828282 |
| Mau2 | 17,7491 | 19,5549 | 0,13978 | 0,439 | 0,828282 |

| | | | | | |
|---|---|---|---|---|---|
| Use1 | 59,1034 | 53,1444 | -0,153323 | 0,43915 | 0,828416 |
| Acot6 | 0,888502 | 0,679301 | -0,387325 | 0,4392 | 0,828416 |
| Tmem230 | 7,93283 | 6,90437 | -0,200326 | 0,43935 | 0,828577 |
| Fam120c | 2,1918 | 1,93178 | -0,182187 | 0,4396 | 0,828872 |
| Ap1g2 | 3,74797 | 3,00755 | -0,317521 | 0,43965 | 0,828872 |
| 9430008C03Rik | 19,7741 | 21,8896 | 0,146629 | 0,4397 | 0,828872 |
| Orc4 | 18,8613 | 22,0075 | 0,222563 | 0,4402 | 0,829515 |
| Fadd | 7,64891 | 8,59124 | 0,167612 | 0,4402 | 0,829515 |
| Mgme1 | 12,0809 | 13,4595 | 0,155897 | 0,44025 | 0,829515 |
| Armcx6 | 3,64398 | 4,23942 | 0,218349 | 0,4403 | 0,829515 |
| Smcr8 | 9,63712 | 10,6902 | 0,149617 | 0,44075 | 0,830036 |
| Fbxo22 | 37,2603 | 41,1945 | 0,14481 | 0,44075 | 0,830036 |
| Zfp711 | 3,99689 | 4,53282 | 0,18153 | 0,4409 | 0,830036 |
| Rnf31 | 6,7976 | 7,63271 | 0,167171 | 0,4409 | 0,830036 |
| Tmem29 | 24,21 | 21,6016 | -0,164468 | 0,4409 | 0,830036 |
| Tmem238 | 1,22252 | 1,60336 | 0,391244 | 0,44115 | 0,830329 |
| Fblim1 | 8,61076 | 9,62947 | 0,161316 | 0,44125 | 0,830329 |
| Man2c1 | 14,7521 | 16,3749 | 0,150567 | 0,44125 | 0,830329 |
| Gas8 | 28,7908 | 25,8774 | -0,153915 | 0,44135 | 0,830396 |
| Foxred2 | 3,59862 | 3,1784 | -0,179145 | 0,44145 | 0,830462 |
| Myo1d | 1,35587 | 1,57773 | 0,218628 | 0,44155 | 0,830501 |
| Lrp2 | 2,50251 | 2,78157 | 0,152527 | 0,4416 | 0,830501 |
| Snord15b | 62,5802 | 67,7896 | 0,115357 | 0,44215 | 0,831413 |
| Rnf113a1 | 6,83671 | 5,82146 | -0,23192 | 0,44225 | 0,831451 |
| 2900026A02Rik | 1,64271 | 1,42051 | -0,209664 | 0,4423 | 0,831451 |
| Otulin | 22,8973 | 20,4896 | -0,160287 | 0,4424 | 0,831518 |
| Mbtps1 | 77,1993 | 84,9958 | 0,138805 | 0,44255 | 0,831528 |
| Polr3f | 15,7027 | 17,3685 | 0,14546 | 0,4426 | 0,831528 |
| Pin1rt1 | 4,9891 | 4,18127 | -0,254838 | 0,4426 | 0,831528 |
| Serpine2 | 35,4468 | 39,1319 | 0,142689 | 0,4428 | 0,831754 |
| 2700099C18Rik | 25,3031 | 28,5898 | 0,176185 | 0,44285 | 0,831754 |
| Npm1 | 693,127 | 628,349 | -0,141555 | 0,4434 | 0,832665 |
| Prpf38a | 72,4399 | 65,667 | -0,141616 | 0,44355 | 0,832816 |
| Isyna1 | 82,2688 | 74,7552 | -0,138171 | 0,4438 | 0,832816 |
| Naf1 | 11,7485 | 10,5532 | -0,154792 | 0,4438 | 0,832816 |
| Rpph1 | 30,7053 | 36,6029 | 0,253469 | 0,44385 | 0,832816 |
| Relb | 0,871012 | 1,08728 | 0,319953 | 0,44395 | 0,832816 |
| Rnf32 | 1,02622 | 0,775753 | -0,40367 | 0,44395 | 0,832816 |
| Srl | 9,09809 | 10,1025 | 0,151077 | 0,444 | 0,832816 |
| Cabin1 | 15,4795 | 17,0692 | 0,141041 | 0,444 | 0,832816 |
| Zfp180 | 14,7233 | 16,274 | 0,14447 | 0,44415 | 0,832854 |
| Slc25a44 | 10,5452 | 9,48014 | -0,153604 | 0,44415 | 0,832854 |
| Hsd17b11 | 13,0241 | 14,6344 | 0,168173 | 0,4444 | 0,832865 |
| Cmc2 | 20,276 | 18,1066 | -0,163252 | 0,4444 | 0,832865 |
| Zbtb8os | 10,7405 | 9,06223 | -0,24512 | 0,44445 | 0,832865 |
| Vipas39 | 26,8513 | 24,298 | -0,144157 | 0,4445 | 0,832865 |
| Chd1 | 29,3072 | 32,1905 | 0,135376 | 0,44465 | 0,832865 |
| 5430416N02Rik | 53,1314 | 59,5067 | 0,163487 | 0,44475 | 0,832865 |
| Calm1 | 116,545 | 106,038 | -0,136309 | 0,44475 | 0,832865 |
| B3galt6 | 5,09494 | 5,74846 | 0,17411 | 0,4449 | 0,832865 |

| | | | | | |
|---|---|---|---|---|---|
| Tldc1 | 4,30861 | 4,93369 | 0,195445 | 0,44495 | 0,832865 |
| Mn1 | 3,28638 | 2,91169 | -0,174642 | 0,44495 | 0,832865 |
| Ero1lb | 0,810443 | 0,970801 | 0,260464 | 0,445 | 0,832865 |
| Farsa | 67,0068 | 60,8399 | -0,139291 | 0,445 | 0,832865 |
| Papss1 | 31,1931 | 28,2614 | -0,142393 | 0,445 | 0,832865 |
| Gnb4 | 14,2967 | 15,8707 | 0,150689 | 0,4452 | 0,83309 |
| Zfp708 | 2,99029 | 3,51949 | 0,235081 | 0,44525 | 0,83309 |
| Slirp | 255,224 | 200,107 | -0,350987 | 0,44535 | 0,833156 |
| Rnf223 | 1,2649 | 1,61325 | 0,350948 | 0,44555 | 0,833381 |
| Zdhhc24 | 3,31092 | 3,83865 | 0,213366 | 0,4456 | 0,83338 |
| Fbxw11 | 39,7338 | 43,7039 | 0,137396 | 0,44585 | 0,833652 |
| Cks2 | 7,70509 | 9,16265 | 0,249953 | 0,44605 | 0,833652 |
| Ighmbp2 | 4,03803 | 4,52398 | 0,163941 | 0,44605 | 0,833652 |
| Chpf2 | 5,15652 | 4,56568 | -0,175569 | 0,4461 | 0,833652 |
| Katnal2 | 3,08741 | 3,6514 | 0,242055 | 0,44615 | 0,833652 |
| Ext1 | 33,7748 | 37,2033 | 0,139484 | 0,44615 | 0,833652 |
| Mecr | 22,7436 | 20,3439 | -0,160862 | 0,4462 | 0,833652 |
| Lix1l | 27,2897 | 24,38 | -0,162658 | 0,44645 | 0,833941 |
| Marf1 | 14,8566 | 16,3768 | 0,140553 | 0,44655 | 0,833941 |
| Mrfap1 | 147,03 | 133,79 | -0,136145 | 0,44655 | 0,833941 |
| Dad1 | 125,136 | 113,07 | -0,146282 | 0,4474 | 0,835407 |
| Hes6 | 34,8742 | 38,6483 | 0,148244 | 0,44755 | 0,835509 |
| Gm10516 | 0,876084 | 1,28243 | 0,54974 | 0,4476 | 0,835509 |
| Tbrg1 | 84,641 | 76,877 | -0,138805 | 0,44765 | 0,835509 |
| Tert | 4,05178 | 4,59817 | 0,182503 | 0,44775 | 0,835574 |
| Smarcb1 | 135,294 | 123,191 | -0,135198 | 0,44805 | 0,835871 |
| Safb2 | 35,8714 | 39,4677 | 0,137835 | 0,4481 | 0,835871 |
| Ep400 | 26,4373 | 29,0406 | 0,135494 | 0,44815 | 0,835871 |
| Lpcat3 | 16,5875 | 20,3447 | 0,29456 | 0,44825 | 0,835871 |
| Aco2 | 75,8111 | 68,5856 | -0,144503 | 0,44825 | 0,835871 |
| Prkcz | 1,03537 | 1,22931 | 0,247697 | 0,4483 | 0,835871 |
| Miip | 21,157 | 23,4803 | 0,15032 | 0,4484 | 0,835936 |
| Kdelc2 | 5,57963 | 4,94946 | -0,172899 | 0,4487 | 0,836374 |
| Zscan18 | 5,95778 | 5,30167 | -0,168327 | 0,4488 | 0,836439 |
| Apitd1 | 40,3014 | 36,113 | -0,15831 | 0,449 | 0,836568 |
| Dusp27 | 0,649075 | 0,527245 | -0,299912 | 0,449 | 0,836568 |
| Snapc4 | 12,7837 | 14,1179 | 0,14322 | 0,4493 | 0,837006 |
| Slc25a42 | 0,675488 | 0,834758 | 0,305427 | 0,4497 | 0,837479 |
| Fkbp10 | 22,6616 | 26,1245 | 0,205152 | 0,4497 | 0,837479 |
| Cers6 | 5,30497 | 5,95843 | 0,167587 | 0,44975 | 0,837479 |
| Ankrd10 | 68,4591 | 75,392 | 0,139168 | 0,44985 | 0,837544 |
| Rpap3 | 28,0292 | 25,3111 | -0,147157 | 0,44995 | 0,837608 |
| Klhl3 | 0,629562 | 0,477809 | -0,397915 | 0,45015 | 0,83783 |
| Hist2h4 | 1,65816 | 2,54659 | 0,618986 | 0,4502 | 0,83783 |
| Pprc1 | 33,3067 | 36,6279 | 0,137131 | 0,45065 | 0,838468 |
| Adam17 | 22,8653 | 25,3415 | 0,148339 | 0,45075 | 0,838468 |
| Tmem86a | 2,4016 | 2,85118 | 0,247566 | 0,45095 | 0,838468 |
| Gpx8 | 76,0116 | 83,5878 | 0,137073 | 0,45095 | 0,838468 |
| Camkv | 9,311 | 8,32591 | -0,161329 | 0,45095 | 0,838468 |
| Oip5 | 17,8965 | 14,1657 | -0,337281 | 0,45095 | 0,838468 |

| | | | | | |
|---|---|---|---|---|---|
| B930041F14Rik | 2,00726 | 2,39072 | 0,252215 | 0,451 | 0,838468 |
| Rmdn2 | 2,97249 | 3,50325 | 0,237024 | 0,4512 | 0,838568 |
| Irak4 | 0,816227 | 1,00038 | 0,293499 | 0,45125 | 0,838568 |
| Simc1 | 10,8529 | 12,9031 | 0,249649 | 0,45125 | 0,838568 |
| Hjurp | 10,9196 | 13,6387 | 0,320786 | 0,45155 | 0,838947 |
| Pard6b | 3,19706 | 2,78028 | -0,201514 | 0,4516 | 0,838947 |
| Rbm17 | 113,099 | 102,84 | -0,137183 | 0,45165 | 0,838947 |
| Trappc11 | 16,4265 | 18,1094 | 0,140714 | 0,45185 | 0,839075 |
| Tmem70 | 33,8721 | 30,5912 | -0,146982 | 0,45185 | 0,839075 |
| Ncapd2 | 127,999 | 116,502 | -0,135774 | 0,452 | 0,839082 |
| Gm15706 | 1,371 | 1,7988 | 0,391813 | 0,45205 | 0,839082 |
| Fam216a | 48,8422 | 44,0793 | -0,148027 | 0,45205 | 0,839082 |
| Terc | 2,83087 | 1,93557 | -0,548486 | 0,45235 | 0,839489 |
| Slc25a38 | 17,5536 | 19,6347 | 0,161641 | 0,4524 | 0,839489 |
| Mcc | 3,62731 | 4,03057 | 0,152081 | 0,4529 | 0,840295 |
| Yae1d1 | 6,42819 | 5,72167 | -0,167974 | 0,45315 | 0,840487 |
| 1700008J07Rik | 3,53141 | 3,0461 | -0,213282 | 0,45315 | 0,840487 |
| Polr2c | 24,2521 | 21,525 | -0,172096 | 0,4532 | 0,840487 |
| Hist1h2bm | 1,75718 | 1,15371 | -0,606987 | 0,45335 | 0,840644 |
| Wdr37 | 11,147 | 10,0708 | -0,146475 | 0,45355 | 0,840755 |
| Smo | 55,6762 | 50,7028 | -0,134994 | 0,45375 | 0,840755 |
| Enthd2 | 8,49611 | 9,50408 | 0,161745 | 0,4538 | 0,840755 |
| Fut11 | 9,45432 | 10,5036 | 0,151833 | 0,4538 | 0,840755 |
| Lrrc4b | 0,724895 | 0,57497 | -0,334285 | 0,45385 | 0,840755 |
| Inpp5b | 9,29169 | 10,3152 | 0,150753 | 0,4539 | 0,840755 |
| Map3k1 | 6,49146 | 5,86608 | -0,146146 | 0,45395 | 0,840755 |
| Rtn4ip1 | 6,56872 | 7,37074 | 0,166198 | 0,454 | 0,840755 |
| Nrp2 | 32,2159 | 29,3042 | -0,136663 | 0,454 | 0,840755 |
| Tlk2 | 24,0446 | 26,3719 | 0,13329 | 0,45415 | 0,840883 |
| Rbm22 | 69,6845 | 63,3586 | -0,137297 | 0,4542 | 0,840883 |
| Eif4a2 | 113,759 | 127,279 | 0,162006 | 0,45445 | 0,841224 |
| Nlgn2 | 11,0287 | 9,97474 | -0,144911 | 0,45485 | 0,841843 |
| Magohb | 42,0663 | 37,6116 | -0,161487 | 0,4555 | 0,842808 |
| BC026585 | 2,72956 | 2,25444 | -0,2759 | 0,4556 | 0,842808 |
| Pld6 | 0,957704 | 0,74476 | -0,362803 | 0,45565 | 0,842808 |
| Aspm | 12,3264 | 11,2106 | -0,136889 | 0,4557 | 0,842808 |
| Lrrc61 | 7,17416 | 6,38239 | -0,168714 | 0,4557 | 0,842808 |
| Mfap1a | 5,96838 | 6,68709 | 0,164039 | 0,4558 | 0,842872 |
| 2610015P09Rik | 4,38438 | 4,93506 | 0,170695 | 0,45595 | 0,842877 |
| Igsf9b | 0,861628 | 1,03369 | 0,262668 | 0,456 | 0,842877 |
| Smarce1 | 68,4627 | 62,3725 | -0,134407 | 0,456 | 0,842877 |
| Exoc5 | 18,2322 | 20,4318 | 0,164326 | 0,4561 | 0,842912 |
| Elac2 | 29,7371 | 26,8689 | -0,146331 | 0,45615 | 0,842912 |
| Trim11 | 10,572 | 11,8245 | 0,161531 | 0,4564 | 0,843252 |
| Gm10865 | 0,308752 | 1,34257 | 2,12047 | 0,45655 | 0,84335 |
| Emd | 78,1347 | 71,1556 | -0,134986 | 0,4566 | 0,84335 |
| Zcchc10 | 19,7544 | 17,6026 | -0,166387 | 0,45665 | 0,84335 |
| Rnft1 | 5,17678 | 4,10708 | -0,333941 | 0,4569 | 0,84369 |
| Gm19705 | 0,697938 | 0,129453 | -2,43068 | 0,457 | 0,843753 |
| Etohi1 | 3,52499 | 4,07922 | 0,210676 | 0,4571 | 0,843797 |

| | | | | | |
|---|---|---|---|---|---|
| Anks3 | 14,2602 | 15,8191 | 0,149674 | 0,45725 | 0,843797 |
| Tysnd1 | 33,001 | 29,0866 | -0,182158 | 0,45725 | 0,843797 |
| Ube3b | 25,6783 | 28,1296 | 0,131538 | 0,4573 | 0,843797 |
| Kctd10 | 28,0655 | 25,5315 | -0,13652 | 0,45745 | 0,843797 |
| Mir199a-1 | 158,881 | 249,844 | 0,653083 | 0,45755 | 0,843797 |
| Lrrc56 | 5,2055 | 6,05114 | 0,217172 | 0,45755 | 0,843797 |
| Zfyve26 | 4,11508 | 3,70879 | -0,14997 | 0,45755 | 0,843797 |
| Sp2 | 11,0172 | 12,295 | 0,15832 | 0,45785 | 0,844171 |
| Kif5a | 0,75133 | 0,626248 | -0,262711 | 0,4579 | 0,844171 |
| Slc18a2 | 0,900902 | 0,747234 | -0,269809 | 0,45795 | 0,844171 |
| Dpp8 | 36,4503 | 39,9504 | 0,132278 | 0,4581 | 0,844297 |
| Arhgdia | 234,672 | 213,342 | -0,137478 | 0,45815 | 0,844297 |
| Prep | 76,505 | 69,7333 | -0,133706 | 0,4585 | 0,844645 |
| Leng8 | 32,7264 | 35,9287 | 0,134685 | 0,45855 | 0,844645 |
| Ctsd | 56,555 | 51,4202 | -0,137318 | 0,4586 | 0,844645 |
| Hyls1 | 13,7133 | 16,2886 | 0,248282 | 0,4587 | 0,844645 |
| Cdipt | 24,2463 | 21,9462 | -0,143791 | 0,4587 | 0,844645 |
| Abcf3 | 20,8982 | 18,9617 | -0,140289 | 0,4588 | 0,844645 |
| Mgat1 | 16,5022 | 14,9143 | -0,145958 | 0,4588 | 0,844645 |
| Jmjd4 | 11,6645 | 12,8768 | 0,142649 | 0,45895 | 0,844712 |
| Sdr39u1 | 23,0719 | 25,721 | 0,156815 | 0,45905 | 0,844712 |
| Setd8 | 48,309 | 44,0062 | -0,134586 | 0,45905 | 0,844712 |
| Spryd7 | 9,08791 | 10,2702 | 0,176448 | 0,4591 | 0,844712 |
| C230091D08Rik | 5,97704 | 6,70432 | 0,16566 | 0,4593 | 0,844959 |
| Mocs2 | 21,1971 | 19,1223 | -0,148612 | 0,4596 | 0,845336 |
| Sdhaf2 | 20,0012 | 18,1296 | -0,14174 | 0,4597 | 0,845336 |
| Fdx1 | 24,237 | 21,4591 | -0,17562 | 0,45975 | 0,845336 |
| C2cd3 | 15,2807 | 13,9351 | -0,132988 | 0,4598 | 0,845336 |
| Gm10336 | 6,97135 | 7,76282 | 0,155144 | 0,4599 | 0,845336 |
| Utp23 | 6,36384 | 5,65477 | -0,170429 | 0,4599 | 0,845336 |
| Cd59b | 0,800094 | 1,12977 | 0,497786 | 0,46 | 0,845398 |
| Mdm2 | 25,8592 | 28,3962 | 0,135024 | 0,46025 | 0,845737 |
| Otud5 | 30,3034 | 27,608 | -0,134396 | 0,46065 | 0,8462 |
| Smim4 | 27,6265 | 23,5704 | -0,229075 | 0,46065 | 0,8462 |
| Gan | 4,0085 | 4,58625 | 0,194253 | 0,4607 | 0,8462 |
| Tars2 | 27,8376 | 30,607 | 0,136824 | 0,4609 | 0,846417 |
| Tpst1 | 37,483 | 41,2061 | 0,136623 | 0,46095 | 0,846417 |
| 2310015A10Rik | 1,49712 | 1,29033 | -0,214452 | 0,46115 | 0,846575 |
| Klhl2 | 8,88218 | 9,8762 | 0,153042 | 0,4612 | 0,846575 |
| Acvr2b | 25,6906 | 28,4804 | 0,148728 | 0,4613 | 0,846575 |
| Ctdsp2 | 50,6791 | 46,2164 | -0,132986 | 0,4613 | 0,846575 |
| 1700012D14Rik | 0,822893 | 1,23202 | 0,582255 | 0,4617 | 0,847158 |
| Mir466i | 6,99585 | 16,7937 | 1,26335 | 0,46175 | 0,847158 |
| Pif1 | 9,95824 | 8,97708 | -0,149645 | 0,46205 | 0,847587 |
| Zfp759 | 1,27592 | 1,07246 | -0,250626 | 0,4623 | 0,847872 |
| Ddx10 | 24,1691 | 26,5514 | 0,135626 | 0,46235 | 0,847872 |
| Ltv1 | 73,0614 | 66,631 | -0,132916 | 0,46245 | 0,847872 |
| Ube2j2 | 7,49637 | 6,54571 | -0,195641 | 0,46255 | 0,847872 |
| Dhx16 | 25,6972 | 28,1872 | 0,133427 | 0,4626 | 0,847872 |
| Rab4b | 16,3735 | 18,453 | 0,172492 | 0,46275 | 0,847872 |

| | | | | | |
|---|---|---|---|---|---|
| Atp5b | 1043,64 | 942,715 | -0,146728 | 0,46275 | 0,847872 |
| Ten1 | 21,354 | 17,4259 | -0,293277 | 0,46275 | 0,847872 |
| Nudt16l1 | 55,7795 | 48,4292 | -0,203858 | 0,4628 | 0,847872 |
| Cops7b | 35,423 | 32,2335 | -0,136124 | 0,4629 | 0,847934 |
| Arhgap9 | 0,701937 | 1,41441 | 1,01079 | 0,463 | 0,847966 |
| Usp45 | 6,42663 | 7,11366 | 0,14653 | 0,46305 | 0,847967 |
| Ctsc | 5,08448 | 4,45963 | -0,189177 | 0,46315 | 0,848029 |
| Mtx3 | 9,64003 | 10,6067 | 0,13787 | 0,46325 | 0,848091 |
| Myl7 | 1,09882 | 1,57703 | 0,521245 | 0,46335 | 0,848094 |
| Exoc6b | 8,44342 | 9,32969 | 0,144001 | 0,4634 | 0,848094 |
| Cep70 | 4,79873 | 5,46439 | 0,18741 | 0,46345 | 0,848094 |
| Zfp932 | 4,84315 | 5,55138 | 0,1969 | 0,46375 | 0,848374 |
| Ugdh | 28,1299 | 30,9287 | 0,136842 | 0,4638 | 0,848374 |
| Cxxc4 | 2,32818 | 2,04431 | -0,187586 | 0,46385 | 0,848374 |
| Atp6v1g2 | 2,24504 | 2,68566 | 0,258541 | 0,46395 | 0,848374 |
| Tbc1d25 | 4,61515 | 5,25364 | 0,186941 | 0,46395 | 0,848374 |
| Dok3 | 2,11557 | 2,5327 | 0,259633 | 0,464 | 0,848374 |
| Sec63 | 27,2526 | 29,8708 | 0,132346 | 0,4641 | 0,848436 |
| Impdh2 | 19,7018 | 17,7141 | -0,153425 | 0,4643 | 0,848651 |
| Ndufa7 | 260,223 | 236,072 | -0,140526 | 0,46435 | 0,848651 |
| Gpd1l | 8,64833 | 7,81403 | -0,146354 | 0,4645 | 0,84878 |
| Impa1 | 38,9012 | 35,4572 | -0,133736 | 0,4646 | 0,84878 |
| Sgce | 19,537 | 17,566 | -0,153423 | 0,46475 | 0,84878 |
| Zfp53 | 2,86367 | 3,29128 | 0,200785 | 0,46485 | 0,84878 |
| Gfra4 | 1,26959 | 1,5782 | 0,313916 | 0,4649 | 0,84878 |
| Zfp566 | 12,4039 | 13,8877 | 0,16301 | 0,4649 | 0,84878 |
| Slc25a3 | 357,515 | 326,24 | -0,132071 | 0,4649 | 0,84878 |
| Wbp2 | 19,9527 | 22,129 | 0,14935 | 0,46495 | 0,84878 |
| Usp1 | 42,8527 | 39,1346 | -0,13094 | 0,46505 | 0,848783 |
| Trafd1 | 15,5663 | 14,0658 | -0,146239 | 0,4651 | 0,848783 |
| Recql4 | 11,8674 | 13,1684 | 0,15008 | 0,46515 | 0,848783 |
| Srebf2 | 57,9305 | 63,4507 | 0,131312 | 0,4653 | 0,848936 |
| Pacrg | 0,569414 | 0,785244 | 0,463663 | 0,4654 | 0,848998 |
| Tchp | 11,9887 | 13,2824 | 0,14784 | 0,4655 | 0,849059 |
| Gm20300 | 1,056 | 1,25394 | 0,247855 | 0,46565 | 0,849185 |
| Me2 | 45,1994 | 41,1769 | -0,134469 | 0,46575 | 0,849185 |
| Slc10a7 | 5,7223 | 5,09498 | -0,16752 | 0,46585 | 0,849185 |
| Paxip1 | 25,7309 | 28,206 | 0,1325 | 0,4659 | 0,849185 |
| Ywhaz | 53,2222 | 48,5276 | -0,133222 | 0,4659 | 0,849185 |
| Tnfrsf4 | 1,01325 | 0,752936 | -0,428397 | 0,46615 | 0,849491 |
| Camsap1 | 12,3296 | 13,5203 | 0,133 | 0,4662 | 0,849491 |
| Osbpl11 | 14,2832 | 15,6906 | 0,13558 | 0,46635 | 0,849643 |
| Tlk1 | 39,5173 | 43,2562 | 0,130425 | 0,46645 | 0,849705 |
| Pomgnt1 | 25,7704 | 28,268 | 0,133457 | 0,4666 | 0,849857 |
| Rtca | 57,4735 | 52,2761 | -0,136747 | 0,46695 | 0,850164 |
| Hist1h1c | 4,62471 | 5,5935 | 0,274391 | 0,46705 | 0,850164 |
| Map1s | 15,5197 | 17,0904 | 0,139085 | 0,46705 | 0,850164 |
| Lemd2 | 14,7789 | 13,3457 | -0,147164 | 0,4671 | 0,850164 |
| Lrrc75a | 8,33019 | 7,36932 | -0,176818 | 0,4671 | 0,850164 |
| Gm12942 | 2,10014 | 1,68504 | -0,317705 | 0,46725 | 0,850289 |

| | | | | | |
|---|---|---|---|---|---|
| B330016D10Rik | 1,61157 | 1,33831 | -0,268052 | 0,46735 | 0,850289 |
| Armc6 | 9,51897 | 7,61465 | -0,322027 | 0,4674 | 0,850289 |
| Tmem104 | 1,89232 | 2,16528 | 0,194393 | 0,4675 | 0,850289 |
| Eif1 | 114,42 | 104,36 | -0,132765 | 0,4675 | 0,850289 |
| Cacnb1 | 2,86121 | 3,33933 | 0,222933 | 0,46775 | 0,85042 |
| Ccnyl1 | 8,64293 | 7,74761 | -0,15777 | 0,4679 | 0,85042 |
| Polr3a | 12,1267 | 13,3621 | 0,139957 | 0,468 | 0,85042 |
| Samd8 | 3,28718 | 3,66677 | 0,157657 | 0,46805 | 0,85042 |
| Acvr2a | 11,2624 | 9,69215 | -0,216622 | 0,4681 | 0,85042 |
| Prkg1 | 2,04628 | 1,54703 | -0,403498 | 0,4681 | 0,85042 |
| Flrt3 | 20,5821 | 18,6584 | -0,141563 | 0,46815 | 0,85042 |
| Slmo2 | 69,5431 | 63,3682 | -0,134147 | 0,46825 | 0,85042 |
| Syt13 | 4,0242 | 4,5337 | 0,171989 | 0,4683 | 0,85042 |
| Rab7 | 67,5448 | 60,7268 | -0,15351 | 0,4683 | 0,85042 |
| Exosc5 | 36,3658 | 32,7697 | -0,15022 | 0,46835 | 0,85042 |
| Adprhl2 | 31,1901 | 28,1894 | -0,145936 | 0,4684 | 0,85042 |
| Cln6 | 8,00714 | 9,02957 | 0,173371 | 0,4685 | 0,85042 |
| Ccni | 55,8969 | 51,0628 | -0,130494 | 0,4685 | 0,85042 |
| Nek7 | 14,8996 | 16,4183 | 0,140032 | 0,4686 | 0,850481 |
| Ccnd3 | 61,1482 | 53,7697 | -0,185518 | 0,46875 | 0,850574 |
| Golm1 | 78,4398 | 71,6337 | -0,130947 | 0,4688 | 0,850574 |
| Taf9b | 4,12338 | 3,6015 | -0,195227 | 0,46885 | 0,850574 |
| Afap1 | 4,27943 | 4,75046 | 0,150648 | 0,4691 | 0,850907 |
| Acp2 | 14,8324 | 16,3578 | 0,141226 | 0,46935 | 0,850972 |
| Dsel | 3,02906 | 3,3998 | 0,166579 | 0,4694 | 0,850972 |
| 2010320M18Rik | 17,0475 | 14,865 | -0,197638 | 0,4694 | 0,850972 |
| Rdh12 | 1,01608 | 0,791731 | -0,359938 | 0,46945 | 0,850972 |
| Wnk4 | 1,8426 | 1,59481 | -0,208359 | 0,4695 | 0,850972 |
| Cmc1 | 11,6142 | 9,09569 | -0,35264 | 0,46955 | 0,850972 |
| Mmachc | 19,897 | 17,9709 | -0,146881 | 0,4696 | 0,850972 |
| 3830406C13Rik | 2,74128 | 2,36219 | -0,214723 | 0,4698 | 0,851005 |
| Bcl2 | 4,3466 | 3,86328 | -0,170062 | 0,46985 | 0,851005 |
| Evl | 61,1645 | 53,963 | -0,180725 | 0,4699 | 0,851005 |
| Kif1b | 20,9116 | 22,9075 | 0,131517 | 0,46995 | 0,851005 |
| Hsd17b12 | 84,23 | 92,2162 | 0,130686 | 0,46995 | 0,851005 |
| Zfp629 | 23,0362 | 21,0753 | -0,128348 | 0,47025 | 0,851099 |
| Dag1 | 27,5875 | 25,2012 | -0,130521 | 0,47025 | 0,851099 |
| Camkmt | 3,62907 | 3,08258 | -0,235459 | 0,47025 | 0,851099 |
| Reep1 | 6,25537 | 6,94839 | 0,151585 | 0,47035 | 0,851099 |
| Ints2 | 20,3173 | 22,2432 | 0,130655 | 0,4704 | 0,851099 |
| Lsm5 | 7,56143 | 6,13309 | -0,302046 | 0,4704 | 0,851099 |
| Rbm28 | 39,2907 | 43,0262 | 0,131031 | 0,47065 | 0,851281 |
| Pkmyt1 | 21,1629 | 19,0726 | -0,150035 | 0,47065 | 0,851281 |
| 5330426P16Rik | 1,2145 | 0,97686 | -0,314142 | 0,4707 | 0,851281 |
| Mb21d2 | 13,1136 | 11,87 | -0,14374 | 0,4708 | 0,851342 |
| Qars | 91,955 | 84,0586 | -0,129531 | 0,4711 | 0,851557 |
| Ccdc122 | 0,547308 | 0,718396 | 0,392427 | 0,47115 | 0,851557 |
| Slc25a37 | 11,1019 | 10,0494 | -0,143694 | 0,4712 | 0,851557 |
| Slc25a13 | 37,0994 | 40,5772 | 0,129274 | 0,47125 | 0,851557 |
| 2610008E11Rik | 15,1623 | 16,6857 | 0,138123 | 0,4713 | 0,851557 |

| | | | | | |
|---|---|---|---|---|---|
| Mppe1 | 5,78358 | 7,63235 | 0,400165 | 0,47135 | 0,851557 |
| Rps6kb2 | 19,2867 | 17,3219 | -0,155012 | 0,4714 | 0,851557 |
| Rpn2 | 159,453 | 174,52 | 0,130258 | 0,47145 | 0,851557 |
| Fam210b | 3,10606 | 2,72862 | -0,186914 | 0,47155 | 0,851618 |
| Amacr | 5,38278 | 4,7536 | -0,179331 | 0,4717 | 0,851741 |
| Prdm4 | 29,6764 | 32,4704 | 0,12981 | 0,4718 | 0,851741 |
| Lrr1 | 9,89493 | 8,73976 | -0,179096 | 0,47185 | 0,851741 |
| Dpf1 | 9,10392 | 10,1542 | 0,157513 | 0,4719 | 0,851741 |
| Them4 | 3,21237 | 2,73476 | -0,232224 | 0,47195 | 0,851741 |
| Mir6907 | 189,793 | 507,27 | 1,41833 | 0,4721 | 0,851833 |
| Zscan21 | 45,6561 | 41,6869 | -0,131213 | 0,47215 | 0,851833 |
| Trib2 | 14,5583 | 13,2431 | -0,1366 | 0,4722 | 0,851833 |
| Spg7 | 26,509 | 29,094 | 0,134241 | 0,4724 | 0,852044 |
| Pcsk1 | 1,03389 | 1,24487 | 0,267914 | 0,47245 | 0,852044 |
| Dpp3 | 73,0592 | 66,8248 | -0,128684 | 0,4727 | 0,85211 |
| Pcdhgc4 | 1,60905 | 2,8675 | 0,83359 | 0,47295 | 0,85211 |
| Tirap | 5,34433 | 4,79838 | -0,15546 | 0,47295 | 0,85211 |
| Amigo2 | 1,81148 | 2,15029 | 0,247364 | 0,47305 | 0,85211 |
| Rdh14 | 6,27428 | 7,18884 | 0,19631 | 0,47305 | 0,85211 |
| Arfip2 | 20,4183 | 18,5728 | -0,136669 | 0,47305 | 0,85211 |
| Xbp1 | 43,231 | 47,3546 | 0,131441 | 0,4731 | 0,85211 |
| Rps10 | 59,9632 | 53,7501 | -0,157811 | 0,4732 | 0,85211 |
| Thyn1 | 103,196 | 89,4206 | -0,20671 | 0,4732 | 0,85211 |
| Zxdc | 4,13374 | 4,62725 | 0,162707 | 0,47325 | 0,85211 |
| Rps17 | 6,65438 | 5,32106 | -0,32259 | 0,47325 | 0,85211 |
| Zfp263 | 12,4573 | 13,742 | 0,141607 | 0,4733 | 0,85211 |
| Slc25a24 | 17,6907 | 16,0784 | -0,137871 | 0,47335 | 0,85211 |
| Mir3062 | 67,5054 | 11,8485 | -2,5103 | 0,4735 | 0,852231 |
| Mtfmt | 5,31728 | 6,03848 | 0,183496 | 0,47355 | 0,852231 |
| Mir6944 | 19,0993 | 35,4494 | 0,892239 | 0,4737 | 0,852261 |
| Asb8 | 14,826 | 13,3635 | -0,149835 | 0,4737 | 0,852261 |
| Lrrc3 | 0,725337 | 0,863175 | 0,251003 | 0,47425 | 0,853012 |
| Triqk | 1,85462 | 1,51983 | -0,287213 | 0,47425 | 0,853012 |
| Zfp2 | 7,17714 | 6,42444 | -0,159838 | 0,4746 | 0,853521 |
| Bmp7 | 8,35489 | 7,53568 | -0,148885 | 0,4747 | 0,853582 |
| Clic1 | 111,464 | 101,74 | -0,131689 | 0,4748 | 0,853642 |
| Cyp2d22 | 1,72301 | 1,46315 | -0,235862 | 0,47495 | 0,853732 |
| Mef2c | 0,84889 | 0,98536 | 0,215073 | 0,47505 | 0,853732 |
| Ube2q2 | 10,498 | 9,46204 | -0,149898 | 0,47505 | 0,853732 |
| Ddx17 | 254,577 | 280,596 | 0,140392 | 0,47525 | 0,853884 |
| Id1 | 118,678 | 108,352 | -0,131327 | 0,47535 | 0,853884 |
| Srsf10 | 70,9975 | 77,6212 | 0,128682 | 0,4755 | 0,853884 |
| E130308A19Rik | 10,7078 | 9,69348 | -0,143569 | 0,4756 | 0,853884 |
| Psmd1 | 124,071 | 113,369 | -0,130143 | 0,47565 | 0,853884 |
| Vopp1 | 40,4229 | 36,856 | -0,133276 | 0,47565 | 0,853884 |
| Rnf138 | 11,9696 | 10,8325 | -0,144002 | 0,4757 | 0,853884 |
| Ints1 | 19,96 | 21,8393 | 0,129814 | 0,47575 | 0,853884 |
| Csnk1g2 | 96,4375 | 105,428 | 0,128588 | 0,4758 | 0,853884 |
| Rbm42 | 62,1627 | 56,6958 | -0,132807 | 0,4758 | 0,853884 |
| Cc2d1a | 7,54221 | 8,38697 | 0,153164 | 0,4759 | 0,853914 |

| | | | | | |
|---|---|---|---|---|---|
| Ccnk | 35,5793 | 32,4503 | -0,132806 | 0,47595 | 0,853914 |
| Pcgf6 | 33,2858 | 30,321 | -0,13459 | 0,47615 | 0,854154 |
| Tbc1d5 | 6,69087 | 6,05823 | -0,143297 | 0,47625 | 0,854214 |
| Itsn2 | 2,58222 | 2,92099 | 0,177848 | 0,47635 | 0,854273 |
| Cep78 | 33,8909 | 37,1091 | 0,130878 | 0,4766 | 0,854602 |
| Qtrtd1 | 27,2283 | 29,846 | 0,132429 | 0,4767 | 0,854662 |
| Abhd3 | 0,56933 | 0,741049 | 0,380305 | 0,47685 | 0,854812 |
| Hist4h4 | 1,92783 | 2,70443 | 0,488347 | 0,477 | 0,854961 |
| Eed | 54,5126 | 59,6691 | 0,130396 | 0,4772 | 0,855081 |
| 1110059E24Rik | 45,8883 | 41,717 | -0,13749 | 0,4772 | 0,855081 |
| Plac8 | 1,23665 | 0,868704 | -0,509504 | 0,47735 | 0,85523 |
| Nedd1 | 15,709 | 17,2591 | 0,135771 | 0,47745 | 0,85523 |
| BC017158 | 13,0107 | 14,3563 | 0,141981 | 0,47755 | 0,85523 |
| Lztr1 | 27,2993 | 24,9229 | -0,131395 | 0,47775 | 0,85523 |
| Pard6a | 6,02565 | 5,20942 | -0,209993 | 0,4779 | 0,85523 |
| Topbp1 | 59,2709 | 64,7735 | 0,128079 | 0,47795 | 0,85523 |
| Foxc1 | 38,6912 | 35,4222 | -0,127353 | 0,47805 | 0,85523 |
| Pex26 | 3,71753 | 3,30211 | -0,170954 | 0,47805 | 0,85523 |
| Slc39a1 | 13,9183 | 15,3979 | 0,145752 | 0,4781 | 0,85523 |
| Pdss1 | 10,9692 | 8,75507 | -0,325265 | 0,47815 | 0,85523 |
| Tfcp2 | 11,5985 | 12,8038 | 0,142633 | 0,47825 | 0,85523 |
| Ube2q1 | 51,094 | 55,8507 | 0,128422 | 0,47825 | 0,85523 |
| P4ha1 | 24,5789 | 26,9123 | 0,130844 | 0,4784 | 0,85523 |
| Tbck | 3,80485 | 4,29853 | 0,176005 | 0,4785 | 0,85523 |
| Etv3 | 6,68997 | 6,06012 | -0,142653 | 0,4785 | 0,85523 |
| Pet112 | 8,693 | 7,7644 | -0,16298 | 0,47855 | 0,85523 |
| Ppp1cc | 5,58993 | 4,93908 | -0,178588 | 0,47855 | 0,85523 |
| Bod1 | 72,1402 | 66,0178 | -0,127948 | 0,4786 | 0,85523 |
| Aurkb | 68,2733 | 62,4942 | -0,1276 | 0,4788 | 0,85523 |
| Idnk | 9,69497 | 8,55746 | -0,180054 | 0,47885 | 0,85523 |
| Gm3604 | 1,19982 | 1,42092 | 0,244005 | 0,47895 | 0,85523 |
| Micu1 | 16,9319 | 18,6431 | 0,138903 | 0,47905 | 0,85523 |
| Slain2 | 19,0397 | 20,8103 | 0,128291 | 0,47905 | 0,85523 |
| Mrpl3 | 68,9018 | 63,0417 | -0,128234 | 0,47905 | 0,85523 |
| Rfc2 | 107,824 | 98,5868 | -0,129212 | 0,47905 | 0,85523 |
| Ptp4a2 | 90,3173 | 82,9152 | -0,123366 | 0,4791 | 0,85523 |
| Wipf1 | 10,8946 | 9,90893 | -0,136812 | 0,4791 | 0,85523 |
| Auh | 11,7452 | 10,4124 | -0,17377 | 0,47915 | 0,85523 |
| Erh | 1,44697 | 1,88831 | 0,384057 | 0,47925 | 0,855289 |
| Pla2g12a | 11,65 | 13,0696 | 0,165888 | 0,4796 | 0,855795 |
| Ergic2 | 38,4295 | 43,1618 | 0,167539 | 0,4798 | 0,856033 |
| Shcbp1 | 50,9184 | 46,5374 | -0,129796 | 0,4799 | 0,856092 |
| Pttg1 | 62,206 | 68,8828 | 0,147089 | 0,48005 | 0,856211 |
| Gpr137b-ps | 2,75868 | 3,16556 | 0,198482 | 0,4801 | 0,856211 |
| Paics | 258,798 | 233,488 | -0,148473 | 0,4803 | 0,856448 |
| Mmp11 | 10,1898 | 9,14771 | -0,155638 | 0,48045 | 0,856597 |
| Pxylp1 | 8,46309 | 9,38717 | 0,149505 | 0,4808 | 0,856863 |
| Cdk2ap1 | 26,1353 | 23,5491 | -0,15033 | 0,4808 | 0,856863 |
| Dph5 | 15,8378 | 14,231 | -0,154331 | 0,4808 | 0,856863 |
| Nckipsd | 3,81939 | 4,33319 | 0,182087 | 0,481 | 0,857071 |

| | | | | | |
|---|---|---|---|---|---|
| Bop1 | 59,8994 | 54,5569 | -0,134781 | 0,48105 | 0,857071 |
| Mfhas1 | 2,97591 | 3,322 | 0,15872 | 0,48125 | 0,857308 |
| Rnh1 | 29,6309 | 26,9406 | -0,137322 | 0,48145 | 0,857545 |
| Kif14 | 6,68729 | 7,33119 | 0,132625 | 0,48155 | 0,857604 |
| Tnrc6b | 5,70981 | 6,24373 | 0,128967 | 0,4818 | 0,857808 |
| Gpt | 4,28724 | 3,71431 | -0,206955 | 0,4818 | 0,857808 |
| Mcfd2 | 33,154 | 30,2302 | -0,133191 | 0,4819 | 0,857808 |
| Socs4 | 10,2987 | 11,4517 | 0,153105 | 0,48195 | 0,857808 |
| Snw1 | 99,4939 | 89,1885 | -0,15775 | 0,4821 | 0,857808 |
| Gm20604 | 18,7006 | 16,6223 | -0,169966 | 0,48215 | 0,857808 |
| Kif3b | 9,7021 | 10,67 | 0,137189 | 0,48225 | 0,857808 |
| Rbbp6 | 32,9496 | 36,211 | 0,136168 | 0,48225 | 0,857808 |
| Mospd1 | 8,34603 | 7,43644 | -0,166479 | 0,4823 | 0,857808 |
| Scly | 5,57068 | 6,28854 | 0,174872 | 0,4824 | 0,857808 |
| Atp6v0d1 | 35,8047 | 32,5402 | -0,137926 | 0,4824 | 0,857808 |
| Mad2l1 | 100,609 | 92,0921 | -0,127607 | 0,48255 | 0,857837 |
| Mrps30 | 41,1629 | 37,4753 | -0,135404 | 0,48255 | 0,857837 |
| Fancg | 8,63542 | 9,57701 | 0,149309 | 0,4828 | 0,858044 |
| B3glct | 13,1064 | 14,3566 | 0,131439 | 0,4828 | 0,858044 |
| Reep6 | 1,09385 | 0,977726 | -0,161914 | 0,483 | 0,85828 |
| Pgm2 | 45,7047 | 41,7866 | -0,1293 | 0,4832 | 0,858456 |
| Flnb | 22,3316 | 20,4751 | -0,125218 | 0,4833 | 0,858456 |
| Crcp | 34,7668 | 31,6014 | -0,137723 | 0,4833 | 0,858456 |
| Rnft2 | 2,99697 | 3,37132 | 0,169808 | 0,4834 | 0,858515 |
| Ecd | 33,7251 | 30,8754 | -0,127363 | 0,4836 | 0,858571 |
| Stk16 | 24,694 | 22,5196 | -0,132976 | 0,4836 | 0,858571 |
| Creb3 | 13,8536 | 10,1944 | -0,44248 | 0,48365 | 0,858571 |
| Bbs2 | 8,47074 | 7,62078 | -0,152549 | 0,4837 | 0,858571 |
| Chsy1 | 25,1102 | 27,4525 | 0,128661 | 0,48385 | 0,858571 |
| Itch | 24,5675 | 26,7972 | 0,125331 | 0,48385 | 0,858571 |
| 4921533I20Rik | 1,38209 | 1,769 | 0,356078 | 0,4839 | 0,858571 |
| Dcp1a | 15,5798 | 17,0696 | 0,131751 | 0,48425 | 0,858983 |
| Nt5dc2 | 336,195 | 307,465 | -0,128876 | 0,4844 | 0,858983 |
| Myeov2 | 189,469 | 172,101 | -0,138707 | 0,4844 | 0,858983 |
| Sprtn | 7,06362 | 6,27225 | -0,171423 | 0,4844 | 0,858983 |
| Gga1 | 20,3703 | 22,3038 | 0,130821 | 0,4847 | 0,859005 |
| Klhdc8b | 2,48627 | 3,2445 | 0,384016 | 0,4848 | 0,859005 |
| Mbtd1 | 34,196 | 31,3988 | -0,123117 | 0,4849 | 0,859005 |
| Nelfe | 65,0215 | 59,4462 | -0,129333 | 0,48495 | 0,859005 |
| Pde12 | 12,1424 | 11,0389 | -0,137449 | 0,48495 | 0,859005 |
| Kcnh2 | 4,58815 | 4,10333 | -0,161116 | 0,48495 | 0,859005 |
| Pnpla6 | 6,40517 | 7,08734 | 0,146007 | 0,485 | 0,859005 |
| 1700066M21Rik | 3,29052 | 2,88264 | -0,190926 | 0,4851 | 0,859005 |
| Snhg18 | 9,7319 | 8,60689 | -0,177229 | 0,48515 | 0,859005 |
| Pmm2 | 23,2732 | 21,1466 | -0,138247 | 0,4852 | 0,859005 |
| Rnf41 | 13,9658 | 15,2782 | 0,129575 | 0,48525 | 0,859005 |
| Ckap2 | 31,0646 | 28,3889 | -0,129943 | 0,48535 | 0,859005 |
| Nusap1 | 42,0023 | 38,1715 | -0,137974 | 0,48535 | 0,859005 |
| Fam102b | 6,90734 | 6,27434 | -0,138666 | 0,48535 | 0,859005 |
| Cops3 | 129,858 | 119,104 | -0,124706 | 0,4855 | 0,859152 |

| | | | | | |
|---|---|---|---|---|---|
| Mogs | 49,3089 | 45,2055 | -0,125348 | 0,4857 | 0,859388 |
| Ippk | 9,27242 | 10,261 | 0,14616 | 0,4859 | 0,859444 |
| Fmnl3 | 15,7021 | 17,1843 | 0,130128 | 0,4859 | 0,859444 |
| Ncapg | 40,8989 | 45,3548 | 0,149194 | 0,486 | 0,859444 |
| Ndst1 | 24,8734 | 22,7714 | -0,127384 | 0,486 | 0,859444 |
| Med6 | 77,2969 | 70,4653 | -0,133496 | 0,48625 | 0,859649 |
| Hist1h2ag | 1,67973 | 1,13568 | -0,564677 | 0,48625 | 0,859649 |
| Ncapg2 | 17,3641 | 18,9603 | 0,126874 | 0,48665 | 0,860202 |
| Smpd2 | 21,5283 | 19,4503 | -0,146441 | 0,4869 | 0,860202 |
| Atg2a | 5,42503 | 5,97391 | 0,139045 | 0,487 | 0,860202 |
| Zfp512 | 29,1916 | 26,7344 | -0,12686 | 0,48705 | 0,860202 |
| Arl16 | 11,9443 | 10,1531 | -0,234406 | 0,48705 | 0,860202 |
| Gm12238 | 41,2858 | 47,9945 | 0,217222 | 0,48725 | 0,860202 |
| Cdc73 | 27,7542 | 30,3699 | 0,129936 | 0,48725 | 0,860202 |
| Rnf20 | 43,3664 | 47,3099 | 0,125562 | 0,4873 | 0,860202 |
| Tpp2 | 36,956 | 40,2813 | 0,1243 | 0,4873 | 0,860202 |
| Gart | 81,1953 | 74,4911 | -0,124328 | 0,4873 | 0,860202 |
| Dnajc11 | 66,6255 | 61,0793 | -0,125391 | 0,4873 | 0,860202 |
| Cep135 | 9,88109 | 10,8549 | 0,135603 | 0,48745 | 0,860283 |
| Tbc1d31 | 12,439 | 13,6688 | 0,136006 | 0,4875 | 0,860283 |
| Calr | 592,432 | 650,01 | 0,133812 | 0,4876 | 0,860283 |
| Lysmd3 | 5,84405 | 6,52589 | 0,159205 | 0,4877 | 0,860283 |
| Stx6 | 29,3102 | 32,0959 | 0,13099 | 0,4877 | 0,860283 |
| 1700048O20Rik | 6,83435 | 6,03528 | -0,179383 | 0,48775 | 0,860283 |
| Zxdb | 0,677133 | 0,574157 | -0,237994 | 0,48785 | 0,860283 |
| Hinfp | 12,8069 | 14,1897 | 0,147923 | 0,4879 | 0,860283 |
| Hnrnpa3 | 16,0706 | 17,5609 | 0,127946 | 0,48795 | 0,860283 |
| Cd164 | 77,1987 | 84,2138 | 0,125481 | 0,4882 | 0,860606 |
| Ino80 | 10,0331 | 10,9762 | 0,129612 | 0,4883 | 0,860662 |
| Gm6251 | 5,61767 | 7,32737 | 0,383325 | 0,48845 | 0,860662 |
| Guk1 | 69,384 | 63,2448 | -0,133657 | 0,4885 | 0,860662 |
| Oaz2 | 23,1339 | 21,0305 | -0,137524 | 0,4885 | 0,860662 |
| Tm2d3 | 27,6214 | 24,7534 | -0,158156 | 0,48865 | 0,860808 |
| Tomm20 | 18,1227 | 16,5766 | -0,128651 | 0,4888 | 0,860891 |
| Mir1668 | 23,5178 | 39,14 | 0,734894 | 0,48895 | 0,860891 |
| Sertad2 | 4,80231 | 5,30468 | 0,143535 | 0,489 | 0,860891 |
| Mok | 3,21357 | 2,75804 | -0,220537 | 0,489 | 0,860891 |
| Arid2 | 28,7634 | 31,3272 | 0,123181 | 0,48905 | 0,860891 |
| Ptdss2 | 38,3313 | 35,0774 | -0,12798 | 0,4891 | 0,860891 |
| Cacna1h | 0,949585 | 1,0821 | 0,18846 | 0,4892 | 0,860949 |
| Setd3 | 24,3946 | 22,2955 | -0,129807 | 0,4894 | 0,861005 |
| Diexf | 11,0167 | 10,0629 | -0,130649 | 0,48945 | 0,861005 |
| Rab9 | 30,56 | 27,7739 | -0,137913 | 0,48945 | 0,861005 |
| Ncbp2 | 124,269 | 114,094 | -0,12325 | 0,4895 | 0,861005 |
| Ezh2 | 132,015 | 121,104 | -0,124452 | 0,4896 | 0,861062 |
| Phactr4 | 21,3126 | 23,2599 | 0,126143 | 0,4898 | 0,861178 |
| Ydjc | 3,79037 | 3,23212 | -0,229858 | 0,4898 | 0,861178 |
| 2300009A05Rik | 7,08731 | 5,91223 | -0,261536 | 0,48995 | 0,861324 |
| Bnip3 | 13,6464 | 15,1457 | 0,150385 | 0,4901 | 0,861439 |
| Lancl1 | 10,4943 | 11,5003 | 0,132055 | 0,49015 | 0,861439 |

| | | | | | |
|---|---|---|---|---|---|
| Tmem81 | 2,33928 | 1,98018 | -0,240434 | 0,4903 | 0,861555 |
| Pfdn2 | 181,775 | 166,268 | -0,128644 | 0,49035 | 0,861555 |
| Zfp568 | 24,6155 | 28,4914 | 0,122513 | 0,49055 | 0,861788 |
| Hexb | 6,07824 | 4,69263 | -0,373256 | 0,4907 | 0,861933 |
| Zfp358 | 29,6892 | 27,0557 | -0,134008 | 0,49085 | 0,862076 |
| G3bp2 | 114,233 | 124,405 | 0,123065 | 0,49095 | 0,862076 |
| Btf3l4 | 3,308 | 3,74338 | 0,178379 | 0,49105 | 0,862076 |
| Noc2l | 123,558 | 113,332 | -0,124637 | 0,49105 | 0,862076 |
| Kansl1l | 4,07384 | 4,94912 | 0,280783 | 0,4913 | 0,862397 |
| Lrrc4 | 2,03239 | 1,08723 | -0,902518 | 0,4917 | 0,862951 |
| Srsf2 | 391,779 | 358,715 | -0,127201 | 0,49175 | 0,862951 |
| Glt25d1 | 38,4432 | 35,275 | -0,124085 | 0,4924 | 0,863635 |
| Dgkd | 22,8327 | 24,8817 | 0,123983 | 0,49255 | 0,863635 |
| Alkbh3 | 12,6936 | 11,2356 | -0,176029 | 0,49255 | 0,863635 |
| Cmtr2 | 4,91219 | 4,40154 | -0,15836 | 0,4926 | 0,863635 |
| Ctsa | 22,6594 | 20,5348 | -0,142041 | 0,49265 | 0,863635 |
| Psmb5 | 73,6606 | 66,7241 | -0,142686 | 0,49265 | 0,863635 |
| Gm15401 | 0,544238 | 1,18195 | 1,11886 | 0,4927 | 0,863635 |
| Bmp2k | 2,86027 | 2,52374 | -0,180588 | 0,4927 | 0,863635 |
| Amer1 | 10,4772 | 11,4315 | 0,125763 | 0,49275 | 0,863635 |
| Tmem234 | 75,4881 | 68,8402 | -0,132997 | 0,49285 | 0,863635 |
| Nucb2 | 8,5987 | 9,64324 | 0,1654 | 0,4929 | 0,863635 |
| Ctage5 | 38,0545 | 41,4285 | 0,122557 | 0,49295 | 0,863635 |
| Mnt | 7,67576 | 6,9759 | -0,137929 | 0,49305 | 0,863635 |
| Paip2 | 220,737 | 202,82 | -0,122127 | 0,4931 | 0,863635 |
| Zfp846 | 6,77353 | 7,53848 | 0,154364 | 0,49315 | 0,863635 |
| Pold4 | 5,24878 | 4,44681 | -0,239211 | 0,49335 | 0,863868 |
| Commd2 | 17,1862 | 15,6446 | -0,135589 | 0,4935 | 0,864012 |
| Vegfa | 23,262 | 25,3915 | 0,126371 | 0,4936 | 0,864069 |
| Samd14 | 19,5535 | 21,3893 | 0,129463 | 0,49385 | 0,864389 |
| Vstm4 | 0,627406 | 0,773233 | 0,301503 | 0,494 | 0,864473 |
| Tsc2 | 21,1618 | 23,82 | 0,170712 | 0,49405 | 0,864473 |
| Fnip2 | 4,71436 | 5,18626 | 0,137632 | 0,4941 | 0,864473 |
| Phkb | 14,1402 | 15,4671 | 0,129404 | 0,4943 | 0,864705 |
| Tut1 | 18,9981 | 20,8225 | 0,132283 | 0,4944 | 0,864731 |
| Dcaf8 | 52,1261 | 56,7103 | 0,121604 | 0,49445 | 0,864731 |
| Lig4 | 6,67544 | 7,36673 | 0,142161 | 0,49475 | 0,864951 |
| 1810043G02Rik | 12,5483 | 11,2727 | -0,154664 | 0,4948 | 0,864951 |
| Rrp36 | 36,283 | 32,9959 | -0,137008 | 0,49485 | 0,864951 |
| Timm10 | 56,7924 | 51,3302 | -0,145889 | 0,49505 | 0,864951 |
| Ankrd16 | 10,1786 | 11,2925 | 0,149819 | 0,4951 | 0,864951 |
| Cd247 | 1,14781 | 0,473087 | -1,27871 | 0,4951 | 0,864951 |
| Mir3070b | 41,7122 | 101,543 | 1,28355 | 0,49515 | 0,864951 |
| Aoc2 | 4,83204 | 5,44494 | 0,172285 | 0,4952 | 0,864951 |
| 1110037F02Rik | 22,0519 | 24,0309 | 0,123983 | 0,49525 | 0,864951 |
| Bend6 | 0,790765 | 0,634481 | -0,317672 | 0,49525 | 0,864951 |
| Trf | 6,2998 | 5,60434 | -0,168763 | 0,49535 | 0,865008 |
| 03/06/16 | 23,6715 | 25,7448 | 0,12113 | 0,4956 | 0,865327 |
| Hypk | 375,36 | 344,326 | -0,124501 | 0,4957 | 0,865384 |
| Mbd1 | 16,9574 | 18,5894 | 0,132564 | 0,4959 | 0,865554 |

| | | | | | |
|---|---|---|---|---|---|
| Grin3b | 0,55104 | 0,741481 | 0,428252 | 0,496 | 0,865554 |
| Gtf2h4 | 35,211 | 32,1068 | -0,133146 | 0,496 | 0,865554 |
| B3galt4 | 1,11916 | 0,894814 | -0,322755 | 0,49615 | 0,865698 |
| Cnot3 | 33,5104 | 36,8469 | 0,136935 | 0,49645 | 0,866043 |
| Siah1b | 20,8624 | 18,9276 | -0,140418 | 0,4965 | 0,866043 |
| Pip5k1b | 1,67999 | 1,29727 | -0,372971 | 0,49655 | 0,866043 |
| Gm6787 | 1,1235 | 0,888155 | -0,339116 | 0,4967 | 0,866086 |
| Dcaf4 | 12,0405 | 10,8728 | -0,147169 | 0,4968 | 0,866086 |
| Syt11 | 12,8931 | 14,1025 | 0,129354 | 0,4969 | 0,866086 |
| Bckdk | 27,3493 | 25,0898 | -0,124402 | 0,497 | 0,866086 |
| Tspan12 | 10,86 | 9,83051 | -0,143687 | 0,49705 | 0,866086 |
| Prr5l | 4,86109 | 5,4017 | 0,152133 | 0,49715 | 0,866086 |
| Alg5 | 17,4464 | 19,3291 | 0,147847 | 0,4972 | 0,866086 |
| Mcoln1 | 14,1253 | 15,6313 | 0,146154 | 0,4972 | 0,866086 |
| Chmp4b | 118,324 | 108,636 | -0,123239 | 0,4972 | 0,866086 |
| Wdr82 | 50,5621 | 46,4585 | -0,122113 | 0,49725 | 0,866086 |
| Tubgcp6 | 13,3836 | 14,6548 | 0,130908 | 0,4976 | 0,866578 |
| D10Jhu81e | 32,4005 | 29,5075 | -0,134938 | 0,4977 | 0,866634 |
| Tmem127 | 14,6868 | 16,0372 | 0,126904 | 0,49785 | 0,866747 |
| Abcb7 | 8,83526 | 9,6775 | 0,131362 | 0,4979 | 0,866747 |
| Sh3gl1 | 59,0982 | 52,6561 | -0,166512 | 0,49805 | 0,866891 |
| Hspbap1 | 5,96916 | 5,40677 | -0,142762 | 0,4983 | 0,867147 |
| Zfp169 | 1,85422 | 2,11878 | 0,192425 | 0,4984 | 0,867147 |
| Lrrc8a | 8,31318 | 9,13128 | 0,135417 | 0,4984 | 0,867147 |
| Slc26a1 | 0,393356 | 0,665135 | 0,75781 | 0,4988 | 0,867695 |
| Snhg6 | 162,712 | 147,933 | -0,137372 | 0,49885 | 0,867695 |
| C1qtnf4 | 1,4707 | 1,18729 | -0,308834 | 0,499 | 0,867776 |
| Prickle3 | 1,61329 | 1,99803 | 0,308577 | 0,49905 | 0,867776 |
| Tmem256 | 121,819 | 110,384 | -0,142199 | 0,4991 | 0,867776 |
| Tspan3 | 173,715 | 159,722 | -0,121161 | 0,4992 | 0,867833 |
| Slc25a19 | 6,82028 | 7,60209 | 0,156565 | 0,4996 | 0,868021 |
| Ush1c | 2,40877 | 2,06511 | -0,222081 | 0,4996 | 0,868021 |
| Garem | 5,55763 | 6,13409 | 0,142377 | 0,49965 | 0,868021 |
| Smim1 | 0,32883 | 0,710594 | 1,11168 | 0,4998 | 0,868021 |
| Ndufa3 | 391,185 | 357,301 | -0,130714 | 0,4999 | 0,868021 |
| Cdkn2d | 13,5185 | 12,0821 | -0,162067 | 0,49995 | 0,868021 |
| Zfp354a | 2,88788 | 2,55117 | -0,178851 | 0,49995 | 0,868021 |
| Lamp1 | 92,1066 | 84,5871 | -0,122866 | 0,50005 | 0,868021 |
| Srp9 | 35,848 | 32,6909 | -0,133003 | 0,50005 | 0,868021 |
| Itpripl2 | 2,98605 | 3,30746 | 0,147487 | 0,5001 | 0,868021 |
| Snrpf | 76,7535 | 69,8889 | -0,135169 | 0,5001 | 0,868021 |
| Vangl2 | 25,1019 | 27,4493 | 0,128973 | 0,50015 | 0,868021 |
| Troap | 16,0384 | 17,623 | 0,135924 | 0,50025 | 0,868021 |
| Clk3 | 36,7607 | 40,0591 | 0,123968 | 0,50035 | 0,868021 |
| Gphn | 8,05631 | 7,28989 | -0,144222 | 0,50045 | 0,868021 |
| Gmppa | 16,8918 | 15,2491 | -0,1476 | 0,50045 | 0,868021 |
| Mir6363 | 8,58631 | 0,00347602 | -11,2704 | 0,5005 | 0,868021 |
| Ncbp1 | 93,7641 | 86,3334 | -0,119117 | 0,5006 | 0,868021 |
| 9330102E08Rik | 0,761579 | 0,638165 | -0,255065 | 0,5006 | 0,868021 |
| Dynlt3 | 8,11876 | 7,27958 | -0,157405 | 0,5008 | 0,868021 |

| | | | | | |
|---|---|---|---|---|---|
| Zfp668 | 9,7345 | 8,8312 | -0,140497 | 0,50085 | 0,868021 |
| Dcaf10 | 6,99499 | 7,65253 | 0,129615 | 0,50095 | 0,868021 |
| Arih1 | 28,2114 | 30,6831 | 0,121169 | 0,50095 | 0,868021 |
| Sufu | 23,3747 | 21,4591 | -0,123359 | 0,50095 | 0,868021 |
| Irf2bp1 | 28,2542 | 25,9032 | -0,125334 | 0,501 | 0,868021 |
| Tmed8 | 4,17306 | 4,59461 | 0,138835 | 0,5013 | 0,868393 |
| Ccnh | 29,2011 | 26,6524 | -0,131759 | 0,50135 | 0,868393 |
| Lamtor1 | 47,6057 | 43,2547 | -0,138276 | 0,5016 | 0,868709 |
| Rab30 | 2,94787 | 3,40146 | 0,206482 | 0,50195 | 0,869049 |
| Ccdc103 | 0,712635 | 0,558238 | -0,352283 | 0,50195 | 0,869049 |
| Hemk1 | 4,52554 | 5,12506 | 0,179479 | 0,502 | 0,869049 |
| Glo1 | 30,2841 | 33,0691 | 0,126925 | 0,50245 | 0,869594 |
| Mt1 | 1,72307 | 1,23169 | -0,484345 | 0,50245 | 0,869594 |
| Fos | 6,85421 | 7,72444 | 0,172441 | 0,50275 | 0,869847 |
| Cbll1 | 9,96211 | 10,916 | 0,13192 | 0,5028 | 0,869847 |
| Ppa1 | 242,931 | 223,55 | -0,119949 | 0,5028 | 0,869847 |
| Fbxw4 | 8,28373 | 9,63623 | 0,218188 | 0,5032 | 0,870422 |
| Zmpste24 | 18,0259 | 19,6761 | 0,126372 | 0,5033 | 0,870478 |
| Sept1 | 5,58147 | 4,86642 | -0,197783 | 0,5035 | 0,870675 |
| Tssk6 | 2,44686 | 3,71008 | 0,600518 | 0,50355 | 0,870675 |
| Cog4 | 32,2583 | 29,618 | -0,123197 | 0,5039 | 0,87094 |
| Hfe2 | 0,679045 | 0,847217 | 0,319224 | 0,50395 | 0,87094 |
| Rcor3 | 6,87078 | 7,72846 | 0,169706 | 0,50395 | 0,87094 |
| Pmp22 | 8,52345 | 7,59451 | -0,166481 | 0,504 | 0,87094 |
| Aurka | 82,8305 | 76,1884 | -0,120591 | 0,50415 | 0,87094 |
| Ing1 | 30,3673 | 27,8566 | -0,1245 | 0,50425 | 0,87094 |
| Rmrp | 9,07131 | 6,68694 | -0,439966 | 0,50425 | 0,87094 |
| Morc2a | 32,2456 | 35,0015 | 0,118313 | 0,5044 | 0,87094 |
| Mrpl53 | 93,5229 | 84,3503 | -0,148927 | 0,5044 | 0,87094 |
| Zfp105 | 25,0138 | 27,3823 | 0,130521 | 0,50445 | 0,87094 |
| Ube2f | 32,8236 | 29,9466 | -0,132339 | 0,50445 | 0,87094 |
| Snrnp200 | 74,6903 | 68,7046 | -0,120515 | 0,5047 | 0,871254 |
| Ap2b1 | 46,7194 | 50,7034 | 0,118062 | 0,5048 | 0,871303 |
| Cdh4 | 1,62263 | 1,43568 | -0,176603 | 0,5049 | 0,871303 |
| Gm16039 | 5,34605 | 4,73617 | -0,174754 | 0,505 | 0,871303 |
| Tha1 | 2,12233 | 1,79991 | -0,237722 | 0,505 | 0,871303 |
| Mir194-1 | 106,706 | 259,761 | 1,28355 | 0,5052 | 0,871531 |
| Numb | 12,6028 | 13,7764 | 0,128457 | 0,5054 | 0,871759 |
| Hvcn1 | 7,80186 | 6,73258 | -0,212658 | 0,5055 | 0,871814 |
| Gm11974 | 37,1554 | 41,4199 | 0,156753 | 0,5056 | 0,871869 |
| Cnih1 | 88,251 | 81,1052 | -0,121819 | 0,50585 | 0,872183 |
| Gm5434 | 2,94281 | 2,51054 | -0,229197 | 0,50605 | 0,872411 |
| Ccnb2 | 86,1786 | 79,2759 | -0,120448 | 0,50625 | 0,8726 |
| Macf1 | 19,5951 | 21,4873 | 0,132987 | 0,5065 | 0,8726 |
| Col4a3bp | 8,9594 | 9,78695 | 0,127456 | 0,5065 | 0,8726 |
| Pacs1 | 18,6373 | 20,2974 | 0,1231 | 0,5065 | 0,8726 |
| Msl2 | 20,7201 | 22,5386 | 0,121365 | 0,5065 | 0,8726 |
| A630072M18Rik | 1,52821 | 1,18166 | -0,371024 | 0,5066 | 0,872655 |
| Lmbr1l | 10,9975 | 12,1517 | 0,143987 | 0,5068 | 0,872766 |
| Prps1l3 | 41,1513 | 37,8323 | -0,121318 | 0,5068 | 0,872766 |

| | | | | | |
|---|---|---|---|---|---|
| Aff1 | 4,74285 | 5,18753 | 0,129294 | 0,5072 | 0,873306 |
| Snx10 | 8,74094 | 9,66255 | 0,144616 | 0,50725 | 0,873306 |
| Zc3h12c | 2,73736 | 3,04594 | 0,154102 | 0,5076 | 0,873674 |
| Setdb2 | 6,84314 | 7,57432 | 0,146459 | 0,5076 | 0,873674 |
| Vps4a | 34,0899 | 31,2192 | -0,126911 | 0,50775 | 0,873815 |
| Noc4l | 41,9111 | 38,508 | -0,122172 | 0,5079 | 0,873893 |
| Gm3414 | 1,25325 | 1,84077 | 0,554638 | 0,508 | 0,873894 |
| 0610037L13Rik | 32,2703 | 29,4937 | -0,129801 | 0,508 | 0,873893 |
| Thg1l | 10,3491 | 9,44884 | -0,131292 | 0,50815 | 0,874034 |
| Lrrc58 | 23,8812 | 25,938 | 0,119191 | 0,50865 | 0,874777 |
| Madd | 4,66962 | 5,1309 | 0,135908 | 0,50875 | 0,874824 |
| Nudt16 | 2,27449 | 1,92685 | -0,239295 | 0,5089 | 0,874824 |
| Ap1ar | 45,3443 | 49,2734 | 0,119888 | 0,50895 | 0,874824 |
| Rab1 | 78,4842 | 72,3529 | -0,117351 | 0,50895 | 0,874824 |
| Ankrd28 | 20,6995 | 22,4908 | 0,119738 | 0,5096 | 0,875776 |
| Srp68 | 69,8542 | 64,2607 | -0,120411 | 0,50975 | 0,875776 |
| Gcsh | 90,1699 | 82,9443 | -0,120503 | 0,5098 | 0,875776 |
| Mettl8 | 12,8525 | 14,1778 | 0,141587 | 0,50985 | 0,875776 |
| Fam173a | 30,307 | 27,182 | -0,156998 | 0,50985 | 0,875776 |
| Lrsam1 | 2,28219 | 2,83341 | 0,312122 | 0,51 | 0,875776 |
| Wdr74 | 68,3722 | 62,8354 | -0,121831 | 0,51 | 0,875776 |
| Mir6357 | 56,6505 | 14,6268 | -1,95347 | 0,51005 | 0,875776 |
| Med26 | 7,26324 | 8,03091 | 0,144952 | 0,51025 | 0,875908 |
| Vps29 | 78,7081 | 72,1671 | -0,125171 | 0,5103 | 0,875908 |
| Desi2 | 13,1623 | 12,0485 | -0,127562 | 0,51035 | 0,875908 |
| Glb1 | 8,10365 | 9,19739 | 0,182652 | 0,5104 | 0,875908 |
| Acad8 | 12,2694 | 13,4756 | 0,135281 | 0,5106 | 0,876017 |
| Cep76 | 13,2711 | 14,4747 | 0,125243 | 0,5106 | 0,876017 |
| Vdac3 | 42,0514 | 38,4832 | -0,127926 | 0,5108 | 0,876243 |
| Adat1 | 10,1531 | 11,1407 | 0,133923 | 0,5109 | 0,876266 |
| Med1 | 24,7818 | 27,1612 | 0,132267 | 0,51095 | 0,876266 |
| Epg5 | 2,99565 | 2,72066 | -0,138909 | 0,51115 | 0,87646 |
| Psmd8 | 147,956 | 136,266 | -0,118739 | 0,5112 | 0,87646 |
| Rtl1 | 0,529877 | 0,633052 | 0,256666 | 0,51155 | 0,876709 |
| Rab11fip2 | 3,47687 | 3,85689 | 0,149647 | 0,51155 | 0,876709 |
| Atpaf1 | 2,81101 | 2,44731 | -0,199895 | 0,51155 | 0,876709 |
| Nup35 | 4,53829 | 4,04458 | -0,166161 | 0,5117 | 0,876786 |
| Wdr19 | 4,12868 | 3,71432 | -0,152585 | 0,51175 | 0,876786 |
| Txndc15 | 28,2906 | 25,82 | -0,131838 | 0,5118 | 0,876786 |
| Dhdds | 17,8313 | 16,3179 | -0,127958 | 0,5119 | 0,87684 |
| Pias2 | 35,2423 | 38,7025 | 0,135119 | 0,51205 | 0,87698 |
| Btbd1 | 18,6245 | 17,0443 | -0,127906 | 0,5122 | 0,876994 |
| Dnajb3 | 4,25424 | 3,6083 | -0,237581 | 0,5123 | 0,876994 |
| Map7d2 | 2,97909 | 3,34218 | 0,165918 | 0,51235 | 0,876994 |
| Hiatl1 | 17,8052 | 16,3255 | -0,125175 | 0,51235 | 0,876994 |
| Ptpn9 | 27,0162 | 29,3484 | 0,119458 | 0,5124 | 0,876994 |
| Smad1 | 13,4101 | 14,6489 | 0,127472 | 0,51275 | 0,877418 |
| Cops6 | 166,967 | 153,773 | -0,118754 | 0,5129 | 0,877418 |
| Zbtb5 | 10,9413 | 11,9509 | 0,12733 | 0,513 | 0,877418 |
| Obsl1 | 18,7102 | 20,3064 | 0,118112 | 0,513 | 0,877418 |

| | | | | | |
|---|---|---|---|---|---|
| Ankrd27 | 21,2165 | 23,1644 | 0,126723 | 0,51315 | 0,877418 |
| Zfp131 | 30,6418 | 33,3095 | 0,120435 | 0,5132 | 0,877418 |
| Snx29 | 2,43773 | 2,12049 | -0,20114 | 0,51335 | 0,877418 |
| Gtf2ird2 | 4,50252 | 5,00186 | 0,151734 | 0,51345 | 0,877418 |
| Tspan15 | 1,99442 | 1,74638 | -0,191605 | 0,51345 | 0,877418 |
| Pcgf1 | 7,84128 | 6,78116 | -0,209555 | 0,51355 | 0,877418 |
| Renbp | 6,37672 | 5,62093 | -0,182006 | 0,5136 | 0,877418 |
| 4933408B17Rik | 0,763115 | 0,618211 | -0,303802 | 0,5136 | 0,877418 |
| Coa4 | 10,805 | 9,44793 | -0,193634 | 0,51365 | 0,877418 |
| Ubac1 | 14,6271 | 13,3801 | -0,128552 | 0,51375 | 0,877418 |
| Cd9 | 3,85761 | 4,47948 | 0,215625 | 0,5138 | 0,877418 |
| Pdzk1 | 4,46637 | 3,95997 | -0,173613 | 0,51395 | 0,877418 |
| Rfx5 | 7,2446 | 7,95718 | 0,135351 | 0,514 | 0,877418 |
| Piga | 12,259 | 13,3884 | 0,127143 | 0,51405 | 0,877418 |
| Pak1ip1 | 52,9363 | 57,5569 | 0,120731 | 0,51415 | 0,877418 |
| Mrps28 | 51,2728 | 46,6504 | -0,136304 | 0,5142 | 0,877418 |
| Lamb2 | 8,88668 | 9,70665 | 0,12733 | 0,51425 | 0,877418 |
| Cenpc1 | 17,432 | 19,0139 | 0,125321 | 0,5143 | 0,877418 |
| Acad9 | 18,5538 | 17,0591 | -0,121176 | 0,5143 | 0,877418 |
| Tceanc2 | 4,70718 | 4,27738 | -0,138135 | 0,51435 | 0,877418 |
| Anln | 24,4556 | 27,6407 | 0,176631 | 0,5144 | 0,877418 |
| 9330133O14Rik | 3,23472 | 2,856 | -0,179643 | 0,5146 | 0,877418 |
| Panx1 | 28,5419 | 26,1217 | -0,127835 | 0,51465 | 0,877418 |
| Snx21 | 5,66379 | 4,81407 | -0,234509 | 0,51465 | 0,877418 |
| Txndc11 | 12,2797 | 14,708 | 0,260319 | 0,5147 | 0,877418 |
| Cdc40 | 25,5461 | 27,7708 | 0,12047 | 0,5147 | 0,877418 |
| Cdk7 | 15,5263 | 14,2927 | -0,119438 | 0,51485 | 0,877557 |
| Hdac5 | 13,242 | 12,0959 | -0,130611 | 0,515 | 0,877665 |
| Slc22a23 | 3,48064 | 3,1449 | -0,146341 | 0,51505 | 0,877665 |
| Zfp493 | 2,05619 | 2,3457 | 0,190049 | 0,51515 | 0,877719 |
| Parn | 19,2786 | 17,6446 | -0,127779 | 0,51535 | 0,877943 |
| Chmp6 | 38,2855 | 35,0773 | -0,126263 | 0,5155 | 0,878041 |
| Setmar | 4,08791 | 4,66024 | 0,189039 | 0,51565 | 0,878041 |
| Lclat1 | 6,79001 | 7,45079 | 0,13398 | 0,5157 | 0,878041 |
| Fgf13 | 6,74423 | 7,51338 | 0,155808 | 0,51575 | 0,878041 |
| Olfml2b | 3,95731 | 3,51877 | -0,169451 | 0,51575 | 0,878041 |
| Prdm15 | 6,43195 | 7,03136 | 0,128547 | 0,5159 | 0,878117 |
| Wbscr22 | 59,8971 | 54,7691 | -0,129123 | 0,516 | 0,878117 |
| 03/05/16 | 51,6679 | 46,7578 | -0,144061 | 0,516 | 0,878117 |
| Rwdd1 | 28,0967 | 25,522 | -0,138661 | 0,5162 | 0,878341 |
| Sbsn | 3,39649 | 4,15059 | 0,289274 | 0,5163 | 0,878363 |
| Erich1 | 5,93342 | 6,70719 | 0,176844 | 0,51635 | 0,878363 |
| Has2os | 1,31661 | 5,32707 | 2,01651 | 0,51665 | 0,878474 |
| Ndst2 | 10,9543 | 11,975 | 0,128528 | 0,5167 | 0,878474 |
| Rxrb | 32,3455 | 29,7748 | -0,119471 | 0,51675 | 0,878474 |
| Mrpl41 | 15,0595 | 13,398 | -0,168657 | 0,5168 | 0,878474 |
| Psmd3 | 129,586 | 119,45 | -0,117505 | 0,51685 | 0,878474 |
| Ndufc1 | 136,843 | 124,504 | -0,136331 | 0,517 | 0,878474 |
| Slc45a3 | 1,78868 | 1,55549 | -0,201527 | 0,51705 | 0,878474 |
| Fbxo46 | 4,84531 | 5,39885 | 0,156063 | 0,5171 | 0,878474 |

| | | | | | |
|---|---|---|---|---|---|
| Fam193a | 17,6453 | 19,1519 | 0,118204 | 0,5171 | 0,878474 |
| Cluh | 30,2499 | 32,7635 | 0,115157 | 0,5171 | 0,878474 |
| Gdi2 | 165,463 | 149,002 | -0,151182 | 0,5175 | 0,879037 |
| Tbl3 | 54,6835 | 50,391 | -0,117938 | 0,51785 | 0,879484 |
| Hnrnpk | 270,052 | 248,532 | -0,119804 | 0,5179 | 0,879484 |
| 2310009A05Rik | 25,9122 | 22,819 | -0,183396 | 0,5182 | 0,879697 |
| Vps18 | 8,29962 | 9,09821 | 0,132538 | 0,5183 | 0,879697 |
| Pggt1b | 12,3664 | 13,5379 | 0,13058 | 0,5183 | 0,879697 |
| 1810055G02Rik | 22,1393 | 24,1891 | 0,127747 | 0,5183 | 0,879697 |
| Lrig1 | 27,8883 | 25,6836 | -0,118815 | 0,5184 | 0,879719 |
| Mir2861 | 41,4193 | 99,4279 | 1,26335 | 0,51845 | 0,879719 |
| Cab39 | 38,4007 | 35,4289 | -0,116208 | 0,51875 | 0,880094 |
| E130309F12Rik | 0,70594 | 0,589478 | -0,260106 | 0,51885 | 0,880094 |
| Mir697 | 40,8397 | 94,1534 | 1,20504 | 0,5191 | 0,880094 |
| A230072C01Rik | 3,11228 | 3,65587 | 0,232243 | 0,5192 | 0,880094 |
| Ephb2 | 15,2537 | 16,5607 | 0,11861 | 0,5192 | 0,880094 |
| Thoc1 | 33,5355 | 38,0226 | 0,18117 | 0,5193 | 0,880094 |
| Nfatc3 | 27,9628 | 30,321 | 0,116809 | 0,51935 | 0,880094 |
| Igf2bp2 | 112,286 | 103,668 | -0,115209 | 0,5194 | 0,880094 |
| Cdr2l | 8,87352 | 8,07882 | -0,135363 | 0,51945 | 0,880094 |
| Mir7650 | 402,738 | 1774,18 | 2,13924 | 0,51955 | 0,880094 |
| Snx4 | 23,5787 | 21,6013 | -0,126367 | 0,51955 | 0,880094 |
| Zfp106 | 49,5122 | 53,7769 | 0,119201 | 0,5196 | 0,880094 |
| Lrrc45 | 17,9624 | 19,6193 | 0,12729 | 0,51965 | 0,880094 |
| Unc45a | 15,9551 | 17,414 | 0,126234 | 0,51965 | 0,880094 |
| Yars | 50,82 | 46,6521 | -0,123453 | 0,5197 | 0,880094 |
| Znhit3 | 35,7943 | 30,5423 | -0,22892 | 0,51985 | 0,8802 |
| Gm8363 | 2,58222 | 2,24013 | -0,205029 | 0,5199 | 0,8802 |
| A730008H23Rik | 30,4827 | 38,5021 | 0,336947 | 0,52 | 0,880253 |
| Gon4l | 11,0789 | 12,2897 | 0,14963 | 0,52015 | 0,880391 |
| Zwilch | 32,6905 | 35,4521 | 0,117002 | 0,52035 | 0,880613 |
| Calcoco1 | 9,30142 | 10,224 | 0,13643 | 0,5206 | 0,880804 |
| Snrpa | 104,761 | 96,2631 | -0,122043 | 0,5206 | 0,880804 |
| Fstl3 | 0,966085 | 1,54578 | 0,678109 | 0,52075 | 0,880942 |
| Msl3l2 | 5,036 | 5,91258 | 0,231509 | 0,5209 | 0,880943 |
| Lsm11 | 4,67932 | 4,26981 | -0,132129 | 0,52095 | 0,880943 |
| Cabyr | 0,6119 | 0,478611 | -0,354443 | 0,521 | 0,880943 |
| Arhgap20os | 1,18837 | 1,53155 | 0,366007 | 0,52115 | 0,880943 |
| Lect1 | 1,31322 | 1,5912 | 0,277006 | 0,52115 | 0,880943 |
| Arhgef9 | 2,73924 | 3,03897 | 0,149807 | 0,5212 | 0,880943 |
| Snip1 | 23,2692 | 25,3518 | 0,123668 | 0,52125 | 0,880943 |
| Lrwd1 | 38,0721 | 35,0395 | -0,119751 | 0,5213 | 0,880943 |
| Ctif | 4,35938 | 4,78234 | 0,133593 | 0,52155 | 0,881186 |
| Sdccag3 | 45,5708 | 49,4671 | 0,118362 | 0,52165 | 0,881186 |
| Hook2 | 4,39162 | 3,9151 | -0,165702 | 0,52165 | 0,881186 |
| 4833420G17Rik | 30,1266 | 27,7557 | -0,118255 | 0,5218 | 0,881229 |
| Szrd1 | 53,5767 | 49,5222 | -0,113528 | 0,5219 | 0,881229 |
| Tmem161b | 7,97355 | 8,80332 | 0,142824 | 0,52195 | 0,881229 |
| Itpa | 47,836 | 43,9214 | -0,123172 | 0,52195 | 0,881229 |
| Ttyh3 | 42,3822 | 39,1402 | -0,114809 | 0,52205 | 0,881281 |

| | | | | | |
|---|---|---|---|---|---|
| Brd7 | 73,6408 | 79,7658 | 0,115265 | 0,5223 | 0,881587 |
| Mir6418 | 19,771 | 49,5208 | 1,32465 | 0,5224 | 0,88164 |
| B3gat3 | 26,9584 | 24,6864 | -0,127016 | 0,5225 | 0,881672 |
| Adam12 | 0,939699 | 1,06419 | 0,179491 | 0,5226 | 0,881672 |
| Zfp39 | 2,78856 | 3,12052 | 0,162262 | 0,52285 | 0,881672 |
| Elp3 | 77,5476 | 71,6789 | -0,113532 | 0,5229 | 0,881672 |
| Rnf7 | 88,0392 | 81,0809 | -0,118783 | 0,5229 | 0,881672 |
| Meaf6 | 42,1065 | 38,6465 | -0,123706 | 0,5229 | 0,881672 |
| 1110007C09Rik | 5,87225 | 5,02037 | -0,226122 | 0,5229 | 0,881672 |
| Actr8 | 27,9664 | 30,457 | 0,123081 | 0,523 | 0,881725 |
| C130046K22Rik | 2,57423 | 2,90852 | 0,176146 | 0,52315 | 0,88183 |
| Tm6sf2 | 0,784803 | 0,606617 | -0,371544 | 0,5232 | 0,88183 |
| Setd6 | 12,8423 | 14,1689 | 0,14183 | 0,52355 | 0,882009 |
| Anapc16 | 43,2545 | 47,8695 | 0,146254 | 0,52365 | 0,882009 |
| Lpar2 | 5,90457 | 5,39384 | -0,130518 | 0,52365 | 0,882009 |
| Kdm5a | 11,8083 | 12,9107 | 0,128766 | 0,5238 | 0,882009 |
| Mnat1 | 24,7914 | 26,9605 | 0,121009 | 0,5239 | 0,882009 |
| Ccar1 | 67,4461 | 73,1058 | 0,116251 | 0,5239 | 0,882009 |
| Ssr3 | 184,563 | 170,192 | -0,116949 | 0,52395 | 0,882009 |
| 2410016O06Rik | 20,0327 | 18,3587 | -0,125891 | 0,524 | 0,882009 |
| Oplah | 0,959091 | 1,11697 | 0,219853 | 0,52405 | 0,882009 |
| Ift88 | 11,1266 | 12,1635 | 0,128553 | 0,52405 | 0,882009 |
| Bloc1s1 | 47,0249 | 42,3977 | -0,149439 | 0,5241 | 0,882009 |
| Relt | 2,63627 | 2,99027 | 0,181781 | 0,5242 | 0,882009 |
| Zfp91 | 30,0148 | 32,5261 | 0,115925 | 0,5242 | 0,882009 |
| Xrcc6 | 52,4029 | 48,3915 | -0,114893 | 0,5243 | 0,88203 |
| Tnfaip8 | 8,34383 | 7,5952 | -0,135622 | 0,52435 | 0,88203 |
| Glyr1 | 109,755 | 101,421 | -0,113938 | 0,5246 | 0,882335 |
| Vps33a | 14,9339 | 13,7133 | -0,123015 | 0,52475 | 0,88244 |
| Rab14 | 52,1083 | 48,1379 | -0,114341 | 0,5248 | 0,88244 |
| Snord104 | 3190,98 | 3955,1 | 0,309716 | 0,5251 | 0,882734 |
| Il6st | 18,0026 | 16,5914 | -0,117773 | 0,52515 | 0,882734 |
| Herpud1 | 14,4324 | 15,8467 | 0,134868 | 0,52525 | 0,882734 |
| Fam64a | 94,7569 | 87,5318 | -0,114424 | 0,52525 | 0,882734 |
| Zfp316 | 3,66053 | 3,33403 | -0,134787 | 0,5255 | 0,88296 |
| Ino80e | 44,5641 | 40,3161 | -0,144526 | 0,5257 | 0,88296 |
| Plrg1 | 94,4176 | 87,2866 | -0,113296 | 0,52575 | 0,88296 |
| Eps8 | 3,57555 | 3,21487 | -0,153403 | 0,52575 | 0,88296 |
| P2rx4 | 0,792684 | 0,63916 | -0,310569 | 0,5258 | 0,88296 |
| Atrip | 16,0102 | 17,4681 | 0,125729 | 0,5259 | 0,88296 |
| Eif4ebp2 | 65,7833 | 60,7774 | -0,114188 | 0,52595 | 0,88296 |
| Bend5 | 9,51971 | 8,59021 | -0,148224 | 0,52615 | 0,88296 |
| Rbl1 | 17,7406 | 19,3088 | 0,122202 | 0,5262 | 0,88296 |
| Hirip3 | 65,5138 | 60,539 | -0,113934 | 0,5262 | 0,88296 |
| Rundc3a | 3,24901 | 3,66833 | 0,175122 | 0,52625 | 0,88296 |
| Zfyve27 | 4,65116 | 5,10438 | 0,134147 | 0,52635 | 0,88296 |
| Imp3 | 69,0411 | 63,353 | -0,124044 | 0,52675 | 0,88296 |
| Slc25a11 | 45,1514 | 48,9682 | 0,117076 | 0,5269 | 0,88296 |
| Lrch4 | 10,6646 | 12,1234 | 0,184967 | 0,52695 | 0,88296 |
| Kdm2a | 26,1145 | 28,2965 | 0,115776 | 0,52695 | 0,88296 |

| | | | | | |
|---|---|---|---|---|---|
| Fbxo38 | 21,149 | 22,9395 | 0,117244 | 0,527 | 0,88296 |
| Rsl24d1 | 30,9209 | 28,389 | -0,123248 | 0,527 | 0,88296 |
| Golim4 | 27,12 | 25,0547 | -0,114274 | 0,5272 | 0,88296 |
| Rhbdd3 | 10,7761 | 11,9241 | 0,14604 | 0,52725 | 0,88296 |
| Eny2 | 44,6272 | 40,021 | -0,157168 | 0,52725 | 0,88296 |
| Atic | 105,058 | 97,0995 | -0,113648 | 0,5273 | 0,88296 |
| Nudt19 | 21,1226 | 19,3333 | -0,127703 | 0,5273 | 0,88296 |
| Fyco1 | 4,6383 | 5,05886 | 0,125217 | 0,52735 | 0,88296 |
| Nus1 | 27,8155 | 30,1243 | 0,115036 | 0,5274 | 0,88296 |
| Chpf | 17,2401 | 15,8469 | -0,121568 | 0,5274 | 0,88296 |
| Phf12 | 25,5539 | 27,6881 | 0,11572 | 0,52745 | 0,88296 |
| Plau | 0,810434 | 0,663499 | -0,288601 | 0,52745 | 0,88296 |
| Nup155 | 33,8132 | 36,6029 | 0,11437 | 0,52755 | 0,88296 |
| Syncrip | 75,3478 | 82,3859 | 0,128832 | 0,5277 | 0,88296 |
| Daam1 | 9,58886 | 8,815 | -0,1214 | 0,5277 | 0,88296 |
| 4933421O10Rik | 0,868571 | 0,741918 | -0,227384 | 0,52775 | 0,88296 |
| Ap4s1 | 19,8658 | 15,6262 | -0,346316 | 0,5278 | 0,88296 |
| Osgepl1 | 11,6459 | 10,5394 | -0,14404 | 0,5279 | 0,88296 |
| Znhit1 | 39,5849 | 33,9694 | -0,220713 | 0,5279 | 0,88296 |
| Kcnf1 | 0,542696 | 0,638808 | 0,23524 | 0,528 | 0,88296 |
| Dusp22 | 8,19073 | 9,22991 | 0,172324 | 0,528 | 0,88296 |
| Taf8 | 17,9178 | 19,5037 | 0,122354 | 0,528 | 0,88296 |
| Klc3 | 0,494811 | 0,96103 | 0,957703 | 0,52815 | 0,882967 |
| Vgll4 | 24,4991 | 22,3081 | -0,135163 | 0,5282 | 0,882967 |
| Gtpbp10 | 6,55059 | 7,23568 | 0,143503 | 0,52835 | 0,882967 |
| Mmgt2 | 10,9821 | 9,94114 | -0,143672 | 0,5284 | 0,882967 |
| Cct6a | 477,796 | 440,258 | -0,118046 | 0,52845 | 0,882967 |
| Ccdc53 | 23,5715 | 21,367 | -0,141662 | 0,52855 | 0,882967 |
| Cisd1 | 59,9158 | 54,978 | -0,124081 | 0,52865 | 0,882967 |
| Olfml3 | 22,4298 | 17,0372 | -0,396725 | 0,52865 | 0,882967 |
| Cnppd1 | 12,0993 | 11,06 | -0,12957 | 0,52875 | 0,882967 |
| Gtf2h2 | 32,5571 | 35,4163 | 0,121441 | 0,52885 | 0,882967 |
| Rhobtb1 | 4,77787 | 4,32805 | -0,142652 | 0,52885 | 0,882967 |
| Hyal2 | 53,3021 | 49,2161 | -0,115064 | 0,5291 | 0,882967 |
| Mthfd1 | 70,3668 | 65,0566 | -0,1132 | 0,5292 | 0,882967 |
| Tbrg4 | 37,7467 | 40,8599 | 0,114338 | 0,5293 | 0,882967 |
| Psmd2 | 148,09 | 137,012 | -0,112173 | 0,5293 | 0,882967 |
| Sf3a2 | 68,8496 | 63,5083 | -0,116502 | 0,5293 | 0,882967 |
| Mtif2 | 19,7109 | 18,1063 | -0,122498 | 0,5293 | 0,882967 |
| Insig1 | 25,1544 | 27,2838 | 0,117231 | 0,52935 | 0,882967 |
| Mir8094 | 112,587 | 168,172 | 0,578897 | 0,5294 | 0,882967 |
| Slc9a8 | 7,85433 | 8,58212 | 0,127845 | 0,5294 | 0,882967 |
| Cyb5r3 | 79,8163 | 86,4253 | 0,11477 | 0,52945 | 0,882967 |
| Fam19a5 | 5,20617 | 4,65993 | -0,159913 | 0,5296 | 0,883071 |
| Iqgap3 | 12,1078 | 11,1616 | -0,117394 | 0,52965 | 0,883071 |
| Gna11 | 31,4226 | 28,9931 | -0,116094 | 0,53 | 0,883403 |
| Rab21 | 12,4743 | 11,3157 | -0,140638 | 0,53005 | 0,883403 |
| Eif4e2 | 72,081 | 66,3032 | -0,120541 | 0,5302 | 0,883403 |
| Mrps10 | 20,016 | 18,0669 | -0,147804 | 0,5302 | 0,883403 |
| Ppp1ca | 465,895 | 428,505 | -0,120694 | 0,53035 | 0,883403 |

| | | | | | |
|---|---|---|---|---|---|
| Mir1907 | 19,7536 | 76,3 | 1,94957 | 0,5304 | 0,883403 |
| Rab24 | 29,6193 | 32,2149 | 0,121192 | 0,5304 | 0,883403 |
| Iba57 | 3,64403 | 3,25032 | -0,164953 | 0,5304 | 0,883403 |
| Rcor2 | 106,602 | 115,305 | 0,113222 | 0,5306 | 0,883622 |
| Prmt1 | 560,766 | 517,295 | -0,116411 | 0,5308 | 0,88384 |
| Nubp2 | 28,6415 | 26,2061 | -0,128206 | 0,53095 | 0,883975 |
| Nenf | 38,2221 | 34,5857 | -0,144231 | 0,53105 | 0,884027 |
| Gnaq | 18,9131 | 20,4619 | 0,11356 | 0,5313 | 0,884328 |
| Mrpl21 | 27,1412 | 24,6733 | -0,137536 | 0,53145 | 0,884463 |
| Snord65 | 441,255 | 1389,28 | 1,65465 | 0,53185 | 0,885014 |
| Myo5a | 10,7779 | 11,665 | 0,114113 | 0,532 | 0,885085 |
| Ech1 | 23,5845 | 21,4077 | -0,139704 | 0,53205 | 0,885085 |
| Klhl25 | 10,5658 | 11,551 | 0,128606 | 0,5321 | 0,885085 |
| Fkbp8 | 109,744 | 101,491 | -0,112803 | 0,53235 | 0,885355 |
| Mboat1 | 0,68926 | 0,825961 | 0,261024 | 0,5324 | 0,885355 |
| Dnaja4 | 1,54053 | 1,76637 | 0,197358 | 0,53275 | 0,885498 |
| Pih1d1 | 57,8667 | 51,7242 | -0,161893 | 0,53275 | 0,885498 |
| Rer1 | 91,2166 | 84,2853 | -0,114016 | 0,5328 | 0,885498 |
| Gm1943 | 10,7194 | 9,53834 | -0,168418 | 0,5328 | 0,885498 |
| Ndufb11 | 89,1385 | 79,0087 | -0,174039 | 0,5329 | 0,885498 |
| Nhej1 | 7,96537 | 6,95266 | -0,196176 | 0,5329 | 0,885498 |
| Pex5l | 0,733359 | 0,607034 | -0,272743 | 0,5331 | 0,88562 |
| Flnc | 5,82631 | 6,34427 | 0,122872 | 0,53315 | 0,88562 |
| Fbxl6 | 28,1192 | 25,8303 | -0,122489 | 0,5332 | 0,88562 |
| Zfp207 | 200,313 | 185,302 | -0,112381 | 0,53325 | 0,88562 |
| Plcd3 | 0,567445 | 0,682944 | 0,267287 | 0,5334 | 0,885628 |
| Tuba1a | 219,045 | 236,897 | 0,113033 | 0,53355 | 0,885628 |
| Map3k11 | 16,3554 | 15,072 | -0,117893 | 0,53355 | 0,885628 |
| Dusp1 | 7,47802 | 6,7049 | -0,15744 | 0,53355 | 0,885628 |
| Timm8a1 | 3,99274 | 3,44438 | -0,213132 | 0,5336 | 0,885628 |
| Opn3 | 6,94086 | 5,6198 | -0,304597 | 0,53385 | 0,885929 |
| Nsmf | 17,5493 | 19,0881 | 0,121263 | 0,5341 | 0,886229 |
| Ehd1 | 41,7615 | 45,1328 | 0,112003 | 0,5343 | 0,886446 |
| Fn3krp | 7,32919 | 8,11219 | 0,146439 | 0,5344 | 0,886466 |
| Zfp335 | 9,86767 | 10,7279 | 0,120586 | 0,53445 | 0,886466 |
| Anxa9 | 1,03003 | 1,4436 | 0,486975 | 0,5346 | 0,886532 |
| Grasp | 2,70815 | 3,1036 | 0,196633 | 0,53485 | 0,886532 |
| Get4 | 27,9045 | 25,6793 | -0,119892 | 0,53485 | 0,886532 |
| Wsb2 | 9,219 | 10,1244 | 0,135149 | 0,5349 | 0,886532 |
| Rbbp9 | 33,7202 | 31,1069 | -0,116378 | 0,53505 | 0,886532 |
| Lrba | 2,57001 | 2,33615 | -0,137641 | 0,53505 | 0,886532 |
| Slc30a3 | 0,372361 | 0,843324 | 1,17939 | 0,5351 | 0,886532 |
| Pla2g4a | 2,59367 | 2,28584 | -0,182267 | 0,53515 | 0,886532 |
| Ell3 | 0,704406 | 0,876643 | 0,315582 | 0,5352 | 0,886532 |
| Slc2a13 | 0,7181 | 0,622941 | -0,205089 | 0,5352 | 0,886532 |
| Unc13b | 2,41959 | 2,67667 | 0,145677 | 0,53525 | 0,886532 |
| Rin3 | 1,92825 | 1,70311 | -0,179117 | 0,5354 | 0,886602 |
| Cpeb4 | 1,56854 | 1,75004 | 0,15797 | 0,5355 | 0,886602 |
| Tmem170 | 4,50239 | 3,97772 | -0,178748 | 0,5355 | 0,886602 |
| Znhit2 | 15,4394 | 13,9601 | -0,145313 | 0,53565 | 0,886661 |

| | | | | | |
|---|---|---|---|---|---|
| Trappc8 | 14,5945 | 15,833 | 0,117505 | 0,5357 | 0,886661 |
| Carm1 | 84,7609 | 77,46 | -0,129947 | 0,5358 | 0,886661 |
| Btbd8 | 0,817372 | 1,04645 | 0,356441 | 0,5359 | 0,886661 |
| Bend4 | 8,56885 | 7,90015 | -0,117222 | 0,53595 | 0,886661 |
| Pigu | 29,638 | 27,2517 | -0,1211 | 0,53595 | 0,886661 |
| Mir6992 | 8,27596 | 56,2767 | 2,76554 | 0,53605 | 0,886712 |
| Mir6340 | 9,63451 | 52,1985 | 2,43773 | 0,5362 | 0,88677 |
| Kat2a | 50,0524 | 54,051 | 0,110883 | 0,53625 | 0,88677 |
| Usp6nl | 15,4237 | 16,6669 | 0,111843 | 0,5364 | 0,88677 |
| Kctd5 | 53,2794 | 57,6844 | 0,114606 | 0,53645 | 0,88677 |
| Rsrp1 | 92,4481 | 99,9857 | 0,113078 | 0,53645 | 0,88677 |
| Smad3 | 40,7819 | 37,7326 | -0,112116 | 0,5365 | 0,88677 |
| Spin1 | 33,7248 | 31,3113 | -0,107127 | 0,53675 | 0,887069 |
| Gmnn | 122,405 | 113,29 | -0,111642 | 0,5372 | 0,887527 |
| Zscan25 | 9,09598 | 8,22512 | -0,145192 | 0,53725 | 0,887527 |
| Ascc2 | 23,4747 | 25,4684 | 0,117599 | 0,53735 | 0,887527 |
| Arsk | 2,94033 | 3,29172 | 0,162864 | 0,53745 | 0,887527 |
| Prpsap1 | 65,2073 | 60,3247 | -0,112284 | 0,5375 | 0,887527 |
| 6430706D22Rik | 95,8365 | 105,686 | 0,141143 | 0,5376 | 0,887527 |
| Zcchc6 | 13,9061 | 15,0828 | 0,117185 | 0,5376 | 0,887527 |
| Cpt2 | 17,0373 | 18,545 | 0,122335 | 0,53765 | 0,887527 |
| Mta2 | 108,557 | 100,57 | -0,110259 | 0,53765 | 0,887527 |
| Sra1 | 60,9007 | 56,0302 | -0,120254 | 0,53775 | 0,887578 |
| Lats2 | 9,09201 | 9,9584 | 0,131315 | 0,53795 | 0,887794 |
| Csnk2b | 143,539 | 133,005 | -0,109964 | 0,53805 | 0,887845 |
| Mbnl1 | 5,39634 | 4,9085 | -0,1367 | 0,5382 | 0,887978 |
| Pum2 | 50,767 | 54,8345 | 0,111192 | 0,53845 | 0,888048 |
| Smg5 | 42,2986 | 45,6767 | 0,110849 | 0,53845 | 0,888048 |
| Rbbp4 | 176,716 | 163,358 | -0,113389 | 0,53845 | 0,888048 |
| Nmnat1 | 10,1692 | 9,02311 | -0,172513 | 0,5386 | 0,888149 |
| Pknox1 | 10,2012 | 11,0937 | 0,121012 | 0,53865 | 0,888149 |
| Trnp1 | 0,690444 | 0,867028 | 0,328554 | 0,53875 | 0,888187 |
| Mir704 | 411,54 | 538,145 | 0,386963 | 0,5389 | 0,888187 |
| Def6 | 0,605819 | 0,741731 | 0,292009 | 0,5389 | 0,888187 |
| Hspa2 | 1,62143 | 1,866 | 0,202686 | 0,53895 | 0,888187 |
| Atg13 | 12,4168 | 11,4103 | -0,121958 | 0,5391 | 0,88832 |
| Cul9 | 0,686139 | 0,59989 | -0,193803 | 0,53925 | 0,888422 |
| 3110007F17Rik | 0,8558 | 0,640896 | -0,417184 | 0,5393 | 0,888422 |
| Calml4 | 1,37366 | 2,22372 | 0,694949 | 0,53945 | 0,888491 |
| Mad1l1 | 22,0444 | 20,2952 | -0,119272 | 0,5395 | 0,888491 |
| Ftx | 5,29975 | 4,81962 | -0,137005 | 0,53955 | 0,888491 |
| Oxsm | 2,12167 | 1,81898 | -0,22207 | 0,53985 | 0,888871 |
| Stim1 | 2,90218 | 3,23962 | 0,158687 | 0,5401 | 0,889169 |
| Slc25a10 | 11,1679 | 10,2789 | -0,119672 | 0,5405 | 0,889355 |
| Prkab2 | 4,67777 | 5,14435 | 0,137167 | 0,54065 | 0,889355 |
| Zbtb1 | 10,9681 | 11,921 | 0,120195 | 0,54075 | 0,889355 |
| Strn3 | 32,922 | 35,6718 | 0,115731 | 0,5408 | 0,889355 |
| Gtf2e2 | 77,7061 | 71,9053 | -0,111929 | 0,541 | 0,889355 |
| Map7 | 1,20792 | 1,01719 | -0,247946 | 0,541 | 0,889355 |
| Anks1 | 8,59242 | 9,32123 | 0,117457 | 0,54105 | 0,889355 |

| | | | | | |
|---|---|---|---|---|---|
| Vhl | 19,7148 | 21,3912 | 0,117739 | 0,54115 | 0,889355 |
| Camsap2 | 15,5818 | 14,3942 | -0,114377 | 0,54115 | 0,889355 |
| Iscu | 14,2852 | 12,7674 | -0,16205 | 0,54115 | 0,889355 |
| Phc1 | 29,9563 | 32,378 | 0,112154 | 0,5412 | 0,889355 |
| Usp3 | 38,3688 | 35,4732 | -0,113204 | 0,5413 | 0,889355 |
| Tpk1 | 2,60616 | 2,9484 | 0,178003 | 0,5414 | 0,889355 |
| Tm2d2 | 37,2913 | 34,3009 | -0,120595 | 0,5414 | 0,889355 |
| Sri | 20,762 | 19,121 | -0,118784 | 0,54145 | 0,889355 |
| U2af1l4 | 40,7141 | 37,2125 | -0,129741 | 0,54145 | 0,889355 |
| Rab33b | 3,89119 | 3,49502 | -0,15491 | 0,54145 | 0,889355 |
| Ifit2 | 2,25338 | 2,00666 | -0,167295 | 0,54155 | 0,889355 |
| Ss18 | 47,8294 | 51,5625 | 0,108424 | 0,5416 | 0,889355 |
| Dgkz | 27,0143 | 24,9739 | -0,113305 | 0,5416 | 0,889355 |
| Ccdc104 | 31,7611 | 29,2214 | -0,120233 | 0,54175 | 0,889488 |
| Ssr1 | 36,6524 | 33,9534 | -0,110352 | 0,54205 | 0,889713 |
| Thap3 | 14,5555 | 13,083 | -0,15387 | 0,54205 | 0,889713 |
| Serpinh1 | 495,283 | 536,686 | 0,115824 | 0,5421 | 0,889713 |
| Ubac2 | 27,2362 | 29,5296 | 0,116637 | 0,54225 | 0,889713 |
| Cdk10 | 16,5423 | 18,1141 | 0,130954 | 0,5424 | 0,889713 |
| Lman1 | 57,4981 | 62,3708 | 0,117356 | 0,5424 | 0,889713 |
| Fam131a | 0,882198 | 0,576902 | -0,612775 | 0,5424 | 0,889713 |
| Sesn1 | 9,8646 | 10,7954 | 0,130087 | 0,5426 | 0,889713 |
| Cpsf4l | 0,952009 | 0,758013 | -0,328753 | 0,5426 | 0,889713 |
| Golph3l | 14,3937 | 13,2378 | -0,120782 | 0,54265 | 0,889713 |
| Cnp | 5,80771 | 5,23009 | -0,151133 | 0,54265 | 0,889713 |
| Eps15l1 | 43,4596 | 46,955 | 0,111605 | 0,54305 | 0,889996 |
| Tmco1 | 13,4413 | 12,3896 | -0,117538 | 0,54305 | 0,889996 |
| Ergic3 | 85,855 | 79,4836 | -0,111245 | 0,5431 | 0,889996 |
| Vldlr | 2,12792 | 1,92797 | -0,142364 | 0,5431 | 0,889996 |
| Fau | 5,62531 | 6,55532 | 0,220733 | 0,54325 | 0,890128 |
| Map4k5 | 15,1371 | 13,3371 | -0,182639 | 0,5434 | 0,890228 |
| Hic2 | 10,7494 | 9,92252 | -0,115478 | 0,54345 | 0,890228 |
| BC002163 | 35,9188 | 32,1417 | -0,160293 | 0,5436 | 0,89036 |
| Ift46 | 21,579 | 19,9526 | -0,11305 | 0,54385 | 0,890515 |
| Tcta | 2,72203 | 3,12336 | 0,198417 | 0,54395 | 0,890515 |
| Eif4a3 | 116,237 | 107,565 | -0,111862 | 0,54395 | 0,890515 |
| Nlgn3 | 2,89718 | 3,22027 | 0,152532 | 0,54405 | 0,890515 |
| Trim33 | 14,447 | 16,4547 | 0,18773 | 0,5442 | 0,890515 |
| Sepn1 | 31,0046 | 33,501 | 0,11172 | 0,5442 | 0,890515 |
| Gm13315 | 6,63999 | 7,39564 | 0,155495 | 0,54425 | 0,890515 |
| Cyp20a1 | 21,3026 | 23,2518 | 0,126312 | 0,54425 | 0,890515 |
| Gpx1 | 244,266 | 226,444 | -0,109297 | 0,5445 | 0,890611 |
| Dnaja1 | 17,4918 | 15,1204 | -0,210178 | 0,5445 | 0,890611 |
| Shroom2 | 3,43268 | 3,1122 | -0,1414 | 0,5447 | 0,890611 |
| Heatr6 | 9,00515 | 9,76343 | 0,116637 | 0,54485 | 0,890611 |
| Ube2j1 | 16,4724 | 15,1747 | -0,118385 | 0,54495 | 0,890611 |
| Eif2s2 | 51,8717 | 48,0669 | -0,109906 | 0,545 | 0,890611 |
| Zfp952 | 8,49104 | 7,74876 | -0,131976 | 0,54505 | 0,890611 |
| Snord72 | 341,515 | 316,935 | -0,107761 | 0,5451 | 0,890611 |
| Fam109a | 22,3607 | 20,5845 | -0,119407 | 0,54515 | 0,890611 |

| | | | | | |
|---|---|---|---|---|---|
| Bin3 | 17,5131 | 13,6667 | -0,357775 | 0,54515 | 0,890611 |
| Scarna8 | 17,7536 | 7,20312 | -1,30142 | 0,5452 | 0,890611 |
| Camk2g | 20,3122 | 21,9518 | 0,11199 | 0,54525 | 0,890611 |
| Coch | 0,959654 | 1,12963 | 0,235259 | 0,5453 | 0,890611 |
| Timm22 | 19,0451 | 17,5698 | -0,11632 | 0,5453 | 0,890611 |
| Mrpl17 | 10,75 | 9,95638 | -0,110646 | 0,54535 | 0,890611 |
| E2f3 | 27,0205 | 29,1864 | 0,111241 | 0,54545 | 0,890661 |
| Ssbp1 | 48,2526 | 44,2485 | -0,124978 | 0,54575 | 0,891037 |
| Zdhhc4 | 18,4521 | 16,8178 | -0,133801 | 0,5459 | 0,891137 |
| Tnpo3 | 77,1084 | 71,4953 | -0,109041 | 0,54595 | 0,891137 |
| Trmt10a | 9,25735 | 8,50452 | -0,122369 | 0,54605 | 0,891187 |
| Mtif3 | 4,43829 | 5,13638 | 0,210747 | 0,5462 | 0,891305 |
| Stx5a | 45,2159 | 41,8546 | -0,111444 | 0,5463 | 0,891305 |
| B4galt4 | 7,97156 | 8,7603 | 0,136118 | 0,54635 | 0,891305 |
| Ddx42 | 59,0904 | 63,698 | 0,108326 | 0,5464 | 0,891305 |
| B4galt2 | 25,9826 | 24,0065 | -0,11412 | 0,5468 | 0,89178 |
| Tnfrsf22 | 2,60968 | 2,9577 | 0,180602 | 0,54685 | 0,89178 |
| Erp29 | 91,5574 | 84,7712 | -0,111102 | 0,5469 | 0,89178 |
| Pgls | 102,51 | 94,842 | -0,11217 | 0,54705 | 0,891879 |
| Lrrc3b | 0,600098 | 0,751172 | 0,323946 | 0,5471 | 0,891879 |
| Depdc5 | 12,1312 | 13,2543 | 0,127735 | 0,54725 | 0,891883 |
| Mrpl50 | 60,9976 | 56,5048 | -0,110379 | 0,54725 | 0,891883 |
| Gm9079 | 0,804612 | 0,649468 | -0,309034 | 0,54735 | 0,891883 |
| Dnmbp | 8,45419 | 7,80042 | -0,116114 | 0,54745 | 0,891883 |
| Asf1a | 45,3589 | 41,7596 | -0,119277 | 0,54745 | 0,891883 |
| Lig3 | 26,9479 | 24,9418 | -0,111611 | 0,54755 | 0,891933 |
| Acot8 | 6,43659 | 7,49469 | 0,219572 | 0,54765 | 0,891983 |
| Mir7089 | 631,103 | 185,859 | -1,76367 | 0,54785 | 0,892195 |
| Ik | 112,15 | 104,048 | -0,108183 | 0,54795 | 0,892245 |
| Vimp | 49,5614 | 45,6911 | -0,117305 | 0,5482 | 0,892538 |
| Rgs19 | 35,4403 | 32,4166 | -0,128658 | 0,54845 | 0,892687 |
| 2810403D21Rik | 0,971134 | 0,748365 | -0,375928 | 0,54845 | 0,892687 |
| Ppp4r2 | 13,7678 | 14,918 | 0,115751 | 0,5485 | 0,892687 |
| Eif3c | 215,285 | 199,351 | -0,110934 | 0,54875 | 0,89298 |
| Zdhhc5 | 26,5244 | 29,231 | 0,140179 | 0,5489 | 0,893111 |
| Vars2 | 9,23464 | 10,0123 | 0,116648 | 0,5492 | 0,893196 |
| Lime1 | 20,4171 | 22,1476 | 0,117368 | 0,54925 | 0,893196 |
| Aak1 | 5,51773 | 5,95092 | 0,109039 | 0,54925 | 0,893196 |
| Rpusd3 | 11,1717 | 12,3873 | 0,149007 | 0,5493 | 0,893196 |
| Poc1a | 16,3121 | 14,9289 | -0,127832 | 0,5493 | 0,893196 |
| Arsg | 0,93816 | 1,18404 | 0,335808 | 0,5495 | 0,893294 |
| Ranbp17 | 3,76302 | 4,13171 | 0,134849 | 0,5495 | 0,893294 |
| Tsr1 | 53,9689 | 58,3443 | 0,112462 | 0,54965 | 0,893329 |
| Tmed2 | 16,9262 | 15,5382 | -0,123443 | 0,5497 | 0,893329 |
| Mrps25 | 13,2298 | 11,9392 | -0,148084 | 0,54975 | 0,893329 |
| Tbc1d15 | 17,648 | 19,094 | 0,113614 | 0,5498 | 0,893329 |
| 6330418K02Rik | 6,46657 | 8,74078 | 0,43476 | 0,5499 | 0,893329 |
| Fbxl12os | 0,738722 | 0,879118 | 0,251024 | 0,5501 | 0,893329 |
| Atad3a | 41,4111 | 38,3929 | -0,109179 | 0,5501 | 0,893329 |
| Tab1 | 17,1126 | 15,7717 | -0,117719 | 0,55025 | 0,893329 |

| | | | | | |
|---|---|---|---|---|---|
| Ctdspl2 | 12,7853 | 13,8111 | 0,11134 | 0,5503 | 0,893329 |
| Snord49a | 3472,61 | 946,15 | -1,87588 | 0,55035 | 0,893329 |
| Tmed10 | 21,4188 | 19,8239 | -0,111632 | 0,5504 | 0,893329 |
| Pkig | 54,8823 | 50,6623 | -0,115429 | 0,55045 | 0,893329 |
| Alg6 | 7,68306 | 7,02502 | -0,129178 | 0,55045 | 0,893329 |
| Matn3 | 2,81892 | 3,1616 | 0,16551 | 0,5505 | 0,893329 |
| Ptrh1 | 3,41502 | 2,88266 | -0,244493 | 0,55065 | 0,893329 |
| Tab2 | 48,4406 | 52,1854 | 0,10743 | 0,5507 | 0,893329 |
| Kifc5b | 5,70298 | 6,29585 | 0,142686 | 0,55075 | 0,893329 |
| Afg3l1 | 25,8716 | 24,0063 | -0,107957 | 0,5508 | 0,893329 |
| Aldoa | 531,73 | 574,836 | 0,112455 | 0,5509 | 0,893329 |
| Gimap4 | 1,01187 | 0,786057 | -0,364322 | 0,551 | 0,893329 |
| Ick | 11,1482 | 10,3165 | -0,11186 | 0,5511 | 0,893329 |
| Gm15471 | 0,404571 | 1,14791 | 1,50454 | 0,55115 | 0,893329 |
| Magee1 | 5,59001 | 5,0988 | -0,132692 | 0,55125 | 0,893329 |
| Dffb | 5,81822 | 5,22728 | -0,154517 | 0,5514 | 0,893329 |
| Tm2d1 | 8,81432 | 7,82101 | -0,172494 | 0,5514 | 0,893329 |
| Ankrd13d | 1,84324 | 2,11294 | 0,197009 | 0,55145 | 0,893329 |
| Tcp11l1 | 1,82026 | 2,05777 | 0,176942 | 0,5515 | 0,893329 |
| Map3k14 | 0,788231 | 0,678297 | -0,2167 | 0,5515 | 0,893329 |
| Nup188 | 47,1033 | 50,7003 | 0,106167 | 0,5516 | 0,893329 |
| Psat1 | 144,865 | 134,438 | -0,107769 | 0,55165 | 0,893329 |
| Rnpepl1 | 12,7383 | 13,8302 | 0,11865 | 0,55175 | 0,893329 |
| Pop5 | 39,3218 | 36,0834 | -0,123997 | 0,55175 | 0,893329 |
| Spen | 10,9965 | 12,0782 | 0,135364 | 0,55185 | 0,893347 |
| Tpm3 | 67,5059 | 72,7677 | 0,108285 | 0,5519 | 0,893347 |
| Ccdc163 | 15,9376 | 14,4889 | -0,137478 | 0,55205 | 0,893477 |
| Capn5 | 1,19597 | 1,3579 | 0,183188 | 0,55215 | 0,893494 |
| Exoc2 | 16,3843 | 15,1865 | -0,10952 | 0,5522 | 0,893494 |
| Tmem205 | 9,61461 | 10,9932 | 0,19331 | 0,55245 | 0,893673 |
| Wac | 19,7171 | 21,2704 | 0,109397 | 0,55245 | 0,893673 |
| Prpf19 | 70,7423 | 65,6318 | -0,108177 | 0,5526 | 0,89369 |
| Klhl7 | 26,0916 | 22,6614 | -0,203345 | 0,5526 | 0,89369 |
| Ccdc126 | 1,60462 | 1,38792 | -0,209305 | 0,5528 | 0,893901 |
| Fzr1 | 49,1589 | 45,5841 | -0,108921 | 0,55295 | 0,894031 |
| Cmtm7 | 33,2473 | 30,5502 | -0,122054 | 0,5531 | 0,894161 |
| Nucks1 | 85,9 | 79,7617 | -0,106962 | 0,55325 | 0,894291 |
| Mrpl52 | 221,798 | 204,661 | -0,11601 | 0,5534 | 0,894359 |
| Kazn | 7,90385 | 8,60072 | 0,121903 | 0,5535 | 0,894359 |
| Poc1b | 26,129 | 28,4725 | 0,123921 | 0,5536 | 0,894359 |
| Nr4a1 | 2,38344 | 2,69682 | 0,178213 | 0,55365 | 0,894359 |
| Wasl | 16,1757 | 17,5907 | 0,120983 | 0,55375 | 0,894359 |
| Prkaca | 59,2985 | 55,1028 | -0,10587 | 0,5538 | 0,894359 |
| 2210013O21Rik | 32,6352 | 29,7797 | -0,132099 | 0,5538 | 0,894359 |
| Uchl3 | 52,8254 | 48,6454 | -0,118928 | 0,55385 | 0,894359 |
| Bend3 | 20,617 | 22,2079 | 0,107238 | 0,55395 | 0,894395 |
| Sec22c | 4,65464 | 5,07327 | 0,124248 | 0,5542 | 0,894395 |
| Fbxw8 | 39,1255 | 36,393 | -0,104447 | 0,55425 | 0,894395 |
| Nbn | 25,7393 | 27,797 | 0,110957 | 0,5543 | 0,894395 |
| Ercc1 | 32,8734 | 28,7474 | -0,193487 | 0,5543 | 0,894395 |

| | | | | | |
|---|---|---|---|---|---|
| Fam178a | 19,109 | 20,5735 | 0,106534 | 0,55435 | 0,894395 |
| Cdca7l | 34,0256 | 36,7025 | 0,109255 | 0,55445 | 0,894395 |
| Mrpl19 | 19,9898 | 18,3656 | -0,122261 | 0,55445 | 0,894395 |
| Magt1 | 24,3123 | 26,1605 | 0,105707 | 0,5545 | 0,894395 |
| Mir6516 | 39,789 | 98,5403 | 1,30835 | 0,5547 | 0,894574 |
| Zfp740 | 55,1468 | 59,3984 | 0,107146 | 0,55475 | 0,894574 |
| Noa1 | 23,9329 | 22,1065 | -0,114527 | 0,55485 | 0,894608 |
| Mir6958 | 79,829 | 339,798 | 2,08969 | 0,55495 | 0,894608 |
| Itfg2 | 14,3727 | 15,634 | 0,121358 | 0,555 | 0,894608 |
| Golga3 | 12,967 | 14,014 | 0,112022 | 0,55505 | 0,894608 |
| Hdac7 | 15,8507 | 17,2343 | 0,120737 | 0,55525 | 0,894817 |
| Pnkp | 12,0963 | 13,2417 | 0,130524 | 0,55545 | 0,895027 |
| Nkapl | 1,19767 | 1,43981 | 0,26565 | 0,55565 | 0,895125 |
| Aldh6a1 | 4,57814 | 5,16005 | 0,172626 | 0,55565 | 0,895125 |
| Adck4 | 8,25096 | 7,50252 | -0,137187 | 0,55575 | 0,895142 |
| Ulk4 | 1,06347 | 1,21287 | 0,189645 | 0,5558 | 0,895142 |
| Srp14 | 136,765 | 126,77 | -0,109489 | 0,55595 | 0,895239 |
| Rabep1 | 18,7551 | 21,3582 | 0,187507 | 0,556 | 0,895239 |
| Rangrf | 4,13752 | 3,42334 | -0,273365 | 0,5561 | 0,895272 |
| Psmc2 | 155,88 | 145,009 | -0,104292 | 0,55625 | 0,895272 |
| Ddr1 | 22,6658 | 21,0047 | -0,109808 | 0,55625 | 0,895272 |
| Mkl1 | 7,18802 | 7,97915 | 0,15064 | 0,5563 | 0,895272 |
| Tnip2 | 0,582143 | 0,721422 | 0,30947 | 0,55655 | 0,895499 |
| Mir6998 | 150,005 | 182,583 | 0,283546 | 0,5566 | 0,895499 |
| Nbas | 8,63166 | 7,98464 | -0,112409 | 0,55665 | 0,895499 |
| Dyrk1a | 20,1983 | 21,7299 | 0,105452 | 0,5568 | 0,895549 |
| Trh | 2,32454 | 2,69369 | 0,21264 | 0,5569 | 0,895549 |
| Gnpnat1 | 8,49706 | 7,72004 | -0,138354 | 0,5569 | 0,895549 |
| Gm2a | 1,11395 | 0,975377 | -0,191648 | 0,557 | 0,895549 |
| Fgd1 | 14,6485 | 16,3229 | 0,156141 | 0,55705 | 0,895549 |
| Srd5a3 | 10,2274 | 9,27667 | -0,140755 | 0,5571 | 0,895549 |
| Nelfb | 42,1443 | 45,3428 | 0,105535 | 0,5573 | 0,895644 |
| Malsu1 | 31,2216 | 28,4959 | -0,131792 | 0,55745 | 0,895644 |
| Fbxl15 | 3,13137 | 2,20058 | -0,508914 | 0,5575 | 0,895644 |
| Polm | 4,12157 | 3,70903 | -0,152149 | 0,5576 | 0,895644 |
| Ddx52 | 46,0724 | 42,8189 | -0,105656 | 0,5577 | 0,895644 |
| Trim46 | 1,61305 | 1,4085 | -0,19563 | 0,5577 | 0,895644 |
| Rngtt | 14,6261 | 15,7829 | 0,109823 | 0,5579 | 0,895644 |
| Gpr89 | 18,4482 | 16,939 | -0,123125 | 0,5579 | 0,895644 |
| Prpf39 | 22,6692 | 27,594 | 0,283619 | 0,558 | 0,895644 |
| Tbcd | 21,7224 | 20,0706 | -0,114098 | 0,55805 | 0,895644 |
| Slc27a4 | 10,1365 | 9,34778 | -0,116871 | 0,5581 | 0,895644 |
| Aebp2 | 38,0307 | 42,8292 | 0,171429 | 0,5582 | 0,895644 |
| 2310061J03Rik | 1,52528 | 0,946105 | -0,689006 | 0,55825 | 0,895644 |
| Nans | 48,9284 | 52,7679 | 0,108989 | 0,5584 | 0,895644 |
| Crtc1 | 5,6191 | 5,16676 | -0,121079 | 0,5584 | 0,895644 |
| Armc9 | 12,5261 | 11,4907 | -0,124463 | 0,5584 | 0,895644 |
| Mir7068 | 168,211 | 108,288 | -0,635394 | 0,55855 | 0,895644 |
| Tmed9 | 140,75 | 130,728 | -0,106571 | 0,55865 | 0,895644 |
| Gjb6 | 0,789418 | 0,945564 | 0,260384 | 0,55875 | 0,895644 |

| | | | | | |
|---|---|---|---|---|---|
| Gtpbp8 | 13,7985 | 12,4764 | -0,145314 | 0,55895 | 0,895644 |
| Snx6 | 97,2894 | 90,4813 | -0,104663 | 0,55905 | 0,895644 |
| Spdl1 | 26,9123 | 28,9873 | 0,107156 | 0,5592 | 0,895644 |
| BC004004 | 10,1421 | 9,31237 | -0,123131 | 0,5592 | 0,895644 |
| Sec16a | 18,4722 | 19,872 | 0,105386 | 0,5594 | 0,895644 |
| Stom | 3,74953 | 3,36264 | -0,157117 | 0,5594 | 0,895644 |
| Mms19 | 21,9929 | 24,0317 | 0,127904 | 0,5595 | 0,895644 |
| Stx1b | 3,20024 | 3,64714 | 0,188585 | 0,55955 | 0,895644 |
| Snord1b | 411,569 | 108,363 | -1,92526 | 0,55965 | 0,895644 |
| Cry2 | 6,9806 | 6,41171 | -0,122641 | 0,5597 | 0,895644 |
| Cstf1 | 41,2501 | 38,2348 | -0,10951 | 0,5598 | 0,895644 |
| Dusp12 | 22,9397 | 21,0747 | -0,122335 | 0,55985 | 0,895644 |
| Zc3h10 | 10,8104 | 9,87195 | -0,131011 | 0,55995 | 0,895644 |
| Cenpq | 38,7866 | 35,8245 | -0,11461 | 0,56005 | 0,895644 |
| 1110004E09Rik | 30,0739 | 27,6919 | -0,119049 | 0,56005 | 0,895644 |
| Kansl2 | 9,03514 | 8,29559 | -0,123203 | 0,56005 | 0,895644 |
| Nfkbib | 13,2562 | 12,1455 | -0,12624 | 0,5601 | 0,895644 |
| Map2k1 | 40,9313 | 44,4147 | 0,117831 | 0,56015 | 0,895644 |
| 2610301B20Rik | 10,6385 | 9,71641 | -0,130798 | 0,56025 | 0,895644 |
| Cmtm6 | 12,6379 | 13,6592 | 0,112112 | 0,56035 | 0,895644 |
| Tubb5 | 1352,74 | 1235,14 | -0,131211 | 0,56035 | 0,895644 |
| Plxnd1 | 4,73014 | 5,13269 | 0,117832 | 0,5604 | 0,895644 |
| Gm10653 | 0,719684 | 0,405013 | -0,829395 | 0,5604 | 0,895644 |
| Polh | 10,2801 | 9,42969 | -0,124578 | 0,56045 | 0,895644 |
| Kif1c | 14,5179 | 15,635 | 0,106946 | 0,56055 | 0,895644 |
| Dlg3 | 33,797 | 36,3519 | 0,105137 | 0,56055 | 0,895644 |
| Ubox5 | 10,0388 | 9,03478 | -0,152021 | 0,5606 | 0,895644 |
| Taok2 | 26,5991 | 28,6271 | 0,106005 | 0,56065 | 0,895644 |
| Tiparp | 13,3762 | 12,3788 | -0,111798 | 0,56065 | 0,895644 |
| Smap2 | 14,96 | 13,8093 | -0,115473 | 0,56065 | 0,895644 |
| Smim18 | 0,686724 | 0,111127 | -2,62753 | 0,56065 | 0,895644 |
| Hpdl | 1,65375 | 1,92904 | 0,222145 | 0,56085 | 0,895708 |
| Papd5 | 14,1944 | 15,3051 | 0,108686 | 0,5609 | 0,895708 |
| Nrm | 35,0554 | 32,4047 | -0,113432 | 0,5609 | 0,895708 |
| Tceal1 | 6,94096 | 6,15664 | -0,172993 | 0,561 | 0,895757 |
| Sgpp1 | 3,18976 | 3,54251 | 0,151323 | 0,5612 | 0,895964 |
| Tbce | 20,5873 | 23,6456 | 0,199817 | 0,5614 | 0,896029 |
| Zfp37 | 7,37902 | 8,03314 | 0,122535 | 0,56145 | 0,896029 |
| Mrps35 | 56,3232 | 52,029 | -0,114413 | 0,56145 | 0,896029 |
| BC022687 | 3,82966 | 4,30558 | 0,168992 | 0,56165 | 0,896142 |
| Asf1b | 42,1051 | 38,9783 | -0,111323 | 0,5617 | 0,896142 |
| Rps27l | 302,845 | 280,891 | -0,108574 | 0,56175 | 0,896142 |
| Pde4d | 2,55919 | 2,80151 | 0,130521 | 0,5618 | 0,896142 |
| Polg2 | 7,12136 | 6,39254 | -0,155763 | 0,5619 | 0,89619 |
| Wdr7 | 5,23949 | 5,66705 | 0,113171 | 0,5621 | 0,896222 |
| Ist1 | 73,6216 | 68,4395 | -0,1053 | 0,5621 | 0,896222 |
| Gng10 | 23,6614 | 21,6687 | -0,126922 | 0,56215 | 0,896222 |
| Mcts1 | 46,8153 | 43,1791 | -0,116645 | 0,5622 | 0,896222 |
| Gabpb1 | 46,3168 | 49,9983 | 0,110341 | 0,5624 | 0,896287 |
| Rab40c | 16,1681 | 14,9363 | -0,114327 | 0,5624 | 0,896287 |

| | | | | | |
|---|---|---|---|---|---|
| Nucb1 | 39,0343 | 42,1418 | 0,110512 | 0,56245 | 0,896287 |
| Dnajc18 | 5,76049 | 6,25857 | 0,11964 | 0,5626 | 0,896414 |
| Prkd1 | 9,02564 | 9,78985 | 0,117256 | 0,5628 | 0,896622 |
| Tmem14a | 2,46349 | 2,87078 | 0,220742 | 0,56315 | 0,897068 |
| Usmg5 | 7,82762 | 4,29407 | -0,86623 | 0,56335 | 0,897275 |
| Rps27a | 9,08071 | 10,2276 | 0,171589 | 0,5635 | 0,897307 |
| Ube2w | 11,4919 | 10,5828 | -0,118898 | 0,56355 | 0,897307 |
| Exo5 | 25,3219 | 23,3935 | -0,114276 | 0,56365 | 0,897307 |
| 4931428F04Rik | 7,59372 | 6,95478 | -0,126801 | 0,56365 | 0,897307 |
| Ciz1 | 25,7703 | 27,7833 | 0,108509 | 0,56385 | 0,89734 |
| Zfp322a | 15,3082 | 16,5006 | 0,108216 | 0,5639 | 0,89734 |
| Cat | 52,6999 | 56,644 | 0,104123 | 0,56395 | 0,89734 |
| Rpl7 | 439,48 | 409,186 | -0,103039 | 0,5641 | 0,89734 |
| Aass | 0,777666 | 0,898923 | 0,209046 | 0,56415 | 0,89734 |
| Atxn3 | 7,39228 | 8,18502 | 0,146967 | 0,56415 | 0,89734 |
| Enpp4 | 0,754723 | 0,864957 | 0,19668 | 0,56425 | 0,89734 |
| Yy1 | 50,0565 | 53,8038 | 0,104151 | 0,5643 | 0,89734 |
| Gtf2h1 | 28,9137 | 26,8573 | -0,106441 | 0,5643 | 0,89734 |
| Fam188b | 5,50928 | 4,9974 | -0,140687 | 0,5644 | 0,897342 |
| Ctu1 | 8,52312 | 7,79459 | -0,128907 | 0,5647 | 0,897342 |
| Dctn1 | 33,7568 | 31,4304 | -0,103019 | 0,5648 | 0,897342 |
| Ppp2r1b | 61,1959 | 65,8727 | 0,106246 | 0,56485 | 0,897342 |
| 4930413G21Rik | 5,19508 | 4,61023 | -0,172307 | 0,5649 | 0,897342 |
| Psmd12 | 134,54 | 125,179 | -0,104043 | 0,56495 | 0,897342 |
| Prmt3 | 32,3972 | 30,1003 | -0,106091 | 0,565 | 0,897342 |
| Tprgl | 31,9925 | 29,5716 | -0,113523 | 0,56505 | 0,897342 |
| Pus7l | 8,84324 | 9,62534 | 0,122263 | 0,5651 | 0,897342 |
| Armcx2 | 59,7039 | 55,638 | -0,101755 | 0,5651 | 0,897342 |
| Fam175b | 21,5991 | 20,0236 | -0,109267 | 0,5651 | 0,897342 |
| Brap | 13,4048 | 14,467 | 0,110022 | 0,5652 | 0,897342 |
| Pip5k1a | 22,0391 | 23,8191 | 0,112053 | 0,56535 | 0,897342 |
| Adi1 | 23,2147 | 25,1409 | 0,114996 | 0,56555 | 0,897342 |
| Fastk | 32,2051 | 29,8277 | -0,110635 | 0,5656 | 0,897342 |
| Zfp191 | 36,8136 | 34,2671 | -0,103414 | 0,56575 | 0,897342 |
| Usp27x | 2,2141 | 2,48411 | 0,166011 | 0,56585 | 0,897342 |
| Sel1l | 17,1355 | 18,4024 | 0,102909 | 0,56585 | 0,897342 |
| D6Wsu163e | 8,85607 | 8,08828 | -0,130834 | 0,56595 | 0,897342 |
| Zfp653 | 8,12198 | 11,0286 | 0,441347 | 0,566 | 0,897342 |
| Ropn1l | 1,4411 | 1,15708 | -0,316682 | 0,566 | 0,897342 |
| Bex4 | 15,8706 | 14,2353 | -0,156889 | 0,56615 | 0,897342 |
| 9130023H24Rik | 2,84266 | 3,31574 | 0,222087 | 0,5662 | 0,897342 |
| Klhl10 | 0,724998 | 0,594065 | -0,287355 | 0,5662 | 0,897342 |
| Slc16a10 | 1,81997 | 1,62072 | -0,167276 | 0,56625 | 0,897342 |
| Tnfsf9 | 3,14989 | 3,74292 | 0,248866 | 0,5663 | 0,897342 |
| Dbnl | 47,9774 | 44,5785 | -0,106008 | 0,5663 | 0,897342 |
| Mettl18 | 6,94822 | 6,23003 | -0,157404 | 0,5663 | 0,897342 |
| Prmt7 | 44,8525 | 41,6632 | -0,106415 | 0,56635 | 0,897342 |
| Snx16 | 3,86649 | 3,47533 | -0,153875 | 0,5664 | 0,897342 |
| Ndufb3 | 97,9274 | 90,2456 | -0,117857 | 0,5666 | 0,897548 |
| Hsp90aa1 | 121,556 | 130,661 | 0,10421 | 0,5667 | 0,897596 |

| | | | | | |
|---|---|---|---|---|---|
| Tmx1 | 47,5052 | 52,2894 | 0,138434 | 0,5668 | 0,897643 |
| Mul1 | 9,5612 | 8,80715 | -0,118517 | 0,567 | 0,897707 |
| Ccdc59 | 49,9612 | 46,17 | -0,11385 | 0,56705 | 0,897707 |
| Mterfd1 | 31,1976 | 27,6988 | -0,171607 | 0,56705 | 0,897707 |
| Suclg1 | 103,319 | 96,1308 | -0,104034 | 0,56725 | 0,897913 |
| Ppme1 | 44,6283 | 47,9599 | 0,103871 | 0,56745 | 0,898118 |
| Nprl2 | 12,5037 | 11,3908 | -0,13449 | 0,56755 | 0,898166 |
| Kif2a | 28,4979 | 30,6512 | 0,10509 | 0,5677 | 0,898292 |
| Epo | 1,5385 | 1,21296 | -0,342995 | 0,5678 | 0,89834 |
| 9330182L06Rik | 0,670891 | 0,771214 | 0,201052 | 0,5679 | 0,898355 |
| 2010015L04Rik | 4,23892 | 3,80729 | -0,15493 | 0,56795 | 0,898355 |
| H1f0 | 221,366 | 206,01 | -0,103721 | 0,56815 | 0,898561 |
| Sgol1 | 20,2371 | 21,7872 | 0,106478 | 0,5683 | 0,898687 |
| Cand2 | 35,9717 | 40,5059 | 0,171269 | 0,56845 | 0,898766 |
| Cdadc1 | 12,5216 | 11,4705 | -0,126495 | 0,56855 | 0,898766 |
| Snap47 | 48,2444 | 51,8604 | 0,104271 | 0,56865 | 0,898766 |
| Sil1 | 7,5626 | 6,83296 | -0,146371 | 0,56865 | 0,898766 |
| Gtf2h5 | 82,2804 | 76,5208 | -0,104697 | 0,5687 | 0,898766 |
| Camk2n1 | 2,5591 | 2,29614 | -0,156427 | 0,56885 | 0,898892 |
| Hmmr | 27,9759 | 30,0544 | 0,103388 | 0,5694 | 0,899286 |
| Mesdc2 | 30,3123 | 28,2225 | -0,103058 | 0,5694 | 0,899286 |
| Tdp1 | 37,8155 | 35,0986 | -0,107565 | 0,5695 | 0,899286 |
| Zc3h3 | 1,11814 | 0,97474 | -0,198012 | 0,5695 | 0,899286 |
| Csnk1e | 75,1174 | 80,5544 | 0,100817 | 0,56955 | 0,899286 |
| Zfp524 | 3,66309 | 3,17569 | -0,205989 | 0,56965 | 0,899286 |
| Ppp1r3f | 1,59178 | 1,97195 | 0,308987 | 0,5697 | 0,899286 |
| Znrd1as | 6,60476 | 7,4466 | 0,173077 | 0,5697 | 0,899286 |
| Cog3 | 16,0491 | 14,9076 | -0,106441 | 0,56975 | 0,899286 |
| Pkp4 | 30,1461 | 32,3647 | 0,10245 | 0,5698 | 0,899286 |
| Wdr73 | 26,2252 | 24,2986 | -0,110085 | 0,57015 | 0,899727 |
| Brca1 | 20,357 | 21,88 | 0,104094 | 0,57075 | 0,900453 |
| Ola1 | 73,7689 | 68,4735 | -0,107468 | 0,57075 | 0,900453 |
| Mfap1b | 2,3743 | 2,6996 | 0,185244 | 0,571 | 0,900626 |
| Clcn4-2 | 11,5798 | 10,7205 | -0,111239 | 0,571 | 0,900626 |
| 2610507B11Rik | 43,027 | 40,0931 | -0,101889 | 0,5711 | 0,900673 |
| Zbed4 | 2,931 | 2,67538 | -0,13165 | 0,57125 | 0,900766 |
| Naa20 | 25,2972 | 27,4895 | 0,119905 | 0,5713 | 0,900767 |
| Sgol2 | 11,7295 | 10,8684 | -0,110003 | 0,5714 | 0,900813 |
| Spag5 | 56,5685 | 60,7306 | 0,102423 | 0,57155 | 0,900875 |
| Col4a5 | 6,18266 | 5,70914 | -0,114954 | 0,5716 | 0,900875 |
| Igfbpl1 | 14,3399 | 15,4938 | 0,111657 | 0,57165 | 0,900875 |
| Gm6524 | 7,27793 | 8,05109 | 0,145656 | 0,5718 | 0,90089 |
| Dhrs7b | 11,8149 | 12,9222 | 0,129238 | 0,5718 | 0,90089 |
| Uevld | 2,42086 | 2,68052 | 0,146992 | 0,57205 | 0,901012 |
| Tle2 | 1,59062 | 1,80891 | 0,185538 | 0,5721 | 0,901012 |
| Dnajc8 | 130,555 | 121,68 | -0,101563 | 0,5723 | 0,901012 |
| Obfc1 | 27,6766 | 25,6485 | -0,109792 | 0,5723 | 0,901012 |
| Cpsf4 | 45,67 | 42,5377 | -0,102507 | 0,57235 | 0,901012 |
| Sdha | 134,611 | 125,376 | -0,102535 | 0,5724 | 0,901012 |
| Sept9 | 110,193 | 102,724 | -0,101257 | 0,5725 | 0,901012 |

| | | | | | |
|---|---|---|---|---|---|
| Nme7 | 25,951 | 28,4077 | 0,130491 | 0,57255 | 0,901012 |
| Trit1 | 15,0648 | 16,3213 | 0,115581 | 0,57255 | 0,901012 |
| Vezf1 | 43,2533 | 40,3195 | -0,101335 | 0,5726 | 0,901012 |
| Aif1l | 43,9889 | 41,0073 | -0,101259 | 0,57265 | 0,901012 |
| Gstt2 | 1,2447 | 1,48526 | 0,254909 | 0,5728 | 0,901027 |
| Ccng1 | 17,8091 | 16,5369 | -0,106924 | 0,5728 | 0,901027 |
| Faf1 | 23,0874 | 21,4909 | -0,103379 | 0,5729 | 0,901072 |
| Rap1a | 10,9637 | 11,9053 | 0,118871 | 0,573 | 0,901072 |
| Poll | 11,716 | 12,7226 | 0,118909 | 0,5732 | 0,901072 |
| 0610030E20Rik | 11,4666 | 10,6199 | -0,11067 | 0,5732 | 0,901072 |
| Ska1 | 13,7073 | 12,6598 | -0,114691 | 0,5732 | 0,901072 |
| Primpol | 3,68687 | 3,17572 | -0,215313 | 0,57325 | 0,901072 |
| Sppl3 | 35,205 | 37,8432 | 0,104251 | 0,5735 | 0,901227 |
| Ubxn2b | 6,60916 | 7,15769 | 0,115027 | 0,57355 | 0,901227 |
| Fam69a | 13,8137 | 12,4577 | -0,149063 | 0,5736 | 0,901227 |
| Coq5 | 25,2254 | 23,3699 | -0,110223 | 0,5737 | 0,901227 |
| Hmga2-ps1 | 37,0712 | 34,002 | -0,124682 | 0,5737 | 0,901227 |
| BC030336 | 9,94064 | 10,7225 | 0,109227 | 0,57385 | 0,901353 |
| Rev1 | 16,4607 | 13,9671 | -0,236998 | 0,57395 | 0,901399 |
| Gm17066 | 11,0447 | 13,1086 | 0,247162 | 0,5742 | 0,901579 |
| Ssr4 | 66,0498 | 61,0551 | -0,113442 | 0,5743 | 0,901579 |
| Hnrnpa0 | 296,834 | 276,182 | -0,104036 | 0,57435 | 0,901579 |
| Tmem45a | 0,935263 | 1,08576 | 0,215266 | 0,5745 | 0,901579 |
| AW209491 | 14,6823 | 13,5044 | -0,120648 | 0,57455 | 0,901579 |
| Trmt11 | 12,15 | 13,1815 | 0,117565 | 0,57465 | 0,901579 |
| Hapln1 | 3,83367 | 3,50206 | -0,130522 | 0,5747 | 0,901579 |
| Gm15413 | 0,112923 | 0,64611 | 2,51645 | 0,57495 | 0,901579 |
| Iqcg | 0,93496 | 1,38964 | 0,571734 | 0,575 | 0,901579 |
| Polr1b | 27,0238 | 29,0203 | 0,102833 | 0,57505 | 0,901579 |
| Pard3 | 26,5991 | 24,7304 | -0,105093 | 0,57515 | 0,901579 |
| Hnrnpm | 383,933 | 356,741 | -0,105977 | 0,57515 | 0,901579 |
| Ubxn11 | 1,3369 | 1,5742 | 0,235722 | 0,5752 | 0,901579 |
| Ahcyl1 | 65,6779 | 70,3896 | 0,0999551 | 0,5752 | 0,901579 |
| Pex14 | 25,6395 | 23,7739 | -0,108993 | 0,5752 | 0,901579 |
| Lurap1 | 1,2499 | 1,4871 | 0,250691 | 0,57525 | 0,901579 |
| Camk1 | 16,5956 | 18,4509 | 0,152893 | 0,57535 | 0,901579 |
| Plxna1 | 22,1358 | 23,7561 | 0,101919 | 0,57535 | 0,901579 |
| Tmem175 | 3,20191 | 3,79397 | 0,244778 | 0,5754 | 0,901579 |
| Wdr26 | 29,5214 | 31,6468 | 0,100299 | 0,5755 | 0,901626 |
| E2f7 | 12,9127 | 13,8753 | 0,103733 | 0,57575 | 0,901908 |
| Ttf1 | 7,66919 | 8,3056 | 0,11501 | 0,5759 | 0,902032 |
| Mycbpap | 0,745549 | 0,87014 | 0,222943 | 0,576 | 0,902079 |
| Got1 | 39,5289 | 42,4545 | 0,103011 | 0,5762 | 0,902282 |
| Gzf1 | 16,3501 | 15,1811 | -0,10703 | 0,57645 | 0,902563 |
| Usf1 | 35,8371 | 33,2492 | -0,108135 | 0,57655 | 0,90261 |
| Sdad1 | 34,8317 | 37,3719 | 0,101553 | 0,5767 | 0,902734 |
| Amt | 25,6579 | 23,6438 | -0,117941 | 0,57685 | 0,902859 |
| Gm12060 | 1,5793 | 1,95076 | 0,304745 | 0,5771 | 0,90314 |
| Slc29a2 | 6,88143 | 6,28352 | -0,131136 | 0,57745 | 0,90348 |
| Pim2 | 2,15718 | 2,45109 | 0,184281 | 0,57765 | 0,90348 |

| | | | | | |
|---|---|---|---|---|---|
| Ubtd2 | 56,3526 | 52,5866 | -0,099786 | 0,57765 | 0,90348 |
| Ncstn | 37,951 | 43,9757 | 0,212566 | 0,57775 | 0,90348 |
| Tmem204 | 0,761375 | 1,28056 | 0,750095 | 0,5778 | 0,90348 |
| Pdcd10 | 6,54648 | 5,92156 | -0,144741 | 0,5778 | 0,90348 |
| Phf20 | 12,7384 | 11,8664 | -0,102299 | 0,57795 | 0,90348 |
| Sbk1 | 45,0134 | 41,9815 | -0,100603 | 0,5781 | 0,90348 |
| Rspry1 | 15,375 | 14,2808 | -0,106508 | 0,57815 | 0,90348 |
| Mettl13 | 19,5747 | 18,1586 | -0,108341 | 0,57825 | 0,90348 |
| Trp53bp2 | 18,217 | 19,57 | 0,103355 | 0,5783 | 0,90348 |
| Rfwd3 | 21,286 | 22,8625 | 0,103076 | 0,5784 | 0,90348 |
| Chek1 | 29,5405 | 27,5099 | -0,102741 | 0,5784 | 0,90348 |
| Tfb2m | 2,09294 | 1,84699 | -0,180354 | 0,5785 | 0,90348 |
| Bap1 | 57,8826 | 54,0198 | -0,0996414 | 0,57875 | 0,90348 |
| Ptges3l | 1,84369 | 2,38525 | 0,371544 | 0,579 | 0,90348 |
| Amn1 | 11,6138 | 12,8359 | 0,144346 | 0,57905 | 0,90348 |
| Zfp280b | 16,3967 | 17,6017 | 0,102308 | 0,57905 | 0,90348 |
| Dph6 | 11,6563 | 10,7632 | -0,114994 | 0,5793 | 0,90348 |
| Eif2s1 | 58,6162 | 54,7028 | -0,0996863 | 0,57935 | 0,90348 |
| Rap1gds1 | 34,4012 | 32,0804 | -0,100767 | 0,57935 | 0,90348 |
| Mpi | 28,2915 | 26,1996 | -0,110824 | 0,57935 | 0,90348 |
| Phka1 | 4,20543 | 4,56584 | 0,118628 | 0,5794 | 0,90348 |
| Zcrb1 | 62,9895 | 58,3965 | -0,109231 | 0,5795 | 0,90348 |
| Dnajc25 | 9,55497 | 8,72237 | -0,131532 | 0,57965 | 0,90348 |
| Glis1 | 1,41108 | 1,23832 | -0,188411 | 0,57965 | 0,90348 |
| Ccdc8 | 38,6238 | 41,4921 | 0,103347 | 0,5797 | 0,90348 |
| Mfn2 | 14,8593 | 15,9633 | 0,103397 | 0,5799 | 0,90348 |
| Atp1b3 | 91,526 | 85,3258 | -0,1012 | 0,58005 | 0,90348 |
| Hist1h4i | 15,4433 | 17,6504 | 0,192724 | 0,58015 | 0,90348 |
| Nuf2 | 48,665 | 45,3627 | -0,101381 | 0,58015 | 0,90348 |
| Rrp8 | 28,1239 | 31,4444 | 0,161009 | 0,58025 | 0,90348 |
| Kdelr1 | 78,3074 | 72,9691 | -0,101864 | 0,5803 | 0,90348 |
| Kcnj13 | 0,246448 | 0,7878 | 1,67655 | 0,58035 | 0,90348 |
| Trim24 | 21,3196 | 19,8824 | -0,100691 | 0,58035 | 0,90348 |
| Acot2 | 1,67186 | 1,45542 | -0,200022 | 0,58035 | 0,90348 |
| Pcid2 | 24,4966 | 26,3527 | 0,105373 | 0,5804 | 0,90348 |
| Gria4 | 0,83958 | 0,986537 | 0,232707 | 0,58045 | 0,90348 |
| Zfp84 | 4,63772 | 5,02099 | 0,114556 | 0,58045 | 0,90348 |
| Rmdn1 | 2,13083 | 2,42382 | 0,18587 | 0,58055 | 0,90348 |
| Cinp | 27,9218 | 25,9325 | -0,106631 | 0,58055 | 0,90348 |
| Ccnf | 41,8157 | 39,0422 | -0,0990072 | 0,58065 | 0,90348 |
| Deb1 | 46,2678 | 50,5434 | 0,127514 | 0,58075 | 0,90348 |
| Atg7 | 7,80469 | 7,19785 | -0,116775 | 0,5808 | 0,90348 |
| Magee2 | 1,97739 | 1,74295 | -0,182072 | 0,58085 | 0,90348 |
| Bambi-ps1 | 2,22927 | 1,89869 | -0,231566 | 0,58085 | 0,90348 |
| Luc7l2 | 62,7125 | 67,3554 | 0,103041 | 0,5809 | 0,90348 |
| Cstf2 | 40,8207 | 43,7877 | 0,101224 | 0,58095 | 0,90348 |
| Slc2a10 | 1,26354 | 1,10987 | -0,187077 | 0,581 | 0,90348 |
| Nap1l3 | 1,37836 | 1,56822 | 0,186175 | 0,58105 | 0,90348 |
| Synj2bp | 4,47817 | 4,82645 | 0,108054 | 0,58105 | 0,90348 |
| Rapgef1 | 30,0988 | 28,0956 | -0,0993609 | 0,58105 | 0,90348 |

| | | | | | |
|---|---|---|---|---|---|
| Ndufb10 | 149,616 | 139,145 | -0,104672 | 0,58105 | 0,90348 |
| Dlst | 91,2655 | 85,1975 | -0,0992588 | 0,58115 | 0,903494 |
| Ndufa4l2 | 0,826217 | 2,29939 | 1,47666 | 0,5812 | 0,903494 |
| Cbr1 | 52,9796 | 57,1392 | 0,109044 | 0,5813 | 0,90354 |
| Msh3 | 20,0516 | 22,5998 | 0,172592 | 0,58155 | 0,903787 |
| Fbln1 | 46,5044 | 43,3955 | -0,0998208 | 0,5816 | 0,903787 |
| Fbxo33 | 7,56802 | 6,96325 | -0,120155 | 0,5818 | 0,90383 |
| Fam222b | 9,13551 | 9,85849 | 0,109882 | 0,58185 | 0,90383 |
| Tmem14c | 96,93 | 90,2997 | -0,102223 | 0,5819 | 0,90383 |
| Fan1 | 5,40855 | 4,95452 | -0,126495 | 0,582 | 0,90383 |
| Pomt2 | 7,24321 | 7,96628 | 0,137277 | 0,58205 | 0,90383 |
| Anapc13 | 174,899 | 161,778 | -0,1125 | 0,58205 | 0,90383 |
| Mnd1-ps | 1,7691 | 1,54172 | -0,198477 | 0,5822 | 0,903921 |
| Ptpla | 12,5409 | 11,3082 | -0,149277 | 0,58225 | 0,903921 |
| C920006O11Rik | 2,02942 | 1,74315 | -0,219372 | 0,58235 | 0,903967 |
| Akirin2 | 73,6386 | 66,8996 | -0,138464 | 0,5826 | 0,904246 |
| LOC102633315 | 0,306702 | 0,600916 | 0,970327 | 0,583 | 0,904757 |
| Asnsd1 | 49,7437 | 46,1675 | -0,107638 | 0,5834 | 0,905168 |
| Tmem87a | 16,1014 | 17,4084 | 0,112603 | 0,58355 | 0,905168 |
| Park7 | 282,632 | 264,362 | -0,0964085 | 0,5837 | 0,905168 |
| Sel1l3 | 0,579184 | 0,66635 | 0,202257 | 0,58375 | 0,905168 |
| Sub1 | 96,3265 | 89,9968 | -0,0980587 | 0,5838 | 0,905168 |
| Nampt | 29,6737 | 31,8596 | 0,102543 | 0,58385 | 0,905168 |
| Psmg2 | 59,9191 | 55,6202 | -0,107408 | 0,58385 | 0,905168 |
| N4bp3 | 5,13387 | 5,67041 | 0,143407 | 0,5839 | 0,905168 |
| Pter | 4,09023 | 3,73192 | -0,132265 | 0,5839 | 0,905168 |
| Noxred1 | 0,56079 | 0,702303 | 0,324634 | 0,58405 | 0,905292 |
| 1500015A07Rik | 2,5515 | 2,88459 | 0,17702 | 0,5842 | 0,905365 |
| Ngrn | 20,1686 | 21,8909 | 0,118217 | 0,5843 | 0,905365 |
| Txndc12 | 59,6498 | 54,3101 | -0,135297 | 0,5844 | 0,905365 |
| Nop58 | 255,592 | 238,933 | -0,0972351 | 0,58445 | 0,905365 |
| Hddc3 | 9,15352 | 8,1597 | -0,165811 | 0,58445 | 0,905365 |
| Mien1 | 35,6087 | 30,1878 | -0,238263 | 0,5851 | 0,906121 |
| Grk5 | 4,05928 | 4,45895 | 0,135478 | 0,58515 | 0,906121 |
| Kmt2c | 4,23496 | 4,5415 | 0,100819 | 0,58515 | 0,906121 |
| Accs | 4,50769 | 4,91077 | 0,123559 | 0,58525 | 0,906166 |
| Zfp771 | 24,3369 | 22,4202 | -0,118346 | 0,5854 | 0,906289 |
| Tada2a | 31,0514 | 33,3193 | 0,1017 | 0,5857 | 0,906612 |
| Itgb1bp1 | 12,7119 | 10,9332 | -0,217457 | 0,58575 | 0,906612 |
| Rab12 | 19,6552 | 18,2249 | -0,109003 | 0,58585 | 0,906657 |
| Sgta | 86,1271 | 80,4393 | -0,0985672 | 0,5861 | 0,906842 |
| 9230110C19Rik | 8,0899 | 7,33831 | -0,140674 | 0,58625 | 0,906842 |
| Fbxo11 | 30,1196 | 34,5865 | 0,199504 | 0,5863 | 0,906842 |
| Qprt | 4,91843 | 5,53497 | 0,170377 | 0,5865 | 0,906842 |
| Cep72 | 7,95802 | 8,59727 | 0,111469 | 0,5867 | 0,906842 |
| Dnajc17 | 11,2395 | 13,0825 | 0,219061 | 0,58675 | 0,906842 |
| Ptch2 | 5,02459 | 5,47 | 0,122535 | 0,5868 | 0,906842 |
| Rps23 | 1,78483 | 1,38375 | -0,3672 | 0,5868 | 0,906842 |
| Pcdhac2 | 1,11509 | 0,515344 | -1,11355 | 0,5868 | 0,906842 |
| Ppp2r2a | 39,73 | 42,4727 | 0,0963052 | 0,58685 | 0,906842 |

| | | | | | |
|---|---|---|---|---|---|
| Jag2 | 0,982081 | 0,861187 | -0,189516 | 0,58685 | 0,906842 |
| Gm14326 | 0,673118 | 0,792922 | 0,23632 | 0,58695 | 0,906842 |
| Tnfrsf10b | 2,17957 | 1,951 | -0,159832 | 0,58695 | 0,906842 |
| Cmas | 38,4883 | 35,7902 | -0,104855 | 0,587 | 0,906842 |
| Vwa8 | 5,28969 | 5,74988 | 0,12035 | 0,58705 | 0,906842 |
| Mthfsd | 17,191 | 18,6017 | 0,113779 | 0,5871 | 0,906842 |
| Chn1 | 18,9651 | 20,3568 | 0,102164 | 0,5872 | 0,906887 |
| Prox1 | 2,32608 | 2,56442 | 0,140732 | 0,5874 | 0,906977 |
| Ube2v1 | 9,72901 | 8,92382 | -0,124632 | 0,5874 | 0,906977 |
| Gpn1 | 53,0278 | 49,4036 | -0,102133 | 0,58765 | 0,907254 |
| Uba1 | 195,272 | 181,942 | -0,102004 | 0,58785 | 0,907454 |
| Angptl2 | 14,3013 | 15,4281 | 0,109418 | 0,58805 | 0,907653 |
| Chuk | 24,8784 | 26,6337 | 0,0983564 | 0,58825 | 0,907743 |
| Tcp1 | 372,258 | 346,828 | -0,102083 | 0,58825 | 0,907743 |
| Mta3 | 21,0903 | 22,7813 | 0,111273 | 0,58835 | 0,907789 |
| Ilf3 | 124,984 | 116,688 | -0,0990777 | 0,58855 | 0,907892 |
| Ufsp2 | 32,2965 | 29,9598 | -0,108347 | 0,58865 | 0,907892 |
| Kif26a | 5,3851 | 4,98859 | -0,11034 | 0,58865 | 0,907892 |
| Traf6 | 10,3456 | 11,1122 | 0,103127 | 0,5887 | 0,907892 |
| Cers5 | 37,0925 | 39,7869 | 0,101165 | 0,58885 | 0,907918 |
| Stxbp4 | 13,5638 | 14,5427 | 0,100531 | 0,58895 | 0,907918 |
| Ggact | 4,31782 | 3,79573 | -0,185926 | 0,58895 | 0,907918 |
| Ermard | 7,59931 | 8,37134 | 0,139591 | 0,589 | 0,907918 |
| Slc12a4 | 8,60019 | 9,2598 | 0,106612 | 0,5894 | 0,908252 |
| Malt1 | 3,47977 | 4,27229 | 0,296018 | 0,58945 | 0,908252 |
| Slc38a1 | 43,0162 | 45,9983 | 0,0966984 | 0,5895 | 0,908252 |
| Alyref | 129,886 | 121,405 | -0,097421 | 0,5895 | 0,908252 |
| Polr2d | 136,104 | 127,135 | -0,0983388 | 0,5896 | 0,908265 |
| Chmp1a | 24,1819 | 22,4571 | -0,106753 | 0,58965 | 0,908265 |
| Ctnnd1 | 54,9133 | 58,707 | 0,0963766 | 0,5899 | 0,908522 |
| Med4 | 60,4616 | 56,4251 | -0,0996822 | 0,59 | 0,908522 |
| Zfp273 | 1,67135 | 1,92062 | 0,200554 | 0,59005 | 0,908522 |
| Dennd1a | 6,8276 | 7,95883 | 0,221177 | 0,5901 | 0,908522 |
| Dnajb12 | 19,6757 | 21,2339 | 0,109955 | 0,5903 | 0,908612 |
| Kcnmb4 | 17,2421 | 15,7528 | -0,130324 | 0,5903 | 0,908612 |
| Rnf157 | 4,3106 | 3,96576 | -0,12029 | 0,5905 | 0,908637 |
| Phax | 65,7252 | 59,4389 | -0,145039 | 0,5905 | 0,908637 |
| Map7d1 | 48,3121 | 45,2041 | -0,0959329 | 0,59055 | 0,908637 |
| Sugp1 | 24,3065 | 22,6204 | -0,103717 | 0,5906 | 0,908637 |
| Atp6v1d | 39,6843 | 36,9352 | -0,10357 | 0,5909 | 0,908758 |
| Api5 | 64,6213 | 60,4507 | -0,0962499 | 0,59105 | 0,908758 |
| Plekhm2 | 7,04688 | 6,51812 | -0,112528 | 0,59105 | 0,908758 |
| Tufm | 7,51214 | 6,83118 | -0,137088 | 0,5913 | 0,908758 |
| Klhl26 | 6,35397 | 5,82173 | -0,12621 | 0,59135 | 0,908758 |
| 0610007P14Rik | 53,4199 | 49,673 | -0,104916 | 0,59145 | 0,908758 |
| D930048N14Rik | 0,656719 | 0,836774 | 0,349563 | 0,5915 | 0,908758 |
| LOC101056043 | 8,73623 | 7,70573 | -0,181079 | 0,59155 | 0,908758 |
| Tnks2 | 72,5977 | 77,623 | 0,0965592 | 0,5916 | 0,908758 |
| Raet1c,Raet1e | 0,969512 | 0,515454 | -0,911416 | 0,59175 | 0,908758 |
| Serinc4 | 0,975122 | 1,41 | 0,532045 | 0,5918 | 0,908758 |

| | | | | | |
|---|---|---|---|---|---|
| Rad18 | 16,8694 | 18,13 | 0,103972 | 0,59185 | 0,908758 |
| Iws1 | 17,6618 | 18,8774 | 0,0960279 | 0,5919 | 0,908758 |
| Calu | 151,237 | 141,36 | -0,0974449 | 0,5919 | 0,908758 |
| Znfx1 | 2,71475 | 3,23475 | 0,252834 | 0,59195 | 0,908758 |
| Apod | 1,04073 | 0,877121 | -0,246751 | 0,5922 | 0,908758 |
| F2r | 50,946 | 54,4503 | 0,0959715 | 0,59225 | 0,908758 |
| Ywhae | 742,498 | 689,689 | -0,106442 | 0,59225 | 0,908758 |
| Amotl1 | 24,9751 | 23,3617 | -0,0963458 | 0,59235 | 0,908758 |
| Gm10857 | 2,50759 | 2,79082 | 0,15439 | 0,59245 | 0,908758 |
| Rab5c | 79,2666 | 74,0732 | -0,0977602 | 0,59245 | 0,908758 |
| Eif2ak3 | 6,02291 | 5,56618 | -0,113773 | 0,5925 | 0,908758 |
| Phf6 | 42,1767 | 39,408 | -0,0979606 | 0,59255 | 0,908758 |
| Ifrd1 | 49,9558 | 46,6445 | -0,0989452 | 0,5926 | 0,908758 |
| Lzts3 | 2,22723 | 2,45853 | 0,142544 | 0,59265 | 0,908758 |
| 1700037H04Rik | 41,752 | 33,6384 | -0,311737 | 0,59265 | 0,908758 |
| Dpm1 | 15,3137 | 14,2001 | -0,108916 | 0,59295 | 0,908758 |
| Spidr | 15,1446 | 16,2865 | 0,104871 | 0,59315 | 0,908758 |
| Pdia3 | 414,632 | 445,221 | 0,10269 | 0,5932 | 0,908758 |
| Zc3h12b | 1,40954 | 1,2494 | -0,173993 | 0,5932 | 0,908758 |
| Eno1b | 1,3729 | 2,73187 | 0,992665 | 0,59325 | 0,908758 |
| Hist1h4c | 1,82449 | 1,27994 | -0,511414 | 0,59335 | 0,908758 |
| 9430020K01Rik | 3,33079 | 3,06963 | -0,117801 | 0,59365 | 0,908758 |
| Tada3 | 18,7542 | 20,1987 | 0,107045 | 0,59375 | 0,908758 |
| Dffa | 17,4351 | 15,8394 | -0,138478 | 0,59375 | 0,908758 |
| Star | 1,79842 | 1,99891 | 0,152484 | 0,5938 | 0,908758 |
| Tbc1d24 | 3,38777 | 3,69809 | 0,126441 | 0,5938 | 0,908758 |
| Fam160b1 | 8,93318 | 9,58478 | 0,101572 | 0,5938 | 0,908758 |
| Ctso | 0,747517 | 0,857676 | 0,198325 | 0,59385 | 0,908758 |
| Smim12 | 36,8482 | 34,1201 | -0,110974 | 0,59385 | 0,908758 |
| Leprot | 20,9111 | 19,3948 | -0,108597 | 0,59395 | 0,908758 |
| Mgrn1 | 21,561 | 23,6331 | 0,132389 | 0,594 | 0,908758 |
| Bend7 | 0,588395 | 0,680173 | 0,209115 | 0,59415 | 0,908758 |
| Fam13a | 0,877851 | 0,993228 | 0,178149 | 0,5942 | 0,908758 |
| Qrsl1 | 14,9714 | 16,1993 | 0,113718 | 0,5942 | 0,908758 |
| Maf | 1,1478 | 1,01339 | -0,179682 | 0,5943 | 0,908758 |
| Rplp2 | 956,54 | 894,719 | -0,0963911 | 0,59435 | 0,908758 |
| Stag3 | 1,1438 | 1,28588 | 0,16892 | 0,5944 | 0,908758 |
| Bbs10 | 3,73107 | 3,37643 | -0,144087 | 0,5944 | 0,908758 |
| Sfxn5 | 2,45778 | 2,71506 | 0,143628 | 0,59455 | 0,908758 |
| Plcd4 | 0,959823 | 0,269671 | -1,83157 | 0,59455 | 0,908758 |
| Abhd10 | 21,6838 | 23,5857 | 0,121296 | 0,5946 | 0,908758 |
| Catsper2 | 0,87009 | 1,01435 | 0,22132 | 0,59465 | 0,908758 |
| Dennd5b | 6,34251 | 5,91736 | -0,100101 | 0,59465 | 0,908758 |
| Csk | 55,1183 | 51,5519 | -0,0965036 | 0,59475 | 0,908758 |
| Prelid1 | 141,459 | 132,004 | -0,0997941 | 0,59475 | 0,908758 |
| 9430091E24Rik | 1,11391 | 1,44163 | 0,372072 | 0,5948 | 0,908758 |
| Sh2b2 | 8,46417 | 7,80211 | -0,117503 | 0,5948 | 0,908758 |
| Ndufab1 | 7,75084 | 6,89337 | -0,169144 | 0,5949 | 0,908758 |
| Ap3m1 | 12,9642 | 12,0787 | -0,102067 | 0,59495 | 0,908758 |
| Adnp | 68,8473 | 73,5734 | 0,0957843 | 0,595 | 0,908758 |

| | | | | | |
|---|---|---|---|---|---|
| Nptn | 34,1163 | 36,5071 | 0,0977161 | 0,5952 | 0,908955 |
| Zfp784 | 2,91921 | 3,26228 | 0,1603 | 0,5953 | 0,908955 |
| Sytl4 | 1,40953 | 1,58228 | 0,166792 | 0,5954 | 0,908955 |
| Rassf5 | 0,783323 | 0,89713 | 0,195709 | 0,59555 | 0,908955 |
| Scamp4 | 36,557 | 34,0082 | -0,104264 | 0,5956 | 0,908955 |
| Snord2 | 1142,98 | 2052,25 | 0,844408 | 0,59575 | 0,908955 |
| Snap23 | 40,4626 | 43,329 | 0,0987432 | 0,59575 | 0,908955 |
| Itpkb | 2,80856 | 2,55925 | -0,134106 | 0,5958 | 0,908955 |
| Zfp772 | 4,37092 | 3,98517 | -0,133295 | 0,59585 | 0,908955 |
| Naa40 | 40,4132 | 37,7495 | -0,0983678 | 0,59595 | 0,908955 |
| Gm15455 | 3,37683 | 3,71467 | 0,137566 | 0,596 | 0,908955 |
| Zkscan17 | 28,5764 | 30,5935 | 0,0983982 | 0,59605 | 0,908955 |
| Uap1 | 33,2738 | 35,6094 | 0,0978709 | 0,59605 | 0,908955 |
| Dtx2 | 12,0214 | 12,9603 | 0,108492 | 0,59645 | 0,909424 |
| Sephs2 | 38,2914 | 41,0077 | 0,0988734 | 0,59665 | 0,909424 |
| Exosc7 | 78,4884 | 73,2818 | -0,0990258 | 0,59665 | 0,909424 |
| Tmem38a | 4,13871 | 3,73129 | -0,149507 | 0,59675 | 0,909424 |
| Krcc1 | 21,7578 | 23,4295 | 0,106796 | 0,5968 | 0,909424 |
| Dtl | 34,7469 | 32,507 | -0,0961321 | 0,5969 | 0,909424 |
| Orai3 | 5,7836 | 6,34698 | 0,134103 | 0,59695 | 0,909424 |
| Bambi | 6,716 | 7,23581 | 0,107552 | 0,59695 | 0,909424 |
| Bsdc1 | 15,2188 | 16,3747 | 0,105611 | 0,59705 | 0,909424 |
| Nr1d2 | 4,06456 | 4,41005 | 0,117695 | 0,59715 | 0,909424 |
| Extl2 | 10,701 | 9,91801 | -0,109626 | 0,59725 | 0,909424 |
| Ap1g1 | 29,5804 | 31,58 | 0,0943681 | 0,59735 | 0,909424 |
| Tmem59 | 82,5329 | 77,1806 | -0,0967302 | 0,59735 | 0,909424 |
| 1500012F01Rik | 133,757 | 123,462 | -0,115553 | 0,59735 | 0,909424 |
| Tnfrsf18 | 1,0234 | 0,809713 | -0,337892 | 0,5976 | 0,909697 |
| Ndc1 | 39,8737 | 42,6294 | 0,0964126 | 0,5977 | 0,909701 |
| Trmt61a | 16,9402 | 15,7427 | -0,105772 | 0,59785 | 0,909701 |
| Pcbp1 | 333,211 | 311,977 | -0,0949984 | 0,59795 | 0,909701 |
| Dnttip1 | 35,595 | 33,0953 | -0,105051 | 0,598 | 0,909701 |
| Arhgap39 | 9,40646 | 10,0978 | 0,102317 | 0,59805 | 0,909701 |
| Naglu | 2,52003 | 2,8157 | 0,16005 | 0,59815 | 0,909701 |
| Mpg | 15,0209 | 16,7445 | 0,156722 | 0,59835 | 0,909701 |
| Yme1l1 | 39,6723 | 42,4079 | 0,0961985 | 0,5984 | 0,909701 |
| Zhx2 | 5,29049 | 4,88143 | -0,116098 | 0,59845 | 0,909701 |
| Rogdi | 15,9038 | 14,6426 | -0,119203 | 0,59845 | 0,909701 |
| Ppp3r1 | 45,8087 | 48,9381 | 0,0953344 | 0,5985 | 0,909701 |
| Araf | 37,2179 | 40,0238 | 0,104861 | 0,59855 | 0,909701 |
| Gcn1l1 | 24,3476 | 26,0093 | 0,0952471 | 0,59865 | 0,909701 |
| Dusp18 | 5,97408 | 5,52946 | -0,111576 | 0,59865 | 0,909701 |
| Folr1 | 0,763335 | 0,924273 | 0,276003 | 0,5988 | 0,909701 |
| Polg | 18,5525 | 20,2716 | 0,127849 | 0,5988 | 0,909701 |
| Kat6a | 22,3951 | 23,9205 | 0,0950641 | 0,5989 | 0,909701 |
| Whamm | 2,0108 | 2,24363 | 0,158065 | 0,59895 | 0,909701 |
| Vps41 | 26,295 | 24,57 | -0,0978907 | 0,59895 | 0,909701 |
| Hip1r | 2,29469 | 2,52161 | 0,136044 | 0,59905 | 0,909713 |
| 2310057M21Rik | 9,77595 | 10,5987 | 0,116574 | 0,5991 | 0,909713 |
| Pigb | 5,01361 | 5,69596 | 0,18409 | 0,59925 | 0,909725 |

| | | | | | |
|---|---|---|---|---|---|
| Nmd3 | 63,4925 | 59,2915 | -0,0987605 | 0,59925 | 0,909725 |
| Rmdn3 | 19,9983 | 21,4926 | 0,103964 | 0,59955 | 0,910041 |
| Cnot4 | 13,9314 | 14,9726 | 0,103982 | 0,5996 | 0,910041 |
| 4930414N06Rik | 0,251372 | 0,673558 | 1,42198 | 0,5998 | 0,910237 |
| Zak | 4,07821 | 4,43811 | 0,122008 | 0,6002 | 0,910629 |
| Abhd13 | 11,8625 | 11,0844 | -0,0978815 | 0,6002 | 0,910629 |
| Dennd5a | 30,1595 | 32,1985 | 0,0943827 | 0,6004 | 0,910824 |
| Jagn1 | 43,8268 | 40,755 | -0,104835 | 0,6008 | 0,911292 |
| Mllt1 | 44,022 | 41,2035 | -0,0954593 | 0,60085 | 0,911292 |
| Ppp1r15b | 30,7788 | 32,8435 | 0,0936696 | 0,601 | 0,911411 |
| Cd68 | 2,70974 | 2,32539 | -0,220683 | 0,6013 | 0,911758 |
| Tor1aip1 | 17,1929 | 19,4464 | 0,17769 | 0,60145 | 0,911878 |
| Nbr1 | 34,8125 | 37,2094 | 0,0960619 | 0,60165 | 0,912042 |
| Mir6912 | 103,972 | 289,467 | 1,4772 | 0,6017 | 0,912042 |
| Sec23ip | 28,2828 | 30,2109 | 0,0951431 | 0,6021 | 0,912418 |
| Cct4 | 363,009 | 339,824 | -0,0952168 | 0,6021 | 0,912418 |
| Gm2518 | 0,608768 | 0,945622 | 0,635372 | 0,6024 | 0,912418 |
| Ovgp1 | 1,94024 | 2,17713 | 0,166188 | 0,60255 | 0,912418 |
| Ago4 | 1,00915 | 1,12096 | 0,151589 | 0,60255 | 0,912418 |
| Chd9 | 4,6339 | 4,97295 | 0,101874 | 0,60275 | 0,912418 |
| Tmem2 | 19,084 | 17,8634 | -0,0953632 | 0,6028 | 0,912418 |
| Etfdh | 20,3456 | 18,9458 | -0,102842 | 0,60285 | 0,912418 |
| 8030462N17Rik | 10,3315 | 11,1175 | 0,105787 | 0,6029 | 0,912418 |
| Pot1b | 10,3858 | 9,6582 | -0,104791 | 0,603 | 0,912418 |
| Chka | 22,9305 | 24,5405 | 0,0978939 | 0,60305 | 0,912418 |
| H2afz | 14,8914 | 13,6173 | -0,129045 | 0,60305 | 0,912418 |
| Ankrd17 | 25,4629 | 27,1894 | 0,0946474 | 0,60315 | 0,912418 |
| Zranb2 | 86,4525 | 92,1691 | 0,0923761 | 0,60315 | 0,912418 |
| Ugp2 | 35,9926 | 39,4917 | 0,133849 | 0,6032 | 0,912418 |
| Mtf1 | 3,14144 | 3,45527 | 0,137374 | 0,60325 | 0,912418 |
| Ninj1 | 6,70346 | 6,01438 | -0,156491 | 0,60325 | 0,912418 |
| Alad | 11,8179 | 11,024 | -0,100333 | 0,6033 | 0,912418 |
| Vaultrc5 | 1,91133 | 1,27741 | -0,581358 | 0,6033 | 0,912418 |
| C330018D20Rik | 6,84329 | 6,27625 | -0,124787 | 0,60345 | 0,912538 |
| Anapc2 | 56,9733 | 53,332 | -0,0952827 | 0,60365 | 0,912732 |
| Actr6 | 16,085 | 17,3707 | 0,110945 | 0,6038 | 0,912852 |
| Tipin | 148,77 | 139,425 | -0,0935985 | 0,6039 | 0,912895 |
| Zfp599 | 4,16579 | 5,55 | 0,413895 | 0,60405 | 0,913014 |
| Tm9sf1 | 27,3904 | 25,6087 | -0,0970318 | 0,60425 | 0,913102 |
| 1700030K09Rik | 6,82511 | 5,27603 | -0,371398 | 0,60425 | 0,913102 |
| Ints10 | 38,5366 | 41,1323 | 0,0940398 | 0,6046 | 0,913383 |
| Zbtb48 | 10,4112 | 11,2472 | 0,111427 | 0,60465 | 0,913383 |
| Camk2d | 1,44934 | 1,3024 | -0,154228 | 0,60465 | 0,913383 |
| Zfand3 | 47,1906 | 50,389 | 0,094609 | 0,605 | 0,913537 |
| Hexa | 19,5309 | 18,1389 | -0,10667 | 0,605 | 0,913537 |
| Pdik1l | 5,76902 | 5,34005 | -0,111475 | 0,60505 | 0,913537 |
| Sfr1 | 216,394 | 202,975 | -0,0923598 | 0,6051 | 0,913537 |
| Fto | 43,297 | 40,634 | -0,0915776 | 0,6052 | 0,913537 |
| Csrp2bp | 30,1058 | 28,2008 | -0,0943064 | 0,6052 | 0,913537 |
| Zfp704 | 9,08129 | 9,68496 | 0,0928494 | 0,60525 | 0,913537 |

| | | | | | |
|---|---|---|---|---|---|
| Ing4 | 58,0134 | 54,352 | -0,0940553 | 0,6055 | 0,913806 |
| Slc12a2 | 16,5596 | 17,689 | 0,0951902 | 0,6058 | 0,913916 |
| Akr1c19 | 0,919255 | 0,747715 | -0,297978 | 0,6058 | 0,913916 |
| Arl6ip6 | 27,0737 | 25,2975 | -0,0978983 | 0,60595 | 0,913916 |
| Bola3 | 44,1966 | 40,5883 | -0,122871 | 0,60595 | 0,913916 |
| Awat2 | 0,863436 | 1,02682 | 0,250017 | 0,606 | 0,913916 |
| Zfp286 | 7,52193 | 6,95906 | -0,11221 | 0,606 | 0,913916 |
| Mrpl37 | 67,052 | 71,6376 | 0,0954364 | 0,6064 | 0,91395 |
| Gabra3 | 1,51761 | 1,36052 | -0,157645 | 0,6064 | 0,91395 |
| Rpl19 | 4,15585 | 4,7733 | 0,199844 | 0,60645 | 0,91395 |
| Cdan1 | 7,20815 | 7,73292 | 0,101384 | 0,60645 | 0,91395 |
| Luzp1 | 27,395 | 29,2069 | 0,0923942 | 0,60645 | 0,91395 |
| Htatsf1 | 55,0243 | 51,5351 | -0,0945122 | 0,60645 | 0,91395 |
| Lgals4 | 0,535254 | 0,665408 | 0,314015 | 0,60665 | 0,914123 |
| Mfsd7b | 7,26481 | 6,73662 | -0,1089 | 0,60675 | 0,914123 |
| Snapc3 | 27,4511 | 29,5126 | 0,10447 | 0,6068 | 0,914123 |
| Zc3h4 | 30,6637 | 32,7273 | 0,0939641 | 0,60685 | 0,914123 |
| Mthfd2 | 54,3957 | 50,9722 | -0,0937809 | 0,60695 | 0,914166 |
| Slc30a1 | 2,84891 | 3,10556 | 0,124444 | 0,6071 | 0,914221 |
| Zfr2 | 0,571033 | 0,663572 | 0,216678 | 0,60715 | 0,914221 |
| Gchfr | 4,94074 | 4,27086 | -0,210201 | 0,6072 | 0,914221 |
| Yipf3 | 19,4776 | 18,1263 | -0,103731 | 0,6075 | 0,914426 |
| Lrrc27 | 2,26258 | 2,57193 | 0,18488 | 0,60755 | 0,914426 |
| Itpr1 | 8,13308 | 8,67839 | 0,0936251 | 0,60755 | 0,914426 |
| Rps29 | 1,52365 | 2,06745 | 0,440318 | 0,60765 | 0,914437 |
| Mir7030 | 683,03 | 502,879 | -0,441738 | 0,6077 | 0,914437 |
| Eid2 | 30,532 | 28,3898 | -0,10495 | 0,60805 | 0,914824 |
| Stx2 | 8,18091 | 7,56391 | -0,113128 | 0,6081 | 0,914824 |
| Suclg2 | 13,4954 | 12,5918 | -0,0999891 | 0,6082 | 0,914867 |
| Cd276 | 25,022 | 26,7272 | 0,0951131 | 0,60865 | 0,915052 |
| Inppl1 | 33,186 | 35,421 | 0,0940299 | 0,60865 | 0,915052 |
| Vwa9 | 32,3947 | 30,3837 | -0,0924642 | 0,60875 | 0,915052 |
| Ppcs | 9,87899 | 10,7571 | 0,122859 | 0,6088 | 0,915052 |
| Polr1d | 103,432 | 96,7305 | -0,0966334 | 0,6088 | 0,915052 |
| Fut4 | 3,46785 | 3,7799 | 0,124306 | 0,60895 | 0,915052 |
| Dip2b | 8,89793 | 8,32936 | -0,0952634 | 0,60905 | 0,915052 |
| Znrf1 | 14,9596 | 15,9399 | 0,0915722 | 0,6091 | 0,915052 |
| Pde1b | 2,31712 | 2,55918 | 0,143346 | 0,6092 | 0,915052 |
| Agpat1 | 48,8825 | 52,0986 | 0,0919255 | 0,6092 | 0,915052 |
| Sept3 | 23,015 | 21,5798 | -0,0928932 | 0,6092 | 0,915052 |
| Gins2 | 90,4307 | 80,6644 | -0,164881 | 0,6092 | 0,915052 |
| Nkrf | 8,83877 | 9,51719 | 0,106689 | 0,60925 | 0,915052 |
| Dnlz | 18,7647 | 16,8308 | -0,156915 | 0,6095 | 0,91532 |
| Hist3h2ba | 3,46863 | 4,20585 | 0,278035 | 0,60965 | 0,915374 |
| Nvl | 45,6544 | 48,6836 | 0,0926838 | 0,6097 | 0,915374 |
| Ticrr | 11,1407 | 12,0607 | 0,114475 | 0,60975 | 0,915374 |
| Gatad1 | 37,8064 | 40,3411 | 0,0936187 | 0,61015 | 0,915868 |
| Cbx5 | 99,4037 | 104,931 | 0,0964239 | 0,6105 | 0,916254 |
| Bcl7b | 15,7136 | 14,5447 | -0,111522 | 0,61055 | 0,916254 |
| Hif1an | 24,5417 | 26,1709 | 0,092732 | 0,61065 | 0,916264 |

| | | | | | |
|---|---|---|---|---|---|
| Nit1 | 25,8798 | 23,2387 | -0,1553 | 0,6107 | 0,916264 |
| Map3k13 | 0,646453 | 0,574578 | -0,170042 | 0,6108 | 0,916307 |
| Wdr12 | 21,7039 | 20,2003 | -0,103573 | 0,61095 | 0,916318 |
| Ddit3 | 10,781 | 7,47587 | -0,528171 | 0,61095 | 0,916318 |
| D19Bwg1357e | 57,7403 | 54,1738 | -0,0919828 | 0,61115 | 0,916318 |
| Cnpy2 | 123,574 | 115,852 | -0,0930833 | 0,6112 | 0,916318 |
| Bcap29 | 10,4392 | 9,60784 | -0,119721 | 0,6112 | 0,916318 |
| Sh3d21 | 1,67107 | 1,46754 | -0,187373 | 0,61135 | 0,916318 |
| Snora33 | 237,15 | 163,036 | -0,540606 | 0,61135 | 0,916318 |
| Neil2 | 0,859937 | 1,00795 | 0,229121 | 0,61145 | 0,916318 |
| Napg | 9,27529 | 9,96908 | 0,104068 | 0,61145 | 0,916318 |
| Hbp1 | 29,6655 | 27,5067 | -0,109004 | 0,6116 | 0,916318 |
| Mcm9 | 5,42877 | 6,27219 | 0,208342 | 0,6118 | 0,916318 |
| Chic2 | 31,0494 | 33,395 | 0,105065 | 0,61185 | 0,916318 |
| Mcl1 | 59,8307 | 56,1498 | -0,0916052 | 0,61185 | 0,916318 |
| Eprs | 108,949 | 102,172 | -0,0926553 | 0,61185 | 0,916318 |
| Pou4f1 | 0,564279 | 0,650613 | 0,205393 | 0,61195 | 0,916318 |
| Rchy1 | 25,2318 | 23,2684 | -0,116871 | 0,61195 | 0,916318 |
| Gm4013 | 1,75339 | 2,29074 | 0,385666 | 0,61225 | 0,916489 |
| Brk1 | 129,063 | 120,95 | -0,0936751 | 0,6123 | 0,916489 |
| Kdm1a | 98,257 | 104,667 | 0,0911748 | 0,61235 | 0,916489 |
| Ubtf | 63,8435 | 59,964 | -0,0904453 | 0,61235 | 0,916489 |
| Mrpl15 | 34,2301 | 31,9142 | -0,101068 | 0,61255 | 0,916585 |
| Brcc3 | 34,6231 | 37,6284 | 0,120086 | 0,6126 | 0,916585 |
| Sgsm3 | 6,93974 | 7,77481 | 0,163926 | 0,6127 | 0,916585 |
| Rbm39 | 186,671 | 199,072 | 0,0927907 | 0,6127 | 0,916585 |
| Tmem246 | 4,33535 | 4,72329 | 0,123645 | 0,6128 | 0,916628 |
| Hspb9 | 0,738511 | 0,960613 | 0,379337 | 0,613 | 0,91682 |
| Ubl5 | 51,8213 | 48,3299 | -0,100628 | 0,61315 | 0,916905 |
| Scaper | 2,31605 | 2,52759 | 0,126095 | 0,6132 | 0,916905 |
| Phf7 | 5,62181 | 6,15366 | 0,130411 | 0,61335 | 0,917023 |
| Pfkm | 28,6877 | 25,6499 | -0,161476 | 0,6136 | 0,917289 |
| Dcp2 | 29,2744 | 31,1817 | 0,0910582 | 0,6137 | 0,917332 |
| Tfrc | 44,9816 | 47,9183 | 0,0912424 | 0,6139 | 0,917385 |
| Ctdp1 | 19,1777 | 17,9549 | -0,0950563 | 0,6139 | 0,917385 |
| Itgb5 | 44,5916 | 41,8537 | -0,0914192 | 0,61395 | 0,917385 |
| Rbm3os | 2,07634 | 3,0165 | 0,538831 | 0,6142 | 0,917652 |
| Tmem57 | 21,9252 | 23,3956 | 0,0936521 | 0,6143 | 0,917694 |
| Derl2 | 10,7887 | 12,1386 | 0,170079 | 0,6145 | 0,917702 |
| Man2c1os | 1,79925 | 1,06573 | -0,755562 | 0,61465 | 0,917702 |
| Zglp1 | 1,59537 | 1,88816 | 0,243087 | 0,61475 | 0,917702 |
| Nadsyn1 | 3,50716 | 3,85343 | 0,135837 | 0,61485 | 0,917702 |
| A730017L22Rik | 1,17842 | 1,03 | -0,194211 | 0,6149 | 0,917702 |
| Frrs1l | 0,634898 | 0,726703 | 0,19484 | 0,615 | 0,917702 |
| Golga2 | 13,7316 | 14,6711 | 0,0954716 | 0,615 | 0,917702 |
| Nipsnap3b | 24,9717 | 23,2606 | -0,102406 | 0,615 | 0,917702 |
| Top2a | 206,789 | 220,94 | 0,0954945 | 0,61505 | 0,917702 |
| Yif1a | 18,0582 | 16,6396 | -0,118029 | 0,61505 | 0,917702 |
| Rabgef1 | 8,1836 | 8,84004 | 0,111317 | 0,6151 | 0,917702 |
| Hk1 | 7,98273 | 8,56348 | 0,101315 | 0,61525 | 0,917702 |

| | | | | | |
|---|---|---|---|---|---|
| Tep1 | 1,84999 | 2,00652 | 0,11718 | 0,6154 | 0,917702 |
| Capzb | 182,279 | 171,327 | -0,0893966 | 0,61545 | 0,917702 |
| Umps | 66,3147 | 62,2855 | -0,0904321 | 0,61545 | 0,917702 |
| Pxk | 5,21517 | 4,79091 | -0,122415 | 0,61545 | 0,917702 |
| 2700029M09Rik | 76,3088 | 71,4838 | -0,0942339 | 0,6156 | 0,917787 |
| Herpud2 | 11,6111 | 12,4727 | 0,103271 | 0,61565 | 0,917787 |
| Vapa | 92,1853 | 86,4881 | -0,0920345 | 0,6158 | 0,91785 |
| Zw10 | 34,9567 | 37,2558 | 0,0918955 | 0,61595 | 0,91785 |
| Osbp | 15,008 | 16,0323 | 0,0952488 | 0,616 | 0,91785 |
| Hectd1 | 25,9887 | 27,6604 | 0,0899366 | 0,61605 | 0,91785 |
| Ap5s1 | 7,17557 | 4,28426 | -0,744049 | 0,61605 | 0,91785 |
| Dusp11 | 25,4407 | 27,1008 | 0,0911982 | 0,61615 | 0,91786 |
| Lrrc40 | 46,2372 | 43,3886 | -0,091738 | 0,6162 | 0,91786 |
| Sdc3 | 19,3786 | 18,168 | -0,0930683 | 0,6163 | 0,917903 |
| Gabpa | 23,9323 | 21,7754 | -0,136261 | 0,61645 | 0,917944 |
| Hist1h2ak | 1,38962 | 1,77211 | 0,35078 | 0,61665 | 0,917944 |
| Ints9 | 30,1958 | 28,3287 | -0,0920852 | 0,61665 | 0,917944 |
| Tbkbp1 | 6,98657 | 6,47182 | -0,110413 | 0,61665 | 0,917944 |
| Man1a | 1,73219 | 1,56053 | -0,150559 | 0,6168 | 0,917944 |
| Snora41 | 159,154 | 213,293 | 0,422418 | 0,6169 | 0,917944 |
| Cisd2 | 24,4921 | 22,9481 | -0,09394 | 0,6169 | 0,917944 |
| Ccdc101 | 26,0375 | 24,1646 | -0,107696 | 0,6169 | 0,917944 |
| Chtf18 | 19,7347 | 21,0731 | 0,0946685 | 0,617 | 0,917954 |
| Zfp385a | 21,1284 | 22,5889 | 0,0964307 | 0,61705 | 0,917954 |
| Gdf1 | 5,9945 | 4,27829 | -0,486606 | 0,61715 | 0,917964 |
| Map1lc3b | 30,041 | 28,076 | -0,0975933 | 0,6172 | 0,917964 |
| Maf1 | 33,5278 | 31,3634 | -0,0962768 | 0,6176 | 0,918453 |
| Garnl3 | 1,64474 | 1,47359 | -0,158526 | 0,61775 | 0,918463 |
| Necab3 | 0,802464 | 0,662993 | -0,275442 | 0,61775 | 0,918463 |
| Tctn2 | 8,59223 | 9,25351 | 0,106968 | 0,61805 | 0,918696 |
| Stk11ip | 14,4971 | 13,5899 | -0,0932375 | 0,61805 | 0,918696 |
| Pigt | 22,5443 | 21,0926 | -0,0960235 | 0,61815 | 0,918738 |
| Ift57 | 7,70624 | 7,12026 | -0,114097 | 0,61825 | 0,91878 |
| Cd81 | 82,7305 | 88,2499 | 0,0931753 | 0,61835 | 0,918822 |
| Zmat1 | 1,07712 | 1,2095 | 0,167224 | 0,61855 | 0,918907 |
| Pusl1 | 17,2575 | 14,6356 | -0,237745 | 0,61855 | 0,918907 |
| Cep250 | 14,6044 | 15,5429 | 0,0898435 | 0,6189 | 0,91932 |
| Atp6v1g1 | 21,8611 | 23,6203 | 0,111656 | 0,61905 | 0,919437 |
| Frs2 | 12,7964 | 13,6363 | 0,0917212 | 0,61915 | 0,919446 |
| Zrsr1 | 2,74987 | 3,31042 | 0,267649 | 0,6192 | 0,919446 |
| Epc1 | 24,0841 | 22,1327 | -0,121903 | 0,6193 | 0,919488 |
| Casc3 | 39,6259 | 43,4697 | 0,133563 | 0,6195 | 0,919624 |
| Rnf214 | 14,5588 | 15,8553 | 0,123074 | 0,61955 | 0,919624 |
| Arl14ep | 14,5992 | 15,6115 | 0,0967182 | 0,6197 | 0,919624 |
| Tma16 | 20,1705 | 18,6511 | -0,112988 | 0,6197 | 0,919624 |
| Ciapin1 | 22,5776 | 21,2149 | -0,0898141 | 0,61975 | 0,919624 |
| Dync1h1 | 41,7718 | 39,1601 | -0,0931473 | 0,6199 | 0,919741 |
| BC053749 | 7,42121 | 8,02568 | 0,112969 | 0,6201 | 0,919825 |
| Phlpp1 | 4,64313 | 4,32008 | -0,104041 | 0,6201 | 0,919825 |
| Zfp128 | 2,69797 | 2,4605 | -0,132922 | 0,62025 | 0,919941 |

| | | | | | |
|---|---|---|---|---|---|
| Ksr1 | 2,22316 | 2,43654 | 0,132218 | 0,62035 | 0,919983 |
| Csad | 9,81292 | 9,09385 | -0,109791 | 0,62055 | 0,920076 |
| Rnf40 | 28,6921 | 30,5764 | 0,0917668 | 0,6206 | 0,920076 |
| Ssh3 | 0,97034 | 1,10702 | 0,190124 | 0,62065 | 0,920076 |
| Arfgap2 | 25,7245 | 24,1486 | -0,0912028 | 0,6207 | 0,920076 |
| Nqo2 | 6,37092 | 5,77163 | -0,142523 | 0,62085 | 0,92016 |
| Fbxo44 | 5,8464 | 5,3184 | -0,136555 | 0,6209 | 0,92016 |
| Chst14 | 15,17 | 14,1154 | -0,103953 | 0,62105 | 0,920244 |
| Mbd4 | 11,057 | 11,8362 | 0,0982558 | 0,6211 | 0,920244 |
| AW011738 | 1,41277 | 1,58409 | 0,165128 | 0,62135 | 0,920295 |
| Ddx25 | 16,278 | 15,134 | -0,105132 | 0,62135 | 0,920295 |
| Mitd1 | 19,839 | 18,3787 | -0,110305 | 0,62135 | 0,920295 |
| Myl12a | 101,701 | 108,069 | 0,0876247 | 0,6215 | 0,920347 |
| H2afx | 186,323 | 175,29 | -0,08806 | 0,62155 | 0,920347 |
| Rbm8a | 61,4085 | 57,7084 | -0,0896582 | 0,6216 | 0,920347 |
| St3gal4 | 10,2248 | 8,85073 | -0,208208 | 0,6218 | 0,920472 |
| Crot | 19,1449 | 17,9263 | -0,0948834 | 0,62185 | 0,920472 |
| Rnf113a2 | 11,9856 | 13,2698 | 0,14685 | 0,6219 | 0,920472 |
| 9930012K11Rik | 1,9214 | 2,15946 | 0,168513 | 0,6221 | 0,92063 |
| Fh1 | 107,792 | 101,365 | -0,0886847 | 0,62215 | 0,92063 |
| Lgi2 | 3,55051 | 3,92332 | 0,144049 | 0,62245 | 0,920968 |
| Ska3 | 32,9801 | 35,1085 | 0,0902257 | 0,62255 | 0,921009 |
| Bri3bp | 15,4282 | 14,4909 | -0,090425 | 0,6227 | 0,921019 |
| Atad3aos | 0,939504 | 0,738213 | -0,347863 | 0,6227 | 0,921019 |
| Ccdc167 | 15,1838 | 17,6891 | 0,220326 | 0,62285 | 0,921028 |
| Fez2 | 11,5867 | 10,6318 | -0,124087 | 0,62285 | 0,921028 |
| Arpc3 | 159,444 | 149,748 | -0,0905121 | 0,62305 | 0,921153 |
| Ccdc120 | 4,11916 | 3,79614 | -0,117815 | 0,6231 | 0,921153 |
| Mob2 | 25,8138 | 24,0712 | -0,10083 | 0,62315 | 0,921153 |
| Cactin | 30,2597 | 28,4182 | -0,0905835 | 0,62365 | 0,921786 |
| Cep104 | 11,2317 | 12,0237 | 0,0983074 | 0,624 | 0,921929 |
| Myef2 | 43,7292 | 46,5698 | 0,0907966 | 0,624 | 0,921929 |
| Dhx29 | 26,9501 | 25,3661 | -0,0873905 | 0,62405 | 0,921929 |
| Drg1 | 140,01 | 124,347 | -0,171151 | 0,6241 | 0,921929 |
| Sipa1l2 | 5,96599 | 5,57169 | -0,0986487 | 0,62415 | 0,921929 |
| Ttc27 | 21,6505 | 20,2976 | -0,0930916 | 0,62425 | 0,921929 |
| Rpp38 | 16,048 | 14,7659 | -0,120129 | 0,62425 | 0,921929 |
| Vpreb3 | 1,50505 | 1,86646 | 0,310494 | 0,6246 | 0,92234 |
| Plekha6 | 1,32256 | 1,20701 | -0,131903 | 0,62475 | 0,922455 |
| Hps5 | 8,58342 | 8,03335 | -0,0955508 | 0,62495 | 0,922645 |
| Rpl38 | 17,4038 | 14,2083 | -0,29267 | 0,62505 | 0,922686 |
| Tmpo | 207,106 | 194,429 | -0,0911215 | 0,62515 | 0,922727 |
| Gm5113 | 2,74987 | 3,02593 | 0,138016 | 0,62545 | 0,922976 |
| Gpr173 | 2,40258 | 2,17915 | -0,140815 | 0,62555 | 0,922976 |
| Cct2 | 433,677 | 406,901 | -0,091942 | 0,6256 | 0,922976 |
| Adrm1 | 7,44983 | 6,79679 | -0,132353 | 0,62565 | 0,922976 |
| Rb1 | 5,70018 | 5,01155 | -0,185751 | 0,6257 | 0,922976 |
| Mpst | 23,448 | 25,1443 | 0,100765 | 0,62575 | 0,922976 |
| Baz1a | 10,9078 | 10,2443 | -0,090539 | 0,6259 | 0,923059 |
| Elmod3 | 4,77792 | 4,37314 | -0,127713 | 0,62595 | 0,923059 |

| | | | | | |
|---|---|---|---|---|---|
| Snhg9 | 25,2816 | 19,3876 | -0,38295 | 0,62605 | 0,9231 |
| Rsl1d1 | 239,402 | 225,359 | -0,087214 | 0,62615 | 0,923118 |
| Fgd6 | 3,88845 | 4,16883 | 0,100448 | 0,62635 | 0,923118 |
| Tbl2 | 13,7791 | 14,6967 | 0,0930079 | 0,62635 | 0,923118 |
| Hnrnpf | 35,6184 | 33,4485 | -0,090678 | 0,62635 | 0,923118 |
| 1110002L01Rik | 5,68825 | 6,37351 | 0,164103 | 0,6268 | 0,923536 |
| Lrrc16a | 2,64337 | 2,87101 | 0,119181 | 0,6268 | 0,923536 |
| Poglut1 | 44,1041 | 41,4905 | -0,0881315 | 0,62685 | 0,923536 |
| Gm5176 | 1,82383 | 1,4478 | -0,333111 | 0,62695 | 0,923578 |
| Fam73b | 10,8625 | 11,6171 | 0,0969012 | 0,6271 | 0,923693 |
| Sestd1 | 7,66444 | 7,18961 | -0,0922663 | 0,62735 | 0,923751 |
| Zcchc4 | 6,93421 | 7,51722 | 0,116467 | 0,6274 | 0,923751 |
| Insig2 | 10,2404 | 11,0047 | 0,103857 | 0,62745 | 0,923751 |
| Smarcd1 | 74,8137 | 70,4047 | -0,0876301 | 0,62745 | 0,923751 |
| Brix1 | 47,1236 | 50,1181 | 0,0888833 | 0,6275 | 0,923751 |
| Mtor | 13,0675 | 12,2841 | -0,0891922 | 0,62765 | 0,923834 |
| Spa17 | 4,2411 | 4,85194 | 0,194122 | 0,6277 | 0,923834 |
| Cryzl1 | 18,2026 | 16,9388 | -0,103817 | 0,6283 | 0,924587 |
| Gps1 | 12,067 | 11,2027 | -0,107213 | 0,6284 | 0,924587 |
| Mcm3 | 178,755 | 168,248 | -0,0874001 | 0,62845 | 0,924587 |
| Ncoa3 | 9,64943 | 10,2791 | 0,0912026 | 0,6285 | 0,924587 |
| Lrrc1 | 7,29887 | 7,86163 | 0,107155 | 0,6286 | 0,924596 |
| Zfp189 | 6,50204 | 7,03076 | 0,112788 | 0,62865 | 0,924596 |
| Ttc21b | 10,0643 | 9,42284 | -0,0950151 | 0,6289 | 0,924857 |
| Ankrd54 | 17,425 | 16,2591 | -0,0999187 | 0,629 | 0,924866 |
| Zfp580 | 11,6006 | 10,6391 | -0,124819 | 0,62905 | 0,924866 |
| Ywhab | 193,882 | 182,28 | -0,0890245 | 0,6292 | 0,924874 |
| Smim7 | 36,4594 | 34,2705 | -0,0893224 | 0,6292 | 0,924874 |
| Mon2 | 10,1282 | 10,781 | 0,0901049 | 0,6294 | 0,925062 |
| Cd99l2 | 12,1786 | 11,4114 | -0,0938747 | 0,62995 | 0,925456 |
| Ctnna1 | 133,135 | 141,353 | 0,0864154 | 0,63 | 0,925456 |
| Pcif1 | 18,3978 | 19,652 | 0,0951406 | 0,63005 | 0,925456 |
| Clic6 | 0,61632 | 0,702703 | 0,189234 | 0,6302 | 0,925456 |
| Srsf11 | 118,004 | 125,251 | 0,0859896 | 0,63025 | 0,925456 |
| Tnfaip1 | 44,9687 | 42,3424 | -0,0868179 | 0,63025 | 0,925456 |
| Prpf8 | 114,589 | 107,598 | -0,0908266 | 0,63035 | 0,925456 |
| Arfgap3 | 7,67383 | 7,116 | -0,10888 | 0,63045 | 0,925456 |
| D17H6S53E | 4,37656 | 4,78922 | 0,129993 | 0,6305 | 0,925456 |
| Saal1 | 15,4589 | 14,5068 | -0,0917092 | 0,6305 | 0,925456 |
| Srp72 | 64,0774 | 60,3587 | -0,0862555 | 0,63055 | 0,925456 |
| Aga | 7,01008 | 6,38479 | -0,134791 | 0,6307 | 0,925456 |
| Wdr31 | 3,38829 | 3,71042 | 0,131024 | 0,63075 | 0,925456 |
| Arfip1 | 18,5929 | 19,822 | 0,0923473 | 0,63075 | 0,925456 |
| Rltpr | 0,829108 | 0,610987 | -0,440418 | 0,63075 | 0,925456 |
| Rbak | 7,98193 | 8,56567 | 0,101828 | 0,63085 | 0,925497 |
| Cyhr1 | 28,7563 | 26,8501 | -0,098952 | 0,63095 | 0,925538 |
| Brwd1 | 45,6481 | 49,0188 | 0,102782 | 0,63145 | 0,926149 |
| Mier1 | 19,776 | 21,0262 | 0,0884392 | 0,6316 | 0,926149 |
| Trmu | 12,9614 | 12,0168 | -0,109168 | 0,6316 | 0,926149 |
| Rdh13 | 17,8661 | 19,0372 | 0,0915982 | 0,6318 | 0,926149 |

| | | | | | |
|---|---|---|---|---|---|
| Wdr3 | 40,6262 | 43,1737 | 0,0877442 | 0,6318 | 0,926149 |
| Pole3 | 119,022 | 111,515 | -0,0939861 | 0,6318 | 0,926149 |
| Trim65 | 0,576226 | 0,667077 | 0,21122 | 0,63195 | 0,926206 |
| Psmd4 | 175,827 | 165,756 | -0,0850917 | 0,632 | 0,926206 |
| Rtn3 | 117,438 | 126,074 | 0,102373 | 0,6321 | 0,926206 |
| Fastkd5 | 11,3313 | 10,3201 | -0,134849 | 0,6322 | 0,926206 |
| Mipol1 | 6,51288 | 4,85777 | -0,423002 | 0,6322 | 0,926206 |
| Lpcat1 | 45,4892 | 48,4382 | 0,0906221 | 0,63235 | 0,926255 |
| Rheb | 78,657 | 74,0693 | -0,0867006 | 0,6324 | 0,926255 |
| Mex3b | 32,7685 | 34,7929 | 0,0864821 | 0,63245 | 0,926255 |
| Tram1l1 | 7,71608 | 7,13172 | -0,113618 | 0,6329 | 0,926743 |
| 5530601H04Rik | 4,18274 | 3,86438 | -0,114211 | 0,63325 | 0,926743 |
| Stac | 0,706028 | 0,606067 | -0,22025 | 0,63325 | 0,926743 |
| Cadps | 1,22366 | 1,34489 | 0,136286 | 0,6334 | 0,926743 |
| Mks1 | 8,52575 | 9,17006 | 0,105105 | 0,6334 | 0,926743 |
| Kank2 | 16,1377 | 17,1417 | 0,0870748 | 0,63345 | 0,926743 |
| Twistnb | 17,49 | 16,3705 | -0,0954314 | 0,63345 | 0,926743 |
| Mtdh | 55,0497 | 58,4276 | 0,0859149 | 0,6335 | 0,926743 |
| Sh3bp2 | 7,60118 | 7,05806 | -0,106952 | 0,6335 | 0,926743 |
| Spats2 | 58,9897 | 62,6118 | 0,0859712 | 0,63355 | 0,926743 |
| Zfp157 | 8,24596 | 8,80296 | 0,0943014 | 0,6336 | 0,926743 |
| U2af1 | 46,7079 | 49,9424 | 0,0965994 | 0,63365 | 0,926743 |
| Mrpl57 | 10,6569 | 9,9121 | -0,104524 | 0,63375 | 0,926784 |
| Elp6 | 15,4861 | 14,4564 | -0,0992631 | 0,6342 | 0,927336 |
| Lonrf1 | 3,23357 | 3,51063 | 0,118605 | 0,63465 | 0,927598 |
| Kif11 | 56,3195 | 59,7415 | 0,0851007 | 0,63465 | 0,927598 |
| Sdccag8 | 6,10061 | 5,30829 | -0,200706 | 0,63465 | 0,927598 |
| Ccdc125 | 1,41117 | 1,96454 | 0,477305 | 0,6348 | 0,927598 |
| B230208H11Rik | 0,878556 | 0,723405 | -0,280331 | 0,6349 | 0,927598 |
| Prdx1 | 294,771 | 277,841 | -0,0853327 | 0,635 | 0,927598 |
| B230219D22Rik | 44,337 | 47,0631 | 0,0860859 | 0,63505 | 0,927598 |
| Ambra1 | 14,8203 | 15,7438 | 0,0872132 | 0,6352 | 0,927598 |
| Ube2k | 35,7149 | 33,6631 | -0,0853586 | 0,6352 | 0,927598 |
| Mto1 | 10,8163 | 10,0806 | -0,101626 | 0,6352 | 0,927598 |
| Mrpl49 | 56,0397 | 51,5507 | -0,120458 | 0,6352 | 0,927598 |
| G6pdx | 25,8312 | 23,9023 | -0,111967 | 0,6353 | 0,927598 |
| Poldip2 | 74,5755 | 70,3641 | -0,0838623 | 0,63545 | 0,927598 |
| Zic2 | 20,3012 | 19,0497 | -0,0917972 | 0,63545 | 0,927598 |
| Fam73a | 10,228 | 10,8784 | 0,0889415 | 0,6355 | 0,927598 |
| Uqcc1 | 23,0717 | 21,6897 | -0,0891184 | 0,63565 | 0,927598 |
| Ppp3cb | 29,4682 | 31,7863 | 0,109247 | 0,6359 | 0,927598 |
| Slc52a2 | 8,74531 | 9,39047 | 0,102689 | 0,6359 | 0,927598 |
| Pdha1 | 70,4624 | 66,3944 | -0,0857926 | 0,6359 | 0,927598 |
| Ly6g5b | 0,902379 | 0,734683 | -0,296611 | 0,6361 | 0,927598 |
| Pitpna | 27,2442 | 25,5848 | -0,0906619 | 0,6362 | 0,927598 |
| Nrde2 | 8,15913 | 7,62949 | -0,0968279 | 0,6362 | 0,927598 |
| Slc6a6 | 22,7144 | 21,4075 | -0,08549 | 0,6363 | 0,927598 |
| Psme3 | 129,804 | 122,487 | -0,0837045 | 0,63645 | 0,927598 |
| Arnt2 | 14,4264 | 13,5866 | -0,0865284 | 0,63655 | 0,927598 |
| Bloc1s4 | 9,4353 | 10,2587 | 0,120702 | 0,6366 | 0,927598 |

| | | | | | |
|---|---|---|---|---|---|
| Cdh13 | 1,29504 | 1,14552 | -0,176983 | 0,6366 | 0,927598 |
| Mir343 | 155,372 | 478,256 | 1,62205 | 0,63665 | 0,927598 |
| Mir6244 | 14,1302 | 34,8226 | 1,30124 | 0,63675 | 0,927598 |
| Dpysl3 | 18,2862 | 19,4219 | 0,0869301 | 0,6368 | 0,927598 |
| Zfp646 | 16,7915 | 17,8175 | 0,0855667 | 0,6368 | 0,927598 |
| Ckap4 | 65,8456 | 62,0905 | -0,0847154 | 0,63685 | 0,927598 |
| Neo1 | 35,3552 | 33,3424 | -0,0845666 | 0,6369 | 0,927598 |
| Xrcc1 | 65,3581 | 61,5946 | -0,0855618 | 0,63705 | 0,927598 |
| Ap1s2 | 7,87364 | 7,30809 | -0,107536 | 0,6371 | 0,927598 |
| 4930452B06Rik | 4,9796 | 4,59363 | -0,116395 | 0,63715 | 0,927598 |
| Ugt8a | 0,663958 | 0,753998 | 0,183467 | 0,63735 | 0,927598 |
| Rabepk | 14,6063 | 15,664 | 0,100857 | 0,63735 | 0,927598 |
| Dhx33 | 27,8763 | 29,5878 | 0,085963 | 0,63735 | 0,927598 |
| Lrpprc | 39,2826 | 37,0632 | -0,0839038 | 0,63735 | 0,927598 |
| Ppp2ca | 265,542 | 250,326 | -0,0851319 | 0,63735 | 0,927598 |
| Ccdc134 | 14,9063 | 15,9498 | 0,0976161 | 0,6375 | 0,927598 |
| Ints3 | 45,1363 | 47,8537 | 0,0843419 | 0,6375 | 0,927598 |
| Kcne3 | 1,02222 | 1,22142 | 0,256857 | 0,6376 | 0,927598 |
| Gins3 | 18,6569 | 17,4787 | -0,0941147 | 0,63765 | 0,927598 |
| Gapdh | 1,12849 | 1,31095 | 0,216221 | 0,63775 | 0,927598 |
| Fam184b | 0,75607 | 0,851423 | 0,171357 | 0,6378 | 0,927598 |
| Clec16a | 3,95802 | 4,24419 | 0,100708 | 0,63785 | 0,927598 |
| Mir1291 | 48,3671 | 111,788 | 1,20867 | 0,6381 | 0,927751 |
| Rnu73b | 1048,58 | 403,934 | -1,37625 | 0,6381 | 0,927751 |
| Nsun2 | 78,5849 | 74,092 | -0,0849349 | 0,63825 | 0,927864 |
| Pde7a | 24,0669 | 25,5347 | 0,0854086 | 0,63845 | 0,928017 |
| Edc3 | 25,1836 | 26,7725 | 0,0882671 | 0,6385 | 0,928017 |
| Tmem132a | 79,4697 | 84,2225 | 0,0838017 | 0,6387 | 0,928203 |
| Thnsl1 | 3,51405 | 3,2434 | -0,115627 | 0,63885 | 0,928283 |
| Mapk14 | 46,8498 | 44,1827 | -0,0845616 | 0,6389 | 0,928283 |
| Ppp1r2 | 11,2131 | 12,5771 | 0,165621 | 0,6391 | 0,928331 |
| Hnrnpul1 | 82,3989 | 77,7205 | -0,0843296 | 0,63915 | 0,928331 |
| Atf1 | 15,0197 | 14,0407 | -0,0972389 | 0,63915 | 0,928331 |
| Stau2 | 10,7058 | 10,0066 | -0,0974465 | 0,6393 | 0,928443 |
| Ubap2l | 101,619 | 95,4718 | -0,0900231 | 0,63945 | 0,928474 |
| Ndufaf1 | 20,9501 | 22,4167 | 0,0976154 | 0,6395 | 0,928474 |
| Mrpl44 | 35,4519 | 33,2323 | -0,0932777 | 0,6396 | 0,928474 |
| Mir7079 | 577,343 | 1416,12 | 1,29444 | 0,63965 | 0,928474 |
| Snhg5 | 123,249 | 130,865 | 0,0865025 | 0,63975 | 0,928474 |
| Cep44 | 21,2836 | 19,8751 | -0,0987855 | 0,6398 | 0,928474 |
| Ddx51 | 17,0591 | 16,054 | -0,0876079 | 0,63985 | 0,928474 |
| Pex12 | 6,80233 | 6,27392 | -0,116662 | 0,6399 | 0,928474 |
| Rnf13 | 14,4176 | 15,4735 | 0,10197 | 0,6401 | 0,928626 |
| Xrcc2 | 11,8157 | 12,5926 | 0,0918636 | 0,64015 | 0,928627 |
| Zbtb17 | 16,4464 | 14,4155 | -0,190155 | 0,6403 | 0,928739 |
| Ubtd1 | 6,94881 | 7,57016 | 0,123559 | 0,64055 | 0,928754 |
| Mns1 | 39,5008 | 42,9018 | 0,119153 | 0,64055 | 0,928754 |
| Cenpt | 24,1988 | 25,8199 | 0,0935488 | 0,6406 | 0,928754 |
| Vars | 96,5671 | 91,0888 | -0,0842576 | 0,6406 | 0,928754 |
| Txnl1 | 96,2207 | 90,8289 | -0,0831964 | 0,6409 | 0,929084 |

| | | | | | |
|---|---|---|---|---|---|
| Snd1 | 70,6284 | 66,5872 | -0,085005 | 0,641 | 0,929092 |
| Map2k5 | 18,6435 | 17,4809 | -0,0928925 | 0,64105 | 0,929092 |
| 4930432K21Rik | 0,89906 | 0,784792 | -0,196106 | 0,6412 | 0,929204 |
| Phospho2 | 11,4036 | 10,6234 | -0,102236 | 0,6413 | 0,929211 |
| Vdac1 | 202,127 | 190,805 | -0,0831598 | 0,64135 | 0,929211 |
| Lins | 6,25486 | 6,6778 | 0,0943949 | 0,64145 | 0,929251 |
| Agpat5 | 19,3578 | 20,5517 | 0,0863406 | 0,64195 | 0,929708 |
| Nt5c3b | 61,7022 | 56,9478 | -0,115683 | 0,64195 | 0,929708 |
| Ackr1 | 1,32059 | 1,53901 | 0,220813 | 0,6422 | 0,929708 |
| Nln | 10,2131 | 10,8751 | 0,0906146 | 0,6422 | 0,929708 |
| Mcm10 | 46,6207 | 49,373 | 0,0827521 | 0,6422 | 0,929708 |
| Hist1h2an | 1,16712 | 0,856594 | -0,446265 | 0,6422 | 0,929708 |
| Aifm2 | 2,15438 | 1,1631 | -0,889293 | 0,6424 | 0,929755 |
| Cdkal1 | 7,34705 | 7,90788 | 0,106126 | 0,64245 | 0,929755 |
| Tfdp1 | 59,6367 | 56,3355 | -0,082156 | 0,64245 | 0,929755 |
| Gm15417 | 0,807328 | 1,02889 | 0,34986 | 0,6426 | 0,929802 |
| Macrod2 | 2,17227 | 2,65114 | 0,287408 | 0,6427 | 0,929802 |
| Nap1l5 | 5,6754 | 6,29704 | 0,149952 | 0,6427 | 0,929802 |
| Marc2 | 11,1966 | 10,4062 | -0,105622 | 0,64305 | 0,930204 |
| Gm10791 | 0,6324 | 0,220787 | -1,51818 | 0,64335 | 0,930533 |
| Esam | 1,40407 | 1,22733 | -0,194089 | 0,64345 | 0,930572 |
| Chmp3 | 27,8225 | 26,1947 | -0,0869763 | 0,6438 | 0,930904 |
| Tmem106b | 5,97428 | 5,59569 | -0,0944486 | 0,64395 | 0,930904 |
| Khsrp | 64,2105 | 67,9907 | 0,0825279 | 0,6441 | 0,930904 |
| Rps4l | 73,4803 | 65,6173 | -0,163282 | 0,64415 | 0,930904 |
| Aamp | 118,208 | 111,577 | -0,0832926 | 0,64425 | 0,930904 |
| Dis3l2 | 13,8632 | 13,0144 | -0,0911552 | 0,6443 | 0,930904 |
| Fbxo34 | 7,25489 | 7,79967 | 0,10446 | 0,64435 | 0,930904 |
| Cdc42se2 | 53,933 | 50,9214 | -0,0828959 | 0,6444 | 0,930904 |
| Stk4 | 21,6558 | 20,4264 | -0,0843173 | 0,6445 | 0,930904 |
| Phyhd1 | 1,31581 | 1,51617 | 0,204479 | 0,64455 | 0,930904 |
| Zmynd11 | 28,2433 | 26,6269 | -0,085024 | 0,64455 | 0,930904 |
| Srsf7 | 266,561 | 251,058 | -0,0864462 | 0,64455 | 0,930904 |
| A330035P11Rik | 1,25599 | 1,40599 | 0,162761 | 0,6447 | 0,931016 |
| Ccdc115 | 17,2843 | 16,144 | -0,0984631 | 0,64495 | 0,931272 |
| Atxn7l3 | 42,6904 | 45,4512 | 0,0904077 | 0,64505 | 0,931311 |
| Ralgps1 | 1,22687 | 1,44376 | 0,234844 | 0,6452 | 0,931318 |
| Slc38a9 | 1,4642 | 1,35441 | -0,112456 | 0,6452 | 0,931318 |
| Irgq | 4,93797 | 4,60986 | -0,0991941 | 0,6455 | 0,931581 |
| Tln1 | 19,2188 | 18,122 | -0,0847761 | 0,6456 | 0,931581 |
| Med18 | 7,47715 | 6,87528 | -0,121069 | 0,6456 | 0,931581 |
| Pcbd2 | 16,8083 | 18,4966 | 0,138083 | 0,6457 | 0,931621 |
| Ttll3 | 1,62026 | 2,33336 | 0,526184 | 0,64605 | 0,932021 |
| Herc1 | 11,1166 | 11,7863 | 0,0844048 | 0,64615 | 0,932028 |
| Iars2 | 31,9407 | 29,6689 | -0,106444 | 0,6462 | 0,932028 |
| Ccdc173 | 2,56247 | 2,29801 | -0,157154 | 0,64675 | 0,932664 |
| Zfp622 | 17,0732 | 16,0336 | -0,0906364 | 0,64695 | 0,932664 |
| Nsd1 | 28,3581 | 30,0341 | 0,0828429 | 0,6471 | 0,932664 |
| Cntln | 5,20152 | 4,85407 | -0,0997379 | 0,6471 | 0,932664 |
| Hpcal1 | 15,4126 | 14,3733 | -0,100719 | 0,6471 | 0,932664 |

| | | | | | |
|---|---|---|---|---|---|
| Slk | 20,2371 | 21,4542 | 0,0842611 | 0,64715 | 0,932664 |
| Egfl7 | 14,5803 | 13,5111 | -0,109877 | 0,64715 | 0,932664 |
| Fam134a | 19,438 | 20,6968 | 0,0905274 | 0,64725 | 0,932673 |
| Ube2d2a | 23,2655 | 21,8726 | -0,0890662 | 0,64765 | 0,932673 |
| Cd151 | 23,2183 | 21,7991 | -0,0909984 | 0,64765 | 0,932673 |
| Prx | 2,26259 | 2,08325 | -0,11914 | 0,64765 | 0,932673 |
| Slc25a40 | 2,82359 | 2,57793 | -0,131317 | 0,6477 | 0,932673 |
| Gm5617 | 6,65056 | 5,91511 | -0,169072 | 0,6477 | 0,932673 |
| Ppp1r14c | 0,589918 | 0,683883 | 0,213236 | 0,64775 | 0,932673 |
| Cblb | 12,5848 | 11,8654 | -0,0849311 | 0,64795 | 0,932673 |
| Ankrd37 | 5,07308 | 3,89595 | -0,380885 | 0,64795 | 0,932673 |
| Mir1894 | 166,31 | 270,194 | 0,700126 | 0,648 | 0,932673 |
| Znhit6 | 17,8502 | 16,7776 | -0,0894086 | 0,6482 | 0,932673 |
| Cops2 | 66,7923 | 70,7701 | 0,0834575 | 0,6483 | 0,932673 |
| Rnf114 | 41,021 | 43,6318 | 0,0890179 | 0,64835 | 0,932673 |
| Mkks | 15,1721 | 14,1202 | -0,103664 | 0,6484 | 0,932673 |
| Tmcc1 | 7,96638 | 8,46741 | 0,087997 | 0,6485 | 0,932673 |
| 1700020L24Rik | 0,840873 | 1,02822 | 0,290186 | 0,64855 | 0,932673 |
| Pgam1 | 14,3048 | 15,2953 | 0,0965859 | 0,64875 | 0,932673 |
| Pibf1 | 7,39244 | 8,78992 | 0,249798 | 0,6488 | 0,932673 |
| Tmem47 | 16,8419 | 17,8719 | 0,0856384 | 0,6488 | 0,932673 |
| Zfp74 | 7,29793 | 7,80718 | 0,0973152 | 0,64895 | 0,932673 |
| Eif1b | 117,802 | 124,816 | 0,0834348 | 0,6491 | 0,932673 |
| Mir6990 | 37,4593 | 81,7667 | 1,12619 | 0,64915 | 0,932673 |
| Tsfm | 39,7401 | 37,33 | -0,0902566 | 0,64915 | 0,932673 |
| Kif15 | 37,0239 | 39,2197 | 0,0831195 | 0,6492 | 0,932673 |
| Gm3258 | 36,4268 | 33,1634 | -0,13541 | 0,64925 | 0,932673 |
| Gspt2 | 8,10802 | 8,70307 | 0,102174 | 0,64935 | 0,932673 |
| Fbxw5 | 20,6429 | 19,3945 | -0,0899969 | 0,64945 | 0,932673 |
| Cep63 | 17,0437 | 18,1137 | 0,0878417 | 0,64955 | 0,932673 |
| Rab5a | 24,2572 | 22,837 | -0,0870392 | 0,6496 | 0,932673 |
| Slc25a20 | 14,1231 | 13,1854 | -0,0991244 | 0,6496 | 0,932673 |
| 1700123O20Rik | 33,7659 | 28,3892 | -0,250225 | 0,64965 | 0,932673 |
| Atp7a | 13,7293 | 14,6791 | 0,0819444 | 0,64975 | 0,932673 |
| Sephs1 | 44,7493 | 42,2447 | -0,0830956 | 0,64975 | 0,932673 |
| Rnf165 | 5,39149 | 5,74234 | 0,090956 | 0,6498 | 0,932673 |
| Ppfia3 | 0,935101 | 1,0351 | 0,146582 | 0,65005 | 0,932673 |
| Dhx38 | 30,7995 | 29,0748 | -0,0831377 | 0,65015 | 0,932673 |
| Zfp868 | 11,5191 | 10,779 | -0,0958108 | 0,65015 | 0,932673 |
| Chst8 | 1,17847 | 1,04495 | -0,173486 | 0,6502 | 0,932673 |
| Gatad2a | 38,0199 | 40,1763 | 0,0795902 | 0,65025 | 0,932673 |
| Ghr | 8,37462 | 7,72164 | -0,117116 | 0,6503 | 0,932673 |
| Ppm1m | 27,6532 | 29,3955 | 0,0881492 | 0,6505 | 0,932673 |
| Acox3 | 2,65891 | 2,8856 | 0,118038 | 0,65055 | 0,932673 |
| Ranbp3 | 75,2007 | 71,0843 | -0,0812151 | 0,65055 | 0,932673 |
| Rfxap | 14,07 | 13,1834 | -0,0939007 | 0,65065 | 0,932673 |
| Gmds | 13,5495 | 12,616 | -0,102985 | 0,65065 | 0,932673 |
| Foxred1 | 25,5428 | 24,0924 | -0,0843416 | 0,65075 | 0,932673 |
| Polr3h | 13,9132 | 11,9518 | -0,219222 | 0,6508 | 0,932673 |
| Slc25a51 | 22,1914 | 23,4898 | 0,082034 | 0,65095 | 0,932673 |

| | | | | | |
|---|---|---|---|---|---|
| Blcap | 16,8581 | 13,9879 | -0,269263 | 0,65105 | 0,932673 |
| Sec11a | 111,331 | 105,093 | -0,0831885 | 0,6511 | 0,932673 |
| 2310003H01Rik | 9,50625 | 8,89497 | -0,0958866 | 0,65135 | 0,932673 |
| Arhgap23 | 1,8248 | 1,677 | -0,121855 | 0,65135 | 0,932673 |
| Gm10638 | 0,478741 | 0,766776 | 0,679561 | 0,6514 | 0,932673 |
| Pdlim2 | 1,02266 | 1,17865 | 0,204803 | 0,6515 | 0,932673 |
| Papolg | 9,74148 | 10,3744 | 0,0908096 | 0,6515 | 0,932673 |
| Pink1 | 12,5938 | 13,4507 | 0,0949657 | 0,65155 | 0,932673 |
| Timp2 | 18,3008 | 19,3895 | 0,0833648 | 0,6516 | 0,932673 |
| Srpr | 38,1159 | 40,6795 | 0,0939097 | 0,6517 | 0,932673 |
| Nob1 | 63,9366 | 60,4162 | -0,081708 | 0,6518 | 0,932673 |
| Sema3b | 0,870607 | 0,776577 | -0,164893 | 0,65185 | 0,932673 |
| Ddx19a | 34,6024 | 32,6471 | -0,0839197 | 0,6519 | 0,932673 |
| Arpc5l | 74,7321 | 70,4994 | -0,0841168 | 0,65195 | 0,932673 |
| Rab3ip | 13,2001 | 14,0603 | 0,0910877 | 0,652 | 0,932673 |
| Rpl27 | 4,60071 | 4,0618 | -0,179738 | 0,6521 | 0,932673 |
| Syap1 | 21,1693 | 24,8842 | 0,233257 | 0,6522 | 0,932673 |
| Bcl2l13 | 26,3899 | 24,972 | -0,0796767 | 0,6523 | 0,932673 |
| Efhd2 | 7,74193 | 7,20015 | -0,104665 | 0,65235 | 0,932673 |
| Rnf135 | 1,25683 | 1,10438 | -0,18655 | 0,65235 | 0,932673 |
| Pde6d | 40,3098 | 37,8414 | -0,0911657 | 0,65245 | 0,932673 |
| Phkg2 | 18,5874 | 17,3285 | -0,101178 | 0,6525 | 0,932673 |
| Mir1949 | 812,252 | 346,077 | -1,23083 | 0,6525 | 0,932673 |
| BC024978 | 4,29188 | 5,3318 | 0,313014 | 0,65255 | 0,932673 |
| Gpr135 | 0,648479 | 0,776468 | 0,259866 | 0,65255 | 0,932673 |
| Nxt1 | 74,7011 | 70,4617 | -0,084291 | 0,65255 | 0,932673 |
| Zfp954 | 5,42797 | 5,00069 | -0,118286 | 0,65275 | 0,932673 |
| Trim28 | 401,161 | 377,215 | -0,0887945 | 0,6528 | 0,932673 |
| Picalm | 55,4885 | 58,6769 | 0,0806056 | 0,65285 | 0,932673 |
| Men1 | 38,0416 | 35,9195 | -0,0828083 | 0,6529 | 0,932673 |
| Ccdc86 | 24,4117 | 23,0441 | -0,0831699 | 0,6529 | 0,932673 |
| Anp32a | 332,133 | 351,852 | 0,0832056 | 0,65305 | 0,932751 |
| Tsr3 | 31,8696 | 29,9091 | -0,0915984 | 0,6531 | 0,932751 |
| Usp33 | 22,2539 | 23,5679 | 0,0827693 | 0,65335 | 0,932874 |
| Osbpl9 | 20,8189 | 23,5027 | 0,174936 | 0,6534 | 0,932874 |
| 2700060E02Rik | 71,4355 | 62,9872 | -0,181582 | 0,65345 | 0,932874 |
| Zbtb45 | 9,40756 | 8,75876 | -0,103094 | 0,65355 | 0,932874 |
| Adora2b | 0,756518 | 0,642488 | -0,235705 | 0,65355 | 0,932874 |
| Smim19 | 8,47778 | 10,1461 | 0,259174 | 0,65365 | 0,932881 |
| Blzf1 | 6,88188 | 7,34303 | 0,0935728 | 0,6537 | 0,932881 |
| Dpcd | 21,5285 | 23,2563 | 0,111373 | 0,6538 | 0,932887 |
| Pfdn4 | 103,346 | 97,5704 | -0,0829729 | 0,65385 | 0,932887 |
| Rap1b | 67,2448 | 63,6057 | -0,0802659 | 0,65395 | 0,932926 |
| Zfp830 | 11,1098 | 10,4512 | -0,0881622 | 0,65465 | 0,93373 |
| Sept2 | 74,5116 | 66,4813 | -0,0804641 | 0,6548 | 0,93373 |
| Ipo8 | 19,3801 | 18,3376 | -0,0797764 | 0,6549 | 0,93373 |
| Cog2 | 16,6614 | 15,6785 | -0,0877248 | 0,6549 | 0,93373 |
| Usb1 | 14,6434 | 13,6931 | -0,0968062 | 0,6549 | 0,93373 |
| Topors | 32,7009 | 34,6194 | 0,0822526 | 0,65495 | 0,93373 |
| Exoc3 | 23,6674 | 22,3837 | -0,0804515 | 0,6555 | 0,93428 |

| | | | | | |
|---|---|---|---|---|---|
| Cdk5rap3 | 31,1208 | 29,3198 | -0,0860039 | 0,6556 | 0,93428 |
| Dohh | 34,8972 | 32,8599 | -0,0867829 | 0,6556 | 0,93428 |
| 1600012H06Rik | 8,34661 | 7,02332 | -0,249038 | 0,65565 | 0,93428 |
| Tcf25 | 72,6009 | 68,6997 | -0,0796849 | 0,6557 | 0,93428 |
| Zfp688 | 11,2963 | 10,3894 | -0,120742 | 0,6561 | 0,934746 |
| Gm13375 | 3,36946 | 3,03939 | -0,148733 | 0,6562 | 0,934758 |
| E130307A14Rik | 8,96492 | 8,22287 | -0,124648 | 0,65635 | 0,934758 |
| Mir6905 | 207,945 | 153,099 | -0,441738 | 0,65635 | 0,934758 |
| Acaa1a | 16,8046 | 15,7391 | -0,0945006 | 0,6564 | 0,934758 |
| Sf3a3 | 150,34 | 142,264 | -0,0796658 | 0,6565 | 0,934797 |
| Wbscr16 | 19,5417 | 18,3976 | -0,0870439 | 0,6566 | 0,934803 |
| Spag1 | 2,86227 | 2,62538 | -0,124632 | 0,65665 | 0,934803 |
| Capza1 | 7,64725 | 7,15106 | -0,0967838 | 0,65675 | 0,934841 |
| Pithd1 | 37,7988 | 35,6147 | -0,0858642 | 0,65705 | 0,935132 |
| B3gntl1 | 3,72689 | 4,42928 | 0,249101 | 0,6571 | 0,935132 |
| Scap | 20,7568 | 21,955 | 0,0809662 | 0,65725 | 0,935242 |
| Zfp595 | 1,1631 | 1,05003 | -0,147547 | 0,65735 | 0,93528 |
| Mri1 | 12,7731 | 13,6343 | 0,0941364 | 0,6576 | 0,935508 |
| C1qbp | 320,801 | 303,152 | -0,0816367 | 0,65775 | 0,935508 |
| Trpc2 | 1,96218 | 2,40501 | 0,293587 | 0,65785 | 0,935508 |
| Snora74a | 15,2613 | 7,52063 | -1,02095 | 0,6579 | 0,935508 |
| Fbxo7 | 8,67398 | 9,31845 | 0,103396 | 0,65815 | 0,935508 |
| Tpp1 | 13,0221 | 13,8198 | 0,0857722 | 0,65815 | 0,935508 |
| Prr13 | 15,1342 | 14,0612 | -0,106095 | 0,6582 | 0,935508 |
| Rabep2 | 8,32118 | 8,9193 | 0,100142 | 0,6584 | 0,935508 |
| 1700025G04Rik | 6,71819 | 6,33615 | -0,084465 | 0,6585 | 0,935508 |
| Palb2 | 9,60351 | 10,244 | 0,0931402 | 0,65855 | 0,935508 |
| Gss | 8,01033 | 7,43417 | -0,107689 | 0,65855 | 0,935508 |
| Rnf145 | 58,331 | 61,5954 | 0,0785606 | 0,6587 | 0,935508 |
| Anapc5 | 157,222 | 148,878 | -0,0786728 | 0,6587 | 0,935508 |
| Tor1aip2 | 64,9357 | 60,6924 | -0,0974966 | 0,6587 | 0,935508 |
| Atp6v0b | 47,521 | 44,7141 | -0,0878353 | 0,65875 | 0,935508 |
| Enkd1 | 27,5271 | 25,8155 | -0,092614 | 0,65875 | 0,935508 |
| Fchsd1 | 1,38597 | 1,26036 | -0,137065 | 0,65875 | 0,935508 |
| Rpia | 16,9672 | 15,9054 | -0,0932347 | 0,6589 | 0,935552 |
| Snord49b | 1742,97 | 621,153 | -1,48853 | 0,65895 | 0,935552 |
| Fam219aos | 1,41222 | 1,09214 | -0,370807 | 0,659 | 0,935552 |
| Shoc2 | 16,7433 | 18,2105 | 0,121187 | 0,65935 | 0,935842 |
| Prr7 | 5,00752 | 4,56084 | -0,134796 | 0,65935 | 0,935842 |
| LOC100504703 | 15,5061 | 17,5985 | 0,182615 | 0,65955 | 0,935854 |
| Fam220a | 16,5881 | 15,5919 | -0,0893508 | 0,65955 | 0,935854 |
| Rnf216 | 22,6513 | 23,9498 | 0,0804175 | 0,65965 | 0,935854 |
| Gstz1 | 17,3481 | 15,7737 | -0,137251 | 0,65965 | 0,935854 |
| Secisbp2 | 12,1723 | 11,4622 | -0,0867124 | 0,6598 | 0,93586 |
| Rbm4 | 11,8485 | 9,99985 | -0,244731 | 0,6598 | 0,93586 |
| Rabl6 | 49,3487 | 46,7618 | -0,0776837 | 0,65995 | 0,935877 |
| Ubqln2 | 43,9314 | 41,5833 | -0,0792476 | 0,66 | 0,935877 |
| Arhgap1 | 18,597 | 17,5467 | -0,0838697 | 0,6601 | 0,935877 |
| Rccd1 | 8,66277 | 9,2799 | 0,0992805 | 0,6602 | 0,935877 |
| Prkci | 35,4866 | 33,5963 | -0,0789721 | 0,6602 | 0,935877 |

| | | | | | |
|---|---|---|---|---|---|
| Npl | 4,27106 | 3,88181 | -0,137867 | 0,66025 | 0,935877 |
| Grik2 | 0,796183 | 0,882098 | 0,147839 | 0,66035 | 0,935915 |
| Bag3 | 15,6272 | 16,6108 | 0,0880619 | 0,66075 | 0,936379 |
| Cir1 | 1,74548 | 1,9459 | 0,156811 | 0,6609 | 0,936455 |
| Dis3 | 46,2626 | 49,1014 | 0,0859171 | 0,66095 | 0,936455 |
| Hps4 | 12,5714 | 13,3662 | 0,0884451 | 0,66105 | 0,936467 |
| Ccdc32 | 10,7616 | 10,0288 | -0,101735 | 0,66115 | 0,936467 |
| Ppp1r18 | 23,8435 | 25,2527 | 0,0828426 | 0,66125 | 0,936467 |
| Ganab | 123,764 | 130,693 | 0,0785885 | 0,66125 | 0,936467 |
| Cadm3 | 3,08708 | 2,86315 | -0,108641 | 0,66145 | 0,936581 |
| Uba2 | 164,843 | 156,232 | -0,0774064 | 0,66155 | 0,936581 |
| Pgrmc1 | 116,469 | 110,279 | -0,0787839 | 0,66155 | 0,936581 |
| Cdca3 | 122,246 | 115,661 | -0,0798889 | 0,6617 | 0,936653 |
| Cpox | 9,89318 | 10,5222 | 0,0889359 | 0,66185 | 0,936653 |
| Chst10 | 5,32443 | 5,70316 | 0,099134 | 0,66195 | 0,936653 |
| Tusc1 | 1,83514 | 1,61905 | -0,180739 | 0,662 | 0,936653 |
| Trabd | 24,3677 | 22,9762 | -0,0848265 | 0,66215 | 0,936653 |
| Snord23 | 79,0195 | 46,1941 | -0,774499 | 0,66225 | 0,936653 |
| AI837181 | 22,7369 | 21,3828 | -0,0885841 | 0,66235 | 0,936653 |
| Pitpnb | 61,4618 | 58,2008 | -0,0786516 | 0,6624 | 0,936653 |
| Ipo11 | 39,3185 | 37,2226 | -0,0790297 | 0,66245 | 0,936653 |
| Wdr53 | 11,5819 | 12,5122 | 0,111467 | 0,6625 | 0,936653 |
| Zdhhc16 | 26,2857 | 23,8575 | -0,139833 | 0,6625 | 0,936653 |
| Ggps1 | 16,3604 | 17,3689 | 0,0862989 | 0,66255 | 0,936653 |
| Snord83b | 572,801 | 226,093 | -1,34112 | 0,66255 | 0,936653 |
| Klf12 | 6,4339 | 6,83177 | 0,0865658 | 0,66265 | 0,936691 |
| Rnf115 | 34,6135 | 32,7151 | -0,0813767 | 0,6628 | 0,936735 |
| Tpx2 | 93,3655 | 88,4529 | -0,0779803 | 0,66285 | 0,936735 |
| Gle1 | 25,9782 | 24,5662 | -0,0806292 | 0,6629 | 0,936735 |
| Dedd2 | 2,76314 | 3,04478 | 0,14003 | 0,6633 | 0,936872 |
| Ipo9 | 69,8562 | 66,1363 | -0,0789457 | 0,66335 | 0,936872 |
| Snx19 | 13,1556 | 13,9217 | 0,0816625 | 0,66345 | 0,936872 |
| Colgalt2 | 1,18248 | 1,07142 | -0,142284 | 0,66345 | 0,936872 |
| Tbcb | 82,4883 | 77,9994 | -0,0807251 | 0,6635 | 0,936872 |
| Lsg1 | 49,4798 | 46,8822 | -0,0777979 | 0,66355 | 0,936872 |
| Snora28 | 112,804 | 360,768 | 1,67725 | 0,6636 | 0,936872 |
| Ssbp3 | 87,4802 | 92,2782 | 0,0770341 | 0,66365 | 0,936872 |
| Trmt1l | 21,1166 | 22,3643 | 0,0828196 | 0,6638 | 0,936872 |
| Kif22 | 90,6358 | 85,8589 | -0,0781138 | 0,6638 | 0,936872 |
| Mapre1 | 45,9535 | 43,5238 | -0,0783718 | 0,6638 | 0,936872 |
| Mir1905 | 351,581 | 778,692 | 1,14719 | 0,66395 | 0,936948 |
| Gas5 | 183,798 | 194,193 | 0,0793647 | 0,664 | 0,936948 |
| Cnst | 9,13557 | 9,69056 | 0,0850856 | 0,66415 | 0,937024 |
| Unc50 | 28,9287 | 27,2429 | -0,0866196 | 0,6642 | 0,937024 |
| Snora61 | 449,214 | 473,075 | 0,0746662 | 0,6643 | 0,937062 |
| Dgcr14 | 15,4118 | 16,3521 | 0,0854397 | 0,66445 | 0,93717 |
| Uimc1 | 14,9017 | 15,7975 | 0,0842201 | 0,6648 | 0,93729 |
| Slc6a15 | 1,15328 | 1,27328 | 0,14281 | 0,66485 | 0,93729 |
| Nat6 | 7,81004 | 8,38441 | 0,102379 | 0,665 | 0,93729 |
| Dhx9 | 219,123 | 231,963 | 0,0821519 | 0,665 | 0,93729 |

| | | | | | |
|---|---|---|---|---|---|
| Wdr70 | 21,9797 | 20,7136 | -0,0855941 | 0,66505 | 0,93729 |
| Ercc8 | 17,3931 | 16,3624 | -0,0881296 | 0,66515 | 0,93729 |
| Raver1-fdx1l | 0,607062 | 1,18367 | 0,963346 | 0,66525 | 0,93729 |
| D2hgdh | 2,37264 | 2,17119 | -0,128005 | 0,6654 | 0,93729 |
| Idi1 | 2,02114 | 1,84192 | -0,133957 | 0,6654 | 0,93729 |
| Rps15 | 1389,34 | 1132,58 | -0,294791 | 0,6654 | 0,93729 |
| Zfp142 | 7,27126 | 8,39251 | 0,206897 | 0,66545 | 0,93729 |
| 1110032F04Rik | 1,79462 | 1,62622 | -0,14215 | 0,66555 | 0,93729 |
| Nr3c1 | 1,92809 | 2,07593 | 0,106591 | 0,6656 | 0,93729 |
| H2-T24 | 1,64445 | 1,8348 | 0,158016 | 0,66565 | 0,93729 |
| Skiv2l | 24,5123 | 25,9035 | 0,0796442 | 0,66565 | 0,93729 |
| Rab11fip3 | 15,8757 | 14,9945 | -0,0823899 | 0,6658 | 0,93729 |
| Rgs16 | 4,54768 | 4,20321 | -0,11364 | 0,6658 | 0,93729 |
| 1700003F12Rik | 1,14262 | 0,867445 | -0,397497 | 0,66585 | 0,93729 |
| Tgfbrap1 | 7,29996 | 6,87562 | -0,0863988 | 0,66615 | 0,937517 |
| Hltf | 6,68766 | 7,11424 | 0,0892074 | 0,6662 | 0,937517 |
| Usp15 | 36,9781 | 39,0296 | 0,077897 | 0,66625 | 0,937517 |
| Anapc4 | 58,2938 | 55,2412 | -0,0775991 | 0,6664 | 0,937517 |
| Ccdc130 | 4,62027 | 5,00412 | 0,11514 | 0,66645 | 0,937517 |
| Edrf1 | 13,056 | 13,7983 | 0,0797757 | 0,66645 | 0,937517 |
| Ppapdc1b | 9,85625 | 9,14556 | -0,107967 | 0,66655 | 0,937555 |
| Zfp709 | 1,5072 | 1,64739 | 0,128315 | 0,66675 | 0,937598 |
| Rbm7 | 47,9784 | 45,4518 | -0,0780465 | 0,66675 | 0,937598 |
| Dazap2 | 80,1387 | 75,9124 | -0,0781628 | 0,6668 | 0,937598 |
| Pik3ip1 | 2,97136 | 5,03856 | 0,761888 | 0,66705 | 0,93762 |
| Galnt10 | 9,26651 | 9,82412 | 0,0843024 | 0,66705 | 0,93762 |
| Mcm3ap | 21,8863 | 23,163 | 0,0817957 | 0,6671 | 0,93762 |
| Cep162 | 6,17408 | 6,55007 | 0,0852855 | 0,6672 | 0,93762 |
| Slc16a6 | 3,41224 | 3,69752 | 0,115839 | 0,66735 | 0,93762 |
| Snord100 | 2477 | 1866,39 | -0,408348 | 0,66735 | 0,93762 |
| Scarna3a | 13,6796 | 22,8583 | 0,740689 | 0,6674 | 0,93762 |
| Keap1 | 15,6806 | 14,7701 | -0,0863016 | 0,6674 | 0,93762 |
| Tcn2 | 8,65544 | 9,27491 | 0,0997271 | 0,66775 | 0,937906 |
| Khdrbs1 | 128,749 | 121,882 | -0,0790773 | 0,66775 | 0,937906 |
| Tnpo1 | 44,5129 | 46,9301 | 0,0762903 | 0,66785 | 0,937944 |
| Pex3 | 11,8976 | 12,8492 | 0,110999 | 0,66825 | 0,938235 |
| Smyd3 | 3,08369 | 2,8547 | -0,111321 | 0,66825 | 0,938235 |
| Vamp4 | 8,02543 | 8,57539 | 0,0956245 | 0,6683 | 0,938235 |
| Sars | 47,8524 | 45,3223 | -0,0783723 | 0,66835 | 0,938235 |
| Rbp4 | 4,56947 | 5,0837 | 0,153851 | 0,66845 | 0,93824 |
| Pfas | 25,5873 | 26,9951 | 0,0772729 | 0,6685 | 0,93824 |
| Frem2 | 0,983637 | 1,05866 | 0,106041 | 0,6686 | 0,938278 |
| Hsph1 | 140,927 | 148,865 | 0,0790609 | 0,6687 | 0,938289 |
| Gm16023 | 1,26817 | 1,63817 | 0,36933 | 0,6688 | 0,938289 |
| Mcm8 | 19,7592 | 20,8653 | 0,0785778 | 0,66885 | 0,938289 |
| Gnpda2 | 5,5422 | 5,99298 | 0,112814 | 0,6689 | 0,938289 |
| Zfp747 | 1,38473 | 1,53205 | 0,145856 | 0,66915 | 0,938504 |
| Xpo5 | 64,5369 | 68,0296 | 0,0760394 | 0,6692 | 0,938504 |
| Selo | 6,01354 | 7,04235 | 0,227843 | 0,6693 | 0,938542 |
| Pomc | 1,93858 | 2,20112 | 0,183235 | 0,66945 | 0,938547 |

| | | | | | |
|---|---|---|---|---|---|
| Nup98 | 31,5981 | 33,377 | 0,079017 | 0,66945 | 0,938547 |
| Gtdc1 | 7,2113 | 7,71226 | 0,0968935 | 0,6696 | 0,938557 |
| Pigq | 29,4595 | 31,0769 | 0,0771118 | 0,66965 | 0,938557 |
| Tprkb | 26,1141 | 27,6308 | 0,0814468 | 0,66975 | 0,938557 |
| Dync1li2 | 36,3809 | 34,4839 | -0,0772612 | 0,66975 | 0,938557 |
| Zfp266 | 34,6928 | 38,7823 | 0,0771657 | 0,66985 | 0,938561 |
| C330006A16Rik | 12,4248 | 11,6653 | -0,0910001 | 0,66995 | 0,938561 |
| Car11 | 2,73578 | 2,46777 | -0,148741 | 0,67005 | 0,938561 |
| Gnb1l | 2,94245 | 2,72785 | -0,109257 | 0,6701 | 0,938561 |
| Dnajc28 | 1,81733 | 1,98612 | 0,128129 | 0,67015 | 0,938561 |
| Ccnl2 | 67,1823 | 70,899 | 0,0776842 | 0,6704 | 0,938561 |
| Gfod2 | 12,7342 | 13,5735 | 0,0920867 | 0,67045 | 0,938561 |
| Chtf8 | 118,313 | 112,249 | -0,0759102 | 0,67055 | 0,938561 |
| Fhit | 0,912695 | 1,21231 | 0,409556 | 0,67065 | 0,938561 |
| Anapc15 | 28,4239 | 26,594 | -0,0960044 | 0,67065 | 0,938561 |
| Wbscr27 | 2,53661 | 2,32256 | -0,127185 | 0,67065 | 0,938561 |
| Rbm4b | 17,3455 | 18,4558 | 0,0895147 | 0,67075 | 0,938561 |
| Ubfd1 | 46,3803 | 48,8964 | 0,0762169 | 0,67075 | 0,938561 |
| Airn | 16,0078 | 21,8281 | 0,447415 | 0,67085 | 0,938561 |
| Map4k3 | 12,6117 | 11,9037 | -0,0833578 | 0,67085 | 0,938561 |
| Rfc4 | 96,5542 | 84,4207 | -0,193742 | 0,6711 | 0,938776 |
| Adora2a | 0,957453 | 0,849158 | -0,17317 | 0,67115 | 0,938776 |
| Gm15760 | 1,05191 | 1,16206 | 0,143676 | 0,6713 | 0,938851 |
| Naa16 | 10,5822 | 11,2134 | 0,0835787 | 0,67135 | 0,938851 |
| Rprd1a | 13,7253 | 14,5051 | 0,0797309 | 0,6715 | 0,938856 |
| Dtd1 | 13,6707 | 12,7503 | -0,10056 | 0,6715 | 0,938856 |
| Slc25a29 | 2,08487 | 1,88547 | -0,145034 | 0,6716 | 0,938893 |
| Lyrm9 | 1,53968 | 1,41488 | -0,121943 | 0,6718 | 0,939038 |
| Tusc3 | 37,2225 | 39,4558 | 0,084062 | 0,67185 | 0,939038 |
| Naprt1 | 0,939902 | 0,820341 | -0,196287 | 0,67205 | 0,93916 |
| Taf15 | 152,359 | 160,666 | 0,0765916 | 0,6722 | 0,93916 |
| Ccdc34os | 0,271441 | 0,717626 | 1,40259 | 0,6723 | 0,93916 |
| Fkbp15 | 7,99679 | 8,47649 | 0,0840456 | 0,6724 | 0,93916 |
| Clptm1l | 50,5062 | 53,3273 | 0,0784152 | 0,6724 | 0,93916 |
| Ehbp1l1 | 1,2603 | 1,37676 | 0,127507 | 0,67245 | 0,93916 |
| Drp2 | 1,05294 | 1,14392 | 0,119558 | 0,67245 | 0,93916 |
| Gfer | 24,074 | 22,726 | -0,0831358 | 0,67265 | 0,939305 |
| Pi4kb | 18,8042 | 17,1821 | -0,130148 | 0,6727 | 0,939305 |
| 2410002F23Rik | 45,6608 | 43,2456 | -0,0784029 | 0,67285 | 0,939315 |
| Chid1 | 7,70574 | 8,23737 | 0,0962499 | 0,6729 | 0,939315 |
| Dnaja2 | 126,403 | 119,876 | -0,0764888 | 0,67295 | 0,939315 |
| Mtfp1 | 6,61885 | 7,1776 | 0,116922 | 0,673 | 0,939315 |
| 2810001G20Rik | 17,7258 | 16,4504 | -0,107726 | 0,6732 | 0,939395 |
| Chfr | 23,6597 | 24,9625 | 0,0773325 | 0,67325 | 0,939395 |
| Isca1 | 12,2373 | 11,4585 | -0,0948644 | 0,6733 | 0,939395 |
| Kdm5c | 27,6113 | 26,2101 | -0,0751396 | 0,67335 | 0,939395 |
| Cenpl | 19,6313 | 20,8247 | 0,085144 | 0,67345 | 0,939432 |
| Hectd3 | 5,59517 | 5,95144 | 0,0890584 | 0,6736 | 0,939539 |
| Dph2 | 11,8742 | 11,1662 | -0,0886855 | 0,6737 | 0,939576 |
| Trim71 | 10,4944 | 11,1037 | 0,0814172 | 0,67385 | 0,939683 |

| | | | | | |
|---|---|---|---|---|---|
| 4933439C10Rik | 4,13091 | 5,1569 | 0,320044 | 0,67395 | 0,939721 |
| Ifnar1 | 12,8693 | 13,5692 | 0,0763987 | 0,67415 | 0,939833 |
| Ogdh | 30,7344 | 29,1407 | -0,0768175 | 0,67425 | 0,939833 |
| Utp11l | 90,3007 | 85,4999 | -0,0788142 | 0,67425 | 0,939833 |
| Loxl2 | 19,1306 | 18,1405 | -0,0766624 | 0,67445 | 0,940009 |
| Zfp511 | 22,2598 | 19,3308 | -0,203537 | 0,67465 | 0,940186 |
| Zkscan14 | 2,67262 | 2,43442 | -0,134675 | 0,6748 | 0,940293 |
| Patl1 | 17,9067 | 18,9077 | 0,0784751 | 0,6749 | 0,94033 |
| Kif7 | 8,76792 | 7,93791 | -0,143476 | 0,675 | 0,940367 |
| Cept1 | 14,6154 | 15,4794 | 0,0828658 | 0,67515 | 0,940372 |
| Ctnnb1 | 439,975 | 415,437 | -0,0827935 | 0,67515 | 0,940372 |
| Cdh5 | 1,9681 | 1,80602 | -0,123986 | 0,67535 | 0,940548 |
| Lyrm5 | 9,14997 | 9,84309 | 0,105344 | 0,67595 | 0,941179 |
| Zfand6 | 46,8072 | 44,3257 | -0,0785871 | 0,67595 | 0,94118 |
| Zfp869 | 18,3675 | 19,4057 | 0,0793219 | 0,6763 | 0,941462 |
| Lipe | 2,95613 | 2,68787 | -0,137251 | 0,6763 | 0,941462 |
| Fuz | 5,34302 | 6,66299 | 0,318514 | 0,6764 | 0,941499 |
| Tfpt | 22,9722 | 21,5366 | -0,0930992 | 0,67655 | 0,941502 |
| Akap13 | 7,57886 | 8,00244 | 0,0784595 | 0,6766 | 0,941502 |
| Napepld | 3,34584 | 3,10579 | -0,107404 | 0,67685 | 0,941502 |
| Nup62 | 87,9601 | 82,7703 | -0,0877368 | 0,6769 | 0,941502 |
| Mcts2 | 45,9014 | 40,697 | -0,173615 | 0,6769 | 0,941502 |
| 2810006K23Rik | 14,6549 | 13,7555 | -0,0913795 | 0,677 | 0,941502 |
| Nsfl1c | 64,0555 | 60,7578 | -0,0762544 | 0,67715 | 0,941502 |
| Gin1 | 4,36825 | 4,03246 | -0,115397 | 0,6772 | 0,941502 |
| 1110006O24Rik | 1,66235 | 1,40908 | -0,23847 | 0,67745 | 0,941502 |
| Dok1 | 6,02028 | 5,58194 | -0,109064 | 0,6775 | 0,941502 |
| Cpne1 | 50,9267 | 47,29 | -0,106889 | 0,67755 | 0,941502 |
| Etnk1 | 20,2097 | 21,3071 | 0,076285 | 0,6777 | 0,941502 |
| Hspb1 | 1,14437 | 0,954279 | -0,262071 | 0,6777 | 0,941502 |
| Ttc37 | 11,4034 | 12,0372 | 0,0780377 | 0,67785 | 0,941502 |
| Mycbp | 21,7547 | 23,0899 | 0,0859358 | 0,6779 | 0,941502 |
| Csrnp2 | 8,80921 | 9,32944 | 0,0827783 | 0,6779 | 0,941502 |
| Cnnm2 | 1,7594 | 1,91889 | 0,125189 | 0,678 | 0,941502 |
| Nsf | 20,3354 | 19,2683 | -0,0777644 | 0,678 | 0,941502 |
| Psph | 16,4633 | 15,4778 | -0,0890546 | 0,67805 | 0,941502 |
| Cpt1a | 0,824723 | 0,910449 | 0,142668 | 0,6781 | 0,941502 |
| Fastkd3 | 18,6834 | 19,761 | 0,0808946 | 0,6782 | 0,941502 |
| Mcph1 | 10,6749 | 11,3087 | 0,0832056 | 0,67835 | 0,941502 |
| Sdf2l1 | 38,7868 | 41,0984 | 0,0835194 | 0,6784 | 0,941502 |
| Hn1l | 132,87 | 126,056 | -0,075957 | 0,67845 | 0,941502 |
| Kctd13 | 8,10457 | 8,69769 | 0,101896 | 0,67855 | 0,941502 |
| Lcmt1 | 32,7007 | 30,8826 | -0,0825294 | 0,67855 | 0,941502 |
| Pqbp1 | 93,1198 | 88,3193 | -0,0763593 | 0,6787 | 0,941502 |
| Gstm4 | 6,75507 | 6,25383 | -0,111231 | 0,67875 | 0,941502 |
| Alg10b | 33,0825 | 34,8412 | 0,0747279 | 0,6788 | 0,941502 |
| Arglu1 | 114,117 | 108,269 | -0,0758879 | 0,6788 | 0,941502 |
| Senp2 | 25,0157 | 26,3324 | 0,0740022 | 0,6789 | 0,941502 |
| Eif4e | 47,9199 | 45,4923 | -0,0750014 | 0,6789 | 0,941502 |
| Zfp341 | 2,86467 | 2,6465 | -0,114285 | 0,6789 | 0,941502 |

| | | | | | |
|---|---|---|---|---|---|
| Hdac3 | 86,7749 | 82,4187 | -0,0743061 | 0,67905 | 0,941502 |
| Ercc5 | 12,9834 | 12,2866 | -0,0795786 | 0,67905 | 0,941502 |
| Cask | 10,6426 | 10,1017 | -0,0752615 | 0,67915 | 0,941502 |
| Rad54b | 9,96312 | 10,6327 | 0,0938372 | 0,67925 | 0,941502 |
| Efna4 | 31,1415 | 29,4565 | -0,0802534 | 0,6793 | 0,941502 |
| Limd2 | 77,286 | 73,3817 | -0,074787 | 0,6796 | 0,941502 |
| Ap1b1 | 24,2344 | 22,9915 | -0,0759533 | 0,67965 | 0,941502 |
| Scg5 | 3,6532 | 3,31268 | -0,141162 | 0,67965 | 0,941502 |
| Ngf | 2,69917 | 2,4139 | -0,161148 | 0,67965 | 0,941502 |
| Pcnxl3 | 18,8514 | 19,8344 | 0,0733331 | 0,67975 | 0,941502 |
| Dcaf13 | 61,8593 | 58,7009 | -0,0756067 | 0,6798 | 0,941502 |
| Rhbdl1 | 3,71716 | 4,06548 | 0,129224 | 0,67995 | 0,941502 |
| Tbc1d22b | 10,8824 | 11,5308 | 0,0834859 | 0,67995 | 0,941502 |
| Uhrf1bp1 | 9,84167 | 9,33212 | -0,0766972 | 0,68 | 0,941502 |
| Itfg3 | 9,83875 | 10,4446 | 0,0862149 | 0,6802 | 0,941502 |
| D5Ertd579e | 12,7864 | 13,4814 | 0,0763626 | 0,6802 | 0,941502 |
| Fahd2a | 6,6907 | 6,14537 | -0,122656 | 0,6802 | 0,941502 |
| Iffo1 | 6,09873 | 6,5162 | 0,0955229 | 0,6803 | 0,941502 |
| Senp8 | 6,97396 | 6,56927 | -0,086246 | 0,6803 | 0,941502 |
| Wdr77 | 71,9727 | 67,5509 | -0,0914747 | 0,6804 | 0,941502 |
| Fam53c | 21,8143 | 23,0172 | 0,0774364 | 0,68045 | 0,941502 |
| Dlat | 25,2647 | 26,5999 | 0,0743008 | 0,68055 | 0,941502 |
| Slmo1 | 6,36146 | 5,92363 | -0,102877 | 0,68065 | 0,941502 |
| Rasal2 | 4,83581 | 4,49684 | -0,104847 | 0,68075 | 0,941502 |
| Nle1 | 22,2299 | 20,9821 | -0,0833408 | 0,6809 | 0,941502 |
| Rnasek | 31,4868 | 33,7293 | 0,0992581 | 0,68095 | 0,941502 |
| 3425401B19Rik | 0,66127 | 0,727586 | 0,137879 | 0,6811 | 0,941502 |
| Sae1 | 238,295 | 226,45 | -0,0735534 | 0,6811 | 0,941502 |
| Cwf19l1 | 9,97457 | 9,41182 | -0,0837813 | 0,68115 | 0,941502 |
| C330007P06Rik | 7,71833 | 8,20113 | 0,087534 | 0,6812 | 0,941502 |
| BC030867 | 15,1514 | 16,0335 | 0,0816447 | 0,68135 | 0,941502 |
| Prokr1 | 0,636679 | 0,568257 | -0,164022 | 0,68135 | 0,941502 |
| Sft2d2 | 4,13295 | 3,88044 | -0,0909508 | 0,6814 | 0,941502 |
| Cox17 | 98,8038 | 105,016 | 0,087969 | 0,68145 | 0,941502 |
| Wdr45b | 41,0565 | 38,9504 | -0,0759729 | 0,6817 | 0,941502 |
| Tmco6 | 9,34215 | 10,5102 | 0,169961 | 0,68175 | 0,941502 |
| Nxpe3 | 1,68627 | 1,56826 | -0,104673 | 0,68175 | 0,941502 |
| Opa3 | 19,2998 | 18,2394 | -0,081527 | 0,6818 | 0,941502 |
| Crlf2 | 2,2457 | 2,49853 | 0,153916 | 0,6819 | 0,941502 |
| Tmc4 | 1,45676 | 1,96817 | 0,43409 | 0,6821 | 0,941502 |
| Clpx | 19,8575 | 18,8204 | -0,0773851 | 0,6821 | 0,941502 |
| Bcl2l15 | 0,785324 | 0,297518 | -1,40031 | 0,6821 | 0,941502 |
| Gmeb1 | 10,8689 | 11,4472 | 0,0747854 | 0,68215 | 0,941502 |
| Mlxipl | 0,679817 | 0,757591 | 0,156274 | 0,6822 | 0,941502 |
| Bid | 9,03033 | 9,65591 | 0,0966325 | 0,6822 | 0,941502 |
| Lsm8 | 191,421 | 181,718 | -0,0750442 | 0,6822 | 0,941502 |
| 9530068E07Rik | 85,4455 | 89,8557 | 0,0726056 | 0,68255 | 0,941819 |
| Dclre1a | 6,15919 | 6,5491 | 0,0885563 | 0,6826 | 0,941819 |
| Tcaim | 2,5426 | 2,7565 | 0,116528 | 0,68265 | 0,941819 |
| Sys1 | 17,6709 | 16,6286 | -0,0877051 | 0,68295 | 0,942132 |

| | | | | | |
|---|---|---|---|---|---|
| Becn1 | 55,5071 | 58,4225 | 0,0738523 | 0,68315 | 0,942242 |
| Srrm1 | 89,1837 | 93,8778 | 0,0740032 | 0,6832 | 0,942242 |
| Syf2 | 44,8767 | 42,4693 | -0,0795478 | 0,68325 | 0,942242 |
| Rslcan18 | 0,807017 | 0,947446 | 0,231444 | 0,68335 | 0,942278 |
| Sdsl | 4,18532 | 3,81473 | -0,133757 | 0,68355 | 0,942323 |
| Tmem63b | 14,0609 | 14,8699 | 0,0807002 | 0,6836 | 0,942323 |
| Raet1e | 2,05687 | 2,42434 | 0,23714 | 0,68365 | 0,942323 |
| Ino80dos | 1,78546 | 2,68832 | 0,590411 | 0,68375 | 0,942323 |
| Phlda3 | 13,4029 | 12,5542 | -0,094372 | 0,68375 | 0,942323 |
| Rspo1 | 0,924971 | 0,807714 | -0,195564 | 0,68405 | 0,942458 |
| Dnajb1 | 40,4757 | 42,6422 | 0,075226 | 0,68425 | 0,942458 |
| Hspa12b | 5,20922 | 6,75793 | 0,375514 | 0,6843 | 0,942458 |
| Rrp9 | 50,6721 | 53,423 | 0,0762686 | 0,6843 | 0,942458 |
| Pygo2 | 32,4808 | 34,2149 | 0,0750363 | 0,6843 | 0,942458 |
| 1600002K03Rik | 7,35648 | 7,60678 | 0,04827 | 0,68445 | 0,942458 |
| Brf2 | 13,7667 | 12,8984 | -0,0939952 | 0,68445 | 0,942458 |
| Hyal1 | 2,54082 | 2,77023 | 0,124713 | 0,6846 | 0,942458 |
| Tmc6 | 6,30119 | 6,75615 | 0,100578 | 0,6846 | 0,942458 |
| Ltbr | 15,5862 | 14,6772 | -0,0866969 | 0,68465 | 0,942458 |
| Acn9 | 21,1616 | 19,4757 | -0,119773 | 0,6847 | 0,942458 |
| Gltscr1 | 7,09979 | 7,49847 | 0,0788197 | 0,68475 | 0,942458 |
| Ubqln1 | 70,2537 | 66,8698 | -0,0712203 | 0,68485 | 0,942458 |
| Tnfrsf21 | 11,9048 | 12,5653 | 0,0778951 | 0,68495 | 0,942458 |
| Hsd17b10 | 158,252 | 150,332 | -0,074071 | 0,68495 | 0,942458 |
| Pdxp | 15,6654 | 16,5989 | 0,0835112 | 0,68515 | 0,942613 |
| Zfp202 | 6,65284 | 7,05782 | 0,0852525 | 0,6853 | 0,942613 |
| Eif3l | 212,763 | 202,035 | -0,0746418 | 0,6853 | 0,942613 |
| Prmt5 | 111,483 | 106,074 | -0,0717488 | 0,6854 | 0,942613 |
| Sik2 | 5,73544 | 6,81427 | 0,248655 | 0,6856 | 0,942613 |
| Rnf150 | 1,81409 | 1,69961 | -0,0940392 | 0,6856 | 0,942613 |
| Zfp874b | 1,70069 | 1,56349 | -0,121346 | 0,6856 | 0,942613 |
| Gpalpp1 | 11,6576 | 11,049 | -0,0773654 | 0,68565 | 0,942613 |
| Pgap2 | 18,1373 | 19,1795 | 0,0806014 | 0,6859 | 0,942694 |
| 1700020I14Rik | 36,2761 | 38,2865 | 0,0778133 | 0,686 | 0,942694 |
| Trnt1 | 31,5907 | 33,7896 | 0,0970775 | 0,68605 | 0,942694 |
| Larp1 | 36,1505 | 38,0243 | 0,0729045 | 0,68605 | 0,942694 |
| Aph1b | 2,29013 | 2,46621 | 0,106866 | 0,68615 | 0,942694 |
| Mterfd3 | 13,7469 | 12,8888 | -0,0929921 | 0,68615 | 0,942694 |
| Wdr11 | 19,9841 | 18,9742 | -0,0748086 | 0,6863 | 0,942799 |
| Them6 | 2,01163 | 1,81061 | -0,151894 | 0,6864 | 0,942835 |
| Cherp | 50,677 | 53,3554 | 0,0743029 | 0,68665 | 0,94288 |
| Rcl1 | 58,0901 | 55,2111 | -0,0733326 | 0,68665 | 0,94288 |
| Arhgef39 | 15,3891 | 14,4675 | -0,0890934 | 0,6867 | 0,94288 |
| Zfp820 | 2,28686 | 2,47068 | 0,111538 | 0,68675 | 0,94288 |
| Zfp108 | 1,11769 | 1,23658 | 0,14584 | 0,6868 | 0,94288 |
| Abi2 | 20,2933 | 21,3396 | 0,0725272 | 0,68715 | 0,94326 |
| Sin3b | 79,4858 | 74,8535 | -0,0866267 | 0,68725 | 0,943296 |
| Stam2 | 15,7574 | 16,6142 | 0,0763836 | 0,68735 | 0,943332 |
| Aldoart1 | 15,0883 | 14,2422 | -0,0832616 | 0,68765 | 0,943643 |
| Kifc3 | 3,05715 | 2,67572 | -0,19226 | 0,68785 | 0,943784 |

| | | | | | |
|---|---|---|---|---|---|
| D4Ertd617e | 0,306001 | 0,652414 | 1,09225 | 0,6879 | 0,943784 |
| Hexim1 | 12,7775 | 12,0862 | -0,0802432 | 0,688 | 0,94382 |
| Mex3a | 50,3524 | 52,9195 | 0,0717394 | 0,6881 | 0,943857 |
| Parp16 | 4,92057 | 5,27962 | 0,101609 | 0,6882 | 0,943872 |
| Wwp2 | 17,8389 | 18,7688 | 0,0733085 | 0,68845 | 0,943872 |
| Ccdc107 | 13,7102 | 12,6923 | -0,111303 | 0,6885 | 0,943872 |
| Pja1 | 70,8117 | 74,4687 | 0,0726479 | 0,68855 | 0,943872 |
| Chchd7 | 79,7419 | 84,7086 | 0,0871706 | 0,6886 | 0,943872 |
| Hmgcs1 | 139,151 | 132,482 | -0,0708548 | 0,6886 | 0,943872 |
| B3gnt5 | 18,2757 | 17,3409 | -0,0757521 | 0,68865 | 0,943872 |
| Strn4 | 48,0959 | 50,9125 | 0,0821053 | 0,6887 | 0,943872 |
| Arf3 | 36,6122 | 38,493 | 0,0722716 | 0,6892 | 0,944392 |
| Plekhh3 | 0,957789 | 0,862382 | -0,151381 | 0,68925 | 0,944392 |
| Ctcf | 70,1913 | 66,757 | -0,0723743 | 0,6893 | 0,944392 |
| Bora | 29,1579 | 30,7235 | 0,0754591 | 0,68945 | 0,944496 |
| Cep83os | 1,33905 | 1,45842 | 0,123196 | 0,6896 | 0,944536 |
| Syp | 1,47072 | 1,33693 | -0,137598 | 0,68965 | 0,944536 |
| Ccdc82 | 1,47942 | 1,36184 | -0,119469 | 0,6897 | 0,944536 |
| Aig1 | 5,70379 | 6,22123 | 0,125281 | 0,6898 | 0,944572 |
| Emc4 | 41,7865 | 38,7231 | -0,109842 | 0,6899 | 0,944609 |
| Nfkbiz | 2,72614 | 2,93559 | 0,106792 | 0,69065 | 0,945343 |
| Hyi | 4,84156 | 3,61073 | -0,423184 | 0,69075 | 0,945343 |
| Eif2d | 38,0783 | 36,1555 | -0,0747542 | 0,69085 | 0,945343 |
| Apba3 | 16,7648 | 15,8322 | -0,0825769 | 0,69085 | 0,945343 |
| AI597479 | 20,3233 | 21,4163 | 0,0755699 | 0,6911 | 0,945343 |
| Smchd1 | 32,1241 | 30,5558 | -0,07221 | 0,6912 | 0,945343 |
| Nxpe4 | 2,04864 | 1,88386 | -0,120976 | 0,6912 | 0,945343 |
| BC048403 | 2,76985 | 2,56641 | -0,110057 | 0,69125 | 0,945343 |
| Snord55 | 928,445 | 361,458 | -1,36099 | 0,69125 | 0,945343 |
| Dzip1 | 17,0907 | 17,9901 | 0,073995 | 0,6913 | 0,945343 |
| Tmem134 | 23,3015 | 24,6467 | 0,0809714 | 0,69135 | 0,945343 |
| Pank2 | 27,1278 | 29,7125 | 0,131295 | 0,6914 | 0,945343 |
| Larp7 | 66,1741 | 69,5348 | 0,0714681 | 0,69145 | 0,945343 |
| Kif18b | 19,1314 | 20,1459 | 0,074549 | 0,69155 | 0,945343 |
| Sh3bgrl3 | 72,3375 | 68,5156 | -0,0783117 | 0,69155 | 0,945343 |
| Nck1 | 30,3018 | 28,7154 | -0,0775801 | 0,6917 | 0,945343 |
| Itfg1 | 27,5458 | 28,9857 | 0,0735079 | 0,6918 | 0,945343 |
| P2ry1 | 0,918304 | 1,00988 | 0,137138 | 0,6919 | 0,945343 |
| Lrp1 | 23,8596 | 25,0773 | 0,0718126 | 0,69215 | 0,945343 |
| D330050I16Rik | 0,81581 | 1,39907 | 0,778159 | 0,69225 | 0,945343 |
| Cerk | 6,18122 | 6,5498 | 0,0835588 | 0,69235 | 0,945343 |
| 1600023N17Rik | 0,643949 | 0,770932 | 0,259657 | 0,6924 | 0,945343 |
| Tyw1 | 14,0248 | 14,7979 | 0,077413 | 0,6924 | 0,945343 |
| Hspa13 | 18,1606 | 19,1187 | 0,0741721 | 0,6924 | 0,945343 |
| Dcaf15 | 37,7277 | 39,6978 | 0,073432 | 0,6924 | 0,945343 |
| Mapk1ip1 | 7,73492 | 8,30542 | 0,102665 | 0,6925 | 0,945343 |
| Drosha | 62,3851 | 65,4543 | 0,0692875 | 0,6925 | 0,945343 |
| Cebpz | 46,5588 | 44,2702 | -0,0727155 | 0,6925 | 0,945343 |
| Peli1 | 24,5234 | 25,7827 | 0,0722425 | 0,6926 | 0,94535 |
| Wdr4 | 18,3859 | 17,464 | -0,0742204 | 0,69275 | 0,94535 |

| | | | | | |
|---|---|---|---|---|---|
| Neil3 | 12,1384 | 11,445 | -0,0848648 | 0,6928 | 0,94535 |
| Gm20337 | 1,09199 | 0,939915 | -0,216363 | 0,6928 | 0,94535 |
| Prr3 | 20,1561 | 18,8995 | -0,0928672 | 0,693 | 0,94549 |
| E130201H02Rik | 10,569 | 9,76299 | -0,114449 | 0,69305 | 0,94549 |
| Snord17 | 9,07039 | 16,8985 | 0,897661 | 0,6933 | 0,94563 |
| Snhg12 | 72,5274 | 68,4424 | -0,0836354 | 0,6933 | 0,94563 |
| Ccng2 | 18,1741 | 17,2155 | -0,0781821 | 0,6934 | 0,945666 |
| Rom1 | 3,92864 | 3,60181 | -0,12531 | 0,69355 | 0,945738 |
| Zfp687 | 22,6369 | 24,1204 | 0,0915783 | 0,6936 | 0,945738 |
| 4930429B21Rik | 0,440874 | 0,665717 | 0,594542 | 0,69375 | 0,945809 |
| Tbp | 34,2403 | 36,0616 | 0,0747646 | 0,6938 | 0,945809 |
| Paqr4 | 2,04735 | 2,3967 | 0,227292 | 0,6941 | 0,946033 |
| 2700097O09Rik | 27,4671 | 25,9596 | -0,0814343 | 0,6943 | 0,946033 |
| Cc2d2a | 5,62135 | 5,94491 | 0,0807397 | 0,69435 | 0,946033 |
| AI507597 | 0,982212 | 1,2374 | 0,3332 | 0,69445 | 0,946033 |
| Dtwd2 | 2,56717 | 2,37464 | -0,112469 | 0,6945 | 0,946033 |
| Zswim8 | 25,9296 | 27,2324 | 0,0707243 | 0,69455 | 0,946033 |
| BC029722 | 5,26528 | 5,70436 | 0,115554 | 0,6946 | 0,946033 |
| Pcgf5 | 5,2481 | 5,67139 | 0,111908 | 0,69465 | 0,946033 |
| Dnajb9 | 8,74306 | 9,33334 | 0,0942551 | 0,69485 | 0,946033 |
| Elovl4 | 7,81702 | 7,36391 | -0,0861455 | 0,6949 | 0,946033 |
| Ncs1 | 20,6846 | 19,6746 | -0,0722202 | 0,69515 | 0,946033 |
| Dcun1d2 | 10,955 | 10,1659 | -0,107854 | 0,69515 | 0,946033 |
| Pole4 | 23,7359 | 22,4724 | -0,0789174 | 0,69535 | 0,946033 |
| Sh2d4a | 0,643652 | 0,569224 | -0,177284 | 0,69535 | 0,946033 |
| Seh1l | 74,638 | 70,8727 | -0,0746819 | 0,6954 | 0,946033 |
| Ezh1 | 6,33073 | 5,9726 | -0,0840133 | 0,69545 | 0,946033 |
| St5 | 10,4927 | 11,0844 | 0,0791476 | 0,6956 | 0,946033 |
| Clint1 | 61,5053 | 58,5903 | -0,0700497 | 0,6956 | 0,946033 |
| Gtf3c5 | 29,7254 | 28,2734 | -0,0722492 | 0,6956 | 0,946033 |
| Cndp2 | 37,9767 | 39,9235 | 0,0721235 | 0,69565 | 0,946033 |
| Mib2 | 6,26424 | 6,6351 | 0,0829778 | 0,6957 | 0,946033 |
| Nubp1 | 33,0321 | 31,1302 | -0,08555 | 0,6957 | 0,946033 |
| Pdzd7 | 0,506819 | 0,728888 | 0,524226 | 0,69585 | 0,946033 |
| Asb6 | 18,6431 | 17,6437 | -0,079494 | 0,69585 | 0,946033 |
| Prdm5 | 13,6049 | 12,8559 | -0,0816934 | 0,6959 | 0,946033 |
| Mdm1 | 21,1752 | 22,266 | 0,0724655 | 0,696 | 0,946033 |
| Srek1ip1 | 18,5492 | 17,5381 | -0,0808618 | 0,696 | 0,946033 |
| Sphk2 | 12,8557 | 12,1739 | -0,0786104 | 0,6961 | 0,946033 |
| Smu1 | 71,8746 | 67,8142 | -0,0838939 | 0,6962 | 0,946033 |
| Spag16 | 2,35768 | 2,18943 | -0,106814 | 0,69625 | 0,946033 |
| Taf10 | 81,293 | 73,1565 | -0,152146 | 0,69625 | 0,946033 |
| Gskip | 8,89607 | 8,39935 | -0,0828904 | 0,69645 | 0,946204 |
| Atl1 | 1,65936 | 1,18506 | -0,485664 | 0,69655 | 0,946214 |
| Arid1b | 8,76346 | 9,21886 | 0,0730882 | 0,69665 | 0,946214 |
| Azi2 | 16,8865 | 17,8944 | 0,0836367 | 0,6968 | 0,946214 |
| 9030624J02Rik | 18,2335 | 17,3171 | -0,0743948 | 0,6968 | 0,946214 |
| Ddx49 | 29,7091 | 27,4828 | -0,112377 | 0,69685 | 0,946214 |
| Alg1 | 9,38457 | 10,1359 | 0,111105 | 0,6969 | 0,946214 |
| Bloc1s5 | 15,5307 | 16,44 | 0,0820866 | 0,697 | 0,946218 |

| | | | | | |
|---|---|---|---|---|---|
| Mrto4 | 51,4221 | 48,8283 | -0,0746709 | 0,69705 | 0,946218 |
| Slc44a5 | 0,898979 | 0,818168 | -0,13589 | 0,69715 | 0,946221 |
| Caskin1 | 0,796009 | 0,57487 | -0,469547 | 0,6972 | 0,946221 |
| Dnase1 | 5,14511 | 6,57667 | 0,354155 | 0,69735 | 0,946224 |
| Nat9 | 6,65982 | 7,21506 | 0,115529 | 0,69735 | 0,946224 |
| Lyst | 1,18144 | 1,10806 | -0,092517 | 0,6975 | 0,946328 |
| Ralgapa2 | 2,64039 | 2,79928 | 0,0843078 | 0,69765 | 0,946431 |
| Megf8 | 14,8018 | 14,0953 | -0,0705609 | 0,69795 | 0,946738 |
| Prkrir | 73,3653 | 69,8817 | -0,0701829 | 0,69805 | 0,946774 |
| Glod4 | 57,3318 | 54,545 | -0,0718904 | 0,6983 | 0,946916 |
| Fam219b | 8,66861 | 9,19053 | 0,0843473 | 0,6984 | 0,946916 |
| Ehd4 | 6,1792 | 5,8193 | -0,0865744 | 0,6984 | 0,946916 |
| Nkap | 10,6699 | 11,2522 | 0,0766538 | 0,69845 | 0,946916 |
| Ahctf1 | 31,6632 | 33,2436 | 0,0702662 | 0,6986 | 0,947019 |
| Fam175a | 10,3108 | 10,9066 | 0,0810364 | 0,69875 | 0,94709 |
| Gstk1 | 16,7903 | 17,9393 | 0,0954953 | 0,6988 | 0,94709 |
| Epdr1 | 12,1829 | 12,8814 | 0,0804313 | 0,6989 | 0,947125 |
| Tctn3 | 6,51372 | 6,11708 | -0,0906383 | 0,6991 | 0,947206 |
| Dcaf5 | 11,6023 | 12,192 | 0,0715172 | 0,69915 | 0,947206 |
| Gtf2ird1 | 20,9314 | 22,0212 | 0,0732227 | 0,69925 | 0,947206 |
| Snx1 | 67,9517 | 64,7585 | -0,0694397 | 0,6993 | 0,947206 |
| Rad54l | 30,4503 | 28,3667 | -0,102257 | 0,6994 | 0,947206 |
| 2410021H03Rik | 0,772432 | 2,26967 | 1,555 | 0,69945 | 0,947206 |
| Supt6 | 40,3806 | 38,4962 | -0,0689448 | 0,6995 | 0,947206 |
| Slco4a1 | 0,706182 | 0,631003 | -0,162394 | 0,69955 | 0,947206 |
| Snora68 | 425,462 | 670,159 | 0,655474 | 0,69965 | 0,947241 |
| Stap2 | 2,56741 | 2,33725 | -0,135502 | 0,6999 | 0,947274 |
| Nek1 | 9,55875 | 10,0621 | 0,074034 | 0,69995 | 0,947274 |
| Heatr1 | 28,5287 | 27,1878 | -0,0694532 | 0,7003 | 0,947274 |
| Cdh1 | 1,39103 | 1,28162 | -0,118189 | 0,7004 | 0,947274 |
| Hsd3b7 | 3,27666 | 3,55469 | 0,117495 | 0,7007 | 0,947274 |
| Gmeb2 | 12,7593 | 13,4551 | 0,0766078 | 0,70075 | 0,947274 |
| Ctbs | 2,1282 | 1,93294 | -0,138839 | 0,7008 | 0,947274 |
| Akip1 | 13,0297 | 12,1615 | -0,0994823 | 0,7009 | 0,947274 |
| LOC106740 | 13,3341 | 15,8978 | 0,253709 | 0,701 | 0,947274 |
| Fam168b | 93,1201 | 88,7762 | -0,068919 | 0,701 | 0,947274 |
| Fbxo6 | 2,84029 | 2,57265 | -0,142786 | 0,70105 | 0,947274 |
| Pxmp4 | 4,49021 | 4,86271 | 0,114976 | 0,7011 | 0,947274 |
| Ric8b | 11,7897 | 12,4048 | 0,07337 | 0,7011 | 0,947274 |
| Rhot1 | 22,2765 | 23,369 | 0,0690747 | 0,7011 | 0,947274 |
| Tmsb15b2 | 4,22974 | 5,16782 | 0,288989 | 0,7012 | 0,947274 |
| Msantd4 | 31,2326 | 29,7284 | -0,0712084 | 0,70135 | 0,947274 |
| Prkar1a | 111,297 | 106,069 | -0,0694113 | 0,7014 | 0,947274 |
| Hspa4 | 176,452 | 168,161 | -0,0694272 | 0,7014 | 0,947274 |
| Bmper | 2,60345 | 2,79153 | 0,100632 | 0,70145 | 0,947274 |
| Zfp276 | 6,22989 | 6,81345 | 0,129178 | 0,7016 | 0,947274 |
| F420014N23Rik | 0,764303 | 1,14911 | 0,588303 | 0,7017 | 0,947274 |
| Pex1 | 5,6714 | 6,0752 | 0,0992271 | 0,702 | 0,947274 |
| Papd4 | 17,7563 | 16,806 | -0,0793579 | 0,70205 | 0,947274 |
| Swsap1 | 8,4837 | 7,95382 | -0,0930471 | 0,70205 | 0,947274 |

| | | | | | |
|---|---|---|---|---|---|
| 2310045N01Rik | 33,3719 | 31,276 | -0,0935777 | 0,70205 | 0,947274 |
| 1110008L16Rik | 8,32754 | 6,99101 | -0,252388 | 0,70205 | 0,947274 |
| Tuba1c | 9,42693 | 8,85894 | -0,0896547 | 0,7022 | 0,947274 |
| Cabp1 | 2,78611 | 3,06522 | 0,137742 | 0,70225 | 0,947274 |
| Rrbp1 | 39,3936 | 41,329 | 0,0691949 | 0,70225 | 0,947274 |
| Msto1 | 32,6872 | 30,1413 | -0,116981 | 0,70225 | 0,947274 |
| Aars2 | 13,2501 | 13,9564 | 0,0749228 | 0,70255 | 0,947274 |
| Toe1 | 25,7405 | 24,1633 | -0,0912213 | 0,70255 | 0,947274 |
| Pex11b | 25,6156 | 24,0146 | -0,0931131 | 0,7026 | 0,947274 |
| Gripap1 | 12,7358 | 12,0599 | -0,078673 | 0,70265 | 0,947274 |
| Mrpl1 | 15,2386 | 14,3688 | -0,0847865 | 0,70265 | 0,947274 |
| Fam45a | 6,96089 | 6,52766 | -0,0927049 | 0,70265 | 0,947274 |
| Xpo6 | 42,5632 | 44,6374 | 0,0686486 | 0,7027 | 0,947274 |
| Exosc4 | 23,6718 | 22,456 | -0,0760682 | 0,70285 | 0,947274 |
| Dexi | 12,8732 | 13,7069 | 0,090529 | 0,7029 | 0,947274 |
| Mrps24 | 35,443 | 33,5397 | -0,0796308 | 0,70295 | 0,947274 |
| Ripk2 | 6,2841 | 6,72778 | 0,0984257 | 0,703 | 0,947274 |
| Kctd18 | 6,03392 | 6,41905 | 0,089263 | 0,703 | 0,947274 |
| Socs6 | 16,1392 | 17,0046 | 0,0753627 | 0,703 | 0,947274 |
| Kif4 | 26,9696 | 28,29 | 0,0689597 | 0,70315 | 0,947274 |
| Scamp5 | 2,37318 | 2,19557 | -0,112222 | 0,70325 | 0,947274 |
| Insl6 | 1,80882 | 1,54593 | -0,22657 | 0,70325 | 0,947274 |
| Dnajc7 | 121,345 | 127,243 | 0,0684704 | 0,7033 | 0,947274 |
| Zfyve20 | 15,7303 | 14,9851 | -0,0700186 | 0,7033 | 0,947274 |
| Ptcd3 | 62,5849 | 59,6222 | -0,0699657 | 0,70335 | 0,947274 |
| Asns | 39,5192 | 41,5338 | 0,0717318 | 0,7035 | 0,947274 |
| 4933411K20Rik | 7,62139 | 8,02943 | 0,0752419 | 0,7036 | 0,947274 |
| Wdr75 | 55,396 | 52,8161 | -0,0688036 | 0,70375 | 0,947274 |
| Hs6st1 | 14,2062 | 14,9525 | 0,0738702 | 0,7038 | 0,947274 |
| Dbp | 2,13055 | 1,71691 | -0,311408 | 0,704 | 0,947274 |
| Pld3 | 10,4811 | 11,1223 | 0,0856623 | 0,70405 | 0,947274 |
| Kctd20 | 51,2809 | 48,9172 | -0,0680803 | 0,7041 | 0,947274 |
| Mllt11 | 9,35713 | 7,87225 | -0,249291 | 0,70415 | 0,947274 |
| Btbd10 | 9,74426 | 10,3118 | 0,081676 | 0,7042 | 0,947274 |
| Smarca5 | 29,1758 | 27,8281 | -0,0682347 | 0,70425 | 0,947274 |
| Pms2 | 17,3104 | 18,1993 | 0,0722469 | 0,7043 | 0,947274 |
| Tceb3 | 20,1168 | 19,1711 | -0,0694675 | 0,7043 | 0,947274 |
| Pan2 | 24,6029 | 25,8258 | 0,0699837 | 0,70435 | 0,947274 |
| Tada2b | 6,45418 | 7,21047 | 0,159858 | 0,7044 | 0,947274 |
| Taf2 | 28,2842 | 29,6858 | 0,069775 | 0,7044 | 0,947274 |
| Etnk2 | 1,27081 | 1,40103 | 0,140737 | 0,70465 | 0,947446 |
| Zranb1 | 23,7113 | 24,9425 | 0,073032 | 0,70475 | 0,947446 |
| Abhd6 | 4,17467 | 3,87083 | -0,109018 | 0,70475 | 0,947446 |
| AU022252 | 14,9242 | 15,7756 | 0,0800404 | 0,70485 | 0,947481 |
| Foxm1 | 42,8869 | 44,9653 | 0,0682726 | 0,70495 | 0,947517 |
| Neurl1a | 1,439 | 1,55939 | 0,115922 | 0,7052 | 0,947654 |
| Nisch | 128,422 | 134,877 | 0,0707559 | 0,7052 | 0,947654 |
| Snord88c | 193,193 | 107,607 | -0,844273 | 0,70545 | 0,947859 |
| Paf1 | 56,4199 | 59,221 | 0,0699061 | 0,7055 | 0,947859 |
| Cox16 | 11,2273 | 10,5891 | -0,0844226 | 0,7056 | 0,947862 |

| | | | | | |
|---|---|---|---|---|---|
| Snord16a | 1049,52 | 2008,9 | 0,936681 | 0,70565 | 0,947862 |
| Map2k3os | 1,37552 | 1,55179 | 0,17395 | 0,70605 | 0,9482 |
| D16Ertd472e | 8,38059 | 8,80317 | 0,0709728 | 0,70605 | 0,9482 |
| Edc4 | 27,7086 | 29,072 | 0,0692965 | 0,70615 | 0,948203 |
| Ptcd1 | 8,0668 | 8,95872 | 0,151297 | 0,7062 | 0,948203 |
| Ilvbl | 7,92904 | 8,42292 | 0,0871733 | 0,70635 | 0,948279 |
| Tmem44 | 4,98858 | 4,70753 | -0,0836581 | 0,70645 | 0,948279 |
| Timmdc1 | 17,5418 | 18,4879 | 0,0757804 | 0,7065 | 0,948279 |
| 4930565N06Rik | 0,495408 | 0,845657 | 0,771455 | 0,7066 | 0,948279 |
| Cdca7 | 118,684 | 124,313 | 0,0668438 | 0,7067 | 0,948279 |
| 1700009P17Rik | 1,98755 | 1,75987 | -0,17552 | 0,7067 | 0,948279 |
| Mon1b | 5,90748 | 5,58851 | -0,0800796 | 0,7068 | 0,948314 |
| Daglb | 5,9913 | 6,35536 | 0,0851039 | 0,70705 | 0,948323 |
| Nup107 | 57,9509 | 55,2934 | -0,0677221 | 0,7071 | 0,948323 |
| Fbxw9 | 6,68027 | 5,38226 | -0,311695 | 0,7071 | 0,948323 |
| Atf4 | 213,656 | 223,981 | 0,0680885 | 0,7072 | 0,948323 |
| 0610009L18Rik | 1,24407 | 1,47803 | 0,248612 | 0,70725 | 0,948323 |
| Nthl1 | 9,75788 | 9,09714 | -0,101154 | 0,70725 | 0,948323 |
| 1600002H07Rik | 19,0774 | 20,0667 | 0,0729437 | 0,70755 | 0,948445 |
| Slc48a1 | 4,95222 | 4,34838 | -0,1876 | 0,70755 | 0,948445 |
| Mthfd2l | 11,3862 | 10,7493 | -0,0830471 | 0,7077 | 0,948445 |
| Aktip | 11,0558 | 11,8266 | 0,0972313 | 0,7078 | 0,948445 |
| AW046200 | 0,275582 | 0,598412 | 1,11866 | 0,7079 | 0,948445 |
| Zfp386 | 9,97568 | 10,4975 | 0,0735577 | 0,708 | 0,948445 |
| Apc2 | 0,678107 | 0,338713 | -1,00145 | 0,7081 | 0,948445 |
| Kalrn | 3,09097 | 3,27187 | 0,0820598 | 0,7082 | 0,948445 |
| Cenpa | 143,473 | 150,28 | 0,066873 | 0,70825 | 0,948445 |
| Chmp1b | 21,8808 | 20,4768 | -0,0956758 | 0,70825 | 0,948445 |
| Ghdc | 2,86273 | 2,64541 | -0,113899 | 0,70825 | 0,948445 |
| Sp3 | 38,9818 | 40,869 | 0,0682088 | 0,7083 | 0,948445 |
| Zfp398 | 3,28323 | 3,06617 | -0,0986779 | 0,70835 | 0,948445 |
| Haghl | 16,62 | 15,3802 | -0,111842 | 0,7084 | 0,948445 |
| Atxn7l1 | 6,79757 | 6,34898 | -0,0984931 | 0,70845 | 0,948445 |
| Kif20a | 94,6266 | 87,64 | -0,110656 | 0,70865 | 0,948613 |
| Ercc6 | 5,73105 | 6,03009 | 0,0733795 | 0,7088 | 0,948622 |
| Timm10b | 30,6355 | 29,0054 | -0,0788809 | 0,70885 | 0,948622 |
| Gins1 | 66,32 | 63,0677 | -0,0725429 | 0,709 | 0,948622 |
| Svip | 0,869216 | 0,786259 | -0,14471 | 0,70905 | 0,948622 |
| Tpcn2 | 4,02828 | 4,30218 | 0,0949056 | 0,7091 | 0,948622 |
| Naga | 7,48326 | 7,96617 | 0,0902197 | 0,7091 | 0,948622 |
| Cacfd1 | 46,0244 | 43,8692 | -0,0691882 | 0,7092 | 0,948625 |
| Abcd3 | 20,363 | 19,4 | -0,069895 | 0,70925 | 0,948625 |
| Apmap | 20,4047 | 19,3993 | -0,0728945 | 0,70945 | 0,948793 |
| Chmp5 | 64,0201 | 67,1243 | 0,0683095 | 0,7097 | 0,948802 |
| Tmem129 | 15,0531 | 15,8607 | 0,0753969 | 0,70975 | 0,948802 |
| Rexo1 | 19,049 | 19,9759 | 0,0685497 | 0,7098 | 0,948802 |
| Sirt4 | 3,89451 | 3,58934 | -0,117723 | 0,70985 | 0,948802 |
| 6330408A02Rik | 4,87216 | 3,89032 | -0,324673 | 0,70985 | 0,948802 |
| Wdr91 | 8,31347 | 7,85043 | -0,0826794 | 0,7099 | 0,948802 |
| 1700105P06Rik | 0,890365 | 0,426943 | -1,06035 | 0,7101 | 0,94897 |

| | | | | | |
|---|---|---|---|---|---|
| Slc35b3 | 9,17127 | 9,73205 | 0,0856216 | 0,7102 | 0,948973 |
| Eef2 | 856,898 | 812,629 | -0,0765276 | 0,71025 | 0,948973 |
| Dus3l | 24,1536 | 22,9739 | -0,0722389 | 0,7104 | 0,949074 |
| Prkacb | 29,7182 | 28,386 | -0,0661648 | 0,7105 | 0,949109 |
| Dimt1 | 4,94756 | 5,277 | 0,0930006 | 0,7108 | 0,949379 |
| Prpf18 | 19,4406 | 20,4183 | 0,0707919 | 0,71085 | 0,949379 |
| Lekr1 | 0,853644 | 0,763858 | -0,160332 | 0,71105 | 0,949547 |
| Ubb | 164,262 | 152,281 | -0,109259 | 0,7113 | 0,949782 |
| Wasf1 | 22,6991 | 23,8559 | 0,0717086 | 0,7114 | 0,949785 |
| 4930579G24Rik | 11,3295 | 10,7154 | -0,0803956 | 0,71145 | 0,949785 |
| Ccdc24 | 6,75293 | 7,40866 | 0,133698 | 0,71165 | 0,949921 |
| Snora43 | 41,3939 | 26,6578 | -0,634862 | 0,7117 | 0,949921 |
| AK010878 | 25,3766 | 22,8451 | -0,151611 | 0,7118 | 0,949955 |
| Eif2b3 | 35,9514 | 37,7957 | 0,0721739 | 0,71195 | 0,950057 |
| Zbtb46 | 8,06145 | 7,58561 | -0,0877738 | 0,7121 | 0,950158 |
| Pmpca | 57,3201 | 60,2693 | 0,072382 | 0,7124 | 0,950264 |
| Zcchc14 | 18,284 | 17,4224 | -0,0696369 | 0,7124 | 0,950264 |
| Ddx28 | 14,7034 | 13,9196 | -0,0790332 | 0,71245 | 0,950264 |
| Nynrin | 27,0216 | 28,2722 | 0,0652667 | 0,7125 | 0,950264 |
| Lman2 | 44,2877 | 42,3162 | -0,065698 | 0,71255 | 0,950264 |
| Fam204a | 16,4976 | 17,3809 | 0,0752449 | 0,71265 | 0,950299 |
| Lipt1 | 3,07302 | 3,34103 | 0,120637 | 0,71275 | 0,950333 |
| Chn1os3 | 0,934218 | 0,665444 | -0,489442 | 0,71285 | 0,950368 |
| Snord52 | 1932,7 | 2300,95 | 0,251606 | 0,71295 | 0,950402 |
| Wdr90 | 14,7824 | 15,8108 | 0,097026 | 0,7132 | 0,950605 |
| Tfip11 | 25,1119 | 23,8555 | -0,074047 | 0,71325 | 0,950605 |
| Coro1c | 57,5027 | 60,249 | 0,0673074 | 0,71335 | 0,950607 |
| Ube2t | 27,0217 | 25,5413 | -0,0812868 | 0,7134 | 0,950607 |
| Ccdc91 | 13,2479 | 12,5667 | -0,0761575 | 0,7136 | 0,950775 |
| Snrnp70 | 349,49 | 333,494 | -0,0675881 | 0,71375 | 0,950876 |
| Snora69 | 27,0521 | 18,8288 | -0,522798 | 0,714 | 0,950904 |
| Fam120aos | 5,21397 | 5,61794 | 0,107658 | 0,71405 | 0,950904 |
| Zbtb43 | 7,07441 | 7,45154 | 0,0749284 | 0,7142 | 0,950904 |
| Rhog | 24,406 | 25,7633 | 0,0780806 | 0,7143 | 0,950904 |
| Utp18 | 40,4973 | 38,6844 | -0,0660723 | 0,7144 | 0,950904 |
| Zfp46 | 8,70289 | 9,14912 | 0,0721392 | 0,71445 | 0,950904 |
| Gga3 | 17,728 | 18,604 | 0,0695826 | 0,7145 | 0,950904 |
| Letm1 | 22,8655 | 21,8179 | -0,0676623 | 0,7146 | 0,950904 |
| Trim12a | 0,539899 | 0,628963 | 0,220287 | 0,71465 | 0,950904 |
| Rsg1 | 1,41567 | 1,62216 | 0,196427 | 0,7147 | 0,950904 |
| Mir7027 | 546,387 | 578,522 | 0,0824492 | 0,7147 | 0,950904 |
| Zfp689 | 6,09577 | 6,43905 | 0,0790384 | 0,7147 | 0,950904 |
| Cox8a | 174,4 | 183,026 | 0,0696494 | 0,715 | 0,950904 |
| Ddx56 | 38,7346 | 37,0118 | -0,0656339 | 0,715 | 0,950904 |
| Mzt1 | 28,5273 | 27,1521 | -0,0712793 | 0,71515 | 0,950904 |
| Spata6 | 7,74905 | 8,21352 | 0,0839802 | 0,7152 | 0,950904 |
| Dlgap5 | 37,6432 | 39,4248 | 0,0667147 | 0,7152 | 0,950904 |
| Jkamp | 17,8127 | 18,7456 | 0,0736527 | 0,71535 | 0,950904 |
| Dclre1b | 9,8976 | 8,57818 | -0,206406 | 0,71535 | 0,950904 |
| 2510003E04Rik | 19,3896 | 18,454 | -0,0713473 | 0,71545 | 0,950904 |

| | | | | | |
|---|---|---|---|---|---|
| Adck1 | 11,1265 | 10,5102 | -0,0822109 | 0,71545 | 0,950904 |
| Mtfr1 | 28,2428 | 29,5868 | 0,0670703 | 0,7155 | 0,950904 |
| Srm | 160,076 | 152,881 | -0,0663494 | 0,7155 | 0,950904 |
| Ndufs1 | 73,8439 | 70,4804 | -0,0672556 | 0,71555 | 0,950904 |
| Nfatc2ip | 7,99573 | 8,42776 | 0,0759199 | 0,7157 | 0,950941 |
| Etfa | 47,9764 | 45,7325 | -0,069104 | 0,7158 | 0,950941 |
| Prr15 | 0,66579 | 0,503832 | -0,402125 | 0,7158 | 0,950941 |
| Ube2n | 15,1607 | 14,4438 | -0,0698876 | 0,7159 | 0,950975 |
| Gpsm2 | 27,5098 | 29,0617 | 0,0791725 | 0,716 | 0,951009 |
| Pbk | 62,0923 | 65,0616 | 0,0673921 | 0,71625 | 0,951243 |
| Gadd45gip1 | 35,8562 | 33,4965 | -0,0982115 | 0,71635 | 0,951245 |
| Usp37 | 14,1171 | 15,4287 | 0,128168 | 0,7164 | 0,951245 |
| Napb | 1,50278 | 1,62513 | 0,112924 | 0,71655 | 0,951254 |
| E130311K13Rik | 1,42222 | 1,57833 | 0,150248 | 0,7166 | 0,951254 |
| Fam114a2 | 36,3671 | 38,051 | 0,0653031 | 0,7167 | 0,951254 |
| Ier3ip1 | 53,4075 | 50,9654 | -0,0675231 | 0,7168 | 0,951254 |
| Zmat3 | 6,65075 | 6,33651 | -0,0698282 | 0,7169 | 0,951254 |
| Cyth2 | 42,4618 | 40,5708 | -0,0657252 | 0,71695 | 0,951254 |
| Narfl | 9,41591 | 8,80205 | -0,0972603 | 0,71695 | 0,951254 |
| Nek2 | 29,3485 | 30,7357 | 0,0666304 | 0,717 | 0,951254 |
| Stat5b | 7,0439 | 6,69892 | -0,0724474 | 0,7173 | 0,951455 |
| Ctsz | 45,6012 | 41,8019 | -0,125502 | 0,7173 | 0,951455 |
| Nudt3 | 55,0177 | 57,5714 | 0,0654577 | 0,7176 | 0,951755 |
| Homer2 | 1,66532 | 1,76259 | 0,0819052 | 0,71785 | 0,951956 |
| Cmip | 14,6967 | 15,4127 | 0,0686284 | 0,7179 | 0,951956 |
| E2f5 | 61,6898 | 64,6339 | 0,0672606 | 0,71815 | 0,952157 |
| Immp2l | 1,18121 | 1,46669 | 0,312296 | 0,7182 | 0,952157 |
| Atg5 | 29,5386 | 28,18 | -0,0679272 | 0,7183 | 0,952191 |
| Prpf3 | 42,1195 | 40,2397 | -0,0658663 | 0,7187 | 0,952524 |
| Mapkapk5 | 31,8042 | 30,3424 | -0,0678834 | 0,7187 | 0,952524 |
| Hist1h3a | 0,97476 | 1,21698 | 0,320191 | 0,7188 | 0,952559 |
| Arrdc2 | 1,9907 | 2,16036 | 0,117992 | 0,71895 | 0,952659 |
| Trpc4ap | 60,0251 | 57,3792 | -0,0650397 | 0,7191 | 0,952727 |
| Psen1 | 26,6281 | 25,437 | -0,0660249 | 0,71915 | 0,952727 |
| Cebpd | 0,726953 | 0,959749 | 0,400795 | 0,7197 | 0,95293 |
| Slc30a5 | 31,6235 | 30,2363 | -0,0647155 | 0,7197 | 0,95293 |
| Riok3 | 24,1165 | 23,0333 | -0,0663035 | 0,7197 | 0,95293 |
| Ss18l1 | 3,94195 | 3,71794 | -0,0844075 | 0,7198 | 0,95293 |
| Ltn1 | 14,7039 | 15,3795 | 0,0648069 | 0,71985 | 0,95293 |
| Srpk1 | 61,7949 | 64,6473 | 0,0651037 | 0,71995 | 0,95293 |
| Supt20 | 34,4697 | 36,1038 | 0,066819 | 0,72005 | 0,95293 |
| Xpnpep3 | 5,02922 | 5,33657 | 0,0855776 | 0,7201 | 0,95293 |
| Polb | 43,2331 | 41,0135 | -0,0760383 | 0,7201 | 0,95293 |
| 9130221H12Rik | 9,93043 | 8,60388 | -0,20687 | 0,72015 | 0,95293 |
| Zfp583 | 2,70969 | 2,51958 | -0,10494 | 0,7203 | 0,95293 |
| Mvk | 24,9357 | 26,1661 | 0,069485 | 0,72055 | 0,95293 |
| Snord4a | 79,2725 | 96,4891 | 0,283546 | 0,72065 | 0,95293 |
| 0610040B10Rik | 3,85624 | 4,41756 | 0,196056 | 0,7207 | 0,95293 |
| Sh3d19 | 4,44128 | 4,21303 | -0,0761164 | 0,7207 | 0,95293 |
| Tfap4 | 28,5264 | 29,9267 | 0,069135 | 0,7208 | 0,95293 |

| | | | | | |
|---|---|---|---|---|---|
| Slc27a1 | 3,94825 | 3,70518 | -0,091668 | 0,7211 | 0,95293 |
| Clk4 | 36,6645 | 38,4338 | 0,0679924 | 0,72115 | 0,95293 |
| Zfp383 | 10,4701 | 9,93176 | -0,0761559 | 0,72125 | 0,95293 |
| Sun2 | 11,3191 | 11,8875 | 0,07069 | 0,7213 | 0,95293 |
| Tiprl | 16,3944 | 17,1696 | 0,0666501 | 0,7213 | 0,95293 |
| Acadsb | 10,9451 | 11,4945 | 0,0706635 | 0,72155 | 0,95293 |
| Elmo2 | 20,0926 | 21,041 | 0,0665337 | 0,72155 | 0,95293 |
| Ankrd49 | 20,6568 | 19,6168 | -0,0745299 | 0,72155 | 0,95293 |
| Vps26b | 17,3821 | 16,4225 | -0,0819308 | 0,72155 | 0,95293 |
| Mff | 55,8887 | 53,4003 | -0,0657092 | 0,72165 | 0,95293 |
| Nupl2 | 7,16108 | 6,77543 | -0,0798638 | 0,72165 | 0,95293 |
| Cbx6 | 8,6097 | 9,05438 | 0,0726526 | 0,72195 | 0,95293 |
| Tinf2 | 36,7992 | 38,5631 | 0,0675441 | 0,72195 | 0,95293 |
| Pde4dip | 8,93394 | 9,35503 | 0,0664456 | 0,72195 | 0,95293 |
| Zfp518b | 20,793 | 21,737 | 0,0640546 | 0,72195 | 0,95293 |
| Pde3b | 13,0856 | 13,6993 | 0,0661242 | 0,722 | 0,95293 |
| Ate1 | 10,8066 | 11,3406 | 0,0695801 | 0,72205 | 0,95293 |
| Bysl | 28,6412 | 26,8971 | -0,0906406 | 0,72205 | 0,95293 |
| D2Wsu81e | 47,3255 | 45,1888 | -0,0666517 | 0,7221 | 0,95293 |
| Lmbr1 | 2,67728 | 2,84076 | 0,085513 | 0,72215 | 0,95293 |
| Ddx55 | 9,11102 | 9,60779 | 0,0765921 | 0,72215 | 0,95293 |
| U2af2 | 107,059 | 102,492 | -0,0628918 | 0,7222 | 0,95293 |
| St13 | 268,797 | 257,183 | -0,0637216 | 0,7222 | 0,95293 |
| Mfsd3 | 2,22605 | 2,62281 | 0,23663 | 0,72235 | 0,953029 |
| Itpkc | 7,24289 | 7,64201 | 0,0773872 | 0,72245 | 0,953031 |
| Vdac2 | 143,15 | 136,98 | -0,063563 | 0,7225 | 0,953031 |
| Xpot | 22,7422 | 23,7867 | 0,0647826 | 0,72265 | 0,953131 |
| 4930563E22Rik | 0,701514 | 0,575392 | -0,285926 | 0,7229 | 0,953343 |
| Knop1 | 62,1028 | 59,1947 | -0,0691896 | 0,723 | 0,953343 |
| Zfp707 | 4,06753 | 4,36632 | 0,102266 | 0,72305 | 0,953343 |
| 2310068J16Rik | 0,877078 | 1,44549 | 0,720777 | 0,7232 | 0,953343 |
| 4930402H24Rik | 3,40133 | 3,20733 | -0,0847271 | 0,7232 | 0,953343 |
| Tsn | 82,6921 | 79,1619 | -0,0629431 | 0,72345 | 0,953343 |
| Pts | 23,4855 | 22,2443 | -0,0783339 | 0,7235 | 0,953343 |
| Ssrp1 | 302,382 | 288,825 | -0,0661779 | 0,7236 | 0,953343 |
| Rfx1 | 7,2498 | 7,64073 | 0,0757698 | 0,72365 | 0,953343 |
| 4932702P03Rik | 0,759597 | 1,51596 | 0,99693 | 0,7237 | 0,953343 |
| 1700037C18Rik | 1,82712 | 2,03463 | 0,155199 | 0,72375 | 0,953343 |
| Zbtb39 | 12,4032 | 12,9995 | 0,0677349 | 0,7238 | 0,953343 |
| Lgi4 | 1,01019 | 0,924288 | -0,12821 | 0,7239 | 0,953343 |
| Elmod2 | 7,15186 | 7,51118 | 0,0707209 | 0,72395 | 0,953343 |
| Hmcn1 | 7,50126 | 7,17068 | -0,0650225 | 0,72395 | 0,953343 |
| Hars2 | 16,3394 | 17,1531 | 0,0701208 | 0,724 | 0,953343 |
| Atp6v1f | 57,0388 | 60,055 | 0,0743389 | 0,72415 | 0,953345 |
| Tigd2 | 5,05165 | 4,75673 | -0,0867844 | 0,72415 | 0,953345 |
| 3110052M02Rik | 1,94998 | 1,82004 | -0,0994899 | 0,72425 | 0,953379 |
| Pisd-ps1 | 4,43833 | 4,93822 | 0,153975 | 0,7244 | 0,953384 |
| A230103J11Rik | 1,88089 | 2,60539 | 0,470085 | 0,72455 | 0,953384 |
| Uprt | 2,64326 | 2,90385 | 0,135651 | 0,72455 | 0,953384 |
| Sesn3 | 13,863 | 13,1558 | -0,0755321 | 0,7246 | 0,953384 |

| | | | | | |
|---|---|---|---|---|---|
| Klhdc3 | 64,2689 | 59,4232 | -0,113095 | 0,72465 | 0,953384 |
| Dnajb2 | 10,1322 | 9,58172 | -0,0805942 | 0,7247 | 0,953384 |
| Sh2b1 | 34,0222 | 35,5722 | 0,0642758 | 0,725 | 0,953584 |
| Idh3a | 45,0595 | 43,0892 | -0,0645043 | 0,725 | 0,953584 |
| Spryd4 | 18,7728 | 19,7737 | 0,074941 | 0,72515 | 0,953683 |
| Wdr8 | 16,8917 | 16,0332 | -0,0752584 | 0,72525 | 0,95369 |
| 4930414L22Rik | 2,65298 | 3,83556 | 0,531826 | 0,72535 | 0,95369 |
| Rbck1 | 28,9859 | 27,7036 | -0,0652758 | 0,7255 | 0,95369 |
| Gns | 20,8065 | 19,8848 | -0,0653677 | 0,7255 | 0,95369 |
| Fbxo16 | 6,93021 | 6,45958 | -0,101458 | 0,72555 | 0,95369 |
| Rwdd2a | 0,95181 | 1,0791 | 0,181077 | 0,72565 | 0,95369 |
| Thnsl2 | 1,70868 | 1,5667 | -0,125155 | 0,7257 | 0,95369 |
| Mdh2 | 352,371 | 337,229 | -0,0633677 | 0,72575 | 0,95369 |
| Nrbf2 | 4,30437 | 4,00337 | -0,104588 | 0,72585 | 0,953724 |
| Ap4m1 | 20,9239 | 18,9643 | -0,141866 | 0,72635 | 0,954283 |
| Ppcdc | 5,16855 | 4,87119 | -0,0854839 | 0,72645 | 0,954315 |
| Klf4 | 1,07565 | 1,17174 | 0,123437 | 0,72675 | 0,954315 |
| Cct8 | 431,292 | 411,009 | -0,069495 | 0,72685 | 0,954315 |
| Timeless | 40,319 | 42,1011 | 0,0623985 | 0,72695 | 0,954315 |
| 9430015G10Rik | 38,7091 | 40,4198 | 0,0623866 | 0,727 | 0,954315 |
| Snx2 | 37,4192 | 35,7359 | -0,0664048 | 0,727 | 0,954315 |
| Actg1 | 8,54485 | 9,06143 | 0,0846844 | 0,7272 | 0,954315 |
| Polr1a | 33,4426 | 34,9202 | 0,0623719 | 0,72725 | 0,954315 |
| Cdc5l | 29,9837 | 31,3787 | 0,0656066 | 0,7275 | 0,954315 |
| Dynlrb1 | 211,197 | 202,199 | -0,0628121 | 0,72755 | 0,954315 |
| Asic1 | 1,29056 | 1,39289 | 0,11009 | 0,72765 | 0,954315 |
| Sec24a | 4,8358 | 5,07304 | 0,0690941 | 0,7278 | 0,954315 |
| Utp6 | 52,4212 | 50,1934 | -0,0626549 | 0,7279 | 0,954315 |
| Unk | 17,7858 | 18,6094 | 0,0653059 | 0,72815 | 0,954315 |
| Nr2c1 | 9,35236 | 8,89854 | -0,0717614 | 0,72815 | 0,954315 |
| Ctbp2 | 58,1103 | 60,7184 | 0,0633402 | 0,7282 | 0,954315 |
| Lrig2 | 5,7441 | 6,02371 | 0,0685714 | 0,72825 | 0,954315 |
| Atp6ap1 | 39,7177 | 41,5095 | 0,0636578 | 0,72825 | 0,954315 |
| Ltbp3 | 1,00646 | 1,20048 | 0,254329 | 0,7284 | 0,954315 |
| Med17 | 17,569 | 18,3714 | 0,0644275 | 0,7284 | 0,954315 |
| AY358078 | 4,49019 | 4,83089 | 0,105511 | 0,7285 | 0,954315 |
| Raver2 | 5,85521 | 5,18855 | -0,174389 | 0,7285 | 0,954315 |
| Aen | 24,3833 | 23,2798 | -0,0668119 | 0,72855 | 0,954315 |
| Hmga1 | 32,205 | 28,5553 | -0,173526 | 0,7286 | 0,954315 |
| Rftn2 | 6,24115 | 5,91748 | -0,0768302 | 0,7287 | 0,954315 |
| Mavs | 9,62431 | 11,3228 | 0,234477 | 0,7288 | 0,954315 |
| Tsta3 | 35,8943 | 34,2321 | -0,068403 | 0,7288 | 0,954315 |
| Rpl35 | 18,3264 | 16,9538 | -0,112318 | 0,72905 | 0,954315 |
| Tmem117 | 0,935009 | 1,02387 | 0,130977 | 0,7291 | 0,954315 |
| Pgbd1 | 3,8282 | 4,06326 | 0,0859727 | 0,7292 | 0,954315 |
| Pabpc4l | 1,94141 | 2,06796 | 0,0911056 | 0,72925 | 0,954315 |
| Nudt17 | 3,02869 | 2,48998 | -0,282556 | 0,72925 | 0,954315 |
| Dnph1 | 38,3085 | 36,4963 | -0,0699147 | 0,7294 | 0,954315 |
| Lmnb1 | 260,661 | 249,216 | -0,0647769 | 0,72945 | 0,954315 |
| Tnfsf12 | 1,20777 | 1,35223 | 0,162996 | 0,7295 | 0,954315 |

| | | | | | |
|---|---|---|---|---|---|
| Klhl18 | 5,18066 | 5,45562 | 0,0746076 | 0,7295 | 0,954315 |
| Leprotl1 | 18,0375 | 17,209 | -0,0678374 | 0,72955 | 0,954315 |
| Nipa2 | 12,9589 | 14,436 | 0,15572 | 0,72975 | 0,954315 |
| Pgam2 | 0,851255 | 0,720932 | -0,239729 | 0,72985 | 0,954315 |
| E230016M11Rik | 0,71095 | 0,901751 | 0,342981 | 0,7299 | 0,954315 |
| Slc35d1 | 5,613 | 5,30794 | -0,0806198 | 0,72995 | 0,954315 |
| Spata5l1 | 11,2901 | 11,9777 | 0,0852926 | 0,73005 | 0,954315 |
| B4galt3 | 20,8687 | 21,8982 | 0,0694674 | 0,73005 | 0,954315 |
| Depdc1b | 14,2828 | 14,992 | 0,0699163 | 0,73015 | 0,954315 |
| BC030307 | 1,72933 | 1,88281 | 0,122668 | 0,7302 | 0,954315 |
| Zfp746 | 16,369 | 17,1306 | 0,0656075 | 0,73035 | 0,954315 |
| Gstm5 | 184,26 | 175,92 | -0,066826 | 0,73035 | 0,954315 |
| Ppil2 | 34,8799 | 32,9848 | -0,0805927 | 0,73035 | 0,954315 |
| 4930515G01Rik | 0,966568 | 1,09656 | 0,182046 | 0,7304 | 0,954315 |
| Rab43 | 4,25921 | 4,50087 | 0,079619 | 0,73045 | 0,954315 |
| Rbm5 | 65,3179 | 68,1743 | 0,0617492 | 0,7305 | 0,954315 |
| Brd9 | 30,631 | 32,0283 | 0,0643538 | 0,73055 | 0,954315 |
| Pcdhgc5 | 0,565802 | 0,952275 | 0,75108 | 0,7308 | 0,954315 |
| Tmem115 | 15,3121 | 14,5836 | -0,0703218 | 0,73085 | 0,954315 |
| Tardbp | 127,317 | 121,652 | -0,0656671 | 0,7309 | 0,954315 |
| BC055324 | 25,8722 | 27,2831 | 0,0766072 | 0,731 | 0,954315 |
| Snhg7 | 13,089 | 12,1996 | -0,101523 | 0,731 | 0,954315 |
| Hspa9 | 212,919 | 203,652 | -0,0642007 | 0,73105 | 0,954315 |
| Rragc | 24,2247 | 23,1617 | -0,0647407 | 0,7311 | 0,954315 |
| Fbxo18 | 35,2523 | 36,7999 | 0,0619843 | 0,73115 | 0,954315 |
| Sfpq | 406,475 | 425,976 | 0,0676047 | 0,7312 | 0,954315 |
| Krt10 | 3,39772 | 3,16102 | -0,104173 | 0,7312 | 0,954315 |
| Rasl12 | 1,83247 | 1,98068 | 0,112205 | 0,73125 | 0,954315 |
| Ubc | 42,9413 | 44,8329 | 0,0621922 | 0,7313 | 0,954315 |
| Mrpl10 | 52,2863 | 50,0528 | -0,0629808 | 0,7313 | 0,954315 |
| Pecr | 15,3568 | 14,5238 | -0,0804496 | 0,73135 | 0,954315 |
| Flywch1 | 47,1309 | 49,2147 | 0,0624158 | 0,7314 | 0,954315 |
| Rbm12 | 39,6891 | 42,2683 | 0,0908351 | 0,7315 | 0,954315 |
| Slc25a1 | 51,1836 | 48,988 | -0,0632536 | 0,7316 | 0,954315 |
| Cdip1 | 37,0749 | 35,2461 | -0,0729757 | 0,7316 | 0,954315 |
| Pafah2 | 3,76997 | 3,5439 | -0,0892145 | 0,73175 | 0,954315 |
| Ap3d1 | 31,0369 | 29,7438 | -0,0613964 | 0,73195 | 0,954315 |
| Prr22 | 1,98367 | 1,80268 | -0,138027 | 0,73195 | 0,954315 |
| 2210404O09Rik | 0,944725 | 0,852229 | -0,148653 | 0,73195 | 0,954315 |
| Ttf2 | 26,5586 | 25,4405 | -0,0620513 | 0,732 | 0,954315 |
| Hnrnpr | 28,6332 | 27,4567 | -0,0605335 | 0,73205 | 0,954315 |
| Csl | 22,0654 | 21,0643 | -0,0669918 | 0,73215 | 0,954315 |
| Reps1 | 35,6908 | 34,1607 | -0,0632152 | 0,7322 | 0,954315 |
| Mmgt1 | 22,9965 | 24,0049 | 0,061917 | 0,73235 | 0,954315 |
| Mblac2 | 0,886396 | 0,813676 | -0,123498 | 0,73235 | 0,954315 |
| Eif2a | 63,3777 | 58,9784 | -0,103789 | 0,7324 | 0,954315 |
| Mettl10 | 38,0086 | 36,2026 | -0,0702319 | 0,7325 | 0,954316 |
| Bin1 | 42,974 | 41,1806 | -0,0615012 | 0,73255 | 0,954316 |
| Slc31a2 | 3,88897 | 3,62362 | -0,101958 | 0,73265 | 0,95435 |
| Tmem167b | 17,7356 | 16,9379 | -0,0663934 | 0,7329 | 0,954429 |

| | | | | | |
|---|---|---|---|---|---|
| Usp5 | 75,8909 | 72,7639 | -0,0607049 | 0,73305 | 0,954429 |
| Usp4 | 49,4129 | 47,3511 | -0,0614902 | 0,73305 | 0,954429 |
| Zfand2a | 9,19162 | 9,65215 | 0,0705304 | 0,7331 | 0,954429 |
| Ttc26 | 7,87738 | 8,27214 | 0,0705449 | 0,73315 | 0,954429 |
| Mrps5 | 28,3554 | 27,0647 | -0,0672103 | 0,73325 | 0,954429 |
| Tex264 | 9,89138 | 10,5149 | 0,0881909 | 0,7333 | 0,954429 |
| Fam169a | 8,56192 | 8,07135 | -0,0851233 | 0,73345 | 0,954429 |
| Cog5 | 12,6643 | 11,7102 | -0,113009 | 0,73345 | 0,954429 |
| Fbxo27 | 0,909446 | 1,01001 | 0,151309 | 0,7335 | 0,954429 |
| Nabp1 | 1,95026 | 1,81331 | -0,10504 | 0,73365 | 0,954429 |
| Ifi30 | 15,6699 | 14,7699 | -0,085337 | 0,73375 | 0,954429 |
| Wdtc1 | 10,5394 | 10,0603 | -0,0671241 | 0,7338 | 0,954429 |
| Ctnnbl1 | 41,2699 | 43,0748 | 0,0617547 | 0,73385 | 0,954429 |
| Zfp319 | 6,56108 | 6,24539 | -0,0711425 | 0,7339 | 0,954429 |
| Gstp1 | 39,1185 | 37,1178 | -0,0757375 | 0,734 | 0,954429 |
| Mt2 | 1,13418 | 0,940877 | -0,269569 | 0,734 | 0,954429 |
| Ptger1 | 4,65783 | 5,18179 | 0,153794 | 0,73405 | 0,954429 |
| Bcl7a | 36,4639 | 38,041 | 0,0610877 | 0,73415 | 0,954462 |
| Epn1 | 50,7262 | 48,5603 | -0,0629545 | 0,73425 | 0,954495 |
| Ctdnep1 | 49,3827 | 46,306 | -0,0928057 | 0,7344 | 0,954562 |
| Rrm2b | 1,42868 | 1,33146 | -0,10168 | 0,73445 | 0,954562 |
| Babam1 | 55,1862 | 52,7893 | -0,0640605 | 0,73455 | 0,954595 |
| Mtmr10 | 5,26599 | 5,52893 | 0,0702971 | 0,73465 | 0,954628 |
| Slc11a2 | 15,2619 | 14,5812 | -0,0658228 | 0,7348 | 0,954695 |
| Gltpd1 | 6,82854 | 7,20398 | 0,0772159 | 0,73485 | 0,954695 |
| Smyd5 | 49,3388 | 51,5041 | 0,061966 | 0,73495 | 0,954728 |
| Kdm4a | 31,7065 | 30,3816 | -0,061583 | 0,73515 | 0,954891 |
| 2410006H16Rik | 187,597 | 196,421 | 0,0663114 | 0,7353 | 0,954989 |
| Lss | 28,3789 | 29,6166 | 0,0615856 | 0,7355 | 0,955056 |
| Gna12 | 34,4802 | 32,9592 | -0,0650856 | 0,7355 | 0,955056 |
| C78339 | 13,2192 | 13,9729 | 0,0800046 | 0,7356 | 0,955089 |
| 1810026B05Rik | 6,24007 | 5,58926 | -0,158903 | 0,73575 | 0,955161 |
| Snora16a | 121,223 | 83,8203 | -0,532297 | 0,73585 | 0,955161 |
| Zfp526 | 3,66017 | 3,44896 | -0,0857473 | 0,7359 | 0,955161 |
| Cic | 20,9129 | 20,0464 | -0,0610509 | 0,7362 | 0,955161 |
| Hoga1 | 0,732196 | 0,828804 | 0,178801 | 0,73635 | 0,955161 |
| Wibg | 51,6814 | 49,3993 | -0,0651546 | 0,73635 | 0,955161 |
| Golga5 | 17,753 | 16,9519 | -0,0666162 | 0,73635 | 0,955161 |
| Fam122a | 10,2483 | 10,8678 | 0,0846691 | 0,7364 | 0,955161 |
| 1810030O07Rik | 15,6752 | 16,4164 | 0,0666549 | 0,7364 | 0,955161 |
| Tbc1d9b | 13,5546 | 14,146 | 0,0616197 | 0,7364 | 0,955161 |
| Slc18b1 | 2,46503 | 2,63978 | 0,0988095 | 0,7367 | 0,955453 |
| Rsph3b | 0,657251 | 0,74729 | 0,185225 | 0,7368 | 0,955456 |
| Zfp109 | 2,28375 | 2,1256 | -0,10353 | 0,7369 | 0,955456 |
| Klhdc4 | 37,3431 | 39,0077 | 0,0629146 | 0,73695 | 0,955456 |
| Ccdc93 | 6,51217 | 6,81187 | 0,0649117 | 0,737 | 0,955456 |
| Ino80c | 14,5654 | 13,8757 | -0,0699873 | 0,73715 | 0,955554 |
| Stat2 | 3,9349 | 4,14834 | 0,0762086 | 0,7373 | 0,955597 |
| Stk40 | 23,1479 | 22,148 | -0,063704 | 0,73745 | 0,955597 |
| Gprasp2 | 10,0744 | 9,62065 | -0,0664869 | 0,7375 | 0,955597 |

| | | | | | |
|---|---|---|---|---|---|
| Zfp606 | 10,6346 | 10,1298 | -0,0701513 | 0,7375 | 0,955597 |
| Trp53inp2 | 7,02762 | 7,37284 | 0,0691843 | 0,73765 | 0,955597 |
| Jpx | 0,946947 | 1,02564 | 0,115175 | 0,73775 | 0,955597 |
| Fam160a2 | 6,17761 | 6,44927 | 0,062087 | 0,73785 | 0,955597 |
| Capn7 | 17,3528 | 18,1289 | 0,0631257 | 0,73795 | 0,955597 |
| Mat2b | 50,5767 | 52,7608 | 0,0609956 | 0,738 | 0,955597 |
| Hdac1 | 32,1503 | 30,77 | -0,0633105 | 0,7381 | 0,955597 |
| Zfp458 | 0,636359 | 0,577778 | -0,139326 | 0,73815 | 0,955597 |
| Pvrl3 | 26,1856 | 25,0897 | -0,0616789 | 0,7382 | 0,955597 |
| Acap3 | 6,36429 | 6,0558 | -0,0716823 | 0,73825 | 0,955597 |
| Pyroxd1 | 11,2071 | 10,3283 | -0,117815 | 0,7383 | 0,955597 |
| Gtf2e1 | 25,6581 | 24,5638 | -0,0628803 | 0,73835 | 0,955597 |
| Krtcap3 | 2,29528 | 2,25739 | -0,0240166 | 0,73845 | 0,955597 |
| Cdk9 | 24,3395 | 23,3145 | -0,0620711 | 0,73845 | 0,955597 |
| Hbs1l | 40,9561 | 39,3221 | -0,0587372 | 0,73855 | 0,955599 |
| Rbm15b | 44,2901 | 42,4482 | -0,0612783 | 0,7386 | 0,955599 |
| Pskh1 | 22,2101 | 23,1945 | 0,0625669 | 0,7387 | 0,955632 |
| Ipo13 | 24,1683 | 23,1576 | -0,061634 | 0,73895 | 0,955859 |
| 2210416O15Rik | 0,815186 | 1,30902 | 0,683287 | 0,73905 | 0,955861 |
| Cyb5d1 | 6,06927 | 6,57789 | 0,116103 | 0,7392 | 0,955861 |
| Rfc1 | 54,768 | 52,5863 | -0,0586454 | 0,7392 | 0,955861 |
| Zfp111 | 3,2431 | 3,40605 | 0,0707268 | 0,73925 | 0,955861 |
| BC027231 | 24,206 | 23,1715 | -0,0630134 | 0,73945 | 0,956004 |
| Atp13a2 | 9,71948 | 9,28436 | -0,0660771 | 0,73965 | 0,956004 |
| Cenpk | 32,2367 | 30,2333 | -0,0925653 | 0,73965 | 0,956004 |
| Sugp2 | 18,4353 | 19,3295 | 0,068332 | 0,7398 | 0,956004 |
| Rundc1 | 7,72523 | 7,33806 | -0,0741777 | 0,73995 | 0,956004 |
| Rbbp5 | 24,8365 | 23,797 | -0,0616831 | 0,74 | 0,956004 |
| Anxa7 | 8,93625 | 8,49856 | -0,0724508 | 0,74015 | 0,956004 |
| Magoh | 438,382 | 420,621 | -0,0596665 | 0,7404 | 0,956004 |
| Tcte2 | 1,36777 | 2,47721 | 0,856892 | 0,74055 | 0,956004 |
| Rnf170 | 1,99909 | 2,13251 | 0,0932129 | 0,74055 | 0,956004 |
| Brip1 | 4,01498 | 4,21605 | 0,0705013 | 0,74055 | 0,956004 |
| Bcam | 4,79956 | 4,52449 | -0,0851465 | 0,74055 | 0,956004 |
| Adss | 71,4423 | 68,5817 | -0,0589565 | 0,7406 | 0,956004 |
| Pus1 | 38,4335 | 36,7557 | -0,0643953 | 0,7406 | 0,956004 |
| Zfp119a | 2,14718 | 2,31714 | 0,109905 | 0,74065 | 0,956004 |
| Pcnxl4 | 7,08554 | 6,75546 | -0,0688238 | 0,74075 | 0,956004 |
| Mmab | 13,5441 | 14,1846 | 0,0666553 | 0,7409 | 0,956004 |
| Slc7a1 | 8,36891 | 8,00766 | -0,0636597 | 0,7409 | 0,956004 |
| Ddx59 | 8,90166 | 9,37513 | 0,0747634 | 0,74095 | 0,956004 |
| Atp1a1 | 120,992 | 126,185 | 0,0606285 | 0,74095 | 0,956004 |
| Srsf6 | 197,569 | 189,228 | -0,0622281 | 0,741 | 0,956004 |
| Mfn1 | 15,7939 | 15,1261 | -0,0623321 | 0,741 | 0,956004 |
| Zfp68 | 15,5872 | 14,9191 | -0,0631934 | 0,74125 | 0,956058 |
| Iars | 65,3119 | 62,724 | -0,058327 | 0,74145 | 0,956058 |
| Calm3 | 128,203 | 123,14 | -0,0581289 | 0,7415 | 0,956058 |
| Cnpy3 | 31,3191 | 29,9799 | -0,063044 | 0,7415 | 0,956058 |
| Clk1 | 71,6614 | 74,7947 | 0,0617399 | 0,74155 | 0,956058 |
| Aars | 69,5464 | 66,7356 | -0,0595184 | 0,74155 | 0,956058 |

| | | | | | |
|---|---|---|---|---|---|
| Pomp | 324,596 | 311,586 | -0,0590161 | 0,7416 | 0,956058 |
| Pip4k2c | 11,307 | 12,3365 | 0,125717 | 0,74175 | 0,956058 |
| Zfp647 | 13,0497 | 12,4463 | -0,0683023 | 0,7421 | 0,956058 |
| Plp2 | 10,8697 | 11,6779 | 0,103478 | 0,7423 | 0,956058 |
| Tmem144 | 1,61235 | 1,74757 | 0,116184 | 0,74235 | 0,956058 |
| Ppard | 6,61189 | 6,28077 | -0,0741204 | 0,74235 | 0,956058 |
| Gnas | 1105,26 | 1157,39 | 0,0664941 | 0,7424 | 0,956058 |
| Dck | 46,2265 | 44,4107 | -0,0578123 | 0,7424 | 0,956058 |
| Casp2 | 37,4672 | 35,9256 | -0,0606149 | 0,74245 | 0,956058 |
| Vwce | 1,10591 | 1,19448 | 0,111143 | 0,74255 | 0,956058 |
| Zc3h18 | 41,5911 | 43,2943 | 0,0579044 | 0,74255 | 0,956058 |
| Fbxo42 | 11,902 | 12,4246 | 0,0619938 | 0,7427 | 0,956058 |
| Elp2 | 89,4658 | 85,9086 | -0,058533 | 0,74285 | 0,956058 |
| Plxnb2 | 49,8531 | 47,8274 | -0,0598457 | 0,7429 | 0,956058 |
| Prss36 | 2,07562 | 1,93768 | -0,0992112 | 0,7429 | 0,956058 |
| 5730408K05Rik | 12,9064 | 9,7695 | -0,401725 | 0,743 | 0,956058 |
| Rbm34 | 14,3579 | 15,0126 | 0,0643315 | 0,7431 | 0,956058 |
| 2010012O05Rik | 5,82677 | 5,50157 | -0,0828552 | 0,7431 | 0,956058 |
| Rwdd3 | 2,34589 | 2,16175 | -0,117937 | 0,7431 | 0,956058 |
| Rpl28 | 3,34313 | 2,94329 | -0,183771 | 0,7431 | 0,956058 |
| Cwc25 | 11,6976 | 11,1822 | -0,0650067 | 0,74315 | 0,956058 |
| Alg8 | 31,7602 | 30,4032 | -0,0629989 | 0,74325 | 0,956058 |
| Naa35 | 48,5758 | 46,6394 | -0,0586884 | 0,7434 | 0,956058 |
| Npr2 | 4,33123 | 4,57404 | 0,078693 | 0,74355 | 0,956058 |
| Smim8 | 15,3303 | 14,4638 | -0,0839464 | 0,74355 | 0,956058 |
| Slx1b | 5,67382 | 5,96963 | 0,0733231 | 0,7436 | 0,956058 |
| Taf5l | 22,9169 | 21,9467 | -0,0624103 | 0,7436 | 0,956058 |
| Urb2 | 12,9468 | 13,5027 | 0,0606525 | 0,74365 | 0,956058 |
| Rai14 | 37,6109 | 36,082 | -0,0598728 | 0,74365 | 0,956058 |
| Ppp4c | 206,355 | 198,245 | -0,057848 | 0,74415 | 0,956509 |
| Tmem184c | 14,6893 | 14,0357 | -0,0656586 | 0,74415 | 0,956509 |
| Cog8 | 16,1777 | 16,9432 | 0,0667016 | 0,74425 | 0,956541 |
| Akap8l | 29,7748 | 31,0666 | 0,0612704 | 0,7444 | 0,956581 |
| Ccdc36 | 0,754635 | 0,827121 | 0,132319 | 0,7448 | 0,956581 |
| Pno1 | 79,6879 | 76,5879 | -0,0572439 | 0,74495 | 0,956581 |
| 1700102P08Rik | 0,797766 | 0,693322 | -0,20244 | 0,74495 | 0,956581 |
| AA413626 | 0,698726 | 0,673743 | -0,0525274 | 0,745 | 0,956581 |
| Lbh | 17,0147 | 16,2756 | -0,064074 | 0,745 | 0,956581 |
| Alg2 | 35,6182 | 34,1781 | -0,0595426 | 0,7451 | 0,956581 |
| AU040320 | 8,80784 | 9,7256 | 0,142999 | 0,74515 | 0,956581 |
| Mdn1 | 15,4495 | 16,1082 | 0,0602335 | 0,74515 | 0,956581 |
| Dapl1 | 4,3247 | 3,89351 | -0,151528 | 0,74515 | 0,956581 |
| Zfp791 | 0,71954 | 0,647766 | -0,151603 | 0,74515 | 0,956581 |
| Cox7c | 1,31533 | 1,597 | 0,279943 | 0,74525 | 0,956581 |
| Kpna4 | 44,195 | 42,4232 | -0,0590305 | 0,74525 | 0,956581 |
| Mettl6 | 33,1654 | 34,6252 | 0,0621442 | 0,74535 | 0,956613 |
| Rnf168 | 18,4844 | 17,7231 | -0,0606754 | 0,74545 | 0,956646 |
| Agbl5 | 10,202 | 10,8348 | 0,0868156 | 0,7458 | 0,956779 |
| Rbms1 | 47,5493 | 45,6519 | -0,0587504 | 0,7458 | 0,956779 |
| Nmt1 | 91,9596 | 88,3633 | -0,0575526 | 0,74585 | 0,956779 |

| | | | | | |
|---|---|---|---|---|---|
| Lin9 | 9,47144 | 9,9451 | 0,0704018 | 0,746 | 0,956779 |
| Pars2 | 5,45498 | 5,15724 | -0,0809753 | 0,746 | 0,956779 |
| Maoa | 17,0064 | 17,7394 | 0,0608792 | 0,74605 | 0,956779 |
| Zbtb42 | 1,56381 | 1,67701 | 0,100825 | 0,74615 | 0,956779 |
| Mthfr | 8,22112 | 8,58789 | 0,0629691 | 0,74615 | 0,956779 |
| Dchs1 | 19,3628 | 18,5927 | -0,0585484 | 0,74625 | 0,956811 |
| Pip5k1c | 25,5838 | 26,6424 | 0,0584903 | 0,74635 | 0,956844 |
| Rtn4 | 65,6849 | 68,5477 | 0,0615456 | 0,74645 | 0,956876 |
| Nup160 | 26,9601 | 28,0721 | 0,0583114 | 0,74655 | 0,956909 |
| Maml1 | 15,7084 | 15,0724 | -0,0596295 | 0,7469 | 0,957199 |
| Ggnbp2 | 34,3473 | 35,7995 | 0,0597433 | 0,747 | 0,957199 |
| Tgif2 | 23,8395 | 22,8469 | -0,0613592 | 0,747 | 0,957199 |
| Os9 | 14,9803 | 14,3395 | -0,0630689 | 0,74735 | 0,957552 |
| Hnrnph1 | 571,321 | 597,028 | 0,0634968 | 0,7475 | 0,957585 |
| Mbtps2 | 6,95523 | 7,33961 | 0,0776062 | 0,74755 | 0,957585 |
| Zfp809 | 3,84156 | 5,19552 | 0,435575 | 0,7476 | 0,957585 |
| Iqck | 0,394422 | 0,857439 | 1,1203 | 0,74785 | 0,957809 |
| Fhl3 | 5,73077 | 6,08998 | 0,0877079 | 0,74795 | 0,957842 |
| Ccdc71 | 18,3713 | 17,5826 | -0,0633044 | 0,7483 | 0,958194 |
| Sharpin | 11,5167 | 10,9396 | -0,0741705 | 0,7484 | 0,958227 |
| Ndufaf6 | 15,5366 | 14,6984 | -0,0800087 | 0,7488 | 0,958612 |
| Eftud1 | 11,482 | 11,9952 | 0,0630846 | 0,74885 | 0,958612 |
| Cobl | 0,766608 | 0,711169 | -0,108298 | 0,7492 | 0,958713 |
| Hoxd3os1 | 0,728101 | 1,78285 | 1,29197 | 0,74935 | 0,958713 |
| Gsg2 | 12,6703 | 13,2456 | 0,0640641 | 0,74935 | 0,958713 |
| Acaca | 6,35174 | 6,08181 | -0,0626527 | 0,7495 | 0,958713 |
| Pold3 | 53,1582 | 51,1035 | -0,0568706 | 0,74955 | 0,958713 |
| Nek3 | 3,78615 | 4,03858 | 0,0931184 | 0,74965 | 0,958713 |
| Uck1 | 9,83634 | 9,13532 | -0,106666 | 0,74975 | 0,958713 |
| Tug1 | 37,1194 | 38,6233 | 0,0572996 | 0,74985 | 0,958713 |
| St7 | 10,6806 | 10,1522 | -0,0731973 | 0,7499 | 0,958713 |
| Fam134c | 25,9616 | 27,0312 | 0,0582431 | 0,75 | 0,958713 |
| Psmd9 | 21,2292 | 20,3232 | -0,0629269 | 0,75005 | 0,958713 |
| Mir7060 | 14,9454 | 24,2912 | 0,700728 | 0,7501 | 0,958713 |
| Fam188a | 15,4667 | 14,7804 | -0,0654766 | 0,7501 | 0,958713 |
| Rps16 | 1,79238 | 1,56203 | -0,198455 | 0,7501 | 0,958713 |
| Prickle4 | 2,18489 | 1,61566 | -0,435441 | 0,7501 | 0,958713 |
| 2900008C10Rik | 1,81419 | 2,58025 | 0,508185 | 0,7502 | 0,958713 |
| Mapk8 | 18,0997 | 17,3722 | -0,0591819 | 0,7502 | 0,958713 |
| Rnmt | 21,2751 | 20,4521 | -0,0569191 | 0,7506 | 0,959046 |
| Tram1 | 21,9648 | 21,0676 | -0,0601673 | 0,75075 | 0,959046 |
| Cst3 | 104,186 | 108,626 | 0,0602113 | 0,751 | 0,959046 |
| Cebpzos | 9,63378 | 8,16875 | -0,237987 | 0,75115 | 0,959046 |
| Ube2c | 212,937 | 221,508 | 0,0569316 | 0,7512 | 0,959046 |
| Zmym1 | 15,501 | 16,1655 | 0,0605615 | 0,75125 | 0,959046 |
| Trerf1 | 2,76449 | 2,92437 | 0,0811089 | 0,75135 | 0,959046 |
| Taf9 | 59,5739 | 56,451 | -0,0776804 | 0,75135 | 0,959046 |
| Asb7 | 6,06762 | 5,79721 | -0,0657736 | 0,7516 | 0,959046 |
| Slc12a5 | 0,954209 | 1,021 | 0,0976105 | 0,75165 | 0,959046 |
| Zfp281 | 15,8145 | 15,7937 | -0,0589648 | 0,7517 | 0,959046 |

| | | | | | |
|---|---|---|---|---|---|
| Trmt61b | 13,562 | 12,5539 | -0,111434 | 0,75175 | 0,959046 |
| Trp53bp1 | 38,2317 | 39,8687 | 0,0604876 | 0,7518 | 0,959046 |
| Gramd1c | 0,611341 | 0,556523 | -0,135535 | 0,7519 | 0,959046 |
| Scaf1 | 39,7998 | 41,4053 | 0,0570553 | 0,75195 | 0,959046 |
| Gm1821 | 82,4535 | 66,9367 | -0,300784 | 0,75205 | 0,959046 |
| Tmppe | 2,44475 | 2,28901 | -0,0949618 | 0,7522 | 0,959046 |
| Gm11696 | 0,776333 | 1,04864 | 0,433776 | 0,75225 | 0,959046 |
| Tyw3 | 4,58294 | 4,27251 | -0,101191 | 0,75225 | 0,959046 |
| Ccdc97 | 17,1947 | 17,9454 | 0,0616507 | 0,75235 | 0,959046 |
| M6pr | 39,7453 | 41,3837 | 0,0582764 | 0,75235 | 0,959046 |
| Fam168a | 12,3688 | 11,8754 | -0,0587293 | 0,75235 | 0,959046 |
| Ormdl2 | 13,405 | 12,3069 | -0,123309 | 0,75245 | 0,959046 |
| Tceanc | 2,50223 | 2,36137 | -0,0835892 | 0,7526 | 0,959046 |
| Rpf1 | 64,931 | 62,3157 | -0,0593116 | 0,75265 | 0,959046 |
| 3830408C21Rik | 1,28333 | 1,17202 | -0,13089 | 0,75265 | 0,959046 |
| Ercc2 | 16,3891 | 17,0903 | 0,0604436 | 0,7527 | 0,959046 |
| Tmem183a | 35,7646 | 34,3562 | -0,0579617 | 0,75275 | 0,959046 |
| Zfp867 | 5,21204 | 5,47438 | 0,0708471 | 0,7529 | 0,959046 |
| AV051173 | 0,966904 | 1,65629 | 0,776509 | 0,753 | 0,959046 |
| Setdb1 | 41,5554 | 43,5975 | 0,0692093 | 0,7531 | 0,959046 |
| Phldb2 | 23,2804 | 24,1998 | 0,0558802 | 0,75315 | 0,959046 |
| Rnf10 | 120,19 | 125,004 | 0,0566543 | 0,7532 | 0,959046 |
| Arhgdig | 2,43132 | 2,65914 | 0,129224 | 0,75325 | 0,959046 |
| Kdm2b | 19,4135 | 18,6129 | -0,0607592 | 0,75325 | 0,959046 |
| Prmt6 | 20,2793 | 19,4418 | -0,0608499 | 0,7533 | 0,959046 |
| Snord57 | 3644,94 | 3213,46 | -0,181767 | 0,7533 | 0,959046 |
| Rraga | 68,9785 | 66,3119 | -0,0568785 | 0,7534 | 0,959046 |
| Bcas3 | 5,83235 | 5,5442 | -0,0730988 | 0,75355 | 0,959046 |
| Tmem131 | 24,7146 | 25,7096 | 0,0569478 | 0,7536 | 0,959046 |
| Snord69 | 6275,17 | 5128,66 | -0,291072 | 0,75365 | 0,959046 |
| Pdlim7 | 41,2404 | 39,3043 | -0,0693707 | 0,75375 | 0,959046 |
| Tmem177 | 5,41106 | 5,13972 | -0,0742216 | 0,75375 | 0,959046 |
| Doc2a | 7,66916 | 7,11188 | -0,108837 | 0,75375 | 0,959046 |
| Unc119b | 35,5765 | 37,0705 | 0,0593442 | 0,75385 | 0,959078 |
| Hscb | 13,4913 | 12,694 | -0,0878883 | 0,75395 | 0,959111 |
| Fermt3 | 0,828103 | 0,927015 | 0,162783 | 0,7541 | 0,959206 |
| Vps4b | 29,2037 | 30,398 | 0,057824 | 0,7542 | 0,959207 |
| Stau1 | 37,9989 | 39,536 | 0,0572094 | 0,75425 | 0,959207 |
| Arrdc1 | 1,53454 | 1,66943 | 0,12155 | 0,7545 | 0,959239 |
| Rbm18 | 30,4818 | 31,8808 | 0,0647358 | 0,7545 | 0,959239 |
| Sart3 | 53,943 | 51,8391 | -0,0573963 | 0,7545 | 0,959239 |
| Dmc1 | 2,08146 | 2,22963 | 0,0992104 | 0,75475 | 0,959242 |
| Col5a2 | 17,1757 | 16,514 | -0,0566781 | 0,7549 | 0,959242 |
| Zfp873 | 5,55277 | 5,2551 | -0,0794879 | 0,7549 | 0,959242 |
| Cnn2 | 74,3572 | 55,3336 | -0,426316 | 0,7549 | 0,959242 |
| 0610009B22Rik | 36,4902 | 34,782 | -0,0691657 | 0,755 | 0,959242 |
| A230056P14Rik | 4,97466 | 5,4753 | 0,138339 | 0,7551 | 0,959242 |
| Cpxm1 | 22,2124 | 23,1693 | 0,060844 | 0,7551 | 0,959242 |
| Gnl2 | 57,0874 | 59,3138 | 0,0551956 | 0,7551 | 0,959242 |
| Tmbim6 | 84,194 | 87,5276 | 0,0560197 | 0,75565 | 0,959816 |

| | | | | | |
|---|---|---|---|---|---|
| Ppp6c | 46,8245 | 48,7856 | 0,0591929 | 0,7558 | 0,959816 |
| 4930473A02Rik | 0,339266 | 0,628026 | 0,888408 | 0,75585 | 0,959816 |
| Srfbp1 | 34,3262 | 35,8028 | 0,0607626 | 0,75595 | 0,959816 |
| Dand5 | 1,81759 | 1,98706 | 0,128603 | 0,75605 | 0,959816 |
| Cldn3 | 1,66317 | 1,82918 | 0,137261 | 0,75615 | 0,959816 |
| Rint1 | 10,2884 | 11,5111 | 0,162009 | 0,7562 | 0,959816 |
| Golga7 | 37,2906 | 38,8349 | 0,0585393 | 0,75625 | 0,959816 |
| Zdhhc3 | 16,3642 | 15,7217 | -0,0577888 | 0,75625 | 0,959816 |
| Gltp | 20,7899 | 21,7332 | 0,0640159 | 0,75635 | 0,959816 |
| Efcab7 | 3,61192 | 3,85526 | 0,0940623 | 0,7564 | 0,959816 |
| Taf1c | 7,05439 | 6,72231 | -0,0695632 | 0,75645 | 0,959816 |
| Sap25 | 3,48166 | 4,18939 | 0,266965 | 0,75655 | 0,959848 |
| Ddx54 | 35,3859 | 34,0259 | -0,0565423 | 0,7567 | 0,959912 |
| Cntd1 | 0,806566 | 0,556926 | -0,534307 | 0,75675 | 0,959912 |
| Yipf1 | 18,4716 | 17,6164 | -0,0683891 | 0,75695 | 0,960071 |
| Fuca1 | 7,94625 | 8,33815 | 0,0694518 | 0,75705 | 0,960071 |
| Wbp4 | 21,2374 | 20,3169 | -0,0639277 | 0,7571 | 0,960071 |
| Cdca5 | 49,7158 | 47,7648 | -0,0577581 | 0,7575 | 0,960388 |
| Ttc8 | 13,9876 | 13,3752 | -0,0645797 | 0,7575 | 0,960388 |
| Pgam5 | 53,8994 | 51,8525 | -0,0558567 | 0,75765 | 0,960389 |
| Mfsd5 | 22,5146 | 21,5708 | -0,0617795 | 0,7577 | 0,960389 |
| Acyp1 | 30,4242 | 32,0329 | 0,0743366 | 0,75775 | 0,960389 |
| Ddx24 | 57,9288 | 55,7463 | -0,0554057 | 0,7578 | 0,960389 |
| Spr | 27,6636 | 26,4622 | -0,0640543 | 0,7579 | 0,960421 |
| Tacc2 | 21,0586 | 20,2512 | -0,0564001 | 0,7581 | 0,960548 |
| Tmem123 | 43,7601 | 42,1342 | -0,0546226 | 0,75815 | 0,960548 |
| Lamtor5 | 22,3221 | 23,4798 | 0,0729468 | 0,75825 | 0,960548 |
| Nop14 | 36,5799 | 38,0504 | 0,056861 | 0,7583 | 0,960548 |
| Thrap3 | 51,7229 | 49,7739 | -0,0554147 | 0,7585 | 0,960707 |
| Phka2 | 7,54985 | 7,22055 | -0,0643385 | 0,75865 | 0,960739 |
| Fancf | 4,23375 | 3,97737 | -0,0901209 | 0,7587 | 0,960739 |
| Atg101 | 25,0679 | 26,2126 | 0,064424 | 0,75875 | 0,960739 |
| Zfp775 | 3,34744 | 3,17511 | -0,0762516 | 0,75885 | 0,960771 |
| Fkbp5 | 33,3261 | 34,6321 | 0,055456 | 0,7593 | 0,961151 |
| Tanc1 | 15,1766 | 15,7757 | 0,0558552 | 0,7594 | 0,961151 |
| Reep4 | 17,6267 | 16,8513 | -0,0649072 | 0,7595 | 0,961151 |
| Snord34 | 3289,9 | 4516,65 | 0,457209 | 0,75965 | 0,961151 |
| Ankrd42 | 2,74255 | 2,911 | 0,0859952 | 0,75965 | 0,961151 |
| Rrs1 | 59,1682 | 56,9612 | -0,0548422 | 0,7597 | 0,961151 |
| Arhgap32 | 6,63097 | 6,37331 | -0,0571773 | 0,75975 | 0,961151 |
| Commd5 | 20,6798 | 21,717 | 0,0705976 | 0,7598 | 0,961151 |
| Ubqln4 | 76,0491 | 79,0144 | 0,0551833 | 0,75985 | 0,961151 |
| Abhd11os | 2,80971 | 2,20029 | -0,352727 | 0,7599 | 0,961151 |
| Snord89 | 117,426 | 148,306 | 0,336834 | 0,76005 | 0,961246 |
| Rufy1 | 10,9188 | 11,4076 | 0,0631743 | 0,76035 | 0,961311 |
| Ufd1l | 45,9936 | 44,2386 | -0,0561282 | 0,7604 | 0,961311 |
| Xpa | 11,3797 | 12,0524 | 0,0828646 | 0,76045 | 0,961311 |
| Gyk | 2,75272 | 2,90489 | 0,0776233 | 0,76055 | 0,961311 |
| Ogfod3 | 12,8155 | 11,6513 | -0,137402 | 0,76065 | 0,961311 |
| Ccdc80 | 7,11324 | 6,80086 | -0,0647891 | 0,7607 | 0,961311 |

| | | | | | |
|---|---|---|---|---|---|
| Trap1 | 68,8306 | 71,5061 | 0,0550157 | 0,76075 | 0,961311 |
| Ccdc85c | 10,1262 | 9,62179 | -0,0737164 | 0,76075 | 0,961311 |
| Slc26a6 | 2,42004 | 2,57574 | 0,0899517 | 0,7608 | 0,961311 |
| Pfdn1 | 116,824 | 112,426 | -0,0553694 | 0,76095 | 0,961311 |
| Gsk3a | 51,1124 | 53,2003 | 0,0577613 | 0,761 | 0,961311 |
| Khk | 3,23783 | 2,9941 | -0,112904 | 0,76105 | 0,961311 |
| Zfp939 | 2,03902 | 1,89607 | -0,104864 | 0,7611 | 0,961311 |
| Dhx37 | 14,8647 | 15,4646 | 0,057086 | 0,76125 | 0,961311 |
| Amdhd2 | 11,7986 | 11,2078 | -0,0741122 | 0,76125 | 0,961311 |
| Cyth1 | 8,00991 | 8,37609 | 0,0644917 | 0,7613 | 0,961311 |
| Slc4a1ap | 22,2871 | 23,2205 | 0,0591899 | 0,76145 | 0,961374 |
| Rnf169 | 9,94941 | 10,3304 | 0,0542182 | 0,7615 | 0,961374 |
| Ddx31 | 10,8438 | 11,3178 | 0,0617279 | 0,7617 | 0,961438 |
| Gpld1 | 7,1496 | 7,46485 | 0,0622504 | 0,76185 | 0,961438 |
| Eme1 | 33,6928 | 30,7 | -0,134203 | 0,76185 | 0,961438 |
| Gm5595 | 2,31722 | 2,17932 | -0,0885124 | 0,7619 | 0,961438 |
| 1700007L15Rik | 3,52976 | 3,84534 | 0,123543 | 0,76195 | 0,961438 |
| Cln8 | 4,58137 | 4,78477 | 0,0626711 | 0,762 | 0,961438 |
| Strada | 18,2005 | 18,9507 | 0,0582731 | 0,7622 | 0,961501 |
| Zwint | 54,8504 | 52,8129 | -0,0546114 | 0,76225 | 0,961501 |
| Skiv2l2 | 67,0866 | 64,5053 | -0,0566076 | 0,7623 | 0,961501 |
| Rpl23a | 0,802964 | 0,964156 | 0,263932 | 0,76235 | 0,961501 |
| Serinc1 | 64,3426 | 66,8197 | 0,0544993 | 0,76255 | 0,961564 |
| Syngap1 | 9,74953 | 10,1661 | 0,0603655 | 0,76265 | 0,961564 |
| Srd5a1 | 6,0547 | 5,75943 | -0,0721283 | 0,76265 | 0,961564 |
| Cep85l | 3,45408 | 3,66224 | 0,084423 | 0,7627 | 0,961564 |
| Nxph3 | 2,6802 | 2,85773 | 0,092533 | 0,76285 | 0,961659 |
| Chmp2a | 47,4475 | 45,4894 | -0,0608045 | 0,763 | 0,961753 |
| Bnip1 | 33,7039 | 35,1776 | 0,0617437 | 0,76315 | 0,961817 |
| Bcl9 | 24,5764 | 25,5159 | 0,0541244 | 0,76325 | 0,961817 |
| Slc24a5 | 1,44757 | 1,90544 | 0,396488 | 0,76335 | 0,961817 |
| Ddx27 | 51,5918 | 49,6336 | -0,0558263 | 0,7634 | 0,961817 |
| Phf1 | 3,34475 | 3,15582 | -0,0838829 | 0,7635 | 0,961817 |
| Vegfb | 38,5564 | 36,9798 | -0,0602353 | 0,76375 | 0,961817 |
| Crocc | 8,25482 | 8,58536 | 0,0566421 | 0,7638 | 0,961817 |
| Tnfaip3 | 0,647338 | 0,699674 | 0,112163 | 0,7639 | 0,961817 |
| BC005764 | 28,283 | 19,3203 | -0,549816 | 0,7639 | 0,961817 |
| 2900011O08Rik | 0,876273 | 0,97526 | 0,154406 | 0,76395 | 0,961817 |
| Rnf25 | 28,4744 | 29,6862 | 0,0601261 | 0,76395 | 0,961817 |
| Spata13 | 15,4728 | 14,9043 | -0,0539997 | 0,76395 | 0,961817 |
| Ciao1 | 21,0528 | 21,88 | 0,0555974 | 0,7642 | 0,96188 |
| Zbtb2 | 8,26964 | 7,91281 | -0,0636346 | 0,76425 | 0,96188 |
| Ccdc88c | 6,79915 | 6,52303 | -0,0598119 | 0,7644 | 0,96188 |
| Smarca4 | 86,92 | 83,6993 | -0,0544725 | 0,76445 | 0,96188 |
| Daxx | 32,728 | 31,4907 | -0,0555977 | 0,76445 | 0,96188 |
| Ftsj3 | 95,8478 | 92,3475 | -0,0536728 | 0,7645 | 0,96188 |
| Ufsp1 | 6,96634 | 6,5436 | -0,0903159 | 0,76475 | 0,96188 |
| Ppp2r2d | 45,5777 | 43,8428 | -0,0559872 | 0,7648 | 0,96188 |
| Eci2 | 41,634 | 40,0019 | -0,0576932 | 0,7648 | 0,96188 |
| Snhg10 | 3,41054 | 3,79444 | 0,153888 | 0,765 | 0,96188 |

| | | | | | |
|---|---|---|---|---|---|
| Utp20 | 21,1523 | 21,9545 | 0,0536988 | 0,7651 | 0,96188 |
| Gemin4 | 26,8084 | 25,7912 | -0,0558067 | 0,7651 | 0,96188 |
| Mief1 | 11,0537 | 10,6168 | -0,0581711 | 0,7651 | 0,96188 |
| Sdf2 | 34,1644 | 32,7935 | -0,0590846 | 0,76515 | 0,96188 |
| Dcaf17 | 9,19442 | 9,7385 | 0,0829408 | 0,7652 | 0,96188 |
| Sp3os | 4,85004 | 3,76459 | -0,365506 | 0,7652 | 0,96188 |
| Ghitm | 68,9496 | 71,4742 | 0,0518805 | 0,76535 | 0,961943 |
| Gm16596 | 0,576582 | 1,01336 | 0,813547 | 0,7654 | 0,961943 |
| Apoa2 | 2,4926 | 2,8161 | 0,176046 | 0,76555 | 0,961943 |
| Commd10 | 33,9534 | 32,6012 | -0,0586286 | 0,7656 | 0,961943 |
| Cdk5rap1 | 6,60032 | 6,95463 | 0,0754378 | 0,7657 | 0,961943 |
| Nek9 | 37,0414 | 38,4081 | 0,0522735 | 0,7657 | 0,961943 |
| Arhgap18 | 1,1916 | 1,11116 | -0,100837 | 0,7658 | 0,961974 |
| Tmem126a | 57,7286 | 55,3647 | -0,0603182 | 0,76615 | 0,962006 |
| Morn1 | 1,59621 | 1,46478 | -0,123963 | 0,76615 | 0,962006 |
| Trex1 | 10,6541 | 11,2572 | 0,0794454 | 0,76625 | 0,962006 |
| Zfyve19 | 14,9894 | 15,7412 | 0,0706021 | 0,7663 | 0,962006 |
| Ttpal | 14,3813 | 13,8284 | -0,0565679 | 0,7663 | 0,962006 |
| Ccnt1 | 18,1293 | 18,9107 | 0,0608773 | 0,76635 | 0,962006 |
| Med29 | 7,45497 | 7,09633 | -0,0711301 | 0,76635 | 0,962006 |
| Tax1bp3 | 0,846802 | 0,756611 | -0,162472 | 0,76645 | 0,962037 |
| Tead2 | 272,25 | 262,35 | -0,0534407 | 0,7666 | 0,962131 |
| Cox7a2 | 294,189 | 283,603 | -0,0528729 | 0,7667 | 0,962163 |
| Txnrd2 | 12,1037 | 11,1487 | -0,118568 | 0,76685 | 0,962163 |
| Parp6 | 9,69534 | 10,1332 | 0,0637247 | 0,767 | 0,962163 |
| Mapk11 | 12,0712 | 12,5918 | 0,0609157 | 0,767 | 0,962163 |
| C87436 | 7,32433 | 7,66096 | 0,0648295 | 0,76705 | 0,962163 |
| Nfkbia | 11,6909 | 13,6886 | 0,227588 | 0,76715 | 0,962163 |
| Ankzf1 | 14,447 | 13,6655 | -0,0802354 | 0,76715 | 0,962163 |
| Arpc5 | 41,9481 | 40,3649 | -0,055503 | 0,76725 | 0,962163 |
| Taok3 | 1,67675 | 1,57937 | -0,0863173 | 0,7673 | 0,962163 |
| Sik3 | 8,767 | 9,11715 | 0,0564994 | 0,76745 | 0,962225 |
| Chd8 | 29,0562 | 30,1488 | 0,0532561 | 0,7675 | 0,962225 |
| Gm11837 | 0,547022 | 0,73898 | 0,433936 | 0,76765 | 0,962319 |
| Pkd2 | 16,0921 | 15,5084 | -0,0533028 | 0,76775 | 0,962351 |
| Mrpl48 | 15,6768 | 13,5253 | -0,212968 | 0,76805 | 0,962382 |
| Mat2a | 133,001 | 128,123 | -0,0539159 | 0,7681 | 0,962382 |
| Clp1 | 26,4825 | 25,4437 | -0,0577271 | 0,7681 | 0,962382 |
| Akap8 | 67,7096 | 70,2031 | 0,0521746 | 0,76815 | 0,962382 |
| Zfyve21 | 18,1586 | 17,3648 | -0,0644893 | 0,76815 | 0,962382 |
| Vkorc1l1 | 21,2886 | 22,0919 | 0,0534377 | 0,7683 | 0,962476 |
| Dnajb6 | 7,99127 | 8,43825 | 0,0785182 | 0,76855 | 0,962507 |
| Sh3pxd2b | 20,9854 | 20,2044 | -0,054716 | 0,76855 | 0,962507 |
| Mzt2 | 26,7838 | 25,6062 | -0,0648682 | 0,76855 | 0,962507 |
| Lck | 5,75661 | 6,06672 | 0,0756977 | 0,76865 | 0,962538 |
| Ngef | 0,584429 | 0,640344 | 0,131819 | 0,76885 | 0,962632 |
| Actr3 | 48,4542 | 50,2562 | 0,0526787 | 0,7689 | 0,962632 |
| Limk2 | 13,6775 | 15,3925 | 0,170424 | 0,76895 | 0,962632 |
| Jrk | 3,9538 | 4,13584 | 0,0649408 | 0,7695 | 0,963007 |
| Gpatch3 | 5,65306 | 5,3626 | -0,0760982 | 0,7696 | 0,963007 |

| | | | | | |
|---|---|---|---|---|---|
| Pcsk7 | 8,29181 | 8,87182 | 0,0975424 | 0,76975 | 0,963007 |
| Cpsf1 | 39,4597 | 41,0031 | 0,0553536 | 0,76975 | 0,963007 |
| Fnbp4 | 33,4493 | 34,6973 | 0,0528449 | 0,76975 | 0,963007 |
| Ppp6r3 | 50,3052 | 52,1706 | 0,0525306 | 0,7699 | 0,963007 |
| Arrdc4 | 11,7511 | 12,374 | 0,0745181 | 0,76995 | 0,963007 |
| Toporsos | 34,3544 | 27,452 | -0,323584 | 0,76995 | 0,963007 |
| Maz | 139,953 | 134,928 | -0,0527582 | 0,77005 | 0,963007 |
| Tril | 16,4722 | 17,1089 | 0,0547185 | 0,77015 | 0,963007 |
| Snapc5 | 43,4 | 41,5502 | -0,0628404 | 0,77015 | 0,963007 |
| Slc29a4 | 5,16161 | 4,90882 | -0,0724438 | 0,77015 | 0,963007 |
| A830082N09Rik | 0,911574 | 0,965225 | 0,0825051 | 0,77025 | 0,963038 |
| Glcci1 | 9,00769 | 8,67165 | -0,0548505 | 0,7705 | 0,963162 |
| Amz2 | 29,8641 | 31,0112 | 0,0543769 | 0,7706 | 0,963162 |
| 2810442I21Rik | 3,37645 | 3,17118 | -0,0904869 | 0,7706 | 0,963162 |
| Cep95 | 13,9453 | 13,3888 | -0,0587493 | 0,77065 | 0,963162 |
| Izumo4 | 1,04499 | 1,1797 | 0,174934 | 0,7708 | 0,963162 |
| Bche | 3,88993 | 4,0621 | 0,0624819 | 0,7708 | 0,963162 |
| Gdap10 | 1,85981 | 2,25223 | 0,276202 | 0,77095 | 0,963224 |
| Sertad1 | 7,58834 | 7,16418 | -0,0829825 | 0,771 | 0,963224 |
| Cdc42bpg | 2,05056 | 1,94728 | -0,0745582 | 0,7712 | 0,963254 |
| BC065397 | 2,70562 | 3,30284 | 0,287749 | 0,77125 | 0,963254 |
| Usp20 | 3,41741 | 3,2543 | -0,0705581 | 0,77125 | 0,963254 |
| Mir7052 | 349,271 | 419,309 | 0,263666 | 0,7714 | 0,963254 |
| Med19 | 37,2758 | 34,9402 | -0,093353 | 0,7714 | 0,963254 |
| Trappc1 | 98,0361 | 94,3581 | -0,0551668 | 0,7715 | 0,963254 |
| Ccdc142 | 3,13647 | 2,83493 | -0,145829 | 0,77155 | 0,963254 |
| Hmgb2 | 29,7562 | 28,6544 | -0,0544301 | 0,77165 | 0,963254 |
| Lrrc41 | 34,1167 | 32,4296 | -0,07317 | 0,77195 | 0,963254 |
| Rsph9 | 1,93669 | 1,75633 | -0,14103 | 0,77195 | 0,963254 |
| St7l | 4,57001 | 4,76411 | 0,0600083 | 0,772 | 0,963254 |
| Atmin | 8,64776 | 8,99273 | 0,0564326 | 0,772 | 0,963254 |
| Epb4.1 | 74,7666 | 77,5936 | 0,053545 | 0,772 | 0,963254 |
| Cad | 53,0596 | 48,5902 | -0,12695 | 0,77235 | 0,963378 |
| Mars | 62,3117 | 60,031 | -0,0537958 | 0,77255 | 0,963378 |
| Vegfc | 19,6208 | 18,8366 | -0,0588393 | 0,77255 | 0,963378 |
| Snora81 | 38,9019 | 65,8433 | 0,759196 | 0,77265 | 0,963378 |
| Chordc1 | 68,3356 | 70,8699 | 0,0525358 | 0,77265 | 0,963378 |
| Rhoj | 1,73554 | 1,6188 | -0,100465 | 0,77265 | 0,963378 |
| Fv1 | 13,0248 | 13,6527 | 0,0679229 | 0,7727 | 0,963378 |
| Scamp3 | 38,4995 | 40,0406 | 0,0566234 | 0,77275 | 0,963378 |
| Thap1 | 6,55735 | 6,87905 | 0,0690953 | 0,77285 | 0,963378 |
| Kdelr3 | 5,15016 | 2,51126 | -1,0362 | 0,77285 | 0,963378 |
| Nudt5 | 74,8655 | 72,1827 | -0,0526463 | 0,77295 | 0,963409 |
| Arpp19 | 52,8282 | 50,9666 | -0,0517584 | 0,77315 | 0,963439 |
| Fabp3 | 1,68961 | 1,90313 | 0,171684 | 0,77325 | 0,963439 |
| Dcaf11 | 21,2463 | 22,0719 | 0,054998 | 0,7733 | 0,963439 |
| Rps6kc1 | 8,7764 | 8,42833 | -0,0583824 | 0,7733 | 0,963439 |
| Polr2m | 269,184 | 259,438 | -0,0532001 | 0,77335 | 0,963439 |
| Pum1 | 43,048 | 44,6164 | 0,0516275 | 0,77355 | 0,963595 |
| Tspan33 | 2,00946 | 1,87555 | -0,0994915 | 0,7737 | 0,963688 |

| | | | | | |
|---|---|---|---|---|---|
| Zhx3 | 2,81317 | 2,42622 | -0,213488 | 0,7739 | 0,963844 |
| Adck5 | 9,83655 | 11,0865 | 0,172576 | 0,774 | 0,963875 |
| Rab35 | 40,7199 | 42,2024 | 0,0515926 | 0,77435 | 0,964029 |
| Hdgfrp2 | 83,3441 | 86,4023 | 0,0519897 | 0,7744 | 0,964029 |
| Tigd3 | 2,58107 | 2,74435 | 0,0884951 | 0,77445 | 0,964029 |
| Erlin2 | 23,6935 | 22,8373 | -0,0531046 | 0,77455 | 0,964029 |
| Spsb1 | 3,93031 | 3,73943 | -0,0718213 | 0,77455 | 0,964029 |
| 3110021A11Rik | 0,992948 | 1,13239 | 0,189586 | 0,7747 | 0,964029 |
| Invs | 4,07559 | 4,46318 | 0,131064 | 0,7748 | 0,964029 |
| Tex10 | 42,256 | 40,6575 | -0,0556359 | 0,7748 | 0,964029 |
| Fdx1l | 51,1967 | 45,7182 | -0,16328 | 0,7748 | 0,964029 |
| Rttn | 5,39425 | 5,18742 | -0,0564058 | 0,7751 | 0,96409 |
| Atp8a1 | 3,03192 | 2,90483 | -0,0617791 | 0,7751 | 0,96409 |
| Fbxw2 | 21,5394 | 20,7223 | -0,0557974 | 0,7753 | 0,96409 |
| Pogk | 20,3851 | 19,5527 | -0,060149 | 0,7753 | 0,96409 |
| Coa6 | 24,2693 | 23,12 | -0,069996 | 0,7753 | 0,96409 |
| 2210015D19Rik | 6,44382 | 5,97795 | -0,108265 | 0,7753 | 0,96409 |
| Zfp146 | 92,9707 | 89,6813 | -0,0519687 | 0,7756 | 0,964276 |
| Hmox2 | 51,8236 | 48,413 | -0,0982147 | 0,7756 | 0,964276 |
| Rab7l1 | 1,46898 | 1,6093 | 0,131619 | 0,7759 | 0,964429 |
| Ankrd32 | 12,379 | 12,8758 | 0,056773 | 0,7759 | 0,964429 |
| Zfp346 | 12,1705 | 12,6481 | 0,055537 | 0,776 | 0,964429 |
| Gnl3 | 138,941 | 133,324 | -0,0595335 | 0,7761 | 0,964429 |
| Aqr | 39,7976 | 41,2272 | 0,0509134 | 0,7762 | 0,964429 |
| Dclre1c | 3,36545 | 3,74856 | 0,155536 | 0,77635 | 0,964429 |
| Celf1 | 84,4837 | 87,6175 | 0,0525459 | 0,7764 | 0,964429 |
| Il20rb | 1,1832 | 1,27212 | 0,104545 | 0,77645 | 0,964429 |
| H60b | 1,29574 | 1,06006 | -0,289633 | 0,77645 | 0,964429 |
| Man1a2 | 16,1705 | 16,7582 | 0,0515029 | 0,7765 | 0,964429 |
| Csnk1a1 | 180,763 | 174,574 | -0,0502576 | 0,77655 | 0,964429 |
| Gprc5c | 14,5817 | 15,2123 | 0,0610773 | 0,7767 | 0,964522 |
| Tmem33 | 17,7812 | 17,1403 | -0,0529602 | 0,7768 | 0,964552 |
| Mea1 | 86,6001 | 78,9345 | -0,133713 | 0,77695 | 0,964645 |
| Aida | 14,2118 | 14,7711 | 0,0556913 | 0,7772 | 0,964706 |
| Dus2 | 9,78542 | 9,34574 | -0,0663236 | 0,77725 | 0,964706 |
| Tatdn2 | 25,7653 | 26,7383 | 0,0534775 | 0,7773 | 0,964706 |
| Mon1a | 4,54881 | 4,30826 | -0,078381 | 0,7773 | 0,964706 |
| Gak | 22,2654 | 23,0469 | 0,049772 | 0,7775 | 0,964828 |
| Tsen2 | 17,1101 | 16,4102 | -0,0602513 | 0,77765 | 0,964828 |
| Klf11 | 4,6736 | 4,88806 | 0,0647296 | 0,7777 | 0,964828 |
| Ap3s1 | 10,8973 | 10,3891 | -0,0689036 | 0,7778 | 0,964828 |
| Tmem138 | 27,412 | 26,2521 | -0,062373 | 0,77785 | 0,964828 |
| Tyw5 | 17,344 | 18,4091 | 0,0859838 | 0,778 | 0,964828 |
| Arhgap35 | 35,5034 | 36,7878 | 0,0512681 | 0,778 | 0,964828 |
| Lace1 | 2,2523 | 2,12152 | -0,0862961 | 0,778 | 0,964828 |
| 4931414P19Rik | 6,6342 | 6,3345 | -0,0666919 | 0,77835 | 0,96485 |
| Pus10 | 21,861 | 20,8054 | -0,0713986 | 0,77835 | 0,96485 |
| Rps12 | 7,29141 | 6,70543 | -0,120866 | 0,77835 | 0,96485 |
| Zbtb49 | 3,70395 | 3,89549 | 0,0727439 | 0,7784 | 0,96485 |
| 9330020H09Rik | 1,15758 | 1,3172 | 0,18636 | 0,7785 | 0,96485 |

| | | | | | |
|---|---|---|---|---|---|
| Wasf2 | 17,5916 | 18,2482 | 0,0528699 | 0,7788 | 0,96485 |
| Asun | 37,327 | 38,6643 | 0,0507856 | 0,77895 | 0,96485 |
| Trim13 | 4,06347 | 4,33104 | 0,0920022 | 0,779 | 0,96485 |
| Sfxn3 | 2,13471 | 2,01278 | -0,0848451 | 0,77905 | 0,96485 |
| Cradd | 3,61174 | 3,40746 | -0,0839955 | 0,7792 | 0,96485 |
| Cnot2 | 78,9375 | 76,2145 | -0,0506459 | 0,7793 | 0,96485 |
| Glce | 14,641 | 15,1843 | 0,0525671 | 0,77935 | 0,96485 |
| Orc5 | 32,2082 | 33,6931 | 0,0650261 | 0,77965 | 0,96485 |
| Naa25 | 33,2601 | 34,443 | 0,0504207 | 0,77975 | 0,96485 |
| Chchd5 | 15,2503 | 14,5747 | -0,065375 | 0,7798 | 0,96485 |
| Phtf1 | 27,2822 | 28,6836 | 0,0722663 | 0,77985 | 0,96485 |
| Ubxn4 | 27,9311 | 26,9672 | -0,0506633 | 0,77985 | 0,96485 |
| Fam78b | 0,821433 | 0,769183 | -0,0948164 | 0,7799 | 0,96485 |
| AV039307 | 1,00676 | 1,54419 | 0,617138 | 0,78 | 0,96485 |
| Rhno1 | 76,2163 | 73,6039 | -0,0503169 | 0,78 | 0,96485 |
| Gusb | 28,3907 | 27,3743 | -0,0525961 | 0,78005 | 0,96485 |
| Slc38a10 | 28,2763 | 27,3028 | -0,0505424 | 0,7801 | 0,96485 |
| Gm5089 | 1,47134 | 1,11744 | -0,396928 | 0,7801 | 0,96485 |
| Tmem237 | 16,8618 | 17,5678 | 0,0591784 | 0,78015 | 0,96485 |
| Snora21 | 65,7019 | 42,19 | -0,639032 | 0,7802 | 0,96485 |
| Dusp14 | 5,81779 | 6,13976 | 0,0777105 | 0,78025 | 0,96485 |
| Tead1 | 17,6523 | 18,2845 | 0,0507641 | 0,78035 | 0,96485 |
| Cdc6 | 24,2118 | 23,3929 | -0,0496378 | 0,78035 | 0,96485 |
| Snora31 | 199,816 | 140,762 | -0,505417 | 0,78045 | 0,96485 |
| Tmem198 | 0,934725 | 1,01342 | 0,116617 | 0,78055 | 0,96485 |
| 1810032O08Rik | 22,3956 | 21,4255 | -0,0638912 | 0,78055 | 0,96485 |
| 4930594C11Rik | 2,33293 | 2,15778 | -0,112598 | 0,78055 | 0,96485 |
| Zfp40 | 2,29149 | 2,41756 | 0,0772652 | 0,7806 | 0,96485 |
| Agfg2 | 8,29761 | 8,67358 | 0,0639317 | 0,7806 | 0,96485 |
| Gorasp2 | 70,6322 | 68,1903 | -0,0507583 | 0,78065 | 0,96485 |
| Ccdc57 | 3,60052 | 3,78043 | 0,0703428 | 0,7808 | 0,964942 |
| Ube2b | 59,7599 | 61,887 | 0,050458 | 0,7809 | 0,964973 |
| Wdr33 | 45,0012 | 47,295 | 0,0717229 | 0,7811 | 0,965127 |
| Arl13b | 10,0634 | 10,4596 | 0,0557151 | 0,7812 | 0,965127 |
| Tmem243 | 7,96413 | 9,5228 | 0,257869 | 0,78125 | 0,965127 |
| Dnal1 | 3,8982 | 4,06063 | 0,0588936 | 0,7815 | 0,965248 |
| Tubg1 | 57,8207 | 55,8039 | -0,0512199 | 0,7815 | 0,965248 |
| Efna2 | 18,1539 | 9,81269 | -0,887562 | 0,7816 | 0,965248 |
| Zfp287 | 3,36039 | 3,51252 | 0,0638778 | 0,78175 | 0,965248 |
| Gm10785 | 1,11438 | 1,31532 | 0,239173 | 0,7818 | 0,965248 |
| Gtpbp4 | 79,1152 | 81,9165 | 0,0501983 | 0,7818 | 0,965248 |
| Cecr6 | 0,748853 | 0,701227 | -0,0948015 | 0,7819 | 0,965279 |
| Acy1 | 21,5004 | 20,6485 | -0,0583283 | 0,7822 | 0,965462 |
| Rnf141 | 3,92835 | 4,20874 | 0,0994665 | 0,7823 | 0,965462 |
| Mef2d | 6,71885 | 7,02169 | 0,0636029 | 0,7824 | 0,965462 |
| Tfam | 37,5437 | 36,2822 | -0,0493071 | 0,7824 | 0,965462 |
| Gapdhs | 0,769905 | 0,851346 | 0,145066 | 0,78245 | 0,965462 |
| Mrps2 | 44,1618 | 42,6185 | -0,0513187 | 0,7825 | 0,965462 |
| Pex13 | 14,3539 | 13,6095 | -0,0768313 | 0,7826 | 0,965493 |
| 1810062O18Rik | 0,76212 | 1,4718 | 0,949491 | 0,78285 | 0,965553 |

| | | | | | |
|---|---|---|---|---|---|
| BC017643 | 14,9199 | 14,1509 | -0,0763441 | 0,78285 | 0,965553 |
| Dvl3 | 21,0739 | 21,8524 | 0,0523388 | 0,7829 | 0,965553 |
| Decr1 | 22,0873 | 21,2984 | -0,0524738 | 0,78295 | 0,965553 |
| Nars | 103,504 | 99,9809 | -0,0499627 | 0,7831 | 0,965645 |
| Pigyl | 18,273 | 14,6045 | -0,323299 | 0,78325 | 0,965734 |
| Pofut2 | 31,7144 | 32,878 | 0,0519845 | 0,78335 | 0,965734 |
| Arhgef7 | 25,7594 | 24,8878 | -0,049659 | 0,78345 | 0,965734 |
| Faf2 | 29,6821 | 30,7172 | 0,0494531 | 0,7836 | 0,965734 |
| Bak1 | 38,5904 | 37,2297 | -0,0517888 | 0,78365 | 0,965734 |
| Vps37b | 14,8213 | 15,3962 | 0,0549061 | 0,7837 | 0,965734 |
| Mrm1 | 3,64684 | 3,8385 | 0,0738953 | 0,7838 | 0,965734 |
| Utp14a | 40,0517 | 38,7064 | -0,04929 | 0,78385 | 0,965734 |
| 2500004C02Rik | 7,16545 | 6,82711 | -0,0697822 | 0,7839 | 0,965734 |
| Kifc1 | 18,457 | 17,785 | -0,0535037 | 0,78395 | 0,965734 |
| Dear1 | 0,714096 | 0,378154 | -0,917145 | 0,784 | 0,965734 |
| Aes | 62,9474 | 65,1973 | 0,0506667 | 0,78425 | 0,965886 |
| Trmt44 | 5,00171 | 5,21634 | 0,060616 | 0,7844 | 0,965886 |
| Ap2m1 | 10,1014 | 9,68323 | -0,0609891 | 0,78445 | 0,965886 |
| Zfp330 | 29,7958 | 30,9236 | 0,0535976 | 0,7845 | 0,965886 |
| AI429214 | 1,34164 | 1,24561 | -0,107142 | 0,7845 | 0,965886 |
| Hnrnpa1 | 68,1775 | 65,8075 | -0,0510439 | 0,7846 | 0,965916 |
| Grb2 | 28,9266 | 27,9111 | -0,0515592 | 0,7847 | 0,965946 |
| Trappc5 | 26,8143 | 25,8473 | -0,0529907 | 0,7848 | 0,965946 |
| Col25a1 | 1,30934 | 1,24393 | -0,0739377 | 0,78485 | 0,965946 |
| Pphln1 | 39,0808 | 40,4355 | 0,0491654 | 0,7851 | 0,96615 |
| Gm10433 | 0,847168 | 0,92414 | 0,125463 | 0,7853 | 0,96615 |
| Hspa4l | 4,01783 | 3,84316 | -0,0641219 | 0,78535 | 0,96615 |
| Traf2 | 11,3636 | 11,797 | 0,0539912 | 0,7855 | 0,96615 |
| Snora5c | 156,745 | 106,76 | -0,554045 | 0,78565 | 0,96615 |
| Aldh4a1 | 8,61624 | 8,95711 | 0,0559739 | 0,7857 | 0,96615 |
| Wiz | 35,329 | 34,1751 | -0,0479084 | 0,7857 | 0,96615 |
| Proser2 | 1,2217 | 1,29861 | 0,0880718 | 0,78575 | 0,96615 |
| Gemin6 | 27,7218 | 26,5989 | -0,0596565 | 0,78575 | 0,96615 |
| Vps37d | 5,72891 | 6,02987 | 0,0738651 | 0,78615 | 0,96615 |
| Sav1 | 42,2322 | 44,1097 | 0,0627528 | 0,78615 | 0,96615 |
| Pola1 | 20,1808 | 20,8747 | 0,0487701 | 0,78615 | 0,96615 |
| Kank3 | 2,15592 | 2,03178 | -0,0855607 | 0,7862 | 0,96615 |
| Crem | 4,28286 | 4,05701 | -0,0781572 | 0,7863 | 0,96615 |
| Tmem253 | 1,27397 | 2,09715 | 0,719098 | 0,7864 | 0,96615 |
| Zfp35 | 21,321 | 20,5707 | -0,0516828 | 0,7864 | 0,96615 |
| Txndc17 | 67,2115 | 69,5911 | 0,0501951 | 0,7865 | 0,96615 |
| Rab2b | 8,86573 | 9,73185 | 0,134475 | 0,78665 | 0,96615 |
| Trappc6a | 27,6606 | 28,8982 | 0,0631432 | 0,78665 | 0,96615 |
| Ddx21 | 96,3734 | 93,2038 | -0,0482457 | 0,7867 | 0,96615 |
| Snora23 | 23,3309 | 38,6876 | 0,729631 | 0,78695 | 0,96615 |
| Tfe3 | 12,1617 | 12,6217 | 0,0535527 | 0,78695 | 0,96615 |
| 2810403A07Rik | 46,1124 | 47,755 | 0,0504977 | 0,78695 | 0,96615 |
| 2700038G22Rik | 17,7925 | 15,5357 | -0,19568 | 0,78695 | 0,96615 |
| Rpe | 18,4994 | 19,3685 | 0,0662352 | 0,78705 | 0,96615 |
| Ppp1r13b | 0,763642 | 0,815125 | 0,0941239 | 0,7871 | 0,96615 |

| | | | | | |
|---|---|---|---|---|---|
| Safb | 76,4267 | 79,0066 | 0,0478957 | 0,7871 | 0,96615 |
| Ada | 3,69384 | 3,48709 | -0,0830961 | 0,78715 | 0,96615 |
| Wtap | 36,9481 | 38,3104 | 0,0522363 | 0,7872 | 0,96615 |
| Tmsb15b1 | 6,03056 | 5,12692 | -0,2342 | 0,7873 | 0,96618 |
| Rnasel | 1,80879 | 1,90191 | 0,0724199 | 0,78765 | 0,9663 |
| Gfpt2 | 9,76097 | 9,39267 | -0,0554886 | 0,78765 | 0,9663 |
| Usp13 | 4,17207 | 3,99842 | -0,0613313 | 0,78775 | 0,9663 |
| C2cd2 | 5,46416 | 5,67054 | 0,0534867 | 0,7878 | 0,9663 |
| Samd10 | 3,46961 | 3,28619 | -0,0783555 | 0,78785 | 0,9663 |
| Zfp251 | 14,4827 | 13,9573 | -0,0533181 | 0,78795 | 0,9663 |
| Tmem248 | 29,8083 | 30,8173 | 0,0480262 | 0,788 | 0,9663 |
| Slc25a26 | 4,63546 | 3,98581 | -0,217839 | 0,788 | 0,9663 |
| 5730559C18Rik | 9,75561 | 9,38834 | -0,0553627 | 0,78825 | 0,966514 |
| Lcmt2 | 4,08916 | 4,3767 | 0,0980395 | 0,78845 | 0,966544 |
| Kctd2 | 19,6247 | 20,3771 | 0,0542793 | 0,7885 | 0,966544 |
| Vps11 | 15,7146 | 16,2948 | 0,0523002 | 0,7885 | 0,966544 |
| Scd1 | 29,6074 | 30,6129 | 0,0481819 | 0,7887 | 0,966634 |
| Mcat | 21,7898 | 22,609 | 0,0532438 | 0,78875 | 0,966634 |
| Klhl5 | 12,0198 | 11,5772 | -0,0541248 | 0,7888 | 0,966634 |
| Galk2 | 11,3634 | 11,8064 | 0,0551657 | 0,7889 | 0,966662 |
| Magi3 | 6,45599 | 6,69781 | 0,0530495 | 0,7891 | 0,966662 |
| Hcn3 | 0,794206 | 0,852529 | 0,102235 | 0,78925 | 0,966662 |
| Ppp2r5e | 25,3224 | 26,2035 | 0,0493418 | 0,78925 | 0,966662 |
| Gtf2i | 137,35 | 132,731 | -0,0493529 | 0,78925 | 0,966662 |
| Msra | 15,4305 | 16,1121 | 0,062361 | 0,7893 | 0,966662 |
| Hsp90ab1 | 2091,84 | 2003 | -0,0626108 | 0,78935 | 0,966662 |
| Pcdhga6 | 0,954935 | 0,639709 | -0,577986 | 0,7896 | 0,966876 |
| Rpn1 | 91,0495 | 94,1154 | 0,04778 | 0,78975 | 0,966905 |
| Lrif1 | 5,37577 | 5,65783 | 0,0737765 | 0,78985 | 0,966905 |
| Nras | 55,2407 | 53,455 | -0,0474075 | 0,78985 | 0,966905 |
| Pnma1 | 0,676659 | 0,736245 | 0,121757 | 0,79 | 0,966996 |
| Sft2d3 | 4,09932 | 3,90564 | -0,0698253 | 0,79015 | 0,967055 |
| Gm10636 | 0,601083 | 0,656555 | 0,127352 | 0,7902 | 0,967055 |
| Usp44 | 4,05788 | 3,86224 | -0,0712877 | 0,7903 | 0,967055 |
| Gramd3 | 10,514 | 10,1012 | -0,0577811 | 0,79035 | 0,967055 |
| Nav1 | 7,70957 | 7,98015 | 0,0497665 | 0,79045 | 0,967086 |
| Polr3e | 21,8529 | 22,5947 | 0,0481637 | 0,7906 | 0,967145 |
| Rnf8 | 41,8407 | 40,3984 | -0,050609 | 0,79065 | 0,967145 |
| Kbtbd2 | 20,2194 | 20,9283 | 0,0497191 | 0,79075 | 0,967145 |
| Lsm12 | 23,4206 | 24,351 | 0,0562008 | 0,7908 | 0,967145 |
| Scml2 | 2,15085 | 2,25664 | 0,0692675 | 0,7909 | 0,967171 |
| Bub3 | 112,21 | 108,649 | -0,0465191 | 0,7911 | 0,967171 |
| Anapc1 | 48,1659 | 46,5953 | -0,0478304 | 0,79115 | 0,967171 |
| Rbm45 | 32,2203 | 33,373 | 0,0507107 | 0,7912 | 0,967171 |
| Gm3716 | 0,787057 | 0,847112 | 0,106085 | 0,7913 | 0,967171 |
| Lrrc14 | 15,0871 | 14,5251 | -0,0547666 | 0,7913 | 0,967171 |
| Gpatch1 | 9,19749 | 9,55445 | 0,0549323 | 0,79135 | 0,967171 |
| Phyhipl | 6,7949 | 7,12406 | 0,0682462 | 0,7915 | 0,967171 |
| Smad2 | 13,1325 | 13,5748 | 0,0477895 | 0,7915 | 0,967171 |
| Dync1li1 | 37,6017 | 38,8502 | 0,0471226 | 0,79175 | 0,967199 |

| | | | | | |
|---|---|---|---|---|---|
| Ythdf1 | 59,0434 | 57,1844 | -0,0461555 | 0,7918 | 0,967199 |
| Napa | 50,3908 | 48,7424 | -0,0479825 | 0,7918 | 0,967199 |
| Gm15545 | 7,9416 | 8,61676 | 0,117715 | 0,79185 | 0,967199 |
| Eif3h | 192,408 | 186,153 | -0,0476735 | 0,7919 | 0,967199 |
| Wdr48 | 31,2917 | 30,104 | -0,0558244 | 0,792 | 0,967229 |
| Coq3 | 6,42377 | 6,14981 | -0,0628788 | 0,79215 | 0,967289 |
| Ccdc166 | 2,8202 | 2,96996 | 0,0746435 | 0,7922 | 0,967289 |
| Snx14 | 13,7729 | 13,2898 | -0,0515224 | 0,7923 | 0,967319 |
| 1190007I07Rik | 13,6355 | 12,8886 | -0,0812783 | 0,7926 | 0,967468 |
| Wdr20 | 7,4022 | 7,09573 | -0,0610015 | 0,7927 | 0,967468 |
| Akirin1 | 13,5796 | 14,0996 | 0,0542109 | 0,79275 | 0,967468 |
| Mrpl11 | 37,5972 | 36,3729 | -0,04776 | 0,79275 | 0,967468 |
| Dis3l | 10,1749 | 9,80945 | -0,0527679 | 0,7928 | 0,967468 |
| Erlin1 | 16,0143 | 15,4466 | -0,0520703 | 0,7929 | 0,967468 |
| Gmps | 52,688 | 54,4068 | 0,0463136 | 0,79295 | 0,967468 |
| Supt16 | 122,666 | 126,771 | 0,047498 | 0,79325 | 0,967554 |
| Nmb | 6,53038 | 6,962 | 0,0923349 | 0,7934 | 0,967554 |
| Sox8 | 1,58572 | 1,4935 | -0,0864411 | 0,79345 | 0,967554 |
| Zmym6 | 7,14257 | 7,41284 | 0,0535839 | 0,7935 | 0,967554 |
| Lipa | 7,77846 | 8,09258 | 0,0571141 | 0,79355 | 0,967554 |
| Gmppb | 11,3787 | 10,6477 | -0,0957928 | 0,7936 | 0,967554 |
| Luc7l | 88,5268 | 91,8258 | 0,052785 | 0,7937 | 0,967554 |
| B3gat2 | 0,96541 | 1,12002 | 0,214314 | 0,79375 | 0,967554 |
| Ephb6 | 3,2014 | 3,3563 | 0,0681653 | 0,79375 | 0,967554 |
| Zfp133-ps | 1,06202 | 1,17682 | 0,148075 | 0,7938 | 0,967554 |
| Zfp617 | 4,36724 | 4,17312 | -0,0655958 | 0,79385 | 0,967554 |
| Ndel1 | 19,7367 | 19,0445 | -0,0515058 | 0,79405 | 0,967706 |
| Rnf217 | 8,14176 | 8,46065 | 0,0554268 | 0,7943 | 0,967919 |
| 4933430I17Rik | 1,50059 | 1,39736 | -0,102818 | 0,79475 | 0,968375 |
| Zfp628 | 5,26801 | 5,05378 | -0,0598977 | 0,79495 | 0,968434 |
| Mlx | 20,0904 | 19,1373 | -0,0701224 | 0,79495 | 0,968434 |
| Tspyl1 | 20,4701 | 19,769 | -0,05028 | 0,79505 | 0,968434 |
| Exog | 12,7067 | 12,268 | -0,0506863 | 0,7951 | 0,968434 |
| Ankle1 | 11,7992 | 11,3506 | -0,0559123 | 0,7953 | 0,968484 |
| 9530027J09Rik | 0,961866 | 0,705515 | -0,447159 | 0,79535 | 0,968484 |
| Yrdc | 24,9142 | 25,8843 | 0,0551142 | 0,79555 | 0,968484 |
| Tmem74 | 0,637402 | 0,707714 | 0,150962 | 0,7956 | 0,968484 |
| Crip1 | 5,19618 | 4,75099 | -0,129222 | 0,79565 | 0,968484 |
| Stx4a | 22,8674 | 22,0125 | -0,0549672 | 0,79585 | 0,968484 |
| Qk | 63,6832 | 65,744 | 0,0459479 | 0,79595 | 0,968484 |
| Gtf2b | 60,3123 | 58,3016 | -0,0489158 | 0,796 | 0,968484 |
| Fbrsl1 | 7,78807 | 8,12191 | 0,0605549 | 0,79605 | 0,968484 |
| Prmt10 | 13,2551 | 13,7443 | 0,0522841 | 0,7961 | 0,968484 |
| Ehmt1 | 48,4144 | 46,8646 | -0,0469373 | 0,7961 | 0,968484 |
| Nup93 | 78,3038 | 80,8335 | 0,0458693 | 0,79615 | 0,968484 |
| 4930483K19Rik | 1,07427 | 1,45705 | 0,439685 | 0,7962 | 0,968484 |
| Gm4432 | 0,426946 | 0,590242 | 0,467254 | 0,79635 | 0,968484 |
| Pdcl3 | 43,5964 | 42,1152 | -0,0498679 | 0,7964 | 0,968484 |
| Cdc42se1 | 31,5288 | 33,3705 | 0,0819056 | 0,7965 | 0,968484 |
| E430025E21Rik | 29,341 | 28,4195 | -0,0460328 | 0,7965 | 0,968484 |

| | | | | | |
|---|---|---|---|---|---|
| Ddx39b | 576,092 | 556,922 | -0,0488246 | 0,7965 | 0,968484 |
| Mynn | 6,50093 | 6,73772 | 0,0516132 | 0,79665 | 0,968574 |
| Ccsap | 4,34496 | 4,54855 | 0,0660662 | 0,79675 | 0,968603 |
| Mtr | 15,2216 | 15,7417 | 0,0484727 | 0,7969 | 0,968603 |
| Kat8 | 26,4936 | 25,5649 | -0,0514773 | 0,7969 | 0,968603 |
| Tmem69 | 8,7305 | 8,37945 | -0,0592096 | 0,7971 | 0,968754 |
| Suv420h2 | 34,893 | 36,0773 | 0,0481565 | 0,7975 | 0,969148 |
| Poli | 5,94705 | 6,21541 | 0,0636766 | 0,79765 | 0,969239 |
| Mast3 | 4,0879 | 4,25353 | 0,0572994 | 0,79775 | 0,969269 |
| Lgalsl | 8,19072 | 8,49394 | 0,0524435 | 0,79805 | 0,969541 |
| Rab18 | 31,8591 | 32,9083 | 0,0467429 | 0,79835 | 0,969688 |
| Nipa1 | 5,77078 | 6,1166 | 0,0839653 | 0,7984 | 0,969688 |
| Rnf185 | 16,9233 | 15,0579 | -0,168493 | 0,7984 | 0,969688 |
| Ccdc61 | 12,2234 | 12,7035 | 0,0555787 | 0,79855 | 0,969688 |
| Blvrb | 3,28789 | 3,52691 | 0,101244 | 0,79865 | 0,969688 |
| Slc25a15 | 10,2615 | 10,6378 | 0,0519547 | 0,79865 | 0,969688 |
| 5730480H06Rik | 1,53996 | 1,66625 | 0,113712 | 0,7987 | 0,969688 |
| Urod | 67,8794 | 70,1801 | 0,0480881 | 0,7988 | 0,969693 |
| Pi4k2b | 8,09759 | 8,40624 | 0,0539682 | 0,7991 | 0,969693 |
| Mterf1a | 1,74796 | 1,62412 | -0,106014 | 0,7991 | 0,969693 |
| Gfra3 | 10,3125 | 9,91063 | -0,057352 | 0,7994 | 0,969693 |
| Ccdc89 | 1,13653 | 1,06118 | -0,0989682 | 0,79945 | 0,969693 |
| C8g | 2,73819 | 3,26866 | 0,255476 | 0,7995 | 0,969693 |
| Gm20605,Lrch4 | 5,16681 | 5,50161 | 0,0905811 | 0,7995 | 0,969693 |
| Tmem11 | 49,1954 | 47,5424 | -0,0493102 | 0,7995 | 0,969693 |
| Pdrg1 | 44,0951 | 45,6519 | 0,0500568 | 0,79955 | 0,969693 |
| Enpp5 | 0,6875 | 0,633745 | -0,117458 | 0,79965 | 0,969693 |
| Cbx7 | 0,613907 | 0,662663 | 0,110256 | 0,7998 | 0,969693 |
| Nrbp1 | 58,9381 | 67,0754 | 0,186582 | 0,79985 | 0,969693 |
| Vta1 | 38,1437 | 39,5071 | 0,0506638 | 0,79985 | 0,969693 |
| Bms1 | 54,8942 | 56,6469 | 0,0453444 | 0,7999 | 0,969693 |
| Lars | 46,587 | 48,0454 | 0,0444732 | 0,80005 | 0,969693 |
| Mapk3 | 43,1429 | 44,5754 | 0,0471245 | 0,80015 | 0,969693 |
| Ager | 3,26222 | 3,45998 | 0,0849073 | 0,8002 | 0,969693 |
| Ntan1 | 34,2713 | 32,0172 | -0,0981567 | 0,8002 | 0,969693 |
| Zfp943 | 5,89656 | 6,14904 | 0,060487 | 0,80025 | 0,969693 |
| Ldhal6b | 0,500453 | 0,71043 | 0,505456 | 0,8003 | 0,969693 |
| 6330416G13Rik | 4,98481 | 4,79092 | -0,0572342 | 0,8004 | 0,969693 |
| Atp6v1a | 31,5147 | 30,5277 | -0,0459066 | 0,8005 | 0,969693 |
| Fyn | 23,1379 | 23,888 | 0,0460284 | 0,8006 | 0,969693 |
| Anxa6 | 19,7081 | 19,0337 | -0,0502296 | 0,8006 | 0,969693 |
| Limk1 | 10,8725 | 10,4959 | -0,0508595 | 0,8006 | 0,969693 |
| Cxx1a | 2,95923 | 2,78317 | -0,0884939 | 0,8007 | 0,969693 |
| Lag3 | 1,00177 | 1,08095 | 0,10975 | 0,8008 | 0,969693 |
| Yipf4 | 14,6048 | 15,1479 | 0,0526692 | 0,8012 | 0,969693 |
| Rbfa | 17,0386 | 17,7194 | 0,0565201 | 0,80125 | 0,969693 |
| Stab1 | 0,621079 | 0,421951 | -0,557702 | 0,80125 | 0,969693 |
| Mios | 13,5005 | 13,9532 | 0,0475885 | 0,8013 | 0,969693 |
| Brox | 8,06917 | 8,37031 | 0,0528615 | 0,80135 | 0,969693 |
| Pole | 25,9422 | 26,7892 | 0,0463511 | 0,80135 | 0,969693 |

| | | | | | |
|---|---|---|---|---|---|
| Psmb1 | 242,425 | 234,847 | -0,045821 | 0,80135 | 0,969693 |
| Lrrc49 | 16,2686 | 15,7203 | -0,0494592 | 0,80135 | 0,969693 |
| Npepl1 | 12,604 | 13,0825 | 0,0537514 | 0,8015 | 0,969748 |
| Scoc | 13,3505 | 13,8727 | 0,0553528 | 0,8017 | 0,969748 |
| Igf2os | 4,8767 | 3,96796 | -0,297507 | 0,80175 | 0,969748 |
| Syt12 | 0,634279 | 0,555731 | -0,190729 | 0,8019 | 0,969748 |
| Rad21 | 133,26 | 129,109 | -0,0456527 | 0,80195 | 0,969748 |
| Rasip1 | 1,72146 | 1,81928 | 0,0797293 | 0,802 | 0,969748 |
| Vamp2 | 13,011 | 13,5037 | 0,05363 | 0,802 | 0,969748 |
| Snora3 | 194,467 | 131,145 | -0,56836 | 0,802 | 0,969748 |
| Ppp6r1 | 39,4042 | 40,6336 | 0,0443214 | 0,80225 | 0,969866 |
| Trappc12 | 12,5706 | 13,0177 | 0,0504175 | 0,80235 | 0,969866 |
| Nova1 | 2,62167 | 2,72782 | 0,057263 | 0,8024 | 0,969866 |
| Cwc15 | 123,735 | 119,973 | -0,0445469 | 0,8024 | 0,969866 |
| Sars2 | 18,3132 | 17,691 | -0,049861 | 0,80265 | 0,970046 |
| Mfsd1 | 26,3059 | 27,173 | 0,0467869 | 0,8027 | 0,970046 |
| Nploc4 | 31,3184 | 32,3229 | 0,0455457 | 0,80285 | 0,970074 |
| Phactr2 | 10,0119 | 10,3349 | 0,0457986 | 0,80295 | 0,970074 |
| Scarb1 | 13,5215 | 13,0488 | -0,051339 | 0,80295 | 0,970074 |
| Gabbr1 | 7,98028 | 8,26177 | 0,0500122 | 0,8032 | 0,970253 |
| Tomm34 | 19,9656 | 19,2778 | -0,0505801 | 0,80325 | 0,970253 |
| Nars2 | 7,37224 | 7,64598 | 0,0525988 | 0,80335 | 0,970283 |
| Herc6 | 1,22308 | 1,16103 | -0,0751206 | 0,80365 | 0,970461 |
| Dbr1 | 20,6701 | 19,9912 | -0,0481784 | 0,80375 | 0,970461 |
| Pla2g15 | 4,88521 | 4,67826 | -0,0624475 | 0,80375 | 0,970461 |
| Ube2h | 11,7364 | 12,1402 | 0,0488033 | 0,8038 | 0,970461 |
| Amd1,Amd2 | 4,46624 | 5,16669 | 0,210179 | 0,8039 | 0,97049 |
| Kctd1 | 8,32499 | 7,9813 | -0,0608238 | 0,80425 | 0,970782 |
| Aasdhppt | 17,6573 | 18,2712 | 0,0493081 | 0,80445 | 0,970782 |
| Hmbox1 | 11,131 | 11,5286 | 0,050642 | 0,80455 | 0,970782 |
| Eif4g1 | 283,721 | 293,366 | 0,048227 | 0,80465 | 0,970782 |
| Anp32e | 242,883 | 235,282 | -0,0458719 | 0,8047 | 0,970782 |
| Ctnnal1 | 18,3725 | 17,7909 | -0,0464031 | 0,8047 | 0,970782 |
| Ikbkg | 5,99765 | 6,31471 | 0,0743187 | 0,8048 | 0,970782 |
| Dgcr8 | 22,3828 | 23,4192 | 0,0653014 | 0,8048 | 0,970782 |
| Igsf6 | 0,90222 | 1,26138 | 0,483455 | 0,8049 | 0,970782 |
| Cep128 | 5,69165 | 5,48592 | -0,0531125 | 0,805 | 0,970782 |
| Atp6v1h | 37,4848 | 36,3156 | -0,0457175 | 0,80505 | 0,970782 |
| Ribc1 | 2,93894 | 2,77313 | -0,0837801 | 0,80505 | 0,970782 |
| Snora34 | 111,946 | 154,209 | 0,462091 | 0,80535 | 0,970961 |
| Usp24 | 14,2681 | 14,7161 | 0,0446084 | 0,80535 | 0,970961 |
| Haus2 | 20,9153 | 20,1312 | -0,0551232 | 0,8055 | 0,971051 |
| Taf1a | 12,8977 | 13,3551 | 0,0502733 | 0,8058 | 0,971106 |
| Tm9sf2 | 46,5101 | 45,0815 | -0,0450067 | 0,8058 | 0,971106 |
| Mtrf1 | 5,77839 | 6,04235 | 0,0644426 | 0,80585 | 0,971106 |
| Fubp1 | 105,717 | 109,179 | 0,0464888 | 0,80595 | 0,971106 |
| Ppp2r3c | 12,1993 | 10,9762 | -0,152421 | 0,80595 | 0,971106 |
| Nacc1 | 40,1956 | 41,4598 | 0,044675 | 0,806 | 0,971106 |
| Zfp3 | 2,2769 | 2,41308 | 0,0838067 | 0,80625 | 0,971316 |
| Pik3r4 | 13,1527 | 12,733 | -0,0467778 | 0,80645 | 0,971466 |

| | | | | | |
|---|---|---|---|---|---|
| Fam49b | 25,5561 | 24,7829 | -0,0443225 | 0,80655 | 0,971495 |
| Klc1 | 51,233 | 53,0502 | 0,0502858 | 0,80685 | 0,971665 |
| Rbm14-rbm4 | 7,28088 | 8,2158 | 0,174289 | 0,8069 | 0,971665 |
| Zscan12 | 8,43375 | 8,71481 | 0,0472942 | 0,80705 | 0,971665 |
| Lin54 | 9,24951 | 8,9462 | -0,0481016 | 0,80705 | 0,971665 |
| Cchcr1 | 9,80102 | 10,1604 | 0,0519475 | 0,8071 | 0,971665 |
| Acads | 35,1456 | 36,2818 | 0,045902 | 0,8072 | 0,971665 |
| 1700001G11Rik | 0,616449 | 0,84003 | 0,446458 | 0,80735 | 0,971665 |
| Nox4 | 0,984099 | 1,04507 | 0,0867196 | 0,80735 | 0,971665 |
| Hnf1b | 3,01952 | 3,16144 | 0,0662595 | 0,8074 | 0,971665 |
| Gpn3 | 42,6971 | 44,1181 | 0,047235 | 0,8075 | 0,971665 |
| A130010J15Rik | 5,97714 | 6,20853 | 0,0547956 | 0,80755 | 0,971665 |
| Rqcd1 | 67,2041 | 70,0623 | 0,0600894 | 0,8076 | 0,971665 |
| Mecp2 | 9,76528 | 10,0699 | 0,0443169 | 0,80785 | 0,971844 |
| Urm1 | 18,5028 | 17,8923 | -0,0484052 | 0,8079 | 0,971844 |
| Bdh1 | 6,11084 | 6,35525 | 0,0565767 | 0,8082 | 0,971989 |
| Scarna6 | 3,81798 | 2,61329 | -0,546944 | 0,80825 | 0,971989 |
| Esyt2 | 11,952 | 12,3284 | 0,0447356 | 0,8083 | 0,971989 |
| Grk6 | 51,8019 | 53,4059 | 0,043995 | 0,80835 | 0,971989 |
| Prune | 14,1796 | 13,7163 | -0,0479176 | 0,8084 | 0,971989 |
| B130034C11Rik | 1,57624 | 1,69013 | 0,100641 | 0,8086 | 0,972139 |
| Gcc1 | 5,03431 | 4,85187 | -0,0532522 | 0,80875 | 0,972228 |
| Snx12 | 38,3462 | 39,7901 | 0,0533269 | 0,809 | 0,972346 |
| Ddx50 | 42,8042 | 41,5117 | -0,0442346 | 0,809 | 0,972346 |
| P4hb | 361,084 | 349,883 | -0,0454606 | 0,8092 | 0,972372 |
| Snupn | 14,7148 | 14,1511 | -0,0563529 | 0,80925 | 0,972372 |
| Nagk | 3,97943 | 3,76854 | -0,0785567 | 0,8093 | 0,972372 |
| Ppp1r3e | 0,634236 | 0,691023 | 0,123713 | 0,8094 | 0,972372 |
| Gpsm3 | 3,28339 | 3,4724 | 0,0807458 | 0,8094 | 0,972372 |
| Ccdc117 | 29,804 | 28,8916 | -0,0448542 | 0,8095 | 0,972401 |
| Foxd1 | 13,2396 | 13,7099 | 0,0503509 | 0,80975 | 0,97261 |
| Arpc4 | 58,6097 | 56,8477 | -0,044038 | 0,80985 | 0,972637 |
| Irf3 | 23,375 | 24,1411 | 0,0465211 | 0,80995 | 0,972637 |
| Zik1 | 17,3241 | 16,7741 | -0,0465413 | 0,81 | 0,972637 |
| Atp5d | 235,613 | 196,08 | -0,264976 | 0,8102 | 0,97275 |
| 2610203C20Rik | 0,981932 | 1,04829 | 0,0943449 | 0,8103 | 0,97275 |
| Myl6b | 4,25603 | 4,50516 | 0,0820681 | 0,8104 | 0,97275 |
| Dnaja3 | 30,9753 | 31,9516 | 0,0447722 | 0,8104 | 0,97275 |
| Spon2 | 0,784977 | 0,8464 | 0,108689 | 0,8106 | 0,97275 |
| Ebna1bp2 | 58,6331 | 56,9268 | -0,0426056 | 0,8106 | 0,97275 |
| Zbtb22 | 17,6389 | 17,0791 | -0,046522 | 0,81065 | 0,97275 |
| Arf1 | 173,509 | 168,42 | -0,0429468 | 0,8108 | 0,97275 |
| Zfp608 | 19,121 | 19,703 | 0,0432582 | 0,81105 | 0,97275 |
| Zfp1 | 13,2384 | 12,7648 | -0,0525536 | 0,81105 | 0,97275 |
| Cgrrf1 | 15,1361 | 15,7483 | 0,0572009 | 0,8111 | 0,97275 |
| Spcs1 | 67,5927 | 62,6715 | -0,109059 | 0,8111 | 0,97275 |
| Zcchc12 | 0,842736 | 0,904166 | 0,101507 | 0,8112 | 0,97275 |
| Atf7 | 3,31901 | 3,47396 | 0,0658268 | 0,8112 | 0,97275 |
| Cc2d1b | 13,3465 | 12,9256 | -0,0462381 | 0,81155 | 0,97275 |
| 5430405H02Rik | 2,03468 | 1,7016 | -0,257915 | 0,81155 | 0,97275 |

| | | | | | |
|---|---|---|---|---|---|
| Nup214 | 14,8355 | 15,287 | 0,0432527 | 0,8116 | 0,97275 |
| Cryl1 | 3,93773 | 4,1517 | 0,0763384 | 0,8117 | 0,97275 |
| Hist1h3c | 2,8837 | 3,17739 | 0,139921 | 0,8118 | 0,97275 |
| Rnaset2b | 1,1692 | 1,27857 | 0,129017 | 0,81185 | 0,97275 |
| Taf4a | 11,0318 | 10,6854 | -0,046021 | 0,812 | 0,97275 |
| Gm13157 | 2,32222 | 2,20974 | -0,071627 | 0,812 | 0,97275 |
| Fam129b | 13,8398 | 13,3699 | -0,0498294 | 0,81205 | 0,97275 |
| 1700048M11Rik | 1,25864 | 0,842213 | -0,579614 | 0,81215 | 0,97275 |
| Phf2 | 20,7726 | 21,3987 | 0,0428459 | 0,8123 | 0,97275 |
| Mtpn | 59,3793 | 57,6569 | -0,0424648 | 0,8125 | 0,97275 |
| Acat3 | 0,597174 | 0,853206 | 0,514742 | 0,81265 | 0,97275 |
| Pdxk | 6,41997 | 6,20484 | -0,0491734 | 0,81265 | 0,97275 |
| Ubald1 | 16,731 | 17,3295 | 0,0507046 | 0,8127 | 0,97275 |
| Lgals8 | 2,35942 | 2,24256 | -0,073281 | 0,8127 | 0,97275 |
| Epm2a | 1,13302 | 1,0362 | -0,128867 | 0,8127 | 0,97275 |
| Ttll1 | 7,29647 | 7,01063 | -0,057655 | 0,81275 | 0,97275 |
| Ppil6 | 0,944533 | 1,03623 | 0,133675 | 0,8128 | 0,97275 |
| Abcf1 | 84,5165 | 87,0249 | 0,0421944 | 0,8128 | 0,97275 |
| Pbxip1 | 5,79872 | 6,00099 | 0,0494655 | 0,8129 | 0,97275 |
| Dpagt1 | 23,0413 | 22,305 | -0,0468544 | 0,8129 | 0,97275 |
| Cenpo | 7,0949 | 6,75091 | -0,0717006 | 0,8129 | 0,97275 |
| Rce1 | 24,2518 | 23,436 | -0,0493681 | 0,81305 | 0,972839 |
| Klhl8 | 3,01348 | 3,14938 | 0,0636369 | 0,81335 | 0,973045 |
| Fdps | 48,1332 | 49,7299 | 0,0470805 | 0,8134 | 0,973045 |
| Maea | 102,448 | 105,503 | 0,0423988 | 0,81345 | 0,973045 |
| Rad51b | 3,03108 | 3,18427 | 0,0711331 | 0,81355 | 0,973074 |
| Elavl1 | 37,0234 | 35,9377 | -0,0429394 | 0,81365 | 0,973103 |
| Cog7 | 11,1902 | 11,5678 | 0,0478874 | 0,8139 | 0,97322 |
| Smpdl3b | 3,82503 | 3,64254 | -0,0705275 | 0,8139 | 0,97322 |
| Cpne3 | 8,32506 | 8,59243 | 0,0456069 | 0,81405 | 0,973239 |
| Zfp551 | 2,03081 | 1,92663 | -0,0759734 | 0,81415 | 0,973239 |
| Amigo3 | 2,24826 | 2,50764 | 0,15752 | 0,81435 | 0,973239 |
| Pkm | 773,092 | 747,758 | -0,0480678 | 0,81435 | 0,973239 |
| Mapkapk2 | 57,312 | 55,6434 | -0,0426277 | 0,8144 | 0,973239 |
| Gm14325 | 1,448 | 1,37837 | -0,0710934 | 0,81445 | 0,973239 |
| 2310035C23Rik | 7,22199 | 7,45862 | 0,0465117 | 0,81455 | 0,973239 |
| Acly | 150,241 | 154,705 | 0,0422334 | 0,8146 | 0,973239 |
| Noc3l | 19,4809 | 20,0553 | 0,0419281 | 0,8146 | 0,973239 |
| Zfp667 | 2,45876 | 2,35408 | -0,0627651 | 0,81475 | 0,973239 |
| Ptrh2 | 25,2847 | 22,2299 | -0,185762 | 0,81475 | 0,973239 |
| Rabif | 20,8123 | 20,1526 | -0,0464691 | 0,81485 | 0,973268 |
| Zbtb11 | 7,07613 | 7,31499 | 0,0478946 | 0,81495 | 0,973297 |
| Zdhhc15 | 6,52941 | 6,31569 | -0,0480107 | 0,8151 | 0,973385 |
| Mir5121 | 1888,09 | 2038,35 | 0,110472 | 0,81535 | 0,973549 |
| Supt5 | 75,3008 | 77,4713 | 0,0409969 | 0,81555 | 0,973549 |
| Thoc7 | 209,101 | 215,246 | 0,0417876 | 0,8157 | 0,973549 |
| Rbm19 | 21,851 | 21,2202 | -0,0422615 | 0,81585 | 0,973549 |
| Cdc20 | 106,769 | 103,472 | -0,0452594 | 0,8159 | 0,973549 |
| Nrp | 1,82935 | 2,75275 | 0,589542 | 0,81595 | 0,973549 |
| Apbb1 | 24,3996 | 23,6755 | -0,0434598 | 0,81595 | 0,973549 |

| | | | | | |
|---|---|---|---|---|---|
| Lonp1 | 55,7446 | 54,1303 | -0,0423949 | 0,816 | 0,973549 |
| Ddb2 | 3,72572 | 4,18872 | 0,16899 | 0,81605 | 0,973549 |
| Dnd1 | 1,40092 | 1,15697 | -0,276021 | 0,8161 | 0,973549 |
| Gas2l1 | 7,02529 | 7,28542 | 0,0524548 | 0,81625 | 0,973549 |
| Arrdc3 | 24,5464 | 23,8283 | -0,0428338 | 0,81625 | 0,973549 |
| Elmo1 | 2,20587 | 2,11515 | -0,0605872 | 0,81625 | 0,973549 |
| Pak2 | 34,4389 | 35,4515 | 0,0418101 | 0,8163 | 0,973549 |
| Zfp960 | 0,712859 | 0,663184 | -0,104208 | 0,81645 | 0,973633 |
| Xndc1 | 17,1159 | 16,5558 | -0,0479988 | 0,8166 | 0,973633 |
| Nek4 | 13,4578 | 13,9045 | 0,0471174 | 0,81665 | 0,973633 |
| Cep164 | 11,5787 | 11,2326 | -0,0437833 | 0,8167 | 0,973633 |
| Dhrs1 | 30,1138 | 29,171 | -0,0458899 | 0,81675 | 0,973633 |
| 4930444P10Rik | 0,75427 | 0,854756 | 0,180431 | 0,8169 | 0,973722 |
| Bbip1 | 16,577 | 15,5753 | -0,0899275 | 0,8171 | 0,97387 |
| Sbds | 38,0768 | 39,2884 | 0,0451924 | 0,8173 | 0,973982 |
| Numa1 | 49,7601 | 48,3645 | -0,0410399 | 0,8175 | 0,973982 |
| Pdcl | 21,095 | 21,7588 | 0,0446982 | 0,81755 | 0,973982 |
| Ganc | 1,43148 | 1,56395 | 0,127688 | 0,8176 | 0,973982 |
| Adat3 | 5,18241 | 5,7096 | 0,139768 | 0,81765 | 0,973982 |
| Tmc7 | 3,18946 | 3,06857 | -0,0557448 | 0,81765 | 0,973982 |
| Sike1 | 18,7376 | 19,3435 | 0,045909 | 0,81785 | 0,974099 |
| 4921536K21Rik | 1,11907 | 1,05507 | -0,0849636 | 0,8179 | 0,974099 |
| AA465934 | 5,14435 | 6,12866 | 0,252584 | 0,818 | 0,974128 |
| Dtna | 0,872753 | 0,824885 | -0,081381 | 0,8182 | 0,974245 |
| Gtf3a | 45,2012 | 46,932 | 0,0542109 | 0,81825 | 0,974245 |
| Fibp | 36,4818 | 37,6606 | 0,0458777 | 0,81835 | 0,974273 |
| Trp53i13 | 8,84902 | 9,2211 | 0,0594215 | 0,8185 | 0,97431 |
| Crk | 43,1202 | 41,9101 | -0,0410682 | 0,8188 | 0,97431 |
| Rgs12 | 15,6332 | 15,1729 | -0,0431159 | 0,8188 | 0,97431 |
| Ikbip | 11,9272 | 11,4594 | -0,0577204 | 0,81905 | 0,97431 |
| Sspn | 0,876782 | 0,92403 | 0,0757219 | 0,8191 | 0,97431 |
| Matk | 0,684623 | 0,634605 | -0,109452 | 0,8191 | 0,97431 |
| Cutc | 6,79335 | 6,49176 | -0,0655138 | 0,81915 | 0,97431 |
| Tbcc | 15,1653 | 14,6593 | -0,0489612 | 0,8192 | 0,97431 |
| Eid3 | 1,38312 | 2,17915 | 0,655843 | 0,81925 | 0,97431 |
| Ift52 | 29,095 | 28,2193 | -0,0440887 | 0,81945 | 0,97431 |
| Gtf2h3 | 40,6766 | 41,8409 | 0,0407138 | 0,8195 | 0,97431 |
| Mmadhc | 108,96 | 105,908 | -0,0409845 | 0,8195 | 0,97431 |
| Dpf2 | 41,6983 | 40,5485 | -0,0403416 | 0,81975 | 0,97431 |
| Emc8 | 20,8267 | 20,2306 | -0,0418898 | 0,81975 | 0,97431 |
| Zscan29 | 7,18963 | 6,92029 | -0,0550835 | 0,8198 | 0,97431 |
| Myo1b | 33,2023 | 34,1753 | 0,0416712 | 0,81985 | 0,97431 |
| BC005624 | 17,1369 | 16,5445 | -0,0507491 | 0,81985 | 0,97431 |
| 4933417G07Rik | 4,74025 | 5,39658 | 0,187083 | 0,8199 | 0,97431 |
| Clcn5 | 13,3578 | 12,9573 | -0,0439135 | 0,8199 | 0,97431 |
| Cd320 | 20,9745 | 20,3265 | -0,045273 | 0,8199 | 0,97431 |
| Iqcc | 10,1055 | 10,4609 | 0,0498616 | 0,82005 | 0,974398 |
| Kiz | 12,0046 | 11,5958 | -0,0499764 | 0,8204 | 0,974603 |
| Dmap1 | 31,0313 | 31,9797 | 0,0434359 | 0,82045 | 0,974603 |
| Mpp1 | 11,0769 | 10,012 | -0,145827 | 0,82045 | 0,974603 |

| | | | | | |
|---|---|---|---|---|---|
| Dnttip2 | 63,118 | 64,9853 | 0,0420621 | 0,82065 | 0,974688 |
| Col11a2 | 1,22043 | 1,27783 | 0,0663043 | 0,8207 | 0,974688 |
| Ncl | 308,676 | 298,176 | -0,0499307 | 0,82075 | 0,974688 |
| Siah1a | 16,5239 | 15,9984 | -0,0466318 | 0,821 | 0,974895 |
| Dtwd1 | 12,5591 | 12,0931 | -0,0545561 | 0,82135 | 0,975189 |
| Zfp282 | 10,161 | 9,85763 | -0,0437252 | 0,8214 | 0,975189 |
| Ccdc132 | 16,3295 | 16,8101 | 0,0418428 | 0,8215 | 0,975218 |
| Fbxo28 | 14,5468 | 14,9716 | 0,041528 | 0,8216 | 0,975246 |
| Map1lc3a | 23,9563 | 24,8051 | 0,0502283 | 0,82175 | 0,975334 |
| Sf3a1 | 66,9807 | 68,8938 | 0,0406291 | 0,8219 | 0,975422 |
| Txnrd1 | 114,162 | 111,025 | -0,0402046 | 0,82205 | 0,975466 |
| Fbxl12 | 9,28508 | 9,96951 | 0,102609 | 0,8221 | 0,975466 |
| Fam171b | 4,10354 | 4,26117 | 0,0543827 | 0,8223 | 0,975466 |
| Snx25 | 8,32755 | 8,05181 | -0,0485794 | 0,82245 | 0,975466 |
| Taf1b | 16,6063 | 17,1507 | 0,0465337 | 0,8225 | 0,975466 |
| Rtel1 | 24,1449 | 23,4769 | -0,040475 | 0,82255 | 0,975466 |
| Zfp212 | 15,4406 | 15,9323 | 0,0452278 | 0,8226 | 0,975466 |
| Jmjd8 | 28,516 | 30,0899 | 0,0775063 | 0,82275 | 0,975466 |
| Tk2 | 12,0498 | 11,6567 | -0,0478514 | 0,82275 | 0,975466 |
| Gria3 | 1,0095 | 0,950859 | -0,0863435 | 0,82275 | 0,975466 |
| Snx22 | 2,05682 | 2,6491 | 0,365085 | 0,8228 | 0,975466 |
| Tmem181b-ps | 3,04476 | 3,16098 | 0,0540468 | 0,82285 | 0,975466 |
| Pus7 | 26,8357 | 27,6144 | 0,041268 | 0,82295 | 0,975495 |
| Minpp1 | 35,7836 | 36,8031 | 0,0405271 | 0,82315 | 0,975577 |
| Shc4 | 0,751023 | 0,911292 | 0,279056 | 0,8232 | 0,975577 |
| Rprd1b | 24,2316 | 23,5463 | -0,0413904 | 0,8233 | 0,975577 |
| Zfp661 | 4,81676 | 4,63948 | -0,0540985 | 0,82335 | 0,975577 |
| Tstd1 | 1,44828 | 1,61306 | 0,155463 | 0,8234 | 0,975577 |
| Lactb2 | 8,64065 | 8,32873 | -0,0530427 | 0,8236 | 0,975688 |
| Zbtb25 | 10,9957 | 11,3982 | 0,0518719 | 0,8238 | 0,975688 |
| Eif4ebp3 | 14,3985 | 13,865 | -0,054466 | 0,8238 | 0,975688 |
| 4930404N11Rik | 1,64606 | 1,42263 | -0,210453 | 0,8238 | 0,975688 |
| Sass6 | 11,0964 | 11,4864 | 0,0498471 | 0,82395 | 0,975688 |
| Fkbp1a | 202,696 | 197,156 | -0,0399757 | 0,82395 | 0,975688 |
| Spsb3 | 18,0946 | 18,6835 | 0,0462022 | 0,82425 | 0,975846 |
| Emc7 | 34,8942 | 33,8025 | -0,0458579 | 0,82435 | 0,975846 |
| C230052I12Rik | 6,30837 | 5,93195 | -0,0887612 | 0,8244 | 0,975846 |
| Tmem55a | 41,798 | 42,9913 | 0,0406101 | 0,8245 | 0,975846 |
| Ift140 | 10,2946 | 9,99687 | -0,042345 | 0,8245 | 0,975846 |
| D130020L05Rik | 0,743126 | 0,977397 | 0,395338 | 0,82455 | 0,975846 |
| Galnt1 | 42,5169 | 41,363 | -0,0396977 | 0,8248 | 0,975846 |
| Ift22 | 33,5472 | 32,4784 | -0,0467102 | 0,82485 | 0,975846 |
| Wif1 | 1,22555 | 1,15531 | -0,0851501 | 0,82485 | 0,975846 |
| Iqgap2 | 14,9868 | 14,5641 | -0,0412722 | 0,825 | 0,975846 |
| Coq10b | 8,18386 | 7,87826 | -0,0549053 | 0,825 | 0,975846 |
| Ip6k2 | 26,7406 | 27,5429 | 0,0426467 | 0,82515 | 0,975846 |
| Pfkfb4 | 2,6527 | 2,76662 | 0,0606664 | 0,8252 | 0,975846 |
| Inpp5f | 14,3873 | 14,8187 | 0,0426235 | 0,82525 | 0,975846 |
| Casp8 | 7,93297 | 7,65768 | -0,0509533 | 0,82525 | 0,975846 |
| Tdrd3 | 22,2726 | 22,9379 | 0,0424647 | 0,8253 | 0,975846 |

| | | | | | |
|---|---|---|---|---|---|
| Snx32 | 1,2313 | 1,31861 | 0,0988342 | 0,8254 | 0,975874 |
| 1700012D01Rik | 0,992641 | 0,893053 | -0,152526 | 0,8255 | 0,975902 |
| Ccdc127 | 25,7721 | 25,0442 | -0,0413332 | 0,8256 | 0,975928 |
| Als2 | 8,68981 | 8,96128 | 0,0443803 | 0,82575 | 0,975928 |
| Ppp4r1l-ps | 2,17357 | 2,08709 | -0,058574 | 0,82575 | 0,975928 |
| Pgk1 | 19,6926 | 20,3073 | 0,0443418 | 0,8261 | 0,976252 |
| Rab26os | 51,4128 | 48,9059 | -0,0721194 | 0,8262 | 0,97628 |
| Mthfd1l | 24,8366 | 25,5091 | 0,038547 | 0,8263 | 0,976308 |
| Bcs1l | 11,5925 | 11,1918 | -0,0507502 | 0,8264 | 0,976336 |
| Gm9776 | 1,34054 | 1,26589 | -0,0826516 | 0,82655 | 0,976393 |
| Pafah1b2 | 73,2319 | 71,2984 | -0,0386033 | 0,8266 | 0,976393 |
| Gamt | 6,78827 | 7,54905 | 0,153251 | 0,8268 | 0,976534 |
| Pih1d2 | 2,38814 | 2,88121 | 0,270785 | 0,82705 | 0,976534 |
| Pdss2 | 9,17742 | 9,51199 | 0,0516582 | 0,82705 | 0,976534 |
| Sap18 | 9,21061 | 9,49521 | 0,0439034 | 0,8271 | 0,976534 |
| D230025D16Rik | 11,9798 | 11,6189 | -0,0441297 | 0,8271 | 0,976534 |
| Cpd | 13,598 | 13,9716 | 0,0390984 | 0,8272 | 0,976562 |
| Mfge8 | 32,9409 | 32,0267 | -0,0406059 | 0,8274 | 0,976708 |
| Sfi1 | 4,72626 | 4,95825 | 0,0691313 | 0,8276 | 0,976734 |
| Cdv3 | 59,1501 | 57,5693 | -0,0390818 | 0,8276 | 0,976734 |
| Hecw2 | 0,905088 | 1,00074 | 0,144933 | 0,82765 | 0,976734 |
| Mark3 | 48,5956 | 49,9165 | 0,0386911 | 0,8278 | 0,976748 |
| Camta2 | 3,70735 | 3,84521 | 0,0526748 | 0,82785 | 0,976748 |
| Mtmr14 | 16,4847 | 16,9771 | 0,042469 | 0,82795 | 0,976748 |
| Pcdhga11 | 2,66436 | 2,24534 | -0,246852 | 0,8281 | 0,976748 |
| Ppp2r4 | 85,8071 | 88,0819 | 0,03775 | 0,82815 | 0,976748 |
| Med27 | 18,4041 | 17,7969 | -0,0484035 | 0,82815 | 0,976748 |
| Idua | 2,53592 | 2,69599 | 0,0883023 | 0,8284 | 0,976748 |
| Gpc6 | 12,5734 | 12,2312 | -0,0398088 | 0,8284 | 0,976748 |
| Pi4ka | 16,5197 | 17,0401 | 0,0447487 | 0,82845 | 0,976748 |
| Serinc3 | 28,6782 | 29,4703 | 0,0393056 | 0,82845 | 0,976748 |
| Erf | 21,2567 | 21,8518 | 0,0398383 | 0,8285 | 0,976748 |
| Ginm1 | 14,1534 | 14,6244 | 0,047221 | 0,82885 | 0,976847 |
| Taf5 | 11,9987 | 11,652 | -0,0422985 | 0,8289 | 0,976847 |
| Dhx15 | 208,451 | 214,348 | 0,0402478 | 0,82905 | 0,976847 |
| Bnip3l | 36,8981 | 35,9241 | -0,0385928 | 0,82905 | 0,976847 |
| Pcyox1 | 24,7672 | 24,1066 | -0,039002 | 0,82905 | 0,976847 |
| Tmem222 | 38,8157 | 37,7158 | -0,0414702 | 0,82925 | 0,976847 |
| Rad9a | 25,2055 | 27,726 | 0,137503 | 0,8293 | 0,976847 |
| Usp21 | 37,9196 | 36,8812 | -0,0400549 | 0,8293 | 0,976847 |
| Commd7 | 10,8007 | 10,4709 | -0,0447416 | 0,8293 | 0,976847 |
| Ints8 | 37,3393 | 38,6654 | 0,0503514 | 0,8294 | 0,976847 |
| Gng13 | 4,1472 | 5,27813 | 0,347888 | 0,82965 | 0,976847 |
| Alkbh1 | 32,8852 | 35,0272 | 0,091036 | 0,82965 | 0,976847 |
| Ncam1 | 28,5445 | 29,3297 | 0,0391494 | 0,82965 | 0,976847 |
| Taldo1 | 106,793 | 103,943 | -0,0390185 | 0,82965 | 0,976847 |
| Eri1 | 17,9073 | 17,4292 | -0,0390448 | 0,8298 | 0,976934 |
| Abtb1 | 8,66644 | 8,98688 | 0,0523811 | 0,8299 | 0,976962 |
| Fxr1 | 95,1048 | 97,7715 | 0,0398964 | 0,8302 | 0,977084 |
| Cox7a1 | 5,91666 | 5,4564 | -0,116834 | 0,8302 | 0,977084 |

| | | | | | |
|---|---|---|---|---|---|
| Atp6v0c | 10,2953 | 10,6997 | 0,0555848 | 0,83025 | 0,977084 |
| Zfp365 | 0,64683 | 0,683655 | 0,0798804 | 0,83045 | 0,977084 |
| Ccs | 20,2369 | 20,9264 | 0,048341 | 0,83045 | 0,977084 |
| Cd47 | 3,43359 | 3,58795 | 0,0634418 | 0,83085 | 0,977084 |
| Pfkl | 109,187 | 112,115 | 0,0381778 | 0,8309 | 0,977084 |
| Snx8 | 11,0711 | 10,7275 | -0,0454852 | 0,8311 | 0,977084 |
| 1700054A03Rik | 0,760216 | 0,846334 | 0,154819 | 0,83115 | 0,977084 |
| Ccdc85b | 5,88056 | 6,06824 | 0,0453243 | 0,8312 | 0,977084 |
| Fbxo5 | 51,7618 | 53,1945 | 0,0393868 | 0,8312 | 0,977084 |
| Slc31a1 | 25,3246 | 26,0254 | 0,0393807 | 0,83125 | 0,977084 |
| Kdm1b | 4,34295 | 3,79698 | -0,193825 | 0,83125 | 0,977084 |
| Armcx5 | 5,26389 | 5,79075 | 0,137622 | 0,8313 | 0,977084 |
| Txlna | 54,4886 | 55,8994 | 0,0368805 | 0,8313 | 0,977084 |
| Spcs2 | 59,7071 | 61,3282 | 0,0386494 | 0,8314 | 0,977084 |
| Trim2 | 8,75938 | 8,51674 | -0,0405275 | 0,83155 | 0,977084 |
| Gnpat | 74,633 | 76,7062 | 0,0395298 | 0,83165 | 0,977084 |
| Rps6kb1 | 19,5901 | 19,0162 | -0,0429007 | 0,83165 | 0,977084 |
| Mcf2l | 1,07504 | 1,12295 | 0,0629088 | 0,8319 | 0,977084 |
| Tagln | 8,49033 | 7,82568 | -0,117604 | 0,832 | 0,977084 |
| Rabl3 | 16,6927 | 16,1892 | -0,0441853 | 0,8322 | 0,977084 |
| Stard4 | 17,1552 | 16,6892 | -0,0397333 | 0,83225 | 0,977084 |
| Fdxacb1 | 4,95962 | 4,77062 | -0,0560521 | 0,8323 | 0,977084 |
| Cdkn2aip | 13,7088 | 13,3243 | -0,0410365 | 0,8324 | 0,977084 |
| Smyd4 | 4,47029 | 4,63111 | 0,0509894 | 0,83245 | 0,977084 |
| Kcnk5 | 6,57486 | 6,36635 | -0,0464922 | 0,83245 | 0,977084 |
| Ccny | 36,8264 | 35,8529 | -0,0386505 | 0,8325 | 0,977084 |
| Eif3b | 210,317 | 204,686 | -0,0391495 | 0,8325 | 0,977084 |
| Tmem192 | 9,5183 | 9,91245 | 0,0585374 | 0,83255 | 0,977084 |
| Lims1 | 27,5216 | 26,8125 | -0,037658 | 0,83255 | 0,977084 |
| Tmem184b | 18,6478 | 19,1823 | 0,040772 | 0,83265 | 0,977084 |
| Ccdc58 | 90,4846 | 88,0341 | -0,0396098 | 0,83265 | 0,977084 |
| Adam9 | 13,2719 | 13,6551 | 0,041073 | 0,83275 | 0,977084 |
| 4930528A17Rik | 0,475142 | 0,759225 | 0,676169 | 0,8328 | 0,977084 |
| Msh5 | 0,703529 | 0,661194 | -0,0895356 | 0,83285 | 0,977084 |
| Nfyc | 14,5088 | 14,0648 | -0,0448371 | 0,8329 | 0,977084 |
| Cdr2 | 20,3827 | 20,9682 | 0,0408606 | 0,833 | 0,977084 |
| Nrd1 | 62,0424 | 63,9437 | 0,0435463 | 0,83305 | 0,977084 |
| Wdr78 | 3,25385 | 3,1348 | -0,0537749 | 0,83305 | 0,977084 |
| Nespas | 1,16632 | 2,14235 | 0,877232 | 0,83325 | 0,977229 |
| Tssk4 | 0,722186 | 0,663209 | -0,122906 | 0,8336 | 0,977427 |
| Nat10 | 31,9078 | 32,7679 | 0,038374 | 0,83365 | 0,977427 |
| Mpp2 | 5,48707 | 5,66686 | 0,046514 | 0,8337 | 0,977427 |
| Zranb3 | 12,7323 | 13,0924 | 0,0402355 | 0,83375 | 0,977427 |
| Hmg20b | 38,0787 | 37,0265 | -0,0404245 | 0,8338 | 0,977427 |
| Slc39a9 | 16,4788 | 16,0394 | -0,0389855 | 0,83425 | 0,977586 |
| Polr2l | 4,63013 | 4,82251 | 0,0587322 | 0,83445 | 0,977586 |
| Gm1976 | 5,42394 | 5,75847 | 0,0863452 | 0,8345 | 0,977586 |
| Hlcs | 5,86392 | 5,69255 | -0,0427904 | 0,8345 | 0,977586 |
| Twf2 | 22,3883 | 21,7285 | -0,0431522 | 0,83485 | 0,977586 |
| Cdk17 | 12,7586 | 12,4017 | -0,0409369 | 0,8349 | 0,977586 |

| | | | | | |
|---|---|---|---|---|---|
| Crbn | 12,4633 | 11,9595 | -0,0595325 | 0,83495 | 0,977586 |
| Senp3 | 65,3935 | 63,6843 | -0,0382092 | 0,8351 | 0,977586 |
| Lin52 | 5,99351 | 5,78054 | -0,0521965 | 0,8351 | 0,977586 |
| Wdr5 | 68,1379 | 69,9529 | 0,0379278 | 0,83515 | 0,977586 |
| 6230400D17Rik | 1,00623 | 1,30077 | 0,370409 | 0,8352 | 0,977586 |
| Tst | 5,55715 | 5,31201 | -0,0650864 | 0,83525 | 0,977586 |
| Mir1982 | 367,704 | 529,611 | 0,526389 | 0,8354 | 0,977586 |
| Hnrnpul2 | 86,1647 | 88,4852 | 0,0383397 | 0,8355 | 0,977586 |
| Cbx3 | 14,4658 | 14,0239 | -0,0447627 | 0,83555 | 0,977586 |
| Dut | 103,143 | 100,514 | -0,0372501 | 0,8356 | 0,977586 |
| Apoe | 8,66907 | 8,34311 | -0,0552918 | 0,8356 | 0,977586 |
| Sec23b | 42,5421 | 41,4687 | -0,0368684 | 0,8357 | 0,977586 |
| B3galnt2 | 25,7608 | 24,9786 | -0,0444872 | 0,8358 | 0,977586 |
| Snora70 | 352,994 | 384,387 | 0,122916 | 0,83585 | 0,977586 |
| Ppapdc3 | 1,96507 | 1,86737 | -0,0735728 | 0,83585 | 0,977586 |
| Ska2 | 71,6945 | 64,7041 | -0,148004 | 0,83585 | 0,977586 |
| Mia | 0,838055 | 0,942908 | 0,170073 | 0,8359 | 0,977586 |
| Rmnd5a | 26,1313 | 26,8335 | 0,0382566 | 0,8359 | 0,977586 |
| Sfxn4 | 1,86512 | 1,95351 | 0,0667979 | 0,83605 | 0,977586 |
| Zfp944 | 5,31706 | 5,5089 | 0,0511353 | 0,83605 | 0,977586 |
| Tia1 | 44,0568 | 42,9558 | -0,036512 | 0,83615 | 0,977586 |
| Bcdin3d | 10,352 | 9,98309 | -0,0523512 | 0,83615 | 0,977586 |
| Snord33 | 1294,3 | 1109,59 | -0,222143 | 0,83625 | 0,977586 |
| Ccdc77 | 14,8901 | 15,7698 | 0,0828095 | 0,83635 | 0,977586 |
| Akr7a5 | 27,4731 | 26,6872 | -0,0418743 | 0,83635 | 0,977586 |
| 5830417I10Rik | 3,34033 | 3,44169 | 0,043127 | 0,8364 | 0,977586 |
| Zfand4 | 2,13748 | 2,04798 | -0,0617049 | 0,83645 | 0,977586 |
| Ly6g6d | 1,79924 | 1,95864 | 0,122464 | 0,8366 | 0,977672 |
| Uri1 | 28,116 | 28,8504 | 0,0371979 | 0,8367 | 0,9777 |
| Mettl2 | 30,6695 | 29,8571 | -0,0387318 | 0,83705 | 0,97802 |
| Tbl1x | 62,4643 | 60,8405 | -0,0379998 | 0,8372 | 0,978106 |
| Vrk1 | 34,3861 | 33,5254 | -0,0365707 | 0,8374 | 0,978107 |
| Vac14 | 21,8343 | 21,2608 | -0,0384012 | 0,8374 | 0,978107 |
| Hgsnat | 11,0009 | 10,681 | -0,0425727 | 0,83765 | 0,978107 |
| 1110012L19Rik | 16,9093 | 16,3418 | -0,0492486 | 0,83775 | 0,978107 |
| Slc2a4rg-ps | 4,23286 | 4,08321 | -0,0519287 | 0,83785 | 0,978107 |
| Morn2 | 10,7319 | 11,9578 | 0,15604 | 0,83805 | 0,978107 |
| Ehd2 | 3,93291 | 4,07595 | 0,0515394 | 0,83805 | 0,978107 |
| Paqr3 | 5,03134 | 4,69455 | -0,0999557 | 0,83805 | 0,978107 |
| Rdm1 | 11,5162 | 11,9285 | 0,0507527 | 0,83815 | 0,978107 |
| Zfp770 | 6,65585 | 6,8652 | 0,0446787 | 0,83815 | 0,978107 |
| Yars2 | 38,0243 | 39,071 | 0,039175 | 0,8382 | 0,978107 |
| Gm16617 | 3,49576 | 4,80782 | 0,459776 | 0,83835 | 0,978107 |
| Tsen15 | 25,385 | 24,6244 | -0,0438865 | 0,83835 | 0,978107 |
| Ptgr2 | 7,34949 | 7,12181 | -0,0454017 | 0,83845 | 0,978107 |
| 1110054M08Rik | 2,26026 | 2,91488 | 0,366947 | 0,8385 | 0,978107 |
| Cbr4 | 13,0273 | 13,5052 | 0,0519838 | 0,8385 | 0,978107 |
| Zfp956 | 3,86106 | 4,05153 | 0,0694679 | 0,83855 | 0,978107 |
| Ints5 | 26,575 | 25,8673 | -0,0389392 | 0,83865 | 0,978107 |
| Fam53b | 7,14811 | 6,9527 | -0,0399887 | 0,83865 | 0,978107 |

| | | | | | |
|---|---|---|---|---|---|
| Zdhhc12 | 13,1705 | 13,8101 | 0,068419 | 0,8391 | 0,978359 |
| Gm11517 | 1,67296 | 2,48569 | 0,571249 | 0,83915 | 0,978359 |
| Farsb | 72,7854 | 70,9396 | -0,0370582 | 0,8393 | 0,978359 |
| Rnf38 | 28,2722 | 28,9913 | 0,0362391 | 0,83945 | 0,978359 |
| 1700109H08Rik | 1,55825 | 1,9303 | 0,308898 | 0,83955 | 0,978359 |
| Crnkl1 | 28,4187 | 29,144 | 0,0363558 | 0,8396 | 0,978359 |
| Plscr1 | 11,3739 | 11,0232 | -0,0451837 | 0,83965 | 0,978359 |
| 1110008P14Rik | 19,2352 | 18,5185 | -0,0547849 | 0,83965 | 0,978359 |
| Fam107b | 23,8446 | 24,4659 | 0,037114 | 0,8397 | 0,978359 |
| Hnrnpa2b1 | 923,314 | 897,587 | -0,0407688 | 0,8401 | 0,978359 |
| Patz1 | 49,9055 | 52,7393 | 0,0796802 | 0,8402 | 0,978359 |
| Nap1l4 | 185,848 | 181,203 | -0,0365179 | 0,8402 | 0,978359 |
| Ttll12 | 27,0117 | 26,3328 | -0,0367246 | 0,8405 | 0,978359 |
| Exosc3 | 59,036 | 57,4682 | -0,0388294 | 0,84055 | 0,978359 |
| Uba52 | 1,48128 | 1,63657 | 0,143839 | 0,8406 | 0,978359 |
| Haus8 | 22,1826 | 21,562 | -0,0409376 | 0,8407 | 0,978359 |
| Nedd4 | 687,188 | 665,74 | -0,0457442 | 0,8408 | 0,978359 |
| Snord12 | 4975,78 | 3658,08 | -0,443836 | 0,84085 | 0,978359 |
| Cdk3-ps | 0,741812 | 0,684625 | -0,11574 | 0,8409 | 0,978359 |
| Taf6 | 56,8193 | 55,2792 | -0,0396445 | 0,84095 | 0,978359 |
| Gfm2 | 20,4171 | 19,8459 | -0,0409389 | 0,84105 | 0,978359 |
| Masp1 | 1,62274 | 1,54816 | -0,0678783 | 0,84105 | 0,978359 |
| Npff | 2,69921 | 2,447 | -0,141527 | 0,84105 | 0,978359 |
| Hilpda | 20,3503 | 19,6887 | -0,0476846 | 0,8411 | 0,978359 |
| Rpusd2 | 4,74314 | 4,60049 | -0,0440548 | 0,8412 | 0,978359 |
| Smg8 | 16,8725 | 19,0468 | 0,174878 | 0,8413 | 0,978359 |
| Opa1 | 21,0586 | 21,5913 | 0,0360419 | 0,8413 | 0,978359 |
| Pcdhga7 | 2,35118 | 1,97686 | -0,250178 | 0,84135 | 0,978359 |
| Eef1e1 | 76,3334 | 78,3042 | 0,0367745 | 0,8414 | 0,978359 |
| Tfdp2 | 22,1785 | 21,6486 | -0,0348926 | 0,8415 | 0,978359 |
| Ptplad1 | 42,4015 | 43,4666 | 0,0357925 | 0,84155 | 0,978359 |
| Atg4b | 33,8124 | 32,9534 | -0,037124 | 0,84165 | 0,978359 |
| Fndc4 | 31,1424 | 26,9363 | -0,20933 | 0,84175 | 0,978359 |
| Dsn1 | 22,5026 | 23,1158 | 0,0387889 | 0,8418 | 0,978359 |
| Nop2 | 75,6025 | 73,7427 | -0,0359341 | 0,84195 | 0,978359 |
| Ube2g2 | 35,1218 | 34,2424 | -0,0365858 | 0,842 | 0,978359 |
| Gstm7 | 7,14095 | 7,42129 | 0,0555541 | 0,84205 | 0,978359 |
| Akap5 | 0,858395 | 0,823778 | -0,0593865 | 0,8421 | 0,978359 |
| E130102H24Rik | 1,69558 | 1,83826 | 0,116565 | 0,84215 | 0,978359 |
| Psmc6 | 206,602 | 211,802 | 0,0358613 | 0,84215 | 0,978359 |
| Gdpd1 | 5,31079 | 3,97324 | -0,418612 | 0,84215 | 0,978359 |
| Srsf9 | 173,016 | 177,304 | 0,0353152 | 0,84225 | 0,978359 |
| Cdc25c | 28,0821 | 27,3169 | -0,0398595 | 0,84225 | 0,978359 |
| Slc25a17 | 43,5441 | 44,7084 | 0,0380698 | 0,84245 | 0,978359 |
| Kpna1 | 34,5451 | 35,4072 | 0,0355604 | 0,84245 | 0,978359 |
| Fnta | 56,81 | 55,4689 | -0,0344652 | 0,84245 | 0,978359 |
| Pcbp2 | 225,636 | 219,944 | -0,0368589 | 0,84245 | 0,978359 |
| 1700065D16Rik | 0,714164 | 1,00988 | 0,499857 | 0,8426 | 0,978444 |
| Pomk | 6,22719 | 6,04244 | -0,0434499 | 0,8427 | 0,978472 |
| Wnt6 | 0,881934 | 0,830845 | -0,0860911 | 0,84295 | 0,978612 |

| | | | | | |
|---|---|---|---|---|---|
| Gnb1 | 210,099 | 204,99 | -0,0355111 | 0,843 | 0,978612 |
| Gm19557 | 0,52332 | 0,620108 | 0,244828 | 0,84305 | 0,978612 |
| Rpgrip1l | 3,71317 | 3,60539 | -0,0424958 | 0,84315 | 0,97864 |
| E2f4 | 71,2127 | 69,4877 | -0,0353755 | 0,8434 | 0,978842 |
| Etaa1 | 14,1812 | 14,5594 | 0,0379727 | 0,84375 | 0,978951 |
| Smndc1 | 36,0372 | 35,1637 | -0,0353977 | 0,84375 | 0,978951 |
| Trmt10c | 29,5384 | 30,3216 | 0,0377527 | 0,8438 | 0,978951 |
| Nadk | 48,1247 | 46,9287 | -0,0363076 | 0,8438 | 0,978951 |
| Rnf103 | 6,15604 | 5,96555 | -0,0453469 | 0,8444 | 0,979308 |
| Map3k12 | 6,3989 | 5,6934 | -0,168534 | 0,8445 | 0,979308 |
| Txnl4a | 121,852 | 118,691 | -0,0379184 | 0,8446 | 0,979308 |
| Zfyve1 | 8,40613 | 8,64217 | 0,039952 | 0,8447 | 0,979308 |
| Aasdh | 4,31034 | 4,45311 | 0,0470122 | 0,84475 | 0,979308 |
| Lrrc47 | 30,6629 | 31,4061 | 0,0345472 | 0,8448 | 0,979308 |
| Irak2 | 1,90314 | 1,98179 | 0,0584217 | 0,84485 | 0,979308 |
| Tmem241 | 1,23842 | 1,18269 | -0,066428 | 0,8449 | 0,979308 |
| Xpo7 | 16,0215 | 15,6394 | -0,03482 | 0,845 | 0,979308 |
| 4930581F22Rik | 0,878488 | 0,724109 | -0,278816 | 0,845 | 0,979308 |
| Pxmp2 | 3,86386 | 4,05015 | 0,067931 | 0,8451 | 0,979308 |
| Tubgcp3 | 28,9849 | 29,7084 | 0,0355693 | 0,8451 | 0,979308 |
| Abhd17a | 44,0151 | 42,8609 | -0,0383368 | 0,8451 | 0,979308 |
| Wdr89 | 5,85654 | 6,50892 | 0,15237 | 0,8453 | 0,979387 |
| Krr1 | 24,4277 | 25,0299 | 0,0351324 | 0,84535 | 0,979387 |
| Elf1 | 4,38761 | 4,63332 | 0,0786122 | 0,84545 | 0,979387 |
| Snord87 | 350,233 | 409,486 | 0,225501 | 0,8455 | 0,979387 |
| Dnajc4 | 6,17665 | 6,46715 | 0,0663053 | 0,84555 | 0,979387 |
| Kri1 | 21,858 | 22,5775 | 0,0467214 | 0,84575 | 0,979412 |
| Rbm48 | 6,96731 | 6,67118 | -0,0626606 | 0,84575 | 0,979412 |
| Gm6642 | 0,741952 | 1,01456 | 0,451459 | 0,8458 | 0,979412 |
| 3110001I22Rik | 7,64014 | 8,10606 | 0,0854021 | 0,846 | 0,979555 |
| Dido1 | 25,797 | 26,4333 | 0,0351527 | 0,8461 | 0,979579 |
| Fads1 | 76,6285 | 74,8267 | -0,034327 | 0,8462 | 0,979579 |
| Gtf2f2 | 48,1401 | 46,9447 | -0,0362776 | 0,84625 | 0,979579 |
| Plaa | 26,5776 | 27,2229 | 0,0346109 | 0,84635 | 0,979606 |
| Csnk1d | 81,9701 | 83,971 | 0,0347937 | 0,84675 | 0,979886 |
| BC068157 | 2,71704 | 2,62657 | -0,0488553 | 0,8468 | 0,979886 |
| Scrn2 | 15,6694 | 15,2059 | -0,0433171 | 0,84695 | 0,979886 |
| Zfp712 | 1,40493 | 1,46581 | 0,0612028 | 0,84705 | 0,979886 |
| Zfp697 | 3,35972 | 3,25795 | -0,0443763 | 0,84705 | 0,979886 |
| N6amt1 | 9,60212 | 9,30569 | -0,0452389 | 0,84705 | 0,979886 |
| Arhgef11 | 10,314 | 10,5811 | 0,0368859 | 0,8473 | 0,979995 |
| Bax | 146,587 | 150,246 | 0,0355725 | 0,8473 | 0,979995 |
| Hagh | 10,6638 | 11,0346 | 0,0493152 | 0,8474 | 0,979995 |
| Rnf5 | 41,0751 | 40,0041 | -0,0381166 | 0,84745 | 0,979995 |
| 3110062M04Rik | 4,1209 | 4,26117 | 0,0482913 | 0,84765 | 0,980138 |
| Aebp1 | 2,1353 | 1,96203 | -0,122093 | 0,8479 | 0,980339 |
| Lpin2 | 15,3889 | 15,7644 | 0,0347762 | 0,84805 | 0,980424 |
| Tnrc6c | 9,30568 | 9,53598 | 0,0352703 | 0,84815 | 0,980451 |
| Trmt2a | 28,7797 | 27,8098 | -0,0494549 | 0,84835 | 0,980594 |
| Stmn2 | 1,48368 | 1,4111 | -0,0723641 | 0,84865 | 0,980781 |

| | | | | | |
|---|---|---|---|---|---|
| Rfng | 10,0647 | 9,77154 | -0,0426456 | 0,84875 | 0,980781 |
| Pcdhgb2 | 1,15954 | 1,42733 | 0,299771 | 0,849 | 0,980781 |
| Ilk | 53,1742 | 51,1659 | -0,0555441 | 0,84905 | 0,980781 |
| Arhgap27 | 0,595063 | 0,624891 | 0,0705607 | 0,8491 | 0,980781 |
| Tvp23a | 8,0157 | 8,3913 | 0,0660655 | 0,8491 | 0,980781 |
| Tmem191c | 1,63305 | 1,72505 | 0,079073 | 0,84915 | 0,980781 |
| Leng1 | 8,66762 | 8,13775 | -0,0910053 | 0,84915 | 0,980781 |
| Smim15 | 35,3251 | 36,1927 | 0,0350065 | 0,8492 | 0,980781 |
| Xylt1 | 1,06172 | 1,0126 | -0,068338 | 0,8494 | 0,980924 |
| Usp16 | 22,8821 | 25,2623 | 0,142768 | 0,8496 | 0,981036 |
| Gng3 | 2,20396 | 2,02996 | -0,118642 | 0,84965 | 0,981036 |
| Mettl17 | 13,6494 | 14,0629 | 0,0430509 | 0,84985 | 0,981148 |
| Smim24 | 1,9486 | 1,79792 | -0,116107 | 0,8499 | 0,981148 |
| Pigw | 8,3801 | 8,84286 | 0,0775457 | 0,85 | 0,981175 |
| Ulk1 | 16,7361 | 16,3323 | -0,0352342 | 0,85025 | 0,981351 |
| Cstb | 71,46 | 73,4571 | 0,0397658 | 0,8506 | 0,981351 |
| Ppp1r12c | 25,2121 | 25,8136 | 0,0340104 | 0,8506 | 0,981351 |
| Tcea1 | 13,2816 | 12,9429 | -0,0372681 | 0,8507 | 0,981351 |
| 4930487H11Rik | 0,643675 | 0,919842 | 0,515054 | 0,85075 | 0,981351 |
| Sf3b1 | 106,224 | 108,846 | 0,0351801 | 0,85075 | 0,981351 |
| 4930577N17Rik | 0,580428 | 0,766715 | 0,401573 | 0,8508 | 0,981351 |
| Emc3 | 33,4247 | 32,601 | -0,0359964 | 0,8508 | 0,981351 |
| Gm20324 | 1,04941 | 1,4232 | 0,439563 | 0,851 | 0,981351 |
| Osbpl3 | 2,82278 | 2,74074 | -0,0425489 | 0,8511 | 0,981351 |
| Tmem79 | 1,29225 | 1,23263 | -0,0681482 | 0,8511 | 0,981351 |
| Snrnp27 | 99,1655 | 96,7139 | -0,0361151 | 0,85115 | 0,981351 |
| Ntf3 | 3,57512 | 3,41799 | -0,0648408 | 0,8512 | 0,981351 |
| Uhrf2 | 61,7068 | 63,168 | 0,0337648 | 0,8513 | 0,981351 |
| Txndc9 | 16,0374 | 15,643 | -0,0359261 | 0,8513 | 0,981351 |
| E2f6 | 48,3209 | 49,4535 | 0,033425 | 0,85145 | 0,981429 |
| Tial1 | 33,6666 | 34,451 | 0,0332295 | 0,85155 | 0,981429 |
| Zcchc11 | 34,7819 | 35,6175 | 0,0342461 | 0,8516 | 0,981429 |
| Tctex1d2 | 19,8439 | 19,1544 | -0,0510234 | 0,8517 | 0,981429 |
| Tagap1 | 7,42816 | 7,21161 | -0,0426842 | 0,85175 | 0,981429 |
| Rad9b | 2,372 | 2,47636 | 0,0621184 | 0,85195 | 0,981483 |
| Abcb6 | 18,3272 | 18,7925 | 0,0361695 | 0,85195 | 0,981483 |
| Cdk16 | 67,4926 | 69,0661 | 0,0332494 | 0,8521 | 0,981503 |
| Ftsj2 | 9,60979 | 9,93754 | 0,0483836 | 0,85225 | 0,981503 |
| Bud13 | 20,2923 | 19,7639 | -0,0380618 | 0,85225 | 0,981503 |
| Prkab1 | 30,2279 | 30,9885 | 0,0358554 | 0,85235 | 0,981503 |
| Atxn7l2 | 10,6089 | 10,3219 | -0,0395741 | 0,85235 | 0,981503 |
| Dhx30 | 45,241 | 46,2959 | 0,0332565 | 0,8526 | 0,98155 |
| Zfp94 | 3,30491 | 3,42577 | 0,0518168 | 0,85275 | 0,98155 |
| 6330549D23Rik | 7,92422 | 8,29626 | 0,066193 | 0,8528 | 0,98155 |
| Cdkn1b | 11,049 | 11,3464 | 0,03832 | 0,8528 | 0,98155 |
| 4931440P22Rik | 0,543008 | 0,663719 | 0,289599 | 0,85285 | 0,98155 |
| Ttc9c | 27,6398 | 27,0034 | -0,033604 | 0,85285 | 0,98155 |
| 4930513N10Rik | 1,41726 | 1,87696 | 0,405292 | 0,85295 | 0,981577 |
| Ypel5 | 4,78344 | 4,93607 | 0,0453152 | 0,8532 | 0,981735 |
| Slc22a5 | 2,15138 | 2,23329 | 0,053911 | 0,85335 | 0,981735 |

| | | | | | |
|---|---|---|---|---|---|
| Homez | 6,39189 | 6,56104 | 0,0376835 | 0,8534 | 0,981735 |
| Nsun5 | 16,6454 | 15,6574 | -0,0882783 | 0,85345 | 0,981735 |
| Entpd7 | 1,80912 | 1,92034 | 0,0860779 | 0,8537 | 0,981735 |
| Ppm1a | 37,7911 | 38,6795 | 0,0335207 | 0,85375 | 0,981735 |
| Bbs9 | 5,32004 | 5,16737 | -0,0420091 | 0,85375 | 0,981735 |
| Clcc1 | 19,7507 | 18,889 | -0,0643575 | 0,8538 | 0,981735 |
| Clip4 | 1,51788 | 1,58607 | 0,0634036 | 0,8539 | 0,981735 |
| Rdh11 | 25,3852 | 26,055 | 0,0375717 | 0,854 | 0,981735 |
| E130012A19Rik | 23,8068 | 24,3952 | 0,0352203 | 0,8541 | 0,981735 |
| Dph1 | 17,3209 | 16,7115 | -0,0516719 | 0,8541 | 0,981735 |
| 1810026J23Rik | 24,2051 | 23,0472 | -0,0707202 | 0,85415 | 0,981735 |
| 4632427E13Rik | 2,28586 | 2,43899 | 0,0935502 | 0,8543 | 0,981735 |
| Cirh1a | 64,6377 | 63,1584 | -0,0334014 | 0,85435 | 0,981735 |
| Ogfod1 | 6,38074 | 6,54407 | 0,0364637 | 0,85455 | 0,981735 |
| Afg3l2 | 30,9436 | 31,6771 | 0,0337961 | 0,85455 | 0,981735 |
| Ppp2cb | 62,6035 | 61,121 | -0,034577 | 0,8546 | 0,981735 |
| Echdc1 | 6,33064 | 5,96988 | -0,0846501 | 0,8547 | 0,981735 |
| Gm166 | 2,11619 | 2,26019 | 0,0949799 | 0,8548 | 0,981735 |
| Sumf1 | 5,02232 | 4,86349 | -0,0463608 | 0,85485 | 0,981735 |
| Tmem218 | 9,39193 | 9,07219 | -0,0499707 | 0,85485 | 0,981735 |
| Agk | 4,8283 | 4,66682 | -0,0490748 | 0,8549 | 0,981735 |
| Tango6 | 7,50042 | 7,69975 | 0,03784 | 0,855 | 0,981735 |
| Tmem18 | 12,3082 | 11,9938 | -0,037335 | 0,855 | 0,981735 |
| Fus | 408,544 | 399,391 | -0,0326872 | 0,8551 | 0,981755 |
| Eefsec | 16,5986 | 16,1949 | -0,0355291 | 0,8552 | 0,981755 |
| Thap4 | 24,5622 | 23,9634 | -0,0356073 | 0,85535 | 0,981755 |
| Ppap2c | 15,8174 | 15,3818 | -0,0402898 | 0,85535 | 0,981755 |
| Slc27a3 | 19,6983 | 19,2105 | -0,0361715 | 0,8554 | 0,981755 |
| Cdc14a | 7,5003 | 7,70103 | 0,0381034 | 0,8556 | 0,981896 |
| BC031181 | 41,1281 | 42,2115 | 0,0375096 | 0,8557 | 0,981923 |
| Ilkap | 81,6249 | 83,5865 | 0,0342606 | 0,8559 | 0,982065 |
| Nacc2 | 0,915319 | 0,949874 | 0,0534612 | 0,8561 | 0,982169 |
| Bcat2 | 23,5126 | 24,1192 | 0,0367513 | 0,85625 | 0,982169 |
| Cpsf3 | 63,7118 | 62,319 | -0,0318889 | 0,85625 | 0,982169 |
| Grik5 | 12,9708 | 12,6503 | -0,0360901 | 0,85635 | 0,982169 |
| Neurl2 | 2,11228 | 2,46756 | 0,224285 | 0,8564 | 0,982169 |
| Npc2 | 55,2559 | 54,0417 | -0,0320537 | 0,85645 | 0,982169 |
| Ptpn1 | 26,2504 | 26,8672 | 0,0335034 | 0,85675 | 0,98236 |
| Nudcd3 | 29,5394 | 28,8623 | -0,0334523 | 0,8568 | 0,98236 |
| Rps6ka2 | 7,83857 | 8,04026 | 0,0366523 | 0,85685 | 0,98236 |
| Rpgr | 14,254 | 14,6178 | 0,0363621 | 0,857 | 0,98236 |
| Trub1 | 10,6962 | 10,9632 | 0,0355814 | 0,857 | 0,98236 |
| Gid8 | 20,1358 | 20,6086 | 0,0334899 | 0,85725 | 0,982529 |
| Dgcr2 | 58,7363 | 60,0964 | 0,0330263 | 0,8573 | 0,982529 |
| Fsbp | 1,58078 | 1,68511 | 0,0922079 | 0,85745 | 0,982582 |
| Fhl2 | 4,43526 | 4,26604 | -0,0561191 | 0,8575 | 0,982582 |
| Polr3d | 33,3019 | 34,0823 | 0,0334147 | 0,85775 | 0,982632 |
| Ermp1 | 11,1513 | 10,8898 | -0,0342338 | 0,85775 | 0,982632 |
| Mlst8 | 17,874 | 17,4531 | -0,0343776 | 0,8578 | 0,982632 |
| F3 | 1,47825 | 1,54981 | 0,0682019 | 0,85785 | 0,982632 |

| | | | | | |
|---|---|---|---|---|---|
| Gm12338 | 33,0532 | 30,911 | -0,0966657 | 0,8582 | 0,982813 |
| Tbc1d14 | 19,8378 | 20,3103 | 0,0339662 | 0,85855 | 0,982813 |
| Rexo4 | 33,4222 | 32,6558 | -0,0334706 | 0,8586 | 0,982813 |
| Ccdc38 | 0,597258 | 0,670002 | 0,165813 | 0,8587 | 0,982813 |
| 1600014C10Rik | 5,81092 | 5,97927 | 0,0412031 | 0,8587 | 0,982813 |
| Ranbp10 | 12,2835 | 12,5723 | 0,0335263 | 0,8587 | 0,982813 |
| Npepps | 34,8436 | 35,621 | 0,031835 | 0,8587 | 0,982813 |
| Hmces | 38,261 | 39,1885 | 0,0345559 | 0,8588 | 0,982813 |
| Atp2b1 | 26,3262 | 25,7477 | -0,0320554 | 0,8588 | 0,982813 |
| Myo10 | 8,64136 | 8,43772 | -0,0344051 | 0,85905 | 0,982813 |
| 3110056K07Rik | 7,4996 | 7,12639 | -0,073644 | 0,8591 | 0,982813 |
| Ovca2 | 16,997 | 17,5224 | 0,0439153 | 0,85915 | 0,982813 |
| Plekhf1 | 3,15623 | 3,02999 | -0,0588898 | 0,85915 | 0,982813 |
| Snora62 | 25,1683 | 30,9713 | 0,299323 | 0,85925 | 0,982813 |
| Cystm1 | 5,33155 | 5,61457 | 0,0746197 | 0,8594 | 0,982813 |
| AU019823 | 17,189 | 16,6276 | -0,047898 | 0,8594 | 0,982813 |
| Polr1e | 15,2159 | 14,7811 | -0,0418277 | 0,8595 | 0,982813 |
| Tmem185b | 10,0399 | 10,3132 | 0,0387479 | 0,85955 | 0,982813 |
| Cldn25 | 36,0882 | 36,9104 | 0,032497 | 0,8596 | 0,982813 |
| Ankrd61 | 0,968688 | 1,25736 | 0,376298 | 0,8597 | 0,982813 |
| Zbtb6 | 5,64846 | 5,79964 | 0,0381059 | 0,8598 | 0,982813 |
| Sirt6 | 19,9403 | 20,4485 | 0,0363097 | 0,85985 | 0,982813 |
| Zdhhc17 | 14,6426 | 14,981 | 0,0329702 | 0,85985 | 0,982813 |
| Ywhah | 64,3779 | 65,8492 | 0,0326017 | 0,85985 | 0,982813 |
| Cstf3 | 43,2746 | 42,082 | -0,0403175 | 0,86 | 0,982813 |
| Snx17 | 63,6472 | 57,7685 | -0,139815 | 0,86 | 0,982813 |
| Pradc1 | 12,8516 | 10,6391 | -0,272565 | 0,86035 | 0,983023 |
| Eid2b | 6,12779 | 5,95337 | -0,0416584 | 0,86045 | 0,983023 |
| 1110059G10Rik | 4,90885 | 5,05266 | 0,0416558 | 0,8605 | 0,983023 |
| Manbal | 32,8844 | 32,0566 | -0,0367797 | 0,86065 | 0,983023 |
| Shmt2 | 78,8494 | 77,1242 | -0,0319161 | 0,86075 | 0,983023 |
| Ncdn | 15,2615 | 15,6229 | 0,0337659 | 0,8608 | 0,983023 |
| Tbc1d20 | 19,6927 | 20,1926 | 0,0361701 | 0,86085 | 0,983023 |
| Zfp58 | 5,66982 | 5,50574 | -0,0423671 | 0,86085 | 0,983023 |
| Vps36 | 47,9117 | 46,781 | -0,0344561 | 0,86095 | 0,983023 |
| Gm2382 | 4,24943 | 4,05843 | -0,0663479 | 0,86095 | 0,983023 |
| Gpcpd1 | 6,29851 | 6,1325 | -0,0385343 | 0,8615 | 0,983381 |
| Zfp639 | 28,5802 | 27,9106 | -0,0342011 | 0,86155 | 0,983381 |
| Recql5 | 10,3158 | 10,5642 | 0,0343314 | 0,86175 | 0,983381 |
| Fam104a | 12,9965 | 12,5462 | -0,0508753 | 0,86175 | 0,983381 |
| Msmp | 2,49652 | 3,37633 | 0,435538 | 0,8618 | 0,983381 |
| Med9 | 15,6014 | 15,2368 | -0,034108 | 0,8618 | 0,983381 |
| Cyp4f41-ps | 1,22563 | 1,15081 | -0,0908678 | 0,8618 | 0,983381 |
| Rpl11 | 9,0452 | 8,68189 | -0,0591426 | 0,86205 | 0,983518 |
| Zfyve9 | 21,7661 | 22,3097 | 0,0355853 | 0,8621 | 0,983518 |
| Lrrc39 | 0,963814 | 1,08206 | 0,166959 | 0,86215 | 0,983518 |
| Fam96b | 44,9476 | 43,7929 | -0,037546 | 0,8623 | 0,983602 |
| Tspan6 | 111,005 | 108,591 | -0,0317193 | 0,86245 | 0,983655 |
| Trappc9 | 5,40095 | 5,58456 | 0,0482312 | 0,8625 | 0,983655 |
| Clhc1 | 28,7941 | 28,0817 | -0,0361442 | 0,86275 | 0,983761 |

| | | | | | |
|---|---|---|---|---|---|
| Ahsa2 | 35,5222 | 36,8909 | 0,0545452 | 0,86285 | 0,983761 |
| Exoc7 | 19,2527 | 19,7069 | 0,0336425 | 0,86285 | 0,983761 |
| Racgap1 | 61,7426 | 60,4124 | -0,0314224 | 0,8629 | 0,983761 |
| Zmiz1 | 33,1152 | 32,4137 | -0,0308919 | 0,86325 | 0,983981 |
| Guf1 | 20,7235 | 21,1916 | 0,0322222 | 0,86335 | 0,983981 |
| Mob3a | 13,0062 | 12,6971 | -0,0346997 | 0,86335 | 0,983981 |
| Rela | 12,5395 | 12,2326 | -0,0357439 | 0,8634 | 0,983981 |
| Cox11 | 11,6236 | 11,2318 | -0,0494787 | 0,86355 | 0,984004 |
| Hebp2 | 10,5042 | 10,7794 | 0,0373208 | 0,8636 | 0,984004 |
| Wisp3 | 0,633919 | 0,74721 | 0,237216 | 0,86365 | 0,984004 |
| Ptpn13 | 17,5386 | 17,1771 | -0,0300445 | 0,86395 | 0,984045 |
| Ube2d1 | 54,6977 | 55,9406 | 0,0324158 | 0,8642 | 0,984045 |
| Ppp2r5c | 48,8505 | 47,7818 | -0,031913 | 0,86425 | 0,984045 |
| Ston1 | 8,11615 | 7,9078 | -0,0375197 | 0,8644 | 0,984045 |
| Trip4 | 8,88604 | 9,33214 | 0,0706683 | 0,86445 | 0,984045 |
| Fam71e1 | 0,90296 | 0,836207 | -0,110801 | 0,86445 | 0,984045 |
| Polr3b | 15,9441 | 16,3091 | 0,0326536 | 0,8647 | 0,984045 |
| Sh3bgr | 1,03052 | 1,09413 | 0,0864053 | 0,8648 | 0,984045 |
| Ppp4r1 | 37,7673 | 36,9682 | -0,0308559 | 0,8648 | 0,984045 |
| Mut | 12,3873 | 12,6717 | 0,0327403 | 0,86495 | 0,984045 |
| Mtfr1l | 39,1581 | 38,318 | -0,0312898 | 0,86495 | 0,984045 |
| Copg2 | 30,9056 | 28,6608 | -0,10879 | 0,86495 | 0,984045 |
| Mrgbp | 19,3404 | 19,85 | 0,0375192 | 0,865 | 0,984045 |
| Nfkb1 | 13,6296 | 13,3296 | -0,0321042 | 0,86505 | 0,984045 |
| Gemin2 | 29,9105 | 29,2059 | -0,0343927 | 0,86505 | 0,984045 |
| Lrrc57 | 12,6977 | 13,2688 | 0,0634761 | 0,8652 | 0,984045 |
| Ccdc137 | 17,1652 | 16,72 | -0,0379113 | 0,8652 | 0,984045 |
| Zfp87 | 6,32958 | 6,50353 | 0,0391128 | 0,86525 | 0,984045 |
| 2810013P06Rik | 7,02021 | 7,22607 | 0,0416971 | 0,8653 | 0,984045 |
| Slc35g2 | 2,95029 | 2,83697 | -0,0565054 | 0,8653 | 0,984045 |
| Tdg | 0,693551 | 0,629934 | -0,1388 | 0,8654 | 0,984045 |
| Spryd3 | 5,09524 | 5,24502 | 0,0418002 | 0,86545 | 0,984045 |
| Zc3h15 | 80,4209 | 82,1775 | 0,0311728 | 0,86545 | 0,984045 |
| Ppan | 64,7888 | 63,3748 | -0,0318356 | 0,86555 | 0,984071 |
| Nsl1 | 21,1666 | 20,7126 | -0,0312809 | 0,86585 | 0,984291 |
| Txnl4b | 7,02468 | 6,81991 | -0,0426795 | 0,8659 | 0,984291 |
| 2310022B05Rik | 29,6557 | 30,2895 | 0,0305125 | 0,86605 | 0,984291 |
| Mtg1 | 9,91867 | 9,63845 | -0,0413465 | 0,86605 | 0,984291 |
| Tmem68 | 9,05095 | 9,27981 | 0,036027 | 0,86615 | 0,984317 |
| Elk3 | 13,7188 | 13,4167 | -0,0321216 | 0,86635 | 0,984396 |
| Zfp513 | 15,069 | 17,7789 | 0,238582 | 0,86645 | 0,984396 |
| Hdhd3 | 3,75477 | 3,91111 | 0,0588528 | 0,86645 | 0,984396 |
| Ptpn2 | 54,1103 | 55,3384 | 0,0323771 | 0,8667 | 0,984415 |
| Zfp763 | 6,40508 | 6,57248 | 0,0372208 | 0,86675 | 0,984415 |
| Fbxo17 | 3,92083 | 3,7924 | -0,0480474 | 0,8668 | 0,984415 |
| Rab3a | 1,95559 | 2,07589 | 0,0861281 | 0,86685 | 0,984415 |
| Mxd3 | 7,40863 | 6,79233 | -0,125301 | 0,86685 | 0,984415 |
| Ergic1 | 85,3903 | 83,6704 | -0,0293556 | 0,867 | 0,984498 |
| Gm6297 | 0,814894 | 0,530582 | -0,619036 | 0,86715 | 0,984544 |
| Cbfa2t2 | 10,1622 | 10,3833 | 0,031052 | 0,8673 | 0,984544 |

| | | | | | |
|---|---|---|---|---|---|
| Clip3 | 40,1536 | 41,0089 | 0,0304083 | 0,86735 | 0,984544 |
| Hira | 34,3668 | 33,6643 | -0,0297973 | 0,86735 | 0,984544 |
| Cyb5 | 116,958 | 119,494 | 0,0309436 | 0,8676 | 0,984544 |
| Nol6 | 25,7207 | 25,1923 | -0,0299511 | 0,8676 | 0,984544 |
| Pcdhgb7 | 3,02798 | 2,67651 | -0,177999 | 0,8678 | 0,984544 |
| Snord47 | 5048,99 | 6584,25 | 0,383025 | 0,86805 | 0,984544 |
| Trio | 11,0272 | 11,2618 | 0,0303694 | 0,86805 | 0,984544 |
| Cacul1 | 19,8863 | 20,3134 | 0,0306512 | 0,86825 | 0,984544 |
| Fam173b | 12,9527 | 13,3365 | 0,0421356 | 0,8683 | 0,984544 |
| Rrp12 | 21,9852 | 21,5154 | -0,0311603 | 0,86835 | 0,984544 |
| Prepl | 20,5674 | 20,1285 | -0,0311218 | 0,86845 | 0,984544 |
| Plekhn1 | 2,89796 | 3,00094 | 0,0503748 | 0,8685 | 0,984544 |
| Rfwd2 | 5,12621 | 5,00226 | -0,0353117 | 0,86865 | 0,984544 |
| Dleu2 | 0,907379 | 0,827672 | -0,132646 | 0,8687 | 0,984544 |
| Hexdc | 9,90674 | 9,50314 | -0,0600054 | 0,86895 | 0,984544 |
| Supt4a | 31,6665 | 32,5323 | 0,038916 | 0,869 | 0,984544 |
| Zbtb24 | 8,2948 | 8,09029 | -0,0360158 | 0,86915 | 0,984544 |
| Zfp664 | 89,6012 | 87,7546 | -0,030043 | 0,8692 | 0,984544 |
| Bad | 19,8688 | 19,202 | -0,0492466 | 0,86925 | 0,984544 |
| Ripk1 | 4,45536 | 4,57365 | 0,0378065 | 0,8694 | 0,984544 |
| Ercc4 | 2,72384 | 2,7975 | 0,0384973 | 0,86945 | 0,984544 |
| Mpnd | 59,3431 | 57,5452 | -0,044383 | 0,86955 | 0,984544 |
| Ubap1 | 34,9274 | 35,6905 | 0,0311786 | 0,86965 | 0,984544 |
| Rsc1a1 | 12,4504 | 11,8313 | -0,0735861 | 0,86965 | 0,984544 |
| Pdgfra | 48,9601 | 47,9716 | -0,0294272 | 0,86975 | 0,984544 |
| Mybbp1a | 169,195 | 172,664 | 0,0292844 | 0,8698 | 0,984544 |
| Dusp5 | 4,64631 | 4,81811 | 0,0523807 | 0,86985 | 0,984544 |
| Champ1 | 30,4817 | 31,1637 | 0,0319248 | 0,86985 | 0,984544 |
| Fsd1 | 13,7478 | 13,4108 | -0,0358046 | 0,86985 | 0,984544 |
| Ankrd23 | 3,08818 | 2,98212 | -0,0504216 | 0,86995 | 0,984544 |
| Chga | 1,13663 | 1,19258 | 0,0693163 | 0,8702 | 0,984544 |
| Map2k3 | 20,3313 | 20,7857 | 0,0318854 | 0,8702 | 0,984544 |
| Mre11a | 30,1017 | 29,4794 | -0,0301377 | 0,8702 | 0,984544 |
| Yaf2 | 27,5643 | 26,9825 | -0,0307779 | 0,8702 | 0,984544 |
| 9430016H08Rik | 29,7925 | 28,9801 | -0,0398856 | 0,8702 | 0,984544 |
| Grsf1 | 78,5791 | 77,0044 | -0,0292045 | 0,87025 | 0,984544 |
| Poc5 | 12,7008 | 12,4133 | -0,0330364 | 0,8703 | 0,984544 |
| 1810013L24Rik | 10,4785 | 10,7166 | 0,0324162 | 0,87035 | 0,984544 |
| Rnf219 | 27,8312 | 27,2643 | -0,0296885 | 0,87055 | 0,984544 |
| Slc36a1 | 2,84537 | 2,77139 | -0,0380036 | 0,87055 | 0,984544 |
| Polr3c | 32,8784 | 32,1748 | -0,031207 | 0,8707 | 0,984544 |
| Rps15a-ps6 | 1,2925 | 0,966835 | -0,418824 | 0,87075 | 0,984544 |
| Ddx47 | 79,0343 | 77,4444 | -0,0293176 | 0,8708 | 0,984544 |
| Ythdf2 | 31,881 | 32,556 | 0,0302291 | 0,871 | 0,984544 |
| Baz2b | 12,4613 | 12,7139 | 0,028953 | 0,871 | 0,984544 |
| Ubr5 | 42,1908 | 41,3675 | -0,0284304 | 0,871 | 0,984544 |
| Pcmt1 | 33,3094 | 32,5821 | -0,0318508 | 0,871 | 0,984544 |
| Nop16 | 62,5546 | 61,2858 | -0,0295612 | 0,8711 | 0,984544 |
| Cacng4 | 4,36564 | 4,20117 | -0,0554028 | 0,8713 | 0,984544 |
| Tbc1d17 | 16,3457 | 15,5146 | -0,075286 | 0,87155 | 0,984544 |

| | | | | | |
|---|---|---|---|---|---|
| Apeh | 26,9539 | 27,5256 | 0,0302814 | 0,8717 | 0,984544 |
| Murc | 2,74332 | 2,6477 | -0,0511855 | 0,87175 | 0,984544 |
| Mycn | 73,5756 | 72,0862 | -0,0295055 | 0,87185 | 0,984544 |
| Actr1b | 80,4411 | 82,0523 | 0,0286115 | 0,87195 | 0,984544 |
| Smg9 | 9,38633 | 9,60814 | 0,0336963 | 0,872 | 0,984544 |
| Scrib | 33,6565 | 34,3443 | 0,0291856 | 0,872 | 0,984544 |
| Iffo2 | 5,81826 | 5,68036 | -0,0346063 | 0,87205 | 0,984544 |
| Ggcx | 7,58189 | 7,76719 | 0,0348363 | 0,87215 | 0,984544 |
| Acp5 | 0,696367 | 1,01447 | 0,542807 | 0,8722 | 0,984544 |
| Rbms3 | 4,80822 | 4,91587 | 0,031946 | 0,8722 | 0,984544 |
| Mir130c | 185,605 | 188,462 | 0,0220391 | 0,8722 | 0,984544 |
| Hdac6 | 16,6702 | 16,3164 | -0,0309462 | 0,8722 | 0,984544 |
| Cnep1r1 | 32,9282 | 33,6417 | 0,030923 | 0,8723 | 0,984544 |
| Sigmar1 | 50,5202 | 49,512 | -0,029081 | 0,8724 | 0,984544 |
| Nelfa | 49,2175 | 48,2495 | -0,0286591 | 0,8725 | 0,984544 |
| Wdr54 | 5,58262 | 5,39322 | -0,0497948 | 0,8725 | 0,984544 |
| Cyb5b | 86,7644 | 85,0659 | -0,0285226 | 0,8726 | 0,984544 |
| Ube2e2 | 12,8183 | 12,5027 | -0,0359732 | 0,87265 | 0,984544 |
| Tead3 | 7,24483 | 7,06439 | -0,036386 | 0,87265 | 0,984544 |
| Micu3 | 2,23504 | 2,16246 | -0,0476261 | 0,87275 | 0,984544 |
| Polr2e | 169,651 | 147,976 | -0,197212 | 0,8728 | 0,984544 |
| D830031N03Rik | 8,23873 | 8,67159 | 0,0738758 | 0,87295 | 0,984544 |
| Ivns1abp | 228,059 | 223,572 | -0,0286676 | 0,87295 | 0,984544 |
| Zfp213 | 11,9492 | 11,6804 | -0,0328317 | 0,87295 | 0,984544 |
| Dapp1 | 0,856409 | 0,819878 | -0,0628898 | 0,87295 | 0,984544 |
| Fahd1 | 9,06369 | 9,32753 | 0,041396 | 0,87305 | 0,98457 |
| Derl3 | 0,802715 | 0,854035 | 0,0894066 | 0,87345 | 0,984802 |
| 9130019O22Rik | 1,30506 | 1,26001 | -0,0506763 | 0,87365 | 0,984802 |
| Trub2 | 24,0928 | 24,6035 | 0,0302636 | 0,8738 | 0,984802 |
| Polr2g | 141,236 | 144,08 | 0,0287644 | 0,8738 | 0,984802 |
| 9930104L06Rik | 10,586 | 10,8339 | 0,0333964 | 0,8739 | 0,984802 |
| Gm5801 | 25,8358 | 25,234 | -0,0340038 | 0,8739 | 0,984802 |
| Ypel2 | 1,15596 | 0,787889 | -0,553027 | 0,8739 | 0,984802 |
| Cluap1 | 16,8792 | 16,4973 | -0,0330151 | 0,87395 | 0,984802 |
| Orc2 | 45,573 | 44,6647 | -0,0290431 | 0,874 | 0,984802 |
| Gar1 | 82,9909 | 84,6803 | 0,0290724 | 0,87405 | 0,984802 |
| Grpel2 | 10,4122 | 10,6456 | 0,0319836 | 0,8741 | 0,984802 |
| Zfp523 | 20,6124 | 21,0524 | 0,0304768 | 0,87425 | 0,984884 |
| Ccdc138 | 5,11781 | 5,25953 | 0,0394074 | 0,8744 | 0,984967 |
| Gdap2 | 7,36652 | 7,18896 | -0,0351998 | 0,8745 | 0,984993 |
| Fam122b | 3,99373 | 3,89343 | -0,036696 | 0,87465 | 0,985076 |
| Prmt2 | 12,4905 | 12,2039 | -0,0334884 | 0,8749 | 0,98516 |
| 0610031J06Rik | 24,3894 | 23,8506 | -0,0322319 | 0,87505 | 0,98516 |
| Thap2 | 8,11663 | 8,30191 | 0,0325623 | 0,8751 | 0,98516 |
| D10Wsu102e | 40,9268 | 40,14 | -0,0280076 | 0,87515 | 0,98516 |
| Bcap31 | 79,0395 | 77,419 | -0,0298873 | 0,87535 | 0,98516 |
| Gbas | 31,1022 | 30,4543 | -0,0303721 | 0,87535 | 0,98516 |
| Zc4h2 | 21,3058 | 20,8571 | -0,0307085 | 0,8754 | 0,98516 |
| Adal | 12,5769 | 12,2426 | -0,0388725 | 0,87545 | 0,98516 |
| Rab3gap2 | 9,05582 | 8,76321 | -0,0473871 | 0,87545 | 0,98516 |

| | | | | | |
|---|---|---|---|---|---|
| Mir6935 | 2660,93 | 3242,69 | 0,285262 | 0,8755 | 0,98516 |
| Brd8 | 46,9779 | 45,1476 | -0,0573308 | 0,87565 | 0,98516 |
| Supt7l | 13,5306 | 13,2386 | -0,0314753 | 0,87575 | 0,98516 |
| Iah1 | 28,7464 | 27,3686 | -0,0708566 | 0,87575 | 0,98516 |
| Vps52 | 22,3532 | 22,8143 | 0,0294606 | 0,8758 | 0,98516 |
| Sppl2b | 10,1592 | 9,93522 | -0,0321692 | 0,876 | 0,985294 |
| Psme4 | 50,8296 | 51,8451 | 0,028539 | 0,87615 | 0,985294 |
| Phf23 | 79,2579 | 80,8337 | 0,0284021 | 0,87615 | 0,985294 |
| Ahdc1 | 12,0515 | 12,2877 | 0,0279932 | 0,87625 | 0,98532 |
| Rgl2 | 19,1352 | 18,7277 | -0,0310546 | 0,87635 | 0,985346 |
| Ccdc25 | 37,526 | 36,788 | -0,0286566 | 0,87665 | 0,985361 |
| Ipp | 10,3837 | 10,1359 | -0,0348389 | 0,87665 | 0,985361 |
| Flot1 | 11,6658 | 11,9566 | 0,0355168 | 0,87695 | 0,985361 |
| Fam63a | 7,93751 | 7,74275 | -0,035841 | 0,87695 | 0,985361 |
| Snapin | 16,3006 | 15,0175 | -0,118279 | 0,877 | 0,985361 |
| Reln | 1,99904 | 2,04702 | 0,034217 | 0,87705 | 0,985361 |
| C530005A16Rik | 0,601169 | 0,685113 | 0,188572 | 0,87715 | 0,985361 |
| Gpr161 | 7,40357 | 7,21419 | -0,0373851 | 0,87715 | 0,985361 |
| Fam133b | 31,0224 | 30,4049 | -0,0290099 | 0,87725 | 0,985361 |
| Ufl1 | 9,74378 | 9,95192 | 0,0304921 | 0,8773 | 0,985361 |
| Zfp57 | 29,8496 | 29,2581 | -0,0288774 | 0,8775 | 0,985361 |
| Tmem109 | 24,0066 | 23,5033 | -0,0305692 | 0,87755 | 0,985361 |
| Tmem66 | 35,6023 | 34,8858 | -0,0293323 | 0,8776 | 0,985361 |
| Brf1 | 21,8241 | 21,3068 | -0,0346131 | 0,8776 | 0,985361 |
| Hiat1 | 32,758 | 32,1248 | -0,0281557 | 0,87765 | 0,985361 |
| Htra2 | 38,8884 | 38,076 | -0,0304611 | 0,87765 | 0,985361 |
| Dph7 | 9,91758 | 10,222 | 0,0436218 | 0,87775 | 0,985361 |
| Snora17 | 66,0448 | 55,5293 | -0,250196 | 0,8778 | 0,985361 |
| Clstn1 | 34,352 | 33,7117 | -0,0271426 | 0,87805 | 0,985361 |
| 3632451O06Rik | 4,66721 | 4,54057 | -0,0396874 | 0,8781 | 0,985361 |
| Nt5c2 | 17,7867 | 17,4279 | -0,0294038 | 0,8782 | 0,985361 |
| Gtpbp2 | 13,5219 | 13,2392 | -0,0304755 | 0,87825 | 0,985361 |
| Sec22b | 41,841 | 41,0305 | -0,02822 | 0,8783 | 0,985361 |
| Jmjd6 | 67,1063 | 65,7723 | -0,0289689 | 0,8783 | 0,985361 |
| Chmp7 | 36,6905 | 36,0052 | -0,0272008 | 0,8784 | 0,985361 |
| Tmbim4 | 31,0115 | 31,7667 | 0,0347097 | 0,8785 | 0,985361 |
| Zfp692 | 20,8802 | 21,3219 | 0,0301977 | 0,8786 | 0,985361 |
| Ap4b1 | 10,3702 | 11,0934 | 0,0972614 | 0,87865 | 0,985361 |
| Afmid | 1,76762 | 1,63253 | -0,114697 | 0,8787 | 0,985361 |
| Mak16 | 82,9047 | 84,6437 | 0,0299491 | 0,87885 | 0,985361 |
| Pex5 | 18,6172 | 18,9865 | 0,0283365 | 0,879 | 0,985361 |
| 2310022A10Rik | 16,2825 | 16,6272 | 0,0302272 | 0,8792 | 0,985361 |
| Gm19710 | 4,92247 | 3,45967 | -0,508747 | 0,8792 | 0,985361 |
| Tor2a | 22,0924 | 22,5885 | 0,0320389 | 0,87925 | 0,985361 |
| Ubl7 | 40,1839 | 39,3765 | -0,029284 | 0,87925 | 0,985361 |
| Mms22l | 19,5232 | 19,9066 | 0,0280546 | 0,87935 | 0,985361 |
| Paip1 | 25,6347 | 25,1537 | -0,0273268 | 0,8794 | 0,985361 |
| Mreg | 1,87151 | 1,80427 | -0,0527871 | 0,8795 | 0,985361 |
| Mapkap1 | 27,1425 | 26,6462 | -0,0266219 | 0,87955 | 0,985361 |
| Ept1 | 17,3433 | 17,6804 | 0,0277704 | 0,87965 | 0,985361 |

| | | | | | |
|---|---|---|---|---|---|
| Cgref1 | 2,71499 | 2,2827 | -0,250209 | 0,87965 | 0,985361 |
| Tymp | 0,691639 | 0,525914 | -0,395193 | 0,8797 | 0,985361 |
| Taf3 | 7,47312 | 7,31364 | -0,0311204 | 0,8798 | 0,985361 |
| Dhx34 | 6,64876 | 6,49891 | -0,0328869 | 0,8799 | 0,985361 |
| Ppp5c | 78,4636 | 79,963 | 0,0273088 | 0,87995 | 0,985361 |
| Rhot2 | 35,0193 | 35,9433 | 0,0375711 | 0,88 | 0,985361 |
| Mpp6 | 52,0166 | 51,0025 | -0,0284046 | 0,88005 | 0,985361 |
| 1700010I14Rik | 2,2537 | 2,1774 | -0,0496874 | 0,88005 | 0,985361 |
| Pomt1 | 11,1359 | 10,8504 | -0,0374813 | 0,88025 | 0,985499 |
| Rnf24 | 4,64845 | 5,01735 | 0,110177 | 0,8804 | 0,985581 |
| Saysd1 | 3,46726 | 3,56883 | 0,0416527 | 0,8807 | 0,985745 |
| 9530082P21Rik | 17,5301 | 17,1782 | -0,0292595 | 0,8807 | 0,985745 |
| Riok2 | 29,5334 | 28,984 | -0,0270883 | 0,8808 | 0,985771 |
| H1fx | 96,7169 | 94,8196 | -0,0285834 | 0,88095 | 0,985852 |
| Ifnar2 | 13,6627 | 14,006 | 0,0358022 | 0,8812 | 0,985912 |
| Gsn | 0,987572 | 0,948747 | -0,0578613 | 0,8813 | 0,985912 |
| Ccdc96 | 1,43152 | 1,47913 | 0,0472069 | 0,88135 | 0,985912 |
| Faim | 27,475 | 26,856 | -0,032875 | 0,8814 | 0,985912 |
| Caml | 27,882 | 27,3104 | -0,0298847 | 0,8816 | 0,985912 |
| Dnal4 | 26,8738 | 26,3165 | -0,0302319 | 0,8816 | 0,985912 |
| Aaas | 61,1976 | 60,0625 | -0,027009 | 0,88175 | 0,985912 |
| Cntf | 3,43472 | 3,93254 | 0,195267 | 0,88185 | 0,985912 |
| Mfap3l | 1,5658 | 1,60932 | 0,0395527 | 0,8819 | 0,985912 |
| Tgds | 12,2391 | 12,502 | 0,0306563 | 0,88195 | 0,985912 |
| Pigm | 10,1421 | 9,9505 | -0,0275147 | 0,882 | 0,985912 |
| Ndufs7 | 92,0493 | 73,2079 | -0,330407 | 0,88205 | 0,985912 |
| Chac2 | 3,03691 | 2,85476 | -0,0892312 | 0,8822 | 0,985912 |
| Hist2h2bb | 5,49305 | 5,24969 | -0,0653748 | 0,8823 | 0,985912 |
| Wdr60 | 5,80977 | 5,94189 | 0,0324421 | 0,8826 | 0,985912 |
| Zkscan3 | 38,8822 | 38,1971 | -0,0256478 | 0,8826 | 0,985912 |
| Tmem50a | 54,9045 | 53,8082 | -0,0290984 | 0,8827 | 0,985912 |
| 2700081O15Rik | 28,6913 | 29,6921 | 0,049468 | 0,8829 | 0,985912 |
| Fam72a | 4,92727 | 4,7951 | -0,0392267 | 0,8831 | 0,985912 |
| Glmn | 17,9924 | 18,4822 | 0,0387526 | 0,88315 | 0,985912 |
| Ctu2 | 18,7425 | 18,1873 | -0,0433811 | 0,88315 | 0,985912 |
| Upk1a | 0,734693 | 0,691356 | -0,0877133 | 0,8833 | 0,985912 |
| Gm6981 | 0,625249 | 0,719864 | 0,203293 | 0,88335 | 0,985912 |
| Actb | 925,601 | 944,593 | 0,0293016 | 0,8834 | 0,985912 |
| Mgmt | 6,68071 | 6,46232 | -0,0479494 | 0,8834 | 0,985912 |
| Pabpc1 | 272,386 | 267,295 | -0,027218 | 0,8837 | 0,985912 |
| Tmem223 | 26,6015 | 25,9823 | -0,0339778 | 0,8837 | 0,985912 |
| Zfp30 | 3,14678 | 3,23957 | 0,0419263 | 0,88375 | 0,985912 |
| Tmem214 | 25,1734 | 25,6443 | 0,0267369 | 0,88375 | 0,985912 |
| Actr10 | 24,2047 | 23,7653 | -0,0264261 | 0,88375 | 0,985912 |
| Gpn2 | 20,9596 | 20,5086 | -0,0313805 | 0,8838 | 0,985912 |
| Smim20 | 23,8002 | 23,2638 | -0,032889 | 0,8838 | 0,985912 |
| D430042O09Rik | 6,4384 | 6,30736 | -0,0296651 | 0,88395 | 0,985912 |
| Zfp691 | 8,00363 | 7,80638 | -0,0360004 | 0,884 | 0,985912 |
| Baz1b | 80,6659 | 82,2109 | 0,0273698 | 0,88405 | 0,985912 |
| Sympk | 61,6438 | 62,7889 | 0,0265535 | 0,88425 | 0,985912 |

| | | | | | |
|---|---|---|---|---|---|
| Ttc33 | 17,6788 | 17,3087 | -0,0305244 | 0,8843 | 0,985912 |
| Ebp | 15,7913 | 15,4557 | -0,0309935 | 0,8844 | 0,985912 |
| Calhm2 | 1,17691 | 1,13018 | -0,0584519 | 0,88445 | 0,985912 |
| Yeats2 | 36,7458 | 37,4292 | 0,0265856 | 0,8845 | 0,985912 |
| Zc3h8 | 9,51281 | 9,27726 | -0,0361733 | 0,8845 | 0,985912 |
| Gid4 | 16,6744 | 16,3657 | -0,0269601 | 0,88455 | 0,985912 |
| Klf6 | 7,04195 | 6,89577 | -0,0302637 | 0,8846 | 0,985912 |
| Pcdhga3 | 2,41019 | 2,67904 | 0,152575 | 0,88465 | 0,985912 |
| Bbs7 | 8,88897 | 8,69799 | -0,0313347 | 0,8848 | 0,985912 |
| Eno1,Eno1b | 1,45592 | 1,86869 | 0,360095 | 0,88485 | 0,985912 |
| Mettl20 | 5,9595 | 6,11892 | 0,0380856 | 0,88485 | 0,985912 |
| Nudt14 | 20,6858 | 21,2082 | 0,0359828 | 0,88485 | 0,985912 |
| Bst2 | 1,3719 | 1,45673 | 0,0865598 | 0,8849 | 0,985912 |
| Zkscan5 | 9,72271 | 9,53332 | -0,0283802 | 0,8849 | 0,985912 |
| Nelfcd | 60,9005 | 59,5477 | -0,0324082 | 0,88495 | 0,985912 |
| Pelo | 22,7903 | 23,4027 | 0,0382552 | 0,885 | 0,985912 |
| Mettl14 | 25,7133 | 26,1874 | 0,0263553 | 0,8853 | 0,986069 |
| Ccdc90b | 30,6814 | 30,0957 | -0,0278043 | 0,8854 | 0,986069 |
| Rabggtb | 107,398 | 105,561 | -0,0248912 | 0,88565 | 0,986069 |
| Ptdss1 | 20,0867 | 19,6331 | -0,0329494 | 0,88565 | 0,986069 |
| Slc33a1 | 8,98011 | 9,17495 | 0,0309673 | 0,88585 | 0,986069 |
| Dalrd3 | 16,9014 | 15,6537 | -0,110634 | 0,8859 | 0,986069 |
| Tmem259 | 45,2767 | 52,2208 | 0,205856 | 0,88595 | 0,986069 |
| Ap2a1 | 46,0697 | 45,2514 | -0,0258556 | 0,88595 | 0,986069 |
| Pcyox1l | 12,4143 | 12,6923 | 0,0319525 | 0,886 | 0,986069 |
| Ddost | 129,775 | 132,174 | 0,0264244 | 0,8861 | 0,986069 |
| Kctd6 | 10,3161 | 10,5615 | 0,0339271 | 0,88625 | 0,986069 |
| Cxx1b | 7,50574 | 7,7197 | 0,0405515 | 0,88645 | 0,986069 |
| Ccnc | 16,3649 | 16,8094 | 0,0386632 | 0,88655 | 0,986069 |
| Fam162a | 236,812 | 241,076 | 0,0257483 | 0,88665 | 0,986069 |
| Mettl22 | 8,46753 | 8,2495 | -0,0376334 | 0,88665 | 0,986069 |
| Cnot1 | 59,2502 | 60,325 | 0,025935 | 0,8868 | 0,986069 |
| Zfp354c | 13,2852 | 13,051 | -0,0256605 | 0,8868 | 0,986069 |
| Nfs1 | 13,7161 | 13,3321 | -0,0409669 | 0,88685 | 0,986069 |
| Hs1bp3 | 9,0699 | 8,87439 | -0,031438 | 0,8869 | 0,986069 |
| Spata7 | 8,06077 | 8,25155 | 0,0337482 | 0,887 | 0,986069 |
| Specc1l | 23,8332 | 24,272 | 0,0263226 | 0,88705 | 0,986069 |
| Gm7120 | 5,55107 | 5,70954 | 0,0406066 | 0,88715 | 0,986069 |
| Tmem19 | 8,50179 | 8,68805 | 0,0312655 | 0,88715 | 0,986069 |
| Sf3b4 | 27,3022 | 27,8399 | 0,0281382 | 0,88715 | 0,986069 |
| Cebpg | 22,8327 | 23,2277 | 0,0247461 | 0,88765 | 0,986069 |
| Cr1l | 8,90262 | 9,12439 | 0,0354972 | 0,8877 | 0,986069 |
| Nhlrc1 | 1,32159 | 1,27476 | -0,052053 | 0,8878 | 0,986069 |
| Npcd | 1,09454 | 1,16689 | 0,092339 | 0,8879 | 0,986069 |
| Fam129c | 1,66932 | 1,72443 | 0,046859 | 0,8879 | 0,986069 |
| Ints4 | 31,3462 | 31,8962 | 0,0250944 | 0,88805 | 0,986069 |
| Cks1b | 250,491 | 246,138 | -0,0252918 | 0,88805 | 0,986069 |
| Xpr1 | 6,10233 | 5,98809 | -0,0272654 | 0,88805 | 0,986069 |
| Ppm1e | 1,17525 | 0,890629 | -0,400072 | 0,8881 | 0,986069 |
| Adar | 13,885 | 14,1394 | 0,0261912 | 0,88825 | 0,986069 |

| | | | | | |
|---|---|---|---|---|---|
| Hint3 | 7,36607 | 7,60971 | 0,0469459 | 0,8883 | 0,986069 |
| Lpar4 | 3,47043 | 3,55122 | 0,0332013 | 0,8883 | 0,986069 |
| Vbp1 | 64,2217 | 65,4146 | 0,0265499 | 0,8883 | 0,986069 |
| Dynlt1a | 0,660913 | 0,698083 | 0,078938 | 0,88835 | 0,986069 |
| Mocs1 | 12,0012 | 11,7671 | -0,0284237 | 0,88835 | 0,986069 |
| Cyp2j6 | 2,13894 | 2,07913 | -0,0409191 | 0,88835 | 0,986069 |
| Rad52 | 14,2284 | 14,5383 | 0,0310787 | 0,8884 | 0,986069 |
| Itsn1 | 38,1881 | 38,8886 | 0,0262216 | 0,8884 | 0,986069 |
| Pmvk | 35,0723 | 34,4043 | -0,0277423 | 0,88845 | 0,986069 |
| Diablo | 46,2281 | 47,0761 | 0,026224 | 0,88855 | 0,986069 |
| Trappc6b | 19,5906 | 20,02 | 0,0312827 | 0,8886 | 0,986069 |
| Iglon5 | 1,77101 | 1,82875 | 0,0462865 | 0,8889 | 0,986317 |
| Adam8 | 1,64064 | 1,58971 | -0,0454958 | 0,88905 | 0,986368 |
| Terf2 | 26,572 | 27,0553 | 0,026006 | 0,8891 | 0,986368 |
| Mbd6 | 15,9913 | 16,546 | 0,0491891 | 0,88935 | 0,986544 |
| Fitm2 | 5,05738 | 4,92014 | -0,0396909 | 0,88945 | 0,986544 |
| 4930579G18Rik | 5,33686 | 6,1795 | 0,211498 | 0,8896 | 0,986544 |
| Msl1 | 44,7401 | 45,7792 | 0,0331247 | 0,88975 | 0,986544 |
| Arih2 | 15,9864 | 16,2839 | 0,0266039 | 0,8898 | 0,986544 |
| 1700086O06Rik | 4,19373 | 4,48552 | 0,0970397 | 0,88995 | 0,986544 |
| Acad11 | 6,76415 | 6,62438 | -0,0301242 | 0,88995 | 0,986544 |
| Ring1 | 12,8471 | 12,5809 | -0,0302119 | 0,89005 | 0,986544 |
| Mir762 | 855,463 | 706,912 | -0,275176 | 0,89005 | 0,986544 |
| Ncoa5 | 47,4548 | 46,6774 | -0,0238282 | 0,8901 | 0,986544 |
| Nsmaf | 24,7163 | 24,2919 | -0,0249901 | 0,89035 | 0,986544 |
| Rnf208 | 3,2904 | 3,18288 | -0,0479333 | 0,89035 | 0,986544 |
| Utp15 | 28,0813 | 28,5752 | 0,0251524 | 0,89045 | 0,986544 |
| Gm3086 | 1,08098 | 1,32357 | 0,292097 | 0,89055 | 0,986544 |
| Fam65a | 6,98408 | 7,12376 | 0,0285698 | 0,8906 | 0,986544 |
| Spcs3 | 37,5986 | 38,2666 | 0,0254075 | 0,89075 | 0,986544 |
| Gfm1 | 39,585 | 38,8304 | -0,0277654 | 0,89075 | 0,986544 |
| Eng | 1,56716 | 1,67723 | 0,0979298 | 0,8908 | 0,986544 |
| Nfkbil1 | 8,96624 | 9,18542 | 0,0348431 | 0,8909 | 0,986544 |
| Stat1 | 3,37628 | 3,45202 | 0,0320027 | 0,8909 | 0,986544 |
| Snx24 | 16,7756 | 17,1055 | 0,0280915 | 0,8909 | 0,986544 |
| Akt1 | 115,64 | 113,715 | -0,0242233 | 0,89095 | 0,986544 |
| Impact | 19,994 | 19,6438 | -0,0254967 | 0,89115 | 0,98662 |
| Zfp444 | 8,86724 | 8,70414 | -0,0267845 | 0,89125 | 0,98662 |
| Manba | 0,819845 | 0,79264 | -0,0486851 | 0,89125 | 0,98662 |
| Ndufs5 | 26,9251 | 26,2291 | -0,0377806 | 0,89135 | 0,986646 |
| Commd6 | 38,891 | 39,6829 | 0,0290844 | 0,89165 | 0,986748 |
| Efnb1 | 17,5596 | 17,2426 | -0,0262781 | 0,8917 | 0,986748 |
| Clcn7 | 11,4398 | 11,2331 | -0,0263076 | 0,89175 | 0,986748 |
| Med16 | 20,2793 | 16,7521 | -0,275662 | 0,89175 | 0,986748 |
| Pnpla7 | 2,57393 | 2,63491 | 0,0337842 | 0,89215 | 0,986786 |
| Lyar | 120,264 | 118,292 | -0,0238594 | 0,89215 | 0,986786 |
| Gm10069 | 1,93666 | 1,87327 | -0,0480086 | 0,89215 | 0,986786 |
| Bri3 | 19,1754 | 18,5709 | -0,0462128 | 0,89235 | 0,986786 |
| Mis12 | 34,6439 | 35,3767 | 0,0301967 | 0,8924 | 0,986786 |
| Macrod1 | 0,863943 | 0,817186 | -0,0802715 | 0,89245 | 0,986786 |

| | | | | | |
|---|---|---|---|---|---|
| Litaf | 26,8026 | 26,3107 | -0,0267241 | 0,8925 | 0,986786 |
| 4931406C07Rik | 8,02276 | 8,53031 | 0,0884997 | 0,89255 | 0,986786 |
| Mdc1 | 31,6481 | 32,1722 | 0,0236937 | 0,8926 | 0,986786 |
| Ybey | 3,07965 | 2,90071 | -0,0863624 | 0,89275 | 0,986786 |
| Alg11 | 10,1326 | 10,316 | 0,0258796 | 0,8928 | 0,986786 |
| Dna2 | 23,3035 | 22,8944 | -0,0255566 | 0,8929 | 0,986786 |
| Anapc11 | 16,2079 | 15,9154 | -0,0262751 | 0,89295 | 0,986786 |
| Snapc2 | 25,15 | 24,6827 | -0,0270557 | 0,89305 | 0,986786 |
| Dhx40 | 25,8497 | 27,5048 | 0,089538 | 0,8931 | 0,986786 |
| Zfp773 | 1,26756 | 1,22844 | -0,0452307 | 0,8932 | 0,986786 |
| Zfp69 | 2,74421 | 2,82341 | 0,0410489 | 0,89325 | 0,986786 |
| Gm6654 | 3,11554 | 3,46809 | 0,154662 | 0,8933 | 0,986786 |
| Phykpl | 4,18722 | 5,61192 | 0,422502 | 0,89335 | 0,986786 |
| Rmnd5b | 11,5991 | 11,1121 | -0,0618814 | 0,89335 | 0,986786 |
| Pex6 | 19,2431 | 18,9124 | -0,0250058 | 0,8934 | 0,986786 |
| Ift172 | 11,277 | 10,2179 | -0,142275 | 0,8936 | 0,986837 |
| Clip1 | 21,3631 | 21,7256 | 0,0242757 | 0,89365 | 0,986837 |
| Pus3 | 24,7935 | 25,3122 | 0,0298721 | 0,8937 | 0,986837 |
| Slu7 | 18,346 | 18,0185 | -0,0259867 | 0,8938 | 0,986837 |
| Sin3a | 42,4813 | 43,1933 | 0,0239782 | 0,89395 | 0,986837 |
| Prom1 | 29,6637 | 29,1622 | -0,0246009 | 0,894 | 0,986837 |
| Slc22a15 | 0,820035 | 0,843749 | 0,041127 | 0,89415 | 0,986837 |
| Snrpn | 22,8092 | 22,3897 | -0,0267824 | 0,89415 | 0,986837 |
| Foxk1 | 10,444 | 10,264 | -0,0250859 | 0,8942 | 0,986837 |
| Frs3 | 2,49831 | 2,57528 | 0,0437763 | 0,89445 | 0,986837 |
| Slc25a47 | 5,18342 | 5,04786 | -0,0382339 | 0,89445 | 0,986837 |
| Cenph | 55,2203 | 54,2295 | -0,0261219 | 0,8945 | 0,986837 |
| Cdca2 | 37,1511 | 37,7868 | 0,0244791 | 0,89455 | 0,986837 |
| Gm5124 | 5,98757 | 6,12746 | 0,0333169 | 0,8946 | 0,986837 |
| Ecsit | 19,3084 | 18,2438 | -0,0818269 | 0,8946 | 0,986837 |
| Gps2 | 34,7255 | 34,0949 | -0,026438 | 0,89495 | 0,98693 |
| Htt | 10,0407 | 10,2099 | 0,0241121 | 0,895 | 0,98693 |
| Ppm1g | 179,28 | 172,235 | -0,0578391 | 0,89505 | 0,98693 |
| Mettl4 | 9,62682 | 9,82755 | 0,029772 | 0,89515 | 0,98693 |
| Cog6 | 16,5517 | 16,2595 | -0,0256987 | 0,89525 | 0,98693 |
| Nae1 | 58,4585 | 59,6185 | 0,0283462 | 0,8953 | 0,98693 |
| Bace1 | 14,6612 | 14,9197 | 0,0252092 | 0,8953 | 0,98693 |
| Cxxc1 | 39,5259 | 38,8892 | -0,0234277 | 0,8953 | 0,98693 |
| 1700109K24Rik | 2,90934 | 3,42689 | 0,236205 | 0,8955 | 0,987066 |
| Bloc1s6 | 17,9861 | 17,6865 | -0,0242367 | 0,8958 | 0,987252 |
| Map4k1 | 3,64993 | 3,56318 | -0,0347053 | 0,8959 | 0,987252 |
| Trp53rk | 1,83439 | 1,7863 | -0,0383261 | 0,8959 | 0,987252 |
| Haus1 | 42,7145 | 41,944 | -0,0262621 | 0,8962 | 0,987468 |
| Snrnp25 | 60,1894 | 59,1302 | -0,0256135 | 0,89625 | 0,987468 |
| Hipk1 | 14,7548 | 15,1874 | 0,0416903 | 0,8965 | 0,987476 |
| Aste1 | 5,80493 | 5,9337 | 0,0316549 | 0,89655 | 0,987476 |
| Btbd2 | 28,2865 | 27,8113 | -0,0244394 | 0,89665 | 0,987476 |
| Dvl1 | 16,1489 | 15,8706 | -0,025085 | 0,89665 | 0,987476 |
| Dcaf12 | 26,4473 | 26,8758 | 0,023185 | 0,89675 | 0,987476 |
| Sac3d1 | 26,1638 | 25,6812 | -0,0268589 | 0,89685 | 0,987476 |

| | | | | | |
|---|---|---|---|---|---|
| Thap6 | 5,60587 | 5,43117 | -0,0456744 | 0,89685 | 0,987476 |
| Dnase2a | 1,46979 | 1,52238 | 0,0507125 | 0,89695 | 0,987476 |
| Prrt2 | 2,87414 | 2,95114 | 0,0381457 | 0,89695 | 0,987476 |
| Trim45 | 5,02093 | 4,91791 | -0,0299085 | 0,8972 | 0,987523 |
| Rgs5 | 2,77087 | 2,85664 | 0,0439812 | 0,89725 | 0,987523 |
| Degs1 | 32,9659 | 32,4217 | -0,0240142 | 0,8973 | 0,987523 |
| Hsdl1 | 7,00872 | 6,87601 | -0,0275788 | 0,8973 | 0,987523 |
| Ctns | 5,37714 | 5,26787 | -0,0296192 | 0,8974 | 0,987548 |
| Pcdhga4 | 1,48101 | 1,28921 | -0,200103 | 0,8979 | 0,987598 |
| Trim68 | 4,50661 | 4,39659 | -0,0356568 | 0,89795 | 0,987598 |
| Tcerg1 | 67,2403 | 68,3231 | 0,0230479 | 0,89805 | 0,987598 |
| Pced1a | 10,581 | 10,9252 | 0,0461858 | 0,89815 | 0,987598 |
| Rab11b | 6,8379 | 6,95795 | 0,0251087 | 0,8982 | 0,987598 |
| Gm3219 | 17,4729 | 17,0173 | -0,038113 | 0,8982 | 0,987598 |
| Pdcd7 | 12,7418 | 12,974 | 0,0260608 | 0,89825 | 0,987598 |
| Lsm14a | 93,4249 | 91,8854 | -0,0239723 | 0,89825 | 0,987598 |
| Vps33b | 9,97428 | 9,78469 | -0,0276874 | 0,8983 | 0,987598 |
| Snn | 52,7096 | 51,7953 | -0,0252429 | 0,89835 | 0,987598 |
| Mansc4 | 0,978512 | 1,02785 | 0,0709641 | 0,89845 | 0,987598 |
| Cep83 | 24,6099 | 25,0316 | 0,0245164 | 0,89845 | 0,987598 |
| Ankra2 | 7,62461 | 7,81553 | 0,0356809 | 0,89855 | 0,987598 |
| Btbd6 | 11,4735 | 11,8443 | 0,0458927 | 0,8986 | 0,987598 |
| Anapc10 | 9,07868 | 8,91303 | -0,0265664 | 0,8986 | 0,987598 |
| Dact1 | 50,5168 | 51,3354 | 0,0231916 | 0,89885 | 0,987649 |
| Snord22 | 798,311 | 715,892 | -0,157208 | 0,89885 | 0,987649 |
| Rps19-ps3 | 6,00329 | 5,74638 | -0,0631 | 0,89895 | 0,987649 |
| Prkaa1 | 11,6074 | 11,4041 | -0,0255034 | 0,89905 | 0,987649 |
| F8a | 1,70456 | 1,65644 | -0,0413151 | 0,89935 | 0,987649 |
| Frg1 | 67,3111 | 68,4613 | 0,0244457 | 0,89945 | 0,987649 |
| Abcb10 | 9,47677 | 9,31225 | -0,0252653 | 0,89945 | 0,987649 |
| Dnajc30 | 14,8806 | 14,2725 | -0,0601944 | 0,8995 | 0,987649 |
| Wdr13 | 8,08954 | 7,94779 | -0,025503 | 0,8996 | 0,987649 |
| Mpv17 | 20,0662 | 17,474 | -0,199559 | 0,8997 | 0,987649 |
| Fig4 | 10,0873 | 10,2739 | 0,0264377 | 0,8998 | 0,987649 |
| Ddx18 | 28,3399 | 27,8518 | -0,0250633 | 0,8998 | 0,987649 |
| Cap1 | 78,6172 | 79,8433 | 0,0223266 | 0,89995 | 0,987649 |
| Tarbp2 | 13,4978 | 13,2299 | -0,0289285 | 0,89995 | 0,987649 |
| Hspa1l | 3,56022 | 3,4754 | -0,0347907 | 0,90005 | 0,987649 |
| Ankrd39 | 4,18393 | 4,29357 | 0,0373201 | 0,90035 | 0,987649 |
| Gm16515 | 9,95341 | 10,1257 | 0,0247635 | 0,9004 | 0,987649 |
| Gys1 | 20,2196 | 19,9005 | -0,0229491 | 0,9004 | 0,987649 |
| Aptx | 5,96619 | 5,86472 | -0,0247476 | 0,90045 | 0,987649 |
| Adat2 | 7,34979 | 7,10385 | -0,0491025 | 0,9005 | 0,987649 |
| Pwwp2b | 1,20336 | 1,24329 | 0,0471031 | 0,9006 | 0,987649 |
| Wdr18 | 45,8747 | 41,1138 | -0,158076 | 0,9006 | 0,987649 |
| Mfsd12 | 1,88439 | 1,81745 | -0,0521778 | 0,9007 | 0,987649 |
| 1110034G24Rik | 4,53806 | 4,65557 | 0,0368809 | 0,9008 | 0,987649 |
| Gm16861 | 0,801525 | 0,64943 | -0,303573 | 0,90085 | 0,987649 |
| Stx16 | 22,857 | 22,4956 | -0,0229879 | 0,90095 | 0,987649 |
| Sh3bp5l | 16,558 | 16,8285 | 0,0233696 | 0,9012 | 0,987649 |

| | | | | | |
|---|---|---|---|---|---|
| Atraid | 25,7989 | 21,5411 | -0,260218 | 0,90125 | 0,987649 |
| Gpt2 | 9,62731 | 9,79862 | 0,0254451 | 0,90135 | 0,987649 |
| Papola | 92,6949 | 94,1519 | 0,0225012 | 0,90155 | 0,987649 |
| Ebpl | 8,16964 | 8,00022 | -0,0302324 | 0,90155 | 0,987649 |
| Atp6v1b2 | 54,5316 | 55,3742 | 0,0221221 | 0,9017 | 0,987649 |
| Rpp21 | 38,5272 | 37,6973 | -0,0314159 | 0,9017 | 0,987649 |
| Helq | 8,95134 | 9,1083 | 0,0250784 | 0,90175 | 0,987649 |
| Pkd2l2 | 1,85527 | 1,9889 | 0,100344 | 0,9018 | 0,987649 |
| Eif4a1 | 108,344 | 110,007 | 0,0219745 | 0,90185 | 0,987649 |
| Med21 | 68,8781 | 67,7183 | -0,0245011 | 0,90185 | 0,987649 |
| Fkbp3 | 500,65 | 509,978 | 0,0266339 | 0,9019 | 0,987649 |
| Usp49 | 8,28806 | 8,15265 | -0,0237655 | 0,90195 | 0,987649 |
| Aph1a | 3,97637 | 3,74967 | -0,084688 | 0,902 | 0,987649 |
| Pdcd11 | 27,9481 | 28,3749 | 0,0218614 | 0,9021 | 0,987649 |
| Msh2 | 78,0128 | 76,8654 | -0,0213774 | 0,9021 | 0,987649 |
| Kif24 | 5,29791 | 5,39551 | 0,0263373 | 0,90215 | 0,987649 |
| Ttc5 | 24,7811 | 25,1887 | 0,0235385 | 0,9023 | 0,987649 |
| Ttc19 | 8,45112 | 8,95407 | 0,0834005 | 0,90235 | 0,987649 |
| Eif2ak4 | 13,8339 | 14,0531 | 0,0226771 | 0,90235 | 0,987649 |
| C130036L24Rik | 1,62719 | 1,68728 | 0,0523155 | 0,9024 | 0,987649 |
| Mdp1 | 36,0477 | 35,4558 | -0,0238864 | 0,9025 | 0,987649 |
| Hnrnpll | 78,7704 | 79,9583 | 0,021594 | 0,9026 | 0,987649 |
| Mapk1 | 93,204 | 91,5614 | -0,0256528 | 0,9026 | 0,987649 |
| Bves | 3,27408 | 3,36684 | 0,0403082 | 0,90265 | 0,987649 |
| Inha | 1,52239 | 1,47044 | -0,0500906 | 0,90265 | 0,987649 |
| Pmpcb | 72,357 | 71,246 | -0,0223233 | 0,90275 | 0,987674 |
| Flad1 | 7,57235 | 7,42302 | -0,0287363 | 0,903 | 0,987834 |
| Rasl2-9 | 6,19184 | 6,36107 | 0,0389008 | 0,90305 | 0,987834 |
| Arid3a | 27,0317 | 29,6908 | 0,135362 | 0,90325 | 0,987968 |
| Kank1 | 13,3624 | 13,1546 | -0,0226092 | 0,9034 | 0,988001 |
| Gtpbp6 | 16,0452 | 16,3671 | 0,0286495 | 0,90345 | 0,988001 |
| Ammecr1 | 7,21928 | 7,35502 | 0,0268733 | 0,90355 | 0,988001 |
| Dazap1 | 83,5242 | 91,2549 | 0,127707 | 0,90365 | 0,988001 |
| Slc39a14 | 8,67867 | 8,53749 | -0,0236626 | 0,90395 | 0,988001 |
| Tmem108 | 1,32639 | 1,29224 | -0,0376348 | 0,90395 | 0,988001 |
| Pop7 | 29,2027 | 29,7594 | 0,0272423 | 0,904 | 0,988001 |
| Parpbp | 10,1685 | 10,341 | 0,02427 | 0,904 | 0,988001 |
| Inpp4a | 2,81976 | 2,8779 | 0,0294427 | 0,90405 | 0,988001 |
| Slc30a7 | 6,766 | 6,65723 | -0,023383 | 0,90405 | 0,988001 |
| Thumpd1 | 55,1875 | 54,3678 | -0,0215899 | 0,90425 | 0,988106 |
| Nsun6 | 6,53545 | 6,40721 | -0,0285894 | 0,9043 | 0,988106 |
| Msh6 | 69,2629 | 67,9942 | -0,0266699 | 0,90455 | 0,988295 |
| Lrpap1 | 19,367 | 19,0561 | -0,0233479 | 0,90475 | 0,988324 |
| Rrn3 | 39,8096 | 40,4138 | 0,0217323 | 0,9048 | 0,988324 |
| Nudcd1 | 22,2022 | 22,7304 | 0,0339203 | 0,90485 | 0,988324 |
| Atg12 | 30,3923 | 29,9386 | -0,0216996 | 0,9052 | 0,988324 |
| Tmem59l | 2,62874 | 2,70347 | 0,0404412 | 0,90525 | 0,988324 |
| 2210408I21Rik | 3,25102 | 3,33827 | 0,0382084 | 0,90535 | 0,988324 |
| Samd4b | 27,0528 | 26,6544 | -0,021405 | 0,90535 | 0,988324 |
| Zfp384 | 20,2532 | 19,9445 | -0,0221538 | 0,90545 | 0,988324 |

| | | | | | |
|---|---|---|---|---|---|
| Srsf4 | 83,2594 | 84,5649 | 0,0224456 | 0,9055 | 0,988324 |
| Parp2 | 46,2074 | 45,5157 | -0,0217595 | 0,9055 | 0,988324 |
| Ttc9 | 1,06288 | 1,02629 | -0,0505355 | 0,9055 | 0,988324 |
| Rnls | 1,48866 | 1,53907 | 0,0480482 | 0,9057 | 0,988324 |
| Agpat6 | 49,7782 | 49,0462 | -0,0213728 | 0,9057 | 0,988324 |
| Fam221a | 1,10703 | 1,07114 | -0,0475488 | 0,9058 | 0,988324 |
| Gpx4 | 16,7322 | 11,7061 | -0,515368 | 0,9059 | 0,988324 |
| Bloc1s2 | 9,09965 | 8,88426 | -0,034559 | 0,906 | 0,988324 |
| Mlf2 | 171,698 | 174,176 | 0,0206747 | 0,90605 | 0,988324 |
| Cers2 | 46,8457 | 45,555 | -0,0403094 | 0,90605 | 0,988324 |
| Cdk11b | 78,4877 | 79,6325 | 0,0208911 | 0,9061 | 0,988324 |
| Dnajc2 | 60,5766 | 59,7027 | -0,0209642 | 0,9062 | 0,988324 |
| 1810009A15Rik | 148,109 | 145,872 | -0,021951 | 0,90625 | 0,988324 |
| Neil1 | 2,87206 | 2,96016 | 0,0435932 | 0,9063 | 0,988324 |
| Man2b1 | 8,2197 | 8,35437 | 0,0234445 | 0,90635 | 0,988324 |
| Crat | 5,61202 | 5,75822 | 0,0371033 | 0,90645 | 0,988324 |
| Rtkn | 18,2813 | 18,5775 | 0,0231835 | 0,9067 | 0,988324 |
| 2510002D24Rik | 7,09205 | 6,92852 | -0,0336557 | 0,9067 | 0,988324 |
| Zfp955a | 3,79578 | 3,87038 | 0,028078 | 0,9068 | 0,988324 |
| Gba2 | 16,3543 | 16,8286 | 0,0412458 | 0,9069 | 0,988324 |
| Rab23 | 13,0089 | 12,8159 | -0,0215611 | 0,9069 | 0,988324 |
| Gabpb2 | 16,5105 | 16,8284 | 0,027516 | 0,90705 | 0,988324 |
| Lmnb2 | 49,635 | 50,3575 | 0,0208493 | 0,90705 | 0,988324 |
| Caprin1 | 241,288 | 245,294 | 0,0237552 | 0,90715 | 0,988324 |
| Sirt7 | 10,8938 | 10,6859 | -0,0277879 | 0,90715 | 0,988324 |
| Zfp934 | 2,66864 | 2,60283 | -0,0360278 | 0,9072 | 0,988324 |
| Bzw1 | 198,529 | 201,556 | 0,0218286 | 0,90735 | 0,988323 |
| Pias1 | 10,7991 | 10,6351 | -0,0220739 | 0,90735 | 0,988324 |
| Endog | 8,00956 | 8,4661 | 0,0799751 | 0,9076 | 0,988507 |
| Gm12359 | 1,3716 | 1,73612 | 0,340002 | 0,90775 | 0,988507 |
| Scpep1 | 44,3559 | 43,7289 | -0,0205363 | 0,90775 | 0,988507 |
| Mmd | 24,3308 | 24,7007 | 0,021767 | 0,90805 | 0,988549 |
| 9130401M01Rik | 43,3272 | 42,6589 | -0,0224244 | 0,9081 | 0,988549 |
| Synj1 | 2,71642 | 2,60449 | -0,0607046 | 0,9082 | 0,988549 |
| Tceb2 | 197,318 | 182,342 | -0,113877 | 0,90825 | 0,988549 |
| Ostc | 175,525 | 178,059 | 0,0206845 | 0,9084 | 0,988549 |
| E2f1 | 23,3205 | 23,6725 | 0,0216112 | 0,9085 | 0,988549 |
| Bmp1 | 25,5958 | 25,2168 | -0,0215205 | 0,9085 | 0,988549 |
| Palm | 33,8867 | 29,7356 | -0,188531 | 0,90855 | 0,988549 |
| Pinx1 | 36,1114 | 35,5416 | -0,0229462 | 0,9087 | 0,988549 |
| Wdhd1 | 40,5554 | 41,1491 | 0,0209666 | 0,90885 | 0,988549 |
| Prpf4 | 26,9658 | 26,5812 | -0,0207211 | 0,90885 | 0,988549 |
| Med11 | 34,6488 | 34,0781 | -0,0239618 | 0,9089 | 0,988549 |
| Mrps31 | 30,7442 | 30,2648 | -0,022673 | 0,909 | 0,988549 |
| Gpatch2 | 10,4116 | 10,2551 | -0,0218553 | 0,90915 | 0,988549 |
| Yap1 | 85,5252 | 86,7243 | 0,0200856 | 0,90945 | 0,988549 |
| Igf2bp1 | 52,028 | 51,2636 | -0,0213521 | 0,90945 | 0,988549 |
| Smim13 | 5,55555 | 5,46315 | -0,0241964 | 0,9095 | 0,988549 |
| Nubpl | 9,39884 | 9,19871 | -0,0310512 | 0,9095 | 0,988549 |
| Gimap9 | 1,5157 | 1,56778 | 0,0487418 | 0,90955 | 0,988549 |

| | | | | | |
|---|---|---|---|---|---|
| Alkbh2 | 10,7542 | 10,982 | 0,0302486 | 0,90955 | 0,988549 |
| Csnk1g3 | 19,0351 | 19,3135 | 0,0209525 | 0,90975 | 0,988549 |
| Tmem55b | 17,106 | 17,7841 | 0,0560791 | 0,9098 | 0,988549 |
| Ercc3 | 32,5564 | 33,0515 | 0,0217728 | 0,9099 | 0,988549 |
| Smarcc2 | 39,7985 | 40,3489 | 0,0198168 | 0,90995 | 0,988549 |
| Flii | 26,282 | 25,6003 | -0,0379135 | 0,90995 | 0,988549 |
| Rad51c | 6,21502 | 5,46375 | -0,185868 | 0,90995 | 0,988549 |
| A630066F11Rik | 1,47949 | 1,62317 | 0,133716 | 0,91 | 0,988549 |
| Cd2bp2 | 54,228 | 53,4988 | -0,019533 | 0,91 | 0,988549 |
| Paxbp1 | 30,3663 | 29,8912 | -0,0227521 | 0,91005 | 0,988549 |
| Maged1 | 284,346 | 288,637 | 0,0216057 | 0,9101 | 0,988549 |
| Nudt22 | 7,58948 | 7,40564 | -0,0353763 | 0,91035 | 0,988737 |
| Selm | 8,72516 | 8,52284 | -0,0338469 | 0,91095 | 0,989044 |
| Map10 | 2,50522 | 2,55683 | 0,0294193 | 0,911 | 0,989044 |
| Shq1 | 12,9758 | 13,1958 | 0,0242514 | 0,91105 | 0,989044 |
| Ung | 35,9937 | 35,4713 | -0,021093 | 0,91115 | 0,989044 |
| Appbp2 | 45,0207 | 44,4018 | -0,01997 | 0,91125 | 0,989044 |
| Gm15441 | 0,721903 | 0,654512 | -0,141385 | 0,91135 | 0,989044 |
| 4930507D05Rik | 0,754506 | 0,827951 | 0,134012 | 0,9116 | 0,989044 |
| Trim37 | 22,5588 | 21,5416 | -0,0665617 | 0,9116 | 0,989044 |
| 1110020A21Rik | 1,03901 | 0,896634 | -0,212612 | 0,9117 | 0,989044 |
| Smpd4 | 35,69 | 35,2072 | -0,0196476 | 0,91175 | 0,989044 |
| Wdr41 | 9,24687 | 9,39959 | 0,0236316 | 0,912 | 0,989044 |
| Dctn4 | 27,9921 | 27,6147 | -0,0195856 | 0,91205 | 0,989044 |
| Cln3 | 4,53435 | 4,44801 | -0,0277356 | 0,91205 | 0,989044 |
| Zfp12 | 11,3382 | 11,1754 | -0,0208623 | 0,9123 | 0,989044 |
| Tra2a | 193,579 | 196,305 | 0,0201756 | 0,91235 | 0,989044 |
| Dpp7 | 6,26561 | 6,39681 | 0,0298977 | 0,91245 | 0,989044 |
| Smarca2 | 18,6868 | 19,0004 | 0,0240111 | 0,91255 | 0,989044 |
| Zbed5 | 5,14447 | 5,04674 | -0,0276713 | 0,9126 | 0,989044 |
| Actn4 | 91,9862 | 90,7497 | -0,0195257 | 0,91265 | 0,989044 |
| Tbc1d22a | 6,60146 | 6,70865 | 0,0232359 | 0,9127 | 0,989044 |
| Stard3 | 14,6026 | 14,831 | 0,0223965 | 0,91275 | 0,989044 |
| 2210016L21Rik | 30,2107 | 29,7499 | -0,0221762 | 0,91275 | 0,989044 |
| Gm561 | 33,3202 | 34,0147 | 0,0297592 | 0,9128 | 0,989044 |
| Kdelc1 | 50,5775 | 49,8972 | -0,0195383 | 0,9128 | 0,989044 |
| Cirbp | 46,1327 | 53,2824 | 0,20787 | 0,9129 | 0,989044 |
| Ror2 | 27,5454 | 27,9221 | 0,0195953 | 0,9129 | 0,989044 |
| Eps15 | 8,33151 | 8,21016 | -0,0211671 | 0,9129 | 0,989044 |
| 2810410L24Rik | 5,06972 | 4,96615 | -0,0297782 | 0,9129 | 0,989044 |
| Gm15421 | 58,7418 | 57,3567 | -0,0344241 | 0,9129 | 0,989044 |
| Plod3 | 22,1092 | 22,4976 | 0,0251272 | 0,91305 | 0,989044 |
| Qrich1 | 62,0164 | 61,1796 | -0,0195994 | 0,91305 | 0,989044 |
| Add1 | 51,2336 | 51,93 | 0,0194803 | 0,9131 | 0,989044 |
| Atp5sl | 18,1612 | 17,8681 | -0,0234779 | 0,9132 | 0,989069 |
| A330069E16Rik | 1,05868 | 1,01112 | -0,0663134 | 0,9136 | 0,989389 |
| Fam134b | 0,674708 | 0,653234 | -0,0466644 | 0,91365 | 0,989389 |
| Lin37 | 10,5828 | 10,7917 | 0,0281949 | 0,91435 | 0,989697 |
| 2410127L17Rik | 0,670222 | 0,646639 | -0,0516797 | 0,91435 | 0,989697 |
| 8430429K09Rik | 5,97418 | 6,75458 | 0,177127 | 0,9144 | 0,989697 |

| | | | | | |
|---|---|---|---|---|---|
| Sp1 | 26,2478 | 25,9031 | -0,0190721 | 0,91455 | 0,989697 |
| Pcdhga9 | 1,01335 | 0,878887 | -0,205383 | 0,9146 | 0,989697 |
| Emc9 | 0,705743 | 0,606413 | -0,218843 | 0,9146 | 0,989697 |
| Gnai2 | 157,441 | 159,515 | 0,0188852 | 0,91465 | 0,989697 |
| Nceh1 | 1,29836 | 1,26806 | -0,0340688 | 0,91465 | 0,989697 |
| Tspan14 | 41,8214 | 42,377 | 0,0190399 | 0,91475 | 0,989697 |
| Mrpl32 | 60,3585 | 59,461 | -0,021613 | 0,91485 | 0,989697 |
| Srprb | 40,4095 | 40,9791 | 0,0201946 | 0,9149 | 0,989697 |
| Efcab11 | 5,51535 | 5,61909 | 0,0268836 | 0,91495 | 0,989697 |
| Tspan5 | 16,0709 | 15,8372 | -0,0211296 | 0,91505 | 0,989697 |
| Tmem168 | 7,79657 | 7,91158 | 0,021126 | 0,91515 | 0,989697 |
| Commd1 | 150,325 | 148,11 | -0,0214162 | 0,9152 | 0,989697 |
| Ell2 | 1,08391 | 1,05618 | -0,0373815 | 0,9152 | 0,989697 |
| Fbxl22 | 0,887918 | 0,798654 | -0,152857 | 0,9153 | 0,989697 |
| 2610002J02Rik | 40,7834 | 41,3925 | 0,0213857 | 0,9154 | 0,989697 |
| Hnrnpl | 272,806 | 276,537 | 0,0195937 | 0,9154 | 0,989697 |
| Cct7 | 265,199 | 268,885 | 0,0199104 | 0,9155 | 0,989721 |
| Zbtb7b | 0,619369 | 0,638794 | 0,0445535 | 0,91565 | 0,9898 |
| Snx9 | 15,7673 | 16,0112 | 0,0221527 | 0,91575 | 0,989825 |
| Cdkn2aipnl | 52,7674 | 52,06 | -0,0194723 | 0,9159 | 0,98987 |
| Tbc1d19 | 12,1308 | 12,3258 | 0,02301 | 0,91605 | 0,98987 |
| Naif1 | 1,22907 | 1,28654 | 0,0659355 | 0,9161 | 0,98987 |
| Usp28 | 24,1134 | 24,4355 | 0,0191424 | 0,9161 | 0,98987 |
| Pcyt2 | 28,902 | 28,4937 | -0,0205233 | 0,91625 | 0,989919 |
| Rpl30 | 64,7772 | 67,2308 | 0,0536364 | 0,9163 | 0,989919 |
| Rps6ka4 | 1,17892 | 1,20822 | 0,0354261 | 0,9166 | 0,989992 |
| Nol9 | 36,7863 | 37,292 | 0,0196978 | 0,91665 | 0,989992 |
| Mettl23 | 15,4523 | 15,1722 | -0,0263887 | 0,91685 | 0,989992 |
| Cggbp1 | 32,4504 | 32,8812 | 0,019023 | 0,91695 | 0,989992 |
| Gaa | 8,46789 | 8,34521 | -0,0210551 | 0,91695 | 0,989992 |
| Stambp | 4,87523 | 4,96869 | 0,0273938 | 0,9171 | 0,989992 |
| Cul1 | 114,588 | 116,097 | 0,0188728 | 0,91715 | 0,989992 |
| Phf14 | 35,2825 | 35,7369 | 0,0184605 | 0,91715 | 0,989992 |
| Ttll4 | 41,2545 | 41,7986 | 0,0189017 | 0,91735 | 0,989992 |
| Inpp5a | 12,016 | 11,8283 | -0,0227154 | 0,91735 | 0,989992 |
| Adam10 | 37,604 | 37,1141 | -0,0189219 | 0,9174 | 0,989992 |
| Timp1 | 6,87195 | 7,26447 | 0,0801374 | 0,9175 | 0,989992 |
| Rpusd4 | 25,9777 | 25,6289 | -0,019505 | 0,91775 | 0,989992 |
| Csnk2a1 | 28,7405 | 28,3506 | -0,019705 | 0,91775 | 0,989992 |
| Sdk2 | 2,05352 | 2,0178 | -0,0253149 | 0,9178 | 0,989992 |
| Ccdc73 | 0,553592 | 0,605624 | 0,129601 | 0,91785 | 0,989992 |
| Gcdh | 22,4623 | 22,8164 | 0,0225639 | 0,91785 | 0,989992 |
| Ahcy | 17,729 | 17,4702 | -0,0212112 | 0,9179 | 0,989992 |
| Sec61a1 | 102,199 | 103,486 | 0,0180507 | 0,91805 | 0,989992 |
| Rnps1 | 51,3744 | 52,0576 | 0,0190595 | 0,9181 | 0,989992 |
| Mum1 | 36,5583 | 39,7522 | 0,120836 | 0,9182 | 0,989992 |
| Grip1 | 6,57314 | 6,70104 | 0,0278019 | 0,91825 | 0,989992 |
| Sepsecs | 6,1677 | 6,28232 | 0,026565 | 0,91825 | 0,989992 |
| Pi4k2a | 21,5617 | 21,8335 | 0,0180727 | 0,91825 | 0,989992 |
| Sart1 | 32,7103 | 33,1354 | 0,0186292 | 0,9183 | 0,989992 |

| | | | | | |
|---|---|---|---|---|---|
| Farp2 | 2,85327 | 2,72884 | -0,0643296 | 0,91845 | 0,989992 |
| Baiap2 | 6,51176 | 6,39949 | -0,0250918 | 0,9185 | 0,989992 |
| Stk24 | 41,7862 | 42,3367 | 0,018881 | 0,9188 | 0,989992 |
| Abca1 | 2,4294 | 2,46615 | 0,0216606 | 0,9189 | 0,989992 |
| Snx5 | 183,998 | 186,276 | 0,0177524 | 0,9189 | 0,989992 |
| Klhl17 | 17,2138 | 17,4465 | 0,0193759 | 0,919 | 0,989992 |
| Fads2 | 114,374 | 112,987 | -0,0176043 | 0,91905 | 0,989992 |
| Xkr5 | 7,02905 | 7,14211 | 0,0230215 | 0,9191 | 0,989992 |
| Ncln | 35,498 | 35,9563 | 0,018506 | 0,9192 | 0,989992 |
| Pcdhga12 | 1,59695 | 1,44251 | -0,146742 | 0,9192 | 0,989992 |
| Sox18 | 0,89701 | 0,927152 | 0,0476819 | 0,91925 | 0,989992 |
| Agpat3 | 7,52027 | 7,63307 | 0,0214787 | 0,91925 | 0,989992 |
| Ikbkap | 49,4382 | 48,75 | -0,0202241 | 0,9193 | 0,989992 |
| Las1l | 50,8239 | 50,1796 | -0,0184052 | 0,9196 | 0,990202 |
| Zfp426 | 14,5221 | 14,7169 | 0,0192294 | 0,91965 | 0,990202 |
| Eif4enif1 | 47,3742 | 46,1721 | -0,0370793 | 0,9198 | 0,990251 |
| Aacs | 17,573 | 17,8086 | 0,0192205 | 0,91985 | 0,990251 |
| Wtip | 22,6184 | 22,3087 | -0,0198932 | 0,92 | 0,99033 |
| Ublcp1 | 11,6279 | 11,8033 | 0,0216007 | 0,92055 | 0,990449 |
| Iqgap1 | 13,4428 | 13,6119 | 0,0180331 | 0,92055 | 0,990449 |
| Otud6b | 20,3548 | 20,0884 | -0,0190071 | 0,92055 | 0,990449 |
| Acin1 | 131,202 | 129,436 | -0,01955 | 0,92065 | 0,990449 |
| Clk2 | 46,2804 | 46,8601 | 0,0179589 | 0,92075 | 0,990449 |
| Fancc | 10,3268 | 10,598 | 0,037399 | 0,9209 | 0,990449 |
| Ttc30b | 2,33368 | 2,28512 | -0,0303355 | 0,92095 | 0,990449 |
| Ube2cbp | 4,96517 | 5,05687 | 0,0264007 | 0,92105 | 0,990449 |
| Srr | 6,88444 | 7,14758 | 0,0541156 | 0,92125 | 0,990449 |
| Twsg1 | 32,637 | 33,038 | 0,0176195 | 0,9213 | 0,990449 |
| Ptpra | 30,8888 | 30,5046 | -0,0180568 | 0,9214 | 0,990449 |
| B630019K06Rik | 2,12773 | 2,08154 | -0,0316701 | 0,9214 | 0,990449 |
| 1700019G17Rik | 0,991955 | 0,963864 | -0,0414443 | 0,92155 | 0,990449 |
| Atpaf2 | 22,925 | 23,2399 | 0,0196803 | 0,9216 | 0,990449 |
| Dstn | 113,656 | 115,093 | 0,0181226 | 0,9216 | 0,990449 |
| Pigk | 12,3937 | 12,2389 | -0,0181287 | 0,92165 | 0,990449 |
| Ube2a | 32,4596 | 32,0254 | -0,0194264 | 0,9217 | 0,990449 |
| Bcl10 | 22,6833 | 22,9953 | 0,0197057 | 0,92175 | 0,990449 |
| Nrg4 | 1,16347 | 1,26376 | 0,119283 | 0,9218 | 0,990449 |
| Lcat | 1,50416 | 1,55142 | 0,0446352 | 0,9218 | 0,990449 |
| Gm6578 | 2,08327 | 2,02121 | -0,0436302 | 0,92185 | 0,990449 |
| 1700092M07Rik | 7,89895 | 8,12598 | 0,0408803 | 0,9219 | 0,990449 |
| Zzef1 | 3,55113 | 3,59897 | 0,0193051 | 0,92205 | 0,990449 |
| Dtd2 | 5,18896 | 5,10236 | -0,024281 | 0,9221 | 0,990449 |
| Scn8a | 1,02921 | 1,0461 | 0,0234908 | 0,9222 | 0,990449 |
| Kctd9 | 12,5215 | 12,3557 | -0,0192269 | 0,9222 | 0,990449 |
| Arpc2 | 127,87 | 126,3 | -0,0178133 | 0,9223 | 0,990449 |
| Nt5dc1 | 4,01387 | 4,08398 | 0,024983 | 0,92235 | 0,990449 |
| Pcgf2 | 29,9267 | 30,3141 | 0,018557 | 0,92235 | 0,990449 |
| Gm16062 | 4,33879 | 4,92597 | 0,183115 | 0,92285 | 0,990845 |
| Mcm2 | 148,721 | 146,987 | -0,0169124 | 0,9229 | 0,990845 |
| E130317F20Rik | 3,83974 | 5,11609 | 0,414033 | 0,92295 | 0,990845 |

| | | | | | |
|---|---|---|---|---|---|
| Tacc1 | 5,11764 | 5,18605 | 0,0191587 | 0,92315 | 0,990976 |
| Slc23a2 | 4,58347 | 4,52107 | -0,0197777 | 0,9235 | 0,991025 |
| Arsb | 4,58817 | 4,51842 | -0,0220993 | 0,9235 | 0,991025 |
| Vill | 3,75241 | 3,68829 | -0,0248648 | 0,9235 | 0,991025 |
| Dcbld1 | 6,86073 | 6,75966 | -0,0214097 | 0,9238 | 0,991025 |
| Dpp9 | 28,1721 | 28,5231 | 0,0178623 | 0,92385 | 0,991025 |
| Gpr137 | 10,3247 | 10,1155 | -0,0295391 | 0,92395 | 0,991025 |
| Zswim5 | 5,50632 | 5,57804 | 0,0186711 | 0,92405 | 0,991025 |
| Zfp811 | 3,71045 | 3,77019 | 0,0230414 | 0,92415 | 0,991025 |
| Hmox1 | 14,0319 | 13,8261 | -0,0213146 | 0,92415 | 0,991025 |
| Rps7 | 8,89635 | 8,72563 | -0,0279538 | 0,92415 | 0,991025 |
| Xxylt1 | 16,5239 | 16,1519 | -0,0328575 | 0,92415 | 0,991025 |
| Ubxn8 | 5,43746 | 5,25683 | -0,048739 | 0,92415 | 0,991025 |
| 4932416H05Rik | 1,77316 | 1,86021 | 0,0691434 | 0,9242 | 0,991025 |
| Zfp961 | 16,1342 | 15,9283 | -0,0185365 | 0,92435 | 0,991074 |
| AI413582 | 2,72512 | 2,81583 | 0,0472379 | 0,9244 | 0,991074 |
| Nfatc4 | 22,9379 | 22,6704 | -0,016929 | 0,92455 | 0,991152 |
| Vti1a | 7,56088 | 7,66632 | 0,0199784 | 0,92475 | 0,991153 |
| Mta1 | 89,9847 | 88,9418 | -0,0168185 | 0,925 | 0,991153 |
| Cmtm8 | 4,45107 | 4,35433 | -0,031702 | 0,92505 | 0,991153 |
| Chst12 | 8,5855 | 8,72686 | 0,0235609 | 0,9251 | 0,991153 |
| Chd4 | 157,561 | 159,535 | 0,0179597 | 0,9252 | 0,991153 |
| Llgl1 | 54,207 | 55,0036 | 0,0210461 | 0,92525 | 0,991153 |
| Zhx1 | 14,0736 | 14,2429 | 0,0172498 | 0,92535 | 0,991153 |
| Zc3h13 | 22,2524 | 22,5158 | 0,0169775 | 0,92535 | 0,991153 |
| Ptbp1 | 228,063 | 220,572 | -0,0481853 | 0,92555 | 0,991153 |
| D030028A08Rik | 1,72597 | 1,79298 | 0,0549529 | 0,92565 | 0,991153 |
| Chaf1a | 74,1157 | 74,9565 | 0,0162744 | 0,92565 | 0,991153 |
| Pnrc1 | 23,4882 | 23,1851 | -0,0187377 | 0,9257 | 0,991153 |
| Gnai3 | 100,126 | 101,26 | 0,0162436 | 0,9258 | 0,991153 |
| Abhd16a | 14,5507 | 14,7481 | 0,0194409 | 0,92585 | 0,991153 |
| Capn10 | 14,7058 | 14,8964 | 0,0185796 | 0,92585 | 0,991153 |
| Melk | 43,9506 | 43,4451 | -0,0166869 | 0,92595 | 0,991153 |
| Urgcp | 5,10789 | 5,03145 | -0,0217524 | 0,92595 | 0,991153 |
| Ube4a | 16,7635 | 16,5726 | -0,0165218 | 0,9261 | 0,991153 |
| Aup1 | 47,2688 | 46,6612 | -0,0186658 | 0,92615 | 0,991153 |
| Nup88 | 87,877 | 89,1358 | 0,0205193 | 0,9263 | 0,991153 |
| Slc30a6 | 10,8069 | 10,9585 | 0,0200962 | 0,9263 | 0,991153 |
| Rpl9 | 7,61078 | 7,447 | -0,0313839 | 0,9263 | 0,991153 |
| Nf2 | 38,523 | 39,0312 | 0,0189109 | 0,92635 | 0,991153 |
| Mfap3 | 22,6025 | 22,8625 | 0,016503 | 0,92645 | 0,991153 |
| Ddb1 | 343,714 | 339,363 | -0,0183792 | 0,9265 | 0,991153 |
| Snora44 | 162,437 | 145,679 | -0,157081 | 0,9266 | 0,991153 |
| Leo1 | 45,8442 | 46,3769 | 0,0166684 | 0,92675 | 0,991153 |
| Flot2 | 20,4482 | 20,7065 | 0,0181112 | 0,9268 | 0,991153 |
| Gm12992 | 1,04118 | 1,12808 | 0,115656 | 0,9272 | 0,991153 |
| Pex19 | 59,3906 | 60,0426 | 0,0157522 | 0,9274 | 0,991153 |
| 1700094D03Rik | 2,89189 | 2,65895 | -0,121157 | 0,92745 | 0,991153 |
| Terf1 | 24,0269 | 23,7431 | -0,0171441 | 0,9275 | 0,991153 |
| Zfp13 | 12,1079 | 11,9588 | -0,0178744 | 0,9275 | 0,991153 |

| | | | | | |
|---|---|---|---|---|---|
| Uchl4 | 26,0375 | 25,3829 | -0,0367348 | 0,9275 | 0,991153 |
| Tmub2 | 13,8759 | 13,4609 | -0,0438032 | 0,92755 | 0,991153 |
| Pdzd11 | 63,8674 | 63,1118 | -0,0171692 | 0,92765 | 0,991153 |
| Pgm3 | 7,66037 | 7,52985 | -0,0247934 | 0,9277 | 0,991153 |
| Rac1 | 131,356 | 129,906 | -0,0160195 | 0,9278 | 0,991153 |
| R74862 | 3,7397 | 3,57309 | -0,0657495 | 0,928 | 0,991153 |
| Tmem67 | 5,33671 | 5,41518 | 0,0210577 | 0,9282 | 0,991153 |
| Vps9d1 | 9,23797 | 9,36573 | 0,0198165 | 0,9282 | 0,991153 |
| Tmem38b | 2,07289 | 2,11315 | 0,0277575 | 0,92835 | 0,991153 |
| Zkscan16 | 1,32105 | 1,34878 | 0,0299645 | 0,9284 | 0,991153 |
| Tcof1 | 52,2798 | 52,869 | 0,0161698 | 0,9284 | 0,991153 |
| Brd3 | 59,1408 | 58,4764 | -0,0163001 | 0,92845 | 0,991153 |
| Taco1 | 19,8579 | 19,6008 | -0,0188014 | 0,92845 | 0,991153 |
| Pdhx | 24,2533 | 23,9633 | -0,017354 | 0,9285 | 0,991153 |
| Dcp1b | 3,37067 | 3,2837 | -0,0377135 | 0,92855 | 0,991153 |
| Pptc7 | 8,12156 | 8,02077 | -0,018016 | 0,92865 | 0,991153 |
| Galnt2 | 26,8674 | 26,578 | -0,0156252 | 0,9287 | 0,991153 |
| Snord32a | 2366,44 | 2074,05 | -0,190263 | 0,9287 | 0,991153 |
| Scx | 0,719491 | 0,847725 | 0,23662 | 0,92875 | 0,991153 |
| Rassf7 | 13,313 | 13,5062 | 0,0207891 | 0,9289 | 0,991153 |
| Tdp2 | 50,6926 | 50,1123 | -0,0166094 | 0,929 | 0,991153 |
| Polr2b | 82,3094 | 83,2367 | 0,0161615 | 0,92905 | 0,991153 |
| Ddx6 | 69,7915 | 70,5682 | 0,0159677 | 0,92905 | 0,991153 |
| Trim59 | 59,1622 | 59,8183 | 0,0159115 | 0,9292 | 0,991153 |
| Zfp874a | 3,2817 | 3,22913 | -0,0232949 | 0,9292 | 0,991153 |
| Vps53 | 16,6277 | 16,4257 | -0,0176307 | 0,9293 | 0,991153 |
| Fam21 | 25,8227 | 25,5245 | -0,0167601 | 0,92935 | 0,991153 |
| Gm11944 | 3,83399 | 4,65304 | 0,27933 | 0,9294 | 0,991153 |
| Ppp6r2 | 5,41731 | 5,49403 | 0,0202888 | 0,92945 | 0,991153 |
| Zfp275 | 17,9684 | 17,7716 | -0,0158844 | 0,92945 | 0,991153 |
| Fam206a | 16,4358 | 16,8899 | 0,0393221 | 0,9296 | 0,991153 |
| Med9os | 0,961592 | 0,902392 | -0,0916694 | 0,9296 | 0,991153 |
| Mob3c | 2,00887 | 1,97117 | -0,0273267 | 0,92965 | 0,991153 |
| Degs2 | 5,65244 | 5,43806 | -0,0557823 | 0,9301 | 0,99155 |
| Zfp334 | 23,2588 | 23,5141 | 0,0157464 | 0,9304 | 0,991705 |
| Tmem39a | 25,2864 | 25,001 | -0,0163738 | 0,9304 | 0,991705 |
| Rrp1 | 144,539 | 146,093 | 0,0154214 | 0,93055 | 0,991783 |
| Rpain | 14,5874 | 15,3401 | 0,0725822 | 0,9307 | 0,991831 |
| Glul | 75,5044 | 74,6818 | -0,0158041 | 0,93075 | 0,991831 |
| Med15 | 15,8828 | 16,0668 | 0,0166126 | 0,93085 | 0,991855 |
| Zfp438 | 1,58444 | 1,61503 | 0,0275949 | 0,9311 | 0,991928 |
| Pikfyve | 4,75651 | 4,81481 | 0,0175739 | 0,9311 | 0,991928 |
| Usp14 | 43,2288 | 43,8569 | 0,0208095 | 0,93115 | 0,991928 |
| Tekt1 | 2,68534 | 2,73933 | 0,0287199 | 0,93135 | 0,991947 |
| 4933404O12Rik | 3,91332 | 3,96683 | 0,0195948 | 0,93135 | 0,991947 |
| Raf1 | 77,1749 | 76,3003 | -0,0164443 | 0,9314 | 0,991947 |
| Nkain1 | 77,5981 | 76,7575 | -0,0157135 | 0,9315 | 0,991971 |
| Aldh16a1 | 20,2602 | 20,5649 | 0,0215311 | 0,93165 | 0,992048 |
| Vasn | 8,37839 | 8,23893 | -0,0242172 | 0,93195 | 0,99228 |
| Cdc42ep4 | 56,9974 | 57,6448 | 0,0162952 | 0,93205 | 0,99228 |

| | | | | | |
|---|---|---|---|---|---|
| Rwdd4a | 20,7284 | 20,4883 | -0,0168076 | 0,9321 | 0,99228 |
| Prkcsh | 98,9443 | 99,9793 | 0,015013 | 0,9323 | 0,992289 |
| Gtf3c3 | 13,9656 | 14,1312 | 0,0170072 | 0,9324 | 0,992289 |
| Sh3glb2 | 18,2985 | 18,5227 | 0,017572 | 0,9325 | 0,992289 |
| Specc1 | 5,48315 | 5,42103 | -0,0164399 | 0,93255 | 0,992289 |
| Gpr62 | 1,23065 | 1,04661 | -0,233696 | 0,93255 | 0,992289 |
| St6galnac6 | 9,38986 | 9,26439 | -0,0194086 | 0,9326 | 0,992289 |
| Abracl | 105,376 | 104,242 | -0,0156109 | 0,93265 | 0,992289 |
| Tmem141 | 7,31696 | 7,45121 | 0,0262292 | 0,93295 | 0,99241 |
| Mxi1 | 8,38958 | 8,48696 | 0,0166492 | 0,93295 | 0,99241 |
| Alg13 | 15,762 | 15,5707 | -0,0176144 | 0,933 | 0,99241 |
| Ap2a2 | 34,1639 | 33,8128 | -0,0149036 | 0,9331 | 0,99241 |
| Kctd15 | 39,9393 | 40,3705 | 0,0154905 | 0,93315 | 0,99241 |
| Rimkla | 1,06565 | 1,0869 | 0,028487 | 0,93325 | 0,992434 |
| Rab10os | 4,83866 | 5,15668 | 0,0918335 | 0,93335 | 0,992458 |
| Oxld1 | 5,88264 | 5,77037 | -0,0277991 | 0,93345 | 0,992482 |
| Pak1 | 17,8405 | 17,6456 | -0,0158509 | 0,9336 | 0,992509 |
| Phactr1 | 1,34631 | 1,37125 | 0,0264837 | 0,9339 | 0,992509 |
| Gsr | 23,7903 | 24,0613 | 0,0163462 | 0,934 | 0,992509 |
| Snord35a | 1379,66 | 1322,1 | -0,0614843 | 0,93405 | 0,992509 |
| Edem2 | 8,69864 | 8,81228 | 0,0187267 | 0,9341 | 0,992509 |
| 1700120C14Rik | 1,43338 | 1,58051 | 0,140962 | 0,93435 | 0,992509 |
| Copb1 | 66,1257 | 65,4829 | -0,014091 | 0,93445 | 0,992509 |
| Mfsd11 | 5,5412 | 5,61665 | 0,0195105 | 0,9345 | 0,992509 |
| Bloc1s3 | 6,01197 | 5,92255 | -0,0216196 | 0,93485 | 0,992509 |
| Gpaa1 | 34,6239 | 34,2602 | -0,0152363 | 0,93495 | 0,992509 |
| Mrpl24 | 88,7227 | 87,6435 | -0,0176557 | 0,93495 | 0,992509 |
| Psen2 | 2,13104 | 2,17166 | 0,02724 | 0,93505 | 0,992509 |
| Gm10677 | 1,1004 | 1,23205 | 0,163024 | 0,9351 | 0,992509 |
| Lrp10 | 9,61031 | 9,71993 | 0,0163625 | 0,9351 | 0,992509 |
| Tbc1d7 | 22,258 | 22,51 | 0,0162407 | 0,9351 | 0,992509 |
| Zfp110 | 13,9751 | 13,8152 | -0,0166004 | 0,93525 | 0,992509 |
| Cd97 | 1,63542 | 1,74747 | 0,0956011 | 0,9353 | 0,992509 |
| Tomt | 6,07162 | 5,8894 | -0,0439601 | 0,9353 | 0,992509 |
| Dcaf7 | 41,5261 | 41,1202 | -0,0141731 | 0,93545 | 0,992509 |
| Tbpl1 | 34,7376 | 34,3763 | -0,0150849 | 0,93545 | 0,992509 |
| Etohd2 | 0,822893 | 0,874229 | 0,0873064 | 0,9355 | 0,992509 |
| Rbm10 | 22,8918 | 23,2036 | 0,0195205 | 0,93555 | 0,992509 |
| Pck2 | 6,69637 | 6,77934 | 0,0177641 | 0,93555 | 0,992509 |
| Ccar2 | 64,333 | 63,6643 | -0,0150736 | 0,93555 | 0,992509 |
| Slc10a3-ubl4,Ubl4 | 4,71872 | 4,493 | -0,0707143 | 0,93575 | 0,992509 |
| Snora65 | 290,96 | 302,342 | 0,0553628 | 0,93585 | 0,992509 |
| Isoc2a | 14,6295 | 14,8295 | 0,0195885 | 0,936 | 0,992509 |
| Gigyf2 | 19,6563 | 19,4584 | -0,0145944 | 0,93605 | 0,992509 |
| Stx12 | 38,5408 | 38,1464 | -0,0148395 | 0,9362 | 0,992509 |
| Mlh1 | 22,3066 | 22,5524 | 0,015813 | 0,93635 | 0,992509 |
| Usp30 | 12,0176 | 12,1542 | 0,0163021 | 0,93645 | 0,992509 |
| Nap1l1 | 194,446 | 192,266 | -0,0162647 | 0,93645 | 0,992509 |
| Pacsin3 | 9,70755 | 9,83496 | 0,018812 | 0,9366 | 0,992509 |
| Nlk | 12,3878 | 12,5213 | 0,0154654 | 0,9367 | 0,992509 |

| | | | | | |
|---|---|---|---|---|---|
| Dolpp1 | 16,025 | 15,7892 | -0,0213857 | 0,9367 | 0,992509 |
| Wdr92 | 12,2459 | 12,1108 | -0,0160064 | 0,93675 | 0,992509 |
| Pcdhga10 | 1,40181 | 1,52498 | 0,121497 | 0,9368 | 0,992509 |
| Gt(ROSA)26Sor | 5,57546 | 5,28584 | -0,0769595 | 0,93685 | 0,992509 |
| Cks1brt | 1,85214 | 1,88701 | 0,0269039 | 0,9369 | 0,992509 |
| Cdk1 | 104,393 | 103,332 | -0,0147423 | 0,9369 | 0,992509 |
| Tbc1d4 | 3,45618 | 3,499 | 0,0177633 | 0,937 | 0,992509 |
| Rab38 | 1,00785 | 1,03158 | 0,0335708 | 0,93705 | 0,992509 |
| 6430562O15Rik | 0,557795 | 0,617533 | 0,146783 | 0,9371 | 0,992509 |
| Crls1 | 22,3917 | 22,1388 | -0,0163888 | 0,93715 | 0,992509 |
| Tmem158 | 2,81857 | 2,77173 | -0,024175 | 0,9375 | 0,992509 |
| Cmtm4 | 15,1298 | 15,2782 | 0,014089 | 0,93755 | 0,992509 |
| Eif2b4 | 31,1322 | 33,1118 | 0,0889346 | 0,93765 | 0,992509 |
| Rptor | 12,9064 | 12,7761 | -0,014643 | 0,93765 | 0,992509 |
| Zscan22 | 8,83283 | 8,93474 | 0,0165503 | 0,9377 | 0,992509 |
| Zbtb26 | 6,28087 | 6,35204 | 0,0162558 | 0,9377 | 0,992509 |
| Pwp2 | 21,6921 | 21,9068 | 0,0142141 | 0,9377 | 0,992509 |
| Snora7a | 268,565 | 311,751 | 0,215124 | 0,9379 | 0,992509 |
| BC031361 | 0,920947 | 0,991355 | 0,106283 | 0,93795 | 0,992509 |
| Tmem251 | 19,9014 | 20,3637 | 0,0331276 | 0,938 | 0,992509 |
| Rala | 38,5355 | 38,1596 | -0,0141427 | 0,9381 | 0,992509 |
| Cnot11 | 27,2481 | 26,9714 | -0,0147282 | 0,9381 | 0,992509 |
| Trmt6 | 34,7453 | 34,397 | -0,0145345 | 0,9382 | 0,992509 |
| Ppp2r5d | 71,1317 | 70,1769 | -0,0194967 | 0,9382 | 0,992509 |
| Aldh7a1 | 44,5826 | 45,1686 | 0,0188412 | 0,9383 | 0,992509 |
| 2410131K14Rik | 13,2088 | 13,0628 | -0,0160363 | 0,9383 | 0,992509 |
| Tom1l1 | 12,954 | 12,7569 | -0,0221238 | 0,9383 | 0,992509 |
| Kat5 | 36,3013 | 35,9324 | -0,0147343 | 0,93835 | 0,992509 |
| Chrm4 | 0,725569 | 0,717993 | -0,0151422 | 0,93835 | 0,992509 |
| Sirt3 | 11,2543 | 11,4944 | 0,0304496 | 0,9389 | 0,99259 |
| Dnmt1 | 106,923 | 105,878 | -0,0141571 | 0,9389 | 0,99259 |
| Med23 | 20,3836 | 20,5844 | 0,0141413 | 0,93895 | 0,99259 |
| Kif13b | 2,45493 | 2,42259 | -0,0191346 | 0,939 | 0,99259 |
| Bhlhb9 | 37,6372 | 37,2689 | -0,0141844 | 0,9391 | 0,99259 |
| Hhat | 3,34479 | 3,39328 | 0,0207644 | 0,93915 | 0,99259 |
| Hmg20a | 30,4707 | 30,7474 | 0,0130393 | 0,93915 | 0,99259 |
| Neu1 | 4,23739 | 4,18596 | -0,0176158 | 0,93915 | 0,99259 |
| Preb | 21,108 | 22,0133 | 0,0605828 | 0,9392 | 0,99259 |
| Klf13 | 6,8058 | 6,8769 | 0,0149922 | 0,9392 | 0,99259 |
| Nudt7 | 6,44127 | 6,55476 | 0,0251987 | 0,9394 | 0,992662 |
| L3hypdh | 2,95275 | 2,90303 | -0,0245009 | 0,93945 | 0,992662 |
| Sat1 | 5,0772 | 5,15728 | 0,0225783 | 0,9395 | 0,992662 |
| 2700046G09Rik | 1,70944 | 1,67383 | -0,0303677 | 0,93965 | 0,992723 |
| Plcg1 | 42,931 | 42,5156 | -0,0140288 | 0,93975 | 0,992723 |
| Wdr83 | 28,9342 | 29,3771 | 0,0219163 | 0,93985 | 0,992723 |
| Tsg101 | 36,436 | 36,0908 | -0,0137324 | 0,9399 | 0,992723 |
| 2810021J22Rik | 3,91775 | 3,87148 | -0,0171404 | 0,93995 | 0,992723 |
| Tmem121 | 6,53569 | 6,62961 | 0,0205851 | 0,9401 | 0,992723 |
| Far2 | 1,85386 | 1,74119 | -0,0904562 | 0,9401 | 0,992723 |
| Nom1 | 20,1861 | 19,9943 | -0,0137692 | 0,9403 | 0,992756 |

| | | | | | |
|---|---|---|---|---|---|
| Sco2 | 4,09083 | 3,82879 | -0,0955051 | 0,9404 | 0,992756 |
| Ddx20 | 28,7467 | 28,4813 | -0,0133802 | 0,94045 | 0,992756 |
| Cenpu | 11,6 | 11,7538 | 0,0190034 | 0,9406 | 0,992756 |
| Exd2 | 11,2959 | 11,1848 | -0,0142583 | 0,94065 | 0,992756 |
| 9230105E05Rik | 0,839153 | 0,854133 | 0,0255263 | 0,94075 | 0,992756 |
| Vps16 | 30,4172 | 30,7503 | 0,0157146 | 0,94075 | 0,992756 |
| Slc25a22 | 18,7847 | 18,5974 | -0,0144563 | 0,94075 | 0,992756 |
| Rpl36a | 1,55545 | 1,63265 | 0,0698835 | 0,9412 | 0,992865 |
| Trim7 | 4,19253 | 4,26027 | 0,0231264 | 0,9412 | 0,992865 |
| Usp54 | 8,62474 | 8,77466 | 0,0248615 | 0,94125 | 0,992865 |
| Med20 | 23,2334 | 23,4672 | 0,0144495 | 0,9413 | 0,992865 |
| Ttc4 | 38,5324 | 38,8991 | 0,0136644 | 0,9413 | 0,992865 |
| Txlng | 7,15776 | 6,84779 | -0,06387 | 0,94135 | 0,992865 |
| Lnx1 | 2,27441 | 2,39028 | 0,071682 | 0,94145 | 0,992865 |
| Prkd3 | 43,728 | 44,1135 | 0,0126614 | 0,9415 | 0,992865 |
| Dnajc5 | 44,0663 | 44,4766 | 0,0133711 | 0,94155 | 0,992865 |
| Lrrc14b | 2,31459 | 2,2264 | -0,0560468 | 0,94165 | 0,992889 |
| Atf7ip | 21,1801 | 21,3855 | 0,0139245 | 0,9419 | 0,993071 |
| Hist1h3e | 1,45463 | 1,40506 | -0,0500207 | 0,94205 | 0,993147 |
| Zdhhc6 | 32,3609 | 32,0592 | -0,0135125 | 0,94245 | 0,993183 |
| Gsta4 | 124,072 | 122,904 | -0,013643 | 0,94245 | 0,993183 |
| Ppil4 | 25,9694 | 25,7137 | -0,0142722 | 0,9426 | 0,993183 |
| Trip13 | 44,5537 | 44,9652 | 0,0132643 | 0,9427 | 0,993183 |
| Lat2 | 2,25102 | 2,21142 | -0,0256111 | 0,9427 | 0,993183 |
| Usp19 | 46,0145 | 46,4394 | 0,0132626 | 0,94285 | 0,993183 |
| Nup205 | 54,2545 | 53,7725 | -0,0128742 | 0,94305 | 0,993183 |
| Tmem194b | 6,48163 | 6,55512 | 0,0162642 | 0,9431 | 0,993183 |
| Lmf2 | 14,3823 | 14,2435 | -0,0139894 | 0,9433 | 0,993183 |
| Tlcd1 | 7,06154 | 7,15567 | 0,0191034 | 0,94335 | 0,993183 |
| H2afy | 222,881 | 224,869 | 0,012813 | 0,94355 | 0,993183 |
| Lmo4 | 99,1565 | 100,076 | 0,0133123 | 0,94365 | 0,993183 |
| Usp40 | 15,0156 | 15,1519 | 0,0130362 | 0,94365 | 0,993183 |
| Hnrnpd | 110,012 | 111,055 | 0,0136115 | 0,9437 | 0,993183 |
| Prr14 | 22,7149 | 22,9267 | 0,0133906 | 0,9439 | 0,993183 |
| Gm15708 | 0,899093 | 0,952321 | 0,0829777 | 0,94405 | 0,993183 |
| Ado | 15,2028 | 15,352 | 0,0140918 | 0,94405 | 0,993183 |
| Pan3 | 11,8416 | 11,9545 | 0,0136847 | 0,94405 | 0,993183 |
| Surf6 | 30,8103 | 30,5254 | -0,0134035 | 0,94405 | 0,993183 |
| Kptn | 9,0582 | 8,95549 | -0,0164524 | 0,94425 | 0,993183 |
| Tjp2 | 24,6879 | 24,467 | -0,0129651 | 0,9444 | 0,993183 |
| Bsg | 264,361 | 254,52 | -0,0547301 | 0,94445 | 0,993183 |
| Smad6 | 8,12462 | 8,03865 | -0,015348 | 0,9445 | 0,993183 |
| 2310011J03Rik | 20,6063 | 17,9076 | -0,202514 | 0,9447 | 0,993183 |
| Pcdhgc3 | 45,8699 | 45,3033 | -0,0179304 | 0,9449 | 0,993183 |
| Dos | 3,93932 | 4,67982 | 0,24851 | 0,945 | 0,993183 |
| Dak | 14,6826 | 14,8268 | 0,0140942 | 0,94515 | 0,993183 |
| Emc10 | 35,0615 | 35,3857 | 0,0132806 | 0,94515 | 0,993183 |
| Vma21 | 5,92201 | 5,98103 | 0,0143072 | 0,94525 | 0,993183 |
| Coq2 | 23,7615 | 23,9856 | 0,0135414 | 0,94525 | 0,993183 |
| Srrt | 109,873 | 108,907 | -0,0127433 | 0,94565 | 0,993183 |

| | | | | | |
|---|---|---|---|---|---|
| Zfp710 | 12,1003 | 11,9878 | -0,0134833 | 0,94565 | 0,993183 |
| Sbno2 | 5,61714 | 6,48985 | 0,20835 | 0,9458 | 0,993183 |
| Abca7 | 1,47526 | 1,65243 | 0,16362 | 0,9458 | 0,993183 |
| Pcdhgb4 | 1,76642 | 1,87788 | 0,0882708 | 0,9458 | 0,993183 |
| Irf8 | 0,967503 | 0,984366 | 0,0249295 | 0,9458 | 0,993183 |
| Nxf1 | 129,179 | 130,272 | 0,0121599 | 0,94595 | 0,993183 |
| Cox6b2 | 10,5016 | 10,6859 | 0,0251085 | 0,94605 | 0,993183 |
| Map3k10 | 9,90555 | 9,99601 | 0,0131158 | 0,94605 | 0,993183 |
| Tmem229b | 0,885376 | 0,900486 | 0,0244133 | 0,94615 | 0,993183 |
| B9d1 | 25,4578 | 25,7454 | 0,0162079 | 0,94625 | 0,993183 |
| Senp1 | 22,2115 | 22,4517 | 0,0155218 | 0,9463 | 0,993183 |
| Kcng1 | 1,56424 | 1,59053 | 0,024049 | 0,94635 | 0,993183 |
| Deaf1 | 21,3746 | 21,5923 | 0,0146177 | 0,94635 | 0,993183 |
| Slc36a4 | 8,14226 | 8,23767 | 0,0168062 | 0,9465 | 0,993183 |
| Adam1a | 0,931147 | 0,913611 | -0,0274288 | 0,9465 | 0,993183 |
| Dpy19l1 | 21,9143 | 21,7284 | -0,0122952 | 0,94665 | 0,993183 |
| Psma3 | 10,8763 | 10,7326 | -0,0191948 | 0,9468 | 0,993183 |
| Pom121 | 33,1797 | 33,4693 | 0,0125397 | 0,94685 | 0,993183 |
| 2810429I04Rik | 5,48281 | 5,39018 | -0,0245819 | 0,94685 | 0,993183 |
| Fbxl2 | 1,29594 | 1,37411 | 0,0844983 | 0,9469 | 0,993183 |
| Tenm3 | 23,6291 | 23,8164 | 0,0113874 | 0,9469 | 0,993183 |
| Bpnt1 | 21,7658 | 21,5723 | -0,0128848 | 0,9469 | 0,993183 |
| Pcdhgb8 | 0,697643 | 0,623432 | -0,162259 | 0,94695 | 0,993183 |
| Incenp | 74,7996 | 75,4377 | 0,012255 | 0,947 | 0,993183 |
| Sf1 | 122,958 | 123,964 | 0,0117479 | 0,947 | 0,993183 |
| Gpatch11 | 4,20764 | 4,16326 | -0,0152995 | 0,9472 | 0,993183 |
| Zfp810 | 7,49222 | 7,73923 | 0,0467982 | 0,94725 | 0,993183 |
| Slc25a46 | 13,9715 | 14,0981 | 0,0130167 | 0,94725 | 0,993183 |
| Hdlbp | 124,721 | 123,649 | -0,0124591 | 0,9473 | 0,993183 |
| Fam35a | 3,78452 | 3,74075 | -0,0167824 | 0,9473 | 0,993183 |
| 4921507L20Rik | 0,67106 | 0,70178 | 0,0645765 | 0,94735 | 0,993183 |
| D17Wsu104e | 71,3707 | 71,9818 | 0,012301 | 0,94745 | 0,993183 |
| Pkn1 | 10,6201 | 10,509 | -0,0151779 | 0,9475 | 0,993183 |
| Fra10ac1 | 23,2157 | 23,4405 | 0,0139035 | 0,94775 | 0,993183 |
| Trib1 | 9,12932 | 9,04752 | -0,0129854 | 0,9478 | 0,993183 |
| Igflr1 | 5,48757 | 5,41814 | -0,0183686 | 0,9479 | 0,993183 |
| Pafah1b1 | 54,0469 | 54,4564 | 0,0108892 | 0,94795 | 0,993183 |
| Slc9a6 | 6,2491 | 6,30533 | 0,0129229 | 0,9482 | 0,993183 |
| Azin1 | 60,5466 | 60,0455 | -0,01199 | 0,94825 | 0,993183 |
| D15Ertd621e | 30,5831 | 30,8306 | 0,0116285 | 0,94835 | 0,993183 |
| Kif21b | 6,79501 | 6,73539 | -0,0127135 | 0,94835 | 0,993183 |
| Kcnn3 | 1,05705 | 1,04381 | -0,0181819 | 0,9484 | 0,993183 |
| Apbb3 | 8,76472 | 8,95646 | 0,0312196 | 0,9485 | 0,993183 |
| Zfp11 | 2,82647 | 2,86085 | 0,0174468 | 0,9485 | 0,993183 |
| Vamp1 | 0,773059 | 0,785898 | 0,0237636 | 0,94855 | 0,993183 |
| Cdh24 | 12,7427 | 12,8644 | 0,0137082 | 0,94865 | 0,993183 |
| Exosc6 | 46,7194 | 47,1301 | 0,0126252 | 0,94875 | 0,993183 |
| Elavl2 | 4,39242 | 4,44081 | 0,0158043 | 0,9488 | 0,993183 |
| Cox18 | 13,954 | 14,1025 | 0,0152672 | 0,9488 | 0,993183 |
| Atxn1l | 12,2176 | 12,312 | 0,0111044 | 0,9488 | 0,993183 |

| | | | | | |
|---|---|---|---|---|---|
| Aip | 18,8535 | 18,6649 | -0,0145011 | 0,9488 | 0,993183 |
| Pcdhga1 | 0,58101 | 0,653528 | 0,169685 | 0,9489 | 0,993183 |
| Gm9833 | 32,5075 | 32,7905 | 0,0125052 | 0,949 | 0,993183 |
| Sdc1 | 66,5938 | 67,1373 | 0,0117257 | 0,949 | 0,993183 |
| Taf7 | 13,0998 | 12,7633 | -0,0375372 | 0,94925 | 0,993183 |
| Fdft1 | 28,267 | 27,95 | -0,016273 | 0,9494 | 0,993183 |
| Rpl34,Rpl34-ps1 | 47,6192 | 46,3663 | -0,0384651 | 0,94955 | 0,993183 |
| Mttp | 3,02314 | 2,99132 | -0,0152673 | 0,9496 | 0,993183 |
| Arfrp1 | 25,0769 | 25,3594 | 0,0161564 | 0,94965 | 0,993183 |
| Gsk3b | 21,2112 | 21,3864 | 0,011869 | 0,94975 | 0,993183 |
| Dda1 | 26,9869 | 26,7195 | -0,0143664 | 0,94975 | 0,993183 |
| Wash | 12,5851 | 12,4823 | -0,0118378 | 0,9498 | 0,993183 |
| Eif4g2 | 329,987 | 326,849 | -0,0137866 | 0,95005 | 0,993183 |
| Nrf1 | 28,7027 | 28,4549 | -0,0125094 | 0,9501 | 0,993183 |
| 2310010J17Rik | 5,54002 | 5,66967 | 0,0333742 | 0,95015 | 0,993183 |
| Gemin5 | 29,9874 | 29,7645 | -0,0107648 | 0,95015 | 0,993183 |
| Pddc1 | 20,8299 | 20,64 | -0,0132114 | 0,95015 | 0,993183 |
| Otub1 | 83,6232 | 84,3172 | 0,0119243 | 0,95025 | 0,993183 |
| Slc25a39 | 91,7364 | 92,4754 | 0,011576 | 0,95025 | 0,993183 |
| Ahrr | 0,708555 | 0,697719 | -0,0222335 | 0,9503 | 0,993183 |
| Slc35f5 | 11,5965 | 11,7058 | 0,0135427 | 0,95035 | 0,993183 |
| Klhl9 | 40,1682 | 39,8588 | -0,011156 | 0,95035 | 0,993183 |
| Grn | 15,7319 | 15,5935 | -0,0127471 | 0,95035 | 0,993183 |
| Got2 | 55,6243 | 56,0816 | 0,011812 | 0,9504 | 0,993183 |
| Angpt2 | 1,37523 | 1,3356 | -0,042181 | 0,95045 | 0,993183 |
| Poldip3 | 52,8364 | 53,2612 | 0,0115545 | 0,9505 | 0,993183 |
| Zfp14 | 1,0765 | 1,09397 | 0,0232186 | 0,95055 | 0,993183 |
| Ipo5 | 177,442 | 178,892 | 0,0117419 | 0,9506 | 0,993183 |
| Cox20 | 92,9416 | 92,0978 | -0,0131582 | 0,9506 | 0,993183 |
| Haus5 | 13,8693 | 13,7426 | -0,0132443 | 0,9507 | 0,993206 |
| Mcee | 19,9378 | 20,1605 | 0,0160233 | 0,9513 | 0,993588 |
| Fam111a | 16,698 | 16,8395 | 0,0121744 | 0,9513 | 0,993588 |
| Cpsf2 | 34,578 | 34,8343 | 0,0106545 | 0,9516 | 0,993588 |
| Oxsr1 | 22,6026 | 22,7702 | 0,010662 | 0,95165 | 0,993588 |
| Mkrn2 | 13,4085 | 13,729 | 0,0340764 | 0,9517 | 0,993588 |
| Asah1 | 31,3915 | 31,6459 | 0,011641 | 0,9517 | 0,993588 |
| Dolk | 7,72165 | 7,64125 | -0,0151014 | 0,9517 | 0,993588 |
| Rnmtl1 | 13,101 | 12,9734 | -0,0141177 | 0,95175 | 0,993588 |
| Katnbl1 | 12,4551 | 12,5596 | 0,0120509 | 0,9518 | 0,993588 |
| Capza2 | 95,3211 | 96,1011 | 0,0117584 | 0,95205 | 0,993588 |
| Ubap2 | 51,9428 | 51,548 | -0,0110075 | 0,9521 | 0,993588 |
| Gm13889 | 0,814348 | 0,790679 | -0,0425534 | 0,9521 | 0,993588 |
| Sf3b3 | 164,865 | 163,63 | -0,0108426 | 0,95215 | 0,993588 |
| Tmx2 | 13,0021 | 12,6828 | -0,0358737 | 0,95215 | 0,993588 |
| 9330151L19Rik | 2,20981 | 2,27163 | 0,0398042 | 0,95235 | 0,993716 |
| Suv39h2 | 24,688 | 24,4837 | -0,0119875 | 0,95275 | 0,99399 |
| Usf2 | 20,7872 | 20,9484 | 0,0111459 | 0,95295 | 0,99399 |
| Slc37a4 | 7,10268 | 7,03694 | -0,0134143 | 0,95295 | 0,99399 |
| Rbm14 | 54,717 | 55,1625 | 0,0116986 | 0,953 | 0,99399 |
| Pdk3 | 44,3028 | 44,6317 | 0,0106725 | 0,953 | 0,99399 |

| | | | | | |
|---|---|---|---|---|---|
| Pabpn1 | 202,938 | 204,478 | 0,0109061 | 0,95325 | 0,994061 |
| Heatr2 | 14,3056 | 14,4205 | 0,0115355 | 0,9533 | 0,994061 |
| Tmem86b | 3,18781 | 3,14976 | -0,0173246 | 0,95335 | 0,994061 |
| Zfp219 | 44,6456 | 45,1495 | 0,0161939 | 0,9535 | 0,994061 |
| Ercc6l | 22,2489 | 22,0834 | -0,010772 | 0,9536 | 0,994061 |
| Zeb2os | 3,20746 | 3,33774 | 0,0574392 | 0,95375 | 0,994061 |
| Kcmf1 | 33,806 | 34,045 | 0,0101645 | 0,95385 | 0,994061 |
| Slc35b2 | 28,1798 | 27,9603 | -0,0112849 | 0,954 | 0,994061 |
| Acyp2 | 5,25453 | 5,18413 | -0,0194593 | 0,954 | 0,994061 |
| Vmp1 | 21,3566 | 20,6858 | -0,046044 | 0,95415 | 0,994061 |
| Nudt13 | 3,89176 | 3,93233 | 0,014961 | 0,9542 | 0,994061 |
| Trim35 | 61,24 | 60,8304 | -0,00968107 | 0,9542 | 0,994061 |
| Rfxank | 6,69159 | 6,76768 | 0,0163115 | 0,9543 | 0,994061 |
| Zdhhc8 | 7,94666 | 7,88403 | -0,0114163 | 0,9543 | 0,994061 |
| Nme3 | 23,4074 | 22,8667 | -0,0337162 | 0,95435 | 0,994061 |
| Ascc1 | 22,1412 | 22,4191 | 0,0179981 | 0,9544 | 0,994061 |
| Traf7 | 60,5319 | 60,9593 | 0,01015 | 0,95455 | 0,994061 |
| Zfp958 | 4,68608 | 4,73373 | 0,0145947 | 0,9546 | 0,994061 |
| Usp7 | 54,8687 | 55,257 | 0,0101747 | 0,9547 | 0,994061 |
| Pxn | 16,3697 | 16,2515 | -0,0104559 | 0,95475 | 0,994061 |
| Slc5a6 | 3,99496 | 3,60323 | -0,148893 | 0,95475 | 0,994061 |
| Smarcc1 | 97,2148 | 96,5379 | -0,0100794 | 0,95485 | 0,994061 |
| Rnaseh1 | 19,8804 | 19,7191 | -0,011751 | 0,95485 | 0,994061 |
| Oraov1 | 10,3781 | 10,2877 | -0,0126108 | 0,95495 | 0,994084 |
| Anxa11 | 1,73328 | 1,75731 | 0,0198675 | 0,9551 | 0,99416 |
| Arnt | 15,0033 | 15,1139 | 0,0105964 | 0,95535 | 0,994178 |
| Tmed1 | 40 | 39,4981 | -0,0182156 | 0,95535 | 0,994178 |
| Midn | 77,3398 | 74,8208 | -0,0477708 | 0,95535 | 0,994178 |
| L3mbtl2 | 19,1078 | 19,2487 | 0,0106001 | 0,9556 | 0,994323 |
| Dhps | 33,2797 | 32,9747 | -0,0132853 | 0,95575 | 0,994323 |
| Pqlc1 | 1,7789 | 1,75575 | -0,0188953 | 0,95575 | 0,994323 |
| Gnl1 | 31,1017 | 31,3199 | 0,0100857 | 0,9558 | 0,994323 |
| Stk35 | 11,8739 | 11,7879 | -0,0104871 | 0,95615 | 0,994606 |
| Ctsf | 5,85199 | 5,91067 | 0,0143939 | 0,95635 | 0,994606 |
| Nop56 | 140,67 | 139,446 | -0,012607 | 0,95645 | 0,994606 |
| Srsf5 | 147,989 | 146,989 | -0,00977729 | 0,95655 | 0,994606 |
| Golph3 | 49,299 | 49,6384 | 0,00990026 | 0,9566 | 0,994606 |
| Stim2 | 8,65455 | 8,71969 | 0,0108178 | 0,95675 | 0,994606 |
| Dbf4 | 69,5704 | 70,0427 | 0,00976165 | 0,95675 | 0,994606 |
| Tpi1 | 322,206 | 319,958 | -0,0100995 | 0,95685 | 0,994606 |
| Sorbs3 | 12,1052 | 12,2129 | 0,0127769 | 0,957 | 0,994606 |
| Vezt | 7,87897 | 7,81915 | -0,0109943 | 0,957 | 0,994606 |
| Pmch | 0,691721 | 0,680419 | -0,0237668 | 0,9572 | 0,994606 |
| Cse1l | 134,576 | 133,644 | -0,0100181 | 0,95725 | 0,994606 |
| Fam167a | 0,955732 | 0,943281 | -0,0189193 | 0,9574 | 0,994606 |
| Cwc27 | 19,9708 | 20,1291 | 0,0113901 | 0,95745 | 0,994606 |
| Arel1 | 13,2021 | 13,2944 | 0,0100423 | 0,95765 | 0,994606 |
| Bag6 | 92,6902 | 92,0689 | -0,00970263 | 0,95785 | 0,994606 |
| Tpra1 | 10,3481 | 10,2581 | -0,0126033 | 0,9579 | 0,994606 |
| Ppp1r16a | 18,1824 | 18,0534 | -0,0102728 | 0,95795 | 0,994606 |

| | | | | | |
|---|---|---|---|---|---|
| Hk1os | 1,02963 | 0,969619 | -0,0866397 | 0,958 | 0,994606 |
| B2m | 44,6088 | 44,9308 | 0,010378 | 0,95815 | 0,994606 |
| Gm20748 | 1,48957 | 1,47803 | -0,0112205 | 0,95815 | 0,994606 |
| Irak1 | 20,1043 | 19,9706 | -0,00962851 | 0,9583 | 0,994606 |
| Dusp19 | 9,12886 | 9,04701 | -0,0129927 | 0,9583 | 0,994606 |
| Pgs1 | 24,6723 | 24,8447 | 0,0100492 | 0,95845 | 0,994606 |
| R3hdm4 | 40,6558 | 38,2134 | -0,0893845 | 0,95845 | 0,994606 |
| Gm6525 | 2,0261 | 2,18151 | 0,106619 | 0,9589 | 0,994606 |
| Cox15 | 13,6542 | 13,7552 | 0,0106343 | 0,95905 | 0,994606 |
| Sms | 16,1793 | 16,0624 | -0,0104605 | 0,95905 | 0,994606 |
| Eif3m | 86,4816 | 85,9207 | -0,00938769 | 0,9592 | 0,994606 |
| Clasp1 | 29,969 | 29,7779 | -0,00923027 | 0,9593 | 0,994606 |
| Gmcl1 | 18,9411 | 19,0642 | 0,0093429 | 0,95935 | 0,994606 |
| Chd3os | 10,5823 | 10,4949 | -0,01196 | 0,95935 | 0,994606 |
| Bmpr1a | 39,7262 | 39,4711 | -0,00929539 | 0,9594 | 0,994606 |
| Ccdc43 | 33,8984 | 33,6796 | -0,00934123 | 0,9594 | 0,994606 |
| BC068281 | 5,32614 | 5,27936 | -0,0127255 | 0,9594 | 0,994606 |
| Wbp11 | 89,9381 | 90,4934 | 0,00888008 | 0,95955 | 0,994606 |
| Zfp418 | 1,70184 | 1,68459 | -0,0147039 | 0,95955 | 0,994606 |
| Ccm2 | 35,4084 | 35,6477 | 0,00971995 | 0,9596 | 0,994606 |
| Dnajc19 | 7,22162 | 6,94318 | -0,0567264 | 0,9596 | 0,994606 |
| Med7 | 18,7501 | 18,5991 | -0,0116627 | 0,9598 | 0,994606 |
| Mettl3 | 31,9359 | 31,5802 | -0,0161585 | 0,9601 | 0,994606 |
| Zfp72 | 2,68201 | 2,71374 | 0,0169666 | 0,96015 | 0,994606 |
| Stx7 | 12,218 | 12,3124 | 0,0111041 | 0,9602 | 0,994606 |
| 2410015M20Rik | 83,9497 | 84,5077 | 0,00955835 | 0,9604 | 0,994606 |
| Msantd2 | 27,9753 | 27,7975 | -0,00919642 | 0,96045 | 0,994606 |
| Snap29 | 8,67631 | 8,55798 | -0,0198114 | 0,9605 | 0,994606 |
| Shank3 | 1,34803 | 1,33575 | -0,0131942 | 0,96055 | 0,994606 |
| Zfp938 | 6,44016 | 6,49965 | 0,0132646 | 0,9606 | 0,994606 |
| Spata2 | 6,78976 | 6,68423 | -0,0226003 | 0,9606 | 0,994606 |
| Vash2 | 26,7574 | 26,5847 | -0,0093378 | 0,9607 | 0,994606 |
| Fam207a | 25,9702 | 26,1423 | 0,00952826 | 0,9609 | 0,994606 |
| 1700113A16Rik | 5,51268 | 5,44988 | -0,0165304 | 0,96095 | 0,994606 |
| Mtbp | 24,7802 | 24,6209 | -0,00930134 | 0,96105 | 0,994606 |
| Dfna5 | 4,35654 | 4,31628 | -0,0133954 | 0,9611 | 0,994606 |
| Snora52 | 133,197 | 139,42 | 0,0658704 | 0,9612 | 0,994606 |
| Dnajc1 | 8,02437 | 8,11523 | 0,0162432 | 0,9612 | 0,994606 |
| Cand1 | 54,6611 | 54,315 | -0,00916355 | 0,9613 | 0,994606 |
| 1110032A03Rik | 7,36267 | 7,29905 | -0,0125205 | 0,9613 | 0,994606 |
| Fkrp | 16,8878 | 16,6362 | -0,0216625 | 0,96135 | 0,994606 |
| Mmaa | 2,38125 | 2,40419 | 0,0138336 | 0,9614 | 0,994606 |
| Uvrag | 5,6635 | 5,70354 | 0,0101629 | 0,9614 | 0,994606 |
| Zfp768 | 26,6062 | 26,4281 | -0,00968992 | 0,9614 | 0,994606 |
| Stk11 | 60,278 | 63,413 | 0,0731472 | 0,9615 | 0,994606 |
| Mtpap | 21,25 | 21,1124 | -0,00936898 | 0,9615 | 0,994606 |
| Isg20l2 | 40,1925 | 39,9384 | -0,00914839 | 0,96155 | 0,994606 |
| Pttg1ip | 22,7051 | 22,857 | 0,00961961 | 0,96165 | 0,994606 |
| Nfil3 | 15,7558 | 15,8667 | 0,0101159 | 0,9617 | 0,994606 |
| Arhgef2 | 35,1495 | 35,3561 | 0,00845577 | 0,9618 | 0,994606 |

| | | | | | |
|---|---|---|---|---|---|
| Crb3 | 0,840251 | 0,836676 | -0,00615008 | 0,9622 | 0,994606 |
| Mvd | 36,587 | 36,3629 | -0,00886417 | 0,9622 | 0,994606 |
| Dyrk2 | 16,5141 | 16,4027 | -0,00977038 | 0,9622 | 0,994606 |
| Gtf3c2 | 59,95 | 60,8452 | 0,0213834 | 0,96225 | 0,994606 |
| Nfyb | 43,9945 | 43,7273 | -0,00878916 | 0,96235 | 0,994606 |
| Cul4a | 46,9067 | 47,1726 | 0,00815602 | 0,96245 | 0,994606 |
| Med24 | 35,5219 | 35,7321 | 0,00851135 | 0,9625 | 0,994606 |
| Csde1 | 86,8579 | 86,3637 | -0,00823099 | 0,9625 | 0,994606 |
| Lrrc46 | 0,740094 | 0,732329 | -0,0152164 | 0,9626 | 0,994606 |
| Asb3 | 8,02301 | 8,07775 | 0,00981 | 0,96275 | 0,994606 |
| Unc119 | 51,8682 | 52,217 | 0,00966849 | 0,96275 | 0,994606 |
| Rsu1 | 30,2977 | 30,0926 | -0,00979749 | 0,96275 | 0,994606 |
| Atl3 | 7,53152 | 7,57875 | 0,00901942 | 0,96285 | 0,994606 |
| Zfp235 | 2,71255 | 2,68823 | -0,0129956 | 0,963 | 0,994606 |
| Syne4 | 9,15551 | 9,23025 | 0,0117293 | 0,96305 | 0,994606 |
| Psmd5 | 59,8677 | 59,5216 | -0,00836533 | 0,96305 | 0,994606 |
| Derl1 | 21,5667 | 21,4363 | -0,00875059 | 0,96305 | 0,994606 |
| Nhlrc3 | 5,74451 | 5,69891 | -0,0114979 | 0,96305 | 0,994606 |
| Fam50a | 41,5045 | 41,7545 | 0,00866358 | 0,9631 | 0,994606 |
| Ccdc74a | 0,876995 | 0,856195 | -0,0346292 | 0,96315 | 0,994606 |
| Csnk2a2 | 21,092 | 20,9643 | -0,0087591 | 0,9632 | 0,994606 |
| N6amt2 | 54,3793 | 54,0391 | -0,00905435 | 0,9632 | 0,994606 |
| 2900076A07Rik | 5,9247 | 5,97899 | 0,0131601 | 0,9633 | 0,994606 |
| Tstd2 | 25,9696 | 25,8143 | -0,00865161 | 0,9633 | 0,994606 |
| Heatr5a | 9,94139 | 9,88185 | -0,0086674 | 0,96335 | 0,994606 |
| Cml1 | 1,17031 | 1,18644 | 0,019753 | 0,9634 | 0,994606 |
| Zfp28 | 2,49557 | 2,51656 | 0,0120784 | 0,9635 | 0,994606 |
| Oard1 | 65,6371 | 66,0004 | 0,00796169 | 0,9636 | 0,994606 |
| 1810037I17Rik | 24,6019 | 24,4211 | -0,0106379 | 0,96365 | 0,994606 |
| Fam3a | 8,84034 | 8,90521 | 0,0105475 | 0,9639 | 0,994606 |
| Mob4 | 28,9033 | 29,0779 | 0,00868955 | 0,96395 | 0,994606 |
| AI450353 | 2,1092 | 2,15043 | 0,027924 | 0,964 | 0,994606 |
| Pip5kl1 | 0,705233 | 0,719558 | 0,0290102 | 0,9642 | 0,994606 |
| Slc4a8 | 1,22681 | 1,2359 | 0,0106481 | 0,96425 | 0,994606 |
| Tmem64 | 7,74096 | 7,69052 | -0,00942993 | 0,96425 | 0,994606 |
| Pak3 | 0,636655 | 0,643142 | 0,0146248 | 0,9645 | 0,994606 |
| Dgat1 | 15,4804 | 15,586 | 0,00980551 | 0,9645 | 0,994606 |
| Atg4a | 1,91941 | 1,80479 | -0,0888285 | 0,9645 | 0,994606 |
| Map3k7 | 26,9936 | 27,1522 | 0,00845642 | 0,96455 | 0,994606 |
| Nsdhl | 39,0412 | 39,2699 | 0,00842541 | 0,96455 | 0,994606 |
| Zeb2 | 9,09679 | 9,14938 | 0,00831676 | 0,96455 | 0,994606 |
| Kat7 | 26,2817 | 26,4351 | 0,00839524 | 0,9646 | 0,994606 |
| Atf5 | 15,4994 | 15,7653 | 0,0245413 | 0,9649 | 0,994708 |
| Plekhb2 | 7,88501 | 7,83617 | -0,00896342 | 0,9649 | 0,994708 |
| Phpt1 | 141,907 | 141,134 | -0,00787989 | 0,965 | 0,994708 |
| Usp11 | 31,7217 | 31,5552 | -0,00759023 | 0,96515 | 0,994708 |
| Sec62 | 25,6467 | 25,5059 | -0,00794506 | 0,9652 | 0,994708 |
| Surf1 | 24,7373 | 25,1992 | 0,0266892 | 0,96545 | 0,994708 |
| Rbmx2 | 12,483 | 12,5636 | 0,00928117 | 0,9655 | 0,994708 |
| Fundc1 | 21,158 | 21,2834 | 0,00852452 | 0,9655 | 0,994708 |

| | | | | | |
|---|---|---|---|---|---|
| Syde2 | 1,77748 | 1,79185 | 0,0116202 | 0,96555 | 0,994708 |
| Qpctl | 5,26303 | 5,22106 | -0,01155 | 0,96575 | 0,994708 |
| Tubgcp4 | 21,9824 | 21,6749 | -0,020319 | 0,96575 | 0,994708 |
| Desi1 | 30,9643 | 31,1366 | 0,00800245 | 0,9659 | 0,994708 |
| Immt | 109,692 | 109,092 | -0,00791609 | 0,966 | 0,994708 |
| Fam98b | 75,651 | 75,2231 | -0,00818283 | 0,966 | 0,994708 |
| Sec24b | 23,0574 | 23,1828 | 0,00782478 | 0,9661 | 0,994708 |
| Cysltr1 | 0,896119 | 0,907688 | 0,0185058 | 0,96615 | 0,994708 |
| Pcdhga2 | 1,06736 | 1,00819 | -0,0822855 | 0,9662 | 0,994708 |
| Ggnbp1 | 1,65382 | 1,71984 | 0,0564767 | 0,96625 | 0,994708 |
| Wdfy3 | 11,9946 | 12,0597 | 0,0078069 | 0,96625 | 0,994708 |
| Ogg1 | 12,5732 | 12,4546 | -0,0136678 | 0,96625 | 0,994708 |
| Cox10 | 13,5888 | 13,6726 | 0,00886785 | 0,9666 | 0,994988 |
| 0610040J01Rik | 2,99102 | 3,01697 | 0,0124651 | 0,9669 | 0,994996 |
| Foxn2 | 8,26124 | 8,30836 | 0,00820691 | 0,96695 | 0,994996 |
| Mettl25 | 5,61282 | 5,57414 | -0,00997797 | 0,967 | 0,994996 |
| Tm7sf2 | 11,1472 | 11,0231 | -0,0161573 | 0,967 | 0,994996 |
| Mpp7 | 3,3736 | 3,39938 | 0,010982 | 0,9671 | 0,994996 |
| Ric8 | 13,1744 | 13,0922 | -0,00903015 | 0,9671 | 0,994996 |
| Atp8b2 | 16,8911 | 16,7966 | -0,00809898 | 0,96715 | 0,994996 |
| Atp6v0a2 | 13,6743 | 13,7439 | 0,00733388 | 0,9674 | 0,994996 |
| 4930520O04Rik | 0,809047 | 0,779609 | -0,0534724 | 0,96745 | 0,994996 |
| Myl6 | 55,9496 | 56,3014 | 0,00904132 | 0,9676 | 0,994996 |
| Tube1 | 9,01569 | 8,96028 | -0,00889339 | 0,9677 | 0,994996 |
| Gla | 5,58367 | 5,54687 | -0,00953884 | 0,968 | 0,994996 |
| Mgarp | 2,65512 | 2,68301 | 0,0150741 | 0,96805 | 0,994996 |
| Dhx35 | 14,3524 | 14,4294 | 0,00772537 | 0,96805 | 0,994996 |
| Xrn2 | 76,3617 | 75,971 | -0,00739934 | 0,9681 | 0,994996 |
| Serhl | 6,1692 | 6,26516 | 0,0222679 | 0,9682 | 0,994996 |
| Gnptg | 19,0505 | 18,9077 | -0,0108509 | 0,9682 | 0,994996 |
| Crebzf | 26,6503 | 26,5067 | -0,00779644 | 0,96825 | 0,994996 |
| Bcl7c | 27,353 | 27,5248 | 0,00903548 | 0,9683 | 0,994996 |
| Pcdhgb5 | 1,3973 | 1,33606 | -0,0646533 | 0,9683 | 0,994996 |
| Mras | 2,49267 | 2,47602 | -0,00967368 | 0,9684 | 0,994996 |
| Pcdhgb6 | 4,13371 | 4,2429 | 0,0376113 | 0,9685 | 0,994996 |
| 2310033P09Rik | 32,5898 | 32,4067 | -0,00813053 | 0,9686 | 0,994996 |
| Enah | 28,3033 | 28,1588 | -0,00738729 | 0,96865 | 0,994996 |
| Zfp324 | 4,67495 | 4,64687 | -0,00869372 | 0,9687 | 0,994996 |
| Cflar | 2,51385 | 2,52897 | 0,00865364 | 0,96875 | 0,994996 |
| Rnd3 | 44,5469 | 44,7866 | 0,0077423 | 0,96885 | 0,994996 |
| Dnajc21 | 28,1796 | 28,0335 | -0,00749591 | 0,96885 | 0,994996 |
| Fanci | 13,9431 | 13,8306 | -0,0116846 | 0,969 | 0,994996 |
| Sucla2 | 51,9487 | 51,7025 | -0,00685335 | 0,9691 | 0,994996 |
| Cstf2t | 27,1516 | 27,0152 | -0,00727057 | 0,9691 | 0,994996 |
| Kif2c | 53,7606 | 53,4976 | -0,00707355 | 0,96915 | 0,994996 |
| Scyl3 | 8,29229 | 8,21809 | -0,0129671 | 0,96925 | 0,994996 |
| Surf2 | 44,9561 | 44,3503 | -0,0195743 | 0,96925 | 0,994996 |
| Brms1 | 37,8552 | 38,0572 | 0,00767694 | 0,96935 | 0,994996 |
| Shisa2 | 10,5037 | 10,5609 | 0,00784289 | 0,9694 | 0,994996 |
| Bivm | 12,1104 | 12,0504 | -0,0071622 | 0,9695 | 0,995014 |

| | | | | | |
|---|---|---|---|---|---|
| Chek2 | 17,2205 | 17,1316 | -0,00747244 | 0,9696 | 0,995014 |
| Fbxo45 | 17,139 | 17,0524 | -0,00731008 | 0,96965 | 0,995014 |
| Hsf1 | 38,784 | 38,9726 | 0,00699818 | 0,9699 | 0,995106 |
| Pcdhgb1 | 0,658709 | 0,618979 | -0,0897512 | 0,97 | 0,995106 |
| Ift122 | 9,90178 | 9,95632 | 0,00792512 | 0,97005 | 0,995106 |
| Ppm1b | 21,6234 | 21,728 | 0,00696216 | 0,97005 | 0,995106 |
| Cdkl3 | 4,97952 | 4,94403 | -0,0103206 | 0,97025 | 0,995232 |
| Zfp940 | 6,28336 | 6,2463 | -0,00853386 | 0,97045 | 0,995248 |
| Nsg1 | 23,1389 | 22,9863 | -0,00954837 | 0,97045 | 0,995248 |
| Bbx | 9,56617 | 9,52218 | -0,00664927 | 0,9705 | 0,995248 |
| Kdelr2 | 77,9651 | 77,6438 | -0,0059569 | 0,97075 | 0,995248 |
| Vrk2 | 8,28149 | 8,35083 | 0,012029 | 0,9708 | 0,995248 |
| Dgkq | 5,49728 | 5,53775 | 0,0105828 | 0,9709 | 0,995248 |
| Zrsr2 | 23,1403 | 23,0265 | -0,00711065 | 0,9709 | 0,995248 |
| Wdr43 | 72,6921 | 72,3671 | -0,00646504 | 0,97095 | 0,995248 |
| Tti2 | 14,1028 | 14,2629 | 0,016289 | 0,971 | 0,995248 |
| Kpna2 | 10,7574 | 10,698 | -0,00798689 | 0,9711 | 0,995248 |
| Mterf1b | 1,01127 | 1,00386 | -0,0106168 | 0,97135 | 0,995248 |
| Ap5z1 | 3,25301 | 3,27216 | 0,00846915 | 0,9714 | 0,995248 |
| Cdc45 | 62,4669 | 62,7304 | 0,00607339 | 0,97145 | 0,995248 |
| Hist1h2bj | 5,59898 | 5,69843 | 0,025402 | 0,97155 | 0,995248 |
| B4galnt1 | 1,85901 | 1,87506 | 0,0123996 | 0,9717 | 0,995248 |
| Ubp1 | 49,6159 | 49,409 | -0,00602949 | 0,9717 | 0,995248 |
| Gm16973 | 10,6824 | 10,5453 | -0,0186328 | 0,9718 | 0,995248 |
| Laptm4a | 166,147 | 165,412 | -0,00639585 | 0,97185 | 0,995248 |
| Ebag9 | 18,154 | 18,0635 | -0,00720716 | 0,97195 | 0,995248 |
| Exosc9 | 68,766 | 68,1517 | -0,0129454 | 0,97195 | 0,995248 |
| Glt8d1 | 19,2598 | 18,998 | -0,0197503 | 0,972 | 0,995248 |
| Ift74 | 30,3712 | 30,5166 | 0,00688944 | 0,97205 | 0,995248 |
| Med13l | 8,15488 | 8,11526 | -0,00702754 | 0,97205 | 0,995248 |
| Lcp1 | 0,680746 | 0,704008 | 0,0484767 | 0,9723 | 0,995323 |
| Ap3b1 | 20,231 | 20,3219 | 0,0064654 | 0,9724 | 0,995323 |
| 0610009O20Rik | 32,0902 | 32,2343 | 0,00646317 | 0,97245 | 0,995323 |
| Ddt | 87,6765 | 87,2612 | -0,00684952 | 0,9725 | 0,995323 |
| BC003331 | 11,0245 | 10,8238 | -0,0264996 | 0,9727 | 0,995323 |
| Utp3 | 87,7256 | 88,12 | 0,00647117 | 0,97275 | 0,995323 |
| Vamp3 | 79,2725 | 78,9231 | -0,00637313 | 0,97275 | 0,995323 |
| Spata24 | 10,3607 | 10,2826 | -0,0109136 | 0,97285 | 0,995323 |
| Ewsr1 | 195,917 | 196,791 | 0,0064198 | 0,9729 | 0,995323 |
| Smarcal1 | 10,0217 | 10,0662 | 0,00639813 | 0,9729 | 0,995323 |
| Prc1 | 54,7149 | 54,4856 | -0,0060572 | 0,97305 | 0,995398 |
| Cercam | 7,99335 | 7,95228 | -0,00743155 | 0,97335 | 0,995406 |
| Impdh1 | 23,7537 | 23,8645 | 0,00671592 | 0,9734 | 0,995406 |
| Gprc5b | 10,1119 | 10,2397 | 0,0181106 | 0,9735 | 0,995406 |
| Ociad1 | 144,004 | 144,594 | 0,00589957 | 0,9735 | 0,995406 |
| Rffl | 1,68966 | 1,72655 | 0,0311575 | 0,9737 | 0,995406 |
| Sfrp4 | 0,831167 | 0,839385 | 0,0141949 | 0,97385 | 0,995406 |
| Hnrnpu | 530,914 | 533,249 | 0,00632936 | 0,97385 | 0,995406 |
| Ccpg1 | 6,02769 | 6,06755 | 0,00950824 | 0,97395 | 0,995406 |
| Alg12 | 7,37876 | 7,33975 | -0,00764842 | 0,97395 | 0,995406 |

| | | | | | |
|---|---|---|---|---|---|
| Snurf | 1,2692 | 1,29514 | 0,0291881 | 0,974 | 0,995406 |
| Qtrt1 | 31,0984 | 30,9612 | -0,00637941 | 0,97405 | 0,995406 |
| Nup54 | 32,6937 | 32,5547 | -0,00614933 | 0,9741 | 0,995406 |
| Rnf215 | 10,4689 | 10,3948 | -0,010258 | 0,9742 | 0,995406 |
| Hmgb3 | 44,5078 | 44,3219 | -0,00603615 | 0,97425 | 0,995406 |
| Fbxo31 | 5,79672 | 5,82564 | 0,00717993 | 0,9743 | 0,995406 |
| Hmgxb4 | 22,423 | 22,3344 | -0,00570957 | 0,9743 | 0,995406 |
| Mthfs | 2,30935 | 2,30449 | -0,00303454 | 0,9744 | 0,995429 |
| Tbc1d13 | 23,1364 | 23,2323 | 0,00596379 | 0,97465 | 0,995445 |
| Ube2s | 20,415 | 20,5131 | 0,00691675 | 0,975 | 0,995445 |
| Clptm1 | 28,2768 | 28,3901 | 0,00577303 | 0,975 | 0,995445 |
| Mettl24 | 1,10136 | 1,08784 | -0,0178223 | 0,975 | 0,995445 |
| Nup133 | 31,7887 | 31,6663 | -0,00556547 | 0,97505 | 0,995445 |
| Dus4l | 10,4492 | 10,3952 | -0,00748712 | 0,97505 | 0,995445 |
| Sec24c | 56,1154 | 56,3336 | 0,00559959 | 0,97515 | 0,995445 |
| Polrmt | 11,8969 | 11,3054 | -0,0735778 | 0,97515 | 0,995445 |
| Tex9 | 15,5903 | 15,501 | -0,00828588 | 0,9752 | 0,995445 |
| Pigf | 31,582 | 31,8489 | 0,0121406 | 0,97535 | 0,995445 |
| Slc25a53 | 1,66461 | 1,64896 | -0,0136333 | 0,9754 | 0,995445 |
| Paqr8 | 3,67889 | 3,6968 | 0,00700558 | 0,9755 | 0,995445 |
| Comt | 13,337 | 13,4419 | 0,011303 | 0,97555 | 0,995445 |
| Mapk12 | 22,6313 | 22,5306 | -0,00643299 | 0,97555 | 0,995445 |
| Katnb1 | 11,3223 | 11,3764 | 0,00687105 | 0,97585 | 0,995445 |
| Ndc80 | 33,1044 | 33,2422 | 0,00599076 | 0,97585 | 0,995445 |
| Ppp1r8 | 96,2955 | 95,9224 | -0,00560087 | 0,97585 | 0,995445 |
| Slc17a5 | 2,55433 | 2,57034 | 0,00901416 | 0,976 | 0,995445 |
| Echdc2 | 12,8768 | 12,813 | -0,00716818 | 0,976 | 0,995445 |
| Kras | 26,2992 | 26,2041 | -0,00522798 | 0,9761 | 0,995445 |
| Sash1 | 7,8866 | 7,85327 | -0,00611021 | 0,97615 | 0,995445 |
| Plekha2 | 1,58319 | 1,57528 | -0,00722705 | 0,97615 | 0,995445 |
| Itgb1 | 105,713 | 105,319 | -0,0053845 | 0,9762 | 0,995445 |
| Snora64 | 561,507 | 537,94 | -0,0618599 | 0,97645 | 0,99562 |
| Zfp870 | 2,95768 | 2,97197 | 0,00695401 | 0,97665 | 0,995684 |
| Usp10 | 68,8371 | 69,0616 | 0,00469731 | 0,97675 | 0,995684 |
| Traip | 15,1633 | 15,2261 | 0,00596838 | 0,97685 | 0,995684 |
| Fam203a | 12,4703 | 12,5222 | 0,00599235 | 0,9769 | 0,995684 |
| Cldn4 | 0,957986 | 0,951029 | -0,0105148 | 0,9769 | 0,995684 |
| Ankfy1 | 11,6679 | 11,7115 | 0,00537599 | 0,97705 | 0,995724 |
| Ankrd40 | 36,9368 | 36,8097 | -0,00497334 | 0,9771 | 0,995724 |
| 2810408A11Rik | 8,81518 | 8,77302 | -0,00691647 | 0,9772 | 0,995724 |
| Ubiad1 | 9,26184 | 9,22098 | -0,0063797 | 0,97725 | 0,995724 |
| Pdcd4 | 26,265 | 26,4331 | 0,00920346 | 0,9774 | 0,995759 |
| Pgap3 | 3,72952 | 3,7468 | 0,00666894 | 0,9775 | 0,995759 |
| Inpp5e | 11,0771 | 11,1604 | 0,0108155 | 0,9776 | 0,995759 |
| Hccs | 20,0082 | 20,0835 | 0,00542061 | 0,97765 | 0,995759 |
| Spsb2 | 2,20742 | 2,3294 | 0,0775981 | 0,97775 | 0,995759 |
| Xpc | 12,7239 | 12,6702 | -0,00610639 | 0,97775 | 0,995759 |
| 5830415F09Rik | 5,29406 | 5,31928 | 0,00685622 | 0,978 | 0,995811 |
| Rmi1 | 3,43753 | 3,35501 | -0,0350542 | 0,978 | 0,995811 |
| Dr1 | 29,8405 | 29,7347 | -0,00512154 | 0,9783 | 0,995811 |

| | | | | | |
|---|---|---|---|---|---|
| Ube4b | 44,4324 | 44,5921 | 0,00517551 | 0,97835 | 0,995811 |
| Trmt10b | 9,55525 | 9,51619 | -0,00590972 | 0,97835 | 0,995811 |
| Sult4a1 | 0,870826 | 0,877178 | 0,0104855 | 0,97845 | 0,995811 |
| Prox2 | 0,831227 | 0,836288 | 0,00875693 | 0,97885 | 0,995811 |
| Large | 9,69826 | 9,66434 | -0,0050549 | 0,97885 | 0,995811 |
| Comtd1 | 1,94608 | 1,99605 | 0,0365716 | 0,97905 | 0,995811 |
| Gm7102 | 1,50599 | 1,5095 | 0,00335445 | 0,97905 | 0,995811 |
| Rnf139 | 10,6064 | 10,6489 | 0,00577394 | 0,97915 | 0,995811 |
| Myo5b | 1,34258 | 1,3358 | -0,00730092 | 0,97915 | 0,995811 |
| Rnf167 | 26,5404 | 26,4442 | -0,00524137 | 0,9792 | 0,995811 |
| Myo1a | 0,68991 | 0,696594 | 0,0139103 | 0,97925 | 0,995811 |
| Top1 | 136,934 | 137,412 | 0,00502808 | 0,9793 | 0,995811 |
| Stac2 | 0,702278 | 0,699115 | -0,00651287 | 0,9793 | 0,995811 |
| Trim12c | 0,655376 | 0,660312 | 0,0108245 | 0,9794 | 0,995811 |
| Grina | 6,35539 | 6,38393 | 0,00646406 | 0,9794 | 0,995811 |
| Klhdc9 | 6,60928 | 6,57458 | -0,00759503 | 0,9794 | 0,995811 |
| Apbb2 | 3,7431 | 3,75762 | 0,00558745 | 0,9795 | 0,995811 |
| Exo1 | 28,6435 | 28,5505 | -0,00469311 | 0,97965 | 0,995811 |
| Sco1 | 8,14719 | 8,17695 | 0,00526058 | 0,97995 | 0,995811 |
| Ints7 | 38,1629 | 38,033 | -0,00492159 | 0,97995 | 0,995811 |
| Ppp2r5b | 7,21127 | 7,18133 | -0,00600259 | 0,98 | 0,995811 |
| Hmha1 | 7,23679 | 6,88192 | -0,0725394 | 0,98015 | 0,995811 |
| Ddx23 | 63,8523 | 63,6552 | -0,00446115 | 0,9802 | 0,995811 |
| Scnn1a | 0,698909 | 0,702413 | 0,00721421 | 0,98045 | 0,995811 |
| Ccdc39 | 0,78704 | 0,771854 | -0,0281079 | 0,98045 | 0,995811 |
| Trappc3 | 47,5082 | 47,3422 | -0,00505089 | 0,98055 | 0,995811 |
| Ccdc9 | 28,914 | 29,0087 | 0,00471856 | 0,98065 | 0,995811 |
| Begain | 1,76069 | 1,77208 | 0,00930157 | 0,9807 | 0,995811 |
| Harbi1 | 7,52369 | 7,55244 | 0,00550273 | 0,981 | 0,995811 |
| Ing5 | 9,99595 | 10,0286 | 0,00471026 | 0,981 | 0,995811 |
| Etf1 | 129,609 | 129,227 | -0,00426567 | 0,9812 | 0,995811 |
| Gm5803 | 2,59187 | 2,58962 | -0,00125153 | 0,9814 | 0,995811 |
| Tmco3 | 5,74905 | 5,77559 | 0,00664421 | 0,9817 | 0,995811 |
| Pot1a | 15,6763 | 15,6284 | -0,00440981 | 0,9817 | 0,995811 |
| Top2b | 69,4567 | 69,2447 | -0,00440913 | 0,98175 | 0,995811 |
| Nasp | 466,046 | 464,573 | -0,00456647 | 0,9819 | 0,995811 |
| 1110065P20Rik | 25,0603 | 25,2131 | 0,00876549 | 0,982 | 0,995811 |
| Gtpbp1 | 34,4408 | 34,5332 | 0,00386539 | 0,982 | 0,995811 |
| Eif5 | 125,592 | 125,228 | -0,00418731 | 0,982 | 0,995811 |
| Dynll2 | 119,739 | 119,427 | -0,00375525 | 0,98205 | 0,995811 |
| Cpne2 | 9,044 | 9,07361 | 0,00471439 | 0,9821 | 0,995811 |
| Man1b1 | 37,4636 | 37,5776 | 0,00438151 | 0,98215 | 0,995811 |
| Pigs | 20,8771 | 20,9371 | 0,00414451 | 0,9822 | 0,995811 |
| Tlr2 | 1,15204 | 1,16137 | 0,0116349 | 0,98225 | 0,995811 |
| Eif2ak2 | 5,61539 | 5,63209 | 0,00428324 | 0,98235 | 0,995811 |
| Riok1 | 23,7932 | 23,7209 | -0,00438772 | 0,98235 | 0,995811 |
| Rars2 | 29,7317 | 29,6415 | -0,00438477 | 0,9824 | 0,995811 |
| Hexim2 | 2,70931 | 2,69446 | -0,00792497 | 0,98245 | 0,995811 |
| A730098P11Rik | 2,82638 | 2,80265 | -0,0121638 | 0,98255 | 0,995811 |
| Aifm1 | 39,8322 | 39,9396 | 0,00388373 | 0,9826 | 0,995811 |

| | | | | | |
|---|---|---|---|---|---|
| Dpysl2 | 15,6215 | 15,5766 | -0,00415261 | 0,9826 | 0,995811 |
| Gm10658 | 1,48213 | 1,52269 | 0,0389461 | 0,9827 | 0,995811 |
| Dars | 106,681 | 106,412 | -0,00363965 | 0,98275 | 0,995811 |
| Cd59a | 1,23586 | 1,23873 | 0,00334103 | 0,9829 | 0,995811 |
| Map3k3 | 20,9879 | 20,9237 | -0,00441972 | 0,9829 | 0,995811 |
| Zfp651 | 3,44454 | 3,43277 | -0,00493624 | 0,98295 | 0,995811 |
| Rps25 | 15,9272 | 15,9997 | 0,00655141 | 0,983 | 0,995811 |
| Acap2 | 12,2346 | 12,1854 | -0,00581819 | 0,983 | 0,995811 |
| Tradd | 2,12068 | 2,13933 | 0,0126278 | 0,98305 | 0,995811 |
| Yipf2 | 8,54219 | 8,44434 | -0,0166208 | 0,98315 | 0,995811 |
| B4galt7 | 10,582 | 10,5419 | -0,00548203 | 0,9832 | 0,995811 |
| Ccdc151 | 1,40334 | 1,39561 | -0,00797433 | 0,9832 | 0,995811 |
| Trpm4 | 0,871997 | 0,905454 | 0,0543167 | 0,9834 | 0,995811 |
| Rad17 | 20,4608 | 20,5174 | 0,00398362 | 0,9836 | 0,995811 |
| Trappc2l | 33,6143 | 33,707 | 0,00397139 | 0,984 | 0,995811 |
| Fam53a | 29,0941 | 29,0143 | -0,00396215 | 0,98405 | 0,995811 |
| Srrd | 16,4786 | 16,5924 | 0,00993603 | 0,9841 | 0,995811 |
| Nde1 | 114,974 | 114,694 | -0,00351739 | 0,9841 | 0,995811 |
| Dctn2 | 102,058 | 101,767 | -0,0041066 | 0,98415 | 0,995811 |
| Trim32 | 24,2107 | 24,275 | 0,00382898 | 0,9842 | 0,995811 |
| Ndrg1 | 0,995525 | 0,990836 | -0,00681141 | 0,9842 | 0,995811 |
| Pik3c3 | 14,88 | 14,9181 | 0,00368765 | 0,9844 | 0,995811 |
| Chpt1 | 6,80736 | 6,78241 | -0,00529773 | 0,9845 | 0,995811 |
| Eml5 | 1,06313 | 1,05217 | -0,014947 | 0,9845 | 0,995811 |
| D130017N08Rik | 2,20438 | 2,21509 | 0,00699676 | 0,9846 | 0,995811 |
| Tmem176b | 15,7911 | 15,8432 | 0,00475335 | 0,9847 | 0,995811 |
| Rrp15 | 79,3405 | 79,5425 | 0,00366881 | 0,9847 | 0,995811 |
| Egfl8 | 1,14122 | 1,1189 | -0,0284974 | 0,98475 | 0,995811 |
| Phyh | 16,0581 | 16,0107 | -0,00426342 | 0,98485 | 0,995811 |
| Pcdhga8 | 0,733505 | 0,710553 | -0,0458645 | 0,98485 | 0,995811 |
| Plgrkt | 20,8086 | 20,8826 | 0,00511943 | 0,9849 | 0,995811 |
| Efemp1 | 1,92936 | 1,93007 | 0,000531232 | 0,9851 | 0,995811 |
| Rtf1 | 40,3225 | 40,2296 | -0,00332891 | 0,98515 | 0,995811 |
| Mtrr | 10,6376 | 10,6661 | 0,00385902 | 0,98525 | 0,995811 |
| Mospd3 | 5,95715 | 5,93526 | -0,00531118 | 0,9853 | 0,995811 |
| Pten | 7,92947 | 7,91095 | -0,00337505 | 0,98535 | 0,995811 |
| Gtf3c4 | 27,3824 | 27,4417 | 0,00312026 | 0,9854 | 0,995811 |
| Cib2 | 6,02517 | 5,99752 | -0,00663707 | 0,9854 | 0,995811 |
| Fbxo4 | 0,942818 | 0,945689 | 0,00438558 | 0,98545 | 0,995811 |
| AI464131 | 0,917128 | 0,913938 | -0,00502698 | 0,98545 | 0,995811 |
| Ppat | 44,7468 | 44,5444 | -0,00653916 | 0,9855 | 0,995811 |
| Cdc23 | 55,9243 | 55,6746 | -0,00645725 | 0,98555 | 0,995811 |
| Hap1 | 3,92961 | 3,9173 | -0,00452809 | 0,98565 | 0,995811 |
| 2610524H06Rik | 19,7094 | 19,7622 | 0,0038575 | 0,9857 | 0,995811 |
| Ddx5 | 231,049 | 231,606 | 0,00347732 | 0,9858 | 0,995811 |
| Tpm4 | 274,641 | 275,267 | 0,00328319 | 0,9858 | 0,995811 |
| Pdgfa | 12,7816 | 12,7436 | -0,00429368 | 0,98585 | 0,995811 |
| Tbc1d23 | 8,22018 | 8,19977 | -0,00358637 | 0,9859 | 0,995811 |
| Rel | 1,62509 | 1,6334 | 0,00735253 | 0,9862 | 0,995811 |
| Stil | 12,0511 | 12,079 | 0,00334208 | 0,98625 | 0,995811 |

| | | | | | |
|---|---|---|---|---|---|
| Zfp248 | 5,09805 | 5,08391 | -0,00400724 | 0,9863 | 0,995811 |
| Acadvl | 16,6431 | 16,6841 | 0,00355278 | 0,98635 | 0,995811 |
| Zc3h14 | 47,7426 | 47,8544 | 0,00337471 | 0,9864 | 0,995811 |
| Tgfbr1 | 43,0273 | 42,9409 | -0,00290022 | 0,9864 | 0,995811 |
| Pfkfb3 | 11,1776 | 11,205 | 0,00353197 | 0,9865 | 0,995811 |
| Cnot10 | 39,1737 | 39,2558 | 0,00302301 | 0,9865 | 0,995811 |
| Snx33 | 4,43877 | 4,45099 | 0,00396609 | 0,98655 | 0,995811 |
| Pam | 10,8595 | 10,8367 | -0,00302912 | 0,9866 | 0,995811 |
| 1200014J11Rik | 18,0708 | 18,0362 | -0,00276416 | 0,9867 | 0,995811 |
| Orc3 | 17,4816 | 17,5376 | 0,00461419 | 0,9868 | 0,995811 |
| Tmem128 | 55,1878 | 55,0724 | -0,00302071 | 0,98695 | 0,995811 |
| Btd | 8,96428 | 8,93835 | -0,00417911 | 0,98695 | 0,995811 |
| Tox4 | 18,7843 | 18,8413 | 0,0043756 | 0,987 | 0,995811 |
| Dhrs11 | 3,1837 | 3,18137 | -0,00105895 | 0,987 | 0,995811 |
| Exoc1 | 21,9925 | 22,0354 | 0,00281576 | 0,98725 | 0,995811 |
| Fzd3 | 11,1524 | 11,1306 | -0,00283212 | 0,98725 | 0,995811 |
| Spef1 | 4,12811 | 4,06785 | -0,0212153 | 0,98725 | 0,995811 |
| Gopc | 12,6804 | 12,7071 | 0,00303789 | 0,9873 | 0,995811 |
| Cpsf7 | 43,0332 | 42,947 | -0,00289096 | 0,98735 | 0,995811 |
| Taf13 | 17,5685 | 17,5256 | -0,00353378 | 0,98735 | 0,995811 |
| Gm16740 | 1,85687 | 1,83948 | -0,0135751 | 0,98745 | 0,995834 |
| Mtmr2 | 30,2882 | 30,3444 | 0,00267126 | 0,9877 | 0,995891 |
| Enox1 | 1,11018 | 1,10708 | -0,00403504 | 0,98775 | 0,995891 |
| Gm10509 | 2,34847 | 2,34114 | -0,00451139 | 0,9878 | 0,995891 |
| Pcdhga5 | 1,56277 | 1,58516 | 0,0205238 | 0,9879 | 0,995891 |
| BC023829 | 19,2484 | 19,2915 | 0,0032264 | 0,98795 | 0,995891 |
| 4930481A15Rik | 2,68584 | 2,71631 | 0,0162756 | 0,98805 | 0,995891 |
| Rilpl1 | 17,9442 | 17,9052 | -0,00314402 | 0,98805 | 0,995891 |
| Pigp | 25,9151 | 26,2535 | 0,0187195 | 0,98825 | 0,995948 |
| Hoxa11 | 44,8269 | 45,0019 | 0,00562096 | 0,9884 | 0,995948 |
| Tmed7 | 35,6972 | 35,7582 | 0,0024659 | 0,98845 | 0,995948 |
| Fam161b | 1,67554 | 1,67694 | 0,00121214 | 0,9885 | 0,995948 |
| Trmt12 | 4,51179 | 4,52086 | 0,00289857 | 0,98855 | 0,995948 |
| Papolb | 1,41115 | 1,40484 | -0,00646235 | 0,9886 | 0,995948 |
| 4933434E20Rik | 19,1106 | 19,0029 | -0,00815223 | 0,98865 | 0,995948 |
| Pqlc2 | 2,55005 | 2,54655 | -0,00197971 | 0,98895 | 0,996144 |
| Lzic | 39,6089 | 39,5401 | -0,0025075 | 0,989 | 0,996144 |
| Gbf1 | 16,6239 | 16,6522 | 0,00245838 | 0,9891 | 0,996167 |
| Eya3 | 28,3321 | 28,3868 | 0,00278075 | 0,9893 | 0,99629 |
| Dph3 | 21,3156 | 21,2734 | -0,00286116 | 0,98955 | 0,996458 |
| Suv39h1 | 60,4197 | 60,3304 | -0,00213399 | 0,98965 | 0,996458 |
| Btbd19 | 2,22972 | 2,2347 | 0,00321984 | 0,9897 | 0,996458 |
| Rnf2 | 56,1558 | 56,0639 | -0,00236271 | 0,98985 | 0,996531 |
| Cib1 | 39,8479 | 39,778 | -0,00253302 | 0,99 | 0,996604 |
| Ncaph | 64,071 | 63,9726 | -0,00221681 | 0,99015 | 0,996649 |
| Thumpd2 | 7,24438 | 7,22735 | -0,00339488 | 0,9902 | 0,996649 |
| Amfr | 29,9574 | 29,9119 | -0,00218896 | 0,9903 | 0,996666 |
| Kit | 5,35178 | 5,35977 | 0,00215232 | 0,9904 | 0,996666 |
| Lacc1 | 2,80462 | 2,79884 | -0,00297486 | 0,99045 | 0,996666 |
| Fubp3 | 37,9915 | 37,9263 | -0,00247974 | 0,99065 | 0,996789 |

| | | | | | |
|---|---|---|---|---|---|
| Med25 | 45,2323 | 45,2918 | 0,00189626 | 0,9908 | 0,996861 |
| 4930549G23Rik | 4,38846 | 4,34012 | -0,0159795 | 0,99105 | 0,997035 |
| Tgs1 | 22,9943 | 23,025 | 0,00192678 | 0,99125 | 0,997079 |
| Ccbl1 | 4,09633 | 4,09901 | 0,000943597 | 0,99125 | 0,99708 |
| Proser1 | 15,9065 | 15,8843 | -0,00201488 | 0,9914 | 0,997152 |
| Cyfip1 | 30,2422 | 30,1973 | -0,00214376 | 0,9916 | 0,997275 |
| Rnf4 | 78,9951 | 79,1016 | 0,00194216 | 0,9918 | 0,997342 |
| Mettl5 | 16,393 | 16,2405 | -0,013491 | 0,99185 | 0,997342 |
| 4930404I05Rik | 2,50623 | 2,53736 | 0,0178096 | 0,9919 | 0,997342 |
| Tnfrsf1a | 16,3331 | 16,3115 | -0,0019114 | 0,99215 | 0,997488 |
| Csf2ra | 5,29496 | 5,29389 | -0,00029127 | 0,9922 | 0,997488 |
| Pcsk5 | 6,56226 | 6,57094 | 0,00190695 | 0,9923 | 0,99751 |
| Sptssa | 76,6598 | 76,7622 | 0,00192611 | 0,9924 | 0,997532 |
| AA987161 | 1,79958 | 1,80293 | 0,00268966 | 0,99255 | 0,997605 |
| Acox1 | 10,9549 | 10,971 | 0,00212557 | 0,9928 | 0,9977 |
| Ppifos | 1,91078 | 1,90397 | -0,0051507 | 0,9928 | 0,9977 |
| Tspyl5 | 2,18318 | 2,18821 | 0,00331591 | 0,9929 | 0,997711 |
| Cul2 | 27,4198 | 27,4543 | 0,00181256 | 0,99305 | 0,997711 |
| Tmem164 | 34,4006 | 34,438 | 0,00156884 | 0,9931 | 0,997711 |
| Abce1 | 88,3899 | 88,284 | -0,00172917 | 0,99315 | 0,997711 |
| Acsl3 | 41,3309 | 41,39 | 0,00206165 | 0,9932 | 0,997711 |
| LOC101669761 | 2,58255 | 2,59577 | 0,00736428 | 0,99335 | 0,997784 |
| Dhrs13 | 8,01494 | 8,02402 | 0,00163263 | 0,9935 | 0,997856 |
| Eepd1 | 3,74691 | 3,74586 | -0,000403371 | 0,99385 | 0,997912 |
| Slc2a4 | 4,43774 | 4,44338 | 0,00183004 | 0,994 | 0,997912 |
| Surf4 | 92,9268 | 92,8352 | -0,00142345 | 0,99405 | 0,997912 |
| Prcc | 44,7886 | 44,8363 | 0,00153436 | 0,99415 | 0,997912 |
| Raver1 | 71,6485 | 71,585 | -0,00127912 | 0,99415 | 0,997912 |
| Prkar2a | 36,7493 | 36,7118 | -0,00147389 | 0,99425 | 0,997912 |
| Fam76a | 5,42743 | 5,42152 | -0,00157256 | 0,99425 | 0,997912 |
| Slc7a6os | 47,0086 | 46,9635 | -0,00138375 | 0,9943 | 0,997912 |
| Smc1a | 93,6529 | 93,5733 | -0,00122641 | 0,9944 | 0,997912 |
| Fam60a | 4,70032 | 4,69742 | -0,000891875 | 0,99445 | 0,997912 |
| Vps35 | 69,5175 | 69,5789 | 0,00127387 | 0,99465 | 0,997912 |
| Wipi2 | 33,3872 | 33,3617 | -0,00110358 | 0,99465 | 0,997912 |
| Lmln | 2,66476 | 2,66195 | -0,00151965 | 0,99465 | 0,997912 |
| Gmip | 3,24628 | 3,24534 | -0,000414776 | 0,99485 | 0,997912 |
| Bicd2 | 14,5645 | 14,5503 | -0,00141116 | 0,99485 | 0,997912 |
| Pigh | 9,98554 | 9,97186 | -0,00197747 | 0,9949 | 0,997912 |
| Tgoln1,Tgoln2 | 13,8834 | 13,8961 | 0,00132515 | 0,99505 | 0,997912 |
| Mxd1 | 2,65718 | 2,65542 | -0,000956139 | 0,99505 | 0,997912 |
| Tkt | 215,472 | 215,318 | -0,00103107 | 0,99515 | 0,997912 |
| Galns | 2,60325 | 2,60542 | 0,00120499 | 0,9952 | 0,997912 |
| Xrcc3 | 12,9281 | 12,9461 | 0,00201112 | 0,9953 | 0,997912 |
| 4933431E20Rik | 3,6628 | 3,66903 | 0,00245056 | 0,99535 | 0,997912 |
| Zfp865 | 10,071 | 10,0782 | 0,0010395 | 0,9955 | 0,997912 |
| Nt5m | 16,8469 | 16,8505 | 0,000309546 | 0,9955 | 0,997912 |
| Exoc8 | 2,43173 | 2,43197 | 0,000145401 | 0,9955 | 0,997912 |
| Tpcn1 | 12,8305 | 12,819 | -0,00129908 | 0,9957 | 0,997973 |
| E530011L22Rik | 0,86537 | 0,862826 | -0,0042465 | 0,9958 | 0,997973 |

| | | | | | |
|---|---|---|---|---|---|
| Nup85 | 105,939 | 105,873 | -0,000901912 | 0,9959 | 0,997973 |
| 2310061I04Rik | 16,813 | 16,8054 | -0,000651656 | 0,99595 | 0,997973 |
| Mrps6 | 76,8981 | 76,838 | -0,00112822 | 0,99595 | 0,997973 |
| Dennd6a | 14,0818 | 14,0716 | -0,00104429 | 0,9962 | 0,998012 |
| Prr11 | 17,8637 | 17,9035 | 0,00320785 | 0,99625 | 0,998012 |
| Hcfc1r1 | 9,84789 | 9,83191 | -0,00234391 | 0,99625 | 0,998012 |
| Abhd5 | 8,21028 | 8,2052 | -0,000893606 | 0,9963 | 0,998012 |
| Lsm10 | 21,4512 | 21,4672 | 0,00108147 | 0,99655 | 0,998178 |
| Tatdn1 | 9,35339 | 9,35067 | -0,000419118 | 0,99665 | 0,998178 |
| Alkbh6 | 28,9021 | 28,9306 | 0,0014172 | 0,9967 | 0,998178 |
| Pnpt1 | 34,5471 | 34,531 | -0,000674009 | 0,9969 | 0,998223 |
| Abhd4 | 12,5448 | 12,5374 | -0,000850398 | 0,9969 | 0,998223 |
| Ctc1 | 32,8195 | 32,8331 | 0,000597007 | 0,99745 | 0,998696 |
| Naa10 | 82,4606 | 82,4246 | -0,000630022 | 0,9977 | 0,998868 |
| Cars | 29,453 | 29,4382 | -0,00072351 | 0,9978 | 0,99889 |
| Timm17a | 58,6222 | 58,651 | 0,000709529 | 0,99815 | 0,999135 |
| Tubb4b | 112,964 | 112,997 | 0,000419108 | 0,9982 | 0,999135 |
| Igsf10 | 2,8252 | 2,82581 | 0,000313104 | 0,9983 | 0,999157 |
| Srbd1 | 20,2307 | 20,2364 | 0,000406878 | 0,9984 | 0,999179 |
| Ift81 | 17,3972 | 17,3909 | -0,000528152 | 0,9986 | 0,999245 |
| Gorasp1 | 11,6848 | 11,6772 | -0,000929135 | 0,99865 | 0,999245 |
| Arl5b | 18,8716 | 18,8718 | 1,87E-005 | 0,9987 | 0,999245 |
| Rnf126 | 53,375 | 53,4362 | 0,00165324 | 0,99905 | 0,999518 |
| Cpsf3l | 42,4907 | 42,4817 | -0,000306552 | 0,9993 | 0,999662 |
| Stip1 | 265,421 | 265,379 | -0,000229507 | 0,99935 | 0,999662 |
| Dguok | 19,7526 | 19,7551 | 0,000182446 | 0,9995 | 0,999734 |
| Hcfc1 | 65,4338 | 65,4351 | 2,87E-005 | 0,99965 | 0,999778 |
| Vmac | 3,67029 | 3,67042 | 5,18E-005 | 0,9997 | 0,999778 |
| Rnpc3 | 13,6709 | 13,6703 | -5,59E-005 | 0,99995 | 0,99995 |
| Snord19 | 887,74 | 0 | -inf | 1 | 1 |
| Snord99 | 543,631 | 0 | -inf | 1 | 1 |
| Mir7687 | 306,216 | 0 | -inf | 1 | 1 |
| Mir3620 | 293,627 | 0 | -inf | 1 | 1 |
| Scarna2 | 223,873 | 0 | -inf | 1 | 1 |
| Mir130a | 196,743 | 0 | -inf | 1 | 1 |
| Mir7024 | 194,332 | 0 | -inf | 1 | 1 |
| Mir707 | 163,51 | 0 | -inf | 1 | 1 |
| Mir7073 | 150,005 | 0 | -inf | 1 | 1 |
| Snord71 | 150,005 | 0 | -inf | 1 | 1 |
| Mir3097 | 106,706 | 0 | -inf | 1 | 1 |
| Snord73a | 96,2356 | 0 | -inf | 1 | 1 |
| Mir7578 | 63,1648 | 0 | -inf | 1 | 1 |
| Mir1943 | 60,887 | 0 | -inf | 1 | 1 |
| Mir5135 | 50,6042 | 0 | -inf | 1 | 1 |
| Mir3091 | 47,9564 | 0 | -inf | 1 | 1 |
| Mir19b-1 | 46,6574 | 0 | -inf | 1 | 1 |
| Mir3058 | 37,4588 | 0 | -inf | 1 | 1 |
| Mir5113 | 32,3348 | 0 | -inf | 1 | 1 |
| Mir20a | 23,5178 | 0 | -inf | 1 | 1 |
| Mir3101 | 22,0448 | 0 | -inf | 1 | 1 |

| | | | | | |
|---|---|---|---|---|---|
| Mir709 | 22,0448 | 0 | -inf | 1 | 1 |
| Mirlet7d | 21,1897 | 0 | -inf | 1 | 1 |
| Mir7064 | 20,8561 | 0 | -inf | 1 | 1 |
| Mir341 | 19,0993 | 0 | -inf | 1 | 1 |
| Mir1955 | 17,3755 | 0 | -inf | 1 | 1 |
| Snora2b | 14,738 | 0 | -inf | 1 | 1 |
| Mir433 | 10,3562 | 0 | -inf | 1 | 1 |
| Mir3102 | 10,1523 | 0 | -inf | 1 | 1 |
| Mir6239 | 9,73437 | 0 | -inf | 1 | 1 |
| Mir701 | 8,27596 | 0 | -inf | 1 | 1 |
| Mir8098 | 6,6604 | 0 | -inf | 1 | 1 |
| Scarna3b | 4,82429 | 0 | -inf | 1 | 1 |
| Mir3473g | 3,32781 | 0 | -inf | 1 | 1 |
| Pf4 | 0,874987 | 0 | -inf | 1 | 1 |
| Lbx1 | 0,707258 | 0 | -inf | 1 | 1 |
| Nnmt | 0,688395 | 0 | -inf | 1 | 1 |
| Mir5131 | 0,635206 | 0 | -inf | 1 | 1 |
| Mir431 | 0,620775 | 0 | -inf | 1 | 1 |
| Clic3 | 0,61999 | 0 | -inf | 1 | 1 |
| 1700018A04Rik | 0,58304 | 0 | -inf | 1 | 1 |
| Hpgd | 0,552563 | 0 | -inf | 1 | 1 |
| Hist1h3b | 0,508896 | 0 | -inf | 1 | 1 |
| Scarletltr | 0,505143 | 0 | -inf | 1 | 1 |
| Gm11346 | 0,494164 | 0 | -inf | 1 | 1 |
| Hbb-b1,Hbb-bs | 0,471815 | 0 | -inf | 1 | 1 |
| Guca1a | 0,46715 | 0 | -inf | 1 | 1 |
| Hist3h2bb-ps | 0,442971 | 0 | -inf | 1 | 1 |
| Adig | 0,408562 | 0 | -inf | 1 | 1 |
| Nupr1 | 0,37842 | 0 | -inf | 1 | 1 |
| Zic4 | 0,373671 | 0 | -inf | 1 | 1 |
| AA467197 | 0,369568 | 0 | -inf | 1 | 1 |
| AU021063 | 0,34697 | 0 | -inf | 1 | 1 |
| Ifnk | 0,344279 | 0 | -inf | 1 | 1 |
| Tmem163 | 0,337554 | 0 | -inf | 1 | 1 |
| Gm12830 | 0,336398 | 0 | -inf | 1 | 1 |
| Trim55 | 0,335737 | 0 | -inf | 1 | 1 |
| Npy | 0,318916 | 0 | -inf | 1 | 1 |
| Gm6607 | 0,312501 | 0 | -inf | 1 | 1 |
| Cidea | 0,304089 | 0 | -inf | 1 | 1 |
| Hamp | 0,304074 | 0 | -inf | 1 | 1 |
| Wnt2 | 0,304018 | 0 | -inf | 1 | 1 |
| Shh | 0,286782 | 0 | -inf | 1 | 1 |
| 1700019D03Rik | 0,277566 | 0 | -inf | 1 | 1 |
| 1700108J01Rik | 0,276435 | 0 | -inf | 1 | 1 |
| Gm15104 | 0,275238 | 0 | -inf | 1 | 1 |
| Gm21057 | 0,27268 | 0 | -inf | 1 | 1 |
| Ctla2b | 0,270137 | 0 | -inf | 1 | 1 |
| Lhx3 | 0,265387 | 0 | -inf | 1 | 1 |
| Spink3 | 0,265157 | 0 | -inf | 1 | 1 |
| Ascl1 | 0,264608 | 0 | -inf | 1 | 1 |

| | | | | | |
|---|---|---|---|---|---|
| Gm6042 | 0,261866 | 0 | -inf | 1 | 1 |
| Dpt | 0,260624 | 0 | -inf | 1 | 1 |
| Sstr4 | 0,253574 | 0 | -inf | 1 | 1 |
| Tdo2 | 0,243736 | 0 | -inf | 1 | 1 |
| Lgals7 | 0,241775 | 0 | -inf | 1 | 1 |
| 1700092C10Rik | 0,24151 | 0 | -inf | 1 | 1 |
| Insc | 0,229551 | 0 | -inf | 1 | 1 |
| Gstp2 | 0,226137 | 0 | -inf | 1 | 1 |
| 1700018G05Rik | 0,225929 | 0 | -inf | 1 | 1 |
| Cntnap4 | 0,223735 | 0 | -inf | 1 | 1 |
| Kcnmb2 | 0,222106 | 0 | -inf | 1 | 1 |
| Gm15350 | 0,221907 | 0 | -inf | 1 | 1 |
| Tmem217 | 0,22028 | 0 | -inf | 1 | 1 |
| Uts2 | 0,218502 | 0 | -inf | 1 | 1 |
| Cbr2 | 0,214036 | 0 | -inf | 1 | 1 |
| C330013F16Rik | 0,209302 | 0 | -inf | 1 | 1 |
| 1700026F02Rik | 0,207942 | 0 | -inf | 1 | 1 |
| 1810013A23Rik | 0,206903 | 0 | -inf | 1 | 1 |
| Inmt | 0,205129 | 0 | -inf | 1 | 1 |
| Npffr2 | 0,201868 | 0 | -inf | 1 | 1 |
| Grip1os2 | 0,196317 | 0 | -inf | 1 | 1 |
| Efcab9 | 0,19572 | 0 | -inf | 1 | 1 |
| 9530026P05Rik | 0,19321 | 0 | -inf | 1 | 1 |
| Spem1 | 0,193102 | 0 | -inf | 1 | 1 |
| Asb12 | 0,191605 | 0 | -inf | 1 | 1 |
| Reg4 | 0,188822 | 0 | -inf | 1 | 1 |
| Rab11fip4os2 | 0,188164 | 0 | -inf | 1 | 1 |
| 2610528J11Rik | 0,185976 | 0 | -inf | 1 | 1 |
| Ppp2r2b | 0,185241 | 0 | -inf | 1 | 1 |
| 9430069I07Rik | 0,181325 | 0 | -inf | 1 | 1 |
| 2010005H15Rik | 0,178421 | 0 | -inf | 1 | 1 |
| Esp5 | 0,177223 | 0 | -inf | 1 | 1 |
| Prima1 | 0,176782 | 0 | -inf | 1 | 1 |
| Gpx6 | 0,173131 | 0 | -inf | 1 | 1 |
| 1810012K16Rik | 0,171031 | 0 | -inf | 1 | 1 |
| Mospd4 | 0,170278 | 0 | -inf | 1 | 1 |
| Gm4971 | 0,166503 | 0 | -inf | 1 | 1 |
| Slitrk6 | 0,164009 | 0 | -inf | 1 | 1 |
| Dynlrb2 | 0,1638 | 0 | -inf | 1 | 1 |
| 1600010M07Rik | 0,16047 | 0 | -inf | 1 | 1 |
| 1700001D01Rik | 0,160134 | 0 | -inf | 1 | 1 |
| Otor | 0,154762 | 0 | -inf | 1 | 1 |
| 03/11/16 | 0,15173 | 0 | -inf | 1 | 1 |
| Ankrd60 | 0,151729 | 0 | -inf | 1 | 1 |
| Gm4984 | 0,150124 | 0 | -inf | 1 | 1 |
| Mc2r | 0,15001 | 0 | -inf | 1 | 1 |
| Gnat3 | 0,143398 | 0 | -inf | 1 | 1 |
| Mybph | 0,143187 | 0 | -inf | 1 | 1 |
| Ttc36 | 0,142584 | 0 | -inf | 1 | 1 |
| Gata5os | 0,142361 | 0 | -inf | 1 | 1 |

| | | | | | |
|---|---|---|---|---|---|
| Gm15217 | 0,140668 | 0 | -inf | 1 | 1 |
| Slco1a5 | 0,140668 | 0 | -inf | 1 | 1 |
| Myoz1 | 0,140211 | 0 | -inf | 1 | 1 |
| Gm4489 | 0,139479 | 0 | -inf | 1 | 1 |
| Gata5 | 0,137996 | 0 | -inf | 1 | 1 |
| Gm14393 | 0,135384 | 0 | -inf | 1 | 1 |
| Gm16894 | 0,13479 | 0 | -inf | 1 | 1 |
| Gm4566 | 0,134376 | 0 | -inf | 1 | 1 |
| Gm6213 | 0,1332 | 0 | -inf | 1 | 1 |
| Wnt10b | 0,132699 | 0 | -inf | 1 | 1 |
| Npffr1 | 0,132333 | 0 | -inf | 1 | 1 |
| Gpr52 | 0,131088 | 0 | -inf | 1 | 1 |
| 4930547E08Rik | 0,130787 | 0 | -inf | 1 | 1 |
| 4933417E11Rik | 0,130056 | 0 | -inf | 1 | 1 |
| Gtsf1 | 0,127633 | 0 | -inf | 1 | 1 |
| Hist1h2bn | 0,127328 | 0 | -inf | 1 | 1 |
| Slc32a1 | 0,126543 | 0 | -inf | 1 | 1 |
| 2310007B03Rik | 0,124496 | 0 | -inf | 1 | 1 |
| 4930452G13Rik | 0,124147 | 0 | -inf | 1 | 1 |
| Scgb1c1 | 0,12405 | 0 | -inf | 1 | 1 |
| 1010001N08Rik | 0,123782 | 0 | -inf | 1 | 1 |
| 1700006F04Rik | 0,122995 | 0 | -inf | 1 | 1 |
| 4930429F24Rik | 0,122443 | 0 | -inf | 1 | 1 |
| Iigp1 | 0,12228 | 0 | -inf | 1 | 1 |
| Clec18a | 0,121924 | 0 | -inf | 1 | 1 |
| Olfr90 | 0,121005 | 0 | -inf | 1 | 1 |
| E130215H24Rik | 0,120427 | 0 | -inf | 1 | 1 |
| Defb48 | 0,119933 | 0 | -inf | 1 | 1 |
| 8430423G03Rik | 0,117109 | 0 | -inf | 1 | 1 |
| Spink4 | 0,116528 | 0 | -inf | 1 | 1 |
| Gm17359 | 0,115382 | 0 | -inf | 1 | 1 |
| Hist1h4a | 0,115155 | 0 | -inf | 1 | 1 |
| Slc51a | 0,114052 | 0 | -inf | 1 | 1 |
| Tomm20l | 0,112438 | 0 | -inf | 1 | 1 |
| Inhba | 0,110161 | 0 | -inf | 1 | 1 |
| Tsga13 | 0,108737 | 0 | -inf | 1 | 1 |
| Epyc | 0,105133 | 0 | -inf | 1 | 1 |
| 4930474N09Rik | 0,104821 | 0 | -inf | 1 | 1 |
| Ppy | 0,104724 | 0 | -inf | 1 | 1 |
| Gm6880 | 0,104198 | 0 | -inf | 1 | 1 |
| Agr2 | 0,103973 | 0 | -inf | 1 | 1 |
| 4930521E06Rik | 0,103962 | 0 | -inf | 1 | 1 |
| Gm9920 | 0,103741 | 0 | -inf | 1 | 1 |
| Gm13446 | 0,103677 | 0 | -inf | 1 | 1 |
| Lyz2 | 0,101658 | 0 | -inf | 1 | 1 |
| Dppa5a | 0,101324 | 0 | -inf | 1 | 1 |
| BC107364 | 0,100711 | 0 | -inf | 1 | 1 |
| Gm16701 | 0,0997009 | 0 | -inf | 1 | 1 |
| 4930483J18Rik | 0,0990481 | 0 | -inf | 1 | 1 |
| Krt13 | 0,0979758 | 0 | -inf | 1 | 1 |

| | | | | | |
|---|---|---|---|---|---|
| Ccdc152 | 0,0973221 | 0 | -inf | 1 | 1 |
| A230065H16Rik | 0,0972975 | 0 | -inf | 1 | 1 |
| Sncb | 0,0957859 | 0 | -inf | 1 | 1 |
| Sfta2 | 0,0957088 | 0 | -inf | 1 | 1 |
| Upk3bl | 0,0951703 | 0 | -inf | 1 | 1 |
| Nmu | 0,0943057 | 0 | -inf | 1 | 1 |
| Rptoros | 0,0940949 | 0 | -inf | 1 | 1 |
| E530001F21Rik | 0,0938471 | 0 | -inf | 1 | 1 |
| Efcab10 | 0,0935793 | 0 | -inf | 1 | 1 |
| Tnfrsf13b | 0,0933015 | 0 | -inf | 1 | 1 |
| Clrn1 | 0,0931587 | 0 | -inf | 1 | 1 |
| Pdzk1ip1 | 0,0929332 | 0 | -inf | 1 | 1 |
| Ube2d2b | 0,0926287 | 0 | -inf | 1 | 1 |
| 2310015B20Rik | 0,0922064 | 0 | -inf | 1 | 1 |
| BC039966 | 0,0908935 | 0 | -inf | 1 | 1 |
| Gm6994 | 0,090422 | 0 | -inf | 1 | 1 |
| Gm17660 | 0,0903254 | 0 | -inf | 1 | 1 |
| Fezf1 | 0,0903235 | 0 | -inf | 1 | 1 |
| H2afb1 | 0,090044 | 0 | -inf | 1 | 1 |
| Bach2os | 0,0891542 | 0 | -inf | 1 | 1 |
| Nts | 0,0888471 | 0 | -inf | 1 | 1 |
| Gm15008 | 0,0886898 | 0 | -inf | 1 | 1 |
| Olfr545 | 0,0885066 | 0 | -inf | 1 | 1 |
| 9030404E10Rik | 0,0877144 | 0 | -inf | 1 | 1 |
| Hist1h2ad | 0,0873238 | 0 | -inf | 1 | 1 |
| Otud6a | 0,0864118 | 0 | -inf | 1 | 1 |
| 2610207O16Rik | 0,0857487 | 0 | -inf | 1 | 1 |
| Ntrk1 | 0,0854757 | 0 | -inf | 1 | 1 |
| Myf6 | 0,0844377 | 0 | -inf | 1 | 1 |
| Gm16336 | 0,0843124 | 0 | -inf | 1 | 1 |
| Gm5166 | 0,0842704 | 0 | -inf | 1 | 1 |
| Gm13272,Gm13285 | 0,0842585 | 0 | -inf | 1 | 1 |
| Ccl6 | 0,0827001 | 0 | -inf | 1 | 1 |
| Fitm1 | 0,0816519 | 0 | -inf | 1 | 1 |
| 4931440L10Rik | 0,0813124 | 0 | -inf | 1 | 1 |
| Bpifb1 | 0,0807324 | 0 | -inf | 1 | 1 |
| Gm9866 | 0,0807197 | 0 | -inf | 1 | 1 |
| Prss33 | 0,0805936 | 0 | -inf | 1 | 1 |
| 5031414D18Rik | 0,0804689 | 0 | -inf | 1 | 1 |
| 9130015A21Rik | 0,0800686 | 0 | -inf | 1 | 1 |
| Aqp8 | 0,0795683 | 0 | -inf | 1 | 1 |
| Nox1 | 0,0791602 | 0 | -inf | 1 | 1 |
| Pcdhb3 | 0,0791581 | 0 | -inf | 1 | 1 |
| Prl3b1 | 0,0789748 | 0 | -inf | 1 | 1 |
| Agtr2 | 0,0789509 | 0 | -inf | 1 | 1 |
| Vsx1 | 0,0777601 | 0 | -inf | 1 | 1 |
| Gm5065 | 0,0771265 | 0 | -inf | 1 | 1 |
| 9230112D13Rik | 0,0767875 | 0 | -inf | 1 | 1 |
| Nppb | 0,0765967 | 0 | -inf | 1 | 1 |
| Prr15l | 0,0761253 | 0 | -inf | 1 | 1 |

| | | | | | |
|---|---|---|---|---|---|
| Mill2 | 0,0759001 | 0 | -inf | 1 | 1 |
| Asb17os | 0,0757304 | 0 | -inf | 1 | 1 |
| Gm14207 | 0,0745931 | 0 | -inf | 1 | 1 |
| Dydc2 | 0,0744957 | 0 | -inf | 1 | 1 |
| Fam183b | 0,0739472 | 0 | -inf | 1 | 1 |
| Gm16063 | 0,0737068 | 0 | -inf | 1 | 1 |
| Mrap | 0,0733625 | 0 | -inf | 1 | 1 |
| 1700104L18Rik | 0,0733613 | 0 | -inf | 1 | 1 |
| Pp2d1 | 0,0733117 | 0 | -inf | 1 | 1 |
| Higd1c | 0,0729721 | 0 | -inf | 1 | 1 |
| Serpinf2 | 0,072699 | 0 | -inf | 1 | 1 |
| 2310002D06Rik | 0,0724921 | 0 | -inf | 1 | 1 |
| Adh4 | 0,0720652 | 0 | -inf | 1 | 1 |
| Myl10 | 0,071722 | 0 | -inf | 1 | 1 |
| Arhgap40 | 0,0717061 | 0 | -inf | 1 | 1 |
| F11 | 0,0715509 | 0 | -inf | 1 | 1 |
| 8430426J06Rik | 0,0712028 | 0 | -inf | 1 | 1 |
| Cxcl3 | 0,0710193 | 0 | -inf | 1 | 1 |
| Vax2 | 0,0709246 | 0 | -inf | 1 | 1 |
| 4930525D18Rik | 0,0695942 | 0 | -inf | 1 | 1 |
| AA543401 | 0,06944 | 0 | -inf | 1 | 1 |
| 1600029O15Rik | 0,0686554 | 0 | -inf | 1 | 1 |
| Prss48 | 0,0683798 | 0 | -inf | 1 | 1 |
| Erp27 | 0,0680146 | 0 | -inf | 1 | 1 |
| A330070K13Rik | 0,0673763 | 0 | -inf | 1 | 1 |
| Sctr | 0,0672994 | 0 | -inf | 1 | 1 |
| AI314278 | 0,0672954 | 0 | -inf | 1 | 1 |
| Tmprss5 | 0,0671682 | 0 | -inf | 1 | 1 |
| Rrh | 0,0670994 | 0 | -inf | 1 | 1 |
| Fbp2 | 0,0669555 | 0 | -inf | 1 | 1 |
| Tsks | 0,0669053 | 0 | -inf | 1 | 1 |
| Aspdh | 0,0668543 | 0 | -inf | 1 | 1 |
| Sox14 | 0,0665907 | 0 | -inf | 1 | 1 |
| Foxe3 | 0,0664799 | 0 | -inf | 1 | 1 |
| Gm11747 | 0,0663212 | 0 | -inf | 1 | 1 |
| C330022C24Rik | 0,0662196 | 0 | -inf | 1 | 1 |
| Gm9961 | 0,0655883 | 0 | -inf | 1 | 1 |
| Kcnmb4os1 | 0,0644647 | 0 | -inf | 1 | 1 |
| Tnp2 | 0,0636928 | 0 | -inf | 1 | 1 |
| Csrp3 | 0,0635885 | 0 | -inf | 1 | 1 |
| Bfsp2 | 0,063481 | 0 | -inf | 1 | 1 |
| Tmprss15 | 0,0633704 | 0 | -inf | 1 | 1 |
| Hic1 | 0,0626162 | 0 | -inf | 1 | 1 |
| Fam229a | 0,0622534 | 0 | -inf | 1 | 1 |
| Plet1os | 0,0619909 | 0 | -inf | 1 | 1 |
| Tnfsf15 | 0,0619312 | 0 | -inf | 1 | 1 |
| Ptcra | 0,0612741 | 0 | -inf | 1 | 1 |
| Cfc1 | 0,0610828 | 0 | -inf | 1 | 1 |
| Sphkap | 0,0600261 | 0 | -inf | 1 | 1 |
| Tm4sf1 | 0,060004 | 0 | -inf | 1 | 1 |

| | | | | | |
|---|---|---|---|---|---|
| Plekhd1os | 0,0599677 | 0 | -inf | 1 | 1 |
| Cyp1a1 | 0,0597918 | 0 | -inf | 1 | 1 |
| 1500009C09Rik | 0,05976 | 0 | -inf | 1 | 1 |
| Ldlrad1 | 0,059322 | 0 | -inf | 1 | 1 |
| Cpne4 | 0,0590651 | 0 | -inf | 1 | 1 |
| Slfn3 | 0,0589455 | 0 | -inf | 1 | 1 |
| Nipal4 | 0,0587952 | 0 | -inf | 1 | 1 |
| Pkd2l1 | 0,0587662 | 0 | -inf | 1 | 1 |
| Dmr | 0,0587047 | 0 | -inf | 1 | 1 |
| Mlc1 | 0,0583178 | 0 | -inf | 1 | 1 |
| BC021767 | 0,0578353 | 0 | -inf | 1 | 1 |
| Snai3 | 0,0576523 | 0 | -inf | 1 | 1 |
| Ltc4s | 0,0573901 | 0 | -inf | 1 | 1 |
| 4930567J20Rik | 0,0572368 | 0 | -inf | 1 | 1 |
| 1700060C20Rik | 0,0570962 | 0 | -inf | 1 | 1 |
| Nmur1 | 0,0570117 | 0 | -inf | 1 | 1 |
| Serpinb13 | 0,0569126 | 0 | -inf | 1 | 1 |
| Ccl19 | 0,0560166 | 0 | -inf | 1 | 1 |
| Hmx1 | 0,0559675 | 0 | -inf | 1 | 1 |
| Cldn22 | 0,0557928 | 0 | -inf | 1 | 1 |
| 4930467K11Rik | 0,0556928 | 0 | -inf | 1 | 1 |
| 3110039I08Rik | 0,0551614 | 0 | -inf | 1 | 1 |
| Hbq1a | 0,0550945 | 0 | -inf | 1 | 1 |
| 4930558C23Rik | 0,054537 | 0 | -inf | 1 | 1 |
| Higd1b | 0,054537 | 0 | -inf | 1 | 1 |
| Tmc1 | 0,0544434 | 0 | -inf | 1 | 1 |
| Khdc3 | 0,0538788 | 0 | -inf | 1 | 1 |
| C030018K13Rik | 0,0534875 | 0 | -inf | 1 | 1 |
| Tex12 | 0,0533604 | 0 | -inf | 1 | 1 |
| Prom2 | 0,0527417 | 0 | -inf | 1 | 1 |
| Gm648 | 0,0527229 | 0 | -inf | 1 | 1 |
| Treml4 | 0,0526973 | 0 | -inf | 1 | 1 |
| C4bp-ps1 | 0,0525539 | 0 | -inf | 1 | 1 |
| Gm11567 | 0,0525344 | 0 | -inf | 1 | 1 |
| Rdh9 | 0,0524428 | 0 | -inf | 1 | 1 |
| Slc18a3 | 0,052317 | 0 | -inf | 1 | 1 |
| 1700019A02Rik | 0,0520989 | 0 | -inf | 1 | 1 |
| Gm5084 | 0,0514322 | 0 | -inf | 1 | 1 |
| Crygb | 0,0513703 | 0 | -inf | 1 | 1 |
| Wdr38 | 0,0508415 | 0 | -inf | 1 | 1 |
| Ppp1r27 | 0,0506211 | 0 | -inf | 1 | 1 |
| Cryge | 0,0503148 | 0 | -inf | 1 | 1 |
| 4930511E03Rik | 0,0502401 | 0 | -inf | 1 | 1 |
| Cd209g | 0,05003 | 0 | -inf | 1 | 1 |
| 1700064M15Rik | 0,0499725 | 0 | -inf | 1 | 1 |
| Prps1l1 | 0,04958 | 0 | -inf | 1 | 1 |
| Opn5 | 0,0492892 | 0 | -inf | 1 | 1 |
| 2010106C02Rik | 0,0486488 | 0 | -inf | 1 | 1 |
| Spata32 | 0,0485423 | 0 | -inf | 1 | 1 |
| Pbp2 | 0,0484251 | 0 | -inf | 1 | 1 |

| | | | | | |
|---|---|---|---|---|---|
| 1700049L16Rik | 0,0480144 | 0 | -inf | 1 | 1 |
| Tspan32 | 0,0480138 | 0 | -inf | 1 | 1 |
| Lce1k | 0,0480129 | 0 | -inf | 1 | 1 |
| Gp9 | 0,0476189 | 0 | -inf | 1 | 1 |
| Hrg | 0,0476071 | 0 | -inf | 1 | 1 |
| A330048O09Rik | 0,0475981 | 0 | -inf | 1 | 1 |
| Ctnna3 | 0,0473151 | 0 | -inf | 1 | 1 |
| Adam18 | 0,0472355 | 0 | -inf | 1 | 1 |
| Hsd17b3 | 0,0470825 | 0 | -inf | 1 | 1 |
| Prf1 | 0,0468731 | 0 | -inf | 1 | 1 |
| Gm16157 | 0,0467407 | 0 | -inf | 1 | 1 |
| Mctp1 | 0,0465217 | 0 | -inf | 1 | 1 |
| Ccnb3 | 0,0465141 | 0 | -inf | 1 | 1 |
| Scin | 0,0463772 | 0 | -inf | 1 | 1 |
| Gm10863 | 0,0462982 | 0 | -inf | 1 | 1 |
| Gimap5 | 0,0459163 | 0 | -inf | 1 | 1 |
| Gjd3 | 0,045663 | 0 | -inf | 1 | 1 |
| Slc38a8 | 0,0455716 | 0 | -inf | 1 | 1 |
| Gm10637 | 0,0455192 | 0 | -inf | 1 | 1 |
| Gpr25 | 0,0454718 | 0 | -inf | 1 | 1 |
| 1810008I18Rik | 0,0452524 | 0 | -inf | 1 | 1 |
| Pnlip | 0,0450585 | 0 | -inf | 1 | 1 |
| Il27ra | 0,044903 | 0 | -inf | 1 | 1 |
| Iapp | 0,0448838 | 0 | -inf | 1 | 1 |
| 8030443G20Rik | 0,0448337 | 0 | -inf | 1 | 1 |
| Nrxn1 | 0,0445699 | 0 | -inf | 1 | 1 |
| Gm17745 | 0,0444005 | 0 | -inf | 1 | 1 |
| Kcne2 | 0,0443725 | 0 | -inf | 1 | 1 |
| 1600002D24Rik | 0,0442314 | 0 | -inf | 1 | 1 |
| Gckr | 0,0441156 | 0 | -inf | 1 | 1 |
| 1700034J05Rik | 0,0438754 | 0 | -inf | 1 | 1 |
| 2410137M14Rik | 0,0437288 | 0 | -inf | 1 | 1 |
| H2-Eb2 | 0,0433436 | 0 | -inf | 1 | 1 |
| Trim30e-ps1 | 0,0432869 | 0 | -inf | 1 | 1 |
| Ly6f | 0,0429491 | 0 | -inf | 1 | 1 |
| Gm20172 | 0,0429176 | 0 | -inf | 1 | 1 |
| C130080G10Rik | 0,04288 | 0 | -inf | 1 | 1 |
| Dazl | 0,0426744 | 0 | -inf | 1 | 1 |
| Rxfp4 | 0,0423405 | 0 | -inf | 1 | 1 |
| Gsx1 | 0,0421652 | 0 | -inf | 1 | 1 |
| Fezf2 | 0,0419779 | 0 | -inf | 1 | 1 |
| Fmo2 | 0,04176 | 0 | -inf | 1 | 1 |
| Sult2a7 | 0,0417096 | 0 | -inf | 1 | 1 |
| Tas2r134 | 0,0417096 | 0 | -inf | 1 | 1 |
| Padi2 | 0,0414148 | 0 | -inf | 1 | 1 |
| Rem1 | 0,0413224 | 0 | -inf | 1 | 1 |
| D430036J16Rik | 0,0412314 | 0 | -inf | 1 | 1 |
| LOC101055769 | 0,0409633 | 0 | -inf | 1 | 1 |
| Ccl24 | 0,0406235 | 0 | -inf | 1 | 1 |
| Krt75 | 0,0404457 | 0 | -inf | 1 | 1 |

| | | | | | |
|---|---|---|---|---|---|
| Cd300lg | 0,0404434 | 0 | -inf | 1 | 1 |
| 1700049E15Rik | 0,0403135 | 0 | -inf | 1 | 1 |
| Cabp2 | 0,0402872 | 0 | -inf | 1 | 1 |
| Clvs1 | 0,040074 | 0 | -inf | 1 | 1 |
| Fgf20 | 0,0400498 | 0 | -inf | 1 | 1 |
| Nat8 | 0,0397044 | 0 | -inf | 1 | 1 |
| Fgf5 | 0,0394851 | 0 | -inf | 1 | 1 |
| Ccdc110 | 0,0392322 | 0 | -inf | 1 | 1 |
| Tm4sf20 | 0,0392173 | 0 | -inf | 1 | 1 |
| Idi2 | 0,0391607 | 0 | -inf | 1 | 1 |
| C330046G13Rik | 0,0390058 | 0 | -inf | 1 | 1 |
| Kcnv2 | 0,0389988 | 0 | -inf | 1 | 1 |
| Adh7 | 0,0389236 | 0 | -inf | 1 | 1 |
| Gm10790 | 0,0387467 | 0 | -inf | 1 | 1 |
| Reg3g | 0,0386632 | 0 | -inf | 1 | 1 |
| Cyp4x1 | 0,0385934 | 0 | -inf | 1 | 1 |
| Olfr657 | 0,0382339 | 0 | -inf | 1 | 1 |
| Slc25a54 | 0,0381884 | 0 | -inf | 1 | 1 |
| 4930542D17Rik | 0,0380627 | 0 | -inf | 1 | 1 |
| Ndst4 | 0,0377247 | 0 | -inf | 1 | 1 |
| 2310007L24Rik | 0,0376719 | 0 | -inf | 1 | 1 |
| Dppa4 | 0,0375422 | 0 | -inf | 1 | 1 |
| Gm3985 | 0,0375229 | 0 | -inf | 1 | 1 |
| 4930433N12Rik | 0,0374164 | 0 | -inf | 1 | 1 |
| 5430435G22Rik | 0,0373935 | 0 | -inf | 1 | 1 |
| 1700057H15Rik | 0,0372485 | 0 | -inf | 1 | 1 |
| Abcc12 | 0,0370787 | 0 | -inf | 1 | 1 |
| Nat3 | 0,037008 | 0 | -inf | 1 | 1 |
| Dnajb8 | 0,0368214 | 0 | -inf | 1 | 1 |
| Sftpc | 0,0367914 | 0 | -inf | 1 | 1 |
| Matn1 | 0,0367715 | 0 | -inf | 1 | 1 |
| Nxph1 | 0,0366119 | 0 | -inf | 1 | 1 |
| Csn3 | 0,036571 | 0 | -inf | 1 | 1 |
| Gm12718 | 0,0364868 | 0 | -inf | 1 | 1 |
| Ifi202b | 0,0363473 | 0 | -inf | 1 | 1 |
| Chrm2 | 0,036089 | 0 | -inf | 1 | 1 |
| Ahsg | 0,0359809 | 0 | -inf | 1 | 1 |
| Pcdhb18 | 0,03587 | 0 | -inf | 1 | 1 |
| 1700086L19Rik | 0,0358333 | 0 | -inf | 1 | 1 |
| Slc26a5 | 0,0357548 | 0 | -inf | 1 | 1 |
| Pcdhb14 | 0,0357516 | 0 | -inf | 1 | 1 |
| Slc34a3 | 0,0357379 | 0 | -inf | 1 | 1 |
| Gjb4 | 0,035517 | 0 | -inf | 1 | 1 |
| Gria2 | 0,0351577 | 0 | -inf | 1 | 1 |
| Gbp6 | 0,0351474 | 0 | -inf | 1 | 1 |
| Olfr49 | 0,0349938 | 0 | -inf | 1 | 1 |
| Gm8096 | 0,0349886 | 0 | -inf | 1 | 1 |
| Tssk3 | 0,0348251 | 0 | -inf | 1 | 1 |
| Atp13a5 | 0,0347822 | 0 | -inf | 1 | 1 |
| 1700123K08Rik | 0,0342881 | 0 | -inf | 1 | 1 |

| | | | | | |
|---|---|---|---|---|---|
| Lhcgr | 0,0339926 | 0 | -inf | 1 | 1 |
| Arrdc5 | 0,0339259 | 0 | -inf | 1 | 1 |
| 3110039M20Rik | 0,0339047 | 0 | -inf | 1 | 1 |
| Chrng | 0,0336693 | 0 | -inf | 1 | 1 |
| Mro | 0,0332738 | 0 | -inf | 1 | 1 |
| 4930557J02Rik | 0,0331861 | 0 | -inf | 1 | 1 |
| E030025P04Rik | 0,0330719 | 0 | -inf | 1 | 1 |
| 4930546K05Rik | 0,0330411 | 0 | -inf | 1 | 1 |
| St18 | 0,0329936 | 0 | -inf | 1 | 1 |
| 4921534H16Rik | 0,0329216 | 0 | -inf | 1 | 1 |
| Ankrd22 | 0,0328667 | 0 | -inf | 1 | 1 |
| Gm1322 | 0,0328094 | 0 | -inf | 1 | 1 |
| Tmem51os1 | 0,0327821 | 0 | -inf | 1 | 1 |
| 4932438H23Rik | 0,0326957 | 0 | -inf | 1 | 1 |
| Lhx8 | 0,0326059 | 0 | -inf | 1 | 1 |
| LOC101056149 | 0,0325872 | 0 | -inf | 1 | 1 |
| Pebp4 | 0,0325505 | 0 | -inf | 1 | 1 |
| Tex21 | 0,0325306 | 0 | -inf | 1 | 1 |
| 9530026F06Rik | 0,0320321 | 0 | -inf | 1 | 1 |
| Hs3st2 | 0,0320015 | 0 | -inf | 1 | 1 |
| 3010033K07Rik | 0,0319393 | 0 | -inf | 1 | 1 |
| Myadml2 | 0,0319038 | 0 | -inf | 1 | 1 |
| Olfr725 | 0,0319038 | 0 | -inf | 1 | 1 |
| 1700010B08Rik | 0,031847 | 0 | -inf | 1 | 1 |
| Nr5a1 | 0,0316134 | 0 | -inf | 1 | 1 |
| Chrna10 | 0,0316076 | 0 | -inf | 1 | 1 |
| Slc11a1 | 0,0314834 | 0 | -inf | 1 | 1 |
| Cxcr1 | 0,0314184 | 0 | -inf | 1 | 1 |
| Icam4 | 0,0313487 | 0 | -inf | 1 | 1 |
| Bbox1 | 0,0309527 | 0 | -inf | 1 | 1 |
| Lypd3 | 0,030931 | 0 | -inf | 1 | 1 |
| Olfr99 | 0,0308657 | 0 | -inf | 1 | 1 |
| Gm2115 | 0,0307073 | 0 | -inf | 1 | 1 |
| Gm20110 | 0,0305712 | 0 | -inf | 1 | 1 |
| Pcdh20 | 0,0303658 | 0 | -inf | 1 | 1 |
| Cxcl13 | 0,0301721 | 0 | -inf | 1 | 1 |
| Olfr71 | 0,0299839 | 0 | -inf | 1 | 1 |
| Emx1 | 0,0298296 | 0 | -inf | 1 | 1 |
| Trap1a | 0,0297989 | 0 | -inf | 1 | 1 |
| Ccdc144b | 0,0296694 | 0 | -inf | 1 | 1 |
| Gm5091 | 0,0296461 | 0 | -inf | 1 | 1 |
| Car1 | 0,0294682 | 0 | -inf | 1 | 1 |
| Mlkl | 0,0294237 | 0 | -inf | 1 | 1 |
| Lrrc4c | 0,0293519 | 0 | -inf | 1 | 1 |
| Speer5-ps1 | 0,0287699 | 0 | -inf | 1 | 1 |
| Akr1c21 | 0,0287611 | 0 | -inf | 1 | 1 |
| Slc13a3 | 0,0287138 | 0 | -inf | 1 | 1 |
| 4833424O15Rik | 0,0286959 | 0 | -inf | 1 | 1 |
| Cps1 | 0,0285239 | 0 | -inf | 1 | 1 |
| Clec1a | 0,0283014 | 0 | -inf | 1 | 1 |

| | | | | | |
|---|---|---|---|---|---|
| D030040B21Rik | 0,0282438 | 0 | -inf | 1 | 1 |
| Gm11437 | 0,0281999 | 0 | -inf | 1 | 1 |
| Pparg | 0,0281607 | 0 | -inf | 1 | 1 |
| Bpi | 0,0279852 | 0 | -inf | 1 | 1 |
| Gm44 | 0,0279318 | 0 | -inf | 1 | 1 |
| 4930528D03Rik | 0,0279275 | 0 | -inf | 1 | 1 |
| 4930568G15Rik | 0,0277559 | 0 | -inf | 1 | 1 |
| Lhfpl5 | 0,0277133 | 0 | -inf | 1 | 1 |
| 5730488B01Rik | 0,0274372 | 0 | -inf | 1 | 1 |
| 03/04/16 | 0,0273046 | 0 | -inf | 1 | 1 |
| Gm20125 | 0,027124 | 0 | -inf | 1 | 1 |
| Sbk3 | 0,0270857 | 0 | -inf | 1 | 1 |
| A930003A15Rik | 0,027038 | 0 | -inf | 1 | 1 |
| Gm805 | 0,0270124 | 0 | -inf | 1 | 1 |
| Trim43a | 0,0270078 | 0 | -inf | 1 | 1 |
| Lair1 | 0,0268112 | 0 | -inf | 1 | 1 |
| 1700026D08Rik | 0,0265183 | 0 | -inf | 1 | 1 |
| AU022793 | 0,0264884 | 0 | -inf | 1 | 1 |
| Me3 | 0,0264504 | 0 | -inf | 1 | 1 |
| Slc6a20a | 0,0260809 | 0 | -inf | 1 | 1 |
| Pcdhb11 | 0,025844 | 0 | -inf | 1 | 1 |
| Fpr1 | 0,0258311 | 0 | -inf | 1 | 1 |
| Pcdhb1 | 0,0256592 | 0 | -inf | 1 | 1 |
| Spatc1l | 0,0256477 | 0 | -inf | 1 | 1 |
| B230312C02Rik | 0,0255344 | 0 | -inf | 1 | 1 |
| Rprml | 0,0253896 | 0 | -inf | 1 | 1 |
| Tgtp1 | 0,0253801 | 0 | -inf | 1 | 1 |
| 1700113H08Rik | 0,0253775 | 0 | -inf | 1 | 1 |
| 8430436N08Rik | 0,0252442 | 0 | -inf | 1 | 1 |
| 1700051A21Rik | 0,0250476 | 0 | -inf | 1 | 1 |
| E030003E18Rik | 0,025015 | 0 | -inf | 1 | 1 |
| 4933433F19Rik | 0,0249866 | 0 | -inf | 1 | 1 |
| 6030440G07Rik | 0,0247899 | 0 | -inf | 1 | 1 |
| Fam26d | 0,0247053 | 0 | -inf | 1 | 1 |
| Rxfp1 | 0,0245748 | 0 | -inf | 1 | 1 |
| Gm14327 | 0,0245688 | 0 | -inf | 1 | 1 |
| Slc22a28 | 0,0244871 | 0 | -inf | 1 | 1 |
| Odf1 | 0,0244597 | 0 | -inf | 1 | 1 |
| Epb4.2 | 0,0244335 | 0 | -inf | 1 | 1 |
| Gm6878 | 0,0242981 | 0 | -inf | 1 | 1 |
| 2310005A03Rik | 0,0242899 | 0 | -inf | 1 | 1 |
| Rnf183 | 0,0240875 | 0 | -inf | 1 | 1 |
| 4931440F15Rik | 0,0238045 | 0 | -inf | 1 | 1 |
| Rgs6 | 0,0237923 | 0 | -inf | 1 | 1 |
| Nmur2 | 0,0237572 | 0 | -inf | 1 | 1 |
| Mfsd6l | 0,0237482 | 0 | -inf | 1 | 1 |
| Mcemp1 | 0,0237481 | 0 | -inf | 1 | 1 |
| A930016O22Rik | 0,0237473 | 0 | -inf | 1 | 1 |
| Pirt | 0,0235286 | 0 | -inf | 1 | 1 |
| Clec4n | 0,0234429 | 0 | -inf | 1 | 1 |

| | | | | | |
|---|---|---|---|---|---|
| Slamf9 | 0,02342 | 0 | -inf | 1 | 1 |
| Fnd3c2 | 0,0231178 | 0 | -inf | 1 | 1 |
| Bpifb4 | 0,0230648 | 0 | -inf | 1 | 1 |
| Prn | 0,0227499 | 0 | -inf | 1 | 1 |
| 2410141K09Rik | 0,0224592 | 0 | -inf | 1 | 1 |
| 2310005E17Rik | 0,0224422 | 0 | -inf | 1 | 1 |
| Lgi3 | 0,0222087 | 0 | -inf | 1 | 1 |
| Catsper4 | 0,0221652 | 0 | -inf | 1 | 1 |
| Itih3 | 0,0221241 | 0 | -inf | 1 | 1 |
| 4930513D17Rik | 0,0220824 | 0 | -inf | 1 | 1 |
| Tgm5 | 0,0218924 | 0 | -inf | 1 | 1 |
| Pglyrp4 | 0,0217825 | 0 | -inf | 1 | 1 |
| Vgf | 0,0216873 | 0 | -inf | 1 | 1 |
| Ceacam20 | 0,0214828 | 0 | -inf | 1 | 1 |
| Fgf6 | 0,021438 | 0 | -inf | 1 | 1 |
| Slco1b2 | 0,0213827 | 0 | -inf | 1 | 1 |
| Olfr658 | 0,021348 | 0 | -inf | 1 | 1 |
| Gad2 | 0,0213283 | 0 | -inf | 1 | 1 |
| Gfap | 0,021257 | 0 | -inf | 1 | 1 |
| Als2cr12 | 0,0210678 | 0 | -inf | 1 | 1 |
| Pik3c2g | 0,020969 | 0 | -inf | 1 | 1 |
| Cxcl11 | 0,0209141 | 0 | -inf | 1 | 1 |
| Tcp10a | 0,0208875 | 0 | -inf | 1 | 1 |
| BC021785 | 0,0208179 | 0 | -inf | 1 | 1 |
| BC089597 | 0,0207712 | 0 | -inf | 1 | 1 |
| Eps8l3 | 0,0206871 | 0 | -inf | 1 | 1 |
| Gm20751 | 0,0206738 | 0 | -inf | 1 | 1 |
| 8430422H06Rik | 0,0205965 | 0 | -inf | 1 | 1 |
| Ifi44l | 0,0203681 | 0 | -inf | 1 | 1 |
| Arx | 0,0201757 | 0 | -inf | 1 | 1 |
| Hnf4g | 0,0200827 | 0 | -inf | 1 | 1 |
| Clec4a2 | 0,0200586 | 0 | -inf | 1 | 1 |
| Chst13 | 0,0199758 | 0 | -inf | 1 | 1 |
| Foxs1 | 0,0199081 | 0 | -inf | 1 | 1 |
| Celf6 | 0,0198921 | 0 | -inf | 1 | 1 |
| Gm9 | 0,0198115 | 0 | -inf | 1 | 1 |
| BC051537 | 0,0197439 | 0 | -inf | 1 | 1 |
| 4933401D09Rik | 0,0196595 | 0 | -inf | 1 | 1 |
| Gm11541 | 0,0195701 | 0 | -inf | 1 | 1 |
| Gm11780 | 0,0195172 | 0 | -inf | 1 | 1 |
| Unc13c | 0,0194071 | 0 | -inf | 1 | 1 |
| 4930413F20Rik | 0,0193662 | 0 | -inf | 1 | 1 |
| Vmn2r2 | 0,0193319 | 0 | -inf | 1 | 1 |
| Aanat | 0,0191777 | 0 | -inf | 1 | 1 |
| Gm14139 | 0,0189666 | 0 | -inf | 1 | 1 |
| C130071C03Rik | 0,0189332 | 0 | -inf | 1 | 1 |
| Sstr5 | 0,0185411 | 0 | -inf | 1 | 1 |
| Fut2 | 0,0185126 | 0 | -inf | 1 | 1 |
| Acot3 | 0,0184499 | 0 | -inf | 1 | 1 |
| Slc6a7 | 0,0184168 | 0 | -inf | 1 | 1 |

| | | | | | |
|---|---|---|---|---|---|
| Lcp2 | 0,0183875 | 0 | -inf | 1 | 1 |
| Uroc1 | 0,018306 | 0 | -inf | 1 | 1 |
| Prph2 | 0,0181535 | 0 | -inf | 1 | 1 |
| Gm5082 | 0,0179447 | 0 | -inf | 1 | 1 |
| Cntnap5b | 0,017932 | 0 | -inf | 1 | 1 |
| Acot12 | 0,0179064 | 0 | -inf | 1 | 1 |
| Gm15941 | 0,0178206 | 0 | -inf | 1 | 1 |
| 2210019I11Rik | 0,0178015 | 0 | -inf | 1 | 1 |
| LOC100862268 | 0,0177114 | 0 | -inf | 1 | 1 |
| AU022751 | 0,0176674 | 0 | -inf | 1 | 1 |
| 4931423N10Rik | 0,0176466 | 0 | -inf | 1 | 1 |
| Syt10 | 0,0176143 | 0 | -inf | 1 | 1 |
| Cyp2g1 | 0,0175288 | 0 | -inf | 1 | 1 |
| Mmp21 | 0,0174759 | 0 | -inf | 1 | 1 |
| Gje1 | 0,0173121 | 0 | -inf | 1 | 1 |
| Zfp648 | 0,0172774 | 0 | -inf | 1 | 1 |
| Il5ra | 0,017262 | 0 | -inf | 1 | 1 |
| Arl11 | 0,0172444 | 0 | -inf | 1 | 1 |
| Brinp3 | 0,0172437 | 0 | -inf | 1 | 1 |
| Lingo2 | 0,0172181 | 0 | -inf | 1 | 1 |
| B230214G05Rik | 0,0171239 | 0 | -inf | 1 | 1 |
| 9130409I23Rik | 0,0170632 | 0 | -inf | 1 | 1 |
| Il17re | 0,0170593 | 0 | -inf | 1 | 1 |
| Lgsn | 0,017013 | 0 | -inf | 1 | 1 |
| Gpr26 | 0,0169853 | 0 | -inf | 1 | 1 |
| Zp1 | 0,0167955 | 0 | -inf | 1 | 1 |
| Pcdha7 | 0,0167682 | 0 | -inf | 1 | 1 |
| Gm7538 | 0,0167592 | 0 | -inf | 1 | 1 |
| Grap2 | 0,0167503 | 0 | -inf | 1 | 1 |
| Dppa2 | 0,0166408 | 0 | -inf | 1 | 1 |
| Il5 | 0,0165702 | 0 | -inf | 1 | 1 |
| Cml2 | 0,0165616 | 0 | -inf | 1 | 1 |
| A530046M15Rik | 0,0165205 | 0 | -inf | 1 | 1 |
| Nkx2-2os | 0,0164809 | 0 | -inf | 1 | 1 |
| F12 | 0,0164514 | 0 | -inf | 1 | 1 |
| Kcns2 | 0,0163475 | 0 | -inf | 1 | 1 |
| Pinc | 0,0162524 | 0 | -inf | 1 | 1 |
| LOC100861615 | 0,0161752 | 0 | -inf | 1 | 1 |
| Chrna9 | 0,0160673 | 0 | -inf | 1 | 1 |
| Slco1a4 | 0,0159741 | 0 | -inf | 1 | 1 |
| Rnf224 | 0,0159081 | 0 | -inf | 1 | 1 |
| Slc45a2 | 0,0158607 | 0 | -inf | 1 | 1 |
| Angptl3 | 0,0157417 | 0 | -inf | 1 | 1 |
| Slc23a3 | 0,0155907 | 0 | -inf | 1 | 1 |
| Cyp2d34 | 0,0155418 | 0 | -inf | 1 | 1 |
| Vstm2a | 0,0154187 | 0 | -inf | 1 | 1 |
| Kcnj1 | 0,0153739 | 0 | -inf | 1 | 1 |
| Gm5039 | 0,0153363 | 0 | -inf | 1 | 1 |
| 2810047C21Rik1 | 0,0153044 | 0 | -inf | 1 | 1 |
| Slc25a31 | 0,0151653 | 0 | -inf | 1 | 1 |

| | | | | | |
|---|---|---|---|---|---|
| Gsdma2 | 0,0150741 | 0 | -inf | 1 | 1 |
| 4930432J09Rik | 0,0150536 | 0 | -inf | 1 | 1 |
| 4932418E24Rik | 0,0149768 | 0 | -inf | 1 | 1 |
| Crp | 0,0147512 | 0 | -inf | 1 | 1 |
| Gm20758 | 0,014695 | 0 | -inf | 1 | 1 |
| 4933400C23Rik | 0,0146531 | 0 | -inf | 1 | 1 |
| Rab11fip4os1 | 0,0145765 | 0 | -inf | 1 | 1 |
| Trim58 | 0,0145398 | 0 | -inf | 1 | 1 |
| 4930480K15Rik | 0,0144888 | 0 | -inf | 1 | 1 |
| Pkib | 0,0144549 | 0 | -inf | 1 | 1 |
| 6330415B21Rik | 0,0143462 | 0 | -inf | 1 | 1 |
| 2700070H01Rik | 0,014292 | 0 | -inf | 1 | 1 |
| Col6a6 | 0,0142625 | 0 | -inf | 1 | 1 |
| Npsr1 | 0,0142023 | 0 | -inf | 1 | 1 |
| Cntnap3 | 0,0141757 | 0 | -inf | 1 | 1 |
| Lrrc30 | 0,0141726 | 0 | -inf | 1 | 1 |
| Cuzd1 | 0,0140863 | 0 | -inf | 1 | 1 |
| Trim43b | 0,0140452 | 0 | -inf | 1 | 1 |
| Pla2g2d | 0,0140316 | 0 | -inf | 1 | 1 |
| Cdh7 | 0,0139971 | 0 | -inf | 1 | 1 |
| G630090E17Rik | 0,0139655 | 0 | -inf | 1 | 1 |
| Gpa33 | 0,0138368 | 0 | -inf | 1 | 1 |
| Ttll10 | 0,0137447 | 0 | -inf | 1 | 1 |
| Ace2 | 0,013712 | 0 | -inf | 1 | 1 |
| Slc6a14 | 0,0136545 | 0 | -inf | 1 | 1 |
| Olfr70 | 0,0134568 | 0 | -inf | 1 | 1 |
| Rpe65 | 0,0134462 | 0 | -inf | 1 | 1 |
| Atp1a4 | 0,013434 | 0 | -inf | 1 | 1 |
| Olfr78 | 0,0133891 | 0 | -inf | 1 | 1 |
| Fam43b | 0,0133634 | 0 | -inf | 1 | 1 |
| Pglyrp2 | 0,0133493 | 0 | -inf | 1 | 1 |
| Nt5c1b | 0,0133485 | 0 | -inf | 1 | 1 |
| Heatr9 | 0,0133404 | 0 | -inf | 1 | 1 |
| Astl | 0,0132714 | 0 | -inf | 1 | 1 |
| 2010109I03Rik | 0,0130947 | 0 | -inf | 1 | 1 |
| Pla2g2f | 0,0129445 | 0 | -inf | 1 | 1 |
| Gm12709 | 0,0129031 | 0 | -inf | 1 | 1 |
| Akr1c14 | 0,0128983 | 0 | -inf | 1 | 1 |
| Mybpc2 | 0,012888 | 0 | -inf | 1 | 1 |
| Sh2d2a | 0,0127456 | 0 | -inf | 1 | 1 |
| Cybb | 0,0127168 | 0 | -inf | 1 | 1 |
| Col28a1 | 0,0126537 | 0 | -inf | 1 | 1 |
| Aldob | 0,0126005 | 0 | -inf | 1 | 1 |
| Gna15 | 0,0125432 | 0 | -inf | 1 | 1 |
| Fbxo40 | 0,0124914 | 0 | -inf | 1 | 1 |
| Bin2 | 0,012423 | 0 | -inf | 1 | 1 |
| Vmn2r118 | 0,0122583 | 0 | -inf | 1 | 1 |
| C86187 | 0,0121357 | 0 | -inf | 1 | 1 |
| Upk3b | 0,0120297 | 0 | -inf | 1 | 1 |
| Gm10768 | 0,0119839 | 0 | -inf | 1 | 1 |

| | | | | | |
|---|---|---|---|---|---|
| Gm13752 | 0,0119839 | 0 | -inf | 1 | 1 |
| 4930442L01Rik | 0,011932 | 0 | -inf | 1 | 1 |
| Dyrk4 | 0,0117915 | 0 | -inf | 1 | 1 |
| Cdsn | 0,0116836 | 0 | -inf | 1 | 1 |
| C7 | 0,011539 | 0 | -inf | 1 | 1 |
| Hephl1 | 0,0114318 | 0 | -inf | 1 | 1 |
| Tmprss3 | 0,0114184 | 0 | -inf | 1 | 1 |
| Inhbe | 0,0114129 | 0 | -inf | 1 | 1 |
| Hspb7 | 0,0112735 | 0 | -inf | 1 | 1 |
| Hcrtr1 | 0,011249 | 0 | -inf | 1 | 1 |
| Rph3a | 0,0112298 | 0 | -inf | 1 | 1 |
| Gm14685 | 0,0111526 | 0 | -inf | 1 | 1 |
| Slc22a14 | 0,0110523 | 0 | -inf | 1 | 1 |
| Gm9159 | 0,0110413 | 0 | -inf | 1 | 1 |
| Aldh8a1 | 0,0108084 | 0 | -inf | 1 | 1 |
| Gpr119 | 0,0106723 | 0 | -inf | 1 | 1 |
| 2310015D24Rik | 0,0106543 | 0 | -inf | 1 | 1 |
| Ces1g | 0,01057 | 0 | -inf | 1 | 1 |
| Nxpe5 | 0,0104546 | 0 | -inf | 1 | 1 |
| Barhl2 | 0,0104447 | 0 | -inf | 1 | 1 |
| Npy5r | 0,0104447 | 0 | -inf | 1 | 1 |
| Il2ra | 0,0104349 | 0 | -inf | 1 | 1 |
| Apol6 | 0,0104348 | 0 | -inf | 1 | 1 |
| AU022754 | 0,0104279 | 0 | -inf | 1 | 1 |
| Htr1a | 0,0104029 | 0 | -inf | 1 | 1 |
| Cd180 | 0,0103857 | 0 | -inf | 1 | 1 |
| Padi6 | 0,0103515 | 0 | -inf | 1 | 1 |
| Slc16a14 | 0,0102743 | 0 | -inf | 1 | 1 |
| Itgbl1 | 0,0102361 | 0 | -inf | 1 | 1 |
| A730020M07Rik | 0,0101606 | 0 | -inf | 1 | 1 |
| Cdh19 | 0,0101422 | 0 | -inf | 1 | 1 |
| Slc47a2 | 0,0101419 | 0 | -inf | 1 | 1 |
| 4932415M13Rik | 0,0101102 | 0 | -inf | 1 | 1 |
| Gm1110 | 0,0101047 | 0 | -inf | 1 | 1 |
| Zfp936 | 0,00997762 | 0 | -inf | 1 | 1 |
| Otop3 | 0,00993616 | 0 | -inf | 1 | 1 |
| Sgk2 | 0,00987136 | 0 | -inf | 1 | 1 |
| B230206H07Rik | 0,00984761 | 0 | -inf | 1 | 1 |
| Gm20139 | 0,00962866 | 0 | -inf | 1 | 1 |
| Pabpc6 | 0,00962223 | 0 | -inf | 1 | 1 |
| Atp6v0d2 | 0,00958608 | 0 | -inf | 1 | 1 |
| Atp8b4 | 0,00956689 | 0 | -inf | 1 | 1 |
| Gm5531 | 0,00948704 | 0 | -inf | 1 | 1 |
| Ankrd63 | 0,00946466 | 0 | -inf | 1 | 1 |
| Musk | 0,00925556 | 0 | -inf | 1 | 1 |
| Gpr149 | 0,00922898 | 0 | -inf | 1 | 1 |
| Vmn2r83 | 0,00921723 | 0 | -inf | 1 | 1 |
| Aknad1 | 0,0091865 | 0 | -inf | 1 | 1 |
| Zfp36l3 | 0,00914836 | 0 | -inf | 1 | 1 |
| Klrb1c | 0,00909375 | 0 | -inf | 1 | 1 |

| | | | | | |
|---|---|---|---|---|---|
| Il1rl1 | 0,00905628 | 0 | -inf | 1 | 1 |
| Slc5a12 | 0,00900637 | 0 | -inf | 1 | 1 |
| Krt78 | 0,00894897 | 0 | -inf | 1 | 1 |
| Adam7 | 0,00885576 | 0 | -inf | 1 | 1 |
| Best3 | 0,00876279 | 0 | -inf | 1 | 1 |
| Arhgap27os3 | 0,00872118 | 0 | -inf | 1 | 1 |
| Wdr27 | 0,0086256 | 0 | -inf | 1 | 1 |
| Adh6-ps1 | 0,0085255 | 0 | -inf | 1 | 1 |
| Nlrp4c | 0,0084766 | 0 | -inf | 1 | 1 |
| Kcnj16 | 0,0084482 | 0 | -inf | 1 | 1 |
| Lrrc2 | 0,00837589 | 0 | -inf | 1 | 1 |
| Dgkb | 0,00825411 | 0 | -inf | 1 | 1 |
| Mgat4c | 0,00817428 | 0 | -inf | 1 | 1 |
| Ccr5 | 0,00809641 | 0 | -inf | 1 | 1 |
| Ern2 | 0,00795898 | 0 | -inf | 1 | 1 |
| Pcdha2 | 0,00793453 | 0 | -inf | 1 | 1 |
| Gm10220 | 0,00792749 | 0 | -inf | 1 | 1 |
| Abca13 | 0,00783995 | 0 | -inf | 1 | 1 |
| Adamtsl5 | 0,00779273 | 0 | -inf | 1 | 1 |
| Pcdhb12 | 0,00778743 | 0 | -inf | 1 | 1 |
| Tmem151a | 0,0077266 | 0 | -inf | 1 | 1 |
| Slfn4 | 0,00769984 | 0 | -inf | 1 | 1 |
| Tubal3 | 0,00762048 | 0 | -inf | 1 | 1 |
| Gprin3 | 0,00760995 | 0 | -inf | 1 | 1 |
| Insm2 | 0,00760733 | 0 | -inf | 1 | 1 |
| Pkhd1 | 0,00746784 | 0 | -inf | 1 | 1 |
| Svop | 0,00739049 | 0 | -inf | 1 | 1 |
| Cdhr2 | 0,00736814 | 0 | -inf | 1 | 1 |
| Ccdc129 | 0,00715533 | 0 | -inf | 1 | 1 |
| Atp1b4 | 0,00710557 | 0 | -inf | 1 | 1 |
| Olfm3 | 0,00707759 | 0 | -inf | 1 | 1 |
| Ush2a | 0,00702534 | 0 | -inf | 1 | 1 |
| Gm4776 | 0,00697398 | 0 | -inf | 1 | 1 |
| 2010002M12Rik | 0,00697198 | 0 | -inf | 1 | 1 |
| Gm11190 | 0,00686556 | 0 | -inf | 1 | 1 |
| Slc6a3 | 0,00677689 | 0 | -inf | 1 | 1 |
| Chrnb3 | 0,00665288 | 0 | -inf | 1 | 1 |
| Xdh | 0,00655447 | 0 | -inf | 1 | 1 |
| 4930407I10Rik | 0,00646794 | 0 | -inf | 1 | 1 |
| Frmpd4 | 0,00639576 | 0 | -inf | 1 | 1 |
| Abcb11 | 0,00616185 | 0 | -inf | 1 | 1 |
| Gm5177 | 0,00615427 | 0 | -inf | 1 | 1 |
| Kcng4 | 0,00611839 | 0 | -inf | 1 | 1 |
| Grin2b | 0,00603711 | 0 | -inf | 1 | 1 |
| Cntnap5c | 0,00593388 | 0 | -inf | 1 | 1 |
| Apob | 0,00582627 | 0 | -inf | 1 | 1 |
| Gm13242 | 0,00582104 | 0 | -inf | 1 | 1 |
| Slc22a6 | 0,00576621 | 0 | -inf | 1 | 1 |
| Rab44 | 0,00564944 | 0 | -inf | 1 | 1 |
| Cilp | 0,00558077 | 0 | -inf | 1 | 1 |

| | | | | | |
|---|---|---|---|---|---|
| Zfp804a | 0,00553869 | 0 | -inf | 1 | 1 |
| Ghsr | 0,00521127 | 0 | -inf | 1 | 1 |
| Cacna2d4 | 0,00516257 | 0 | -inf | 1 | 1 |
| Scn11a | 0,00513323 | 0 | -inf | 1 | 1 |
| Mug1 | 0,00494913 | 0 | -inf | 1 | 1 |
| Slc6a2 | 0,00492507 | 0 | -inf | 1 | 1 |
| Aox2 | 0,00482031 | 0 | -inf | 1 | 1 |
| Tnfsf10 | 0,00464946 | 0 | -inf | 1 | 1 |
| Dlx1as | 0,00422161 | 0 | -inf | 1 | 1 |
| Slfn5 | 0,00380959 | 0 | -inf | 1 | 1 |
| Myh7 | 0,00376726 | 0 | -inf | 1 | 1 |
| Hbb-bs | 0,00199039 | 0 | -inf | 1 | 1 |
| ChkbCpt1b | 0,000144244 | 0 | -inf | 1 | 1 |
| DXBay18,Gm14685 | 9,56E-005 | 0 | -inf | 1 | 1 |
| Esp6-esp5 | 3,79E-005 | 0 | -inf | 1 | 1 |
| Duxbl3 | 5,31E-006 | 0 | -inf | 1 | 1 |
| 1700001C19Rik | 0 | 0,0749329 | inf | 1 | 1 |
| 1700001J03Rik | 0 | 0,222449 | inf | 1 | 1 |
| 1700003D09Rik | 0 | 0,179588 | inf | 1 | 1 |
| 1700008I05Rik | 0 | 0,0541508 | inf | 1 | 1 |
| 1700009J07Rik | 0 | 0,113343 | inf | 1 | 1 |
| 1700012B09Rik | 0 | 0,144254 | inf | 1 | 1 |
| 1700017G19Rik | 0 | 0,0571311 | inf | 1 | 1 |
| 1700017J07Rik | 0 | 0,358985 | inf | 1 | 1 |
| 1700019B21Rik | 0 | 0,0783828 | inf | 1 | 1 |
| 1700023F06Rik | 0 | 0,112512 | inf | 1 | 1 |
| 1700023L04Rik | 0 | 0,118087 | inf | 1 | 1 |
| 1700028I16Rik | 0 | 0,0499557 | inf | 1 | 1 |
| 1700029J03Rik | 0 | 0,0458535 | inf | 1 | 1 |
| 1700039E22Rik | 0 | 0,395576 | inf | 1 | 1 |
| 1700040L02Rik | 0 | 0,0259425 | inf | 1 | 1 |
| 1700047A11Rik | 0 | 0,064481 | inf | 1 | 1 |
| 1700066B17Rik | 0 | 0,260313 | inf | 1 | 1 |
| 1700100L14Rik | 0 | 0,0561175 | inf | 1 | 1 |
| 1700120E14Rik | 0 | 0,0155236 | inf | 1 | 1 |
| 1810006J02Rik | 0 | 0,0287003 | inf | 1 | 1 |
| 2010009K17Rik | 0 | 0,186039 | inf | 1 | 1 |
| 2310001H17Rik | 0 | 0,185799 | inf | 1 | 1 |
| 2310005G13Rik | 0 | 0,0429749 | inf | 1 | 1 |
| 2610203C22Rik | 0 | 0,040886 | inf | 1 | 1 |
| 3830417A13Rik | 0 | 0,0480188 | inf | 1 | 1 |
| 4921511I17Rik | 0 | 0,115871 | inf | 1 | 1 |
| 4921530L21Rik | 0 | 0,0447083 | inf | 1 | 1 |
| 4930417O22Rik | 0 | 0,0980749 | inf | 1 | 1 |
| 4930429D17Rik | 0 | 0,0299736 | inf | 1 | 1 |
| 4930430A15Rik | 0 | 0,0520962 | inf | 1 | 1 |
| 4930431P03Rik | 0 | 0,0903625 | inf | 1 | 1 |
| 4930444M15Rik | 0 | 0,0585454 | inf | 1 | 1 |
| 4930449E01Rik | 0 | 0,174775 | inf | 1 | 1 |
| 4930461G14Rik | 0 | 0,04657 | inf | 1 | 1 |

| | | | | | |
|---|---|---|---|---|---|
| 4930471M09Rik | 0 | 0,0590849 | inf | 1 | 1 |
| 4930483O08Rik | 0 | 0,0649503 | inf | 1 | 1 |
| 4930488L21Rik | 0 | 0,016179 | inf | 1 | 1 |
| 4930511A02Rik | 0 | 0,0309933 | inf | 1 | 1 |
| 4930511M06Rik | 0 | 0,0742969 | inf | 1 | 1 |
| 4930515L03Rik | 0 | 0,151462 | inf | 1 | 1 |
| 4930517E11Rik | 0 | 0,0339723 | inf | 1 | 1 |
| 4930524O08Rik | 0 | 0,0215113 | inf | 1 | 1 |
| 4930529C04Rik | 0 | 0,0255863 | inf | 1 | 1 |
| 4930542C21Rik | 0 | 0,107298 | inf | 1 | 1 |
| 4930545E07Rik | 0 | 0,064684 | inf | 1 | 1 |
| 4930547E14Rik | 0 | 0,123366 | inf | 1 | 1 |
| 4930563E18Rik | 0 | 0,0397697 | inf | 1 | 1 |
| 4930571K23Rik | 0 | 0,0218382 | inf | 1 | 1 |
| 4930590L20Rik | 0 | 0,0555618 | inf | 1 | 1 |
| 4930593A02Rik | 0 | 0,115973 | inf | 1 | 1 |
| 4931402G19Rik | 0 | 0,029425 | inf | 1 | 1 |
| 4931420L22Rik | 0 | 0,07467 | inf | 1 | 1 |
| 4931428L18Rik | 0 | 0,0956456 | inf | 1 | 1 |
| 4931429L15Rik | 0 | 0,0378545 | inf | 1 | 1 |
| 4931440J10Rik | 0 | 0,0619536 | inf | 1 | 1 |
| 4932413F04Rik | 0 | 0,0209487 | inf | 1 | 1 |
| 4933406G16Rik | 0 | 0,0351175 | inf | 1 | 1 |
| 4933406I18Rik | 0 | 0,060801 | inf | 1 | 1 |
| 4933406J10Rik | 0 | 0,054497 | inf | 1 | 1 |
| 4933406M09Rik | 0 | 0,0685177 | inf | 1 | 1 |
| 4933411E08Rik | 0 | 0,028375 | inf | 1 | 1 |
| 4933424G06Rik | 0 | 0,1333 | inf | 1 | 1 |
| 4933427D06Rik | 0 | 0,0221967 | inf | 1 | 1 |
| 4933427E13Rik | 0 | 0,0330198 | inf | 1 | 1 |
| 4933427I22Rik | 0 | 0,0228489 | inf | 1 | 1 |
| 4933430H16Rik | 0 | 0,0434056 | inf | 1 | 1 |
| 5031426D15Rik | 0 | 0,0411967 | inf | 1 | 1 |
| 5330413P13Rik | 0 | 0,0359023 | inf | 1 | 1 |
| 5430428K19Rik | 0 | 0,0383206 | inf | 1 | 1 |
| 5730403I07Rik | 0 | 0,0146824 | inf | 1 | 1 |
| 5730420D15Rik | 0 | 0,218172 | inf | 1 | 1 |
| 5730435O14Rik | 0 | 0,103055 | inf | 1 | 1 |
| 5830403L16Rik | 0 | 0,0293535 | inf | 1 | 1 |
| 5830416P10Rik | 0 | 0,0334613 | inf | 1 | 1 |
| 5830418P13Rik | 0 | 0,0215616 | inf | 1 | 1 |
| 5830473C10Rik | 0 | 0,0340635 | inf | 1 | 1 |
| 6030466F02Rik | 0 | 0,0346036 | inf | 1 | 1 |
| 6330410L21Rik | 0 | 0,00775404 | inf | 1 | 1 |
| 8030423F21Rik | 0 | 0,11679 | inf | 1 | 1 |
| 9430019J16Rik | 0 | 0,0587064 | inf | 1 | 1 |
| 9530003J23Rik | 0 | 0,0419334 | inf | 1 | 1 |
| 9530059O14Rik | 0 | 0,0326013 | inf | 1 | 1 |
| 9830166K06Rik | 0 | 0,0211136 | inf | 1 | 1 |
| 9930111J21Rik1 | 0 | 0,0116323 | inf | 1 | 1 |

| | | | | | |
|---|---|---|---|---|---|
| A230009B12Rik | 0 | 0,21899 | inf | 1 | 1 |
| A530088E08Rik | 0 | 0,0168603 | inf | 1 | 1 |
| A630019I02Rik | 0 | 0,0934138 | inf | 1 | 1 |
| A730036I17Rik | 0 | 0,0131814 | inf | 1 | 1 |
| A930001A20Rik | 0 | 0,0366274 | inf | 1 | 1 |
| A930019D19Rik | 0 | 0,169345 | inf | 1 | 1 |
| AA387883 | 0 | 0,0288559 | inf | 1 | 1 |
| AI317395 | 0 | 0,0280353 | inf | 1 | 1 |
| AI593442 | 0 | 0,00493205 | inf | 1 | 1 |
| AU018091 | 0 | 0,0262409 | inf | 1 | 1 |
| Abca14 | 0 | 0,0246503 | inf | 1 | 1 |
| Abca8a | 0 | 0,00786452 | inf | 1 | 1 |
| Abcb5 | 0 | 0,0219157 | inf | 1 | 1 |
| Acer1 | 0 | 0,0673536 | inf | 1 | 1 |
| Acsm2 | 0 | 0,00672243 | inf | 1 | 1 |
| Acss2os | 0 | 0,107993 | inf | 1 | 1 |
| Actl11 | 0 | 0,0224665 | inf | 1 | 1 |
| Actl7b | 0 | 0,0349732 | inf | 1 | 1 |
| Actl9 | 0 | 0,0230254 | inf | 1 | 1 |
| Actrt3 | 0 | 0,104672 | inf | 1 | 1 |
| Adam26a | 0 | 0,0186107 | inf | 1 | 1 |
| Adamts13 | 0 | 0,161441 | inf | 1 | 1 |
| Afm | 0 | 0,0147551 | inf | 1 | 1 |
| Afp | 0 | 0,0226156 | inf | 1 | 1 |
| Aif1 | 0 | 0,0399767 | inf | 1 | 1 |
| Akp3 | 0 | 0,0576728 | inf | 1 | 1 |
| Akr1d1 | 0 | 0,0341184 | inf | 1 | 1 |
| Aldh3a1 | 0 | 0,0305964 | inf | 1 | 1 |
| Alox15 | 0 | 0,0192591 | inf | 1 | 1 |
| Als2cr11 | 0 | 0,0139205 | inf | 1 | 1 |
| Ambp | 0 | 0,208549 | inf | 1 | 1 |
| Apol10b | 0 | 0,0451595 | inf | 1 | 1 |
| Aqp12 | 0 | 0,0964595 | inf | 1 | 1 |
| Aqp5 | 0 | 0,122939 | inf | 1 | 1 |
| Areg | 0 | 0,0269594 | inf | 1 | 1 |
| Ascl5 | 0 | 0,0738952 | inf | 1 | 1 |
| Aspa | 0 | 0,0201235 | inf | 1 | 1 |
| Asphd1 | 0 | 0,391823 | inf | 1 | 1 |
| Atp10b | 0 | 0,00800145 | inf | 1 | 1 |
| Atp12a | 0 | 0,00715901 | inf | 1 | 1 |
| Atp6v1e2 | 0 | 0,0531363 | inf | 1 | 1 |
| Atp6v1g3 | 0 | 0,232492 | inf | 1 | 1 |
| BB031773 | 0 | 0,0120754 | inf | 1 | 1 |
| BB283400 | 0 | 0,0220961 | inf | 1 | 1 |
| BC016579 | 0 | 0,0642293 | inf | 1 | 1 |
| BC027072 | 0 | 0,00886355 | inf | 1 | 1 |
| BC033916 | 0 | 0,0705837 | inf | 1 | 1 |
| Banf2 | 0 | 0,207569 | inf | 1 | 1 |
| Bhmt | 0 | 0,0227973 | inf | 1 | 1 |
| Bpifc | 0 | 0,0400778 | inf | 1 | 1 |

| | | | | | |
|---|---|---|---|---|---|
| Bsx | 0 | 0,0399767 | inf | 1 | 1 |
| Btc | 0 | 0,0259684 | inf | 1 | 1 |
| Btg4 | 0 | 0,0726954 | inf | 1 | 1 |
| C030029H02Rik | 0 | 0,0129276 | inf | 1 | 1 |
| C030034L19Rik | 0 | 0,0145216 | inf | 1 | 1 |
| C130026I21Rik | 0 | 0,050576 | inf | 1 | 1 |
| C330011F03Rik | 0 | 0,0125 | inf | 1 | 1 |
| C3ar1 | 0 | 0,00655063 | inf | 1 | 1 |
| C86695 | 0 | 0,0159386 | inf | 1 | 1 |
| Cabp4 | 0 | 0,103895 | inf | 1 | 1 |
| Cacna1f | 0 | 0,0163878 | inf | 1 | 1 |
| Cacng1 | 0 | 0,0268514 | inf | 1 | 1 |
| Cacng6 | 0 | 0,0463287 | inf | 1 | 1 |
| Calcoco2 | 0 | 0,0146985 | inf | 1 | 1 |
| Calr4 | 0 | 0,122981 | inf | 1 | 1 |
| Capn9 | 0 | 0,0400401 | inf | 1 | 1 |
| Catsper3 | 0 | 0,0269571 | inf | 1 | 1 |
| Catsperg2 | 0 | 0,011341 | inf | 1 | 1 |
| Ccdc175 | 0 | 0,0177703 | inf | 1 | 1 |
| Ccdc183 | 0 | 0,0279833 | inf | 1 | 1 |
| Ccdc87 | 0 | 0,036262 | inf | 1 | 1 |
| Ccl12 | 0 | 0,208991 | inf | 1 | 1 |
| Ccl17 | 0 | 0,173057 | inf | 1 | 1 |
| Ccl2 | 0 | 0,0445587 | inf | 1 | 1 |
| Cd19 | 0 | 0,0190693 | inf | 1 | 1 |
| Cd3e | 0 | 0,125416 | inf | 1 | 1 |
| Cd52 | 0 | 0,232056 | inf | 1 | 1 |
| Cdrt4 | 0 | 0,206738 | inf | 1 | 1 |
| Ceacam10 | 0 | 0,100739 | inf | 1 | 1 |
| Ceacam12 | 0 | 0,0508835 | inf | 1 | 1 |
| Ceacam18 | 0 | 0,0154788 | inf | 1 | 1 |
| Ceacam19 | 0 | 0,013482 | inf | 1 | 1 |
| Ces2g | 0 | 0,0843316 | inf | 1 | 1 |
| Ces5a | 0 | 0,0569639 | inf | 1 | 1 |
| Chp2 | 0 | 0,0554799 | inf | 1 | 1 |
| Chrm3 | 0 | 0,0234101 | inf | 1 | 1 |
| Chrne | 0 | 0,0193401 | inf | 1 | 1 |
| Cistr-act | 0 | 0,363049 | inf | 1 | 1 |
| Ckm | 0 | 0,0351465 | inf | 1 | 1 |
| Clcnkb | 0 | 0,0199263 | inf | 1 | 1 |
| Clec2h | 0 | 0,02176 | inf | 1 | 1 |
| Clec2i | 0 | 0,0208192 | inf | 1 | 1 |
| Clec7a | 0 | 0,0722921 | inf | 1 | 1 |
| Clec9a | 0 | 0,00978544 | inf | 1 | 1 |
| Clic5 | 0 | 0,0122463 | inf | 1 | 1 |
| Cma1 | 0 | 0,0488099 | inf | 1 | 1 |
| Cmtm1 | 0 | 0,0459636 | inf | 1 | 1 |
| Cmya5 | 0 | 0,0174586 | inf | 1 | 1 |
| Cnga3 | 0 | 0,0126998 | inf | 1 | 1 |
| Cngb1 | 0 | 0,00448052 | inf | 1 | 1 |

| | | | | | |
|---|---|---|---|---|---|
| Cntn5 | 0 | 0,0125107 | inf | 1 | 1 |
| Col6a5 | 0 | 0,00763089 | inf | 1 | 1 |
| Cplx4 | 0 | 0,0721326 | inf | 1 | 1 |
| Crisp2 | 0 | 0,0430059 | inf | 1 | 1 |
| Cst6 | 0 | 0,00715901 | inf | 1 | 1 |
| Cst7 | 0 | 0,0544274 | inf | 1 | 1 |
| Ctse | 0 | 0,0444286 | inf | 1 | 1 |
| Cx3cr1 | 0 | 0,00755841 | inf | 1 | 1 |
| Cxcr3 | 0 | 0,0494211 | inf | 1 | 1 |
| Cyct | 0 | 0,190007 | inf | 1 | 1 |
| Cyp11a1 | 0 | 0,0713056 | inf | 1 | 1 |
| Cyp1a2 | 0 | 0,0252148 | inf | 1 | 1 |
| Cyp21a1 | 0 | 0,0468145 | inf | 1 | 1 |
| Cyp2b19 | 0 | 0,0170866 | inf | 1 | 1 |
| Cyp2b23 | 0 | 0,0271231 | inf | 1 | 1 |
| Cyp2b9 | 0 | 0,0160339 | inf | 1 | 1 |
| Cyp2c44 | 0 | 0,0457239 | inf | 1 | 1 |
| Cyp2c55 | 0 | 0,0715338 | inf | 1 | 1 |
| Cyp2j11 | 0 | 0,0246026 | inf | 1 | 1 |
| Cyp4f18 | 0 | 0,0852387 | inf | 1 | 1 |
| Cyp8b1 | 0 | 0,0487544 | inf | 1 | 1 |
| D14Ertd670e | 0 | 0,117899 | inf | 1 | 1 |
| D16Ertd519e | 0 | 0,209663 | inf | 1 | 1 |
| D530049I02Rik | 0 | 0,122655 | inf | 1 | 1 |
| D630029K05Rik | 0 | 0,0277807 | inf | 1 | 1 |
| D630041G03Rik | 0 | 0,0175954 | inf | 1 | 1 |
| Dbhos | 0 | 0,0140591 | inf | 1 | 1 |
| Ddx60 | 0 | 0,0212683 | inf | 1 | 1 |
| Defb37 | 0 | 0,180342 | inf | 1 | 1 |
| Defb40 | 0 | 0,923859 | inf | 1 | 1 |
| Defb47 | 0 | 0,820767 | inf | 1 | 1 |
| Dlx4 | 0 | 0,0165994 | inf | 1 | 1 |
| Dmrt1 | 0 | 0,0207927 | inf | 1 | 1 |
| Dnmt3aos | 0 | 0,0720786 | inf | 1 | 1 |
| Dnmt3l | 0 | 0,116786 | inf | 1 | 1 |
| Dpp10 | 0 | 0,00961653 | inf | 1 | 1 |
| Duox2 | 0 | 0,0123333 | inf | 1 | 1 |
| E130114P18Rik | 0 | 0,0414434 | inf | 1 | 1 |
| E230025N22Rik | 0 | 0,0469786 | inf | 1 | 1 |
| E430016F16Rik | 0 | 0,0294858 | inf | 1 | 1 |
| Ear14 | 0 | 0,161746 | inf | 1 | 1 |
| Ear4 | 0 | 0,00315687 | inf | 1 | 1 |
| Efcab8 | 0 | 0,0889194 | inf | 1 | 1 |
| Eif4e1b | 0 | 0,027723 | inf | 1 | 1 |
| Elovl3 | 0 | 0,050821 | inf | 1 | 1 |
| Emr4 | 0 | 0,0470073 | inf | 1 | 1 |
| Enpp6 | 0 | 0,018752 | inf | 1 | 1 |
| Enpp7 | 0 | 0,0365029 | inf | 1 | 1 |
| Ephx3 | 0 | 0,0192417 | inf | 1 | 1 |
| Epsti1 | 0 | 0,0234476 | inf | 1 | 1 |

| | | | | | |
|---|---|---|---|---|---|
| Eras | 0 | 0,0614536 | inf | 1 | 1 |
| Erich5 | 0 | 0,0442966 | inf | 1 | 1 |
| Ermn | 0 | 0,00808707 | inf | 1 | 1 |
| Erv3 | 0 | 0,00845142 | inf | 1 | 1 |
| Etnppl | 0 | 0,0173711 | inf | 1 | 1 |
| F2 | 0 | 0,0300726 | inf | 1 | 1 |
| Fam135b | 0 | 0,0192635 | inf | 1 | 1 |
| Fam150b | 0 | 0,0844813 | inf | 1 | 1 |
| Fam19a3 | 0 | 0,0364362 | inf | 1 | 1 |
| Fam71d | 0 | 0,0276292 | inf | 1 | 1 |
| Fas | 0 | 0,185393 | inf | 1 | 1 |
| Fbxw13 | 0 | 0,033226 | inf | 1 | 1 |
| Fbxw19 | 0 | 0,0338334 | inf | 1 | 1 |
| Fcgr1 | 0 | 0,0290933 | inf | 1 | 1 |
| Fcgr2b | 0 | 0,134009 | inf | 1 | 1 |
| Fer1l4 | 0 | 0,0390335 | inf | 1 | 1 |
| Fetub | 0 | 0,0236116 | inf | 1 | 1 |
| Fgf21 | 0 | 0,0572842 | inf | 1 | 1 |
| Fkbp6 | 0 | 0,0378983 | inf | 1 | 1 |
| Flt3 | 0 | 0,0123299 | inf | 1 | 1 |
| Fmo6 | 0 | 0,0255478 | inf | 1 | 1 |
| Fmr1nb | 0 | 0,219498 | inf | 1 | 1 |
| Fosl1 | 0 | 0,10244 | inf | 1 | 1 |
| Foxh1 | 0 | 0,165597 | inf | 1 | 1 |
| Frem3 | 0 | 0,0132553 | inf | 1 | 1 |
| Fsd2 | 0 | 0,0193541 | inf | 1 | 1 |
| Fstl5 | 0 | 0,0196221 | inf | 1 | 1 |
| Fut7 | 0 | 0,0274067 | inf | 1 | 1 |
| G630055G22Rik | 0 | 0,0237807 | inf | 1 | 1 |
| Gbp11 | 0 | 0,0107128 | inf | 1 | 1 |
| Gcsam | 0 | 0,0275086 | inf | 1 | 1 |
| Gdpd4 | 0 | 0,0188509 | inf | 1 | 1 |
| Gfi1b | 0 | 0,0280171 | inf | 1 | 1 |
| Gipc3 | 0 | 0,10381 | inf | 1 | 1 |
| Gjc2 | 0 | 0,352366 | inf | 1 | 1 |
| Gjc3 | 0 | 0,0128549 | inf | 1 | 1 |
| Gjd2 | 0 | 0,0159113 | inf | 1 | 1 |
| Gm10373 | 0 | 0,0578107 | inf | 1 | 1 |
| Gm10400 | 0 | 0,00738772 | inf | 1 | 1 |
| Gm10440 | 0 | 0,0167109 | inf | 1 | 1 |
| Gm1045 | 0 | 0,0766881 | inf | 1 | 1 |
| Gm10510 | 0 | 0,164015 | inf | 1 | 1 |
| Gm10640 | 0 | 0,189984 | inf | 1 | 1 |
| Gm10845 | 0 | 0,0168445 | inf | 1 | 1 |
| Gm10872 | 0 | 0,0554918 | inf | 1 | 1 |
| Gm11186 | 0 | 0,142332 | inf | 1 | 1 |
| Gm1123 | 0 | 0,098808 | inf | 1 | 1 |
| Gm11413 | 0 | 0,0765408 | inf | 1 | 1 |
| Gm11529 | 0 | 0,0144297 | inf | 1 | 1 |
| Gm11545 | 0 | 0,0107773 | inf | 1 | 1 |

| | | | | | |
|---|---|---|---|---|---|
| Gm11570 | 0 | 0,0159008 | inf | 1 | 1 |
| Gm11762 | 0 | 0,0283561 | inf | 1 | 1 |
| Gm12216 | 0 | 0,11809 | inf | 1 | 1 |
| Gm12633 | 0 | 0,0286423 | inf | 1 | 1 |
| Gm13247 | 0 | 0,0154805 | inf | 1 | 1 |
| Gm13271 | 0 | 0,155501 | inf | 1 | 1 |
| Gm13286 | 0 | 0,123366 | inf | 1 | 1 |
| Gm13288 | 0 | 0,0160857 | inf | 1 | 1 |
| Gm13539 | 0 | 0,136857 | inf | 1 | 1 |
| Gm136 | 0 | 0,0244082 | inf | 1 | 1 |
| Gm13944 | 0 | 0,147117 | inf | 1 | 1 |
| Gm14446 | 0 | 0,0266745 | inf | 1 | 1 |
| Gm14496 | 0 | 0,0181427 | inf | 1 | 1 |
| Gm15133 | 0 | 0,0677739 | inf | 1 | 1 |
| Gm15816 | 0 | 0,0570161 | inf | 1 | 1 |
| Gm16130 | 0 | 0,0103371 | inf | 1 | 1 |
| Gm16432 | 0 | 0,0183857 | inf | 1 | 1 |
| Gm1661 | 0 | 0,0348587 | inf | 1 | 1 |
| Gm1965 | 0 | 0,0315582 | inf | 1 | 1 |
| Gm19990 | 0 | 0,0418509 | inf | 1 | 1 |
| Gm20594 | 0 | 0,612476 | inf | 1 | 1 |
| Gm20750 | 0 | 0,0128549 | inf | 1 | 1 |
| Gm20755 | 0 | 0,144785 | inf | 1 | 1 |
| Gm21319 | 0 | 0,0161692 | inf | 1 | 1 |
| Gm2176 | 0 | 0,0278186 | inf | 1 | 1 |
| Gm2721 | 0 | 0,0195658 | inf | 1 | 1 |
| Gm3317 | 0 | 0,0348599 | inf | 1 | 1 |
| Gm4265 | 0 | 0,181114 | inf | 1 | 1 |
| Gm4371 | 0 | 0,0190787 | inf | 1 | 1 |
| Gm4559 | 0 | 0,107479 | inf | 1 | 1 |
| Gm5142 | 0 | 0,0635642 | inf | 1 | 1 |
| Gm5475 | 0 | 0,0201147 | inf | 1 | 1 |
| Gm5591 | 0 | 0,0394465 | inf | 1 | 1 |
| Gm5800 | 0 | 0,197351 | inf | 1 | 1 |
| Gm6086 | 0 | 0,0110409 | inf | 1 | 1 |
| Gm6289 | 0 | 0,0842808 | inf | 1 | 1 |
| Gm6377 | 0 | 0,0206583 | inf | 1 | 1 |
| Gm6904 | 0 | 0,115956 | inf | 1 | 1 |
| Gm7616 | 0 | 0,414224 | inf | 1 | 1 |
| Gm8298 | 0 | 0,0424774 | inf | 1 | 1 |
| Gm9767 | 0 | 0,0624714 | inf | 1 | 1 |
| Gpr101 | 0 | 0,00515268 | inf | 1 | 1 |
| Gpr128 | 0 | 0,194518 | inf | 1 | 1 |
| Gpr139 | 0 | 0,0510339 | inf | 1 | 1 |
| Gpr141 | 0 | 0,0346703 | inf | 1 | 1 |
| Gpr158 | 0 | 0,00613562 | inf | 1 | 1 |
| Gpr33 | 0 | 0,0359784 | inf | 1 | 1 |
| Gpr84 | 0 | 0,0398287 | inf | 1 | 1 |
| Gramd2 | 0 | 0,0365657 | inf | 1 | 1 |
| Grid2 | 0 | 0,0150932 | inf | 1 | 1 |

| | | | | | |
|---|---|---|---|---|---|
| Grifin | 0 | 0,0800713 | inf | 1 | 1 |
| Grm2 | 0 | 0,0136463 | inf | 1 | 1 |
| Gstt4 | 0 | 0,0353418 | inf | 1 | 1 |
| Gtf2a1l | 0 | 0,0379147 | inf | 1 | 1 |
| Gykl1 | 0 | 0,0156778 | inf | 1 | 1 |
| H2-Q2 | 0 | 0,0299715 | inf | 1 | 1 |
| H60c | 0 | 0,0325776 | inf | 1 | 1 |
| Haao | 0 | 0,449886 | inf | 1 | 1 |
| Hand1 | 0 | 0,154552 | inf | 1 | 1 |
| Hbq1b | 0 | 0,0493093 | inf | 1 | 1 |
| Hcar2 | 0 | 0,01539 | inf | 1 | 1 |
| Helt | 0 | 0,0201993 | inf | 1 | 1 |
| Hes2 | 0 | 0,0231547 | inf | 1 | 1 |
| Hist1h2af | 0 | 0,161215 | inf | 1 | 1 |
| Hist2h3b | 0 | 0,0900126 | inf | 1 | 1 |
| Hmx2 | 0 | 0,0366207 | inf | 1 | 1 |
| Hnf4a | 0 | 0,0102462 | inf | 1 | 1 |
| Hrh4 | 0 | 0,0319298 | inf | 1 | 1 |
| Hsd17b1 | 0 | 0,124685 | inf | 1 | 1 |
| Hsd3b6 | 0 | 0,133174 | inf | 1 | 1 |
| Htr4 | 0 | 0,121123 | inf | 1 | 1 |
| I830012O16Rik | 0 | 0,0470565 | inf | 1 | 1 |
| Icos | 0 | 0,0930796 | inf | 1 | 1 |
| Ifi27 | 0 | 0,0475583 | inf | 1 | 1 |
| Igll1 | 0 | 0,127814 | inf | 1 | 1 |
| Il10 | 0 | 0,0243417 | inf | 1 | 1 |
| Il11 | 0 | 0,0913175 | inf | 1 | 1 |
| Il12a | 0 | 0,0251072 | inf | 1 | 1 |
| Il12rb1 | 0 | 0,015953 | inf | 1 | 1 |
| Il17c | 0 | 0,2138 | inf | 1 | 1 |
| Il17f | 0 | 0,0550812 | inf | 1 | 1 |
| Il18r1 | 0 | 0,00678422 | inf | 1 | 1 |
| Il1bos | 0 | 0,0357067 | inf | 1 | 1 |
| Il1f5 | 0 | 0,0821142 | inf | 1 | 1 |
| Il22 | 0 | 0,0482019 | inf | 1 | 1 |
| Il25 | 0 | 0,0990972 | inf | 1 | 1 |
| Il7 | 0 | 0,0374836 | inf | 1 | 1 |
| Il9r | 0 | 0,0706515 | inf | 1 | 1 |
| Ins1 | 0 | 0,0522877 | inf | 1 | 1 |
| Insl3 | 0 | 0,111173 | inf | 1 | 1 |
| Iqch | 0 | 0,0606826 | inf | 1 | 1 |
| Iqcj | 0 | 0,0309598 | inf | 1 | 1 |
| Irg1 | 0 | 0,0178535 | inf | 1 | 1 |
| Itgam | 0 | 0,0190161 | inf | 1 | 1 |
| Itgax | 0 | 0,0111392 | inf | 1 | 1 |
| Kcnd3os | 0 | 0,030276 | inf | 1 | 1 |
| Kcnk4 | 0 | 0,0278005 | inf | 1 | 1 |
| Kcnmb3 | 0 | 0,164968 | inf | 1 | 1 |
| Kctd16 | 0 | 0,0231047 | inf | 1 | 1 |
| Khdc1a | 0 | 0,0243772 | inf | 1 | 1 |

| | | | | | |
|---|---|---|---|---|---|
| Kif2b | 0 | 0,0206015 | inf | 1 | 1 |
| Kiss1 | 0 | 0,275586 | inf | 1 | 1 |
| Klhl30 | 0 | 0,0302292 | inf | 1 | 1 |
| Klk1 | 0 | 0,0399173 | inf | 1 | 1 |
| Klk11 | 0 | 0,0253062 | inf | 1 | 1 |
| Klrb1f | 0 | 0,0198662 | inf | 1 | 1 |
| Kng1 | 0 | 0,0812652 | inf | 1 | 1 |
| Kng2 | 0 | 0,0190478 | inf | 1 | 1 |
| Krt23 | 0 | 0,0519754 | inf | 1 | 1 |
| Krt25 | 0 | 0,0370763 | inf | 1 | 1 |
| Krt27 | 0 | 0,0921621 | inf | 1 | 1 |
| Krt4 | 0 | 0,013906 | inf | 1 | 1 |
| Krt42 | 0 | 0,0199737 | inf | 1 | 1 |
| Krt71 | 0 | 0,0432108 | inf | 1 | 1 |
| Krt80 | 0 | 0,0542198 | inf | 1 | 1 |
| Krt81 | 0 | 0,0257488 | inf | 1 | 1 |
| Krt83 | 0 | 0,194489 | inf | 1 | 1 |
| Krt84 | 0 | 0,0719572 | inf | 1 | 1 |
| Krtap3-1 | 0 | 0,20118 | inf | 1 | 1 |
| Krtap3-3 | 0 | 0,322438 | inf | 1 | 1 |
| Krtap9-3 | 0 | 0,0766881 | inf | 1 | 1 |
| L3mbtl4 | 0 | 0,00974869 | inf | 1 | 1 |
| LOC101056236 | 0 | 0,0599431 | inf | 1 | 1 |
| LOC102308570 | 0 | 0,0557701 | inf | 1 | 1 |
| LOC102632423 | 0 | 0,125346 | inf | 1 | 1 |
| LOC102634431 | 0 | 0,0430035 | inf | 1 | 1 |
| LOC381967 | 0 | 0,303428 | inf | 1 | 1 |
| Lactbl1 | 0 | 0,0306568 | inf | 1 | 1 |
| Lce1b | 0 | 0,096076 | inf | 1 | 1 |
| Lce1i | 0 | 0,0807625 | inf | 1 | 1 |
| Lctl | 0 | 0,0479727 | inf | 1 | 1 |
| Ldhc | 0 | 0,0242537 | inf | 1 | 1 |
| Leap2 | 0 | 0,145259 | inf | 1 | 1 |
| Lgals12 | 0 | 0,157031 | inf | 1 | 1 |
| Lhb | 0 | 0,175894 | inf | 1 | 1 |
| Lif | 0 | 0,0405367 | inf | 1 | 1 |
| Lipc | 0 | 0,0248034 | inf | 1 | 1 |
| Lman1l | 0 | 0,0277642 | inf | 1 | 1 |
| Lrrc74 | 0 | 0,0243912 | inf | 1 | 1 |
| Ltb4r2 | 0 | 0,0561839 | inf | 1 | 1 |
| Magea5 | 0 | 0,062638 | inf | 1 | 1 |
| Mep1a | 0 | 0,0255195 | inf | 1 | 1 |
| Mepe | 0 | 0,0575948 | inf | 1 | 1 |
| Mettl7a2 | 0 | 2,93E-006 | inf | 1 | 1 |
| Miox | 0 | 0,0318203 | inf | 1 | 1 |
| Mir1892 | 0 | 138,786 | inf | 1 | 1 |
| Mir1903 | 0 | 25,1449 | inf | 1 | 1 |
| Mir1932 | 0 | 24,0437 | inf | 1 | 1 |
| Mir1966 | 0 | 16,6312 | inf | 1 | 1 |
| Mir2139 | 0 | 27,5454 | inf | 1 | 1 |

| | | | | | |
|---|---|---|---|---|---|
| Mir25 | 0 | 53,8034 | inf | 1 | 1 |
| Mir3072 | 0 | 57,2407 | inf | 1 | 1 |
| Mir345 | 0 | 35,4494 | inf | 1 | 1 |
| Mir599 | 0 | 42,575 | inf | 1 | 1 |
| Mir6361 | 0 | 22,328 | inf | 1 | 1 |
| Mir6403 | 0 | 7,93725 | inf | 1 | 1 |
| Mir6939 | 0 | 100,362 | inf | 1 | 1 |
| Mir7075 | 0 | 60,9897 | inf | 1 | 1 |
| Mirlet7i | 0 | 50,6452 | inf | 1 | 1 |
| Mmp12 | 0 | 0,00788269 | inf | 1 | 1 |
| Mmp27 | 0 | 0,0184743 | inf | 1 | 1 |
| Mmp7 | 0 | 0,0795976 | inf | 1 | 1 |
| Morc1 | 0 | 0,0377017 | inf | 1 | 1 |
| Moxd2 | 0 | 0,025702 | inf | 1 | 1 |
| Mpo | 0 | 0,052535 | inf | 1 | 1 |
| Mrgprb5 | 0 | 0,0556368 | inf | 1 | 1 |
| Mrgprh | 0 | 0,0530374 | inf | 1 | 1 |
| Ms4a10 | 0 | 0,0528723 | inf | 1 | 1 |
| Ms4a13 | 0 | 0,0363471 | inf | 1 | 1 |
| Ms4a7 | 0 | 0,024948 | inf | 1 | 1 |
| Mslnl | 0 | 0,0351177 | inf | 1 | 1 |
| Msmb | 0 | 0,134262 | inf | 1 | 1 |
| Msr1 | 0 | 0,067399 | inf | 1 | 1 |
| Mtnr1b | 0 | 0,0667182 | inf | 1 | 1 |
| Muc13 | 0 | 0,0194031 | inf | 1 | 1 |
| Muc5ac | 0 | 0,00512743 | inf | 1 | 1 |
| Muc5b | 0 | 0,00181753 | inf | 1 | 1 |
| Mug2 | 0 | 0,00980709 | inf | 1 | 1 |
| Mx1 | 0 | 0,0512156 | inf | 1 | 1 |
| Mycs | 0 | 0,0248608 | inf | 1 | 1 |
| Myh4 | 0 | 0,0164945 | inf | 1 | 1 |
| Mylk2 | 0 | 0,025287 | inf | 1 | 1 |
| Myo1f | 0 | 0,0117285 | inf | 1 | 1 |
| Myo3b | 0 | 0,113589 | inf | 1 | 1 |
| Nanos2 | 0 | 0,0344635 | inf | 1 | 1 |
| Ncf4 | 0 | 0,0680999 | inf | 1 | 1 |
| Nek10 | 0 | 0,0135378 | inf | 1 | 1 |
| Nek5 | 0 | 0,0243861 | inf | 1 | 1 |
| Neto1 | 0 | 0,0127931 | inf | 1 | 1 |
| Neurod2 | 0 | 0,0145117 | inf | 1 | 1 |
| Ninj2 | 0 | 0,0434051 | inf | 1 | 1 |
| Nkx1-1 | 0 | 0,0380361 | inf | 1 | 1 |
| Nlrp4f | 0 | 0,0128085 | inf | 1 | 1 |
| Nlrp4g | 0 | 0,0316548 | inf | 1 | 1 |
| Npc1l1 | 0 | 0,0122852 | inf | 1 | 1 |
| Nphs1 | 0 | 0,0155993 | inf | 1 | 1 |
| Nphs2 | 0 | 0,0146566 | inf | 1 | 1 |
| Nrap | 0 | 0,0596634 | inf | 1 | 1 |
| Nt5c1a | 0 | 0,047344 | inf | 1 | 1 |
| Nxf3 | 0 | 0,00967007 | inf | 1 | 1 |

| | | | | | |
|---|---|---|---|---|---|
| Oas1h | 0 | 0,0358379 | inf | 1 | 1 |
| Oas3 | 0 | 0,00594259 | inf | 1 | 1 |
| Odf3l1 | 0 | 0,0485871 | inf | 1 | 1 |
| Ofcc1 | 0 | 0,0102698 | inf | 1 | 1 |
| Oit1 | 0 | 0,128102 | inf | 1 | 1 |
| Olfr1168 | 0 | 0,0579077 | inf | 1 | 1 |
| Olfr1188 | 0 | 0,0557097 | inf | 1 | 1 |
| Olfr1198 | 0 | 0,117738 | inf | 1 | 1 |
| Olfr1220 | 0 | 0,0517789 | inf | 1 | 1 |
| Olfr1249 | 0 | 0,0565235 | inf | 1 | 1 |
| Olfr1270 | 0 | 0,0522877 | inf | 1 | 1 |
| Olfr1347 | 0 | 0,0562992 | inf | 1 | 1 |
| Olfr1349 | 0 | 0,0925155 | inf | 1 | 1 |
| Olfr1366 | 0 | 0,0315582 | inf | 1 | 1 |
| Olfr1392 | 0 | 0,0517789 | inf | 1 | 1 |
| Olfr1396 | 0 | 0,0547776 | inf | 1 | 1 |
| Olfr1414 | 0 | 0,086421 | inf | 1 | 1 |
| Olfr1417 | 0 | 0,114414 | inf | 1 | 1 |
| Olfr1420 | 0 | 0,0369483 | inf | 1 | 1 |
| Olfr1474 | 0 | 0,0574388 | inf | 1 | 1 |
| Olfr218 | 0 | 0,0576723 | inf | 1 | 1 |
| Olfr267 | 0 | 0,0576723 | inf | 1 | 1 |
| Olfr370 | 0 | 0,0477156 | inf | 1 | 1 |
| Olfr38 | 0 | 0,0567495 | inf | 1 | 1 |
| Olfr412 | 0 | 0,0579077 | inf | 1 | 1 |
| Olfr444 | 0 | 0,0583843 | inf | 1 | 1 |
| Olfr456 | 0 | 0,0316327 | inf | 1 | 1 |
| Olfr457 | 0 | 0,0576723 | inf | 1 | 1 |
| Olfr469 | 0 | 0,0574388 | inf | 1 | 1 |
| Olfr5 | 0 | 0,0506693 | inf | 1 | 1 |
| Olfr521 | 0 | 0,0558559 | inf | 1 | 1 |
| Olfr655 | 0 | 0,0742031 | inf | 1 | 1 |
| Olfr698 | 0 | 0,058145 | inf | 1 | 1 |
| Olfr726 | 0 | 0,0704054 | inf | 1 | 1 |
| Olfr750 | 0 | 0,0458645 | inf | 1 | 1 |
| Olfr767 | 0 | 0,0586255 | inf | 1 | 1 |
| Olfr920 | 0 | 0,0242384 | inf | 1 | 1 |
| Olfr94 | 0 | 0,0444747 | inf | 1 | 1 |
| Olfr95 | 0 | 0,173723 | inf | 1 | 1 |
| Olfr960 | 0 | 0,143638 | inf | 1 | 1 |
| Olfr995 | 0 | 0,0572072 | inf | 1 | 1 |
| Onecut3 | 0 | 0,0189801 | inf | 1 | 1 |
| Orm2 | 0 | 0,0470602 | inf | 1 | 1 |
| Ovch2 | 0 | 0,0247457 | inf | 1 | 1 |
| Oxct2a | 0 | 0,026585 | inf | 1 | 1 |
| Oxt | 0 | 0,079833 | inf | 1 | 1 |
| Pabpc2 | 0 | 0,0112614 | inf | 1 | 1 |
| Patl2 | 0 | 0,012029 | inf | 1 | 1 |
| Pbld1 | 0 | 0,0979013 | inf | 1 | 1 |
| Pcdha11 | 0 | 0,020603 | inf | 1 | 1 |

| | | | | | |
|---|---|---|---|---|---|
| Pcdha9 | 0 | 0,0227174 | inf | 1 | 1 |
| Pck1 | 0 | 0,0111214 | inf | 1 | 1 |
| Pdcd1lg2 | 0 | 0,0780832 | inf | 1 | 1 |
| Pde6h | 0 | 0,303095 | inf | 1 | 1 |
| Pdx1 | 0 | 0,0393004 | inf | 1 | 1 |
| Pfn3 | 0 | 0,0786624 | inf | 1 | 1 |
| Pglyrp1 | 0 | 0,234691 | inf | 1 | 1 |
| Pifo | 0 | 0,0604517 | inf | 1 | 1 |
| Pik3cg | 0 | 0,0556698 | inf | 1 | 1 |
| Pirb | 0 | 0,06561 | inf | 1 | 1 |
| Piwil1 | 0 | 0,0299332 | inf | 1 | 1 |
| Pla2g10 | 0 | 0,0697137 | inf | 1 | 1 |
| Pla2g1b | 0 | 0,244605 | inf | 1 | 1 |
| Pla2g2e | 0 | 0,297406 | inf | 1 | 1 |
| Pla2g5 | 0 | 0,165602 | inf | 1 | 1 |
| Pldi | 0 | 0,065581 | inf | 1 | 1 |
| Plxdc1 | 0 | 0,0158655 | inf | 1 | 1 |
| Pou4f2 | 0 | 0,0141498 | inf | 1 | 1 |
| Ppbp | 0 | 0,0485871 | inf | 1 | 1 |
| Ppp1r42 | 0 | 0,20881 | inf | 1 | 1 |
| Prlhr | 0 | 0,0311129 | inf | 1 | 1 |
| Prr23a | 0 | 0,0313737 | inf | 1 | 1 |
| Psg25 | 0 | 0,0297329 | inf | 1 | 1 |
| Psmb9 | 0 | 0,550156 | inf | 1 | 1 |
| Ptchd3 | 0 | 0,00884723 | inf | 1 | 1 |
| Ptger3 | 0 | 0,0295752 | inf | 1 | 1 |
| Pthlh | 0 | 0,102541 | inf | 1 | 1 |
| Ptprc | 0 | 0,0161782 | inf | 1 | 1 |
| Pydc3 | 0 | 0,0305876 | inf | 1 | 1 |
| Qrfpr | 0 | 0,552063 | inf | 1 | 1 |
| R3hdml | 0 | 0,0478228 | inf | 1 | 1 |
| Rac2 | 0 | 0,0189441 | inf | 1 | 1 |
| Ramp1 | 0 | 0,0295895 | inf | 1 | 1 |
| Rasal3 | 0 | 0,0476821 | inf | 1 | 1 |
| Rasgef1a | 0 | 0,0228851 | inf | 1 | 1 |
| Rasgrf1 | 0 | 0,0423089 | inf | 1 | 1 |
| Rbm46os | 0 | 0,0139437 | inf | 1 | 1 |
| Rbmxl2 | 0 | 0,188988 | inf | 1 | 1 |
| Rbp3 | 0 | 0,00839172 | inf | 1 | 1 |
| Rdh7 | 0 | 0,030607 | inf | 1 | 1 |
| Ren1 | 0 | 0,0349084 | inf | 1 | 1 |
| Ren2 | 0 | 3,33E-005 | inf | 1 | 1 |
| Rfpl4 | 0 | 0,0577903 | inf | 1 | 1 |
| Rgs18 | 0 | 0,0162083 | inf | 1 | 1 |
| Rhag | 0 | 0,045555 | inf | 1 | 1 |
| Rnase1 | 0 | 0,111128 | inf | 1 | 1 |
| Rnase6 | 0 | 0,0309395 | inf | 1 | 1 |
| Rnu12 | 0 | 2,76288 | inf | 1 | 1 |
| Rp1l1 | 0 | 0,00649318 | inf | 1 | 1 |
| Rptn | 0 | 0,0107972 | inf | 1 | 1 |

| | | | | | |
|---|---|---|---|---|---|
| S100a14 | 0 | 0,190881 | inf | 1 | 1 |
| S100g | 0 | 0,0972274 | inf | 1 | 1 |
| Selp | 0 | 0,0394831 | inf | 1 | 1 |
| Sept14 | 0 | 0,0180638 | inf | 1 | 1 |
| Serpina12 | 0 | 0,0583797 | inf | 1 | 1 |
| Serpinb6c | 0 | 0,151468 | inf | 1 | 1 |
| Serpinb6d | 0 | 0,067399 | inf | 1 | 1 |
| Siglec15 | 0 | 0,0325145 | inf | 1 | 1 |
| Siglech | 0 | 0,0388168 | inf | 1 | 1 |
| Six3 | 0 | 0,0244895 | inf | 1 | 1 |
| Six6 | 0 | 0,0142989 | inf | 1 | 1 |
| Skor2 | 0 | 0,0144919 | inf | 1 | 1 |
| Slamf1 | 0 | 0,0342695 | inf | 1 | 1 |
| Slc17a2 | 0 | 0,0307312 | inf | 1 | 1 |
| Slc17a3 | 0 | 0,0107785 | inf | 1 | 1 |
| Slc1a7 | 0 | 0,0303911 | inf | 1 | 1 |
| Slc22a12 | 0 | 0,0261833 | inf | 1 | 1 |
| Slc22a2 | 0 | 0,0416102 | inf | 1 | 1 |
| Slc25a41 | 0 | 0,0262471 | inf | 1 | 1 |
| Slc34a1 | 0 | 0,0443824 | inf | 1 | 1 |
| Slc46a2 | 0 | 0,0229324 | inf | 1 | 1 |
| Slc4a9 | 0 | 0,0503848 | inf | 1 | 1 |
| Slc5a1 | 0 | 0,120086 | inf | 1 | 1 |
| Slc5a10 | 0 | 0,0682215 | inf | 1 | 1 |
| Slc5a11 | 0 | 0,0649391 | inf | 1 | 1 |
| Slc5a2 | 0 | 0,0172571 | inf | 1 | 1 |
| Slc5a4b | 0 | 0,0278265 | inf | 1 | 1 |
| Slc5a9 | 0 | 0,026937 | inf | 1 | 1 |
| Slc6a18 | 0 | 0,0555182 | inf | 1 | 1 |
| Slc6a19 | 0 | 0,0670522 | inf | 1 | 1 |
| Slc6a19os | 0 | 0,0319778 | inf | 1 | 1 |
| Slc7a13 | 0 | 0,0441063 | inf | 1 | 1 |
| Slc9a2 | 0 | 0,0258819 | inf | 1 | 1 |
| Slco6b1 | 0 | 0,0187418 | inf | 1 | 1 |
| Slfn2 | 0 | 0,0183731 | inf | 1 | 1 |
| Slfn8 | 0 | 0,0295674 | inf | 1 | 1 |
| Slpi | 0 | 0,0399173 | inf | 1 | 1 |
| Slurp1 | 0 | 0,132589 | inf | 1 | 1 |
| Smpx | 0 | 0,126224 | inf | 1 | 1 |
| Snora24 | 0 | 14,5425 | inf | 1 | 1 |
| Snora26 | 0 | 34,8395 | inf | 1 | 1 |
| Snord7 | 0 | 147,509 | inf | 1 | 1 |
| Sntg1 | 0 | 0,0125357 | inf | 1 | 1 |
| Soat2 | 0 | 0,0779717 | inf | 1 | 1 |
| Sp100 | 0 | 0,0274562 | inf | 1 | 1 |
| Spag11b | 0 | 0,121662 | inf | 1 | 1 |
| Spata16 | 0 | 0,0265008 | inf | 1 | 1 |
| Spata3 | 0 | 0,234716 | inf | 1 | 1 |
| Spata31d1a | 0 | 0,017036 | inf | 1 | 1 |
| Spatc1 | 0 | 0,0382936 | inf | 1 | 1 |

| | | | | | |
|---|---|---|---|---|---|
| Speer4b | 0 | 0,0240074 | inf | 1 | 1 |
| Spi1 | 0 | 0,0237596 | inf | 1 | 1 |
| Strc | 0 | 0,00766483 | inf | 1 | 1 |
| Stx19 | 0 | 0,211545 | inf | 1 | 1 |
| Svs1 | 0 | 0,10787 | inf | 1 | 1 |
| Sycp1-ps1 | 0 | 0,0184426 | inf | 1 | 1 |
| Taar7d | 0 | 0,0307268 | inf | 1 | 1 |
| Tas1r2 | 0 | 0,0188774 | inf | 1 | 1 |
| Tas2r135 | 0 | 0,0558559 | inf | 1 | 1 |
| Tas2r143 | 0 | 0,181468 | inf | 1 | 1 |
| Tbata | 0 | 0,0377237 | inf | 1 | 1 |
| Tbx19 | 0 | 0,017316 | inf | 1 | 1 |
| Tc2n | 0 | 0,00837273 | inf | 1 | 1 |
| Tcte1 | 0 | 0,0130327 | inf | 1 | 1 |
| Tepp | 0 | 0,093336 | inf | 1 | 1 |
| Tex35 | 0 | 0,177284 | inf | 1 | 1 |
| Tfap2e | 0 | 0,14845 | inf | 1 | 1 |
| Tff2 | 0 | 0,0730898 | inf | 1 | 1 |
| Them5 | 0 | 0,0512408 | inf | 1 | 1 |
| Tifab | 0 | 0,0249101 | inf | 1 | 1 |
| Tigd4 | 0 | 0,0639516 | inf | 1 | 1 |
| Tktl2 | 0 | 0,00690506 | inf | 1 | 1 |
| Tlr13 | 0 | 0,0111713 | inf | 1 | 1 |
| Tlx3 | 0 | 0,0201993 | inf | 1 | 1 |
| Tm4sf5 | 0 | 0,0695836 | inf | 1 | 1 |
| Tmc5 | 0 | 0,0394334 | inf | 1 | 1 |
| Tmem150cos | 0 | 0,0236132 | inf | 1 | 1 |
| Tmem154 | 0 | 0,0678182 | inf | 1 | 1 |
| Tmem221 | 0 | 0,0519754 | inf | 1 | 1 |
| Tmem52b | 0 | 0,0619991 | inf | 1 | 1 |
| Tmem8c | 0 | 0,422326 | inf | 1 | 1 |
| Tmprss4 | 0 | 0,0130028 | inf | 1 | 1 |
| Tnf | 0 | 0,346676 | inf | 1 | 1 |
| Tnfrsf8 | 0 | 0,101221 | inf | 1 | 1 |
| Tnfrsf9 | 0 | 0,0685418 | inf | 1 | 1 |
| Tnip3 | 0 | 0,00752449 | inf | 1 | 1 |
| Tnmd | 0 | 0,0239078 | inf | 1 | 1 |
| Tnn | 0 | 0,0123796 | inf | 1 | 1 |
| Tnxb | 0 | 0,00908977 | inf | 1 | 1 |
| Toporsl | 0 | 0,0366292 | inf | 1 | 1 |
| Tpo | 0 | 0,013828 | inf | 1 | 1 |
| Tppp2 | 0 | 0,301143 | inf | 1 | 1 |
| Tprg | 0 | 0,351162 | inf | 1 | 1 |
| Tpsg1 | 0 | 0,0529606 | inf | 1 | 1 |
| Traf1 | 0 | 0,0327145 | inf | 1 | 1 |
| Trcg1 | 0 | 0,0170455 | inf | 1 | 1 |
| Treh | 0 | 0,015407 | inf | 1 | 1 |
| Trim14 | 0 | 0,142471 | inf | 1 | 1 |
| Trim54 | 0 | 0,0201993 | inf | 1 | 1 |
| Tshb | 0 | 0,227959 | inf | 1 | 1 |

| | | | | | |
|---|---|---|---|---|---|
| Tspan2os | 0 | 0,110263 | inf | 1 | 1 |
| Txk | 0 | 0,164628 | inf | 1 | 1 |
| Ubash3a | 0 | 0,00975802 | inf | 1 | 1 |
| Ucp1 | 0 | 0,0777428 | inf | 1 | 1 |
| Ucp3 | 0 | 0,011954 | inf | 1 | 1 |
| Ugt2b5 | 0 | 0,0251999 | inf | 1 | 1 |
| Usp50 | 0 | 0,0449399 | inf | 1 | 1 |
| Usp9y | 0 | 0,00679971 | inf | 1 | 1 |
| Utf1 | 0 | 0,0561183 | inf | 1 | 1 |
| Uts2r | 0 | 0,0283939 | inf | 1 | 1 |
| Vmn1r197 | 0 | 0,122834 | inf | 1 | 1 |
| Vmn1r221 | 0 | 0,0579077 | inf | 1 | 1 |
| Vmn1r26 | 0 | 0,104319 | inf | 1 | 1 |
| Vmn1r49 | 0 | 0,0583843 | inf | 1 | 1 |
| Vmn1r50 | 0 | 0,0367962 | inf | 1 | 1 |
| Vmn1r90 | 0 | 0,0247518 | inf | 1 | 1 |
| Vmn2r109 | 0 | 0,0179362 | inf | 1 | 1 |
| Vmn2r120 | 0 | 0,0359634 | inf | 1 | 1 |
| Vmn2r18 | 0 | 0,0318145 | inf | 1 | 1 |
| Vmn2r23 | 0 | 0,0228685 | inf | 1 | 1 |
| Vmn2r28 | 0 | 0,016757 | inf | 1 | 1 |
| Vmn2r3 | 0 | 0,0172598 | inf | 1 | 1 |
| Vmn2r90 | 0 | 0,0178236 | inf | 1 | 1 |
| Vwde | 0 | 0,0071078 | inf | 1 | 1 |
| Wdr95 | 0 | 0,0422354 | inf | 1 | 1 |
| Wfdc10 | 0 | 0,439 | inf | 1 | 1 |
| Wfdc8 | 0 | 0,01815 | inf | 1 | 1 |
| Yipf7 | 0 | 0,0419334 | inf | 1 | 1 |
| Zbp1 | 0 | 0,0300726 | inf | 1 | 1 |
| Zcchc16 | 0 | 0,0202296 | inf | 1 | 1 |
| Zdhhc19 | 0 | 0,0242537 | inf | 1 | 1 |
| Zfp42 | 0 | 0,0266516 | inf | 1 | 1 |
| Zfp541 | 0 | 0,00976882 | inf | 1 | 1 |
| Zfp831 | 0 | 0,00443317 | inf | 1 | 1 |
| Zp3 | 0 | 0,114723 | inf | 1 | 1 |
| H2-Q6,H2-Q8 | 0,00459447 | 1,52503 | 8,37472 | 1 | 1 |
| H2-L | 0,000976763 | 0,107704 | 6,78484 | 1 | 1 |
| Acnat1 | 0,00454964 | 0,383511 | 6,39737 | 1 | 1 |
| Slc4a1 | 0,00507097 | 0,261215 | 5,68683 | 1 | 1 |
| 4933416I08Rik | 0,00115572 | 0,0509137 | 5,46119 | 1 | 1 |
| Oas1b | 0,0135081 | 0,467827 | 5,11408 | 1 | 1 |
| 1700023E05Rik | 0,0145468 | 0,443366 | 4,92972 | 1 | 1 |
| Crhr1 | 0,0198187 | 0,551618 | 4,79874 | 1 | 1 |
| Syne3 | 0,00590668 | 0,145528 | 4,6228 | 1 | 1 |
| Il22ra1 | 0,0142055 | 0,347614 | 4,61297 | 1 | 1 |
| Adcy8 | 0,00608837 | 0,14389 | 4,56277 | 1 | 1 |
| Btnl9 | 0,00975582 | 0,228945 | 4,5526 | 1 | 1 |
| Gm16880 | 0,022149 | 0,477007 | 4,4287 | 1 | 1 |
| Psmb8 | 0,028366 | 0,608274 | 4,42249 | 1 | 1 |
| Entpd2 | 0,0127169 | 0,253028 | 4,31448 | 1 | 1 |

| | | | | | |
|---|---|---|---|---|---|
| Ccdc147 | 0,00701637 | 0,137556 | 4,29315 | 1 | 1 |
| Glp1r | 0,0226528 | 0,431384 | 4,25121 | 1 | 1 |
| Ibsp | 0,0125862 | 0,228775 | 4,18402 | 1 | 1 |
| Crygs | 0,0437261 | 0,763573 | 4,1262 | 1 | 1 |
| Oas1a | 0,0130869 | 0,212603 | 4,02196 | 1 | 1 |
| Vrtn | 0,00761476 | 0,122834 | 4,01176 | 1 | 1 |
| Slc14a2 | 0,0321757 | 0,516877 | 4,00578 | 1 | 1 |
| Dok7 | 0,0253665 | 0,405533 | 3,99882 | 1 | 1 |
| A230028O05Rik | 0,00851233 | 0,134004 | 3,97658 | 1 | 1 |
| Card11 | 0,00563932 | 0,0871295 | 3,94957 | 1 | 1 |
| Tnfsf12Tnfsf13 | 0,00807344 | 0,124307 | 3,94459 | 1 | 1 |
| Sox15 | 0,0309474 | 0,46948 | 3,92317 | 1 | 1 |
| Nodal | 0,0116605 | 0,175237 | 3,90961 | 1 | 1 |
| Pou5f1 | 0,0367272 | 0,539455 | 3,87658 | 1 | 1 |
| Nxph4 | 0,0196499 | 0,284234 | 3,85448 | 1 | 1 |
| Syt15 | 0,0128842 | 0,181186 | 3,81379 | 1 | 1 |
| Agbl1 | 0,00696317 | 0,0976385 | 3,80963 | 1 | 1 |
| Fgf1 | 0,0334482 | 0,465691 | 3,79937 | 1 | 1 |
| C920025E04Rik | 0,0266405 | 0,358984 | 3,75222 | 1 | 1 |
| Sidt1 | 0,00543629 | 0,0696961 | 3,68038 | 1 | 1 |
| Dnah9 | 0,00537821 | 0,0689242 | 3,67981 | 1 | 1 |
| B230216N24Rik | 0,0226265 | 0,284956 | 3,65465 | 1 | 1 |
| Grin2a | 0,0153471 | 0,190041 | 3,63027 | 1 | 1 |
| Spta1 | 0,0035246 | 0,0423044 | 3,58528 | 1 | 1 |
| Abcg3 | 0,00778743 | 0,0925131 | 3,57044 | 1 | 1 |
| C230037L18Rik | 0,0346462 | 0,41132 | 3,56949 | 1 | 1 |
| Cacna1i | 0,0129224 | 0,152615 | 3,56195 | 1 | 1 |
| Egr4 | 0,0304906 | 0,347854 | 3,51205 | 1 | 1 |
| Gm6277 | 0,0361877 | 0,407114 | 3,49186 | 1 | 1 |
| Spata21 | 0,00954456 | 0,103784 | 3,44277 | 1 | 1 |
| 4930590J08Rik | 0,00905835 | 0,096843 | 3,41833 | 1 | 1 |
| Ablim3 | 0,0164826 | 0,175005 | 3,40838 | 1 | 1 |
| Gm5148 | 0,0435969 | 0,460257 | 3,40014 | 1 | 1 |
| Lrp2bp | 0,00898793 | 0,0946439 | 3,39645 | 1 | 1 |
| Sstr1 | 0,00788682 | 0,0830491 | 3,39645 | 1 | 1 |
| Ptger2 | 0,0471388 | 0,484115 | 3,36036 | 1 | 1 |
| Armc4 | 0,0088314 | 0,0901536 | 3,35167 | 1 | 1 |
| Kcnn4 | 0,0126043 | 0,128082 | 3,34509 | 1 | 1 |
| Runx3 | 0,00597734 | 0,060727 | 3,34476 | 1 | 1 |
| 4930578M01Rik | 0,0411163 | 0,417189 | 3,34292 | 1 | 1 |
| B020014A21Rik | 0,0166696 | 0,166457 | 3,31986 | 1 | 1 |
| Cd163 | 0,0173488 | 0,171384 | 3,30433 | 1 | 1 |
| A930003O13Rik | 0,0165456 | 0,162329 | 3,29441 | 1 | 1 |
| Gm5796 | 0,0179423 | 0,172284 | 3,26335 | 1 | 1 |
| Bmp15 | 0,00770844 | 0,0728095 | 3,23962 | 1 | 1 |
| Syndig1l | 0,0125948 | 0,11891 | 3,23897 | 1 | 1 |
| Hus1b | 0,0294347 | 0,271345 | 3,20454 | 1 | 1 |
| Kcnh7 | 0,0624304 | 0,562973 | 3,17275 | 1 | 1 |
| E130310I04Rik | 0,0485432 | 0,437194 | 3,17093 | 1 | 1 |
| Catip | 0,0457118 | 0,409137 | 3,16194 | 1 | 1 |

| | | | | | |
|---|---|---|---|---|---|
| Nostrin | 0,0137482 | 0,122941 | 3,16066 | 1 | 1 |
| Gm8801 | 0,0132522 | 0,118506 | 3,16066 | 1 | 1 |
| Hsf3 | 0,0124651 | 0,111468 | 3,16066 | 1 | 1 |
| Prkag3 | 0,008386 | 0,0749907 | 3,16066 | 1 | 1 |
| Tmprss13 | 0,00724948 | 0,0632976 | 3,1262 | 1 | 1 |
| Shisa9 | 0,012583 | 0,109189 | 3,11728 | 1 | 1 |
| 9430021M05Rik | 0,0375421 | 0,324548 | 3,11185 | 1 | 1 |
| Shisa6 | 0,0612657 | 0,507749 | 3,05096 | 1 | 1 |
| Hist1h4h | 0,0953993 | 0,784969 | 3,04059 | 1 | 1 |
| BC051628 | 0,0187241 | 0,154067 | 3,04059 | 1 | 1 |
| C4b | 0,00424715 | 0,0349466 | 3,04059 | 1 | 1 |
| Htr2a | 0,0552334 | 0,451382 | 3,03074 | 1 | 1 |
| Rbm44 | 0,00792575 | 0,0644331 | 3,02318 | 1 | 1 |
| 9430018G01Rik | 0,0187566 | 0,152406 | 3,02245 | 1 | 1 |
| Gbp3 | 0,0212276 | 0,170585 | 3,00648 | 1 | 1 |
| Nuggc | 0,0093304 | 0,0748038 | 3,0031 | 1 | 1 |
| Plk5 | 0,011587 | 0,092773 | 3,0012 | 1 | 1 |
| 4930519F16Rik | 0,0529554 | 0,416407 | 2,97514 | 1 | 1 |
| Gpbar1 | 0,0560767 | 0,435474 | 2,95711 | 1 | 1 |
| Tfpi2 | 0,0221664 | 0,171702 | 2,95346 | 1 | 1 |
| 4933407L21Rik | 0,0233456 | 0,180349 | 2,94957 | 1 | 1 |
| Rhcg | 0,0116852 | 0,0902706 | 2,94957 | 1 | 1 |
| Entpd8 | 0,0104845 | 0,0809945 | 2,94957 | 1 | 1 |
| Ptprt | 0,0743961 | 0,573904 | 2,94751 | 1 | 1 |
| Comp | 0,0626 | 0,480484 | 2,94026 | 1 | 1 |
| 5430440P10Rik | 0,0355971 | 0,270008 | 2,92317 | 1 | 1 |
| Pdxk-ps | 0,0270394 | 0,205097 | 2,92317 | 1 | 1 |
| Gm13003 | 0,0107056 | 0,0812034 | 2,92317 | 1 | 1 |
| Ccr9 | 0,00534458 | 0,0401854 | 2,91052 | 1 | 1 |
| Slc35g3 | 0,0340623 | 0,255949 | 2,90961 | 1 | 1 |
| Arhgef37 | 0,0145468 | 0,109307 | 2,90961 | 1 | 1 |
| Mmel1 | 0,0211527 | 0,158091 | 2,90184 | 1 | 1 |
| Lgals6 | 0,0116074 | 0,0858876 | 2,8874 | 1 | 1 |
| Parp10 | 0,054417 | 0,400994 | 2,88145 | 1 | 1 |
| Kndc1 | 0,0184241 | 0,135278 | 2,87626 | 1 | 1 |
| D730005E14Rik | 0,0129524 | 0,095044 | 2,87538 | 1 | 1 |
| 9630001P10Rik | 0,00845355 | 0,0618571 | 2,87131 | 1 | 1 |
| Mroh6 | 0,0160807 | 0,116512 | 2,85708 | 1 | 1 |
| Vwa3a | 0,0167007 | 0,120887 | 2,85568 | 1 | 1 |
| Gm5820 | 0,033341 | 0,240247 | 2,84915 | 1 | 1 |
| Rp1 | 0,0101671 | 0,0732408 | 2,84873 | 1 | 1 |
| Ptgir | 0,00923488 | 0,0665056 | 2,84831 | 1 | 1 |
| Ptprh | 0,00757474 | 0,05455 | 2,84831 | 1 | 1 |
| H2-T10 | 0,0544069 | 0,389697 | 2,84049 | 1 | 1 |
| Psors1c2 | 0,068869 | 0,492901 | 2,83937 | 1 | 1 |
| Gypa | 0,058546 | 0,413862 | 2,82151 | 1 | 1 |
| Fam166a | 0,0241853 | 0,169753 | 2,81124 | 1 | 1 |
| Gabra1 | 0,00491601 | 0,0344665 | 2,80963 | 1 | 1 |
| 9230102K24Rik | 0,0526023 | 0,368731 | 2,80937 | 1 | 1 |
| Styk1 | 0,0572577 | 0,39846 | 2,79889 | 1 | 1 |

| | | | | | |
|---|---|---|---|---|---|
| Adam21 | 0,0269605 | 0,18762 | 2,79889 | 1 | 1 |
| Calb2 | 0,0415186 | 0,287996 | 2,79422 | 1 | 1 |
| Inhbc | 0,0249868 | 0,172548 | 2,78776 | 1 | 1 |
| Gpr87 | 0,0301435 | 0,205533 | 2,76946 | 1 | 1 |
| Hist1h3i | 0,0789875 | 0,537117 | 2,76554 | 1 | 1 |
| Gm17746 | 0,0241912 | 0,164501 | 2,76554 | 1 | 1 |
| Kbtbd12 | 0,0119904 | 0,0815351 | 2,76554 | 1 | 1 |
| Cpne6 | 0,0160569 | 0,106899 | 2,73499 | 1 | 1 |
| 1700022I11Rik | 0,00820704 | 0,0546351 | 2,73489 | 1 | 1 |
| Rasgrf2 | 0,00512595 | 0,034124 | 2,73489 | 1 | 1 |
| Myl4 | 0,0649134 | 0,431935 | 2,73422 | 1 | 1 |
| Tmem184a | 0,0440946 | 0,292029 | 2,72744 | 1 | 1 |
| Trpc5 | 0,0128841 | 0,0849396 | 2,72085 | 1 | 1 |
| C1ra | 0,017231 | 0,112837 | 2,71116 | 1 | 1 |
| Bcl2l14 | 0,0580648 | 0,378602 | 2,70494 | 1 | 1 |
| Tdrd6 | 0,00743709 | 0,0483974 | 2,70212 | 1 | 1 |
| Rorc | 0,044318 | 0,288125 | 2,70073 | 1 | 1 |
| Igfbp1 | 0,0387107 | 0,25167 | 2,70073 | 1 | 1 |
| Gm11985 | 0,108529 | 0,696969 | 2,68302 | 1 | 1 |
| Kctd4 | 0,0181229 | 0,114513 | 2,65963 | 1 | 1 |
| Mrap2 | 0,0127585 | 0,0803518 | 2,65487 | 1 | 1 |
| Gdf15 | 0,0250434 | 0,157697 | 2,65465 | 1 | 1 |
| Lta | 0,0186924 | 0,117705 | 2,65465 | 1 | 1 |
| Kcnj3 | 0,0149412 | 0,0940843 | 2,65465 | 1 | 1 |
| Kcnj5 | 0,00513478 | 0,0323335 | 2,65465 | 1 | 1 |
| Fgfbp1 | 0,0290736 | 0,182419 | 2,64948 | 1 | 1 |
| Noto | 0,041352 | 0,259423 | 2,64928 | 1 | 1 |
| Serpina3g | 0,0159169 | 0,0998553 | 2,64928 | 1 | 1 |
| Capn8 | 0,00998568 | 0,0621073 | 2,63683 | 1 | 1 |
| 4930563D23Rik | 0,0178088 | 0,110262 | 2,63027 | 1 | 1 |
| Gpnmb | 0,00804263 | 0,0491616 | 2,61179 | 1 | 1 |
| Gm10125 | 0,0168876 | 0,102776 | 2,60547 | 1 | 1 |
| Cdkl2 | 0,0153188 | 0,0931353 | 2,60402 | 1 | 1 |
| Pla1a | 0,0287816 | 0,171963 | 2,57888 | 1 | 1 |
| Cd300e | 0,104146 | 0,618178 | 2,56941 | 1 | 1 |
| 4931408D14Rik | 0,0101228 | 0,0596309 | 2,55845 | 1 | 1 |
| Chrnd | 0,0184511 | 0,108556 | 2,55666 | 1 | 1 |
| Bcl2l10 | 0,0215217 | 0,124694 | 2,53453 | 1 | 1 |
| Actl7a | 0,0169525 | 0,0982211 | 2,53453 | 1 | 1 |
| BC006965 | 0,0126078 | 0,0730479 | 2,53453 | 1 | 1 |
| Sash3 | 0,00925983 | 0,0536504 | 2,53453 | 1 | 1 |
| Phf11d | 0,00887929 | 0,0514456 | 2,53453 | 1 | 1 |
| Slc12a3 | 0,00712412 | 0,0412763 | 2,53453 | 1 | 1 |
| Fhad1 | 0,00512403 | 0,029688 | 2,53453 | 1 | 1 |
| Adamts2 | 0,0533231 | 0,308283 | 2,53142 | 1 | 1 |
| Gm5086 | 0,119663 | 0,689803 | 2,52721 | 1 | 1 |
| Col4a3 | 0,0335736 | 0,193069 | 2,52372 | 1 | 1 |
| Phkg1 | 0,0134741 | 0,0773784 | 2,52175 | 1 | 1 |
| Lgals9 | 0,0224212 | 0,128465 | 2,51843 | 1 | 1 |
| Pou6f2 | 0,030283 | 0,173493 | 2,5183 | 1 | 1 |

| | | | | | |
|---|---|---|---|---|---|
| Colec11 | 0,0245165 | 0,140457 | 2,5183 | 1 | 1 |
| Hcls1 | 0,0159345 | 0,0912896 | 2,5183 | 1 | 1 |
| Kdm4d | 0,0117596 | 0,0673717 | 2,5183 | 1 | 1 |
| Luzp2 | 0,0301442 | 0,171709 | 2,51001 | 1 | 1 |
| Erich6 | 0,022138 | 0,125929 | 2,50801 | 1 | 1 |
| Casc1 | 0,0437284 | 0,247757 | 2,50229 | 1 | 1 |
| Wdr93 | 0,0244803 | 0,138701 | 2,50228 | 1 | 1 |
| Gm5741 | 0,17534 | 0,978894 | 2,481 | 1 | 1 |
| Gm6938 | 0,0388678 | 0,216993 | 2,481 | 1 | 1 |
| 2210011C24Rik | 0,0313487 | 0,175015 | 2,481 | 1 | 1 |
| 2310002F09Rik | 0,00937804 | 0,0523561 | 2,481 | 1 | 1 |
| Fam179a | 0,00668034 | 0,0372953 | 2,481 | 1 | 1 |
| Omp | 0,0116932 | 0,0652752 | 2,48086 | 1 | 1 |
| 1700013H16Rik | 0,0283382 | 0,157792 | 2,4772 | 1 | 1 |
| Snx31 | 0,0348115 | 0,19312 | 2,47186 | 1 | 1 |
| Vdr | 0,0265524 | 0,145566 | 2,45476 | 1 | 1 |
| Pcp2 | 0,0765139 | 0,417317 | 2,44735 | 1 | 1 |
| Pate4 | 0,0542812 | 0,295118 | 2,44277 | 1 | 1 |
| 4933427G17Rik | 0,0262518 | 0,142726 | 2,44277 | 1 | 1 |
| Parp14 | 0,0330183 | 0,178809 | 2,43708 | 1 | 1 |
| Spdef | 0,0492892 | 0,266705 | 2,4359 | 1 | 1 |
| Dpy19l2 | 0,0225462 | 0,121764 | 2,43314 | 1 | 1 |
| Igfbp6 | 0,0266405 | 0,142407 | 2,41833 | 1 | 1 |
| Srms | 0,0123368 | 0,0659465 | 2,41833 | 1 | 1 |
| Foxr2 | 0,00962223 | 0,0514357 | 2,41833 | 1 | 1 |
| Gpr1 | 0,027666 | 0,14742 | 2,41375 | 1 | 1 |
| Ryr2 | 0,0617582 | 0,328747 | 2,41228 | 1 | 1 |
| Pgr | 0,0959863 | 0,503035 | 2,38976 | 1 | 1 |
| Hemgn | 0,0241897 | 0,126525 | 2,38696 | 1 | 1 |
| Hspb3 | 0,126483 | 0,655817 | 2,37435 | 1 | 1 |
| 1700011L22Rik | 0,0373447 | 0,193618 | 2,37423 | 1 | 1 |
| Gja6 | 0,0159432 | 0,0826594 | 2,37423 | 1 | 1 |
| 1700001J11Rik | 0,0127673 | 0,0661933 | 2,37423 | 1 | 1 |
| Slc25a34 | 0,0116647 | 0,0604769 | 2,37423 | 1 | 1 |
| Adra2b | 0,050922 | 0,263267 | 2,37016 | 1 | 1 |
| Zpld1 | 0,02136 | 0,109774 | 2,36156 | 1 | 1 |
| Krt35 | 0,029036 | 0,147496 | 2,34476 | 1 | 1 |
| Rad21l | 0,0151466 | 0,0769414 | 2,34476 | 1 | 1 |
| Slc24a4 | 0,0116729 | 0,0592955 | 2,34476 | 1 | 1 |
| P2rx5 | 0,010575 | 0,0537188 | 2,34476 | 1 | 1 |
| Slco1c1 | 0,00741411 | 0,037662 | 2,34476 | 1 | 1 |
| Kcna7 | 0,00669861 | 0,0340274 | 2,34476 | 1 | 1 |
| Treml2 | 0,0540488 | 0,274419 | 2,34405 | 1 | 1 |
| C1ql2 | 0,0321883 | 0,1633 | 2,34292 | 1 | 1 |
| Slc35f3 | 0,0282986 | 0,143122 | 2,33844 | 1 | 1 |
| Ache | 0,0253893 | 0,128164 | 2,3357 | 1 | 1 |
| Gm21284 | 0,0144112 | 0,0727472 | 2,3357 | 1 | 1 |
| Chit1 | 0,0210054 | 0,105671 | 2,33075 | 1 | 1 |
| Fgr | 0,0838187 | 0,418077 | 2,31843 | 1 | 1 |
| Nrgn | 0,0404743 | 0,201329 | 2,31448 | 1 | 1 |

| | | | | | |
|---|---|---|---|---|---|
| Mdh1b | 0,029248 | 0,145487 | 2,31448 | 1 | 1 |
| Csmd1 | 0,00723942 | 0,0359602 | 2,31245 | 1 | 1 |
| Rgn | 0,0693576 | 0,343045 | 2,30627 | 1 | 1 |
| H2-T23 | 0,0594023 | 0,293393 | 2,30424 | 1 | 1 |
| Slc6a12 | 0,0373691 | 0,183155 | 2,29315 | 1 | 1 |
| A330023F24Rik | 0,023454 | 0,114374 | 2,28585 | 1 | 1 |
| Hist1h2ac | 0,0778711 | 0,379133 | 2,28355 | 1 | 1 |
| Nps | 0,0335946 | 0,163563 | 2,28355 | 1 | 1 |
| Gm13582 | 0,0089916 | 0,0437777 | 2,28355 | 1 | 1 |
| Rsph4a | 0,00639537 | 0,0311373 | 2,28355 | 1 | 1 |
| A630075F10Rik | 0,073617 | 0,353806 | 2,26485 | 1 | 1 |
| Tnfaip8l2 | 0,0314868 | 0,15117 | 2,26335 | 1 | 1 |
| Adrb1 | 0,117381 | 0,562696 | 2,26115 | 1 | 1 |
| A330093E20Rik | 0,0247661 | 0,118704 | 2,26093 | 1 | 1 |
| Galr2 | 0,0459957 | 0,219007 | 2,25141 | 1 | 1 |
| Chrm1 | 0,00482227 | 0,0229496 | 2,25068 | 1 | 1 |
| Sardh | 0,106881 | 0,508652 | 2,25067 | 1 | 1 |
| Gm5424 | 0,15303 | 0,727407 | 2,24895 | 1 | 1 |
| Adra1b | 0,0146874 | 0,069319 | 2,23867 | 1 | 1 |
| Tap2 | 0,08914 | 0,420095 | 2,23657 | 1 | 1 |
| Gm7444 | 0,00774108 | 0,0359197 | 2,21417 | 1 | 1 |
| Aph1c | 0,0439563 | 0,203571 | 2,21139 | 1 | 1 |
| Adrb3 | 0,033468 | 0,154607 | 2,20775 | 1 | 1 |
| Glyatl3 | 0,0761477 | 0,351246 | 2,20561 | 1 | 1 |
| Tnfrsf13c | 0,059863 | 0,27613 | 2,20561 | 1 | 1 |
| Abcc2 | 0,0140756 | 0,0647736 | 2,20221 | 1 | 1 |
| Igfn1 | 0,0249029 | 0,114273 | 2,1981 | 1 | 1 |
| 1700112J05Rik | 0,112248 | 0,511058 | 2,18679 | 1 | 1 |
| Cdh23 | 0,00670159 | 0,0304218 | 2,18253 | 1 | 1 |
| Tlr3 | 0,0687906 | 0,311429 | 2,17862 | 1 | 1 |
| Gm11549 | 0,103729 | 0,46606 | 2,16769 | 1 | 1 |
| Hr | 0,0675006 | 0,302423 | 2,16359 | 1 | 1 |
| Tex11 | 0,0122528 | 0,0543675 | 2,14964 | 1 | 1 |
| Tbr1 | 0,0405172 | 0,179309 | 2,14584 | 1 | 1 |
| Tspan8 | 0,0245708 | 0,108569 | 2,1436 | 1 | 1 |
| Serpina3f | 0,0157249 | 0,0694695 | 2,14333 | 1 | 1 |
| S100a4 | 0,19596 | 0,865648 | 2,14322 | 1 | 1 |
| Ifi27l2a | 0,111169 | 0,491086 | 2,14322 | 1 | 1 |
| Gpr142 | 0,0323321 | 0,142826 | 2,14322 | 1 | 1 |
| 6530411M01Rik | 0,0284778 | 0,1258 | 2,14322 | 1 | 1 |
| A530016L24Rik | 0,0118853 | 0,0525033 | 2,14322 | 1 | 1 |
| Scn7a | 0,0040546 | 0,0179111 | 2,14322 | 1 | 1 |
| Slc6a4 | 0,0200329 | 0,0880457 | 2,13588 | 1 | 1 |
| Nkg7 | 0,119837 | 0,525628 | 2,13297 | 1 | 1 |
| Cdcp2 | 0,136144 | 0,594971 | 2,12768 | 1 | 1 |
| Grm5 | 0,00267633 | 0,0116926 | 2,12727 | 1 | 1 |
| Adm | 0,0865646 | 0,378022 | 2,12662 | 1 | 1 |
| Slc22a18 | 0,016945 | 0,0739945 | 2,12656 | 1 | 1 |
| Npas2 | 0,0564323 | 0,24639 | 2,12635 | 1 | 1 |
| 6430531B16Rik | 0,0690853 | 0,301603 | 2,1262 | 1 | 1 |

| | | | | | |
|---|---|---|---|---|---|
| Srgn | 0,0300247 | 0,131078 | 2,1262 | 1 | 1 |
| Ctxn3 | 0,0173804 | 0,0758768 | 2,1262 | 1 | 1 |
| C330024D21Rik | 0,0145084 | 0,0633388 | 2,1262 | 1 | 1 |
| Alox5 | 0,00840199 | 0,0366802 | 2,1262 | 1 | 1 |
| Tmc2 | 0,00731688 | 0,031943 | 2,1262 | 1 | 1 |
| Slc10a5 | 0,00592591 | 0,0258705 | 2,1262 | 1 | 1 |
| Cd5 | 0,0782647 | 0,34079 | 2,12245 | 1 | 1 |
| Zfp92 | 0,05786 | 0,250593 | 2,11471 | 1 | 1 |
| Runx2 | 0,0364062 | 0,157039 | 2,10887 | 1 | 1 |
| Oasl1 | 0,0265331 | 0,114192 | 2,1056 | 1 | 1 |
| Hpd | 0,120014 | 0,511655 | 2,09197 | 1 | 1 |
| Cd6 | 0,0106519 | 0,0453484 | 2,08995 | 1 | 1 |
| Gm14858 | 0,0186004 | 0,0791741 | 2,08969 | 1 | 1 |
| Gm10432 | 0,0111317 | 0,0465081 | 2,06281 | 1 | 1 |
| A330076C08Rik | 0,104868 | 0,436713 | 2,05812 | 1 | 1 |
| Synpr | 0,0463319 | 0,192773 | 2,05683 | 1 | 1 |
| Pstpip1 | 0,0618621 | 0,255601 | 2,04677 | 1 | 1 |
| Ace | 0,0385107 | 0,158977 | 2,04549 | 1 | 1 |
| Snx20 | 0,0537519 | 0,221868 | 2,04531 | 1 | 1 |
| Gm6537 | 0,0602164 | 0,248389 | 2,04438 | 1 | 1 |
| Bmp8a | 0,0204267 | 0,0840537 | 2,04086 | 1 | 1 |
| Gm5083 | 0,161134 | 0,662394 | 2,03942 | 1 | 1 |
| Crtam | 0,0714887 | 0,292983 | 2,03503 | 1 | 1 |
| Duox1 | 0,0429604 | 0,175973 | 2,03428 | 1 | 1 |
| Lrrc7 | 0,0240649 | 0,0980724 | 2,02691 | 1 | 1 |
| Ttll6 | 0,0362272 | 0,147618 | 2,02673 | 1 | 1 |
| Umodl1 | 0,128049 | 0,521533 | 2,02606 | 1 | 1 |
| Plet1 | 0,1164 | 0,472674 | 2,02176 | 1 | 1 |
| Cfi | 0,0353746 | 0,14257 | 2,01088 | 1 | 1 |
| Ntsr1 | 0,0296854 | 0,119383 | 2,00777 | 1 | 1 |
| Gm6559 | 0,048981 | 0,196837 | 2,0067 | 1 | 1 |
| Scn10a | 0,0117514 | 0,0472246 | 2,0067 | 1 | 1 |
| Klhl33 | 0,0315284 | 0,126385 | 2,0031 | 1 | 1 |
| Slc13a5 | 0,0143807 | 0,0576466 | 2,0031 | 1 | 1 |
| Mef2b | 0,144111 | 0,573599 | 1,99287 | 1 | 1 |
| Nebl | 0,0367029 | 0,145979 | 1,9918 | 1 | 1 |
| Ccdc11 | 0,0462999 | 0,183634 | 1,98775 | 1 | 1 |
| 4930428G15Rik | 0,121029 | 0,47866 | 1,98364 | 1 | 1 |
| Gch1 | 0,114034 | 0,450995 | 1,98364 | 1 | 1 |
| Padi4 | 0,0723757 | 0,284842 | 1,97658 | 1 | 1 |
| Nfix | 0,0179269 | 0,0705299 | 1,97611 | 1 | 1 |
| Drd5 | 0,0477004 | 0,186956 | 1,97063 | 1 | 1 |
| Kcna2 | 0,107233 | 0,41705 | 1,95947 | 1 | 1 |
| Ccdc33 | 0,0122943 | 0,0476171 | 1,95349 | 1 | 1 |
| Ccdc105 | 0,0190918 | 0,073943 | 1,95346 | 1 | 1 |
| Ocstamp | 0,0164983 | 0,0638983 | 1,95346 | 1 | 1 |
| Gm16532 | 0,0145544 | 0,0563696 | 1,95346 | 1 | 1 |
| Kctd19 | 0,0180395 | 0,069786 | 1,95178 | 1 | 1 |
| 1700012B07Rik | 0,018504 | 0,0714766 | 1,94963 | 1 | 1 |
| Gm3458 | 0,0402146 | 0,155335 | 1,94959 | 1 | 1 |

| | | | | | |
|---|---|---|---|---|---|
| Serpina3a | 0,0220379 | 0,0851247 | 1,94959 | 1 | 1 |
| 4930474M22Rik | 0,107764 | 0,41625 | 1,94957 | 1 | 1 |
| 1810010D01Rik | 0,0301479 | 0,116449 | 1,94957 | 1 | 1 |
| Tekt4 | 0,0165951 | 0,0641001 | 1,94957 | 1 | 1 |
| Nobox | 0,0130482 | 0,0503997 | 1,94957 | 1 | 1 |
| Asb14 | 0,0121692 | 0,0470045 | 1,94957 | 1 | 1 |
| Eef1a2 | 0,0119644 | 0,0462135 | 1,94957 | 1 | 1 |
| Slc22a22 | 0,0101841 | 0,039337 | 1,94957 | 1 | 1 |
| Pramel1 | 0,00985617 | 0,0380703 | 1,94957 | 1 | 1 |
| Ifi44 | 0,0081503 | 0,0314813 | 1,94957 | 1 | 1 |
| Ush1g | 0,00739793 | 0,0285752 | 1,94957 | 1 | 1 |
| Htr7 | 0,139222 | 0,5365 | 1,94619 | 1 | 1 |
| Hapln4 | 0,0322521 | 0,123465 | 1,93664 | 1 | 1 |
| P2rx1 | 0,0523284 | 0,200179 | 1,93562 | 1 | 1 |
| Cnksr1 | 0,0608303 | 0,231385 | 1,92743 | 1 | 1 |
| Hgfac | 0,0384317 | 0,146156 | 1,92713 | 1 | 1 |
| Sowahd | 0,0421971 | 0,160035 | 1,92317 | 1 | 1 |
| Tnk1 | 0,0537058 | 0,203628 | 1,92279 | 1 | 1 |
| Gm6083 | 0,107929 | 0,40767 | 1,91732 | 1 | 1 |
| Gal | 0,191956 | 0,723266 | 1,91375 | 1 | 1 |
| Fam159b | 0,0784041 | 0,29525 | 1,91294 | 1 | 1 |
| Gm10778 | 0,00974937 | 0,0366291 | 1,90961 | 1 | 1 |
| 1700084C01Rik | 0,118093 | 0,440137 | 1,89803 | 1 | 1 |
| Ckmt1 | 0,0686065 | 0,255142 | 1,89488 | 1 | 1 |
| Ankrd1 | 0,018517 | 0,0687372 | 1,89224 | 1 | 1 |
| Cd46 | 0,122819 | 0,454791 | 1,88867 | 1 | 1 |
| Cyp2t4 | 0,122092 | 0,452006 | 1,88837 | 1 | 1 |
| Gm10390 | 0,0192078 | 0,0708243 | 1,88255 | 1 | 1 |
| Abcg8 | 0,0205891 | 0,0757634 | 1,87962 | 1 | 1 |
| Dlec1 | 0,0634184 | 0,233167 | 1,87839 | 1 | 1 |
| Muc2 | 0,0457994 | 0,168033 | 1,87535 | 1 | 1 |
| Ciita | 0,0253548 | 0,0929446 | 1,87411 | 1 | 1 |
| Rad51ap2 | 0,0806282 | 0,295186 | 1,87227 | 1 | 1 |
| Htr1d | 0,00850614 | 0,0310682 | 1,86886 | 1 | 1 |
| Sell | 0,0109671 | 0,0400542 | 1,86877 | 1 | 1 |
| 1700007B14Rik | 0,0119815 | 0,0437574 | 1,86872 | 1 | 1 |
| Gpha2 | 0,0573901 | 0,209563 | 1,86851 | 1 | 1 |
| A530006G24Rik | 0,0195548 | 0,0714053 | 1,86851 | 1 | 1 |
| Rhov | 0,0151675 | 0,0553847 | 1,86851 | 1 | 1 |
| P2ry6 | 0,0128654 | 0,0469785 | 1,86851 | 1 | 1 |
| Spata18 | 0,0126294 | 0,046117 | 1,86851 | 1 | 1 |
| Slc44a4 | 0,0102935 | 0,0375872 | 1,86851 | 1 | 1 |
| Elfn2 | 0,00473233 | 0,0172803 | 1,86851 | 1 | 1 |
| Ccdc79 | 0,0346439 | 0,124669 | 1,84743 | 1 | 1 |
| Gsg1 | 0,143378 | 0,514358 | 1,84295 | 1 | 1 |
| Gldn | 0,0164189 | 0,0589004 | 1,84292 | 1 | 1 |
| Tmem171 | 0,144751 | 0,517501 | 1,83799 | 1 | 1 |
| Lrfn5 | 0,0144644 | 0,0514872 | 1,83171 | 1 | 1 |
| Nos1 | 0,0470621 | 0,166952 | 1,82679 | 1 | 1 |
| AW495222 | 0,0499844 | 0,177038 | 1,82451 | 1 | 1 |

| | | | | | |
|---|---|---|---|---|---|
| Tnfrsf11a | 0,0257214 | 0,0910128 | 1,8231 | 1 | 1 |
| Kcnq3 | 0,119982 | 0,421736 | 1,81352 | 1 | 1 |
| Slc17a9 | 0,04351 | 0,152722 | 1,81149 | 1 | 1 |
| Lama3 | 0,13136 | 0,460826 | 1,8107 | 1 | 1 |
| 4933429K18Rik | 0,0543481 | 0,190519 | 1,80963 | 1 | 1 |
| Gpr65 | 0,0226849 | 0,0795227 | 1,80963 | 1 | 1 |
| LOC100504608 | 0,233572 | 0,818645 | 1,80937 | 1 | 1 |
| Myt1l | 0,0115653 | 0,040218 | 1,79804 | 1 | 1 |
| 2010109A12Rik | 0,232111 | 0,806805 | 1,79741 | 1 | 1 |
| Creg2 | 0,129472 | 0,449082 | 1,79434 | 1 | 1 |
| Isg15 | 0,209454 | 0,725687 | 1,79272 | 1 | 1 |
| Efcab4a | 0,0779637 | 0,270118 | 1,79272 | 1 | 1 |
| C2cd4b | 0,193167 | 0,668002 | 1,79 | 1 | 1 |
| Nlrc5 | 0,0226719 | 0,0783677 | 1,78935 | 1 | 1 |
| Gm16675 | 0,111143 | 0,383326 | 1,78615 | 1 | 1 |
| Egr3 | 0,0968769 | 0,333482 | 1,78338 | 1 | 1 |
| Grid2ip | 0,0291913 | 0,100321 | 1,78101 | 1 | 1 |
| Phldb3 | 0,0351212 | 0,120657 | 1,78049 | 1 | 1 |
| Dmrtb1 | 0,109911 | 0,376773 | 1,77736 | 1 | 1 |
| Neurl1b | 0,157403 | 0,536323 | 1,76863 | 1 | 1 |
| 1700016L04Rik | 0,0624309 | 0,212266 | 1,76554 | 1 | 1 |
| Cxcr6 | 0,0693064 | 0,23442 | 1,75803 | 1 | 1 |
| Unc13d | 0,0848779 | 0,286666 | 1,75591 | 1 | 1 |
| Acvr1c | 0,00602442 | 0,0201378 | 1,74101 | 1 | 1 |
| Acaa1b | 0,164494 | 0,548831 | 1,73833 | 1 | 1 |
| Hspb2 | 0,0454477 | 0,151548 | 1,73749 | 1 | 1 |
| AI197445 | 0,0440098 | 0,146515 | 1,73515 | 1 | 1 |
| Htr6 | 0,0249611 | 0,0830843 | 1,73489 | 1 | 1 |
| BC030870 | 0,0226706 | 0,0754602 | 1,73489 | 1 | 1 |
| Chil1 | 0,0194647 | 0,0647891 | 1,73489 | 1 | 1 |
| Clcnka | 0,0131208 | 0,0436734 | 1,73489 | 1 | 1 |
| Diras2 | 0,0069752 | 0,0232173 | 1,73489 | 1 | 1 |
| Pkd1l2 | 0,00402468 | 0,0133963 | 1,73489 | 1 | 1 |
| Arg2 | 0,0726884 | 0,241859 | 1,73437 | 1 | 1 |
| Zfhx2os | 0,0892425 | 0,29612 | 1,73038 | 1 | 1 |
| Stmn1-rs1 | 0,168762 | 0,559797 | 1,72991 | 1 | 1 |
| Upb1 | 0,0460861 | 0,152871 | 1,72991 | 1 | 1 |
| Oas1c | 0,0800019 | 0,264384 | 1,72453 | 1 | 1 |
| Abcc3 | 0,146917 | 0,485512 | 1,72451 | 1 | 1 |
| Stxbp5l | 0,0119694 | 0,0395036 | 1,72264 | 1 | 1 |
| 5830444B04Rik | 0,0990836 | 0,326832 | 1,72183 | 1 | 1 |
| Mcoln2 | 0,111933 | 0,368743 | 1,71998 | 1 | 1 |
| Ackr4 | 0,026742 | 0,0876248 | 1,71223 | 1 | 1 |
| Ano9 | 0,0411754 | 0,134763 | 1,71056 | 1 | 1 |
| Cdkn2a | 0,0341112 | 0,110998 | 1,70221 | 1 | 1 |
| Il17rb | 0,0511687 | 0,166411 | 1,70142 | 1 | 1 |
| Tmprss11d | 0,0313845 | 0,10194 | 1,69959 | 1 | 1 |
| Barhl1 | 0,0362779 | 0,117275 | 1,69274 | 1 | 1 |
| Mfi2 | 0,0336636 | 0,108812 | 1,69257 | 1 | 1 |
| Ppm1j | 0,124408 | 0,39848 | 1,67943 | 1 | 1 |

| | | | | | |
|---|---|---|---|---|---|
| Lrrc38 | 0,0862807 | 0,276295 | 1,6791 | 1 | 1 |
| 1700016H13Rik | 0,0873769 | 0,276063 | 1,65967 | 1 | 1 |
| Vax1 | 0,116875 | 0,36866 | 1,65732 | 1 | 1 |
| Kl | 0,0324985 | 0,102409 | 1,65589 | 1 | 1 |
| Fam46d | 0,0157783 | 0,0496812 | 1,65475 | 1 | 1 |
| Sftpb | 0,0171065 | 0,0538627 | 1,65474 | 1 | 1 |
| Gprc5d | 0,0791988 | 0,249361 | 1,65468 | 1 | 1 |
| Zfp652os | 0,132358 | 0,416727 | 1,65465 | 1 | 1 |
| Lgals2 | 0,0567984 | 0,178829 | 1,65465 | 1 | 1 |
| Wfdc13 | 0,0506625 | 0,15951 | 1,65465 | 1 | 1 |
| Tac4 | 0,0211297 | 0,0665263 | 1,65465 | 1 | 1 |
| C6 | 0,0167148 | 0,0526261 | 1,65465 | 1 | 1 |
| 9030617O03Rik | 0,0145709 | 0,045876 | 1,65465 | 1 | 1 |
| 4933425B07Rik | 0,0127981 | 0,0402946 | 1,65465 | 1 | 1 |
| BC049762 | 0,160165 | 0,504096 | 1,65414 | 1 | 1 |
| Ubap1l | 0,0736745 | 0,231858 | 1,654 | 1 | 1 |
| Bspry | 0,103767 | 0,325827 | 1,65076 | 1 | 1 |
| N28178 | 0,0250388 | 0,0786219 | 1,65076 | 1 | 1 |
| Gm17365 | 0,0513866 | 0,161188 | 1,64928 | 1 | 1 |
| Clca5 | 0,015553 | 0,0487861 | 1,64928 | 1 | 1 |
| Nkpd1 | 0,151804 | 0,475813 | 1,64819 | 1 | 1 |
| Ovol1 | 0,175158 | 0,548706 | 1,64738 | 1 | 1 |
| Foxo6 | 0,162927 | 0,510336 | 1,64722 | 1 | 1 |
| Stat4 | 0,171085 | 0,533344 | 1,64036 | 1 | 1 |
| Necab1 | 0,105087 | 0,326905 | 1,63728 | 1 | 1 |
| Ptgdr | 0,04663 | 0,144842 | 1,63515 | 1 | 1 |
| Gpx2 | 0,143597 | 0,445939 | 1,63482 | 1 | 1 |
| Ccdc17 | 0,115451 | 0,357623 | 1,63115 | 1 | 1 |
| Klk7 | 0,051816 | 0,159684 | 1,62375 | 1 | 1 |
| Abcc6 | 0,133832 | 0,411385 | 1,62006 | 1 | 1 |
| Ldlrad4 | 0,152791 | 0,469232 | 1,61874 | 1 | 1 |
| F10 | 0,0413282 | 0,12674 | 1,61667 | 1 | 1 |
| LOC102632430 | 0,0369664 | 0,112898 | 1,61074 | 1 | 1 |
| Krtap17-1 | 0,078848 | 0,239931 | 1,60547 | 1 | 1 |
| Nags | 0,0230648 | 0,0701851 | 1,60547 | 1 | 1 |
| Nmbr | 0,119319 | 0,362833 | 1,60449 | 1 | 1 |
| Gata1 | 0,030002 | 0,0909967 | 1,60075 | 1 | 1 |
| Serpina3h | 0,16843 | 0,510456 | 1,59964 | 1 | 1 |
| Igfbp7 | 0,183825 | 0,550425 | 1,58221 | 1 | 1 |
| Irgm2 | 0,136443 | 0,408346 | 1,58149 | 1 | 1 |
| Dennd2d | 0,0725011 | 0,21662 | 1,57909 | 1 | 1 |
| Tmem102 | 0,076408 | 0,227936 | 1,57683 | 1 | 1 |
| Cd28 | 0,0160747 | 0,0479154 | 1,57569 | 1 | 1 |
| Sptlc3 | 0,0743799 | 0,221599 | 1,57497 | 1 | 1 |
| Myo15 | 0,0245277 | 0,0730754 | 1,57497 | 1 | 1 |
| Ccdc67 | 0,167906 | 0,498147 | 1,56891 | 1 | 1 |
| Trpc6 | 0,160594 | 0,473943 | 1,56129 | 1 | 1 |
| Msh4 | 0,00951134 | 0,028017 | 1,55858 | 1 | 1 |
| 4932441J04Rik | 0,0553728 | 0,163072 | 1,55826 | 1 | 1 |
| Sugct | 0,038033 | 0,112007 | 1,55826 | 1 | 1 |

| | | | | | |
|---|---|---|---|---|---|
| Arr3 | 0,025026 | 0,0737012 | 1,55826 | 1 | 1 |
| D830030K20Rik | 0,0239082 | 0,0704093 | 1,55826 | 1 | 1 |
| 4930428O21Rik | 0,0188184 | 0,05542 | 1,55826 | 1 | 1 |
| Asb10 | 0,0136732 | 0,0402674 | 1,55826 | 1 | 1 |
| Apoa5 | 0,0136538 | 0,0402103 | 1,55826 | 1 | 1 |
| 1700073E17Rik | 0,0131537 | 0,0387374 | 1,55826 | 1 | 1 |
| Kel | 0,0125022 | 0,0368189 | 1,55826 | 1 | 1 |
| Kcnk10 | 0,169092 | 0,497723 | 1,55754 | 1 | 1 |
| Nfe2 | 0,0742851 | 0,218655 | 1,55751 | 1 | 1 |
| Ankef1 | 0,0606712 | 0,178478 | 1,55666 | 1 | 1 |
| Wdr16 | 0,0586172 | 0,1714 | 1,54797 | 1 | 1 |
| Dscam | 0,0350033 | 0,102085 | 1,54422 | 1 | 1 |
| Dnah11 | 0,132383 | 0,38586 | 1,54336 | 1 | 1 |
| Tnfrsf14 | 0,191914 | 0,558554 | 1,54124 | 1 | 1 |
| Mmp24 | 0,0142948 | 0,0415785 | 1,54035 | 1 | 1 |
| Gabbr2 | 0,0286647 | 0,0831476 | 1,5364 | 1 | 1 |
| Lrit3 | 0,0239287 | 0,0693202 | 1,53453 | 1 | 1 |
| Iqca | 0,014663 | 0,0424778 | 1,53453 | 1 | 1 |
| Myom3 | 0,0161362 | 0,0466553 | 1,53174 | 1 | 1 |
| Gabra5 | 0,0942022 | 0,271944 | 1,52948 | 1 | 1 |
| Pcsk2os1 | 0,0198185 | 0,0571697 | 1,5284 | 1 | 1 |
| 2010300C02Rik | 0,0861303 | 0,248138 | 1,52655 | 1 | 1 |
| D030018L15Rik | 0,040084 | 0,115415 | 1,52574 | 1 | 1 |
| Gm19299 | 0,0827673 | 0,23825 | 1,52534 | 1 | 1 |
| 1700001L19Rik | 0,0729791 | 0,209382 | 1,52059 | 1 | 1 |
| Csta1 | 0,022877 | 0,0655321 | 1,5183 | 1 | 1 |
| Dnase1l3 | 0,0165409 | 0,0473819 | 1,5183 | 1 | 1 |
| Ncf1 | 0,0552269 | 0,158153 | 1,51788 | 1 | 1 |
| Myrip | 0,0238601 | 0,0681902 | 1,51497 | 1 | 1 |
| Styxl1 | 0,0394906 | 0,112698 | 1,51289 | 1 | 1 |
| A4galt | 0,0386407 | 0,110241 | 1,51246 | 1 | 1 |
| E030011O05Rik | 0,130556 | 0,372377 | 1,5121 | 1 | 1 |
| Gpr123 | 0,0160259 | 0,0457099 | 1,5121 | 1 | 1 |
| Hpx | 0,10901 | 0,310293 | 1,50917 | 1 | 1 |
| Atp6v1b1 | 0,0383721 | 0,10898 | 1,50594 | 1 | 1 |
| Gpr64 | 0,01953 | 0,0554285 | 1,50494 | 1 | 1 |
| Baiap3 | 0,0332051 | 0,0936978 | 1,49661 | 1 | 1 |
| Xlr3b | 0,143196 | 0,403988 | 1,49633 | 1 | 1 |
| Cpb1 | 0,135557 | 0,381508 | 1,49282 | 1 | 1 |
| Dntt | 0,0427814 | 0,119979 | 1,48773 | 1 | 1 |
| Ddo | 0,0339844 | 0,0953015 | 1,48763 | 1 | 1 |
| Psd4 | 0,0534822 | 0,149967 | 1,48752 | 1 | 1 |
| Kcnc1 | 0,0288695 | 0,0808816 | 1,48626 | 1 | 1 |
| Rasl10a | 0,137085 | 0,383777 | 1,4852 | 1 | 1 |
| Gm10046 | 0,254091 | 0,711138 | 1,48478 | 1 | 1 |
| Dlx1 | 0,0155393 | 0,0434758 | 1,48429 | 1 | 1 |
| Gucy2c | 0,0417489 | 0,116771 | 1,48387 | 1 | 1 |
| Plekha4 | 0,0739967 | 0,206794 | 1,48266 | 1 | 1 |
| Ccdc42b | 0,0615023 | 0,17173 | 1,48143 | 1 | 1 |
| Serpina6 | 0,035037 | 0,0978031 | 1,481 | 1 | 1 |

| | | | | | |
|---|---|---|---|---|---|
| Akap3 | 0,0298927 | 0,0834433 | 1,481 | 1 | 1 |
| LOC100504039 | 0,0489856 | 0,136509 | 1,47856 | 1 | 1 |
| Fxyd4 | 0,0831762 | 0,231777 | 1,4785 | 1 | 1 |
| Arsj | 0,0940047 | 0,261941 | 1,47844 | 1 | 1 |
| Myh6 | 0,00496918 | 0,013844 | 1,47818 | 1 | 1 |
| Fndc1 | 0,109824 | 0,305812 | 1,47744 | 1 | 1 |
| Sun3 | 0,0322204 | 0,0897175 | 1,47741 | 1 | 1 |
| U90926 | 0,0908935 | 0,253055 | 1,4772 | 1 | 1 |
| A630076J17Rik | 0,0625638 | 0,174183 | 1,4772 | 1 | 1 |
| 4930469G21Rik | 0,0385397 | 0,107298 | 1,4772 | 1 | 1 |
| Dupd1 | 0,0321523 | 0,0895145 | 1,4772 | 1 | 1 |
| Prss30 | 0,0228498 | 0,0636157 | 1,4772 | 1 | 1 |
| Smtnl1 | 0,0185646 | 0,0516853 | 1,4772 | 1 | 1 |
| Ccdc64b | 0,0168248 | 0,0468418 | 1,4772 | 1 | 1 |
| Nctc1 | 0,0151972 | 0,0423103 | 1,4772 | 1 | 1 |
| Fa2h | 0,0126334 | 0,0351723 | 1,4772 | 1 | 1 |
| Icam5 | 0,010405 | 0,0289685 | 1,4772 | 1 | 1 |
| F630042J09Rik | 0,00976526 | 0,0271873 | 1,4772 | 1 | 1 |
| 1700071K01Rik | 0,0346589 | 0,096491 | 1,47717 | 1 | 1 |
| Gabrr2 | 0,0809356 | 0,224427 | 1,4714 | 1 | 1 |
| Zfp872 | 0,0449657 | 0,124197 | 1,46573 | 1 | 1 |
| Tcp11 | 0,111767 | 0,308538 | 1,46495 | 1 | 1 |
| Mrps36 | 0,271992 | 0,747837 | 1,45916 | 1 | 1 |
| Htr3a | 0,0710365 | 0,194848 | 1,45572 | 1 | 1 |
| Cldn20 | 0,144986 | 0,397661 | 1,45562 | 1 | 1 |
| Nek11 | 0,0273091 | 0,0749019 | 1,45562 | 1 | 1 |
| Zpbp2 | 0,0929577 | 0,254917 | 1,45538 | 1 | 1 |
| 4930465K10Rik | 0,221282 | 0,60681 | 1,45536 | 1 | 1 |
| Adm2 | 0,239545 | 0,655844 | 1,45306 | 1 | 1 |
| Jazf1 | 0,0637263 | 0,174421 | 1,45262 | 1 | 1 |
| Atp6v1c2 | 0,150089 | 0,410218 | 1,45058 | 1 | 1 |
| Lpxn | 0,178069 | 0,486222 | 1,44918 | 1 | 1 |
| Thpo | 0,177055 | 0,483303 | 1,44873 | 1 | 1 |
| Arid3c | 0,0349583 | 0,0952403 | 1,44594 | 1 | 1 |
| 5730416F02Rik | 0,0837874 | 0,228245 | 1,44577 | 1 | 1 |
| Ido1 | 0,0377079 | 0,10272 | 1,44577 | 1 | 1 |
| Lgi1 | 0,0125632 | 0,0342232 | 1,44577 | 1 | 1 |
| Capn3 | 0,129811 | 0,353386 | 1,44483 | 1 | 1 |
| Il1r1 | 0,136307 | 0,371067 | 1,44482 | 1 | 1 |
| Lonrf3 | 0,185037 | 0,502711 | 1,44192 | 1 | 1 |
| Gm13032 | 0,0775037 | 0,210518 | 1,4416 | 1 | 1 |
| Slc26a4 | 0,0607396 | 0,164858 | 1,44051 | 1 | 1 |
| Gucy2e | 0,0584599 | 0,158348 | 1,43758 | 1 | 1 |
| Gpr21 | 0,180222 | 0,48754 | 1,43575 | 1 | 1 |
| Prnd | 0,0168325 | 0,0454983 | 1,43456 | 1 | 1 |
| Arhgap36 | 0,207861 | 0,559651 | 1,42891 | 1 | 1 |
| Kcnd3 | 0,12237 | 0,329077 | 1,42717 | 1 | 1 |
| Gm7173 | 0,0723139 | 0,19326 | 1,4182 | 1 | 1 |
| Gpr179 | 0,0491841 | 0,131444 | 1,41819 | 1 | 1 |
| Optc | 0,142208 | 0,377508 | 1,40851 | 1 | 1 |

| | | | | | |
|---|---|---|---|---|---|
| Mip | 0,20236 | 0,534378 | 1,40094 | 1 | 1 |
| Thsd7b | 0,0350603 | 0,0925363 | 1,40018 | 1 | 1 |
| Padi1 | 0,0162255 | 0,0427249 | 1,39681 | 1 | 1 |
| Mbd3l2 | 0,0261115 | 0,0687461 | 1,39659 | 1 | 1 |
| Myo1g | 0,0164175 | 0,043224 | 1,39659 | 1 | 1 |
| Misp | 0,0329403 | 0,0867066 | 1,39629 | 1 | 1 |
| Cftr | 0,0905042 | 0,23782 | 1,39382 | 1 | 1 |
| Barx1 | 0,25107 | 0,659687 | 1,39369 | 1 | 1 |
| Slain1os | 0,0782052 | 0,205398 | 1,39309 | 1 | 1 |
| Atp8b5 | 0,0528836 | 0,138879 | 1,39293 | 1 | 1 |
| Tcf24 | 0,0311102 | 0,0814899 | 1,38923 | 1 | 1 |
| Car5a | 0,0570917 | 0,149502 | 1,38881 | 1 | 1 |
| 1700056E22Rik | 0,276066 | 0,721418 | 1,38582 | 1 | 1 |
| Gm4925 | 0,0801425 | 0,209274 | 1,38475 | 1 | 1 |
| 4933405L10Rik | 0,0867913 | 0,226261 | 1,38237 | 1 | 1 |
| 4930509E16Rik | 0,0808998 | 0,210902 | 1,38237 | 1 | 1 |
| Pou5f2 | 0,0672526 | 0,17485 | 1,37845 | 1 | 1 |
| Scgb3a1 | 0,0538822 | 0,140001 | 1,37756 | 1 | 1 |
| Gsx2 | 0,0848221 | 0,220326 | 1,37713 | 1 | 1 |
| AI463170 | 0,172049 | 0,446002 | 1,37423 | 1 | 1 |
| Gm15412 | 0,0648091 | 0,168005 | 1,37423 | 1 | 1 |
| Ptf1a | 0,0443328 | 0,114924 | 1,37423 | 1 | 1 |
| Nanog | 0,0496814 | 0,128568 | 1,37176 | 1 | 1 |
| Srpk3 | 0,0704521 | 0,181985 | 1,36911 | 1 | 1 |
| Anxa8 | 0,0493528 | 0,127426 | 1,36845 | 1 | 1 |
| Slc47a1 | 0,141099 | 0,360169 | 1,35196 | 1 | 1 |
| Gm9047 | 0,180649 | 0,460422 | 1,34977 | 1 | 1 |
| Bhlha9 | 0,0796093 | 0,202901 | 1,34977 | 1 | 1 |
| Abca12 | 0,00983483 | 0,0250661 | 1,34977 | 1 | 1 |
| Chmp4c | 0,184065 | 0,468393 | 1,3475 | 1 | 1 |
| Oca2 | 0,0325859 | 0,0827926 | 1,34525 | 1 | 1 |
| Kcnip2 | 0,158112 | 0,400483 | 1,34079 | 1 | 1 |
| Mroh2a | 0,00739354 | 0,0187251 | 1,34064 | 1 | 1 |
| Cldn19 | 0,0252697 | 0,0639583 | 1,33972 | 1 | 1 |
| Pcdhb2 | 0,0541024 | 0,136752 | 1,33779 | 1 | 1 |
| Acap1 | 0,111816 | 0,281047 | 1,32968 | 1 | 1 |
| Cdcp1 | 0,148482 | 0,372582 | 1,32727 | 1 | 1 |
| Egfem1 | 0,0220025 | 0,0551551 | 1,32582 | 1 | 1 |
| H2-DMb1 | 0,0471696 | 0,118229 | 1,32565 | 1 | 1 |
| 9530053A07Rik | 0,00660064 | 0,0165442 | 1,32565 | 1 | 1 |
| Cdhr3 | 0,0425823 | 0,106689 | 1,32508 | 1 | 1 |
| Mkln1os | 0,0869927 | 0,217892 | 1,32465 | 1 | 1 |
| Trank1 | 0,0225553 | 0,0564101 | 1,32249 | 1 | 1 |
| Tnfaip8l3 | 0,0613188 | 0,152907 | 1,31825 | 1 | 1 |
| Ankfn1 | 0,108015 | 0,269263 | 1,31779 | 1 | 1 |
| Morn3 | 0,100942 | 0,250809 | 1,31307 | 1 | 1 |
| Fam186b | 0,024466 | 0,0607904 | 1,31307 | 1 | 1 |
| Zap70 | 0,0105649 | 0,0262273 | 1,31179 | 1 | 1 |
| Atp2a3 | 0,0873895 | 0,216536 | 1,30908 | 1 | 1 |
| Eomes | 0,0251006 | 0,0621253 | 1,30746 | 1 | 1 |

| | | | | | |
|---|---|---|---|---|---|
| Rab42 | 0,178943 | 0,442528 | 1,30627 | 1 | 1 |
| Plekhd1 | 0,0857844 | 0,212146 | 1,30627 | 1 | 1 |
| 4930550C14Rik | 0,175929 | 0,433492 | 1,30101 | 1 | 1 |
| Slc45a1 | 0,253933 | 0,62512 | 1,29969 | 1 | 1 |
| Pdzd9 | 0,228481 | 0,562235 | 1,2991 | 1 | 1 |
| Lyl1 | 0,136197 | 0,334821 | 1,29769 | 1 | 1 |
| 4931409K22Rik | 0,0117209 | 0,0287975 | 1,29686 | 1 | 1 |
| Slc9a7 | 0,19333 | 0,474992 | 1,29684 | 1 | 1 |
| Rit2 | 0,178386 | 0,43824 | 1,29672 | 1 | 1 |
| 0610040F04Rik | 0,0188738 | 0,0462486 | 1,29303 | 1 | 1 |
| Trp53cor1 | 0,162874 | 0,398989 | 1,29259 | 1 | 1 |
| P2rx2 | 0,130021 | 0,318168 | 1,29104 | 1 | 1 |
| Arhgef6 | 0,184832 | 0,450835 | 1,28638 | 1 | 1 |
| Muc6 | 0,0321644 | 0,078414 | 1,28565 | 1 | 1 |
| Gm10267 | 0,155247 | 0,377927 | 1,28355 | 1 | 1 |
| Tyrobp | 0,0608777 | 0,148199 | 1,28355 | 1 | 1 |
| Gm16998 | 0,0214798 | 0,0522896 | 1,28355 | 1 | 1 |
| Acpt | 0,0205198 | 0,0499527 | 1,28355 | 1 | 1 |
| Il23a | 0,0190808 | 0,0464495 | 1,28355 | 1 | 1 |
| Hmx3 | 0,0189495 | 0,04613 | 1,28355 | 1 | 1 |
| 4921506M07Rik | 0,0172849 | 0,0420779 | 1,28355 | 1 | 1 |
| Kis2 | 0,0120626 | 0,0293648 | 1,28355 | 1 | 1 |
| Adam32 | 0,00982541 | 0,0239186 | 1,28355 | 1 | 1 |
| Fam115e | 0,00758377 | 0,0184617 | 1,28355 | 1 | 1 |
| Dlx6os1 | 0,00545101 | 0,0132698 | 1,28355 | 1 | 1 |
| Abhd1 | 0,0427876 | 0,104129 | 1,2831 | 1 | 1 |
| Cnih3 | 0,0628573 | 0,152181 | 1,27563 | 1 | 1 |
| Igfals | 0,0916984 | 0,221688 | 1,27357 | 1 | 1 |
| Iqub | 0,0147791 | 0,035694 | 1,27212 | 1 | 1 |
| Trp53tg5 | 0,109832 | 0,26468 | 1,26895 | 1 | 1 |
| Olfr1372-ps1 | 0,148772 | 0,357896 | 1,26644 | 1 | 1 |
| Aqp9 | 0,0118347 | 0,0284104 | 1,26339 | 1 | 1 |
| Fabp4 | 0,0661429 | 0,158777 | 1,26335 | 1 | 1 |
| 6030468B19Rik | 0,0330719 | 0,0793899 | 1,26335 | 1 | 1 |
| Fcgr3 | 0,0251566 | 0,0603891 | 1,26335 | 1 | 1 |
| Plxna4os1 | 0,0169531 | 0,0406964 | 1,26335 | 1 | 1 |
| Tacr2 | 0,0140726 | 0,0337816 | 1,26335 | 1 | 1 |
| Micalcl | 0,0131806 | 0,0316404 | 1,26335 | 1 | 1 |
| BC048546 | 0,00643196 | 0,0154401 | 1,26335 | 1 | 1 |
| Mettl7a2Higd1c | 0,0232169 | 0,0557289 | 1,26325 | 1 | 1 |
| Brinp2 | 0,107735 | 0,258546 | 1,26293 | 1 | 1 |
| Erich3 | 0,1259 | 0,301391 | 1,25935 | 1 | 1 |
| Spo11 | 0,135922 | 0,324907 | 1,25725 | 1 | 1 |
| Slc16a9 | 0,0758315 | 0,18123 | 1,25695 | 1 | 1 |
| Wdfy4 | 0,0408304 | 0,0975218 | 1,25608 | 1 | 1 |
| Vstm5 | 0,200696 | 0,47916 | 1,25549 | 1 | 1 |
| Inpp5j | 0,0425504 | 0,101315 | 1,25161 | 1 | 1 |
| A230050P20Rik | 0,234815 | 0,558672 | 1,25048 | 1 | 1 |
| Foxa3 | 0,0750184 | 0,177955 | 1,2462 | 1 | 1 |
| Lrrc23 | 0,097887 | 0,231664 | 1,24284 | 1 | 1 |

| | | | | | |
|---|---|---|---|---|---|
| Tlr12 | 0,156202 | 0,368506 | 1,23827 | 1 | 1 |
| F8 | 0,235451 | 0,554932 | 1,23688 | 1 | 1 |
| Nxpe2 | 0,0300318 | 0,070765 | 1,23655 | 1 | 1 |
| Dnah8 | 0,0227903 | 0,0536537 | 1,23526 | 1 | 1 |
| Capn11 | 0,225839 | 0,529485 | 1,2293 | 1 | 1 |
| Capsl | 0,205969 | 0,482506 | 1,22812 | 1 | 1 |
| Nyx | 0,0487458 | 0,113989 | 1,22554 | 1 | 1 |
| Sim2 | 0,0587924 | 0,137453 | 1,22523 | 1 | 1 |
| Foxp3 | 0,0945846 | 0,221091 | 1,22496 | 1 | 1 |
| Gipr | 0,132645 | 0,309916 | 1,22431 | 1 | 1 |
| A630012P03Rik | 0,126908 | 0,296167 | 1,22262 | 1 | 1 |
| Ildr1 | 0,0449922 | 0,10472 | 1,21879 | 1 | 1 |
| Efhc2 | 0,0990125 | 0,230275 | 1,21768 | 1 | 1 |
| Tbx10 | 0,0203969 | 0,0470777 | 1,20669 | 1 | 1 |
| Nckap1l | 0,0466744 | 0,107625 | 1,20532 | 1 | 1 |
| Gm10406 | 0,254002 | 0,585421 | 1,20463 | 1 | 1 |
| Cysltr2 | 0,0902537 | 0,20772 | 1,20258 | 1 | 1 |
| Gsap | 0,123664 | 0,28373 | 1,1981 | 1 | 1 |
| Slc5a7 | 0,158286 | 0,362915 | 1,19709 | 1 | 1 |
| Ltbp2 | 0,0946263 | 0,215029 | 1,18421 | 1 | 1 |
| Myh11 | 0,0113997 | 0,0259027 | 1,18411 | 1 | 1 |
| A730043L09Rik | 0,0797683 | 0,181155 | 1,18334 | 1 | 1 |
| Cmah | 0,00805108 | 0,0182739 | 1,18253 | 1 | 1 |
| Gm7904 | 0,0634499 | 0,14382 | 1,18058 | 1 | 1 |
| Klk9 | 0,052018 | 0,117908 | 1,18058 | 1 | 1 |
| Irs3 | 0,047767 | 0,107987 | 1,17677 | 1 | 1 |
| Klf1 | 0,228738 | 0,516869 | 1,1761 | 1 | 1 |
| Ncam2 | 0,0151569 | 0,0341165 | 1,1705 | 1 | 1 |
| Wfikkn2 | 0,123531 | 0,278011 | 1,17028 | 1 | 1 |
| Chd5 | 0,122672 | 0,275997 | 1,16984 | 1 | 1 |
| Elovl7 | 0,104462 | 0,234777 | 1,16832 | 1 | 1 |
| 2900005J15Rik | 0,256006 | 0,575121 | 1,16769 | 1 | 1 |
| Rab3c | 0,202164 | 0,453207 | 1,16464 | 1 | 1 |
| Trim10 | 0,0465557 | 0,104269 | 1,16328 | 1 | 1 |
| Cd164l2 | 0,223615 | 0,500358 | 1,16194 | 1 | 1 |
| Mir143hg | 0,0955693 | 0,213419 | 1,15907 | 1 | 1 |
| Gm996 | 0,049193 | 0,109456 | 1,15383 | 1 | 1 |
| Rlbp1 | 0,099127 | 0,220525 | 1,15359 | 1 | 1 |
| Hrh1 | 0,0388487 | 0,0863401 | 1,15216 | 1 | 1 |
| Gm5126 | 0,210772 | 0,468428 | 1,15215 | 1 | 1 |
| Grm8 | 0,129588 | 0,287863 | 1,15145 | 1 | 1 |
| Spata17 | 0,26124 | 0,57994 | 1,15053 | 1 | 1 |
| Adcyap1 | 0,115967 | 0,257262 | 1,14953 | 1 | 1 |
| Cnpy1 | 0,200218 | 0,442734 | 1,14487 | 1 | 1 |
| Scnn1b | 0,0232021 | 0,0512499 | 1,1433 | 1 | 1 |
| Trpv3 | 0,0258543 | 0,0571055 | 1,14322 | 1 | 1 |
| 4931429I11Rik | 0,0201852 | 0,0445838 | 1,14322 | 1 | 1 |
| Vwc2 | 0,0171341 | 0,0378448 | 1,14322 | 1 | 1 |
| Insl5 | 0,153538 | 0,338782 | 1,14176 | 1 | 1 |
| Pom121l2 | 0,0356574 | 0,0786619 | 1,14146 | 1 | 1 |

| | | | | | |
|---|---|---|---|---|---|
| Rpgrip1 | 0,0855131 | 0,188631 | 1,14135 | 1 | 1 |
| Slc52a3 | 0,0263536 | 0,0579142 | 1,13592 | 1 | 1 |
| Gm19583 | 0,043841 | 0,096342 | 1,13588 | 1 | 1 |
| Galnt6 | 0,00927655 | 0,0203835 | 1,13574 | 1 | 1 |
| Glra4 | 0,12679 | 0,278363 | 1,13453 | 1 | 1 |
| Prkag2os1 | 0,12089 | 0,264835 | 1,1314 | 1 | 1 |
| Fam19a1 | 0,0311747 | 0,0682948 | 1,1314 | 1 | 1 |
| Itih4 | 0,0155161 | 0,0338697 | 1,12623 | 1 | 1 |
| 4930415F15Rik | 0,0802842 | 0,175247 | 1,1262 | 1 | 1 |
| 4933433C11Rik | 0,0359515 | 0,0784761 | 1,1262 | 1 | 1 |
| Prss56 | 0,0245553 | 0,0536 | 1,1262 | 1 | 1 |
| E330013P04Rik | 0,116563 | 0,254354 | 1,12572 | 1 | 1 |
| Sytl5 | 0,0424733 | 0,0926027 | 1,1245 | 1 | 1 |
| Agtr1b | 0,146712 | 0,319009 | 1,12062 | 1 | 1 |
| Fam83f | 0,258084 | 0,558762 | 1,11439 | 1 | 1 |
| Abcc8 | 0,220375 | 0,476991 | 1,114 | 1 | 1 |
| Acpp | 0,100613 | 0,21771 | 1,11359 | 1 | 1 |
| Cyp2u1 | 0,099708 | 0,215744 | 1,11354 | 1 | 1 |
| Flrt1 | 0,0759258 | 0,164168 | 1,11251 | 1 | 1 |
| Tdrd12 | 0,0767491 | 0,165834 | 1,11152 | 1 | 1 |
| Slc17a7 | 0,0397492 | 0,0858871 | 1,11152 | 1 | 1 |
| Sec14l4 | 0,0477693 | 0,103068 | 1,10944 | 1 | 1 |
| Gm3558 | 0,107156 | 0,230908 | 1,10761 | 1 | 1 |
| Syn2 | 0,0940343 | 0,202428 | 1,10615 | 1 | 1 |
| Gja4 | 0,201615 | 0,433203 | 1,10344 | 1 | 1 |
| Srpx2 | 0,0820831 | 0,176125 | 1,10145 | 1 | 1 |
| Zc3hav1 | 0,230616 | 0,493722 | 1,0982 | 1 | 1 |
| Avpr2 | 0,0771653 | 0,164989 | 1,09635 | 1 | 1 |
| P2rx7 | 0,0526431 | 0,112421 | 1,09459 | 1 | 1 |
| Gm19897 | 0,0841917 | 0,179561 | 1,09272 | 1 | 1 |
| Mroh8 | 0,193866 | 0,413224 | 1,09186 | 1 | 1 |
| E330011O21Rik | 0,155402 | 0,330741 | 1,08969 | 1 | 1 |
| H2-Aa | 0,0668321 | 0,142238 | 1,08969 | 1 | 1 |
| Pigz | 0,0291971 | 0,0621398 | 1,08969 | 1 | 1 |
| Kif12 | 0,278652 | 0,592789 | 1,08905 | 1 | 1 |
| Sptb | 0,171327 | 0,363881 | 1,08671 | 1 | 1 |
| 4921504A21Rik | 0,0701608 | 0,148893 | 1,08554 | 1 | 1 |
| BC030499 | 0,0430097 | 0,0912693 | 1,08547 | 1 | 1 |
| Xkrx | 0,0273341 | 0,05787 | 1,08211 | 1 | 1 |
| Hist1h3d | 0,413069 | 0,872673 | 1,07906 | 1 | 1 |
| Lrrc71 | 0,0811223 | 0,171215 | 1,07764 | 1 | 1 |
| Plin1 | 0,0307524 | 0,0647887 | 1,07504 | 1 | 1 |
| 6030408B16Rik | 0,0288959 | 0,0608615 | 1,07467 | 1 | 1 |
| Cckbr | 0,0223535 | 0,0470816 | 1,07467 | 1 | 1 |
| Mfsd7c | 0,180705 | 0,38025 | 1,07331 | 1 | 1 |
| Gm3002 | 0,0420358 | 0,0884356 | 1,07301 | 1 | 1 |
| Trim66 | 0,0283873 | 0,0597026 | 1,07255 | 1 | 1 |
| Mst1 | 0,0317556 | 0,0666545 | 1,06969 | 1 | 1 |
| Bcmo1 | 0,031141 | 0,0653645 | 1,06969 | 1 | 1 |
| C230024C17Rik | 0,0211029 | 0,0442946 | 1,06969 | 1 | 1 |

| | | | | | |
|---|---|---|---|---|---|
| Fmo4 | 0,111375 | 0,233171 | 1,06596 | 1 | 1 |
| 4933402D24Rik | 0,169473 | 0,354669 | 1,06542 | 1 | 1 |
| Gm4262 | 0,159338 | 0,333152 | 1,06409 | 1 | 1 |
| Apobec1 | 0,0753235 | 0,157327 | 1,06259 | 1 | 1 |
| L1td1 | 0,226832 | 0,471708 | 1,05627 | 1 | 1 |
| Scarf1 | 0,243479 | 0,506072 | 1,05555 | 1 | 1 |
| Ggt5 | 0,0353668 | 0,0733886 | 1,05316 | 1 | 1 |
| Ccdc3 | 0,0712588 | 0,147789 | 1,0524 | 1 | 1 |
| 4930478L05Rik | 0,0567091 | 0,117611 | 1,05238 | 1 | 1 |
| Ascl4 | 0,083786 | 0,173765 | 1,05236 | 1 | 1 |
| Gm101 | 0,0346115 | 0,0717814 | 1,05236 | 1 | 1 |
| Spib | 0,04212 | 0,0867158 | 1,04179 | 1 | 1 |
| Zdhhc23 | 0,0820792 | 0,168842 | 1,04059 | 1 | 1 |
| Ngb | 0,128708 | 0,264703 | 1,04028 | 1 | 1 |
| Gm10474 | 0,129066 | 0,265256 | 1,03927 | 1 | 1 |
| Paqr5 | 0,0931735 | 0,191421 | 1,03876 | 1 | 1 |
| Ttc39d | 0,0560126 | 0,114438 | 1,03074 | 1 | 1 |
| Krt86 | 0,0493823 | 0,100619 | 1,02683 | 1 | 1 |
| Klk13 | 0,262371 | 0,534553 | 1,02673 | 1 | 1 |
| Ptpn18 | 0,207174 | 0,421673 | 1,02528 | 1 | 1 |
| Rnf151 | 0,244901 | 0,498333 | 1,02491 | 1 | 1 |
| Mfsd6 | 0,113175 | 0,230266 | 1,02475 | 1 | 1 |
| Lrp1b | 0,00795248 | 0,0161626 | 1,02318 | 1 | 1 |
| Drd4 | 0,257885 | 0,523858 | 1,02245 | 1 | 1 |
| Pax5 | 0,0658513 | 0,133588 | 1,02051 | 1 | 1 |
| Gm14164 | 0,036874 | 0,074804 | 1,02051 | 1 | 1 |
| 4930539E08Rik | 0,112916 | 0,228858 | 1,0192 | 1 | 1 |
| Ephx2 | 0,280719 | 0,568599 | 1,01828 | 1 | 1 |
| Cd79a | 0,0846124 | 0,171347 | 1,01798 | 1 | 1 |
| Serpine3 | 0,182758 | 0,369906 | 1,01722 | 1 | 1 |
| Tuba8 | 0,100916 | 0,204172 | 1,01663 | 1 | 1 |
| Nfatc2 | 0,232732 | 0,469177 | 1,01146 | 1 | 1 |
| Bcl3 | 0,0489568 | 0,0986807 | 1,01126 | 1 | 1 |
| Gm16982 | 0,15107 | 0,304497 | 1,01121 | 1 | 1 |
| Cbln4 | 0,224993 | 0,453457 | 1,01109 | 1 | 1 |
| Frmpd1 | 0,0460242 | 0,0927545 | 1,01102 | 1 | 1 |
| Tnfrsf11b | 0,0753281 | 0,151467 | 1,00775 | 1 | 1 |
| Plek2 | 0,130888 | 0,26234 | 1,0031 | 1 | 1 |
| Aifm3 | 0,118352 | 0,237144 | 1,00267 | 1 | 1 |
| Gm17455 | 0,200292 | 0,400931 | 1,00125 | 1 | 1 |
| Ankrd33b | 0,248305 | 0,497012 | 1,00117 | 1 | 1 |
| Ovol3 | 0,0489597 | 0,0979496 | 1,00044 | 1 | 1 |
| Hormad2 | 0,147499 | 0,294862 | 0,999337 | 1 | 1 |
| En2 | 0,0987514 | 0,197344 | 0,998842 | 1 | 1 |
| Matn4 | 0,049967 | 0,099807 | 0,998165 | 1 | 1 |
| Klhl35 | 0,0582877 | 0,116425 | 0,998137 | 1 | 1 |
| Rph3al | 0,287389 | 0,574012 | 0,998074 | 1 | 1 |
| Bmp8b | 0,0880791 | 0,175826 | 0,997281 | 1 | 1 |
| Ppp2r2c | 0,191079 | 0,380752 | 0,994684 | 1 | 1 |
| Lrrc63 | 0,0499441 | 0,0995144 | 0,994591 | 1 | 1 |

| | | | | | |
|---|---|---|---|---|---|
| Rhd | 0,116664 | 0,232346 | 0,993914 | 1 | 1 |
| Gm12253 | 0,0956522 | 0,190319 | 0,992552 | 1 | 1 |
| Gucy1b2 | 0,0446418 | 0,0888235 | 0,992548 | 1 | 1 |
| Tex14 | 0,0412258 | 0,081949 | 0,991178 | 1 | 1 |
| Htra3 | 0,287139 | 0,570163 | 0,989624 | 1 | 1 |
| 4931403E22Rik | 0,267075 | 0,53032 | 0,989619 | 1 | 1 |
| 5330417C22Rik | 0,221388 | 0,439466 | 0,989177 | 1 | 1 |
| Gm14634 | 0,204934 | 0,406728 | 0,988905 | 1 | 1 |
| Prelp | 0,0879326 | 0,174518 | 0,988905 | 1 | 1 |
| Plbd1 | 0,0987959 | 0,196009 | 0,988399 | 1 | 1 |
| Vwf | 0,0354186 | 0,0702068 | 0,987105 | 1 | 1 |
| Kcnip1 | 0,185764 | 0,367791 | 0,985418 | 1 | 1 |
| E130112N10Rik | 0,127256 | 0,251643 | 0,983643 | 1 | 1 |
| Pabpc5 | 0,0944578 | 0,186693 | 0,982923 | 1 | 1 |
| C030016D13Rik | 0,127838 | 0,252655 | 0,982849 | 1 | 1 |
| Hba-a2 | 0,00681439 | 0,0134641 | 0,982459 | 1 | 1 |
| Galr3 | 0,0494778 | 0,097722 | 0,981901 | 1 | 1 |
| Lypd1 | 0,0958772 | 0,188569 | 0,975829 | 1 | 1 |
| F2rl3 | 0,139607 | 0,274333 | 0,974559 | 1 | 1 |
| Doc2b | 0,156734 | 0,307925 | 0,974261 | 1 | 1 |
| Mall | 0,0866877 | 0,170007 | 0,971698 | 1 | 1 |
| Ky | 0,16119 | 0,31607 | 0,971479 | 1 | 1 |
| Thy1 | 0,143008 | 0,280336 | 0,971063 | 1 | 1 |
| LOC100503676 | 0,212006 | 0,415467 | 0,970632 | 1 | 1 |
| Gpr20 | 0,128698 | 0,251676 | 0,967573 | 1 | 1 |
| Cd27 | 0,137268 | 0,268274 | 0,966708 | 1 | 1 |
| Gpr4 | 0,30009 | 0,586146 | 0,965867 | 1 | 1 |
| E130018N17Rik | 0,139609 | 0,271241 | 0,958185 | 1 | 1 |
| Calb1 | 0,0713634 | 0,138607 | 0,957746 | 1 | 1 |
| Slc2a9 | 0,127044 | 0,246658 | 0,957188 | 1 | 1 |
| Mogat1 | 0,0576945 | 0,111726 | 0,953458 | 1 | 1 |
| 4831440E17Rik | 0,0258486 | 0,0500561 | 0,953458 | 1 | 1 |
| Ccin | 0,0437287 | 0,084618 | 0,952385 | 1 | 1 |
| Spaca4 | 0,201354 | 0,389469 | 0,951777 | 1 | 1 |
| M1ap | 0,0551635 | 0,1067 | 0,951777 | 1 | 1 |
| Faim2 | 0,00496565 | 0,00960293 | 0,951493 | 1 | 1 |
| Dclk3 | 0,11495 | 0,222158 | 0,950583 | 1 | 1 |
| Park2 | 0,084991 | 0,164214 | 0,950192 | 1 | 1 |
| Trp73 | 0,00516751 | 0,0099809 | 0,949701 | 1 | 1 |
| Ucma | 0,0379223 | 0,0732445 | 0,949676 | 1 | 1 |
| Efcab6 | 0,0100673 | 0,0194433 | 0,949604 | 1 | 1 |
| Xaf1 | 0,0208984 | 0,0403619 | 0,9496 | 1 | 1 |
| Kif17 | 0,0153095 | 0,0295674 | 0,94958 | 1 | 1 |
| Rpl3l | 0,0527854 | 0,101945 | 0,949578 | 1 | 1 |
| Shc3 | 0,0661811 | 0,127815 | 0,949569 | 1 | 1 |
| 4933417O13Rik | 0,0500113 | 0,0965866 | 0,949569 | 1 | 1 |
| C1ql4 | 0,042959 | 0,0829665 | 0,949569 | 1 | 1 |
| Spata25 | 0,0380622 | 0,0735094 | 0,949569 | 1 | 1 |
| Gtsf1l | 0,0326007 | 0,0629617 | 0,949569 | 1 | 1 |
| 4833419F23Rik | 0,0325526 | 0,0628687 | 0,949569 | 1 | 1 |

| | | | | | |
|---|---|---|---|---|---|
| Pitpnm2os1 | 0,0311716 | 0,0602016 | 0,949569 | 1 | 1 |
| Gm5150 | 0,0310834 | 0,0600312 | 0,949569 | 1 | 1 |
| Olfr61 | 0,0301067 | 0,058145 | 0,949569 | 1 | 1 |
| 3830403N18Rik | 0,0270076 | 0,0521596 | 0,949569 | 1 | 1 |
| Wbscr25 | 0,0235579 | 0,0454972 | 0,949569 | 1 | 1 |
| Gp5 | 0,0220936 | 0,0426693 | 0,949569 | 1 | 1 |
| Gm867 | 0,0215217 | 0,0415648 | 0,949569 | 1 | 1 |
| Alox12b | 0,0205678 | 0,0397225 | 0,949569 | 1 | 1 |
| Gbp2 | 0,019581 | 0,0378167 | 0,949569 | 1 | 1 |
| Gm13939 | 0,0192474 | 0,0371724 | 0,949569 | 1 | 1 |
| Gm13277,Gm13285 | 0,0175185 | 0,0338334 | 0,949569 | 1 | 1 |
| Cacng2 | 0,0164282 | 0,0317277 | 0,949569 | 1 | 1 |
| Gm8994 | 0,0160981 | 0,0310902 | 0,949569 | 1 | 1 |
| 4930455D15Rik | 0,0156079 | 0,0301434 | 0,949569 | 1 | 1 |
| Tekt3 | 0,0143572 | 0,027728 | 0,949569 | 1 | 1 |
| Tex19.1 | 0,0143572 | 0,027728 | 0,949569 | 1 | 1 |
| Ces2f | 0,0136123 | 0,0262894 | 0,949569 | 1 | 1 |
| Kcnk12 | 0,0126078 | 0,0243493 | 0,949569 | 1 | 1 |
| Ces1a | 0,012416 | 0,0239789 | 0,949569 | 1 | 1 |
| Dbh | 0,0106056 | 0,0204825 | 0,949569 | 1 | 1 |
| 4930463O16Rik | 0,010371 | 0,0200295 | 0,949569 | 1 | 1 |
| Mlph | 0,0102983 | 0,0198891 | 0,949569 | 1 | 1 |
| Scel | 0,00773549 | 0,0149395 | 0,949569 | 1 | 1 |
| Nr2e1 | 0,00727581 | 0,0140517 | 0,949569 | 1 | 1 |
| Grm3 | 0,00661418 | 0,0127739 | 0,949569 | 1 | 1 |
| Pnpla1 | 0,00515038 | 0,0099469 | 0,949569 | 1 | 1 |
| D630023F18Rik | 0,0291944 | 0,0563541 | 0,948829 | 1 | 1 |
| Kcnc4 | 0,0887467 | 0,171257 | 0,9484 | 1 | 1 |
| Bnc1 | 0,214316 | 0,411409 | 0,940834 | 1 | 1 |
| Avil | 0,0900382 | 0,172321 | 0,936493 | 1 | 1 |
| Cyp2j9 | 0,213664 | 0,408848 | 0,936219 | 1 | 1 |
| Gata4 | 0,0953978 | 0,182479 | 0,935702 | 1 | 1 |
| Gsc | 0,330753 | 0,632366 | 0,935006 | 1 | 1 |
| Ryr3 | 0,228298 | 0,436451 | 0,934902 | 1 | 1 |
| Gm13139,Gm13251 | 0,108823 | 0,208042 | 0,934894 | 1 | 1 |
| Prss44 | 0,0530693 | 0,101346 | 0,933339 | 1 | 1 |
| Tssk2 | 0,0802532 | 0,153156 | 0,932369 | 1 | 1 |
| Wee2 | 0,0354289 | 0,0676128 | 0,932369 | 1 | 1 |
| Sh2d4b | 0,233256 | 0,444788 | 0,931202 | 1 | 1 |
| 2010107G12Rik | 0,160679 | 0,305866 | 0,928721 | 1 | 1 |
| Capn12 | 0,181093 | 0,34418 | 0,926434 | 1 | 1 |
| 1700018L02Rik | 0,217876 | 0,412879 | 0,922215 | 1 | 1 |
| Gm6225 | 0,156501 | 0,296225 | 0,920521 | 1 | 1 |
| Mustn1 | 0,217515 | 0,410855 | 0,917512 | 1 | 1 |
| Best2 | 0,0288444 | 0,0544778 | 0,917376 | 1 | 1 |
| Gm6484 | 0,231208 | 0,43666 | 0,91732 | 1 | 1 |
| Aldh3b1 | 0,031292 | 0,0590982 | 0,91732 | 1 | 1 |
| Lingo4 | 0,0185993 | 0,0351268 | 0,91732 | 1 | 1 |
| Bcl6 | 0,28332 | 0,534819 | 0,91662 | 1 | 1 |
| Tmem239 | 0,20325 | 0,38291 | 0,913748 | 1 | 1 |

| | | | | | |
|---|---|---|---|---|---|
| Dlx2 | 0,0764349 | 0,143917 | 0,912935 | 1 | 1 |
| Kcnt1 | 0,035932 | 0,0675847 | 0,911427 | 1 | 1 |
| Hsbp1l1 | 0,0761183 | 0,142861 | 0,908291 | 1 | 1 |
| Serpina3i | 0,0871949 | 0,16328 | 0,905028 | 1 | 1 |
| Cdhr5 | 0,149977 | 0,280517 | 0,903345 | 1 | 1 |
| Zyg11a | 0,203433 | 0,378938 | 0,897408 | 1 | 1 |
| Inpp5d | 0,0750208 | 0,139653 | 0,896484 | 1 | 1 |
| Gpr114 | 0,0253124 | 0,047105 | 0,896038 | 1 | 1 |
| Prrt3 | 0,166519 | 0,309297 | 0,893302 | 1 | 1 |
| Asprv1 | 0,153559 | 0,285163 | 0,89299 | 1 | 1 |
| Cacna1s | 0,00965594 | 0,0179263 | 0,892587 | 1 | 1 |
| Ugt1a6b | 0,0138363 | 0,0256815 | 0,892265 | 1 | 1 |
| Stard6 | 0,181596 | 0,337056 | 0,892254 | 1 | 1 |
| D030047H15Rik | 0,012395 | 0,0230058 | 0,892242 | 1 | 1 |
| Lkaaear1 | 0,0564542 | 0,104782 | 0,892239 | 1 | 1 |
| B230119M05Rik | 0,0372965 | 0,0692244 | 0,892239 | 1 | 1 |
| Esm1 | 0,0304424 | 0,0565029 | 0,892239 | 1 | 1 |
| Ccdc154 | 0,0262416 | 0,0487059 | 0,892239 | 1 | 1 |
| Cyp2w1 | 0,0220986 | 0,0410163 | 0,892239 | 1 | 1 |
| BC049635 | 0,0199071 | 0,0369486 | 0,892239 | 1 | 1 |
| Prodh2 | 0,0179201 | 0,0332606 | 0,892239 | 1 | 1 |
| A530065N20Rik | 0,0163862 | 0,0304136 | 0,892239 | 1 | 1 |
| Lrrc6 | 0,0153751 | 0,0285369 | 0,892239 | 1 | 1 |
| Serpinc1 | 0,0147701 | 0,0274141 | 0,892239 | 1 | 1 |
| Ptk6 | 0,0138833 | 0,0257682 | 0,892239 | 1 | 1 |
| Rgsl1 | 0,00841245 | 0,015614 | 0,892239 | 1 | 1 |
| Myh2 | 0,00491669 | 0,00912565 | 0,892239 | 1 | 1 |
| Kcnma1 | 0,105472 | 0,195494 | 0,89026 | 1 | 1 |
| Cr2 | 0,11285 | 0,209131 | 0,889998 | 1 | 1 |
| Tecta | 0,106711 | 0,197635 | 0,889123 | 1 | 1 |
| Obscn | 0,00580276 | 0,0107331 | 0,887259 | 1 | 1 |
| Esrp2 | 0,195195 | 0,361001 | 0,887087 | 1 | 1 |
| Csgalnact1 | 0,10494 | 0,193994 | 0,886449 | 1 | 1 |
| A730085K08Rik | 0,304389 | 0,562169 | 0,885089 | 1 | 1 |
| Slc6a11 | 0,0188975 | 0,0347292 | 0,877954 | 1 | 1 |
| A430088P11Rik | 0,0339081 | 0,0622665 | 0,876825 | 1 | 1 |
| a | 0,375349 | 0,688203 | 0,874604 | 1 | 1 |
| Il2rg | 0,231719 | 0,424474 | 0,873301 | 1 | 1 |
| Kcnk3 | 0,0716782 | 0,131299 | 0,873248 | 1 | 1 |
| Camk1d | 0,0850838 | 0,155495 | 0,869908 | 1 | 1 |
| Als2cl | 0,187811 | 0,342988 | 0,868881 | 1 | 1 |
| Galnt13 | 0,189574 | 0,346137 | 0,868582 | 1 | 1 |
| Ovol2 | 0,0436823 | 0,079755 | 0,868525 | 1 | 1 |
| 0610043K17Rik | 0,212926 | 0,388755 | 0,868508 | 1 | 1 |
| 1700027J07Rik | 0,102982 | 0,188022 | 0,868508 | 1 | 1 |
| AI467606 | 0,0257157 | 0,0469511 | 0,868508 | 1 | 1 |
| Olfr558 | 0,0162107 | 0,029597 | 0,868508 | 1 | 1 |
| Hepacam | 0,0152832 | 0,0279037 | 0,868508 | 1 | 1 |
| Slc28a3 | 0,0101094 | 0,0184575 | 0,868508 | 1 | 1 |
| Strip2 | 0,189141 | 0,345051 | 0,867344 | 1 | 1 |

| | | | | | |
|---|---|---|---|---|---|
| Lca5l | 0,250208 | 0,456307 | 0,866881 | 1 | 1 |
| H2-M10.2 | 0,250516 | 0,455496 | 0,862539 | 1 | 1 |
| Sorcs1 | 0,101454 | 0,184356 | 0,861668 | 1 | 1 |
| Cd14 | 0,366327 | 0,66471 | 0,859591 | 1 | 1 |
| 2610206C17Rik | 0,196534 | 0,356484 | 0,85906 | 1 | 1 |
| Apobec2 | 0,272933 | 0,494523 | 0,85749 | 1 | 1 |
| F630111L10Rik | 0,0438443 | 0,079425 | 0,857204 | 1 | 1 |
| Trpv6 | 0,147093 | 0,266281 | 0,856221 | 1 | 1 |
| Mpz | 0,170052 | 0,307684 | 0,855478 | 1 | 1 |
| Prune2 | 0,244232 | 0,44107 | 0,852754 | 1 | 1 |
| Sdr42e1 | 0,304585 | 0,54853 | 0,848726 | 1 | 1 |
| Aipl1 | 0,223763 | 0,402962 | 0,848674 | 1 | 1 |
| Oprl1 | 0,0991074 | 0,178366 | 0,84778 | 1 | 1 |
| Gm10814 | 0,0403012 | 0,0724675 | 0,84651 | 1 | 1 |
| 4930455C13Rik | 0,313938 | 0,564087 | 0,845438 | 1 | 1 |
| Otog | 0,0132933 | 0,0238833 | 0,845304 | 1 | 1 |
| Dennd3 | 0,236993 | 0,425271 | 0,84354 | 1 | 1 |
| Ldb3 | 0,0531576 | 0,0953104 | 0,842358 | 1 | 1 |
| Amh | 0,348688 | 0,624997 | 0,841911 | 1 | 1 |
| Ptchd1 | 0,0151186 | 0,0270885 | 0,841352 | 1 | 1 |
| Flt1 | 0,164718 | 0,295094 | 0,841178 | 1 | 1 |
| Pla2g2c | 0,196742 | 0,351342 | 0,836569 | 1 | 1 |
| 4930415O20Rik | 0,422171 | 0,753841 | 0,836432 | 1 | 1 |
| Dpcr1 | 0,17584 | 0,313627 | 0,834785 | 1 | 1 |
| Ankrd7 | 0,103386 | 0,184006 | 0,831706 | 1 | 1 |
| Rax | 0,0695473 | 0,123779 | 0,831706 | 1 | 1 |
| Ccr4 | 0,0394453 | 0,0702043 | 0,831706 | 1 | 1 |
| Map3k8 | 0,240968 | 0,428642 | 0,830928 | 1 | 1 |
| Ssmem1 | 0,130444 | 0,231912 | 0,83015 | 1 | 1 |
| 4933416M07Rik | 0,2017 | 0,356899 | 0,823302 | 1 | 1 |
| Hist1h2bl | 0,479062 | 0,847632 | 0,823225 | 1 | 1 |
| 4933428G20Rik | 0,0684544 | 0,120994 | 0,821723 | 1 | 1 |
| Mylk4 | 0,0417972 | 0,0736781 | 0,817829 | 1 | 1 |
| Npb | 0,482779 | 0,850082 | 0,816239 | 1 | 1 |
| Cd200 | 0,186972 | 0,32896 | 0,815087 | 1 | 1 |
| Opn1sw | 0,0761569 | 0,133759 | 0,812588 | 1 | 1 |
| Jakmip1 | 0,0632043 | 0,110846 | 0,810464 | 1 | 1 |
| C430002E04Rik | 0,0448847 | 0,0786724 | 0,809634 | 1 | 1 |
| Tlx2 | 0,235 | 0,411462 | 0,808099 | 1 | 1 |
| Dsc3 | 0,0944715 | 0,165393 | 0,807945 | 1 | 1 |
| Cpeb1 | 0,0504286 | 0,0882854 | 0,807932 | 1 | 1 |
| Elf5 | 0,038005 | 0,0665008 | 0,807185 | 1 | 1 |
| Lmod1 | 0,118598 | 0,207156 | 0,804634 | 1 | 1 |
| 6030443J06Rik | 0,258347 | 0,449776 | 0,799897 | 1 | 1 |
| Gm11744 | 0,292908 | 0,509832 | 0,799573 | 1 | 1 |
| Ptprcap | 0,396751 | 0,690254 | 0,798894 | 1 | 1 |
| Klk10 | 0,10179 | 0,176965 | 0,797876 | 1 | 1 |
| Asic3 | 0,0459428 | 0,0798473 | 0,797406 | 1 | 1 |
| Kcnj14 | 0,31887 | 0,550528 | 0,787847 | 1 | 1 |
| Efhb | 0,234249 | 0,404395 | 0,787719 | 1 | 1 |

| | | | | | |
|---|---|---|---|---|---|
| D630039A03Rik | 0,141189 | 0,243671 | 0,787305 | 1 | 1 |
| Ubxn10 | 0,341983 | 0,589453 | 0,785452 | 1 | 1 |
| Ccdc170 | 0,0827608 | 0,142552 | 0,784471 | 1 | 1 |
| Adcy9 | 0,24005 | 0,413267 | 0,783739 | 1 | 1 |
| Lrrc15 | 0,271681 | 0,467265 | 0,782327 | 1 | 1 |
| Duxbl1,Duxbl2 | 0,0808152 | 0,138829 | 0,780606 | 1 | 1 |
| Vstm2l | 0,341027 | 0,585101 | 0,778799 | 1 | 1 |
| 2900041M22Rik | 0,257337 | 0,441431 | 0,77853 | 1 | 1 |
| Rell2 | 0,197553 | 0,338821 | 0,778282 | 1 | 1 |
| Ppm1n | 0,24889 | 0,426592 | 0,777351 | 1 | 1 |
| Tunar | 0,215161 | 0,368473 | 0,776138 | 1 | 1 |
| Slc35d3 | 0,0539349 | 0,09233 | 0,775579 | 1 | 1 |
| Ttc16 | 0,134846 | 0,230767 | 0,775124 | 1 | 1 |
| Flt3l | 0,157658 | 0,269741 | 0,774781 | 1 | 1 |
| Sspo | 0,0690102 | 0,118006 | 0,773979 | 1 | 1 |
| Gm16796 | 0,0812013 | 0,138712 | 0,772514 | 1 | 1 |
| Pde7b | 0,342967 | 0,585779 | 0,772287 | 1 | 1 |
| Oscar | 0,0847821 | 0,144752 | 0,77175 | 1 | 1 |
| Gm7104 | 0,0674725 | 0,115168 | 0,771372 | 1 | 1 |
| AI854517 | 0,0372728 | 0,0635984 | 0,770867 | 1 | 1 |
| Tmc3 | 0,144979 | 0,246593 | 0,766289 | 1 | 1 |
| 4933433G15Rik | 0,0928903 | 0,157914 | 0,76554 | 1 | 1 |
| Nfe2l3 | 0,338914 | 0,575628 | 0,76422 | 1 | 1 |
| Tenm2 | 0,206291 | 0,3501 | 0,763086 | 1 | 1 |
| Gcgr | 0,18997 | 0,321796 | 0,760372 | 1 | 1 |
| Slc16a11 | 0,0851998 | 0,144265 | 0,759802 | 1 | 1 |
| 1700019B03Rik | 0,0469233 | 0,0794533 | 0,759802 | 1 | 1 |
| Tmem25 | 0,318281 | 0,538331 | 0,758193 | 1 | 1 |
| Gm9899 | 0,0984498 | 0,166269 | 0,756057 | 1 | 1 |
| Myh13 | 0,0569029 | 0,0960918 | 0,755912 | 1 | 1 |
| Cpvl | 0,0567466 | 0,0956694 | 0,753525 | 1 | 1 |
| Gm10941 | 0,374826 | 0,631373 | 0,75227 | 1 | 1 |
| Gnb3 | 0,0712902 | 0,120021 | 0,751515 | 1 | 1 |
| Scn5a | 0,104948 | 0,176596 | 0,75078 | 1 | 1 |
| Ifitm10 | 0,0263876 | 0,0443239 | 0,748225 | 1 | 1 |
| Chad | 0,292724 | 0,490955 | 0,746052 | 1 | 1 |
| Gm1653 | 0,0589154 | 0,0987608 | 0,745294 | 1 | 1 |
| D10Bwg1379e | 0,0876629 | 0,146647 | 0,742314 | 1 | 1 |
| Syna | 0,0694925 | 0,116235 | 0,742114 | 1 | 1 |
| Gm2373 | 0,0162482 | 0,0271644 | 0,741433 | 1 | 1 |
| Gpr160 | 0,0685126 | 0,11449 | 0,740774 | 1 | 1 |
| 1100001G20Rik | 0,217697 | 0,363765 | 0,740689 | 1 | 1 |
| Fam167b | 0,0688418 | 0,115033 | 0,740689 | 1 | 1 |
| 4933406C10Rik | 0,0356497 | 0,0595696 | 0,740689 | 1 | 1 |
| Akap14 | 0,0320802 | 0,0536052 | 0,740689 | 1 | 1 |
| Spata31d1d | 0,0281224 | 0,0469917 | 0,740689 | 1 | 1 |
| Gbp5 | 0,0180843 | 0,0302185 | 0,740689 | 1 | 1 |
| Tas1r1 | 0,288087 | 0,480998 | 0,739526 | 1 | 1 |
| Ak8 | 0,116908 | 0,195067 | 0,738594 | 1 | 1 |
| 1810034E14Rik | 0,255036 | 0,425481 | 0,738396 | 1 | 1 |

| | | | | | |
|---|---|---|---|---|---|
| Bpifb5 | 0,0758921 | 0,126544 | 0,737611 | 1 | 1 |
| Serpind1 | 0,0548154 | 0,0913999 | 0,737611 | 1 | 1 |
| Piwil2 | 0,023095 | 0,0385088 | 0,737611 | 1 | 1 |
| Elmod1 | 0,245433 | 0,409209 | 0,73751 | 1 | 1 |
| Gimap1 | 0,123182 | 0,205115 | 0,735644 | 1 | 1 |
| Scpep1os | 0,309662 | 0,51557 | 0,735473 | 1 | 1 |
| Napsa | 0,136024 | 0,226381 | 0,734894 | 1 | 1 |
| Unc45b | 0,0172671 | 0,0287373 | 0,734894 | 1 | 1 |
| Pi15 | 0,00860403 | 0,0143195 | 0,734894 | 1 | 1 |
| Ifih1 | 0,174924 | 0,290539 | 0,732005 | 1 | 1 |
| Nav3 | 0,0990265 | 0,164438 | 0,731659 | 1 | 1 |
| Mak | 0,335937 | 0,557003 | 0,729492 | 1 | 1 |
| Ncf2 | 0,229132 | 0,379796 | 0,729046 | 1 | 1 |
| Lrguk | 0,340156 | 0,563694 | 0,728718 | 1 | 1 |
| Sptbn4 | 0,0386319 | 0,0637789 | 0,723285 | 1 | 1 |
| Gp1bb | 0,165574 | 0,273076 | 0,721825 | 1 | 1 |
| Ggn | 0,319429 | 0,526794 | 0,721743 | 1 | 1 |
| Lrriq3 | 0,0833092 | 0,137306 | 0,720847 | 1 | 1 |
| Zmynd15 | 0,140951 | 0,231553 | 0,716154 | 1 | 1 |
| Rfx8 | 0,276003 | 0,453254 | 0,715636 | 1 | 1 |
| 5730405O15Rik | 0,18144 | 0,297963 | 0,715636 | 1 | 1 |
| Gdf5 | 0,0587046 | 0,0964051 | 0,715636 | 1 | 1 |
| Irf7 | 0,106744 | 0,174895 | 0,712333 | 1 | 1 |
| Rtn4rl1 | 0,244898 | 0,400878 | 0,71098 | 1 | 1 |
| Inhbb | 0,0523524 | 0,0856425 | 0,710072 | 1 | 1 |
| 1700034H15Rik | 0,151117 | 0,247063 | 0,709206 | 1 | 1 |
| Arhgef19 | 0,325547 | 0,531964 | 0,708462 | 1 | 1 |
| Tdrd5 | 0,0621965 | 0,101167 | 0,701831 | 1 | 1 |
| Prkaa2 | 0,338847 | 0,551015 | 0,701458 | 1 | 1 |
| 4930502E18Rik | 0,197103 | 0,32043 | 0,701061 | 1 | 1 |
| Ucn2 | 0,0828502 | 0,134459 | 0,698583 | 1 | 1 |
| Cntnap1 | 0,123119 | 0,199714 | 0,697886 | 1 | 1 |
| 1700028K03Rik | 0,3269 | 0,529989 | 0,697113 | 1 | 1 |
| Wdr96 | 0,244789 | 0,396591 | 0,696114 | 1 | 1 |
| Cacna1b | 0,172871 | 0,279725 | 0,694312 | 1 | 1 |
| Fbxl21 | 0,293186 | 0,47384 | 0,692582 | 1 | 1 |
| 2900052N01Rik | 0,380566 | 0,614688 | 0,691709 | 1 | 1 |
| Cnnm1 | 0,10138 | 0,163376 | 0,688424 | 1 | 1 |
| Cds1 | 0,243589 | 0,392337 | 0,687642 | 1 | 1 |
| Sox7 | 0,182072 | 0,293072 | 0,686749 | 1 | 1 |
| Cdnf | 0,129907 | 0,20882 | 0,684776 | 1 | 1 |
| Ccdc149 | 0,306519 | 0,492439 | 0,683965 | 1 | 1 |
| Samd9l | 0,0609932 | 0,0978633 | 0,68212 | 1 | 1 |
| Aire | 0,0151558 | 0,0243147 | 0,681959 | 1 | 1 |
| Syndig1 | 0,279708 | 0,448288 | 0,680505 | 1 | 1 |
| Trpm5 | 0,0207503 | 0,0332077 | 0,678386 | 1 | 1 |
| Birc7 | 0,0819795 | 0,131095 | 0,677282 | 1 | 1 |
| Mx2 | 0,0360411 | 0,0576342 | 0,677282 | 1 | 1 |
| Fndc9 | 0,314277 | 0,50102 | 0,672829 | 1 | 1 |
| 9430037G07Rik | 0,167219 | 0,266355 | 0,671613 | 1 | 1 |

| | | | | | |
|---|---|---|---|---|---|
| Htr5b | 0,0726622 | 0,115646 | 0,670432 | 1 | 1 |
| Mctp2 | 0,0468371 | 0,0745436 | 0,670432 | 1 | 1 |
| Npas4 | 0,0425618 | 0,0677392 | 0,670432 | 1 | 1 |
| Isoc2b | 0,362979 | 0,576997 | 0,668676 | 1 | 1 |
| Poln | 0,270763 | 0,429366 | 0,665178 | 1 | 1 |
| BC037032 | 0,262232 | 0,415079 | 0,662542 | 1 | 1 |
| Xlr3a | 0,182274 | 0,288512 | 0,662522 | 1 | 1 |
| Trim34a | 0,177912 | 0,281496 | 0,661946 | 1 | 1 |
| Kcna1 | 0,0342498 | 0,0541441 | 0,660707 | 1 | 1 |
| Fmnl1 | 0,014264 | 0,0225389 | 0,660038 | 1 | 1 |
| 2010310C07Rik | 0,0794739 | 0,125543 | 0,659629 | 1 | 1 |
| Adap2 | 0,036571 | 0,0577703 | 0,659629 | 1 | 1 |
| A930011G23Rik | 0,0161726 | 0,0255474 | 0,659629 | 1 | 1 |
| Cdk5r2 | 0,192212 | 0,303444 | 0,658732 | 1 | 1 |
| Ntf5 | 0,111943 | 0,176697 | 0,658517 | 1 | 1 |
| Vipr1 | 0,149195 | 0,235324 | 0,657451 | 1 | 1 |
| Trpa1 | 0,242911 | 0,383133 | 0,657414 | 1 | 1 |
| Akr1b7 | 0,091015 | 0,143545 | 0,657324 | 1 | 1 |
| Ecm2 | 0,127824 | 0,201294 | 0,655148 | 1 | 1 |
| Tldc2 | 0,101324 | 0,159507 | 0,654655 | 1 | 1 |
| Thrsp | 0,0528494 | 0,0831975 | 0,654655 | 1 | 1 |
| C2 | 0,0361285 | 0,0568749 | 0,654655 | 1 | 1 |
| Olfr856-ps1 | 0,0306514 | 0,0482526 | 0,654655 | 1 | 1 |
| 2410012M07Rik | 0,0273768 | 0,0430977 | 0,654655 | 1 | 1 |
| 1110028F11Rik | 0,0176237 | 0,0277439 | 0,654655 | 1 | 1 |
| Nwd2 | 0,00529008 | 0,00832786 | 0,654655 | 1 | 1 |
| Cd74 | 0,0432531 | 0,0680902 | 0,654641 | 1 | 1 |
| Gpr75 | 0,144892 | 0,228049 | 0,654368 | 1 | 1 |
| Lpin3 | 0,29506 | 0,463436 | 0,651361 | 1 | 1 |
| AI661453 | 0,119211 | 0,187205 | 0,651095 | 1 | 1 |
| Rab27b | 0,117419 | 0,184135 | 0,649106 | 1 | 1 |
| Cxcl10 | 0,121376 | 0,19001 | 0,646594 | 1 | 1 |
| Zfp85os | 0,380405 | 0,594964 | 0,645268 | 1 | 1 |
| Tnk2os | 0,394851 | 0,617426 | 0,644958 | 1 | 1 |
| Fbxo15 | 0,131501 | 0,205488 | 0,643975 | 1 | 1 |
| Carns1 | 0,318294 | 0,496971 | 0,642803 | 1 | 1 |
| Gm10536 | 0,0682507 | 0,106551 | 0,642625 | 1 | 1 |
| Pou6f1 | 0,367332 | 0,572956 | 0,64134 | 1 | 1 |
| Baat | 0,354047 | 0,551283 | 0,638853 | 1 | 1 |
| 1700028J19Rik | 0,289989 | 0,45081 | 0,636522 | 1 | 1 |
| A230072E10Rik | 0,0509101 | 0,0790577 | 0,634953 | 1 | 1 |
| Gm11201 | 0,0457371 | 0,0709767 | 0,633982 | 1 | 1 |
| Tctex1d1 | 0,243773 | 0,378176 | 0,633518 | 1 | 1 |
| Gm5878 | 0,0263706 | 0,0408511 | 0,631445 | 1 | 1 |
| Pcdha5 | 0,0158233 | 0,0244802 | 0,629562 | 1 | 1 |
| Dcaf12l2 | 0,32554 | 0,502498 | 0,626285 | 1 | 1 |
| Dnajb7 | 0,162623 | 0,250582 | 0,623751 | 1 | 1 |
| Prdm14 | 0,066847 | 0,103003 | 0,623751 | 1 | 1 |
| Pcdhb15 | 0,0308177 | 0,0474863 | 0,623751 | 1 | 1 |
| Has1 | 0,150986 | 0,232255 | 0,6213 | 1 | 1 |

| | | | | | |
|---|---|---|---|---|---|
| 4931431C16Rik | 0,0863572 | 0,132758 | 0,620414 | 1 | 1 |
| 1700020A23Rik | 0,381602 | 0,586629 | 0,620379 | 1 | 1 |
| C130030K03Rik | 0,0372547 | 0,0571465 | 0,617244 | 1 | 1 |
| A230056J06Rik | 0,0247397 | 0,0379492 | 0,617244 | 1 | 1 |
| Cbln3 | 0,205074 | 0,314453 | 0,6167 | 1 | 1 |
| Tbc1d10c | 0,249765 | 0,382504 | 0,614902 | 1 | 1 |
| F730043M19Rik | 0,31892 | 0,488303 | 0,614584 | 1 | 1 |
| Ltb | 0,188553 | 0,288618 | 0,61419 | 1 | 1 |
| Stpg1 | 0,131663 | 0,201516 | 0,61404 | 1 | 1 |
| Mamstr | 0,239101 | 0,365814 | 0,61349 | 1 | 1 |
| Ptk2b | 0,101172 | 0,154683 | 0,6125 | 1 | 1 |
| Ehf | 0,0661623 | 0,101067 | 0,611225 | 1 | 1 |
| Chdh | 0,0556365 | 0,0848534 | 0,608941 | 1 | 1 |
| Creb3l1 | 0,225833 | 0,344169 | 0,60786 | 1 | 1 |
| Cnbd2 | 0,379878 | 0,57761 | 0,604559 | 1 | 1 |
| Lrrc29 | 0,407745 | 0,619411 | 0,603232 | 1 | 1 |
| Ihh | 0,103875 | 0,157756 | 0,602849 | 1 | 1 |
| Plch2 | 0,0559549 | 0,0848978 | 0,601464 | 1 | 1 |
| Gm20199 | 0,336447 | 0,509768 | 0,599461 | 1 | 1 |
| Syt5 | 0,287682 | 0,43563 | 0,598625 | 1 | 1 |
| 4833417C18Rik | 0,345747 | 0,523503 | 0,598481 | 1 | 1 |
| Ralyl | 0,0712869 | 0,107828 | 0,597026 | 1 | 1 |
| 4833411C07Rik | 0,137934 | 0,208578 | 0,59661 | 1 | 1 |
| Spock1 | 0,128964 | 0,194988 | 0,596422 | 1 | 1 |
| Trim15 | 0,235889 | 0,356294 | 0,59496 | 1 | 1 |
| 1700019L03Rik | 0,238119 | 0,359522 | 0,594396 | 1 | 1 |
| Hgd | 0,0827202 | 0,124859 | 0,59399 | 1 | 1 |
| Faah | 0,161644 | 0,243595 | 0,591667 | 1 | 1 |
| Epgn | 0,097309 | 0,146166 | 0,58696 | 1 | 1 |
| Btbd18 | 0,191002 | 0,286802 | 0,586471 | 1 | 1 |
| Coro2a | 0,314407 | 0,471966 | 0,586048 | 1 | 1 |
| Adamts8 | 0,093592 | 0,140333 | 0,584397 | 1 | 1 |
| Gpr156 | 0,399337 | 0,598536 | 0,583834 | 1 | 1 |
| Tagap | 0,121274 | 0,181699 | 0,583274 | 1 | 1 |
| Fbxo41 | 0,11265 | 0,16856 | 0,581416 | 1 | 1 |
| 9530051G07Rik | 0,117744 | 0,176031 | 0,580172 | 1 | 1 |
| Mfrp | 0,113411 | 0,169523 | 0,579921 | 1 | 1 |
| Fam228a | 0,191711 | 0,286521 | 0,579709 | 1 | 1 |
| Scd3 | 0,109187 | 0,163174 | 0,579608 | 1 | 1 |
| Arl5c | 0,33787 | 0,504473 | 0,578306 | 1 | 1 |
| Slc41a2 | 0,0401728 | 0,0599535 | 0,577625 | 1 | 1 |
| L3mbtl1 | 0,0778598 | 0,11597 | 0,574798 | 1 | 1 |
| Padi3 | 0,109123 | 0,162473 | 0,574247 | 1 | 1 |
| Slc10a4 | 0,432566 | 0,643839 | 0,573777 | 1 | 1 |
| Sh3gl2 | 0,41017 | 0,609765 | 0,572032 | 1 | 1 |
| Gmfg | 0,392333 | 0,583117 | 0,571708 | 1 | 1 |
| Prkcg | 0,151122 | 0,224601 | 0,571653 | 1 | 1 |
| Zmynd10 | 0,1831 | 0,271772 | 0,569763 | 1 | 1 |
| Car5b | 0,0878464 | 0,130281 | 0,568569 | 1 | 1 |
| 1700003M02Rik | 0,386147 | 0,571509 | 0,565628 | 1 | 1 |

| | | | | | |
|---|---|---|---|---|---|
| Fam166b | 0,367953 | 0,544164 | 0,564518 | 1 | 1 |
| Paqr7 | 0,10344 | 0,152517 | 0,560175 | 1 | 1 |
| Cd101 | 0,0190922 | 0,0281321 | 0,559232 | 1 | 1 |
| Guca1b | 0,453348 | 0,667935 | 0,559087 | 1 | 1 |
| Mir22hg | 0,28034 | 0,412851 | 0,558447 | 1 | 1 |
| Cartpt | 0,0457279 | 0,0673374 | 0,558332 | 1 | 1 |
| Boll | 0,0349039 | 0,0513983 | 0,558332 | 1 | 1 |
| Amy1 | 0,0198089 | 0,0291698 | 0,55833 | 1 | 1 |
| Cabp5 | 0,112544 | 0,16572 | 0,558262 | 1 | 1 |
| Ccl7 | 0,0971586 | 0,143065 | 0,558262 | 1 | 1 |
| 1700125H20Rik | 0,03749 | 0,0552039 | 0,558262 | 1 | 1 |
| Cyp4f39 | 0,0359219 | 0,0528948 | 0,558262 | 1 | 1 |
| 5430427M07Rik | 0,035336 | 0,0520321 | 0,558262 | 1 | 1 |
| Olfr544 | 0,0295854 | 0,0435643 | 0,558262 | 1 | 1 |
| Clec2d | 0,0286187 | 0,0421409 | 0,558262 | 1 | 1 |
| Klk14 | 0,0271153 | 0,0399272 | 0,558262 | 1 | 1 |
| Gm15408 | 0,0246 | 0,0362233 | 0,558262 | 1 | 1 |
| Shbg | 0,0240274 | 0,0353802 | 0,558262 | 1 | 1 |
| 1700123L14Rik | 0,0192829 | 0,0283939 | 0,558262 | 1 | 1 |
| Slfnl1 | 0,0185527 | 0,0273187 | 0,558262 | 1 | 1 |
| Serpinb9b | 0,0167566 | 0,024674 | 0,558262 | 1 | 1 |
| Hapln2 | 0,0156159 | 0,0229944 | 0,558262 | 1 | 1 |
| Gabrr1 | 0,0141207 | 0,0207927 | 0,558262 | 1 | 1 |
| A330050F15Rik | 0,013817 | 0,0203454 | 0,558262 | 1 | 1 |
| Aicda | 0,0131806 | 0,0194084 | 0,558262 | 1 | 1 |
| En1 | 0,0123001 | 0,0181118 | 0,558262 | 1 | 1 |
| Nxph2 | 0,0121706 | 0,0179211 | 0,558262 | 1 | 1 |
| Epx | 0,0116225 | 0,0171141 | 0,558262 | 1 | 1 |
| Cnga2 | 0,0101994 | 0,0150186 | 0,558262 | 1 | 1 |
| Tmprss6 | 0,00967375 | 0,0142445 | 0,558262 | 1 | 1 |
| Grm6 | 0,00700223 | 0,0103107 | 0,558262 | 1 | 1 |
| Pramef12 | 0,0579753 | 0,0852945 | 0,557015 | 1 | 1 |
| Rai2 | 0,167786 | 0,24681 | 0,55678 | 1 | 1 |
| C1qtnf2 | 0,101178 | 0,148819 | 0,556661 | 1 | 1 |
| Creb3l3 | 0,0490077 | 0,0720835 | 0,556661 | 1 | 1 |
| Smok4a | 0,0345231 | 0,0507564 | 0,556026 | 1 | 1 |
| Dmrtc1a | 0,0336872 | 0,0495227 | 0,555887 | 1 | 1 |
| Fam83c | 0,310486 | 0,456263 | 0,555338 | 1 | 1 |
| Snca | 0,332347 | 0,487882 | 0,553842 | 1 | 1 |
| F13a1 | 0,131626 | 0,193114 | 0,55301 | 1 | 1 |
| Ptpn20 | 0,0629716 | 0,0923296 | 0,552092 | 1 | 1 |
| Cutal | 0,0934095 | 0,136924 | 0,551737 | 1 | 1 |
| Tmod1 | 0,052947 | 0,0775096 | 0,549827 | 1 | 1 |
| Trpm2 | 0,0174839 | 0,0255838 | 0,549204 | 1 | 1 |
| Unc80 | 0,0197332 | 0,0288732 | 0,549105 | 1 | 1 |
| 4932435O22Rik | 0,0822462 | 0,120341 | 0,549104 | 1 | 1 |
| Adarb2 | 0,0397459 | 0,058135 | 0,548602 | 1 | 1 |
| Tcap | 0,127714 | 0,186523 | 0,546438 | 1 | 1 |
| Ascl2 | 0,0674858 | 0,0985613 | 0,546438 | 1 | 1 |
| Fscn3 | 0,0599118 | 0,0874997 | 0,546438 | 1 | 1 |

| | | | | | |
|---|---|---|---|---|---|
| Upp2 | 0,0475339 | 0,0694221 | 0,546438 | 1 | 1 |
| Rbfox1 | 0,136942 | 0,199523 | 0,542985 | 1 | 1 |
| Slco4c1 | 0,242682 | 0,353584 | 0,542984 | 1 | 1 |
| Dscaml1 | 0,00989604 | 0,0144009 | 0,541238 | 1 | 1 |
| Gm8787 | 0,0986999 | 0,14355 | 0,540427 | 1 | 1 |
| Nod1 | 0,24002 | 0,349064 | 0,540334 | 1 | 1 |
| Stk33 | 0,172896 | 0,251346 | 0,539768 | 1 | 1 |
| Itgb7 | 0,193964 | 0,281643 | 0,538077 | 1 | 1 |
| Rnf125 | 0,476405 | 0,690956 | 0,536405 | 1 | 1 |
| Zfp459 | 0,355872 | 0,516134 | 0,53639 | 1 | 1 |
| Styx | 0,310769 | 0,450481 | 0,535624 | 1 | 1 |
| Gsdmd | 0,342455 | 0,495545 | 0,533103 | 1 | 1 |
| Gsto2 | 0,283472 | 0,410093 | 0,532747 | 1 | 1 |
| Wisp1 | 0,14233 | 0,205856 | 0,532401 | 1 | 1 |
| Klhl41 | 0,165171 | 0,238596 | 0,530609 | 1 | 1 |
| Gm8773 | 0,424457 | 0,612131 | 0,528222 | 1 | 1 |
| Cilp2 | 0,125483 | 0,18052 | 0,524668 | 1 | 1 |
| Slc24a2 | 0,0560228 | 0,0805818 | 0,52444 | 1 | 1 |
| Gm4944 | 0,403343 | 0,579719 | 0,523349 | 1 | 1 |
| Fam131c | 0,255439 | 0,366775 | 0,521918 | 1 | 1 |
| Foxa1 | 0,418224 | 0,59951 | 0,519508 | 1 | 1 |
| Fam46b | 0,171806 | 0,245939 | 0,517523 | 1 | 1 |
| Dnajc6 | 0,134493 | 0,192215 | 0,51519 | 1 | 1 |
| Hid1 | 0,322819 | 0,461206 | 0,514685 | 1 | 1 |
| Cd55 | 0,15 | 0,213953 | 0,512332 | 1 | 1 |
| Galr1 | 0,0547297 | 0,0780511 | 0,512097 | 1 | 1 |
| Sdcbp2 | 0,439367 | 0,626063 | 0,510882 | 1 | 1 |
| Fbxo47 | 0,285227 | 0,405853 | 0,508846 | 1 | 1 |
| Anks1b | 0,0487988 | 0,0693462 | 0,506971 | 1 | 1 |
| Tmem232 | 0,0456433 | 0,0648511 | 0,506729 | 1 | 1 |
| Pld4 | 0,0613042 | 0,0870983 | 0,50666 | 1 | 1 |
| P2ry13 | 0,0373463 | 0,0530016 | 0,505071 | 1 | 1 |
| 4930524B15Rik | 0,0494104 | 0,0701064 | 0,504731 | 1 | 1 |
| Hck | 0,274988 | 0,390061 | 0,504331 | 1 | 1 |
| Fam83e | 0,107748 | 0,152698 | 0,503014 | 1 | 1 |
| Dbil5 | 0,489839 | 0,693135 | 0,500828 | 1 | 1 |
| Adh6a | 0,102953 | 0,145645 | 0,500467 | 1 | 1 |
| Pabpc1l | 0,0985343 | 0,139394 | 0,500467 | 1 | 1 |
| 4933408J17Rik | 0,103624 | 0,146375 | 0,498313 | 1 | 1 |
| Cd160 | 0,167977 | 0,237217 | 0,497945 | 1 | 1 |
| Tomm6os | 0,395233 | 0,5574 | 0,496008 | 1 | 1 |
| Tas1r3 | 0,184802 | 0,260357 | 0,494513 | 1 | 1 |
| Trib3 | 0,386923 | 0,54492 | 0,493999 | 1 | 1 |
| Cyth4 | 0,0558662 | 0,0786669 | 0,493782 | 1 | 1 |
| Smc1b | 0,0376762 | 0,0530531 | 0,493782 | 1 | 1 |
| Cyb5r2 | 0,161713 | 0,227579 | 0,49293 | 1 | 1 |
| Tmem252 | 0,0735526 | 0,10331 | 0,490125 | 1 | 1 |
| Gltpd2 | 0,159856 | 0,224123 | 0,487515 | 1 | 1 |
| Cyp2d26 | 0,0554118 | 0,0776889 | 0,487515 | 1 | 1 |
| Becn2 | 0,0503776 | 0,0706308 | 0,487515 | 1 | 1 |

| | | | | | |
|---|---|---|---|---|---|
| Adam1b | 0,0297943 | 0,0417724 | 0,487515 | 1 | 1 |
| Sez6l | 0,113347 | 0,158791 | 0,48638 | 1 | 1 |
| Rho | 0,134416 | 0,188271 | 0,486108 | 1 | 1 |
| Sertm1 | 0,197391 | 0,276475 | 0,486088 | 1 | 1 |
| 9230116L04Rik | 0,0589028 | 0,0824301 | 0,484835 | 1 | 1 |
| Grin2d | 0,251757 | 0,352189 | 0,484315 | 1 | 1 |
| 5430402O13Rik | 0,119768 | 0,167159 | 0,48098 | 1 | 1 |
| P2ry14 | 0,141324 | 0,197124 | 0,480095 | 1 | 1 |
| AY074887 | 0,549235 | 0,766002 | 0,479926 | 1 | 1 |
| Sult1a1 | 0,483064 | 0,673611 | 0,479704 | 1 | 1 |
| Pla2g16 | 0,408652 | 0,569746 | 0,479446 | 1 | 1 |
| 4930447N08Rik | 0,0871766 | 0,121498 | 0,478919 | 1 | 1 |
| Aspn | 0,0542638 | 0,0755448 | 0,477342 | 1 | 1 |
| Crygn | 0,144522 | 0,20118 | 0,477202 | 1 | 1 |
| Nanp | 0,049368 | 0,0687223 | 0,477202 | 1 | 1 |
| Gm16853 | 0,255062 | 0,354864 | 0,47642 | 1 | 1 |
| Mmp9 | 0,105713 | 0,147042 | 0,476078 | 1 | 1 |
| 1700001L05Rik | 0,344788 | 0,47763 | 0,470183 | 1 | 1 |
| Pianp | 0,240372 | 0,332775 | 0,46928 | 1 | 1 |
| Slc3a1 | 0,278313 | 0,385096 | 0,46851 | 1 | 1 |
| Hpn | 0,255882 | 0,353474 | 0,466126 | 1 | 1 |
| Grk1 | 0,0444061 | 0,0613358 | 0,465971 | 1 | 1 |
| Slc25a2 | 0,153124 | 0,211402 | 0,465287 | 1 | 1 |
| AA986860 | 0,0946973 | 0,130637 | 0,46417 | 1 | 1 |
| 4933427E11Rik | 0,107676 | 0,148377 | 0,462566 | 1 | 1 |
| Gm20554 | 0,331109 | 0,456236 | 0,462474 | 1 | 1 |
| Hist1h2bh | 0,597591 | 0,823279 | 0,462222 | 1 | 1 |
| Fgf11 | 0,23033 | 0,316826 | 0,45999 | 1 | 1 |
| Ccdc65 | 0,209048 | 0,287543 | 0,459945 | 1 | 1 |
| Efcab4b | 0,310959 | 0,427185 | 0,458135 | 1 | 1 |
| Gm14124 | 0,0567511 | 0,0779138 | 0,457231 | 1 | 1 |
| E130008D07Rik | 0,384636 | 0,527529 | 0,455758 | 1 | 1 |
| D130043K22Rik | 0,0629025 | 0,0862507 | 0,45542 | 1 | 1 |
| Apba1 | 0,416438 | 0,570944 | 0,455247 | 1 | 1 |
| 4632428C04Rik | 0,0947126 | 0,129783 | 0,454468 | 1 | 1 |
| Jakmip3 | 0,0341107 | 0,0466901 | 0,452891 | 1 | 1 |
| Cemip | 0,12413 | 0,169776 | 0,451784 | 1 | 1 |
| Irf6 | 0,403971 | 0,552012 | 0,450446 | 1 | 1 |
| 4732416N19Rik | 0,632507 | 0,864072 | 0,45007 | 1 | 1 |
| Tfeb | 0,208314 | 0,284359 | 0,448948 | 1 | 1 |
| Dnah7b | 0,241169 | 0,329146 | 0,448686 | 1 | 1 |
| Tgoln2 | 11,2174 | 15,2819 | 0,446085 | 1 | 1 |
| Gfi1 | 0,0208246 | 0,0283651 | 0,445824 | 1 | 1 |
| Lst1 | 0,240263 | 0,327249 | 0,445775 | 1 | 1 |
| 2900060B14Rik | 0,129624 | 0,176553 | 0,445775 | 1 | 1 |
| Gm19395 | 0,112208 | 0,152832 | 0,445775 | 1 | 1 |
| Mc5r | 0,0531765 | 0,0724289 | 0,445775 | 1 | 1 |
| Ribc2 | 0,0351819 | 0,0479193 | 0,445775 | 1 | 1 |
| Pcdhb7 | 0,0308562 | 0,0420276 | 0,445775 | 1 | 1 |
| Tspan10 | 0,027389 | 0,037305 | 0,445775 | 1 | 1 |

| | | | | | |
|---|---|---|---|---|---|
| Fam83a | 0,0270832 | 0,0368885 | 0,445775 | 1 | 1 |
| Dnah10 | 0,0184125 | 0,0250787 | 0,445775 | 1 | 1 |
| Slfn14 | 0,0145553 | 0,019825 | 0,445775 | 1 | 1 |
| 1700110C19Rik | 0,311066 | 0,423253 | 0,444296 | 1 | 1 |
| Ank1 | 0,108056 | 0,146629 | 0,440395 | 1 | 1 |
| Ccdc88b | 0,0610916 | 0,0828921 | 0,440261 | 1 | 1 |
| Osmr | 0,141525 | 0,19177 | 0,438321 | 1 | 1 |
| Hivep3 | 0,047002 | 0,063688 | 0,4383 | 1 | 1 |
| Nupr1l | 0,464393 | 0,628434 | 0,436417 | 1 | 1 |
| Crhr2 | 0,0360831 | 0,0488174 | 0,436072 | 1 | 1 |
| Tal1 | 0,157589 | 0,21319 | 0,435973 | 1 | 1 |
| 4933424G05Rik | 0,0848747 | 0,114657 | 0,433915 | 1 | 1 |
| Hydin | 0,0177648 | 0,0239915 | 0,433504 | 1 | 1 |
| Nrg2 | 0,108553 | 0,146553 | 0,433021 | 1 | 1 |
| Ncan | 0,358393 | 0,483688 | 0,432538 | 1 | 1 |
| Gm3696 | 0,448309 | 0,604797 | 0,431959 | 1 | 1 |
| Aqp4 | 0,0479947 | 0,0646704 | 0,43023 | 1 | 1 |
| Prss8 | 0,242825 | 0,326778 | 0,428393 | 1 | 1 |
| C4a | 0,215718 | 0,289907 | 0,426447 | 1 | 1 |
| Vmo1 | 0,20303 | 0,272832 | 0,426314 | 1 | 1 |
| Gdf7 | 0,0795145 | 0,106851 | 0,426314 | 1 | 1 |
| Pla2g4f | 0,0325301 | 0,0437138 | 0,426314 | 1 | 1 |
| Syt17 | 0,180267 | 0,24224 | 0,426303 | 1 | 1 |
| Ppp1r2-ps3 | 0,133079 | 0,178829 | 0,426297 | 1 | 1 |
| Kank4os | 0,193553 | 0,26006 | 0,426115 | 1 | 1 |
| Hrasls | 0,132071 | 0,177448 | 0,426087 | 1 | 1 |
| A230073K19Rik | 0,290311 | 0,389931 | 0,425617 | 1 | 1 |
| Exd1 | 0,433831 | 0,582686 | 0,425586 | 1 | 1 |
| Frmd3 | 0,0676389 | 0,090725 | 0,423647 | 1 | 1 |
| Hsf2bp | 0,386262 | 0,517629 | 0,422342 | 1 | 1 |
| Islr2 | 0,0711315 | 0,0953183 | 0,422264 | 1 | 1 |
| Mt3 | 0,655505 | 0,878077 | 0,421742 | 1 | 1 |
| 4732471J01Rik | 0,376337 | 0,503699 | 0,420538 | 1 | 1 |
| Hyal3 | 0,206644 | 0,276497 | 0,420113 | 1 | 1 |
| Nlrx1 | 0,434297 | 0,580513 | 0,418646 | 1 | 1 |
| Mei1 | 0,032724 | 0,0437285 | 0,418226 | 1 | 1 |
| Pzp | 0,0269035 | 0,0359502 | 0,418205 | 1 | 1 |
| Abcg1 | 0,228339 | 0,305044 | 0,417838 | 1 | 1 |
| Slc7a10 | 0,155029 | 0,20674 | 0,415278 | 1 | 1 |
| Susd2 | 0,370791 | 0,494335 | 0,414882 | 1 | 1 |
| Aqp7 | 0,0724148 | 0,0965087 | 0,414375 | 1 | 1 |
| Trim29 | 0,11585 | 0,154318 | 0,413639 | 1 | 1 |
| 9030624G23Rik | 0,397541 | 0,528873 | 0,411818 | 1 | 1 |
| Mrc1 | 0,0737595 | 0,0980546 | 0,410757 | 1 | 1 |
| Lemd1 | 0,289872 | 0,38475 | 0,408506 | 1 | 1 |
| Stox1 | 0,235138 | 0,311878 | 0,407476 | 1 | 1 |
| Smim6 | 0,339276 | 0,449978 | 0,407397 | 1 | 1 |
| Gm16938 | 0,0896493 | 0,118834 | 0,406585 | 1 | 1 |
| Gm2694 | 0,622071 | 0,824566 | 0,406555 | 1 | 1 |
| Ephx1 | 0,193144 | 0,255825 | 0,40548 | 1 | 1 |

| | | | | | |
|---|---|---|---|---|---|
| Fry | 0,28702 | 0,380026 | 0,404943 | 1 | 1 |
| Acbd4 | 0,459728 | 0,608337 | 0,404091 | 1 | 1 |
| Otoa | 0,0339794 | 0,0448514 | 0,400493 | 1 | 1 |
| Cpa4 | 0,141638 | 0,186826 | 0,399487 | 1 | 1 |
| Clvs2 | 0,241703 | 0,318722 | 0,399063 | 1 | 1 |
| 1700101E01Rik | 0,100387 | 0,13234 | 0,39868 | 1 | 1 |
| Serpinb6b | 0,0440977 | 0,0581341 | 0,39868 | 1 | 1 |
| Pcdhb6 | 0,0345068 | 0,0454904 | 0,39868 | 1 | 1 |
| Hsf5 | 0,0184578 | 0,0243329 | 0,39868 | 1 | 1 |
| Sult2b1 | 0,540775 | 0,712637 | 0,398138 | 1 | 1 |
| Ppp1r3fos | 0,27191 | 0,357959 | 0,396664 | 1 | 1 |
| Zfp114 | 0,250706 | 0,32915 | 0,39275 | 1 | 1 |
| 4933417D19Rik | 0,298204 | 0,390813 | 0,390181 | 1 | 1 |
| 4933411K16Rik | 0,201322 | 0,263542 | 0,388524 | 1 | 1 |
| Ttc21a | 0,17965 | 0,235118 | 0,388192 | 1 | 1 |
| Kif9 | 0,247873 | 0,324393 | 0,388139 | 1 | 1 |
| Iqsec3 | 0,0365254 | 0,0477805 | 0,387522 | 1 | 1 |
| Atp4a | 0,0847162 | 0,110765 | 0,386797 | 1 | 1 |
| Kcnt2 | 0,261567 | 0,341952 | 0,38661 | 1 | 1 |
| Gm9199 | 0,335304 | 0,437843 | 0,384946 | 1 | 1 |
| Ankrd53 | 0,419506 | 0,547347 | 0,383765 | 1 | 1 |
| Eqtn | 0,196455 | 0,25632 | 0,383752 | 1 | 1 |
| Ido2 | 0,106333 | 0,138612 | 0,382469 | 1 | 1 |
| Fcer2a | 0,0713457 | 0,0928765 | 0,380486 | 1 | 1 |
| Mrpl23 | 0,673837 | 0,876163 | 0,3788 | 1 | 1 |
| 1700026L06Rik | 0,545422 | 0,70919 | 0,3788 | 1 | 1 |
| Tmprss9 | 0,0783748 | 0,101908 | 0,3788 | 1 | 1 |
| Atp2b2 | 0,0729468 | 0,0948449 | 0,378725 | 1 | 1 |
| Wdr52 | 0,0367014 | 0,0477152 | 0,378615 | 1 | 1 |
| Ano3 | 0,0248241 | 0,0322525 | 0,377666 | 1 | 1 |
| Aox1 | 0,200712 | 0,260418 | 0,375703 | 1 | 1 |
| Smc2os | 0,225161 | 0,291549 | 0,372783 | 1 | 1 |
| 4833412C05Rik | 0,1081 | 0,139972 | 0,372783 | 1 | 1 |
| 4930453L07Rik | 0,0769392 | 0,0996246 | 0,372783 | 1 | 1 |
| Slc25a43 | 0,351156 | 0,45464 | 0,372613 | 1 | 1 |
| Il33 | 0,159532 | 0,206435 | 0,371844 | 1 | 1 |
| Pla2r1 | 0,156069 | 0,201822 | 0,370904 | 1 | 1 |
| Cyp27b1 | 0,163947 | 0,211986 | 0,370739 | 1 | 1 |
| Gm16845 | 0,43238 | 0,558533 | 0,369345 | 1 | 1 |
| Ccdc146 | 0,0272262 | 0,0351492 | 0,368495 | 1 | 1 |
| Mos | 0,193716 | 0,249846 | 0,367096 | 1 | 1 |
| Prg4 | 0,232212 | 0,299489 | 0,367057 | 1 | 1 |
| C1rl | 0,192366 | 0,247919 | 0,366012 | 1 | 1 |
| Grm1 | 0,00986297 | 0,0126999 | 0,364724 | 1 | 1 |
| Calcr | 0,0202365 | 0,0260554 | 0,364622 | 1 | 1 |
| 4933405D12Rik | 0,117223 | 0,150929 | 0,364606 | 1 | 1 |
| Vsig2 | 0,0684417 | 0,0881208 | 0,364606 | 1 | 1 |
| Spag6 | 0,0292286 | 0,0376328 | 0,364606 | 1 | 1 |
| Il10ra | 0,0201019 | 0,0258819 | 0,364606 | 1 | 1 |
| Gm6623 | 0,471926 | 0,607313 | 0,363878 | 1 | 1 |

| | | | | | |
|---|---|---|---|---|---|
| Paqr9 | 0,277209 | 0,356644 | 0,363507 | 1 | 1 |
| Il6ra | 0,243897 | 0,31357 | 0,362517 | 1 | 1 |
| Psma8 | 0,246035 | 0,31563 | 0,359367 | 1 | 1 |
| Zfa-ps | 0,312543 | 0,400706 | 0,35849 | 1 | 1 |
| Ppp1r1b | 0,119609 | 0,153146 | 0,356581 | 1 | 1 |
| B4galnt2 | 0,150401 | 0,192562 | 0,356508 | 1 | 1 |
| Gabre | 0,371496 | 0,474907 | 0,354298 | 1 | 1 |
| Ptgdr2 | 0,0804238 | 0,10276 | 0,353583 | 1 | 1 |
| Hhipl2 | 0,193063 | 0,246638 | 0,353323 | 1 | 1 |
| Kcnb2 | 0,266466 | 0,340364 | 0,353127 | 1 | 1 |
| Pld5 | 0,311744 | 0,39814 | 0,352914 | 1 | 1 |
| Pycard | 0,355123 | 0,453242 | 0,351965 | 1 | 1 |
| Fam221b | 0,0697585 | 0,089015 | 0,35168 | 1 | 1 |
| Slc7a4 | 0,120552 | 0,153724 | 0,350693 | 1 | 1 |
| 4931403G20Rik | 0,279858 | 0,355757 | 0,346195 | 1 | 1 |
| Fam71f2 | 0,246445 | 0,31314 | 0,345544 | 1 | 1 |
| Gja3 | 0,132974 | 0,168942 | 0,345384 | 1 | 1 |
| Odf3b | 0,105446 | 0,13391 | 0,344764 | 1 | 1 |
| Pdyn | 0,0892975 | 0,113333 | 0,34388 | 1 | 1 |
| Zbtb4 | 0,380028 | 0,482242 | 0,343651 | 1 | 1 |
| 1810020O05Rik | 0,25972 | 0,329237 | 0,342172 | 1 | 1 |
| Mb21d1 | 0,331891 | 0,42042 | 0,341122 | 1 | 1 |
| Panx2 | 0,0407707 | 0,0516209 | 0,340423 | 1 | 1 |
| Rs1 | 0,0232066 | 0,0293826 | 0,340423 | 1 | 1 |
| Cdh10 | 0,304978 | 0,385824 | 0,339239 | 1 | 1 |
| Slc23a1 | 0,278347 | 0,35193 | 0,338402 | 1 | 1 |
| Liph | 0,108672 | 0,137165 | 0,335931 | 1 | 1 |
| Gpr35 | 0,0545924 | 0,0688973 | 0,33575 | 1 | 1 |
| Zbtb7c | 0,127049 | 0,160328 | 0,335635 | 1 | 1 |
| Asb16 | 0,269884 | 0,34037 | 0,334762 | 1 | 1 |
| F2rl1 | 0,123028 | 0,155147 | 0,334657 | 1 | 1 |
| Ptchd4 | 0,0284238 | 0,0358419 | 0,334549 | 1 | 1 |
| Rxrg | 0,103824 | 0,13091 | 0,334439 | 1 | 1 |
| Ccdc19 | 0,518276 | 0,652995 | 0,333351 | 1 | 1 |
| Prkcq | 0,297617 | 0,374923 | 0,333138 | 1 | 1 |
| Rasd2 | 0,164845 | 0,207654 | 0,333069 | 1 | 1 |
| 4930443O20Rik | 0,0889358 | 0,112005 | 0,332727 | 1 | 1 |
| Prss50 | 0,0880827 | 0,110931 | 0,332727 | 1 | 1 |
| Atoh1 | 0,0584262 | 0,0735815 | 0,332727 | 1 | 1 |
| Trim60 | 0,13371 | 0,168351 | 0,332358 | 1 | 1 |
| Sag | 0,100945 | 0,127096 | 0,332358 | 1 | 1 |
| Sgcg | 0,0461585 | 0,0581168 | 0,332358 | 1 | 1 |
| Tap1 | 0,202765 | 0,254901 | 0,330129 | 1 | 1 |
| Rapgef3 | 0,356483 | 0,447206 | 0,327105 | 1 | 1 |
| Gpr115 | 0,0630147 | 0,0790186 | 0,326504 | 1 | 1 |
| Hist1h3g | 0,707521 | 0,886687 | 0,325652 | 1 | 1 |
| Cacnb4 | 0,224671 | 0,281108 | 0,323314 | 1 | 1 |
| Mmp19 | 0,189918 | 0,23761 | 0,323221 | 1 | 1 |
| Tom1 | 0,0661105 | 0,0827059 | 0,323111 | 1 | 1 |
| Il18rap | 0,0546759 | 0,0683273 | 0,321556 | 1 | 1 |

| | | | | | |
|---|---|---|---|---|---|
| Scn1a | 0,133737 | 0,166998 | 0,320424 | 1 | 1 |
| Ttc18 | 0,106107 | 0,132394 | 0,319318 | 1 | 1 |
| Tmem196 | 0,160029 | 0,199609 | 0,318845 | 1 | 1 |
| Trim17 | 0,376956 | 0,470103 | 0,318581 | 1 | 1 |
| Tsx | 0,561155 | 0,699517 | 0,317958 | 1 | 1 |
| Myl1 | 0,440267 | 0,547868 | 0,315451 | 1 | 1 |
| Apon | 0,339836 | 0,422673 | 0,314703 | 1 | 1 |
| Alox8 | 0,0475608 | 0,0590869 | 0,313066 | 1 | 1 |
| Nup62-il4i1 | 0,497825 | 0,618313 | 0,312699 | 1 | 1 |
| Ccdc116 | 0,271547 | 0,336693 | 0,310234 | 1 | 1 |
| 8430408G22Rik | 0,287505 | 0,356421 | 0,309995 | 1 | 1 |
| Lysmd1 | 0,274473 | 0,340247 | 0,309919 | 1 | 1 |
| Tjp3 | 0,244465 | 0,303028 | 0,309821 | 1 | 1 |
| Rhox5 | 0,600734 | 0,743527 | 0,30766 | 1 | 1 |
| Htr2b | 0,370005 | 0,457613 | 0,306582 | 1 | 1 |
| Gm16712 | 0,175343 | 0,216531 | 0,30439 | 1 | 1 |
| Vwa5b2 | 0,217549 | 0,268511 | 0,303643 | 1 | 1 |
| Gm15910 | 0,41819 | 0,51551 | 0,30184 | 1 | 1 |
| 1700096K18Rik | 0,439071 | 0,540949 | 0,301041 | 1 | 1 |
| Mapk4 | 0,47493 | 0,584432 | 0,29932 | 1 | 1 |
| Fam196a | 0,183871 | 0,225589 | 0,295005 | 1 | 1 |
| 6430571L13Rik | 0,294978 | 0,361323 | 0,292681 | 1 | 1 |
| Clec11a | 0,408724 | 0,500568 | 0,292438 | 1 | 1 |
| Noxa1 | 0,111719 | 0,136643 | 0,290533 | 1 | 1 |
| Gm10416 | 0,69784 | 0,853317 | 0,290186 | 1 | 1 |
| Lamb3 | 0,0971365 | 0,118675 | 0,288936 | 1 | 1 |
| Prrg4 | 0,149059 | 0,182082 | 0,288702 | 1 | 1 |
| 4930538K18Rik | 0,208713 | 0,254767 | 0,287664 | 1 | 1 |
| Cacnb2 | 0,313273 | 0,382244 | 0,287071 | 1 | 1 |
| Gipc2 | 0,384477 | 0,468675 | 0,285691 | 1 | 1 |
| Myh14 | 0,253217 | 0,308497 | 0,284887 | 1 | 1 |
| Tspear | 0,367903 | 0,44812 | 0,284557 | 1 | 1 |
| Hist2h2aa1,Hist2h2aa2 | 0,125147 | 0,152336 | 0,283638 | 1 | 1 |
| Lpar5 | 0,0185306 | 0,0225562 | 0,283613 | 1 | 1 |
| Gm20556 | 0,0213342 | 0,0259681 | 0,283571 | 1 | 1 |
| Glod5 | 0,0941791 | 0,114633 | 0,283546 | 1 | 1 |
| Camp | 0,0557919 | 0,0679089 | 0,283546 | 1 | 1 |
| Galnt15 | 0,0515638 | 0,0627626 | 0,283546 | 1 | 1 |
| Resp18 | 0,0435533 | 0,0530123 | 0,283546 | 1 | 1 |
| 4930502E09Rik | 0,0399964 | 0,0486829 | 0,283546 | 1 | 1 |
| Gm3448 | 0,0378011 | 0,0460108 | 0,283546 | 1 | 1 |
| Themis3 | 0,0276516 | 0,0336571 | 0,283546 | 1 | 1 |
| Stap1 | 0,0236462 | 0,0287817 | 0,283546 | 1 | 1 |
| Fcrlb | 0,0204058 | 0,0248376 | 0,283546 | 1 | 1 |
| C2cd4d | 0,0198186 | 0,0241228 | 0,283546 | 1 | 1 |
| Gm11961 | 0,0197653 | 0,0240579 | 0,283546 | 1 | 1 |
| Atp4b | 0,0196245 | 0,0238866 | 0,283546 | 1 | 1 |
| Krt17 | 0,0168617 | 0,0205238 | 0,283546 | 1 | 1 |
| G630071F17Rik | 0,0153098 | 0,0186348 | 0,283546 | 1 | 1 |
| Ccdc121 | 0,013646 | 0,0166097 | 0,283546 | 1 | 1 |

| | | | | | |
|---|---|---|---|---|---|
| Pde6a | 0,0110358 | 0,0134326 | 0,283546 | 1 | 1 |
| 4930552P12Rik | 0,00961536 | 0,0117036 | 0,283546 | 1 | 1 |
| Trim75 | 0,00958191 | 0,0116629 | 0,283546 | 1 | 1 |
| Ifit1 | 0,0090544 | 0,0110209 | 0,283546 | 1 | 1 |
| Gm5134 | 0,00837326 | 0,0101918 | 0,283546 | 1 | 1 |
| Cytip | 0,00796907 | 0,00969982 | 0,283546 | 1 | 1 |
| Tmem178b | 0,00763103 | 0,00928836 | 0,283546 | 1 | 1 |
| Kcnh3 | 0,00652218 | 0,00793868 | 0,283546 | 1 | 1 |
| Nlrc4 | 0,00606451 | 0,00738162 | 0,283546 | 1 | 1 |
| Pkp1 | 0,00514677 | 0,00626456 | 0,283546 | 1 | 1 |
| Sox17 | 0,148203 | 0,180384 | 0,283496 | 1 | 1 |
| Ppp1r32 | 0,1965 | 0,239013 | 0,282555 | 1 | 1 |
| 1700034P13Rik | 0,201538 | 0,245057 | 0,282062 | 1 | 1 |
| Eaf2 | 0,277736 | 0,337686 | 0,281968 | 1 | 1 |
| Gareml | 0,295274 | 0,358768 | 0,280995 | 1 | 1 |
| 2310001K24Rik | 0,172842 | 0,209991 | 0,280871 | 1 | 1 |
| Oacyl | 0,0943045 | 0,1144 | 0,278684 | 1 | 1 |
| E130304I02Rik | 0,448114 | 0,543393 | 0,278131 | 1 | 1 |
| Krt36 | 0,335573 | 0,405893 | 0,274472 | 1 | 1 |
| Nxnl2 | 0,129223 | 0,156267 | 0,274159 | 1 | 1 |
| Xrra1 | 0,102122 | 0,123495 | 0,274159 | 1 | 1 |
| Kif6 | 0,490962 | 0,593636 | 0,273966 | 1 | 1 |
| Myom1 | 0,395001 | 0,477599 | 0,273946 | 1 | 1 |
| Alpk3 | 0,218656 | 0,264262 | 0,273309 | 1 | 1 |
| Pcdhb8 | 0,0476999 | 0,0576016 | 0,272121 | 1 | 1 |
| Gja10 | 0,0775393 | 0,093635 | 0,27212 | 1 | 1 |
| Ccr10 | 0,0669867 | 0,0808919 | 0,27212 | 1 | 1 |
| 6820431F20Rik | 0,319539 | 0,385848 | 0,27204 | 1 | 1 |
| Kcnu1 | 0,0703691 | 0,0848215 | 0,269489 | 1 | 1 |
| Ccdc63 | 0,0431784 | 0,0520285 | 0,268991 | 1 | 1 |
| Mchr1 | 0,040226 | 0,0484696 | 0,268951 | 1 | 1 |
| Chgb | 0,0315732 | 0,0380436 | 0,268951 | 1 | 1 |
| Cplx3 | 0,0300749 | 0,0362381 | 0,268951 | 1 | 1 |
| Gadl1 | 0,430879 | 0,519177 | 0,268945 | 1 | 1 |
| Fam154b | 0,185964 | 0,223403 | 0,264627 | 1 | 1 |
| Gpat2 | 0,141449 | 0,169832 | 0,263833 | 1 | 1 |
| Fam160a1 | 0,123399 | 0,148145 | 0,26367 | 1 | 1 |
| Npw | 0,143803 | 0,172601 | 0,263348 | 1 | 1 |
| Rgs14 | 0,0268635 | 0,0322432 | 0,263348 | 1 | 1 |
| Foxl2 | 0,0249613 | 0,02996 | 0,263348 | 1 | 1 |
| 6430584L05Rik | 0,0196167 | 0,0235451 | 0,263348 | 1 | 1 |
| Fam189a1 | 0,329856 | 0,395086 | 0,260332 | 1 | 1 |
| Catsperg1 | 0,143358 | 0,171706 | 0,260315 | 1 | 1 |
| Tulp2 | 0,214377 | 0,256725 | 0,260069 | 1 | 1 |
| Fmn1 | 0,296663 | 0,35503 | 0,259114 | 1 | 1 |
| Cpxm2 | 0,251444 | 0,300908 | 0,259084 | 1 | 1 |
| Zc3h12a | 0,212808 | 0,254671 | 0,259084 | 1 | 1 |
| Kcnab1 | 0,338793 | 0,404898 | 0,257153 | 1 | 1 |
| Npy1r | 0,0574869 | 0,0686495 | 0,256017 | 1 | 1 |
| Lct | 0,280825 | 0,335171 | 0,255228 | 1 | 1 |

| | | | | | |
|---|---|---|---|---|---|
| Fut9 | 0,258029 | 0,307778 | 0,254356 | 1 | 1 |
| Gpr165 | 0,100412 | 0,119741 | 0,253975 | 1 | 1 |
| Ang | 0,425185 | 0,506846 | 0,253457 | 1 | 1 |
| Klhdc8a | 0,120255 | 0,143155 | 0,251483 | 1 | 1 |
| Slc25a45 | 0,129307 | 0,153913 | 0,25132 | 1 | 1 |
| Lrrn4 | 0,189431 | 0,225142 | 0,249161 | 1 | 1 |
| Mccc1os | 0,15811 | 0,187717 | 0,247634 | 1 | 1 |
| Tfcp2l1 | 0,0990321 | 0,117447 | 0,246041 | 1 | 1 |
| Prss41 | 0,432129 | 0,512466 | 0,245993 | 1 | 1 |
| Gpr116 | 0,467995 | 0,554898 | 0,245729 | 1 | 1 |
| Ccl28 | 0,277698 | 0,329236 | 0,245607 | 1 | 1 |
| Abcb1a | 0,0245253 | 0,0290751 | 0,245513 | 1 | 1 |
| B3gnt9 | 0,312678 | 0,369925 | 0,242553 | 1 | 1 |
| Folr2 | 0,54173 | 0,640285 | 0,24114 | 1 | 1 |
| Cers3 | 0,172172 | 0,203425 | 0,240649 | 1 | 1 |
| Chadl | 0,326367 | 0,385468 | 0,240116 | 1 | 1 |
| Lsm7 | 0,855313 | 1,00929 | 0,238808 | 1 | 1 |
| Zfp296 | 0,170295 | 0,200952 | 0,238808 | 1 | 1 |
| Scn1b | 0,221578 | 0,261087 | 0,236716 | 1 | 1 |
| 1700016K19Rik | 0,132523 | 0,156135 | 0,236547 | 1 | 1 |
| Nutm1 | 0,0269409 | 0,0317409 | 0,236547 | 1 | 1 |
| Gm10789 | 0,199304 | 0,234722 | 0,235984 | 1 | 1 |
| Evi2a | 0,138175 | 0,162593 | 0,234773 | 1 | 1 |
| Pls1 | 0,348062 | 0,40948 | 0,234449 | 1 | 1 |
| Esyt3 | 0,144201 | 0,169614 | 0,234174 | 1 | 1 |
| Scn3b | 0,0782128 | 0,0919332 | 0,233181 | 1 | 1 |
| Gjb5 | 0,330467 | 0,387948 | 0,231358 | 1 | 1 |
| Uba7 | 0,303037 | 0,355612 | 0,230807 | 1 | 1 |
| Ctf2 | 0,326505 | 0,382844 | 0,229649 | 1 | 1 |
| Gm14204 | 0,0439048 | 0,0514806 | 0,229649 | 1 | 1 |
| Rpp25 | 0,45462 | 0,533052 | 0,229617 | 1 | 1 |
| Nptxr | 0,258673 | 0,302954 | 0,227969 | 1 | 1 |
| Agbl2 | 0,117886 | 0,137994 | 0,227219 | 1 | 1 |
| Rgs17 | 0,120532 | 0,141053 | 0,226817 | 1 | 1 |
| Gm3435 | 0,141055 | 0,165046 | 0,22661 | 1 | 1 |
| Chrna5 | 0,255932 | 0,299373 | 0,226181 | 1 | 1 |
| Osbp2 | 0,16659 | 0,194866 | 0,226181 | 1 | 1 |
| Gm6402 | 0,234278 | 0,273836 | 0,225095 | 1 | 1 |
| Tekt5 | 0,055338 | 0,0646643 | 0,224697 | 1 | 1 |
| Grm4 | 0,00963476 | 0,0112511 | 0,22374 | 1 | 1 |
| Gm15328 | 0,128599 | 0,150135 | 0,223383 | 1 | 1 |
| Asb17 | 0,0864706 | 0,100858 | 0,222049 | 1 | 1 |
| Trim69 | 0,0469638 | 0,054778 | 0,222049 | 1 | 1 |
| Slc27a5 | 0,0343283 | 0,0400401 | 0,222049 | 1 | 1 |
| 4732490B19Rik | 0,0325522 | 0,0379685 | 0,222049 | 1 | 1 |
| Btk | 0,0313093 | 0,0365188 | 0,222049 | 1 | 1 |
| Pcdhb10 | 0,0276762 | 0,0322812 | 0,222049 | 1 | 1 |
| Smyd1 | 0,182372 | 0,212429 | 0,22009 | 1 | 1 |
| 1700020D05Rik | 0,332914 | 0,387065 | 0,217428 | 1 | 1 |
| Pln | 0,157766 | 0,183407 | 0,217258 | 1 | 1 |

| | | | | | |
|---|---|---|---|---|---|
| Edn1 | 0,088132 | 0,102453 | 0,217222 | 1 | 1 |
| Slfn10-ps | 0,0474202 | 0,0550129 | 0,21427 | 1 | 1 |
| Gpr82 | 0,0504263 | 0,0585 | 0,214259 | 1 | 1 |
| Aoc3 | 0,104459 | 0,121156 | 0,21393 | 1 | 1 |
| Qrfp | 0,137236 | 0,158992 | 0,212301 | 1 | 1 |
| Lynx1 | 0,242828 | 0,281133 | 0,211317 | 1 | 1 |
| Cd72 | 0,103182 | 0,119362 | 0,210154 | 1 | 1 |
| N4bp2l1 | 0,521784 | 0,602275 | 0,206968 | 1 | 1 |
| Sez6l2 | 0,22534 | 0,26005 | 0,206684 | 1 | 1 |
| Slc25a18 | 0,350427 | 0,404103 | 0,20561 | 1 | 1 |
| Zfp947 | 0,377939 | 0,435609 | 0,204881 | 1 | 1 |
| Sbspon | 0,0893282 | 0,102913 | 0,204245 | 1 | 1 |
| Zpbp | 0,221586 | 0,255271 | 0,204164 | 1 | 1 |
| Plcb2 | 0,0494546 | 0,0569599 | 0,203844 | 1 | 1 |
| Iqcd | 0,557741 | 0,642039 | 0,203067 | 1 | 1 |
| Card6 | 0,197307 | 0,227066 | 0,202668 | 1 | 1 |
| Pstpip2 | 0,505088 | 0,579425 | 0,198089 | 1 | 1 |
| 9130008F23Rik | 0,509888 | 0,584889 | 0,197982 | 1 | 1 |
| 1700028E10Rik | 0,207101 | 0,237522 | 0,197723 | 1 | 1 |
| Henmt1 | 0,184878 | 0,211855 | 0,1965 | 1 | 1 |
| Irf4 | 0,0322675 | 0,0369725 | 0,196373 | 1 | 1 |
| Tmem88b | 0,0758651 | 0,0869256 | 0,196345 | 1 | 1 |
| Scn2a1 | 0,0368288 | 0,0421901 | 0,196071 | 1 | 1 |
| Cytl1 | 0,370205 | 0,424084 | 0,196025 | 1 | 1 |
| Pbld2 | 0,323135 | 0,370141 | 0,195936 | 1 | 1 |
| 1700067K01Rik | 0,233194 | 0,267028 | 0,195464 | 1 | 1 |
| Zbtb20 | 0,153732 | 0,17592 | 0,194498 | 1 | 1 |
| Rnf207 | 0,390626 | 0,44697 | 0,194391 | 1 | 1 |
| Sh3tc1 | 0,123568 | 0,141174 | 0,192168 | 1 | 1 |
| Gnat2 | 0,531242 | 0,606504 | 0,191148 | 1 | 1 |
| A3galt2 | 0,223858 | 0,254947 | 0,187616 | 1 | 1 |
| Gm2897 | 0,445423 | 0,506859 | 0,18641 | 1 | 1 |
| Drc1 | 0,106879 | 0,121593 | 0,186086 | 1 | 1 |
| Gm14379 | 0,243457 | 0,276904 | 0,185717 | 1 | 1 |
| Slc26a8 | 0,117449 | 0,133585 | 0,185717 | 1 | 1 |
| Lin7b | 0,28508 | 0,32343 | 0,182088 | 1 | 1 |
| 1700025F24Rik | 0,211539 | 0,239996 | 0,182088 | 1 | 1 |
| Themis2 | 0,124862 | 0,141659 | 0,182088 | 1 | 1 |
| Ldhd | 0,0723668 | 0,0821019 | 0,182088 | 1 | 1 |
| Apoa1 | 0,354836 | 0,40237 | 0,18137 | 1 | 1 |
| Hecw1 | 0,0300557 | 0,034082 | 0,18137 | 1 | 1 |
| Wnt9a | 0,523967 | 0,593585 | 0,179978 | 1 | 1 |
| Rimbp3 | 0,347918 | 0,394042 | 0,179601 | 1 | 1 |
| Adipoq | 0,105791 | 0,119685 | 0,178026 | 1 | 1 |
| Gpr151 | 0,0761046 | 0,0860997 | 0,178026 | 1 | 1 |
| Dapk2 | 0,352762 | 0,399032 | 0,177809 | 1 | 1 |
| Gpr137c | 0,39404 | 0,445647 | 0,177558 | 1 | 1 |
| Rasef | 0,129965 | 0,146975 | 0,177445 | 1 | 1 |
| Cldn15 | 0,318522 | 0,360054 | 0,176822 | 1 | 1 |
| Hist1h2ab | 0,31567 | 0,356462 | 0,175329 | 1 | 1 |

| | | | | | |
|---|---|---|---|---|---|
| Tbx21 | 0,0404686 | 0,0456981 | 0,175329 | 1 | 1 |
| Vsig10l | 0,34602 | 0,390464 | 0,174334 | 1 | 1 |
| Slitrk4 | 0,319952 | 0,361022 | 0,17423 | 1 | 1 |
| Kcnj11 | 0,0648373 | 0,0731124 | 0,173292 | 1 | 1 |
| Upk1b | 0,111737 | 0,125926 | 0,172473 | 1 | 1 |
| Slc2a12 | 0,331245 | 0,372344 | 0,168736 | 1 | 1 |
| Cetn4 | 0,510467 | 0,573518 | 0,168022 | 1 | 1 |
| Rd3 | 0,0595948 | 0,0668918 | 0,166644 | 1 | 1 |
| Plin5 | 0,118335 | 0,132824 | 0,166638 | 1 | 1 |
| Zfp957 | 0,0540475 | 0,0606646 | 0,166627 | 1 | 1 |
| Klhdc1 | 0,426104 | 0,477162 | 0,163273 | 1 | 1 |
| Nlrp10 | 0,0289614 | 0,0324317 | 0,163273 | 1 | 1 |
| Gm14137 | 0,400668 | 0,447864 | 0,160655 | 1 | 1 |
| Tmem150b | 0,54736 | 0,61129 | 0,159366 | 1 | 1 |
| Magix | 0,0429679 | 0,0479686 | 0,158833 | 1 | 1 |
| Rab3d | 0,508262 | 0,567015 | 0,157815 | 1 | 1 |
| Cntn4 | 0,0460024 | 0,0512775 | 0,156618 | 1 | 1 |
| Glyctk | 0,510241 | 0,568528 | 0,156053 | 1 | 1 |
| Pdzrn4 | 0,158046 | 0,176084 | 0,155925 | 1 | 1 |
| 8430431K14Rik | 0,054538 | 0,0607396 | 0,155376 | 1 | 1 |
| Bcan | 0,11097 | 0,123543 | 0,154847 | 1 | 1 |
| Plcxd3 | 0,250723 | 0,27875 | 0,152879 | 1 | 1 |
| Pxdc1 | 0,475454 | 0,528564 | 0,152775 | 1 | 1 |
| Gm10190 | 0,181126 | 0,201138 | 0,151192 | 1 | 1 |
| 1600014C23Rik | 0,276154 | 0,306397 | 0,149931 | 1 | 1 |
| C3 | 0,017593 | 0,0195197 | 0,149931 | 1 | 1 |
| Pkhd1l1 | 0,00691789 | 0,00767552 | 0,149931 | 1 | 1 |
| 9230112J17Rik | 0,040728 | 0,0451863 | 0,149863 | 1 | 1 |
| Hepacam2 | 0,193806 | 0,214918 | 0,149169 | 1 | 1 |
| Kcnj6 | 0,353084 | 0,391262 | 0,148125 | 1 | 1 |
| Gm10532 | 0,134384 | 0,14873 | 0,146336 | 1 | 1 |
| Cd1d2 | 0,353879 | 0,39165 | 0,146305 | 1 | 1 |
| Myh7b | 0,0675916 | 0,0747555 | 0,145337 | 1 | 1 |
| Gcnt7 | 0,134587 | 0,148672 | 0,143596 | 1 | 1 |
| Scn9a | 0,0120203 | 0,0132766 | 0,143413 | 1 | 1 |
| Ano5 | 0,0155548 | 0,0171792 | 0,143305 | 1 | 1 |
| Igj | 0,060917 | 0,0672749 | 0,143225 | 1 | 1 |
| Fam209 | 0,423816 | 0,467769 | 0,14236 | 1 | 1 |
| Spag17 | 0,17083 | 0,188514 | 0,142111 | 1 | 1 |
| 9230102O04Rik | 0,26278 | 0,289887 | 0,141634 | 1 | 1 |
| Apol7d | 0,640401 | 0,706289 | 0,141284 | 1 | 1 |
| Gm14092 | 0,0453303 | 0,0499839 | 0,140988 | 1 | 1 |
| B3gnt6 | 0,0337411 | 0,0372049 | 0,140988 | 1 | 1 |
| E230008N13Rik | 0,0149646 | 0,0165008 | 0,140988 | 1 | 1 |
| Adgb | 0,0145557 | 0,01605 | 0,140988 | 1 | 1 |
| Atp2b3 | 0,0111525 | 0,0122975 | 0,140988 | 1 | 1 |
| Mal | 0,199755 | 0,219814 | 0,138053 | 1 | 1 |
| Sytl1 | 0,304826 | 0,335428 | 0,138017 | 1 | 1 |
| Ccdc150 | 0,503295 | 0,552831 | 0,135435 | 1 | 1 |
| Srp54c | 2,71493 | 2,9803 | 0,134542 | 1 | 1 |

| | | | | | |
|---|---|---|---|---|---|
| Nrip3 | 0,183035 | 0,200771 | 0,133429 | 1 | 1 |
| 9330162012Rik | 0,256085 | 0,280774 | 0,132784 | 1 | 1 |
| 4930412C18Rik | 0,156899 | 0,171756 | 0,130529 | 1 | 1 |
| 4933433G19Rik | 0,333118 | 0,364163 | 0,12855 | 1 | 1 |
| 0610005C13Rik | 0,586423 | 0,640941 | 0,12825 | 1 | 1 |
| AI182371 | 0,296303 | 0,323541 | 0,126875 | 1 | 1 |
| Trpm6 | 0,035815 | 0,0390285 | 0,123966 | 1 | 1 |
| Rab26 | 0,390673 | 0,425708 | 0,123903 | 1 | 1 |
| Esrrb | 0,0637535 | 0,0694686 | 0,123857 | 1 | 1 |
| Map3k9 | 0,516613 | 0,562275 | 0,122194 | 1 | 1 |
| Ccdc60 | 0,566262 | 0,616214 | 0,121962 | 1 | 1 |
| Tlx1 | 0,287298 | 0,312616 | 0,121842 | 1 | 1 |
| Espn | 0,455555 | 0,495587 | 0,121512 | 1 | 1 |
| C1ql3 | 0,036615 | 0,0398016 | 0,120391 | 1 | 1 |
| Tmem140 | 0,484051 | 0,525644 | 0,118927 | 1 | 1 |
| Ttll9 | 0,52795 | 0,571642 | 0,114711 | 1 | 1 |
| 1700001K19Rik | 0,532318 | 0,576076 | 0,113971 | 1 | 1 |
| BC048644 | 0,0528061 | 0,0570359 | 0,111165 | 1 | 1 |
| Sh3rf3 | 0,168632 | 0,182136 | 0,111137 | 1 | 1 |
| Jak3 | 0,146185 | 0,157653 | 0,108952 | 1 | 1 |
| Slc41a3 | 0,531833 | 0,573171 | 0,107992 | 1 | 1 |
| 1810024B03Rik | 0,219094 | 0,236086 | 0,107763 | 1 | 1 |
| Bfsp1 | 0,40643 | 0,437813 | 0,107307 | 1 | 1 |
| Cd37 | 0,4373 | 0,470887 | 0,106757 | 1 | 1 |
| Hist2h2aa2 | 6,08E-005 | 6,54E-005 | 0,103224 | 1 | 1 |
| Ttll7 | 0,438972 | 0,471164 | 0,1021 | 1 | 1 |
| Slc8a3 | 0,483178 | 0,51811 | 0,100705 | 1 | 1 |
| Eml6 | 0,160518 | 0,172052 | 0,100107 | 1 | 1 |
| Gabrb3 | 0,522659 | 0,560028 | 0,0996281 | 1 | 1 |
| Wbscr17 | 0,387724 | 0,415277 | 0,0990413 | 1 | 1 |
| Klhl6 | 0,36213 | 0,387568 | 0,0979423 | 1 | 1 |
| Gpr182 | 0,415856 | 0,444561 | 0,0962983 | 1 | 1 |
| Kcnrg | 0,0722809 | 0,0771795 | 0,0946036 | 1 | 1 |
| Plekhb1 | 0,176709 | 0,188414 | 0,0925359 | 1 | 1 |
| Adcy10 | 0,0159538 | 0,0170081 | 0,0923197 | 1 | 1 |
| Zfp781 | 0,193514 | 0,206193 | 0,0915576 | 1 | 1 |
| Slc18a1 | 0,222064 | 0,236522 | 0,0909951 | 1 | 1 |
| Kcng2 | 0,111807 | 0,119039 | 0,0904228 | 1 | 1 |
| Kcnq2 | 0,341315 | 0,363241 | 0,0898231 | 1 | 1 |
| Kcns1 | 0,0839876 | 0,0892744 | 0,08807 | 1 | 1 |
| LOC100505025 | 0,146148 | 0,155343 | 0,0880297 | 1 | 1 |
| Extl1 | 0,158473 | 0,168222 | 0,0861327 | 1 | 1 |
| Tchh | 0,249167 | 0,264415 | 0,0856919 | 1 | 1 |
| Caps2 | 0,269849 | 0,286301 | 0,0853836 | 1 | 1 |
| Arhgap6 | 0,379915 | 0,40296 | 0,0849604 | 1 | 1 |
| Gdf9 | 0,277074 | 0,293733 | 0,084232 | 1 | 1 |
| Nol4 | 0,153115 | 0,162299 | 0,0840346 | 1 | 1 |
| Pram1 | 0,552733 | 0,585254 | 0,082478 | 1 | 1 |
| Ptprv | 0,0618572 | 0,0654952 | 0,0824492 | 1 | 1 |
| 0610038B21Rik | 0,353113 | 0,373828 | 0,0822455 | 1 | 1 |

| | | | | | |
|---|---|---|---|---|---|
| Rbfox3 | 0,243376 | 0,257595 | 0,0819204 | 1 | 1 |
| Pde8b | 0,307456 | 0,325324 | 0,0814948 | 1 | 1 |
| 4930500J02Rik | 0,554402 | 0,586503 | 0,0812044 | 1 | 1 |
| Abcg4 | 0,428133 | 0,452572 | 0,0800879 | 1 | 1 |
| 9430083A17Rik | 0,371673 | 0,392843 | 0,0799197 | 1 | 1 |
| Atxn1 | 0,148654 | 0,156857 | 0,0774909 | 1 | 1 |
| 4930405A21Rik | 0,461327 | 0,486533 | 0,0767481 | 1 | 1 |
| Stac3 | 0,402767 | 0,424643 | 0,076306 | 1 | 1 |
| Siglec5 | 0,0231085 | 0,0243391 | 0,0748517 | 1 | 1 |
| Tmem95 | 0,155785 | 0,16406 | 0,0746662 | 1 | 1 |
| 6030407O03Rik | 0,0318994 | 0,0335939 | 0,0746662 | 1 | 1 |
| Sntg2 | 0,0238835 | 0,0251521 | 0,0746662 | 1 | 1 |
| Tmie | 0,0228139 | 0,0240258 | 0,0746662 | 1 | 1 |
| Foxi2 | 0,0213421 | 0,0224757 | 0,0746662 | 1 | 1 |
| A730020E08Rik | 0,150271 | 0,158089 | 0,073166 | 1 | 1 |
| E130218I03Rik | 0,0588528 | 0,0619137 | 0,0731477 | 1 | 1 |
| C330013E15Rik | 0,249258 | 0,262143 | 0,0727107 | 1 | 1 |
| Zan | 0,0268142 | 0,0281925 | 0,0723117 | 1 | 1 |
| Stx11 | 0,12055 | 0,126706 | 0,0718616 | 1 | 1 |
| Fgf23 | 0,046861 | 0,0492542 | 0,0718589 | 1 | 1 |
| Dnah17 | 0,206809 | 0,217291 | 0,0713322 | 1 | 1 |
| Ankmy1 | 0,0602745 | 0,0633184 | 0,0710772 | 1 | 1 |
| Dock3 | 0,255672 | 0,268483 | 0,0705336 | 1 | 1 |
| Golt1a | 0,0403633 | 0,042361 | 0,0696921 | 1 | 1 |
| E330020D12Rik | 0,025206 | 0,0264535 | 0,0696921 | 1 | 1 |
| Adra2c | 0,0220826 | 0,0231755 | 0,0696921 | 1 | 1 |
| Gm9926 | 0,0141031 | 0,0148011 | 0,0696921 | 1 | 1 |
| Nlrc3 | 0,0650686 | 0,0681866 | 0,0675268 | 1 | 1 |
| Gjb3 | 0,268816 | 0,281631 | 0,0671874 | 1 | 1 |
| Cnn1 | 0,254101 | 0,266194 | 0,0670757 | 1 | 1 |
| Pnldc1 | 0,535981 | 0,560705 | 0,0650586 | 1 | 1 |
| Rgcc | 0,511925 | 0,535483 | 0,0649075 | 1 | 1 |
| 1700066B19Rik | 0,142478 | 0,14882 | 0,0628285 | 1 | 1 |
| Sybu | 0,319617 | 0,333678 | 0,0621125 | 1 | 1 |
| Lepr | 0,123131 | 0,128488 | 0,0614354 | 1 | 1 |
| Emx2 | 0,364022 | 0,379479 | 0,0599936 | 1 | 1 |
| D930015M05Rik | 0,499422 | 0,52061 | 0,0599437 | 1 | 1 |
| Map3k15 | 0,401635 | 0,418405 | 0,0590161 | 1 | 1 |
| Arap2 | 0,0609744 | 0,0634588 | 0,0576147 | 1 | 1 |
| Npr1 | 0,470697 | 0,489766 | 0,0572941 | 1 | 1 |
| Serpine1 | 0,288736 | 0,300043 | 0,0554168 | 1 | 1 |
| Hcn2 | 0,350873 | 0,364472 | 0,0548608 | 1 | 1 |
| Rassf10 | 0,128369 | 0,133161 | 0,0528688 | 1 | 1 |
| Sox30 | 0,133206 | 0,137889 | 0,0498561 | 1 | 1 |
| Slc27a2 | 0,323736 | 0,334809 | 0,0485182 | 1 | 1 |
| Tdh | 0,126463 | 0,130739 | 0,0479688 | 1 | 1 |
| Gm1564 | 0,0450281 | 0,0465382 | 0,0475909 | 1 | 1 |
| Nfia | 0,267082 | 0,275976 | 0,0472595 | 1 | 1 |
| Cd300a | 0,213459 | 0,220339 | 0,045765 | 1 | 1 |
| Fgf12 | 0,185778 | 0,191764 | 0,0457502 | 1 | 1 |

| | | | | | |
|---|---|---|---|---|---|
| Kif27 | 0,552305 | 0,56968 | 0,0446885 | 1 | 1 |
| 4930431F12Rik | 0,0528444 | 0,0543975 | 0,0417895 | 1 | 1 |
| Arhgap30 | 0,025392 | 0,0261383 | 0,0417895 | 1 | 1 |
| Dach2 | 0,478127 | 0,491955 | 0,0411325 | 1 | 1 |
| Gpr3 | 0,13919 | 0,14314 | 0,0403779 | 1 | 1 |
| 9030025P20Rik | 0,273803 | 0,281504 | 0,0400152 | 1 | 1 |
| Gm14295 | 0,300927 | 0,3092 | 0,0391251 | 1 | 1 |
| Slc39a4 | 0,0721878 | 0,0741155 | 0,0380204 | 1 | 1 |
| Sprn | 0,0494458 | 0,0507662 | 0,0380204 | 1 | 1 |
| St8sia6 | 0,0442614 | 0,0453888 | 0,0362887 | 1 | 1 |
| Wnt8b | 0,0412334 | 0,0422837 | 0,0362887 | 1 | 1 |
| A330009N23Rik | 0,416613 | 0,427075 | 0,035783 | 1 | 1 |
| Gm5464 | 0,213776 | 0,218961 | 0,0345701 | 1 | 1 |
| Ttc39b | 0,437361 | 0,447879 | 0,0342859 | 1 | 1 |
| Oaz3 | 0,642728 | 0,658149 | 0,0342065 | 1 | 1 |
| Dmrta2 | 0,600891 | 0,615193 | 0,0339359 | 1 | 1 |
| Tmem200c | 0,577068 | 0,590645 | 0,03355 | 1 | 1 |
| Pcdh17 | 0,124566 | 0,127461 | 0,0331482 | 1 | 1 |
| Cdh12 | 0,093468 | 0,0954874 | 0,0308376 | 1 | 1 |
| Fam20a | 0,250447 | 0,255739 | 0,0301691 | 1 | 1 |
| Xkr4 | 0,125244 | 0,127842 | 0,0296217 | 1 | 1 |
| Gm5868 | 0,160375 | 0,163625 | 0,0289427 | 1 | 1 |
| Galntl6 | 0,0462258 | 0,0471518 | 0,0286141 | 1 | 1 |
| Mzf1 | 0,384511 | 0,392201 | 0,0285717 | 1 | 1 |
| Tacr1 | 0,333803 | 0,340162 | 0,0272269 | 1 | 1 |
| Tbc1d2 | 0,176287 | 0,179575 | 0,0266606 | 1 | 1 |
| LOC171588 | 0,209624 | 0,21333 | 0,0252803 | 1 | 1 |
| Tnfaip2 | 0,251171 | 0,25534 | 0,0237514 | 1 | 1 |
| Mir719 | 304,157 | 309,196 | 0,0237074 | 1 | 1 |
| Fam178b | 0,414388 | 0,420925 | 0,0225825 | 1 | 1 |
| Ffar4 | 0,372849 | 0,378118 | 0,0202452 | 1 | 1 |
| Gsta3 | 0,0943137 | 0,0955956 | 0,0194771 | 1 | 1 |
| 4833422C13Rik | 0,361341 | 0,366252 | 0,0194735 | 1 | 1 |
| Rgs20 | 0,503783 | 0,510343 | 0,0186625 | 1 | 1 |
| Zfp174 | 0,512047 | 0,518674 | 0,0185522 | 1 | 1 |
| Rln3 | 0,350614 | 0,355012 | 0,0179832 | 1 | 1 |
| Dmbx1 | 0,0248716 | 0,0251836 | 0,0179832 | 1 | 1 |
| Nwd1 | 0,0126357 | 0,0127942 | 0,0179832 | 1 | 1 |
| Vwa7 | 0,383674 | 0,388217 | 0,0169797 | 1 | 1 |
| AI839979 | 0,255778 | 0,258744 | 0,0166325 | 1 | 1 |
| B3gnt3 | 0,0805218 | 0,0814081 | 0,0157921 | 1 | 1 |
| BC100451 | 0,270873 | 0,273794 | 0,0154737 | 1 | 1 |
| Raet1b | 0,68723 | 0,694637 | 0,0154668 | 1 | 1 |
| Kcnj4 | 0,168223 | 0,17002 | 0,0153251 | 1 | 1 |
| Kcne1 | 0,0592576 | 0,0598735 | 0,0149168 | 1 | 1 |
| 1700029I15Rik | 1,31265 | 1,32619 | 0,0148027 | 1 | 1 |
| Abcb9 | 0,0672975 | 0,0679916 | 0,0148027 | 1 | 1 |
| Phyhip | 0,60296 | 0,608796 | 0,0138978 | 1 | 1 |
| Col20a1 | 0,0929333 | 0,0937924 | 0,0132758 | 1 | 1 |
| Kcnn2 | 0,214781 | 0,216671 | 0,0126423 | 1 | 1 |

| | | | | | |
|---|---|---|---|---|---|
| Wdr17 | 0,430025 | 0,433718 | 0,0123386 | 1 | 1 |
| Dcdc2b | 0,417142 | 0,420524 | 0,0116476 | 1 | 1 |
| Spef2 | 0,0493225 | 0,0497223 | 0,0116476 | 1 | 1 |
| A930017M01Rik | 0,0691379 | 0,0696915 | 0,0115051 | 1 | 1 |
| Ptx4 | 0,0619876 | 0,0624765 | 0,0113347 | 1 | 1 |
| 9430014N10Rik | 0,0861359 | 0,0868108 | 0,0112594 | 1 | 1 |
| Myh3 | 0,0137956 | 0,0139037 | 0,0112594 | 1 | 1 |
| Gpr61 | 0,105212 | 0,106006 | 0,0108398 | 1 | 1 |
| Caln1 | 0,324072 | 0,326349 | 0,0100986 | 1 | 1 |
| Zfp300 | 0,396269 | 0,399024 | 0,00999669 | 1 | 1 |
| Ctsw | 0,432641 | 0,4356 | 0,00983374 | 1 | 1 |
| Shisa7 | 0,263986 | 0,26569 | 0,00928082 | 1 | 1 |
| Ppara | 0,0758902 | 0,0763523 | 0,0087593 | 1 | 1 |
| Apom | 0,727532 | 0,731418 | 0,00768402 | 1 | 1 |
| Fam57b | 0,292939 | 0,294446 | 0,00740171 | 1 | 1 |
| E030024N20Rik | 0,15809 | 0,158821 | 0,00666024 | 1 | 1 |
| Gm960 | 0,162569 | 0,163192 | 0,00552557 | 1 | 1 |
| Hhex | 0,624621 | 0,626999 | 0,00548214 | 1 | 1 |
| Tmod4 | 0,68391 | 0,68601 | 0,00442313 | 1 | 1 |
| Mog | 0,184135 | 0,184663 | 0,00413092 | 1 | 1 |
| Hcn4 | 0,432882 | 0,433997 | 0,00371441 | 1 | 1 |
| Apobr | 0,328098 | 0,328906 | 0,00354888 | 1 | 1 |
| 4930512B01Rik | 0,189644 | 0,189932 | 0,00219175 | 1 | 1 |
| Hnf1a | 0,0734226 | 0,0735243 | 0,00199719 | 1 | 1 |
| Ccr7 | 0,147704 | 0,147832 | 0,00125333 | 1 | 1 |
| Gspt1 | 72,8616 | 0 | 0 | 1 | 1 |
| 2610020C07Rik | 3,97187 | 0 | 0 | 1 | 1 |
| 0610010B08Rik,Gm14308, | 0 | 0 | 0 | 1 | 1 |
| 0610031O16Rik | 0 | 0 | 0 | 1 | 1 |
| 1110015O18Rik | 0 | 0 | 0 | 1 | 1 |
| 1110025L11Rik | 0 | 0 | 0 | 1 | 1 |
| 1110028F18Rik | 0 | 0 | 0 | 1 | 1 |
| 1110036E04Rik | 0 | 0 | 0 | 1 | 1 |
| 1190003K10Rik | 0 | 0 | 0 | 1 | 1 |
| 1500012K07Rik | 0 | 0 | 0 | 1 | 1 |
| 1500015L24Rik | 0 | 0 | 0 | 1 | 1 |
| 1500017E21Rik | 0 | 0 | 0 | 1 | 1 |
| 1600014K23Rik | 0 | 0 | 0 | 1 | 1 |
| 1600015I10Rik | 0 | 0 | 0 | 1 | 1 |
| 1600019K03Rik | 0 | 0 | 0 | 1 | 1 |
| 1600025M17Rik | 0 | 0 | 0 | 1 | 1 |
| 1600029I14Rik | 0 | 0 | 0 | 1 | 1 |
| 1700001C02Rik | 0 | 0 | 0 | 1 | 1 |
| 1700001F09Rik | 0 | 0 | 0 | 1 | 1 |
| 1700003C15Rik | 0 | 0 | 0 | 1 | 1 |
| 1700003E24Rik | 0 | 0 | 0 | 1 | 1 |
| 1700003G13Rik | 0 | 0 | 0 | 1 | 1 |
| 1700003G18Rik | 0 | 0 | 0 | 1 | 1 |
| 1700003H04Rik | 0 | 0 | 0 | 1 | 1 |
| 1700003L19Rik | 0 | 0 | 0 | 1 | 1 |

| | | | | | |
|---|---|---|---|---|---|
| 1700003P14Rik | 0 | 0 | 0 | 1 | 1 |
| 1700006A11Rik | 0 | 0 | 0 | 1 | 1 |
| 1700006E09Rik | 0 | 0 | 0 | 1 | 1 |
| 1700006H21Rik | 0 | 0 | 0 | 1 | 1 |
| 1700007F19Rik | 0 | 0 | 0 | 1 | 1 |
| 1700007K09Rik | 0 | 0 | 0 | 1 | 1 |
| 1700007P06Rik | 0 | 0 | 0 | 1 | 1 |
| 1700008F21Rik | 0 | 0 | 0 | 1 | 1 |
| 1700008K24Rik | 0 | 0 | 0 | 1 | 1 |
| 1700008P02Rik | 0 | 0 | 0 | 1 | 1 |
| 1700009C05Rik | 0 | 0 | 0 | 1 | 1 |
| 1700009N14Rik | 0 | 0 | 0 | 1 | 1 |
| 1700010D01Rik | 0 | 0 | 0 | 1 | 1 |
| 1700010I02Rik | 0 | 0 | 0 | 1 | 1 |
| 1700010J16Rik | 0 | 0 | 0 | 1 | 1 |
| 1700010K23Rik | 0 | 0 | 0 | 1 | 1 |
| 1700011A15Rik | 0 | 0 | 0 | 1 | 1 |
| 1700011B04Rik | 0 | 0 | 0 | 1 | 1 |
| 1700011E24Rik | 0 | 0 | 0 | 1 | 1 |
| 1700011I03Rik | 0 | 0 | 0 | 1 | 1 |
| 1700011M02Rik | 0 | 0 | 0 | 1 | 1 |
| 1700012A03Rik | 0 | 0 | 0 | 1 | 1 |
| 1700012I11Rik | 0 | 0 | 0 | 1 | 1 |
| 1700012L04Rik | 0 | 0 | 0 | 1 | 1 |
| 1700012P22Rik | 0 | 0 | 0 | 1 | 1 |
| 1700013D24Rik | 0 | 0 | 0 | 1 | 1 |
| 1700013F07Rik | 0 | 0 | 0 | 1 | 1 |
| 1700013G24Rik | 0 | 0 | 0 | 1 | 1 |
| 1700015F17Rik | 0 | 0 | 0 | 1 | 1 |
| 1700015G11Rik | 0 | 0 | 0 | 1 | 1 |
| 1700016C15Rik | 0 | 0 | 0 | 1 | 1 |
| 1700016G22Rik | 0 | 0 | 0 | 1 | 1 |
| 1700016L21Rik | 0 | 0 | 0 | 1 | 1 |
| 1700016P04Rik | 0 | 0 | 0 | 1 | 1 |
| 1700017D01Rik | 0 | 0 | 0 | 1 | 1 |
| 1700017N19Rik | 0 | 0 | 0 | 1 | 1 |
| 1700018B08Rik | 0 | 0 | 0 | 1 | 1 |
| 1700018B24Rik | 0 | 0 | 0 | 1 | 1 |
| 1700018C11Rik | 0 | 0 | 0 | 1 | 1 |
| 1700018F24Rik | 0 | 0 | 0 | 1 | 1 |
| 1700019E08Rik | 0 | 0 | 0 | 1 | 1 |
| 1700019G24Rik | 0 | 0 | 0 | 1 | 1 |
| 1700019M22Rik | 0 | 0 | 0 | 1 | 1 |
| 1700019O17Rik | 0 | 0 | 0 | 1 | 1 |
| 1700020G17Rik | 0 | 0 | 0 | 1 | 1 |
| 1700020M21Rik | 0 | 0 | 0 | 1 | 1 |
| 1700020N01Rik | 0 | 0 | 0 | 1 | 1 |
| 1700020N15Rik | 0 | 0 | 0 | 1 | 1 |
| 1700020N18Rik | 0 | 0 | 0 | 1 | 1 |
| 1700021F07Rik | 0 | 0 | 0 | 1 | 1 |

| | | | | | |
|---|---|---|---|---|---|
| 1700021N21Rik | 0 | 0 | 0 | 1 | 1 |
| 1700022A21Rik | 0 | 0 | 0 | 1 | 1 |
| 1700022A22Rik | 0 | 0 | 0 | 1 | 1 |
| 1700022E09Rik | 0 | 0 | 0 | 1 | 1 |
| 1700022H16Rik | 0 | 0 | 0 | 1 | 1 |
| 1700023C21Rik | 0 | 0 | 0 | 1 | 1 |
| 1700023F02Rik | 0 | 0 | 0 | 1 | 1 |
| 1700024B18Rik | 0 | 0 | 0 | 1 | 1 |
| 1700024G13Rik | 0 | 0 | 0 | 1 | 1 |
| 1700024P04Rik | 0 | 0 | 0 | 1 | 1 |
| 1700024P16Rik | 0 | 0 | 0 | 1 | 1 |
| 1700025B11Rik | 0 | 0 | 0 | 1 | 1 |
| 1700025C18Rik | 0 | 0 | 0 | 1 | 1 |
| 1700025F22Rik | 0 | 0 | 0 | 1 | 1 |
| 1700025K24Rik | 0 | 0 | 0 | 1 | 1 |
| 1700025M24Rik | 0 | 0 | 0 | 1 | 1 |
| 1700025N23Rik | 0 | 0 | 0 | 1 | 1 |
| 1700026D11Rik | 0 | 0 | 0 | 1 | 1 |
| 1700027A15Rik | 0 | 0 | 0 | 1 | 1 |
| 1700027F09Rik | 0 | 0 | 0 | 1 | 1 |
| 1700027H10Rik | 0 | 0 | 0 | 1 | 1 |
| 1700027I24Rik | 0 | 0 | 0 | 1 | 1 |
| 1700028B04Rik | 0 | 0 | 0 | 1 | 1 |
| 1700028D13Rik | 0 | 0 | 0 | 1 | 1 |
| 1700028M03Rik | 0 | 0 | 0 | 1 | 1 |
| 1700028P14Rik | 0 | 0 | 0 | 1 | 1 |
| 1700028P15Rik | 0 | 0 | 0 | 1 | 1 |
| 1700029B22Rik | 0 | 0 | 0 | 1 | 1 |
| 1700029F12Rik | 0 | 0 | 0 | 1 | 1 |
| 1700029H14Rik | 0 | 0 | 0 | 1 | 1 |
| 1700029M20Rik | 0 | 0 | 0 | 1 | 1 |
| 1700029N11Rik | 0 | 0 | 0 | 1 | 1 |
| 1700029P11Rik | 0 | 0 | 0 | 1 | 1 |
| 1700030A11Rik | 0 | 0 | 0 | 1 | 1 |
| 1700030F04Rik | 0 | 0 | 0 | 1 | 1 |
| 1700030F18Rik | 0 | 0 | 0 | 1 | 1 |
| 1700030L20Rik | 0 | 0 | 0 | 1 | 1 |
| 1700030M09Rik | 0 | 0 | 0 | 1 | 1 |
| 1700030N03Rik | 0 | 0 | 0 | 1 | 1 |
| 1700030O20Rik | 0 | 0 | 0 | 1 | 1 |
| 1700031A10Rik | 0 | 0 | 0 | 1 | 1 |
| 1700031F05Rik | 0 | 0 | 0 | 1 | 1 |
| 1700031M16Rik | 0 | 0 | 0 | 1 | 1 |
| 1700031P21Rik | 0 | 0 | 0 | 1 | 1 |
| 1700034E13Rik | 0 | 0 | 0 | 1 | 1 |
| 1700034G24Rik | 0 | 0 | 0 | 1 | 1 |
| 1700034I23Rik | 0 | 0 | 0 | 1 | 1 |
| 1700034K08Rik | 0 | 0 | 0 | 1 | 1 |
| 1700034O15Rik | 0 | 0 | 0 | 1 | 1 |
| 1700036G14Rik | 0 | 0 | 0 | 1 | 1 |

| | | | | | |
|---|---|---|---|---|---|
| 1700039E15Rik | 0 | 0 | 0 | 1 | 1 |
| 1700041C23Rik | 0 | 0 | 0 | 1 | 1 |
| 1700041M19Rik | 0 | 0 | 0 | 1 | 1 |
| 1700042B14Rik | 0 | 0 | 0 | 1 | 1 |
| 1700042G07Rik | 0 | 0 | 0 | 1 | 1 |
| 1700042G15Rik | 0 | 0 | 0 | 1 | 1 |
| 1700044C05Rik | 0 | 0 | 0 | 1 | 1 |
| 1700044K03Rik | 0 | 0 | 0 | 1 | 1 |
| 1700045H11Rik | 0 | 0 | 0 | 1 | 1 |
| 1700046C09Rik | 0 | 0 | 0 | 1 | 1 |
| 1700047E10Rik | 0 | 0 | 0 | 1 | 1 |
| 1700047G03Rik | 0 | 0 | 0 | 1 | 1 |
| 1700047L14Rik | 0 | 0 | 0 | 1 | 1 |
| 1700047M11Rik | 0 | 0 | 0 | 1 | 1 |
| 1700049E22Rik | 0 | 0 | 0 | 1 | 1 |
| 1700052I22Rik | 0 | 0 | 0 | 1 | 1 |
| 1700054K19Rik | 0 | 0 | 0 | 1 | 1 |
| 1700054M17Rik | 0 | 0 | 0 | 1 | 1 |
| 1700054O13Rik | 0 | 0 | 0 | 1 | 1 |
| 1700055C04Rik | 0 | 0 | 0 | 1 | 1 |
| 1700055N04Rik | 0 | 0 | 0 | 1 | 1 |
| 1700057G04Rik | 0 | 0 | 0 | 1 | 1 |
| 1700060C16Rik | 0 | 0 | 0 | 1 | 1 |
| 1700061F12Rik | 0 | 0 | 0 | 1 | 1 |
| 1700061I17Rik | 0 | 0 | 0 | 1 | 1 |
| 1700063A18Rik | 0 | 0 | 0 | 1 | 1 |
| 1700063D05Rik | 0 | 0 | 0 | 1 | 1 |
| 1700063O14Rik | 0 | 0 | 0 | 1 | 1 |
| 1700064J06Rik | 0 | 0 | 0 | 1 | 1 |
| 1700065I16Rik | 0 | 0 | 0 | 1 | 1 |
| 1700065J11Rik | 0 | 0 | 0 | 1 | 1 |
| 1700065J18Rik | 0 | 0 | 0 | 1 | 1 |
| 1700065L07Rik | 0 | 0 | 0 | 1 | 1 |
| 1700065O20Rik | 0 | 0 | 0 | 1 | 1 |
| 1700066N21Rik | 0 | 0 | 0 | 1 | 1 |
| 1700066O22Rik | 0 | 0 | 0 | 1 | 1 |
| 1700067G17Rik | 0 | 0 | 0 | 1 | 1 |
| 1700067P10Rik | 0 | 0 | 0 | 1 | 1 |
| 1700069L16Rik | 0 | 0 | 0 | 1 | 1 |
| 1700069P05Rik | 0 | 0 | 0 | 1 | 1 |
| 1700072B07Rik | 0 | 0 | 0 | 1 | 1 |
| 1700072O05Rik | 0 | 0 | 0 | 1 | 1 |
| 1700074H08Rik | 0 | 0 | 0 | 1 | 1 |
| 1700074P13Rik | 0 | 0 | 0 | 1 | 1 |
| 1700080E11Rik | 0 | 0 | 0 | 1 | 1 |
| 1700080N15Rik | 0 | 0 | 0 | 1 | 1 |
| 1700080O16Rik | 0 | 0 | 0 | 1 | 1 |
| 1700081H04Rik | 0 | 0 | 0 | 1 | 1 |
| 1700084F23Rik | 0 | 0 | 0 | 1 | 1 |
| 1700085C21Rik | 0 | 0 | 0 | 1 | 1 |

| | | | | | |
|---|---|---|---|---|---|
| 1700091H14Rik | 0 | 0 | 0 | 1 | 1 |
| 1700092C02Rik | 0 | 0 | 0 | 1 | 1 |
| 1700092E19Rik | 0 | 0 | 0 | 1 | 1 |
| 1700092K14Rik | 0 | 0 | 0 | 1 | 1 |
| 1700093K21Rik | 0 | 0 | 0 | 1 | 1 |
| 1700094M24Rik | 0 | 0 | 0 | 1 | 1 |
| 1700095A21Rik | 0 | 0 | 0 | 1 | 1 |
| 1700097N02Rik | 0 | 0 | 0 | 1 | 1 |
| 1700101O22Rik | 0 | 0 | 0 | 1 | 1 |
| 1700109G15Rik | 0 | 0 | 0 | 1 | 1 |
| 1700109I08Rik | 0 | 0 | 0 | 1 | 1 |
| 1700110I01Rik | 0 | 0 | 0 | 1 | 1 |
| 1700110K17Rik | 0 | 0 | 0 | 1 | 1 |
| 1700111N16Rik | 0 | 0 | 0 | 1 | 1 |
| 1700112H15Rik | 0 | 0 | 0 | 1 | 1 |
| 1700120G07Rik | 0 | 0 | 0 | 1 | 1 |
| 1700121L16Rik | 0 | 0 | 0 | 1 | 1 |
| 1700121N20Rik | 0 | 0 | 0 | 1 | 1 |
| 1700122O11Rik | 0 | 0 | 0 | 1 | 1 |
| 1700123I01Rik | 0 | 0 | 0 | 1 | 1 |
| 1700123O12Rik | 0 | 0 | 0 | 1 | 1 |
| 1700123O21Rik | 0 | 0 | 0 | 1 | 1 |
| 1700125G02Rik | 0 | 0 | 0 | 1 | 1 |
| 1700125G22Rik | 0 | 0 | 0 | 1 | 1 |
| 1700125H03Rik | 0 | 0 | 0 | 1 | 1 |
| 1700126H18Rik | 0 | 0 | 0 | 1 | 1 |
| 1700128A07Rik | 0 | 0 | 0 | 1 | 1 |
| 1700128F08Rik | 0 | 0 | 0 | 1 | 1 |
| 1700129C05Rik | 0 | 0 | 0 | 1 | 1 |
| 1810007C17Rik | 0 | 0 | 0 | 1 | 1 |
| 1810007D17Rik | 0 | 0 | 0 | 1 | 1 |
| 1810009J06Rik | 0 | 0 | 0 | 1 | 1 |
| 1810011H11Rik | 0 | 0 | 0 | 1 | 1 |
| 1810018F18Rik | 0 | 0 | 0 | 1 | 1 |
| 1810046K07Rik | 0 | 0 | 0 | 1 | 1 |
| 1810053B23Rik | 0 | 0 | 0 | 1 | 1 |
| 1810064F22Rik | 0 | 0 | 0 | 1 | 1 |
| 1810065E05Rik | 0 | 0 | 0 | 1 | 1 |
| 2010003K11Rik | 0 | 0 | 0 | 1 | 1 |
| 2010016I18Rik | 0 | 0 | 0 | 1 | 1 |
| 2010106E10Rik | 0 | 0 | 0 | 1 | 1 |
| 2200002J24Rik | 0 | 0 | 0 | 1 | 1 |
| 2210010C04Rik | 0 | 0 | 0 | 1 | 1 |
| 2210407C18Rik | 0 | 0 | 0 | 1 | 1 |
| 2210408F21Rik | 0 | 0 | 0 | 1 | 1 |
| 2210409D07Rik | 0 | 0 | 0 | 1 | 1 |
| 2210409E12Rik | 0 | 0 | 0 | 1 | 1 |
| 2210414B05Rik | 0 | 0 | 0 | 1 | 1 |
| 2210417A02Rik | 0 | 0 | 0 | 1 | 1 |
| 2210420H20Rik | 0 | 0 | 0 | 1 | 1 |

| | | | | | |
|---|---|---|---|---|---|
| 2300002M23Rik | 0 | 0 | 0 | 1 | 1 |
| 2300003K06Rik | 0 | 0 | 0 | 1 | 1 |
| 2300005B03Rik | 0 | 0 | 0 | 1 | 1 |
| 2310002J15Rik | 0 | 0 | 0 | 1 | 1 |
| 2310002L09Rik | 0 | 0 | 0 | 1 | 1 |
| 2310008N11Rik | 0 | 0 | 0 | 1 | 1 |
| 2310016D03Rik | 0 | 0 | 0 | 1 | 1 |
| 2310020H05Rik | 0 | 0 | 0 | 1 | 1 |
| 2310030A07Rik | 0 | 0 | 0 | 1 | 1 |
| 2310034C09Rik | 0 | 0 | 0 | 1 | 1 |
| 2310039L15Rik | 0 | 0 | 0 | 1 | 1 |
| 2310042E22Rik | 0 | 0 | 0 | 1 | 1 |
| 2310043O21Rik | 0 | 0 | 0 | 1 | 1 |
| 2310050C09Rik | 0 | 0 | 0 | 1 | 1 |
| 2310057J18Rik | 0 | 0 | 0 | 1 | 1 |
| 2310057N15Rik | 0 | 0 | 0 | 1 | 1 |
| 2310061N02Rik | 0 | 0 | 0 | 1 | 1 |
| 2310065F04Rik | 0 | 0 | 0 | 1 | 1 |
| 2310069B03Rik | 0 | 0 | 0 | 1 | 1 |
| 2310079G19Rik | 0 | 0 | 0 | 1 | 1 |
| 2310081J21Rik | 0 | 0 | 0 | 1 | 1 |
| 2410003L11Rik | 0 | 0 | 0 | 1 | 1 |
| 2410007B07Rik | 0 | 0 | 0 | 1 | 1 |
| 2410012E07Rik | 0 | 0 | 0 | 1 | 1 |
| 2410017I17Rik | 0 | 0 | 0 | 1 | 1 |
| 2410088K16Rik | 0 | 0 | 0 | 1 | 1 |
| 2410114N07Rik | 0 | 0 | 0 | 1 | 1 |
| 2410124H12Rik | 0 | 0 | 0 | 1 | 1 |
| 2700089I24Rik | 0 | 0 | 0 | 1 | 1 |
| 2810007J24Rik | 0 | 0 | 0 | 1 | 1 |
| 2810049E08Rik | 0 | 0 | 0 | 1 | 1 |
| 2810404M03Rik | 0 | 0 | 0 | 1 | 1 |
| 2810471M01Rik | 0 | 0 | 0 | 1 | 1 |
| 2900055J20Rik | 0 | 0 | 0 | 1 | 1 |
| 2900057B20Rik | 0 | 0 | 0 | 1 | 1 |
| 2900079G21Rik | 0 | 0 | 0 | 1 | 1 |
| 2900092C05Rik | 0 | 0 | 0 | 1 | 1 |
| 3100003L05Rik | 0 | 0 | 0 | 1 | 1 |
| 3110009F21Rik | 0 | 0 | 0 | 1 | 1 |
| 3110015C05Rik | 0 | 0 | 0 | 1 | 1 |
| 3110079O15Rik | 0 | 0 | 0 | 1 | 1 |
| 3110082J24Rik | 0 | 0 | 0 | 1 | 1 |
| 3300005D01Rik | 0 | 0 | 0 | 1 | 1 |
| 4430402I18Rik | 0 | 0 | 0 | 1 | 1 |
| 4833423E24Rik | 0 | 0 | 0 | 1 | 1 |
| 4833427F10Rik | 0 | 0 | 0 | 1 | 1 |
| 4833428L15Rik | 0 | 0 | 0 | 1 | 1 |
| 4921501E09Rik | 0 | 0 | 0 | 1 | 1 |
| 4921504E06Rik | 0 | 0 | 0 | 1 | 1 |
| 4921508D12Rik | 0 | 0 | 0 | 1 | 1 |

| | | | | | |
|---|---|---|---|---|---|
| 4921509C19Rik | 0 | 0 | 0 | 1 | 1 |
| 4921509O07Rik | 0 | 0 | 0 | 1 | 1 |
| 4921511C20Rik | 0 | 0 | 0 | 1 | 1 |
| 4921511H03Rik | 0 | 0 | 0 | 1 | 1 |
| 4921511M17Rik,Gm10057 | 0 | 0 | 0 | 1 | 1 |
| 4921515E04Rik | 0 | 0 | 0 | 1 | 1 |
| 4921524L21Rik | 0 | 0 | 0 | 1 | 1 |
| 4921529L05Rik | 0 | 0 | 0 | 1 | 1 |
| 4921531P14Rik | 0 | 0 | 0 | 1 | 1 |
| 4921539E11Rik | 0 | 0 | 0 | 1 | 1 |
| 4922502D21Rik | 0 | 0 | 0 | 1 | 1 |
| 4922502H24Rik | 0 | 0 | 0 | 1 | 1 |
| 4922502N22Rik | 0 | 0 | 0 | 1 | 1 |
| 4930401C15Rik | 0 | 0 | 0 | 1 | 1 |
| 4930401O10Rik | 0 | 0 | 0 | 1 | 1 |
| 4930401O12Rik | 0 | 0 | 0 | 1 | 1 |
| 4930402F06Rik | 0 | 0 | 0 | 1 | 1 |
| 4930402F11Rik | 0 | 0 | 0 | 1 | 1 |
| 4930402K13Rik | 0 | 0 | 0 | 1 | 1 |
| 4930404A05Rik | 0 | 0 | 0 | 1 | 1 |
| 4930404A10Rik | 0 | 0 | 0 | 1 | 1 |
| 4930405A10Rik | 0 | 0 | 0 | 1 | 1 |
| 4930405D11Rik | 0 | 0 | 0 | 1 | 1 |
| 4930405J17Rik | 0 | 0 | 0 | 1 | 1 |
| 4930405L22Rik | 0 | 0 | 0 | 1 | 1 |
| 4930406D18Rik | 0 | 0 | 0 | 1 | 1 |
| 4930412B13Rik | 0 | 0 | 0 | 1 | 1 |
| 4930412D23Rik | 0 | 0 | 0 | 1 | 1 |
| 4930413E15Rik | 0 | 0 | 0 | 1 | 1 |
| 4930413M19Rik | 0 | 0 | 0 | 1 | 1 |
| 4930415L06Rik | 0 | 0 | 0 | 1 | 1 |
| 4930417O13Rik | 0 | 0 | 0 | 1 | 1 |
| 4930419G24Rik | 0 | 0 | 0 | 1 | 1 |
| 4930423M02Rik | 0 | 0 | 0 | 1 | 1 |
| 4930425K10Rik | 0 | 0 | 0 | 1 | 1 |
| 4930425O10Rik | 0 | 0 | 0 | 1 | 1 |
| 4930428D18Rik | 0 | 0 | 0 | 1 | 1 |
| 4930428E07Rik | 0 | 0 | 0 | 1 | 1 |
| 4930429F11Rik | 0 | 0 | 0 | 1 | 1 |
| 4930430D24Rik | 0 | 0 | 0 | 1 | 1 |
| 4930430F21Rik | 0 | 0 | 0 | 1 | 1 |
| 4930430J02Rik | 0 | 0 | 0 | 1 | 1 |
| 4930432M17Rik | 0 | 0 | 0 | 1 | 1 |
| 4930433B08Rik | 0 | 0 | 0 | 1 | 1 |
| 4930433I11Rik | 0 | 0 | 0 | 1 | 1 |
| 4930434J06Rik | 0 | 0 | 0 | 1 | 1 |
| 4930435E12Rik | 0 | 0 | 0 | 1 | 1 |
| 4930440C22Rik | 0 | 0 | 0 | 1 | 1 |
| 4930440I19Rik | 0 | 0 | 0 | 1 | 1 |
| 4930441J16Rik | 0 | 0 | 0 | 1 | 1 |

| | | | | | |
|---|---|---|---|---|---|
| 4930442J19Rik | 0 | 0 | 0 | 1 | 1 |
| 4930444F02Rik | 0 | 0 | 0 | 1 | 1 |
| 4930444G20Rik | 0 | 0 | 0 | 1 | 1 |
| 4930447A16Rik | 0 | 0 | 0 | 1 | 1 |
| 4930447J18Rik | 0 | 0 | 0 | 1 | 1 |
| 4930447K03Rik | 0 | 0 | 0 | 1 | 1 |
| 4930448C13Rik | 0 | 0 | 0 | 1 | 1 |
| 4930448F12Rik | 0 | 0 | 0 | 1 | 1 |
| 4930448H16Rik | 0 | 0 | 0 | 1 | 1 |
| 4930448I06Rik | 0 | 0 | 0 | 1 | 1 |
| 4930448I18Rik | 0 | 0 | 0 | 1 | 1 |
| 4930448K20Rik | 0 | 0 | 0 | 1 | 1 |
| 4930449E18Rik | 0 | 0 | 0 | 1 | 1 |
| 4930449I24Rik | 0 | 0 | 0 | 1 | 1 |
| 4930451I11Rik | 0 | 0 | 0 | 1 | 1 |
| 4930452A19Rik | 0 | 0 | 0 | 1 | 1 |
| 4930452N14Rik | 0 | 0 | 0 | 1 | 1 |
| 4930453H23Rik | 0 | 0 | 0 | 1 | 1 |
| 4930455B14Rik | 0 | 0 | 0 | 1 | 1 |
| 4930455F16Rik | 0 | 0 | 0 | 1 | 1 |
| 4930455H04Rik | 0 | 0 | 0 | 1 | 1 |
| 4930455J16Rik | 0 | 0 | 0 | 1 | 1 |
| 4930456L15Rik | 0 | 0 | 0 | 1 | 1 |
| 4930459C07Rik | 0 | 0 | 0 | 1 | 1 |
| 4930459L07Rik | 0 | 0 | 0 | 1 | 1 |
| 4930465M20Rik | 0 | 0 | 0 | 1 | 1 |
| 4930467D21Rik | 0 | 0 | 0 | 1 | 1 |
| 4930467E23Rik | 0 | 0 | 0 | 1 | 1 |
| 4930468A15Rik | 0 | 0 | 0 | 1 | 1 |
| 4930470H14Rik | 0 | 0 | 0 | 1 | 1 |
| 4930470P17Rik | 0 | 0 | 0 | 1 | 1 |
| 4930471C04Rik | 0 | 0 | 0 | 1 | 1 |
| 4930471G03Rik | 0 | 0 | 0 | 1 | 1 |
| 4930473O22Rik | 0 | 0 | 0 | 1 | 1 |
| 4930474G06Rik | 0 | 0 | 0 | 1 | 1 |
| 4930474H20Rik | 0 | 0 | 0 | 1 | 1 |
| 4930474N05Rik | 0 | 0 | 0 | 1 | 1 |
| 4930478P22Rik | 0 | 0 | 0 | 1 | 1 |
| 4930480E11Rik | 0 | 0 | 0 | 1 | 1 |
| 4930480G23Rik | 0 | 0 | 0 | 1 | 1 |
| 4930480M12Rik | 0 | 0 | 0 | 1 | 1 |
| 4930482G09Rik | 0 | 0 | 0 | 1 | 1 |
| 4930486F22Rik | 0 | 0 | 0 | 1 | 1 |
| 4930486I03Rik | 0 | 0 | 0 | 1 | 1 |
| 4930486L24Rik | 0 | 0 | 0 | 1 | 1 |
| 4930487D11Rik | 0 | 0 | 0 | 1 | 1 |
| 4930488B22Rik | 0 | 0 | 0 | 1 | 1 |
| 4930500F04Rik | 0 | 0 | 0 | 1 | 1 |
| 4930502A04Rik | 0 | 0 | 0 | 1 | 1 |
| 4930503E14Rik | 0 | 0 | 0 | 1 | 1 |

| | | | | | |
|---|---|---|---|---|---|
| 4930503H13Rik | 0 | 0 | 0 | 1 | 1 |
| 4930503O07Rik | 0 | 0 | 0 | 1 | 1 |
| 4930504O13Rik | 0 | 0 | 0 | 1 | 1 |
| 4930505A04Rik | 0 | 0 | 0 | 1 | 1 |
| 4930505G20Rik | 0 | 0 | 0 | 1 | 1 |
| 4930507D10Rik | 0 | 0 | 0 | 1 | 1 |
| 4930509J09Rik | 0 | 0 | 0 | 1 | 1 |
| 4930509K18Rik | 0 | 0 | 0 | 1 | 1 |
| 4930513O06Rik | 0 | 0 | 0 | 1 | 1 |
| 4930515B02Rik | 0 | 0 | 0 | 1 | 1 |
| 4930515G16Rik | 0 | 0 | 0 | 1 | 1 |
| 4930515L19Rik | 0 | 0 | 0 | 1 | 1 |
| 4930518P08Rik | 0 | 0 | 0 | 1 | 1 |
| 4930519D14Rik | 0 | 0 | 0 | 1 | 1 |
| 4930519F24Rik | 0 | 0 | 0 | 1 | 1 |
| 4930519H02Rik | 0 | 0 | 0 | 1 | 1 |
| 4930520P13Rik | 0 | 0 | 0 | 1 | 1 |
| 4930522H14Rik | 0 | 0 | 0 | 1 | 1 |
| 4930522O17Rik | 0 | 0 | 0 | 1 | 1 |
| 4930523O13Rik | 0 | 0 | 0 | 1 | 1 |
| 4930524N10Rik | 0 | 0 | 0 | 1 | 1 |
| 4930525M21Rik | 0 | 0 | 0 | 1 | 1 |
| 4930526L06Rik | 0 | 0 | 0 | 1 | 1 |
| 4930527F14Rik | 0 | 0 | 0 | 1 | 1 |
| 4930527G23Rik | 0 | 0 | 0 | 1 | 1 |
| 4930528P14Rik | 0 | 0 | 0 | 1 | 1 |
| 4930529K09Rik | 0 | 0 | 0 | 1 | 1 |
| 4930529L06Rik | 0 | 0 | 0 | 1 | 1 |
| 4930532M18Rik | 0 | 0 | 0 | 1 | 1 |
| 4930533B01Rik | 0 | 0 | 0 | 1 | 1 |
| 4930533P14Rik | 0 | 0 | 0 | 1 | 1 |
| 4930539C22Rik | 0 | 0 | 0 | 1 | 1 |
| 4930539M17Rik | 0 | 0 | 0 | 1 | 1 |
| 4930539N22Rik | 0 | 0 | 0 | 1 | 1 |
| 4930540M03Rik | 0 | 0 | 0 | 1 | 1 |
| 4930543E12Rik | 0 | 0 | 0 | 1 | 1 |
| 4930544D05Rik | 0 | 0 | 0 | 1 | 1 |
| 4930544G11Rik | 0 | 0 | 0 | 1 | 1 |
| 4930544M13Rik | 0 | 0 | 0 | 1 | 1 |
| 4930545H06Rik | 0 | 0 | 0 | 1 | 1 |
| 4930545L23Rik | 0 | 0 | 0 | 1 | 1 |
| 4930546C10Rik | 0 | 0 | 0 | 1 | 1 |
| 4930548G14Rik | 0 | 0 | 0 | 1 | 1 |
| 4930548J01Rik | 0 | 0 | 0 | 1 | 1 |
| 4930548K13Rik | 0 | 0 | 0 | 1 | 1 |
| 4930549C01Rik | 0 | 0 | 0 | 1 | 1 |
| 4930550L24Rik | 0 | 0 | 0 | 1 | 1 |
| 4930552N02Rik | 0 | 0 | 0 | 1 | 1 |
| 4930553E22Rik | 0 | 0 | 0 | 1 | 1 |
| 4930554C24Rik | 0 | 0 | 0 | 1 | 1 |

| | | | | | |
|---|---|---|---|---|---|
| 4930555B11Rik | 0 | 0 | 0 | 1 | 1 |
| 4930555G01Rik | 0 | 0 | 0 | 1 | 1 |
| 4930556C24Rik | 0 | 0 | 0 | 1 | 1 |
| 4930556G01Rik | 0 | 0 | 0 | 1 | 1 |
| 4930556J02Rik | 0 | 0 | 0 | 1 | 1 |
| 4930556N09Rik | 0 | 0 | 0 | 1 | 1 |
| 4930557A04Rik | 0 | 0 | 0 | 1 | 1 |
| 4930558G05Rik | 0 | 0 | 0 | 1 | 1 |
| 4930558K02Rik | 0 | 0 | 0 | 1 | 1 |
| 4930562F07Rik | 0 | 0 | 0 | 1 | 1 |
| 4930563F08Rik | 0 | 0 | 0 | 1 | 1 |
| 4930563M20Rik | 0 | 0 | 0 | 1 | 1 |
| 4930564B18Rik | 0 | 0 | 0 | 1 | 1 |
| 4930564D02Rik | 0 | 0 | 0 | 1 | 1 |
| 4930565D16Rik | 0 | 0 | 0 | 1 | 1 |
| 4930567H12Rik | 0 | 0 | 0 | 1 | 1 |
| 4930567H17Rik | 0 | 0 | 0 | 1 | 1 |
| 4930567K20Rik | 0 | 0 | 0 | 1 | 1 |
| 4930568D16Rik | 0 | 0 | 0 | 1 | 1 |
| 4930568E12Rik | 0 | 0 | 0 | 1 | 1 |
| 4930571O06Rik | 0 | 0 | 0 | 1 | 1 |
| 4930572K03Rik | 0 | 0 | 0 | 1 | 1 |
| 4930572O03Rik | 0 | 0 | 0 | 1 | 1 |
| 4930572O13Rik | 0 | 0 | 0 | 1 | 1 |
| 4930573O16Rik | 0 | 0 | 0 | 1 | 1 |
| 4930578C19Rik | 0 | 0 | 0 | 1 | 1 |
| 4930578E11Rik | 0 | 0 | 0 | 1 | 1 |
| 4930578I06Rik | 0 | 0 | 0 | 1 | 1 |
| 4930578N18Rik | 0 | 0 | 0 | 1 | 1 |
| 4930579F01Rik | 0 | 0 | 0 | 1 | 1 |
| 4930583K01Rik | 0 | 0 | 0 | 1 | 1 |
| 4930583P06Rik | 0 | 0 | 0 | 1 | 1 |
| 4930584F24Rik | 0 | 0 | 0 | 1 | 1 |
| 4930591A17Rik | 0 | 0 | 0 | 1 | 1 |
| 4930592A05Rik | 0 | 0 | 0 | 1 | 1 |
| 4930593C16Rik | 0 | 0 | 0 | 1 | 1 |
| 4930595M18Rik | 0 | 0 | 0 | 1 | 1 |
| 4930596D02Rik | 0 | 0 | 0 | 1 | 1 |
| 4930597G03Rik | 0 | 0 | 0 | 1 | 1 |
| 4930598F16Rik | 0 | 0 | 0 | 1 | 1 |
| 4931406B18Rik | 0 | 0 | 0 | 1 | 1 |
| 4931408C20Rik | 0 | 0 | 0 | 1 | 1 |
| 4931412M21 | 0 | 0 | 0 | 1 | 1 |
| 4931417E11Rik | 0 | 0 | 0 | 1 | 1 |
| 4931419H13Rik | 0 | 0 | 0 | 1 | 1 |
| 4931429P17Rik | 0 | 0 | 0 | 1 | 1 |
| 4931431B13Rik | 0 | 0 | 0 | 1 | 1 |
| 4931431F19Rik | 0 | 0 | 0 | 1 | 1 |
| 4932411N23Rik | 0 | 0 | 0 | 1 | 1 |
| 4932412D23Rik | 0 | 0 | 0 | 1 | 1 |

| | | | | | |
|---|---|---|---|---|---|
| 4932414J04Rik | 0 | 0 | 0 | 1 | 1 |
| 4932414N04Rik | 0 | 0 | 0 | 1 | 1 |
| 4932416K20Rik | 0 | 0 | 0 | 1 | 1 |
| 4932429P05Rik | 0 | 0 | 0 | 1 | 1 |
| 4932443I19Rik | 0 | 0 | 0 | 1 | 1 |
| 4933400A11Rik | 0 | 0 | 0 | 1 | 1 |
| 4933400B14Rik | 0 | 0 | 0 | 1 | 1 |
| 4933400L20Rik | 0 | 0 | 0 | 1 | 1 |
| 4933401B06Rik | 0 | 0 | 0 | 1 | 1 |
| 4933401H06Rik | 0 | 0 | 0 | 1 | 1 |
| 4933402C06Rik | 0 | 0 | 0 | 1 | 1 |
| 4933402E13Rik | 0 | 0 | 0 | 1 | 1 |
| 4933402J07Rik | 0 | 0 | 0 | 1 | 1 |
| 4933402J10Rik | 0 | 0 | 0 | 1 | 1 |
| 4933402J15Rik | 0 | 0 | 0 | 1 | 1 |
| 4933402N03Rik | 0 | 0 | 0 | 1 | 1 |
| 4933402N22Rik | 0 | 0 | 0 | 1 | 1 |
| 4933402P03Rik | 0 | 0 | 0 | 1 | 1 |
| 4933404G15Rik | 0 | 0 | 0 | 1 | 1 |
| 4933404K08Rik | 0 | 0 | 0 | 1 | 1 |
| 4933405E24Rik | 0 | 0 | 0 | 1 | 1 |
| 4933405O20Rik | 0 | 0 | 0 | 1 | 1 |
| 4933406D12Rik | 0 | 0 | 0 | 1 | 1 |
| 4933406F09Rik | 0 | 0 | 0 | 1 | 1 |
| 4933406J08Rik | 0 | 0 | 0 | 1 | 1 |
| 4933406K04Rik | 0 | 0 | 0 | 1 | 1 |
| 4933407E24Rik | 0 | 0 | 0 | 1 | 1 |
| 4933407G14Rik | 0 | 0 | 0 | 1 | 1 |
| 4933407I05Rik | 0 | 0 | 0 | 1 | 1 |
| 4933409G03Rik | 0 | 0 | 0 | 1 | 1 |
| 4933409K07Rik | 0 | 0 | 0 | 1 | 1 |
| 4933411G06Rik | 0 | 0 | 0 | 1 | 1 |
| 4933411G11Rik | 0 | 0 | 0 | 1 | 1 |
| 4933412E24Rik | 0 | 0 | 0 | 1 | 1 |
| 4933412O06Rik | 0 | 0 | 0 | 1 | 1 |
| 4933413L06Rik | 0 | 0 | 0 | 1 | 1 |
| 4933415F23Rik | 0 | 0 | 0 | 1 | 1 |
| 4933416C03Rik | 0 | 0 | 0 | 1 | 1 |
| 4933416E03Rik | 0 | 0 | 0 | 1 | 1 |
| 4933416M06Rik | 0 | 0 | 0 | 1 | 1 |
| 4933417A18Rik | 0 | 0 | 0 | 1 | 1 |
| 4933421I07Rik | 0 | 0 | 0 | 1 | 1 |
| 4933425L06Rik | 0 | 0 | 0 | 1 | 1 |
| 4933428C19Rik | 0 | 0 | 0 | 1 | 1 |
| 4933429O19Rik | 0 | 0 | 0 | 1 | 1 |
| 4933430M04Rik | 0 | 0 | 0 | 1 | 1 |
| 4933430N04Rik | 0 | 0 | 0 | 1 | 1 |
| 4933432G23Rik | 0 | 0 | 0 | 1 | 1 |
| 4933432I09Rik | 0 | 0 | 0 | 1 | 1 |
| 4933432K03Rik | 0 | 0 | 0 | 1 | 1 |

| | | | | | |
|---|---|---|---|---|---|
| 4933433G08Rik | 0 | 0 | 0 | 1 | 1 |
| 4933433H22Rik | 0 | 0 | 0 | 1 | 1 |
| 4933434I20Rik | 0 | 0 | 0 | 1 | 1 |
| 4933436E23Rik | 0 | 0 | 0 | 1 | 1 |
| 4933436H12Rik | 0 | 0 | 0 | 1 | 1 |
| 4933436I01Rik | 0 | 0 | 0 | 1 | 1 |
| 4933438B17Rik | 0 | 0 | 0 | 1 | 1 |
| 4933438K21Rik | 0 | 0 | 0 | 1 | 1 |
| 4933440J02Rik | 0 | 0 | 0 | 1 | 1 |
| 5031410I06Rik | 0 | 0 | 0 | 1 | 1 |
| 5031434C07Rik | 0 | 0 | 0 | 1 | 1 |
| 5033403H07Rik | 0 | 0 | 0 | 1 | 1 |
| 5133400J02Rik | 0 | 0 | 0 | 1 | 1 |
| 5330439B14Rik | 0 | 0 | 0 | 1 | 1 |
| 5430401F13Rik | 0 | 0 | 0 | 1 | 1 |
| 5430402E10Rik | 0 | 0 | 0 | 1 | 1 |
| 5430403N17Rik | 0 | 0 | 0 | 1 | 1 |
| 5430416O09Rik | 0 | 0 | 0 | 1 | 1 |
| 5430419D17Rik | 0 | 0 | 0 | 1 | 1 |
| 5430421F17Rik | 0 | 0 | 0 | 1 | 1 |
| 5430425K12Rik | 0 | 0 | 0 | 1 | 1 |
| 5430427O19Rik | 0 | 0 | 0 | 1 | 1 |
| 5430434I15Rik | 0 | 0 | 0 | 1 | 1 |
| 5430437J10Rik | 0 | 0 | 0 | 1 | 1 |
| 5530400C23Rik | 0 | 0 | 0 | 1 | 1 |
| 5530401A14Rik | 0 | 0 | 0 | 1 | 1 |
| 5730412P04Rik | 0 | 0 | 0 | 1 | 1 |
| 5730460C07Rik | 0 | 0 | 0 | 1 | 1 |
| 5830411N06Rik | 0 | 0 | 0 | 1 | 1 |
| 5830416I19Rik | 0 | 0 | 0 | 1 | 1 |
| 5830428M24Rik | 0 | 0 | 0 | 1 | 1 |
| 5830432E09Rik | 0 | 0 | 0 | 1 | 1 |
| 5930438M14Rik | 0 | 0 | 0 | 1 | 1 |
| 6030469F06Rik | 0 | 0 | 0 | 1 | 1 |
| 6030498E09Rik | 0 | 0 | 0 | 1 | 1 |
| 6330407A03Rik | 0 | 0 | 0 | 1 | 1 |
| 6430503K07Rik | 0 | 0 | 0 | 1 | 1 |
| 6430710C18Rik | 0 | 0 | 0 | 1 | 1 |
| 7420426K07Rik | 0 | 0 | 0 | 1 | 1 |
| 7420461P10Rik | 0 | 0 | 0 | 1 | 1 |
| 7420700N18Rik | 0 | 0 | 0 | 1 | 1 |
| 7420701I03Rik | 0 | 0 | 0 | 1 | 1 |
| 7530416G11Rik | 0 | 0 | 0 | 1 | 1 |
| 7630403G23Rik | 0 | 0 | 0 | 1 | 1 |
| 8030411F24Rik | 0 | 0 | 0 | 1 | 1 |
| 8030423J24Rik | 0 | 0 | 0 | 1 | 1 |
| 8030442B05Rik | 0 | 0 | 0 | 1 | 1 |
| 8430437L04Rik | 0 | 0 | 0 | 1 | 1 |
| 9030204H09Rik | 0 | 0 | 0 | 1 | 1 |
| 9030619P08Rik | 0 | 0 | 0 | 1 | 1 |

| | | | | | |
|---|---|---|---|---|---|
| 9130015L21Rik | 0 | 0 | 0 | 1 | 1 |
| 9130204L05Rik | 0 | 0 | 0 | 1 | 1 |
| 9130209A04Rik | 0 | 0 | 0 | 1 | 1 |
| 9130221F21Rik | 0 | 0 | 0 | 1 | 1 |
| 9130227L01Rik | 0 | 0 | 0 | 1 | 1 |
| 9130230L23Rik | 0 | 0 | 0 | 1 | 1 |
| 9230009I02Rik | 0 | 0 | 0 | 1 | 1 |
| 9230104L09Rik | 0 | 0 | 0 | 1 | 1 |
| 9230110F15Rik | 0 | 0 | 0 | 1 | 1 |
| 9330111N05Rik | 0 | 0 | 0 | 1 | 1 |
| 9330158H04Rik | 0 | 0 | 0 | 1 | 1 |
| 9330162B11Rik | 0 | 0 | 0 | 1 | 1 |
| 9330175E14Rik | 0 | 0 | 0 | 1 | 1 |
| 9330175M20Rik | 0 | 0 | 0 | 1 | 1 |
| 9330178D15Rik | 0 | 0 | 0 | 1 | 1 |
| 9330182O14Rik | 0 | 0 | 0 | 1 | 1 |
| 9330188P03Rik | 0 | 0 | 0 | 1 | 1 |
| 9430007A20Rik | 0 | 0 | 0 | 1 | 1 |
| 9430041J12Rik | 0 | 0 | 0 | 1 | 1 |
| 9530002B09Rik | 0 | 0 | 0 | 1 | 1 |
| 9530036O11Rik | 0 | 0 | 0 | 1 | 1 |
| 9530080O11Rik | 0 | 0 | 0 | 1 | 1 |
| 9630013A20Rik | 0 | 0 | 0 | 1 | 1 |
| 9630028H03Rik | 0 | 0 | 0 | 1 | 1 |
| 9830107B12Rik | 0 | 0 | 0 | 1 | 1 |
| 9830132P13Rik | 0 | 0 | 0 | 1 | 1 |
| 9930111H07Rik | 0 | 0 | 0 | 1 | 1 |
| 9930111J21Rik2 | 0 | 0 | 0 | 1 | 1 |
| A130077B15Rik | 0 | 0 | 0 | 1 | 1 |
| A1bg | 0 | 0 | 0 | 1 | 1 |
| A1cf | 0 | 0 | 0 | 1 | 1 |
| A230001M10Rik | 0 | 0 | 0 | 1 | 1 |
| A330049N07Rik | 0 | 0 | 0 | 1 | 1 |
| A430078G23Rik | 0 | 0 | 0 | 1 | 1 |
| A430089I19Rik | 0 | 0 | 0 | 1 | 1 |
| A430107P09Rik | 0 | 0 | 0 | 1 | 1 |
| A4gnt | 0 | 0 | 0 | 1 | 1 |
| A530013C23Rik | 0 | 0 | 0 | 1 | 1 |
| A530032D15Rik | 0 | 0 | 0 | 1 | 1 |
| A530050N04Rik | 0 | 0 | 0 | 1 | 1 |
| A530053G22Rik | 0 | 0 | 0 | 1 | 1 |
| A530058N18Rik | 0 | 0 | 0 | 1 | 1 |
| A530064D06Rik | 0 | 0 | 0 | 1 | 1 |
| A530099J19Rik | 0 | 0 | 0 | 1 | 1 |
| A630010A05Rik | 0 | 0 | 0 | 1 | 1 |
| A630023A22Rik | 0 | 0 | 0 | 1 | 1 |
| A630023P12Rik | 0 | 0 | 0 | 1 | 1 |
| A630033H20Rik | 0 | 0 | 0 | 1 | 1 |
| A630073D07Rik | 0 | 0 | 0 | 1 | 1 |
| A630077J23Rik | 0 | 0 | 0 | 1 | 1 |

| | | | | | |
|---|---|---|---|---|---|
| A630095E13Rik | 0 | 0 | 0 | 1 | 1 |
| A630095N17Rik | 0 | 0 | 0 | 1 | 1 |
| A730006G06Rik | 0 | 0 | 0 | 1 | 1 |
| A730018C14Rik | 0 | 0 | 0 | 1 | 1 |
| A730046J19Rik | 0 | 0 | 0 | 1 | 1 |
| A730082K24Rik | 0 | 0 | 0 | 1 | 1 |
| A730090H04Rik | 0 | 0 | 0 | 1 | 1 |
| A830019L24Rik | 0 | 0 | 0 | 1 | 1 |
| A930006I01Rik | 0 | 0 | 0 | 1 | 1 |
| A930009A15Rik | 0 | 0 | 0 | 1 | 1 |
| A930018P22Rik | 0 | 0 | 0 | 1 | 1 |
| AA536875 | 0 | 0 | 0 | 1 | 1 |
| AA545190 | 0 | 0 | 0 | 1 | 1 |
| AA619741 | 0 | 0 | 0 | 1 | 1 |
| AA792892 | 0 | 0 | 0 | 1 | 1 |
| AB041803 | 0 | 0 | 0 | 1 | 1 |
| AF067061 | 0 | 0 | 0 | 1 | 1 |
| AF067063 | 0 | 0 | 0 | 1 | 1 |
| AF251705 | 0 | 0 | 0 | 1 | 1 |
| AF357359 | 0 | 0 | 0 | 1 | 1 |
| AF357425 | 0 | 0 | 0 | 1 | 1 |
| AF357426 | 0 | 0 | 0 | 1 | 1 |
| AF366264 | 0 | 0 | 0 | 1 | 1 |
| AI115009 | 0 | 0 | 0 | 1 | 1 |
| AI427809 | 0 | 0 | 0 | 1 | 1 |
| AI606473 | 0 | 0 | 0 | 1 | 1 |
| AI607873 | 0 | 0 | 0 | 1 | 1 |
| AI747448 | 0 | 0 | 0 | 1 | 1 |
| AI847159 | 0 | 0 | 0 | 1 | 1 |
| AU015791 | 0 | 0 | 0 | 1 | 1 |
| AU016765 | 0 | 0 | 0 | 1 | 1 |
| AU018829,Gm16367 | 0 | 0 | 0 | 1 | 1 |
| AU019990 | 0 | 0 | 0 | 1 | 1 |
| AU021092 | 0 | 0 | 0 | 1 | 1 |
| AU023762 | 0 | 0 | 0 | 1 | 1 |
| AV320801,Gm15023 | 0 | 0 | 0 | 1 | 1 |
| AW112010 | 0 | 0 | 0 | 1 | 1 |
| AY512915 | 0 | 0 | 0 | 1 | 1 |
| AY761184 | 0 | 0 | 0 | 1 | 1 |
| AY761185 | 0 | 0 | 0 | 1 | 1 |
| Aadac | 0 | 0 | 0 | 1 | 1 |
| Aadacl2 | 0 | 0 | 0 | 1 | 1 |
| Aadacl3 | 0 | 0 | 0 | 1 | 1 |
| Abca15 | 0 | 0 | 0 | 1 | 1 |
| Abca16 | 0 | 0 | 0 | 1 | 1 |
| Abca17 | 0 | 0 | 0 | 1 | 1 |
| Abhd12b | 0 | 0 | 0 | 1 | 1 |
| Abo | 0 | 0 | 0 | 1 | 1 |
| Abra | 0 | 0 | 0 | 1 | 1 |
| Accsl | 0 | 0 | 0 | 1 | 1 |

| | | | | | |
|---|---|---|---|---|---|
| Ace3 | 0 | 0 | 0 | 1 | 1 |
| Acnat2 | 0 | 0 | 0 | 1 | 1 |
| Acot5 | 0 | 0 | 0 | 1 | 1 |
| Acox2 | 0 | 0 | 0 | 1 | 1 |
| Acrv1 | 0 | 0 | 0 | 1 | 1 |
| Acsbg2 | 0 | 0 | 0 | 1 | 1 |
| Acsm1 | 0 | 0 | 0 | 1 | 1 |
| Acsm5 | 0 | 0 | 0 | 1 | 1 |
| Actbl2 | 0 | 0 | 0 | 1 | 1 |
| Actl6b | 0 | 0 | 0 | 1 | 1 |
| Actrt1 | 0 | 0 | 0 | 1 | 1 |
| Actrt2 | 0 | 0 | 0 | 1 | 1 |
| Adam20 | 0 | 0 | 0 | 1 | 1 |
| Adam24 | 0 | 0 | 0 | 1 | 1 |
| Adam25 | 0 | 0 | 0 | 1 | 1 |
| Adam26b | 0 | 0 | 0 | 1 | 1 |
| Adam29 | 0 | 0 | 0 | 1 | 1 |
| Adam3 | 0 | 0 | 0 | 1 | 1 |
| Adam30 | 0 | 0 | 0 | 1 | 1 |
| Adam34 | 0 | 0 | 0 | 1 | 1 |
| Adam39 | 0 | 0 | 0 | 1 | 1 |
| Adam6a | 0 | 0 | 0 | 1 | 1 |
| Adam6b | 0 | 0 | 0 | 1 | 1 |
| Adora3 | 0 | 0 | 0 | 1 | 1 |
| Adtrp | 0 | 0 | 0 | 1 | 1 |
| Agbl4 | 0 | 0 | 0 | 1 | 1 |
| Agmat | 0 | 0 | 0 | 1 | 1 |
| Agmo | 0 | 0 | 0 | 1 | 1 |
| Agr3 | 0 | 0 | 0 | 1 | 1 |
| Agt | 0 | 0 | 0 | 1 | 1 |
| Agxt | 0 | 0 | 0 | 1 | 1 |
| Agxt2 | 0 | 0 | 0 | 1 | 1 |
| Akap4 | 0 | 0 | 0 | 1 | 1 |
| Akr1c20 | 0 | 0 | 0 | 1 | 1 |
| Akr1c6 | 0 | 0 | 0 | 1 | 1 |
| Akr1cl | 0 | 0 | 0 | 1 | 1 |
| Alb | 0 | 0 | 0 | 1 | 1 |
| Allc | 0 | 0 | 0 | 1 | 1 |
| Alox12e | 0 | 0 | 0 | 1 | 1 |
| Alpi | 0 | 0 | 0 | 1 | 1 |
| Alppl2 | 0 | 0 | 0 | 1 | 1 |
| Ambn | 0 | 0 | 0 | 1 | 1 |
| Amelx | 0 | 0 | 0 | 1 | 1 |
| Amer3 | 0 | 0 | 0 | 1 | 1 |
| Ampd1 | 0 | 0 | 0 | 1 | 1 |
| Amtn | 0 | 0 | 0 | 1 | 1 |
| Amy2a4,Amy2a5 | 0 | 0 | 0 | 1 | 1 |
| Amy2a5 | 0 | 0 | 0 | 1 | 1 |
| Amy2b | 0 | 0 | 0 | 1 | 1 |
| Ang2 | 0 | 0 | 0 | 1 | 1 |

| | | | | | |
|---|---|---|---|---|---|
| Ang3,Ang5 | 0 | 0 | 0 | 1 | 1 |
| Ang4 | 0 | 0 | 0 | 1 | 1 |
| Ang5 | 0 | 0 | 0 | 1 | 1 |
| Ang6 | 0 | 0 | 0 | 1 | 1 |
| Ankk1 | 0 | 0 | 0 | 1 | 1 |
| Ankrd2 | 0 | 0 | 0 | 1 | 1 |
| Ankrd33 | 0 | 0 | 0 | 1 | 1 |
| Ankrd34c | 0 | 0 | 0 | 1 | 1 |
| Ankrd36 | 0 | 0 | 0 | 1 | 1 |
| Anks4b | 0 | 0 | 0 | 1 | 1 |
| Ankub1 | 0 | 0 | 0 | 1 | 1 |
| Antxrl | 0 | 0 | 0 | 1 | 1 |
| Anxa10 | 0 | 0 | 0 | 1 | 1 |
| Anxa13 | 0 | 0 | 0 | 1 | 1 |
| Aoah | 0 | 0 | 0 | 1 | 1 |
| Aoc1 | 0 | 0 | 0 | 1 | 1 |
| Apcs | 0 | 0 | 0 | 1 | 1 |
| Apoa4 | 0 | 0 | 0 | 1 | 1 |
| Apobec4 | 0 | 0 | 0 | 1 | 1 |
| Apoc2 | 0 | 0 | 0 | 1 | 1 |
| Apoc3 | 0 | 0 | 0 | 1 | 1 |
| Apoc4 | 0 | 0 | 0 | 1 | 1 |
| Apof | 0 | 0 | 0 | 1 | 1 |
| Apoh | 0 | 0 | 0 | 1 | 1 |
| Apol10a | 0 | 0 | 0 | 1 | 1 |
| Apol11a | 0 | 0 | 0 | 1 | 1 |
| Apol11b | 0 | 0 | 0 | 1 | 1 |
| Apol7a | 0 | 0 | 0 | 1 | 1 |
| Apol7b | 0 | 0 | 0 | 1 | 1 |
| Apol7c | 0 | 0 | 0 | 1 | 1 |
| Apol7e | 0 | 0 | 0 | 1 | 1 |
| Apol9a | 0 | 0 | 0 | 1 | 1 |
| Apol9b | 0 | 0 | 0 | 1 | 1 |
| Apoo-ps | 0 | 0 | 0 | 1 | 1 |
| Aqp2 | 0 | 0 | 0 | 1 | 1 |
| Aqp6 | 0 | 0 | 0 | 1 | 1 |
| Arhgap15 | 0 | 0 | 0 | 1 | 1 |
| Arhgap15os | 0 | 0 | 0 | 1 | 1 |
| Arhgef38 | 0 | 0 | 0 | 1 | 1 |
| Arl14 | 0 | 0 | 0 | 1 | 1 |
| Arl14epl | 0 | 0 | 0 | 1 | 1 |
| Armc12 | 0 | 0 | 0 | 1 | 1 |
| Art2a-ps | 0 | 0 | 0 | 1 | 1 |
| Art2b | 0 | 0 | 0 | 1 | 1 |
| Asb11 | 0 | 0 | 0 | 1 | 1 |
| Asb15 | 0 | 0 | 0 | 1 | 1 |
| Asb18 | 0 | 0 | 0 | 1 | 1 |
| Asb9 | 0 | 0 | 0 | 1 | 1 |
| Ascl3 | 0 | 0 | 0 | 1 | 1 |
| Asic5 | 0 | 0 | 0 | 1 | 1 |

| | | | | | |
|---|---|---|---|---|---|
| Asmt | 0 | 0 | 0 | 1 | 1 |
| Asz1 | 0 | 0 | 0 | 1 | 1 |
| Atoh7 | 0 | 0 | 0 | 1 | 1 |
| Atp13a4 | 0 | 0 | 0 | 1 | 1 |
| Atp6ap1l | 0 | 0 | 0 | 1 | 1 |
| Atp6v0c-ps2 | 0 | 0 | 0 | 1 | 1 |
| Aurkc | 0 | 0 | 0 | 1 | 1 |
| Avp | 0 | 0 | 0 | 1 | 1 |
| Awat1 | 0 | 0 | 0 | 1 | 1 |
| Aym1 | 0 | 0 | 0 | 1 | 1 |
| Azgp1 | 0 | 0 | 0 | 1 | 1 |
| B020004C17Rik | 0 | 0 | 0 | 1 | 1 |
| B020004J07Rik | 0 | 0 | 0 | 1 | 1 |
| B020031M17Rik | 0 | 0 | 0 | 1 | 1 |
| B230112J18Rik | 0 | 0 | 0 | 1 | 1 |
| B230209E15Rik | 0 | 0 | 0 | 1 | 1 |
| B430010I23Rik | 0 | 0 | 0 | 1 | 1 |
| B430306N03Rik | 0 | 0 | 0 | 1 | 1 |
| B930018H19Rik | 0 | 0 | 0 | 1 | 1 |
| B930025P03Rik | 0 | 0 | 0 | 1 | 1 |
| BB014433 | 0 | 0 | 0 | 1 | 1 |
| BB019430 | 0 | 0 | 0 | 1 | 1 |
| BB123696 | 0 | 0 | 0 | 1 | 1 |
| BB287469 | 0 | 0 | 0 | 1 | 1 |
| BC018473 | 0 | 0 | 0 | 1 | 1 |
| BC021614 | 0 | 0 | 0 | 1 | 1 |
| BC024386 | 0 | 0 | 0 | 1 | 1 |
| BC048562 | 0 | 0 | 0 | 1 | 1 |
| BC048602 | 0 | 0 | 0 | 1 | 1 |
| BC048671 | 0 | 0 | 0 | 1 | 1 |
| BC048679 | 0 | 0 | 0 | 1 | 1 |
| BC049352 | 0 | 0 | 0 | 1 | 1 |
| BC049730 | 0 | 0 | 0 | 1 | 1 |
| BC051665 | 0 | 0 | 0 | 1 | 1 |
| BC052688 | 0 | 0 | 0 | 1 | 1 |
| BC053393 | 0 | 0 | 0 | 1 | 1 |
| BC055111 | 0 | 0 | 0 | 1 | 1 |
| BC055402 | 0 | 0 | 0 | 1 | 1 |
| BC061195 | 0 | 0 | 0 | 1 | 1 |
| BC061212 | 0 | 0 | 0 | 1 | 1 |
| BC061237 | 0 | 0 | 0 | 1 | 1 |
| BC080695 | 0 | 0 | 0 | 1 | 1 |
| BC094916 | 0 | 0 | 0 | 1 | 1 |
| BC100530 | 0 | 0 | 0 | 1 | 1 |
| BC117090 | 0 | 0 | 0 | 1 | 1 |
| BC147527 | 0 | 0 | 0 | 1 | 1 |
| Bank1 | 0 | 0 | 0 | 1 | 1 |
| Batf2 | 0 | 0 | 0 | 1 | 1 |
| Bc1 | 0 | 0 | 0 | 1 | 1 |
| Bcas1os2 | 0 | 0 | 0 | 1 | 1 |

| | | | | | |
|---|---|---|---|---|---|
| Bcl2a1a | 0 | 0 | 0 | 1 | 1 |
| Bcl2a1b | 0 | 0 | 0 | 1 | 1 |
| Bcl2a1c | 0 | 0 | 0 | 1 | 1 |
| Bcl2a1d | 0 | 0 | 0 | 1 | 1 |
| Bex6 | 0 | 0 | 0 | 1 | 1 |
| Bglap2 | 0 | 0 | 0 | 1 | 1 |
| Bglap3 | 0 | 0 | 0 | 1 | 1 |
| Bhlhe23 | 0 | 0 | 0 | 1 | 1 |
| Blk | 0 | 0 | 0 | 1 | 1 |
| Bmp10 | 0 | 0 | 0 | 1 | 1 |
| Bpifa1 | 0 | 0 | 0 | 1 | 1 |
| Bpifa2 | 0 | 0 | 0 | 1 | 1 |
| Bpifa3 | 0 | 0 | 0 | 1 | 1 |
| Bpifa5 | 0 | 0 | 0 | 1 | 1 |
| Bpifa6 | 0 | 0 | 0 | 1 | 1 |
| Bpifb2 | 0 | 0 | 0 | 1 | 1 |
| Bpifb3 | 0 | 0 | 0 | 1 | 1 |
| Bpifb6 | 0 | 0 | 0 | 1 | 1 |
| Bpifb9a | 0 | 0 | 0 | 1 | 1 |
| Bpifb9b | 0 | 0 | 0 | 1 | 1 |
| Bricd5 | 0 | 0 | 0 | 1 | 1 |
| Brs3 | 0 | 0 | 0 | 1 | 1 |
| Bsnd | 0 | 0 | 0 | 1 | 1 |
| Bsph1 | 0 | 0 | 0 | 1 | 1 |
| Bsph2 | 0 | 0 | 0 | 1 | 1 |
| Btbd16 | 0 | 0 | 0 | 1 | 1 |
| Btla | 0 | 0 | 0 | 1 | 1 |
| Btn1a1 | 0 | 0 | 0 | 1 | 1 |
| Btnl1 | 0 | 0 | 0 | 1 | 1 |
| Btnl10 | 0 | 0 | 0 | 1 | 1 |
| Btnl2 | 0 | 0 | 0 | 1 | 1 |
| Btnl4 | 0 | 0 | 0 | 1 | 1 |
| Btnl5-ps | 0 | 0 | 0 | 1 | 1 |
| Btnl6 | 0 | 0 | 0 | 1 | 1 |
| C030007H22Rik | 0 | 0 | 0 | 1 | 1 |
| C030013G03Rik | 0 | 0 | 0 | 1 | 1 |
| C030023E24Rik | 0 | 0 | 0 | 1 | 1 |
| C130079G13Rik | 0 | 0 | 0 | 1 | 1 |
| C1qa | 0 | 0 | 0 | 1 | 1 |
| C1qb | 0 | 0 | 0 | 1 | 1 |
| C1qc | 0 | 0 | 0 | 1 | 1 |
| C1rb | 0 | 0 | 0 | 1 | 1 |
| C230004F18Rik | 0 | 0 | 0 | 1 | 1 |
| C230029M16 | 0 | 0 | 0 | 1 | 1 |
| C230079O03Rik | 0 | 0 | 0 | 1 | 1 |
| C430002N11Rik | 0 | 0 | 0 | 1 | 1 |
| C4bp | 0 | 0 | 0 | 1 | 1 |
| C530044C16Rik | 0 | 0 | 0 | 1 | 1 |
| C5ar1 | 0 | 0 | 0 | 1 | 1 |
| C5ar2 | 0 | 0 | 0 | 1 | 1 |

| | | | | | |
|---|---|---|---|---|---|
| C630028M04Rik | 0 | 0 | 0 | 1 | 1 |
| C630031E19Rik | 0 | 0 | 0 | 1 | 1 |
| C730002L08Rik | 0 | 0 | 0 | 1 | 1 |
| C730027H18Rik | 0 | 0 | 0 | 1 | 1 |
| C87198 | 0 | 0 | 0 | 1 | 1 |
| C87414 | 0 | 0 | 0 | 1 | 1 |
| C87499 | 0 | 0 | 0 | 1 | 1 |
| C87977 | 0 | 0 | 0 | 1 | 1 |
| C8a | 0 | 0 | 0 | 1 | 1 |
| C8b | 0 | 0 | 0 | 1 | 1 |
| C9 | 0 | 0 | 0 | 1 | 1 |
| C920009B18Rik | 0 | 0 | 0 | 1 | 1 |
| Cabp7 | 0 | 0 | 0 | 1 | 1 |
| Cabs1 | 0 | 0 | 0 | 1 | 1 |
| Cacng3 | 0 | 0 | 0 | 1 | 1 |
| Calcb | 0 | 0 | 0 | 1 | 1 |
| Calhm1 | 0 | 0 | 0 | 1 | 1 |
| Calm4 | 0 | 0 | 0 | 1 | 1 |
| Calm5 | 0 | 0 | 0 | 1 | 1 |
| Calml3 | 0 | 0 | 0 | 1 | 1 |
| Capn13 | 0 | 0 | 0 | 1 | 1 |
| Capza3 | 0 | 0 | 0 | 1 | 1 |
| Car10 | 0 | 0 | 0 | 1 | 1 |
| Card14 | 0 | 0 | 0 | 1 | 1 |
| Casp1 | 0 | 0 | 0 | 1 | 1 |
| Casp12 | 0 | 0 | 0 | 1 | 1 |
| Casp14 | 0 | 0 | 0 | 1 | 1 |
| Catsper1 | 0 | 0 | 0 | 1 | 1 |
| Catsperb | 0 | 0 | 0 | 1 | 1 |
| Cav3 | 0 | 0 | 0 | 1 | 1 |
| Cblc | 0 | 0 | 0 | 1 | 1 |
| Ccdc108 | 0 | 0 | 0 | 1 | 1 |
| Ccdc113 | 0 | 0 | 0 | 1 | 1 |
| Ccdc135 | 0 | 0 | 0 | 1 | 1 |
| Ccdc153 | 0 | 0 | 0 | 1 | 1 |
| Ccdc172 | 0 | 0 | 0 | 1 | 1 |
| Ccdc178 | 0 | 0 | 0 | 1 | 1 |
| Ccdc185 | 0 | 0 | 0 | 1 | 1 |
| Ccdc27 | 0 | 0 | 0 | 1 | 1 |
| Ccdc54 | 0 | 0 | 0 | 1 | 1 |
| Ccdc7 | 0 | 0 | 0 | 1 | 1 |
| Ccdc70 | 0 | 0 | 0 | 1 | 1 |
| Ccdc83 | 0 | 0 | 0 | 1 | 1 |
| Ccer1 | 0 | 0 | 0 | 1 | 1 |
| Cckar | 0 | 0 | 0 | 1 | 1 |
| Ccl1 | 0 | 0 | 0 | 1 | 1 |
| Ccl11 | 0 | 0 | 0 | 1 | 1 |
| Ccl20 | 0 | 0 | 0 | 1 | 1 |
| Ccl21a,Gm1987 | 0 | 0 | 0 | 1 | 1 |
| Ccl21b,Gm13304 | 0 | 0 | 0 | 1 | 1 |

| | | | | | |
|---|---|---|---|---|---|
| Ccl21c | 0 | 0 | 0 | 1 | 1 |
| Ccl22 | 0 | 0 | 0 | 1 | 1 |
| Ccl26 | 0 | 0 | 0 | 1 | 1 |
| Ccl27b | 0 | 0 | 0 | 1 | 1 |
| Ccl3 | 0 | 0 | 0 | 1 | 1 |
| Ccl4 | 0 | 0 | 0 | 1 | 1 |
| Ccl5 | 0 | 0 | 0 | 1 | 1 |
| Ccl8 | 0 | 0 | 0 | 1 | 1 |
| Ccl9 | 0 | 0 | 0 | 1 | 1 |
| Ccnb1ip1 | 0 | 0 | 0 | 1 | 1 |
| Ccr1 | 0 | 0 | 0 | 1 | 1 |
| Ccr1l1 | 0 | 0 | 0 | 1 | 1 |
| Ccr2 | 0 | 0 | 0 | 1 | 1 |
| Ccr3 | 0 | 0 | 0 | 1 | 1 |
| Ccr6 | 0 | 0 | 0 | 1 | 1 |
| Ccr8 | 0 | 0 | 0 | 1 | 1 |
| Ccrl2 | 0 | 0 | 0 | 1 | 1 |
| Cct8l1 | 0 | 0 | 0 | 1 | 1 |
| Cd163l1 | 0 | 0 | 0 | 1 | 1 |
| Cd177 | 0 | 0 | 0 | 1 | 1 |
| Cd200r1 | 0 | 0 | 0 | 1 | 1 |
| Cd200r2 | 0 | 0 | 0 | 1 | 1 |
| Cd200r3 | 0 | 0 | 0 | 1 | 1 |
| Cd200r4 | 0 | 0 | 0 | 1 | 1 |
| Cd207 | 0 | 0 | 0 | 1 | 1 |
| Cd209a | 0 | 0 | 0 | 1 | 1 |
| Cd209b | 0 | 0 | 0 | 1 | 1 |
| Cd209c | 0 | 0 | 0 | 1 | 1 |
| Cd209d | 0 | 0 | 0 | 1 | 1 |
| Cd209e | 0 | 0 | 0 | 1 | 1 |
| Cd209f | 0 | 0 | 0 | 1 | 1 |
| Cd226 | 0 | 0 | 0 | 1 | 1 |
| Cd244 | 0 | 0 | 0 | 1 | 1 |
| Cd300c | 0 | 0 | 0 | 1 | 1 |
| Cd300lb | 0 | 0 | 0 | 1 | 1 |
| Cd300ld | 0 | 0 | 0 | 1 | 1 |
| Cd300lh | 0 | 0 | 0 | 1 | 1 |
| Cd33 | 0 | 0 | 0 | 1 | 1 |
| Cd3d | 0 | 0 | 0 | 1 | 1 |
| Cd3g | 0 | 0 | 0 | 1 | 1 |
| Cd40lg | 0 | 0 | 0 | 1 | 1 |
| Cd48 | 0 | 0 | 0 | 1 | 1 |
| Cd53 | 0 | 0 | 0 | 1 | 1 |
| Cd5l | 0 | 0 | 0 | 1 | 1 |
| Cd69 | 0 | 0 | 0 | 1 | 1 |
| Cd7 | 0 | 0 | 0 | 1 | 1 |
| Cd84 | 0 | 0 | 0 | 1 | 1 |
| Cd8b1 | 0 | 0 | 0 | 1 | 1 |
| Cd96 | 0 | 0 | 0 | 1 | 1 |
| Cdc20b | 0 | 0 | 0 | 1 | 1 |

| | | | | | |
|---|---|---|---|---|---|
| Cdh17 | 0 | 0 | 0 | 1 | 1 |
| Cdh9 | 0 | 0 | 0 | 1 | 1 |
| Ceacam-ps1 | 0 | 0 | 0 | 1 | 1 |
| Ceacam11 | 0 | 0 | 0 | 1 | 1 |
| Ceacam13 | 0 | 0 | 0 | 1 | 1 |
| Ceacam14 | 0 | 0 | 0 | 1 | 1 |
| Ceacam15 | 0 | 0 | 0 | 1 | 1 |
| Ceacam16 | 0 | 0 | 0 | 1 | 1 |
| Ceacam3 | 0 | 0 | 0 | 1 | 1 |
| Ceacam5 | 0 | 0 | 0 | 1 | 1 |
| Ceacam9 | 0 | 0 | 0 | 1 | 1 |
| Cebpe | 0 | 0 | 0 | 1 | 1 |
| Cel | 0 | 0 | 0 | 1 | 1 |
| Cela2a | 0 | 0 | 0 | 1 | 1 |
| Cela3b | 0 | 0 | 0 | 1 | 1 |
| Celrr | 0 | 0 | 0 | 1 | 1 |
| Cerkl | 0 | 0 | 0 | 1 | 1 |
| Ces1b | 0 | 0 | 0 | 1 | 1 |
| Ces1c | 0 | 0 | 0 | 1 | 1 |
| Ces1d | 0 | 0 | 0 | 1 | 1 |
| Ces1e | 0 | 0 | 0 | 1 | 1 |
| Ces1f | 0 | 0 | 0 | 1 | 1 |
| Ces2a | 0 | 0 | 0 | 1 | 1 |
| Ces2b | 0 | 0 | 0 | 1 | 1 |
| Ces2c | 0 | 0 | 0 | 1 | 1 |
| Ces2d-ps | 0 | 0 | 0 | 1 | 1 |
| Ces2e | 0 | 0 | 0 | 1 | 1 |
| Ces2h | 0 | 0 | 0 | 1 | 1 |
| Ces3a | 0 | 0 | 0 | 1 | 1 |
| Ces3b | 0 | 0 | 0 | 1 | 1 |
| Ces4a | 0 | 0 | 0 | 1 | 1 |
| Cetn1 | 0 | 0 | 0 | 1 | 1 |
| Cfd | 0 | 0 | 0 | 1 | 1 |
| Cfh | 0 | 0 | 0 | 1 | 1 |
| Cfhr1 | 0 | 0 | 0 | 1 | 1 |
| Cfhr2 | 0 | 0 | 0 | 1 | 1 |
| Cga | 0 | 0 | 0 | 1 | 1 |
| Chat | 0 | 0 | 0 | 1 | 1 |
| Chia1 | 0 | 0 | 0 | 1 | 1 |
| Chil3 | 0 | 0 | 0 | 1 | 1 |
| Chil4 | 0 | 0 | 0 | 1 | 1 |
| Chil6 | 0 | 0 | 0 | 1 | 1 |
| Chrm5 | 0 | 0 | 0 | 1 | 1 |
| Chrna2 | 0 | 0 | 0 | 1 | 1 |
| Chst4 | 0 | 0 | 0 | 1 | 1 |
| Cib3 | 0 | 0 | 0 | 1 | 1 |
| Cib4 | 0 | 0 | 0 | 1 | 1 |
| Cidec | 0 | 0 | 0 | 1 | 1 |
| Ckmt2 | 0 | 0 | 0 | 1 | 1 |
| Clca1 | 0 | 0 | 0 | 1 | 1 |

| | | | | | |
|---|---|---|---|---|---|
| Clca2 | 0 | 0 | 0 | 1 | 1 |
| Clca3 | 0 | 0 | 0 | 1 | 1 |
| Clca4 | 0 | 0 | 0 | 1 | 1 |
| Clca6 | 0 | 0 | 0 | 1 | 1 |
| Cldn13 | 0 | 0 | 0 | 1 | 1 |
| Cldn16 | 0 | 0 | 0 | 1 | 1 |
| Cldn17 | 0 | 0 | 0 | 1 | 1 |
| Cldn18 | 0 | 0 | 0 | 1 | 1 |
| Cldn24 | 0 | 0 | 0 | 1 | 1 |
| Clec10a | 0 | 0 | 0 | 1 | 1 |
| Clec12a | 0 | 0 | 0 | 1 | 1 |
| Clec12b | 0 | 0 | 0 | 1 | 1 |
| Clec2e | 0 | 0 | 0 | 1 | 1 |
| Clec2f | 0 | 0 | 0 | 1 | 1 |
| Clec2g | 0 | 0 | 0 | 1 | 1 |
| Clec4a3 | 0 | 0 | 0 | 1 | 1 |
| Clec4a4 | 0 | 0 | 0 | 1 | 1 |
| Clec4b1 | 0 | 0 | 0 | 1 | 1 |
| Clec4b2 | 0 | 0 | 0 | 1 | 1 |
| Clec4d | 0 | 0 | 0 | 1 | 1 |
| Clec4e | 0 | 0 | 0 | 1 | 1 |
| Clec4f | 0 | 0 | 0 | 1 | 1 |
| Clec4g | 0 | 0 | 0 | 1 | 1 |
| Clec5a | 0 | 0 | 0 | 1 | 1 |
| Clps | 0 | 0 | 0 | 1 | 1 |
| Clpsl2 | 0 | 0 | 0 | 1 | 1 |
| Clrn2 | 0 | 0 | 0 | 1 | 1 |
| Clrn3 | 0 | 0 | 0 | 1 | 1 |
| Cma2 | 0 | 0 | 0 | 1 | 1 |
| Cml3 | 0 | 0 | 0 | 1 | 1 |
| Cml5 | 0 | 0 | 0 | 1 | 1 |
| Cmtm2a | 0 | 0 | 0 | 1 | 1 |
| Cmtm2b | 0 | 0 | 0 | 1 | 1 |
| Cndp1 | 0 | 0 | 0 | 1 | 1 |
| Cnga1 | 0 | 0 | 0 | 1 | 1 |
| Cnga4 | 0 | 0 | 0 | 1 | 1 |
| Cngb3 | 0 | 0 | 0 | 1 | 1 |
| Cntn6 | 0 | 0 | 0 | 1 | 1 |
| Cntnap5a | 0 | 0 | 0 | 1 | 1 |
| Colec10 | 0 | 0 | 0 | 1 | 1 |
| Colq | 0 | 0 | 0 | 1 | 1 |
| Cox6a2 | 0 | 0 | 0 | 1 | 1 |
| Cox7b2 | 0 | 0 | 0 | 1 | 1 |
| Cox8b | 0 | 0 | 0 | 1 | 1 |
| Cox8c | 0 | 0 | 0 | 1 | 1 |
| Cpa3 | 0 | 0 | 0 | 1 | 1 |
| Cpb2 | 0 | 0 | 0 | 1 | 1 |
| Cphx1,Cphx2 | 0 | 0 | 0 | 1 | 1 |
| Cphx2 | 0 | 0 | 0 | 1 | 1 |
| Cpn2 | 0 | 0 | 0 | 1 | 1 |

| | | | | | |
|---|---|---|---|---|---|
| Cpxcr1 | 0 | 0 | 0 | 1 | 1 |
| Crct1 | 0 | 0 | 0 | 1 | 1 |
| Crh | 0 | 0 | 0 | 1 | 1 |
| Crisp1 | 0 | 0 | 0 | 1 | 1 |
| Crisp3 | 0 | 0 | 0 | 1 | 1 |
| Crisp4 | 0 | 0 | 0 | 1 | 1 |
| Crnn | 0 | 0 | 0 | 1 | 1 |
| Crx | 0 | 0 | 0 | 1 | 1 |
| Crxos | 0 | 0 | 0 | 1 | 1 |
| Cryaa | 0 | 0 | 0 | 1 | 1 |
| Cryba2 | 0 | 0 | 0 | 1 | 1 |
| Cryba4 | 0 | 0 | 0 | 1 | 1 |
| Crybb1 | 0 | 0 | 0 | 1 | 1 |
| Crybb2 | 0 | 0 | 0 | 1 | 1 |
| Cryga | 0 | 0 | 0 | 1 | 1 |
| Crygc | 0 | 0 | 0 | 1 | 1 |
| Crygd | 0 | 0 | 0 | 1 | 1 |
| Crygf | 0 | 0 | 0 | 1 | 1 |
| Csf1r | 0 | 0 | 0 | 1 | 1 |
| Csf2 | 0 | 0 | 0 | 1 | 1 |
| Csmd2os | 0 | 0 | 0 | 1 | 1 |
| Csmd3 | 0 | 0 | 0 | 1 | 1 |
| Csn1s1 | 0 | 0 | 0 | 1 | 1 |
| Csn1s2a | 0 | 0 | 0 | 1 | 1 |
| Csn1s2b | 0 | 0 | 0 | 1 | 1 |
| Csn2 | 0 | 0 | 0 | 1 | 1 |
| Csnka2ip | 0 | 0 | 0 | 1 | 1 |
| Csprs | 0 | 0 | 0 | 1 | 1 |
| Cst10 | 0 | 0 | 0 | 1 | 1 |
| Cst11 | 0 | 0 | 0 | 1 | 1 |
| Cst12 | 0 | 0 | 0 | 1 | 1 |
| Cst13 | 0 | 0 | 0 | 1 | 1 |
| Cst8 | 0 | 0 | 0 | 1 | 1 |
| Cst9 | 0 | 0 | 0 | 1 | 1 |
| Cstl1 | 0 | 0 | 0 | 1 | 1 |
| Ctag2 | 0 | 0 | 0 | 1 | 1 |
| Ctcfl | 0 | 0 | 0 | 1 | 1 |
| Ctcflos | 0 | 0 | 0 | 1 | 1 |
| Ctla4 | 0 | 0 | 0 | 1 | 1 |
| Ctrb1 | 0 | 0 | 0 | 1 | 1 |
| Ctrc | 0 | 0 | 0 | 1 | 1 |
| Ctrcos | 0 | 0 | 0 | 1 | 1 |
| Cts3 | 0 | 0 | 0 | 1 | 1 |
| Cts6 | 0 | 0 | 0 | 1 | 1 |
| Cts7 | 0 | 0 | 0 | 1 | 1 |
| Cts8 | 0 | 0 | 0 | 1 | 1 |
| Cts8-ps | 0 | 0 | 0 | 1 | 1 |
| Ctsg | 0 | 0 | 0 | 1 | 1 |
| Ctsj | 0 | 0 | 0 | 1 | 1 |
| Ctsll3 | 0 | 0 | 0 | 1 | 1 |

| | | | | | |
|---|---|---|---|---|---|
| Ctsm | 0 | 0 | 0 | 1 | 1 |
| Ctsq | 0 | 0 | 0 | 1 | 1 |
| Ctsr | 0 | 0 | 0 | 1 | 1 |
| Ctss | 0 | 0 | 0 | 1 | 1 |
| Ctxn2 | 0 | 0 | 0 | 1 | 1 |
| Cwh43 | 0 | 0 | 0 | 1 | 1 |
| Cxcl15 | 0 | 0 | 0 | 1 | 1 |
| Cxcl2 | 0 | 0 | 0 | 1 | 1 |
| Cxcl9 | 0 | 0 | 0 | 1 | 1 |
| Cxcr2 | 0 | 0 | 0 | 1 | 1 |
| Cylc1 | 0 | 0 | 0 | 1 | 1 |
| Cylc2 | 0 | 0 | 0 | 1 | 1 |
| Cyp11b1 | 0 | 0 | 0 | 1 | 1 |
| Cyp11b2 | 0 | 0 | 0 | 1 | 1 |
| Cyp19a1 | 0 | 0 | 0 | 1 | 1 |
| Cyp2a12 | 0 | 0 | 0 | 1 | 1 |
| Cyp2a22 | 0 | 0 | 0 | 1 | 1 |
| Cyp2a4 | 0 | 0 | 0 | 1 | 1 |
| Cyp2a5 | 0 | 0 | 0 | 1 | 1 |
| Cyp2ab1 | 0 | 0 | 0 | 1 | 1 |
| Cyp2b10 | 0 | 0 | 0 | 1 | 1 |
| Cyp2b13 | 0 | 0 | 0 | 1 | 1 |
| Cyp2c29 | 0 | 0 | 0 | 1 | 1 |
| Cyp2c37 | 0 | 0 | 0 | 1 | 1 |
| Cyp2c38 | 0 | 0 | 0 | 1 | 1 |
| Cyp2c39 | 0 | 0 | 0 | 1 | 1 |
| Cyp2c40 | 0 | 0 | 0 | 1 | 1 |
| Cyp2c50 | 0 | 0 | 0 | 1 | 1 |
| Cyp2c53-ps | 0 | 0 | 0 | 1 | 1 |
| Cyp2c54 | 0 | 0 | 0 | 1 | 1 |
| Cyp2c65 | 0 | 0 | 0 | 1 | 1 |
| Cyp2c66 | 0 | 0 | 0 | 1 | 1 |
| Cyp2c67 | 0 | 0 | 0 | 1 | 1 |
| Cyp2c68 | 0 | 0 | 0 | 1 | 1 |
| Cyp2c69 | 0 | 0 | 0 | 1 | 1 |
| Cyp2c70 | 0 | 0 | 0 | 1 | 1 |
| Cyp2d10 | 0 | 0 | 0 | 1 | 1 |
| Cyp2d11 | 0 | 0 | 0 | 1 | 1 |
| Cyp2d12 | 0 | 0 | 0 | 1 | 1 |
| Cyp2d13 | 0 | 0 | 0 | 1 | 1 |
| Cyp2d37-ps | 0 | 0 | 0 | 1 | 1 |
| Cyp2d40 | 0 | 0 | 0 | 1 | 1 |
| Cyp2d9 | 0 | 0 | 0 | 1 | 1 |
| Cyp2e1 | 0 | 0 | 0 | 1 | 1 |
| Cyp2f2 | 0 | 0 | 0 | 1 | 1 |
| Cyp2j12 | 0 | 0 | 0 | 1 | 1 |
| Cyp2j5 | 0 | 0 | 0 | 1 | 1 |
| Cyp2j8 | 0 | 0 | 0 | 1 | 1 |
| Cyp3a11 | 0 | 0 | 0 | 1 | 1 |
| Cyp3a13 | 0 | 0 | 0 | 1 | 1 |

| | | | | | |
|---|---|---|---|---|---|
| Cyp3a16 | 0 | 0 | 0 | 1 | 1 |
| Cyp3a25 | 0 | 0 | 0 | 1 | 1 |
| Cyp3a41a | 0 | 0 | 0 | 1 | 1 |
| Cyp3a44 | 0 | 0 | 0 | 1 | 1 |
| Cyp3a57 | 0 | 0 | 0 | 1 | 1 |
| Cyp3a59 | 0 | 0 | 0 | 1 | 1 |
| Cyp4a10 | 0 | 0 | 0 | 1 | 1 |
| Cyp4a12a | 0 | 0 | 0 | 1 | 1 |
| Cyp4a12b | 0 | 0 | 0 | 1 | 1 |
| Cyp4a14 | 0 | 0 | 0 | 1 | 1 |
| Cyp4a29 | 0 | 0 | 0 | 1 | 1 |
| Cyp4a30b | 0 | 0 | 0 | 1 | 1 |
| Cyp4a31 | 0 | 0 | 0 | 1 | 1 |
| Cyp4a32 | 0 | 0 | 0 | 1 | 1 |
| Cyp4b1 | 0 | 0 | 0 | 1 | 1 |
| Cyp4b1-ps2 | 0 | 0 | 0 | 1 | 1 |
| Cyp4f14 | 0 | 0 | 0 | 1 | 1 |
| Cyp4f40 | 0 | 0 | 0 | 1 | 1 |
| Cyp7a1 | 0 | 0 | 0 | 1 | 1 |
| Cypt1,Cypt8 | 0 | 0 | 0 | 1 | 1 |
| Cypt10,Cypt4 | 0 | 0 | 0 | 1 | 1 |
| Cypt12 | 0 | 0 | 0 | 1 | 1 |
| Cypt14 | 0 | 0 | 0 | 1 | 1 |
| Cypt15 | 0 | 0 | 0 | 1 | 1 |
| Cypt2 | 0 | 0 | 0 | 1 | 1 |
| Cypt3 | 0 | 0 | 0 | 1 | 1 |
| Cypt4 | 0 | 0 | 0 | 1 | 1 |
| Cypt7 | 0 | 0 | 0 | 1 | 1 |
| Cypt8 | 0 | 0 | 0 | 1 | 1 |
| D030024E09Rik | 0 | 0 | 0 | 1 | 1 |
| D030025E07Rik | 0 | 0 | 0 | 1 | 1 |
| D130009I18Rik | 0 | 0 | 0 | 1 | 1 |
| D130058E03 | 0 | 0 | 0 | 1 | 1 |
| D17Ertd648e | 0 | 0 | 0 | 1 | 1 |
| D3Bwg0562e | 0 | 0 | 0 | 1 | 1 |
| D5Ertd577e,Gm3286 | 0 | 0 | 0 | 1 | 1 |
| D630010B17Rik | 0 | 0 | 0 | 1 | 1 |
| D630013N20Rik | 0 | 0 | 0 | 1 | 1 |
| D630024D03Rik | 0 | 0 | 0 | 1 | 1 |
| D6Ertd474e | 0 | 0 | 0 | 1 | 1 |
| D730001G18Rik | 0 | 0 | 0 | 1 | 1 |
| D730045A05Rik | 0 | 0 | 0 | 1 | 1 |
| D730048I06Rik | 0 | 0 | 0 | 1 | 1 |
| D730050B12Rik | 0 | 0 | 0 | 1 | 1 |
| D7Ertd143e | 0 | 0 | 0 | 1 | 1 |
| D830005E20Rik | 0 | 0 | 0 | 1 | 1 |
| D830013O20Rik | 0 | 0 | 0 | 1 | 1 |
| D830015G02Rik | 0 | 0 | 0 | 1 | 1 |
| D830032E09Rik | 0 | 0 | 0 | 1 | 1 |
| D930032P07Rik | 0 | 0 | 0 | 1 | 1 |

| | | | | | |
|---|---|---|---|---|---|
| DQ267100 | 0 | 0 | 0 | 1 | 1 |
| DQ267101 | 0 | 0 | 0 | 1 | 1 |
| Daf2 | 0 | 0 | 0 | 1 | 1 |
| Dao | 0 | 0 | 0 | 1 | 1 |
| Dbpht2 | 0 | 0 | 0 | 1 | 1 |
| Dcdc2c | 0 | 0 | 0 | 1 | 1 |
| Dcpp1 | 0 | 0 | 0 | 1 | 1 |
| Dcpp2 | 0 | 0 | 0 | 1 | 1 |
| Dcstamp | 0 | 0 | 0 | 1 | 1 |
| Ddi1 | 0 | 0 | 0 | 1 | 1 |
| Defa-ps1 | 0 | 0 | 0 | 1 | 1 |
| Defa-ps12 | 0 | 0 | 0 | 1 | 1 |
| Defa-ps13 | 0 | 0 | 0 | 1 | 1 |
| Defa-rs1 | 0 | 0 | 0 | 1 | 1 |
| Defa-rs7,Defa23 | 0 | 0 | 0 | 1 | 1 |
| Defa17 | 0 | 0 | 0 | 1 | 1 |
| Defa2 | 0 | 0 | 0 | 1 | 1 |
| Defa20 | 0 | 0 | 0 | 1 | 1 |
| Defa21 | 0 | 0 | 0 | 1 | 1 |
| Defa22 | 0 | 0 | 0 | 1 | 1 |
| Defa23 | 0 | 0 | 0 | 1 | 1 |
| Defa24,Defa6 | 0 | 0 | 0 | 1 | 1 |
| Defa25 | 0 | 0 | 0 | 1 | 1 |
| Defa26 | 0 | 0 | 0 | 1 | 1 |
| Defa3 | 0 | 0 | 0 | 1 | 1 |
| Defa4 | 0 | 0 | 0 | 1 | 1 |
| Defa5 | 0 | 0 | 0 | 1 | 1 |
| Defa6 | 0 | 0 | 0 | 1 | 1 |
| Defb1 | 0 | 0 | 0 | 1 | 1 |
| Defb10 | 0 | 0 | 0 | 1 | 1 |
| Defb11 | 0 | 0 | 0 | 1 | 1 |
| Defb12 | 0 | 0 | 0 | 1 | 1 |
| Defb13 | 0 | 0 | 0 | 1 | 1 |
| Defb14 | 0 | 0 | 0 | 1 | 1 |
| Defb15 | 0 | 0 | 0 | 1 | 1 |
| Defb18 | 0 | 0 | 0 | 1 | 1 |
| Defb19 | 0 | 0 | 0 | 1 | 1 |
| Defb2 | 0 | 0 | 0 | 1 | 1 |
| Defb20 | 0 | 0 | 0 | 1 | 1 |
| Defb21 | 0 | 0 | 0 | 1 | 1 |
| Defb22 | 0 | 0 | 0 | 1 | 1 |
| Defb23 | 0 | 0 | 0 | 1 | 1 |
| Defb25 | 0 | 0 | 0 | 1 | 1 |
| Defb26 | 0 | 0 | 0 | 1 | 1 |
| Defb28 | 0 | 0 | 0 | 1 | 1 |
| Defb29 | 0 | 0 | 0 | 1 | 1 |
| Defb3 | 0 | 0 | 0 | 1 | 1 |
| Defb30 | 0 | 0 | 0 | 1 | 1 |
| Defb33 | 0 | 0 | 0 | 1 | 1 |
| Defb34 | 0 | 0 | 0 | 1 | 1 |

| | | | | | |
|---|---|---|---|---|---|
| Defb35 | 0 | 0 | 0 | 1 | 1 |
| Defb36 | 0 | 0 | 0 | 1 | 1 |
| Defb38 | 0 | 0 | 0 | 1 | 1 |
| Defb39 | 0 | 0 | 0 | 1 | 1 |
| Defb4 | 0 | 0 | 0 | 1 | 1 |
| Defb41 | 0 | 0 | 0 | 1 | 1 |
| Defb42 | 0 | 0 | 0 | 1 | 1 |
| Defb43 | 0 | 0 | 0 | 1 | 1 |
| Defb44-ps | 0 | 0 | 0 | 1 | 1 |
| Defb45 | 0 | 0 | 0 | 1 | 1 |
| Defb46 | 0 | 0 | 0 | 1 | 1 |
| Defb5 | 0 | 0 | 0 | 1 | 1 |
| Defb50 | 0 | 0 | 0 | 1 | 1 |
| Defb6 | 0 | 0 | 0 | 1 | 1 |
| Defb7 | 0 | 0 | 0 | 1 | 1 |
| Defb8 | 0 | 0 | 0 | 1 | 1 |
| Defb9 | 0 | 0 | 0 | 1 | 1 |
| Dgat2l6 | 0 | 0 | 0 | 1 | 1 |
| Dgkeos | 0 | 0 | 0 | 1 | 1 |
| Dhrs7c | 0 | 0 | 0 | 1 | 1 |
| Dhrs9 | 0 | 0 | 0 | 1 | 1 |
| Dhrsx | 0 | 0 | 0 | 1 | 1 |
| Dio1 | 0 | 0 | 0 | 1 | 1 |
| Dkk4 | 0 | 0 | 0 | 1 | 1 |
| Dlx6as2 | 0 | 0 | 0 | 1 | 1 |
| Dmbt1 | 0 | 0 | 0 | 1 | 1 |
| Dmkn | 0 | 0 | 0 | 1 | 1 |
| Dmp1 | 0 | 0 | 0 | 1 | 1 |
| Dmrt3 | 0 | 0 | 0 | 1 | 1 |
| Dmrtc1b | 0 | 0 | 0 | 1 | 1 |
| Dmrtc1c1 | 0 | 0 | 0 | 1 | 1 |
| Dmrtc1c2 | 0 | 0 | 0 | 1 | 1 |
| Dmrtc2 | 0 | 0 | 0 | 1 | 1 |
| Dnajc5b | 0 | 0 | 0 | 1 | 1 |
| Dnajc5g | 0 | 0 | 0 | 1 | 1 |
| Dnase2b | 0 | 0 | 0 | 1 | 1 |
| Doxl2 | 0 | 0 | 0 | 1 | 1 |
| Dpep2 | 0 | 0 | 0 | 1 | 1 |
| Dpep3 | 0 | 0 | 0 | 1 | 1 |
| Dppa1 | 0 | 0 | 0 | 1 | 1 |
| Dppa3 | 0 | 0 | 0 | 1 | 1 |
| Drd3 | 0 | 0 | 0 | 1 | 1 |
| Dreh | 0 | 0 | 0 | 1 | 1 |
| Dsc1 | 0 | 0 | 0 | 1 | 1 |
| Dsg1a | 0 | 0 | 0 | 1 | 1 |
| Dsg1b | 0 | 0 | 0 | 1 | 1 |
| Dsg1c | 0 | 0 | 0 | 1 | 1 |
| Dsg3 | 0 | 0 | 0 | 1 | 1 |
| Dsg4 | 0 | 0 | 0 | 1 | 1 |
| Dspp | 0 | 0 | 0 | 1 | 1 |

| | | | | | |
|---|---|---|---|---|---|
| Dthd1 | 0 | 0 | 0 | 1 | 1 |
| Duoxa1 | 0 | 0 | 0 | 1 | 1 |
| Duoxa2 | 0 | 0 | 0 | 1 | 1 |
| Dusp21 | 0 | 0 | 0 | 1 | 1 |
| Dux | 0 | 0 | 0 | 1 | 1 |
| Duxbl2 | 0 | 0 | 0 | 1 | 1 |
| Dydc1 | 0 | 0 | 0 | 1 | 1 |
| Dynap | 0 | 0 | 0 | 1 | 1 |
| Dynlt1b | 0 | 0 | 0 | 1 | 1 |
| Dynlt1c,Dynlt1f | 0 | 0 | 0 | 1 | 1 |
| Dynlt1f | 0 | 0 | 0 | 1 | 1 |
| Dytn | 0 | 0 | 0 | 1 | 1 |
| E030002O03Rik | 0 | 0 | 0 | 1 | 1 |
| E030019B06Rik | 0 | 0 | 0 | 1 | 1 |
| E030019B13Rik | 0 | 0 | 0 | 1 | 1 |
| E030044B06Rik | 0 | 0 | 0 | 1 | 1 |
| E230016K23Rik | 0 | 0 | 0 | 1 | 1 |
| E230019M04Rik | 0 | 0 | 0 | 1 | 1 |
| E230029C05Rik | 0 | 0 | 0 | 1 | 1 |
| E330012B07Rik | 0 | 0 | 0 | 1 | 1 |
| E330014E10Rik,Gm3286 | 0 | 0 | 0 | 1 | 1 |
| E330017A01Rik | 0 | 0 | 0 | 1 | 1 |
| E330017L17Rik | 0 | 0 | 0 | 1 | 1 |
| E330021D16Rik | 0 | 0 | 0 | 1 | 1 |
| E330023G01Rik | 0 | 0 | 0 | 1 | 1 |
| E330034G19Rik | 0 | 0 | 0 | 1 | 1 |
| Ear1 | 0 | 0 | 0 | 1 | 1 |
| Ear10 | 0 | 0 | 0 | 1 | 1 |
| Ear2,Ear3 | 0 | 0 | 0 | 1 | 1 |
| Ear3 | 0 | 0 | 0 | 1 | 1 |
| Ear6,Ear7 | 0 | 0 | 0 | 1 | 1 |
| Ear7 | 0 | 0 | 0 | 1 | 1 |
| Eddm3b | 0 | 0 | 0 | 1 | 1 |
| Edn2 | 0 | 0 | 0 | 1 | 1 |
| Efcab3 | 0 | 0 | 0 | 1 | 1 |
| Eftud2 | 0 | 125,145 | 0 | 1 | 1 |
| Egfbp2 | 0 | 0 | 0 | 1 | 1 |
| Elane | 0 | 0 | 0 | 1 | 1 |
| Emr1 | 0 | 0 | 0 | 1 | 1 |
| Enam | 0 | 0 | 0 | 1 | 1 |
| Endou | 0 | 0 | 0 | 1 | 1 |
| Enpep | 0 | 0 | 0 | 1 | 1 |
| Enthd1 | 0 | 0 | 0 | 1 | 1 |
| Entpd4 | 0 | 0 | 0 | 1 | 1 |
| Epha6 | 0 | 0 | 0 | 1 | 1 |
| Eppin | 0 | 0 | 0 | 1 | 1 |
| Erbb4 | 0 | 0 | 0 | 1 | 1 |
| Erdr1 | 0 | 0 | 0 | 1 | 1 |
| Esp1 | 0 | 0 | 0 | 1 | 1 |
| Esp15 | 0 | 0 | 0 | 1 | 1 |

| | | | | | |
|---|---|---|---|---|---|
| Esp16 | 0 | 0 | 0 | 1 | 1 |
| Esp18 | 0 | 0 | 0 | 1 | 1 |
| Esp23 | 0 | 0 | 0 | 1 | 1 |
| Esp24 | 0 | 0 | 0 | 1 | 1 |
| Esp3 | 0 | 0 | 0 | 1 | 1 |
| Esp31 | 0 | 0 | 0 | 1 | 1 |
| Esp34 | 0 | 0 | 0 | 1 | 1 |
| Esp36 | 0 | 0 | 0 | 1 | 1 |
| Esp38 | 0 | 0 | 0 | 1 | 1 |
| Esp4 | 0 | 0 | 0 | 1 | 1 |
| Esp6 | 0 | 0 | 0 | 1 | 1 |
| Esp8 | 0 | 0 | 0 | 1 | 1 |
| Esx1 | 0 | 0 | 0 | 1 | 1 |
| Etd | 0 | 0 | 0 | 1 | 1 |
| F13b | 0 | 0 | 0 | 1 | 1 |
| F630028O10Rik | 0 | 0 | 0 | 1 | 1 |
| F630206G17Rik | 0 | 0 | 0 | 1 | 1 |
| F730035M05Rik | 0 | 0 | 0 | 1 | 1 |
| F830002L21Rik | 0 | 0 | 0 | 1 | 1 |
| F830016B08Rik | 0 | 0 | 0 | 1 | 1 |
| F830045P16Rik | 0 | 0 | 0 | 1 | 1 |
| F9 | 0 | 0 | 0 | 1 | 1 |
| Fabp1 | 0 | 0 | 0 | 1 | 1 |
| Fabp12 | 0 | 0 | 0 | 1 | 1 |
| Fabp2 | 0 | 0 | 0 | 1 | 1 |
| Fabp6 | 0 | 0 | 0 | 1 | 1 |
| Fabp9 | 0 | 0 | 0 | 1 | 1 |
| Fam122c | 0 | 0 | 0 | 1 | 1 |
| Fam151a | 0 | 0 | 0 | 1 | 1 |
| Fam154a | 0 | 0 | 0 | 1 | 1 |
| Fam170a | 0 | 0 | 0 | 1 | 1 |
| Fam170b | 0 | 0 | 0 | 1 | 1 |
| Fam177a | 0 | 0 | 0 | 1 | 1 |
| Fam180a | 0 | 0 | 0 | 1 | 1 |
| Fam187a | 0 | 0 | 0 | 1 | 1 |
| Fam216b | 0 | 0 | 0 | 1 | 1 |
| Fam227b | 0 | 0 | 0 | 1 | 1 |
| Fam24a | 0 | 0 | 0 | 1 | 1 |
| Fam25c | 0 | 0 | 0 | 1 | 1 |
| Fam3b | 0 | 0 | 0 | 1 | 1 |
| Fam47c | 0 | 0 | 0 | 1 | 1 |
| Fam47e | 0 | 0 | 0 | 1 | 1 |
| Fam50b | 0 | 0 | 0 | 1 | 1 |
| Fam71a | 0 | 0 | 0 | 1 | 1 |
| Fam71b | 0 | 0 | 0 | 1 | 1 |
| Fam71e2 | 0 | 0 | 0 | 1 | 1 |
| Fam71f1 | 0 | 0 | 0 | 1 | 1 |
| Fasl | 0 | 0 | 0 | 1 | 1 |
| Fate1 | 0 | 0 | 0 | 1 | 1 |
| Fbll1 | 0 | 0 | 0 | 1 | 1 |

| | | | | | |
|---|---|---|---|---|---|
| Fbxo39 | 0 | 0 | 0 | 1 | 1 |
| Fbxw14 | 0 | 0 | 0 | 1 | 1 |
| Fbxw15 | 0 | 0 | 0 | 1 | 1 |
| Fbxw16 | 0 | 0 | 0 | 1 | 1 |
| Fbxw18 | 0 | 0 | 0 | 1 | 1 |
| Fbxw20 | 0 | 0 | 0 | 1 | 1 |
| Fbxw21 | 0 | 0 | 0 | 1 | 1 |
| Fbxw22 | 0 | 0 | 0 | 1 | 1 |
| Fbxw24 | 0 | 0 | 0 | 1 | 1 |
| Fbxw26 | 0 | 0 | 0 | 1 | 1 |
| Fbxw28 | 0 | 0 | 0 | 1 | 1 |
| Fcamr | 0 | 0 | 0 | 1 | 1 |
| Fcer1a | 0 | 0 | 0 | 1 | 1 |
| Fcgbp | 0 | 0 | 0 | 1 | 1 |
| Fcgr4 | 0 | 0 | 0 | 1 | 1 |
| Fcna | 0 | 0 | 0 | 1 | 1 |
| Fcnb | 0 | 0 | 0 | 1 | 1 |
| Fcrl1 | 0 | 0 | 0 | 1 | 1 |
| Fcrl5 | 0 | 0 | 0 | 1 | 1 |
| Fcrla | 0 | 0 | 0 | 1 | 1 |
| Fcrls | 0 | 0 | 0 | 1 | 1 |
| Ffar1 | 0 | 0 | 0 | 1 | 1 |
| Ffar2 | 0 | 0 | 0 | 1 | 1 |
| Ffar3 | 0 | 0 | 0 | 1 | 1 |
| Fga | 0 | 0 | 0 | 1 | 1 |
| Fgb | 0 | 0 | 0 | 1 | 1 |
| Fgf16 | 0 | 0 | 0 | 1 | 1 |
| Fgf22 | 0 | 0 | 0 | 1 | 1 |
| Fhad1os1 | 0 | 0 | 0 | 1 | 1 |
| Fhl5 | 0 | 0 | 0 | 1 | 1 |
| Figla | 0 | 0 | 0 | 1 | 1 |
| Flg2 | 0 | 0 | 0 | 1 | 1 |
| Fmo3 | 0 | 0 | 0 | 1 | 1 |
| Fmo9 | 0 | 0 | 0 | 1 | 1 |
| Folr4 | 0 | 0 | 0 | 1 | 1 |
| Foxb2 | 0 | 0 | 0 | 1 | 1 |
| Foxi1 | 0 | 0 | 0 | 1 | 1 |
| Foxi3 | 0 | 0 | 0 | 1 | 1 |
| Foxr1 | 0 | 0 | 0 | 1 | 1 |
| Fpr-rs3 | 0 | 0 | 0 | 1 | 1 |
| Fpr-rs4 | 0 | 0 | 0 | 1 | 1 |
| Fpr-rs6 | 0 | 0 | 0 | 1 | 1 |
| Fpr2 | 0 | 0 | 0 | 1 | 1 |
| Fpr3 | 0 | 0 | 0 | 1 | 1 |
| Frmpd1os | 0 | 0 | 0 | 1 | 1 |
| Fscb | 0 | 0 | 0 | 1 | 1 |
| Fshb | 0 | 0 | 0 | 1 | 1 |
| Fshr | 0 | 0 | 0 | 1 | 1 |
| Ftcd | 0 | 0 | 0 | 1 | 1 |
| Fthl17 | 0 | 0 | 0 | 1 | 1 |

| | | | | | |
|---|---|---|---|---|---|
| Ftmt | 0 | 0 | 0 | 1 | 1 |
| Fut1 | 0 | 0 | 0 | 1 | 1 |
| Fut4-ps1 | 0 | 0 | 0 | 1 | 1 |
| Fyb | 0 | 0 | 0 | 1 | 1 |
| G630093K05Rik | 0 | 0 | 0 | 1 | 1 |
| G6pc | 0 | 0 | 0 | 1 | 1 |
| G730013B05Rik | 0 | 0 | 0 | 1 | 1 |
| Gabra6 | 0 | 0 | 0 | 1 | 1 |
| Gabrb1 | 0 | 0 | 0 | 1 | 1 |
| Gabrd | 0 | 0 | 0 | 1 | 1 |
| Gabrg2 | 0 | 0 | 0 | 1 | 1 |
| Gabrg3 | 0 | 0 | 0 | 1 | 1 |
| Gal3st2 | 0 | 0 | 0 | 1 | 1 |
| Galntl5 | 0 | 0 | 0 | 1 | 1 |
| Galp | 0 | 0 | 0 | 1 | 1 |
| Gapt | 0 | 0 | 0 | 1 | 1 |
| Gas2l2 | 0 | 0 | 0 | 1 | 1 |
| Gast | 0 | 0 | 0 | 1 | 1 |
| Gbgt1 | 0 | 0 | 0 | 1 | 1 |
| Gbp10 | 0 | 0 | 0 | 1 | 1 |
| Gbp2b | 0 | 0 | 0 | 1 | 1 |
| Gc | 0 | 0 | 0 | 1 | 1 |
| Gcg | 0 | 0 | 0 | 1 | 1 |
| Gcm1 | 0 | 0 | 0 | 1 | 1 |
| Gcm2 | 0 | 0 | 0 | 1 | 1 |
| Gcnt3 | 0 | 0 | 0 | 1 | 1 |
| Gdf2 | 0 | 0 | 0 | 1 | 1 |
| Gdf3 | 0 | 0 | 0 | 1 | 1 |
| Gfral | 0 | 0 | 0 | 1 | 1 |
| Gfy | 0 | 0 | 0 | 1 | 1 |
| Ggt6 | 0 | 0 | 0 | 1 | 1 |
| Gh | 0 | 0 | 0 | 1 | 1 |
| Ghrh | 0 | 0 | 0 | 1 | 1 |
| Ghrhr | 0 | 0 | 0 | 1 | 1 |
| Gif | 0 | 0 | 0 | 1 | 1 |
| Gimap3 | 0 | 0 | 0 | 1 | 1 |
| Gimap7 | 0 | 0 | 0 | 1 | 1 |
| Gip | 0 | 0 | 0 | 1 | 1 |
| Gjb1 | 0 | 0 | 0 | 1 | 1 |
| Gjd4 | 0 | 0 | 0 | 1 | 1 |
| Gkn1 | 0 | 0 | 0 | 1 | 1 |
| Gkn3 | 0 | 0 | 0 | 1 | 1 |
| Glb1l3 | 0 | 0 | 0 | 1 | 1 |
| Gldnos | 0 | 0 | 0 | 1 | 1 |
| Glipr1 | 0 | 0 | 0 | 1 | 1 |
| Glipr1l1 | 0 | 0 | 0 | 1 | 1 |
| Glipr1l2 | 0 | 0 | 0 | 1 | 1 |
| Glra2 | 0 | 0 | 0 | 1 | 1 |
| Glra3 | 0 | 0 | 0 | 1 | 1 |
| Glrp1 | 0 | 0 | 0 | 1 | 1 |

| | | | | | |
|---|---|---|---|---|---|
| Glt6d1 | 0 | 0 | 0 | 1 | 1 |
| Glyat | 0 | 0 | 0 | 1 | 1 |
| Glycam1 | 0 | 0 | 0 | 1 | 1 |
| Gm10007 | 0 | 0 | 0 | 1 | 1 |
| Gm10024 | 0 | 0 | 0 | 1 | 1 |
| Gm10052 | 0 | 0 | 0 | 1 | 1 |
| Gm10057,Samt1 | 0 | 0 | 0 | 1 | 1 |
| Gm10058,Gm10096,Gm10 | 0 | 0 | 0 | 1 | 1 |
| Gm10081 | 0 | 0 | 0 | 1 | 1 |
| Gm10094,Sap18 | 0 | 0 | 0 | 1 | 1 |
| Gm10096,Gm10486,Gm10 | 0 | 0 | 0 | 1 | 1 |
| Gm10100 | 0 | 0 | 0 | 1 | 1 |
| Gm10104 | 0 | 0 | 0 | 1 | 1 |
| Gm10142 | 0 | 0 | 0 | 1 | 1 |
| Gm10147,Gm14632 | 0 | 0 | 0 | 1 | 1 |
| Gm10228 | 0 | 0 | 0 | 1 | 1 |
| Gm10229 | 0 | 0 | 0 | 1 | 1 |
| Gm10230 | 0 | 0 | 0 | 1 | 1 |
| Gm10248 | 0 | 0 | 0 | 1 | 1 |
| Gm10272 | 0 | 0 | 0 | 1 | 1 |
| Gm10280 | 0 | 0 | 0 | 1 | 1 |
| Gm10318 | 0 | 0 | 0 | 1 | 1 |
| Gm10319 | 0 | 0 | 0 | 1 | 1 |
| Gm10324 | 0 | 0 | 0 | 1 | 1 |
| Gm10334 | 0 | 0 | 0 | 1 | 1 |
| Gm10352,Rbmy | 0 | 0 | 0 | 1 | 1 |
| Gm10354 | 0 | 0 | 0 | 1 | 1 |
| Gm10375 | 0 | 0 | 0 | 1 | 1 |
| Gm10377 | 0 | 0 | 0 | 1 | 1 |
| Gm10408 | 0 | 0 | 0 | 1 | 1 |
| Gm10409,Gm3020 | 0 | 0 | 0 | 1 | 1 |
| Gm10413 | 0 | 0 | 0 | 1 | 1 |
| Gm10415 | 0 | 0 | 0 | 1 | 1 |
| Gm10436 | 0 | 0 | 0 | 1 | 1 |
| Gm10439 | 0 | 0 | 0 | 1 | 1 |
| Gm10445 | 0 | 0 | 0 | 1 | 1 |
| Gm10466 | 0 | 0 | 0 | 1 | 1 |
| Gm10471 | 0 | 0 | 0 | 1 | 1 |
| Gm10486 | 0 | 0 | 0 | 1 | 1 |
| Gm10487 | 0 | 0 | 0 | 1 | 1 |
| Gm10488 | 0 | 0 | 0 | 1 | 1 |
| Gm10494 | 0 | 0 | 0 | 1 | 1 |
| Gm10512 | 0 | 0 | 0 | 1 | 1 |
| Gm10538 | 0 | 0 | 0 | 1 | 1 |
| Gm10556 | 0 | 0 | 0 | 1 | 1 |
| Gm10591 | 0 | 0 | 0 | 1 | 1 |
| Gm10619 | 0 | 0 | 0 | 1 | 1 |
| Gm10635 | 0 | 0 | 0 | 1 | 1 |
| Gm10639 | 0 | 0 | 0 | 1 | 1 |
| Gm10649 | 0 | 0 | 0 | 1 | 1 |

| | | | | | |
|---|---|---|---|---|---|
| Gm10662,Gm5891 | 0 | 0 | 0 | 1 | 1 |
| Gm10665 | 0 | 0 | 0 | 1 | 1 |
| Gm10666 | 0 | 0 | 0 | 1 | 1 |
| Gm10670 | 0 | 0 | 0 | 1 | 1 |
| Gm10681,Hsd3b4 | 0 | 0 | 0 | 1 | 1 |
| Gm10696 | 0 | 0 | 0 | 1 | 1 |
| Gm10714 | 0 | 0 | 0 | 1 | 1 |
| Gm10731 | 0 | 0 | 0 | 1 | 1 |
| Gm10745 | 0 | 0 | 0 | 1 | 1 |
| Gm10754 | 0 | 0 | 0 | 1 | 1 |
| Gm10782 | 0 | 0 | 0 | 1 | 1 |
| Gm10787 | 0 | 0 | 0 | 1 | 1 |
| Gm10804 | 0 | 0 | 0 | 1 | 1 |
| Gm10823 | 0 | 0 | 0 | 1 | 1 |
| Gm10825 | 0 | 0 | 0 | 1 | 1 |
| Gm10921 | 0 | 0 | 0 | 1 | 1 |
| Gm10922 | 0 | 0 | 0 | 1 | 1 |
| Gm11127 | 0 | 0 | 0 | 1 | 1 |
| Gm11128 | 0 | 0 | 0 | 1 | 1 |
| Gm11166 | 0 | 0 | 0 | 1 | 1 |
| Gm11213 | 0 | 0 | 0 | 1 | 1 |
| Gm11237,LOC101055863 | 0 | 0 | 0 | 1 | 1 |
| Gm11240 | 0 | 0 | 0 | 1 | 1 |
| Gm11351 | 0 | 0 | 0 | 1 | 1 |
| Gm1140,Gm14692 | 0 | 0 | 0 | 1 | 1 |
| Gm11426 | 0 | 0 | 0 | 1 | 1 |
| Gm11487 | 0 | 0 | 0 | 1 | 1 |
| Gm11544 | 0 | 0 | 0 | 1 | 1 |
| Gm11548 | 0 | 0 | 0 | 1 | 1 |
| Gm11554 | 0 | 0 | 0 | 1 | 1 |
| Gm11559 | 0 | 0 | 0 | 1 | 1 |
| Gm11562 | 0 | 0 | 0 | 1 | 1 |
| Gm11563 | 0 | 0 | 0 | 1 | 1 |
| Gm11564 | 0 | 0 | 0 | 1 | 1 |
| Gm11565 | 0 | 0 | 0 | 1 | 1 |
| Gm11568 | 0 | 0 | 0 | 1 | 1 |
| Gm11569 | 0 | 0 | 0 | 1 | 1 |
| Gm11595 | 0 | 0 | 0 | 1 | 1 |
| Gm11596 | 0 | 0 | 0 | 1 | 1 |
| Gm11651 | 0 | 0 | 0 | 1 | 1 |
| Gm11710 | 0 | 0 | 0 | 1 | 1 |
| Gm11757,Gm11758,Gm13 | 0 | 0 | 0 | 1 | 1 |
| Gm11758 | 0 | 0 | 0 | 1 | 1 |
| Gm11937 | 0 | 0 | 0 | 1 | 1 |
| Gm11938 | 0 | 0 | 0 | 1 | 1 |
| Gm11978 | 0 | 0 | 0 | 1 | 1 |
| Gm11981 | 0 | 0 | 0 | 1 | 1 |
| Gm12 | 0 | 0 | 0 | 1 | 1 |
| Gm12130 | 0 | 0 | 0 | 1 | 1 |
| Gm12159 | 0 | 0 | 0 | 1 | 1 |

| | | | | | |
|---|---|---|---|---|---|
| Gm12169 | 0 | 0 | 0 | 1 | 1 |
| Gm12171 | 0 | 0 | 0 | 1 | 1 |
| Gm12185 | 0 | 0 | 0 | 1 | 1 |
| Gm12298 | 0 | 0 | 0 | 1 | 1 |
| Gm12409 | 0 | 0 | 0 | 1 | 1 |
| Gm12429 | 0 | 0 | 0 | 1 | 1 |
| Gm12505 | 0 | 0 | 0 | 1 | 1 |
| Gm12530 | 0 | 0 | 0 | 1 | 1 |
| Gm12603 | 0 | 0 | 0 | 1 | 1 |
| Gm12669 | 0 | 0 | 0 | 1 | 1 |
| Gm12695 | 0 | 0 | 0 | 1 | 1 |
| Gm12789 | 0 | 0 | 0 | 1 | 1 |
| Gm12794 | 0 | 0 | 0 | 1 | 1 |
| Gm12886 | 0 | 0 | 0 | 1 | 1 |
| Gm12887 | 0 | 0 | 0 | 1 | 1 |
| Gm12888 | 0 | 0 | 0 | 1 | 1 |
| Gm13011 | 0 | 0 | 0 | 1 | 1 |
| Gm13023 | 0 | 0 | 0 | 1 | 1 |
| Gm13031 | 0 | 0 | 0 | 1 | 1 |
| Gm13040 | 0 | 0 | 0 | 1 | 1 |
| Gm13043 | 0 | 0 | 0 | 1 | 1 |
| Gm13051 | 0 | 0 | 0 | 1 | 1 |
| Gm13078 | 0 | 0 | 0 | 1 | 1 |
| Gm13083 | 0 | 0 | 0 | 1 | 1 |
| Gm13084 | 0 | 0 | 0 | 1 | 1 |
| Gm13088 | 0 | 0 | 0 | 1 | 1 |
| Gm13102 | 0 | 0 | 0 | 1 | 1 |
| Gm13103 | 0 | 0 | 0 | 1 | 1 |
| Gm13119 | 0 | 0 | 0 | 1 | 1 |
| Gm13124 | 0 | 0 | 0 | 1 | 1 |
| Gm13125 | 0 | 0 | 0 | 1 | 1 |
| Gm13128 | 0 | 0 | 0 | 1 | 1 |
| Gm13177 | 0 | 0 | 0 | 1 | 1 |
| Gm13178 | 0 | 0 | 0 | 1 | 1 |
| Gm13238 | 0 | 0 | 0 | 1 | 1 |
| Gm13275,Gm13287 | 0 | 0 | 0 | 1 | 1 |
| Gm13276 | 0 | 0 | 0 | 1 | 1 |
| Gm13278,Gm13279,Gm13 | 0 | 0 | 0 | 1 | 1 |
| Gm13279 | 0 | 0 | 0 | 1 | 1 |
| Gm13285,Gm13288,Gm13 | 0 | 0 | 0 | 1 | 1 |
| Gm13287 | 0 | 0 | 0 | 1 | 1 |
| Gm13290 | 0 | 0 | 0 | 1 | 1 |
| Gm13293 | 0 | 0 | 0 | 1 | 1 |
| Gm13298 | 0 | 0 | 0 | 1 | 1 |
| Gm13304 | 0 | 0 | 0 | 1 | 1 |
| Gm13305,Gm2002 | 0 | 0 | 0 | 1 | 1 |
| Gm13306 | 0 | 0 | 0 | 1 | 1 |
| Gm13308,Gm21586 | 0 | 0 | 0 | 1 | 1 |
| Gm13363 | 0 | 0 | 0 | 1 | 1 |
| Gm13490 | 0 | 0 | 0 | 1 | 1 |

| | | | | | |
|---|---|---|---|---|---|
| Gm13497 | 0 | 0 | 0 | 1 | 1 |
| Gm13544 | 0 | 0 | 0 | 1 | 1 |
| Gm13546 | 0 | 0 | 0 | 1 | 1 |
| Gm13580 | 0 | 0 | 0 | 1 | 1 |
| Gm13629 | 0 | 0 | 0 | 1 | 1 |
| Gm13749 | 0 | 0 | 0 | 1 | 1 |
| Gm13769 | 0 | 0 | 0 | 1 | 1 |
| Gm13807 | 0 | 0 | 0 | 1 | 1 |
| Gm13871 | 0 | 0 | 0 | 1 | 1 |
| Gm14015 | 0 | 0 | 0 | 1 | 1 |
| Gm14085 | 0 | 0 | 0 | 1 | 1 |
| Gm14151 | 0 | 0 | 0 | 1 | 1 |
| Gm14169 | 0 | 0 | 0 | 1 | 1 |
| Gm14288 | 0 | 0 | 0 | 1 | 1 |
| Gm14306,Gm14436 | 0 | 0 | 0 | 1 | 1 |
| Gm14308,OTTMUSG0000( | 0 | 0 | 0 | 1 | 1 |
| Gm14345,Gm14351 | 0 | 0 | 0 | 1 | 1 |
| Gm14346,Gm3706 | 0 | 0 | 0 | 1 | 1 |
| Gm14347 | 0 | 0 | 0 | 1 | 1 |
| Gm14351,Gm2913,Gm293 | 0 | 0 | 0 | 1 | 1 |
| Gm14374 | 0 | 0 | 0 | 1 | 1 |
| Gm14391 | 0 | 0 | 0 | 1 | 1 |
| Gm14405 | 0 | 0 | 0 | 1 | 1 |
| Gm14430,OTTMUSG0000( | 0 | 0 | 0 | 1 | 1 |
| Gm14431 | 0 | 0 | 0 | 1 | 1 |
| Gm14436 | 0 | 0 | 0 | 1 | 1 |
| Gm14440 | 0 | 0 | 0 | 1 | 1 |
| Gm14458 | 0 | 0 | 0 | 1 | 1 |
| Gm14459 | 0 | 0 | 0 | 1 | 1 |
| Gm14461 | 0 | 0 | 0 | 1 | 1 |
| Gm14474 | 0 | 0 | 0 | 1 | 1 |
| Gm14475,Gm14478,Gm14 | 0 | 0 | 0 | 1 | 1 |
| Gm14477,Gm14478,Gm14 | 0 | 0 | 0 | 1 | 1 |
| Gm14478 | 0 | 0 | 0 | 1 | 1 |
| Gm14479,Gm14483 | 0 | 0 | 0 | 1 | 1 |
| Gm14482 | 0 | 0 | 0 | 1 | 1 |
| Gm14483 | 0 | 0 | 0 | 1 | 1 |
| Gm14484 | 0 | 0 | 0 | 1 | 1 |
| Gm14499 | 0 | 0 | 0 | 1 | 1 |
| Gm14501 | 0 | 0 | 0 | 1 | 1 |
| Gm14511 | 0 | 0 | 0 | 1 | 1 |
| Gm14525 | 0 | 0 | 0 | 1 | 1 |
| Gm14548 | 0 | 0 | 0 | 1 | 1 |
| Gm14625 | 0 | 0 | 0 | 1 | 1 |
| Gm14632 | 0 | 0 | 0 | 1 | 1 |
| Gm14635 | 0 | 0 | 0 | 1 | 1 |
| Gm14692 | 0 | 0 | 0 | 1 | 1 |
| Gm14718 | 0 | 0 | 0 | 1 | 1 |
| Gm14725 | 0 | 0 | 0 | 1 | 1 |
| Gm14743 | 0 | 0 | 0 | 1 | 1 |

| | | | | | |
|---|---|---|---|---|---|
| Gm14744 | 0 | 0 | 0 | 1 | 1 |
| Gm14781 | 0 | 0 | 0 | 1 | 1 |
| Gm14812 | 0 | 0 | 0 | 1 | 1 |
| Gm14819 | 0 | 0 | 0 | 1 | 1 |
| Gm14850,Gm21498 | 0 | 0 | 0 | 1 | 1 |
| Gm14851 | 0 | 0 | 0 | 1 | 1 |
| Gm14920 | 0 | 0 | 0 | 1 | 1 |
| Gm15023 | 0 | 0 | 0 | 1 | 1 |
| Gm15056 | 0 | 0 | 0 | 1 | 1 |
| Gm15080 | 0 | 0 | 0 | 1 | 1 |
| Gm15085 | 0 | 0 | 0 | 1 | 1 |
| Gm15091 | 0 | 0 | 0 | 1 | 1 |
| Gm15093 | 0 | 0 | 0 | 1 | 1 |
| Gm15097 | 0 | 0 | 0 | 1 | 1 |
| Gm15107 | 0 | 0 | 0 | 1 | 1 |
| Gm15114 | 0 | 0 | 0 | 1 | 1 |
| Gm15127 | 0 | 0 | 0 | 1 | 1 |
| Gm15140 | 0 | 0 | 0 | 1 | 1 |
| Gm1527 | 0 | 0 | 0 | 1 | 1 |
| Gm15284 | 0 | 0 | 0 | 1 | 1 |
| Gm15292 | 0 | 0 | 0 | 1 | 1 |
| Gm15293 | 0 | 0 | 0 | 1 | 1 |
| Gm15299 | 0 | 0 | 0 | 1 | 1 |
| Gm15308 | 0 | 0 | 0 | 1 | 1 |
| Gm15315 | 0 | 0 | 0 | 1 | 1 |
| Gm15319 | 0 | 0 | 0 | 1 | 1 |
| Gm15386 | 0 | 0 | 0 | 1 | 1 |
| Gm1553 | 0 | 0 | 0 | 1 | 1 |
| Gm156 | 0 | 0 | 0 | 1 | 1 |
| Gm15645 | 0 | 0 | 0 | 1 | 1 |
| Gm15679 | 0 | 0 | 0 | 1 | 1 |
| Gm15698 | 0 | 0 | 0 | 1 | 1 |
| Gm15850 | 0 | 0 | 0 | 1 | 1 |
| Gm1587 | 0 | 0 | 0 | 1 | 1 |
| Gm15881 | 0 | 0 | 0 | 1 | 1 |
| Gm15915 | 0 | 0 | 0 | 1 | 1 |
| Gm15987 | 0 | 0 | 0 | 1 | 1 |
| Gm15997 | 0 | 0 | 0 | 1 | 1 |
| Gm1604b | 0 | 0 | 0 | 1 | 1 |
| Gm16287 | 0 | 0 | 0 | 1 | 1 |
| Gm16291 | 0 | 0 | 0 | 1 | 1 |
| Gm16294 | 0 | 0 | 0 | 1 | 1 |
| Gm16367 | 0 | 0 | 0 | 1 | 1 |
| Gm16381 | 0 | 0 | 0 | 1 | 1 |
| Gm16390 | 0 | 0 | 0 | 1 | 1 |
| Gm16404 | 0 | 0 | 0 | 1 | 1 |
| Gm16405 | 0 | 0 | 0 | 1 | 1 |
| Gm16442 | 0 | 0 | 0 | 1 | 1 |
| Gm16445 | 0 | 0 | 0 | 1 | 1 |
| Gm16451 | 0 | 0 | 0 | 1 | 1 |

| | | | | | |
|---|---|---|---|---|---|
| Gm1647 | 0 | 0 | 0 | 1 | 1 |
| Gm16497 | 0 | 0 | 0 | 1 | 1 |
| Gm16501 | 0 | 0 | 0 | 1 | 1 |
| Gm16548 | 0 | 0 | 0 | 1 | 1 |
| Gm16677 | 0 | 0 | 0 | 1 | 1 |
| Gm16833 | 0 | 0 | 0 | 1 | 1 |
| Gm17019 | 0 | 0 | 0 | 1 | 1 |
| Gm1715 | 0 | 0 | 0 | 1 | 1 |
| Gm1720 | 0 | 0 | 0 | 1 | 1 |
| Gm17252 | 0 | 0 | 0 | 1 | 1 |
| Gm17644 | 0 | 0 | 0 | 1 | 1 |
| Gm17677 | 0 | 0 | 0 | 1 | 1 |
| Gm17689 | 0 | 0 | 0 | 1 | 1 |
| Gm17727 | 0 | 0 | 0 | 1 | 1 |
| Gm17751 | 0 | 0 | 0 | 1 | 1 |
| Gm17757,Gm18853 | 0 | 0 | 0 | 1 | 1 |
| Gm17821 | 0 | 0 | 0 | 1 | 1 |
| Gm17830 | 0 | 0 | 0 | 1 | 1 |
| Gm18409 | 0 | 0 | 0 | 1 | 1 |
| Gm18853 | 0 | 0 | 0 | 1 | 1 |
| Gm19276 | 0 | 0 | 0 | 1 | 1 |
| Gm19277 | 0 | 0 | 0 | 1 | 1 |
| Gm19303 | 0 | 0 | 0 | 1 | 1 |
| Gm19345 | 0 | 0 | 0 | 1 | 1 |
| Gm19402 | 0 | 0 | 0 | 1 | 1 |
| Gm19510 | 0 | 0 | 0 | 1 | 1 |
| Gm19589 | 0 | 0 | 0 | 1 | 1 |
| Gm19619 | 0 | 0 | 0 | 1 | 1 |
| Gm1966 | 0 | 0 | 0 | 1 | 1 |
| Gm19668 | 0 | 0 | 0 | 1 | 1 |
| Gm1968 | 0 | 0 | 0 | 1 | 1 |
| Gm19689 | 0 | 0 | 0 | 1 | 1 |
| Gm19757 | 0 | 0 | 0 | 1 | 1 |
| Gm19782 | 0 | 0 | 0 | 1 | 1 |
| Gm19784 | 0 | 0 | 0 | 1 | 1 |
| Gm1979 | 0 | 0 | 0 | 1 | 1 |
| Gm1987 | 0 | 0 | 0 | 1 | 1 |
| Gm1993 | 0 | 0 | 0 | 1 | 1 |
| Gm2002 | 0 | 0 | 0 | 1 | 1 |
| Gm20063 | 0 | 0 | 0 | 1 | 1 |
| Gm20098 | 0 | 0 | 0 | 1 | 1 |
| Gm2012 | 0 | 0 | 0 | 1 | 1 |
| Gm2016 | 0 | 0 | 0 | 1 | 1 |
| Gm20187 | 0 | 0 | 0 | 1 | 1 |
| Gm2022 | 0 | 0 | 0 | 1 | 1 |
| Gm20268 | 0 | 0 | 0 | 1 | 1 |
| Gm2030 | 0 | 0 | 0 | 1 | 1 |
| Gm20356 | 0 | 0 | 0 | 1 | 1 |
| Gm20362 | 0 | 0 | 0 | 1 | 1 |
| Gm2042 | 0 | 0 | 0 | 1 | 1 |

| | | | | | |
|---|---|---|---|---|---|
| Gm20611 | 0 | 0 | 0 | 1 | 1 |
| Gm20735 | 0 | 0 | 0 | 1 | 1 |
| Gm20736 | 0 | 0 | 0 | 1 | 1 |
| Gm20738,Gm20823 | 0 | 0 | 0 | 1 | 1 |
| Gm20740 | 0 | 0 | 0 | 1 | 1 |
| Gm20741 | 0 | 0 | 0 | 1 | 1 |
| Gm20743 | 0 | 0 | 0 | 1 | 1 |
| Gm20744 | 0 | 0 | 0 | 1 | 1 |
| Gm20745 | 0 | 0 | 0 | 1 | 1 |
| Gm20747 | 0 | 0 | 0 | 1 | 1 |
| Gm20752 | 0 | 0 | 0 | 1 | 1 |
| Gm20753 | 0 | 0 | 0 | 1 | 1 |
| Gm20754 | 0 | 0 | 0 | 1 | 1 |
| Gm20756 | 0 | 0 | 0 | 1 | 1 |
| Gm20757 | 0 | 0 | 0 | 1 | 1 |
| Gm20759 | 0 | 0 | 0 | 1 | 1 |
| Gm20765,Gm21304 | 0 | 0 | 0 | 1 | 1 |
| Gm20767 | 0 | 0 | 0 | 1 | 1 |
| Gm20806 | 0 | 0 | 0 | 1 | 1 |
| Gm20809 | 0 | 0 | 0 | 1 | 1 |
| Gm20815 | 0 | 0 | 0 | 1 | 1 |
| Gm20816,Gm20823,Gm2( | 0 | 0 | 0 | 1 | 1 |
| Gm20822 | 0 | 0 | 0 | 1 | 1 |
| Gm20823 | 0 | 0 | 0 | 1 | 1 |
| Gm20826 | 0 | 0 | 0 | 1 | 1 |
| Gm2083,Mup19 | 0 | 0 | 0 | 1 | 1 |
| Gm20831 | 0 | 0 | 0 | 1 | 1 |
| Gm20854 | 0 | 0 | 0 | 1 | 1 |
| Gm20857 | 0 | 0 | 0 | 1 | 1 |
| Gm20865 | 0 | 0 | 0 | 1 | 1 |
| Gm20867 | 0 | 0 | 0 | 1 | 1 |
| Gm2087 | 0 | 0 | 0 | 1 | 1 |
| Gm20871 | 0 | 0 | 0 | 1 | 1 |
| Gm20877 | 0 | 0 | 0 | 1 | 1 |
| Gm20878 | 0 | 0 | 0 | 1 | 1 |
| Gm20917 | 0 | 0 | 0 | 1 | 1 |
| Gm21002 | 0 | 0 | 0 | 1 | 1 |
| Gm2109 | 0 | 0 | 0 | 1 | 1 |
| Gm21119 | 0 | 0 | 0 | 1 | 1 |
| Gm21221 | 0 | 0 | 0 | 1 | 1 |
| Gm21269 | 0 | 0 | 0 | 1 | 1 |
| Gm21276 | 0 | 0 | 0 | 1 | 1 |
| Gm21283 | 0 | 0 | 0 | 1 | 1 |
| Gm21293,Gm21312 | 0 | 0 | 0 | 1 | 1 |
| Gm21304,Gm21312 | 0 | 0 | 0 | 1 | 1 |
| Gm21312 | 0 | 0 | 0 | 1 | 1 |
| Gm21498 | 0 | 0 | 0 | 1 | 1 |
| Gm21541 | 0 | 0 | 0 | 1 | 1 |
| Gm21586 | 0 | 0 | 0 | 1 | 1 |
| Gm21637 | 0 | 0 | 0 | 1 | 1 |

| | | | | | |
|---|---|---|---|---|---|
| Gm21671 | 0 | 0 | 0 | 1 | 1 |
| Gm21677,Rbmy | 0 | 0 | 0 | 1 | 1 |
| Gm21693 | 0 | 0 | 0 | 1 | 1 |
| Gm21704 | 0 | 0 | 0 | 1 | 1 |
| Gm21943 | 0 | 0 | 0 | 1 | 1 |
| Gm21944 | 0 | 0 | 0 | 1 | 1 |
| Gm21949 | 0 | 0 | 0 | 1 | 1 |
| Gm21950 | 0 | 0 | 0 | 1 | 1 |
| Gm21951 | 0 | 0 | 0 | 1 | 1 |
| Gm2381 | 0 | 0 | 0 | 1 | 1 |
| Gm2447 | 0 | 0 | 0 | 1 | 1 |
| Gm2506 | 0 | 0 | 0 | 1 | 1 |
| Gm2516 | 0 | 0 | 0 | 1 | 1 |
| Gm2663 | 0 | 0 | 0 | 1 | 1 |
| Gm2696 | 0 | 0 | 0 | 1 | 1 |
| Gm2799,Gmcl1l | 0 | 0 | 0 | 1 | 1 |
| Gm2825 | 0 | 0 | 0 | 1 | 1 |
| Gm2837 | 0 | 0 | 0 | 1 | 1 |
| Gm2848 | 0 | 0 | 0 | 1 | 1 |
| Gm2863 | 0 | 0 | 0 | 1 | 1 |
| Gm2913 | 0 | 0 | 0 | 1 | 1 |
| Gm2927 | 0 | 0 | 0 | 1 | 1 |
| Gm2933 | 0 | 0 | 0 | 1 | 1 |
| Gm3020 | 0 | 0 | 0 | 1 | 1 |
| Gm3139 | 0 | 0 | 0 | 1 | 1 |
| Gm3143 | 0 | 0 | 0 | 1 | 1 |
| Gm3238 | 0 | 0 | 0 | 1 | 1 |
| Gm3259 | 0 | 0 | 0 | 1 | 1 |
| Gm3264 | 0 | 0 | 0 | 1 | 1 |
| Gm3279 | 0 | 0 | 0 | 1 | 1 |
| Gm3285 | 0 | 0 | 0 | 1 | 1 |
| Gm3286 | 0 | 0 | 0 | 1 | 1 |
| Gm3336 | 0 | 0 | 0 | 1 | 1 |
| Gm3376,Rbmy | 0 | 0 | 0 | 1 | 1 |
| Gm3383 | 0 | 0 | 0 | 1 | 1 |
| Gm3402,Gm3415 | 0 | 0 | 0 | 1 | 1 |
| Gm3404 | 0 | 0 | 0 | 1 | 1 |
| Gm3409 | 0 | 0 | 0 | 1 | 1 |
| Gm3415 | 0 | 0 | 0 | 1 | 1 |
| Gm3417 | 0 | 0 | 0 | 1 | 1 |
| Gm3428 | 0 | 0 | 0 | 1 | 1 |
| Gm3434 | 0 | 0 | 0 | 1 | 1 |
| Gm3488 | 0 | 0 | 0 | 1 | 1 |
| Gm3500 | 0 | 0 | 0 | 1 | 1 |
| Gm362 | 0 | 0 | 0 | 1 | 1 |
| Gm364 | 0 | 0 | 0 | 1 | 1 |
| Gm3646 | 0 | 0 | 0 | 1 | 1 |
| Gm3701 | 0 | 0 | 0 | 1 | 1 |
| Gm3706 | 0 | 0 | 0 | 1 | 1 |
| Gm3750 | 0 | 0 | 0 | 1 | 1 |

| | | | | | |
|---|---|---|---|---|---|
| Gm3763 | 0 | 0 | 0 | 1 | 1 |
| Gm3776 | 0 | 0 | 0 | 1 | 1 |
| Gm382 | 0 | 0 | 0 | 1 | 1 |
| Gm3893 | 0 | 0 | 0 | 1 | 1 |
| Gm4027 | 0 | 0 | 0 | 1 | 1 |
| Gm4070,Gvin1 | 0 | 0 | 0 | 1 | 1 |
| Gm41 | 0 | 0 | 0 | 1 | 1 |
| Gm4133 | 0 | 0 | 0 | 1 | 1 |
| Gm4141 | 0 | 0 | 0 | 1 | 1 |
| Gm4175 | 0 | 0 | 0 | 1 | 1 |
| Gm4177 | 0 | 0 | 0 | 1 | 1 |
| Gm4201 | 0 | 0 | 0 | 1 | 1 |
| Gm4216 | 0 | 0 | 0 | 1 | 1 |
| Gm4224 | 0 | 0 | 0 | 1 | 1 |
| Gm4251 | 0 | 0 | 0 | 1 | 1 |
| Gm4278 | 0 | 0 | 0 | 1 | 1 |
| Gm428 | 0 | 0 | 0 | 1 | 1 |
| Gm4297 | 0 | 0 | 0 | 1 | 1 |
| Gm4301 | 0 | 0 | 0 | 1 | 1 |
| Gm4302 | 0 | 0 | 0 | 1 | 1 |
| Gm4303 | 0 | 0 | 0 | 1 | 1 |
| Gm4307 | 0 | 0 | 0 | 1 | 1 |
| Gm4312 | 0 | 0 | 0 | 1 | 1 |
| Gm4340 | 0 | 0 | 0 | 1 | 1 |
| Gm436 | 0 | 0 | 0 | 1 | 1 |
| Gm438 | 0 | 0 | 0 | 1 | 1 |
| Gm4461 | 0 | 0 | 0 | 1 | 1 |
| Gm4477 | 0 | 0 | 0 | 1 | 1 |
| Gm4498 | 0 | 0 | 0 | 1 | 1 |
| Gm4541 | 0 | 0 | 0 | 1 | 1 |
| Gm4567,Gm6902 | 0 | 0 | 0 | 1 | 1 |
| Gm4598 | 0 | 0 | 0 | 1 | 1 |
| Gm4710 | 0 | 0 | 0 | 1 | 1 |
| Gm4719 | 0 | 0 | 0 | 1 | 1 |
| Gm4736 | 0 | 0 | 0 | 1 | 1 |
| Gm4745 | 0 | 0 | 0 | 1 | 1 |
| Gm4759 | 0 | 0 | 0 | 1 | 1 |
| Gm4763 | 0 | 0 | 0 | 1 | 1 |
| Gm4788 | 0 | 0 | 0 | 1 | 1 |
| Gm4791 | 0 | 0 | 0 | 1 | 1 |
| Gm4792 | 0 | 0 | 0 | 1 | 1 |
| Gm4794 | 0 | 0 | 0 | 1 | 1 |
| Gm4814 | 0 | 0 | 0 | 1 | 1 |
| Gm4827 | 0 | 0 | 0 | 1 | 1 |
| Gm4836 | 0 | 0 | 0 | 1 | 1 |
| Gm4841 | 0 | 0 | 0 | 1 | 1 |
| Gm4846 | 0 | 0 | 0 | 1 | 1 |
| Gm4847 | 0 | 0 | 0 | 1 | 1 |
| Gm4850 | 0 | 0 | 0 | 1 | 1 |
| Gm4858 | 0 | 0 | 0 | 1 | 1 |

| | | | | | |
|---|---|---|---|---|---|
| Gm4861 | 0 | 0 | 0 | 1 | 1 |
| Gm4871 | 0 | 0 | 0 | 1 | 1 |
| Gm4872 | 0 | 0 | 0 | 1 | 1 |
| Gm4884 | 0 | 0 | 0 | 1 | 1 |
| Gm4894 | 0 | 0 | 0 | 1 | 1 |
| Gm4906 | 0 | 0 | 0 | 1 | 1 |
| Gm4922 | 0 | 0 | 0 | 1 | 1 |
| Gm4926 | 0 | 0 | 0 | 1 | 1 |
| Gm4937 | 0 | 0 | 0 | 1 | 1 |
| Gm4951 | 0 | 0 | 0 | 1 | 1 |
| Gm4952 | 0 | 0 | 0 | 1 | 1 |
| Gm4956 | 0 | 0 | 0 | 1 | 1 |
| Gm4981 | 0 | 0 | 0 | 1 | 1 |
| Gm5 | 0 | 0 | 0 | 1 | 1 |
| Gm5071 | 0 | 0 | 0 | 1 | 1 |
| Gm5072 | 0 | 0 | 0 | 1 | 1 |
| Gm5087 | 0 | 0 | 0 | 1 | 1 |
| Gm5095 | 0 | 0 | 0 | 1 | 1 |
| Gm5108 | 0 | 0 | 0 | 1 | 1 |
| Gm5111 | 0 | 0 | 0 | 1 | 1 |
| Gm5114 | 0 | 0 | 0 | 1 | 1 |
| Gm5122 | 0 | 0 | 0 | 1 | 1 |
| Gm5132 | 0 | 0 | 0 | 1 | 1 |
| Gm5136 | 0 | 0 | 0 | 1 | 1 |
| Gm5168 | 0 | 0 | 0 | 1 | 1 |
| Gm5169 | 0 | 0 | 0 | 1 | 1 |
| Gm525 | 0 | 0 | 0 | 1 | 1 |
| Gm5294 | 0 | 0 | 0 | 1 | 1 |
| Gm5334 | 0 | 0 | 0 | 1 | 1 |
| Gm5346 | 0 | 0 | 0 | 1 | 1 |
| Gm5347 | 0 | 0 | 0 | 1 | 1 |
| Gm5382 | 0 | 0 | 0 | 1 | 1 |
| Gm5409 | 0 | 0 | 0 | 1 | 1 |
| Gm5414 | 0 | 0 | 0 | 1 | 1 |
| Gm5415 | 0 | 0 | 0 | 1 | 1 |
| Gm5416 | 0 | 0 | 0 | 1 | 1 |
| Gm5420 | 0 | 0 | 0 | 1 | 1 |
| Gm5431 | 0 | 0 | 0 | 1 | 1 |
| Gm5441 | 0 | 0 | 0 | 1 | 1 |
| Gm5458 | 0 | 0 | 0 | 1 | 1 |
| Gm5460 | 0 | 0 | 0 | 1 | 1 |
| Gm5476 | 0 | 0 | 0 | 1 | 1 |
| Gm5477 | 0 | 0 | 0 | 1 | 1 |
| Gm5478 | 0 | 0 | 0 | 1 | 1 |
| Gm5483 | 0 | 0 | 0 | 1 | 1 |
| Gm5485 | 0 | 0 | 0 | 1 | 1 |
| Gm5535 | 0 | 0 | 0 | 1 | 1 |
| Gm5538 | 0 | 0 | 0 | 1 | 1 |
| Gm5547 | 0 | 0 | 0 | 1 | 1 |
| Gm5549 | 0 | 0 | 0 | 1 | 1 |

| | | | | | |
|---|---|---|---|---|---|
| Gm5592 | 0 | 0 | 0 | 1 | 1 |
| Gm5615 | 0 | 0 | 0 | 1 | 1 |
| Gm5627 | 0 | 0 | 0 | 1 | 1 |
| Gm5634 | 0 | 0 | 0 | 1 | 1 |
| Gm5635 | 0 | 0 | 0 | 1 | 1 |
| Gm5640 | 0 | 0 | 0 | 1 | 1 |
| Gm5643 | 0 | 0 | 0 | 1 | 1 |
| Gm5662 | 0 | 0 | 0 | 1 | 1 |
| Gm5712 | 0 | 0 | 0 | 1 | 1 |
| Gm572 | 0 | 0 | 0 | 1 | 1 |
| Gm5725 | 0 | 0 | 0 | 1 | 1 |
| Gm5726 | 0 | 0 | 0 | 1 | 1 |
| Gm5728 | 0 | 0 | 0 | 1 | 1 |
| Gm5766 | 0 | 0 | 0 | 1 | 1 |
| Gm5771 | 0 | 0 | 0 | 1 | 1 |
| Gm5795 | 0 | 0 | 0 | 1 | 1 |
| Gm5797 | 0 | 0 | 0 | 1 | 1 |
| Gm5833 | 0 | 0 | 0 | 1 | 1 |
| Gm5860 | 0 | 0 | 0 | 1 | 1 |
| Gm5862 | 0 | 0 | 0 | 1 | 1 |
| Gm5885 | 0 | 0 | 0 | 1 | 1 |
| Gm5886 | 0 | 0 | 0 | 1 | 1 |
| Gm5891 | 0 | 0 | 0 | 1 | 1 |
| Gm5893 | 0 | 0 | 0 | 1 | 1 |
| Gm590 | 0 | 0 | 0 | 1 | 1 |
| Gm5901 | 0 | 0 | 0 | 1 | 1 |
| Gm5916 | 0 | 0 | 0 | 1 | 1 |
| Gm5925 | 0 | 0 | 0 | 1 | 1 |
| Gm5934 | 0 | 0 | 0 | 1 | 1 |
| Gm5935 | 0 | 0 | 0 | 1 | 1 |
| Gm5936 | 0 | 0 | 0 | 1 | 1 |
| Gm5938 | 0 | 0 | 0 | 1 | 1 |
| Gm5941 | 0 | 0 | 0 | 1 | 1 |
| Gm595 | 0 | 0 | 0 | 1 | 1 |
| Gm597 | 0 | 0 | 0 | 1 | 1 |
| Gm6026 | 0 | 0 | 0 | 1 | 1 |
| Gm6034 | 0 | 0 | 0 | 1 | 1 |
| Gm6040 | 0 | 0 | 0 | 1 | 1 |
| Gm609 | 0 | 0 | 0 | 1 | 1 |
| Gm6116 | 0 | 0 | 0 | 1 | 1 |
| Gm6121 | 0 | 0 | 0 | 1 | 1 |
| Gm614 | 0 | 0 | 0 | 1 | 1 |
| Gm6150 | 0 | 0 | 0 | 1 | 1 |
| Gm6164 | 0 | 0 | 0 | 1 | 1 |
| Gm6249 | 0 | 0 | 0 | 1 | 1 |
| Gm6260 | 0 | 0 | 0 | 1 | 1 |
| Gm6268 | 0 | 0 | 0 | 1 | 1 |
| Gm6300 | 0 | 0 | 0 | 1 | 1 |
| Gm6307 | 0 | 0 | 0 | 1 | 1 |
| Gm6313 | 0 | 0 | 0 | 1 | 1 |

| | | | | | |
|---|---|---|---|---|---|
| Gm6329 | 0 | 0 | 0 | 1 | 1 |
| Gm6367 | 0 | 0 | 0 | 1 | 1 |
| Gm6370 | 0 | 0 | 0 | 1 | 1 |
| Gm6406 | 0 | 0 | 0 | 1 | 1 |
| Gm6408 | 0 | 0 | 0 | 1 | 1 |
| Gm6416 | 0 | 0 | 0 | 1 | 1 |
| Gm6432 | 0 | 0 | 0 | 1 | 1 |
| Gm6455 | 0 | 0 | 0 | 1 | 1 |
| Gm6460 | 0 | 0 | 0 | 1 | 1 |
| Gm6567 | 0 | 0 | 0 | 1 | 1 |
| Gm6588 | 0 | 0 | 0 | 1 | 1 |
| Gm6592 | 0 | 0 | 0 | 1 | 1 |
| Gm6602 | 0 | 0 | 0 | 1 | 1 |
| Gm6614 | 0 | 0 | 0 | 1 | 1 |
| Gm6634 | 0 | 0 | 0 | 1 | 1 |
| Gm6639 | 0 | 0 | 0 | 1 | 1 |
| Gm6696 | 0 | 0 | 0 | 1 | 1 |
| Gm6710 | 0 | 0 | 0 | 1 | 1 |
| Gm6760 | 0 | 0 | 0 | 1 | 1 |
| Gm6763 | 0 | 0 | 0 | 1 | 1 |
| Gm6792 | 0 | 0 | 0 | 1 | 1 |
| Gm6793 | 0 | 0 | 0 | 1 | 1 |
| Gm6812 | 0 | 0 | 0 | 1 | 1 |
| Gm6890 | 0 | 0 | 0 | 1 | 1 |
| Gm6902 | 0 | 0 | 0 | 1 | 1 |
| Gm6927 | 0 | 0 | 0 | 1 | 1 |
| Gm6936 | 0 | 0 | 0 | 1 | 1 |
| Gm694 | 0 | 0 | 0 | 1 | 1 |
| Gm7030 | 0 | 0 | 0 | 1 | 1 |
| Gm7056 | 0 | 0 | 0 | 1 | 1 |
| Gm7073 | 0 | 0 | 0 | 1 | 1 |
| Gm7157 | 0 | 0 | 0 | 1 | 1 |
| Gm7168 | 0 | 0 | 0 | 1 | 1 |
| Gm7257 | 0 | 0 | 0 | 1 | 1 |
| Gm7271 | 0 | 0 | 0 | 1 | 1 |
| Gm732 | 0 | 0 | 0 | 1 | 1 |
| Gm7325 | 0 | 0 | 0 | 1 | 1 |
| Gm7337 | 0 | 0 | 0 | 1 | 1 |
| Gm7361 | 0 | 0 | 0 | 1 | 1 |
| Gm7367 | 0 | 0 | 0 | 1 | 1 |
| Gm7534 | 0 | 0 | 0 | 1 | 1 |
| Gm7550 | 0 | 0 | 0 | 1 | 1 |
| Gm7609 | 0 | 0 | 0 | 1 | 1 |
| Gm765 | 0 | 0 | 0 | 1 | 1 |
| Gm766 | 0 | 0 | 0 | 1 | 1 |
| Gm7714 | 0 | 0 | 0 | 1 | 1 |
| Gm773 | 0 | 0 | 0 | 1 | 1 |
| Gm7788 | 0 | 0 | 0 | 1 | 1 |
| Gm7849,Gm7861 | 0 | 0 | 0 | 1 | 1 |
| Gm7854 | 0 | 0 | 0 | 1 | 1 |

| | | | | | |
|---|---|---|---|---|---|
| Gm7861 | 0 | 0 | 0 | 1 | 1 |
| Gm7903 | 0 | 0 | 0 | 1 | 1 |
| Gm7977 | 0 | 0 | 0 | 1 | 1 |
| Gm7978 | 0 | 0 | 0 | 1 | 1 |
| Gm806 | 0 | 0 | 0 | 1 | 1 |
| Gm813 | 0 | 0 | 0 | 1 | 1 |
| Gm815 | 0 | 0 | 0 | 1 | 1 |
| Gm8179 | 0 | 0 | 0 | 1 | 1 |
| Gm8221 | 0 | 0 | 0 | 1 | 1 |
| Gm826 | 0 | 0 | 0 | 1 | 1 |
| Gm8267 | 0 | 0 | 0 | 1 | 1 |
| Gm829 | 0 | 0 | 0 | 1 | 1 |
| Gm8300 | 0 | 0 | 0 | 1 | 1 |
| Gm833 | 0 | 0 | 0 | 1 | 1 |
| Gm8369 | 0 | 0 | 0 | 1 | 1 |
| Gm839 | 0 | 0 | 0 | 1 | 1 |
| Gm8439 | 0 | 0 | 0 | 1 | 1 |
| Gm8453 | 0 | 0 | 0 | 1 | 1 |
| Gm853 | 0 | 0 | 0 | 1 | 1 |
| Gm8579 | 0 | 0 | 0 | 1 | 1 |
| Gm8633 | 0 | 0 | 0 | 1 | 1 |
| Gm8653 | 0 | 0 | 0 | 1 | 1 |
| Gm8693 | 0 | 0 | 0 | 1 | 1 |
| Gm8709 | 0 | 0 | 0 | 1 | 1 |
| Gm8765 | 0 | 0 | 0 | 1 | 1 |
| Gm8817 | 0 | 0 | 0 | 1 | 1 |
| Gm884 | 0 | 0 | 0 | 1 | 1 |
| Gm8882 | 0 | 0 | 0 | 1 | 1 |
| Gm8883 | 0 | 0 | 0 | 1 | 1 |
| Gm8884 | 0 | 0 | 0 | 1 | 1 |
| Gm8979 | 0 | 0 | 0 | 1 | 1 |
| Gm8989 | 0 | 0 | 0 | 1 | 1 |
| Gm904 | 0 | 0 | 0 | 1 | 1 |
| Gm906 | 0 | 0 | 0 | 1 | 1 |
| Gm9112 | 0 | 0 | 0 | 1 | 1 |
| Gm9125 | 0 | 0 | 0 | 1 | 1 |
| Gm9268 | 0 | 0 | 0 | 1 | 1 |
| Gm933 | 0 | 0 | 0 | 1 | 1 |
| Gm9376 | 0 | 0 | 0 | 1 | 1 |
| Gm94 | 0 | 0 | 0 | 1 | 1 |
| Gm9513 | 0 | 0 | 0 | 1 | 1 |
| Gm9573 | 0 | 0 | 0 | 1 | 1 |
| Gm9696 | 0 | 0 | 0 | 1 | 1 |
| Gm9731 | 0 | 0 | 0 | 1 | 1 |
| Gm9733 | 0 | 0 | 0 | 1 | 1 |
| Gm9758 | 0 | 0 | 0 | 1 | 1 |
| Gm9839 | 0 | 0 | 0 | 1 | 1 |
| Gm9962 | 0 | 0 | 0 | 1 | 1 |
| Gm9992 | 0 | 0 | 0 | 1 | 1 |
| Gm9994 | 0 | 0 | 0 | 1 | 1 |

| | | | | | |
|---|---|---|---|---|---|
| Gm9999 | 0 | 0 | 0 | 1 | 1 |
| Gmcl1l | 0 | 0 | 0 | 1 | 1 |
| Gml | 0 | 0 | 0 | 1 | 1 |
| Gngt1 | 0 | 0 | 0 | 1 | 1 |
| Gnrhr | 0 | 0 | 0 | 1 | 1 |
| Golga7b | 0 | 0 | 0 | 1 | 1 |
| Gp2 | 0 | 0 | 0 | 1 | 1 |
| Gp49a | 0 | 0 | 0 | 1 | 1 |
| Gp6 | 0 | 0 | 0 | 1 | 1 |
| Gphb5 | 0 | 0 | 0 | 1 | 1 |
| Gpr110 | 0 | 0 | 0 | 1 | 1 |
| Gpr113 | 0 | 0 | 0 | 1 | 1 |
| Gpr132 | 0 | 0 | 0 | 1 | 1 |
| Gpr143 | 0 | 0 | 0 | 1 | 1 |
| Gpr152 | 0 | 0 | 0 | 1 | 1 |
| Gpr174 | 0 | 0 | 0 | 1 | 1 |
| Gpr31b | 0 | 0 | 0 | 1 | 1 |
| Gpr37l1 | 0 | 0 | 0 | 1 | 1 |
| Gpr6 | 0 | 0 | 0 | 1 | 1 |
| Gprc6a | 0 | 0 | 0 | 1 | 1 |
| Gpx5 | 0 | 0 | 0 | 1 | 1 |
| Grin1os | 0 | 0 | 0 | 1 | 1 |
| Grm7 | 0 | 0 | 0 | 1 | 1 |
| Grp | 0 | 0 | 0 | 1 | 1 |
| Grxcr1 | 0 | 0 | 0 | 1 | 1 |
| Grxcr2 | 0 | 0 | 0 | 1 | 1 |
| Gsc2 | 0 | 0 | 0 | 1 | 1 |
| Gsdma | 0 | 0 | 0 | 1 | 1 |
| Gsdma3 | 0 | 0 | 0 | 1 | 1 |
| Gsdmc | 0 | 0 | 0 | 1 | 1 |
| Gsdmc2 | 0 | 0 | 0 | 1 | 1 |
| Gsdmc3 | 0 | 0 | 0 | 1 | 1 |
| Gsdmcl-ps | 0 | 0 | 0 | 1 | 1 |
| Gsdmcl1 | 0 | 0 | 0 | 1 | 1 |
| Gsdmcl2 | 0 | 0 | 0 | 1 | 1 |
| Gsta1 | 0 | 0 | 0 | 1 | 1 |
| Gsta2 | 0 | 0 | 0 | 1 | 1 |
| Gstm3 | 0 | 0 | 0 | 1 | 1 |
| Guca2a | 0 | 0 | 0 | 1 | 1 |
| Guca2b | 0 | 0 | 0 | 1 | 1 |
| Gucy2d | 0 | 0 | 0 | 1 | 1 |
| Gucy2f | 0 | 0 | 0 | 1 | 1 |
| Gvin1 | 0 | 0 | 0 | 1 | 1 |
| Gys2 | 0 | 0 | 0 | 1 | 1 |
| Gzma | 0 | 0 | 0 | 1 | 1 |
| Gzmb | 0 | 0 | 0 | 1 | 1 |
| Gzmc | 0 | 0 | 0 | 1 | 1 |
| Gzmd | 0 | 0 | 0 | 1 | 1 |
| Gzme | 0 | 0 | 0 | 1 | 1 |
| Gzmf | 0 | 0 | 0 | 1 | 1 |

| | | | | | |
|---|---|---|---|---|---|
| Gzmg | 0 | 0 | 0 | 1 | 1 |
| Gzmn | 0 | 0 | 0 | 1 | 1 |
| H1fnt | 0 | 0 | 0 | 1 | 1 |
| H1foo | 0 | 0 | 0 | 1 | 1 |
| H2-DMb2 | 0 | 0 | 0 | 1 | 1 |
| H2-Ea-ps | 0 | 0 | 0 | 1 | 1 |
| H2-M1 | 0 | 0 | 0 | 1 | 1 |
| H2-M10.1 | 0 | 0 | 0 | 1 | 1 |
| H2-M10.3 | 0 | 0 | 0 | 1 | 1 |
| H2-M10.4 | 0 | 0 | 0 | 1 | 1 |
| H2-M10.5 | 0 | 0 | 0 | 1 | 1 |
| H2-M10.6 | 0 | 0 | 0 | 1 | 1 |
| H2-M11 | 0 | 0 | 0 | 1 | 1 |
| H2-M2 | 0 | 0 | 0 | 1 | 1 |
| H2-M9 | 0 | 0 | 0 | 1 | 1 |
| H2-Ob | 0 | 0 | 0 | 1 | 1 |
| H2-Q1 | 0 | 0 | 0 | 1 | 1 |
| H2-Q9 | 0 | 0 | 0 | 1 | 1 |
| H2-T3 | 0 | 0 | 0 | 1 | 1 |
| H2afb2 | 0 | 0 | 0 | 1 | 1 |
| H2afb3 | 0 | 0 | 0 | 1 | 1 |
| H2bfm | 0 | 0 | 0 | 1 | 1 |
| Habp2 | 0 | 0 | 0 | 1 | 1 |
| Hamp2 | 0 | 0 | 0 | 1 | 1 |
| Hao1 | 0 | 0 | 0 | 1 | 1 |
| Hao2 | 0 | 0 | 0 | 1 | 1 |
| Havcr1 | 0 | 0 | 0 | 1 | 1 |
| Havcr2 | 0 | 0 | 0 | 1 | 1 |
| Hbb-bh2 | 0 | 0 | 0 | 1 | 1 |
| Hc | 0 | 0 | 0 | 1 | 1 |
| Hcrt | 0 | 0 | 0 | 1 | 1 |
| Hdgfl1 | 0 | 0 | 0 | 1 | 1 |
| Hdhd1a | 0 | 0 | 0 | 1 | 1 |
| Hemt1 | 0 | 0 | 0 | 1 | 1 |
| Hhatl | 0 | 0 | 0 | 1 | 1 |
| Hhla1 | 0 | 0 | 0 | 1 | 1 |
| Hils1 | 0 | 0 | 0 | 1 | 1 |
| Hist1h2aa | 0 | 0 | 0 | 1 | 1 |
| Hist1h2ao,Hist1h2ap | 0 | 0 | 0 | 1 | 1 |
| Hist1h2ap | 0 | 0 | 0 | 1 | 1 |
| Hist1h2bq | 0 | 0 | 0 | 1 | 1 |
| Hist1h2br | 0 | 0 | 0 | 1 | 1 |
| Hist1h4m,Hist1h4n | 0 | 0 | 0 | 1 | 1 |
| Hist1h4n | 0 | 0 | 0 | 1 | 1 |
| Hist2h3c1,Hist2h3c2 | 0 | 0 | 0 | 1 | 1 |
| Hist2h3c2 | 0 | 0 | 0 | 1 | 1 |
| Hmgb4 | 0 | 0 | 0 | 1 | 1 |
| Hnf4aos | 0 | 0 | 0 | 1 | 1 |
| Hormad1 | 0 | 0 | 0 | 1 | 1 |
| Hrasls5 | 0 | 0 | 0 | 1 | 1 |

| | | | | | |
|---|---|---|---|---|---|
| Hrct1 | 0 | 0 | 0 | 1 | 1 |
| Hrh2 | 0 | 0 | 0 | 1 | 1 |
| Hrnr | 0 | 0 | 0 | 1 | 1 |
| Hsd11b1 | 0 | 0 | 0 | 1 | 1 |
| Hsd17b13 | 0 | 0 | 0 | 1 | 1 |
| Hsd17b2 | 0 | 0 | 0 | 1 | 1 |
| Hsd17b6 | 0 | 0 | 0 | 1 | 1 |
| Hsd3b1 | 0 | 0 | 0 | 1 | 1 |
| Hsd3b2 | 0 | 0 | 0 | 1 | 1 |
| Hsd3b4 | 0 | 0 | 0 | 1 | 1 |
| Hsd3b5 | 0 | 0 | 0 | 1 | 1 |
| Hsf4 | 0 | 0 | 0 | 1 | 1 |
| Hsfy2 | 0 | 0 | 0 | 1 | 1 |
| Hsh2d | 0 | 0 | 0 | 1 | 1 |
| Htr2c | 0 | 0 | 0 | 1 | 1 |
| Htr3b | 0 | 0 | 0 | 1 | 1 |
| Htr5a | 0 | 0 | 0 | 1 | 1 |
| Hyal4 | 0 | 0 | 0 | 1 | 1 |
| Hyal5 | 0 | 0 | 0 | 1 | 1 |
| Hyal6 | 0 | 0 | 0 | 1 | 1 |
| I730028E13Rik | 0 | 0 | 0 | 1 | 1 |
| I730030J21Rik | 0 | 0 | 0 | 1 | 1 |
| I830077J02Rik | 0 | 0 | 0 | 1 | 1 |
| Ifi203 | 0 | 0 | 0 | 1 | 1 |
| Ifi205 | 0 | 0 | 0 | 1 | 1 |
| Ifi27l2b | 0 | 0 | 0 | 1 | 1 |
| Ifi47 | 0 | 0 | 0 | 1 | 1 |
| Ifitm6 | 0 | 0 | 0 | 1 | 1 |
| Ifltd1 | 0 | 0 | 0 | 1 | 1 |
| Ifna1 | 0 | 0 | 0 | 1 | 1 |
| Ifna11 | 0 | 0 | 0 | 1 | 1 |
| Ifna12 | 0 | 0 | 0 | 1 | 1 |
| Ifna13 | 0 | 0 | 0 | 1 | 1 |
| Ifna14 | 0 | 0 | 0 | 1 | 1 |
| Ifna15 | 0 | 0 | 0 | 1 | 1 |
| Ifna16 | 0 | 0 | 0 | 1 | 1 |
| Ifna2 | 0 | 0 | 0 | 1 | 1 |
| Ifna4 | 0 | 0 | 0 | 1 | 1 |
| Ifna5 | 0 | 0 | 0 | 1 | 1 |
| Ifna6 | 0 | 0 | 0 | 1 | 1 |
| Ifna7 | 0 | 0 | 0 | 1 | 1 |
| Ifna9 | 0 | 0 | 0 | 1 | 1 |
| Ifnab | 0 | 0 | 0 | 1 | 1 |
| Ifnb1 | 0 | 0 | 0 | 1 | 1 |
| Ifne | 0 | 0 | 0 | 1 | 1 |
| Ifng | 0 | 0 | 0 | 1 | 1 |
| Ifnl2 | 0 | 0 | 0 | 1 | 1 |
| Ifnl3 | 0 | 0 | 0 | 1 | 1 |
| Ifnz | 0 | 0 | 0 | 1 | 1 |
| Igbp1b | 0 | 0 | 0 | 1 | 1 |

| | | | | | |
|---|---|---|---|---|---|
| Igfl3 | 0 | 0 | 0 | 1 | 1 |
| Il11ra2 | 0 | 0 | 0 | 1 | 1 |
| Il12b | 0 | 0 | 0 | 1 | 1 |
| Il13 | 0 | 0 | 0 | 1 | 1 |
| Il13ra2 | 0 | 0 | 0 | 1 | 1 |
| Il17a | 0 | 0 | 0 | 1 | 1 |
| Il1b | 0 | 0 | 0 | 1 | 1 |
| Il1f10 | 0 | 0 | 0 | 1 | 1 |
| Il1f6 | 0 | 0 | 0 | 1 | 1 |
| Il1f8 | 0 | 0 | 0 | 1 | 1 |
| Il1f9 | 0 | 0 | 0 | 1 | 1 |
| Il1rapl1 | 0 | 0 | 0 | 1 | 1 |
| Il1rapl2 | 0 | 0 | 0 | 1 | 1 |
| Il1rn | 0 | 0 | 0 | 1 | 1 |
| Il2 | 0 | 0 | 0 | 1 | 1 |
| Il20 | 0 | 0 | 0 | 1 | 1 |
| Il20ra | 0 | 0 | 0 | 1 | 1 |
| Il21 | 0 | 0 | 0 | 1 | 1 |
| Il21r | 0 | 0 | 0 | 1 | 1 |
| Il22ra2 | 0 | 0 | 0 | 1 | 1 |
| Il23r | 0 | 0 | 0 | 1 | 1 |
| Il24 | 0 | 0 | 0 | 1 | 1 |
| Il27 | 0 | 0 | 0 | 1 | 1 |
| Il2rb | 0 | 0 | 0 | 1 | 1 |
| Il3 | 0 | 0 | 0 | 1 | 1 |
| Il31 | 0 | 0 | 0 | 1 | 1 |
| Il34 | 0 | 0 | 0 | 1 | 1 |
| Il4 | 0 | 0 | 0 | 1 | 1 |
| Il4i1 | 0 | 0 | 0 | 1 | 1 |
| Il6 | 0 | 0 | 0 | 1 | 1 |
| Il7r | 0 | 0 | 0 | 1 | 1 |
| Il9 | 0 | 0 | 0 | 1 | 1 |
| Iltifb | 0 | 0 | 0 | 1 | 1 |
| Impg1 | 0 | 0 | 0 | 1 | 1 |
| Ins2 | 0 | 0 | 0 | 1 | 1 |
| Ip6k3 | 0 | 0 | 0 | 1 | 1 |
| Ipw | 0 | 0 | 0 | 1 | 1 |
| Iqcf1 | 0 | 0 | 0 | 1 | 1 |
| Iqcf3 | 0 | 0 | 0 | 1 | 1 |
| Iqcf4 | 0 | 0 | 0 | 1 | 1 |
| Iqcf5 | 0 | 0 | 0 | 1 | 1 |
| Iqcf6 | 0 | 0 | 0 | 1 | 1 |
| Irgc1 | 0 | 0 | 0 | 1 | 1 |
| Isx | 0 | 0 | 0 | 1 | 1 |
| Itgad | 0 | 0 | 0 | 1 | 1 |
| Itgb6 | 0 | 0 | 0 | 1 | 1 |
| Itih1 | 0 | 0 | 0 | 1 | 1 |
| Itk | 0 | 0 | 0 | 1 | 1 |
| Itln1 | 0 | 0 | 0 | 1 | 1 |
| Ivl | 0 | 0 | 0 | 1 | 1 |

| | | | | | |
|---|---|---|---|---|---|
| Iyd | 0 | 0 | 0 | 1 | 1 |
| Izumo2 | 0 | 0 | 0 | 1 | 1 |
| Izumo3 | 0 | 0 | 0 | 1 | 1 |
| Kap | 0 | 0 | 0 | 1 | 1 |
| Kcna10 | 0 | 0 | 0 | 1 | 1 |
| Kcna4 | 0 | 0 | 0 | 1 | 1 |
| Kcnk15 | 0 | 0 | 0 | 1 | 1 |
| Kcnk18 | 0 | 0 | 0 | 1 | 1 |
| Kcnk7 | 0 | 0 | 0 | 1 | 1 |
| Kcnk9 | 0 | 0 | 0 | 1 | 1 |
| Keg1 | 0 | 0 | 0 | 1 | 1 |
| Kera | 0 | 0 | 0 | 1 | 1 |
| Khdc1b | 0 | 0 | 0 | 1 | 1 |
| Khdc1c | 0 | 0 | 0 | 1 | 1 |
| Kir3dl1 | 0 | 0 | 0 | 1 | 1 |
| Kir3dl2 | 0 | 0 | 0 | 1 | 1 |
| Klf17 | 0 | 0 | 0 | 1 | 1 |
| Klhdc7b | 0 | 0 | 0 | 1 | 1 |
| Klhl1 | 0 | 0 | 0 | 1 | 1 |
| Klhl31 | 0 | 0 | 0 | 1 | 1 |
| Klhl38 | 0 | 0 | 0 | 1 | 1 |
| Klk12 | 0 | 0 | 0 | 1 | 1 |
| Klk1b1 | 0 | 0 | 0 | 1 | 1 |
| Klk1b11 | 0 | 0 | 0 | 1 | 1 |
| Klk1b16 | 0 | 0 | 0 | 1 | 1 |
| Klk1b21 | 0 | 0 | 0 | 1 | 1 |
| Klk1b22 | 0 | 0 | 0 | 1 | 1 |
| Klk1b24 | 0 | 0 | 0 | 1 | 1 |
| Klk1b26 | 0 | 0 | 0 | 1 | 1 |
| Klk1b27 | 0 | 0 | 0 | 1 | 1 |
| Klk1b3 | 0 | 0 | 0 | 1 | 1 |
| Klk1b4 | 0 | 0 | 0 | 1 | 1 |
| Klk1b5 | 0 | 0 | 0 | 1 | 1 |
| Klk1b7-ps | 0 | 0 | 0 | 1 | 1 |
| Klk1b8 | 0 | 0 | 0 | 1 | 1 |
| Klk1b9 | 0 | 0 | 0 | 1 | 1 |
| Klk4 | 0 | 0 | 0 | 1 | 1 |
| Klk5 | 0 | 0 | 0 | 1 | 1 |
| Klk6 | 0 | 0 | 0 | 1 | 1 |
| Klkb1 | 0 | 0 | 0 | 1 | 1 |
| Klra1 | 0 | 0 | 0 | 1 | 1 |
| Klra10 | 0 | 0 | 0 | 1 | 1 |
| Klra12 | 0 | 0 | 0 | 1 | 1 |
| Klra13-ps | 0 | 0 | 0 | 1 | 1 |
| Klra14-ps | 0 | 0 | 0 | 1 | 1 |
| Klra15 | 0 | 0 | 0 | 1 | 1 |
| Klra17 | 0 | 0 | 0 | 1 | 1 |
| Klra18,Klra33,Klra4 | 0 | 0 | 0 | 1 | 1 |
| Klra19,Klra6 | 0 | 0 | 0 | 1 | 1 |
| Klra2 | 0 | 0 | 0 | 1 | 1 |

| | | | | | |
|---|---|---|---|---|---|
| Klra21 | 0 | 0 | 0 | 1 | 1 |
| Klra22 | 0 | 0 | 0 | 1 | 1 |
| Klra23 | 0 | 0 | 0 | 1 | 1 |
| Klra3 | 0 | 0 | 0 | 1 | 1 |
| Klra33 | 0 | 0 | 0 | 1 | 1 |
| Klra4 | 0 | 0 | 0 | 1 | 1 |
| Klra5 | 0 | 0 | 0 | 1 | 1 |
| Klra6 | 0 | 0 | 0 | 1 | 1 |
| Klra7 | 0 | 0 | 0 | 1 | 1 |
| Klra8 | 0 | 0 | 0 | 1 | 1 |
| Klra9 | 0 | 0 | 0 | 1 | 1 |
| Klrb1 | 0 | 0 | 0 | 1 | 1 |
| Klrb1-ps1 | 0 | 0 | 0 | 1 | 1 |
| Klrb1a | 0 | 0 | 0 | 1 | 1 |
| Klrb1b | 0 | 0 | 0 | 1 | 1 |
| Klrc1 | 0 | 0 | 0 | 1 | 1 |
| Klrc2 | 0 | 0 | 0 | 1 | 1 |
| Klrc3 | 0 | 0 | 0 | 1 | 1 |
| Klrd1 | 0 | 0 | 0 | 1 | 1 |
| Klre1 | 0 | 0 | 0 | 1 | 1 |
| Klrg1 | 0 | 0 | 0 | 1 | 1 |
| Klri1 | 0 | 0 | 0 | 1 | 1 |
| Klri2 | 0 | 0 | 0 | 1 | 1 |
| Klrk1 | 0 | 0 | 0 | 1 | 1 |
| Kmo | 0 | 0 | 0 | 1 | 1 |
| Kncn | 0 | 0 | 0 | 1 | 1 |
| Kpna7 | 0 | 0 | 0 | 1 | 1 |
| Kprp | 0 | 0 | 0 | 1 | 1 |
| Krt12 | 0 | 0 | 0 | 1 | 1 |
| Krt16 | 0 | 0 | 0 | 1 | 1 |
| Krt24 | 0 | 0 | 0 | 1 | 1 |
| Krt26 | 0 | 0 | 0 | 1 | 1 |
| Krt31 | 0 | 0 | 0 | 1 | 1 |
| Krt32 | 0 | 0 | 0 | 1 | 1 |
| Krt33a | 0 | 0 | 0 | 1 | 1 |
| Krt33b | 0 | 0 | 0 | 1 | 1 |
| Krt34 | 0 | 0 | 0 | 1 | 1 |
| Krt40 | 0 | 0 | 0 | 1 | 1 |
| Krt6a | 0 | 0 | 0 | 1 | 1 |
| Krt6b | 0 | 0 | 0 | 1 | 1 |
| Krt72 | 0 | 0 | 0 | 1 | 1 |
| Krt73 | 0 | 0 | 0 | 1 | 1 |
| Krt74 | 0 | 0 | 0 | 1 | 1 |
| Krt76 | 0 | 0 | 0 | 1 | 1 |
| Krt77 | 0 | 0 | 0 | 1 | 1 |
| Krt85 | 0 | 0 | 0 | 1 | 1 |
| Krtap1-3 | 0 | 0 | 0 | 1 | 1 |
| Krtap1-4 | 0 | 0 | 0 | 1 | 1 |
| Krtap1-5 | 0 | 0 | 0 | 1 | 1 |
| Krtap10-10 | 0 | 0 | 0 | 1 | 1 |

| | | | | | |
|---|---|---|---|---|---|
| Krtap10-4 | 0 | 0 | 0 | 1 | 1 |
| Krtap11-1 | 0 | 0 | 0 | 1 | 1 |
| Krtap12-1 | 0 | 0 | 0 | 1 | 1 |
| Krtap13 | 0 | 0 | 0 | 1 | 1 |
| Krtap13-1 | 0 | 0 | 0 | 1 | 1 |
| Krtap14 | 0 | 0 | 0 | 1 | 1 |
| Krtap15 | 0 | 0 | 0 | 1 | 1 |
| Krtap16-1 | 0 | 0 | 0 | 1 | 1 |
| Krtap16-3 | 0 | 0 | 0 | 1 | 1 |
| Krtap19-1 | 0 | 0 | 0 | 1 | 1 |
| Krtap19-3 | 0 | 0 | 0 | 1 | 1 |
| Krtap19-4 | 0 | 0 | 0 | 1 | 1 |
| Krtap19-5 | 0 | 0 | 0 | 1 | 1 |
| Krtap19-9b | 0 | 0 | 0 | 1 | 1 |
| Krtap2-4 | 0 | 0 | 0 | 1 | 1 |
| Krtap20-2 | 0 | 0 | 0 | 1 | 1 |
| Krtap21-1 | 0 | 0 | 0 | 1 | 1 |
| Krtap22-2 | 0 | 0 | 0 | 1 | 1 |
| Krtap24-1 | 0 | 0 | 0 | 1 | 1 |
| Krtap27-1 | 0 | 0 | 0 | 1 | 1 |
| Krtap3-2 | 0 | 0 | 0 | 1 | 1 |
| Krtap31-1 | 0 | 0 | 0 | 1 | 1 |
| Krtap31-2 | 0 | 0 | 0 | 1 | 1 |
| Krtap4-1 | 0 | 0 | 0 | 1 | 1 |
| Krtap4-13 | 0 | 0 | 0 | 1 | 1 |
| Krtap4-16 | 0 | 0 | 0 | 1 | 1 |
| Krtap4-2 | 0 | 0 | 0 | 1 | 1 |
| Krtap4-6 | 0 | 0 | 0 | 1 | 1 |
| Krtap4-7 | 0 | 0 | 0 | 1 | 1 |
| Krtap4-8 | 0 | 0 | 0 | 1 | 1 |
| Krtap4-9 | 0 | 0 | 0 | 1 | 1 |
| Krtap5-1 | 0 | 0 | 0 | 1 | 1 |
| Krtap5-2 | 0 | 0 | 0 | 1 | 1 |
| Krtap5-3 | 0 | 0 | 0 | 1 | 1 |
| Krtap5-4 | 0 | 0 | 0 | 1 | 1 |
| Krtap5-5 | 0 | 0 | 0 | 1 | 1 |
| Krtap6-1 | 0 | 0 | 0 | 1 | 1 |
| Krtap6-2 | 0 | 0 | 0 | 1 | 1 |
| Krtap6-5 | 0 | 0 | 0 | 1 | 1 |
| Krtap7-1 | 0 | 0 | 0 | 1 | 1 |
| Krtap8-1 | 0 | 0 | 0 | 1 | 1 |
| Krtap9-1 | 0 | 0 | 0 | 1 | 1 |
| Krtap9-5 | 0 | 0 | 0 | 1 | 1 |
| Krtdap | 0 | 0 | 0 | 1 | 1 |
| LOC100038947 | 0 | 0 | 0 | 1 | 1 |
| LOC100040786 | 0 | 0 | 0 | 1 | 1 |
| LOC100043315 | 0 | 0 | 0 | 1 | 1 |
| LOC100048884 | 0 | 0 | 0 | 1 | 1 |
| LOC100502896 | 0 | 0 | 0 | 1 | 1 |
| LOC100503280 | 0 | 0 | 0 | 1 | 1 |

| | | | | | |
|---|---|---|---|---|---|
| LOC100861978 | 0 | 0 | 0 | 1 | 1 |
| LOC100862015 | 0 | 0 | 0 | 1 | 1 |
| LOC101055863 | 0 | 0 | 0 | 1 | 1 |
| LOC101056136 | 0 | 0 | 0 | 1 | 1 |
| LOC101243624 | 0 | 0 | 0 | 1 | 1 |
| LOC102631757 | 0 | 0 | 0 | 1 | 1 |
| LOC102633035 | 0 | 0 | 0 | 1 | 1 |
| LOC102634753 | 0 | 0 | 0 | 1 | 1 |
| LOC666331 | 0 | 0 | 0 | 1 | 1 |
| Lalba | 0 | 0 | 0 | 1 | 1 |
| Lao1 | 0 | 0 | 0 | 1 | 1 |
| Lce1a1 | 0 | 0 | 0 | 1 | 1 |
| Lce1a2 | 0 | 0 | 0 | 1 | 1 |
| Lce1c | 0 | 0 | 0 | 1 | 1 |
| Lce1d | 0 | 0 | 0 | 1 | 1 |
| Lce1e | 0 | 0 | 0 | 1 | 1 |
| Lce1f | 0 | 0 | 0 | 1 | 1 |
| Lce1g | 0 | 0 | 0 | 1 | 1 |
| Lce1h | 0 | 0 | 0 | 1 | 1 |
| Lce1j | 0 | 0 | 0 | 1 | 1 |
| Lce1l | 0 | 0 | 0 | 1 | 1 |
| Lce1m | 0 | 0 | 0 | 1 | 1 |
| Lce3a | 0 | 0 | 0 | 1 | 1 |
| Lce3b | 0 | 0 | 0 | 1 | 1 |
| Lce3c | 0 | 0 | 0 | 1 | 1 |
| Lce3d | 0 | 0 | 0 | 1 | 1 |
| Lce3e | 0 | 0 | 0 | 1 | 1 |
| Lce3f | 0 | 0 | 0 | 1 | 1 |
| Lce6a | 0 | 0 | 0 | 1 | 1 |
| Lcn10 | 0 | 0 | 0 | 1 | 1 |
| Lcn11 | 0 | 0 | 0 | 1 | 1 |
| Lcn12 | 0 | 0 | 0 | 1 | 1 |
| Lcn2 | 0 | 0 | 0 | 1 | 1 |
| Lcn3 | 0 | 0 | 0 | 1 | 1 |
| Lcn4 | 0 | 0 | 0 | 1 | 1 |
| Lcn5 | 0 | 0 | 0 | 1 | 1 |
| Lcn6 | 0 | 0 | 0 | 1 | 1 |
| Lcn8 | 0 | 0 | 0 | 1 | 1 |
| Lcn9 | 0 | 0 | 0 | 1 | 1 |
| Ldlrad2 | 0 | 0 | 0 | 1 | 1 |
| Lect2 | 0 | 0 | 0 | 1 | 1 |
| Lelp1 | 0 | 0 | 0 | 1 | 1 |
| Lep | 0 | 0 | 0 | 1 | 1 |
| Lilra5 | 0 | 0 | 0 | 1 | 1 |
| Lilra6 | 0 | 0 | 0 | 1 | 1 |
| Lilrb4 | 0 | 0 | 0 | 1 | 1 |
| Lim2 | 0 | 0 | 0 | 1 | 1 |
| Lincrna-cox2 | 0 | 0 | 0 | 1 | 1 |
| Lipf | 0 | 0 | 0 | 1 | 1 |
| Lipi | 0 | 0 | 0 | 1 | 1 |

| | | | | | |
|---|---|---|---|---|---|
| Lipk | 0 | 0 | 0 | 1 | 1 |
| Lipm | 0 | 0 | 0 | 1 | 1 |
| Lipn | 0 | 0 | 0 | 1 | 1 |
| Lmod2 | 0 | 0 | 0 | 1 | 1 |
| Lmod3 | 0 | 0 | 0 | 1 | 1 |
| Loxhd1 | 0 | 0 | 0 | 1 | 1 |
| Lpcat2b | 0 | 0 | 0 | 1 | 1 |
| Lrcol1 | 0 | 0 | 0 | 1 | 1 |
| Lrg1 | 0 | 0 | 0 | 1 | 1 |
| Lrit1 | 0 | 0 | 0 | 1 | 1 |
| Lrit2 | 0 | 0 | 0 | 1 | 1 |
| Lrrc25 | 0 | 0 | 0 | 1 | 1 |
| Lrrc26 | 0 | 0 | 0 | 1 | 1 |
| Lrrc43 | 0 | 0 | 0 | 1 | 1 |
| Lrrc52 | 0 | 0 | 0 | 1 | 1 |
| Lrrc66 | 0 | 0 | 0 | 1 | 1 |
| Lrrc69 | 0 | 0 | 0 | 1 | 1 |
| Lrrc72 | 0 | 0 | 0 | 1 | 1 |
| Lrriq4 | 0 | 0 | 0 | 1 | 1 |
| Lrtm2 | 0 | 0 | 0 | 1 | 1 |
| Lsmem1 | 0 | 0 | 0 | 1 | 1 |
| Ltb4r1 | 0 | 0 | 0 | 1 | 1 |
| Ltf | 0 | 0 | 0 | 1 | 1 |
| Luzp4 | 0 | 0 | 0 | 1 | 1 |
| Ly6c2 | 0 | 0 | 0 | 1 | 1 |
| Ly6d | 0 | 0 | 0 | 1 | 1 |
| Ly6g5c | 0 | 0 | 0 | 1 | 1 |
| Ly6g6f | 0 | 0 | 0 | 1 | 1 |
| Ly6i | 0 | 0 | 0 | 1 | 1 |
| Ly6k | 0 | 0 | 0 | 1 | 1 |
| Ly86 | 0 | 0 | 0 | 1 | 1 |
| Ly9 | 0 | 0 | 0 | 1 | 1 |
| Lyg1 | 0 | 0 | 0 | 1 | 1 |
| Lyg2 | 0 | 0 | 0 | 1 | 1 |
| Lypd4 | 0 | 0 | 0 | 1 | 1 |
| Lypd5 | 0 | 0 | 0 | 1 | 1 |
| Lypd8 | 0 | 0 | 0 | 1 | 1 |
| Lyrm7os | 0 | 0 | 0 | 1 | 1 |
| Lyve1 | 0 | 0 | 0 | 1 | 1 |
| Lyzl1 | 0 | 0 | 0 | 1 | 1 |
| Lyzl4 | 0 | 0 | 0 | 1 | 1 |
| Lyzl4os | 0 | 0 | 0 | 1 | 1 |
| Lyzl6 | 0 | 0 | 0 | 1 | 1 |
| Magea1 | 0 | 0 | 0 | 1 | 1 |
| Magea10 | 0 | 0 | 0 | 1 | 1 |
| Magea2 | 0 | 0 | 0 | 1 | 1 |
| Magea3 | 0 | 0 | 0 | 1 | 1 |
| Magea4 | 0 | 0 | 0 | 1 | 1 |
| Magea6 | 0 | 0 | 0 | 1 | 1 |
| Magea8 | 0 | 0 | 0 | 1 | 1 |

| | | | | | |
|---|---|---|---|---|---|
| Mageb1 | 0 | 0 | 0 | 1 | 1 |
| Mageb16 | 0 | 0 | 0 | 1 | 1 |
| Mageb16-ps1 | 0 | 0 | 0 | 1 | 1 |
| Mageb18 | 0 | 0 | 0 | 1 | 1 |
| Mageb2 | 0 | 0 | 0 | 1 | 1 |
| Mageb4 | 0 | 0 | 0 | 1 | 1 |
| Mageb5 | 0 | 0 | 0 | 1 | 1 |
| Manr | 0 | 0 | 0 | 1 | 1 |
| Map3k19 | 0 | 0 | 0 | 1 | 1 |
| Marco | 0 | 0 | 0 | 1 | 1 |
| Mat1a | 0 | 0 | 0 | 1 | 1 |
| Mbd3l1 | 0 | 0 | 0 | 1 | 1 |
| Mbl1 | 0 | 0 | 0 | 1 | 1 |
| Mbl2 | 0 | 0 | 0 | 1 | 1 |
| Mc3r | 0 | 0 | 0 | 1 | 1 |
| Mcidas | 0 | 0 | 0 | 1 | 1 |
| Mcpt-ps1 | 0 | 0 | 0 | 1 | 1 |
| Mcpt1 | 0 | 0 | 0 | 1 | 1 |
| Mcpt2 | 0 | 0 | 0 | 1 | 1 |
| Mcpt4 | 0 | 0 | 0 | 1 | 1 |
| Mcpt8 | 0 | 0 | 0 | 1 | 1 |
| Mcpt9 | 0 | 0 | 0 | 1 | 1 |
| Mefv | 0 | 0 | 0 | 1 | 1 |
| Meig1 | 0 | 0 | 0 | 1 | 1 |
| Meiob | 0 | 0 | 0 | 1 | 1 |
| Mep1b | 0 | 0 | 0 | 1 | 1 |
| Mettl11b | 0 | 0 | 0 | 1 | 1 |
| Mettl21e | 0 | 0 | 0 | 1 | 1 |
| Mettl7a3 | 0 | 0 | 0 | 1 | 1 |
| Mettl7b | 0 | 0 | 0 | 1 | 1 |
| Mgl2 | 0 | 0 | 0 | 1 | 1 |
| Mill1 | 0 | 0 | 0 | 1 | 1 |
| Mir100 | 0 | 0 | 0 | 1 | 1 |
| Mir101a | 0 | 0 | 0 | 1 | 1 |
| Mir101b | 0 | 0 | 0 | 1 | 1 |
| Mir101c | 0 | 0 | 0 | 1 | 1 |
| Mir103-1 | 0 | 0 | 0 | 1 | 1 |
| Mir103-2 | 0 | 0 | 0 | 1 | 1 |
| Mir105 | 0 | 0 | 0 | 1 | 1 |
| Mir106a | 0 | 0 | 0 | 1 | 1 |
| Mir106b | 0 | 0 | 0 | 1 | 1 |
| Mir107 | 0 | 0 | 0 | 1 | 1 |
| Mir10a | 0 | 0 | 0 | 1 | 1 |
| Mir10b | 0 | 0 | 0 | 1 | 1 |
| Mir1187 | 0 | 0 | 0 | 1 | 1 |
| Mir1190 | 0 | 0 | 0 | 1 | 1 |
| Mir1191 | 0 | 0 | 0 | 1 | 1 |
| Mir1191b | 0 | 0 | 0 | 1 | 1 |
| Mir1192 | 0 | 0 | 0 | 1 | 1 |
| Mir1193 | 0 | 0 | 0 | 1 | 1 |

| | | | | | |
|---|---|---|---|---|---|
| Mir1195 | 0 | 0 | 0 | 1 | 1 |
| Mir1197 | 0 | 0 | 0 | 1 | 1 |
| Mir1198 | 0 | 0 | 0 | 1 | 1 |
| Mir1199 | 0 | 0 | 0 | 1 | 1 |
| Mir1224 | 0 | 0 | 0 | 1 | 1 |
| Mir122a | 0 | 0 | 0 | 1 | 1 |
| Mir1231 | 0 | 0 | 0 | 1 | 1 |
| Mir1247 | 0 | 0 | 0 | 1 | 1 |
| Mir1249 | 0 | 0 | 0 | 1 | 1 |
| Mir124a-1 | 0 | 0 | 0 | 1 | 1 |
| Mir124a-2 | 0 | 0 | 0 | 1 | 1 |
| Mir124a-3 | 0 | 0 | 0 | 1 | 1 |
| Mir1251 | 0 | 0 | 0 | 1 | 1 |
| Mir1258 | 0 | 0 | 0 | 1 | 1 |
| Mir125a | 0 | 0 | 0 | 1 | 1 |
| Mir125b-1 | 0 | 0 | 0 | 1 | 1 |
| Mir125b-2 | 0 | 0 | 0 | 1 | 1 |
| Mir126 | 0 | 0 | 0 | 1 | 1 |
| Mir1264 | 0 | 0 | 0 | 1 | 1 |
| Mir126b | 0 | 0 | 0 | 1 | 1 |
| Mir127 | 0 | 0 | 0 | 1 | 1 |
| Mir128-1 | 0 | 0 | 0 | 1 | 1 |
| Mir128-2 | 0 | 0 | 0 | 1 | 1 |
| Mir129-1 | 0 | 0 | 0 | 1 | 1 |
| Mir129-2 | 0 | 0 | 0 | 1 | 1 |
| Mir1298 | 0 | 0 | 0 | 1 | 1 |
| Mir129b | 0 | 0 | 0 | 1 | 1 |
| Mir1306 | 0 | 0 | 0 | 1 | 1 |
| Mir130b | 0 | 0 | 0 | 1 | 1 |
| Mir132 | 0 | 0 | 0 | 1 | 1 |
| Mir133a-1 | 0 | 0 | 0 | 1 | 1 |
| Mir133a-2 | 0 | 0 | 0 | 1 | 1 |
| Mir133b | 0 | 0 | 0 | 1 | 1 |
| Mir133c | 0 | 0 | 0 | 1 | 1 |
| Mir134 | 0 | 0 | 0 | 1 | 1 |
| Mir135a-1 | 0 | 0 | 0 | 1 | 1 |
| Mir135a-2 | 0 | 0 | 0 | 1 | 1 |
| Mir135b | 0 | 0 | 0 | 1 | 1 |
| Mir136 | 0 | 0 | 0 | 1 | 1 |
| Mir137 | 0 | 0 | 0 | 1 | 1 |
| Mir138-1 | 0 | 0 | 0 | 1 | 1 |
| Mir138-2 | 0 | 0 | 0 | 1 | 1 |
| Mir139 | 0 | 0 | 0 | 1 | 1 |
| Mir140 | 0 | 0 | 0 | 1 | 1 |
| Mir141 | 0 | 0 | 0 | 1 | 1 |
| Mir142 | 0 | 0 | 0 | 1 | 1 |
| Mir142b | 0 | 0 | 0 | 1 | 1 |
| Mir143 | 0 | 0 | 0 | 1 | 1 |
| Mir144 | 0 | 0 | 0 | 1 | 1 |
| Mir145 | 0 | 0 | 0 | 1 | 1 |

| | | | | | |
|---|---|---|---|---|---|
| Mir145b | 0 | 0 | 0 | 1 | 1 |
| Mir146 | 0 | 0 | 0 | 1 | 1 |
| Mir146b | 0 | 0 | 0 | 1 | 1 |
| Mir147 | 0 | 0 | 0 | 1 | 1 |
| Mir148a | 0 | 0 | 0 | 1 | 1 |
| Mir148b | 0 | 0 | 0 | 1 | 1 |
| Mir149 | 0 | 0 | 0 | 1 | 1 |
| Mir152 | 0 | 0 | 0 | 1 | 1 |
| Mir153 | 0 | 0 | 0 | 1 | 1 |
| Mir154 | 0 | 0 | 0 | 1 | 1 |
| Mir155 | 0 | 0 | 0 | 1 | 1 |
| Mir15a | 0 | 0 | 0 | 1 | 1 |
| Mir15b | 0 | 0 | 0 | 1 | 1 |
| Mir16-1 | 0 | 0 | 0 | 1 | 1 |
| Mir16-2 | 0 | 0 | 0 | 1 | 1 |
| Mir17 | 0 | 0 | 0 | 1 | 1 |
| Mir18 | 0 | 0 | 0 | 1 | 1 |
| Mir181a-1 | 0 | 0 | 0 | 1 | 1 |
| Mir181a-2 | 0 | 0 | 0 | 1 | 1 |
| Mir181b-1 | 0 | 0 | 0 | 1 | 1 |
| Mir181b-2 | 0 | 0 | 0 | 1 | 1 |
| Mir181c | 0 | 0 | 0 | 1 | 1 |
| Mir181d | 0 | 0 | 0 | 1 | 1 |
| Mir182 | 0 | 0 | 0 | 1 | 1 |
| Mir183 | 0 | 0 | 0 | 1 | 1 |
| Mir184 | 0 | 0 | 0 | 1 | 1 |
| Mir1843 | 0 | 0 | 0 | 1 | 1 |
| Mir1843b | 0 | 0 | 0 | 1 | 1 |
| Mir185 | 0 | 0 | 0 | 1 | 1 |
| Mir186 | 0 | 0 | 0 | 1 | 1 |
| Mir187 | 0 | 0 | 0 | 1 | 1 |
| Mir188 | 0 | 0 | 0 | 1 | 1 |
| Mir1893 | 0 | 0 | 0 | 1 | 1 |
| Mir1896 | 0 | 0 | 0 | 1 | 1 |
| Mir1897 | 0 | 0 | 0 | 1 | 1 |
| Mir1898 | 0 | 0 | 0 | 1 | 1 |
| Mir1899 | 0 | 0 | 0 | 1 | 1 |
| Mir18b | 0 | 0 | 0 | 1 | 1 |
| Mir190 | 0 | 0 | 0 | 1 | 1 |
| Mir1900 | 0 | 0 | 0 | 1 | 1 |
| Mir1902 | 0 | 0 | 0 | 1 | 1 |
| Mir1904 | 0 | 0 | 0 | 1 | 1 |
| Mir1906-1 | 0 | 0 | 0 | 1 | 1 |
| Mir1906-2 | 0 | 0 | 0 | 1 | 1 |
| Mir190b | 0 | 0 | 0 | 1 | 1 |
| Mir191 | 0 | 0 | 0 | 1 | 1 |
| Mir1912 | 0 | 0 | 0 | 1 | 1 |
| Mir192 | 0 | 0 | 0 | 1 | 1 |
| Mir1928 | 0 | 0 | 0 | 1 | 1 |
| Mir1929 | 0 | 0 | 0 | 1 | 1 |

| | | | | | |
|---|---|---|---|---|---|
| Mir193 | 0 | 0 | 0 | 1 | 1 |
| Mir1930 | 0 | 0 | 0 | 1 | 1 |
| Mir1931 | 0 | 0 | 0 | 1 | 1 |
| Mir1933 | 0 | 0 | 0 | 1 | 1 |
| Mir1936 | 0 | 0 | 0 | 1 | 1 |
| Mir193b | 0 | 0 | 0 | 1 | 1 |
| Mir194-2 | 0 | 0 | 0 | 1 | 1 |
| Mir1940 | 0 | 0 | 0 | 1 | 1 |
| Mir1941 | 0 | 0 | 0 | 1 | 1 |
| Mir1942 | 0 | 0 | 0 | 1 | 1 |
| Mir1945 | 0 | 0 | 0 | 1 | 1 |
| Mir1946a | 0 | 0 | 0 | 1 | 1 |
| Mir1946b | 0 | 0 | 0 | 1 | 1 |
| Mir1947 | 0 | 0 | 0 | 1 | 1 |
| Mir1948 | 0 | 0 | 0 | 1 | 1 |
| Mir195 | 0 | 0 | 0 | 1 | 1 |
| Mir1950 | 0 | 0 | 0 | 1 | 1 |
| Mir1951 | 0 | 0 | 0 | 1 | 1 |
| Mir1952 | 0 | 0 | 0 | 1 | 1 |
| Mir1953 | 0 | 0 | 0 | 1 | 1 |
| Mir1954 | 0 | 0 | 0 | 1 | 1 |
| Mir1957 | 0 | 0 | 0 | 1 | 1 |
| Mir1958 | 0 | 0 | 0 | 1 | 1 |
| Mir195b | 0 | 0 | 0 | 1 | 1 |
| Mir1960 | 0 | 0 | 0 | 1 | 1 |
| Mir1961 | 0 | 0 | 0 | 1 | 1 |
| Mir1962 | 0 | 0 | 0 | 1 | 1 |
| Mir1963 | 0 | 0 | 0 | 1 | 1 |
| Mir1964 | 0 | 0 | 0 | 1 | 1 |
| Mir1967 | 0 | 0 | 0 | 1 | 1 |
| Mir1968 | 0 | 0 | 0 | 1 | 1 |
| Mir1969 | 0 | 0 | 0 | 1 | 1 |
| Mir196a-2 | 0 | 0 | 0 | 1 | 1 |
| Mir196b | 0 | 0 | 0 | 1 | 1 |
| Mir1970 | 0 | 0 | 0 | 1 | 1 |
| Mir1971 | 0 | 0 | 0 | 1 | 1 |
| Mir1981 | 0 | 0 | 0 | 1 | 1 |
| Mir1983 | 0 | 0 | 0 | 1 | 1 |
| Mir199a-2 | 0 | 0 | 0 | 1 | 1 |
| Mir19a | 0 | 0 | 0 | 1 | 1 |
| Mir19b-2 | 0 | 0 | 0 | 1 | 1 |
| Mir1a-1 | 0 | 0 | 0 | 1 | 1 |
| Mir1a-2 | 0 | 0 | 0 | 1 | 1 |
| Mir200a | 0 | 0 | 0 | 1 | 1 |
| Mir200b | 0 | 0 | 0 | 1 | 1 |
| Mir200c | 0 | 0 | 0 | 1 | 1 |
| Mir201 | 0 | 0 | 0 | 1 | 1 |
| Mir202 | 0 | 0 | 0 | 1 | 1 |
| Mir203 | 0 | 0 | 0 | 1 | 1 |
| Mir204 | 0 | 0 | 0 | 1 | 1 |

| | | | | | |
|---|---|---|---|---|---|
| Mir205 | 0 | 0 | 0 | 1 | 1 |
| Mir206 | 0 | 0 | 0 | 1 | 1 |
| Mir207 | 0 | 0 | 0 | 1 | 1 |
| Mir208a | 0 | 0 | 0 | 1 | 1 |
| Mir208b | 0 | 0 | 0 | 1 | 1 |
| Mir20b | 0 | 0 | 0 | 1 | 1 |
| Mir21 | 0 | 0 | 0 | 1 | 1 |
| Mir210 | 0 | 0 | 0 | 1 | 1 |
| Mir211 | 0 | 0 | 0 | 1 | 1 |
| Mir212 | 0 | 0 | 0 | 1 | 1 |
| Mir2136 | 0 | 0 | 0 | 1 | 1 |
| Mir2137 | 0 | 0 | 0 | 1 | 1 |
| Mir214 | 0 | 0 | 0 | 1 | 1 |
| Mir216a | 0 | 0 | 0 | 1 | 1 |
| Mir216b | 0 | 0 | 0 | 1 | 1 |
| Mir216c | 0 | 0 | 0 | 1 | 1 |
| Mir217 | 0 | 0 | 0 | 1 | 1 |
| Mir218-1 | 0 | 0 | 0 | 1 | 1 |
| Mir218-2 | 0 | 0 | 0 | 1 | 1 |
| Mir219-2 | 0 | 0 | 0 | 1 | 1 |
| Mir219c | 0 | 0 | 0 | 1 | 1 |
| Mir21b | 0 | 0 | 0 | 1 | 1 |
| Mir21c | 0 | 0 | 0 | 1 | 1 |
| Mir22 | 0 | 0 | 0 | 1 | 1 |
| Mir221 | 0 | 0 | 0 | 1 | 1 |
| Mir222 | 0 | 0 | 0 | 1 | 1 |
| Mir223 | 0 | 0 | 0 | 1 | 1 |
| Mir23b | 0 | 0 | 0 | 1 | 1 |
| Mir24-1 | 0 | 0 | 0 | 1 | 1 |
| Mir24-2 | 0 | 0 | 0 | 1 | 1 |
| Mir26a-1 | 0 | 0 | 0 | 1 | 1 |
| Mir26a-2 | 0 | 0 | 0 | 1 | 1 |
| Mir26b | 0 | 0 | 0 | 1 | 1 |
| Mir27a | 0 | 0 | 0 | 1 | 1 |
| Mir27b | 0 | 0 | 0 | 1 | 1 |
| Mir28 | 0 | 0 | 0 | 1 | 1 |
| Mir28b | 0 | 0 | 0 | 1 | 1 |
| Mir28c | 0 | 0 | 0 | 1 | 1 |
| Mir290 | 0 | 0 | 0 | 1 | 1 |
| Mir290b | 0 | 0 | 0 | 1 | 1 |
| Mir291a | 0 | 0 | 0 | 1 | 1 |
| Mir291b | 0 | 0 | 0 | 1 | 1 |
| Mir292 | 0 | 0 | 0 | 1 | 1 |
| Mir292b | 0 | 0 | 0 | 1 | 1 |
| Mir293 | 0 | 0 | 0 | 1 | 1 |
| Mir294 | 0 | 0 | 0 | 1 | 1 |
| Mir295 | 0 | 0 | 0 | 1 | 1 |
| Mir296 | 0 | 0 | 0 | 1 | 1 |
| Mir297-1 | 0 | 0 | 0 | 1 | 1 |
| Mir297-2 | 0 | 0 | 0 | 1 | 1 |

| | | | | | |
|---|---|---|---|---|---|
| Mir297a-3,Mir297a-4 | 0 | 0 | 0 | 1 | 1 |
| Mir297a-4 | 0 | 0 | 0 | 1 | 1 |
| Mir297b | 0 | 0 | 0 | 1 | 1 |
| Mir297c | 0 | 0 | 0 | 1 | 1 |
| Mir298 | 0 | 0 | 0 | 1 | 1 |
| Mir299 | 0 | 0 | 0 | 1 | 1 |
| Mir299b | 0 | 0 | 0 | 1 | 1 |
| Mir29a | 0 | 0 | 0 | 1 | 1 |
| Mir29b-1 | 0 | 0 | 0 | 1 | 1 |
| Mir29b-2 | 0 | 0 | 0 | 1 | 1 |
| Mir29c | 0 | 0 | 0 | 1 | 1 |
| Mir300 | 0 | 0 | 0 | 1 | 1 |
| Mir301 | 0 | 0 | 0 | 1 | 1 |
| Mir301b | 0 | 0 | 0 | 1 | 1 |
| Mir302a | 0 | 0 | 0 | 1 | 1 |
| Mir302b | 0 | 0 | 0 | 1 | 1 |
| Mir302c | 0 | 0 | 0 | 1 | 1 |
| Mir302d | 0 | 0 | 0 | 1 | 1 |
| Mir3057 | 0 | 0 | 0 | 1 | 1 |
| Mir3059 | 0 | 0 | 0 | 1 | 1 |
| Mir3060 | 0 | 0 | 0 | 1 | 1 |
| Mir3061 | 0 | 0 | 0 | 1 | 1 |
| Mir3063 | 0 | 0 | 0 | 1 | 1 |
| Mir3065 | 0 | 0 | 0 | 1 | 1 |
| Mir3066 | 0 | 0 | 0 | 1 | 1 |
| Mir3067 | 0 | 0 | 0 | 1 | 1 |
| Mir3068 | 0 | 0 | 0 | 1 | 1 |
| Mir3069 | 0 | 0 | 0 | 1 | 1 |
| Mir3070a | 0 | 0 | 0 | 1 | 1 |
| Mir3071 | 0 | 0 | 0 | 1 | 1 |
| Mir3073 | 0 | 0 | 0 | 1 | 1 |
| Mir3073b | 0 | 0 | 0 | 1 | 1 |
| Mir3074-1 | 0 | 0 | 0 | 1 | 1 |
| Mir3074-2 | 0 | 0 | 0 | 1 | 1 |
| Mir3075 | 0 | 0 | 0 | 1 | 1 |
| Mir3076 | 0 | 0 | 0 | 1 | 1 |
| Mir3077 | 0 | 0 | 0 | 1 | 1 |
| Mir3078 | 0 | 0 | 0 | 1 | 1 |
| Mir3079 | 0 | 0 | 0 | 1 | 1 |
| Mir3081 | 0 | 0 | 0 | 1 | 1 |
| Mir3082 | 0 | 0 | 0 | 1 | 1 |
| Mir3083 | 0 | 0 | 0 | 1 | 1 |
| Mir3084 | 0 | 0 | 0 | 1 | 1 |
| Mir3084-2 | 0 | 0 | 0 | 1 | 1 |
| Mir3085 | 0 | 0 | 0 | 1 | 1 |
| Mir3086 | 0 | 0 | 0 | 1 | 1 |
| Mir3087 | 0 | 0 | 0 | 1 | 1 |
| Mir3088 | 0 | 0 | 0 | 1 | 1 |
| Mir3089 | 0 | 0 | 0 | 1 | 1 |
| Mir3092 | 0 | 0 | 0 | 1 | 1 |

| | | | | | |
|---|---|---|---|---|---|
| Mir3093 | 0 | 0 | 0 | 1 | 1 |
| Mir3094 | 0 | 0 | 0 | 1 | 1 |
| Mir3095 | 0 | 0 | 0 | 1 | 1 |
| Mir3098 | 0 | 0 | 0 | 1 | 1 |
| Mir3099 | 0 | 0 | 0 | 1 | 1 |
| Mir30a | 0 | 0 | 0 | 1 | 1 |
| Mir30b | 0 | 0 | 0 | 1 | 1 |
| Mir30c-1 | 0 | 0 | 0 | 1 | 1 |
| Mir30c-2 | 0 | 0 | 0 | 1 | 1 |
| Mir30d | 0 | 0 | 0 | 1 | 1 |
| Mir30f | 0 | 0 | 0 | 1 | 1 |
| Mir31 | 0 | 0 | 0 | 1 | 1 |
| Mir3103 | 0 | 0 | 0 | 1 | 1 |
| Mir3104 | 0 | 0 | 0 | 1 | 1 |
| Mir3106 | 0 | 0 | 0 | 1 | 1 |
| Mir3107 | 0 | 0 | 0 | 1 | 1 |
| Mir3108 | 0 | 0 | 0 | 1 | 1 |
| Mir3109 | 0 | 0 | 0 | 1 | 1 |
| Mir3110 | 0 | 0 | 0 | 1 | 1 |
| Mir3112 | 0 | 0 | 0 | 1 | 1 |
| Mir32 | 0 | 0 | 0 | 1 | 1 |
| Mir320 | 0 | 0 | 0 | 1 | 1 |
| Mir322 | 0 | 0 | 0 | 1 | 1 |
| Mir323 | 0 | 0 | 0 | 1 | 1 |
| Mir324 | 0 | 0 | 0 | 1 | 1 |
| Mir325 | 0 | 0 | 0 | 1 | 1 |
| Mir326 | 0 | 0 | 0 | 1 | 1 |
| Mir328 | 0 | 0 | 0 | 1 | 1 |
| Mir329 | 0 | 0 | 0 | 1 | 1 |
| Mir33 | 0 | 0 | 0 | 1 | 1 |
| Mir330 | 0 | 0 | 0 | 1 | 1 |
| Mir331 | 0 | 0 | 0 | 1 | 1 |
| Mir335 | 0 | 0 | 0 | 1 | 1 |
| Mir337 | 0 | 0 | 0 | 1 | 1 |
| Mir338 | 0 | 0 | 0 | 1 | 1 |
| Mir339 | 0 | 0 | 0 | 1 | 1 |
| Mir340 | 0 | 0 | 0 | 1 | 1 |
| Mir344 | 0 | 0 | 0 | 1 | 1 |
| Mir344-2 | 0 | 0 | 0 | 1 | 1 |
| Mir344b | 0 | 0 | 0 | 1 | 1 |
| Mir344c | 0 | 0 | 0 | 1 | 1 |
| Mir344d-1 | 0 | 0 | 0 | 1 | 1 |
| Mir344d-2 | 0 | 0 | 0 | 1 | 1 |
| Mir344d-3 | 0 | 0 | 0 | 1 | 1 |
| Mir344e | 0 | 0 | 0 | 1 | 1 |
| Mir344f | 0 | 0 | 0 | 1 | 1 |
| Mir344g | 0 | 0 | 0 | 1 | 1 |
| Mir344h-2 | 0 | 0 | 0 | 1 | 1 |
| Mir344i | 0 | 0 | 0 | 1 | 1 |
| Mir346 | 0 | 0 | 0 | 1 | 1 |

| | | | | | |
|---|---|---|---|---|---|
| Mir3470a | 0 | 0 | 0 | 1 | 1 |
| Mir3470b | 0 | 0 | 0 | 1 | 1 |
| Mir3471-1 | 0 | 0 | 0 | 1 | 1 |
| Mir3473 | 0 | 0 | 0 | 1 | 1 |
| Mir3473c | 0 | 0 | 0 | 1 | 1 |
| Mir3473d | 0 | 0 | 0 | 1 | 1 |
| Mir3473e | 0 | 0 | 0 | 1 | 1 |
| Mir3473f | 0 | 0 | 0 | 1 | 1 |
| Mir3474 | 0 | 0 | 0 | 1 | 1 |
| Mir3475 | 0 | 0 | 0 | 1 | 1 |
| Mir34a | 0 | 0 | 0 | 1 | 1 |
| Mir34b | 0 | 0 | 0 | 1 | 1 |
| Mir34c | 0 | 0 | 0 | 1 | 1 |
| Mir350 | 0 | 0 | 0 | 1 | 1 |
| Mir351 | 0 | 0 | 0 | 1 | 1 |
| Mir3544 | 0 | 0 | 0 | 1 | 1 |
| Mir3547 | 0 | 0 | 0 | 1 | 1 |
| Mir3569 | 0 | 0 | 0 | 1 | 1 |
| Mir3572 | 0 | 0 | 0 | 1 | 1 |
| Mir362 | 0 | 0 | 0 | 1 | 1 |
| Mir363 | 0 | 0 | 0 | 1 | 1 |
| Mir365-1 | 0 | 0 | 0 | 1 | 1 |
| Mir365-2 | 0 | 0 | 0 | 1 | 1 |
| Mir367 | 0 | 0 | 0 | 1 | 1 |
| Mir369 | 0 | 0 | 0 | 1 | 1 |
| Mir370 | 0 | 0 | 0 | 1 | 1 |
| Mir374 | 0 | 0 | 0 | 1 | 1 |
| Mir374c | 0 | 0 | 0 | 1 | 1 |
| Mir375 | 0 | 0 | 0 | 1 | 1 |
| Mir376a | 0 | 0 | 0 | 1 | 1 |
| Mir376b | 0 | 0 | 0 | 1 | 1 |
| Mir376c | 0 | 0 | 0 | 1 | 1 |
| Mir377 | 0 | 0 | 0 | 1 | 1 |
| Mir378 | 0 | 0 | 0 | 1 | 1 |
| Mir378c | 0 | 0 | 0 | 1 | 1 |
| Mir379 | 0 | 0 | 0 | 1 | 1 |
| Mir380 | 0 | 0 | 0 | 1 | 1 |
| Mir381 | 0 | 0 | 0 | 1 | 1 |
| Mir382 | 0 | 0 | 0 | 1 | 1 |
| Mir383 | 0 | 0 | 0 | 1 | 1 |
| Mir384 | 0 | 0 | 0 | 1 | 1 |
| Mir3962 | 0 | 0 | 0 | 1 | 1 |
| Mir3963 | 0 | 0 | 0 | 1 | 1 |
| Mir3964 | 0 | 0 | 0 | 1 | 1 |
| Mir3965 | 0 | 0 | 0 | 1 | 1 |
| Mir3966 | 0 | 0 | 0 | 1 | 1 |
| Mir3967 | 0 | 0 | 0 | 1 | 1 |
| Mir3968 | 0 | 0 | 0 | 1 | 1 |
| Mir3969 | 0 | 0 | 0 | 1 | 1 |
| Mir3970 | 0 | 0 | 0 | 1 | 1 |

| | | | | | |
|---|---|---|---|---|---|
| Mir3971 | 0 | 0 | 0 | 1 | 1 |
| Mir409 | 0 | 0 | 0 | 1 | 1 |
| Mir410 | 0 | 0 | 0 | 1 | 1 |
| Mir411 | 0 | 0 | 0 | 1 | 1 |
| Mir412 | 0 | 0 | 0 | 1 | 1 |
| Mir421 | 0 | 0 | 0 | 1 | 1 |
| Mir423 | 0 | 0 | 0 | 1 | 1 |
| Mir425 | 0 | 0 | 0 | 1 | 1 |
| Mir429 | 0 | 0 | 0 | 1 | 1 |
| Mir432 | 0 | 0 | 0 | 1 | 1 |
| Mir434 | 0 | 0 | 0 | 1 | 1 |
| Mir448 | 0 | 0 | 0 | 1 | 1 |
| Mir449a | 0 | 0 | 0 | 1 | 1 |
| Mir449b | 0 | 0 | 0 | 1 | 1 |
| Mir449c | 0 | 0 | 0 | 1 | 1 |
| Mir450-1 | 0 | 0 | 0 | 1 | 1 |
| Mir450-2 | 0 | 0 | 0 | 1 | 1 |
| Mir450b | 0 | 0 | 0 | 1 | 1 |
| Mir451 | 0 | 0 | 0 | 1 | 1 |
| Mir452 | 0 | 0 | 0 | 1 | 1 |
| Mir453 | 0 | 0 | 0 | 1 | 1 |
| Mir455 | 0 | 0 | 0 | 1 | 1 |
| Mir463 | 0 | 0 | 0 | 1 | 1 |
| Mir465 | 0 | 0 | 0 | 1 | 1 |
| Mir465b-1 | 0 | 0 | 0 | 1 | 1 |
| Mir465c-1 | 0 | 0 | 0 | 1 | 1 |
| Mir465d | 0 | 0 | 0 | 1 | 1 |
| Mir466 | 0 | 0 | 0 | 1 | 1 |
| Mir4660 | 0 | 0 | 0 | 1 | 1 |
| Mir466b-2,Mir466b-3 | 0 | 0 | 0 | 1 | 1 |
| Mir466b-3 | 0 | 0 | 0 | 1 | 1 |
| Mir466d | 0 | 0 | 0 | 1 | 1 |
| Mir466f-1 | 0 | 0 | 0 | 1 | 1 |
| Mir466f-2 | 0 | 0 | 0 | 1 | 1 |
| Mir466f-3 | 0 | 0 | 0 | 1 | 1 |
| Mir466g | 0 | 0 | 0 | 1 | 1 |
| Mir466h | 0 | 0 | 0 | 1 | 1 |
| Mir466n | 0 | 0 | 0 | 1 | 1 |
| Mir466p | 0 | 0 | 0 | 1 | 1 |
| Mir467a-1 | 0 | 0 | 0 | 1 | 1 |
| Mir467a-10 | 0 | 0 | 0 | 1 | 1 |
| Mir467a-2 | 0 | 0 | 0 | 1 | 1 |
| Mir467a-3 | 0 | 0 | 0 | 1 | 1 |
| Mir467a-5 | 0 | 0 | 0 | 1 | 1 |
| Mir467a-7 | 0 | 0 | 0 | 1 | 1 |
| Mir467a-8 | 0 | 0 | 0 | 1 | 1 |
| Mir467b | 0 | 0 | 0 | 1 | 1 |
| Mir467c | 0 | 0 | 0 | 1 | 1 |
| Mir467d | 0 | 0 | 0 | 1 | 1 |
| Mir467e | 0 | 0 | 0 | 1 | 1 |

| | | | | | |
|---|---|---|---|---|---|
| Mir467f | 0 | 0 | 0 | 1 | 1 |
| Mir468 | 0 | 0 | 0 | 1 | 1 |
| Mir470 | 0 | 0 | 0 | 1 | 1 |
| Mir471 | 0 | 0 | 0 | 1 | 1 |
| Mir483 | 0 | 0 | 0 | 1 | 1 |
| Mir484 | 0 | 0 | 0 | 1 | 1 |
| Mir485 | 0 | 0 | 0 | 1 | 1 |
| Mir486 | 0 | 0 | 0 | 1 | 1 |
| Mir487b | 0 | 0 | 0 | 1 | 1 |
| Mir488 | 0 | 0 | 0 | 1 | 1 |
| Mir489 | 0 | 0 | 0 | 1 | 1 |
| Mir490 | 0 | 0 | 0 | 1 | 1 |
| Mir491 | 0 | 0 | 0 | 1 | 1 |
| Mir493 | 0 | 0 | 0 | 1 | 1 |
| Mir494 | 0 | 0 | 0 | 1 | 1 |
| Mir495 | 0 | 0 | 0 | 1 | 1 |
| Mir496 | 0 | 0 | 0 | 1 | 1 |
| Mir496b | 0 | 0 | 0 | 1 | 1 |
| Mir497 | 0 | 0 | 0 | 1 | 1 |
| Mir497b | 0 | 0 | 0 | 1 | 1 |
| Mir499 | 0 | 0 | 0 | 1 | 1 |
| Mir500 | 0 | 0 | 0 | 1 | 1 |
| Mir501 | 0 | 0 | 0 | 1 | 1 |
| Mir503 | 0 | 0 | 0 | 1 | 1 |
| Mir504 | 0 | 0 | 0 | 1 | 1 |
| Mir5046 | 0 | 0 | 0 | 1 | 1 |
| Mir505 | 0 | 0 | 0 | 1 | 1 |
| Mir509 | 0 | 0 | 0 | 1 | 1 |
| Mir5098 | 0 | 0 | 0 | 1 | 1 |
| Mir5100 | 0 | 0 | 0 | 1 | 1 |
| Mir5101 | 0 | 0 | 0 | 1 | 1 |
| Mir5103 | 0 | 0 | 0 | 1 | 1 |
| Mir5104 | 0 | 0 | 0 | 1 | 1 |
| Mir5106 | 0 | 0 | 0 | 1 | 1 |
| Mir5107 | 0 | 0 | 0 | 1 | 1 |
| Mir5108 | 0 | 0 | 0 | 1 | 1 |
| Mir511 | 0 | 0 | 0 | 1 | 1 |
| Mir5112 | 0 | 0 | 0 | 1 | 1 |
| Mir5114 | 0 | 0 | 0 | 1 | 1 |
| Mir5116 | 0 | 0 | 0 | 1 | 1 |
| Mir5119 | 0 | 0 | 0 | 1 | 1 |
| Mir5120 | 0 | 0 | 0 | 1 | 1 |
| Mir5122 | 0 | 0 | 0 | 1 | 1 |
| Mir5123 | 0 | 0 | 0 | 1 | 1 |
| Mir5124 | 0 | 0 | 0 | 1 | 1 |
| Mir5125 | 0 | 0 | 0 | 1 | 1 |
| Mir5126 | 0 | 0 | 0 | 1 | 1 |
| Mir5127 | 0 | 0 | 0 | 1 | 1 |
| Mir5128 | 0 | 0 | 0 | 1 | 1 |
| Mir5129 | 0 | 0 | 0 | 1 | 1 |

| | | | | | |
|---|---|---|---|---|---|
| Mir5130 | 0 | 0 | 0 | 1 | 1 |
| Mir5132 | 0 | 0 | 0 | 1 | 1 |
| Mir5134 | 0 | 0 | 0 | 1 | 1 |
| Mir5136 | 0 | 0 | 0 | 1 | 1 |
| Mir532 | 0 | 0 | 0 | 1 | 1 |
| Mir539 | 0 | 0 | 0 | 1 | 1 |
| Mir540 | 0 | 0 | 0 | 1 | 1 |
| Mir541 | 0 | 0 | 0 | 1 | 1 |
| Mir542 | 0 | 0 | 0 | 1 | 1 |
| Mir543 | 0 | 0 | 0 | 1 | 1 |
| Mir544 | 0 | 0 | 0 | 1 | 1 |
| Mir547 | 0 | 0 | 0 | 1 | 1 |
| Mir551b | 0 | 0 | 0 | 1 | 1 |
| Mir5615-1 | 0 | 0 | 0 | 1 | 1 |
| Mir5615-2 | 0 | 0 | 0 | 1 | 1 |
| Mir5616 | 0 | 0 | 0 | 1 | 1 |
| Mir5617 | 0 | 0 | 0 | 1 | 1 |
| Mir5618 | 0 | 0 | 0 | 1 | 1 |
| Mir5619 | 0 | 0 | 0 | 1 | 1 |
| Mir5620 | 0 | 0 | 0 | 1 | 1 |
| Mir5621 | 0 | 0 | 0 | 1 | 1 |
| Mir5622 | 0 | 0 | 0 | 1 | 1 |
| Mir5623 | 0 | 0 | 0 | 1 | 1 |
| Mir5624 | 0 | 0 | 0 | 1 | 1 |
| Mir5625 | 0 | 0 | 0 | 1 | 1 |
| Mir5626 | 0 | 0 | 0 | 1 | 1 |
| Mir5627 | 0 | 0 | 0 | 1 | 1 |
| Mir568 | 0 | 0 | 0 | 1 | 1 |
| Mir5709 | 0 | 0 | 0 | 1 | 1 |
| Mir5710 | 0 | 0 | 0 | 1 | 1 |
| Mir574 | 0 | 0 | 0 | 1 | 1 |
| Mir582 | 0 | 0 | 0 | 1 | 1 |
| Mir592 | 0 | 0 | 0 | 1 | 1 |
| Mir598 | 0 | 0 | 0 | 1 | 1 |
| Mir615 | 0 | 0 | 0 | 1 | 1 |
| Mir6237 | 0 | 0 | 0 | 1 | 1 |
| Mir6238 | 0 | 0 | 0 | 1 | 1 |
| Mir6241 | 0 | 0 | 0 | 1 | 1 |
| Mir6335 | 0 | 0 | 0 | 1 | 1 |
| Mir6336 | 0 | 0 | 0 | 1 | 1 |
| Mir6337 | 0 | 0 | 0 | 1 | 1 |
| Mir6338 | 0 | 0 | 0 | 1 | 1 |
| Mir6339 | 0 | 0 | 0 | 1 | 1 |
| Mir6341 | 0 | 0 | 0 | 1 | 1 |
| Mir6342 | 0 | 0 | 0 | 1 | 1 |
| Mir6343 | 0 | 0 | 0 | 1 | 1 |
| Mir6344 | 0 | 0 | 0 | 1 | 1 |
| Mir6345 | 0 | 0 | 0 | 1 | 1 |
| Mir6348 | 0 | 0 | 0 | 1 | 1 |
| Mir6349 | 0 | 0 | 0 | 1 | 1 |

| | | | | | |
|---|---|---|---|---|---|
| Mir6350 | 0 | 0 | 0 | 1 | 1 |
| Mir6353 | 0 | 0 | 0 | 1 | 1 |
| Mir6354 | 0 | 0 | 0 | 1 | 1 |
| Mir6355 | 0 | 0 | 0 | 1 | 1 |
| Mir6356 | 0 | 0 | 0 | 1 | 1 |
| Mir6358 | 0 | 0 | 0 | 1 | 1 |
| Mir6359 | 0 | 0 | 0 | 1 | 1 |
| Mir6360 | 0 | 0 | 0 | 1 | 1 |
| Mir6362 | 0 | 0 | 0 | 1 | 1 |
| Mir6364 | 0 | 0 | 0 | 1 | 1 |
| Mir6365 | 0 | 0 | 0 | 1 | 1 |
| Mir6366 | 0 | 0 | 0 | 1 | 1 |
| Mir6367 | 0 | 0 | 0 | 1 | 1 |
| Mir6368 | 0 | 0 | 0 | 1 | 1 |
| Mir6369 | 0 | 0 | 0 | 1 | 1 |
| Mir6370 | 0 | 0 | 0 | 1 | 1 |
| Mir6372 | 0 | 0 | 0 | 1 | 1 |
| Mir6373 | 0 | 0 | 0 | 1 | 1 |
| Mir6374 | 0 | 0 | 0 | 1 | 1 |
| Mir6375 | 0 | 0 | 0 | 1 | 1 |
| Mir6376 | 0 | 0 | 0 | 1 | 1 |
| Mir6378 | 0 | 0 | 0 | 1 | 1 |
| Mir6380 | 0 | 0 | 0 | 1 | 1 |
| Mir6381 | 0 | 0 | 0 | 1 | 1 |
| Mir6382 | 0 | 0 | 0 | 1 | 1 |
| Mir6383 | 0 | 0 | 0 | 1 | 1 |
| Mir6384 | 0 | 0 | 0 | 1 | 1 |
| Mir6386 | 0 | 0 | 0 | 1 | 1 |
| Mir6388 | 0 | 0 | 0 | 1 | 1 |
| Mir6389 | 0 | 0 | 0 | 1 | 1 |
| Mir6390 | 0 | 0 | 0 | 1 | 1 |
| Mir6391 | 0 | 0 | 0 | 1 | 1 |
| Mir6392 | 0 | 0 | 0 | 1 | 1 |
| Mir6393 | 0 | 0 | 0 | 1 | 1 |
| Mir6394 | 0 | 0 | 0 | 1 | 1 |
| Mir6396 | 0 | 0 | 0 | 1 | 1 |
| Mir6397 | 0 | 0 | 0 | 1 | 1 |
| Mir6398 | 0 | 0 | 0 | 1 | 1 |
| Mir6399 | 0 | 0 | 0 | 1 | 1 |
| Mir6400 | 0 | 0 | 0 | 1 | 1 |
| Mir6401 | 0 | 0 | 0 | 1 | 1 |
| Mir6402 | 0 | 0 | 0 | 1 | 1 |
| Mir6404 | 0 | 0 | 0 | 1 | 1 |
| Mir6405 | 0 | 0 | 0 | 1 | 1 |
| Mir6408 | 0 | 0 | 0 | 1 | 1 |
| Mir6409 | 0 | 0 | 0 | 1 | 1 |
| Mir6410 | 0 | 0 | 0 | 1 | 1 |
| Mir6411 | 0 | 0 | 0 | 1 | 1 |
| Mir6412 | 0 | 0 | 0 | 1 | 1 |
| Mir6413 | 0 | 0 | 0 | 1 | 1 |

| | | | | | |
|---|---|---|---|---|---|
| Mir6414 | 0 | 0 | 0 | 1 | 1 |
| Mir6415 | 0 | 0 | 0 | 1 | 1 |
| Mir6416 | 0 | 0 | 0 | 1 | 1 |
| Mir6417 | 0 | 0 | 0 | 1 | 1 |
| Mir6419 | 0 | 0 | 0 | 1 | 1 |
| Mir6420 | 0 | 0 | 0 | 1 | 1 |
| Mir653 | 0 | 0 | 0 | 1 | 1 |
| Mir6537 | 0 | 0 | 0 | 1 | 1 |
| Mir6539 | 0 | 0 | 0 | 1 | 1 |
| Mir654 | 0 | 0 | 0 | 1 | 1 |
| Mir6540 | 0 | 0 | 0 | 1 | 1 |
| Mir6541 | 0 | 0 | 0 | 1 | 1 |
| Mir6546 | 0 | 0 | 0 | 1 | 1 |
| Mir664 | 0 | 0 | 0 | 1 | 1 |
| Mir665 | 0 | 0 | 0 | 1 | 1 |
| Mir666 | 0 | 0 | 0 | 1 | 1 |
| Mir667 | 0 | 0 | 0 | 1 | 1 |
| Mir668 | 0 | 0 | 0 | 1 | 1 |
| Mir669a-1 | 0 | 0 | 0 | 1 | 1 |
| Mir669a-2 | 0 | 0 | 0 | 1 | 1 |
| Mir669a-3 | 0 | 0 | 0 | 1 | 1 |
| Mir669a-4,Mir669p-1 | 0 | 0 | 0 | 1 | 1 |
| Mir669b | 0 | 0 | 0 | 1 | 1 |
| Mir669c | 0 | 0 | 0 | 1 | 1 |
| Mir669e | 0 | 0 | 0 | 1 | 1 |
| Mir669g | 0 | 0 | 0 | 1 | 1 |
| Mir669h | 0 | 0 | 0 | 1 | 1 |
| Mir669i | 0 | 0 | 0 | 1 | 1 |
| Mir669j | 0 | 0 | 0 | 1 | 1 |
| Mir669k | 0 | 0 | 0 | 1 | 1 |
| Mir669m-1 | 0 | 0 | 0 | 1 | 1 |
| Mir669m-2 | 0 | 0 | 0 | 1 | 1 |
| Mir669p-1 | 0 | 0 | 0 | 1 | 1 |
| Mir670 | 0 | 0 | 0 | 1 | 1 |
| Mir671 | 0 | 0 | 0 | 1 | 1 |
| Mir6715 | 0 | 0 | 0 | 1 | 1 |
| Mir672 | 0 | 0 | 0 | 1 | 1 |
| Mir673 | 0 | 0 | 0 | 1 | 1 |
| Mir674 | 0 | 0 | 0 | 1 | 1 |
| Mir675 | 0 | 0 | 0 | 1 | 1 |
| Mir676 | 0 | 0 | 0 | 1 | 1 |
| Mir6769b | 0 | 0 | 0 | 1 | 1 |
| Mir679 | 0 | 0 | 0 | 1 | 1 |
| Mir680-2 | 0 | 0 | 0 | 1 | 1 |
| Mir680-3 | 0 | 0 | 0 | 1 | 1 |
| Mir683-1 | 0 | 0 | 0 | 1 | 1 |
| Mir684-1 | 0 | 0 | 0 | 1 | 1 |
| Mir684-2 | 0 | 0 | 0 | 1 | 1 |
| Mir687 | 0 | 0 | 0 | 1 | 1 |
| Mir688 | 0 | 0 | 0 | 1 | 1 |

| | | | | | |
|---|---|---|---|---|---|
| Mir6896 | 0 | 0 | 0 | 1 | 1 |
| Mir6897 | 0 | 0 | 0 | 1 | 1 |
| Mir6898 | 0 | 0 | 0 | 1 | 1 |
| Mir6899 | 0 | 0 | 0 | 1 | 1 |
| Mir690 | 0 | 0 | 0 | 1 | 1 |
| Mir6900 | 0 | 0 | 0 | 1 | 1 |
| Mir6901 | 0 | 0 | 0 | 1 | 1 |
| Mir6902 | 0 | 0 | 0 | 1 | 1 |
| Mir6903 | 0 | 0 | 0 | 1 | 1 |
| Mir6906 | 0 | 0 | 0 | 1 | 1 |
| Mir6908 | 0 | 0 | 0 | 1 | 1 |
| Mir6909 | 0 | 0 | 0 | 1 | 1 |
| Mir691 | 0 | 0 | 0 | 1 | 1 |
| Mir6911 | 0 | 0 | 0 | 1 | 1 |
| Mir6913 | 0 | 0 | 0 | 1 | 1 |
| Mir6914 | 0 | 0 | 0 | 1 | 1 |
| Mir6916 | 0 | 0 | 0 | 1 | 1 |
| Mir6917 | 0 | 0 | 0 | 1 | 1 |
| Mir6918 | 0 | 0 | 0 | 1 | 1 |
| Mir6919 | 0 | 0 | 0 | 1 | 1 |
| Mir692-1 | 0 | 0 | 0 | 1 | 1 |
| Mir692-2b | 0 | 0 | 0 | 1 | 1 |
| Mir6922 | 0 | 0 | 0 | 1 | 1 |
| Mir6923 | 0 | 0 | 0 | 1 | 1 |
| Mir6924 | 0 | 0 | 0 | 1 | 1 |
| Mir6925 | 0 | 0 | 0 | 1 | 1 |
| Mir6926 | 0 | 0 | 0 | 1 | 1 |
| Mir6927 | 0 | 0 | 0 | 1 | 1 |
| Mir6928 | 0 | 0 | 0 | 1 | 1 |
| Mir6929 | 0 | 0 | 0 | 1 | 1 |
| Mir693 | 0 | 0 | 0 | 1 | 1 |
| Mir6930 | 0 | 0 | 0 | 1 | 1 |
| Mir6931 | 0 | 0 | 0 | 1 | 1 |
| Mir6932 | 0 | 0 | 0 | 1 | 1 |
| Mir6933 | 0 | 0 | 0 | 1 | 1 |
| Mir6934 | 0 | 0 | 0 | 1 | 1 |
| Mir6936 | 0 | 0 | 0 | 1 | 1 |
| Mir6937 | 0 | 0 | 0 | 1 | 1 |
| Mir6938 | 0 | 0 | 0 | 1 | 1 |
| Mir694 | 0 | 0 | 0 | 1 | 1 |
| Mir6940 | 0 | 0 | 0 | 1 | 1 |
| Mir6941 | 0 | 0 | 0 | 1 | 1 |
| Mir6942 | 0 | 0 | 0 | 1 | 1 |
| Mir6946 | 0 | 0 | 0 | 1 | 1 |
| Mir6947 | 0 | 0 | 0 | 1 | 1 |
| Mir6949 | 0 | 0 | 0 | 1 | 1 |
| Mir695 | 0 | 0 | 0 | 1 | 1 |
| Mir6950 | 0 | 0 | 0 | 1 | 1 |
| Mir6951 | 0 | 0 | 0 | 1 | 1 |
| Mir6952 | 0 | 0 | 0 | 1 | 1 |

| | | | | | |
|---|---|---|---|---|---|
| Mir6953 | 0 | 0 | 0 | 1 | 1 |
| Mir6954 | 0 | 0 | 0 | 1 | 1 |
| Mir6956 | 0 | 0 | 0 | 1 | 1 |
| Mir6957 | 0 | 0 | 0 | 1 | 1 |
| Mir6959 | 0 | 0 | 0 | 1 | 1 |
| Mir6960 | 0 | 0 | 0 | 1 | 1 |
| Mir6961 | 0 | 0 | 0 | 1 | 1 |
| Mir6962 | 0 | 0 | 0 | 1 | 1 |
| Mir6964 | 0 | 0 | 0 | 1 | 1 |
| Mir6965 | 0 | 0 | 0 | 1 | 1 |
| Mir6966 | 0 | 0 | 0 | 1 | 1 |
| Mir6968 | 0 | 0 | 0 | 1 | 1 |
| Mir6969 | 0 | 0 | 0 | 1 | 1 |
| Mir6971 | 0 | 0 | 0 | 1 | 1 |
| Mir6973a | 0 | 0 | 0 | 1 | 1 |
| Mir6974 | 0 | 0 | 0 | 1 | 1 |
| Mir6975 | 0 | 0 | 0 | 1 | 1 |
| Mir6978 | 0 | 0 | 0 | 1 | 1 |
| Mir6979 | 0 | 0 | 0 | 1 | 1 |
| Mir698 | 0 | 0 | 0 | 1 | 1 |
| Mir6980 | 0 | 0 | 0 | 1 | 1 |
| Mir6981 | 0 | 0 | 0 | 1 | 1 |
| Mir6982 | 0 | 0 | 0 | 1 | 1 |
| Mir6983 | 0 | 0 | 0 | 1 | 1 |
| Mir6984 | 0 | 0 | 0 | 1 | 1 |
| Mir6985 | 0 | 0 | 0 | 1 | 1 |
| Mir6986 | 0 | 0 | 0 | 1 | 1 |
| Mir6987 | 0 | 0 | 0 | 1 | 1 |
| Mir6988 | 0 | 0 | 0 | 1 | 1 |
| Mir6989 | 0 | 0 | 0 | 1 | 1 |
| Mir6993 | 0 | 0 | 0 | 1 | 1 |
| Mir6994 | 0 | 0 | 0 | 1 | 1 |
| Mir6995 | 0 | 0 | 0 | 1 | 1 |
| Mir6996 | 0 | 0 | 0 | 1 | 1 |
| Mir6997 | 0 | 0 | 0 | 1 | 1 |
| Mir6999 | 0 | 0 | 0 | 1 | 1 |
| Mir7-2 | 0 | 0 | 0 | 1 | 1 |
| Mir700 | 0 | 0 | 0 | 1 | 1 |
| Mir7000 | 0 | 0 | 0 | 1 | 1 |
| Mir7001 | 0 | 0 | 0 | 1 | 1 |
| Mir7002 | 0 | 0 | 0 | 1 | 1 |
| Mir7003 | 0 | 0 | 0 | 1 | 1 |
| Mir7004 | 0 | 0 | 0 | 1 | 1 |
| Mir7007 | 0 | 0 | 0 | 1 | 1 |
| Mir7008 | 0 | 0 | 0 | 1 | 1 |
| Mir7009 | 0 | 0 | 0 | 1 | 1 |
| Mir7010 | 0 | 0 | 0 | 1 | 1 |
| Mir7011 | 0 | 0 | 0 | 1 | 1 |
| Mir7012 | 0 | 0 | 0 | 1 | 1 |
| Mir7015 | 0 | 0 | 0 | 1 | 1 |

| | | | | | |
|---|---|---|---|---|---|
| Mir7016 | 0 | 0 | 0 | 1 | 1 |
| Mir7017 | 0 | 0 | 0 | 1 | 1 |
| Mir7018 | 0 | 0 | 0 | 1 | 1 |
| Mir7019 | 0 | 0 | 0 | 1 | 1 |
| Mir702 | 0 | 0 | 0 | 1 | 1 |
| Mir7020 | 0 | 0 | 0 | 1 | 1 |
| Mir7021 | 0 | 0 | 0 | 1 | 1 |
| Mir7022 | 0 | 0 | 0 | 1 | 1 |
| Mir7023 | 0 | 0 | 0 | 1 | 1 |
| Mir7025 | 0 | 0 | 0 | 1 | 1 |
| Mir7026 | 0 | 0 | 0 | 1 | 1 |
| Mir7028 | 0 | 0 | 0 | 1 | 1 |
| Mir7029 | 0 | 0 | 0 | 1 | 1 |
| Mir7031 | 0 | 0 | 0 | 1 | 1 |
| Mir7032 | 0 | 0 | 0 | 1 | 1 |
| Mir7034 | 0 | 0 | 0 | 1 | 1 |
| Mir7036 | 0 | 0 | 0 | 1 | 1 |
| Mir7036b | 0 | 0 | 0 | 1 | 1 |
| Mir7037 | 0 | 0 | 0 | 1 | 1 |
| Mir7038 | 0 | 0 | 0 | 1 | 1 |
| Mir7039 | 0 | 0 | 0 | 1 | 1 |
| Mir7040 | 0 | 0 | 0 | 1 | 1 |
| Mir7041 | 0 | 0 | 0 | 1 | 1 |
| Mir7042 | 0 | 0 | 0 | 1 | 1 |
| Mir7043 | 0 | 0 | 0 | 1 | 1 |
| Mir7044 | 0 | 0 | 0 | 1 | 1 |
| Mir7045 | 0 | 0 | 0 | 1 | 1 |
| Mir7046 | 0 | 0 | 0 | 1 | 1 |
| Mir7047 | 0 | 0 | 0 | 1 | 1 |
| Mir7048 | 0 | 0 | 0 | 1 | 1 |
| Mir7049 | 0 | 0 | 0 | 1 | 1 |
| Mir705 | 0 | 0 | 0 | 1 | 1 |
| Mir7050 | 0 | 0 | 0 | 1 | 1 |
| Mir7051 | 0 | 0 | 0 | 1 | 1 |
| Mir7053 | 0 | 0 | 0 | 1 | 1 |
| Mir7054 | 0 | 0 | 0 | 1 | 1 |
| Mir7055 | 0 | 0 | 0 | 1 | 1 |
| Mir7056 | 0 | 0 | 0 | 1 | 1 |
| Mir7057 | 0 | 0 | 0 | 1 | 1 |
| Mir7058 | 0 | 0 | 0 | 1 | 1 |
| Mir7059 | 0 | 0 | 0 | 1 | 1 |
| Mir706 | 0 | 0 | 0 | 1 | 1 |
| Mir7061 | 0 | 0 | 0 | 1 | 1 |
| Mir7062 | 0 | 0 | 0 | 1 | 1 |
| Mir7063 | 0 | 0 | 0 | 1 | 1 |
| Mir7065 | 0 | 0 | 0 | 1 | 1 |
| Mir7066 | 0 | 0 | 0 | 1 | 1 |
| Mir7069 | 0 | 0 | 0 | 1 | 1 |
| Mir7070 | 0 | 0 | 0 | 1 | 1 |
| Mir7071 | 0 | 0 | 0 | 1 | 1 |

| | | | | | |
|---|---|---|---|---|---|
| Mir7072 | 0 | 0 | 0 | 1 | 1 |
| Mir7074 | 0 | 0 | 0 | 1 | 1 |
| Mir7076 | 0 | 0 | 0 | 1 | 1 |
| Mir7077 | 0 | 0 | 0 | 1 | 1 |
| Mir708 | 0 | 0 | 0 | 1 | 1 |
| Mir7080 | 0 | 0 | 0 | 1 | 1 |
| Mir7081 | 0 | 0 | 0 | 1 | 1 |
| Mir7083 | 0 | 0 | 0 | 1 | 1 |
| Mir7084 | 0 | 0 | 0 | 1 | 1 |
| Mir7085 | 0 | 0 | 0 | 1 | 1 |
| Mir7086 | 0 | 0 | 0 | 1 | 1 |
| Mir7087 | 0 | 0 | 0 | 1 | 1 |
| Mir7088 | 0 | 0 | 0 | 1 | 1 |
| Mir7090 | 0 | 0 | 0 | 1 | 1 |
| Mir7091 | 0 | 0 | 0 | 1 | 1 |
| Mir7092 | 0 | 0 | 0 | 1 | 1 |
| Mir7093 | 0 | 0 | 0 | 1 | 1 |
| Mir7094-1 | 0 | 0 | 0 | 1 | 1 |
| Mir7094-2 | 0 | 0 | 0 | 1 | 1 |
| Mir710 | 0 | 0 | 0 | 1 | 1 |
| Mir711 | 0 | 0 | 0 | 1 | 1 |
| Mir7117 | 0 | 0 | 0 | 1 | 1 |
| Mir7118 | 0 | 0 | 0 | 1 | 1 |
| Mir7119 | 0 | 0 | 0 | 1 | 1 |
| Mir713 | 0 | 0 | 0 | 1 | 1 |
| Mir718 | 0 | 0 | 0 | 1 | 1 |
| Mir721 | 0 | 0 | 0 | 1 | 1 |
| Mir7210 | 0 | 0 | 0 | 1 | 1 |
| Mir7211 | 0 | 0 | 0 | 1 | 1 |
| Mir7212 | 0 | 0 | 0 | 1 | 1 |
| Mir7213 | 0 | 0 | 0 | 1 | 1 |
| Mir7214 | 0 | 0 | 0 | 1 | 1 |
| Mir7215 | 0 | 0 | 0 | 1 | 1 |
| Mir7216 | 0 | 0 | 0 | 1 | 1 |
| Mir7217 | 0 | 0 | 0 | 1 | 1 |
| Mir7218 | 0 | 0 | 0 | 1 | 1 |
| Mir7219 | 0 | 0 | 0 | 1 | 1 |
| Mir7220 | 0 | 0 | 0 | 1 | 1 |
| Mir7221 | 0 | 0 | 0 | 1 | 1 |
| Mir7222 | 0 | 0 | 0 | 1 | 1 |
| Mir7223 | 0 | 0 | 0 | 1 | 1 |
| Mir7224 | 0 | 0 | 0 | 1 | 1 |
| Mir7225 | 0 | 0 | 0 | 1 | 1 |
| Mir7226 | 0 | 0 | 0 | 1 | 1 |
| Mir7227 | 0 | 0 | 0 | 1 | 1 |
| Mir7228 | 0 | 0 | 0 | 1 | 1 |
| Mir7229 | 0 | 0 | 0 | 1 | 1 |
| Mir7230 | 0 | 0 | 0 | 1 | 1 |
| Mir7231 | 0 | 0 | 0 | 1 | 1 |
| Mir7232 | 0 | 0 | 0 | 1 | 1 |

| | | | | | |
|---|---|---|---|---|---|
| Mir7233 | 0 | 0 | 0 | 1 | 1 |
| Mir7234 | 0 | 0 | 0 | 1 | 1 |
| Mir7235 | 0 | 0 | 0 | 1 | 1 |
| Mir7236 | 0 | 0 | 0 | 1 | 1 |
| Mir7237 | 0 | 0 | 0 | 1 | 1 |
| Mir7238 | 0 | 0 | 0 | 1 | 1 |
| Mir7239 | 0 | 0 | 0 | 1 | 1 |
| Mir7240 | 0 | 0 | 0 | 1 | 1 |
| Mir7241 | 0 | 0 | 0 | 1 | 1 |
| Mir7242 | 0 | 0 | 0 | 1 | 1 |
| Mir7243 | 0 | 0 | 0 | 1 | 1 |
| Mir741 | 0 | 0 | 0 | 1 | 1 |
| Mir742 | 0 | 0 | 0 | 1 | 1 |
| Mir743 | 0 | 0 | 0 | 1 | 1 |
| Mir743b | 0 | 0 | 0 | 1 | 1 |
| Mir744 | 0 | 0 | 0 | 1 | 1 |
| Mir758 | 0 | 0 | 0 | 1 | 1 |
| Mir759 | 0 | 0 | 0 | 1 | 1 |
| Mir760 | 0 | 0 | 0 | 1 | 1 |
| Mir761 | 0 | 0 | 0 | 1 | 1 |
| Mir764 | 0 | 0 | 0 | 1 | 1 |
| Mir7646 | 0 | 0 | 0 | 1 | 1 |
| Mir7647 | 0 | 0 | 0 | 1 | 1 |
| Mir7648 | 0 | 0 | 0 | 1 | 1 |
| Mir7649 | 0 | 0 | 0 | 1 | 1 |
| Mir7652 | 0 | 0 | 0 | 1 | 1 |
| Mir7653 | 0 | 0 | 0 | 1 | 1 |
| Mir7654 | 0 | 0 | 0 | 1 | 1 |
| Mir7655 | 0 | 0 | 0 | 1 | 1 |
| Mir7656 | 0 | 0 | 0 | 1 | 1 |
| Mir7657 | 0 | 0 | 0 | 1 | 1 |
| Mir7658 | 0 | 0 | 0 | 1 | 1 |
| Mir7661 | 0 | 0 | 0 | 1 | 1 |
| Mir7662 | 0 | 0 | 0 | 1 | 1 |
| Mir7663 | 0 | 0 | 0 | 1 | 1 |
| Mir7665 | 0 | 0 | 0 | 1 | 1 |
| Mir7666 | 0 | 0 | 0 | 1 | 1 |
| Mir7667 | 0 | 0 | 0 | 1 | 1 |
| Mir7668 | 0 | 0 | 0 | 1 | 1 |
| Mir7669 | 0 | 0 | 0 | 1 | 1 |
| Mir767 | 0 | 0 | 0 | 1 | 1 |
| Mir7670 | 0 | 0 | 0 | 1 | 1 |
| Mir7671 | 0 | 0 | 0 | 1 | 1 |
| Mir7672 | 0 | 0 | 0 | 1 | 1 |
| Mir7673 | 0 | 0 | 0 | 1 | 1 |
| Mir7674 | 0 | 0 | 0 | 1 | 1 |
| Mir7675 | 0 | 0 | 0 | 1 | 1 |
| Mir7676-1 | 0 | 0 | 0 | 1 | 1 |
| Mir7676-2 | 0 | 0 | 0 | 1 | 1 |
| Mir7677 | 0 | 0 | 0 | 1 | 1 |

| | | | | | |
|---|---|---|---|---|---|
| Mir7678 | 0 | 0 | 0 | 1 | 1 |
| Mir7679 | 0 | 0 | 0 | 1 | 1 |
| Mir7680 | 0 | 0 | 0 | 1 | 1 |
| Mir7681 | 0 | 0 | 0 | 1 | 1 |
| Mir7682 | 0 | 0 | 0 | 1 | 1 |
| Mir7684 | 0 | 0 | 0 | 1 | 1 |
| Mir7685 | 0 | 0 | 0 | 1 | 1 |
| Mir7686 | 0 | 0 | 0 | 1 | 1 |
| Mir770 | 0 | 0 | 0 | 1 | 1 |
| Mir7b | 0 | 0 | 0 | 1 | 1 |
| Mir802 | 0 | 0 | 0 | 1 | 1 |
| Mir804 | 0 | 0 | 0 | 1 | 1 |
| Mir8092 | 0 | 0 | 0 | 1 | 1 |
| Mir8095 | 0 | 0 | 0 | 1 | 1 |
| Mir8096 | 0 | 0 | 0 | 1 | 1 |
| Mir8099-1 | 0 | 0 | 0 | 1 | 1 |
| Mir8099-2 | 0 | 0 | 0 | 1 | 1 |
| Mir8100 | 0 | 0 | 0 | 1 | 1 |
| Mir8101 | 0 | 0 | 0 | 1 | 1 |
| Mir8103 | 0 | 0 | 0 | 1 | 1 |
| Mir8104 | 0 | 0 | 0 | 1 | 1 |
| Mir8105 | 0 | 0 | 0 | 1 | 1 |
| Mir8106 | 0 | 0 | 0 | 1 | 1 |
| Mir8107 | 0 | 0 | 0 | 1 | 1 |
| Mir8108 | 0 | 0 | 0 | 1 | 1 |
| Mir8109 | 0 | 0 | 0 | 1 | 1 |
| Mir8110 | 0 | 0 | 0 | 1 | 1 |
| Mir8111 | 0 | 0 | 0 | 1 | 1 |
| Mir8112 | 0 | 0 | 0 | 1 | 1 |
| Mir8113 | 0 | 0 | 0 | 1 | 1 |
| Mir8114 | 0 | 0 | 0 | 1 | 1 |
| Mir8115 | 0 | 0 | 0 | 1 | 1 |
| Mir8116 | 0 | 0 | 0 | 1 | 1 |
| Mir8119 | 0 | 0 | 0 | 1 | 1 |
| Mir8120 | 0 | 0 | 0 | 1 | 1 |
| Mir871 | 0 | 0 | 0 | 1 | 1 |
| Mir872 | 0 | 0 | 0 | 1 | 1 |
| Mir873 | 0 | 0 | 0 | 1 | 1 |
| Mir874 | 0 | 0 | 0 | 1 | 1 |
| Mir875 | 0 | 0 | 0 | 1 | 1 |
| Mir876 | 0 | 0 | 0 | 1 | 1 |
| Mir877 | 0 | 0 | 0 | 1 | 1 |
| Mir878 | 0 | 0 | 0 | 1 | 1 |
| Mir879 | 0 | 0 | 0 | 1 | 1 |
| Mir880 | 0 | 0 | 0 | 1 | 1 |
| Mir881 | 0 | 0 | 0 | 1 | 1 |
| Mir882 | 0 | 0 | 0 | 1 | 1 |
| Mir883a | 0 | 0 | 0 | 1 | 1 |
| Mir883b | 0 | 0 | 0 | 1 | 1 |
| Mir9-1 | 0 | 0 | 0 | 1 | 1 |

| | | | | | |
|---|---|---|---|---|---|
| Mir9-2 | 0 | 0 | 0 | 1 | 1 |
| Mir9-3 | 0 | 0 | 0 | 1 | 1 |
| Mir92-1 | 0 | 0 | 0 | 1 | 1 |
| Mir92-2 | 0 | 0 | 0 | 1 | 1 |
| Mir92b | 0 | 0 | 0 | 1 | 1 |
| Mir93 | 0 | 0 | 0 | 1 | 1 |
| Mir96 | 0 | 0 | 0 | 1 | 1 |
| Mir98 | 0 | 0 | 0 | 1 | 1 |
| Mir99a | 0 | 0 | 0 | 1 | 1 |
| Mir99b | 0 | 0 | 0 | 1 | 1 |
| Mirlet7a-1 | 0 | 0 | 0 | 1 | 1 |
| Mirlet7a-2 | 0 | 0 | 0 | 1 | 1 |
| Mirlet7b | 0 | 0 | 0 | 1 | 1 |
| Mirlet7bhg | 0 | 0 | 0 | 1 | 1 |
| Mirlet7c-1 | 0 | 0 | 0 | 1 | 1 |
| Mirlet7c-2 | 0 | 0 | 0 | 1 | 1 |
| Mirlet7e | 0 | 0 | 0 | 1 | 1 |
| Mirlet7f-1 | 0 | 0 | 0 | 1 | 1 |
| Mirlet7f-2 | 0 | 0 | 0 | 1 | 1 |
| Mirlet7g | 0 | 0 | 0 | 1 | 1 |
| Mirlet7j | 0 | 0 | 0 | 1 | 1 |
| Mirlet7k | 0 | 0 | 0 | 1 | 1 |
| Mlana | 0 | 0 | 0 | 1 | 1 |
| Mlip | 0 | 0 | 0 | 1 | 1 |
| Mmp10 | 0 | 0 | 0 | 1 | 1 |
| Mmp13 | 0 | 0 | 0 | 1 | 1 |
| Mmp1a | 0 | 0 | 0 | 1 | 1 |
| Mmp1b | 0 | 0 | 0 | 1 | 1 |
| Mmp20 | 0 | 0 | 0 | 1 | 1 |
| Mmp3 | 0 | 0 | 0 | 1 | 1 |
| Mmp8 | 0 | 0 | 0 | 1 | 1 |
| Mocos | 0 | 0 | 0 | 1 | 1 |
| Mov10l1 | 0 | 0 | 0 | 1 | 1 |
| Mptx1 | 0 | 0 | 0 | 1 | 1 |
| Mptx2 | 0 | 0 | 0 | 1 | 1 |
| Mrgpra1 | 0 | 0 | 0 | 1 | 1 |
| Mrgpra2a | 0 | 0 | 0 | 1 | 1 |
| Mrgpra2b | 0 | 0 | 0 | 1 | 1 |
| Mrgpra3 | 0 | 0 | 0 | 1 | 1 |
| Mrgpra4 | 0 | 0 | 0 | 1 | 1 |
| Mrgpra6 | 0 | 0 | 0 | 1 | 1 |
| Mrgpra9 | 0 | 0 | 0 | 1 | 1 |
| Mrgprb1 | 0 | 0 | 0 | 1 | 1 |
| Mrgprb2 | 0 | 0 | 0 | 1 | 1 |
| Mrgprb3 | 0 | 0 | 0 | 1 | 1 |
| Mrgprb4 | 0 | 0 | 0 | 1 | 1 |
| Mrgprb8 | 0 | 0 | 0 | 1 | 1 |
| Mrgprd | 0 | 0 | 0 | 1 | 1 |
| Mrgprg | 0 | 0 | 0 | 1 | 1 |
| Mrgprx1 | 0 | 0 | 0 | 1 | 1 |

| | | | | | |
|---|---|---|---|---|---|
| Mrgprx2 | 0 | 0 | 0 | 1 | 1 |
| Mroh4 | 0 | 0 | 0 | 1 | 1 |
| Mroh5 | 0 | 0 | 0 | 1 | 1 |
| Mroh9 | 0 | 0 | 0 | 1 | 1 |
| Ms4a1 | 0 | 0 | 0 | 1 | 1 |
| Ms4a15 | 0 | 0 | 0 | 1 | 1 |
| Ms4a18 | 0 | 0 | 0 | 1 | 1 |
| Ms4a2 | 0 | 0 | 0 | 1 | 1 |
| Ms4a3 | 0 | 0 | 0 | 1 | 1 |
| Ms4a4b | 0 | 0 | 0 | 1 | 1 |
| Ms4a4c | 0 | 0 | 0 | 1 | 1 |
| Ms4a5 | 0 | 0 | 0 | 1 | 1 |
| Ms4a6b | 0 | 0 | 0 | 1 | 1 |
| Ms4a6c | 0 | 0 | 0 | 1 | 1 |
| Ms4a6d | 0 | 0 | 0 | 1 | 1 |
| Ms4a8a | 0 | 0 | 0 | 1 | 1 |
| Msln | 0 | 0 | 0 | 1 | 1 |
| Mstn | 0 | 0 | 0 | 1 | 1 |
| Mt4 | 0 | 0 | 0 | 1 | 1 |
| Muc15 | 0 | 0 | 0 | 1 | 1 |
| Muc19 | 0 | 0 | 0 | 1 | 1 |
| Muc20 | 0 | 0 | 0 | 1 | 1 |
| Muc4 | 0 | 0 | 0 | 1 | 1 |
| Mug-ps1 | 0 | 0 | 0 | 1 | 1 |
| Mup1 | 0 | 0 | 0 | 1 | 1 |
| Mup10 | 0 | 0 | 0 | 1 | 1 |
| Mup11 | 0 | 0 | 0 | 1 | 1 |
| Mup12 | 0 | 0 | 0 | 1 | 1 |
| Mup13 | 0 | 0 | 0 | 1 | 1 |
| Mup14 | 0 | 0 | 0 | 1 | 1 |
| Mup15 | 0 | 0 | 0 | 1 | 1 |
| Mup16 | 0 | 0 | 0 | 1 | 1 |
| Mup17 | 0 | 0 | 0 | 1 | 1 |
| Mup19 | 0 | 0 | 0 | 1 | 1 |
| Mup2 | 0 | 0 | 0 | 1 | 1 |
| Mup20 | 0 | 0 | 0 | 1 | 1 |
| Mup21 | 0 | 0 | 0 | 1 | 1 |
| Mup3 | 0 | 0 | 0 | 1 | 1 |
| Mup4 | 0 | 0 | 0 | 1 | 1 |
| Mup5 | 0 | 0 | 0 | 1 | 1 |
| Mup6 | 0 | 0 | 0 | 1 | 1 |
| Mup7 | 0 | 0 | 0 | 1 | 1 |
| Mup8 | 0 | 0 | 0 | 1 | 1 |
| Mup9 | 0 | 0 | 0 | 1 | 1 |
| Mybpc3 | 0 | 0 | 0 | 1 | 1 |
| Myh15 | 0 | 0 | 0 | 1 | 1 |
| Myo3a | 0 | 0 | 0 | 1 | 1 |
| Myoc | 0 | 0 | 0 | 1 | 1 |
| Myot | 0 | 0 | 0 | 1 | 1 |
| Myoz2 | 0 | 0 | 0 | 1 | 1 |

| | | | | | |
|---|---|---|---|---|---|
| Myoz3 | 0 | 0 | 0 | 1 | 1 |
| Naa11 | 0 | 0 | 0 | 1 | 1 |
| Naip1 | 0 | 0 | 0 | 1 | 1 |
| Naip6 | 0 | 0 | 0 | 1 | 1 |
| Naip7 | 0 | 0 | 0 | 1 | 1 |
| Nccrp1 | 0 | 0 | 0 | 1 | 1 |
| Ncr1 | 0 | 0 | 0 | 1 | 1 |
| Nell1os | 0 | 0 | 0 | 1 | 1 |
| Neu2 | 0 | 0 | 0 | 1 | 1 |
| Neu4 | 0 | 0 | 0 | 1 | 1 |
| Neurod6 | 0 | 0 | 0 | 1 | 1 |
| Ngp | 0 | 0 | 0 | 1 | 1 |
| Nipsnap3a | 0 | 0 | 0 | 1 | 1 |
| Nkain2 | 0 | 0 | 0 | 1 | 1 |
| Nkx2-1 | 0 | 0 | 0 | 1 | 1 |
| Nkx2-5 | 0 | 0 | 0 | 1 | 1 |
| Nkx6-3 | 0 | 0 | 0 | 1 | 1 |
| Nlrp12 | 0 | 0 | 0 | 1 | 1 |
| Nlrp14 | 0 | 0 | 0 | 1 | 1 |
| Nlrp1a | 0 | 0 | 0 | 1 | 1 |
| Nlrp1b | 0 | 0 | 0 | 1 | 1 |
| Nlrp1c-ps | 0 | 0 | 0 | 1 | 1 |
| Nlrp2 | 0 | 0 | 0 | 1 | 1 |
| Nlrp4a | 0 | 0 | 0 | 1 | 1 |
| Nlrp4b | 0 | 0 | 0 | 1 | 1 |
| Nlrp4e | 0 | 0 | 0 | 1 | 1 |
| Nlrp5 | 0 | 0 | 0 | 1 | 1 |
| Nlrp9a | 0 | 0 | 0 | 1 | 1 |
| Nlrp9b | 0 | 0 | 0 | 1 | 1 |
| Nlrp9c | 0 | 0 | 0 | 1 | 1 |
| Nme8 | 0 | 0 | 0 | 1 | 1 |
| Nme9 | 0 | 0 | 0 | 1 | 1 |
| Nmrk2 | 0 | 0 | 0 | 1 | 1 |
| Nms | 0 | 0 | 0 | 1 | 1 |
| Nov | 0 | 0 | 0 | 1 | 1 |
| Nox3 | 0 | 0 | 0 | 1 | 1 |
| Nphs1os | 0 | 0 | 0 | 1 | 1 |
| Npvf | 0 | 0 | 0 | 1 | 1 |
| Npy2r | 0 | 0 | 0 | 1 | 1 |
| Npy4r | 0 | 0 | 0 | 1 | 1 |
| Npy6r | 0 | 0 | 0 | 1 | 1 |
| Nr0b2 | 0 | 0 | 0 | 1 | 1 |
| Nr1h4 | 0 | 0 | 0 | 1 | 1 |
| Nr1i2 | 0 | 0 | 0 | 1 | 1 |
| Nr2e3 | 0 | 0 | 0 | 1 | 1 |
| Nrg3 | 0 | 0 | 0 | 1 | 1 |
| Nrg3os | 0 | 0 | 0 | 1 | 1 |
| Nrl | 0 | 0 | 0 | 1 | 1 |
| Ntsr2 | 0 | 0 | 0 | 1 | 1 |
| Nutf2-ps1 | 0 | 0 | 0 | 1 | 1 |

| | | | | | |
|---|---|---|---|---|---|
| Nxf7 | 0 | 0 | 0 | 1 | 1 |
| Nxnl1 | 0 | 0 | 0 | 1 | 1 |
| Nyap2 | 0 | 0 | 0 | 1 | 1 |
| OTTMUSG00000016609 | 0 | 0 | 0 | 1 | 1 |
| Oas1d | 0 | 0 | 0 | 1 | 1 |
| Oas1e | 0 | 0 | 0 | 1 | 1 |
| Oas1f | 0 | 0 | 0 | 1 | 1 |
| Oas1g | 0 | 0 | 0 | 1 | 1 |
| Oas2 | 0 | 0 | 0 | 1 | 1 |
| Oaz1-ps | 0 | 0 | 0 | 1 | 1 |
| Obox1 | 0 | 0 | 0 | 1 | 1 |
| Obox2 | 0 | 0 | 0 | 1 | 1 |
| Obox3 | 0 | 0 | 0 | 1 | 1 |
| Obox5 | 0 | 0 | 0 | 1 | 1 |
| Obox6 | 0 | 0 | 0 | 1 | 1 |
| Obp1a | 0 | 0 | 0 | 1 | 1 |
| Obp2a | 0 | 0 | 0 | 1 | 1 |
| Obp2b | 0 | 0 | 0 | 1 | 1 |
| Oc90 | 0 | 0 | 0 | 1 | 1 |
| Ocm | 0 | 0 | 0 | 1 | 1 |
| Odam | 0 | 0 | 0 | 1 | 1 |
| Odf3 | 0 | 0 | 0 | 1 | 1 |
| Odf4 | 0 | 0 | 0 | 1 | 1 |
| Olah | 0 | 0 | 0 | 1 | 1 |
| Olfr1 | 0 | 0 | 0 | 1 | 1 |
| Olfr10 | 0 | 0 | 0 | 1 | 1 |
| Olfr100,Olfr102 | 0 | 0 | 0 | 1 | 1 |
| Olfr1000 | 0 | 0 | 0 | 1 | 1 |
| Olfr1002 | 0 | 0 | 0 | 1 | 1 |
| Olfr1006 | 0 | 0 | 0 | 1 | 1 |
| Olfr1008 | 0 | 0 | 0 | 1 | 1 |
| Olfr1009 | 0 | 0 | 0 | 1 | 1 |
| Olfr101 | 0 | 0 | 0 | 1 | 1 |
| Olfr1010 | 0 | 0 | 0 | 1 | 1 |
| Olfr1012 | 0 | 0 | 0 | 1 | 1 |
| Olfr1013 | 0 | 0 | 0 | 1 | 1 |
| Olfr1014 | 0 | 0 | 0 | 1 | 1 |
| Olfr1015 | 0 | 0 | 0 | 1 | 1 |
| Olfr1016 | 0 | 0 | 0 | 1 | 1 |
| Olfr1018 | 0 | 0 | 0 | 1 | 1 |
| Olfr1019 | 0 | 0 | 0 | 1 | 1 |
| Olfr102 | 0 | 0 | 0 | 1 | 1 |
| Olfr1020 | 0 | 0 | 0 | 1 | 1 |
| Olfr1022 | 0 | 0 | 0 | 1 | 1 |
| Olfr1023 | 0 | 0 | 0 | 1 | 1 |
| Olfr1024 | 0 | 0 | 0 | 1 | 1 |
| Olfr1026 | 0 | 0 | 0 | 1 | 1 |
| Olfr1028 | 0 | 0 | 0 | 1 | 1 |
| Olfr1029 | 0 | 0 | 0 | 1 | 1 |
| Olfr103 | 0 | 0 | 0 | 1 | 1 |

| | | | | | |
|---|---|---|---|---|---|
| Olfr1030 | 0 | 0 | 0 | 1 | 1 |
| Olfr1031 | 0 | 0 | 0 | 1 | 1 |
| Olfr1032 | 0 | 0 | 0 | 1 | 1 |
| Olfr1033 | 0 | 0 | 0 | 1 | 1 |
| Olfr1034 | 0 | 0 | 0 | 1 | 1 |
| Olfr1036 | 0 | 0 | 0 | 1 | 1 |
| Olfr1037 | 0 | 0 | 0 | 1 | 1 |
| Olfr1038-ps | 0 | 0 | 0 | 1 | 1 |
| Olfr1039 | 0 | 0 | 0 | 1 | 1 |
| Olfr1040 | 0 | 0 | 0 | 1 | 1 |
| Olfr1042 | 0 | 0 | 0 | 1 | 1 |
| Olfr1043 | 0 | 0 | 0 | 1 | 1 |
| Olfr1044 | 0 | 0 | 0 | 1 | 1 |
| Olfr1045 | 0 | 0 | 0 | 1 | 1 |
| Olfr1046 | 0 | 0 | 0 | 1 | 1 |
| Olfr1047 | 0 | 0 | 0 | 1 | 1 |
| Olfr1048 | 0 | 0 | 0 | 1 | 1 |
| Olfr1049 | 0 | 0 | 0 | 1 | 1 |
| Olfr1051 | 0 | 0 | 0 | 1 | 1 |
| Olfr1052 | 0 | 0 | 0 | 1 | 1 |
| Olfr1053 | 0 | 0 | 0 | 1 | 1 |
| Olfr1054 | 0 | 0 | 0 | 1 | 1 |
| Olfr1055 | 0 | 0 | 0 | 1 | 1 |
| Olfr1056 | 0 | 0 | 0 | 1 | 1 |
| Olfr1057 | 0 | 0 | 0 | 1 | 1 |
| Olfr1058 | 0 | 0 | 0 | 1 | 1 |
| Olfr1061 | 0 | 0 | 0 | 1 | 1 |
| Olfr1062 | 0 | 0 | 0 | 1 | 1 |
| Olfr1065 | 0 | 0 | 0 | 1 | 1 |
| Olfr1066 | 0 | 0 | 0 | 1 | 1 |
| Olfr107 | 0 | 0 | 0 | 1 | 1 |
| Olfr1076 | 0 | 0 | 0 | 1 | 1 |
| Olfr1077-ps1 | 0 | 0 | 0 | 1 | 1 |
| Olfr1079 | 0 | 0 | 0 | 1 | 1 |
| Olfr108 | 0 | 0 | 0 | 1 | 1 |
| Olfr1080 | 0 | 0 | 0 | 1 | 1 |
| Olfr1082 | 0 | 0 | 0 | 1 | 1 |
| Olfr1084 | 0 | 0 | 0 | 1 | 1 |
| Olfr1085 | 0 | 0 | 0 | 1 | 1 |
| Olfr1086 | 0 | 0 | 0 | 1 | 1 |
| Olfr1087 | 0 | 0 | 0 | 1 | 1 |
| Olfr1089 | 0 | 0 | 0 | 1 | 1 |
| Olfr109 | 0 | 0 | 0 | 1 | 1 |
| Olfr1090 | 0 | 0 | 0 | 1 | 1 |
| Olfr1093 | 0 | 0 | 0 | 1 | 1 |
| Olfr1094 | 0 | 0 | 0 | 1 | 1 |
| Olfr1095 | 0 | 0 | 0 | 1 | 1 |
| Olfr1097 | 0 | 0 | 0 | 1 | 1 |
| Olfr1098 | 0 | 0 | 0 | 1 | 1 |
| Olfr1099 | 0 | 0 | 0 | 1 | 1 |

| | | | | | |
|---|---|---|---|---|---|
| Olfr11 | 0 | 0 | 0 | 1 | 1 |
| Olfr110 | 0 | 0 | 0 | 1 | 1 |
| Olfr1100 | 0 | 0 | 0 | 1 | 1 |
| Olfr1101 | 0 | 0 | 0 | 1 | 1 |
| Olfr1102 | 0 | 0 | 0 | 1 | 1 |
| Olfr1104 | 0 | 0 | 0 | 1 | 1 |
| Olfr1105 | 0 | 0 | 0 | 1 | 1 |
| Olfr1106 | 0 | 0 | 0 | 1 | 1 |
| Olfr1107 | 0 | 0 | 0 | 1 | 1 |
| Olfr1109 | 0 | 0 | 0 | 1 | 1 |
| Olfr111 | 0 | 0 | 0 | 1 | 1 |
| Olfr1110 | 0 | 0 | 0 | 1 | 1 |
| Olfr1111 | 0 | 0 | 0 | 1 | 1 |
| Olfr1112 | 0 | 0 | 0 | 1 | 1 |
| Olfr1113 | 0 | 0 | 0 | 1 | 1 |
| Olfr1115 | 0 | 0 | 0 | 1 | 1 |
| Olfr1116-ps | 0 | 0 | 0 | 1 | 1 |
| Olfr1118 | 0 | 0 | 0 | 1 | 1 |
| Olfr112 | 0 | 0 | 0 | 1 | 1 |
| Olfr1120 | 0 | 0 | 0 | 1 | 1 |
| Olfr1121 | 0 | 0 | 0 | 1 | 1 |
| Olfr1122 | 0 | 0 | 0 | 1 | 1 |
| Olfr1123 | 0 | 0 | 0 | 1 | 1 |
| Olfr1124 | 0 | 0 | 0 | 1 | 1 |
| Olfr1126 | 0 | 0 | 0 | 1 | 1 |
| Olfr1128 | 0 | 0 | 0 | 1 | 1 |
| Olfr1129 | 0 | 0 | 0 | 1 | 1 |
| Olfr113 | 0 | 0 | 0 | 1 | 1 |
| Olfr1130 | 0 | 0 | 0 | 1 | 1 |
| Olfr1131 | 0 | 0 | 0 | 1 | 1 |
| Olfr1132 | 0 | 0 | 0 | 1 | 1 |
| Olfr1133 | 0 | 0 | 0 | 1 | 1 |
| Olfr1134 | 0 | 0 | 0 | 1 | 1 |
| Olfr1135 | 0 | 0 | 0 | 1 | 1 |
| Olfr1136 | 0 | 0 | 0 | 1 | 1 |
| Olfr1137 | 0 | 0 | 0 | 1 | 1 |
| Olfr1138 | 0 | 0 | 0 | 1 | 1 |
| Olfr114 | 0 | 0 | 0 | 1 | 1 |
| Olfr1140 | 0 | 0 | 0 | 1 | 1 |
| Olfr1141 | 0 | 0 | 0 | 1 | 1 |
| Olfr1143 | 0 | 0 | 0 | 1 | 1 |
| Olfr1145 | 0 | 0 | 0 | 1 | 1 |
| Olfr1148 | 0 | 0 | 0 | 1 | 1 |
| Olfr115 | 0 | 0 | 0 | 1 | 1 |
| Olfr1151 | 0 | 0 | 0 | 1 | 1 |
| Olfr1152 | 0 | 0 | 0 | 1 | 1 |
| Olfr1153 | 0 | 0 | 0 | 1 | 1 |
| Olfr1154 | 0 | 0 | 0 | 1 | 1 |
| Olfr1155 | 0 | 0 | 0 | 1 | 1 |
| Olfr1156 | 0 | 0 | 0 | 1 | 1 |

| | | | | | |
|---|---|---|---|---|---|
| Olfr1157 | 0 | 0 | 0 | 1 | 1 |
| Olfr1158 | 0 | 0 | 0 | 1 | 1 |
| Olfr116 | 0 | 0 | 0 | 1 | 1 |
| Olfr1160 | 0 | 0 | 0 | 1 | 1 |
| Olfr1161 | 0 | 0 | 0 | 1 | 1 |
| Olfr1162 | 0 | 0 | 0 | 1 | 1 |
| Olfr1163 | 0 | 0 | 0 | 1 | 1 |
| Olfr1164 | 0 | 0 | 0 | 1 | 1 |
| Olfr1166 | 0 | 0 | 0 | 1 | 1 |
| Olfr1167 | 0 | 0 | 0 | 1 | 1 |
| Olfr117 | 0 | 0 | 0 | 1 | 1 |
| Olfr1170 | 0 | 0 | 0 | 1 | 1 |
| Olfr1173 | 0 | 0 | 0 | 1 | 1 |
| Olfr1176 | 0 | 0 | 0 | 1 | 1 |
| Olfr1178 | 0 | 0 | 0 | 1 | 1 |
| Olfr1179 | 0 | 0 | 0 | 1 | 1 |
| Olfr118 | 0 | 0 | 0 | 1 | 1 |
| Olfr1180 | 0 | 0 | 0 | 1 | 1 |
| Olfr1181 | 0 | 0 | 0 | 1 | 1 |
| Olfr1182 | 0 | 0 | 0 | 1 | 1 |
| Olfr1183 | 0 | 0 | 0 | 1 | 1 |
| Olfr1184 | 0 | 0 | 0 | 1 | 1 |
| Olfr1186 | 0 | 0 | 0 | 1 | 1 |
| Olfr1189 | 0 | 0 | 0 | 1 | 1 |
| Olfr119 | 0 | 0 | 0 | 1 | 1 |
| Olfr1193 | 0 | 0 | 0 | 1 | 1 |
| Olfr1195 | 0 | 0 | 0 | 1 | 1 |
| Olfr1196 | 0 | 0 | 0 | 1 | 1 |
| Olfr1197 | 0 | 0 | 0 | 1 | 1 |
| Olfr1199 | 0 | 0 | 0 | 1 | 1 |
| Olfr12 | 0 | 0 | 0 | 1 | 1 |
| Olfr120 | 0 | 0 | 0 | 1 | 1 |
| Olfr1200 | 0 | 0 | 0 | 1 | 1 |
| Olfr1201 | 0 | 0 | 0 | 1 | 1 |
| Olfr1202 | 0 | 0 | 0 | 1 | 1 |
| Olfr1204 | 0 | 0 | 0 | 1 | 1 |
| Olfr1205 | 0 | 0 | 0 | 1 | 1 |
| Olfr1206 | 0 | 0 | 0 | 1 | 1 |
| Olfr1208 | 0 | 0 | 0 | 1 | 1 |
| Olfr1209 | 0 | 0 | 0 | 1 | 1 |
| Olfr121 | 0 | 0 | 0 | 1 | 1 |
| Olfr1211 | 0 | 0 | 0 | 1 | 1 |
| Olfr1212 | 0 | 0 | 0 | 1 | 1 |
| Olfr1213 | 0 | 0 | 0 | 1 | 1 |
| Olfr1214 | 0 | 0 | 0 | 1 | 1 |
| Olfr1215 | 0 | 0 | 0 | 1 | 1 |
| Olfr1216 | 0 | 0 | 0 | 1 | 1 |
| Olfr1217 | 0 | 0 | 0 | 1 | 1 |
| Olfr1218 | 0 | 0 | 0 | 1 | 1 |
| Olfr1219 | 0 | 0 | 0 | 1 | 1 |

| | | | | | |
|---|---|---|---|---|---|
| Olfr122 | 0 | 0 | 0 | 1 | 1 |
| Olfr1221 | 0 | 0 | 0 | 1 | 1 |
| Olfr1222 | 0 | 0 | 0 | 1 | 1 |
| Olfr1223 | 0 | 0 | 0 | 1 | 1 |
| Olfr1225 | 0 | 0 | 0 | 1 | 1 |
| Olfr1226 | 0 | 0 | 0 | 1 | 1 |
| Olfr1228 | 0 | 0 | 0 | 1 | 1 |
| Olfr1229 | 0 | 0 | 0 | 1 | 1 |
| Olfr123 | 0 | 0 | 0 | 1 | 1 |
| Olfr1230 | 0 | 0 | 0 | 1 | 1 |
| Olfr1231 | 0 | 0 | 0 | 1 | 1 |
| Olfr1232 | 0 | 0 | 0 | 1 | 1 |
| Olfr1233 | 0 | 0 | 0 | 1 | 1 |
| Olfr1234 | 0 | 0 | 0 | 1 | 1 |
| Olfr1238 | 0 | 0 | 0 | 1 | 1 |
| Olfr1239 | 0 | 0 | 0 | 1 | 1 |
| Olfr124 | 0 | 0 | 0 | 1 | 1 |
| Olfr1240 | 0 | 0 | 0 | 1 | 1 |
| Olfr1241 | 0 | 0 | 0 | 1 | 1 |
| Olfr1242 | 0 | 0 | 0 | 1 | 1 |
| Olfr1243 | 0 | 0 | 0 | 1 | 1 |
| Olfr1245 | 0 | 0 | 0 | 1 | 1 |
| Olfr1246 | 0 | 0 | 0 | 1 | 1 |
| Olfr1247 | 0 | 0 | 0 | 1 | 1 |
| Olfr1248 | 0 | 0 | 0 | 1 | 1 |
| Olfr125 | 0 | 0 | 0 | 1 | 1 |
| Olfr1250 | 0 | 0 | 0 | 1 | 1 |
| Olfr1251 | 0 | 0 | 0 | 1 | 1 |
| Olfr1252 | 0 | 0 | 0 | 1 | 1 |
| Olfr1253 | 0 | 0 | 0 | 1 | 1 |
| Olfr1254 | 0 | 0 | 0 | 1 | 1 |
| Olfr1255 | 0 | 0 | 0 | 1 | 1 |
| Olfr1256 | 0 | 0 | 0 | 1 | 1 |
| Olfr1257 | 0 | 0 | 0 | 1 | 1 |
| Olfr1258 | 0 | 0 | 0 | 1 | 1 |
| Olfr1259 | 0 | 0 | 0 | 1 | 1 |
| Olfr126 | 0 | 0 | 0 | 1 | 1 |
| Olfr1260 | 0 | 0 | 0 | 1 | 1 |
| Olfr1261 | 0 | 0 | 0 | 1 | 1 |
| Olfr1262 | 0 | 0 | 0 | 1 | 1 |
| Olfr1263 | 0 | 0 | 0 | 1 | 1 |
| Olfr1264 | 0 | 0 | 0 | 1 | 1 |
| Olfr1265 | 0 | 0 | 0 | 1 | 1 |
| Olfr1269 | 0 | 0 | 0 | 1 | 1 |
| Olfr127 | 0 | 0 | 0 | 1 | 1 |
| Olfr1271 | 0 | 0 | 0 | 1 | 1 |
| Olfr1272 | 0 | 0 | 0 | 1 | 1 |
| Olfr1273-ps | 0 | 0 | 0 | 1 | 1 |
| Olfr1274-ps | 0 | 0 | 0 | 1 | 1 |
| Olfr1275 | 0 | 0 | 0 | 1 | 1 |

| | | | | | |
|---|---|---|---|---|---|
| Olfr1276 | 0 | 0 | 0 | 1 | 1 |
| Olfr1277 | 0 | 0 | 0 | 1 | 1 |
| Olfr1278 | 0 | 0 | 0 | 1 | 1 |
| Olfr1279 | 0 | 0 | 0 | 1 | 1 |
| Olfr128 | 0 | 0 | 0 | 1 | 1 |
| Olfr1280 | 0 | 0 | 0 | 1 | 1 |
| Olfr1281 | 0 | 0 | 0 | 1 | 1 |
| Olfr1282 | 0 | 0 | 0 | 1 | 1 |
| Olfr1283 | 0 | 0 | 0 | 1 | 1 |
| Olfr1284 | 0 | 0 | 0 | 1 | 1 |
| Olfr1286 | 0 | 0 | 0 | 1 | 1 |
| Olfr1287 | 0 | 0 | 0 | 1 | 1 |
| Olfr1288 | 0 | 0 | 0 | 1 | 1 |
| Olfr1289 | 0 | 0 | 0 | 1 | 1 |
| Olfr129 | 0 | 0 | 0 | 1 | 1 |
| Olfr1290 | 0 | 0 | 0 | 1 | 1 |
| Olfr1294 | 0 | 0 | 0 | 1 | 1 |
| Olfr1295 | 0 | 0 | 0 | 1 | 1 |
| Olfr1297 | 0 | 0 | 0 | 1 | 1 |
| Olfr1298 | 0 | 0 | 0 | 1 | 1 |
| Olfr1299 | 0 | 0 | 0 | 1 | 1 |
| Olfr13 | 0 | 0 | 0 | 1 | 1 |
| Olfr130 | 0 | 0 | 0 | 1 | 1 |
| Olfr1300-ps1 | 0 | 0 | 0 | 1 | 1 |
| Olfr1301 | 0 | 0 | 0 | 1 | 1 |
| Olfr1302 | 0 | 0 | 0 | 1 | 1 |
| Olfr1303 | 0 | 0 | 0 | 1 | 1 |
| Olfr1305 | 0 | 0 | 0 | 1 | 1 |
| Olfr1306 | 0 | 0 | 0 | 1 | 1 |
| Olfr1307 | 0 | 0 | 0 | 1 | 1 |
| Olfr1308 | 0 | 0 | 0 | 1 | 1 |
| Olfr1309 | 0 | 0 | 0 | 1 | 1 |
| Olfr131 | 0 | 0 | 0 | 1 | 1 |
| Olfr1310 | 0 | 0 | 0 | 1 | 1 |
| Olfr1311 | 0 | 0 | 0 | 1 | 1 |
| Olfr1312 | 0 | 0 | 0 | 1 | 1 |
| Olfr1313 | 0 | 0 | 0 | 1 | 1 |
| Olfr1314 | 0 | 0 | 0 | 1 | 1 |
| Olfr1316 | 0 | 0 | 0 | 1 | 1 |
| Olfr1317 | 0 | 0 | 0 | 1 | 1 |
| Olfr1318 | 0 | 0 | 0 | 1 | 1 |
| Olfr132 | 0 | 0 | 0 | 1 | 1 |
| Olfr1320 | 0 | 0 | 0 | 1 | 1 |
| Olfr1321 | 0 | 0 | 0 | 1 | 1 |
| Olfr1322 | 0 | 0 | 0 | 1 | 1 |
| Olfr1323 | 0 | 0 | 0 | 1 | 1 |
| Olfr1324 | 0 | 0 | 0 | 1 | 1 |
| Olfr1325 | 0 | 0 | 0 | 1 | 1 |
| Olfr1328 | 0 | 0 | 0 | 1 | 1 |
| Olfr1329 | 0 | 0 | 0 | 1 | 1 |

| | | | | | |
|---|---|---|---|---|---|
| Olfr133 | 0 | 0 | 0 | 1 | 1 |
| Olfr1330 | 0 | 0 | 0 | 1 | 1 |
| Olfr1331 | 0 | 0 | 0 | 1 | 1 |
| Olfr1333 | 0 | 0 | 0 | 1 | 1 |
| Olfr1335 | 0 | 0 | 0 | 1 | 1 |
| Olfr1336 | 0 | 0 | 0 | 1 | 1 |
| Olfr1337 | 0 | 0 | 0 | 1 | 1 |
| Olfr1338 | 0 | 0 | 0 | 1 | 1 |
| Olfr1339 | 0 | 0 | 0 | 1 | 1 |
| Olfr134 | 0 | 0 | 0 | 1 | 1 |
| Olfr1340 | 0 | 0 | 0 | 1 | 1 |
| Olfr1341 | 0 | 0 | 0 | 1 | 1 |
| Olfr1342 | 0 | 0 | 0 | 1 | 1 |
| Olfr1344 | 0 | 0 | 0 | 1 | 1 |
| Olfr1346 | 0 | 0 | 0 | 1 | 1 |
| Olfr1348 | 0 | 0 | 0 | 1 | 1 |
| Olfr135 | 0 | 0 | 0 | 1 | 1 |
| Olfr1350 | 0 | 0 | 0 | 1 | 1 |
| Olfr1351 | 0 | 0 | 0 | 1 | 1 |
| Olfr1352 | 0 | 0 | 0 | 1 | 1 |
| Olfr1353 | 0 | 0 | 0 | 1 | 1 |
| Olfr1354 | 0 | 0 | 0 | 1 | 1 |
| Olfr1355 | 0 | 0 | 0 | 1 | 1 |
| Olfr1356 | 0 | 0 | 0 | 1 | 1 |
| Olfr1357 | 0 | 0 | 0 | 1 | 1 |
| Olfr1359,Olfr1360 | 0 | 0 | 0 | 1 | 1 |
| Olfr136 | 0 | 0 | 0 | 1 | 1 |
| Olfr1360 | 0 | 0 | 0 | 1 | 1 |
| Olfr1361 | 0 | 0 | 0 | 1 | 1 |
| Olfr1362 | 0 | 0 | 0 | 1 | 1 |
| Olfr1364 | 0 | 0 | 0 | 1 | 1 |
| Olfr1367 | 0 | 0 | 0 | 1 | 1 |
| Olfr1368 | 0 | 0 | 0 | 1 | 1 |
| Olfr137 | 0 | 0 | 0 | 1 | 1 |
| Olfr1370 | 0 | 0 | 0 | 1 | 1 |
| Olfr1371 | 0 | 0 | 0 | 1 | 1 |
| Olfr1373 | 0 | 0 | 0 | 1 | 1 |
| Olfr1377 | 0 | 0 | 0 | 1 | 1 |
| Olfr1378 | 0 | 0 | 0 | 1 | 1 |
| Olfr138 | 0 | 0 | 0 | 1 | 1 |
| Olfr1380 | 0 | 0 | 0 | 1 | 1 |
| Olfr1381 | 0 | 0 | 0 | 1 | 1 |
| Olfr1382 | 0 | 0 | 0 | 1 | 1 |
| Olfr1383 | 0 | 0 | 0 | 1 | 1 |
| Olfr1384 | 0 | 0 | 0 | 1 | 1 |
| Olfr1385 | 0 | 0 | 0 | 1 | 1 |
| Olfr1386 | 0 | 0 | 0 | 1 | 1 |
| Olfr1387 | 0 | 0 | 0 | 1 | 1 |
| Olfr1388 | 0 | 0 | 0 | 1 | 1 |
| Olfr1389 | 0 | 0 | 0 | 1 | 1 |

| | | | | | |
|---|---|---|---|---|---|
| Olfr139 | 0 | 0 | 0 | 1 | 1 |
| Olfr1390 | 0 | 0 | 0 | 1 | 1 |
| Olfr1391 | 0 | 0 | 0 | 1 | 1 |
| Olfr1393 | 0 | 0 | 0 | 1 | 1 |
| Olfr1394 | 0 | 0 | 0 | 1 | 1 |
| Olfr1395 | 0 | 0 | 0 | 1 | 1 |
| Olfr140 | 0 | 0 | 0 | 1 | 1 |
| Olfr1402 | 0 | 0 | 0 | 1 | 1 |
| Olfr1404 | 0 | 0 | 0 | 1 | 1 |
| Olfr1406 | 0 | 0 | 0 | 1 | 1 |
| Olfr1408 | 0 | 0 | 0 | 1 | 1 |
| Olfr141 | 0 | 0 | 0 | 1 | 1 |
| Olfr1410 | 0 | 0 | 0 | 1 | 1 |
| Olfr1411 | 0 | 0 | 0 | 1 | 1 |
| Olfr1412 | 0 | 0 | 0 | 1 | 1 |
| Olfr1413 | 0 | 0 | 0 | 1 | 1 |
| Olfr1415 | 0 | 0 | 0 | 1 | 1 |
| Olfr1416 | 0 | 0 | 0 | 1 | 1 |
| Olfr1418 | 0 | 0 | 0 | 1 | 1 |
| Olfr1419 | 0 | 0 | 0 | 1 | 1 |
| Olfr142 | 0 | 0 | 0 | 1 | 1 |
| Olfr1423 | 0 | 0 | 0 | 1 | 1 |
| Olfr1424 | 0 | 0 | 0 | 1 | 1 |
| Olfr1425 | 0 | 0 | 0 | 1 | 1 |
| Olfr1426 | 0 | 0 | 0 | 1 | 1 |
| Olfr1427 | 0 | 0 | 0 | 1 | 1 |
| Olfr1428 | 0 | 0 | 0 | 1 | 1 |
| Olfr143 | 0 | 0 | 0 | 1 | 1 |
| Olfr1431 | 0 | 0 | 0 | 1 | 1 |
| Olfr1433,Olfr1434 | 0 | 0 | 0 | 1 | 1 |
| Olfr1434 | 0 | 0 | 0 | 1 | 1 |
| Olfr1436 | 0 | 0 | 0 | 1 | 1 |
| Olfr1437 | 0 | 0 | 0 | 1 | 1 |
| Olfr1440 | 0 | 0 | 0 | 1 | 1 |
| Olfr1441 | 0 | 0 | 0 | 1 | 1 |
| Olfr1442 | 0 | 0 | 0 | 1 | 1 |
| Olfr1443 | 0 | 0 | 0 | 1 | 1 |
| Olfr1444 | 0 | 0 | 0 | 1 | 1 |
| Olfr1445 | 0 | 0 | 0 | 1 | 1 |
| Olfr1446 | 0 | 0 | 0 | 1 | 1 |
| Olfr1447 | 0 | 0 | 0 | 1 | 1 |
| Olfr1448 | 0 | 0 | 0 | 1 | 1 |
| Olfr1449 | 0 | 0 | 0 | 1 | 1 |
| Olfr145 | 0 | 0 | 0 | 1 | 1 |
| Olfr1450 | 0 | 0 | 0 | 1 | 1 |
| Olfr1451 | 0 | 0 | 0 | 1 | 1 |
| Olfr1453 | 0 | 0 | 0 | 1 | 1 |
| Olfr1454 | 0 | 0 | 0 | 1 | 1 |
| Olfr1457 | 0 | 0 | 0 | 1 | 1 |
| Olfr1459 | 0 | 0 | 0 | 1 | 1 |

| | | | | | |
|---|---|---|---|---|---|
| Olfr146 | 0 | 0 | 0 | 1 | 1 |
| Olfr1461 | 0 | 0 | 0 | 1 | 1 |
| Olfr1462 | 0 | 0 | 0 | 1 | 1 |
| Olfr1463 | 0 | 0 | 0 | 1 | 1 |
| Olfr1465 | 0 | 0 | 0 | 1 | 1 |
| Olfr1466 | 0 | 0 | 0 | 1 | 1 |
| Olfr1467 | 0 | 0 | 0 | 1 | 1 |
| Olfr1469 | 0 | 0 | 0 | 1 | 1 |
| Olfr147 | 0 | 0 | 0 | 1 | 1 |
| Olfr1471 | 0 | 0 | 0 | 1 | 1 |
| Olfr1472 | 0 | 0 | 0 | 1 | 1 |
| Olfr1475 | 0 | 0 | 0 | 1 | 1 |
| Olfr1477 | 0 | 0 | 0 | 1 | 1 |
| Olfr148 | 0 | 0 | 0 | 1 | 1 |
| Olfr1480 | 0 | 0 | 0 | 1 | 1 |
| Olfr1484 | 0 | 0 | 0 | 1 | 1 |
| Olfr1487 | 0 | 0 | 0 | 1 | 1 |
| Olfr1489 | 0 | 0 | 0 | 1 | 1 |
| Olfr149 | 0 | 0 | 0 | 1 | 1 |
| Olfr1490 | 0 | 0 | 0 | 1 | 1 |
| Olfr1491 | 0 | 0 | 0 | 1 | 1 |
| Olfr1494 | 0 | 0 | 0 | 1 | 1 |
| Olfr1495 | 0 | 0 | 0 | 1 | 1 |
| Olfr1496 | 0 | 0 | 0 | 1 | 1 |
| Olfr1497 | 0 | 0 | 0 | 1 | 1 |
| Olfr1499 | 0 | 0 | 0 | 1 | 1 |
| Olfr15 | 0 | 0 | 0 | 1 | 1 |
| Olfr150 | 0 | 0 | 0 | 1 | 1 |
| Olfr1500 | 0 | 0 | 0 | 1 | 1 |
| Olfr1501 | 0 | 0 | 0 | 1 | 1 |
| Olfr1502 | 0 | 0 | 0 | 1 | 1 |
| Olfr1504 | 0 | 0 | 0 | 1 | 1 |
| Olfr1505 | 0 | 0 | 0 | 1 | 1 |
| Olfr1507 | 0 | 0 | 0 | 1 | 1 |
| Olfr1508 | 0 | 0 | 0 | 1 | 1 |
| Olfr1509 | 0 | 0 | 0 | 1 | 1 |
| Olfr151 | 0 | 0 | 0 | 1 | 1 |
| Olfr1510 | 0 | 0 | 0 | 1 | 1 |
| Olfr1511 | 0 | 0 | 0 | 1 | 1 |
| Olfr1512 | 0 | 0 | 0 | 1 | 1 |
| Olfr1513 | 0 | 0 | 0 | 1 | 1 |
| Olfr152 | 0 | 0 | 0 | 1 | 1 |
| Olfr153 | 0 | 0 | 0 | 1 | 1 |
| Olfr1532-ps1 | 0 | 0 | 0 | 1 | 1 |
| Olfr1535 | 0 | 0 | 0 | 1 | 1 |
| Olfr1537 | 0 | 0 | 0 | 1 | 1 |
| Olfr154 | 0 | 0 | 0 | 1 | 1 |
| Olfr155 | 0 | 0 | 0 | 1 | 1 |
| Olfr156 | 0 | 0 | 0 | 1 | 1 |
| Olfr157 | 0 | 0 | 0 | 1 | 1 |

| | | | | | |
|---|---|---|---|---|---|
| Olfr159 | 0 | 0 | 0 | 1 | 1 |
| Olfr16 | 0 | 0 | 0 | 1 | 1 |
| Olfr160 | 0 | 0 | 0 | 1 | 1 |
| Olfr161 | 0 | 0 | 0 | 1 | 1 |
| Olfr164 | 0 | 0 | 0 | 1 | 1 |
| Olfr165 | 0 | 0 | 0 | 1 | 1 |
| Olfr166 | 0 | 0 | 0 | 1 | 1 |
| Olfr167 | 0 | 0 | 0 | 1 | 1 |
| Olfr168 | 0 | 0 | 0 | 1 | 1 |
| Olfr169 | 0 | 0 | 0 | 1 | 1 |
| Olfr17 | 0 | 0 | 0 | 1 | 1 |
| Olfr170 | 0 | 0 | 0 | 1 | 1 |
| Olfr171 | 0 | 0 | 0 | 1 | 1 |
| Olfr172 | 0 | 0 | 0 | 1 | 1 |
| Olfr173 | 0 | 0 | 0 | 1 | 1 |
| Olfr175-ps1 | 0 | 0 | 0 | 1 | 1 |
| Olfr176 | 0 | 0 | 0 | 1 | 1 |
| Olfr177 | 0 | 0 | 0 | 1 | 1 |
| Olfr178 | 0 | 0 | 0 | 1 | 1 |
| Olfr18 | 0 | 0 | 0 | 1 | 1 |
| Olfr180 | 0 | 0 | 0 | 1 | 1 |
| Olfr181 | 0 | 0 | 0 | 1 | 1 |
| Olfr183 | 0 | 0 | 0 | 1 | 1 |
| Olfr186 | 0 | 0 | 0 | 1 | 1 |
| Olfr187 | 0 | 0 | 0 | 1 | 1 |
| Olfr19 | 0 | 0 | 0 | 1 | 1 |
| Olfr190 | 0 | 0 | 0 | 1 | 1 |
| Olfr191 | 0 | 0 | 0 | 1 | 1 |
| Olfr192 | 0 | 0 | 0 | 1 | 1 |
| Olfr193 | 0 | 0 | 0 | 1 | 1 |
| Olfr194 | 0 | 0 | 0 | 1 | 1 |
| Olfr195 | 0 | 0 | 0 | 1 | 1 |
| Olfr196 | 0 | 0 | 0 | 1 | 1 |
| Olfr197 | 0 | 0 | 0 | 1 | 1 |
| Olfr198 | 0 | 0 | 0 | 1 | 1 |
| Olfr199 | 0 | 0 | 0 | 1 | 1 |
| Olfr2 | 0 | 0 | 0 | 1 | 1 |
| Olfr20 | 0 | 0 | 0 | 1 | 1 |
| Olfr201 | 0 | 0 | 0 | 1 | 1 |
| Olfr202 | 0 | 0 | 0 | 1 | 1 |
| Olfr203 | 0 | 0 | 0 | 1 | 1 |
| Olfr204 | 0 | 0 | 0 | 1 | 1 |
| Olfr205 | 0 | 0 | 0 | 1 | 1 |
| Olfr206 | 0 | 0 | 0 | 1 | 1 |
| Olfr207 | 0 | 0 | 0 | 1 | 1 |
| Olfr211 | 0 | 0 | 0 | 1 | 1 |
| Olfr212 | 0 | 0 | 0 | 1 | 1 |
| Olfr213 | 0 | 0 | 0 | 1 | 1 |
| Olfr214 | 0 | 0 | 0 | 1 | 1 |
| Olfr215 | 0 | 0 | 0 | 1 | 1 |

| | | | | | |
|---|---|---|---|---|---|
| Olfr220 | 0 | 0 | 0 | 1 | 1 |
| Olfr221 | 0 | 0 | 0 | 1 | 1 |
| Olfr222 | 0 | 0 | 0 | 1 | 1 |
| Olfr223 | 0 | 0 | 0 | 1 | 1 |
| Olfr224 | 0 | 0 | 0 | 1 | 1 |
| Olfr225 | 0 | 0 | 0 | 1 | 1 |
| Olfr228 | 0 | 0 | 0 | 1 | 1 |
| Olfr229 | 0 | 0 | 0 | 1 | 1 |
| Olfr23 | 0 | 0 | 0 | 1 | 1 |
| Olfr231 | 0 | 0 | 0 | 1 | 1 |
| Olfr235 | 0 | 0 | 0 | 1 | 1 |
| Olfr237-ps1 | 0 | 0 | 0 | 1 | 1 |
| Olfr239 | 0 | 0 | 0 | 1 | 1 |
| Olfr24 | 0 | 0 | 0 | 1 | 1 |
| Olfr242 | 0 | 0 | 0 | 1 | 1 |
| Olfr243 | 0 | 0 | 0 | 1 | 1 |
| Olfr247 | 0 | 0 | 0 | 1 | 1 |
| Olfr248 | 0 | 0 | 0 | 1 | 1 |
| Olfr25 | 0 | 0 | 0 | 1 | 1 |
| Olfr259 | 0 | 0 | 0 | 1 | 1 |
| Olfr26 | 0 | 0 | 0 | 1 | 1 |
| Olfr262 | 0 | 0 | 0 | 1 | 1 |
| Olfr263 | 0 | 0 | 0 | 1 | 1 |
| Olfr266 | 0 | 0 | 0 | 1 | 1 |
| Olfr27 | 0 | 0 | 0 | 1 | 1 |
| Olfr270 | 0 | 0 | 0 | 1 | 1 |
| Olfr272 | 0 | 0 | 0 | 1 | 1 |
| Olfr273 | 0 | 0 | 0 | 1 | 1 |
| Olfr275 | 0 | 0 | 0 | 1 | 1 |
| Olfr279 | 0 | 0 | 0 | 1 | 1 |
| Olfr281 | 0 | 0 | 0 | 1 | 1 |
| Olfr282 | 0 | 0 | 0 | 1 | 1 |
| Olfr283 | 0 | 0 | 0 | 1 | 1 |
| Olfr284 | 0 | 0 | 0 | 1 | 1 |
| Olfr285 | 0 | 0 | 0 | 1 | 1 |
| Olfr286 | 0 | 0 | 0 | 1 | 1 |
| Olfr287 | 0 | 0 | 0 | 1 | 1 |
| Olfr288 | 0 | 0 | 0 | 1 | 1 |
| Olfr29-ps1 | 0 | 0 | 0 | 1 | 1 |
| Olfr290 | 0 | 0 | 0 | 1 | 1 |
| Olfr291 | 0 | 0 | 0 | 1 | 1 |
| Olfr292 | 0 | 0 | 0 | 1 | 1 |
| Olfr293 | 0 | 0 | 0 | 1 | 1 |
| Olfr294 | 0 | 0 | 0 | 1 | 1 |
| Olfr295 | 0 | 0 | 0 | 1 | 1 |
| Olfr297 | 0 | 0 | 0 | 1 | 1 |
| Olfr298 | 0 | 0 | 0 | 1 | 1 |
| Olfr299 | 0 | 0 | 0 | 1 | 1 |
| Olfr3 | 0 | 0 | 0 | 1 | 1 |
| Olfr30 | 0 | 0 | 0 | 1 | 1 |

| | | | | | |
|---|---|---|---|---|---|
| Olfr301 | 0 | 0 | 0 | 1 | 1 |
| Olfr303 | 0 | 0 | 0 | 1 | 1 |
| Olfr304 | 0 | 0 | 0 | 1 | 1 |
| Olfr305 | 0 | 0 | 0 | 1 | 1 |
| Olfr307 | 0 | 0 | 0 | 1 | 1 |
| Olfr308 | 0 | 0 | 0 | 1 | 1 |
| Olfr309 | 0 | 0 | 0 | 1 | 1 |
| Olfr31 | 0 | 0 | 0 | 1 | 1 |
| Olfr310 | 0 | 0 | 0 | 1 | 1 |
| Olfr311 | 0 | 0 | 0 | 1 | 1 |
| Olfr312 | 0 | 0 | 0 | 1 | 1 |
| Olfr313 | 0 | 0 | 0 | 1 | 1 |
| Olfr314 | 0 | 0 | 0 | 1 | 1 |
| Olfr315 | 0 | 0 | 0 | 1 | 1 |
| Olfr316 | 0 | 0 | 0 | 1 | 1 |
| Olfr317 | 0 | 0 | 0 | 1 | 1 |
| Olfr318 | 0 | 0 | 0 | 1 | 1 |
| Olfr319 | 0 | 0 | 0 | 1 | 1 |
| Olfr32 | 0 | 0 | 0 | 1 | 1 |
| Olfr320 | 0 | 0 | 0 | 1 | 1 |
| Olfr322 | 0 | 0 | 0 | 1 | 1 |
| Olfr323 | 0 | 0 | 0 | 1 | 1 |
| Olfr324 | 0 | 0 | 0 | 1 | 1 |
| Olfr325 | 0 | 0 | 0 | 1 | 1 |
| Olfr328 | 0 | 0 | 0 | 1 | 1 |
| Olfr329-ps | 0 | 0 | 0 | 1 | 1 |
| Olfr33 | 0 | 0 | 0 | 1 | 1 |
| Olfr330 | 0 | 0 | 0 | 1 | 1 |
| Olfr331 | 0 | 0 | 0 | 1 | 1 |
| Olfr332 | 0 | 0 | 0 | 1 | 1 |
| Olfr338 | 0 | 0 | 0 | 1 | 1 |
| Olfr339 | 0 | 0 | 0 | 1 | 1 |
| Olfr340 | 0 | 0 | 0 | 1 | 1 |
| Olfr341 | 0 | 0 | 0 | 1 | 1 |
| Olfr342 | 0 | 0 | 0 | 1 | 1 |
| Olfr344 | 0 | 0 | 0 | 1 | 1 |
| Olfr345 | 0 | 0 | 0 | 1 | 1 |
| Olfr346 | 0 | 0 | 0 | 1 | 1 |
| Olfr347 | 0 | 0 | 0 | 1 | 1 |
| Olfr348 | 0 | 0 | 0 | 1 | 1 |
| Olfr350 | 0 | 0 | 0 | 1 | 1 |
| Olfr351 | 0 | 0 | 0 | 1 | 1 |
| Olfr352 | 0 | 0 | 0 | 1 | 1 |
| Olfr353 | 0 | 0 | 0 | 1 | 1 |
| Olfr354 | 0 | 0 | 0 | 1 | 1 |
| Olfr355 | 0 | 0 | 0 | 1 | 1 |
| Olfr356 | 0 | 0 | 0 | 1 | 1 |
| Olfr357 | 0 | 0 | 0 | 1 | 1 |
| Olfr358 | 0 | 0 | 0 | 1 | 1 |
| Olfr360 | 0 | 0 | 0 | 1 | 1 |

| | | | | | |
|---|---|---|---|---|---|
| Olfr361 | 0 | 0 | 0 | 1 | 1 |
| Olfr362 | 0 | 0 | 0 | 1 | 1 |
| Olfr365 | 0 | 0 | 0 | 1 | 1 |
| Olfr366 | 0 | 0 | 0 | 1 | 1 |
| Olfr367-ps | 0 | 0 | 0 | 1 | 1 |
| Olfr368 | 0 | 0 | 0 | 1 | 1 |
| Olfr371 | 0 | 0 | 0 | 1 | 1 |
| Olfr372 | 0 | 0 | 0 | 1 | 1 |
| Olfr373 | 0 | 0 | 0 | 1 | 1 |
| Olfr374 | 0 | 0 | 0 | 1 | 1 |
| Olfr376 | 0 | 0 | 0 | 1 | 1 |
| Olfr378 | 0 | 0 | 0 | 1 | 1 |
| Olfr380 | 0 | 0 | 0 | 1 | 1 |
| Olfr381 | 0 | 0 | 0 | 1 | 1 |
| Olfr382 | 0 | 0 | 0 | 1 | 1 |
| Olfr384 | 0 | 0 | 0 | 1 | 1 |
| Olfr385 | 0 | 0 | 0 | 1 | 1 |
| Olfr389 | 0 | 0 | 0 | 1 | 1 |
| Olfr39 | 0 | 0 | 0 | 1 | 1 |
| Olfr390 | 0 | 0 | 0 | 1 | 1 |
| Olfr391-ps | 0 | 0 | 0 | 1 | 1 |
| Olfr392 | 0 | 0 | 0 | 1 | 1 |
| Olfr393 | 0 | 0 | 0 | 1 | 1 |
| Olfr394 | 0 | 0 | 0 | 1 | 1 |
| Olfr395 | 0 | 0 | 0 | 1 | 1 |
| Olfr397 | 0 | 0 | 0 | 1 | 1 |
| Olfr398 | 0 | 0 | 0 | 1 | 1 |
| Olfr399 | 0 | 0 | 0 | 1 | 1 |
| Olfr401 | 0 | 0 | 0 | 1 | 1 |
| Olfr402 | 0 | 0 | 0 | 1 | 1 |
| Olfr403 | 0 | 0 | 0 | 1 | 1 |
| Olfr406 | 0 | 0 | 0 | 1 | 1 |
| Olfr410 | 0 | 0 | 0 | 1 | 1 |
| Olfr411 | 0 | 0 | 0 | 1 | 1 |
| Olfr414 | 0 | 0 | 0 | 1 | 1 |
| Olfr417 | 0 | 0 | 0 | 1 | 1 |
| Olfr418-ps1 | 0 | 0 | 0 | 1 | 1 |
| Olfr419 | 0 | 0 | 0 | 1 | 1 |
| Olfr420 | 0 | 0 | 0 | 1 | 1 |
| Olfr421-ps1 | 0 | 0 | 0 | 1 | 1 |
| Olfr424 | 0 | 0 | 0 | 1 | 1 |
| Olfr426 | 0 | 0 | 0 | 1 | 1 |
| Olfr427 | 0 | 0 | 0 | 1 | 1 |
| Olfr429 | 0 | 0 | 0 | 1 | 1 |
| Olfr43 | 0 | 0 | 0 | 1 | 1 |
| Olfr430 | 0 | 0 | 0 | 1 | 1 |
| Olfr432 | 0 | 0 | 0 | 1 | 1 |
| Olfr433 | 0 | 0 | 0 | 1 | 1 |
| Olfr434 | 0 | 0 | 0 | 1 | 1 |
| Olfr435 | 0 | 0 | 0 | 1 | 1 |

| | | | | | |
|---|---|---|---|---|---|
| Olfr437 | 0 | 0 | 0 | 1 | 1 |
| Olfr44 | 0 | 0 | 0 | 1 | 1 |
| Olfr441 | 0 | 0 | 0 | 1 | 1 |
| Olfr446 | 0 | 0 | 0 | 1 | 1 |
| Olfr447 | 0 | 0 | 0 | 1 | 1 |
| Olfr448 | 0 | 0 | 0 | 1 | 1 |
| Olfr449 | 0 | 0 | 0 | 1 | 1 |
| Olfr45 | 0 | 0 | 0 | 1 | 1 |
| Olfr450 | 0 | 0 | 0 | 1 | 1 |
| Olfr452 | 0 | 0 | 0 | 1 | 1 |
| Olfr453 | 0 | 0 | 0 | 1 | 1 |
| Olfr455 | 0 | 0 | 0 | 1 | 1 |
| Olfr458 | 0 | 0 | 0 | 1 | 1 |
| Olfr459 | 0 | 0 | 0 | 1 | 1 |
| Olfr46 | 0 | 0 | 0 | 1 | 1 |
| Olfr460 | 0 | 0 | 0 | 1 | 1 |
| Olfr461 | 0 | 0 | 0 | 1 | 1 |
| Olfr462 | 0 | 0 | 0 | 1 | 1 |
| Olfr463 | 0 | 0 | 0 | 1 | 1 |
| Olfr464 | 0 | 0 | 0 | 1 | 1 |
| Olfr466 | 0 | 0 | 0 | 1 | 1 |
| Olfr467 | 0 | 0 | 0 | 1 | 1 |
| Olfr47 | 0 | 0 | 0 | 1 | 1 |
| Olfr470 | 0 | 0 | 0 | 1 | 1 |
| Olfr472 | 0 | 0 | 0 | 1 | 1 |
| Olfr473 | 0 | 0 | 0 | 1 | 1 |
| Olfr474 | 0 | 0 | 0 | 1 | 1 |
| Olfr476 | 0 | 0 | 0 | 1 | 1 |
| Olfr477 | 0 | 0 | 0 | 1 | 1 |
| Olfr478 | 0 | 0 | 0 | 1 | 1 |
| Olfr479 | 0 | 0 | 0 | 1 | 1 |
| Olfr48 | 0 | 0 | 0 | 1 | 1 |
| Olfr480 | 0 | 0 | 0 | 1 | 1 |
| Olfr481 | 0 | 0 | 0 | 1 | 1 |
| Olfr482 | 0 | 0 | 0 | 1 | 1 |
| Olfr483 | 0 | 0 | 0 | 1 | 1 |
| Olfr484 | 0 | 0 | 0 | 1 | 1 |
| Olfr485 | 0 | 0 | 0 | 1 | 1 |
| Olfr486 | 0 | 0 | 0 | 1 | 1 |
| Olfr487 | 0 | 0 | 0 | 1 | 1 |
| Olfr488 | 0 | 0 | 0 | 1 | 1 |
| Olfr490 | 0 | 0 | 0 | 1 | 1 |
| Olfr491 | 0 | 0 | 0 | 1 | 1 |
| Olfr492 | 0 | 0 | 0 | 1 | 1 |
| Olfr493 | 0 | 0 | 0 | 1 | 1 |
| Olfr494 | 0 | 0 | 0 | 1 | 1 |
| Olfr495 | 0 | 0 | 0 | 1 | 1 |
| Olfr497 | 0 | 0 | 0 | 1 | 1 |
| Olfr498 | 0 | 0 | 0 | 1 | 1 |
| Olfr50 | 0 | 0 | 0 | 1 | 1 |

| | | | | | |
|---|---|---|---|---|---|
| Olfr502 | 0 | 0 | 0 | 1 | 1 |
| Olfr503 | 0 | 0 | 0 | 1 | 1 |
| Olfr504 | 0 | 0 | 0 | 1 | 1 |
| Olfr506 | 0 | 0 | 0 | 1 | 1 |
| Olfr507 | 0 | 0 | 0 | 1 | 1 |
| Olfr508 | 0 | 0 | 0 | 1 | 1 |
| Olfr509 | 0 | 0 | 0 | 1 | 1 |
| Olfr51 | 0 | 0 | 0 | 1 | 1 |
| Olfr510 | 0 | 0 | 0 | 1 | 1 |
| Olfr512 | 0 | 0 | 0 | 1 | 1 |
| Olfr513 | 0 | 0 | 0 | 1 | 1 |
| Olfr514 | 0 | 0 | 0 | 1 | 1 |
| Olfr516 | 0 | 0 | 0 | 1 | 1 |
| Olfr517 | 0 | 0 | 0 | 1 | 1 |
| Olfr518 | 0 | 0 | 0 | 1 | 1 |
| Olfr519 | 0 | 0 | 0 | 1 | 1 |
| Olfr52 | 0 | 0 | 0 | 1 | 1 |
| Olfr520 | 0 | 0 | 0 | 1 | 1 |
| Olfr522 | 0 | 0 | 0 | 1 | 1 |
| Olfr523 | 0 | 0 | 0 | 1 | 1 |
| Olfr524 | 0 | 0 | 0 | 1 | 1 |
| Olfr525 | 0 | 0 | 0 | 1 | 1 |
| Olfr527 | 0 | 0 | 0 | 1 | 1 |
| Olfr53 | 0 | 0 | 0 | 1 | 1 |
| Olfr530 | 0 | 0 | 0 | 1 | 1 |
| Olfr531 | 0 | 0 | 0 | 1 | 1 |
| Olfr532 | 0 | 0 | 0 | 1 | 1 |
| Olfr533 | 0 | 0 | 0 | 1 | 1 |
| Olfr535 | 0 | 0 | 0 | 1 | 1 |
| Olfr536 | 0 | 0 | 0 | 1 | 1 |
| Olfr538 | 0 | 0 | 0 | 1 | 1 |
| Olfr539 | 0 | 0 | 0 | 1 | 1 |
| Olfr54 | 0 | 0 | 0 | 1 | 1 |
| Olfr541 | 0 | 0 | 0 | 1 | 1 |
| Olfr547 | 0 | 0 | 0 | 1 | 1 |
| Olfr549 | 0 | 0 | 0 | 1 | 1 |
| Olfr55 | 0 | 0 | 0 | 1 | 1 |
| Olfr550 | 0 | 0 | 0 | 1 | 1 |
| Olfr551 | 0 | 0 | 0 | 1 | 1 |
| Olfr552 | 0 | 0 | 0 | 1 | 1 |
| Olfr553 | 0 | 0 | 0 | 1 | 1 |
| Olfr554 | 0 | 0 | 0 | 1 | 1 |
| Olfr555 | 0 | 0 | 0 | 1 | 1 |
| Olfr556 | 0 | 0 | 0 | 1 | 1 |
| Olfr557 | 0 | 0 | 0 | 1 | 1 |
| Olfr559 | 0 | 0 | 0 | 1 | 1 |
| Olfr56 | 0 | 0 | 0 | 1 | 1 |
| Olfr560 | 0 | 0 | 0 | 1 | 1 |
| Olfr561 | 0 | 0 | 0 | 1 | 1 |
| Olfr564 | 0 | 0 | 0 | 1 | 1 |

| | | | | | |
|---|---|---|---|---|---|
| Olfr566 | 0 | 0 | 0 | 1 | 1 |
| Olfr568 | 0 | 0 | 0 | 1 | 1 |
| Olfr569 | 0 | 0 | 0 | 1 | 1 |
| Olfr57 | 0 | 0 | 0 | 1 | 1 |
| Olfr570 | 0 | 0 | 0 | 1 | 1 |
| Olfr571 | 0 | 0 | 0 | 1 | 1 |
| Olfr572 | 0 | 0 | 0 | 1 | 1 |
| Olfr574 | 0 | 0 | 0 | 1 | 1 |
| Olfr575 | 0 | 0 | 0 | 1 | 1 |
| Olfr576 | 0 | 0 | 0 | 1 | 1 |
| Olfr577 | 0 | 0 | 0 | 1 | 1 |
| Olfr578 | 0 | 0 | 0 | 1 | 1 |
| Olfr58 | 0 | 0 | 0 | 1 | 1 |
| Olfr582 | 0 | 0 | 0 | 1 | 1 |
| Olfr583 | 0 | 0 | 0 | 1 | 1 |
| Olfr584 | 0 | 0 | 0 | 1 | 1 |
| Olfr585 | 0 | 0 | 0 | 1 | 1 |
| Olfr586 | 0 | 0 | 0 | 1 | 1 |
| Olfr589 | 0 | 0 | 0 | 1 | 1 |
| Olfr59 | 0 | 0 | 0 | 1 | 1 |
| Olfr591 | 0 | 0 | 0 | 1 | 1 |
| Olfr592 | 0 | 0 | 0 | 1 | 1 |
| Olfr593 | 0 | 0 | 0 | 1 | 1 |
| Olfr594 | 0 | 0 | 0 | 1 | 1 |
| Olfr596 | 0 | 0 | 0 | 1 | 1 |
| Olfr597 | 0 | 0 | 0 | 1 | 1 |
| Olfr598 | 0 | 0 | 0 | 1 | 1 |
| Olfr599 | 0 | 0 | 0 | 1 | 1 |
| Olfr6 | 0 | 0 | 0 | 1 | 1 |
| Olfr60 | 0 | 0 | 0 | 1 | 1 |
| Olfr600 | 0 | 0 | 0 | 1 | 1 |
| Olfr601 | 0 | 0 | 0 | 1 | 1 |
| Olfr603 | 0 | 0 | 0 | 1 | 1 |
| Olfr605 | 0 | 0 | 0 | 1 | 1 |
| Olfr606 | 0 | 0 | 0 | 1 | 1 |
| Olfr608 | 0 | 0 | 0 | 1 | 1 |
| Olfr609 | 0 | 0 | 0 | 1 | 1 |
| Olfr610 | 0 | 0 | 0 | 1 | 1 |
| Olfr611 | 0 | 0 | 0 | 1 | 1 |
| Olfr612 | 0 | 0 | 0 | 1 | 1 |
| Olfr613 | 0 | 0 | 0 | 1 | 1 |
| Olfr615 | 0 | 0 | 0 | 1 | 1 |
| Olfr616 | 0 | 0 | 0 | 1 | 1 |
| Olfr617 | 0 | 0 | 0 | 1 | 1 |
| Olfr618 | 0 | 0 | 0 | 1 | 1 |
| Olfr619 | 0 | 0 | 0 | 1 | 1 |
| Olfr62 | 0 | 0 | 0 | 1 | 1 |
| Olfr620 | 0 | 0 | 0 | 1 | 1 |
| Olfr622 | 0 | 0 | 0 | 1 | 1 |
| Olfr623 | 0 | 0 | 0 | 1 | 1 |

| | | | | | |
|---|---|---|---|---|---|
| Olfr624 | 0 | 0 | 0 | 1 | 1 |
| Olfr628 | 0 | 0 | 0 | 1 | 1 |
| Olfr629 | 0 | 0 | 0 | 1 | 1 |
| Olfr63 | 0 | 0 | 0 | 1 | 1 |
| Olfr630 | 0 | 0 | 0 | 1 | 1 |
| Olfr631 | 0 | 0 | 0 | 1 | 1 |
| Olfr632 | 0 | 0 | 0 | 1 | 1 |
| Olfr633 | 0 | 0 | 0 | 1 | 1 |
| Olfr635 | 0 | 0 | 0 | 1 | 1 |
| Olfr638 | 0 | 0 | 0 | 1 | 1 |
| Olfr639 | 0 | 0 | 0 | 1 | 1 |
| Olfr64 | 0 | 0 | 0 | 1 | 1 |
| Olfr640 | 0 | 0 | 0 | 1 | 1 |
| Olfr641 | 0 | 0 | 0 | 1 | 1 |
| Olfr642 | 0 | 0 | 0 | 1 | 1 |
| Olfr643 | 0 | 0 | 0 | 1 | 1 |
| Olfr644 | 0 | 0 | 0 | 1 | 1 |
| Olfr645 | 0 | 0 | 0 | 1 | 1 |
| Olfr646 | 0 | 0 | 0 | 1 | 1 |
| Olfr648 | 0 | 0 | 0 | 1 | 1 |
| Olfr649 | 0 | 0 | 0 | 1 | 1 |
| Olfr65 | 0 | 0 | 0 | 1 | 1 |
| Olfr651 | 0 | 0 | 0 | 1 | 1 |
| Olfr652 | 0 | 0 | 0 | 1 | 1 |
| Olfr653 | 0 | 0 | 0 | 1 | 1 |
| Olfr654 | 0 | 0 | 0 | 1 | 1 |
| Olfr656 | 0 | 0 | 0 | 1 | 1 |
| Olfr659 | 0 | 0 | 0 | 1 | 1 |
| Olfr66 | 0 | 0 | 0 | 1 | 1 |
| Olfr661 | 0 | 0 | 0 | 1 | 1 |
| Olfr663 | 0 | 0 | 0 | 1 | 1 |
| Olfr665 | 0 | 0 | 0 | 1 | 1 |
| Olfr666 | 0 | 0 | 0 | 1 | 1 |
| Olfr667 | 0 | 0 | 0 | 1 | 1 |
| Olfr668 | 0 | 0 | 0 | 1 | 1 |
| Olfr669 | 0 | 0 | 0 | 1 | 1 |
| Olfr67 | 0 | 0 | 0 | 1 | 1 |
| Olfr670 | 0 | 0 | 0 | 1 | 1 |
| Olfr671 | 0 | 0 | 0 | 1 | 1 |
| Olfr672 | 0 | 0 | 0 | 1 | 1 |
| Olfr675 | 0 | 0 | 0 | 1 | 1 |
| Olfr676 | 0 | 0 | 0 | 1 | 1 |
| Olfr677 | 0 | 0 | 0 | 1 | 1 |
| Olfr678 | 0 | 0 | 0 | 1 | 1 |
| Olfr679 | 0 | 0 | 0 | 1 | 1 |
| Olfr68 | 0 | 0 | 0 | 1 | 1 |
| Olfr681 | 0 | 0 | 0 | 1 | 1 |
| Olfr683 | 0 | 0 | 0 | 1 | 1 |
| Olfr684 | 0 | 0 | 0 | 1 | 1 |
| Olfr685 | 0 | 0 | 0 | 1 | 1 |

| | | | | | |
|---|---|---|---|---|---|
| Olfr686 | 0 | 0 | 0 | 1 | 1 |
| Olfr688 | 0 | 0 | 0 | 1 | 1 |
| Olfr689 | 0 | 0 | 0 | 1 | 1 |
| Olfr69 | 0 | 0 | 0 | 1 | 1 |
| Olfr690 | 0 | 0 | 0 | 1 | 1 |
| Olfr691 | 0 | 0 | 0 | 1 | 1 |
| Olfr692 | 0 | 0 | 0 | 1 | 1 |
| Olfr693 | 0 | 0 | 0 | 1 | 1 |
| Olfr694 | 0 | 0 | 0 | 1 | 1 |
| Olfr695 | 0 | 0 | 0 | 1 | 1 |
| Olfr697 | 0 | 0 | 0 | 1 | 1 |
| Olfr699 | 0 | 0 | 0 | 1 | 1 |
| Olfr700 | 0 | 0 | 0 | 1 | 1 |
| Olfr701 | 0 | 0 | 0 | 1 | 1 |
| Olfr702 | 0 | 0 | 0 | 1 | 1 |
| Olfr703 | 0 | 0 | 0 | 1 | 1 |
| Olfr704 | 0 | 0 | 0 | 1 | 1 |
| Olfr705 | 0 | 0 | 0 | 1 | 1 |
| Olfr706 | 0 | 0 | 0 | 1 | 1 |
| Olfr707 | 0 | 0 | 0 | 1 | 1 |
| Olfr710 | 0 | 0 | 0 | 1 | 1 |
| Olfr711 | 0 | 0 | 0 | 1 | 1 |
| Olfr713 | 0 | 0 | 0 | 1 | 1 |
| Olfr714 | 0 | 0 | 0 | 1 | 1 |
| Olfr715 | 0 | 0 | 0 | 1 | 1 |
| Olfr716 | 0 | 0 | 0 | 1 | 1 |
| Olfr720 | 0 | 0 | 0 | 1 | 1 |
| Olfr722 | 0 | 0 | 0 | 1 | 1 |
| Olfr723 | 0 | 0 | 0 | 1 | 1 |
| Olfr724 | 0 | 0 | 0 | 1 | 1 |
| Olfr727 | 0 | 0 | 0 | 1 | 1 |
| Olfr728 | 0 | 0 | 0 | 1 | 1 |
| Olfr729 | 0 | 0 | 0 | 1 | 1 |
| Olfr73 | 0 | 0 | 0 | 1 | 1 |
| Olfr730 | 0 | 0 | 0 | 1 | 1 |
| Olfr731 | 0 | 0 | 0 | 1 | 1 |
| Olfr732 | 0 | 0 | 0 | 1 | 1 |
| Olfr733 | 0 | 0 | 0 | 1 | 1 |
| Olfr734 | 0 | 0 | 0 | 1 | 1 |
| Olfr735 | 0 | 0 | 0 | 1 | 1 |
| Olfr736 | 0 | 0 | 0 | 1 | 1 |
| Olfr738 | 0 | 0 | 0 | 1 | 1 |
| Olfr739 | 0 | 0 | 0 | 1 | 1 |
| Olfr74 | 0 | 0 | 0 | 1 | 1 |
| Olfr740 | 0 | 0 | 0 | 1 | 1 |
| Olfr741 | 0 | 0 | 0 | 1 | 1 |
| Olfr742 | 0 | 0 | 0 | 1 | 1 |
| Olfr743 | 0 | 0 | 0 | 1 | 1 |
| Olfr744 | 0 | 0 | 0 | 1 | 1 |
| Olfr745 | 0 | 0 | 0 | 1 | 1 |

| | | | | | |
|---|---|---|---|---|---|
| Olfr746 | 0 | 0 | 0 | 1 | 1 |
| Olfr747 | 0 | 0 | 0 | 1 | 1 |
| Olfr748 | 0 | 0 | 0 | 1 | 1 |
| Olfr749 | 0 | 0 | 0 | 1 | 1 |
| Olfr75-ps1 | 0 | 0 | 0 | 1 | 1 |
| Olfr76 | 0 | 0 | 0 | 1 | 1 |
| Olfr761 | 0 | 0 | 0 | 1 | 1 |
| Olfr763 | 0 | 0 | 0 | 1 | 1 |
| Olfr765 | 0 | 0 | 0 | 1 | 1 |
| Olfr768 | 0 | 0 | 0 | 1 | 1 |
| Olfr769 | 0 | 0 | 0 | 1 | 1 |
| Olfr77 | 0 | 0 | 0 | 1 | 1 |
| Olfr770 | 0 | 0 | 0 | 1 | 1 |
| Olfr771 | 0 | 0 | 0 | 1 | 1 |
| Olfr772 | 0 | 0 | 0 | 1 | 1 |
| Olfr773 | 0 | 0 | 0 | 1 | 1 |
| Olfr774 | 0 | 0 | 0 | 1 | 1 |
| Olfr775 | 0 | 0 | 0 | 1 | 1 |
| Olfr776 | 0 | 0 | 0 | 1 | 1 |
| Olfr777 | 0 | 0 | 0 | 1 | 1 |
| Olfr780 | 0 | 0 | 0 | 1 | 1 |
| Olfr781 | 0 | 0 | 0 | 1 | 1 |
| Olfr782 | 0 | 0 | 0 | 1 | 1 |
| Olfr784 | 0 | 0 | 0 | 1 | 1 |
| Olfr786 | 0 | 0 | 0 | 1 | 1 |
| Olfr787 | 0 | 0 | 0 | 1 | 1 |
| Olfr788 | 0 | 0 | 0 | 1 | 1 |
| Olfr790 | 0 | 0 | 0 | 1 | 1 |
| Olfr791 | 0 | 0 | 0 | 1 | 1 |
| Olfr792 | 0 | 0 | 0 | 1 | 1 |
| Olfr794 | 0 | 0 | 0 | 1 | 1 |
| Olfr796 | 0 | 0 | 0 | 1 | 1 |
| Olfr798 | 0 | 0 | 0 | 1 | 1 |
| Olfr799 | 0 | 0 | 0 | 1 | 1 |
| Olfr8 | 0 | 0 | 0 | 1 | 1 |
| Olfr800 | 0 | 0 | 0 | 1 | 1 |
| Olfr801 | 0 | 0 | 0 | 1 | 1 |
| Olfr802 | 0 | 0 | 0 | 1 | 1 |
| Olfr803 | 0 | 0 | 0 | 1 | 1 |
| Olfr804 | 0 | 0 | 0 | 1 | 1 |
| Olfr805 | 0 | 0 | 0 | 1 | 1 |
| Olfr806 | 0 | 0 | 0 | 1 | 1 |
| Olfr807 | 0 | 0 | 0 | 1 | 1 |
| Olfr808 | 0 | 0 | 0 | 1 | 1 |
| Olfr809 | 0 | 0 | 0 | 1 | 1 |
| Olfr810 | 0 | 0 | 0 | 1 | 1 |
| Olfr811 | 0 | 0 | 0 | 1 | 1 |
| Olfr812 | 0 | 0 | 0 | 1 | 1 |
| Olfr813 | 0 | 0 | 0 | 1 | 1 |
| Olfr814 | 0 | 0 | 0 | 1 | 1 |

| | | | | | |
|---|---|---|---|---|---|
| Olfr815 | 0 | 0 | 0 | 1 | 1 |
| Olfr816 | 0 | 0 | 0 | 1 | 1 |
| Olfr818 | 0 | 0 | 0 | 1 | 1 |
| Olfr819 | 0 | 0 | 0 | 1 | 1 |
| Olfr820 | 0 | 0 | 0 | 1 | 1 |
| Olfr821 | 0 | 0 | 0 | 1 | 1 |
| Olfr822 | 0 | 0 | 0 | 1 | 1 |
| Olfr823 | 0 | 0 | 0 | 1 | 1 |
| Olfr824 | 0 | 0 | 0 | 1 | 1 |
| Olfr825 | 0 | 0 | 0 | 1 | 1 |
| Olfr826 | 0 | 0 | 0 | 1 | 1 |
| Olfr827 | 0 | 0 | 0 | 1 | 1 |
| Olfr828 | 0 | 0 | 0 | 1 | 1 |
| Olfr829 | 0 | 0 | 0 | 1 | 1 |
| Olfr830 | 0 | 0 | 0 | 1 | 1 |
| Olfr832 | 0 | 0 | 0 | 1 | 1 |
| Olfr834 | 0 | 0 | 0 | 1 | 1 |
| Olfr835 | 0 | 0 | 0 | 1 | 1 |
| Olfr836 | 0 | 0 | 0 | 1 | 1 |
| Olfr837 | 0 | 0 | 0 | 1 | 1 |
| Olfr843 | 0 | 0 | 0 | 1 | 1 |
| Olfr845 | 0 | 0 | 0 | 1 | 1 |
| Olfr846 | 0 | 0 | 0 | 1 | 1 |
| Olfr847 | 0 | 0 | 0 | 1 | 1 |
| Olfr849 | 0 | 0 | 0 | 1 | 1 |
| Olfr850 | 0 | 0 | 0 | 1 | 1 |
| Olfr851 | 0 | 0 | 0 | 1 | 1 |
| Olfr853 | 0 | 0 | 0 | 1 | 1 |
| Olfr854 | 0 | 0 | 0 | 1 | 1 |
| Olfr855 | 0 | 0 | 0 | 1 | 1 |
| Olfr857 | 0 | 0 | 0 | 1 | 1 |
| Olfr859 | 0 | 0 | 0 | 1 | 1 |
| Olfr860 | 0 | 0 | 0 | 1 | 1 |
| Olfr862 | 0 | 0 | 0 | 1 | 1 |
| Olfr866 | 0 | 0 | 0 | 1 | 1 |
| Olfr867 | 0 | 0 | 0 | 1 | 1 |
| Olfr868 | 0 | 0 | 0 | 1 | 1 |
| Olfr869 | 0 | 0 | 0 | 1 | 1 |
| Olfr870 | 0 | 0 | 0 | 1 | 1 |
| Olfr871 | 0 | 0 | 0 | 1 | 1 |
| Olfr872 | 0 | 0 | 0 | 1 | 1 |
| Olfr873 | 0 | 0 | 0 | 1 | 1 |
| Olfr874 | 0 | 0 | 0 | 1 | 1 |
| Olfr875 | 0 | 0 | 0 | 1 | 1 |
| Olfr876 | 0 | 0 | 0 | 1 | 1 |
| Olfr877 | 0 | 0 | 0 | 1 | 1 |
| Olfr878 | 0 | 0 | 0 | 1 | 1 |
| Olfr881 | 0 | 0 | 0 | 1 | 1 |
| Olfr883 | 0 | 0 | 0 | 1 | 1 |
| Olfr884 | 0 | 0 | 0 | 1 | 1 |

| | | | | | |
|---|---|---|---|---|---|
| Olfr885 | 0 | 0 | 0 | 1 | 1 |
| Olfr887 | 0 | 0 | 0 | 1 | 1 |
| Olfr888 | 0 | 0 | 0 | 1 | 1 |
| Olfr889 | 0 | 0 | 0 | 1 | 1 |
| Olfr890 | 0 | 0 | 0 | 1 | 1 |
| Olfr891 | 0 | 0 | 0 | 1 | 1 |
| Olfr893 | 0 | 0 | 0 | 1 | 1 |
| Olfr894 | 0 | 0 | 0 | 1 | 1 |
| Olfr895 | 0 | 0 | 0 | 1 | 1 |
| Olfr898 | 0 | 0 | 0 | 1 | 1 |
| Olfr899 | 0 | 0 | 0 | 1 | 1 |
| Olfr9 | 0 | 0 | 0 | 1 | 1 |
| Olfr900 | 0 | 0 | 0 | 1 | 1 |
| Olfr901 | 0 | 0 | 0 | 1 | 1 |
| Olfr902 | 0 | 0 | 0 | 1 | 1 |
| Olfr904 | 0 | 0 | 0 | 1 | 1 |
| Olfr905 | 0 | 0 | 0 | 1 | 1 |
| Olfr906 | 0 | 0 | 0 | 1 | 1 |
| Olfr907 | 0 | 0 | 0 | 1 | 1 |
| Olfr908 | 0 | 0 | 0 | 1 | 1 |
| Olfr910 | 0 | 0 | 0 | 1 | 1 |
| Olfr911-ps1 | 0 | 0 | 0 | 1 | 1 |
| Olfr912 | 0 | 0 | 0 | 1 | 1 |
| Olfr913 | 0 | 0 | 0 | 1 | 1 |
| Olfr914 | 0 | 0 | 0 | 1 | 1 |
| Olfr915 | 0 | 0 | 0 | 1 | 1 |
| Olfr916 | 0 | 0 | 0 | 1 | 1 |
| Olfr917 | 0 | 0 | 0 | 1 | 1 |
| Olfr918 | 0 | 0 | 0 | 1 | 1 |
| Olfr919 | 0 | 0 | 0 | 1 | 1 |
| Olfr92 | 0 | 0 | 0 | 1 | 1 |
| Olfr921 | 0 | 0 | 0 | 1 | 1 |
| Olfr922 | 0 | 0 | 0 | 1 | 1 |
| Olfr923 | 0 | 0 | 0 | 1 | 1 |
| Olfr924 | 0 | 0 | 0 | 1 | 1 |
| Olfr926 | 0 | 0 | 0 | 1 | 1 |
| Olfr93 | 0 | 0 | 0 | 1 | 1 |
| Olfr930 | 0 | 0 | 0 | 1 | 1 |
| Olfr933 | 0 | 0 | 0 | 1 | 1 |
| Olfr934 | 0 | 0 | 0 | 1 | 1 |
| Olfr935 | 0 | 0 | 0 | 1 | 1 |
| Olfr936 | 0 | 0 | 0 | 1 | 1 |
| Olfr937 | 0 | 0 | 0 | 1 | 1 |
| Olfr938 | 0 | 0 | 0 | 1 | 1 |
| Olfr943 | 0 | 0 | 0 | 1 | 1 |
| Olfr944 | 0 | 0 | 0 | 1 | 1 |
| Olfr945 | 0 | 0 | 0 | 1 | 1 |
| Olfr947-ps1 | 0 | 0 | 0 | 1 | 1 |
| Olfr948 | 0 | 0 | 0 | 1 | 1 |
| Olfr951 | 0 | 0 | 0 | 1 | 1 |

| | | | | | |
|---|---|---|---|---|---|
| Olfr952 | 0 | 0 | 0 | 1 | 1 |
| Olfr954 | 0 | 0 | 0 | 1 | 1 |
| Olfr955 | 0 | 0 | 0 | 1 | 1 |
| Olfr957 | 0 | 0 | 0 | 1 | 1 |
| Olfr958 | 0 | 0 | 0 | 1 | 1 |
| Olfr959 | 0 | 0 | 0 | 1 | 1 |
| Olfr96 | 0 | 0 | 0 | 1 | 1 |
| Olfr961 | 0 | 0 | 0 | 1 | 1 |
| Olfr963 | 0 | 0 | 0 | 1 | 1 |
| Olfr965 | 0 | 0 | 0 | 1 | 1 |
| Olfr967 | 0 | 0 | 0 | 1 | 1 |
| Olfr968 | 0 | 0 | 0 | 1 | 1 |
| Olfr969 | 0 | 0 | 0 | 1 | 1 |
| Olfr97 | 0 | 0 | 0 | 1 | 1 |
| Olfr970 | 0 | 0 | 0 | 1 | 1 |
| Olfr971 | 0 | 0 | 0 | 1 | 1 |
| Olfr972 | 0 | 0 | 0 | 1 | 1 |
| Olfr974 | 0 | 0 | 0 | 1 | 1 |
| Olfr975 | 0 | 0 | 0 | 1 | 1 |
| Olfr976 | 0 | 0 | 0 | 1 | 1 |
| Olfr978 | 0 | 0 | 0 | 1 | 1 |
| Olfr979 | 0 | 0 | 0 | 1 | 1 |
| Olfr98 | 0 | 0 | 0 | 1 | 1 |
| Olfr980 | 0 | 0 | 0 | 1 | 1 |
| Olfr981 | 0 | 0 | 0 | 1 | 1 |
| Olfr982 | 0 | 0 | 0 | 1 | 1 |
| Olfr983 | 0 | 0 | 0 | 1 | 1 |
| Olfr984 | 0 | 0 | 0 | 1 | 1 |
| Olfr985 | 0 | 0 | 0 | 1 | 1 |
| Olfr986 | 0 | 0 | 0 | 1 | 1 |
| Olfr987 | 0 | 0 | 0 | 1 | 1 |
| Olfr988 | 0 | 0 | 0 | 1 | 1 |
| Olfr992 | 0 | 0 | 0 | 1 | 1 |
| Olfr993 | 0 | 0 | 0 | 1 | 1 |
| Olfr994 | 0 | 0 | 0 | 1 | 1 |
| Olfr996 | 0 | 0 | 0 | 1 | 1 |
| Olfr998 | 0 | 0 | 0 | 1 | 1 |
| Olr1 | 0 | 0 | 0 | 1 | 1 |
| Omt2a | 0 | 0 | 0 | 1 | 1 |
| Omt2b | 0 | 0 | 0 | 1 | 1 |
| Oog1 | 0 | 0 | 0 | 1 | 1 |
| Oog2 | 0 | 0 | 0 | 1 | 1 |
| Oog3 | 0 | 0 | 0 | 1 | 1 |
| Oog4 | 0 | 0 | 0 | 1 | 1 |
| Oosp1 | 0 | 0 | 0 | 1 | 1 |
| Oosp2 | 0 | 0 | 0 | 1 | 1 |
| Oosp3 | 0 | 0 | 0 | 1 | 1 |
| Opalin | 0 | 0 | 0 | 1 | 1 |
| Opn1mw | 0 | 0 | 0 | 1 | 1 |
| Opn4 | 0 | 0 | 0 | 1 | 1 |

| | | | | | |
|---|---|---|---|---|---|
| Oprm1 | 0 | 0 | 0 | 1 | 1 |
| Orm1 | 0 | 0 | 0 | 1 | 1 |
| Orm3 | 0 | 0 | 0 | 1 | 1 |
| Osm | 0 | 0 | 0 | 1 | 1 |
| Ostn | 0 | 0 | 0 | 1 | 1 |
| Otc | 0 | 0 | 0 | 1 | 1 |
| Otogl | 0 | 0 | 0 | 1 | 1 |
| Otol1 | 0 | 0 | 0 | 1 | 1 |
| Otos | 0 | 0 | 0 | 1 | 1 |
| Otp | 0 | 0 | 0 | 1 | 1 |
| Otx2os1 | 0 | 0 | 0 | 1 | 1 |
| Oxgr1 | 0 | 0 | 0 | 1 | 1 |
| P2ry10 | 0 | 0 | 0 | 1 | 1 |
| P2ry12 | 0 | 0 | 0 | 1 | 1 |
| P2ry4 | 0 | 0 | 0 | 1 | 1 |
| Pabpn1l | 0 | 0 | 0 | 1 | 1 |
| Pah | 0 | 0 | 0 | 1 | 1 |
| Panx3 | 0 | 0 | 0 | 1 | 1 |
| Papl | 0 | 0 | 0 | 1 | 1 |
| Paqr6 | 0 | 0 | 0 | 1 | 1 |
| Pax6os1 | 0 | 0 | 0 | 1 | 1 |
| Pbsn | 0 | 0 | 0 | 1 | 1 |
| Pcdha1 | 0 | 0 | 0 | 1 | 1 |
| Pcdha10 | 0 | 0 | 0 | 1 | 1 |
| Pcdha12 | 0 | 0 | 0 | 1 | 1 |
| Pcdha3 | 0 | 0 | 0 | 1 | 1 |
| Pcdha4 | 0 | 0 | 0 | 1 | 1 |
| Pcdha4-g | 0 | 0 | 0 | 1 | 1 |
| Pcdha6 | 0 | 0 | 0 | 1 | 1 |
| Pcdha8 | 0 | 0 | 0 | 1 | 1 |
| Pcdhb13 | 0 | 0 | 0 | 1 | 1 |
| Pcdhb9 | 0 | 0 | 0 | 1 | 1 |
| Pcsk2os2 | 0 | 0 | 0 | 1 | 1 |
| Pdc | 0 | 0 | 0 | 1 | 1 |
| Pdcd1 | 0 | 0 | 0 | 1 | 1 |
| Pdcl2 | 0 | 0 | 0 | 1 | 1 |
| Pde6b | 0 | 0 | 0 | 1 | 1 |
| Pde6c | 0 | 0 | 0 | 1 | 1 |
| Pdha2 | 0 | 0 | 0 | 1 | 1 |
| Pdilt | 0 | 0 | 0 | 1 | 1 |
| Pdzd3 | 0 | 0 | 0 | 1 | 1 |
| Pea15b | 0 | 0 | 0 | 1 | 1 |
| Peril | 0 | 0 | 0 | 1 | 1 |
| Pet2 | 0 | 0 | 0 | 1 | 1 |
| Pfpl | 0 | 0 | 0 | 1 | 1 |
| Pga5 | 0 | 0 | 0 | 1 | 1 |
| Pgc | 0 | 0 | 0 | 1 | 1 |
| Pgk2 | 0 | 0 | 0 | 1 | 1 |
| Pgr15l | 0 | 0 | 0 | 1 | 1 |
| Phf11a | 0 | 0 | 0 | 1 | 1 |

| | | | | | |
|---|---|---|---|---|---|
| Phf11b | 0 | 0 | 0 | 1 | 1 |
| Phgr1 | 0 | 0 | 0 | 1 | 1 |
| Phox2b | 0 | 0 | 0 | 1 | 1 |
| Pigr | 0 | 0 | 0 | 1 | 1 |
| Pih1d3 | 0 | 0 | 0 | 1 | 1 |
| Pilrb1 | 0 | 0 | 0 | 1 | 1 |
| Pilrb2 | 0 | 0 | 0 | 1 | 1 |
| Pinlyp | 0 | 0 | 0 | 1 | 1 |
| Pip | 0 | 0 | 0 | 1 | 1 |
| Pira1 | 0 | 0 | 0 | 1 | 1 |
| Pira11 | 0 | 0 | 0 | 1 | 1 |
| Pira2 | 0 | 0 | 0 | 1 | 1 |
| Pira4,Pira7 | 0 | 0 | 0 | 1 | 1 |
| Pira6 | 0 | 0 | 0 | 1 | 1 |
| Pira7 | 0 | 0 | 0 | 1 | 1 |
| Pisd-ps3 | 0 | 0 | 0 | 1 | 1 |
| Pitx3 | 0 | 0 | 0 | 1 | 1 |
| Pla2g10os | 0 | 0 | 0 | 1 | 1 |
| Pla2g12b | 0 | 0 | 0 | 1 | 1 |
| Pla2g2a | 0 | 0 | 0 | 1 | 1 |
| Pla2g4e | 0 | 0 | 0 | 1 | 1 |
| Plac8l1 | 0 | 0 | 0 | 1 | 1 |
| Plac9a,Plac9b | 0 | 0 | 0 | 1 | 1 |
| Plac9b | 0 | 0 | 0 | 1 | 1 |
| Plcz1 | 0 | 0 | 0 | 1 | 1 |
| Plscr5 | 0 | 0 | 0 | 1 | 1 |
| Plxnb3 | 0 | 0 | 0 | 1 | 1 |
| Pmis2 | 0 | 0 | 0 | 1 | 1 |
| Pmp2 | 0 | 0 | 0 | 1 | 1 |
| Pnma5 | 0 | 0 | 0 | 1 | 1 |
| Pnmt | 0 | 0 | 0 | 1 | 1 |
| Pnpla5 | 0 | 0 | 0 | 1 | 1 |
| Podnl1 | 0 | 0 | 0 | 1 | 1 |
| Pom121l12 | 0 | 0 | 0 | 1 | 1 |
| Poteg | 0 | 0 | 0 | 1 | 1 |
| Pou1f1 | 0 | 0 | 0 | 1 | 1 |
| Pou2af1 | 0 | 0 | 0 | 1 | 1 |
| Pou2f3 | 0 | 0 | 0 | 1 | 1 |
| Pou4f3 | 0 | 0 | 0 | 1 | 1 |
| Ppp1r1c | 0 | 0 | 0 | 1 | 1 |
| Ppp1r2-ps7 | 0 | 0 | 0 | 1 | 1 |
| Ppp1r2-ps9 | 0 | 0 | 0 | 1 | 1 |
| Ppp1r3a | 0 | 0 | 0 | 1 | 1 |
| Ppp2r2cos | 0 | 0 | 0 | 1 | 1 |
| Ppp3r2 | 0 | 0 | 0 | 1 | 1 |
| Prame | 0 | 0 | 0 | 1 | 1 |
| Pramef17 | 0 | 0 | 0 | 1 | 1 |
| Pramef25 | 0 | 0 | 0 | 1 | 1 |
| Pramef6 | 0 | 0 | 0 | 1 | 1 |
| Pramel3 | 0 | 0 | 0 | 1 | 1 |

| | | | | | |
|---|---|---|---|---|---|
| Pramel4 | 0 | 0 | 0 | 1 | 1 |
| Pramel5 | 0 | 0 | 0 | 1 | 1 |
| Pramel6 | 0 | 0 | 0 | 1 | 1 |
| Pramel7 | 0 | 0 | 0 | 1 | 1 |
| Prb1 | 0 | 0 | 0 | 1 | 1 |
| Prg3 | 0 | 0 | 0 | 1 | 1 |
| Prh1 | 0 | 0 | 0 | 1 | 1 |
| Prl | 0 | 0 | 0 | 1 | 1 |
| Prl2a1 | 0 | 0 | 0 | 1 | 1 |
| Prl2b1 | 0 | 0 | 0 | 1 | 1 |
| Prl2c1 | 0 | 0 | 0 | 1 | 1 |
| Prl2c2 | 0 | 0 | 0 | 1 | 1 |
| Prl2c3,Prl2c4 | 0 | 0 | 0 | 1 | 1 |
| Prl2c4 | 0 | 0 | 0 | 1 | 1 |
| Prl2c5 | 0 | 0 | 0 | 1 | 1 |
| Prl3a1 | 0 | 0 | 0 | 1 | 1 |
| Prl3c1 | 0 | 0 | 0 | 1 | 1 |
| Prl3d1 | 0 | 0 | 0 | 1 | 1 |
| Prl3d2 | 0 | 0 | 0 | 1 | 1 |
| Prl3d3 | 0 | 0 | 0 | 1 | 1 |
| Prl4a1 | 0 | 0 | 0 | 1 | 1 |
| Prl5a1 | 0 | 0 | 0 | 1 | 1 |
| Prl6a1 | 0 | 0 | 0 | 1 | 1 |
| Prl7a1 | 0 | 0 | 0 | 1 | 1 |
| Prl7a2 | 0 | 0 | 0 | 1 | 1 |
| Prl7b1 | 0 | 0 | 0 | 1 | 1 |
| Prl7c1 | 0 | 0 | 0 | 1 | 1 |
| Prl7d1 | 0 | 0 | 0 | 1 | 1 |
| Prl8a1 | 0 | 0 | 0 | 1 | 1 |
| Prl8a2 | 0 | 0 | 0 | 1 | 1 |
| Prl8a6 | 0 | 0 | 0 | 1 | 1 |
| Prl8a8 | 0 | 0 | 0 | 1 | 1 |
| Prl8a9 | 0 | 0 | 0 | 1 | 1 |
| Prlh | 0 | 0 | 0 | 1 | 1 |
| Prm1 | 0 | 0 | 0 | 1 | 1 |
| Prm2 | 0 | 0 | 0 | 1 | 1 |
| Prm3 | 0 | 0 | 0 | 1 | 1 |
| Prok1 | 0 | 0 | 0 | 1 | 1 |
| Prok2 | 0 | 0 | 0 | 1 | 1 |
| Prol1 | 0 | 0 | 0 | 1 | 1 |
| Prop1 | 0 | 0 | 0 | 1 | 1 |
| Prp2 | 0 | 0 | 0 | 1 | 1 |
| Prpmp5 | 0 | 0 | 0 | 1 | 1 |
| Prr27 | 0 | 0 | 0 | 1 | 1 |
| Prr30 | 0 | 0 | 0 | 1 | 1 |
| Prr32 | 0 | 0 | 0 | 1 | 1 |
| Prrxl1 | 0 | 0 | 0 | 1 | 1 |
| Prss1 | 0 | 0 | 0 | 1 | 1 |
| Prss2 | 0 | 0 | 0 | 1 | 1 |
| Prss21 | 0 | 0 | 0 | 1 | 1 |

| | | | | | |
|---|---|---|---|---|---|
| Prss27 | 0 | 0 | 0 | 1 | 1 |
| Prss28 | 0 | 0 | 0 | 1 | 1 |
| Prss29 | 0 | 0 | 0 | 1 | 1 |
| Prss3 | 0 | 0 | 0 | 1 | 1 |
| Prss32 | 0 | 0 | 0 | 1 | 1 |
| Prss34 | 0 | 0 | 0 | 1 | 1 |
| Prss37 | 0 | 0 | 0 | 1 | 1 |
| Prss39 | 0 | 0 | 0 | 1 | 1 |
| Prss40 | 0 | 0 | 0 | 1 | 1 |
| Prss43 | 0 | 0 | 0 | 1 | 1 |
| Prss45 | 0 | 0 | 0 | 1 | 1 |
| Prss46 | 0 | 0 | 0 | 1 | 1 |
| Prss51 | 0 | 0 | 0 | 1 | 1 |
| Prss52 | 0 | 0 | 0 | 1 | 1 |
| Prss54 | 0 | 0 | 0 | 1 | 1 |
| Prss55 | 0 | 0 | 0 | 1 | 1 |
| Prss57 | 0 | 0 | 0 | 1 | 1 |
| Prss58 | 0 | 0 | 0 | 1 | 1 |
| Psapl1 | 0 | 0 | 0 | 1 | 1 |
| Psca | 0 | 0 | 0 | 1 | 1 |
| Psg-ps1 | 0 | 0 | 0 | 1 | 1 |
| Psg16 | 0 | 0 | 0 | 1 | 1 |
| Psg17 | 0 | 0 | 0 | 1 | 1 |
| Psg18 | 0 | 0 | 0 | 1 | 1 |
| Psg19 | 0 | 0 | 0 | 1 | 1 |
| Psg20 | 0 | 0 | 0 | 1 | 1 |
| Psg21 | 0 | 0 | 0 | 1 | 1 |
| Psg22 | 0 | 0 | 0 | 1 | 1 |
| Psg23 | 0 | 0 | 0 | 1 | 1 |
| Psg26 | 0 | 0 | 0 | 1 | 1 |
| Psg27 | 0 | 0 | 0 | 1 | 1 |
| Psg28 | 0 | 0 | 0 | 1 | 1 |
| Psg29 | 0 | 0 | 0 | 1 | 1 |
| Ptafr | 0 | 0 | 0 | 1 | 1 |
| Ptgs2os | 0 | 0 | 0 | 1 | 1 |
| Pth | 0 | 0 | 0 | 1 | 1 |
| Pth2 | 0 | 0 | 0 | 1 | 1 |
| Ptprtos | 0 | 0 | 0 | 1 | 1 |
| Pvalb | 0 | 0 | 0 | 1 | 1 |
| Pydc4 | 0 | 0 | 0 | 1 | 1 |
| Pyhin1 | 0 | 0 | 0 | 1 | 1 |
| Pyy | 0 | 0 | 0 | 1 | 1 |
| Rag1 | 0 | 0 | 0 | 1 | 1 |
| Rag2 | 0 | 0 | 0 | 1 | 1 |
| Rasal1 | 0 | 0 | 0 | 1 | 1 |
| Rbakdn | 0 | 0 | 0 | 1 | 1 |
| Rbbp8nl | 0 | 0 | 0 | 1 | 1 |
| Rbm31y | 0 | 0 | 0 | 1 | 1 |
| Rbmy | 0 | 0 | 0 | 1 | 1 |
| Rbpjl | 0 | 0 | 0 | 1 | 1 |

| | | | | | |
|---|---|---|---|---|---|
| Rcvrn | 0 | 0 | 0 | 1 | 1 |
| Rd3l | 0 | 0 | 0 | 1 | 1 |
| Rdh16 | 0 | 0 | 0 | 1 | 1 |
| Rdh18-ps | 0 | 0 | 0 | 1 | 1 |
| Rdh19 | 0 | 0 | 0 | 1 | 1 |
| Rdh8 | 0 | 0 | 0 | 1 | 1 |
| Reg1 | 0 | 0 | 0 | 1 | 1 |
| Reg2 | 0 | 0 | 0 | 1 | 1 |
| Reg3a | 0 | 0 | 0 | 1 | 1 |
| Reg3b | 0 | 0 | 0 | 1 | 1 |
| Reg3d | 0 | 0 | 0 | 1 | 1 |
| Rergl | 0 | 0 | 0 | 1 | 1 |
| Retn | 0 | 0 | 0 | 1 | 1 |
| Retnla | 0 | 0 | 0 | 1 | 1 |
| Retnlb | 0 | 0 | 0 | 1 | 1 |
| Retnlg | 0 | 0 | 0 | 1 | 1 |
| Rex2 | 0 | 0 | 0 | 1 | 1 |
| Rfpl4b | 0 | 0 | 0 | 1 | 1 |
| Rfx6 | 0 | 0 | 0 | 1 | 1 |
| Rgr | 0 | 0 | 0 | 1 | 1 |
| Rgs1 | 0 | 0 | 0 | 1 | 1 |
| Rgs13 | 0 | 0 | 0 | 1 | 1 |
| Rgs21 | 0 | 0 | 0 | 1 | 1 |
| Rhbdl2 | 0 | 0 | 0 | 1 | 1 |
| Rhox1 | 0 | 0 | 0 | 1 | 1 |
| Rhox10 | 0 | 0 | 0 | 1 | 1 |
| Rhox11 | 0 | 0 | 0 | 1 | 1 |
| Rhox12 | 0 | 0 | 0 | 1 | 1 |
| Rhox13 | 0 | 0 | 0 | 1 | 1 |
| Rhox2a | 0 | 0 | 0 | 1 | 1 |
| Rhox2b | 0 | 0 | 0 | 1 | 1 |
| Rhox2c | 0 | 0 | 0 | 1 | 1 |
| Rhox2d | 0 | 0 | 0 | 1 | 1 |
| Rhox2e | 0 | 0 | 0 | 1 | 1 |
| Rhox2f | 0 | 0 | 0 | 1 | 1 |
| Rhox2g | 0 | 0 | 0 | 1 | 1 |
| Rhox2h | 0 | 0 | 0 | 1 | 1 |
| Rhox3a,Rhox3e | 0 | 0 | 0 | 1 | 1 |
| Rhox3c | 0 | 0 | 0 | 1 | 1 |
| Rhox3e | 0 | 0 | 0 | 1 | 1 |
| Rhox3f | 0 | 0 | 0 | 1 | 1 |
| Rhox3g | 0 | 0 | 0 | 1 | 1 |
| Rhox3h | 0 | 0 | 0 | 1 | 1 |
| Rhox4a | 0 | 0 | 0 | 1 | 1 |
| Rhox4b | 0 | 0 | 0 | 1 | 1 |
| Rhox4c | 0 | 0 | 0 | 1 | 1 |
| Rhox4d | 0 | 0 | 0 | 1 | 1 |
| Rhox4e | 0 | 0 | 0 | 1 | 1 |
| Rhox4f | 0 | 0 | 0 | 1 | 1 |
| Rhox4g | 0 | 0 | 0 | 1 | 1 |

| | | | | | |
|---|---|---|---|---|---|
| Rhox6 | 0 | 0 | 0 | 1 | 1 |
| Rhox7 | 0 | 0 | 0 | 1 | 1 |
| Rhox8 | 0 | 0 | 0 | 1 | 1 |
| Rhox9 | 0 | 0 | 0 | 1 | 1 |
| Rn4.5s | 0 | 0 | 0 | 1 | 1 |
| Rnase10 | 0 | 0 | 0 | 1 | 1 |
| Rnase11 | 0 | 0 | 0 | 1 | 1 |
| Rnase12 | 0 | 0 | 0 | 1 | 1 |
| Rnase13 | 0 | 0 | 0 | 1 | 1 |
| Rnase2a | 0 | 0 | 0 | 1 | 1 |
| Rnase2b | 0 | 0 | 0 | 1 | 1 |
| Rnase9 | 0 | 0 | 0 | 1 | 1 |
| Rnf133 | 0 | 0 | 0 | 1 | 1 |
| Rnf148 | 0 | 0 | 0 | 1 | 1 |
| Rnf186 | 0 | 0 | 0 | 1 | 1 |
| Rnf222 | 0 | 0 | 0 | 1 | 1 |
| Rnu6 | 0 | 0 | 0 | 1 | 1 |
| Rnu7 | 0 | 0 | 0 | 1 | 1 |
| Robo3 | 0 | 0 | 0 | 1 | 1 |
| Ropn1 | 0 | 0 | 0 | 1 | 1 |
| Ros1 | 0 | 0 | 0 | 1 | 1 |
| Rpl10l | 0 | 0 | 0 | 1 | 1 |
| Rpl26 | 0 | 0 | 0 | 1 | 1 |
| Rpl39l | 0 | 0 | 0 | 1 | 1 |
| Rprl1 | 0 | 0 | 0 | 1 | 1 |
| Rprl2 | 0 | 0 | 0 | 1 | 1 |
| Rprl3 | 0 | 0 | 0 | 1 | 1 |
| Rps27rt | 0 | 0 | 0 | 1 | 1 |
| Rtdr1 | 0 | 0 | 0 | 1 | 1 |
| Rtp1 | 0 | 0 | 0 | 1 | 1 |
| Rtp2 | 0 | 0 | 0 | 1 | 1 |
| Rtp3 | 0 | 0 | 0 | 1 | 1 |
| S100a2 | 0 | 0 | 0 | 1 | 1 |
| S100a5 | 0 | 0 | 0 | 1 | 1 |
| S100a7a | 0 | 0 | 0 | 1 | 1 |
| S100a8 | 0 | 0 | 0 | 1 | 1 |
| S100a9 | 0 | 0 | 0 | 1 | 1 |
| S100z | 0 | 0 | 0 | 1 | 1 |
| Saa4 | 0 | 0 | 0 | 1 | 1 |
| Samd7 | 0 | 0 | 0 | 1 | 1 |
| Samt1 | 0 | 0 | 0 | 1 | 1 |
| Samt2 | 0 | 0 | 0 | 1 | 1 |
| Samt3 | 0 | 0 | 0 | 1 | 1 |
| Samt4 | 0 | 0 | 0 | 1 | 1 |
| Satl1 | 0 | 0 | 0 | 1 | 1 |
| Sbp | 0 | 0 | 0 | 1 | 1 |
| Sbpl | 0 | 0 | 0 | 1 | 1 |
| Scarna17 | 0 | 0 | 0 | 1 | 1 |
| Scarna9 | 0 | 0 | 0 | 1 | 1 |
| Scgb1a1 | 0 | 0 | 0 | 1 | 1 |

| | | | | | |
|---|---|---|---|---|---|
| Scgb1b19 | 0 | 0 | 0 | 1 | 1 |
| Scgb1b2 | 0 | 0 | 0 | 1 | 1 |
| Scgb1b20 | 0 | 0 | 0 | 1 | 1 |
| Scgb1b24 | 0 | 0 | 0 | 1 | 1 |
| Scgb1b27 | 0 | 0 | 0 | 1 | 1 |
| Scgb1b29 | 0 | 0 | 0 | 1 | 1 |
| Scgb1b3 | 0 | 0 | 0 | 1 | 1 |
| Scgb1b30 | 0 | 0 | 0 | 1 | 1 |
| Scgb1b7 | 0 | 0 | 0 | 1 | 1 |
| Scgb2b12 | 0 | 0 | 0 | 1 | 1 |
| Scgb2b17 | 0 | 0 | 0 | 1 | 1 |
| Scgb2b19 | 0 | 0 | 0 | 1 | 1 |
| Scgb2b2 | 0 | 0 | 0 | 1 | 1 |
| Scgb2b20 | 0 | 0 | 0 | 1 | 1 |
| Scgb2b23-ps | 0 | 0 | 0 | 1 | 1 |
| Scgb2b24 | 0 | 0 | 0 | 1 | 1 |
| Scgb2b26 | 0 | 0 | 0 | 1 | 1 |
| Scgb2b27 | 0 | 0 | 0 | 1 | 1 |
| Scgb2b3 | 0 | 0 | 0 | 1 | 1 |
| Scgb2b7 | 0 | 0 | 0 | 1 | 1 |
| Scgb3a2 | 0 | 0 | 0 | 1 | 1 |
| Scimp | 0 | 0 | 0 | 1 | 1 |
| Scnn1g | 0 | 0 | 0 | 1 | 1 |
| Scp2d1 | 0 | 0 | 0 | 1 | 1 |
| Scrg1 | 0 | 0 | 0 | 1 | 1 |
| Sdr16c5 | 0 | 0 | 0 | 1 | 1 |
| Sdr16c6 | 0 | 0 | 0 | 1 | 1 |
| Sdr9c7 | 0 | 0 | 0 | 1 | 1 |
| Sebox | 0 | 0 | 0 | 1 | 1 |
| Sec14l3 | 0 | 0 | 0 | 1 | 1 |
| Sectm1a | 0 | 0 | 0 | 1 | 1 |
| Sectm1b | 0 | 0 | 0 | 1 | 1 |
| Sel1l2 | 0 | 0 | 0 | 1 | 1 |
| Sele | 0 | 0 | 0 | 1 | 1 |
| Sept12 | 0 | 0 | 0 | 1 | 1 |
| Serpina10 | 0 | 0 | 0 | 1 | 1 |
| Serpina11 | 0 | 0 | 0 | 1 | 1 |
| Serpina1a | 0 | 0 | 0 | 1 | 1 |
| Serpina1b | 0 | 0 | 0 | 1 | 1 |
| Serpina1c | 0 | 0 | 0 | 1 | 1 |
| Serpina1d | 0 | 0 | 0 | 1 | 1 |
| Serpina1e | 0 | 0 | 0 | 1 | 1 |
| Serpina1f | 0 | 0 | 0 | 1 | 1 |
| Serpina3c | 0 | 0 | 0 | 1 | 1 |
| Serpina3j | 0 | 0 | 0 | 1 | 1 |
| Serpina3k | 0 | 0 | 0 | 1 | 1 |
| Serpina3n | 0 | 0 | 0 | 1 | 1 |
| Serpina4-ps1 | 0 | 0 | 0 | 1 | 1 |
| Serpina5 | 0 | 0 | 0 | 1 | 1 |
| Serpina7 | 0 | 0 | 0 | 1 | 1 |

| | | | | | |
|---|---|---|---|---|---|
| Serpina9 | 0 | 0 | 0 | 1 | 1 |
| Serpinb10 | 0 | 0 | 0 | 1 | 1 |
| Serpinb11 | 0 | 0 | 0 | 1 | 1 |
| Serpinb12 | 0 | 0 | 0 | 1 | 1 |
| Serpinb1b | 0 | 0 | 0 | 1 | 1 |
| Serpinb1c | 0 | 0 | 0 | 1 | 1 |
| Serpinb2 | 0 | 0 | 0 | 1 | 1 |
| Serpinb3a | 0 | 0 | 0 | 1 | 1 |
| Serpinb3b | 0 | 0 | 0 | 1 | 1 |
| Serpinb3c | 0 | 0 | 0 | 1 | 1 |
| Serpinb3d | 0 | 0 | 0 | 1 | 1 |
| Serpinb5 | 0 | 0 | 0 | 1 | 1 |
| Serpinb6e | 0 | 0 | 0 | 1 | 1 |
| Serpinb7 | 0 | 0 | 0 | 1 | 1 |
| Serpinb8 | 0 | 0 | 0 | 1 | 1 |
| Serpinb9c | 0 | 0 | 0 | 1 | 1 |
| Serpinb9d | 0 | 0 | 0 | 1 | 1 |
| Serpinb9e | 0 | 0 | 0 | 1 | 1 |
| Serpinb9f | 0 | 0 | 0 | 1 | 1 |
| Serpinb9g | 0 | 0 | 0 | 1 | 1 |
| Serpini2 | 0 | 0 | 0 | 1 | 1 |
| Sftpa1 | 0 | 0 | 0 | 1 | 1 |
| Sftpd | 0 | 0 | 0 | 1 | 1 |
| Sgca | 0 | 0 | 0 | 1 | 1 |
| Sh2d1a | 0 | 0 | 0 | 1 | 1 |
| Sh2d1b1 | 0 | 0 | 0 | 1 | 1 |
| Sh2d1b2 | 0 | 0 | 0 | 1 | 1 |
| Siah3 | 0 | 0 | 0 | 1 | 1 |
| Siglec1 | 0 | 0 | 0 | 1 | 1 |
| Sirpb1a | 0 | 0 | 0 | 1 | 1 |
| Sirpb1b | 0 | 0 | 0 | 1 | 1 |
| Sis | 0 | 0 | 0 | 1 | 1 |
| Sit1 | 0 | 0 | 0 | 1 | 1 |
| Six3os1 | 0 | 0 | 0 | 1 | 1 |
| Skint1 | 0 | 0 | 0 | 1 | 1 |
| Skint10 | 0 | 0 | 0 | 1 | 1 |
| Skint11 | 0 | 0 | 0 | 1 | 1 |
| Skint2 | 0 | 0 | 0 | 1 | 1 |
| Skint3 | 0 | 0 | 0 | 1 | 1 |
| Skint4 | 0 | 0 | 0 | 1 | 1 |
| Skint5 | 0 | 0 | 0 | 1 | 1 |
| Skint6 | 0 | 0 | 0 | 1 | 1 |
| Skint7 | 0 | 0 | 0 | 1 | 1 |
| Skint8 | 0 | 0 | 0 | 1 | 1 |
| Skint9 | 0 | 0 | 0 | 1 | 1 |
| Sla | 0 | 0 | 0 | 1 | 1 |
| Slamf6 | 0 | 0 | 0 | 1 | 1 |
| Slamf7 | 0 | 0 | 0 | 1 | 1 |
| Slc10a2 | 0 | 0 | 0 | 1 | 1 |
| Slc10a6 | 0 | 0 | 0 | 1 | 1 |

| | | | | | |
|---|---|---|---|---|---|
| Slc13a1 | 0 | 0 | 0 | 1 | 1 |
| Slc13a2 | 0 | 0 | 0 | 1 | 1 |
| Slc13a2os | 0 | 0 | 0 | 1 | 1 |
| Slc15a1 | 0 | 0 | 0 | 1 | 1 |
| Slc15a5 | 0 | 0 | 0 | 1 | 1 |
| Slc17a1 | 0 | 0 | 0 | 1 | 1 |
| Slc17a4 | 0 | 0 | 0 | 1 | 1 |
| Slc17a6 | 0 | 0 | 0 | 1 | 1 |
| Slc22a13 | 0 | 0 | 0 | 1 | 1 |
| Slc22a16 | 0 | 0 | 0 | 1 | 1 |
| Slc22a19 | 0 | 0 | 0 | 1 | 1 |
| Slc22a20 | 0 | 0 | 0 | 1 | 1 |
| Slc22a26 | 0 | 0 | 0 | 1 | 1 |
| Slc22a27 | 0 | 0 | 0 | 1 | 1 |
| Slc22a29 | 0 | 0 | 0 | 1 | 1 |
| Slc22a8 | 0 | 0 | 0 | 1 | 1 |
| Slc24a1 | 0 | 0 | 0 | 1 | 1 |
| Slc26a3 | 0 | 0 | 0 | 1 | 1 |
| Slc26a7 | 0 | 0 | 0 | 1 | 1 |
| Slc26a9 | 0 | 0 | 0 | 1 | 1 |
| Slc28a1 | 0 | 0 | 0 | 1 | 1 |
| Slc2a7 | 0 | 0 | 0 | 1 | 1 |
| Slc30a8 | 0 | 0 | 0 | 1 | 1 |
| Slc34a2 | 0 | 0 | 0 | 1 | 1 |
| Slc35f4 | 0 | 0 | 0 | 1 | 1 |
| Slc36a1os | 0 | 0 | 0 | 1 | 1 |
| Slc36a3 | 0 | 0 | 0 | 1 | 1 |
| Slc38a11 | 0 | 0 | 0 | 1 | 1 |
| Slc39a12 | 0 | 0 | 0 | 1 | 1 |
| Slc51b | 0 | 0 | 0 | 1 | 1 |
| Slc5a4a | 0 | 0 | 0 | 1 | 1 |
| Slc6a20b | 0 | 0 | 0 | 1 | 1 |
| Slc6a5 | 0 | 0 | 0 | 1 | 1 |
| Slc7a12 | 0 | 0 | 0 | 1 | 1 |
| Slc7a15 | 0 | 0 | 0 | 1 | 1 |
| Slc9a3 | 0 | 0 | 0 | 1 | 1 |
| Slc9a4 | 0 | 0 | 0 | 1 | 1 |
| Slc9c1 | 0 | 0 | 0 | 1 | 1 |
| Slco1a1 | 0 | 0 | 0 | 1 | 1 |
| Slco1a6 | 0 | 0 | 0 | 1 | 1 |
| Slco6c1 | 0 | 0 | 0 | 1 | 1 |
| Slco6d1 | 0 | 0 | 0 | 1 | 1 |
| Slfn1 | 0 | 0 | 0 | 1 | 1 |
| Slfn5os | 0 | 0 | 0 | 1 | 1 |
| Sln | 0 | 0 | 0 | 1 | 1 |
| Slx | 0 | 0 | 0 | 1 | 1 |
| Slxl1 | 0 | 0 | 0 | 1 | 1 |
| Sly | 0 | 0 | 0 | 1 | 1 |
| Smco1 | 0 | 0 | 0 | 1 | 1 |
| Smco2 | 0 | 0 | 0 | 1 | 1 |

| | | | | | |
|---|---|---|---|---|---|
| Smcp | 0 | 0 | 0 | 1 | 1 |
| Smgc | 0 | 0 | 0 | 1 | 1 |
| Smim23 | 0 | 0 | 0 | 1 | 1 |
| Smim9 | 0 | 0 | 0 | 1 | 1 |
| Smlr1 | 0 | 0 | 0 | 1 | 1 |
| Smok2a | 0 | 0 | 0 | 1 | 1 |
| Smok2b | 0 | 0 | 0 | 1 | 1 |
| Smok3a | 0 | 0 | 0 | 1 | 1 |
| Smok3b | 0 | 0 | 0 | 1 | 1 |
| Smr2 | 0 | 0 | 0 | 1 | 1 |
| Smr3a | 0 | 0 | 0 | 1 | 1 |
| Snora19 | 0 | 0 | 0 | 1 | 1 |
| Snora20 | 0 | 0 | 0 | 1 | 1 |
| Snora35 | 0 | 0 | 0 | 1 | 1 |
| Snora36b | 0 | 0 | 0 | 1 | 1 |
| Snora47 | 0 | 0 | 0 | 1 | 1 |
| Snord11 | 0 | 0 | 0 | 1 | 1 |
| Snord110 | 0 | 0 | 0 | 1 | 1 |
| Snord111 | 0 | 0 | 0 | 1 | 1 |
| Snord116 | 0 | 0 | 0 | 1 | 1 |
| Snord116l1 | 0 | 0 | 0 | 1 | 1 |
| Snord116l2 | 0 | 0 | 0 | 1 | 1 |
| Snord118 | 0 | 0 | 0 | 1 | 1 |
| Snord123 | 0 | 0 | 0 | 1 | 1 |
| Snord14a | 0 | 0 | 0 | 1 | 1 |
| Snord14c | 0 | 0 | 0 | 1 | 1 |
| Snord14d | 0 | 0 | 0 | 1 | 1 |
| Snord1a | 0 | 0 | 0 | 1 | 1 |
| Snord1c | 0 | 0 | 0 | 1 | 1 |
| Snord38a | 0 | 0 | 0 | 1 | 1 |
| Snord42a | 0 | 0 | 0 | 1 | 1 |
| Snord43 | 0 | 0 | 0 | 1 | 1 |
| Snord45b | 0 | 0 | 0 | 1 | 1 |
| Snord58b | 0 | 0 | 0 | 1 | 1 |
| Snord61 | 0 | 0 | 0 | 1 | 1 |
| Snord64 | 0 | 0 | 0 | 1 | 1 |
| Snord68 | 0 | 0 | 0 | 1 | 1 |
| Snord70 | 0 | 0 | 0 | 1 | 1 |
| Snord8 | 0 | 0 | 0 | 1 | 1 |
| Snord90 | 0 | 0 | 0 | 1 | 1 |
| Snord91a | 0 | 0 | 0 | 1 | 1 |
| Snord93 | 0 | 0 | 0 | 1 | 1 |
| Snord98 | 0 | 0 | 0 | 1 | 1 |
| Sntn | 0 | 0 | 0 | 1 | 1 |
| Sohlh1 | 0 | 0 | 0 | 1 | 1 |
| Sohlh2 | 0 | 0 | 0 | 1 | 1 |
| Sorbs2os | 0 | 0 | 0 | 1 | 1 |
| Sp110 | 0 | 0 | 0 | 1 | 1 |
| Sp140 | 0 | 0 | 0 | 1 | 1 |
| Spaca3 | 0 | 0 | 0 | 1 | 1 |

| | | | | | |
|---|---|---|---|---|---|
| Spaca5 | 0 | 0 | 0 | 1 | 1 |
| Spaca7 | 0 | 0 | 0 | 1 | 1 |
| Spag11a | 0 | 0 | 0 | 1 | 1 |
| Spam1 | 0 | 0 | 0 | 1 | 1 |
| Spata19 | 0 | 0 | 0 | 1 | 1 |
| Spata20 | 0 | 0 | 0 | 1 | 1 |
| Spata22 | 0 | 0 | 0 | 1 | 1 |
| Spata31 | 0 | 0 | 0 | 1 | 1 |
| Spata31d1b | 0 | 0 | 0 | 1 | 1 |
| Spata31d1c | 0 | 0 | 0 | 1 | 1 |
| Spata4 | 0 | 0 | 0 | 1 | 1 |
| Spdyb | 0 | 0 | 0 | 1 | 1 |
| Speer1-ps1 | 0 | 0 | 0 | 1 | 1 |
| Speer2 | 0 | 0 | 0 | 1 | 1 |
| Speer3 | 0 | 0 | 0 | 1 | 1 |
| Speer4a | 0 | 0 | 0 | 1 | 1 |
| Speer4c | 0 | 0 | 0 | 1 | 1 |
| Speer4d | 0 | 0 | 0 | 1 | 1 |
| Speer4e | 0 | 0 | 0 | 1 | 1 |
| Speer4f | 0 | 0 | 0 | 1 | 1 |
| Speer6-ps1 | 0 | 0 | 0 | 1 | 1 |
| Speer7-ps1 | 0 | 0 | 0 | 1 | 1 |
| Speer8-ps1 | 0 | 0 | 0 | 1 | 1 |
| Speer9-ps1 | 0 | 0 | 0 | 1 | 1 |
| Spert | 0 | 0 | 0 | 1 | 1 |
| Spesp1 | 0 | 0 | 0 | 1 | 1 |
| Spic | 0 | 0 | 0 | 1 | 1 |
| Spin2d | 0 | 0 | 0 | 1 | 1 |
| Spink11 | 0 | 0 | 0 | 1 | 1 |
| Spink12 | 0 | 0 | 0 | 1 | 1 |
| Spink13 | 0 | 0 | 0 | 1 | 1 |
| Spink14 | 0 | 0 | 0 | 1 | 1 |
| Spink5 | 0 | 0 | 0 | 1 | 1 |
| Spink6 | 0 | 0 | 0 | 1 | 1 |
| Spink7 | 0 | 0 | 0 | 1 | 1 |
| Spinkl | 0 | 0 | 0 | 1 | 1 |
| Spint3 | 0 | 0 | 0 | 1 | 1 |
| Spint4 | 0 | 0 | 0 | 1 | 1 |
| Spint5 | 0 | 0 | 0 | 1 | 1 |
| Spn-ps | 0 | 0 | 0 | 1 | 1 |
| Spns3 | 0 | 0 | 0 | 1 | 1 |
| Spp2 | 0 | 0 | 0 | 1 | 1 |
| Sppl2c | 0 | 0 | 0 | 1 | 1 |
| Sprr1a | 0 | 0 | 0 | 1 | 1 |
| Sprr1b | 0 | 0 | 0 | 1 | 1 |
| Sprr2a2 | 0 | 0 | 0 | 1 | 1 |
| Sprr2b | 0 | 0 | 0 | 1 | 1 |
| Sprr2d | 0 | 0 | 0 | 1 | 1 |
| Sprr2e | 0 | 0 | 0 | 1 | 1 |
| Sprr2f | 0 | 0 | 0 | 1 | 1 |

| | | | | | |
|---|---|---|---|---|---|
| Sprr2g | 0 | 0 | 0 | 1 | 1 |
| Sprr2h | 0 | 0 | 0 | 1 | 1 |
| Sprr2i | 0 | 0 | 0 | 1 | 1 |
| Sprr2j-ps | 0 | 0 | 0 | 1 | 1 |
| Sprr2k | 0 | 0 | 0 | 1 | 1 |
| Sprr3 | 0 | 0 | 0 | 1 | 1 |
| Sprr4 | 0 | 0 | 0 | 1 | 1 |
| Spry3 | 0 | 0 | 0 | 1 | 1 |
| Sptssb | 0 | 0 | 0 | 1 | 1 |
| Spz1 | 0 | 0 | 0 | 1 | 1 |
| Srp54b,Srp54c | 0 | 0 | 0 | 1 | 1 |
| Sry | 0 | 0 | 0 | 1 | 1 |
| Sstr3 | 0 | 0 | 0 | 1 | 1 |
| Ssty1 | 0 | 0 | 0 | 1 | 1 |
| Ssty2 | 0 | 0 | 0 | 1 | 1 |
| Ssu2 | 0 | 0 | 0 | 1 | 1 |
| Ssx9 | 0 | 0 | 0 | 1 | 1 |
| Ssxb1 | 0 | 0 | 0 | 1 | 1 |
| Ssxb10 | 0 | 0 | 0 | 1 | 1 |
| Ssxb2 | 0 | 0 | 0 | 1 | 1 |
| Ssxb3 | 0 | 0 | 0 | 1 | 1 |
| Ssxb5 | 0 | 0 | 0 | 1 | 1 |
| Ssxb6 | 0 | 0 | 0 | 1 | 1 |
| Ssxb8 | 0 | 0 | 0 | 1 | 1 |
| Ssxb9 | 0 | 0 | 0 | 1 | 1 |
| St6galnac1 | 0 | 0 | 0 | 1 | 1 |
| St8sia3os | 0 | 0 | 0 | 1 | 1 |
| Stfa1 | 0 | 0 | 0 | 1 | 1 |
| Stfa2 | 0 | 0 | 0 | 1 | 1 |
| Stfa2l1 | 0 | 0 | 0 | 1 | 1 |
| Stfa3 | 0 | 0 | 0 | 1 | 1 |
| Stmnd1 | 0 | 0 | 0 | 1 | 1 |
| Stoml3 | 0 | 0 | 0 | 1 | 1 |
| Stpg2 | 0 | 0 | 0 | 1 | 1 |
| Stra8 | 0 | 0 | 0 | 1 | 1 |
| Sucnr1 | 0 | 0 | 0 | 1 | 1 |
| Sult1b1 | 0 | 0 | 0 | 1 | 1 |
| Sult1c1 | 0 | 0 | 0 | 1 | 1 |
| Sult1c2 | 0 | 0 | 0 | 1 | 1 |
| Sult1d1 | 0 | 0 | 0 | 1 | 1 |
| Sult1e1 | 0 | 0 | 0 | 1 | 1 |
| Sult2a1 | 0 | 0 | 0 | 1 | 1 |
| Sult2a2 | 0 | 0 | 0 | 1 | 1 |
| Sult2a3 | 0 | 0 | 0 | 1 | 1 |
| Sult2a4 | 0 | 0 | 0 | 1 | 1 |
| Sult2a5 | 0 | 0 | 0 | 1 | 1 |
| Sult2a6 | 0 | 0 | 0 | 1 | 1 |
| Sult3a1 | 0 | 0 | 0 | 1 | 1 |
| Sun5 | 0 | 0 | 0 | 1 | 1 |
| Sva | 0 | 0 | 0 | 1 | 1 |

| | | | | | |
|---|---|---|---|---|---|
| Sval1 | 0 | 0 | 0 | 1 | 1 |
| Sval2 | 0 | 0 | 0 | 1 | 1 |
| Sval3 | 0 | 0 | 0 | 1 | 1 |
| Svs2 | 0 | 0 | 0 | 1 | 1 |
| Svs3a | 0 | 0 | 0 | 1 | 1 |
| Svs3b | 0 | 0 | 0 | 1 | 1 |
| Svs4 | 0 | 0 | 0 | 1 | 1 |
| Svs5 | 0 | 0 | 0 | 1 | 1 |
| Svs6 | 0 | 0 | 0 | 1 | 1 |
| Syce1 | 0 | 0 | 0 | 1 | 1 |
| Syce3 | 0 | 0 | 0 | 1 | 1 |
| Sycn | 0 | 0 | 0 | 1 | 1 |
| Synb | 0 | 0 | 0 | 1 | 1 |
| Syt8 | 0 | 0 | 0 | 1 | 1 |
| Taar1 | 0 | 0 | 0 | 1 | 1 |
| Taar3 | 0 | 0 | 0 | 1 | 1 |
| Taar5 | 0 | 0 | 0 | 1 | 1 |
| Taar6 | 0 | 0 | 0 | 1 | 1 |
| Taar7a | 0 | 0 | 0 | 1 | 1 |
| Taar7b | 0 | 0 | 0 | 1 | 1 |
| Taar7e | 0 | 0 | 0 | 1 | 1 |
| Taar7f | 0 | 0 | 0 | 1 | 1 |
| Taar8a | 0 | 0 | 0 | 1 | 1 |
| Taar8b | 0 | 0 | 0 | 1 | 1 |
| Taar8c | 0 | 0 | 0 | 1 | 1 |
| Taar9 | 0 | 0 | 0 | 1 | 1 |
| Tac2 | 0 | 0 | 0 | 1 | 1 |
| Taf7l | 0 | 0 | 0 | 1 | 1 |
| Tarm1 | 0 | 0 | 0 | 1 | 1 |
| Tas2r102 | 0 | 0 | 0 | 1 | 1 |
| Tas2r103 | 0 | 0 | 0 | 1 | 1 |
| Tas2r104 | 0 | 0 | 0 | 1 | 1 |
| Tas2r105 | 0 | 0 | 0 | 1 | 1 |
| Tas2r106 | 0 | 0 | 0 | 1 | 1 |
| Tas2r107 | 0 | 0 | 0 | 1 | 1 |
| Tas2r108 | 0 | 0 | 0 | 1 | 1 |
| Tas2r109 | 0 | 0 | 0 | 1 | 1 |
| Tas2r110 | 0 | 0 | 0 | 1 | 1 |
| Tas2r113 | 0 | 0 | 0 | 1 | 1 |
| Tas2r114 | 0 | 0 | 0 | 1 | 1 |
| Tas2r115 | 0 | 0 | 0 | 1 | 1 |
| Tas2r116 | 0 | 0 | 0 | 1 | 1 |
| Tas2r117 | 0 | 0 | 0 | 1 | 1 |
| Tas2r118 | 0 | 0 | 0 | 1 | 1 |
| Tas2r119 | 0 | 0 | 0 | 1 | 1 |
| Tas2r120 | 0 | 0 | 0 | 1 | 1 |
| Tas2r121 | 0 | 0 | 0 | 1 | 1 |
| Tas2r122 | 0 | 0 | 0 | 1 | 1 |
| Tas2r123 | 0 | 0 | 0 | 1 | 1 |
| Tas2r124 | 0 | 0 | 0 | 1 | 1 |

| | | | | | |
|---|---|---|---|---|---|
| Tas2r125 | 0 | 0 | 0 | 1 | 1 |
| Tas2r126 | 0 | 0 | 0 | 1 | 1 |
| Tas2r129 | 0 | 0 | 0 | 1 | 1 |
| Tas2r130 | 0 | 0 | 0 | 1 | 1 |
| Tas2r131 | 0 | 0 | 0 | 1 | 1 |
| Tas2r136 | 0 | 0 | 0 | 1 | 1 |
| Tas2r138 | 0 | 0 | 0 | 1 | 1 |
| Tas2r139 | 0 | 0 | 0 | 1 | 1 |
| Tas2r140 | 0 | 0 | 0 | 1 | 1 |
| Tas2r144 | 0 | 0 | 0 | 1 | 1 |
| Tat | 0 | 0 | 0 | 1 | 1 |
| Tbc1d21 | 0 | 0 | 0 | 1 | 1 |
| Tbpl2 | 0 | 0 | 0 | 1 | 1 |
| Tcf23 | 0 | 0 | 0 | 1 | 1 |
| Tchhl1 | 0 | 0 | 0 | 1 | 1 |
| Tcl1 | 0 | 0 | 0 | 1 | 1 |
| Tcl1b1 | 0 | 0 | 0 | 1 | 1 |
| Tcl1b2 | 0 | 0 | 0 | 1 | 1 |
| Tcl1b3 | 0 | 0 | 0 | 1 | 1 |
| Tcl1b4 | 0 | 0 | 0 | 1 | 1 |
| Tcl1b5 | 0 | 0 | 0 | 1 | 1 |
| Tcp10b | 0 | 0 | 0 | 1 | 1 |
| Tcp10c | 0 | 0 | 0 | 1 | 1 |
| Tcstv1 | 0 | 0 | 0 | 1 | 1 |
| Tcstv3 | 0 | 0 | 0 | 1 | 1 |
| Tcte3 | 0 | 0 | 0 | 1 | 1 |
| Tctex1d4 | 0 | 0 | 0 | 1 | 1 |
| Tdgf1 | 0 | 0 | 0 | 1 | 1 |
| Tdpoz1 | 0 | 0 | 0 | 1 | 1 |
| Tdpoz2 | 0 | 0 | 0 | 1 | 1 |
| Tdpoz3 | 0 | 0 | 0 | 1 | 1 |
| Tdpoz4 | 0 | 0 | 0 | 1 | 1 |
| Tdpoz5 | 0 | 0 | 0 | 1 | 1 |
| Tectb | 0 | 0 | 0 | 1 | 1 |
| Teddm1 | 0 | 0 | 0 | 1 | 1 |
| Tenm1 | 0 | 0 | 0 | 1 | 1 |
| Tespa1 | 0 | 0 | 0 | 1 | 1 |
| Tex101 | 0 | 0 | 0 | 1 | 1 |
| Tex13 | 0 | 0 | 0 | 1 | 1 |
| Tex13a | 0 | 0 | 0 | 1 | 1 |
| Tex19.2 | 0 | 0 | 0 | 1 | 1 |
| Tex24 | 0 | 0 | 0 | 1 | 1 |
| Tex28 | 0 | 0 | 0 | 1 | 1 |
| Tex33 | 0 | 0 | 0 | 1 | 1 |
| Tex36 | 0 | 0 | 0 | 1 | 1 |
| Tex37 | 0 | 0 | 0 | 1 | 1 |
| Tex43 | 0 | 0 | 0 | 1 | 1 |
| Tfap2d | 0 | 0 | 0 | 1 | 1 |
| Tff1 | 0 | 0 | 0 | 1 | 1 |
| Tgif2lx1,Tgif2lx2 | 0 | 0 | 0 | 1 | 1 |

| | | | | | |
|---|---|---|---|---|---|
| Tgif2lx2 | 0 | 0 | 0 | 1 | 1 |
| Tgm6 | 0 | 0 | 0 | 1 | 1 |
| Tgm7 | 0 | 0 | 0 | 1 | 1 |
| Tgtp2 | 0 | 0 | 0 | 1 | 1 |
| Theg | 0 | 0 | 0 | 1 | 1 |
| Themis | 0 | 0 | 0 | 1 | 1 |
| Tigit | 0 | 0 | 0 | 1 | 1 |
| Timd2 | 0 | 0 | 0 | 1 | 1 |
| Timd4 | 0 | 0 | 0 | 1 | 1 |
| Timm8a2 | 0 | 0 | 0 | 1 | 1 |
| Tinag | 0 | 0 | 0 | 1 | 1 |
| Tlr1 | 0 | 0 | 0 | 1 | 1 |
| Tlr11 | 0 | 0 | 0 | 1 | 1 |
| Tlr8 | 0 | 0 | 0 | 1 | 1 |
| Tlr9 | 0 | 0 | 0 | 1 | 1 |
| Tm4sf19 | 0 | 0 | 0 | 1 | 1 |
| Tmbim7 | 0 | 0 | 0 | 1 | 1 |
| Tmco2 | 0 | 0 | 0 | 1 | 1 |
| Tmco5 | 0 | 0 | 0 | 1 | 1 |
| Tmco5b | 0 | 0 | 0 | 1 | 1 |
| Tmed11 | 0 | 0 | 0 | 1 | 1 |
| Tmed6 | 0 | 0 | 0 | 1 | 1 |
| Tmem179 | 0 | 0 | 0 | 1 | 1 |
| Tmem182 | 0 | 0 | 0 | 1 | 1 |
| Tmem190 | 0 | 0 | 0 | 1 | 1 |
| Tmem202 | 0 | 0 | 0 | 1 | 1 |
| Tmem207 | 0 | 0 | 0 | 1 | 1 |
| Tmem210 | 0 | 0 | 0 | 1 | 1 |
| Tmem212 | 0 | 0 | 0 | 1 | 1 |
| Tmem213 | 0 | 0 | 0 | 1 | 1 |
| Tmem225 | 0 | 0 | 0 | 1 | 1 |
| Tmem233 | 0 | 0 | 0 | 1 | 1 |
| Tmem235 | 0 | 0 | 0 | 1 | 1 |
| Tmem247 | 0 | 0 | 0 | 1 | 1 |
| Tmem254a,Tmem254b,Tn | 0 | 0 | 0 | 1 | 1 |
| Tmem254b | 0 | 0 | 0 | 1 | 1 |
| Tmem254c | 0 | 0 | 0 | 1 | 1 |
| Tmem255b | 0 | 0 | 0 | 1 | 1 |
| Tmem27 | 0 | 0 | 0 | 1 | 1 |
| Tmem45b | 0 | 0 | 0 | 1 | 1 |
| Tmem52 | 0 | 0 | 0 | 1 | 1 |
| Tmem63c | 0 | 0 | 0 | 1 | 1 |
| Tmem71 | 0 | 0 | 0 | 1 | 1 |
| Tmem72 | 0 | 0 | 0 | 1 | 1 |
| Tmem92 | 0 | 0 | 0 | 1 | 1 |
| Tmevpg1 | 0 | 0 | 0 | 1 | 1 |
| Tmlhe | 0 | 0 | 0 | 1 | 1 |
| Tmprss11a | 0 | 0 | 0 | 1 | 1 |
| Tmprss11bnl | 0 | 0 | 0 | 1 | 1 |
| Tmprss11c | 0 | 0 | 0 | 1 | 1 |

| | | | | | |
|---|---|---|---|---|---|
| Tmprss11e | 0 | 0 | 0 | 1 | 1 |
| Tmprss11f | 0 | 0 | 0 | 1 | 1 |
| Tmprss11g | 0 | 0 | 0 | 1 | 1 |
| Tmprss12 | 0 | 0 | 0 | 1 | 1 |
| Tmsb15a | 0 | 0 | 0 | 1 | 1 |
| Tnfsf14 | 0 | 0 | 0 | 1 | 1 |
| Tnfsf18 | 0 | 0 | 0 | 1 | 1 |
| Tnfsf8 | 0 | 0 | 0 | 1 | 1 |
| Tnni3k | 0 | 0 | 0 | 1 | 1 |
| Tnp1 | 0 | 0 | 0 | 1 | 1 |
| Tnr | 0 | 0 | 0 | 1 | 1 |
| Topaz1 | 0 | 0 | 0 | 1 | 1 |
| Tpbpa | 0 | 0 | 0 | 1 | 1 |
| Tpbpb | 0 | 0 | 0 | 1 | 1 |
| Tph2 | 0 | 0 | 0 | 1 | 1 |
| Tpsab1 | 0 | 0 | 0 | 1 | 1 |
| Tpsb2 | 0 | 0 | 0 | 1 | 1 |
| Tpte | 0 | 0 | 0 | 1 | 1 |
| Trat1 | 0 | 0 | 0 | 1 | 1 |
| Trdn | 0 | 0 | 0 | 1 | 1 |
| Trem1 | 0 | 0 | 0 | 1 | 1 |
| Trem2 | 0 | 0 | 0 | 1 | 1 |
| Trem3 | 0 | 0 | 0 | 1 | 1 |
| Trex2 | 0 | 0 | 0 | 1 | 1 |
| Trhr | 0 | 0 | 0 | 1 | 1 |
| Trhr2 | 0 | 0 | 0 | 1 | 1 |
| Trim30b | 0 | 0 | 0 | 1 | 1 |
| Trim31 | 0 | 0 | 0 | 1 | 1 |
| Trim34b | 0 | 0 | 0 | 1 | 1 |
| Trim40 | 0 | 0 | 0 | 1 | 1 |
| Trim42 | 0 | 0 | 0 | 1 | 1 |
| Trim50 | 0 | 0 | 0 | 1 | 1 |
| Trim52 | 0 | 0 | 0 | 1 | 1 |
| Triml1 | 0 | 0 | 0 | 1 | 1 |
| Triml2 | 0 | 0 | 0 | 1 | 1 |
| Trpc5os | 0 | 0 | 0 | 1 | 1 |
| Trpc7 | 0 | 0 | 0 | 1 | 1 |
| Trpd52l3 | 0 | 0 | 0 | 1 | 1 |
| Trpm8 | 0 | 0 | 0 | 1 | 1 |
| Trpv1 | 0 | 0 | 0 | 1 | 1 |
| Trpv5 | 0 | 0 | 0 | 1 | 1 |
| Try10 | 0 | 0 | 0 | 1 | 1 |
| Try4 | 0 | 0 | 0 | 1 | 1 |
| Try5 | 0 | 0 | 0 | 1 | 1 |
| Tsga8 | 0 | 0 | 0 | 1 | 1 |
| Tsix | 0 | 0 | 0 | 1 | 1 |
| Tspo2 | 0 | 0 | 0 | 1 | 1 |
| Tspy-ps | 0 | 0 | 0 | 1 | 1 |
| Tssk5 | 0 | 0 | 0 | 1 | 1 |
| Ttc22 | 0 | 0 | 0 | 1 | 1 |

| | | | | | |
|---|---|---|---|---|---|
| Ttc23l | 0 | 0 | 0 | 1 | 1 |
| Ttc24 | 0 | 0 | 0 | 1 | 1 |
| Ttc29 | 0 | 0 | 0 | 1 | 1 |
| Ttll2 | 0 | 0 | 0 | 1 | 1 |
| Ttll8 | 0 | 0 | 0 | 1 | 1 |
| Ttr | 0 | 0 | 0 | 1 | 1 |
| Tuba3b | 0 | 0 | 0 | 1 | 1 |
| Txndc8 | 0 | 0 | 0 | 1 | 1 |
| Tyr | 0 | 0 | 0 | 1 | 1 |
| Tyrp1 | 0 | 0 | 0 | 1 | 1 |
| Uba1y | 0 | 0 | 0 | 1 | 1 |
| Ubd | 0 | 0 | 0 | 1 | 1 |
| Ube2dnl1 | 0 | 0 | 0 | 1 | 1 |
| Ube2dnl2 | 0 | 0 | 0 | 1 | 1 |
| Ubqln3 | 0 | 0 | 0 | 1 | 1 |
| Ubqlnl | 0 | 0 | 0 | 1 | 1 |
| Ubtfl1 | 0 | 0 | 0 | 1 | 1 |
| Uchl1os | 0 | 0 | 0 | 1 | 1 |
| Ucn | 0 | 0 | 0 | 1 | 1 |
| Ucn3 | 0 | 0 | 0 | 1 | 1 |
| Ugt1a1 | 0 | 0 | 0 | 1 | 1 |
| Ugt1a10 | 0 | 0 | 0 | 1 | 1 |
| Ugt1a2 | 0 | 0 | 0 | 1 | 1 |
| Ugt1a5 | 0 | 0 | 0 | 1 | 1 |
| Ugt1a6a | 0 | 0 | 0 | 1 | 1 |
| Ugt1a7c | 0 | 0 | 0 | 1 | 1 |
| Ugt1a9 | 0 | 0 | 0 | 1 | 1 |
| Ugt2a1 | 0 | 0 | 0 | 1 | 1 |
| Ugt2a2 | 0 | 0 | 0 | 1 | 1 |
| Ugt2a3 | 0 | 0 | 0 | 1 | 1 |
| Ugt2b1 | 0 | 0 | 0 | 1 | 1 |
| Ugt2b34 | 0 | 0 | 0 | 1 | 1 |
| Ugt2b35 | 0 | 0 | 0 | 1 | 1 |
| Ugt2b36 | 0 | 0 | 0 | 1 | 1 |
| Ugt2b37 | 0 | 0 | 0 | 1 | 1 |
| Ugt2b38 | 0 | 0 | 0 | 1 | 1 |
| Ugt3a1 | 0 | 0 | 0 | 1 | 1 |
| Ugt3a2 | 0 | 0 | 0 | 1 | 1 |
| Umod | 0 | 0 | 0 | 1 | 1 |
| Unc93a | 0 | 0 | 0 | 1 | 1 |
| Uox | 0 | 0 | 0 | 1 | 1 |
| Upk2 | 0 | 0 | 0 | 1 | 1 |
| Usp17la | 0 | 0 | 0 | 1 | 1 |
| Usp17lb | 0 | 0 | 0 | 1 | 1 |
| Usp17lc | 0 | 0 | 0 | 1 | 1 |
| Usp17ld | 0 | 0 | 0 | 1 | 1 |
| Usp17le | 0 | 0 | 0 | 1 | 1 |
| Usp26 | 0 | 0 | 0 | 1 | 1 |
| Uts2b | 0 | 0 | 0 | 1 | 1 |
| V1ra8 | 0 | 0 | 0 | 1 | 1 |

| | | | | | |
|---|---|---|---|---|---|
| V1rd18,V1rd19 | 0 | 0 | 0 | 1 | 1 |
| V1rd19 | 0 | 0 | 0 | 1 | 1 |
| Vamp7 | 0 | 0 | 0 | 1 | 1 |
| Vav1 | 0 | 0 | 0 | 1 | 1 |
| Vax2os | 0 | 0 | 0 | 1 | 1 |
| Vgll1 | 0 | 0 | 0 | 1 | 1 |
| Vip | 0 | 0 | 0 | 1 | 1 |
| Vmn1r-ps103 | 0 | 0 | 0 | 1 | 1 |
| Vmn1r-ps79 | 0 | 0 | 0 | 1 | 1 |
| Vmn1r1 | 0 | 0 | 0 | 1 | 1 |
| Vmn1r10 | 0 | 0 | 0 | 1 | 1 |
| Vmn1r100,Vmn1r148 | 0 | 0 | 0 | 1 | 1 |
| Vmn1r101 | 0 | 0 | 0 | 1 | 1 |
| Vmn1r103 | 0 | 0 | 0 | 1 | 1 |
| Vmn1r107 | 0 | 0 | 0 | 1 | 1 |
| Vmn1r11 | 0 | 0 | 0 | 1 | 1 |
| Vmn1r112 | 0 | 0 | 0 | 1 | 1 |
| Vmn1r113 | 0 | 0 | 0 | 1 | 1 |
| Vmn1r114 | 0 | 0 | 0 | 1 | 1 |
| Vmn1r115 | 0 | 0 | 0 | 1 | 1 |
| Vmn1r116 | 0 | 0 | 0 | 1 | 1 |
| Vmn1r117 | 0 | 0 | 0 | 1 | 1 |
| Vmn1r118 | 0 | 0 | 0 | 1 | 1 |
| Vmn1r119 | 0 | 0 | 0 | 1 | 1 |
| Vmn1r12 | 0 | 0 | 0 | 1 | 1 |
| Vmn1r120 | 0 | 0 | 0 | 1 | 1 |
| Vmn1r121 | 0 | 0 | 0 | 1 | 1 |
| Vmn1r122 | 0 | 0 | 0 | 1 | 1 |
| Vmn1r123 | 0 | 0 | 0 | 1 | 1 |
| Vmn1r124 | 0 | 0 | 0 | 1 | 1 |
| Vmn1r125 | 0 | 0 | 0 | 1 | 1 |
| Vmn1r126 | 0 | 0 | 0 | 1 | 1 |
| Vmn1r127 | 0 | 0 | 0 | 1 | 1 |
| Vmn1r128 | 0 | 0 | 0 | 1 | 1 |
| Vmn1r129 | 0 | 0 | 0 | 1 | 1 |
| Vmn1r13 | 0 | 0 | 0 | 1 | 1 |
| Vmn1r130 | 0 | 0 | 0 | 1 | 1 |
| Vmn1r131 | 0 | 0 | 0 | 1 | 1 |
| Vmn1r132,Vmn1r139 | 0 | 0 | 0 | 1 | 1 |
| Vmn1r138 | 0 | 0 | 0 | 1 | 1 |
| Vmn1r139 | 0 | 0 | 0 | 1 | 1 |
| Vmn1r14 | 0 | 0 | 0 | 1 | 1 |
| Vmn1r142 | 0 | 0 | 0 | 1 | 1 |
| Vmn1r143 | 0 | 0 | 0 | 1 | 1 |
| Vmn1r148 | 0 | 0 | 0 | 1 | 1 |
| Vmn1r15 | 0 | 0 | 0 | 1 | 1 |
| Vmn1r151 | 0 | 0 | 0 | 1 | 1 |
| Vmn1r152 | 0 | 0 | 0 | 1 | 1 |
| Vmn1r155 | 0 | 0 | 0 | 1 | 1 |
| Vmn1r157 | 0 | 0 | 0 | 1 | 1 |

| | | | | | |
|---|---|---|---|---|---|
| Vmn1r158 | 0 | 0 | 0 | 1 | 1 |
| Vmn1r159 | 0 | 0 | 0 | 1 | 1 |
| Vmn1r16 | 0 | 0 | 0 | 1 | 1 |
| Vmn1r160 | 0 | 0 | 0 | 1 | 1 |
| Vmn1r163 | 0 | 0 | 0 | 1 | 1 |
| Vmn1r165 | 0 | 0 | 0 | 1 | 1 |
| Vmn1r166 | 0 | 0 | 0 | 1 | 1 |
| Vmn1r167 | 0 | 0 | 0 | 1 | 1 |
| Vmn1r168 | 0 | 0 | 0 | 1 | 1 |
| Vmn1r169 | 0 | 0 | 0 | 1 | 1 |
| Vmn1r17 | 0 | 0 | 0 | 1 | 1 |
| Vmn1r170 | 0 | 0 | 0 | 1 | 1 |
| Vmn1r171 | 0 | 0 | 0 | 1 | 1 |
| Vmn1r172 | 0 | 0 | 0 | 1 | 1 |
| Vmn1r173 | 0 | 0 | 0 | 1 | 1 |
| Vmn1r174 | 0 | 0 | 0 | 1 | 1 |
| Vmn1r175 | 0 | 0 | 0 | 1 | 1 |
| Vmn1r176 | 0 | 0 | 0 | 1 | 1 |
| Vmn1r177 | 0 | 0 | 0 | 1 | 1 |
| Vmn1r178 | 0 | 0 | 0 | 1 | 1 |
| Vmn1r179 | 0 | 0 | 0 | 1 | 1 |
| Vmn1r18 | 0 | 0 | 0 | 1 | 1 |
| Vmn1r180 | 0 | 0 | 0 | 1 | 1 |
| Vmn1r181 | 0 | 0 | 0 | 1 | 1 |
| Vmn1r183 | 0 | 0 | 0 | 1 | 1 |
| Vmn1r184 | 0 | 0 | 0 | 1 | 1 |
| Vmn1r185 | 0 | 0 | 0 | 1 | 1 |
| Vmn1r186,Vmn1r62 | 0 | 0 | 0 | 1 | 1 |
| Vmn1r187,Vmn1r63 | 0 | 0 | 0 | 1 | 1 |
| Vmn1r188 | 0 | 0 | 0 | 1 | 1 |
| Vmn1r189 | 0 | 0 | 0 | 1 | 1 |
| Vmn1r19 | 0 | 0 | 0 | 1 | 1 |
| Vmn1r191 | 0 | 0 | 0 | 1 | 1 |
| Vmn1r192 | 0 | 0 | 0 | 1 | 1 |
| Vmn1r193 | 0 | 0 | 0 | 1 | 1 |
| Vmn1r194 | 0 | 0 | 0 | 1 | 1 |
| Vmn1r195 | 0 | 0 | 0 | 1 | 1 |
| Vmn1r196 | 0 | 0 | 0 | 1 | 1 |
| Vmn1r198 | 0 | 0 | 0 | 1 | 1 |
| Vmn1r199 | 0 | 0 | 0 | 1 | 1 |
| Vmn1r2 | 0 | 0 | 0 | 1 | 1 |
| Vmn1r20 | 0 | 0 | 0 | 1 | 1 |
| Vmn1r200 | 0 | 0 | 0 | 1 | 1 |
| Vmn1r201 | 0 | 0 | 0 | 1 | 1 |
| Vmn1r202 | 0 | 0 | 0 | 1 | 1 |
| Vmn1r203 | 0 | 0 | 0 | 1 | 1 |
| Vmn1r204 | 0 | 0 | 0 | 1 | 1 |
| Vmn1r205 | 0 | 0 | 0 | 1 | 1 |
| Vmn1r206 | 0 | 0 | 0 | 1 | 1 |
| Vmn1r207-ps | 0 | 0 | 0 | 1 | 1 |

| | | | | | |
|---|---|---|---|---|---|
| Vmn1r208 | 0 | 0 | 0 | 1 | 1 |
| Vmn1r209 | 0 | 0 | 0 | 1 | 1 |
| Vmn1r21 | 0 | 0 | 0 | 1 | 1 |
| Vmn1r210 | 0 | 0 | 0 | 1 | 1 |
| Vmn1r211 | 0 | 0 | 0 | 1 | 1 |
| Vmn1r212 | 0 | 0 | 0 | 1 | 1 |
| Vmn1r213 | 0 | 0 | 0 | 1 | 1 |
| Vmn1r214 | 0 | 0 | 0 | 1 | 1 |
| Vmn1r215 | 0 | 0 | 0 | 1 | 1 |
| Vmn1r216 | 0 | 0 | 0 | 1 | 1 |
| Vmn1r217 | 0 | 0 | 0 | 1 | 1 |
| Vmn1r218 | 0 | 0 | 0 | 1 | 1 |
| Vmn1r219 | 0 | 0 | 0 | 1 | 1 |
| Vmn1r22 | 0 | 0 | 0 | 1 | 1 |
| Vmn1r220 | 0 | 0 | 0 | 1 | 1 |
| Vmn1r222 | 0 | 0 | 0 | 1 | 1 |
| Vmn1r223 | 0 | 0 | 0 | 1 | 1 |
| Vmn1r224 | 0 | 0 | 0 | 1 | 1 |
| Vmn1r225 | 0 | 0 | 0 | 1 | 1 |
| Vmn1r226 | 0 | 0 | 0 | 1 | 1 |
| Vmn1r227 | 0 | 0 | 0 | 1 | 1 |
| Vmn1r228 | 0 | 0 | 0 | 1 | 1 |
| Vmn1r229 | 0 | 0 | 0 | 1 | 1 |
| Vmn1r23 | 0 | 0 | 0 | 1 | 1 |
| Vmn1r230 | 0 | 0 | 0 | 1 | 1 |
| Vmn1r231 | 0 | 0 | 0 | 1 | 1 |
| Vmn1r232 | 0 | 0 | 0 | 1 | 1 |
| Vmn1r233 | 0 | 0 | 0 | 1 | 1 |
| Vmn1r234 | 0 | 0 | 0 | 1 | 1 |
| Vmn1r235 | 0 | 0 | 0 | 1 | 1 |
| Vmn1r236 | 0 | 0 | 0 | 1 | 1 |
| Vmn1r237 | 0 | 0 | 0 | 1 | 1 |
| Vmn1r238 | 0 | 0 | 0 | 1 | 1 |
| Vmn1r24 | 0 | 0 | 0 | 1 | 1 |
| Vmn1r25 | 0 | 0 | 0 | 1 | 1 |
| Vmn1r27 | 0 | 0 | 0 | 1 | 1 |
| Vmn1r28 | 0 | 0 | 0 | 1 | 1 |
| Vmn1r29 | 0 | 0 | 0 | 1 | 1 |
| Vmn1r3 | 0 | 0 | 0 | 1 | 1 |
| Vmn1r30 | 0 | 0 | 0 | 1 | 1 |
| Vmn1r31 | 0 | 0 | 0 | 1 | 1 |
| Vmn1r33 | 0 | 0 | 0 | 1 | 1 |
| Vmn1r34 | 0 | 0 | 0 | 1 | 1 |
| Vmn1r35 | 0 | 0 | 0 | 1 | 1 |
| Vmn1r36 | 0 | 0 | 0 | 1 | 1 |
| Vmn1r37 | 0 | 0 | 0 | 1 | 1 |
| Vmn1r38 | 0 | 0 | 0 | 1 | 1 |
| Vmn1r39 | 0 | 0 | 0 | 1 | 1 |
| Vmn1r4 | 0 | 0 | 0 | 1 | 1 |
| Vmn1r40 | 0 | 0 | 0 | 1 | 1 |

| | | | | | |
|---|---|---|---|---|---|
| Vmn1r41 | 0 | 0 | 0 | 1 | 1 |
| Vmn1r42 | 0 | 0 | 0 | 1 | 1 |
| Vmn1r43 | 0 | 0 | 0 | 1 | 1 |
| Vmn1r44 | 0 | 0 | 0 | 1 | 1 |
| Vmn1r45 | 0 | 0 | 0 | 1 | 1 |
| Vmn1r46 | 0 | 0 | 0 | 1 | 1 |
| Vmn1r47 | 0 | 0 | 0 | 1 | 1 |
| Vmn1r48 | 0 | 0 | 0 | 1 | 1 |
| Vmn1r5 | 0 | 0 | 0 | 1 | 1 |
| Vmn1r51 | 0 | 0 | 0 | 1 | 1 |
| Vmn1r52 | 0 | 0 | 0 | 1 | 1 |
| Vmn1r54 | 0 | 0 | 0 | 1 | 1 |
| Vmn1r55 | 0 | 0 | 0 | 1 | 1 |
| Vmn1r56 | 0 | 0 | 0 | 1 | 1 |
| Vmn1r57 | 0 | 0 | 0 | 1 | 1 |
| Vmn1r58 | 0 | 0 | 0 | 1 | 1 |
| Vmn1r59 | 0 | 0 | 0 | 1 | 1 |
| Vmn1r6 | 0 | 0 | 0 | 1 | 1 |
| Vmn1r60 | 0 | 0 | 0 | 1 | 1 |
| Vmn1r61 | 0 | 0 | 0 | 1 | 1 |
| Vmn1r62 | 0 | 0 | 0 | 1 | 1 |
| Vmn1r63 | 0 | 0 | 0 | 1 | 1 |
| Vmn1r64 | 0 | 0 | 0 | 1 | 1 |
| Vmn1r65 | 0 | 0 | 0 | 1 | 1 |
| Vmn1r66 | 0 | 0 | 0 | 1 | 1 |
| Vmn1r67 | 0 | 0 | 0 | 1 | 1 |
| Vmn1r68 | 0 | 0 | 0 | 1 | 1 |
| Vmn1r69 | 0 | 0 | 0 | 1 | 1 |
| Vmn1r7 | 0 | 0 | 0 | 1 | 1 |
| Vmn1r70 | 0 | 0 | 0 | 1 | 1 |
| Vmn1r71 | 0 | 0 | 0 | 1 | 1 |
| Vmn1r72 | 0 | 0 | 0 | 1 | 1 |
| Vmn1r73 | 0 | 0 | 0 | 1 | 1 |
| Vmn1r74 | 0 | 0 | 0 | 1 | 1 |
| Vmn1r75 | 0 | 0 | 0 | 1 | 1 |
| Vmn1r76 | 0 | 0 | 0 | 1 | 1 |
| Vmn1r77 | 0 | 0 | 0 | 1 | 1 |
| Vmn1r78 | 0 | 0 | 0 | 1 | 1 |
| Vmn1r79 | 0 | 0 | 0 | 1 | 1 |
| Vmn1r8 | 0 | 0 | 0 | 1 | 1 |
| Vmn1r80 | 0 | 0 | 0 | 1 | 1 |
| Vmn1r81 | 0 | 0 | 0 | 1 | 1 |
| Vmn1r82 | 0 | 0 | 0 | 1 | 1 |
| Vmn1r83 | 0 | 0 | 0 | 1 | 1 |
| Vmn1r84 | 0 | 0 | 0 | 1 | 1 |
| Vmn1r85 | 0 | 0 | 0 | 1 | 1 |
| Vmn1r86 | 0 | 0 | 0 | 1 | 1 |
| Vmn1r87 | 0 | 0 | 0 | 1 | 1 |
| Vmn1r88 | 0 | 0 | 0 | 1 | 1 |
| Vmn1r89 | 0 | 0 | 0 | 1 | 1 |

| | | | | | |
|---|---|---|---|---|---|
| Vmn1r9 | 0 | 0 | 0 | 1 | 1 |
| Vmn1r91 | 0 | 0 | 0 | 1 | 1 |
| Vmn1r93 | 0 | 0 | 0 | 1 | 1 |
| Vmn1r94 | 0 | 0 | 0 | 1 | 1 |
| Vmn2r-ps11 | 0 | 0 | 0 | 1 | 1 |
| Vmn2r-ps129 | 0 | 0 | 0 | 1 | 1 |
| Vmn2r-ps159 | 0 | 0 | 0 | 1 | 1 |
| Vmn2r-ps54 | 0 | 0 | 0 | 1 | 1 |
| Vmn2r-ps60 | 0 | 0 | 0 | 1 | 1 |
| Vmn2r10 | 0 | 0 | 0 | 1 | 1 |
| Vmn2r100 | 0 | 0 | 0 | 1 | 1 |
| Vmn2r101 | 0 | 0 | 0 | 1 | 1 |
| Vmn2r102 | 0 | 0 | 0 | 1 | 1 |
| Vmn2r103 | 0 | 0 | 0 | 1 | 1 |
| Vmn2r104 | 0 | 0 | 0 | 1 | 1 |
| Vmn2r105 | 0 | 0 | 0 | 1 | 1 |
| Vmn2r106 | 0 | 0 | 0 | 1 | 1 |
| Vmn2r107 | 0 | 0 | 0 | 1 | 1 |
| Vmn2r108 | 0 | 0 | 0 | 1 | 1 |
| Vmn2r11 | 0 | 0 | 0 | 1 | 1 |
| Vmn2r110 | 0 | 0 | 0 | 1 | 1 |
| Vmn2r111 | 0 | 0 | 0 | 1 | 1 |
| Vmn2r112 | 0 | 0 | 0 | 1 | 1 |
| Vmn2r113 | 0 | 0 | 0 | 1 | 1 |
| Vmn2r114 | 0 | 0 | 0 | 1 | 1 |
| Vmn2r115 | 0 | 0 | 0 | 1 | 1 |
| Vmn2r116 | 0 | 0 | 0 | 1 | 1 |
| Vmn2r117 | 0 | 0 | 0 | 1 | 1 |
| Vmn2r12 | 0 | 0 | 0 | 1 | 1 |
| Vmn2r121 | 0 | 0 | 0 | 1 | 1 |
| Vmn2r122 | 0 | 0 | 0 | 1 | 1 |
| Vmn2r123 | 0 | 0 | 0 | 1 | 1 |
| Vmn2r124 | 0 | 0 | 0 | 1 | 1 |
| Vmn2r13 | 0 | 0 | 0 | 1 | 1 |
| Vmn2r14 | 0 | 0 | 0 | 1 | 1 |
| Vmn2r15 | 0 | 0 | 0 | 1 | 1 |
| Vmn2r16 | 0 | 0 | 0 | 1 | 1 |
| Vmn2r17 | 0 | 0 | 0 | 1 | 1 |
| Vmn2r19 | 0 | 0 | 0 | 1 | 1 |
| Vmn2r20 | 0 | 0 | 0 | 1 | 1 |
| Vmn2r21 | 0 | 0 | 0 | 1 | 1 |
| Vmn2r22 | 0 | 0 | 0 | 1 | 1 |
| Vmn2r24 | 0 | 0 | 0 | 1 | 1 |
| Vmn2r25 | 0 | 0 | 0 | 1 | 1 |
| Vmn2r26 | 0 | 0 | 0 | 1 | 1 |
| Vmn2r27 | 0 | 0 | 0 | 1 | 1 |
| Vmn2r30 | 0 | 0 | 0 | 1 | 1 |
| Vmn2r31 | 0 | 0 | 0 | 1 | 1 |
| Vmn2r32 | 0 | 0 | 0 | 1 | 1 |
| Vmn2r33 | 0 | 0 | 0 | 1 | 1 |

| | | | | | |
|---|---|---|---|---|---|
| Vmn2r34 | 0 | 0 | 0 | 1 | 1 |
| Vmn2r35 | 0 | 0 | 0 | 1 | 1 |
| Vmn2r36 | 0 | 0 | 0 | 1 | 1 |
| Vmn2r37 | 0 | 0 | 0 | 1 | 1 |
| Vmn2r38 | 0 | 0 | 0 | 1 | 1 |
| Vmn2r39 | 0 | 0 | 0 | 1 | 1 |
| Vmn2r4 | 0 | 0 | 0 | 1 | 1 |
| Vmn2r40 | 0 | 0 | 0 | 1 | 1 |
| Vmn2r41 | 0 | 0 | 0 | 1 | 1 |
| Vmn2r42 | 0 | 0 | 0 | 1 | 1 |
| Vmn2r43 | 0 | 0 | 0 | 1 | 1 |
| Vmn2r44 | 0 | 0 | 0 | 1 | 1 |
| Vmn2r45 | 0 | 0 | 0 | 1 | 1 |
| Vmn2r46 | 0 | 0 | 0 | 1 | 1 |
| Vmn2r47 | 0 | 0 | 0 | 1 | 1 |
| Vmn2r48 | 0 | 0 | 0 | 1 | 1 |
| Vmn2r49 | 0 | 0 | 0 | 1 | 1 |
| Vmn2r5 | 0 | 0 | 0 | 1 | 1 |
| Vmn2r50 | 0 | 0 | 0 | 1 | 1 |
| Vmn2r51 | 0 | 0 | 0 | 1 | 1 |
| Vmn2r52 | 0 | 0 | 0 | 1 | 1 |
| Vmn2r53 | 0 | 0 | 0 | 1 | 1 |
| Vmn2r54 | 0 | 0 | 0 | 1 | 1 |
| Vmn2r55 | 0 | 0 | 0 | 1 | 1 |
| Vmn2r56 | 0 | 0 | 0 | 1 | 1 |
| Vmn2r58 | 0 | 0 | 0 | 1 | 1 |
| Vmn2r59 | 0 | 0 | 0 | 1 | 1 |
| Vmn2r6 | 0 | 0 | 0 | 1 | 1 |
| Vmn2r60 | 0 | 0 | 0 | 1 | 1 |
| Vmn2r61 | 0 | 0 | 0 | 1 | 1 |
| Vmn2r62 | 0 | 0 | 0 | 1 | 1 |
| Vmn2r63 | 0 | 0 | 0 | 1 | 1 |
| Vmn2r65 | 0 | 0 | 0 | 1 | 1 |
| Vmn2r66 | 0 | 0 | 0 | 1 | 1 |
| Vmn2r67 | 0 | 0 | 0 | 1 | 1 |
| Vmn2r68 | 0 | 0 | 0 | 1 | 1 |
| Vmn2r69 | 0 | 0 | 0 | 1 | 1 |
| Vmn2r7 | 0 | 0 | 0 | 1 | 1 |
| Vmn2r70 | 0 | 0 | 0 | 1 | 1 |
| Vmn2r71 | 0 | 0 | 0 | 1 | 1 |
| Vmn2r72 | 0 | 0 | 0 | 1 | 1 |
| Vmn2r73 | 0 | 0 | 0 | 1 | 1 |
| Vmn2r74 | 0 | 0 | 0 | 1 | 1 |
| Vmn2r75 | 0 | 0 | 0 | 1 | 1 |
| Vmn2r76 | 0 | 0 | 0 | 1 | 1 |
| Vmn2r77 | 0 | 0 | 0 | 1 | 1 |
| Vmn2r78 | 0 | 0 | 0 | 1 | 1 |
| Vmn2r79 | 0 | 0 | 0 | 1 | 1 |
| Vmn2r8 | 0 | 0 | 0 | 1 | 1 |
| Vmn2r80 | 0 | 0 | 0 | 1 | 1 |

| | | | | | |
|---|---|---|---|---|---|
| Vmn2r81 | 0 | 0 | 0 | 1 | 1 |
| Vmn2r82 | 0 | 0 | 0 | 1 | 1 |
| Vmn2r84 | 0 | 0 | 0 | 1 | 1 |
| Vmn2r85 | 0 | 0 | 0 | 1 | 1 |
| Vmn2r86 | 0 | 0 | 0 | 1 | 1 |
| Vmn2r88 | 0 | 0 | 0 | 1 | 1 |
| Vmn2r89 | 0 | 0 | 0 | 1 | 1 |
| Vmn2r9 | 0 | 0 | 0 | 1 | 1 |
| Vmn2r91 | 0 | 0 | 0 | 1 | 1 |
| Vmn2r92 | 0 | 0 | 0 | 1 | 1 |
| Vmn2r93 | 0 | 0 | 0 | 1 | 1 |
| Vmn2r94 | 0 | 0 | 0 | 1 | 1 |
| Vmn2r95 | 0 | 0 | 0 | 1 | 1 |
| Vmn2r96 | 0 | 0 | 0 | 1 | 1 |
| Vmn2r97 | 0 | 0 | 0 | 1 | 1 |
| Vmn2r98 | 0 | 0 | 0 | 1 | 1 |
| Vmn2r99 | 0 | 0 | 0 | 1 | 1 |
| Vnn3 | 0 | 0 | 0 | 1 | 1 |
| Vpreb2 | 0 | 0 | 0 | 1 | 1 |
| Vsig1 | 0 | 0 | 0 | 1 | 1 |
| Vsig4 | 0 | 0 | 0 | 1 | 1 |
| Vsx2 | 0 | 0 | 0 | 1 | 1 |
| Vtcn1 | 0 | 0 | 0 | 1 | 1 |
| Vwc2l | 0 | 0 | 0 | 1 | 1 |
| Wap | 0 | 0 | 0 | 1 | 1 |
| Wbscr28 | 0 | 0 | 0 | 1 | 1 |
| Wdr63 | 0 | 0 | 0 | 1 | 1 |
| Wdr64 | 0 | 0 | 0 | 1 | 1 |
| Wfdc11 | 0 | 0 | 0 | 1 | 1 |
| Wfdc12 | 0 | 0 | 0 | 1 | 1 |
| Wfdc15a | 0 | 0 | 0 | 1 | 1 |
| Wfdc15b | 0 | 0 | 0 | 1 | 1 |
| Wfdc16 | 0 | 0 | 0 | 1 | 1 |
| Wfdc17 | 0 | 0 | 0 | 1 | 1 |
| Wfdc18 | 0 | 0 | 0 | 1 | 1 |
| Wfdc5 | 0 | 0 | 0 | 1 | 1 |
| Wfdc6a | 0 | 0 | 0 | 1 | 1 |
| Wfdc6b | 0 | 0 | 0 | 1 | 1 |
| Wfdc9 | 0 | 0 | 0 | 1 | 1 |
| Xcl1 | 0 | 0 | 0 | 1 | 1 |
| Xcr1 | 0 | 0 | 0 | 1 | 1 |
| Xirp1 | 0 | 0 | 0 | 1 | 1 |
| Xirp2 | 0 | 0 | 0 | 1 | 1 |
| Xkr9 | 0 | 0 | 0 | 1 | 1 |
| Xlr3c | 0 | 0 | 0 | 1 | 1 |
| Xlr4c | 0 | 0 | 0 | 1 | 1 |
| Xlr5a | 0 | 0 | 0 | 1 | 1 |
| Xlr5b | 0 | 0 | 0 | 1 | 1 |
| Xlr5c | 0 | 0 | 0 | 1 | 1 |
| Zar1l | 0 | 0 | 0 | 1 | 1 |

| | | | | | |
|---|---|---|---|---|---|
| Zbbx | 0 | 0 | 0 | 1 | 1 |
| Zc2hc1b | 0 | 0 | 0 | 1 | 1 |
| Zcchc13 | 0 | 0 | 0 | 1 | 1 |
| Zcchc5 | 0 | 0 | 0 | 1 | 1 |
| Zdhhc11 | 0 | 0 | 0 | 1 | 1 |
| Zdhhc22 | 0 | 0 | 0 | 1 | 1 |
| Zdhhc25 | 0 | 0 | 0 | 1 | 1 |
| Zf12 | 0 | 0 | 0 | 1 | 1 |
| Zfp352 | 0 | 0 | 0 | 1 | 1 |
| Zfp474 | 0 | 0 | 0 | 1 | 1 |
| Zfp534 | 0 | 0 | 0 | 1 | 1 |
| Zfp600 | 0 | 0 | 0 | 1 | 1 |
| Zfp616 | 0 | 0 | 0 | 1 | 1 |
| Zfp735 | 0 | 0 | 0 | 1 | 1 |
| Zfp804b | 0 | 0 | 0 | 1 | 1 |
| Zfp91Cntf | 0 | 0 | 0 | 1 | 1 |
| Zfy1 | 0 | 0 | 0 | 1 | 1 |
| Zfy2 | 0 | 0 | 0 | 1 | 1 |
| Zg16 | 0 | 0 | 0 | 1 | 1 |
| Zim3 | 0 | 0 | 0 | 1 | 1 |
| Znrf4 | 0 | 0 | 0 | 1 | 1 |
| Zp2 | 0 | 0 | 0 | 1 | 1 |
| Zp3r | 0 | 0 | 0 | 1 | 1 |
| Zp4-ps | 0 | 0 | 0 | 1 | 1 |
| Zscan4a | 0 | 0 | 0 | 1 | 1 |
| Zscan4b | 0 | 0 | 0 | 1 | 1 |
| Zscan4c | 0 | 0 | 0 | 1 | 1 |
| Zscan4d | 0 | 0 | 0 | 1 | 1 |
| Zscan4e | 0 | 0 | 0 | 1 | 1 |
| Zscan4f | 0 | 0 | 0 | 1 | 1 |
| Zscan5b | 0 | 0 | 0 | 1 | 1 |
| Zswim2 | 0 | 0 | 0 | 1 | 1 |
| Sp9 | 0,186263 | 0,186168 | -0,000733083 | 1 | 1 |
| Il12rb2 | 0,557166 | 0,556516 | -0,00168405 | 1 | 1 |
| Ccna1 | 0,149613 | 0,14942 | -0,00186269 | 1 | 1 |
| Rin1 | 0,467636 | 0,466984 | -0,00201136 | 1 | 1 |
| Znf41-ps | 0,296147 | 0,295234 | -0,0044564 | 1 | 1 |
| Tinagl1 | 0,23288 | 0,23156 | -0,00820287 | 1 | 1 |
| Btg3 | 0,000708577 | 0,000704227 | -0,0088848 | 1 | 1 |
| Qrich2 | 0,24966 | 0,248065 | -0,00924716 | 1 | 1 |
| Pyroxd2 | 0,461452 | 0,45828 | -0,00995362 | 1 | 1 |
| 5830454E08Rik | 0,459826 | 0,456529 | -0,0103813 | 1 | 1 |
| Magi2 | 0,304513 | 0,302092 | -0,0115186 | 1 | 1 |
| Mettl21c | 0,312149 | 0,309406 | -0,0127332 | 1 | 1 |
| Akr1b3 | 0,000266365 | 0,000263965 | -0,0130599 | 1 | 1 |
| Adssl1 | 0,30893 | 0,306056 | -0,0134854 | 1 | 1 |
| 4930562C15Rik | 0,0641875 | 0,0635411 | -0,014603 | 1 | 1 |
| Tusc5 | 0,139374 | 0,137854 | -0,0158221 | 1 | 1 |
| Ncmap | 0,221313 | 0,218707 | -0,0170835 | 1 | 1 |
| Gm14322 | 0,150449 | 0,148372 | -0,0200566 | 1 | 1 |

| | | | | | |
|---|---|---|---|---|---|
| Fscn2 | 0,182797 | 0,18025 | -0,0202498 | 1 | 1 |
| Tssk1 | 0,182237 | 0,17955 | -0,0214295 | 1 | 1 |
| Adck3 | 0,452833 | 0,446022 | -0,0218666 | 1 | 1 |
| Pate2 | 0,560728 | 0,552063 | -0,0224689 | 1 | 1 |
| Itga4 | 0,529569 | 0,521383 | -0,0224755 | 1 | 1 |
| Mucl1 | 1,59E-006 | 1,56E-006 | -0,0227987 | 1 | 1 |
| Frs3os | 0,516458 | 0,507925 | -0,024035 | 1 | 1 |
| Zfp97 | 0,578318 | 0,568296 | -0,0252194 | 1 | 1 |
| Tbxas1 | 0,370257 | 0,363514 | -0,0265153 | 1 | 1 |
| Ptgds | 0,144283 | 0,141637 | -0,0267003 | 1 | 1 |
| EU599041 | 0,0708456 | 0,0695465 | -0,0267003 | 1 | 1 |
| Ankar | 0,0198479 | 0,019484 | -0,0267003 | 1 | 1 |
| Mc4r | 0,14948 | 0,146729 | -0,0268067 | 1 | 1 |
| Ptpro | 0,0327848 | 0,0321442 | -0,0284703 | 1 | 1 |
| Lims2 | 0,084582 | 0,0829087 | -0,0288272 | 1 | 1 |
| Trpm3 | 0,0886172 | 0,0868291 | -0,0294074 | 1 | 1 |
| Gm7457 | 0,0334884 | 0,0327858 | -0,0305898 | 1 | 1 |
| Tmem181c-ps | 0,147682 | 0,14452 | -0,0312277 | 1 | 1 |
| Grid1 | 0,161036 | 0,157533 | -0,0317269 | 1 | 1 |
| Gas6 | 0,512383 | 0,500345 | -0,0342998 | 1 | 1 |
| Ak5 | 0,406543 | 0,396653 | -0,0355329 | 1 | 1 |
| Fam198a | 0,348642 | 0,340109 | -0,0357504 | 1 | 1 |
| Phospho1 | 0,302135 | 0,294695 | -0,0359727 | 1 | 1 |
| Rnf144b | 0,537029 | 0,523789 | -0,0360118 | 1 | 1 |
| Pik3r5 | 0,289715 | 0,282505 | -0,0363576 | 1 | 1 |
| Dnah2 | 0,198049 | 0,192902 | -0,0379924 | 1 | 1 |
| Lrrc19 | 0,335559 | 0,326619 | -0,0389545 | 1 | 1 |
| Epn3 | 0,1564 | 0,152106 | -0,0401591 | 1 | 1 |
| Ushbp1 | 0,127997 | 0,124462 | -0,0404028 | 1 | 1 |
| Cntn1 | 0,0400365 | 0,038921 | -0,0407669 | 1 | 1 |
| 3632454L22Rik | 0,0132221 | 0,0128356 | -0,0428026 | 1 | 1 |
| Chac1 | 0,335644 | 0,325719 | -0,0433045 | 1 | 1 |
| B230217O12Rik | 0,396598 | 0,384848 | -0,043388 | 1 | 1 |
| Ccdc162 | 0,0454305 | 0,0439883 | -0,0465424 | 1 | 1 |
| Syne1 | 0,305322 | 0,295535 | -0,0470021 | 1 | 1 |
| Tmem169 | 0,421355 | 0,407802 | -0,0471687 | 1 | 1 |
| Tmprss7 | 0,335358 | 0,32447 | -0,0476147 | 1 | 1 |
| Cyp24a1 | 0,0142579 | 0,0137682 | -0,0504203 | 1 | 1 |
| Mir6481 | 16,5519 | 15,9833 | -0,0504313 | 1 | 1 |
| Prap1 | 0,105444 | 0,101822 | -0,0504313 | 1 | 1 |
| Gm9895 | 0,095817 | 0,0925254 | -0,0504313 | 1 | 1 |
| Prss38 | 0,042586 | 0,0411231 | -0,0504313 | 1 | 1 |
| Bdkrb1 | 0,0388682 | 0,037533 | -0,0504313 | 1 | 1 |
| Gm5105 | 0,0301688 | 0,0291325 | -0,0504313 | 1 | 1 |
| Sec14l5 | 0,0206642 | 0,0199544 | -0,0504313 | 1 | 1 |
| Slc2a2 | 0,0186292 | 0,0179893 | -0,0504313 | 1 | 1 |
| B020018J22Rik | 0,0145998 | 0,0140983 | -0,0504313 | 1 | 1 |
| Gabrp | 0,0143293 | 0,013837 | -0,0504313 | 1 | 1 |
| Gm15348 | 0,0127722 | 0,0123334 | -0,0504313 | 1 | 1 |
| Slc12a1 | 0,00992723 | 0,00958621 | -0,0504313 | 1 | 1 |

| | | | | | |
|---|---|---|---|---|---|
| Csf2rb | 0,00966387 | 0,00933189 | -0,0504313 | 1 | 1 |
| Treml1 | 0,0474936 | 0,0458618 | -0,0504402 | 1 | 1 |
| Gmnc | 0,0111571 | 0,0107738 | -0,0504408 | 1 | 1 |
| Cd8a | 0,0152151 | 0,014692 | -0,05047 | 1 | 1 |
| Sdpr | 0,0642491 | 0,0619736 | -0,0520227 | 1 | 1 |
| Fam84a | 0,174733 | 0,16836 | -0,053601 | 1 | 1 |
| Spag8 | 0,313125 | 0,301555 | -0,0543165 | 1 | 1 |
| Cxcl16 | 0,359666 | 0,346302 | -0,0546258 | 1 | 1 |
| Hist1h2ae | 0,925677 | 0,891106 | -0,054911 | 1 | 1 |
| Nrsn2 | 0,241175 | 0,232055 | -0,0556084 | 1 | 1 |
| Gm14023 | 0,401657 | 0,385893 | -0,0577622 | 1 | 1 |
| Mast1 | 0,215017 | 0,206371 | -0,0592089 | 1 | 1 |
| Hipk4 | 0,545213 | 0,522419 | -0,0616128 | 1 | 1 |
| Tmco4 | 0,559558 | 0,534665 | -0,065653 | 1 | 1 |
| Ptpre | 0,0922857 | 0,0881292 | -0,0664878 | 1 | 1 |
| Trim47 | 0,0871732 | 0,0832369 | -0,0666612 | 1 | 1 |
| Bcas1 | 0,0631096 | 0,0602577 | -0,0667146 | 1 | 1 |
| Abcb4 | 0,0636562 | 0,0607046 | -0,068496 | 1 | 1 |
| Actl10 | 0,151817 | 0,144774 | -0,0685253 | 1 | 1 |
| Stk32b | 0,346341 | 0,329925 | -0,0700547 | 1 | 1 |
| Acmsd | 0,0351035 | 0,033437 | -0,0701673 | 1 | 1 |
| Slc4a11 | 0,152984 | 0,145717 | -0,0702092 | 1 | 1 |
| Sumo2 | 0,586088 | 0,558117 | -0,0705501 | 1 | 1 |
| Gm9855 | 5,1623 | 4,91413 | -0,0710804 | 1 | 1 |
| Nbeal2 | 0,292728 | 0,278513 | -0,0718176 | 1 | 1 |
| Zxda | 0,431216 | 0,410258 | -0,0718787 | 1 | 1 |
| Bdnf | 0,353925 | 0,336694 | -0,0720049 | 1 | 1 |
| Hrc | 0,222041 | 0,211129 | -0,0727 | 1 | 1 |
| Hist1h3f | 0,417631 | 0,397062 | -0,0728654 | 1 | 1 |
| Hs6st3 | 0,046191 | 0,043916 | -0,0728654 | 1 | 1 |
| Col4a4 | 0,0287936 | 0,0273755 | -0,0728654 | 1 | 1 |
| Plekhg6 | 0,0259261 | 0,0246492 | -0,0728654 | 1 | 1 |
| Gm19461 | 0,025097 | 0,0238609 | -0,0728654 | 1 | 1 |
| Hcar1 | 0,0236878 | 0,0225211 | -0,0728654 | 1 | 1 |
| Asgr1 | 0,154269 | 0,14667 | -0,072876 | 1 | 1 |
| Rspo4 | 0,545198 | 0,51826 | -0,0731035 | 1 | 1 |
| Proca1 | 0,554241 | 0,526524 | -0,074014 | 1 | 1 |
| Trpv4 | 0,197271 | 0,187355 | -0,0744007 | 1 | 1 |
| Usp18 | 0,56419 | 0,535471 | -0,0753736 | 1 | 1 |
| Atp2a1 | 0,240386 | 0,228128 | -0,0755057 | 1 | 1 |
| Grik3 | 0,36423 | 0,345619 | -0,0756668 | 1 | 1 |
| Syt16 | 0,220741 | 0,209366 | -0,0763317 | 1 | 1 |
| Ccdc37 | 0,424348 | 0,402168 | -0,0774517 | 1 | 1 |
| Fgl1 | 0,641748 | 0,607947 | -0,0780607 | 1 | 1 |
| Spn | 0,303754 | 0,287363 | -0,0800275 | 1 | 1 |
| 5730508B09Rik | 0,395647 | 0,374283 | -0,0800851 | 1 | 1 |
| Gm4975 | 0,0782904 | 0,0740629 | -0,0800851 | 1 | 1 |
| 2310043L19Rik | 0,206245 | 0,194757 | -0,0826796 | 1 | 1 |
| Slc9b1 | 0,0499201 | 0,0471397 | -0,0826796 | 1 | 1 |
| Rnf17 | 0,438709 | 0,413818 | -0,0842694 | 1 | 1 |

| | | | | | |
|---|---|---|---|---|---|
| Gm10767 | 0,602834 | 0,568608 | -0,0843273 | 1 | 1 |
| A930015D03Rik | 0,590203 | 0,556628 | -0,084496 | 1 | 1 |
| Pnliprp1 | 0,231589 | 0,218392 | -0,0846483 | 1 | 1 |
| Hpgds | 0,108134 | 0,101959 | -0,0848294 | 1 | 1 |
| Mir6991 | 103,972 | 97,9815 | -0,0856185 | 1 | 1 |
| Zfp879 | 0,572907 | 0,539673 | -0,0862152 | 1 | 1 |
| Wfdc3 | 0,212322 | 0,199732 | -0,0881851 | 1 | 1 |
| Krt14 | 0,0898295 | 0,0845031 | -0,0881851 | 1 | 1 |
| B3gnt4 | 0,560614 | 0,527221 | -0,0885999 | 1 | 1 |
| Rgs7bp | 0,100862 | 0,0947595 | -0,0900395 | 1 | 1 |
| Ksr2 | 0,153451 | 0,144128 | -0,0904266 | 1 | 1 |
| Gpr39 | 0,568442 | 0,533824 | -0,0906498 | 1 | 1 |
| Smim5 | 0,447193 | 0,419867 | -0,0909661 | 1 | 1 |
| Gja8 | 0,202673 | 0,190171 | -0,0918557 | 1 | 1 |
| Lrrc24 | 0,634186 | 0,594775 | -0,0925621 | 1 | 1 |
| Tmem145 | 0,236038 | 0,221365 | -0,0925928 | 1 | 1 |
| 3110070M22Rik | 0,521606 | 0,489075 | -0,0929059 | 1 | 1 |
| Sarm1 | 0,412505 | 0,386702 | -0,0931877 | 1 | 1 |
| Pde3a | 0,586784 | 0,549866 | -0,0937496 | 1 | 1 |
| Kcnj8 | 0,0916006 | 0,0858117 | -0,0941822 | 1 | 1 |
| Drd2 | 0,174064 | 0,16301 | -0,0946616 | 1 | 1 |
| Gbp4 | 0,0599664 | 0,0560987 | -0,0961864 | 1 | 1 |
| Cyp4v3 | 0,447049 | 0,417758 | -0,0977644 | 1 | 1 |
| 6430548M08Rik | 0,541144 | 0,50521 | -0,0991306 | 1 | 1 |
| Gm10451 | 0,549547 | 0,512871 | -0,0996474 | 1 | 1 |
| Nron | 0,116173 | 0,108368 | -0,100337 | 1 | 1 |
| Slc12a8 | 0,326192 | 0,304251 | -0,100457 | 1 | 1 |
| 1700061G19Rik | 0,200633 | 0,187017 | -0,101389 | 1 | 1 |
| Zfp641 | 0,265902 | 0,247759 | -0,101958 | 1 | 1 |
| Tmem200a | 0,107717 | 0,100346 | -0,102271 | 1 | 1 |
| 4921507P07Rik | 0,320594 | 0,298391 | -0,103544 | 1 | 1 |
| Mapt | 0,438187 | 0,407813 | -0,103641 | 1 | 1 |
| Syt3 | 0,511406 | 0,475815 | -0,104066 | 1 | 1 |
| Gm4787 | 0,453521 | 0,421803 | -0,104601 | 1 | 1 |
| 1700001P01Rik | 0,504937 | 0,469215 | -0,105855 | 1 | 1 |
| Cdc42ep2 | 0,430077 | 0,399626 | -0,105945 | 1 | 1 |
| A330102I10Rik | 0,134328 | 0,12476 | -0,1066 | 1 | 1 |
| 6330409D20Rik | 0,0316266 | 0,0293531 | -0,107626 | 1 | 1 |
| Kcnk16 | 0,128466 | 0,11922 | -0,107761 | 1 | 1 |
| Tnni2 | 0,0624287 | 0,0579356 | -0,107761 | 1 | 1 |
| Fxyd7 | 0,06098 | 0,0565911 | -0,107761 | 1 | 1 |
| Il17b | 0,0592306 | 0,0549676 | -0,107761 | 1 | 1 |
| Gkn2 | 0,0504444 | 0,0468138 | -0,107761 | 1 | 1 |
| Sox5os3 | 0,0412925 | 0,0383206 | -0,107761 | 1 | 1 |
| 5033406O09Rik | 0,0379083 | 0,0351799 | -0,107761 | 1 | 1 |
| Gm6756 | 0,034026 | 0,0315771 | -0,107761 | 1 | 1 |
| Nkx2-4 | 0,0335713 | 0,0311551 | -0,107761 | 1 | 1 |
| Sbk2 | 0,0309143 | 0,0286893 | -0,107761 | 1 | 1 |
| Acsm4 | 0,0296615 | 0,0275267 | -0,107761 | 1 | 1 |
| Mael | 0,0215388 | 0,0199886 | -0,107761 | 1 | 1 |

| | | | | | |
|---|---|---|---|---|---|
| 5330411J11Rik | 0,0200729 | 0,0186282 | -0,107761 | 1 | 1 |
| Nepn | 0,0178537 | 0,0165687 | -0,107761 | 1 | 1 |
| Cyp17a1 | 0,0176573 | 0,0163865 | -0,107761 | 1 | 1 |
| Serpina3b | 0,0154904 | 0,0143755 | -0,107761 | 1 | 1 |
| Was | 0,0152937 | 0,014193 | -0,107761 | 1 | 1 |
| Gm19424 | 0,014211 | 0,0131882 | -0,107761 | 1 | 1 |
| 2010001E11Rik | 0,01389 | 0,0128903 | -0,107761 | 1 | 1 |
| Foxd4 | 0,0135008 | 0,0125291 | -0,107761 | 1 | 1 |
| AY512931 | 0,0131927 | 0,0122432 | -0,107761 | 1 | 1 |
| Scg2 | 0,0126721 | 0,0117601 | -0,107761 | 1 | 1 |
| Krt2 | 0,0119988 | 0,0111352 | -0,107761 | 1 | 1 |
| Cd4 | 0,00999833 | 0,00927872 | -0,107761 | 1 | 1 |
| Stk31 | 0,00952708 | 0,0088414 | -0,107761 | 1 | 1 |
| Gpr88 | 0,00890486 | 0,00826396 | -0,107761 | 1 | 1 |
| Mroh2b | 0,00575114 | 0,00533721 | -0,107761 | 1 | 1 |
| Ndst3 | 0,00563016 | 0,00522494 | -0,107761 | 1 | 1 |
| Spt1 | 0,0665978 | 0,0618045 | -0,107764 | 1 | 1 |
| Cnfn | 0,0886617 | 0,0822802 | -0,107766 | 1 | 1 |
| Adam33 | 0,0601527 | 0,0558231 | -0,107768 | 1 | 1 |
| St8sia5 | 0,0161364 | 0,0149749 | -0,107775 | 1 | 1 |
| 4930451C15Rik | 0,014966 | 0,0138886 | -0,10779 | 1 | 1 |
| Proc | 0,0211511 | 0,0196267 | -0,107912 | 1 | 1 |
| Mpl | 0,0115748 | 0,010739 | -0,108132 | 1 | 1 |
| Gm12657 | 0,289886 | 0,26894 | -0,108199 | 1 | 1 |
| 4732491K20Rik | 0,465436 | 0,431729 | -0,108455 | 1 | 1 |
| Mag | 0,213074 | 0,197255 | -0,111291 | 1 | 1 |
| Zfp456 | 0,403634 | 0,373635 | -0,111417 | 1 | 1 |
| Entpd3 | 0,0856511 | 0,0791752 | -0,113422 | 1 | 1 |
| Fibcd1 | 0,220423 | 0,20365 | -0,114182 | 1 | 1 |
| Lax1 | 0,0565714 | 0,0522447 | -0,114789 | 1 | 1 |
| Hivep2 | 0,571068 | 0,527377 | -0,114829 | 1 | 1 |
| Gm16863 | 0,126279 | 0,116568 | -0,115444 | 1 | 1 |
| Ttc30a2 | 0,141196 | 0,130259 | -0,116318 | 1 | 1 |
| Cd274 | 0,182439 | 0,168105 | -0,118047 | 1 | 1 |
| Efhc1 | 0,458294 | 0,421916 | -0,119317 | 1 | 1 |
| Fbxo24 | 0,582509 | 0,535676 | -0,12092 | 1 | 1 |
| Pllp | 0,18633 | 0,171297 | -0,121355 | 1 | 1 |
| Otud7a | 0,168279 | 0,154509 | -0,123165 | 1 | 1 |
| Vsnl1 | 0,503987 | 0,46257 | -0,123714 | 1 | 1 |
| 4933431G14Rik | 0,110147 | 0,101034 | -0,12459 | 1 | 1 |
| Hist2h2ac | 0,443385 | 0,406418 | -0,125595 | 1 | 1 |
| 4930525G20Rik | 0,398298 | 0,364992 | -0,125984 | 1 | 1 |
| Dram1 | 0,203625 | 0,186597 | -0,125988 | 1 | 1 |
| Pcdh12 | 0,22096 | 0,202455 | -0,126188 | 1 | 1 |
| Gm15787 | 0,498409 | 0,456369 | -0,127129 | 1 | 1 |
| Kiss1r | 0,10671 | 0,0976881 | -0,127445 | 1 | 1 |
| Gm2011 | 0,14941 | 0,136395 | -0,131492 | 1 | 1 |
| Sds | 0,0918259 | 0,0838267 | -0,131492 | 1 | 1 |
| Grk4 | 0,472562 | 0,431162 | -0,132272 | 1 | 1 |
| Csmd2 | 0,0741495 | 0,0675635 | -0,134193 | 1 | 1 |

| | | | | | |
|---|---|---|---|---|---|
| Gm973 | 0,126685 | 0,115389 | -0,134731 | 1 | 1 |
| Disc1 | 0,218676 | 0,198953 | -0,136369 | 1 | 1 |
| Grap | 0,622645 | 0,566368 | -0,136669 | 1 | 1 |
| Efcab5 | 0,0912609 | 0,0829465 | -0,137817 | 1 | 1 |
| Akap6 | 0,314714 | 0,28602 | -0,137923 | 1 | 1 |
| Gm13283 | 0,251776 | 0,228621 | -0,139188 | 1 | 1 |
| H2-Oa | 0,180863 | 0,164229 | -0,139188 | 1 | 1 |
| F7 | 0,0923446 | 0,0838517 | -0,139188 | 1 | 1 |
| Coro6 | 0,139949 | 0,127078 | -0,139191 | 1 | 1 |
| 4833418N02Rik | 0,319877 | 0,290232 | -0,140311 | 1 | 1 |
| Psmb11 | 0,052456 | 0,0475946 | -0,140311 | 1 | 1 |
| Fzd5 | 0,236265 | 0,214063 | -0,142367 | 1 | 1 |
| Mitf | 0,582075 | 0,527113 | -0,143094 | 1 | 1 |
| Atp6v0a4 | 0,0856022 | 0,0774541 | -0,144306 | 1 | 1 |
| Tmem178 | 0,522334 | 0,472493 | -0,144681 | 1 | 1 |
| Siglecg | 0,108707 | 0,0982018 | -0,146617 | 1 | 1 |
| Nr3c2 | 0,0485714 | 0,0438452 | -0,147689 | 1 | 1 |
| Medag | 0,132922 | 0,119803 | -0,149918 | 1 | 1 |
| Gm12250 | 0,318053 | 0,286649 | -0,149985 | 1 | 1 |
| Mylk3 | 0,188339 | 0,169474 | -0,152267 | 1 | 1 |
| Hhipl1 | 0,359038 | 0,322511 | -0,154788 | 1 | 1 |
| Gper1 | 0,582464 | 0,522145 | -0,157718 | 1 | 1 |
| Sycp1 | 0,0625847 | 0,0560896 | -0,158076 | 1 | 1 |
| Ttc9b | 0,135742 | 0,121606 | -0,158648 | 1 | 1 |
| B930092H01Rik | 0,0384076 | 0,034408 | -0,158648 | 1 | 1 |
| Slc4a5 | 0,122565 | 0,109627 | -0,160948 | 1 | 1 |
| Tff3 | 0,54194 | 0,484375 | -0,162007 | 1 | 1 |
| Rbm46 | 0,295172 | 0,263439 | -0,164087 | 1 | 1 |
| Crip3 | 0,120107 | 0,10715 | -0,164686 | 1 | 1 |
| Hsd3b3 | 0,0371233 | 0,0330952 | -0,165706 | 1 | 1 |
| Ccdc62 | 0,46805 | 0,416765 | -0,167428 | 1 | 1 |
| Fhl4 | 0,476309 | 0,423951 | -0,168001 | 1 | 1 |
| Jsrp1 | 0,52094 | 0,463549 | -0,168396 | 1 | 1 |
| Armc2 | 0,100458 | 0,0893211 | -0,169524 | 1 | 1 |
| 4932411E22Rik | 0,124786 | 0,110889 | -0,170341 | 1 | 1 |
| Adamts16 | 0,114601 | 0,101827 | -0,170504 | 1 | 1 |
| Nr0b1 | 0,362885 | 0,322349 | -0,170888 | 1 | 1 |
| Nacad | 0,0886872 | 0,0787187 | -0,17202 | 1 | 1 |
| Itpka | 0,278177 | 0,246842 | -0,172414 | 1 | 1 |
| Shank2 | 0,459113 | 0,407352 | -0,172572 | 1 | 1 |
| Cdkn2c | 0,467049 | 0,414366 | -0,172666 | 1 | 1 |
| 6820408C15Rik | 0,165168 | 0,146479 | -0,173239 | 1 | 1 |
| Dkkl1 | 0,259991 | 0,230252 | -0,175243 | 1 | 1 |
| Cdkn2b | 0,178367 | 0,157925 | -0,17561 | 1 | 1 |
| Ccdc68 | 0,269949 | 0,238804 | -0,176863 | 1 | 1 |
| Pdgfb | 0,302189 | 0,26727 | -0,177149 | 1 | 1 |
| Ccm2l | 0,235276 | 0,208058 | -0,177366 | 1 | 1 |
| Mfsd2b | 0,262592 | 0,232159 | -0,177707 | 1 | 1 |
| Zfp433 | 0,291192 | 0,256766 | -0,181516 | 1 | 1 |
| Rplp2-ps1 | 506,575 | 446,446 | -0,182291 | 1 | 1 |

| | | | | | |
|---|---|---|---|---|---|
| Raet1a,Raet1b | 0,27437 | 0,241618 | -0,183395 | 1 | 1 |
| Nptx1 | 0,215498 | 0,189395 | -0,186282 | 1 | 1 |
| Tbc1d22bos | 0,290848 | 0,255545 | -0,186686 | 1 | 1 |
| Ankrd29 | 0,429018 | 0,376768 | -0,18736 | 1 | 1 |
| Srp54a | 0,459545 | 0,403566 | -0,187403 | 1 | 1 |
| Pkdrej | 0,0843929 | 0,0741009 | -0,187631 | 1 | 1 |
| Zfp575 | 0,292517 | 0,256725 | -0,188294 | 1 | 1 |
| Klb | 0,294171 | 0,25794 | -0,189621 | 1 | 1 |
| Ttn | 0,00812101 | 0,00711194 | -0,191415 | 1 | 1 |
| Fam26e | 0,166259 | 0,145598 | -0,191447 | 1 | 1 |
| Sp7 | 0,513642 | 0,449662 | -0,191923 | 1 | 1 |
| Synpo2l | 0,149261 | 0,130638 | -0,192258 | 1 | 1 |
| G6pc2 | 0,453198 | 0,39665 | -0,192274 | 1 | 1 |
| Exoc3l | 0,628065 | 0,549525 | -0,192729 | 1 | 1 |
| H2-K2 | 0,404518 | 0,353512 | -0,194447 | 1 | 1 |
| Gm16523 | 0,32045 | 0,27962 | -0,196634 | 1 | 1 |
| Gm6644 | 11,3923 | 9,93053 | -0,198117 | 1 | 1 |
| Prss16 | 0,138547 | 0,12063 | -0,199787 | 1 | 1 |
| Mab21l3 | 0,584178 | 0,508347 | -0,200594 | 1 | 1 |
| Ocel1 | 0,271971 | 0,236385 | -0,202315 | 1 | 1 |
| Alms1-ps2 | 0,0826584 | 0,0718291 | -0,202594 | 1 | 1 |
| Pik3r6 | 0,052803 | 0,0458654 | -0,203215 | 1 | 1 |
| C130074G19Rik | 0,457329 | 0,396419 | -0,206206 | 1 | 1 |
| Cdkl4 | 0,250937 | 0,217459 | -0,206579 | 1 | 1 |
| Kcnj12 | 0,0293421 | 0,0254066 | -0,207765 | 1 | 1 |
| Prr9 | 0,126583 | 0,1096 | -0,207836 | 1 | 1 |
| Dlk2 | 0,102315 | 0,0885834 | -0,207913 | 1 | 1 |
| Slc30a10 | 0,500003 | 0,432805 | -0,20822 | 1 | 1 |
| AA414768 | 0,391884 | 0,339111 | -0,208672 | 1 | 1 |
| 1700007J10Rik | 0,538954 | 0,466104 | -0,209508 | 1 | 1 |
| Rps15a-ps4 | 0,488802 | 0,422227 | -0,21123 | 1 | 1 |
| Cass4 | 0,168032 | 0,145037 | -0,212311 | 1 | 1 |
| E130309D14Rik | 0,0424265 | 0,036539 | -0,215529 | 1 | 1 |
| Gm16325 | 0,0542611 | 0,046703 | -0,216401 | 1 | 1 |
| Agap2 | 0,318043 | 0,27334 | -0,218524 | 1 | 1 |
| Lbp | 0,605695 | 0,52055 | -0,218557 | 1 | 1 |
| Gem | 0,598885 | 0,514636 | -0,218726 | 1 | 1 |
| Cd80 | 0,195138 | 0,167596 | -0,219506 | 1 | 1 |
| Gnmt | 0,299334 | 0,257018 | -0,219885 | 1 | 1 |
| Kcnh6 | 0,101341 | 0,0869521 | -0,22092 | 1 | 1 |
| Cacna2d3 | 0,172735 | 0,148149 | -0,221512 | 1 | 1 |
| Esr2 | 0,0589738 | 0,0505397 | -0,222658 | 1 | 1 |
| Zfp78 | 0,314938 | 0,26966 | -0,223927 | 1 | 1 |
| Hist1h4j | 1,18099 | 1,01085 | -0,224421 | 1 | 1 |
| Grpr | 0,0571395 | 0,0489078 | -0,224421 | 1 | 1 |
| Megf11 | 0,36161 | 0,309175 | -0,226009 | 1 | 1 |
| Galnt14 | 0,290251 | 0,248137 | -0,22616 | 1 | 1 |
| Aldh1a7 | 0,286354 | 0,244739 | -0,226556 | 1 | 1 |
| Prr19 | 0,327459 | 0,279497 | -0,228479 | 1 | 1 |
| Rab9b | 0,053023 | 0,0452485 | -0,228749 | 1 | 1 |

| | | | | | |
|---|---|---|---|---|---|
| Prdm16 | 0,506325 | 0,431356 | -0,231187 | 1 | 1 |
| Zfp354b | 0,285572 | 0,243181 | -0,231828 | 1 | 1 |
| Sncg | 0,273718 | 0,23302 | -0,232234 | 1 | 1 |
| Tmem91 | 0,18925 | 0,161104 | -0,232298 | 1 | 1 |
| Calca | 0,287619 | 0,244809 | -0,232505 | 1 | 1 |
| Cyyr1 | 0,57334 | 0,487832 | -0,233006 | 1 | 1 |
| Parvg | 0,0741411 | 0,0630274 | -0,234294 | 1 | 1 |
| Rpl34-ps1 | 760,043 | 645,678 | -0,235266 | 1 | 1 |
| Itgb4 | 0,276128 | 0,23441 | -0,236302 | 1 | 1 |
| Ms4a4d | 0,249178 | 0,211477 | -0,236675 | 1 | 1 |
| Chrnb1 | 0,294434 | 0,249723 | -0,237614 | 1 | 1 |
| Trim72 | 0,0861857 | 0,0730157 | -0,239242 | 1 | 1 |
| C130050O18Rik | 0,181853 | 0,154023 | -0,239628 | 1 | 1 |
| S1pr4 | 0,0868162 | 0,073526 | -0,239708 | 1 | 1 |
| Clnk | 0,0687554 | 0,0582301 | -0,239708 | 1 | 1 |
| B830017H08Rik | 0,144513 | 0,122349 | -0,240198 | 1 | 1 |
| Lrrn4cl | 0,278985 | 0,236038 | -0,241164 | 1 | 1 |
| Ly6g6e | 0,547556 | 0,463086 | -0,241727 | 1 | 1 |
| Bco2 | 0,131701 | 0,111257 | -0,243369 | 1 | 1 |
| Plin4 | 0,0592936 | 0,0500001 | -0,245946 | 1 | 1 |
| Opcml | 0,0820842 | 0,0692101 | -0,246122 | 1 | 1 |
| Wdr20rt | 0,144233 | 0,121578 | -0,24652 | 1 | 1 |
| Syn3 | 0,418218 | 0,351742 | -0,24974 | 1 | 1 |
| Prss12 | 0,206883 | 0,173995 | -0,249772 | 1 | 1 |
| Mmrn2 | 0,356173 | 0,299366 | -0,250665 | 1 | 1 |
| Tm4sf4 | 0,298662 | 0,251023 | -0,250694 | 1 | 1 |
| Gm15612 | 0,364799 | 0,306248 | -0,252398 | 1 | 1 |
| Samd15 | 0,402307 | 0,337272 | -0,254386 | 1 | 1 |
| Gpr146 | 0,264625 | 0,22173 | -0,255147 | 1 | 1 |
| Sh2d5 | 0,446948 | 0,374357 | -0,25569 | 1 | 1 |
| Itgb3 | 0,155344 | 0,130108 | -0,255757 | 1 | 1 |
| Gm10684 | 0,0160275 | 0,013414 | -0,256812 | 1 | 1 |
| 4933413J09Rik | 0,118506 | 0,0991083 | -0,257886 | 1 | 1 |
| Fam124a | 0,373692 | 0,312332 | -0,258771 | 1 | 1 |
| Otof | 0,0075967 | 0,00634807 | -0,259055 | 1 | 1 |
| Fndc7 | 0,0109298 | 0,00913255 | -0,259183 | 1 | 1 |
| Mb | 0,0666731 | 0,0557067 | -0,259254 | 1 | 1 |
| Rab17 | 0,0375941 | 0,0314103 | -0,259267 | 1 | 1 |
| Foxg1 | 0,0185717 | 0,0155166 | -0,259288 | 1 | 1 |
| Hist1h1t | 0,191874 | 0,160308 | -0,259311 | 1 | 1 |
| Tmem89 | 0,152521 | 0,127429 | -0,259311 | 1 | 1 |
| Tex29 | 0,0892166 | 0,0745393 | -0,259311 | 1 | 1 |
| Rfpl3s | 0,0866372 | 0,0723843 | -0,259311 | 1 | 1 |
| Clec4a1 | 0,0752486 | 0,0628693 | -0,259311 | 1 | 1 |
| Tktl1 | 0,0449436 | 0,0375498 | -0,259311 | 1 | 1 |
| Gm3230 | 0,0269828 | 0,0225438 | -0,259311 | 1 | 1 |
| Kcnh8 | 0,012477 | 0,0104244 | -0,259311 | 1 | 1 |
| Sorcs3 | 0,448332 | 0,374333 | -0,260242 | 1 | 1 |
| Tacstd2 | 0,127144 | 0,106153 | -0,260317 | 1 | 1 |
| 4930564C03Rik | 0,0521433 | 0,0435294 | -0,260493 | 1 | 1 |

| | | | | | |
|---|---|---|---|---|---|
| C130060C02Rik | 0,54327 | 0,453367 | -0,260989 | 1 | 1 |
| Slc30a2 | 0,16653 | 0,138972 | -0,260989 | 1 | 1 |
| A430035B10Rik | 0,251163 | 0,209396 | -0,262389 | 1 | 1 |
| Gm12522 | 0,212573 | 0,177152 | -0,262971 | 1 | 1 |
| AF529169 | 0,176181 | 0,146785 | -0,263355 | 1 | 1 |
| Phtf1os | 0,607165 | 0,505848 | -0,263386 | 1 | 1 |
| Klf2 | 0,421829 | 0,351231 | -0,264237 | 1 | 1 |
| Hs3st4 | 0,0602827 | 0,0501773 | -0,264709 | 1 | 1 |
| Bsn | 0,320422 | 0,266665 | -0,264945 | 1 | 1 |
| Pnma3 | 0,234433 | 0,195091 | -0,265027 | 1 | 1 |
| 9030625G05Rik | 0,137152 | 0,11413 | -0,265106 | 1 | 1 |
| Cxcl5 | 0,0796282 | 0,0662616 | -0,265106 | 1 | 1 |
| Atp8b3 | 0,287016 | 0,238813 | -0,265249 | 1 | 1 |
| BC021891 | 0,104351 | 0,0868097 | -0,265511 | 1 | 1 |
| Unc13a | 0,0291147 | 0,0242102 | -0,266134 | 1 | 1 |
| Wipf3 | 0,406615 | 0,337475 | -0,268883 | 1 | 1 |
| Gpr155 | 0,307019 | 0,254357 | -0,271473 | 1 | 1 |
| Samsn1 | 0,220617 | 0,182646 | -0,272491 | 1 | 1 |
| Stat5a | 0,383283 | 0,317083 | -0,27355 | 1 | 1 |
| Ly6c1 | 0,102483 | 0,0846682 | -0,275494 | 1 | 1 |
| Tnfrsf26 | 0,0888615 | 0,0733903 | -0,275969 | 1 | 1 |
| Myo1h | 0,0684662 | 0,0565091 | -0,276909 | 1 | 1 |
| Fam228b | 0,412761 | 0,340464 | -0,277805 | 1 | 1 |
| Gpr18 | 0,164317 | 0,135498 | -0,278211 | 1 | 1 |
| Crb1 | 0,0936601 | 0,0771316 | -0,280111 | 1 | 1 |
| Sqrdl | 0,403383 | 0,332123 | -0,280434 | 1 | 1 |
| 1700019N19Rik | 0,162806 | 0,133812 | -0,282941 | 1 | 1 |
| Pcnxl2 | 0,0166466 | 0,0136821 | -0,282941 | 1 | 1 |
| Plscr4 | 0,197858 | 0,162618 | -0,282975 | 1 | 1 |
| 9530091C08Rik | 0,0719083 | 0,0590363 | -0,284556 | 1 | 1 |
| Slc46a3 | 0,170791 | 0,140194 | -0,284805 | 1 | 1 |
| Mboat4 | 0,0799951 | 0,0656262 | -0,285639 | 1 | 1 |
| Actg2 | 0,169758 | 0,139075 | -0,287623 | 1 | 1 |
| Gm13212 | 0,129623 | 0,106194 | -0,287623 | 1 | 1 |
| Cacng8 | 0,587727 | 0,481448 | -0,287766 | 1 | 1 |
| Mvp | 0,490797 | 0,40186 | -0,288434 | 1 | 1 |
| Rgs11 | 0,206313 | 0,168909 | -0,288588 | 1 | 1 |
| Pfn4 | 0,633424 | 0,517998 | -0,290226 | 1 | 1 |
| Nos3 | 0,285488 | 0,233394 | -0,290664 | 1 | 1 |
| Ankrd55 | 0,305633 | 0,249502 | -0,292748 | 1 | 1 |
| Psd | 0,320735 | 0,261573 | -0,294168 | 1 | 1 |
| Vwa5b1 | 0,399573 | 0,32516 | -0,297311 | 1 | 1 |
| Adam5 | 0,108034 | 0,0877917 | -0,299325 | 1 | 1 |
| Aqp3 | 0,173116 | 0,140634 | -0,299797 | 1 | 1 |
| Filip1 | 0,0674219 | 0,0547713 | -0,299797 | 1 | 1 |
| Gm11149 | 0,257318 | 0,20895 | -0,300392 | 1 | 1 |
| Tmem132cos | 0,554781 | 0,450348 | -0,300879 | 1 | 1 |
| Morn5 | 0,136037 | 0,110388 | -0,301417 | 1 | 1 |
| Krt28 | 0,0514513 | 0,0417504 | -0,301417 | 1 | 1 |
| Rhoh | 0,0278029 | 0,0225608 | -0,301417 | 1 | 1 |

| | | | | | |
|---|---|---|---|---|---|
| Nlrp3 | 0,0173213 | 0,0140554 | -0,301417 | 1 | 1 |
| Tfr2 | 0,0203336 | 0,0164992 | -0,301465 | 1 | 1 |
| Smco3 | 0,446554 | 0,361923 | -0,303154 | 1 | 1 |
| Lrrc48 | 0,609439 | 0,493899 | -0,303264 | 1 | 1 |
| Ap5b1 | 0,306623 | 0,247939 | -0,306479 | 1 | 1 |
| Ppp1r14d | 0,38862 | 0,314106 | -0,307111 | 1 | 1 |
| Rap1gap2 | 0,242069 | 0,195432 | -0,308754 | 1 | 1 |
| Acoxl | 0,141967 | 0,114556 | -0,309502 | 1 | 1 |
| Ikzf1 | 0,242688 | 0,195732 | -0,310221 | 1 | 1 |
| Milr1 | 0,0629932 | 0,0507008 | -0,313188 | 1 | 1 |
| Car7 | 0,114649 | 0,0922726 | -0,313249 | 1 | 1 |
| Cpne7 | 0,525456 | 0,422829 | -0,313498 | 1 | 1 |
| Krt15 | 0,346995 | 0,278865 | -0,315347 | 1 | 1 |
| Cct6b | 0,132545 | 0,106402 | -0,316951 | 1 | 1 |
| Prkcb | 0,2203 | 0,176657 | -0,318524 | 1 | 1 |
| Dlgap3 | 0,392902 | 0,314704 | -0,320174 | 1 | 1 |
| Camk2b | 0,127855 | 0,102394 | -0,320376 | 1 | 1 |
| Plcxd2 | 0,563657 | 0,451083 | -0,321423 | 1 | 1 |
| Xlr4b | 0,159991 | 0,12802 | -0,321614 | 1 | 1 |
| Tbx20 | 0,196528 | 0,157157 | -0,322534 | 1 | 1 |
| Fgl2 | 0,244452 | 0,195136 | -0,325069 | 1 | 1 |
| Gucy1b3 | 0,440164 | 0,351162 | -0,325905 | 1 | 1 |
| A230057D06Rik | 0,294464 | 0,234866 | -0,326249 | 1 | 1 |
| Dzank1 | 0,467889 | 0,373158 | -0,326381 | 1 | 1 |
| Gm20319 | 0,360122 | 0,286566 | -0,329619 | 1 | 1 |
| Slc38a5 | 0,355642 | 0,282946 | -0,329899 | 1 | 1 |
| Gm10389 | 0,168914 | 0,134185 | -0,332063 | 1 | 1 |
| A930012L18Rik | 0,275146 | 0,21849 | -0,332629 | 1 | 1 |
| Kcnmb1 | 0,128232 | 0,101717 | -0,334197 | 1 | 1 |
| Slc9a9 | 0,193066 | 0,153052 | -0,335069 | 1 | 1 |
| Myo18b | 0,0611671 | 0,0484496 | -0,336272 | 1 | 1 |
| Tdrd9 | 0,333684 | 0,263999 | -0,337949 | 1 | 1 |
| 4931406H21Rik | 0,110748 | 0,0876062 | -0,338175 | 1 | 1 |
| Itgae | 0,276206 | 0,218407 | -0,338721 | 1 | 1 |
| Ipcef1 | 0,0190195 | 0,0150223 | -0,340371 | 1 | 1 |
| Gm6568 | 0,690355 | 0,545105 | -0,340806 | 1 | 1 |
| Spata9 | 0,334064 | 0,26372 | -0,341118 | 1 | 1 |
| Spdya | 0,460185 | 0,363027 | -0,342137 | 1 | 1 |
| Astn2 | 0,072973 | 0,0575465 | -0,342634 | 1 | 1 |
| Slc22a7 | 0,0488927 | 0,0384844 | -0,345345 | 1 | 1 |
| Gpr55 | 0,0375761 | 0,0295769 | -0,345345 | 1 | 1 |
| Gucy2g | 0,0239223 | 0,0188297 | -0,345345 | 1 | 1 |
| Rdh1 | 0,0239158 | 0,0188246 | -0,345345 | 1 | 1 |
| Sycp2 | 0,0160777 | 0,0126551 | -0,345345 | 1 | 1 |
| Bace2 | 0,207761 | 0,163441 | -0,346152 | 1 | 1 |
| Zfyve28 | 0,307428 | 0,241823 | -0,346293 | 1 | 1 |
| Pde1a | 0,062598 | 0,0492263 | -0,346686 | 1 | 1 |
| 4930579K19Rik | 0,698587 | 0,547435 | -0,351753 | 1 | 1 |
| Casp4 | 0,0885613 | 0,0693199 | -0,353406 | 1 | 1 |
| Grin2c | 0,542143 | 0,424 | -0,354609 | 1 | 1 |

| | | | | | |
|---|---|---|---|---|---|
| Mdga2 | 0,0595688 | 0,0465719 | -0,355098 | 1 | 1 |
| Dlx3 | 0,182455 | 0,142619 | -0,355371 | 1 | 1 |
| Shank1 | 0,294336 | 0,229784 | -0,357185 | 1 | 1 |
| Kdf1 | 0,366735 | 0,286029 | -0,358578 | 1 | 1 |
| Phf19 | 0,104645 | 0,0815589 | -0,35959 | 1 | 1 |
| Scn3a | 0,137167 | 0,106887 | -0,359846 | 1 | 1 |
| Gm17769 | 0,45683 | 0,355544 | -0,361628 | 1 | 1 |
| Gm1141 | 0,343464 | 0,266961 | -0,36353 | 1 | 1 |
| Slc7a8 | 0,35734 | 0,277542 | -0,364589 | 1 | 1 |
| Ceacam2 | 0,00953286 | 0,00740403 | -0,364598 | 1 | 1 |
| Pmfbp1 | 0,0387995 | 0,0301228 | -0,365181 | 1 | 1 |
| Csf2rb2 | 0,0588983 | 0,0457265 | -0,365194 | 1 | 1 |
| Pgpep1l | 0,339167 | 0,262414 | -0,370152 | 1 | 1 |
| A930001C03Rik | 0,264327 | 0,204457 | -0,370521 | 1 | 1 |
| Mcf2 | 0,174748 | 0,13516 | -0,370611 | 1 | 1 |
| Kcnv1 | 0,0660819 | 0,0510833 | -0,371403 | 1 | 1 |
| G6b | 0,253054 | 0,195594 | -0,37158 | 1 | 1 |
| Amd2 | 7,5601 | 5,8396 | -0,372536 | 1 | 1 |
| Rab39 | 0,503623 | 0,388817 | -0,373251 | 1 | 1 |
| Mir546 | 506,551 | 390,718 | -0,374578 | 1 | 1 |
| Fam163a | 0,29164 | 0,224512 | -0,377398 | 1 | 1 |
| Amz1 | 0,249749 | 0,192023 | -0,379198 | 1 | 1 |
| B930059L03Rik | 0,158682 | 0,121709 | -0,382709 | 1 | 1 |
| Slc39a5 | 0,106493 | 0,0816165 | -0,383822 | 1 | 1 |
| 9230116N13Rik | 0,33549 | 0,256583 | -0,386843 | 1 | 1 |
| Tescl | 0,716758 | 0,548031 | -0,387229 | 1 | 1 |
| Wisp2 | 0,13408 | 0,102449 | -0,388186 | 1 | 1 |
| A530072M11Rik | 0,208052 | 0,158866 | -0,389137 | 1 | 1 |
| Bmp6 | 0,563574 | 0,430304 | -0,389251 | 1 | 1 |
| Masp2 | 0,336557 | 0,256692 | -0,390813 | 1 | 1 |
| Syce1l | 0,0512369 | 0,039072 | -0,391046 | 1 | 1 |
| Catsperd | 0,475305 | 0,36206 | -0,392627 | 1 | 1 |
| Xk | 0,440746 | 0,335346 | -0,394295 | 1 | 1 |
| Mapk10 | 0,158637 | 0,120595 | -0,395567 | 1 | 1 |
| Gm5544 | 0,244542 | 0,185861 | -0,395858 | 1 | 1 |
| Scgn | 0,139189 | 0,105789 | -0,395858 | 1 | 1 |
| Gm7134 | 0,182128 | 0,138352 | -0,396609 | 1 | 1 |
| Rtn4r | 0,652941 | 0,495806 | -0,397178 | 1 | 1 |
| Slc39a2 | 0,314147 | 0,238444 | -0,397789 | 1 | 1 |
| Rab25 | 0,247754 | 0,188022 | -0,398003 | 1 | 1 |
| Neb | 0,0287292 | 0,0217956 | -0,398484 | 1 | 1 |
| Smpd5 | 0,149309 | 0,11311 | -0,400569 | 1 | 1 |
| Eno2 | 0,534432 | 0,404825 | -0,400706 | 1 | 1 |
| Foxq1 | 0,363664 | 0,275297 | -0,401614 | 1 | 1 |
| Vmn1r32 | 0,107036 | 0,0810266 | -0,401626 | 1 | 1 |
| Fam129a | 0,235934 | 0,178595 | -0,401695 | 1 | 1 |
| Pcdh10 | 0,138052 | 0,104443 | -0,402486 | 1 | 1 |
| Gm5779 | 0,311441 | 0,235558 | -0,402875 | 1 | 1 |
| Fam217a | 0,0999296 | 0,0755817 | -0,402875 | 1 | 1 |
| Syt4 | 0,0592227 | 0,0447931 | -0,402875 | 1 | 1 |

| | | | | | |
|---|---|---|---|---|---|
| Ttc25 | 0,4962 | 0,374086 | -0,407551 | 1 | 1 |
| Ermap | 0,0472079 | 0,0355448 | -0,409392 | 1 | 1 |
| Gm15772 | 401,086 | 301,725 | -0,410677 | 1 | 1 |
| Gm1995 | 0,312216 | 0,234814 | -0,411027 | 1 | 1 |
| Gm10548 | 0,371881 | 0,279644 | -0,411252 | 1 | 1 |
| 0610039K10Rik | 0,420609 | 0,316234 | -0,411487 | 1 | 1 |
| 2610005L07Rik | 0,067321 | 0,0505716 | -0,412731 | 1 | 1 |
| Krt222 | 0,247528 | 0,185687 | -0,41472 | 1 | 1 |
| Fbxw10 | 0,223369 | 0,16729 | -0,417077 | 1 | 1 |
| Brdt | 0,20719 | 0,155022 | -0,418482 | 1 | 1 |
| Dmrta1 | 0,334503 | 0,249877 | -0,420799 | 1 | 1 |
| Tbc1d30 | 0,13265 | 0,0990738 | -0,421054 | 1 | 1 |
| Aqp1 | 0,185314 | 0,138252 | -0,422671 | 1 | 1 |
| Ccdc155 | 0,247109 | 0,184003 | -0,425419 | 1 | 1 |
| Grin1 | 0,0986016 | 0,0732351 | -0,429075 | 1 | 1 |
| Tnnt3 | 0,544187 | 0,404099 | -0,429395 | 1 | 1 |
| Zfp455 | 0,56601 | 0,419417 | -0,432441 | 1 | 1 |
| Oxct2b | 0,0702089 | 0,0519523 | -0,434466 | 1 | 1 |
| Amdhd1 | 0,0490545 | 0,0362987 | -0,434466 | 1 | 1 |
| Dnah1 | 0,505707 | 0,374128 | -0,434768 | 1 | 1 |
| Slc14a1 | 0,0551976 | 0,0407944 | -0,436236 | 1 | 1 |
| 6430550D23Rik | 0,346136 | 0,255698 | -0,436902 | 1 | 1 |
| Trim38 | 0,142425 | 0,105112 | -0,438277 | 1 | 1 |
| Il1a | 0,0979528 | 0,0722493 | -0,439104 | 1 | 1 |
| Slc16a5 | 0,0850661 | 0,0627438 | -0,439111 | 1 | 1 |
| Gm5069 | 0,541716 | 0,398867 | -0,441629 | 1 | 1 |
| Gm11468 | 0,322591 | 0,237507 | -0,441738 | 1 | 1 |
| Spata45 | 0,258063 | 0,189998 | -0,441738 | 1 | 1 |
| Atp5l | 0,187081 | 0,137738 | -0,441738 | 1 | 1 |
| Cldnd2 | 0,140313 | 0,103305 | -0,441738 | 1 | 1 |
| A330040F15Rik | 0,128181 | 0,0943731 | -0,441738 | 1 | 1 |
| 4933432I03Rik | 0,0674559 | 0,0496642 | -0,441738 | 1 | 1 |
| Kcne4 | 0,0511822 | 0,0376827 | -0,441738 | 1 | 1 |
| Csf3 | 0,0498792 | 0,0367234 | -0,441738 | 1 | 1 |
| Mfap5 | 0,0479353 | 0,0352922 | -0,441738 | 1 | 1 |
| Prss42 | 0,0427906 | 0,0315044 | -0,441738 | 1 | 1 |
| Serpina3m | 0,039732 | 0,0292526 | -0,441738 | 1 | 1 |
| Ereg | 0,0294051 | 0,0216494 | -0,441738 | 1 | 1 |
| Otop2 | 0,0271282 | 0,0199731 | -0,441738 | 1 | 1 |
| Adamdec1 | 0,0261822 | 0,0192766 | -0,441738 | 1 | 1 |
| Vmn2r87 | 0,0246736 | 0,0181659 | -0,441738 | 1 | 1 |
| Cyp4f37 | 0,0237122 | 0,017458 | -0,441738 | 1 | 1 |
| Kcnh4 | 0,0161482 | 0,011889 | -0,441738 | 1 | 1 |
| Mroh7 | 0,0139571 | 0,0102759 | -0,441738 | 1 | 1 |
| D830026I12Rik | 0,0424069 | 0,0312202 | -0,441821 | 1 | 1 |
| Slc4a10 | 0,0111159 | 0,0081823 | -0,442052 | 1 | 1 |
| Tnnc2 | 0,386472 | 0,284369 | -0,442603 | 1 | 1 |
| Scd4 | 0,065761 | 0,0483874 | -0,442603 | 1 | 1 |
| Marveld3 | 0,179906 | 0,132363 | -0,442743 | 1 | 1 |
| 4930479D17Rik | 0,136158 | 0,100091 | -0,443974 | 1 | 1 |

| | | | | | |
|---|---|---|---|---|---|
| 4833427G06Rik | 0,127812 | 0,0939555 | -0,443974 | 1 | 1 |
| Bst1 | 0,0342317 | 0,0251639 | -0,443974 | 1 | 1 |
| Mc1r | 0,0210992 | 0,0155102 | -0,443974 | 1 | 1 |
| Col19a1 | 0,00705961 | 0,00518957 | -0,443974 | 1 | 1 |
| Lrtm1 | 0,0377358 | 0,0277026 | -0,445912 | 1 | 1 |
| Gabra4 | 0,456722 | 0,335216 | -0,446225 | 1 | 1 |
| Gm128 | 0,236014 | 0,173182 | -0,446582 | 1 | 1 |
| A430105I19Rik | 0,0695849 | 0,0510498 | -0,446869 | 1 | 1 |
| Lrrc8e | 0,0614624 | 0,0450735 | -0,447427 | 1 | 1 |
| Irak3 | 0,469102 | 0,343955 | -0,447681 | 1 | 1 |
| Nppa | 0,244525 | 0,179117 | -0,449078 | 1 | 1 |
| Prph | 0,0976488 | 0,0715272 | -0,449111 | 1 | 1 |
| Mir686 | 2985,03 | 2186,45 | -0,449157 | 1 | 1 |
| Gprin2 | 0,120259 | 0,0880719 | -0,449389 | 1 | 1 |
| 4930570G19Rik | 0,321433 | 0,235215 | -0,450542 | 1 | 1 |
| Hist1h2ah | 0,73167 | 0,535361 | -0,450683 | 1 | 1 |
| Pcdhb20 | 0,0823491 | 0,0602457 | -0,450895 | 1 | 1 |
| Fank1 | 0,0536837 | 0,0392084 | -0,453321 | 1 | 1 |
| Asgr2 | 0,0813526 | 0,0594067 | -0,453562 | 1 | 1 |
| Pth2r | 0,0429956 | 0,031397 | -0,453562 | 1 | 1 |
| A830009L08Rik | 0,0365042 | 0,0266567 | -0,453562 | 1 | 1 |
| Pla2g4d | 0,0308308 | 0,0225138 | -0,453562 | 1 | 1 |
| Cadm2 | 0,0101665 | 0,00742399 | -0,453562 | 1 | 1 |
| AI854703 | 0,517016 | 0,377308 | -0,454467 | 1 | 1 |
| Runx1 | 0,321976 | 0,234807 | -0,455477 | 1 | 1 |
| Enpp1 | 0,616001 | 0,449097 | -0,455904 | 1 | 1 |
| Cubn | 0,0381405 | 0,0277877 | -0,456877 | 1 | 1 |
| Slc37a1 | 0,31067 | 0,22549 | -0,462319 | 1 | 1 |
| Ccdc148 | 0,115776 | 0,0840038 | -0,462808 | 1 | 1 |
| Nr5a2 | 0,198278 | 0,143435 | -0,467127 | 1 | 1 |
| AI118078 | 0,17434 | 0,125987 | -0,46863 | 1 | 1 |
| Clec3b | 0,657024 | 0,474312 | -0,470109 | 1 | 1 |
| Tex16 | 0,34635 | 0,24993 | -0,470711 | 1 | 1 |
| 1700102H20Rik | 0,479712 | 0,346126 | -0,470869 | 1 | 1 |
| Epas1 | 0,475558 | 0,343078 | -0,471084 | 1 | 1 |
| Impg2 | 0,0759447 | 0,0547596 | -0,471838 | 1 | 1 |
| Gm6682 | 0,365102 | 0,263145 | -0,472443 | 1 | 1 |
| Tmem130 | 0,149661 | 0,107865 | -0,472474 | 1 | 1 |
| Mmrn1 | 0,504558 | 0,362311 | -0,477794 | 1 | 1 |
| Acad12 | 0,508076 | 0,364749 | -0,47814 | 1 | 1 |
| Acta1 | 0,239351 | 0,171684 | -0,479374 | 1 | 1 |
| Gm10584 | 0,474133 | 0,338967 | -0,48415 | 1 | 1 |
| Rab32 | 0,567919 | 0,404066 | -0,491094 | 1 | 1 |
| F2rl2 | 0,0932039 | 0,0663105 | -0,491154 | 1 | 1 |
| Zfp366 | 0,0851636 | 0,0604984 | -0,49334 | 1 | 1 |
| St8sia4 | 0,490601 | 0,348481 | -0,49347 | 1 | 1 |
| Nup62cl | 0,521606 | 0,370436 | -0,493736 | 1 | 1 |
| Gm14827 | 0,304948 | 0,21646 | -0,494461 | 1 | 1 |
| Nppc | 0,358269 | 0,253951 | -0,496493 | 1 | 1 |
| Trhde | 0,117531 | 0,0831621 | -0,499039 | 1 | 1 |

| | | | | | |
|---|---|---|---|---|---|
| Trim67 | 0,0552189 | 0,0390344 | -0,500418 | 1 | 1 |
| Lin7a | 0,3004 | 0,212197 | -0,501477 | 1 | 1 |
| Usp43 | 0,265721 | 0,187685 | -0,5016 | 1 | 1 |
| Sh3rf2 | 0,0439073 | 0,0309428 | -0,504857 | 1 | 1 |
| Kcnb1 | 0,328236 | 0,231183 | -0,5057 | 1 | 1 |
| Gabrq | 0,0612163 | 0,0431008 | -0,506199 | 1 | 1 |
| Tmem211 | 0,107744 | 0,0758082 | -0,507183 | 1 | 1 |
| AU015228 | 0,191315 | 0,134544 | -0,507879 | 1 | 1 |
| Adam4 | 0,576292 | 0,405107 | -0,508495 | 1 | 1 |
| Dnaic2 | 0,171913 | 0,120697 | -0,510296 | 1 | 1 |
| Cdh22 | 0,101807 | 0,0713848 | -0,512144 | 1 | 1 |
| Mtus2 | 0,125687 | 0,0881132 | -0,512403 | 1 | 1 |
| Aim1l | 0,11726 | 0,0820195 | -0,515666 | 1 | 1 |
| Wbp2nl | 0,123571 | 0,0863659 | -0,516807 | 1 | 1 |
| Adad2 | 0,0798484 | 0,0558074 | -0,516807 | 1 | 1 |
| Slc36a2 | 0,113731 | 0,0793383 | -0,519539 | 1 | 1 |
| Ttc34 | 0,112631 | 0,0784337 | -0,522062 | 1 | 1 |
| 4930596I21Rik | 0,451622 | 0,314339 | -0,522798 | 1 | 1 |
| 2410076I21Rik | 0,170278 | 0,118517 | -0,522798 | 1 | 1 |
| Fndc8 | 0,384397 | 0,267155 | -0,524923 | 1 | 1 |
| Abhd15 | 0,134314 | 0,093263 | -0,526236 | 1 | 1 |
| Tmem132b | 0,294536 | 0,204461 | -0,526622 | 1 | 1 |
| Rps6kl1 | 0,413707 | 0,287053 | -0,52729 | 1 | 1 |
| B430319G15Rik | 0,509716 | 0,353638 | -0,527421 | 1 | 1 |
| Acot4 | 0,138315 | 0,0959506 | -0,527593 | 1 | 1 |
| Eps8l2 | 0,426155 | 0,295534 | -0,528056 | 1 | 1 |
| Snord45c | 485,297 | 336,468 | -0,528397 | 1 | 1 |
| Kcnq5 | 0,118236 | 0,081914 | -0,529491 | 1 | 1 |
| 4930506M07Rik | 0,549004 | 0,380036 | -0,530681 | 1 | 1 |
| A630001G21Rik | 0,08031 | 0,0555497 | -0,531801 | 1 | 1 |
| Tcerg1l | 0,150784 | 0,104048 | -0,535234 | 1 | 1 |
| Gm9054 | 0,522897 | 0,360505 | -0,53651 | 1 | 1 |
| Evi2a-evi2b,Evi2b | 0,0175449 | 0,0120759 | -0,538921 | 1 | 1 |
| Arpp21 | 0,172624 | 0,118792 | -0,539197 | 1 | 1 |
| Cav2 | 0,195727 | 0,134579 | -0,540383 | 1 | 1 |
| Ldoc1 | 0,1035 | 0,0711528 | -0,540633 | 1 | 1 |
| Cpne8 | 0,179458 | 0,123355 | -0,540823 | 1 | 1 |
| Scn4a | 0,0376893 | 0,0258993 | -0,541239 | 1 | 1 |
| Hesx1 | 0,132824 | 0,0912375 | -0,541813 | 1 | 1 |
| Aldh1a1 | 0,0677928 | 0,0465674 | -0,541813 | 1 | 1 |
| Gpr126 | 0,059003 | 0,0405296 | -0,541813 | 1 | 1 |
| Pnoc | 0,0412129 | 0,028276 | -0,543515 | 1 | 1 |
| Pcdhb16 | 0,0399401 | 0,0273935 | -0,544006 | 1 | 1 |
| Tsnaxip1 | 0,109145 | 0,0747067 | -0,546942 | 1 | 1 |
| Tppp | 0,231671 | 0,158484 | -0,547742 | 1 | 1 |
| Hs3st6 | 0,322537 | 0,220373 | -0,549519 | 1 | 1 |
| Tnfsf13 | 0,407627 | 0,278254 | -0,550847 | 1 | 1 |
| Eppk1 | 0,0129813 | 0,00885237 | -0,552296 | 1 | 1 |
| Ect2l | 0,0730369 | 0,0497492 | -0,553952 | 1 | 1 |
| Cacna1e | 0,0402282 | 0,0273984 | -0,554118 | 1 | 1 |

| | | | | | |
|---|---|---|---|---|---|
| Hist1h2bg | 0,391856 | 0,266863 | -0,554225 | 1 | 1 |
| 1700015E13Rik | 0,200167 | 0,136318 | -0,554225 | 1 | 1 |
| 1700007G11Rik | 0,155077 | 0,105611 | -0,554225 | 1 | 1 |
| Izumo1 | 0,0790432 | 0,0538302 | -0,554225 | 1 | 1 |
| Asb5 | 0,0740994 | 0,0504634 | -0,554225 | 1 | 1 |
| Slc22a30 | 0,0568248 | 0,038699 | -0,554225 | 1 | 1 |
| Procr | 0,435235 | 0,296036 | -0,55602 | 1 | 1 |
| 3200001D21Rik | 0,423171 | 0,287476 | -0,557803 | 1 | 1 |
| Icam1 | 0,393806 | 0,267446 | -0,558235 | 1 | 1 |
| Lrrc75b | 0,504519 | 0,342538 | -0,558645 | 1 | 1 |
| Nudt15 | 0,406534 | 0,275439 | -0,561646 | 1 | 1 |
| Mir678 | 92,2872 | 62,5105 | -0,56203 | 1 | 1 |
| Abcc9 | 0,143495 | 0,0971785 | -0,56229 | 1 | 1 |
| A330076H08Rik | 0,271642 | 0,183654 | -0,564717 | 1 | 1 |
| Lhx1 | 0,283392 | 0,191476 | -0,565636 | 1 | 1 |
| Ppl | 0,0807303 | 0,0545365 | -0,565887 | 1 | 1 |
| Atcayos | 0,514957 | 0,347018 | -0,569443 | 1 | 1 |
| Mtag2 | 0,2118 | 0,142568 | -0,57105 | 1 | 1 |
| Cxcl1 | 0,107178 | 0,0720119 | -0,573703 | 1 | 1 |
| Gpr15 | 0,073525 | 0,0494007 | -0,573703 | 1 | 1 |
| Itgb2 | 0,0300409 | 0,0201842 | -0,573703 | 1 | 1 |
| 2610037D02Rik | 0,0232999 | 0,015655 | -0,573703 | 1 | 1 |
| Epha10 | 0,0773823 | 0,0519429 | -0,575077 | 1 | 1 |
| Ttbk1 | 0,14393 | 0,0964195 | -0,577967 | 1 | 1 |
| Kbtbd13 | 0,601076 | 0,402164 | -0,579763 | 1 | 1 |
| 1700112E06Rik | 0,463501 | 0,310108 | -0,579803 | 1 | 1 |
| Ube2u | 0,24958 | 0,166983 | -0,579803 | 1 | 1 |
| Apold1 | 0,415733 | 0,27754 | -0,58296 | 1 | 1 |
| Adamts5 | 0,240154 | 0,160317 | -0,583034 | 1 | 1 |
| Bnipl | 0,123745 | 0,0825564 | -0,583914 | 1 | 1 |
| Pkp3 | 0,198538 | 0,132342 | -0,585139 | 1 | 1 |
| Mzb1 | 0,239316 | 0,159471 | -0,585625 | 1 | 1 |
| Fgf7 | 0,0680723 | 0,0453607 | -0,585625 | 1 | 1 |
| Slc10a1 | 0,0743851 | 0,0495551 | -0,58598 | 1 | 1 |
| Gm11992 | 0,142972 | 0,0947887 | -0,592943 | 1 | 1 |
| Cyp2r1 | 0,544445 | 0,360067 | -0,596519 | 1 | 1 |
| Trpm1 | 0,0834254 | 0,0550154 | -0,600652 | 1 | 1 |
| Cd79b | 0,440532 | 0,290377 | -0,60132 | 1 | 1 |
| C1s2 | 0,0593832 | 0,0391425 | -0,60132 | 1 | 1 |
| Ppfia2 | 0,0234236 | 0,0154397 | -0,60132 | 1 | 1 |
| Fam169b | 0,48666 | 0,320629 | -0,602011 | 1 | 1 |
| Th | 0,135634 | 0,089359 | -0,60204 | 1 | 1 |
| Slc5a8 | 0,0682152 | 0,0449262 | -0,602535 | 1 | 1 |
| Lrrtm2 | 0,0848741 | 0,0556852 | -0,60803 | 1 | 1 |
| B230319C09Rik | 0,225297 | 0,147642 | -0,609721 | 1 | 1 |
| Gm19466 | 0,131663 | 0,086269 | -0,609938 | 1 | 1 |
| Mas1 | 0,121865 | 0,0798492 | -0,609938 | 1 | 1 |
| Gpr68 | 0,480619 | 0,314733 | -0,610767 | 1 | 1 |
| Ttll13 | 0,153608 | 0,100571 | -0,61105 | 1 | 1 |
| Nfasc | 0,506327 | 0,330808 | -0,614075 | 1 | 1 |

| | | | | | |
|---|---|---|---|---|---|
| Npas3 | 0,346018 | 0,225631 | -0,616884 | 1 | 1 |
| 5730522E02Rik | 0,336868 | 0,219328 | -0,619095 | 1 | 1 |
| Fbxl13 | 0,0562005 | 0,0365787 | -0,61958 | 1 | 1 |
| Cldn23 | 0,0794122 | 0,051685 | -0,619617 | 1 | 1 |
| Hrk | 0,0986898 | 0,0641612 | -0,6212 | 1 | 1 |
| Lpo | 0,0935431 | 0,0608152 | -0,6212 | 1 | 1 |
| Cpeb2 | 0,394594 | 0,256084 | -0,623754 | 1 | 1 |
| Chrnb2 | 0,427979 | 0,27771 | -0,623963 | 1 | 1 |
| Galnt3 | 0,0921935 | 0,0598169 | -0,624114 | 1 | 1 |
| Pcdh15 | 0,00464729 | 0,00300947 | -0,626877 | 1 | 1 |
| Arhgap25 | 0,148886 | 0,0962571 | -0,629246 | 1 | 1 |
| Il16 | 0,167775 | 0,108459 | -0,629385 | 1 | 1 |
| Mpeg1 | 0,164646 | 0,106123 | -0,63363 | 1 | 1 |
| Mmd2 | 0,119814 | 0,0771687 | -0,634711 | 1 | 1 |
| Tmem8 | 0,417841 | 0,2691 | -0,63481 | 1 | 1 |
| Aloxe3 | 0,128018 | 0,082444 | -0,634862 | 1 | 1 |
| Zfp112 | 0,617808 | 0,397733 | -0,635359 | 1 | 1 |
| Prg2 | 0,104281 | 0,0671325 | -0,635394 | 1 | 1 |
| Htr1b | 0,0690853 | 0,0444747 | -0,635394 | 1 | 1 |
| 1700016D06Rik | 0,0624313 | 0,0401911 | -0,635394 | 1 | 1 |
| Car6 | 0,0567509 | 0,0365343 | -0,635394 | 1 | 1 |
| Rbm11 | 0,0264991 | 0,0170592 | -0,635394 | 1 | 1 |
| Kirrel2 | 0,0226577 | 0,0145863 | -0,635394 | 1 | 1 |
| D930020B18Rik | 0,0205136 | 0,0132059 | -0,635394 | 1 | 1 |
| Ptgs2 | 0,0167763 | 0,0108 | -0,635394 | 1 | 1 |
| Dfnb59 | 0,66338 | 0,426515 | -0,637241 | 1 | 1 |
| Tgm1 | 0,117458 | 0,0754781 | -0,638013 | 1 | 1 |
| Fgf14 | 0,259452 | 0,166469 | -0,640217 | 1 | 1 |
| Gm16793 | 0,0631895 | 0,0404977 | -0,641845 | 1 | 1 |
| Aatk | 0,288221 | 0,184583 | -0,642906 | 1 | 1 |
| Ifi204 | 0,317942 | 0,203552 | -0,643372 | 1 | 1 |
| Ccdc169 | 0,0639192 | 0,0409035 | -0,644025 | 1 | 1 |
| Gria1 | 0,0726294 | 0,0463567 | -0,647775 | 1 | 1 |
| Gab3 | 0,25676 | 0,163876 | -0,647818 | 1 | 1 |
| Vmn1r53 | 0,118467 | 0,0755848 | -0,64832 | 1 | 1 |
| Tubb1 | 0,0604187 | 0,0385485 | -0,64832 | 1 | 1 |
| Htra4 | 0,0550095 | 0,0350973 | -0,64832 | 1 | 1 |
| Rnf138rt1 | 0,212767 | 0,135478 | -0,65122 | 1 | 1 |
| Rgag1 | 0,0534894 | 0,0340589 | -0,65122 | 1 | 1 |
| Sema6b | 0,255729 | 0,162673 | -0,652639 | 1 | 1 |
| 2610028H24Rik | 0,110453 | 0,0702508 | -0,652845 | 1 | 1 |
| Gm5468 | 0,144492 | 0,0918873 | -0,653048 | 1 | 1 |
| Ddn | 0,223153 | 0,141813 | -0,654044 | 1 | 1 |
| Rinl | 0,378489 | 0,240453 | -0,654497 | 1 | 1 |
| Them7 | 0,24509 | 0,155463 | -0,656735 | 1 | 1 |
| Egfl6 | 0,250423 | 0,158628 | -0,65872 | 1 | 1 |
| 4921525O09Rik | 0,13237 | 0,0838204 | -0,659201 | 1 | 1 |
| 4930592I03Rik | 0,550473 | 0,47962 | -0,659406 | 1 | 1 |
| Pcdhb4 | 0,0744196 | 0,0471124 | -0,659577 | 1 | 1 |
| Pcdhac1 | 0,110311 | 0,0698283 | -0,659691 | 1 | 1 |

| | | | | | |
|---|---|---|---|---|---|
| Pde8a | 0,323841 | 0,204875 | -0,660539 | 1 | 1 |
| Ephx4 | 0,191968 | 0,121145 | -0,66413 | 1 | 1 |
| Gm8580 | 0,375495 | 0,236387 | -0,667642 | 1 | 1 |
| Gm17801 | 0,201387 | 0,126774 | -0,667707 | 1 | 1 |
| Acr | 0,130127 | 0,0818994 | -0,668 | 1 | 1 |
| Efcab12 | 0,536181 | 0,337397 | -0,668272 | 1 | 1 |
| Krtap26-1 | 0,333099 | 0,209294 | -0,670422 | 1 | 1 |
| Pcsk1n | 0,5088 | 0,319544 | -0,671084 | 1 | 1 |
| Ntn5 | 0,160339 | 0,100287 | -0,676999 | 1 | 1 |
| Casz1 | 0,120894 | 0,0755704 | -0,677846 | 1 | 1 |
| Tmtc1 | 0,240589 | 0,150113 | -0,680527 | 1 | 1 |
| Slc15a3 | 0,230029 | 0,143488 | -0,680886 | 1 | 1 |
| Pde6g | 0,346735 | 0,215626 | -0,685307 | 1 | 1 |
| Col6a4 | 0,0736578 | 0,0457775 | -0,686198 | 1 | 1 |
| Oprd1 | 0,404078 | 0,250844 | -0,687843 | 1 | 1 |
| Ica1l | 0,187364 | 0,116247 | -0,688652 | 1 | 1 |
| Rsph6a | 0,0942298 | 0,0584209 | -0,689699 | 1 | 1 |
| C1qtnf5 | 0,11384 | 0,07046 | -0,692129 | 1 | 1 |
| 5033404E19Rik | 0,17171 | 0,106235 | -0,692723 | 1 | 1 |
| Srd5a2 | 0,133406 | 0,0825365 | -0,692723 | 1 | 1 |
| Krt9 | 0,0364964 | 0,0225798 | -0,692723 | 1 | 1 |
| Lhx4 | 0,0835814 | 0,051685 | -0,693438 | 1 | 1 |
| A230108P19Rik | 0,059956 | 0,0370755 | -0,693438 | 1 | 1 |
| Col8a2 | 0,10444 | 0,0645659 | -0,693837 | 1 | 1 |
| Wnt3 | 0,418144 | 0,258391 | -0,694446 | 1 | 1 |
| Pkd1l3 | 0,0776574 | 0,047918 | -0,696555 | 1 | 1 |
| Tfec | 0,1381 | 0,0851577 | -0,697501 | 1 | 1 |
| 1810010H24Rik | 0,612379 | 0,377242 | -0,698931 | 1 | 1 |
| Lrrd1 | 0,066158 | 0,0407536 | -0,698987 | 1 | 1 |
| Trim30d | 0,0449347 | 0,0276555 | -0,700262 | 1 | 1 |
| Gm14005 | 0,301989 | 0,185787 | -0,700848 | 1 | 1 |
| Susd3 | 0,433555 | 0,266723 | -0,700873 | 1 | 1 |
| 6720483E21Rik | 0,246518 | 0,1515 | -0,70238 | 1 | 1 |
| Fbxo2 | 0,169775 | 0,104173 | -0,704637 | 1 | 1 |
| Mapk15 | 0,307088 | 0,188159 | -0,7067 | 1 | 1 |
| Dtx1 | 0,34041 | 0,208427 | -0,707732 | 1 | 1 |
| Pde4c | 0,263029 | 0,160944 | -0,708661 | 1 | 1 |
| Nkx2-3 | 0,178948 | 0,1091 | -0,713894 | 1 | 1 |
| Hist1h2bf | 0,194159 | 0,118164 | -0,716454 | 1 | 1 |
| Hcst | 0,185188 | 0,112704 | -0,716454 | 1 | 1 |
| Lrrc10 | 0,0720206 | 0,0438311 | -0,716454 | 1 | 1 |
| Tuba3a | 0,0675504 | 0,0411106 | -0,716454 | 1 | 1 |
| Fam150a | 0,0673948 | 0,0410159 | -0,716454 | 1 | 1 |
| Ly6g6c | 0,058302 | 0,0354821 | -0,716454 | 1 | 1 |
| Tex22 | 0,0414643 | 0,0252348 | -0,716454 | 1 | 1 |
| Plscr2 | 0,0260272 | 0,0158399 | -0,716454 | 1 | 1 |
| Gsdmc4 | 0,0234451 | 0,0142685 | -0,716454 | 1 | 1 |
| Htr1f | 0,0191223 | 0,0116376 | -0,716454 | 1 | 1 |
| Aox4 | 0,0184886 | 0,011252 | -0,716454 | 1 | 1 |
| Lrat | 0,0171339 | 0,0104275 | -0,716454 | 1 | 1 |

| | | | | | |
|---|---|---|---|---|---|
| Clec3a | 0,0161691 | 0,00984035 | -0,716454 | 1 | 1 |
| Slitrk3 | 0,0124934 | 0,0076034 | -0,716454 | 1 | 1 |
| Zc3h12d | 0,0113337 | 0,00689761 | -0,716454 | 1 | 1 |
| Naip5 | 0,00866292 | 0,00527218 | -0,716454 | 1 | 1 |
| D7Ertd443e | 0,0595343 | 0,0362313 | -0,716485 | 1 | 1 |
| Sytl3 | 0,0339037 | 0,020633 | -0,716488 | 1 | 1 |
| Hs3st5 | 0,0161605 | 0,0098341 | -0,716609 | 1 | 1 |
| Omg | 0,100075 | 0,0608799 | -0,71704 | 1 | 1 |
| Lamp3 | 0,419773 | 0,255232 | -0,717798 | 1 | 1 |
| Gpr37 | 0,0995651 | 0,0604258 | -0,720476 | 1 | 1 |
| Itga11 | 0,117859 | 0,0713366 | -0,724352 | 1 | 1 |
| Espnl | 0,0791173 | 0,0478667 | -0,724971 | 1 | 1 |
| Pcdhb5 | 0,111246 | 0,0672576 | -0,725988 | 1 | 1 |
| Wscd2 | 0,52297 | 0,315722 | -0,728072 | 1 | 1 |
| Mageb3 | 0,292553 | 0,176469 | -0,729282 | 1 | 1 |
| Gm6498 | 0,401704 | 0,242031 | -0,730938 | 1 | 1 |
| Gpx2-ps1 | 0,196186 | 0,118196 | -0,731049 | 1 | 1 |
| Igsf11 | 0,611662 | 0,368258 | -0,732016 | 1 | 1 |
| Celf5 | 0,19308 | 0,116033 | -0,734665 | 1 | 1 |
| Ptger4 | 0,532739 | 0,319553 | -0,737373 | 1 | 1 |
| Dlgap1 | 0,206616 | 0,123662 | -0,740552 | 1 | 1 |
| D5Ertd605e | 0,322556 | 0,193005 | -0,740916 | 1 | 1 |
| Sync | 0,167545 | 0,100197 | -0,741706 | 1 | 1 |
| Ankrd35 | 0,0604363 | 0,0360455 | -0,745597 | 1 | 1 |
| Dcst1 | 0,131971 | 0,0785576 | -0,748394 | 1 | 1 |
| Pcdhb22 | 0,249113 | 0,147664 | -0,754486 | 1 | 1 |
| Thsd7a | 0,206723 | 0,122282 | -0,757493 | 1 | 1 |
| Zar1 | 0,283216 | 0,167474 | -0,757962 | 1 | 1 |
| Samd5 | 0,457972 | 0,270552 | -0,759351 | 1 | 1 |
| Fam83b | 0,0619836 | 0,0365959 | -0,760205 | 1 | 1 |
| Msantd1 | 0,427588 | 0,252258 | -0,761319 | 1 | 1 |
| Slc5a5 | 0,285568 | 0,168461 | -0,76142 | 1 | 1 |
| 1700030C10Rik | 0,422147 | 0,248955 | -0,761861 | 1 | 1 |
| Plcb1 | 0,422135 | 0,248795 | -0,76275 | 1 | 1 |
| Marc1 | 0,0980601 | 0,0577657 | -0,763453 | 1 | 1 |
| Arhgap33os | 0,0591349 | 0,0348303 | -0,763666 | 1 | 1 |
| Tnnt2 | 0,138625 | 0,0815897 | -0,764728 | 1 | 1 |
| 1810041L15Rik | 0,114469 | 0,0673639 | -0,764913 | 1 | 1 |
| Fbxl8 | 0,219926 | 0,129198 | -0,767434 | 1 | 1 |
| Clec14a | 0,431902 | 0,253544 | -0,768467 | 1 | 1 |
| Mei4 | 0,21347 | 0,125315 | -0,768479 | 1 | 1 |
| Trp63 | 0,258803 | 0,151805 | -0,769642 | 1 | 1 |
| BC048609 | 0,550054 | 0,322209 | -0,771577 | 1 | 1 |
| 4930426L09Rik | 0,142881 | 0,0836962 | -0,771577 | 1 | 1 |
| 4930524O05Rik | 0,133805 | 0,0783234 | -0,772621 | 1 | 1 |
| Irx4 | 0,182083 | 0,10654 | -0,773205 | 1 | 1 |
| AB124611 | 0,513726 | 0,30046 | -0,773828 | 1 | 1 |
| Shcbp1l | 0,114416 | 0,0668315 | -0,775694 | 1 | 1 |
| D330050G23Rik | 0,139861 | 0,0816915 | -0,775736 | 1 | 1 |
| Pvrl4 | 0,0653073 | 0,0381432 | -0,775817 | 1 | 1 |

| | | | | | |
|---|---|---|---|---|---|
| Pak7 | 0,182033 | 0,106306 | -0,775974 | 1 | 1 |
| Arhgap8 | 0,536644 | 0,312984 | -0,777879 | 1 | 1 |
| Eva1c | 0,0813582 | 0,0474479 | -0,777943 | 1 | 1 |
| Enkur | 0,0733481 | 0,0427762 | -0,777951 | 1 | 1 |
| Scml4 | 0,0482155 | 0,028119 | -0,777951 | 1 | 1 |
| Mndal | 0,0428297 | 0,024978 | -0,777951 | 1 | 1 |
| Cd86 | 0,0312556 | 0,0182281 | -0,777951 | 1 | 1 |
| Mypn | 0,0144748 | 0,00844165 | -0,777951 | 1 | 1 |
| Cadps2 | 0,447035 | 0,260336 | -0,78001 | 1 | 1 |
| Cideb | 0,246442 | 0,14338 | -0,781407 | 1 | 1 |
| Pde2a | 0,551858 | 0,320795 | -0,782643 | 1 | 1 |
| Zfp488 | 0,111909 | 0,0650387 | -0,782962 | 1 | 1 |
| Gpr176 | 0,206138 | 0,119544 | -0,786069 | 1 | 1 |
| Apol8 | 0,356436 | 0,206631 | -0,786589 | 1 | 1 |
| Hpcal4 | 0,0538549 | 0,0311995 | -0,787557 | 1 | 1 |
| Oprk1 | 0,0846308 | 0,0489952 | -0,78854 | 1 | 1 |
| Trim43c | 0,249024 | 0,143987 | -0,790351 | 1 | 1 |
| Ppargc1b | 0,222199 | 0,128375 | -0,791489 | 1 | 1 |
| Vipr2 | 0,584823 | 0,337653 | -0,792461 | 1 | 1 |
| 1700034F02Rik | 0,155475 | 0,0896676 | -0,794029 | 1 | 1 |
| B130006D01Rik | 0,227506 | 0,131128 | -0,79493 | 1 | 1 |
| Adamts18 | 0,327431 | 0,188561 | -0,796157 | 1 | 1 |
| Srcin1 | 0,22156 | 0,127387 | -0,798483 | 1 | 1 |
| Arhgef33 | 0,138287 | 0,0789254 | -0,809102 | 1 | 1 |
| Btn2a2 | 0,568495 | 0,324117 | -0,810631 | 1 | 1 |
| Rims1 | 0,29219 | 0,165899 | -0,816602 | 1 | 1 |
| Rps13 | 0,750303 | 0,425778 | -0,817374 | 1 | 1 |
| 1110046J04Rik | 0,509503 | 0,289001 | -0,818017 | 1 | 1 |
| Hmga1-rs1 | 45,6252 | 25,8754 | -0,818252 | 1 | 1 |
| BC037704 | 0,374829 | 0,212519 | -0,81864 | 1 | 1 |
| Pilra | 0,0232003 | 0,0131062 | -0,823894 | 1 | 1 |
| 2610028E06Rik | 0,0284578 | 0,0160676 | -0,824671 | 1 | 1 |
| Slc17a8 | 0,0223683 | 0,0126294 | -0,824671 | 1 | 1 |
| Chst5 | 0,213944 | 0,120255 | -0,831135 | 1 | 1 |
| Plxna4 | 0,346362 | 0,194372 | -0,833461 | 1 | 1 |
| Ccdc158 | 0,216234 | 0,121307 | -0,833934 | 1 | 1 |
| Ar | 0,0886766 | 0,0496512 | -0,836727 | 1 | 1 |
| Krt20 | 0,138787 | 0,0777077 | -0,836738 | 1 | 1 |
| 1700003M07Rik | 0,0696964 | 0,0390227 | -0,836771 | 1 | 1 |
| Aqp11 | 0,411384 | 0,230017 | -0,838744 | 1 | 1 |
| Sgip1 | 0,39908 | 0,223087 | -0,83907 | 1 | 1 |
| Ccdc81 | 0,0614518 | 0,0342281 | -0,844273 | 1 | 1 |
| Adora1 | 0,064945 | 0,0361474 | -0,845329 | 1 | 1 |
| Kcnq1 | 0,0671349 | 0,0373461 | -0,846103 | 1 | 1 |
| 6720468P15Rik | 0,109883 | 0,061025 | -0,848503 | 1 | 1 |
| Mrgprf | 0,0962434 | 0,0533918 | -0,850069 | 1 | 1 |
| Cybrd1 | 0,572923 | 0,317426 | -0,851921 | 1 | 1 |
| Disp2 | 0,0857024 | 0,0473736 | -0,855251 | 1 | 1 |
| Pi16 | 0,17472 | 0,0965582 | -0,855578 | 1 | 1 |
| Fam212b | 0,293397 | 0,162056 | -0,856356 | 1 | 1 |

| | | | | | |
|---|---|---|---|---|---|
| Abhd16b | 0,0981976 | 0,0541853 | -0,857786 | 1 | 1 |
| Tcam1 | 0,163431 | 0,0901565 | -0,85818 | 1 | 1 |
| Selenbp2 | 0,0978963 | 0,0539732 | -0,859012 | 1 | 1 |
| Trpv2 | 0,0573082 | 0,0315957 | -0,859012 | 1 | 1 |
| Pde11a | 0,135417 | 0,0745369 | -0,861385 | 1 | 1 |
| Ctnna2 | 0,421953 | 0,231878 | -0,863712 | 1 | 1 |
| Tmem240 | 0,191119 | 0,104801 | -0,866816 | 1 | 1 |
| Il1rl2 | 0,145062 | 0,0794814 | -0,867984 | 1 | 1 |
| Kcnc2 | 0,115349 | 0,0631532 | -0,869082 | 1 | 1 |
| Myh1 | 0,0774185 | 0,0423172 | -0,871433 | 1 | 1 |
| Lingo3 | 0,270277 | 0,147268 | -0,875999 | 1 | 1 |
| Cldn14 | 0,446873 | 0,242734 | -0,880488 | 1 | 1 |
| Arid5a | 0,386051 | 0,20897 | -0,8855 | 1 | 1 |
| Col15a1 | 0,508379 | 0,274387 | -0,88969 | 1 | 1 |
| Gfod1 | 0,150513 | 0,0811822 | -0,890649 | 1 | 1 |
| Cobll1 | 0,283664 | 0,152837 | -0,892187 | 1 | 1 |
| Gm13710 | 0,398219 | 0,214252 | -0,894255 | 1 | 1 |
| C130060K24Rik | 0,242363 | 0,130238 | -0,896021 | 1 | 1 |
| Ccdc69 | 0,312823 | 0,167498 | -0,9012 | 1 | 1 |
| Mr1 | 0,246717 | 0,13205 | -0,90178 | 1 | 1 |
| Prr33 | 0,38113 | 0,203984 | -0,901825 | 1 | 1 |
| Larp6 | 0,228389 | 0,122225 | -0,901957 | 1 | 1 |
| Myct1 | 0,132037 | 0,0704224 | -0,906836 | 1 | 1 |
| Ano2 | 0,145371 | 0,0775281 | -0,906953 | 1 | 1 |
| Eltd1 | 0,362328 | 0,193224 | -0,907019 | 1 | 1 |
| Ackr2 | 0,0296124 | 0,0157847 | -0,907668 | 1 | 1 |
| Upk3a | 0,0885233 | 0,0471866 | -0,90768 | 1 | 1 |
| Cers1 | 1,291 | 0,687547 | -0,908955 | 1 | 1 |
| Tmem30c | 0,264082 | 0,140576 | -0,909632 | 1 | 1 |
| Gpc5 | 0,127353 | 0,0676381 | -0,912926 | 1 | 1 |
| Vmn2r29 | 0,187405 | 0,0994798 | -0,913686 | 1 | 1 |
| Marcksl1-ps4 | 0,155889 | 0,0827017 | -0,914531 | 1 | 1 |
| Atp8b1 | 0,0209738 | 0,0111269 | -0,914531 | 1 | 1 |
| Ddx4 | 0,195773 | 0,103796 | -0,915435 | 1 | 1 |
| 03/01/16 | 0,151365 | 0,080181 | -0,916696 | 1 | 1 |
| Pcdh9 | 0,150527 | 0,0797146 | -0,917102 | 1 | 1 |
| Mtl5 | 0,154059 | 0,0815551 | -0,917636 | 1 | 1 |
| Spata2l | 0,190448 | 0,100758 | -0,918505 | 1 | 1 |
| Ap1m2 | 0,337333 | 0,178467 | -0,918514 | 1 | 1 |
| 4930404H11Rik | 0,413278 | 0,218534 | -0,919251 | 1 | 1 |
| LOC102634101 | 0,283407 | 0,149861 | -0,919251 | 1 | 1 |
| Npas1 | 0,208699 | 0,110314 | -0,919809 | 1 | 1 |
| Kcnk6 | 0,205901 | 0,108819 | -0,920025 | 1 | 1 |
| Ifit3 | 0,089819 | 0,0474499 | -0,920615 | 1 | 1 |
| P2rx6 | 0,0247106 | 0,0130118 | -0,925306 | 1 | 1 |
| Gm10012 | 0,302628 | 0,159351 | -0,925334 | 1 | 1 |
| Cldn8 | 0,236507 | 0,124535 | -0,925334 | 1 | 1 |
| Atp5g2 | 0,229988 | 0,121102 | -0,925334 | 1 | 1 |
| C330024C12Rik | 0,0854753 | 0,0450078 | -0,925334 | 1 | 1 |
| Dnajb13 | 0,0542525 | 0,0285671 | -0,925334 | 1 | 1 |

| | | | | | |
|---|---|---|---|---|---|
| Fam92b | 0,0517189 | 0,0272331 | -0,925334 | 1 | 1 |
| Tnfsf11 | 0,0501659 | 0,0264153 | -0,925334 | 1 | 1 |
| Srrm4os | 0,0484562 | 0,025515 | -0,925334 | 1 | 1 |
| Gabrr3 | 0,0406895 | 0,0214254 | -0,925334 | 1 | 1 |
| Myod1 | 0,0308748 | 0,0162574 | -0,925334 | 1 | 1 |
| LOC102635087 | 0,0273449 | 0,0143987 | -0,925334 | 1 | 1 |
| Zfp457 | 0,0266441 | 0,0140297 | -0,925334 | 1 | 1 |
| Klhl40 | 0,0226315 | 0,0119168 | -0,925334 | 1 | 1 |
| Cpne9 | 0,0197839 | 0,0104174 | -0,925334 | 1 | 1 |
| Fam65c | 0,0186538 | 0,00982233 | -0,925334 | 1 | 1 |
| Cnr2 | 0,0139343 | 0,0073372 | -0,925334 | 1 | 1 |
| 1700119H24Rik | 0,26673 | 0,14044 | -0,925426 | 1 | 1 |
| Sod3 | 0,293198 | 0,154256 | -0,926551 | 1 | 1 |
| Sult6b1 | 0,153589 | 0,0807443 | -0,927639 | 1 | 1 |
| Adamtsl4 | 0,279778 | 0,146416 | -0,934208 | 1 | 1 |
| G6bos | 0,0536466 | 0,0280592 | -0,935016 | 1 | 1 |
| 2610316D01Rik | 0,214856 | 0,11208 | -0,938847 | 1 | 1 |
| 9030612E09Rik | 0,332839 | 0,173563 | -0,939361 | 1 | 1 |
| Bcl6b | 0,394387 | 0,205586 | -0,939866 | 1 | 1 |
| Fads6 | 0,54806 | 0,285092 | -0,942905 | 1 | 1 |
| Cbfa2t3 | 0,32172 | 0,16718 | -0,944405 | 1 | 1 |
| Tmem40 | 0,501772 | 0,260215 | -0,947326 | 1 | 1 |
| A430090L17Rik | 0,0707474 | 0,0366604 | -0,948454 | 1 | 1 |
| Spns2 | 0,389171 | 0,200833 | -0,954408 | 1 | 1 |
| Rapsn | 0,205147 | 0,105834 | -0,954855 | 1 | 1 |
| Adra1d | 0,365264 | 0,187982 | -0,958342 | 1 | 1 |
| Vmn2r1 | 0,0944955 | 0,0485353 | -0,961211 | 1 | 1 |
| Metrnl | 0,498067 | 0,254979 | -0,965961 | 1 | 1 |
| Asb2 | 0,163937 | 0,0837338 | -0,969263 | 1 | 1 |
| D030025P21Rik | 0,0481814 | 0,0246027 | -0,969657 | 1 | 1 |
| Mfsd7a | 0,346565 | 0,176659 | -0,97216 | 1 | 1 |
| Tmem255a | 0,465179 | 0,2367 | -0,974727 | 1 | 1 |
| 4933403O08Rik | 0,319029 | 0,161712 | -0,980264 | 1 | 1 |
| Rbm47 | 0,0374397 | 0,0189704 | -0,980818 | 1 | 1 |
| Alpk2 | 0,0238643 | 0,0120866 | -0,981442 | 1 | 1 |
| Batf | 0,302966 | 0,153282 | -0,982965 | 1 | 1 |
| Fmo1 | 0,293882 | 0,148662 | -0,983199 | 1 | 1 |
| 2410018L13Rik | 0,526482 | 0,266094 | -0,984449 | 1 | 1 |
| Pcdhb17 | 0,11407 | 0,0576278 | -0,985083 | 1 | 1 |
| Pou3f1 | 0,15799 | 0,0797934 | -0,985494 | 1 | 1 |
| Adap1 | 0,289202 | 0,145837 | -0,987722 | 1 | 1 |
| Cryab | 0,238653 | 0,120262 | -0,988736 | 1 | 1 |
| Dio3os | 0,148555 | 0,0748593 | -0,988741 | 1 | 1 |
| Samd12 | 0,420679 | 0,211659 | -0,99098 | 1 | 1 |
| Mir8097 | 68,657 | 34,5105 | -0,992373 | 1 | 1 |
| Foxj1 | 0,54347 | 0,272393 | -0,996512 | 1 | 1 |
| Pdgfrl | 0,619562 | 0,309761 | -1,0001 | 1 | 1 |
| Abi3 | 0,175873 | 0,0878587 | -1,00127 | 1 | 1 |
| C2cd4a | 0,140361 | 0,0700992 | -1,00167 | 1 | 1 |
| Kcnk13 | 0,0563607 | 0,0281452 | -1,0018 | 1 | 1 |

| | | | | | |
|---|---|---|---|---|---|
| Gdf6 | 0,0740424 | 0,0369703 | -1,00199 | 1 | 1 |
| Dlx6 | 0,160433 | 0,0800309 | -1,00335 | 1 | 1 |
| Tmem116 | 0,131834 | 0,0657594 | -1,00346 | 1 | 1 |
| Rapgef4 | 0,166167 | 0,0828151 | -1,00467 | 1 | 1 |
| Zdhhc14 | 0,369373 | 0,183686 | -1,00783 | 1 | 1 |
| Ankrd66 | 0,241618 | 0,120074 | -1,0088 | 1 | 1 |
| Ccdc13 | 0,110792 | 0,055059 | -1,0088 | 1 | 1 |
| Dhrs2 | 0,333585 | 0,165281 | -1,01313 | 1 | 1 |
| Zfp536 | 0,394444 | 0,194604 | -1,01928 | 1 | 1 |
| Il19 | 0,132787 | 0,0655055 | -1,01943 | 1 | 1 |
| Madcam1 | 0,0894352 | 0,0441195 | -1,01943 | 1 | 1 |
| Plg | 0,0426319 | 0,0210308 | -1,01943 | 1 | 1 |
| Cd22 | 0,051829 | 0,0255298 | -1,02158 | 1 | 1 |
| Gm20257 | 0,267031 | 0,131436 | -1,02265 | 1 | 1 |
| 4921511C10Rik | 0,0688761 | 0,0338066 | -1,0267 | 1 | 1 |
| 5031425F14Rik | 0,0605552 | 0,0297224 | -1,0267 | 1 | 1 |
| Txndc2 | 0,0569355 | 0,0279449 | -1,02674 | 1 | 1 |
| Slc13a4 | 0,0679535 | 0,0333021 | -1,02894 | 1 | 1 |
| Tbx1 | 0,509944 | 0,249651 | -1,03043 | 1 | 1 |
| Prss53 | 0,344659 | 0,168473 | -1,03266 | 1 | 1 |
| A930011O12Rik | 0,296201 | 0,144746 | -1,03306 | 1 | 1 |
| Gm13154 | 0,104963 | 0,0511338 | -1,03753 | 1 | 1 |
| Spink8 | 0,231978 | 0,112944 | -1,03838 | 1 | 1 |
| Nol3 | 0,245886 | 0,119697 | -1,03861 | 1 | 1 |
| Gpr45 | 0,0719429 | 0,0349257 | -1,04256 | 1 | 1 |
| Pnpla3 | 0,0768131 | 0,0372554 | -1,0439 | 1 | 1 |
| Alk | 0,059754 | 0,0289815 | -1,0439 | 1 | 1 |
| Cdkl1 | 0,348652 | 0,168846 | -1,04608 | 1 | 1 |
| C1qtnf9 | 0,149922 | 0,0724171 | -1,04981 | 1 | 1 |
| Syt1 | 0,0756822 | 0,0365472 | -1,05019 | 1 | 1 |
| Ptprr | 0,0270247 | 0,013048 | -1,05044 | 1 | 1 |
| AU040972 | 0,173309 | 0,0832723 | -1,05744 | 1 | 1 |
| Elfn1 | 0,223446 | 0,1073 | -1,05827 | 1 | 1 |
| Col10a1 | 0,0493937 | 0,0237141 | -1,05858 | 1 | 1 |
| Rbp2 | 0,479422 | 0,229266 | -1,06427 | 1 | 1 |
| Bhlha15 | 0,146495 | 0,0700111 | -1,0652 | 1 | 1 |
| Dll4 | 0,335037 | 0,159832 | -1,06777 | 1 | 1 |
| Abcg5 | 0,173892 | 0,082686 | -1,07247 | 1 | 1 |
| 1700001K23Rik | 0,513937 | 0,244312 | -1,07287 | 1 | 1 |
| Neurl3 | 0,0949191 | 0,0451026 | -1,07349 | 1 | 1 |
| Prrt4 | 0,415792 | 0,196611 | -1,08052 | 1 | 1 |
| Tbrg3 | 0,431154 | 0,20287 | -1,08765 | 1 | 1 |
| Zfp345 | 0,10455 | 0,0491755 | -1,08819 | 1 | 1 |
| Pin4 | 0,757209 | 0,355012 | -1,09283 | 1 | 1 |
| Unc5cl | 0,0690486 | 0,0323425 | -1,09418 | 1 | 1 |
| Sntb1 | 0,561687 | 0,262974 | -1,09485 | 1 | 1 |
| Samd11 | 0,107218 | 0,0500746 | -1,0984 | 1 | 1 |
| Slc44a3 | 0,179423 | 0,0836483 | -1,10095 | 1 | 1 |
| Gpr97 | 0,18486 | 0,0861344 | -1,10177 | 1 | 1 |
| Lpar3 | 0,401058 | 0,186643 | -1,10353 | 1 | 1 |

| | | | | | |
|---|---|---|---|---|---|
| Dnah5 | 0,0193872 | 0,00900438 | -1,10641 | 1 | 1 |
| Gpr171 | 0,0286061 | 0,0132778 | -1,10731 | 1 | 1 |
| Gpr34 | 0,13651 | 0,0633503 | -1,10758 | 1 | 1 |
| Dcpp3 | 0,142386 | 0,0660691 | -1,10776 | 1 | 1 |
| 4930529M08Rik | 0,127804 | 0,0593028 | -1,10776 | 1 | 1 |
| Tnfrsf17 | 0,0950812 | 0,044119 | -1,10776 | 1 | 1 |
| Tas2r137 | 0,0724432 | 0,0336146 | -1,10776 | 1 | 1 |
| Siglece | 0,0702859 | 0,0326136 | -1,10776 | 1 | 1 |
| Zfp572 | 0,0646642 | 0,0300051 | -1,10776 | 1 | 1 |
| Cd2 | 0,0614998 | 0,0285367 | -1,10776 | 1 | 1 |
| Odf3l2 | 0,0553735 | 0,025694 | -1,10776 | 1 | 1 |
| Vgll2 | 0,0458082 | 0,0212557 | -1,10776 | 1 | 1 |
| Arl13a | 0,0375878 | 0,0174413 | -1,10776 | 1 | 1 |
| Tmem174 | 0,0282277 | 0,013098 | -1,10776 | 1 | 1 |
| Sgcz | 0,0264578 | 0,0122768 | -1,10776 | 1 | 1 |
| A330033J07Rik | 0,0221326 | 0,0102698 | -1,10776 | 1 | 1 |
| A630020A06 | 0,0174023 | 0,0080749 | -1,10776 | 1 | 1 |
| Adam28 | 0,0237316 | 0,0110112 | -1,10784 | 1 | 1 |
| Naip2 | 0,0126236 | 0,00585519 | -1,10834 | 1 | 1 |
| Cd300lf | 0,0348109 | 0,0161442 | -1,10852 | 1 | 1 |
| Baiap2l2 | 0,573475 | 0,265952 | -1,10856 | 1 | 1 |
| G530011O06Rik | 0,275288 | 0,12755 | -1,10988 | 1 | 1 |
| Arhgef16 | 0,400548 | 0,185166 | -1,11315 | 1 | 1 |
| Aldh1a3 | 0,314949 | 0,145596 | -1,11315 | 1 | 1 |
| Crybb3 | 0,209616 | 0,0968346 | -1,11415 | 1 | 1 |
| 2510049J12Rik | 0,228228 | 0,105385 | -1,11481 | 1 | 1 |
| Traf3ip3 | 0,0352492 | 0,0162649 | -1,11583 | 1 | 1 |
| Lhx2 | 0,561043 | 0,258773 | -1,11642 | 1 | 1 |
| Slc16a8 | 0,233357 | 0,107278 | -1,12118 | 1 | 1 |
| Sypl2 | 0,148887 | 0,068407 | -1,122 | 1 | 1 |
| C2cd4c | 0,121443 | 0,0557886 | -1,12223 | 1 | 1 |
| Tmigd1 | 0,100738 | 0,0461697 | -1,1256 | 1 | 1 |
| Gm5622 | 0,0850097 | 0,038961 | -1,1256 | 1 | 1 |
| Arl9 | 0,468486 | 0,213761 | -1,13201 | 1 | 1 |
| Trappc3l | 0,370508 | 0,168806 | -1,13414 | 1 | 1 |
| Hist1h3h | 0,538114 | 0,244312 | -1,13919 | 1 | 1 |
| Dgki | 0,104673 | 0,0473743 | -1,14371 | 1 | 1 |
| 4930524C18Rik | 0,111533 | 0,0503401 | -1,14769 | 1 | 1 |
| Gm4980 | 0,125875 | 0,0568121 | -1,14773 | 1 | 1 |
| Ccdc176 | 0,411853 | 0,185261 | -1,15257 | 1 | 1 |
| Ptpn7 | 0,17097 | 0,0767932 | -1,1547 | 1 | 1 |
| Celsr3 | 0,3075 | 0,138079 | -1,1551 | 1 | 1 |
| B430212C06Rik | 0,294837 | 0,132333 | -1,15575 | 1 | 1 |
| P2ry2 | 0,174834 | 0,0784154 | -1,15678 | 1 | 1 |
| Amhr2 | 0,109031 | 0,0488383 | -1,15865 | 1 | 1 |
| Selplg | 0,0395918 | 0,0177344 | -1,15865 | 1 | 1 |
| Hcrtr2 | 0,0269638 | 0,012078 | -1,15865 | 1 | 1 |
| Ggt1 | 0,18403 | 0,0824321 | -1,15866 | 1 | 1 |
| Redrum | 0,165299 | 0,0740423 | -1,15866 | 1 | 1 |
| B3galt5 | 0,15756 | 0,0703169 | -1,16396 | 1 | 1 |

| | | | | | |
|---|---|---|---|---|---|
| Dlg2 | 0,0464911 | 0,0207389 | -1,16462 | 1 | 1 |
| Dio2 | 0,0878323 | 0,039075 | -1,16851 | 1 | 1 |
| Laptm5 | 0,240929 | 0,107104 | -1,16959 | 1 | 1 |
| Dnah7a | 0,0279619 | 0,0124182 | -1,17101 | 1 | 1 |
| Kif4-ps | 0,0903001 | 0,0400968 | -1,17124 | 1 | 1 |
| Il31ra | 0,232908 | 0,103356 | -1,17214 | 1 | 1 |
| Lrrc73 | 0,654622 | 0,290071 | -1,17425 | 1 | 1 |
| Gm7334 | 0,239094 | 0,105894 | -1,17496 | 1 | 1 |
| 4933400F21Rik | 0,053342 | 0,0236203 | -1,17524 | 1 | 1 |
| Nxf2 | 0,117178 | 0,0518352 | -1,17669 | 1 | 1 |
| Hkdc1 | 0,162084 | 0,0716643 | -1,17742 | 1 | 1 |
| Kcnj10 | 0,211447 | 0,0933526 | -1,17954 | 1 | 1 |
| Proz | 0,182626 | 0,0805316 | -1,18126 | 1 | 1 |
| Nat1 | 0,223044 | 0,0980293 | -1,18604 | 1 | 1 |
| 1700042O10Rik | 0,350928 | 0,153987 | -1,18837 | 1 | 1 |
| Pcdhb19 | 0,105344 | 0,0461558 | -1,19053 | 1 | 1 |
| Myo16 | 0,421791 | 0,184352 | -1,19406 | 1 | 1 |
| 9330179D12Rik | 0,426984 | 0,186328 | -1,19634 | 1 | 1 |
| Grin3a | 0,177659 | 0,0775079 | -1,19669 | 1 | 1 |
| Spink10 | 0,436205 | 0,18978 | -1,20068 | 1 | 1 |
| Dpyd | 0,580656 | 0,252218 | -1,20301 | 1 | 1 |
| Pof1b | 0,0769461 | 0,0333796 | -1,20488 | 1 | 1 |
| Faim3 | 0,125202 | 0,0539138 | -1,21553 | 1 | 1 |
| Zfp750 | 0,0530226 | 0,0228323 | -1,21553 | 1 | 1 |
| Barx2 | 0,288904 | 0,124181 | -1,21815 | 1 | 1 |
| Gm13483 | 0,310817 | 0,133564 | -1,21854 | 1 | 1 |
| Klhl14 | 0,436244 | 0,187368 | -1,21926 | 1 | 1 |
| 9630028B13Rik | 0,322488 | 0,138398 | -1,22042 | 1 | 1 |
| 3300002I08Rik | 0,672945 | 0,287579 | -1,22653 | 1 | 1 |
| Kcnq4 | 0,379455 | 0,161942 | -1,22845 | 1 | 1 |
| 4933422H20Rik | 0,130678 | 0,0556782 | -1,23083 | 1 | 1 |
| Wnt9b | 0,0609556 | 0,0259445 | -1,23233 | 1 | 1 |
| Lrfn2 | 0,511409 | 0,217644 | -1,23251 | 1 | 1 |
| Fmo5 | 0,0908978 | 0,0385867 | -1,23614 | 1 | 1 |
| Rilp | 0,148121 | 0,0627232 | -1,23971 | 1 | 1 |
| Slc7a9 | 0,0846002 | 0,0358115 | -1,24024 | 1 | 1 |
| Tll1 | 0,583181 | 0,245385 | -1,2489 | 1 | 1 |
| Rgs22 | 0,18749 | 0,0787918 | -1,25069 | 1 | 1 |
| Gsg1l | 0,372371 | 0,156432 | -1,25121 | 1 | 1 |
| A330032B11Rik | 0,0594093 | 0,0248178 | -1,25931 | 1 | 1 |
| Gm15880 | 0,0380176 | 0,0158816 | -1,25931 | 1 | 1 |
| Casr | 0,0235306 | 0,00982974 | -1,25931 | 1 | 1 |
| Gm15713 | 0,0759406 | 0,0317157 | -1,25967 | 1 | 1 |
| Lmx1b | 0,666005 | 0,278028 | -1,2603 | 1 | 1 |
| Hcn1 | 0,039847 | 0,0165679 | -1,26608 | 1 | 1 |
| 9330159M07Rik | 0,21791 | 0,0904944 | -1,26783 | 1 | 1 |
| Cym | 0,387612 | 0,160775 | -1,26957 | 1 | 1 |
| Klhl4 | 0,179786 | 0,0744034 | -1,27284 | 1 | 1 |
| 1810062G17Rik | 0,420583 | 0,174025 | -1,2731 | 1 | 1 |
| Ctsk | 0,423389 | 0,17507 | -1,27405 | 1 | 1 |

| | | | | | |
|---|---|---|---|---|---|
| Plekhs1 | 0,0770299 | 0,0318145 | -1,27573 | 1 | 1 |
| Tnfrsf23 | 0,054564 | 0,0225357 | -1,27573 | 1 | 1 |
| Nckap5 | 0,398473 | 0,164408 | -1,2772 | 1 | 1 |
| Sphk1 | 0,234334 | 0,0963083 | -1,28283 | 1 | 1 |
| Gucy1a3 | 0,271666 | 0,11164 | -1,28297 | 1 | 1 |
| 9330117O12Rik | 0,302996 | 0,124395 | -1,28436 | 1 | 1 |
| Card9 | 0,0929579 | 0,0381303 | -1,28564 | 1 | 1 |
| 5730422E09Rik | 0,0865598 | 0,0355059 | -1,28564 | 1 | 1 |
| Sla2 | 0,134716 | 0,0552488 | -1,2859 | 1 | 1 |
| Grhl2 | 0,236954 | 0,0971475 | -1,28636 | 1 | 1 |
| Tmem215 | 0,295734 | 0,121218 | -1,28669 | 1 | 1 |
| 5031434O11Rik | 0,128626 | 0,0521872 | -1,30142 | 1 | 1 |
| Tesc | 0,0895859 | 0,0363475 | -1,30142 | 1 | 1 |
| 2310034O05Rik | 0,0732166 | 0,029706 | -1,30142 | 1 | 1 |
| Ch25h | 0,0575913 | 0,0233664 | -1,30142 | 1 | 1 |
| Adprhl1 | 0,0569465 | 0,0231047 | -1,30142 | 1 | 1 |
| Rtp4 | 0,0482943 | 0,0195943 | -1,30142 | 1 | 1 |
| Tspan1 | 0,0374166 | 0,0151809 | -1,30142 | 1 | 1 |
| 4930519G04Rik | 0,0359908 | 0,0146025 | -1,30142 | 1 | 1 |
| Dnah6 | 0,00531176 | 0,00215513 | -1,30142 | 1 | 1 |
| Pak6 | 0,0191189 | 0,0077567 | -1,30148 | 1 | 1 |
| Ltk | 0,0325674 | 0,0132126 | -1,30152 | 1 | 1 |
| Cdh16 | 0,0242318 | 0,00981691 | -1,30356 | 1 | 1 |
| Adad1 | 0,172072 | 0,0695909 | -1,30604 | 1 | 1 |
| Shpk | 0,174694 | 0,0703677 | -1,31185 | 1 | 1 |
| Ogn | 0,193333 | 0,0776274 | -1,31645 | 1 | 1 |
| Klrg2 | 0,129187 | 0,0516357 | -1,32302 | 1 | 1 |
| Mrvi1 | 0,169687 | 0,0678057 | -1,32339 | 1 | 1 |
| Kazald1 | 0,204917 | 0,0818126 | -1,32465 | 1 | 1 |
| Eci3 | 0,294223 | 0,117446 | -1,32492 | 1 | 1 |
| Gdap1l1 | 0,49025 | 0,195092 | -1,32937 | 1 | 1 |
| Ccdc78 | 0,613736 | 0,242976 | -1,3368 | 1 | 1 |
| Aox3 | 0,0751994 | 0,0296047 | -1,34489 | 1 | 1 |
| Itga2b | 0,202277 | 0,0795426 | -1,34653 | 1 | 1 |
| Xkr6 | 0,367037 | 0,143956 | -1,35029 | 1 | 1 |
| A830052D11Rik | 0,339053 | 0,132868 | -1,35152 | 1 | 1 |
| Adamtsl1 | 0,17206 | 0,0673639 | -1,35287 | 1 | 1 |
| Nrn1l | 0,270525 | 0,105874 | -1,35341 | 1 | 1 |
| Gm10421 | 0,170458 | 0,0667116 | -1,35341 | 1 | 1 |
| Mybphl | 0,101449 | 0,0397037 | -1,35341 | 1 | 1 |
| Gm20597 | 0,0205896 | 0,00805808 | -1,35341 | 1 | 1 |
| Tlr7 | 0,030279 | 0,0118486 | -1,3536 | 1 | 1 |
| Rasgef1c | 0,0674798 | 0,0262988 | -1,35946 | 1 | 1 |
| Phex | 0,122204 | 0,0476221 | -1,35959 | 1 | 1 |
| Sycp3 | 0,526193 | 0,204544 | -1,36318 | 1 | 1 |
| Slc22a4 | 0,289257 | 0,112031 | -1,36845 | 1 | 1 |
| 1700109G14Rik | 0,185505 | 0,0716531 | -1,37236 | 1 | 1 |
| Vmn2r57 | 0,0461441 | 0,0178236 | -1,37236 | 1 | 1 |
| Slc28a2 | 0,0297817 | 0,0115034 | -1,37236 | 1 | 1 |
| Dusp13 | 0,127241 | 0,0491477 | -1,37237 | 1 | 1 |

| | | | | | |
|---|---|---|---|---|---|
| Nefh | 0,0658104 | 0,0253993 | -1,37353 | 1 | 1 |
| Spats1 | 0,145498 | 0,0561183 | -1,37446 | 1 | 1 |
| A430093F15Rik | 0,0727875 | 0,028074 | -1,37446 | 1 | 1 |
| Armc3 | 0,0534748 | 0,0206248 | -1,37448 | 1 | 1 |
| Sh2d7 | 0,222577 | 0,0858118 | -1,37506 | 1 | 1 |
| A330074K22Rik | 0,46819 | 0,180163 | -1,37779 | 1 | 1 |
| Grem2 | 0,125603 | 0,0480799 | -1,38536 | 1 | 1 |
| Fam124b | 0,0792479 | 0,0302763 | -1,38819 | 1 | 1 |
| Tex38 | 0,295982 | 0,113007 | -1,3891 | 1 | 1 |
| Synpo2 | 0,458994 | 0,174797 | -1,39279 | 1 | 1 |
| Dnaaf1 | 0,0839772 | 0,0319446 | -1,39442 | 1 | 1 |
| Eif3j2 | 4,41895 | 1,67844 | -1,39658 | 1 | 1 |
| Ahr | 0,434385 | 0,164905 | -1,39734 | 1 | 1 |
| Slco2a1 | 0,0852082 | 0,032133 | -1,40694 | 1 | 1 |
| Pcdh11x | 0,0961745 | 0,0362045 | -1,40948 | 1 | 1 |
| Sigirr | 0,400138 | 0,150194 | -1,41367 | 1 | 1 |
| Fam19a2 | 0,202571 | 0,0759153 | -1,41596 | 1 | 1 |
| Kynu | 0,153499 | 0,0574814 | -1,41706 | 1 | 1 |
| Mpp4 | 0,187147 | 0,0699961 | -1,41882 | 1 | 1 |
| 4933413G19Rik | 0,421239 | 0,157238 | -1,42169 | 1 | 1 |
| 4931430N09Rik | 0,0736624 | 0,0273443 | -1,42969 | 1 | 1 |
| Lrriq1 | 0,154787 | 0,0573669 | -1,43199 | 1 | 1 |
| Nipal2 | 0,0918155 | 0,0339788 | -1,4341 | 1 | 1 |
| Rassf6 | 0,248833 | 0,0917607 | -1,43923 | 1 | 1 |
| Fev | 0,0906824 | 0,0333823 | -1,44174 | 1 | 1 |
| Dhh | 0,0534538 | 0,0196776 | -1,44174 | 1 | 1 |
| Gabra2 | 0,052843 | 0,0194528 | -1,44174 | 1 | 1 |
| Nkx2-2 | 0,201129 | 0,0740212 | -1,44211 | 1 | 1 |
| Myh8 | 0,0808432 | 0,0297253 | -1,44344 | 1 | 1 |
| Gk2 | 0,0887856 | 0,0326335 | -1,44397 | 1 | 1 |
| 4631405J19Rik | 0,0710627 | 0,0261194 | -1,44397 | 1 | 1 |
| Cdh18 | 0,0287331 | 0,0105609 | -1,44397 | 1 | 1 |
| Gm6815 | 0,0227855 | 0,0083749 | -1,44397 | 1 | 1 |
| Tmem236 | 0,0211297 | 0,0077663 | -1,44397 | 1 | 1 |
| Stab2 | 0,0184089 | 0,00676627 | -1,44397 | 1 | 1 |
| Cela1 | 0,549554 | 0,201038 | -1,4508 | 1 | 1 |
| Ppp1r3d | 0,344861 | 0,126015 | -1,45242 | 1 | 1 |
| Pcsk2 | 0,0418685 | 0,015287 | -1,45356 | 1 | 1 |
| Edaradd | 0,367151 | 0,133984 | -1,45432 | 1 | 1 |
| Bcas3os1 | 0,0398974 | 0,0145069 | -1,45956 | 1 | 1 |
| BC061194 | 0,0808973 | 0,0294143 | -1,45957 | 1 | 1 |
| Tlr6 | 0,0488559 | 0,017764 | -1,45957 | 1 | 1 |
| Hrh3 | 0,0483915 | 0,0175952 | -1,45957 | 1 | 1 |
| Gpr111 | 0,0380185 | 0,0138236 | -1,45957 | 1 | 1 |
| Gm2061 | 0,149253 | 0,0541404 | -1,46298 | 1 | 1 |
| Sv2c | 0,318015 | 0,114952 | -1,46807 | 1 | 1 |
| Itgal | 0,108953 | 0,0392738 | -1,47207 | 1 | 1 |
| Glra1 | 0,0535475 | 0,0192678 | -1,47463 | 1 | 1 |
| Spaca1 | 0,359121 | 0,128822 | -1,47909 | 1 | 1 |
| Otx2 | 0,115352 | 0,0413749 | -1,47922 | 1 | 1 |

| | | | | | |
|---|---|---|---|---|---|
| Gal3st1 | 0,431672 | 0,154808 | -1,47946 | 1 | 1 |
| Adhfe1 | 0,093244 | 0,0332307 | -1,48849 | 1 | 1 |
| Mansc1 | 0,283955 | 0,100943 | -1,49212 | 1 | 1 |
| Akr1c12 | 0,382058 | 0,135125 | -1,49949 | 1 | 1 |
| Gpr12 | 0,280617 | 0,0992227 | -1,49986 | 1 | 1 |
| S100b | 0,165397 | 0,0584249 | -1,50128 | 1 | 1 |
| 1700094J05Rik | 0,183821 | 0,0648935 | -1,50216 | 1 | 1 |
| Reps2 | 0,0872011 | 0,030783 | -1,50221 | 1 | 1 |
| Zmat4 | 0,383405 | 0,1353 | -1,50271 | 1 | 1 |
| Gm8234 | 0,211626 | 0,0746791 | -1,50274 | 1 | 1 |
| Gm17762 | 0,421379 | 0,148505 | -1,50461 | 1 | 1 |
| Rnaset2a,Rnaset2b | 3,78532 | 1,33121 | -1,50768 | 1 | 1 |
| A230070E04Rik | 0,179798 | 0,0631826 | -1,50878 | 1 | 1 |
| Dpys | 0,0534551 | 0,0187749 | -1,50953 | 1 | 1 |
| Ppp1r16b | 0,357592 | 0,125555 | -1,50999 | 1 | 1 |
| Taar4 | 0,181939 | 0,0638679 | -1,5103 | 1 | 1 |
| Tnfrsf25 | 0,226992 | 0,0795604 | -1,51252 | 1 | 1 |
| Cyp2s1 | 0,467729 | 0,163813 | -1,51362 | 1 | 1 |
| Gpr22 | 0,0628151 | 0,0219513 | -1,51681 | 1 | 1 |
| Taar2 | 0,189053 | 0,0657463 | -1,52381 | 1 | 1 |
| Atp2c2 | 0,0916716 | 0,0318689 | -1,52432 | 1 | 1 |
| Scrt1 | 0,0727159 | 0,0252791 | -1,52432 | 1 | 1 |
| Scn4b | 0,606339 | 0,210439 | -1,52672 | 1 | 1 |
| Lancl3 | 0,199708 | 0,0692038 | -1,52897 | 1 | 1 |
| Nos2 | 0,0614579 | 0,0212558 | -1,53174 | 1 | 1 |
| Nalcn | 0,0739221 | 0,0255661 | -1,53177 | 1 | 1 |
| Unc79 | 0,0554503 | 0,0191092 | -1,53693 | 1 | 1 |
| Xlr4a | 0,125557 | 0,0431229 | -1,54181 | 1 | 1 |
| Cacna2d2 | 0,565629 | 0,194243 | -1,54199 | 1 | 1 |
| Tlr4 | 0,0555564 | 0,019052 | -1,54401 | 1 | 1 |
| Frat1 | 0,422768 | 0,143957 | -1,55423 | 1 | 1 |
| Xlr | 0,0951454 | 0,032397 | -1,55427 | 1 | 1 |
| Ccser1 | 0,230305 | 0,077976 | -1,56245 | 1 | 1 |
| Bcas3os2 | 0,703496 | 0,236336 | -1,5737 | 1 | 1 |
| Vpreb1 | 0,283146 | 0,0951216 | -1,5737 | 1 | 1 |
| Rbp7 | 0,204735 | 0,0687795 | -1,5737 | 1 | 1 |
| Gpihbp1 | 0,13465 | 0,045235 | -1,5737 | 1 | 1 |
| Olfr91 | 0,108636 | 0,0364958 | -1,5737 | 1 | 1 |
| Nhlrc4 | 0,0824456 | 0,0276972 | -1,5737 | 1 | 1 |
| Lyz1 | 0,0752574 | 0,0252824 | -1,5737 | 1 | 1 |
| Fgg | 0,060537 | 0,0203371 | -1,5737 | 1 | 1 |
| Pdia2 | 0,0526358 | 0,0176827 | -1,5737 | 1 | 1 |
| Zbtb32 | 0,0508265 | 0,0170749 | -1,5737 | 1 | 1 |
| 4921517D22Rik | 0,0271779 | 0,00913029 | -1,5737 | 1 | 1 |
| Morc2b | 0,0227749 | 0,00765112 | -1,5737 | 1 | 1 |
| Exoc3l4 | 0,0807859 | 0,0271387 | -1,57375 | 1 | 1 |
| Igsf23 | 0,237799 | 0,0793161 | -1,58406 | 1 | 1 |
| Thrb | 0,0579951 | 0,0193181 | -1,58598 | 1 | 1 |
| Khdrbs2 | 0,139511 | 0,0464495 | -1,58664 | 1 | 1 |
| 1700084J12Rik | 0,553932 | 0,182579 | -1,60119 | 1 | 1 |

| | | | | | |
|---|---|---|---|---|---|
| Mia2 | 0,147921 | 0,0487269 | -1,60204 | 1 | 1 |
| Cryba1 | 0,712789 | 0,233711 | -1,60875 | 1 | 1 |
| 1300017J02Rik | 0,102404 | 0,0335043 | -1,61186 | 1 | 1 |
| 2610027K06Rik | 0,275914 | 0,0901524 | -1,61378 | 1 | 1 |
| Chrna1 | 0,0949804 | 0,0310224 | -1,61432 | 1 | 1 |
| Ptprq | 0,0359518 | 0,0117413 | -1,61448 | 1 | 1 |
| Ulbp1 | 0,0809857 | 0,0263545 | -1,61962 | 1 | 1 |
| Gda | 0,0977243 | 0,0317668 | -1,6212 | 1 | 1 |
| Dnali1 | 0,201192 | 0,0653755 | -1,62175 | 1 | 1 |
| Slc1a1 | 0,0848621 | 0,0274579 | -1,6279 | 1 | 1 |
| C1qtnf3 | 0,538873 | 0,174164 | -1,6295 | 1 | 1 |
| 5330434G04Rik | 0,183714 | 0,0592988 | -1,63139 | 1 | 1 |
| Tnfsf13b | 0,311484 | 0,100309 | -1,63471 | 1 | 1 |
| B3galt2 | 0,106392 | 0,034262 | -1,63471 | 1 | 1 |
| Acbd7 | 0,399474 | 0,128584 | -1,63539 | 1 | 1 |
| Gm711 | 0,0672928 | 0,0216604 | -1,63539 | 1 | 1 |
| 4921513I03Rik | 0,096598 | 0,0310222 | -1,63869 | 1 | 1 |
| 9530052E02Rik | 0,129657 | 0,0415202 | -1,64282 | 1 | 1 |
| Sostdc1 | 0,226074 | 0,0723657 | -1,64342 | 1 | 1 |
| Angptl7 | 0,0904209 | 0,0288748 | -1,64684 | 1 | 1 |
| Evx2 | 0,397023 | 0,126235 | -1,65311 | 1 | 1 |
| C1s1 | 0,237828 | 0,0751912 | -1,66128 | 1 | 1 |
| Hlf | 0,463917 | 0,145857 | -1,66931 | 1 | 1 |
| E030013I19Rik | 0,258351 | 0,0810286 | -1,67283 | 1 | 1 |
| A230020J21Rik | 0,0944904 | 0,0295033 | -1,67929 | 1 | 1 |
| Gm19522 | 0,0372855 | 0,0115385 | -1,69215 | 1 | 1 |
| Hist1h2ba | 0,380308 | 0,117646 | -1,69272 | 1 | 1 |
| Dok6 | 0,109488 | 0,0338693 | -1,69272 | 1 | 1 |
| Tbx5 | 0,0464092 | 0,0143563 | -1,69272 | 1 | 1 |
| Irx6 | 0,0398508 | 0,0123276 | -1,69272 | 1 | 1 |
| Ppef1 | 0,0380498 | 0,0117704 | -1,69272 | 1 | 1 |
| Slc6a1 | 0,0336493 | 0,010404 | -1,69344 | 1 | 1 |
| Kcna5 | 0,598254 | 0,184644 | -1,69601 | 1 | 1 |
| Art4 | 0,143361 | 0,044134 | -1,69969 | 1 | 1 |
| 4930548H24Rik | 0,142983 | 0,0438669 | -1,70464 | 1 | 1 |
| Amn | 0,118075 | 0,0362251 | -1,70464 | 1 | 1 |
| Gm2762 | 0,0651509 | 0,0199357 | -1,70843 | 1 | 1 |
| Skor1 | 0,0409557 | 0,0125217 | -1,70963 | 1 | 1 |
| Zfp385c | 0,0625657 | 0,0190643 | -1,7145 | 1 | 1 |
| Gm4907 | 0,571355 | 0,173981 | -1,71546 | 1 | 1 |
| Ecel1 | 0,0330414 | 0,0100544 | -1,71645 | 1 | 1 |
| Ranbp3l | 0,022 | 0,00669449 | -1,71645 | 1 | 1 |
| 4933440M02Rik | 0,0598825 | 0,018219 | -1,71669 | 1 | 1 |
| Creb5 | 0,43697 | 0,132399 | -1,72264 | 1 | 1 |
| Krt1 | 0,181698 | 0,0548537 | -1,72788 | 1 | 1 |
| Ube2ql1 | 0,600124 | 0,18071 | -1,73158 | 1 | 1 |
| Hbb-bt | 0,542806 | 0,162877 | -1,73665 | 1 | 1 |
| D630032N06Rik | 0,533655 | 0,159142 | -1,7456 | 1 | 1 |
| Vnn1 | 0,10519 | 0,0313686 | -1,7456 | 1 | 1 |
| Cdk15 | 0,424019 | 0,126432 | -1,74577 | 1 | 1 |

| | | | | | |
|---|---|---|---|---|---|
| Slc22a3 | 0,163818 | 0,0488464 | -1,74577 | 1 | 1 |
| Cacng5 | 0,391507 | 0,116643 | -1,74694 | 1 | 1 |
| Pon1 | 0,178851 | 0,0530077 | -1,75449 | 1 | 1 |
| Cdh8 | 0,0308507 | 0,00904777 | -1,76967 | 1 | 1 |
| 1700096J18Rik | 0,732662 | 0,214588 | -1,77158 | 1 | 1 |
| Nlrp6 | 0,0432832 | 0,0126771 | -1,77158 | 1 | 1 |
| Ccdc85a | 0,0802553 | 0,0234416 | -1,77552 | 1 | 1 |
| 9430060I03Rik | 0,14472 | 0,042258 | -1,77597 | 1 | 1 |
| Tac1 | 0,175858 | 0,0512798 | -1,77795 | 1 | 1 |
| Csf3r | 0,0431957 | 0,0125954 | -1,77799 | 1 | 1 |
| Kcnj2 | 0,178518 | 0,0520403 | -1,77837 | 1 | 1 |
| Gdpd2 | 0,323069 | 0,0936429 | -1,7866 | 1 | 1 |
| Bai1 | 0,458838 | 0,132247 | -1,79475 | 1 | 1 |
| Dcc | 0,183919 | 0,0526557 | -1,80441 | 1 | 1 |
| Fam46a | 0,351052 | 0,100497 | -1,80453 | 1 | 1 |
| Fam26f | 0,338672 | 0,0969528 | -1,80453 | 1 | 1 |
| Stk32a | 0,257912 | 0,0735221 | -1,81063 | 1 | 1 |
| Lum | 0,479093 | 0,136491 | -1,81149 | 1 | 1 |
| Dusp8 | 0,175443 | 0,0499104 | -1,81359 | 1 | 1 |
| Mafa | 0,215298 | 0,0612432 | -1,81371 | 1 | 1 |
| Cxcl14 | 0,649809 | 0,184672 | -1,81505 | 1 | 1 |
| E130006D01Rik | 0,531886 | 0,150833 | -1,81817 | 1 | 1 |
| Dock2 | 0,276439 | 0,0781896 | -1,82191 | 1 | 1 |
| Folh1 | 0,248328 | 0,0701197 | -1,82436 | 1 | 1 |
| Myl3 | 0,136915 | 0,0386519 | -1,82467 | 1 | 1 |
| Palmd | 0,0433883 | 0,0122487 | -1,82467 | 1 | 1 |
| 4833403I15Rik | 0,0315159 | 0,00889711 | -1,82467 | 1 | 1 |
| Macc1 | 0,0263069 | 0,00742658 | -1,82467 | 1 | 1 |
| Pitpnm3 | 0,029616 | 0,00835761 | -1,82521 | 1 | 1 |
| 1700007K13Rik | 0,569046 | 0,160351 | -1,82731 | 1 | 1 |
| Grip2 | 0,100094 | 0,0281035 | -1,83254 | 1 | 1 |
| Cspg5 | 0,41471 | 0,11609 | -1,83686 | 1 | 1 |
| Gm13547 | 0,359593 | 0,100145 | -1,84427 | 1 | 1 |
| Gm8909 | 0,13255 | 0,0369145 | -1,84427 | 1 | 1 |
| A330041J22Rik | 0,093246 | 0,0259686 | -1,84427 | 1 | 1 |
| BC035044 | 0,0634679 | 0,0176755 | -1,84427 | 1 | 1 |
| Lox | 0,20156 | 0,0560674 | -1,84598 | 1 | 1 |
| Cys1 | 0,284539 | 0,078704 | -1,85412 | 1 | 1 |
| Rorb | 0,122879 | 0,0337399 | -1,86471 | 1 | 1 |
| Papln | 0,0476083 | 0,0130649 | -1,86552 | 1 | 1 |
| Ebi3 | 0,505323 | 0,138259 | -1,86983 | 1 | 1 |
| Atp1a3 | 0,0989983 | 0,0270828 | -1,87003 | 1 | 1 |
| Ppp1r17 | 0,304299 | 0,0832383 | -1,87017 | 1 | 1 |
| Kcnj9 | 0,064223 | 0,0174982 | -1,87588 | 1 | 1 |
| Slc22a13b-ps | 0,20361 | 0,0554653 | -1,87615 | 1 | 1 |
| Rnf152 | 0,470882 | 0,127895 | -1,8804 | 1 | 1 |
| 03/10/16 | 0,172332 | 0,0467331 | -1,88268 | 1 | 1 |
| Gm10364 | 0,316828 | 0,0858266 | -1,8842 | 1 | 1 |
| Bmx | 0,182903 | 0,0495448 | -1,88428 | 1 | 1 |
| BC030500 | 0,556261 | 0,148581 | -1,90452 | 1 | 1 |

| | | | | | |
|---|---|---|---|---|---|
| Gm1631 | 0,365431 | 0,097395 | -1,90768 | 1 | 1 |
| Gm9871 | 0,11777 | 0,0313883 | -1,90768 | 1 | 1 |
| Rnf112 | 0,0565216 | 0,0150621 | -1,90788 | 1 | 1 |
| Rxfp2 | 0,238379 | 0,063294 | -1,91312 | 1 | 1 |
| Kctd8 | 0,0974244 | 0,0258426 | -1,91453 | 1 | 1 |
| Gm4832 | 0,362573 | 0,0954579 | -1,92533 | 1 | 1 |
| Erich4 | 0,144314 | 0,0379949 | -1,92533 | 1 | 1 |
| Maats1 | 0,0795682 | 0,0209487 | -1,92533 | 1 | 1 |
| Dusp15 | 0,0349288 | 0,00919602 | -1,92533 | 1 | 1 |
| A930041C12Rik | 0,306088 | 0,0805522 | -1,92595 | 1 | 1 |
| 9130019P16Rik | 0,142482 | 0,037434 | -1,92836 | 1 | 1 |
| Gimap8 | 0,207685 | 0,0544757 | -1,93071 | 1 | 1 |
| Diras1 | 0,113071 | 0,0296297 | -1,93211 | 1 | 1 |
| Omd | 0,238461 | 0,0621926 | -1,93894 | 1 | 1 |
| Nlgn1 | 0,0603953 | 0,0156885 | -1,94473 | 1 | 1 |
| 2900092D14Rik | 0,522697 | 0,13577 | -1,94482 | 1 | 1 |
| Steap4 | 0,168346 | 0,0434905 | -1,95266 | 1 | 1 |
| Agrp | 0,586004 | 0,149851 | -1,96738 | 1 | 1 |
| Ccno | 0,0996195 | 0,0254561 | -1,96842 | 1 | 1 |
| Trpc4 | 0,168994 | 0,0430318 | -1,9735 | 1 | 1 |
| Ak7 | 0,188719 | 0,0480356 | -1,97406 | 1 | 1 |
| Ano7 | 0,10247 | 0,0260337 | -1,97675 | 1 | 1 |
| 4930426D05Rik | 0,421432 | 0,106766 | -1,98084 | 1 | 1 |
| Plcl1 | 0,23091 | 0,0581187 | -1,99026 | 1 | 1 |
| Pdk4 | 0,262329 | 0,0655013 | -2,00178 | 1 | 1 |
| Fbln5 | 0,551828 | 0,137245 | -2,00746 | 1 | 1 |
| Fam65b | 0,110537 | 0,0273452 | -2,01517 | 1 | 1 |
| Gdap1 | 0,343396 | 0,0845063 | -2,02274 | 1 | 1 |
| Wt1os | 0,0566213 | 0,0138958 | -2,0267 | 1 | 1 |
| Trim30a | 0,0486347 | 0,0119357 | -2,0267 | 1 | 1 |
| Abca6 | 0,17352 | 0,0422669 | -2,0375 | 1 | 1 |
| Krt79 | 0,0582102 | 0,0141705 | -2,03838 | 1 | 1 |
| Fbxo43 | 0,0287633 | 0,00700204 | -2,03838 | 1 | 1 |
| Sfrp5 | 0,103107 | 0,0250959 | -2,03861 | 1 | 1 |
| 2610034M16Rik | 0,0440477 | 0,0107211 | -2,03861 | 1 | 1 |
| Cpz | 0,325884 | 0,0784726 | -2,0541 | 1 | 1 |
| Frmpd3 | 0,279503 | 0,0672296 | -2,05569 | 1 | 1 |
| Myo7b | 0,0909631 | 0,0217278 | -2,06574 | 1 | 1 |
| Itga2 | 0,318998 | 0,0758284 | -2,07274 | 1 | 1 |
| Saa2 | 0,719015 | 0,1709 | -2,07287 | 1 | 1 |
| Chst9 | 0,193829 | 0,0460508 | -2,07349 | 1 | 1 |
| Prkg2 | 0,548313 | 0,129547 | -2,08152 | 1 | 1 |
| Col5a3 | 0,419472 | 0,0989334 | -2,08405 | 1 | 1 |
| Olfr543 | 0,513064 | 0,120895 | -2,08538 | 1 | 1 |
| Angpt4 | 0,258575 | 0,0606893 | -2,09107 | 1 | 1 |
| Popdc2 | 0,272691 | 0,0636033 | -2,10009 | 1 | 1 |
| 4933408N05Rik | 0,132927 | 0,0308872 | -2,10555 | 1 | 1 |
| Bglap | 0,380832 | 0,088486 | -2,10564 | 1 | 1 |
| Gm5523 | 0,39809 | 0,0924222 | -2,10678 | 1 | 1 |
| Cd70 | 0,181221 | 0,0420445 | -2,10776 | 1 | 1 |

| | | | | | |
|---|---|---|---|---|---|
| E030018B13Rik | 0,142564 | 0,0330758 | -2,10776 | 1 | 1 |
| Hist1h4f | 0,757201 | 0,173518 | -2,1256 | 1 | 1 |
| 1500015O10Rik | 0,171136 | 0,039217 | -2,1256 | 1 | 1 |
| Foxe1 | 0,128353 | 0,0294129 | -2,1256 | 1 | 1 |
| Rassf4 | 0,531009 | 0,120232 | -2,14292 | 1 | 1 |
| Cplx1 | 0,189741 | 0,0428195 | -2,14769 | 1 | 1 |
| Gm10578 | 0,14117 | 0,0318583 | -2,14769 | 1 | 1 |
| Vit | 0,0769717 | 0,0173681 | -2,14789 | 1 | 1 |
| Tmem54 | 0,272197 | 0,0609524 | -2,15889 | 1 | 1 |
| Paupar | 0,116415 | 0,0259687 | -2,16443 | 1 | 1 |
| Mgst2 | 0,625915 | 0,139346 | -2,1673 | 1 | 1 |
| Lsamp | 0,39307 | 0,0870859 | -2,17427 | 1 | 1 |
| 2810433D01Rik | 0,450825 | 0,0990703 | -2,18604 | 1 | 1 |
| Gm10549 | 0,174598 | 0,0383621 | -2,18628 | 1 | 1 |
| Clec1b | 0,54381 | 0,118556 | -2,19754 | 1 | 1 |
| Fam107a | 0,427055 | 0,0928228 | -2,20187 | 1 | 1 |
| Il1r2 | 0,108706 | 0,0235304 | -2,20784 | 1 | 1 |
| Osgin1 | 0,0705323 | 0,0152673 | -2,20784 | 1 | 1 |
| Aldh3b2 | 0,0633241 | 0,0137071 | -2,20784 | 1 | 1 |
| Daw1 | 0,107853 | 0,0233452 | -2,20786 | 1 | 1 |
| Gm12295 | 0,131931 | 0,0285177 | -2,20986 | 1 | 1 |
| Gm7008 | 1,00641 | 0,215965 | -2,22036 | 1 | 1 |
| Nr1i3 | 0,121453 | 0,0260131 | -2,22308 | 1 | 1 |
| Alox12 | 0,117066 | 0,0248968 | -2,23328 | 1 | 1 |
| A730090N16Rik | 0,49277 | 0,104356 | -2,23939 | 1 | 1 |
| Gbp8 | 0,0903655 | 0,0188748 | -2,25931 | 1 | 1 |
| Tmem56 | 0,2219 | 0,0461201 | -2,26644 | 1 | 1 |
| 2610100L16Rik | 0,451394 | 0,0925785 | -2,28564 | 1 | 1 |
| 1700101I11Rik | 0,217259 | 0,0445587 | -2,28564 | 1 | 1 |
| Cstad | 0,213361 | 0,0437591 | -2,28564 | 1 | 1 |
| Zmynd12 | 0,100532 | 0,0206185 | -2,28564 | 1 | 1 |
| Klf14 | 0,455999 | 0,0923269 | -2,30421 | 1 | 1 |
| Dcn | 0,480592 | 0,0971712 | -2,30621 | 1 | 1 |
| Sowahb | 0,429233 | 0,0866107 | -2,30914 | 1 | 1 |
| Dner | 0,240619 | 0,048308 | -2,31641 | 1 | 1 |
| Pcdhb21 | 0,107611 | 0,0213914 | -2,33073 | 1 | 1 |
| A730056A06Rik | 0,461507 | 0,0917225 | -2,33101 | 1 | 1 |
| Tbx3os2 | 0,543218 | 0,107174 | -2,34157 | 1 | 1 |
| Nsun7 | 0,0926353 | 0,0181272 | -2,35341 | 1 | 1 |
| Ap3b2 | 0,518376 | 0,101183 | -2,35703 | 1 | 1 |
| Tslp | 0,147745 | 0,0288338 | -2,35727 | 1 | 1 |
| Hist2h2ab | 0,588257 | 0,11463 | -2,35946 | 1 | 1 |
| Cntn3 | 0,522395 | 0,101647 | -2,36157 | 1 | 1 |
| D830046C22Rik | 0,101693 | 0,019769 | -2,36291 | 1 | 1 |
| Galnt9 | 0,102806 | 0,0199663 | -2,36428 | 1 | 1 |
| Tecrl | 0,205052 | 0,0392463 | -2,38536 | 1 | 1 |
| Rspo2 | 0,376278 | 0,0713968 | -2,39787 | 1 | 1 |
| Wnt7b | 0,364592 | 0,0691776 | -2,39791 | 1 | 1 |
| Slc16a12 | 0,0773659 | 0,0146344 | -2,40234 | 1 | 1 |
| Emx2os | 0,0588553 | 0,0111329 | -2,40234 | 1 | 1 |

| | | | | | |
|---|---|---|---|---|---|
| Dct | 0,413942 | 0,0781683 | -2,40477 | 1 | 1 |
| Anxa1 | 0,358277 | 0,0675683 | -2,40666 | 1 | 1 |
| Tmem125 | 0,136195 | 0,025671 | -2,40747 | 1 | 1 |
| C730036E19Rik | 0,0926981 | 0,0174724 | -2,40747 | 1 | 1 |
| 9430076C15Rik | 0,203226 | 0,037982 | -2,4197 | 1 | 1 |
| Mtnr1a | 0,509825 | 0,0947162 | -2,42832 | 1 | 1 |
| Wfdc1 | 0,149766 | 0,0277975 | -2,42969 | 1 | 1 |
| BC051019 | 0,0839837 | 0,0154342 | -2,44397 | 1 | 1 |
| Foxn4 | 0,0527708 | 0,00969804 | -2,44397 | 1 | 1 |
| Ptpn22 | 0,057337 | 0,0105127 | -2,44733 | 1 | 1 |
| Acan | 0,0529499 | 0,00970321 | -2,44809 | 1 | 1 |
| Veph1 | 0,0844784 | 0,0154509 | -2,4509 | 1 | 1 |
| Mal2 | 0,116217 | 0,0208672 | -2,47752 | 1 | 1 |
| Myog | 0,306377 | 0,0543786 | -2,4942 | 1 | 1 |
| CK137956 | 0,271924 | 0,0479861 | -2,50251 | 1 | 1 |
| Slitrk1 | 0,339165 | 0,0596549 | -2,50728 | 1 | 1 |
| Ferd3l | 0,224418 | 0,0393897 | -2,5103 | 1 | 1 |
| Pxt1 | 0,188445 | 0,0330758 | -2,5103 | 1 | 1 |
| Phxr4 | 0,129957 | 0,02281 | -2,5103 | 1 | 1 |
| Gm19434 | 0,06454 | 0,011328 | -2,5103 | 1 | 1 |
| Kcng3 | 0,0484869 | 0,00851041 | -2,5103 | 1 | 1 |
| Lhx1os | 0,731171 | 0,127734 | -2,51707 | 1 | 1 |
| Mnda | 0,212231 | 0,0370763 | -2,51707 | 1 | 1 |
| Cp | 0,494869 | 0,0861774 | -2,52167 | 1 | 1 |
| Map3k7cl | 0,170174 | 0,029394 | -2,53342 | 1 | 1 |
| Sez6 | 0,062816 | 0,0107835 | -2,54231 | 1 | 1 |
| Fam110c | 0,328333 | 0,0562753 | -2,54458 | 1 | 1 |
| Kremen2 | 0,24685 | 0,042067 | -2,55288 | 1 | 1 |
| Tceal7 | 0,531348 | 0,090465 | -2,55423 | 1 | 1 |
| Lhfpl1 | 0,432404 | 0,0713056 | -2,60029 | 1 | 1 |
| Epha8 | 0,568667 | 0,0937595 | -2,60055 | 1 | 1 |
| Cyp2j13 | 0,0623853 | 0,0102069 | -2,61166 | 1 | 1 |
| Nkx2-6 | 0,274771 | 0,0447083 | -2,61962 | 1 | 1 |
| Bdkrb2 | 0,223967 | 0,0363478 | -2,62334 | 1 | 1 |
| Ano4 | 0,109403 | 0,0177177 | -2,62639 | 1 | 1 |
| Foxl2os | 0,074553 | 0,0120643 | -2,62753 | 1 | 1 |
| Oit3 | 0,0663074 | 0,0106787 | -2,63444 | 1 | 1 |
| Unc5a | 0,562841 | 0,0878514 | -2,67959 | 1 | 1 |
| Ntng1 | 0,310548 | 0,0483791 | -2,68236 | 1 | 1 |
| Lgr6 | 0,463587 | 0,0719726 | -2,68732 | 1 | 1 |
| Fat2 | 0,08291 | 0,0127532 | -2,70068 | 1 | 1 |
| Tbx15 | 0,396752 | 0,0605149 | -2,71288 | 1 | 1 |
| Gm15663 | 0,455245 | 0,0693875 | -2,71389 | 1 | 1 |
| Gm15179 | 0,373524 | 0,0568309 | -2,71645 | 1 | 1 |
| Cldn2 | 0,160702 | 0,0242444 | -2,72867 | 1 | 1 |
| Gpr17 | 0,0568184 | 0,00853133 | -2,73552 | 1 | 1 |
| Ntrk2 | 0,538673 | 0,0807711 | -2,7375 | 1 | 1 |
| Pax7 | 0,35172 | 0,052714 | -2,73817 | 1 | 1 |
| 4933422A05Rik | 0,157756 | 0,0232853 | -2,76021 | 1 | 1 |
| Tacr3 | 0,0510495 | 0,00753506 | -2,76021 | 1 | 1 |

| | | | | | |
|---|---|---|---|---|---|
| Xpnpep2 | 0,055169 | 0,00814212 | -2,76038 | 1 | 1 |
| BB557941 | 0,204662 | 0,0299715 | -2,77158 | 1 | 1 |
| Casq2 | 0,239616 | 0,0348374 | -2,78202 | 1 | 1 |
| Nrxn3 | 0,189419 | 0,0273228 | -2,7934 | 1 | 1 |
| Gm6583 | 0,145257 | 0,0206916 | -2,81149 | 1 | 1 |
| Cpa1 | 0,138311 | 0,019523 | -2,82467 | 1 | 1 |
| Fam155a | 0,400657 | 0,0563648 | -2,8295 | 1 | 1 |
| Stmn4 | 0,480798 | 0,067497 | -2,83254 | 1 | 1 |
| Scrt2 | 0,241173 | 0,0336256 | -2,84244 | 1 | 1 |
| Glp2r | 0,379561 | 0,0527039 | -2,84835 | 1 | 1 |
| Ccdc184 | 0,56176 | 0,0779292 | -2,84972 | 1 | 1 |
| Hpca | 0,14453 | 0,0198996 | -2,86056 | 1 | 1 |
| 1810019D21Rik | 0,16662 | 0,022867 | -2,86522 | 1 | 1 |
| Gja5 | 0,066954 | 0,00918443 | -2,86591 | 1 | 1 |
| Slamf8 | 0,170222 | 0,0231822 | -2,87633 | 1 | 1 |
| Gbx1 | 0,409352 | 0,0555986 | -2,88022 | 1 | 1 |
| AI662270 | 0,169247 | 0,0222796 | -2,92533 | 1 | 1 |
| Cxcr5 | 0,0845185 | 0,011126 | -2,92533 | 1 | 1 |
| Lamp5 | 0,18715 | 0,0243076 | -2,94471 | 1 | 1 |
| Srrm3 | 0,111163 | 0,014391 | -2,94944 | 1 | 1 |
| Gnat1 | 0,211656 | 0,0267712 | -2,98297 | 1 | 1 |
| Pax4 | 0,576021 | 0,0723534 | -2,99299 | 1 | 1 |
| Fbp1 | 0,336929 | 0,0420779 | -3,00131 | 1 | 1 |
| Xkr7 | 0,139505 | 0,0169775 | -3,03861 | 1 | 1 |
| Rsad2 | 0,123396 | 0,0149817 | -3,04202 | 1 | 1 |
| Adamtsl3 | 0,161796 | 0,0195983 | -3,04538 | 1 | 1 |
| Prr16 | 0,502967 | 0,0609093 | -3,04573 | 1 | 1 |
| St8sia3 | 0,13076 | 0,0158348 | -3,04575 | 1 | 1 |
| Tnfsf4 | 0,25381 | 0,0302936 | -3,06667 | 1 | 1 |
| Syt6 | 0,169654 | 0,0198211 | -3,09748 | 1 | 1 |
| Podn | 0,489418 | 0,0563411 | -3,11881 | 1 | 1 |
| Hdc | 0,106659 | 0,0122208 | -3,1256 | 1 | 1 |
| Kcnd2 | 0,0539194 | 0,00617799 | -3,1256 | 1 | 1 |
| Avpr1a | 0,2869 | 0,0326203 | -3,1367 | 1 | 1 |
| Lrrc36 | 0,252343 | 0,0285252 | -3,14508 | 1 | 1 |
| Nrsn1 | 0,124668 | 0,0138773 | -3,1673 | 1 | 1 |
| Gm16897 | 0,11194 | 0,0124357 | -3,17017 | 1 | 1 |
| Negr1 | 0,340665 | 0,0377945 | -3,17211 | 1 | 1 |
| Brinp1 | 0,244769 | 0,0270239 | -3,17911 | 1 | 1 |
| 4930438E09Rik | 0,154112 | 0,0468956 | -3,19944 | 1 | 1 |
| Hnmt | 0,344674 | 0,0374124 | -3,20364 | 1 | 1 |
| Slc22a1 | 0,276279 | 0,0299885 | -3,20364 | 1 | 1 |
| Popdc3 | 0,150486 | 0,016287 | -3,20784 | 1 | 1 |
| Cmtm5 | 0,367264 | 0,0391026 | -3,23148 | 1 | 1 |
| Shox2 | 0,578972 | 0,061197 | -3,24196 | 1 | 1 |
| Neurod1 | 0,57777 | 0,0606004 | -3,2531 | 1 | 1 |
| Dcx | 0,123193 | 0,0127598 | -3,27124 | 1 | 1 |
| Cdr1 | 0,12335 | 0,0127192 | -3,27769 | 1 | 1 |
| Slc7a14 | 0,0310853 | 0,00310617 | -3,32302 | 1 | 1 |
| Isl2 | 0,590986 | 0,0584165 | -3,33868 | 1 | 1 |

| | | | | | |
|---|---|---|---|---|---|
| Gata6 | 0,445657 | 0,0427051 | -3,38345 | 1 | 1 |
| Bean1 | 0,0954625 | 0,00902767 | -3,40251 | 1 | 1 |
| Rufy4 | 0,135594 | 0,0127511 | -3,4106 | 1 | 1 |
| Mobp | 0,190557 | 0,0178616 | -3,41529 | 1 | 1 |
| Myt1 | 0,33523 | 0,0302857 | -3,46844 | 1 | 1 |
| Lamc3 | 0,0521822 | 0,00468475 | -3,47752 | 1 | 1 |
| Fap | 0,569993 | 0,0510752 | -3,48025 | 1 | 1 |
| 3110035E14Rik | 0,48145 | 0,0428053 | -3,49152 | 1 | 1 |
| Rab40b | 0,288435 | 0,0256247 | -3,49264 | 1 | 1 |
| Sec16b | 0,427284 | 0,0361562 | -3,56288 | 1 | 1 |
| Chrdl2 | 0,41376 | 0,0334014 | -3,63081 | 1 | 1 |
| 4930503B20Rik | 0,171699 | 0,01377 | -3,64028 | 1 | 1 |
| Cpa5 | 0,309952 | 0,024197 | -3,67914 | 1 | 1 |
| Pappa2 | 0,205373 | 0,0152355 | -3,75274 | 1 | 1 |
| Kif19a | 0,224785 | 0,0166149 | -3,75799 | 1 | 1 |
| Evi2b | 0,121689 | 0,00884973 | -3,78142 | 1 | 1 |
| Hsd17b14 | 0,31898 | 0,0231047 | -3,78721 | 1 | 1 |
| Gabrg1 | 0,133542 | 0,00931963 | -3,84088 | 1 | 1 |
| Syt2 | 0,314177 | 0,0218063 | -3,84876 | 1 | 1 |
| Dnm3 | 0,526524 | 0,0355069 | -3,89033 | 1 | 1 |
| Gpr183 | 0,158356 | 0,00979722 | -4,01465 | 1 | 1 |
| Igsf5 | 0,500939 | 0,0308639 | -4,02064 | 1 | 1 |
| Gpr150 | 0,225485 | 0,0138129 | -4,02894 | 1 | 1 |
| Lama2 | 0,596187 | 0,0364853 | -4,03038 | 1 | 1 |
| 2210039B01Rik | 0,540863 | 0,0321251 | -4,07349 | 1 | 1 |
| Insm1 | 0,415792 | 0,024623 | -4,07779 | 1 | 1 |
| Cdh26 | 0,166474 | 0,00984953 | -4,0791 | 1 | 1 |
| Hal | 0,179519 | 0,010503 | -4,09526 | 1 | 1 |
| Lrrc10b | 0,405852 | 0,0227243 | -4,15865 | 1 | 1 |
| 6430411K18Rik | 0,224581 | 0,0120461 | -4,2206 | 1 | 1 |
| Hk3 | 0,309052 | 0,0158205 | -4,28798 | 1 | 1 |
| Gm5607 | 0,362416 | 0,0175047 | -4,37183 | 1 | 1 |
| Hmgcs2 | 0,306579 | 0,0137743 | -4,4762 | 1 | 1 |
| Rxfp3 | 0,478661 | 0,021029 | -4,50855 | 1 | 1 |
| 1700106J16Rik | 0,617231 | 0,0264281 | -4,54566 | 1 | 1 |
| Baalc | 0,225054 | 0,00712857 | -4,98052 | 1 | 1 |
| Sec1 | 0,386125 | 0,0118379 | -5,02758 | 1 | 1 |
| 2810055G20Rik | 0,300374 | 0,00910241 | -5,04437 | 1 | 1 |
| 3110099E03Rik | 0,363377 | 0,0109938 | -5,04671 | 1 | 1 |
| Insrr | 0,184413 | 0,00547669 | -5,07349 | 1 | 1 |
| Nr4a3 | 0,572059 | 0,0159671 | -5,16299 | 1 | 1 |
| 1700095B10Rik | 0,000375854 | 1,01E-005 | -5,22419 | 1 | 1 |
| Vgll3 | 0,576791 | 0,0136114 | -5,40517 | 1 | 1 |
| Corin | 0,304787 | 0,00595654 | -5,67718 | 1 | 1 |
| Fendrr | 0,633945 | 0,0123276 | -5,6844 | 1 | 1 |
| Gm5512 | 0,0358756 | 0,000443293 | -6,3386 | 1 | 1 |
| Ass1 | 0,244334 | 0,000124364 | -10,9401 | 1 | 1 |
| G630025P09Rik | 0,0468128 | 4,91E-006 | -13,2175 | 1 | 1 |
