## Supplementary material for "Epha1 is a cell surface marker for neuromesodermal progenitors and their early mesoderm derivatives": Table S2 containing datasets

| Gene | $Tail^{Tot}$ | $Tail^{Prog}$ | $Log_2$fold_change | p_value | q_value |
|---|---|---|---|---|---|
| 2610528J11Rik | 0,840903 | 0 | -inf | 5,00E-005 | 0,001041 |
| AI646519 | 0,929593 | 0 | -inf | 5,00E-005 | 0,001041 |
| Apoa4 | 0,797064 | 0 | -inf | 5,00E-005 | 0,001041 |
| Apoc2 | 0,904617 | 0 | -inf | 5,00E-005 | 0,001041 |
| Aqp8 | 1,19393 | 0 | -inf | 5,00E-005 | 0,001041 |
| Bin2 | 2,77716 | 0 | -inf | 5,00E-005 | 0,001041 |
| C1qa | 3,47272 | 0 | -inf | 5,00E-005 | 0,001041 |
| C1qb | 6,54157 | 0 | -inf | 5,00E-005 | 0,001041 |
| C1qc | 4,51354 | 0 | -inf | 5,00E-005 | 0,001041 |
| C330013F16Rik | 1,15267 | 0 | -inf | 5,00E-005 | 0,001041 |
| Cbr2 | 1,29722 | 0 | -inf | 5,00E-005 | 0,001041 |
| Cldn13 | 3,21867 | 0 | -inf | 5,00E-005 | 0,001041 |
| Csf1r | 2,25752 | 0 | -inf | 5,00E-005 | 0,001041 |
| Ctla2b | 1,57788 | 0 | -inf | 5,00E-005 | 0,001041 |
| Ctsj | 1,91581 | 0 | -inf | 5,00E-005 | 0,001041 |
| Ctss | 2,77271 | 0 | -inf | 5,00E-005 | 0,001041 |
| Epb4.2 | 6,00408 | 0 | -inf | 5,00E-005 | 0,001041 |
| Fcrls | 1,03021 | 0 | -inf | 5,00E-005 | 0,001041 |
| Gp9 | 2,37035 | 0 | -inf | 5,00E-005 | 0,001041 |
| Grap2 | 1,25904 | 0 | -inf | 5,00E-005 | 0,001041 |
| Gsdmc2 | 0,61308 | 0 | -inf | 5,00E-005 | 0,001041 |
| Gzmc | 0,843881 | 0 | -inf | 5,00E-005 | 0,001041 |
| Gzmd | 2,03608 | 0 | -inf | 5,00E-005 | 0,001041 |
| Gzmf | 1,89884 | 0 | -inf | 5,00E-005 | 0,001041 |
| Gzmg | 1,86218 | 0 | -inf | 5,00E-005 | 0,001041 |
| Hbb-b1,Hbb-bs | 31,3241 | 0 | -inf | 5,00E-005 | 0,001041 |
| Hpgd | 1,08255 | 0 | -inf | 5,00E-005 | 0,001041 |
| I830077J02Rik | 0,72781 | 0 | -inf | 5,00E-005 | 0,001041 |
| Icam4 | 3,54386 | 0 | -inf | 5,00E-005 | 0,001041 |
| Il17re | 0,705281 | 0 | -inf | 5,00E-005 | 0,001041 |
| Lcp2 | 0,860397 | 0 | -inf | 5,00E-005 | 0,001041 |
| Ly86 | 1,76944 | 0 | -inf | 5,00E-005 | 0,001041 |
| Lyz2 | 2,00627 | 0 | -inf | 5,00E-005 | 0,001041 |
| Mrap | 1,78004 | 0 | -inf | 5,00E-005 | 0,001041 |
| Ms4a6b | 1,03008 | 0 | -inf | 5,00E-005 | 0,001041 |
| Ms4a6c | 1,2146 | 0 | -inf | 5,00E-005 | 0,001041 |
| Ms4a6d | 0,760855 | 0 | -inf | 5,00E-005 | 0,001041 |
| Pdzk1ip1 | 8,7018 | 0 | -inf | 5,00E-005 | 0,001041 |
| Pf4 | 25,9456 | 0 | -inf | 5,00E-005 | 0,001041 |
| Prl2c3,Prl2c4 | 0,739859 | 0 | -inf | 5,00E-005 | 0,001041 |
| Prl2c5 | 1,99492 | 0 | -inf | 5,00E-005 | 0,001041 |
| Prl3b1 | 5,16345 | 0 | -inf | 5,00E-005 | 0,001041 |
| Prl3d1 | 1,39213 | 0 | -inf | 5,00E-005 | 0,001041 |
| Prl4a1 | 1,1337 | 0 | -inf | 5,00E-005 | 0,001041 |
| Prl7a1 | 4,96935 | 0 | -inf | 5,00E-005 | 0,001041 |
| Prl8a2 | 9,59576 | 0 | -inf | 5,00E-005 | 0,001041 |
| Prom2 | 1,85961 | 0 | -inf | 5,00E-005 | 0,001041 |

| | | | | | |
|---|---|---|---|---|---|
| S100a8 | 3,21077 | 0 | -inf | 5,00E-005 | 0,001041 |
| S100a9 | 1,15791 | 0 | -inf | 5,00E-005 | 0,001041 |
| Shh | 9,67731 | 0 | -inf | 5,00E-005 | 0,001041 |
| Spp1 | 1,84072 | 0 | -inf | 5,00E-005 | 0,001041 |
| Spp2 | 0,959266 | 0 | -inf | 5,00E-005 | 0,001041 |
| Tm4sf1 | 0,728656 | 0 | -inf | 5,00E-005 | 0,001041 |
| Trap1a | 0,683527 | 0 | -inf | 5,00E-005 | 0,001041 |
| Tspan32 | 3,55922 | 0 | -inf | 5,00E-005 | 0,001041 |
| Ttr | 1,52373 | 0 | -inf | 5,00E-005 | 0,001041 |
| Upk3bl | 1,02296 | 0 | -inf | 5,00E-005 | 0,001041 |
| Vav1 | 0,559907 | 0 | -inf | 5,00E-005 | 0,001041 |
| Esr1 | 0,138129 | 3,94949 | 4,83757 | 5,00E-005 | 0,001041 |
| Fos | 0,322515 | 7,75633 | 4,58793 | 5,00E-005 | 0,001041 |
| Hspa1b | 0,315665 | 7,05413 | 4,482 | 5,00E-005 | 0,001041 |
| Urad | 1,67447 | 30,3311 | 4,17902 | 5,00E-005 | 0,001041 |
| Egr1 | 0,559422 | 6,80263 | 3,60408 | 5,00E-005 | 0,001041 |
| Rpph1 | 7,75479 | 36,7519 | 2,24466 | 5,00E-005 | 0,001041 |
| Tgm3 | 0,835319 | 3,11744 | 1,89996 | 5,00E-005 | 0,001041 |
| Kcnh5 | 0,592289 | 2,09512 | 1,82266 | 5,00E-005 | 0,001041 |
| Gm1943 | 3,72182 | 9,57799 | 1,36371 | 5,00E-005 | 0,001041 |
| Arl4d | 83,9505 | 193,23 | 1,20271 | 5,00E-005 | 0,001041 |
| Hes3 | 3,46843 | 7,66182 | 1,1434 | 5,00E-005 | 0,001041 |
| Hoxc9 | 16,3129 | 35,5662 | 1,1245 | 5,00E-005 | 0,001041 |
| Tpt1 | 20,8952 | 44,7163 | 1,09763 | 5,00E-005 | 0,001041 |
| Cyr61 | 21,2938 | 45,4629 | 1,09425 | 5,00E-005 | 0,001041 |
| 2610307P16Rik | 2,79723 | 5,91566 | 1,08054 | 5,00E-005 | 0,001041 |
| Tmem181b-ps | 1,50167 | 3,17445 | 1,07993 | 5,00E-005 | 0,001041 |
| Hoxc10 | 95,538 | 200,664 | 1,07064 | 5,00E-005 | 0,001041 |
| Cyp26a1 | 54,9184 | 114,744 | 1,06306 | 5,00E-005 | 0,001041 |
| Oasl2 | 1,85777 | 3,85829 | 1,05439 | 5,00E-005 | 0,001041 |
| Cdx4 | 66,567 | 135,87 | 1,02935 | 5,00E-005 | 0,001041 |
| Nkx1-2 | 13,7981 | 28,1324 | 1,02777 | 5,00E-005 | 0,001041 |
| Uap1l1 | 11,25 | 22,904 | 1,02567 | 5,00E-005 | 0,001041 |
| Tg | 1,25263 | 2,53702 | 1,01818 | 5,00E-005 | 0,001041 |
| Sox3 | 3,72198 | 7,31317 | 0,974427 | 5,00E-005 | 0,001041 |
| Lin28a | 48,2616 | 94,5552 | 0,970281 | 5,00E-005 | 0,001041 |
| Map6 | 3,61552 | 7,01695 | 0,956641 | 5,00E-005 | 0,001041 |
| Dmpk | 6,13448 | 11,7744 | 0,940637 | 5,00E-005 | 0,001041 |
| Fam131b | 8,19245 | 15,5854 | 0,927831 | 5,00E-005 | 0,001041 |
| Soga3 | 4,17009 | 7,89412 | 0,920701 | 5,00E-005 | 0,001041 |
| Arhgap24 | 7,36589 | 13,7983 | 0,905562 | 5,00E-005 | 0,001041 |
| Tcea3 | 9,51681 | 17,688 | 0,894219 | 5,00E-005 | 0,001041 |
| Kcnq1ot1 | 0,435962 | 0,809292 | 0,892458 | 5,00E-005 | 0,001041 |
| 2610528A11Rik | 77,1884 | 142,836 | 0,887903 | 5,00E-005 | 0,001041 |
| Smtnl2 | 7,11847 | 13,1307 | 0,883301 | 5,00E-005 | 0,001041 |
| Amph | 7,20448 | 13,1899 | 0,872463 | 5,00E-005 | 0,001041 |
| Dgkk | 1,72685 | 3,12034 | 0,853562 | 5,00E-005 | 0,001041 |
| Jun | 17,6694 | 31,8849 | 0,851619 | 5,00E-005 | 0,001041 |

| | | | | | |
|---|---|---|---|---|---|
| Zbtb16 | 6,71267 | 12,0679 | 0,846218 | 5,00E-005 | 0,001041 |
| Adamtsl2 | 23,5511 | 41,7364 | 0,825509 | 5,00E-005 | 0,001041 |
| Pmaip1 | 27,2147 | 48,1286 | 0,822506 | 5,00E-005 | 0,001041 |
| Kcp | 4,31059 | 7,60197 | 0,818489 | 5,00E-005 | 0,001041 |
| Rbm15 | 10,4601 | 18,3741 | 0,812783 | 5,00E-005 | 0,001041 |
| Cacna1a | 3,63771 | 6,37237 | 0,808797 | 5,00E-005 | 0,001041 |
| Acta2 | 8,59394 | 15,0099 | 0,804522 | 5,00E-005 | 0,001041 |
| Spry4 | 20,1928 | 35,0073 | 0,793815 | 5,00E-005 | 0,001041 |
| Msgn1 | 215,275 | 372,536 | 0,791201 | 5,00E-005 | 0,001041 |
| Fgf3 | 35,0492 | 60,5281 | 0,788223 | 5,00E-005 | 0,001041 |
| Tmsb10 | 256,676 | 438,543 | 0,772773 | 5,00E-005 | 0,001041 |
| Hoxb1 | 51,6144 | 88,0201 | 0,770059 | 5,00E-005 | 0,001041 |
| Cdhr1 | 3,12608 | 5,29498 | 0,760271 | 5,00E-005 | 0,001041 |
| Gbx2 | 21,6404 | 35,8839 | 0,729609 | 5,00E-005 | 0,001041 |
| Snhg1 | 312,476 | 516,741 | 0,725698 | 5,00E-005 | 0,001041 |
| Cdx2 | 54,7004 | 90,065 | 0,719415 | 5,00E-005 | 0,001041 |
| Tbx6 | 121,583 | 199,842 | 0,716915 | 5,00E-005 | 0,001041 |
| Nkd2 | 60,6478 | 99,6428 | 0,716311 | 5,00E-005 | 0,001041 |
| Mif | 171,801 | 279,625 | 0,70275 | 5,00E-005 | 0,001041 |
| Peli3 | 29,1668 | 47,3315 | 0,698475 | 5,00E-005 | 0,001041 |
| Crabp1 | 159,682 | 256,987 | 0,686494 | 5,00E-005 | 0,001041 |
| Fgf17 | 62,8964 | 101,108 | 0,684851 | 5,00E-005 | 0,001041 |
| AW551984 | 8,84736 | 14,1212 | 0,67454 | 5,00E-005 | 0,001041 |
| Hoxa10 | 68,1954 | 108,107 | 0,664718 | 5,00E-005 | 0,001041 |
| Chst7 | 19,828 | 31,219 | 0,654884 | 5,00E-005 | 0,001041 |
| Rspo3 | 71,7764 | 112,982 | 0,654509 | 5,00E-005 | 0,001041 |
| Rftn1 | 31,4232 | 48,7823 | 0,634528 | 5,00E-005 | 0,001041 |
| Ube2d3 | 82,585 | 53,0823 | -0,63765 | 5,00E-005 | 0,001041 |
| Cdkn1c | 55,244 | 35,4401 | -0,640434 | 5,00E-005 | 0,001041 |
| Igfbp3 | 40,7376 | 26,0396 | -0,645656 | 5,00E-005 | 0,001041 |
| Tspan9 | 23,6877 | 15,0475 | -0,654612 | 5,00E-005 | 0,001041 |
| Iqgap1 | 21,5537 | 13,6695 | -0,656976 | 5,00E-005 | 0,001041 |
| Tram1 | 33,3668 | 21,1578 | -0,657222 | 5,00E-005 | 0,001041 |
| Tmed5 | 17,3646 | 10,9915 | -0,659756 | 5,00E-005 | 0,001041 |
| Mfge8 | 51,2543 | 32,164 | -0,672228 | 5,00E-005 | 0,001041 |
| Tfrc | 76,9452 | 48,1213 | -0,677155 | 5,00E-005 | 0,001041 |
| Ctsd | 82,5642 | 51,6352 | -0,677164 | 5,00E-005 | 0,001041 |
| Tacc1 | 8,37461 | 5,20802 | -0,685288 | 5,00E-005 | 0,001041 |
| Lamp2 | 48,1432 | 29,9299 | -0,685747 | 5,00E-005 | 0,001041 |
| Hoxb13 | 84,2068 | 51,9235 | -0,697549 | 5,00E-005 | 0,001041 |
| Arrb1 | 12,4517 | 7,60124 | -0,712039 | 5,00E-005 | 0,001041 |
| Col2a1 | 37,8493 | 23,0585 | -0,714965 | 5,00E-005 | 0,001041 |
| Zfp760 | 12,8369 | 7,77377 | -0,723607 | 5,00E-005 | 0,001041 |
| Map4k2 | 9,29472 | 5,61804 | -0,726346 | 5,00E-005 | 0,001041 |
| Ucp2 | 15,0703 | 9,0977 | -0,728138 | 5,00E-005 | 0,001041 |
| Creg1 | 29,7709 | 17,9407 | -0,730671 | 5,00E-005 | 0,001041 |
| Nudt4 | 86,6442 | 52,1134 | -0,733449 | 5,00E-005 | 0,001041 |
| Mmp14 | 49,7395 | 29,9155 | -0,7335 | 5,00E-005 | 0,001041 |

| | | | | | |
|---|---|---|---|---|---|
| Sh3bp5 | 20,497 | 12,3226 | -0,734105 | 5,00E-005 | 0,001041 |
| Vamp2 | 22,5963 | 13,5618 | -0,736534 | 5,00E-005 | 0,001041 |
| Sema3f | 18,2523 | 10,9415 | -0,738259 | 5,00E-005 | 0,001041 |
| Lgr4 | 24,2536 | 14,5318 | -0,738979 | 5,00E-005 | 0,001041 |
| Plec | 4,34366 | 2,59572 | -0,742775 | 5,00E-005 | 0,001041 |
| Dsc2 | 9,84272 | 5,74136 | -0,777664 | 5,00E-005 | 0,001041 |
| Kit | 9,29509 | 5,38262 | -0,788159 | 5,00E-005 | 0,001041 |
| Foxo1 | 6,26765 | 3,61326 | -0,794623 | 5,00E-005 | 0,001041 |
| Dsp | 13,1718 | 7,5856 | -0,796119 | 5,00E-005 | 0,001041 |
| Igfbp2 | 235,544 | 135,106 | -0,801906 | 5,00E-005 | 0,001041 |
| Ston2 | 4,22727 | 2,41694 | -0,806544 | 5,00E-005 | 0,001041 |
| Deptor | 3,99956 | 2,28063 | -0,810407 | 5,00E-005 | 0,001041 |
| Lrp5 | 16,2188 | 9,22974 | -0,813302 | 5,00E-005 | 0,001041 |
| Galnt16 | 6,649 | 3,78171 | -0,814101 | 5,00E-005 | 0,001041 |
| Nedd9 | 8,32242 | 4,73236 | -0,814445 | 5,00E-005 | 0,001041 |
| Ctsb | 33,0539 | 18,741 | -0,818619 | 5,00E-005 | 0,001041 |
| Itm2b | 62,5224 | 35,312 | -0,824213 | 5,00E-005 | 0,001041 |
| Ptprd | 5,65812 | 3,17921 | -0,831654 | 5,00E-005 | 0,001041 |
| Gprc5c | 27,2523 | 15,2747 | -0,835236 | 5,00E-005 | 0,001041 |
| Dram2 | 14,1163 | 7,89924 | -0,837582 | 5,00E-005 | 0,001041 |
| Kif26a | 8,95548 | 5,00965 | -0,838062 | 5,00E-005 | 0,001041 |
| Txnip | 26,8674 | 14,9727 | -0,843527 | 5,00E-005 | 0,001041 |
| Pvrl2 | 45,7721 | 25,2588 | -0,857684 | 5,00E-005 | 0,001041 |
| Slc30a1 | 5,65597 | 3,11887 | -0,858751 | 5,00E-005 | 0,001041 |
| Serinc3 | 53,6872 | 29,5953 | -0,859209 | 5,00E-005 | 0,001041 |
| Itpkb | 4,66775 | 2,56996 | -0,860979 | 5,00E-005 | 0,001041 |
| Snhg4 | 27,5399 | 15,1585 | -0,861402 | 5,00E-005 | 0,001041 |
| Cmtm3 | 35,7575 | 19,5866 | -0,868379 | 5,00E-005 | 0,001041 |
| Anxa3 | 19,2645 | 10,5319 | -0,871172 | 5,00E-005 | 0,001041 |
| Myo1c | 6,5982 | 3,60579 | -0,871756 | 5,00E-005 | 0,001041 |
| Acsl5 | 18,153 | 9,9103 | -0,873206 | 5,00E-005 | 0,001041 |
| Slc38a4 | 32,6617 | 17,8143 | -0,874561 | 5,00E-005 | 0,001041 |
| Col26a1 | 31,5485 | 17,1477 | -0,87956 | 5,00E-005 | 0,001041 |
| Ackr3 | 11,7328 | 6,37653 | -0,879698 | 5,00E-005 | 0,001041 |
| Shroom2 | 5,75773 | 3,12542 | -0,881452 | 5,00E-005 | 0,001041 |
| Lars2 | 101,361 | 54,9182 | -0,884147 | 5,00E-005 | 0,001041 |
| Sema3e | 5,24492 | 2,83447 | -0,887841 | 5,00E-005 | 0,001041 |
| Ptprb | 2,06289 | 1,1117 | -0,891901 | 5,00E-005 | 0,001041 |
| Hmbs | 45,8706 | 24,6721 | -0,894691 | 5,00E-005 | 0,001041 |
| 1190002N15Rik | 12,1512 | 6,52743 | -0,896506 | 5,00E-005 | 0,001041 |
| Sipa1 | 11,4065 | 6,1093 | -0,90078 | 5,00E-005 | 0,001041 |
| Efna5 | 5,64716 | 3,01459 | -0,905566 | 5,00E-005 | 0,001041 |
| Ctsc | 8,43406 | 4,47854 | -0,913199 | 5,00E-005 | 0,001041 |
| Adcy7 | 3,25428 | 1,71879 | -0,92095 | 5,00E-005 | 0,001041 |
| Ets1 | 17,7093 | 9,35079 | -0,921345 | 5,00E-005 | 0,001041 |
| Atp2b4 | 3,90874 | 2,06156 | -0,92297 | 5,00E-005 | 0,001041 |
| Islr | 17,5872 | 9,24098 | -0,928412 | 5,00E-005 | 0,001041 |
| Plch1 | 12,4426 | 6,53766 | -0,928446 | 5,00E-005 | 0,001041 |

| | | | | | |
|---|---|---|---|---|---|
| F11r | 33,7896 | 17,6898 | -0,933657 | 5,00E-005 | 0,001041 |
| Fras1 | 5,83207 | 3,04844 | -0,935937 | 5,00E-005 | 0,001041 |
| Hs6st2 | 7,02924 | 3,66476 | -0,939649 | 5,00E-005 | 0,001041 |
| Mtus1 | 4,57867 | 2,38276 | -0,942295 | 5,00E-005 | 0,001041 |
| Fkbp14 | 6,94152 | 3,60315 | -0,945994 | 5,00E-005 | 0,001041 |
| Flrt2 | 4,9843 | 2,58574 | -0,946815 | 5,00E-005 | 0,001041 |
| Gypc | 18,0007 | 9,23342 | -0,963117 | 5,00E-005 | 0,001041 |
| Plekhg1 | 3,45561 | 1,77239 | -0,96324 | 5,00E-005 | 0,001041 |
| Myo6 | 2,50883 | 1,28526 | -0,964949 | 5,00E-005 | 0,001041 |
| Tfpi | 27,072 | 13,8277 | -0,969234 | 5,00E-005 | 0,001041 |
| Trf | 11,0504 | 5,62863 | -0,973249 | 5,00E-005 | 0,001041 |
| Sox5 | 2,46191 | 1,25313 | -0,97424 | 5,00E-005 | 0,001041 |
| Mgat5 | 9,9853 | 5,0807 | -0,97478 | 5,00E-005 | 0,001041 |
| Arap3 | 7,9052 | 4,01282 | -0,978185 | 5,00E-005 | 0,001041 |
| Gcnt1 | 7,32177 | 3,71509 | -0,978797 | 5,00E-005 | 0,001041 |
| Ntn1 | 27,4023 | 13,8502 | -0,984395 | 5,00E-005 | 0,001041 |
| Cd151 | 43,4396 | 21,8917 | -0,988625 | 5,00E-005 | 0,001041 |
| Sec24d | 3,2285 | 1,62688 | -0,988758 | 5,00E-005 | 0,001041 |
| 2900026A02Rik | 2,8307 | 1,42635 | -0,988827 | 5,00E-005 | 0,001041 |
| Plk2 | 23,5116 | 11,7907 | -0,995727 | 5,00E-005 | 0,001041 |
| Lama5 | 15,0787 | 7,53626 | -1,00059 | 5,00E-005 | 0,001041 |
| Clu | 10,0241 | 5,00652 | -1,0016 | 5,00E-005 | 0,001041 |
| St8sia1 | 2,60787 | 1,29448 | -1,01049 | 5,00E-005 | 0,001041 |
| Josd2 | 25,27 | 12,516 | -1,01365 | 5,00E-005 | 0,001041 |
| Arxes2 | 22,3061 | 11,0304 | -1,01595 | 5,00E-005 | 0,001041 |
| Tmprss2 | 8,83308 | 4,35589 | -1,01995 | 5,00E-005 | 0,001041 |
| Adam12 | 2,17122 | 1,0687 | -1,02264 | 5,00E-005 | 0,001041 |
| Gclm | 11,6024 | 5,70905 | -1,0231 | 5,00E-005 | 0,001041 |
| Col4a1 | 91,9529 | 45,1802 | -1,0252 | 5,00E-005 | 0,001041 |
| Spaca6 | 13,6229 | 6,69329 | -1,02524 | 5,00E-005 | 0,001041 |
| Sema3a | 7,60739 | 3,72435 | -1,03041 | 5,00E-005 | 0,001041 |
| Fzd4 | 4,68915 | 2,29466 | -1,03105 | 5,00E-005 | 0,001041 |
| Elf4 | 3,06678 | 1,49855 | -1,03316 | 5,00E-005 | 0,001041 |
| Mid1 | 3,64168 | 1,77499 | -1,03679 | 5,00E-005 | 0,001041 |
| Itga9 | 5,97326 | 2,91049 | -1,03726 | 5,00E-005 | 0,001041 |
| Efnb2 | 34,5534 | 16,8026 | -1,04015 | 5,00E-005 | 0,001041 |
| Thsd4 | 6,47466 | 3,13286 | -1,04732 | 5,00E-005 | 0,001041 |
| Col4a2 | 52,0722 | 25,1936 | -1,04746 | 5,00E-005 | 0,001041 |
| Cdc42ep1 | 9,70448 | 4,68595 | -1,05031 | 5,00E-005 | 0,001041 |
| Grina | 13,3192 | 6,41094 | -1,0549 | 5,00E-005 | 0,001041 |
| Foxp4 | 15,6689 | 7,50567 | -1,06185 | 5,00E-005 | 0,001041 |
| Abca1 | 5,20023 | 2,47632 | -1,07038 | 5,00E-005 | 0,001041 |
| Ocln | 3,97841 | 1,89381 | -1,0709 | 5,00E-005 | 0,001041 |
| Igf2 | 408,187 | 194,244 | -1,07136 | 5,00E-005 | 0,001041 |
| Cpeb4 | 3,70037 | 1,7578 | -1,0739 | 5,00E-005 | 0,001041 |
| Dmd | 0,699495 | 0,331442 | -1,07756 | 5,00E-005 | 0,001041 |
| Trp53inp1 | 4,58881 | 2,17133 | -1,07954 | 5,00E-005 | 0,001041 |
| Ostm1 | 3,98241 | 1,88426 | -1,07965 | 5,00E-005 | 0,001041 |

| | | | | | |
|---|---|---|---|---|---|
| Slc9a3r2 | 6,56889 | 3,1072 | -1,08003 | 5,00E-005 | 0,001041 |
| Vangl1 | 7,18979 | 3,37905 | -1,08933 | 5,00E-005 | 0,001041 |
| Krt19 | 19,0698 | 8,89322 | -1,10051 | 5,00E-005 | 0,001041 |
| Col8a1 | 4,2903 | 1,99781 | -1,10266 | 5,00E-005 | 0,001041 |
| Wnk4 | 3,4735 | 1,60151 | -1,11696 | 5,00E-005 | 0,001041 |
| Tspan13 | 26,1823 | 12,0661 | -1,11763 | 5,00E-005 | 0,001041 |
| Ddit4l | 3,49082 | 1,60773 | -1,11854 | 5,00E-005 | 0,001041 |
| Wnt16 | 13,2415 | 6,06302 | -1,12696 | 5,00E-005 | 0,001041 |
| Sp6 | 3,08593 | 1,41259 | -1,12736 | 5,00E-005 | 0,001041 |
| Aplnr | 34,2828 | 15,585 | -1,13733 | 5,00E-005 | 0,001041 |
| Itm2a | 11,5344 | 5,23265 | -1,14033 | 5,00E-005 | 0,001041 |
| Mbnl1 | 10,9022 | 4,92957 | -1,14509 | 5,00E-005 | 0,001041 |
| Klf3 | 7,73612 | 3,49292 | -1,14718 | 5,00E-005 | 0,001041 |
| Gmpr | 23,6163 | 10,615 | -1,15368 | 5,00E-005 | 0,001041 |
| Mfng | 6,55185 | 2,92881 | -1,16159 | 5,00E-005 | 0,001041 |
| Timp3 | 29,2686 | 13,0741 | -1,16264 | 5,00E-005 | 0,001041 |
| Mme | 2,65382 | 1,18445 | -1,16386 | 5,00E-005 | 0,001041 |
| Epor | 10,8845 | 4,85689 | -1,16417 | 5,00E-005 | 0,001041 |
| Dag1 | 56,8755 | 25,3091 | -1,16815 | 5,00E-005 | 0,001041 |
| Dancr | 29,7314 | 13,1906 | -1,17248 | 5,00E-005 | 0,001041 |
| Sat1 | 11,7014 | 5,17961 | -1,17576 | 5,00E-005 | 0,001041 |
| Axl | 7,56918 | 3,34539 | -1,17796 | 5,00E-005 | 0,001041 |
| Eya1 | 6,73446 | 2,97129 | -1,18047 | 5,00E-005 | 0,001041 |
| Vsig10 | 3,05795 | 1,34717 | -1,18263 | 5,00E-005 | 0,001041 |
| Gpr56 | 9,0306 | 3,97275 | -1,18468 | 5,00E-005 | 0,001041 |
| Shank3 | 3,06019 | 1,34151 | -1,18977 | 5,00E-005 | 0,001041 |
| Crip2 | 16,2326 | 7,11196 | -1,19057 | 5,00E-005 | 0,001041 |
| Klhdc7a | 1,7027 | 0,745559 | -1,19143 | 5,00E-005 | 0,001041 |
| Emp1 | 49,9416 | 21,8447 | -1,19296 | 5,00E-005 | 0,001041 |
| Ehd2 | 9,3697 | 4,09358 | -1,19464 | 5,00E-005 | 0,001041 |
| Sft2d2 | 8,94793 | 3,89687 | -1,19924 | 5,00E-005 | 0,001041 |
| Gdpd5 | 2,84888 | 1,23291 | -1,20833 | 5,00E-005 | 0,001041 |
| Ptp4a3 | 11,6122 | 5,00724 | -1,21355 | 5,00E-005 | 0,001041 |
| Met | 17,2694 | 7,43846 | -1,21514 | 5,00E-005 | 0,001041 |
| Slc6a9 | 8,66163 | 3,72888 | -1,2159 | 5,00E-005 | 0,001041 |
| Ecm1 | 6,29363 | 2,70352 | -1,21905 | 5,00E-005 | 0,001041 |
| Vamp8 | 20,9739 | 8,92699 | -1,23235 | 5,00E-005 | 0,001041 |
| Angpt1 | 5,12681 | 2,18021 | -1,23359 | 5,00E-005 | 0,001041 |
| Plat | 5,29598 | 2,25108 | -1,23428 | 5,00E-005 | 0,001041 |
| Bmp2 | 5,09976 | 2,16755 | -1,23437 | 5,00E-005 | 0,001041 |
| Apoe | 19,7214 | 8,37976 | -1,23478 | 5,00E-005 | 0,001041 |
| Myocd | 2,45567 | 1,03652 | -1,24437 | 5,00E-005 | 0,001041 |
| Pla2g4a | 5,44082 | 2,29541 | -1,24508 | 5,00E-005 | 0,001041 |
| 5430417L22Rik | 7,93194 | 3,33775 | -1,2488 | 5,00E-005 | 0,001041 |
| Itga4 | 1,24806 | 0,523654 | -1,253 | 5,00E-005 | 0,001041 |
| Cdc42ep3 | 4,10377 | 1,72147 | -1,25331 | 5,00E-005 | 0,001041 |
| Frk | 1,44027 | 0,60201 | -1,25848 | 5,00E-005 | 0,001041 |
| Cd44 | 2,7051 | 1,12688 | -1,26335 | 5,00E-005 | 0,001041 |

| | | | | | |
|---|---|---|---|---|---|
| Bmp7 | 18,2713 | 7,5682 | -1,27156 | 5,00E-005 | 0,001041 |
| Pear1 | 2,46283 | 1,01779 | -1,27487 | 5,00E-005 | 0,001041 |
| Ggh | 8,68472 | 3,56849 | -1,28317 | 5,00E-005 | 0,001041 |
| Nell2 | 3,39804 | 1,39251 | -1,28702 | 5,00E-005 | 0,001041 |
| Coro2b | 2,84181 | 1,16275 | -1,28926 | 5,00E-005 | 0,001041 |
| Ikzf2 | 1,20801 | 0,494223 | -1,2894 | 5,00E-005 | 0,001041 |
| Sparc | 91,2083 | 37,2887 | -1,29043 | 5,00E-005 | 0,001041 |
| Slc16a10 | 3,99733 | 1,62754 | -1,29634 | 5,00E-005 | 0,001041 |
| Mbnl2 | 7,03848 | 2,85832 | -1,3001 | 5,00E-005 | 0,001041 |
| H2afz | 33,703 | 13,677 | -1,30113 | 5,00E-005 | 0,001041 |
| Megf6 | 10,2921 | 4,10286 | -1,32683 | 5,00E-005 | 0,001041 |
| Dmrt2 | 19,5824 | 7,79864 | -1,32826 | 5,00E-005 | 0,001041 |
| Tgfbr2 | 4,50064 | 1,78726 | -1,33238 | 5,00E-005 | 0,001041 |
| Grb7 | 5,05336 | 2,00133 | -1,33628 | 5,00E-005 | 0,001041 |
| Lyn | 4,36066 | 1,71777 | -1,34401 | 5,00E-005 | 0,001041 |
| Foxp2 | 1,15777 | 0,45333 | -1,35271 | 5,00E-005 | 0,001041 |
| Cpm | 8,09305 | 3,16699 | -1,35357 | 5,00E-005 | 0,001041 |
| Tmem51 | 3,99524 | 1,56282 | -1,35413 | 5,00E-005 | 0,001041 |
| Dhrs11 | 8,17107 | 3,19522 | -1,35461 | 5,00E-005 | 0,001041 |
| Abcb10 | 23,9166 | 9,35196 | -1,35467 | 5,00E-005 | 0,001041 |
| Mef2c | 2,53082 | 0,989609 | -1,35467 | 5,00E-005 | 0,001041 |
| Cer1 | 57,712 | 22,5412 | -1,35631 | 5,00E-005 | 0,001041 |
| Ctsh | 4,88531 | 1,88641 | -1,37281 | 5,00E-005 | 0,001041 |
| Cgn | 3,62379 | 1,39529 | -1,37694 | 5,00E-005 | 0,001041 |
| Pon2 | 20,7255 | 7,96484 | -1,37969 | 5,00E-005 | 0,001041 |
| Sox6 | 2,53822 | 0,975281 | -1,37993 | 5,00E-005 | 0,001041 |
| Dusp3 | 4,4014 | 1,68821 | -1,38247 | 5,00E-005 | 0,001041 |
| Ramp2 | 35,746 | 13,6578 | -1,38806 | 5,00E-005 | 0,001041 |
| Edil3 | 2,56237 | 0,978675 | -1,38858 | 5,00E-005 | 0,001041 |
| Piezo2 | 1,73767 | 0,663253 | -1,38952 | 5,00E-005 | 0,001041 |
| Cd38 | 5,04203 | 1,89817 | -1,4094 | 5,00E-005 | 0,001041 |
| Snhg7 | 32,6283 | 12,2482 | -1,41355 | 5,00E-005 | 0,001041 |
| Rab26os | 130,917 | 49,097 | -1,41494 | 5,00E-005 | 0,001041 |
| Alcam | 3,52845 | 1,32313 | -1,41508 | 5,00E-005 | 0,001041 |
| Arhgef28 | 1,79799 | 0,671276 | -1,42141 | 5,00E-005 | 0,001041 |
| 03/02/16 | 13,7997 | 5,14898 | -1,42228 | 5,00E-005 | 0,001041 |
| Arhgap23 | 4,52036 | 1,68427 | -1,42432 | 5,00E-005 | 0,001041 |
| Slc29a1 | 21,5659 | 8,02628 | -1,42595 | 5,00E-005 | 0,001041 |
| Sh2d3c | 3,90868 | 1,44417 | -1,43644 | 5,00E-005 | 0,001041 |
| Crim1 | 3,73826 | 1,38075 | -1,43691 | 5,00E-005 | 0,001041 |
| Rapgef5 | 2,45864 | 0,906063 | -1,44018 | 5,00E-005 | 0,001041 |
| Prkch | 4,21425 | 1,54957 | -1,44341 | 5,00E-005 | 0,001041 |
| Pgm5 | 7,61286 | 2,79357 | -1,44633 | 5,00E-005 | 0,001041 |
| Mest | 616,452 | 225,489 | -1,45093 | 5,00E-005 | 0,001041 |
| Atp8a1 | 7,98596 | 2,91721 | -1,45288 | 5,00E-005 | 0,001041 |
| Sdc4 | 8,87214 | 3,24059 | -1,45303 | 5,00E-005 | 0,001041 |
| Cdh3 | 36,6805 | 13,3266 | -1,4607 | 5,00E-005 | 0,001041 |
| Fst | 2,93956 | 1,06408 | -1,46599 | 5,00E-005 | 0,001041 |

| | | | | | |
|---|---|---|---|---|---|
| Sox9 | 6,20156 | 2,24006 | -1,4691 | 5,00E-005 | 0,001041 |
| Cpt1a | 2,54048 | 0,914274 | -1,4744 | 5,00E-005 | 0,001041 |
| Cpox | 29,4026 | 10,5666 | -1,47643 | 5,00E-005 | 0,001041 |
| Arhgap29 | 14,1721 | 5,09259 | -1,47658 | 5,00E-005 | 0,001041 |
| Pdgfa | 35,8966 | 12,7969 | -1,48805 | 5,00E-005 | 0,001041 |
| Egfl7 | 38,136 | 13,5681 | -1,49094 | 5,00E-005 | 0,001041 |
| Manba | 2,24176 | 0,79608 | -1,49365 | 5,00E-005 | 0,001041 |
| Man1a | 4,41447 | 1,56706 | -1,49418 | 5,00E-005 | 0,001041 |
| Fli1 | 6,17165 | 2,18777 | -1,4962 | 5,00E-005 | 0,001041 |
| Sepp1 | 44,6563 | 15,7857 | -1,50024 | 5,00E-005 | 0,001041 |
| Enpp2 | 2,32784 | 0,818899 | -1,50724 | 5,00E-005 | 0,001041 |
| Ostf1 | 8,69937 | 3,03999 | -1,51684 | 5,00E-005 | 0,001041 |
| Zfp185 | 3,23509 | 1,12653 | -1,52192 | 5,00E-005 | 0,001041 |
| Pdgfc | 7,11283 | 2,47204 | -1,52472 | 5,00E-005 | 0,001041 |
| Arrdc1 | 4,83795 | 1,67617 | -1,52922 | 5,00E-005 | 0,001041 |
| Fermt1 | 5,55205 | 1,9197 | -1,53214 | 5,00E-005 | 0,001041 |
| Slc1a5 | 33,1448 | 11,4432 | -1,53429 | 5,00E-005 | 0,001041 |
| Fgfr2 | 21,5647 | 7,44153 | -1,535 | 5,00E-005 | 0,001041 |
| Fam210b | 7,97383 | 2,74025 | -1,54097 | 5,00E-005 | 0,001041 |
| Meox2 | 4,33129 | 1,48573 | -1,54363 | 5,00E-005 | 0,001041 |
| Scube3 | 9,70944 | 3,3066 | -1,55404 | 5,00E-005 | 0,001041 |
| Ehbp1l1 | 4,06809 | 1,38249 | -1,55709 | 5,00E-005 | 0,001041 |
| Leprel1 | 6,19915 | 2,10232 | -1,56009 | 5,00E-005 | 0,001041 |
| Slco3a1 | 3,25359 | 1,10234 | -1,56146 | 5,00E-005 | 0,001041 |
| Gm20554 | 1,3545 | 0,458176 | -1,56379 | 5,00E-005 | 0,001041 |
| Htra1 | 4,30363 | 1,44672 | -1,57277 | 5,00E-005 | 0,001041 |
| Arhgef15 | 4,63468 | 1,55619 | -1,57445 | 5,00E-005 | 0,001041 |
| Tgfbr3 | 2,37768 | 0,793676 | -1,58294 | 5,00E-005 | 0,001041 |
| Nr2f2 | 9,19744 | 3,06922 | -1,58336 | 5,00E-005 | 0,001041 |
| Col3a1 | 4,19661 | 1,39908 | -1,58474 | 5,00E-005 | 0,001041 |
| Jup | 5,9118 | 1,9598 | -1,59289 | 5,00E-005 | 0,001041 |
| Itga6 | 16,5027 | 5,45575 | -1,59686 | 5,00E-005 | 0,001041 |
| Sel1l3 | 2,02436 | 0,669102 | -1,59717 | 5,00E-005 | 0,001041 |
| Stard8 | 2,2055 | 0,725178 | -1,6047 | 5,00E-005 | 0,001041 |
| Calcrl | 4,75093 | 1,55908 | -1,60751 | 5,00E-005 | 0,001041 |
| Maf | 3,13801 | 1,01766 | -1,62459 | 5,00E-005 | 0,001041 |
| Afap1l1 | 15,6405 | 5,0642 | -1,62688 | 5,00E-005 | 0,001041 |
| Sema3g | 3,13483 | 1,01392 | -1,62844 | 5,00E-005 | 0,001041 |
| Sertad4 | 4,53211 | 1,46418 | -1,63009 | 5,00E-005 | 0,001041 |
| Fech | 23,1529 | 7,45145 | -1,6356 | 5,00E-005 | 0,001041 |
| Nrp1 | 12,792 | 4,11512 | -1,63623 | 5,00E-005 | 0,001041 |
| Adam15 | 8,53883 | 2,7274 | -1,64651 | 5,00E-005 | 0,001041 |
| Rhoj | 5,12064 | 1,62588 | -1,6551 | 5,00E-005 | 0,001041 |
| Enpp5 | 2,01427 | 0,63628 | -1,66253 | 5,00E-005 | 0,001041 |
| Oxtr | 2,93407 | 0,923794 | -1,66726 | 5,00E-005 | 0,001041 |
| Eya2 | 8,36484 | 2,61858 | -1,67555 | 5,00E-005 | 0,001041 |
| Spint1 | 11,2199 | 3,47976 | -1,689 | 5,00E-005 | 0,001041 |
| Tns4 | 1,42071 | 0,43989 | -1,6914 | 5,00E-005 | 0,001041 |

| | | | | | |
|---|---|---|---|---|---|
| Adamts15 | 6,29357 | 1,94765 | -1,69214 | 5,00E-005 | 0,001041 |
| Rnase4 | 6,4512 | 1,98553 | -1,70005 | 5,00E-005 | 0,001041 |
| Fzd8 | 2,40075 | 0,737522 | -1,70273 | 5,00E-005 | 0,001041 |
| F3 | 5,07718 | 1,5564 | -1,70582 | 5,00E-005 | 0,001041 |
| Gyltl1b | 6,41573 | 1,95715 | -1,71286 | 5,00E-005 | 0,001041 |
| Tgfb2 | 7,08257 | 2,1507 | -1,71947 | 5,00E-005 | 0,001041 |
| Pag1 | 1,38058 | 0,413697 | -1,73863 | 5,00E-005 | 0,001041 |
| Npr1 | 1,64531 | 0,491869 | -1,74202 | 5,00E-005 | 0,001041 |
| Ppp1r13b | 2,74125 | 0,818607 | -1,74359 | 5,00E-005 | 0,001041 |
| She | 3,01185 | 0,891842 | -1,75579 | 5,00E-005 | 0,001041 |
| Fmod | 7,21746 | 2,13651 | -1,75623 | 5,00E-005 | 0,001041 |
| Tspan18 | 11,1659 | 3,30507 | -1,75635 | 5,00E-005 | 0,001041 |
| Card10 | 3,73243 | 1,09477 | -1,76949 | 5,00E-005 | 0,001041 |
| Dpp4 | 1,4271 | 0,416942 | -1,77517 | 5,00E-005 | 0,001041 |
| Cmpk2 | 2,00532 | 0,585818 | -1,77531 | 5,00E-005 | 0,001041 |
| Fry | 1,3175 | 0,381562 | -1,78781 | 5,00E-005 | 0,001041 |
| Cyp1b1 | 1,41754 | 0,405588 | -1,8053 | 5,00E-005 | 0,001041 |
| Tgfb1 | 7,47103 | 2,13632 | -1,80618 | 5,00E-005 | 0,001041 |
| Nt5e | 2,68463 | 0,767554 | -1,80638 | 5,00E-005 | 0,001041 |
| Cd36 | 6,85855 | 1,95411 | -1,81139 | 5,00E-005 | 0,001041 |
| Vav3 | 3,22774 | 0,913506 | -1,82104 | 5,00E-005 | 0,001041 |
| Cd109 | 4,78171 | 1,35263 | -1,82177 | 5,00E-005 | 0,001041 |
| Abcg4 | 1,60695 | 0,454542 | -1,82184 | 5,00E-005 | 0,001041 |
| Bmp3 | 2,22134 | 0,625776 | -1,82771 | 5,00E-005 | 0,001041 |
| Gpr124 | 6,51584 | 1,83387 | -1,82906 | 5,00E-005 | 0,001041 |
| Runx1t1 | 0,78174 | 0,219907 | -1,8298 | 5,00E-005 | 0,001041 |
| Car8 | 1,39141 | 0,391193 | -1,8306 | 5,00E-005 | 0,001041 |
| Mir17hg | 54,9216 | 15,3977 | -1,83465 | 5,00E-005 | 0,001041 |
| Uncx | 7,34326 | 2,05855 | -1,83479 | 5,00E-005 | 0,001041 |
| 4930506M07Rik | 1,36219 | 0,381626 | -1,8357 | 5,00E-005 | 0,001041 |
| Plxnc1 | 0,99973 | 0,279927 | -1,83649 | 5,00E-005 | 0,001041 |
| Spint2 | 39,6556 | 11,059 | -1,8423 | 5,00E-005 | 0,001041 |
| Erg | 4,4384 | 1,23631 | -1,844 | 5,00E-005 | 0,001041 |
| Arhgef19 | 1,92087 | 0,534257 | -1,84615 | 5,00E-005 | 0,001041 |
| Nid1 | 18,0653 | 5,01946 | -1,84762 | 5,00E-005 | 0,001041 |
| Pnpo | 23,7524 | 6,54144 | -1,86039 | 5,00E-005 | 0,001041 |
| Sh2d4a | 2,11834 | 0,571615 | -1,88982 | 5,00E-005 | 0,001041 |
| Gng11 | 27,0089 | 7,26916 | -1,89358 | 5,00E-005 | 0,001041 |
| Rap1gap2 | 0,732636 | 0,196262 | -1,90032 | 5,00E-005 | 0,001041 |
| Asb4 | 7,42946 | 1,96229 | -1,92072 | 5,00E-005 | 0,001041 |
| Tnc | 0,954022 | 0,251074 | -1,92591 | 5,00E-005 | 0,001041 |
| Parm1 | 6,92189 | 1,77418 | -1,96401 | 5,00E-005 | 0,001041 |
| Samd3 | 7,58746 | 1,94421 | -1,96443 | 5,00E-005 | 0,001041 |
| Rasip1 | 7,20556 | 1,82749 | -1,97925 | 5,00E-005 | 0,001041 |
| Anxa2 | 44,7967 | 11,3073 | -1,98614 | 5,00E-005 | 0,001041 |
| Fam134b | 2,60252 | 0,656092 | -1,98794 | 5,00E-005 | 0,001041 |
| Podxl | 24,1136 | 6,07884 | -1,98798 | 5,00E-005 | 0,001041 |
| Pcdh1 | 5,98565 | 1,50464 | -1,99209 | 5,00E-005 | 0,001041 |

| | | | | | |
|---|---|---|---|---|---|
| Rab3il1 | 8,47579 | 2,1221 | -1,99786 | 5,00E-005 | 0,001041 |
| Cd9 | 18,0719 | 4,49809 | -2,00637 | 5,00E-005 | 0,001041 |
| Fzd5 | 0,863454 | 0,214873 | -2,00664 | 5,00E-005 | 0,001041 |
| Lrrc32 | 1,55916 | 0,387646 | -2,00796 | 5,00E-005 | 0,001041 |
| Abca9 | 0,991236 | 0,245011 | -2,01638 | 5,00E-005 | 0,001041 |
| Krt8 | 12,3694 | 3,04395 | -2,02276 | 5,00E-005 | 0,001041 |
| Slc25a21 | 2,78413 | 0,682308 | -2,02873 | 5,00E-005 | 0,001041 |
| Fermt3 | 3,80874 | 0,931102 | -2,0323 | 5,00E-005 | 0,001041 |
| Sfn | 7,27694 | 1,7759 | -2,03478 | 5,00E-005 | 0,001041 |
| Cd59a | 5,1661 | 1,24419 | -2,05387 | 5,00E-005 | 0,001041 |
| Scn4b | 0,887917 | 0,211297 | -2,07115 | 5,00E-005 | 0,001041 |
| Ptgfr | 1,97343 | 0,465689 | -2,08326 | 5,00E-005 | 0,001041 |
| S1pr1 | 7,49439 | 1,74194 | -2,10511 | 5,00E-005 | 0,001041 |
| Ebf3 | 1,69084 | 0,391591 | -2,11032 | 5,00E-005 | 0,001041 |
| Bnc1 | 1,78426 | 0,413063 | -2,11089 | 5,00E-005 | 0,001041 |
| Capn1 | 12,3528 | 2,78507 | -2,14906 | 5,00E-005 | 0,001041 |
| Plet1 | 2,10761 | 0,47449 | -2,15116 | 5,00E-005 | 0,001041 |
| Enpp1 | 2,01722 | 0,451066 | -2,16096 | 5,00E-005 | 0,001041 |
| Col18a1 | 88,2889 | 19,5891 | -2,17218 | 5,00E-005 | 0,001041 |
| Zc3hav1 | 2,23906 | 0,496095 | -2,17421 | 5,00E-005 | 0,001041 |
| Atp2a3 | 0,981728 | 0,217512 | -2,17423 | 5,00E-005 | 0,001041 |
| Plxnd1 | 23,3995 | 5,15498 | -2,18244 | 5,00E-005 | 0,001041 |
| Arhgap18 | 5,07009 | 1,11602 | -2,18365 | 5,00E-005 | 0,001041 |
| Gas6 | 2,28497 | 0,502609 | -2,18467 | 5,00E-005 | 0,001041 |
| Cntnap2 | 5,24191 | 1,14219 | -2,19828 | 5,00E-005 | 0,001041 |
| Bgn | 11,947 | 2,60113 | -2,19943 | 5,00E-005 | 0,001041 |
| Rab27b | 0,849506 | 0,184927 | -2,19967 | 5,00E-005 | 0,001041 |
| Ramp3 | 4,08083 | 0,88143 | -2,21095 | 5,00E-005 | 0,001041 |
| Fxyd5 | 6,66089 | 1,43179 | -2,2179 | 5,00E-005 | 0,001041 |
| Kctd12b | 2,00906 | 0,430499 | -2,22244 | 5,00E-005 | 0,001041 |
| Dsg2 | 3,48944 | 0,747042 | -2,22373 | 5,00E-005 | 0,001041 |
| Ecscr | 18,532 | 3,94267 | -2,23277 | 5,00E-005 | 0,001041 |
| Slc7a11 | 1,36229 | 0,289633 | -2,23374 | 5,00E-005 | 0,001041 |
| Sost | 15,393 | 3,24584 | -2,24561 | 5,00E-005 | 0,001041 |
| Steap3 | 4,90086 | 1,02822 | -2,25289 | 5,00E-005 | 0,001041 |
| Fam198b | 2,43182 | 0,506256 | -2,2641 | 5,00E-005 | 0,001041 |
| Tspan33 | 9,11344 | 1,88347 | -2,2746 | 5,00E-005 | 0,001041 |
| Hlx | 2,87976 | 0,594192 | -2,27695 | 5,00E-005 | 0,001041 |
| Enpp3 | 2,28989 | 0,470716 | -2,28235 | 5,00E-005 | 0,001041 |
| Igfbp5 | 79,5105 | 15,9826 | -2,31465 | 5,00E-005 | 0,001041 |
| Krt18 | 20,5827 | 4,10878 | -2,32465 | 5,00E-005 | 0,001041 |
| Rapgef3 | 2,25744 | 0,449292 | -2,32896 | 5,00E-005 | 0,001041 |
| Cobl | 3,61022 | 0,714157 | -2,33777 | 5,00E-005 | 0,001041 |
| Tbx18 | 1,3234 | 0,259072 | -2,35283 | 5,00E-005 | 0,001041 |
| Myzap | 3,78147 | 0,7329 | -2,36726 | 5,00E-005 | 0,001041 |
| Cldn3 | 9,5421 | 1,83632 | -2,37749 | 5,00E-005 | 0,001041 |
| Mir6236 | 10173,9 | 1953,37 | -2,38084 | 5,00E-005 | 0,001041 |
| Arhgap27 | 3,29646 | 0,627562 | -2,39309 | 5,00E-005 | 0,001041 |

| | | | | | |
|---|---|---|---|---|---|
| Plau | 3,56769 | 0,666524 | -2,42026 | 5,00E-005 | 0,001041 |
| Acvrl1 | 5,63116 | 1,04015 | -2,43665 | 5,00E-005 | 0,001041 |
| Ell2 | 5,74381 | 1,06067 | -2,43704 | 5,00E-005 | 0,001041 |
| Sorcs2 | 2,30228 | 0,424032 | -2,44082 | 5,00E-005 | 0,001041 |
| Add2 | 6,64018 | 1,22242 | -2,44149 | 5,00E-005 | 0,001041 |
| L1cam | 0,968996 | 0,178221 | -2,44283 | 5,00E-005 | 0,001041 |
| Ptrf | 5,96307 | 1,09166 | -2,44953 | 5,00E-005 | 0,001041 |
| Abcg1 | 1,67554 | 0,306441 | -2,45095 | 5,00E-005 | 0,001041 |
| Ube2l6 | 13,5702 | 2,46866 | -2,45864 | 5,00E-005 | 0,001041 |
| Itih5 | 1,88561 | 0,339147 | -2,47505 | 5,00E-005 | 0,001041 |
| Chl1 | 3,35543 | 0,600018 | -2,48342 | 5,00E-005 | 0,001041 |
| Rgs3 | 3,2055 | 0,572461 | -2,4853 | 5,00E-005 | 0,001041 |
| Slc25a37 | 56,525 | 10,094 | -2,48539 | 5,00E-005 | 0,001041 |
| Plek | 4,34032 | 0,772789 | -2,48965 | 5,00E-005 | 0,001041 |
| Apold1 | 1,57627 | 0,278722 | -2,49962 | 5,00E-005 | 0,001041 |
| Foxa1 | 3,43381 | 0,602171 | -2,51157 | 5,00E-005 | 0,001041 |
| Krt7 | 7,72347 | 1,3467 | -2,51982 | 5,00E-005 | 0,001041 |
| Fzd6 | 6,82809 | 1,17582 | -2,53781 | 5,00E-005 | 0,001041 |
| Cthrc1 | 17,6999 | 3,04568 | -2,53891 | 5,00E-005 | 0,001041 |
| Foxa2 | 5,45041 | 0,934775 | -2,54367 | 5,00E-005 | 0,001041 |
| Vwa1 | 2,31693 | 0,396515 | -2,54677 | 5,00E-005 | 0,001041 |
| Marveld2 | 3,59806 | 0,61425 | -2,55032 | 5,00E-005 | 0,001041 |
| Eng | 9,97348 | 1,68462 | -2,56568 | 5,00E-005 | 0,001041 |
| Treml2 | 1,65278 | 0,275554 | -2,58449 | 5,00E-005 | 0,001041 |
| Npr3 | 2,18018 | 0,35921 | -2,60155 | 5,00E-005 | 0,001041 |
| Apln | 7,21342 | 1,18092 | -2,61077 | 5,00E-005 | 0,001041 |
| Epcam | 36,1945 | 5,91418 | -2,61352 | 5,00E-005 | 0,001041 |
| Lama4 | 4,53189 | 0,737452 | -2,61949 | 5,00E-005 | 0,001041 |
| Prss23 | 3,72316 | 0,601247 | -2,6305 | 5,00E-005 | 0,001041 |
| Exoc3l | 3,51378 | 0,551943 | -2,67043 | 5,00E-005 | 0,001041 |
| Adcy4 | 2,04297 | 0,320708 | -2,67134 | 5,00E-005 | 0,001041 |
| Cxcl12 | 122,551 | 19,1838 | -2,67542 | 5,00E-005 | 0,001041 |
| Cldn1 | 1,79791 | 0,278572 | -2,6902 | 5,00E-005 | 0,001041 |
| Flt4 | 4,80766 | 0,728901 | -2,72154 | 5,00E-005 | 0,001041 |
| Acss1 | 4,76465 | 0,718171 | -2,72997 | 5,00E-005 | 0,001041 |
| Adamts18 | 1,25858 | 0,189326 | -2,73286 | 5,00E-005 | 0,001041 |
| St14 | 4,78798 | 0,720138 | -2,73307 | 5,00E-005 | 0,001041 |
| Lamc2 | 10,0937 | 1,50776 | -2,74298 | 5,00E-005 | 0,001041 |
| Cldn7 | 18,0614 | 2,64905 | -2,76936 | 5,00E-005 | 0,001041 |
| Coro1a | 7,53907 | 1,10522 | -2,77006 | 5,00E-005 | 0,001041 |
| Frem2 | 7,4527 | 1,06303 | -2,80958 | 5,00E-005 | 0,001041 |
| Thbd | 1,3613 | 0,193827 | -2,81214 | 5,00E-005 | 0,001041 |
| C130074G19Rik | 2,79794 | 0,398308 | -2,81241 | 5,00E-005 | 0,001041 |
| Neat1 | 5,69831 | 0,807074 | -2,81976 | 5,00E-005 | 0,001041 |
| Scarf1 | 3,60063 | 0,50829 | -2,82453 | 5,00E-005 | 0,001041 |
| Irf6 | 3,93522 | 0,554356 | -2,82756 | 5,00E-005 | 0,001041 |
| Eps8l2 | 2,13099 | 0,296684 | -2,84452 | 5,00E-005 | 0,001041 |
| Gimap4 | 5,69839 | 0,789416 | -2,8517 | 5,00E-005 | 0,001041 |

| | | | | | |
|---|---|---|---|---|---|
| Cdcp1 | 2,78207 | 0,374105 | -2,89464 | 5,00E-005 | 0,001041 |
| Esrp1 | 7,29902 | 0,978035 | -2,89974 | 5,00E-005 | 0,001041 |
| Cav1 | 2,89793 | 0,38605 | -2,90816 | 5,00E-005 | 0,001041 |
| Car2 | 60,1523 | 7,94988 | -2,91962 | 5,00E-005 | 0,001041 |
| Esrp2 | 2,7505 | 0,362653 | -2,92303 | 5,00E-005 | 0,001041 |
| Erbb3 | 3,5073 | 0,461506 | -2,92594 | 5,00E-005 | 0,001041 |
| Thsd1 | 4,56195 | 0,59584 | -2,93666 | 5,00E-005 | 0,001041 |
| Uba7 | 2,77543 | 0,357171 | -2,95802 | 5,00E-005 | 0,001041 |
| Pcdh17 | 1,00146 | 0,128005 | -2,96783 | 5,00E-005 | 0,001041 |
| Xk | 2,69237 | 0,33684 | -2,99874 | 5,00E-005 | 0,001041 |
| Pkdcc | 3,60038 | 0,449926 | -3,00039 | 5,00E-005 | 0,001041 |
| Upk1a | 5,56035 | 0,694448 | -3,00124 | 5,00E-005 | 0,001041 |
| Foxd2 | 2,98492 | 0,370611 | -3,00971 | 5,00E-005 | 0,001041 |
| Jag2 | 6,99764 | 0,864672 | -3,01664 | 5,00E-005 | 0,001041 |
| Galnt18 | 5,37091 | 0,661273 | -3,02185 | 5,00E-005 | 0,001041 |
| Sh3tc1 | 1,15405 | 0,141827 | -3,02451 | 5,00E-005 | 0,001041 |
| Nkx3-1 | 12,8943 | 1,5799 | -3,02884 | 5,00E-005 | 0,001041 |
| Cldn6 | 75,3138 | 9,16666 | -3,03844 | 5,00E-005 | 0,001041 |
| Wnt4 | 3,871 | 0,470983 | -3,03896 | 5,00E-005 | 0,001041 |
| Acpp | 1,80582 | 0,218528 | -3,04677 | 5,00E-005 | 0,001041 |
| Bcam | 37,6256 | 4,54395 | -3,0497 | 5,00E-005 | 0,001041 |
| Npnt | 8,97566 | 1,07193 | -3,06581 | 5,00E-005 | 0,001041 |
| Cldn5 | 16,7053 | 1,99166 | -3,06826 | 5,00E-005 | 0,001041 |
| Cited4 | 13,9675 | 1,66364 | -3,06966 | 5,00E-005 | 0,001041 |
| Pde2a | 2,71905 | 0,322169 | -3,07721 | 5,00E-005 | 0,001041 |
| Dnm3os | 0,892727 | 0,104174 | -3,09923 | 5,00E-005 | 0,001041 |
| Adra2b | 2,28223 | 0,26442 | -3,10954 | 5,00E-005 | 0,001041 |
| Gpr116 | 4,8147 | 0,557222 | -3,11112 | 5,00E-005 | 0,001041 |
| Blvrb | 30,7954 | 3,54235 | -3,11993 | 5,00E-005 | 0,001041 |
| Tmem184a | 2,55032 | 0,293304 | -3,12021 | 5,00E-005 | 0,001041 |
| Foxd2os | 2,48728 | 0,28099 | -3,14598 | 5,00E-005 | 0,001041 |
| Slc30a10 | 3,85595 | 0,43464 | -3,1492 | 5,00E-005 | 0,001041 |
| Kcne3 | 10,9284 | 1,22679 | -3,15513 | 5,00E-005 | 0,001041 |
| Gjb2 | 2,8619 | 0,310228 | -3,20557 | 5,00E-005 | 0,001041 |
| Clec14a | 2,35058 | 0,2547 | -3,20615 | 5,00E-005 | 0,001041 |
| Epas1 | 3,18323 | 0,344637 | -3,20734 | 5,00E-005 | 0,001041 |
| Mrc1 | 0,938755 | 0,0985052 | -3,25248 | 5,00E-005 | 0,001041 |
| Sox18 | 8,93692 | 0,931472 | -3,26219 | 5,00E-005 | 0,001041 |
| Sox17 | 1,74292 | 0,181182 | -3,26599 | 5,00E-005 | 0,001041 |
| 2610203C20Rik | 10,4152 | 1,05285 | -3,30632 | 5,00E-005 | 0,001041 |
| Hapln1 | 35,1492 | 3,51727 | -3,32097 | 5,00E-005 | 0,001041 |
| Icam1 | 2,69754 | 0,268642 | -3,32788 | 5,00E-005 | 0,001041 |
| Plvap | 30,5025 | 2,97928 | -3,35589 | 5,00E-005 | 0,001041 |
| Inpp5d | 1,45528 | 0,140258 | -3,37514 | 5,00E-005 | 0,001041 |
| Rassf4 | 1,25341 | 0,12071 | -3,37625 | 5,00E-005 | 0,001041 |
| Kdr | 15,035 | 1,4108 | -3,41374 | 5,00E-005 | 0,001041 |
| Nos3 | 2,50887 | 0,234439 | -3,41976 | 5,00E-005 | 0,001041 |
| Ikzf1 | 2,10887 | 0,196536 | -3,4236 | 5,00E-005 | 0,001041 |

| | | | | | |
|---|---|---|---|---|---|
| Esam | 13,3663 | 1,23253 | -3,43891 | 5,00E-005 | 0,001041 |
| Ahnak | 2,91095 | 0,26638 | -3,44993 | 5,00E-005 | 0,001041 |
| Sox7 | 3,24626 | 0,294333 | -3,46326 | 5,00E-005 | 0,001041 |
| Gja4 | 4,82579 | 0,434913 | -3,47196 | 5,00E-005 | 0,001041 |
| Slc39a8 | 17,5895 | 1,57141 | -3,48458 | 5,00E-005 | 0,001041 |
| Tnfaip2 | 2,98658 | 0,256473 | -3,54162 | 5,00E-005 | 0,001041 |
| Mcam | 13,4132 | 1,14424 | -3,5512 | 5,00E-005 | 0,001041 |
| Cobll1 | 1,83288 | 0,15343 | -3,57846 | 5,00E-005 | 0,001041 |
| Lyl1 | 4,11445 | 0,336184 | -3,61338 | 5,00E-005 | 0,001041 |
| Mgst3 | 61,2423 | 4,98583 | -3,61862 | 5,00E-005 | 0,001041 |
| Klhl6 | 4,99163 | 0,389399 | -3,68019 | 5,00E-005 | 0,001041 |
| Slc40a1 | 2,98965 | 0,232923 | -3,68205 | 5,00E-005 | 0,001041 |
| Mfsd2b | 3,09676 | 0,233177 | -3,73127 | 5,00E-005 | 0,001041 |
| Tie1 | 11,0077 | 0,808569 | -3,767 | 5,00E-005 | 0,001041 |
| Flt1 | 4,09607 | 0,296429 | -3,78848 | 5,00E-005 | 0,001041 |
| Mmrn1 | 5,03428 | 0,364 | -3,78977 | 5,00E-005 | 0,001041 |
| Tgm2 | 5,17983 | 0,365984 | -3,82305 | 5,00E-005 | 0,001041 |
| Cd34 | 39,7694 | 2,80175 | -3,82726 | 5,00E-005 | 0,001041 |
| Cdh5 | 26,0734 | 1,81388 | -3,84543 | 5,00E-005 | 0,001041 |
| Gimap6 | 8,90137 | 0,590586 | -3,91381 | 5,00E-005 | 0,001041 |
| Wnt6 | 12,6314 | 0,834462 | -3,92003 | 5,00E-005 | 0,001041 |
| Tfap2a | 3,05227 | 0,200685 | -3,92688 | 5,00E-005 | 0,001041 |
| Icam2 | 10,7811 | 0,695943 | -3,9534 | 5,00E-005 | 0,001041 |
| Pdgfb | 4,21402 | 0,268421 | -3,97263 | 5,00E-005 | 0,001041 |
| Eltd1 | 3,09493 | 0,19401 | -3,99571 | 5,00E-005 | 0,001041 |
| Wnt3 | 4,17253 | 0,259578 | -4,00668 | 5,00E-005 | 0,001041 |
| Tek | 6,24248 | 0,383663 | -4,02421 | 5,00E-005 | 0,001041 |
| Bcl6b | 3,44754 | 0,206468 | -4,06158 | 5,00E-005 | 0,001041 |
| Pcdh12 | 3,43126 | 0,20332 | -4,07691 | 5,00E-005 | 0,001041 |
| Emcn | 32,6012 | 1,88113 | -4,11525 | 5,00E-005 | 0,001041 |
| Klf1 | 9,26025 | 0,519365 | -4,15623 | 5,00E-005 | 0,001041 |
| Moxd1 | 6,52459 | 0,361944 | -4,17205 | 5,00E-005 | 0,001041 |
| Trim29 | 2,84496 | 0,155043 | -4,19767 | 5,00E-005 | 0,001041 |
| Sptb | 6,7191 | 0,365427 | -4,20061 | 5,00E-005 | 0,001041 |
| Col15a1 | 5,08659 | 0,275626 | -4,20591 | 5,00E-005 | 0,001041 |
| Mmrn2 | 5,63261 | 0,30064 | -4,2277 | 5,00E-005 | 0,001041 |
| Grhl2 | 1,85238 | 0,0975044 | -4,24777 | 5,00E-005 | 0,001041 |
| Pecam1 | 19,6499 | 1,01202 | -4,27921 | 5,00E-005 | 0,001041 |
| Itga3 | 14,4334 | 0,735947 | -4,29367 | 5,00E-005 | 0,001041 |
| Arhgdib | 9,06655 | 0,461337 | -4,29666 | 5,00E-005 | 0,001041 |
| Cd93 | 8,82681 | 0,444893 | -4,31036 | 5,00E-005 | 0,001041 |
| Pklr | 12,7617 | 0,624638 | -4,35265 | 5,00E-005 | 0,001041 |
| Pax1 | 9,30799 | 0,455033 | -4,35443 | 5,00E-005 | 0,001041 |
| Trp63 | 3,15452 | 0,152401 | -4,37147 | 5,00E-005 | 0,001041 |
| Cdh1 | 27,2994 | 1,28688 | -4,40692 | 5,00E-005 | 0,001041 |
| Gjb3 | 6,30313 | 0,282913 | -4,47764 | 5,00E-005 | 0,001041 |
| Cldn4 | 21,6399 | 0,955035 | -4,50199 | 5,00E-005 | 0,001041 |
| Dll4 | 3,67342 | 0,160462 | -4,51682 | 5,00E-005 | 0,001041 |

| | | | | | |
|---|---|---|---|---|---|
| Thbs1 | 12,332 | 0,466837 | -4,72335 | 5,00E-005 | 0,001041 |
| Itgb3 | 3,50503 | 0,130712 | -4,74496 | 5,00E-005 | 0,001041 |
| Lmx1a | 4,8108 | 0,164263 | -4,8722 | 5,00E-005 | 0,001041 |
| Tinagl1 | 7,30411 | 0,232648 | -4,97249 | 5,00E-005 | 0,001041 |
| Perp | 26,6703 | 0,806859 | -5,04677 | 5,00E-005 | 0,001041 |
| Tal1 | 7,47688 | 0,214101 | -5,12607 | 5,00E-005 | 0,001041 |
| Vwf | 2,50169 | 0,0704498 | -5,15016 | 5,00E-005 | 0,001041 |
| Spns2 | 7,59709 | 0,201706 | -5,23512 | 5,00E-005 | 0,001041 |
| Ank1 | 5,72032 | 0,14725 | -5,27975 | 5,00E-005 | 0,001041 |
| Sema3d | 2,87969 | 0,0579 | -5,63621 | 5,00E-005 | 0,001041 |
| Alas2 | 52,2908 | 0,961067 | -5,76578 | 5,00E-005 | 0,001041 |
| Gypa | 24,3304 | 0,415827 | -5,87063 | 5,00E-005 | 0,001041 |
| Mt1 | 74,0058 | 1,2372 | -5,90249 | 5,00E-005 | 0,001041 |
| Noto | 19,5729 | 0,260627 | -6,23072 | 5,00E-005 | 0,001041 |
| Hba-x | 4627,75 | 50,5338 | -6,51692 | 5,00E-005 | 0,001041 |
| Hba-a1,Hba-a2 | 339,078 | 3,65138 | -6,53703 | 5,00E-005 | 0,001041 |
| Hbb-bh1 | 7541,24 | 54,6533 | -7,10835 | 5,00E-005 | 0,001041 |
| Hbb-y | 12373,9 | 84,7314 | -7,19019 | 5,00E-005 | 0,001041 |
| Slc4a1 | 55,6685 | 0,262468 | -7,72857 | 5,00E-005 | 0,001041 |
| Smim22 | 1,26886 | 0 | -inf | 0,0001 | 0,001968 |
| Hspa1a | 0,139231 | 4,45362 | 4,99943 | 0,0001 | 0,001968 |
| Gzmk | 4,68087 | 10,6562 | 1,18684 | 0,0001 | 0,001968 |
| Dgkg | 1,5297 | 2,78923 | 0,866617 | 0,0001 | 0,001968 |
| Fam222a | 4,82307 | 8,51271 | 0,819666 | 0,0001 | 0,001968 |
| Cap2 | 3,71046 | 6,41736 | 0,790381 | 0,0001 | 0,001968 |
| Aldoart2 | 10,508 | 17,8466 | 0,764167 | 0,0001 | 0,001968 |
| Svep1 | 2,53141 | 4,29394 | 0,76236 | 0,0001 | 0,001968 |
| Tpm1 | 992,172 | 1682,66 | 0,762079 | 0,0001 | 0,001968 |
| Pde9a | 17,2136 | 27,6252 | 0,682433 | 0,0001 | 0,001968 |
| Mogat2 | 39,933 | 64,0411 | 0,681417 | 0,0001 | 0,001968 |
| Greb1 | 70,1639 | 108,405 | 0,627636 | 0,0001 | 0,001968 |
| BC030336 | 17,3333 | 10,7683 | -0,686751 | 0,0001 | 0,001968 |
| Mmp15 | 10,2582 | 6,06766 | -0,75756 | 0,0001 | 0,001968 |
| Pcsk6 | 2,54486 | 1,20062 | -1,0838 | 0,0001 | 0,001968 |
| Rin3 | 3,63196 | 1,71069 | -1,08617 | 0,0001 | 0,001968 |
| Kank3 | 4,55358 | 2,03988 | -1,15852 | 0,0001 | 0,001968 |
| Bmp5 | 3,53356 | 1,56545 | -1,17454 | 0,0001 | 0,001968 |
| Gpm6b | 2,60546 | 1,15151 | -1,17801 | 0,0001 | 0,001968 |
| Pou6f1 | 1,45738 | 0,575462 | -1,34059 | 0,0001 | 0,001968 |
| Sorbs2 | 1,01421 | 0,388075 | -1,38595 | 0,0001 | 0,001968 |
| Rin1 | 1,31607 | 0,468918 | -1,48882 | 0,0001 | 0,001968 |
| Sh3gl2 | 1,80188 | 0,612375 | -1,55702 | 0,0001 | 0,001968 |
| Cbln4 | 1,68355 | 0,455272 | -1,8867 | 0,0001 | 0,001968 |
| Hhex | 2,56166 | 0,629745 | -2,02424 | 0,0001 | 0,001968 |
| Fam46c | 2,44536 | 0,555466 | -2,13827 | 0,0001 | 0,001968 |
| Grap | 2,65635 | 0,568879 | -2,22325 | 0,0001 | 0,001968 |
| Ppp1r16b | 0,768013 | 0,126104 | -2,60651 | 0,0001 | 0,001968 |
| Jak3 | 1,02749 | 0,158338 | -2,69804 | 0,0001 | 0,001968 |

| | | | | | |
|---|---|---|---|---|---|
| Gjb5 | 2,53305 | 0,389663 | -2,70058 | 0,0001 | 0,001968 |
| Klf2 | 2,36189 | 0,352745 | -2,74325 | 0,0001 | 0,001968 |
| C130021I20Rik | 1,31509 | 0,156135 | -3,0743 | 0,0001 | 0,001968 |
| Snca | 5,4152 | 0,489909 | -3,46643 | 0,0001 | 0,001968 |
| Slc38a5 | 3,96996 | 0,284174 | -3,80428 | 0,0001 | 0,001968 |
| F2rl1 | 2,36385 | 0,155757 | -3,92377 | 0,0001 | 0,001968 |
| Ctla2a | 7,62323 | 0,425791 | -4,16219 | 0,0001 | 0,001968 |
| Cxcl14 | 10,6849 | 0,185385 | -5,8489 | 0,0001 | 0,001968 |
| Nptx2 | 13,0529 | 20,7997 | 0,672195 | 0,00015 | 0,002856 |
| Rarg | 48,2068 | 72,9123 | 0,596926 | 0,00015 | 0,002856 |
| Arl6ip1 | 200,233 | 136,263 | -0,555292 | 0,00015 | 0,002856 |
| Kidins220 | 11,7056 | 7,75824 | -0,593399 | 0,00015 | 0,002856 |
| Tmed7 | 55,5388 | 35,9113 | -0,629058 | 0,00015 | 0,002856 |
| Mocs1 | 19,2177 | 11,8156 | -0,70174 | 0,00015 | 0,002856 |
| Reln | 3,3978 | 2,05544 | -0,725155 | 0,00015 | 0,002856 |
| Slc22a23 | 5,3116 | 3,15839 | -0,74996 | 0,00015 | 0,002856 |
| Sgk3 | 3,71059 | 1,96901 | -0,914178 | 0,00015 | 0,002856 |
| Pdk2 | 3,91343 | 1,84 | -1,08873 | 0,00015 | 0,002856 |
| Slc38a3 | 4,65532 | 2,159 | -1,10851 | 0,00015 | 0,002856 |
| Il4ra | 1,61 | 0,701234 | -1,19909 | 0,00015 | 0,002856 |
| Hhip | 0,904188 | 0,388903 | -1,21721 | 0,00015 | 0,002856 |
| Fam109b | 2,96997 | 1,22238 | -1,28076 | 0,00015 | 0,002856 |
| Fam78a | 1,61085 | 0,646943 | -1,31611 | 0,00015 | 0,002856 |
| Lmna | 3,17912 | 1,09939 | -1,53192 | 0,00015 | 0,002856 |
| Kif12 | 2,0546 | 0,595305 | -1,78715 | 0,00015 | 0,002856 |
| Trpc6 | 1,88042 | 0,475933 | -1,98222 | 0,00015 | 0,002856 |
| Dok4 | 7,35651 | 1,83844 | -2,00054 | 0,00015 | 0,002856 |
| Ednrb | 0,907326 | 0,160486 | -2,49918 | 0,00015 | 0,002856 |
| Cd55 | 1,48388 | 0,21487 | -2,78783 | 0,00015 | 0,002856 |
| Nfe2 | 4,53991 | 0,219654 | -4,36936 | 0,00015 | 0,002856 |
| Spta1 | 5,29264 | 0,0424936 | -6,9606 | 0,00015 | 0,002856 |
| Col4a5 | 9,13071 | 5,73329 | -0,671365 | 0,0002 | 0,003692 |
| Mbd2 | 15,9795 | 9,84014 | -0,699471 | 0,0002 | 0,003692 |
| Srgap1 | 4,54539 | 2,79658 | -0,700741 | 0,0002 | 0,003692 |
| Ank3 | 7,04541 | 4,28389 | -0,717762 | 0,0002 | 0,003692 |
| Ids | 5,34061 | 3,22391 | -0,728193 | 0,0002 | 0,003692 |
| Abcc4 | 3,95625 | 2,23602 | -0,823201 | 0,0002 | 0,003692 |
| Susd1 | 4,00425 | 2,20377 | -0,861561 | 0,0002 | 0,003692 |
| C630043F03Rik | 5,69663 | 3,09828 | -0,878642 | 0,0002 | 0,003692 |
| Tmcc3 | 1,91318 | 0,909621 | -1,07264 | 0,0002 | 0,003692 |
| Grik1 | 3,04478 | 1,42359 | -1,09681 | 0,0002 | 0,003692 |
| Nrros | 2,63107 | 1,17736 | -1,16009 | 0,0002 | 0,003692 |
| Cyb561d2 | 4,60737 | 2,01428 | -1,19368 | 0,0002 | 0,003692 |
| 2010315B03Rik | 2,81865 | 1,14329 | -1,30181 | 0,0002 | 0,003692 |
| Nova2 | 0,763218 | 0,304827 | -1,3241 | 0,0002 | 0,003692 |
| Svip | 1,98018 | 0,789474 | -1,32666 | 0,0002 | 0,003692 |
| Ndrg1 | 2,64261 | 0,994549 | -1,40985 | 0,0002 | 0,003692 |
| Dapl1 | 12,079 | 3,91034 | -1,62713 | 0,0002 | 0,003692 |

| | | | | | |
|---|---|---|---|---|---|
| Wipf3 | 1,07255 | 0,338843 | -1,66235 | 0,0002 | 0,003692 |
| Pde8a | 2,01802 | 0,205717 | -3,29421 | 0,0002 | 0,003692 |
| Arhgap25 | 0,959917 | 0,0966782 | -3,31165 | 0,0002 | 0,003692 |
| Rhd | 12,3935 | 0,233362 | -5,73087 | 0,0002 | 0,003692 |
| Mt2 | 64,0742 | 0,944912 | -6,08342 | 0,0002 | 0,003692 |
| Hoxc11 | 38,9655 | 61,499 | 0,658365 | 0,00025 | 0,004437 |
| Ankrd26 | 11,8986 | 18,3587 | 0,625671 | 0,00025 | 0,004437 |
| Lss | 19,4986 | 29,7418 | 0,609122 | 0,00025 | 0,004437 |
| Lhpp | 50,0534 | 75,4356 | 0,591777 | 0,00025 | 0,004437 |
| Pea15a | 82,8692 | 123,895 | 0,580214 | 0,00025 | 0,004437 |
| Fzd2 | 28,6891 | 18,9574 | -0,597745 | 0,00025 | 0,004437 |
| Marcks | 106,85 | 70,4075 | -0,601788 | 0,00025 | 0,004437 |
| Slc12a4 | 14,1445 | 9,29968 | -0,604985 | 0,00025 | 0,004437 |
| Tmem14c | 138,956 | 90,6848 | -0,615692 | 0,00025 | 0,004437 |
| Lrp10 | 15,167 | 9,7619 | -0,635702 | 0,00025 | 0,004437 |
| Ppap2b | 23,419 | 14,8305 | -0,659107 | 0,00025 | 0,004437 |
| Sema3c | 7,60304 | 4,65527 | -0,707711 | 0,00025 | 0,004437 |
| Rdh10 | 11,0565 | 6,61137 | -0,741878 | 0,00025 | 0,004437 |
| Snhg3 | 128,914 | 76,0503 | -0,761387 | 0,00025 | 0,004437 |
| Tcf7l2 | 4,34203 | 2,51305 | -0,788932 | 0,00025 | 0,004437 |
| 1700066M21Rik | 5,22068 | 2,89476 | -0,850795 | 0,00025 | 0,004437 |
| Llgl2 | 3,48457 | 1,79199 | -0,959417 | 0,00025 | 0,004437 |
| Capn5 | 2,70599 | 1,3636 | -0,98873 | 0,00025 | 0,004437 |
| Abca3 | 2,04822 | 0,955777 | -1,09963 | 0,00025 | 0,004437 |
| Slc18a2 | 1,65285 | 0,750295 | -1,13943 | 0,00025 | 0,004437 |
| Unc5c | 0,622549 | 0,252219 | -1,30351 | 0,00025 | 0,004437 |
| Tor4a | 1,86189 | 0,751723 | -1,30849 | 0,00025 | 0,004437 |
| Emx2 | 1,44406 | 0,381074 | -1,92198 | 0,00025 | 0,004437 |
| Olfml2a | 1,842 | 0,484784 | -1,92586 | 0,00025 | 0,004437 |
| Ahr | 0,680498 | 0,165544 | -2,03937 | 0,00025 | 0,004437 |
| Htatip2 | 2,46544 | 0,502072 | -2,29588 | 0,00025 | 0,004437 |
| Ly6h | 4,09138 | 0,627565 | -2,70475 | 0,00025 | 0,004437 |
| Arhgef16 | 1,63487 | 0,185909 | -3,13651 | 0,00025 | 0,004437 |
| Ppl | 1,08595 | 0,0547704 | -4,30942 | 0,00025 | 0,004437 |
| Rtl1 | 0,150789 | 0,635861 | 2,07618 | 0,0003 | 0,005214 |
| Dhtkd1 | 3,68889 | 6,19933 | 0,748926 | 0,0003 | 0,005214 |
| Crmp1 | 11,3473 | 17,7968 | 0,649271 | 0,0003 | 0,005214 |
| App | 54,222 | 36,3519 | -0,576845 | 0,0003 | 0,005214 |
| Mmp11 | 14,7434 | 9,18722 | -0,682373 | 0,0003 | 0,005214 |
| Fam43a | 8,5521 | 5,06804 | -0,754851 | 0,0003 | 0,005214 |
| Rnf128 | 8,55168 | 5,02166 | -0,754851 | 0,0003 | 0,005214 |
| Eda | 2,3602 | 1,25783 | -0,754851 | 0,0003 | 0,005214 |
| Ttyh1 | 1,67692 | 0,760679 | -0,754851 | 0,0003 | 0,005214 |
| Bst2 | 5,94138 | 1,46243 | -0,754851 | 0,0003 | 0,005214 |
| Mpeg1 | 0,832425 | 0,106553 | -0,754851 | 0,0003 | 0,005214 |
| Ushbp1 | 1,16817 | 0,125019 | -0,754851 | 0,0003 | 0,005214 |
| Mrvi1 | 0,645083 | 0,0680711 | -0,754851 | 0,0003 | 0,005214 |
| Rapgef4 | 0,875243 | 0,0831942 | -0,754851 | 0,0003 | 0,005214 |

| | | | | | |
|---|---|---|---|---|---|
| Tbx22 | 2,52592 | 0,187534 | -0,754851 | 0,0003 | 0,005214 |
| Krt5 | 3,47328 | 0,149071 | -0,754851 | 0,0003 | 0,005214 |
| Hsph1 | 99,4626 | 149,504 | -0,754851 | 0,00035 | 0,005974 |
| Crabp2 | 286,788 | 428,463 | -0,754851 | 0,00035 | 0,005974 |
| Sppl2a | 19,1459 | 12,9161 | -0,754851 | 0,00035 | 0,005974 |
| Man2a1 | 28,0977 | 18,7251 | -0,754851 | 0,00035 | 0,005974 |
| Slc15a2 | 5,80476 | 3,37515 | -0,754851 | 0,00035 | 0,005974 |
| Vat1 | 8,7725 | 5,06751 | -0,754851 | 0,00035 | 0,005974 |
| Frrs1 | 6,94864 | 3,78313 | -0,754851 | 0,00035 | 0,005974 |
| Gsn | 2,24456 | 0,952953 | -0,754851 | 0,00035 | 0,005974 |
| Ifnlr1 | 1,61107 | 0,678657 | -0,754851 | 0,00035 | 0,005974 |
| Ptges | 1,1023 | 0,350553 | -0,754851 | 0,00035 | 0,005974 |
| Sqrdl | 2,03182 | 0,333566 | -0,754851 | 0,00035 | 0,005974 |
| St8sia4 | 2,39185 | 0,350136 | -0,754851 | 0,00035 | 0,005974 |
| Kcnk6 | 1,11098 | 0,109242 | -0,754851 | 0,00035 | 0,005974 |
| Tmem56 | 1,40666 | 0,0463264 | -0,754851 | 0,00035 | 0,005974 |
| Dusp5 | 1,99767 | 4,83782 | -0,754851 | 0,0004 | 0,006665 |
| Kcnip3 | 2,73567 | 5,05275 | -0,754851 | 0,0004 | 0,006665 |
| Sept5 | 8,87393 | 14,4609 | -0,754851 | 0,0004 | 0,006665 |
| Hoxc4 | 24,7902 | 37,4639 | -0,754851 | 0,0004 | 0,006665 |
| Rps2 | 162,12 | 238,216 | -0,754851 | 0,0004 | 0,006665 |
| Serpine2 | 57,9307 | 39,3013 | -0,754851 | 0,0004 | 0,006665 |
| Casd1 | 33,9322 | 22,581 | -0,754851 | 0,0004 | 0,006665 |
| Syt13 | 7,44192 | 4,55202 | -0,754851 | 0,0004 | 0,006665 |
| Tpst2 | 15,5034 | 9,47941 | -0,754851 | 0,0004 | 0,006665 |
| Wwc1 | 5,91985 | 3,49191 | -0,754851 | 0,0004 | 0,006665 |
| Stk17b | 3,46457 | 1,87606 | -0,754851 | 0,0004 | 0,006665 |
| Dact2 | 3,49114 | 1,76561 | -0,754851 | 0,0004 | 0,006665 |
| Sfmbt2 | 0,984948 | 0,469394 | -0,754851 | 0,0004 | 0,006665 |
| Crb3 | 3,35818 | 0,840129 | -0,754851 | 0,0004 | 0,006665 |
| 5730508B09Rik | 2,28045 | 0,375869 | -0,754851 | 0,0004 | 0,006665 |
| C1qtnf3 | 1,24427 | 0,174929 | -0,754851 | 0,0004 | 0,006665 |
| Nckap1l | 0,876249 | 0,108107 | -0,754851 | 0,0004 | 0,006665 |
| Cav2 | 1,54298 | 0,135119 | -0,754851 | 0,0004 | 0,006665 |
| Hemgn | 20,2871 | 0,127135 | -0,754851 | 0,0004 | 0,006665 |
| Spink3 | 1,24779 | 0 | -0,754851 | 0,00045 | 0,007406 |
| Sh3bgrl | 83,7974 | 57,6116 | -0,754851 | 0,00045 | 0,007406 |
| Ccnb1 | 40,2596 | 27,3377 | -0,754851 | 0,00045 | 0,007406 |
| Tmem106b | 8,87578 | 5,61964 | -0,754851 | 0,00045 | 0,007406 |
| Celf2 | 1,39349 | 0,720698 | -0,754851 | 0,00045 | 0,007406 |
| Spon1 | 1,45703 | 0,725037 | -0,754851 | 0,00045 | 0,007406 |
| Rinl | 1,14404 | 0,241402 | -0,754851 | 0,00045 | 0,007406 |
| Hpn | 1,95422 | 0,355116 | -0,754851 | 0,00045 | 0,007406 |
| Plek2 | 2,63625 | 0,263331 | -0,754851 | 0,00045 | 0,007406 |
| Wnt7b | 4,27094 | 0,0694586 | -0,754851 | 0,00045 | 0,007406 |
| Selt | 45,0776 | 30,6903 | -0,754851 | 0,0005 | 0,008169 |
| Megf9 | 8,37522 | 5,13169 | -0,754851 | 0,0005 | 0,008169 |
| Fam63b | 2,5657 | 1,50948 | -0,754851 | 0,0005 | 0,008169 |

| | | | | | |
|---|---|---|---|---|---|
| Unc13d | 0,956378 | 0,287916 | -0,754851 | 0,0005 | 0,008169 |
| Vtn | 2,51677 | 0,330936 | -0,754851 | 0,0005 | 0,008169 |
| Klhl4 | 1,33173 | 0,0747624 | -0,754851 | 0,0005 | 0,008169 |
| Sall3 | 15,8192 | 23,3948 | -0,754851 | 0,00055 | 0,008824 |
| Adam10 | 54,7349 | 37,2733 | -0,754851 | 0,00055 | 0,008824 |
| Fth1 | 393,957 | 266,915 | -0,754851 | 0,00055 | 0,008824 |
| Ggta1 | 12,1243 | 7,84439 | -0,754851 | 0,00055 | 0,008824 |
| Dysf | 4,87737 | 3,00024 | -0,754851 | 0,00055 | 0,008824 |
| Plxdc2 | 1,96032 | 1,09004 | -0,754851 | 0,00055 | 0,008824 |
| 2810008D09Rik | 32,6068 | 17,9539 | -0,754851 | 0,00055 | 0,008824 |
| Colgalt2 | 2,1657 | 1,07615 | -0,754851 | 0,00055 | 0,008824 |
| Lhx6 | 1,36304 | 0,557333 | -0,754851 | 0,00055 | 0,008824 |
| Nmnat3 | 2,64814 | 1,07781 | -0,754851 | 0,00055 | 0,008824 |
| Penk | 3,88118 | 1,48413 | -0,754851 | 0,00055 | 0,008824 |
| Qpct | 2,2508 | 0,776337 | -0,754851 | 0,00055 | 0,008824 |
| AI661453 | 0,811502 | 0,187959 | -0,754851 | 0,00055 | 0,008824 |
| Ap1m2 | 2,95774 | 0,179179 | -0,754851 | 0,00055 | 0,008824 |
| Wnt7a | 2,09156 | 0,0884663 | -0,754851 | 0,00055 | 0,008824 |
| Rassf9 | 2,16183 | 4,28651 | -0,754851 | 0,0006 | 0,009478 |
| Dll3 | 114,4 | 169,774 | -0,754851 | 0,0006 | 0,009478 |
| Csrp2 | 285,376 | 417,377 | -0,754851 | 0,0006 | 0,009478 |
| Rgs2 | 6,18354 | 3,63935 | -0,754851 | 0,0006 | 0,009478 |
| Mafb | 2,61584 | 1,34615 | -0,754851 | 0,0006 | 0,009478 |
| Sim1 | 0,980272 | 0,47469 | -0,754851 | 0,0006 | 0,009478 |
| Pisd-ps1 | 11,6769 | 4,96055 | -0,754851 | 0,0006 | 0,009478 |
| Btbd11 | 1,05355 | 0,433152 | -0,754851 | 0,0006 | 0,009478 |
| Als2cl | 0,919409 | 0,344586 | -0,754851 | 0,0006 | 0,009478 |
| Ccm2l | 1,65415 | 0,208989 | -0,754851 | 0,0006 | 0,009478 |
| Tbx1 | 1,98968 | 0,250892 | -0,754851 | 0,0006 | 0,009478 |
| Col8a2 | 1,16762 | 0,064877 | -0,754851 | 0,0006 | 0,009478 |
| Cldn8 | 3,57842 | 0,124949 | -0,754851 | 0,0006 | 0,009478 |
| Rnu11 | 0 | 90,3337 | -0,754851 | 0,00065 | 0,010077 |
| Evx1 | 22,1177 | 38,9754 | -0,754851 | 0,00065 | 0,010077 |
| Lama1 | 43,6799 | 64,4841 | -0,754851 | 0,00065 | 0,010077 |
| Tmem123 | 61,3445 | 42,314 | -0,754851 | 0,00065 | 0,010077 |
| Cadm1 | 56,0447 | 38,2236 | -0,754851 | 0,00065 | 0,010077 |
| Rnpc3 | 20,8591 | 13,729 | -0,754851 | 0,00065 | 0,010077 |
| E2f2 | 5,30494 | 3,27264 | -0,754851 | 0,00065 | 0,010077 |
| Fhdc1 | 4,14787 | 2,54389 | -0,754851 | 0,00065 | 0,010077 |
| St6galnac3 | 6,26085 | 3,83698 | -0,754851 | 0,00065 | 0,010077 |
| Kcnk1 | 8,02633 | 4,68678 | -0,754851 | 0,00065 | 0,010077 |
| Fam13a | 2,00226 | 0,997408 | -0,754851 | 0,00065 | 0,010077 |
| Skida1 | 1,25156 | 0,533947 | -0,754851 | 0,00065 | 0,010077 |
| Arid5a | 0,647233 | 0,20983 | -0,754851 | 0,00065 | 0,010077 |
| Trib3 | 1,77215 | 0,546992 | -0,754851 | 0,00065 | 0,010077 |
| Sp9 | 1,27072 | 0,187001 | -0,754851 | 0,00065 | 0,010077 |
| Apoa1 | 4,05894 | 0,404013 | -0,754851 | 0,00065 | 0,010077 |
| Nefm | 1,96745 | 3,77214 | -0,754851 | 0,0007 | 0,010716 |

| | | | | | |
|---|---|---|---|---|---|
| Igsf21 | 3,84251 | 6,90696 | -0,754851 | 0,0007 | 0,010716 |
| Hoxc6 | 37,5081 | 55,8343 | -0,754851 | 0,0007 | 0,010716 |
| Slc7a1 | 11,8253 | 8,04188 | -0,754851 | 0,0007 | 0,010716 |
| Kctd12 | 8,11803 | 5,4755 | -0,754851 | 0,0007 | 0,010716 |
| Arl4c | 10,26 | 6,63651 | -0,754851 | 0,0007 | 0,010716 |
| Rfesd | 8,41127 | 4,96468 | -0,754851 | 0,0007 | 0,010716 |
| Ppapdc2 | 2,92424 | 1,475 | -0,754851 | 0,0007 | 0,010716 |
| Sema3b | 1,71049 | 0,77984 | -0,754851 | 0,0007 | 0,010716 |
| Zbtb7b | 1,54886 | 0,641581 | -0,754851 | 0,0007 | 0,010716 |
| Lrmp | 1,2634 | 0,25692 | -0,754851 | 0,0007 | 0,010716 |
| Plod2 | 37,135 | 25,1787 | -0,754851 | 0,00075 | 0,011377 |
| Vill | 6,28216 | 3,70419 | -0,754851 | 0,00075 | 0,011377 |
| Evpl | 1,96406 | 1,10498 | -0,754851 | 0,00075 | 0,011377 |
| Snap91 | 3,63213 | 2,01342 | -0,754851 | 0,00075 | 0,011377 |
| Tspan15 | 3,2205 | 1,75392 | -0,754851 | 0,00075 | 0,011377 |
| Mafk | 3,54624 | 1,82855 | -0,754851 | 0,00075 | 0,011377 |
| Pqlc3 | 3,63206 | 1,63332 | -0,754851 | 0,00075 | 0,011377 |
| Chrnb1 | 1,57381 | 0,25084 | -0,754851 | 0,00075 | 0,011377 |
| Pik3c2b | 11,495 | 16,9991 | -0,754851 | 0,0008 | 0,01204 |
| Tmem30a | 44,6966 | 30,6288 | -0,754851 | 0,0008 | 0,01204 |
| Uaca | 8,18736 | 5,31059 | -0,754851 | 0,0008 | 0,01204 |
| Rasgrp2 | 2,0211 | 0,766063 | -0,754851 | 0,0008 | 0,01204 |
| Gpr155 | 0,756722 | 0,255475 | -0,754851 | 0,0008 | 0,01204 |
| Procr | 2,21382 | 0,297266 | -0,754851 | 0,0008 | 0,01204 |
| Kcnj2 | 0,85897 | 0,0522439 | -0,754851 | 0,0008 | 0,01204 |
| Actg1 | 5,40843 | 9,10047 | -0,754851 | 0,00085 | 0,012608 |
| Rps10 | 32,8431 | 53,971 | 0,716592 | 0,00085 | 0,012608 |
| Aldoa | 395,651 | 577,243 | 0,54495 | 0,00085 | 0,012608 |
| Daam1 | 13,0496 | 8,85224 | -0,559889 | 0,00085 | 0,012608 |
| Bmp2k | 4,47849 | 2,53426 | -0,82145 | 0,00085 | 0,012608 |
| Pard3b | 1,50594 | 0,847608 | -0,8292 | 0,00085 | 0,012608 |
| Zbtb4 | 0,999847 | 0,484326 | -1,04573 | 0,00085 | 0,012608 |
| Vamp1 | 1,78081 | 0,789268 | -1,17395 | 0,00085 | 0,012608 |
| Adora2a | 1,9611 | 0,852547 | -1,20181 | 0,00085 | 0,012608 |
| Dusp2 | 3,32882 | 1,39918 | -1,25043 | 0,00085 | 0,012608 |
| Lmcd1 | 2,6922 | 1,0881 | -1,30697 | 0,00085 | 0,012608 |
| Usp43 | 0,73244 | 0,188462 | -1,95844 | 0,00085 | 0,012608 |
| Mapk13 | 2,69538 | 0,317062 | -3,08765 | 0,00085 | 0,012608 |
| Fgf8 | 39,5986 | 59,6221 | 0,590398 | 0,0009 | 0,013232 |
| Ier2 | 27,9165 | 41,6566 | 0,577428 | 0,0009 | 0,013232 |
| Dusp4 | 90,2551 | 131,748 | 0,545705 | 0,0009 | 0,013232 |
| Cd164 | 122,082 | 84,5803 | -0,529456 | 0,0009 | 0,013232 |
| Ptpru | 8,52381 | 5,72219 | -0,57493 | 0,0009 | 0,013232 |
| Foxd3 | 1,31297 | 0,363653 | -1,8522 | 0,0009 | 0,013232 |
| Fam129a | 0,773625 | 0,179402 | -2,10844 | 0,0009 | 0,013232 |
| Gucy1a3 | 0,576137 | 0,112119 | -2,36138 | 0,0009 | 0,013232 |
| Glis3 | 2,19834 | 3,73997 | 0,766614 | 0,00095 | 0,0138 |
| Nrp2 | 42,8338 | 29,4285 | -0,541538 | 0,00095 | 0,0138 |

| | | | | | |
|---|---|---|---|---|---|
| Ormdl1 | 26,8917 | 17,7853 | -0,596471 | 0,00095 | 0,0138 |
| Slc30a4 | 8,44397 | 5,55891 | -0,603119 | 0,00095 | 0,0138 |
| Rbm3 | 37,633 | 24,355 | -0,627781 | 0,00095 | 0,0138 |
| Prkcdbp | 13,8546 | 7,81747 | -0,825589 | 0,00095 | 0,0138 |
| Synpo | 1,01127 | 0,435139 | -1,21662 | 0,00095 | 0,0138 |
| Slc39a2 | 1,32583 | 0,239424 | -2,46926 | 0,00095 | 0,0138 |
| Vamp5 | 1,75088 | 0,241336 | -2,85897 | 0,00095 | 0,0138 |
| Ntf5 | 2,12683 | 0,17733 | -3,5842 | 0,00095 | 0,0138 |
| Ache | 2,4626 | 0,12863 | -4,25888 | 0,00095 | 0,0138 |
| Pam | 15,9149 | 10,8821 | -0,548422 | 0,001 | 0,014432 |
| Abca8b | 0,829436 | 0,368261 | -1,1714 | 0,001 | 0,014432 |
| Fam160a1 | 0,615553 | 0,148771 | -2,04879 | 0,001 | 0,014432 |
| Ptpn7 | 1,61368 | 0,0771267 | -4,38698 | 0,001 | 0,014432 |
| Itga2b | 3,40945 | 0,079832 | -5,41643 | 0,001 | 0,014432 |
| Laptm5 | 4,67982 | 0,107569 | -5,44311 | 0,001 | 0,014432 |
| Wfikkn1 | 1,69096 | 3,31904 | 0,97292 | 0,00105 | 0,014992 |
| Snph | 3,03647 | 4,92322 | 0,697206 | 0,00105 | 0,014992 |
| Tll2 | 4,86038 | 7,71669 | 0,666911 | 0,00105 | 0,014992 |
| Hoxd11 | 82,8255 | 119,085 | 0,523849 | 0,00105 | 0,014992 |
| Chpf2 | 7,226 | 4,58542 | -0,656143 | 0,00105 | 0,014992 |
| Prrg3 | 5,29837 | 3,34873 | -0,661936 | 0,00105 | 0,014992 |
| Camk2n1 | 3,99571 | 2,30588 | -0,793135 | 0,00105 | 0,014992 |
| Plaur | 2,00158 | 0,395983 | -2,33762 | 0,00105 | 0,014992 |
| Dlx2 | 1,23169 | 0,144466 | -3,09183 | 0,00105 | 0,014992 |
| Samsn1 | 1,6664 | 0,183351 | -3,18406 | 0,00105 | 0,014992 |
| Rnf165 | 3,7549 | 5,76682 | 0,619001 | 0,0011 | 0,015622 |
| Epha7 | 4,91153 | 3,02185 | -0,700742 | 0,0011 | 0,015622 |
| Nabp1 | 3,38042 | 1,82059 | -0,8928 | 0,0011 | 0,015622 |
| Creg2 | 0,947016 | 0,450931 | -1,07048 | 0,0011 | 0,015622 |
| Gm15055 | 9,55256 | 3,42873 | -1,47821 | 0,0011 | 0,015622 |
| Nr1h3 | 1,15039 | 2,70241 | 1,23212 | 0,00115 | 0,016212 |
| Tubb3 | 3,00058 | 5,68056 | 0,920792 | 0,00115 | 0,016212 |
| Cited1 | 114,673 | 165,25 | 0,527126 | 0,00115 | 0,016212 |
| 2810410L24Rik | 8,65484 | 4,98802 | -0,79504 | 0,00115 | 0,016212 |
| Nrbp2 | 1,93015 | 0,812018 | -1,24913 | 0,00115 | 0,016212 |
| Mmp28 | 1,5107 | 0,477862 | -1,66055 | 0,00115 | 0,016212 |
| Kcnn4 | 2,94842 | 0,128691 | -4,51796 | 0,00115 | 0,016212 |
| Ermp1 | 15,9613 | 10,936 | -0,545483 | 0,0012 | 0,016793 |
| Slc1a3 | 6,58895 | 4,255 | -0,63089 | 0,0012 | 0,016793 |
| Chrdl1 | 4,07938 | 2,50114 | -0,705766 | 0,0012 | 0,016793 |
| Card6 | 0,708709 | 0,22806 | -1,63578 | 0,0012 | 0,016793 |
| Dapk2 | 1,81918 | 0,400817 | -2,18227 | 0,0012 | 0,016793 |
| Tec | 0,812302 | 0,171172 | -2,24657 | 0,0012 | 0,016793 |
| Ncf1 | 1,32858 | 0,158743 | -3,06512 | 0,0012 | 0,016793 |
| Fgf15 | 36,4045 | 52,4956 | 0,528078 | 0,00125 | 0,017347 |
| Slc7a5 | 41,6309 | 29,1142 | -0,515931 | 0,00125 | 0,017347 |
| Pald1 | 19,8078 | 13,4188 | -0,56181 | 0,00125 | 0,017347 |
| Scamp2 | 18,1259 | 11,867 | -0,6111 | 0,00125 | 0,017347 |

| | | | | | |
|---|---|---|---|---|---|
| Ccdc82 | 2,43567 | 1,36758 | -0,832701 | 0,00125 | 0,017347 |
| Sncaip | 2,14701 | 1,14189 | -0,910907 | 0,00125 | 0,017347 |
| Rtn4r | 1,75157 | 0,497932 | -1,81463 | 0,00125 | 0,017347 |
| 8430408G22Rik | 3,2904 | 0,357832 | -3,20091 | 0,00125 | 0,017347 |
| Malat1 | 52,8418 | 76,5589 | 0,534889 | 0,0013 | 0,017892 |
| Tmem47 | 25,7675 | 17,9479 | -0,521741 | 0,0013 | 0,017892 |
| Pmp22 | 12,2888 | 7,62675 | -0,6882 | 0,0013 | 0,017892 |
| Robo4 | 4,64787 | 2,84013 | -0,710613 | 0,0013 | 0,017892 |
| Cbx7 | 1,58523 | 0,665695 | -1,25176 | 0,0013 | 0,017892 |
| Runx1 | 0,565342 | 0,235707 | -1,26213 | 0,0013 | 0,017892 |
| Kdf1 | 1,49451 | 0,287189 | -2,3796 | 0,0013 | 0,017892 |
| Itga2 | 0,724171 | 0,0761599 | -3,24923 | 0,0013 | 0,017892 |
| Cdh20 | 2,63835 | 4,36804 | 0,72735 | 0,00135 | 0,018428 |
| Lcor | 2,18506 | 3,59142 | 0,716878 | 0,00135 | 0,018428 |
| Ccdc136 | 7,48359 | 11,3748 | 0,604043 | 0,00135 | 0,018428 |
| Oat | 96,0046 | 136,559 | 0,508348 | 0,00135 | 0,018428 |
| Ppic | 89,8668 | 62,447 | -0,525158 | 0,00135 | 0,018428 |
| Kcnf1 | 1,28861 | 0,641472 | -1,00636 | 0,00135 | 0,018428 |
| Cox6b2 | 22,0164 | 10,7291 | -1,03705 | 0,00135 | 0,018428 |
| Tnfaip8l1 | 1,93709 | 0,804081 | -1,26848 | 0,00135 | 0,018428 |
| Ccdc171 | 1,40409 | 2,48691 | 0,824721 | 0,0014 | 0,018956 |
| Sp5 | 27,7379 | 40,0071 | 0,528398 | 0,0014 | 0,018956 |
| Hspa13 | 27,4526 | 19,2002 | -0,51582 | 0,0014 | 0,018956 |
| Asah1 | 45,8841 | 31,7818 | -0,529793 | 0,0014 | 0,018956 |
| Zfp943 | 9,61478 | 6,17528 | -0,638749 | 0,0014 | 0,018956 |
| F13a1 | 0,720142 | 0,193955 | -1,89256 | 0,0014 | 0,018956 |
| Epn3 | 0,64843 | 0,152828 | -2,08504 | 0,0014 | 0,018956 |
| Nxpe2 | 1,31129 | 0,0711009 | -4,20498 | 0,0014 | 0,018956 |
| Sct | 14,1281 | 26,003 | 0,88011 | 0,00145 | 0,019436 |
| Fstl4 | 1,78823 | 3,19503 | 0,837302 | 0,00145 | 0,019436 |
| Smad7 | 4,2547 | 6,66613 | 0,647792 | 0,00145 | 0,019436 |
| Bcap29 | 15,0143 | 9,64898 | -0,637885 | 0,00145 | 0,019436 |
| Lpar4 | 5,59545 | 3,56643 | -0,649773 | 0,00145 | 0,019436 |
| 1810011O10Rik | 15,2875 | 9,46573 | -0,691568 | 0,00145 | 0,019436 |
| C430049B03Rik | 5,32469 | 3,22674 | -0,722619 | 0,00145 | 0,019436 |
| Rhbdf2 | 2,55047 | 1,40203 | -0,863248 | 0,00145 | 0,019436 |
| Ror1 | 1,77015 | 0,898681 | -0,97799 | 0,00145 | 0,019436 |
| Coro2a | 1,06404 | 0,47386 | -1,16702 | 0,00145 | 0,019436 |
| Vps13a | 4,58651 | 3,11571 | -0,557834 | 0,0015 | 0,020066 |
| Vma21 | 9,00175 | 6,00692 | -0,583579 | 0,0015 | 0,020066 |
| Elk3 | 19,4078 | 13,4749 | -0,52636 | 0,00155 | 0,020632 |
| Pros1 | 14,8991 | 10,1485 | -0,55396 | 0,00155 | 0,020632 |
| Lypd6b | 13,9841 | 9,11628 | -0,617268 | 0,00155 | 0,020632 |
| Fes | 3,31934 | 1,79453 | -0,88729 | 0,00155 | 0,020632 |
| Inhbb | 0,591844 | 0,0860256 | -2,78238 | 0,00155 | 0,020632 |
| Cpne2 | 5,77584 | 9,11217 | 0,657764 | 0,0016 | 0,02115 |
| D430041D05Rik | 1,55087 | 2,42838 | 0,646916 | 0,0016 | 0,02115 |
| Ets2 | 106,306 | 151,641 | 0,512438 | 0,0016 | 0,02115 |

| | | | | | |
|---|---|---|---|---|---|
| Ppp1r9a | 6,59095 | 4,58766 | -0,522728 | 0,0016 | 0,02115 |
| Ebf1 | 1,77794 | 0,990931 | -0,843346 | 0,0016 | 0,02115 |
| Cd200 | 1,14926 | 0,330388 | -1,79847 | 0,0016 | 0,02115 |
| F2rl3 | 1,60729 | 0,275501 | -2,54451 | 0,0016 | 0,02115 |
| Klhdc4 | 26,9574 | 39,1738 | 0,539205 | 0,00165 | 0,021682 |
| Gsto1 | 87,5096 | 124,425 | 0,507767 | 0,00165 | 0,021682 |
| Clcn3 | 15,298 | 10,5752 | -0,532653 | 0,00165 | 0,021682 |
| D7Ertd715e | 15,6161 | 10,2946 | -0,601143 | 0,00165 | 0,021682 |
| Astn1 | 1,37506 | 0,772228 | -0,832391 | 0,00165 | 0,021682 |
| Pltp | 18,4632 | 7,79313 | -1,24438 | 0,00165 | 0,021682 |
| Trim71 | 7,64446 | 11,1499 | 0,544539 | 0,0017 | 0,022208 |
| Nrarp | 67,7711 | 95,2835 | 0,491557 | 0,0017 | 0,022208 |
| Mbnl3 | 3,03858 | 1,97107 | -0,624421 | 0,0017 | 0,022208 |
| Pxdc1 | 1,66799 | 0,5308 | -1,65187 | 0,0017 | 0,022208 |
| Tbc1d2 | 0,61351 | 0,180386 | -1,766 | 0,0017 | 0,022208 |
| Ccdc68 | 1,8239 | 0,239779 | -2,92725 | 0,0017 | 0,022208 |
| Hoxb2 | 33,0469 | 48,1584 | 0,543271 | 0,00175 | 0,022794 |
| Lrrc8c | 5,3018 | 3,22328 | -0,717955 | 0,00175 | 0,022794 |
| 2610305D13Rik | 1,798 | 0,812462 | -1,14602 | 0,00175 | 0,022794 |
| Zfp61 | 7,46487 | 11,4316 | 0,614838 | 0,0018 | 0,023309 |
| Lamp1 | 119,532 | 84,9499 | -0,492713 | 0,0018 | 0,023309 |
| Abcb7 | 14,0663 | 9,71846 | -0,533447 | 0,0018 | 0,023309 |
| Ell3 | 2,27161 | 0,88021 | -1,3678 | 0,0018 | 0,023309 |
| Ampd3 | 0,687186 | 0,201864 | -1,76732 | 0,0018 | 0,023309 |
| Aqp3 | 4,40815 | 0,141204 | -4,96432 | 0,0018 | 0,023309 |
| Ccnjl | 30,9484 | 43,45 | 0,489495 | 0,00185 | 0,023887 |
| Fstl1 | 122,52 | 86,9093 | -0,495441 | 0,00185 | 0,023887 |
| Pik3r1 | 7,33322 | 5,04173 | -0,540528 | 0,00185 | 0,023887 |
| Chd3 | 38,1924 | 53,8817 | 0,496507 | 0,0019 | 0,024486 |
| Mgat4a | 3,35472 | 2,03597 | -0,720477 | 0,0019 | 0,024486 |
| Hoxd10 | 85,4526 | 121,058 | 0,502505 | 0,00195 | 0,025082 |
| Nrk | 1,38058 | 0,768175 | -0,845764 | 0,00195 | 0,025082 |
| Sptbn2 | 5,89281 | 4,01292 | -0,554305 | 0,002 | 0,025602 |
| Mdfic | 4,93081 | 3,0478 | -0,694055 | 0,002 | 0,025602 |
| Il10rb | 3,69705 | 1,88116 | -0,974747 | 0,002 | 0,025602 |
| Slc35d2 | 1,62561 | 0,671102 | -1,27638 | 0,002 | 0,025602 |
| Hpgds | 0,736782 | 0,102376 | -2,84737 | 0,002 | 0,025602 |
| Exosc6 | 32,6267 | 47,3288 | 0,536666 | 0,00205 | 0,026166 |
| Sec62 | 36,6499 | 25,6129 | -0,516937 | 0,00205 | 0,026166 |
| Gm5141 | 3,77701 | 2,21544 | -0,76965 | 0,00205 | 0,026166 |
| Mgst1 | 31,5514 | 20,4343 | -0,626707 | 0,0021 | 0,026677 |
| Cd47 | 6,17665 | 3,60296 | -0,777642 | 0,0021 | 0,026677 |
| 0610040J01Rik | 5,43003 | 3,02997 | -0,841657 | 0,0021 | 0,026677 |
| Ano1 | 1,73736 | 0,949548 | -0,87158 | 0,0021 | 0,026677 |
| Trim16 | 1,95472 | 1,04503 | -0,903413 | 0,0021 | 0,026677 |
| 2410006H16Rik | 136,17 | 197,253 | 0,534642 | 0,00215 | 0,027183 |
| Mvd | 25,3826 | 36,5175 | 0,52475 | 0,00215 | 0,027183 |
| Sh3bp4 | 21,5555 | 30,6711 | 0,508822 | 0,00215 | 0,027183 |

| | | | | | |
|---|---|---|---|---|---|
| Twsg1 | 46,7039 | 33,1779 | -0,493321 | 0,00215 | 0,027183 |
| Fam169b | 0,991003 | 0,321969 | -1,62197 | 0,00215 | 0,027183 |
| Slc22a21 | 1,44503 | 2,84472 | 0,977192 | 0,0022 | 0,027581 |
| Dtx4 | 4,6351 | 6,8473 | 0,562934 | 0,0022 | 0,027581 |
| Lmo2 | 116,845 | 163,634 | 0,485883 | 0,0022 | 0,027581 |
| Ece1 | 21,8232 | 15,4145 | -0,501571 | 0,0022 | 0,027581 |
| Hiatl1 | 23,2702 | 16,3952 | -0,505211 | 0,0022 | 0,027581 |
| Scara3 | 15,7971 | 10,9659 | -0,526637 | 0,0022 | 0,027581 |
| Zfp944 | 8,62218 | 5,53268 | -0,640074 | 0,0022 | 0,027581 |
| Tmem100 | 1,73818 | 0,574627 | -1,59688 | 0,0022 | 0,027581 |
| Myf5 | 1,28598 | 0,379375 | -1,76117 | 0,0022 | 0,027581 |
| Rnf43 | 3,40725 | 2,07294 | -0,716931 | 0,00225 | 0,028128 |
| Il13ra1 | 2,21252 | 1,23307 | -0,843434 | 0,00225 | 0,028128 |
| Ldlrad4 | 1,29815 | 0,47121 | -1,46201 | 0,00225 | 0,028128 |
| Meox1 | 152,687 | 108,925 | -0,487241 | 0,0023 | 0,028673 |
| Tmem59l | 5,12802 | 2,71475 | -0,917584 | 0,0023 | 0,028673 |
| Gimap1 | 1,90153 | 0,2061 | -3,20574 | 0,0023 | 0,028673 |
| Nod2 | 1,34283 | 2,35381 | 0,809724 | 0,00235 | 0,029133 |
| Hoxd9 | 23,2593 | 33,3516 | 0,51995 | 0,00235 | 0,029133 |
| Gas1 | 30,8918 | 21,5958 | -0,516473 | 0,00235 | 0,029133 |
| Dock9 | 4,04792 | 2,69352 | -0,58769 | 0,00235 | 0,029133 |
| Zfp9 | 2,57248 | 1,4589 | -0,818282 | 0,00235 | 0,029133 |
| Sim2 | 0,637229 | 0,137979 | -2,20736 | 0,00235 | 0,029133 |
| Hist1h1b | 0,551715 | 3,81759 | 2,79067 | 0,0024 | 0,029616 |
| Arc | 0,941787 | 1,89591 | 1,00942 | 0,0024 | 0,029616 |
| Rpl36 | 26,9668 | 46,7252 | 0,79302 | 0,0024 | 0,029616 |
| Kat6b | 7,18272 | 10,1738 | 0,502254 | 0,0024 | 0,029616 |
| Myd88 | 2,69735 | 1,28036 | -1,07499 | 0,0024 | 0,029616 |
| Tspan12 | 14,7612 | 9,87275 | -0,580282 | 0,00245 | 0,030121 |
| Fam132a | 12,2843 | 7,45307 | -0,720909 | 0,00245 | 0,030121 |
| Srcrb4d | 1,7831 | 0,810843 | -1,13689 | 0,00245 | 0,030121 |
| Galnt14 | 0,941605 | 0,249139 | -1,91817 | 0,00245 | 0,030121 |
| B230323A14Rik | 6,76672 | 10,8893 | 0,686388 | 0,0025 | 0,030651 |
| T | 271,096 | 388,818 | 0,520294 | 0,0025 | 0,030651 |
| Lamb3 | 0,568195 | 0,11928 | -2,25204 | 0,0025 | 0,030651 |
| Serinc1 | 93,0437 | 67,105 | -0,471488 | 0,00255 | 0,031179 |
| Itfg1 | 40,7326 | 29,109 | -0,484719 | 0,00255 | 0,031179 |
| Arl6ip5 | 30,2108 | 20,6573 | -0,548407 | 0,00255 | 0,031179 |
| Plekha7 | 4,9579 | 3,16115 | -0,649279 | 0,0026 | 0,031761 |
| Lgmn | 35,7312 | 24,9164 | -0,520086 | 0,00265 | 0,032166 |
| Nt5c3 | 28,9959 | 19,8125 | -0,549434 | 0,00265 | 0,032166 |
| Col4a6 | 3,55961 | 2,31067 | -0,623408 | 0,00265 | 0,032166 |
| Stom | 5,44841 | 3,37697 | -0,690108 | 0,00265 | 0,032166 |
| Ctnnd2 | 1,5063 | 0,856178 | -0,815027 | 0,00265 | 0,032166 |
| Gpx2 | 2,34556 | 0,447791 | -2,38903 | 0,00265 | 0,032166 |
| Gt(ROSA)26Sor | 31,2042 | 5,30847 | -2,55537 | 0,00265 | 0,032166 |
| St3gal5 | 14,9731 | 10,0702 | -0,572283 | 0,0027 | 0,032743 |
| Mmp17 | 8,8752 | 12,9804 | 0,548487 | 0,00275 | 0,033229 |

| | | | | | |
|---|---|---|---|---|---|
| Dynlt3 | 11,1866 | 7,31082 | -0,613669 | 0,00275 | 0,033229 |
| Ildr1 | 0,634535 | 0,105157 | -2,59315 | 0,00275 | 0,033229 |
| Chst5 | 2,01091 | 0,120793 | -4,05724 | 0,00275 | 0,033229 |
| Hspa8 | 219,791 | 309,211 | 0,492458 | 0,0028 | 0,03359 |
| Fabp7 | 133,393 | 187,58 | 0,491823 | 0,0028 | 0,03359 |
| Vcl | 41,5624 | 29,8053 | -0,479708 | 0,0028 | 0,03359 |
| Atp9a | 16,439 | 11,5931 | -0,503854 | 0,0028 | 0,03359 |
| Cdon | 10,6977 | 7,54301 | -0,504094 | 0,0028 | 0,03359 |
| Sumf1 | 7,6684 | 4,88468 | -0,65066 | 0,0028 | 0,03359 |
| Plac8 | 4,25912 | 0,872403 | -2,28749 | 0,0028 | 0,03359 |
| Cyth4 | 1,36401 | 0,0789595 | -4,1106 | 0,0028 | 0,03359 |
| Nim1k | 2,24087 | 3,60795 | 0,687122 | 0,00285 | 0,034129 |
| Entpd1 | 1,74809 | 0,93618 | -0,900917 | 0,00285 | 0,034129 |
| Kcnab3 | 12,2332 | 17,7262 | 0,535076 | 0,0029 | 0,034665 |
| Hypk | 246,879 | 345,783 | 0,486063 | 0,0029 | 0,034665 |
| Wdfy1 | 4,66425 | 3,0467 | -0,614397 | 0,00295 | 0,035137 |
| Gm11538 | 9,24834 | 5,87458 | -0,654708 | 0,00295 | 0,035137 |
| Syk | 2,70161 | 1,65541 | -0,706635 | 0,00295 | 0,035137 |
| Slc7a8 | 0,743563 | 0,278736 | -1,41556 | 0,00295 | 0,035137 |
| Tbx3 | 3,73618 | 5,61567 | 0,587896 | 0,003 | 0,035574 |
| Snrpf | 48,2351 | 70,1836 | 0,54105 | 0,003 | 0,035574 |
| Kmt2d | 7,8524 | 11,0346 | 0,490833 | 0,003 | 0,035574 |
| Dll1 | 127,782 | 178,414 | 0,481548 | 0,003 | 0,035574 |
| Fgfbp1 | 3,48947 | 0,183267 | -4,25099 | 0,003 | 0,035574 |
| Rhpn2 | 8,86424 | 13,2838 | 0,583596 | 0,00305 | 0,036007 |
| Csrp1 | 42,6849 | 30,0992 | -0,504003 | 0,00305 | 0,036007 |
| Arl4a | 11,879 | 8,25324 | -0,525385 | 0,00305 | 0,036007 |
| Morc4 | 3,84901 | 2,42497 | -0,66652 | 0,00305 | 0,036007 |
| Edn1 | 1,1511 | 0,102868 | -3,48414 | 0,00305 | 0,036007 |
| Rab6b | 1,50134 | 2,48948 | 0,729598 | 0,0031 | 0,036533 |
| Foxp1 | 13,5791 | 9,72678 | -0,481358 | 0,0031 | 0,036533 |
| Garnl3 | 2,6076 | 1,47989 | -0,817228 | 0,00315 | 0,037024 |
| 1500015A07Rik | 5,15732 | 2,89668 | -0,832221 | 0,00315 | 0,037024 |
| Gpr97 | 2,2315 | 0,0864922 | -4,6893 | 0,00315 | 0,037024 |
| Pgap1 | 2,33458 | 1,54632 | -0,594321 | 0,0032 | 0,037579 |
| Ubash3b | 2,78202 | 1,59885 | -0,7991 | 0,00325 | 0,037999 |
| Slc8a2 | 1,47282 | 0,783551 | -0,910483 | 0,00325 | 0,037999 |
| Magi2 | 0,628899 | 0,303316 | -1,05201 | 0,00325 | 0,037999 |
| Cyp7b1 | 1,19788 | 0,40519 | -1,56381 | 0,00325 | 0,037999 |
| Entpd3 | 0,758219 | 0,0794633 | -3,25425 | 0,00325 | 0,037999 |
| Ephb3 | 12,2763 | 8,63105 | -0,508265 | 0,0033 | 0,038416 |
| Meis1 | 16,2388 | 11,2507 | -0,529427 | 0,0033 | 0,038416 |
| Tcea2 | 7,1367 | 3,8483 | -0,891034 | 0,0033 | 0,038416 |
| Hsd3b6 | 5,02199 | 0,133866 | -5,2294 | 0,0033 | 0,038416 |
| Trim10 | 6,52754 | 0,104811 | -5,96068 | 0,0033 | 0,038416 |
| Flnc | 4,44862 | 6,37087 | 0,51813 | 0,00335 | 0,038694 |
| Smug1 | 16,1297 | 22,8219 | 0,500704 | 0,00335 | 0,038694 |
| Igfbp4 | 171,83 | 124,592 | -0,463767 | 0,00335 | 0,038694 |

| | | | | | |
|---|---|---|---|---|---|
| Atl3 | 10,7322 | 7,61167 | -0,495662 | 0,00335 | 0,038694 |
| Cept1 | 22,0958 | 15,5459 | -0,507232 | 0,00335 | 0,038694 |
| Cd68 | 5,15216 | 2,33526 | -1,1416 | 0,00335 | 0,038694 |
| Six2 | 1,90351 | 0,840134 | -1,17997 | 0,00335 | 0,038694 |
| Il2rg | 1,4509 | 0,426395 | -1,76669 | 0,00335 | 0,038694 |
| Itgal | 0,804855 | 0,0394678 | -4,34998 | 0,00335 | 0,038694 |
| Lzts1 | 2,32577 | 4,31085 | 0,890263 | 0,0034 | 0,039069 |
| Nkd1 | 26,0847 | 36,359 | 0,479106 | 0,0034 | 0,039069 |
| Prkce | 2,54549 | 1,62027 | -0,651705 | 0,0034 | 0,039069 |
| Cmtm8 | 8,06883 | 4,37159 | -0,8842 | 0,0034 | 0,039069 |
| Gimap9 | 3,36907 | 1,57448 | -1,09748 | 0,0034 | 0,039069 |
| Cdc42ep2 | 1,61035 | 0,40108 | -2,00541 | 0,0034 | 0,039069 |
| Scara5 | 8,94858 | 12,6657 | 0,501201 | 0,00345 | 0,039508 |
| Tmed10 | 27,8097 | 19,9089 | -0,482173 | 0,00345 | 0,039508 |
| Mgat1 | 21,0834 | 14,9776 | -0,493295 | 0,00345 | 0,039508 |
| Slc25a45 | 1,43221 | 0,15462 | -3,21144 | 0,00345 | 0,039508 |
| Hoxd12 | 67,3934 | 92,6039 | 0,458466 | 0,0035 | 0,040012 |
| Hs6st1 | 21,0329 | 15,0167 | -0,486076 | 0,0035 | 0,040012 |
| Cep170b | 5,79047 | 4,02171 | -0,525874 | 0,00355 | 0,04048 |
| 2900056M20Rik | 2,65647 | 1,5805 | -0,749127 | 0,00355 | 0,04048 |
| Hck | 1,22295 | 0,391822 | -1,64209 | 0,00355 | 0,04048 |
| Sntb2 | 5,10213 | 7,4766 | 0,551284 | 0,0036 | 0,04091 |
| Ptprm | 6,89002 | 4,7759 | -0,528736 | 0,0036 | 0,04091 |
| Egfl6 | 0,759127 | 0,159252 | -2,25303 | 0,0036 | 0,04091 |
| Slc22a4 | 1,17164 | 0,112443 | -3,38127 | 0,0036 | 0,04091 |
| Snai2 | 7,41016 | 4,64916 | -0,672535 | 0,00365 | 0,041373 |
| Mtmr1 | 3,10169 | 1,94455 | -0,67362 | 0,00365 | 0,041373 |
| Lrrc8b | 3,75139 | 2,23433 | -0,747582 | 0,00365 | 0,041373 |
| Aldoc | 2,82189 | 5,04432 | 0,838001 | 0,0037 | 0,041798 |
| Tssc1 | 15,6016 | 22,6614 | 0,538538 | 0,0037 | 0,041798 |
| Tmx3 | 28,5216 | 20,5367 | -0,473851 | 0,0037 | 0,041798 |
| Pcdh7 | 4,1318 | 2,75619 | -0,584097 | 0,0037 | 0,041798 |
| 5730457N03Rik | 27,1116 | 42,4299 | 0,64617 | 0,00375 | 0,042149 |
| Slc8a1 | 0,603081 | 0,937995 | 0,637229 | 0,00375 | 0,042149 |
| Dlc1 | 27,2008 | 37,2971 | 0,455414 | 0,00375 | 0,042149 |
| B3gnt1 | 16,9156 | 11,5177 | -0,554501 | 0,00375 | 0,042149 |
| Ano9 | 0,704382 | 0,135409 | -2,37903 | 0,00375 | 0,042149 |
| Lcp1 | 8,73663 | 0,707003 | -3,62729 | 0,00375 | 0,042149 |
| Gli2 | 6,40268 | 4,4378 | -0,52883 | 0,0038 | 0,042675 |
| Zfp36l1 | 38,4525 | 27,7708 | -0,469507 | 0,00385 | 0,043092 |
| Bmi1 | 12,9431 | 9,06917 | -0,513136 | 0,00385 | 0,043092 |
| Itsn2 | 4,50593 | 2,93371 | -0,619097 | 0,00385 | 0,043092 |
| Lgals9 | 1,85514 | 0,129016 | -3,8459 | 0,00385 | 0,043092 |
| C230091D08Rik | 9,82483 | 6,73284 | -0,545216 | 0,0039 | 0,043615 |
| Wnt3a | 31,5563 | 43,5171 | 0,463654 | 0,00395 | 0,04399 |
| Slc23a2 | 6,54442 | 4,53979 | -0,527641 | 0,00395 | 0,04399 |
| Cdc42bpg | 3,02216 | 1,95571 | -0,627886 | 0,00395 | 0,04399 |
| Bbs10 | 5,38328 | 3,39085 | -0,666839 | 0,00395 | 0,04399 |

| | | | | | |
|---|---|---|---|---|---|
| Frmd7 | 2,36634 | 1,42164 | -0,735101 | 0,00395 | 0,04399 |
| AY358078 | 2,63732 | 4,85089 | 0,879174 | 0,004 | 0,044362 |
| Eogt | 49,0851 | 67,7067 | 0,464015 | 0,004 | 0,044362 |
| Cd24a | 277,645 | 201,746 | -0,460704 | 0,004 | 0,044362 |
| Tgfbr1 | 59,4414 | 43,1246 | -0,462959 | 0,004 | 0,044362 |
| Tfap2c | 1,16321 | 0,502681 | -1,21039 | 0,004 | 0,044362 |
| Alpl | 13,5575 | 19,3711 | 0,514812 | 0,00405 | 0,044731 |
| Grhpr | 28,8601 | 41,0839 | 0,509496 | 0,00405 | 0,044731 |
| Axin2 | 23,6779 | 32,5786 | 0,460385 | 0,00405 | 0,044731 |
| Exoc6 | 7,87911 | 5,19678 | -0,600416 | 0,00405 | 0,044731 |
| Enpp4 | 1,57065 | 0,868438 | -0,854864 | 0,00405 | 0,044731 |
| Lix1 | 5,09279 | 7,63042 | 0,583306 | 0,0041 | 0,045171 |
| Sh3bp1 | 20,2564 | 28,6635 | 0,500837 | 0,0041 | 0,045171 |
| Gngt2 | 4,94562 | 0,445551 | -3,47249 | 0,0041 | 0,045171 |
| Ddx43 | 0,814754 | 1,76805 | 1,11772 | 0,00415 | 0,045609 |
| Sgk1 | 11,5674 | 7,79297 | -0,56982 | 0,00415 | 0,045609 |
| Mall | 1,07116 | 0,170663 | -2,64995 | 0,00415 | 0,045609 |
| Pgf | 2,93825 | 1,27813 | -1,20092 | 0,0042 | 0,046121 |
| Rassf2 | 4,7687 | 3,15848 | -0,594368 | 0,00425 | 0,046517 |
| Gdf10 | 5,47352 | 3,43844 | -0,670715 | 0,00425 | 0,046517 |
| Inha | 3,12257 | 1,47716 | -1,07991 | 0,00425 | 0,046517 |
| Gm12942 | 3,70395 | 1,69234 | -1,13004 | 0,00425 | 0,046517 |
| Slc2a6 | 1,33576 | 2,56834 | 0,943168 | 0,0043 | 0,046834 |
| Phyhipl | 4,25765 | 7,15468 | 0,748829 | 0,0043 | 0,046834 |
| Pask | 13,1426 | 18,1771 | 0,467867 | 0,0043 | 0,046834 |
| Prkar2b | 18,8073 | 13,424 | -0,486474 | 0,0043 | 0,046834 |
| Cyyr1 | 1,3491 | 0,489847 | -1,46159 | 0,0043 | 0,046834 |
| Dhrs7 | 2,25515 | 0,814923 | -1,46849 | 0,0043 | 0,046834 |
| Ptprf | 67,2056 | 49,1248 | -0,45213 | 0,0044 | 0,047806 |
| Tpd52 | 10,6655 | 7,22478 | -0,561931 | 0,0044 | 0,047806 |
| Kcnj15 | 1,77111 | 1,05148 | -0,752233 | 0,0044 | 0,047806 |
| Cacna1c | 1,18152 | 1,89435 | 0,681054 | 0,00445 | 0,048232 |
| Grn | 22,0918 | 15,6605 | -0,496381 | 0,00445 | 0,048232 |
| Efcab4a | 1,3336 | 0,271285 | -2,29745 | 0,00445 | 0,048232 |
| Zfp54 | 1,59925 | 0,743051 | -1,10586 | 0,0045 | 0,048695 |
| Srpx2 | 0,679312 | 0,176801 | -1,94194 | 0,0045 | 0,048695 |
| Ppef2 | 1,86733 | 3,33005 | 0,834567 | 0,00455 | 0,049156 |
| Fkbp9 | 81,9114 | 59,8701 | -0,452228 | 0,00455 | 0,049156 |
| Rplp2 | 656,281 | 898,544 | 0,453277 | 0,0046 | 0,049576 |
| Dab2 | 9,64141 | 6,75713 | -0,512834 | 0,0046 | 0,049576 |
| Ddx58 | 3,48979 | 2,2878 | -0,609176 | 0,0046 | 0,049576 |
| Rpl10 | 3,17474 | 7,13658 | 1,16859 | 0,00465 | 0,049954 |
| Tmsb4x | 626,381 | 459,028 | -0,44846 | 0,00465 | 0,049954 |
| Slc39a10 | 38,9192 | 28,4951 | -0,449766 | 0,00465 | 0,049954 |
| Cyb561 | 3,98265 | 2,36179 | -0,75385 | 0,00465 | 0,049954 |
| E130201H02Rik | 5,36244 | 9,80395 | 0,870474 | 0,00475 | 0,050743 |
| Fdps | 35,5455 | 49,9356 | 0,490402 | 0,00475 | 0,050743 |
| Atp6ap2 | 40,7943 | 29,6783 | -0,458959 | 0,00475 | 0,050743 |

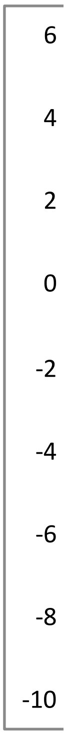

| | | | | | |
|---|---|---|---|---|---|
| Nup210 | 8,63449 | 6,18389 | -0,481595 | 0,00475 | 0,050743 |
| Zfp958 | 7,37374 | 4,75379 | -0,633317 | 0,00475 | 0,050743 |
| Zdhhc24 | 6,15574 | 3,85554 | -0,675 | 0,00475 | 0,050743 |
| Mylk3 | 0,675426 | 0,170232 | -1,9883 | 0,00475 | 0,050743 |
| Sept8 | 61,6218 | 84,5524 | 0,456405 | 0,0048 | 0,051154 |
| Zdhhc2 | 6,0355 | 3,81742 | -0,660877 | 0,0048 | 0,051154 |
| Myct1 | 1,53346 | 0,0707493 | -4,43794 | 0,0048 | 0,051154 |
| Hist1h1d | 0,765661 | 3,14028 | 2,03612 | 0,00485 | 0,051564 |
| Ahcy | 12,272 | 17,5421 | 0,515448 | 0,00485 | 0,051564 |
| S1pr3 | 44,3497 | 60,4884 | 0,447734 | 0,00485 | 0,051564 |
| Loxl4 | 0,705976 | 1,25924 | 0,834862 | 0,0049 | 0,051848 |
| 5430416N02Rik | 40,7637 | 59,7562 | 0,551803 | 0,0049 | 0,051848 |
| Slc8b1 | 4,41836 | 2,75881 | -0,679464 | 0,0049 | 0,051848 |
| Tmem150c | 1,16339 | 0,527571 | -1,1409 | 0,0049 | 0,051848 |
| Fcho1 | 0,859689 | 0,338523 | -1,34456 | 0,0049 | 0,051848 |
| Tfpi2 | 1,67981 | 0,172517 | -3,28349 | 0,0049 | 0,051848 |
| Bhlhe40 | 2,74734 | 4,35877 | 0,665885 | 0,00495 | 0,052253 |
| Sun2 | 16,6671 | 11,9393 | -0,481285 | 0,00495 | 0,052253 |
| Pls1 | 0,935442 | 0,41132 | -1,18539 | 0,00495 | 0,052253 |
| Cecr2 | 18,0145 | 24,6437 | 0,452066 | 0,005 | 0,052697 |
| Slc14a1 | 0,915573 | 0,0409775 | -4,48177 | 0,005 | 0,052697 |
| Gbe1 | 5,81037 | 8,58444 | 0,563093 | 0,00505 | 0,053099 |
| Utp3 | 64,8335 | 88,4928 | 0,448821 | 0,00505 | 0,053099 |
| Gpx1 | 312,131 | 227,392 | -0,45697 | 0,00505 | 0,053099 |
| Dlg4 | 10,8471 | 15,3886 | 0,50455 | 0,0051 | 0,053289 |
| Trip10 | 27,0529 | 37,1569 | 0,457845 | 0,0051 | 0,053289 |
| Tril | 12,5221 | 17,1818 | 0,456401 | 0,0051 | 0,053289 |
| Col5a1 | 16,4474 | 11,9611 | -0,459516 | 0,0051 | 0,053289 |
| B4galt5 | 16,4512 | 11,885 | -0,469055 | 0,0051 | 0,053289 |
| Yipf6 | 7,65728 | 5,39523 | -0,505149 | 0,0051 | 0,053289 |
| C1qtnf6 | 6,54468 | 4,16121 | -0,653319 | 0,0051 | 0,053289 |
| Tll1 | 0,604012 | 0,246437 | -1,29336 | 0,0051 | 0,053289 |
| 6330403K07Rik | 55,1865 | 76,0938 | 0,463464 | 0,00515 | 0,053769 |
| Flrt3 | 25,9322 | 18,7375 | -0,468818 | 0,0052 | 0,054249 |
| Ehd4 | 8,51968 | 5,84371 | -0,543914 | 0,00525 | 0,054728 |
| Ctso | 1,58345 | 0,861483 | -0,878174 | 0,0053 | 0,05512 |
| Nrn1 | 2,16237 | 0,966026 | -1,16248 | 0,0053 | 0,05512 |
| H2-T10 | 0,983887 | 0,391508 | -1,32945 | 0,0053 | 0,05512 |
| Cdk18 | 3,45183 | 5,31648 | 0,623107 | 0,00535 | 0,055554 |
| Asah2 | 1,74764 | 1,03019 | -0,762497 | 0,00535 | 0,055554 |
| Mfsd2a | 7,08176 | 10,6222 | 0,584898 | 0,0054 | 0,055942 |
| Sema6d | 5,95252 | 4,14876 | -0,520819 | 0,0054 | 0,055942 |
| Ppm1j | 1,24226 | 0,400121 | -1,63446 | 0,0054 | 0,055942 |
| Ccpg1os | 8,57293 | 14,6913 | 0,777106 | 0,00545 | 0,056373 |
| Cxx1c | 21,1867 | 30,8937 | 0,544157 | 0,00545 | 0,056373 |
| Notch4 | 14,0463 | 19,1128 | 0,44435 | 0,0055 | 0,056715 |
| Chrd | 2,95713 | 1,71655 | -0,784687 | 0,0055 | 0,056715 |
| Hexb | 12,2298 | 4,712 | -1,37599 | 0,0055 | 0,056715 |

| | | | | | |
|---|---|---|---|---|---|
| Samd5 | 0,842542 | 0,271696 | -1,63276 | 0,0055 | 0,056715 |
| Apitd1 | 24,919 | 36,2678 | 0,541441 | 0,00555 | 0,057098 |
| Slc35g1 | 7,05107 | 4,78687 | -0,558758 | 0,00555 | 0,057098 |
| Ncald | 1,68783 | 0,919436 | -0,876351 | 0,00555 | 0,057098 |
| Irx4 | 0,901405 | 0,107093 | -3,07331 | 0,0056 | 0,057568 |
| Rpl12 | 8,67548 | 15,6392 | 0,850155 | 0,00565 | 0,057904 |
| Nrxn2 | 1,77611 | 2,6821 | 0,594642 | 0,00565 | 0,057904 |
| Med12 | 17,8509 | 24,264 | 0,442822 | 0,00565 | 0,057904 |
| Prss22 | 2,40541 | 0,992613 | -1,27698 | 0,00565 | 0,057904 |
| Fgd2 | 1,34655 | 0,625266 | -1,10673 | 0,0057 | 0,058238 |
| Prkcq | 0,887282 | 0,376452 | -1,23693 | 0,0057 | 0,058238 |
| C230035I16Rik | 4,02828 | 1,03927 | -1,95459 | 0,0057 | 0,058238 |
| Prss12 | 0,721683 | 0,174607 | -2,04725 | 0,0057 | 0,058238 |
| Nup62cl | 1,76811 | 0,371927 | -2,24911 | 0,00575 | 0,058704 |
| Nphp4 | 2,04929 | 3,15028 | 0,620353 | 0,0058 | 0,059034 |
| Sap30l | 15,7041 | 22,4858 | 0,517874 | 0,0058 | 0,059034 |
| Fam102a | 6,2446 | 4,28373 | -0,543742 | 0,0058 | 0,059034 |
| Marveld3 | 0,807609 | 0,132909 | -2,60321 | 0,0058 | 0,059034 |
| Fasn | 60,3036 | 83,7877 | 0,474494 | 0,00585 | 0,059317 |
| Ifitm2 | 253,663 | 345,427 | 0,445466 | 0,00585 | 0,059317 |
| Ogfrl1 | 6,51826 | 4,50532 | -0,532858 | 0,00585 | 0,059317 |
| Mxd4 | 12,8179 | 8,15988 | -0,651537 | 0,00585 | 0,059317 |
| Trdmt1 | 3,54768 | 2,11903 | -0,74347 | 0,00585 | 0,059317 |
| Dact1 | 37,8867 | 51,5536 | 0,444381 | 0,0059 | 0,059733 |
| G2e3 | 19,8129 | 14,5763 | -0,442815 | 0,0059 | 0,059733 |
| Lima1 | 29,7863 | 22,0049 | -0,436824 | 0,006 | 0,060654 |
| Tm9sf3 | 70,6054 | 51,859 | -0,445183 | 0,006 | 0,060654 |
| Gm10560 | 3,71497 | 1,41256 | -1,39504 | 0,00605 | 0,061113 |
| Manf | 68,8903 | 93,4446 | 0,439811 | 0,0061 | 0,061525 |
| Aph1b | 3,73202 | 2,47648 | -0,591664 | 0,0061 | 0,061525 |
| Sgsm1 | 4,18158 | 6,13046 | 0,551947 | 0,00615 | 0,061842 |
| Ypel3 | 35,0628 | 49,3237 | 0,49234 | 0,00615 | 0,061842 |
| Gm3604 | 2,50308 | 1,42693 | -0,810785 | 0,00615 | 0,061842 |
| P2rx4 | 1,52873 | 0,641865 | -1,25199 | 0,00615 | 0,061842 |
| Shf | 43,6299 | 59,8509 | 0,456057 | 0,0062 | 0,062158 |
| Reep4 | 24,0082 | 16,9239 | -0,504468 | 0,0062 | 0,062158 |
| Lmbrd1 | 5,62443 | 3,90668 | -0,525764 | 0,0062 | 0,062158 |
| Dio3 | 1,33421 | 0,49309 | -1,43607 | 0,0062 | 0,062158 |
| Rab32 | 1,10732 | 0,40584 | -1,44809 | 0,00625 | 0,062612 |
| 2700086A05Rik | 59,2956 | 116,383 | 0,972879 | 0,0063 | 0,062925 |
| Ankrd45 | 1,88731 | 3,01663 | 0,676606 | 0,0063 | 0,062925 |
| Alg6 | 10,1064 | 7,0552 | -0,518516 | 0,0063 | 0,062925 |
| Pou3f2 | 0,960248 | 0,541899 | -0,825383 | 0,0063 | 0,062925 |
| H19 | 961,794 | 655,743 | -0,552598 | 0,00635 | 0,063235 |
| 2610524H06Rik | 29,1528 | 19,8464 | -0,554758 | 0,00635 | 0,063235 |
| Lama3 | 0,773105 | 0,462736 | -0,740475 | 0,00635 | 0,063235 |
| Ptplad2 | 1,08977 | 0,374868 | -1,53956 | 0,00635 | 0,063235 |
| Mmp2 | 56,4796 | 76,2945 | 0,433849 | 0,0064 | 0,063638 |

| | | | | | |
|---|---|---|---|---|---|
| Abca5 | 2,62692 | 1,79131 | -0,552359 | 0,0064 | 0,063638 |
| Zfp932 | 8,4672 | 5,57559 | -0,602761 | 0,00645 | 0,06404 |
| Ptx3 | 3,15722 | 1,71395 | -0,881333 | 0,00645 | 0,06404 |
| Nuak1 | 9,53922 | 13,171 | 0,465417 | 0,0065 | 0,064345 |
| Ano6 | 18,9594 | 14,0536 | -0,431971 | 0,0065 | 0,064345 |
| Tmem208 | 54,8307 | 38,668 | -0,503842 | 0,0065 | 0,064345 |
| Nt5dc3 | 0,790927 | 0,410946 | -0,944595 | 0,0065 | 0,064345 |
| Drd1a | 1,26222 | 2,15085 | 0,768938 | 0,00655 | 0,064696 |
| Glrb | 1,05806 | 0,473722 | -1,15931 | 0,00655 | 0,064696 |
| Tmprss13 | 0,681539 | 0,0635605 | -3,42259 | 0,00655 | 0,064696 |
| Hist1h1c | 2,48762 | 5,61852 | 1,17543 | 0,0066 | 0,064998 |
| Spry2 | 32,1981 | 43,637 | 0,438577 | 0,0066 | 0,064998 |
| C1galt1c1 | 13,1976 | 8,70941 | -0,599629 | 0,0066 | 0,064998 |
| Pianp | 0,975662 | 0,334198 | -1,54568 | 0,0066 | 0,064998 |
| Tmem33 | 23,1585 | 17,2142 | -0,427946 | 0,00665 | 0,065394 |
| Slco4a1 | 1,28922 | 0,633847 | -1,02429 | 0,00665 | 0,065394 |
| Rps21 | 373,848 | 509,423 | 0,446413 | 0,0067 | 0,065837 |
| E130012A19Rik | 17,702 | 24,4998 | 0,468857 | 0,00675 | 0,066183 |
| Gpc4 | 25,1896 | 18,2297 | -0,466535 | 0,00675 | 0,066183 |
| Arid5b | 1,00848 | 0,521341 | -0,951877 | 0,00675 | 0,066183 |
| Prdm1 | 0,838988 | 0,434816 | -0,948245 | 0,0068 | 0,066624 |
| Pvrl1 | 5,14597 | 3,53483 | -0,541804 | 0,00685 | 0,067016 |
| Arhgef3 | 3,93587 | 2,56464 | -0,617929 | 0,00685 | 0,067016 |
| Ypel4 | 4,21389 | 1,68489 | -1,3225 | 0,0069 | 0,067455 |
| Cep70 | 8,09597 | 5,4881 | -0,560897 | 0,00695 | 0,067845 |
| 5330426P16Rik | 2,0863 | 0,981345 | -1,08811 | 0,00695 | 0,067845 |
| Ing2 | 9,61647 | 13,6186 | 0,501997 | 0,007 | 0,068084 |
| Lnp | 10,1555 | 7,49829 | -0,437632 | 0,007 | 0,068084 |
| Eps15 | 11,3204 | 8,24538 | -0,457271 | 0,007 | 0,068084 |
| Mgat5b | 2,13 | 1,29105 | -0,722312 | 0,007 | 0,068084 |
| Sema7a | 2,67199 | 1,59364 | -0,745588 | 0,007 | 0,068084 |
| Mansc1 | 0,785009 | 0,101353 | -2,95333 | 0,00705 | 0,068521 |
| Rab3b | 1,12662 | 1,96351 | 0,801435 | 0,0071 | 0,068907 |
| Foxj1 | 0,782916 | 0,273534 | -1,51714 | 0,0071 | 0,068907 |
| Hoxb4 | 20,358 | 27,8813 | 0,453702 | 0,0072 | 0,069826 |
| 2610002M06Rik | 3,60756 | 2,4642 | -0,549901 | 0,0073 | 0,070694 |
| 1190002F15Rik | 18,3221 | 10,7986 | -0,762747 | 0,0073 | 0,070694 |
| Ripk3 | 14,034 | 19,641 | 0,484944 | 0,00735 | 0,071075 |
| Tug1 | 52,2521 | 38,7874 | -0,429902 | 0,00735 | 0,071075 |
| Egr2 | 0,062207 | 0,925784 | 3,89554 | 0,0074 | 0,071455 |
| Tap1 | 0,713451 | 0,255989 | -1,47874 | 0,0074 | 0,071455 |
| Fam212a | 29,0575 | 41,0623 | 0,498904 | 0,00745 | 0,071834 |
| Tm9sf2 | 61,2251 | 45,2748 | -0,435417 | 0,00745 | 0,071834 |
| Itm2c | 73,5491 | 54,5141 | -0,432078 | 0,0075 | 0,072212 |
| Tnfrsf18 | 2,41642 | 0,812954 | -1,57163 | 0,0075 | 0,072212 |
| Dtx1 | 0,581712 | 0,209324 | -1,47456 | 0,00755 | 0,072641 |
| Hist1h4i | 8,65363 | 17,7275 | 1,03461 | 0,0076 | 0,072964 |
| Ppt1 | 28,2687 | 20,6198 | -0,455179 | 0,0076 | 0,072964 |

| | | | | | |
|---|---|---|---|---|---|
| Mfap3l | 2,4319 | 1,61615 | -0,589518 | 0,0076 | 0,072964 |
| Dclk2 | 4,72742 | 3,19963 | -0,563149 | 0,00765 | 0,073339 |
| Arhgap6 | 0,80629 | 0,404756 | -0,994247 | 0,00765 | 0,073339 |
| Nox4 | 1,78918 | 1,04948 | -0,769625 | 0,0077 | 0,073765 |
| Lpin3 | 0,96064 | 0,465452 | -1,04536 | 0,00775 | 0,074191 |
| Tdrd7 | 4,3344 | 2,87112 | -0,594221 | 0,0078 | 0,074563 |
| Arsi | 1,98129 | 0,912843 | -1,118 | 0,0078 | 0,074563 |
| Zfp821 | 26,5014 | 36,2736 | 0,452847 | 0,00785 | 0,074827 |
| Ndrg3 | 20,574 | 27,9172 | 0,440331 | 0,00785 | 0,074827 |
| Ttyh2 | 1,45643 | 0,714488 | -1,02745 | 0,00785 | 0,074827 |
| P2rx1 | 0,74909 | 0,201129 | -1,89702 | 0,00785 | 0,074827 |
| Sema5b | 8,65134 | 11,921 | 0,462514 | 0,0079 | 0,075196 |
| Chchd10 | 2,99164 | 1,19649 | -1,32212 | 0,0079 | 0,075196 |
| Rpl9 | 3,81742 | 7,47819 | 0,970091 | 0,008 | 0,07561 |
| Chn1 | 15,0907 | 20,4436 | 0,437984 | 0,008 | 0,07561 |
| Kif26b | 32,7433 | 44,2366 | 0,434043 | 0,008 | 0,07561 |
| Cox5a | 215,762 | 290,295 | 0,428082 | 0,008 | 0,07561 |
| Nog | 5,87843 | 3,70508 | -0,665926 | 0,008 | 0,07561 |
| Ica1 | 4,0222 | 2,29969 | -0,806546 | 0,008 | 0,07561 |
| Spo11 | 1,05445 | 0,326274 | -1,69234 | 0,008 | 0,07561 |
| Arhgap8 | 1,08478 | 0,314272 | -1,78732 | 0,00795 | 0,07561 |
| Hcls1 | 2,22572 | 0,091681 | -4,6015 | 0,008 | 0,07561 |
| Tacstd2 | 3,3213 | 0,106608 | -4,96135 | 0,008 | 0,07561 |
| Grik3 | 0,622557 | 0,347119 | -0,842777 | 0,00805 | 0,076028 |
| Car3 | 20,0582 | 28,2526 | 0,494192 | 0,0081 | 0,076285 |
| Dock5 | 5,03667 | 3,67249 | -0,455711 | 0,0081 | 0,076285 |
| Ginm1 | 20,9486 | 14,687 | -0,512313 | 0,0081 | 0,076285 |
| Marc2 | 14,9424 | 10,4503 | -0,515873 | 0,0081 | 0,076285 |
| Dbndd2 | 0,725531 | 1,32381 | 0,867586 | 0,00815 | 0,076648 |
| Gstp1 | 25,5522 | 37,2765 | 0,544818 | 0,00815 | 0,076648 |
| Mtrf1l | 5,93946 | 8,65703 | 0,543539 | 0,0082 | 0,076956 |
| Zgrf1 | 7,54778 | 10,2729 | 0,444723 | 0,0082 | 0,076956 |
| Pkp2 | 3,98792 | 2,55821 | -0,640498 | 0,0082 | 0,076956 |
| Fjx1 | 21,4155 | 15,5034 | -0,466065 | 0,00825 | 0,077316 |
| Ssh3 | 1,99233 | 1,11184 | -0,841509 | 0,00825 | 0,077316 |
| Gfpt2 | 6,5713 | 9,4327 | 0,521492 | 0,0083 | 0,077567 |
| Foxb1 | 35,7651 | 48,1422 | 0,428752 | 0,0083 | 0,077567 |
| Lipa | 11,4567 | 8,12735 | -0,495332 | 0,0083 | 0,077567 |
| Pdlim3 | 2,53315 | 1,27369 | -0,991917 | 0,0083 | 0,077567 |
| Rgs5 | 1,5675 | 2,86868 | 0,871924 | 0,0084 | 0,078228 |
| Cox8a | 134,872 | 183,801 | 0,446554 | 0,0084 | 0,078228 |
| Naa60 | 44,3896 | 59,4615 | 0,421733 | 0,0084 | 0,078228 |
| B4galt4 | 12,4518 | 8,79812 | -0,501082 | 0,0084 | 0,078228 |
| Vwa5a | 5,8941 | 4,08549 | -0,528761 | 0,0084 | 0,078228 |
| Scamp5 | 3,50255 | 2,20472 | -0,667815 | 0,00845 | 0,078639 |
| Itpripl2 | 4,69939 | 3,32173 | -0,500541 | 0,0085 | 0,07894 |
| 2310015A10Rik | 2,05375 | 1,29586 | -0,66435 | 0,0085 | 0,07894 |
| Pcyt1b | 1,45116 | 0,850706 | -0,770477 | 0,0085 | 0,07894 |

| | | | | | |
|---|---|---|---|---|---|
| Ciart | 4,1952 | 7,3458 | 0,808179 | 0,00855 | 0,079239 |
| Fam220a | 11,119 | 15,6597 | 0,494033 | 0,00855 | 0,079239 |
| Foxl1 | 3,34055 | 2,10156 | -0,668626 | 0,00855 | 0,079239 |
| Gmnn | 83,8678 | 113,763 | 0,439849 | 0,0086 | 0,079427 |
| Hif3a | 3,02006 | 2,07987 | -0,538081 | 0,0086 | 0,079427 |
| Shisa5 | 11,8503 | 8,11672 | -0,545957 | 0,0086 | 0,079427 |
| C030034I22Rik | 1,85168 | 1,01313 | -0,87001 | 0,0086 | 0,079427 |
| Rab37 | 1,28597 | 0,574196 | -1,16325 | 0,0086 | 0,079427 |
| Slc38a9 | 2,00247 | 1,3602 | -0,55796 | 0,00865 | 0,079833 |
| Gm2a | 1,67437 | 0,979603 | -0,773346 | 0,0087 | 0,080074 |
| Fgd3 | 1,0329 | 0,497025 | -1,05531 | 0,0087 | 0,080074 |
| Mir22hg | 1,18265 | 0,414835 | -1,51141 | 0,0087 | 0,080074 |
| Prrg4 | 0,614083 | 0,182839 | -1,74786 | 0,0087 | 0,080074 |
| Lyar | 88,8704 | 118,793 | 0,41867 | 0,00875 | 0,080368 |
| Tm2d3 | 35,6884 | 24,8574 | -0,521776 | 0,00875 | 0,080368 |
| Folr1 | 2,08963 | 0,928367 | -1,17048 | 0,00875 | 0,080368 |
| Rpl35 | 9,09284 | 17,0259 | 0,904926 | 0,0088 | 0,080716 |
| Rplp1 | 1138,83 | 1525,7 | 0,421919 | 0,0088 | 0,080716 |
| Epha1 | 34,2307 | 56,4128 | 0,720731 | 0,00885 | 0,081063 |
| Etv5 | 25,1348 | 33,832 | 0,428703 | 0,00885 | 0,081063 |
| Prcp | 16,2386 | 11,7359 | -0,468493 | 0,0089 | 0,08141 |
| Lrrc16a | 4,20386 | 2,8832 | -0,544044 | 0,0089 | 0,08141 |
| Cln5 | 17,3488 | 12,4713 | -0,476225 | 0,00895 | 0,081644 |
| Arhgap10 | 20,5454 | 27,7715 | 0,434793 | 0,00895 | 0,081644 |
| Elmo3 | 1,63148 | 0,900057 | -0,85809 | 0,00895 | 0,081644 |
| Lbp | 1,14538 | 0,522693 | -1,13179 | 0,00895 | 0,081644 |
| Leo1 | 34,4652 | 46,5733 | 0,434361 | 0,009 | 0,082044 |
| Zdhhc1 | 4,38785 | 2,70027 | -0,700412 | 0,00905 | 0,082443 |
| Grhl1 | 1,18052 | 0,605151 | -0,96405 | 0,0091 | 0,082842 |
| Stip1 | 198,461 | 266,512 | 0,425345 | 0,00915 | 0,083128 |
| Sft2d3 | 5,91354 | 3,92178 | -0,592511 | 0,00915 | 0,083128 |
| D930048N14Rik | 2,08926 | 0,840298 | -1,31402 | 0,00915 | 0,083128 |
| Lhx9 | 0,233727 | 0,680241 | 1,54122 | 0,0092 | 0,083468 |
| Abhd13 | 15,0432 | 11,1317 | -0,43443 | 0,0092 | 0,083468 |
| Hdac1 | 22,5812 | 30,8998 | 0,452474 | 0,00925 | 0,083638 |
| Pigt | 28,7338 | 21,1831 | -0,439835 | 0,00925 | 0,083638 |
| Inf2 | 2,62318 | 1,7154 | -0,612775 | 0,00925 | 0,083638 |
| Robo2 | 0,866246 | 0,517983 | -0,741873 | 0,00925 | 0,083638 |
| Trim12c | 1,23007 | 0,663185 | -0,891252 | 0,00925 | 0,083638 |
| Fgl2 | 0,592853 | 0,195897 | -1,59758 | 0,0093 | 0,084033 |
| Tcfl5 | 1,0764 | 2,37099 | 1,13927 | 0,00935 | 0,084371 |
| Atp6v0b | 61,8291 | 44,9037 | -0,461451 | 0,00935 | 0,084371 |
| 2700049A03Rik | 8,75161 | 11,8365 | 0,43562 | 0,0094 | 0,084651 |
| Yme1l1 | 56,6201 | 42,5894 | -0,410819 | 0,0094 | 0,084651 |
| Alg10b | 46,5407 | 34,989 | -0,411592 | 0,0094 | 0,084651 |
| Snx13 | 6,07627 | 4,38275 | -0,47135 | 0,00945 | 0,084986 |
| Pla2g16 | 1,10936 | 0,572192 | -0,955157 | 0,00945 | 0,084986 |
| Hoxc5 | 14,7946 | 20,2369 | 0,451915 | 0,0095 | 0,085379 |

| | | | | | |
|---|---|---|---|---|---|
| Kalrn | 2,30112 | 3,28586 | 0,513937 | 0,00955 | 0,08577 |
| Hmga2-ps1 | 23,636 | 34,1442 | 0,530652 | 0,0096 | 0,086162 |
| Fam163b | 1,65782 | 2,71854 | 0,713544 | 0,00965 | 0,086378 |
| Egln3 | 19,0043 | 25,7421 | 0,437803 | 0,00965 | 0,086378 |
| 2210018M11Rik | 18,5263 | 24,7971 | 0,420599 | 0,00965 | 0,086378 |
| Flt3l | 1,55879 | 0,27074 | -2,52545 | 0,00965 | 0,086378 |
| Emilin1 | 25,6783 | 19,0405 | -0,431475 | 0,0097 | 0,086652 |
| Fbxo6 | 4,60507 | 2,58427 | -0,833467 | 0,0097 | 0,086652 |
| Gm4944 | 1,04949 | 0,582345 | -0,849741 | 0,0097 | 0,086652 |
| Jakmip2 | 1,3497 | 2,22195 | 0,719191 | 0,00985 | 0,087816 |
| Nxpe4 | 3,01319 | 1,89152 | -0,671745 | 0,00985 | 0,087816 |
| Gm7694 | 1,26923 | 0,709286 | -0,839514 | 0,00985 | 0,087816 |
| Cyp27a1 | 6,5694 | 9,71512 | 0,56447 | 0,0099 | 0,088085 |
| Pbxip1 | 8,3853 | 6,02681 | -0,476469 | 0,0099 | 0,088085 |
| 9330151L19Rik | 5,94881 | 2,28095 | -1,38296 | 0,0099 | 0,088085 |
| Degs1 | 43,727 | 32,5595 | -0,425447 | 0,01005 | 0,089301 |
| Map7 | 2,08255 | 1,02146 | -1,02772 | 0,01005 | 0,089301 |
| Izumo4 | 2,86397 | 1,1846 | -1,27362 | 0,0101 | 0,089685 |
| Parvg | 0,804253 | 0,0633352 | -3,66657 | 0,01015 | 0,09007 |
| Ust | 5,89373 | 4,15081 | -0,505787 | 0,0102 | 0,090453 |
| Zdhhc20 | 26,8101 | 20,0169 | -0,421558 | 0,01025 | 0,090836 |
| E330009J07Rik | 10,5586 | 14,8812 | 0,495067 | 0,01035 | 0,091662 |
| Ccnl1 | 65,8298 | 87,5781 | 0,41183 | 0,0104 | 0,092043 |
| Wnt11 | 5,7618 | 3,86717 | -0,57524 | 0,01045 | 0,092425 |
| Sar1b | 47,0305 | 34,45 | -0,449092 | 0,0105 | 0,092683 |
| Cttnbp2 | 1,42365 | 0,874804 | -0,702559 | 0,0105 | 0,092683 |
| Glrx | 5,01352 | 2,88319 | -0,79816 | 0,0105 | 0,092683 |
| Scd1 | 23,1217 | 30,7415 | 0,410937 | 0,01055 | 0,09294 |
| Cdkn1b | 15,8401 | 11,3956 | -0,475111 | 0,01055 | 0,09294 |
| Pdik1l | 7,4835 | 5,36282 | -0,480721 | 0,01055 | 0,09294 |
| Heyl | 3,26127 | 2,20483 | -0,564767 | 0,0106 | 0,093258 |
| Tmem229b | 1,55634 | 0,90438 | -0,783155 | 0,0106 | 0,093258 |
| Sox13 | 11,111 | 15,2005 | 0,45213 | 0,01065 | 0,093636 |
| Mfsd7c | 0,806239 | 0,381884 | -1,07807 | 0,0107 | 0,094014 |
| Tdrkh | 7,22676 | 10,2243 | 0,500576 | 0,01075 | 0,094329 |
| Fuca1 | 11,7796 | 8,37354 | -0,492385 | 0,01075 | 0,094329 |
| Gnal | 9,43193 | 14,1513 | 0,585312 | 0,0108 | 0,094581 |
| Maml3 | 13,072 | 17,4985 | 0,420754 | 0,0108 | 0,094581 |
| Nrep | 167,842 | 126,712 | -0,405554 | 0,0108 | 0,094581 |
| Gm13826 | 177,858 | 237,152 | 0,415089 | 0,01085 | 0,094833 |
| Pdcd10 | 8,77505 | 5,94658 | -0,561345 | 0,01085 | 0,094833 |
| Adap1 | 0,641993 | 0,146503 | -2,13162 | 0,01085 | 0,094833 |
| Parp3 | 0,743309 | 1,42989 | 0,943869 | 0,0109 | 0,095208 |
| Rgs4 | 0,62071 | 1,25284 | 1,01321 | 0,01095 | 0,095457 |
| P4ha2 | 8,37163 | 11,8709 | 0,503853 | 0,01095 | 0,095457 |
| Ccdc126 | 2,45475 | 1,39385 | -0,816499 | 0,01095 | 0,095457 |
| Nr1d1 | 2,27709 | 3,58917 | 0,656459 | 0,011 | 0,095768 |
| Rasgrp3 | 8,68656 | 6,33308 | -0,455878 | 0,011 | 0,095768 |

| | | | | | |
|---|---|---|---|---|---|
| Lgals3 | 1,55697 | 0,616054 | -1,33761 | 0,0111 | 0,096576 |
| Itgb1 | 141,183 | 105,765 | -0,4167 | 0,0112 | 0,097382 |
| Cacna1g | 1,23481 | 1,83951 | 0,575031 | 0,01125 | 0,097627 |
| Rrp15 | 59,8987 | 79,8759 | 0,415236 | 0,01125 | 0,097627 |
| Erlec1 | 12,2291 | 8,94031 | -0,451923 | 0,01125 | 0,097627 |
| Scg5 | 1,7131 | 3,32759 | 0,957865 | 0,0113 | 0,097933 |
| Slc35a3 | 8,51962 | 6,19034 | -0,46077 | 0,0113 | 0,097933 |
| H2-Q10 | 1,78168 | 3,23996 | 0,862738 | 0,01135 | 0,098303 |
| Bex2 | 72,529 | 97,6139 | 0,428529 | 0,0114 | 0,098672 |
| Tsen2 | 11,6661 | 16,4806 | 0,498444 | 0,01145 | 0,098913 |
| Chst11 | 12,1649 | 9,0594 | -0,42524 | 0,01145 | 0,098913 |
| Fcer1g | 4,36972 | 0,37694 | -3,53513 | 0,01145 | 0,098913 |
| Lrrtm3 | 1,08242 | 0,588694 | -0,878674 | 0,0115 | 0,09928 |
| 2610020H08Rik | 9,98415 | 14,1452 | 0,502597 | 0,0116 | 0,09995 |
| Plekha8 | 8,86065 | 6,60131 | -0,424661 | 0,0116 | 0,09995 |
| Stat3 | 7,72655 | 5,57013 | -0,472112 | 0,0116 | 0,09995 |
| Tmc8 | 1,25856 | 0,644275 | -0,966029 | 0,01165 | 0,100252 |
| 2610016A17Rik | 6,07467 | 2,02929 | -1,58183 | 0,01165 | 0,100252 |
| Tmem168 | 10,8476 | 7,94534 | -0,44919 | 0,0117 | 0,100617 |
| Mtm1 | 0,891017 | 0,416963 | -1,09553 | 0,01175 | 0,100917 |
| Gca | 0,645074 | 0,255761 | -1,33467 | 0,01175 | 0,100917 |
| Lrp2 | 2,02906 | 2,79291 | 0,460962 | 0,0118 | 0,101281 |
| Ddr1 | 28,0804 | 21,0934 | -0,412773 | 0,01185 | 0,101385 |
| Zfp526 | 5,05659 | 3,46357 | -0,545905 | 0,01185 | 0,101385 |
| Ghr | 11,3368 | 7,75481 | -0,547855 | 0,01185 | 0,101385 |
| Nr1h5 | 4,02435 | 2,58036 | -0,641182 | 0,01185 | 0,101385 |
| Zxdb | 1,00178 | 0,576742 | -0,796573 | 0,01185 | 0,101385 |
| Tox | 0,842948 | 0,376835 | -1,16151 | 0,01195 | 0,102175 |
| Stambpl1 | 2,69623 | 4,33224 | 0,684171 | 0,012 | 0,102341 |
| Pgbd5 | 3,55365 | 5,24592 | 0,561895 | 0,012 | 0,102341 |
| Cycs | 2,18965 | 1,31201 | -0,738924 | 0,012 | 0,102341 |
| Grin2c | 0,798926 | 0,425733 | -0,908112 | 0,012 | 0,102341 |
| Ap5b1 | 0,641499 | 0,248985 | -1,36539 | 0,01205 | 0,102702 |
| Dut | 76,5622 | 100,937 | 0,398746 | 0,0121 | 0,103062 |
| Arl5a | 37,202 | 28,2113 | -0,399108 | 0,01215 | 0,103356 |
| Zfp770 | 9,46398 | 6,89422 | -0,45706 | 0,01215 | 0,103356 |
| Itgav | 17,8933 | 13,5026 | -0,406182 | 0,0122 | 0,10365 |
| Tmem186 | 14,9923 | 10,9714 | -0,450475 | 0,0122 | 0,10365 |
| Slc41a3 | 1,21455 | 0,575466 | -1,07762 | 0,0123 | 0,104433 |
| Rps27a | 6,18163 | 10,2717 | 0,732611 | 0,0124 | 0,105082 |
| Zdhhc21 | 6,46543 | 4,81445 | -0,425375 | 0,0124 | 0,105082 |
| Mesp2 | 16,2431 | 11,6214 | -0,483041 | 0,0124 | 0,105082 |
| Metrnl | 0,729564 | 0,256038 | -1,51067 | 0,01245 | 0,105438 |
| Upp1 | 2,01891 | 0,117461 | -4,10333 | 0,01255 | 0,106218 |
| Snai1 | 49,2695 | 65,3118 | 0,406647 | 0,0126 | 0,106372 |
| Bhlha9 | 1,40446 | 0,203656 | -2,78581 | 0,0126 | 0,106372 |
| Gata1 | 6,39672 | 0,0914398 | -6,12837 | 0,0126 | 0,106372 |
| Ermap | 8,47175 | 0,0357179 | -7,88987 | 0,0126 | 0,106372 |

| | | | | | |
|---|---|---|---|---|---|
| Hspg2 | 22,7744 | 17,1921 | -0,405664 | 0,01265 | 0,106391 |
| Cnih4 | 16,2142 | 11,9139 | -0,444612 | 0,01265 | 0,106391 |
| Tuba1c | 12,5983 | 8,89671 | -0,501883 | 0,01265 | 0,106391 |
| Tlr12 | 0,815004 | 0,370133 | -1,13876 | 0,01265 | 0,106391 |
| Ifi35 | 1,75639 | 0,727676 | -1,27124 | 0,01265 | 0,106391 |
| Abi3 | 1,32095 | 0,0882686 | -3,90353 | 0,01265 | 0,106391 |
| D10Wsu102e | 30,5303 | 40,3113 | 0,400944 | 0,0127 | 0,106677 |
| Arhgef6 | 0,855772 | 0,452935 | -0,917922 | 0,0127 | 0,106677 |
| Ptar1 | 10,9412 | 15,1933 | 0,473667 | 0,0128 | 0,10718 |
| Wdr70 | 15,318 | 20,802 | 0,441496 | 0,0128 | 0,10718 |
| Mcm10 | 37,6293 | 49,5804 | 0,397911 | 0,0128 | 0,10718 |
| S100a10 | 67,7697 | 49,1107 | -0,464605 | 0,0128 | 0,10718 |
| Gm9958 | 4,2154 | 2,49402 | -0,757196 | 0,0128 | 0,10718 |
| Rpl30 | 31,2479 | 67,509 | 1,11132 | 0,01285 | 0,107531 |
| Rnf103 | 8,39242 | 5,99065 | -0,486373 | 0,0129 | 0,107882 |
| Epm2aip1 | 27,3152 | 20,7393 | -0,397337 | 0,01295 | 0,108233 |
| Cst3 | 144,755 | 109,094 | -0,408046 | 0,013 | 0,108447 |
| Tmem9 | 47,5148 | 35,5906 | -0,416878 | 0,013 | 0,108447 |
| Mxra8 | 10,4374 | 7,3378 | -0,508346 | 0,013 | 0,108447 |
| Ablim2 | 0,573696 | 1,07303 | 0,903331 | 0,01305 | 0,108796 |
| Jmjd7-pla2g4b | 1,17784 | 3,82949 | 1,70101 | 0,0131 | 0,109077 |
| Leprot | 26,4483 | 19,4783 | -0,441312 | 0,0131 | 0,109077 |
| Zyx | 71,6115 | 104 | 0,538323 | 0,01315 | 0,109289 |
| Zfp36l2 | 21,8793 | 28,9731 | 0,405149 | 0,01315 | 0,109289 |
| Fam3c | 17,2114 | 12,6297 | -0,446541 | 0,01315 | 0,109289 |
| Ppargc1a | 0,960757 | 0,576083 | -0,737896 | 0,0132 | 0,109636 |
| Btg2 | 59,5536 | 78,4044 | 0,396745 | 0,01325 | 0,109983 |
| Catsper2 | 0,434457 | 1,01874 | 1,2295 | 0,0133 | 0,110193 |
| Krba1 | 10,8267 | 14,3443 | 0,405888 | 0,0133 | 0,110193 |
| Pvrl4 | 1,57512 | 0,0382953 | -5,36215 | 0,0133 | 0,110193 |
| Nr3c1 | 2,9866 | 2,08479 | -0,518603 | 0,01335 | 0,110334 |
| Tmem178 | 1,19024 | 0,474544 | -1,32664 | 0,01335 | 0,110334 |
| Acap1 | 0,745921 | 0,282331 | -1,40164 | 0,01335 | 0,110334 |
| Zfp366 | 0,771958 | 0,060793 | -3,66655 | 0,01335 | 0,110334 |
| Hsp90aa1 | 99,7923 | 131,213 | 0,394907 | 0,0134 | 0,11061 |
| Ltbp1 | 14,4704 | 10,8745 | -0,412158 | 0,0134 | 0,11061 |
| U2af1 | 36,8684 | 50,1574 | 0,444078 | 0,01345 | 0,110817 |
| Tgoln1,Tgoln2 | 19,5965 | 13,9556 | -0,489756 | 0,01345 | 0,110817 |
| Tph1 | 2,12944 | 0,602519 | -1,8214 | 0,01345 | 0,110817 |
| Wdr86 | 8,21507 | 5,49319 | -0,580629 | 0,0135 | 0,111092 |
| Crip1 | 10,0834 | 4,77073 | -1,0797 | 0,0135 | 0,111092 |
| Slc35c1 | 4,657 | 3,11482 | -0,580252 | 0,01355 | 0,111435 |
| Filip1l | 4,9339 | 2,81081 | -0,811743 | 0,0137 | 0,1126 |
| Slain1 | 4,38616 | 6,36627 | 0,537492 | 0,01375 | 0,112664 |
| Tmem165 | 80,2365 | 56,3443 | -0,509989 | 0,01375 | 0,112664 |
| Amigo1 | 1,81631 | 1,18669 | -0,614066 | 0,01375 | 0,112664 |
| Cebpa | 1,53841 | 0,847043 | -0,860935 | 0,01375 | 0,112664 |
| Serpine1 | 0,712173 | 0,301211 | -1,24145 | 0,01375 | 0,112664 |

| | | | | | |
|---|---|---|---|---|---|
| Bet1l | 12,3415 | 8,58094 | -0,524312 | 0,01385 | 0,113206 |
| Dock8 | 1,60942 | 1,06825 | -0,591295 | 0,01385 | 0,113206 |
| Dlx5 | 4,26782 | 2,45634 | -0,796986 | 0,01385 | 0,113206 |
| Mfsd6 | 0,558814 | 0,231179 | -1,27336 | 0,01385 | 0,113206 |
| Lat | 1,33367 | 0,224447 | -2,57095 | 0,0139 | 0,113545 |
| Manea | 16,3697 | 12,3279 | -0,409098 | 0,01395 | 0,113884 |
| Klhl3 | 1,11506 | 0,479646 | -1,21708 | 0,014 | 0,114153 |
| Alox5ap | 2,0772 | 0,660346 | -1,65334 | 0,014 | 0,114153 |
| Ifi30 | 21,52 | 14,833 | -0,536861 | 0,01405 | 0,114351 |
| Gatm | 1,96912 | 1,10131 | -0,838332 | 0,01405 | 0,114351 |
| Gm4285 | 1,66013 | 0,672226 | -1,30428 | 0,01405 | 0,114351 |
| Cks2 | 5,31546 | 9,2032 | 0,791941 | 0,01415 | 0,115025 |
| Atrnl1 | 11,3441 | 8,55872 | -0,40647 | 0,01415 | 0,115025 |
| Fut11 | 14,1811 | 10,5486 | -0,426918 | 0,0142 | 0,115361 |
| Pik3r6 | 0,576096 | 0,046075 | -3,64425 | 0,01425 | 0,115697 |
| Il1rap | 1,70295 | 1,05572 | -0,689805 | 0,0143 | 0,116032 |
| Redrum | 1,81595 | 0,074328 | -4,61067 | 0,01435 | 0,116367 |
| D2hgdh | 3,34347 | 2,1805 | -0,616684 | 0,0144 | 0,116702 |
| Igfbpl1 | 11,4764 | 15,5595 | 0,439128 | 0,01445 | 0,116895 |
| Dennd3 | 0,775203 | 0,427185 | -0,859714 | 0,01445 | 0,116895 |
| Slc6a13 | 1,17451 | 0,0773072 | -3,92531 | 0,01445 | 0,116895 |
| Aspm | 14,7759 | 11,2574 | -0,39237 | 0,0145 | 0,117086 |
| Dapp1 | 1,45546 | 0,823131 | -0,822283 | 0,0145 | 0,117086 |
| Il15 | 1,22719 | 0,380092 | -1,69094 | 0,0145 | 0,117086 |
| Fsd1 | 9,52049 | 13,4682 | 0,500447 | 0,01455 | 0,117207 |
| Myl12b | 79,5039 | 105,556 | 0,408911 | 0,01455 | 0,117207 |
| Slc38a10 | 36,0441 | 27,419 | -0,394589 | 0,01455 | 0,117207 |
| Ggt1 | 1,22101 | 0,0827502 | -3,88317 | 0,01455 | 0,117207 |
| Dync2h1 | 5,95421 | 7,8625 | 0,401077 | 0,0146 | 0,117539 |
| Gm13889 | 1,86833 | 0,794188 | -1,2342 | 0,01465 | 0,11787 |
| Ifitm1 | 377,762 | 492,918 | 0,38387 | 0,0147 | 0,118201 |
| Ercc2 | 22,8681 | 17,1652 | -0,413847 | 0,01475 | 0,118532 |
| Erlin2 | 30,1565 | 22,9342 | -0,39497 | 0,0149 | 0,119594 |
| A830082N09Rik | 1,47213 | 0,969249 | -0,602966 | 0,0149 | 0,119594 |
| Foxa3 | 0,70984 | 0,178634 | -1,99049 | 0,01495 | 0,119923 |
| Kdelc2 | 6,92115 | 4,97068 | -0,477567 | 0,015 | 0,12018 |
| Hmgcll1 | 2,52536 | 1,63042 | -0,631249 | 0,015 | 0,12018 |
| Rps7 | 5,43489 | 8,76223 | 0,689047 | 0,01505 | 0,120292 |
| Clcn4-2 | 14,3262 | 10,7659 | -0,412188 | 0,01505 | 0,120292 |
| Ttpa | 2,76566 | 1,75124 | -0,659242 | 0,01505 | 0,120292 |
| Megf11 | 0,588068 | 0,31045 | -0,921622 | 0,01505 | 0,120292 |
| Pnmal2 | 0,831807 | 1,39189 | 0,742726 | 0,0151 | 0,120547 |
| Siae | 2,43456 | 1,53426 | -0,666121 | 0,0151 | 0,120547 |
| Snrpa | 72,7255 | 96,6751 | 0,410683 | 0,01515 | 0,120658 |
| Lef1 | 96,5332 | 127,926 | 0,406216 | 0,01515 | 0,120658 |
| Eid1 | 73,7822 | 56,5027 | -0,384954 | 0,01515 | 0,120658 |
| Cbx8 | 4,52128 | 2,69068 | -0,74876 | 0,01515 | 0,120658 |
| Rpl29 | 44,9316 | 61,4121 | 0,450792 | 0,0152 | 0,12084 |

| | | | | | |
|---|---|---|---|---|---|
| Tapt1 | 24,2254 | 18,3701 | -0,399165 | 0,0152 | 0,12084 |
| Ube2v2 | 3,16769 | 2,21294 | -0,51747 | 0,0152 | 0,12084 |
| Hsd17b11 | 20,1402 | 14,6966 | -0,454593 | 0,01525 | 0,121166 |
| Efhd2 | 10,1953 | 7,23142 | -0,495555 | 0,0153 | 0,121347 |
| Lgals8 | 3,52449 | 2,25249 | -0,645894 | 0,0153 | 0,121347 |
| Ripply3 | 5,76347 | 3,61707 | -0,672117 | 0,0153 | 0,121347 |
| 6330419J24Rik | 4,80804 | 6,84557 | 0,509723 | 0,01535 | 0,121527 |
| Arhgef5 | 6,45465 | 4,74133 | -0,445048 | 0,01535 | 0,121527 |
| Bnipl | 0,98453 | 0,0829484 | -3,56915 | 0,01535 | 0,121527 |
| Trim9 | 20,8041 | 32,1768 | 0,629154 | 0,0155 | 0,122642 |
| Mbd5 | 3,77217 | 5,13398 | 0,444685 | 0,01555 | 0,122892 |
| Rhbdd1 | 7,30194 | 5,23177 | -0,480982 | 0,01555 | 0,122892 |
| Itga7 | 0,282278 | 0,59894 | 1,08529 | 0,0156 | 0,123069 |
| Zfp758 | 4,61838 | 3,12655 | -0,562815 | 0,0156 | 0,123069 |
| Tifa | 1,52516 | 0,783264 | -0,961388 | 0,0156 | 0,123069 |
| AA987161 | 2,67138 | 1,81067 | -0,561064 | 0,01565 | 0,12339 |
| Pld4 | 0,947607 | 0,0875225 | -3,43656 | 0,01575 | 0,124106 |
| D630039A03Rik | 0,615677 | 0,244807 | -1,33053 | 0,0158 | 0,124426 |
| Nudc | 109,746 | 143,442 | 0,386304 | 0,01585 | 0,1246 |
| Alg11 | 13,6526 | 10,3592 | -0,398273 | 0,01585 | 0,1246 |
| E130309D14Rik | 1,19057 | 0,0367024 | -5,01963 | 0,01585 | 0,1246 |
| Cux2 | 2,4801 | 1,67678 | -0,564703 | 0,0159 | 0,124919 |
| Zfp105 | 20,6732 | 27,4986 | 0,411598 | 0,01595 | 0,125165 |
| Ndn | 49,5859 | 65,0622 | 0,39189 | 0,01595 | 0,125165 |
| Krt39 | 0,755743 | 1,70107 | 1,17048 | 0,016 | 0,125484 |
| Ngf | 1,11565 | 2,42439 | 1,11974 | 0,01615 | 0,126437 |
| Mira | 71,8986 | 95,4205 | 0,408336 | 0,01615 | 0,126437 |
| Acvr2a | 14,6126 | 9,73331 | -0,586212 | 0,01615 | 0,126437 |
| Zfp451 | 20,9345 | 28,0454 | 0,421885 | 0,0162 | 0,126755 |
| Rplp0 | 494,463 | 655,242 | 0,406165 | 0,01625 | 0,127071 |
| Smc6 | 79,4482 | 104,195 | 0,391197 | 0,0163 | 0,127388 |
| Hfe | 1,30637 | 0,546872 | -1,25629 | 0,0164 | 0,128094 |
| Tmem60 | 30,2749 | 21,5301 | -0,49177 | 0,01645 | 0,128185 |
| Anpep | 3,908 | 2,66706 | -0,551178 | 0,01645 | 0,128185 |
| Prokr1 | 1,0081 | 0,570767 | -0,82067 | 0,01645 | 0,128185 |
| Tmem8 | 0,617694 | 0,270191 | -1,19291 | 0,01645 | 0,128185 |
| Pnliprp2 | 1,46658 | 2,65104 | 0,854106 | 0,0165 | 0,12835 |
| Rec8 | 1,96361 | 1,07743 | -0,865919 | 0,0165 | 0,12835 |
| Trim47 | 0,660747 | 0,0835938 | -2,98263 | 0,0165 | 0,12835 |
| Zbtb26 | 8,58082 | 6,37896 | -0,427795 | 0,01655 | 0,12859 |
| Atf3 | 2,25223 | 1,26996 | -0,826568 | 0,01655 | 0,12859 |
| Atp6v0e | 52,0725 | 38,1113 | -0,450305 | 0,0166 | 0,128903 |
| Otud3 | 5,10381 | 7,60171 | 0,574749 | 0,01665 | 0,129066 |
| Tspan2 | 9,69998 | 7,18297 | -0,433401 | 0,01665 | 0,129066 |
| Lysmd3 | 9,02844 | 6,55321 | -0,462275 | 0,01665 | 0,129066 |
| Scamp1 | 20,3948 | 15,4783 | -0,397952 | 0,0167 | 0,129304 |
| Nceh1 | 1,98354 | 1,2736 | -0,639158 | 0,0167 | 0,129304 |
| Dbndd1 | 3,6488 | 5,73435 | 0,652208 | 0,01675 | 0,129541 |

| | | | | | |
|---|---|---|---|---|---|
| Sgol2 | 14,4416 | 10,9146 | -0,403976 | 0,01675 | 0,129541 |
| Ndufs5 | 16,8033 | 26,3352 | 0,648247 | 0,0168 | 0,129853 |
| Sept4 | 1,42958 | 0,627525 | -1,18785 | 0,01685 | 0,130164 |
| Dct | 0,885537 | 0,0784286 | -3,4971 | 0,0169 | 0,130475 |
| 8430419L09Rik | 10,7778 | 8,04855 | -0,421259 | 0,01695 | 0,13071 |
| Tmem37 | 2,32998 | 0,991414 | -1,23275 | 0,01695 | 0,13071 |
| Furin | 21,6264 | 16,5066 | -0,389748 | 0,017 | 0,130944 |
| Zfp119a | 3,69316 | 2,32719 | -0,666271 | 0,017 | 0,130944 |
| Mycl | 23,2876 | 30,453 | 0,38702 | 0,0171 | 0,131638 |
| Ctps | 82,4378 | 107,191 | 0,378806 | 0,01715 | 0,131871 |
| Cdc42ep4 | 44,5323 | 57,8893 | 0,378445 | 0,01715 | 0,131871 |
| Atp11c | 17,0139 | 13,019 | -0,38609 | 0,0172 | 0,13218 |
| Ssc5d | 3,99735 | 2,79043 | -0,518558 | 0,0173 | 0,132872 |
| Xpo4 | 19,0941 | 25,0922 | 0,394114 | 0,0174 | 0,133487 |
| Slc10a7 | 7,0734 | 5,11654 | -0,467234 | 0,0174 | 0,133487 |
| Rtn4 | 90,6119 | 68,8413 | -0,396425 | 0,01745 | 0,133717 |
| Ccdc141 | 1,2718 | 0,806248 | -0,657576 | 0,01745 | 0,133717 |
| Gm10033 | 5,2531 | 3,27405 | -0,682092 | 0,0175 | 0,134023 |
| Fat1 | 46,8223 | 62,9554 | 0,427135 | 0,01755 | 0,134252 |
| Bmf | 13,7267 | 10,3765 | -0,403668 | 0,01755 | 0,134252 |
| Stap2 | 3,83086 | 2,3475 | -0,706547 | 0,01765 | 0,13494 |
| Etv6 | 17,778 | 23,1791 | 0,38273 | 0,0177 | 0,135245 |
| Hist1h2bj | 1,73539 | 5,72269 | 1,72143 | 0,01785 | 0,135847 |
| Tceal6 | 4,2684 | 6,85684 | 0,683848 | 0,01785 | 0,135847 |
| Spata5l1 | 8,27305 | 12,0285 | 0,539966 | 0,0178 | 0,135847 |
| Pdlim4 | 46,305 | 61,3837 | 0,406688 | 0,01785 | 0,135847 |
| Col9a1 | 8,66613 | 6,30641 | -0,45857 | 0,01785 | 0,135847 |
| Stim1 | 4,6434 | 3,25354 | -0,513173 | 0,01785 | 0,135847 |
| Cpz | 0,745716 | 0,0788547 | -3,24136 | 0,0178 | 0,135847 |
| Lrp1 | 19,2563 | 25,1836 | 0,387156 | 0,0179 | 0,136073 |
| Mvp | 0,872075 | 0,403564 | -1,11166 | 0,0179 | 0,136073 |
| Tdp1 | 26,7561 | 35,2486 | 0,397698 | 0,01795 | 0,136375 |
| Mbtps2 | 10,377 | 7,37047 | -0,493559 | 0,018 | 0,136677 |
| 2010010A06Rik | 3,81146 | 7,09629 | 0,89672 | 0,01805 | 0,136901 |
| Ldlrap1 | 8,99697 | 6,45859 | -0,478219 | 0,01805 | 0,136901 |
| Ccdc40 | 1,69611 | 2,50064 | 0,560067 | 0,0181 | 0,137125 |
| Dpf3 | 2,70591 | 1,46013 | -0,890019 | 0,0181 | 0,137125 |
| Hk2 | 28,9659 | 37,7423 | 0,381828 | 0,01815 | 0,137348 |
| Sgpp2 | 4,93709 | 1,08658 | -2,18387 | 0,01815 | 0,137348 |
| Pinx1 | 26,6411 | 35,6904 | 0,421883 | 0,0182 | 0,137415 |
| Mex3b | 26,7402 | 34,9382 | 0,385796 | 0,0182 | 0,137415 |
| Slc16a6 | 5,39167 | 3,71292 | -0,538178 | 0,0182 | 0,137415 |
| Il1r1 | 0,697785 | 0,372652 | -0,904953 | 0,0182 | 0,137415 |
| Galnt2 | 34,8295 | 26,6914 | -0,383934 | 0,01835 | 0,138313 |
| Camsap3 | 5,24986 | 3,78255 | -0,472918 | 0,01835 | 0,138313 |
| Prss44 | 0,721102 | 0,10178 | -2,82474 | 0,01835 | 0,138313 |
| Hipk3 | 9,35867 | 7,13266 | -0,391864 | 0,01845 | 0,13891 |
| Tnik | 0,703698 | 0,390844 | -0,848365 | 0,01845 | 0,13891 |

| | | | | | |
|---|---|---|---|---|---|
| Kcnk2 | 0,913532 | 0,481823 | -0,922952 | 0,0185 | 0,139208 |
| Neu1 | 5,85024 | 4,2039 | -0,476767 | 0,01855 | 0,139506 |
| Pnck | 1,85307 | 0,959738 | -0,949204 | 0,0186 | 0,139803 |
| Ryr1 | 0,405322 | 0,610777 | 0,591578 | 0,01865 | 0,140022 |
| Ttc9c | 35,2151 | 27,1186 | -0,376911 | 0,01865 | 0,140022 |
| Ccl25 | 3,79715 | 6,23119 | 0,714589 | 0,0187 | 0,14024 |
| Jph1 | 1,28487 | 1,9611 | 0,61004 | 0,0187 | 0,14024 |
| Rps19 | 78,0789 | 104,481 | 0,420237 | 0,0188 | 0,140831 |
| Sae1 | 175,421 | 227,408 | 0,374464 | 0,0188 | 0,140831 |
| Ddt | 64,9283 | 87,6297 | 0,432574 | 0,01885 | 0,141127 |
| 4933426M11Rik | 8,70468 | 6,54573 | -0,411238 | 0,01895 | 0,141796 |
| Pglyrp3 | 1,17054 | 2,46751 | 1,07588 | 0,019 | 0,142091 |
| Hivep2 | 0,818877 | 0,529614 | -0,628706 | 0,0191 | 0,142759 |
| Hdgfrp2 | 66,6278 | 86,7722 | 0,381109 | 0,01915 | 0,143052 |
| Nap1l2 | 1,1533 | 1,91625 | 0,732523 | 0,0192 | 0,143266 |
| Aldh1a2 | 38,977 | 29,7768 | -0,388438 | 0,0192 | 0,143266 |
| Arhgef26 | 2,85477 | 1,98732 | -0,522544 | 0,01925 | 0,143559 |
| Gabra3 | 2,11568 | 1,36626 | -0,63089 | 0,0194 | 0,144597 |
| 2610018G03Rik | 14,7578 | 19,89 | 0,430559 | 0,01945 | 0,144888 |
| Nphp1 | 24,0666 | 31,6249 | 0,394025 | 0,0195 | 0,14518 |
| Ncf2 | 0,829048 | 0,381392 | -1,12018 | 0,01955 | 0,145471 |
| Capg | 9,30776 | 6,24968 | -0,574652 | 0,0196 | 0,145681 |
| Gpr114 | 0,739688 | 0,0472922 | -3,96724 | 0,0196 | 0,145681 |
| Prdx3 | 76,631 | 58,7524 | -0,383281 | 0,01965 | 0,14589 |
| Fgd5 | 4,07965 | 2,92759 | -0,478733 | 0,01965 | 0,14589 |
| Lpar2 | 7,26313 | 5,4171 | -0,423072 | 0,0197 | 0,146099 |
| Entpd6 | 12,5189 | 9,21123 | -0,442637 | 0,0197 | 0,146099 |
| Map7d2 | 2,30478 | 3,35663 | 0,542386 | 0,01985 | 0,146561 |
| Pgk1 | 15,1081 | 20,3942 | 0,432834 | 0,01985 | 0,146561 |
| Pfn2 | 34,0517 | 25,9232 | -0,393479 | 0,0198 | 0,146561 |
| Tmem19 | 11,7622 | 8,72529 | -0,430885 | 0,0198 | 0,146561 |
| Ctdspl | 6,33142 | 4,6738 | -0,437934 | 0,0198 | 0,146561 |
| Depdc1a | 4,31482 | 2,98011 | -0,533933 | 0,01985 | 0,146561 |
| Nkx2-9 | 1,17102 | 0,382428 | -1,6145 | 0,01985 | 0,146561 |
| Tmem255a | 3,57846 | 0,23774 | -3,91188 | 0,01985 | 0,146561 |
| Prrx2 | 6,0245 | 9,29033 | 0,624889 | 0,01995 | 0,146975 |
| Mat2a | 167,859 | 128,662 | -0,383664 | 0,01995 | 0,146975 |
| Anxa6 | 25,1038 | 19,1157 | -0,393149 | 0,01995 | 0,146975 |
| Xpr1 | 7,90261 | 6,01348 | -0,39413 | 0,01995 | 0,146975 |
| Cacna1h | 1,58442 | 1,08684 | -0,543807 | 0,02005 | 0,14763 |
| Nrgn | 0,92962 | 0,202145 | -2,20125 | 0,0201 | 0,147917 |
| Gtf2b | 44,5645 | 58,5505 | 0,393787 | 0,0202 | 0,148407 |
| Klhl42 | 3,59246 | 2,61434 | -0,458525 | 0,0202 | 0,148407 |
| B3gnt3 | 0,68918 | 0,0818045 | -3,07463 | 0,0202 | 0,148407 |
| Aldoart1 | 10,4885 | 14,3013 | 0,447335 | 0,0203 | 0,148896 |
| Sort1 | 22,0245 | 16,9963 | -0,373886 | 0,0203 | 0,148896 |
| Cmpk1 | 38,4358 | 29,3989 | -0,38669 | 0,0203 | 0,148896 |
| Gm4349 | 0,375233 | 1,28437 | 1,77521 | 0,02045 | 0,149832 |

| | | | | | |
|---|---|---|---|---|---|
| Mfsd1 | 35,5481 | 27,2905 | -0,381372 | 0,02045 | 0,149832 |
| Rnd1 | 1,03945 | 1,8507 | 0,832248 | 0,0205 | 0,150116 |
| Ptpn5 | 0,590029 | 1,09174 | 0,887776 | 0,0206 | 0,150683 |
| Gm10069 | 1,1006 | 1,8815 | 0,773595 | 0,0206 | 0,150683 |
| Upk1b | 0,730116 | 0,126548 | -2,52844 | 0,02065 | 0,150884 |
| Milr1 | 0,715863 | 0,0509641 | -3,81213 | 0,02065 | 0,150884 |
| Zmpste24 | 25,7451 | 19,7596 | -0,381745 | 0,0207 | 0,151167 |
| Tomm6 | 102,43 | 133,417 | 0,381298 | 0,02075 | 0,151366 |
| Tuba4a | 6,38202 | 1,57739 | -2,01647 | 0,02075 | 0,151366 |
| Ephb1 | 1,86352 | 2,7107 | 0,540638 | 0,0208 | 0,151648 |
| Rps26 | 631,768 | 816,13 | 0,369403 | 0,02085 | 0,15193 |
| B2m | 59,9792 | 45,1237 | -0,410576 | 0,0209 | 0,152211 |
| Hist1h2bb | 0,600508 | 4,03174 | 2,74715 | 0,02095 | 0,152326 |
| Hoxc8 | 19,0849 | 25,1629 | 0,398865 | 0,02095 | 0,152326 |
| Mgat4b | 20,8476 | 15,7766 | -0,402092 | 0,02095 | 0,152326 |
| Atp6v1f | 44,2782 | 60,3111 | 0,445827 | 0,021 | 0,152607 |
| Hes7 | 90,5009 | 117,408 | 0,375526 | 0,0211 | 0,153001 |
| Rcan3 | 4,54158 | 3,297 | -0,462043 | 0,0211 | 0,153001 |
| Myom1 | 0,810199 | 0,479608 | -0,75642 | 0,0211 | 0,153001 |
| Barx2 | 0,694064 | 0,124794 | -2,47552 | 0,0211 | 0,153001 |
| Hoxa2 | 14,3357 | 19,4026 | 0,436631 | 0,02115 | 0,15328 |
| Gstm1 | 6,7132 | 9,89781 | 0,560109 | 0,02135 | 0,154646 |
| Rbm24 | 5,24532 | 7,23214 | 0,463392 | 0,0214 | 0,15484 |
| Focad | 8,89187 | 11,6033 | 0,38398 | 0,0214 | 0,15484 |
| Mirlet7d | 51,2551 | 0 | -inf | 0,02145 | 0,155034 |
| Pttg1ip | 30,0926 | 22,9541 | -0,39066 | 0,02145 | 0,155034 |
| Cdkl5 | 1,82899 | 2,81273 | 0,620926 | 0,0215 | 0,155311 |
| Pigm | 12,9718 | 9,9927 | -0,376438 | 0,02155 | 0,155588 |
| Qpctl | 7,48942 | 5,24401 | -0,514183 | 0,0216 | 0,155865 |
| Hs3st3b1 | 6,26792 | 4,6889 | -0,418737 | 0,0217 | 0,156502 |
| Vegfa | 33,2656 | 25,4984 | -0,383623 | 0,02175 | 0,156778 |
| Tmem9b | 10,2088 | 7,21346 | -0,501054 | 0,0219 | 0,157689 |
| Cldn14 | 0,756931 | 0,243794 | -1,6345 | 0,0219 | 0,157689 |
| 4933421O10Rik | 1,24225 | 0,744976 | -0,737687 | 0,02195 | 0,157964 |
| Rab30 | 2,11964 | 3,41596 | 0,688471 | 0,022 | 0,157984 |
| Dym | 16,5479 | 21,776 | 0,396093 | 0,022 | 0,157984 |
| Fam171a2 | 12,9943 | 9,79081 | -0,40838 | 0,022 | 0,157984 |
| Ube2q2 | 12,7094 | 9,50305 | -0,419431 | 0,022 | 0,157984 |
| Ccnf | 50,4029 | 39,2066 | -0,362413 | 0,02205 | 0,158003 |
| Slc30a5 | 39,1864 | 30,3655 | -0,36792 | 0,02205 | 0,158003 |
| Colec12 | 27,23 | 20,9778 | -0,376329 | 0,02205 | 0,158003 |
| Ablim1 | 10,0382 | 7,65241 | -0,391509 | 0,02205 | 0,158003 |
| Slc31a2 | 5,46483 | 3,6394 | -0,586474 | 0,02215 | 0,15855 |
| Mst1 | 0,571619 | 0,0669527 | -3,09384 | 0,02215 | 0,15855 |
| Pld1 | 0,97018 | 0,590393 | -0,716578 | 0,0222 | 0,158823 |
| Tmem2 | 23,0117 | 17,9387 | -0,359294 | 0,02225 | 0,158925 |
| Zfp933 | 5,29325 | 3,82524 | -0,468605 | 0,02225 | 0,158925 |
| Pla2g15 | 6,58708 | 4,69872 | -0,487371 | 0,02225 | 0,158925 |

| | | | | | |
|---|---|---|---|---|---|
| Pcgf2 | 23,2624 | 30,4481 | 0,388353 | 0,0223 | 0,159112 |
| Pwwp2a | 24,2679 | 31,4539 | 0,374189 | 0,0223 | 0,159112 |
| Nktr | 32,2626 | 41,3628 | 0,358472 | 0,02235 | 0,159299 |
| Krt36 | 1,0172 | 0,407624 | -1,3193 | 0,02235 | 0,159299 |
| Myl2 | 2,82808 | 5,57844 | 0,980036 | 0,0224 | 0,159485 |
| Lgr5 | 0,576579 | 0,301175 | -0,936916 | 0,0224 | 0,159485 |
| Sult4a1 | 0,404814 | 0,881045 | 1,12196 | 0,02245 | 0,159671 |
| Dusp9 | 43,508 | 33,631 | -0,371488 | 0,02245 | 0,159671 |
| Rest | 13,4087 | 17,4196 | 0,377541 | 0,02255 | 0,160297 |
| Stmn1 | 30,2635 | 40,3574 | 0,415255 | 0,0226 | 0,160396 |
| St13 | 200,744 | 258,27 | 0,363522 | 0,0226 | 0,160396 |
| Gab1 | 9,58412 | 7,23396 | -0,40586 | 0,0226 | 0,160396 |
| Lenep | 5,7935 | 3,04436 | -0,928296 | 0,02265 | 0,160666 |
| Hey2 | 10,4156 | 13,9537 | 0,421896 | 0,02275 | 0,161033 |
| Cd81 | 114,809 | 88,6258 | -0,373431 | 0,02275 | 0,161033 |
| Alad | 14,4204 | 11,0717 | -0,381243 | 0,02275 | 0,161033 |
| 4632428N05Rik | 1,3103 | 0,793847 | -0,722959 | 0,02275 | 0,161033 |
| Ifitm5 | 7,20809 | 11,5521 | 0,680464 | 0,0228 | 0,161131 |
| Rad50 | 13,8307 | 17,9649 | 0,377311 | 0,0228 | 0,161131 |
| Ptges3 | 50,9375 | 39,2904 | -0,374549 | 0,0228 | 0,161131 |
| Lhx5 | 0,621753 | 1,15668 | 0,89557 | 0,02285 | 0,161229 |
| Tmem136 | 9,13912 | 6,77999 | -0,430772 | 0,02285 | 0,161229 |
| Gipc2 | 1,26199 | 0,470684 | -1,42287 | 0,02285 | 0,161229 |
| Tspan3 | 206,109 | 160,405 | -0,36169 | 0,0229 | 0,161411 |
| Il6st | 21,5045 | 16,6617 | -0,3681 | 0,0229 | 0,161411 |
| Nat2 | 3,20064 | 1,83586 | -0,801908 | 0,023 | 0,161945 |
| Adm | 1,10681 | 0,379348 | -1,54481 | 0,023 | 0,161945 |
| Tpi1 | 248,998 | 321,293 | 0,367754 | 0,02305 | 0,162041 |
| Sh3tc2 | 2,66229 | 1,83392 | -0,537733 | 0,02305 | 0,162041 |
| Slc25a40 | 3,80339 | 2,58927 | -0,554741 | 0,02305 | 0,162041 |
| Rps18 | 225,316 | 292,383 | 0,375911 | 0,02315 | 0,162573 |
| Zmiz2 | 17,4898 | 13,3822 | -0,386196 | 0,02315 | 0,162573 |
| Ptbp3 | 25,4543 | 19,8107 | -0,361633 | 0,0232 | 0,162753 |
| Ifngr1 | 15,2167 | 11,3217 | -0,42656 | 0,0232 | 0,162753 |
| Sox1 | 1,03809 | 1,6351 | 0,655455 | 0,02325 | 0,163018 |
| Cers4 | 6,63262 | 4,87241 | -0,444941 | 0,02335 | 0,163634 |
| Bhlhe22 | 0,426183 | 0,823203 | 0,949775 | 0,0234 | 0,163726 |
| Mgme1 | 10,1774 | 13,5172 | 0,409431 | 0,0234 | 0,163726 |
| Tmem35 | 4,44154 | 2,8972 | -0,616401 | 0,0234 | 0,163726 |
| Pitpnm2 | 4,99644 | 3,74804 | -0,414765 | 0,02345 | 0,163905 |
| Gpr146 | 1,24659 | 0,222653 | -2,48511 | 0,02345 | 0,163905 |
| Gata2 | 0,993507 | 0,545664 | -0,864517 | 0,0235 | 0,164168 |
| Trim7 | 2,62914 | 4,27883 | 0,702625 | 0,02355 | 0,16426 |
| Glt25d1 | 45,6759 | 35,4265 | -0,366607 | 0,02355 | 0,16426 |
| Rab25 | 1,54949 | 0,188648 | -3,03803 | 0,02355 | 0,16426 |
| Fbxo21 | 39,8649 | 51,0077 | 0,355597 | 0,0236 | 0,164523 |
| Acsl6 | 5,62257 | 7,78297 | 0,469091 | 0,0237 | 0,164705 |
| Spata5 | 16,77 | 21,8152 | 0,37945 | 0,0237 | 0,164705 |

| | | | | | |
|---|---|---|---|---|---|
| Laptm4a | 213,855 | 166,122 | -0,364392 | 0,0237 | 0,164705 |
| Tmem68 | 12,5174 | 9,31914 | -0,425666 | 0,02365 | 0,164705 |
| Pde3a | 0,948327 | 0,552272 | -0,780007 | 0,0237 | 0,164705 |
| Cdkn2b | 0,995749 | 0,15845 | -2,65175 | 0,0237 | 0,164705 |
| Cops4 | 78,8771 | 101,035 | 0,357181 | 0,02375 | 0,164966 |
| Klhl32 | 2,47434 | 3,73965 | 0,595859 | 0,0238 | 0,165228 |
| Mir6910 | 0 | 291,278 | inf | 0,02385 | 0,165489 |
| Prr18 | 1,51696 | 2,33267 | 0,620804 | 0,0239 | 0,16575 |
| Mir3070b | 0 | 101,881 | inf | 0,02395 | 0,165924 |
| Myo18a | 8,33992 | 6,38953 | -0,384324 | 0,02395 | 0,165924 |
| Nme2 | 232,75 | 299,435 | 0,363457 | 0,024 | 0,166098 |
| Diap2 | 1,58682 | 1,11241 | -0,512451 | 0,024 | 0,166098 |
| Kif18a | 14,3956 | 10,8895 | -0,402685 | 0,02405 | 0,166272 |
| Cr1l | 12,6915 | 9,1632 | -0,46994 | 0,02405 | 0,166272 |
| Col7a1 | 0,736596 | 1,08453 | 0,558129 | 0,0242 | 0,16662 |
| Xylb | 3,75263 | 5,16911 | 0,462014 | 0,0242 | 0,16662 |
| Blnk | 24,2974 | 31,7139 | 0,384312 | 0,0242 | 0,16662 |
| Ndufa10 | 88,6529 | 113,983 | 0,362584 | 0,0242 | 0,16662 |
| Agpat3 | 10,1986 | 7,66599 | -0,411825 | 0,0242 | 0,16662 |
| Nrtn | 3,52822 | 1,92025 | -0,877646 | 0,02415 | 0,16662 |
| Ifi204 | 0,63185 | 0,204609 | -1,62672 | 0,0242 | 0,16662 |
| Tmem40 | 0,882021 | 0,261279 | -1,75522 | 0,02415 | 0,16662 |
| Slc6a6 | 27,667 | 21,4986 | -0,363926 | 0,0243 | 0,167222 |
| Zfp40 | 3,45385 | 2,42802 | -0,508427 | 0,02435 | 0,16748 |
| Slc35d1 | 7,34679 | 5,33075 | -0,462776 | 0,02445 | 0,168081 |
| Sorl1 | 4,70889 | 3,52401 | -0,418169 | 0,02455 | 0,168682 |
| Sptbn1 | 41,9052 | 32,7166 | -0,357107 | 0,02465 | 0,169108 |
| Sgcb | 8,14322 | 6,04328 | -0,430269 | 0,02465 | 0,169108 |
| Th | 0,697782 | 0,0897586 | -2,95865 | 0,02465 | 0,169108 |
| Ik | 81,4721 | 104,491 | 0,358997 | 0,0247 | 0,169277 |
| Nostrin | 0,657214 | 0,123549 | -2,41128 | 0,0247 | 0,169277 |
| Cyb561d1 | 2,32055 | 1,56172 | -0,571327 | 0,02475 | 0,169533 |
| Clic6 | 1,19286 | 0,705872 | -0,756949 | 0,0248 | 0,169702 |
| Pdgfrl | 0,881955 | 0,311031 | -1,50365 | 0,0248 | 0,169702 |
| Lrrc75a | 5,22172 | 7,40054 | 0,503105 | 0,0249 | 0,170212 |
| Nsdhl | 30,5177 | 39,4345 | 0,369808 | 0,0249 | 0,170212 |
| Tma16 | 13,5063 | 18,7304 | 0,471754 | 0,025 | 0,170808 |
| Snord37 | 0 | 1457,09 | inf | 0,0251 | 0,171403 |
| Fosb | 0,034792 | 0,949739 | 4,77071 | 0,02515 | 0,171569 |
| Zfp52 | 1,51208 | 0,916686 | -0,722035 | 0,02515 | 0,171569 |
| Tdrd3 | 17,591 | 23,0365 | 0,389084 | 0,0252 | 0,171823 |
| Ssh2 | 3,84037 | 2,89281 | -0,408776 | 0,02525 | 0,172076 |
| Plcd3 | 1,20219 | 0,685695 | -0,810024 | 0,0253 | 0,172329 |
| G530011O06Rik | 1,28976 | 0,128121 | -3,33153 | 0,0255 | 0,173603 |
| Nolc1 | 83,8024 | 107,372 | 0,35756 | 0,02555 | 0,173854 |
| Efcc1 | 2,15495 | 3,2368 | 0,586915 | 0,0256 | 0,174106 |
| Tes | 17,1705 | 13,0583 | -0,394966 | 0,02565 | 0,174269 |
| Rusc2 | 1,39518 | 0,903738 | -0,626478 | 0,02565 | 0,174269 |

| | | | | | |
|---|---|---|---|---|---|
| B3gat3 | 32,5783 | 24,7928 | -0,393991 | 0,0257 | 0,174343 |
| Parvb | 9,08837 | 6,81629 | -0,415036 | 0,0257 | 0,174343 |
| Mypop | 2,8354 | 1,78489 | -0,667715 | 0,0257 | 0,174343 |
| Tnfaip6 | 1,97556 | 1,06797 | -0,887386 | 0,02575 | 0,174416 |
| Sncg | 3,68856 | 0,233946 | -3,97881 | 0,02575 | 0,174416 |
| Mpo | 1,61276 | 0,0527438 | -4,93438 | 0,02575 | 0,174416 |
| Tcf7 | 28,2474 | 36,5553 | 0,371962 | 0,0258 | 0,174578 |
| Etv4 | 65,4389 | 84,1181 | 0,362268 | 0,0258 | 0,174578 |
| Dlgap2 | 1,4157 | 2,09649 | 0,566465 | 0,02585 | 0,174739 |
| Tada2a | 25,7381 | 33,4619 | 0,378614 | 0,02585 | 0,174739 |
| Pclo | 0,529857 | 0,756358 | 0,513465 | 0,0259 | 0,174989 |
| Atrip | 13,3822 | 17,5423 | 0,390519 | 0,02605 | 0,175824 |
| Mccc1 | 13,0511 | 17,0363 | 0,384441 | 0,02605 | 0,175824 |
| Krt14 | 1,08168 | 0,084839 | -3,6724 | 0,0261 | 0,176073 |
| Slc44a2 | 24,1036 | 18,7302 | -0,36388 | 0,02615 | 0,176321 |
| Mdh2 | 265,464 | 338,658 | 0,351312 | 0,0262 | 0,17648 |
| Sorcs3 | 0,660594 | 0,376047 | -0,81285 | 0,0262 | 0,17648 |
| Pdk3 | 34,8557 | 44,8207 | 0,362773 | 0,02635 | 0,177401 |
| Psme1 | 43,4457 | 57,0012 | 0,391777 | 0,0264 | 0,177559 |
| Gnpda2 | 8,48483 | 6,01856 | -0,495466 | 0,0264 | 0,177559 |
| Rnps1 | 40,7099 | 52,2787 | 0,360841 | 0,02645 | 0,177806 |
| Bend5 | 6,13125 | 8,62601 | 0,492511 | 0,0265 | 0,177963 |
| Gm13375 | 4,82273 | 3,05247 | -0,659875 | 0,0265 | 0,177963 |
| Tagln2 | 132,454 | 103,696 | -0,35314 | 0,02655 | 0,17803 |
| Fam213a | 20,1596 | 14,9732 | -0,429087 | 0,02655 | 0,17803 |
| Pfkfb1 | 3,54385 | 2,22446 | -0,671865 | 0,02655 | 0,17803 |
| Amotl2 | 33,1661 | 42,3568 | 0,352883 | 0,0266 | 0,178187 |
| Itpr3 | 2,04698 | 1,48369 | -0,464307 | 0,0266 | 0,178187 |
| Cyp4v3 | 0,823048 | 0,419469 | -0,972411 | 0,02665 | 0,178432 |
| Hist1h2bc | 7,17133 | 4,1104 | -0,802961 | 0,0267 | 0,178677 |
| Rnf8 | 31,5556 | 40,5705 | 0,362536 | 0,02685 | 0,179232 |
| Slc19a1 | 16,8452 | 12,7703 | -0,399545 | 0,0268 | 0,179232 |
| Zbtb6 | 7,70611 | 5,82421 | -0,403939 | 0,02685 | 0,179232 |
| Trim25 | 3,90394 | 2,86257 | -0,447617 | 0,02685 | 0,179232 |
| Bmpr1b | 3,54142 | 2,56559 | -0,465036 | 0,02685 | 0,179232 |
| Exo1 | 22,4216 | 28,6711 | 0,354708 | 0,0269 | 0,179477 |
| Tesk2 | 7,08284 | 11,1857 | 0,65925 | 0,02695 | 0,17972 |
| Srxn1 | 7,87716 | 5,77811 | -0,447077 | 0,027 | 0,179874 |
| Zfp120 | 3,7676 | 2,70177 | -0,479743 | 0,027 | 0,179874 |
| Hoxd1 | 19,4748 | 25,6217 | 0,395757 | 0,02705 | 0,180118 |
| Peg10 | 72,2687 | 56,3018 | -0,360192 | 0,0271 | 0,180361 |
| Cpa6 | 1,76018 | 2,84885 | 0,694656 | 0,0272 | 0,180846 |
| Calu | 181,758 | 141,962 | -0,356513 | 0,0272 | 0,180846 |
| Cacng4 | 2,3978 | 4,22004 | 0,815542 | 0,02725 | 0,180999 |
| Rprm | 6,02823 | 8,73532 | 0,535126 | 0,02725 | 0,180999 |
| Tprkb | 21,3222 | 27,7484 | 0,380052 | 0,02735 | 0,181482 |
| Mamdc2 | 0,731048 | 0,385116 | -0,924675 | 0,02735 | 0,181482 |
| Tnfrsf14 | 1,6492 | 0,560874 | -1,55602 | 0,02745 | 0,182055 |

| | | | | | |
|---|---|---|---|---|---|
| Pcyox1 | 30,8699 | 24,2087 | -0,350675 | 0,0275 | 0,182297 |
| Gli1 | 17,2168 | 13,1874 | -0,384659 | 0,0276 | 0,182869 |
| H2-Q7,H2-Q9 | 5,82386 | 9,10438 | 0,644585 | 0,0277 | 0,182987 |
| Rbm17 | 80,784 | 103,276 | 0,354362 | 0,0277 | 0,182987 |
| Map3k1 | 7,67044 | 5,8907 | -0,38087 | 0,0277 | 0,182987 |
| Scarb2 | 14,7626 | 11,1061 | -0,410596 | 0,02765 | 0,182987 |
| Cflar | 3,40469 | 2,53961 | -0,422916 | 0,02765 | 0,182987 |
| St6galnac2 | 2,22022 | 1,32659 | -0,742978 | 0,0277 | 0,182987 |
| Cbl | 14,3531 | 18,4481 | 0,362109 | 0,0278 | 0,183376 |
| Csnk2b | 104,409 | 133,574 | 0,35539 | 0,0278 | 0,183376 |
| Ppm1h | 1,13172 | 0,748236 | -0,596958 | 0,0278 | 0,183376 |
| A630007B06Rik | 17,1315 | 22,2152 | 0,374898 | 0,028 | 0,184513 |
| Cldn26 | 1,64455 | 0,795727 | -1,04735 | 0,028 | 0,184513 |
| Net1 | 22,7437 | 17,679 | -0,363432 | 0,02805 | 0,184751 |
| Cep112 | 3,38915 | 4,69939 | 0,471548 | 0,0281 | 0,184989 |
| Junb | 3,96229 | 5,81255 | 0,552835 | 0,02815 | 0,185045 |
| Parp6 | 7,5645 | 10,1774 | 0,428052 | 0,02815 | 0,185045 |
| Esyt1 | 10,3269 | 7,83396 | -0,398591 | 0,02815 | 0,185045 |
| Rnf34 | 38,2753 | 49,2292 | 0,363103 | 0,0282 | 0,185101 |
| Emc10 | 45,7683 | 35,5356 | -0,365082 | 0,0282 | 0,185101 |
| Sash3 | 0,77114 | 0,053929 | -3,83786 | 0,0282 | 0,185101 |
| Rasl2-9 | 4,01447 | 6,38895 | 0,67037 | 0,02825 | 0,185156 |
| Pomgnt2 | 4,52249 | 3,08676 | -0,551024 | 0,02825 | 0,185156 |
| Abhd3 | 1,43526 | 0,744601 | -0,94677 | 0,02825 | 0,185156 |
| Acat2 | 52,7331 | 67,3936 | 0,353902 | 0,0283 | 0,185302 |
| Fam122b | 5,30697 | 3,91014 | -0,44067 | 0,0283 | 0,185302 |
| Styk1 | 0,760939 | 0,400262 | -0,926836 | 0,0284 | 0,185866 |
| Ccdc174 | 14,7516 | 19,644 | 0,413218 | 0,0285 | 0,186338 |
| Gm5176 | 3,66157 | 1,45419 | -1,33225 | 0,0285 | 0,186338 |
| Lmo1 | 1,61823 | 2,98317 | 0,882431 | 0,02855 | 0,186391 |
| Pcnp | 26,1262 | 20,1255 | -0,376472 | 0,02855 | 0,186391 |
| Foxh1 | 0,799685 | 0,166285 | -2,26578 | 0,02855 | 0,186391 |
| Snord82 | 0 | 304,351 | inf | 0,0286 | 0,186626 |
| Lgalsl | 11,2288 | 8,5302 | -0,396548 | 0,02865 | 0,186861 |
| Rab11fip4 | 7,71657 | 5,71099 | -0,43422 | 0,0287 | 0,187096 |
| Tmem127 | 20,6429 | 16,1051 | -0,358125 | 0,0288 | 0,187565 |
| Bmp6 | 0,891018 | 0,432312 | -1,04338 | 0,0288 | 0,187565 |
| Rps20 | 541,884 | 688,835 | 0,346174 | 0,02885 | 0,187708 |
| Trove2 | 9,55445 | 7,45489 | -0,357986 | 0,02885 | 0,187708 |
| Nmrk1 | 1,25806 | 0,59426 | -1,08203 | 0,0289 | 0,187941 |
| Cdc45 | 49,2777 | 62,9988 | 0,354389 | 0,02895 | 0,188175 |
| Slc25a16 | 7,10764 | 5,23659 | -0,440745 | 0,029 | 0,188408 |
| Rbpms2 | 14,4787 | 19,0957 | 0,399318 | 0,02915 | 0,189291 |
| Odf2 | 60,4354 | 77,0418 | 0,350248 | 0,0292 | 0,189431 |
| Pm20d1 | 4,33178 | 3,10204 | -0,481741 | 0,0292 | 0,189431 |
| Ybx1 | 169,868 | 216,791 | 0,351886 | 0,0293 | 0,189896 |
| Ldb2 | 6,13376 | 4,36521 | -0,490718 | 0,0293 | 0,189896 |
| C030046E11Rik | 13,6815 | 17,502 | 0,355294 | 0,0295 | 0,191007 |

| | | | | | |
|---|---|---|---|---|---|
| Tspan6 | 138,375 | 109,048 | -0,343618 | 0,0295 | 0,191007 |
| Gpi1 | 58,3012 | 74,1073 | 0,34609 | 0,0296 | 0,191561 |
| Crybg3 | 3,84835 | 2,50071 | -0,621903 | 0,02985 | 0,193086 |
| Tcp11l2 | 1,96295 | 1,2023 | -0,707226 | 0,0299 | 0,193316 |
| Tnfrsf10b | 2,88343 | 1,95915 | -0,55756 | 0,02995 | 0,193545 |
| Lpin1 | 1,70117 | 2,40772 | 0,501143 | 0,03005 | 0,194098 |
| Tmem170 | 5,81443 | 3,99484 | -0,541499 | 0,03015 | 0,19465 |
| Rcor2 | 90,7618 | 115,792 | 0,351373 | 0,0302 | 0,194784 |
| Zbtbd6 | 1,94133 | 1,2274 | -0,661434 | 0,0302 | 0,194784 |
| Bcl7c | 20,3378 | 27,6417 | 0,442683 | 0,03025 | 0,194919 |
| Tmem184b | 24,6835 | 19,265 | -0,357565 | 0,03025 | 0,194919 |
| Gtf2h3 | 32,9082 | 42,0168 | 0,35252 | 0,0303 | 0,195053 |
| Rps11 | 780,509 | 994,603 | 0,349706 | 0,0303 | 0,195053 |
| Eps8 | 4,43881 | 3,22818 | -0,459453 | 0,03035 | 0,19528 |
| Rpl11 | 5,1146 | 8,71954 | 0,769632 | 0,03045 | 0,19583 |
| 0610030E20Rik | 13,7833 | 10,6654 | -0,369972 | 0,0305 | 0,196057 |
| Meg3 | 64,8082 | 88,7456 | 0,4535 | 0,03055 | 0,196284 |
| Tjp3 | 1,51881 | 0,304406 | -2,31888 | 0,03065 | 0,196832 |
| Ppp1r9b | 24,6095 | 19,3095 | -0,349905 | 0,03075 | 0,197284 |
| Mfsd7b | 8,84515 | 6,76559 | -0,386671 | 0,03075 | 0,197284 |
| Znf41-ps | 0,593157 | 0,296436 | -1,00069 | 0,0308 | 0,19751 |
| Cpsf6 | 62,0553 | 48,7617 | -0,347806 | 0,03085 | 0,197736 |
| Col1a2 | 11,7673 | 15,1079 | 0,360523 | 0,031 | 0,198507 |
| Got1 | 33,3312 | 42,6342 | 0,355139 | 0,031 | 0,198507 |
| AI846148 | 23,4778 | 30,5728 | 0,380952 | 0,03105 | 0,198637 |
| Hyal1 | 4,13303 | 2,78203 | -0,571061 | 0,03105 | 0,198637 |
| Dusp6 | 108,494 | 137,56 | 0,342441 | 0,03115 | 0,198991 |
| Mfap3 | 29,0812 | 22,959 | -0,341026 | 0,03115 | 0,198991 |
| Zfp367 | 8,95229 | 6,77273 | -0,40252 | 0,03115 | 0,198991 |
| Smim1 | 3,61698 | 0,713771 | -2,34125 | 0,0312 | 0,199215 |
| Art5 | 6,42164 | 9,05738 | 0,496152 | 0,03125 | 0,199439 |
| Chrna7 | 3,59848 | 2,36684 | -0,604427 | 0,0313 | 0,199663 |
| Hist1h1a | 0,229712 | 1,92352 | 3,06585 | 0,03135 | 0,199791 |
| Ism2 | 1,71339 | 1,04166 | -0,71797 | 0,03135 | 0,199791 |
| B4galt1 | 20,5472 | 16,1042 | -0,351498 | 0,0314 | 0,200014 |
| Cnksr2 | 3,13632 | 4,20973 | 0,424653 | 0,03145 | 0,200142 |
| Dlg3 | 28,6365 | 36,5067 | 0,350306 | 0,03145 | 0,200142 |
| Mir142b | 3,53706 | 0 | -inf | 0,03155 | 0,200587 |
| Acot7 | 34,535 | 44,7214 | 0,372909 | 0,03155 | 0,200587 |
| Kctd13 | 6,20193 | 8,73589 | 0,494239 | 0,03175 | 0,201763 |
| Aacs | 13,7931 | 17,8838 | 0,374708 | 0,0318 | 0,201888 |
| Map1b | 10,3749 | 8,11554 | -0,354343 | 0,0318 | 0,201888 |
| Anks6 | 2,04023 | 2,95923 | 0,536489 | 0,0319 | 0,202235 |
| Cd99l2 | 14,7651 | 11,4598 | -0,365608 | 0,0319 | 0,202235 |
| Leprel4 | 18,2719 | 13,8497 | -0,399772 | 0,0319 | 0,202235 |
| Glce | 19,4984 | 15,2493 | -0,354618 | 0,03195 | 0,202456 |
| Piwil4 | 0,469573 | 0,90621 | 0,948495 | 0,03205 | 0,202897 |
| Mphosph6 | 36,5902 | 47,6843 | 0,382058 | 0,03205 | 0,202897 |

| | | | | | |
|---|---|---|---|---|---|
| Kat8 | 19,1532 | 25,6758 | 0,422819 | 0,0321 | 0,203118 |
| Gm17066 | 20,8354 | 13,1654 | -0,662282 | 0,03215 | 0,203338 |
| Slc16a1 | 60,2197 | 47,4336 | -0,344328 | 0,03225 | 0,203874 |
| H2-Bl | 1,10989 | 2,18094 | 0,974529 | 0,03235 | 0,204216 |
| Srsf4 | 66,7422 | 84,9261 | 0,347608 | 0,03235 | 0,204216 |
| Cdh10 | 0,726781 | 0,387406 | -0,907675 | 0,03235 | 0,204216 |
| Cfb | 0,878126 | 1,464 | 0,737413 | 0,03245 | 0,204557 |
| Ppp4r4 | 2,47404 | 3,48947 | 0,496137 | 0,03245 | 0,204557 |
| Srfbp1 | 27,7215 | 35,9543 | 0,375159 | 0,03245 | 0,204557 |
| Zfp703 | 15,4452 | 11,8927 | -0,377083 | 0,0325 | 0,204776 |
| Gm266 | 3,97636 | 2,4327 | -0,708891 | 0,03255 | 0,204994 |
| Tmx1 | 74,1796 | 52,5134 | -0,498338 | 0,03265 | 0,205527 |
| 5430421N21Rik | 1,32442 | 2,25909 | 0,770374 | 0,0327 | 0,205552 |
| Snhg5 | 103,063 | 131,428 | 0,350755 | 0,0327 | 0,205552 |
| Mcm4 | 93,5313 | 118,757 | 0,344492 | 0,0327 | 0,205552 |
| Lrrc8d | 9,39736 | 7,18834 | -0,386597 | 0,03275 | 0,205769 |
| Timp2 | 24,8086 | 19,4743 | -0,349272 | 0,0328 | 0,205986 |
| Sf3b4 | 21,4802 | 27,9618 | 0,380445 | 0,03285 | 0,206204 |
| Zfp459 | 0,912727 | 0,518335 | -0,816299 | 0,0329 | 0,20642 |
| Acaca | 4,73385 | 6,10698 | 0,367447 | 0,03295 | 0,206637 |
| Tmem204 | 5,52577 | 1,28638 | -2,10285 | 0,033 | 0,206854 |
| Gm12070 | 0,64025 | 1,29401 | 1,01514 | 0,0331 | 0,207383 |
| Uqcc2 | 105,906 | 138,431 | 0,386391 | 0,0332 | 0,207718 |
| Cd1d1 | 3,75298 | 2,44117 | -0,620464 | 0,0332 | 0,207718 |
| Lmx1b | 0,962274 | 0,279423 | -1,784 | 0,0332 | 0,207718 |
| Zbtb8a | 3,0868 | 4,49862 | 0,543369 | 0,03325 | 0,207836 |
| Tspan14 | 53,9457 | 42,5552 | -0,342172 | 0,03325 | 0,207836 |
| Zcchc24 | 15,1836 | 11,8463 | -0,358082 | 0,0333 | 0,207857 |
| Apela | 79,9064 | 62,1308 | -0,363003 | 0,0333 | 0,207857 |
| Cd82 | 3,49575 | 2,26747 | -0,624516 | 0,0333 | 0,207857 |
| Chchd5 | 10,6104 | 14,6374 | 0,464185 | 0,03335 | 0,207975 |
| Cldn11 | 0,859206 | 0,376902 | -1,18881 | 0,03335 | 0,207975 |
| Kcns3 | 0,62605 | 1,09813 | 0,810701 | 0,0334 | 0,208092 |
| Cecr5 | 14,5903 | 19,1797 | 0,394574 | 0,0334 | 0,208092 |
| Eif3h | 147,694 | 186,934 | 0,339915 | 0,03345 | 0,208307 |
| Dcbld1 | 9,05134 | 6,78878 | -0,414979 | 0,03355 | 0,208832 |
| Il17rd | 41,3149 | 52,5059 | 0,345817 | 0,0336 | 0,208949 |
| Cldn25 | 47,1296 | 37,0677 | -0,346473 | 0,0336 | 0,208949 |
| Arpc5 | 51,7767 | 40,5374 | -0,353049 | 0,03365 | 0,209162 |
| Apcdd1 | 29,9524 | 23,4673 | -0,35202 | 0,0338 | 0,209997 |
| Gpr27 | 0,758728 | 1,65663 | 1,12659 | 0,03385 | 0,21021 |
| Bbs4 | 8,32285 | 11,0437 | 0,408073 | 0,03395 | 0,210635 |
| Herc3 | 3,6568 | 2,42986 | -0,589708 | 0,03395 | 0,210635 |
| Slc39a1 | 20,028 | 15,4636 | -0,37314 | 0,03405 | 0,211157 |
| Dcaf12l1 | 1,11435 | 0,676009 | -0,721087 | 0,0341 | 0,211369 |
| Shroom4 | 6,01752 | 7,85381 | 0,384225 | 0,03415 | 0,211581 |
| Spon2 | 1,46767 | 0,849959 | -0,788064 | 0,0342 | 0,211793 |
| Tro | 24,3025 | 30,8367 | 0,343542 | 0,03425 | 0,212004 |

| | | | | | |
|---|---|---|---|---|---|
| Parp8 | 8,89614 | 11,6376 | 0,387542 | 0,0343 | 0,212019 |
| BC017643 | 20,2255 | 14,2103 | -0,509237 | 0,0343 | 0,212019 |
| Rhox5 | 1,93671 | 0,746801 | -1,37481 | 0,0343 | 0,212019 |
| Pax9 | 13,2556 | 10,0647 | -0,397299 | 0,03435 | 0,21223 |
| Wscd1 | 4,16993 | 2,91909 | -0,514506 | 0,0344 | 0,212441 |
| Zfp407 | 5,68202 | 7,33496 | 0,368384 | 0,03445 | 0,212651 |
| Kat2b | 0,999087 | 0,607797 | -0,71702 | 0,0345 | 0,212861 |
| Haus7 | 49,7381 | 63,3572 | 0,349157 | 0,03455 | 0,212875 |
| Cish | 1,73348 | 1,02296 | -0,76092 | 0,03455 | 0,212875 |
| Pamr1 | 0,757692 | 0,382539 | -0,986005 | 0,03455 | 0,212875 |
| Ppm1k | 2,18269 | 2,99933 | 0,458535 | 0,0346 | 0,213085 |
| Wasf3 | 9,68105 | 12,7055 | 0,392215 | 0,03465 | 0,213295 |
| 2700097O09Rik | 19,6191 | 26,0721 | 0,410247 | 0,0348 | 0,213922 |
| Nup93 | 64,4418 | 81,176 | 0,333057 | 0,0348 | 0,213922 |
| Creb3l2 | 12,3801 | 9,50647 | -0,381036 | 0,0348 | 0,213922 |
| Gng4 | 0,803351 | 0,41894 | -0,939289 | 0,03485 | 0,214131 |
| Sart1 | 26,3096 | 33,2762 | 0,338904 | 0,035 | 0,214363 |
| Ylpm1 | 25,7678 | 32,5371 | 0,336516 | 0,03495 | 0,214363 |
| Naa30 | 16,3292 | 12,818 | -0,349281 | 0,035 | 0,214363 |
| Tpbg | 10,2406 | 7,8407 | -0,385251 | 0,035 | 0,214363 |
| Slc36a1 | 3,75665 | 2,7832 | -0,432699 | 0,03495 | 0,214363 |
| Tuft1 | 3,40146 | 2,30125 | -0,563734 | 0,03495 | 0,214363 |
| Zfp759 | 1,67413 | 1,07702 | -0,636368 | 0,035 | 0,214363 |
| Atg16l1 | 14,8925 | 19,0405 | 0,354492 | 0,03525 | 0,215795 |
| Zfp143 | 15,2385 | 19,5036 | 0,356018 | 0,0353 | 0,215805 |
| Kcnn3 | 1,48328 | 1,04821 | -0,500861 | 0,0353 | 0,215805 |
| Oxsm | 2,86348 | 1,82674 | -0,6485 | 0,0353 | 0,215805 |
| Zfp184 | 11,4893 | 14,9421 | 0,379088 | 0,03535 | 0,215913 |
| Cdk5rap2 | 20,9371 | 26,4867 | 0,339209 | 0,03535 | 0,215913 |
| Cox7a1 | 1,60301 | 5,48042 | 1,77351 | 0,03545 | 0,215932 |
| Hoxa11 | 25,0691 | 45,1914 | 0,850137 | 0,03545 | 0,215932 |
| Morn4 | 20,1944 | 26,1777 | 0,374386 | 0,03545 | 0,215932 |
| Ube4a | 21,096 | 16,643 | -0,342054 | 0,03545 | 0,215932 |
| Tpcn1 | 16,44 | 12,8741 | -0,35273 | 0,0354 | 0,215932 |
| Prr13 | 19,1856 | 14,1223 | -0,442045 | 0,03545 | 0,215932 |
| Zfp423 | 18,3045 | 23,1948 | 0,3416 | 0,0355 | 0,216138 |
| Kars | 135,249 | 170,647 | 0,335398 | 0,03555 | 0,216343 |
| Mtnr1a | 0,733684 | 0,0950927 | -2,94775 | 0,03565 | 0,216853 |
| Col5a2 | 21,0133 | 16,585 | -0,341418 | 0,0357 | 0,21696 |
| N4bp3 | 7,87986 | 5,69491 | -0,468497 | 0,0357 | 0,21696 |
| Psmd4 | 132,185 | 166,457 | 0,332591 | 0,03575 | 0,216968 |
| Rcsd1 | 9,90992 | 7,43538 | -0,414467 | 0,03575 | 0,216968 |
| Mtcp1 | 11,7719 | 8,22428 | -0,51739 | 0,03575 | 0,216968 |
| Car11 | 1,44646 | 2,47836 | 0,776857 | 0,0358 | 0,217074 |
| Tm9sf1 | 32,7265 | 25,7198 | -0,347577 | 0,0358 | 0,217074 |
| Dock6 | 16,7618 | 21,1654 | 0,336535 | 0,03595 | 0,21749 |
| Kat7 | 21,0646 | 26,5482 | 0,33379 | 0,03595 | 0,21749 |
| Slc2a4rg-ps | 5,64427 | 4,10008 | -0,461133 | 0,03595 | 0,21749 |

| | | | | | |
|---|---|---|---|---|---|
| Cyp2j6 | 2,98373 | 2,08776 | -0,515159 | 0,03595 | 0,21749 |
| Serpinb9 | 0,749053 | 0,408257 | -0,87559 | 0,03595 | 0,21749 |
| Cspg4 | 1,04027 | 1,47269 | 0,501502 | 0,036 | 0,217596 |
| Sostdc1 | 1,25958 | 0,0727092 | -4,11466 | 0,036 | 0,217596 |
| Zfp819 | 0,415568 | 0,838539 | 1,01279 | 0,03615 | 0,218404 |
| Ccdc57 | 2,71667 | 3,79675 | 0,482923 | 0,0363 | 0,219013 |
| Atoh8 | 4,85254 | 6,68877 | 0,463002 | 0,0363 | 0,219013 |
| Tmem18 | 15,5281 | 12,0446 | -0,366491 | 0,0363 | 0,219013 |
| Tnfaip8 | 9,96543 | 7,62787 | -0,385651 | 0,03635 | 0,219215 |
| Gm5803 | 1,44764 | 2,60088 | 0,845304 | 0,0364 | 0,219418 |
| Zfp60 | 17,6154 | 13,8247 | -0,349593 | 0,03665 | 0,220825 |
| Zfc3h1 | 18,3531 | 14,531 | -0,336889 | 0,0367 | 0,221027 |
| Tpp1 | 17,8071 | 13,8793 | -0,35952 | 0,03675 | 0,221228 |
| Nsun3 | 2,62231 | 4,11506 | 0,650076 | 0,0368 | 0,22143 |
| Fbn2 | 60,896 | 47,8187 | -0,348772 | 0,03685 | 0,221531 |
| Galns | 3,79188 | 2,61644 | -0,535308 | 0,03685 | 0,221531 |
| Yipf5 | 25,8771 | 20,2004 | -0,357295 | 0,0369 | 0,221732 |
| Dbi | 298,806 | 376,459 | 0,33328 | 0,03695 | 0,221932 |
| Eif1 | 82,8589 | 104,813 | 0,339083 | 0,037 | 0,222033 |
| Hfm1 | 1,35787 | 0,740694 | -0,874392 | 0,037 | 0,222033 |
| Rabac1 | 24,3027 | 17,7877 | -0,450237 | 0,03705 | 0,222233 |
| Ppp2r2c | 0,683685 | 0,382455 | -0,838041 | 0,03715 | 0,222733 |
| Efcab1 | 1,76864 | 0,84319 | -1,06871 | 0,03725 | 0,223233 |
| Cln8 | 6,2152 | 4,805 | -0,371266 | 0,0373 | 0,223432 |
| Gna14 | 1,9123 | 2,76663 | 0,532822 | 0,0374 | 0,22373 |
| Dnajb1 | 33,7955 | 42,8255 | 0,341638 | 0,0374 | 0,22373 |
| Ppp1r12a | 32,0395 | 40,3775 | 0,333701 | 0,0374 | 0,22373 |
| Erp29 | 107,37 | 85,13 | -0,334855 | 0,0375 | 0,224128 |
| Aqp11 | 0,788985 | 0,230854 | -1,77302 | 0,0375 | 0,224128 |
| Anxa7 | 11,1879 | 8,53424 | -0,390601 | 0,03755 | 0,224326 |
| Dnmt3b | 18,215 | 23,1133 | 0,343597 | 0,0376 | 0,224424 |
| Fchsd1 | 1,88007 | 1,26596 | -0,570554 | 0,0376 | 0,224424 |
| Mrpl15 | 24,8297 | 32,0494 | 0,368229 | 0,03765 | 0,224623 |
| Sgpp1 | 4,86901 | 3,55768 | -0,452688 | 0,0377 | 0,224821 |
| Ier3 | 2,39221 | 3,96447 | 0,728787 | 0,03775 | 0,225018 |
| Lefty1 | 0,727163 | 1,40628 | 0,951535 | 0,0379 | 0,225812 |
| Gstt1 | 0,674954 | 1,57068 | 1,21853 | 0,03795 | 0,226 |
| Gpr133 | 0,47276 | 0,769124 | 0,702109 | 0,03805 | 0,226 |
| Fam161a | 1,56794 | 2,48516 | 0,66447 | 0,03805 | 0,226 |
| Nyap1 | 4,14393 | 5,58282 | 0,429996 | 0,03805 | 0,226 |
| Zfp566 | 10,4228 | 13,9477 | 0,420281 | 0,038 | 0,226 |
| Zfp322a | 20,9267 | 16,5708 | -0,336703 | 0,038 | 0,226 |
| BC020402 | 2,21386 | 1,39304 | -0,668332 | 0,03805 | 0,226 |
| Sash1 | 10,0368 | 7,88678 | -0,347793 | 0,0381 | 0,226096 |
| Mpzl2 | 16,1885 | 12,6027 | -0,361232 | 0,0381 | 0,226096 |
| Hdx | 2,18236 | 3,05565 | 0,485591 | 0,0382 | 0,226288 |
| Gstk1 | 13,0051 | 18,016 | 0,470208 | 0,0382 | 0,226288 |
| Helz | 10,2735 | 12,9649 | 0,335677 | 0,0382 | 0,226288 |

| | | | | | |
|---|---|---|---|---|---|
| Fkbp10 | 35,9871 | 26,2363 | -0,455914 | 0,0382 | 0,226288 |
| Acvr1b | 26,1936 | 20,6612 | -0,34229 | 0,0383 | 0,226679 |
| 1600012H06Rik | 13,7865 | 7,05303 | -0,966938 | 0,0383 | 0,226679 |
| Drosha | 52,3569 | 65,7332 | 0,328242 | 0,0384 | 0,22717 |
| Zfp397 | 26,0433 | 20,6682 | -0,3335 | 0,0385 | 0,227661 |
| Cdx1 | 5,41404 | 7,5525 | 0,480249 | 0,03855 | 0,227856 |
| Nelfa | 38,4242 | 48,4569 | 0,334686 | 0,0386 | 0,228051 |
| Tmem167 | 12,8516 | 10,2544 | -0,325701 | 0,0387 | 0,228339 |
| Tmem69 | 11,1122 | 8,41494 | -0,401114 | 0,0387 | 0,228339 |
| Hps3 | 5,53689 | 4,08963 | -0,437108 | 0,0387 | 0,228339 |
| Dnajc27 | 8,01602 | 10,3087 | 0,362907 | 0,03875 | 0,228432 |
| Rab8b | 31,9763 | 40,1391 | 0,328004 | 0,03875 | 0,228432 |
| Sema5a | 11,7863 | 9,36368 | -0,331967 | 0,0388 | 0,228626 |
| Osbpl10 | 5,55358 | 7,69637 | 0,47076 | 0,03885 | 0,228719 |
| Uchl1 | 40,4429 | 51,7184 | 0,354793 | 0,03885 | 0,228719 |
| Cyp26c1 | 1,17696 | 2,0473 | 0,798659 | 0,0389 | 0,228811 |
| Snrpn | 17,3655 | 22,4843 | 0,372692 | 0,0389 | 0,228811 |
| Lactb2 | 11,2454 | 8,36302 | -0,42724 | 0,03895 | 0,229005 |
| Cep78 | 29,5318 | 37,2651 | 0,335554 | 0,039 | 0,229097 |
| Myadm | 25,0292 | 19,7524 | -0,341584 | 0,039 | 0,229097 |
| Lrrfip2 | 14,2988 | 18,207 | 0,348605 | 0,0391 | 0,229584 |
| Atad1 | 61,2433 | 48,8541 | -0,326071 | 0,0393 | 0,230555 |
| Slc18b1 | 3,77734 | 2,65104 | -0,510811 | 0,0393 | 0,230555 |
| Gcnt2 | 9,95308 | 12,7434 | 0,356535 | 0,0395 | 0,231615 |
| Sall1 | 126,572 | 161,197 | 0,348876 | 0,03955 | 0,231615 |
| Frmd4b | 12,7117 | 16,0892 | 0,339929 | 0,03955 | 0,231615 |
| H3f3a | 28,7736 | 21,7667 | -0,402623 | 0,03955 | 0,231615 |
| Mrto4 | 38,3726 | 49,0351 | 0,353739 | 0,03965 | 0,231998 |
| Ttc7 | 1,32331 | 0,861856 | -0,618637 | 0,03965 | 0,231998 |
| Nptn | 46,1671 | 36,6641 | -0,332495 | 0,0397 | 0,232087 |
| Loxl1 | 0,69889 | 0,375433 | -0,89651 | 0,0397 | 0,232087 |
| Dock7 | 38,6401 | 48,637 | 0,331957 | 0,0398 | 0,23257 |
| Fip1l1 | 41,8874 | 52,8666 | 0,33584 | 0,03985 | 0,23276 |
| Ppp1r3b | 1,33458 | 0,865913 | -0,624094 | 0,0399 | 0,23295 |
| Fam129b | 17,3554 | 13,4261 | -0,370343 | 0,03995 | 0,233141 |
| Pcolce | 25,2823 | 19,3811 | -0,383479 | 0,04 | 0,23333 |
| Pmvk | 26,6773 | 34,5524 | 0,373174 | 0,04005 | 0,23352 |
| Pkm | 575,253 | 750,895 | 0,384415 | 0,0401 | 0,233608 |
| Fam45a | 8,77431 | 6,55546 | -0,420589 | 0,0401 | 0,233608 |
| Zfp109 | 3,10081 | 2,1343 | -0,538879 | 0,04015 | 0,233797 |
| Cdipt | 28,1227 | 22,0401 | -0,351608 | 0,0402 | 0,233986 |
| Rasl11a | 8,52635 | 11,9252 | 0,484009 | 0,04025 | 0,234074 |
| Smek2 | 28,7463 | 22,8672 | -0,330097 | 0,04025 | 0,234074 |
| Cdk11b | 63,5999 | 79,9742 | 0,330511 | 0,0403 | 0,234161 |
| Elmo1 | 2,89495 | 2,12439 | -0,446488 | 0,0403 | 0,234161 |
| Cep131 | 17,2308 | 21,775 | 0,337689 | 0,04035 | 0,234349 |
| Mical2 | 0,851663 | 1,23994 | 0,541914 | 0,0404 | 0,234436 |
| Ppip5k1 | 2,21827 | 3,16515 | 0,512842 | 0,0404 | 0,234436 |

| | | | | | |
|---|---|---|---|---|---|
| Adamts12 | 2,26606 | 1,64853 | -0,459005 | 0,04045 | 0,234624 |
| Usf1 | 26,0847 | 33,3906 | 0,356236 | 0,0406 | 0,23529 |
| Tppp3 | 8,10833 | 5,4348 | -0,577178 | 0,0406 | 0,23529 |
| Fam50a | 32,8234 | 41,9309 | 0,353289 | 0,04065 | 0,235478 |
| Gm5124 | 4,37608 | 6,15374 | 0,491824 | 0,04085 | 0,236022 |
| Thumpd3 | 39,8105 | 51,1765 | 0,36233 | 0,04085 | 0,236022 |
| Sfrp1 | 9,36724 | 11,9816 | 0,355126 | 0,0408 | 0,236022 |
| Pop1 | 17,6511 | 22,3397 | 0,339849 | 0,04085 | 0,236022 |
| Dixdc1 | 15,7026 | 19,7966 | 0,334253 | 0,0408 | 0,236022 |
| Idi1 | 2,7146 | 1,84972 | -0,553428 | 0,0408 | 0,236022 |
| Slc25a14 | 6,8112 | 4,71685 | -0,530086 | 0,0409 | 0,236209 |
| Anxa4 | 4,96563 | 6,89467 | 0,473503 | 0,041 | 0,23648 |
| Gdf11 | 52,8438 | 66,4881 | 0,33136 | 0,041 | 0,23648 |
| Dnaja4 | 2,58461 | 1,77387 | -0,54305 | 0,041 | 0,23648 |
| Ddhd1 | 4,6487 | 6,04493 | 0,378899 | 0,04105 | 0,236666 |
| Ldha | 866,792 | 1127,13 | 0,378896 | 0,04115 | 0,236936 |
| Flnb | 25,7807 | 20,5612 | -0,326363 | 0,04115 | 0,236936 |
| Apom | 1,99188 | 0,734629 | -1,43904 | 0,04115 | 0,236936 |
| Ncapg2 | 23,8562 | 19,0396 | -0,325363 | 0,0412 | 0,237122 |
| Tjap1 | 28,4433 | 35,8318 | 0,333152 | 0,0413 | 0,237391 |
| Rassf3 | 11,7855 | 9,11511 | -0,370679 | 0,0413 | 0,237391 |
| Zfand4 | 2,94877 | 2,05647 | -0,519944 | 0,0413 | 0,237391 |
| Agpat9 | 7,06123 | 9,31009 | 0,398876 | 0,04135 | 0,237474 |
| Rasgrp1 | 1,21838 | 0,805969 | -0,59617 | 0,04135 | 0,237474 |
| Hoxa1 | 13,7145 | 17,619 | 0,361427 | 0,04145 | 0,237946 |
| Ftl1 | 240,226 | 301,447 | 0,327508 | 0,0415 | 0,238028 |
| Zfpm1 | 8,92408 | 6,82991 | -0,385837 | 0,0415 | 0,238028 |
| Shmt1 | 42,4163 | 53,2832 | 0,32906 | 0,0416 | 0,238193 |
| Zadh2 | 15,3469 | 12,0017 | -0,354705 | 0,04155 | 0,238193 |
| H2-D1 | 16,5588 | 12,5921 | -0,395087 | 0,0416 | 0,238193 |
| Srd5a3 | 12,4646 | 9,31587 | -0,42007 | 0,0416 | 0,238193 |
| Gmfb | 54,0632 | 43,0896 | -0,327307 | 0,04165 | 0,238275 |
| Caskin2 | 10,4147 | 8,14721 | -0,354238 | 0,04165 | 0,238275 |
| Zfp229 | 5,37706 | 4,07487 | -0,400063 | 0,04175 | 0,238745 |
| Bccip | 92,6717 | 118,135 | 0,350239 | 0,04185 | 0,239214 |
| Mpzl1 | 93,0961 | 74,6477 | -0,318623 | 0,04195 | 0,239684 |
| Ngfrap1 | 235,396 | 295,284 | 0,327015 | 0,042 | 0,239867 |
| Vstm2b | 25,1392 | 32,3498 | 0,363815 | 0,04205 | 0,239947 |
| Ssr3 | 214,935 | 170,918 | -0,330598 | 0,04205 | 0,239947 |
| Zbtb37 | 2,83625 | 3,8794 | 0,451849 | 0,0421 | 0,240027 |
| Mid2 | 1,60626 | 1,12645 | -0,511918 | 0,0421 | 0,240027 |
| Irak2 | 2,86993 | 1,99031 | -0,528021 | 0,04225 | 0,240677 |
| Tspan4 | 11,6953 | 6,94017 | -0,752882 | 0,04225 | 0,240677 |
| Slc12a2 | 22,3114 | 17,7642 | -0,328806 | 0,0423 | 0,240859 |
| Kdm7a | 15,9783 | 19,9945 | 0,323483 | 0,04235 | 0,240939 |
| Srsf3 | 126,841 | 101,499 | -0,321558 | 0,04235 | 0,240939 |
| Arhgef25 | 43,806 | 54,8049 | 0,323177 | 0,0424 | 0,241121 |
| Aven | 32,1178 | 41,6369 | 0,374493 | 0,0425 | 0,241587 |

| | | | | | |
|---|---|---|---|---|---|
| Pax6 | 0,522018 | 0,937049 | 0,844026 | 0,0426 | 0,242052 |
| D630033O11Rik | 0,735679 | 2,21751 | 1,59179 | 0,04275 | 0,242595 |
| Dnajb4 | 10,9894 | 8,40244 | -0,387233 | 0,04275 | 0,242595 |
| Stat5a | 0,59147 | 0,318465 | -0,893171 | 0,04275 | 0,242595 |
| Kirrel3 | 2,24073 | 3,12643 | 0,480549 | 0,04285 | 0,243059 |
| Anapc13 | 126,014 | 162,468 | 0,366573 | 0,0429 | 0,24324 |
| Polr2d | 101,8 | 127,674 | 0,32673 | 0,04295 | 0,24342 |
| Zbtb44 | 5,09271 | 6,45241 | 0,341405 | 0,043 | 0,2436 |
| Gpc6 | 15,4344 | 12,2822 | -0,329579 | 0,0431 | 0,244063 |
| Zfand6 | 55,9206 | 44,5158 | -0,329063 | 0,04315 | 0,244139 |
| Ctbs | 2,98813 | 1,94127 | -0,622247 | 0,04315 | 0,244139 |
| Aco1 | 28,7975 | 36,1056 | 0,326278 | 0,04325 | 0,244394 |
| Man2b2 | 4,54883 | 3,33547 | -0,447604 | 0,04325 | 0,244394 |
| Pfdn5 | 150,562 | 189,644 | 0,332935 | 0,04325 | 0,244395 |
| Mir150 | 0 | 487,817 | inf | 0,0434 | 0,244725 |
| Zfp647 | 9,54974 | 12,4992 | 0,388299 | 0,0434 | 0,244725 |
| Khdrbs3 | 35,1093 | 44,2788 | 0,334763 | 0,0434 | 0,244725 |
| Srm | 122,758 | 153,538 | 0,32278 | 0,0434 | 0,244725 |
| Cdh2 | 52,8617 | 65,8713 | 0,317426 | 0,0434 | 0,244725 |
| AI450353 | 4,33017 | 2,15958 | -1,00367 | 0,04345 | 0,244903 |
| Pmf1 | 71,9688 | 90,6808 | 0,333425 | 0,04355 | 0,245363 |
| Armcx3 | 1,56723 | 1,0207 | -0,618654 | 0,04365 | 0,245823 |
| Bzrap1 | 1,10232 | 0,763817 | -0,529244 | 0,04375 | 0,246179 |
| Spag8 | 0,753009 | 0,302673 | -1,31491 | 0,04375 | 0,246179 |
| Ahsa1 | 157,556 | 197,137 | 0,323331 | 0,0438 | 0,246253 |
| Gbp9 | 1,06187 | 0,655467 | -0,696009 | 0,0438 | 0,246253 |
| Emb | 18,5039 | 14,4247 | -0,359288 | 0,04385 | 0,24643 |
| Cep152 | 7,35785 | 9,37791 | 0,349984 | 0,0439 | 0,246608 |
| Rln1 | 1,9453 | 3,73309 | 0,940374 | 0,04395 | 0,246785 |
| Bms1 | 45,604 | 56,887 | 0,318938 | 0,044 | 0,246962 |
| Mfsd8 | 2,782 | 1,92482 | -0,531396 | 0,04405 | 0,247139 |
| 4930451G09Rik | 0,873946 | 1,48808 | 0,767836 | 0,04425 | 0,248157 |
| Slc35e3 | 5,94683 | 4,48122 | -0,40823 | 0,0443 | 0,248333 |
| Mlf1 | 10,0209 | 13,8099 | 0,46269 | 0,04435 | 0,248405 |
| Rgcc | 1,39253 | 0,537489 | -1,3734 | 0,04435 | 0,248405 |
| C2cd2l | 3,34124 | 2,45489 | -0,444724 | 0,0445 | 0,24914 |
| A930006K02Rik | 1,76424 | 0,700889 | -1,33179 | 0,0446 | 0,249596 |
| Dzip1l | 13,0161 | 16,4704 | 0,339581 | 0,04465 | 0,249666 |
| Ung | 28,2053 | 35,6254 | 0,336941 | 0,04465 | 0,249666 |
| Rhbg | 14,0555 | 18,2116 | 0,373726 | 0,0447 | 0,249737 |
| C1qbp | 243,754 | 304,453 | 0,320793 | 0,0447 | 0,249737 |
| Pigk | 15,4683 | 12,2907 | -0,331752 | 0,0448 | 0,250191 |
| Awat2 | 0,502021 | 1,0313 | 1,03865 | 0,04505 | 0,251167 |
| Tmem128 | 70,0413 | 55,3048 | -0,3408 | 0,04505 | 0,251167 |
| Lrrn1 | 14,05 | 11,0539 | -0,346004 | 0,045 | 0,251167 |
| Nrcam | 1,13529 | 0,794697 | -0,514579 | 0,04505 | 0,251167 |
| Zfp606 | 13,0537 | 10,1726 | -0,359768 | 0,0451 | 0,251341 |
| Ciz1 | 22,143 | 27,9036 | 0,3336 | 0,04525 | 0,251653 |

| | | | | | |
|---|---|---|---|---|---|
| Pqbp1 | 70,5871 | 88,6927 | 0,329411 | 0,0452 | 0,251653 |
| Notch2 | 16,8657 | 13,475 | -0,32381 | 0,04525 | 0,251653 |
| Myo10 | 10,6555 | 8,47318 | -0,330617 | 0,04525 | 0,251653 |
| Iba57 | 4,43854 | 3,26411 | -0,443395 | 0,04525 | 0,251653 |
| Zfp455 | 0,785922 | 0,421257 | -0,899687 | 0,0453 | 0,251826 |
| 2610507I01Rik | 5,64454 | 4,17524 | -0,434997 | 0,04535 | 0,251999 |
| Hoxa5 | 46,4259 | 66,3346 | 0,51483 | 0,0454 | 0,252067 |
| Csdc2 | 2,84623 | 4,04387 | 0,506686 | 0,0454 | 0,252067 |
| Mboat1 | 1,32917 | 0,829541 | -0,680138 | 0,0455 | 0,252517 |
| 1500012F01Rik | 94,6147 | 123,989 | 0,390075 | 0,04565 | 0,25272 |
| Atxn2l | 65,958 | 82,4894 | 0,322663 | 0,0456 | 0,25272 |
| Mtdh | 73,0567 | 58,674 | -0,316294 | 0,04565 | 0,25272 |
| Atrn | 8,92958 | 7,09906 | -0,330964 | 0,04565 | 0,25272 |
| Cyba | 49,6617 | 38,1665 | -0,379828 | 0,04565 | 0,25272 |
| Slc39a3 | 3,85575 | 2,80737 | -0,457794 | 0,04565 | 0,25272 |
| Fn3k | 0,830538 | 1,69872 | 1,03233 | 0,0457 | 0,252892 |
| Mmgt2 | 13,1626 | 9,98336 | -0,398848 | 0,0459 | 0,253894 |
| Rps27,Rps27rt | 3,734 | 6,53981 | 0,808526 | 0,04595 | 0,254065 |
| Gm11974 | 30,9861 | 41,5959 | 0,42482 | 0,046 | 0,254131 |
| 9530068E07Rik | 112,586 | 90,2412 | -0,319167 | 0,046 | 0,254131 |
| Fermt2 | 65,0448 | 81,0763 | 0,317847 | 0,0462 | 0,255131 |
| 2410004P03Rik | 0,377011 | 0,810399 | 1,10402 | 0,0463 | 0,255367 |
| Pfkl | 90,1806 | 112,584 | 0,320117 | 0,0463 | 0,255367 |
| Apoo | 2,0843 | 1,08843 | -0,937309 | 0,0463 | 0,255367 |
| B230219D22Rik | 59,0321 | 47,2645 | -0,320743 | 0,04635 | 0,255432 |
| Zfp160 | 14,2561 | 11,2327 | -0,343871 | 0,04635 | 0,255432 |
| Gm20748 | 3,59541 | 1,48487 | -1,27582 | 0,04645 | 0,255877 |
| Plekhh3 | 1,37963 | 0,865933 | -0,67195 | 0,0465 | 0,256047 |
| 3930402G23Rik | 0,425458 | 0,787998 | 0,889175 | 0,04655 | 0,256217 |
| St6gal1 | 17,7264 | 14,0518 | -0,335152 | 0,04665 | 0,256239 |
| Kremen1 | 11,2662 | 8,89581 | -0,340797 | 0,04665 | 0,256239 |
| Lrig2 | 7,66245 | 6,04943 | -0,341008 | 0,04665 | 0,256239 |
| Cmtm6 | 17,4403 | 13,7168 | -0,346477 | 0,0466 | 0,256239 |
| Tm6sf1 | 9,24438 | 5,03417 | -0,876823 | 0,0466 | 0,256239 |
| B130034C11Rik | 2,79777 | 1,69707 | -0,721226 | 0,0467 | 0,256409 |
| Rabl3 | 12,5658 | 16,2575 | 0,371611 | 0,0468 | 0,256852 |
| Fads6 | 0,886697 | 0,286041 | -1,63222 | 0,04685 | 0,257021 |
| Doc2g | 5,40427 | 7,63827 | 0,499145 | 0,0469 | 0,25719 |
| 03/06/16 | 32,2298 | 25,8549 | -0,317956 | 0,04695 | 0,257358 |
| Epha5 | 0,424045 | 0,649307 | 0,614684 | 0,0471 | 0,257441 |
| Atp13a2 | 11,7949 | 9,32469 | -0,339031 | 0,04705 | 0,257441 |
| Suox | 3,98533 | 2,79991 | -0,509319 | 0,0471 | 0,257441 |
| Gm20300 | 1,8624 | 1,25946 | -0,564355 | 0,0471 | 0,257441 |
| Dagla | 0,726213 | 0,459677 | -0,659772 | 0,04705 | 0,257441 |
| Acad12 | 0,738948 | 0,366183 | -1,01291 | 0,04705 | 0,257441 |
| Acss3 | 0,759017 | 0,370842 | -1,03333 | 0,04705 | 0,257441 |
| Mrps21 | 120,048 | 154,759 | 0,366409 | 0,04725 | 0,25805 |
| Dxo | 41,0964 | 51,7066 | 0,331335 | 0,04725 | 0,25805 |

| | | | | | |
|---|---|---|---|---|---|
| Apba2 | 0,921206 | 1,42208 | 0,626403 | 0,04735 | 0,258174 |
| Psmd2 | 110,041 | 137,593 | 0,322365 | 0,04735 | 0,258174 |
| Msh2 | 61,9149 | 77,1913 | 0,318151 | 0,04735 | 0,258174 |
| Tmem42 | 7,58239 | 5,11921 | -0,566732 | 0,0473 | 0,258174 |
| Smarca5-ps | 3,77058 | 5,01576 | 0,411683 | 0,04745 | 0,258509 |
| Elmo2 | 16,9083 | 21,1319 | 0,32169 | 0,04745 | 0,258509 |
| Rorc | 0,60527 | 0,28936 | -1,06471 | 0,0475 | 0,258676 |
| Whrn | 6,69377 | 8,64613 | 0,369236 | 0,0477 | 0,259342 |
| Mcm8 | 16,628 | 20,9538 | 0,333601 | 0,0477 | 0,259342 |
| Cbx1 | 122,102 | 151,699 | 0,31312 | 0,04765 | 0,259342 |
| Tor3a | 2,56382 | 1,79008 | -0,518273 | 0,0477 | 0,259342 |
| Ocel1 | 0,633908 | 0,237336 | -1,41734 | 0,04775 | 0,259508 |
| Rras | 14,8146 | 19,9842 | 0,431839 | 0,0481 | 0,261092 |
| Asph | 29,0928 | 22,679 | -0,359303 | 0,0481 | 0,261092 |
| Gm4532 | 10,8291 | 7,19757 | -0,589325 | 0,0481 | 0,261092 |
| Prtg | 5,48695 | 6,90859 | 0,332388 | 0,04815 | 0,261257 |
| Thnsl2 | 2,42132 | 1,57341 | -0,621904 | 0,04825 | 0,261694 |
| Maged2 | 118,442 | 146,72 | 0,30889 | 0,04835 | 0,26213 |
| Usp28 | 19,6483 | 24,5396 | 0,320701 | 0,0484 | 0,262188 |
| Tmem30b | 1,86208 | 1,2443 | -0,581586 | 0,0484 | 0,262188 |
| Fzd7 | 43,4833 | 35,0218 | -0,312209 | 0,04845 | 0,262352 |
| Zswim5 | 4,32021 | 5,60161 | 0,374739 | 0,04855 | 0,262469 |
| Ssr1 | 42,4341 | 34,0979 | -0,315541 | 0,04855 | 0,262469 |
| Epb4.1l1 | 3,85795 | 2,95289 | -0,385709 | 0,04855 | 0,262469 |
| Mcoln2 | 0,719269 | 0,370354 | -0,957629 | 0,0485 | 0,262469 |
| Plac1 | 4,42958 | 2,80571 | -0,658803 | 0,04865 | 0,262903 |
| Egf | 0,635116 | 0,98302 | 0,6302 | 0,04915 | 0,265283 |
| Rora | 3,61965 | 5,10916 | 0,497237 | 0,04915 | 0,265283 |
| Dkk3 | 2,55199 | 1,76886 | -0,5288 | 0,04915 | 0,265283 |
| Pou2f2 | 0,666773 | 1,06428 | 0,674606 | 0,0492 | 0,265446 |
| Plag1 | 6,11557 | 7,83933 | 0,358244 | 0,0493 | 0,265878 |
| Id1 | 86,7404 | 108,801 | 0,326909 | 0,0494 | 0,265988 |
| Foxo4 | 56,8956 | 70,6558 | 0,31249 | 0,0494 | 0,265988 |
| Spcs3 | 47,8031 | 38,4299 | -0,314875 | 0,0494 | 0,265988 |
| Esm1 | 1,50528 | 0,056691 | -4,73077 | 0,04935 | 0,265988 |
| Mcm6 | 152,943 | 191,185 | 0,321973 | 0,04945 | 0,26615 |
| Nfic | 2,16977 | 1,59714 | -0,442054 | 0,0495 | 0,266312 |
| A230072C01Rik | 5,39693 | 3,67166 | -0,555709 | 0,04955 | 0,266473 |
| Gstm7 | 5,20979 | 7,45357 | 0,516706 | 0,0496 | 0,266528 |
| Gm16702 | 3,41376 | 4,87896 | 0,515211 | 0,0496 | 0,266528 |
| Foxn3 | 8,67589 | 11,2638 | 0,376612 | 0,04965 | 0,266689 |
| 4930447C04Rik | 2,08217 | 1,29603 | -0,683988 | 0,04975 | 0,267119 |
| Fam73a | 13,6628 | 10,9249 | -0,322632 | 0,0498 | 0,267173 |
| Gramd1c | 0,925065 | 0,558893 | -0,726983 | 0,0498 | 0,267173 |
| Bcat1 | 14,0064 | 17,4371 | 0,316075 | 0,04985 | 0,267334 |
| Setbp1 | 11,2016 | 13,9525 | 0,316811 | 0,0499 | 0,267494 |
| Vegfb | 29,1054 | 37,1396 | 0,351671 | 0,04995 | 0,267548 |
| Ctnnbl1 | 34,7408 | 43,2578 | 0,316332 | 0,04995 | 0,267548 |

| | | | | | |
|---|---|---|---|---|---|
| Lrp11 | 1,38713 | 0,898857 | -0,625937 | 0,05005 | 0,267976 |
| Utp14a | 31,1315 | 38,8703 | 0,320294 | 0,0501 | 0,268029 |
| Tmed4 | 48,5819 | 38,7824 | -0,325015 | 0,0501 | 0,268029 |
| Mpv17l2 | 9,54736 | 6,42843 | -0,570637 | 0,05015 | 0,268189 |
| Slc1a4 | 21,928 | 27,3656 | 0,319595 | 0,05025 | 0,268509 |
| Flywch1 | 39,6816 | 49,4222 | 0,316689 | 0,05025 | 0,268509 |
| Clspn | 30,1398 | 37,3591 | 0,309789 | 0,0503 | 0,268669 |
| Hoxb9 | 80,1829 | 100,027 | 0,319022 | 0,05035 | 0,268828 |
| Gabrb2 | 0,833679 | 1,17524 | 0,495396 | 0,0504 | 0,26888 |
| Mmgt1 | 30,0592 | 24,1074 | -0,318332 | 0,0504 | 0,26888 |
| Cpne5 | 0,68277 | 1,05187 | 0,623488 | 0,0505 | 0,269306 |
| Taf1d | 72,1659 | 102,936 | 0,512352 | 0,05075 | 0,270424 |
| Scmh1 | 10,5311 | 13,3719 | 0,344544 | 0,05075 | 0,270424 |
| Fbln7 | 2,47286 | 3,4624 | 0,485591 | 0,05085 | 0,270849 |
| Tmub1 | 8,5402 | 6,09261 | -0,487209 | 0,0509 | 0,271007 |
| Esyt2 | 15,4723 | 12,3813 | -0,321525 | 0,05095 | 0,271165 |
| Eef1e1 | 62,4747 | 78,6372 | 0,331939 | 0,05105 | 0,271373 |
| Adamts10 | 16,0816 | 20,1216 | 0,323341 | 0,05105 | 0,271373 |
| Micu3 | 3,04652 | 2,1716 | -0,488403 | 0,05105 | 0,271373 |
| Tmem64 | 9,79076 | 7,72288 | -0,342282 | 0,0511 | 0,271531 |
| A630089N07Rik | 4,73669 | 6,44638 | 0,444608 | 0,05125 | 0,27222 |
| Zfp14 | 0,69082 | 1,09882 | 0,669577 | 0,0514 | 0,272691 |
| Mki67 | 92,43 | 72,7705 | -0,345007 | 0,0514 | 0,272691 |
| Klk8 | 11,4891 | 8,45499 | -0,442393 | 0,0514 | 0,272691 |
| Gm15421 | 37,567 | 57,5972 | 0,61653 | 0,05145 | 0,272848 |
| Hyal2 | 61,4028 | 49,4237 | -0,313103 | 0,0515 | 0,273005 |
| Whsc1l1 | 17,5679 | 22,6079 | 0,36388 | 0,0516 | 0,273427 |
| Zbtb21 | 5,21463 | 6,65065 | 0,35093 | 0,0517 | 0,273787 |
| Greb1l | 62,2592 | 77,9449 | 0,324168 | 0,05175 | 0,273787 |
| Trak2 | 10,8799 | 8,67671 | -0,326452 | 0,05175 | 0,273787 |
| Eif2ak2 | 7,26216 | 5,65627 | -0,360548 | 0,05175 | 0,273787 |
| Reep5 | 24,1061 | 19,2409 | -0,325223 | 0,0519 | 0,274255 |
| Cpn1 | 15,1568 | 11,6357 | -0,381399 | 0,0519 | 0,274255 |
| Fam3a | 11,848 | 8,94319 | -0,405779 | 0,0519 | 0,274255 |
| Elavl3 | 0,344856 | 0,574756 | 0,736957 | 0,05215 | 0,275467 |
| Cpsf3 | 50,4235 | 62,5851 | 0,311722 | 0,0522 | 0,275623 |
| Rnf139 | 13,719 | 10,6943 | -0,359337 | 0,05225 | 0,275778 |
| Fam184a | 1,3924 | 1,99582 | 0,51941 | 0,05235 | 0,276087 |
| Katna1 | 32,0786 | 40,2069 | 0,325832 | 0,05235 | 0,276087 |
| Mfsd12 | 3,03096 | 1,82472 | -0,732101 | 0,0524 | 0,276242 |
| Clasrp | 26,1308 | 32,8234 | 0,328969 | 0,05245 | 0,276397 |
| Zfp658 | 3,26845 | 4,39748 | 0,42807 | 0,0525 | 0,276442 |
| Bmpr1a | 49,1192 | 39,6391 | -0,309362 | 0,0525 | 0,276442 |
| Zbtb40 | 4,31681 | 5,48887 | 0,346545 | 0,05265 | 0,276905 |
| Nucks1 | 99,8381 | 80,0991 | -0,317805 | 0,05265 | 0,276905 |
| Slc9a6 | 8,039 | 6,33217 | -0,344316 | 0,05265 | 0,276905 |
| Trp53bp1 | 31,6732 | 40,0377 | 0,338095 | 0,0528 | 0,277148 |
| Zfp521 | 16,1562 | 20,0452 | 0,311165 | 0,0528 | 0,277148 |

| | | | | | |
|---|---|---|---|---|---|
| Oxct1 | 67,7339 | 54,6277 | -0,310247 | 0,05275 | 0,277148 |
| Tgfb3 | 3,3409 | 2,41237 | -0,469787 | 0,05275 | 0,277148 |
| Apbb1ip | 1,19608 | 0,723463 | -0,725321 | 0,0528 | 0,277148 |
| Tmem203 | 9,67238 | 6,69468 | -0,530856 | 0,05285 | 0,277302 |
| Ffar4 | 0,886185 | 0,379578 | -1,22321 | 0,0529 | 0,277455 |
| Fmr1 | 39,0778 | 31,6074 | -0,30609 | 0,05295 | 0,277608 |
| Soat1 | 12,44 | 9,91714 | -0,326986 | 0,053 | 0,277762 |
| Fam21 | 20,6361 | 25,6331 | 0,312835 | 0,05325 | 0,278962 |
| Dzip1 | 14,4427 | 18,0681 | 0,323104 | 0,0533 | 0,279115 |
| Dync2li1 | 12,9632 | 17,0486 | 0,395231 | 0,05335 | 0,279158 |
| Unc50 | 34,6713 | 27,3581 | -0,341772 | 0,05335 | 0,279158 |
| Col9a2 | 1,85168 | 1,2151 | -0,607757 | 0,0534 | 0,279201 |
| Wnt9a | 0,97546 | 0,596122 | -0,710476 | 0,0534 | 0,279201 |
| Stard10 | 9,26627 | 6,62378 | -0,484335 | 0,0535 | 0,279614 |
| Actn3 | 2,46886 | 3,45993 | 0,486901 | 0,0536 | 0,279809 |
| Dph3 | 16,5286 | 21,3652 | 0,370296 | 0,0536 | 0,279809 |
| Naga | 10,5695 | 7,99994 | -0,401843 | 0,0536 | 0,279809 |
| Cyp4f15 | 0,74253 | 1,27428 | 0,779156 | 0,05365 | 0,27996 |
| Mir2861 | 0 | 99,8726 | inf | 0,0538 | 0,280087 |
| Ubc | 36,2629 | 45,0271 | 0,3123 | 0,0537 | 0,280087 |
| Lypla1 | 37,1437 | 29,833 | -0,316211 | 0,05375 | 0,280087 |
| Tmem206 | 5,12911 | 3,39441 | -0,595547 | 0,0538 | 0,280087 |
| Lurap1l | 1,02572 | 0,524373 | -0,967969 | 0,0538 | 0,280087 |
| 2810405F15Rik | 0,959769 | 0,443816 | -1,11273 | 0,0538 | 0,280087 |
| Gm16576 | 0,935693 | 0,622271 | -0,588492 | 0,05385 | 0,280238 |
| Phc1 | 26,2382 | 32,5163 | 0,309495 | 0,0539 | 0,28028 |
| Chn2 | 1,691 | 1,08904 | -0,634824 | 0,0539 | 0,28028 |
| Stab1 | 5,28304 | 0,423866 | -3,63969 | 0,05395 | 0,280431 |
| Vps53 | 13,0816 | 16,4954 | 0,334524 | 0,054 | 0,280582 |
| Usp35 | 1,34114 | 1,93855 | 0,531517 | 0,05405 | 0,280623 |
| Exoc8 | 3,17483 | 2,44236 | -0,378405 | 0,05405 | 0,280623 |
| Tubb2a-ps2 | 61,9191 | 78,857 | 0,348856 | 0,0541 | 0,280774 |
| Edn3 | 4,59262 | 3,39371 | -0,436456 | 0,0542 | 0,281184 |
| Tcta | 4,5657 | 3,1366 | -0,541637 | 0,05425 | 0,281334 |
| Slit1 | 0,692701 | 1,03198 | 0,575105 | 0,0543 | 0,281484 |
| Nabp2 | 59,6188 | 74,8308 | 0,327866 | 0,05435 | 0,281634 |
| Snord66 | 0 | 1452,68 | inf | 0,0544 | 0,281784 |
| Trappc6a | 21,8851 | 29,0197 | 0,407081 | 0,05445 | 0,281933 |
| 4930539J05Rik | 2,16727 | 4,49356 | 1,05198 | 0,0545 | 0,282083 |
| Fbxo25 | 11,4685 | 14,8615 | 0,373904 | 0,05475 | 0,283267 |
| Txndc15 | 32,784 | 25,9311 | -0,338308 | 0,05485 | 0,283675 |
| Akr1c19 | 1,48245 | 0,750587 | -0,981894 | 0,0549 | 0,283714 |
| Ly96 | 2,37467 | 0,93206 | -1,34923 | 0,0549 | 0,283714 |
| Hmgcs1 | 107,414 | 133,036 | 0,30864 | 0,05495 | 0,283753 |
| Edem3 | 16,0891 | 12,974 | -0,310457 | 0,05495 | 0,283753 |
| Zfp937 | 1,82364 | 1,34277 | -0,441603 | 0,05505 | 0,284159 |
| Ddx25 | 11,7075 | 15,1977 | 0,376419 | 0,0551 | 0,284198 |
| Herc2 | 12,1402 | 15,0598 | 0,31091 | 0,0551 | 0,284198 |

| | | | | | |
|---|---|---|---|---|---|
| Ercc3 | 26,6967 | 33,1922 | 0,314182 | 0,05515 | 0,284236 |
| Dcun1d1 | 4,81487 | 3,69717 | -0,381074 | 0,05515 | 0,284236 |
| Abca4 | 1,17725 | 0,830772 | -0,502896 | 0,0552 | 0,284384 |
| Cdc6 | 18,9794 | 23,4918 | 0,307722 | 0,05535 | 0,284937 |
| Ttc28 | 6,11763 | 4,90966 | -0,317349 | 0,05535 | 0,284937 |
| Susd3 | 0,749639 | 0,267949 | -1,48424 | 0,0554 | 0,285085 |
| Mcm3 | 136,409 | 168,957 | 0,308714 | 0,0555 | 0,28549 |
| Srgn | 2,58958 | 0,131622 | -4,29824 | 0,0557 | 0,285968 |
| Atp2c2 | 1,00197 | 0,0320013 | -4,96856 | 0,0557 | 0,285968 |
| Art4 | 2,08528 | 0,0443173 | -5,55622 | 0,0557 | 0,285968 |
| Pkhd1l1 | 0,987771 | 0,0077074 | -7,00179 | 0,0557 | 0,285968 |
| Rhag | 7,99024 | 0,0457442 | -7,4485 | 0,0557 | 0,285968 |
| Mospd1 | 9,75623 | 7,46662 | -0,385869 | 0,05585 | 0,286628 |
| 2610035D17Rik | 2,34056 | 3,45586 | 0,562192 | 0,056 | 0,287288 |
| Per3 | 3,73757 | 4,77933 | 0,354706 | 0,0561 | 0,28769 |
| Gucy1a2 | 4,48831 | 5,59162 | 0,317095 | 0,0562 | 0,287982 |
| Ank | 5,64669 | 4,30421 | -0,391656 | 0,0562 | 0,287982 |
| Dleu7 | 0,939568 | 0,408659 | | 0,05625 | 0,288128 |
| Depdc7 | 3,32188 | 2,21763 | -0,582981 | 0,0563 | 0,288273 |
| Pus7 | 22,3601 | 27,7316 | 0,310602 | 0,05635 | 0,288308 |
| Gmps | 44,228 | 54,6348 | 0,30486 | 0,05635 | 0,288308 |
| Hinfp | 11,1174 | 14,2497 | 0,358111 | 0,05645 | 0,288599 |
| Slc43a3 | 31,2726 | 25,1126 | -0,31649 | 0,05645 | 0,288599 |
| Trit1 | 12,8387 | 16,3915 | 0,352454 | 0,0566 | 0,289145 |
| Yipf4 | 19,2416 | 15,2122 | -0,339002 | 0,0566 | 0,289145 |
| Rps6ka5 | 4,34597 | 5,6161 | 0,369891 | 0,05665 | 0,289289 |
| Draxin | 2,66967 | 3,51493 | 0,396831 | 0,0568 | 0,289834 |
| Itgb8 | 5,85448 | 7,60811 | 0,377996 | 0,0568 | 0,289834 |
| Gtf2f2 | 37,865 | 47,1429 | 0,316175 | 0,0569 | 0,290012 |
| Zswim6 | 6,35465 | 5,00162 | -0,345417 | 0,0569 | 0,290012 |
| Rnft2 | 4,44902 | 3,38577 | -0,394002 | 0,0569 | 0,290012 |
| Wfdc2 | 13,3077 | 9,22562 | -0,528545 | 0,05695 | 0,290156 |
| Mnx1 | 6,77532 | 9,21628 | 0,443896 | 0,057 | 0,29019 |
| Ctbp2 | 49,3709 | 60,979 | 0,304651 | 0,057 | 0,29019 |
| Gstm6 | 1,66694 | 2,77341 | 0,734457 | 0,05705 | 0,290223 |
| Serpinb1a | 1,46441 | 0,870454 | -0,750479 | 0,05705 | 0,290223 |
| Stac2 | 0,391207 | 0,701973 | 0,843482 | 0,0571 | 0,290367 |
| Pam16 | 0,340346 | 1,89367 | 2,47611 | 0,05715 | 0,290511 |
| Acp1 | 15,6828 | 12,4505 | -0,332981 | 0,05725 | 0,290908 |
| 4930563E22Rik | 1,26479 | 0,577747 | -1,13039 | 0,0573 | 0,291052 |
| Pgm2 | 33,8783 | 41,9604 | 0,308666 | 0,05745 | 0,291703 |
| Fkbp7 | 7,70309 | 5,21236 | -0,563502 | 0,0576 | 0,292353 |
| Zfp711 | 5,86666 | 4,55195 | -0,366055 | 0,0577 | 0,292528 |
| Gm14378 | 2,57683 | 1,50215 | -0,778567 | 0,0577 | 0,292528 |
| 1810019D21Rik | 1,38842 | 0,0229431 | -5,91923 | 0,0577 | 0,292528 |
| Aldh9a1 | 14,443 | 11,387 | -0,342983 | 0,0578 | 0,292923 |
| Cdca7 | 101,438 | 124,84 | 0,299484 | 0,05785 | 0,292955 |
| Lypd6 | 12,8109 | 10,1318 | -0,338476 | 0,05785 | 0,292955 |

| | | | | | |
|---|---|---|---|---|---|
| 9430020K01Rik | 3,96881 | 3,0825 | -0,364608 | 0,0579 | 0,292986 |
| Ghdc | 3,77683 | 2,65637 | -0,507724 | 0,0579 | 0,292986 |
| Emp2 | 3,35481 | 2,46733 | -0,443283 | 0,058 | 0,29327 |
| Cysltr1 | 1,42847 | 0,911631 | -0,647951 | 0,058 | 0,29327 |
| Pop4 | 75,0005 | 60,359 | -0,313331 | 0,05815 | 0,293806 |
| Gm14005 | 0,845693 | 0,186533 | -2,18071 | 0,05815 | 0,293806 |
| Sec11c | 18,3972 | 13,41 | -0,456178 | 0,0582 | 0,293947 |
| Prodh | 7,17592 | 5,3839 | -0,414511 | 0,0583 | 0,294341 |
| Klhdc1 | 0,916095 | 0,479296 | -0,934579 | 0,05835 | 0,294482 |
| Lingo1 | 3,85507 | 5,11972 | 0,40931 | 0,0584 | 0,294623 |
| Fgf4 | 6,74425 | 8,72151 | 0,37092 | 0,05845 | 0,294683 |
| Lrrfip1 | 24,7587 | 30,731 | 0,311759 | 0,0585 | 0,294683 |
| Clip3 | 33,3635 | 41,1873 | 0,303929 | 0,0585 | 0,294683 |
| A730017C20Rik | 1,55203 | 0,942297 | -0,719905 | 0,0585 | 0,294683 |
| Lap3 | 65,1083 | 80,1921 | 0,300618 | 0,05855 | 0,294713 |
| Camk2a | 0,856237 | 0,544553 | -0,652937 | 0,05855 | 0,294713 |
| Serf1 | 84,0519 | 105,677 | 0,330305 | 0,0586 | 0,294853 |
| Tctex1d2 | 13,6261 | 19,2362 | 0,497449 | 0,0587 | 0,295245 |
| Trps1 | 11,8845 | 14,6537 | 0,302182 | 0,05875 | 0,295275 |
| Ran | 117,558 | 144,78 | 0,300486 | 0,05875 | 0,295275 |
| Cntrob | 17,7038 | 21,8863 | 0,30597 | 0,0588 | 0,295415 |
| Cdc5l | 25,4108 | 31,5113 | 0,310428 | 0,05885 | 0,295555 |
| Higd2a | 90,0965 | 112,754 | 0,32363 | 0,0589 | 0,295695 |
| Pick1 | 13,9749 | 10,8895 | -0,359907 | 0,05895 | 0,295835 |
| Plagl1 | 70,9961 | 57,4978 | -0,304232 | 0,05905 | 0,296225 |
| Pnma1 | 0,395487 | 0,739396 | 0,902715 | 0,05915 | 0,296504 |
| Zcchc7 | 10,7262 | 13,6683 | 0,349691 | 0,05915 | 0,296504 |
| Elf3 | 5,4291 | 7,29794 | 0,426778 | 0,0592 | 0,296644 |
| Rpl13 | 54,6234 | 68,8291 | 0,333499 | 0,05925 | 0,296783 |
| Dse | 14,4274 | 11,5449 | -0,321557 | 0,05935 | 0,297061 |
| Rrm2b | 1,88136 | 1,33723 | -0,492528 | 0,05935 | 0,297061 |
| Adam9 | 17,0585 | 13,7133 | -0,31492 | 0,0594 | 0,297089 |
| Mkrn3 | 2,40769 | 1,6426 | -0,551674 | 0,0594 | 0,297089 |
| Hoxb6 | 25,0293 | 31,7442 | 0,342877 | 0,05945 | 0,297228 |
| Slc25a1 | 39,649 | 49,197 | 0,311284 | 0,05955 | 0,297505 |
| Etnk1 | 26,3783 | 21,3995 | -0,301774 | 0,05955 | 0,297505 |
| Mfap4 | 53,9768 | 66,8179 | 0,307894 | 0,05965 | 0,297893 |
| Trim59 | 73,827 | 60,0733 | -0,297425 | 0,0597 | 0,298032 |
| Col17a1 | 0,908167 | 0,608921 | -0,576702 | 0,0598 | 0,298419 |
| Tango6 | 6,08359 | 7,73192 | 0,345902 | 0,05985 | 0,298446 |
| Rrp12 | 17,4529 | 21,608 | 0,308097 | 0,05985 | 0,298446 |
| Ccdc91 | 9,88058 | 12,6207 | 0,353122 | 0,05995 | 0,298833 |
| Mre11a | 23,9404 | 29,6029 | 0,306291 | 0,06 | 0,298971 |
| Ddx59 | 7,24062 | 9,41465 | 0,378795 | 0,06025 | 0,299324 |
| Cdk2ap1 | 18,3599 | 23,6487 | 0,365202 | 0,06015 | 0,299324 |
| Eif3c | 161,601 | 200,189 | 0,308925 | 0,06025 | 0,299324 |
| Ttf2 | 20,6516 | 25,5489 | 0,307006 | 0,0602 | 0,299324 |
| Cep250 | 12,6795 | 15,6091 | 0,299897 | 0,0601 | 0,299324 |

| | | | | | |
|---|---|---|---|---|---|
| Plin2 | 34,4035 | 27,5742 | -0,319236 | 0,0602 | 0,299324 |
| Lrif1 | 7,75511 | 5,68194 | -0,448765 | 0,0602 | 0,299324 |
| Tfeb | 0,58291 | 0,285498 | -1,02979 | 0,06025 | 0,299324 |
| Hebp1 | 31,0167 | 24,1316 | -0,362127 | 0,0603 | 0,299461 |
| Eif3l | 164,897 | 202,892 | 0,299144 | 0,0604 | 0,299735 |
| Gulp1 | 9,28675 | 7,24783 | -0,357624 | 0,0604 | 0,299735 |
| Klc4 | 14,8608 | 18,726 | 0,333531 | 0,0605 | 0,299897 |
| Rsrc2 | 78,7007 | 96,8689 | 0,299657 | 0,0605 | 0,299897 |
| Mzt1 | 33,8577 | 27,268 | -0,312278 | 0,0605 | 0,299897 |
| Slc25a42 | 1,29258 | 0,838419 | -0,624514 | 0,0606 | 0,300281 |
| Dennd2a | 29,6396 | 36,4637 | 0,298936 | 0,06065 | 0,300306 |
| Itgb5 | 51,6356 | 42,0312 | -0,296906 | 0,06065 | 0,300306 |
| B3gnt5 | 21,4991 | 17,4144 | -0,303998 | 0,0607 | 0,300443 |
| Gabpb1 | 40,404 | 50,2118 | 0,313528 | 0,06075 | 0,300579 |
| Lyrm9 | 1,96662 | 1,42087 | -0,46894 | 0,06085 | 0,300962 |
| Cercam | 10,2221 | 7,98637 | -0,356086 | 0,06095 | 0,301345 |
| Pnma2 | 0,532401 | 0,828939 | 0,638753 | 0,061 | 0,301481 |
| Nr4a2 | 1,30846 | 1,91939 | 0,552779 | 0,06125 | 0,302381 |
| Xpo1 | 160,488 | 129,125 | -0,313695 | 0,06125 | 0,302381 |
| Leprotl1 | 21,576 | 17,2836 | -0,320021 | 0,06125 | 0,302381 |
| Masp1 | 2,27185 | 1,55459 | -0,547336 | 0,06135 | 0,302763 |
| Bpgm | 36,9091 | 29,7105 | -0,313003 | 0,0614 | 0,302898 |
| Raet1e | 4,26122 | 2,4346 | -0,807582 | 0,06145 | 0,303032 |
| Gm2061 | 1,68284 | 0,0543541 | -4,95237 | 0,0615 | 0,303167 |
| Psmf1 | 12,9927 | 16,2374 | 0,321613 | 0,06175 | 0,304287 |
| Socs5 | 4,3868 | 3,36106 | -0,384251 | 0,0618 | 0,304421 |
| Cct2 | 329,31 | 408,618 | 0,311308 | 0,06185 | 0,304555 |
| Pdia5 | 33,6062 | 41,6898 | 0,310968 | 0,06195 | 0,304823 |
| Eda2r | 3,11427 | 2,21866 | -0,489204 | 0,06195 | 0,304823 |
| Gm6251 | 2,76069 | 7,36117 | 1,41491 | 0,06205 | 0,305203 |
| Zgpat | 24,0743 | 32,7141 | 0,442422 | 0,06215 | 0,305358 |
| Cbr1 | 45,9211 | 57,3818 | 0,321435 | 0,06215 | 0,305358 |
| Mllt6 | 22,9925 | 28,2315 | 0,29614 | 0,06215 | 0,305358 |
| Tango2 | 6,91394 | 4,9944 | -0,469197 | 0,06225 | 0,305736 |
| Slc31a1 | 32,1635 | 26,1355 | -0,299416 | 0,0623 | 0,30587 |
| Pygo1 | 2,76554 | 3,99182 | 0,529489 | 0,06245 | 0,306493 |
| Rpl7a | 72,4914 | 90,2137 | 0,315537 | 0,06255 | 0,306646 |
| Eps15l1 | 38,1785 | 47,154 | 0,30462 | 0,06255 | 0,306646 |
| Pigh | 13,738 | 10,0141 | -0,456141 | 0,06255 | 0,306646 |
| Tbc1d7 | 17,565 | 22,6054 | 0,363965 | 0,06295 | 0,308381 |
| Rab21 | 14,5587 | 11,3638 | -0,357446 | 0,06295 | 0,308381 |
| Zdhhc15 | 7,97119 | 6,34281 | -0,329674 | 0,06305 | 0,308758 |
| Apex1 | 192,434 | 239,931 | 0,318255 | 0,06315 | 0,309021 |
| Smyd2 | 71,3818 | 87,878 | 0,299946 | 0,06315 | 0,309021 |
| Wbp1l | 22,2545 | 18,037 | -0,303138 | 0,0632 | 0,309153 |
| Sec22b | 33,2711 | 41,2036 | 0,308501 | 0,06325 | 0,309284 |
| Sf3a3 | 116,547 | 142,869 | 0,293778 | 0,0633 | 0,309415 |
| Tmem66 | 43,3235 | 35,0337 | -0,306406 | 0,06335 | 0,309546 |

| | | | | | |
|---|---|---|---|---|---|
| 1700092M07Rik | 4,16667 | 8,16024 | 0,969714 | 0,06345 | 0,309826 |
| Hnrnpf | 27,1254 | 33,5916 | 0,308458 | 0,0635 | 0,309826 |
| Map3k9 | 0,790555 | 0,564658 | -0,48549 | 0,0635 | 0,309826 |
| Ankrd9 | 1,68982 | 1,10753 | -0,609521 | 0,0635 | 0,309826 |
| Calr3 | 1,38274 | 0,794109 | -0,800125 | 0,0636 | 0,310201 |
| BC026585 | 3,47608 | 2,26421 | -0,618453 | 0,06365 | 0,310331 |
| Snora43 | 231,295 | 26,7685 | -3,11112 | 0,06385 | 0,311193 |
| Celf4 | 1,37036 | 1,94087 | 0,502147 | 0,0639 | 0,31121 |
| Trim36 | 4,22823 | 5,43966 | 0,363462 | 0,0639 | 0,31121 |
| Dok2 | 4,03381 | 2,79223 | -0,530728 | 0,064 | 0,311583 |
| Nek4 | 11,0519 | 13,9631 | 0,337318 | 0,06405 | 0,3116 |
| Man1a2 | 20,682 | 16,8291 | -0,297411 | 0,06405 | 0,3116 |
| Adrb2 | 1,49358 | 0,937369 | -0,672082 | 0,0642 | 0,312216 |
| Zic5 | 2,82299 | 3,92735 | 0,476332 | 0,06425 | 0,312345 |
| Rnf41 | 12,3869 | 15,343 | 0,308758 | 0,06445 | 0,31309 |
| 09/15/16 | 173,629 | 141,467 | -0,295541 | 0,06445 | 0,31309 |
| Rab11a | 52,8882 | 42,9929 | -0,298847 | 0,06455 | 0,313462 |
| Zfp35 | 16,6312 | 20,6581 | 0,312812 | 0,0646 | 0,31359 |
| Pecr | 11,007 | 14,5859 | 0,406144 | 0,06465 | 0,313719 |
| Tbl1xr1 | 9,57386 | 7,76976 | -0,301232 | 0,0648 | 0,314219 |
| Slc2a3 | 269,896 | 215,861 | -0,322303 | 0,0648 | 0,314219 |
| Impad1 | 45,1189 | 36,7927 | -0,29431 | 0,06485 | 0,314347 |
| Zc3hav1l | 21,031 | 17,1159 | -0,297179 | 0,0649 | 0,314476 |
| Sfxn3 | 2,82775 | 2,02159 | -0,484167 | 0,06495 | 0,314604 |
| Cyp4f17 | 2,39344 | 3,51113 | 0,552847 | 0,0651 | 0,314774 |
| Ahi1 | 8,73294 | 10,9525 | 0,326718 | 0,06505 | 0,314774 |
| Fhod1 | 9,51427 | 7,54001 | -0,335526 | 0,06515 | 0,314774 |
| Znf512b | 10,5036 | 8,0888 | -0,376891 | 0,06515 | 0,314774 |
| 6430573F11Rik | 1,04105 | 0,569607 | -0,869999 | 0,0651 | 0,314774 |
| Fcgrt | 1,03027 | 0,545222 | -0,918105 | 0,06515 | 0,314774 |
| Tstd1 | 3,39599 | 1,62072 | -1,0672 | 0,06505 | 0,314774 |
| Nup85 | 86,7305 | 106,327 | 0,293899 | 0,0652 | 0,314902 |
| 3830406C13Rik | 3,32967 | 2,37196 | -0,489301 | 0,06525 | 0,315029 |
| Eepd1 | 2,74007 | 3,76177 | 0,457203 | 0,06545 | 0,315653 |
| Actr8 | 24,6795 | 30,5871 | 0,309613 | 0,06545 | 0,315653 |
| A930015D03Rik | 2,08975 | 0,559015 | -1,90237 | 0,06545 | 0,315653 |
| Crebbp | 8,78156 | 10,8312 | 0,302647 | 0,0655 | 0,31578 |
| Ccdc71l | 6,60645 | 5,16505 | -0,355093 | 0,06555 | 0,315793 |
| Kank4 | 1,42186 | 0,98266 | -0,533019 | 0,06555 | 0,315793 |
| 2610015P09Rik | 3,79737 | 4,9562 | 0,384234 | 0,0656 | 0,31592 |
| Cd2ap | 16,3188 | 13,2556 | -0,299932 | 0,06565 | 0,316047 |
| Hmga1 | 12,5397 | 28,6861 | 1,19385 | 0,0658 | 0,316213 |
| Kcna3 | 1,29532 | 2,00139 | 0,627698 | 0,0658 | 0,316213 |
| Uap1 | 28,9549 | 35,7611 | 0,304583 | 0,06585 | 0,316213 |
| Slc12a6 | 7,15576 | 5,34822 | -0,420044 | 0,06585 | 0,316213 |
| Cnnm2 | 2,67607 | 1,92704 | -0,473728 | 0,06585 | 0,316213 |
| Atp1b1 | 14,4516 | 10,3122 | -0,486874 | 0,06575 | 0,316213 |
| Camk2d | 1,83677 | 1,30799 | -0,489819 | 0,06575 | 0,316213 |

| | | | | | |
|---|---|---|---|---|---|
| Coprs | 45,4167 | 57,2805 | 0,334824 | 0,0659 | 0,316339 |
| Tanc2 | 3,76266 | 4,68661 | 0,316789 | 0,06595 | 0,316465 |
| Adamts20 | 6,88162 | 8,84271 | 0,36174 | 0,066 | 0,316478 |
| Elf1 | 7,0465 | 4,65288 | -0,598781 | 0,066 | 0,316478 |
| Rpl23 | 119,635 | 146,925 | 0,296441 | 0,06605 | 0,316491 |
| Ccng2 | 21,5451 | 17,2882 | -0,317569 | 0,06605 | 0,316491 |
| Fem1c | 18,9808 | 23,5659 | 0,312161 | 0,0661 | 0,316617 |
| Pdzk1 | 5,3519 | 3,97671 | -0,428474 | 0,06615 | 0,316743 |
| A230077H06Rik | 0,922567 | 1,41296 | 0,614995 | 0,0664 | 0,317031 |
| S1pr5 | 7,02753 | 9,08554 | 0,370554 | 0,06635 | 0,317031 |
| Prpf3 | 32,8598 | 40,4095 | 0,298369 | 0,0664 | 0,317031 |
| Sgce | 22,2526 | 17,6407 | -0,335065 | 0,0664 | 0,317031 |
| Agpat4 | 16,8639 | 13,317 | -0,340664 | 0,0664 | 0,317031 |
| Slc25a32 | 4,82283 | 3,63118 | -0,409441 | 0,06635 | 0,317031 |
| Irf8 | 1,49873 | 0,98843 | -0,600531 | 0,06635 | 0,317031 |
| Igtp | 2,12206 | 1,38965 | -0,610746 | 0,06635 | 0,317031 |
| 0610031J06Rik | 30,0209 | 23,9514 | -0,325854 | 0,06645 | 0,317156 |
| Trpm7 | 30,6108 | 25,6097 | -0,291989 | 0,0665 | 0,317282 |
| Ttll9 | 0,250783 | 0,573923 | 1,19442 | 0,06655 | 0,317306 |
| Fert2 | 8,9125 | 11,2197 | 0,332138 | 0,0666 | 0,317306 |
| Stx17 | 4,15216 | 3,23182 | -0,361514 | 0,0666 | 0,317306 |
| 2210404O09Rik | 1,38211 | 0,85601 | -0,691172 | 0,0666 | 0,317306 |
| Nudt5 | 59,1253 | 72,4898 | 0,294001 | 0,06665 | 0,317431 |
| Acbd6 | 29,4095 | 36,999 | 0,331207 | 0,0667 | 0,317556 |
| Tuba1a | 194,167 | 237,901 | 0,293059 | 0,06685 | 0,318043 |
| Lgi2 | 5,40857 | 3,93984 | -0,457113 | 0,06685 | 0,318043 |
| Mir219-1 | 4,27098 | 71,513 | 4,06557 | 0,067 | 0,318077 |
| Klhdc10 | 29,2254 | 23,8753 | -0,291702 | 0,06695 | 0,318077 |
| Tmem120a | 15,8692 | 12,1618 | -0,383865 | 0,0669 | 0,318077 |
| Map4k5 | 18,6227 | 13,3943 | -0,475441 | 0,06695 | 0,318077 |
| Tlr2 | 1,72842 | 1,16619 | -0,567647 | 0,067 | 0,318077 |
| Cyp2r1 | 0,860875 | 0,361468 | -1,25194 | 0,067 | 0,318077 |
| Cdkn3 | 14,0861 | 10,1231 | -0,47662 | 0,06705 | 0,318202 |
| Ankrd27 | 18,6467 | 23,2616 | 0,319032 | 0,0671 | 0,318326 |
| Pink1 | 10,6725 | 13,5074 | 0,339845 | 0,06715 | 0,318337 |
| Espn | 1,03441 | 0,497669 | -1,05555 | 0,06715 | 0,318337 |
| Wnt5b | 26,0092 | 36,4113 | 0,485365 | 0,0673 | 0,318371 |
| Nanos1 | 2,23348 | 2,99938 | 0,425367 | 0,0673 | 0,318371 |
| Pnrc1 | 18,6153 | 23,2862 | 0,322985 | 0,06725 | 0,318371 |
| Atg2b | 6,15399 | 7,6102 | 0,306412 | 0,0673 | 0,318371 |
| Stx2 | 9,65062 | 7,59646 | -0,345294 | 0,06725 | 0,318371 |
| Igsf1 | 2,52518 | 1,66223 | -0,603269 | 0,0673 | 0,318371 |
| Mical3 | 6,55593 | 8,35347 | 0,349575 | 0,06735 | 0,318494 |
| Srl | 8,14641 | 10,1457 | 0,316632 | 0,0674 | 0,318506 |
| Dhx8 | 32,225 | 39,5184 | 0,294345 | 0,0674 | 0,318506 |
| Commd10 | 26,2946 | 32,7415 | 0,316353 | 0,0675 | 0,318753 |
| Ppfibp1 | 51,3591 | 62,8001 | 0,290148 | 0,0675 | 0,318753 |
| Gnai1 | 13,7116 | 10,9742 | -0,321284 | 0,06755 | 0,318876 |

| | | | | | |
|---|---|---|---|---|---|
| Zfp748 | 6,42284 | 5,07074 | -0,341014 | 0,06765 | 0,319123 |
| 1700102H20Rik | 1,56189 | 0,347409 | -2,16858 | 0,06765 | 0,319123 |
| Ninl | 16,0434 | 19,6785 | 0,294636 | 0,0677 | 0,319133 |
| St8sia2 | 3,66488 | 2,8228 | -0,376639 | 0,0677 | 0,319133 |
| Fgf2 | 0,39764 | 1,32252 | 1,73375 | 0,06785 | 0,319165 |
| Dmtf1 | 36,6129 | 45,0226 | 0,298298 | 0,06785 | 0,319165 |
| Idh1 | 70,8263 | 86,4324 | 0,287288 | 0,0678 | 0,319165 |
| Col4a3bp | 12,1384 | 9,8291 | -0,304449 | 0,0678 | 0,319165 |
| Smpd1 | 13,6552 | 10,786 | -0,340295 | 0,0678 | 0,319165 |
| 2410018L13Rik | 0,669398 | 0,267309 | -1,32435 | 0,06785 | 0,319165 |
| Hoxa7 | 20,9127 | 26,1597 | 0,322971 | 0,06795 | 0,319298 |
| Ube2d2a | 27,1981 | 21,9672 | -0,308156 | 0,06795 | 0,319298 |
| Slc27a1 | 4,95408 | 3,72101 | -0,412923 | 0,06795 | 0,319298 |
| R3hcc1l | 11,2417 | 14,0273 | 0,319378 | 0,06805 | 0,319543 |
| Dtna | 1,21104 | 0,828202 | -0,548197 | 0,06805 | 0,319543 |
| Snrpa1 | 106,145 | 129,921 | 0,291602 | 0,0682 | 0,320135 |
| Gucy1b3 | 0,617393 | 0,352723 | -0,807653 | 0,0683 | 0,320492 |
| Pax8 | 0,513083 | 0,870695 | 0,762976 | 0,0684 | 0,320849 |
| Atf4 | 184,168 | 224,933 | 0,288474 | 0,06845 | 0,320858 |
| Gfra2 | 1,14301 | 0,746215 | -0,615179 | 0,06845 | 0,320858 |
| Gm15455 | 2,75756 | 3,73043 | 0,435947 | 0,06855 | 0,321214 |
| Rpa2 | 74,5356 | 91,4253 | 0,294664 | 0,06865 | 0,32157 |
| 4632427E13Rik | 1,18401 | 2,44865 | 1,0483 | 0,06875 | 0,321587 |
| Cdk2 | 64,3033 | 78,4502 | 0,286883 | 0,06875 | 0,321587 |
| Tcf15 | 83,9316 | 67,8276 | -0,307342 | 0,0687 | 0,321587 |
| Fuca2 | 6,88607 | 5,40696 | -0,348863 | 0,06875 | 0,321587 |
| Slc37a2 | 1,56969 | 1,11041 | -0,49939 | 0,0688 | 0,321709 |
| Tpm3 | 59,6362 | 73,0786 | 0,29326 | 0,0689 | 0,322064 |
| Psme2 | 9,02451 | 12,5755 | 0,478692 | 0,06895 | 0,322185 |
| Rpl19 | 2,94012 | 4,79397 | 0,705344 | 0,06905 | 0,322314 |
| Usp25 | 12,3684 | 9,95548 | -0,313098 | 0,06905 | 0,322314 |
| Matn3 | 4,2427 | 3,17512 | -0,418175 | 0,06905 | 0,322314 |
| Nasp | 372,106 | 466,546 | 0,326305 | 0,0691 | 0,322322 |
| Eftud1 | 9,65668 | 12,0453 | 0,318871 | 0,0691 | 0,322322 |
| Cdc26 | 38,9178 | 49,2769 | 0,340481 | 0,06915 | 0,32233 |
| Bambi | 9,06931 | 7,26671 | -0,319689 | 0,06915 | 0,32233 |
| Ccdc130 | 3,68333 | 5,0261 | 0,448427 | 0,0692 | 0,322451 |
| Dirc2 | 5,06518 | 3,94792 | -0,359518 | 0,06935 | 0,323037 |
| Cwc15 | 98,3482 | 120,478 | 0,292802 | 0,0694 | 0,323157 |
| Egfr | 2,01672 | 1,47539 | -0,450912 | 0,0695 | 0,32351 |
| 1500011B03Rik | 11,2819 | 8,11739 | -0,474916 | 0,06955 | 0,32363 |
| Stx16 | 27,8073 | 22,5923 | -0,299631 | 0,06965 | 0,323983 |
| Maff | 5,73803 | 7,6406 | 0,41313 | 0,0697 | 0,324103 |
| Hcn4 | 0,708171 | 0,435862 | -0,700227 | 0,06975 | 0,324222 |
| Bnip1 | 27,9743 | 35,3263 | 0,336642 | 0,0699 | 0,324243 |
| Itpk1 | 10,1243 | 12,6978 | 0,326761 | 0,0699 | 0,324243 |
| Heatr5b | 6,09965 | 7,60732 | 0,318662 | 0,06985 | 0,324243 |
| Zbtb33 | 17,9593 | 14,5139 | -0,3073 | 0,0699 | 0,324243 |

| | | | | | |
|---|---|---|---|---|---|
| Fam83h | 2,58887 | 1,93202 | -0,422213 | 0,0698 | 0,324243 |
| Cib2 | 8,85787 | 6,02256 | -0,556584 | 0,0699 | 0,324243 |
| Mettl21a | 7,52216 | 5,80574 | -0,373666 | 0,06995 | 0,324362 |
| Vopp1 | 30,1747 | 37,0148 | 0,294762 | 0,07015 | 0,324614 |
| Rragd | 27,5043 | 22,4577 | -0,292449 | 0,07015 | 0,324614 |
| Scarb1 | 16,3952 | 13,1042 | -0,32324 | 0,07015 | 0,324614 |
| Gpr180 | 10,7217 | 8,47951 | -0,338478 | 0,0701 | 0,324614 |
| Dclk1 | 5,34799 | 4,21728 | -0,342686 | 0,0701 | 0,324614 |
| Tmed3 | 17,9924 | 13,9291 | -0,369285 | 0,07015 | 0,324614 |
| Adamts3 | 2,53995 | 3,24859 | 0,355011 | 0,07025 | 0,324852 |
| Pot1b | 12,1014 | 9,69949 | -0,319196 | 0,07025 | 0,324852 |
| Tbcb | 63,6411 | 78,3291 | 0,299591 | 0,07035 | 0,325089 |
| Kmt2a | 12,4815 | 15,2744 | 0,291323 | 0,07035 | 0,325089 |
| Atg16l2 | 4,92603 | 6,55995 | 0,41326 | 0,0704 | 0,325095 |
| Agrn | 54,135 | 44,2849 | -0,289748 | 0,0704 | 0,325095 |
| Phpt1 | 114,464 | 141,734 | 0,308296 | 0,07065 | 0,326024 |
| Fut10 | 20,4682 | 25,2768 | 0,304427 | 0,07065 | 0,326024 |
| Pole3 | 90,0886 | 111,984 | 0,313881 | 0,07075 | 0,326373 |
| Sparcl1 | 98,6167 | 120,56 | 0,289843 | 0,0711 | 0,327761 |
| Etohi1 | 5,48318 | 4,09648 | -0,420627 | 0,0711 | 0,327761 |
| Gnptab | 6,569 | 5,22074 | -0,331419 | 0,07115 | 0,327879 |
| Bag1 | 90,8023 | 110,805 | 0,287221 | 0,0712 | 0,327996 |
| Cdkn2c | 0,971453 | 0,416186 | -1,22292 | 0,07125 | 0,328113 |
| Fam120b | 10,9091 | 13,4996 | 0,307382 | 0,07135 | 0,328348 |
| Trpc3 | 1,32846 | 0,8981 | -0,564811 | 0,07135 | 0,328348 |
| Luc7l | 73,6726 | 92,2202 | 0,323953 | 0,0714 | 0,328465 |
| Cacnb1 | 2,36086 | 3,35431 | 0,506702 | 0,07145 | 0,328468 |
| Ilvbl | 10,8289 | 8,45925 | -0,356279 | 0,07145 | 0,328468 |
| Dusp7 | 13,1856 | 10,5449 | -0,32242 | 0,0715 | 0,328585 |
| Hist1h2ak | 0,570501 | 1,7792 | 1,64093 | 0,0716 | 0,328819 |
| Ccdc102a | 10,2337 | 7,97439 | -0,359885 | 0,0716 | 0,328819 |
| Arid3b | 95,1638 | 116,358 | 0,290084 | 0,07165 | 0,328935 |
| Shc2 | 8,62192 | 10,7746 | 0,321547 | 0,0717 | 0,328939 |
| Mut | 15,7545 | 12,7251 | -0,308086 | 0,0717 | 0,328939 |
| Saa3 | 0,579224 | 1,83592 | 1,66431 | 0,0718 | 0,328946 |
| Gpr157 | 0,545492 | 0,824227 | 0,595483 | 0,07175 | 0,328946 |
| Bdh2 | 3,06964 | 1,9273 | -0,671484 | 0,0718 | 0,328946 |
| Prnd | 2,74789 | 0,0456817 | -5,91056 | 0,0718 | 0,328946 |
| Apba1 | 0,377651 | 0,573388 | 0,602457 | 0,07185 | 0,329062 |
| Tbccd1 | 12,405 | 15,5092 | 0,322199 | 0,072 | 0,329073 |
| Pfdn2 | 135,516 | 166,979 | 0,301205 | 0,072 | 0,329073 |
| Atp5e | 451,691 | 551,791 | 0,288787 | 0,0719 | 0,329073 |
| Psmc2 | 119,455 | 145,62 | 0,28574 | 0,072 | 0,329073 |
| Zak | 5,80241 | 4,45708 | -0,380553 | 0,07195 | 0,329073 |
| Fam89a | 4,28407 | 2,89695 | -0,564444 | 0,072 | 0,329073 |
| Ddx27 | 40,8076 | 49,844 | 0,288581 | 0,0722 | 0,329874 |
| Ndufb10 | 113,215 | 139,731 | 0,303584 | 0,07225 | 0,329989 |
| Fam204a | 13,8729 | 17,4544 | 0,331328 | 0,0723 | 0,330105 |

| | | | | | |
|---|---|---|---|---|---|
| Aldh1l2 | 0,587325 | 0,860067 | 0,55029 | 0,0724 | 0,33011 |
| Ube2v1 | 6,87243 | 8,96157 | 0,382933 | 0,0724 | 0,33011 |
| Fam185a | 7,14542 | 9,05066 | 0,341005 | 0,07235 | 0,33011 |
| Shb | 9,33898 | 11,7751 | 0,334406 | 0,0724 | 0,33011 |
| Rnf32 | 1,39001 | 0,779385 | -0,834691 | 0,07245 | 0,330113 |
| Vax1 | 0,961007 | 0,370082 | -1,3767 | 0,07245 | 0,330113 |
| Chka | 19,9537 | 24,6452 | 0,304646 | 0,0725 | 0,330228 |
| Zfp410 | 34,2986 | 41,982 | 0,291619 | 0,07255 | 0,330344 |
| C330021F23Rik | 1,25301 | 1,99085 | 0,667988 | 0,0726 | 0,330346 |
| Rpl6 | 49,703 | 61,1604 | 0,299265 | 0,0726 | 0,330346 |
| Rilpl2 | 7,98789 | 10,6936 | 0,420864 | 0,07265 | 0,330461 |
| Poc1a | 11,7765 | 14,9926 | 0,348339 | 0,0727 | 0,330466 |
| Wrap53 | 18,3858 | 22,9839 | 0,32203 | 0,07275 | 0,330466 |
| Eif4a3 | 88,6346 | 108,022 | 0,285388 | 0,07275 | 0,330466 |
| Kras | 32,1162 | 26,3163 | -0,287344 | 0,07275 | 0,330466 |
| Epo | 0,382077 | 1,21809 | 1,67268 | 0,0729 | 0,330586 |
| Hint2 | 19,3773 | 26,2029 | 0,435362 | 0,07285 | 0,330586 |
| Ndufaf6 | 11,1587 | 14,7589 | 0,403411 | 0,0729 | 0,330586 |
| Rhno1 | 60,2761 | 73,9204 | 0,294387 | 0,0729 | 0,330586 |
| Fxyd3 | 5,81641 | 0,609354 | -3,25478 | 0,07285 | 0,330586 |
| Gprc5a | 12,8152 | 10,0371 | -0,352512 | 0,073 | 0,330927 |
| Bcl6 | 0,866527 | 0,537321 | -0,689461 | 0,07305 | 0,331041 |
| Zfp764 | 3,26356 | 2,34666 | -0,475839 | 0,07315 | 0,331382 |
| BC030867 | 12,8911 | 16,1026 | 0,320915 | 0,0732 | 0,331384 |
| Tfap4 | 24,41 | 30,0552 | 0,300144 | 0,0732 | 0,331384 |
| Zfp429 | 2,228 | 1,48247 | -0,587744 | 0,0733 | 0,331724 |
| 2310033P09Rik | 25,9752 | 32,5422 | 0,325175 | 0,07335 | 0,331838 |
| Ctsl | 168,321 | 138,303 | -0,283389 | 0,0734 | 0,33184 |
| 1700123M08Rik | 0,721044 | 0,236968 | -1,6054 | 0,0734 | 0,33184 |
| Tnip2 | 1,21982 | 0,724544 | -0,751525 | 0,07345 | 0,331953 |
| Gm14420 | 1,62455 | 1,15984 | -0,486119 | 0,07355 | 0,332293 |
| Gxylt1 | 22,8094 | 18,7377 | -0,283685 | 0,0736 | 0,332406 |
| Mum1l1 | 14,2697 | 17,5295 | 0,296832 | 0,07365 | 0,332407 |
| Hmces | 31,7095 | 39,3562 | 0,311679 | 0,07365 | 0,332408 |
| Atxn7 | 9,3058 | 11,4345 | 0,297195 | 0,074 | 0,333762 |
| Mcf2l | 1,54513 | 1,12774 | -0,454295 | 0,074 | 0,333762 |
| Tmbim4 | 40,6814 | 31,9032 | -0,35067 | 0,07405 | 0,333875 |
| Ddx18 | 22,7437 | 27,9711 | 0,298471 | 0,0741 | 0,333987 |
| Chst1 | 1,13728 | 0,715974 | -0,667612 | 0,07415 | 0,3341 |
| Vps8 | 5,32123 | 6,67428 | 0,326853 | 0,07435 | 0,334663 |
| Etv3 | 7,58745 | 6,08581 | -0,318166 | 0,07435 | 0,334663 |
| Dnajc15 | 61,0356 | 48,2348 | -0,339579 | 0,0743 | 0,334663 |
| Fbln2 | 18,7042 | 22,8699 | 0,290091 | 0,07445 | 0,335 |
| Slmo1 | 4,46031 | 5,94917 | 0,415544 | 0,07455 | 0,335111 |
| Dnmt3a | 48,528 | 59,4143 | 0,291995 | 0,07455 | 0,335111 |
| Nbl1 | 4,96998 | 3,59479 | -0,467331 | 0,0745 | 0,335111 |
| Cpd | 17,0883 | 14,0311 | -0,284384 | 0,0746 | 0,335223 |
| Ift57 | 5,52663 | 7,15045 | 0,371634 | 0,07475 | 0,335671 |

| | | | | | |
|---|---|---|---|---|---|
| Daam2 | 16,0967 | 13,1619 | -0,290397 | 0,07475 | 0,335671 |
| Pigu | 33,9279 | 27,367 | -0,310033 | 0,0748 | 0,335671 |
| C1galt1 | 12,7334 | 10,0412 | -0,342682 | 0,0748 | 0,335671 |
| Dpysl3 | 15,951 | 19,5049 | 0,290189 | 0,07485 | 0,335782 |
| Mark1 | 7,06714 | 8,80607 | 0,317372 | 0,0749 | 0,335893 |
| Ddx19a | 26,7817 | 32,7869 | 0,291873 | 0,07495 | 0,335893 |
| Nap1l4 | 149,215 | 181,971 | 0,286311 | 0,07495 | 0,335893 |
| Atp13a3 | 30,0515 | 24,7814 | -0,278181 | 0,075 | 0,336004 |
| Cog4 | 24,2457 | 29,7439 | 0,294865 | 0,07505 | 0,336115 |
| Dcakd | 132,041 | 160,38 | 0,280502 | 0,07515 | 0,336338 |
| Tarsl2 | 4,93189 | 3,77261 | -0,386576 | 0,07515 | 0,336338 |
| Gatsl3 | 3,97694 | 5,54014 | 0,478261 | 0,07535 | 0,336893 |
| Hpcal1 | 11,2334 | 14,4347 | 0,361749 | 0,0754 | 0,336893 |
| Wnk3 | 3,64522 | 4,5381 | 0,316082 | 0,0754 | 0,336893 |
| Ift81 | 14,1051 | 17,4647 | 0,308223 | 0,0753 | 0,336893 |
| Cox20 | 116,668 | 92,5029 | -0,334837 | 0,0754 | 0,336893 |
| Gnl1 | 25,6862 | 31,4541 | 0,292256 | 0,07545 | 0,337003 |
| Cux1 | 29,6776 | 38,2587 | 0,366411 | 0,0757 | 0,337668 |
| Cct7 | 220,035 | 270,024 | 0,295359 | 0,07565 | 0,337668 |
| Sfxn2 | 11,2485 | 8,9885 | -0,323579 | 0,07565 | 0,337668 |
| Vkorc1 | 17,3461 | 12,7663 | -0,442269 | 0,0757 | 0,337668 |
| Rrp9 | 43,7981 | 53,6512 | 0,292742 | 0,07575 | 0,337779 |
| B930003M22Rik | 1,39434 | 0,742905 | -0,908331 | 0,0758 | 0,337889 |
| Hist2h2be | 2,49428 | 1,75464 | -0,507452 | 0,07585 | 0,337999 |
| Plekha5 | 13,3628 | 16,3422 | 0,290375 | 0,07605 | 0,338321 |
| Lims1 | 32,7412 | 26,927 | -0,282055 | 0,0761 | 0,338321 |
| Hiat1 | 39,377 | 32,261 | -0,287559 | 0,07595 | 0,338321 |
| Arl8b | 36,0075 | 29,4972 | -0,287717 | 0,0761 | 0,338321 |
| Fgf18 | 31,2884 | 25,1394 | -0,315676 | 0,076 | 0,338321 |
| Shisa2 | 13,2168 | 10,6069 | -0,317368 | 0,0761 | 0,338321 |
| Chmp2b | 23,076 | 18,5143 | -0,317752 | 0,07615 | 0,338321 |
| Tapbp | 6,90862 | 5,3503 | -0,368777 | 0,07615 | 0,338321 |
| Tlcd2 | 4,30214 | 2,96722 | -0,535942 | 0,0761 | 0,338321 |
| Zim1 | 2,2414 | 1,60727 | -0,479781 | 0,07625 | 0,338652 |
| Klc2 | 18,0751 | 22,209 | 0,297142 | 0,0763 | 0,338762 |
| Rpl28 | 1,25679 | 2,95635 | 1,23407 | 0,0764 | 0,339093 |
| 4632415L05Rik | 6,4244 | 5,06674 | -0,342504 | 0,0765 | 0,339424 |
| Uhrf1 | 116,155 | 141,886 | 0,28868 | 0,0766 | 0,339755 |
| Gm10416 | 1,98648 | 0,857064 | -1,21274 | 0,0767 | 0,340086 |
| Pla2g7 | 27,6202 | 22,2763 | -0,310214 | 0,07675 | 0,340195 |
| Gapdh | 0,727892 | 1,31688 | 0,85533 | 0,0769 | 0,340516 |
| Ccdc96 | 1,0263 | 1,48551 | 0,533506 | 0,07695 | 0,340516 |
| 9330159F19Rik | 1,0633 | 1,46275 | 0,460142 | 0,07695 | 0,340516 |
| Phf14 | 29,4446 | 35,889 | 0,285539 | 0,07695 | 0,340516 |
| Gtpbp8 | 16,2946 | 12,5281 | -0,379225 | 0,07685 | 0,340516 |
| Ctsw | 0,96177 | 0,437668 | -1,13586 | 0,07715 | 0,341176 |
| Krtap17-1 | 1,06955 | 0,24073 | -2,15152 | 0,07715 | 0,341176 |
| Zfp94 | 2,51995 | 3,44028 | 0,449133 | 0,07725 | 0,341387 |

| | | | | | |
|---|---|---|---|---|---|
| Narf | 13,1772 | 10,7034 | -0,299974 | 0,0773 | 0,341387 |
| Nuak2 | 5,55997 | 4,26762 | -0,381643 | 0,07725 | 0,341387 |
| Etv1 | 0,960392 | 0,632974 | -0,601477 | 0,0773 | 0,341387 |
| Slc38a1 | 56,2466 | 46,1924 | -0,284112 | 0,07735 | 0,341495 |
| C2cd4b | 0,268226 | 0,671071 | 1,32302 | 0,0775 | 0,341706 |
| Ttc14 | 16,8317 | 13,7964 | -0,286887 | 0,0775 | 0,341706 |
| Zfp111 | 4,29689 | 3,42055 | -0,329066 | 0,0775 | 0,341706 |
| Sertad3 | 6,93769 | 4,97346 | -0,480205 | 0,07745 | 0,341706 |
| Rpl14-ps1 | 774,991 | 948,362 | 0,29126 | 0,0776 | 0,342034 |
| Tns3 | 9,35648 | 7,63655 | -0,293045 | 0,0777 | 0,342362 |
| Mir684-2 | 59,789 | 0 | -inf | 0,078 | 0,343344 |
| Tmc6 | 8,77018 | 6,78546 | -0,370159 | 0,078 | 0,343344 |
| Tnfsf9 | 5,57404 | 3,7591 | -0,568338 | 0,078 | 0,343344 |
| Syt9 | 2,05669 | 2,82357 | 0,457198 | 0,0781 | 0,343445 |
| Rbpj | 8,92888 | 7,23544 | -0,303398 | 0,0781 | 0,343445 |
| Syngr1 | 1,28008 | 0,490278 | -1,38456 | 0,07805 | 0,343445 |
| Sfr1 | 167,881 | 203,831 | 0,279938 | 0,07815 | 0,343552 |
| Szt2 | 7,10789 | 8,67802 | 0,287944 | 0,0782 | 0,343658 |
| Lrp8 | 13,4633 | 16,3837 | 0,283228 | 0,07825 | 0,343765 |
| Zmynd8 | 38,844 | 47,4285 | 0,288062 | 0,0783 | 0,343872 |
| Nfkbiz | 2,18694 | 2,94837 | 0,431009 | 0,07835 | 0,343972 |
| Mlxip | 15,0147 | 18,2839 | 0,284196 | 0,0784 | 0,343972 |
| F2r | 66,4422 | 54,6835 | -0,280996 | 0,0784 | 0,343972 |
| Acp5 | 26,2221 | 1,01861 | -4,68612 | 0,07845 | 0,344078 |
| Fxyd6 | 61,0703 | 50,0839 | -0,286125 | 0,07855 | 0,344404 |
| Sgms1 | 5,3288 | 4,14048 | -0,364014 | 0,0786 | 0,34451 |
| Fastkd3 | 15,9962 | 19,8457 | 0,311101 | 0,07865 | 0,344616 |
| Slc38a2 | 99,8936 | 82,2612 | -0,28018 | 0,0787 | 0,344722 |
| Rmdn1 | 3,46031 | 2,43407 | -0,507526 | 0,0788 | 0,345047 |
| Gm12191 | 333,5 | 267,033 | -0,32067 | 0,07885 | 0,345153 |
| Pias4 | 23,1538 | 28,6591 | 0,307741 | 0,0789 | 0,345251 |
| Slit2 | 13,8737 | 11,3898 | -0,284612 | 0,07895 | 0,345251 |
| 2810442I21Rik | 4,47679 | 3,18502 | -0,49116 | 0,07895 | 0,345251 |
| Mir1934 | 0 | 72,3912 | inf | 0,0796 | 0,346732 |
| Mir345 | 0 | 35,5675 | inf | 0,0796 | 0,346732 |
| Bloc1s1 | 32,7107 | 42,5822 | 0,380486 | 0,0794 | 0,346732 |
| Nfkbib | 9,60905 | 12,1964 | 0,343989 | 0,07935 | 0,346732 |
| Nhlrc2 | 11,7886 | 14,5043 | 0,299088 | 0,0795 | 0,346732 |
| Ints4 | 26,2719 | 32,0328 | 0,286029 | 0,0795 | 0,346732 |
| Fam120a | 54,0688 | 44,6079 | -0,277496 | 0,0796 | 0,346732 |
| Spast | 20,1408 | 16,549 | -0,283372 | 0,0795 | 0,346732 |
| Peli2 | 11,7974 | 9,63503 | -0,292109 | 0,0796 | 0,346732 |
| Wwtr1 | 13,8084 | 11,2598 | -0,294365 | 0,0794 | 0,346732 |
| Zdhhc8 | 9,78724 | 7,91848 | -0,30568 | 0,07955 | 0,346732 |
| Ndufb4 | 18,7512 | 12,8598 | -0,544108 | 0,0794 | 0,346732 |
| Msi1 | 39,1615 | 48,098 | 0,29654 | 0,0797 | 0,347055 |
| Spire2 | 1,59023 | 2,30914 | 0,538117 | 0,0798 | 0,347108 |
| Nfam1 | 3,30374 | 4,28994 | 0,376856 | 0,0798 | 0,347108 |

| | | | | | |
|---|---|---|---|---|---|
| Tagln3 | 16,3029 | 20,8383 | 0,35411 | 0,07995 | 0,347108 |
| Snx12 | 31,46 | 39,9577 | 0,344957 | 0,07985 | 0,347108 |
| S100pbp | 27,1156 | 33,5794 | 0,308454 | 0,08 | 0,347108 |
| Dynlrb1 | 166,806 | 203,065 | 0,283772 | 0,07985 | 0,347108 |
| Stt3a | 98,7264 | 81,4209 | -0,278037 | 0,08 | 0,347108 |
| Mis18bp1 | 33,5306 | 27,5528 | -0,28328 | 0,08 | 0,347108 |
| Tmtc3 | 15,0025 | 12,3265 | -0,283437 | 0,0799 | 0,347108 |
| Fbxw2 | 25,5647 | 20,8104 | -0,296848 | 0,08005 | 0,347108 |
| Slc17a5 | 3,4562 | 2,58085 | -0,42134 | 0,0799 | 0,347108 |
| Zfp874b | 2,1583 | 1,5701 | -0,459039 | 0,08005 | 0,347108 |
| Crlf1 | 2,98889 | 2,03099 | -0,557426 | 0,07995 | 0,347108 |
| Zcchc8 | 27,413 | 33,3422 | 0,282493 | 0,0801 | 0,347213 |
| S100a6 | 18,8269 | 24,7232 | 0,393075 | 0,0802 | 0,347533 |
| Ttc21b | 7,64152 | 9,46273 | 0,308397 | 0,08025 | 0,347637 |
| Mtr | 12,9407 | 15,8083 | 0,288768 | 0,0803 | 0,347741 |
| Commd3 | 54,3037 | 43,6438 | -0,315274 | 0,08045 | 0,348278 |
| Tmco1 | 15,2698 | 12,442 | -0,295455 | 0,0806 | 0,348588 |
| Mageh1 | 25,3962 | 20,2787 | -0,324648 | 0,0806 | 0,348588 |
| Birc3 | 1,58289 | 1,07417 | -0,559339 | 0,0806 | 0,348588 |
| Ndufc2 | 167,45 | 137,516 | -0,284136 | 0,0807 | 0,348908 |
| Klhl11 | 6,70942 | 8,61989 | 0,361481 | 0,08085 | 0,349208 |
| Ruvbl1 | 17,6074 | 21,9642 | 0,318973 | 0,0809 | 0,349208 |
| Ppil1 | 79,4535 | 96,7631 | 0,284347 | 0,08085 | 0,349208 |
| Umps | 51,6366 | 62,5482 | 0,276573 | 0,08085 | 0,349208 |
| Hmgxb3 | 17,3949 | 21,0556 | 0,27554 | 0,0809 | 0,349208 |
| Bod1l | 11,6734 | 14,1768 | 0,280309 | 0,08115 | 0,350174 |
| Zfp106 | 44,3763 | 54,0023 | 0,283233 | 0,0813 | 0,350708 |
| Asap2 | 12,0936 | 15,0646 | 0,316921 | 0,08135 | 0,35081 |
| Mical1 | 30,7233 | 37,3241 | 0,280776 | 0,08145 | 0,351128 |
| 1810058I24Rik | 51,2079 | 62,7692 | 0,29369 | 0,08155 | 0,351446 |
| Alyref | 100,422 | 121,922 | 0,27988 | 0,0816 | 0,351548 |
| Atp6v0a2 | 16,7803 | 13,8029 | -0,281797 | 0,0817 | 0,351639 |
| Asb13 | 10,3892 | 8,14599 | -0,350918 | 0,08165 | 0,351639 |
| 4921507P07Rik | 0,597609 | 0,299683 | -0,995766 | 0,0817 | 0,351639 |
| Mpdu1 | 38,9761 | 31,5018 | -0,307158 | 0,08175 | 0,351741 |
| Rab4a | 8,19441 | 6,09817 | -0,426263 | 0,0818 | 0,351843 |
| Prrt1 | 0,255809 | 0,808457 | 1,66011 | 0,082 | 0,352476 |
| Ripk1 | 5,79012 | 4,59275 | -0,334235 | 0,082 | 0,352476 |
| Rpl32 | 705,922 | 926,632 | 0,392487 | 0,0821 | 0,352566 |
| Jmjd6 | 54,3932 | 66,0498 | 0,280129 | 0,08205 | 0,352566 |
| Samd8 | 4,59973 | 3,68235 | -0,320922 | 0,0821 | 0,352566 |
| Fam192a | 35,377 | 43,2467 | 0,289779 | 0,0822 | 0,352768 |
| Rsad1 | 3,65092 | 2,80753 | -0,378956 | 0,0822 | 0,352768 |
| Ifnar1 | 16,5816 | 13,6264 | -0,283185 | 0,08225 | 0,35287 |
| Mcoln3 | 5,75903 | 4,23291 | -0,444176 | 0,0825 | 0,353829 |
| Cpne3 | 10,5715 | 8,62897 | -0,292921 | 0,08255 | 0,353929 |
| Snrpc | 40,7591 | 51,1298 | 0,32704 | 0,08265 | 0,354219 |
| Gad1 | 22,0509 | 27,1454 | 0,29987 | 0,0827 | 0,354219 |

| | | | | | |
|---|---|---|---|---|---|
| D2Wsu81e | 36,9991 | 45,3827 | 0,29465 | 0,0827 | 0,354219 |
| Spred2 | 34,0563 | 41,2961 | 0,278083 | 0,08275 | 0,354219 |
| Rrp1b | 31,8212 | 38,5459 | 0,276591 | 0,08275 | 0,354219 |
| Slc25a33 | 9,29583 | 11,8151 | 0,345978 | 0,0829 | 0,354293 |
| Osbpl8 | 6,22159 | 5,05136 | -0,300613 | 0,0829 | 0,354293 |
| Hexa | 22,5745 | 18,2159 | -0,309495 | 0,0828 | 0,354293 |
| Cped1 | 3,06049 | 2,35222 | -0,379739 | 0,08285 | 0,354293 |
| Prss8 | 5,01156 | 0,327962 | -3,93366 | 0,08285 | 0,354293 |
| Cd14 | 0,292854 | 0,667659 | 1,18893 | 0,08315 | 0,354503 |
| AA415398 | 0,69373 | 1,17249 | 0,757128 | 0,0834 | 0,354503 |
| Dqx1 | 1,76961 | 2,44634 | 0,467197 | 0,0832 | 0,354503 |
| Vcan | 82,815 | 102,837 | 0,3124 | 0,083 | 0,354503 |
| Ipo5 | 146,401 | 179,648 | 0,295251 | 0,08315 | 0,354503 |
| Vps72 | 68,7216 | 83,5965 | 0,282678 | 0,08325 | 0,354503 |
| Mllt3 | 50,2688 | 60,9986 | 0,279112 | 0,0832 | 0,354503 |
| Ythdf2 | 26,9779 | 32,6935 | 0,277228 | 0,08305 | 0,354503 |
| Ptms | 232,431 | 281,64 | 0,277046 | 0,08335 | 0,354503 |
| Actn1 | 72,9839 | 88,2969 | 0,274785 | 0,08325 | 0,354503 |
| Crls1 | 27,5666 | 22,233 | -0,310215 | 0,08305 | 0,354503 |
| Hsd11b2 | 10,4712 | 8,16015 | -0,359755 | 0,0834 | 0,354503 |
| Ncf4 | 1,5245 | 0,0684536 | -4,47706 | 0,0834 | 0,354503 |
| Gp5 | 1,01198 | 0,0428909 | -4,56036 | 0,0834 | 0,354503 |
| Btk | 0,972876 | 0,0367084 | -4,72807 | 0,0834 | 0,354503 |
| Ctse | 1,78867 | 0,0446593 | -5,32378 | 0,0834 | 0,354503 |
| Kel | 8,18285 | 0,0370101 | -7,78854 | 0,0834 | 0,354503 |
| Asb1 | 4,7814 | 3,82224 | -0,323012 | 0,0837 | 0,355665 |
| Mdm4 | 23,7803 | 28,9112 | 0,281859 | 0,08385 | 0,356189 |
| AW822252 | 0,438602 | 0,978853 | 1,15818 | 0,08405 | 0,356925 |
| Zbtb46 | 5,88344 | 7,61782 | 0,372719 | 0,0843 | 0,357518 |
| Rhod | 14,9328 | 19,061 | 0,352131 | 0,08435 | 0,357518 |
| Dtnbp1 | 43,9476 | 53,7468 | 0,290396 | 0,08425 | 0,357518 |
| Ogt | 75,8554 | 62,7253 | -0,274204 | 0,08435 | 0,357518 |
| B3gnt2 | 11,0457 | 8,80843 | -0,326527 | 0,08435 | 0,357518 |
| Cenpp | 31,3385 | 24,9816 | -0,327073 | 0,08435 | 0,357518 |
| 2300009A05Rik | 8,93977 | 5,93735 | -0,590419 | 0,0845 | 0,35804 |
| Tars2 | 25,2153 | 30,7368 | 0,285667 | 0,08465 | 0,358221 |
| Slc25a12 | 9,32024 | 7,41738 | -0,329458 | 0,0846 | 0,358221 |
| Ngly1 | 4,33089 | 3,1096 | -0,477938 | 0,0846 | 0,358221 |
| Vsig2 | 0,858453 | 0,0885784 | -3,27671 | 0,08465 | 0,358221 |
| Isoc2b | 1,18971 | 0,579471 | -1,03781 | 0,0848 | 0,358742 |
| Ssr2 | 157,877 | 130,304 | -0,276919 | 0,0849 | 0,359052 |
| Pgm1 | 17,7473 | 21,8849 | 0,302337 | 0,08495 | 0,359134 |
| Igf2bp2 | 86,0211 | 104,113 | 0,275389 | 0,085 | 0,359134 |
| 1700048O20Rik | 7,92289 | 6,06088 | -0,386501 | 0,085 | 0,359134 |
| Znrd1 | 65,5824 | 81,1173 | 0,306703 | 0,0851 | 0,359411 |
| Sapcd2 | 11,0953 | 13,6657 | 0,300609 | 0,0852 | 0,359411 |
| Arf6 | 36,8219 | 30,3214 | -0,280228 | 0,0852 | 0,359411 |
| Pnpla6 | 8,81049 | 7,11767 | -0,307817 | 0,0852 | 0,359411 |

| | | | | | |
|---|---|---|---|---|---|
| Plcg2 | 1,94769 | 1,42686 | -0,448923 | 0,08515 | 0,359411 |
| Snapc1 | 13,2847 | 16,5372 | 0,315942 | 0,08525 | 0,359508 |
| Pcna | 111,996 | 135,852 | 0,278586 | 0,08535 | 0,359784 |
| Tomm20 | 20,1835 | 16,6469 | -0,277928 | 0,08545 | 0,359784 |
| Tmx4 | 9,77499 | 7,96034 | -0,296265 | 0,08545 | 0,359784 |
| Wdr13 | 9,85592 | 7,98236 | -0,304174 | 0,08545 | 0,359784 |
| B3galt1 | 0,923747 | 0,636037 | -0,538388 | 0,0854 | 0,359784 |
| 3110021A11Rik | 2,13929 | 1,13642 | -0,912634 | 0,08585 | 0,361354 |
| Bscl2 | 13,2963 | 10,407 | -0,353465 | 0,08595 | 0,361661 |
| Mcm5 | 100,459 | 121,512 | 0,274496 | 0,086 | 0,361758 |
| Cct4 | 280,786 | 341,261 | 0,281405 | 0,08605 | 0,361854 |
| Ncam1 | 35,583 | 29,452 | -0,272821 | 0,08625 | 0,362581 |
| Slc26a11 | 6,1912 | 7,87104 | 0,346336 | 0,0863 | 0,362677 |
| Klhl36 | 1,57615 | 1,02729 | -0,617563 | 0,08635 | 0,362773 |
| Utp14b | 9,02539 | 13,8777 | 0,620707 | 0,08645 | 0,363079 |
| Emid1 | 5,90245 | 4,43601 | -0,41205 | 0,0866 | 0,363595 |
| Dfna5 | 3,18835 | 4,33439 | 0,443022 | 0,08665 | 0,363691 |
| Zfp942 | 11,4534 | 9,26521 | -0,305877 | 0,08675 | 0,363996 |
| Galnt1 | 50,1802 | 41,5398 | -0,272624 | 0,0868 | 0,364091 |
| Ubap2l | 78,085 | 95,8786 | 0,296162 | 0,0869 | 0,364282 |
| Shkbp1 | 10,2188 | 8,08133 | -0,338557 | 0,0869 | 0,364282 |
| B3galnt2 | 31,237 | 25,0854 | -0,316409 | 0,08695 | 0,364377 |
| Pfdn1 | 93,1796 | 112,901 | 0,276967 | 0,087 | 0,364473 |
| Dvl2 | 30,8532 | 37,4011 | 0,277661 | 0,0871 | 0,364663 |
| Spata9 | 0,7317 | 0,264971 | -1,46542 | 0,0871 | 0,364663 |
| Nufip2 | 12,0141 | 14,5423 | 0,275519 | 0,08715 | 0,364758 |
| 4930562F07Rik | 0,572924 | 0 | -inf | 0,08725 | 0,364948 |
| Nfil3 | 12,7606 | 15,9329 | 0,320317 | 0,08725 | 0,364948 |
| Tcp11l1 | 1,49241 | 2,0665 | 0,469543 | 0,0873 | 0,365024 |
| Cadm4 | 5,81978 | 4,41 | -0,400185 | 0,08735 | 0,365024 |
| Gp1ba | 1,06439 | 0,678612 | -0,649371 | 0,08735 | 0,365024 |
| Rap1b | 77,1506 | 63,8772 | -0,272375 | 0,08745 | 0,365099 |
| Tnnt1 | 3,52235 | 2,28159 | -0,626498 | 0,08745 | 0,365099 |
| Msx1os | 3,50233 | 2,05625 | -0,768298 | 0,08745 | 0,365099 |
| Adcy3 | 4,24006 | 6,12375 | 0,530331 | 0,08755 | 0,365402 |
| Zfp326 | 43,5688 | 52,6315 | 0,27263 | 0,0876 | 0,365497 |
| Uros | 12,9956 | 10,2426 | -0,34345 | 0,0877 | 0,3658 |
| Polr2f | 170,394 | 207,836 | 0,286569 | 0,0878 | 0,366103 |
| Ttyh3 | 47,4322 | 39,309 | -0,271007 | 0,088 | 0,366822 |
| Cbx3 | 17,5933 | 14,0834 | -0,321029 | 0,08805 | 0,366916 |
| Aagab | 27,4854 | 33,4405 | 0,282932 | 0,08815 | 0,367104 |
| Ube2e2 | 16,0504 | 12,5559 | -0,354248 | 0,08815 | 0,367104 |
| Kank1 | 16,0521 | 13,2092 | -0,281216 | 0,0883 | 0,367614 |
| Tspan31 | 74,4175 | 61,3851 | -0,277753 | 0,0884 | 0,367801 |
| Pdlim1 | 7,20812 | 5,38854 | -0,419728 | 0,0884 | 0,367801 |
| Tcof1 | 43,9341 | 53,0937 | 0,273202 | 0,08845 | 0,367895 |
| Slx4ip | 5,72774 | 8,52838 | 0,574306 | 0,08865 | 0,368612 |
| 1810009A15Rik | 119,769 | 146,486 | 0,290509 | 0,0887 | 0,368705 |

| | | | | | |
|---|---|---|---|---|---|
| Paqr8 | 4,71533 | 3,71248 | -0,344976 | 0,0888 | 0,368891 |
| Zfp579 | 8,82255 | 6,90148 | -0,354289 | 0,0888 | 0,368891 |
| Osgep | 19,2085 | 23,8452 | 0,31195 | 0,08895 | 0,3694 |
| Bmp4 | 9,89405 | 7,76708 | -0,349188 | 0,0891 | 0,369908 |
| Cdk19 | 6,93855 | 5,61856 | -0,304433 | 0,08915 | 0,37 |
| Rbm4b | 14,9122 | 18,5337 | 0,313662 | 0,08925 | 0,370185 |
| Fbxo18 | 30,5546 | 36,9566 | 0,27444 | 0,08925 | 0,370185 |
| Col11a2 | 1,7215 | 1,28332 | -0,423787 | 0,08935 | 0,370485 |
| Tmtc2 | 2,19888 | 1,68305 | -0,385688 | 0,08945 | 0,370785 |
| Wdr62 | 6,19416 | 7,66673 | 0,307702 | 0,0895 | 0,370877 |
| Fcrl6 | 0,941953 | 0,536324 | -0,812551 | 0,08955 | 0,370969 |
| Ears2 | 13,0608 | 16,3547 | 0,324463 | 0,08965 | 0,37113 |
| Map3k4 | 17,6512 | 21,3495 | 0,274431 | 0,0897 | 0,37113 |
| Arhgap19 | 13,5035 | 11,1148 | -0,280854 | 0,08965 | 0,37113 |
| Cd164l2 | 1,12777 | 0,502126 | -1,16735 | 0,0897 | 0,37113 |
| Snrnp48 | 28,3443 | 34,7126 | 0,292398 | 0,0899 | 0,371613 |
| H1fx | 78,5726 | 95,2348 | 0,277463 | 0,08985 | 0,371613 |
| 3110056K07Rik | 10,7909 | 7,1553 | -0,592737 | 0,0899 | 0,371613 |
| Thap1 | 5,33991 | 6,90818 | 0,371491 | 0,09005 | 0,371864 |
| Kdm1a | 87,0955 | 105,11 | 0,271232 | 0,0901 | 0,371864 |
| Pcdh19 | 24,7773 | 29,8731 | 0,269827 | 0,0901 | 0,371864 |
| Dgke | 5,26829 | 4,20545 | -0,325073 | 0,09 | 0,371864 |
| Fam167b | 0,831594 | 0,11563 | -2,84636 | 0,09005 | 0,371864 |
| Rassf1 | 19,0996 | 23,5943 | 0,304893 | 0,0902 | 0,372024 |
| Fam124b | 0,599108 | 0,0303771 | -4,30176 | 0,09025 | 0,372024 |
| Anxa1 | 1,35718 | 0,0677933 | -4,32333 | 0,09025 | 0,372024 |
| Kremen2 | 3,87787 | 0,0422071 | -6,52164 | 0,09025 | 0,372024 |
| 2810002D19Rik | 8,37885 | 6,32899 | -0,404778 | 0,09035 | 0,372321 |
| Ak4 | 13,6127 | 16,5054 | 0,277991 | 0,09065 | 0,373188 |
| Mtmr4 | 28,1688 | 33,9985 | 0,271375 | 0,0907 | 0,373188 |
| N4bp2 | 24,1982 | 29,1749 | 0,269829 | 0,0907 | 0,373188 |
| Ube2g1 | 32,593 | 26,997 | -0,271763 | 0,09065 | 0,373188 |
| Zfp945 | 4,53865 | 3,64963 | -0,314514 | 0,0907 | 0,373188 |
| Cirh1a | 52,4517 | 63,4255 | 0,274074 | 0,09085 | 0,373575 |
| Adh1 | 0,686401 | 0,294352 | -1,22151 | 0,09085 | 0,373575 |
| Bloc1s2 | 6,41447 | 8,92312 | 0,476218 | 0,09095 | 0,373756 |
| Atad3aos | 1,60987 | 0,741589 | -1,11825 | 0,09095 | 0,373756 |
| Ttc30a1 | 4,89559 | 3,76248 | -0,379797 | 0,091 | 0,373847 |
| Man2b1 | 10,3 | 8,3905 | -0,295819 | 0,09155 | 0,375991 |
| Fam96b | 34,834 | 43,9774 | 0,336267 | 0,0917 | 0,376376 |
| Pon3 | 8,72179 | 6,73223 | -0,37354 | 0,0917 | 0,376376 |
| Zfp846 | 5,98443 | 7,57036 | 0,339147 | 0,092 | 0,376566 |
| Ppp1r15a | 9,08698 | 11,4121 | 0,328694 | 0,092 | 0,376566 |
| Mybbp1a | 142,576 | 173,398 | 0,282358 | 0,09185 | 0,376566 |
| Abcg2 | 47,9666 | 39,6938 | -0,273117 | 0,09195 | 0,376566 |
| 5530601H04Rik | 4,94826 | 3,88056 | -0,350657 | 0,09195 | 0,376566 |
| Pisd-ps2 | 4,31192 | 3,05927 | -0,495144 | 0,0918 | 0,376566 |
| Dnase2a | 2,28271 | 1,52882 | -0,578324 | 0,092 | 0,376566 |

| | | | | | |
|---|---|---|---|---|---|
| B4galnt3 | 1,06703 | 0,695598 | -0,617281 | 0,09195 | 0,376566 |
| Sytl1 | 0,657063 | 0,336822 | -0,964046 | 0,09185 | 0,376566 |
| Zfp948 | 9,03413 | 7,17276 | -0,332856 | 0,09205 | 0,376655 |
| Elovl5 | 60,0439 | 49,8447 | -0,268579 | 0,0921 | 0,376745 |
| Polr2k | 2,76327 | 4,56391 | 0,723895 | 0,09225 | 0,376986 |
| Cep95 | 10,8852 | 13,4463 | 0,304839 | 0,0923 | 0,376986 |
| Ranbp1 | 478,365 | 577,735 | 0,272296 | 0,0922 | 0,376986 |
| Gart | 62,1243 | 74,8059 | 0,267996 | 0,09225 | 0,376986 |
| Gtpbp2 | 16,3202 | 13,2965 | -0,295618 | 0,0923 | 0,376986 |
| Kif3a | 13,8738 | 16,8001 | 0,276105 | 0,0924 | 0,377164 |
| Sltm | 55,0055 | 66,0258 | 0,263452 | 0,0924 | 0,377164 |
| Gnas | 934,614 | 1162,36 | 0,314611 | 0,09255 | 0,377661 |
| Peg12 | 20,2755 | 24,7257 | 0,286276 | 0,09265 | 0,377953 |
| Dzip3 | 22,921 | 19,0102 | -0,269898 | 0,09275 | 0,378246 |
| Cry1 | 12,2608 | 15,0414 | 0,294888 | 0,09295 | 0,378715 |
| Wipf1 | 12,1416 | 9,95184 | -0,286922 | 0,09295 | 0,378715 |
| Stim2 | 10,7394 | 8,75742 | -0,294333 | 0,0929 | 0,378715 |
| Asb17os | 1,85528 | 0 | -inf | 0,09315 | 0,378782 |
| Mtmr11 | 7,54769 | 9,44896 | 0,324121 | 0,09315 | 0,378782 |
| Vps37b | 12,5387 | 15,4622 | 0,30236 | 0,0932 | 0,378782 |
| Desi1 | 25,8568 | 31,2704 | 0,274252 | 0,0931 | 0,378782 |
| Phb2 | 210,081 | 252,729 | 0,266652 | 0,0932 | 0,378782 |
| Maea | 88,1082 | 105,95 | 0,266032 | 0,09325 | 0,378782 |
| Sox12 | 24,5365 | 20,2887 | -0,274252 | 0,0932 | 0,378782 |
| Reep3 | 12,722 | 10,5034 | -0,276477 | 0,0931 | 0,378782 |
| Ammecr1 | 9,20515 | 7,38707 | -0,317439 | 0,09325 | 0,378782 |
| Gba | 14,1836 | 11,2908 | -0,329077 | 0,09315 | 0,378782 |
| Slc1a2 | 0,575745 | 0,849331 | 0,560897 | 0,09335 | 0,378958 |
| Cep85 | 15,5338 | 18,8763 | 0,28116 | 0,09335 | 0,378958 |
| Thumpd2 | 5,57197 | 7,25756 | 0,381298 | 0,09355 | 0,379654 |
| Atp6ap1 | 50,3592 | 41,6871 | -0,272654 | 0,0936 | 0,379742 |
| Exo5 | 19,1696 | 23,493 | 0,29341 | 0,09365 | 0,37983 |
| Fam134a | 25,2877 | 20,7863 | -0,282802 | 0,09385 | 0,38041 |
| Irf2 | 8,23241 | 6,48373 | -0,344491 | 0,09385 | 0,38041 |
| Hottip | 7,33197 | 5,7588 | -0,348433 | 0,0939 | 0,380497 |
| Map1a | 0,714689 | 1,01483 | 0,50585 | 0,09405 | 0,380874 |
| Rab11fip1 | 6,59429 | 8,05485 | 0,288639 | 0,09405 | 0,380874 |
| Eefsec | 13,2394 | 16,2636 | 0,296808 | 0,0941 | 0,380961 |
| Rbm8a | 48,0822 | 57,9529 | 0,269378 | 0,0942 | 0,38125 |
| Ttll5 | 6,16103 | 7,59883 | 0,302606 | 0,09425 | 0,381337 |
| Sla | 1,29633 | 0 | -inf | 0,0945 | 0,381771 |
| Pld3 | 13,9298 | 11,1695 | -0,318609 | 0,0945 | 0,381771 |
| Tbc1d8b | 1,83532 | 1,38888 | -0,402107 | 0,09445 | 0,381771 |
| Fam214a | 2,02744 | 1,51151 | -0,42367 | 0,0945 | 0,381771 |
| Hddc3 | 11,1875 | 8,1946 | -0,449147 | 0,0944 | 0,381771 |
| Zfp358 | 22,3108 | 27,1715 | 0,284352 | 0,09455 | 0,381857 |
| Fkbpl | 8,82373 | 11,5657 | 0,390388 | 0,0946 | 0,381944 |
| Slc4a1ap | 19,081 | 23,3198 | 0,289422 | 0,09465 | 0,38203 |

| | | | | | |
|---|---|---|---|---|---|
| Peg13 | 4,45775 | 3,13123 | -0,509586 | 0,0947 | 0,382117 |
| 2010015L04Rik | 2,84963 | 3,82412 | 0,424354 | 0,09475 | 0,382174 |
| Pdap1 | 113,433 | 136,157 | 0,263428 | 0,0948 | 0,382174 |
| Arsg | 2,06821 | 1,18906 | -0,798555 | 0,0948 | 0,382174 |
| Yae1d1 | 7,19299 | 5,74593 | -0,324052 | 0,0949 | 0,382347 |
| Slc35a2 | 10,3937 | 8,2136 | -0,339629 | 0,0949 | 0,382347 |
| Sap30 | 45,2838 | 55,2437 | 0,286816 | 0,09495 | 0,382433 |
| Pin1 | 21,7854 | 26,2975 | 0,271565 | 0,0951 | 0,382662 |
| Mthfd1 | 54,3432 | 65,3326 | 0,265702 | 0,09505 | 0,382662 |
| Rfc1 | 43,9531 | 52,8071 | 0,264766 | 0,09515 | 0,382662 |
| Rab14 | 58,1175 | 48,3469 | -0,265549 | 0,0951 | 0,382662 |
| Nagpa | 5,56671 | 4,23616 | -0,394068 | 0,09515 | 0,382662 |
| Cdk12 | 17,9968 | 21,7704 | 0,274624 | 0,0952 | 0,382748 |
| Sdk2 | 1,562 | 2,02637 | 0,375507 | 0,0953 | 0,382804 |
| Pde10a | 2,59283 | 3,24688 | 0,324525 | 0,0953 | 0,382804 |
| Setd1b | 10,0895 | 12,1747 | 0,271033 | 0,0953 | 0,382804 |
| Lphn2 | 56,4563 | 46,9607 | -0,265681 | 0,09535 | 0,38289 |
| Ctc1 | 27,2945 | 32,9745 | 0,272739 | 0,0954 | 0,382976 |
| Sbds | 32,3072 | 39,4547 | 0,288341 | 0,0956 | 0,383548 |
| Stoml2 | 75,7625 | 91,1531 | 0,266808 | 0,0956 | 0,383548 |
| Ednra | 7,47614 | 5,99442 | -0,318674 | 0,09565 | 0,383633 |
| Vcam1 | 28,4414 | 34,1984 | 0,265933 | 0,0957 | 0,383719 |
| Mrs2 | 15,1853 | 18,6728 | 0,298258 | 0,0958 | 0,384005 |
| Pwwp2b | 1,80785 | 1,24863 | -0,533932 | 0,09595 | 0,38449 |
| Mettl1 | 34,4611 | 42,7572 | 0,311199 | 0,096 | 0,384576 |
| H2afy | 188,199 | 225,827 | 0,262957 | 0,09615 | 0,385061 |
| Zfp389 | 0,818794 | 1,55786 | 0,927996 | 0,09625 | 0,385115 |
| Bdh1 | 5,02977 | 6,38226 | 0,343575 | 0,09625 | 0,385115 |
| Mrpl11 | 30,3334 | 36,5282 | 0,268104 | 0,0962 | 0,385115 |
| 1110038B12Rik | 209,334 | 254,024 | 0,279159 | 0,09635 | 0,3854 |
| Psmd9 | 16,7773 | 20,4102 | 0,282776 | 0,09645 | 0,385685 |
| Dkk2 | 1,23089 | 1,7052 | 0,470235 | 0,09655 | 0,385738 |
| Tulp4 | 3,86998 | 4,73277 | 0,290358 | 0,0965 | 0,385738 |
| Prmt5 | 88,8201 | 106,524 | 0,262216 | 0,09655 | 0,385738 |
| Fdft1 | 20,4193 | 28,0678 | 0,458985 | 0,0966 | 0,385823 |
| Gm17296 | 8,11256 | 9,88358 | 0,284878 | 0,0967 | 0,386107 |
| Pspc1 | 56,2801 | 67,6964 | 0,266455 | 0,0968 | 0,38639 |
| Zfp287 | 2,76757 | 3,52749 | 0,350026 | 0,09695 | 0,386874 |
| Oxr1 | 11,6736 | 9,55361 | -0,289137 | 0,097 | 0,386958 |
| Cyp2j9 | 0,751948 | 0,410483 | -0,87331 | 0,0971 | 0,387241 |
| Eef1d | 141,615 | 170,152 | 0,264852 | 0,0973 | 0,387923 |
| Dennd2c | 2,88788 | 2,24227 | -0,365047 | 0,0974 | 0,388206 |
| Maged1 | 238,902 | 289,858 | 0,278929 | 0,09745 | 0,388289 |
| Immt | 91,2693 | 109,558 | 0,263497 | 0,0975 | 0,388373 |
| Laptm4b | 84,059 | 100,777 | 0,261699 | 0,09755 | 0,388456 |
| Derl1 | 26,0339 | 21,5275 | -0,274213 | 0,09765 | 0,388738 |
| Clic1 | 122,844 | 102,17 | -0,265849 | 0,0977 | 0,388822 |
| Ccdc15 | 2,68235 | 3,50277 | 0,384998 | 0,09775 | 0,388905 |

| | | | | | |
|---|---|---|---|---|---|
| 5730408K05Rik | 30,5228 | 9,80775 | -1,63789 | 0,0978 | 0,388988 |
| Rcl1 | 46,0202 | 55,4468 | 0,268836 | 0,09795 | 0,389469 |
| Prdm2 | 19,6074 | 23,5567 | 0,264744 | 0,098 | 0,389551 |
| Lor | 1,26671 | 1,91164 | 0,593723 | 0,09815 | 0,389684 |
| Rps6ka2 | 6,61366 | 8,07425 | 0,28788 | 0,09815 | 0,389684 |
| Zfp281 | 19,1224 | 15,8599 | -0,270515 | 0,0981 | 0,389684 |
| Ier3ip1 | 62,017 | 51,1835 | -0,276984 | 0,0981 | 0,389684 |
| Iqcb1 | 31,4777 | 37,9772 | 0,270803 | 0,09825 | 0,389965 |
| Polr2b | 69,5302 | 83,5922 | 0,265727 | 0,0983 | 0,390048 |
| Zfp90 | 11,9741 | 14,733 | 0,299134 | 0,09835 | 0,39013 |
| 4933404O12Rik | 5,07001 | 3,98524 | -0,34732 | 0,0984 | 0,390213 |
| Rhou | 18,4711 | 15,2109 | -0,280169 | 0,0985 | 0,39046 |
| 2510039O18Rik | 21,5839 | 17,7646 | -0,280949 | 0,09855 | 0,39046 |
| Tmppe | 3,11935 | 2,29867 | -0,440447 | 0,09855 | 0,39046 |
| Strada | 15,5865 | 19,0315 | 0,288092 | 0,0988 | 0,391335 |
| Kctd18 | 5,05085 | 6,44703 | 0,35211 | 0,09895 | 0,391813 |
| Il18 | 0,670627 | 1,39475 | 1,05642 | 0,09905 | 0,391976 |
| Pqlc1 | 2,51791 | 1,76344 | -0,513835 | 0,09905 | 0,391976 |
| Serpina3h | 0,888139 | 0,512972 | -0,791905 | 0,0991 | 0,392058 |
| Rufy3 | 11,6365 | 14,097 | 0,276736 | 0,09915 | 0,39214 |
| Dedd2 | 4,1556 | 3,05736 | -0,442769 | 0,0992 | 0,392222 |
| Gltp | 17,6811 | 21,827 | 0,303902 | 0,09935 | 0,392582 |
| Gm53 | 67,6726 | 83,0866 | 0,296044 | 0,09935 | 0,392582 |
| Cks1brt | 1,31995 | 1,89528 | 0,521921 | 0,0994 | 0,392595 |
| Bcl2l11 | 23,6433 | 28,6614 | 0,277674 | 0,09945 | 0,392595 |
| Anp32a | 293,123 | 353,348 | 0,269585 | 0,0995 | 0,392595 |
| Crtap | 19,1842 | 15,4601 | -0,311369 | 0,09945 | 0,392595 |
| Fam131c | 0,749254 | 0,368123 | -1,02527 | 0,0995 | 0,392595 |
| Nfxl1 | 20,0667 | 24,1493 | 0,267182 | 0,09955 | 0,392676 |
| 1600002H07Rik | 16,4721 | 20,1529 | 0,290961 | 0,0996 | 0,392757 |
| Fgf9 | 4,67713 | 5,84335 | 0,32117 | 0,09965 | 0,392839 |
| Rimbp2 | 1,98267 | 2,60072 | 0,391464 | 0,09985 | 0,393163 |
| Lrp6 | 19,504 | 16,2729 | -0,261302 | 0,09985 | 0,393163 |
| Fam101b | 22,3566 | 18,4989 | -0,273263 | 0,09985 | 0,393163 |
| Klk15 | 3,51318 | 2,1403 | -0,714961 | 0,0998 | 0,393163 |
| Tubb2b | 55,6273 | 66,8267 | 0,264632 | 0,0999 | 0,393244 |
| Fbxo27 | 1,55214 | 1,0142 | -0,613919 | 0,1 | 0,393522 |
| Jade1 | 20,5764 | 17,0735 | -0,26923 | 0,10005 | 0,393603 |
| Ddx11 | 15,897 | 19,1614 | 0,269446 | 0,10015 | 0,393764 |
| Larp7 | 58,2493 | 69,8297 | 0,261598 | 0,10015 | 0,393764 |
| Fam134c | 32,6755 | 27,1472 | -0,267406 | 0,1002 | 0,393845 |
| Mfap1b | 1,8992 | 2,71151 | 0,513707 | 0,10025 | 0,393926 |
| Map2k3os | 0,918823 | 1,55867 | 0,762456 | 0,1004 | 0,393936 |
| Cenpt | 21,1846 | 25,9316 | 0,291695 | 0,10035 | 0,393936 |
| Cfl2 | 54,1199 | 45,1257 | -0,262212 | 0,1004 | 0,393936 |
| Map3k11 | 18,2963 | 15,1358 | -0,273585 | 0,10035 | 0,393936 |
| Pcsk1 | 1,80013 | 1,25032 | -0,525802 | 0,10035 | 0,393936 |
| Hsdl1 | 8,58142 | 6,9057 | -0,313428 | 0,1005 | 0,393981 |

| | | | | | |
|---|---|---|---|---|---|
| Ttc39a | 2,05188 | 1,43583 | -0,515063 | 0,1005 | 0,393981 |
| Rfx4 | 1,2343 | 0,851756 | -0,535179 | 0,1005 | 0,393981 |
| Rab38 | 1,63847 | 1,03588 | -0,661494 | 0,10055 | 0,394062 |
| Scaf11 | 46,2012 | 55,4814 | 0,264074 | 0,1007 | 0,394534 |
| Crcp | 25,9341 | 31,7365 | 0,291292 | 0,1008 | 0,39481 |
| Mphosph8 | 43,417 | 52,5491 | 0,275407 | 0,10085 | 0,39489 |
| Amfr | 36,0596 | 30,0397 | -0,263514 | 0,1009 | 0,39497 |
| Kcnc3 | 1,87122 | 2,4355 | 0,380237 | 0,10095 | 0,39505 |
| Mob1b | 9,69444 | 7,82164 | -0,309685 | 0,101 | 0,39513 |
| Tmem11 | 39,2943 | 47,747 | 0,281091 | 0,10115 | 0,395601 |
| 6330403A02Rik | 1,1742 | 0,863561 | -0,443315 | 0,1014 | 0,396463 |
| Rps25 | 10,4548 | 16,0689 | 0,620099 | 0,1017 | 0,397403 |
| Dynll1 | 89,1217 | 106,77 | 0,260658 | 0,1017 | 0,397403 |
| Irs2 | 8,32824 | 10,1678 | 0,287929 | 0,1018 | 0,397445 |
| Nxt1 | 58,4068 | 70,7614 | 0,276826 | 0,1018 | 0,397445 |
| Bri3bp | 17,4956 | 14,5522 | -0,26575 | 0,10175 | 0,397445 |
| Mdn1 | 13,4552 | 16,1751 | 0,265614 | 0,1019 | 0,39772 |
| Vdac1 | 159,886 | 191,615 | 0,261169 | 0,102 | 0,397994 |
| Bche | 5,03141 | 4,07926 | -0,302656 | 0,10205 | 0,398072 |
| Slit3 | 17,457 | 20,954 | 0,263416 | 0,10225 | 0,398504 |
| Pcsk5 | 8,01848 | 6,59899 | -0,281082 | 0,10225 | 0,398504 |
| Nlrx1 | 0,878747 | 0,582893 | -0,592219 | 0,1022 | 0,398504 |
| Rab2a | 51,2139 | 42,5623 | -0,266958 | 0,1024 | 0,398856 |
| Trp53inp2 | 9,09285 | 7,40442 | -0,296345 | 0,1024 | 0,398856 |
| Hmgb2 | 23,8185 | 28,7755 | 0,272756 | 0,1028 | 0,399058 |
| Eif2b1 | 42,8071 | 51,4462 | 0,265214 | 0,1025 | 0,399058 |
| Prdx6 | 125,261 | 149,997 | 0,259999 | 0,1026 | 0,399058 |
| Hip1r | 3,24099 | 2,53245 | -0,355903 | 0,10275 | 0,399058 |
| 2610316D01Rik | 0,783285 | 0,112453 | -2,80022 | 0,1029 | 0,399058 |
| Marcksl1-ps4 | 1,03049 | 0,0830716 | -3,63284 | 0,1029 | 0,399058 |
| Cd52 | 4,37737 | 0,233094 | -4,23108 | 0,1029 | 0,399058 |
| Dbx2 | 0,639108 | 0,0337193 | -4,24441 | 0,1029 | 0,399058 |
| 6030468B19Rik | 2,89241 | 0,079745 | -5,18073 | 0,1029 | 0,399058 |
| Tlr4 | 0,728201 | 0,0191373 | -5,24988 | 0,1029 | 0,399058 |
| Alox12 | 1,04939 | 0,0250082 | -5,391 | 0,1029 | 0,399058 |
| Tubb1 | 3,37454 | 0,0387209 | -6,44543 | 0,1029 | 0,399058 |
| 1300017J02Rik | 3,10649 | 0,0336541 | -6,52836 | 0,1029 | 0,399058 |
| Hbb-bt | 83,4572 | 0,163606 | -8,99467 | 0,1029 | 0,399058 |
| Fcgr3 | 1,55613 | 0,0606592 | -4,68109 | 0,1029 | 0,399059 |
| Pbdc1 | 29,9515 | 36,2898 | 0,276935 | 0,10295 | 0,399099 |
| Mprip | 38,8853 | 46,5453 | 0,259411 | 0,103 | 0,399099 |
| Elmod2 | 9,18716 | 7,54329 | -0,284426 | 0,103 | 0,399099 |
| Tnnc1 | 0,593232 | 1,37738 | 1,21526 | 0,10315 | 0,399333 |
| Med4 | 46,7847 | 56,6648 | 0,276418 | 0,10315 | 0,399333 |
| Pip4k2a | 7,13032 | 5,72622 | -0,316383 | 0,10315 | 0,399333 |
| Hist4h4 | 1,10865 | 2,71614 | 1,29276 | 0,10325 | 0,399604 |
| Pdia4 | 134,804 | 161,527 | 0,260913 | 0,10335 | 0,399644 |
| B3galt4 | 1,45295 | 0,89866 | -0,693139 | 0,10335 | 0,399644 |

| | | | | | |
|---|---|---|---|---|---|
| Kcnj6 | 0,660463 | 0,393013 | -0,748898 | 0,10335 | 0,399644 |
| Zfp652 | 11,6493 | 14,3097 | 0,296755 | 0,1035 | 0,399992 |
| Mmadhc | 88,6302 | 106,358 | 0,26306 | 0,1035 | 0,399992 |
| Wbscr27 | 3,12084 | 2,33206 | -0,420328 | 0,10355 | 0,40007 |
| Eef1g | 92,6271 | 111,11 | 0,262483 | 0,1036 | 0,400148 |
| Cort | 0,462923 | 1,60543 | 1,79412 | 0,1037 | 0,400149 |
| Rps6ka1 | 28,7208 | 34,4254 | 0,261378 | 0,10375 | 0,400149 |
| Sulf1 | 65,1377 | 78,0433 | 0,260783 | 0,1037 | 0,400149 |
| Las1l | 42,083 | 50,3913 | 0,259937 | 0,10365 | 0,400149 |
| Trp53 | 87,852 | 105,098 | 0,25859 | 0,10375 | 0,400149 |
| Amdhd2 | 8,7829 | 11,2535 | 0,357599 | 0,10425 | 0,401151 |
| Clstn2 | 4,40528 | 5,49154 | 0,317978 | 0,1042 | 0,401151 |
| Gstcd | 21,883 | 26,4469 | 0,273286 | 0,1042 | 0,401151 |
| Iqgap2 | 12,1594 | 14,6247 | 0,266332 | 0,10425 | 0,401151 |
| Nmral1 | 80,2762 | 96,0878 | 0,259381 | 0,1041 | 0,401151 |
| Slc30a7 | 8,05201 | 6,68582 | -0,268245 | 0,10405 | 0,401151 |
| Robo1 | 9,54926 | 7,92653 | -0,2687 | 0,10415 | 0,401151 |
| Tom1l2 | 5,74998 | 4,603 | -0,320983 | 0,1042 | 0,401151 |
| Cda | 2,43684 | 1,40297 | -0,796524 | 0,1044 | 0,401612 |
| Mapk7 | 32,7625 | 39,2327 | 0,260014 | 0,10445 | 0,401689 |
| Nova1 | 2,17158 | 2,73981 | 0,335333 | 0,1045 | 0,401766 |
| Rpl36a | 0,477541 | 1,6392 | 1,77929 | 0,1046 | 0,402035 |
| Msc | 1,88984 | 1,25519 | -0,590364 | 0,10465 | 0,402111 |
| Rnf19b | 15,7003 | 19,0721 | 0,280675 | 0,1049 | 0,402685 |
| Banp | 8,98263 | 10,8681 | 0,274886 | 0,10495 | 0,402685 |
| Tnfrsf19 | 54,9334 | 65,4654 | 0,253049 | 0,10495 | 0,402685 |
| Zfp518a | 11,2564 | 9,35825 | -0,26644 | 0,1049 | 0,402685 |
| Slc50a1 | 12,4916 | 9,54209 | -0,388584 | 0,1049 | 0,402685 |
| Rab12 | 22,3223 | 18,3021 | -0,28648 | 0,105 | 0,402761 |
| Cep19 | 14,2798 | 17,6983 | 0,309632 | 0,10505 | 0,402837 |
| Hist2h4 | 0,427113 | 2,55815 | 2,58241 | 0,1051 | 0,402913 |
| Gsta4 | 102,617 | 123,42 | 0,266306 | 0,10515 | 0,40299 |
| Raet1a,Raet1b | 1,02728 | 0,242715 | -2,08149 | 0,1052 | 0,403066 |
| Lypd2 | 0,677908 | 1,88087 | 1,47224 | 0,1053 | 0,403333 |
| Eif3g | 175,711 | 209,713 | 0,255215 | 0,10535 | 0,403409 |
| Tbc1d8 | 1,95702 | 1,28942 | -0,601929 | 0,10545 | 0,403676 |
| Tvp23b | 11,0696 | 8,74581 | -0,339943 | 0,1055 | 0,403752 |
| Scube2 | 1,00137 | 1,4075 | 0,491152 | 0,1057 | 0,404286 |
| Alx3 | 4,15964 | 5,46762 | 0,394452 | 0,1057 | 0,404286 |
| Ubap1 | 29,909 | 35,842 | 0,26107 | 0,10585 | 0,404628 |
| Oaf | 29,2364 | 24,2363 | -0,270597 | 0,10585 | 0,404628 |
| Adamts6 | 3,97616 | 4,9585 | 0,318528 | 0,106 | 0,404698 |
| Commd1 | 122,389 | 148,735 | 0,281278 | 0,10605 | 0,404698 |
| Gm15800 | 4,66376 | 5,60054 | 0,264071 | 0,106 | 0,404698 |
| Gars | 98,4943 | 117,743 | 0,257526 | 0,106 | 0,404698 |
| Fam159a | 0,724348 | 0,100283 | -2,8526 | 0,10605 | 0,404698 |
| Dlx6 | 1,15401 | 0,0803888 | -3,84352 | 0,106 | 0,404698 |
| Pcgf1 | 4,82334 | 6,81108 | 0,497853 | 0,1062 | 0,405155 |

| | | | | | |
|---|---|---|---|---|---|
| Dnajc9 | 90,9917 | 108,692 | 0,256434 | 0,10635 | 0,405611 |
| Cdc25b | 11,4922 | 4,96398 | -1,21109 | 0,10645 | 0,405877 |
| Pgrmc1 | 132,333 | 110,749 | -0,256873 | 0,1065 | 0,405952 |
| Cpeb3 | 0,951673 | 0,689961 | -0,463951 | 0,10655 | 0,406026 |
| Gramd3 | 12,4657 | 10,1435 | -0,297414 | 0,10665 | 0,40606 |
| Stard5 | 4,65662 | 3,56071 | -0,38712 | 0,10665 | 0,40606 |
| Rffl | 3,86285 | 1,73394 | -1,15561 | 0,1066 | 0,40606 |
| Gprin1 | 1,03309 | 1,43258 | 0,471646 | 0,1068 | 0,406515 |
| Plekhg2 | 32,3313 | 38,6528 | 0,257642 | 0,1069 | 0,406664 |
| Rnf144b | 0,771354 | 0,525991 | -0,552355 | 0,1069 | 0,406664 |
| 4930444P10Rik | 0,173081 | 0,85822 | 2,3099 | 0,10695 | 0,406739 |
| Snrnp200 | 57,5733 | 68,9939 | 0,261069 | 0,107 | 0,406813 |
| Gm14305 | 0,743596 | 1,23845 | 0,735939 | 0,1071 | 0,406962 |
| Kcnh2 | 3,26117 | 4,12075 | 0,337515 | 0,1071 | 0,406962 |
| Zfp456 | 0,585149 | 0,375227 | -0,641038 | 0,10715 | 0,407036 |
| Kdm6b | 17,4085 | 20,8062 | 0,257226 | 0,1072 | 0,40711 |
| Klc1 | 43,6238 | 53,2779 | 0,288422 | 0,1073 | 0,407217 |
| Gins4 | 48,0336 | 57,8459 | 0,26817 | 0,10735 | 0,407217 |
| Slc35b2 | 33,8408 | 28,0803 | -0,269207 | 0,10735 | 0,407217 |
| B3galnt1 | 6,98723 | 5,46714 | -0,353935 | 0,1073 | 0,407217 |
| Cnot3 | 30,3052 | 37,0079 | 0,288269 | 0,1074 | 0,407291 |
| Tmem59 | 92,7862 | 77,5093 | -0,259541 | 0,10745 | 0,407365 |
| Ubl7 | 32,6302 | 39,5491 | 0,277441 | 0,10755 | 0,407397 |
| Ppa1 | 188,168 | 224,496 | 0,254668 | 0,10755 | 0,407397 |
| Gm16675 | 0,780646 | 0,384958 | -1,01997 | 0,1075 | 0,407397 |
| Fam20b | 26,914 | 22,5209 | -0,257089 | 0,1076 | 0,407471 |
| Aff3 | 1,75929 | 1,34124 | -0,391426 | 0,1077 | 0,407504 |
| Fosl2 | 1,10893 | 0,812157 | -0,449339 | 0,1077 | 0,407504 |
| Anxa11 | 2,45064 | 1,76443 | -0,473953 | 0,10765 | 0,407504 |
| Comp | 0,799137 | 0,482619 | -0,727557 | 0,1078 | 0,407767 |
| Tbkbp1 | 8,04784 | 6,49947 | -0,30828 | 0,108 | 0,408407 |
| Slbp | 55,1762 | 66,3095 | 0,265169 | 0,10805 | 0,408481 |
| Paf1 | 49,6485 | 59,4759 | 0,260555 | 0,1081 | 0,408554 |
| Slc35f5 | 14,3628 | 11,7556 | -0,288989 | 0,10815 | 0,408628 |
| Vash1 | 23,6332 | 19,7711 | -0,257421 | 0,1082 | 0,408701 |
| Pold3 | 42,9301 | 51,3188 | 0,257496 | 0,1083 | 0,408848 |
| C330027C09Rik | 40,2142 | 33,6761 | -0,255982 | 0,1083 | 0,408848 |
| Rab1 | 86,6931 | 72,6645 | -0,254668 | 0,10835 | 0,408921 |
| Hipk2 | 19,048 | 22,9313 | 0,267683 | 0,1085 | 0,409256 |
| Aldh5a1 | 5,97317 | 4,85255 | -0,299754 | 0,1085 | 0,409256 |
| Rwdd1 | 20,6047 | 25,6303 | 0,314874 | 0,1086 | 0,40936 |
| Rbmxl1 | 23,6742 | 28,7842 | 0,28196 | 0,10865 | 0,40936 |
| Ccnt2 | 31,8535 | 38,0503 | 0,256455 | 0,10865 | 0,40936 |
| Rbm33 | 14,785 | 17,6599 | 0,256339 | 0,10865 | 0,40936 |
| BC029214 | 2,86439 | 1,8241 | -0,651046 | 0,1087 | 0,409433 |
| 1700030J22Rik | 1,13819 | 1,60737 | 0,497954 | 0,10875 | 0,409505 |
| Hmcn1 | 6,00463 | 7,20072 | 0,262067 | 0,1089 | 0,409955 |
| Irx5 | 0,508932 | 0,837583 | 0,718758 | 0,10895 | 0,410028 |

| | | | | | |
|---|---|---|---|---|---|
| Gcsh | 69,6009 | 83,2955 | 0,259132 | 0,109 | 0,4101 |
| Smpdl3a | 9,3099 | 7,33435 | -0,344097 | 0,10905 | 0,410173 |
| Cbx6 | 11,0536 | 9,09302 | -0,281683 | 0,10935 | 0,411186 |
| Nufip1 | 12,7983 | 15,4359 | 0,270335 | 0,1094 | 0,411258 |
| Scand1 | 24,23 | 30,7314 | 0,342918 | 0,10945 | 0,41133 |
| Inca1 | 0,508084 | 0,948305 | 0,900285 | 0,10955 | 0,41159 |
| Lrrc49 | 13,0212 | 15,7874 | 0,277912 | 0,1096 | 0,411662 |
| Slc39a13 | 6,79477 | 5,32221 | -0,352397 | 0,10965 | 0,411734 |
| Ebna1bp2 | 47,8904 | 57,1683 | 0,255477 | 0,10985 | 0,412369 |
| Dpf2 | 34,1172 | 40,7222 | 0,255315 | 0,1099 | 0,412441 |
| Phactr1 | 1,84437 | 1,37737 | -0,421214 | 0,10995 | 0,412513 |
| Cdh11 | 54,6311 | 45,7578 | -0,255705 | 0,11 | 0,412585 |
| Snrpd3 | 553,905 | 659,893 | 0,252594 | 0,11005 | 0,412657 |
| Zscan10 | 1,48473 | 2,0573 | 0,470551 | 0,1101 | 0,412728 |
| Mst1r | 0,337444 | 0,673839 | 0,997756 | 0,11015 | 0,4128 |
| Efnb3 | 15,9364 | 13,1354 | -0,278861 | 0,1102 | 0,412872 |
| Fam57b | 0,550219 | 0,295829 | -0,89524 | 0,1103 | 0,41313 |
| Psd2 | 0,775995 | 1,08826 | 0,487902 | 0,11055 | 0,41379 |
| Steap2 | 3,90726 | 3,22275 | -0,277866 | 0,1106 | 0,41379 |
| Bahd1 | 7,69855 | 6,29914 | -0,289431 | 0,1106 | 0,41379 |
| Fbxo46 | 6,79351 | 5,42183 | -0,325377 | 0,11055 | 0,41379 |
| Ckb | 18,6796 | 22,9827 | 0,299084 | 0,1108 | 0,414101 |
| Qrsl1 | 13,26 | 16,2685 | 0,294999 | 0,1109 | 0,414101 |
| Mocs2 | 15,684 | 19,2042 | 0,292129 | 0,11085 | 0,414101 |
| Ccnd3 | 68,5986 | 54 | -0,345218 | 0,11085 | 0,414101 |
| Fhl2 | 5,74155 | 4,28394 | -0,422503 | 0,1108 | 0,414101 |
| Fads3 | 2,31774 | 1,71879 | -0,431329 | 0,1109 | 0,414101 |
| Myb | 0,723083 | 0,467768 | -0,62837 | 0,11075 | 0,414101 |
| Zfp202 | 5,74724 | 7,08814 | 0,302539 | 0,11105 | 0,414545 |
| Nrbf2 | 2,95435 | 4,0199 | 0,444317 | 0,1111 | 0,414616 |
| Gys1 | 16,6363 | 19,9858 | 0,26464 | 0,1112 | 0,414873 |
| Acly | 129,559 | 155,354 | 0,261949 | 0,11135 | 0,415317 |
| B630005N14Rik | 12,5784 | 10,3872 | -0,276143 | 0,1114 | 0,415388 |
| Nob1 | 50,642 | 60,6735 | 0,260732 | 0,11155 | 0,415831 |
| Rnf113a2 | 9,80916 | 13,3259 | 0,442028 | 0,1117 | 0,416274 |
| Zfp101 | 21,0761 | 17,4667 | -0,271003 | 0,11185 | 0,416601 |
| Tmem106c | 11,8334 | 9,36062 | -0,338187 | 0,11185 | 0,416601 |
| Otx1 | 1,40461 | 1,95255 | 0,475195 | 0,11195 | 0,416857 |
| Mfsd9 | 1,11459 | 0,761941 | -0,548762 | 0,112 | 0,416927 |
| Snora15 | 15,3727 | 20,0991 | 0,386759 | 0,1121 | 0,417183 |
| Sema4g | 18,6593 | 22,3243 | 0,25872 | 0,1122 | 0,417323 |
| Fam162a | 202,46 | 242,076 | 0,257826 | 0,1122 | 0,417323 |
| Ptprg | 16,9318 | 14,2059 | -0,253245 | 0,1123 | 0,417462 |
| Sppl2b | 12,1696 | 9,97758 | -0,286517 | 0,1123 | 0,417462 |
| Gas8 | 21,3969 | 25,9883 | 0,280463 | 0,11235 | 0,417532 |
| Hist2h2bb | 0,245556 | 5,27211 | 4,42426 | 0,11255 | 0,417927 |
| Med8 | 36,3224 | 43,4525 | 0,258577 | 0,1125 | 0,417927 |
| Plagl2 | 36,1124 | 42,9533 | 0,250273 | 0,11255 | 0,417927 |

| | | | | | |
|---|---|---|---|---|---|
| Tbcc | 11,898 | 14,7216 | 0,307208 | 0,11265 | 0,418019 |
| Ugcg | 35,1758 | 29,4592 | -0,255867 | 0,11265 | 0,418019 |
| Pptc7 | 9,76292 | 8,0549 | -0,277447 | 0,1127 | 0,418019 |
| Dpy19l4 | 14,1947 | 11,685 | -0,280697 | 0,1127 | 0,418019 |
| Bri3 | 12,8839 | 18,6516 | 0,533727 | 0,1129 | 0,418645 |
| Rcbtb1 | 36,0806 | 43,0404 | 0,254469 | 0,11295 | 0,418667 |
| Ubxn1 | 136,697 | 163,016 | 0,254034 | 0,113 | 0,418667 |
| Nr2f1 | 1,55696 | 1,05216 | -0,565384 | 0,113 | 0,418667 |
| Mcm2 | 123,663 | 147,612 | 0,255392 | 0,1131 | 0,418921 |
| Rbp4 | 7,10832 | 5,10521 | -0,477538 | 0,11325 | 0,419361 |
| Tmem159 | 7,62221 | 5,81334 | -0,390841 | 0,11385 | 0,421466 |
| 4732456N10Rik | 0,74172 | 1,12488 | 0,600819 | 0,1139 | 0,421534 |
| Zc3h18 | 36,5598 | 43,4786 | 0,250045 | 0,11415 | 0,422342 |
| Casc4 | 21,3003 | 17,8363 | -0,256061 | 0,1142 | 0,42241 |
| Tmem74b | 0,668651 | 1,15423 | 0,787611 | 0,1143 | 0,422663 |
| Sox8 | 1,06546 | 1,50015 | 0,493633 | 0,11475 | 0,423809 |
| Dnajc21 | 23,382 | 28,1523 | 0,267851 | 0,11475 | 0,423809 |
| Zswim4 | 17,5154 | 20,9326 | 0,257133 | 0,11475 | 0,423809 |
| Rps9 | 732,359 | 874,55 | 0,25599 | 0,11475 | 0,423809 |
| AW549877 | 26,5774 | 22,2829 | -0,254263 | 0,1148 | 0,423809 |
| Adcy9 | 0,581082 | 0,41508 | -0,485353 | 0,1148 | 0,423809 |
| Oard1 | 55,1646 | 66,2788 | 0,264803 | 0,11485 | 0,423876 |
| Cyp26b1 | 1,95545 | 1,48744 | -0,394672 | 0,11495 | 0,424128 |
| Pgap3 | 4,86337 | 3,76281 | -0,370146 | 0,11505 | 0,42438 |
| Gabrb3 | 0,378137 | 0,562475 | 0,572882 | 0,1151 | 0,424447 |
| Naa10 | 69,0078 | 82,7738 | 0,262414 | 0,11515 | 0,424514 |
| Mthfd1l | 21,4274 | 25,6153 | 0,257548 | 0,11565 | 0,42624 |
| Ago2 | 21,8475 | 26,0271 | 0,252543 | 0,1157 | 0,426307 |
| Cfdp1 | 159,85 | 189,946 | 0,248872 | 0,11585 | 0,426624 |
| Arf5 | 99,5067 | 83,0986 | -0,259969 | 0,11585 | 0,426624 |
| Utp23 | 7,09342 | 5,67863 | -0,320939 | 0,1159 | 0,426691 |
| Gpr125 | 102,702 | 122,607 | 0,255581 | 0,11595 | 0,426757 |
| Spa17 | 3,17885 | 4,87168 | 0,615915 | 0,116 | 0,426824 |
| Rbp1 | 31,4591 | 37,596 | 0,2571 | 0,1161 | 0,427074 |
| Adam17 | 30,6568 | 25,449 | -0,268595 | 0,11625 | 0,427508 |
| Ndufv3 | 85,059 | 106,217 | 0,320483 | 0,1163 | 0,427574 |
| Ift22 | 26,3823 | 32,6151 | 0,305966 | 0,11665 | 0,428625 |
| Casp6 | 30,9372 | 37,3061 | 0,27007 | 0,11665 | 0,428625 |
| Mutyh | 11,2237 | 13,9637 | 0,315126 | 0,1168 | 0,428823 |
| Snrpd2 | 188,652 | 226,224 | 0,262028 | 0,11675 | 0,428823 |
| Ints10 | 34,629 | 41,307 | 0,254408 | 0,1168 | 0,428823 |
| Sik1 | 9,63137 | 11,6274 | 0,271714 | 0,11685 | 0,428889 |
| Klf6 | 8,44152 | 6,92525 | -0,285637 | 0,11705 | 0,429387 |
| Zfp808 | 3,44394 | 2,64627 | -0,380101 | 0,11705 | 0,429387 |
| Tlr5 | 1,06339 | 0,7304 | -0,541916 | 0,11715 | 0,429636 |
| Zdhhc18 | 23,4895 | 19,5471 | -0,265064 | 0,11725 | 0,429884 |
| Bod1 | 55,4647 | 66,3044 | 0,257533 | 0,1173 | 0,42995 |
| Ccdc42 | 0,921969 | 1,51228 | 0,713934 | 0,11745 | 0,429975 |

| | | | | | |
|---|---|---|---|---|---|
| Dis3 | 41,006 | 49,3104 | 0,266057 | 0,1175 | 0,429975 |
| Rpl41 | 330,967 | 394,893 | 0,254774 | 0,1175 | 0,429975 |
| Dnajc11 | 51,5858 | 61,3386 | 0,249819 | 0,1175 | 0,429975 |
| Acer3 | 7,81966 | 6,39142 | -0,290969 | 0,11735 | 0,429975 |
| Gpm6a | 1,24749 | 0,866476 | -0,525797 | 0,11745 | 0,429975 |
| Tlcd1 | 9,3945 | 7,1854 | -0,386748 | 0,1176 | 0,430223 |
| 1700007L15Rik | 2,46352 | 3,86209 | 0,648659 | 0,1177 | 0,430236 |
| Zw10 | 31,3934 | 37,4162 | 0,253199 | 0,11765 | 0,430236 |
| Mxra7 | 11,5478 | 9,21335 | -0,325816 | 0,1177 | 0,430236 |
| Polr2g | 121,266 | 144,694 | 0,25483 | 0,11785 | 0,430549 |
| Slc35f2 | 4,18221 | 3,21777 | -0,378202 | 0,11785 | 0,430549 |
| Hirip3 | 51,0484 | 60,8013 | 0,252235 | 0,11795 | 0,430796 |
| Gm16907 | 2,24982 | 5,04115 | 1,16394 | 0,11815 | 0,43082 |
| Klhl5 | 9,58774 | 11,6265 | 0,278157 | 0,11815 | 0,43082 |
| Cntrl | 15,1353 | 18,1161 | 0,259355 | 0,1181 | 0,43082 |
| Bmpr2 | 15,103 | 12,7097 | -0,24891 | 0,1181 | 0,43082 |
| Tmem241 | 1,64428 | 1,18768 | -0,469303 | 0,1181 | 0,43082 |
| Ccdc160 | 1,80896 | 1,19262 | -0,60103 | 0,11805 | 0,43082 |
| Col13a1 | 20,1759 | 24,1399 | 0,258784 | 0,11825 | 0,43095 |
| Tmeff1 | 71,5728 | 60,1657 | -0,25047 | 0,11825 | 0,43095 |
| Pcyt2 | 23,7215 | 28,6161 | 0,270632 | 0,1183 | 0,431015 |
| Tsga10 | 1,70859 | 2,39167 | 0,485212 | 0,1186 | 0,431167 |
| Fn1 | 298,741 | 382,055 | 0,354887 | 0,1185 | 0,431167 |
| Acyp1 | 25,3052 | 32,1689 | 0,34623 | 0,11845 | 0,431167 |
| Hn1l | 106,474 | 126,59 | 0,249652 | 0,11845 | 0,431167 |
| Rfc2 | 83,5781 | 99,0064 | 0,244398 | 0,1186 | 0,431167 |
| Ralgps2 | 17,725 | 14,8466 | -0,255648 | 0,1186 | 0,431167 |
| Slc16a13 | 7,0331 | 5,60766 | -0,326762 | 0,1186 | 0,431167 |
| Def6 | 1,14202 | 0,745005 | -0,61627 | 0,11855 | 0,431167 |
| Stxbp6 | 7,64047 | 6,25412 | -0,288855 | 0,11865 | 0,431232 |
| Tbrg4 | 34,3644 | 41,0345 | 0,255923 | 0,1188 | 0,431542 |
| Fam199x | 7,93004 | 6,61196 | -0,262249 | 0,1188 | 0,431542 |
| Ccdc58 | 73,266 | 88,4045 | 0,270975 | 0,1191 | 0,432397 |
| Ikbkb | 16,1952 | 19,4021 | 0,260646 | 0,1191 | 0,432397 |
| Uba1 | 152,967 | 182,715 | 0,256379 | 0,1192 | 0,432642 |
| Ccdc47 | 28,9393 | 24,2694 | -0,253896 | 0,11925 | 0,432706 |
| Srsf9 | 149,752 | 178,06 | 0,249793 | 0,11965 | 0,433804 |
| Fitm2 | 6,45979 | 4,94086 | -0,386723 | 0,11965 | 0,433804 |
| Plcd1 | 2,1252 | 1,54831 | -0,456907 | 0,11965 | 0,433804 |
| Prlr | 0,333312 | 0,60154 | 0,851788 | 0,11975 | 0,433995 |
| Slc43a1 | 23,5279 | 19,5448 | -0,26759 | 0,1198 | 0,433995 |
| Psen2 | 2,98804 | 2,18102 | -0,454197 | 0,1198 | 0,433995 |
| Tmem176b | 19,877 | 15,9105 | -0,321121 | 0,11995 | 0,434421 |
| Tnfaip3 | 0,989857 | 0,702594 | -0,494528 | 0,12005 | 0,434547 |
| Moap1 | 1,55926 | 0,473454 | -1,71956 | 0,12005 | 0,434547 |
| Auh | 8,19176 | 10,457 | 0,352225 | 0,12015 | 0,434791 |
| Kctd5 | 48,8177 | 57,9287 | 0,246872 | 0,1202 | 0,434855 |
| Ggt7 | 1,91124 | 2,57167 | 0,428199 | 0,12025 | 0,434918 |

| | | | | | |
|---|---|---|---|---|---|
| Os9 | 17,244 | 14,4004 | -0,259982 | 0,12035 | 0,435162 |
| Tigd5 | 1,23615 | 0,906852 | -0,44691 | 0,12065 | 0,436128 |
| Otud1 | 0,67732 | 1,02874 | 0,602965 | 0,12075 | 0,436254 |
| Itpripl1 | 5,97087 | 4,82968 | -0,306012 | 0,12075 | 0,436254 |
| Rnf13 | 19,0626 | 15,5393 | -0,294825 | 0,1208 | 0,436317 |
| Parp10 | 0,64036 | 0,402817 | -0,668757 | 0,12085 | 0,436379 |
| Dmc1 | 1,60274 | 2,23903 | 0,48233 | 0,1209 | 0,436442 |
| Gspt2 | 7,04989 | 8,7413 | 0,310247 | 0,121 | 0,436567 |
| Ywhah | 55,5564 | 66,1339 | 0,251437 | 0,121 | 0,436567 |
| Npepl1 | 16,025 | 13,1392 | -0,286447 | 0,12105 | 0,436629 |
| Adora2b | 1,04677 | 0,645063 | -0,698432 | 0,1211 | 0,436692 |
| Ccm2 | 29,8413 | 35,7979 | 0,262565 | 0,1212 | 0,436935 |
| Sssca1 | 59,0308 | 71,5395 | 0,277272 | 0,12125 | 0,436997 |
| Sod1 | 212,732 | 252,965 | 0,249895 | 0,1213 | 0,437059 |
| Prkag2 | 34,6185 | 41,2627 | 0,253295 | 0,1214 | 0,437066 |
| Cast | 7,40122 | 5,96472 | -0,311309 | 0,1214 | 0,437066 |
| Rasgef1b | 2,61017 | 1,96904 | -0,406653 | 0,1214 | 0,437066 |
| Prosc | 15,9141 | 19,4037 | 0,286026 | 0,1216 | 0,437668 |
| 1110001J03Rik | 44,0337 | 34,188 | -0,365118 | 0,1217 | 0,43791 |
| Zfp618 | 7,32571 | 9,01202 | 0,298882 | 0,12175 | 0,437972 |
| Pik3cd | 2,62346 | 2,05807 | -0,35018 | 0,1218 | 0,438034 |
| Gxylt2 | 2,69716 | 3,65293 | 0,437611 | 0,1219 | 0,438158 |
| Acvr2b | 23,7167 | 28,5999 | 0,270109 | 0,1219 | 0,438158 |
| Prdx4 | 165,619 | 139,293 | -0,249748 | 0,12195 | 0,43822 |
| Gatad2b | 19,8243 | 23,9805 | 0,27459 | 0,1221 | 0,438641 |
| 2500004C02Rik | 8,63195 | 6,8564 | -0,332235 | 0,12215 | 0,438703 |
| Mgat2 | 33,7806 | 28,369 | -0,251881 | 0,1222 | 0,438764 |
| Zfp532 | 21,3718 | 26,1582 | 0,29155 | 0,12225 | 0,438826 |
| Camk4 | 1,17043 | 0,925615 | -0,33856 | 0,1223 | 0,438888 |
| Kdm8 | 8,94137 | 11,0315 | 0,303059 | 0,1225 | 0,439369 |
| Zfp820 | 3,17933 | 2,48141 | -0,357567 | 0,1225 | 0,439369 |
| Slc16a4 | 1,53107 | 1,0447 | -0,551455 | 0,1226 | 0,43961 |
| Tmem214 | 30,6451 | 25,7529 | -0,25092 | 0,1227 | 0,439615 |
| Cds2 | 16,5262 | 13,3834 | -0,304316 | 0,1227 | 0,439615 |
| Gm12504 | 1,12166 | 0,729907 | -0,61985 | 0,12265 | 0,439615 |
| Crtc2 | 17,7594 | 21,3948 | 0,268678 | 0,12285 | 0,440034 |
| Zfp617 | 5,28836 | 4,19031 | -0,335762 | 0,1229 | 0,440095 |
| Clhc1 | 23,0327 | 28,2 | 0,292013 | 0,12305 | 0,440161 |
| Hnrnpm | 299,184 | 358,265 | 0,259992 | 0,12305 | 0,440161 |
| Prmt3 | 25,336 | 30,2298 | 0,254784 | 0,12305 | 0,440161 |
| Pcmtd1 | 15,3502 | 18,2863 | 0,252513 | 0,123 | 0,440161 |
| Dpysl4 | 7,74177 | 9,53619 | 0,300749 | 0,12315 | 0,4404 |
| Arhgef40 | 23,5554 | 29,8563 | 0,34198 | 0,1232 | 0,440461 |
| Polr1d | 81,1953 | 97,1374 | 0,258631 | 0,12325 | 0,440522 |
| Magel2 | 1,01093 | 1,36437 | 0,432557 | 0,1233 | 0,440583 |
| Ube2o | 16,1635 | 19,2384 | 0,251247 | 0,1236 | 0,441301 |
| Ube4b | 37,8125 | 44,781 | 0,244021 | 0,1236 | 0,441301 |
| Fam84b | 4,41543 | 3,50636 | -0,332578 | 0,12355 | 0,441301 |

| | | | | | |
|---|---|---|---|---|---|
| Gins1 | 52,7043 | 63,3365 | 0,265117 | 0,1237 | 0,441305 |
| Acsf2 | 5,5192 | 4,3763 | -0,334747 | 0,1237 | 0,441305 |
| Them6 | 2,61935 | 1,81818 | -0,526713 | 0,12365 | 0,441305 |
| Mns1 | 33,6305 | 43,0829 | 0,357343 | 0,12375 | 0,441365 |
| Ckap4 | 73,851 | 62,3551 | -0,24411 | 0,1239 | 0,441782 |
| Lrrc29 | 1,00583 | 0,621843 | -0,693763 | 0,12395 | 0,441843 |
| Dedd | 16,6675 | 20,7983 | 0,319426 | 0,12415 | 0,44232 |
| Fam208a | 10,6114 | 8,51009 | -0,318372 | 0,12415 | 0,44232 |
| Rbmx2 | 10,194 | 12,6178 | 0,307748 | 0,12425 | 0,44244 |
| Lman2 | 50,3707 | 42,4955 | -0,245275 | 0,12425 | 0,44244 |
| A930013F10Rik | 3,24227 | 4,99051 | 0,622182 | 0,12435 | 0,442678 |
| Rpl8 | 1448,73 | 1749,87 | 0,272464 | 0,12445 | 0,442916 |
| Snx25 | 9,87092 | 8,08577 | -0,2878 | 0,1245 | 0,442976 |
| Mir7007 | 42,5458 | 0 | -inf | 0,1246 | 0,443214 |
| Sec14l2 | 4,34057 | 5,51681 | 0,345949 | 0,12465 | 0,443274 |
| Zfp442 | 2,30858 | 1,70204 | -0,439744 | 0,1247 | 0,443334 |
| Synrg | 3,92615 | 3,2121 | -0,289601 | 0,12475 | 0,443393 |
| Mpp6 | 43,1291 | 51,2189 | 0,248014 | 0,1249 | 0,443573 |
| Foxk2 | 36,4423 | 43,1014 | 0,24212 | 0,12485 | 0,443573 |
| Chrnb4 | 1,29663 | 0,926006 | -0,485678 | 0,1249 | 0,443573 |
| Hoxa13 | 16,7143 | 20,5424 | 0,297524 | 0,125 | 0,44381 |
| Htra3 | 0,885774 | 0,572374 | -0,629982 | 0,12505 | 0,443869 |
| Abcf1 | 73,7835 | 87,398 | 0,244302 | 0,1253 | 0,444639 |
| Zcchc9 | 21,2685 | 25,8167 | 0,279582 | 0,1254 | 0,444875 |
| Rnf169 | 8,70079 | 10,3744 | 0,253809 | 0,1255 | 0,445112 |
| Pa2g4 | 181,219 | 215,133 | 0,247493 | 0,1256 | 0,44523 |
| Chaf1a | 63,4426 | 75,2757 | 0,246734 | 0,1256 | 0,44523 |
| Hist1h2bp | 0,676651 | 1,76745 | 1,38519 | 0,12575 | 0,445467 |
| Gpn1 | 41,6803 | 49,614 | 0,25138 | 0,1258 | 0,445467 |
| Ptpn21 | 5,45275 | 4,48424 | -0,28212 | 0,1258 | 0,445467 |
| Chmp4c | 0,813723 | 0,470352 | -0,790796 | 0,1258 | 0,445467 |
| Tmem198 | 0,648611 | 1,01787 | 0,650135 | 0,12585 | 0,445526 |
| Mettl20 | 7,82045 | 6,14492 | -0,347858 | 0,1259 | 0,445585 |
| Aak1 | 5,03153 | 5,97615 | 0,248218 | 0,12595 | 0,445644 |
| Fnip2 | 4,30644 | 5,20795 | 0,274217 | 0,12615 | 0,446233 |
| Mdfi | 12,0464 | 9,60061 | -0,327409 | 0,1264 | 0,446881 |
| Zfp467 | 6,84046 | 5,16853 | -0,404339 | 0,1264 | 0,446881 |
| Nsun2 | 62,936 | 74,4053 | 0,24152 | 0,12645 | 0,446939 |
| Ube2s | 16,4769 | 20,6001 | 0,322214 | 0,1265 | 0,446998 |
| Stxbp4 | 12,3032 | 14,6039 | 0,247322 | 0,1266 | 0,447233 |
| Tmem209 | 22,8447 | 19,1888 | -0,251599 | 0,12685 | 0,447998 |
| Ubxn10 | 0,894172 | 0,592112 | -0,594682 | 0,12695 | 0,448232 |
| Rpl14 | 202,42 | 239,653 | 0,243597 | 0,12705 | 0,448349 |
| Mturn | 1,43604 | 1,078 | -0,413739 | 0,12705 | 0,448349 |
| Tkt | 181,674 | 216,223 | 0,251164 | 0,1271 | 0,448407 |
| Dyrk2 | 13,5514 | 16,4703 | 0,281424 | 0,12715 | 0,448465 |
| Erlin1 | 18,5681 | 15,5126 | -0,259384 | 0,12735 | 0,449052 |
| C1qtnf1 | 1,03776 | 1,39726 | 0,429126 | 0,1275 | 0,449106 |

| | | | | | |
|---|---|---|---|---|---|
| Nf1 | 7,64737 | 9,08653 | 0,248765 | 0,1275 | 0,449106 |
| Ankrd32 | 15,4635 | 12,9308 | -0,258061 | 0,1274 | 0,449106 |
| Pla2g4b | 6,30112 | 4,41017 | -0,514776 | 0,1275 | 0,449106 |
| Fat3 | 3,88819 | 4,63687 | 0,25405 | 0,1276 | 0,44934 |
| Peg3 | 61,6885 | 73,313 | 0,249069 | 0,1278 | 0,449689 |
| 4931428F04Rik | 8,5195 | 6,98465 | -0,286581 | 0,1278 | 0,449689 |
| Gm13157 | 2,93534 | 2,21946 | -0,40332 | 0,1278 | 0,449689 |
| Kptn | 7,11628 | 8,99349 | 0,337757 | 0,12785 | 0,449743 |
| Dnajc25 | 6,96667 | 8,75972 | 0,330416 | 0,12795 | 0,449743 |
| Zfp64 | 9,91898 | 12,0495 | 0,280707 | 0,1279 | 0,449743 |
| Fam173a | 34,0979 | 27,2992 | -0,320823 | 0,12795 | 0,449743 |
| Scarna13 | 5,52065 | 0 | -inf | 0,12815 | 0,449794 |
| Cwc27 | 16,689 | 20,2137 | 0,276429 | 0,12815 | 0,449794 |
| Mapkbp1 | 5,5538 | 6,67107 | 0,264443 | 0,1282 | 0,449794 |
| Rabggtb | 89,5063 | 106,016 | 0,244218 | 0,1281 | 0,449794 |
| Psmb2 | 220,037 | 259,74 | 0,23932 | 0,1282 | 0,449794 |
| Hdgfrp3 | 12,2596 | 10,1551 | -0,27171 | 0,1282 | 0,449794 |
| Slc2a12 | 1,27438 | 0,37394 | -1,76892 | 0,1281 | 0,449794 |
| Nde1 | 97,1354 | 115,178 | 0,2458 | 0,12825 | 0,449851 |
| 1810013L24Rik | 12,9068 | 10,7614 | -0,262265 | 0,1284 | 0,450259 |
| Gng7 | 1,43402 | 1,9236 | 0,423748 | 0,12855 | 0,450606 |
| Socs2 | 15,4241 | 18,5974 | 0,269914 | 0,1286 | 0,450606 |
| Bag3 | 13,8425 | 16,6827 | 0,26925 | 0,1286 | 0,450606 |
| Plekhg4 | 1,37757 | 1,82349 | 0,404582 | 0,1287 | 0,450659 |
| Parp2 | 38,4061 | 45,7112 | 0,251213 | 0,12875 | 0,450659 |
| Srgap3 | 1,97492 | 1,585 | -0,31731 | 0,12875 | 0,450659 |
| Shisa6 | 0,698957 | 0,51005 | -0,454567 | 0,12875 | 0,450659 |
| Cttn | 62,431 | 73,8994 | 0,243301 | 0,12895 | 0,451241 |
| Tmem57 | 27,8744 | 23,4954 | -0,246562 | 0,1291 | 0,451647 |
| Pigq | 37,0177 | 31,2106 | -0,246179 | 0,1292 | 0,451761 |
| Pmepa1 | 5,52472 | 4,50284 | -0,295066 | 0,1292 | 0,451761 |
| Ric3 | 5,83638 | 7,36522 | 0,335655 | 0,12925 | 0,451817 |
| Slc25a19 | 6,13967 | 7,63417 | 0,314309 | 0,1293 | 0,451874 |
| Tbrg1 | 65,0678 | 77,2052 | 0,246755 | 0,12935 | 0,45193 |
| Med9 | 12,7068 | 15,3023 | 0,268141 | 0,12955 | 0,451977 |
| Mcat | 18,8772 | 22,7049 | 0,266352 | 0,12945 | 0,451977 |
| Hsf1 | 32,8321 | 39,1435 | 0,253666 | 0,1295 | 0,451977 |
| Ormdl3 | 33,7467 | 28,2858 | -0,254671 | 0,1294 | 0,451977 |
| Sh2b3 | 7,98122 | 6,40855 | -0,31661 | 0,1296 | 0,451977 |
| Rdh14 | 9,19311 | 7,22022 | -0,34851 | 0,1296 | 0,451977 |
| Pdgfd | 1,64443 | 1,10995 | -0,567098 | 0,1296 | 0,451977 |
| Pds5a | 61,1463 | 72,1528 | 0,238791 | 0,12965 | 0,452033 |
| Mpc2 | 19,2162 | 15,188 | -0,339396 | 0,1297 | 0,452089 |
| Nme7 | 22,6361 | 28,527 | 0,333703 | 0,12985 | 0,452494 |
| Trim28 | 314,637 | 378,83 | 0,267861 | 0,1301 | 0,453247 |
| Neil3 | 13,998 | 11,4952 | -0,284181 | 0,13015 | 0,453303 |
| Zscan2 | 6,70719 | 8,34694 | 0,315539 | 0,1303 | 0,453707 |
| 8430427H17Rik | 11,2409 | 13,3562 | 0,248747 | 0,13035 | 0,453763 |

| | | | | | |
|---|---|---|---|---|---|
| Lmo7 | 4,13065 | 3,37217 | -0,292692 | 0,13045 | 0,453993 |
| Xpo5 | 57,8769 | 68,3185 | 0,239288 | 0,1305 | 0,454049 |
| Tenm4 | 31,5063 | 37,4731 | 0,250213 | 0,13065 | 0,454272 |
| Chaf1b | 41,2019 | 48,9917 | 0,249827 | 0,1306 | 0,454272 |
| Skap2 | 65,1015 | 77,0615 | 0,243318 | 0,1307 | 0,454272 |
| Rsrp1 | 118,811 | 100,412 | -0,242744 | 0,1307 | 0,454272 |
| Mamstr | 0,638769 | 0,367233 | -0,798597 | 0,13075 | 0,454327 |
| Gm3716 | 0,559088 | 0,850706 | 0,605585 | 0,1314 | 0,456158 |
| Stk36 | 1,74996 | 2,23598 | 0,35359 | 0,1315 | 0,456158 |
| Cchcr1 | 8,34325 | 10,2042 | 0,290482 | 0,13155 | 0,456158 |
| Tpm2 | 12,8948 | 15,6929 | 0,283319 | 0,1314 | 0,456158 |
| Sec14l1 | 25,3037 | 30,2548 | 0,25782 | 0,13145 | 0,456158 |
| Gpatch4 | 35,2461 | 41,9857 | 0,252433 | 0,13155 | 0,456158 |
| Rps6 | 15,3868 | 12,4289 | -0,307987 | 0,13135 | 0,456158 |
| Trabd2b | 0,721949 | 0,525337 | -0,458653 | 0,13155 | 0,456158 |
| Smarcb1 | 104,775 | 123,725 | 0,239848 | 0,1317 | 0,45656 |
| Dennd4b | 7,24911 | 8,70501 | 0,264043 | 0,1318 | 0,456788 |
| Ttbk2 | 7,60687 | 9,02537 | 0,246684 | 0,13185 | 0,456843 |
| Rsph3a | 1,87109 | 2,58267 | 0,464985 | 0,13195 | 0,456953 |
| Eea1 | 4,39783 | 3,63976 | -0,27295 | 0,13195 | 0,456953 |
| Wdr34 | 7,32868 | 9,17168 | 0,323633 | 0,132 | 0,457007 |
| Nusap1 | 45,9897 | 38,3334 | -0,262711 | 0,13205 | 0,457062 |
| Fnta | 47,1129 | 55,7064 | 0,241721 | 0,1323 | 0,457809 |
| Hdlbp | 104,2 | 124,168 | 0,252948 | 0,1325 | 0,458382 |
| Adnp2 | 13,9732 | 16,6301 | 0,251136 | 0,13265 | 0,458664 |
| Cops2 | 83,7594 | 71,0721 | -0,236968 | 0,13265 | 0,458664 |
| Smim13 | 6,65873 | 5,4871 | -0,279202 | 0,1327 | 0,458718 |
| Acot11 | 1,21537 | 0,913378 | -0,412116 | 0,1329 | 0,45929 |
| Zbed3 | 18,1394 | 15,0368 | -0,270626 | 0,133 | 0,459517 |
| Ctnna1 | 119,805 | 141,953 | 0,244723 | 0,13305 | 0,459571 |
| Abhd10 | 19,168 | 23,6849 | 0,305271 | 0,13315 | 0,459798 |
| Dad1 | 135,394 | 113,551 | -0,253824 | 0,13325 | 0,459905 |
| Tceanc | 3,01288 | 2,37156 | -0,345306 | 0,13325 | 0,459905 |
| Rnf39 | 0,800621 | 1,29069 | 0,688955 | 0,1333 | 0,459959 |
| Rpl7 | 348,08 | 410,902 | 0,239379 | 0,1334 | 0,460185 |
| Rpap3 | 21,2769 | 25,4192 | 0,256633 | 0,1335 | 0,460411 |
| Cmtm4 | 18,1203 | 15,343 | -0,240021 | 0,13365 | 0,460691 |
| Gm5577 | 8,44719 | 6,59282 | -0,357575 | 0,13365 | 0,460691 |
| Chic2 | 27,6812 | 33,538 | 0,276887 | 0,1337 | 0,460744 |
| Nxn | 62,4661 | 73,5483 | 0,235619 | 0,13375 | 0,460798 |
| G6pdx | 19,0155 | 24,0036 | 0,336076 | 0,1338 | 0,460851 |
| Rasa3 | 14,7327 | 17,4797 | 0,246654 | 0,1341 | 0,461409 |
| Eif1b | 105,963 | 125,347 | 0,242365 | 0,134 | 0,461409 |
| Eif3f | 142,084 | 167,334 | 0,23599 | 0,13405 | 0,461409 |
| 1110032F04Rik | 2,2117 | 1,63334 | -0,437328 | 0,1341 | 0,461409 |
| Ugdh | 26,1298 | 31,0605 | 0,249383 | 0,1342 | 0,461634 |
| Naaa | 43,4212 | 51,3708 | 0,242551 | 0,1343 | 0,461859 |
| Klhl15 | 2,31599 | 3,06416 | 0,403861 | 0,1345 | 0,46194 |

| | | | | | |
|---|---|---|---|---|---|
| Scly | 4,99493 | 6,31613 | 0,338576 | 0,1346 | 0,46194 |
| Npm3-ps1 | 38,5016 | 46,2862 | 0,265665 | 0,1346 | 0,46194 |
| Snx5 | 158,309 | 187,068 | 0,240823 | 0,1344 | 0,46194 |
| Unc5b | 31,9334 | 27,0715 | -0,238291 | 0,1346 | 0,46194 |
| Mef2a | 12,694 | 10,6996 | -0,246582 | 0,13445 | 0,46194 |
| Tmem129 | 19,1615 | 15,9293 | -0,266525 | 0,13455 | 0,46194 |
| Sesn2 | 6,16408 | 4,92717 | -0,323127 | 0,1346 | 0,46194 |
| Lipg | 3,46115 | 4,3175 | 0,318946 | 0,1347 | 0,462165 |
| Gm13315 | 5,84282 | 7,42589 | 0,3459 | 0,135 | 0,462363 |
| Fbxl18 | 6,09502 | 7,51184 | 0,301535 | 0,1349 | 0,462363 |
| Nphp3 | 4,5624 | 5,56048 | 0,285418 | 0,135 | 0,462363 |
| Ddx39b | 468,899 | 559,302 | 0,254351 | 0,1349 | 0,462363 |
| Saal1 | 12,2491 | 14,5685 | 0,250181 | 0,1349 | 0,462363 |
| Ubr4 | 17,3706 | 20,5847 | 0,244925 | 0,135 | 0,462363 |
| Crispld2 | 0,907715 | 0,643039 | -0,497331 | 0,1348 | 0,462363 |
| Lrrtm4 | 0,551051 | 0,802841 | 0,54293 | 0,1352 | 0,462574 |
| Lta4h | 47,7616 | 56,4287 | 0,240579 | 0,13515 | 0,462574 |
| Chst14 | 17,1343 | 14,1761 | -0,273419 | 0,1352 | 0,462574 |
| Hap1 | 4,91798 | 3,93406 | -0,322049 | 0,1352 | 0,462574 |
| Timm10b | 23,8077 | 29,1285 | 0,291001 | 0,13525 | 0,462626 |
| Fgd4 | 5,51628 | 6,80168 | 0,302195 | 0,13535 | 0,462784 |
| Nmt1 | 75,3104 | 88,7379 | 0,236702 | 0,1354 | 0,462784 |
| Rian | 81,4275 | 95,8563 | 0,235358 | 0,1354 | 0,462784 |
| Actn4 | 107,511 | 91,1367 | -0,238378 | 0,13545 | 0,462837 |
| Cldn9 | 13,7791 | 16,9089 | 0,295295 | 0,13555 | 0,462876 |
| Nsmce2 | 47,6228 | 56,8227 | 0,254817 | 0,1356 | 0,462876 |
| Cbll1 | 13,0696 | 10,9626 | -0,25363 | 0,1356 | 0,462876 |
| Rasa2 | 8,62408 | 7,21128 | -0,258116 | 0,1355 | 0,462876 |
| Pank4 | 23,9072 | 28,3988 | 0,248383 | 0,13575 | 0,463151 |
| Itsn1 | 33,1407 | 39,053 | 0,23683 | 0,13575 | 0,463151 |
| Ufd1l | 37,531 | 44,4266 | 0,243341 | 0,1358 | 0,463203 |
| Rab3d | 0,822714 | 0,569509 | -0,530672 | 0,1359 | 0,463426 |
| Dpysl5 | 40,1501 | 47,3872 | 0,239092 | 0,13595 | 0,463479 |
| Ppp1r15b | 38,8638 | 32,9821 | -0,236745 | 0,13615 | 0,463924 |
| Nfe2l3 | 0,885347 | 0,57802 | -0,615124 | 0,13615 | 0,463924 |
| Pcyox1l | 15,4607 | 12,7476 | -0,278381 | 0,13625 | 0,464146 |
| Ptdss2 | 41,5667 | 35,2268 | -0,238756 | 0,13635 | 0,464368 |
| Hist1h2ai | 0,339314 | 1,42817 | 2,07347 | 0,1365 | 0,464761 |
| Sepw1 | 299,79 | 353,038 | 0,235874 | 0,1366 | 0,464864 |
| St5 | 13,3386 | 11,1323 | -0,260865 | 0,1366 | 0,464864 |
| Prcc | 38,0609 | 45,0284 | 0,242525 | 0,13665 | 0,464916 |
| Katnbl1 | 15,1345 | 12,6131 | -0,262917 | 0,1368 | 0,46519 |
| Rnf151 | 0,938913 | 0,500717 | -0,906995 | 0,1368 | 0,46519 |
| Atp5s | 5,32798 | 4,02733 | -0,403767 | 0,13685 | 0,465241 |
| Ccdc25 | 31,135 | 36,9441 | 0,246806 | 0,13695 | 0,465463 |
| Slc48a1 | 6,77659 | 4,36695 | -0,633932 | 0,13705 | 0,465684 |
| Mnd1-ps | 1,06637 | 1,548 | 0,537694 | 0,13715 | 0,465787 |
| Pvr | 2,55415 | 1,92626 | -0,407039 | 0,13715 | 0,465787 |

| | | | | | |
|---|---|---|---|---|---|
| Poc1b | 23,4482 | 28,5929 | 0,286182 | 0,13725 | 0,466008 |
| Tgfbi | 12,5694 | 10,3774 | -0,276472 | 0,1373 | 0,46606 |
| Crlf2 | 3,49625 | 2,50897 | -0,478712 | 0,13735 | 0,466111 |
| Tubg1 | 47,282 | 56,0447 | 0,245286 | 0,13765 | 0,466892 |
| Pygb | 24,9455 | 21,1146 | -0,240542 | 0,13765 | 0,466892 |
| Vav2 | 14,1556 | 16,8673 | 0,252854 | 0,1377 | 0,466943 |
| Egflam | 5,46226 | 6,59252 | 0,271332 | 0,1378 | 0,467096 |
| Ptpn2 | 46,467 | 55,5738 | 0,258197 | 0,13785 | 0,467096 |
| Map4k4 | 94,5122 | 112,136 | 0,246681 | 0,13785 | 0,467096 |
| Thap7 | 12,1727 | 9,61073 | -0,340935 | 0,1379 | 0,467147 |
| Dhx29 | 21,5762 | 25,4737 | 0,239568 | 0,1381 | 0,467706 |
| Frrs1l | 0,497854 | 0,72997 | 0,552115 | 0,1383 | 0,467909 |
| Zfp7 | 15,7122 | 18,7811 | 0,257394 | 0,1383 | 0,467909 |
| Ctns | 6,53202 | 5,28999 | -0,304263 | 0,1383 | 0,467909 |
| Ttc39b | 0,60858 | 0,449843 | -0,436024 | 0,13825 | 0,467909 |
| Gm6623 | 0,372451 | 0,609901 | 0,711524 | 0,13835 | 0,46796 |
| Mrpl42 | 76,3662 | 91,2981 | 0,25765 | 0,13855 | 0,46845 |
| Rad51 | 47,7868 | 56,4169 | 0,239514 | 0,1386 | 0,46845 |
| Yif1b | 32,7901 | 27,1023 | -0,274845 | 0,1386 | 0,46845 |
| Syngap1 | 8,53436 | 10,2098 | 0,258593 | 0,1387 | 0,468669 |
| Adi1 | 21,0397 | 25,2486 | 0,263093 | 0,13905 | 0,469733 |
| 2310067B10Rik | 13,0329 | 10,9647 | -0,249292 | 0,1391 | 0,469783 |
| Ncor2 | 19,5606 | 23,0228 | 0,235114 | 0,13925 | 0,470052 |
| Mdga1 | 3,00023 | 2,44923 | -0,292745 | 0,13925 | 0,470052 |
| Ipo4 | 41,2838 | 48,5938 | 0,235198 | 0,1393 | 0,470102 |
| Camkv | 6,87272 | 8,36154 | 0,282888 | 0,13935 | 0,470134 |
| Rrs1 | 48,5013 | 57,203 | 0,238067 | 0,1394 | 0,470134 |
| Ipo9 | 56,4517 | 66,4193 | 0,234585 | 0,13945 | 0,470134 |
| Ppox | 24,8155 | 20,7122 | -0,260758 | 0,13945 | 0,470134 |
| Usp6nl | 14,1383 | 16,7381 | 0,243532 | 0,1396 | 0,470521 |
| AK010878 | 15,3131 | 22,9437 | 0,58333 | 0,1397 | 0,47067 |
| Tmem43 | 29,1353 | 24,6565 | -0,2408 | 0,13975 | 0,47067 |
| Msrb3 | 7,67567 | 6,31474 | -0,281571 | 0,13975 | 0,47067 |
| Ddx42 | 54,3994 | 63,9708 | 0,233823 | 0,1399 | 0,471057 |
| Fam71e1 | 1,53311 | 0,839852 | -0,868253 | 0,14015 | 0,471779 |
| Hk1 | 10,269 | 8,59962 | -0,255951 | 0,1402 | 0,471829 |
| 2810004N23Rik | 62,5042 | 73,6915 | 0,237543 | 0,1403 | 0,472047 |
| Zmynd19 | 36,9086 | 44,0225 | 0,254284 | 0,14035 | 0,472096 |
| Zfp964 | 1,81914 | 2,52221 | 0,471433 | 0,14065 | 0,472748 |
| Ndufaf2 | 57,2082 | 68,895 | 0,268177 | 0,14065 | 0,472748 |
| Stk10 | 2,16793 | 1,69935 | -0,351336 | 0,14065 | 0,472748 |
| Eif3k | 267,627 | 314,871 | 0,234537 | 0,1407 | 0,472797 |
| Ly6g6e | 0,894542 | 0,465223 | -0,943229 | 0,14075 | 0,472846 |
| Cpa2 | 10,4031 | 8,23417 | -0,337317 | 0,1408 | 0,472896 |
| AF357399 | 94,9343 | 0 | -inf | 0,1409 | 0,472903 |
| Zfp668 | 7,33052 | 8,86947 | 0,274932 | 0,141 | 0,472903 |
| Mtrr | 8,92439 | 10,7117 | 0,263365 | 0,141 | 0,472903 |
| Smn1 | 55,4615 | 65,6598 | 0,243523 | 0,14105 | 0,472903 |

| | | | | | |
|---|---|---|---|---|---|
| Mast2 | 23,2101 | 27,3642 | 0,237538 | 0,14085 | 0,472903 |
| Snrnp40 | 108,058 | 127,008 | 0,233115 | 0,141 | 0,472903 |
| Supt5 | 66,3211 | 77,8032 | 0,230361 | 0,14105 | 0,472903 |
| Stc2 | 10,9015 | 13,3626 | 0,293676 | 0,1412 | 0,473169 |
| Hist1h2ab | 1,25959 | 0,357649 | -1,81634 | 0,1412 | 0,473169 |
| Ovgp1 | 1,60704 | 2,1865 | 0,444218 | 0,14145 | 0,473294 |
| Purg | 17,0593 | 20,6534 | 0,275823 | 0,14145 | 0,473294 |
| Tubgcp2 | 27,2572 | 32,6652 | 0,261116 | 0,14135 | 0,473294 |
| Traf3ip1 | 10,4315 | 12,4811 | 0,2588 | 0,1414 | 0,473294 |
| Telo2 | 14,5656 | 17,3198 | 0,249859 | 0,1414 | 0,473294 |
| Dusp27 | 0,762679 | 0,529498 | -0,526451 | 0,1414 | 0,473294 |
| Dnajc8 | 103,924 | 122,198 | 0,233689 | 0,14155 | 0,47351 |
| Ifitm7 | 0,927642 | 2,29697 | 1,30809 | 0,14165 | 0,473587 |
| Rfxap | 10,9021 | 13,2398 | 0,280281 | 0,1417 | 0,473587 |
| Slc25a25 | 9,616 | 11,6514 | 0,276997 | 0,1417 | 0,473587 |
| Utp11l | 72,3521 | 85,8613 | 0,246974 | 0,14175 | 0,473587 |
| Golga3 | 11,8959 | 14,0736 | 0,242533 | 0,14175 | 0,473587 |
| Necap2 | 18,6325 | 15,4795 | -0,267467 | 0,14185 | 0,473802 |
| Ddi2 | 26,1059 | 31,4349 | 0,267993 | 0,1419 | 0,473851 |
| Prep | 59,4887 | 70,0301 | 0,235358 | 0,14195 | 0,4739 |
| Zfp395 | 18,8699 | 22,2654 | 0,238717 | 0,14215 | 0,474449 |
| Mphosph10 | 49,2885 | 58,0787 | 0,23676 | 0,14235 | 0,474761 |
| Mplkip | 10,6879 | 8,82315 | -0,276617 | 0,14235 | 0,474761 |
| Rgmb | 8,42703 | 6,8829 | -0,292008 | 0,1423 | 0,474761 |
| Hbp1 | 33,5662 | 27,6221 | -0,281185 | 0,14245 | 0,474975 |
| Tmem63b | 17,738 | 14,9334 | -0,248304 | 0,14265 | 0,475524 |
| Asrgl1 | 10,2551 | 12,4401 | 0,278653 | 0,1428 | 0,475905 |
| Itga1 | 6,60605 | 8,36736 | 0,340985 | 0,1429 | 0,47612 |
| Col27a1 | 3,31803 | 3,99848 | 0,269127 | 0,14305 | 0,47643 |
| Pim3 | 15,0071 | 17,9455 | 0,25798 | 0,1431 | 0,47643 |
| Cmbl | 1,19557 | 0,667827 | -0,840147 | 0,1431 | 0,47643 |
| AW146154 | 2,00832 | 1,52175 | -0,400262 | 0,14325 | 0,47681 |
| Map6d1 | 0,865779 | 1,20102 | 0,472188 | 0,1435 | 0,477025 |
| Bcl7b | 11,9699 | 14,6073 | 0,287279 | 0,14355 | 0,477025 |
| Acad8 | 11,2789 | 13,5346 | 0,263034 | 0,1434 | 0,477025 |
| Ift46 | 16,8899 | 20,0362 | 0,246451 | 0,1434 | 0,477025 |
| Gemin5 | 25,3921 | 29,8896 | 0,235262 | 0,14355 | 0,477025 |
| Stt3b | 82,0213 | 69,7998 | -0,232775 | 0,1436 | 0,477025 |
| Psap | 125,839 | 106,956 | -0,234562 | 0,1435 | 0,477025 |
| Ssfa2 | 7,5162 | 6,27132 | -0,261235 | 0,1436 | 0,477025 |
| Sh3d19 | 5,09813 | 4,23055 | -0,269122 | 0,1437 | 0,477238 |
| Rapgef2 | 4,10776 | 4,95709 | 0,271142 | 0,1439 | 0,477665 |
| Ndufb3 | 75,2005 | 90,6312 | 0,269265 | 0,1439 | 0,477665 |
| Bloc1s6 | 14,9527 | 17,7624 | 0,248422 | 0,14395 | 0,477712 |
| E2f1 | 20,0454 | 23,7748 | 0,246159 | 0,14415 | 0,478257 |
| Safb | 67,5049 | 79,3469 | 0,233182 | 0,14425 | 0,47847 |
| Mef2b | 0,283249 | 0,576083 | 1,0242 | 0,1444 | 0,478587 |
| Nars2 | 6,3413 | 7,67856 | 0,276057 | 0,1445 | 0,478587 |

| | | | | | |
|---|---|---|---|---|---|
| Pkd2 | 18,3514 | 15,5756 | -0,236596 | 0,14435 | 0,478587 |
| Dnajc18 | 7,52907 | 6,28502 | -0,260556 | 0,1445 | 0,478587 |
| Hspa12a | 2,29041 | 1,82875 | -0,324747 | 0,14445 | 0,478587 |
| 1810014B01Rik | 2,13213 | 1,49323 | -0,51386 | 0,1445 | 0,478587 |
| Lrrk1 | 3,78874 | 3,13546 | -0,273042 | 0,1446 | 0,478681 |
| Micall2 | 3,252 | 2,55367 | -0,348756 | 0,1446 | 0,478681 |
| Spns1 | 25,5327 | 21,5285 | -0,246102 | 0,14465 | 0,478728 |
| Aste1 | 4,77143 | 5,95897 | 0,32064 | 0,14485 | 0,479152 |
| Nampt | 27,1718 | 31,9932 | 0,235657 | 0,14485 | 0,479152 |
| Tmem107 | 14,6948 | 18,2193 | 0,310163 | 0,1449 | 0,479199 |
| Snhg6 | 123,552 | 148,56 | 0,265936 | 0,14495 | 0,479246 |
| Zfp382 | 2,98765 | 3,88342 | 0,378317 | 0,145 | 0,479293 |
| Emc7 | 40,6678 | 33,948 | -0,26056 | 0,14505 | 0,479339 |
| Gnpda1 | 8,85435 | 10,7796 | 0,28384 | 0,14515 | 0,479551 |
| Mus81 | 9,39092 | 11,4822 | 0,290058 | 0,1455 | 0,480233 |
| Ccdc104 | 24,5324 | 29,3474 | 0,258547 | 0,1455 | 0,480233 |
| Safb2 | 33,665 | 39,6387 | 0,23566 | 0,1455 | 0,480233 |
| Tln1 | 21,3569 | 18,1987 | -0,230865 | 0,1454 | 0,480233 |
| Rnf17 | 0,595281 | 0,415613 | -0,51833 | 0,1456 | 0,480444 |
| Mnat1 | 22,8503 | 27,074 | 0,244696 | 0,1457 | 0,480655 |
| Pja1 | 63,6009 | 74,7853 | 0,233708 | 0,1458 | 0,480867 |
| Diap1 | 15,7026 | 13,2474 | -0,2453 | 0,146 | 0,481407 |
| Ttll4 | 35,7496 | 41,9767 | 0,231658 | 0,14605 | 0,481453 |
| Commd6 | 32,9409 | 39,8503 | 0,274711 | 0,1461 | 0,481499 |
| Ranbp6 | 12,5655 | 10,6073 | -0,244409 | 0,1462 | 0,48171 |
| Lrrc9 | 0,429638 | 0,654766 | 0,607859 | 0,1467 | 0,481905 |
| Osbpl6 | 3,46623 | 4,23991 | 0,290668 | 0,1468 | 0,481905 |
| Timm50 | 132,973 | 156,197 | 0,232234 | 0,14665 | 0,481905 |
| Psmc4 | 174,21 | 204,595 | 0,231948 | 0,1468 | 0,481905 |
| Dhx33 | 25,3237 | 29,7141 | 0,230663 | 0,14645 | 0,481905 |
| Ptk2 | 49,7581 | 58,2785 | 0,228032 | 0,14645 | 0,481905 |
| B3glct | 16,983 | 14,4178 | -0,236237 | 0,1464 | 0,481905 |
| Flot2 | 24,6885 | 20,7955 | -0,247567 | 0,14675 | 0,481905 |
| Gpam | 10,6491 | 8,91243 | -0,256847 | 0,1465 | 0,481905 |
| Sema4b | 6,0492 | 4,96926 | -0,283712 | 0,14655 | 0,481905 |
| Arxes1 | 12,4516 | 10,0844 | -0,304211 | 0,1468 | 0,481905 |
| Mpp7 | 4,26772 | 3,41357 | -0,322186 | 0,14645 | 0,481905 |
| Mpped2 | 4,07463 | 3,16872 | -0,362767 | 0,14665 | 0,481905 |
| Ifitm3 | 15,8523 | 12,0855 | -0,391413 | 0,14665 | 0,481905 |
| Sgms2 | 0,945205 | 0,699226 | -0,434869 | 0,1466 | 0,481905 |
| Rragc | 27,4805 | 23,2616 | -0,240457 | 0,14685 | 0,481951 |
| Atp5j2 | 84,8602 | 102,729 | 0,275687 | 0,147 | 0,482088 |
| Gnpat | 65,6799 | 77,0309 | 0,229986 | 0,147 | 0,482088 |
| Nucb1 | 50,0091 | 42,3223 | -0,240772 | 0,14695 | 0,482088 |
| Ndst1 | 26,8774 | 22,8678 | -0,233075 | 0,14705 | 0,482134 |
| Akna | 5,0475 | 6,07683 | 0,267748 | 0,14715 | 0,482343 |
| Srsf1 | 224,472 | 188,045 | -0,255459 | 0,14725 | 0,482552 |
| Mapk12 | 18,9231 | 22,6258 | 0,257821 | 0,14735 | 0,482689 |

| | | | | | |
|---|---|---|---|---|---|
| Ints3 | 40,8977 | 48,0572 | 0,232733 | 0,1474 | 0,482689 |
| Fez1 | 3,14975 | 2,24769 | -0,486793 | 0,1474 | 0,482689 |
| Ubl3 | 12,1291 | 10,0778 | -0,267296 | 0,1476 | 0,483225 |
| Rabep2 | 7,33492 | 8,95718 | 0,288263 | 0,1477 | 0,483243 |
| Akr7a5 | 22,4289 | 26,8008 | 0,256915 | 0,14775 | 0,483243 |
| Hdac2 | 155,87 | 182,861 | 0,230407 | 0,14775 | 0,483243 |
| Urod | 83,0141 | 70,4774 | -0,236197 | 0,14775 | 0,483243 |
| Rps12 | 4,35504 | 6,73357 | 0,628685 | 0,14795 | 0,483751 |
| Gfod2 | 11,2286 | 13,6317 | 0,279791 | 0,14805 | 0,483751 |
| Hps4 | 11,2174 | 13,4237 | 0,259039 | 0,14805 | 0,483751 |
| Zranb2 | 108,4 | 92,5631 | -0,227853 | 0,14805 | 0,483751 |
| Gnl2 | 50,7841 | 59,567 | 0,230136 | 0,1481 | 0,483796 |
| Cd40 | 8,23597 | 10,2383 | 0,313972 | 0,14825 | 0,484167 |
| Thoc5 | 31,2993 | 36,8948 | 0,237285 | 0,1485 | 0,484865 |
| Snord95 | 154,95 | 235,053 | 0,601182 | 0,14875 | 0,485207 |
| Trpc1 | 1,09814 | 1,50107 | 0,450924 | 0,14875 | 0,485207 |
| Fam171a1 | 21,6423 | 18,3693 | -0,236553 | 0,1487 | 0,485207 |
| Ltbr | 17,6818 | 14,7399 | -0,262543 | 0,1487 | 0,485207 |
| Zfp771 | 18,5149 | 22,515 | 0,282202 | 0,1488 | 0,485252 |
| Pank3 | 20,4301 | 17,4315 | -0,229004 | 0,1489 | 0,485341 |
| Edem2 | 10,756 | 8,8502 | -0,281358 | 0,1489 | 0,485341 |
| Arsb | 5,53981 | 4,53795 | -0,287794 | 0,14895 | 0,485386 |
| Ago3 | 2,35778 | 2,87633 | 0,286802 | 0,149 | 0,485431 |
| Zfp663 | 0,499232 | 0,740722 | 0,569222 | 0,1493 | 0,486289 |
| Xrcc6 | 41,2722 | 48,5957 | 0,235659 | 0,1494 | 0,486496 |
| Zfp236 | 7,93694 | 9,3395 | 0,234762 | 0,14945 | 0,486541 |
| Fkbp4 | 309,381 | 365,595 | 0,240863 | 0,1495 | 0,486585 |
| Pkd1 | 7,60341 | 9,72968 | 0,355745 | 0,14955 | 0,486629 |
| Dpp9 | 24,3467 | 28,6454 | 0,234579 | 0,1496 | 0,486673 |
| Qtrtd1 | 25,4393 | 29,9736 | 0,236634 | 0,14975 | 0,487043 |
| Acsl4 | 16,423 | 13,9332 | -0,237193 | 0,15005 | 0,4879 |
| Rbm42 | 48,3474 | 56,9347 | 0,235869 | 0,1501 | 0,487944 |
| Lphn1 | 16,3485 | 19,1886 | 0,23109 | 0,15015 | 0,487988 |
| Dusp26 | 2,25938 | 3,05594 | 0,435689 | 0,15045 | 0,488487 |
| Rbm20 | 2,2322 | 2,74814 | 0,299987 | 0,1504 | 0,488487 |
| Ddx31 | 9,48691 | 11,366 | 0,260721 | 0,1504 | 0,488487 |
| Parp1 | 124,956 | 147,171 | 0,236076 | 0,15045 | 0,488487 |
| Ubn2 | 11,041 | 12,9232 | 0,227089 | 0,1506 | 0,488856 |
| Zrsr2 | 19,5385 | 23,1255 | 0,243167 | 0,15085 | 0,489192 |
| Ccdc34 | 41,499 | 48,7526 | 0,232401 | 0,15075 | 0,489192 |
| Rmdn3 | 25,5906 | 21,5843 | -0,245633 | 0,1508 | 0,489192 |
| Hgsnat | 12,8671 | 10,726 | -0,262579 | 0,15085 | 0,489192 |
| Pms2 | 15,4264 | 18,2764 | 0,24458 | 0,15095 | 0,489279 |
| Gtf2f1 | 52,0932 | 61,1923 | 0,232254 | 0,15095 | 0,489279 |
| Sult1a1 | 0,350182 | 0,676467 | 0,949916 | 0,151 | 0,489322 |
| Prdx6b | 2,50673 | 3,22066 | 0,361551 | 0,1512 | 0,489495 |
| Gpatch2 | 8,64853 | 10,2987 | 0,251933 | 0,1511 | 0,489495 |
| Vash2 | 22,7489 | 26,6977 | 0,230921 | 0,1512 | 0,489495 |

| | | | | | |
|---|---|---|---|---|---|
| Hsd3b7 | 4,64753 | 3,56996 | -0,380557 | 0,1512 | 0,489495 |
| Zfp87 | 7,9872 | 6,5306 | -0,290474 | 0,15125 | 0,489539 |
| Bhlhe41 | 0,797274 | 0,590308 | -0,433607 | 0,1513 | 0,489582 |
| Specc1 | 6,47797 | 5,44393 | -0,250894 | 0,1514 | 0,489787 |
| Rtca | 44,6172 | 52,4994 | 0,234703 | 0,1516 | 0,490078 |
| Pard6g | 36,1662 | 42,4446 | 0,23094 | 0,1516 | 0,490078 |
| Brd7 | 68,461 | 80,1046 | 0,226603 | 0,1516 | 0,490078 |
| Upf2 | 17,5141 | 20,5652 | 0,231691 | 0,15165 | 0,490121 |
| Hspa9 | 173,878 | 204,516 | 0,234142 | 0,15175 | 0,490326 |
| 4930500J02Rik | 0,225677 | 0,589036 | 1,3841 | 0,15195 | 0,490735 |
| Gm8615 | 1,36441 | 1,9056 | 0,481972 | 0,15195 | 0,490735 |
| Rnft1 | 6,15945 | 4,12436 | -0,578631 | 0,152 | 0,490777 |
| Pycard | 0,73572 | 0,455285 | -0,692386 | 0,15205 | 0,49082 |
| Fam193a | 16,3493 | 19,2338 | 0,234413 | 0,1521 | 0,490863 |
| Myl6 | 46,7661 | 56,5447 | 0,273929 | 0,1522 | 0,490916 |
| Ppp5c | 68,5822 | 80,3058 | 0,227669 | 0,15225 | 0,490916 |
| H3f3b | 244,624 | 208,108 | -0,233229 | 0,1523 | 0,490916 |
| Lhfpl2 | 19,3791 | 16,4823 | -0,233584 | 0,15225 | 0,490916 |
| Bik | 3,28463 | 2,2458 | -0,548502 | 0,1523 | 0,490916 |
| Phf8 | 8,99534 | 10,6062 | 0,237655 | 0,15235 | 0,490959 |
| Zfyve20 | 12,7818 | 15,0487 | 0,235544 | 0,15245 | 0,491044 |
| Ccdc107 | 16,1972 | 12,746 | -0,345699 | 0,15245 | 0,491044 |
| Ehmt1 | 40,1004 | 47,0618 | 0,230942 | 0,15255 | 0,491129 |
| Acadm | 16,0387 | 13,3582 | -0,263841 | 0,15255 | 0,491129 |
| Ptk7 | 288,246 | 345,889 | 0,26301 | 0,1526 | 0,491172 |
| Zfp108 | 0,878045 | 1,24197 | 0,500267 | 0,1529 | 0,491782 |
| Zfp27 | 5,73747 | 6,95739 | 0,278133 | 0,15285 | 0,491782 |
| Eif4ebp1 | 43,9307 | 52,2868 | 0,251218 | 0,1529 | 0,491782 |
| Bmp1 | 29,6452 | 25,3248 | -0,227252 | 0,15305 | 0,492146 |
| Derl3 | 0,481221 | 0,857719 | 0,833806 | 0,15315 | 0,492349 |
| 03/09/16 | 4,99916 | 3,90548 | -0,356185 | 0,1532 | 0,492392 |
| Ripply2 | 33,5008 | 41,9617 | 0,32488 | 0,1535 | 0,492409 |
| Dhfr | 23,9557 | 29,121 | 0,281689 | 0,1534 | 0,492409 |
| Zfp800 | 5,79331 | 6,99318 | 0,271559 | 0,1534 | 0,492409 |
| Nup54 | 27,6801 | 32,6923 | 0,240105 | 0,15325 | 0,492409 |
| Zfp329 | 9,81478 | 8,33504 | -0,235767 | 0,1535 | 0,492409 |
| Ralgds | 9,80637 | 8,17151 | -0,263116 | 0,15345 | 0,492409 |
| Epb4.1l4a | 1,97629 | 1,51614 | -0,382389 | 0,1533 | 0,492409 |
| Prox2 | 1,19208 | 0,839905 | -0,50518 | 0,1535 | 0,492409 |
| Prdm4 | 27,8262 | 32,6089 | 0,228823 | 0,15355 | 0,492451 |
| Fbxl14 | 23,1212 | 28,271 | 0,290108 | 0,1536 | 0,492493 |
| Tnk1 | 0,704508 | 0,204476 | -1,78468 | 0,1537 | 0,492695 |
| Parp4 | 2,21594 | 1,78282 | -0,31376 | 0,1538 | 0,492898 |
| Ube2f | 25,1255 | 30,0747 | 0,259394 | 0,15385 | 0,49294 |
| 1200014J11Rik | 15,442 | 18,1131 | 0,230173 | 0,1539 | 0,492982 |
| Soga1 | 15,8813 | 18,5766 | 0,226156 | 0,1541 | 0,493504 |
| Hist1h2ag | 2,34086 | 1,14064 | -1,0372 | 0,15415 | 0,493546 |
| Gzmm | 0,661891 | 1,11419 | 0,751324 | 0,15435 | 0,494068 |

| | | | | | |
|---|---|---|---|---|---|
| Timm17a | 49,6548 | 58,9002 | 0,246339 | 0,15445 | 0,49427 |
| Ptplad1 | 51,1183 | 43,6538 | -0,227731 | 0,15455 | 0,494471 |
| Tnip1 | 7,5286 | 9,13292 | 0,278694 | 0,1548 | 0,495153 |
| 2410016O06Rik | 15,5075 | 18,436 | 0,249562 | 0,1549 | 0,495235 |
| Fut8 | 16,3215 | 13,6956 | -0,253068 | 0,1549 | 0,495235 |
| Tomm40 | 82,071 | 96,0516 | 0,226937 | 0,155 | 0,495436 |
| Myo19 | 11,8239 | 14,8924 | 0,332875 | 0,15505 | 0,495478 |
| Ikbkap | 41,0093 | 48,9541 | 0,255479 | 0,1551 | 0,495519 |
| Amer2 | 1,30523 | 1,792 | 0,457267 | 0,15515 | 0,49556 |
| Cep97 | 10,2659 | 13,5337 | 0,398691 | 0,1552 | 0,495602 |
| P2rx3 | 1,15531 | 1,52074 | 0,3965 | 0,15535 | 0,495844 |
| Mov10 | 9,31761 | 11,1058 | 0,253283 | 0,15535 | 0,495844 |
| Slc25a43 | 0,855185 | 0,456749 | -0,904834 | 0,1555 | 0,496204 |
| D8Ertd82e | 12,2078 | 14,3882 | 0,23708 | 0,15585 | 0,496846 |
| Ep400 | 24,8908 | 29,165 | 0,228624 | 0,15585 | 0,496846 |
| Paxip1 | 24,2198 | 28,3267 | 0,225976 | 0,1558 | 0,496846 |
| Luc7l3 | 97,15 | 83,0821 | -0,225677 | 0,15585 | 0,496846 |
| Mpc1 | 1,18697 | 2,31169 | 0,961662 | 0,15595 | 0,497046 |
| Rtkn | 15,7221 | 18,658 | 0,247002 | 0,156 | 0,497087 |
| St7l | 3,97838 | 4,78467 | 0,266238 | 0,15615 | 0,497091 |
| 2310022B05Rik | 25,9108 | 30,4192 | 0,231432 | 0,1561 | 0,497091 |
| Rnf20 | 40,6848 | 47,5109 | 0,223769 | 0,1561 | 0,497091 |
| Lrig1 | 30,3256 | 25,792 | -0,233613 | 0,15615 | 0,497091 |
| Prmt10 | 11,5842 | 13,8035 | 0,252881 | 0,1563 | 0,497136 |
| Ppme1 | 41,0813 | 48,1625 | 0,229429 | 0,15635 | 0,497136 |
| Elp3 | 61,637 | 71,9815 | 0,223829 | 0,15635 | 0,497136 |
| Hnrnpul1 | 66,8601 | 78,0477 | 0,22321 | 0,1563 | 0,497136 |
| Eno2 | 0,64425 | 0,406598 | -0,664017 | 0,15635 | 0,497136 |
| Rdm1 | 9,50131 | 11,9806 | 0,334498 | 0,15665 | 0,497853 |
| Rtcb | 115,365 | 134,7 | 0,22354 | 0,15665 | 0,497853 |
| Tfcp2 | 10,8154 | 12,8591 | 0,249708 | 0,1567 | 0,497893 |
| Cct8 | 346,878 | 412,757 | 0,250866 | 0,157 | 0,498728 |
| Cables2 | 10,1299 | 12,1113 | 0,257731 | 0,1571 | 0,498927 |
| Atg5 | 23,9673 | 28,2979 | 0,239626 | 0,15715 | 0,498967 |
| BC029722 | 7,32102 | 5,72935 | -0,353672 | 0,1572 | 0,499007 |
| Ikzf3 | 0,892247 | 0,642237 | -0,474338 | 0,15725 | 0,499047 |
| Rp2h | 13,6883 | 16,9927 | 0,31197 | 0,1573 | 0,499088 |
| Tpra1 | 12,5192 | 10,3032 | -0,281049 | 0,15735 | 0,499128 |
| Prdm5 | 10,7325 | 12,9103 | 0,266537 | 0,15745 | 0,499326 |
| Rrad | 2,26675 | 3,08244 | 0,443446 | 0,15755 | 0,499407 |
| Gas2l1 | 8,92188 | 7,31696 | -0,286103 | 0,15755 | 0,499407 |
| Gtse1 | 31,7967 | 37,3512 | 0,232279 | 0,15765 | 0,499605 |
| Zfp212 | 13,4851 | 15,9994 | 0,246655 | 0,15785 | 0,500002 |
| Xylt1 | 1,39201 | 1,01683 | -0,453094 | 0,15785 | 0,500002 |
| Mthfd2l | 8,91322 | 10,7942 | 0,276234 | 0,15795 | 0,5002 |
| Efr3b | 3,91953 | 4,70232 | 0,262689 | 0,15805 | 0,500398 |
| Shq1 | 10,9586 | 13,2518 | 0,274126 | 0,15835 | 0,500676 |
| Atpaf2 | 19,6468 | 23,3392 | 0,248462 | 0,1582 | 0,500676 |

| | | | | | |
|---|---|---|---|---|---|
| Exoc1 | 18,8615 | 22,1301 | 0,230568 | 0,1583 | 0,500676 |
| Limd2 | 63,1453 | 73,6943 | 0,222878 | 0,1584 | 0,500676 |
| Wrb | 19,9994 | 16,8778 | -0,244832 | 0,1584 | 0,500676 |
| Letmd1 | 10,0065 | 8,27515 | -0,274079 | 0,1583 | 0,500676 |
| Dusp23 | 5,20337 | 3,96925 | -0,39058 | 0,1583 | 0,500676 |
| Iqsec1 | 9,79477 | 8,30534 | -0,237972 | 0,15845 | 0,500715 |
| Drg2 | 48,788 | 57,3008 | 0,232029 | 0,1585 | 0,500755 |
| Ric8b | 10,5563 | 12,4581 | 0,238981 | 0,15855 | 0,500795 |
| Camsap1 | 11,6093 | 13,5773 | 0,225915 | 0,1586 | 0,500834 |
| Serpini1 | 0,927815 | 0,648856 | -0,515938 | 0,1587 | 0,501032 |
| 4930523C07Rik | 0,861559 | 0,609938 | -0,498287 | 0,1588 | 0,501229 |
| Stac3 | 0,724126 | 0,426407 | -0,764011 | 0,15885 | 0,501268 |
| Fndc3b | 10,1484 | 8,6254 | -0,234584 | 0,1589 | 0,501308 |
| Slc33a1 | 11,0817 | 9,21369 | -0,266333 | 0,15895 | 0,501347 |
| Mak16 | 71,6311 | 85,0043 | 0,24695 | 0,159 | 0,501387 |
| Qtrt1 | 26,1277 | 31,096 | 0,251151 | 0,15935 | 0,501898 |
| Atf6 | 7,66726 | 6,50671 | -0,236782 | 0,15935 | 0,501898 |
| Tenc1 | 3,06106 | 2,46716 | -0,311175 | 0,1592 | 0,501898 |
| Mrps33 | 4,69844 | 3,45362 | -0,444071 | 0,15925 | 0,501898 |
| Csf2rb | 2,53958 | 0,00938036 | -8,08073 | 0,15935 | 0,501898 |
| Gpr162 | 4,08551 | 3,22728 | -0,340196 | 0,1594 | 0,501937 |
| Aadat | 2,56584 | 1,87627 | -0,451567 | 0,15945 | 0,501976 |
| Zufsp | 6,46567 | 7,92028 | 0,292752 | 0,15955 | 0,502055 |
| Lmf2 | 16,9521 | 14,3053 | -0,244919 | 0,15955 | 0,502055 |
| Nans | 45,1447 | 52,9929 | 0,231244 | 0,15965 | 0,502133 |
| 5330417C22Rik | 0,661665 | 0,441412 | -0,583976 | 0,15965 | 0,502133 |
| Znrf2 | 6,82822 | 5,58291 | -0,290493 | 0,1597 | 0,502172 |
| Pde4b | 10,0102 | 11,9046 | 0,250056 | 0,15975 | 0,502211 |
| Mrps6 | 60,4632 | 77,1631 | 0,351855 | 0,16 | 0,502406 |
| Top3a | 13,6273 | 16,0899 | 0,239663 | 0,16 | 0,502406 |
| Smarca1 | 11,7645 | 13,8551 | 0,235974 | 0,15995 | 0,502406 |
| Nhlrc3 | 7,25116 | 5,72367 | -0,341271 | 0,15995 | 0,502406 |
| 1600027J07Rik | 2,49289 | 1,26147 | -0,982713 | 0,15985 | 0,502406 |
| Glb1 | 11,5839 | 9,23628 | -0,326737 | 0,1601 | 0,502602 |
| Mapk11 | 10,5549 | 12,6456 | 0,260728 | 0,16015 | 0,502641 |
| Elmsan1 | 7,95819 | 9,34552 | 0,231834 | 0,1603 | 0,502914 |
| Nol10 | 31,9034 | 37,253 | 0,223648 | 0,16035 | 0,502914 |
| Ppifos | 3,47081 | 1,91224 | -0,860013 | 0,16035 | 0,502914 |
| Kif5c | 9,3553 | 10,9768 | 0,230601 | 0,1604 | 0,502953 |
| Spice1 | 11,1488 | 13,185 | 0,242004 | 0,16055 | 0,503305 |
| Prickle1 | 43,9915 | 51,3512 | 0,223171 | 0,16065 | 0,503501 |
| H2-Q8 | 1,62067 | 4,00659 | 1,30579 | 0,16115 | 0,503571 |
| Rsph3b | 0,436443 | 0,750285 | 0,781644 | 0,16085 | 0,503571 |
| Bola3 | 32,9182 | 40,7621 | 0,308341 | 0,16095 | 0,503571 |
| Trmt5 | 22,3051 | 26,4614 | 0,246519 | 0,16095 | 0,503571 |
| Fibp | 31,8818 | 37,8214 | 0,246471 | 0,16115 | 0,503571 |
| Mapk1ip1l | 43,3213 | 50,5373 | 0,222273 | 0,16095 | 0,503571 |
| Rgma | 20,4696 | 17,4028 | -0,234164 | 0,16105 | 0,503571 |

| | | | | | |
|---|---|---|---|---|---|
| Rspry1 | 16,8728 | 14,3413 | -0,234527 | 0,16095 | 0,503571 |
| Mospd3 | 7,56691 | 5,9613 | -0,344079 | 0,1608 | 0,503571 |
| Madcam1 | 0,720872 | 0,0442664 | -4,02546 | 0,1612 | 0,503571 |
| 5033406O09Rik | 0,98631 | 0,0352971 | -4,80442 | 0,1612 | 0,503571 |
| Tyrobp | 5,93494 | 0,148692 | -5,31883 | 0,1612 | 0,503571 |
| Mal2 | 0,838195 | 0,0209367 | -5,32318 | 0,1612 | 0,503571 |
| Rac2 | 0,96377 | 0,0190071 | -5,66408 | 0,1612 | 0,503571 |
| Tcf20 | 17,3782 | 20,2881 | 0,223358 | 0,16125 | 0,503609 |
| Sf1 | 106,461 | 124,5 | 0,22582 | 0,1614 | 0,50396 |
| Clns1a | 46,067 | 53,8086 | 0,224104 | 0,16145 | 0,503999 |
| Dgkh | 1,15452 | 1,51603 | 0,393001 | 0,1615 | 0,504037 |
| Arhgap31 | 8,9896 | 10,5369 | 0,229119 | 0,1617 | 0,504425 |
| Ajuba | 31,7593 | 27,1766 | -0,224814 | 0,1617 | 0,504425 |
| Dnajc7 | 109,494 | 127,784 | 0,222862 | 0,1619 | 0,504932 |
| Snord99 | 5843,43 | 0 | -inf | 0,1621 | 0,505202 |
| Zfp46 | 7,74664 | 9,18825 | 0,246219 | 0,16205 | 0,505202 |
| Comt | 17,9631 | 13,4993 | -0,412151 | 0,1621 | 0,505202 |
| Rps5 | 1261,38 | 1489,77 | 0,240087 | 0,1622 | 0,505395 |
| Ldhb | 41,369 | 48,8526 | 0,239887 | 0,1625 | 0,506132 |
| Park7 | 227,405 | 265,479 | 0,223336 | 0,16255 | 0,506132 |
| Slc39a9 | 18,8251 | 16,1064 | -0,225028 | 0,16255 | 0,506132 |
| Tmem108 | 1,6956 | 1,29786 | -0,385665 | 0,1626 | 0,50617 |
| Sf3b3 | 140,004 | 164,325 | 0,231092 | 0,163 | 0,506825 |
| Ero1l | 12,6958 | 10,7658 | -0,237901 | 0,163 | 0,506825 |
| Abcb8 | 16,7931 | 14,207 | -0,241273 | 0,16285 | 0,506825 |
| Aim2 | 1,69517 | 1,24647 | -0,443584 | 0,16295 | 0,506825 |
| Ctf2 | 0,825038 | 0,384688 | -1,10077 | 0,16295 | 0,506825 |
| Tmem263 | 11,0374 | 9,30591 | -0,246178 | 0,16305 | 0,506862 |
| 1700021F05Rik | 24,6333 | 29,8283 | 0,276069 | 0,16325 | 0,507285 |
| Rundc3b | 1,42587 | 1,06823 | -0,416627 | 0,1633 | 0,507285 |
| Tssk4 | 1,09709 | 0,666024 | -0,720041 | 0,1633 | 0,507285 |
| Tmem80 | 18,3144 | 12,5488 | -0,545425 | 0,16335 | 0,507323 |
| Srpk1 | 55,5679 | 64,9217 | 0,22445 | 0,16345 | 0,507397 |
| Ang | 1,42518 | 0,508957 | -1,48553 | 0,16345 | 0,507397 |
| Jak2 | 9,67191 | 8,18323 | -0,241132 | 0,16355 | 0,50759 |
| Fam172a | 6,6385 | 5,51783 | -0,266757 | 0,1637 | 0,507702 |
| Nat6 | 10,2733 | 8,41984 | -0,287029 | 0,1637 | 0,507702 |
| Pitx1 | 0,891597 | 0,595054 | -0,58337 | 0,1637 | 0,507702 |
| Rcn3 | 49,5419 | 42,1181 | -0,234208 | 0,16385 | 0,508049 |
| Dyx1c1 | 1,1082 | 1,57855 | 0,510385 | 0,1639 | 0,508086 |
| Pomp | 268,372 | 312,9 | 0,221468 | 0,16405 | 0,508315 |
| Clstn1 | 39,446 | 33,8561 | -0,220463 | 0,16405 | 0,508315 |
| Dab1 | 1,3331 | 0,833319 | -0,677847 | 0,1641 | 0,508352 |
| Psmb8 | 1,04595 | 0,610844 | -0,775942 | 0,16415 | 0,508389 |
| Scnn1a | 0,485205 | 0,705377 | 0,539802 | 0,1644 | 0,508457 |
| Polr2l | 3,75285 | 4,84371 | 0,368125 | 0,1644 | 0,508457 |
| Prkd1 | 8,27215 | 9,83114 | 0,249097 | 0,16435 | 0,508457 |
| Arhgap42 | 8,89781 | 7,53439 | -0,239958 | 0,1643 | 0,508457 |

| | | | | | |
|---|---|---|---|---|---|
| Man1c1 | 8,48201 | 7,15506 | -0,245441 | 0,16435 | 0,508457 |
| Nexn | 1,8389 | 1,36379 | -0,431228 | 0,16435 | 0,508457 |
| Golt1b | 36,8532 | 31,4826 | -0,227236 | 0,16445 | 0,508494 |
| Ndufaf4 | 23,9885 | 20,5189 | -0,225389 | 0,16455 | 0,508568 |
| Chac2 | 4,59943 | 2,86666 | -0,682084 | 0,16455 | 0,508568 |
| Cd276 | 31,384 | 26,8419 | -0,22554 | 0,1649 | 0,509531 |
| Atp11b | 22,3134 | 19,1418 | -0,221179 | 0,165 | 0,509723 |
| Fam69b | 11,1501 | 9,08397 | -0,295658 | 0,1651 | 0,509796 |
| 2810403D21Rik | 1,29339 | 0,751253 | -0,78379 | 0,1651 | 0,509796 |
| Rnf114 | 36,9859 | 43,8182 | 0,244557 | 0,16515 | 0,509832 |
| Cct3 | 316,155 | 370,634 | 0,229364 | 0,1652 | 0,509869 |
| Eif3b | 175,669 | 205,55 | 0,226631 | 0,1653 | 0,509942 |
| Tmem229a | 3,04451 | 2,47397 | -0,299381 | 0,1653 | 0,509942 |
| Myh10 | 69,3964 | 81,5136 | 0,23218 | 0,16535 | 0,509979 |
| Mad1l1 | 17,2827 | 20,3818 | 0,237951 | 0,1656 | 0,510632 |
| Fcho2 | 15,3111 | 13,07 | -0,228313 | 0,16565 | 0,510668 |
| Lrrc27 | 1,88525 | 2,58279 | 0,454175 | 0,16575 | 0,510859 |
| Fnbp1 | 13,4347 | 15,8739 | 0,240694 | 0,1658 | 0,510895 |
| Smc3 | 57,4646 | 67,0032 | 0,221557 | 0,16585 | 0,510931 |
| Stag2 | 48,7466 | 41,8247 | -0,220947 | 0,16595 | 0,511121 |
| Adprm | 11,5511 | 14,0716 | 0,284755 | 0,1662 | 0,511656 |
| Tshz1 | 7,99723 | 6,77486 | -0,239309 | 0,1662 | 0,511656 |
| Fgl1 | 1,05973 | 0,610231 | -0,796267 | 0,1663 | 0,511846 |
| Dpcd | 18,7996 | 23,3557 | 0,313077 | 0,1667 | 0,512287 |
| Hoxc12 | 86,7109 | 101,614 | 0,228819 | 0,16675 | 0,512287 |
| Tle4 | 29,0698 | 33,8756 | 0,220726 | 0,1666 | 0,512287 |
| C2cd3 | 12,0214 | 13,9942 | 0,219226 | 0,16665 | 0,512287 |
| Wdhd1 | 35,5199 | 41,324 | 0,218353 | 0,16665 | 0,512287 |
| Vbp1 | 76,6434 | 65,6924 | -0,222434 | 0,16665 | 0,512287 |
| Slitrk2 | 0,815452 | 0,623702 | -0,386744 | 0,16675 | 0,512287 |
| LOC100504703 | 24,5917 | 17,6721 | -0,476696 | 0,16655 | 0,512287 |
| Spopl | 6,07235 | 7,38657 | 0,282649 | 0,1671 | 0,512302 |
| Isyna1 | 64,4287 | 75,0747 | 0,220623 | 0,167 | 0,512302 |
| Mcm7 | 99,0576 | 115,265 | 0,218617 | 0,16695 | 0,512302 |
| Slc11a2 | 17,2367 | 14,6438 | -0,235194 | 0,16695 | 0,512302 |
| Atp10d | 2,3094 | 1,8682 | -0,305869 | 0,1671 | 0,512302 |
| Sytl4 | 2,06497 | 1,58939 | -0,37765 | 0,1671 | 0,512302 |
| Lad1 | 1,90042 | 1,43315 | -0,407133 | 0,16695 | 0,512302 |
| Znfx1 | 4,37212 | 3,24853 | -0,428549 | 0,1671 | 0,512302 |
| Chst8 | 1,45775 | 1,04944 | -0,474126 | 0,1671 | 0,512302 |
| Nup214 | 13,1542 | 15,352 | 0,222907 | 0,1672 | 0,512373 |
| Actn2 | 0,606972 | 0,876989 | 0,530929 | 0,1672 | 0,512374 |
| Taf10 | 50,7637 | 73,4673 | 0,533304 | 0,1673 | 0,512562 |
| Aldh1b1 | 6,97222 | 8,49779 | 0,285469 | 0,1676 | 0,512694 |
| Ptgs1 | 6,1617 | 7,48385 | 0,280452 | 0,16765 | 0,512694 |
| Nup107 | 47,6148 | 55,5271 | 0,221784 | 0,16765 | 0,512694 |
| Xrcc1 | 53,0599 | 61,8576 | 0,221329 | 0,16755 | 0,512694 |
| Faf2 | 35,9085 | 30,8488 | -0,219113 | 0,16745 | 0,512694 |

| | | | | | |
|---|---|---|---|---|---|
| Apmap | 22,9879 | 19,4818 | -0,238749 | 0,16755 | 0,512694 |
| Fam76a | 6,61235 | 5,4448 | -0,280285 | 0,1675 | 0,512694 |
| 1700120K04Rik | 2,16511 | 1,089 | -0,991431 | 0,16755 | 0,512694 |
| Igf1 | 1,27288 | 0,888554 | -0,51856 | 0,1677 | 0,51273 |
| Spin4 | 2,58155 | 3,19572 | 0,307901 | 0,16775 | 0,512766 |
| Fancd2 | 13,0167 | 15,2411 | 0,227611 | 0,16795 | 0,51326 |
| Bcorl1 | 8,71447 | 10,2273 | 0,230939 | 0,1681 | 0,5136 |
| D8Ertd738e | 159,769 | 186,884 | 0,22616 | 0,1682 | 0,513789 |
| Magt1 | 30,6154 | 26,2724 | -0,220711 | 0,1683 | 0,513859 |
| Larp4 | 17,2794 | 14,7931 | -0,224127 | 0,1683 | 0,513859 |
| 4921524J17Rik | 8,24 | 10,1307 | 0,298022 | 0,16855 | 0,514305 |
| Ncoa6 | 17,3816 | 20,2821 | 0,22265 | 0,1686 | 0,514305 |
| Snrpb | 354,158 | 412,171 | 0,21885 | 0,1686 | 0,514305 |
| Dnajb14 | 4,61897 | 3,6092 | -0,355894 | 0,16855 | 0,514305 |
| Pou3f3os | 0,986836 | 0,250189 | -1,97979 | 0,1687 | 0,514493 |
| Nr4a1 | 2,0718 | 2,70839 | 0,386548 | 0,1688 | 0,514516 |
| Ccs | 17,2468 | 21,0154 | 0,285115 | 0,16885 | 0,514516 |
| Cnot8 | 28,9949 | 34,0088 | 0,230108 | 0,1689 | 0,514516 |
| Adh5 | 140,997 | 163,713 | 0,215511 | 0,1688 | 0,514516 |
| Tmem62 | 0,740455 | 0,495655 | -0,579077 | 0,1689 | 0,514516 |
| Nup188 | 43,7252 | 50,9171 | 0,219686 | 0,169 | 0,514703 |
| 1110038F14Rik | 27,6902 | 33,0489 | 0,255227 | 0,1693 | 0,51483 |
| Pknox2 | 20,2986 | 23,7717 | 0,227864 | 0,16925 | 0,51483 |
| Map2k6 | 40,9681 | 47,8968 | 0,225428 | 0,16935 | 0,51483 |
| Zfp598 | 29,9811 | 35,013 | 0,223841 | 0,16925 | 0,51483 |
| Ankhd1 | 20,056 | 23,3429 | 0,218948 | 0,1692 | 0,51483 |
| Psmc5 | 214,996 | 249,789 | 0,216403 | 0,16915 | 0,51483 |
| Nudt16 | 2,63512 | 1,93536 | -0,445263 | 0,1692 | 0,51483 |
| Pex11g | 3,19119 | 2,27978 | -0,485197 | 0,16935 | 0,51483 |
| Crocc | 7,34366 | 8,622 | 0,231524 | 0,1695 | 0,514853 |
| Gar1 | 72,8768 | 85,0431 | 0,222734 | 0,16945 | 0,514853 |
| Sgpl1 | 49,8475 | 42,7899 | -0,220249 | 0,16955 | 0,514853 |
| Pofut2 | 38,6313 | 33,0191 | -0,226472 | 0,16945 | 0,514853 |
| Rab39 | 0,601215 | 0,390313 | -0,623251 | 0,16955 | 0,514853 |
| Rtn1 | 5,80964 | 7,32565 | 0,334508 | 0,1696 | 0,514887 |
| Zfp777 | 13,7523 | 16,2375 | 0,239651 | 0,1697 | 0,515074 |
| Mfsd10 | 25,5822 | 21,6858 | -0,238387 | 0,1698 | 0,51526 |
| Snord67 | 0 | 16,1786 | inf | 0,16995 | 0,515282 |
| Apol7d | 0,359117 | 0,709603 | 0,982559 | 0,17 | 0,515282 |
| Fam114a2 | 32,7843 | 38,2141 | 0,221101 | 0,17 | 0,515282 |
| Mkrn1 | 28,8781 | 33,6461 | 0,220462 | 0,16995 | 0,515282 |
| Lamc1 | 48,9019 | 41,9847 | -0,220026 | 0,1699 | 0,515282 |
| Srf | 27,0746 | 31,519 | 0,219281 | 0,17005 | 0,515317 |
| Txlnb | 0,822571 | 1,10238 | 0,42241 | 0,1705 | 0,515512 |
| Map2k5 | 14,8476 | 17,5567 | 0,241797 | 0,17045 | 0,515512 |
| Gle1 | 21,0972 | 24,6716 | 0,225804 | 0,17025 | 0,515512 |
| Napa | 41,9862 | 48,9498 | 0,221387 | 0,1703 | 0,515512 |
| Vcp | 47,0943 | 54,7895 | 0,218345 | 0,17045 | 0,515512 |

| | | | | | |
|---|---|---|---|---|---|
| Prpf31 | 44,2087 | 51,3889 | 0,217126 | 0,1702 | 0,515512 |
| Cachd1 | 48,3423 | 56,105 | 0,214844 | 0,17045 | 0,515512 |
| Lemd2 | 15,8896 | 13,4029 | -0,245538 | 0,1705 | 0,515512 |
| Sh3kbp1 | 4,74552 | 3,93566 | -0,269963 | 0,17035 | 0,515512 |
| Vsig8 | 0,94658 | 0,592043 | -0,677023 | 0,17025 | 0,515512 |
| Ccdc124 | 68,9684 | 81,0042 | 0,232061 | 0,17055 | 0,515546 |
| Tmem230 | 8,60759 | 6,9337 | -0,311985 | 0,17095 | 0,516522 |
| Ndufaf3 | 10,5915 | 8,36651 | -0,340215 | 0,17095 | 0,516522 |
| Tlk2 | 22,8167 | 26,4852 | 0,215095 | 0,17105 | 0,516707 |
| Zfp420 | 3,41754 | 4,21017 | 0,300921 | 0,1711 | 0,516741 |
| Gtf2e1 | 21,0723 | 24,6693 | 0,227371 | 0,1713 | 0,516878 |
| Ralbp1 | 24,6249 | 28,6953 | 0,220695 | 0,1713 | 0,516878 |
| Tbl1x | 52,5464 | 61,0948 | 0,217458 | 0,1713 | 0,516878 |
| Phlda1 | 14,4354 | 12,0407 | -0,261689 | 0,17125 | 0,516878 |
| Blvra | 11,2093 | 8,96318 | -0,322617 | 0,17135 | 0,516912 |
| Itgb4 | 0,669006 | 0,235407 | -1,50686 | 0,17145 | 0,517097 |
| Nell1 | 2,47586 | 1,91969 | -0,367055 | 0,1715 | 0,517131 |
| Psmb1 | 202,727 | 235,84 | 0,218274 | 0,1717 | 0,517618 |
| Cib1 | 33,5402 | 39,9452 | 0,252132 | 0,17185 | 0,517836 |
| Prpf6 | 40,9094 | 47,6008 | 0,218554 | 0,17185 | 0,517836 |
| Rundc3a | 2,903 | 3,68389 | 0,343687 | 0,172 | 0,518171 |
| Ccdc101 | 20,2027 | 24,2661 | 0,264392 | 0,1721 | 0,518273 |
| Sub1 | 77,7262 | 90,3796 | 0,217596 | 0,17215 | 0,518273 |
| Pdlim5 | 28,1108 | 24,1006 | -0,222057 | 0,17215 | 0,518273 |
| Kcnk5 | 7,65838 | 6,3929 | -0,26057 | 0,1722 | 0,518306 |
| Gjb6 | 1,35379 | 0,949713 | -0,511444 | 0,1723 | 0,518491 |
| Stac | 0,894216 | 0,608565 | -0,555211 | 0,17235 | 0,518524 |
| Mir717 | 13,3252 | 32,1326 | 1,26988 | 0,1726 | 0,518809 |
| Pdpr | 10,2739 | 12,2748 | 0,25672 | 0,1726 | 0,518809 |
| Sclt1 | 11,8437 | 13,9693 | 0,238134 | 0,1726 | 0,518809 |
| Anapc11 | 13,5861 | 15,9842 | 0,23452 | 0,17255 | 0,518809 |
| Cth | 20,4574 | 24,2017 | 0,242485 | 0,17265 | 0,518843 |
| Firre | 17,7934 | 15,2443 | -0,223075 | 0,1727 | 0,518877 |
| Lrrtm1 | 0,815044 | 0,139391 | -2,54774 | 0,1728 | 0,51906 |
| Nme5 | 0,880388 | 1,53805 | 0,804887 | 0,17295 | 0,519078 |
| Tmbim6 | 102,223 | 87,9019 | -0,217749 | 0,17295 | 0,519078 |
| Snx14 | 15,7082 | 13,3465 | -0,235053 | 0,1729 | 0,519078 |
| Sri | 22,61 | 19,2031 | -0,23562 | 0,173 | 0,519078 |
| Mtfr2 | 9,1068 | 7,53115 | -0,274074 | 0,173 | 0,519078 |
| 2900009J06Rik | 1,45409 | 2,82348 | 0,957357 | 0,17335 | 0,519312 |
| Lrrc20 | 7,08775 | 8,55077 | 0,270728 | 0,17325 | 0,519312 |
| Depdc1b | 12,7177 | 15,0569 | 0,24358 | 0,17335 | 0,519312 |
| Aplp2 | 113,127 | 97,242 | -0,218296 | 0,17325 | 0,519312 |
| Cox6a1 | 288,815 | 247,954 | -0,220075 | 0,1732 | 0,519312 |
| Zfp97 | 0,835224 | 0,570623 | -0,549627 | 0,17325 | 0,519312 |
| 2310034G01Rik | 1,72348 | 1,0091 | -0,772264 | 0,1733 | 0,519312 |
| Hmgn2 | 36,7736 | 43,327 | 0,236598 | 0,1734 | 0,519345 |
| Lats1 | 11,0642 | 12,8972 | 0,221167 | 0,1735 | 0,519412 |

| | | | | | |
|---|---|---|---|---|---|
| Cltb | 9,06998 | 7,3409 | -0,305141 | 0,1735 | 0,519412 |
| 1700019G17Rik | 1,41343 | 0,967734 | -0,546514 | 0,17355 | 0,519445 |
| Mrpl21 | 20,4679 | 24,7796 | 0,275786 | 0,1736 | 0,519478 |
| Rab20 | 0,925553 | 1,65454 | 0,838042 | 0,17365 | 0,519512 |
| Fam98a | 29,8351 | 25,5859 | -0,221661 | 0,1737 | 0,519545 |
| Smtn | 15,1429 | 12,8666 | -0,235011 | 0,1738 | 0,519728 |
| Pip5k1b | 1,99362 | 1,30289 | -0,613676 | 0,17415 | 0,520658 |
| Rbm43 | 4,9467 | 3,92747 | -0,332868 | 0,17425 | 0,52084 |
| Lmtk3 | 5,81494 | 6,93423 | 0,253971 | 0,1743 | 0,520873 |
| Ndnl2 | 29,4006 | 34,8074 | 0,243552 | 0,17445 | 0,521005 |
| Basp1 | 73,2507 | 85,1568 | 0,217279 | 0,1745 | 0,521005 |
| Lrrc8a | 10,801 | 9,16929 | -0,236279 | 0,1745 | 0,521005 |
| Acat1 | 7,09312 | 5,90675 | -0,264056 | 0,1744 | 0,521005 |
| Zfp747 | 2,03158 | 1,5385 | -0,401082 | 0,17475 | 0,521635 |
| Tshr | 0,497538 | 0,699825 | 0,492188 | 0,1749 | 0,521966 |
| Spag1 | 3,3009 | 2,63673 | -0,32411 | 0,175 | 0,522148 |
| B230217C12Rik | 8,08267 | 10,1431 | 0,327593 | 0,1751 | 0,52233 |
| Meaf6 | 32,9215 | 38,8127 | 0,237499 | 0,1753 | 0,522809 |
| Zfp563 | 1,76513 | 1,41871 | -0,315194 | 0,17535 | 0,522842 |
| Emc3 | 38,2603 | 32,7391 | -0,224833 | 0,1754 | 0,522874 |
| Fam122a | 8,83223 | 10,9141 | 0,305342 | 0,176 | 0,523475 |
| Zc3h12c | 2,51772 | 3,05896 | 0,280924 | 0,17615 | 0,523475 |
| Tsen15 | 20,6413 | 24,729 | 0,260667 | 0,17605 | 0,523475 |
| Recql4 | 11,1934 | 13,2247 | 0,240582 | 0,17615 | 0,523475 |
| Per1 | 7,19668 | 8,49489 | 0,239264 | 0,1761 | 0,523475 |
| Polr2j | 123,951 | 145,401 | 0,230275 | 0,1758 | 0,523475 |
| Pafah1b3 | 73,8164 | 86,572 | 0,229959 | 0,17615 | 0,523475 |
| Rab34 | 44,9193 | 52,4632 | 0,223971 | 0,17615 | 0,523475 |
| Sec23ip | 26,133 | 30,3385 | 0,215276 | 0,17575 | 0,523475 |
| Atxn10 | 106,707 | 123,862 | 0,215075 | 0,1759 | 0,523475 |
| Rbm6 | 36,9415 | 42,776 | 0,211558 | 0,17585 | 0,523475 |
| Cers6 | 7,13928 | 5,98333 | -0,254829 | 0,1758 | 0,523475 |
| Ftx | 5,79193 | 4,84019 | -0,25898 | 0,1757 | 0,523475 |
| Lcat | 2,20852 | 1,55775 | -0,503614 | 0,17565 | 0,523475 |
| Phlpp2 | 5,55381 | 6,52489 | 0,232476 | 0,1763 | 0,523688 |
| Prss41 | 0,919351 | 0,514522 | -0,837385 | 0,1763 | 0,523688 |
| Aldh7a1 | 37,2004 | 45,361 | 0,286135 | 0,1764 | 0,523752 |
| Taf12 | 26,786 | 31,7105 | 0,24348 | 0,1764 | 0,523752 |
| Commd5 | 17,9818 | 21,8086 | 0,27836 | 0,17655 | 0,523796 |
| Ddb1 | 287,082 | 340,798 | 0,247453 | 0,17655 | 0,523796 |
| Mylip | 40,6058 | 47,1563 | 0,215765 | 0,1766 | 0,523796 |
| Echs1 | 76,7513 | 89,0975 | 0,215193 | 0,1765 | 0,523796 |
| Glcci1 | 10,1945 | 8,70882 | -0,22725 | 0,17665 | 0,523796 |
| Zfp3 | 3,19616 | 2,42317 | -0,39944 | 0,17665 | 0,523796 |
| Pno1 | 65,9717 | 76,9117 | 0,221355 | 0,1767 | 0,523828 |
| Cetn4 | 1,01784 | 0,576263 | -0,820713 | 0,17695 | 0,524453 |
| Rnf11 | 32,4459 | 27,6673 | -0,229856 | 0,17715 | 0,52493 |
| Usf2 | 17,8863 | 21,0398 | 0,234268 | 0,1772 | 0,524961 |

| | | | | | |
|---|---|---|---|---|---|
| Klf11 | 4,05847 | 4,90965 | 0,274682 | 0,1774 | 0,525363 |
| Impdh2 | 14,8809 | 17,7891 | 0,257531 | 0,17755 | 0,525363 |
| Ly75 | 8,89386 | 10,4445 | 0,231861 | 0,17745 | 0,525363 |
| Usp39 | 55,1453 | 63,996 | 0,214743 | 0,17765 | 0,525363 |
| Hexim1 | 14,2796 | 12,1369 | -0,234555 | 0,17755 | 0,525363 |
| Ppp3ca | 21,61 | 18,3392 | -0,236766 | 0,17765 | 0,525363 |
| Gaa | 9,91062 | 8,38059 | -0,241923 | 0,1776 | 0,525363 |
| Batf3 | 2,19119 | 1,319 | -0,732274 | 0,1775 | 0,525363 |
| Tle3 | 33,7839 | 39,1922 | 0,214233 | 0,1779 | 0,525986 |
| Unc93b1 | 1,8116 | 1,33787 | -0,437323 | 0,178 | 0,526165 |
| Atp7a | 17,0922 | 14,7406 | -0,213541 | 0,1782 | 0,52664 |
| A630072M18Rik | 1,83063 | 1,18667 | -0,625421 | 0,1783 | 0,526819 |
| Trim61 | 0,614573 | 0,351392 | -0,806502 | 0,17835 | 0,52685 |
| Rpl37 | 73,9549 | 86,5808 | 0,227401 | 0,1784 | 0,526881 |
| Kctd17 | 22,9077 | 19,2853 | -0,248333 | 0,17845 | 0,526912 |
| Ccdc134 | 13,4635 | 16,0177 | 0,250611 | 0,17855 | 0,527037 |
| Elk1 | 13,0476 | 15,312 | 0,230881 | 0,17865 | 0,527037 |
| Pign | 9,95941 | 8,52303 | -0,224694 | 0,17865 | 0,527037 |
| Abhd11 | 17,4937 | 14,2506 | -0,295818 | 0,1786 | 0,527037 |
| Cep63 | 15,4928 | 18,1914 | 0,231651 | 0,1788 | 0,527162 |
| Zpr1 | 21,6187 | 25,2438 | 0,22365 | 0,17875 | 0,527162 |
| Loxl2 | 21,1526 | 18,2167 | -0,215575 | 0,17885 | 0,527162 |
| Zfp69 | 3,61244 | 2,83572 | -0,349261 | 0,17885 | 0,527162 |
| Sppl3 | 44,168 | 38,0073 | -0,216725 | 0,17905 | 0,527635 |
| Slc38a6 | 5,79032 | 6,97469 | 0,268486 | 0,1791 | 0,527666 |
| Tef | 9,83992 | 11,553 | 0,231545 | 0,17915 | 0,527697 |
| Phf1 | 3,99439 | 3,16911 | -0,333896 | 0,1792 | 0,527728 |
| Trmt44 | 4,3325 | 5,2381 | 0,273844 | 0,1793 | 0,527906 |
| Rab11fip3 | 12,8455 | 15,0582 | 0,22928 | 0,17945 | 0,528231 |
| Gpr19 | 3,50589 | 4,49551 | 0,358705 | 0,17955 | 0,528385 |
| Prelid2 | 20,8563 | 25,6176 | 0,296652 | 0,1797 | 0,528385 |
| Mrps18b | 17,8639 | 21,6379 | 0,276507 | 0,1797 | 0,528385 |
| Gnl3 | 112,084 | 133,891 | 0,256478 | 0,1797 | 0,528385 |
| Tmem219 | 8,25005 | 6,3748 | -0,372025 | 0,17965 | 0,528385 |
| Ddx24 | 48,3167 | 55,9836 | 0,212485 | 0,1798 | 0,528563 |
| Nvl | 42,0585 | 48,8894 | 0,217125 | 0,17985 | 0,528594 |
| Rpl21 | 3,1071 | 4,00557 | 0,366437 | 0,1804 | 0,529022 |
| Lzts3 | 1,96946 | 2,46918 | 0,326238 | 0,18045 | 0,529022 |
| Cdc14a | 6,52536 | 7,73421 | 0,245197 | 0,1801 | 0,529022 |
| Nfyb | 37,7521 | 43,9148 | 0,21815 | 0,18055 | 0,529022 |
| Trap1 | 61,8021 | 71,8101 | 0,216531 | 0,18025 | 0,529022 |
| Mapre2 | 23,4136 | 27,202 | 0,216363 | 0,1805 | 0,529022 |
| Ei24 | 74,6668 | 64,4979 | -0,211213 | 0,1804 | 0,529022 |
| Prom1 | 34,0413 | 29,287 | -0,217024 | 0,18055 | 0,529022 |
| Abcb6 | 22,0221 | 18,8725 | -0,222671 | 0,1805 | 0,529022 |
| Cisd2 | 26,8924 | 23,0454 | -0,222717 | 0,18055 | 0,529022 |
| Btd | 10,8346 | 8,9766 | -0,271408 | 0,1805 | 0,529022 |
| Tmem104 | 2,68186 | 2,17481 | -0,302341 | 0,1805 | 0,529022 |

| | | | | | |
|---|---|---|---|---|---|
| Mob3c | 2,54413 | 1,97954 | -0,362007 | 0,18035 | 0,529022 |
| Zfp595 | 1,38699 | 1,05477 | -0,395032 | 0,18015 | 0,529022 |
| 9930014A18Rik | 1,46476 | 2,02522 | 0,467417 | 0,18065 | 0,529083 |
| Rab24 | 27,6116 | 32,3532 | 0,228629 | 0,18065 | 0,529083 |
| Gabarapl2 | 8,49039 | 6,58681 | -0,366251 | 0,1808 | 0,52929 |
| Tnfrsf1a | 19,3212 | 16,3812 | -0,238147 | 0,1808 | 0,529291 |
| Rims4 | 0,377205 | 0,812047 | 1,10621 | 0,181 | 0,529528 |
| Dpy19l1 | 25,3243 | 21,821 | -0,214807 | 0,181 | 0,529528 |
| Apoa2 | 4,47941 | 2,82773 | -0,663667 | 0,181 | 0,529528 |
| Chsy1 | 31,9939 | 27,5683 | -0,214786 | 0,18115 | 0,529851 |
| Myef2 | 54,3069 | 46,7671 | -0,21564 | 0,1813 | 0,530174 |
| Rpap1 | 18,516 | 21,5152 | 0,216581 | 0,18135 | 0,530204 |
| Tmem86a | 3,71953 | 2,86338 | -0,377405 | 0,1814 | 0,530234 |
| Copg1 | 52,9992 | 62,6384 | 0,241078 | 0,18155 | 0,530239 |
| Rilpl1 | 15,2613 | 17,9822 | 0,236696 | 0,1816 | 0,530239 |
| Pak1 | 15,1256 | 17,7211 | 0,228483 | 0,18155 | 0,530239 |
| Ints9 | 24,4204 | 28,4475 | 0,22021 | 0,1816 | 0,530239 |
| Cul4a | 40,8651 | 47,3717 | 0,213157 | 0,18155 | 0,530239 |
| Aplp1 | 5,92801 | 4,83371 | -0,294418 | 0,18165 | 0,530269 |
| Cdk10 | 15,2807 | 18,1932 | 0,251691 | 0,1817 | 0,530299 |
| Pla2g3 | 0,677724 | 0,968504 | 0,515059 | 0,1823 | 0,530835 |
| H2-Ab1 | 4,30334 | 5,61275 | 0,383252 | 0,1823 | 0,530835 |
| AA388235 | 2,31389 | 2,88241 | 0,316954 | 0,1824 | 0,530835 |
| Zbtb38 | 3,04589 | 3,71939 | 0,288203 | 0,1824 | 0,530835 |
| Mrps5 | 22,9903 | 27,179 | 0,241467 | 0,1824 | 0,530835 |
| Ndufa7 | 202,493 | 237,061 | 0,22739 | 0,18215 | 0,530835 |
| Sept6 | 30,442 | 35,4143 | 0,21827 | 0,18235 | 0,530835 |
| Hspd1 | 155,336 | 180,441 | 0,216137 | 0,1821 | 0,530835 |
| Gls | 31,244 | 27,0386 | -0,208558 | 0,18215 | 0,530835 |
| Phf6 | 45,7864 | 39,5756 | -0,210309 | 0,1824 | 0,530835 |
| Irx2 | 13,1914 | 11,1174 | -0,246782 | 0,18235 | 0,530835 |
| Zfp58 | 6,7575 | 5,5293 | -0,289394 | 0,1824 | 0,530835 |
| Usp2 | 0,791684 | 0,560772 | -0,497511 | 0,1824 | 0,530835 |
| Ddx56 | 32,0139 | 37,1687 | 0,215388 | 0,18245 | 0,530865 |
| Engase | 5,90865 | 7,02574 | 0,249821 | 0,18275 | 0,531249 |
| Rab9 | 23,6396 | 27,8922 | 0,238657 | 0,18285 | 0,531249 |
| Nbea | 4,84598 | 5,71288 | 0,237428 | 0,18285 | 0,531249 |
| Zranb3 | 11,2404 | 13,1485 | 0,226202 | 0,18285 | 0,531249 |
| Med25 | 39,1254 | 45,4882 | 0,217387 | 0,18275 | 0,531249 |
| Traf4 | 31,8572 | 37,0247 | 0,21687 | 0,1829 | 0,531249 |
| Sec24c | 48,9229 | 56,5744 | 0,209641 | 0,18285 | 0,531249 |
| Fam219a | 1,43886 | 1,00008 | -0,524808 | 0,1829 | 0,531249 |
| Klhl2 | 8,36387 | 9,91815 | 0,2459 | 0,18305 | 0,531483 |
| Ssx2ip | 17,0947 | 14,6586 | -0,221803 | 0,1831 | 0,531483 |
| Naprt1 | 1,20465 | 0,823919 | -0,548042 | 0,1831 | 0,531483 |
| Wdr37 | 11,8375 | 10,1132 | -0,227128 | 0,1833 | 0,531948 |
| Cyth2 | 35,1156 | 40,7441 | 0,214481 | 0,18335 | 0,531978 |
| Pxylp1 | 11,1403 | 9,42719 | -0,240889 | 0,18345 | 0,532152 |

| | | | | | |
|---|---|---|---|---|---|
| Mettl3 | 24,6472 | 31,7149 | 0,363739 | 0,1835 | 0,532182 |
| Scarna10 | 77,664 | 0 | -inf | 0,1837 | 0,532646 |
| Clip1 | 18,8673 | 21,8175 | 0,209595 | 0,18385 | 0,532966 |
| Ube2cbp | 4,0242 | 5,0783 | 0,335645 | 0,184 | 0,533285 |
| Tpr | 59,632 | 69,7519 | 0,226144 | 0,18405 | 0,533314 |
| Clgn | 0,48523 | 0,744645 | 0,617885 | 0,18455 | 0,533404 |
| Fndc5 | 0,618542 | 0,888983 | 0,523285 | 0,1845 | 0,533404 |
| Snrpg | 153,335 | 181,353 | 0,242112 | 0,18435 | 0,533404 |
| Zfp346 | 10,8346 | 12,7018 | 0,229387 | 0,1845 | 0,533404 |
| Rhbdd2 | 13,5307 | 15,7905 | 0,222821 | 0,1846 | 0,533404 |
| Zfp280b | 15,1937 | 17,6762 | 0,218338 | 0,1842 | 0,533404 |
| Sept3 | 18,668 | 21,6722 | 0,215276 | 0,1845 | 0,533404 |
| Arl14ep | 18,3302 | 15,678 | -0,225479 | 0,1846 | 0,533404 |
| Celsr2 | 2,8487 | 2,40238 | -0,24584 | 0,18425 | 0,533404 |
| Rnf26 | 14,446 | 12,1744 | -0,246813 | 0,1843 | 0,533404 |
| Gpcpd1 | 7,33601 | 6,15866 | -0,252378 | 0,1846 | 0,533404 |
| 3110009E18Rik | 8,0381 | 5,62178 | -0,515827 | 0,1846 | 0,533404 |
| Grem1 | 0,750982 | 0,45252 | -0,730795 | 0,1844 | 0,533404 |
| Gan | 3,72112 | 4,60508 | 0,307489 | 0,18475 | 0,533607 |
| Carhsp1 | 38,085 | 44,1755 | 0,214022 | 0,18475 | 0,533607 |
| Pde4a | 1,58177 | 1,99029 | 0,331438 | 0,185 | 0,533983 |
| Pgrmc2 | 36,4938 | 31,4481 | -0,214679 | 0,185 | 0,533983 |
| Alg5 | 23,0069 | 19,4115 | -0,24515 | 0,18495 | 0,533983 |
| Dld | 94,6637 | 81,7635 | -0,211353 | 0,1851 | 0,534041 |
| Lst1 | 1,31208 | 0,328712 | -1,99696 | 0,1851 | 0,534041 |
| Tex261 | 30,9451 | 26,3224 | -0,23342 | 0,18535 | 0,534301 |
| Tmem260 | 6,97812 | 5,911 | -0,239438 | 0,18525 | 0,534301 |
| Dsel | 4,09008 | 3,41418 | -0,26059 | 0,18535 | 0,534301 |
| Prtn3 | 1,22945 | 0,463456 | -1,4075 | 0,18535 | 0,534301 |
| Pex26 | 4,02746 | 3,31638 | -0,280264 | 0,18545 | 0,534474 |
| Adamts17 | 5,14661 | 6,08374 | 0,241337 | 0,18565 | 0,534848 |
| Angpt2 | 2,17316 | 1,3413 | -0,696162 | 0,1857 | 0,534848 |
| Glyatl3 | 0,752255 | 0,352772 | -1,09249 | 0,1857 | 0,534848 |
| Ankrd10 | 65,3448 | 75,7197 | 0,212597 | 0,18575 | 0,534877 |
| Mob1a | 62,3292 | 53,8121 | -0,211979 | 0,18595 | 0,535337 |
| Brca1 | 18,9826 | 21,972 | 0,210986 | 0,18645 | 0,536199 |
| Dapk1 | 16,5699 | 19,1788 | 0,21095 | 0,18645 | 0,536199 |
| Denr | 60,471 | 69,9842 | 0,210787 | 0,18635 | 0,536199 |
| Pcbp4 | 151,439 | 175,129 | 0,209678 | 0,18635 | 0,536199 |
| Tet2 | 1,92144 | 1,5875 | -0,275431 | 0,1864 | 0,536199 |
| Fryl | 9,60689 | 11,1448 | 0,214224 | 0,1866 | 0,536515 |
| Aff2 | 2,08721 | 2,58098 | 0,306344 | 0,18665 | 0,536543 |
| Nudt8 | 14,0056 | 17,4 | 0,313085 | 0,18675 | 0,5366 |
| Ezh2 | 105,414 | 121,622 | 0,206342 | 0,18675 | 0,5366 |
| 2310014L17Rik | 1,08039 | 1,43201 | 0,406482 | 0,1868 | 0,536628 |
| BC064078 | 4,34672 | 5,84498 | 0,427273 | 0,187 | 0,536654 |
| 9130401M01Rik | 36,6085 | 42,8405 | 0,226794 | 0,18685 | 0,536654 |
| Tspyl2 | 16,9734 | 19,8361 | 0,224853 | 0,18705 | 0,536654 |

| | | | | | |
|---|---|---|---|---|---|
| Fam13b | 20,2623 | 23,6385 | 0,222342 | 0,18705 | 0,536654 |
| Creld2 | 44,7451 | 52,1829 | 0,221847 | 0,187 | 0,536654 |
| Tmem147 | 65,7024 | 56,0341 | -0,229642 | 0,187 | 0,536654 |
| Hnrnpab | 491,902 | 581,073 | 0,240349 | 0,1871 | 0,536683 |
| St6gal2 | 0,896241 | 1,25638 | 0,487312 | 0,1874 | 0,537197 |
| Asphd2 | 5,2052 | 6,52658 | 0,326372 | 0,1874 | 0,537197 |
| Prkdc | 4,76945 | 5,56219 | 0,221832 | 0,18735 | 0,537197 |
| Snora75 | 63,1729 | 162,587 | 1,36383 | 0,1875 | 0,537253 |
| Ubald1 | 14,54 | 17,4043 | 0,259419 | 0,1875 | 0,537253 |
| Cox4i2 | 9,05397 | 14,6208 | 0,691397 | 0,18775 | 0,537739 |
| Nop58 | 207,247 | 239,951 | 0,211391 | 0,18775 | 0,537739 |
| Gusb | 31,9617 | 27,4905 | -0,217409 | 0,1878 | 0,537767 |
| Sap130 | 25,6253 | 29,6701 | 0,211439 | 0,18785 | 0,537795 |
| Srek1 | 36,8977 | 42,6975 | 0,21062 | 0,18795 | 0,537966 |
| Ogfr | 59,9081 | 69,246 | 0,208979 | 0,1882 | 0,538566 |
| Prdm10 | 4,97757 | 5,9775 | 0,2641 | 0,18825 | 0,538594 |
| Cmklr1 | 3,02574 | 3,76709 | 0,316161 | 0,1884 | 0,538908 |
| Dr1 | 34,5468 | 29,8603 | -0,210325 | 0,1886 | 0,539365 |
| Dnajb5 | 9,43498 | 11,2409 | 0,252671 | 0,18865 | 0,539392 |
| Usp3 | 30,6856 | 35,624 | 0,215289 | 0,1888 | 0,53959 |
| Rab33b | 4,28051 | 3,50996 | -0,286327 | 0,1888 | 0,53959 |
| Prmt1 | 446,794 | 519,502 | 0,21752 | 0,1889 | 0,53976 |
| Neurl4 | 27,0864 | 31,2947 | 0,208348 | 0,18895 | 0,539788 |
| Flywch2 | 19,8358 | 24,3891 | 0,298125 | 0,1893 | 0,540095 |
| Usp38 | 13,1624 | 11,287 | -0,221763 | 0,1891 | 0,540095 |
| Clpb | 9,78533 | 8,34795 | -0,229199 | 0,1892 | 0,540095 |
| Pars2 | 6,33906 | 5,17899 | -0,291598 | 0,18915 | 0,540095 |
| Zfp709 | 2,09256 | 1,65448 | -0,33889 | 0,18925 | 0,540095 |
| Dusp10 | 0,753039 | 0,511511 | -0,55796 | 0,1893 | 0,540095 |
| Znrd1as | 5,77432 | 7,4792 | 0,373233 | 0,18935 | 0,540122 |
| Cep85l | 2,93752 | 3,67794 | 0,3243 | 0,1895 | 0,540435 |
| Huwe1 | 61,328 | 71,9553 | 0,230554 | 0,18965 | 0,540747 |
| Dguok | 16,3879 | 19,8397 | 0,275764 | 0,1899 | 0,541114 |
| 1110057K04Rik | 23,1757 | 27,1684 | 0,229316 | 0,1899 | 0,541114 |
| Ap3m2 | 1,02644 | 0,7541 | -0,444814 | 0,18985 | 0,541114 |
| Clmn | 2,57761 | 3,03085 | 0,233685 | 0,19005 | 0,541426 |
| Ube2e1 | 65,8546 | 56,3867 | -0,223929 | 0,1902 | 0,541738 |
| Eif3d | 158,221 | 182,705 | 0,207579 | 0,1903 | 0,541791 |
| Tead4 | 2,38043 | 1,87401 | -0,345091 | 0,1903 | 0,541791 |
| Acd | 17,4129 | 20,8334 | 0,258745 | 0,19045 | 0,541899 |
| Lrrc42 | 48,2898 | 56,017 | 0,214148 | 0,19045 | 0,541899 |
| Atxn2 | 23,5488 | 27,2825 | 0,212321 | 0,1905 | 0,541899 |
| Dhrs13 | 9,77606 | 8,05825 | -0,278786 | 0,1905 | 0,541899 |
| Cdc37 | 117,218 | 135,485 | 0,208932 | 0,19055 | 0,541926 |
| Rab10 | 42,2118 | 36,4048 | -0,213515 | 0,19065 | 0,542095 |
| Gstm5 | 151,131 | 176,661 | 0,225184 | 0,1907 | 0,542122 |
| Mir6904 | 0 | 207,151 | inf | 0,19085 | 0,542371 |
| Fam193b | 27,7716 | 32,1231 | 0,21 | 0,19095 | 0,542371 |

| | | | | | |
|---|---|---|---|---|---|
| Ankle1 | 13,5379 | 11,4006 | -0,247894 | 0,1909 | 0,542371 |
| D630003M21Rik | 1,32743 | 1,01662 | -0,38485 | 0,19095 | 0,542371 |
| Lbx2 | 1,83437 | 2,76478 | 0,591881 | 0,191 | 0,542398 |
| Csnk1e | 70,0996 | 80,8983 | 0,206702 | 0,19105 | 0,542425 |
| Lipt2 | 12,1507 | 9,91224 | -0,29376 | 0,1911 | 0,542451 |
| Brcc3 | 31,4272 | 37,7894 | 0,265967 | 0,19155 | 0,542744 |
| Nrip1 | 5,15886 | 6,1152 | 0,245348 | 0,1916 | 0,542744 |
| Snx24 | 14,5099 | 17,178 | 0,243521 | 0,19145 | 0,542744 |
| Rnf219 | 23,6105 | 27,3795 | 0,21367 | 0,19145 | 0,542744 |
| Kctd3 | 22,1893 | 25,7104 | 0,212487 | 0,1916 | 0,542744 |
| Trrap | 22,8546 | 26,4164 | 0,208949 | 0,19165 | 0,542744 |
| Qrich1 | 53,1957 | 61,4393 | 0,20785 | 0,19155 | 0,542744 |
| Lars | 41,7964 | 48,249 | 0,20712 | 0,19165 | 0,542744 |
| Mpp5 | 16,6396 | 14,3647 | -0,212092 | 0,1914 | 0,542744 |
| Adamts1 | 4,84 | 4,05012 | -0,257044 | 0,19135 | 0,542744 |
| Orai2 | 1,61608 | 1,25446 | -0,365432 | 0,19125 | 0,542744 |
| Dtd1 | 10,5014 | 12,8045 | 0,28607 | 0,1918 | 0,542824 |
| Nkain1 | 66,7906 | 77,0809 | 0,206729 | 0,1918 | 0,542824 |
| Serinc5 | 35,5507 | 30,7903 | -0,207402 | 0,19175 | 0,542824 |
| Vps36 | 40,2364 | 46,9775 | 0,223466 | 0,19205 | 0,542842 |
| Palld | 22,2137 | 25,7084 | 0,210788 | 0,19205 | 0,542842 |
| Ubqln4 | 68,8646 | 79,3532 | 0,204524 | 0,1919 | 0,542842 |
| 1700020I14Rik | 44,7131 | 38,4477 | -0,217801 | 0,19195 | 0,542842 |
| Syt7 | 1,13439 | 0,892539 | -0,345928 | 0,192 | 0,542842 |
| Gm11549 | 0,705551 | 0,468104 | -0,591921 | 0,192 | 0,542842 |
| Rer1 | 97,8677 | 84,6399 | -0,209495 | 0,1921 | 0,542868 |
| Hes1 | 32,6443 | 38,1347 | 0,224277 | 0,19225 | 0,543063 |
| Nfx1 | 42,1803 | 48,6572 | 0,206085 | 0,19225 | 0,543063 |
| 1600014C10Rik | 4,98219 | 6,00487 | 0,269353 | 0,1925 | 0,543539 |
| Ssr4 | 71,9844 | 61,3174 | -0,231389 | 0,1925 | 0,543539 |
| Sept2 | 79,4278 | 66,8266 | -0,205222 | 0,1927 | 0,543989 |
| Psme2b | 36,3269 | 42,7436 | 0,234668 | 0,19275 | 0,544015 |
| Akap1 | 17,5009 | 15,0676 | -0,215981 | 0,1928 | 0,544041 |
| Dhx38 | 25,2424 | 29,1977 | 0,210009 | 0,1929 | 0,54412 |
| 4930427A07Rik | 11,8635 | 10,1202 | -0,229291 | 0,19295 | 0,54412 |
| 4930404N11Rik | 3,13427 | 1,4293 | -1,13282 | 0,19295 | 0,54412 |
| Crb2 | 20,4566 | 23,967 | 0,228479 | 0,1931 | 0,544313 |
| Elp2 | 74,864 | 86,2724 | 0,204627 | 0,1931 | 0,544313 |
| Hotair | 27,9769 | 24,0249 | -0,219703 | 0,1932 | 0,54448 |
| Agtrap | 13,3642 | 15,5703 | 0,220421 | 0,1933 | 0,544647 |
| Psmd8 | 118,54 | 136,843 | 0,207148 | 0,19335 | 0,544673 |
| Dnajc24 | 17,5202 | 20,953 | 0,258133 | 0,19345 | 0,54484 |
| Xrn1 | 9,14977 | 10,6702 | 0,221783 | 0,1935 | 0,544866 |
| Mir8116 | 67,0634 | 0 | -inf | 0,19355 | 0,544892 |
| Zfp384 | 17,2205 | 20,0305 | 0,218067 | 0,1936 | 0,544918 |
| Csgalnact2 | 6,71438 | 5,63464 | -0,252931 | 0,19385 | 0,545507 |
| Dhx16 | 24,4813 | 28,3069 | 0,209471 | 0,19395 | 0,545558 |
| Bnip3l | 41,67 | 36,0755 | -0,207989 | 0,19395 | 0,545558 |

| | | | | | |
|---|---|---|---|---|---|
| Pomt2 | 9,64056 | 8,00049 | -0,26903 | 0,1941 | 0,545865 |
| Ticam1 | 2,49808 | 3,13638 | 0,328283 | 0,19425 | 0,546172 |
| Thap4 | 20,6111 | 24,0647 | 0,223498 | 0,19435 | 0,546339 |
| Zwint | 45,8706 | 53,0348 | 0,20937 | 0,1944 | 0,546364 |
| Phf10 | 86,8689 | 102,102 | 0,2331 | 0,1946 | 0,546811 |
| Raly | 252,669 | 291,949 | 0,208468 | 0,19465 | 0,546837 |
| Nop56 | 117,575 | 140,043 | 0,252284 | 0,19485 | 0,546849 |
| Ubl5 | 41,3632 | 48,5345 | 0,230662 | 0,19485 | 0,546849 |
| Sfswap | 34,7969 | 40,1964 | 0,208107 | 0,1949 | 0,546849 |
| Strbp | 9,14694 | 10,545 | 0,205191 | 0,1947 | 0,546849 |
| Sepn1 | 38,8899 | 33,6456 | -0,208978 | 0,19475 | 0,546849 |
| Zfp938 | 7,94486 | 6,52689 | -0,283626 | 0,1949 | 0,546849 |
| Zfp30 | 2,58012 | 3,25373 | 0,334652 | 0,19515 | 0,547091 |
| Ssrp1 | 249,956 | 290,041 | 0,21458 | 0,19515 | 0,547091 |
| Hyou1 | 88,7417 | 102,545 | 0,20857 | 0,1951 | 0,547091 |
| Prkaca | 47,93 | 55,3384 | 0,207352 | 0,19515 | 0,547091 |
| Aimp1 | 81,1669 | 94,0257 | 0,212164 | 0,19535 | 0,547448 |
| Igdcc4 | 4,82219 | 4,09084 | -0,23729 | 0,1954 | 0,547448 |
| Ap1s2 | 8,73478 | 7,33993 | -0,251004 | 0,1954 | 0,547448 |
| Psma3 | 8,47095 | 10,7785 | 0,347561 | 0,19555 | 0,547524 |
| Ece2 | 20,3057 | 24,5067 | 0,271289 | 0,19555 | 0,547524 |
| Tmem109 | 27,5896 | 23,6043 | -0,225075 | 0,1955 | 0,547524 |
| Snord8 | 37,3897 | 0 | -inf | 0,1957 | 0,547586 |
| Csnk2a2 | 18,1804 | 21,0549 | 0,211773 | 0,1958 | 0,547586 |
| Dlg5 | 20,9704 | 24,1996 | 0,206626 | 0,1958 | 0,547586 |
| Scaf8 | 29,3543 | 33,8706 | 0,206458 | 0,1959 | 0,547586 |
| Pole | 23,3487 | 26,9027 | 0,204408 | 0,19585 | 0,547586 |
| Pax3 | 22,6988 | 19,6022 | -0,211602 | 0,1959 | 0,547586 |
| Tcn2 | 11,1458 | 9,31522 | -0,258841 | 0,19575 | 0,547586 |
| N6amt1 | 11,2525 | 9,34594 | -0,267839 | 0,19565 | 0,547586 |
| H2-T9 | 0,326684 | 2,14564 | 2,71544 | 0,19595 | 0,547611 |
| Zfhx3 | 1,24656 | 1,04404 | -0,255782 | 0,19625 | 0,548335 |
| Cep120 | 11,6021 | 9,95307 | -0,22117 | 0,1963 | 0,54836 |
| Tox2 | 2,97877 | 2,25694 | -0,400346 | 0,19635 | 0,548385 |
| Dock4 | 4,18525 | 3,55383 | -0,235938 | 0,19645 | 0,548435 |
| Tapbpl | 1,77375 | 1,29006 | -0,459365 | 0,19645 | 0,548435 |
| Glod4 | 47,4295 | 54,7749 | 0,207731 | 0,19655 | 0,548599 |
| Mir7091 | 186,653 | 0 | -inf | 0,1969 | 0,54867 |
| Snord11 | 91,2968 | 0 | -inf | 0,19675 | 0,54867 |
| Pip5kl1 | 0,424545 | 0,722704 | 0,767489 | 0,19685 | 0,54867 |
| Igsf9b | 0,773992 | 1,03786 | 0,423225 | 0,19695 | 0,54867 |
| Psmg4 | 39,3006 | 47,7688 | 0,281516 | 0,1968 | 0,54867 |
| Dcps | 55,9326 | 66,2788 | 0,244858 | 0,19705 | 0,54867 |
| Ascc2 | 22,0252 | 25,577 | 0,215693 | 0,1971 | 0,54867 |
| Nbn | 24,047 | 27,9159 | 0,215228 | 0,1971 | 0,54867 |
| Smg6 | 16,1195 | 18,6487 | 0,210263 | 0,19715 | 0,54867 |
| Eif3i | 211,645 | 244,18 | 0,206293 | 0,19695 | 0,54867 |
| Hat1 | 111,247 | 128,09 | 0,203386 | 0,19705 | 0,54867 |

| | | | | | |
|---|---|---|---|---|---|
| Sdf4 | 30,8953 | 26,7804 | -0,206209 | 0,19715 | 0,54867 |
| Slc2a1 | 128,451 | 111,268 | -0,207178 | 0,19675 | 0,54867 |
| Ctxn1 | 37,1645 | 31,6793 | -0,230388 | 0,19675 | 0,54867 |
| Arhgef39 | 17,2351 | 14,5301 | -0,246307 | 0,1973 | 0,548973 |
| Gm2694 | 0,33422 | 0,828638 | 1,30994 | 0,1975 | 0,548983 |
| 4933430I17Rik | 0,993417 | 1,40332 | 0,498376 | 0,19745 | 0,548983 |
| Llgl1 | 46,5628 | 55,2376 | 0,246471 | 0,19755 | 0,548983 |
| Thap11 | 33,8626 | 39,2657 | 0,213579 | 0,19755 | 0,548983 |
| Gtpbp4 | 71,4084 | 82,2611 | 0,204116 | 0,1975 | 0,548983 |
| Tmem50a | 62,903 | 54,0349 | -0,219237 | 0,19745 | 0,548983 |
| Alg14 | 19,4256 | 15,9988 | -0,279994 | 0,19765 | 0,549146 |
| Snrpd1 | 195,827 | 225,727 | 0,205 | 0,1977 | 0,549171 |
| Nudt19 | 16,5261 | 19,4143 | 0,232376 | 0,19775 | 0,549196 |
| Afap1l2 | 1,73224 | 2,16861 | 0,324132 | 0,19785 | 0,54936 |
| Ntf3 | 2,59325 | 3,43248 | 0,404491 | 0,19795 | 0,549409 |
| Lmbr1 | 3,43194 | 2,85287 | -0,266608 | 0,19795 | 0,549409 |
| Fau | 4,88578 | 6,58305 | 0,43017 | 0,19825 | 0,549671 |
| Cep290 | 5,13468 | 6,09621 | 0,24764 | 0,19815 | 0,549671 |
| Gas5 | 168,633 | 195,044 | 0,209907 | 0,19825 | 0,549671 |
| Pgm2l1 | 4,75989 | 4,06504 | -0,227657 | 0,19815 | 0,549671 |
| Eva1a | 2,5474 | 1,91713 | -0,410081 | 0,1982 | 0,549671 |
| Cyld | 5,48088 | 6,38514 | 0,220309 | 0,1983 | 0,549695 |
| Crkl | 34,7611 | 39,9555 | 0,200919 | 0,19835 | 0,54972 |
| Extl2 | 11,7154 | 9,96005 | -0,234176 | 0,1984 | 0,549744 |
| Fanca | 14,4747 | 17,1887 | 0,247925 | 0,19845 | 0,549769 |
| Nt5m | 14,1128 | 16,9226 | 0,261939 | 0,1986 | 0,549843 |
| Kiz | 9,78922 | 11,6442 | 0,250344 | 0,19855 | 0,549843 |
| Cdca5 | 41,5355 | 47,9671 | 0,207703 | 0,1986 | 0,549843 |
| Cpsf3l | 36,5573 | 42,6636 | 0,222846 | 0,19885 | 0,549851 |
| Sra1 | 48,2828 | 56,2682 | 0,220811 | 0,19875 | 0,549851 |
| Cherp | 46,2153 | 53,5862 | 0,21349 | 0,1987 | 0,549851 |
| Pnn | 105,892 | 122,254 | 0,207288 | 0,19885 | 0,549851 |
| Eed | 51,9491 | 59,924 | 0,206036 | 0,19865 | 0,549851 |
| Kat2a | 47,1413 | 54,2863 | 0,203595 | 0,1988 | 0,549851 |
| Usp29 | 0,476155 | 0,62014 | 0,381164 | 0,19895 | 0,5499 |
| Spag7 | 73,562 | 85,0457 | 0,209277 | 0,19895 | 0,5499 |
| Aard | 2,41145 | 3,21755 | 0,41606 | 0,1996 | 0,549959 |
| BC025920 | 2,27879 | 2,96293 | 0,378755 | 0,1998 | 0,549959 |
| 2310057M21Rik | 8,93267 | 10,6447 | 0,252976 | 0,1999 | 0,549959 |
| Tonsl | 10,8663 | 12,6643 | 0,220916 | 0,1994 | 0,549959 |
| Wdr55 | 33,8355 | 39,3942 | 0,219444 | 0,19985 | 0,549959 |
| Cpsf1 | 35,5278 | 41,1783 | 0,212936 | 0,19995 | 0,549959 |
| Ube2g2 | 29,6716 | 34,3884 | 0,212837 | 0,1994 | 0,549959 |
| Snap47 | 44,9689 | 52,0802 | 0,211808 | 0,19915 | 0,549959 |
| Kctd15 | 35,1072 | 40,5438 | 0,207716 | 0,19975 | 0,549959 |
| Map2k2 | 59,3066 | 68,4412 | 0,206672 | 0,1995 | 0,549959 |
| Zscan21 | 36,2834 | 41,8631 | 0,206371 | 0,19975 | 0,549959 |
| Edc4 | 25,3232 | 29,1973 | 0,205379 | 0,19965 | 0,549959 |

| | | | | | |
|---|---|---|---|---|---|
| Ddx47 | 67,498 | 77,7725 | 0,204416 | 0,2 | 0,549959 |
| Atp9b | 15,7641 | 13,6604 | -0,206645 | 0,19995 | 0,549959 |
| Pofut1 | 14,6087 | 12,6231 | -0,210766 | 0,19955 | 0,549959 |
| Pxn | 18,8908 | 16,321 | -0,210954 | 0,19975 | 0,549959 |
| Golph3l | 15,5224 | 13,2934 | -0,223647 | 0,1993 | 0,549959 |
| Slc38a7 | 6,78032 | 5,67985 | -0,255499 | 0,2 | 0,549959 |
| Nudt11 | 7,2811 | 6,06067 | -0,264678 | 0,19905 | 0,549959 |
| 1700020L24Rik | 1,637 | 1,03273 | -0,664584 | 0,19975 | 0,549959 |
| Gm9895 | 0,862616 | 0,0930059 | -3,21332 | 0,19985 | 0,549959 |
| Prap1 | 1,11906 | 0,102351 | -3,4507 | 0,19975 | 0,549959 |
| Vgll3 | 0,634338 | 0,013682 | -5,5349 | 0,1995 | 0,549959 |
| Ppbp | 6,71752 | 0,0488394 | -7,10374 | 0,1995 | 0,549959 |
| Afp | 14,7422 | 0,0227331 | -9,34094 | 0,1995 | 0,549959 |
| Mterf1b | 1,49014 | 1,00843 | -0,563336 | 0,20015 | 0,550145 |
| Syt5 | 0,708275 | 0,437439 | -0,695227 | 0,20015 | 0,550145 |
| Hfe2 | 0,56706 | 0,850736 | 0,58521 | 0,20025 | 0,550307 |
| Gpsm2 | 24,4896 | 29,1863 | 0,253121 | 0,2004 | 0,550606 |
| Lsm10 | 17,7419 | 21,5607 | 0,281248 | 0,2006 | 0,550613 |
| Ptpdc1 | 4,18738 | 4,98398 | 0,251251 | 0,2006 | 0,550613 |
| Lactb | 10,9035 | 12,9348 | 0,246459 | 0,20055 | 0,550613 |
| Stau2 | 8,486 | 10,0485 | 0,243818 | 0,20045 | 0,550613 |
| Slc43a2 | 1,72139 | 1,40674 | -0,291227 | 0,20065 | 0,550613 |
| Dnajb3 | 4,83494 | 3,62354 | -0,416101 | 0,20065 | 0,550613 |
| Fgfr4 | 0,770661 | 0,543718 | -0,503237 | 0,2008 | 0,550912 |
| Nop16 | 53,2953 | 61,5462 | 0,207661 | 0,20095 | 0,55121 |
| Lrrk2 | 2,42729 | 2,88131 | 0,24738 | 0,20105 | 0,551371 |
| Ppp1r13l | 3,43569 | 4,19543 | 0,288218 | 0,2011 | 0,551395 |
| Ppp1r14b | 61,8973 | 72,0398 | 0,218917 | 0,20125 | 0,551693 |
| Dcaf13 | 50,9909 | 58,9499 | 0,20925 | 0,20145 | 0,552128 |
| Hes5 | 2,45286 | 3,28059 | 0,419491 | 0,20155 | 0,552176 |
| Rab3ip | 12,0328 | 14,1195 | 0,230721 | 0,20155 | 0,552176 |
| Slc25a4 | 331,126 | 382,112 | 0,206616 | 0,2017 | 0,552474 |
| Hook1 | 6,04727 | 5,10611 | -0,244061 | 0,20185 | 0,552771 |
| Alkbh2 | 8,89631 | 11,0276 | 0,309844 | 0,20195 | 0,552932 |
| AK129341 | 4,3598 | 5,23945 | 0,265155 | 0,2022 | 0,553389 |
| Mbp | 3,42852 | 2,83405 | -0,274724 | 0,2022 | 0,553389 |
| Apip | 32,1789 | 37,9751 | 0,238939 | 0,2026 | 0,554144 |
| Cenpn | 27,7197 | 32,248 | 0,218299 | 0,2026 | 0,554144 |
| Kif13b | 2,9326 | 2,43287 | -0,26952 | 0,2026 | 0,554144 |
| Xpa | 9,73484 | 12,1034 | 0,314186 | 0,2027 | 0,554304 |
| Mir7022 | 842,732 | 0 | -inf | 0,203 | 0,554784 |
| Noxo1 | 1,01289 | 1,38457 | 0,450968 | 0,203 | 0,554784 |
| Ccpg1 | 7,54797 | 6,09395 | -0,308711 | 0,20295 | 0,554784 |
| Slc35a5 | 3,46629 | 2,86101 | -0,276867 | 0,2031 | 0,554943 |
| Dopey2 | 2,12366 | 2,81439 | 0,406268 | 0,2034 | 0,555422 |
| Tmem87a | 20,5048 | 17,4843 | -0,229902 | 0,20335 | 0,555422 |
| Ly6a | 0,999477 | 0,6001 | -0,73597 | 0,2034 | 0,555422 |
| Csnk1g2 | 92,0094 | 105,878 | 0,202554 | 0,2035 | 0,555582 |

| | | | | | |
|---|---|---|---|---|---|
| Sf3a1 | 60,0929 | 69,1933 | 0,203437 | 0,20365 | 0,555764 |
| Erc2 | 2,56156 | 2,1232 | -0,270786 | 0,20365 | 0,555764 |
| Mir3535 | 26,36 | 57,3242 | 1,1208 | 0,20375 | 0,55581 |
| Fign | 3,34721 | 2,85469 | -0,229626 | 0,20375 | 0,55581 |
| Zbtb9 | 13,0691 | 15,2567 | 0,223286 | 0,20395 | 0,556129 |
| Pcbp1 | 271,708 | 313,312 | 0,20554 | 0,20395 | 0,556129 |
| Nelfb | 39,5042 | 45,5374 | 0,205047 | 0,204 | 0,556152 |
| 9630033F20Rik | 4,66698 | 5,57229 | 0,255782 | 0,20405 | 0,556175 |
| Isg20 | 0,856439 | 1,3312 | 0,636305 | 0,20425 | 0,556493 |
| Uty | 4,80891 | 4,05164 | -0,247203 | 0,20425 | 0,556493 |
| Hsp90ab1 | 1591,6 | 2011,49 | 0,337786 | 0,2043 | 0,556516 |
| Tnnt2 | 0,738877 | 0,0818631 | -3,17405 | 0,2044 | 0,556674 |
| Ccdc159 | 1,35163 | 2,08166 | 0,623033 | 0,20475 | 0,55672 |
| Dnaaf3 | 3,08371 | 3,8773 | 0,330389 | 0,2046 | 0,55672 |
| 4930453N24Rik | 25,5698 | 30,02 | 0,231482 | 0,20475 | 0,55672 |
| Btf3 | 124,55 | 143,562 | 0,204953 | 0,2045 | 0,55672 |
| Sdc1 | 77,6339 | 67,4222 | -0,203463 | 0,2046 | 0,55672 |
| Abcd3 | 22,5096 | 19,4824 | -0,208368 | 0,20475 | 0,55672 |
| Ddah1 | 8,92762 | 7,60123 | -0,232042 | 0,20465 | 0,55672 |
| Jpx | 1,34263 | 1,02992 | -0,382526 | 0,2046 | 0,55672 |
| Kansl1 | 30,9833 | 35,6142 | 0,200964 | 0,2048 | 0,556743 |
| Bop1 | 46,9376 | 54,7898 | 0,223165 | 0,20485 | 0,556765 |
| Tada3 | 17,3114 | 20,2852 | 0,228702 | 0,2049 | 0,556788 |
| Sdad1 | 32,619 | 37,5305 | 0,202352 | 0,20495 | 0,556811 |
| Vldlr | 2,31476 | 1,93586 | -0,257886 | 0,205 | 0,556833 |
| Mir6973b | 77,3442 | 0 | -inf | 0,2051 | 0,556992 |
| Angptl4 | 7,82901 | 9,40077 | 0,26395 | 0,20515 | 0,557014 |
| Adsl | 26,2567 | 30,2405 | 0,2038 | 0,2052 | 0,557037 |
| Rpl5 | 9,2076 | 11,942 | 0,375144 | 0,20525 | 0,55706 |
| Rasd1 | 1,11235 | 2,21738 | 0,995253 | 0,2053 | 0,557082 |
| Ppp1r8 | 83,7234 | 96,3316 | 0,202379 | 0,2054 | 0,557127 |
| Igflr1 | 6,89794 | 5,44256 | -0,341879 | 0,2054 | 0,557127 |
| Zfp955a | 4,66296 | 3,8867 | -0,2627 | 0,2056 | 0,557556 |
| Exosc2 | 62,6098 | 72,2127 | 0,205864 | 0,20565 | 0,557579 |
| Slc20a1 | 50,8616 | 44,1864 | -0,202974 | 0,2057 | 0,557601 |
| Cox10 | 11,7594 | 13,732 | 0,223726 | 0,20585 | 0,557895 |
| Syngr2 | 23,2801 | 19,8094 | -0,232912 | 0,20605 | 0,558324 |
| Zscan20 | 3,50071 | 4,16161 | 0,249494 | 0,2061 | 0,558346 |
| Myo5c | 1,28934 | 1,03152 | -0,321856 | 0,20635 | 0,55891 |
| Pigx | 53,4019 | 45,9123 | -0,218011 | 0,2064 | 0,558932 |
| Tfam | 31,6741 | 36,4348 | 0,202012 | 0,2065 | 0,55909 |
| Xpc | 10,9504 | 12,7243 | 0,216597 | 0,20655 | 0,559112 |
| Ccnyl1 | 9,19188 | 7,7811 | -0,240385 | 0,20675 | 0,55954 |
| Zdhhc5 | 35,0209 | 29,3559 | -0,254563 | 0,20685 | 0,559697 |
| Ndst2 | 13,9862 | 12,0269 | -0,217737 | 0,20695 | 0,559854 |
| Taf15 | 140,236 | 161,359 | 0,202412 | 0,207 | 0,559876 |
| Sall4 | 64,8152 | 74,6462 | 0,203736 | 0,20725 | 0,560439 |
| Zbtb18 | 24,5632 | 21,2715 | -0,207581 | 0,2073 | 0,560461 |

| | | | | | |
|---|---|---|---|---|---|
| Arntl2 | 1,30872 | 1,71342 | 0,388725 | 0,20735 | 0,560482 |
| Pycr2 | 50,705 | 58,4683 | 0,205525 | 0,2075 | 0,560661 |
| Chpt1 | 8,38319 | 6,81148 | -0,299531 | 0,2075 | 0,560661 |
| 2700046G09Rik | 2,25759 | 1,68079 | -0,425646 | 0,20765 | 0,560953 |
| Mcmdc2 | 0,942449 | 0,668927 | -0,494565 | 0,2079 | 0,561515 |
| Smim7 | 39,6198 | 34,4135 | -0,203249 | 0,2081 | 0,561941 |
| Tead2 | 229,144 | 263,465 | 0,201353 | 0,20815 | 0,561963 |
| 3110001I22Rik | 5,76793 | 8,14027 | 0,497023 | 0,2084 | 0,562411 |
| Rbm28 | 37,6453 | 43,2096 | 0,198884 | 0,2084 | 0,562411 |
| Pxmp2 | 3,0505 | 4,06762 | 0,41514 | 0,20855 | 0,562475 |
| Fam69a | 15,0373 | 12,5121 | -0,265214 | 0,20855 | 0,562475 |
| Naglu | 3,52623 | 2,82773 | -0,318485 | 0,20855 | 0,562475 |
| Tecpr2 | 4,93349 | 5,738 | 0,217939 | 0,20865 | 0,562631 |
| Anapc16 | 40,4904 | 48,0712 | 0,247595 | 0,2088 | 0,562808 |
| Gm16386 | 3,19591 | 2,6351 | -0,278369 | 0,2088 | 0,562808 |
| Fbxo22 | 35,8289 | 41,3717 | 0,20752 | 0,20885 | 0,562829 |
| Ptprz1 | 3,82603 | 4,46923 | 0,224179 | 0,2089 | 0,562851 |
| Eif3m | 74,809 | 86,2846 | 0,205891 | 0,209 | 0,562893 |
| Srd5a1 | 6,93022 | 5,78396 | -0,260843 | 0,209 | 0,562893 |
| Bend3 | 19,4055 | 22,3016 | 0,200681 | 0,20915 | 0,563184 |
| Ikbke | 0,714629 | 0,96966 | 0,440284 | 0,20925 | 0,563339 |
| Fam174a | 4,65322 | 3,74957 | -0,311503 | 0,2094 | 0,56363 |
| 2310045N01Rik | 25,8476 | 31,4084 | 0,281121 | 0,2097 | 0,56389 |
| Taco1 | 16,6106 | 19,6846 | 0,24496 | 0,20975 | 0,56389 |
| Noa1 | 19,1086 | 22,1994 | 0,2163 | 0,2097 | 0,56389 |
| Eef1b2 | 328,469 | 379,327 | 0,207683 | 0,20975 | 0,56389 |
| Peli1 | 22,451 | 25,8933 | 0,205803 | 0,20965 | 0,56389 |
| Mtpn | 66,3825 | 57,9033 | -0,197157 | 0,20965 | 0,56389 |
| Gm10636 | 0,438322 | 0,659242 | 0,588819 | 0,2099 | 0,563953 |
| Aarsd1 | 55,82 | 64,3906 | 0,206069 | 0,20985 | 0,563953 |
| Atg9a | 9,53314 | 8,1639 | -0,223693 | 0,2099 | 0,563953 |
| 4930519F09Rik | 2,6971 | 3,5065 | 0,378624 | 0,2102 | 0,563987 |
| Ssbp2 | 6,1878 | 7,18591 | 0,215746 | 0,21005 | 0,563987 |
| Nup133 | 27,6919 | 31,8007 | 0,199592 | 0,2101 | 0,563987 |
| Cdc25a | 45,002 | 51,5871 | 0,197023 | 0,21005 | 0,563987 |
| Sdcbp | 77,3063 | 67,3936 | -0,197975 | 0,21015 | 0,563987 |
| Ireb2 | 46,1439 | 40,1418 | -0,201034 | 0,21025 | 0,563987 |
| D17Wsu104e | 83,7213 | 72,2874 | -0,21185 | 0,21025 | 0,563987 |
| Gch1 | 0,665968 | 0,452868 | -0,556362 | 0,2102 | 0,563987 |
| Pttg1 | 59,0792 | 69,1686 | 0,227467 | 0,2103 | 0,564007 |
| Supt3 | 10,9045 | 13,1244 | 0,267334 | 0,2104 | 0,564049 |
| Acot10 | 6,07081 | 4,86318 | -0,319988 | 0,2104 | 0,564049 |
| Slc25a17 | 51,8844 | 44,8981 | -0,208645 | 0,2105 | 0,564204 |
| Dnajb11 | 70,0034 | 80,4752 | 0,20112 | 0,21055 | 0,564225 |
| Smad9 | 0,45413 | 0,613286 | 0,433455 | 0,2106 | 0,564246 |
| Mthfd2 | 44,5002 | 51,1906 | 0,202066 | 0,21075 | 0,564421 |
| Mir196a-1 | 1495,08 | 1105,14 | -0,435991 | 0,21075 | 0,564421 |
| Utrn | 9,36489 | 8,13843 | -0,202511 | 0,2108 | 0,564442 |

| | | | | | |
|---|---|---|---|---|---|
| Qsox1 | 23,3101 | 26,9629 | 0,210022 | 0,2111 | 0,565019 |
| Bfar | 17,4474 | 14,8877 | -0,228892 | 0,2111 | 0,565019 |
| Ywhaz | 55,9339 | 48,7344 | -0,198784 | 0,21125 | 0,565235 |
| Sypl | 19,9505 | 17,3214 | -0,203866 | 0,21135 | 0,565235 |
| Rnasel | 2,34151 | 1,91022 | -0,293695 | 0,21135 | 0,565235 |
| Synj2 | 3,42662 | 2,64507 | -0,373481 | 0,2113 | 0,565235 |
| Lrfn1 | 7,35261 | 8,75911 | 0,252529 | 0,2114 | 0,565256 |
| Akap12 | 6,31248 | 5,4162 | -0,220926 | 0,21145 | 0,565277 |
| Pebp1 | 76,7088 | 88,3285 | 0,203487 | 0,2115 | 0,565297 |
| Rtf1 | 35,2524 | 40,4013 | 0,196682 | 0,21185 | 0,566119 |
| D19Bwg1357e | 47,453 | 54,4031 | 0,197188 | 0,2119 | 0,56614 |
| Dcaf17 | 8,02241 | 9,77978 | 0,285767 | 0,21195 | 0,56616 |
| Ebpl | 9,59876 | 8,03448 | -0,256643 | 0,2121 | 0,566448 |
| Ighmbp2 | 3,84917 | 4,54325 | 0,239176 | 0,2123 | 0,566662 |
| Vegfc | 16,171 | 18,9185 | 0,226393 | 0,21235 | 0,566662 |
| Lsm3 | 248,931 | 286,001 | 0,200277 | 0,21235 | 0,566662 |
| Hsd17b12 | 106,33 | 92,6068 | -0,19936 | 0,21235 | 0,566662 |
| Bcs1l | 9,43251 | 11,2408 | 0,253035 | 0,21255 | 0,56697 |
| Urb1 | 9,27175 | 10,6686 | 0,202459 | 0,21255 | 0,56697 |
| Irx3 | 2,17511 | 2,76534 | 0,34637 | 0,21265 | 0,567123 |
| Mir199a-1 | 0 | 250,944 | inf | 0,21275 | 0,567184 |
| Snrnp27 | 84,0732 | 97,1223 | 0,208156 | 0,2128 | 0,567184 |
| B230208H11Rik | 1,15906 | 0,726711 | -0,673503 | 0,2128 | 0,567184 |
| Gm1976 | 7,70301 | 5,78347 | -0,413487 | 0,2129 | 0,567337 |
| Tbx2 | 4,49275 | 3,73505 | -0,26647 | 0,213 | 0,56749 |
| Mdc1 | 28,2375 | 32,3084 | 0,194299 | 0,21315 | 0,567777 |
| Pms1 | 9,80103 | 11,4521 | 0,2246 | 0,2132 | 0,567797 |
| Tnks | 19,8216 | 22,7655 | 0,199779 | 0,2133 | 0,56795 |
| 4930503L19Rik | 23,9035 | 27,5675 | 0,205751 | 0,21345 | 0,568236 |
| Murc | 2,05276 | 2,65906 | 0,373353 | 0,2136 | 0,568316 |
| Rufy1 | 9,75191 | 11,4568 | 0,232448 | 0,21365 | 0,568316 |
| Tcp1 | 300,312 | 348,298 | 0,21386 | 0,21365 | 0,568316 |
| Agps | 24,792 | 21,619 | -0,197574 | 0,21355 | 0,568316 |
| Atm | 11,0052 | 12,6372 | 0,199496 | 0,2137 | 0,568336 |
| Rad52 | 12,326 | 14,6015 | 0,244413 | 0,21375 | 0,568356 |
| Pop7 | 25,0084 | 29,8855 | 0,257032 | 0,21385 | 0,568508 |
| Shroom3 | 16,4443 | 14,3201 | -0,199538 | 0,2139 | 0,568528 |
| Ddx39 | 96,5845 | 111,148 | 0,202615 | 0,214 | 0,568681 |
| Polb | 34,9614 | 41,1855 | 0,236372 | 0,2141 | 0,56872 |
| Tial1 | 39,6924 | 34,5987 | -0,198146 | 0,2141 | 0,56872 |
| Gper1 | 1,12493 | 0,524228 | -1,10157 | 0,21415 | 0,56874 |
| Mkks | 11,8888 | 14,1808 | 0,254331 | 0,2142 | 0,56876 |
| Zfp949 | 3,29742 | 3,9872 | 0,274037 | 0,2145 | 0,569443 |
| Syf2 | 36,7174 | 42,6484 | 0,21603 | 0,2147 | 0,569861 |
| C2cd2 | 6,61307 | 5,69472 | -0,215695 | 0,21475 | 0,569881 |
| Rce1 | 27,4222 | 23,5365 | -0,220441 | 0,21495 | 0,570298 |
| Rbm10 | 19,4023 | 23,3019 | 0,264221 | 0,2151 | 0,570321 |
| Gtf3c6 | 66,6428 | 76,8774 | 0,20611 | 0,21525 | 0,570321 |

| | | | | | |
|---|---|---|---|---|---|
| Uck2 | 84,7242 | 97,2318 | 0,198654 | 0,21525 | 0,570321 |
| Mlf2 | 152,728 | 174,914 | 0,195674 | 0,2153 | 0,570321 |
| Tmed2 | 18,2755 | 15,6045 | -0,227944 | 0,21515 | 0,570321 |
| Dnase1l1 | 2,70433 | 2,0499 | -0,39972 | 0,21525 | 0,570321 |
| Rbks | 1,56423 | 1,02629 | -0,608014 | 0,21525 | 0,570321 |
| Tm6sf2 | 0,960831 | 0,609356 | -0,656998 | 0,2153 | 0,570321 |
| Gm11985 | 0,355896 | 0,699588 | 0,975049 | 0,21555 | 0,570418 |
| Poc5 | 10,6951 | 12,4661 | 0,22106 | 0,21555 | 0,570418 |
| Zfp330 | 26,7498 | 31,055 | 0,215299 | 0,21545 | 0,570418 |
| Polh | 11,1347 | 9,47004 | -0,233622 | 0,21545 | 0,570418 |
| Agtpbp1 | 5,20746 | 4,38238 | -0,248863 | 0,2155 | 0,570418 |
| AI429214 | 0,893793 | 1,2508 | 0,484839 | 0,2157 | 0,570476 |
| Ehd3 | 5,64784 | 6,66748 | 0,239443 | 0,2157 | 0,570476 |
| Tmod1 | 1,92026 | 0,0777677 | -4,62598 | 0,21565 | 0,570476 |
| Agtr1a | 0,959091 | 0,673161 | -0,510717 | 0,21585 | 0,57076 |
| 1300002K09Rik | 13,0805 | 15,1485 | 0,211762 | 0,21595 | 0,570817 |
| Hsd17b7 | 12,0992 | 13,9719 | 0,207617 | 0,216 | 0,570817 |
| Abhd5 | 9,67392 | 8,23998 | -0,231459 | 0,216 | 0,570817 |
| Grip1 | 5,44533 | 6,73007 | 0,305602 | 0,2166 | 0,571725 |
| Stk19 | 10,2007 | 12,4576 | 0,288357 | 0,2165 | 0,571725 |
| Dgcr14 | 14,1686 | 16,422 | 0,212937 | 0,21655 | 0,571725 |
| Exosc7 | 63,7637 | 73,5897 | 0,206769 | 0,2166 | 0,571725 |
| Grcc10 | 170,731 | 196,672 | 0,20406 | 0,2165 | 0,571725 |
| 1810044D09Rik | 6,88542 | 4,84747 | -0,506312 | 0,2165 | 0,571725 |
| 2610034B18Rik | 8,228 | 6,87365 | -0,259466 | 0,21675 | 0,572008 |
| Mir1839 | 212,556 | 0 | -inf | 0,2169 | 0,572178 |
| Dna2 | 20,0334 | 22,9917 | 0,198703 | 0,2169 | 0,572178 |
| Ccdc90b | 26,0731 | 30,2243 | 0,213146 | 0,217 | 0,572328 |
| Exog | 10,6245 | 12,321 | 0,213727 | 0,21715 | 0,572611 |
| 3110045C21Rik | 6,9953 | 9,22984 | 0,399919 | 0,21755 | 0,572987 |
| Drp2 | 0,921296 | 1,14888 | 0,318495 | 0,2174 | 0,572987 |
| Brms1 | 32,9369 | 38,2173 | 0,21452 | 0,21755 | 0,572987 |
| Lzic | 34,4707 | 39,7088 | 0,204086 | 0,2175 | 0,572987 |
| Gtf2ird1 | 19,2019 | 22,1155 | 0,20381 | 0,21755 | 0,572987 |
| Pdcd2l | 23,8562 | 20,2808 | -0,234249 | 0,2174 | 0,572987 |
| Cxxc5 | 33,4892 | 38,4871 | 0,200676 | 0,2176 | 0,573006 |
| Utp18 | 33,7886 | 38,8497 | 0,201365 | 0,21765 | 0,573025 |
| Zc4h2 | 18,0548 | 20,9454 | 0,214247 | 0,21805 | 0,573419 |
| Pde4dip | 8,12963 | 9,39423 | 0,208585 | 0,21805 | 0,573419 |
| Usp47 | 66,26 | 75,9858 | 0,197591 | 0,218 | 0,573419 |
| Mcl1 | 64,5701 | 56,3876 | -0,195487 | 0,21795 | 0,573419 |
| Cbfb | 64,0371 | 55,8659 | -0,196941 | 0,21785 | 0,573419 |
| Ap1g2 | 4,11129 | 3,0204 | -0,444853 | 0,2181 | 0,573419 |
| B230217O12Rik | 0,615619 | 0,386447 | -0,671769 | 0,2181 | 0,573419 |
| Arid4a | 8,58031 | 9,92081 | 0,209428 | 0,21825 | 0,573475 |
| Fnbp4 | 30,435 | 34,8458 | 0,195252 | 0,2182 | 0,573475 |
| Nr6a1 | 2,00534 | 1,65288 | -0,278867 | 0,21825 | 0,573475 |
| Tbc1d4 | 4,1264 | 3,51358 | -0,231943 | 0,21835 | 0,573625 |

| | | | | | |
|---|---|---|---|---|---|
| Pced1b | 2,28481 | 1,65536 | -0,464931 | 0,2184 | 0,573643 |
| Utp6 | 44,0946 | 50,4068 | 0,193014 | 0,2186 | 0,574056 |
| Mir1898 | 15,9071 | 0 | -inf | 0,21865 | 0,574074 |
| Malsu1 | 23,9228 | 28,6194 | 0,258607 | 0,21915 | 0,574165 |
| Col22a1 | 2,40912 | 2,87404 | 0,254576 | 0,2192 | 0,574165 |
| Gins3 | 15,0938 | 17,5533 | 0,21779 | 0,2188 | 0,574165 |
| Sirt2 | 19,0071 | 22,103 | 0,217702 | 0,21875 | 0,574165 |
| Eci2 | 34,6973 | 40,1691 | 0,211262 | 0,21905 | 0,574165 |
| Pfdn4 | 85,0361 | 97,9845 | 0,20448 | 0,2192 | 0,574165 |
| Noc3l | 17,5027 | 20,141 | 0,202561 | 0,21885 | 0,574165 |
| Thop1 | 47,6901 | 54,7052 | 0,197989 | 0,219 | 0,574165 |
| C1d | 22,9744 | 19,9149 | -0,206176 | 0,2191 | 0,574165 |
| Zfp654 | 6,04607 | 5,16128 | -0,228268 | 0,2189 | 0,574165 |
| 2810408I11Rik | 17,1885 | 13,9098 | -0,305344 | 0,2191 | 0,574165 |
| Mir677 | 656,515 | 160,733 | -2,03016 | 0,2191 | 0,574165 |
| Mylk | 1,9253 | 1,58832 | -0,277582 | 0,21925 | 0,574183 |
| Rnf25 | 25,6873 | 29,8161 | 0,215036 | 0,21935 | 0,574333 |
| Zfr2 | 0,900387 | 0,66654 | -0,433853 | 0,21945 | 0,574482 |
| Prpf40b | 23,8529 | 27,4038 | 0,200213 | 0,21965 | 0,574893 |
| Atn1 | 23,0768 | 26,4947 | 0,199261 | 0,2197 | 0,574911 |
| Dnpep | 34,1761 | 29,5259 | -0,211009 | 0,2198 | 0,57506 |
| Ankrd49 | 22,9421 | 19,6996 | -0,219831 | 0,21985 | 0,575078 |
| Exosc10 | 40,1901 | 46,0765 | 0,197193 | 0,2199 | 0,575097 |
| Smc5 | 31,265 | 35,7841 | 0,194768 | 0,21995 | 0,575115 |
| Tchp | 11,4663 | 13,3409 | 0,218452 | 0,2203 | 0,575355 |
| Cct5 | 407,939 | 469,827 | 0,203778 | 0,2203 | 0,575355 |
| Lsm8 | 159,136 | 182,485 | 0,197515 | 0,22025 | 0,575355 |
| Cops6 | 134,796 | 154,422 | 0,196103 | 0,2203 | 0,575355 |
| Foxc2 | 45,6638 | 39,8532 | -0,196356 | 0,2201 | 0,575355 |
| Chst12 | 10,456 | 8,76469 | -0,254559 | 0,22025 | 0,575355 |
| Paox | 1,96961 | 1,47334 | -0,418823 | 0,22045 | 0,575634 |
| Gm10677 | 0,063478 | 1,23751 | 4,28503 | 0,2208 | 0,575873 |
| Mto1 | 8,5563 | 10,1237 | 0,242685 | 0,2206 | 0,575873 |
| Shc1 | 25,1533 | 28,9086 | 0,200752 | 0,22075 | 0,575873 |
| St3gal6 | 13,1526 | 11,0666 | -0,249135 | 0,22065 | 0,575873 |
| Zfp873 | 6,349 | 5,27767 | -0,266628 | 0,22075 | 0,575873 |
| Fam105a | 1,86801 | 1,46286 | -0,352709 | 0,2208 | 0,575873 |
| Rrp8 | 25,4435 | 31,5785 | 0,311642 | 0,22095 | 0,576075 |
| Nrf1 | 24,7889 | 28,5775 | 0,205186 | 0,22105 | 0,576075 |
| Sfxn1 | 78,409 | 89,7297 | 0,194566 | 0,22105 | 0,576075 |
| Nos1ap | 0,672353 | 0,418455 | -0,684145 | 0,22105 | 0,576075 |
| Mir6993 | 71,4612 | 0 | -inf | 0,2213 | 0,576295 |
| Mkln1 | 30,5996 | 35,0087 | 0,194202 | 0,2213 | 0,576295 |
| Pigc | 6,26536 | 5,29061 | -0,243964 | 0,22135 | 0,576295 |
| Snx32 | 1,8304 | 1,32432 | -0,466904 | 0,2213 | 0,576295 |
| Dio3os | 0,661231 | 0,0751086 | -3,13811 | 0,22135 | 0,576295 |
| Rnf223 | 1,11484 | 1,62029 | 0,539419 | 0,2215 | 0,576402 |
| Tspyl5 | 1,77675 | 2,19739 | 0,306548 | 0,22165 | 0,576402 |

| | | | | | |
|---|---|---|---|---|---|
| Zkscan17 | 26,7286 | 30,7245 | 0,201007 | 0,22165 | 0,576402 |
| Zfp217 | 13,5937 | 15,5816 | 0,1969 | 0,22155 | 0,576402 |
| Taf1 | 18,5085 | 21,1672 | 0,193641 | 0,2216 | 0,576402 |
| 1700001L05Rik | 0,628758 | 0,479576 | -0,390746 | 0,22145 | 0,576402 |
| Ngef | 0,443976 | 0,642944 | 0,534212 | 0,2217 | 0,57642 |
| Dusp16 | 6,02552 | 5,12379 | -0,233876 | 0,2218 | 0,576568 |
| Gpr135 | 0,475422 | 0,779731 | 0,713768 | 0,22275 | 0,57657 |
| 3110021N24Rik | 2,43134 | 3,89691 | 0,680581 | 0,2227 | 0,57657 |
| D1Pas1 | 1,10609 | 1,43524 | 0,375817 | 0,22205 | 0,57657 |
| Zfp438 | 1,25688 | 1,62176 | 0,367713 | 0,2219 | 0,57657 |
| Acacb | 0,493174 | 0,632503 | 0,358976 | 0,22235 | 0,57657 |
| Kri1 | 18,6988 | 22,6756 | 0,278198 | 0,2225 | 0,57657 |
| S100a11 | 43,2101 | 51,6759 | 0,258124 | 0,22235 | 0,57657 |
| Zfp369 | 5,56173 | 6,60642 | 0,248335 | 0,22185 | 0,57657 |
| Cstb | 63,1052 | 73,7659 | 0,225194 | 0,22255 | 0,57657 |
| Zfp623 | 12,6931 | 14,7875 | 0,220325 | 0,2224 | 0,57657 |
| Osgin2 | 12,9708 | 15,0767 | 0,217053 | 0,22275 | 0,57657 |
| Cacnb3 | 14,9746 | 17,3901 | 0,21575 | 0,22245 | 0,57657 |
| Ppp1r7 | 22,707 | 26,0467 | 0,197967 | 0,2222 | 0,57657 |
| Bax | 131,559 | 150,89 | 0,197791 | 0,2224 | 0,57657 |
| Samhd1 | 24,821 | 28,4602 | 0,197382 | 0,2224 | 0,57657 |
| Sgta | 70,5681 | 80,7826 | 0,195028 | 0,2223 | 0,57657 |
| Prr12 | 16,3147 | 18,6514 | 0,193109 | 0,2225 | 0,57657 |
| Lamb1 | 60,0172 | 52,5253 | -0,192364 | 0,2223 | 0,57657 |
| Tmtc4 | 7,61684 | 6,48518 | -0,232047 | 0,22265 | 0,57657 |
| Slc44a1 | 13,0417 | 11,0811 | -0,235028 | 0,22275 | 0,57657 |
| Stard13 | 3,59643 | 3,03401 | -0,245341 | 0,2221 | 0,57657 |
| Tmem88 | 10,1882 | 7,72061 | -0,400115 | 0,2221 | 0,57657 |
| 4930432K21Rik | 0,547355 | 0,788145 | 0,525983 | 0,2228 | 0,576588 |
| Vrk1 | 29,3991 | 33,6674 | 0,195579 | 0,22285 | 0,576605 |
| Mrpl23 | 0,375916 | 0,87992 | 1,22696 | 0,22295 | 0,576753 |
| Lca5 | 10,017 | 11,5933 | 0,210853 | 0,22315 | 0,576935 |
| Tstd2 | 22,5562 | 25,9239 | 0,200757 | 0,22315 | 0,576935 |
| Rhoq | 19,0202 | 16,1343 | -0,237395 | 0,22315 | 0,576935 |
| Gm867 | 0,606974 | 0,0417806 | -3,86073 | 0,2233 | 0,577211 |
| Appbp2 | 38,9209 | 44,5886 | 0,196131 | 0,22365 | 0,578004 |
| Slc2a13 | 0,802481 | 0,625616 | -0,35919 | 0,22375 | 0,578151 |
| Trib2 | 15,3046 | 13,2997 | -0,202567 | 0,2238 | 0,578168 |
| Cmss1 | 22,834 | 27,4625 | 0,266283 | 0,22395 | 0,578332 |
| Tmem41b | 4,36295 | 3,64684 | -0,258659 | 0,22395 | 0,578332 |
| Mir6352 | 0 | 7,76137 | inf | 0,2242 | 0,578642 |
| Btbd7 | 18,535 | 21,2235 | 0,195417 | 0,2242 | 0,578642 |
| Mboat2 | 22,0396 | 19,1385 | -0,203622 | 0,2242 | 0,578642 |
| Pi4k2a | 19,1198 | 21,9275 | 0,197673 | 0,22425 | 0,578659 |
| Dpy30 | 130,96 | 150,393 | 0,199608 | 0,22435 | 0,578806 |
| Desi2 | 13,9843 | 12,0998 | -0,208823 | 0,22445 | 0,578952 |
| D030028A08Rik | 2,68575 | 1,80065 | -0,57681 | 0,22455 | 0,579098 |
| AW554918 | 5,94736 | 6,91759 | 0,218019 | 0,2246 | 0,579115 |

| | | | | | |
|---|---|---|---|---|---|
| Gm10046 | 1,32763 | 0,714247 | -0,894355 | 0,22465 | 0,579132 |
| Alas1 | 12,5919 | 14,6042 | 0,213892 | 0,22475 | 0,579278 |
| Snord88a | 160,27 | 0 | -inf | 0,22485 | 0,579313 |
| Lpin2 | 13,8428 | 15,8316 | 0,193674 | 0,22485 | 0,579313 |
| Mir1943 | 147,224 | 0 | -inf | 0,225 | 0,579381 |
| Fastkd2 | 17,3116 | 14,9714 | -0,209523 | 0,22495 | 0,579381 |
| 2900076A07Rik | 7,6948 | 6,00366 | -0,358043 | 0,22505 | 0,579381 |
| 2610203C22Rik | 0,665235 | 0,0410984 | -4,01671 | 0,22505 | 0,579381 |
| Snord92 | 15,9071 | 0 | -inf | 0,22565 | 0,579398 |
| Hist1h3a | 0,536638 | 1,22263 | 1,18797 | 0,2255 | 0,579398 |
| Ppp1r14a | 15,1148 | 19,0665 | 0,335074 | 0,22545 | 0,579398 |
| Rybp | 6,64522 | 7,76408 | 0,224498 | 0,22575 | 0,579398 |
| Ppp1r16a | 15,6866 | 18,1307 | 0,208906 | 0,2257 | 0,579398 |
| Lig1 | 75,3944 | 86,3928 | 0,196454 | 0,2255 | 0,579398 |
| Nsmce4a | 105,036 | 120,167 | 0,194159 | 0,22555 | 0,579398 |
| Micu2 | 46,9363 | 53,6813 | 0,193716 | 0,2256 | 0,579398 |
| Smyd5 | 45,2303 | 51,7248 | 0,193567 | 0,22535 | 0,579398 |
| Arhgap35 | 32,3256 | 36,9418 | 0,19258 | 0,22575 | 0,579398 |
| Plxna2 | 14,866 | 16,9774 | 0,191593 | 0,22545 | 0,579398 |
| Mogs | 51,8649 | 45,3968 | -0,192168 | 0,2252 | 0,579398 |
| Col23a1 | 8,4514 | 7,31151 | -0,209021 | 0,2255 | 0,579398 |
| Twist1 | 26,3468 | 22,5902 | -0,221927 | 0,22565 | 0,579398 |
| Casp8 | 9,06493 | 7,69064 | -0,237193 | 0,2255 | 0,579398 |
| Nipal3 | 3,38892 | 2,8425 | -0,253666 | 0,2254 | 0,579398 |
| Fam78b | 0,590486 | 0,772504 | 0,387639 | 0,2259 | 0,579449 |
| Tcirg1 | 8,84937 | 10,3664 | 0,228263 | 0,2259 | 0,579449 |
| Dmxl1 | 7,17576 | 6,25687 | -0,197691 | 0,2259 | 0,579449 |
| Ctsz | 53,0598 | 41,9799 | -0,337921 | 0,22595 | 0,579466 |
| Mtfr1l | 33,5152 | 38,4814 | 0,199343 | 0,226 | 0,579483 |
| Btbd9 | 5,46622 | 4,70373 | -0,216739 | 0,22605 | 0,5795 |
| Hnrnpll | 70,3243 | 80,2998 | 0,191374 | 0,22615 | 0,579645 |
| Mms19 | 20,4619 | 24,1331 | 0,238075 | 0,2262 | 0,579662 |
| Mir7033 | 0 | 172,01 | inf | 0,2263 | 0,579807 |
| Caprin1 | 211,192 | 246,336 | 0,222074 | 0,22635 | 0,579824 |
| Ddx21 | 81,7967 | 93,5991 | 0,194452 | 0,2264 | 0,579841 |
| Ttc30b | 2,88714 | 2,29566 | -0,330732 | 0,2265 | 0,579986 |
| Snord85 | 0 | 162,606 | inf | 0,2266 | 0,58002 |
| Slc29a2 | 7,51964 | 6,31049 | -0,252913 | 0,2266 | 0,58002 |
| Dpf1 | 8,62969 | 10,1975 | 0,240835 | 0,22675 | 0,58007 |
| Dnajc2 | 52,3395 | 59,9547 | 0,195971 | 0,22675 | 0,58007 |
| Got2 | 49,1811 | 56,322 | 0,195596 | 0,2267 | 0,58007 |
| Golgb1 | 12,2799 | 14,0928 | 0,198664 | 0,22685 | 0,580215 |
| 5930412G12Rik | 1,75354 | 2,24958 | 0,359384 | 0,2269 | 0,580232 |
| Snord71 | 80,508 | 0 | -inf | 0,2273 | 0,580306 |
| Pvt1 | 3,4155 | 4,68991 | 0,457464 | 0,2274 | 0,580306 |
| Sacs | 1,47448 | 1,86625 | 0,33994 | 0,22705 | 0,580306 |
| Gm20604 | 13,9296 | 16,6923 | 0,261026 | 0,2272 | 0,580306 |
| Zranb1 | 21,7191 | 25,0489 | 0,205786 | 0,22705 | 0,580306 |

| | | | | | |
|---|---|---|---|---|---|
| Pus1 | 32,0278 | 36,9148 | 0,204874 | 0,2274 | 0,580306 |
| Slu7 | 15,7397 | 18,0952 | 0,201196 | 0,22735 | 0,580306 |
| Wars | 49,3501 | 56,7236 | 0,200896 | 0,22745 | 0,580306 |
| Traf7 | 53,4511 | 61,2201 | 0,195784 | 0,22745 | 0,580306 |
| Mga | 16,3562 | 18,7082 | 0,193835 | 0,2271 | 0,580306 |
| Kif22 | 75,4584 | 86,2256 | 0,192434 | 0,2273 | 0,580306 |
| Ptpn18 | 0,687529 | 0,423425 | -0,699314 | 0,2272 | 0,580306 |
| Zfp444 | 7,53017 | 8,74102 | 0,21512 | 0,2275 | 0,580323 |
| Mir1938 | 6,48835 | 0 | -inf | 0,22755 | 0,580339 |
| Mir6963 | 0 | 88,3598 | inf | 0,2276 | 0,580356 |
| Rock2 | 23,944 | 27,3111 | 0,189819 | 0,22765 | 0,580373 |
| Spag5 | 53,4538 | 60,9907 | 0,190297 | 0,22775 | 0,580517 |
| Foxn2 | 7,19813 | 8,34352 | 0,213035 | 0,22805 | 0,580839 |
| Ncln | 41,2962 | 36,1108 | -0,193579 | 0,228 | 0,580839 |
| Zbtb7a | 3,92381 | 3,31891 | -0,241542 | 0,22795 | 0,580839 |
| 5730480H06Rik | 2,28687 | 1,67357 | -0,450446 | 0,22805 | 0,580839 |
| Unc119 | 45,5467 | 52,4401 | 0,203326 | 0,2281 | 0,580855 |
| Mrpl13 | 145,373 | 166,329 | 0,194275 | 0,22825 | 0,581127 |
| Cdo1 | 3,70993 | 2,89408 | -0,358289 | 0,22835 | 0,58127 |
| Cdk6 | 9,09268 | 10,664 | 0,229973 | 0,22855 | 0,581669 |
| Mir6357 | 0 | 14,6762 | inf | 0,22865 | 0,581767 |
| Leng9 | 0,637432 | 0,945488 | 0,568788 | 0,22885 | 0,581767 |
| Zfp341 | 2,15028 | 2,65818 | 0,305914 | 0,22885 | 0,581767 |
| Mrps17 | 43,0743 | 49,7342 | 0,207411 | 0,2287 | 0,581767 |
| Rlim | 16,3733 | 14,3149 | -0,19383 | 0,2288 | 0,581767 |
| Leng8 | 41,3024 | 36,0874 | -0,194732 | 0,22885 | 0,581767 |
| Diexf | 8,77441 | 10,1054 | 0,203754 | 0,229 | 0,581882 |
| Raf1 | 66,6377 | 76,6259 | 0,201494 | 0,22905 | 0,581882 |
| Magoh | 369,806 | 422,391 | 0,191809 | 0,2291 | 0,581882 |
| Sec22a | 6,81027 | 5,81124 | -0,228866 | 0,22895 | 0,581882 |
| Scml2 | 2,73786 | 2,26666 | -0,272481 | 0,2292 | 0,581882 |
| Mir6991 | 320,852 | 98,2219 | -1,70779 | 0,2292 | 0,581882 |
| Gja5 | 2,50043 | 0,0092154 | -8,08392 | 0,22915 | 0,581882 |
| Mrpl19 | 15,8012 | 18,4438 | 0,223098 | 0,2294 | 0,58228 |
| Mir106b | 16,9489 | 0 | -inf | 0,2296 | 0,582345 |
| Psenen | 0,99504 | 1,6942 | 0,767778 | 0,2295 | 0,582345 |
| C030039L03Rik | 1,14324 | 1,43348 | 0,326395 | 0,22955 | 0,582345 |
| Mbtd1 | 36,0129 | 31,5323 | -0,19168 | 0,2296 | 0,582345 |
| Cnppd1 | 12,9124 | 11,1072 | -0,217259 | 0,22965 | 0,582361 |
| 2010308F09Rik | 0,41956 | 0,981008 | 1,22539 | 0,22985 | 0,582647 |
| Rnf135 | 1,5012 | 1,10924 | -0,436548 | 0,22985 | 0,582647 |
| Gp1bb | 0,805599 | 0,274416 | -1,5537 | 0,2299 | 0,582663 |
| Gltscr1 | 6,51028 | 7,53121 | 0,21016 | 0,22995 | 0,58268 |
| Nelfcd | 50,9306 | 59,801 | 0,231637 | 0,23 | 0,582696 |
| Mir26b | 14,0742 | 0 | -inf | 0,2305 | 0,583064 |
| Efhd1 | 0,617452 | 0,912067 | 0,562812 | 0,2303 | 0,583064 |
| Brsk1 | 11,5991 | 13,5191 | 0,22098 | 0,23075 | 0,583064 |
| Mta3 | 19,6528 | 22,8769 | 0,219155 | 0,23055 | 0,583064 |

| | | | | | |
|---|---|---|---|---|---|
| Pgs1 | 21,6019 | 24,9512 | 0,207954 | 0,23035 | 0,583064 |
| Atp5k | 506,798 | 582,897 | 0,201829 | 0,2303 | 0,583064 |
| Tatdn2 | 23,3747 | 26,8519 | 0,200076 | 0,2306 | 0,583064 |
| Isy1 | 48,8653 | 55,9551 | 0,195459 | 0,2307 | 0,583064 |
| Rsl1d1 | 198,036 | 226,31 | 0,192539 | 0,2303 | 0,583064 |
| Zfp608 | 17,3268 | 19,7871 | 0,191559 | 0,2308 | 0,583064 |
| Mesdc2 | 32,3749 | 28,3418 | -0,191949 | 0,2306 | 0,583064 |
| Gpr107 | 20,6783 | 18,0284 | -0,197849 | 0,23045 | 0,583064 |
| Dennd1c | 1,44876 | 1,09842 | -0,399386 | 0,23065 | 0,583064 |
| Klf7 | 2,5103 | 1,86235 | -0,430735 | 0,2304 | 0,583064 |
| Slc24a5 | 4,17903 | 1,91388 | -1,12667 | 0,2308 | 0,583064 |
| H2afy3 | 21,184 | 24,94 | 0,235483 | 0,23105 | 0,583365 |
| Mob4 | 33,4489 | 29,2024 | -0,195869 | 0,23105 | 0,583365 |
| Prpsap2 | 8,25461 | 6,84251 | -0,270676 | 0,231 | 0,583365 |
| Adcy5 | 1,53338 | 1,87841 | 0,292793 | 0,2312 | 0,583634 |
| Bysl | 22,1575 | 27,013 | 0,285855 | 0,2313 | 0,583776 |
| Anapc2 | 46,9969 | 53,5602 | 0,188598 | 0,2314 | 0,583918 |
| Gm1821 | 38,5675 | 67,1851 | 0,800754 | 0,23175 | 0,583919 |
| Ccdc36 | 0,600338 | 0,83059 | 0,468361 | 0,2316 | 0,583919 |
| Prdm15 | 6,11616 | 7,06161 | 0,207373 | 0,2317 | 0,583919 |
| Syde1 | 21,4942 | 24,6729 | 0,198979 | 0,2315 | 0,583919 |
| Atad5 | 16,9222 | 14,818 | -0,191572 | 0,23155 | 0,583919 |
| Lpar6 | 41,4047 | 35,5945 | -0,21814 | 0,23155 | 0,583919 |
| C1qtnf4 | 1,67248 | 1,19233 | -0,488207 | 0,2317 | 0,583919 |
| Snhg10 | 5,50035 | 3,81122 | -0,529268 | 0,23175 | 0,583919 |
| Mir6939 | 0 | 100,863 | inf | 0,23185 | 0,583967 |
| Mir6966 | 124,709 | 0 | -inf | 0,2319 | 0,583967 |
| Deaf1 | 18,3556 | 21,6848 | 0,240467 | 0,2319 | 0,583967 |
| BC003965 | 15,5349 | 13,454 | -0,207478 | 0,23195 | 0,583983 |
| Ilkap | 73,4403 | 83,9423 | 0,192826 | 0,232 | 0,583998 |
| Mir8091 | 8,19311 | 30,4971 | 1,89619 | 0,23225 | 0,584 |
| Smarcd3 | 29,2253 | 33,6741 | 0,204423 | 0,23235 | 0,584 |
| Rbm19 | 18,6311 | 21,3095 | 0,193782 | 0,23205 | 0,584 |
| Hbs1l | 34,549 | 39,4894 | 0,192823 | 0,23215 | 0,584 |
| Dnajc10 | 66,4459 | 58,2568 | -0,189755 | 0,2322 | 0,584 |
| Smim15 | 41,6818 | 36,3473 | -0,19757 | 0,23235 | 0,584 |
| Nlk | 14,447 | 12,5749 | -0,200229 | 0,23225 | 0,584 |
| Tab3 | 6,23772 | 5,38892 | -0,211022 | 0,2323 | 0,584 |
| Mir6976 | 182,594 | 0 | -inf | 0,23245 | 0,584141 |
| Smim3 | 13,5074 | 15,7608 | 0,222591 | 0,2325 | 0,584157 |
| Habp4 | 6,84941 | 5,76477 | -0,248716 | 0,23255 | 0,584173 |
| 2700029M09Rik | 62,6691 | 71,7857 | 0,195941 | 0,23275 | 0,584565 |
| Rai14 | 31,8338 | 36,236 | 0,186862 | 0,23305 | 0,585099 |
| Tsen54 | 12,0278 | 10,2174 | -0,235342 | 0,23305 | 0,585099 |
| Mir6922 | 63,8043 | 0 | -inf | 0,23315 | 0,58524 |
| Stat6 | 4,98736 | 4,05439 | -0,29879 | 0,2332 | 0,585255 |
| Axin1 | 29,9384 | 34,1195 | 0,188601 | 0,23325 | 0,585271 |
| Aaas | 52,7856 | 60,3195 | 0,192479 | 0,2333 | 0,585287 |

| | | | | | |
|---|---|---|---|---|---|
| Prkd3 | 38,8463 | 44,3005 | 0,189544 | 0,23335 | 0,585302 |
| Atxn7l1 | 5,23235 | 6,37691 | 0,285398 | 0,23345 | 0,585333 |
| Ppp1r10 | 23,1757 | 26,4211 | 0,189078 | 0,23345 | 0,585333 |
| Hoxa6 | 40,6444 | 61,2862 | 0,592506 | 0,23365 | 0,58563 |
| Rcn1 | 109,514 | 96,1202 | -0,188199 | 0,2337 | 0,58563 |
| Tmem140 | 1,24219 | 0,528201 | -1,23373 | 0,2337 | 0,58563 |
| Cpq | 4,45623 | 3,6154 | -0,301669 | 0,23395 | 0,586147 |
| Mir615 | 9,54071 | 0 | -inf | 0,23405 | 0,586224 |
| Mir6938 | 91,2968 | 0 | -inf | 0,2342 | 0,586224 |
| St3gal3 | 15,8193 | 18,3027 | 0,210367 | 0,2342 | 0,586224 |
| Clk2 | 41,2408 | 47,0643 | 0,190563 | 0,23415 | 0,586224 |
| Lman1 | 71,7592 | 62,6348 | -0,1962 | 0,2341 | 0,586224 |
| Mir7036b | 91,2968 | 0 | -inf | 0,2344 | 0,5863 |
| Phyh | 13,6492 | 16,0799 | 0,236438 | 0,2344 | 0,5863 |
| Dennd6b | 6,75708 | 7,83737 | 0,21397 | 0,2343 | 0,5863 |
| Ripk4 | 2,48396 | 2,0187 | -0,299213 | 0,23445 | 0,5863 |
| Spaca4 | 1,09993 | 0,391393 | -1,49072 | 0,23445 | 0,5863 |
| Mir7019 | 46,7846 | 0 | -inf | 0,2345 | 0,586316 |
| Homer1 | 14,4675 | 16,834 | 0,218564 | 0,23465 | 0,586581 |
| Cdkn2d | 14,4602 | 12,1327 | -0,253189 | 0,2347 | 0,586596 |
| Mir6406 | 0 | 6,74081 | inf | 0,235 | 0,587017 |
| Hnrnph1 | 517,799 | 599,599 | 0,211607 | 0,235 | 0,587017 |
| Dmwd | 35,6535 | 40,6939 | 0,190769 | 0,23495 | 0,587017 |
| Trak1 | 6,57225 | 7,61584 | 0,212617 | 0,2351 | 0,587157 |
| Tm9sf4 | 43,5498 | 49,5665 | 0,186697 | 0,23515 | 0,587172 |
| Ercc8 | 14,1169 | 16,4307 | 0,21897 | 0,23535 | 0,587482 |
| Mib1 | 23,5094 | 26,8583 | 0,192133 | 0,23545 | 0,587482 |
| Bcap31 | 88,9135 | 77,7465 | -0,193624 | 0,2354 | 0,587482 |
| Peg3os | 28,4704 | 8,21485 | -1,79316 | 0,23545 | 0,587482 |
| Mir215 | 12,3267 | 0 | -inf | 0,23555 | 0,587622 |
| Lrr1 | 10,5023 | 8,77632 | -0,259015 | 0,2356 | 0,587637 |
| Gemin6 | 22,7905 | 26,7119 | 0,22905 | 0,23575 | 0,587682 |
| Aplf | 5,434 | 4,57323 | -0,248802 | 0,2357 | 0,587682 |
| Exoc3l4 | 0,876403 | 0,0272293 | -5,00836 | 0,23575 | 0,587682 |
| Inpp4b | 1,33391 | 1,67237 | 0,326231 | 0,23585 | 0,587822 |
| Clec1b | 3,8419 | 0,119172 | -5,01071 | 0,2359 | 0,587837 |
| Fam64a | 76,9777 | 87,9004 | 0,19143 | 0,23615 | 0,58835 |
| Olfml2b | 4,24876 | 3,53372 | -0,265853 | 0,23625 | 0,588489 |
| Cdc123 | 87,5123 | 99,7286 | 0,18852 | 0,23635 | 0,588629 |
| Mir7069 | 208,597 | 0 | -inf | 0,2365 | 0,588783 |
| Bcar3 | 5,94513 | 6,97169 | 0,229799 | 0,2365 | 0,588783 |
| Psat1 | 118,308 | 135,005 | 0,190463 | 0,23655 | 0,588798 |
| Fam168a | 13,645 | 11,9264 | -0,194217 | 0,23675 | 0,588966 |
| Dolk | 9,08622 | 7,67383 | -0,243733 | 0,2367 | 0,588966 |
| Uevld | 3,21731 | 2,69245 | -0,256934 | 0,23675 | 0,588966 |
| Mir7012 | 80,508 | 0 | -inf | 0,2368 | 0,588981 |
| Hist1h4k | 0,635317 | 1,56827 | 1,30362 | 0,237 | 0,58904 |
| Snord15a | 12,8477 | 18,5631 | 0,530926 | 0,2369 | 0,58904 |

| | | | | | |
|---|---|---|---|---|---|
| Phax | 48,9131 | 59,6913 | 0,2873 | 0,237 | 0,58904 |
| Capn2 | 54,508 | 47,8467 | -0,188048 | 0,23695 | 0,58904 |
| C530008M17Rik | 14,6241 | 16,6733 | 0,189194 | 0,23715 | 0,589303 |
| Treml1 | 1,27162 | 0,0461002 | -4,78574 | 0,2372 | 0,589318 |
| Luzp1 | 25,7536 | 29,3303 | 0,187615 | 0,23755 | 0,590078 |
| Ptprn | 0,799692 | 1,04456 | 0,385378 | 0,2376 | 0,590092 |
| Txndc12 | 64,7358 | 54,5413 | -0,247215 | 0,23775 | 0,590355 |
| Mpl | 0,623289 | 0,010775 | -5,85414 | 0,23785 | 0,590494 |
| Mir6386 | 5,44879 | 0 | -inf | 0,238 | 0,590537 |
| Map1lc3a | 21,1936 | 24,914 | 0,233326 | 0,23795 | 0,590537 |
| Spock2 | 52,8513 | 60,2157 | 0,1882 | 0,238 | 0,590537 |
| Slc4a2 | 14,6926 | 12,8291 | -0,195665 | 0,2381 | 0,590675 |
| Mir7024 | 208,597 | 0 | -inf | 0,2382 | 0,590814 |
| A830010M20Rik | 1,43921 | 1,73543 | 0,27002 | 0,23825 | 0,590828 |
| Ftsj2 | 8,3133 | 9,98023 | 0,263651 | 0,2383 | 0,590843 |
| Polr1a | 30,7799 | 35,0679 | 0,188163 | 0,2385 | 0,591229 |
| Casc5 | 14,3613 | 12,5638 | -0,192916 | 0,23855 | 0,591243 |
| Gm12238 | 0 | 48,189 | inf | 0,2387 | 0,591286 |
| Ccdc92 | 2,70342 | 2,13817 | -0,338408 | 0,23865 | 0,591286 |
| Pkp3 | 1,14355 | 0,132783 | -3,10638 | 0,2387 | 0,591286 |
| Ccdc30 | 0,981974 | 1,3082 | 0,413822 | 0,23875 | 0,5913 |
| Vps33b | 8,38999 | 9,82609 | 0,227949 | 0,2388 | 0,591314 |
| Sacm1l | 21,3784 | 18,6746 | -0,195082 | 0,23885 | 0,591329 |
| Ppp6c | 42,8389 | 48,9961 | 0,193745 | 0,23925 | 0,592099 |
| Aldh6a1 | 6,31491 | 5,18188 | -0,285285 | 0,23925 | 0,592099 |
| Edf1 | 108,097 | 124,01 | 0,19814 | 0,2393 | 0,592114 |
| Stk39 | 20,2217 | 23,1361 | 0,194242 | 0,23945 | 0,592375 |
| Fmnl2 | 20,805 | 18,2536 | -0,188755 | 0,2395 | 0,592389 |
| Txn1 | 178,965 | 203,467 | 0,185112 | 0,2396 | 0,592527 |
| Mir6965 | 80,508 | 0 | -inf | 0,23965 | 0,592541 |
| Epm2a | 0,64293 | 1,04101 | 0,695257 | 0,24 | 0,592639 |
| Zfp941 | 0,897878 | 1,15424 | 0,362349 | 0,23995 | 0,592639 |
| Haus8 | 18,6677 | 21,6554 | 0,214181 | 0,23985 | 0,592639 |
| AI314180 | 23,0938 | 26,3221 | 0,188771 | 0,23975 | 0,592639 |
| Tmed9 | 149,283 | 131,288 | -0,185323 | 0,24 | 0,592639 |
| Fhod3 | 3,9943 | 3,12055 | -0,356144 | 0,2399 | 0,592639 |
| H2-T24 | 2,38195 | 1,84243 | -0,37053 | 0,23985 | 0,592639 |
| Khdrbs1 | 106,992 | 122,402 | 0,194131 | 0,24005 | 0,592653 |
| Lrch3 | 18,1458 | 20,7291 | 0,19202 | 0,2401 | 0,592666 |
| Abhd6 | 4,75486 | 3,88713 | -0,290697 | 0,2402 | 0,592804 |
| Wbp11 | 79,9375 | 90,8835 | 0,185147 | 0,24025 | 0,592818 |
| Hdac10 | 6,62851 | 7,8208 | 0,238631 | 0,24075 | 0,593942 |
| Mir6970 | 602,515 | 0 | -inf | 0,2409 | 0,593983 |
| Rpp21 | 31,4156 | 37,8553 | 0,269012 | 0,2409 | 0,593983 |
| Kifc3 | 3,7898 | 2,68689 | -0,496188 | 0,2409 | 0,593983 |
| Cdca2 | 43,1687 | 37,9465 | -0,186016 | 0,24105 | 0,594243 |
| Med18 | 8,28748 | 6,90409 | -0,263481 | 0,2411 | 0,594257 |
| Mir6912 | 0 | 290,431 | inf | 0,2423 | 0,594623 |

| | | | | | |
|---|---|---|---|---|---|
| Mir6920 | 0 | 140,46 | inf | 0,2424 | 0,594623 |
| Mir7052 | 0 | 421,177 | inf | 0,242 | 0,594623 |
| Mir1940 | 23,9013 | 0 | -inf | 0,24195 | 0,594623 |
| Mir7078 | 80,508 | 0 | -inf | 0,24175 | 0,594623 |
| Pus7l | 8,28871 | 9,66612 | 0,221789 | 0,2424 | 0,594623 |
| 0610010K14Rik | 66,8263 | 77,1234 | 0,206753 | 0,24245 | 0,594623 |
| Imp3 | 55,2358 | 63,6237 | 0,203961 | 0,24215 | 0,594623 |
| Uimc1 | 13,7919 | 15,8651 | 0,20203 | 0,24135 | 0,594623 |
| Trim32 | 21,3556 | 24,3783 | 0,190981 | 0,24245 | 0,594623 |
| Slc25a11 | 43,0973 | 49,1765 | 0,190371 | 0,2415 | 0,594623 |
| Usp48 | 28,1182 | 32,0189 | 0,187418 | 0,24165 | 0,594623 |
| Rere | 26,4617 | 30,1241 | 0,187015 | 0,24225 | 0,594623 |
| Ptn | 54,4639 | 47,8054 | -0,188127 | 0,2416 | 0,594623 |
| Ephb2 | 18,9634 | 16,6307 | -0,189374 | 0,2421 | 0,594623 |
| Rpl15 | 50,2935 | 44,034 | -0,191753 | 0,2413 | 0,594623 |
| Eif2s3y | 22,3058 | 19,3267 | -0,206823 | 0,2423 | 0,594623 |
| Slc44a5 | 1,05719 | 0,821647 | -0,363649 | 0,2414 | 0,594623 |
| Tfr2 | 0,68826 | 0,0165543 | -5,37767 | 0,24175 | 0,594623 |
| Aif1 | 1,82625 | 0,0401099 | -5,50878 | 0,2422 | 0,594623 |
| Hbq1b | 2,43665 | 0,0494735 | -5,6221 | 0,2422 | 0,594623 |
| Was | 0,798833 | 0,0142402 | -5,80985 | 0,2422 | 0,594623 |
| Fam65c | 0,665504 | 0,00985504 | -6,07744 | 0,2422 | 0,594623 |
| Rgs18 | 1,19109 | 0,0162623 | -6,19461 | 0,2422 | 0,594623 |
| Spi1 | 2,17641 | 0,0238387 | -6,5125 | 0,2422 | 0,594623 |
| Erv3 | 0,899233 | 0,00847956 | -6,72856 | 0,2422 | 0,594623 |
| Cx3cr1 | 2,84131 | 0,00758358 | -8,54946 | 0,2422 | 0,594623 |
| Vps16 | 26,602 | 30,8827 | 0,215261 | 0,24275 | 0,59525 |
| Topors | 30,4511 | 34,7652 | 0,191151 | 0,24285 | 0,595276 |
| D17H6S53E | 5,80626 | 4,80905 | -0,271859 | 0,24285 | 0,595276 |
| Mir1907 | 0 | 76,6962 | inf | 0,24305 | 0,595657 |
| Tmed1 | 49,7404 | 39,6648 | -0,326559 | 0,2431 | 0,595671 |
| Gpr183 | 0,983779 | 0,00982984 | -6,64502 | 0,2432 | 0,595807 |
| Mir6395 | 55,3561 | 0 | -inf | 0,2434 | 0,595873 |
| Gm13152 | 2,63445 | 3,43776 | 0,383969 | 0,24345 | 0,595873 |
| Pgp | 29,7488 | 34,5746 | 0,216883 | 0,24335 | 0,595873 |
| Cxxc1 | 34,2844 | 39,0569 | 0,188026 | 0,2434 | 0,595873 |
| Cdc42bpa | 7,49022 | 6,53511 | -0,196796 | 0,24345 | 0,595873 |
| Tsfm | 32,4809 | 37,4883 | 0,206848 | 0,2435 | 0,595886 |
| Nemf | 24,5862 | 28,0537 | 0,19034 | 0,24355 | 0,595899 |
| Wwp1 | 13,394 | 15,2722 | 0,189322 | 0,24365 | 0,596035 |
| Sdf2l1 | 35,7909 | 41,2733 | 0,205616 | 0,24375 | 0,596061 |
| Slc27a4 | 10,8154 | 9,3883 | -0,204154 | 0,24375 | 0,596061 |
| Gtf2h4 | 27,9863 | 32,2423 | 0,204232 | 0,24385 | 0,596197 |
| Gm14325 | 1,7139 | 1,38414 | -0,308291 | 0,24395 | 0,596332 |
| Cdc14b | 19,7079 | 22,4285 | 0,186558 | 0,2441 | 0,596385 |
| Cnep1r1 | 38,7291 | 33,7847 | -0,197048 | 0,24415 | 0,596385 |
| Gla | 6,56267 | 5,57044 | -0,236491 | 0,24415 | 0,596385 |
| H2-T22,H2-T9 | 1,70555 | 9,14E-005 | -14,1877 | 0,2441 | 0,596385 |

| | | | | | |
|---|---|---|---|---|---|
| Hecw2 | 1,39415 | 1,00475 | -0,472555 | 0,24425 | 0,596411 |
| Gfi1b | 1,92158 | 0,028163 | -6,09235 | 0,24425 | 0,596411 |
| Rab18 | 37,5672 | 33,0487 | -0,184879 | 0,2443 | 0,596424 |
| Nfatc2 | 0,621296 | 0,471171 | -0,39903 | 0,24435 | 0,596437 |
| Mir6945 | 0 | 422,161 | inf | 0,2445 | 0,596529 |
| Wdr90 | 13,2163 | 15,8792 | 0,264817 | 0,2447 | 0,596529 |
| Lrrc28 | 7,04481 | 8,27577 | 0,232334 | 0,2446 | 0,596529 |
| Fam98b | 66,4058 | 75,5432 | 0,185994 | 0,24465 | 0,596529 |
| Bag6 | 81,4384 | 92,4643 | 0,183187 | 0,2447 | 0,596529 |
| Lrrc3 | 1,10558 | 0,866864 | -0,350922 | 0,24465 | 0,596529 |
| Nr2c2ap | 17,782 | 13,5763 | -0,389331 | 0,24455 | 0,596529 |
| Rpl31-ps12 | 0,337932 | 1,284 | 1,92584 | 0,24475 | 0,596542 |
| Aco2 | 78,8374 | 68,8763 | -0,194872 | 0,24485 | 0,596677 |
| Prpsap1 | 53,2976 | 60,5847 | 0,184884 | 0,24495 | 0,596812 |
| Mir6387 | 0 | 10,5166 | inf | 0,2452 | 0,596998 |
| Mir6944 | 0 | 35,5675 | inf | 0,24525 | 0,596998 |
| Mir7068 | 0 | 108,851 | inf | 0,24525 | 0,596998 |
| Snora47 | 2,71792 | 0 | -inf | 0,24515 | 0,596998 |
| Haghl | 19,11 | 15,4458 | -0,307113 | 0,2452 | 0,596998 |
| Hmgb1-rs17 | 0,581115 | 0,889388 | 0,613988 | 0,24535 | 0,597133 |
| Mir6998 | 0 | 183,191 | inf | 0,24585 | 0,597154 |
| Snx11 | 14,7216 | 16,9658 | 0,204692 | 0,2458 | 0,597154 |
| Cpt2 | 16,1729 | 18,6245 | 0,203619 | 0,2457 | 0,597154 |
| Ocrl | 11,8129 | 13,502 | 0,192801 | 0,24545 | 0,597154 |
| Vdac2 | 120,893 | 137,561 | 0,186347 | 0,24585 | 0,597154 |
| Spats2l | 3,29395 | 2,74803 | -0,261422 | 0,24585 | 0,597154 |
| Pfkfb4 | 3,35418 | 2,77847 | -0,271667 | 0,24555 | 0,597154 |
| Fgfbp3 | 3,08242 | 2,42283 | -0,34737 | 0,24565 | 0,597154 |
| Phf11c | 1,36673 | 0,978349 | -0,482304 | 0,2458 | 0,597154 |
| Gimap8 | 0,652591 | 0,0546571 | -3,5777 | 0,24585 | 0,597154 |
| Tmem54 | 3,12473 | 0,0611563 | -5,67509 | 0,24585 | 0,597154 |
| Mir681 | 0 | 30,8987 | inf | 0,2462 | 0,597691 |
| 2900011O08Rik | 0,639655 | 0,979414 | 0,614626 | 0,2462 | 0,597691 |
| Rpusd4 | 22,3239 | 25,7375 | 0,205284 | 0,24625 | 0,597691 |
| Ppm1n | 0,676162 | 0,428457 | -0,658219 | 0,24625 | 0,597691 |
| Mir6915 | 0 | 169,226 | inf | 0,2468 | 0,598074 |
| Prr3 | 16,156 | 18,981 | 0,232485 | 0,24665 | 0,598074 |
| Psmb3 | 59,98 | 69,2348 | 0,207016 | 0,2469 | 0,598074 |
| Dlat | 23,4295 | 26,713 | 0,189214 | 0,24675 | 0,598074 |
| Ncapd2 | 102,767 | 116,999 | 0,187118 | 0,24655 | 0,598074 |
| Ltv1 | 58,8496 | 66,9165 | 0,185329 | 0,2467 | 0,598074 |
| Setd5 | 51,6219 | 58,6907 | 0,185149 | 0,24675 | 0,598074 |
| Bptf | 20,5331 | 23,3192 | 0,183564 | 0,24685 | 0,598074 |
| E130102H24Rik | 2,7205 | 1,8457 | -0,559708 | 0,2469 | 0,598074 |
| 1700096K18Rik | 0,84164 | 0,543197 | -0,631729 | 0,24665 | 0,598074 |
| Apoc1 | 2,44096 | 0,236072 | -3,37015 | 0,24665 | 0,598074 |
| Cdk20 | 3,84028 | 4,7285 | 0,300169 | 0,24705 | 0,59822 |
| 2310022A10Rik | 14,4767 | 16,6986 | 0,205996 | 0,24705 | 0,59822 |

| | | | | | |
|---|---|---|---|---|---|
| Nupl2 | 5,78007 | 6,80365 | 0,235222 | 0,2471 | 0,598233 |
| Mir1b | 0 | 22,6046 | inf | 0,2472 | 0,598366 |
| Crnkl1 | 25,6609 | 29,2674 | 0,189722 | 0,24725 | 0,598379 |
| Exosc1 | 25,9502 | 29,8452 | 0,201754 | 0,24735 | 0,598404 |
| Tfap2b | 1,37017 | 0,0519374 | -4,72144 | 0,24735 | 0,598404 |
| Pdss2 | 8,04444 | 9,55291 | 0,247948 | 0,24745 | 0,598538 |
| Zfand1 | 3,79922 | 3,00661 | -0,337568 | 0,2475 | 0,59855 |
| Vprbp | 21,3217 | 24,2269 | 0,184288 | 0,24755 | 0,598563 |
| Mir6955 | 90,2793 | 0 | -inf | 0,2476 | 0,598576 |
| Ptprc | 0,858581 | 0,0162323 | -5,72501 | 0,24765 | 0,598588 |
| Hsd17b10 | 132,919 | 150,965 | 0,18367 | 0,24785 | 0,598963 |
| Thada | 6,53415 | 7,48832 | 0,196644 | 0,24795 | 0,599097 |
| Mir6943 | 0 | 230,789 | inf | 0,248 | 0,599109 |
| Ado | 17,5805 | 15,417 | -0,189454 | 0,2482 | 0,599484 |
| Mir7036 | 250,608 | 0 | -inf | 0,24845 | 0,599979 |
| Naaladl1 | 1,50479 | 1,14708 | -0,391598 | 0,2486 | 0,600233 |
| Srek1ip1 | 20,3727 | 17,6124 | -0,210044 | 0,249 | 0,60109 |
| Ttc8 | 11,5887 | 13,4326 | 0,21302 | 0,24915 | 0,601127 |
| Scpep1 | 38,5466 | 43,9123 | 0,188022 | 0,2491 | 0,601127 |
| Cul1 | 102,566 | 116,588 | 0,184867 | 0,24915 | 0,601127 |
| Snapc3 | 25,6845 | 29,6394 | 0,206618 | 0,2492 | 0,601139 |
| Nubpl | 7,59486 | 9,23688 | 0,282382 | 0,24925 | 0,601151 |
| Smpd3 | 1,50619 | 1,8345 | 0,284483 | 0,2493 | 0,601163 |
| Hspb11 | 26,7788 | 22,0607 | -0,279615 | 0,24935 | 0,601175 |
| Sept9 | 90,6277 | 103,16 | 0,186855 | 0,2495 | 0,601428 |
| Tbxa2r | 0,96015 | 0,674024 | -0,510459 | 0,2497 | 0,601802 |
| Kbtbd7 | 6,07471 | 5,224 | -0,217661 | 0,2499 | 0,602175 |
| Gstt2 | 2,0205 | 1,49143 | -0,438017 | 0,24995 | 0,602187 |
| Rtbdn | 0,552365 | 0,808734 | 0,550045 | 0,25005 | 0,602319 |
| Mir199b | 0 | 9,71834 | inf | 0,2505 | 0,602667 |
| Hoxa4 | 8,81663 | 14,7032 | 0,737831 | 0,25055 | 0,602667 |
| Etnk2 | 1,06342 | 1,40676 | 0,403675 | 0,2504 | 0,602667 |
| Stx18 | 9,31085 | 11,5935 | 0,316333 | 0,2503 | 0,602667 |
| Zfp318 | 8,3965 | 9,80191 | 0,223275 | 0,25035 | 0,602667 |
| Trmt1l | 19,6214 | 22,4606 | 0,194966 | 0,25055 | 0,602667 |
| Gjc1 | 74,1502 | 84,242 | 0,184089 | 0,2503 | 0,602667 |
| Cbx2 | 42,4555 | 48,1484 | 0,181536 | 0,2506 | 0,602667 |
| Zfp182 | 2,21859 | 1,85661 | -0,256972 | 0,2506 | 0,602667 |
| Nat9 | 5,85559 | 7,24508 | 0,307186 | 0,2507 | 0,602676 |
| Eno3 | 32,8673 | 37,7044 | 0,198083 | 0,25105 | 0,602676 |
| Pim1 | 31,7847 | 36,4148 | 0,196192 | 0,2511 | 0,602676 |
| Mdm1 | 19,6297 | 22,3622 | 0,188024 | 0,25065 | 0,602676 |
| Ap1ar | 43,4851 | 49,4838 | 0,186437 | 0,2511 | 0,602676 |
| Tceal8 | 59,6203 | 67,8168 | 0,185839 | 0,25095 | 0,602676 |
| Rpl7l1 | 64,7718 | 73,5202 | 0,182775 | 0,2508 | 0,602676 |
| Rrbp1 | 47,2625 | 41,5052 | -0,187404 | 0,25105 | 0,602676 |
| Zcchc3 | 21,5243 | 18,8458 | -0,191723 | 0,25095 | 0,602676 |
| Ppp3cb | 37,1886 | 31,9221 | -0,220303 | 0,25095 | 0,602676 |

| | | | | | |
|---|---|---|---|---|---|
| Ggact | 4,86474 | 3,81236 | -0,35168 | 0,25095 | 0,602676 |
| Ccdc86 | 20,2713 | 23,1427 | 0,191122 | 0,25115 | 0,602688 |
| Mir7063 | 30,1564 | 0 | -inf | 0,25125 | 0,60282 |
| Polr3g | 11,33 | 9,80471 | -0,208596 | 0,25145 | 0,603191 |
| Mvk | 22,7923 | 26,2767 | 0,205233 | 0,2515 | 0,603203 |
| Mrpl44 | 29,0347 | 33,3722 | 0,200869 | 0,2516 | 0,603334 |
| Ttc39c | 3,5151 | 4,26009 | 0,277315 | 0,2517 | 0,603369 |
| Utp20 | 19,4004 | 22,0458 | 0,184418 | 0,25175 | 0,603369 |
| Prkaa1 | 13,0609 | 11,4527 | -0,18957 | 0,25175 | 0,603369 |
| Mir1188 | 0 | 26,9632 | inf | 0,2523 | 0,603674 |
| Mir6385 | 0 | 11,888 | inf | 0,2526 | 0,603674 |
| Mir7031 | 250,608 | 0 | -inf | 0,25225 | 0,603674 |
| Satb2 | 2,16997 | 2,58598 | 0,253035 | 0,2526 | 0,603674 |
| 4833439L19Rik | 52,1565 | 60,913 | 0,223903 | 0,25225 | 0,603674 |
| Kif16b | 5,94099 | 6,83739 | 0,202743 | 0,252 | 0,603674 |
| Epb4.1l2 | 99,7692 | 113,548 | 0,18663 | 0,25255 | 0,603674 |
| Numa1 | 42,7306 | 48,571 | 0,184823 | 0,252 | 0,603674 |
| Htt | 9,0313 | 10,2531 | 0,183053 | 0,2525 | 0,603674 |
| Psmc3 | 157,714 | 178,733 | 0,180492 | 0,2524 | 0,603674 |
| Ptov1 | 119,031 | 134,832 | 0,17983 | 0,25225 | 0,603674 |
| Emc1 | 11,8087 | 10,3654 | -0,188077 | 0,2521 | 0,603674 |
| Usp20 | 3,87139 | 3,2685 | -0,244221 | 0,25235 | 0,603674 |
| Tox3 | 3,37737 | 2,79265 | -0,274267 | 0,2526 | 0,603674 |
| Gm14403 | 1,85956 | 1,50076 | -0,309269 | 0,25215 | 0,603674 |
| Caly | 0,760826 | 0,447924 | -0,764311 | 0,2523 | 0,603674 |
| Mrps18a | 51,5663 | 59,248 | 0,200338 | 0,25265 | 0,603686 |
| Igdcc3 | 38,9546 | 44,2425 | 0,18364 | 0,2527 | 0,603697 |
| Tubb4a | 9,78135 | 11,4041 | 0,221444 | 0,25285 | 0,603948 |
| Lrpap1 | 21,8642 | 19,135 | -0,192358 | 0,2529 | 0,603959 |
| Mir6905 | 0 | 153,894 | inf | 0,25295 | 0,603971 |
| Prrc2c | 47,0689 | 53,6989 | 0,190118 | 0,2531 | 0,604113 |
| Dgkz | 28,5858 | 25,0805 | -0,188732 | 0,2531 | 0,604113 |
| Tmem184c | 16,1765 | 14,096 | -0,198616 | 0,25315 | 0,604124 |
| Tbck | 3,62161 | 4,31602 | 0,253068 | 0,2532 | 0,604136 |
| Snord123 | 23,6631 | 0 | -inf | 0,25335 | 0,604386 |
| Kdm5d | 7,38802 | 8,4774 | 0,198435 | 0,2534 | 0,604397 |
| Plxnb2 | 54,4439 | 48,0317 | -0,180786 | 0,25345 | 0,604409 |
| Cryab | 0,846946 | 0,120664 | -2,81128 | 0,2535 | 0,60442 |
| Tmsb15b2 | 1,90484 | 5,19027 | 1,44614 | 0,2538 | 0,604919 |
| Eif6 | 63,5637 | 72,2793 | 0,185379 | 0,2538 | 0,604919 |
| Ogdhl | 12,2428 | 14,0609 | 0,19975 | 0,25395 | 0,605169 |
| Mir7062 | 63,8043 | 0 | -inf | 0,25405 | 0,605299 |
| Fpgs | 17,1838 | 20,3891 | 0,246749 | 0,25425 | 0,605344 |
| Dscr3 | 22,1921 | 25,3754 | 0,193381 | 0,25415 | 0,605344 |
| Sirpa | 12,179 | 10,6163 | -0,198119 | 0,25425 | 0,605344 |
| Snora17 | 152,974 | 55,7412 | -1,45647 | 0,25425 | 0,605344 |
| Akr1e1 | 15,5582 | 18,0619 | 0,215269 | 0,25435 | 0,605474 |
| Ccdc151 | 1,06018 | 1,40154 | 0,402706 | 0,2544 | 0,605486 |

| | | | | | |
|---|---|---|---|---|---|
| Acaa1a | 13,6195 | 15,8061 | 0,214805 | 0,25455 | 0,605735 |
| Zfp622 | 14,0219 | 16,1011 | 0,199473 | 0,2546 | 0,605746 |
| Cdk5rap3 | 25,6591 | 29,4459 | 0,198594 | 0,2547 | 0,605876 |
| Gm684 | 1,03995 | 0,683557 | -0,605386 | 0,25475 | 0,605887 |
| Lmln | 3,18823 | 2,67309 | -0,254247 | 0,2549 | 0,606028 |
| Rfx8 | 0,678972 | 0,45494 | -0,577678 | 0,2549 | 0,606028 |
| Exph5 | 0,784288 | 0,638243 | -0,297277 | 0,25505 | 0,606277 |
| Twf1 | 32,0557 | 36,4719 | 0,186208 | 0,25535 | 0,606666 |
| Rbm22 | 55,9883 | 63,6323 | 0,184635 | 0,2553 | 0,606666 |
| Spryd3 | 6,25378 | 5,26725 | -0,24768 | 0,25535 | 0,606666 |
| 4921536K21Rik | 0,792094 | 1,0597 | 0,419917 | 0,2557 | 0,607066 |
| Taf6l | 17,8568 | 21,0087 | 0,234514 | 0,2556 | 0,607066 |
| Tsku | 14,2361 | 16,3858 | 0,202891 | 0,2557 | 0,607066 |
| Arsa | 5,09532 | 4,31649 | -0,239316 | 0,25565 | 0,607066 |
| Nxf1 | 114,869 | 130,833 | 0,187738 | 0,2558 | 0,607196 |
| Map2k1 | 38,7518 | 44,6028 | 0,202869 | 0,25595 | 0,607271 |
| Timm44 | 47,0357 | 53,5446 | 0,186985 | 0,2559 | 0,607271 |
| Ankrd11 | 39,3822 | 44,5664 | 0,178412 | 0,25615 | 0,607271 |
| Pdzd8 | 20,2921 | 17,8975 | -0,181157 | 0,25615 | 0,607271 |
| Pcf11 | 19,1366 | 16,8614 | -0,182617 | 0,25605 | 0,607271 |
| Tmem132d | 0,651105 | 0,488683 | -0,41399 | 0,2561 | 0,607271 |
| Cdkn2a | 0,759438 | 0,111367 | -2,76961 | 0,256 | 0,607271 |
| Fbxo3 | 39,2396 | 45,5683 | 0,21572 | 0,25635 | 0,607638 |
| Pura | 5,13658 | 6,24234 | 0,281281 | 0,2564 | 0,607648 |
| Paics | 202,055 | 234,478 | 0,214702 | 0,25655 | 0,607681 |
| Sars | 40,2118 | 45,5139 | 0,178687 | 0,25655 | 0,607681 |
| Lifr | 6,80465 | 5,84168 | -0,220136 | 0,25655 | 0,607681 |
| D5Ertd579e | 15,3471 | 13,5385 | -0,1809 | 0,2566 | 0,607691 |
| Arhgap39 | 8,84606 | 10,1412 | 0,197121 | 0,25685 | 0,608079 |
| Irf2bpl | 8,01084 | 6,94216 | -0,206568 | 0,2569 | 0,608079 |
| Slc35f1 | 1,11101 | 0,883543 | -0,330495 | 0,2569 | 0,608079 |
| Cggbp1 | 37,4415 | 33,0224 | -0,181194 | 0,25695 | 0,608089 |
| Ralgapa2 | 2,42096 | 2,81104 | 0,215523 | 0,2571 | 0,608229 |
| B9d2 | 10,6375 | 8,7285 | -0,285358 | 0,2571 | 0,608229 |
| H2-M5 | 1,12128 | 1,59874 | 0,511785 | 0,2572 | 0,60825 |
| Wdr46 | 37,1366 | 42,1679 | 0,183302 | 0,2572 | 0,60825 |
| 1810021B22Rik | 0,513083 | 1,02435 | 0,997452 | 0,2573 | 0,608292 |
| H2-Q5 | 0,878469 | 1,29045 | 0,55481 | 0,2574 | 0,608292 |
| Dis3l2 | 11,3623 | 13,0693 | 0,20193 | 0,2574 | 0,608292 |
| Dap3 | 16,9865 | 19,3139 | 0,185247 | 0,25735 | 0,608292 |
| Zfp879 | 0,764231 | 0,541852 | -0,496111 | 0,25745 | 0,608303 |
| Triap1 | 32,0095 | 36,9806 | 0,20827 | 0,25755 | 0,608432 |
| Cnot10 | 34,7194 | 39,4255 | 0,183389 | 0,2576 | 0,608442 |
| Katnal2 | 2,92524 | 3,66696 | 0,32603 | 0,25765 | 0,608453 |
| Clcn1 | 0,632455 | 0,833011 | 0,397373 | 0,2579 | 0,608537 |
| Cbwd1 | 16,4655 | 18,9824 | 0,205212 | 0,258 | 0,608537 |
| Mrps30 | 32,9283 | 37,6323 | 0,192646 | 0,25775 | 0,608537 |
| Rdh13 | 16,733 | 19,1183 | 0,192257 | 0,2578 | 0,608537 |

| | | | | | |
|---|---|---|---|---|---|
| Ift80 | 16,3376 | 18,5873 | 0,186118 | 0,25805 | 0,608537 |
| Chtf8 | 99,3059 | 112,726 | 0,182865 | 0,25795 | 0,608537 |
| Cald1 | 47,2056 | 41,6585 | -0,180345 | 0,25795 | 0,608537 |
| Rbm38 | 22,9594 | 19,9358 | -0,203718 | 0,25805 | 0,608537 |
| Gem | 0,738329 | 0,516968 | -0,51419 | 0,25815 | 0,608666 |
| Gid4 | 14,4288 | 16,4359 | 0,187895 | 0,25825 | 0,608794 |
| Polr1e | 12,4 | 14,8437 | 0,259506 | 0,25835 | 0,608922 |
| Sp4 | 8,59864 | 9,76644 | 0,183724 | 0,25855 | 0,609286 |
| Apex2 | 16,802 | 19,3547 | 0,204052 | 0,2589 | 0,610003 |
| Tet3 | 11,3345 | 9,99921 | -0,180839 | 0,25895 | 0,610014 |
| Cnot4 | 13,1234 | 15,0366 | 0,19633 | 0,259 | 0,610024 |
| Nek1 | 8,80363 | 10,1048 | 0,198872 | 0,25915 | 0,610162 |
| Cript | 39,4971 | 34,3404 | -0,201838 | 0,25915 | 0,610162 |
| Mkx | 2,01859 | 2,47711 | 0,29531 | 0,25945 | 0,610653 |
| Mir703 | 2658,67 | 2173,95 | -0,290389 | 0,25945 | 0,610653 |
| Tmem185b | 11,9523 | 10,3575 | -0,206611 | 0,25955 | 0,610781 |
| Pex5 | 16,6968 | 19,0686 | 0,191627 | 0,2596 | 0,610791 |
| H2-Eb1 | 0,472816 | 0,721166 | 0,609051 | 0,2601 | 0,61102 |
| 2310047M10Rik | 4,69713 | 5,66358 | 0,269935 | 0,2601 | 0,61102 |
| Adam23 | 3,503 | 4,14704 | 0,243492 | 0,26 | 0,61102 |
| Sh2b2 | 6,72216 | 7,83571 | 0,221139 | 0,2602 | 0,61102 |
| Tex10 | 35,6028 | 40,831 | 0,197674 | 0,2602 | 0,61102 |
| Arpc5l | 62,2236 | 70,7986 | 0,186259 | 0,2602 | 0,61102 |
| Rps24 | 130,069 | 147,692 | 0,183324 | 0,2602 | 0,61102 |
| Rrm2 | 203,527 | 230,714 | 0,180889 | 0,25985 | 0,61102 |
| Prpf4 | 23,5655 | 26,6938 | 0,179829 | 0,25995 | 0,61102 |
| Tspan7 | 31,2566 | 27,2995 | -0,195289 | 0,25975 | 0,61102 |
| BC048403 | 3,11745 | 2,57721 | -0,274554 | 0,2602 | 0,61102 |
| Med11 | 29,4334 | 34,2226 | 0,217493 | 0,26035 | 0,611157 |
| 1500011K16Rik | 11,9657 | 9,73493 | -0,297666 | 0,26035 | 0,611157 |
| Srrm2 | 142,87 | 165,4 | 0,211257 | 0,2605 | 0,611402 |
| Has2 | 32,8915 | 37,2214 | 0,178417 | 0,26055 | 0,611412 |
| Klhl22 | 24,9692 | 28,4329 | 0,187409 | 0,2607 | 0,611656 |
| Wasf1 | 20,9861 | 23,9598 | 0,191187 | 0,26075 | 0,611666 |
| Zfp953 | 1,37432 | 1,10768 | -0,311174 | 0,26085 | 0,611794 |
| Snx15 | 9,67485 | 8,1555 | -0,246466 | 0,26115 | 0,61239 |
| F2rl2 | 1,00838 | 0,0665664 | -3,92111 | 0,2613 | 0,612634 |
| 1700010I14Rik | 1,70082 | 2,1865 | 0,362388 | 0,2615 | 0,612995 |
| Bbc3 | 3,65509 | 4,45985 | 0,287091 | 0,26225 | 0,614388 |
| Zfyve19 | 13,4363 | 15,8081 | 0,234529 | 0,2626 | 0,614388 |
| Rnf216 | 21,2151 | 24,0527 | 0,181106 | 0,2622 | 0,614388 |
| Timeless | 37,3253 | 42,2817 | 0,179881 | 0,2626 | 0,614388 |
| Thrap3 | 44,1861 | 49,9837 | 0,177864 | 0,26235 | 0,614388 |
| Sfrp2 | 38,0694 | 33,4864 | -0,185057 | 0,2623 | 0,614388 |
| Tmem256 | 127,963 | 110,847 | -0,207151 | 0,2626 | 0,614388 |
| Ddx17 | 325,463 | 281,802 | -0,207812 | 0,2625 | 0,614388 |
| Gyg | 17,0607 | 14,7255 | -0,21236 | 0,2625 | 0,614388 |
| Grik2 | 1,11118 | 0,885763 | -0,327101 | 0,2625 | 0,614388 |

| | | | | | |
|---|---|---|---|---|---|
| Fam219aos | 1,9739 | 1,09672 | -0,847856 | 0,2626 | 0,614388 |
| Chtf18 | 18,5566 | 21,1633 | 0,189631 | 0,2627 | 0,614515 |
| Rufy2 | 7,31176 | 9,04816 | 0,307406 | 0,2628 | 0,614534 |
| Tmbim1 | 8,33826 | 6,71885 | -0,311532 | 0,2628 | 0,614534 |
| Swi5 | 384,846 | 435,773 | 0,179297 | 0,26285 | 0,614543 |
| Dck | 39,3868 | 44,6004 | 0,179347 | 0,2629 | 0,614552 |
| Zfp91 | 36,9601 | 32,6647 | -0,178236 | 0,263 | 0,614571 |
| Agbl3 | 1,44429 | 0,985752 | -0,551065 | 0,263 | 0,614571 |
| Pdrg1 | 40,0372 | 45,848 | 0,195517 | 0,2632 | 0,614716 |
| Tsr1 | 51,4235 | 58,5909 | 0,188249 | 0,26315 | 0,614716 |
| Bag4 | 12,2833 | 13,967 | 0,185316 | 0,2632 | 0,614716 |
| Sec63 | 33,9273 | 29,9998 | -0,177493 | 0,2633 | 0,614842 |
| Hadh | 75,2353 | 85,2098 | 0,179608 | 0,26345 | 0,615085 |
| Lamtor2 | 72,1369 | 83,0651 | 0,203505 | 0,26385 | 0,615911 |
| Gpatch8 | 19,9149 | 22,5087 | 0,176635 | 0,2639 | 0,61592 |
| Nsfl1c | 53,7971 | 61,0136 | 0,181604 | 0,26395 | 0,615929 |
| Jph3 | 1,15546 | 1,44602 | 0,323627 | 0,26405 | 0,616055 |
| Mtx3 | 12,102 | 10,6521 | -0,184105 | 0,2642 | 0,616298 |
| Syt12 | 0,916588 | 0,558186 | -0,715528 | 0,26425 | 0,616307 |
| Tinf2 | 33,9691 | 38,7268 | 0,18911 | 0,2643 | 0,616316 |
| Fbxw17 | 7,2492 | 8,59173 | 0,245126 | 0,26435 | 0,616325 |
| Zfp426 | 12,9614 | 14,7791 | 0,189337 | 0,2645 | 0,616567 |
| Rras2 | 29,6314 | 33,6398 | 0,18304 | 0,2646 | 0,616585 |
| Ptgfrn | 60,4389 | 53,4307 | -0,177807 | 0,2646 | 0,616585 |
| Idh2 | 144,98 | 163,747 | 0,175616 | 0,26495 | 0,617293 |
| Csl | 18,4337 | 21,1528 | 0,198503 | 0,265 | 0,617302 |
| Rps14 | 637,558 | 722,333 | 0,180108 | 0,2651 | 0,617319 |
| Mir6363 | 72,2716 | 0,00297683 | -14,5674 | 0,2651 | 0,617319 |
| Suclg1 | 85,3245 | 96,5401 | 0,178169 | 0,26515 | 0,617328 |
| Rimkla | 0,863111 | 1,09169 | 0,338943 | 0,26535 | 0,61748 |
| 2810006K23Rik | 11,849 | 13,8136 | 0,221325 | 0,2654 | 0,61748 |
| Tbc1d31 | 12,0032 | 13,7275 | 0,193649 | 0,2654 | 0,61748 |
| Wwox | 2,6896 | 2,15722 | -0,318216 | 0,26535 | 0,61748 |
| Stxbp1 | 8,39993 | 9,66685 | 0,202669 | 0,26545 | 0,617489 |
| Iqcc | 8,99816 | 10,5051 | 0,223383 | 0,26555 | 0,617614 |
| Tulp3 | 29,7631 | 33,7297 | 0,180497 | 0,26565 | 0,617739 |
| Gpr153 | 5,97852 | 6,92613 | 0,21226 | 0,2659 | 0,617783 |
| Rnf146 | 22,8808 | 26,2619 | 0,198833 | 0,2659 | 0,617783 |
| Prkcd | 20,2892 | 23,1036 | 0,187411 | 0,2659 | 0,617783 |
| Ager | 4,34976 | 3,47552 | -0,32371 | 0,26585 | 0,617783 |
| Sycp3 | 0,89546 | 0,205225 | -2,12542 | 0,2658 | 0,617783 |
| 2010107G23Rik | 1,41126 | 1,84835 | 0,389256 | 0,26625 | 0,618056 |
| Hax1 | 19,0054 | 22,3338 | 0,232822 | 0,2667 | 0,618056 |
| Cenpm | 16,1413 | 18,8852 | 0,226495 | 0,26645 | 0,618056 |
| Tpgs1 | 21,0059 | 24,4259 | 0,217616 | 0,26675 | 0,618056 |
| Wbp4 | 17,6537 | 20,4029 | 0,208803 | 0,26675 | 0,618056 |
| Sfpq | 371,656 | 427,812 | 0,20301 | 0,26625 | 0,618056 |
| Rbm14 | 48,1935 | 55,401 | 0,201075 | 0,26685 | 0,618056 |

| | | | | | |
|---|---|---|---|---|---|
| Fam53b | 6,08085 | 6,98207 | 0,199382 | 0,26635 | 0,618056 |
| Bcl10 | 20,1772 | 23,0931 | 0,19474 | 0,2668 | 0,618056 |
| Mrpl39 | 40,8735 | 46,386 | 0,182521 | 0,2666 | 0,618056 |
| Emd | 63,1107 | 71,4595 | 0,179242 | 0,26685 | 0,618056 |
| Nop2 | 65,469 | 74,0554 | 0,177793 | 0,26655 | 0,618056 |
| D17Wsu92e | 53,7409 | 60,6119 | 0,173581 | 0,2664 | 0,618056 |
| Ect2 | 44,6482 | 39,5669 | -0,174309 | 0,26685 | 0,618056 |
| Gpr161 | 8,52737 | 7,24409 | -0,235297 | 0,26615 | 0,618056 |
| Tmem63a | 2,66434 | 2,18917 | -0,283392 | 0,26655 | 0,618056 |
| AW011738 | 1,9918 | 1,59065 | -0,32446 | 0,26625 | 0,618056 |
| Gm20337 | 1,37586 | 0,94389 | -0,543643 | 0,26685 | 0,618056 |
| Pir | 4,67344 | 5,68119 | 0,281707 | 0,26705 | 0,618305 |
| Sh3pxd2b | 17,955 | 20,291 | 0,176454 | 0,26705 | 0,618305 |
| Enkd1 | 22,4331 | 25,9246 | 0,208693 | 0,2672 | 0,618438 |
| Pdcl | 24,7726 | 21,8515 | -0,181016 | 0,2672 | 0,618438 |
| Ppapdc1a | 0,455476 | 0,763442 | 0,745142 | 0,26795 | 0,619012 |
| Ube2j2 | 5,42513 | 6,57414 | 0,277144 | 0,2675 | 0,619012 |
| Camkk1 | 2,2771 | 2,75268 | 0,273637 | 0,26795 | 0,619012 |
| Zfp607 | 3,05363 | 3,62904 | 0,249061 | 0,2679 | 0,619012 |
| Rps19bp1 | 43,0465 | 49,6036 | 0,204548 | 0,26785 | 0,619012 |
| Gsg2 | 11,5926 | 13,3014 | 0,198368 | 0,26805 | 0,619012 |
| Anp32b | 321,16 | 363,15 | 0,177275 | 0,2677 | 0,619012 |
| Paip2 | 180,433 | 203,678 | 0,174825 | 0,268 | 0,619012 |
| Capza2 | 108,977 | 96,5144 | -0,175205 | 0,26765 | 0,619012 |
| Lclat1 | 8,58455 | 7,48252 | -0,198218 | 0,2679 | 0,619012 |
| Casp7 | 8,31254 | 7,1006 | -0,227349 | 0,26805 | 0,619012 |
| Sprtn | 7,42464 | 6,29915 | -0,237163 | 0,268 | 0,619012 |
| Ctsf | 7,03973 | 5,93564 | -0,246117 | 0,26775 | 0,619012 |
| Rps6kb1 | 21,9124 | 19,0981 | -0,198321 | 0,2681 | 0,61902 |
| Notch3 | 31,1954 | 35,2212 | 0,175111 | 0,2682 | 0,619144 |
| Ddx1 | 130,053 | 147,076 | 0,177468 | 0,2687 | 0,620191 |
| Lancl1 | 10,1485 | 11,5482 | 0,186392 | 0,269 | 0,620455 |
| Daxx | 27,9246 | 31,6244 | 0,179503 | 0,269 | 0,620455 |
| Hmgn5 | 27,0451 | 23,7083 | -0,189978 | 0,269 | 0,620455 |
| D330023K18Rik | 3,998 | 2,7154 | -0,558113 | 0,26895 | 0,620455 |
| Kntc1 | 18,0469 | 20,4193 | 0,178177 | 0,26915 | 0,620694 |
| Capn6 | 12,2569 | 13,9962 | 0,191441 | 0,2692 | 0,620702 |
| Gm13034 | 0,73477 | 1,10786 | 0,592416 | 0,26945 | 0,620751 |
| Msh6 | 58,4863 | 68,2815 | 0,223395 | 0,2695 | 0,620751 |
| Hnrnph3 | 63,8293 | 74,1031 | 0,215315 | 0,26935 | 0,620751 |
| Mrps2 | 37,7736 | 42,8002 | 0,18024 | 0,26935 | 0,620751 |
| Add3 | 42,5114 | 37,5616 | -0,178591 | 0,2695 | 0,620751 |
| Cdk17 | 14,2073 | 12,4556 | -0,189846 | 0,2695 | 0,620751 |
| Gpt2 | 8,56519 | 9,83961 | 0,200117 | 0,2696 | 0,620874 |
| Pomgnt1 | 32,1571 | 28,3889 | -0,179809 | 0,26965 | 0,620883 |
| Eef2 | 697,504 | 816,071 | 0,226493 | 0,2698 | 0,621014 |
| Mink1 | 13,3589 | 15,166 | 0,183043 | 0,2698 | 0,621014 |
| 9530082P21Rik | 15,1375 | 17,2514 | 0,188586 | 0,26995 | 0,621145 |

| | | | | | |
|---|---|---|---|---|---|
| Cdh13 | 1,4926 | 1,15025 | -0,375877 | 0,26995 | 0,621145 |
| Zfp458 | 0,769742 | 0,580185 | -0,407862 | 0,27005 | 0,621268 |
| Gfra4 | 1,15319 | 1,58476 | 0,458628 | 0,27035 | 0,621745 |
| Cdca7l | 32,4884 | 36,859 | 0,182094 | 0,27035 | 0,621745 |
| Myrfl | 1,00704 | 1,28341 | 0,34987 | 0,27045 | 0,62176 |
| Efcab14 | 14,936 | 13,1188 | -0,187155 | 0,27045 | 0,621761 |
| Tra2b | 69,6697 | 61,7385 | -0,174362 | 0,27055 | 0,621883 |
| Polq | 7,90466 | 8,97511 | 0,183226 | 0,2706 | 0,621891 |
| Mir5133 | 47,7757 | 345,622 | 2,85484 | 0,27075 | 0,622129 |
| Slc35f6 | 5,66683 | 4,81931 | -0,233714 | 0,2708 | 0,622137 |
| Dido1 | 30,1543 | 26,5444 | -0,183958 | 0,27085 | 0,622145 |
| Stamos | 1,06242 | 1,62098 | 0,609514 | 0,2717 | 0,623692 |
| Ppp1r35 | 22,4954 | 26,1824 | 0,218967 | 0,27185 | 0,623692 |
| Caprin2 | 11,1091 | 12,6779 | 0,190564 | 0,27165 | 0,623692 |
| Rps3a1 | 137,846 | 155,791 | 0,176558 | 0,27185 | 0,623692 |
| Taf9b | 4,34102 | 3,61689 | -0,263286 | 0,2716 | 0,623692 |
| Zfp707 | 5,29913 | 4,38533 | -0,273067 | 0,2718 | 0,623692 |
| Proser2 | 1,61641 | 1,30407 | -0,30977 | 0,2718 | 0,623692 |
| Pdxp | 14,4855 | 16,6702 | 0,202667 | 0,27195 | 0,623814 |
| Srbd1 | 17,9192 | 20,3222 | 0,181552 | 0,272 | 0,623822 |
| Sec61g | 0,301221 | 0,596655 | 0,986077 | 0,27205 | 0,623829 |
| Primpol | 4,02631 | 3,18923 | -0,336251 | 0,2722 | 0,624066 |
| Kat6a | 21,2712 | 24,0211 | 0,175403 | 0,27225 | 0,624074 |
| Ptp4a1 | 3,71654 | 3,15241 | -0,237506 | 0,2723 | 0,624081 |
| Ccdc88c | 7,46634 | 6,5506 | -0,188774 | 0,2724 | 0,624203 |
| Gpr4 | 0,795095 | 0,588631 | -0,433764 | 0,27245 | 0,624211 |
| Rnf157 | 3,40355 | 3,98267 | 0,226698 | 0,27255 | 0,624333 |
| Spats2 | 55,6528 | 62,88 | 0,176146 | 0,27265 | 0,624455 |
| Nedd8 | 306,062 | 345,628 | 0,175398 | 0,27285 | 0,624485 |
| Cul4b | 48,5206 | 43,0353 | -0,173078 | 0,27275 | 0,624485 |
| Gbp7 | 1,49426 | 1,23469 | -0,275288 | 0,2728 | 0,624485 |
| Fah | 2,24118 | 1,72495 | -0,377707 | 0,27285 | 0,624485 |
| Adamts19 | 1,73269 | 1,42333 | -0,28375 | 0,27315 | 0,624957 |
| Kansl3 | 29,0433 | 32,7438 | 0,173015 | 0,27315 | 0,624958 |
| Pfn1 | 536,924 | 606,724 | 0,176324 | 0,2733 | 0,625194 |
| Hint1 | 510,385 | 575,837 | 0,174073 | 0,2736 | 0,625773 |
| Ccdc51 | 5,17453 | 6,10032 | 0,237457 | 0,2737 | 0,625894 |
| Ccl27a | 8,26571 | 12,1993 | 0,561584 | 0,27375 | 0,625902 |
| Rps19-ps3 | 3,97306 | 5,77194 | 0,538805 | 0,27385 | 0,626023 |
| Rpusd3 | 10,5342 | 12,4408 | 0,240005 | 0,2739 | 0,62603 |
| Fzd10 | 52,6413 | 46,6818 | -0,173335 | 0,274 | 0,626152 |
| Etv2 | 0,799389 | 1,20109 | 0,587371 | 0,27415 | 0,626173 |
| Zfp189 | 6,06062 | 7,06061 | 0,220328 | 0,2741 | 0,626173 |
| Ift140 | 8,76702 | 10,0391 | 0,195476 | 0,27415 | 0,626173 |
| Scn8a | 0,88261 | 1,05071 | 0,251521 | 0,2743 | 0,626409 |
| Ldoc1l | 6,21721 | 5,39612 | -0,204344 | 0,2745 | 0,626759 |
| Nfya | 36,7722 | 41,419 | 0,171678 | 0,27465 | 0,626994 |
| Ick | 9,15252 | 10,3608 | 0,178901 | 0,2748 | 0,627229 |

| | | | | | |
|---|---|---|---|---|---|
| 2410089E03Rik | 8,03429 | 9,08958 | 0,178042 | 0,27495 | 0,627464 |
| Zfhx2 | 1,27068 | 1,05937 | -0,262395 | 0,2751 | 0,6277 |
| Eapp | 30,2469 | 34,7201 | 0,198982 | 0,27515 | 0,627706 |
| Zscan22 | 7,75789 | 8,97197 | 0,209762 | 0,2754 | 0,62817 |
| 9330182L06Rik | 0,978372 | 0,774643 | -0,336853 | 0,2755 | 0,62829 |
| Atp6v0c | 8,97551 | 10,7433 | 0,259366 | 0,2756 | 0,628411 |
| Insig1 | 24,2097 | 27,3994 | 0,178559 | 0,2758 | 0,62876 |
| Hspa5 | 551,346 | 633,619 | 0,200658 | 0,27595 | 0,628995 |
| Ppapdc1b | 10,8359 | 9,18489 | -0,238482 | 0,276 | 0,629001 |
| Ift20 | 39,6081 | 34,2267 | -0,210674 | 0,27615 | 0,629236 |
| Ilf3 | 104,019 | 117,184 | 0,171936 | 0,27635 | 0,629584 |
| Cramp1l | 12,7939 | 14,4462 | 0,175228 | 0,27645 | 0,629597 |
| Taldo1 | 92,4484 | 104,386 | 0,175214 | 0,27645 | 0,629597 |
| Dars2 | 14,1546 | 16,0951 | 0,185352 | 0,27655 | 0,629611 |
| Morn1 | 1,918 | 1,47064 | -0,38316 | 0,27655 | 0,629611 |
| Txlna | 49,8527 | 56,1385 | 0,171317 | 0,2766 | 0,629617 |
| Nrip2 | 0,955097 | 1,28147 | 0,424085 | 0,27685 | 0,629864 |
| Cog1 | 14,696 | 17,2218 | 0,228818 | 0,27685 | 0,629864 |
| Nelfe | 52,8071 | 59,699 | 0,176975 | 0,27685 | 0,629864 |
| Enc1 | 27,1916 | 30,7085 | 0,175475 | 0,27695 | 0,629877 |
| Aqr | 36,7344 | 41,4008 | 0,172526 | 0,27695 | 0,629877 |
| Kctd10 | 28,9558 | 25,6383 | -0,17555 | 0,277 | 0,629884 |
| Cmc2 | 15,7234 | 18,1838 | 0,209743 | 0,2772 | 0,630232 |
| Hebp2 | 9,28174 | 10,8253 | 0,221938 | 0,2773 | 0,630264 |
| Ift74 | 27,0236 | 30,6467 | 0,181513 | 0,27735 | 0,630264 |
| Nfatc3 | 27,0083 | 30,4486 | 0,172971 | 0,27745 | 0,630264 |
| Arhgef1 | 23,4903 | 20,7266 | -0,180583 | 0,27745 | 0,630264 |
| Rtkn2 | 2,56154 | 2,13486 | -0,26287 | 0,27745 | 0,630264 |
| Epha2 | 24,922 | 22,0717 | -0,175217 | 0,27775 | 0,630838 |
| Camta1 | 8,45309 | 9,92558 | 0,231672 | 0,27795 | 0,631078 |
| Ino80d | 5,08687 | 5,76578 | 0,180737 | 0,27795 | 0,631078 |
| Snora70 | 267,139 | 386,066 | 0,531257 | 0,27885 | 0,631625 |
| Rab19 | 2,96831 | 3,74324 | 0,334647 | 0,2789 | 0,631625 |
| Camk1 | 15,4925 | 18,5278 | 0,258116 | 0,27865 | 0,631625 |
| Dctn2 | 87,5563 | 102,204 | 0,223167 | 0,27855 | 0,631625 |
| Msx2 | 7,75327 | 9,03978 | 0,221483 | 0,27845 | 0,631625 |
| Dnm2 | 28,1346 | 32,755 | 0,219371 | 0,27825 | 0,631625 |
| Senp1 | 19,4599 | 22,5474 | 0,212458 | 0,2788 | 0,631625 |
| Schip1 | 24,1092 | 27,6648 | 0,19847 | 0,2787 | 0,631625 |
| Phb | 18,039 | 20,676 | 0,196837 | 0,27885 | 0,631625 |
| Irs1 | 9,99015 | 11,2968 | 0,177342 | 0,2789 | 0,631625 |
| Zfp516 | 15,1207 | 17,6437 | 0,173115 | 0,27875 | 0,631625 |
| Cab39 | 40,1064 | 35,5813 | -0,172714 | 0,2788 | 0,631625 |
| Agap1 | 5,88747 | 5,18765 | -0,182566 | 0,27865 | 0,631625 |
| Dip2a | 6,8175 | 5,98559 | -0,187748 | 0,2785 | 0,631625 |
| Bambi-ps1 | 2,51794 | 1,90643 | -0,40137 | 0,27865 | 0,631625 |
| Dnajc17 | 10,1107 | 13,1388 | 0,377951 | 0,27935 | 0,632109 |
| Nup210l | 0,749002 | 0,933 | 0,316908 | 0,27935 | 0,632109 |

| | | | | | |
|---|---|---|---|---|---|
| Gtf2h5 | 67,9303 | 76,8427 | 0,177853 | 0,2793 | 0,632109 |
| Slc29a3 | 2,24394 | 1,88649 | -0,250328 | 0,2793 | 0,632109 |
| Zkscan4 | 1,33279 | 1,02809 | -0,374485 | 0,2793 | 0,632109 |
| Rps6ka3 | 23,1423 | 20,521 | -0,173428 | 0,27945 | 0,632228 |
| Cldn12 | 22,7728 | 20,1308 | -0,177907 | 0,27955 | 0,63224 |
| Galnt5 | 0,743112 | 0,557421 | -0,414813 | 0,27955 | 0,63224 |
| Lcmt1 | 27,1992 | 31,0138 | 0,189351 | 0,27965 | 0,632252 |
| Rbbp4 | 144,788 | 164,056 | 0,180248 | 0,27965 | 0,632252 |
| C330006A16Rik | 13,4772 | 11,7154 | -0,20212 | 0,27975 | 0,632371 |
| Amh | 0,409344 | 0,627729 | 0,61683 | 0,27995 | 0,632426 |
| Ranbp17 | 3,57709 | 4,14892 | 0,213952 | 0,28015 | 0,632426 |
| Hmbox1 | 10,0935 | 11,5771 | 0,197843 | 0,2801 | 0,632426 |
| E4f1 | 14,0241 | 15,9985 | 0,190032 | 0,2799 | 0,632426 |
| Cep135 | 9,60269 | 10,9013 | 0,18299 | 0,2802 | 0,632426 |
| Ptpn13 | 15,3075 | 17,2497 | 0,172341 | 0,2802 | 0,632426 |
| Spc25 | 83,3653 | 73,8245 | -0,175348 | 0,2802 | 0,632426 |
| Sptlc1 | 24,0987 | 21,2007 | -0,184849 | 0,28005 | 0,632426 |
| Ficd | 1,25328 | 0,985064 | -0,34742 | 0,27985 | 0,632426 |
| Trim62 | 7,19735 | 6,21927 | -0,210721 | 0,2803 | 0,632545 |
| Smurf1 | 14,4157 | 16,2977 | 0,177031 | 0,2804 | 0,632663 |
| Lgi4 | 0,700805 | 0,928294 | 0,405568 | 0,28055 | 0,632895 |
| Mrpl45 | 26,209 | 29,6822 | 0,179535 | 0,2807 | 0,632913 |
| Imp4 | 37,079 | 41,9135 | 0,17681 | 0,2807 | 0,632913 |
| Nup205 | 47,928 | 53,9989 | 0,17206 | 0,2807 | 0,632913 |
| Htatsf1 | 45,9113 | 51,7549 | 0,172848 | 0,2808 | 0,633032 |
| Gpd1l | 8,95827 | 7,8471 | -0,191062 | 0,2809 | 0,63315 |
| Tk1 | 30,0262 | 34,4865 | 0,199811 | 0,281 | 0,633269 |
| 9930012K11Rik | 2,69848 | 2,16858 | -0,315395 | 0,28105 | 0,633275 |
| Lrrc45 | 17,2906 | 19,7041 | 0,188509 | 0,2812 | 0,633506 |
| Ywhaq | 46,2189 | 40,9827 | -0,173467 | 0,28135 | 0,633737 |
| Tst | 6,57705 | 5,33343 | -0,302375 | 0,28145 | 0,633855 |
| Vps4a | 27,6239 | 31,3528 | 0,182673 | 0,28165 | 0,633879 |
| Atp1a2 | 20,7528 | 23,3996 | 0,173178 | 0,2816 | 0,633879 |
| Ruvbl2 | 105,766 | 118,953 | 0,169511 | 0,28155 | 0,633879 |
| Gnpnat1 | 9,00912 | 7,75306 | -0,21662 | 0,28165 | 0,633879 |
| 1600023N17Rik | 0,463714 | 0,774189 | 0,739452 | 0,282 | 0,63393 |
| Tekt1 | 2,15326 | 2,75083 | 0,353348 | 0,2821 | 0,63393 |
| 2210013O21Rik | 25,7566 | 29,906 | 0,215489 | 0,2819 | 0,63393 |
| Uchl3 | 42,7039 | 48,8453 | 0,193853 | 0,2821 | 0,63393 |
| Mrpl9 | 44,6224 | 50,5016 | 0,17856 | 0,282 | 0,63393 |
| Gtf3c1 | 32,542 | 36,6548 | 0,171698 | 0,28205 | 0,63393 |
| Shmt2 | 68,765 | 77,4504 | 0,171597 | 0,282 | 0,63393 |
| Slc35c2 | 23,3244 | 20,4956 | -0,186531 | 0,282 | 0,63393 |
| Thap2 | 9,56144 | 8,33701 | -0,197699 | 0,2821 | 0,63393 |
| Tti1 | 18,8033 | 21,2127 | 0,173941 | 0,2822 | 0,633942 |
| Mtss1l | 21,4699 | 24,1873 | 0,171937 | 0,2822 | 0,633942 |
| Ush1c | 1,63719 | 2,07358 | 0,340904 | 0,2825 | 0,6341 |
| Sh3gl3 | 5,77937 | 6,89116 | 0,253833 | 0,28245 | 0,6341 |

| | | | | | |
|---|---|---|---|---|---|
| Helq | 7,98589 | 9,14755 | 0,195932 | 0,28265 | 0,6341 |
| Abat | 7,6145 | 8,67078 | 0,187412 | 0,28265 | 0,6341 |
| Mta1 | 79,3588 | 89,3256 | 0,170683 | 0,2825 | 0,6341 |
| Pan3 | 13,622 | 12,0056 | -0,182226 | 0,28265 | 0,6341 |
| Rhobtb1 | 5,02907 | 4,34602 | -0,210597 | 0,28265 | 0,6341 |
| Tex15 | 0,95022 | 0,784721 | -0,276082 | 0,28255 | 0,6341 |
| Mier3 | 12,3118 | 10,8815 | -0,178168 | 0,2827 | 0,634106 |
| Ctrl | 1,22871 | 1,78067 | 0,535279 | 0,2828 | 0,634117 |
| Slc35e1 | 18,9454 | 16,8314 | -0,170692 | 0,2828 | 0,634117 |
| Narg2 | 12,9954 | 14,7544 | 0,183146 | 0,28285 | 0,634123 |
| Adat2 | 5,44535 | 7,13375 | 0,389635 | 0,28295 | 0,63414 |
| Dicer1 | 39,8544 | 45,0123 | 0,175582 | 0,283 | 0,63414 |
| Zfp85 | 2,95566 | 2,37966 | -0,312727 | 0,283 | 0,63414 |
| Nhs | 2,35624 | 2,77013 | 0,233468 | 0,2831 | 0,634152 |
| Zfp146 | 79,9662 | 90,0617 | 0,171523 | 0,2831 | 0,634152 |
| Chst15 | 13,7253 | 12,1035 | -0,181416 | 0,28315 | 0,634157 |
| Nfkbie | 2,1843 | 2,69123 | 0,301095 | 0,2838 | 0,634231 |
| Ubox5 | 7,57808 | 9,07307 | 0,259759 | 0,2834 | 0,634231 |
| Nfatc1 | 3,47837 | 4,0643 | 0,224595 | 0,2835 | 0,634231 |
| Ncl | 256,569 | 299,435 | 0,222898 | 0,28375 | 0,634231 |
| Hdac4 | 4,85129 | 5,62066 | 0,212373 | 0,2838 | 0,634231 |
| Acp6 | 12,2256 | 14,14 | 0,209878 | 0,2836 | 0,634231 |
| Spg11 | 6,64146 | 7,65394 | 0,204702 | 0,2837 | 0,634231 |
| Elp4 | 10,359 | 11,7852 | 0,186086 | 0,28325 | 0,634231 |
| BC018507 | 28,7824 | 32,4083 | 0,171173 | 0,28375 | 0,634231 |
| Tln2 | 19,138 | 21,5334 | 0,17014 | 0,2834 | 0,634231 |
| Tmem131 | 29,0348 | 25,8194 | -0,169326 | 0,28355 | 0,634231 |
| Arg1 | 18,5398 | 16,0638 | -0,206813 | 0,2837 | 0,634231 |
| Ppp1r36 | 1,92878 | 1,45971 | -0,402006 | 0,2834 | 0,634231 |
| Glipr2 | 21,2466 | 24,1837 | 0,186801 | 0,28385 | 0,634237 |
| Adam19 | 27,1193 | 30,4985 | 0,169422 | 0,2839 | 0,634243 |
| BC027231 | 20,5486 | 23,2704 | 0,17946 | 0,284 | 0,63436 |
| Zfpm2 | 0,554707 | 0,709321 | 0,354713 | 0,28405 | 0,634366 |
| Mras | 2,07854 | 2,48685 | 0,258748 | 0,2845 | 0,634963 |
| Rrn3 | 36,015 | 40,5833 | 0,172288 | 0,28465 | 0,634963 |
| Cand1 | 48,4177 | 54,5446 | 0,171903 | 0,2846 | 0,634963 |
| Melk | 38,7634 | 43,6283 | 0,170569 | 0,28465 | 0,634963 |
| Zfp704 | 10,9607 | 9,72567 | -0,172476 | 0,2846 | 0,634963 |
| Gpalpp1 | 12,5397 | 11,0965 | -0,176403 | 0,2846 | 0,634963 |
| Pigs | 23,8573 | 21,0269 | -0,18219 | 0,28445 | 0,634963 |
| Gm715 | 0,488989 | 0,769044 | 0,653265 | 0,28495 | 0,635108 |
| Ppp1r3c | 0,986533 | 1,26851 | 0,362696 | 0,285 | 0,635108 |
| Xndc1 | 14,5567 | 16,6284 | 0,191963 | 0,2849 | 0,635108 |
| Pacs1 | 18,0544 | 20,3845 | 0,175123 | 0,28485 | 0,635108 |
| Tubb6 | 53,5851 | 60,4053 | 0,172844 | 0,285 | 0,635108 |
| Pqlc2 | 3,14468 | 2,55774 | -0,298039 | 0,28495 | 0,635108 |
| C1ql1 | 4,4771 | 5,51198 | 0,300008 | 0,28505 | 0,635113 |
| Opa3 | 20,8352 | 18,317 | -0,185834 | 0,2851 | 0,635119 |

| | | | | | |
|---|---|---|---|---|---|
| Nkiras1 | 1,71084 | 2,48571 | 0,538957 | 0,2852 | 0,635235 |
| Bcl2l12 | 30,1871 | 25,8306 | -0,224852 | 0,28525 | 0,635241 |
| Btbd3 | 9,22894 | 8,11328 | -0,18588 | 0,2854 | 0,635469 |
| Psma5 | 29,5154 | 33,7185 | 0,19207 | 0,2859 | 0,63637 |
| Itfg3 | 11,9832 | 10,4889 | -0,192153 | 0,2859 | 0,63637 |
| Plekhm1 | 4,82817 | 4,18954 | -0,204682 | 0,286 | 0,636487 |
| Ap2b1 | 45,3513 | 50,9179 | 0,167029 | 0,28615 | 0,636715 |
| Hs2st1 | 28,4229 | 25,2457 | -0,171014 | 0,28625 | 0,636831 |
| Tnfrsf21 | 14,3323 | 12,6187 | -0,183706 | 0,2864 | 0,636958 |
| Arl15 | 12,1774 | 10,6778 | -0,189589 | 0,28645 | 0,636958 |
| Spock3 | 1,66925 | 1,34852 | -0,307819 | 0,28645 | 0,636958 |
| Rpp25l | 42,5836 | 48,7496 | 0,195091 | 0,28665 | 0,637085 |
| Fam133b | 27,003 | 30,5334 | 0,177264 | 0,28665 | 0,637085 |
| Uhrf2 | 56,5189 | 63,4388 | 0,166631 | 0,2866 | 0,637085 |
| Tet1 | 3,90346 | 4,41143 | 0,176493 | 0,2868 | 0,637206 |
| Prmt7 | 37,0976 | 41,8392 | 0,173529 | 0,2868 | 0,637206 |
| Pi4k2b | 9,69751 | 8,44168 | -0,200084 | 0,287 | 0,637545 |
| Kcna6 | 1,60837 | 1,90517 | 0,244321 | 0,2873 | 0,637681 |
| C87436 | 6,65004 | 7,69331 | 0,210243 | 0,2873 | 0,637681 |
| Vti1a | 6,67811 | 7,69851 | 0,205139 | 0,28725 | 0,637681 |
| Vac14 | 18,8309 | 21,3518 | 0,181252 | 0,28725 | 0,637681 |
| Gm7120 | 6,84454 | 5,73365 | -0,255499 | 0,28725 | 0,637681 |
| Creld1 | 11,982 | 13,7782 | 0,201527 | 0,28745 | 0,637908 |
| Ppp1r21 | 10,0197 | 11,4423 | 0,191534 | 0,2877 | 0,638145 |
| Exoc3 | 19,9881 | 22,4785 | 0,169405 | 0,28765 | 0,638145 |
| Slc12a9 | 6,61271 | 5,70987 | -0,211784 | 0,2877 | 0,638145 |
| Ubap2 | 46,0341 | 51,7708 | 0,169435 | 0,28775 | 0,63815 |
| Pigf | 40,2396 | 31,9836 | -0,331286 | 0,2879 | 0,638377 |
| Chd3os | 9,0397 | 10,5405 | 0,221591 | 0,28805 | 0,638497 |
| Pja2 | 28,1782 | 31,7106 | 0,170387 | 0,28805 | 0,638497 |
| Gm4961 | 1,06234 | 0,798411 | -0,412048 | 0,2882 | 0,638724 |
| Cdip1 | 30,893 | 35,3969 | 0,196341 | 0,2883 | 0,63884 |
| Casp3 | 69,4287 | 78,1773 | 0,171217 | 0,28835 | 0,638844 |
| 5930430L01Rik | 0,747357 | 0,571584 | -0,386831 | 0,2884 | 0,638849 |
| Ss18 | 46,0472 | 51,7846 | 0,169411 | 0,28845 | 0,638854 |
| Tmem120b | 3,46347 | 2,81076 | -0,301258 | 0,2885 | 0,638859 |
| Fxr1 | 87,1127 | 98,1895 | 0,172687 | 0,28855 | 0,638864 |
| Zfp74 | 6,82671 | 7,84119 | 0,199882 | 0,28905 | 0,639446 |
| Spop | 37,0858 | 41,7631 | 0,171362 | 0,289 | 0,639446 |
| Map4k3 | 13,5538 | 11,9546 | -0,181134 | 0,28895 | 0,639446 |
| Ifngr2 | 23,8481 | 21,0268 | -0,181641 | 0,2891 | 0,639446 |
| Zfp930 | 15,2013 | 13,3637 | -0,18588 | 0,2891 | 0,639446 |
| 2700038G22Rik | 23,8144 | 15,5995 | -0,610335 | 0,28895 | 0,639446 |
| Golga4 | 17,3718 | 19,5335 | 0,169204 | 0,28925 | 0,639672 |
| Ddx54 | 30,3966 | 34,1703 | 0,168833 | 0,28935 | 0,639681 |
| Trib1 | 10,329 | 9,08593 | -0,184993 | 0,28935 | 0,639681 |
| Tbc1d2b | 5,47872 | 4,79482 | -0,192362 | 0,2895 | 0,639801 |
| Pgam2 | 1,13835 | 0,724462 | -0,651959 | 0,2895 | 0,639801 |

| | | | | | |
|---|---|---|---|---|---|
| Wdr41 | 8,23011 | 9,4398 | 0,197845 | 0,2897 | 0,640137 |
| Unkl | 2,10291 | 2,63991 | 0,328106 | 0,2898 | 0,640147 |
| Smg1 | 17,6511 | 19,8994 | 0,172965 | 0,2898 | 0,640147 |
| Plscr1 | 9,58837 | 11,0698 | 0,207274 | 0,28995 | 0,640372 |
| Nutf2 | 13,4165 | 15,4012 | 0,199031 | 0,2901 | 0,640386 |
| Prr14 | 20,2867 | 23,0242 | 0,18262 | 0,2901 | 0,640386 |
| Map7d1 | 40,3729 | 45,398 | 0,169244 | 0,2901 | 0,640386 |
| Wdr3 | 38,4813 | 43,3566 | 0,172092 | 0,29015 | 0,640391 |
| E2f8 | 15,7317 | 17,7551 | 0,174562 | 0,29025 | 0,6404 |
| Disp1 | 6,84738 | 5,99834 | -0,190988 | 0,29025 | 0,6404 |
| Lrrc55 | 2,05981 | 2,50795 | 0,283997 | 0,2903 | 0,640404 |
| Naa25 | 30,7161 | 34,5879 | 0,171274 | 0,2905 | 0,64074 |
| Slc39a7 | 98,162 | 87,4499 | -0,166707 | 0,29055 | 0,640744 |
| Tsen34 | 35,7735 | 40,5822 | 0,181958 | 0,2906 | 0,640749 |
| Bex4 | 11,8916 | 14,2957 | 0,265635 | 0,29065 | 0,640753 |
| Slc26a2 | 6,55241 | 5,70173 | -0,200627 | 0,2907 | 0,640758 |
| Pde6d | 33,272 | 38,0016 | 0,191752 | 0,291 | 0,641208 |
| Zfp768 | 23,4274 | 26,5406 | 0,180007 | 0,291 | 0,641208 |
| Ptpmt1 | 27,2674 | 23,8247 | -0,194721 | 0,2911 | 0,641322 |
| Ccnd2 | 31,4111 | 27,921 | -0,169923 | 0,29115 | 0,641327 |
| Fbrsl1 | 6,99415 | 8,15689 | 0,221871 | 0,29135 | 0,64145 |
| Tfpt | 18,7364 | 21,6268 | 0,206971 | 0,2913 | 0,64145 |
| Zbtb1 | 10,5487 | 11,9711 | 0,1825 | 0,29135 | 0,64145 |
| Tm2d2 | 39,136 | 34,4462 | -0,184151 | 0,2915 | 0,641675 |
| Bex1 | 46,9773 | 53,5954 | 0,190148 | 0,2916 | 0,641789 |
| Dctpp1 | 178,751 | 201,003 | 0,169271 | 0,29165 | 0,641793 |
| BC039771 | 0,953992 | 1,25941 | 0,400696 | 0,29185 | 0,642022 |
| St6galnac4 | 15,1124 | 17,0604 | 0,174919 | 0,29185 | 0,642022 |
| Mier1 | 23,7513 | 21,115 | -0,169732 | 0,2919 | 0,642026 |
| Ppfia1 | 21,2692 | 23,9251 | 0,169756 | 0,29195 | 0,642031 |
| L3mbtl3 | 16,6663 | 18,8317 | 0,176232 | 0,2921 | 0,642153 |
| Phf23 | 72,185 | 81,1769 | 0,16937 | 0,29215 | 0,642153 |
| Gm5595 | 2,63474 | 2,18874 | -0,267559 | 0,29215 | 0,642153 |
| Gtf2h1 | 23,9058 | 26,972 | 0,174103 | 0,2924 | 0,642597 |
| D930028M14Rik | 0,908061 | 1,36496 | 0,588 | 0,29335 | 0,643838 |
| 1810043G02Rik | 9,74076 | 11,3206 | 0,216849 | 0,2932 | 0,643838 |
| Marcksl1 | 417,126 | 470,206 | 0,172809 | 0,2931 | 0,643838 |
| Rpl13a | 526,541 | 593,277 | 0,172161 | 0,29335 | 0,643838 |
| Cox6b1 | 411,655 | 462,614 | 0,168372 | 0,29325 | 0,643838 |
| Ski | 43,023 | 48,2643 | 0,165848 | 0,2933 | 0,643838 |
| Fam102b | 7,16485 | 6,30089 | -0,185382 | 0,29335 | 0,643838 |
| 2700089E24Rik | 24,248 | 21,2844 | -0,188069 | 0,29335 | 0,643838 |
| Jund | 20,4029 | 23,0648 | 0,17692 | 0,29345 | 0,643952 |
| Als2 | 7,88536 | 8,99848 | 0,190505 | 0,29355 | 0,64396 |
| Spdl1 | 25,7779 | 29,1109 | 0,175424 | 0,29355 | 0,64396 |
| Klhl23 | 14,3314 | 12,7017 | -0,174153 | 0,29385 | 0,6443 |
| Bbs5 | 9,15991 | 7,7709 | -0,237251 | 0,29385 | 0,6443 |
| Trim13 | 5,36207 | 4,34858 | -0,302244 | 0,29385 | 0,6443 |

| | | | | | |
|---|---|---|---|---|---|
| Cstf2 | 39,0548 | 43,9719 | 0,171081 | 0,29405 | 0,644633 |
| Hdac3 | 73,7312 | 82,7713 | 0,166856 | 0,29425 | 0,644966 |
| Casq1 | 0,7302 | 0,999588 | 0,453043 | 0,29455 | 0,645215 |
| Msh3 | 18,7469 | 22,6961 | 0,275789 | 0,2946 | 0,645215 |
| Anks3 | 13,9434 | 15,887 | 0,18826 | 0,2945 | 0,645215 |
| Siva1 | 73,3011 | 83,131 | 0,181552 | 0,29475 | 0,645215 |
| Zfp57 | 25,9335 | 29,3843 | 0,180231 | 0,29475 | 0,645215 |
| 9430008C03Rik | 19,466 | 21,9862 | 0,175637 | 0,29465 | 0,645215 |
| Kdelr1 | 82,289 | 73,2805 | -0,167269 | 0,2947 | 0,645215 |
| Ankra2 | 9,30759 | 7,84794 | -0,246093 | 0,29475 | 0,645215 |
| Zfp850 | 5,18875 | 5,89847 | 0,184954 | 0,2951 | 0,645876 |
| Snhg9 | 8,40213 | 19,4638 | 1,21197 | 0,29535 | 0,646317 |
| Pex19 | 53,7863 | 60,2975 | 0,164861 | 0,29545 | 0,64643 |
| Lemd1 | 0,668913 | 0,386235 | -0,792341 | 0,29555 | 0,646543 |
| Ppp1r37 | 24,9284 | 28,0664 | 0,171056 | 0,29565 | 0,64655 |
| Cebpz | 39,5574 | 44,459 | 0,168528 | 0,29565 | 0,64655 |
| Rps4l | 51,4251 | 65,897 | 0,357739 | 0,29575 | 0,646663 |
| Zc3h3 | 0,766772 | 0,978973 | 0,35247 | 0,2959 | 0,646885 |
| Thoc7 | 192,835 | 216,161 | 0,164739 | 0,29595 | 0,646888 |
| Smc4 | 119,216 | 106,066 | -0,168617 | 0,296 | 0,646892 |
| Ssh1 | 5,77596 | 6,68219 | 0,21026 | 0,2962 | 0,647223 |
| Gpbp1 | 62,1515 | 55,4172 | -0,165455 | 0,29635 | 0,647445 |
| Snhg8 | 41,4977 | 36,2045 | -0,196863 | 0,2965 | 0,647667 |
| Gse1 | 21,7296 | 24,9282 | 0,198118 | 0,2968 | 0,648216 |
| Gdpgp1 | 2,08167 | 1,71377 | -0,28057 | 0,29715 | 0,648875 |
| Epc2 | 15,5569 | 17,5066 | 0,170348 | 0,2972 | 0,648878 |
| Matn2 | 1,3467 | 1,66025 | 0,301974 | 0,29795 | 0,648928 |
| Sorbs1 | 5,71823 | 6,51776 | 0,188807 | 0,29775 | 0,648928 |
| Ufsp2 | 26,4451 | 30,0887 | 0,18622 | 0,2974 | 0,648928 |
| Ttc1 | 31,1143 | 35,312 | 0,18258 | 0,29775 | 0,648928 |
| Zer1 | 10,3302 | 11,682 | 0,177415 | 0,29765 | 0,648928 |
| Ccar2 | 56,7977 | 63,9354 | 0,170784 | 0,298 | 0,648928 |
| Usp11 | 28,205 | 31,6909 | 0,168117 | 0,29765 | 0,648928 |
| Fxr2 | 71,2805 | 80,0034 | 0,166554 | 0,29735 | 0,648928 |
| Mcmbp | 36,3522 | 40,7752 | 0,165646 | 0,2976 | 0,648928 |
| Cnot6 | 83,3028 | 74,2094 | -0,166762 | 0,2978 | 0,648928 |
| Fam73b | 13,2195 | 11,6658 | -0,180382 | 0,2977 | 0,648928 |
| Pts | 25,7711 | 22,3403 | -0,206102 | 0,2973 | 0,648928 |
| Pdp1 | 3,23697 | 2,75895 | -0,230527 | 0,2978 | 0,648928 |
| Susd2 | 0,667478 | 0,496474 | -0,427003 | 0,2979 | 0,648928 |
| Ly6g6d | 2,86393 | 1,96694 | -0,542044 | 0,29785 | 0,648928 |
| Rltpr | 1,08145 | 0,613574 | -0,817654 | 0,298 | 0,648928 |
| Uba2 | 139,923 | 156,896 | 0,165172 | 0,29805 | 0,648931 |
| Pafah2 | 3,00238 | 3,55918 | 0,245441 | 0,2983 | 0,64937 |
| Sh3bgrl3 | 60,531 | 68,8079 | 0,184901 | 0,29855 | 0,649808 |
| Alkbh5 | 24,4622 | 27,4526 | 0,16639 | 0,2986 | 0,649811 |
| Sptlc2 | 20,217 | 17,9451 | -0,171978 | 0,2988 | 0,650141 |
| Ptch1 | 37,1865 | 33,1572 | -0,165457 | 0,299 | 0,65047 |

| | | | | | |
|---|---|---|---|---|---|
| Frmd4a | 28,073 | 31,4916 | 0,165784 | 0,2992 | 0,650799 |
| Erap1 | 9,93776 | 8,74665 | -0,184191 | 0,29925 | 0,650802 |
| Zfp51 | 5,74177 | 4,90809 | -0,226333 | 0,2993 | 0,650805 |
| Sox2ot | 0,78801 | 0,302377 | -1,38187 | 0,2994 | 0,650916 |
| Nupl1 | 43,0944 | 48,3543 | 0,166142 | 0,2995 | 0,651028 |
| Gm6524 | 6,83339 | 8,0847 | 0,242593 | 0,29965 | 0,651142 |
| Rrp36 | 29,1225 | 33,1348 | 0,186214 | 0,29965 | 0,651142 |
| Atp5g1 | 3,7383 | 2,85405 | -0,389369 | 0,2999 | 0,651579 |
| Mrpl34 | 82,1021 | 93,1478 | 0,182102 | 0,29995 | 0,651582 |
| Rpl34,Rpl34-ps1 | 32,6241 | 46,5655 | 0,513323 | 0,3001 | 0,651802 |
| Trappc12 | 14,8179 | 13,0723 | -0,180829 | 0,30015 | 0,651804 |
| Sapcd1 | 1,59693 | 2,39668 | 0,585737 | 0,30025 | 0,651842 |
| Emg1 | 145,546 | 166,243 | 0,191812 | 0,3008 | 0,651842 |
| Pcbp2 | 195,632 | 220,894 | 0,175211 | 0,30085 | 0,651842 |
| Arih2 | 14,4944 | 16,3542 | 0,174161 | 0,30075 | 0,651842 |
| Dnmt1 | 94,4992 | 106,327 | 0,170134 | 0,30085 | 0,651842 |
| E2f6 | 44,2851 | 49,665 | 0,165406 | 0,30065 | 0,651842 |
| Lasp1 | 43,52 | 48,7663 | 0,164204 | 0,3005 | 0,651842 |
| Ppp4c | 177,94 | 199,083 | 0,161983 | 0,3007 | 0,651842 |
| Armcx4 | 29,2735 | 26,1444 | -0,163095 | 0,3005 | 0,651842 |
| Zfp354a | 3,018 | 2,56188 | -0,236386 | 0,30045 | 0,651842 |
| Aim1 | 0,802602 | 0,655543 | -0,291993 | 0,30045 | 0,651842 |
| Cers2 | 56,1253 | 45,751 | -0,294849 | 0,30085 | 0,651842 |
| Atp6v1g2 | 3,37576 | 2,6962 | -0,324282 | 0,3007 | 0,651842 |
| Capsl | 0,797692 | 0,484541 | -0,719214 | 0,30035 | 0,651842 |
| Cep170 | 21,1 | 23,659 | 0,165141 | 0,3009 | 0,651844 |
| Hjurp | 10,4428 | 13,6979 | 0,39144 | 0,301 | 0,651955 |
| Stard3nl | 35,3269 | 40,0367 | 0,180553 | 0,3011 | 0,652066 |
| Spred3 | 5,28996 | 6,07297 | 0,199146 | 0,30125 | 0,652182 |
| Phf21a | 13,998 | 15,7131 | 0,166744 | 0,3013 | 0,652182 |
| Tnni1 | 12,5488 | 10,5894 | -0,24493 | 0,3013 | 0,652182 |
| Gng3 | 1,26673 | 2,03908 | 0,68681 | 0,30135 | 0,652185 |
| Myrf | 3,29952 | 3,80293 | 0,204856 | 0,3014 | 0,652187 |
| Ttc37 | 10,7198 | 12,088 | 0,173298 | 0,3015 | 0,652298 |
| Rmnd1 | 2,4519 | 3,77889 | 0,624063 | 0,30165 | 0,652517 |
| Errfi1 | 17,5329 | 15,4916 | -0,178577 | 0,30175 | 0,652522 |
| Fam65a | 8,15718 | 7,15445 | -0,189231 | 0,30175 | 0,652522 |
| Terf2 | 24,1622 | 27,1715 | 0,169341 | 0,3019 | 0,652741 |
| Clcf1 | 3,4134 | 2,56364 | -0,413015 | 0,30205 | 0,65296 |
| Prnp | 28,2607 | 24,8846 | -0,183544 | 0,30215 | 0,65307 |
| Det1 | 7,73718 | 8,94844 | 0,209829 | 0,3026 | 0,653622 |
| Kirrel | 25,1926 | 22,4992 | -0,163125 | 0,30265 | 0,653622 |
| Mettl7a1 | 6,38941 | 5,40676 | -0,240917 | 0,30265 | 0,653622 |
| 2810468N07Rik | 2,83317 | 2,23116 | -0,344624 | 0,30265 | 0,653622 |
| Tmem74 | 1,00834 | 0,710758 | -0,504546 | 0,3025 | 0,653622 |
| Armc10 | 31,3042 | 27,7882 | -0,171885 | 0,3028 | 0,653841 |
| Mylpf | 1,79435 | 2,51231 | 0,485548 | 0,303 | 0,654061 |
| 2610035F20Rik | 7,28494 | 8,7271 | 0,260586 | 0,303 | 0,654061 |

| | | | | | |
|---|---|---|---|---|---|
| Acbd4 | 0,837931 | 0,610829 | -0,456064 | 0,30305 | 0,654063 |
| Tgif2 | 20,3183 | 22,9437 | 0,175319 | 0,30315 | 0,654068 |
| Endov | 9,44454 | 10,6528 | 0,173682 | 0,30315 | 0,654068 |
| Me2 | 36,835 | 41,3528 | 0,166908 | 0,3033 | 0,654286 |
| Arhgef9 | 3,53663 | 3,05185 | -0,212689 | 0,30345 | 0,654504 |
| Canx | 220,684 | 197,791 | -0,173942 | 0,3037 | 0,654728 |
| Tesk1 | 9,99049 | 8,77902 | -0,186495 | 0,30375 | 0,654728 |
| Gap43 | 8,02449 | 6,74287 | -0,251047 | 0,30375 | 0,654728 |
| Atg10 | 2,89473 | 2,32122 | -0,318549 | 0,3037 | 0,654728 |
| Sp2 | 10,7302 | 12,3477 | 0,20256 | 0,3038 | 0,65473 |
| Tmod2 | 3,41016 | 2,99604 | -0,186782 | 0,30385 | 0,654733 |
| Rpa1 | 72,599 | 81,2077 | 0,161666 | 0,3039 | 0,654735 |
| Bid | 8,36253 | 9,69633 | 0,2135 | 0,3041 | 0,654957 |
| Armc9 | 10,0513 | 11,5384 | 0,199067 | 0,30415 | 0,654957 |
| Hoxb3 | 55,5351 | 62,1961 | 0,163424 | 0,30415 | 0,654957 |
| Csnk2a1 | 31,9441 | 28,4701 | -0,166099 | 0,30435 | 0,655282 |
| Dkk1 | 26,1474 | 23,1833 | -0,173584 | 0,3044 | 0,655284 |
| Zfp518b | 19,4849 | 21,8297 | 0,163936 | 0,3045 | 0,655393 |
| Gemin4 | 23,0601 | 25,8994 | 0,167522 | 0,3047 | 0,655718 |
| Clasp2 | 15,7741 | 17,726 | 0,168313 | 0,3049 | 0,656043 |
| Doc2a | 5,72096 | 7,14236 | 0,320142 | 0,30505 | 0,65626 |
| Usp44 | 3,26078 | 3,87825 | 0,25019 | 0,3052 | 0,656477 |
| Orc2 | 40,0524 | 44,8547 | 0,163371 | 0,30525 | 0,656479 |
| Dpp6 | 0,421657 | 0,549676 | 0,38251 | 0,3053 | 0,656481 |
| Agpat5 | 23,1629 | 20,6387 | -0,16646 | 0,30555 | 0,656913 |
| Mettl6 | 30,8667 | 34,7726 | 0,171901 | 0,3059 | 0,656926 |
| Gbas | 27,1722 | 30,5851 | 0,170695 | 0,3057 | 0,656926 |
| Auts2 | 19,9592 | 22,3573 | 0,163695 | 0,30585 | 0,656926 |
| Tiparp | 14,0179 | 12,4307 | -0,17337 | 0,3058 | 0,656926 |
| Adck2 | 5,20819 | 4,49619 | -0,212077 | 0,30575 | 0,656926 |
| Nmi | 4,52875 | 3,67856 | -0,299971 | 0,3058 | 0,656926 |
| A930005H10Rik | 6,15209 | 4,82591 | -0,350276 | 0,3059 | 0,656926 |
| Alkbh8 | 15,1679 | 17,2063 | 0,181916 | 0,3062 | 0,657464 |
| Heca | 6,70949 | 7,67081 | 0,193177 | 0,3063 | 0,657573 |
| Pbx3 | 7,0687 | 6,12844 | -0,205923 | 0,30635 | 0,657575 |
| Angptl6 | 1,02976 | 1,40921 | 0,452571 | 0,30645 | 0,657578 |
| Uso1 | 42,3589 | 47,414 | 0,162648 | 0,30645 | 0,657578 |
| 1700020A23Rik | 0,276599 | 0,589219 | 1,09101 | 0,30675 | 0,658116 |
| Slc9a5 | 7,76297 | 8,78128 | 0,177823 | 0,3072 | 0,658553 |
| Smad2 | 12,1651 | 13,6326 | 0,164314 | 0,3072 | 0,658553 |
| Eml4 | 37,7365 | 42,2018 | 0,161346 | 0,3072 | 0,658553 |
| Vps26a | 22,8116 | 20,2841 | -0,169421 | 0,3072 | 0,658553 |
| Rnf215 | 12,6052 | 10,4398 | -0,271928 | 0,3071 | 0,658553 |
| Zfand3 | 45,2379 | 50,6045 | 0,161733 | 0,30725 | 0,658555 |
| Apool | 34,0979 | 38,833 | 0,187603 | 0,3073 | 0,658556 |
| Rims2 | 1,0419 | 1,25522 | 0,268715 | 0,30735 | 0,658558 |
| Tbc1d12 | 3,91251 | 4,52543 | 0,20996 | 0,3074 | 0,658559 |
| Echdc2 | 11,0291 | 12,8677 | 0,222443 | 0,30755 | 0,65867 |

| | | | | | |
|---|---|---|---|---|---|
| Stx6 | 28,7279 | 32,2337 | 0,166115 | 0,30755 | 0,65867 |
| Psmc1 | 100,766 | 112,867 | 0,163614 | 0,3076 | 0,658671 |
| Mtcl1 | 5,42327 | 6,1249 | 0,175524 | 0,3077 | 0,65878 |
| Mrpl33 | 150,867 | 170,477 | 0,176302 | 0,3078 | 0,658783 |
| Eif4e2 | 59,1119 | 66,5836 | 0,171719 | 0,3078 | 0,658783 |
| Snhg12 | 59,9942 | 68,7368 | 0,196259 | 0,3079 | 0,65879 |
| Copa | 67,6736 | 77,3272 | 0,192384 | 0,30805 | 0,65879 |
| Hnrnpd | 98,9258 | 111,533 | 0,173048 | 0,30795 | 0,65879 |
| Nt5dc2 | 274,79 | 308,768 | 0,168195 | 0,30805 | 0,65879 |
| Metrn | 41,498 | 36,7337 | -0,175937 | 0,30805 | 0,65879 |
| P2ry1 | 1,25408 | 1,01408 | -0,306458 | 0,3083 | 0,659219 |
| M6pr | 46,5725 | 41,5615 | -0,164233 | 0,3084 | 0,659328 |
| Vim | 226,083 | 253,181 | 0,163316 | 0,30845 | 0,659329 |
| Ints6 | 12,4954 | 14,0307 | 0,167186 | 0,30855 | 0,659437 |
| Prim2 | 30,0569 | 33,8427 | 0,171146 | 0,30875 | 0,659759 |
| Acox3 | 3,39295 | 2,89806 | -0,227453 | 0,3088 | 0,659761 |
| Osbpl7 | 3,3714 | 3,9416 | 0,225433 | 0,30885 | 0,659762 |
| Cxadr | 36,3432 | 32,1376 | -0,17742 | 0,30895 | 0,65987 |
| Bbs7 | 7,59395 | 8,73527 | 0,202002 | 0,30925 | 0,660405 |
| F5 | 1,2502 | 1,05094 | -0,250477 | 0,3093 | 0,660407 |
| Tmem189 | 21,6934 | 19,0824 | -0,185015 | 0,3095 | 0,660728 |
| Sac3d1 | 22,6935 | 25,7895 | 0,184507 | 0,3097 | 0,660733 |
| Lonp1 | 48,5848 | 54,3604 | 0,162052 | 0,3097 | 0,660733 |
| Itga5 | 50,5364 | 56,4834 | 0,160503 | 0,3096 | 0,660733 |
| Cln3 | 5,25195 | 4,46724 | -0,233468 | 0,3097 | 0,660733 |
| Cct6a | 391,858 | 442,121 | 0,17411 | 0,3098 | 0,660736 |
| Rab23 | 11,4153 | 12,8701 | 0,173058 | 0,30985 | 0,660736 |
| 2010107E04Rik | 511,605 | 573,294 | 0,164245 | 0,30985 | 0,660736 |
| Msrb2 | 9,41571 | 11,0881 | 0,235872 | 0,30995 | 0,660739 |
| Mettl16 | 28,1278 | 31,5728 | 0,166686 | 0,30995 | 0,660739 |
| Rbfa | 15,4851 | 17,7955 | 0,20063 | 0,31035 | 0,661064 |
| 2610318N02Rik | 18,9766 | 21,6971 | 0,193283 | 0,31025 | 0,661064 |
| Rrm1 | 124,726 | 139,877 | 0,16539 | 0,3103 | 0,661064 |
| Lepre1 | 15,4002 | 13,6248 | -0,176718 | 0,31035 | 0,661064 |
| Tgm4 | 3,38606 | 2,84272 | -0,252338 | 0,31025 | 0,661064 |
| Tmem17 | 10,9304 | 9,35687 | -0,224251 | 0,3105 | 0,661278 |
| Myg1 | 35,2907 | 39,7772 | 0,172655 | 0,31065 | 0,661492 |
| Aamp | 100,232 | 112,044 | 0,160723 | 0,3108 | 0,661601 |
| Hspb1 | 1,41467 | 0,958591 | -0,561483 | 0,3108 | 0,661601 |
| Gyk | 3,39651 | 2,91746 | -0,219338 | 0,31095 | 0,661815 |
| Stam2 | 14,8298 | 16,6849 | 0,170042 | 0,3111 | 0,661819 |
| Cactin | 25,421 | 28,5396 | 0,166941 | 0,3111 | 0,661819 |
| Nup43 | 64,3602 | 72,1977 | 0,165785 | 0,31115 | 0,661819 |
| Maob | 2,22021 | 1,7996 | -0,303015 | 0,31115 | 0,661819 |
| Epc1 | 19,1367 | 22,2285 | 0,216066 | 0,3112 | 0,66182 |
| Birc6 | 14,568 | 16,3331 | 0,164997 | 0,31135 | 0,662034 |
| Ddx50 | 37,1624 | 41,6884 | 0,165803 | 0,31145 | 0,662141 |
| Baz1b | 73,7017 | 82,5583 | 0,163715 | 0,31165 | 0,662461 |

| | | | | | |
|---|---|---|---|---|---|
| Sap25 | 2,32317 | 4,20771 | 0,856938 | 0,31175 | 0,662464 |
| Wdr83os | 39,742 | 45,5332 | 0,196255 | 0,31185 | 0,662464 |
| Taf11 | 47,0843 | 53,0551 | 0,172245 | 0,31185 | 0,662464 |
| Wdr26 | 35,5275 | 31,7821 | -0,160723 | 0,3118 | 0,662464 |
| A2m | 1,15162 | 1,39578 | 0,277408 | 0,3119 | 0,662465 |
| Syne4 | 7,86294 | 9,27084 | 0,237631 | 0,31215 | 0,662787 |
| Metap1d | 11,4359 | 13,2966 | 0,217495 | 0,31225 | 0,662787 |
| C330007P06Rik | 9,3643 | 8,2363 | -0,185175 | 0,3122 | 0,662787 |
| Zfp697 | 3,77282 | 3,27191 | -0,20551 | 0,31225 | 0,662787 |
| Kpna6 | 22,9048 | 25,5885 | 0,15985 | 0,31245 | 0,663107 |
| Gpatch11 | 3,6172 | 4,18118 | 0,209036 | 0,3126 | 0,663109 |
| Ints5 | 23,1447 | 25,9779 | 0,166604 | 0,3126 | 0,663109 |
| Neo1 | 37,3504 | 33,4841 | -0,15765 | 0,3126 | 0,663109 |
| Irf1 | 9,42646 | 10,8144 | 0,198167 | 0,31285 | 0,663323 |
| Snapc5 | 36,4872 | 41,7251 | 0,193526 | 0,31285 | 0,663323 |
| Stam | 19,8203 | 22,2315 | 0,165627 | 0,31285 | 0,663323 |
| Cks1b | 220,919 | 247,187 | 0,162086 | 0,3129 | 0,663324 |
| Gcfc2 | 8,73243 | 10,8267 | 0,310137 | 0,31305 | 0,663537 |
| Rpl18a | 52,1734 | 46,4002 | -0,169183 | 0,31325 | 0,66375 |
| Mon1b | 6,38191 | 5,61236 | -0,185382 | 0,31325 | 0,66375 |
| Tprgl | 33,4215 | 29,6968 | -0,170467 | 0,31335 | 0,663857 |
| Tbca | 184,269 | 207,126 | 0,168695 | 0,3135 | 0,663859 |
| Trpc4ap | 51,5894 | 57,6222 | 0,15955 | 0,3135 | 0,663859 |
| Rbbp6 | 40,9276 | 36,3632 | -0,170598 | 0,3135 | 0,663859 |
| Srsf11 | 112,408 | 125,786 | 0,162225 | 0,31355 | 0,66386 |
| Cep68 | 27,6912 | 30,9711 | 0,161495 | 0,3136 | 0,66386 |
| Zfp260 | 41,7305 | 37,363 | -0,159492 | 0,31365 | 0,663861 |
| Tssc4 | 40,7363 | 36,1297 | -0,17313 | 0,3137 | 0,663862 |
| Rap1gds1 | 28,7971 | 32,2176 | 0,161925 | 0,3138 | 0,663863 |
| Ssna1 | 67,3863 | 59,7112 | -0,174454 | 0,3138 | 0,663863 |
| Vps28 | 46,7725 | 41,1461 | -0,184905 | 0,31385 | 0,663864 |
| Fam72a | 4,06693 | 4,81603 | 0,243903 | 0,31405 | 0,663867 |
| Dus4l | 9,01211 | 10,4386 | 0,211991 | 0,31405 | 0,663867 |
| Ppm1a | 34,7064 | 38,8438 | 0,162483 | 0,31405 | 0,663867 |
| Tns1 | 3,32302 | 2,92693 | -0,183105 | 0,31405 | 0,663867 |
| Diablo | 42,2796 | 47,2783 | 0,161219 | 0,3145 | 0,664713 |
| Golim4 | 28,121 | 25,1621 | -0,16039 | 0,31455 | 0,664713 |
| Brd4 | 37,4343 | 42,1608 | 0,171543 | 0,3146 | 0,664714 |
| Gm10857 | 2,3398 | 2,80268 | 0,26042 | 0,31485 | 0,664718 |
| Mad2l2 | 15,96 | 18,3905 | 0,204503 | 0,3147 | 0,664718 |
| Cuta | 64,8356 | 73,3303 | 0,177623 | 0,31475 | 0,664718 |
| Cspp1 | 14,0397 | 15,7813 | 0,168702 | 0,315 | 0,664718 |
| Tead3 | 8,18195 | 7,09432 | -0,20578 | 0,31495 | 0,664718 |
| Ypel5 | 5,74012 | 4,95745 | -0,211483 | 0,31495 | 0,664718 |
| Grrp1 | 11,3213 | 9,65637 | -0,229481 | 0,315 | 0,664718 |
| Crbn | 14,3082 | 12,0097 | -0,252636 | 0,31495 | 0,664718 |
| Wdr65 | 0,779289 | 0,977075 | 0,326312 | 0,3151 | 0,664824 |
| Gdpd3 | 1,28103 | 0,887348 | -0,529738 | 0,31515 | 0,664825 |

| | | | | | |
|---|---|---|---|---|---|
| Lhfpl3 | 0,954126 | 1,21154 | 0,344596 | 0,3153 | 0,664931 |
| Hcfc1 | 58,7104 | 65,7155 | 0,162617 | 0,31525 | 0,664931 |
| Upf3a | 25,8169 | 29,2525 | 0,180244 | 0,3155 | 0,665143 |
| Iars | 56,3722 | 62,9911 | 0,160163 | 0,3155 | 0,665143 |
| Fgd1 | 14,0207 | 16,3925 | 0,22548 | 0,3156 | 0,665144 |
| Nsf | 17,2478 | 19,3504 | 0,165952 | 0,31555 | 0,665144 |
| Barx1 | 0,42979 | 0,662637 | 0,624588 | 0,31585 | 0,665566 |
| Jkamp | 21,3151 | 18,8273 | -0,17905 | 0,316 | 0,665672 |
| Angel1 | 5,97498 | 5,21139 | -0,197267 | 0,31595 | 0,665672 |
| 2200002D01Rik | 3,5164 | 6,47693 | 0,881212 | 0,3166 | 0,666726 |
| Ahdc1 | 10,9992 | 12,3402 | 0,165962 | 0,31655 | 0,666726 |
| Xrcc2 | 11,1745 | 12,6454 | 0,178411 | 0,3169 | 0,667147 |
| Stag3 | 1,56107 | 1,2915 | -0,273489 | 0,3169 | 0,667147 |
| Ppp2r5d | 60,4409 | 70,4737 | 0,22156 | 0,3171 | 0,667253 |
| Nsg2 | 20,6894 | 23,3159 | 0,172421 | 0,31705 | 0,667253 |
| Rrp1 | 131,462 | 146,71 | 0,158318 | 0,3171 | 0,667253 |
| Cndp2 | 35,7819 | 40,0929 | 0,16412 | 0,3173 | 0,667464 |
| Layn | 1,02912 | 0,762925 | -0,431801 | 0,3173 | 0,667464 |
| Gm5129 | 0,192888 | 0,980607 | 2,34591 | 0,31745 | 0,667674 |
| Suco | 18,4301 | 16,4691 | -0,162304 | 0,31755 | 0,667779 |
| Pitpnm1 | 4,07485 | 4,98247 | 0,290114 | 0,31775 | 0,668095 |
| Zkscan3 | 34,3214 | 38,3629 | 0,160601 | 0,3179 | 0,668095 |
| Sec31a | 35,7439 | 39,9168 | 0,159298 | 0,3178 | 0,668095 |
| 1110006O24Rik | 2,01557 | 1,4156 | -0,509772 | 0,31785 | 0,668095 |
| Taf3 | 6,48611 | 7,34494 | 0,179398 | 0,318 | 0,6682 |
| Acadl | 30,4332 | 34,1751 | 0,167297 | 0,31805 | 0,6682 |
| Rbbp8 | 23,0295 | 25,7926 | 0,163474 | 0,3181 | 0,6682 |
| Bmyc | 25,7522 | 29,581 | 0,199976 | 0,3183 | 0,66841 |
| Picalm | 65,7003 | 58,9254 | -0,157012 | 0,3183 | 0,66841 |
| Jarid2 | 28,3668 | 31,6578 | 0,158356 | 0,3184 | 0,668515 |
| Abcc10 | 2,33643 | 2,72357 | 0,221195 | 0,31885 | 0,668619 |
| Atp5j | 204,546 | 238,131 | 0,219329 | 0,31875 | 0,668619 |
| Cdt1 | 41,4233 | 46,3876 | 0,163297 | 0,3192 | 0,668619 |
| Sart3 | 46,63 | 52,0605 | 0,158928 | 0,31915 | 0,668619 |
| Prpf4b | 64,8942 | 72,4484 | 0,158863 | 0,3185 | 0,668619 |
| Icmt | 22,8694 | 20,4758 | -0,159497 | 0,3192 | 0,668619 |
| Klhl24 | 7,7678 | 6,89043 | -0,172913 | 0,31895 | 0,668619 |
| Fam219b | 10,4709 | 9,22985 | -0,182008 | 0,31855 | 0,668619 |
| Insig2 | 12,548 | 11,0519 | -0,183173 | 0,319 | 0,668619 |
| Gm2027 | 1,89709 | 1,55459 | -0,287254 | 0,3192 | 0,668619 |
| Amigo2 | 2,63899 | 2,15976 | -0,28912 | 0,3191 | 0,668619 |
| Ggct | 4,36051 | 3,56126 | -0,292107 | 0,3187 | 0,668619 |
| Nipa1 | 7,53423 | 6,14373 | -0,294345 | 0,3188 | 0,668619 |
| Cenpb | 42,3899 | 33,1988 | -0,352589 | 0,3191 | 0,668619 |
| 2700069I18Rik | 1,0658 | 0,074019 | -3,84789 | 0,31875 | 0,668619 |
| Ccdc117 | 25,9626 | 29,0148 | 0,160355 | 0,31935 | 0,668723 |
| Copz1 | 115,936 | 129,467 | 0,159254 | 0,3194 | 0,668723 |
| Plin3 | 22,4893 | 19,9725 | -0,171223 | 0,31945 | 0,668723 |

| | | | | | |
|---|---|---|---|---|---|
| Surf4 | 105,32 | 93,2331 | -0,175862 | 0,3193 | 0,668723 |
| Kcnj13 | 0,087878 | 0,791324 | 3,1707 | 0,3196 | 0,668932 |
| Snip1 | 22,6517 | 25,4589 | 0,16855 | 0,31965 | 0,668932 |
| Bud31 | 147,247 | 167,705 | 0,18769 | 0,3199 | 0,669351 |
| Ptrhd1 | 5,27513 | 6,59605 | 0,322395 | 0,3203 | 0,669873 |
| Dtwd2 | 1,99004 | 2,38475 | 0,261039 | 0,3202 | 0,669873 |
| Zfp574 | 9,87158 | 11,1179 | 0,17153 | 0,3203 | 0,669873 |
| Slc7a7 | 5,12399 | 6,95989 | 0,441798 | 0,3218 | 0,670074 |
| Foxn1 | 0,522565 | 0,685698 | 0,391963 | 0,32105 | 0,670074 |
| Car15 | 1,91863 | 2,49989 | 0,381789 | 0,3217 | 0,670074 |
| Letm2 | 2,11698 | 2,59217 | 0,292153 | 0,32165 | 0,670074 |
| Rad54l | 24,1047 | 28,4876 | 0,241018 | 0,32135 | 0,670074 |
| Eif4e3 | 7,28664 | 8,3596 | 0,19818 | 0,32185 | 0,670074 |
| Ctgf | 10,7123 | 12,1964 | 0,187187 | 0,32075 | 0,670074 |
| Ftsj1 | 8,61726 | 9,76476 | 0,180355 | 0,32185 | 0,670074 |
| Mrpl4 | 30,518 | 34,4644 | 0,175449 | 0,32105 | 0,670074 |
| Trmt6 | 30,902 | 34,5442 | 0,160742 | 0,32085 | 0,670074 |
| Kcmf1 | 30,5888 | 34,1898 | 0,160563 | 0,32175 | 0,670074 |
| Pacsin2 | 22,9829 | 25,6802 | 0,160095 | 0,3217 | 0,670074 |
| Ccdc50 | 38,5319 | 34,5237 | -0,158463 | 0,3208 | 0,670074 |
| Osbp | 18,0174 | 16,1004 | -0,162296 | 0,32185 | 0,670074 |
| Ism1 | 25,0384 | 22,3398 | -0,164526 | 0,3207 | 0,670074 |
| Nadk2 | 21,0529 | 18,6123 | -0,177758 | 0,32075 | 0,670074 |
| Bcl9l | 5,72387 | 5,0423 | -0,182911 | 0,32125 | 0,670074 |
| Gcc1 | 5,56471 | 4,87234 | -0,191692 | 0,32095 | 0,670074 |
| Kdsr | 3,88882 | 3,3996 | -0,193965 | 0,32095 | 0,670074 |
| Tmem135 | 5,76078 | 5,02559 | -0,196972 | 0,3215 | 0,670074 |
| Fam120c | 2,23765 | 1,94002 | -0,205916 | 0,321 | 0,670074 |
| Atxn3 | 9,55677 | 8,22025 | -0,217341 | 0,32125 | 0,670074 |
| Cdh4 | 1,69415 | 1,44166 | -0,232831 | 0,32065 | 0,670074 |
| Herc6 | 1,39873 | 1,166 | -0,262545 | 0,3214 | 0,670074 |
| Derl2 | 14,7964 | 12,1916 | -0,279367 | 0,32155 | 0,670074 |
| Ano8 | 8,85836 | 7,24314 | -0,290425 | 0,32055 | 0,670074 |
| Lrrc15 | 0,592693 | 0,469163 | -0,337196 | 0,32175 | 0,670074 |
| Ntn4 | 1,00874 | 0,77294 | -0,384127 | 0,3217 | 0,670074 |
| Gpr182 | 0,651597 | 0,446467 | -0,545426 | 0,32155 | 0,670074 |
| Ifit2 | 2,38541 | 2,01533 | -0,24322 | 0,322 | 0,670281 |
| Spata24 | 12,4299 | 10,3261 | -0,267524 | 0,32205 | 0,670281 |
| Atp1b2 | 6,24653 | 5,42672 | -0,202975 | 0,3222 | 0,670489 |
| Ctps2 | 13,7253 | 17,4342 | 0,345079 | 0,32245 | 0,670692 |
| Il17d | 3,7674 | 4,61159 | 0,291694 | 0,32295 | 0,670692 |
| Gins2 | 66,3139 | 81,0084 | 0,28876 | 0,3229 | 0,670692 |
| Mrps31 | 26,9274 | 30,3905 | 0,174549 | 0,3226 | 0,670692 |
| Wbscr16 | 16,397 | 18,4753 | 0,172163 | 0,3226 | 0,670692 |
| Pdk1 | 16,4598 | 18,4375 | 0,163691 | 0,3224 | 0,670692 |
| Zfp592 | 11,8781 | 13,2835 | 0,161335 | 0,32265 | 0,670692 |
| Thoc3 | 43,6651 | 48,7676 | 0,159442 | 0,3225 | 0,670692 |
| Rars | 93,9597 | 104,733 | 0,156598 | 0,3229 | 0,670692 |

| | | | | | |
|---|---|---|---|---|---|
| Cant1 | 24,3187 | 21,6779 | -0,165843 | 0,3229 | 0,670692 |
| Zkscan8 | 10,6184 | 9,41368 | -0,173733 | 0,32295 | 0,670692 |
| St7 | 11,691 | 10,1954 | -0,197471 | 0,3228 | 0,670692 |
| Hint3 | 9,38469 | 7,64262 | -0,296241 | 0,3224 | 0,670692 |
| B3gat2 | 0,615692 | 1,12474 | 0,869307 | 0,3234 | 0,671313 |
| S100a13 | 5,89509 | 7,33958 | 0,316182 | 0,32335 | 0,671313 |
| Palb2 | 9,02125 | 10,2877 | 0,189522 | 0,32345 | 0,671313 |
| Prepl | 18,0022 | 20,2141 | 0,167188 | 0,3235 | 0,671313 |
| Snora33 | 295,507 | 163,684 | -0,852277 | 0,3234 | 0,671313 |
| Vhl | 19,1104 | 21,4819 | 0,168767 | 0,3236 | 0,671416 |
| Arhgap17 | 16,0863 | 18,0453 | 0,165796 | 0,32385 | 0,671621 |
| Sf3b2 | 153,674 | 171,491 | 0,158254 | 0,3239 | 0,671621 |
| Arel1 | 14,9256 | 13,3506 | -0,160882 | 0,3238 | 0,671621 |
| Dgcr8 | 27,3683 | 23,5188 | -0,218688 | 0,3238 | 0,671621 |
| Hmgcr | 82,9481 | 92,6018 | 0,158832 | 0,3241 | 0,671932 |
| Chd1l | 12,1403 | 13,6815 | 0,172418 | 0,3243 | 0,672242 |
| Uqcrh | 361,34 | 403,648 | 0,159741 | 0,3244 | 0,672345 |
| Thoc2 | 23,285 | 20,9048 | -0,155561 | 0,32445 | 0,672345 |
| Tubb5 | 1062,68 | 1240,37 | 0,223058 | 0,32465 | 0,672655 |
| Eif3j1,Eif3j2 | 2,19517 | 4,70514 | 1,09991 | 0,32475 | 0,672757 |
| Jdp2 | 8,44741 | 9,80637 | 0,215211 | 0,3248 | 0,672757 |
| Steap1 | 7,19458 | 6,02387 | -0,256221 | 0,32515 | 0,673377 |
| Spata17 | 0,361959 | 0,582088 | 0,685413 | 0,3255 | 0,673893 |
| Pcmtd2 | 19,6637 | 17,5813 | -0,161493 | 0,32545 | 0,673893 |
| Gm7102 | 1,16741 | 1,51575 | 0,376722 | 0,326 | 0,673989 |
| Baz2a | 21,7212 | 24,2157 | 0,15684 | 0,32575 | 0,673989 |
| U2af2 | 92,3324 | 102,93 | 0,156753 | 0,32565 | 0,673989 |
| Lrpprc | 33,4115 | 37,2197 | 0,155725 | 0,32595 | 0,673989 |
| Dock1 | 20,6071 | 22,9495 | 0,155325 | 0,3259 | 0,673989 |
| Ppm1l | 3,8202 | 3,30321 | -0,209778 | 0,32595 | 0,673989 |
| Lyrm5 | 11,4436 | 9,88465 | -0,211273 | 0,3258 | 0,673989 |
| F930015N05Rik | 1,74573 | 1,36738 | -0,352414 | 0,32575 | 0,673989 |
| Fam196b | 0,958727 | 0,733494 | -0,386336 | 0,3257 | 0,673989 |
| Kctd20 | 43,9945 | 49,1241 | 0,159108 | 0,32615 | 0,674195 |
| Eif5a2 | 9,13231 | 10,2843 | 0,171388 | 0,3263 | 0,6744 |
| Btf3l4 | 4,37024 | 3,75857 | -0,217529 | 0,32645 | 0,674606 |
| Ubb | 133,097 | 152,947 | 0,200546 | 0,3268 | 0,675016 |
| Rif1 | 34,3949 | 38,3444 | 0,156821 | 0,3268 | 0,675016 |
| Ankrd12 | 2,41301 | 2,11345 | -0,191235 | 0,3268 | 0,675016 |
| Tceal5 | 2,40648 | 1,81783 | -0,40471 | 0,3269 | 0,675118 |
| Mt3 | 1,47083 | 0,882188 | -0,737473 | 0,327 | 0,675221 |
| Uqcrfs1 | 118,957 | 132,65 | 0,157187 | 0,3271 | 0,675323 |
| Noc4l | 34,5754 | 38,6713 | 0,161518 | 0,3273 | 0,675422 |
| Gcc2 | 11,7263 | 13,0979 | 0,159579 | 0,32725 | 0,675422 |
| Zswim8 | 24,5544 | 27,3486 | 0,155483 | 0,3272 | 0,675422 |
| Nus1 | 33,6577 | 30,2532 | -0,153851 | 0,32735 | 0,675422 |
| 0610007P14Rik | 55,9347 | 49,8846 | -0,165149 | 0,32755 | 0,67573 |
| Coil | 17,8601 | 20,0687 | 0,168207 | 0,32815 | 0,676863 |

| | | | | | |
|---|---|---|---|---|---|
| C030037D09Rik | 2,49413 | 3,17337 | 0,347479 | 0,32875 | 0,677677 |
| Glt1d1 | 2,61231 | 3,21412 | 0,299094 | 0,32865 | 0,677677 |
| Zfp110 | 12,3522 | 13,8735 | 0,167566 | 0,3288 | 0,677677 |
| Pdzrn3 | 28,7167 | 32,032 | 0,157626 | 0,3289 | 0,677677 |
| Zfp462 | 15,401 | 17,1492 | 0,155112 | 0,32885 | 0,677677 |
| Lfng | 69,2588 | 62,1352 | -0,156587 | 0,32865 | 0,677677 |
| Pgpep1 | 6,42832 | 5,67603 | -0,179559 | 0,3288 | 0,677677 |
| Gm9776 | 1,59498 | 1,27128 | -0,327258 | 0,32895 | 0,677677 |
| Zfp746 | 15,3498 | 17,2027 | 0,164417 | 0,32905 | 0,677778 |
| Rpl4 | 1004,47 | 1137,43 | 0,179334 | 0,32935 | 0,677977 |
| Dhx9 | 207,054 | 232,952 | 0,170031 | 0,3292 | 0,677977 |
| Znhit6 | 14,9895 | 16,849 | 0,16871 | 0,32925 | 0,677977 |
| Rae1 | 56,7719 | 63,3433 | 0,158014 | 0,3293 | 0,677977 |
| Slc35a1 | 43,8355 | 39,2533 | -0,159284 | 0,3294 | 0,677977 |
| Sirt3 | 9,19036 | 11,5441 | 0,32896 | 0,32955 | 0,678178 |
| Crtc1 | 5,86402 | 5,189 | -0,176434 | 0,32965 | 0,678178 |
| Ddrgk1 | 27,4742 | 24,1565 | -0,185663 | 0,3296 | 0,678178 |
| Ddit4 | 89,7947 | 100,026 | 0,155666 | 0,32985 | 0,678485 |
| Hoxb5 | 9,91234 | 11,3476 | 0,195086 | 0,33 | 0,678689 |
| Stx7 | 13,97 | 12,3647 | -0,176107 | 0,3301 | 0,67879 |
| Prrx1 | 6,09966 | 6,92575 | 0,183242 | 0,3302 | 0,678891 |
| Ncdn | 13,9934 | 15,6901 | 0,165113 | 0,3303 | 0,678991 |
| Rgl1 | 8,4763 | 7,49329 | -0,177834 | 0,33035 | 0,678991 |
| Acsl3 | 46,9044 | 41,5659 | -0,174322 | 0,3306 | 0,6794 |
| Spcs1 | 81,1025 | 62,9371 | -0,365837 | 0,3308 | 0,679707 |
| Zfp637 | 26,4793 | 30,0504 | 0,182519 | 0,331 | 0,679908 |
| Tmc4 | 3,33181 | 1,9765 | -0,753362 | 0,33095 | 0,679908 |
| Pigw | 11,2069 | 8,88009 | -0,33574 | 0,33115 | 0,680112 |
| Itgb3bp | 7,33527 | 6,40055 | -0,196654 | 0,33135 | 0,680418 |
| Yars | 41,6978 | 46,851 | 0,168109 | 0,3315 | 0,680517 |
| Lmo4 | 90,2036 | 100,501 | 0,155952 | 0,33145 | 0,680517 |
| Rpl22 | 155,526 | 173,287 | 0,156008 | 0,3318 | 0,680924 |
| Sec16a | 17,936 | 19,9565 | 0,154005 | 0,33175 | 0,680924 |
| Mlxipl | 0,598125 | 0,760989 | 0,347428 | 0,33225 | 0,681743 |
| Tex264 | 12,2201 | 10,5589 | -0,210795 | 0,33235 | 0,681843 |
| Tomm70a | 89,8537 | 80,8961 | -0,151507 | 0,33285 | 0,682764 |
| Cyb5r3 | 78,0171 | 86,7916 | 0,153765 | 0,33295 | 0,682864 |
| Afap1 | 4,21812 | 4,77066 | 0,177588 | 0,33315 | 0,683056 |
| 2610002J02Rik | 36,8685 | 41,5722 | 0,173228 | 0,33335 | 0,683056 |
| Pagr1a | 41,5168 | 46,4609 | 0,16232 | 0,33325 | 0,683056 |
| Ube2d1 | 50,2952 | 56,1793 | 0,159617 | 0,33325 | 0,683056 |
| Nsmaf | 27,213 | 24,3951 | -0,1577 | 0,3333 | 0,683056 |
| Sirt5 | 5,09174 | 4,21555 | -0,272438 | 0,3331 | 0,683056 |
| AU015836 | 0,616014 | 0,796459 | 0,370636 | 0,33365 | 0,683566 |
| Lrrn3 | 3,90663 | 3,35342 | -0,220292 | 0,33375 | 0,683666 |
| Pygl | 15,6076 | 17,5279 | 0,167411 | 0,33385 | 0,683766 |
| Tpk1 | 2,47525 | 2,9608 | 0,258415 | 0,33405 | 0,683861 |
| Vdac3 | 34,432 | 38,6471 | 0,166607 | 0,334 | 0,683861 |

| | | | | | |
|---|---|---|---|---|---|
| Cluh | 29,5339 | 32,9036 | 0,155874 | 0,33395 | 0,683861 |
| Tgif1 | 71,1155 | 79,3996 | 0,158969 | 0,33415 | 0,683961 |
| Tubgcp6 | 13,111 | 14,7182 | 0,166827 | 0,3343 | 0,684163 |
| Pard6a | 4,33947 | 5,23141 | 0,269681 | 0,33445 | 0,684365 |
| Phldb1 | 19,3368 | 21,5134 | 0,153888 | 0,33455 | 0,684465 |
| Wibg | 44,2576 | 49,6093 | 0,164686 | 0,33475 | 0,684769 |
| 1110004E09Rik | 24,5444 | 27,8089 | 0,180154 | 0,33505 | 0,684964 |
| Stxbp2 | 10,907 | 9,61339 | -0,182141 | 0,33495 | 0,684964 |
| Ildr2 | 2,62214 | 2,29613 | -0,191536 | 0,33505 | 0,684964 |
| Dyrk1b | 6,09488 | 5,26931 | -0,209985 | 0,335 | 0,684964 |
| H2-Ke6 | 20,581 | 17,8729 | -0,203538 | 0,33525 | 0,685268 |
| Ppp2r1b | 59,2461 | 66,1524 | 0,159073 | 0,33535 | 0,685365 |
| Ncaph | 57,741 | 64,2463 | 0,154019 | 0,3354 | 0,685365 |
| Dennd1a | 6,20166 | 7,9928 | 0,366047 | 0,3356 | 0,685459 |
| Glrx3 | 28,1528 | 31,5463 | 0,164194 | 0,3356 | 0,685459 |
| Scrn3 | 3,53074 | 3,02157 | -0,224671 | 0,3355 | 0,685459 |
| Ahsa2 | 29,8408 | 37,0474 | 0,312086 | 0,3366 | 0,685613 |
| Acn9 | 16,372 | 19,5609 | 0,256747 | 0,3365 | 0,685613 |
| Lonrf2 | 1,42366 | 1,68336 | 0,241743 | 0,33655 | 0,685613 |
| Mir682 | 8338,57 | 9620,29 | 0,206279 | 0,3358 | 0,685613 |
| Siah2 | 6,864 | 7,87098 | 0,197494 | 0,336 | 0,685613 |
| Dnal4 | 23,4054 | 26,4318 | 0,175429 | 0,336 | 0,685613 |
| Fat4 | 3,78222 | 4,23329 | 0,162548 | 0,3365 | 0,685613 |
| Dctd | 41,5429 | 46,4596 | 0,161377 | 0,33595 | 0,685613 |
| Sf3a2 | 57,1651 | 63,7884 | 0,158159 | 0,3361 | 0,685613 |
| Cfl1 | 549,344 | 612,12 | 0,156104 | 0,3366 | 0,685613 |
| Ppp1cb | 132,762 | 119,103 | -0,156634 | 0,3361 | 0,685613 |
| Slc25a46 | 15,8153 | 14,1579 | -0,159711 | 0,3366 | 0,685613 |
| Nfe2l2 | 19,5396 | 17,3992 | -0,167376 | 0,33665 | 0,685613 |
| Iars2 | 33,9776 | 29,7932 | -0,189598 | 0,3365 | 0,685613 |
| Lyst | 1,28257 | 1,11275 | -0,204909 | 0,3363 | 0,685613 |
| Lrrc51 | 4,45258 | 3,54489 | -0,3289 | 0,3361 | 0,685613 |
| Dnase1l2 | 2,66154 | 2,11035 | -0,33478 | 0,33665 | 0,685613 |
| Traf3ip2 | 1,66064 | 1,25371 | -0,405534 | 0,3359 | 0,685613 |
| Gapdhs | 1,15358 | 0,855132 | -0,431897 | 0,33615 | 0,685613 |
| Dnaaf2 | 15,1372 | 17,2551 | 0,188927 | 0,3368 | 0,685704 |
| Nme1 | 226,16 | 251,073 | 0,150761 | 0,3369 | 0,685704 |
| Lpl | 17,6552 | 15,803 | -0,159897 | 0,3369 | 0,685704 |
| Sox2 | 21,785 | 19,414 | -0,166243 | 0,33675 | 0,685704 |
| Wdr60 | 5,23752 | 5,96789 | 0,188338 | 0,33705 | 0,6858 |
| Med26 | 7,08977 | 8,06532 | 0,185992 | 0,337 | 0,6858 |
| Aif1l | 37,0175 | 41,1811 | 0,153773 | 0,33725 | 0,685894 |
| Fam49b | 27,6935 | 24,8902 | -0,15397 | 0,33715 | 0,685894 |
| Eno1b | 4,21184 | 2,74606 | -0,61709 | 0,33725 | 0,685894 |
| Slc10a4 | 0,465846 | 0,646783 | 0,473428 | 0,3378 | 0,68607 |
| A730008H23Rik | 28,3814 | 38,666 | 0,446122 | 0,33785 | 0,68607 |
| Mrpl54 | 103,802 | 116,482 | 0,166274 | 0,33775 | 0,68607 |
| Prkra | 46,9292 | 52,3607 | 0,157998 | 0,33745 | 0,68607 |

| | | | | | |
|---|---|---|---|---|---|
| Brpf1 | 18,4062 | 20,5078 | 0,155985 | 0,3377 | 0,68607 |
| Angel2 | 29,4048 | 32,7557 | 0,155695 | 0,33765 | 0,68607 |
| Pvrl3 | 28,1723 | 25,1973 | -0,161009 | 0,3374 | 0,68607 |
| Pggt1b | 15,2956 | 13,5956 | -0,169974 | 0,33775 | 0,68607 |
| Wsb2 | 11,5311 | 10,1671 | -0,181616 | 0,3378 | 0,68607 |
| Mcur1 | 5,71617 | 5,03784 | -0,182244 | 0,33765 | 0,68607 |
| BC089491 | 2,33075 | 2,92619 | 0,328233 | 0,338 | 0,686163 |
| Plbd2 | 17,9661 | 16,0727 | -0,160667 | 0,338 | 0,686163 |
| Mdk | 1233,15 | 1102,28 | -0,161853 | 0,33805 | 0,686163 |
| Parp16 | 4,57722 | 5,30204 | 0,212075 | 0,33825 | 0,686361 |
| Prex2 | 6,64028 | 5,95224 | -0,157811 | 0,3382 | 0,686361 |
| Ncan | 0,59685 | 0,485698 | -0,297309 | 0,3384 | 0,686561 |
| Pcid2 | 23,6567 | 26,4642 | 0,161793 | 0,3386 | 0,686755 |
| Naa50 | 76,2452 | 68,6419 | -0,151557 | 0,33855 | 0,686755 |
| H6pd | 5,6915 | 5,02541 | -0,179565 | 0,33865 | 0,686755 |
| Cntfr | 11,9174 | 13,6478 | 0,195593 | 0,33905 | 0,687358 |
| Hbegf | 3,58869 | 3,02923 | -0,244504 | 0,33905 | 0,687358 |
| Cyb5b | 76,8368 | 85,4283 | 0,152916 | 0,3392 | 0,687454 |
| Pcbd1 | 17,8563 | 15,3614 | -0,217126 | 0,33915 | 0,687454 |
| Mpnd | 49,4618 | 57,7921 | 0,224556 | 0,33935 | 0,687653 |
| Trim41 | 21,6598 | 24,1203 | 0,155229 | 0,3396 | 0,688056 |
| Ppp4r1l-ps | 2,44905 | 2,096 | -0,224584 | 0,33975 | 0,688255 |
| Cdpf1 | 11,463 | 13,4553 | 0,231198 | 0,33985 | 0,688353 |
| Dhx35 | 12,9328 | 14,4918 | 0,164198 | 0,34035 | 0,689262 |
| Zfp688 | 8,86546 | 10,4336 | 0,234965 | 0,34065 | 0,689657 |
| Mob3b | 3,30727 | 3,76241 | 0,186017 | 0,34065 | 0,689657 |
| G6pc2 | 0,566213 | 0,398406 | -0,507105 | 0,3407 | 0,689657 |
| Rbm15b | 38,3717 | 42,6269 | 0,151724 | 0,34095 | 0,690058 |
| D11Wsu47e | 13,9283 | 15,7135 | 0,17399 | 0,34125 | 0,690558 |
| Psmb6 | 242,381 | 269,418 | 0,15257 | 0,3413 | 0,690558 |
| Ccdc157 | 5,9001 | 6,80484 | 0,205823 | 0,3415 | 0,690649 |
| Skp2 | 52,703 | 58,9235 | 0,160959 | 0,3414 | 0,690649 |
| Wfs1 | 3,56239 | 3,06942 | -0,214881 | 0,34145 | 0,690649 |
| Cpt1b | 1,27887 | 1,82422 | 0,512413 | 0,3417 | 0,690712 |
| Map3k13 | 0,466887 | 0,577101 | 0,305751 | 0,34195 | 0,690712 |
| Tgfb1i1 | 6,37031 | 7,20927 | 0,178491 | 0,34205 | 0,690712 |
| Anxa5 | 44,7102 | 49,8399 | 0,156697 | 0,34185 | 0,690712 |
| Srp14 | 114,312 | 127,312 | 0,155392 | 0,34185 | 0,690712 |
| Xiap | 24,6731 | 22,21 | -0,151728 | 0,34205 | 0,690712 |
| Frmd8 | 21,7913 | 19,5093 | -0,159587 | 0,3421 | 0,690712 |
| Pif1 | 10,132 | 9,01612 | -0,168342 | 0,342 | 0,690712 |
| Katnal1 | 3,75612 | 3,30019 | -0,186694 | 0,342 | 0,690712 |
| Rps6ka4 | 1,48411 | 1,2133 | -0,290661 | 0,34185 | 0,690712 |
| Ankdd1b | 0,985617 | 0,786576 | -0,325441 | 0,342 | 0,690712 |
| Wdr33 | 41,091 | 47,4966 | 0,209002 | 0,3422 | 0,6908 |
| Zbtb48 | 9,96924 | 11,2959 | 0,180242 | 0,3423 | 0,6908 |
| Dolpp1 | 18,3489 | 15,8574 | -0,210538 | 0,34225 | 0,6908 |
| Map3k6 | 0,828752 | 0,667855 | -0,311406 | 0,34235 | 0,6908 |

| | | | | | |
|---|---|---|---|---|---|
| Ncoa3 | 9,26331 | 10,3227 | 0,156227 | 0,3426 | 0,691095 |
| Brox | 9,48429 | 8,40599 | -0,174123 | 0,34255 | 0,691095 |
| Rab7l1 | 2,09882 | 1,61579 | -0,37734 | 0,3427 | 0,691193 |
| 1110008P14Rik | 15,7988 | 18,5968 | 0,235238 | 0,3428 | 0,69129 |
| Actr1b | 74,1193 | 82,3994 | 0,152785 | 0,3429 | 0,691387 |
| Cyp4f13 | 0,541997 | 0,758772 | 0,485383 | 0,34315 | 0,691464 |
| Ankrd42 | 2,46809 | 2,92309 | 0,244098 | 0,343 | 0,691464 |
| Rhot2 | 31,2414 | 36,0973 | 0,208429 | 0,34325 | 0,691464 |
| Fn3krp | 7,08479 | 8,14703 | 0,201549 | 0,34325 | 0,691464 |
| Dbnl | 40,2066 | 44,7686 | 0,155056 | 0,3433 | 0,691464 |
| Wdfy3 | 10,8913 | 12,1107 | 0,153106 | 0,3433 | 0,691464 |
| Prkar2a | 33,2368 | 36,8675 | 0,149568 | 0,34325 | 0,691464 |
| Ccsap | 3,9273 | 4,56839 | 0,218147 | 0,34345 | 0,691662 |
| Zfp142 | 6,36112 | 8,42846 | 0,405989 | 0,3437 | 0,691748 |
| Rnaseh1 | 17,4628 | 19,8034 | 0,181461 | 0,34355 | 0,691748 |
| Tdrd1 | 1,41078 | 1,18741 | -0,24868 | 0,3436 | 0,691748 |
| Atp8a2 | 1,19025 | 0,97278 | -0,291082 | 0,3437 | 0,691748 |
| 2210016F16Rik | 8,85248 | 10,0202 | 0,178757 | 0,344 | 0,692032 |
| Vta1 | 35,4129 | 39,6737 | 0,163907 | 0,34405 | 0,692032 |
| Nxt2 | 4,77012 | 4,10214 | -0,217647 | 0,34395 | 0,692032 |
| Nudt10 | 5,15823 | 4,37198 | -0,23859 | 0,3441 | 0,692032 |
| Evi5l | 4,06187 | 3,36218 | -0,272748 | 0,34395 | 0,692032 |
| Mir7-1 | 25,4517 | 81,7025 | 1,68262 | 0,34455 | 0,692104 |
| Nup88 | 78,4524 | 89,515 | 0,190312 | 0,34425 | 0,692104 |
| Gltscr1l | 3,71526 | 4,19939 | 0,176717 | 0,34435 | 0,692104 |
| Dbr1 | 17,8848 | 20,0753 | 0,166686 | 0,34445 | 0,692104 |
| Rin2 | 11,4653 | 12,8255 | 0,161736 | 0,34455 | 0,692104 |
| Uqcc1 | 19,507 | 21,7808 | 0,159067 | 0,34435 | 0,692104 |
| Dnaja2 | 108,454 | 120,386 | 0,150592 | 0,3443 | 0,692104 |
| Fancf | 4,74549 | 3,9943 | -0,248615 | 0,3444 | 0,692104 |
| Mgea5 | 66,9645 | 74,4123 | 0,152145 | 0,34465 | 0,692201 |
| Simc1 | 10,6742 | 12,9574 | 0,279643 | 0,34475 | 0,692294 |
| Fam126b | 2,63447 | 2,32009 | -0,18333 | 0,3448 | 0,692294 |
| Rad54b | 9,30732 | 10,679 | 0,198339 | 0,34495 | 0,692383 |
| Samd10 | 3,91779 | 3,30107 | -0,247105 | 0,3449 | 0,692383 |
| AI464131 | 1,1165 | 0,91774 | -0,282829 | 0,345 | 0,692383 |
| Lsm4 | 118,094 | 131,232 | 0,152187 | 0,3451 | 0,69248 |
| Ier5l | 8,62941 | 9,95193 | 0,205714 | 0,3452 | 0,692573 |
| Cluap1 | 14,5977 | 16,5674 | 0,182608 | 0,34525 | 0,692573 |
| AI506816 | 14,4362 | 23,0566 | 0,675483 | 0,34545 | 0,692766 |
| Cbr3 | 2,14473 | 2,71373 | 0,339484 | 0,34545 | 0,692766 |
| Nudt1 | 32,419 | 36,6566 | 0,177231 | 0,34575 | 0,693045 |
| Nipsnap1 | 20,8935 | 23,4285 | 0,165212 | 0,3459 | 0,693045 |
| Osbpl11 | 14,1352 | 15,7573 | 0,156728 | 0,3459 | 0,693045 |
| E2f4 | 62,8366 | 69,7834 | 0,15128 | 0,34565 | 0,693045 |
| Stx3 | 8,15834 | 7,20844 | -0,178589 | 0,3459 | 0,693045 |
| Mesp1 | 8,09379 | 6,79727 | -0,25186 | 0,34575 | 0,693045 |
| Galt | 6,51321 | 7,50787 | 0,205035 | 0,3465 | 0,693204 |

| | | | | | |
|---|---|---|---|---|---|
| Dscc1 | 14,2817 | 16,3471 | 0,194865 | 0,3465 | 0,693204 |
| Taf1a | 11,9022 | 13,4134 | 0,172444 | 0,34645 | 0,693204 |
| Trappc1 | 84,6771 | 94,7577 | 0,162271 | 0,34645 | 0,693204 |
| Poldip3 | 48,2741 | 53,4877 | 0,147956 | 0,34655 | 0,693204 |
| Npat | 10,3121 | 9,27215 | -0,153357 | 0,34655 | 0,693204 |
| Fam160a2 | 7,21552 | 6,47706 | -0,155765 | 0,34625 | 0,693204 |
| Gde1 | 25,6452 | 22,8075 | -0,169181 | 0,34625 | 0,693204 |
| Ptdss1 | 22,5183 | 19,717 | -0,191653 | 0,34625 | 0,693204 |
| Rfx2 | 1,45527 | 1,19652 | -0,282444 | 0,3465 | 0,693204 |
| Pnldc1 | 0,785159 | 0,563213 | -0,479305 | 0,3462 | 0,693204 |
| Gm9833 | 29,4703 | 32,9291 | 0,1601 | 0,34695 | 0,6939 |
| Blzf1 | 6,54936 | 7,37429 | 0,17115 | 0,34725 | 0,694085 |
| Dnttip1 | 29,5367 | 33,2346 | 0,170174 | 0,34715 | 0,694085 |
| Ccnd1 | 123,251 | 137,028 | 0,152871 | 0,3472 | 0,694085 |
| Shfm1 | 560,506 | 621,929 | 0,150018 | 0,34715 | 0,694085 |
| 2210016L21Rik | 26,5706 | 29,8762 | 0,169168 | 0,3474 | 0,694165 |
| Glo1 | 29,7425 | 33,2099 | 0,159086 | 0,34755 | 0,694165 |
| Mrpl28 | 84,5156 | 93,8559 | 0,151231 | 0,3475 | 0,694165 |
| Gli3 | 13,6224 | 15,1161 | 0,150107 | 0,3474 | 0,694165 |
| Klf10 | 10,7834 | 9,58378 | -0,17014 | 0,34745 | 0,694165 |
| Tatdn3 | 5,25693 | 6,25982 | 0,2519 | 0,34815 | 0,695052 |
| Zcchc4 | 6,54307 | 7,54917 | 0,206349 | 0,3481 | 0,695052 |
| Prune | 12,2791 | 13,7749 | 0,16583 | 0,3481 | 0,695052 |
| Qprt | 4,62655 | 5,55883 | 0,264844 | 0,34845 | 0,695091 |
| Wdr72 | 1,0341 | 1,2384 | 0,260106 | 0,3488 | 0,695091 |
| Ptpn6 | 5,07937 | 5,88633 | 0,212719 | 0,34855 | 0,695091 |
| Gpsm1 | 22,4984 | 25,7378 | 0,194068 | 0,34885 | 0,695091 |
| Sec61b | 102,041 | 114,536 | 0,166653 | 0,3488 | 0,695091 |
| Sh3bp5l | 15,0907 | 16,8993 | 0,163309 | 0,34845 | 0,695091 |
| Clpx | 16,9372 | 18,9005 | 0,158235 | 0,3486 | 0,695091 |
| Mettl14 | 23,594 | 26,2995 | 0,156615 | 0,3485 | 0,695091 |
| Mcm3ap | 20,87 | 23,261 | 0,156482 | 0,3484 | 0,695091 |
| Mphosph9 | 9,76106 | 10,8591 | 0,153799 | 0,34855 | 0,695091 |
| Otub1 | 76,3678 | 84,6788 | 0,149037 | 0,34895 | 0,695091 |
| Rnf130 | 26,3651 | 23,1185 | -0,189584 | 0,3486 | 0,695091 |
| Tmem216 | 21,5877 | 18,9053 | -0,191421 | 0,34885 | 0,695091 |
| Fam120aos | 6,65002 | 5,64301 | -0,236894 | 0,3488 | 0,695091 |
| Nipa2 | 18,2221 | 14,4979 | -0,329842 | 0,34895 | 0,695091 |
| Kdm2b | 16,7045 | 18,6934 | 0,162293 | 0,3492 | 0,695278 |
| Atic | 87,9721 | 97,509 | 0,148489 | 0,3492 | 0,695278 |
| Ccdc88a | 15,236 | 16,8754 | 0,147437 | 0,34915 | 0,695278 |
| Rhebl1 | 10,9361 | 12,6067 | 0,205101 | 0,3494 | 0,695557 |
| Lsm2 | 100,294 | 111,597 | 0,154051 | 0,34955 | 0,695557 |
| Noc2l | 102,79 | 113,817 | 0,147009 | 0,3496 | 0,695557 |
| Top2b | 77,1619 | 69,5394 | -0,150059 | 0,3496 | 0,695557 |
| Eif2ak3 | 6,29257 | 5,59028 | -0,170729 | 0,3495 | 0,695557 |
| Ibtk | 20,4193 | 22,6467 | 0,149369 | 0,3498 | 0,695851 |
| 1700052K11Rik | 3,49702 | 4,09425 | 0,227471 | 0,35015 | 0,696025 |

| | | | | | |
|---|---|---|---|---|---|
| Abhd14a | 23,4315 | 26,521 | 0,178687 | 0,3502 | 0,696025 |
| Depdc5 | 11,7995 | 13,3102 | 0,173812 | 0,35005 | 0,696025 |
| Adpgk | 37,6615 | 41,8255 | 0,151292 | 0,3502 | 0,696025 |
| Tbc1d16 | 19,656 | 17,7358 | -0,148307 | 0,35005 | 0,696025 |
| Kbtbd8 | 7,14053 | 6,34459 | -0,170504 | 0,35 | 0,696025 |
| Bloc1s5 | 14,6088 | 16,5095 | 0,176452 | 0,3504 | 0,696319 |
| Slc25a24 | 18,0062 | 16,1476 | -0,15718 | 0,35055 | 0,696514 |
| 1700001O22Rik | 0,529937 | 0,771952 | 0,542691 | 0,35065 | 0,6966 |
| Eif2b2 | 46,2891 | 51,4218 | 0,151707 | 0,3507 | 0,6966 |
| E130309F12Rik | 0,748901 | 0,592132 | -0,338856 | 0,35075 | 0,6966 |
| Magohb | 33,2288 | 37,7686 | 0,184753 | 0,35095 | 0,696786 |
| Mrpl10 | 45,1533 | 50,2661 | 0,154753 | 0,351 | 0,696786 |
| Fam161b | 2,00885 | 1,68459 | -0,253974 | 0,3509 | 0,696786 |
| Tfe3 | 11,2859 | 12,6757 | 0,167547 | 0,35115 | 0,69698 |
| Ggcx | 6,90287 | 7,80086 | 0,176436 | 0,3518 | 0,697537 |
| Orc1 | 13,9737 | 15,6277 | 0,161389 | 0,35175 | 0,697537 |
| Polr1b | 26,2782 | 29,145 | 0,149383 | 0,35165 | 0,697537 |
| Paip1 | 28,0044 | 25,2605 | -0,148769 | 0,3519 | 0,697537 |
| Ccdc85b | 6,85252 | 6,09414 | -0,169211 | 0,35185 | 0,697537 |
| 9830147E19Rik | 3,38898 | 2,96334 | -0,193629 | 0,35165 | 0,697537 |
| 0610040B10Rik | 5,9955 | 4,43563 | -0,43474 | 0,3517 | 0,697537 |
| Nanos3 | 1,87932 | 1,32503 | -0,504192 | 0,35185 | 0,697537 |
| Pcsk4 | 0,720317 | 0,438974 | -0,714496 | 0,35165 | 0,697537 |
| Aatf | 24,0276 | 26,8838 | 0,162045 | 0,3522 | 0,697614 |
| Atg12 | 27,0282 | 30,0659 | 0,153661 | 0,3521 | 0,697614 |
| Gab2 | 2,26258 | 1,96783 | -0,201364 | 0,35205 | 0,697614 |
| Tgfa | 1,88559 | 1,59723 | -0,239448 | 0,35215 | 0,697614 |
| Tiam2 | 2,65214 | 2,15872 | -0,29698 | 0,3522 | 0,697614 |
| BC005624 | 14,6205 | 16,6153 | 0,184517 | 0,3524 | 0,697907 |
| Tsc2 | 20,5542 | 23,9221 | 0,21891 | 0,35255 | 0,697997 |
| Zfp935 | 7,13607 | 6,24023 | -0,193529 | 0,35255 | 0,697997 |
| Med30 | 49,9944 | 55,9912 | 0,163434 | 0,3528 | 0,698181 |
| Nop9 | 19,9846 | 22,1678 | 0,149574 | 0,3527 | 0,698181 |
| Rnd3 | 40,551 | 44,9772 | 0,149456 | 0,3528 | 0,698181 |
| Gpank1 | 8,48511 | 9,83522 | 0,213024 | 0,3534 | 0,698541 |
| Gpn2 | 18,1946 | 20,5975 | 0,178956 | 0,3534 | 0,698541 |
| Dhrs1 | 26,1421 | 29,2943 | 0,164249 | 0,3533 | 0,698541 |
| Fam207a | 23,5864 | 26,2523 | 0,15449 | 0,3534 | 0,698541 |
| Pan2 | 23,3773 | 25,9367 | 0,149887 | 0,3531 | 0,698541 |
| Dnajc4 | 7,8479 | 6,49491 | -0,272995 | 0,3533 | 0,698541 |
| 1700026L06Rik | 1,14112 | 0,712231 | -0,680039 | 0,35335 | 0,698541 |
| A930001C03Rik | 0,713991 | 0,205081 | -1,79971 | 0,35305 | 0,698541 |
| Clic4 | 115,584 | 128,222 | 0,1497 | 0,3536 | 0,69873 |
| Mef2d | 8,03108 | 7,05228 | -0,187504 | 0,35355 | 0,69873 |
| Bak1 | 33,5579 | 37,3887 | 0,155951 | 0,3537 | 0,69882 |
| Mab21l1 | 1,23764 | 0,639949 | -0,95156 | 0,35375 | 0,69882 |
| 2210408I21Rik | 2,79397 | 3,35281 | 0,263053 | 0,35415 | 0,6993 |
| Eya3 | 25,7237 | 28,5083 | 0,14828 | 0,3541 | 0,6993 |

| | | | | | |
|---|---|---|---|---|---|
| Cd2bp2 | 48,6213 | 53,7267 | 0,14405 | 0,3541 | 0,6993 |
| Zfand2b | 9,09855 | 7,79834 | -0,222469 | 0,35425 | 0,699394 |
| Txnl4a | 106,991 | 119,195 | 0,155832 | 0,35435 | 0,699488 |
| Zbtb24 | 7,15601 | 8,12509 | 0,183229 | 0,3545 | 0,699676 |
| 1700025G04Rik | 5,72097 | 6,36302 | 0,153453 | 0,35455 | 0,699676 |
| Kif1b | 20,7252 | 23,0045 | 0,150528 | 0,35465 | 0,69977 |
| Trim26 | 15,6571 | 17,4608 | 0,157302 | 0,355 | 0,700146 |
| Jag1 | 12,5898 | 14,0009 | 0,153261 | 0,35505 | 0,700146 |
| Ska3 | 31,7569 | 35,2599 | 0,150958 | 0,35505 | 0,700146 |
| Ooep | 1,12774 | 0,800831 | -0,493867 | 0,35505 | 0,700146 |
| Epha4 | 47,169 | 52,2192 | 0,14674 | 0,35515 | 0,70024 |
| Tmem126b | 25,0311 | 21,927 | -0,191009 | 0,3553 | 0,700324 |
| Abcb1b | 1,7147 | 1,44768 | -0,24422 | 0,35535 | 0,700324 |
| Sult5a1 | 1,0159 | 0,710992 | -0,514855 | 0,35535 | 0,700324 |
| Slc25a10 | 11,5285 | 10,3229 | -0,159356 | 0,35555 | 0,700507 |
| 2810428I15Rik | 41,8296 | 36,8127 | -0,184319 | 0,3556 | 0,700507 |
| 5730409E04Rik | 2,73358 | 2,31946 | -0,237003 | 0,3556 | 0,700507 |
| Vps13d | 3,3406 | 3,72884 | 0,158617 | 0,35585 | 0,700896 |
| Dtx2 | 11,5751 | 13,0161 | 0,169271 | 0,35595 | 0,70099 |
| Adamts9 | 20,6419 | 18,6154 | -0,149079 | 0,35605 | 0,701083 |
| Miat | 4,72444 | 5,27468 | 0,15894 | 0,3562 | 0,701167 |
| R3hcc1 | 30,5022 | 33,9814 | 0,155831 | 0,3562 | 0,701167 |
| Cln6 | 10,3291 | 9,06785 | -0,187887 | 0,35625 | 0,701167 |
| Hook3 | 9,87541 | 8,92359 | -0,146215 | 0,35645 | 0,701457 |
| Lrch4 | 10,1722 | 12,1756 | 0,259362 | 0,35665 | 0,701748 |
| E030030I06Rik | 2,63367 | 3,31833 | 0,333381 | 0,35695 | 0,701821 |
| Aurkaip1 | 46,7568 | 52,1555 | 0,157645 | 0,3569 | 0,701821 |
| Polr1c | 80,3416 | 89,0626 | 0,148673 | 0,35685 | 0,701821 |
| Brd3 | 53,0942 | 58,7271 | 0,145473 | 0,3569 | 0,701821 |
| Pde3b | 15,2815 | 13,7574 | -0,151574 | 0,35685 | 0,701821 |
| Zbtb3 | 2,64579 | 3,16 | 0,256226 | 0,35715 | 0,701988 |
| Zfp286 | 6,15829 | 6,98922 | 0,182602 | 0,3571 | 0,701988 |
| Stk38 | 38,3919 | 42,5294 | 0,147656 | 0,35735 | 0,701988 |
| Ube2k | 30,5444 | 33,8066 | 0,146394 | 0,3573 | 0,701988 |
| Atp1b3 | 94,9138 | 85,6924 | -0,14745 | 0,3573 | 0,701988 |
| Magi3 | 7,50722 | 6,72612 | -0,158505 | 0,35725 | 0,701988 |
| Tmem160 | 18,3796 | 21,1189 | 0,200427 | 0,35755 | 0,702278 |
| Ndufv1 | 73,6061 | 81,5755 | 0,148312 | 0,35775 | 0,702356 |
| Srrt | 98,8532 | 109,374 | 0,145909 | 0,35775 | 0,702356 |
| Zfp354c | 14,5263 | 13,1068 | -0,148354 | 0,3578 | 0,702356 |
| Trpt1 | 5,11176 | 4,02573 | -0,344572 | 0,35765 | 0,702356 |
| Raver1-fdx1l | 0,189548 | 1,18818 | 2,64812 | 0,35835 | 0,702492 |
| Il18bp | 5,14881 | 6,08973 | 0,242138 | 0,35815 | 0,702492 |
| Tufm | 5,91437 | 6,86091 | 0,214175 | 0,35835 | 0,702492 |
| Setd6 | 12,5594 | 14,2307 | 0,180232 | 0,35845 | 0,702492 |
| Edc3 | 24,193 | 26,8858 | 0,152255 | 0,35835 | 0,702492 |
| Mfn2 | 14,4326 | 16,0311 | 0,151542 | 0,3584 | 0,702492 |
| Pelp1 | 34,9462 | 38,7011 | 0,147239 | 0,3581 | 0,702492 |

| | | | | | |
|---|---|---|---|---|---|
| Heatr1 | 24,667 | 27,3015 | 0,146394 | 0,3584 | 0,702492 |
| Chchd2 | 102,821 | 92,5334 | -0,152082 | 0,3585 | 0,702492 |
| Abca2 | 4,8732 | 4,36307 | -0,159528 | 0,35835 | 0,702492 |
| Prickle3 | 2,54479 | 2,00692 | -0,342563 | 0,3585 | 0,702492 |
| Gm9079 | 0,878108 | 0,652138 | -0,429221 | 0,35825 | 0,702492 |
| Tacc2 | 18,3242 | 20,3384 | 0,150456 | 0,3587 | 0,702575 |
| Pogz | 18,8133 | 20,8418 | 0,147732 | 0,35865 | 0,702575 |
| Nifk | 71,2502 | 78,813 | 0,145538 | 0,3587 | 0,702575 |
| Pdcl3 | 38,0525 | 42,2921 | 0,152396 | 0,3589 | 0,702853 |
| Zfp507 | 16,6146 | 18,4002 | 0,147276 | 0,359 | 0,702853 |
| 2810403A07Rik | 53,156 | 47,9617 | -0,148349 | 0,35895 | 0,702853 |
| Mmaa | 2,03216 | 2,41463 | 0,248789 | 0,35925 | 0,70324 |
| Ston1 | 7,02554 | 7,94178 | 0,176853 | 0,3595 | 0,70342 |
| Snx1 | 58,7578 | 65,0329 | 0,14639 | 0,3594 | 0,70342 |
| Bub3 | 98,7025 | 109,109 | 0,144612 | 0,35945 | 0,70342 |
| Tsc22d2 | 5,6418 | 6,28244 | 0,155171 | 0,3597 | 0,703708 |
| Enox2 | 3,5391 | 3,08511 | -0,19806 | 0,35985 | 0,703899 |
| Nup62 | 73,8313 | 83,1206 | 0,170974 | 0,36 | 0,704089 |
| Trim24 | 18,0136 | 19,967 | 0,148527 | 0,36075 | 0,705349 |
| B4galt6 | 8,82687 | 7,91935 | -0,156519 | 0,36075 | 0,705349 |
| Nisch | 121,789 | 135,452 | 0,153395 | 0,3609 | 0,705436 |
| Polr3a | 12,0734 | 13,419 | 0,152441 | 0,3609 | 0,705436 |
| Mapk9 | 14,4558 | 16,0446 | 0,150443 | 0,3613 | 0,706115 |
| Senp7 | 12,4925 | 14,45 | 0,210005 | 0,3615 | 0,706196 |
| Gne | 7,1563 | 6,29888 | -0,184117 | 0,36145 | 0,706196 |
| Ppapdc3 | 2,28752 | 1,87515 | -0,286775 | 0,3614 | 0,706196 |
| 1700084E18Rik | 1,82731 | 2,77517 | 0,602852 | 0,3616 | 0,706288 |
| Eif5b | 35,485 | 39,5883 | 0,157863 | 0,36195 | 0,706868 |
| Gng5 | 90,3278 | 101,088 | 0,162374 | 0,36205 | 0,70696 |
| Chst2 | 5,73212 | 5,11793 | -0,163507 | 0,36225 | 0,707247 |
| Rab11b | 6,25303 | 6,98716 | 0,160152 | 0,3628 | 0,707489 |
| Tbl3 | 45,6885 | 50,6025 | 0,147377 | 0,36285 | 0,707489 |
| Nol6 | 22,8824 | 25,2996 | 0,144876 | 0,36285 | 0,707489 |
| Topbp1 | 58,8607 | 65,0518 | 0,144283 | 0,36245 | 0,707489 |
| Serf2 | 22,9845 | 20,6451 | -0,154861 | 0,3627 | 0,707489 |
| Slc19a2 | 7,89488 | 7,01702 | -0,170059 | 0,3627 | 0,707489 |
| Tctn3 | 6,98824 | 6,14339 | -0,185893 | 0,3625 | 0,707489 |
| Entpd7 | 2,47797 | 1,92845 | -0,361718 | 0,36285 | 0,707489 |
| 4833417C18Rik | 0,759284 | 0,525459 | -0,53106 | 0,36275 | 0,707489 |
| Hltf | 8,00013 | 7,14427 | -0,163236 | 0,3631 | 0,70777 |
| 2510002D24Rik | 8,14872 | 6,95784 | -0,227935 | 0,36305 | 0,70777 |
| Zfp558 | 3,05339 | 3,5104 | 0,201224 | 0,3633 | 0,70785 |
| Epb4.1l3 | 81,5722 | 90,1658 | 0,144502 | 0,3632 | 0,70785 |
| Daglb | 7,2233 | 6,38247 | -0,178542 | 0,36325 | 0,70785 |
| Msra | 14,1446 | 16,178 | 0,193782 | 0,3635 | 0,708021 |
| E130308A19Rik | 8,69582 | 9,73419 | 0,162738 | 0,3636 | 0,708021 |
| Slc25a3 | 295,741 | 327,629 | 0,147729 | 0,36355 | 0,708021 |
| Cyp51 | 70,2439 | 77,6361 | 0,144356 | 0,3635 | 0,708021 |

| | | | | | |
|---|---|---|---|---|---|
| 2610301B20Rik | 11,0442 | 9,75846 | -0,178559 | 0,36375 | 0,70821 |
| Sec31b | 0,765712 | 0,938724 | 0,2939 | 0,3639 | 0,708399 |
| Tomm7 | 77,0241 | 93,8258 | 0,284675 | 0,36425 | 0,708662 |
| Mlh1 | 20,3986 | 22,6501 | 0,151052 | 0,3643 | 0,708662 |
| Ccny | 39,7791 | 36,0056 | -0,143788 | 0,3641 | 0,708662 |
| Id3 | 127,815 | 115,314 | -0,148494 | 0,36425 | 0,708662 |
| Fam83f | 0,73399 | 0,561058 | -0,387611 | 0,36425 | 0,708662 |
| Gnb2l1 | 1069,64 | 1197,45 | 0,162845 | 0,3644 | 0,708753 |
| Spidr | 14,676 | 16,3554 | 0,156304 | 0,3646 | 0,708815 |
| Map3k3 | 18,9126 | 21,0125 | 0,1519 | 0,36475 | 0,708815 |
| E2f7 | 12,5564 | 13,9356 | 0,150347 | 0,3647 | 0,708815 |
| Tusc3 | 44,2936 | 39,624 | -0,160722 | 0,3646 | 0,708815 |
| Cyb5r1 | 25,9686 | 23,2186 | -0,161488 | 0,3647 | 0,708815 |
| A530054K11Rik | 2,74979 | 2,4044 | -0,193645 | 0,3647 | 0,708815 |
| Gtf2ird2 | 4,4106 | 5,0236 | 0,187745 | 0,36485 | 0,7089 |
| Klhl9 | 44,2031 | 40,0276 | -0,14315 | 0,3649 | 0,7089 |
| Eif4a1 | 100,067 | 110,474 | 0,14274 | 0,36515 | 0,708973 |
| Setdb2 | 6,74365 | 7,60652 | 0,173707 | 0,3651 | 0,708974 |
| Calm1 | 117,756 | 106,489 | -0,145098 | 0,36505 | 0,708974 |
| Abhd16a | 16,6478 | 14,8123 | -0,168536 | 0,365 | 0,708974 |
| Herc1 | 10,6798 | 11,8354 | 0,148226 | 0,36535 | 0,709259 |
| Mzt2 | 22,726 | 25,7151 | 0,178276 | 0,3657 | 0,709423 |
| Pcnt | 13,6496 | 15,0722 | 0,143029 | 0,3656 | 0,709423 |
| Eaf1 | 36,252 | 39,9839 | 0,141359 | 0,36555 | 0,709423 |
| Cotl1 | 77,8236 | 70,3612 | -0,145429 | 0,36555 | 0,709423 |
| Zfp946 | 11,423 | 10,177 | -0,166637 | 0,3657 | 0,709423 |
| Rgs16 | 4,8837 | 4,22081 | -0,210455 | 0,3664 | 0,710672 |
| Lrrn2 | 0,710572 | 0,556546 | -0,352482 | 0,36645 | 0,710672 |
| Ppia | 1,01267 | 1,48261 | 0,54998 | 0,3667 | 0,71095 |
| Timm10 | 45,7289 | 51,5518 | 0,172915 | 0,3667 | 0,71095 |
| Npm2 | 1,22389 | 1,65599 | 0,436217 | 0,36705 | 0,711519 |
| Chchd6 | 13,5598 | 11,7807 | -0,202904 | 0,3671 | 0,711519 |
| D030056L22Rik | 38,9565 | 35,0746 | -0,151436 | 0,3673 | 0,7117 |
| Gm10509 | 2,81812 | 2,35119 | -0,261346 | 0,3673 | 0,7117 |
| Hmgb3 | 40,0784 | 44,5134 | 0,151415 | 0,36745 | 0,711888 |
| Nsun6 | 5,65548 | 6,43442 | 0,186159 | 0,3677 | 0,712062 |
| Mapk8ip3 | 17,9333 | 19,9883 | 0,156517 | 0,3677 | 0,712062 |
| Rbmx | 104,236 | 94,5456 | -0,140774 | 0,3677 | 0,712062 |
| Anks1 | 8,44074 | 9,36102 | 0,149296 | 0,36785 | 0,71225 |
| Usp8 | 24,5463 | 27,1231 | 0,144014 | 0,36795 | 0,71234 |
| 8030462N17Rik | 9,98804 | 11,1652 | 0,160737 | 0,36805 | 0,712424 |
| Mdh1 | 133,017 | 147,026 | 0,14446 | 0,3681 | 0,712424 |
| Trim56 | 4,03461 | 4,59963 | 0,189091 | 0,3685 | 0,712566 |
| Klf16 | 11,8449 | 13,2485 | 0,161564 | 0,3683 | 0,712566 |
| Mrpl32 | 53,4203 | 59,7096 | 0,160574 | 0,3683 | 0,712566 |
| Brd9 | 29,0225 | 32,1654 | 0,148335 | 0,36845 | 0,712566 |
| Abr | 18,1564 | 20,115 | 0,147799 | 0,3685 | 0,712566 |
| Golm1 | 79,3532 | 71,9381 | -0,141533 | 0,36845 | 0,712566 |

| | | | | | |
|---|---|---|---|---|---|
| Gtf2a1 | 9,73338 | 8,76355 | -0,151427 | 0,3686 | 0,712566 |
| Rdh5 | 1,72646 | 1,33505 | -0,370924 | 0,36855 | 0,712566 |
| Srr | 5,24583 | 7,17869 | 0,452549 | 0,36875 | 0,712753 |
| Gabbr1 | 9,26798 | 8,2979 | -0,15951 | 0,36895 | 0,713037 |
| Kcnip4 | 1,10373 | 1,43058 | 0,37421 | 0,3694 | 0,7137 |
| Hes6 | 34,8587 | 38,8122 | 0,154991 | 0,36935 | 0,7137 |
| Rab8a | 51,5078 | 56,9012 | 0,143669 | 0,36955 | 0,713886 |
| Morc3 | 21,132 | 23,3547 | 0,144285 | 0,36985 | 0,714363 |
| Kifc2 | 0,90652 | 0,728118 | -0,316167 | 0,37 | 0,714549 |
| 6330418K02Rik | 5,75147 | 8,77704 | 0,609804 | 0,3703 | 0,715025 |
| Cnot1 | 54,8049 | 60,5778 | 0,144484 | 0,37065 | 0,715598 |
| Rps28 | 2,08863 | 3,44089 | 0,720225 | 0,3708 | 0,71568 |
| Tsr3 | 26,7652 | 30,0343 | 0,166251 | 0,37075 | 0,71568 |
| Crtac1 | 0,984068 | 0,782778 | -0,330156 | 0,37115 | 0,716253 |
| Prob1 | 1,15315 | 1,36605 | 0,244429 | 0,3713 | 0,716335 |
| Polr3c | 29,0296 | 32,3107 | 0,154485 | 0,3713 | 0,716335 |
| Mtmr3 | 17,8317 | 19,7055 | 0,144155 | 0,3714 | 0,716418 |
| Ubr2 | 15,8109 | 17,4582 | 0,142983 | 0,37145 | 0,716418 |
| Rbms2 | 13,17 | 11,8914 | -0,147339 | 0,3717 | 0,716693 |
| Cd302 | 9,43742 | 8,18853 | -0,204789 | 0,3717 | 0,716693 |
| Cops7b | 29,182 | 32,3705 | 0,149603 | 0,3718 | 0,716775 |
| Prr5l | 6,10762 | 5,42474 | -0,171055 | 0,37185 | 0,716775 |
| Arrdc4 | 10,8904 | 12,4272 | 0,190439 | 0,37215 | 0,71725 |
| Mir6935 | 1047,71 | 3255,1 | 1,63546 | 0,37225 | 0,717339 |
| Hist1h2bk | 1,31972 | 2,01189 | 0,608311 | 0,37235 | 0,717422 |
| Mrps9 | 41,0285 | 45,5139 | 0,149681 | 0,3724 | 0,717422 |
| Amt | 21,1386 | 23,7434 | 0,167646 | 0,37255 | 0,717504 |
| Tmem132a | 93,341 | 84,5851 | -0,142107 | 0,3725 | 0,717504 |
| Cby1 | 23,0957 | 25,9718 | 0,169321 | 0,37275 | 0,717785 |
| Iqce | 6,37295 | 7,15209 | 0,166403 | 0,3732 | 0,718135 |
| Tmem134 | 22,1177 | 24,7543 | 0,162483 | 0,37315 | 0,718135 |
| Actr2 | 77,5729 | 70,2576 | -0,142898 | 0,3732 | 0,718135 |
| Tiam1 | 14,4374 | 12,9994 | -0,151364 | 0,373 | 0,718135 |
| Eef2k | 4,03527 | 3,59098 | -0,168285 | 0,3732 | 0,718135 |
| Mir6244 | 3,50817 | 35,0034 | 3,31871 | 0,3735 | 0,718469 |
| Tbc1d10a | 10,5891 | 11,9708 | 0,17694 | 0,3736 | 0,718469 |
| Dbt | 15,6685 | 17,4094 | 0,152001 | 0,37375 | 0,718469 |
| Bora | 27,8818 | 30,8554 | 0,146203 | 0,37375 | 0,718469 |
| Abcf2 | 76,8856 | 84,8198 | 0,141687 | 0,37365 | 0,718469 |
| Syn3 | 0,608921 | 0,353279 | -0,785449 | 0,3735 | 0,718469 |
| Phtf1os | 1,1306 | 0,508027 | -1,15411 | 0,3736 | 0,718469 |
| Pomc | 1,72284 | 2,21012 | 0,359338 | 0,37385 | 0,718558 |
| T2 | 1,65807 | 2,20211 | 0,409379 | 0,37405 | 0,718735 |
| 6720416L17Rik | 14,6103 | 16,3789 | 0,164852 | 0,374 | 0,718735 |
| Rasl12 | 1,64812 | 1,98956 | 0,271624 | 0,3744 | 0,719304 |
| Wipi1 | 7,42141 | 6,467 | -0,198598 | 0,37455 | 0,719489 |
| Ppp2r3a | 6,77778 | 7,53079 | 0,151988 | 0,37465 | 0,719578 |
| Klhl10 | 0,789178 | 0,59657 | -0,403659 | 0,37485 | 0,719859 |

| | | | | | |
|---|---|---|---|---|---|
| Acy3 | 2,5069 | 3,03603 | 0,276281 | 0,37505 | 0,720028 |
| Snrnp70 | 303,217 | 334,922 | 0,143474 | 0,3751 | 0,720028 |
| Dcbld2 | 55,659 | 50,444 | -0,14193 | 0,375 | 0,720028 |
| Ppp1ca | 388,203 | 430,331 | 0,148637 | 0,3752 | 0,720117 |
| Zfp772 | 3,46692 | 4,00236 | 0,207197 | 0,37535 | 0,720198 |
| Il20rb | 1,5571 | 1,2773 | -0,285766 | 0,3753 | 0,720198 |
| Mgam | 0,609418 | 0,732916 | 0,266216 | 0,3755 | 0,720279 |
| Rnf150 | 1,92994 | 1,70683 | -0,177234 | 0,3755 | 0,720279 |
| Akap9 | 10,5856 | 11,6937 | 0,143625 | 0,37565 | 0,720352 |
| Ctdspl2 | 14,8284 | 13,8695 | -0,144249 | 0,3757 | 0,720352 |
| Lsr | 34,7435 | 31,3286 | -0,149259 | 0,3756 | 0,720352 |
| Man1b1 | 41,611 | 37,7381 | -0,140945 | 0,376 | 0,720824 |
| Wdr61 | 82,4322 | 91,0658 | 0,143701 | 0,3761 | 0,720912 |
| Hist1h3c | 2,25189 | 3,19027 | 0,502545 | 0,3765 | 0,721472 |
| Ezh1 | 6,73384 | 5,99819 | -0,166903 | 0,3765 | 0,721472 |
| Nif3l1 | 33,2943 | 36,8268 | 0,145482 | 0,3766 | 0,72156 |
| Ripk2 | 5,87412 | 6,75645 | 0,201893 | 0,3769 | 0,72161 |
| Phf21b | 3,20887 | 3,67889 | 0,197206 | 0,37695 | 0,72161 |
| Sap30bp | 29,3918 | 32,5926 | 0,149134 | 0,37695 | 0,72161 |
| Skil | 15,1467 | 16,7388 | 0,14419 | 0,37695 | 0,72161 |
| Cep192 | 19,9773 | 21,9973 | 0,138962 | 0,37685 | 0,72161 |
| Asb7 | 6,50804 | 5,82192 | -0,160728 | 0,3769 | 0,72161 |
| Dgat2 | 8,62968 | 9,75522 | 0,176867 | 0,3772 | 0,721667 |
| Hars | 60,9924 | 67,2654 | 0,141237 | 0,3772 | 0,721667 |
| Timm22 | 19,5914 | 17,6442 | -0,151023 | 0,3772 | 0,721667 |
| Slc7a2 | 1,12337 | 0,966499 | -0,216989 | 0,37705 | 0,721667 |
| Lpxn | 0,680986 | 0,488543 | -0,479138 | 0,37725 | 0,721667 |
| Anapc15 | 23,462 | 26,7052 | 0,186797 | 0,3774 | 0,721725 |
| Psmd1 | 103,204 | 113,849 | 0,141624 | 0,37735 | 0,721725 |
| Zc3h15 | 74,8805 | 82,5256 | 0,140251 | 0,37745 | 0,721725 |
| Zfp783 | 4,58147 | 3,97807 | -0,20374 | 0,37755 | 0,721725 |
| Zbtb8os | 10,8774 | 9,09954 | -0,25747 | 0,3775 | 0,721725 |
| Surf6 | 27,7302 | 30,655 | 0,144663 | 0,3779 | 0,722291 |
| Smagp | 2,85964 | 2,28238 | -0,325297 | 0,378 | 0,722378 |
| 2610507B11Rik | 44,4382 | 40,2638 | -0,142316 | 0,3781 | 0,722466 |
| Ctu1 | 6,91734 | 7,82836 | 0,178492 | 0,3783 | 0,722642 |
| Serp1 | 102,939 | 91,3716 | -0,171965 | 0,37825 | 0,722642 |
| Xpo6 | 40,6704 | 44,8281 | 0,140424 | 0,3784 | 0,722729 |
| Rccd1 | 8,24531 | 9,31934 | 0,176653 | 0,3787 | 0,723064 |
| Rars2 | 26,8031 | 29,7681 | 0,151368 | 0,3788 | 0,723064 |
| Guk1 | 57,2229 | 63,5103 | 0,150398 | 0,37865 | 0,723064 |
| Spg20 | 27,9651 | 30,8693 | 0,142544 | 0,37885 | 0,723064 |
| Ufl1 | 11,0994 | 9,99459 | -0,151263 | 0,3788 | 0,723064 |
| Fam101a | 1,54802 | 1,22696 | -0,335343 | 0,3789 | 0,723064 |
| Scnm1 | 32,7151 | 36,8024 | 0,169843 | 0,37915 | 0,723216 |
| Armc8 | 12,389 | 13,7398 | 0,149296 | 0,3792 | 0,723216 |
| Zkscan1 | 9,72083 | 10,7426 | 0,144187 | 0,3792 | 0,723216 |
| Espl1 | 25,9342 | 28,5704 | 0,139663 | 0,3792 | 0,723216 |

| | | | | | |
|---|---|---|---|---|---|
| Myl6b | 5,43721 | 4,52411 | -0,265231 | 0,37925 | 0,723216 |
| AI987944 | 4,13781 | 3,59438 | -0,203125 | 0,37935 | 0,723303 |
| Smpd4 | 32,1147 | 35,3571 | 0,138764 | 0,37945 | 0,723391 |
| Fam136a | 56,4525 | 62,2095 | 0,140098 | 0,37955 | 0,723478 |
| Slc15a4 | 12,0551 | 13,4439 | 0,157306 | 0,3797 | 0,723653 |
| Fbxl17 | 3,02358 | 3,35249 | 0,148973 | 0,37975 | 0,723653 |
| Pycr1 | 4,35568 | 4,96147 | 0,18787 | 0,38 | 0,723717 |
| Smu1 | 60,5678 | 68,101 | 0,169126 | 0,37995 | 0,723717 |
| Snapc4 | 12,8246 | 14,1788 | 0,144822 | 0,38 | 0,723717 |
| Tmem234 | 76,756 | 69,131 | -0,150947 | 0,37995 | 0,723717 |
| Hspa2 | 1,55932 | 1,87361 | 0,264901 | 0,3804 | 0,72405 |
| Ascc1 | 18,9927 | 22,5141 | 0,245383 | 0,3804 | 0,72405 |
| Aebp2 | 36,6694 | 43,0095 | 0,230077 | 0,38045 | 0,72405 |
| Aprt | 112,729 | 102,073 | -0,143265 | 0,3805 | 0,72405 |
| Dusp22 | 10,6957 | 9,26901 | -0,20654 | 0,38025 | 0,72405 |
| Rom1 | 4,29892 | 3,61692 | -0,249212 | 0,3805 | 0,72405 |
| Fiz1 | 24,6496 | 27,2453 | 0,144439 | 0,3809 | 0,724065 |
| Zfp609 | 18,8998 | 20,799 | 0,138144 | 0,38075 | 0,724065 |
| Fbxl19 | 53,1981 | 58,5395 | 0,138036 | 0,38095 | 0,724065 |
| Atp8b2 | 18,612 | 16,8684 | -0,141915 | 0,38105 | 0,724065 |
| Mast4 | 5,39843 | 4,87365 | -0,147538 | 0,38105 | 0,724065 |
| Ap1s1 | 51,308 | 46,3169 | -0,147644 | 0,381 | 0,724065 |
| Reck | 9,50772 | 8,55268 | -0,152723 | 0,3809 | 0,724065 |
| Usp45 | 7,94248 | 7,14322 | -0,153015 | 0,38105 | 0,724065 |
| Ppap2c | 17,3208 | 15,446 | -0,165279 | 0,38075 | 0,724065 |
| Trmt2b | 3,24013 | 2,57827 | -0,329648 | 0,381 | 0,724065 |
| Cdca4 | 57,2651 | 63,1592 | 0,141336 | 0,38115 | 0,724145 |
| Acsf3 | 8,07204 | 7,12304 | -0,180441 | 0,3812 | 0,724145 |
| Ift122 | 8,97523 | 9,99936 | 0,155886 | 0,3816 | 0,724184 |
| Stk40 | 20,1354 | 22,2423 | 0,143571 | 0,38145 | 0,724184 |
| Champ1 | 28,3613 | 31,2955 | 0,142031 | 0,3816 | 0,724184 |
| Stau1 | 36,0733 | 39,7058 | 0,13842 | 0,38155 | 0,724184 |
| Tmem101 | 22,1695 | 19,8543 | -0,159125 | 0,3815 | 0,724184 |
| Stat4 | 0,684236 | 0,53552 | -0,353556 | 0,38155 | 0,724184 |
| Gm5512 | 1,95434 | 0,0004169 | -12,1947 | 0,38145 | 0,724184 |
| Wdfy2 | 7,50482 | 8,52346 | 0,183621 | 0,38205 | 0,724318 |
| Polr2i | 55,2468 | 61,6696 | 0,158667 | 0,3818 | 0,724318 |
| Mfn1 | 13,7395 | 15,1906 | 0,144847 | 0,3818 | 0,724318 |
| Rab35 | 38,411 | 42,3824 | 0,141944 | 0,38195 | 0,724318 |
| Eif4g3 | 53,7314 | 59,2016 | 0,139869 | 0,3819 | 0,724318 |
| Olig3 | 3,21484 | 2,72623 | -0,237838 | 0,38195 | 0,724318 |
| Ogfod2 | 4,37816 | 3,69478 | -0,244837 | 0,38205 | 0,724318 |
| Rnf141 | 3,45046 | 4,22626 | 0,292593 | 0,38295 | 0,724356 |
| Pin1rt1 | 3,46177 | 4,19781 | 0,278128 | 0,3823 | 0,724356 |
| Itga10 | 1,20987 | 1,42037 | 0,231409 | 0,38235 | 0,724356 |
| Gm3258 | 28,3799 | 33,3047 | 0,230853 | 0,38305 | 0,724356 |
| Car12 | 2,02293 | 2,35498 | 0,219264 | 0,3831 | 0,724356 |
| Foxred1 | 21,8515 | 24,1942 | 0,146929 | 0,3825 | 0,724356 |

| | | | | | |
|---|---|---|---|---|---|
| Gga3 | 16,8892 | 18,6843 | 0,145726 | 0,3829 | 0,724356 |
| Pdhb | 45,7208 | 50,5579 | 0,145088 | 0,38295 | 0,724356 |
| Dkc1 | 97,9715 | 108,281 | 0,14435 | 0,383 | 0,724356 |
| Erc1 | 14,8175 | 16,3591 | 0,142792 | 0,38275 | 0,724356 |
| Ing4 | 49,4563 | 54,586 | 0,142377 | 0,3829 | 0,724356 |
| Rell1 | 35,2067 | 38,8168 | 0,140834 | 0,38255 | 0,724356 |
| 2700094K13Rik | 304,919 | 335,972 | 0,139916 | 0,3826 | 0,724356 |
| Glul | 68,1938 | 75,0008 | 0,137266 | 0,3824 | 0,724356 |
| Ppp6r2 | 6,24687 | 5,51745 | -0,179134 | 0,3825 | 0,724356 |
| Slc37a4 | 8,03765 | 7,06594 | -0,185892 | 0,38265 | 0,724356 |
| Slc10a3 | 7,07228 | 5,58141 | -0,341546 | 0,3825 | 0,724356 |
| Fsbp | 2,17727 | 1,69285 | -0,363073 | 0,3822 | 0,724356 |
| Tdg | 0,977122 | 0,632762 | -0,626877 | 0,3827 | 0,724356 |
| Arntl | 3,11709 | 3,60853 | 0,211214 | 0,38335 | 0,724419 |
| Mrpl47 | 37,2731 | 41,7114 | 0,162304 | 0,3833 | 0,724419 |
| BC005561 | 8,63185 | 9,5759 | 0,149739 | 0,3833 | 0,724419 |
| Rwdd2a | 1,42433 | 1,08374 | -0,394259 | 0,38325 | 0,724419 |
| Syt14 | 3,65187 | 4,30004 | 0,235715 | 0,38345 | 0,724505 |
| Ppp3cc | 3,16682 | 3,72621 | 0,234672 | 0,38385 | 0,725159 |
| Rnf125 | 0,487234 | 0,693748 | 0,509796 | 0,38415 | 0,725224 |
| D930007P13Rik | 2,50002 | 3,03633 | 0,280392 | 0,38525 | 0,725224 |
| Cyb5d1 | 5,45645 | 6,60601 | 0,275816 | 0,38445 | 0,725224 |
| Bves | 2,82691 | 3,38105 | 0,258247 | 0,3846 | 0,725224 |
| Klhl8 | 2,73266 | 3,16308 | 0,211026 | 0,38495 | 0,725224 |
| Fam188b | 4,38257 | 5,01776 | 0,195265 | 0,3847 | 0,725224 |
| Zfp775 | 2,79127 | 3,18855 | 0,191978 | 0,38475 | 0,725224 |
| Gfra3 | 8,79327 | 9,95366 | 0,178826 | 0,38495 | 0,725224 |
| Ndufc1 | 111,603 | 125,024 | 0,16383 | 0,3848 | 0,725224 |
| Fuk | 7,0262 | 7,83789 | 0,15772 | 0,3852 | 0,725224 |
| Acbd3 | 21,0321 | 23,2183 | 0,142665 | 0,38445 | 0,725224 |
| Nom1 | 18,1959 | 20,0794 | 0,142106 | 0,3848 | 0,725224 |
| Egln1 | 22,8612 | 25,2267 | 0,142053 | 0,38405 | 0,725224 |
| Sde2 | 25,2608 | 27,8716 | 0,141898 | 0,38525 | 0,725224 |
| Ncapd3 | 19,4808 | 21,4448 | 0,138574 | 0,3849 | 0,725224 |
| Rad54l2 | 21,0783 | 23,1843 | 0,13739 | 0,38475 | 0,725224 |
| Eif2b5 | 66,177 | 72,7084 | 0,135792 | 0,38465 | 0,725224 |
| Ankfy1 | 12,9483 | 11,7611 | -0,138746 | 0,3853 | 0,725224 |
| Msmo1 | 78,6267 | 71,3723 | -0,139655 | 0,38495 | 0,725224 |
| Rnaseh2c | 78,8843 | 70,9363 | -0,153215 | 0,3842 | 0,725224 |
| Aldh4a1 | 10,0254 | 8,99496 | -0,156471 | 0,3852 | 0,725224 |
| Hibch | 10,4898 | 9,25584 | -0,180549 | 0,38525 | 0,725224 |
| Fam20c | 3,6705 | 3,1852 | -0,204595 | 0,38535 | 0,725224 |
| Mamld1 | 0,600956 | 0,470438 | -0,353256 | 0,38435 | 0,725224 |
| Ghrl | 1,38385 | 0,880156 | -0,65286 | 0,3848 | 0,725224 |
| Hist1h4b | 1,08278 | 0,566622 | -0,934285 | 0,38495 | 0,725224 |
| Slc18a1 | 0,739533 | 0,237527 | -1,63852 | 0,3851 | 0,725224 |
| Spata7 | 7,28541 | 8,28699 | 0,185838 | 0,38545 | 0,725302 |
| Hapln3 | 3,31876 | 2,75186 | -0,270235 | 0,3855 | 0,725302 |

| | | | | | |
|---|---|---|---|---|---|
| Shroom1 | 0,524496 | 0,66547 | 0,343441 | 0,3856 | 0,725388 |
| Socs7 | 10,6366 | 11,7507 | 0,143716 | 0,38575 | 0,725568 |
| Cinp | 23,4676 | 26,0419 | 0,150165 | 0,38605 | 0,725692 |
| Top3b | 23,4841 | 25,9035 | 0,141462 | 0,386 | 0,725692 |
| Arid1a | 55,1612 | 60,8219 | 0,140937 | 0,38615 | 0,725692 |
| Zic3 | 44,1495 | 40,172 | -0,136204 | 0,38595 | 0,725692 |
| Pcyt1a | 14,9498 | 13,5435 | -0,142523 | 0,3859 | 0,725692 |
| Cd320 | 22,6517 | 20,4123 | -0,150182 | 0,3862 | 0,725692 |
| Arpc1b | 22,7223 | 20,3873 | -0,156442 | 0,3862 | 0,725692 |
| Abcd2 | 1,40905 | 1,20752 | -0,22268 | 0,38625 | 0,725692 |
| Kctd11 | 5,8121 | 6,58209 | 0,179487 | 0,3864 | 0,725761 |
| Fance | 10,3186 | 11,6172 | 0,171018 | 0,38645 | 0,725761 |
| 4930402H24Rik | 3,65031 | 3,2213 | -0,180374 | 0,38635 | 0,725761 |
| Esf1 | 32,6963 | 36,0051 | 0,139075 | 0,3866 | 0,725839 |
| Setd7 | 18,5345 | 16,8559 | -0,136967 | 0,3866 | 0,725839 |
| Klhl12 | 20,1815 | 22,2861 | 0,143109 | 0,3868 | 0,726113 |
| Bet1 | 28,0925 | 25,2431 | -0,154298 | 0,38695 | 0,726292 |
| Babam1 | 48,0377 | 53,0119 | 0,142149 | 0,38715 | 0,726565 |
| Eno1,Eno1b | 0,283088 | 1,87644 | 2,72868 | 0,38825 | 0,726996 |
| Rbm12b1 | 2,35827 | 2,73928 | 0,216066 | 0,38755 | 0,726996 |
| Irgm1 | 3,88327 | 4,48876 | 0,209045 | 0,38785 | 0,726996 |
| Cox16 | 9,42916 | 10,6337 | 0,173447 | 0,38755 | 0,726996 |
| Gpatch1 | 8,58009 | 9,59466 | 0,161238 | 0,38785 | 0,726996 |
| Lrrc14 | 13,0726 | 14,5876 | 0,158195 | 0,388 | 0,726996 |
| Mybl2 | 10,4811 | 11,6282 | 0,149849 | 0,38825 | 0,726996 |
| Ankrd13b | 14,3199 | 15,8699 | 0,148276 | 0,3881 | 0,726996 |
| Lrrc47 | 28,517 | 31,5405 | 0,145387 | 0,3876 | 0,726996 |
| Acad9 | 15,5139 | 17,1321 | 0,14314 | 0,38825 | 0,726996 |
| Ubtf | 54,6925 | 60,2188 | 0,138872 | 0,38825 | 0,726996 |
| Psmd12 | 114,359 | 125,707 | 0,13649 | 0,38775 | 0,726996 |
| Morc2a | 31,9905 | 35,1509 | 0,135916 | 0,38795 | 0,726996 |
| Ppib | 584,079 | 529,875 | -0,140512 | 0,38815 | 0,726996 |
| Dach1 | 7,82245 | 7,04081 | -0,151879 | 0,3878 | 0,726996 |
| Cryl1 | 4,92361 | 4,1702 | -0,239602 | 0,38825 | 0,726996 |
| Ncapg | 50,7229 | 45,547 | -0,155282 | 0,3887 | 0,727431 |
| Inpp5k | 6,14644 | 5,41276 | -0,183385 | 0,38865 | 0,727431 |
| Slc22a5 | 2,61415 | 2,24294 | -0,22095 | 0,3887 | 0,727431 |
| 2410076I21Rik | 1,01965 | 0,118912 | -3,10011 | 0,38865 | 0,727431 |
| Cep104 | 10,8748 | 12,0746 | 0,150983 | 0,3888 | 0,727517 |
| Fam89b | 45,5859 | 41,1506 | -0,147673 | 0,3889 | 0,727602 |
| Bin1 | 37,5375 | 41,3556 | 0,139751 | 0,389 | 0,727687 |
| Erh | 1,40228 | 1,89655 | 0,435602 | 0,38935 | 0,728036 |
| Pdhx | 21,7402 | 24,0654 | 0,146596 | 0,38925 | 0,728036 |
| Rb1 | 6,22205 | 5,03288 | -0,306006 | 0,38935 | 0,728036 |
| Nkx6-1 | 0,819011 | 0,61418 | -0,41522 | 0,38945 | 0,728121 |
| 1700009P17Rik | 1,34815 | 1,76678 | 0,390137 | 0,3907 | 0,7282 |
| Cox17 | 94,1665 | 105,455 | 0,163348 | 0,39065 | 0,7282 |
| Naa20 | 24,6682 | 27,6068 | 0,162374 | 0,39005 | 0,7282 |

| | | | | | |
|---|---|---|---|---|---|
| Ell | 8,21354 | 9,17634 | 0,159916 | 0,39035 | 0,7282 |
| Tank | 15,3383 | 17,0754 | 0,154781 | 0,39025 | 0,7282 |
| Fam160b2 | 8,80205 | 9,77899 | 0,151845 | 0,38965 | 0,7282 |
| Trappc4 | 50,8685 | 56,2763 | 0,145755 | 0,39015 | 0,7282 |
| Dvl3 | 19,8563 | 21,9468 | 0,144417 | 0,39035 | 0,7282 |
| Lmnb1 | 226,872 | 250,28 | 0,141666 | 0,39075 | 0,7282 |
| Plekho1 | 65,5822 | 72,2425 | 0,139542 | 0,39075 | 0,7282 |
| Zfp334 | 21,4534 | 23,6131 | 0,138383 | 0,3903 | 0,7282 |
| 1110008F13Rik | 103,86 | 94,4489 | -0,137039 | 0,38975 | 0,7282 |
| Wbp5 | 161,178 | 146,432 | -0,138428 | 0,39045 | 0,7282 |
| Ccng1 | 18,3378 | 16,6074 | -0,142992 | 0,39055 | 0,7282 |
| Sec22c | 5,68288 | 5,09496 | -0,157554 | 0,39075 | 0,7282 |
| Kctd6 | 11,983 | 10,6069 | -0,17599 | 0,38995 | 0,7282 |
| Cgrrf1 | 17,8719 | 15,8144 | -0,176449 | 0,3901 | 0,7282 |
| BC068157 | 3,00465 | 2,63801 | -0,187745 | 0,38975 | 0,7282 |
| Notum | 7,15296 | 6,27229 | -0,189549 | 0,3906 | 0,7282 |
| Col14a1 | 0,726152 | 0,604435 | -0,264683 | 0,3899 | 0,7282 |
| Tmem86b | 3,80338 | 3,16297 | -0,266 | 0,3908 | 0,7282 |
| Snta1 | 1,36915 | 1,10246 | -0,312559 | 0,39075 | 0,7282 |
| Prkg1 | 2,06969 | 1,55358 | -0,413812 | 0,38975 | 0,7282 |
| Cd247 | 1,32875 | 0,47524 | -1,48334 | 0,39 | 0,7282 |
| Gdap2 | 6,41803 | 7,21956 | 0,169779 | 0,39105 | 0,728564 |
| 2310036O22Rik | 162,617 | 178,774 | 0,136656 | 0,3913 | 0,728725 |
| Rnf170 | 2,47699 | 2,14184 | -0,209742 | 0,39125 | 0,728725 |
| Fam86 | 6,49379 | 5,44987 | -0,25284 | 0,3912 | 0,728725 |
| Ythdf1 | 52,2458 | 57,4283 | 0,136448 | 0,39175 | 0,729462 |
| Crispld1 | 7,68936 | 8,56196 | 0,155077 | 0,3919 | 0,729639 |
| Mrpl24 | 78,2283 | 88,0187 | 0,17012 | 0,392 | 0,729724 |
| Sorbs3 | 13,7482 | 12,2652 | -0,164677 | 0,39225 | 0,730088 |
| Camk2g | 20,0085 | 22,0461 | 0,139915 | 0,39245 | 0,730256 |
| Wdr44 | 1,54969 | 1,31483 | -0,23711 | 0,39245 | 0,730256 |
| Nln | 9,84659 | 10,9208 | 0,149375 | 0,3927 | 0,730611 |
| Gigyf2 | 21,489 | 19,5415 | -0,13706 | 0,39275 | 0,730611 |
| Lin37 | 9,38194 | 10,8381 | 0,208154 | 0,39305 | 0,730736 |
| Chmp7 | 32,8719 | 36,1592 | 0,137508 | 0,393 | 0,730736 |
| Atp6v1b2 | 50,5664 | 55,61 | 0,137164 | 0,39315 | 0,730736 |
| Psmc6 | 193,554 | 212,696 | 0,136057 | 0,393 | 0,730736 |
| Ptma | 194,328 | 213,407 | 0,135113 | 0,3932 | 0,730736 |
| Gapvd1 | 26,06 | 28,6076 | 0,134565 | 0,3931 | 0,730736 |
| Sipa1l2 | 6,21349 | 5,59539 | -0,151166 | 0,39295 | 0,730736 |
| Ap3b1 | 18,531 | 20,4081 | 0,139202 | 0,3936 | 0,730768 |
| Edrf1 | 12,5905 | 13,8569 | 0,138274 | 0,39335 | 0,730768 |
| Ccp110 | 28,2122 | 30,9993 | 0,135917 | 0,39335 | 0,730768 |
| Dchs1 | 16,9971 | 18,671 | 0,13551 | 0,3936 | 0,730768 |
| Slc52a2 | 10,5398 | 9,43137 | -0,160308 | 0,39345 | 0,730768 |
| BC068281 | 6,04647 | 5,30219 | -0,189505 | 0,3935 | 0,730768 |
| S100g | 2,70096 | 0,097546 | -4,79125 | 0,3934 | 0,730768 |
| Rpl27 | 3,23962 | 4,07922 | 0,332466 | 0,3939 | 0,731021 |

| | | | | | |
|---|---|---|---|---|---|
| Wnt2b | 0,678082 | 0,832231 | 0,295524 | 0,39385 | 0,731021 |
| Txndc17 | 63,4775 | 69,8837 | 0,138709 | 0,3938 | 0,731021 |
| Rita1 | 7,62313 | 8,70397 | 0,19129 | 0,39405 | 0,731096 |
| Scamp4 | 37,8255 | 34,1538 | -0,147313 | 0,394 | 0,731096 |
| Wars2 | 3,6463 | 3,21127 | -0,183291 | 0,39425 | 0,731365 |
| 4930481A15Rik | 1,45026 | 2,72795 | 0,911508 | 0,3944 | 0,731542 |
| Supt6 | 35,2107 | 38,6589 | 0,134788 | 0,3945 | 0,731626 |
| Hspa4 | 152,834 | 168,873 | 0,143978 | 0,3946 | 0,73171 |
| Tmem87b | 15,0348 | 13,6278 | -0,141753 | 0,3948 | 0,731776 |
| B4galt7 | 11,9423 | 10,5862 | -0,173899 | 0,3947 | 0,731776 |
| Tram1l1 | 8,08263 | 7,16183 | -0,174498 | 0,3948 | 0,731776 |
| Nfyc | 12,6354 | 14,1261 | 0,1609 | 0,395 | 0,732045 |
| 4930558J18Rik | 1,54068 | 1,86522 | 0,275775 | 0,3962 | 0,73305 |
| Abhd14b | 5,07249 | 5,91207 | 0,220968 | 0,39615 | 0,73305 |
| 5730559C18Rik | 8,44684 | 9,42827 | 0,15858 | 0,3959 | 0,73305 |
| Pank1 | 5,29792 | 5,86558 | 0,146848 | 0,39605 | 0,73305 |
| Mtpap | 19,1918 | 21,2044 | 0,143875 | 0,3958 | 0,73305 |
| Gltscr2 | 51,1058 | 56,275 | 0,139006 | 0,396 | 0,73305 |
| Dync1i2 | 66,045 | 72,4656 | 0,133847 | 0,39575 | 0,73305 |
| Tnrc6b | 6,90638 | 6,26992 | -0,139483 | 0,396 | 0,73305 |
| Ndufa13 | 65,9145 | 59,706 | -0,14272 | 0,3961 | 0,73305 |
| Stk35 | 13,0872 | 11,8379 | -0,144741 | 0,3962 | 0,73305 |
| Tm7sf3 | 15,2176 | 13,7559 | -0,145695 | 0,3961 | 0,73305 |
| Gcnt4 | 1,1693 | 0,991182 | -0,238418 | 0,3958 | 0,73305 |
| Spire1 | 16,4753 | 18,0856 | 0,134536 | 0,3963 | 0,733124 |
| Cnih1 | 89,585 | 81,4493 | -0,137354 | 0,39635 | 0,733124 |
| Uqcr10 | 271,777 | 299,406 | 0,139682 | 0,3965 | 0,733199 |
| Add1 | 57,2602 | 52,1517 | -0,134817 | 0,39645 | 0,733199 |
| Prss36 | 2,27523 | 1,94586 | -0,225604 | 0,3967 | 0,733467 |
| Rnf6 | 12,0815 | 13,3964 | 0,149044 | 0,3968 | 0,733532 |
| Car14 | 105,728 | 116,446 | 0,139303 | 0,3969 | 0,733532 |
| Sh3bgrl2 | 8,98407 | 8,06682 | -0,155368 | 0,39685 | 0,733532 |
| Rpl24 | 0,872556 | 1,29731 | 0,572204 | 0,3972 | 0,733561 |
| Hddc2 | 56,4527 | 62,8407 | 0,154656 | 0,3973 | 0,733561 |
| Psmb7 | 258,952 | 284,203 | 0,134235 | 0,3973 | 0,733561 |
| Chmp4b | 99,5391 | 109,099 | 0,132296 | 0,39715 | 0,733561 |
| Ankle2 | 24,3984 | 22,2187 | -0,135008 | 0,3973 | 0,733561 |
| Slc25a44 | 10,5629 | 9,5203 | -0,149922 | 0,3972 | 0,733561 |
| Cx3cl1 | 4,26846 | 3,74016 | -0,190619 | 0,3972 | 0,733561 |
| Tanc1 | 17,4013 | 15,8419 | -0,135451 | 0,3974 | 0,733645 |
| Bcl2l2 | 11,8153 | 10,65 | -0,149806 | 0,3975 | 0,733728 |
| Sel1l | 20,3069 | 18,4817 | -0,135873 | 0,3978 | 0,734171 |
| Abhd17b | 16,2376 | 14,6315 | -0,150264 | 0,39785 | 0,734171 |
| Hs1bp3 | 7,97124 | 8,91195 | 0,160936 | 0,39795 | 0,734254 |
| Tarbp2 | 11,8007 | 13,2863 | 0,171066 | 0,39815 | 0,734522 |
| Dnaja3 | 29,1668 | 32,0884 | 0,137725 | 0,39865 | 0,735223 |
| Jmjd1c | 12,2829 | 13,4999 | 0,136297 | 0,39875 | 0,735223 |
| Yipf1 | 19,8057 | 17,6921 | -0,162813 | 0,39865 | 0,735223 |

| | | | | | |
|---|---|---|---|---|---|
| Ninj1 | 7,05487 | 6,04039 | -0,223979 | 0,39875 | 0,735223 |
| Gm6194 | 0,53401 | 0,721649 | 0,434431 | 0,39895 | 0,735324 |
| Trnp1 | 0,653258 | 0,870462 | 0,414129 | 0,39905 | 0,735324 |
| Nucb2 | 8,53496 | 9,6848 | 0,182338 | 0,3991 | 0,735324 |
| 4931414P19Rik | 5,62164 | 6,36145 | 0,178365 | 0,3993 | 0,735324 |
| Asap3 | 3,60239 | 4,04207 | 0,166143 | 0,39915 | 0,735324 |
| Tube1 | 8,04003 | 8,99876 | 0,162526 | 0,39905 | 0,735324 |
| Zfp282 | 8,95549 | 9,89942 | 0,144572 | 0,3993 | 0,735324 |
| Ptpn9 | 26,7934 | 29,4725 | 0,137492 | 0,3993 | 0,735324 |
| Sirt4 | 4,24368 | 3,60508 | -0,235285 | 0,39905 | 0,735324 |
| 2310039H08Rik | 9,55077 | 11,2086 | 0,230911 | 0,3996 | 0,735351 |
| Cc2d2a | 5,3622 | 5,96949 | 0,154782 | 0,39945 | 0,735351 |
| Alms1 | 6,61055 | 7,29074 | 0,141293 | 0,3996 | 0,735351 |
| Baz1a | 9,3305 | 10,2875 | 0,140869 | 0,39965 | 0,735351 |
| H2afy2 | 134,248 | 147,316 | 0,134009 | 0,3996 | 0,735351 |
| Entpd5 | 3,5022 | 3,11097 | -0,170897 | 0,39965 | 0,735351 |
| C78339 | 15,8031 | 14,0331 | -0,171373 | 0,3997 | 0,735351 |
| Ipp | 9,05579 | 10,1796 | 0,168765 | 0,40015 | 0,735636 |
| Ngdn | 37,621 | 41,6021 | 0,145115 | 0,40025 | 0,735636 |
| Bap1 | 49,4859 | 54,2514 | 0,132644 | 0,4002 | 0,735636 |
| BC004004 | 10,4261 | 9,35175 | -0,156897 | 0,4001 | 0,735636 |
| Slc39a11 | 7,21077 | 6,41124 | -0,16955 | 0,40025 | 0,735636 |
| Hipk1 | 17,9422 | 15,2518 | -0,234374 | 0,40015 | 0,735636 |
| Fam103a1 | 4,66242 | 3,95253 | -0,238305 | 0,40035 | 0,735636 |
| Zfp931 | 2,18364 | 1,80604 | -0,273902 | 0,40035 | 0,735636 |
| Rubie | 0,922893 | 0,683749 | -0,432696 | 0,4003 | 0,735636 |
| Nhp2 | 99,8784 | 110,747 | 0,149022 | 0,40055 | 0,7357 |
| Wdr92 | 10,9797 | 12,1627 | 0,14763 | 0,4005 | 0,7357 |
| Ncbp2 | 104,423 | 114,579 | 0,133906 | 0,40055 | 0,7357 |
| Irak4 | 1,22023 | 1,00483 | -0,280211 | 0,40065 | 0,735782 |
| Atg4b | 30,099 | 33,093 | 0,13681 | 0,40085 | 0,736048 |
| Fam213b | 13,4496 | 15,5047 | 0,205138 | 0,4011 | 0,736195 |
| Calr | 588,363 | 652,81 | 0,149957 | 0,40115 | 0,736195 |
| Dpm2 | 43,5849 | 39,165 | -0,154264 | 0,401 | 0,736195 |
| Gm5089 | 2,04226 | 1,12246 | -0,863506 | 0,40115 | 0,736195 |
| Csnk1g1 | 10,145 | 11,1516 | 0,136489 | 0,4013 | 0,736268 |
| Dlg1 | 53,9018 | 59,1793 | 0,134759 | 0,4013 | 0,736268 |
| Efr3a | 14,799 | 16,2848 | 0,13802 | 0,40155 | 0,736625 |
| Tiprl | 15,6735 | 17,2437 | 0,137741 | 0,40185 | 0,73667 |
| Blmh | 101,427 | 111,119 | 0,131663 | 0,40185 | 0,73667 |
| Mex3c | 28,8571 | 26,2959 | -0,134091 | 0,40165 | 0,73667 |
| Kif3c | 5,41959 | 4,79336 | -0,177147 | 0,40185 | 0,73667 |
| Glmn | 21,1113 | 18,5601 | -0,185811 | 0,40175 | 0,73667 |
| Pycrl | 44,0822 | 48,6689 | 0,142805 | 0,402 | 0,736743 |
| Zc3h7b | 53,5394 | 58,7599 | 0,134231 | 0,402 | 0,736743 |
| Ifrd1 | 42,5796 | 46,8433 | 0,13768 | 0,4022 | 0,736907 |
| Nop14 | 34,8119 | 38,212 | 0,134448 | 0,40215 | 0,736907 |
| Ctf1 | 5,44806 | 4,63462 | -0,233293 | 0,4026 | 0,737539 |

| | | | | | |
|---|---|---|---|---|---|
| Bbs2 | 6,82003 | 7,65208 | 0,166075 | 0,4029 | 0,737987 |
| Lefty2 | 1,26932 | 1,53548 | 0,274642 | 0,403 | 0,738021 |
| L3mbtl2 | 17,5653 | 19,3313 | 0,138205 | 0,40305 | 0,738021 |
| Antxr2 | 12,0136 | 10,8652 | -0,144952 | 0,4032 | 0,738021 |
| Tmem119 | 24,3387 | 21,9906 | -0,146367 | 0,40325 | 0,738021 |
| Snora16a | 167,18 | 84,1952 | -0,989595 | 0,4032 | 0,738021 |
| Ppp2r1a | 129,328 | 141,676 | 0,131568 | 0,40325 | 0,738022 |
| Uckl1 | 10,6414 | 12,9294 | 0,280971 | 0,4034 | 0,738084 |
| Acot13 | 30,2966 | 34,1641 | 0,173327 | 0,40335 | 0,738084 |
| Arrdc3 | 21,783 | 23,9299 | 0,135612 | 0,40345 | 0,738084 |
| Gls2 | 3,43029 | 2,67658 | -0,357938 | 0,40365 | 0,738349 |
| Tmem143 | 5,55807 | 6,31151 | 0,183402 | 0,40385 | 0,738393 |
| Dusp12 | 18,9437 | 21,1642 | 0,159908 | 0,40395 | 0,738393 |
| Taf6 | 50,0523 | 55,5147 | 0,149433 | 0,40385 | 0,738393 |
| Zbtb5 | 10,8712 | 12,0011 | 0,142647 | 0,4039 | 0,738393 |
| Atp6v1g1 | 26,5214 | 23,7203 | -0,161036 | 0,4038 | 0,738393 |
| Clta | 162,003 | 177,464 | 0,13151 | 0,4042 | 0,738546 |
| Cited2 | 80,1703 | 73,1165 | -0,132869 | 0,4041 | 0,738546 |
| Clcc1 | 22,5337 | 18,97 | -0,248365 | 0,4042 | 0,738546 |
| Tbp | 32,8487 | 36,2157 | 0,140782 | 0,4043 | 0,738628 |
| Znhit2 | 12,4107 | 14,02 | 0,175909 | 0,4045 | 0,738792 |
| Thumpd1 | 49,8263 | 54,598 | 0,131938 | 0,40445 | 0,738792 |
| Smim11 | 55,2495 | 62,1518 | 0,169836 | 0,4049 | 0,73932 |
| Rhbdd3 | 13,4128 | 11,9755 | -0,163521 | 0,40485 | 0,73932 |
| Ttc27 | 18,4666 | 20,3828 | 0,142437 | 0,4051 | 0,739382 |
| Rfng | 10,9915 | 9,81398 | -0,163472 | 0,4051 | 0,739382 |
| Med31 | 15,3318 | 13,4454 | -0,189417 | 0,40505 | 0,739382 |
| Kctd2 | 18,439 | 20,4634 | 0,150287 | 0,40525 | 0,739415 |
| Rab40c | 13,5299 | 15 | 0,148808 | 0,40545 | 0,739415 |
| D430019H16Rik | 6,67259 | 7,38089 | 0,145547 | 0,4054 | 0,739415 |
| Psmd7 | 151,094 | 165,495 | 0,131339 | 0,4052 | 0,739415 |
| Ech1 | 24,0284 | 21,4995 | -0,160442 | 0,4053 | 0,739415 |
| Evi2a | 0,68072 | 0,163268 | -2,05982 | 0,4054 | 0,739415 |
| Rgp1 | 21,1957 | 23,8156 | 0,168137 | 0,40565 | 0,739669 |
| Ube2q1 | 51,2219 | 56,09 | 0,130982 | 0,4057 | 0,739669 |
| Akr1a1 | 236,424 | 259,269 | 0,133072 | 0,40645 | 0,740935 |
| Ccnt1 | 17,0846 | 18,9905 | 0,152574 | 0,40655 | 0,741017 |
| Cdkn1a | 12,4085 | 13,8634 | 0,159942 | 0,4067 | 0,741088 |
| Galk2 | 10,6679 | 11,8564 | 0,152397 | 0,40665 | 0,741088 |
| Rusc1 | 4,5771 | 5,27472 | 0,20466 | 0,40715 | 0,741503 |
| Pdlim7 | 35,375 | 39,4726 | 0,158121 | 0,4071 | 0,741503 |
| Ube2i | 12,5403 | 13,8981 | 0,148308 | 0,407 | 0,741503 |
| Nras | 58,7659 | 53,6821 | -0,130539 | 0,40705 | 0,741503 |
| Alg1 | 11,6997 | 10,1801 | -0,200722 | 0,40735 | 0,741767 |
| Ptges2 | 9,09025 | 8,06541 | -0,172572 | 0,40755 | 0,741828 |
| Fam221a | 1,3305 | 1,07567 | -0,306733 | 0,40745 | 0,741828 |
| Gm10451 | 0,773217 | 0,514809 | -0,586836 | 0,4075 | 0,741828 |
| Tceal3 | 2,34577 | 2,87393 | 0,292965 | 0,4079 | 0,74204 |

| | | | | | |
|---|---|---|---|---|---|
| Pcp4 | 30,8713 | 34,759 | 0,171119 | 0,40775 | 0,74204 |
| Hnrnpa2b1 | 806,823 | 901,428 | 0,15996 | 0,408 | 0,74204 |
| Mapk6 | 60,3415 | 66,1086 | 0,131688 | 0,4079 | 0,74204 |
| Pes1 | 33,8967 | 37,1266 | 0,131306 | 0,40795 | 0,74204 |
| Pkia | 3,61027 | 3,18276 | -0,18183 | 0,4078 | 0,74204 |
| Ehhadh | 1,32055 | 1,56756 | 0,247384 | 0,4081 | 0,742121 |
| Rslcan18 | 0,704945 | 0,95162 | 0,432876 | 0,40845 | 0,742444 |
| Snrnp35 | 18,28 | 20,5476 | 0,168704 | 0,4085 | 0,742444 |
| Gpn3 | 40,2086 | 44,3061 | 0,140002 | 0,4085 | 0,742444 |
| Ak3 | 31,5132 | 28,7304 | -0,133377 | 0,4085 | 0,742444 |
| Zfp422 | 46,168 | 42,1174 | -0,132474 | 0,4086 | 0,742525 |
| Ap1m1 | 48,1717 | 52,9091 | 0,135331 | 0,4088 | 0,742687 |
| Fbxo32 | 0,617338 | 0,518076 | -0,252896 | 0,4088 | 0,742687 |
| Trappc3 | 43,0742 | 47,5426 | 0,142396 | 0,4089 | 0,742767 |
| Dgcr2 | 66,1548 | 60,3553 | -0,132367 | 0,409 | 0,742838 |
| Pld6 | 0,969306 | 0,748028 | -0,373861 | 0,40905 | 0,742838 |
| Tmem8b | 2,2358 | 2,54681 | 0,187897 | 0,4093 | 0,742969 |
| Epb4.1l4b | 3,65049 | 4,15823 | 0,187879 | 0,4093 | 0,742969 |
| Bcl2 | 3,45812 | 3,8798 | 0,165996 | 0,4094 | 0,742969 |
| Rap2b | 7,40242 | 8,20099 | 0,147801 | 0,4094 | 0,742969 |
| Psmd6 | 137,629 | 150,682 | 0,13072 | 0,4093 | 0,742969 |
| Gm16515 | 9,20049 | 10,1689 | 0,144384 | 0,4097 | 0,743301 |
| Ift52 | 25,7124 | 28,3386 | 0,140305 | 0,40975 | 0,743301 |
| Kif21a | 3,30716 | 2,95067 | -0,16455 | 0,40965 | 0,743301 |
| Tnnt3 | 0,702515 | 0,405954 | -0,791213 | 0,41005 | 0,743744 |
| Fkbp3 | 460,382 | 512,144 | 0,153718 | 0,4102 | 0,743915 |
| Nmb | 8,33469 | 6,99061 | -0,253711 | 0,41055 | 0,744449 |
| Cnksr3 | 18,2619 | 20,0939 | 0,137924 | 0,41065 | 0,744509 |
| Slain2 | 19,064 | 20,8991 | 0,132593 | 0,41075 | 0,744509 |
| Zfp871 | 18,5229 | 16,9299 | -0,129737 | 0,41075 | 0,744509 |
| Cnnm3 | 11,0547 | 10,0366 | -0,139391 | 0,411 | 0,744861 |
| Tmem150b | 0,235293 | 0,613681 | 1,38303 | 0,41125 | 0,745202 |
| Cpsf4l | 1,01428 | 0,76113 | -0,414244 | 0,4113 | 0,745202 |
| Clcn6 | 8,58071 | 9,54867 | 0,154203 | 0,4116 | 0,745321 |
| Farsb | 64,7474 | 71,2406 | 0,137878 | 0,4117 | 0,745321 |
| Hspa14 | 62,1078 | 68,2056 | 0,135114 | 0,41155 | 0,745321 |
| Gja1 | 375,854 | 340,012 | -0,144586 | 0,4117 | 0,745321 |
| Akap7 | 6,42925 | 5,72798 | -0,166623 | 0,4116 | 0,745321 |
| 2810029C07Rik | 2,27027 | 1,90227 | -0,25514 | 0,4117 | 0,745321 |
| Kcne1l | 6,93568 | 6,02128 | -0,203968 | 0,41185 | 0,745491 |
| N6amt2 | 49,023 | 54,27 | 0,146695 | 0,41235 | 0,745689 |
| Ntrk3 | 1,3279 | 1,55435 | 0,227167 | 0,41215 | 0,74569 |
| Med12l | 2,76348 | 3,1882 | 0,206256 | 0,41215 | 0,74569 |
| Ift43 | 30,4033 | 34,0166 | 0,162011 | 0,41225 | 0,74569 |
| Atad2 | 48,583 | 53,1481 | 0,129566 | 0,4122 | 0,74569 |
| Atp6v0e2 | 7,92236 | 6,98634 | -0,181393 | 0,4122 | 0,74569 |
| Uprt | 3,53393 | 2,91684 | -0,276868 | 0,4123 | 0,74569 |
| Prox1 | 2,25474 | 2,5754 | 0,191831 | 0,41285 | 0,746291 |

| | | | | | |
|---|---|---|---|---|---|
| Aga | 7,39156 | 6,41252 | -0,204989 | 0,4128 | 0,746291 |
| Mpv17l | 0,609769 | 0,486029 | -0,327221 | 0,4128 | 0,746291 |
| Anapc10 | 8,02472 | 8,95089 | 0,15758 | 0,413 | 0,746349 |
| Capn15 | 14,7871 | 16,2941 | 0,140014 | 0,41305 | 0,746349 |
| Wasl | 19,5483 | 17,6667 | -0,146012 | 0,413 | 0,746349 |
| Dap | 22,7957 | 25,2497 | 0,147502 | 0,41365 | 0,747221 |
| Cops8 | 72,1211 | 78,9458 | 0,130442 | 0,4137 | 0,747221 |
| Capn3 | 0,692543 | 0,35489 | -0,964532 | 0,4136 | 0,747221 |
| Pdzd11 | 57,5969 | 63,383 | 0,138105 | 0,41395 | 0,74747 |
| Ccbe1 | 1,04975 | 0,898056 | -0,225161 | 0,4139 | 0,74747 |
| Hist1h2be | 0,331198 | 0,595131 | 0,845516 | 0,4145 | 0,747512 |
| Lonrf1 | 3,11741 | 3,52579 | 0,1776 | 0,41455 | 0,747512 |
| Clec16a | 3,83417 | 4,26237 | 0,152741 | 0,4144 | 0,747512 |
| Pidd1 | 9,52227 | 10,5775 | 0,15162 | 0,4144 | 0,747512 |
| Fabp5 | 48,1976 | 53,2121 | 0,142792 | 0,41425 | 0,747512 |
| Nfat5 | 10,4402 | 11,4867 | 0,137812 | 0,41465 | 0,747512 |
| Med28 | 16,2065 | 17,7866 | 0,13422 | 0,4141 | 0,747512 |
| Ppp1r12c | 23,6323 | 25,9238 | 0,133517 | 0,4146 | 0,747512 |
| Cdca3 | 106,105 | 116,148 | 0,130471 | 0,4147 | 0,747512 |
| Cul7 | 35,7034 | 39,0675 | 0,129908 | 0,41435 | 0,747512 |
| Ptpn11 | 56,3177 | 61,6187 | 0,129778 | 0,41415 | 0,747512 |
| Tsc22d1 | 128,307 | 116,522 | -0,139004 | 0,41445 | 0,747512 |
| Wdr45 | 8,3452 | 7,32891 | -0,187348 | 0,4144 | 0,747512 |
| Ints12 | 12,8149 | 11,5684 | -0,147639 | 0,41485 | 0,74758 |
| Kdm4c | 4,69944 | 4,19345 | -0,16435 | 0,4148 | 0,74758 |
| Dus2 | 8,33287 | 9,38567 | 0,171646 | 0,415 | 0,747628 |
| Uqcr11 | 164,925 | 181,863 | 0,141044 | 0,4151 | 0,747628 |
| Nf2 | 35,5576 | 39,1951 | 0,140516 | 0,415 | 0,747628 |
| Tmem164 | 37,9426 | 34,5838 | -0,133725 | 0,4151 | 0,747628 |
| Palm3 | 7,54128 | 8,44388 | 0,163097 | 0,4155 | 0,747743 |
| Pgls | 86,6798 | 95,2463 | 0,135967 | 0,4154 | 0,747743 |
| Usp19 | 42,6537 | 46,6388 | 0,128861 | 0,41545 | 0,747743 |
| Mfsd5 | 23,9331 | 21,6622 | -0,143833 | 0,4155 | 0,747743 |
| Tm2d1 | 9,08289 | 7,85466 | -0,209602 | 0,41545 | 0,747743 |
| Adam11 | 1,01074 | 0,85987 | -0,233221 | 0,4155 | 0,747743 |
| Wdr53 | 11,0362 | 12,5671 | 0,187416 | 0,41585 | 0,748129 |
| Faim | 24,1332 | 26,9718 | 0,160434 | 0,41605 | 0,748129 |
| Evc2 | 15,4165 | 16,9209 | 0,134328 | 0,41595 | 0,748129 |
| Dync1li1 | 35,5941 | 39,014 | 0,132351 | 0,416 | 0,748129 |
| Ttc4 | 35,6477 | 39,065 | 0,132066 | 0,41595 | 0,748129 |
| Kcnab2 | 0,983603 | 0,81789 | -0,266169 | 0,416 | 0,748129 |
| Dpm3 | 35,5971 | 31,2104 | -0,189733 | 0,41635 | 0,748567 |
| Syn1 | 0,678333 | 1,14462 | 0,754807 | 0,41655 | 0,748815 |
| Orc6 | 78,0577 | 85,4094 | 0,129854 | 0,4166 | 0,748815 |
| Eid2 | 25,7206 | 28,5115 | 0,148622 | 0,41675 | 0,748984 |
| Gale | 15,4911 | 17,2364 | 0,15402 | 0,4169 | 0,749041 |
| Siah1b | 17,1356 | 19,009 | 0,149683 | 0,41695 | 0,749041 |
| Ndufaf7 | 11,7745 | 10,5748 | -0,155039 | 0,41695 | 0,749042 |

| | | | | | |
|---|---|---|---|---|---|
| Plcxd1 | 3,34144 | 3,97307 | 0,249786 | 0,41735 | 0,749055 |
| Zc3h8 | 8,22188 | 9,31781 | 0,180522 | 0,41735 | 0,749055 |
| Atp5b | 851,981 | 946,7 | 0,152086 | 0,4173 | 0,749055 |
| Trafd1 | 12,7735 | 14,1257 | 0,145171 | 0,4172 | 0,749055 |
| Ddx46 | 47,5371 | 51,9375 | 0,12772 | 0,4172 | 0,749055 |
| Tbc1d23 | 9,11163 | 8,23474 | -0,145986 | 0,41705 | 0,749055 |
| Fam83g | 1,15742 | 0,985831 | -0,231502 | 0,41715 | 0,749055 |
| Parva | 19,9178 | 18,1842 | -0,131377 | 0,4176 | 0,749403 |
| Pip5k1a | 21,7754 | 23,9193 | 0,135478 | 0,41785 | 0,749449 |
| Specc1l | 22,2776 | 24,3747 | 0,129787 | 0,41785 | 0,749449 |
| Tnpo3 | 65,6751 | 71,7984 | 0,128607 | 0,4178 | 0,749449 |
| Klhl26 | 6,54942 | 5,84671 | -0,163743 | 0,41785 | 0,749449 |
| 4933427D14Rik | 10,4227 | 11,4695 | 0,13807 | 0,41795 | 0,749528 |
| 4931440P22Rik | 0,262729 | 0,666644 | 1,34334 | 0,41825 | 0,749664 |
| Zfp511 | 15,1013 | 19,4162 | 0,362588 | 0,41815 | 0,749664 |
| Pet112 | 6,94447 | 7,79731 | 0,167111 | 0,41825 | 0,749664 |
| Extl3 | 34,1868 | 31,28 | -0,128202 | 0,41825 | 0,749664 |
| Paip2b | 43,3237 | 47,4015 | 0,129776 | 0,41835 | 0,749731 |
| Brd2 | 103,342 | 113,229 | 0,131806 | 0,4184 | 0,749732 |
| Ufc1 | 61,8241 | 68,0931 | 0,139339 | 0,4186 | 0,749788 |
| Ankrd44 | 1,50894 | 1,31576 | -0,197633 | 0,4185 | 0,749788 |
| Cbx4 | 0,892838 | 0,752419 | -0,246862 | 0,41855 | 0,749788 |
| Rtn4ip1 | 6,6273 | 7,40205 | 0,159505 | 0,4189 | 0,750125 |
| Fam126a | 11,1144 | 9,0685 | -0,293496 | 0,41885 | 0,750125 |
| BC051142 | 0,910098 | 0,709029 | -0,360177 | 0,419 | 0,750203 |
| Mpi | 23,8588 | 26,3106 | 0,141123 | 0,41925 | 0,750439 |
| Psmd5 | 54,6262 | 59,7725 | 0,129889 | 0,4192 | 0,750439 |
| Sptssa | 84,4031 | 77,0868 | -0,130813 | 0,4193 | 0,750439 |
| Porcn | 3,67187 | 4,23986 | 0,207504 | 0,41955 | 0,750685 |
| Hnrnpa1 | 72,4558 | 66,0931 | -0,1326 | 0,4195 | 0,750685 |
| Timm23 | 2,41047 | 1,95425 | -0,3027 | 0,41965 | 0,750763 |
| Hmmr | 33,067 | 30,1836 | -0,131629 | 0,41975 | 0,750842 |
| Hells | 89,4501 | 97,7496 | 0,128007 | 0,41985 | 0,75092 |
| Sult2b1 | 0,977906 | 0,715265 | -0,451217 | 0,4201 | 0,751267 |
| Gtf2a2 | 88,4049 | 80,5185 | -0,134806 | 0,4203 | 0,751323 |
| Ascc3 | 8,18675 | 7,44815 | -0,136409 | 0,4202 | 0,751323 |
| Slc22a15 | 0,983641 | 0,847322 | -0,215221 | 0,4203 | 0,751323 |
| Zfp82 | 5,6189 | 6,35959 | 0,178647 | 0,4205 | 0,75158 |
| Cend1 | 0,794647 | 1,02349 | 0,365111 | 0,42075 | 0,751826 |
| Ikbip | 13,04 | 11,5082 | -0,180285 | 0,42075 | 0,751826 |
| Serpinf1 | 11,8408 | 13,2666 | 0,164032 | 0,4209 | 0,751993 |
| Pcsk9 | 14,1337 | 15,5505 | 0,137821 | 0,42125 | 0,752305 |
| Map4 | 35,4647 | 38,7279 | 0,126991 | 0,42125 | 0,752305 |
| Ptpra | 33,4665 | 30,6347 | -0,127554 | 0,4213 | 0,752305 |
| Ap1s3 | 1,22074 | 1,0173 | -0,26302 | 0,4213 | 0,752305 |
| Fv1 | 12,2129 | 13,7106 | 0,166884 | 0,4215 | 0,752361 |
| Zfp790 | 5,45843 | 6,05925 | 0,150653 | 0,42145 | 0,752361 |
| Fdxacb1 | 5,47941 | 4,79089 | -0,193727 | 0,4214 | 0,752361 |

| | | | | | |
|---|---|---|---|---|---|
| Mtap | 58,6387 | 64,0558 | 0,127475 | 0,4216 | 0,752428 |
| Lrrc58 | 23,8496 | 26,0479 | 0,127205 | 0,42165 | 0,752428 |
| Kdm3a | 32,4234 | 35,4473 | 0,128639 | 0,4218 | 0,752494 |
| Aph1a | 5,19861 | 3,76595 | -0,465113 | 0,4218 | 0,752494 |
| Cabin1 | 15,683 | 17,1425 | 0,128382 | 0,42195 | 0,752661 |
| Acin1 | 117,151 | 129,993 | 0,150069 | 0,42225 | 0,752672 |
| Syncrip | 74,8001 | 82,7384 | 0,145517 | 0,42225 | 0,752672 |
| Med13l | 7,4334 | 8,1497 | 0,132725 | 0,4223 | 0,752672 |
| Eif2d | 33,1416 | 36,309 | 0,131684 | 0,4222 | 0,752672 |
| Smc1a | 85,8973 | 93,9663 | 0,12953 | 0,4221 | 0,752672 |
| E430025E21Rik | 26,1098 | 28,5399 | 0,12839 | 0,42225 | 0,752672 |
| Zfp639 | 30,8355 | 28,0283 | -0,137708 | 0,42235 | 0,752672 |
| Uckl1os | 0,955154 | 2,44896 | 1,35837 | 0,42265 | 0,752805 |
| Wrn | 5,87216 | 6,46583 | 0,138946 | 0,4226 | 0,752805 |
| Psmd13 | 116,518 | 127,444 | 0,129308 | 0,4225 | 0,752805 |
| Lsm14b | 46,6288 | 50,9517 | 0,127908 | 0,4226 | 0,752805 |
| Lipe | 2,28336 | 2,69957 | 0,241573 | 0,42275 | 0,752826 |
| Rnf10 | 114,856 | 125,54 | 0,12832 | 0,42295 | 0,752826 |
| Ubr7 | 35,887 | 39,1871 | 0,12692 | 0,42295 | 0,752826 |
| Ndnf | 0,948927 | 0,801144 | -0,244236 | 0,423 | 0,752826 |
| Ahrr | 0,835415 | 0,700603 | -0,253896 | 0,42285 | 0,752826 |
| Dcxr | 4,21113 | 3,48515 | -0,272988 | 0,42295 | 0,752826 |
| Hnrnpk | 227,716 | 249,592 | 0,132333 | 0,4231 | 0,75287 |
| Srsf5 | 135,078 | 147,619 | 0,128089 | 0,4232 | 0,75287 |
| Cs | 121,704 | 133,002 | 0,128066 | 0,42315 | 0,75287 |
| Ubac1 | 14,7714 | 13,4362 | -0,136675 | 0,42325 | 0,75287 |
| Mks1 | 8,25753 | 9,20959 | 0,157427 | 0,42335 | 0,752948 |
| Wbscr22 | 49,8467 | 55,0027 | 0,142006 | 0,42355 | 0,753203 |
| Rbm5 | 62,7016 | 68,4686 | 0,12694 | 0,42395 | 0,753502 |
| Rab43 | 5,03974 | 4,51961 | -0,157151 | 0,4239 | 0,753502 |
| Adck4 | 8,44488 | 7,53393 | -0,164675 | 0,4239 | 0,753502 |
| Ccbl1 | 4,72504 | 4,11603 | -0,199074 | 0,4239 | 0,753502 |
| Cdk5r1 | 1,83342 | 1,58473 | -0,210297 | 0,424 | 0,753502 |
| Asun | 35,546 | 38,8284 | 0,127422 | 0,4241 | 0,75358 |
| Kif11 | 65,5467 | 59,9962 | -0,127652 | 0,4243 | 0,753823 |
| Pex12 | 7,12763 | 6,29997 | -0,178079 | 0,42435 | 0,753823 |
| Sugt1 | 52,4014 | 57,3091 | 0,129161 | 0,42445 | 0,753901 |
| Fbxl6 | 28,5629 | 25,9415 | -0,138881 | 0,42455 | 0,753978 |
| Prkrip1 | 17,7556 | 19,6854 | 0,14885 | 0,4247 | 0,754033 |
| Pkig | 46,2307 | 50,8776 | 0,138178 | 0,4247 | 0,754033 |
| Marveld1 | 4,16248 | 3,68171 | -0,177065 | 0,42475 | 0,754033 |
| Abcc1 | 9,98981 | 10,9555 | 0,133129 | 0,4249 | 0,754099 |
| Nub1 | 22,4188 | 24,5102 | 0,128672 | 0,4249 | 0,754099 |
| Mir6948 | 312,896 | 1466,5 | 2,22862 | 0,4252 | 0,75423 |
| Wrnip1 | 39,5928 | 36,193 | -0,129525 | 0,4251 | 0,75423 |
| Klhl29 | 2,28235 | 2,03244 | -0,167309 | 0,4251 | 0,75423 |
| Car9 | 0,785959 | 0,990889 | 0,33427 | 0,4252 | 0,754231 |
| Cdyl | 10,5376 | 11,6239 | 0,141551 | 0,4255 | 0,754451 |

| | | | | | |
|---|---|---|---|---|---|
| Supt20 | 33,0547 | 36,2578 | 0,133436 | 0,4254 | 0,754451 |
| Abhd17c | 31,072 | 28,3697 | -0,131267 | 0,4255 | 0,754451 |
| Piezo1 | 14,9112 | 13,3936 | -0,154854 | 0,42555 | 0,754451 |
| Eif2ak4 | 12,8911 | 14,112 | 0,130544 | 0,42565 | 0,754528 |
| 6030419C18Rik | 1,10741 | 1,34879 | 0,284476 | 0,426 | 0,754636 |
| Hyls1 | 13,974 | 16,3586 | 0,227297 | 0,42605 | 0,754636 |
| Cep76 | 13,2507 | 14,5358 | 0,133543 | 0,42605 | 0,754636 |
| Cblb | 10,8809 | 11,9164 | 0,131147 | 0,426 | 0,754636 |
| Pola1 | 19,1934 | 20,9639 | 0,127297 | 0,4258 | 0,754636 |
| Fnip1 | 5,96446 | 5,40305 | -0,142619 | 0,426 | 0,754636 |
| Elp6 | 13,1065 | 14,518 | 0,14756 | 0,42625 | 0,754879 |
| Akap10 | 12,9236 | 14,1986 | 0,135747 | 0,4263 | 0,754879 |
| Kcng1 | 1,33682 | 1,59744 | 0,256959 | 0,4279 | 0,755 |
| Rqcd1 | 62,3376 | 70,3616 | 0,174685 | 0,4269 | 0,755 |
| Gps1 | 10,0789 | 11,2513 | 0,158759 | 0,4277 | 0,755 |
| Timmdc1 | 16,8384 | 18,567 | 0,140987 | 0,4277 | 0,755 |
| Bpnt1 | 19,6924 | 21,6643 | 0,137679 | 0,4278 | 0,755 |
| Dennd5b | 5,40916 | 5,94196 | 0,135535 | 0,42775 | 0,755 |
| Ciao1 | 20,0746 | 21,9728 | 0,130348 | 0,42745 | 0,755 |
| Nono | 118,256 | 129,343 | 0,129278 | 0,4276 | 0,755 |
| Mrpl50 | 51,9054 | 56,7465 | 0,128648 | 0,42735 | 0,755 |
| Lsm14a | 84,4644 | 92,2747 | 0,127592 | 0,42755 | 0,755 |
| Prex1 | 15,3587 | 16,7744 | 0,12721 | 0,4273 | 0,755 |
| Nol11 | 37,4179 | 40,8167 | 0,125431 | 0,42705 | 0,755 |
| Xrn2 | 69,9521 | 76,293 | 0,125183 | 0,42765 | 0,755 |
| Actl6a | 102,337 | 111,461 | 0,123215 | 0,42795 | 0,755 |
| Tmem248 | 33,7859 | 30,9481 | -0,126571 | 0,4268 | 0,755 |
| Atp6v0a1 | 18,4446 | 16,8226 | -0,132794 | 0,42725 | 0,755 |
| Erbb2 | 21,9682 | 20,0275 | -0,133431 | 0,42735 | 0,755 |
| Paxbp1 | 33,1111 | 30,0196 | -0,141408 | 0,42705 | 0,755 |
| Chm | 7,01939 | 6,36171 | -0,141932 | 0,4278 | 0,755 |
| Ube2w | 11,7476 | 10,6281 | -0,144485 | 0,42795 | 0,755 |
| Pla2g6 | 6,232 | 5,56778 | -0,162594 | 0,42675 | 0,755 |
| 03/03/16 | 8,59591 | 7,61229 | -0,17532 | 0,42705 | 0,755 |
| Armcx6 | 4,86523 | 4,25739 | -0,192538 | 0,42645 | 0,755 |
| Zbtb42 | 1,95832 | 1,6839 | -0,217805 | 0,4269 | 0,755 |
| Wdr5b | 3,84935 | 3,30986 | -0,217843 | 0,42765 | 0,755 |
| E330033B04Rik | 0,964936 | 0,828577 | -0,219798 | 0,42705 | 0,755 |
| Zfp119b | 2,88122 | 2,45216 | -0,232622 | 0,4269 | 0,755 |
| Neil1 | 3,55507 | 2,97248 | -0,25821 | 0,4276 | 0,755 |
| Dhx36 | 37,9654 | 34,8196 | -0,124784 | 0,42815 | 0,755042 |
| Ndufaf1 | 24,8665 | 22,512 | -0,143509 | 0,4282 | 0,755042 |
| D130017N08Rik | 2,63911 | 2,22426 | -0,246722 | 0,4281 | 0,755042 |
| Txnrd2 | 13,5205 | 11,1959 | -0,27218 | 0,42805 | 0,755042 |
| Mrfap1 | 123,091 | 134,357 | 0,126344 | 0,42875 | 0,755701 |
| Cdc37l1 | 20,0215 | 18,2318 | -0,135089 | 0,42875 | 0,755701 |
| Taok3 | 1,8249 | 1,58636 | -0,202096 | 0,4288 | 0,755701 |
| Acss2 | 2,73906 | 2,37372 | -0,206533 | 0,42875 | 0,755701 |

| | | | | | |
|---|---|---|---|---|---|
| Hspb6 | 2,48753 | 2,977 | 0,259147 | 0,429 | 0,755919 |
| Mrps7 | 57,0624 | 62,4539 | 0,130252 | 0,42915 | 0,755919 |
| Cdc40 | 25,5231 | 27,8899 | 0,127941 | 0,42915 | 0,755919 |
| Psen1 | 27,917 | 25,5446 | -0,128128 | 0,42915 | 0,755919 |
| Abt1 | 6,13009 | 6,78202 | 0,145806 | 0,42935 | 0,755961 |
| Tmcc1 | 7,72434 | 8,50313 | 0,138582 | 0,4294 | 0,755961 |
| Nup37 | 50,2732 | 55,0291 | 0,130407 | 0,42925 | 0,755961 |
| Ppp2r2a | 39,1141 | 42,6578 | 0,12512 | 0,4293 | 0,755961 |
| Knstrn | 33,7808 | 30,9225 | -0,127543 | 0,42955 | 0,756126 |
| Hdac11 | 14,4581 | 13,1013 | -0,142165 | 0,4298 | 0,756466 |
| Ccdc114 | 2,53484 | 2,92632 | 0,207192 | 0,43005 | 0,756806 |
| Dpp8 | 36,8091 | 40,1212 | 0,124305 | 0,43015 | 0,756883 |
| Gm14326 | 0,974156 | 0,796311 | -0,290821 | 0,43025 | 0,756959 |
| Zfp827 | 6,94443 | 7,61574 | 0,133128 | 0,43045 | 0,757211 |
| Sar1a | 92,2386 | 84,5892 | -0,124898 | 0,4306 | 0,757375 |
| Gng8 | 0,868298 | 1,19222 | 0,457383 | 0,4312 | 0,75744 |
| Ankrd34a | 0,999185 | 1,20633 | 0,271801 | 0,4308 | 0,75744 |
| Prdm11 | 2,54054 | 2,97571 | 0,228103 | 0,43115 | 0,75744 |
| Cenpo | 5,84721 | 6,77961 | 0,213455 | 0,4313 | 0,75744 |
| 2700054A10Rik | 2,05183 | 2,33964 | 0,189373 | 0,43155 | 0,75744 |
| Gtpbp6 | 14,5184 | 16,4364 | 0,179014 | 0,43075 | 0,75744 |
| Zfp691 | 6,9501 | 7,83886 | 0,173612 | 0,43155 | 0,75744 |
| Asb3 | 7,27618 | 8,11152 | 0,156791 | 0,43105 | 0,75744 |
| H2afj | 11,0126 | 12,2709 | 0,156087 | 0,43135 | 0,75744 |
| Ofd1 | 5,7972 | 6,41652 | 0,146434 | 0,4313 | 0,75744 |
| Exosc4 | 20,4525 | 22,5523 | 0,140998 | 0,43145 | 0,75744 |
| Nck2 | 14,8977 | 16,4044 | 0,138993 | 0,43115 | 0,75744 |
| Frg1 | 62,7616 | 68,7531 | 0,131542 | 0,43095 | 0,75744 |
| Tbpl1 | 31,568 | 34,524 | 0,129136 | 0,4314 | 0,75744 |
| Psma2 | 183,233 | 199,565 | 0,123173 | 0,4316 | 0,75744 |
| A830018L16Rik | 1,3582 | 1,14033 | -0,252251 | 0,43145 | 0,75744 |
| 0610009L18Rik | 2,03996 | 1,48487 | -0,45821 | 0,4312 | 0,75744 |
| 4933411K20Rik | 8,88098 | 8,06417 | -0,139193 | 0,43175 | 0,757603 |
| Dph6 | 9,7623 | 10,8093 | 0,146986 | 0,43185 | 0,757679 |
| Tmem82 | 0,679487 | 0,882359 | 0,376921 | 0,43195 | 0,757755 |
| Fam13c | 3,81156 | 4,43751 | 0,219367 | 0,4322 | 0,757895 |
| Trip13 | 41,3552 | 45,1573 | 0,126892 | 0,4322 | 0,757895 |
| Alg9 | 18,6004 | 16,9586 | -0,133319 | 0,4321 | 0,757895 |
| Smarcc2 | 37,1347 | 40,5232 | 0,125979 | 0,43235 | 0,75796 |
| Phf20 | 13,0323 | 11,9167 | -0,12911 | 0,43235 | 0,75796 |
| Zfp830 | 9,52309 | 10,4956 | 0,140283 | 0,43255 | 0,758199 |
| Angptl2 | 17,0658 | 15,4945 | -0,139356 | 0,4326 | 0,758199 |
| Ankib1 | 9,06202 | 8,26395 | -0,133002 | 0,43275 | 0,758362 |
| Sh3bgr | 0,842637 | 1,09905 | 0,383274 | 0,4329 | 0,758526 |
| Myeov2 | 157,221 | 172,833 | 0,136581 | 0,4332 | 0,758653 |
| Gm15446 | 0,900722 | 1,14654 | 0,348133 | 0,43315 | 0,758654 |
| Itgb2l | 0,958822 | 1,15747 | 0,271639 | 0,43305 | 0,758654 |
| H2afv | 99,4234 | 108,336 | 0,123853 | 0,4331 | 0,758654 |

| | | | | | |
|---|---|---|---|---|---|
| Ptgr2 | 7,9274 | 7,15188 | -0,148527 | 0,43335 | 0,758817 |
| Skp1a | 249,337 | 272,067 | 0,125865 | 0,43345 | 0,758881 |
| Tfb2m | 2,1739 | 1,85498 | -0,228886 | 0,4335 | 0,758881 |
| Thg1l | 8,60847 | 9,48918 | 0,140528 | 0,4336 | 0,758944 |
| Chchd4 | 47,2819 | 51,9309 | 0,135305 | 0,43365 | 0,758944 |
| Cenpv | 22,0663 | 24,55 | 0,153882 | 0,43375 | 0,75902 |
| Fchsd2 | 13,2117 | 14,4616 | 0,130418 | 0,43395 | 0,75927 |
| 0610012G03Rik | 16,8442 | 18,6877 | 0,149832 | 0,43405 | 0,75931 |
| Foxo3 | 9,30601 | 10,2809 | 0,143726 | 0,4342 | 0,75931 |
| Polr2m | 238,562 | 260,553 | 0,127208 | 0,4341 | 0,75931 |
| Pde4d | 3,12654 | 2,81357 | -0,152166 | 0,43415 | 0,75931 |
| Pbrm1 | 43,8692 | 40,1745 | -0,126929 | 0,4343 | 0,759386 |
| Mpdz | 28,7718 | 31,3509 | 0,123851 | 0,43445 | 0,759549 |
| 4930483K19Rik | 0,575529 | 1,4627 | 1,34567 | 0,43475 | 0,759652 |
| Tefm | 15,2111 | 16,9754 | 0,158319 | 0,43485 | 0,759652 |
| Litaf | 24,0837 | 26,424 | 0,133792 | 0,43475 | 0,759652 |
| Dock11 | 12,5984 | 11,5246 | -0,128531 | 0,4348 | 0,759652 |
| Gm20939 | 1,17509 | 0,97011 | -0,276548 | 0,4348 | 0,759652 |
| Trpm4 | 2,07885 | 0,909352 | -1,19287 | 0,43485 | 0,759652 |
| Ranbp2 | 44,741 | 40,9469 | -0,127842 | 0,43495 | 0,759727 |
| Nr2f6 | 31,3723 | 28,6706 | -0,129922 | 0,43515 | 0,759977 |
| Kctd14 | 0,511699 | 0,651717 | 0,348951 | 0,43555 | 0,760278 |
| Akap8l | 28,456 | 31,2013 | 0,132875 | 0,43555 | 0,760278 |
| Ppp4r1 | 34,0915 | 37,1259 | 0,123013 | 0,43555 | 0,760278 |
| Trim65 | 0,819601 | 0,67019 | -0,290351 | 0,43545 | 0,760278 |
| Vps25 | 83,586 | 91,2261 | 0,126186 | 0,4357 | 0,760441 |
| Mycbp | 20,9845 | 23,188 | 0,144054 | 0,43595 | 0,760678 |
| Apbb2 | 4,19336 | 3,77358 | -0,152175 | 0,43595 | 0,760678 |
| Echdc3 | 0,8172 | 1,03213 | 0,33686 | 0,43625 | 0,76078 |
| Tox4 | 16,7295 | 18,9213 | 0,177614 | 0,4362 | 0,76078 |
| Zfp692 | 19,4349 | 21,4154 | 0,140001 | 0,43635 | 0,76078 |
| Pcnxl3 | 21,6734 | 19,9187 | -0,121798 | 0,4361 | 0,76078 |
| Abi3bp | 0,828545 | 0,689191 | -0,265677 | 0,43615 | 0,76078 |
| Mir3064 | 2952,93 | 805,241 | -1,87466 | 0,4363 | 0,76078 |
| A730098P11Rik | 2,01186 | 2,81452 | 0,484359 | 0,4367 | 0,761068 |
| Usp31 | 6,28083 | 6,87006 | 0,129368 | 0,4368 | 0,761068 |
| Ftsj3 | 85,1129 | 92,736 | 0,123752 | 0,43665 | 0,761068 |
| Ubfd1 | 45,0905 | 49,1044 | 0,12303 | 0,43675 | 0,761068 |
| Uba6 | 14,2256 | 13,0145 | -0,128372 | 0,43665 | 0,761068 |
| Gpr50 | 0,985872 | 1,22287 | 0,310801 | 0,437 | 0,761107 |
| Tmem50b | 23,4996 | 25,8093 | 0,135255 | 0,437 | 0,761107 |
| Nbr1 | 34,255 | 37,3669 | 0,125446 | 0,43695 | 0,761107 |
| Ryk | 53,0406 | 48,6879 | -0,123533 | 0,43705 | 0,761107 |
| Exosc9 | 56,6037 | 68,44 | 0,273943 | 0,43725 | 0,761146 |
| Myl12a | 99,7355 | 108,527 | 0,121877 | 0,4372 | 0,761146 |
| Dbn1 | 129,371 | 118,537 | -0,126187 | 0,4373 | 0,761146 |
| Pex2 | 29,4389 | 26,8799 | -0,131194 | 0,4373 | 0,761146 |
| Bckdhb | 21,9925 | 24,2828 | 0,142924 | 0,43755 | 0,761482 |

| | | | | | |
|---|---|---|---|---|---|
| Slc25a22 | 17,0048 | 18,6778 | 0,135384 | 0,4377 | 0,761544 |
| Puf60 | 118,279 | 128,696 | 0,12178 | 0,4377 | 0,761544 |
| Fam189a2 | 0,805795 | 0,984488 | 0,28896 | 0,4378 | 0,761619 |
| Ergic3 | 87,0875 | 79,8254 | -0,125617 | 0,43805 | 0,761955 |
| Ccdc6 | 15,6882 | 17,1274 | 0,126627 | 0,4384 | 0,762365 |
| 5031425E22Rik | 6,95137 | 6,22312 | -0,159659 | 0,4384 | 0,762365 |
| Fam229b | 4,28879 | 5,17943 | 0,272224 | 0,43865 | 0,762664 |
| Tshz3 | 1,27716 | 1,47106 | 0,203915 | 0,4388 | 0,762664 |
| Igfbp7 | 2,53667 | 0,552523 | -2,19883 | 0,43875 | 0,762664 |
| Rnu73b | 2196,42 | 405,54 | -2,43724 | 0,4388 | 0,762664 |
| Hsf2 | 26,3643 | 28,7898 | 0,12697 | 0,43905 | 0,762999 |
| Klf12 | 6,23442 | 6,86079 | 0,13812 | 0,4393 | 0,763334 |
| Prdm9 | 1,09072 | 1,28479 | 0,236254 | 0,43955 | 0,763446 |
| Pus3 | 22,7944 | 25,42 | 0,157288 | 0,4396 | 0,763446 |
| Hdhd2 | 18,0874 | 19,8892 | 0,137004 | 0,43945 | 0,763446 |
| Amot | 30,0208 | 32,6818 | 0,122526 | 0,43965 | 0,763446 |
| Prkx | 6,55092 | 5,88235 | -0,155307 | 0,43955 | 0,763446 |
| Snupn | 12,7103 | 14,2117 | 0,161086 | 0,43985 | 0,763496 |
| Lrp3 | 18,1096 | 19,7648 | 0,126174 | 0,4398 | 0,763496 |
| Nfe2l1 | 64,1748 | 58,8739 | -0,124381 | 0,43975 | 0,763496 |
| G6pd2 | 3,83481 | 4,43707 | 0,210454 | 0,44025 | 0,763781 |
| Rbbp5 | 21,8907 | 23,899 | 0,126634 | 0,4401 | 0,763781 |
| Slc35b3 | 10,8731 | 9,77397 | -0,153751 | 0,44025 | 0,763781 |
| Ppih | 24,759 | 21,9529 | -0,173546 | 0,44025 | 0,763781 |
| Ero1lb | 1,14144 | 0,975173 | -0,227124 | 0,4403 | 0,763781 |
| Mir1901 | 31,1465 | 226,35 | 2,86142 | 0,44055 | 0,764016 |
| Tmod4 | 0,496711 | 0,688753 | 0,471578 | 0,44055 | 0,764016 |
| Smcr8 | 9,79354 | 10,7349 | 0,132404 | 0,4407 | 0,764177 |
| Oma1 | 10,4655 | 9,38708 | -0,156897 | 0,4408 | 0,764252 |
| Vasp | 60,655 | 66,0944 | 0,123901 | 0,44105 | 0,764573 |
| Lphn3 | 1,93249 | 1,7118 | -0,174944 | 0,4411 | 0,764573 |
| Prelid1 | 121,379 | 132,564 | 0,127173 | 0,44145 | 0,764882 |
| Snx27 | 24,1337 | 26,3044 | 0,124258 | 0,4414 | 0,764882 |
| Fancb | 9,34326 | 8,4428 | -0,146205 | 0,44135 | 0,764882 |
| Stmn2 | 1,70274 | 1,4171 | -0,264914 | 0,44165 | 0,76513 |
| Mgrn1 | 21,145 | 23,7354 | 0,16672 | 0,44175 | 0,765204 |
| Ndufb7 | 141,943 | 155,381 | 0,130499 | 0,4419 | 0,765352 |
| Radil | 1,33292 | 1,11158 | -0,261973 | 0,44195 | 0,765352 |
| Wee1 | 27,3611 | 29,8023 | 0,1233 | 0,44215 | 0,765363 |
| Clasp1 | 27,4561 | 29,9021 | 0,123119 | 0,4423 | 0,765363 |
| Pnrc2 | 84,9689 | 78,1454 | -0,120774 | 0,4421 | 0,765363 |
| Arhgap5 | 36,2648 | 33,3086 | -0,122677 | 0,44225 | 0,765363 |
| Hdac6 | 17,8983 | 16,3863 | -0,127335 | 0,44225 | 0,765363 |
| Slc9a8 | 9,46321 | 8,61916 | -0,134782 | 0,44225 | 0,765363 |
| Ikzf5 | 6,16896 | 6,81313 | 0,143291 | 0,44255 | 0,765696 |
| Adam1a | 1,09235 | 0,91756 | -0,251567 | 0,44265 | 0,76577 |
| Chp1 | 26,769 | 24,4962 | -0,128008 | 0,44275 | 0,765844 |
| Trmt13 | 5,30063 | 6,38017 | 0,26743 | 0,44315 | 0,765916 |

| | | | | | |
|---|---|---|---|---|---|
| Psph | 13,9841 | 15,5421 | 0,152396 | 0,44295 | 0,765916 |
| Pex7 | 11,2121 | 12,4366 | 0,149538 | 0,44325 | 0,765916 |
| Btg1 | 11,9971 | 13,1096 | 0,127939 | 0,4431 | 0,765916 |
| Ddx51 | 14,7871 | 16,1224 | 0,124733 | 0,44325 | 0,765916 |
| Arhgap32 | 5,8742 | 6,40006 | 0,123694 | 0,44305 | 0,765916 |
| Polr2a | 70,4857 | 76,7675 | 0,123164 | 0,44315 | 0,765916 |
| Ssbp4 | 15,1435 | 13,6039 | -0,154684 | 0,443 | 0,765916 |
| Ccdc138 | 4,70015 | 5,28149 | 0,168238 | 0,44345 | 0,765939 |
| Pias3 | 11,8453 | 13,0647 | 0,141362 | 0,44355 | 0,765939 |
| Fktn | 15,0563 | 13,7513 | -0,130798 | 0,44335 | 0,765939 |
| Ppp1r1a | 34,0121 | 30,9302 | -0,137031 | 0,44355 | 0,765939 |
| Efcab2 | 4,06088 | 3,58229 | -0,180913 | 0,44345 | 0,765939 |
| Rnf113a1 | 5,03046 | 5,84577 | 0,216702 | 0,4438 | 0,766074 |
| Prmt6 | 17,8297 | 19,5247 | 0,131015 | 0,4438 | 0,766074 |
| Aktip | 13,4435 | 11,8769 | -0,178753 | 0,44375 | 0,766074 |
| Wdr83 | 25,7828 | 29,5032 | 0,194463 | 0,44395 | 0,76611 |
| 9430038I01Rik | 7,36605 | 8,42327 | 0,19349 | 0,4439 | 0,76611 |
| Snap23 | 39,9008 | 43,5138 | 0,125055 | 0,44405 | 0,76611 |
| Zfyve27 | 5,64639 | 5,12658 | -0,13933 | 0,44405 | 0,76611 |
| Ptgr1 | 6,62403 | 7,42312 | 0,164316 | 0,4442 | 0,766269 |
| Lyrm4 | 6,29034 | 5,51012 | -0,191054 | 0,44435 | 0,766429 |
| Elavl4 | 1,01787 | 1,1924 | 0,228313 | 0,44455 | 0,766452 |
| 2700099C18Rik | 25,7603 | 28,7125 | 0,156531 | 0,44465 | 0,766452 |
| Sugp1 | 20,7602 | 22,7177 | 0,129996 | 0,4446 | 0,766452 |
| Rnf19a | 15,9882 | 14,6378 | -0,127315 | 0,4445 | 0,766452 |
| Slc25a36 | 19,1724 | 17,5254 | -0,129583 | 0,44445 | 0,766452 |
| A330021E22Rik | 1,04326 | 1,23652 | 0,24519 | 0,44485 | 0,7665 |
| Zmym2 | 30,9602 | 33,6537 | 0,120351 | 0,44485 | 0,7665 |
| Nab1 | 3,72504 | 3,34882 | -0,153602 | 0,44475 | 0,7665 |
| D130040H23Rik | 2,16289 | 1,83228 | -0,239313 | 0,44495 | 0,766574 |
| Intu | 4,69424 | 5,16576 | 0,13809 | 0,44505 | 0,766647 |
| Elavl2 | 3,99075 | 4,4596 | 0,160255 | 0,44515 | 0,766721 |
| Gnb5 | 5,85535 | 5,20994 | -0,16849 | 0,44535 | 0,766966 |
| Nadk | 43,3069 | 47,1271 | 0,12196 | 0,44545 | 0,76704 |
| Mis12 | 31,9191 | 35,5267 | 0,154485 | 0,44555 | 0,767113 |
| Cep83 | 23,0327 | 25,1372 | 0,12614 | 0,4458 | 0,767221 |
| Uqcrc1 | 136,196 | 125,201 | -0,121441 | 0,44585 | 0,767221 |
| Zdhhc13 | 24,8094 | 22,6771 | -0,129646 | 0,4458 | 0,767221 |
| Gadd45g | 18,0556 | 16,1711 | -0,159028 | 0,4459 | 0,767221 |
| Nfu1 | 19,8192 | 17,7308 | -0,160641 | 0,44585 | 0,767221 |
| Clptm1 | 31,0609 | 28,5112 | -0,123575 | 0,446 | 0,767295 |
| Frmd6 | 15,9976 | 14,6549 | -0,126466 | 0,44615 | 0,767355 |
| Galnt10 | 10,7897 | 9,86618 | -0,129089 | 0,44615 | 0,767355 |
| Api5 | 66,0896 | 60,7074 | -0,12255 | 0,4463 | 0,767514 |
| Sytl2 | 0,614638 | 0,753095 | 0,293095 | 0,44645 | 0,767575 |
| Akap17b | 2,75337 | 2,46397 | -0,160217 | 0,44645 | 0,767575 |
| Ret | 1,60655 | 1,80575 | 0,16863 | 0,4466 | 0,767622 |
| Pik3r3 | 16,2816 | 14,942 | -0,123864 | 0,44665 | 0,767622 |

| | | | | | |
|---|---|---|---|---|---|
| Large | 10,6539 | 9,70519 | -0,134555 | 0,44655 | 0,767622 |
| Prokr2 | 0,841106 | 0,987069 | 0,230864 | 0,4471 | 0,767765 |
| Ccndbp1 | 18,4356 | 20,3644 | 0,143559 | 0,4472 | 0,767765 |
| Bud13 | 18,0924 | 19,8484 | 0,133645 | 0,4471 | 0,767765 |
| Ints1 | 20,1624 | 21,9332 | 0,121447 | 0,44725 | 0,767765 |
| Tor1a | 29,613 | 26,9828 | -0,134193 | 0,4471 | 0,767765 |
| Rnf166 | 12,8448 | 11,5885 | -0,14849 | 0,4469 | 0,767765 |
| 2810408A11Rik | 9,85944 | 8,81049 | -0,162284 | 0,44715 | 0,767765 |
| Rpp38 | 16,6096 | 14,8278 | -0,163706 | 0,44695 | 0,767765 |
| Gsdmd | 0,655333 | 0,497801 | -0,39666 | 0,44705 | 0,767765 |
| Serpinh1 | 592,83 | 538,972 | -0,137408 | 0,4478 | 0,768413 |
| Relt | 3,41199 | 3,00278 | -0,184316 | 0,44775 | 0,768413 |
| Nudt6 | 2,63985 | 2,16759 | -0,284369 | 0,4477 | 0,768413 |
| Cisd3 | 6,37024 | 7,49581 | 0,234737 | 0,4491 | 0,768473 |
| Zfp493 | 2,0531 | 2,35568 | 0,198341 | 0,4481 | 0,768473 |
| C330018D20Rik | 5,62732 | 6,3035 | 0,163705 | 0,44805 | 0,768473 |
| Zfp839 | 4,30939 | 4,81169 | 0,15906 | 0,44885 | 0,768473 |
| Tbc1d22a | 6,0739 | 6,73704 | 0,149492 | 0,44895 | 0,768473 |
| Ift27 | 25,499 | 28,2104 | 0,145786 | 0,4491 | 0,768473 |
| Mrpl52 | 187,746 | 205,523 | 0,130522 | 0,4487 | 0,768473 |
| Rcan1 | 18,9555 | 20,7432 | 0,13002 | 0,4489 | 0,768473 |
| Nin | 11,0514 | 12,0845 | 0,128924 | 0,4491 | 0,768473 |
| Fhl1 | 35,8903 | 39,1285 | 0,124628 | 0,44825 | 0,768473 |
| Fbxo5 | 49,0407 | 53,4228 | 0,123476 | 0,4483 | 0,768473 |
| Cdc34 | 74,1962 | 80,823 | 0,123422 | 0,44795 | 0,768473 |
| Txnrd1 | 102,487 | 111,496 | 0,121538 | 0,44835 | 0,768473 |
| Dot1l | 18,138 | 19,7317 | 0,121501 | 0,4486 | 0,768473 |
| Senp3 | 58,788 | 63,953 | 0,121492 | 0,44885 | 0,768473 |
| Igf1r | 28,6519 | 31,1559 | 0,120875 | 0,4483 | 0,768473 |
| Poldip2 | 64,9893 | 70,6613 | 0,120719 | 0,4489 | 0,768473 |
| Lamb2 | 10,6342 | 9,74838 | -0,125475 | 0,4488 | 0,768473 |
| Fastk | 32,7076 | 29,9556 | -0,126799 | 0,4489 | 0,768473 |
| Yes1 | 21,4342 | 19,6252 | -0,127205 | 0,44895 | 0,768473 |
| Plekhn1 | 3,45868 | 3,01386 | -0,198611 | 0,44855 | 0,768473 |
| Rgs10 | 6,78439 | 5,81161 | -0,223282 | 0,4489 | 0,768473 |
| Plce1 | 2,75861 | 3,03942 | 0,139852 | 0,44955 | 0,768652 |
| Taf8 | 17,9142 | 19,5876 | 0,12884 | 0,44945 | 0,768652 |
| Rfc3 | 67,9115 | 73,979 | 0,123461 | 0,44955 | 0,768652 |
| Cers5 | 43,6037 | 39,96 | -0,125895 | 0,4495 | 0,768652 |
| Lsm12 | 26,9752 | 24,4558 | -0,141461 | 0,4495 | 0,768652 |
| Map4k1 | 4,05718 | 3,57772 | -0,18144 | 0,4495 | 0,768652 |
| Rpa3 | 74,0499 | 67,5056 | -0,13349 | 0,44975 | 0,768896 |
| Tmem28 | 1,18701 | 1,4741 | 0,312505 | 0,45 | 0,769028 |
| Ccdc28a | 3,19964 | 3,77678 | 0,239247 | 0,45 | 0,769028 |
| Susd5 | 1,82286 | 1,58907 | -0,198021 | 0,45 | 0,769028 |
| Mir1668 | 9,62355 | 39,3026 | 2,02998 | 0,4512 | 0,76921 |
| 1700109K24Rik | 1,16858 | 3,44107 | 1,5581 | 0,4511 | 0,76921 |
| Zfp85os | 0,436452 | 0,597444 | 0,45298 | 0,45055 | 0,76921 |

| | | | | | |
|---|---|---|---|---|---|
| Tmem121 | 5,86103 | 6,6576 | 0,183848 | 0,4507 | 0,76921 |
| Smyd4 | 4,1651 | 4,65163 | 0,159386 | 0,451 | 0,76921 |
| Gm13498 | 8,11695 | 9,03692 | 0,154893 | 0,45035 | 0,76921 |
| Pmm1 | 14,846 | 16,4939 | 0,15186 | 0,45095 | 0,76921 |
| Mrps35 | 47,7351 | 52,2514 | 0,13042 | 0,4506 | 0,76921 |
| Ndufa3 | 327,808 | 358,816 | 0,130394 | 0,45075 | 0,76921 |
| Bag5 | 21,7859 | 23,8231 | 0,128972 | 0,45025 | 0,76921 |
| Kat5 | 33,0038 | 36,0874 | 0,128861 | 0,45055 | 0,76921 |
| Agap3 | 31,0933 | 33,871 | 0,123447 | 0,45075 | 0,76921 |
| Zfp280d | 23,3415 | 25,3916 | 0,121457 | 0,4512 | 0,76921 |
| Kif5b | 48,5706 | 52,8237 | 0,121102 | 0,45075 | 0,76921 |
| Plrg1 | 80,645 | 87,6557 | 0,120262 | 0,45115 | 0,76921 |
| Ube2z | 42,4526 | 46,1267 | 0,119747 | 0,4509 | 0,76921 |
| Napg | 10,9944 | 10,0114 | -0,135126 | 0,45025 | 0,76921 |
| Cyb561a3 | 6,66521 | 6,00424 | -0,150668 | 0,4508 | 0,76921 |
| Zfp791 | 0,80954 | 0,650519 | -0,315511 | 0,4505 | 0,76921 |
| Id2 | 63,8129 | 58,5192 | -0,124939 | 0,45135 | 0,769269 |
| Tagap1 | 8,06929 | 7,24203 | -0,156049 | 0,45135 | 0,769269 |
| Mir7089 | 51,65 | 186,824 | 1,85484 | 0,4521 | 0,769638 |
| Fabp3 | 1,41215 | 1,91037 | 0,435957 | 0,45195 | 0,769638 |
| Alkbh1 | 29,1194 | 35,1787 | 0,272722 | 0,45205 | 0,769638 |
| Asf1b | 35,7905 | 39,1455 | 0,129273 | 0,45215 | 0,769638 |
| Cetn2 | 49,2859 | 53,8507 | 0,127789 | 0,452 | 0,769638 |
| Atf2 | 26,5647 | 28,9448 | 0,123794 | 0,45215 | 0,769638 |
| Med24 | 32,9508 | 35,8854 | 0,123082 | 0,4522 | 0,769638 |
| Pmpcb | 65,7939 | 71,552 | 0,12104 | 0,45205 | 0,769638 |
| Gng2 | 36,3252 | 39,4535 | 0,119184 | 0,4519 | 0,769638 |
| Synj2bp | 5,29028 | 4,84711 | -0,126219 | 0,45205 | 0,769638 |
| Trmt10a | 9,36913 | 8,54058 | -0,133581 | 0,45165 | 0,769638 |
| Snord15b | 42,3927 | 68,0712 | 0,683229 | 0,4529 | 0,769641 |
| Rgs9bp | 0,655788 | 0,765226 | 0,222656 | 0,4525 | 0,769641 |
| Zcchc18 | 6,06451 | 6,81127 | 0,167533 | 0,4526 | 0,769641 |
| Zmat5 | 25,2594 | 28,1208 | 0,154819 | 0,4527 | 0,769641 |
| Dusp18 | 5,02982 | 5,55278 | 0,142705 | 0,4525 | 0,769641 |
| Papd4 | 15,3371 | 16,8782 | 0,138139 | 0,45295 | 0,769641 |
| Parg | 23,3436 | 25,3778 | 0,120539 | 0,4527 | 0,769641 |
| Kif20b | 26,5928 | 24,4751 | -0,11972 | 0,4528 | 0,769641 |
| Bok | 28,0545 | 25,5347 | -0,135777 | 0,45285 | 0,769641 |
| Ankrd6 | 4,03506 | 3,62523 | -0,154519 | 0,4526 | 0,769641 |
| Ifnar2 | 15,7264 | 14,0652 | -0,16106 | 0,45275 | 0,769641 |
| Mttp | 3,3655 | 3,00379 | -0,164034 | 0,4526 | 0,769641 |
| Sv2a | 0,993302 | 0,84211 | -0,238223 | 0,45245 | 0,769641 |
| Blcap | 18,0828 | 14,047 | -0,36435 | 0,45315 | 0,769883 |
| Tmem200c | 0,480059 | 0,593297 | 0,305542 | 0,4535 | 0,770379 |
| Cdk2ap2 | 24,892 | 27,6205 | 0,150056 | 0,45385 | 0,770765 |
| Vamp3 | 72,9241 | 79,2579 | 0,12016 | 0,4538 | 0,770765 |
| Gtdc1 | 8,58875 | 7,74472 | -0,149234 | 0,4539 | 0,770765 |
| H2-Q4 | 10,2225 | 11,3492 | 0,150838 | 0,4541 | 0,770909 |

| | | | | | |
|---|---|---|---|---|---|
| Exoc6b | 8,57036 | 9,36977 | 0,128657 | 0,45405 | 0,770909 |
| 1500004A13Rik | 1,70763 | 2,13835 | 0,324503 | 0,45425 | 0,770967 |
| Lect1 | 1,93579 | 1,59805 | -0,276618 | 0,45425 | 0,770967 |
| Wdr24 | 12,0285 | 13,1805 | 0,13195 | 0,4546 | 0,771313 |
| Lsg1 | 43,3225 | 47,0812 | 0,120034 | 0,45475 | 0,771313 |
| Tra2a | 181,598 | 197,142 | 0,118491 | 0,45475 | 0,771313 |
| Senp2 | 24,3717 | 26,4447 | 0,117772 | 0,45455 | 0,771313 |
| Fzr1 | 49,7747 | 45,7806 | -0,120675 | 0,45475 | 0,771313 |
| Bace1 | 16,3126 | 14,9828 | -0,122684 | 0,4548 | 0,771313 |
| Elk4 | 5,55145 | 6,14308 | 0,146099 | 0,4551 | 0,77143 |
| Tnrc6c | 8,79975 | 9,57682 | 0,122084 | 0,45505 | 0,77143 |
| Nip7 | 31,8087 | 34,5607 | 0,119708 | 0,45505 | 0,77143 |
| Pex11a | 2,60493 | 2,09782 | -0,312355 | 0,4551 | 0,77143 |
| 1700056E22Rik | 0,504561 | 0,724444 | 0,521846 | 0,45545 | 0,771926 |
| Sms | 14,7681 | 16,1299 | 0,12725 | 0,45555 | 0,771997 |
| Cnrip1 | 6,20008 | 7,02846 | 0,180922 | 0,4557 | 0,772056 |
| Xkr8 | 1,72867 | 1,50778 | -0,197238 | 0,4557 | 0,772056 |
| Pithd1 | 32,7114 | 35,7667 | 0,128821 | 0,4559 | 0,772186 |
| Wnt5a | 172,019 | 187,593 | 0,125034 | 0,4559 | 0,772186 |
| Grwd1 | 56,6058 | 61,5399 | 0,120572 | 0,45595 | 0,772186 |
| Pacsin1 | 1,07415 | 1,24443 | 0,212295 | 0,45625 | 0,7724 |
| Cdca8 | 106,207 | 115,313 | 0,118671 | 0,45615 | 0,7724 |
| Ccar1 | 67,6826 | 73,4185 | 0,117358 | 0,4562 | 0,7724 |
| Bbs12 | 12,0973 | 11,0076 | -0,136178 | 0,45645 | 0,772641 |
| U2af1l4 | 33,9146 | 37,3726 | 0,140075 | 0,45675 | 0,772939 |
| Dcp1a | 15,6949 | 17,1425 | 0,127276 | 0,45675 | 0,772939 |
| Fancm | 11,9771 | 13,0072 | 0,119034 | 0,4568 | 0,772939 |
| Psma7 | 266,273 | 289,161 | 0,118964 | 0,4569 | 0,773011 |
| Zfp940 | 5,6356 | 6,27197 | 0,154348 | 0,4572 | 0,773322 |
| Tbc1d20 | 18,5151 | 20,2785 | 0,131252 | 0,45715 | 0,773322 |
| Xrcc4 | 5,90242 | 6,68807 | 0,180282 | 0,4573 | 0,773394 |
| Cxx1b | 6,79607 | 7,75328 | 0,190105 | 0,45745 | 0,77355 |
| Mapk1ip1 | 7,40666 | 8,34156 | 0,171495 | 0,45755 | 0,773621 |
| Atp6v1d | 33,9632 | 37,092 | 0,127136 | 0,45825 | 0,774706 |
| Plcg1 | 39,2848 | 42,6982 | 0,120204 | 0,4584 | 0,774764 |
| Npl | 4,51235 | 3,89817 | -0,211081 | 0,45835 | 0,774764 |
| Dmap1 | 29,3377 | 32,1174 | 0,130599 | 0,4587 | 0,775075 |
| Car13 | 22,3548 | 24,4095 | 0,126856 | 0,45865 | 0,775075 |
| Mir6921 | 58,0081 | 201,935 | 1,79956 | 0,45915 | 0,775247 |
| Calml4 | 1,17831 | 2,23236 | 0,921849 | 0,45915 | 0,775247 |
| Msx1 | 9,85304 | 11,0594 | 0,166632 | 0,459 | 0,775247 |
| Mrpl46 | 40,3684 | 44,1992 | 0,130794 | 0,4591 | 0,775247 |
| Wdr43 | 66,9789 | 72,675 | 0,117751 | 0,4589 | 0,775247 |
| Tmc7 | 3,43675 | 3,0818 | -0,157271 | 0,45895 | 0,775247 |
| Chid1 | 9,14126 | 8,27206 | -0,144146 | 0,45925 | 0,775318 |
| Rrp7a | 21,6614 | 23,7859 | 0,134981 | 0,45935 | 0,775389 |
| Hvcn1 | 7,85965 | 6,76073 | -0,217285 | 0,4595 | 0,775544 |
| Zdhhc3 | 14,5534 | 15,7882 | 0,117495 | 0,4596 | 0,775601 |

| | | | | | |
|---|---|---|---|---|---|
| Ptpn1 | 24,8758 | 26,9811 | 0,117208 | 0,45965 | 0,775601 |
| Hip1 | 29,1696 | 31,623 | 0,116506 | 0,45975 | 0,775672 |
| Brms1l | 16,8276 | 15,3805 | -0,129725 | 0,45985 | 0,775743 |
| Osbpl5 | 2,19286 | 1,92531 | -0,187722 | 0,45995 | 0,775814 |
| Htra2 | 35,0471 | 38,2404 | 0,1258 | 0,4601 | 0,775871 |
| Pet100 | 15,1482 | 13,5934 | -0,156238 | 0,4601 | 0,775871 |
| Bphl | 10,7321 | 12,017 | 0,163139 | 0,4602 | 0,775942 |
| Rwdd3 | 1,81794 | 2,1711 | 0,256118 | 0,4607 | 0,776197 |
| Ppdpf | 29,0198 | 32,1785 | 0,149061 | 0,46055 | 0,776197 |
| Vps39 | 12,0147 | 13,0969 | 0,12442 | 0,4605 | 0,776197 |
| Hspe1 | 133,556 | 145,264 | 0,121229 | 0,4607 | 0,776197 |
| Pnkd | 19,4136 | 16,7374 | -0,213993 | 0,4605 | 0,776197 |
| Scarna8 | 14,3145 | 7,22711 | -0,985992 | 0,4606 | 0,776197 |
| 4921533I20Rik | 1,40553 | 1,77718 | 0,338473 | 0,4611 | 0,776479 |
| Pdgfrb | 11,8104 | 10,8382 | -0,12393 | 0,461 | 0,776479 |
| Cnpy4 | 35,2424 | 32,3156 | -0,12508 | 0,46105 | 0,776479 |
| Zdhhc12 | 16,2713 | 13,8692 | -0,230446 | 0,4611 | 0,776479 |
| Tpgs2 | 9,4053 | 10,4759 | 0,155526 | 0,46125 | 0,776634 |
| E130307A14Rik | 7,29032 | 8,2578 | 0,179776 | 0,4614 | 0,776691 |
| Jtb | 35,5924 | 32,459 | -0,132952 | 0,4614 | 0,776691 |
| Snapc2 | 22,6372 | 24,7886 | 0,130978 | 0,46155 | 0,776707 |
| Asns | 38,3489 | 41,7081 | 0,121144 | 0,4616 | 0,776707 |
| Gas2l3 | 4,83514 | 4,40814 | -0,133387 | 0,46165 | 0,776707 |
| Slc35g2 | 3,32177 | 2,84877 | -0,221612 | 0,4616 | 0,776707 |
| Dnd1 | 1,84854 | 1,16195 | -0,66984 | 0,4617 | 0,776707 |
| Mir378b | 19,0814 | 121,347 | 2,6689 | 0,46215 | 0,777242 |
| Ticam2 | 0,99442 | 1,17022 | 0,234848 | 0,4626 | 0,777242 |
| Ptpn3 | 1,70822 | 1,91377 | 0,163923 | 0,4623 | 0,777242 |
| Rassf8 | 12,741 | 14,0611 | 0,142236 | 0,46255 | 0,777242 |
| Traf2 | 10,8148 | 11,8473 | 0,131552 | 0,46245 | 0,777242 |
| Kdelr2 | 84,5216 | 77,9729 | -0,116348 | 0,4623 | 0,777242 |
| Ccnh | 29,178 | 26,7659 | -0,124484 | 0,4626 | 0,777242 |
| Ifrd2 | 26,17 | 23,9574 | -0,127443 | 0,46255 | 0,777242 |
| Rep15 | 4,19918 | 3,55418 | -0,240594 | 0,4623 | 0,777242 |
| Akr1c18 | 0,856174 | 0,638042 | -0,424252 | 0,4626 | 0,777242 |
| Csk | 47,7244 | 51,7737 | 0,117492 | 0,46275 | 0,777299 |
| Tceal1 | 7,09896 | 6,18319 | -0,199256 | 0,46275 | 0,777299 |
| Usp13 | 3,62298 | 4,01563 | 0,14845 | 0,463 | 0,777621 |
| Triobp | 14,7065 | 16,1689 | 0,136765 | 0,4632 | 0,777664 |
| Eprs | 94,5775 | 102,602 | 0,117488 | 0,46315 | 0,777664 |
| Klhdc8b | 4,06201 | 3,25813 | -0,31815 | 0,4632 | 0,777664 |
| Msantd2 | 25,6592 | 27,9167 | 0,121655 | 0,46335 | 0,777818 |
| Cox7c | 1,0634 | 1,60265 | 0,591776 | 0,4635 | 0,777847 |
| Fam115c | 3,59013 | 3,9893 | 0,1521 | 0,46355 | 0,777847 |
| Bdp1 | 10,8397 | 11,7642 | 0,118072 | 0,4636 | 0,777847 |
| Qser1 | 20,7117 | 22,4325 | 0,115143 | 0,4636 | 0,777847 |
| Tipin | 129,036 | 140,011 | 0,117761 | 0,46375 | 0,778001 |
| Efcab7 | 3,3679 | 3,87213 | 0,201277 | 0,46405 | 0,778309 |

| | | | | | |
|---|---|---|---|---|---|
| Klhl25 | 10,6087 | 11,6005 | 0,12894 | 0,46405 | 0,778309 |
| Lipo1 | 3,05983 | 2,68959 | -0,186068 | 0,46425 | 0,778546 |
| 1500009L16Rik | 10,8657 | 12,1593 | 0,16227 | 0,46455 | 0,778715 |
| Ext2 | 34,8977 | 32,1829 | -0,116839 | 0,4647 | 0,778715 |
| Ykt6 | 67,2495 | 61,9924 | -0,117432 | 0,46455 | 0,778715 |
| Mtg2 | 13,0256 | 11,7508 | -0,148584 | 0,46455 | 0,778715 |
| Slitrk5 | 3,50356 | 3,15223 | -0,152447 | 0,4647 | 0,778715 |
| Rabgap1l | 4,2461 | 3,81348 | -0,155033 | 0,4647 | 0,778715 |
| Mapk8ip1 | 15,3199 | 16,7145 | 0,125691 | 0,46585 | 0,78053 |
| Tgfbrap1 | 7,54654 | 6,90523 | -0,128125 | 0,4659 | 0,78053 |
| Nlrp5-ps | 0,619161 | 0,759994 | 0,295674 | 0,46615 | 0,780543 |
| Ccdc166 | 2,61456 | 2,98274 | 0,190069 | 0,46605 | 0,780543 |
| Zmynd11 | 24,6672 | 26,7422 | 0,116522 | 0,4662 | 0,780543 |
| Iqgap3 | 12,1782 | 11,2096 | -0,119565 | 0,4661 | 0,780543 |
| Fgfr3 | 3,15365 | 2,81675 | -0,162992 | 0,4662 | 0,780543 |
| 2310010J17Rik | 4,37687 | 5,69398 | 0,379537 | 0,4669 | 0,781168 |
| Mapre3 | 4,35538 | 4,93434 | 0,180058 | 0,46735 | 0,781168 |
| 1600020E01Rik | 27,5948 | 30,812 | 0,159094 | 0,4671 | 0,781168 |
| Hilpda | 17,7497 | 19,7721 | 0,155675 | 0,46725 | 0,781168 |
| Ubald2 | 9,44967 | 10,5256 | 0,155567 | 0,46735 | 0,781168 |
| Bola2 | 194,613 | 213,794 | 0,135614 | 0,46745 | 0,781168 |
| Cyp39a1 | 9,79587 | 10,7329 | 0,131796 | 0,46745 | 0,781168 |
| Secisbp2 | 10,507 | 11,5111 | 0,131669 | 0,4667 | 0,781168 |
| Mdm2 | 26,2685 | 28,5172 | 0,118495 | 0,46745 | 0,781168 |
| Tnks1bp1 | 12,9638 | 14,0712 | 0,118261 | 0,46705 | 0,781168 |
| Fbxo38 | 21,2372 | 23,0367 | 0,117344 | 0,46685 | 0,781168 |
| Dhx30 | 42,9157 | 46,493 | 0,115505 | 0,4667 | 0,781168 |
| Ash2l | 50,3377 | 54,4546 | 0,113415 | 0,4672 | 0,781168 |
| Pom121 | 36,6406 | 33,612 | -0,124469 | 0,46705 | 0,781168 |
| Pgbd1 | 4,56154 | 4,08013 | -0,160905 | 0,4668 | 0,781168 |
| Usp5 | 67,5336 | 73,0729 | 0,113731 | 0,46755 | 0,781238 |
| Arf3 | 41,8956 | 38,6579 | -0,116037 | 0,46785 | 0,781641 |
| 4930515G01Rik | 0,854097 | 1,10135 | 0,366804 | 0,468 | 0,781696 |
| Grsf1 | 71,523 | 77,3329 | 0,112675 | 0,46795 | 0,781696 |
| Nynrin | 26,2145 | 28,391 | 0,115066 | 0,46815 | 0,781849 |
| Bcl2l13 | 23,1534 | 25,0778 | 0,115185 | 0,46825 | 0,781918 |
| Akap5 | 0,948182 | 0,82723 | -0,196875 | 0,4684 | 0,782071 |
| Ndufa12 | 73,0396 | 80,167 | 0,134329 | 0,46855 | 0,782224 |
| Cdk14 | 5,00205 | 5,50014 | 0,13695 | 0,46905 | 0,782289 |
| Zcchc17 | 30,4853 | 33,2908 | 0,127007 | 0,4691 | 0,782289 |
| Rpl22l1 | 233,692 | 253,685 | 0,118426 | 0,4692 | 0,782289 |
| Rfx3 | 10,5531 | 11,4443 | 0,116957 | 0,4693 | 0,782289 |
| Utp15 | 26,4732 | 28,6957 | 0,116301 | 0,46935 | 0,782289 |
| Copb2 | 88,149 | 95,5487 | 0,116292 | 0,469 | 0,782289 |
| Pola2 | 44,1081 | 47,799 | 0,115936 | 0,46885 | 0,782289 |
| Psmd14 | 112,614 | 121,833 | 0,113529 | 0,46915 | 0,782289 |
| Scube1 | 5,74436 | 5,26168 | -0,126623 | 0,4692 | 0,782289 |
| Cdadc1 | 12,7054 | 11,5193 | -0,14139 | 0,46905 | 0,782289 |

| | | | | | |
|---|---|---|---|---|---|
| Has3 | 1,21435 | 1,06632 | -0,187537 | 0,4693 | 0,782289 |
| 1110007C09Rik | 5,92021 | 5,04211 | -0,231621 | 0,4691 | 0,782289 |
| Cntd1 | 1,14467 | 0,559272 | -1,03331 | 0,4691 | 0,782289 |
| Spr | 29,1358 | 26,576 | -0,132669 | 0,46945 | 0,782358 |
| Muc1 | 0,824692 | 1,0028 | 0,282112 | 0,4696 | 0,782413 |
| Cog6 | 14,9761 | 16,3297 | 0,124835 | 0,46955 | 0,782413 |
| D6Wsu163e | 8,9909 | 8,12254 | -0,146534 | 0,4698 | 0,782649 |
| Coq5 | 21,5162 | 23,4704 | 0,125418 | 0,4699 | 0,782703 |
| Pafah1b1 | 59,2186 | 54,6889 | -0,114804 | 0,46995 | 0,782703 |
| Mir1894 | 18,0879 | 271,215 | 3,90634 | 0,47075 | 0,783241 |
| Trmt2a | 24,817 | 27,9288 | 0,170423 | 0,47055 | 0,783241 |
| Sdha | 116,266 | 125,908 | 0,114942 | 0,4708 | 0,783241 |
| Lin7c | 45,6409 | 42,1754 | -0,113928 | 0,47045 | 0,783241 |
| Zfp84 | 5,52506 | 5,04249 | -0,131856 | 0,4704 | 0,783241 |
| L2hgdh | 8,60642 | 7,82145 | -0,137977 | 0,4707 | 0,783241 |
| Smg9 | 10,6374 | 9,64955 | -0,140611 | 0,4708 | 0,783241 |
| Igsf10 | 3,22262 | 2,83757 | -0,183579 | 0,47065 | 0,783241 |
| Spdya | 0,596423 | 0,364763 | -0,709379 | 0,4705 | 0,783241 |
| Baiap2 | 7,17393 | 6,42647 | -0,158738 | 0,471 | 0,783476 |
| Mthfs | 1,82683 | 2,31371 | 0,340868 | 0,47115 | 0,78353 |
| Apbb1 | 21,8787 | 23,7754 | 0,11994 | 0,47115 | 0,78353 |
| Elac1 | 3,24132 | 2,91784 | -0,151679 | 0,4713 | 0,783585 |
| Setd4 | 4,44826 | 3,90549 | -0,18774 | 0,47125 | 0,783585 |
| Zfp560 | 3,30756 | 2,97339 | -0,153658 | 0,4714 | 0,783653 |
| Hscb | 11,2171 | 12,7468 | 0,184431 | 0,47155 | 0,783791 |
| Mrps27 | 37,3132 | 40,5345 | 0,119466 | 0,4716 | 0,783791 |
| Rpl39 | 343,224 | 372,572 | 0,118371 | 0,4719 | 0,784066 |
| Arhgdia | 197,457 | 214,255 | 0,117793 | 0,4719 | 0,784066 |
| Gpaa1 | 37,3676 | 34,4061 | -0,119122 | 0,47195 | 0,784066 |
| Zfp580 | 11,9732 | 10,6854 | -0,164164 | 0,472 | 0,784066 |
| Dnajb6 | 7,51505 | 8,47422 | 0,173298 | 0,47225 | 0,784286 |
| Pprc1 | 33,9813 | 36,7832 | 0,114303 | 0,47225 | 0,784286 |
| Coq2 | 22,0448 | 24,0861 | 0,12776 | 0,47235 | 0,784355 |
| Zfp62 | 27,3935 | 29,6899 | 0,11614 | 0,47255 | 0,784394 |
| Ncoa5 | 43,2859 | 46,8762 | 0,114957 | 0,47245 | 0,784394 |
| Capzb | 158,978 | 172,052 | 0,11402 | 0,47255 | 0,784394 |
| BC023829 | 21,1426 | 19,3741 | -0,126029 | 0,4727 | 0,784546 |
| Fam111a | 18,3667 | 16,9124 | -0,11901 | 0,4729 | 0,784683 |
| Abhd2 | 7,76158 | 7,04912 | -0,138908 | 0,4729 | 0,784683 |
| Rad18 | 16,6948 | 18,2066 | 0,125066 | 0,473 | 0,784751 |
| Lbr | 72,553 | 67,139 | -0,111883 | 0,47315 | 0,784805 |
| Gm1673 | 49,9652 | 44,5702 | -0,164844 | 0,4731 | 0,784805 |
| AU022252 | 14,4416 | 15,8424 | 0,133564 | 0,47345 | 0,785205 |
| Cystm1 | 4,67885 | 5,63874 | 0,269219 | 0,4736 | 0,78523 |
| Fam203a | 11,4275 | 12,576 | 0,138175 | 0,47365 | 0,78523 |
| Rabl2 | 22,4817 | 24,5878 | 0,129192 | 0,4737 | 0,78523 |
| Sfmbt1 | 5,6028 | 6,10139 | 0,122989 | 0,47365 | 0,78523 |
| Snrpe | 229,248 | 211,064 | -0,119231 | 0,47385 | 0,785382 |

| | | | | | |
|---|---|---|---|---|---|
| Arl2 | 47,0845 | 51,4908 | 0,129063 | 0,47405 | 0,785616 |
| Ltbp4 | 3,04117 | 3,36145 | 0,144456 | 0,47415 | 0,785684 |
| Slc25a30 | 4,29604 | 4,88111 | 0,184202 | 0,4744 | 0,785874 |
| Ndufs3 | 21,1202 | 23,351 | 0,144858 | 0,47445 | 0,785874 |
| Eif4g1 | 268,505 | 294,608 | 0,133849 | 0,4744 | 0,785874 |
| Pfkfb2 | 4,29943 | 3,83494 | -0,164943 | 0,4745 | 0,785874 |
| Tctn2 | 10,1946 | 9,29279 | -0,133625 | 0,4746 | 0,785942 |
| Psmg1 | 59,6906 | 64,9917 | 0,122751 | 0,47485 | 0,786259 |
| Trmt1 | 32,1084 | 34,8617 | 0,11869 | 0,47505 | 0,786395 |
| Rpgrip1l | 3,97524 | 3,62087 | -0,134704 | 0,47505 | 0,786395 |
| Agl | 3,80167 | 3,48193 | -0,126748 | 0,47525 | 0,786629 |
| Gm15545 | 10,3885 | 8,65401 | -0,263548 | 0,47565 | 0,787193 |
| 3425401B19Rik | 0,627044 | 0,730766 | 0,220841 | 0,4758 | 0,787329 |
| Nhej1 | 6,02101 | 6,98137 | 0,213504 | 0,47585 | 0,787329 |
| Rbm7 | 42,0406 | 45,6434 | 0,118623 | 0,47605 | 0,787563 |
| Pdcd5 | 38,1619 | 42,1934 | 0,144887 | 0,4762 | 0,787572 |
| 0610037L13Rik | 27,186 | 29,6201 | 0,123712 | 0,47635 | 0,787572 |
| Vps11 | 15,041 | 16,3651 | 0,121723 | 0,47635 | 0,787572 |
| Fbxl20 | 6,15222 | 6,68438 | 0,119686 | 0,47635 | 0,787572 |
| Srp19 | 83,2595 | 76,6281 | -0,119742 | 0,4763 | 0,787572 |
| Mir7082 | 9,54071 | 34,51 | 1,85484 | 0,4768 | 0,787716 |
| Rel | 1,40782 | 1,64053 | 0,220701 | 0,47685 | 0,787716 |
| Tmem242 | 74,5489 | 80,8692 | 0,117403 | 0,4768 | 0,787716 |
| Csrp2bp | 26,1638 | 28,3213 | 0,114314 | 0,4766 | 0,787716 |
| Vars | 84,5358 | 91,4783 | 0,113866 | 0,47655 | 0,787716 |
| 03/07/16 | 40,9403 | 44,3 | 0,113782 | 0,47685 | 0,787716 |
| Dpysl2 | 16,9732 | 15,6421 | -0,117821 | 0,47675 | 0,787716 |
| Ccdc94 | 8,84901 | 9,915 | 0,164098 | 0,477 | 0,787867 |
| Sdccag8 | 4,42738 | 5,33113 | 0,267986 | 0,4773 | 0,788265 |
| Mir6907 | 330,34 | 509,741 | 0,625811 | 0,4774 | 0,788332 |
| Pram1 | 0,726085 | 0,587746 | -0,304945 | 0,4776 | 0,788565 |
| Recql5 | 9,72276 | 10,6094 | 0,12591 | 0,4778 | 0,788798 |
| LOC102634401 | 5,06072 | 6,64132 | 0,392126 | 0,478 | 0,788829 |
| Hgf | 0,523073 | 0,655699 | 0,326022 | 0,47805 | 0,788829 |
| Pibf1 | 7,07124 | 8,82683 | 0,319932 | 0,47835 | 0,788829 |
| Eif2s3x | 4,51112 | 4,9854 | 0,144223 | 0,47805 | 0,788829 |
| Lrwd1 | 32,4438 | 35,1869 | 0,117097 | 0,47825 | 0,788829 |
| Snrpb2 | 104,418 | 112,939 | 0,113172 | 0,47825 | 0,788829 |
| Wbp2 | 24,2364 | 22,2248 | -0,125006 | 0,47815 | 0,788829 |
| Twist2 | 8,55714 | 7,58648 | -0,173697 | 0,47825 | 0,788829 |
| A330009N23Rik | 0,653373 | 0,428996 | -0,606943 | 0,4783 | 0,788829 |
| Slc45a1 | 0,504498 | 0,627773 | 0,315395 | 0,47865 | 0,789226 |
| Dip2c | 4,87879 | 5,31279 | 0,122946 | 0,47875 | 0,789294 |
| Trnt1 | 37,603 | 33,9321 | -0,148193 | 0,47885 | 0,789361 |
| Cttnbp2nl | 18,9276 | 20,496 | 0,114848 | 0,479 | 0,789414 |
| Dera | 37,4761 | 34,5152 | -0,118739 | 0,47895 | 0,789414 |
| Sec23b | 38,4856 | 41,6452 | 0,113834 | 0,4791 | 0,789481 |
| Pknox1 | 10,25 | 11,1411 | 0,120266 | 0,4792 | 0,789549 |

| | | | | | |
|---|---|---|---|---|---|
| Ephx2 | 0,448187 | 0,571048 | 0,349509 | 0,4793 | 0,789586 |
| Abce1 | 95,8563 | 88,6611 | -0,112573 | 0,4794 | 0,789586 |
| Slc6a8 | 12,8645 | 11,8375 | -0,120031 | 0,4794 | 0,789586 |
| Spag4 | 3,763 | 4,32228 | 0,199911 | 0,47965 | 0,789788 |
| Elavl1 | 33,4133 | 36,0926 | 0,111277 | 0,4797 | 0,789788 |
| Cep350 | 7,1194 | 6,57382 | -0,115024 | 0,4796 | 0,789788 |
| Oser1 | 1,85236 | 2,18812 | 0,240333 | 0,4799 | 0,789893 |
| Shcbp1 | 50,6318 | 46,7362 | -0,115505 | 0,4799 | 0,789893 |
| Sigmar1 | 53,9006 | 49,7244 | -0,116346 | 0,48 | 0,789893 |
| Ndufb9 | 149,276 | 137,699 | -0,116469 | 0,48 | 0,789893 |
| Med10 | 61,8012 | 67,2547 | 0,122 | 0,4801 | 0,789915 |
| Prc1 | 59,1111 | 54,7194 | -0,111379 | 0,4802 | 0,789915 |
| Plod1 | 15,4101 | 14,156 | -0,122464 | 0,48025 | 0,789915 |
| Fmn2 | 1,10006 | 0,966872 | -0,186182 | 0,4802 | 0,789915 |
| Fgfr1 | 84,5596 | 91,5678 | 0,114871 | 0,48055 | 0,789923 |
| Mrpl12 | 129,666 | 140,292 | 0,113635 | 0,4804 | 0,789923 |
| Efnb1 | 18,8162 | 17,3167 | -0,119815 | 0,48055 | 0,789923 |
| Sc5d | 21,5721 | 19,8241 | -0,121909 | 0,4805 | 0,789923 |
| Sdf2 | 35,8863 | 32,9343 | -0,123843 | 0,48055 | 0,789923 |
| R3hdm1 | 22,4542 | 24,3054 | 0,114296 | 0,48095 | 0,790483 |
| Gm6654 | 2,04797 | 3,48302 | 0,766142 | 0,48105 | 0,790511 |
| Rims3 | 0,604721 | 0,825766 | 0,449464 | 0,48145 | 0,790511 |
| Frmd5 | 1,45916 | 1,66409 | 0,189596 | 0,4814 | 0,790511 |
| Zfp473 | 2,2744 | 2,56327 | 0,172499 | 0,48205 | 0,790511 |
| Rnf182 | 3,2546 | 3,631 | 0,157885 | 0,4811 | 0,790511 |
| Snw1 | 81,1454 | 89,5681 | 0,142476 | 0,48125 | 0,790511 |
| Ttc3 | 86,9567 | 94,5165 | 0,120269 | 0,48215 | 0,790511 |
| Rab31 | 14,849 | 16,1241 | 0,118854 | 0,48115 | 0,790511 |
| Dpy19l3 | 7,14043 | 7,74618 | 0,117474 | 0,482 | 0,790511 |
| Fam195b | 65,9245 | 71,4461 | 0,116041 | 0,48175 | 0,790511 |
| Fbxo28 | 13,8744 | 15,0346 | 0,115859 | 0,48155 | 0,790511 |
| Sh2b1 | 32,9905 | 35,7261 | 0,114929 | 0,482 | 0,790511 |
| Ppp2r2d | 40,6994 | 44,0319 | 0,113541 | 0,48165 | 0,790511 |
| Ash1l | 12,0007 | 12,9822 | 0,113418 | 0,4818 | 0,790511 |
| Ttll12 | 24,447 | 26,4442 | 0,113295 | 0,48215 | 0,790511 |
| Ngfr | 28,8995 | 31,2421 | 0,112444 | 0,48205 | 0,790511 |
| Rnf4 | 73,554 | 79,4378 | 0,111022 | 0,48145 | 0,790511 |
| Msl3 | 15,5576 | 14,3204 | -0,119544 | 0,48175 | 0,790511 |
| Nme6 | 26,901 | 24,462 | -0,137121 | 0,4816 | 0,790511 |
| Immp1l | 10,2155 | 9,03069 | -0,177855 | 0,48165 | 0,790511 |
| Rabif | 18,5896 | 20,2384 | 0,122599 | 0,4825 | 0,790987 |
| Scai | 7,19668 | 6,63834 | -0,116509 | 0,48295 | 0,791628 |
| Mir466i | 5,72546 | 16,8688 | 1,5589 | 0,4831 | 0,79168 |
| Tmem194b | 7,25106 | 6,58325 | -0,139391 | 0,48305 | 0,79168 |
| Mir6418 | 7,07412 | 49,7331 | 2,81358 | 0,48325 | 0,791828 |
| Clptm1l | 57,9269 | 53,5562 | -0,11318 | 0,4834 | 0,791977 |
| Plekhg3 | 12,3156 | 13,3472 | 0,11605 | 0,4835 | 0,792044 |
| Mecr | 18,5999 | 20,4313 | 0,135489 | 0,48395 | 0,792684 |

| | | | | | |
|---|---|---|---|---|---|
| Pus10 | 23,2585 | 20,8936 | -0,154698 | 0,4842 | 0,792996 |
| Rnf180 | 0,624955 | 0,740382 | 0,244519 | 0,4846 | 0,793523 |
| Son | 87,1669 | 94,6891 | 0,119418 | 0,48465 | 0,793523 |
| Top1mt | 3,73205 | 3,27166 | -0,189942 | 0,4847 | 0,793523 |
| Mettl24 | 0,875691 | 1,09266 | 0,319354 | 0,4848 | 0,793574 |
| Strn3 | 38,8266 | 35,8233 | -0,116148 | 0,48485 | 0,793574 |
| Tert | 4,16516 | 4,6181 | 0,14893 | 0,48495 | 0,79361 |
| Hadhb | 6,64654 | 5,96403 | -0,156316 | 0,485 | 0,79361 |
| Fam214b | 3,77343 | 3,37688 | -0,160184 | 0,48505 | 0,79361 |
| Rps17 | 4,27621 | 5,34406 | 0,321604 | 0,48535 | 0,793712 |
| Chmp2a | 41,8471 | 45,6814 | 0,126479 | 0,4853 | 0,793712 |
| Taf5l | 20,3294 | 22,0407 | 0,116606 | 0,48525 | 0,793712 |
| Rexo1 | 18,5801 | 20,0619 | 0,1107 | 0,4852 | 0,793712 |
| Trim8 | 14,918 | 16,2068 | 0,119549 | 0,4856 | 0,793927 |
| Snora3 | 287,682 | 131,701 | -1,12721 | 0,48555 | 0,793927 |
| Aldh16a1 | 18,4811 | 20,6523 | 0,160252 | 0,48595 | 0,794064 |
| Ak1 | 6,16125 | 6,84588 | 0,152013 | 0,4861 | 0,794064 |
| Trmu | 10,9131 | 12,068 | 0,145128 | 0,4859 | 0,794064 |
| Pi4ka | 15,6748 | 17,1129 | 0,126642 | 0,486 | 0,794064 |
| Ilf2 | 200,619 | 217,116 | 0,114006 | 0,4861 | 0,794064 |
| Fbxo31 | 6,40389 | 5,85066 | -0,130348 | 0,48595 | 0,794064 |
| Ppp2r5a | 12,4909 | 11,3122 | -0,143001 | 0,4858 | 0,794064 |
| Cdcp2 | 0,479424 | 0,59748 | 0,317588 | 0,4863 | 0,794182 |
| Naf1 | 9,70046 | 10,5994 | 0,127857 | 0,48635 | 0,794182 |
| 2810417H13Rik | 104,791 | 96,1914 | -0,123533 | 0,4863 | 0,794182 |
| Rabep1 | 18,8219 | 21,4489 | 0,188493 | 0,48655 | 0,794349 |
| Aptx | 5,39787 | 5,88946 | 0,125743 | 0,48665 | 0,794349 |
| Ttc5 | 23,2802 | 25,2964 | 0,119826 | 0,4867 | 0,794349 |
| Ccdc9 | 26,8593 | 29,1326 | 0,117214 | 0,48675 | 0,794349 |
| Anapc1 | 43,3299 | 46,7921 | 0,110903 | 0,4867 | 0,794349 |
| Phlda2 | 3,33204 | 2,72147 | -0,292017 | 0,48685 | 0,794416 |
| Pskh1 | 21,5046 | 23,2923 | 0,115204 | 0,4872 | 0,794793 |
| Pcgf5 | 6,41896 | 5,69617 | -0,172347 | 0,4872 | 0,794793 |
| Psma6 | 250,225 | 273,311 | 0,127318 | 0,48735 | 0,794812 |
| Mycbp2 | 9,61488 | 10,3864 | 0,111353 | 0,48745 | 0,794812 |
| Rbms3 | 5,37971 | 4,93748 | -0,123754 | 0,48745 | 0,794812 |
| Zfp667 | 2,65544 | 2,36423 | -0,167577 | 0,48745 | 0,794812 |
| Fkbp2 | 63,1049 | 57,8021 | -0,12663 | 0,48755 | 0,794878 |
| Ppp2r5b | 6,53976 | 7,2125 | 0,141261 | 0,48795 | 0,795239 |
| Smoc1 | 87,026 | 93,8919 | 0,109555 | 0,48795 | 0,79524 |
| Copb1 | 60,9567 | 65,762 | 0,109469 | 0,4879 | 0,79524 |
| Arhgef18 | 8,08641 | 8,78122 | 0,118922 | 0,48805 | 0,795305 |
| Tyw5 | 16,1877 | 18,4879 | 0,191685 | 0,48815 | 0,795371 |
| Cdk4 | 704,938 | 765,453 | 0,118816 | 0,4883 | 0,795406 |
| Cdc16 | 90,5686 | 97,7029 | 0,109391 | 0,48835 | 0,795406 |
| Acad10 | 1,01158 | 0,865791 | -0,224526 | 0,48835 | 0,795406 |
| Xab2 | 32,0983 | 34,7226 | 0,113379 | 0,48855 | 0,795441 |
| Sbf2 | 16,7128 | 18,038 | 0,110088 | 0,48855 | 0,795441 |

| | | | | | |
|---|---|---|---|---|---|
| Rtn2 | 6,81296 | 6,03239 | -0,175551 | 0,4885 | 0,795441 |
| Sh3glb1 | 29,1825 | 31,638 | 0,116554 | 0,48865 | 0,795492 |
| Cyth3 | 10,0789 | 9,27091 | -0,120549 | 0,4887 | 0,795492 |
| Zwilch | 32,9466 | 35,6041 | 0,111915 | 0,489 | 0,795868 |
| Armcx2 | 60,2918 | 55,8748 | -0,109765 | 0,48905 | 0,795868 |
| Amz2 | 28,8021 | 31,143 | 0,112735 | 0,48925 | 0,79608 |
| Sin3a | 40,1955 | 43,3765 | 0,10988 | 0,4893 | 0,79608 |
| Sbsn | 3,43499 | 4,16742 | 0,27885 | 0,48955 | 0,796278 |
| Pdcd2 | 31,7654 | 34,6185 | 0,124087 | 0,48955 | 0,796278 |
| Ntmt1 | 39,0718 | 42,4517 | 0,119694 | 0,4896 | 0,796278 |
| Rbm48 | 5,79037 | 6,69927 | 0,210349 | 0,48985 | 0,796394 |
| Tmem222 | 41,0198 | 37,8762 | -0,115031 | 0,48985 | 0,796394 |
| Prps2 | 15,4015 | 14,209 | -0,116266 | 0,48975 | 0,796394 |
| Fam96a | 71,3422 | 77,117 | 0,112293 | 0,49005 | 0,796622 |
| Tbce | 20,4605 | 23,7463 | 0,214862 | 0,49015 | 0,796672 |
| Zeb1 | 17,2865 | 18,6614 | 0,110412 | 0,4902 | 0,796672 |
| Hp | 0,75746 | 0,967672 | 0,353348 | 0,4904 | 0,7969 |
| Wdr5 | 65,1024 | 70,2504 | 0,109796 | 0,49065 | 0,797112 |
| Rpusd2 | 5,06024 | 4,61979 | -0,131378 | 0,49065 | 0,797112 |
| Idnk | 7,70554 | 8,59358 | 0,157363 | 0,49075 | 0,797162 |
| Plxnb1 | 29,7514 | 27,5868 | -0,108982 | 0,4908 | 0,797162 |
| Ccdc14 | 9,55995 | 10,3824 | 0,119061 | 0,491 | 0,79739 |
| Ywhab | 169,55 | 183,061 | 0,110614 | 0,49115 | 0,797409 |
| Fndc3c1 | 34,7599 | 37,494 | 0,109234 | 0,49125 | 0,797409 |
| Zfp825 | 7,28003 | 6,47353 | -0,169391 | 0,49125 | 0,797409 |
| Nme3 | 29,0634 | 22,9615 | -0,339985 | 0,49125 | 0,797409 |
| Adam8 | 1,38247 | 1,5964 | 0,207577 | 0,49135 | 0,797474 |
| Brk1 | 112,502 | 121,459 | 0,11052 | 0,4915 | 0,797621 |
| Mir704 | 215,321 | 540,738 | 1,32844 | 0,49165 | 0,797736 |
| Mir3062 | 10,4489 | 11,888 | 0,186145 | 0,49175 | 0,797736 |
| Zbtb14 | 7,09139 | 7,76463 | 0,13085 | 0,49175 | 0,797736 |
| Npr2 | 4,16064 | 4,59407 | 0,142969 | 0,49205 | 0,797932 |
| Pml | 7,66907 | 8,3307 | 0,119385 | 0,49195 | 0,797932 |
| Fbxo11 | 39,7669 | 34,7365 | -0,195115 | 0,49205 | 0,797932 |
| Exosc3 | 53,2019 | 57,7144 | 0,117453 | 0,4923 | 0,79824 |
| Mrgpre | 0,745649 | 0,871868 | 0,225614 | 0,49245 | 0,798387 |
| Poll | 11,6978 | 12,7764 | 0,127239 | 0,4928 | 0,798438 |
| Smarca4 | 77,6756 | 84,0554 | 0,113878 | 0,4929 | 0,798438 |
| Rapgef6 | 11,5848 | 12,5196 | 0,111949 | 0,4929 | 0,798438 |
| Snord96a | 1123,72 | 1094,04 | -0,0386208 | 0,4929 | 0,798438 |
| Cenpi | 19,1586 | 17,6714 | -0,116576 | 0,4929 | 0,798438 |
| Rhbdf1 | 14,7321 | 13,5338 | -0,12239 | 0,49285 | 0,798438 |
| Snhg18 | 9,59532 | 8,64404 | -0,150624 | 0,4928 | 0,798438 |
| Snora23 | 6,53438 | 38,8621 | 2,57224 | 0,4932 | 0,798731 |
| Rnf40 | 28,4572 | 30,7071 | 0,109779 | 0,4932 | 0,798731 |
| Hnrnpu | 487,488 | 535,501 | 0,135523 | 0,49335 | 0,798764 |
| Ube2m | 136,034 | 125,939 | -0,111243 | 0,4934 | 0,798764 |
| Tmsb15b1 | 7,18412 | 5,14693 | -0,481099 | 0,49335 | 0,798764 |

| | | | | | |
|---|---|---|---|---|---|
| Papss1 | 26,2038 | 28,3823 | 0,115214 | 0,4935 | 0,798829 |
| Hnrnpul2 | 82,2625 | 88,8623 | 0,111335 | 0,49375 | 0,798847 |
| Dctn5 | 49,2392 | 53,1724 | 0,110872 | 0,49375 | 0,798847 |
| Setd1a | 27,1313 | 29,2519 | 0,108576 | 0,49365 | 0,798847 |
| Rnf149 | 29,3365 | 27,1292 | -0,112851 | 0,4937 | 0,798847 |
| Mllt11 | 10,3326 | 7,90544 | -0,386287 | 0,49385 | 0,798912 |
| Dennd5a | 29,9928 | 32,3359 | 0,108519 | 0,49415 | 0,799204 |
| Med1 | 29,6517 | 27,2756 | -0,120506 | 0,4941 | 0,799204 |
| BC037034 | 19,4734 | 17,7947 | -0,130054 | 0,4943 | 0,79935 |
| Parp9 | 2,19475 | 1,88779 | -0,217363 | 0,4944 | 0,799415 |
| Rpsa | 8,72357 | 9,63284 | 0,143042 | 0,4947 | 0,799578 |
| Gatad1 | 37,517 | 40,5131 | 0,110844 | 0,49475 | 0,799578 |
| Prdx1 | 258,735 | 279,016 | 0,108869 | 0,4948 | 0,799578 |
| Atp5c1 | 207,192 | 223,315 | 0,108111 | 0,49475 | 0,799578 |
| Kctd9 | 13,4757 | 12,4088 | -0,119 | 0,49465 | 0,799578 |
| Dusp1 | 6,05597 | 6,73403 | 0,153111 | 0,4951 | 0,799579 |
| Smg5 | 42,582 | 45,8708 | 0,107335 | 0,495 | 0,799579 |
| Ankrd16 | 12,4006 | 11,342 | -0,128739 | 0,49495 | 0,799579 |
| Nsun4 | 6,91238 | 6,31404 | -0,130619 | 0,4949 | 0,799579 |
| Aig1 | 7,118 | 6,24801 | -0,188074 | 0,4951 | 0,799579 |
| Cntn2 | 1,61325 | 1,46012 | -0,143888 | 0,49555 | 0,799822 |
| Asl | 10,0113 | 11,0699 | 0,145017 | 0,4955 | 0,799823 |
| Rbm41 | 5,67221 | 6,18355 | 0,124525 | 0,49545 | 0,799823 |
| Dgcr6 | 35,1876 | 38,2105 | 0,118899 | 0,4954 | 0,799823 |
| 1700048M11Rik | 1,80374 | 0,84587 | -1,09249 | 0,49545 | 0,799823 |
| Myh9 | 84,6481 | 78,2666 | -0,113081 | 0,4957 | 0,799968 |
| Plekha1 | 32,0562 | 34,5997 | 0,110159 | 0,4958 | 0,800033 |
| Gtpbp1 | 32,1527 | 34,6802 | 0,109176 | 0,49595 | 0,800178 |
| Tmf1 | 9,14433 | 9,88981 | 0,113066 | 0,49605 | 0,800243 |
| 9130023H24Rik | 2,83264 | 3,3296 | 0,233199 | 0,4962 | 0,800292 |
| Pacrgl | 9,664 | 10,6951 | 0,146259 | 0,49615 | 0,800292 |
| Chrac1 | 31,6477 | 34,6838 | 0,132162 | 0,49645 | 0,800598 |
| LOC101669761 | 1,13398 | 2,60714 | 1,20107 | 0,4966 | 0,80068 |
| Npepps | 33,1745 | 35,7735 | 0,108816 | 0,49675 | 0,80068 |
| Pcmt1 | 35,4301 | 32,7192 | -0,114836 | 0,49675 | 0,80068 |
| Sike1 | 21,0657 | 19,425 | -0,11698 | 0,4967 | 0,80068 |
| Btbd8 | 1,31748 | 1,0507 | -0,326427 | 0,4968 | 0,80068 |
| Btbd19 | 1,96414 | 2,24401 | 0,192179 | 0,497 | 0,800793 |
| Rexo4 | 30,3779 | 32,7947 | 0,11044 | 0,49705 | 0,800793 |
| Tmem115 | 15,9265 | 14,6459 | -0,120931 | 0,497 | 0,800793 |
| Rbm26 | 33,7144 | 36,8724 | 0,129178 | 0,4975 | 0,801405 |
| Rdx | 119,643 | 110,869 | -0,109878 | 0,49755 | 0,801405 |
| Snord35b | 172,642 | 204,328 | 0,243099 | 0,49765 | 0,80147 |
| Synj1 | 3,17969 | 2,61555 | -0,281772 | 0,49805 | 0,802017 |
| Acsl1 | 6,45088 | 5,89874 | -0,12909 | 0,49825 | 0,802243 |
| Mettl13 | 16,821 | 18,2362 | 0,116545 | 0,4986 | 0,802516 |
| Csnk1d | 78,3103 | 84,3305 | 0,106853 | 0,4985 | 0,802516 |
| 2310003H01Rik | 9,74242 | 8,93388 | -0,124994 | 0,49855 | 0,802516 |

| | | | | | |
|---|---|---|---|---|---|
| Mapkapk5 | 28,1658 | 30,4714 | 0,113512 | 0,49885 | 0,802822 |
| Eid3 | 0,457363 | 2,18879 | 2,25872 | 0,49975 | 0,803219 |
| Serp2 | 3,29087 | 3,89131 | 0,241789 | 0,5003 | 0,803219 |
| Adra2a | 0,813125 | 0,948116 | 0,221587 | 0,5005 | 0,803219 |
| Hmox2 | 42,1682 | 48,6184 | 0,205347 | 0,4998 | 0,803219 |
| Fer1l5 | 1,06695 | 1,20229 | 0,172287 | 0,50005 | 0,803219 |
| Mapkapk3 | 4,48491 | 4,96559 | 0,146885 | 0,4995 | 0,803219 |
| Gorab | 9,22698 | 10,0991 | 0,130303 | 0,4993 | 0,803219 |
| 03/05/16 | 42,9588 | 46,9603 | 0,128488 | 0,5005 | 0,803219 |
| Tk2 | 10,7212 | 11,7067 | 0,126864 | 0,50035 | 0,803219 |
| Hoxa11os | 178,217 | 194,039 | 0,122713 | 0,5004 | 0,803219 |
| Kpnb1 | 199,308 | 216,874 | 0,12186 | 0,5005 | 0,803219 |
| Nbeal1 | 2,72619 | 2,96441 | 0,12086 | 0,49955 | 0,803219 |
| Chd4 | 147,676 | 160,208 | 0,117513 | 0,49955 | 0,803219 |
| Kank2 | 15,9241 | 17,2151 | 0,112461 | 0,4996 | 0,803219 |
| Dst | 9,36593 | 10,1202 | 0,111745 | 0,5007 | 0,803219 |
| Smarcal1 | 9,35867 | 10,109 | 0,111263 | 0,5001 | 0,803219 |
| Anapc7 | 34,5541 | 37,306 | 0,110551 | 0,5006 | 0,803219 |
| Ddx10 | 24,706 | 26,6629 | 0,109973 | 0,4993 | 0,803219 |
| Pphln1 | 37,6595 | 40,6075 | 0,108729 | 0,50025 | 0,803219 |
| N4bp1 | 19,0421 | 20,515 | 0,107486 | 0,5009 | 0,803219 |
| Cox7a2 | 264,517 | 284,806 | 0,106616 | 0,50045 | 0,803219 |
| Apeh | 29,8333 | 27,643 | -0,110009 | 0,5008 | 0,803219 |
| Papolg | 11,2928 | 10,4193 | -0,116146 | 0,5008 | 0,803219 |
| Rsrc1 | 15,2055 | 14,0289 | -0,116195 | 0,49935 | 0,803219 |
| Rnh1 | 29,3424 | 27,0534 | -0,117176 | 0,50025 | 0,803219 |
| Tbc1d19 | 13,4577 | 12,3784 | -0,120602 | 0,4997 | 0,803219 |
| AI837181 | 23,3698 | 21,4756 | -0,121947 | 0,5008 | 0,803219 |
| Slc25a23 | 8,54938 | 7,8328 | -0,126291 | 0,50085 | 0,803219 |
| Myof | 1,06794 | 0,945285 | -0,176013 | 0,50075 | 0,803219 |
| Pld2 | 1,72481 | 1,5183 | -0,183982 | 0,4997 | 0,803219 |
| Mir7079 | 397,992 | 1422,09 | 1,8372 | 0,50105 | 0,803347 |
| Txn2 | 107,984 | 116,342 | 0,107554 | 0,5011 | 0,803347 |
| Pcbd2 | 16,3759 | 18,5751 | 0,181801 | 0,50145 | 0,803794 |
| Mamdc4 | 3,65471 | 4,03148 | 0,141553 | 0,5016 | 0,803794 |
| Mmab | 13,1354 | 14,2455 | 0,117048 | 0,5017 | 0,803794 |
| Eif2s2 | 44,8468 | 48,2717 | 0,106172 | 0,5018 | 0,803794 |
| Ppig | 22,3355 | 20,7376 | -0,10709 | 0,50155 | 0,803794 |
| Cul2 | 29,6981 | 27,5707 | -0,107234 | 0,5018 | 0,803794 |
| Pum2 | 59,3413 | 55,0678 | -0,107827 | 0,5018 | 0,803794 |
| Lck | 5,47995 | 6,09227 | 0,152817 | 0,5023 | 0,804386 |
| Sox4 | 30,2072 | 32,5946 | 0,109739 | 0,5023 | 0,804386 |
| Plp2 | 13,338 | 11,7288 | -0,185478 | 0,50235 | 0,804386 |
| Speg | 12,1852 | 13,5794 | 0,156285 | 0,5027 | 0,804754 |
| Urm1 | 16,5272 | 17,9681 | 0,120596 | 0,50265 | 0,804754 |
| Srrd | 13,6771 | 16,6639 | 0,284969 | 0,5029 | 0,804865 |
| Ccdc18 | 5,36381 | 5,84905 | 0,124945 | 0,5029 | 0,804865 |
| Fmnl3 | 18,6096 | 17,258 | -0,10878 | 0,50295 | 0,804865 |

| | | | | | |
|---|---|---|---|---|---|
| Spata13 | 13,9056 | 14,9673 | 0,106146 | 0,5031 | 0,804912 |
| Eif2ak1 | 37,4002 | 33,8639 | -0,143298 | 0,5031 | 0,804912 |
| Ndufb2 | 104,561 | 95,8458 | -0,125552 | 0,5033 | 0,805135 |
| Ogfod3 | 14,0606 | 11,7025 | -0,264844 | 0,5035 | 0,805359 |
| Foxj3 | 15,7158 | 16,9656 | 0,110402 | 0,5037 | 0,80539 |
| Slc35b4 | 17,4144 | 16,1272 | -0,110789 | 0,5037 | 0,80539 |
| Hook2 | 4,37656 | 3,93212 | -0,154489 | 0,5037 | 0,80539 |
| 3110043O21Rik | 7,94439 | 8,66523 | 0,1253 | 0,50425 | 0,806173 |
| Heatr2 | 13,3725 | 14,4815 | 0,11495 | 0,5044 | 0,806316 |
| Zkscan6 | 6,12268 | 6,76232 | 0,143355 | 0,5048 | 0,80652 |
| Gprasp1 | 29,4685 | 31,8264 | 0,111051 | 0,50495 | 0,80652 |
| Vps52 | 21,2199 | 22,9122 | 0,110698 | 0,5048 | 0,80652 |
| Orc4 | 24,6565 | 22,1021 | -0,157783 | 0,5049 | 0,80652 |
| Il11ra1 | 6,49336 | 5,78078 | -0,167702 | 0,50465 | 0,80652 |
| Msh5 | 0,797748 | 0,664102 | -0,264529 | 0,50495 | 0,80652 |
| Cldn10 | 0,681422 | 0,507203 | -0,425985 | 0,5047 | 0,80652 |
| Mcu | 5,24052 | 5,76922 | 0,138666 | 0,5051 | 0,80655 |
| Tfg | 56,7837 | 61,1197 | 0,106159 | 0,50505 | 0,80655 |
| Tmub2 | 16,3914 | 13,5171 | -0,278155 | 0,50515 | 0,80655 |
| AV051173 | 0,295889 | 1,66275 | 2,49044 | 0,50585 | 0,806555 |
| Myl1 | 0,376385 | 0,550469 | 0,548452 | 0,50535 | 0,806555 |
| Yod1 | 2,52477 | 2,94434 | 0,221792 | 0,50625 | 0,806555 |
| Sfxn4 | 1,71455 | 1,96183 | 0,19437 | 0,5063 | 0,806555 |
| Fbxo48 | 3,06052 | 3,45436 | 0,174642 | 0,50575 | 0,806555 |
| 1110058L19Rik | 19,8928 | 22,1657 | 0,156081 | 0,50565 | 0,806555 |
| Nudt2 | 28,5845 | 31,3384 | 0,132699 | 0,50595 | 0,806555 |
| Ncor1 | 36,9792 | 40,0886 | 0,116479 | 0,50625 | 0,806555 |
| Cdk9 | 21,7226 | 23,4134 | 0,108143 | 0,5063 | 0,806555 |
| Dhx15 | 199,811 | 215,261 | 0,10745 | 0,50615 | 0,806555 |
| Smarcad1 | 32,7486 | 30,4282 | -0,106027 | 0,5061 | 0,806555 |
| Rb1cc1 | 8,1216 | 7,51027 | -0,1129 | 0,50535 | 0,806555 |
| Etaa1 | 15,812 | 14,6209 | -0,112991 | 0,5053 | 0,806555 |
| Cab39l | 17,4948 | 16,1664 | -0,113931 | 0,50585 | 0,806555 |
| Atp6v1c1 | 22,9482 | 21,1892 | -0,115054 | 0,5057 | 0,806555 |
| Mcc | 4,39518 | 4,04781 | -0,118781 | 0,50595 | 0,806555 |
| Pex13 | 15,2391 | 13,6676 | -0,157011 | 0,5054 | 0,806555 |
| Nab2 | 5,33361 | 4,64674 | -0,198895 | 0,5057 | 0,806555 |
| Tex40 | 1,23163 | 0,930465 | -0,40455 | 0,50615 | 0,806555 |
| Ten1 | 14,7202 | 17,5015 | 0,249688 | 0,50665 | 0,806599 |
| Vps9d1 | 8,6156 | 9,40653 | 0,126712 | 0,50665 | 0,806599 |
| Sulf2 | 211,842 | 229,207 | 0,113665 | 0,50665 | 0,806599 |
| Ebf2 | 11,8725 | 12,8063 | 0,10923 | 0,5067 | 0,806599 |
| Usp4 | 44,1782 | 47,5524 | 0,106185 | 0,50675 | 0,806599 |
| Cdyl2 | 1,7833 | 1,61627 | -0,141885 | 0,5065 | 0,806599 |
| Neurog2 | 1,09123 | 0,914974 | -0,254148 | 0,5064 | 0,806599 |
| Igsf3 | 18,4074 | 19,807 | 0,105721 | 0,50685 | 0,806662 |
| Parl | 41,64 | 45,0416 | 0,113288 | 0,50705 | 0,806788 |
| Rnf138 | 11,797 | 10,8791 | -0,116856 | 0,507 | 0,806788 |

| | | | | | |
|---|---|---|---|---|---|
| Ahcyl1 | 65,7065 | 70,6908 | 0,105486 | 0,5072 | 0,806835 |
| Ctnnb1 | 454,573 | 417,208 | -0,123742 | 0,50715 | 0,806835 |
| Eef1a1 | 1240,96 | 1361,05 | 0,133267 | 0,50735 | 0,806961 |
| Cnot2 | 71,2206 | 76,5379 | 0,10388 | 0,5074 | 0,806961 |
| Rap1gap | 1,04904 | 1,21111 | 0,207268 | 0,50755 | 0,807007 |
| Zfp865 | 9,30102 | 10,1214 | 0,121942 | 0,50755 | 0,807008 |
| Grtp1 | 5,17012 | 5,83697 | 0,175021 | 0,50765 | 0,807054 |
| Vkorc1l1 | 23,8862 | 22,1862 | -0,106511 | 0,5077 | 0,807054 |
| Sh3pxd2a | 14,6204 | 15,7376 | 0,106232 | 0,5078 | 0,807117 |
| Gfm1 | 35,8662 | 38,9954 | 0,120678 | 0,5079 | 0,80718 |
| Akr1b10 | 8,30408 | 9,25441 | 0,15632 | 0,50805 | 0,807226 |
| Ccdc39 | 1,26842 | 0,775196 | -0,710402 | 0,50805 | 0,807226 |
| Asf1a | 38,6319 | 41,9376 | 0,118453 | 0,5083 | 0,807431 |
| Rpl10a | 3,21544 | 3,79315 | 0,23838 | 0,50825 | 0,807432 |
| Ccdc115 | 14,8616 | 16,2139 | 0,125639 | 0,5084 | 0,807445 |
| Ccne1 | 43,9697 | 47,3899 | 0,10807 | 0,50855 | 0,807445 |
| Ccna2 | 129,568 | 119,225 | -0,120027 | 0,50855 | 0,807445 |
| Zfp36 | 3,13669 | 2,74044 | -0,194836 | 0,5085 | 0,807445 |
| Dtymk | 159,826 | 172,539 | 0,110416 | 0,5092 | 0,808172 |
| Pard3 | 23,0193 | 24,8339 | 0,109468 | 0,50915 | 0,808172 |
| Atp6v1e1 | 62,3102 | 67,1577 | 0,108085 | 0,50925 | 0,808172 |
| Frs3 | 2,93798 | 2,58671 | -0,183705 | 0,50925 | 0,808172 |
| Wipi2 | 31,1394 | 33,5047 | 0,105622 | 0,50945 | 0,808297 |
| Arfrp1 | 22,9665 | 25,4684 | 0,149175 | 0,50945 | 0,808298 |
| Pank2 | 25,9234 | 29,8395 | 0,202969 | 0,5097 | 0,808406 |
| Jph4 | 1,15096 | 1,00686 | -0,192974 | 0,5096 | 0,808406 |
| Clec2l | 1,00021 | 0,801619 | -0,31932 | 0,50965 | 0,808406 |
| Gga1 | 20,7712 | 22,3999 | 0,108903 | 0,50985 | 0,808452 |
| Aup1 | 50,5463 | 46,8635 | -0,109141 | 0,5098 | 0,808452 |
| Rai1 | 23,1714 | 25,39 | 0,131914 | 0,50995 | 0,808515 |
| Higd1a | 7,23308 | 7,99032 | 0,143642 | 0,5101 | 0,808561 |
| Hdac7 | 15,9273 | 17,3093 | 0,120053 | 0,5101 | 0,808561 |
| Ubiad1 | 10,0802 | 9,26071 | -0,122326 | 0,5102 | 0,808623 |
| H2-M3 | 1,12942 | 1,37112 | 0,279763 | 0,5112 | 0,808878 |
| Tram2 | 4,35279 | 4,93994 | 0,182552 | 0,51135 | 0,808878 |
| Alg13 | 14,335 | 15,6375 | 0,125463 | 0,5107 | 0,808878 |
| Ict1 | 60,4161 | 65,4928 | 0,116403 | 0,51145 | 0,808878 |
| Wdr7 | 5,25674 | 5,69149 | 0,114638 | 0,51105 | 0,808878 |
| Bard1 | 8,25603 | 8,92401 | 0,112243 | 0,51065 | 0,808878 |
| Wdr35 | 10,469 | 11,3142 | 0,112011 | 0,5109 | 0,808878 |
| Raver1 | 66,5571 | 71,8982 | 0,111362 | 0,511 | 0,808878 |
| Usp21 | 34,3632 | 37,0409 | 0,108256 | 0,51055 | 0,808878 |
| Mtx2 | 118,786 | 127,71 | 0,104497 | 0,5114 | 0,808878 |
| Sumf2 | 19,456 | 17,9481 | -0,116379 | 0,51075 | 0,808878 |
| Gm10336 | 8,47079 | 7,79612 | -0,119741 | 0,5114 | 0,808878 |
| Uvssa | 3,61722 | 3,3201 | -0,123655 | 0,51075 | 0,808878 |
| Plekhg5 | 3,63775 | 3,29018 | -0,14488 | 0,5114 | 0,808878 |
| Mertk | 3,088 | 2,77886 | -0,15218 | 0,5113 | 0,808878 |

| | | | | | |
|---|---|---|---|---|---|
| B3gnt8 | 0,996854 | 0,825773 | -0,271637 | 0,51105 | 0,808878 |
| Fbxo36 | 2,00695 | 1,62703 | -0,302765 | 0,51135 | 0,808878 |
| Cdkl1 | 0,765418 | 0,169621 | -2,17393 | 0,51055 | 0,808878 |
| Ap2m1 | 8,8679 | 9,72481 | 0,133077 | 0,5116 | 0,809019 |
| Srebf2 | 68,4405 | 63,7212 | -0,103079 | 0,51195 | 0,809381 |
| Nradd | 27,9332 | 25,6306 | -0,124114 | 0,5119 | 0,809381 |
| Dync1i1 | 2,5067 | 2,82639 | 0,173174 | 0,5124 | 0,809389 |
| Klf9 | 2,72681 | 3,03217 | 0,153135 | 0,51245 | 0,809389 |
| Itpkc | 7,02562 | 7,67449 | 0,127445 | 0,5123 | 0,809389 |
| Ift88 | 11,2721 | 12,2155 | 0,115968 | 0,51235 | 0,809389 |
| Prrc2a | 128,236 | 138,776 | 0,113955 | 0,51245 | 0,809389 |
| Hnrnpdl | 180,057 | 167,312 | -0,105911 | 0,51235 | 0,809389 |
| Aftph | 12,548 | 11,622 | -0,110602 | 0,51245 | 0,809389 |
| Pex10 | 8,34804 | 7,54735 | -0,145467 | 0,5125 | 0,809389 |
| Vat1l | 1,18886 | 1,03263 | -0,203262 | 0,51225 | 0,809389 |
| Frat2 | 0,851797 | 1,0194 | 0,259139 | 0,5127 | 0,809418 |
| Rab15 | 15,0625 | 13,9396 | -0,111772 | 0,5127 | 0,809418 |
| Arhgap4 | 1,8256 | 1,60374 | -0,186931 | 0,51265 | 0,809418 |
| Cwc25 | 10,3521 | 11,2303 | 0,117483 | 0,5128 | 0,80948 |
| Uck1 | 7,99455 | 9,17409 | 0,198549 | 0,51305 | 0,809746 |
| DQ267102 | 505,352 | 493,9 | -0,0330712 | 0,51315 | 0,809746 |
| Ikbkg | 7,07772 | 6,3414 | -0,158484 | 0,51315 | 0,809746 |
| S1pr2 | 15,1607 | 14,0126 | -0,113614 | 0,5133 | 0,809887 |
| Ciapin1 | 19,8019 | 21,3056 | 0,105589 | 0,51375 | 0,810389 |
| Zmat2 | 66,0016 | 70,9288 | 0,103871 | 0,5138 | 0,810389 |
| Fignl1 | 48,731 | 45,2873 | -0,105732 | 0,51375 | 0,810389 |
| Tsr2 | 14,8406 | 16,5031 | 0,153187 | 0,514 | 0,810608 |
| Snx33 | 4,08439 | 4,46957 | 0,130016 | 0,5144 | 0,811048 |
| Ccdc93 | 7,3833 | 6,84148 | -0,109958 | 0,5144 | 0,811048 |
| Ppp1r3g | 0,931363 | 1,10367 | 0,244888 | 0,5147 | 0,811425 |
| Med17 | 17,1124 | 18,4486 | 0,108465 | 0,5149 | 0,811645 |
| Efna3 | 23,8803 | 26,0073 | 0,1231 | 0,51505 | 0,811768 |
| Hmg20b | 34,4054 | 37,186 | 0,112126 | 0,5151 | 0,811768 |
| Cacfd1 | 47,4565 | 44,0569 | -0,107238 | 0,51555 | 0,811807 |
| Slc26a1 | 0,429184 | 0,667927 | 0,638093 | 0,5152 | 0,811808 |
| Jmjd4 | 11,9742 | 12,9306 | 0,11086 | 0,51545 | 0,811808 |
| Ssb | 263,416 | 283,753 | 0,10729 | 0,5155 | 0,811808 |
| Cox7b | 110,291 | 118,633 | 0,105187 | 0,51545 | 0,811808 |
| Srp68 | 60,0768 | 64,5308 | 0,103181 | 0,51525 | 0,811808 |
| Ldlrad3 | 28,6034 | 26,6101 | -0,104213 | 0,5154 | 0,811808 |
| Cox15 | 12,7395 | 13,8136 | 0,11678 | 0,51585 | 0,811993 |
| Gosr2 | 30,7758 | 33,1099 | 0,105465 | 0,5158 | 0,811993 |
| Zfhx4 | 2,42774 | 2,23836 | -0,11717 | 0,51585 | 0,811993 |
| Zfp335 | 9,96234 | 10,7739 | 0,112983 | 0,51605 | 0,812212 |
| Mthfr | 9,29882 | 8,62459 | -0,108593 | 0,5164 | 0,812667 |
| Gm6642 | 0,206332 | 1,01942 | 2,30471 | 0,51685 | 0,812688 |
| E430018J23Rik | 3,40237 | 3,81678 | 0,165813 | 0,51675 | 0,812688 |
| Tasp1 | 8,65158 | 9,454 | 0,127961 | 0,5169 | 0,812688 |

| | | | | | |
|---|---|---|---|---|---|
| Spen | 11,1713 | 12,1288 | 0,118638 | 0,5165 | 0,812688 |
| Mta2 | 94,0447 | 100,997 | 0,102893 | 0,51655 | 0,812688 |
| Tmem201 | 26,4793 | 28,4362 | 0,102865 | 0,51675 | 0,812688 |
| Plcl2 | 5,01414 | 4,58269 | -0,129807 | 0,5169 | 0,812688 |
| Ercc4 | 3,07521 | 2,8096 | -0,130321 | 0,51675 | 0,812688 |
| B3gntl1 | 3,55654 | 4,44865 | 0,322893 | 0,51725 | 0,812884 |
| Piga | 12,4438 | 13,4455 | 0,111698 | 0,51735 | 0,812884 |
| Odc1 | 35,9322 | 38,6626 | 0,105661 | 0,51745 | 0,812884 |
| Creb1 | 20,3586 | 18,9466 | -0,103696 | 0,51745 | 0,812884 |
| Spsb4 | 27,1382 | 25,1315 | -0,110829 | 0,5174 | 0,812884 |
| Tmco3 | 6,51234 | 5,80059 | -0,166977 | 0,51725 | 0,812884 |
| Hist1h1e | 0,84994 | 0,613672 | -0,469895 | 0,5174 | 0,812884 |
| Renbp | 6,3102 | 5,64447 | -0,160848 | 0,51755 | 0,812945 |
| Ndufab1 | 6,03877 | 6,92271 | 0,197081 | 0,51795 | 0,813174 |
| Ncoa7 | 7,99643 | 8,67813 | 0,118027 | 0,518 | 0,813174 |
| Fbxo42 | 11,5893 | 12,4776 | 0,106542 | 0,518 | 0,813174 |
| Rcc1 | 79,7297 | 85,5764 | 0,102095 | 0,51795 | 0,813174 |
| Scrn1 | 1,28707 | 1,07616 | -0,2582 | 0,51785 | 0,813174 |
| Stmn1-rs1 | 0,061048 | 0,561865 | 3,20222 | 0,51815 | 0,813297 |
| Itfg2 | 14,4831 | 15,7003 | 0,116415 | 0,5182 | 0,813297 |
| Timp4 | 3,0724 | 3,54264 | 0,205459 | 0,5184 | 0,81342 |
| Ccdc19 | 0,856011 | 0,655954 | -0,384035 | 0,5184 | 0,81342 |
| Nt5c3b | 51,5265 | 57,19 | 0,150449 | 0,51855 | 0,813464 |
| Reps1 | 31,8899 | 34,306 | 0,10536 | 0,5185 | 0,813464 |
| Gbf1 | 15,5618 | 16,7227 | 0,1038 | 0,51865 | 0,813526 |
| 4930430F08Rik | 5,8874 | 6,8556 | 0,219652 | 0,51925 | 0,814077 |
| Zcchc10 | 16,1502 | 17,676 | 0,130237 | 0,5191 | 0,814077 |
| Tbc1d14 | 18,9447 | 20,3966 | 0,10654 | 0,5195 | 0,814077 |
| Tpst1 | 38,4802 | 41,3811 | 0,104858 | 0,5193 | 0,814077 |
| Slc25a13 | 37,9044 | 40,7486 | 0,104385 | 0,51945 | 0,814077 |
| Tcf25 | 64,2251 | 68,9898 | 0,103245 | 0,5195 | 0,814077 |
| Grin3b | 0,709997 | 0,74464 | 0,0687304 | 0,51925 | 0,814077 |
| Gm15910 | 0,724776 | 0,517627 | -0,485621 | 0,51955 | 0,814077 |
| Cd97 | 2,58498 | 1,75539 | -0,558363 | 0,51955 | 0,814077 |
| Vipas39 | 22,6559 | 24,4018 | 0,107096 | 0,5197 | 0,814217 |
| Acp2 | 17,7035 | 16,4283 | -0,107852 | 0,5198 | 0,814278 |
| Ccdc109b | 0,584408 | 0,755111 | 0,369712 | 0,5202 | 0,814376 |
| Stxbp3b | 3,62758 | 4,18516 | 0,206277 | 0,52035 | 0,814376 |
| Eif2b3 | 35,1351 | 37,9576 | 0,111475 | 0,52035 | 0,814376 |
| Zfp710 | 11,1575 | 12,0386 | 0,109651 | 0,52015 | 0,814376 |
| Rpap2 | 10,2218 | 8,93347 | -0,194361 | 0,52015 | 0,814376 |
| BC030307 | 2,1733 | 1,891 | -0,20074 | 0,52025 | 0,814376 |
| 6230400D17Rik | 2,23687 | 1,306 | -0,776325 | 0,52025 | 0,814376 |
| Angptl1 | 0,763265 | 0,212889 | -1,84208 | 0,52 | 0,814376 |
| Gm2382 | 3,4563 | 4,07605 | 0,237943 | 0,52055 | 0,814403 |
| Nek3 | 3,6278 | 4,05518 | 0,160671 | 0,52055 | 0,814403 |
| Atf7ip | 19,99 | 21,4776 | 0,103558 | 0,52055 | 0,814403 |
| Rab13 | 25,8589 | 28,1315 | 0,121524 | 0,5207 | 0,814413 |

| | | | | | |
|---|---|---|---|---|---|
| Syne2 | 6,52797 | 7,01966 | 0,104766 | 0,52075 | 0,814413 |
| Faf1 | 20,0871 | 21,5824 | 0,103588 | 0,5208 | 0,814413 |
| Naa35 | 43,6107 | 46,8383 | 0,103005 | 0,52065 | 0,814413 |
| Fbxo30 | 16,0343 | 17,2322 | 0,103947 | 0,521 | 0,814535 |
| Ap3m1 | 13,0702 | 12,1303 | -0,10767 | 0,52095 | 0,814535 |
| Stat2 | 4,55866 | 4,16619 | -0,129882 | 0,5211 | 0,814579 |
| Plekhh1 | 4,29356 | 3,85218 | -0,156498 | 0,52115 | 0,814579 |
| Fancd2os | 1,06681 | 0,86469 | -0,303043 | 0,52145 | 0,814857 |
| Dleu2 | 1,13997 | 0,831333 | -0,455495 | 0,5214 | 0,814857 |
| Usp53 | 3,04519 | 3,35696 | 0,140622 | 0,52155 | 0,814901 |
| Tle2 | 2,05998 | 1,81696 | -0,181097 | 0,5216 | 0,814901 |
| Zbtb34 | 5,58113 | 5,16379 | -0,112127 | 0,5219 | 0,815179 |
| Pik3cb | 3,37619 | 3,07882 | -0,133019 | 0,52185 | 0,815179 |
| Nup153 | 52,4153 | 56,264 | 0,102222 | 0,5221 | 0,815396 |
| Cnp | 4,75206 | 5,25227 | 0,144389 | 0,5222 | 0,815457 |
| Nt5c2 | 16,2589 | 17,5019 | 0,10628 | 0,5224 | 0,815674 |
| Wdr31 | 3,34594 | 3,72654 | 0,155424 | 0,52265 | 0,815787 |
| Ranbp9 | 25,1397 | 27,8021 | 0,145222 | 0,5228 | 0,815787 |
| Asnsd1 | 42,9699 | 46,3621 | 0,10962 | 0,5226 | 0,815787 |
| 9030624J02Rik | 16,1569 | 17,3898 | 0,106088 | 0,52285 | 0,815787 |
| R3hdm2 | 19,1743 | 20,5722 | 0,101524 | 0,5229 | 0,815787 |
| Prpf38b | 67,4633 | 72,3262 | 0,100415 | 0,52275 | 0,815787 |
| Ppp1r14c | 0,819089 | 0,68681 | -0,254109 | 0,5229 | 0,815787 |
| Gas2 | 7,84968 | 8,59587 | 0,13101 | 0,52315 | 0,816048 |
| Dcaf5 | 11,3934 | 12,2435 | 0,103815 | 0,5232 | 0,816048 |
| Suv39h1 | 56,5096 | 60,5881 | 0,100537 | 0,52325 | 0,816048 |
| Gm15401 | 0,616652 | 1,18667 | 0,944395 | 0,5236 | 0,816083 |
| Zscan18 | 4,87161 | 5,32503 | 0,128392 | 0,52355 | 0,816083 |
| Zfp955b | 5,69932 | 6,21828 | 0,125724 | 0,5237 | 0,816083 |
| Plcb3 | 15,6242 | 16,8092 | 0,105471 | 0,52355 | 0,816083 |
| P4ha1 | 25,1861 | 27,0257 | 0,101702 | 0,5237 | 0,816083 |
| Qsox2 | 17,9388 | 16,6229 | -0,109911 | 0,5237 | 0,816083 |
| Clmp | 4,86965 | 4,41856 | -0,140242 | 0,52365 | 0,816083 |
| Trim2 | 9,20135 | 8,55274 | -0,105459 | 0,52405 | 0,816438 |
| Aldh1l1 | 0,914738 | 0,780899 | -0,228223 | 0,524 | 0,816438 |
| Ercc1 | 25,3093 | 28,8712 | 0,189962 | 0,52415 | 0,816498 |
| Mir599 | 11,8315 | 42,7961 | 1,85484 | 0,52535 | 0,816939 |
| Tmco6 | 9,0418 | 10,5555 | 0,223315 | 0,5257 | 0,816939 |
| Saysd1 | 3,19931 | 3,58415 | 0,163868 | 0,5256 | 0,816939 |
| Tctn1 | 7,38688 | 8,1589 | 0,14341 | 0,52575 | 0,816939 |
| Hmox1 | 12,725 | 13,8836 | 0,125722 | 0,5258 | 0,816939 |
| Npc1 | 11,3887 | 12,3414 | 0,115893 | 0,52575 | 0,816939 |
| Myl9 | 36,0027 | 38,9819 | 0,114696 | 0,5255 | 0,816939 |
| Mcph1 | 10,499 | 11,3571 | 0,113332 | 0,52485 | 0,816939 |
| Pcolce2 | 23,7556 | 25,6343 | 0,109807 | 0,52545 | 0,816939 |
| Rad51ap1 | 35,3242 | 38,0467 | 0,107113 | 0,52485 | 0,816939 |
| Sp8 | 14,1389 | 15,2084 | 0,105201 | 0,52575 | 0,816939 |
| Zc3hc1 | 36,7918 | 39,5472 | 0,104188 | 0,52455 | 0,816939 |

| | | | | | |
|---|---|---|---|---|---|
| Atad3a | 35,872 | 38,5567 | 0,104123 | 0,52565 | 0,816939 |
| Ecd | 28,8715 | 31,0058 | 0,102892 | 0,5251 | 0,816939 |
| Tbc1d1 | 17,4646 | 18,7452 | 0,102085 | 0,5256 | 0,816939 |
| Sipa1l1 | 18,3248 | 19,6521 | 0,100887 | 0,52545 | 0,816939 |
| Nol8 | 20,1215 | 21,575 | 0,100629 | 0,5247 | 0,816939 |
| Psmb4 | 240,593 | 257,973 | 0,100625 | 0,5259 | 0,816939 |
| Aifm1 | 43,0026 | 40,1078 | -0,10054 | 0,52585 | 0,816939 |
| Afg3l1 | 25,8848 | 24,1087 | -0,10255 | 0,52575 | 0,816939 |
| Zzef1 | 3,90066 | 3,61434 | -0,109987 | 0,525 | 0,816939 |
| Rexo2 | 45,6129 | 42,2403 | -0,110823 | 0,52555 | 0,816939 |
| Prr22 | 2,12929 | 1,81069 | -0,233831 | 0,5251 | 0,816939 |
| Nupr1l | 0,907574 | 0,631462 | -0,52332 | 0,52585 | 0,816939 |
| Gm2518 | 0,576343 | 0,949965 | 0,720946 | 0,5262 | 0,817241 |
| Cenpj | 11,5433 | 12,4466 | 0,108694 | 0,52635 | 0,817241 |
| Mex3d | 17,3402 | 16,102 | -0,106888 | 0,5264 | 0,817241 |
| Lrfn3 | 5,33687 | 4,85526 | -0,136446 | 0,5264 | 0,817241 |
| Gm20605,Lrch4 | 6,35919 | 5,52617 | -0,202564 | 0,52635 | 0,817241 |
| Wdr89 | 4,89371 | 6,53638 | 0,417562 | 0,5267 | 0,817612 |
| Arhgap11a | 30,9515 | 28,8552 | -0,101179 | 0,5268 | 0,817672 |
| Asic1 | 1,23384 | 1,39878 | 0,18101 | 0,52695 | 0,817715 |
| Ube3b | 26,3683 | 28,2496 | 0,0994272 | 0,5269 | 0,817715 |
| Rsbn1l | 8,72245 | 9,4371 | 0,113611 | 0,52705 | 0,817775 |
| Sec24b | 21,6962 | 23,2821 | 0,101779 | 0,52725 | 0,81799 |
| Aasdhppt | 19,7504 | 18,3488 | -0,106195 | 0,5274 | 0,817998 |
| Akt2 | 12,7961 | 11,8372 | -0,112373 | 0,5275 | 0,817998 |
| Mirg | 16,5777 | 15,1741 | -0,127634 | 0,5275 | 0,817998 |
| Crtc3 | 2,39531 | 2,17419 | -0,139737 | 0,52735 | 0,817998 |
| Mettl23 | 13,8535 | 15,2378 | 0,137404 | 0,52775 | 0,818006 |
| Slc25a27 | 7,08761 | 7,73428 | 0,125967 | 0,52765 | 0,818006 |
| Mettl10 | 33,5853 | 36,3559 | 0,114358 | 0,52775 | 0,818006 |
| Jak1 | 28,2615 | 30,4094 | 0,10568 | 0,52775 | 0,818006 |
| Ehbp1 | 7,86613 | 8,47416 | 0,107416 | 0,5279 | 0,818127 |
| Kdm3b | 25,6417 | 27,464 | 0,0990491 | 0,52795 | 0,818127 |
| Pex1 | 5,536 | 6,10106 | 0,140214 | 0,52835 | 0,818332 |
| Sf3b5 | 117,664 | 126,542 | 0,104939 | 0,5283 | 0,818332 |
| Gns | 21,4693 | 19,9699 | -0,104451 | 0,5283 | 0,818332 |
| Mknk1 | 7,43054 | 6,79486 | -0,129024 | 0,52845 | 0,818332 |
| Hs3st3a1 | 2,22478 | 1,96474 | -0,179324 | 0,5284 | 0,818332 |
| Slc2a10 | 1,27397 | 1,1146 | -0,192802 | 0,5283 | 0,818332 |
| Tmem223 | 28,5983 | 26,0981 | -0,131984 | 0,5286 | 0,81847 |
| Tmem39b | 16,4576 | 17,8342 | 0,115895 | 0,5287 | 0,81853 |
| Matr3 | 149,356 | 139,074 | -0,102899 | 0,52885 | 0,818667 |
| 9130011E15Rik | 8,85622 | 9,61219 | 0,118175 | 0,529 | 0,818692 |
| Secisbp2l | 11,1188 | 11,9584 | 0,105024 | 0,52905 | 0,818692 |
| Glyr1 | 95,0456 | 101,853 | 0,0997929 | 0,52905 | 0,818692 |
| Mir7013 | 71,1221 | 242,751 | 1,77111 | 0,52955 | 0,81887 |
| Endog | 6,63869 | 8,50261 | 0,357007 | 0,5299 | 0,81887 |
| 2810429I04Rik | 4,56861 | 5,41187 | 0,244374 | 0,52975 | 0,81887 |

| | | | | | |
|---|---|---|---|---|---|
| Perm1 | 0,932363 | 1,06718 | 0,194835 | 0,5295 | 0,81887 |
| Myo5b | 1,20734 | 1,3415 | 0,152014 | 0,52965 | 0,81887 |
| Haus1 | 38,9408 | 42,123 | 0,113324 | 0,52935 | 0,81887 |
| Fundc1 | 19,8132 | 21,3743 | 0,109419 | 0,5298 | 0,81887 |
| Lmnb2 | 47,1935 | 50,5724 | 0,0997623 | 0,52965 | 0,81887 |
| Sp3 | 44,0445 | 41,0434 | -0,101812 | 0,5297 | 0,81887 |
| Agfg1 | 33,4492 | 31,157 | -0,102415 | 0,5297 | 0,81887 |
| Tecpr1 | 7,23277 | 6,6675 | -0,117402 | 0,52985 | 0,81887 |
| Larp1b | 2,80016 | 2,44908 | -0,193271 | 0,52985 | 0,81887 |
| Lman2l | 19,8715 | 18,4136 | -0,109928 | 0,53 | 0,818929 |
| Ttc38 | 2,4432 | 2,70548 | 0,147114 | 0,5303 | 0,819298 |
| Lsm7 | 0,67264 | 1,01415 | 0,592362 | 0,5305 | 0,819512 |
| Hist1h4d | 2,4482 | 3,26576 | 0,415697 | 0,53085 | 0,819706 |
| Slc9b2 | 2,43315 | 2,74984 | 0,176521 | 0,53075 | 0,819706 |
| Kcnh1 | 0,769106 | 0,863053 | 0,166267 | 0,53125 | 0,819706 |
| Sin3b | 69,1748 | 75,1702 | 0,119915 | 0,5312 | 0,819706 |
| Atp5sl | 16,5269 | 17,9447 | 0,118742 | 0,5309 | 0,819706 |
| Otulin | 18,9661 | 20,5767 | 0,117589 | 0,53115 | 0,819706 |
| Foxd1 | 12,7228 | 13,7689 | 0,114002 | 0,5313 | 0,819706 |
| Zbtb39 | 12,1667 | 13,0553 | 0,101699 | 0,5313 | 0,819706 |
| Gna13 | 18,4462 | 19,7782 | 0,100593 | 0,5308 | 0,819706 |
| Slc4a3 | 6,51915 | 6,00715 | -0,118004 | 0,5313 | 0,819706 |
| Smpdl3b | 4,09358 | 3,6585 | -0,162112 | 0,53115 | 0,819706 |
| Pdzd4 | 2,62505 | 2,90142 | 0,144413 | 0,5315 | 0,81992 |
| Gm15413 | 0,116257 | 0,648628 | 2,48008 | 0,5317 | 0,819994 |
| Necap1 | 26,2605 | 28,2287 | 0,104267 | 0,5318 | 0,819994 |
| Dcaf8 | 53,1456 | 56,9517 | 0,0997887 | 0,53195 | 0,819994 |
| Wtap | 41,4806 | 38,4755 | -0,108497 | 0,53205 | 0,819994 |
| Cyp20a1 | 25,2612 | 23,352 | -0,113378 | 0,5319 | 0,819994 |
| Hlcs | 6,18981 | 5,717 | -0,114636 | 0,5321 | 0,819994 |
| Mvb12a | 18,4221 | 16,8409 | -0,12947 | 0,5319 | 0,819994 |
| Ksr1 | 2,70447 | 2,44698 | -0,144342 | 0,532 | 0,819994 |
| Tmem158 | 3,15952 | 2,78334 | -0,182886 | 0,53205 | 0,819994 |
| Fbxo16 | 5,77454 | 6,48674 | 0,167788 | 0,53225 | 0,820018 |
| Brf2 | 11,8311 | 12,9529 | 0,130683 | 0,5322 | 0,820018 |
| Eya4 | 0,7324 | 0,631968 | -0,21278 | 0,5323 | 0,820018 |
| Podxl2 | 8,63361 | 7,86584 | -0,134362 | 0,5324 | 0,820078 |
| Aen | 21,7393 | 23,3786 | 0,10488 | 0,53255 | 0,820214 |
| Cars | 27,5419 | 29,5642 | 0,102221 | 0,53265 | 0,820274 |
| Pold2 | 36,837 | 39,8003 | 0,111623 | 0,53295 | 0,820434 |
| Fastkd1 | 10,821 | 9,995 | -0,114557 | 0,533 | 0,820434 |
| Arrb2 | 18,0399 | 16,6181 | -0,118438 | 0,53285 | 0,820434 |
| Phospho2 | 11,5867 | 10,6687 | -0,119095 | 0,533 | 0,820434 |
| Prorsd1 | 4,59056 | 5,08657 | 0,148023 | 0,53345 | 0,82056 |
| Grpel1 | 26,523 | 28,7546 | 0,116547 | 0,53335 | 0,82056 |
| Prpf8 | 100,444 | 108,053 | 0,105347 | 0,53345 | 0,82056 |
| Kif3b | 9,96342 | 10,7153 | 0,104963 | 0,53345 | 0,82056 |
| Blm | 31,1946 | 33,4139 | 0,0991514 | 0,53315 | 0,82056 |

| | | | | | |
|---|---|---|---|---|---|
| Fbxl16 | 1,09802 | 0,956066 | -0,199723 | 0,53345 | 0,82056 |
| Ncbp1 | 80,948 | 86,701 | 0,0990533 | 0,5336 | 0,820602 |
| Nt5dc1 | 4,52491 | 4,10175 | -0,141649 | 0,53355 | 0,820602 |
| Pex6 | 17,656 | 18,9927 | 0,10529 | 0,5341 | 0,821276 |
| Ddx49 | 24,7259 | 27,5994 | 0,158615 | 0,5342 | 0,821336 |
| Alkbh6 | 26,7086 | 29,0512 | 0,121293 | 0,5343 | 0,821395 |
| Tspyl4 | 4,75411 | 4,35989 | -0,124884 | 0,5344 | 0,821454 |
| Plxna1 | 25,5246 | 23,8568 | -0,0974855 | 0,5346 | 0,821478 |
| Snx4 | 23,3188 | 21,6929 | -0,104269 | 0,5345 | 0,821478 |
| Decr2 | 8,02946 | 7,32262 | -0,132942 | 0,5346 | 0,821478 |
| Fam115a | 25,5037 | 27,3079 | 0,0986088 | 0,5351 | 0,821945 |
| Dars | 99,8332 | 106,863 | 0,0981748 | 0,5351 | 0,821945 |
| Glis2 | 15,3681 | 14,2888 | -0,105055 | 0,53515 | 0,821945 |
| Cpe | 25,7595 | 23,9363 | -0,105902 | 0,53515 | 0,821945 |
| Mir7650 | 328,694 | 1783,39 | 2,43981 | 0,53545 | 0,822217 |
| Kpna2 | 11,691 | 10,7433 | -0,121955 | 0,53545 | 0,822217 |
| Prpf19 | 61,4396 | 65,9109 | 0,101348 | 0,5357 | 0,822506 |
| Zfp687 | 22,2062 | 24,2218 | 0,125348 | 0,53585 | 0,822536 |
| Csrnp2 | 8,68799 | 9,36926 | 0,108913 | 0,53615 | 0,822536 |
| Pde7a | 23,9549 | 25,6437 | 0,0982824 | 0,53605 | 0,822536 |
| Uqcrq | 182,138 | 169,27 | -0,105706 | 0,53595 | 0,822536 |
| Usp32 | 7,47546 | 6,9447 | -0,106249 | 0,5359 | 0,822536 |
| 9130019O22Rik | 1,42758 | 1,26522 | -0,174178 | 0,53615 | 0,822536 |
| Glt8d1 | 23,2268 | 19,0798 | -0,283742 | 0,536 | 0,822536 |
| Ccdc89 | 0,909071 | 1,06579 | 0,229454 | 0,53635 | 0,822654 |
| Nfkbid | 1,15072 | 0,975138 | -0,238854 | 0,53635 | 0,822654 |
| Zfp13 | 11,0606 | 12,0096 | 0,118748 | 0,53645 | 0,822713 |
| Banf1 | 308,446 | 335,877 | 0,122917 | 0,5368 | 0,822913 |
| Rnf5 | 37,2412 | 40,1737 | 0,10935 | 0,53695 | 0,822913 |
| Ppp3r1 | 52,676 | 49,147 | -0,100042 | 0,53675 | 0,822913 |
| Bach1 | 14,3578 | 13,3909 | -0,100585 | 0,53685 | 0,822913 |
| Akirin1 | 15,2785 | 14,16 | -0,10968 | 0,5369 | 0,822913 |
| Plod3 | 24,5412 | 22,5961 | -0,119131 | 0,5368 | 0,822913 |
| Zkscan5 | 8,8754 | 9,57359 | 0,109248 | 0,53715 | 0,823031 |
| Mrpl43 | 38,735 | 41,7685 | 0,10878 | 0,5371 | 0,823031 |
| Gpr108 | 14,7215 | 15,9193 | 0,112853 | 0,5373 | 0,82313 |
| Rabl6 | 43,8547 | 46,9609 | 0,0987289 | 0,5374 | 0,82313 |
| Rpia | 17,3066 | 15,9731 | -0,115677 | 0,53735 | 0,82313 |
| Asb6 | 16,4078 | 17,7173 | 0,110781 | 0,53755 | 0,823266 |
| Jmy | 9,95076 | 10,6604 | 0,09938 | 0,53775 | 0,823478 |
| Zfp454 | 7,00691 | 7,67583 | 0,131546 | 0,53795 | 0,823595 |
| Cyfip1 | 28,0875 | 30,3248 | 0,110571 | 0,53795 | 0,823595 |
| Fancg | 8,87455 | 9,61829 | 0,116106 | 0,5381 | 0,823636 |
| Upf3b | 18,4071 | 17,0593 | -0,109701 | 0,5381 | 0,823636 |
| Gon4l | 11,2995 | 12,3414 | 0,127249 | 0,53855 | 0,823665 |
| Mfhas1 | 3,0697 | 3,33632 | 0,120162 | 0,53845 | 0,823665 |
| Coro1c | 56,5928 | 60,5047 | 0,0964275 | 0,5385 | 0,823665 |
| Purb | 35,3004 | 32,9863 | -0,0978151 | 0,5383 | 0,823665 |

| | | | | | |
|---|---|---|---|---|---|
| Ap3s2 | 18,0076 | 16,8048 | -0,0997312 | 0,53855 | 0,823665 |
| Sirt1 | 22,8685 | 21,3374 | -0,099979 | 0,53855 | 0,823665 |
| Lmbr1l | 13,2341 | 12,2042 | -0,116885 | 0,5382 | 0,823665 |
| Tmem200b | 0,61963 | 0,782406 | 0,336509 | 0,5389 | 0,823729 |
| Ccdc122 | 0,581253 | 0,721468 | 0,311767 | 0,5388 | 0,823729 |
| Dexi | 12,5896 | 13,766 | 0,12888 | 0,53875 | 0,823729 |
| Taz | 23,1607 | 21,493 | -0,107817 | 0,5389 | 0,823729 |
| Gm5083 | 0,8466 | 0,665234 | -0,34782 | 0,53875 | 0,823729 |
| Slc9a1 | 11,3292 | 10,5353 | -0,104818 | 0,539 | 0,823787 |
| Irak1bp1 | 7,43774 | 8,15593 | 0,132985 | 0,53945 | 0,824287 |
| Ubl4 | 20,7893 | 19,1452 | -0,118856 | 0,53945 | 0,824287 |
| Zfp386 | 11,3402 | 10,5423 | -0,105249 | 0,5396 | 0,824421 |
| Ints7 | 35,6813 | 38,1961 | 0,098256 | 0,53995 | 0,824673 |
| Pacs2 | 19,947 | 21,3461 | 0,0978035 | 0,5399 | 0,824673 |
| Emc6 | 53,3287 | 49,7151 | -0,101229 | 0,53995 | 0,824673 |
| Cep89 | 18,8221 | 20,2638 | 0,106482 | 0,5402 | 0,824867 |
| Eif5a | 636,304 | 684,099 | 0,104489 | 0,5402 | 0,824867 |
| Eif4b | 148,182 | 158,859 | 0,10037 | 0,5404 | 0,825078 |
| Ogg1 | 11,0978 | 12,508 | 0,172584 | 0,54055 | 0,825195 |
| Mknk2 | 47,1096 | 43,9872 | -0,0989372 | 0,5406 | 0,825195 |
| Slc27a3 | 20,7784 | 19,2925 | -0,107046 | 0,54075 | 0,825329 |
| Patz1 | 45,6116 | 52,9649 | 0,215636 | 0,54115 | 0,825379 |
| Tprn | 3,1416 | 3,4846 | 0,149495 | 0,5413 | 0,825379 |
| Fbxl4 | 4,1317 | 4,56961 | 0,145336 | 0,5411 | 0,825379 |
| Tmem29 | 20,0224 | 21,6938 | 0,115666 | 0,5413 | 0,825379 |
| Bach2 | 3,34824 | 3,61818 | 0,111862 | 0,5411 | 0,825379 |
| Gps2 | 31,7283 | 34,2403 | 0,109926 | 0,5413 | 0,825379 |
| Wash | 11,6177 | 12,5361 | 0,109769 | 0,5414 | 0,825379 |
| Mrpl17 | 9,32013 | 9,99839 | 0,101346 | 0,54095 | 0,825379 |
| Arl6ip6 | 27,2468 | 25,4044 | -0,10101 | 0,541 | 0,825379 |
| 9330102E08Rik | 0,740899 | 0,640834 | -0,209326 | 0,54135 | 0,825379 |
| Yeats2 | 35,0871 | 37,5887 | 0,099356 | 0,54165 | 0,825383 |
| Rhot1 | 21,9244 | 23,469 | 0,0982166 | 0,54155 | 0,825383 |
| Bcl9 | 23,9489 | 25,6247 | 0,0975756 | 0,5416 | 0,825383 |
| Abi1 | 23,8696 | 22,1615 | -0,107116 | 0,54155 | 0,825383 |
| Exd2 | 12,0986 | 11,2333 | -0,107052 | 0,54175 | 0,825441 |
| Timm13 | 58,1584 | 62,3186 | 0,0996744 | 0,54185 | 0,8255 |
| Ccdc59 | 43,0517 | 46,3639 | 0,10693 | 0,54195 | 0,825558 |
| Atp2a2 | 129,499 | 138,867 | 0,100764 | 0,5422 | 0,825656 |
| Scaf4 | 23,5894 | 25,2354 | 0,0973095 | 0,5422 | 0,825656 |
| Pgd | 95,801 | 102,434 | 0,0965796 | 0,54215 | 0,825656 |
| Ccdc77 | 13,5917 | 15,8366 | 0,220531 | 0,5426 | 0,826171 |
| Slc45a3 | 1,75869 | 1,5622 | -0,170924 | 0,5427 | 0,82623 |
| Slc8a3 | 0,600787 | 0,520422 | -0,207172 | 0,5431 | 0,826744 |
| Gm11517 | 0,741124 | 2,49662 | 1,75219 | 0,5434 | 0,826864 |
| 1700012D01Rik | 0,651076 | 0,896673 | 0,461756 | 0,54355 | 0,826864 |
| Gng10 | 20,0089 | 21,7608 | 0,121086 | 0,54345 | 0,826864 |
| Bcat2 | 22,4982 | 24,2219 | 0,106505 | 0,54345 | 0,826864 |

| | | | | | |
|---|---|---|---|---|---|
| Rnpep | 37,5448 | 40,206 | 0,0987972 | 0,5435 | 0,826864 |
| Fads2 | 106,125 | 113,463 | 0,0964586 | 0,54345 | 0,826864 |
| Brat1 | 6,74481 | 7,31616 | 0,117309 | 0,54385 | 0,826944 |
| Prps1 | 37,74 | 40,4229 | 0,0990804 | 0,5438 | 0,826944 |
| Fbxo4 | 1,08648 | 0,949767 | -0,194017 | 0,54375 | 0,826944 |
| Reep6 | 1,96646 | 0,982068 | -1,00171 | 0,5437 | 0,826944 |
| Spink2 | 1,53257 | 1,96316 | 0,357228 | 0,54395 | 0,826984 |
| Taf9 | 51,9498 | 56,6927 | 0,126044 | 0,544 | 0,826984 |
| Surf2 | 36,5431 | 44,5395 | 0,285486 | 0,5442 | 0,827006 |
| Cmtr2 | 4,8155 | 4,41998 | -0,123647 | 0,5442 | 0,827006 |
| Emc9 | 1,28058 | 0,608564 | -1,07332 | 0,5442 | 0,827006 |
| E130311K13Rik | 1,84606 | 1,58507 | -0,21991 | 0,5445 | 0,827368 |
| 1700037C18Rik | 1,72371 | 2,04319 | 0,245305 | 0,5446 | 0,827425 |
| Phf7 | 5,61687 | 6,17972 | 0,137775 | 0,54525 | 0,828225 |
| Ola1 | 64,1766 | 68,7652 | 0,0996311 | 0,54525 | 0,828225 |
| 9430015G10Rik | 37,9776 | 40,5935 | 0,0960997 | 0,54545 | 0,82834 |
| Tcf19 | 16,7784 | 15,4767 | -0,116507 | 0,5454 | 0,82834 |
| Reep1 | 7,55182 | 6,97782 | -0,114049 | 0,54585 | 0,828853 |
| Fam117a | 14,6696 | 13,6008 | -0,109138 | 0,546 | 0,828893 |
| Mir25 | 329,354 | 54,0828 | -2,6064 | 0,54595 | 0,828893 |
| Mir194-1 | 573,485 | 260,626 | -1,13777 | 0,54615 | 0,829026 |
| Eif3a | 163,989 | 176,178 | 0,103434 | 0,54645 | 0,829181 |
| Ckap5 | 68,528 | 73,3438 | 0,0979823 | 0,54645 | 0,829181 |
| Fzd3 | 11,951 | 11,1774 | -0,0965465 | 0,5465 | 0,829181 |
| Tep1 | 2,19563 | 2,01519 | -0,123714 | 0,5464 | 0,829181 |
| Sphk2 | 11,343 | 12,2259 | 0,108131 | 0,54665 | 0,829202 |
| Eif2s1 | 51,4102 | 54,9356 | 0,0956853 | 0,5467 | 0,829202 |
| Trim21 | 3,21366 | 2,89605 | -0,15013 | 0,5467 | 0,829202 |
| Rem2 | 1,98768 | 1,7336 | -0,197314 | 0,5469 | 0,829411 |
| Mtap7d3 | 2,28211 | 2,55417 | 0,162487 | 0,54715 | 0,829414 |
| Tsta3 | 31,9337 | 34,3775 | 0,106383 | 0,5471 | 0,829414 |
| Wnk1 | 44,3418 | 47,419 | 0,0967984 | 0,54715 | 0,829414 |
| Sbf1 | 24,383 | 26,0496 | 0,0953819 | 0,54705 | 0,829414 |
| Mrpl49 | 46,9653 | 51,7707 | 0,14054 | 0,54745 | 0,829568 |
| Cenph | 50,7711 | 54,4608 | 0,101211 | 0,5475 | 0,829568 |
| Afg3l2 | 34,017 | 31,8108 | -0,096738 | 0,5474 | 0,829568 |
| Nudt13 | 4,35825 | 3,94915 | -0,142208 | 0,54735 | 0,829568 |
| Bcar1 | 26,4116 | 28,227 | 0,0959077 | 0,54765 | 0,829701 |
| Vasn | 7,43069 | 8,27473 | 0,155216 | 0,5478 | 0,82974 |
| Mbtps1 | 91,2293 | 85,3566 | -0,0959946 | 0,54775 | 0,82974 |
| Ppan | 59,5316 | 63,6481 | 0,0964621 | 0,5479 | 0,829798 |
| Mpzl3 | 1,70958 | 1,42001 | -0,267746 | 0,548 | 0,829855 |
| Slc25a20 | 14,3485 | 13,2401 | -0,115988 | 0,5482 | 0,830064 |
| Rab33a | 0,701336 | 0,88735 | 0,339396 | 0,54845 | 0,830163 |
| Selenbp1 | 3,08961 | 3,47601 | 0,170009 | 0,54845 | 0,830163 |
| Meis2 | 23,3577 | 21,8355 | -0,0972251 | 0,5487 | 0,830163 |
| Txndc16 | 6,82226 | 6,31 | -0,11261 | 0,54865 | 0,830163 |
| Mrps10 | 19,776 | 18,1444 | -0,124227 | 0,5485 | 0,830163 |

| | | | | | |
|---|---|---|---|---|---|
| Ccdc163 | 15,8623 | 14,5517 | -0,124413 | 0,5487 | 0,830163 |
| Mterf1a | 1,90482 | 1,63116 | -0,223763 | 0,54865 | 0,830163 |
| Nr1h2 | 22,6719 | 24,3511 | 0,103084 | 0,5488 | 0,83022 |
| 4930507D05Rik | 0,521982 | 0,831465 | 0,671657 | 0,5491 | 0,83058 |
| Cep83os | 1,3056 | 1,46461 | 0,165804 | 0,54955 | 0,830602 |
| Adar | 13,2808 | 14,1996 | 0,0965163 | 0,5495 | 0,830602 |
| Rnf14 | 30,7053 | 32,824 | 0,0962643 | 0,5492 | 0,830602 |
| Cacul1 | 21,8057 | 20,3995 | -0,0961694 | 0,54945 | 0,830602 |
| Gucd1 | 15,0056 | 13,9552 | -0,104698 | 0,54955 | 0,830602 |
| Stard3 | 16,0649 | 14,8935 | -0,109229 | 0,54955 | 0,830602 |
| Prrg1 | 1,90514 | 1,71062 | -0,155373 | 0,54935 | 0,830602 |
| Wdr73 | 22,7277 | 24,4015 | 0,102521 | 0,5499 | 0,830794 |
| Grb2 | 26,1734 | 28,0312 | 0,0989298 | 0,55 | 0,830794 |
| Gigyf1 | 16,8787 | 18,0661 | 0,098077 | 0,54985 | 0,830794 |
| Cep57 | 38,6678 | 41,3177 | 0,0956265 | 0,54995 | 0,830794 |
| Ncoa2 | 14,1948 | 15,1658 | 0,0954515 | 0,55005 | 0,830794 |
| Gm6297 | 1,60458 | 0,532739 | -1,5907 | 0,5498 | 0,830794 |
| Mir7060 | 5,20505 | 24,3953 | 2,22862 | 0,5502 | 0,830927 |
| Brap | 13,5376 | 14,529 | 0,10197 | 0,5503 | 0,830966 |
| Gnai3 | 108,591 | 101,689 | -0,0947379 | 0,55035 | 0,830966 |
| Zcchc11 | 38,1718 | 35,7694 | -0,0937824 | 0,55055 | 0,830968 |
| 2010204K13Rik | 19,2323 | 17,2629 | -0,155856 | 0,5506 | 0,830968 |
| Irf5 | 0,957076 | 0,810997 | -0,238937 | 0,55045 | 0,830968 |
| Chrna3 | 0,598161 | 0,503554 | -0,248388 | 0,5505 | 0,830968 |
| Eif4a2 | 118,64 | 127,832 | 0,107662 | 0,5507 | 0,831025 |
| Tmem97 | 30,8825 | 33,2228 | 0,105385 | 0,5508 | 0,831082 |
| Ankrd34b | 1,32608 | 1,48988 | 0,168028 | 0,55105 | 0,831084 |
| Rgag4 | 7,18572 | 7,96138 | 0,147884 | 0,55105 | 0,831084 |
| Ranbp3 | 66,9103 | 71,3873 | 0,0934387 | 0,5509 | 0,831084 |
| Slc25a47 | 5,62734 | 5,06928 | -0,150672 | 0,55095 | 0,831084 |
| Dstyk | 8,14545 | 7,59306 | -0,101313 | 0,5514 | 0,831518 |
| Cox5b | 47,5555 | 51,6433 | 0,118969 | 0,5517 | 0,831689 |
| Serbp1 | 139,099 | 149,473 | 0,103777 | 0,5517 | 0,831689 |
| Heg1 | 25,2018 | 26,9519 | 0,0968637 | 0,5516 | 0,831689 |
| Gm15417 | 0,777431 | 1,0334 | 0,410618 | 0,55205 | 0,831935 |
| Map3k2 | 3,68158 | 3,95885 | 0,104757 | 0,552 | 0,831935 |
| Usp14 | 48,0642 | 44,0441 | -0,126016 | 0,552 | 0,831935 |
| Gm5801 | 23,4333 | 25,3423 | 0,112986 | 0,5523 | 0,83203 |
| Fundc2 | 14,8946 | 13,8463 | -0,105281 | 0,5523 | 0,83203 |
| Jagn1 | 44,0303 | 40,9288 | -0,105382 | 0,55225 | 0,83203 |
| Lrrc59 | 105,463 | 113,298 | 0,103388 | 0,55265 | 0,832182 |
| 1110059E24Rik | 39,058 | 41,8952 | 0,101167 | 0,5526 | 0,832182 |
| Polr3d | 32,0169 | 34,2288 | 0,0963785 | 0,5525 | 0,832182 |
| Hdhd3 | 4,46546 | 3,92846 | -0,184844 | 0,5526 | 0,832182 |
| Klhl34 | 2,12392 | 2,38167 | 0,165247 | 0,55275 | 0,83222 |
| 4932416H05Rik | 2,38203 | 1,86803 | -0,350672 | 0,5528 | 0,83222 |
| Kif5a | 0,548862 | 0,62898 | 0,196572 | 0,5529 | 0,832277 |
| Zfp398 | 2,79633 | 3,07957 | 0,139195 | 0,55325 | 0,832598 |

| | | | | | |
|---|---|---|---|---|---|
| Supt4a | 29,963 | 32,6716 | 0,124858 | 0,5533 | 0,832598 |
| Mrpl37 | 67,2602 | 71,9398 | 0,0970376 | 0,5533 | 0,832598 |
| Pcdh18 | 9,13722 | 8,51609 | -0,101565 | 0,55345 | 0,83273 |
| Zmym4 | 26,6906 | 28,6963 | 0,104536 | 0,5536 | 0,832843 |
| Trerf1 | 3,2135 | 2,93654 | -0,130029 | 0,55365 | 0,832843 |
| Snord2 | 361,84 | 2061,16 | 2,51003 | 0,55435 | 0,832865 |
| Zfp93 | 2,10585 | 2,33617 | 0,149742 | 0,5543 | 0,832865 |
| Fra10ac1 | 21,8143 | 23,5407 | 0,10988 | 0,5542 | 0,832865 |
| Iffo2 | 5,29701 | 5,70467 | 0,106965 | 0,5538 | 0,832865 |
| Pias2 | 36,1144 | 38,8679 | 0,106008 | 0,5543 | 0,832865 |
| Twistnb | 15,2791 | 16,4402 | 0,105671 | 0,55425 | 0,832865 |
| Socs6 | 15,9242 | 17,076 | 0,10075 | 0,55425 | 0,832865 |
| Gpx8 | 89,7808 | 83,9419 | -0,0970149 | 0,5539 | 0,832865 |
| Gnao1 | 4,28555 | 3,94379 | -0,119899 | 0,5539 | 0,832865 |
| Rgl3 | 4,42129 | 4,02075 | -0,137002 | 0,55415 | 0,832865 |
| Thpo | 0,648894 | 0,485304 | -0,419095 | 0,55415 | 0,832865 |
| Wls | 121,979 | 130,17 | 0,0937683 | 0,55445 | 0,832903 |
| Coq9 | 16,5069 | 15,273 | -0,112078 | 0,5545 | 0,832903 |
| Orai3 | 6,97415 | 6,37475 | -0,129648 | 0,55465 | 0,833035 |
| Hist1h4c | 0,82956 | 1,28513 | 0,631497 | 0,5548 | 0,833166 |
| Clcn5 | 13,9218 | 13,0118 | -0,097523 | 0,5549 | 0,833204 |
| Gnptg | 20,8176 | 18,9876 | -0,132745 | 0,55495 | 0,833204 |
| Snora69 | 81,7736 | 18,8915 | -2,1139 | 0,5551 | 0,833336 |
| Bckdk | 23,5481 | 25,1962 | 0,0975938 | 0,5552 | 0,833392 |
| Nudt12 | 2,58935 | 2,30859 | -0,165581 | 0,5555 | 0,833749 |
| Gm10865 | 0,569058 | 1,34877 | 1,245 | 0,5557 | 0,833862 |
| Kif23 | 42,0441 | 44,8864 | 0,0943735 | 0,5557 | 0,833862 |
| Ikzf4 | 1,80609 | 2,01137 | 0,155313 | 0,5558 | 0,8339 |
| Endod1 | 8,72961 | 8,12683 | -0,103225 | 0,55585 | 0,8339 |
| Tmem141 | 6,66078 | 7,48262 | 0,167853 | 0,5561 | 0,833994 |
| Tpmt | 4,2482 | 4,65826 | 0,13294 | 0,55605 | 0,833994 |
| Gm10012 | 0,703613 | 0,159882 | -2,13777 | 0,5561 | 0,833994 |
| Gm12338 | 25,1317 | 31,042 | 0,304714 | 0,55625 | 0,834051 |
| Dusp14 | 5,56195 | 6,16507 | 0,148525 | 0,55635 | 0,834051 |
| Zbed6 | 35,8969 | 39,7388 | 0,146688 | 0,55655 | 0,834051 |
| Mrpl22 | 20,7236 | 22,6741 | 0,129774 | 0,5567 | 0,834051 |
| Dtwd1 | 11,117 | 12,1434 | 0,127404 | 0,55665 | 0,834051 |
| Mrpl40 | 50,6621 | 54,3481 | 0,101323 | 0,5565 | 0,834051 |
| Aars | 62,8093 | 67,018 | 0,0935684 | 0,5565 | 0,834051 |
| Sdc2 | 17,1448 | 16,0153 | -0,0983175 | 0,5567 | 0,834051 |
| Asxl3 | 3,22555 | 2,99872 | -0,105198 | 0,5564 | 0,834051 |
| Ccdc55 | 13,334 | 14,3188 | 0,1028 | 0,5568 | 0,834108 |
| Rad21 | 138,391 | 129,66 | -0,0940144 | 0,5569 | 0,834164 |
| Lamtor1 | 40,371 | 43,4383 | 0,105648 | 0,557 | 0,83422 |
| Uchl4 | 22,0602 | 25,4904 | 0,208504 | 0,55735 | 0,834502 |
| Lat2 | 1,94252 | 2,22105 | 0,193311 | 0,5574 | 0,834502 |
| Samd14 | 20,0572 | 21,4808 | 0,0989222 | 0,5575 | 0,834502 |
| Tcf4 | 29,8359 | 27,9668 | -0,0933355 | 0,55745 | 0,834502 |

| | | | | | |
|---|---|---|---|---|---|
| Csf2ra | 5,85966 | 5,31716 | -0,140162 | 0,55735 | 0,834502 |
| Cnot6l | 9,30574 | 8,70794 | -0,09579 | 0,5577 | 0,834707 |
| Cbs | 1,04222 | 1,19932 | 0,202554 | 0,55785 | 0,834801 |
| 1110065P20Rik | 22,4044 | 25,3185 | 0,176414 | 0,55795 | 0,834801 |
| Capn10 | 13,9384 | 14,9588 | 0,101928 | 0,5579 | 0,834801 |
| Mettl17 | 13,0299 | 14,1235 | 0,116267 | 0,5583 | 0,835044 |
| Madd | 4,78458 | 5,15301 | 0,107023 | 0,5583 | 0,835044 |
| Zfp365 | 0,782926 | 0,686675 | -0,18925 | 0,55825 | 0,835044 |
| Rc3h1 | 8,47068 | 9,05089 | 0,0955831 | 0,55845 | 0,835175 |
| Phldb2 | 22,7693 | 24,3028 | 0,0940349 | 0,5586 | 0,835213 |
| Megf8 | 15,1092 | 14,1553 | -0,0940876 | 0,5586 | 0,835213 |
| Chd2 | 13,6415 | 12,6868 | -0,104678 | 0,5587 | 0,835269 |
| Fam175b | 21,534 | 20,1091 | -0,0987715 | 0,55915 | 0,835848 |
| Gm4432 | 0,153616 | 0,592966 | 1,94862 | 0,5594 | 0,836072 |
| Zfyve9 | 20,8256 | 22,403 | 0,105333 | 0,55955 | 0,836072 |
| Gtf2e2 | 67,6629 | 72,2075 | 0,0937837 | 0,55955 | 0,836072 |
| Pex16 | 7,14045 | 6,44831 | -0,147094 | 0,5595 | 0,836072 |
| Vps13c | 3,38589 | 3,15376 | -0,102461 | 0,55965 | 0,836128 |
| Mir7067 | 182,594 | 537,973 | 1,5589 | 0,55995 | 0,836164 |
| Pfkp | 4,38418 | 6,09288 | 0,474819 | 0,56005 | 0,836164 |
| Snord72 | 279,498 | 317,991 | 0,186145 | 0,5604 | 0,836164 |
| 4933407K13Rik | 2,57537 | 2,83517 | 0,138659 | 0,55975 | 0,836164 |
| Ccdc137 | 15,4942 | 16,7908 | 0,115935 | 0,56055 | 0,836164 |
| Sesn3 | 12,2428 | 13,2115 | 0,109864 | 0,56055 | 0,836164 |
| Aspscr1 | 38,2158 | 40,9147 | 0,0984511 | 0,5604 | 0,836164 |
| Tenm3 | 22,4198 | 23,9169 | 0,0932556 | 0,56015 | 0,836164 |
| Fam53c | 21,6734 | 23,1159 | 0,0929649 | 0,55985 | 0,836164 |
| Fnbp1l | 50,8589 | 54,2419 | 0,0929094 | 0,56045 | 0,836164 |
| Ralb | 31,4486 | 29,4192 | -0,0962394 | 0,5605 | 0,836164 |
| Nfkb1 | 14,3284 | 13,3866 | -0,0980865 | 0,56045 | 0,836164 |
| Zmat3 | 6,81293 | 6,3636 | -0,0984325 | 0,5604 | 0,836164 |
| Snx22 | 4,42506 | 2,66052 | -0,73399 | 0,55995 | 0,836164 |
| Rab36 | 2,35259 | 2,62578 | 0,1585 | 0,5607 | 0,836275 |
| Gen1 | 13,9163 | 14,9238 | 0,100847 | 0,56075 | 0,836275 |
| Dyrk3 | 5,64718 | 6,18083 | 0,13027 | 0,56085 | 0,836331 |
| Wdr36 | 38,8485 | 41,4932 | 0,0950188 | 0,5611 | 0,836611 |
| Sco1 | 7,63534 | 8,21225 | 0,105086 | 0,5613 | 0,836722 |
| Wdr20 | 7,73596 | 7,12678 | -0,118329 | 0,5613 | 0,836722 |
| Phyhd1 | 1,77822 | 1,52244 | -0,224047 | 0,5615 | 0,836833 |
| Trh | 2,36782 | 2,70421 | 0,191646 | 0,5615 | 0,836834 |
| Smco4 | 14,4209 | 15,7786 | 0,129804 | 0,56175 | 0,836907 |
| Eci1 | 29,0597 | 31,3016 | 0,107218 | 0,5618 | 0,836907 |
| Senp6 | 42,9425 | 45,7661 | 0,0918727 | 0,56175 | 0,836907 |
| Snord4a | 234,296 | 96,8104 | -1,2751 | 0,56175 | 0,836907 |
| Gldc | 10,8368 | 10,0994 | -0,101675 | 0,562 | 0,837112 |
| Paqr4 | 2,90671 | 2,40734 | -0,27195 | 0,5621 | 0,837168 |
| Zfp651 | 3,72485 | 3,44761 | -0,111588 | 0,5623 | 0,837372 |
| Zfp324 | 4,30528 | 4,66607 | 0,1161 | 0,56245 | 0,837502 |

| | | | | | |
|---|---|---|---|---|---|
| Rabggta | 10,8775 | 11,6999 | 0,105144 | 0,56275 | 0,837762 |
| 5430405H02Rik | 2,40789 | 1,70866 | -0,494907 | 0,5627 | 0,837762 |
| Fam184b | 0,750003 | 0,855128 | 0,189244 | 0,5631 | 0,838096 |
| Vps37a | 10,1146 | 8,90332 | -0,184024 | 0,5631 | 0,838096 |
| 1810032O08Rik | 19,8502 | 21,5177 | 0,116372 | 0,5632 | 0,838133 |
| Unc45a | 16,3429 | 17,4887 | 0,09776 | 0,56325 | 0,838133 |
| Ccnj | 24,2695 | 22,762 | -0,0925116 | 0,56335 | 0,838189 |
| Rps3 | 621,933 | 667,631 | 0,10229 | 0,5637 | 0,838523 |
| 3010026O09Rik | 11,328 | 10,335 | -0,132354 | 0,5637 | 0,838523 |
| Gadd45b | 1,58855 | 1,85516 | 0,223833 | 0,564 | 0,838595 |
| Spsb3 | 17,423 | 18,7633 | 0,106924 | 0,564 | 0,838595 |
| Nfrkb | 11,4998 | 12,3047 | 0,0975994 | 0,5639 | 0,838595 |
| Snord65 | 4266,09 | 1395,5 | -1,61214 | 0,56385 | 0,838595 |
| Hist1h3e | 1,02391 | 1,41123 | 0,462858 | 0,5644 | 0,838724 |
| Terc | 1,41051 | 1,94407 | 0,462858 | 0,5644 | 0,838724 |
| Rfk | 22,3603 | 23,9175 | 0,0971316 | 0,5644 | 0,838724 |
| Donson | 23,9399 | 25,6069 | 0,0971185 | 0,56435 | 0,838724 |
| Def8 | 16,3733 | 15,3056 | -0,0972808 | 0,56415 | 0,838724 |
| Farsa | 57,3569 | 61,0983 | 0,0911634 | 0,5645 | 0,838779 |
| LOC102633315 | 0,146911 | 0,603704 | 2,03889 | 0,5648 | 0,838833 |
| Lrp4 | 13,0138 | 13,8914 | 0,0941539 | 0,56485 | 0,838833 |
| Usp15 | 36,8115 | 39,1945 | 0,0904961 | 0,5647 | 0,838833 |
| Crebrf | 1,06903 | 0,96721 | -0,144408 | 0,56465 | 0,838833 |
| Ggn | 0,68772 | 0,529159 | -0,378119 | 0,56475 | 0,838833 |
| Amacr | 4,36189 | 4,77354 | 0,130106 | 0,5651 | 0,839111 |
| Pcsk7 | 9,9641 | 8,90934 | -0,161421 | 0,5653 | 0,839314 |
| 2700060E02Rik | 54,5567 | 63,2545 | 0,213412 | 0,56545 | 0,839332 |
| Pomt1 | 11,8469 | 10,897 | -0,120578 | 0,56545 | 0,839332 |
| Mpp1 | 12,5015 | 10,0552 | -0,31416 | 0,5655 | 0,839332 |
| Mtmr14 | 15,9013 | 17,0489 | 0,100532 | 0,56575 | 0,839609 |
| Samd4b | 25,1283 | 26,7693 | 0,0912674 | 0,5659 | 0,839739 |
| Ccdc66 | 13,5727 | 15,1827 | 0,161724 | 0,566 | 0,839756 |
| Brpf3 | 13,734 | 14,6567 | 0,0938127 | 0,56605 | 0,839756 |
| Zfand5 | 11,0984 | 10,4089 | -0,0925268 | 0,5661 | 0,839756 |
| Gad1os | 3,22328 | 4,69853 | 0,543678 | 0,56625 | 0,839885 |
| Tpd52l1 | 16,3007 | 14,7565 | -0,143584 | 0,56645 | 0,840089 |
| Casp9 | 10,1353 | 10,8601 | 0,0996561 | 0,5667 | 0,840273 |
| Phf3 | 20,2271 | 21,5577 | 0,0919145 | 0,56665 | 0,840273 |
| Arl2bp | 96,0883 | 90,2902 | -0,0897904 | 0,56695 | 0,840531 |
| Armc1 | 35,1277 | 32,96 | -0,091893 | 0,567 | 0,840531 |
| Ptpla | 12,5252 | 11,3552 | -0,141482 | 0,5673 | 0,840883 |
| Ank2 | 4,91268 | 5,28264 | 0,104747 | 0,56745 | 0,841012 |
| Fancc | 9,12139 | 10,6431 | 0,222596 | 0,5676 | 0,841047 |
| Pdia6 | 379,14 | 405,151 | 0,095729 | 0,56755 | 0,841047 |
| Mir7115 | 161,016 | 582,415 | 1,85484 | 0,5679 | 0,841097 |
| Dhx58 | 0,892852 | 1,03231 | 0,209377 | 0,56785 | 0,841097 |
| Col6a2 | 1,97654 | 2,18556 | 0,145026 | 0,56805 | 0,841097 |
| Aimp2 | 63,8396 | 70,5194 | 0,143569 | 0,5681 | 0,841097 |

| | | | | | |
|---|---|---|---|---|---|
| Nek8 | 6,20551 | 6,72137 | 0,115206 | 0,56785 | 0,841097 |
| Mrps15 | 55,7652 | 59,7197 | 0,0988407 | 0,56815 | 0,841097 |
| Arl3 | 89,0183 | 95,124 | 0,0957067 | 0,5682 | 0,841097 |
| Baiap2l1 | 9,75162 | 8,96065 | -0,122038 | 0,568 | 0,841097 |
| Clybl | 10,3442 | 9,42219 | -0,13469 | 0,56775 | 0,841097 |
| Skap1 | 1,02891 | 1,19147 | 0,211622 | 0,5684 | 0,841111 |
| Trim46 | 1,24995 | 1,4147 | 0,178633 | 0,5685 | 0,841111 |
| Copz2 | 7,66648 | 8,50931 | 0,150478 | 0,56865 | 0,841111 |
| Cdkl3 | 4,51028 | 4,96502 | 0,138583 | 0,56835 | 0,841111 |
| Sufu | 20,2125 | 21,5498 | 0,0924262 | 0,56865 | 0,841111 |
| Map2 | 3,43892 | 3,1799 | -0,112976 | 0,56845 | 0,841111 |
| Rsbn1 | 11,3286 | 10,3321 | -0,132842 | 0,56865 | 0,841111 |
| S100a1 | 7,4307 | 8,44361 | 0,184362 | 0,56875 | 0,841163 |
| Mocs3 | 3,87675 | 4,3165 | 0,155015 | 0,569 | 0,841163 |
| Pdcd7 | 12,1133 | 13,0291 | 0,105146 | 0,569 | 0,841163 |
| Klhl20 | 10,1381 | 9,45628 | -0,100438 | 0,56895 | 0,841163 |
| Mgat3 | 7,79024 | 7,26035 | -0,101629 | 0,5689 | 0,841163 |
| 1110037F02Rik | 22,6757 | 24,1328 | 0,0898474 | 0,5692 | 0,841347 |
| 4930579K19Rik | 0,705713 | 0,549592 | -0,36072 | 0,56925 | 0,841347 |
| Uhmk1 | 12,8412 | 11,9001 | -0,109806 | 0,56945 | 0,841549 |
| Hnrnpr | 25,904 | 27,5732 | 0,0900939 | 0,5697 | 0,841826 |
| Actb | 880,106 | 948,643 | 0,108188 | 0,56985 | 0,841861 |
| Vezt | 8,42462 | 7,85227 | -0,101502 | 0,56985 | 0,841861 |
| Calm3 | 116,206 | 123,662 | 0,0897131 | 0,57005 | 0,842063 |
| Snora68 | 355,898 | 673,249 | 0,919676 | 0,5702 | 0,842192 |
| Fam53a | 27,2673 | 29,1372 | 0,0956942 | 0,5705 | 0,842449 |
| Mon2 | 10,1526 | 10,8259 | 0,0926382 | 0,57045 | 0,842449 |
| Rnf214 | 14,5975 | 15,9228 | 0,125378 | 0,57065 | 0,842558 |
| Acrbp | 6,93658 | 6,27488 | -0,144635 | 0,5707 | 0,842558 |
| Wdr78 | 2,87668 | 3,14853 | 0,130274 | 0,5708 | 0,842613 |
| Iws1 | 17,8317 | 18,9573 | 0,0883065 | 0,57095 | 0,842741 |
| Zfp951 | 1,59887 | 1,79386 | 0,166021 | 0,57115 | 0,842757 |
| Dtx3 | 55,2162 | 59,9285 | 0,118151 | 0,57115 | 0,842757 |
| Tmem98 | 55,2003 | 51,7234 | -0,0938595 | 0,57115 | 0,842757 |
| Metap1 | 37,1875 | 39,6637 | 0,0930027 | 0,57125 | 0,842811 |
| Ppp1cc | 4,53192 | 4,95972 | 0,130137 | 0,57165 | 0,843215 |
| Des | 2,11104 | 1,89925 | -0,152523 | 0,5716 | 0,843215 |
| Ddx23 | 60,0779 | 63,9262 | 0,0895739 | 0,57175 | 0,84327 |
| Fggy | 3,47292 | 3,84924 | 0,148424 | 0,57195 | 0,843278 |
| Polr3gl | 6,94331 | 7,63637 | 0,137264 | 0,57215 | 0,843278 |
| Mrpl14 | 40,8913 | 44,3177 | 0,116089 | 0,57245 | 0,843278 |
| Galk1 | 66,1147 | 70,5856 | 0,0944029 | 0,5722 | 0,843278 |
| Gnl3l | 19,5215 | 20,7761 | 0,0898639 | 0,5723 | 0,843278 |
| Sgol1 | 23,3197 | 21,8798 | -0,0919481 | 0,57185 | 0,843278 |
| Haus3 | 38,353 | 35,9366 | -0,0938832 | 0,57245 | 0,843278 |
| Tmem5 | 28,6897 | 26,7514 | -0,100918 | 0,57225 | 0,843278 |
| Crat | 6,4139 | 5,78343 | -0,149275 | 0,57215 | 0,843278 |
| Slc2a5 | 1,94453 | 1,74167 | -0,15895 | 0,5722 | 0,843278 |

| | | | | | |
|---|---|---|---|---|---|
| Arhgdig | 3,06758 | 2,6703 | -0,200101 | 0,57235 | 0,843278 |
| D330041H03Rik | 2,10915 | 1,71336 | -0,299834 | 0,57265 | 0,84348 |
| Rnaset2b | 1,04627 | 1,28459 | 0,296054 | 0,57285 | 0,843569 |
| Abcd1 | 4,71793 | 5,11247 | 0,115869 | 0,5728 | 0,843569 |
| Ggps1 | 16,2751 | 17,4434 | 0,100014 | 0,5729 | 0,843569 |
| Thyn1 | 80,8904 | 89,7937 | 0,150647 | 0,5733 | 0,843601 |
| Mpst | 23,4995 | 25,2518 | 0,103758 | 0,573 | 0,843601 |
| Rsu1 | 28,2277 | 30,2222 | 0,0984973 | 0,5731 | 0,843601 |
| Hmg20a | 28,9976 | 30,8802 | 0,0907466 | 0,5732 | 0,843601 |
| Foxred2 | 3,46742 | 3,19192 | -0,11944 | 0,5732 | 0,843601 |
| Map3k15 | 0,755999 | 0,420231 | -0,847204 | 0,57325 | 0,843601 |
| Nav2 | 4,61524 | 4,32264 | -0,0944949 | 0,57375 | 0,84417 |
| Smim12 | 31,8779 | 34,2667 | 0,104247 | 0,57395 | 0,844352 |
| Sdhc | 108,409 | 115,442 | 0,0906836 | 0,574 | 0,844352 |
| Snord17 | 4,35104 | 16,9749 | 1,96397 | 0,5743 | 0,844514 |
| Trmt10b | 8,8336 | 9,55607 | 0,113415 | 0,57425 | 0,844514 |
| Slc3a2 | 60,8852 | 64,7648 | 0,0891176 | 0,5743 | 0,844514 |
| Prim1 | 62,7954 | 66,8681 | 0,0906598 | 0,5747 | 0,844916 |
| Narfl | 9,71255 | 8,83966 | -0,13586 | 0,5747 | 0,844916 |
| Traf6 | 10,448 | 11,1597 | 0,0950729 | 0,5748 | 0,844971 |
| 1110002L01Rik | 5,70126 | 6,40079 | 0,16697 | 0,5753 | 0,845148 |
| Grhl3 | 4,14947 | 4,5342 | 0,127919 | 0,5753 | 0,845148 |
| Gas7 | 1,95125 | 2,12161 | 0,12076 | 0,57505 | 0,845148 |
| Bbs9 | 4,7923 | 5,18909 | 0,114764 | 0,5753 | 0,845148 |
| Tubgcp5 | 12,1834 | 13,0106 | 0,0947672 | 0,575 | 0,845148 |
| Rnf187 | 87,9661 | 93,51 | 0,0881743 | 0,5752 | 0,845148 |
| Pcgf3 | 24,4135 | 25,9954 | 0,0905794 | 0,57575 | 0,845623 |
| Arap1 | 3,57218 | 3,28743 | -0,119847 | 0,5757 | 0,845623 |
| Gm6578 | 2,37871 | 2,03009 | -0,228636 | 0,5763 | 0,846338 |
| Foxm1 | 42,4615 | 45,1609 | 0,0889189 | 0,5765 | 0,846538 |
| Gm10638 | 0,455385 | 0,769916 | 0,757615 | 0,5767 | 0,846739 |
| Thtpa | 9,68949 | 8,95444 | -0,113817 | 0,577 | 0,847086 |
| Mir7030 | 279,498 | 505,49 | 0,854844 | 0,57745 | 0,847369 |
| Fzd9 | 1,68652 | 1,90624 | 0,176679 | 0,5776 | 0,847369 |
| Pgap2 | 17,9593 | 19,2624 | 0,101057 | 0,5775 | 0,847369 |
| Cwf19l1 | 8,82668 | 9,45152 | 0,0986745 | 0,5775 | 0,847369 |
| Ttc23 | 21,9078 | 23,41 | 0,0956775 | 0,5777 | 0,847369 |
| Slc7a6os | 44,223 | 47,1661 | 0,0929524 | 0,57735 | 0,847369 |
| Mospd2 | 9,86398 | 9,19479 | -0,101354 | 0,5777 | 0,847369 |
| Sned1 | 0,934654 | 0,830198 | -0,170978 | 0,57765 | 0,847369 |
| Zfr | 47,3518 | 50,2943 | 0,0869768 | 0,5778 | 0,847423 |
| Khnyn | 21,2156 | 22,5871 | 0,0903715 | 0,5779 | 0,847477 |
| Tada1 | 25,0492 | 22,1121 | -0,179926 | 0,57825 | 0,847897 |
| Magi1 | 35,6165 | 33,4199 | -0,0918382 | 0,57845 | 0,847911 |
| Vcpkmt | 15,5592 | 14,3046 | -0,121291 | 0,57835 | 0,847911 |
| Cecr6 | 0,794487 | 0,70414 | -0,174163 | 0,5784 | 0,847911 |
| Nrp | 1,07587 | 2,7635 | 1,36099 | 0,5789 | 0,848063 |
| Snord42b | 1289,25 | 2659,54 | 1,04465 | 0,57955 | 0,848063 |

| | | | | | |
|---|---|---|---|---|---|
| Nkain4 | 0,474742 | 0,632643 | 0,414246 | 0,57905 | 0,848063 |
| Ppat | 39,4727 | 44,733 | 0,180483 | 0,5793 | 0,848063 |
| Tmem218 | 8,26264 | 9,11134 | 0,141061 | 0,57915 | 0,848063 |
| Eva1b | 14,2212 | 15,3627 | 0,111382 | 0,5788 | 0,848063 |
| Ticrr | 11,2396 | 12,1112 | 0,107746 | 0,57945 | 0,848063 |
| Calcoco1 | 9,55562 | 10,2685 | 0,103803 | 0,57975 | 0,848063 |
| Fblim1 | 9,00593 | 9,67158 | 0,102876 | 0,57965 | 0,848063 |
| Proser1 | 14,9666 | 15,9527 | 0,0920526 | 0,5799 | 0,848063 |
| Pcgf6 | 28,5739 | 30,4513 | 0,0918062 | 0,5796 | 0,848063 |
| Zdhhc6 | 30,2218 | 32,1972 | 0,0913424 | 0,5798 | 0,848063 |
| Hsd17b4 | 29,3945 | 31,2995 | 0,0905933 | 0,5797 | 0,848063 |
| Arhgef7 | 23,485 | 24,9946 | 0,0898741 | 0,57985 | 0,848063 |
| Smurf2 | 20,892 | 22,2132 | 0,0884669 | 0,57975 | 0,848063 |
| Wdr45b | 36,7919 | 39,117 | 0,0884071 | 0,5793 | 0,848063 |
| Pnpt1 | 36,9295 | 34,6794 | -0,0906975 | 0,5795 | 0,848063 |
| Coasy | 14,4993 | 13,5164 | -0,101273 | 0,57875 | 0,848063 |
| Pck2 | 7,32613 | 6,80841 | -0,105734 | 0,5791 | 0,848063 |
| Snap29 | 9,82002 | 8,59422 | -0,192359 | 0,5794 | 0,848063 |
| Rmnd5b | 12,8634 | 11,1586 | -0,205119 | 0,57995 | 0,848063 |
| Klf5 | 0,837101 | 0,698703 | -0,260722 | 0,5797 | 0,848063 |
| Lmf1 | 9,25117 | 8,53253 | -0,116662 | 0,5801 | 0,848097 |
| AA474331 | 3,41633 | 2,93969 | -0,216782 | 0,5801 | 0,848097 |
| Snora34 | 73,5182 | 154,923 | 1,07538 | 0,58065 | 0,848114 |
| Mir8094 | 100,72 | 168,862 | 0,745497 | 0,58095 | 0,848114 |
| Ankrd37 | 2,74729 | 3,91176 | 0,509809 | 0,58035 | 0,848114 |
| Lhfpl4 | 1,37191 | 1,51619 | 0,14426 | 0,581 | 0,848114 |
| Zfp719 | 5,11284 | 5,49951 | 0,105179 | 0,58075 | 0,848114 |
| Smap1 | 31,9052 | 34,0487 | 0,0938082 | 0,58035 | 0,848114 |
| Top1 | 129,612 | 137,996 | 0,0904217 | 0,58095 | 0,848114 |
| Nek2 | 29,0143 | 30,8672 | 0,0893119 | 0,58065 | 0,848114 |
| Ak2 | 102,714 | 109,234 | 0,0887983 | 0,5806 | 0,848114 |
| Fh1 | 95,7416 | 101,796 | 0,0884561 | 0,5804 | 0,848114 |
| Spty2d1 | 7,30856 | 6,83627 | -0,0963788 | 0,5807 | 0,848114 |
| Myo1d | 1,7384 | 1,58455 | -0,13369 | 0,58065 | 0,848114 |
| 3010001F23Rik | 1,26445 | 1,02645 | -0,300838 | 0,58095 | 0,848114 |
| Vmp1 | 27,6441 | 20,7744 | -0,412162 | 0,5803 | 0,848114 |
| Dohh | 35,2603 | 32,9991 | -0,0956143 | 0,5812 | 0,848313 |
| Kif27 | 0,497773 | 0,572099 | 0,200776 | 0,58155 | 0,848731 |
| Rbm12 | 38,8643 | 42,4471 | 0,127222 | 0,5817 | 0,848765 |
| 2410015M20Rik | 79,4984 | 84,8661 | 0,0942622 | 0,5817 | 0,848765 |
| Ankzf1 | 12,4307 | 13,7232 | 0,142713 | 0,5818 | 0,848818 |
| Elovl2 | 13,628 | 12,7686 | -0,0939778 | 0,5819 | 0,848871 |
| Arf1 | 159,321 | 169,139 | 0,0862721 | 0,5821 | 0,848885 |
| Prkca | 1,54134 | 1,41524 | -0,123145 | 0,5821 | 0,848885 |
| Tmco4 | 0,622761 | 0,536929 | -0,213948 | 0,582 | 0,848885 |
| Txndc5 | 122,85 | 115,527 | -0,0886793 | 0,5824 | 0,84923 |
| Zfp811 | 3,47867 | 3,78613 | 0,122186 | 0,5828 | 0,84935 |
| Ncoa1 | 2,69746 | 2,90687 | 0,107868 | 0,58275 | 0,84935 |

| | | | | | |
|---|---|---|---|---|---|
| Slk | 22,8847 | 21,545 | -0,0870324 | 0,5828 | 0,84935 |
| Dcaf6 | 6,55775 | 6,08188 | -0,108683 | 0,5826 | 0,84935 |
| Neil2 | 1,17694 | 1,01251 | -0,217101 | 0,58275 | 0,84935 |
| Dffb | 4,79956 | 5,25006 | 0,129433 | 0,583 | 0,849509 |
| Gemin7 | 58,5804 | 62,7557 | 0,0993276 | 0,5831 | 0,849509 |
| Dnajc1 | 9,10297 | 8,15018 | -0,159505 | 0,5831 | 0,849509 |
| Glrx2 | 5,4064 | 5,00844 | -0,110308 | 0,58325 | 0,849635 |
| Oplah | 0,99907 | 1,12155 | 0,166834 | 0,5835 | 0,849721 |
| Mrpl53 | 78,4204 | 84,7008 | 0,111145 | 0,5834 | 0,849721 |
| Ap3d1 | 28,0956 | 29,8706 | 0,0883835 | 0,5835 | 0,849721 |
| Svil | 11,7556 | 11,0531 | -0,0888887 | 0,5836 | 0,849775 |
| Cog8 | 15,8931 | 17,0163 | 0,0985191 | 0,58405 | 0,850317 |
| Emc4 | 42,7871 | 38,8876 | -0,137866 | 0,5841 | 0,850317 |
| Ttk | 30,9039 | 32,8622 | 0,0886405 | 0,58425 | 0,850443 |
| Rps8 | 164,702 | 175,103 | 0,0883394 | 0,5846 | 0,850674 |
| Ergic1 | 89,2103 | 84,0298 | -0,0863093 | 0,58455 | 0,850674 |
| Fdx1 | 23,3369 | 21,5488 | -0,115005 | 0,58455 | 0,850674 |
| Ing1 | 26,3053 | 27,976 | 0,0888338 | 0,58485 | 0,850853 |
| Tm7sf2 | 12,5455 | 11,0694 | -0,180589 | 0,58485 | 0,850853 |
| Cdk7 | 15,2813 | 14,3529 | -0,0904316 | 0,585 | 0,850979 |
| Atp2b1 | 27,4498 | 25,857 | -0,0862418 | 0,5853 | 0,851322 |
| A730017L22Rik | 1,18347 | 1,03464 | -0,193894 | 0,5855 | 0,851428 |
| Smim5 | 0,884547 | 0,421911 | -1,068 | 0,58545 | 0,851428 |
| Nosip | 20,0953 | 21,4486 | 0,0940287 | 0,58575 | 0,851474 |
| Msi2 | 42,1938 | 39,7128 | -0,0874286 | 0,5857 | 0,851474 |
| Gpr89 | 18,2547 | 17,0102 | -0,101867 | 0,58585 | 0,851474 |
| 6720489N17Rik | 8,18323 | 7,59372 | -0,107865 | 0,5857 | 0,851474 |
| Gm14137 | 0,626444 | 0,449771 | -0,477994 | 0,58585 | 0,851474 |
| Dnajc12 | 10,5812 | 11,508 | 0,12114 | 0,58595 | 0,851506 |
| Nmd3 | 56,0203 | 59,5439 | 0,0880045 | 0,586 | 0,851506 |
| Slc10a3-ubl4,Ubl4 | 3,49996 | 4,51192 | 0,366401 | 0,5861 | 0,851559 |
| Lgals1 | 39,5037 | 36,7602 | -0,103841 | 0,58625 | 0,851684 |
| Tubgcp4 | 18,4629 | 21,7661 | 0,23746 | 0,5865 | 0,851783 |
| Med20 | 22,064 | 23,5662 | 0,0950221 | 0,5867 | 0,851783 |
| Mia3 | 25,0731 | 23,6233 | -0,0859299 | 0,58655 | 0,851783 |
| Usp22 | 55,3434 | 52,1199 | -0,086576 | 0,5867 | 0,851783 |
| Tmpo | 208,011 | 195,258 | -0,0912789 | 0,58665 | 0,851783 |
| Retsat | 6,51409 | 5,97422 | -0,124814 | 0,5864 | 0,851783 |
| Cat | 53,5934 | 56,8855 | 0,0860056 | 0,5869 | 0,851888 |
| Gopc | 13,5832 | 12,7614 | -0,0900327 | 0,58685 | 0,851888 |
| Pum1 | 42,255 | 44,807 | 0,0845999 | 0,587 | 0,85194 |
| Ddx41 | 35,5056 | 37,759 | 0,0887722 | 0,5875 | 0,852481 |
| Maml2 | 10,139 | 9,49992 | -0,0939356 | 0,58745 | 0,852481 |
| Gm4013 | 1,73085 | 2,30099 | 0,41077 | 0,58765 | 0,852514 |
| 2610001J05Rik | 25,3242 | 23,7082 | -0,0951289 | 0,58765 | 0,852514 |
| Cyp4f41-ps | 0,956504 | 1,15569 | 0,272911 | 0,58785 | 0,852524 |
| Slc25a29 | 1,67507 | 1,89341 | 0,176765 | 0,5881 | 0,852524 |
| Kifc5b | 5,83716 | 6,32349 | 0,115456 | 0,58855 | 0,852524 |

| | | | | | |
|---|---|---|---|---|---|
| Rn45s | 360,982 | 391,002 | 0,115247 | 0,5883 | 0,852524 |
| Mina | 16,2365 | 17,5779 | 0,114517 | 0,5884 | 0,852524 |
| 5830417I10Rik | 3,22062 | 3,45623 | 0,101864 | 0,5885 | 0,852524 |
| Zc3h14 | 45,1421 | 48,0594 | 0,0903446 | 0,5881 | 0,852524 |
| Chmp5 | 63,4288 | 67,4114 | 0,0878551 | 0,58825 | 0,852524 |
| Mtss1 | 19,992 | 21,2471 | 0,0878447 | 0,58845 | 0,852524 |
| Aldh2 | 52,1478 | 49,1494 | -0,0854305 | 0,5884 | 0,852524 |
| Cep55 | 38,6748 | 36,3867 | -0,0879834 | 0,58815 | 0,852524 |
| Zfp715 | 7,3575 | 6,8676 | -0,0994095 | 0,58775 | 0,852524 |
| Uvrag | 6,16892 | 5,72765 | -0,107073 | 0,5878 | 0,852524 |
| Tor1aip1 | 21,8404 | 19,5292 | -0,161368 | 0,58795 | 0,852524 |
| Slc7a6 | 61,14 | 64,8744 | 0,085533 | 0,589 | 0,85299 |
| Hykk | 1,69634 | 1,5454 | -0,134443 | 0,58895 | 0,85299 |
| Agbl5 | 11,8278 | 10,8815 | -0,120307 | 0,5891 | 0,853043 |
| Brwd1 | 45,6909 | 49,2268 | 0,107536 | 0,5892 | 0,853095 |
| Atp5g3 | 598,436 | 635,371 | 0,0864016 | 0,5893 | 0,853148 |
| Timm8a1 | 3,88736 | 3,45885 | -0,168497 | 0,5894 | 0,8532 |
| Btrc | 9,27044 | 9,87417 | 0,0910218 | 0,5899 | 0,853831 |
| Cand2 | 37,1477 | 40,6793 | 0,131023 | 0,59015 | 0,854008 |
| Mios | 13,1486 | 14,013 | 0,0918632 | 0,5901 | 0,854008 |
| Inpp5a | 11,1045 | 11,8793 | 0,0973109 | 0,5903 | 0,85402 |
| Zc3h13 | 21,1948 | 22,6107 | 0,093299 | 0,59035 | 0,85402 |
| Psmd3 | 113,13 | 119,959 | 0,0845566 | 0,59025 | 0,85402 |
| A430033K04Rik | 2,50038 | 2,74507 | 0,134695 | 0,5907 | 0,854249 |
| Pappa | 1,56978 | 1,69228 | 0,108411 | 0,5906 | 0,854249 |
| Gpx3 | 72,4333 | 76,9745 | 0,0877276 | 0,59065 | 0,854249 |
| Cltc | 80,851 | 89,0372 | 0,139143 | 0,59085 | 0,854261 |
| Cfp | 6,21603 | 6,81134 | 0,131946 | 0,59085 | 0,854261 |
| Mgmt | 7,24776 | 6,49003 | -0,159309 | 0,5909 | 0,854261 |
| Ctsa | 22,1363 | 20,6217 | -0,10225 | 0,591 | 0,854313 |
| Tmem220 | 1,27248 | 1,46846 | 0,206659 | 0,5911 | 0,854365 |
| Rbpms | 15,0584 | 13,8594 | -0,119702 | 0,5912 | 0,854417 |
| Atg14 | 6,2533 | 6,71042 | 0,101785 | 0,5914 | 0,854594 |
| AU040320 | 11,2466 | 9,7673 | -0,203457 | 0,59145 | 0,854594 |
| Ppp1r18 | 23,8728 | 25,3605 | 0,0872136 | 0,5917 | 0,854822 |
| Rsf1 | 12,0924 | 12,8328 | 0,085727 | 0,5918 | 0,854822 |
| 2810013P06Rik | 7,84476 | 7,25682 | -0,11239 | 0,5918 | 0,854822 |
| Papss2 | 0,575418 | 0,661605 | 0,20136 | 0,5921 | 0,854866 |
| Jmjd7 | 14,1796 | 15,8115 | 0,157155 | 0,592 | 0,854866 |
| Thbs2 | 0,820834 | 0,913592 | 0,15446 | 0,59195 | 0,854866 |
| Rnf144a | 22,0208 | 23,3458 | 0,0843006 | 0,59205 | 0,854866 |
| Jam3 | 58,3129 | 54,924 | -0,0863798 | 0,59215 | 0,854866 |
| Rxra | 13,5488 | 12,7537 | -0,0872482 | 0,59225 | 0,854918 |
| Lyrm1 | 1,91012 | 2,22432 | 0,2197 | 0,59235 | 0,854921 |
| Psmd10 | 17,2407 | 18,5235 | 0,103531 | 0,5924 | 0,854921 |
| Ivns1abp | 210,925 | 224,532 | 0,0901864 | 0,5927 | 0,854921 |
| Kif21b | 6,36234 | 6,76407 | 0,0883348 | 0,5926 | 0,854921 |
| Cdc27 | 45,5154 | 42,8758 | -0,0861904 | 0,5925 | 0,854921 |

| | | | | | |
|---|---|---|---|---|---|
| Ccdc127 | 26,8037 | 25,1503 | -0,0918526 | 0,59255 | 0,854921 |
| Mast3 | 4,58306 | 4,27205 | -0,101383 | 0,5927 | 0,854921 |
| Ndufa4l2 | 1,05191 | 2,30903 | 1,13428 | 0,59375 | 0,855143 |
| Necab2 | 7,07603 | 7,66155 | 0,114698 | 0,5932 | 0,855143 |
| Myo7a | 5,13608 | 5,48248 | 0,0941616 | 0,59295 | 0,855143 |
| Strn4 | 47,9128 | 51,1306 | 0,0937778 | 0,59365 | 0,855143 |
| Sccpdh | 33,9394 | 36,1151 | 0,0896419 | 0,5935 | 0,855143 |
| Nipbl | 23,6954 | 22,3465 | -0,0845572 | 0,59325 | 0,855143 |
| Appl1 | 12,7782 | 12,0305 | -0,0869859 | 0,5937 | 0,855143 |
| Chchd1 | 175,901 | 165,587 | -0,0871736 | 0,59365 | 0,855143 |
| Rbms1 | 48,7159 | 45,8478 | -0,0875426 | 0,59315 | 0,855143 |
| Cmas | 38,2749 | 35,9406 | -0,0907842 | 0,5931 | 0,855143 |
| Pard6b | 3,04267 | 2,79188 | -0,124102 | 0,59325 | 0,855143 |
| Cox11 | 12,3778 | 11,2804 | -0,133938 | 0,59375 | 0,855143 |
| Ufsp1 | 7,28779 | 6,5714 | -0,149281 | 0,5935 | 0,855143 |
| Kansl1l | 5,58297 | 4,96999 | -0,167789 | 0,59365 | 0,855143 |
| Rbm3os | 2,04501 | 3,02962 | 0,567033 | 0,59415 | 0,855221 |
| Gulo | 0,910463 | 1,04668 | 0,201141 | 0,59405 | 0,855221 |
| Tamm41 | 31,2804 | 33,4373 | 0,0961966 | 0,5946 | 0,855221 |
| Pak1ip1 | 54,4503 | 57,8015 | 0,0861655 | 0,5946 | 0,855221 |
| Suds3 | 51,9103 | 55,0739 | 0,0853482 | 0,5946 | 0,855221 |
| Rnf145 | 65,5661 | 61,8568 | -0,0840163 | 0,5945 | 0,855221 |
| Metap2 | 39,5107 | 37,2634 | -0,0844871 | 0,5947 | 0,855221 |
| Ktn1 | 66,3024 | 62,4491 | -0,0863804 | 0,59445 | 0,855221 |
| Uggt2 | 7,12028 | 6,67609 | -0,0929298 | 0,59465 | 0,855221 |
| Herpud1 | 17,029 | 15,9153 | -0,0975846 | 0,5944 | 0,855221 |
| Pigg | 6,97631 | 6,49952 | -0,102131 | 0,5947 | 0,855221 |
| Mfsd11 | 6,08537 | 5,64147 | -0,109273 | 0,59415 | 0,855221 |
| Vps37d | 6,6247 | 6,05521 | -0,129677 | 0,594 | 0,855221 |
| Adam22 | 0,807388 | 0,723937 | -0,157399 | 0,5942 | 0,855221 |
| Gmcl1 | 17,9871 | 19,1453 | 0,0900246 | 0,5949 | 0,855417 |
| Shd | 7,91342 | 7,27681 | -0,120996 | 0,59505 | 0,855448 |
| Rnd2 | 8,3744 | 7,47716 | -0,163496 | 0,59505 | 0,855448 |
| Irgq | 4,954 | 4,62929 | -0,0978036 | 0,5953 | 0,855716 |
| Hmgn3 | 38,413 | 41,085 | 0,097015 | 0,59555 | 0,855983 |
| Sgtb | 5,77302 | 6,23171 | 0,110302 | 0,5957 | 0,856046 |
| Irx1 | 18,3013 | 19,5554 | 0,0956192 | 0,5958 | 0,856046 |
| Fbxl3 | 15,6312 | 14,71 | -0,0876304 | 0,5958 | 0,856046 |
| Cenpc1 | 20,3072 | 19,0943 | -0,0888472 | 0,59585 | 0,856046 |
| Mir1949 | 958,75 | 347,515 | -1,46408 | 0,59595 | 0,856098 |
| Grb10 | 307,579 | 330,229 | 0,10251 | 0,5965 | 0,856427 |
| Atxn7l3 | 42,8332 | 45,6462 | 0,0917665 | 0,59645 | 0,856427 |
| Ppt2 | 30,0792 | 32,0331 | 0,0907949 | 0,59645 | 0,856427 |
| Arid2 | 29,6822 | 31,4588 | 0,0838674 | 0,5963 | 0,856427 |
| BC031361 | 1,4714 | 0,995217 | -0,56411 | 0,59635 | 0,856427 |
| Olfm4 | 6,62796 | 6,07714 | -0,125172 | 0,5968 | 0,856766 |
| Phgdh | 12,3084 | 11,4466 | -0,104724 | 0,59705 | 0,857033 |
| Phtf1 | 26,4859 | 28,8055 | 0,121124 | 0,5972 | 0,857044 |

| | | | | | |
|---|---|---|---|---|---|
| Med13 | 13,8836 | 14,7226 | 0,0846493 | 0,59715 | 0,857044 |
| Ube3a | 25,6813 | 24,2307 | -0,0838782 | 0,59725 | 0,857044 |
| Rsl24d1 | 26,7504 | 28,5095 | 0,0918817 | 0,5974 | 0,857075 |
| 3000002C10Rik | 5,79834 | 5,2604 | -0,140467 | 0,59735 | 0,857075 |
| Mir7027 | 266,953 | 580,926 | 1,12177 | 0,5976 | 0,857178 |
| Limd1 | 21,2551 | 22,5255 | 0,0837467 | 0,59755 | 0,857178 |
| Sucla2 | 48,9306 | 51,9228 | 0,0856329 | 0,5978 | 0,857281 |
| 2900097C17Rik | 10,6683 | 10,0148 | -0,0911922 | 0,5978 | 0,857281 |
| Gfm2 | 21,3045 | 19,9307 | -0,0961618 | 0,5979 | 0,857332 |
| Bad | 17,544 | 19,2839 | 0,136422 | 0,598 | 0,857384 |
| Papolb | 1,62637 | 1,4109 | -0,205048 | 0,59815 | 0,857507 |
| Mroh1 | 3,99236 | 4,63994 | 0,216865 | 0,5986 | 0,857702 |
| Slc5a3 | 4,32496 | 4,69066 | 0,117104 | 0,5985 | 0,857702 |
| Cep128 | 5,14265 | 5,50942 | 0,0993883 | 0,59855 | 0,857702 |
| Dhx37 | 14,5954 | 15,5308 | 0,0896239 | 0,59865 | 0,857702 |
| Polr2h | 77,4214 | 82,379 | 0,0895443 | 0,59875 | 0,857702 |
| E2f5 | 61,0033 | 64,9052 | 0,0894474 | 0,5988 | 0,857702 |
| Dlgap5 | 37,3709 | 39,5911 | 0,0832606 | 0,59875 | 0,857702 |
| Gnaz | 1,17568 | 1,03281 | -0,186914 | 0,5986 | 0,857702 |
| Klhdc3 | 53,8115 | 59,6759 | 0,149234 | 0,599 | 0,857805 |
| Wipf2 | 7,29874 | 7,75379 | 0,0872545 | 0,59895 | 0,857805 |
| Taf1c | 7,24268 | 6,75072 | -0,101481 | 0,5992 | 0,857979 |
| Mief2 | 6,08828 | 5,63795 | -0,110865 | 0,59925 | 0,857979 |
| Fam206a | 14,7616 | 16,9622 | 0,200476 | 0,59945 | 0,858174 |
| Slc25a51 | 24,996 | 23,5892 | -0,0835714 | 0,59955 | 0,858225 |
| Snord53 | 181,131 | 1127,04 | 2,63742 | 0,6002 | 0,858379 |
| Gm6525 | 1,026 | 2,1917 | 1,09501 | 0,6002 | 0,858379 |
| Tex26 | 0,858205 | 1,04287 | 0,28116 | 0,6002 | 0,858379 |
| Nhlrc1 | 1,12723 | 1,2798 | 0,183138 | 0,6002 | 0,858379 |
| Plcb4 | 4,70148 | 5,06066 | 0,106212 | 0,6 | 0,858379 |
| Bicc1 | 9,1852 | 9,8117 | 0,0951916 | 0,60025 | 0,858379 |
| Obfc1 | 24,1491 | 25,7569 | 0,0929909 | 0,60025 | 0,858379 |
| Tcf3 | 115,468 | 122,334 | 0,0833398 | 0,59975 | 0,858379 |
| Cdc42se2 | 48,2979 | 51,1381 | 0,0824377 | 0,6003 | 0,858379 |
| Cnot7 | 70,2006 | 65,4641 | -0,100779 | 0,60005 | 0,858379 |
| Ubr1 | 11,0437 | 10,4117 | -0,0850157 | 0,6004 | 0,85843 |
| Antxr1 | 33,2923 | 35,2536 | 0,0825827 | 0,60055 | 0,858553 |
| Gpd1 | 1,12856 | 1,2692 | 0,169433 | 0,60065 | 0,858604 |
| Tyms | 26,4086 | 27,9785 | 0,0833158 | 0,60075 | 0,858614 |
| Lrig3 | 36,5119 | 34,4606 | -0,0834165 | 0,60085 | 0,858614 |
| Tysnd1 | 31,5609 | 29,2117 | -0,111594 | 0,6008 | 0,858614 |
| Dlgap4 | 20,6639 | 21,9542 | 0,0873832 | 0,60095 | 0,858665 |
| Mrpl57 | 10,6628 | 9,95451 | -0,0991658 | 0,60115 | 0,858859 |
| Haus6 | 29,392 | 31,1736 | 0,0848997 | 0,60125 | 0,85891 |
| Rpl18 | 235,691 | 249,869 | 0,0842774 | 0,6014 | 0,859012 |
| Mark3 | 47,3004 | 50,1304 | 0,083835 | 0,60145 | 0,859012 |
| Rpl38 | 11,391 | 14,2685 | 0,324938 | 0,60185 | 0,859288 |
| Dus1l | 34,5564 | 36,6985 | 0,0867697 | 0,60185 | 0,859288 |

| | | | | | |
|---|---|---|---|---|---|
| Rbbp9 | 29,4227 | 31,2396 | 0,0864453 | 0,6019 | 0,859288 |
| Vcpip1 | 12,2772 | 11,5523 | -0,0878071 | 0,6019 | 0,859288 |
| Snord57 | 730,375 | 3227,84 | 2,14386 | 0,60215 | 0,859491 |
| Mdp1 | 33,503 | 35,6067 | 0,0878585 | 0,6022 | 0,859491 |
| Zfp512 | 25,3065 | 26,8481 | 0,085312 | 0,6023 | 0,859491 |
| Pip4k2c | 14,0426 | 12,3889 | -0,180765 | 0,6023 | 0,859491 |
| Lag3 | 0,941195 | 1,08556 | 0,205878 | 0,6025 | 0,859593 |
| Impdh1 | 22,5343 | 23,9656 | 0,0888464 | 0,6025 | 0,859593 |
| Scyl3 | 9,11805 | 8,25296 | -0,143813 | 0,6026 | 0,859644 |
| BC018242 | 4,07523 | 4,42426 | 0,118553 | 0,6029 | 0,859889 |
| Cbfa2t2 | 9,82097 | 10,4272 | 0,0864118 | 0,60285 | 0,859889 |
| Pex11b | 22,267 | 24,1169 | 0,115134 | 0,6031 | 0,85999 |
| Hoxa9 | 57,2673 | 60,626 | 0,0822251 | 0,60305 | 0,85999 |
| Hist1h2ae | 1,18727 | 0,894745 | -0,408095 | 0,6032 | 0,860041 |
| AA543186 | 1,26181 | 1,44491 | 0,195481 | 0,60335 | 0,860071 |
| Adamts7 | 1,93308 | 1,7758 | -0,122431 | 0,6033 | 0,860071 |
| Btbd6 | 10,6201 | 11,8945 | 0,163486 | 0,60345 | 0,860102 |
| Zeb2 | 8,65034 | 9,18793 | 0,086982 | 0,6035 | 0,860102 |
| Stk24 | 40,1485 | 42,5166 | 0,0826803 | 0,6036 | 0,860153 |
| Ensa | 72,8624 | 68,632 | -0,0862914 | 0,6038 | 0,860346 |
| Ap5m1 | 12,8369 | 14,2428 | 0,149934 | 0,604 | 0,860427 |
| Rgs12 | 14,3433 | 15,2371 | 0,0872078 | 0,60405 | 0,860427 |
| Rpl37a | 44,1198 | 40,951 | -0,107526 | 0,60395 | 0,860427 |
| Art1 | 0,98099 | 1,1566 | 0,237579 | 0,6043 | 0,860599 |
| Tmem117 | 0,909618 | 1,02829 | 0,176919 | 0,6043 | 0,860599 |
| Timm21 | 38,4821 | 40,935 | 0,0891492 | 0,60485 | 0,861107 |
| Gpc2 | 34,5327 | 32,5808 | -0,0839395 | 0,60475 | 0,861107 |
| Mtg1 | 10,4442 | 9,67995 | -0,109629 | 0,6048 | 0,861107 |
| Foxc1 | 37,6664 | 35,5729 | -0,0825003 | 0,605 | 0,861138 |
| Acadsb | 12,2878 | 11,5437 | -0,0901217 | 0,60495 | 0,861138 |
| 4930414L22Rik | 2,24472 | 3,85123 | 0,77878 | 0,6056 | 0,861165 |
| Ganc | 1,30668 | 1,57061 | 0,265421 | 0,6058 | 0,861165 |
| Mcm9 | 5,55495 | 6,29878 | 0,181298 | 0,6054 | 0,861165 |
| Zkscan16 | 1,20665 | 1,35458 | 0,16684 | 0,6059 | 0,861165 |
| Nrd1 | 60,1589 | 64,2164 | 0,094165 | 0,60575 | 0,861165 |
| Tmem199 | 40,7786 | 43,3725 | 0,088967 | 0,60525 | 0,861165 |
| Exoc7 | 18,6436 | 19,7912 | 0,0861834 | 0,6056 | 0,861165 |
| Zic2 | 18,0229 | 19,13 | 0,0860068 | 0,60585 | 0,861165 |
| Pdcd11 | 26,9296 | 28,4949 | 0,0815079 | 0,6058 | 0,861165 |
| Tmem167b | 18,063 | 17,01 | -0,0866522 | 0,60595 | 0,861165 |
| Plscr3 | 15,9259 | 14,9668 | -0,0896061 | 0,60605 | 0,861165 |
| Parpbp | 11,0612 | 10,3852 | -0,0909806 | 0,6053 | 0,861165 |
| Cnnm4 | 4,27289 | 3,97925 | -0,102715 | 0,60605 | 0,861165 |
| Sepsecs | 6,83743 | 6,30858 | -0,116139 | 0,60515 | 0,861165 |
| Ndrg4 | 2,13085 | 1,93643 | -0,138031 | 0,6059 | 0,861165 |
| 1810022K09Rik | 67,3445 | 56,2902 | -0,258678 | 0,60605 | 0,861165 |
| Zdhhc16 | 26,48 | 23,9598 | -0,144286 | 0,60625 | 0,861357 |
| Snora41 | 93,391 | 214,208 | 1,19765 | 0,60655 | 0,861527 |

| | | | | | |
|---|---|---|---|---|---|
| A230050P20Rik | 0,437952 | 0,561182 | 0,357695 | 0,60665 | 0,861527 |
| Rnf217 | 7,92599 | 8,49664 | 0,100303 | 0,6068 | 0,861527 |
| Troap | 16,6106 | 17,6989 | 0,0915502 | 0,60685 | 0,861527 |
| Rpp14 | 18,2181 | 19,3961 | 0,0903964 | 0,60665 | 0,861527 |
| Sephs2 | 38,8547 | 41,1825 | 0,0839417 | 0,6066 | 0,861527 |
| Slc25a5 | 158,846 | 168,06 | 0,0813493 | 0,6069 | 0,861527 |
| Ube2r2 | 50,7273 | 53,6503 | 0,0808227 | 0,60695 | 0,861527 |
| Snx9 | 17,1431 | 16,0786 | -0,0924838 | 0,6068 | 0,861527 |
| Med27 | 16,6416 | 17,8716 | 0,102877 | 0,6071 | 0,861557 |
| Chordc1 | 67,2257 | 71,1715 | 0,0822866 | 0,60705 | 0,861557 |
| Mbip | 19,466 | 18,1935 | -0,0975351 | 0,60755 | 0,862013 |
| Bre | 16,7059 | 15,5765 | -0,100986 | 0,60755 | 0,862013 |
| Gpr137b | 2,51978 | 2,29491 | -0,134862 | 0,60775 | 0,862205 |
| Qk | 69,8818 | 66,0234 | -0,0819397 | 0,60795 | 0,862306 |
| Tgds | 13,4184 | 12,5547 | -0,0959823 | 0,60795 | 0,862306 |
| Mrps26 | 39,0648 | 36,6176 | -0,0933319 | 0,60805 | 0,862356 |
| Rsl1 | 2,26701 | 2,5148 | 0,149656 | 0,60815 | 0,862406 |
| Ppie | 60,1065 | 63,835 | 0,0868266 | 0,6086 | 0,862861 |
| Fam166b | 0,726727 | 0,546422 | -0,411399 | 0,60855 | 0,862861 |
| Mettl4 | 9,19855 | 9,86964 | 0,101592 | 0,60875 | 0,862962 |
| Tceanc2 | 4,01615 | 4,29502 | 0,0968532 | 0,6088 | 0,862962 |
| Taf4b | 3,78565 | 4,05902 | 0,100591 | 0,609 | 0,863154 |
| Lsp1 | 7,23537 | 7,82049 | 0,112192 | 0,60945 | 0,863517 |
| Cdv3 | 54,6162 | 57,8114 | 0,0820251 | 0,60935 | 0,863517 |
| Pm20d2 | 1,02988 | 0,932496 | -0,143314 | 0,60945 | 0,863517 |
| Znhit1 | 30,3437 | 34,1142 | 0,168973 | 0,61055 | 0,86359 |
| Avpr1b | 4,23554 | 4,55114 | 0,10368 | 0,61015 | 0,86359 |
| Setdb1 | 40,8378 | 43,7836 | 0,100486 | 0,6099 | 0,86359 |
| Slc25a38 | 18,448 | 19,7186 | 0,0960915 | 0,6103 | 0,86359 |
| Hnrnpc | 188,693 | 200,011 | 0,0840372 | 0,6104 | 0,86359 |
| Frem1 | 9,3689 | 9,92878 | 0,0837368 | 0,60985 | 0,86359 |
| Prrc2b | 53,4656 | 56,6387 | 0,0831789 | 0,6101 | 0,86359 |
| Krr1 | 23,7675 | 25,1355 | 0,080732 | 0,60965 | 0,86359 |
| Tbl2 | 15,6448 | 14,758 | -0,084182 | 0,6106 | 0,86359 |
| 1700017B05Rik | 7,93911 | 7,45978 | -0,0898441 | 0,61055 | 0,86359 |
| Tmem110 | 8,143 | 7,63506 | -0,0929205 | 0,6104 | 0,86359 |
| Homer3 | 34,9805 | 32,6099 | -0,101241 | 0,6105 | 0,86359 |
| Arfgap3 | 7,67292 | 7,14666 | -0,102508 | 0,61 | 0,86359 |
| 3110062M04Rik | 4,64289 | 4,27897 | -0,117759 | 0,61025 | 0,86359 |
| Fstl3 | 1,79814 | 1,55236 | -0,212038 | 0,6103 | 0,86359 |
| 9130221H12Rik | 10,3377 | 8,64144 | -0,258566 | 0,61035 | 0,86359 |
| C920021L13Rik | 0,933767 | 0,732845 | -0,349554 | 0,60985 | 0,86359 |
| AI413582 | 2,39184 | 2,82722 | 0,241261 | 0,6109 | 0,863831 |
| Alx4 | 5,17681 | 5,52676 | 0,0943721 | 0,61085 | 0,863831 |
| Ier5 | 14,4348 | 15,3306 | 0,0868641 | 0,6111 | 0,86391 |
| Kifap3 | 16,3084 | 17,2935 | 0,0846132 | 0,61115 | 0,86391 |
| Ccdc43 | 31,946 | 33,8235 | 0,0823888 | 0,61115 | 0,86391 |
| Phox2a | 0,229193 | 0,682346 | 1,57394 | 0,6116 | 0,864007 |

| | | | | | |
|---|---|---|---|---|---|
| Arhgap20os | 1,28484 | 1,53787 | 0,259344 | 0,6114 | 0,864007 |
| C130036L24Rik | 1,47169 | 1,69402 | 0,202981 | 0,61175 | 0,864007 |
| Col6a1 | 0,830341 | 0,93139 | 0,165682 | 0,6118 | 0,864007 |
| Pik3ca | 12,0903 | 12,7904 | 0,0812189 | 0,6117 | 0,864007 |
| Ipo11 | 39,5261 | 37,3814 | -0,0804835 | 0,6114 | 0,864007 |
| Tmem70 | 32,6037 | 30,7222 | -0,0857521 | 0,61165 | 0,864007 |
| Kctd1 | 8,64264 | 8,01541 | -0,108696 | 0,61155 | 0,864007 |
| Mfsd3 | 3,16741 | 2,63372 | -0,266201 | 0,6118 | 0,864007 |
| Ccdc71 | 16,567 | 17,657 | 0,091933 | 0,612 | 0,864106 |
| Gmeb1 | 10,8546 | 11,4966 | 0,0829089 | 0,61195 | 0,864106 |
| Ankrd52 | 26,692 | 28,2046 | 0,0795269 | 0,6121 | 0,864156 |
| Ppp1r11 | 30,0002 | 31,8649 | 0,0869933 | 0,61235 | 0,864418 |
| Actc1 | 12,1466 | 13,0649 | 0,105149 | 0,6128 | 0,864941 |
| Coro1b | 49,5357 | 46,7841 | -0,0824492 | 0,61285 | 0,864941 |
| Rfx7 | 19,1825 | 20,2743 | 0,0798625 | 0,6131 | 0,865203 |
| Churc1 | 24,6596 | 26,6413 | 0,111514 | 0,61365 | 0,865438 |
| Dclre1a | 6,16016 | 6,57701 | 0,0944643 | 0,61365 | 0,865438 |
| Rabepk | 14,7428 | 15,7304 | 0,0935449 | 0,61375 | 0,865438 |
| Gtf2h2 | 33,5064 | 35,5691 | 0,0861871 | 0,6138 | 0,865438 |
| Tars | 76,4331 | 80,8032 | 0,0802163 | 0,61375 | 0,865438 |
| Kif4 | 26,8797 | 28,4111 | 0,0799378 | 0,61385 | 0,865438 |
| Ptpn12 | 34,5927 | 36,5532 | 0,0795308 | 0,61385 | 0,865438 |
| Pls3 | 36,6645 | 38,7414 | 0,0794926 | 0,61335 | 0,865438 |
| Pim2 | 2,72037 | 2,46167 | -0,144165 | 0,61385 | 0,865438 |
| Fam63a | 7,23442 | 7,7764 | 0,104225 | 0,614 | 0,865559 |
| Ddost | 140,279 | 132,737 | -0,0797262 | 0,61415 | 0,865679 |
| Atp7b | 0,941731 | 1,04377 | 0,148423 | 0,61435 | 0,865757 |
| Zmiz1 | 30,8193 | 32,5518 | 0,0789015 | 0,6143 | 0,865757 |
| Zbtb2 | 8,48388 | 7,94679 | -0,0943528 | 0,6144 | 0,865757 |
| Plk3 | 1,30298 | 1,15556 | -0,173229 | 0,61475 | 0,866159 |
| Zzz3 | 19,2808 | 20,3829 | 0,0801882 | 0,61485 | 0,866209 |
| Herc4 | 24,9596 | 26,4181 | 0,081934 | 0,6151 | 0,866378 |
| Rabgap1 | 23,766 | 25,1368 | 0,0808992 | 0,61505 | 0,866378 |
| Pex3 | 11,8875 | 12,9041 | 0,118381 | 0,6155 | 0,866422 |
| Fbxo9 | 12,884 | 13,7653 | 0,0954616 | 0,61565 | 0,866422 |
| Tmem39a | 23,7044 | 25,1084 | 0,0830132 | 0,6155 | 0,866422 |
| Ndc1 | 40,4634 | 42,8107 | 0,0813547 | 0,6155 | 0,866422 |
| Rheb | 70,4379 | 74,3885 | 0,0787286 | 0,61565 | 0,866422 |
| Racgap1 | 64,1076 | 60,671 | -0,0794894 | 0,6156 | 0,866422 |
| Anln | 30,1699 | 27,758 | -0,120206 | 0,61545 | 0,866422 |
| Fuom | 1,21604 | 1,07891 | -0,172617 | 0,6155 | 0,866422 |
| Gtf3c5 | 26,8285 | 28,3943 | 0,0818357 | 0,61575 | 0,866451 |
| Maneal | 2,12135 | 1,78661 | -0,247758 | 0,6158 | 0,866451 |
| Tmem67 | 5,83037 | 5,43892 | -0,100267 | 0,6159 | 0,8665 |
| Reep2 | 5,74066 | 5,30096 | -0,114963 | 0,616 | 0,86655 |
| Mapre1 | 41,3445 | 43,7094 | 0,0802475 | 0,61615 | 0,86667 |
| 1700071M16Rik | 0,616188 | 0,874503 | 0,505093 | 0,61635 | 0,866677 |
| Ipo7 | 125,126 | 132,395 | 0,0814637 | 0,61635 | 0,866677 |

| | | | | | |
|---|---|---|---|---|---|
| C77370 | 0,813093 | 0,748006 | -0,120371 | 0,61625 | 0,866677 |
| Ttf1 | 7,84651 | 8,34094 | 0,0881603 | 0,61655 | 0,866776 |
| Inppl1 | 33,6766 | 35,5726 | 0,0790203 | 0,61655 | 0,866776 |
| Gm16973 | 8,98995 | 10,5912 | 0,236477 | 0,61685 | 0,866903 |
| Slmo2 | 60,1335 | 63,6409 | 0,0817835 | 0,61675 | 0,866903 |
| Nat10 | 31,1095 | 32,9078 | 0,081074 | 0,6169 | 0,866903 |
| Rnmt | 19,4461 | 20,5389 | 0,0788783 | 0,6168 | 0,866903 |
| Ppil3 | 43,7448 | 46,5753 | 0,0904546 | 0,6171 | 0,867002 |
| Rab5b | 11,0226 | 11,7223 | 0,0887988 | 0,61705 | 0,867002 |
| Bag2 | 21,4761 | 22,8138 | 0,0871754 | 0,61725 | 0,86701 |
| Fbxw8 | 34,6018 | 36,5476 | 0,0789273 | 0,61725 | 0,86701 |
| Keap1 | 15,7465 | 14,8328 | -0,0862426 | 0,6173 | 0,86701 |
| Mettl22 | 7,62281 | 8,28527 | 0,120226 | 0,61755 | 0,867087 |
| Taf2 | 28,1734 | 29,8125 | 0,0815822 | 0,6175 | 0,867087 |
| Itpr2 | 1,12155 | 1,03974 | -0,109269 | 0,6175 | 0,867087 |
| Gria3 | 1,07576 | 0,954751 | -0,172166 | 0,61765 | 0,867137 |
| Zswim1 | 7,1129 | 7,61439 | 0,0982902 | 0,61785 | 0,867326 |
| Ap2a1 | 42,9282 | 45,447 | 0,0822593 | 0,618 | 0,867446 |
| Sh3rf1 | 7,7785 | 7,3171 | -0,0882206 | 0,61815 | 0,867565 |
| Ak6 | 44,0636 | 48,244 | 0,130764 | 0,6188 | 0,867595 |
| 9230110C19Rik | 6,80663 | 7,36915 | 0,114558 | 0,61865 | 0,867595 |
| Gss | 6,93183 | 7,46574 | 0,10705 | 0,6188 | 0,867595 |
| Chchd7 | 79,5229 | 85,0682 | 0,0972486 | 0,6186 | 0,867595 |
| Haus4 | 14,8916 | 15,9218 | 0,0965044 | 0,61875 | 0,867595 |
| Atg7 | 6,77789 | 7,22868 | 0,0928962 | 0,6189 | 0,867595 |
| Klhl21 | 7,07489 | 7,5367 | 0,0912235 | 0,6184 | 0,867595 |
| Fkbp5 | 32,8995 | 34,7804 | 0,0802085 | 0,61825 | 0,867595 |
| Adnp | 69,9323 | 73,8867 | 0,0793567 | 0,61865 | 0,867595 |
| Rmi2 | 4,26898 | 3,97067 | -0,104511 | 0,61895 | 0,867595 |
| Serpinb6a | 22,6872 | 20,9367 | -0,11584 | 0,61865 | 0,867595 |
| Cyp2d22 | 1,63138 | 1,46954 | -0,150733 | 0,61895 | 0,867595 |
| Rnf121 | 27,3339 | 25,7146 | -0,0881074 | 0,6191 | 0,867714 |
| Rfc4 | 73,8162 | 84,778 | 0,199753 | 0,61925 | 0,867721 |
| Akap13 | 8,49319 | 8,03624 | -0,0797867 | 0,6193 | 0,867721 |
| Cd59b | 1,37969 | 1,1342 | -0,28267 | 0,6192 | 0,867721 |
| Ercc5 | 11,645 | 12,339 | 0,0835139 | 0,61965 | 0,86803 |
| Btaf1 | 18,3104 | 19,3574 | 0,0802176 | 0,61965 | 0,86803 |
| Wdr91 | 8,42842 | 7,88335 | -0,0964527 | 0,6198 | 0,868149 |
| Psmg2 | 52,6402 | 55,8581 | 0,0856018 | 0,62005 | 0,868345 |
| Tpd52l2 | 28,7983 | 30,427 | 0,0793671 | 0,62015 | 0,868345 |
| Dnajb12 | 22,6822 | 21,3252 | -0,0889987 | 0,62015 | 0,868345 |
| Eln | 1,62343 | 1,4749 | -0,138428 | 0,6202 | 0,868345 |
| Tomt | 5,00998 | 5,91532 | 0,239653 | 0,62065 | 0,86852 |
| Map9 | 4,0878 | 4,36175 | 0,0935803 | 0,6206 | 0,86852 |
| Ppm1m | 27,8272 | 29,5227 | 0,0853328 | 0,62065 | 0,86852 |
| Rprd1a | 13,7445 | 14,5668 | 0,0838245 | 0,62065 | 0,86852 |
| Fen1 | 76,2983 | 80,599 | 0,0791099 | 0,62055 | 0,86852 |
| Dcaf4 | 10,1964 | 10,9185 | 0,0987198 | 0,62135 | 0,869115 |

| | | | | | |
|---|---|---|---|---|---|
| Ywhae | 650,778 | 692,641 | 0,0899413 | 0,62125 | 0,869115 |
| Ube2e3 | 106,082 | 100,498 | -0,0780171 | 0,62115 | 0,869115 |
| Mterfd1 | 30,5206 | 27,8158 | -0,13388 | 0,6214 | 0,869115 |
| Bsg | 329,392 | 255,596 | -0,365939 | 0,6214 | 0,869115 |
| Mtrf1 | 5,60204 | 6,06797 | 0,115262 | 0,6216 | 0,869219 |
| Cbr4 | 12,5923 | 13,5629 | 0,107122 | 0,6218 | 0,869219 |
| Fis1 | 106,379 | 112,536 | 0,0811764 | 0,62165 | 0,869219 |
| Ly6e | 43,6074 | 46,1001 | 0,0801954 | 0,62175 | 0,869219 |
| Ganab | 138,771 | 131,25 | -0,0803849 | 0,6217 | 0,869219 |
| C230037L18Rik | 0,716139 | 0,412932 | -0,794334 | 0,62195 | 0,869338 |
| Sdhaf2 | 17,1747 | 18,2055 | 0,08409 | 0,6222 | 0,869505 |
| Zfp180 | 15,4401 | 16,343 | 0,0819973 | 0,6222 | 0,869505 |
| Bola1 | 19,9132 | 21,5157 | 0,111664 | 0,62255 | 0,869519 |
| Aip | 17,4106 | 18,7441 | 0,106476 | 0,6226 | 0,869519 |
| Cacna2d1 | 12,4325 | 11,7608 | -0,0801232 | 0,6226 | 0,869519 |
| Ckap2l | 35,5179 | 33,5851 | -0,0807235 | 0,62245 | 0,869519 |
| Leprel2 | 10,8288 | 10,1663 | -0,0910846 | 0,62235 | 0,869519 |
| Acot9 | 7,87107 | 7,27356 | -0,113899 | 0,6225 | 0,869519 |
| Capns2 | 0,68587 | 0,933437 | 0,444617 | 0,6228 | 0,869707 |
| Rspo1 | 0,690267 | 0,811571 | 0,233562 | 0,62295 | 0,869826 |
| Eif2a | 54,0466 | 59,2294 | 0,13211 | 0,62345 | 0,869888 |
| Sh3gl1 | 49,11 | 52,8808 | 0,106727 | 0,62315 | 0,869888 |
| Arpc4 | 53,9658 | 57,0911 | 0,0812199 | 0,62335 | 0,869888 |
| Lrrc40 | 41,2659 | 43,5721 | 0,0784567 | 0,6234 | 0,869888 |
| Ate1 | 12,0534 | 11,3895 | -0,0817379 | 0,62325 | 0,869888 |
| Isoc2a | 16,0164 | 14,894 | -0,104817 | 0,6233 | 0,869888 |
| Slc7a3 | 1,58156 | 1,41647 | -0,159053 | 0,6234 | 0,869888 |
| Uba52 | 1,28613 | 1,64421 | 0,354356 | 0,624 | 0,869908 |
| 1300002E11Rik | 8,72627 | 9,79097 | 0,166087 | 0,6239 | 0,869908 |
| Pdxk | 5,86659 | 6,23122 | 0,0869928 | 0,62405 | 0,869908 |
| Unk | 17,6641 | 18,6899 | 0,081442 | 0,6239 | 0,869908 |
| Smarcc1 | 91,6583 | 96,9503 | 0,0809806 | 0,6239 | 0,869908 |
| Anapc4 | 52,5732 | 55,4749 | 0,0775076 | 0,6237 | 0,869908 |
| Adss | 65,2993 | 68,872 | 0,0768502 | 0,62375 | 0,869908 |
| Dnajc14 | 23,0041 | 21,6539 | -0,0872663 | 0,62395 | 0,869908 |
| Tubg2 | 0,963421 | 0,831777 | -0,21197 | 0,6237 | 0,869908 |
| Src | 23,7174 | 22,4626 | -0,0784193 | 0,62435 | 0,870235 |
| Zfp383 | 9,353 | 9,97392 | 0,0927314 | 0,62465 | 0,870562 |
| Znrf1 | 16,9053 | 16,0089 | -0,0786004 | 0,62485 | 0,870569 |
| Rassf7 | 14,5284 | 13,5624 | -0,0992665 | 0,62475 | 0,870569 |
| Zfp273 | 2,13899 | 1,92877 | -0,149245 | 0,6248 | 0,870569 |
| 0610010F05Rik | 12,1673 | 11,4747 | -0,0845458 | 0,62515 | 0,870805 |
| Bend6 | 0,737048 | 0,637201 | -0,21001 | 0,62515 | 0,870805 |
| Phf13 | 22,5421 | 21,3123 | -0,0809333 | 0,62545 | 0,871132 |
| Trp53i11 | 23,8504 | 25,234 | 0,0813545 | 0,62555 | 0,871181 |
| Erbb2ip | 33,9156 | 32,1233 | -0,0783295 | 0,6257 | 0,871299 |
| Snora30 | 21,6862 | 39,3026 | 0,857847 | 0,62595 | 0,871444 |
| Slc9a3r1 | 49,7096 | 52,4723 | 0,0780338 | 0,626 | 0,871444 |

| | | | | | |
|---|---|---|---|---|---|
| Ubxn2a | 8,8577 | 8,29295 | -0,0950472 | 0,626 | 0,871444 |
| Sept1 | 4,46041 | 4,88745 | 0,131906 | 0,6261 | 0,871493 |
| Zxdc | 4,96546 | 4,64683 | -0,0956789 | 0,6262 | 0,871541 |
| Ssbp1 | 41,7379 | 44,4387 | 0,0904576 | 0,62645 | 0,871686 |
| As3mt | 18,5094 | 19,6952 | 0,0895879 | 0,6265 | 0,871686 |
| Kif14 | 7,7854 | 7,36218 | -0,0806383 | 0,62645 | 0,871686 |
| Pcdhgc4 | 2,04859 | 2,87989 | 0,491378 | 0,62665 | 0,871713 |
| Vrk3 | 15,7949 | 16,8299 | 0,0915614 | 0,62665 | 0,871713 |
| Mtbp | 23,4106 | 24,7244 | 0,0787708 | 0,6269 | 0,871949 |
| Zfp41 | 32,0662 | 33,8431 | 0,0778078 | 0,62695 | 0,871949 |
| Fbn1 | 3,53932 | 3,3298 | -0,088034 | 0,6273 | 0,872345 |
| Parn | 16,7204 | 17,7186 | 0,083655 | 0,6274 | 0,872393 |
| Tbc1d32 | 2,33994 | 2,54199 | 0,119489 | 0,62755 | 0,87242 |
| D3Ertd751e | 6,85458 | 7,39481 | 0,109445 | 0,62755 | 0,87242 |
| Cxxc4 | 1,90615 | 2,05304 | 0,1071 | 0,62775 | 0,872517 |
| Scd2 | 485,399 | 522,254 | 0,105583 | 0,62775 | 0,872517 |
| Sarnp | 12,2392 | 13,2023 | 0,109287 | 0,62785 | 0,872544 |
| Ric8 | 12,3055 | 13,148 | 0,0955391 | 0,6279 | 0,872544 |
| Zfp738 | 4,16278 | 4,45361 | 0,0974288 | 0,6284 | 0,872854 |
| Mapk8 | 16,5288 | 17,446 | 0,077917 | 0,6284 | 0,872854 |
| Msl2 | 21,4502 | 22,6353 | 0,0775807 | 0,62825 | 0,872854 |
| Ttl | 8,11159 | 7,63654 | -0,0870667 | 0,62845 | 0,872854 |
| Trim33 | 17,996 | 16,5243 | -0,123083 | 0,62835 | 0,872854 |
| Fam104a | 11,5367 | 12,6003 | 0,127221 | 0,62855 | 0,872903 |
| 4932702P03Rik | 0,635736 | 1,52101 | 1,25853 | 0,62885 | 0,872956 |
| Stard4 | 15,8224 | 16,7593 | 0,0829963 | 0,62885 | 0,872956 |
| Srgap2 | 13,8986 | 14,6641 | 0,077352 | 0,6288 | 0,872956 |
| Mars2 | 5,85825 | 5,47068 | -0,0987492 | 0,62885 | 0,872956 |
| A430005L14Rik | 35,3599 | 33,2946 | -0,086825 | 0,629 | 0,873074 |
| Lins | 6,31291 | 6,70646 | 0,0872456 | 0,62915 | 0,87317 |
| Ugt8a | 0,849383 | 0,757162 | -0,165813 | 0,6292 | 0,87317 |
| Ethe1 | 5,95194 | 6,48007 | 0,122649 | 0,6295 | 0,873405 |
| Gfer | 21,5095 | 22,8223 | 0,0854715 | 0,6295 | 0,873405 |
| Srsf7 | 266,896 | 252,134 | -0,0820905 | 0,62965 | 0,873523 |
| Arl13b | 9,87993 | 10,5042 | 0,0883975 | 0,6298 | 0,87364 |
| Sod2 | 13,9942 | 14,812 | 0,0819409 | 0,63015 | 0,874035 |
| Prr5 | 12,4901 | 13,326 | 0,0934617 | 0,63025 | 0,874061 |
| Nes | 58,3363 | 61,5564 | 0,0775149 | 0,6303 | 0,874061 |
| Prkcsh | 105,813 | 100,408 | -0,0756351 | 0,63045 | 0,874179 |
| Stk38l | 9,2156 | 9,76204 | 0,0831043 | 0,63065 | 0,874365 |
| Rimklb | 15,7204 | 14,8738 | -0,0798638 | 0,63085 | 0,874461 |
| Sfi1 | 5,47209 | 4,97929 | -0,136153 | 0,6308 | 0,874461 |
| Heatr5a | 9,39158 | 9,924 | 0,0795527 | 0,63095 | 0,874488 |
| Apobec3 | 3,91492 | 3,61081 | -0,116661 | 0,631 | 0,874488 |
| Mtmr10 | 5,22109 | 5,55215 | 0,088694 | 0,63125 | 0,874743 |
| Gpr83 | 1,97418 | 2,17942 | 0,142686 | 0,63135 | 0,874791 |
| Kif6 | 0,523575 | 0,596042 | 0,187017 | 0,63205 | 0,875205 |
| Nudt22 | 6,79825 | 7,43718 | 0,129593 | 0,63235 | 0,875205 |

| | | | | | |
|---|---|---|---|---|---|
| Nup35 | 3,76298 | 4,06147 | 0,110127 | 0,6321 | 0,875205 |
| Smap2 | 13,082 | 13,8688 | 0,084261 | 0,6322 | 0,875205 |
| Rab28 | 29,6285 | 31,3837 | 0,0830279 | 0,6325 | 0,875205 |
| Med6 | 66,8546 | 70,7646 | 0,0820012 | 0,63235 | 0,875205 |
| Fgfr1op2 | 26,8332 | 28,351 | 0,079381 | 0,63185 | 0,875205 |
| Trim35 | 57,9459 | 61,0871 | 0,0761627 | 0,63215 | 0,875205 |
| Nudcd2 | 100,466 | 95,1889 | -0,077845 | 0,63195 | 0,875205 |
| Myc | 25,5699 | 24,2009 | -0,0793818 | 0,6324 | 0,875205 |
| Nnt | 14,4041 | 13,6232 | -0,0804051 | 0,63245 | 0,875205 |
| Crebzf | 28,1484 | 26,6212 | -0,08048 | 0,6319 | 0,875205 |
| Selm | 9,35806 | 8,55977 | -0,128638 | 0,63225 | 0,875205 |
| Xxylt1 | 17,9724 | 16,2209 | -0,147934 | 0,63265 | 0,875322 |
| Ncs1 | 18,7199 | 19,7581 | 0,0778721 | 0,6328 | 0,875348 |
| Klc3 | 1,38245 | 0,964865 | -0,518829 | 0,6328 | 0,875348 |
| Gm6787 | 0,757778 | 0,892127 | 0,235475 | 0,6337 | 0,876058 |
| Cyp4f16 | 5,86312 | 6,30953 | 0,105864 | 0,6341 | 0,876058 |
| Hrsp12 | 23,9121 | 25,538 | 0,094908 | 0,63395 | 0,876058 |
| Dus3l | 21,8051 | 23,0733 | 0,0815628 | 0,63365 | 0,876058 |
| Btbd2 | 26,4575 | 27,9293 | 0,0781048 | 0,6337 | 0,876058 |
| Gtf2i | 126,435 | 133,301 | 0,076282 | 0,63345 | 0,876058 |
| Ago1 | 32,7307 | 34,4719 | 0,0747766 | 0,63385 | 0,876058 |
| Gpc1 | 60,9577 | 57,9142 | -0,0738931 | 0,6338 | 0,876058 |
| Rpl36al | 64,7377 | 60,5994 | -0,0953005 | 0,63355 | 0,876058 |
| Mybl1 | 3,77534 | 3,53275 | -0,0958168 | 0,63385 | 0,876058 |
| B930041F14Rik | 2,6423 | 2,40138 | -0,137932 | 0,6341 | 0,876058 |
| Galnt12 | 1,34447 | 1,19975 | -0,164307 | 0,63405 | 0,876058 |
| Oxa1l | 29,6413 | 27,9215 | -0,0862324 | 0,63425 | 0,876175 |
| Pnp2 | 0,859914 | 1,01986 | 0,246111 | 0,63435 | 0,876223 |
| Zscan29 | 6,4936 | 6,95041 | 0,0980787 | 0,63455 | 0,876318 |
| Pcdhga7 | 2,82403 | 1,98569 | -0,508114 | 0,6345 | 0,876318 |
| Yap1 | 82,6688 | 87,0937 | 0,0752263 | 0,6347 | 0,876434 |
| Taf13 | 16,4618 | 17,6011 | 0,0965393 | 0,63485 | 0,876439 |
| Mff | 50,8407 | 53,6277 | 0,0769953 | 0,6349 | 0,876439 |
| Igf2r | 46,6941 | 44,1947 | -0,0793642 | 0,6349 | 0,876439 |
| Pdlim2 | 1,35114 | 1,18366 | -0,190921 | 0,6351 | 0,876624 |
| Atad2b | 13,7549 | 13,0482 | -0,076094 | 0,63525 | 0,87665 |
| Plekho2 | 2,88854 | 2,69389 | -0,100648 | 0,6352 | 0,87665 |
| Ccdc84 | 9,54571 | 10,4775 | 0,134364 | 0,63575 | 0,87725 |
| Vps29 | 68,5469 | 72,4765 | 0,0804234 | 0,63585 | 0,877297 |
| Ccdc34os | 0,251812 | 0,720322 | 1,5163 | 0,63935 | 0,877325 |
| Gm16062 | 2,71723 | 4,94835 | 0,864809 | 0,63685 | 0,877325 |
| Snord89 | 103,554 | 148,858 | 0,523555 | 0,6372 | 0,877325 |
| Rph3al | 0,494085 | 0,576499 | 0,222561 | 0,6379 | 0,877325 |
| Mkrn2 | 11,8294 | 13,7874 | 0,220976 | 0,64015 | 0,877325 |
| Eif4enif1 | 41,6675 | 46,3682 | 0,154213 | 0,63615 | 0,877325 |
| Msl3l2 | 5,34824 | 5,93804 | 0,150924 | 0,63925 | 0,877325 |
| Begain | 1,61687 | 1,77968 | 0,138413 | 0,6383 | 0,877325 |
| Rrnad1 | 11,6794 | 12,802 | 0,132407 | 0,6381 | 0,877325 |

| | | | | | |
|---|---|---|---|---|---|
| Rhbdl1 | 3,72801 | 4,08375 | 0,131489 | 0,6372 | 0,877325 |
| Napb | 1,49138 | 1,63195 | 0,129951 | 0,6369 | 0,877325 |
| Zcwpw1 | 3,50918 | 3,80421 | 0,116462 | 0,63775 | 0,877325 |
| Mrps23 | 48,1095 | 52,0114 | 0,112508 | 0,63805 | 0,877325 |
| Syvn1 | 25,3783 | 27,2821 | 0,104357 | 0,63935 | 0,877325 |
| B9d1 | 24,1356 | 25,8534 | 0,0991883 | 0,6392 | 0,877325 |
| Armc5 | 7,17091 | 7,67195 | 0,0974376 | 0,6386 | 0,877325 |
| BC053749 | 7,55065 | 8,0604 | 0,09425 | 0,63715 | 0,877325 |
| Zdhhc4 | 15,8268 | 16,8906 | 0,0938504 | 0,6371 | 0,877325 |
| Txnrd3 | 6,63769 | 7,08202 | 0,0934802 | 0,6394 | 0,877325 |
| Stub1 | 84,6158 | 90,0851 | 0,0903615 | 0,63855 | 0,877325 |
| Azi2 | 16,8939 | 17,9704 | 0,0891239 | 0,63885 | 0,877325 |
| Gpatch2l | 7,10229 | 7,54819 | 0,0878466 | 0,63815 | 0,877325 |
| Tmem151b | 5,03573 | 5,3509 | 0,0875786 | 0,6363 | 0,877325 |
| Atp5h | 78,421 | 83,2437 | 0,0861011 | 0,6368 | 0,877325 |
| Epdr1 | 12,1864 | 12,9355 | 0,0860678 | 0,63785 | 0,877325 |
| Mipep | 14,4681 | 15,3268 | 0,0831798 | 0,636 | 0,877325 |
| Nid2 | 70,7214 | 74,9005 | 0,0828274 | 0,6376 | 0,877325 |
| Ndufa11 | 15,518 | 16,4336 | 0,0827092 | 0,6366 | 0,877325 |
| Ddhd2 | 9,3558 | 9,9059 | 0,0824267 | 0,64 | 0,877325 |
| Mxi1 | 8,05027 | 8,52258 | 0,0822535 | 0,6395 | 0,877325 |
| Bcr | 8,80186 | 9,29886 | 0,0792453 | 0,63885 | 0,877325 |
| Rfc5 | 34,5954 | 36,5249 | 0,0783019 | 0,64025 | 0,877325 |
| Etfa | 43,5183 | 45,9248 | 0,0776527 | 0,6402 | 0,877325 |
| Cox6c | 241,62 | 254,947 | 0,0774567 | 0,63905 | 0,877325 |
| Lpar1 | 30,5406 | 32,2237 | 0,0773938 | 0,63685 | 0,877325 |
| Suv420h2 | 34,3476 | 36,2339 | 0,0771281 | 0,63735 | 0,877325 |
| Fam179b | 11,7509 | 12,3938 | 0,0768428 | 0,6404 | 0,877325 |
| Zfp317 | 17,4482 | 18,4002 | 0,076645 | 0,6384 | 0,877325 |
| Anapc5 | 141,791 | 149,509 | 0,076468 | 0,638 | 0,877325 |
| Gga2 | 37,043 | 39,0356 | 0,0755896 | 0,638 | 0,877325 |
| Cdk16 | 65,8494 | 69,3615 | 0,0749667 | 0,63975 | 0,877325 |
| Ist1 | 65,2696 | 68,7359 | 0,074653 | 0,63885 | 0,877325 |
| Map2k4 | 39,6662 | 41,7643 | 0,0743582 | 0,63985 | 0,877325 |
| Wdr75 | 50,3838 | 53,042 | 0,0741769 | 0,6399 | 0,877325 |
| Ep300 | 18,311 | 19,2767 | 0,0741498 | 0,63995 | 0,877325 |
| Ndufb8 | 259,144 | 272,654 | 0,0733138 | 0,63815 | 0,877325 |
| Calm2 | 184,664 | 175,445 | -0,0738842 | 0,638 | 0,877325 |
| Ostc | 188,21 | 178,811 | -0,0739089 | 0,63955 | 0,877325 |
| Ept1 | 18,7125 | 17,7552 | -0,0757588 | 0,63605 | 0,877325 |
| 5031439G07Rik | 13,0283 | 12,3596 | -0,0760137 | 0,6404 | 0,877325 |
| Arnt | 16,0088 | 15,1777 | -0,0769085 | 0,6374 | 0,877325 |
| Sema4d | 11,8139 | 11,1885 | -0,0784696 | 0,64015 | 0,877325 |
| Pex14 | 25,2655 | 23,8745 | -0,0816989 | 0,6395 | 0,877325 |
| Tex30 | 37,7061 | 35,6225 | -0,0820105 | 0,6369 | 0,877325 |
| 1810030O07Rik | 17,4519 | 16,4862 | -0,0821222 | 0,6365 | 0,877325 |
| Lime1 | 23,5495 | 22,2437 | -0,0822974 | 0,63615 | 0,877325 |
| Hspbp1 | 34,2606 | 32,3494 | -0,082812 | 0,6377 | 0,877325 |

| | | | | | |
|---|---|---|---|---|---|
| Enthd2 | 10,1506 | 9,54561 | -0,0886594 | 0,639 | 0,877325 |
| Prmt2 | 13,0344 | 12,256 | -0,0888406 | 0,6387 | 0,877325 |
| Senp8 | 7,01728 | 6,59704 | -0,0890935 | 0,63685 | 0,877325 |
| Dnajb9 | 10,008 | 9,37401 | -0,0944121 | 0,64 | 0,877325 |
| Bicd1 | 1,36096 | 1,2686 | -0,101378 | 0,63985 | 0,877325 |
| Zfp53 | 3,56612 | 3,30566 | -0,109417 | 0,63645 | 0,877325 |
| Plekha6 | 1,30878 | 1,21218 | -0,110609 | 0,64 | 0,877325 |
| Sema4a | 1,00368 | 0,899712 | -0,157766 | 0,63855 | 0,877325 |
| Zfp672 | 10,0073 | 8,84735 | -0,177733 | 0,6373 | 0,877325 |
| Dynlt1a | 0,813446 | 0,701047 | -0,214536 | 0,6403 | 0,877325 |
| Sft2d1 | 1,81629 | 1,50282 | -0,27332 | 0,6393 | 0,877325 |
| 1700124L16Rik | 1,49078 | 1,05259 | -0,502131 | 0,64015 | 0,877325 |
| Dnajc19 | 5,10395 | 6,97287 | 0,450138 | 0,64085 | 0,877432 |
| Serping1 | 3,57156 | 3,90178 | 0,127576 | 0,6407 | 0,877432 |
| Mrpl51 | 21,01 | 22,2936 | 0,0855537 | 0,6406 | 0,877432 |
| Cep164 | 10,6862 | 11,2806 | 0,0780969 | 0,6412 | 0,877432 |
| Por | 42,5197 | 44,7716 | 0,0744538 | 0,64105 | 0,877432 |
| Zfp26 | 9,67403 | 9,18546 | -0,0747648 | 0,6408 | 0,877432 |
| St3gal1 | 11,3164 | 10,7211 | -0,0779576 | 0,64105 | 0,877432 |
| Phtf2 | 10,4732 | 9,90087 | -0,0810754 | 0,6412 | 0,877432 |
| Mb21d2 | 12,6141 | 11,9203 | -0,0816174 | 0,64115 | 0,877432 |
| Isca2 | 36,2405 | 34,091 | -0,0882097 | 0,64115 | 0,877432 |
| Fbxl12os | 0,998843 | 0,882893 | -0,178019 | 0,6407 | 0,877432 |
| Scyl1 | 25,0703 | 26,4902 | 0,0794806 | 0,64135 | 0,877474 |
| Gm6548 | 26,9809 | 28,4839 | 0,0782093 | 0,64205 | 0,877474 |
| Med14 | 19,3553 | 20,3872 | 0,0749346 | 0,64165 | 0,877474 |
| Tcerg1 | 65,1843 | 68,6165 | 0,074031 | 0,64215 | 0,877474 |
| Nup160 | 26,7929 | 28,1912 | 0,0733972 | 0,6416 | 0,877474 |
| Zfp644 | 15,0095 | 14,2509 | -0,0748176 | 0,64155 | 0,877474 |
| Zdhhc17 | 15,874 | 15,046 | -0,0772915 | 0,6421 | 0,877474 |
| Boc | 11,7121 | 11,0926 | -0,0783944 | 0,6414 | 0,877474 |
| Swt1 | 8,11217 | 7,64708 | -0,0851782 | 0,6418 | 0,877474 |
| Zfp763 | 7,03107 | 6,60037 | -0,0911958 | 0,64215 | 0,877474 |
| Nlgn3 | 3,47085 | 3,23414 | -0,101909 | 0,64215 | 0,877474 |
| Nxpe3 | 1,6964 | 1,57502 | -0,107106 | 0,642 | 0,877474 |
| Il3ra | 4,47759 | 4,08363 | -0,132869 | 0,6417 | 0,877474 |
| Ankrd33b | 0,566519 | 0,499233 | -0,182411 | 0,64165 | 0,877474 |
| Eral1 | 16,2451 | 15,3148 | -0,0850773 | 0,6423 | 0,877589 |
| Rit1 | 6,96257 | 6,51395 | -0,0960864 | 0,64275 | 0,878114 |
| Otub2 | 5,44082 | 5,82563 | 0,0985877 | 0,64315 | 0,878549 |
| Ndufb6 | 109,707 | 116,085 | 0,0815237 | 0,6432 | 0,878549 |
| Mrpl18 | 83,2214 | 87,7063 | 0,0757253 | 0,64335 | 0,878574 |
| Yars2 | 41,4082 | 39,2373 | -0,0776922 | 0,64335 | 0,878574 |
| 1700008J07Rik | 2,81101 | 3,05891 | 0,12193 | 0,64355 | 0,878578 |
| Dapk3 | 18,1124 | 19,2138 | 0,0851653 | 0,64345 | 0,878578 |
| Rnf220 | 53,9013 | 56,7558 | 0,0744454 | 0,64355 | 0,878578 |
| 1700054A03Rik | 0,67849 | 0,849428 | 0,324164 | 0,6437 | 0,87865 |
| D830031N03Rik | 7,58301 | 8,70743 | 0,199478 | 0,6438 | 0,87865 |

| | | | | | |
|---|---|---|---|---|---|
| Mapkap1 | 25,3887 | 26,7579 | 0,0757755 | 0,6438 | 0,87865 |
| Snord49a | 2348,88 | 949,3 | -1,30704 | 0,644 | 0,878833 |
| Pcdhgc5 | 0,496856 | 0,956496 | 0,944932 | 0,64425 | 0,878905 |
| Fam216a | 41,8419 | 44,2667 | 0,0812711 | 0,64425 | 0,878905 |
| Ndfip2 | 25,2719 | 23,9433 | -0,0779145 | 0,6442 | 0,878905 |
| Hoxd4 | 113,03 | 120,45 | 0,0917226 | 0,6444 | 0,87902 |
| Tvp23a | 7,64355 | 8,42683 | 0,140747 | 0,6447 | 0,87925 |
| Ddx52 | 40,8664 | 43,0004 | 0,0734345 | 0,64465 | 0,87925 |
| Tmem14a | 3,19852 | 2,88284 | -0,149914 | 0,6456 | 0,880388 |
| Zfp191 | 36,1775 | 34,4121 | -0,072174 | 0,6457 | 0,880412 |
| Ino80 | 11,618 | 11,023 | -0,0758472 | 0,64575 | 0,880412 |
| Cdh24 | 12,2173 | 12,9199 | 0,0806686 | 0,64585 | 0,880459 |
| Mrpl41 | 12,4766 | 13,4539 | 0,108806 | 0,6461 | 0,88053 |
| Fam222b | 9,36596 | 9,90063 | 0,0800933 | 0,6461 | 0,88053 |
| Pnpla8 | 14,8672 | 14,0945 | -0,077001 | 0,64605 | 0,88053 |
| Kctd21 | 1,20999 | 1,09082 | -0,149574 | 0,6464 | 0,880849 |
| Lypla2 | 54,2746 | 57,1349 | 0,0740962 | 0,6466 | 0,881032 |
| Mrps25 | 11,1511 | 11,9903 | 0,104679 | 0,6467 | 0,881078 |
| Zfp959 | 5,65801 | 5,26142 | -0,104841 | 0,6468 | 0,881103 |
| Rps29 | 2,60872 | 2,07604 | -0,32951 | 0,64685 | 0,881103 |
| Fam98c | 6,97491 | 7,68519 | 0,139907 | 0,64695 | 0,881128 |
| Hoxb7 | 49,5997 | 46,9653 | -0,0787363 | 0,647 | 0,881128 |
| Zfp605 | 4,17093 | 3,89264 | -0,0996211 | 0,6473 | 0,881447 |
| Cenpu | 11,0351 | 11,8042 | 0,0971992 | 0,6475 | 0,88145 |
| Mrpl16 | 39,0517 | 41,2711 | 0,0797478 | 0,6475 | 0,88145 |
| Triqk | 1,71754 | 1,52619 | -0,17041 | 0,64745 | 0,88145 |
| Agk | 4,34722 | 4,68648 | 0,108411 | 0,6479 | 0,881793 |
| Socs4 | 10,8186 | 11,5006 | 0,0882026 | 0,64795 | 0,881793 |
| Snord1b | 286,984 | 108,724 | -1,4003 | 0,64795 | 0,881793 |
| Cars2 | 7,40936 | 7,94393 | 0,100504 | 0,64815 | 0,881885 |
| Msantd3 | 30,5835 | 32,3024 | 0,0788848 | 0,64815 | 0,881885 |
| Galnt7 | 28,9647 | 27,5263 | -0,0734811 | 0,64845 | 0,882114 |
| Tecr | 138,397 | 131,493 | -0,0738277 | 0,6484 | 0,882114 |
| Tut1 | 19,8215 | 20,9093 | 0,0770783 | 0,6487 | 0,882364 |
| Epg5 | 2,90137 | 2,73207 | -0,0867396 | 0,6488 | 0,88241 |
| Harbi1 | 7,10551 | 7,58496 | 0,094204 | 0,64945 | 0,882665 |
| Naif1 | 1,07341 | 1,29194 | 0,267335 | 0,64945 | 0,882666 |
| Stc1 | 0,757958 | 0,843944 | 0,155029 | 0,6491 | 0,882666 |
| Tbk1 | 17,0161 | 17,9629 | 0,0781202 | 0,6493 | 0,882666 |
| Atrx | 40,7532 | 38,7686 | -0,072025 | 0,6493 | 0,882666 |
| Dffa | 17,1435 | 15,9056 | -0,108125 | 0,6494 | 0,882666 |
| Gchfr | 4,84782 | 4,28974 | -0,176444 | 0,64945 | 0,882666 |
| Mrpl36 | 39,7328 | 42,1065 | 0,0837128 | 0,64955 | 0,88269 |
| Gpr85 | 0,875093 | 0,780766 | -0,164546 | 0,6496 | 0,88269 |
| Mcts2 | 35,947 | 40,8684 | 0,185114 | 0,6507 | 0,882739 |
| Klf4 | 1,06523 | 1,17678 | 0,143679 | 0,64975 | 0,882739 |
| Camkmt | 2,80881 | 3,09612 | 0,140503 | 0,6503 | 0,882739 |
| Fkbp1b | 4,56226 | 5,02526 | 0,139449 | 0,6505 | 0,882739 |

| | | | | | |
|---|---|---|---|---|---|
| Unc5d | 0,688108 | 0,747562 | 0,119557 | 0,6514 | 0,882739 |
| Fxn | 10,9094 | 11,7204 | 0,103456 | 0,6514 | 0,882739 |
| Zhx2 | 4,60566 | 4,90192 | 0,0899387 | 0,65015 | 0,882739 |
| Zfp1 | 12,0588 | 12,8191 | 0,0882056 | 0,6512 | 0,882739 |
| Fadd | 8,12653 | 8,62797 | 0,0863815 | 0,65005 | 0,882739 |
| Sh3glb2 | 17,5829 | 18,602 | 0,0812885 | 0,6517 | 0,882739 |
| Hdac5 | 11,5107 | 12,1471 | 0,0776377 | 0,6511 | 0,882739 |
| Rcor1 | 19,2265 | 20,2817 | 0,0770819 | 0,6516 | 0,882739 |
| Pnp | 22,5163 | 23,7339 | 0,0759774 | 0,6502 | 0,882739 |
| Brd1 | 23,0302 | 24,2329 | 0,0734413 | 0,65045 | 0,882739 |
| Phf5a | 56,7812 | 59,7415 | 0,0733219 | 0,65145 | 0,882739 |
| Prkci | 32,0752 | 33,7398 | 0,0729919 | 0,6517 | 0,882739 |
| Rcbtb2 | 67,0239 | 70,421 | 0,0713297 | 0,65175 | 0,882739 |
| Ythdc1 | 52,8031 | 55,4779 | 0,0712924 | 0,6516 | 0,882739 |
| Sec61a1 | 109,217 | 103,926 | -0,0716378 | 0,651 | 0,882739 |
| Chmp3 | 27,6719 | 26,3066 | -0,0729979 | 0,6515 | 0,882739 |
| Nek6 | 18,0845 | 17,1625 | -0,0754917 | 0,65005 | 0,882739 |
| Eml3 | 16,4165 | 15,5561 | -0,0776665 | 0,65045 | 0,882739 |
| Mtif2 | 19,1913 | 18,1852 | -0,0776863 | 0,65155 | 0,882739 |
| Exoc5 | 21,8044 | 20,5188 | -0,0876713 | 0,65055 | 0,882739 |
| Slc4a4 | 2,34414 | 2,1994 | -0,0919521 | 0,6509 | 0,882739 |
| Vps45 | 7,40614 | 6,94698 | -0,0923368 | 0,65025 | 0,882739 |
| Nenf | 37,0312 | 34,7328 | -0,0924401 | 0,6512 | 0,882739 |
| Lace1 | 2,32504 | 2,13062 | -0,125988 | 0,65145 | 0,882739 |
| Arhgef4 | 1,16745 | 1,0519 | -0,150373 | 0,65105 | 0,882739 |
| Ntng2 | 0,984452 | 0,824417 | -0,255946 | 0,6512 | 0,882739 |
| Urah | 1,10517 | 0,88785 | -0,315882 | 0,6512 | 0,882739 |
| Snord83b | 573,968 | 227,166 | -1,33722 | 0,65125 | 0,882739 |
| Psd3 | 6,408 | 6,08646 | -0,0742713 | 0,6519 | 0,882853 |
| Rbm45 | 31,799 | 33,5148 | 0,0758178 | 0,65205 | 0,882966 |
| Strip1 | 36,5366 | 38,4556 | 0,0738516 | 0,65245 | 0,883419 |
| Evi5 | 13,7593 | 13,0784 | -0,0732299 | 0,65275 | 0,883646 |
| Lnpep | 8,54475 | 8,10244 | -0,0766814 | 0,6527 | 0,883646 |
| Gm7244 | 5,94298 | 6,57307 | 0,145383 | 0,6533 | 0,883764 |
| Vps37c | 9,81098 | 10,4149 | 0,0861761 | 0,6533 | 0,883764 |
| Ndufa2 | 133,272 | 141,219 | 0,0835581 | 0,65325 | 0,883764 |
| Vimp | 43,4957 | 45,8856 | 0,0771669 | 0,6532 | 0,883764 |
| Jade3 | 9,95714 | 10,4966 | 0,0761219 | 0,6533 | 0,883764 |
| Ndufs2 | 134,619 | 141,374 | 0,0706417 | 0,65325 | 0,883764 |
| Osbpl2 | 12,1215 | 11,4568 | -0,0813607 | 0,65315 | 0,883764 |
| Ndufb5 | 103,481 | 108,772 | 0,0719403 | 0,65345 | 0,883855 |
| Lacc1 | 3,05238 | 2,8111 | -0,1188 | 0,6535 | 0,883855 |
| 1700003E16Rik | 0,651015 | 0,74728 | 0,198957 | 0,65435 | 0,884021 |
| Tle6 | 2,86983 | 3,12919 | 0,124824 | 0,65395 | 0,884021 |
| 2410004N09Rik | 62,1735 | 66,3798 | 0,0944442 | 0,65425 | 0,884021 |
| Ccdc53 | 20,1565 | 21,4573 | 0,0902185 | 0,65425 | 0,884021 |
| Fscn1 | 235,234 | 247,598 | 0,073904 | 0,65415 | 0,884021 |
| Mettl2 | 28,502 | 29,9841 | 0,0731353 | 0,6537 | 0,884021 |

| | | | | | |
|---|---|---|---|---|---|
| Prdx2 | 538,731 | 566,559 | 0,0726624 | 0,6542 | 0,884021 |
| Slc16a3 | 84,2708 | 80,2427 | -0,0706613 | 0,6542 | 0,884021 |
| Mis18a | 39,9453 | 37,8807 | -0,076564 | 0,6542 | 0,884021 |
| Wdtc1 | 10,6598 | 10,1033 | -0,077354 | 0,6543 | 0,884021 |
| Napepld | 3,34126 | 3,11881 | -0,0993953 | 0,6541 | 0,884021 |
| Gm608 | 7,44142 | 7,8248 | 0,0724757 | 0,65445 | 0,884066 |
| Dnajc30 | 12,5403 | 14,3343 | 0,192895 | 0,65495 | 0,884206 |
| Bhmt2 | 0,797348 | 0,907793 | 0,187154 | 0,65485 | 0,884206 |
| 9530077C05Rik | 1,93672 | 2,12396 | 0,133137 | 0,6549 | 0,884206 |
| Fbxo17 | 3,50715 | 3,80912 | 0,119157 | 0,65495 | 0,884206 |
| Cutc | 6,01656 | 6,5189 | 0,115691 | 0,6547 | 0,884206 |
| Patl1 | 18,0464 | 18,9887 | 0,0734317 | 0,6549 | 0,884206 |
| Mxd1 | 2,85296 | 2,66667 | -0,0974233 | 0,65505 | 0,884251 |
| Dcaf7 | 43,395 | 41,2947 | -0,0715731 | 0,65515 | 0,884253 |
| Clk1 | 79,0452 | 75,1146 | -0,0735849 | 0,65525 | 0,884253 |
| Mtmr12 | 9,52523 | 9,02293 | -0,0781585 | 0,6552 | 0,884253 |
| Hspa12b | 8,54766 | 6,78645 | -0,332872 | 0,6557 | 0,884771 |
| Prkar1a | 111,845 | 106,522 | -0,070353 | 0,6558 | 0,884817 |
| Impa2 | 9,83237 | 9,16716 | -0,101064 | 0,65595 | 0,88493 |
| Gmppa | 14,4082 | 15,3131 | 0,0878732 | 0,65635 | 0,88538 |
| 1190005I06Rik | 4,34476 | 4,84092 | 0,156004 | 0,6566 | 0,885451 |
| Trim45 | 4,6449 | 4,93881 | 0,0885171 | 0,6568 | 0,885451 |
| Crk | 44,2191 | 42,0887 | -0,0712355 | 0,6568 | 0,885451 |
| Gabpb2 | 18,0787 | 16,8991 | -0,0973394 | 0,65655 | 0,885451 |
| Rdh12 | 0,917833 | 0,79523 | -0,206858 | 0,65675 | 0,885451 |
| Ccdc73 | 0,82525 | 0,608341 | -0,439951 | 0,65675 | 0,885451 |
| Fhit | 0,937403 | 1,21797 | 0,377736 | 0,65715 | 0,885744 |
| Apba3 | 15,0099 | 15,8994 | 0,0830544 | 0,65715 | 0,885744 |
| Arhgef17 | 15,9273 | 15,1612 | -0,0711139 | 0,65735 | 0,885924 |
| Acad11 | 6,2676 | 6,65239 | 0,0859596 | 0,6579 | 0,886107 |
| Eri3 | 43,7275 | 46,2308 | 0,0803125 | 0,65795 | 0,886107 |
| Phactr2 | 10,9159 | 10,3786 | -0,0728223 | 0,65775 | 0,886107 |
| Xist | 46,4004 | 44,0806 | -0,0739922 | 0,65775 | 0,886107 |
| Zfp868 | 11,4749 | 10,8252 | -0,0840866 | 0,6577 | 0,886107 |
| Hpse | 1,09221 | 0,985725 | -0,147993 | 0,6579 | 0,886107 |
| Cebpb | 0,964409 | 0,836598 | -0,20511 | 0,65785 | 0,886107 |
| Cacng7 | 24,8837 | 26,5311 | 0,0924883 | 0,6581 | 0,886131 |
| G3bp1 | 127,38 | 133,855 | 0,0715288 | 0,65805 | 0,886131 |
| Qdpr | 43,3144 | 45,5934 | 0,0739764 | 0,6585 | 0,886268 |
| Ptcd3 | 57,0109 | 59,8761 | 0,0707418 | 0,6586 | 0,886268 |
| Smo | 53,3921 | 50,9166 | -0,0684895 | 0,65855 | 0,886268 |
| Gphn | 7,7631 | 7,32077 | -0,0846377 | 0,6586 | 0,886268 |
| Ppwd1 | 22,5985 | 21,1026 | -0,0988081 | 0,6583 | 0,886268 |
| 3110052M02Rik | 1,97071 | 1,82774 | -0,108657 | 0,65835 | 0,886268 |
| Ppil6 | 0,881989 | 1,04069 | 0,238707 | 0,6588 | 0,886359 |
| Isg20l2 | 38,2043 | 40,1092 | 0,0701978 | 0,6588 | 0,886359 |
| Svopl | 2,11867 | 1,93203 | -0,133047 | 0,659 | 0,886539 |
| Rbm4 | 8,64454 | 10,042 | 0,216178 | 0,6591 | 0,886584 |

| | | | | | |
|---|---|---|---|---|---|
| Git1 | 25,6118 | 24,3557 | -0,0725476 | 0,65925 | 0,886697 |
| Uhrf1bp1l | 13,0888 | 13,7472 | 0,0707968 | 0,65955 | 0,887011 |
| Ammecr1l | 26,8571 | 25,5876 | -0,0698564 | 0,65975 | 0,88719 |
| Uxs1 | 23,9411 | 25,3328 | 0,0815199 | 0,6599 | 0,887214 |
| Snf8 | 33,3836 | 31,5206 | -0,0828473 | 0,65985 | 0,887214 |
| Wnt8a | 4,38415 | 4,76128 | 0,119054 | 0,66005 | 0,887237 |
| Cog2 | 14,9244 | 15,745 | 0,0772185 | 0,66005 | 0,887237 |
| Sco2 | 2,64657 | 3,84465 | 0,538728 | 0,66025 | 0,887239 |
| Dtd2 | 4,78634 | 5,12415 | 0,0983911 | 0,6603 | 0,887239 |
| Csad | 8,59992 | 9,1331 | 0,0867816 | 0,66045 | 0,887239 |
| Mief1 | 11,2245 | 10,6617 | -0,0742192 | 0,66035 | 0,887239 |
| Stradb | 6,70717 | 6,28845 | -0,0929998 | 0,66045 | 0,887239 |
| Sil1 | 7,36068 | 6,86203 | -0,101204 | 0,66015 | 0,887239 |
| 1700024F13Rik | 1,6673 | 1,96187 | 0,234718 | 0,6607 | 0,887307 |
| Nup98 | 31,9164 | 33,518 | 0,0706391 | 0,6607 | 0,887307 |
| Scaf1 | 39,6661 | 41,5839 | 0,0681153 | 0,6606 | 0,887307 |
| Pde1c | 5,48382 | 5,82238 | 0,0864274 | 0,66085 | 0,88742 |
| Ppil2 | 31,2167 | 33,125 | 0,0856029 | 0,66115 | 0,887733 |
| Zrsr1 | 2,91564 | 3,32426 | 0,189221 | 0,6618 | 0,887737 |
| Ccdc167 | 15,7229 | 17,7648 | 0,176158 | 0,66195 | 0,887737 |
| Idh3b | 73,9378 | 80,539 | 0,123376 | 0,6618 | 0,887737 |
| Rhog | 24,4495 | 25,8746 | 0,0817301 | 0,6617 | 0,887737 |
| Rngtt | 15,0712 | 15,8508 | 0,0727552 | 0,6615 | 0,887737 |
| Zfp740 | 56,8498 | 59,6526 | 0,0694291 | 0,6619 | 0,887737 |
| Usp12 | 13,3112 | 12,6463 | -0,0739264 | 0,66185 | 0,887737 |
| Ebag9 | 19,2017 | 18,1406 | -0,0820126 | 0,66125 | 0,887737 |
| Arhgap33 | 5,69049 | 5,36703 | -0,084427 | 0,6615 | 0,887737 |
| Phlda3 | 13,4197 | 12,6088 | -0,0899203 | 0,66195 | 0,887737 |
| Dok1 | 6,01946 | 5,60588 | -0,102691 | 0,66175 | 0,887737 |
| Ubxn8 | 6,05683 | 5,279 | -0,198298 | 0,66165 | 0,887737 |
| Mrrf | 7,13999 | 8,04846 | 0,172791 | 0,6625 | 0,887762 |
| Zfp597 | 3,63069 | 3,84887 | 0,0841924 | 0,66245 | 0,887762 |
| Ptpn4 | 5,54351 | 5,86649 | 0,081698 | 0,6623 | 0,887762 |
| Ube3c | 30,9151 | 32,468 | 0,0707055 | 0,6622 | 0,887762 |
| Sbk1 | 40,1908 | 42,1596 | 0,0689962 | 0,66245 | 0,887762 |
| Chd8 | 28,8718 | 30,2751 | 0,0684695 | 0,66215 | 0,887762 |
| Sp1 | 27,2649 | 26,0135 | -0,0677837 | 0,66235 | 0,887762 |
| St6galnac6 | 9,87584 | 9,30412 | -0,0860334 | 0,66245 | 0,887762 |
| Cuedc1 | 22,7405 | 24,318 | 0,0967618 | 0,6628 | 0,888053 |
| Plekhf2 | 9,43967 | 9,993 | 0,0821804 | 0,66285 | 0,888053 |
| Gin1 | 4,36933 | 4,04994 | -0,10951 | 0,6631 | 0,888299 |
| Pkn2 | 36,0509 | 37,7787 | 0,0675354 | 0,6632 | 0,888344 |
| Fbxo10 | 10,2374 | 10,9546 | 0,0976868 | 0,6633 | 0,888389 |
| Nlgn2 | 9,51343 | 10,018 | 0,0745558 | 0,66345 | 0,8885 |
| Heatr6 | 9,32011 | 9,80515 | 0,0731926 | 0,66365 | 0,88859 |
| Grik4 | 2,13614 | 1,97695 | -0,111731 | 0,66365 | 0,88859 |
| Tjp2 | 23,4095 | 24,5701 | 0,0698099 | 0,6638 | 0,888702 |
| 5830418K08Rik | 17,3485 | 18,5223 | 0,0944536 | 0,6639 | 0,888747 |

| | | | | | |
|---|---|---|---|---|---|
| Usp40 | 14,4888 | 15,2158 | 0,0706293 | 0,664 | 0,888792 |
| Helz2 | 2,6026 | 2,75505 | 0,0821249 | 0,66465 | 0,888994 |
| Cdc20 | 98,5843 | 103,912 | 0,0759312 | 0,66475 | 0,888994 |
| Rala | 36,4693 | 38,3215 | 0,0714703 | 0,6647 | 0,888994 |
| Farp1 | 29,2491 | 27,8926 | -0,0685109 | 0,6644 | 0,888994 |
| Sdhd | 153,945 | 146,787 | -0,0686849 | 0,6643 | 0,888994 |
| Myo9a | 7,71866 | 7,35848 | -0,0689431 | 0,66475 | 0,888994 |
| Tmem79 | 1,36817 | 1,23781 | -0,144457 | 0,66425 | 0,888994 |
| 9130024F11Rik | 0,944191 | 0,82649 | -0,192081 | 0,66465 | 0,888994 |
| Zfp133-ps | 1,37345 | 1,18219 | -0,216343 | 0,6647 | 0,888994 |
| Tob1 | 10,5371 | 11,166 | 0,0836334 | 0,66505 | 0,889017 |
| Trp53bp2 | 18,7119 | 19,6533 | 0,0708142 | 0,665 | 0,889017 |
| Rad17 | 21,6642 | 20,6046 | -0,0723496 | 0,6651 | 0,889017 |
| Zfp157 | 9,30651 | 8,83947 | -0,0742799 | 0,6651 | 0,889017 |
| Scn2b | 1,81089 | 1,6777 | -0,11021 | 0,6649 | 0,889017 |
| 2310040G24Rik | 1,9296 | 2,27115 | 0,235122 | 0,6652 | 0,889062 |
| Nbas | 8,43196 | 8,01875 | -0,0724907 | 0,6653 | 0,889107 |
| Smdt1 | 133,545 | 140,521 | 0,0734631 | 0,6657 | 0,88933 |
| Chd1 | 30,7989 | 32,3256 | 0,0698003 | 0,66555 | 0,88933 |
| Egln2 | 31,4294 | 29,8755 | -0,073153 | 0,66575 | 0,88933 |
| Pisd | 20,458 | 19,4329 | -0,0741644 | 0,6658 | 0,88933 |
| Fam187b | 1,15346 | 1,00081 | -0,204808 | 0,66575 | 0,88933 |
| S100a3 | 0,664053 | 0,835428 | 0,331218 | 0,6661 | 0,889376 |
| Macrod1 | 0,685331 | 0,820869 | 0,260351 | 0,66605 | 0,889376 |
| Postn | 0,995521 | 1,10115 | 0,145481 | 0,6661 | 0,889376 |
| Cdkn2aipnl | 49,8122 | 52,2827 | 0,0698335 | 0,6661 | 0,889376 |
| Cenpf | 29,605 | 31,0709 | 0,0697232 | 0,6662 | 0,88942 |
| Smarcd2 | 29,6961 | 31,1433 | 0,0686471 | 0,6665 | 0,889643 |
| Pcdhga2 | 1,57917 | 1,01242 | -0,64136 | 0,66645 | 0,889643 |
| 4930429B21Rik | 0,446303 | 0,668605 | 0,58313 | 0,6672 | 0,889955 |
| D430042O09Rik | 6,00625 | 6,33421 | 0,0766999 | 0,6669 | 0,889955 |
| Whsc1 | 58,9084 | 61,894 | 0,0713268 | 0,667 | 0,889955 |
| Fzd1 | 20,3556 | 21,3807 | 0,0708819 | 0,66705 | 0,889955 |
| Epb4.1l5 | 18,5184 | 19,4391 | 0,0700052 | 0,66715 | 0,889955 |
| Med23 | 19,7126 | 20,6718 | 0,0685514 | 0,66715 | 0,889955 |
| Cxcr4 | 42,4133 | 40,392 | -0,0704492 | 0,6672 | 0,889955 |
| Slc16a7 | 0,684035 | 0,772327 | 0,175142 | 0,66735 | 0,890066 |
| Myo9b | 15,8991 | 16,6806 | 0,0692271 | 0,6675 | 0,890155 |
| Stat5b | 7,09772 | 6,72766 | -0,077251 | 0,66755 | 0,890155 |
| 1700120C14Rik | 0,795034 | 1,58772 | 0,99787 | 0,6678 | 0,890311 |
| Mfsd4 | 2,27272 | 2,46091 | 0,114771 | 0,66775 | 0,890311 |
| Hcfc2 | 5,65671 | 6,02984 | 0,0921568 | 0,668 | 0,890467 |
| Nsmf | 18,228 | 19,1705 | 0,0727335 | 0,66805 | 0,890467 |
| Slc24a3 | 2,26087 | 2,43181 | 0,105155 | 0,66865 | 0,890911 |
| Kdm5a | 12,3071 | 12,9647 | 0,0750986 | 0,66855 | 0,890911 |
| Pabpn1 | 215,303 | 205,36 | -0,0682122 | 0,6685 | 0,890911 |
| Ctage5 | 43,6228 | 41,6046 | -0,0683367 | 0,6686 | 0,890911 |
| Pdp2 | 9,65192 | 10,6142 | 0,137109 | 0,669 | 0,891178 |

| | | | | | |
|---|---|---|---|---|---|
| Chmp6 | 33,4819 | 35,2271 | 0,0733042 | 0,66905 | 0,891178 |
| Pten | 8,33995 | 7,94484 | -0,0700204 | 0,669 | 0,891178 |
| Sall2 | 23,6283 | 24,7682 | 0,0679777 | 0,6692 | 0,891288 |
| Wdr59 | 7,17617 | 7,56598 | 0,0763135 | 0,66935 | 0,891311 |
| Tmem175 | 4,24244 | 3,81028 | -0,154998 | 0,66935 | 0,891311 |
| Vps35 | 73,1918 | 69,8759 | -0,0668867 | 0,6695 | 0,891422 |
| Ano10 | 11,752 | 11,133 | -0,0780616 | 0,6698 | 0,891643 |
| Ankrd39 | 4,65564 | 4,31233 | -0,11051 | 0,66975 | 0,891643 |
| Adarb1 | 2,81585 | 2,98722 | 0,0852316 | 0,6699 | 0,891688 |
| Lrrc46 | 0,622345 | 0,735493 | 0,240996 | 0,67 | 0,891732 |
| Cdkal1 | 7,48503 | 7,94071 | 0,0852591 | 0,67035 | 0,891753 |
| Mrps24 | 31,906 | 33,6839 | 0,0782288 | 0,6704 | 0,891753 |
| Gabarap | 113,188 | 118,728 | 0,068937 | 0,67025 | 0,891753 |
| Erp44 | 33,9868 | 32,4051 | -0,0687558 | 0,67025 | 0,891753 |
| Znrf3 | 5,8002 | 5,51091 | -0,0738122 | 0,6705 | 0,891753 |
| Osgepl1 | 11,2347 | 10,5839 | -0,0860943 | 0,67055 | 0,891753 |
| Tubd1 | 4,98158 | 4,63035 | -0,105482 | 0,67045 | 0,891753 |
| Fam195a | 4,49731 | 4,06038 | -0,147447 | 0,67045 | 0,891753 |
| Trim11 | 12,5382 | 11,875 | -0,0783983 | 0,67085 | 0,891975 |
| Klf15 | 1,49564 | 1,35804 | -0,139232 | 0,6708 | 0,891975 |
| Ddx3x | 243,548 | 231,376 | -0,0739637 | 0,67095 | 0,892019 |
| 2900052N01Rik | 0,541822 | 0,617334 | 0,188233 | 0,67215 | 0,892348 |
| Trex1 | 10,5533 | 11,3056 | 0,0993337 | 0,6726 | 0,892348 |
| Eny2 | 37,6676 | 40,1906 | 0,0935313 | 0,67205 | 0,892348 |
| Plgrkt | 19,724 | 20,9721 | 0,088521 | 0,6721 | 0,892348 |
| Dhx34 | 6,16916 | 6,52689 | 0,0813211 | 0,67235 | 0,892348 |
| Tbc1d5 | 5,77652 | 6,08427 | 0,0748837 | 0,67205 | 0,892348 |
| Mbd4 | 11,2884 | 11,8866 | 0,074497 | 0,67245 | 0,892348 |
| Fam189b | 14,1697 | 14,9101 | 0,0734789 | 0,67225 | 0,892348 |
| Decr1 | 20,3606 | 21,3878 | 0,0710113 | 0,67155 | 0,892348 |
| Dip2b | 7,96708 | 8,36424 | 0,0701821 | 0,6716 | 0,892348 |
| Fubp3 | 36,2943 | 38,0884 | 0,0696075 | 0,67155 | 0,892348 |
| Ccnk | 31,0837 | 32,5918 | 0,0683519 | 0,6725 | 0,892348 |
| Phf12 | 26,5349 | 27,8076 | 0,0675865 | 0,67245 | 0,892348 |
| Zfp445 | 48,9814 | 51,3081 | 0,0669548 | 0,6725 | 0,892348 |
| Sqle | 66,0284 | 63,0521 | -0,0665427 | 0,67195 | 0,892348 |
| Phip | 26,1304 | 24,922 | -0,0683122 | 0,67245 | 0,892348 |
| Ccni | 53,7739 | 51,2808 | -0,0684868 | 0,6717 | 0,892348 |
| Cdk8 | 24,5262 | 23,3646 | -0,069999 | 0,6726 | 0,892348 |
| Dhcr7 | 23,1563 | 22,0393 | -0,071328 | 0,6723 | 0,892348 |
| Dcun1d3 | 4,5177 | 4,27895 | -0,0783334 | 0,6725 | 0,892348 |
| Gm4890 | 1,85496 | 1,69872 | -0,126935 | 0,67245 | 0,892348 |
| Rab11fip5 | 1,45151 | 1,565 | 0,108607 | 0,6727 | 0,892393 |
| Usp9x | 17,8573 | 18,7129 | 0,0675198 | 0,67295 | 0,892458 |
| Slc12a7 | 30,5719 | 32,0249 | 0,066988 | 0,67295 | 0,892458 |
| Kazn | 9,11114 | 8,63776 | -0,0769749 | 0,6729 | 0,892458 |
| Fto | 38,947 | 40,8077 | 0,067327 | 0,67345 | 0,893033 |
| Maz | 129,339 | 135,512 | 0,0672603 | 0,67365 | 0,893099 |

| | | | | | |
|---|---|---|---|---|---|
| Zfp503 | 22,3295 | 23,3926 | 0,0670997 | 0,6737 | 0,893099 |
| Gabpa | 23,4584 | 21,8682 | -0,101268 | 0,67365 | 0,893099 |
| Kif20a | 81,3432 | 88,0112 | 0,113665 | 0,67395 | 0,89323 |
| Cenpl | 19,8359 | 20,914 | 0,0763498 | 0,67395 | 0,89323 |
| Zfp799 | 11,8366 | 12,4379 | 0,071488 | 0,674 | 0,89323 |
| Emc2 | 32,3452 | 34,0644 | 0,0747128 | 0,6742 | 0,893407 |
| Ino80e | 38,2531 | 40,4881 | 0,081921 | 0,67485 | 0,893447 |
| Coa7 | 10,5725 | 11,1477 | 0,0764301 | 0,67475 | 0,893447 |
| Macf1 | 20,4832 | 21,5772 | 0,0750653 | 0,6753 | 0,893447 |
| Peo1 | 24,935 | 26,145 | 0,0683626 | 0,67515 | 0,893447 |
| Srrm1 | 89,9196 | 94,2821 | 0,0683483 | 0,67445 | 0,893447 |
| Xbp1 | 45,3829 | 47,5557 | 0,067469 | 0,6752 | 0,893447 |
| Pold1 | 34,1274 | 32,5976 | -0,0661621 | 0,67485 | 0,893447 |
| Eif4e | 47,8362 | 45,6869 | -0,0663236 | 0,67485 | 0,893447 |
| Msn | 79,6565 | 76,041 | -0,067015 | 0,6751 | 0,893447 |
| Rundc1 | 7,80187 | 7,3691 | -0,0823332 | 0,6751 | 0,893447 |
| Dhrs7b | 13,7482 | 12,9786 | -0,0831105 | 0,6751 | 0,893447 |
| Gstz1 | 17,1782 | 15,8418 | -0,116843 | 0,67515 | 0,893447 |
| Gm11110 | 2,94588 | 2,56639 | -0,19896 | 0,675 | 0,893447 |
| Cml1 | 1,37172 | 1,19118 | -0,20359 | 0,67525 | 0,893447 |
| Cebpzos | 9,87837 | 8,20305 | -0,268111 | 0,67455 | 0,893447 |
| AA413626 | 1,72552 | 0,675987 | -1,35197 | 0,67515 | 0,893447 |
| Sirt6 | 21,6284 | 20,5381 | -0,0746204 | 0,6754 | 0,893469 |
| Mblac2 | 0,898459 | 0,81714 | -0,13687 | 0,67545 | 0,893469 |
| Ago4 | 1,04395 | 1,12585 | 0,108959 | 0,6756 | 0,893579 |
| Hprt | 66,7885 | 63,6923 | -0,0684795 | 0,67585 | 0,893821 |
| Zfp599 | 4,53781 | 5,5737 | 0,29664 | 0,6762 | 0,893924 |
| Rad9b | 2,26736 | 2,48706 | 0,13343 | 0,6776 | 0,893924 |
| Oxld1 | 5,28433 | 5,7939 | 0,132814 | 0,6773 | 0,893924 |
| Tigd3 | 2,5338 | 2,75564 | 0,121088 | 0,67625 | 0,893924 |
| Rab3gap2 | 8,15367 | 8,80011 | 0,110071 | 0,6773 | 0,893924 |
| Chmp1b | 19,0821 | 20,565 | 0,107971 | 0,6773 | 0,893924 |
| Crem | 3,78369 | 4,07352 | 0,106481 | 0,6775 | 0,893924 |
| Per2 | 1,44927 | 1,55413 | 0,100777 | 0,6765 | 0,893924 |
| Nprl2 | 10,7634 | 11,4397 | 0,0879068 | 0,67705 | 0,893924 |
| Ddx28 | 13,2285 | 13,978 | 0,0795089 | 0,67705 | 0,893924 |
| Athl1 | 10,2377 | 10,8072 | 0,0780968 | 0,6765 | 0,893924 |
| Atg13 | 10,8927 | 11,4592 | 0,0731408 | 0,67735 | 0,893924 |
| Chfr | 23,9146 | 25,0692 | 0,0680261 | 0,67685 | 0,893924 |
| Supt16 | 121,567 | 127,306 | 0,0665563 | 0,6773 | 0,893924 |
| Smarcd1 | 67,5228 | 70,7071 | 0,0664799 | 0,67755 | 0,893924 |
| Rangap1 | 77,8706 | 81,5301 | 0,0662534 | 0,67735 | 0,893924 |
| Phactr4 | 22,3188 | 23,3612 | 0,0658503 | 0,67725 | 0,893924 |
| Pcm1 | 36,5691 | 34,9399 | -0,0657503 | 0,6767 | 0,893924 |
| Hoxc13 | 70,7627 | 67,5961 | -0,0660478 | 0,6769 | 0,893924 |
| Wwc2 | 14,7082 | 14,0379 | -0,0672932 | 0,67755 | 0,893924 |
| Taok2 | 30,1252 | 28,7508 | -0,0673677 | 0,67745 | 0,893924 |
| Pccb | 23,0842 | 21,9828 | -0,0705292 | 0,6766 | 0,893924 |

| | | | | | |
|---|---|---|---|---|---|
| Lsm11 | 4,52337 | 4,28815 | -0,0770422 | 0,67695 | 0,893924 |
| Slc26a10 | 1,52821 | 1,39012 | -0,136635 | 0,6761 | 0,893924 |
| B330016D10Rik | 1,48014 | 1,34403 | -0,139174 | 0,6763 | 0,893924 |
| Etf1 | 123,93 | 129,778 | 0,066517 | 0,67785 | 0,894054 |
| Slc25a39 | 88,7521 | 92,8716 | 0,0654556 | 0,6781 | 0,894054 |
| Tacc3 | 79,0175 | 82,679 | 0,0653492 | 0,67785 | 0,894054 |
| Acbd5 | 7,76056 | 7,36316 | -0,0758362 | 0,67805 | 0,894054 |
| Plp1 | 2,29763 | 2,13557 | -0,105528 | 0,678 | 0,894054 |
| Crhbp | 1,098 | 0,972738 | -0,174757 | 0,6781 | 0,894054 |
| Pcp4l1 | 3,43239 | 3,73175 | 0,120637 | 0,67865 | 0,894425 |
| Adprhl2 | 26,8966 | 28,3098 | 0,0738752 | 0,67865 | 0,894425 |
| Snord23 | 44,1793 | 46,3479 | 0,0691361 | 0,6788 | 0,894425 |
| Tmem183a | 36,1277 | 34,503 | -0,0663854 | 0,67885 | 0,894425 |
| Cox14 | 37,0913 | 35,0579 | -0,0813407 | 0,6787 | 0,894425 |
| Ccnc | 18,0294 | 16,8822 | -0,0948457 | 0,67875 | 0,894425 |
| Cage1 | 0,800871 | 0,717907 | -0,157773 | 0,67865 | 0,894425 |
| Heatr3 | 37,5214 | 39,3032 | 0,0669364 | 0,67895 | 0,894468 |
| Zbtb12 | 65,4968 | 62,5396 | -0,0666543 | 0,6791 | 0,894578 |
| Cdr2l | 8,55979 | 8,11341 | -0,077267 | 0,67925 | 0,894687 |
| Nap1l5 | 5,77918 | 6,32462 | 0,130116 | 0,6795 | 0,894818 |
| Lrrc56 | 5,63133 | 6,07673 | 0,109817 | 0,67945 | 0,894818 |
| Kif2c | 51,3021 | 53,7212 | 0,0664746 | 0,67955 | 0,894818 |
| H2-Ke2 | 113,971 | 120,194 | 0,0766952 | 0,6797 | 0,894927 |
| Drap1 | 105,591 | 110,659 | 0,0676391 | 0,67985 | 0,895036 |
| Srp72 | 57,9334 | 60,6153 | 0,0652874 | 0,68 | 0,895058 |
| Tradd | 2,36609 | 2,14853 | -0,139155 | 0,67995 | 0,895058 |
| Polr3b | 15,6294 | 16,379 | 0,0675777 | 0,68015 | 0,895145 |
| Pkn1 | 11,151 | 10,5541 | -0,0793708 | 0,6802 | 0,895145 |
| E230016M11Rik | 0,727226 | 0,905967 | 0,317055 | 0,6817 | 0,895198 |
| Fbxw4 | 8,91423 | 9,67661 | 0,118391 | 0,6814 | 0,895198 |
| Casc3 | 40,6906 | 43,6571 | 0,101519 | 0,6819 | 0,895198 |
| Gnb1l | 2,5611 | 2,73916 | 0,0969685 | 0,68095 | 0,895198 |
| Txnl4b | 6,42205 | 6,84879 | 0,0928169 | 0,6812 | 0,895198 |
| Deb1 | 47,8856 | 50,7578 | 0,0840387 | 0,68095 | 0,895198 |
| Rp9 | 32,2407 | 33,9692 | 0,075341 | 0,68175 | 0,895198 |
| Rfx5 | 7,58871 | 7,99162 | 0,0746329 | 0,68175 | 0,895198 |
| Tmem126a | 52,8058 | 55,5962 | 0,0742899 | 0,6807 | 0,895198 |
| Klf13 | 6,57437 | 6,90576 | 0,0709485 | 0,68045 | 0,895198 |
| Tpx2 | 84,7612 | 88,8256 | 0,0675715 | 0,68205 | 0,895198 |
| Zfp553 | 28,3825 | 29,7288 | 0,0668593 | 0,68225 | 0,895198 |
| Gatad2a | 38,5491 | 40,35 | 0,0658708 | 0,68195 | 0,895198 |
| Fkbp1a | 189,33 | 198,005 | 0,0646325 | 0,6807 | 0,895198 |
| Limch1 | 43,4523 | 45,4106 | 0,0635979 | 0,68215 | 0,895198 |
| Dlst | 81,8849 | 85,5605 | 0,0633478 | 0,68225 | 0,895198 |
| Eif3e | 94,2643 | 90,0603 | -0,0658199 | 0,6811 | 0,895198 |
| Mastl | 10,3739 | 9,88102 | -0,0702265 | 0,68125 | 0,895198 |
| Efna4 | 31,0668 | 29,5812 | -0,0706902 | 0,682 | 0,895198 |
| Nrm | 34,1991 | 32,5443 | -0,0715527 | 0,682 | 0,895198 |

| | | | | | |
|---|---|---|---|---|---|
| Pef1 | 26,2623 | 24,972 | -0,0726816 | 0,68205 | 0,895198 |
| Pcca | 14,8979 | 14,1529 | -0,0740167 | 0,682 | 0,895198 |
| Nkrf | 10,0681 | 9,5575 | -0,0750836 | 0,68085 | 0,895198 |
| Plekhm2 | 6,90548 | 6,54604 | -0,0771193 | 0,68095 | 0,895198 |
| Clstn3 | 4,67774 | 4,41722 | -0,0826726 | 0,68105 | 0,895198 |
| 1190007I07Rik | 13,9902 | 12,9413 | -0,112433 | 0,68215 | 0,895198 |
| Ankrd23 | 3,24087 | 2,99495 | -0,113851 | 0,68035 | 0,895198 |
| Ccdc60 | 0,69711 | 0,618699 | -0,172149 | 0,6819 | 0,895198 |
| Tmx2 | 14,6663 | 12,7369 | -0,203494 | 0,68135 | 0,895198 |
| Snora61 | 1044,43 | 474,65 | -1,13777 | 0,68055 | 0,895198 |
| Ogfod1 | 6,90875 | 6,5719 | -0,0721127 | 0,68245 | 0,895372 |
| Dph1 | 15,6271 | 16,7826 | 0,102918 | 0,68295 | 0,895477 |
| Mpp2 | 5,38789 | 5,6911 | 0,0789866 | 0,6832 | 0,895477 |
| Psmb5 | 63,6428 | 67,0087 | 0,0743528 | 0,68315 | 0,895477 |
| Use1 | 50,7522 | 53,369 | 0,07253 | 0,68305 | 0,895477 |
| Cope | 76,9187 | 80,5731 | 0,0669647 | 0,6832 | 0,895477 |
| Elf2 | 16,8343 | 17,6254 | 0,0662508 | 0,683 | 0,895477 |
| Zbtb11 | 7,7224 | 7,34597 | -0,0720976 | 0,68305 | 0,895477 |
| Lig4 | 7,78886 | 7,39815 | -0,0742472 | 0,6829 | 0,895477 |
| Vmac | 3,96423 | 3,68589 | -0,105026 | 0,6826 | 0,895477 |
| Mia | 1,1748 | 0,946704 | -0,311427 | 0,6831 | 0,895477 |
| Ube2n | 15,215 | 14,5057 | -0,0688682 | 0,6833 | 0,89552 |
| Adamts14 | 2,4194 | 2,57245 | 0,0884953 | 0,68385 | 0,895845 |
| Kif24 | 5,14339 | 5,41796 | 0,0750297 | 0,68385 | 0,895845 |
| Sos1 | 9,08638 | 8,67828 | -0,0662976 | 0,6839 | 0,895845 |
| Men1 | 37,7791 | 36,0739 | -0,066632 | 0,6837 | 0,895845 |
| Rwdd4a | 21,5733 | 20,5754 | -0,0683239 | 0,6838 | 0,895845 |
| Smim24 | 2,12249 | 1,80605 | -0,232919 | 0,68395 | 0,895845 |
| Pxk | 4,52152 | 4,81122 | 0,0895955 | 0,68415 | 0,895931 |
| Cops5 | 82,352 | 86,1002 | 0,0642119 | 0,6841 | 0,895931 |
| Lgals3bp | 1,42882 | 1,56779 | 0,133912 | 0,6849 | 0,896363 |
| Zfp593 | 11,8701 | 12,5333 | 0,0784395 | 0,6851 | 0,896363 |
| Rab22a | 19,0261 | 19,9914 | 0,0714016 | 0,6847 | 0,896363 |
| Kdm6a | 18,1076 | 18,9531 | 0,0658381 | 0,6849 | 0,896363 |
| Dgkd | 23,8963 | 24,9881 | 0,0644497 | 0,68485 | 0,896363 |
| Usp46 | 19,474 | 18,5847 | -0,0674308 | 0,68485 | 0,896363 |
| 2810474O19Rik | 12,5519 | 11,9711 | -0,0683535 | 0,68515 | 0,896363 |
| Krcc1 | 24,7175 | 23,5297 | -0,0710493 | 0,68505 | 0,896363 |
| Acsbg1 | 0,990268 | 0,896481 | -0,143546 | 0,6851 | 0,896363 |
| Zfp960 | 0,745795 | 0,665841 | -0,163602 | 0,68505 | 0,896363 |
| A230056P14Rik | 4,85708 | 5,49774 | 0,178748 | 0,6855 | 0,896447 |
| Pih1d1 | 48,6594 | 51,9432 | 0,0942166 | 0,68555 | 0,896447 |
| Idh3g | 72,7402 | 76,1491 | 0,0660753 | 0,6854 | 0,896447 |
| Mepce | 25,2442 | 24,0961 | -0,0671569 | 0,68555 | 0,896447 |
| Il12rb2 | 0,627857 | 0,559006 | -0,167574 | 0,6853 | 0,896447 |
| Zfp292 | 15,2881 | 15,9983 | 0,0655066 | 0,6857 | 0,896556 |
| Fam110b | 3,78476 | 4,06571 | 0,103305 | 0,686 | 0,896661 |
| Riok2 | 27,8024 | 29,1062 | 0,0661191 | 0,686 | 0,896661 |

| | | | | | |
|---|---|---|---|---|---|
| Nars | 96,0144 | 100,401 | 0,0644517 | 0,68605 | 0,896661 |
| Pcnx | 9,17417 | 8,77112 | -0,0648164 | 0,6862 | 0,896661 |
| Rab39b | 2,95626 | 2,76897 | -0,0944248 | 0,68625 | 0,896661 |
| Klhl7 | 24,7278 | 22,7585 | -0,11973 | 0,68625 | 0,896661 |
| Ccdc24 | 8,16839 | 7,43966 | -0,134815 | 0,6859 | 0,896661 |
| 4930473A02Rik | 0,310383 | 0,630702 | 1,02291 | 0,6864 | 0,896681 |
| Ap4e1 | 7,64004 | 7,28726 | -0,0682043 | 0,6864 | 0,896681 |
| Chtop | 76,8228 | 73,4616 | -0,0645449 | 0,6866 | 0,896767 |
| Inpp1 | 4,47996 | 4,16553 | -0,104985 | 0,6866 | 0,896767 |
| Gpsm3 | 3,7894 | 3,48793 | -0,1196 | 0,68675 | 0,896876 |
| Rab5a | 21,8451 | 22,9337 | 0,0701634 | 0,68685 | 0,896896 |
| Ube2b | 59,4395 | 62,1504 | 0,0643432 | 0,6869 | 0,896896 |
| Ghitm | 68,6617 | 71,7772 | 0,0640189 | 0,68715 | 0,897135 |
| Prpf39 | 30,4213 | 27,7129 | -0,134523 | 0,68725 | 0,897178 |
| Me1 | 2,7673 | 2,95404 | 0,0942111 | 0,68765 | 0,897262 |
| Cnr1 | 1,78641 | 1,90447 | 0,0923292 | 0,68755 | 0,897262 |
| Hmgcl | 20,1929 | 21,2897 | 0,0763073 | 0,68765 | 0,897262 |
| Rfwd2 | 4,76914 | 5,02359 | 0,0749897 | 0,6875 | 0,897262 |
| Tyw1 | 14,1585 | 14,8613 | 0,0698986 | 0,6875 | 0,897262 |
| Hspa1l | 3,25254 | 3,49005 | 0,101684 | 0,688 | 0,89728 |
| Tfip11 | 22,8329 | 23,9577 | 0,0693766 | 0,688 | 0,89728 |
| Psma4 | 93,7717 | 98,0233 | 0,0639721 | 0,6878 | 0,89728 |
| Tmem246 | 5,02946 | 4,7439 | -0,0843304 | 0,6879 | 0,89728 |
| Sox21 | 0,950852 | 0,871966 | -0,124948 | 0,6878 | 0,89728 |
| Zscan12 | 8,3301 | 8,75198 | 0,0712759 | 0,6881 | 0,897323 |
| 2900008C10Rik | 1,69253 | 2,59116 | 0,614414 | 0,6885 | 0,897362 |
| Cnbd2 | 0,452449 | 0,579911 | 0,358077 | 0,68845 | 0,897362 |
| Med15 | 15,3958 | 16,137 | 0,067829 | 0,6886 | 0,897362 |
| Wdr4 | 16,7522 | 17,5391 | 0,066223 | 0,68845 | 0,897362 |
| Ankrd50 | 13,6013 | 14,2386 | 0,0660706 | 0,68825 | 0,897362 |
| Dnm1l | 47,0136 | 44,9615 | -0,0643865 | 0,68855 | 0,897362 |
| Diap3 | 8,41651 | 8,00098 | -0,0730455 | 0,68835 | 0,897362 |
| Six4 | 9,68073 | 10,146 | 0,0677203 | 0,68875 | 0,897426 |
| Ndufs1 | 67,6469 | 70,7802 | 0,0653234 | 0,68885 | 0,897426 |
| Sympk | 65,9033 | 63,059 | -0,0636501 | 0,68885 | 0,897426 |
| Lrba | 2,4775 | 2,34603 | -0,0786643 | 0,68905 | 0,897576 |
| Ttll11 | 0,662007 | 0,594903 | -0,154194 | 0,6891 | 0,897576 |
| Il17ra | 7,23126 | 7,60633 | 0,0729535 | 0,6895 | 0,89766 |
| Hn1 | 111,788 | 106,969 | -0,0635632 | 0,68925 | 0,89766 |
| Ralgapb | 14,006 | 14,4255 | -0,0644786 | 0,68945 | 0,89766 |
| Spg7 | 30,5784 | 29,2174 | -0,0656859 | 0,6895 | 0,89766 |
| Col16a1 | 1,67596 | 1,56655 | -0,0973941 | 0,6893 | 0,89766 |
| Gm12992 | 0,831662 | 1,13304 | 0,446124 | 0,69005 | 0,897764 |
| Coa3 | 83,4084 | 87,4024 | 0,0674787 | 0,68995 | 0,897764 |
| Mier2 | 23,3776 | 24,4756 | 0,0662191 | 0,69005 | 0,897764 |
| Pik3r2 | 25,4499 | 26,6391 | 0,0658838 | 0,6898 | 0,897764 |
| Igf2bp1 | 49,1919 | 51,4817 | 0,0656372 | 0,68975 | 0,897764 |
| Ercc6l | 21,2235 | 22,1769 | 0,0633949 | 0,69 | 0,897764 |

| | | | | | |
|---|---|---|---|---|---|
| 4930415O20Rik | 0,948879 | 0,756789 | -0,326332 | 0,68985 | 0,897764 |
| Timm8b | 167,445 | 178,459 | 0,0919007 | 0,6902 | 0,897871 |
| Fam117b | 12,8957 | 13,4989 | 0,0659444 | 0,69035 | 0,897979 |
| Adcy6 | 21,6005 | 20,6668 | -0,0637503 | 0,69055 | 0,898152 |
| 4930414N06Rik | 0,268953 | 0,676601 | 1,33095 | 0,69095 | 0,898213 |
| Zswim7 | 12,291 | 13,1944 | 0,102328 | 0,6908 | 0,898213 |
| Plaa | 26,1696 | 27,3389 | 0,0630626 | 0,6909 | 0,898213 |
| Wdr1 | 83,026 | 79,5159 | -0,0623194 | 0,6908 | 0,898213 |
| Rps16 | 1,82959 | 1,56819 | -0,222419 | 0,691 | 0,898213 |
| Wisp3 | 0,990163 | 0,750678 | -0,399472 | 0,691 | 0,898213 |
| Acap3 | 5,772 | 6,08206 | 0,07549 | 0,69135 | 0,898318 |
| Phc3 | 4,42275 | 4,63055 | 0,0662399 | 0,69125 | 0,898318 |
| Ppid | 76,3532 | 79,7652 | 0,0630724 | 0,69115 | 0,898318 |
| Usp10 | 66,4057 | 69,3572 | 0,0627373 | 0,69135 | 0,898318 |
| Zfp869 | 18,6099 | 19,4881 | 0,0665209 | 0,69165 | 0,898336 |
| Sidt2 | 17,6422 | 16,8811 | -0,0636204 | 0,69155 | 0,898336 |
| Cyb5r4 | 15,1848 | 14,4676 | -0,0698103 | 0,6917 | 0,898336 |
| Actr6 | 18,3594 | 17,4449 | -0,0737114 | 0,6916 | 0,898336 |
| Gprc5b | 11,6061 | 10,2834 | -0,174576 | 0,6916 | 0,898336 |
| Tcaim | 2,58505 | 2,76836 | 0,0988395 | 0,6918 | 0,898356 |
| Cenpa | 144,44 | 150,916 | 0,0632791 | 0,69185 | 0,898356 |
| Ydjc | 3,52732 | 3,24572 | -0,120033 | 0,692 | 0,898464 |
| Mib2 | 6,32318 | 6,66387 | 0,0757113 | 0,69215 | 0,898484 |
| Phlpp1 | 4,1248 | 4,33852 | 0,0728808 | 0,6921 | 0,898484 |
| Neurl2 | 3,25982 | 2,47796 | -0,395637 | 0,69255 | 0,898916 |
| Sec11a | 110,35 | 105,537 | -0,0643434 | 0,69265 | 0,898958 |
| A930004D18Rik | 0,520975 | 0,59743 | 0,197556 | 0,69285 | 0,899043 |
| Pdcd6 | 41,096 | 43,0129 | 0,0657707 | 0,6928 | 0,899043 |
| Elp5 | 61,3861 | 65,4645 | 0,0928013 | 0,693 | 0,899063 |
| Wdr12 | 19,2815 | 20,2854 | 0,0732215 | 0,69295 | 0,899063 |
| Zbtb25 | 10,8218 | 11,4468 | 0,0810075 | 0,6933 | 0,899256 |
| Fam149b | 20,1889 | 21,2374 | 0,0730449 | 0,69325 | 0,899256 |
| Ublcp1 | 12,479 | 11,8538 | -0,0741522 | 0,69335 | 0,899256 |
| Fam217b | 0,620826 | 0,699788 | 0,172728 | 0,6935 | 0,89934 |
| Folr2 | 0,762846 | 0,642869 | -0,246866 | 0,69355 | 0,89934 |
| Tssk6 | 3,06127 | 3,72583 | 0,283433 | 0,69365 | 0,899383 |
| Tuba1b | 104,457 | 109,044 | 0,0620039 | 0,69385 | 0,899468 |
| Zbtb41 | 6,94039 | 6,63033 | -0,0659379 | 0,6938 | 0,899468 |
| 4930528A17Rik | 0,33402 | 0,7628 | 1,19137 | 0,69425 | 0,899637 |
| Traf5 | 1,47229 | 1,61223 | 0,130995 | 0,69425 | 0,899637 |
| Tnk2 | 19,4124 | 20,3176 | 0,0657488 | 0,69415 | 0,899637 |
| Inpp5e | 12,3209 | 11,2078 | -0,136609 | 0,69415 | 0,899637 |
| Kdm5c | 25,1928 | 26,3209 | 0,0631963 | 0,6944 | 0,899657 |
| Fbxo34 | 8,2567 | 7,83227 | -0,0761333 | 0,6944 | 0,899657 |
| Otop1 | 0,771333 | 0,849779 | 0,139733 | 0,6947 | 0,899674 |
| Pot1a | 14,9874 | 15,6948 | 0,0665375 | 0,69475 | 0,899674 |
| Ptplb | 8,06375 | 7,66136 | -0,0738506 | 0,6947 | 0,899674 |
| Eif4ebp3 | 14,7507 | 13,9233 | -0,083291 | 0,69475 | 0,899674 |

| | | | | | |
|---|---|---|---|---|---|
| 1600002K03Rik | 9,07321 | 7,64075 | -0,247899 | 0,69475 | 0,899674 |
| Fez2 | 11,3919 | 10,6779 | -0,0933828 | 0,69485 | 0,899717 |
| Gclc | 14,4376 | 13,774 | -0,0678777 | 0,695 | 0,899737 |
| Rttn | 5,46621 | 5,20939 | -0,0694282 | 0,69495 | 0,899737 |
| Ptbp1 | 189,839 | 221,502 | 0,222541 | 0,69545 | 0,900002 |
| 4933439K11Rik | 1,81528 | 2,09084 | 0,203885 | 0,6961 | 0,900002 |
| Hoga1 | 0,727942 | 0,832339 | 0,193349 | 0,69595 | 0,900002 |
| Cox18 | 13,3861 | 14,1627 | 0,0813545 | 0,69615 | 0,900002 |
| l7Rn6 | 38,8493 | 40,8261 | 0,0716036 | 0,69635 | 0,900002 |
| Elac2 | 25,695 | 26,9842 | 0,070622 | 0,69635 | 0,900002 |
| Thbs3 | 11,7727 | 12,348 | 0,068833 | 0,6957 | 0,900002 |
| Fbxw5 | 18,583 | 19,4782 | 0,067878 | 0,69575 | 0,900002 |
| Vps13b | 4,91165 | 5,13256 | 0,063471 | 0,69555 | 0,900002 |
| Arhgef2 | 33,994 | 35,5073 | 0,0628354 | 0,6955 | 0,900002 |
| Hif1a | 63,5276 | 66,3089 | 0,0618184 | 0,69575 | 0,900002 |
| Fam49a | 7,18445 | 6,84619 | -0,069577 | 0,6957 | 0,900002 |
| Nckipsd | 4,62381 | 4,35196 | -0,0874179 | 0,6963 | 0,900002 |
| Gpr173 | 2,3462 | 2,18855 | -0,100357 | 0,69545 | 0,900002 |
| Prkaa2 | 0,598157 | 0,553396 | -0,112214 | 0,6959 | 0,900002 |
| Zkscan14 | 2,64518 | 2,44498 | -0,113538 | 0,6961 | 0,900002 |
| Chga | 1,32626 | 1,19797 | -0,146782 | 0,69605 | 0,900002 |
| Stk25 | 38,5903 | 36,8844 | -0,0652301 | 0,6965 | 0,900109 |
| 2210416O15Rik | 0,806585 | 1,31414 | 0,704225 | 0,69695 | 0,900233 |
| Pced1a | 10,084 | 10,9722 | 0,121788 | 0,69695 | 0,900233 |
| 2310061I04Rik | 15,9992 | 16,8795 | 0,0772686 | 0,697 | 0,900233 |
| Rmnd5a | 28,1188 | 26,9484 | -0,0613369 | 0,69695 | 0,900233 |
| Fyco1 | 5,32853 | 5,08027 | -0,0688331 | 0,69685 | 0,900233 |
| Gal3st4 | 2,11198 | 1,83553 | -0,202402 | 0,6969 | 0,900233 |
| Cd83 | 16,7406 | 17,5612 | 0,0690422 | 0,6971 | 0,900275 |
| 4930577N17Rik | 0,460515 | 0,769915 | 0,74145 | 0,69745 | 0,900356 |
| Stk3 | 21,9905 | 22,9942 | 0,0643912 | 0,6973 | 0,900356 |
| Dazap2 | 73,0492 | 76,2335 | 0,0615575 | 0,6975 | 0,900356 |
| Nnat | 233,994 | 223,645 | -0,0652558 | 0,69725 | 0,900356 |
| Fam83d | 19,3844 | 18,5012 | -0,067273 | 0,6975 | 0,900356 |
| Adck5 | 9,72503 | 11,134 | 0,195204 | 0,698 | 0,900477 |
| Cad | 43,7304 | 48,7958 | 0,15812 | 0,6978 | 0,900477 |
| 4930452B06Rik | 4,34656 | 4,61276 | 0,0857542 | 0,69815 | 0,900477 |
| Tex2 | 6,66635 | 7,00049 | 0,0705572 | 0,6982 | 0,900477 |
| Pdzd2 | 2,99097 | 3,13383 | 0,0673134 | 0,6982 | 0,900477 |
| Pigo | 9,30191 | 8,87197 | -0,0682736 | 0,6982 | 0,900477 |
| Bcas3 | 5,87097 | 5,56806 | -0,0764236 | 0,6978 | 0,900477 |
| Pnpla7 | 2,80765 | 2,64624 | -0,0854195 | 0,69775 | 0,900477 |
| Ddb2 | 4,89687 | 4,20694 | -0,219086 | 0,6981 | 0,900477 |
| Fdx1l | 39,8243 | 45,9149 | 0,205313 | 0,69875 | 0,900665 |
| C030006K11Rik | 4,38112 | 4,69563 | 0,100019 | 0,69845 | 0,900665 |
| Nprl3 | 11,7635 | 12,5212 | 0,0900595 | 0,6986 | 0,900665 |
| Hsdl2 | 9,38255 | 9,88261 | 0,0749129 | 0,6987 | 0,900665 |
| Chic1 | 6,45833 | 6,76843 | 0,067659 | 0,69865 | 0,900665 |

| | | | | | |
|---|---|---|---|---|---|
| Sept7 | 111,711 | 107,153 | -0,0600932 | 0,6986 | 0,900665 |
| Tmem150a | 8,48185 | 9,49779 | 0,163213 | 0,69895 | 0,900727 |
| Rbm18 | 30,5061 | 32,0165 | 0,069719 | 0,699 | 0,900727 |
| Bsdc1 | 15,6938 | 16,4448 | 0,0674321 | 0,6989 | 0,900727 |
| Zbtb10 | 4,72324 | 4,95197 | 0,0682266 | 0,69915 | 0,900833 |
| Pepd | 22,314 | 23,4113 | 0,0692602 | 0,69925 | 0,900875 |
| Hemk1 | 4,84021 | 5,1475 | 0,0888022 | 0,69965 | 0,90095 |
| Get4 | 24,6665 | 25,79 | 0,0642558 | 0,6998 | 0,90095 |
| Ptpn23 | 18,1818 | 18,982 | 0,0621369 | 0,6996 | 0,90095 |
| Rnf44 | 65,89 | 68,7734 | 0,0617908 | 0,69975 | 0,90095 |
| Nek9 | 36,9678 | 38,5705 | 0,0612268 | 0,69995 | 0,90095 |
| Chmp1a | 23,5977 | 22,5527 | -0,0653434 | 0,69995 | 0,90095 |
| Dpm1 | 14,9766 | 14,2598 | -0,0707537 | 0,69965 | 0,90095 |
| Slc30a6 | 11,5798 | 11,0052 | -0,0734169 | 0,7 | 0,90095 |
| Mapk4 | 0,642903 | 0,58681 | -0,131708 | 0,70005 | 0,90095 |
| Slc25a53 | 1,84515 | 1,65601 | -0,156021 | 0,6998 | 0,90095 |
| Tspan17 | 1,04204 | 0,924356 | -0,172895 | 0,70005 | 0,90095 |
| Mir762 | 354,018 | 709,266 | 1,00251 | 0,7023 | 0,901242 |
| 4921507L20Rik | 0,545192 | 0,704941 | 0,370739 | 0,70185 | 0,901242 |
| 2410127L17Rik | 0,573104 | 0,649547 | 0,180637 | 0,70145 | 0,901242 |
| Tub | 1,99228 | 2,19623 | 0,140605 | 0,70035 | 0,901242 |
| Flii | 23,8635 | 25,7094 | 0,107491 | 0,7019 | 0,901242 |
| Trp53rk | 1,67655 | 1,79383 | 0,0975495 | 0,70145 | 0,901242 |
| Gkap1 | 6,00568 | 6,41501 | 0,0951225 | 0,7022 | 0,901242 |
| Pgam1 | 14,597 | 15,3598 | 0,0734906 | 0,70165 | 0,901242 |
| Sugp2 | 18,4727 | 19,4129 | 0,0716213 | 0,7018 | 0,901242 |
| Rdh11 | 24,9836 | 26,1666 | 0,0667444 | 0,7022 | 0,901242 |
| Nudt9 | 37,8244 | 39,5781 | 0,0653858 | 0,70165 | 0,901242 |
| Inpp5f | 14,2396 | 14,8817 | 0,0636274 | 0,70155 | 0,901242 |
| Eif1a | 54,9873 | 57,403 | 0,0620272 | 0,7016 | 0,901242 |
| Wdr82 | 44,767 | 46,6581 | 0,059692 | 0,7011 | 0,901242 |
| Smad5 | 46,5958 | 44,6605 | -0,0612014 | 0,70075 | 0,901242 |
| Exoc2 | 15,9157 | 15,2513 | -0,0615179 | 0,70225 | 0,901242 |
| 2310035C23Rik | 7,84055 | 7,49032 | -0,0659278 | 0,7015 | 0,901242 |
| Zmym6 | 7,80637 | 7,44462 | -0,0684538 | 0,70215 | 0,901242 |
| Dak | 15,6244 | 14,8889 | -0,0695634 | 0,70185 | 0,901242 |
| Sys1 | 17,5503 | 16,6995 | -0,0716877 | 0,70085 | 0,901242 |
| Fam174b | 6,01371 | 5,67937 | -0,0825249 | 0,70075 | 0,901242 |
| Iqsec2 | 0,956462 | 0,888103 | -0,10698 | 0,70085 | 0,901242 |
| Arrdc2 | 2,34073 | 2,16948 | -0,109613 | 0,70165 | 0,901242 |
| Them4 | 2,97121 | 2,74631 | -0,113558 | 0,7005 | 0,901242 |
| Sobp | 0,849133 | 0,782407 | -0,118071 | 0,70055 | 0,901242 |
| Tspan11 | 0,799754 | 0,732816 | -0,126106 | 0,70145 | 0,901242 |
| Prss35 | 1,00083 | 0,909097 | -0,138686 | 0,70225 | 0,901242 |
| Rab3a | 2,34057 | 2,08511 | -0,166736 | 0,70175 | 0,901242 |
| Lmbrd2 | 8,66917 | 7,67776 | -0,175209 | 0,7022 | 0,901242 |
| Fbxw9 | 6,59817 | 5,4052 | -0,287718 | 0,70195 | 0,901242 |
| Fus | 383,777 | 401,105 | 0,0637127 | 0,7024 | 0,901284 |

| | | | | | |
|---|---|---|---|---|---|
| 2310009B15Rik | 6,0978 | 5,52803 | -0,141521 | 0,7025 | 0,901326 |
| Clock | 6,56466 | 7,04564 | 0,102009 | 0,7027 | 0,901496 |
| Ceacam1 | 0,960653 | 1,08051 | 0,169631 | 0,7033 | 0,901573 |
| Nudt7 | 6,08222 | 6,58351 | 0,114259 | 0,70295 | 0,901573 |
| Msrb1 | 20,1135 | 21,2567 | 0,079749 | 0,70325 | 0,901573 |
| Sestd1 | 6,90893 | 7,22032 | 0,0636014 | 0,7032 | 0,901573 |
| Mapk3 | 42,8852 | 44,7642 | 0,0618644 | 0,70295 | 0,901573 |
| Minos1 | 31,9314 | 30,5971 | -0,0615802 | 0,70315 | 0,901573 |
| Ncstn | 47,9256 | 44,1635 | -0,117943 | 0,7031 | 0,901573 |
| Mbd6 | 18,2882 | 16,6187 | -0,138105 | 0,70315 | 0,901573 |
| B3galt6 | 6,07707 | 5,77272 | -0,0741248 | 0,7034 | 0,901615 |
| Trip11 | 7,04606 | 7,37256 | 0,0653496 | 0,7037 | 0,901759 |
| Tsg101 | 34,668 | 36,2441 | 0,0641395 | 0,70385 | 0,901759 |
| Arpp19 | 53,3408 | 51,1842 | -0,0595406 | 0,7037 | 0,901759 |
| Rnf115 | 34,3045 | 32,8554 | -0,0622713 | 0,70375 | 0,901759 |
| Tmem205 | 11,9112 | 11,0401 | -0,109574 | 0,7038 | 0,901759 |
| Stx1b | 3,39694 | 3,66247 | 0,108579 | 0,7042 | 0,901968 |
| Cog7 | 11,0805 | 11,6163 | 0,0681323 | 0,7043 | 0,901968 |
| Knop1 | 56,8599 | 59,4466 | 0,0641814 | 0,7042 | 0,901968 |
| Slc39a6 | 60,6248 | 58,1471 | -0,0602002 | 0,70435 | 0,901968 |
| Lyrm2 | 30,9993 | 29,5673 | -0,0682326 | 0,7043 | 0,901968 |
| Aasdh | 4,23405 | 4,4721 | 0,0789132 | 0,70465 | 0,902044 |
| Xpnpep3 | 5,07418 | 5,35908 | 0,0788102 | 0,70475 | 0,902044 |
| 6430706D22Rik | 100,641 | 106,142 | 0,0767792 | 0,70495 | 0,902044 |
| Edem1 | 19,7379 | 18,932 | -0,0601436 | 0,70495 | 0,902044 |
| B4galnt4 | 30,7349 | 29,4744 | -0,0604155 | 0,70485 | 0,902044 |
| Pikfyve | 5,05221 | 4,83486 | -0,0634407 | 0,70485 | 0,902044 |
| Crot | 18,8253 | 18,0019 | -0,0645203 | 0,7047 | 0,902044 |
| Snora31 | 211,837 | 141,443 | -0,582732 | 0,7045 | 0,902044 |
| 2610306M01Rik | 4,80736 | 5,20383 | 0,11433 | 0,7051 | 0,902064 |
| Anp32e | 225,982 | 236,285 | 0,0643186 | 0,7051 | 0,902064 |
| Lnx1 | 2,93733 | 2,40041 | -0,291225 | 0,70525 | 0,902169 |
| Hist1h2bm | 0,912324 | 1,15755 | 0,343453 | 0,70545 | 0,902185 |
| Slx1b | 5,69528 | 5,99476 | 0,0739359 | 0,7056 | 0,902185 |
| Kin | 27,2229 | 28,5053 | 0,0664049 | 0,7056 | 0,902185 |
| Lzts2 | 37,4705 | 39,0927 | 0,0611419 | 0,7055 | 0,902185 |
| BC002163 | 34,3239 | 32,2833 | -0,0884245 | 0,7056 | 0,902185 |
| Nme4 | 46,3982 | 44,2202 | -0,069363 | 0,7059 | 0,90246 |
| Ccdc120 | 4,03586 | 3,81261 | -0,0820947 | 0,70595 | 0,90246 |
| Abcf3 | 18,2327 | 19,0426 | 0,0627039 | 0,70615 | 0,90252 |
| Mmd | 25,9039 | 24,8056 | -0,0625049 | 0,7062 | 0,90252 |
| Lin54 | 9,41341 | 8,98456 | -0,0672691 | 0,7061 | 0,90252 |
| Dcun1d4 | 15,6452 | 14,9884 | -0,0618742 | 0,70635 | 0,902626 |
| Gna12 | 31,6582 | 33,0964 | 0,064095 | 0,7065 | 0,902731 |
| Ccdc22 | 15,7752 | 16,583 | 0,0720483 | 0,70685 | 0,902833 |
| Memo1 | 60,6868 | 63,3028 | 0,0608852 | 0,7067 | 0,902833 |
| Cpxm1 | 24,3059 | 23,2682 | -0,062943 | 0,7068 | 0,902833 |
| Snx16 | 3,71369 | 3,49017 | -0,0895585 | 0,70685 | 0,902833 |

| | | | | | |
|---|---|---|---|---|---|
| 2810408M09Rik | 13,4768 | 12,8246 | -0,0715631 | 0,70695 | 0,902875 |
| Pacrg | 0,678945 | 0,78841 | 0,215652 | 0,70825 | 0,903413 |
| Igip | 1,32052 | 1,47067 | 0,155369 | 0,7079 | 0,903413 |
| Crebl2 | 0,944527 | 1,03719 | 0,135016 | 0,70785 | 0,903413 |
| Zfp689 | 6,15032 | 6,46604 | 0,0722214 | 0,70815 | 0,903413 |
| Phrf1 | 34,8113 | 36,2855 | 0,0598395 | 0,7081 | 0,903413 |
| Zc3h4 | 31,5677 | 32,8671 | 0,0581958 | 0,70755 | 0,903413 |
| Ap1g1 | 33,0335 | 31,7143 | -0,0587967 | 0,7076 | 0,903413 |
| Gnb4 | 16,6815 | 15,9383 | -0,0657543 | 0,70825 | 0,903413 |
| Tusc2 | 17,2484 | 16,4037 | -0,0724371 | 0,70775 | 0,903413 |
| Itgb1bp1 | 12,0417 | 10,9796 | -0,133218 | 0,7082 | 0,903413 |
| Raver2 | 5,86716 | 5,21054 | -0,171229 | 0,70815 | 0,903413 |
| Npff | 2,98964 | 2,45723 | -0,282936 | 0,7082 | 0,903413 |
| Snord52 | 4689,56 | 2310,81 | -1,02105 | 0,7081 | 0,903413 |
| Vstm4 | 0,701789 | 0,776776 | 0,14646 | 0,71005 | 0,903654 |
| Pak3 | 0,601423 | 0,645909 | 0,102951 | 0,7104 | 0,903654 |
| Med29 | 6,73904 | 7,12672 | 0,0806954 | 0,70855 | 0,903654 |
| Wdr77 | 64,3707 | 67,8401 | 0,0757337 | 0,70865 | 0,903654 |
| Trappc6b | 19,1057 | 20,1048 | 0,0735421 | 0,70915 | 0,903654 |
| Ccdc80 | 6,50082 | 6,82913 | 0,0710784 | 0,70925 | 0,903654 |
| Gemin2 | 27,9461 | 29,329 | 0,0696811 | 0,71 | 0,903654 |
| Araf | 38,3781 | 40,1966 | 0,066788 | 0,7095 | 0,903654 |
| Scyl2 | 18,626 | 19,45 | 0,062451 | 0,7097 | 0,903654 |
| Hnrnpl | 266,327 | 277,723 | 0,0604453 | 0,7099 | 0,903654 |
| Zfp629 | 20,3006 | 21,1642 | 0,0601064 | 0,71005 | 0,903654 |
| Klhdc2 | 92,423 | 96,3355 | 0,0598154 | 0,70875 | 0,903654 |
| Tomm22 | 104,953 | 109,388 | 0,0597052 | 0,7094 | 0,903654 |
| Asap1 | 22,3303 | 23,2696 | 0,0594491 | 0,7103 | 0,903654 |
| Dctn1 | 30,325 | 31,5651 | 0,0578203 | 0,7098 | 0,903654 |
| Ggnbp2 | 37,4683 | 35,9536 | -0,0595334 | 0,71035 | 0,903654 |
| Dnmbp | 8,1828 | 7,83354 | -0,0629289 | 0,70915 | 0,903654 |
| Nek7 | 17,232 | 16,4878 | -0,0636878 | 0,7089 | 0,903654 |
| Rap1a | 12,5393 | 11,956 | -0,0687202 | 0,7101 | 0,903654 |
| Mllt10 | 17,3662 | 16,4716 | -0,0762984 | 0,70885 | 0,903654 |
| Kcnmb4 | 16,7333 | 15,8217 | -0,0808175 | 0,70925 | 0,903654 |
| Zmat1 | 1,31166 | 1,2147 | -0,110792 | 0,70965 | 0,903654 |
| Nkain3 | 0,958522 | 0,882701 | -0,118887 | 0,7098 | 0,903654 |
| Khk | 3,27447 | 3,00598 | -0,123423 | 0,71035 | 0,903654 |
| Glyctk | 0,626264 | 0,571095 | -0,133041 | 0,7096 | 0,903654 |
| Tulp1 | 0,817063 | 0,72949 | -0,16356 | 0,70905 | 0,903654 |
| Creb3 | 11,9246 | 10,239 | -0,219866 | 0,7104 | 0,903654 |
| AA465934 | 7,69389 | 6,1554 | -0,321861 | 0,7101 | 0,903654 |
| Snord22 | 902,056 | 718,862 | -0,327502 | 0,70995 | 0,903654 |
| Lrrc39 | 1,33089 | 1,08684 | -0,29225 | 0,7106 | 0,903822 |
| Rtfdc1 | 26,7381 | 27,923 | 0,0625589 | 0,7108 | 0,90399 |
| Rac3 | 15,4303 | 16,306 | 0,0796351 | 0,71095 | 0,904095 |
| Rgs7 | 0,821532 | 0,905469 | 0,140349 | 0,7112 | 0,904327 |
| Tbc1d13 | 22,3713 | 23,3308 | 0,0605847 | 0,7116 | 0,904663 |

| | | | | | |
|---|---|---|---|---|---|
| Trim44 | 55,2778 | 57,5671 | 0,0585455 | 0,7116 | 0,904663 |
| Vps33a | 14,3739 | 13,771 | -0,0618188 | 0,7117 | 0,904705 |
| Dnm1 | 2,92388 | 3,09977 | 0,0842808 | 0,7118 | 0,904746 |
| Sik2 | 5,98053 | 6,84312 | 0,194379 | 0,7124 | 0,904961 |
| Tyms-ps | 10,5862 | 11,1934 | 0,0804703 | 0,71215 | 0,904961 |
| Adal | 11,6302 | 12,2943 | 0,0801099 | 0,71255 | 0,904961 |
| Vti1b | 35,3259 | 36,9791 | 0,0659825 | 0,71205 | 0,904961 |
| Cenpw | 26,4427 | 27,6317 | 0,0634535 | 0,71285 | 0,904961 |
| Aar2 | 15,5425 | 16,2202 | 0,0615732 | 0,71265 | 0,904961 |
| Snx6 | 87,2195 | 90,8641 | 0,0590609 | 0,71285 | 0,904961 |
| Rpn1 | 98,4242 | 94,5167 | -0,0584435 | 0,71245 | 0,904961 |
| Irak1 | 20,9091 | 20,0567 | -0,0600416 | 0,7128 | 0,904961 |
| Bckdha | 23,8422 | 22,7846 | -0,0654598 | 0,71275 | 0,904961 |
| Lrrc61 | 6,75571 | 6,41023 | -0,0757319 | 0,71275 | 0,904961 |
| Lyrm7 | 0,862791 | 0,775987 | -0,15298 | 0,7128 | 0,904961 |
| LOC100503676 | 0,573356 | 0,417568 | -0,45742 | 0,7127 | 0,904961 |
| Camk2n2 | 2,06507 | 2,40952 | 0,222555 | 0,713 | 0,905066 |
| Pdcd6ip | 23,156 | 24,1064 | 0,0580337 | 0,71315 | 0,90517 |
| Atpaf1 | 2,2861 | 2,4578 | 0,104483 | 0,7137 | 0,905588 |
| Arhgap26 | 0,77543 | 0,830078 | 0,0982514 | 0,71355 | 0,905588 |
| Twf2 | 20,8091 | 21,8209 | 0,0684972 | 0,71365 | 0,905588 |
| Impact | 18,9165 | 19,7264 | 0,0604866 | 0,71375 | 0,905588 |
| Car4 | 6,59499 | 6,1805 | -0,0936453 | 0,71385 | 0,905629 |
| Usp54 | 8,17816 | 8,81164 | 0,107634 | 0,714 | 0,905733 |
| Tsc22d3 | 24,3091 | 25,4062 | 0,0636839 | 0,7142 | 0,905901 |
| Etohd2 | 1,11907 | 0,87776 | -0,350402 | 0,7143 | 0,905941 |
| Tmem253 | 1,15852 | 2,10568 | 0,862001 | 0,71445 | 0,906015 |
| Eif1ad | 42,813 | 45,8633 | 0,0992924 | 0,7149 | 0,906015 |
| Galnt11 | 16,1072 | 16,8266 | 0,0630332 | 0,71465 | 0,906015 |
| Uchl5 | 32,1331 | 33,4984 | 0,0600314 | 0,71465 | 0,906015 |
| Ddah2 | 137,342 | 143,029 | 0,0585312 | 0,7149 | 0,906015 |
| Lin28b | 16,517 | 15,8408 | -0,0603082 | 0,7145 | 0,906015 |
| Ctdsp1 | 22,1003 | 21,1784 | -0,0614741 | 0,71475 | 0,906015 |
| Nmnat2 | 1,40717 | 1,31322 | -0,0996879 | 0,7148 | 0,906015 |
| 9230105E05Rik | 0,321129 | 0,857713 | 1,41734 | 0,7153 | 0,906178 |
| Pds5b | 33,8057 | 35,1631 | 0,0567919 | 0,7153 | 0,906178 |
| Ddx3y | 9,02686 | 8,63477 | -0,0640656 | 0,7151 | 0,906178 |
| Zfp934 | 2,80027 | 2,61378 | -0,0994254 | 0,7152 | 0,906178 |
| Nmnat1 | 8,47752 | 9,06204 | 0,0961927 | 0,7155 | 0,906214 |
| Zfp81 | 12,4349 | 13,0311 | 0,0675665 | 0,71555 | 0,906214 |
| Mars | 62,8381 | 60,2888 | -0,0597492 | 0,7156 | 0,906214 |
| Aaed1 | 13,311 | 12,549 | -0,0850494 | 0,7156 | 0,906214 |
| Iqcd | 0,566106 | 0,644827 | 0,187838 | 0,7157 | 0,906255 |
| Galnt4 | 6,10183 | 5,65605 | -0,109448 | 0,7158 | 0,906296 |
| Gm15471 | 0,70437 | 1,15245 | 0,710304 | 0,7165 | 0,906495 |
| Carm1 | 74,2574 | 77,7921 | 0,0670884 | 0,71645 | 0,906495 |
| Arid1b | 9,63709 | 9,25774 | -0,0579375 | 0,7164 | 0,906495 |
| Golga7 | 40,6495 | 39,0028 | -0,0596619 | 0,7163 | 0,906495 |

| | | | | | |
|---|---|---|---|---|---|
| Rpusd1 | 8,55005 | 8,09453 | -0,0789846 | 0,7162 | 0,906495 |
| Tusc1 | 1,79154 | 1,62643 | -0,139494 | 0,71605 | 0,906495 |
| 2810032G03Rik | 1,25257 | 1,11367 | -0,16957 | 0,7164 | 0,906495 |
| BC003331 | 12,7343 | 10,8699 | -0,228372 | 0,7161 | 0,906495 |
| Tti2 | 13,0154 | 14,3238 | 0,138194 | 0,717 | 0,906956 |
| Cdc25c | 26,2844 | 27,4343 | 0,0617751 | 0,71695 | 0,906956 |
| Lancl2 | 15,0006 | 14,3661 | -0,0623576 | 0,7171 | 0,906996 |
| Ndufb11 | 74,5351 | 79,3429 | 0,090181 | 0,71735 | 0,907227 |
| Arv1 | 14,1929 | 14,9676 | 0,0766744 | 0,71745 | 0,907245 |
| Stxbp5 | 3,3223 | 3,47453 | 0,0646356 | 0,7175 | 0,907245 |
| Camsap2 | 13,8887 | 14,4548 | 0,0576461 | 0,7176 | 0,907285 |
| Gm16596 | 0,566873 | 1,01773 | 0,844254 | 0,7179 | 0,907493 |
| 1810043H04Rik | 29,6978 | 31,5499 | 0,0872791 | 0,7179 | 0,907493 |
| Pde1b | 2,41189 | 2,56965 | 0,0914103 | 0,71815 | 0,907506 |
| Csde1 | 83,3114 | 86,7284 | 0,0579902 | 0,71825 | 0,907506 |
| Nckap1 | 75,2483 | 72,2894 | -0,0578752 | 0,71825 | 0,907506 |
| Esco2 | 20,6014 | 19,745 | -0,0612492 | 0,7182 | 0,907506 |
| Rab3gap1 | 10,6634 | 10,2155 | -0,0619039 | 0,71825 | 0,907506 |
| Krt10 | 2,96313 | 3,17445 | 0,0993818 | 0,7186 | 0,907795 |
| Timm9 | 47,7555 | 49,9337 | 0,064346 | 0,7187 | 0,907795 |
| Tmod3 | 31,3812 | 30,158 | -0,0573577 | 0,71875 | 0,907795 |
| Zfp446 | 5,03407 | 4,79804 | -0,069281 | 0,7187 | 0,907795 |
| Skiv2l2 | 62,2351 | 64,779 | 0,0577985 | 0,71895 | 0,907961 |
| Snurf | 1,07071 | 1,30061 | 0,280624 | 0,7193 | 0,907992 |
| 4933417G07Rik | 4,51217 | 5,41853 | 0,264081 | 0,71935 | 0,907992 |
| Gcdh | 21,8258 | 22,9128 | 0,0701245 | 0,71935 | 0,907992 |
| Cdr2 | 20,1757 | 21,0573 | 0,0617026 | 0,7194 | 0,907992 |
| H1f0 | 215,505 | 206,89 | -0,0588562 | 0,7193 | 0,907992 |
| Sirt7 | 11,2954 | 10,7321 | -0,0738082 | 0,7192 | 0,907992 |
| Tmem38b | 2,27025 | 2,12219 | -0,0972952 | 0,71945 | 0,907992 |
| Hivep1 | 6,23466 | 6,49765 | 0,0596077 | 0,7196 | 0,90801 |
| Usp36 | 15,5028 | 16,1402 | 0,0581273 | 0,71955 | 0,90801 |
| BC065397 | 2,68045 | 3,31715 | 0,307468 | 0,71985 | 0,908068 |
| Mapk1 | 87,967 | 91,9502 | 0,0638901 | 0,71985 | 0,908068 |
| Tomm40l | 10,0538 | 9,59288 | -0,0677041 | 0,71985 | 0,908068 |
| Alkbh7 | 14,8519 | 15,7083 | 0,0808771 | 0,71995 | 0,908109 |
| Xpo7 | 15,103 | 15,706 | 0,0564807 | 0,7203 | 0,908207 |
| Zc2hc1a | 9,60833 | 9,17785 | -0,0661285 | 0,7201 | 0,908207 |
| Rftn2 | 6,23191 | 5,94282 | -0,0685255 | 0,7203 | 0,908207 |
| Bend7 | 0,743381 | 0,682962 | -0,122296 | 0,72025 | 0,908207 |
| Zc2hc1c | 0,935442 | 1,00819 | 0,10804 | 0,72045 | 0,908311 |
| Mea1 | 71,1523 | 79,2681 | 0,155831 | 0,721 | 0,908833 |
| Plekhm3 | 2,43065 | 2,55104 | 0,0697444 | 0,72095 | 0,908833 |
| Rpe | 20,5011 | 19,4536 | -0,0756649 | 0,7211 | 0,908873 |
| Zfp275 | 18,5604 | 17,847 | -0,056546 | 0,7214 | 0,909143 |
| Cd37 | 0,549865 | 0,472933 | -0,217441 | 0,72145 | 0,909143 |
| Usmg5 | 6,08578 | 4,31238 | -0,496957 | 0,7217 | 0,909372 |
| Cep162 | 6,29455 | 6,57783 | 0,0635088 | 0,7222 | 0,909487 |

| | | | | | |
|---|---|---|---|---|---|
| Nkiras2 | 24,4481 | 25,4961 | 0,0605557 | 0,72215 | 0,909487 |
| Sertad2 | 5,5694 | 5,32694 | -0,0642152 | 0,7219 | 0,909487 |
| Orc3 | 18,6021 | 17,6122 | -0,0788927 | 0,72215 | 0,909487 |
| 1600016N20Rik | 4,241 | 3,90009 | -0,120896 | 0,72195 | 0,909487 |
| Ripply1 | 2,19298 | 1,93073 | -0,183748 | 0,722 | 0,909487 |
| A330035P11Rik | 1,3058 | 1,41209 | 0,112894 | 0,7223 | 0,909528 |
| Fut4 | 3,59941 | 3,79567 | 0,0765952 | 0,72245 | 0,909631 |
| Msx3 | 1,1766 | 1,27978 | 0,121267 | 0,72275 | 0,909666 |
| Zfp472 | 8,7196 | 9,15258 | 0,0699167 | 0,72265 | 0,909666 |
| Rock1 | 15,7921 | 15,1799 | -0,0570444 | 0,7226 | 0,909666 |
| Rnls | 1,69557 | 1,54575 | -0,13346 | 0,72275 | 0,909666 |
| Pdzd7 | 0,559155 | 0,731946 | 0,388489 | 0,72305 | 0,909935 |
| Adc | 1,75915 | 1,90386 | 0,114048 | 0,7231 | 0,909935 |
| Tceb2 | 143,819 | 183,119 | 0,348529 | 0,7237 | 0,909948 |
| Fam129c | 1,60351 | 1,73216 | 0,111336 | 0,72325 | 0,909948 |
| Sord | 6,11282 | 6,44178 | 0,0756213 | 0,7239 | 0,909948 |
| Fam110a | 11,6643 | 12,2491 | 0,0705651 | 0,72395 | 0,909948 |
| Tcf7l1 | 19,1415 | 19,9673 | 0,0609371 | 0,7238 | 0,909948 |
| Spred1 | 28,5416 | 29,7526 | 0,0599493 | 0,7242 | 0,909948 |
| Dync1h1 | 37,732 | 39,3244 | 0,0596345 | 0,72365 | 0,909948 |
| Myo1b | 33,0216 | 34,3194 | 0,0556173 | 0,7238 | 0,909948 |
| Rps15a | 32,963 | 31,7133 | -0,0557596 | 0,72325 | 0,909948 |
| Nup50 | 34,743 | 33,4192 | -0,0560444 | 0,72405 | 0,909948 |
| Srsf6 | 197,885 | 190,045 | -0,0583239 | 0,7242 | 0,909948 |
| Arfip1 | 20,7602 | 19,9075 | -0,0605045 | 0,7242 | 0,909948 |
| Armcx1 | 18,5715 | 17,7922 | -0,0618504 | 0,7242 | 0,909948 |
| Dgat1 | 16,3854 | 15,6534 | -0,065943 | 0,72335 | 0,909948 |
| Accs | 5,17509 | 4,93208 | -0,0693885 | 0,72395 | 0,909948 |
| Hhat | 3,61356 | 3,40817 | -0,0844227 | 0,7241 | 0,909948 |
| Mok | 2,57212 | 2,77004 | 0,106951 | 0,7249 | 0,910007 |
| Pfkm | 24,0744 | 25,7573 | 0,097483 | 0,7245 | 0,910007 |
| Coro7 | 7,91022 | 8,38144 | 0,0834815 | 0,7247 | 0,910007 |
| Bcdin3d | 9,47648 | 10,0258 | 0,0812998 | 0,72505 | 0,910007 |
| 2010012O05Rik | 5,22357 | 5,52519 | 0,0809894 | 0,72485 | 0,910007 |
| Ddx55 | 9,20986 | 9,6484 | 0,0671109 | 0,7251 | 0,910007 |
| 6030458C11Rik | 23,3589 | 24,3613 | 0,060619 | 0,7251 | 0,910007 |
| Taf4a | 10,2951 | 10,7315 | 0,0598868 | 0,72455 | 0,910007 |
| Hsbp1 | 79,5774 | 82,7676 | 0,0567065 | 0,725 | 0,910007 |
| Epn1 | 46,8919 | 48,7693 | 0,0566333 | 0,72465 | 0,910007 |
| Psme3 | 118,318 | 123,008 | 0,0560757 | 0,7251 | 0,910007 |
| Ythdf3 | 42,9812 | 41,368 | -0,0551911 | 0,7244 | 0,910007 |
| Kmt2c | 4,74875 | 4,56045 | -0,0583709 | 0,7252 | 0,910007 |
| Pxdn | 87,0672 | 83,601 | -0,0586092 | 0,7252 | 0,910007 |
| Abhd12 | 24,033 | 23,0443 | -0,0606099 | 0,7253 | 0,910047 |
| Zdhhc7 | 12,381 | 11,8512 | -0,0630957 | 0,72545 | 0,91015 |
| Ube2a | 30,8719 | 32,162 | 0,0590638 | 0,72555 | 0,91019 |
| Eml1 | 18,2078 | 19,4701 | 0,0966978 | 0,7257 | 0,910207 |
| Scrn2 | 14,5569 | 15,269 | 0,0689051 | 0,72565 | 0,910207 |

| | | | | | |
|---|---|---|---|---|---|
| Erich2 | 2,30637 | 2,78534 | 0,272225 | 0,726 | 0,910241 |
| Arl5b | 19,7464 | 18,9517 | -0,0592618 | 0,726 | 0,910241 |
| Rnf167 | 27,725 | 26,5581 | -0,0620386 | 0,7259 | 0,910241 |
| Opn3 | 6,32132 | 5,64282 | -0,163811 | 0,72585 | 0,910241 |
| Cpped1 | 6,86093 | 6,52901 | -0,0715402 | 0,7262 | 0,910407 |
| Ino80c | 14,5527 | 13,9345 | -0,0626273 | 0,7263 | 0,910424 |
| Pax2 | 2,58996 | 2,43349 | -0,0899007 | 0,72635 | 0,910424 |
| Smad1 | 15,3485 | 14,7115 | -0,0611554 | 0,7265 | 0,910527 |
| Ywhag | 117,088 | 121,647 | 0,0551076 | 0,72665 | 0,910544 |
| Gpatch3 | 5,69968 | 5,38562 | -0,0817686 | 0,7266 | 0,910544 |
| Brix1 | 48,4389 | 50,3321 | 0,0553125 | 0,72675 | 0,910561 |
| Chst10 | 6,01757 | 5,72757 | -0,0712585 | 0,7268 | 0,910561 |
| Snn | 54,2928 | 52,0153 | -0,0618241 | 0,72695 | 0,910664 |
| Igsf8 | 26,9974 | 28,1245 | 0,059011 | 0,7271 | 0,910766 |
| Fntb | 14,7046 | 15,3599 | 0,0628956 | 0,7273 | 0,910846 |
| Fgfrl1 | 29,9557 | 28,7992 | -0,056803 | 0,7273 | 0,910846 |
| Ppp2r3d | 5,48317 | 5,77901 | 0,075812 | 0,7276 | 0,910966 |
| Snx3 | 65,2288 | 67,8505 | 0,056851 | 0,7275 | 0,910966 |
| Slc25a26 | 4,7435 | 4,00275 | -0,24496 | 0,72755 | 0,910966 |
| Vrk2 | 7,79233 | 8,38664 | 0,106038 | 0,7277 | 0,911005 |
| Dph5 | 13,6247 | 14,2902 | 0,0688011 | 0,72795 | 0,911233 |
| Star | 1,88786 | 2,00747 | 0,0886199 | 0,7282 | 0,911438 |
| Dhx57 | 15,0277 | 15,6473 | 0,0582921 | 0,72825 | 0,911438 |
| 1110012L19Rik | 15,587 | 16,4098 | 0,074212 | 0,7284 | 0,91154 |
| Yrdc | 24,8468 | 25,994 | 0,0651226 | 0,72855 | 0,911643 |
| Snora7a | 124,701 | 313,1 | 1,32815 | 0,72915 | 0,911881 |
| Ly6g5b | 0,633625 | 0,737969 | 0,219932 | 0,72905 | 0,911881 |
| Brca2 | 9,37622 | 9,74811 | 0,0561165 | 0,72915 | 0,911881 |
| Tia1 | 44,8168 | 43,1415 | -0,0549647 | 0,72905 | 0,911881 |
| Ezr | 64,0734 | 61,6759 | -0,0550183 | 0,729 | 0,911881 |
| Slco2b1 | 0,842067 | 0,781652 | -0,107409 | 0,72885 | 0,911881 |
| Gprasp2 | 10,0797 | 9,6626 | -0,0609731 | 0,7294 | 0,912108 |
| Naca | 122,611 | 127,745 | 0,0591818 | 0,72955 | 0,912211 |
| Rpp30 | 37,8535 | 39,4619 | 0,0600334 | 0,7299 | 0,912563 |
| Hey1 | 37,7328 | 39,2495 | 0,0568543 | 0,73005 | 0,912619 |
| Rab6a | 53,4641 | 55,5902 | 0,0562591 | 0,7301 | 0,912619 |
| Ptp4a2 | 86,4553 | 83,2664 | -0,0542207 | 0,73015 | 0,912619 |
| Brd8 | 42,0613 | 45,3408 | 0,108314 | 0,73035 | 0,912784 |
| Rnasek | 32,1676 | 33,8758 | 0,0746478 | 0,73065 | 0,912988 |
| Thbs4 | 0,86079 | 0,792045 | -0,120078 | 0,73065 | 0,912988 |
| Socs3 | 2,7205 | 2,88986 | 0,0871317 | 0,7312 | 0,913504 |
| Ctdsp2 | 48,2264 | 46,414 | -0,055262 | 0,73115 | 0,913504 |
| Drg1 | 115,745 | 124,874 | 0,109523 | 0,73195 | 0,913752 |
| Smim14 | 5,8347 | 6,15125 | 0,0762201 | 0,7319 | 0,913752 |
| Cdc73 | 29,3644 | 30,5 | 0,0547404 | 0,7317 | 0,913752 |
| Kitl | 30,8436 | 29,7065 | -0,0541908 | 0,73215 | 0,913752 |
| Sqstm1 | 46,8705 | 45,0985 | -0,0556019 | 0,7318 | 0,913752 |
| Capn7 | 18,9412 | 18,2057 | -0,0571369 | 0,73215 | 0,913752 |

| | | | | | |
|---|---|---|---|---|---|
| Zfp277 | 16,042 | 15,3746 | -0,0613091 | 0,73205 | 0,913752 |
| Katnb1 | 11,9578 | 11,4261 | -0,0656265 | 0,73215 | 0,913752 |
| Tbc1d24 | 3,8974 | 3,71389 | -0,0695823 | 0,73155 | 0,913752 |
| Fbxl15 | 2,64936 | 2,20988 | -0,261679 | 0,7319 | 0,913752 |
| Pigp | 33,9573 | 26,3683 | -0,364914 | 0,732 | 0,913752 |
| Gal3st3 | 1,41992 | 1,3139 | -0,111953 | 0,73225 | 0,913791 |
| Hacl1 | 2,45475 | 2,61796 | 0,0928722 | 0,73265 | 0,914028 |
| Yeats4 | 88,6777 | 92,084 | 0,0543793 | 0,73285 | 0,914028 |
| Rfwd3 | 23,8384 | 22,9582 | -0,0542722 | 0,73285 | 0,914028 |
| Grpel2 | 11,1382 | 10,6911 | -0,0591083 | 0,73285 | 0,914028 |
| Hnrnpa0 | 289,109 | 277,362 | -0,0598445 | 0,7326 | 0,914028 |
| Mul1 | 9,23425 | 8,84472 | -0,0621779 | 0,73275 | 0,914028 |
| Ndrg2 | 9,64228 | 10,093 | 0,0659118 | 0,733 | 0,914107 |
| Mcoln1 | 15,0222 | 15,6995 | 0,0636229 | 0,73305 | 0,914107 |
| Fopnl | 33,5639 | 34,9464 | 0,0582332 | 0,7332 | 0,914208 |
| Pcdhgb2 | 1,00187 | 1,43331 | 0,516654 | 0,7334 | 0,914372 |
| Lysmd4 | 5,26576 | 5,5267 | 0,0697754 | 0,7336 | 0,914536 |
| Pfkfb3 | 10,8039 | 11,2511 | 0,058514 | 0,7339 | 0,914825 |
| Ctif | 4,5967 | 4,80311 | 0,0633688 | 0,734 | 0,914864 |
| F8 | 0,519392 | 0,557385 | 0,10185 | 0,73415 | 0,914966 |
| Cacna1d | 0,874215 | 1,01248 | 0,211839 | 0,73425 | 0,915005 |
| Pwp2 | 21,167 | 22,0004 | 0,0557175 | 0,7347 | 0,915481 |
| Rcor3 | 7,36972 | 7,76216 | 0,0748475 | 0,7349 | 0,915645 |
| Mark2 | 20,8676 | 21,6711 | 0,0545056 | 0,7351 | 0,915723 |
| Tyw3 | 4,56518 | 4,29076 | -0,0894368 | 0,73505 | 0,915723 |
| Rbbp7 | 99,8408 | 105,052 | 0,0733968 | 0,7358 | 0,915741 |
| Mterfd2 | 8,59385 | 8,9815 | 0,0636527 | 0,73555 | 0,915741 |
| Sesn1 | 10,3782 | 10,8415 | 0,0630082 | 0,7353 | 0,915741 |
| Synm | 4,50703 | 4,69934 | 0,0602803 | 0,7358 | 0,915741 |
| Obsl1 | 19,638 | 20,3937 | 0,0544713 | 0,7358 | 0,915741 |
| Rps4x | 294,21 | 305,217 | 0,0529869 | 0,7355 | 0,915741 |
| Kifc1 | 18,6095 | 17,861 | -0,0592255 | 0,73565 | 0,915741 |
| Mon1a | 4,58327 | 4,32731 | -0,0829084 | 0,73545 | 0,915741 |
| Cenpk | 32,4228 | 30,3638 | -0,0946558 | 0,7353 | 0,915741 |
| Afmid | 1,91768 | 1,6399 | -0,225753 | 0,7355 | 0,915741 |
| Suz12 | 52,5883 | 50,7022 | -0,0526929 | 0,73605 | 0,915882 |
| D630045J12Rik | 9,34049 | 8,99573 | -0,0542579 | 0,736 | 0,915882 |
| Zbtb45 | 9,21695 | 8,79673 | -0,0673217 | 0,73615 | 0,915921 |
| Smg7 | 33,071 | 34,3259 | 0,0537288 | 0,7364 | 0,916147 |
| Bbip1 | 16,9947 | 15,6411 | -0,119742 | 0,7366 | 0,91631 |
| Gfpt1 | 13,8744 | 14,399 | 0,053534 | 0,7367 | 0,916326 |
| Limk2 | 17,288 | 15,4586 | -0,161354 | 0,73675 | 0,916326 |
| Dhps | 31,4377 | 33,1145 | 0,0749706 | 0,737 | 0,916552 |
| 2410021H03Rik | 0,975643 | 2,27921 | 1,22411 | 0,73715 | 0,916653 |
| Pbx4 | 1,99964 | 2,17165 | 0,119052 | 0,73725 | 0,916692 |
| 1110004F10Rik | 81,9836 | 78,9656 | -0,0541097 | 0,73735 | 0,916731 |
| Pet117 | 1,40127 | 3,06611 | 1,12968 | 0,73785 | 0,917136 |
| Ythdc2 | 6,77917 | 7,05585 | 0,0577113 | 0,73795 | 0,917136 |

| | | | | | |
|---|---|---|---|---|---|
| Clip2 | 8,77791 | 8,4297 | -0,0583962 | 0,7379 | 0,917136 |
| Rad51d | 2,90931 | 2,70736 | -0,103788 | 0,7379 | 0,917136 |
| Mir6972 | 241,524 | 366,382 | 0,601182 | 0,7383 | 0,917137 |
| Mnd1 | 4,24972 | 4,58335 | 0,109035 | 0,7381 | 0,917137 |
| Tmsb15l | 11,5158 | 12,3626 | 0,102365 | 0,7385 | 0,917137 |
| Birc2 | 19,8909 | 20,675 | 0,0557791 | 0,7384 | 0,917137 |
| Bub1 | 72,9313 | 75,738 | 0,0544798 | 0,7383 | 0,917137 |
| Gsk3b | 20,699 | 21,4775 | 0,0532663 | 0,7385 | 0,917137 |
| Fbxw11 | 42,302 | 43,8886 | 0,0531197 | 0,7383 | 0,917137 |
| 1110019D14Rik | 1,99932 | 1,84465 | -0,116162 | 0,7384 | 0,917137 |
| Osbpl9 | 21,8427 | 23,6025 | 0,111787 | 0,73875 | 0,917362 |
| Znhit3 | 28,3309 | 30,6716 | 0,11453 | 0,7389 | 0,917426 |
| Slc35e4 | 1,09471 | 1,17883 | 0,106804 | 0,7392 | 0,917426 |
| Mrgbp | 19,0551 | 19,9347 | 0,0651044 | 0,73935 | 0,917426 |
| Crlf3 | 25,6892 | 26,711 | 0,0562684 | 0,73925 | 0,917426 |
| Vps4b | 31,6731 | 30,5274 | -0,0531523 | 0,73935 | 0,917426 |
| 4932438A13Rik | 5,2156 | 5,02119 | -0,0548039 | 0,7391 | 0,917426 |
| Klhl13 | 6,86422 | 6,55728 | -0,0659984 | 0,739 | 0,917426 |
| Snord88c | 179,629 | 108,166 | -0,731777 | 0,73935 | 0,917426 |
| Gtf3c2 | 54,872 | 61,1036 | 0,155188 | 0,7396 | 0,917496 |
| Zfp385b | 1,43214 | 1,53501 | 0,100072 | 0,7397 | 0,917496 |
| Asxl1 | 34,7358 | 36,014 | 0,0521359 | 0,73965 | 0,917496 |
| Npc2 | 52,3467 | 54,2702 | 0,0520625 | 0,73965 | 0,917496 |
| Snx30 | 4,33611 | 4,15857 | -0,060314 | 0,73975 | 0,917496 |
| Adck1 | 10,0905 | 10,5541 | 0,0648129 | 0,74 | 0,917721 |
| Rcn2 | 183,079 | 176,584 | -0,052117 | 0,74025 | 0,917775 |
| Zcchc14 | 18,1608 | 17,4964 | -0,0537682 | 0,74025 | 0,917775 |
| Mlh3 | 5,10867 | 4,89202 | -0,0625186 | 0,7402 | 0,917775 |
| Rtel1 | 22,7066 | 23,5786 | 0,0543633 | 0,7404 | 0,917876 |
| Mir6481 | 13,3252 | 16,0663 | 0,269881 | 0,7405 | 0,917884 |
| Slc30a3 | 0,764556 | 0,847146 | 0,147987 | 0,74065 | 0,917884 |
| Dgkq | 5,23852 | 5,56202 | 0,086449 | 0,7407 | 0,917884 |
| Cdk13 | 16,6724 | 17,2899 | 0,052466 | 0,74075 | 0,917884 |
| Agpat2 | 10,4864 | 9,98604 | -0,0705294 | 0,7406 | 0,917884 |
| Maml1 | 14,5829 | 15,1367 | 0,0537676 | 0,741 | 0,918109 |
| Rad9a | 24,6244 | 27,8443 | 0,177294 | 0,7414 | 0,918326 |
| Cenpe | 27,1076 | 26,154 | -0,0516672 | 0,7414 | 0,918326 |
| Zbtb49 | 4,11688 | 3,9122 | -0,0735699 | 0,7414 | 0,918326 |
| Stx1a | 5,50484 | 5,22088 | -0,0764087 | 0,74145 | 0,918326 |
| Snora28 | 156,197 | 362,312 | 1,21387 | 0,7418 | 0,91862 |
| Mir6992 | 37,7854 | 56,4939 | 0,580266 | 0,74205 | 0,91862 |
| Dact3 | 6,30695 | 6,60785 | 0,067238 | 0,7421 | 0,91862 |
| Ankrd40 | 35,5784 | 36,9681 | 0,0552776 | 0,742 | 0,91862 |
| Slc35b1 | 69,9622 | 72,6889 | 0,0551588 | 0,74185 | 0,91862 |
| Hira | 32,6081 | 33,8073 | 0,0521035 | 0,74195 | 0,91862 |
| Snord16a | 945,733 | 2018,02 | 1,09343 | 0,7424 | 0,918907 |
| Mir697 | 77,8224 | 94,586 | 0,281441 | 0,74405 | 0,919045 |
| Tbc1d17 | 14,204 | 15,5803 | 0,133425 | 0,74305 | 0,919045 |

| | | | | | |
|---|---|---|---|---|---|
| Zfp418 | 1,57973 | 1,69175 | 0,0988429 | 0,7438 | 0,919045 |
| Cklf | 13,5932 | 14,5383 | 0,0969746 | 0,74335 | 0,919045 |
| Snx29 | 1,99504 | 2,12925 | 0,0939304 | 0,7442 | 0,919045 |
| Pacsin3 | 9,43286 | 9,87591 | 0,0662171 | 0,74455 | 0,919045 |
| Mad2l1bp | 17,4506 | 18,2606 | 0,0654567 | 0,7445 | 0,919045 |
| Zfp263 | 13,2602 | 13,8001 | 0,0575725 | 0,74485 | 0,919045 |
| Arhgef10 | 7,45538 | 7,75487 | 0,05682 | 0,74485 | 0,919045 |
| Ahcyl2 | 8,60923 | 8,95299 | 0,0564857 | 0,7434 | 0,919045 |
| Snrnp25 | 57,1041 | 59,3771 | 0,0563134 | 0,74465 | 0,919045 |
| Kbtbd11 | 5,82995 | 6,06128 | 0,0561388 | 0,74285 | 0,919045 |
| Zfp251 | 13,4832 | 14,017 | 0,0560229 | 0,74415 | 0,919045 |
| Cops3 | 115,339 | 119,609 | 0,0524432 | 0,74365 | 0,919045 |
| Wdr11 | 18,3772 | 19,0551 | 0,0522579 | 0,74455 | 0,919045 |
| Zfp706 | 37,3216 | 38,6792 | 0,0515478 | 0,74355 | 0,919045 |
| Srprb | 42,6631 | 41,1536 | -0,0519697 | 0,74425 | 0,919045 |
| Azin1 | 62,5446 | 60,3009 | -0,0527064 | 0,7433 | 0,919045 |
| Aes | 67,9487 | 65,4784 | -0,0534264 | 0,74365 | 0,919045 |
| Ssu72 | 47,0613 | 45,3405 | -0,0537408 | 0,74485 | 0,919045 |
| Cc2d1b | 13,4987 | 12,9812 | -0,0563876 | 0,7439 | 0,919045 |
| Fam92a | 33,619 | 32,3181 | -0,0569347 | 0,74265 | 0,919045 |
| Rasl11b | 28,1079 | 26,9987 | -0,0580829 | 0,7431 | 0,919045 |
| Ccrn4l | 9,28401 | 8,90048 | -0,0608652 | 0,74475 | 0,919045 |
| Fsd1l | 5,22159 | 5,00247 | -0,0618467 | 0,7428 | 0,919045 |
| Pkmyt1 | 19,9956 | 19,156 | -0,0618878 | 0,74415 | 0,919045 |
| Ppard | 6,60007 | 6,30782 | -0,0653403 | 0,74275 | 0,919045 |
| Zfp882 | 3,4126 | 3,26128 | -0,0654334 | 0,74385 | 0,919045 |
| Zfp780b | 3,28833 | 3,14135 | -0,0659709 | 0,74445 | 0,919045 |
| Zfp870 | 3,12761 | 2,9845 | -0,0675745 | 0,7431 | 0,919045 |
| Naa38 | 80,8996 | 76,8699 | -0,0737149 | 0,7429 | 0,919045 |
| Usp27x | 2,64014 | 2,49484 | -0,0816666 | 0,74425 | 0,919045 |
| Idua | 2,95577 | 2,70729 | -0,126682 | 0,74425 | 0,919045 |
| Oaz3 | 0,755901 | 0,661191 | -0,19313 | 0,74325 | 0,919045 |
| Irf2bp2 | 10,4012 | 10,81 | 0,0556122 | 0,7452 | 0,919114 |
| Chuk | 27,7591 | 26,748 | -0,0535324 | 0,7452 | 0,919114 |
| Atf1 | 14,6933 | 14,1007 | -0,059392 | 0,74525 | 0,919114 |
| Ring1 | 13,1795 | 12,6349 | -0,0608808 | 0,74505 | 0,919114 |
| Dok3 | 2,73083 | 2,54332 | -0,102626 | 0,745 | 0,919114 |
| Amd1,Amd2 | 4,24342 | 5,19767 | 0,292637 | 0,7456 | 0,919307 |
| Slx4 | 9,96509 | 10,3403 | 0,0533173 | 0,7457 | 0,919307 |
| Fbxo33 | 7,30195 | 6,99238 | -0,0624978 | 0,7455 | 0,919307 |
| Lrfn4 | 34,2522 | 32,7033 | -0,06676 | 0,74565 | 0,919307 |
| Fbxl2 | 1,74073 | 1,37999 | -0,335035 | 0,74575 | 0,919307 |
| Pih1d2 | 2,28361 | 2,89366 | 0,34158 | 0,746 | 0,919437 |
| Zeb2os | 2,76188 | 3,35169 | 0,279236 | 0,7461 | 0,919437 |
| Rprd1b | 22,8082 | 23,6466 | 0,0520794 | 0,7462 | 0,919437 |
| Atl2 | 39,591 | 38,1906 | -0,0519539 | 0,746 | 0,919437 |
| Fcf1 | 37,0996 | 35,4673 | -0,0649173 | 0,74615 | 0,919437 |
| BC049715 | 0,727119 | 0,841791 | 0,211269 | 0,74645 | 0,919661 |

| | | | | | |
|---|---|---|---|---|---|
| 9430016H08Rik | 27,7422 | 29,1032 | 0,0690949 | 0,74655 | 0,919699 |
| Has2os | 3,36395 | 5,35252 | 0,670062 | 0,74665 | 0,919706 |
| Gadd45gip1 | 31,9441 | 33,6396 | 0,0746101 | 0,7469 | 0,919706 |
| Slc25a15 | 10,2672 | 10,6834 | 0,0573386 | 0,7467 | 0,919706 |
| Psme4 | 53,9499 | 52,0644 | -0,0513228 | 0,7469 | 0,919706 |
| Lpcat4 | 8,27038 | 7,88555 | -0,0687419 | 0,7468 | 0,919706 |
| Noxred1 | 0,622228 | 0,705495 | 0,181194 | 0,7477 | 0,920352 |
| Pde7b | 0,541725 | 0,588235 | 0,118831 | 0,74765 | 0,920352 |
| Scfd1 | 26,2956 | 27,3173 | 0,0549915 | 0,7476 | 0,920352 |
| Mafg | 19,2201 | 18,4626 | -0,0580076 | 0,7477 | 0,920352 |
| Csf1 | 9,0126 | 9,39443 | 0,0598628 | 0,7484 | 0,920512 |
| Npm1 | 608,257 | 631,016 | 0,0529956 | 0,74845 | 0,920512 |
| Ercc6l2 | 15,9202 | 16,5049 | 0,0520352 | 0,7482 | 0,920512 |
| Heph | 25,3918 | 24,5093 | -0,0510346 | 0,74815 | 0,920512 |
| Lmtk2 | 9,90618 | 9,54987 | -0,0528466 | 0,74815 | 0,920512 |
| Oraov1 | 10,814 | 10,3319 | -0,065786 | 0,74845 | 0,920512 |
| Nol7 | 43,0231 | 40,7613 | -0,0779141 | 0,74825 | 0,920512 |
| Rapgefl1 | 2,07251 | 1,96008 | -0,0804638 | 0,74795 | 0,920512 |
| Ddc | 1,27032 | 1,17082 | -0,117672 | 0,7483 | 0,920512 |
| Plekhf1 | 3,25493 | 3,04248 | -0,0973827 | 0,74855 | 0,92055 |
| Ralgps1 | 1,60582 | 1,44987 | -0,147384 | 0,74865 | 0,920589 |
| AI414108 | 1,69736 | 1,60461 | -0,0810698 | 0,74905 | 0,920996 |
| Col25a1 | 1,31639 | 1,24945 | -0,0752939 | 0,74935 | 0,92128 |
| Kcnd1 | 0,905404 | 0,989036 | 0,127462 | 0,74945 | 0,921318 |
| Nxph3 | 3,0524 | 2,87027 | -0,0887573 | 0,74965 | 0,921479 |
| Samm50 | 72,462 | 75,0591 | 0,0508022 | 0,7503 | 0,922193 |
| Zfp939 | 1,7732 | 1,90395 | 0,102637 | 0,7504 | 0,922232 |
| Ybx3 | 140,057 | 145,018 | 0,0502198 | 0,7506 | 0,922392 |
| Gid8 | 21,4463 | 20,696 | -0,0513766 | 0,75085 | 0,92253 |
| Krtcap3 | 5,44723 | 2,26739 | -1,26449 | 0,7508 | 0,92253 |
| Zmym3 | 17,9292 | 18,7501 | 0,0645917 | 0,751 | 0,922583 |
| Pabpc4 | 21,0789 | 20,2909 | -0,0549688 | 0,7511 | 0,922583 |
| Zfp72 | 2,908 | 2,72545 | -0,0935342 | 0,75105 | 0,922583 |
| Snord55 | 255,971 | 362,959 | 0,503826 | 0,75135 | 0,92272 |
| Elovl6 | 32,0725 | 33,2214 | 0,0507784 | 0,7513 | 0,92272 |
| Ppp1r12b | 2,61834 | 2,76176 | 0,0769355 | 0,75165 | 0,922919 |
| Samd1 | 31,4227 | 32,6152 | 0,05374 | 0,7516 | 0,922919 |
| Foxk1 | 9,93979 | 10,3082 | 0,0525028 | 0,7518 | 0,923018 |
| Gcat | 16,8226 | 17,4996 | 0,0569222 | 0,752 | 0,923179 |
| Gtpbp3 | 11,2176 | 11,6886 | 0,0593345 | 0,7521 | 0,923217 |
| Il15ra | 0,532723 | 0,596236 | 0,162497 | 0,7524 | 0,923403 |
| Rwdd2b | 5,29385 | 5,56458 | 0,0719565 | 0,75235 | 0,923403 |
| AW209491 | 13,0071 | 13,5615 | 0,0602162 | 0,7527 | 0,923403 |
| Senp5 | 20,5175 | 21,241 | 0,0499928 | 0,753 | 0,923403 |
| Akt1 | 110,345 | 114,199 | 0,049532 | 0,7529 | 0,923403 |
| Atp1a1 | 131,231 | 126,719 | -0,0504788 | 0,7531 | 0,923403 |
| Zfx | 10,4003 | 10,0324 | -0,0519568 | 0,75315 | 0,923403 |
| Pole4 | 23,4553 | 22,567 | -0,0556971 | 0,7525 | 0,923403 |

| | | | | | |
|---|---|---|---|---|---|
| Snx10 | 10,1155 | 9,70304 | -0,0600646 | 0,75305 | 0,923403 |
| Tpcn2 | 4,53247 | 4,32085 | -0,0689815 | 0,7527 | 0,923403 |
| Psmb10 | 8,57962 | 8,13786 | -0,0762643 | 0,75255 | 0,923403 |
| Mycbpap | 0,943152 | 0,873998 | -0,10986 | 0,753 | 0,923403 |
| Ccdc38 | 0,782073 | 0,672747 | -0,217239 | 0,75275 | 0,923403 |
| Jmjd8 | 32,2306 | 30,2194 | -0,092957 | 0,75325 | 0,923441 |
| Mtif3 | 4,77926 | 5,15895 | 0,110289 | 0,7535 | 0,92364 |
| Mtmr7 | 4,93363 | 4,29191 | -0,20103 | 0,75355 | 0,92364 |
| Farp2 | 2,40436 | 2,74073 | 0,188906 | 0,75385 | 0,923753 |
| Efs | 10,1164 | 10,514 | 0,0556144 | 0,75385 | 0,923753 |
| Phkb | 14,9867 | 15,5331 | 0,0516583 | 0,75375 | 0,923753 |
| Pex5l | 0,550604 | 0,609656 | 0,146982 | 0,7541 | 0,923928 |
| Rps6ka6 | 31,1778 | 32,2613 | 0,0492811 | 0,7542 | 0,923928 |
| Ubr5 | 40,1744 | 41,5421 | 0,0482977 | 0,7542 | 0,923928 |
| Ndufv2 | 73,8929 | 76,5002 | 0,0500275 | 0,7543 | 0,923942 |
| Adk | 53,1948 | 51,3568 | -0,050729 | 0,75435 | 0,923942 |
| Nfatc4 | 21,9607 | 22,7674 | 0,0520439 | 0,75445 | 0,92398 |
| Iah1 | 25,0031 | 27,484 | 0,136488 | 0,75465 | 0,92414 |
| Acvr1 | 12,1915 | 12,6792 | 0,0565882 | 0,75485 | 0,9243 |
| Pcdhga12 | 1,05919 | 1,44874 | 0,451842 | 0,75585 | 0,924339 |
| Fxyd1 | 0,800286 | 0,936064 | 0,226092 | 0,75575 | 0,924339 |
| Ccdc85c | 9,22638 | 9,66216 | 0,0665815 | 0,7553 | 0,924339 |
| Cwc22 | 13,4605 | 13,9758 | 0,0542007 | 0,75565 | 0,924339 |
| Atp2c1 | 48,1346 | 49,9318 | 0,052884 | 0,7552 | 0,924339 |
| Eif4h | 231,201 | 239,57 | 0,0513004 | 0,75565 | 0,924339 |
| Pgam5 | 50,2841 | 52,0707 | 0,0503694 | 0,75575 | 0,924339 |
| Wwp2 | 18,2019 | 18,8483 | 0,0503408 | 0,7551 | 0,924339 |
| Spag9 | 24,2329 | 25,0815 | 0,0496578 | 0,7558 | 0,924339 |
| Ccnb2 | 82,3702 | 79,6126 | -0,0491263 | 0,7556 | 0,924339 |
| Inpp5b | 10,7659 | 10,3593 | -0,0555393 | 0,7553 | 0,924339 |
| Nudt18 | 1,03165 | 0,963399 | -0,098749 | 0,75515 | 0,924339 |
| Slc1a6 | 1,38512 | 1,28314 | -0,110327 | 0,75545 | 0,924339 |
| Syap1 | 27,3765 | 24,989 | -0,131643 | 0,7557 | 0,924339 |
| G6pc3 | 18,9706 | 18,226 | -0,0577663 | 0,756 | 0,924438 |
| Kbtbd4 | 16,2976 | 16,9251 | 0,0545076 | 0,7561 | 0,924476 |
| Tfb1m | 13,7238 | 14,4667 | 0,0760591 | 0,75655 | 0,924772 |
| Tmem238 | 1,76796 | 1,60991 | -0,135103 | 0,75655 | 0,924772 |
| 4933408B17Rik | 0,683033 | 0,621004 | -0,137352 | 0,75655 | 0,924772 |
| Pdzd9 | 0,253328 | 0,564532 | 1,15605 | 0,7569 | 0,924862 |
| Wdr76 | 12,8814 | 12,4184 | -0,0528107 | 0,7569 | 0,924862 |
| Tsc22d4 | 22,007 | 21,2132 | -0,0529982 | 0,75685 | 0,924862 |
| Tmem139 | 1,31204 | 1,18916 | -0,141868 | 0,7568 | 0,924862 |
| 2410004B18Rik | 26,2981 | 27,4473 | 0,0617011 | 0,757 | 0,924899 |
| Tmem170b | 2,56015 | 2,67435 | 0,0629597 | 0,75725 | 0,924989 |
| BC055324 | 26,3028 | 27,3989 | 0,0589014 | 0,7573 | 0,924989 |
| Sox11 | 33,3374 | 34,4935 | 0,0491833 | 0,75735 | 0,924989 |
| Ankrd28 | 21,8436 | 22,5874 | 0,0483078 | 0,75725 | 0,924989 |
| Stx5a | 40,6259 | 42,0344 | 0,04917 | 0,75745 | 0,925026 |

| | | | | | |
|---|---|---|---|---|---|
| Olfm2 | 1,96704 | 1,82472 | -0,108347 | 0,7577 | 0,925247 |
| Mrpl3 | 61,1575 | 63,309 | 0,0498798 | 0,75825 | 0,925665 |
| Fam76b | 19,6463 | 18,9763 | -0,0500638 | 0,7582 | 0,925665 |
| Abhd8 | 19,6379 | 18,9127 | -0,0542849 | 0,7582 | 0,925665 |
| Snora5c | 62,0491 | 107,157 | 0,788248 | 0,75945 | 0,925668 |
| 4930549G23Rik | 2,66625 | 4,35903 | 0,709197 | 0,7587 | 0,925668 |
| Ttll3 | 1,9007 | 2,34393 | 0,302397 | 0,75965 | 0,925668 |
| Rsg1 | 1,46606 | 1,62777 | 0,150954 | 0,7602 | 0,925668 |
| Ptcd1 | 8,36918 | 8,99578 | 0,104162 | 0,75845 | 0,925668 |
| Pdcd4 | 24,9522 | 26,5465 | 0,0893566 | 0,75885 | 0,925668 |
| Rad23a | 27,9986 | 29,3762 | 0,0692958 | 0,75905 | 0,925668 |
| Gtpbp10 | 6,96782 | 7,2671 | 0,0606713 | 0,75875 | 0,925668 |
| Homer2 | 1,69727 | 1,77007 | 0,060593 | 0,7603 | 0,925668 |
| Ptprj | 5,93022 | 6,17357 | 0,0580206 | 0,7598 | 0,925668 |
| Suv39h2 | 23,7004 | 24,588 | 0,0530433 | 0,76005 | 0,925668 |
| Mtmr2 | 29,4548 | 30,4739 | 0,0490729 | 0,7594 | 0,925668 |
| Ctcf | 64,8438 | 67,0429 | 0,0481157 | 0,7593 | 0,925668 |
| Rab1b | 77,4697 | 74,8913 | -0,0488343 | 0,7604 | 0,925668 |
| Miip | 24,4688 | 23,5815 | -0,0532867 | 0,76015 | 0,925668 |
| P4hb | 364,705 | 351,364 | -0,0537656 | 0,7585 | 0,925668 |
| Taf5 | 12,1464 | 11,7018 | -0,0537999 | 0,76 | 0,925668 |
| Flna | 155,341 | 149,551 | -0,0548039 | 0,7596 | 0,925668 |
| Scoc | 14,5018 | 13,9323 | -0,0577958 | 0,75845 | 0,925668 |
| Tirap | 5,01984 | 4,8186 | -0,0590277 | 0,7603 | 0,925668 |
| Zfp787 | 13,054 | 12,5265 | -0,0595148 | 0,75925 | 0,925668 |
| Osbpl3 | 2,87268 | 2,75265 | -0,0615719 | 0,7595 | 0,925668 |
| Wnk2 | 1,33008 | 1,26331 | -0,0743027 | 0,7593 | 0,925668 |
| Zfp784 | 3,46269 | 3,27628 | -0,0798346 | 0,75975 | 0,925668 |
| Setmar | 4,95073 | 4,68062 | -0,0809425 | 0,75885 | 0,925668 |
| Tdrp | 1,83175 | 1,71924 | -0,0914453 | 0,75885 | 0,925668 |
| Tstd3 | 11,3836 | 10,6041 | -0,102337 | 0,7604 | 0,925668 |
| Tmem179b | 29,3707 | 27,2458 | -0,108342 | 0,7592 | 0,925668 |
| Zglp1 | 2,07179 | 1,89641 | -0,127606 | 0,7594 | 0,925668 |
| 3110007F17Rik | 0,744288 | 0,643805 | -0,209237 | 0,7603 | 0,925668 |
| Atraid | 31,4588 | 21,6328 | -0,540246 | 0,7601 | 0,925668 |
| Pomk | 5,81691 | 6,06854 | 0,0610947 | 0,76095 | 0,926085 |
| Pdf | 33,1555 | 34,3749 | 0,0521069 | 0,7609 | 0,926085 |
| Acads | 35,1823 | 36,4371 | 0,0505606 | 0,76095 | 0,926085 |
| Bzw2 | 124,159 | 128,395 | 0,0483929 | 0,76105 | 0,926122 |
| Trim12a | 0,560172 | 0,632078 | 0,174234 | 0,7618 | 0,926266 |
| Lxn | 29,0525 | 31,2989 | 0,107447 | 0,7619 | 0,926266 |
| Rchy1 | 22,3673 | 23,3675 | 0,0631122 | 0,7619 | 0,926266 |
| Pcbp3 | 10,9453 | 11,4016 | 0,0589237 | 0,7616 | 0,926266 |
| Mrpl35 | 10,6949 | 11,1012 | 0,0537946 | 0,76195 | 0,926266 |
| Slc4a7 | 14,9 | 15,4152 | 0,049045 | 0,76175 | 0,926266 |
| Sema4c | 35,335 | 36,5482 | 0,0487021 | 0,76185 | 0,926266 |
| Xpot | 23,1065 | 23,8876 | 0,0479603 | 0,7618 | 0,926266 |
| Arih1 | 31,8454 | 30,8155 | -0,047429 | 0,76195 | 0,926266 |

| | | | | | |
|---|---|---|---|---|---|
| Ubac2 | 30,7202 | 29,6548 | -0,0509218 | 0,7613 | 0,926266 |
| Tardbp | 126,649 | 122,171 | -0,0519401 | 0,7617 | 0,926266 |
| Gmfg | 0,670143 | 0,585773 | -0,194127 | 0,762 | 0,926266 |
| Ctdp1 | 17,408 | 18,0308 | 0,0507149 | 0,7621 | 0,92628 |
| Apaf1 | 10,3214 | 9,97363 | -0,0494445 | 0,76215 | 0,92628 |
| Gm10658 | 1,06895 | 1,52912 | 0,516504 | 0,76245 | 0,926284 |
| Mgll | 1,54965 | 1,64255 | 0,0839947 | 0,7623 | 0,926284 |
| Mrps34 | 40,5704 | 42,176 | 0,0559915 | 0,76245 | 0,926284 |
| Ephb4 | 71,6308 | 74,1003 | 0,0488995 | 0,7625 | 0,926284 |
| Pde5a | 7,91384 | 7,64124 | -0,0505696 | 0,7624 | 0,926284 |
| Acsm3 | 1,02156 | 1,19793 | 0,229765 | 0,7632 | 0,926451 |
| Cmtr1 | 35,4443 | 36,8273 | 0,0552216 | 0,76295 | 0,926451 |
| Epn2 | 17,9629 | 18,583 | 0,048963 | 0,7632 | 0,926451 |
| Cnn3 | 159,985 | 165,432 | 0,048298 | 0,76285 | 0,926451 |
| Mrps28 | 48,7375 | 46,8488 | -0,0570209 | 0,76285 | 0,926451 |
| Klhl28 | 4,92993 | 4,71052 | -0,0656795 | 0,7633 | 0,926451 |
| Myom2 | 1,32756 | 1,25361 | -0,0826935 | 0,76325 | 0,926451 |
| Mak | 0,607779 | 0,559376 | -0,119729 | 0,76335 | 0,926451 |
| Pcdhgb7 | 3,25366 | 2,68814 | -0,275458 | 0,7629 | 0,926451 |
| Zfp809 | 6,4753 | 5,21825 | -0,311379 | 0,7631 | 0,926451 |
| Tymp | 0,802044 | 0,528167 | -0,602686 | 0,7634 | 0,926451 |
| Mir1957b | 13,2298 | 20,0447 | 0,599434 | 0,7642 | 0,926496 |
| Gemin8 | 4,21148 | 4,44408 | 0,077558 | 0,76385 | 0,926496 |
| Mmp16 | 4,29458 | 4,47798 | 0,0603324 | 0,7639 | 0,926496 |
| Zfp952 | 7,46297 | 7,78141 | 0,0602817 | 0,76405 | 0,926496 |
| Prkd2 | 6,30412 | 6,55807 | 0,0569781 | 0,7639 | 0,926496 |
| Rbak | 8,27534 | 8,60236 | 0,0559141 | 0,76415 | 0,926496 |
| Hars2 | 16,6254 | 17,2279 | 0,0513524 | 0,76415 | 0,926496 |
| Rreb1 | 7,7454 | 8,01963 | 0,0501973 | 0,7636 | 0,926496 |
| Bcor | 30,3646 | 31,376 | 0,0472673 | 0,76375 | 0,926496 |
| Acap2 | 12,8762 | 12,2375 | -0,0733952 | 0,7638 | 0,926496 |
| Hs3st1 | 1,65902 | 1,53999 | -0,107418 | 0,764 | 0,926496 |
| Nrg4 | 1,00965 | 1,26895 | 0,32978 | 0,7643 | 0,926533 |
| Ppp1r3f | 1,79904 | 1,98039 | 0,138559 | 0,7644 | 0,926571 |
| Gskip | 8,09999 | 8,43546 | 0,0585468 | 0,7646 | 0,926729 |
| 1700086O06Rik | 5,12438 | 4,50445 | -0,186028 | 0,7648 | 0,926887 |
| Abhd4 | 13,078 | 12,591 | -0,0547541 | 0,7649 | 0,926925 |
| Tmem106a | 0,937331 | 1,01291 | 0,11187 | 0,7652 | 0,927012 |
| Mcee | 19,3098 | 20,2448 | 0,0682224 | 0,76515 | 0,927012 |
| Cask | 10,4921 | 10,1444 | -0,048619 | 0,76525 | 0,927012 |
| Mif4gd | 8,50821 | 8,10382 | -0,070255 | 0,76515 | 0,927012 |
| AU041133 | 2,88927 | 3,04799 | 0,0771568 | 0,76575 | 0,92703 |
| Itprip | 16,9292 | 16,3586 | -0,0494644 | 0,76545 | 0,92703 |
| Mlec | 137,314 | 132,475 | -0,0517549 | 0,76565 | 0,92703 |
| Pias1 | 11,0749 | 10,6801 | -0,0523633 | 0,7656 | 0,92703 |
| Chml | 8,44147 | 8,12776 | -0,0546364 | 0,7657 | 0,92703 |
| Tmem161b | 9,19702 | 8,8416 | -0,0568578 | 0,7657 | 0,92703 |
| Tnni3 | 2,54932 | 2,33735 | -0,125239 | 0,7657 | 0,92703 |

| | | | | | |
|---|---|---|---|---|---|
| Prkar1b | 0,851544 | 0,918966 | 0,10993 | 0,7661 | 0,927038 |
| Mreg | 1,70192 | 1,81246 | 0,0907932 | 0,7664 | 0,927038 |
| Dnajc28 | 1,89119 | 1,99492 | 0,0770352 | 0,76635 | 0,927038 |
| Tldc1 | 4,73551 | 4,9549 | 0,0653369 | 0,76605 | 0,927038 |
| Coa6 | 22,2049 | 23,2204 | 0,0645103 | 0,7661 | 0,927038 |
| Rft1 | 9,87835 | 10,2792 | 0,0573797 | 0,7659 | 0,927038 |
| Pwp1 | 45,2327 | 46,7551 | 0,0477577 | 0,76625 | 0,927038 |
| Tnrc6a | 30,209 | 29,251 | -0,0464948 | 0,76645 | 0,927038 |
| Glrx5 | 88,1548 | 85,2373 | -0,0485534 | 0,76615 | 0,927038 |
| 4933412E12Rik | 1,72562 | 1,60095 | -0,108186 | 0,76645 | 0,927038 |
| Mir8102 | 16,4088 | 21,5099 | 0,390527 | 0,76745 | 0,927137 |
| Gm11837 | 0,599742 | 0,742538 | 0,308124 | 0,7678 | 0,927137 |
| 1700001G17Rik | 1,18387 | 1,32271 | 0,159996 | 0,76705 | 0,927137 |
| Kti12 | 34,7795 | 36,5293 | 0,0708193 | 0,76785 | 0,927137 |
| Zkscan7 | 5,70378 | 5,96664 | 0,0649985 | 0,7677 | 0,927137 |
| Stra13 | 37,8724 | 39,487 | 0,0602308 | 0,7674 | 0,927137 |
| Rabgef1 | 8,53805 | 8,87847 | 0,0564042 | 0,767 | 0,927137 |
| Bcl2l1 | 20,7364 | 21,524 | 0,0537779 | 0,7675 | 0,927137 |
| Sik3 | 8,84034 | 9,15645 | 0,050687 | 0,76765 | 0,927137 |
| Emilin2 | 12,7751 | 13,2314 | 0,0506251 | 0,7677 | 0,927137 |
| Ewsr1 | 191,011 | 197,635 | 0,0491824 | 0,7675 | 0,927137 |
| Ap1b1 | 22,3372 | 23,0902 | 0,0478326 | 0,76695 | 0,927137 |
| Tubgcp3 | 28,888 | 29,8358 | 0,0465753 | 0,76735 | 0,927137 |
| Cyc1 | 158,672 | 153,659 | -0,0463191 | 0,76725 | 0,927137 |
| Ppp4r2 | 15,5242 | 14,9818 | -0,0512996 | 0,7668 | 0,927137 |
| Mccc2 | 17,1864 | 16,5702 | -0,0526704 | 0,76725 | 0,927137 |
| Wdr81 | 3,37637 | 3,24499 | -0,0572576 | 0,7669 | 0,927137 |
| 3110040N11Rik | 12,1379 | 11,5305 | -0,074069 | 0,7667 | 0,927137 |
| 2210015D19Rik | 6,441 | 6,00298 | -0,101606 | 0,76785 | 0,927137 |
| 9330020H09Rik | 1,17783 | 1,32277 | 0,167434 | 0,7681 | 0,927187 |
| Erich1 | 6,41106 | 6,73471 | 0,0710538 | 0,76805 | 0,927187 |
| Slc35a4 | 34,3264 | 33,1219 | -0,0515344 | 0,768 | 0,927187 |
| Spin2c | 9,72834 | 9,28036 | -0,068013 | 0,76835 | 0,927405 |
| 6330408A02Rik | 3,3581 | 3,90665 | 0,218287 | 0,76885 | 0,92759 |
| Pgm3 | 7,20024 | 7,56161 | 0,07065 | 0,7688 | 0,92759 |
| Thoc1 | 36,3811 | 38,1842 | 0,069786 | 0,7688 | 0,92759 |
| Gramd1a | 37,1269 | 38,3539 | 0,04691 | 0,7686 | 0,92759 |
| Vps18 | 9,47641 | 9,13654 | -0,052692 | 0,76875 | 0,92759 |
| 1700113A16Rik | 5,81409 | 5,47363 | -0,0870555 | 0,76905 | 0,927747 |
| Wdr25 | 2,55086 | 2,69279 | 0,0781172 | 0,76945 | 0,928039 |
| Otud6b | 19,5128 | 20,1744 | 0,0481005 | 0,7694 | 0,928039 |
| Ankrd13c | 16,3216 | 15,7589 | -0,0506103 | 0,7695 | 0,928039 |
| Gamt | 5,6163 | 7,58187 | 0,432934 | 0,7706 | 0,928146 |
| 1700102P08Rik | 0,615853 | 0,696252 | 0,177023 | 0,7703 | 0,928146 |
| Tax1bp3 | 0,683112 | 0,760046 | 0,153964 | 0,7699 | 0,928146 |
| Rab4b | 17,7975 | 18,5334 | 0,0584559 | 0,7707 | 0,928146 |
| Hsp90b1 | 617,503 | 641,907 | 0,0559195 | 0,7704 | 0,928146 |
| Mri1 | 13,1945 | 13,692 | 0,0534014 | 0,77035 | 0,928146 |

| | | | | | |
|---|---|---|---|---|---|
| Aars2 | 13,5418 | 14,0156 | 0,0496147 | 0,77065 | 0,928146 |
| Zfp523 | 20,4312 | 21,1427 | 0,0493883 | 0,7706 | 0,928146 |
| Sf3b1 | 105,694 | 109,309 | 0,0485113 | 0,77035 | 0,928146 |
| Exosc8 | 86,4654 | 89,3216 | 0,0468866 | 0,7703 | 0,928146 |
| Zfp362 | 66,7073 | 64,6048 | -0,0462046 | 0,7702 | 0,928146 |
| Cops7a | 51,3491 | 49,66 | -0,0482524 | 0,77005 | 0,928146 |
| Pstk | 26,1877 | 25,1972 | -0,0556247 | 0,77025 | 0,928146 |
| 0610011F06Rik | 23,8151 | 22,8166 | -0,0617889 | 0,77025 | 0,928146 |
| 2310009A05Rik | 24,1308 | 22,9156 | -0,0745463 | 0,76985 | 0,928146 |
| Tagln | 8,67765 | 7,8586 | -0,143032 | 0,77045 | 0,928146 |
| Gmpr2 | 34,8559 | 36,0701 | 0,0494023 | 0,7708 | 0,928159 |
| Tsc1 | 12,2641 | 12,6679 | 0,046742 | 0,77085 | 0,928159 |
| Polr3e | 21,9526 | 22,6914 | 0,0477575 | 0,77095 | 0,928196 |
| Gdi1 | 47,4157 | 48,918 | 0,0450003 | 0,7711 | 0,928246 |
| Nat8l | 1,61738 | 1,54427 | -0,0667411 | 0,7712 | 0,928246 |
| Zfp235 | 2,83228 | 2,69957 | -0,0692349 | 0,7712 | 0,928246 |
| Nfs1 | 12,7329 | 13,3889 | 0,0724747 | 0,77175 | 0,928574 |
| Tma7 | 58,0378 | 60,2541 | 0,054066 | 0,77165 | 0,928574 |
| Dcaf10 | 7,42797 | 7,68506 | 0,0490888 | 0,77175 | 0,928574 |
| Hras | 22,2516 | 21,492 | -0,0501088 | 0,7717 | 0,928574 |
| Ugp2 | 37,9 | 39,6604 | 0,0655014 | 0,772 | 0,928707 |
| Hunk | 21,9086 | 22,6145 | 0,0457478 | 0,77195 | 0,928707 |
| Morf4l1 | 22,9345 | 22,1511 | -0,0501414 | 0,7721 | 0,928744 |
| Vps51 | 15,8524 | 16,486 | 0,0565398 | 0,7727 | 0,92888 |
| Atg3 | 14,0251 | 14,5565 | 0,0536552 | 0,77265 | 0,92888 |
| Smarce1 | 60,7174 | 62,6401 | 0,0449749 | 0,77235 | 0,92888 |
| Casp8ap2 | 22,2298 | 22,9286 | 0,0446503 | 0,7725 | 0,92888 |
| Mcfd2 | 31,3943 | 30,3596 | -0,0483468 | 0,77265 | 0,92888 |
| Ndufa6 | 140,873 | 136,085 | -0,0498859 | 0,77255 | 0,92888 |
| Ttc32 | 9,25901 | 8,79497 | -0,0741787 | 0,7727 | 0,92888 |
| 4833420G17Rik | 26,9632 | 27,8753 | 0,0479934 | 0,7728 | 0,928917 |
| Socs1 | 2,46083 | 2,6301 | 0,0959709 | 0,7733 | 0,929233 |
| Tmem81 | 1,8627 | 1,98858 | 0,0943409 | 0,7734 | 0,929233 |
| Fyttd1 | 39,9635 | 41,2396 | 0,0453468 | 0,77325 | 0,929233 |
| Aldh18a1 | 40,0302 | 41,3048 | 0,0452194 | 0,7735 | 0,929233 |
| Git2 | 8,66388 | 8,36855 | -0,050035 | 0,7735 | 0,929233 |
| Dpp7 | 6,73243 | 6,42379 | -0,0677019 | 0,77325 | 0,929233 |
| Megf10 | 0,675296 | 0,637881 | -0,0822337 | 0,77355 | 0,929233 |
| Phf20l1 | 12,9258 | 12,5227 | -0,0457104 | 0,77395 | 0,929547 |
| Akap11 | 23,5913 | 22,8549 | -0,0457503 | 0,77395 | 0,929547 |
| Zc3h6 | 1,43355 | 1,50979 | 0,0747561 | 0,7743 | 0,929633 |
| Gk5 | 2,83425 | 2,98212 | 0,0733717 | 0,7743 | 0,929633 |
| Commd2 | 15,1867 | 15,7102 | 0,0488936 | 0,7741 | 0,929633 |
| Mrps22 | 49,2812 | 50,9293 | 0,0474599 | 0,77425 | 0,929633 |
| Ddx19b | 9,53581 | 8,92901 | 0,0462441 | 0,77445 | 0,92973 |
| Ltn1 | 15,9433 | 15,4447 | -0,045842 | 0,77465 | 0,929886 |
| Srsf2 | 348,613 | 360,251 | 0,0473782 | 0,7748 | 0,929983 |
| Nsl1 | 20,1176 | 20,8 | 0,0481239 | 0,7749 | 0,929996 |

| | | | | | |
|---|---|---|---|---|---|
| Gmds | 13,1992 | 12,6698 | -0,0590568 | 0,77495 | 0,929996 |
| Npdc1 | 3,96038 | 3,73927 | -0,0828825 | 0,77505 | 0,930032 |
| Ldhal6b | 0,519903 | 0,713434 | 0,456539 | 0,7757 | 0,930146 |
| Shc4 | 0,742545 | 0,915082 | 0,301423 | 0,7755 | 0,930146 |
| LOC100504608 | 0,705716 | 0,822492 | 0,220914 | 0,776 | 0,930146 |
| Tollip | 24,5429 | 25,3515 | 0,0467636 | 0,7757 | 0,930146 |
| Rbx1 | 54,1032 | 55,884 | 0,0467234 | 0,77605 | 0,930146 |
| Sdccag3 | 48,1015 | 49,6794 | 0,0465663 | 0,77595 | 0,930146 |
| Bcas2 | 83,7523 | 86,4972 | 0,0465249 | 0,7755 | 0,930146 |
| Atp13a1 | 33,4723 | 34,542 | 0,0453859 | 0,77545 | 0,930146 |
| Gnaq | 19,9229 | 20,5496 | 0,0446828 | 0,77585 | 0,930146 |
| Opa1 | 21,0328 | 21,6826 | 0,0438989 | 0,7756 | 0,930146 |
| Cit | 7,08984 | 6,85434 | -0,0487363 | 0,77585 | 0,930146 |
| Fbxo8 | 4,89542 | 4,6983 | -0,0592965 | 0,77575 | 0,930146 |
| Pabpc4l | 2,17238 | 2,07659 | -0,0650614 | 0,7757 | 0,930146 |
| D230030E09Rik | 5,28401 | 4,96141 | -0,090883 | 0,77625 | 0,930303 |
| Tspo | 25,3297 | 24,3536 | -0,0566964 | 0,77705 | 0,931178 |
| Gipc1 | 51,8419 | 50,2165 | -0,045956 | 0,7772 | 0,931274 |
| Uggt1 | 26,5276 | 27,3582 | 0,0444798 | 0,77735 | 0,931287 |
| Acadvl | 17,3356 | 16,7553 | -0,0491186 | 0,77735 | 0,931287 |
| Sdc3 | 17,678 | 18,2451 | 0,0455592 | 0,77745 | 0,931323 |
| Prkab2 | 5,36709 | 5,16701 | -0,0548093 | 0,77775 | 0,931516 |
| Uxt | 5,69337 | 5,25287 | -0,116176 | 0,7777 | 0,931516 |
| 4933431E20Rik | 3,99237 | 3,68449 | -0,115782 | 0,7779 | 0,931612 |
| Fndc3a | 21,2653 | 21,9371 | 0,0448671 | 0,7781 | 0,931768 |
| Nckap5l | 13,8151 | 14,2632 | 0,0460555 | 0,7784 | 0,93202 |
| Rhobtb3 | 42,2765 | 40,9827 | -0,0448404 | 0,77845 | 0,93202 |
| Nr2c2 | 9,22456 | 8,81462 | -0,0655827 | 0,77855 | 0,932056 |
| Trnau1ap | 33,2356 | 34,4212 | 0,050568 | 0,7787 | 0,932152 |
| 6430562O15Rik | 0,444212 | 0,620294 | 0,481705 | 0,779 | 0,932261 |
| Asna1 | 77,4815 | 80,001 | 0,046165 | 0,779 | 0,932261 |
| Spc24 | 54,1646 | 55,9193 | 0,0459937 | 0,779 | 0,932261 |
| Usp24 | 15,2447 | 14,7787 | -0,044789 | 0,7792 | 0,932333 |
| Zfyve26 | 3,85501 | 3,72439 | -0,049732 | 0,7792 | 0,932333 |
| Tomm6os | 0,424278 | 0,559902 | 0,400163 | 0,77935 | 0,932346 |
| Mettl9 | 120,384 | 116,695 | -0,044903 | 0,77935 | 0,932346 |
| Mkl1 | 8,36745 | 8,01273 | -0,0624947 | 0,77985 | 0,93286 |
| Gm16023 | 1,42339 | 1,64482 | 0,2086 | 0,78015 | 0,933136 |
| Ctbp1 | 119,283 | 123,084 | 0,045252 | 0,7804 | 0,933244 |
| Plk1 | 57,8843 | 59,6866 | 0,0442355 | 0,7804 | 0,933244 |
| Mvb12b | 16,6567 | 16,1301 | -0,0463397 | 0,78045 | 0,933244 |
| Ubp1 | 48,1136 | 49,6231 | 0,0445685 | 0,78055 | 0,933256 |
| Wapal | 37,8339 | 39,0107 | 0,0441883 | 0,7806 | 0,933256 |
| Cdc23 | 52,7216 | 55,9132 | 0,084795 | 0,78075 | 0,933341 |
| Ankrd13a | 27,4975 | 28,3624 | 0,044683 | 0,78085 | 0,933341 |
| Rbm12b2 | 7,22944 | 6,96852 | -0,0530316 | 0,78095 | 0,933341 |
| Mss51 | 2,40392 | 2,26516 | -0,0857703 | 0,7809 | 0,933341 |
| Nadsyn1 | 3,69764 | 3,87008 | 0,0657576 | 0,78105 | 0,933377 |

| | | | | | |
|---|---|---|---|---|---|
| Rpain | 13,4438 | 15,4047 | 0,196424 | 0,78135 | 0,933401 |
| Gm8363 | 2,12261 | 2,24966 | 0,0838634 | 0,78125 | 0,933401 |
| Nedd1 | 16,7851 | 17,3329 | 0,0463349 | 0,78135 | 0,933401 |
| Psrc1 | 34,4868 | 35,6095 | 0,0462174 | 0,78125 | 0,933401 |
| Ube2h | 12,6013 | 12,1922 | -0,0476096 | 0,78145 | 0,933437 |
| 4930556M19Rik | 1,13986 | 1,22368 | 0,10237 | 0,78155 | 0,933473 |
| Ppfibp2 | 2,22404 | 2,33425 | 0,0697756 | 0,7824 | 0,934155 |
| Kbtbd2 | 20,3767 | 21,0185 | 0,0447425 | 0,78235 | 0,934155 |
| Vapb | 12,6445 | 13,034 | 0,0437713 | 0,78235 | 0,934155 |
| Btbd10 | 10,7431 | 10,3562 | -0,0529142 | 0,78225 | 0,934155 |
| Riok1 | 23,0689 | 23,8217 | 0,0463269 | 0,7829 | 0,934585 |
| Bzw1 | 196,151 | 202,418 | 0,0453731 | 0,7829 | 0,934585 |
| Gna11 | 28,2331 | 29,1162 | 0,0444354 | 0,78315 | 0,9348 |
| Ttc13 | 23,904 | 23,1873 | -0,0439207 | 0,78325 | 0,934835 |
| Scarna3a | 17,4907 | 22,977 | 0,393603 | 0,78365 | 0,934848 |
| Figf | 0,885349 | 0,955873 | 0,110573 | 0,78395 | 0,934848 |
| Hpdl | 1,81675 | 1,93745 | 0,0928018 | 0,7835 | 0,934848 |
| Med19 | 33,1156 | 35,0896 | 0,0835338 | 0,7841 | 0,934848 |
| Cradd | 3,2426 | 3,42177 | 0,0775908 | 0,78335 | 0,934848 |
| Mtch1 | 108,33 | 111,636 | 0,043375 | 0,7834 | 0,934848 |
| Arcn1 | 72,4094 | 74,5863 | 0,0427348 | 0,78395 | 0,934848 |
| Cyb5 | 123,822 | 119,999 | -0,0452522 | 0,784 | 0,934848 |
| Arfip2 | 19,2751 | 18,652 | -0,0474078 | 0,7839 | 0,934848 |
| D930016D06Rik | 10,847 | 10,4897 | -0,0483264 | 0,7841 | 0,934848 |
| H2-DMa | 1,07046 | 0,989084 | -0,114066 | 0,784 | 0,934848 |
| Tnfsf12 | 1,48095 | 1,35828 | -0,124742 | 0,7838 | 0,934848 |
| Arhgef11 | 10,2944 | 10,6266 | 0,0458235 | 0,78425 | 0,934944 |
| Anxa9 | 1,28925 | 1,45004 | 0,169564 | 0,78485 | 0,934992 |
| Trappc2 | 3,55622 | 3,81941 | 0,103004 | 0,78475 | 0,934992 |
| H13 | 30,4218 | 31,6874 | 0,0588016 | 0,78445 | 0,934992 |
| Atp6v1h | 35,3461 | 36,4703 | 0,0451712 | 0,78475 | 0,934992 |
| Nav1 | 7,77469 | 8,01385 | 0,0437109 | 0,7847 | 0,934992 |
| Kif2a | 31,734 | 30,7826 | -0,0439152 | 0,78445 | 0,934992 |
| Xkr5 | 7,44783 | 7,17316 | -0,0542109 | 0,78455 | 0,934992 |
| 2810454H06Rik | 1,03666 | 0,971265 | -0,0940021 | 0,7848 | 0,934992 |
| Zfyve1 | 8,38945 | 8,6785 | 0,0488699 | 0,7851 | 0,935123 |
| Rpn2 | 170,044 | 175,268 | 0,043651 | 0,78505 | 0,935123 |
| Ankrd61 | 0,89367 | 1,26265 | 0,498646 | 0,78535 | 0,935254 |
| Tmem55a | 44,4867 | 43,173 | -0,0432435 | 0,78535 | 0,935254 |
| Bai2 | 1,11391 | 1,17099 | 0,0721068 | 0,78565 | 0,935528 |
| Ldb1 | 103,461 | 106,547 | 0,0424022 | 0,78615 | 0,93604 |
| Hk1os | 0,637082 | 0,973521 | 0,611734 | 0,78635 | 0,936111 |
| Ppp2ca | 243,981 | 251,392 | 0,0431709 | 0,7863 | 0,936111 |
| Celsr1 | 20,1067 | 20,7066 | 0,0424133 | 0,7865 | 0,936206 |
| Rbm14-rbm4 | 7,38814 | 8,24954 | 0,159103 | 0,7871 | 0,936336 |
| Armc7 | 2,28672 | 2,48867 | 0,122096 | 0,78725 | 0,936336 |
| Dda1 | 25,8262 | 26,8343 | 0,0552429 | 0,7874 | 0,936336 |
| Enoph1 | 21,638 | 22,3379 | 0,0459201 | 0,78735 | 0,936336 |

| | | | | | |
|---|---|---|---|---|---|
| Ndufa4 | 489,293 | 475,24 | -0,0420437 | 0,78715 | 0,936336 |
| Slc22a17 | 18,8004 | 18,1805 | -0,0483717 | 0,787 | 0,936336 |
| Vps54 | 23,6734 | 22,7145 | -0,0596507 | 0,7869 | 0,936336 |
| Zfp708 | 3,70875 | 3,53459 | -0,0693884 | 0,78745 | 0,936336 |
| Ntn3 | 1,72051 | 1,59219 | -0,111824 | 0,7874 | 0,936336 |
| Loxl3 | 1,30646 | 1,16695 | -0,162927 | 0,78745 | 0,936336 |
| Mir130c | 297,82 | 189,44 | -0,652695 | 0,787 | 0,936336 |
| Snord47 | 12306,1 | 6612,41 | -0,896129 | 0,78735 | 0,936336 |
| Spg21 | 19,8308 | 20,4665 | 0,0455248 | 0,7876 | 0,936407 |
| Neurl1a | 1,64846 | 1,56618 | -0,0738625 | 0,78765 | 0,936407 |
| Dctn6 | 44,0565 | 42,6285 | -0,0475374 | 0,7878 | 0,936419 |
| Myo1a | 0,750362 | 0,69972 | -0,10081 | 0,7878 | 0,936419 |
| Gm19557 | 0,509514 | 0,622968 | 0,290036 | 0,788 | 0,936502 |
| Ptges3l | 2,15084 | 2,39518 | 0,155232 | 0,78815 | 0,936502 |
| Mrps16 | 61,5498 | 59,3568 | -0,0523389 | 0,78815 | 0,936502 |
| Zfp874a | 3,38678 | 3,24289 | -0,0626336 | 0,7881 | 0,936502 |
| Psmc3ip | 36,8672 | 38,5475 | 0,0643004 | 0,7885 | 0,936834 |
| Msl1 | 44,1387 | 45,9745 | 0,0587886 | 0,789 | 0,936845 |
| Krtcap2 | 157,357 | 162,449 | 0,0459471 | 0,78935 | 0,936845 |
| Pabpc1 | 260,154 | 268,452 | 0,0452965 | 0,78915 | 0,936845 |
| Zmym1 | 15,7477 | 16,2352 | 0,0439879 | 0,7892 | 0,936845 |
| Bclaf1 | 90,8463 | 93,631 | 0,043558 | 0,7886 | 0,936845 |
| Plxna3 | 11,8836 | 12,2408 | 0,0427211 | 0,78865 | 0,936845 |
| Hif1an | 27,0567 | 26,2817 | -0,0419273 | 0,78915 | 0,936845 |
| Fignl2 | 45,0498 | 43,7452 | -0,0423971 | 0,7892 | 0,936845 |
| Szrd1 | 51,228 | 49,7324 | -0,0427477 | 0,789 | 0,936845 |
| Tbc1d15 | 19,7565 | 19,1754 | -0,0430665 | 0,78905 | 0,936845 |
| Nubp1 | 32,4076 | 31,2619 | -0,0519275 | 0,78935 | 0,936845 |
| Nqo2 | 6,08588 | 5,79625 | -0,0703471 | 0,78925 | 0,936845 |
| Snord49b | 867,95 | 623,221 | -0,477868 | 0,7895 | 0,93694 |
| Brf1 | 22,161 | 21,3977 | -0,0505667 | 0,7897 | 0,937011 |
| Coq10b | 8,23415 | 7,91175 | -0,0576228 | 0,7897 | 0,937011 |
| 4931406P16Rik | 13,2897 | 13,6953 | 0,0433778 | 0,78995 | 0,937141 |
| Ing5 | 10,399 | 10,0718 | -0,0461264 | 0,78995 | 0,937141 |
| Fbxl5 | 37,4284 | 38,5569 | 0,0428532 | 0,7901 | 0,937236 |
| Nkap | 10,9427 | 11,3005 | 0,0464152 | 0,7902 | 0,937247 |
| Caml | 28,3343 | 27,4264 | -0,0469822 | 0,79025 | 0,937247 |
| Mmp25 | 0,840675 | 0,895649 | 0,0913854 | 0,79045 | 0,937258 |
| Fanci | 13,2323 | 13,8881 | 0,0697861 | 0,79075 | 0,937258 |
| Usp49 | 7,93225 | 8,18757 | 0,0457052 | 0,7905 | 0,937258 |
| Igbp1 | 52,4857 | 50,9472 | -0,0429216 | 0,7907 | 0,937258 |
| Dcaf12 | 27,8169 | 26,9913 | -0,0434638 | 0,7904 | 0,937258 |
| Fam173b | 13,9499 | 13,3941 | -0,0586662 | 0,79065 | 0,937258 |
| Mir1895 | 92,2015 | 87,0721 | -0,0825794 | 0,7906 | 0,937258 |
| Ddit3 | 8,95877 | 7,50611 | -0,255236 | 0,79095 | 0,937329 |
| Frs3os | 0,61564 | 0,510109 | -0,271283 | 0,7909 | 0,937329 |
| Ckap2 | 27,6708 | 28,5099 | 0,0430975 | 0,7911 | 0,937399 |
| Amotl1 | 24,1392 | 23,4603 | -0,0411567 | 0,79115 | 0,937399 |

| | | | | | |
|---|---|---|---|---|---|
| Mum1 | 31,6761 | 39,9229 | 0,333825 | 0,79135 | 0,93754 |
| 1700029I15Rik | 1,10666 | 1,33308 | 0,268554 | 0,79175 | 0,93754 |
| Map3k14 | 0,639957 | 0,681215 | 0,0901367 | 0,79185 | 0,93754 |
| Fam171b | 4,11267 | 4,27959 | 0,0573962 | 0,79155 | 0,93754 |
| Urb2 | 13,1546 | 13,5601 | 0,0437993 | 0,7916 | 0,93754 |
| Enah | 27,4702 | 28,2789 | 0,0418607 | 0,79185 | 0,93754 |
| Gatc | 24,4927 | 23,7129 | -0,0466825 | 0,79165 | 0,93754 |
| Pdss1 | 9,50791 | 8,79254 | -0,112848 | 0,79145 | 0,93754 |
| Olfml3 | 19,1375 | 17,111 | -0,161478 | 0,7919 | 0,93754 |
| Riok3 | 23,82 | 23,1307 | -0,0423634 | 0,792 | 0,937575 |
| Rfxank | 7,10067 | 6,7973 | -0,0629932 | 0,79225 | 0,937705 |
| Prkcz | 1,29891 | 1,23447 | -0,0734135 | 0,7922 | 0,937705 |
| Becn1 | 56,9398 | 58,6697 | 0,0431785 | 0,7924 | 0,937717 |
| Tmem144 | 1,86272 | 1,75559 | -0,085456 | 0,79235 | 0,937717 |
| Pnisr | 52,7269 | 54,2445 | 0,040938 | 0,7928 | 0,938107 |
| Pla2g4c | 2,07648 | 1,98123 | -0,067742 | 0,7931 | 0,938379 |
| Guf1 | 20,6653 | 21,2832 | 0,0425052 | 0,7932 | 0,93839 |
| Ociad1 | 149,617 | 145,212 | -0,0431174 | 0,79325 | 0,93839 |
| Pole2 | 26,1143 | 26,9418 | 0,0450052 | 0,7935 | 0,938603 |
| Lpcat1 | 50,0863 | 48,6445 | -0,0421415 | 0,79375 | 0,938815 |
| Scarna6 | 1,96088 | 2,62337 | 0,41992 | 0,79525 | 0,938845 |
| C130083M11Rik | 0,745233 | 0,80429 | 0,110025 | 0,7946 | 0,938845 |
| Cck | 3,63411 | 3,90009 | 0,101905 | 0,79445 | 0,938845 |
| Ino80b | 18,099 | 18,7843 | 0,0536176 | 0,79525 | 0,938845 |
| Tyk2 | 4,64566 | 4,80957 | 0,0500259 | 0,79445 | 0,938845 |
| Baz2b | 12,3961 | 12,7684 | 0,0426962 | 0,794 | 0,938845 |
| Dnttip2 | 63,3875 | 65,26 | 0,0420024 | 0,7945 | 0,938845 |
| Ipmk | 15,0935 | 15,5382 | 0,0418941 | 0,7948 | 0,938845 |
| Apc | 15,3875 | 15,8406 | 0,0418695 | 0,79455 | 0,938845 |
| H2afx | 171,097 | 176,038 | 0,0410748 | 0,79525 | 0,938845 |
| Braf | 11,3031 | 11,6291 | 0,0410227 | 0,7951 | 0,938845 |
| Fads1 | 73,063 | 75,1453 | 0,0405408 | 0,79485 | 0,938845 |
| Fam178a | 21,2547 | 20,6607 | -0,0408944 | 0,79425 | 0,938845 |
| Smek1 | 45,672 | 44,3811 | -0,0413645 | 0,7946 | 0,938845 |
| Pip4k2b | 24,7038 | 24,0013 | -0,0416187 | 0,79455 | 0,938845 |
| Ppp6r3 | 53,9433 | 52,392 | -0,0420963 | 0,79495 | 0,938845 |
| Tmem161a | 20,1597 | 19,5558 | -0,0438802 | 0,79485 | 0,938845 |
| Ebp | 16,0511 | 15,5212 | -0,0484332 | 0,79515 | 0,938845 |
| 6330416G13Rik | 4,99011 | 4,81125 | -0,0526601 | 0,79475 | 0,938845 |
| Cd248 | 4,18839 | 4,01671 | -0,0603822 | 0,79455 | 0,938845 |
| Tmem45a | 1,15831 | 1,09046 | -0,0870866 | 0,79425 | 0,938845 |
| Man2c1 | 15,9547 | 16,4465 | 0,0438017 | 0,79545 | 0,938889 |
| Arpc2 | 123,293 | 126,838 | 0,0408967 | 0,79575 | 0,938889 |
| Ptprk | 32,4742 | 31,5625 | -0,0410834 | 0,7958 | 0,938889 |
| Irf2bp1 | 26,7838 | 26,0151 | -0,0420109 | 0,79575 | 0,938889 |
| Hoxb8 | 16,3949 | 15,8838 | -0,0456946 | 0,7957 | 0,938889 |
| Pctp | 12,1647 | 11,7474 | -0,050356 | 0,7957 | 0,938889 |
| Zc3h10 | 10,2756 | 9,91269 | -0,0518743 | 0,79585 | 0,938889 |

| | | | | | |
|---|---|---|---|---|---|
| Tsacc | 7,86426 | 7,3589 | -0,0958217 | 0,7955 | 0,938889 |
| Sav1 | 42,7113 | 44,2985 | 0,0526406 | 0,7962 | 0,939006 |
| Zfp524 | 3,38317 | 3,18847 | -0,0855121 | 0,79625 | 0,939006 |
| Col9a3 | 1,62635 | 1,52906 | -0,0889849 | 0,7961 | 0,939006 |
| Id4 | 1,35356 | 1,26404 | -0,0987156 | 0,7963 | 0,939006 |
| G0s2 | 2,08455 | 1,91899 | -0,119391 | 0,79615 | 0,939006 |
| 4930599N23Rik | 0,423356 | 0,765481 | 0,854496 | 0,797 | 0,9395 |
| Tjp1 | 39,8056 | 38,7025 | -0,0405456 | 0,79695 | 0,9395 |
| Arsk | 3,43914 | 3,30592 | -0,0569966 | 0,7968 | 0,9395 |
| Glt8d2 | 0,795933 | 0,69717 | -0,191137 | 0,79695 | 0,9395 |
| Strap | 125,082 | 128,679 | 0,0409045 | 0,7973 | 0,93977 |
| Chd9 | 4,85043 | 4,9939 | 0,0420544 | 0,79765 | 0,940076 |
| 2310061J03Rik | 1,14365 | 0,950077 | -0,267533 | 0,7977 | 0,940076 |
| Msantd4 | 30,7084 | 29,8547 | -0,040674 | 0,79795 | 0,940205 |
| Dnal1 | 4,21553 | 4,07765 | -0,0479781 | 0,7979 | 0,940205 |
| Sumo1 | 50,6714 | 49,1688 | -0,0434261 | 0,7981 | 0,940299 |
| 2310068J16Rik | 1,04628 | 1,45092 | 0,471703 | 0,79915 | 0,940303 |
| Cntf | 3,16418 | 3,94951 | 0,319845 | 0,7993 | 0,940303 |
| Syce2 | 22,8579 | 24,2565 | 0,0856808 | 0,7987 | 0,940303 |
| Rab11fip2 | 3,74149 | 3,87367 | 0,0500881 | 0,79885 | 0,940303 |
| Lrch1 | 4,97441 | 5,14702 | 0,0492137 | 0,7992 | 0,940303 |
| Zfand2a | 9,38202 | 9,69262 | 0,0469883 | 0,7988 | 0,940303 |
| Ufm1 | 7,19567 | 7,42991 | 0,0462172 | 0,7986 | 0,940303 |
| B4galt2 | 23,4042 | 24,1112 | 0,0429375 | 0,79915 | 0,940303 |
| Wdr6 | 103,585 | 106,625 | 0,0417239 | 0,7989 | 0,940303 |
| Otud4 | 34,7698 | 35,7689 | 0,0408733 | 0,7983 | 0,940303 |
| Sephs1 | 41,2621 | 42,4244 | 0,0400769 | 0,79855 | 0,940303 |
| Prpf40a | 84,4955 | 82,1917 | -0,0398824 | 0,79925 | 0,940303 |
| BC005537 | 78,1594 | 76,0142 | -0,0401499 | 0,7987 | 0,940303 |
| Acox1 | 11,3901 | 11,0174 | -0,0480062 | 0,7984 | 0,940303 |
| Iglon5 | 1,93696 | 1,83661 | -0,0767421 | 0,79835 | 0,940303 |
| Wif1 | 1,23462 | 1,1604 | -0,0894467 | 0,79925 | 0,940303 |
| Bin3 | 15,0354 | 13,7241 | -0,13165 | 0,79915 | 0,940303 |
| Sos2 | 9,39255 | 9,67573 | 0,0428542 | 0,7997 | 0,940667 |
| Lnx2 | 5,60358 | 5,42094 | -0,0478067 | 0,79975 | 0,940667 |
| Ippk | 9,97421 | 10,3037 | 0,0468885 | 0,79985 | 0,940702 |
| Toe1 | 25,1798 | 24,2655 | -0,053363 | 0,8 | 0,940795 |
| Zfp612 | 4,20365 | 4,06265 | -0,0492223 | 0,8001 | 0,94083 |
| Fam181a | 1,64537 | 1,53632 | -0,0989269 | 0,8003 | 0,940983 |
| Dph7 | 9,84293 | 10,2654 | 0,0606346 | 0,80075 | 0,941346 |
| Lrrc14b | 2,5255 | 2,23618 | -0,175536 | 0,80075 | 0,941346 |
| Rsc1a1 | 12,7326 | 11,8822 | -0,0997305 | 0,80085 | 0,941381 |
| Mir6958 | 245,223 | 341,149 | 0,476308 | 0,8017 | 0,941386 |
| Mir6990 | 60,3133 | 82,1064 | 0,445017 | 0,8014 | 0,941386 |
| Hexim2 | 2,57092 | 2,70602 | 0,0738866 | 0,80135 | 0,941386 |
| Ptprt | 0,550529 | 0,57633 | 0,0660768 | 0,80145 | 0,941386 |
| Rnf31 | 7,41305 | 7,66604 | 0,0484152 | 0,80115 | 0,941386 |
| Nck1 | 27,9722 | 28,8384 | 0,0439979 | 0,80155 | 0,941386 |

| | | | | | |
|---|---|---|---|---|---|
| Sf3b6 | 106,179 | 109,24 | 0,0409979 | 0,80105 | 0,941386 |
| Slc41a1 | 11,8239 | 11,4808 | -0,0424892 | 0,80115 | 0,941386 |
| Cwf19l2 | 11,4893 | 11,1485 | -0,0434481 | 0,8017 | 0,941386 |
| Slc6a15 | 1,34567 | 1,2788 | -0,0735337 | 0,8016 | 0,941386 |
| Npbwr1 | 0,770385 | 0,725108 | -0,0873841 | 0,80115 | 0,941386 |
| Timp1 | 8,16658 | 7,2956 | -0,162705 | 0,8016 | 0,941386 |
| Mtmr6 | 13,0637 | 13,4506 | 0,0421114 | 0,80185 | 0,94148 |
| Plekha3 | 12,068 | 12,4471 | 0,0446277 | 0,80195 | 0,941515 |
| Pop5 | 37,4143 | 36,2373 | -0,0461143 | 0,80225 | 0,941784 |
| Tnfrsf4 | 0,833994 | 0,75611 | -0,141441 | 0,80235 | 0,941819 |
| 1810062O18Rik | 0,866233 | 1,4783 | 0,771112 | 0,8025 | 0,941912 |
| Gm5434 | 2,38327 | 2,52125 | 0,0811963 | 0,80265 | 0,942005 |
| Sdhaf1 | 23,5699 | 24,376 | 0,0485197 | 0,80305 | 0,942061 |
| Mad2l1 | 89,9587 | 92,4843 | 0,0399459 | 0,803 | 0,942061 |
| Dnajc5 | 43,45 | 44,6659 | 0,0398186 | 0,80305 | 0,942061 |
| Dock10 | 0,649888 | 0,61826 | -0,071977 | 0,80295 | 0,942061 |
| Degs2 | 5,99858 | 5,46161 | -0,135294 | 0,803 | 0,942061 |
| Osbpl1a | 16,7859 | 17,3131 | 0,0446088 | 0,8032 | 0,942154 |
| Fam35a | 3,6142 | 3,75699 | 0,0559019 | 0,80345 | 0,942258 |
| 4930503E24Rik | 17,9947 | 18,6064 | 0,048233 | 0,80345 | 0,942258 |
| Stard7 | 54,0945 | 52,6272 | -0,0396725 | 0,8035 | 0,942258 |
| Zfp408 | 9,76942 | 10,0685 | 0,0434998 | 0,8041 | 0,942796 |
| Myo5a | 11,4009 | 11,7143 | 0,0391172 | 0,8041 | 0,942796 |
| Ern1 | 3,6849 | 3,82821 | 0,0550467 | 0,80445 | 0,943124 |
| Dnase1 | 5,82318 | 6,60433 | 0,181605 | 0,80485 | 0,943345 |
| Adprh | 92,4456 | 89,9699 | -0,0391619 | 0,8048 | 0,943345 |
| Stoml1 | 7,23938 | 6,97733 | -0,0531918 | 0,8048 | 0,943345 |
| Hectd2 | 1,78196 | 1,85787 | 0,0601837 | 0,8053 | 0,943376 |
| Nudcd3 | 28,181 | 28,9837 | 0,0405236 | 0,8052 | 0,943376 |
| Cdc7 | 27,4333 | 26,6868 | -0,0398013 | 0,8052 | 0,943376 |
| 5830415F09Rik | 5,58246 | 5,34274 | -0,0633229 | 0,80505 | 0,943376 |
| Ttc19 | 9,89792 | 8,99278 | -0,138358 | 0,80525 | 0,943376 |
| Mgp | 1,9037 | 1,72781 | -0,139863 | 0,805 | 0,943376 |
| Nup62-il4i1 | 0,377157 | 0,620939 | 0,719288 | 0,8074 | 0,943681 |
| Snora64 | 408,461 | 540,261 | 0,403458 | 0,8067 | 0,943681 |
| Syngr4 | 0,716447 | 0,801781 | 0,162349 | 0,80755 | 0,943681 |
| Itgb1bp2 | 1,21782 | 1,31488 | 0,11063 | 0,8061 | 0,943681 |
| Tatdn1 | 8,75625 | 9,38972 | 0,10077 | 0,80745 | 0,943681 |
| Icosl | 1,59425 | 1,67855 | 0,0743339 | 0,80705 | 0,943681 |
| Ypel1 | 10,0458 | 10,5748 | 0,0740438 | 0,80605 | 0,943681 |
| 1110032A03Rik | 7,06399 | 7,33024 | 0,0533777 | 0,8072 | 0,943681 |
| Trmt12 | 4,38731 | 4,53991 | 0,0493278 | 0,80695 | 0,943681 |
| Sap18 | 9,24896 | 9,53521 | 0,0439737 | 0,80705 | 0,943681 |
| Sowahc | 7,38324 | 7,60517 | 0,0427258 | 0,8077 | 0,943681 |
| Naa16 | 10,9384 | 11,2618 | 0,0420342 | 0,80575 | 0,943681 |
| Rnf168 | 17,309 | 17,7977 | 0,040171 | 0,807 | 0,943681 |
| Tgs1 | 22,5 | 23,1234 | 0,0394318 | 0,8065 | 0,943681 |
| Nuf2 | 44,3311 | 45,5536 | 0,0392475 | 0,80775 | 0,943681 |

| | | | | | |
|---|---|---|---|---|---|
| Pou2f1 | 13,6217 | 13,9971 | 0,039227 | 0,8062 | 0,943681 |
| Eif4ebp2 | 59,4097 | 61,0363 | 0,0389698 | 0,8077 | 0,943681 |
| Atp5o | 181,173 | 185,958 | 0,0376134 | 0,80715 | 0,943681 |
| Dusp11 | 27,9413 | 27,217 | -0,0378934 | 0,80775 | 0,943681 |
| Minpp1 | 37,9619 | 36,9598 | -0,038593 | 0,80655 | 0,943681 |
| Akap2 | 15,4627 | 15,0531 | -0,0387359 | 0,8073 | 0,943681 |
| Snd1 | 68,6965 | 66,8711 | -0,0388543 | 0,80685 | 0,943681 |
| Tfdp2 | 22,3364 | 21,7396 | -0,0390687 | 0,80715 | 0,943681 |
| Clcn7 | 11,6067 | 11,2817 | -0,0409817 | 0,8069 | 0,943681 |
| Arfgap1 | 25,592 | 24,8675 | -0,0414298 | 0,80695 | 0,943681 |
| Slc46a1 | 9,06075 | 8,74101 | -0,0518299 | 0,80575 | 0,943681 |
| Rpl31 | 18,1235 | 17,4298 | -0,0563012 | 0,8067 | 0,943681 |
| Cir1 | 2,06041 | 1,95411 | -0,076419 | 0,80745 | 0,943681 |
| Hexdc | 10,0965 | 9,54432 | -0,0811352 | 0,8066 | 0,943681 |
| Cbln1 | 1,43903 | 1,35944 | -0,0820888 | 0,807 | 0,943681 |
| Ltbp3 | 1,32744 | 1,20582 | -0,138634 | 0,8069 | 0,943681 |
| Phkg2 | 16,8404 | 17,4049 | 0,0475697 | 0,80815 | 0,943901 |
| Tspyl1 | 19,2942 | 19,8532 | 0,0411988 | 0,8081 | 0,943901 |
| Snx8 | 11,1175 | 10,7736 | -0,0453256 | 0,8081 | 0,943901 |
| Mark4 | 15,735 | 16,209 | 0,0428166 | 0,80845 | 0,944062 |
| Prpf38a | 64,1836 | 65,9449 | 0,0390557 | 0,8085 | 0,944062 |
| Pkd2l2 | 2,24404 | 1,99762 | -0,16782 | 0,80845 | 0,944062 |
| Tfdp1 | 55,1321 | 56,5774 | 0,0373334 | 0,8086 | 0,944097 |
| Rps15 | 1003,6 | 1137,35 | 0,180496 | 0,80875 | 0,944107 |
| Akap8 | 72,4157 | 70,5046 | -0,0385833 | 0,80875 | 0,944107 |
| Hmgxb4 | 21,8225 | 22,43 | 0,0396129 | 0,80885 | 0,944141 |
| Ndufaf5 | 6,44645 | 6,66935 | 0,0490401 | 0,809 | 0,944234 |
| Slc2a4 | 4,62417 | 4,46259 | -0,0513106 | 0,8092 | 0,944302 |
| Snord69 | 5347,52 | 5149,04 | -0,0545658 | 0,8092 | 0,944302 |
| Snx21 | 4,58323 | 4,83474 | 0,0770752 | 0,80935 | 0,944395 |
| 4930404I05Rik | 1,94245 | 2,54725 | 0,39106 | 0,80955 | 0,944532 |
| Hcfc1r1 | 9,17356 | 9,87319 | 0,106035 | 0,8097 | 0,944532 |
| Ndufa8 | 82,0146 | 84,4463 | 0,0421523 | 0,80965 | 0,944532 |
| Tdp2 | 51,7058 | 50,3242 | -0,0390736 | 0,80975 | 0,944532 |
| Mms22l | 19,454 | 19,9898 | 0,0391998 | 0,80985 | 0,944566 |
| Sgsm3 | 7,45071 | 7,80856 | 0,067678 | 0,81 | 0,944616 |
| Slc12a5 | 0,979616 | 1,02534 | 0,0658182 | 0,8105 | 0,944616 |
| 2810025M15Rik | 32,8771 | 34,3147 | 0,0617462 | 0,8104 | 0,944616 |
| Cpne1 | 45,778 | 47,492 | 0,053029 | 0,8105 | 0,944616 |
| Zfp12 | 10,9171 | 11,2233 | 0,0399118 | 0,8105 | 0,944616 |
| Mllt4 | 56,8606 | 58,4202 | 0,0390366 | 0,8106 | 0,944616 |
| Malt1 | 4,62239 | 4,29035 | -0,107544 | 0,81055 | 0,944616 |
| Igf2os | 4,96376 | 3,98513 | -0,316809 | 0,81035 | 0,944616 |
| 1110020A21Rik | 1,17638 | 0,900598 | -0,385403 | 0,81025 | 0,944616 |
| Aifm2 | 1,56717 | 1,16819 | -0,423883 | 0,8105 | 0,944616 |
| Mixl1 | 3,90627 | 4,07068 | 0,0594765 | 0,81105 | 0,944976 |
| Stx12 | 39,3254 | 38,3087 | -0,0377895 | 0,81105 | 0,944976 |
| Irf3 | 23,5679 | 24,2456 | 0,0409 | 0,8112 | 0,945068 |

| | | | | | |
|---|---|---|---|---|---|
| S100a16 | 11,7418 | 12,2032 | 0,0555957 | 0,8113 | 0,945102 |
| Zfp39 | 3,01782 | 3,13383 | 0,0544196 | 0,81145 | 0,945112 |
| Ubxn7 | 15,9051 | 16,335 | 0,0384789 | 0,81145 | 0,945112 |
| Slc36a4 | 8,55774 | 8,27338 | -0,0487535 | 0,81155 | 0,945146 |
| Iqcg | 1,22519 | 1,39561 | 0,18788 | 0,81165 | 0,94518 |
| Cul3 | 80,6292 | 78,4986 | -0,0386348 | 0,8118 | 0,945273 |
| Gm20324 | 1,03862 | 1,42933 | 0,460679 | 0,8119 | 0,945283 |
| Npcd | 1,29504 | 1,17201 | -0,144013 | 0,81195 | 0,945283 |
| Caap1 | 11,4144 | 11,7788 | 0,0453458 | 0,81215 | 0,945433 |
| Jrkl | 7,42142 | 7,18736 | -0,0462333 | 0,8123 | 0,945525 |
| Slc29a4 | 4,75904 | 4,93014 | 0,0509565 | 0,81245 | 0,945618 |
| Sv2b | 11,4366 | 11,7625 | 0,0405342 | 0,81255 | 0,945628 |
| Swap70 | 18,7487 | 19,2569 | 0,0385859 | 0,8126 | 0,945628 |
| Dcp1b | 3,12285 | 3,29782 | 0,0786499 | 0,81295 | 0,945846 |
| Mrpl30 | 87,3818 | 84,9465 | -0,0407779 | 0,81295 | 0,945846 |
| BC051226 | 2,26125 | 2,10169 | -0,105566 | 0,813 | 0,945846 |
| Hoxd3os1 | 0,993309 | 1,79067 | 0,850183 | 0,81325 | 0,945875 |
| Nsg1 | 22,3345 | 23,0847 | 0,0476645 | 0,81335 | 0,945875 |
| Mfap1a | 6,50383 | 6,71518 | 0,0461353 | 0,8131 | 0,945875 |
| Rfx1 | 7,45274 | 7,67453 | 0,0423069 | 0,8134 | 0,945875 |
| Zfp131 | 32,5778 | 33,4536 | 0,0382696 | 0,81345 | 0,945875 |
| Tmed8 | 4,75345 | 4,61393 | -0,0429783 | 0,8133 | 0,945875 |
| Mir6340 | 34,1384 | 52,3723 | 0,617408 | 0,81425 | 0,945944 |
| Mavs | 10,306 | 11,3715 | 0,141938 | 0,8142 | 0,945944 |
| Lsm5 | 5,7592 | 6,15938 | 0,0969178 | 0,814 | 0,945944 |
| Trappc2l | 32,6725 | 33,8489 | 0,0510326 | 0,8142 | 0,945944 |
| Prdx5 | 57,3025 | 59,0764 | 0,0439844 | 0,81385 | 0,945944 |
| Stk16 | 21,9788 | 22,6167 | 0,0412766 | 0,8141 | 0,945944 |
| Lsm1 | 15,0165 | 15,4466 | 0,0407401 | 0,81445 | 0,945944 |
| Snx19 | 13,6026 | 13,9805 | 0,0395344 | 0,81375 | 0,945944 |
| Rcc2 | 138,605 | 135,074 | -0,0372222 | 0,814 | 0,945944 |
| Ipo8 | 18,8977 | 18,4153 | -0,0373092 | 0,8145 | 0,945944 |
| Apoa1bp | 53,5369 | 52,0428 | -0,0408328 | 0,81395 | 0,945944 |
| Slc6a17 | 1,78082 | 1,71605 | -0,0534544 | 0,81375 | 0,945944 |
| Vwce | 1,25996 | 1,19974 | -0,0706541 | 0,8144 | 0,945944 |
| C77080 | 5,2632 | 4,94115 | -0,0910946 | 0,8145 | 0,945944 |
| Gpbp1l1 | 14,4775 | 14,0765 | -0,0405276 | 0,8147 | 0,946094 |
| Mien1 | 28,2065 | 30,3158 | 0,104043 | 0,8153 | 0,946283 |
| Pemt | 5,18324 | 5,44253 | 0,0704251 | 0,81545 | 0,946283 |
| Cpt1c | 7,53292 | 7,77924 | 0,0464204 | 0,8155 | 0,946283 |
| Golga5 | 16,5543 | 17,0246 | 0,0404134 | 0,81495 | 0,946283 |
| Casp2 | 35,1909 | 36,081 | 0,0360369 | 0,8152 | 0,946283 |
| Cstf1 | 39,4543 | 38,3969 | -0,0391934 | 0,8155 | 0,946283 |
| Smoc2 | 19,2828 | 18,7587 | -0,0397532 | 0,81525 | 0,946283 |
| Hmgb1 | 12,2502 | 11,8792 | -0,0443685 | 0,81525 | 0,946283 |
| Tekt2 | 0,798464 | 0,736624 | -0,1163 | 0,8153 | 0,946283 |
| Zhx1 | 13,9313 | 14,3033 | 0,0380224 | 0,81565 | 0,946293 |
| Pspn | 1,33638 | 1,1731 | -0,18801 | 0,81565 | 0,946293 |

| | | | | | |
|---|---|---|---|---|---|
| Acot8 | 7,16963 | 7,52601 | 0,0699863 | 0,81575 | 0,946303 |
| Fam114a1 | 2,89016 | 2,77694 | -0,057653 | 0,8158 | 0,946303 |
| 4930487H11Rik | 0,58259 | 0,923816 | 0,665125 | 0,81625 | 0,946308 |
| F8a | 1,58209 | 1,66362 | 0,0724903 | 0,8163 | 0,946308 |
| Atmin | 8,78756 | 9,03062 | 0,0393616 | 0,81625 | 0,946308 |
| Snx18 | 16,7034 | 17,1531 | 0,0383212 | 0,81595 | 0,946308 |
| D930015E06Rik | 24,7574 | 24,1363 | -0,0366546 | 0,81595 | 0,946308 |
| Camta2 | 3,99013 | 3,86178 | -0,0471685 | 0,81605 | 0,946308 |
| Kdelr3 | 4,57588 | 2,52237 | -0,859273 | 0,8162 | 0,946308 |
| Zfp213 | 11,3906 | 11,7305 | 0,0424313 | 0,81645 | 0,946317 |
| Sun1 | 29,1103 | 28,3816 | -0,0365751 | 0,81645 | 0,946317 |
| Slc27a6 | 2,15363 | 2,2533 | 0,0652686 | 0,8166 | 0,946327 |
| Srebf1 | 24,9423 | 23,9181 | -0,0604916 | 0,81655 | 0,946327 |
| Shisa3 | 9,70975 | 9,44155 | -0,0404105 | 0,81675 | 0,946419 |
| Adat1 | 10,8739 | 11,1878 | 0,0410543 | 0,81705 | 0,946602 |
| Fem1a | 7,91346 | 7,70103 | -0,039257 | 0,817 | 0,946602 |
| Lsm6 | 20,7093 | 21,2538 | 0,0374376 | 0,81745 | 0,946737 |
| 03/08/16 | 17,0466 | 16,6128 | -0,0371844 | 0,8174 | 0,946737 |
| Ctnnal1 | 18,3396 | 17,8673 | -0,0376403 | 0,81745 | 0,946737 |
| Ap4m1 | 20,1612 | 19,0462 | -0,0820835 | 0,81735 | 0,946737 |
| Sdr39u1 | 26,6258 | 25,8317 | -0,0436834 | 0,81755 | 0,946771 |
| Smchd1 | 29,9281 | 30,6844 | 0,0360053 | 0,81775 | 0,946921 |
| Rraga | 64,9079 | 66,593 | 0,036977 | 0,81795 | 0,94707 |
| Dnajb2 | 9,30372 | 9,62209 | 0,0485432 | 0,8182 | 0,947253 |
| Jam2 | 3,398 | 3,51325 | 0,0481195 | 0,81825 | 0,947253 |
| Nsun5 | 14,6827 | 15,7259 | 0,0990253 | 0,8194 | 0,947354 |
| Trim63 | 1,6439 | 1,73259 | 0,0758107 | 0,81925 | 0,947354 |
| Glb1l | 5,76071 | 6,053 | 0,0714028 | 0,8188 | 0,947354 |
| Glb1l2 | 2,15911 | 2,24722 | 0,0577026 | 0,819 | 0,947354 |
| Zswim3 | 3,59498 | 3,73091 | 0,0535444 | 0,81915 | 0,947354 |
| Rasal2 | 4,35823 | 4,51587 | 0,0512643 | 0,8187 | 0,947354 |
| Eid2b | 5,77877 | 5,97884 | 0,0491029 | 0,8193 | 0,947354 |
| 2700062C07Rik | 15,6778 | 16,2076 | 0,0479501 | 0,8186 | 0,947354 |
| Top2a | 215,501 | 221,875 | 0,0420531 | 0,8191 | 0,947354 |
| Gripap1 | 11,7681 | 12,1117 | 0,0415293 | 0,81935 | 0,947354 |
| Trappc8 | 16,3174 | 15,8998 | -0,0374069 | 0,8194 | 0,947354 |
| Tmcc2 | 10,5062 | 10,2154 | -0,0404975 | 0,8192 | 0,947354 |
| Fam57a | 21,9583 | 21,2336 | -0,0484226 | 0,81925 | 0,947354 |
| Polg2 | 6,67135 | 6,4197 | -0,0554735 | 0,81885 | 0,947354 |
| Efemp1 | 2,03872 | 1,9384 | -0,0727988 | 0,81915 | 0,947354 |
| Ppp1r26 | 5,53517 | 5,69731 | 0,0416524 | 0,81965 | 0,947373 |
| Ptgis | 21,8995 | 22,513 | 0,0398617 | 0,8197 | 0,947373 |
| Cisd1 | 53,7754 | 55,2088 | 0,0379514 | 0,8196 | 0,947373 |
| Actr10 | 23,2528 | 23,8666 | 0,0375888 | 0,8195 | 0,947373 |
| Ppil4 | 25,1822 | 25,8235 | 0,0362808 | 0,81985 | 0,947464 |
| Dennd6a | 14,5345 | 14,1313 | -0,0405843 | 0,82 | 0,947556 |
| Spata1 | 1,57974 | 1,69472 | 0,101363 | 0,82045 | 0,94756 |
| P4ha3 | 0,584645 | 0,627059 | 0,10104 | 0,82035 | 0,94756 |

| | | | | | |
|---|---|---|---|---|---|
| Srsf12 | 1,99572 | 2,08362 | 0,0621824 | 0,8205 | 0,94756 |
| Abi2 | 21,9723 | 21,4289 | -0,0361261 | 0,8204 | 0,94756 |
| 2410002F23Rik | 44,5538 | 43,4304 | -0,0368407 | 0,82025 | 0,94756 |
| Sfrp4 | 0,902839 | 0,842843 | -0,0992048 | 0,8201 | 0,94756 |
| Snord87 | 602,277 | 410,85 | -0,551816 | 0,8205 | 0,94756 |
| Nr1d2 | 4,57082 | 4,42892 | -0,0454982 | 0,8206 | 0,947593 |
| 4930455C13Rik | 0,350708 | 0,566379 | 0,691495 | 0,821 | 0,947785 |
| Rnf208 | 3,04152 | 3,1979 | 0,0723306 | 0,82085 | 0,947785 |
| Thap6 | 5,19772 | 5,45382 | 0,0693883 | 0,82105 | 0,947785 |
| Suclg2 | 12,9985 | 12,6454 | -0,0397378 | 0,82095 | 0,947785 |
| AV039307 | 1,16309 | 1,55119 | 0,415413 | 0,8214 | 0,947847 |
| Foxf2 | 1,02434 | 1,08363 | 0,0811755 | 0,82145 | 0,947847 |
| Nthl1 | 8,79854 | 9,1364 | 0,0543603 | 0,8215 | 0,947847 |
| Npm3 | 39,8761 | 41,0655 | 0,0424054 | 0,82135 | 0,947847 |
| Cstf2t | 26,3963 | 27,129 | 0,039501 | 0,8215 | 0,947847 |
| Hopx | 8,57447 | 8,24759 | -0,0560762 | 0,8216 | 0,947847 |
| Tnfrsf12a | 11,6462 | 10,7989 | -0,108975 | 0,82155 | 0,947847 |
| 1110054M08Rik | 2,22775 | 2,92708 | 0,393874 | 0,8217 | 0,947881 |
| Bbs1 | 5,79208 | 5,95673 | 0,0404405 | 0,8221 | 0,948058 |
| Mbd1 | 18,1804 | 18,6694 | 0,0382869 | 0,82235 | 0,948058 |
| Coa5 | 30,4879 | 31,2622 | 0,0361803 | 0,82225 | 0,948058 |
| Erf | 22,4999 | 21,9448 | -0,036041 | 0,8222 | 0,948058 |
| Abl1 | 45,6501 | 44,5091 | -0,0365164 | 0,82215 | 0,948058 |
| Orc5 | 34,9059 | 33,8388 | -0,044791 | 0,82235 | 0,948058 |
| Dhx32 | 6,47794 | 6,03764 | -0,101551 | 0,822 | 0,948058 |
| Ube2c | 217,088 | 222,467 | 0,0353118 | 0,82245 | 0,948091 |
| Pafah1b2 | 73,3678 | 71,6004 | -0,0351805 | 0,8231 | 0,948104 |
| Zfp653 | 9,99306 | 11,0762 | 0,148469 | 0,82295 | 0,948105 |
| Zfp449 | 5,13609 | 5,301 | 0,0455934 | 0,82265 | 0,948105 |
| Ssbp3 | 90,354 | 92,6745 | 0,036584 | 0,8231 | 0,948105 |
| Gnb1 | 200,714 | 205,86 | 0,0365262 | 0,823 | 0,948105 |
| Fam168b | 91,3748 | 89,1522 | -0,0355268 | 0,82295 | 0,948105 |
| N4bp2l2 | 12,099 | 11,7987 | -0,0362608 | 0,82275 | 0,948105 |
| Micall1 | 7,46078 | 7,26429 | -0,0385032 | 0,8227 | 0,948105 |
| Cnn2 | 69,3019 | 55,5675 | -0,318655 | 0,82275 | 0,948105 |
| Rmdn2 | 3,36713 | 3,51903 | 0,0636612 | 0,82355 | 0,948133 |
| Lyplal1 | 6,09728 | 6,3253 | 0,0529664 | 0,82335 | 0,948133 |
| Stbd1 | 8,11039 | 7,84932 | -0,0472047 | 0,8235 | 0,948133 |
| Wdr54 | 5,64631 | 5,41611 | -0,060052 | 0,82355 | 0,948133 |
| Pik3ip1 | 6,13122 | 5,06005 | -0,277024 | 0,82335 | 0,948133 |
| Ap5s1 | 5,35032 | 4,30233 | -0,314506 | 0,8234 | 0,948133 |
| Aass | 0,858198 | 0,902619 | 0,0728055 | 0,82365 | 0,948166 |
| Zc3h11a | 57,2662 | 59,2384 | 0,048847 | 0,8238 | 0,948242 |
| Rgl2 | 18,3201 | 18,8085 | 0,0379604 | 0,824 | 0,948242 |
| Nol9 | 36,528 | 37,45 | 0,0359631 | 0,8239 | 0,948242 |
| Zfp646 | 18,336 | 17,8929 | -0,0352921 | 0,824 | 0,948242 |
| Lrch2 | 11,3845 | 11,0873 | -0,0381677 | 0,82415 | 0,948333 |
| 1700012D14Rik | 1,11397 | 1,23725 | 0,15142 | 0,82445 | 0,948352 |

| | | | | | |
|---|---|---|---|---|---|
| Olfm1 | 17,0843 | 17,5944 | 0,042448 | 0,8243 | 0,948352 |
| Peak1 | 5,50225 | 5,64423 | 0,0367569 | 0,82445 | 0,948352 |
| Grk6 | 52,3373 | 53,6345 | 0,0353197 | 0,8244 | 0,948352 |
| Lpcat3 | 19,4468 | 20,4312 | 0,0712416 | 0,8247 | 0,948558 |
| Birc5 | 162,546 | 166,818 | 0,0374314 | 0,8249 | 0,948576 |
| Sept11 | 101,377 | 103,97 | 0,0364304 | 0,8248 | 0,948576 |
| Hnrnpa3 | 17,2017 | 17,6363 | 0,0360011 | 0,825 | 0,948576 |
| Naa40 | 37,0287 | 37,9107 | 0,0339643 | 0,825 | 0,948576 |
| Cuedc2 | 31,0164 | 31,8367 | 0,0376594 | 0,8251 | 0,948609 |
| Ino80dos | 2,25724 | 2,69985 | 0,25832 | 0,8252 | 0,948619 |
| Ip6k1 | 32,021 | 31,2407 | -0,0355899 | 0,82525 | 0,948619 |
| Stk4 | 20,0119 | 20,5132 | 0,0356914 | 0,8255 | 0,948661 |
| Scrib | 35,3366 | 34,4924 | -0,0348866 | 0,8254 | 0,948661 |
| Acot2 | 1,53489 | 1,4615 | -0,0706934 | 0,82545 | 0,948661 |
| Adrbk1 | 32,3371 | 33,1518 | 0,0358941 | 0,82565 | 0,948752 |
| Lats2 | 10,2962 | 10,0004 | -0,0420486 | 0,82575 | 0,948785 |
| Cnpy2 | 113,469 | 116,34 | 0,0360498 | 0,82585 | 0,948795 |
| Necab3 | 0,722372 | 0,665596 | -0,118095 | 0,8259 | 0,948795 |
| Atxn7l3b | 89,5188 | 91,7175 | 0,0350061 | 0,82615 | 0,948895 |
| Bloc1s3 | 6,16666 | 5,94781 | -0,0521296 | 0,8262 | 0,948895 |
| Gm16861 | 0,871673 | 0,652122 | -0,418645 | 0,8261 | 0,948895 |
| Gdap10 | 2,00294 | 2,26163 | 0,175248 | 0,82675 | 0,949094 |
| Acer2 | 8,31308 | 8,64937 | 0,0572122 | 0,82645 | 0,949094 |
| Trip6 | 37,4677 | 38,4414 | 0,0370114 | 0,8268 | 0,949094 |
| Cox4i1 | 529,526 | 542,182 | 0,0340761 | 0,82675 | 0,949094 |
| Dhodh | 13,594 | 13,2147 | -0,0408267 | 0,82655 | 0,949094 |
| Zscan25 | 8,53006 | 8,26092 | -0,0462541 | 0,8267 | 0,949094 |
| Txnl1 | 89,0797 | 91,2157 | 0,0341859 | 0,82715 | 0,949333 |
| Camkk2 | 8,72321 | 8,4992 | -0,0375319 | 0,8271 | 0,949333 |
| AI462493 | 28,353 | 29,2218 | 0,0435461 | 0,8273 | 0,949424 |
| Nhp2l1 | 9,46675 | 9,80854 | 0,0511686 | 0,8278 | 0,949427 |
| D230025D16Rik | 11,3576 | 11,6684 | 0,0389557 | 0,8277 | 0,949427 |
| Mrpl2 | 46,8701 | 48,1302 | 0,0382771 | 0,82775 | 0,949427 |
| Pdpn | 26,8317 | 26,1289 | -0,0382902 | 0,82755 | 0,949427 |
| Alkbh4 | 7,35607 | 7,10292 | -0,050523 | 0,8277 | 0,949427 |
| Cxx1a | 2,93513 | 2,7951 | -0,0705222 | 0,8275 | 0,949427 |
| St3gal4 | 9,38689 | 8,88898 | -0,0786293 | 0,8277 | 0,949427 |
| Alyref2 | 2,96259 | 3,10697 | 0,0686448 | 0,82885 | 0,949647 |
| Sh3yl1 | 4,86685 | 5,04994 | 0,0532781 | 0,8285 | 0,949647 |
| Mrps18c | 77,7893 | 80,0073 | 0,0405596 | 0,8288 | 0,949647 |
| Rpgr | 14,294 | 14,6798 | 0,0384231 | 0,8287 | 0,949647 |
| Sec23a | 24,6285 | 25,2459 | 0,0357172 | 0,8288 | 0,949647 |
| Fbf1 | 14,3955 | 14,7522 | 0,0353107 | 0,8283 | 0,949647 |
| Bbx | 9,33434 | 9,56228 | 0,0348056 | 0,82915 | 0,949647 |
| Zyg11b | 13,1419 | 12,8313 | -0,0345114 | 0,82895 | 0,949647 |
| Manbal | 33,0877 | 32,1929 | -0,0395549 | 0,82845 | 0,949647 |
| Mthfsd | 19,2033 | 18,681 | -0,0397839 | 0,8292 | 0,949647 |
| Gpx7 | 26,9845 | 26,238 | -0,040472 | 0,8292 | 0,949647 |

| | | | | | |
|---|---|---|---|---|---|
| Oxnad1 | 12,8019 | 12,4329 | -0,0421925 | 0,8287 | 0,949647 |
| Sass6 | 11,8871 | 11,535 | -0,0433728 | 0,829 | 0,949647 |
| Nsa2 | 10,1273 | 9,71842 | -0,0594592 | 0,8283 | 0,949647 |
| Isl1 | 0,821317 | 0,774189 | -0,0852534 | 0,8281 | 0,949647 |
| Dhdh | 2,75688 | 2,4744 | -0,155957 | 0,82875 | 0,949647 |
| Snora44 | 181,218 | 146,297 | -0,308826 | 0,8292 | 0,949647 |
| Tmem198b | 15,6696 | 16,0913 | 0,0383133 | 0,8295 | 0,949909 |
| Msto1 | 28,995 | 30,2692 | 0,0620464 | 0,8299 | 0,94996 |
| Kpna1 | 34,7476 | 35,5567 | 0,0332094 | 0,8298 | 0,94996 |
| Tubb4b | 116,147 | 113,478 | -0,0335438 | 0,82975 | 0,94996 |
| Arl6ip4 | 52,0296 | 50,7165 | -0,0368758 | 0,8299 | 0,94996 |
| Myl7 | 1,73275 | 1,58392 | -0,12956 | 0,8299 | 0,94996 |
| Sbno1 | 19,4206 | 19,8902 | 0,0344681 | 0,83025 | 0,950117 |
| Tsn | 77,6428 | 79,4981 | 0,0340671 | 0,83025 | 0,950117 |
| Maf1 | 32,3243 | 31,4985 | -0,0373367 | 0,8302 | 0,950117 |
| Rbl2 | 3,49477 | 3,65007 | 0,0627238 | 0,83035 | 0,95015 |
| Zfat | 2,68221 | 2,5941 | -0,0481882 | 0,8305 | 0,95024 |
| Gm11944 | 2,95923 | 4,67227 | 0,658903 | 0,831 | 0,95039 |
| Gm527 | 3,53607 | 3,69758 | 0,0644346 | 0,8312 | 0,95039 |
| Sat2 | 8,64483 | 9,00144 | 0,0583188 | 0,8308 | 0,95039 |
| Map2k3 | 20,3404 | 20,8765 | 0,0375274 | 0,8309 | 0,95039 |
| Cox7a2l | 41,8953 | 42,9813 | 0,0369212 | 0,8311 | 0,95039 |
| Ubxn6 | 25,6788 | 26,3378 | 0,0365531 | 0,8308 | 0,95039 |
| Fam135a | 16,5734 | 16,9711 | 0,0342143 | 0,8312 | 0,95039 |
| Polr3k | 33,5706 | 34,3682 | 0,0338768 | 0,83105 | 0,95039 |
| Preb | 19,9373 | 22,1079 | 0,149093 | 0,83145 | 0,950513 |
| Rhoc | 56,3281 | 54,9208 | -0,0365032 | 0,83145 | 0,950513 |
| Wdpcp | 3,18068 | 3,30278 | 0,0543493 | 0,83165 | 0,950555 |
| Alg8 | 31,2895 | 30,5325 | -0,0353297 | 0,8316 | 0,950555 |
| Adm2 | 0,719024 | 0,658375 | -0,12713 | 0,8317 | 0,950555 |
| Trim68 | 4,26705 | 4,41526 | 0,0492586 | 0,8322 | 0,950557 |
| Aoc2 | 5,30006 | 5,46882 | 0,0452212 | 0,83195 | 0,950557 |
| Zfml | 44,7618 | 45,8036 | 0,0331942 | 0,83215 | 0,950557 |
| Slc16a2 | 8,58944 | 8,36915 | -0,0374829 | 0,832 | 0,950557 |
| Zfp583 | 2,62816 | 2,52996 | -0,0549416 | 0,83185 | 0,950557 |
| Avpi1 | 6,54073 | 6,27412 | -0,0600378 | 0,8322 | 0,950557 |
| Zfp677 | 1,63932 | 1,56925 | -0,0630261 | 0,8321 | 0,950557 |
| Kdm5b | 38,7073 | 39,6094 | 0,0332396 | 0,8323 | 0,95059 |
| Ccbl2 | 3,55981 | 3,69882 | 0,0552628 | 0,8327 | 0,950722 |
| Pdpk1 | 13,096 | 13,4079 | 0,0339571 | 0,8325 | 0,950722 |
| Glg1 | 52,0693 | 50,8641 | -0,0337853 | 0,8326 | 0,950722 |
| Hist3h2a | 12,2104 | 11,6605 | -0,0664783 | 0,83265 | 0,950722 |
| Fbl | 36,6315 | 35,6933 | -0,0374314 | 0,8329 | 0,950869 |
| Zfp207 | 190,56 | 186,095 | -0,0342073 | 0,833 | 0,950902 |
| Pla2g12a | 12,7321 | 13,1247 | 0,0438133 | 0,83315 | 0,950992 |
| Trmt112 | 19,5191 | 20,2305 | 0,051647 | 0,83375 | 0,951108 |
| Hagh | 10,7136 | 11,081 | 0,0486422 | 0,8336 | 0,951108 |
| Atp5a1 | 493,32 | 506,095 | 0,0368839 | 0,8336 | 0,951108 |

| | | | | | |
|---|---|---|---|---|---|
| Zfp113 | 7,56609 | 7,75846 | 0,0362223 | 0,8336 | 0,951108 |
| Hoxd13 | 75,2816 | 77,0339 | 0,0331973 | 0,8334 | 0,951108 |
| Dhrs3 | 20,0809 | 19,5263 | -0,0404048 | 0,83375 | 0,951108 |
| Sertad1 | 7,4599 | 7,19529 | -0,0521032 | 0,83375 | 0,951108 |
| Trim39 | 15,621 | 16,0097 | 0,0354532 | 0,83395 | 0,951255 |
| Serac1 | 3,80172 | 3,93326 | 0,0490705 | 0,83425 | 0,951338 |
| Arhgap12 | 12,3429 | 12,6351 | 0,0337521 | 0,8343 | 0,951338 |
| Btbd1 | 17,5325 | 17,1169 | -0,0346117 | 0,83445 | 0,951338 |
| B4galt3 | 22,5581 | 21,9933 | -0,0365798 | 0,83445 | 0,951338 |
| Mnt | 7,1902 | 7,00572 | -0,0374984 | 0,8344 | 0,951338 |
| Carkd | 31,0973 | 30,2916 | -0,0378692 | 0,83435 | 0,951338 |
| Fig4 | 10,589 | 10,3179 | -0,0374231 | 0,83465 | 0,951485 |
| Ttc33 | 17,8436 | 17,3829 | -0,0377396 | 0,8352 | 0,952031 |
| Rasa1 | 23,6968 | 23,1489 | -0,0337515 | 0,8354 | 0,952129 |
| Ube2j1 | 15,6198 | 15,2392 | -0,0355839 | 0,83545 | 0,952129 |
| Snora65 | 332,235 | 303,711 | -0,129506 | 0,8355 | 0,952129 |
| Tmem44 | 4,59589 | 4,72784 | 0,0408384 | 0,8356 | 0,952162 |
| Gpt | 3,87928 | 3,73027 | -0,0565083 | 0,8359 | 0,952423 |
| Ttc9 | 0,972973 | 1,03062 | 0,0830377 | 0,83645 | 0,952514 |
| Ociad2 | 4,29862 | 4,44802 | 0,0492925 | 0,83635 | 0,952514 |
| Tmem41a | 13,3725 | 13,7728 | 0,0425521 | 0,83655 | 0,952514 |
| Fam149a | 4,1493 | 4,26478 | 0,0396018 | 0,8364 | 0,952514 |
| Trmt11 | 12,9029 | 13,2377 | 0,0369586 | 0,8365 | 0,952514 |
| Pde12 | 11,3675 | 11,0856 | -0,0362332 | 0,8363 | 0,952514 |
| Fgf13 | 7,77297 | 7,54545 | -0,0428585 | 0,83625 | 0,952514 |
| Zfp954 | 5,19361 | 5,02219 | -0,0484227 | 0,83625 | 0,952514 |
| Mir5121 | 1461,38 | 2047,16 | 0,486284 | 0,8375 | 0,952938 |
| Cmc1 | 8,52938 | 9,1342 | 0,0988363 | 0,8373 | 0,952938 |
| D6Ertd527e | 0,76478 | 0,811407 | 0,0853805 | 0,83765 | 0,952938 |
| Hoxa3 | 19,2277 | 20,2663 | 0,075899 | 0,8372 | 0,952938 |
| Rab27a | 1,77366 | 1,84778 | 0,059064 | 0,8377 | 0,952938 |
| Rad51b | 3,07382 | 3,19797 | 0,0571214 | 0,8374 | 0,952938 |
| Rbck1 | 27,1681 | 27,8238 | 0,0344085 | 0,8371 | 0,952938 |
| Morf4l2 | 146,612 | 143,271 | -0,0332572 | 0,8376 | 0,952938 |
| Bivm | 12,3969 | 12,102 | -0,0347338 | 0,8377 | 0,952938 |
| Ngrn | 22,5721 | 21,9842 | -0,0380755 | 0,83785 | 0,952938 |
| Coq3 | 6,36208 | 6,17552 | -0,0429362 | 0,8373 | 0,952938 |
| Prr11 | 19,8876 | 17,9804 | -0,14544 | 0,83775 | 0,952938 |
| Arl1 | 51,6713 | 50,3665 | -0,0368963 | 0,83785 | 0,952939 |
| Apopt1 | 25,1598 | 25,829 | 0,0378711 | 0,83805 | 0,953004 |
| Nudt16l1 | 50,3966 | 48,6352 | -0,0513237 | 0,83805 | 0,953004 |
| Ppfia3 | 0,999466 | 1,03971 | 0,0569493 | 0,83855 | 0,953127 |
| Llph | 17,0666 | 17,5646 | 0,0414907 | 0,83865 | 0,953127 |
| Ttc26 | 8,10348 | 8,30731 | 0,0358384 | 0,83845 | 0,953127 |
| Mtx1 | 33,9124 | 34,761 | 0,0356578 | 0,8387 | 0,953127 |
| Map3k7 | 26,6584 | 27,2684 | 0,0326442 | 0,83855 | 0,953127 |
| Hibadh | 46,6888 | 47,7554 | 0,0325857 | 0,83875 | 0,953127 |
| Tceb1 | 121,87 | 119,143 | -0,0326492 | 0,8384 | 0,953127 |

| | | | | | |
|---|---|---|---|---|---|
| Fgd6 | 4,29542 | 4,18621 | -0,0371551 | 0,8385 | 0,953127 |
| Insl6 | 1,68706 | 1,55264 | -0,119781 | 0,8388 | 0,953127 |
| AI848285 | 0,636472 | 0,681026 | 0,0976119 | 0,8389 | 0,953135 |
| Ptpn14 | 16,1989 | 15,8458 | -0,0317944 | 0,83895 | 0,953135 |
| Mrpl27 | 31,5453 | 32,4701 | 0,0416868 | 0,83905 | 0,953143 |
| Gm3414 | 2,01125 | 1,8485 | -0,121742 | 0,8391 | 0,953143 |
| 8430429K09Rik | 5,52513 | 6,78319 | 0,295956 | 0,84 | 0,953355 |
| Ybey | 2,69015 | 2,91291 | 0,114775 | 0,84 | 0,953355 |
| Fam167a | 0,907586 | 0,947386 | 0,0619169 | 0,83995 | 0,953355 |
| Traip | 14,9291 | 15,2901 | 0,0344677 | 0,8396 | 0,953355 |
| Map2k7 | 28,4699 | 29,1514 | 0,0341291 | 0,83955 | 0,953355 |
| Gsk3a | 54,7061 | 53,431 | -0,0340243 | 0,83985 | 0,953355 |
| Rnf111 | 10,9282 | 10,6712 | -0,0343354 | 0,83995 | 0,953355 |
| Csrnp3 | 0,807193 | 0,779826 | -0,0497603 | 0,83985 | 0,953355 |
| BC028528 | 9,49781 | 8,8107 | -0,10834 | 0,83975 | 0,953355 |
| 1700094D03Rik | 3,09539 | 2,67042 | -0,213053 | 0,8394 | 0,953355 |
| Rev1 | 13,247 | 14,026 | 0,0824391 | 0,84085 | 0,953429 |
| Dusp28 | 5,95215 | 6,15926 | 0,0493459 | 0,84085 | 0,953429 |
| Mcrs1 | 9,3191 | 9,60125 | 0,0430315 | 0,84045 | 0,953429 |
| Flad1 | 7,24479 | 7,45489 | 0,0412438 | 0,84055 | 0,953429 |
| A130010J15Rik | 6,06865 | 6,23555 | 0,0391398 | 0,8407 | 0,953429 |
| Chd6 | 17,2096 | 16,8279 | -0,0323585 | 0,8405 | 0,953429 |
| Sdhb | 130,308 | 127,412 | -0,0324179 | 0,8406 | 0,953429 |
| Lrp12 | 17,4959 | 17,1025 | -0,032809 | 0,8408 | 0,953429 |
| Ajap1 | 4,9433 | 4,79924 | -0,0426664 | 0,8406 | 0,953429 |
| 1700011H14Rik | 2,50799 | 2,38482 | -0,0726553 | 0,84075 | 0,953429 |
| Pcdhga11 | 2,60901 | 2,25508 | -0,210323 | 0,8408 | 0,953429 |
| Gpkow | 16,6154 | 17,004 | 0,0333556 | 0,841 | 0,953438 |
| Kpna3 | 51,2776 | 50,1689 | -0,0315366 | 0,84095 | 0,953438 |
| Rapgef1 | 28,8477 | 28,2153 | -0,0319797 | 0,8412 | 0,953559 |
| Arl6 | 20,0559 | 19,552 | -0,0367103 | 0,84125 | 0,953559 |
| Airn | 19,3788 | 21,9182 | 0,17765 | 0,8415 | 0,9536 |
| Agpat1 | 53,5631 | 52,3242 | -0,0337627 | 0,8414 | 0,9536 |
| Gm11627 | 0,888921 | 0,806795 | -0,139853 | 0,84145 | 0,9536 |
| Pigb | 5,47654 | 5,72015 | 0,0627882 | 0,8416 | 0,953625 |
| Hspb8 | 9,76691 | 10,0525 | 0,0415864 | 0,8419 | 0,953625 |
| Nmt2 | 18,47 | 18,8857 | 0,0321085 | 0,8417 | 0,953625 |
| Naa15 | 59,9318 | 58,6425 | -0,0313751 | 0,84195 | 0,953625 |
| Efna1 | 39,773 | 38,8412 | -0,034202 | 0,84195 | 0,953625 |
| Atp10a | 4,38229 | 4,26962 | -0,0375791 | 0,8418 | 0,953625 |
| Kmt2e | 17,5648 | 18,0559 | 0,0397803 | 0,8421 | 0,953714 |
| Mrpl1 | 14,0481 | 14,4296 | 0,0386555 | 0,8426 | 0,954199 |
| 4732416N19Rik | 0,92643 | 0,867396 | -0,0949911 | 0,8427 | 0,954232 |
| Supt7l | 12,97 | 13,296 | 0,0358122 | 0,8429 | 0,954377 |
| Pi4kb | 16,6142 | 17,2561 | 0,0546855 | 0,8431 | 0,954523 |
| Cyb5d2 | 1,56979 | 1,63719 | 0,0606484 | 0,84325 | 0,954531 |
| Dvl1 | 16,309 | 15,9399 | -0,0330239 | 0,84325 | 0,954531 |
| Pold4 | 4,27488 | 4,46553 | 0,0629475 | 0,84335 | 0,954563 |

| | | | | | |
|---|---|---|---|---|---|
| Psip1 | 187,658 | 192,013 | 0,0331019 | 0,84345 | 0,954596 |
| Wsb1 | 88,8116 | 87,0152 | -0,029481 | 0,84365 | 0,954741 |
| Smyd3 | 2,77606 | 2,86666 | 0,0463311 | 0,84395 | 0,954757 |
| Chst3 | 2,73151 | 2,8081 | 0,0398989 | 0,8439 | 0,954757 |
| Lrrcc1 | 18,1474 | 18,558 | 0,0322791 | 0,8438 | 0,954757 |
| Gm16845 | 0,882971 | 0,561215 | -0,653813 | 0,84385 | 0,954757 |
| Stra6 | 26,9568 | 27,5636 | 0,0321154 | 0,84445 | 0,955161 |
| Dhdds | 16,7739 | 16,3879 | -0,0335864 | 0,84445 | 0,955161 |
| Gtl3 | 50,5197 | 51,6672 | 0,0324036 | 0,84455 | 0,955193 |
| Yif1a | 17,1943 | 16,7118 | -0,0410686 | 0,8447 | 0,955282 |
| Mrpl38 | 42,2814 | 43,2816 | 0,0337306 | 0,84505 | 0,955516 |
| Srpx | 0,604807 | 0,569657 | -0,0863815 | 0,84505 | 0,955516 |
| Akr1c13 | 1,23807 | 1,16468 | -0,0881567 | 0,8452 | 0,955605 |
| Kif18b | 20,6918 | 20,2321 | -0,0324145 | 0,8453 | 0,955637 |
| A330069E16Rik | 0,938019 | 1,01506 | 0,113879 | 0,84545 | 0,95571 |
| Zbtb17 | 13,7999 | 14,4769 | 0,0690957 | 0,8456 | 0,95571 |
| Ppcs | 10,4936 | 10,8023 | 0,0418309 | 0,84555 | 0,95571 |
| Far2 | 1,99144 | 1,74869 | -0,187533 | 0,84565 | 0,95571 |
| Ide | 29,4682 | 30,1203 | 0,0315754 | 0,84575 | 0,955742 |
| 2810001G20Rik | 16,0173 | 16,5202 | 0,0446025 | 0,846 | 0,955758 |
| Tmem132c | 79,9582 | 78,2316 | -0,0314945 | 0,84605 | 0,955758 |
| Emilin3 | 4,23687 | 4,11669 | -0,0415129 | 0,84585 | 0,955758 |
| Nefl | 2,19754 | 2,10826 | -0,0598368 | 0,84605 | 0,955758 |
| Gm6981 | 0,50663 | 0,722889 | 0,512842 | 0,8464 | 0,955805 |
| Dazap1 | 78,7781 | 91,6434 | 0,218236 | 0,8464 | 0,955805 |
| 4930594C11Rik | 2,04884 | 2,16683 | 0,0807752 | 0,84635 | 0,955805 |
| Cpsf4 | 41,8058 | 42,7193 | 0,0311853 | 0,84645 | 0,955805 |
| Uhrf1bp1 | 9,1726 | 9,37189 | 0,0310093 | 0,8462 | 0,955805 |
| Immp2l | 1,34365 | 1,47309 | 0,132681 | 0,8466 | 0,955894 |
| Arpc3 | 147,213 | 150,38 | 0,0307142 | 0,84675 | 0,955983 |
| Actr5 | 8,13168 | 7,91577 | -0,0388239 | 0,84695 | 0,956047 |
| Mblac1 | 2,98124 | 2,85345 | -0,0632039 | 0,84695 | 0,956047 |
| Mtfr1 | 29,0563 | 29,7125 | 0,0322175 | 0,84705 | 0,956079 |
| Mob2 | 23,5992 | 24,1748 | 0,0347661 | 0,8472 | 0,956168 |
| Surf1 | 23,3716 | 25,3069 | 0,114776 | 0,84755 | 0,956352 |
| 5730455P16Rik | 10,3979 | 10,6417 | 0,0334366 | 0,8475 | 0,956352 |
| Rhob | 38,7994 | 39,6454 | 0,031116 | 0,84765 | 0,956352 |
| 4933439C10Rik | 5,61595 | 5,17964 | -0,116678 | 0,8476 | 0,956352 |
| Skiv2l | 25,4693 | 26,0153 | 0,0306007 | 0,8478 | 0,95636 |
| Atf6b | 61,5218 | 60,2536 | -0,0300503 | 0,84775 | 0,95636 |
| Mlx | 19,8194 | 19,219 | -0,0443848 | 0,84795 | 0,956449 |
| Sept10 | 27,0843 | 27,7018 | 0,0325242 | 0,84805 | 0,956481 |
| Fam181b | 12,8255 | 13,1531 | 0,0363891 | 0,8484 | 0,956795 |
| Dctn3 | 30,1749 | 29,4125 | -0,0369177 | 0,8485 | 0,956827 |
| Tmem191c | 1,81973 | 1,73264 | -0,0707573 | 0,8486 | 0,956834 |
| Dclre1b | 9,21405 | 8,61394 | -0,0971623 | 0,84865 | 0,956834 |
| Urgcp | 4,91045 | 5,05279 | 0,0412267 | 0,8488 | 0,956842 |
| Ccdc97 | 17,6328 | 18,0225 | 0,031538 | 0,8488 | 0,956842 |

| | | | | | |
|---|---|---|---|---|---|
| 1700008O03Rik | 0,703343 | 0,768175 | 0,127208 | 0,849 | 0,956857 |
| Cdk5 | 14,2387 | 14,5926 | 0,0354167 | 0,84905 | 0,956857 |
| Taok1 | 12,7714 | 13,0445 | 0,0305223 | 0,8491 | 0,956857 |
| Ldlr | 35,9618 | 35,2017 | -0,0308195 | 0,84895 | 0,956857 |
| Gm16617 | 3,73759 | 4,82864 | 0,369507 | 0,85025 | 0,956919 |
| Mir1905 | 660,572 | 781,714 | 0,242926 | 0,84985 | 0,956919 |
| Ecsit | 16,945 | 18,3238 | 0,112863 | 0,84955 | 0,956919 |
| 3830408C21Rik | 1,11823 | 1,17728 | 0,0742405 | 0,8502 | 0,956919 |
| Sh3d21 | 1,4078 | 1,47416 | 0,0664465 | 0,85005 | 0,956919 |
| Ulk4 | 1,17194 | 1,21789 | 0,0554942 | 0,8493 | 0,956919 |
| Map1s | 16,7863 | 17,1642 | 0,0321176 | 0,85025 | 0,956919 |
| Pcdh8 | 57,3016 | 58,5586 | 0,0313057 | 0,8497 | 0,956919 |
| Rev3l | 10,3748 | 10,5983 | 0,0307449 | 0,85 | 0,956919 |
| Cebpg | 23,8147 | 23,3265 | -0,0298843 | 0,84985 | 0,956919 |
| Larp4b | 22,2745 | 21,8002 | -0,0310514 | 0,8499 | 0,956919 |
| Abracl | 106,991 | 104,681 | -0,0314896 | 0,84955 | 0,956919 |
| Exoc4 | 16,5333 | 16,1729 | -0,0317981 | 0,84985 | 0,956919 |
| 1700021K19Rik | 5,54036 | 5,40741 | -0,0350429 | 0,8503 | 0,956919 |
| Fam58b | 13,6272 | 13,2637 | -0,0390087 | 0,84965 | 0,956919 |
| Grk5 | 4,60531 | 4,47797 | -0,0404526 | 0,84945 | 0,956919 |
| Ccdc103 | 0,523043 | 0,560792 | 0,100537 | 0,8508 | 0,956998 |
| Tnpo2 | 42,5852 | 41,6922 | -0,0305738 | 0,85075 | 0,956998 |
| Bcl11b | 2,27946 | 2,22039 | -0,0378829 | 0,8508 | 0,956998 |
| Polk | 3,78704 | 3,68272 | -0,0403014 | 0,8507 | 0,956998 |
| Tmem192 | 10,2665 | 9,95424 | -0,0445563 | 0,85055 | 0,956998 |
| Zfp810 | 8,4058 | 7,77219 | -0,113064 | 0,8508 | 0,956998 |
| Tor1b | 29,635 | 29,0131 | -0,0305976 | 0,8512 | 0,957287 |
| Nudt14 | 21,8896 | 21,2962 | -0,0396504 | 0,85115 | 0,957287 |
| B230118H07Rik | 9,0105 | 9,24476 | 0,037028 | 0,8513 | 0,957319 |
| Xrcc3 | 12,4246 | 13,002 | 0,0655387 | 0,8514 | 0,95735 |
| Fam132b | 5,144 | 5,00712 | -0,0389081 | 0,8516 | 0,957495 |
| Atxn1l | 12,6346 | 12,3639 | -0,0312491 | 0,8517 | 0,957502 |
| Rassf5 | 0,941252 | 0,901133 | -0,0628406 | 0,85175 | 0,957502 |
| Slco5a1 | 2,12857 | 2,18349 | 0,0367471 | 0,85205 | 0,957525 |
| Ubqln2 | 42,6332 | 41,7618 | -0,0297941 | 0,85185 | 0,957525 |
| Galc | 11,8734 | 11,618 | -0,0313636 | 0,8521 | 0,957525 |
| Ispd | 2,8475 | 2,75486 | -0,0477172 | 0,85195 | 0,957525 |
| Spata2 | 7,04807 | 6,71207 | -0,0704709 | 0,8522 | 0,957525 |
| Prr15 | 0,563316 | 0,505981 | -0,154859 | 0,85215 | 0,957525 |
| Ppp2r3c | 10,2466 | 11,0231 | 0,105375 | 0,8524 | 0,95754 |
| Suv420h1 | 18,0199 | 18,4337 | 0,0327536 | 0,8525 | 0,95754 |
| Mesdc1 | 14,5417 | 14,2288 | -0,0313803 | 0,85245 | 0,95754 |
| Nt5c | 28,85 | 28,0411 | -0,0410295 | 0,85245 | 0,95754 |
| Krt82 | 1,8938 | 1,97279 | 0,0589516 | 0,8527 | 0,957579 |
| Orai1 | 5,11768 | 5,27722 | 0,044287 | 0,8527 | 0,957579 |
| Satb1 | 8,586 | 8,4025 | -0,0311677 | 0,85275 | 0,957579 |
| Ccdc61 | 13,0725 | 12,7581 | -0,0351289 | 0,85285 | 0,957611 |
| Trmt61b | 12,101 | 12,6069 | 0,0590925 | 0,853 | 0,957619 |

| | | | | | |
|---|---|---|---|---|---|
| 1110034G24Rik | 4,84067 | 4,67533 | -0,0501384 | 0,853 | 0,957619 |
| Map3k12 | 5,20701 | 5,71797 | 0,135048 | 0,8545 | 0,957628 |
| Hspb9 | 0,884942 | 0,965027 | 0,124987 | 0,85425 | 0,957628 |
| Rnf185 | 14,0516 | 15,1226 | 0,105971 | 0,8547 | 0,957628 |
| Lrrc18 | 0,558031 | 0,599536 | 0,103501 | 0,85425 | 0,957628 |
| Hcn3 | 0,818733 | 0,856172 | 0,0645078 | 0,8538 | 0,957628 |
| Aspg | 6,79116 | 6,96931 | 0,0373594 | 0,8534 | 0,957628 |
| Ubxn2b | 7,02369 | 7,18809 | 0,0333777 | 0,8538 | 0,957628 |
| D10Jhu81e | 28,9603 | 29,6329 | 0,0331249 | 0,8539 | 0,957628 |
| Selk | 42,0704 | 43,0374 | 0,0327852 | 0,8544 | 0,957628 |
| Avl9 | 7,50767 | 7,66849 | 0,0305773 | 0,85405 | 0,957628 |
| Sipa1l3 | 9,82421 | 10,0342 | 0,0305183 | 0,85325 | 0,957628 |
| Mecp2 | 9,90436 | 10,1123 | 0,0299797 | 0,8542 | 0,957628 |
| Pkp4 | 31,8347 | 32,5025 | 0,0299532 | 0,85435 | 0,957628 |
| Eif5 | 128,349 | 125,763 | -0,029359 | 0,8548 | 0,957628 |
| 4632434I11Rik | 12,9977 | 12,7171 | -0,0314833 | 0,8548 | 0,957628 |
| D3Ertd254e | 6,30127 | 6,16176 | -0,0322988 | 0,8541 | 0,957628 |
| Scp2 | 10,9468 | 10,6893 | -0,0343507 | 0,85375 | 0,957628 |
| Six5 | 8,53089 | 8,32812 | -0,0347061 | 0,85415 | 0,957628 |
| Nhsl2 | 3,21254 | 3,12643 | -0,0391976 | 0,85415 | 0,957628 |
| Ap3s1 | 10,7254 | 10,434 | -0,0397478 | 0,8544 | 0,957628 |
| Thnsl1 | 3,35159 | 3,25748 | -0,0410878 | 0,85445 | 0,957628 |
| Nudt21 | 8,66736 | 8,40114 | -0,0450075 | 0,85455 | 0,957628 |
| Rhpn1 | 3,46392 | 3,35516 | -0,0460273 | 0,85475 | 0,957628 |
| Gnat2 | 0,648221 | 0,608826 | -0,0904565 | 0,85375 | 0,957628 |
| Cebpd | 1,07609 | 0,963725 | -0,159104 | 0,85385 | 0,957628 |
| 3110082I17Rik | 22,193 | 22,738 | 0,0350013 | 0,8552 | 0,957835 |
| Larp1 | 37,4338 | 38,1847 | 0,0286539 | 0,85515 | 0,957835 |
| Snora21 | 44,4306 | 42,361 | -0,0688153 | 0,8552 | 0,957835 |
| Zfp551 | 1,86133 | 1,93457 | 0,0556807 | 0,8554 | 0,957955 |
| Vgll4 | 22,9336 | 22,4 | -0,0339604 | 0,85545 | 0,957955 |
| Lrrc3b | 0,710949 | 0,754489 | 0,0857551 | 0,85575 | 0,95813 |
| L3hypdh | 3,0248 | 2,9157 | -0,0529948 | 0,85575 | 0,95813 |
| Fam208b | 22,5352 | 23,3468 | 0,0510416 | 0,85595 | 0,958201 |
| Tnrc18 | 20,1671 | 19,762 | -0,0292785 | 0,8561 | 0,958201 |
| Stx8 | 3,73995 | 3,64029 | -0,0389622 | 0,856 | 0,958201 |
| 9430091E24Rik | 1,55323 | 1,44757 | -0,10164 | 0,85605 | 0,958201 |
| 1700030K09Rik | 4,86746 | 5,29828 | 0,122354 | 0,85735 | 0,958234 |
| Ppp1r3e | 0,653821 | 0,693853 | 0,0857354 | 0,8577 | 0,958234 |
| 6330549D23Rik | 8,01078 | 8,33174 | 0,0566749 | 0,85725 | 0,958234 |
| Pcx | 11,7621 | 12,1925 | 0,0518569 | 0,85675 | 0,958234 |
| Ccdc28b | 9,00985 | 9,32038 | 0,0488869 | 0,8568 | 0,958234 |
| Gm3219 | 16,5331 | 17,0894 | 0,0477479 | 0,8575 | 0,958234 |
| Agfg2 | 8,49329 | 8,70916 | 0,0362107 | 0,8577 | 0,958234 |
| Ndufa1 | 119,716 | 122,395 | 0,0319377 | 0,857 | 0,958234 |
| Vps41 | 24,1497 | 24,6749 | 0,0310364 | 0,8571 | 0,958234 |
| Rps27l | 276,353 | 282,09 | 0,0296465 | 0,85655 | 0,958234 |
| Mlst8 | 17,177 | 17,5272 | 0,0291156 | 0,85735 | 0,958234 |

| | | | | | |
|---|---|---|---|---|---|
| Mtch2 | 41,1395 | 41,9565 | 0,0283672 | 0,85765 | 0,958234 |
| Tpp2 | 39,6671 | 40,4524 | 0,0282837 | 0,85785 | 0,958234 |
| Ror2 | 27,5006 | 28,0409 | 0,0280733 | 0,85725 | 0,958234 |
| Tbc1d9b | 14,5022 | 14,2073 | -0,0296411 | 0,85625 | 0,958234 |
| Commd7 | 10,7564 | 10,5157 | -0,0326533 | 0,85685 | 0,958234 |
| Sema6c | 6,61458 | 6,46166 | -0,0337464 | 0,8569 | 0,958234 |
| Fbxo7 | 9,59403 | 9,35834 | -0,0358846 | 0,85785 | 0,958234 |
| Iffo1 | 6,71303 | 6,54402 | -0,0367862 | 0,8574 | 0,958234 |
| Zfp661 | 4,79034 | 4,659 | -0,0401073 | 0,85725 | 0,958234 |
| Rasl10b | 1,54292 | 1,48924 | -0,0510863 | 0,85725 | 0,958234 |
| 3110057O12Rik | 1,87588 | 1,80726 | -0,0537625 | 0,8568 | 0,958234 |
| Ntm | 1,88202 | 1,80521 | -0,0601173 | 0,85685 | 0,958234 |
| Hist3h2ba | 4,47446 | 4,22324 | -0,0833647 | 0,8578 | 0,958234 |
| Nqo1 | 1,84855 | 1,77076 | -0,0620279 | 0,85795 | 0,958265 |
| Capza1 | 7,01058 | 7,18122 | 0,0346964 | 0,85815 | 0,958384 |
| Etfdh | 18,6234 | 19,0263 | 0,030873 | 0,8582 | 0,958384 |
| Tbcd | 20,5886 | 20,155 | -0,0307074 | 0,8586 | 0,958671 |
| Fam131a | 0,650404 | 0,579245 | -0,167162 | 0,8586 | 0,958671 |
| Strn | 8,81616 | 9,02281 | 0,0334255 | 0,8588 | 0,958685 |
| Ppp2r5c | 46,9764 | 47,9848 | 0,0306406 | 0,8587 | 0,958685 |
| Lztfl1 | 10,1687 | 9,94326 | -0,0323454 | 0,8589 | 0,958685 |
| Adrbk2 | 4,05009 | 3,95739 | -0,0334047 | 0,8589 | 0,958685 |
| Notch1 | 43,4483 | 44,3435 | 0,0294218 | 0,8594 | 0,959163 |
| 1110051M20Rik | 17,6566 | 18,0565 | 0,0323095 | 0,85965 | 0,959362 |
| Matk | 0,675188 | 0,637333 | -0,0832423 | 0,86015 | 0,95984 |
| Mllt1 | 42,1717 | 41,381 | -0,0273069 | 0,86035 | 0,959983 |
| Capns1 | 82,8926 | 84,5504 | 0,0285689 | 0,8605 | 0,95999 |
| 4930581F22Rik | 0,838302 | 0,727129 | -0,20526 | 0,86045 | 0,95999 |
| Gpr62 | 0,763008 | 1,05103 | 0,462032 | 0,8609 | 0,960036 |
| 5730507C01Rik | 1,11032 | 1,15815 | 0,0608507 | 0,86085 | 0,960036 |
| Golga1 | 9,33453 | 9,13987 | -0,0304039 | 0,8609 | 0,960036 |
| Unc13b | 2,75494 | 2,68796 | -0,0355094 | 0,86065 | 0,960036 |
| Hps6 | 2,71497 | 2,63073 | -0,0454758 | 0,8607 | 0,960036 |
| Zdhhc9 | 13,4865 | 13,2048 | -0,0304593 | 0,86105 | 0,960043 |
| Gm11696 | 1,1974 | 1,05328 | -0,185006 | 0,861 | 0,960043 |
| Amn1 | 13,2412 | 12,8901 | -0,0387744 | 0,86125 | 0,960186 |
| Morn2 | 11,0652 | 12,011 | 0,118336 | 0,8616 | 0,960335 |
| Phka2 | 7,09251 | 7,25126 | 0,0319357 | 0,86155 | 0,960335 |
| Optn | 4,22605 | 4,10396 | -0,0422939 | 0,86155 | 0,960335 |
| Ddx26b | 26,1642 | 26,6637 | 0,0272788 | 0,8619 | 0,960565 |
| Zfp316 | 3,42783 | 3,34819 | -0,0339168 | 0,86195 | 0,960565 |
| Snora81 | 47,7999 | 66,1265 | 0,468222 | 0,8625 | 0,96069 |
| A230103J11Rik | 2,26301 | 2,61709 | 0,20972 | 0,8636 | 0,96069 |
| Caskin1 | 0,504833 | 0,577462 | 0,19392 | 0,8645 | 0,96069 |
| Ift172 | 9,04472 | 10,2615 | 0,182092 | 0,8645 | 0,96069 |
| R74862 | 3,30469 | 3,58805 | 0,118687 | 0,8629 | 0,96069 |
| 4933434E20Rik | 17,8716 | 19,0847 | 0,0947503 | 0,86415 | 0,96069 |
| Zfp773 | 1,1878 | 1,2334 | 0,0543512 | 0,8626 | 0,96069 |

| | | | | | |
|---|---|---|---|---|---|
| Ribc1 | 2,68256 | 2,78419 | 0,0536464 | 0,86275 | 0,96069 |
| Zfp786 | 1,72764 | 1,7879 | 0,0494619 | 0,86425 | 0,96069 |
| Pelo | 22,7572 | 23,5021 | 0,0464665 | 0,8625 | 0,96069 |
| Tom1l1 | 12,4325 | 12,8105 | 0,0432185 | 0,86425 | 0,96069 |
| Hspa4l | 3,76457 | 3,85965 | 0,0359859 | 0,86435 | 0,96069 |
| Recql | 17,3205 | 17,743 | 0,0347767 | 0,86345 | 0,96069 |
| Zfyve21 | 17,031 | 17,4379 | 0,0340605 | 0,86335 | 0,96069 |
| BC017158 | 14,1133 | 14,4186 | 0,0308771 | 0,8634 | 0,96069 |
| Cpsf7 | 42,2705 | 43,1325 | 0,0291231 | 0,8633 | 0,96069 |
| Vangl2 | 27,0197 | 27,566 | 0,0288782 | 0,86315 | 0,96069 |
| Ndfip1 | 143,289 | 146,085 | 0,0278847 | 0,86365 | 0,96069 |
| Atr | 13,7167 | 13,9825 | 0,0276883 | 0,8636 | 0,96069 |
| Pbk | 64,101 | 65,3407 | 0,0276352 | 0,8629 | 0,96069 |
| Ubr3 | 19,6484 | 20,0183 | 0,0269046 | 0,8639 | 0,96069 |
| Poglut1 | 42,4708 | 41,6693 | -0,0274839 | 0,8633 | 0,96069 |
| Rac1 | 132,978 | 130,464 | -0,0275301 | 0,86225 | 0,96069 |
| Adipor2 | 44,3913 | 43,5482 | -0,0276662 | 0,8639 | 0,96069 |
| Pitrm1 | 126,05 | 123,578 | -0,0285811 | 0,86415 | 0,96069 |
| Tmem245 | 7,89556 | 7,74009 | -0,0286897 | 0,86465 | 0,96069 |
| Eri2 | 16,0265 | 15,7043 | -0,029303 | 0,86425 | 0,96069 |
| Snrk | 7,29068 | 7,13772 | -0,03059 | 0,86465 | 0,96069 |
| Kxd1 | 21,1481 | 20,6647 | -0,0333576 | 0,8642 | 0,96069 |
| Zbed4 | 2,75367 | 2,68679 | -0,0354712 | 0,8646 | 0,96069 |
| Tmem180 | 4,05753 | 3,93771 | -0,0432453 | 0,8646 | 0,96069 |
| Esrra | 5,89611 | 5,69131 | -0,0510044 | 0,86265 | 0,96069 |
| Ntan1 | 33,4726 | 32,1551 | -0,057935 | 0,86285 | 0,96069 |
| Ptger1 | 5,43154 | 5,20424 | -0,0616738 | 0,86305 | 0,96069 |
| Kdm1b | 4,20709 | 3,81344 | -0,141729 | 0,86295 | 0,96069 |
| H60b | 1,18864 | 1,06422 | -0,15951 | 0,86395 | 0,96069 |
| Gnai2 | 157,129 | 160,195 | 0,0278783 | 0,86475 | 0,960722 |
| Smim8 | 14,148 | 14,5231 | 0,0377576 | 0,8652 | 0,960798 |
| Ncaph2 | 47,6572 | 48,5767 | 0,0275697 | 0,86515 | 0,960798 |
| Uqcrc2 | 138,53 | 141,16 | 0,0271338 | 0,86505 | 0,960798 |
| Ptbp2 | 63,688 | 62,4874 | -0,0274554 | 0,86505 | 0,960798 |
| Haus2 | 20,7056 | 20,2163 | -0,0344982 | 0,86525 | 0,960798 |
| 2610044O15Rik8 | 6,45809 | 6,28445 | -0,0393207 | 0,86515 | 0,960798 |
| Letm1 | 21,4984 | 21,91 | 0,0273601 | 0,8655 | 0,960916 |
| Smim4 | 24,4991 | 23,6692 | -0,0497144 | 0,8655 | 0,960916 |
| Cadps | 1,31108 | 1,3506 | 0,0428389 | 0,8657 | 0,960929 |
| Spryd7 | 10,5719 | 10,3127 | -0,0358146 | 0,86575 | 0,960929 |
| Nfkb2 | 3,77274 | 3,6705 | -0,0396348 | 0,8658 | 0,960929 |
| Mrm1 | 3,96414 | 3,85481 | -0,040348 | 0,8656 | 0,960929 |
| Prr24 | 4,72443 | 4,86699 | 0,0428898 | 0,86595 | 0,961016 |
| Cdan1 | 7,6082 | 7,76644 | 0,0297 | 0,86655 | 0,961283 |
| Sema6a | 21,816 | 21,4219 | -0,0262978 | 0,8664 | 0,961283 |
| Polr3f | 17,7817 | 17,4438 | -0,0276796 | 0,86635 | 0,961283 |
| Ska1 | 12,9915 | 12,7144 | -0,0311069 | 0,86655 | 0,961283 |
| Pcdhga6 | 0,813427 | 0,642386 | -0,340573 | 0,86645 | 0,961283 |

| | | | | | |
|---|---|---|---|---|---|
| Acat3 | 0,605576 | 0,856856 | 0,500744 | 0,86705 | 0,961598 |
| Comtd1 | 1,74048 | 2,00452 | 0,20377 | 0,867 | 0,961598 |
| Chodl | 3,01287 | 3,10388 | 0,0429314 | 0,86695 | 0,961598 |
| Relb | 1,04707 | 1,0919 | 0,0604775 | 0,86735 | 0,961612 |
| Atp5f1 | 94,9809 | 98,1055 | 0,0466961 | 0,8673 | 0,961612 |
| Dnajc22 | 4,74817 | 4,89365 | 0,0435392 | 0,86735 | 0,961612 |
| Cyhr1 | 27,5574 | 26,964 | -0,0314052 | 0,86725 | 0,961612 |
| Ahctf1 | 33,9852 | 33,3838 | -0,0257594 | 0,8676 | 0,961809 |
| Gorasp1 | 11,3149 | 11,7269 | 0,051589 | 0,86785 | 0,961867 |
| Smim20 | 22,8341 | 23,3631 | 0,0330472 | 0,86805 | 0,961867 |
| Dmtn | 9,04957 | 9,23065 | 0,0285829 | 0,8679 | 0,961867 |
| Wbp1 | 45,1254 | 46,0129 | 0,0280979 | 0,8683 | 0,961867 |
| Bhlhb9 | 36,7193 | 37,4278 | 0,0275696 | 0,8679 | 0,961867 |
| Arfgap2 | 23,8018 | 24,2519 | 0,0270285 | 0,86795 | 0,961867 |
| Ints2 | 22,7588 | 22,3377 | -0,0269446 | 0,8683 | 0,961867 |
| Aggf1 | 29,8409 | 29,2801 | -0,0273701 | 0,86825 | 0,961867 |
| 2700081O15Rik | 30,9204 | 29,8211 | -0,0522238 | 0,86815 | 0,961867 |
| Cstf3 | 41,2899 | 42,2614 | 0,0335492 | 0,8684 | 0,961898 |
| Taf1b | 16,8653 | 17,2231 | 0,0302877 | 0,8685 | 0,961905 |
| Pddc1 | 21,1765 | 20,7295 | -0,0307732 | 0,86855 | 0,961905 |
| Smarca2 | 19,5009 | 19,0814 | -0,0313763 | 0,86865 | 0,961911 |
| Hyi | 4,05697 | 3,62635 | -0,161883 | 0,8687 | 0,961911 |
| 2010111I01Rik | 68,8664 | 70,4271 | 0,0323294 | 0,8688 | 0,961918 |
| Rnf38 | 29,6491 | 29,1156 | -0,0261973 | 0,86885 | 0,961918 |
| Eme2 | 8,55421 | 8,77559 | 0,0368621 | 0,86895 | 0,961949 |
| Gm3086 | 0,991707 | 1,32903 | 0,422383 | 0,86935 | 0,962128 |
| Mir3960 | 261,781 | 297,321 | 0,183657 | 0,86935 | 0,962128 |
| Ube2l3 | 57,9113 | 58,9911 | 0,0266529 | 0,86925 | 0,962128 |
| Rc3h2 | 10,0062 | 9,82306 | -0,0266553 | 0,8694 | 0,962128 |
| Mir1982 | 421,72 | 531,98 | 0,335087 | 0,87035 | 0,962239 |
| D130020L05Rik | 0,823772 | 0,981418 | 0,252623 | 0,87045 | 0,962239 |
| Atf7 | 3,39352 | 3,4887 | 0,039905 | 0,871 | 0,962239 |
| Gltpd1 | 7,07131 | 7,23545 | 0,0331056 | 0,87015 | 0,962239 |
| Dennd1b | 2,59684 | 2,65416 | 0,0314998 | 0,8712 | 0,962239 |
| Hspbap1 | 5,31516 | 5,42929 | 0,030651 | 0,87075 | 0,962239 |
| Ndufa5 | 104,797 | 106,909 | 0,0287911 | 0,87005 | 0,962239 |
| Lonp2 | 20,0964 | 20,4956 | 0,0283775 | 0,87035 | 0,962239 |
| Prps1l3 | 37,2983 | 37,9937 | 0,0266479 | 0,8713 | 0,962239 |
| Arnt2 | 13,3979 | 13,644 | 0,0262665 | 0,87105 | 0,962239 |
| Frs2 | 13,4484 | 13,6946 | 0,0261767 | 0,8711 | 0,962239 |
| Gcn1l1 | 25,6568 | 26,1197 | 0,0258002 | 0,8702 | 0,962239 |
| Zmym5 | 13,8518 | 14,0994 | 0,0255641 | 0,87125 | 0,962239 |
| Rbm27 | 20,5405 | 20,9068 | 0,0255004 | 0,8697 | 0,962239 |
| Ehmt2 | 59,7387 | 60,7928 | 0,0252334 | 0,8699 | 0,962239 |
| Vezf1 | 39,7981 | 40,4906 | 0,0248848 | 0,87125 | 0,962239 |
| Nop10 | 128,143 | 125,813 | -0,0264659 | 0,87025 | 0,962239 |
| Ndufs8 | 83,7285 | 82,1592 | -0,0272958 | 0,8713 | 0,962239 |
| 0610009O20Rik | 33,0038 | 32,3724 | -0,0278665 | 0,86985 | 0,962239 |

| | | | | | |
|---|---|---|---|---|---|
| Scamp3 | 41,0058 | 40,2119 | -0,0282048 | 0,87085 | 0,962239 |
| Ptcd2 | 26,6568 | 26,14 | -0,0282474 | 0,8703 | 0,962239 |
| Nr2c1 | 9,12101 | 8,93697 | -0,0294069 | 0,87125 | 0,962239 |
| Nfib | 3,07058 | 3,00547 | -0,030921 | 0,87035 | 0,962239 |
| 2010320M18Rik | 15,3294 | 14,9277 | -0,0383032 | 0,87085 | 0,962239 |
| Fam227a | 0,609492 | 0,586772 | -0,0548071 | 0,87105 | 0,962239 |
| E2f3 | 29,8647 | 29,3104 | -0,0270296 | 0,8714 | 0,962246 |
| Ovca2 | 18,0264 | 17,597 | -0,0347789 | 0,87145 | 0,962246 |
| Gm12359 | 1,31601 | 1,74312 | 0,405499 | 0,8724 | 0,962338 |
| Polr2e | 128,695 | 148,602 | 0,207498 | 0,87285 | 0,962338 |
| Pusl1 | 14,0235 | 14,6996 | 0,0679304 | 0,87345 | 0,962338 |
| Lrsam1 | 2,71862 | 2,84528 | 0,0656972 | 0,8729 | 0,962338 |
| Trip4 | 9,00978 | 9,37153 | 0,0567926 | 0,87255 | 0,962338 |
| Gadd45a | 9,38882 | 9,65401 | 0,0401832 | 0,8718 | 0,962338 |
| Arl10 | 20,7337 | 21,1855 | 0,0310988 | 0,87225 | 0,962338 |
| Fam175a | 10,7275 | 10,9535 | 0,0300774 | 0,8722 | 0,962338 |
| Tcea1 | 12,7449 | 12,9983 | 0,028409 | 0,8728 | 0,962338 |
| Zfp961 | 15,6951 | 15,9965 | 0,0274404 | 0,8725 | 0,962338 |
| Repin1 | 18,4554 | 18,8028 | 0,0269028 | 0,87345 | 0,962338 |
| Txndc9 | 15,4284 | 15,7092 | 0,0260221 | 0,8734 | 0,962338 |
| Gnb2 | 148,08 | 150,76 | 0,0258767 | 0,8731 | 0,962338 |
| Plk4 | 41,0165 | 41,7346 | 0,0250381 | 0,87235 | 0,962338 |
| Cse1l | 131,925 | 134,213 | 0,0248028 | 0,87325 | 0,962338 |
| Arf4 | 146,581 | 144,024 | -0,0253957 | 0,87265 | 0,962338 |
| A230046K03Rik | 15,2222 | 14,9564 | -0,0254137 | 0,87215 | 0,962338 |
| Dnajc3 | 27,4085 | 26,9274 | -0,0255512 | 0,87355 | 0,962338 |
| Mrpl20 | 177,833 | 174,674 | -0,0258547 | 0,8718 | 0,962338 |
| Pcnxl4 | 6,91795 | 6,78398 | -0,0282131 | 0,87245 | 0,962338 |
| Rps6kc1 | 8,63415 | 8,46343 | -0,0288114 | 0,87305 | 0,962338 |
| Atg101 | 26,8622 | 26,3232 | -0,029241 | 0,8721 | 0,962338 |
| Trim23 | 6,30778 | 6,18114 | -0,0292601 | 0,8734 | 0,962338 |
| Flcn | 10,6768 | 10,46 | -0,0295851 | 0,87345 | 0,962338 |
| Spata6 | 8,43498 | 8,24825 | -0,0322969 | 0,87355 | 0,962338 |
| Camk1g | 3,36697 | 3,27337 | -0,0406743 | 0,873 | 0,962338 |
| Magee2 | 1,8111 | 1,75036 | -0,0492111 | 0,872 | 0,962338 |
| Chn1os3 | 0,757751 | 0,668389 | -0,181037 | 0,87175 | 0,962338 |
| Tomm34 | 18,9949 | 19,3602 | 0,0274821 | 0,87385 | 0,962565 |
| Clpp | 45,194 | 46,0466 | 0,0269617 | 0,8739 | 0,962565 |
| Neu3 | 1,4153 | 1,46112 | 0,0459749 | 0,87405 | 0,962571 |
| Wdr8 | 15,7616 | 16,1003 | 0,0306663 | 0,874 | 0,962571 |
| Ppa2 | 35,4848 | 36,1876 | 0,0282973 | 0,87415 | 0,962602 |
| Zfp169 | 2,18678 | 2,12813 | -0,0392204 | 0,87425 | 0,962633 |
| Stat1 | 3,54421 | 3,46666 | -0,0319212 | 0,8746 | 0,962939 |
| Txndc11 | 14,237 | 14,7714 | 0,0531647 | 0,87485 | 0,962976 |
| Alkbh3 | 11,009 | 11,2825 | 0,0354088 | 0,8748 | 0,962976 |
| Ndor1 | 12,5107 | 12,7423 | 0,0264565 | 0,87485 | 0,962976 |
| Glis1 | 1,2016 | 1,24368 | 0,0496545 | 0,875 | 0,963062 |
| Gm5617 | 6,16755 | 5,94035 | -0,0541504 | 0,8752 | 0,963203 |

| | | | | | |
|---|---|---|---|---|---|
| Prkab1 | 30,5644 | 31,1198 | 0,0259818 | 0,8753 | 0,963234 |
| Pcdhgb1 | 0,475725 | 0,621672 | 0,386025 | 0,87555 | 0,96335 |
| Zfp414 | 19,4644 | 19,0566 | -0,0305493 | 0,8755 | 0,96335 |
| Amigo3 | 2,35411 | 2,5184 | 0,0973244 | 0,87605 | 0,963479 |
| Meis3 | 18,0567 | 18,4126 | 0,0281536 | 0,8759 | 0,963479 |
| Tmem231 | 9,33578 | 9,51918 | 0,0280674 | 0,8761 | 0,963479 |
| Ndc80 | 32,8242 | 33,3834 | 0,024373 | 0,876 | 0,963479 |
| Lhfp | 8,18975 | 8,00529 | -0,0328657 | 0,8761 | 0,963479 |
| 3632451O06Rik | 4,67269 | 4,55986 | -0,0352633 | 0,8759 | 0,963479 |
| Ccdc74a | 0,909564 | 0,85971 | -0,0813253 | 0,87625 | 0,963486 |
| C530005A16Rik | 0,771161 | 0,687797 | -0,16505 | 0,8762 | 0,963486 |
| Mapk14 | 45,1451 | 44,3711 | -0,0249493 | 0,87635 | 0,963516 |
| E130309D02Rik | 23,6725 | 24,118 | 0,0268972 | 0,8765 | 0,963602 |
| 1700003F12Rik | 0,783488 | 0,871383 | 0,153396 | 0,87725 | 0,96361 |
| Hdac8 | 6,52403 | 6,68836 | 0,0358885 | 0,8767 | 0,96361 |
| Rnmtl1 | 12,7583 | 13,029 | 0,0302907 | 0,87725 | 0,96361 |
| Bnip3 | 14,8954 | 15,2081 | 0,0299675 | 0,87675 | 0,96361 |
| Tmem261 | 62,9724 | 64,1685 | 0,027146 | 0,877 | 0,96361 |
| Brwd3 | 7,0098 | 7,13958 | 0,0264661 | 0,877 | 0,96361 |
| Zfp866 | 12,0851 | 12,3085 | 0,0264254 | 0,87705 | 0,96361 |
| Fbrs | 18,1957 | 18,5261 | 0,0259645 | 0,8766 | 0,96361 |
| Dennd4a | 3,62403 | 3,55519 | -0,0276675 | 0,8771 | 0,96361 |
| Cul9 | 0,622163 | 0,602468 | -0,046406 | 0,877 | 0,96361 |
| Acyp2 | 5,39995 | 5,20581 | -0,0528238 | 0,8773 | 0,96361 |
| 1700065D16Rik | 0,845708 | 1,01375 | 0,261466 | 0,87795 | 0,96369 |
| Mgarp | 2,59976 | 2,69422 | 0,0514877 | 0,8778 | 0,96369 |
| Sfxn5 | 2,65786 | 2,72653 | 0,0368039 | 0,8779 | 0,96369 |
| Cdc42ep5 | 13,8759 | 14,2269 | 0,0360374 | 0,8776 | 0,96369 |
| Pak2 | 36,2182 | 35,6022 | -0,0247457 | 0,87765 | 0,96369 |
| Mcts1 | 44,1947 | 43,3627 | -0,0274204 | 0,87775 | 0,96369 |
| Nicn1 | 10,597 | 10,3705 | -0,0311763 | 0,87745 | 0,96369 |
| Fahd2a | 6,34829 | 6,17088 | -0,0408913 | 0,87795 | 0,96369 |
| 9930021J03Rik | 8,42904 | 8,57807 | 0,0252851 | 0,87845 | 0,963922 |
| Max | 40,9571 | 41,6687 | 0,0248494 | 0,8784 | 0,963922 |
| Dctn4 | 28,2171 | 27,7325 | -0,0249929 | 0,87835 | 0,963922 |
| Pogk | 20,0209 | 19,6355 | -0,0280384 | 0,8784 | 0,963922 |
| Gramd4 | 15,6144 | 15,8943 | 0,0256325 | 0,8787 | 0,963959 |
| Phc2 | 42,971 | 42,2524 | -0,0243299 | 0,8786 | 0,963959 |
| Inpp4a | 2,9584 | 2,88985 | -0,0338268 | 0,87865 | 0,963959 |
| Cenpq | 36,6272 | 35,9777 | -0,0258113 | 0,8788 | 0,963989 |
| Tmem258 | 294,278 | 289,495 | -0,023642 | 0,879 | 0,96413 |
| Dbp | 1,59425 | 1,72432 | 0,113145 | 0,87915 | 0,964215 |
| Dennd4c | 4,57493 | 4,48946 | -0,0272055 | 0,8793 | 0,9643 |
| Cdh6 | 10,3465 | 10,1426 | -0,0287103 | 0,87965 | 0,964605 |
| Prickle4 | 1,42967 | 1,62228 | 0,182335 | 0,8811 | 0,964722 |
| Lamtor4 | 14,4597 | 14,8053 | 0,0340768 | 0,88095 | 0,964722 |
| Spryd4 | 19,4466 | 19,8569 | 0,0301211 | 0,8812 | 0,964722 |
| Dbf4 | 69,1947 | 70,3427 | 0,0237397 | 0,8804 | 0,964722 |

| | | | | | |
|---|---|---|---|---|---|
| Hectd1 | 28,2296 | 27,7762 | -0,0233566 | 0,88115 | 0,964722 |
| Prrc1 | 19,2957 | 18,9798 | -0,0238184 | 0,881 | 0,964722 |
| Ddx6 | 72,057 | 70,8675 | -0,0240138 | 0,8811 | 0,964722 |
| Zcchc6 | 15,4115 | 15,147 | -0,024976 | 0,8808 | 0,964722 |
| Stil | 12,3455 | 12,1298 | -0,0254313 | 0,88075 | 0,964722 |
| Arfgef2 | 5,97261 | 5,86437 | -0,0263865 | 0,88015 | 0,964722 |
| Grik5 | 12,9427 | 12,705 | -0,0267452 | 0,87985 | 0,964722 |
| Rnf123 | 12,5163 | 12,2619 | -0,0296235 | 0,88075 | 0,964722 |
| Tor1aip2 | 62,2338 | 60,9507 | -0,030056 | 0,88045 | 0,964722 |
| Slc4a8 | 1,26937 | 1,24107 | -0,0325273 | 0,8804 | 0,964722 |
| C130046K22Rik | 2,99476 | 2,9207 | -0,0361268 | 0,88085 | 0,964722 |
| Fbxl12 | 10,4193 | 10,012 | -0,0575247 | 0,88055 | 0,964722 |
| Eno4 | 0,736643 | 0,705849 | -0,0616066 | 0,8809 | 0,964722 |
| Yipf2 | 9,1251 | 8,48055 | -0,105681 | 0,881 | 0,964722 |
| Arid3a | 33,4081 | 29,8156 | -0,164133 | 0,88035 | 0,964722 |
| BC005764 | 23,7671 | 19,4051 | -0,292528 | 0,88055 | 0,964722 |
| Sdsl | 3,71118 | 3,83146 | 0,0460134 | 0,88135 | 0,964758 |
| Fam109a | 20,3251 | 20,6736 | 0,0245242 | 0,88145 | 0,964758 |
| Acot1 | 4,6961 | 4,56554 | -0,0406771 | 0,88145 | 0,964758 |
| Rbm34 | 15,3421 | 15,0766 | -0,0251831 | 0,88155 | 0,964789 |
| Ip6k2 | 27,1607 | 27,6613 | 0,0263486 | 0,8817 | 0,964874 |
| BC048507 | 37,9793 | 38,803 | 0,0309545 | 0,88205 | 0,964941 |
| Nomo1 | 48,5028 | 47,7334 | -0,0230678 | 0,88205 | 0,964941 |
| Six1 | 17,5685 | 17,2565 | -0,0258465 | 0,8819 | 0,964941 |
| Zfp2 | 6,58594 | 6,45159 | -0,0297328 | 0,88195 | 0,964941 |
| Sec13 | 133,453 | 135,7 | 0,0240865 | 0,88215 | 0,964947 |
| Trpc2 | 2,53271 | 2,4169 | -0,0675229 | 0,8822 | 0,964947 |
| Dcp2 | 31,822 | 31,3106 | -0,0233722 | 0,8823 | 0,964977 |
| Eif2b4 | 29,8892 | 33,2523 | 0,153827 | 0,8824 | 0,965008 |
| Mapk8ip2 | 1,33321 | 1,37293 | 0,0423494 | 0,883 | 0,965166 |
| Coq10a | 4,34427 | 4,45094 | 0,0349946 | 0,8836 | 0,965166 |
| Ccdc12 | 25,0709 | 25,5953 | 0,0298653 | 0,8837 | 0,965166 |
| Praf2 | 20,0686 | 20,4861 | 0,0297034 | 0,88295 | 0,965166 |
| Ccser2 | 6,91734 | 7,03564 | 0,0244635 | 0,8829 | 0,965166 |
| Clint1 | 57,8954 | 58,8386 | 0,0233139 | 0,88265 | 0,965166 |
| Setd3 | 22,7756 | 22,3893 | -0,0246851 | 0,8836 | 0,965166 |
| Etl4 | 11,212 | 11,0194 | -0,0250051 | 0,8834 | 0,965166 |
| Klhl17 | 17,8321 | 17,5229 | -0,0252361 | 0,88365 | 0,965166 |
| Numb | 14,0853 | 13,8349 | -0,0258726 | 0,88335 | 0,965166 |
| D1Ertd622e | 4,40595 | 4,30852 | -0,03226 | 0,88285 | 0,965166 |
| Prr7 | 4,71202 | 4,58042 | -0,040866 | 0,8837 | 0,965166 |
| Ccdc106 | 2,68636 | 2,59948 | -0,0474287 | 0,88355 | 0,965166 |
| Elovl1 | 22,5783 | 21,7801 | -0,0519216 | 0,8837 | 0,965166 |
| Lrrc24 | 0,64421 | 0,597129 | -0,109488 | 0,8833 | 0,965166 |
| Egfl8 | 1,32797 | 1,12355 | -0,241159 | 0,88295 | 0,965166 |
| 1700029J07Rik | 1,62944 | 1,67949 | 0,043648 | 0,88385 | 0,965251 |
| Rps15a-ps6 | 0,735397 | 0,970566 | 0,400304 | 0,88455 | 0,965275 |
| 1700001G11Rik | 0,69141 | 0,84395 | 0,287616 | 0,8845 | 0,965275 |

| | | | | | |
|---|---|---|---|---|---|
| Rint1 | 11,0526 | 11,5602 | 0,0647712 | 0,8847 | 0,965275 |
| Fsip1 | 1,29348 | 1,34446 | 0,0557754 | 0,8851 | 0,965275 |
| Pkn3 | 7,84029 | 8,04217 | 0,0366783 | 0,8843 | 0,965275 |
| Bloc1s4 | 10,0774 | 10,3023 | 0,031847 | 0,88425 | 0,965275 |
| Tomm5 | 35,1388 | 35,8953 | 0,0307305 | 0,8846 | 0,965275 |
| Sars2 | 17,4488 | 17,7677 | 0,026133 | 0,88505 | 0,965275 |
| Rpl3 | 46,2743 | 47,078 | 0,0248419 | 0,8847 | 0,965275 |
| Xrcc5 | 27,8437 | 28,314 | 0,0241647 | 0,8842 | 0,965275 |
| Bnip2 | 26,8227 | 26,4018 | -0,0228229 | 0,8847 | 0,965275 |
| Cetn3 | 122,132 | 120,19 | -0,0231235 | 0,88485 | 0,965275 |
| Ivd | 35,8063 | 35,2297 | -0,0234212 | 0,885 | 0,965275 |
| Cnot11 | 27,5371 | 27,0866 | -0,0237976 | 0,8849 | 0,965275 |
| Zcchc2 | 3,12273 | 3,06081 | -0,0288929 | 0,88485 | 0,965275 |
| Akirin2 | 68,8056 | 67,1836 | -0,0344162 | 0,8842 | 0,965275 |
| 2310030G06Rik | 2,42325 | 2,3375 | -0,0519749 | 0,88405 | 0,965275 |
| Gm5088 | 13,3694 | 13,5978 | 0,0244461 | 0,8857 | 0,965771 |
| B3gnt4 | 0,643003 | 0,529705 | -0,279638 | 0,88565 | 0,965771 |
| 9930104L06Rik | 11,0853 | 10,8812 | -0,0268009 | 0,88595 | 0,965965 |
| Ube2t | 25,1313 | 25,6489 | 0,0294155 | 0,88625 | 0,96611 |
| Arfgef1 | 25,7866 | 25,3771 | -0,0230933 | 0,8863 | 0,96611 |
| Fbxo44 | 5,47517 | 5,34045 | -0,0359419 | 0,8862 | 0,96611 |
| Wnt10a | 1,27383 | 1,32084 | 0,052285 | 0,8867 | 0,966128 |
| Edar | 0,820637 | 0,847517 | 0,0464987 | 0,88675 | 0,966128 |
| Gtf3a | 46,126 | 47,1313 | 0,0311043 | 0,8865 | 0,966128 |
| Gpld1 | 7,3629 | 7,49674 | 0,025989 | 0,88675 | 0,966128 |
| Tmem65 | 13,0753 | 12,8596 | -0,0239989 | 0,88675 | 0,966128 |
| Raet1d | 3,95916 | 3,81159 | -0,0548011 | 0,8866 | 0,966128 |
| Ss18l1 | 3,65866 | 3,73422 | 0,0294951 | 0,88705 | 0,966273 |
| Pitpnb | 57,5512 | 58,45 | 0,0223566 | 0,88705 | 0,966273 |
| Cgref1 | 3,04744 | 2,29242 | -0,410726 | 0,8871 | 0,966273 |
| Man2c1os | 0,917067 | 1,0701 | 0,222646 | 0,88775 | 0,966563 |
| Zfp712 | 1,43207 | 1,47197 | 0,0396496 | 0,88755 | 0,966563 |
| Nacc2 | 0,929269 | 0,953843 | 0,0376564 | 0,8878 | 0,966563 |
| Ankrd54 | 16,0346 | 16,3294 | 0,0262801 | 0,8877 | 0,966563 |
| Zfp385a | 22,3149 | 22,6879 | 0,0239213 | 0,88765 | 0,966563 |
| Vwa2 | 37,4018 | 36,8099 | -0,023012 | 0,8875 | 0,966563 |
| Mir343 | 459,215 | 480,035 | 0,0639714 | 0,88805 | 0,966592 |
| Tmem177 | 5,06092 | 5,16133 | 0,0283436 | 0,8885 | 0,966592 |
| Nrde2 | 7,52424 | 7,66191 | 0,0261565 | 0,8884 | 0,966592 |
| Uqcrb | 82,7719 | 84,2582 | 0,0256758 | 0,88835 | 0,966592 |
| Pmm2 | 20,8727 | 21,2366 | 0,0249352 | 0,8881 | 0,966592 |
| Arhgap1 | 17,3359 | 17,6215 | 0,0235728 | 0,88815 | 0,966592 |
| Tab1 | 16,1151 | 15,8388 | -0,0249526 | 0,88835 | 0,966592 |
| Fastkd5 | 10,5996 | 10,3633 | -0,032527 | 0,88855 | 0,966592 |
| Galm | 4,00263 | 3,90888 | -0,0341931 | 0,88855 | 0,966592 |
| Klf8 | 0,871298 | 0,845845 | -0,0427726 | 0,88825 | 0,966592 |
| Pigl | 8,8737 | 9,05983 | 0,0299492 | 0,88875 | 0,966707 |
| Tspan5 | 15,644 | 15,9042 | 0,0238039 | 0,8888 | 0,966707 |

| | | | | | |
|---|---|---|---|---|---|
| Nrbp1 | 63,2337 | 67,3593 | 0,0911817 | 0,88905 | 0,966797 |
| Sergef | 4,72709 | 4,84869 | 0,0366406 | 0,88905 | 0,966797 |
| Ehd1 | 44,6604 | 45,3253 | 0,021322 | 0,8891 | 0,966797 |
| Ctu2 | 17,7724 | 18,2662 | 0,0395427 | 0,8898 | 0,96685 |
| Aamdc | 9,80207 | 10,0632 | 0,0379332 | 0,8894 | 0,96685 |
| Ipo13 | 22,8882 | 23,2576 | 0,0231004 | 0,8897 | 0,96685 |
| Ralgapa1 | 12,1794 | 12,3729 | 0,022735 | 0,88965 | 0,96685 |
| Hnrnph2 | 45,5086 | 46,2252 | 0,0225434 | 0,88925 | 0,96685 |
| Snx2 | 36,4653 | 35,888 | -0,0230232 | 0,8898 | 0,96685 |
| Trmt61a | 16,0764 | 15,8098 | -0,0241217 | 0,88955 | 0,96685 |
| Mlycd | 2,73177 | 2,65987 | -0,038481 | 0,88945 | 0,96685 |
| 2700046A07Rik | 1,04349 | 1,0049 | -0,0543655 | 0,8893 | 0,96685 |
| Cryz | 10,5899 | 10,8188 | 0,0308594 | 0,88995 | 0,966856 |
| Atp6v0d1 | 32,1731 | 32,6803 | 0,0225681 | 0,8899 | 0,966856 |
| Ttc12 | 1,43765 | 1,47765 | 0,0395905 | 0,8904 | 0,966963 |
| Pik3ap1 | 3,36419 | 3,44297 | 0,0333961 | 0,8902 | 0,966963 |
| Ulk3 | 3,84509 | 3,92889 | 0,0311045 | 0,8907 | 0,966963 |
| Tpm4 | 272,283 | 276,439 | 0,0218541 | 0,89035 | 0,966963 |
| Zfp664 | 86,823 | 88,1299 | 0,0215544 | 0,8907 | 0,966963 |
| Rragb | 10,2043 | 10,018 | -0,0265837 | 0,89065 | 0,966963 |
| Zfp11 | 2,93435 | 2,87282 | -0,0305765 | 0,8906 | 0,966963 |
| Spata33 | 3,37464 | 3,21739 | -0,0688434 | 0,8905 | 0,966963 |
| Tmem243 | 10,2241 | 9,56332 | -0,0963953 | 0,89035 | 0,966963 |
| Tcte2 | 2,00571 | 2,4875 | 0,310583 | 0,89135 | 0,967016 |
| Spag16 | 2,14746 | 2,19869 | 0,034011 | 0,89105 | 0,967016 |
| Cmip | 15,2368 | 15,4787 | 0,022726 | 0,8914 | 0,967016 |
| Cdc42 | 172,488 | 169,87 | -0,0220668 | 0,8913 | 0,967016 |
| Otud5 | 28,1568 | 27,7265 | -0,0222162 | 0,891 | 0,967016 |
| Ulk1 | 16,6667 | 16,4027 | -0,0230321 | 0,8912 | 0,967016 |
| Fam19a4 | 1,66674 | 1,6167 | -0,0439752 | 0,89115 | 0,967016 |
| Ap4s1 | 16,4893 | 15,6919 | -0,0715124 | 0,89135 | 0,967016 |
| Ccdc125 | 2,13368 | 1,9732 | -0,112808 | 0,89115 | 0,967016 |
| Akr1b8 | 12,524 | 12,2765 | -0,0287926 | 0,8915 | 0,967046 |
| Polr2c | 21,2211 | 21,6161 | 0,0266075 | 0,89165 | 0,967087 |
| Ddx20 | 28,1514 | 28,6029 | 0,0229524 | 0,89185 | 0,967087 |
| Gosr1 | 13,1397 | 12,9374 | -0,0223856 | 0,89185 | 0,967087 |
| Gm561 | 34,9359 | 34,1615 | -0,0323366 | 0,8919 | 0,967087 |
| Plekha2 | 1,61973 | 1,58182 | -0,03417 | 0,8919 | 0,967087 |
| 2410131K14Rik | 12,904 | 13,1177 | 0,0237032 | 0,89205 | 0,967093 |
| Lin52 | 5,9292 | 5,80453 | -0,030658 | 0,892 | 0,967093 |
| Hace1 | 12,3682 | 12,566 | 0,0228928 | 0,8922 | 0,967099 |
| Tubb2a | 9,10197 | 8,91537 | -0,0298841 | 0,8922 | 0,967099 |
| Apbb3 | 9,34009 | 8,99478 | -0,0543478 | 0,8924 | 0,967237 |
| Hoxd3 | 47,5571 | 48,8934 | 0,0399804 | 0,8927 | 0,967278 |
| Lbh | 16,0892 | 16,3458 | 0,0228323 | 0,89275 | 0,967278 |
| Tead1 | 18,6341 | 18,3615 | -0,0212626 | 0,89255 | 0,967278 |
| Dstn | 117,315 | 115,584 | -0,021443 | 0,8928 | 0,967278 |
| Hus1 | 13,8621 | 13,6431 | -0,0229772 | 0,8926 | 0,967278 |

| | | | | | |
|---|---|---|---|---|---|
| Pfas | 26,7237 | 27,1094 | 0,0206747 | 0,8929 | 0,967308 |
| Rbm39 | 196,784 | 199,93 | 0,0228827 | 0,89315 | 0,967398 |
| Ap2s1 | 120,313 | 122,18 | 0,0222061 | 0,8931 | 0,967398 |
| Snord33 | 1306,94 | 1114,42 | -0,229892 | 0,8932 | 0,967398 |
| C2cd5 | 11,6541 | 11,4709 | -0,0228594 | 0,8933 | 0,967428 |
| Prrg2 | 6,27644 | 6,42959 | 0,0347806 | 0,89365 | 0,967704 |
| Hmga2 | 167,012 | 169,677 | 0,0228384 | 0,8937 | 0,967704 |
| Ext1 | 37,9276 | 37,3606 | -0,0217303 | 0,89395 | 0,967897 |
| Ercc6 | 5,95776 | 6,05599 | 0,0235934 | 0,89455 | 0,968076 |
| Vwa9 | 30,0684 | 30,5127 | 0,0211644 | 0,89455 | 0,968076 |
| Incenp | 74,7077 | 75,7615 | 0,0202091 | 0,8944 | 0,968076 |
| Scap | 22,3805 | 22,0484 | -0,0215695 | 0,8945 | 0,968076 |
| Arhgef10l | 8,02913 | 7,89852 | -0,0236614 | 0,8944 | 0,968076 |
| Mtf1 | 3,53978 | 3,46986 | -0,0287826 | 0,8943 | 0,968076 |
| Col11a1 | 6,55626 | 6,65843 | 0,0223087 | 0,8948 | 0,968165 |
| Zfp788 | 7,51718 | 7,39285 | -0,0240604 | 0,89475 | 0,968165 |
| Pcdhga5 | 1,79067 | 1,59248 | -0,16922 | 0,89485 | 0,968165 |
| Golph3 | 50,6017 | 49,8515 | -0,0215478 | 0,89495 | 0,968195 |
| Csrnp1 | 1,8584 | 1,80869 | -0,0391135 | 0,8951 | 0,968279 |
| Eps8l1 | 1,34671 | 1,38239 | 0,0377248 | 0,89535 | 0,968392 |
| Zfpl1 | 19,0414 | 18,7104 | -0,025295 | 0,8953 | 0,968392 |
| Gck | 0,737613 | 0,765112 | 0,0528072 | 0,8958 | 0,968776 |
| Aff1 | 5,28772 | 5,20944 | -0,021518 | 0,89585 | 0,968776 |
| Rpl17 | 6,85252 | 7,04815 | 0,0406103 | 0,89645 | 0,968876 |
| Ccz1 | 59,1341 | 60,0171 | 0,0213844 | 0,8962 | 0,968876 |
| Hadha | 39,6306 | 40,2126 | 0,0210319 | 0,8962 | 0,968876 |
| Ppp2r4 | 89,7293 | 88,4556 | -0,0206251 | 0,8962 | 0,968876 |
| Chek2 | 17,4823 | 17,2051 | -0,0230631 | 0,8964 | 0,968876 |
| Fars2 | 7,40748 | 7,2609 | -0,0288349 | 0,89625 | 0,968876 |
| Smox | 28,569 | 27,3196 | -0,0645157 | 0,8964 | 0,968876 |
| Trim37 | 20,5782 | 21,6316 | 0,0720227 | 0,8969 | 0,969206 |
| Prr14l | 7,23944 | 7,34872 | 0,0216148 | 0,89685 | 0,969206 |
| Nle1 | 20,7364 | 21,0719 | 0,0231544 | 0,89705 | 0,969211 |
| Map10 | 2,6216 | 2,56759 | -0,0300315 | 0,897 | 0,969211 |
| Rad51c | 4,70765 | 5,48788 | 0,221244 | 0,89715 | 0,969241 |
| Eme1 | 29,8187 | 30,8305 | 0,0481426 | 0,8974 | 0,969262 |
| Atox1 | 128,431 | 130,54 | 0,0235036 | 0,89735 | 0,969262 |
| Nap1l1 | 190,086 | 193,088 | 0,0226112 | 0,89775 | 0,969262 |
| Cnpy3 | 30,5555 | 30,1077 | -0,0212968 | 0,8975 | 0,969262 |
| Dph2 | 11,4048 | 11,2143 | -0,0243099 | 0,89735 | 0,969262 |
| Wdr19 | 3,80094 | 3,7295 | -0,0273737 | 0,89775 | 0,969262 |
| Scfd2 | 3,32135 | 3,25709 | -0,0281892 | 0,89775 | 0,969262 |
| Cables1 | 3,0009 | 2,93646 | -0,031318 | 0,89755 | 0,969262 |
| Kif13a | 6,87105 | 6,97962 | 0,0226178 | 0,89815 | 0,969357 |
| Cdc42bpb | 26,7828 | 27,1703 | 0,0207251 | 0,8982 | 0,969357 |
| Ppp2cb | 62,3508 | 61,3804 | -0,0226298 | 0,8981 | 0,969357 |
| Myo1e | 6,07815 | 5,97796 | -0,0239779 | 0,898 | 0,969357 |
| Ptrh1 | 2,99687 | 2,89414 | -0,0503231 | 0,89795 | 0,969357 |

| | | | | | |
|---|---|---|---|---|---|
| Gtf3c3 | 13,9702 | 14,1907 | 0,0225977 | 0,8986 | 0,969632 |
| Smad4 | 33,6996 | 34,173 | 0,0201278 | 0,89855 | 0,969632 |
| Snord34 | 4152,57 | 4537,68 | 0,127952 | 0,89935 | 0,969712 |
| Gm15760 | 1,13611 | 1,16703 | 0,0387386 | 0,8994 | 0,969712 |
| Evc | 9,12706 | 9,26689 | 0,021934 | 0,8989 | 0,969712 |
| Trmt10c | 29,9941 | 30,4485 | 0,0216939 | 0,899 | 0,969712 |
| St3gal2 | 21,2406 | 21,55 | 0,0208659 | 0,89925 | 0,969712 |
| Lpgat1 | 32,1621 | 32,6148 | 0,0201637 | 0,89875 | 0,969712 |
| Pdgfra | 47,5101 | 48,1719 | 0,0199581 | 0,89935 | 0,969712 |
| Kif1c | 15,4905 | 15,7022 | 0,0195865 | 0,8991 | 0,969712 |
| Zfp862-ps | 7,84212 | 7,73145 | -0,0205031 | 0,8993 | 0,969712 |
| Bcl11a | 5,07637 | 4,96065 | -0,0332694 | 0,8993 | 0,969712 |
| Map3k10 | 9,88177 | 10,0381 | 0,0226485 | 0,89955 | 0,969717 |
| Oaz2 | 20,8064 | 21,1211 | 0,0216574 | 0,89955 | 0,969717 |
| 1700109H08Rik | 1,72829 | 1,93883 | 0,165839 | 0,90005 | 0,969919 |
| Fuz | 6,32582 | 6,69146 | 0,0810683 | 0,90005 | 0,969919 |
| Cep44 | 19,6352 | 19,9592 | 0,0236104 | 0,90005 | 0,969919 |
| Slc25a28 | 15,2031 | 14,9339 | -0,0257794 | 0,89995 | 0,969919 |
| Foxj2 | 2,96314 | 2,90786 | -0,0271726 | 0,9001 | 0,969919 |
| Nudt17 | 2,34571 | 2,50021 | 0,0920226 | 0,9002 | 0,969949 |
| Lrrc41 | 31,9496 | 32,5695 | 0,0277211 | 0,90085 | 0,970034 |
| Cry2 | 6,33395 | 6,43941 | 0,0238215 | 0,9008 | 0,970034 |
| Rela | 12,0955 | 12,2859 | 0,0225391 | 0,9005 | 0,970034 |
| Nubp2 | 25,9161 | 26,317 | 0,0221493 | 0,90065 | 0,970034 |
| Mysm1 | 7,47619 | 7,58559 | 0,0209584 | 0,90105 | 0,970034 |
| D16Ertd472e | 8,71396 | 8,84082 | 0,020852 | 0,9011 | 0,970034 |
| Tyro3 | 24,1553 | 24,4894 | 0,019817 | 0,9004 | 0,970034 |
| Kdelc1 | 49,4381 | 50,111 | 0,0195024 | 0,9008 | 0,970034 |
| Wdr74 | 64,0172 | 63,1027 | -0,020757 | 0,9011 | 0,970034 |
| Mettl8 | 14,4663 | 14,2392 | -0,0228234 | 0,9009 | 0,970034 |
| Cplx2 | 4,68169 | 4,60446 | -0,0239997 | 0,9009 | 0,970034 |
| 1700049G17Rik | 2,69649 | 2,64723 | -0,0265989 | 0,90115 | 0,970034 |
| Mmp23 | 1,86254 | 1,91988 | 0,0437449 | 0,90125 | 0,970063 |
| Setd8 | 43,5818 | 44,1973 | 0,0202325 | 0,90135 | 0,970093 |
| Dclre1c | 3,62116 | 3,76446 | 0,0559919 | 0,90215 | 0,970329 |
| Thra | 15,1106 | 15,3531 | 0,0229742 | 0,90195 | 0,970329 |
| Zfp28 | 2,57855 | 2,52733 | -0,0289463 | 0,90175 | 0,970329 |
| Zfp128 | 2,52185 | 2,47055 | -0,0296497 | 0,90205 | 0,970329 |
| Pid1 | 2,75911 | 2,69784 | -0,0324006 | 0,9019 | 0,970329 |
| Lrrc34 | 0,839463 | 0,807349 | -0,0562733 | 0,902 | 0,970329 |
| Ggnbp1 | 1,89072 | 1,72778 | -0,130018 | 0,90175 | 0,970329 |
| Pcdhga8 | 0,841602 | 0,713422 | -0,238383 | 0,9021 | 0,970329 |
| Med21 | 66,9998 | 68,0067 | 0,0215193 | 0,90235 | 0,970466 |
| Zfp59 | 3,16455 | 3,22468 | 0,027154 | 0,9025 | 0,970471 |
| Kdm4a | 30,0921 | 30,5115 | 0,0199682 | 0,9025 | 0,970471 |
| Sec24a | 5,0186 | 5,09504 | 0,0218069 | 0,90265 | 0,970554 |
| Prx | 2,13475 | 2,09216 | -0,0290723 | 0,9028 | 0,970637 |
| Gng13 | 4,72868 | 5,30025 | 0,164623 | 0,90295 | 0,970642 |

| | | | | | |
|---|---|---|---|---|---|
| Ppif | 33,2282 | 33,7266 | 0,0214819 | 0,90295 | 0,970642 |
| D4Ertd617e | 0,792923 | 0,655435 | -0,274728 | 0,90315 | 0,970779 |
| C230052I12Rik | 5,76222 | 5,95747 | 0,048075 | 0,90345 | 0,970882 |
| Sh3bp2 | 6,97348 | 7,08817 | 0,0235333 | 0,90385 | 0,970882 |
| Kansl2 | 8,20426 | 8,33101 | 0,0221176 | 0,90355 | 0,970882 |
| Zcrb1 | 57,7781 | 58,6418 | 0,021406 | 0,9039 | 0,970882 |
| Fkbp8 | 100,573 | 101,922 | 0,0192217 | 0,90375 | 0,970882 |
| Arhgap21 | 18,5619 | 18,8042 | 0,0187119 | 0,9039 | 0,970882 |
| Rhoa | 59,8484 | 59,0474 | -0,0194384 | 0,90375 | 0,970882 |
| Tbcel | 11,7237 | 11,5581 | -0,0205174 | 0,9039 | 0,970882 |
| Usp30 | 12,399 | 12,2064 | -0,0225866 | 0,9038 | 0,970882 |
| Rnf2 | 55,5833 | 56,3021 | 0,0185371 | 0,9041 | 0,970941 |
| Rap2a | 10,1154 | 9,96552 | -0,0215386 | 0,90405 | 0,970941 |
| Loh12cr1 | 4,01588 | 3,92437 | -0,0332544 | 0,90435 | 0,971132 |
| Mkl2 | 8,11606 | 8,23916 | 0,0217178 | 0,9046 | 0,971244 |
| LOC106740 | 16,7413 | 15,9662 | -0,068396 | 0,9046 | 0,971244 |
| Ankmy2 | 7,06073 | 6,94175 | -0,0245175 | 0,90475 | 0,971249 |
| Guca1b | 0,69947 | 0,670755 | -0,0604772 | 0,90475 | 0,971249 |
| Slc25a35 | 0,955813 | 0,914573 | -0,0636308 | 0,90485 | 0,971278 |
| Emp3 | 6,37973 | 6,20972 | -0,0389679 | 0,9052 | 0,971498 |
| Crnde | 2,07076 | 2,00449 | -0,0469243 | 0,90515 | 0,971498 |
| Pxmp4 | 4,99239 | 4,88298 | -0,0319695 | 0,9053 | 0,971527 |
| Rab10os | 5,5802 | 5,17759 | -0,108035 | 0,90545 | 0,97161 |
| Pbx1 | 21,1848 | 21,4695 | 0,0192565 | 0,9057 | 0,9718 |
| Gm15708 | 0,875605 | 0,956051 | 0,126808 | 0,90605 | 0,972098 |
| Vamp4 | 8,4751 | 8,61209 | 0,0231321 | 0,9062 | 0,972181 |
| Gpr137 | 9,92753 | 10,1594 | 0,0333084 | 0,90645 | 0,972293 |
| Trappc13 | 24,6961 | 25,0428 | 0,0201095 | 0,90645 | 0,972293 |
| Acaa2 | 20,3083 | 19,9932 | -0,0225595 | 0,90655 | 0,972322 |
| Akip1 | 12,4561 | 12,214 | -0,0283196 | 0,9068 | 0,972512 |
| Otud7b | 11,0237 | 11,1696 | 0,0189732 | 0,9069 | 0,972541 |
| Pnkp | 13,5078 | 13,2972 | -0,0226656 | 0,907 | 0,972571 |
| Vwa8 | 5,86264 | 5,77462 | -0,0218239 | 0,90735 | 0,972712 |
| Ppp1r2 | 12,9371 | 12,6312 | -0,0345226 | 0,90725 | 0,972712 |
| P4htm | 1,68487 | 1,63712 | -0,0414776 | 0,9073 | 0,972712 |
| Nedd4l | 2,96455 | 3,06839 | 0,0496667 | 0,90755 | 0,97277 |
| Ypel2 | 1,0472 | 0,791289 | -0,404255 | 0,9075 | 0,97277 |
| Garem | 6,06982 | 6,16012 | 0,0213055 | 0,9078 | 0,97278 |
| Efemp2 | 54,6336 | 55,3697 | 0,0193082 | 0,90785 | 0,97278 |
| Ppp2r5e | 26,6531 | 26,3162 | -0,0183548 | 0,9077 | 0,97278 |
| Rps23 | 1,46746 | 1,38994 | -0,0782945 | 0,9078 | 0,97278 |
| D15Ertd621e | 31,3579 | 30,9619 | -0,0183307 | 0,90805 | 0,972838 |
| Nfkbia | 14,4281 | 13,7469 | -0,0697815 | 0,90805 | 0,972838 |
| Inip | 19,0774 | 18,8171 | -0,0198179 | 0,9083 | 0,972872 |
| Lrrc4 | 1,20902 | 1,09166 | -0,147306 | 0,9082 | 0,972872 |
| Pcdhgb5 | 1,5007 | 1,34183 | -0,161425 | 0,9083 | 0,972872 |
| Commd8 | 15,9222 | 15,7091 | -0,0194358 | 0,90845 | 0,972954 |
| Pygm | 1,53364 | 1,56995 | 0,0337598 | 0,90865 | 0,973091 |

| | | | | | |
|---|---|---|---|---|---|
| Ccdc181 | 3,12797 | 3,19755 | 0,0317377 | 0,9097 | 0,973104 |
| Arl8a | 16,7683 | 17,018 | 0,0213208 | 0,9093 | 0,973104 |
| Pdia3 | 440,774 | 447,124 | 0,0206366 | 0,90965 | 0,973104 |
| Pitpnc1 | 7,08027 | 7,17819 | 0,0198145 | 0,9089 | 0,973104 |
| Rprd2 | 10,3244 | 10,4591 | 0,0187059 | 0,90875 | 0,973104 |
| Usp42 | 16,8451 | 17,0618 | 0,0184414 | 0,9098 | 0,973104 |
| Pygo2 | 33,939 | 34,3623 | 0,0178821 | 0,9099 | 0,973104 |
| Aurka | 75,5732 | 76,511 | 0,0177931 | 0,9098 | 0,973104 |
| Akt3 | 25,8514 | 25,5249 | -0,0183354 | 0,9097 | 0,973104 |
| Rnaseh2b | 32,2625 | 31,8356 | -0,0192187 | 0,9092 | 0,973104 |
| Ddx5 | 235,794 | 232,6 | -0,0196713 | 0,9099 | 0,973104 |
| Pawr | 17,1884 | 16,9414 | -0,0208803 | 0,9093 | 0,973104 |
| Samd4 | 1,60083 | 1,57199 | -0,0262217 | 0,90965 | 0,973104 |
| Hnf1b | 3,23562 | 3,17513 | -0,0272269 | 0,9098 | 0,973104 |
| Cyfip2 | 4,05209 | 3,95675 | -0,0343481 | 0,90985 | 0,973104 |
| Armc6 | 7,90403 | 7,6463 | -0,0478265 | 0,90955 | 0,973104 |
| Hist1h3g | 0,966519 | 0,891292 | -0,1169 | 0,9091 | 0,973104 |
| Rnpepl1 | 13,7029 | 13,89 | 0,0195594 | 0,9102 | 0,973225 |
| Ankrd46 | 27,1605 | 26,8009 | -0,0192263 | 0,9104 | 0,973225 |
| Itga8 | 4,29099 | 4,22736 | -0,0215519 | 0,91035 | 0,973225 |
| Cdc42se1 | 34,2716 | 33,5122 | -0,0323254 | 0,91035 | 0,973225 |
| Ppap2a | 12,2156 | 11,6885 | -0,0636387 | 0,91015 | 0,973225 |
| Serinc4 | 1,50462 | 1,41633 | -0,0872427 | 0,91045 | 0,973225 |
| Mat2b | 52,3328 | 52,9836 | 0,017829 | 0,91055 | 0,973254 |
| Adcy2 | 0,627984 | 0,643928 | 0,0361705 | 0,9108 | 0,973394 |
| Zfyve16 | 9,83098 | 9,96447 | 0,0194582 | 0,91085 | 0,973394 |
| Qars | 83,3971 | 84,4119 | 0,01745 | 0,9109 | 0,973394 |
| Plekhj1 | 45,2377 | 44,6531 | -0,0187641 | 0,91105 | 0,973399 |
| Ccdc142 | 2,95338 | 2,84737 | -0,0527354 | 0,911 | 0,973399 |
| Zfp280c | 18,7264 | 18,4901 | -0,0183168 | 0,9113 | 0,973588 |
| Mtfmt | 6,17087 | 6,06394 | -0,0252202 | 0,9114 | 0,973617 |
| Pyurf | 4,69959 | 4,63427 | -0,0201938 | 0,91155 | 0,973622 |
| Abtb1 | 9,17323 | 9,02499 | -0,023504 | 0,91155 | 0,973622 |
| Tceb3 | 19,0208 | 19,2521 | 0,0174341 | 0,91175 | 0,97368 |
| Rtn3 | 128,536 | 126,616 | -0,0217091 | 0,91175 | 0,97368 |
| Bahcc1 | 13,408 | 13,2491 | -0,0172039 | 0,912 | 0,973713 |
| Parp12 | 5,27816 | 5,19257 | -0,0235859 | 0,91195 | 0,973713 |
| Zfp219 | 46,1785 | 45,3432 | -0,0263362 | 0,91195 | 0,973713 |
| Itch | 27,2474 | 26,9105 | -0,0179495 | 0,9121 | 0,973742 |
| Gm5113 | 3,09671 | 3,03877 | -0,0272476 | 0,91225 | 0,973825 |
| Ubtd2 | 52,1847 | 52,8099 | 0,0171811 | 0,91235 | 0,973853 |
| Dis3l | 9,71804 | 9,85112 | 0,0196222 | 0,9125 | 0,973936 |
| BC052040 | 7,47234 | 7,58425 | 0,0214462 | 0,91275 | 0,974125 |
| Gm12060 | 1,89144 | 1,95821 | 0,0500538 | 0,9129 | 0,974207 |
| Exosc5 | 32,4621 | 32,9069 | 0,0196354 | 0,9132 | 0,974245 |
| Mmachc | 17,8049 | 18,0476 | 0,0195326 | 0,9131 | 0,974245 |
| Krit1 | 12,2562 | 12,1013 | -0,0183475 | 0,9132 | 0,974245 |
| Supv3l1 | 22,9012 | 22,6074 | -0,0186315 | 0,9133 | 0,974245 |

| | | | | | |
|---|---|---|---|---|---|
| Rnf24 | 5,28154 | 5,0388 | -0,0678777 | 0,91325 | 0,974245 |
| Rpl23a | 0,900493 | 0,968037 | 0,104347 | 0,91355 | 0,974356 |
| Scarf2 | 6,52075 | 6,4279 | -0,0206892 | 0,91355 | 0,974356 |
| Zfp628 | 4,99858 | 5,07583 | 0,0221256 | 0,91375 | 0,974492 |
| Grasp | 3,18259 | 3,11717 | -0,0299639 | 0,91385 | 0,974521 |
| H2-K1 | 13,8149 | 14,0145 | 0,0207039 | 0,9142 | 0,97466 |
| Cc2d1a | 8,54635 | 8,42372 | -0,0208514 | 0,9142 | 0,97466 |
| Tmem259 | 58,1149 | 52,4458 | -0,14808 | 0,9142 | 0,97466 |
| Unc119b | 36,7812 | 37,2266 | 0,0173648 | 0,9143 | 0,974689 |
| Rpp40 | 6,39455 | 6,29415 | -0,022832 | 0,9144 | 0,974718 |
| Gstm4 | 6,17027 | 6,28104 | 0,0256691 | 0,91465 | 0,974751 |
| Stox2 | 11,1481 | 11,0125 | -0,0176585 | 0,9146 | 0,974751 |
| Trim6 | 3,15071 | 3,10072 | -0,023075 | 0,91465 | 0,974751 |
| Fancl | 17,7491 | 18,0358 | 0,0231187 | 0,9152 | 0,974924 |
| Dnajc16 | 8,03934 | 8,14834 | 0,019429 | 0,91525 | 0,974924 |
| Etfb | 76,906 | 77,8995 | 0,0185183 | 0,91495 | 0,974924 |
| Bicd2 | 14,4375 | 14,6125 | 0,0173774 | 0,9151 | 0,974924 |
| Uba3 | 52,5613 | 53,1739 | 0,0167173 | 0,9152 | 0,974924 |
| Olfml1 | 0,557053 | 0,516088 | -0,1102 | 0,9152 | 0,974924 |
| Pth1r | 35,4945 | 35,926 | 0,0174347 | 0,91535 | 0,974953 |
| Haus5 | 13,6186 | 13,8003 | 0,019123 | 0,9157 | 0,975015 |
| Vars2 | 9,92981 | 10,0551 | 0,0180903 | 0,9157 | 0,975015 |
| Ccdc8 | 41,1945 | 41,6684 | 0,016503 | 0,91565 | 0,975015 |
| Insr | 14,166 | 14,0009 | -0,0169179 | 0,91555 | 0,975015 |
| Atp11a | 13,7196 | 13,8857 | 0,0173657 | 0,91585 | 0,975097 |
| Fam107b | 24,2802 | 24,571 | 0,0171764 | 0,916 | 0,975179 |
| Macrod2 | 2,55835 | 2,66247 | 0,0575508 | 0,91635 | 0,975474 |
| Atat1 | 28,7329 | 28,3511 | -0,0192975 | 0,9165 | 0,975556 |
| Slmap | 19,8192 | 19,58 | -0,017518 | 0,9167 | 0,975691 |
| Emc8 | 20,0783 | 20,3163 | 0,0169993 | 0,91705 | 0,975831 |
| Nsd1 | 29,8083 | 30,16 | 0,0169203 | 0,91695 | 0,975831 |
| Ubxn4 | 27,3984 | 27,0827 | -0,0167174 | 0,917 | 0,975831 |
| Toporsos | 25,6026 | 27,569 | 0,106755 | 0,9173 | 0,975917 |
| Glud1 | 39,69 | 39,241 | -0,0164123 | 0,91735 | 0,975917 |
| Rnf7 | 82,3841 | 81,4253 | -0,0168889 | 0,9173 | 0,975917 |
| Gzf1 | 15,4223 | 15,2457 | -0,0166076 | 0,91765 | 0,97608 |
| Snord12 | 4214,61 | 3674,11 | -0,198006 | 0,91765 | 0,97608 |
| Serinc2 | 23,4199 | 23,7135 | 0,0179762 | 0,91795 | 0,976244 |
| Tab2 | 51,8205 | 52,4071 | 0,0162404 | 0,9179 | 0,976244 |
| Rhof | 1,12351 | 1,15345 | 0,0379447 | 0,9182 | 0,976314 |
| Lpcat2 | 1,74875 | 1,78714 | 0,0313286 | 0,9186 | 0,976314 |
| Mynn | 6,68292 | 6,76686 | 0,0180073 | 0,91825 | 0,976314 |
| Lpp | 9,55808 | 9,6774 | 0,0178997 | 0,91815 | 0,976314 |
| Pik3c3 | 14,8001 | 14,982 | 0,0176215 | 0,91845 | 0,976314 |
| Kif15 | 38,9492 | 39,3871 | 0,0161281 | 0,9186 | 0,976314 |
| Psma1 | 193,582 | 195,709 | 0,0157623 | 0,91855 | 0,976314 |
| Arvcf | 10,6831 | 10,5013 | -0,0247648 | 0,91835 | 0,976314 |
| Ppfia4 | 1,22174 | 1,19949 | -0,026506 | 0,9188 | 0,976449 |

| | | | | | |
|---|---|---|---|---|---|
| Fam69c | 0,671314 | 0,691503 | 0,0427487 | 0,91925 | 0,976694 |
| Nol12 | 17,4487 | 17,2382 | -0,0175174 | 0,9192 | 0,976694 |
| 1110008L16Rik | 7,3185 | 7,02023 | -0,0600297 | 0,91925 | 0,976694 |
| Leng1 | 8,41333 | 8,17331 | -0,0417561 | 0,91935 | 0,976723 |
| Cdkn2aip | 13,2223 | 13,3803 | 0,0171374 | 0,91945 | 0,976751 |
| Psmg3 | 40,5533 | 41,0935 | 0,0190899 | 0,9198 | 0,976845 |
| Slc20a2 | 26,8416 | 27,1761 | 0,0178639 | 0,91995 | 0,976845 |
| Pbx2 | 85,6305 | 86,5669 | 0,0156912 | 0,91965 | 0,976845 |
| Ubqln1 | 66,4388 | 67,1552 | 0,0154728 | 0,91975 | 0,976845 |
| Irf9 | 7,74023 | 7,63138 | -0,0204323 | 0,92005 | 0,976845 |
| Ackr1 | 1,59481 | 1,54579 | -0,0450377 | 0,92005 | 0,976845 |
| Abca7 | 2,28291 | 1,65959 | -0,460046 | 0,91995 | 0,976845 |
| Pcdhga1 | 0,564331 | 0,656142 | 0,217467 | 0,92135 | 0,976881 |
| Snora62 | 29,6204 | 31,0744 | 0,0691361 | 0,9208 | 0,976881 |
| Eml5 | 1,01028 | 1,0568 | 0,0649461 | 0,9208 | 0,976881 |
| 1700037H04Rik | 32,4716 | 33,7825 | 0,0570992 | 0,92135 | 0,976881 |
| Syngr3 | 0,731856 | 0,752606 | 0,0403367 | 0,9212 | 0,976881 |
| Rab2b | 9,51081 | 9,77442 | 0,0394418 | 0,92095 | 0,976881 |
| Gmppb | 10,4334 | 10,6931 | 0,0354767 | 0,9214 | 0,976881 |
| AU019823 | 16,4279 | 16,6974 | 0,0234743 | 0,9207 | 0,976881 |
| Elovl4 | 7,29669 | 7,39433 | 0,0191773 | 0,92055 | 0,976881 |
| Pik3r4 | 12,6375 | 12,7871 | 0,0169702 | 0,92045 | 0,976881 |
| Eri1 | 17,3118 | 17,5023 | 0,0157888 | 0,9211 | 0,976881 |
| Ap2a2 | 33,5895 | 33,9566 | 0,0156822 | 0,92035 | 0,976881 |
| Spin1 | 31,1147 | 31,4455 | 0,0152592 | 0,92105 | 0,976881 |
| Arid4b | 19,0662 | 18,8545 | -0,0161049 | 0,9209 | 0,976881 |
| Rxrb | 30,2408 | 29,9009 | -0,0163087 | 0,92105 | 0,976881 |
| Fdxr | 7,5451 | 7,43435 | -0,021334 | 0,921 | 0,976881 |
| Mrpl48 | 14,2278 | 13,5819 | -0,0670265 | 0,921 | 0,976881 |
| Gdpd1 | 4,5907 | 3,99036 | -0,202196 | 0,92115 | 0,976881 |
| Lage3 | 35,6678 | 36,1444 | 0,0191517 | 0,9216 | 0,976991 |
| Creb3l4 | 2,19406 | 2,14851 | -0,030264 | 0,92165 | 0,976991 |
| Abhd11os | 2,04063 | 2,21003 | 0,115053 | 0,92205 | 0,977056 |
| Asxl2 | 7,11473 | 7,19672 | 0,0165304 | 0,92185 | 0,977056 |
| Mid1ip1 | 26,385 | 26,6886 | 0,0165087 | 0,92195 | 0,977056 |
| Phka1 | 4,64257 | 4,58555 | -0,0178297 | 0,92205 | 0,977056 |
| Eif4g2 | 333,228 | 328,239 | -0,0217641 | 0,92215 | 0,977056 |
| Bmper | 2,8465 | 2,80307 | -0,0221833 | 0,9221 | 0,977056 |
| Pcdhgb6 | 4,47785 | 4,26144 | -0,071464 | 0,9223 | 0,977138 |
| Hccs | 19,935 | 20,1683 | 0,0167831 | 0,92245 | 0,977219 |
| Rgs19 | 32,9586 | 32,5559 | -0,0177364 | 0,9226 | 0,977301 |
| Apod | 0,856958 | 0,880711 | 0,0394438 | 0,92285 | 0,977488 |
| Nploc4 | 32,139 | 32,46 | 0,0143399 | 0,9232 | 0,977781 |
| Pter | 3,80349 | 3,74808 | -0,0211701 | 0,9233 | 0,97781 |
| Slc26a6 | 2,541 | 2,58652 | 0,0256146 | 0,92375 | 0,978131 |
| Susd4 | 21,4913 | 21,7302 | 0,0159526 | 0,9237 | 0,978131 |
| Ccdc32 | 9,93904 | 10,072 | 0,0191731 | 0,92395 | 0,978266 |
| Shoc2 | 18,0079 | 18,2881 | 0,0222693 | 0,9242 | 0,978326 |

| | | | | | |
|---|---|---|---|---|---|
| Rnaseh2a | 21,1703 | 21,4128 | 0,0164312 | 0,9243 | 0,978326 |
| Cacybp | 111,814 | 112,965 | 0,0147728 | 0,92425 | 0,978326 |
| Sptan1 | 49,2374 | 48,7034 | -0,0157308 | 0,9243 | 0,978326 |
| Nespas | 1,67968 | 2,15095 | 0,35679 | 0,9246 | 0,978464 |
| 2610008E11Rik | 16,9451 | 16,7563 | -0,0161664 | 0,9246 | 0,978464 |
| Ermard | 8,53185 | 8,40781 | -0,0211298 | 0,92465 | 0,978464 |
| Odf2l | 14,5596 | 14,3932 | -0,0165747 | 0,925 | 0,978732 |
| Fpgt | 4,89855 | 4,83039 | -0,020214 | 0,92505 | 0,978732 |
| Bcl7a | 37,7951 | 38,2038 | 0,0155184 | 0,92515 | 0,978761 |
| Trappc11 | 18,3806 | 18,1859 | -0,015366 | 0,9254 | 0,978948 |
| Thap3 | 12,9517 | 13,1399 | 0,0208194 | 0,92585 | 0,979346 |
| Cep41 | 7,19152 | 7,10288 | -0,0178924 | 0,9261 | 0,979533 |
| 4930565N06Rik | 0,767808 | 0,84919 | 0,145342 | 0,92625 | 0,979537 |
| Pradc1 | 11,6434 | 10,6854 | -0,123875 | 0,9262 | 0,979537 |
| Ska2 | 62,1344 | 64,9806 | 0,0646181 | 0,9264 | 0,979618 |
| Phykpl | 4,79522 | 5,63511 | 0,232847 | 0,92655 | 0,979674 |
| Dgka | 2,21665 | 2,25767 | 0,0264565 | 0,9266 | 0,979674 |
| Mir6516 | 89,7495 | 98,9257 | 0,140442 | 0,9268 | 0,979731 |
| Tmem251 | 20,9114 | 20,4487 | -0,0322784 | 0,9268 | 0,979731 |
| Tmem176a | 12,7619 | 12,5778 | -0,0209585 | 0,9269 | 0,979759 |
| 2410004I01Rik | 0,77363 | 0,808771 | 0,0640868 | 0,9272 | 0,979998 |
| Rab5c | 75,1445 | 74,3889 | -0,0145791 | 0,9273 | 0,980027 |
| Wdr18 | 37,9395 | 41,2874 | 0,122003 | 0,9276 | 0,980087 |
| Taf7 | 12,4075 | 12,8169 | 0,046831 | 0,9276 | 0,980087 |
| Gria4 | 0,967196 | 0,990529 | 0,0343922 | 0,92755 | 0,980087 |
| Rpf2 | 33,9081 | 34,2904 | 0,0161756 | 0,92765 | 0,980087 |
| Gm166 | 2,33627 | 2,26989 | -0,041585 | 0,92775 | 0,980115 |
| Dopey1 | 3,06429 | 3,11512 | 0,023736 | 0,92785 | 0,980143 |
| Msmp | 2,99582 | 3,38968 | 0,1782 | 0,92815 | 0,98015 |
| Ubxn11 | 1,54406 | 1,5809 | 0,0340112 | 0,92805 | 0,98015 |
| Lztr1 | 24,767 | 25,029 | 0,0151845 | 0,92805 | 0,98015 |
| Rasa4 | 1,13394 | 1,11007 | -0,0306843 | 0,9281 | 0,98015 |
| Kctd7 | 1,33629 | 1,35943 | 0,0247697 | 0,9286 | 0,980238 |
| Nfatc2ip | 8,36511 | 8,46454 | 0,0170473 | 0,92845 | 0,980238 |
| Appl2 | 16,0402 | 16,212 | 0,0153741 | 0,92845 | 0,980238 |
| Wiz | 33,9774 | 34,3234 | 0,0146135 | 0,9286 | 0,980238 |
| Spsb2 | 2,7141 | 2,33919 | -0,214468 | 0,9286 | 0,980238 |
| Atp5d | 182,962 | 196,92 | 0,106065 | 0,92895 | 0,980428 |
| Spsb1 | 3,70196 | 3,75503 | 0,0205344 | 0,929 | 0,980428 |
| Srpr | 40,3917 | 40,8525 | 0,0163663 | 0,929 | 0,980428 |
| Spcs2 | 62,1632 | 61,5881 | -0,0134073 | 0,92955 | 0,980931 |
| Snora74a | 6,99963 | 7,5486 | 0,108932 | 0,93005 | 0,980994 |
| Fbln1 | 43,9943 | 43,5809 | -0,0136207 | 0,93 | 0,980994 |
| Mtor | 12,4533 | 12,3361 | -0,0136409 | 0,9298 | 0,980994 |
| Rad1 | 27,7677 | 27,4683 | -0,0156388 | 0,9299 | 0,980994 |
| Tbc1d9 | 2,14085 | 2,11294 | -0,0189356 | 0,9299 | 0,980994 |
| Sp3os | 4,06388 | 3,78017 | -0,104407 | 0,92995 | 0,980994 |
| Rbfox2 | 31,4885 | 31,1838 | -0,01403 | 0,93035 | 0,981155 |

| | | | | | |
|---|---|---|---|---|---|
| Lysmd2 | 4,71314 | 4,63098 | -0,0253721 | 0,93035 | 0,981155 |
| Sharpin | 10,853 | 10,9871 | 0,0177148 | 0,9306 | 0,981317 |
| Cd3eap | 21,3553 | 21,1019 | -0,0172232 | 0,93065 | 0,981317 |
| Syde2 | 1,77247 | 1,79936 | 0,0217209 | 0,93075 | 0,981345 |
| Rad23b | 96,434 | 95,5038 | -0,013984 | 0,9309 | 0,981425 |
| Poli | 6,16514 | 6,24206 | 0,0178868 | 0,9311 | 0,981481 |
| Oip5 | 14,5966 | 14,2265 | -0,0370525 | 0,93105 | 0,981481 |
| Cap1 | 80,965 | 80,1856 | -0,0139543 | 0,9314 | 0,98172 |
| Seh1l | 70,4621 | 71,1711 | 0,0144439 | 0,93155 | 0,981801 |
| Stk11ip | 13,5147 | 13,6484 | 0,0142004 | 0,9317 | 0,981857 |
| E530011L22Rik | 0,956228 | 0,866546 | -0,142079 | 0,93175 | 0,981857 |
| Ptch2 | 5,42852 | 5,49303 | 0,0170431 | 0,93185 | 0,981885 |
| Enox1 | 1,09256 | 1,11159 | 0,0249089 | 0,9324 | 0,98208 |
| Ephb6 | 3,3296 | 3,37043 | 0,0175858 | 0,93245 | 0,98208 |
| Fem1b | 42,7575 | 43,1566 | 0,0134029 | 0,93215 | 0,98208 |
| Hgs | 19,4513 | 19,2607 | -0,0142015 | 0,93235 | 0,98208 |
| Traf3 | 8,37595 | 8,2919 | -0,0145497 | 0,93245 | 0,98208 |
| Hps1 | 4,39247 | 4,33645 | -0,0185206 | 0,93255 | 0,98208 |
| Tspyl3 | 2,42955 | 2,39411 | -0,021197 | 0,93255 | 0,98208 |
| Il17rc | 2,06431 | 2,09904 | 0,0240708 | 0,9327 | 0,982083 |
| Med9os | 0,892379 | 0,906617 | 0,0228371 | 0,93265 | 0,982083 |
| 2810459M11Rik | 0,763277 | 0,746985 | -0,0311287 | 0,9333 | 0,982173 |
| 4930513N10Rik | 1,69391 | 1,88452 | 0,153836 | 0,9333 | 0,982174 |
| Ropn1l | 1,12364 | 1,16217 | 0,0486463 | 0,9333 | 0,982174 |
| Srpk2 | 12,8944 | 13,081 | 0,020729 | 0,93325 | 0,982174 |
| Ccdc112 | 5,62588 | 5,6947 | 0,0175411 | 0,93295 | 0,982174 |
| Hoxd8 | 41,5862 | 41,9875 | 0,0138562 | 0,9329 | 0,982174 |
| Fgfr1op | 20,3974 | 20,2073 | -0,0135055 | 0,93315 | 0,982174 |
| Xrcc6bp1 | 8,43916 | 8,55332 | 0,019386 | 0,9339 | 0,982452 |
| Evl | 53,5279 | 54,1959 | 0,017893 | 0,93395 | 0,982452 |
| Ulk2 | 10,1972 | 10,2945 | 0,0137085 | 0,93395 | 0,982452 |
| Fam210a | 12,5829 | 12,6973 | 0,0130626 | 0,9343 | 0,982452 |
| Dnajc13 | 23,2848 | 23,0796 | -0,012769 | 0,9343 | 0,982452 |
| Cdk1 | 104,727 | 103,769 | -0,0132521 | 0,93405 | 0,982452 |
| Ybx2 | 5,054 | 4,97817 | -0,0218119 | 0,93395 | 0,982452 |
| Map3k5 | 0,814554 | 0,800333 | -0,0254104 | 0,9342 | 0,982452 |
| Actr3b | 1,09184 | 1,06878 | -0,0307935 | 0,9341 | 0,982452 |
| Polrmt | 12,9443 | 11,3542 | -0,189082 | 0,9338 | 0,982452 |
| Mfap2 | 70,8723 | 71,5542 | 0,0138143 | 0,93445 | 0,982455 |
| Ttpal | 14,0184 | 13,8877 | -0,0135212 | 0,93445 | 0,982455 |
| E130317F20Rik | 4,01108 | 5,13691 | 0,356909 | 0,9352 | 0,982486 |
| Cd300e | 0,60392 | 0,620873 | 0,0399398 | 0,9351 | 0,982486 |
| Gstt3 | 0,670349 | 0,688014 | 0,0375245 | 0,9359 | 0,982486 |
| Ntpcr | 5,95723 | 6,05249 | 0,0228866 | 0,93565 | 0,982486 |
| Abcc5 | 23,6428 | 23,9231 | 0,0170017 | 0,93585 | 0,982486 |
| Jade2 | 2,75657 | 2,78738 | 0,0160312 | 0,93545 | 0,982486 |
| Rogdi | 14,5467 | 14,706 | 0,0157039 | 0,9358 | 0,982486 |
| B3gnt7 | 10,0349 | 10,137 | 0,0146147 | 0,93575 | 0,982486 |

| | | | | | |
|---|---|---|---|---|---|
| Plekhh2 | 5,63995 | 5,69666 | 0,0144337 | 0,9357 | 0,982486 |
| Csnk1a1 | 173,696 | 175,315 | 0,0133879 | 0,93525 | 0,982486 |
| Cic | 19,9493 | 20,134 | 0,0132992 | 0,935 | 0,982486 |
| Mtf2 | 41,445 | 41,8065 | 0,0125285 | 0,9356 | 0,982486 |
| Arhgap28 | 25,437 | 25,2122 | -0,0128078 | 0,93525 | 0,982486 |
| Prkag1 | 40,689 | 40,3132 | -0,0133879 | 0,9347 | 0,982486 |
| Chpf | 16,0675 | 15,9146 | -0,0137911 | 0,9353 | 0,982486 |
| Hps5 | 8,14544 | 8,06737 | -0,0138957 | 0,93535 | 0,982486 |
| 0610009B22Rik | 35,3041 | 34,9309 | -0,015332 | 0,935 | 0,982486 |
| Tex9 | 15,7671 | 15,5677 | -0,0183613 | 0,9353 | 0,982486 |
| 4921531C22Rik | 1,13459 | 1,11079 | -0,0305853 | 0,9357 | 0,982486 |
| Ada | 3,4488 | 3,50184 | 0,0220208 | 0,93595 | 0,982487 |
| LOC100503496 | 1,23351 | 1,28603 | 0,060158 | 0,9362 | 0,982672 |
| Nedd4 | 659,479 | 668,562 | 0,0197362 | 0,93655 | 0,982783 |
| Trabd | 22,8515 | 23,0747 | 0,0140257 | 0,93655 | 0,982783 |
| Zfp568 | 27,1457 | 28,5863 | -0,012592 | 0,9366 | 0,982783 |
| Mppe1 | 7,86762 | 7,66465 | -0,0377065 | 0,9365 | 0,982783 |
| Josd1 | 17,0265 | 17,1934 | 0,0140754 | 0,93675 | 0,982863 |
| Kif1a | 1,68321 | 1,70308 | 0,0169319 | 0,93695 | 0,982996 |
| Nrg1 | 0,673374 | 0,688987 | 0,0330678 | 0,93765 | 0,98314 |
| Tnfrsf1b | 2,26003 | 2,2886 | 0,0181268 | 0,9377 | 0,98314 |
| Tmem181a | 3,17444 | 3,21284 | 0,0173446 | 0,9375 | 0,98314 |
| Tmem173 | 10,7351 | 10,8454 | 0,0147349 | 0,93735 | 0,98314 |
| Ambra1 | 15,6628 | 15,8102 | 0,0135217 | 0,9372 | 0,98314 |
| Igsf9 | 42,7966 | 42,4431 | -0,0119666 | 0,9374 | 0,98314 |
| Nsmce1 | 35,2231 | 34,8768 | -0,0142545 | 0,93775 | 0,98314 |
| Parp11 | 8,20706 | 8,12633 | -0,0142616 | 0,9376 | 0,98314 |
| Elof1 | 69,5386 | 68,2045 | -0,0279457 | 0,9374 | 0,98314 |
| Dek | 187,067 | 185,35 | -0,0133063 | 0,93785 | 0,983168 |
| Ccdc132 | 16,7337 | 16,8814 | 0,0126837 | 0,938 | 0,983248 |
| Ogdh | 29,0083 | 29,2666 | 0,0127898 | 0,93875 | 0,983957 |
| Eif1ax | 232,782 | 234,854 | 0,0127831 | 0,939 | 0,984065 |
| Rbm25 | 89,8868 | 89,1212 | -0,0123404 | 0,93895 | 0,984065 |
| 1810026J23Rik | 22,7468 | 23,1456 | 0,0250753 | 0,9406 | 0,984303 |
| Stambp | 4,93183 | 4,99027 | 0,0169952 | 0,94065 | 0,984303 |
| Nae1 | 59,2767 | 59,8723 | 0,0144214 | 0,9404 | 0,984303 |
| Cog3 | 14,8427 | 14,9713 | 0,0124471 | 0,9403 | 0,984303 |
| Zfp68 | 14,8543 | 14,9827 | 0,0124138 | 0,9398 | 0,984303 |
| Tcf12 | 87,6453 | 88,3973 | 0,0123244 | 0,94 | 0,984303 |
| Phf2 | 21,3083 | 21,4899 | 0,0122466 | 0,9399 | 0,984303 |
| Slc37a3 | 40,9527 | 40,6268 | -0,0115282 | 0,94045 | 0,984303 |
| Kpna4 | 42,9589 | 42,6047 | -0,0119456 | 0,9395 | 0,984303 |
| Man2a2 | 16,7651 | 16,6253 | -0,0120762 | 0,9407 | 0,984303 |
| Frzb | 47,0043 | 46,5973 | -0,0125476 | 0,93975 | 0,984303 |
| Fam160b1 | 9,71456 | 9,62619 | -0,0131852 | 0,93975 | 0,984303 |
| Zfp319 | 6,33391 | 6,27244 | -0,0140699 | 0,93985 | 0,984303 |
| Zfp248 | 5,15783 | 5,10604 | -0,014558 | 0,9402 | 0,984303 |
| Usp34 | 22,5134 | 22,2815 | -0,0149349 | 0,9406 | 0,984303 |

| | | | | | |
|---|---|---|---|---|---|
| Romo1 | 105,742 | 104,371 | -0,0188204 | 0,93985 | 0,984303 |
| Lipt1 | 3,40489 | 3,35539 | -0,0211311 | 0,94015 | 0,984303 |
| C920006O11Rik | 1,7861 | 1,751 | -0,0286365 | 0,9396 | 0,984303 |
| 4930506C21Rik | 0,90597 | 0,870882 | -0,0569845 | 0,94055 | 0,984303 |
| Spef1 | 4,33887 | 4,08489 | -0,0870194 | 0,94035 | 0,984303 |
| Calhm2 | 1,15502 | 1,13481 | -0,0254713 | 0,9412 | 0,984749 |
| Coq4 | 11,9638 | 12,0828 | 0,0142755 | 0,94135 | 0,984829 |
| Zc3h12b | 1,23606 | 1,25464 | 0,0215218 | 0,9419 | 0,9849 |
| Nap1l3 | 1,55162 | 1,57474 | 0,0213429 | 0,9424 | 0,9849 |
| Cadm3 | 2,84272 | 2,87549 | 0,016535 | 0,94185 | 0,9849 |
| Cep72 | 8,55201 | 8,63448 | 0,0138456 | 0,94225 | 0,9849 |
| Syt11 | 14,0448 | 14,1615 | 0,0119475 | 0,94245 | 0,9849 |
| Rlf | 16,7685 | 16,6328 | -0,0117236 | 0,9418 | 0,9849 |
| Dlk1 | 26,491 | 26,2728 | -0,0119356 | 0,94235 | 0,9849 |
| 2510003E04Rik | 18,6963 | 18,5321 | -0,0127237 | 0,94215 | 0,9849 |
| Dhrs4 | 28,0059 | 27,7501 | -0,0132365 | 0,9418 | 0,9849 |
| Neto2 | 5,74555 | 5,69211 | -0,0134821 | 0,9417 | 0,9849 |
| Gdi2 | 151,104 | 149,633 | -0,0141176 | 0,94225 | 0,9849 |
| Scaper | 2,56591 | 2,53823 | -0,0156489 | 0,94225 | 0,9849 |
| Fahd1 | 9,47728 | 9,36795 | -0,0167396 | 0,942 | 0,9849 |
| Adam2 | 0,948152 | 0,930004 | -0,0278815 | 0,9422 | 0,9849 |
| Clk3 | 39,8892 | 40,23 | 0,0122723 | 0,94255 | 0,984928 |
| Zbed5 | 5,01145 | 5,06805 | 0,0162036 | 0,94275 | 0,985059 |
| Ndel1 | 18,9559 | 19,1263 | 0,0129094 | 0,94295 | 0,985114 |
| Ost4 | 59,2629 | 58,5149 | -0,0183253 | 0,9429 | 0,985114 |
| Ppm1e | 0,770088 | 0,894477 | 0,216021 | 0,9436 | 0,985436 |
| Zfp513 | 16,7497 | 17,8549 | 0,0921837 | 0,94345 | 0,985436 |
| Gpr63 | 3,2768 | 3,31692 | 0,0175557 | 0,94335 | 0,985436 |
| Setx | 16,1554 | 16,2787 | 0,0109754 | 0,9437 | 0,985436 |
| Isoc1 | 46,9795 | 46,6039 | -0,011583 | 0,94365 | 0,985436 |
| Cdk5rap1 | 7,05902 | 6,98367 | -0,0154814 | 0,94355 | 0,985436 |
| Plekhb2 | 7,80012 | 7,87003 | 0,0128738 | 0,944 | 0,985441 |
| Trub2 | 24,4951 | 24,7072 | 0,0124363 | 0,94395 | 0,985441 |
| Abcd4 | 5,97404 | 5,91278 | -0,0148705 | 0,944 | 0,985441 |
| Nat14 | 1,66908 | 1,64138 | -0,0241482 | 0,94385 | 0,985441 |
| Trappc10 | 12,6566 | 12,5572 | -0,0113809 | 0,9441 | 0,985468 |
| Snord35a | 1176,71 | 1328,31 | 0,174825 | 0,9445 | 0,985475 |
| Chrm4 | 0,660578 | 0,720786 | 0,125842 | 0,9451 | 0,985475 |
| Pcdhga9 | 0,810063 | 0,882791 | 0,124038 | 0,9459 | 0,985475 |
| Ap4b1 | 10,8366 | 11,14 | 0,039836 | 0,94515 | 0,985475 |
| Zfp956 | 4,00741 | 4,06871 | 0,0219012 | 0,9449 | 0,985475 |
| Cyb5rl | 6,35476 | 6,42696 | 0,016299 | 0,94585 | 0,985475 |
| Lrrc16b | 3,36075 | 3,39575 | 0,0149483 | 0,94565 | 0,985475 |
| Nipsnap3b | 23,16 | 23,3615 | 0,0124986 | 0,94535 | 0,985475 |
| Mrc2 | 19,7424 | 19,8986 | 0,0113722 | 0,9453 | 0,985475 |
| Esd | 39,6372 | 39,9495 | 0,011323 | 0,9458 | 0,985475 |
| Yaf2 | 26,8864 | 27,0975 | 0,0112826 | 0,94525 | 0,985475 |
| Smc2 | 60,5021 | 60,968 | 0,0110675 | 0,9458 | 0,985475 |

| | | | | | |
|---|---|---|---|---|---|
| Chek1 | 27,4179 | 27,6277 | 0,0109988 | 0,9457 | 0,985475 |
| Kmt2b | 20,3462 | 20,4993 | 0,0108112 | 0,9456 | 0,985475 |
| Csnk1g3 | 19,2517 | 19,3961 | 0,0107819 | 0,9449 | 0,985475 |
| Zfp266 | 37,027 | 38,9115 | -0,0107187 | 0,94555 | 0,985475 |
| Ppip5k2 | 27,2043 | 27,0004 | -0,0108523 | 0,94525 | 0,985475 |
| Raph1 | 12,0352 | 11,9421 | -0,0111989 | 0,9444 | 0,985475 |
| Pip5k1c | 26,9657 | 26,7572 | -0,0111999 | 0,94525 | 0,985475 |
| 4930579G24Rik | 10,8672 | 10,7617 | -0,0140691 | 0,94455 | 0,985475 |
| Magee1 | 5,17082 | 5,12021 | -0,0141905 | 0,9447 | 0,985475 |
| Col24a1 | 1,23903 | 1,22479 | -0,0166784 | 0,945 | 0,985475 |
| Dalrd3 | 16,2532 | 15,7218 | -0,0479625 | 0,94595 | 0,985475 |
| Cirbp | 58,9385 | 53,5108 | -0,139381 | 0,9447 | 0,985475 |
| Hmha1 | 8,0984 | 6,91165 | -0,228607 | 0,94515 | 0,985475 |
| Mettl15 | 1,64546 | 1,67222 | 0,0232784 | 0,9462 | 0,985634 |
| Mir1291 | 121,414 | 112,305 | -0,112519 | 0,94625 | 0,985634 |
| Zbtb22 | 17,2972 | 17,1529 | -0,0120837 | 0,94635 | 0,985661 |
| Adrm1 | 6,75165 | 6,82532 | 0,0156546 | 0,9467 | 0,985719 |
| Actr3 | 50,1077 | 50,4681 | 0,0103375 | 0,94665 | 0,985719 |
| Papd5 | 15,4856 | 15,3702 | -0,0107844 | 0,94655 | 0,985719 |
| Pmch | 0,741756 | 0,683307 | -0,118412 | 0,9467 | 0,985719 |
| Cnih2 | 3,67412 | 3,77307 | 0,0383398 | 0,94735 | 0,985935 |
| Snx7 | 5,81182 | 5,8757 | 0,0157706 | 0,94735 | 0,985935 |
| Srsf10 | 77,3497 | 77,9557 | 0,0112603 | 0,94735 | 0,985935 |
| Epb4.1 | 77,3662 | 77,9219 | 0,0103248 | 0,94735 | 0,985935 |
| Itpr1 | 8,78202 | 8,7148 | -0,0110861 | 0,9471 | 0,985935 |
| Lcorl | 6,2971 | 6,17294 | -0,0287295 | 0,947 | 0,985935 |
| Zbtb43 | 7,42207 | 7,4833 | 0,0118527 | 0,94765 | 0,986016 |
| Slc39a14 | 8,50846 | 8,57385 | 0,0110444 | 0,94765 | 0,986016 |
| Ebf4 | 1,76229 | 1,7389 | -0,019276 | 0,9476 | 0,986016 |
| Acot6 | 0,665943 | 0,682396 | 0,0352102 | 0,9479 | 0,986079 |
| Iscu | 12,6804 | 12,8214 | 0,0159582 | 0,9482 | 0,986079 |
| Chkb | 20,1992 | 20,3685 | 0,012044 | 0,94815 | 0,986079 |
| Terf1 | 23,667 | 23,8455 | 0,0108405 | 0,94805 | 0,986079 |
| Ccnl2 | 71,7075 | 71,205 | -0,0101451 | 0,94805 | 0,986079 |
| Fbxw7 | 10,4997 | 10,4223 | -0,0106773 | 0,94825 | 0,986079 |
| Xylt2 | 16,9154 | 16,7879 | -0,0109213 | 0,94825 | 0,986079 |
| Dcaf11 | 22,3379 | 22,1664 | -0,0111176 | 0,9483 | 0,986079 |
| Akt1s1 | 42,5527 | 42,8812 | 0,0110928 | 0,94845 | 0,986081 |
| Zfp619 | 1,7326 | 1,71598 | -0,0139057 | 0,94845 | 0,986081 |
| Lamtor3 | 12,2866 | 12,4018 | 0,0134633 | 0,9489 | 0,986089 |
| Pdha1 | 66,1992 | 66,6756 | 0,0103447 | 0,94865 | 0,986089 |
| Arpc1a | 66,318 | 66,7941 | 0,010321 | 0,94865 | 0,986089 |
| Ctr9 | 30,7102 | 30,4975 | -0,0100297 | 0,9487 | 0,986089 |
| Wtip | 22,5807 | 22,4026 | -0,0114202 | 0,94885 | 0,986089 |
| Numbl | 5,30688 | 5,25571 | -0,0139783 | 0,94885 | 0,986089 |
| Far1 | 38,409 | 38,1417 | -0,0100768 | 0,949 | 0,986116 |
| Ints8 | 38,4748 | 38,831 | 0,0132949 | 0,94925 | 0,986222 |
| Mtfp1 | 7,28518 | 7,20875 | -0,0152158 | 0,94925 | 0,986222 |

| | | | | | |
|---|---|---|---|---|---|
| Pcdhgb8 | 0,570647 | 0,625895 | 0,133321 | 0,94985 | 0,986259 |
| Gpx4 | 11,2593 | 11,7559 | 0,062273 | 0,94955 | 0,986259 |
| Ophn1 | 4,4494 | 4,49018 | 0,0131648 | 0,94955 | 0,986259 |
| Tbc1d10b | 14,3423 | 14,4541 | 0,0111974 | 0,9494 | 0,986259 |
| Tob2 | 9,8658 | 9,94048 | 0,0108789 | 0,94995 | 0,986259 |
| Stag1 | 34,4029 | 34,1739 | -0,00963303 | 0,94975 | 0,986259 |
| Trappc5 | 26,1541 | 25,9558 | -0,0109837 | 0,9497 | 0,986259 |
| Sgsm2 | 2,88475 | 2,85742 | -0,0137346 | 0,9499 | 0,986259 |
| A630066F11Rik | 1,7066 | 1,62987 | -0,0663608 | 0,94995 | 0,986259 |
| Smarca5 | 28,1304 | 27,9462 | -0,0094792 | 0,9501 | 0,986339 |
| Mpv17 | 16,4101 | 17,5475 | 0,09668 | 0,95155 | 0,986388 |
| 9530027J09Rik | 0,664232 | 0,708532 | 0,0931452 | 0,95095 | 0,986388 |
| Gm10516 | 1,26111 | 1,28789 | 0,0303065 | 0,95035 | 0,986388 |
| Efcab11 | 5,58606 | 5,64205 | 0,014387 | 0,95135 | 0,986388 |
| Hectd3 | 5,928 | 5,97666 | 0,0117927 | 0,95145 | 0,986388 |
| Tor2a | 22,513 | 22,6841 | 0,0109274 | 0,951 | 0,986388 |
| Rgs9 | 6,0444 | 6,08903 | 0,0106119 | 0,9505 | 0,986388 |
| Dtl | 32,8692 | 32,6455 | -0,00984984 | 0,9511 | 0,986388 |
| Sumo3 | 77,1712 | 76,6392 | -0,00998051 | 0,95065 | 0,986388 |
| Trio | 11,3897 | 11,3095 | -0,0101869 | 0,9515 | 0,986388 |
| Spry1 | 18,5608 | 18,4275 | -0,010395 | 0,9515 | 0,986388 |
| A830080D01Rik | 9,8455 | 9,77418 | -0,0104884 | 0,9505 | 0,986388 |
| Mboat7 | 15,1782 | 15,0678 | -0,010534 | 0,9512 | 0,986388 |
| BC031181 | 42,7155 | 42,3933 | -0,0109223 | 0,9515 | 0,986388 |
| Dtnb | 10,4016 | 10,3109 | -0,0126451 | 0,9505 | 0,986388 |
| Ccdc64 | 1,59023 | 1,56764 | -0,0206419 | 0,9509 | 0,986388 |
| Snapin | 15,4949 | 15,0811 | -0,0390583 | 0,95075 | 0,986388 |
| Mipol1 | 5,04399 | 4,87847 | -0,0481348 | 0,95125 | 0,986388 |
| Rmrp | 6,9525 | 6,71528 | -0,0500844 | 0,95155 | 0,986388 |
| Gm10433 | 0,912489 | 0,928377 | 0,0249046 | 0,9518 | 0,986425 |
| Psmd11 | 31,2427 | 31,4751 | 0,0106897 | 0,95225 | 0,986425 |
| Abl2 | 12,2055 | 12,2887 | 0,00980817 | 0,952 | 0,986425 |
| Uspl1 | 30,1319 | 30,3288 | 0,00939268 | 0,9519 | 0,986425 |
| Cul5 | 16,1233 | 16,015 | -0,00972997 | 0,95205 | 0,986425 |
| Aida | 14,9448 | 14,8346 | -0,0106783 | 0,9519 | 0,986425 |
| Cox19 | 42,5142 | 42,181 | -0,0113519 | 0,95215 | 0,986425 |
| Vil1 | 1,30267 | 1,20866 | -0,108067 | 0,9522 | 0,986425 |
| Dos | 5,52988 | 4,69983 | -0,234639 | 0,952 | 0,986425 |
| B230216G23Rik | 2,53335 | 2,57071 | 0,0211209 | 0,9524 | 0,986479 |
| Lemd3 | 9,65206 | 9,58181 | -0,0105379 | 0,95245 | 0,986479 |
| Pcdhga3 | 2,58757 | 2,69073 | 0,0563965 | 0,9536 | 0,986585 |
| AI507597 | 1,20453 | 1,24233 | 0,0445739 | 0,95335 | 0,986585 |
| BC024978 | 5,23692 | 5,35472 | 0,032091 | 0,954 | 0,986585 |
| Snora78 | 39,9489 | 40,4164 | 0,0167862 | 0,9528 | 0,986585 |
| Tha1 | 1,78757 | 1,80746 | 0,0159601 | 0,9539 | 0,986585 |
| Tmem26 | 1,08912 | 1,10103 | 0,0156842 | 0,95315 | 0,986585 |
| Zkscan2 | 2,58355 | 2,60755 | 0,0133412 | 0,9534 | 0,986585 |
| Dimt1 | 5,25237 | 5,29985 | 0,0129838 | 0,95425 | 0,986585 |

| | | | | | |
|---|---|---|---|---|---|
| Sgsh | 4,31825 | 4,35505 | 0,0122455 | 0,95295 | 0,986585 |
| Zfp239 | 7,71605 | 7,77851 | 0,0116319 | 0,95425 | 0,986585 |
| Clcn2 | 8,30055 | 8,3654 | 0,0112272 | 0,95375 | 0,986585 |
| Smad6 | 8,01338 | 8,07275 | 0,0106499 | 0,9542 | 0,986585 |
| Ndufs4 | 47,503 | 47,8014 | 0,00903453 | 0,9536 | 0,986585 |
| Prkacb | 28,3344 | 28,5068 | 0,00875119 | 0,9542 | 0,986585 |
| Ampd2 | 22,4855 | 22,3383 | -0,00947436 | 0,9536 | 0,986585 |
| Snord100 | 1887,01 | 1874,33 | -0,00972217 | 0,95395 | 0,986585 |
| Med22 | 38,1771 | 37,884 | -0,0111192 | 0,95405 | 0,986585 |
| Tshz2 | 3,93507 | 3,90307 | -0,0117784 | 0,95315 | 0,986585 |
| Slc19a3 | 0,99587 | 0,984557 | -0,0164822 | 0,954 | 0,986585 |
| Cabp1 | 3,1147 | 3,07884 | -0,0167064 | 0,95265 | 0,986585 |
| Polr3h | 12,2222 | 12,0028 | -0,0261375 | 0,95305 | 0,986585 |
| Pcdhgb4 | 1,9796 | 1,88589 | -0,0699634 | 0,954 | 0,986585 |
| AW046200 | 0,654379 | 0,601017 | -0,122719 | 0,95395 | 0,986585 |
| Asb8 | 13,3181 | 13,4212 | 0,0111229 | 0,9544 | 0,986639 |
| Whamm | 2,27498 | 2,25354 | -0,0136639 | 0,95445 | 0,986639 |
| Tbc1d25 | 5,23133 | 5,27652 | 0,012409 | 0,9547 | 0,986669 |
| Ttc17 | 7,04257 | 7,09086 | 0,00985775 | 0,9546 | 0,986669 |
| Jrk | 4,1836 | 4,15296 | -0,0106036 | 0,95465 | 0,986669 |
| Ctnnd1 | 58,5829 | 58,9567 | 0,00917472 | 0,9549 | 0,986723 |
| Nhsl1 | 10,2777 | 10,2114 | -0,00932894 | 0,9549 | 0,986723 |
| Trp53i13 | 9,3394 | 9,26081 | -0,0121914 | 0,955 | 0,98675 |
| Cipc | 12,5254 | 12,6088 | 0,00957157 | 0,9551 | 0,986752 |
| Mitd1 | 18,5969 | 18,4557 | -0,0109959 | 0,95515 | 0,986752 |
| Armcx5 | 5,68791 | 5,81527 | 0,0319467 | 0,95525 | 0,986779 |
| Acy1 | 20,5809 | 20,7368 | 0,010881 | 0,9554 | 0,986858 |
| Prpf18 | 20,3713 | 20,5049 | 0,00942832 | 0,95555 | 0,986936 |
| 1700123O20Rik | 27,9595 | 28,5079 | 0,0280221 | 0,95565 | 0,986939 |
| Trim3 | 7,92398 | 7,86586 | -0,0106209 | 0,9557 | 0,986939 |
| Mrpl55 | 36,9186 | 36,6384 | -0,0109911 | 0,9558 | 0,986966 |
| Oaz1 | 21,778 | 21,6102 | -0,011159 | 0,95605 | 0,987071 |
| Slc2a8 | 6,27949 | 6,22722 | -0,0120586 | 0,95605 | 0,987071 |
| 4930413G21Rik | 4,58063 | 4,63009 | 0,0154921 | 0,95625 | 0,987177 |
| Lias | 26,2553 | 26,4433 | 0,0102921 | 0,9563 | 0,987177 |
| Ppm1f | 23,9953 | 23,8438 | -0,00913832 | 0,9564 | 0,987204 |
| Ctdnep1 | 46,0701 | 46,5049 | 0,0135528 | 0,9565 | 0,987231 |
| Cnbp | 367,184 | 369,535 | 0,00920642 | 0,95665 | 0,987309 |
| Tmem53 | 1,48227 | 1,50792 | 0,0247506 | 0,95725 | 0,987626 |
| Cpsf2 | 34,783 | 34,9822 | 0,00823823 | 0,9574 | 0,987626 |
| Atp6v1a | 30,8468 | 30,6559 | -0,00895916 | 0,9574 | 0,987626 |
| Rpl35a | 166,442 | 165,337 | -0,00960392 | 0,9574 | 0,987626 |
| Col1a1 | 3,52566 | 3,49952 | -0,0107343 | 0,95735 | 0,987626 |
| Cog5 | 11,8926 | 11,7598 | -0,0162003 | 0,95735 | 0,987626 |
| Atxn7l2 | 10,4414 | 10,3656 | -0,0105209 | 0,95765 | 0,987808 |
| Ndufs6 | 77,8325 | 78,3376 | 0,00933274 | 0,95775 | 0,987834 |
| Smg8 | 18,588 | 19,1268 | 0,0412264 | 0,95955 | 0,988074 |
| Usp16 | 24,8241 | 25,3701 | 0,0313854 | 0,95945 | 0,988074 |

| | | | | | |
|---|---|---|---|---|---|
| Sspn | 0,916681 | 0,927937 | 0,0176078 | 0,95875 | 0,988074 |
| Ap5z1 | 3,25931 | 3,28616 | 0,0118357 | 0,9589 | 0,988074 |
| 9330133O14Rik | 2,84552 | 2,86818 | 0,0114424 | 0,95865 | 0,988074 |
| Shisa4 | 13,594 | 13,7 | 0,0112064 | 0,95945 | 0,988074 |
| Fan1 | 4,93922 | 4,97512 | 0,0104467 | 0,9586 | 0,988074 |
| Rhobtb2 | 4,82239 | 4,85694 | 0,0102993 | 0,95895 | 0,988074 |
| Gm5086 | 0,687844 | 0,692608 | 0,00995727 | 0,9602 | 0,988074 |
| Slc35e2 | 8,68082 | 8,73563 | 0,00908052 | 0,95875 | 0,988074 |
| Cmtm7 | 30,4938 | 30,6799 | 0,0087813 | 0,9593 | 0,988074 |
| Dyrk1a | 21,6938 | 21,8221 | 0,00851057 | 0,95945 | 0,988074 |
| Pitpna | 25,5432 | 25,694 | 0,00849047 | 0,96 | 0,988074 |
| Gabarapl1 | 33,0199 | 33,2127 | 0,00839959 | 0,95925 | 0,988074 |
| Zik1 | 16,7519 | 16,8462 | 0,00809391 | 0,9586 | 0,988074 |
| Wac | 21,2476 | 21,3621 | 0,00775 | 0,9595 | 0,988074 |
| Nacc1 | 41,856 | 41,6365 | -0,00758884 | 0,9598 | 0,988074 |
| Khsrp | 68,6408 | 68,2785 | -0,00763463 | 0,95935 | 0,988074 |
| Alg2 | 34,5194 | 34,3238 | -0,00819716 | 0,9601 | 0,988074 |
| Usp33 | 23,8032 | 23,6681 | -0,00821027 | 0,9602 | 0,988074 |
| Aurkb | 63,1226 | 62,7642 | -0,00821546 | 0,96005 | 0,988074 |
| Fam118a | 16,2103 | 16,1103 | -0,00891917 | 0,95935 | 0,988074 |
| Map1lc3b | 28,3777 | 28,1958 | -0,0092784 | 0,9591 | 0,988074 |
| Gatsl2 | 5,08809 | 5,05276 | -0,010054 | 0,9597 | 0,988074 |
| 2810021J22Rik | 3,91707 | 3,88819 | -0,0106747 | 0,9587 | 0,988074 |
| Lamtor5 | 23,7565 | 23,5813 | -0,0106766 | 0,9597 | 0,988074 |
| Vpreb3 | 1,88894 | 1,87427 | -0,0112457 | 0,9585 | 0,988074 |
| Fam60a | 4,75485 | 4,7169 | -0,0115599 | 0,9595 | 0,988074 |
| Arhgap44 | 2,99473 | 2,94808 | -0,0226521 | 0,9595 | 0,988074 |
| Pcdhga4 | 1,36407 | 1,29434 | -0,0757027 | 0,9602 | 0,988074 |
| Midn | 72,9588 | 75,1406 | 0,0425102 | 0,96055 | 0,98808 |
| Clip4 | 1,57716 | 1,59311 | 0,0145122 | 0,9606 | 0,98808 |
| Zfp276 | 6,7756 | 6,84272 | 0,0142207 | 0,96045 | 0,98808 |
| Wdyhv1 | 15,9107 | 15,804 | -0,00970681 | 0,9606 | 0,98808 |
| Tmem138 | 26,5558 | 26,3648 | -0,0104101 | 0,9604 | 0,98808 |
| Copg2 | 29,3233 | 28,7821 | -0,0268761 | 0,96065 | 0,98808 |
| Pcdhga10 | 1,46441 | 1,53174 | 0,0648569 | 0,96155 | 0,988131 |
| Ergic2 | 43,0406 | 43,3464 | 0,0102154 | 0,96095 | 0,988131 |
| 1810037I17Rik | 24,3682 | 24,5253 | 0,00927328 | 0,962 | 0,988131 |
| Limk1 | 10,4771 | 10,5419 | 0,00889117 | 0,961 | 0,988131 |
| Yy1 | 53,7216 | 54,0324 | 0,00832407 | 0,96185 | 0,988131 |
| Zc3h7a | 17,7074 | 17,8088 | 0,00824143 | 0,9616 | 0,988131 |
| Clk4 | 38,3891 | 38,5995 | 0,00788482 | 0,96235 | 0,988131 |
| Igf2bp3 | 63,208 | 63,544 | 0,00764997 | 0,96125 | 0,988131 |
| Gtf3c4 | 27,4158 | 27,5573 | 0,00742847 | 0,96185 | 0,988131 |
| Gorasp2 | 68,1308 | 68,4815 | 0,0074082 | 0,96235 | 0,988131 |
| Dab2ip | 30,15 | 30,3039 | 0,00734548 | 0,9622 | 0,988131 |
| Actr1a | 51,9859 | 52,247 | 0,00722697 | 0,9619 | 0,988131 |
| Tnks2 | 78,3561 | 77,9516 | -0,00746721 | 0,96185 | 0,988131 |
| Zfp148 | 12,8139 | 12,7454 | -0,0077313 | 0,9624 | 0,988131 |

| | | | | | |
|---|---|---|---|---|---|
| 2510009E07Rik | 9,23063 | 9,17867 | -0,00814456 | 0,9609 | 0,988131 |
| Agpat6 | 49,5331 | 49,2532 | -0,0081752 | 0,962 | 0,988131 |
| Rbl1 | 19,5046 | 19,3924 | -0,00832864 | 0,96135 | 0,988131 |
| Nudcd1 | 22,996 | 22,827 | -0,0106446 | 0,96185 | 0,988131 |
| Nagk | 3,82148 | 3,78548 | -0,0136553 | 0,96215 | 0,988131 |
| Rarres1 | 0,735506 | 0,719228 | -0,0322873 | 0,96225 | 0,988131 |
| Mettl5 | 16,9119 | 16,3087 | -0,0524005 | 0,96155 | 0,988131 |
| Palm | 31,7098 | 29,8629 | -0,0865751 | 0,96155 | 0,988131 |
| Slc5a6 | 3,91751 | 3,61836 | -0,114601 | 0,9618 | 0,988131 |
| Fam169a | 8,16601 | 8,10486 | -0,010844 | 0,9626 | 0,98826 |
| Trim27 | 31,8558 | 31,7009 | -0,00703365 | 0,963 | 0,988443 |
| Pyroxd1 | 10,4712 | 10,3722 | -0,0136957 | 0,963 | 0,988443 |
| Ptrh2 | 22,8382 | 22,3237 | -0,0328759 | 0,963 | 0,988443 |
| Med16 | 18,0029 | 16,8244 | -0,0976721 | 0,96315 | 0,988521 |
| Rnf126 | 50,8969 | 53,665 | 0,0764053 | 0,9635 | 0,988579 |
| Atl1 | 1,1436 | 1,1902 | 0,0576247 | 0,96365 | 0,988579 |
| Elmod3 | 4,35795 | 4,39139 | 0,0110309 | 0,96355 | 0,988579 |
| Micu1 | 18,6166 | 18,7229 | 0,00821395 | 0,96365 | 0,988579 |
| Esco1 | 6,42486 | 6,38677 | -0,00857764 | 0,9633 | 0,988579 |
| Plb1 | 1,02575 | 1,00549 | -0,0287816 | 0,9635 | 0,988579 |
| Eml2 | 9,93728 | 9,99871 | 0,00889202 | 0,9639 | 0,988684 |
| Zfp867 | 5,53594 | 5,49765 | -0,0100139 | 0,96385 | 0,988684 |
| Klhl18 | 5,51225 | 5,47872 | -0,008802 | 0,96405 | 0,988762 |
| Cdk3-ps | 0,697003 | 0,687812 | -0,0191501 | 0,96425 | 0,988891 |
| Mrps14 | 23,986 | 24,1224 | 0,00817865 | 0,96445 | 0,988944 |
| Mau2 | 19,7395 | 19,6385 | -0,00740368 | 0,96445 | 0,988944 |
| Atf5 | 15,6144 | 15,8333 | 0,0200884 | 0,96455 | 0,988971 |
| Kdm4b | 15,5335 | 15,4542 | -0,00738405 | 0,9647 | 0,989049 |
| Pou4f1 | 0,646 | 0,653575 | 0,0168174 | 0,9651 | 0,989155 |
| Arhgap22 | 1,84974 | 1,86586 | 0,012521 | 0,9649 | 0,989155 |
| Atf7ip2 | 4,54511 | 4,57659 | 0,00995844 | 0,965 | 0,989155 |
| Rsph9 | 1,79658 | 1,76444 | -0,0260435 | 0,96505 | 0,989155 |
| Smndc1 | 35,1453 | 35,3138 | 0,00689669 | 0,9652 | 0,989182 |
| Aldh3a2 | 10,8543 | 10,9128 | 0,00776131 | 0,96555 | 0,98944 |
| Lgals4 | 0,669127 | 0,66836 | -0,00165553 | 0,9656 | 0,98944 |
| Sbno2 | 5,9689 | 6,51789 | 0,126941 | 0,96625 | 0,989466 |
| Snord32a | 1992,16 | 2083,14 | 0,0644259 | 0,96755 | 0,989466 |
| D330050I16Rik | 1,34733 | 1,40505 | 0,0605211 | 0,96665 | 0,989466 |
| Gstm2 | 1,80499 | 1,83366 | 0,0227324 | 0,9669 | 0,989466 |
| B4galnt1 | 1,86587 | 1,88332 | 0,0134279 | 0,96595 | 0,989466 |
| Ubtd1 | 7,54825 | 7,60184 | 0,0102068 | 0,9659 | 0,989466 |
| 1110059G10Rik | 5,04264 | 5,0743 | 0,00902967 | 0,9663 | 0,989466 |
| Vps26b | 16,3983 | 16,4922 | 0,00823204 | 0,96695 | 0,989466 |
| Nudt3 | 57,5381 | 57,8177 | 0,0069924 | 0,9663 | 0,989466 |
| BC048502 | 1,8132 | 1,80961 | -0,00285933 | 0,9668 | 0,989466 |
| Dhcr24 | 96,6929 | 96,2536 | -0,00656951 | 0,96725 | 0,989466 |
| Adipor1 | 63,7411 | 63,4514 | -0,00657236 | 0,96745 | 0,989466 |
| Uba5 | 26,2728 | 26,15 | -0,00676044 | 0,9675 | 0,989466 |

| | | | | | |
|---|---|---|---|---|---|
| Cd63 | 71,6058 | 71,2686 | -0,00680948 | 0,96655 | 0,989466 |
| Ppm1b | 21,9301 | 21,8215 | -0,00715951 | 0,96745 | 0,989466 |
| Pdxdc1 | 28,2953 | 28,1533 | -0,00725652 | 0,96755 | 0,989466 |
| Zdbf2 | 7,3244 | 7,28754 | -0,00727877 | 0,9665 | 0,989466 |
| Shprh | 10,259 | 10,2065 | -0,00739878 | 0,96665 | 0,989466 |
| Itpa | 44,3383 | 44,1071 | -0,00754079 | 0,9669 | 0,989466 |
| Tmem237 | 17,7359 | 17,6434 | -0,00754212 | 0,96685 | 0,989466 |
| Fbxl7 | 4,0285 | 4,00551 | -0,00826001 | 0,96725 | 0,989466 |
| 4930526I15Rik | 6,60723 | 6,5589 | -0,0105909 | 0,96635 | 0,989466 |
| 3110002H16Rik | 8,72079 | 8,63593 | -0,0141073 | 0,9663 | 0,989466 |
| Mansc4 | 1,04322 | 1,03248 | -0,0149348 | 0,9663 | 0,989466 |
| Aebp1 | 1,99928 | 1,9709 | -0,0206271 | 0,9675 | 0,989466 |
| Efna2 | 10,9165 | 9,85432 | -0,147679 | 0,9673 | 0,989466 |
| Zscan26 | 25,3967 | 25,2837 | -0,00643156 | 0,9677 | 0,989468 |
| Sema4f | 0,69656 | 0,689721 | -0,0142353 | 0,96765 | 0,989468 |
| Rps6kb2 | 17,2971 | 17,3977 | 0,00837067 | 0,96795 | 0,989547 |
| Flot1 | 12,075 | 12,0078 | -0,00804586 | 0,968 | 0,989547 |
| Slfn9 | 4,75269 | 4,72553 | -0,00826872 | 0,96795 | 0,989547 |
| 1810026B05Rik | 5,67224 | 5,61362 | -0,0149869 | 0,9681 | 0,989574 |
| Swsap1 | 8,03625 | 7,98803 | -0,00868263 | 0,96825 | 0,989651 |
| Homez | 6,5565 | 6,5886 | 0,00704605 | 0,96865 | 0,989884 |
| Faxc | 7,1422 | 7,17558 | 0,00672754 | 0,9686 | 0,989884 |
| Snord104 | 4037,67 | 3971,78 | -0,0237394 | 0,9687 | 0,989884 |
| Tada2b | 7,1749 | 7,24149 | 0,0133268 | 0,9691 | 0,990217 |
| Cyth1 | 8,45527 | 8,41199 | -0,00740394 | 0,9693 | 0,990346 |
| Mrps12 | 59,3597 | 59,6578 | 0,00722622 | 0,9694 | 0,990372 |
| Dpp3 | 67,3754 | 67,1084 | -0,00572808 | 0,96955 | 0,990425 |
| Polg | 20,474 | 20,3583 | -0,00817039 | 0,9696 | 0,990425 |
| Dynll2 | 120,456 | 119,942 | -0,00616086 | 0,9697 | 0,990452 |
| Phyhip | 0,60249 | 0,61159 | 0,021629 | 0,97025 | 0,990839 |
| Gm15706 | 1,78801 | 1,80712 | 0,0153395 | 0,9704 | 0,990839 |
| Mbd3 | 126,024 | 126,548 | 0,00598374 | 0,97045 | 0,990839 |
| Xpnpep1 | 46,3464 | 46,1629 | -0,00572456 | 0,97035 | 0,990839 |
| Ncoa4 | 1,71337 | 1,70193 | -0,00966341 | 0,97045 | 0,990839 |
| AI504432 | 2,2436 | 2,25582 | 0,007833 | 0,97075 | 0,990894 |
| Rpf1 | 62,3316 | 62,5808 | 0,00575542 | 0,97075 | 0,990894 |
| Dusp19 | 9,13631 | 9,08558 | -0,0080321 | 0,9708 | 0,990894 |
| Fkrp | 16,8604 | 16,7055 | -0,0133186 | 0,9708 | 0,990894 |
| Zfp655 | 12,1703 | 12,2245 | 0,00641005 | 0,97095 | 0,990895 |
| Ctnnbip1 | 17,0175 | 16,9396 | -0,0066189 | 0,9709 | 0,990895 |
| Chchd3 | 66,1581 | 65,8774 | -0,00613528 | 0,9712 | 0,991026 |
| Lin9 | 10,0343 | 9,98757 | -0,00673914 | 0,9713 | 0,991026 |
| Arhgap9 | 1,4502 | 1,42011 | -0,0302504 | 0,97125 | 0,991026 |
| Atg4a | 1,72767 | 1,81254 | 0,069191 | 0,97165 | 0,991207 |
| Tmem38a | 3,72616 | 3,74695 | 0,00802349 | 0,9716 | 0,991207 |
| Pcif1 | 19,8215 | 19,7369 | -0,00616653 | 0,9717 | 0,991207 |
| Marf1 | 16,5112 | 16,4458 | -0,00572861 | 0,9719 | 0,991259 |
| Selo | 7,15095 | 7,07221 | -0,0159744 | 0,97185 | 0,991259 |

| | | | | | |
|---|---|---|---|---|---|
| Gm10785 | 1,34315 | 1,32102 | -0,0239713 | 0,97205 | 0,991261 |
| Scx | 0,895327 | 0,851484 | -0,0724355 | 0,97205 | 0,991261 |
| Iqck | 0,915287 | 0,861077 | -0,088082 | 0,9724 | 0,991542 |
| Cxcl17 | 1,17061 | 1,17659 | 0,00734211 | 0,97275 | 0,991824 |
| Dpagt1 | 22,4883 | 22,401 | -0,00560903 | 0,97285 | 0,99185 |
| A930024E05Rik | 3,76632 | 3,7889 | 0,00862492 | 0,97305 | 0,991902 |
| Pik3c2a | 16,9822 | 17,0432 | 0,0051736 | 0,973 | 0,991902 |
| Gm19710 | 3,28805 | 3,47527 | 0,0798922 | 0,97345 | 0,992234 |
| Arhgef12 | 14,9754 | 14,9203 | -0,00532458 | 0,9736 | 0,992287 |
| Ankrd13d | 2,13633 | 2,1219 | -0,00977702 | 0,97365 | 0,992287 |
| Tmem55b | 17,7033 | 17,8601 | 0,0127249 | 0,97425 | 0,992376 |
| Yy2 | 7,46451 | 7,51756 | 0,0102164 | 0,97455 | 0,992376 |
| Prrt2 | 2,94759 | 2,96437 | 0,00819207 | 0,97435 | 0,992376 |
| Abtb2 | 1,64096 | 1,64915 | 0,00718545 | 0,9743 | 0,992376 |
| Mn1 | 2,9099 | 2,92378 | 0,00686717 | 0,97465 | 0,992376 |
| Gmip | 3,24426 | 3,25939 | 0,00671079 | 0,97495 | 0,992376 |
| Nit2 | 29,6273 | 29,7396 | 0,00545947 | 0,9742 | 0,992376 |
| Mycn | 72,1163 | 72,3865 | 0,00539451 | 0,97385 | 0,992376 |
| Usp1 | 39,1556 | 39,3013 | 0,00535844 | 0,97435 | 0,992376 |
| Wdr48 | 30,1227 | 30,2318 | 0,00521302 | 0,9745 | 0,992376 |
| Ppp6r1 | 40,6627 | 40,8085 | 0,00516288 | 0,97455 | 0,992376 |
| U2surp | 39,5961 | 39,7272 | 0,00476709 | 0,97485 | 0,992376 |
| Rictor | 9,04476 | 9,01361 | -0,00497775 | 0,97465 | 0,992376 |
| Lig3 | 25,1364 | 25,0483 | -0,00506135 | 0,9745 | 0,992376 |
| Dmxl2 | 3,79539 | 3,78056 | -0,00564728 | 0,9749 | 0,992376 |
| Siah1a | 16,1311 | 16,0668 | -0,00576819 | 0,9749 | 0,992376 |
| 2310011J03Rik | 19,2013 | 17,9831 | -0,094566 | 0,975 | 0,992376 |
| Dmgdh | 2,63728 | 2,65058 | 0,00725792 | 0,9755 | 0,992507 |
| Cep57l1 | 8,25587 | 8,29223 | 0,00633984 | 0,97535 | 0,992507 |
| Nup155 | 36,6298 | 36,7571 | 0,00500236 | 0,97525 | 0,992507 |
| Tlk1 | 43,5829 | 43,4401 | -0,00473309 | 0,97545 | 0,992507 |
| Gmeb2 | 13,5583 | 13,5124 | -0,00488889 | 0,9755 | 0,992507 |
| Invs | 4,44896 | 4,48231 | 0,0107742 | 0,97565 | 0,992584 |
| Tmem194 | 12,4637 | 12,4203 | -0,00503436 | 0,9759 | 0,992763 |
| Setd2 | 26,8869 | 26,9786 | 0,00491191 | 0,97615 | 0,992791 |
| Fyn | 24,0706 | 23,9893 | -0,00488145 | 0,97615 | 0,992791 |
| Lurap1 | 1,49999 | 1,49302 | -0,00671719 | 0,97615 | 0,992791 |
| Kbtbd3 | 2,61119 | 2,62584 | 0,00807616 | 0,97645 | 0,992794 |
| Hdgf | 211,904 | 212,573 | 0,00454309 | 0,97645 | 0,992794 |
| Sec61a2 | 43,2562 | 43,1163 | -0,00467348 | 0,97645 | 0,992794 |
| Impa1 | 35,7344 | 35,6076 | -0,00512903 | 0,97625 | 0,992794 |
| Golga2 | 14,7826 | 14,7335 | -0,00480054 | 0,9766 | 0,99287 |
| Nkapl | 1,43365 | 1,44581 | 0,0121791 | 0,97725 | 0,992927 |
| Lrrc1 | 7,8664 | 7,89519 | 0,00527035 | 0,97705 | 0,992927 |
| Mapkapk2 | 56,0722 | 55,88 | -0,00495302 | 0,97725 | 0,992927 |
| 1810055G02Rik | 24,3834 | 24,2926 | -0,00538197 | 0,977 | 0,992927 |
| Enho | 7,45286 | 7,42146 | -0,0060911 | 0,97695 | 0,992927 |
| Fkbp11 | 15,6213 | 15,5487 | -0,00671817 | 0,9769 | 0,992927 |

| | | | | | |
|---|---|---|---|---|---|
| Mpg | 16,9097 | 16,8161 | -0,00801457 | 0,97725 | 0,992927 |
| Adat3 | 5,8019 | 5,73499 | -0,016734 | 0,97685 | 0,992927 |
| Fubp1 | 109,303 | 109,644 | 0,00448839 | 0,97855 | 0,992964 |
| Snora52 | 136,517 | 140,023 | 0,0365847 | 0,9784 | 0,992965 |
| Dmrta2 | 0,610728 | 0,617877 | 0,0167898 | 0,9775 | 0,992965 |
| Ormdl2 | 12,2755 | 12,3588 | 0,0097613 | 0,97845 | 0,992965 |
| Fam19a5 | 4,66024 | 4,67963 | 0,00599118 | 0,97795 | 0,992965 |
| Col12a1 | 0,747408 | 0,750437 | 0,00583369 | 0,9782 | 0,992965 |
| Cryzl1 | 16,9515 | 17,0103 | 0,00499019 | 0,9777 | 0,992965 |
| Pigv | 10,6478 | 10,6845 | 0,00496242 | 0,97855 | 0,992965 |
| Trub1 | 11,0464 | 11,0107 | -0,00467005 | 0,97815 | 0,992965 |
| Pmpca | 60,7237 | 60,5263 | -0,00469697 | 0,9779 | 0,992965 |
| Ndufa9 | 84,1525 | 83,871 | -0,00483309 | 0,97815 | 0,992965 |
| Brip1 | 4,24838 | 4,23364 | -0,00501443 | 0,97755 | 0,992965 |
| Rara | 9,7882 | 9,75411 | -0,00503362 | 0,97825 | 0,992965 |
| Ilk | 51,6428 | 51,3864 | -0,00718172 | 0,9781 | 0,992965 |
| Zcchc12 | 0,915611 | 0,908055 | -0,0119541 | 0,97785 | 0,992965 |
| Fndc4 | 27,6093 | 27,0508 | -0,0294833 | 0,97805 | 0,992965 |
| Fbxl22 | 0,819989 | 0,801943 | -0,0321058 | 0,9785 | 0,992965 |
| Slc45a4 | 8,54661 | 8,5193 | -0,00461879 | 0,9789 | 0,993144 |
| Gm16039 | 4,7765 | 4,75657 | -0,00603005 | 0,9789 | 0,993144 |
| Gdf1 | 4,36292 | 4,29029 | -0,0242189 | 0,97895 | 0,993144 |
| Zfp963 | 4,229 | 4,2486 | 0,0066737 | 0,97905 | 0,99317 |
| Usp37 | 15,4097 | 15,4934 | 0,00780946 | 0,97915 | 0,993196 |
| A930007I19Rik | 0,882305 | 0,873956 | -0,0137172 | 0,97925 | 0,993222 |
| Ankrd17 | 27,2263 | 27,3033 | 0,00407238 | 0,9794 | 0,993224 |
| AI597479 | 21,5729 | 21,5071 | -0,004402 | 0,97935 | 0,993224 |
| Arglu1 | 108,424 | 108,739 | 0,00418513 | 0,97955 | 0,9933 |
| Fbxo45 | 17,1725 | 17,1251 | -0,00398404 | 0,9799 | 0,993429 |
| Panx1 | 26,3127 | 26,2323 | -0,00441428 | 0,9798 | 0,993429 |
| Gng12 | 28,1296 | 28,0391 | -0,00464782 | 0,97985 | 0,993429 |
| Ankrd24 | 2,89323 | 2,90697 | 0,00683671 | 0,98 | 0,993455 |
| Dsn1 | 23,278 | 23,2142 | -0,00395906 | 0,9802 | 0,993507 |
| Fam188a | 14,8938 | 14,8434 | -0,00489077 | 0,9802 | 0,993507 |
| Mpp3 | 5,18588 | 5,20603 | 0,00559331 | 0,98045 | 0,99351 |
| Ing3 | 9,46916 | 9,49889 | 0,00452342 | 0,9804 | 0,99351 |
| Prickle2 | 3,35524 | 3,36512 | 0,00423889 | 0,9805 | 0,99351 |
| Hmgn1 | 312,241 | 313,088 | 0,00390682 | 0,98045 | 0,99351 |
| Smpd2 | 19,4724 | 19,5335 | 0,00451988 | 0,9807 | 0,993637 |
| Wdr47 | 5,20234 | 5,21795 | 0,00432143 | 0,98095 | 0,99374 |
| Gpr156 | 0,601461 | 0,601199 | -0,000627287 | 0,9809 | 0,99374 |
| Mrps11 | 40,8861 | 40,763 | -0,0043518 | 0,98115 | 0,993867 |
| F420014N23Rik | 1,13216 | 1,15416 | 0,0277675 | 0,9817 | 0,993937 |
| Txlng | 6,80473 | 6,87787 | 0,0154253 | 0,98295 | 0,993937 |
| Dhx40 | 27,3928 | 27,6214 | 0,0119916 | 0,9828 | 0,993937 |
| Lrrc57 | 13,2572 | 13,327 | 0,0075796 | 0,98215 | 0,993937 |
| LOC101056043 | 7,70587 | 7,73904 | 0,00619744 | 0,9831 | 0,993937 |
| Helb | 5,76252 | 5,7868 | 0,00606579 | 0,98345 | 0,993937 |

| | | | | | |
|---|---|---|---|---|---|
| Atg4d | 15,1191 | 15,1774 | 0,0055475 | 0,98325 | 0,993937 |
| Med7 | 18,6193 | 18,6787 | 0,00459909 | 0,98335 | 0,993937 |
| Alg3 | 18,7733 | 18,8321 | 0,00451127 | 0,9823 | 0,993937 |
| Snhg11 | 5,07757 | 5,09114 | 0,00385056 | 0,98235 | 0,993937 |
| Gsr | 24,0998 | 24,1638 | 0,00382654 | 0,98235 | 0,993937 |
| Tsnax | 45,7859 | 45,9049 | 0,00374329 | 0,9817 | 0,993937 |
| Chd7 | 25,5967 | 25,6616 | 0,00365501 | 0,98165 | 0,993937 |
| Tnfaip1 | 42,4162 | 42,522 | 0,00359319 | 0,98225 | 0,993937 |
| Srp9 | 32,7483 | 32,8283 | 0,00352063 | 0,9829 | 0,993937 |
| Tnpo1 | 47,019 | 43,7462 | 0,00330555 | 0,98225 | 0,993937 |
| Rptor | 12,8012 | 12,83 | 0,00324387 | 0,98255 | 0,993937 |
| Zbtb8b | 0,775527 | 0,777121 | 0,00296185 | 0,98155 | 0,993937 |
| Atg2a | 6,01396 | 5,99954 | -0,0034612 | 0,98255 | 0,993937 |
| Upf1 | 38,7923 | 38,697 | -0,00354927 | 0,9834 | 0,993937 |
| Maoa | 17,8605 | 17,815 | -0,00367321 | 0,9827 | 0,993937 |
| Zfp428 | 33,0775 | 32,9923 | -0,00371814 | 0,983 | 0,993937 |
| Usb1 | 13,7873 | 13,7512 | -0,00377922 | 0,9819 | 0,993937 |
| Gpc3 | 514,187 | 512,784 | -0,00393954 | 0,98205 | 0,993937 |
| Ttll1 | 7,06051 | 7,03995 | -0,00420693 | 0,9833 | 0,993937 |
| 9230114K14Rik | 1,16934 | 1,16421 | -0,0063498 | 0,98145 | 0,993937 |
| Rasgrp4 | 1,4108 | 1,40451 | -0,00644844 | 0,9834 | 0,993937 |
| Fam118b | 19,7952 | 19,6868 | -0,00791853 | 0,9827 | 0,993937 |
| Gm13251 | 0,917514 | 0,910721 | -0,01072 | 0,9828 | 0,993937 |
| 4930520O04Rik | 0,795137 | 0,783096 | -0,0220141 | 0,9828 | 0,993937 |
| Cnst | 9,70987 | 9,73299 | 0,00343044 | 0,9836 | 0,993989 |
| Syp | 1,34347 | 1,3426 | -0,000930033 | 0,98365 | 0,993989 |
| Gba2 | 16,8301 | 16,8999 | 0,00596829 | 0,9849 | 0,994024 |
| Polm | 3,71413 | 3,72482 | 0,00414897 | 0,98475 | 0,994024 |
| Thoc6 | 39,9597 | 40,0676 | 0,00389081 | 0,98495 | 0,994024 |
| Yipf3 | 18,1595 | 18,204 | 0,00353171 | 0,9847 | 0,994024 |
| Pramef8 | 6,97598 | 6,99297 | 0,00350996 | 0,9843 | 0,994024 |
| Mex3a | 53,0197 | 53,1451 | 0,0034089 | 0,984 | 0,994024 |
| Gpd2 | 9,54151 | 9,56295 | 0,00323724 | 0,984 | 0,994024 |
| Papola | 94,3425 | 94,5533 | 0,00321991 | 0,98435 | 0,994024 |
| Dcun1d5 | 89,6714 | 89,8679 | 0,00315812 | 0,98435 | 0,994024 |
| Dync1li2 | 34,5565 | 34,631 | 0,00310926 | 0,98435 | 0,994024 |
| Stxbp3a | 9,73201 | 9,70592 | -0,00387317 | 0,98465 | 0,994024 |
| Wdsub1 | 7,14589 | 7,12589 | -0,00404312 | 0,98485 | 0,994024 |
| Alg12 | 7,39371 | 7,3714 | -0,00436023 | 0,9839 | 0,994024 |
| Dcun1d2 | 10,24 | 10,2084 | -0,00445079 | 0,98495 | 0,994024 |
| Gramd1b | 1,23122 | 1,22576 | -0,00641079 | 0,9839 | 0,994024 |
| Snx17 | 58,6024 | 58,0126 | -0,0145943 | 0,98475 | 0,994024 |
| Pigyl | 14,8891 | 14,6685 | -0,021539 | 0,98495 | 0,994024 |
| Smim19 | 10,131 | 10,1899 | 0,00835546 | 0,986 | 0,994037 |
| Slirp | 199,862 | 200,947 | 0,00781114 | 0,9854 | 0,994037 |
| D330045A20Rik | 1,59962 | 1,60687 | 0,00651788 | 0,9856 | 0,994037 |
| Arl16 | 10,1609 | 10,1964 | 0,00503214 | 0,98635 | 0,994037 |
| Trappc9 | 5,59159 | 5,60845 | 0,004344 | 0,98645 | 0,994037 |

| | | | | | |
|---|---|---|---|---|---|
| Pmel | 1,21435 | 1,21727 | 0,003473 | 0,98585 | 0,994037 |
| Timm17b | 22,7481 | 22,801 | 0,00334813 | 0,9854 | 0,994037 |
| Tigd2 | 4,76697 | 4,77715 | 0,00307852 | 0,98645 | 0,994037 |
| Fam32a | 36,1617 | 36,2351 | 0,00292315 | 0,98645 | 0,994037 |
| Uri1 | 28,9163 | 28,9733 | 0,00284266 | 0,98615 | 0,994037 |
| Rap2c | 51,5659 | 51,6591 | 0,00260548 | 0,9861 | 0,994037 |
| Gm15441 | 0,656678 | 0,657287 | 0,00133729 | 0,98635 | 0,994037 |
| Ccdc173 | 2,3064 | 2,30794 | 0,000968243 | 0,98605 | 0,994037 |
| Mterfd3 | 12,9691 | 12,9444 | -0,00275092 | 0,98645 | 0,994037 |
| Ranbp10 | 12,6509 | 12,6263 | -0,00280161 | 0,98555 | 0,994037 |
| Carf | 2,59887 | 2,58676 | -0,00673547 | 0,98545 | 0,994037 |
| Ppm1g | 173,963 | 172,969 | -0,00826593 | 0,9856 | 0,994037 |
| 4931406C07Rik | 8,62194 | 8,56641 | -0,00932211 | 0,9863 | 0,994037 |
| C8g | 3,31571 | 3,28261 | -0,0144765 | 0,98565 | 0,994037 |
| Gm16740 | 1,8896 | 1,8469 | -0,0329754 | 0,98555 | 0,994037 |
| Mob3a | 12,7266 | 12,7517 | 0,00283892 | 0,98665 | 0,994164 |
| 6530402F18Rik | 0,922203 | 0,927199 | 0,00779419 | 0,98685 | 0,994166 |
| Ubn1 | 18,8075 | 18,8413 | 0,00258658 | 0,98685 | 0,994166 |
| Luc7l2 | 67,7784 | 67,6443 | -0,00285876 | 0,98695 | 0,994166 |
| Ndufs7 | 75,4929 | 73,5168 | -0,0382686 | 0,9869 | 0,994166 |
| Set | 76,8951 | 76,7549 | -0,0026342 | 0,98705 | 0,994192 |
| Tle1 | 11,3174 | 11,3453 | 0,00354301 | 0,9874 | 0,994469 |
| Zfp37 | 8,05051 | 8,06736 | 0,00301745 | 0,9875 | 0,994495 |
| Hp1bp3 | 47,8066 | 47,7251 | -0,00246004 | 0,9878 | 0,994722 |
| B630019K06Rik | 2,08728 | 2,09031 | 0,00209384 | 0,98795 | 0,994798 |
| Atg4c | 6,94727 | 6,96071 | 0,0027895 | 0,98855 | 0,995028 |
| Abhd17a | 42,966 | 43,0448 | 0,00264192 | 0,98845 | 0,995028 |
| Amer1 | 11,4982 | 11,479 | -0,00241513 | 0,98845 | 0,995028 |
| Commd4 | 30,2947 | 30,2396 | -0,00262678 | 0,9885 | 0,995028 |
| Nfkbil1 | 9,24526 | 9,22397 | -0,00332694 | 0,98855 | 0,995028 |
| Adcy1 | 1,69949 | 1,69699 | -0,00212329 | 0,9888 | 0,995204 |
| Dand5 | 1,98771 | 1,99546 | 0,00561374 | 0,9889 | 0,99523 |
| Dcaf15 | 39,931 | 39,8695 | -0,00222358 | 0,989 | 0,995256 |
| Dtx3l | 1,79645 | 1,80173 | 0,00423336 | 0,98935 | 0,995284 |
| Kif7 | 7,95069 | 7,97155 | 0,00378182 | 0,9893 | 0,995284 |
| Rab7 | 60,8936 | 60,9853 | 0,00217192 | 0,9892 | 0,995284 |
| Slc30a9 | 38,0156 | 38,0701 | 0,00206659 | 0,9892 | 0,995284 |
| Vapa | 86,7434 | 86,8589 | 0,00191816 | 0,9894 | 0,995284 |
| Tnfrsf22 | 2,96144 | 2,97057 | 0,00444077 | 0,9897 | 0,995336 |
| Lix1l | 24,4449 | 24,4851 | 0,00237598 | 0,9897 | 0,995336 |
| Bend4 | 7,922 | 7,93336 | 0,00206599 | 0,98955 | 0,995336 |
| Usp51 | 2,84555 | 2,83599 | -0,00485893 | 0,98975 | 0,995336 |
| Hic2 | 9,94864 | 9,96456 | 0,00230648 | 0,98995 | 0,995462 |
| Zfp65 | 5,66467 | 5,67647 | 0,00300083 | 0,99015 | 0,995464 |
| Coa4 | 9,49043 | 9,48795 | -0,000377984 | 0,9902 | 0,995464 |
| Tax1bp1 | 62,3919 | 62,3084 | -0,00193045 | 0,9902 | 0,995464 |
| Stx4a | 22,1462 | 22,1076 | -0,00251572 | 0,99025 | 0,995464 |
| Usp7 | 55,4149 | 55,4912 | 0,00198666 | 0,9906 | 0,995567 |

| | | | | | |
|---|---|---|---|---|---|
| Kdm2a | 28,4478 | 28,4163 | -0,00160224 | 0,9905 | 0,995567 |
| Cerk | 6,58589 | 6,5785 | -0,00162038 | 0,99065 | 0,995567 |
| Ttc7b | 10,7608 | 10,7469 | -0,00186898 | 0,99055 | 0,995567 |
| Stk11 | 64,5071 | 63,6866 | -0,0184688 | 0,99075 | 0,995592 |
| Coq6 | 21,5001 | 21,464 | -0,0024198 | 0,991 | 0,995769 |
| Mettl18 | 6,25505 | 6,25533 | 6,52E-005 | 0,99125 | 0,995795 |
| Mettl25 | 5,6004 | 5,59884 | -0,000401564 | 0,9912 | 0,995795 |
| Dnph1 | 36,6885 | 36,6521 | -0,00143323 | 0,99115 | 0,995795 |
| BC022687 | 4,31807 | 4,32399 | 0,00197631 | 0,99165 | 0,996047 |
| Ccne2 | 29,5676 | 29,5145 | -0,00259333 | 0,9916 | 0,996047 |
| Cgnl1 | 28,113 | 28,1456 | 0,0016737 | 0,9919 | 0,996149 |
| Echdc1 | 6,01337 | 5,99529 | -0,00434476 | 0,99185 | 0,996149 |
| Gpr137b-ps | 3,18584 | 3,1796 | -0,00282566 | 0,9923 | 0,996476 |
| Wasf2 | 18,3484 | 18,3268 | -0,00169706 | 0,9924 | 0,996501 |
| Rpl27a | 65,2779 | 65,3392 | 0,00135425 | 0,99285 | 0,996504 |
| Atpif1 | 338,755 | 339,03 | 0,00116772 | 0,99275 | 0,996504 |
| Mtmr9 | 18,3045 | 18,3185 | 0,00110255 | 0,99275 | 0,996504 |
| Ppm1d | 14,6199 | 14,6044 | -0,00152912 | 0,9927 | 0,996504 |
| Cntln | 4,87964 | 4,87445 | -0,00153548 | 0,99285 | 0,996504 |
| Klhdc9 | 6,61693 | 6,60299 | -0,0030439 | 0,99285 | 0,996504 |
| Inadl | 5,49666 | 5,48857 | -0,00212672 | 0,99295 | 0,996529 |
| Oscp1 | 5,63136 | 5,63328 | 0,000491566 | 0,99315 | 0,996655 |
| Fhl3 | 6,11259 | 6,11657 | 0,000939159 | 0,9935 | 0,996708 |
| Tbc1d22b | 11,5898 | 11,5799 | -0,00123244 | 0,99365 | 0,996708 |
| Ccdc23 | 45,8085 | 45,7549 | -0,00169068 | 0,99345 | 0,996708 |
| Lekr1 | 0,76869 | 0,76729 | -0,00263074 | 0,99365 | 0,996708 |
| Mxd3 | 6,84874 | 6,82122 | -0,00580852 | 0,99365 | 0,996708 |
| Igsf6 | 1,2796 | 1,2671 | -0,0141631 | 0,99345 | 0,996708 |
| Clp1 | 25,5691 | 25,5508 | -0,00102819 | 0,99395 | 0,99671 |
| Papd7 | 20,153 | 20,1354 | -0,00125466 | 0,99395 | 0,99671 |
| Ppcdc | 4,9011 | 4,89198 | -0,00268716 | 0,9939 | 0,99671 |
| Lcmt2 | 4,40307 | 4,39474 | -0,0027312 | 0,99375 | 0,99671 |
| Paqr3 | 4,70147 | 4,7156 | 0,00432888 | 0,99455 | 0,996862 |
| Trip12 | 48,7033 | 48,7443 | 0,00121208 | 0,9944 | 0,996862 |
| Bub1b | 82,2652 | 82,3295 | 0,00112728 | 0,99455 | 0,996862 |
| Aff4 | 18,5108 | 18,4982 | -0,000979608 | 0,9943 | 0,996862 |
| Rnf122 | 13,1787 | 13,169 | -0,001068 | 0,99445 | 0,996862 |
| D430020J02Rik | 5,71132 | 5,70292 | -0,00212347 | 0,9945 | 0,996862 |
| Pnpla2 | 4,147 | 4,1419 | -0,00177553 | 0,99465 | 0,996888 |
| Coq7 | 39,0975 | 39,1376 | 0,00147946 | 0,9949 | 0,997064 |
| Adamts4 | 4,15519 | 4,15919 | 0,00138646 | 0,9951 | 0,997189 |
| Serhl | 6,27854 | 6,29164 | 0,00300619 | 0,99525 | 0,99719 |
| Arf2 | 21,2764 | 21,2548 | -0,00146433 | 0,9952 | 0,99719 |
| 1700052N19Rik | 19,2415 | 19,2552 | 0,00102119 | 0,9955 | 0,997291 |
| Rnf181 | 46,3366 | 46,3031 | -0,00104169 | 0,9955 | 0,997291 |
| Coch | 1,13454 | 1,1344 | -0,000176939 | 0,9956 | 0,997292 |
| Commd9 | 15,6998 | 15,6835 | -0,00149908 | 0,99565 | 0,997292 |
| Isca1 | 11,5112 | 11,5075 | -0,000460916 | 0,9958 | 0,997293 |

| | | | | | |
|---|---|---|---|---|---|
| Pak4 | 30,0107 | 29,9914 | -0,0009273 | 0,99575 | 0,997293 |
| Celf1 | 88,0472 | 87,9892 | -0,000951432 | 0,9959 | 0,997318 |
| Idh3a | 43,2582 | 43,2747 | 0,000548856 | 0,9971 | 0,99837 |
| Herpud2 | 12,5286 | 12,5269 | -0,000202635 | 0,9971 | 0,99837 |
| Zhx3 | 2,43663 | 2,43659 | -2,09E-005 | 0,9974 | 0,998596 |
| Terf2ip | 10,4612 | 10,4653 | 0,000564933 | 0,99785 | 0,998896 |
| 4930579G18Rik | 6,21112 | 6,20606 | -0,00117431 | 0,9978 | 0,998896 |
| Fkbp15 | 8,51227 | 8,5128 | 9,15E-005 | 0,9983 | 0,999122 |
| Oxsr1 | 22,8724 | 22,8665 | -0,000373626 | 0,9983 | 0,999122 |
| Prkrir | 70,2021 | 70,1794 | -0,000465339 | 0,99815 | 0,999122 |
| Ptprs | 44,3724 | 44,3654 | -0,000227655 | 0,99855 | 0,999298 |
| Nit1 | 23,3456 | 23,3383 | -0,000451516 | 0,99875 | 0,999423 |
| Pcdhgc3 | 45,4891 | 45,4969 | 0,000246886 | 0,9989 | 0,999498 |
| G3bp2 | 124,913 | 124,932 | 0,000216626 | 0,99905 | 0,999574 |
| Dnaja1 | 15,1826 | 15,1847 | 0,000196218 | 0,9995 | 0,99965 |
| Gak | 23,1435 | 23,1451 | 0,000103159 | 0,9995 | 0,99965 |
| Rangrf | 3,43817 | 3,438 | -7,05E-005 | 0,99935 | 0,99965 |
| Dnlz | 16,9036 | 16,9022 | -0,000118614 | 0,9993 | 0,99965 |
| R3hdm4 | 38,4423 | 38,3771 | -0,00244987 | 0,9995 | 0,99965 |
| Smad3 | 37,8946 | 37,8926 | -7,36E-005 | 0,99965 | 0,999725 |
| Rmi1 | 3,37023 | 3,36955 | -0,000290824 | 0,99985 | 0,99985 |
| 1100001G20Rik | 0 | 0,365654 | inf | 1 | 1 |
| 1700001J03Rik | 0 | 0,223604 | inf | 1 | 1 |
| 1700003D09Rik | 0 | 0,180521 | inf | 1 | 1 |
| 1700009J07Rik | 0 | 0,11372 | inf | 1 | 1 |
| 1700012B07Rik | 0 | 0,0718472 | inf | 1 | 1 |
| 1700016D06Rik | 0 | 0,0403998 | inf | 1 | 1 |
| 1700017G19Rik | 0 | 0,0574278 | inf | 1 | 1 |
| 1700017J07Rik | 0 | 0,360477 | inf | 1 | 1 |
| 1700047A11Rik | 0 | 0,0646957 | inf | 1 | 1 |
| 1700073E17Rik | 0 | 0,0389386 | inf | 1 | 1 |
| 1700094J05Rik | 0 | 0,0652305 | inf | 1 | 1 |
| 1700095B10Rik | 0 | 8,34E-006 | inf | 1 | 1 |
| 1700100L14Rik | 0 | 0,0563044 | inf | 1 | 1 |
| 1700120E14Rik | 0 | 0,0155753 | inf | 1 | 1 |
| 1700123L14Rik | 0 | 0,0285414 | inf | 1 | 1 |
| 2310001K24Rik | 0 | 0,211082 | inf | 1 | 1 |
| 2610028H24Rik | 0 | 0,0705687 | inf | 1 | 1 |
| 4921511C10Rik | 0 | 0,0339821 | inf | 1 | 1 |
| 4921517D22Rik | 0 | 0,00916069 | inf | 1 | 1 |
| 4921530L21Rik | 0 | 0,0449405 | inf | 1 | 1 |
| 4930429D17Rik | 0 | 0,0301292 | inf | 1 | 1 |
| 4930430A15Rik | 0 | 0,0523667 | inf | 1 | 1 |
| 4930455D15Rik | 0 | 0,0303 | inf | 1 | 1 |
| 4930511A02Rik | 0 | 0,0310965 | inf | 1 | 1 |
| 4930511M06Rik | 0 | 0,0746624 | inf | 1 | 1 |
| 4930515L03Rik | 0 | 0,152249 | inf | 1 | 1 |
| 4930517E11Rik | 0 | 0,0341483 | inf | 1 | 1 |

| | | | | | |
|---|---|---|---|---|---|
| 4930524O05Rik | 0 | 0,0785842 | inf | 1 | 1 |
| 4930524O08Rik | 0 | 0,021583 | inf | 1 | 1 |
| 4930542C21Rik | 0 | 0,107655 | inf | 1 | 1 |
| 4930545E07Rik | 0 | 0,0650199 | inf | 1 | 1 |
| 4930547E14Rik | 0 | 0,124007 | inf | 1 | 1 |
| 4930564C03Rik | 0 | 0,0437546 | inf | 1 | 1 |
| 4930571K23Rik | 0 | 0,0219516 | inf | 1 | 1 |
| 4930590L20Rik | 0 | 0,0558278 | inf | 1 | 1 |
| 4930596I21Rik | 0 | 0,315385 | inf | 1 | 1 |
| 4931420L22Rik | 0 | 0,0750578 | inf | 1 | 1 |
| 4931428L18Rik | 0 | 0,0961422 | inf | 1 | 1 |
| 4931430N09Rik | 0 | 0,0274353 | inf | 1 | 1 |
| 4933406G16Rik | 0 | 0,0352999 | inf | 1 | 1 |
| 4933424G06Rik | 0 | 0,13383 | inf | 1 | 1 |
| 4933425B07Rik | 0 | 0,0404748 | inf | 1 | 1 |
| 4933427D06Rik | 0 | 0,0222707 | inf | 1 | 1 |
| 4933427E13Rik | 0 | 0,0331912 | inf | 1 | 1 |
| 5031426D15Rik | 0 | 0,0414019 | inf | 1 | 1 |
| 5730403I07Rik | 0 | 0,0147313 | inf | 1 | 1 |
| 6430584L05Rik | 0 | 0,0236504 | inf | 1 | 1 |
| 9430019J16Rik | 0 | 0,0590113 | inf | 1 | 1 |
| 9530059O14Rik | 0 | 0,0327308 | inf | 1 | 1 |
| 9830166K06Rik | 0 | 0,0211839 | inf | 1 | 1 |
| A330050F15Rik | 0 | 0,0204511 | inf | 1 | 1 |
| A330093E20Rik | 0 | 0,11932 | inf | 1 | 1 |
| A530006G24Rik | 0 | 0,0716431 | inf | 1 | 1 |
| A530088E08Rik | 0 | 0,0169164 | inf | 1 | 1 |
| A630076J17Rik | 0 | 0,174763 | inf | 1 | 1 |
| AA387883 | 0 | 0,0290058 | inf | 1 | 1 |
| Abca14 | 0 | 0,0247783 | inf | 1 | 1 |
| Acsm2 | 0 | 0,00675765 | inf | 1 | 1 |
| Actl11 | 0 | 0,0225831 | inf | 1 | 1 |
| Adam26a | 0 | 0,0187073 | inf | 1 | 1 |
| Akp3 | 0 | 0,0579723 | inf | 1 | 1 |
| Akr1b7 | 0 | 0,144099 | inf | 1 | 1 |
| Akr1d1 | 0 | 0,0342956 | inf | 1 | 1 |
| Ano5 | 0 | 0,017269 | inf | 1 | 1 |
| Aqp12 | 0 | 0,0969604 | inf | 1 | 1 |
| Arid3c | 0 | 0,0956667 | inf | 1 | 1 |
| Arl13a | 0 | 0,0174994 | inf | 1 | 1 |
| Ascl5 | 0 | 0,0741413 | inf | 1 | 1 |
| Aspa | 0 | 0,0201905 | inf | 1 | 1 |
| Atp6v1b1 | 0 | 0,109343 | inf | 1 | 1 |
| Atp6v1e2 | 0 | 0,0534123 | inf | 1 | 1 |
| BC027072 | 0 | 0,00890958 | inf | 1 | 1 |
| Banf2 | 0 | 0,208648 | inf | 1 | 1 |
| Bcl2l10 | 0 | 0,125342 | inf | 1 | 1 |
| Bdkrb1 | 0 | 0,0377279 | inf | 1 | 1 |

| | | | | | |
|---|---|---|---|---|---|
| Bhmt | 0 | 0,0229157 | inf | 1 | 1 |
| Bmp15 | 0 | 0,0731352 | inf | 1 | 1 |
| Bsx | 0 | 0,0401099 | inf | 1 | 1 |
| Btg4 | 0 | 0,0729375 | inf | 1 | 1 |
| C030029H02Rik | 0 | 0,0129706 | inf | 1 | 1 |
| C86695 | 0 | 0,0159917 | inf | 1 | 1 |
| Capn9 | 0 | 0,040248 | inf | 1 | 1 |
| Cartpt | 0 | 0,0676879 | inf | 1 | 1 |
| Catsper3 | 0 | 0,0270469 | inf | 1 | 1 |
| Ccdc81 | 0 | 0,0344059 | inf | 1 | 1 |
| Cdh18 | 0 | 0,0105961 | inf | 1 | 1 |
| Cdh8 | 0 | 0,00907846 | inf | 1 | 1 |
| Cdrt4 | 0 | 0,207426 | inf | 1 | 1 |
| Ceacam12 | 0 | 0,0511326 | inf | 1 | 1 |
| Ceacam19 | 0 | 0,013552 | inf | 1 | 1 |
| Chrne | 0 | 0,0194045 | inf | 1 | 1 |
| Cistr-act | 0 | 0,364557 | inf | 1 | 1 |
| Cldnd2 | 0 | 0,103841 | inf | 1 | 1 |
| Clec2h | 0 | 0,021873 | inf | 1 | 1 |
| Cma1 | 0 | 0,0490634 | inf | 1 | 1 |
| Cmtm1 | 0 | 0,0462023 | inf | 1 | 1 |
| Cnfn | 0 | 0,0825542 | inf | 1 | 1 |
| Cnga3 | 0 | 0,0127662 | inf | 1 | 1 |
| Cntn4 | 0 | 0,0515074 | inf | 1 | 1 |
| Cpne6 | 0 | 0,107343 | inf | 1 | 1 |
| Cyp1a2 | 0 | 0,0253457 | inf | 1 | 1 |
| Cyp2b9 | 0 | 0,0160873 | inf | 1 | 1 |
| Cyp2j11 | 0 | 0,0247304 | inf | 1 | 1 |
| Cyp4f37 | 0 | 0,0175487 | inf | 1 | 1 |
| D14Ertd670e | 0 | 0,118511 | inf | 1 | 1 |
| D530049I02Rik | 0 | 0,123292 | inf | 1 | 1 |
| D630029K05Rik | 0 | 0,0278733 | inf | 1 | 1 |
| Defb37 | 0 | 0,180942 | inf | 1 | 1 |
| Defb40 | 0 | 0,926935 | inf | 1 | 1 |
| Defb47 | 0 | 0,825029 | inf | 1 | 1 |
| Dmrt1 | 0 | 0,0209007 | inf | 1 | 1 |
| Dmrtc1a | 0 | 0,0497891 | inf | 1 | 1 |
| Dnmt3aos | 0 | 0,0724529 | inf | 1 | 1 |
| E130006D01Rik | 0 | 0,151508 | inf | 1 | 1 |
| E430016F16Rik | 0 | 0,0296389 | inf | 1 | 1 |
| Ear14 | 0 | 0,162817 | inf | 1 | 1 |
| Ear4 | 0 | 0,00294717 | inf | 1 | 1 |
| Egr4 | 0 | 0,349054 | inf | 1 | 1 |
| Eif4e1b | 0 | 0,0278673 | inf | 1 | 1 |
| Enpp7 | 0 | 0,0366925 | inf | 1 | 1 |
| Ereg | 0 | 0,0217618 | inf | 1 | 1 |
| Fam135b | 0 | 0,0193636 | inf | 1 | 1 |
| Fam150b | 0 | 0,0847602 | inf | 1 | 1 |

| | | | | | |
|---|---|---|---|---|---|
| Fam166a | 0 | 0,17058 | inf | 1 | 1 |
| Fam71d | 0 | 0,0277734 | inf | 1 | 1 |
| Fbxw13 | 0 | 0,0333985 | inf | 1 | 1 |
| Fbxw19 | 0 | 0,0340091 | inf | 1 | 1 |
| Fetub | 0 | 0,0236904 | inf | 1 | 1 |
| Fmo6 | 0 | 0,0256804 | inf | 1 | 1 |
| Foxd4 | 0 | 0,0125708 | inf | 1 | 1 |
| Frem3 | 0 | 0,0133241 | inf | 1 | 1 |
| Fsd2 | 0 | 0,0194185 | inf | 1 | 1 |
| Fstl5 | 0 | 0,0197241 | inf | 1 | 1 |
| Fut7 | 0 | 0,0275509 | inf | 1 | 1 |
| Fxyd7 | 0 | 0,0567795 | inf | 1 | 1 |
| G630071F17Rik | 0 | 0,0186968 | inf | 1 | 1 |
| Gdpd4 | 0 | 0,0189136 | inf | 1 | 1 |
| Gipc3 | 0 | 0,104349 | inf | 1 | 1 |
| Gjc3 | 0 | 0,0129217 | inf | 1 | 1 |
| Glra1 | 0 | 0,0193679 | inf | 1 | 1 |
| Gm10373 | 0 | 0,0580032 | inf | 1 | 1 |
| Gm10400 | 0 | 0,00741232 | inf | 1 | 1 |
| Gm10440 | 0 | 0,0167977 | inf | 1 | 1 |
| Gm10510 | 0 | 0,164867 | inf | 1 | 1 |
| Gm10532 | 0 | 0,149308 | inf | 1 | 1 |
| Gm10578 | 0 | 0,0320237 | inf | 1 | 1 |
| Gm10684 | 0 | 0,0134592 | inf | 1 | 1 |
| Gm10872 | 0 | 0,0557799 | inf | 1 | 1 |
| Gm11186 | 0 | 0,142969 | inf | 1 | 1 |
| Gm11468 | 0 | 0,23874 | inf | 1 | 1 |
| Gm11529 | 0 | 0,0144777 | inf | 1 | 1 |
| Gm11570 | 0 | 0,0159538 | inf | 1 | 1 |
| Gm11762 | 0 | 0,0285034 | inf | 1 | 1 |
| Gm12633 | 0 | 0,0287911 | inf | 1 | 1 |
| Gm13277,Gm132 | 0 | 0,0340091 | inf | 1 | 1 |
| Gm13286 | 0 | 0,124007 | inf | 1 | 1 |
| Gm13288 | 0 | 0,0161692 | inf | 1 | 1 |
| Gm13944 | 0 | 0,147607 | inf | 1 | 1 |
| Gm14092 | 0 | 0,0501503 | inf | 1 | 1 |
| Gm14496 | 0 | 0,0182369 | inf | 1 | 1 |
| Gm15348 | 0 | 0,0123975 | inf | 1 | 1 |
| Gm15713 | 0 | 0,0318804 | inf | 1 | 1 |
| Gm16130 | 0 | 0,0103715 | inf | 1 | 1 |
| Gm1631 | 0 | 0,0979008 | inf | 1 | 1 |
| Gm16432 | 0 | 0,0184469 | inf | 1 | 1 |
| Gm20319 | 0 | 0,288054 | inf | 1 | 1 |
| Gm20594 | 0 | 0,614515 | inf | 1 | 1 |
| Gm20597 | 0 | 0,00809993 | inf | 1 | 1 |
| Gm21319 | 0 | 0,016223 | inf | 1 | 1 |
| Gm2176 | 0 | 0,0279631 | inf | 1 | 1 |
| Gm2721 | 0 | 0,0196309 | inf | 1 | 1 |

| | | | | | |
|---|---|---|---|---|---|
| Gm3448 | 0 | 0,046164 | inf | 1 | 1 |
| Gm4265 | 0 | 0,182054 | inf | 1 | 1 |
| Gm4371 | 0 | 0,0191422 | inf | 1 | 1 |
| Gm4559 | 0 | 0,108037 | inf | 1 | 1 |
| Gm5142 | 0 | 0,0637759 | inf | 1 | 1 |
| Gm5475 | 0 | 0,0202191 | inf | 1 | 1 |
| Gm5591 | 0 | 0,0396513 | inf | 1 | 1 |
| Gm5800 | 0 | 0,198376 | inf | 1 | 1 |
| Gm6086 | 0 | 0,0110983 | inf | 1 | 1 |
| Gm6289 | 0 | 0,0847185 | inf | 1 | 1 |
| Gm7616 | 0 | 0,416375 | inf | 1 | 1 |
| Gpr139 | 0 | 0,0512989 | inf | 1 | 1 |
| Gpr17 | 0 | 0,00857563 | inf | 1 | 1 |
| Gpr33 | 0 | 0,0361652 | inf | 1 | 1 |
| Gpr88 | 0 | 0,00829148 | inf | 1 | 1 |
| Grm2 | 0 | 0,0137171 | inf | 1 | 1 |
| Grm4 | 0 | 0,0113095 | inf | 1 | 1 |
| Grm5 | 0 | 0,0117416 | inf | 1 | 1 |
| Gstt4 | 0 | 0,0354595 | inf | 1 | 1 |
| Gykl1 | 0 | 0,01573 | inf | 1 | 1 |
| H2-L | 0 | 0,107318 | inf | 1 | 1 |
| H60c | 0 | 0,0327468 | inf | 1 | 1 |
| Hapln2 | 0 | 0,0231138 | inf | 1 | 1 |
| Hcar2 | 0 | 0,0154412 | inf | 1 | 1 |
| Helt | 0 | 0,0202665 | inf | 1 | 1 |
| Hist1h2ba | 0 | 0,118037 | inf | 1 | 1 |
| Hmx2 | 0 | 0,0367427 | inf | 1 | 1 |
| Hrh4 | 0 | 0,0320957 | inf | 1 | 1 |
| Htr1f | 0 | 0,0116764 | inf | 1 | 1 |
| Htr6 | 0 | 0,0834295 | inf | 1 | 1 |
| Ido1 | 0 | 0,103179 | inf | 1 | 1 |
| Il10 | 0 | 0,0244228 | inf | 1 | 1 |
| Il22 | 0 | 0,0484522 | inf | 1 | 1 |
| Il7 | 0 | 0,0376783 | inf | 1 | 1 |
| Iqca | 0 | 0,0426983 | inf | 1 | 1 |
| Irg1 | 0 | 0,0179462 | inf | 1 | 1 |
| Kcnd3os | 0 | 0,0303768 | inf | 1 | 1 |
| Kcne4 | 0 | 0,0378784 | inf | 1 | 1 |
| Kif2b | 0 | 0,0207084 | inf | 1 | 1 |
| Klhl30 | 0 | 0,0303862 | inf | 1 | 1 |
| Krt4 | 0 | 0,0139523 | inf | 1 | 1 |
| Krt71 | 0 | 0,0434352 | inf | 1 | 1 |
| Krt81 | 0 | 0,0258825 | inf | 1 | 1 |
| Krtap3-1 | 0 | 0,20185 | inf | 1 | 1 |
| Krtap3-3 | 0 | 0,324113 | inf | 1 | 1 |
| Krtap9-3 | 0 | 0,0770864 | inf | 1 | 1 |
| L3mbtl4 | 0 | 0,00979931 | inf | 1 | 1 |
| LOC101056236 | 0 | 0,0601427 | inf | 1 | 1 |

| | | | | | |
|---|---|---|---|---|---|
| LOC102308570 | 0 | 0,0559559 | inf | 1 | 1 |
| LOC102632423 | 0 | 0,125764 | inf | 1 | 1 |
| LOC102634431 | 0 | 0,0432269 | inf | 1 | 1 |
| LOC102635087 | 0 | 0,0144466 | inf | 1 | 1 |
| Lactbl1 | 0 | 0,0307589 | inf | 1 | 1 |
| Lce1b | 0 | 0,0965749 | inf | 1 | 1 |
| Lce1i | 0 | 0,0811819 | inf | 1 | 1 |
| Lhb | 0 | 0,176479 | inf | 1 | 1 |
| Lkaaear1 | 0 | 0,105131 | inf | 1 | 1 |
| Lrrc74 | 0 | 0,0245178 | inf | 1 | 1 |
| Magea5 | 0 | 0,0629632 | inf | 1 | 1 |
| Mecom | 0 | 0,0308999 | inf | 1 | 1 |
| Mepe | 0 | 0,0578938 | inf | 1 | 1 |
| Mettl7a2 | 0 | 2,94E-006 | inf | 1 | 1 |
| Miox | 0 | 0,0319262 | inf | 1 | 1 |
| Mir1903 | 0 | 25,1811 | inf | 1 | 1 |
| Mir1932 | 0 | 24,1238 | inf | 1 | 1 |
| Mir2139 | 0 | 27,6885 | inf | 1 | 1 |
| Mir3072 | 0 | 57,5379 | inf | 1 | 1 |
| Mir6361 | 0 | 22,444 | inf | 1 | 1 |
| Mir7075 | 0 | 61,3064 | inf | 1 | 1 |
| Moxd2 | 0 | 0,0258355 | inf | 1 | 1 |
| Mrgprb5 | 0 | 0,0559258 | inf | 1 | 1 |
| Ms4a13 | 0 | 0,0365358 | inf | 1 | 1 |
| Muc5ac | 0 | 0,00515406 | inf | 1 | 1 |
| Mug2 | 0 | 0,00985802 | inf | 1 | 1 |
| Mycs | 0 | 0,0249435 | inf | 1 | 1 |
| Myh2 | 0 | 0,00915604 | inf | 1 | 1 |
| Myh4 | 0 | 0,016563 | inf | 1 | 1 |
| Mylk2 | 0 | 0,0254001 | inf | 1 | 1 |
| Nek10 | 0 | 0,0136081 | inf | 1 | 1 |
| Nepn | 0 | 0,0166239 | inf | 1 | 1 |
| Neurod2 | 0 | 0,014587 | inf | 1 | 1 |
| Ninj2 | 0 | 0,0435496 | inf | 1 | 1 |
| Nkx1-1 | 0 | 0,0382336 | inf | 1 | 1 |
| Nkx2-4 | 0 | 0,0312588 | inf | 1 | 1 |
| Nr2e1 | 0 | 0,0141247 | inf | 1 | 1 |
| Nt5c1a | 0 | 0,0475899 | inf | 1 | 1 |
| Nxf3 | 0 | 0,00970227 | inf | 1 | 1 |
| Oas1h | 0 | 0,0359573 | inf | 1 | 1 |
| Odf3l1 | 0 | 0,0488394 | inf | 1 | 1 |
| Olfr1168 | 0 | 0,0582084 | inf | 1 | 1 |
| Olfr1188 | 0 | 0,055999 | inf | 1 | 1 |
| Olfr1198 | 0 | 0,118349 | inf | 1 | 1 |
| Olfr1220 | 0 | 0,0520477 | inf | 1 | 1 |
| Olfr1249 | 0 | 0,056817 | inf | 1 | 1 |
| Olfr1270 | 0 | 0,0525593 | inf | 1 | 1 |
| Olfr1347 | 0 | 0,0565915 | inf | 1 | 1 |

| | | | | | |
|---|---|---|---|---|---|
| Olfr1366 | 0 | 0,0316633 | inf | 1 | 1 |
| Olfr1392 | 0 | 0,0520477 | inf | 1 | 1 |
| Olfr1396 | 0 | 0,055062 | inf | 1 | 1 |
| Olfr1414 | 0 | 0,0868698 | inf | 1 | 1 |
| Olfr1417 | 0 | 0,115009 | inf | 1 | 1 |
| Olfr1474 | 0 | 0,0577371 | inf | 1 | 1 |
| Olfr218 | 0 | 0,0579718 | inf | 1 | 1 |
| Olfr267 | 0 | 0,0579718 | inf | 1 | 1 |
| Olfr370 | 0 | 0,0479633 | inf | 1 | 1 |
| Olfr38 | 0 | 0,0570442 | inf | 1 | 1 |
| Olfr412 | 0 | 0,0582084 | inf | 1 | 1 |
| Olfr444 | 0 | 0,0586875 | inf | 1 | 1 |
| Olfr457 | 0 | 0,0579718 | inf | 1 | 1 |
| Olfr469 | 0 | 0,0577371 | inf | 1 | 1 |
| Olfr521 | 0 | 0,0561459 | inf | 1 | 1 |
| Olfr544 | 0 | 0,0437905 | inf | 1 | 1 |
| Olfr61 | 0 | 0,0584469 | inf | 1 | 1 |
| Olfr655 | 0 | 0,0744502 | inf | 1 | 1 |
| Olfr698 | 0 | 0,0584469 | inf | 1 | 1 |
| Olfr750 | 0 | 0,0461027 | inf | 1 | 1 |
| Olfr767 | 0 | 0,05893 | inf | 1 | 1 |
| Olfr95 | 0 | 0,174625 | inf | 1 | 1 |
| Olfr960 | 0 | 0,144117 | inf | 1 | 1 |
| Olfr995 | 0 | 0,0575043 | inf | 1 | 1 |
| Onecut1 | 0 | 0,0959016 | inf | 1 | 1 |
| Ovch2 | 0 | 0,0248742 | inf | 1 | 1 |
| Oxt | 0 | 0,0800988 | inf | 1 | 1 |
| Paupar | 0 | 0,0261036 | inf | 1 | 1 |
| Pcdh15 | 0 | 0,0030195 | inf | 1 | 1 |
| Pcdha11 | 0 | 0,0206716 | inf | 1 | 1 |
| Pcdha5 | 0 | 0,0246073 | inf | 1 | 1 |
| Pck1 | 0 | 0,0111584 | inf | 1 | 1 |
| Pfn3 | 0 | 0,0789244 | inf | 1 | 1 |
| Pifo | 0 | 0,060766 | inf | 1 | 1 |
| Pla2g1b | 0 | 0,245875 | inf | 1 | 1 |
| Pou4f2 | 0 | 0,0142233 | inf | 1 | 1 |
| Pramel1 | 0 | 0,038268 | inf | 1 | 1 |
| Prr23a | 0 | 0,0314782 | inf | 1 | 1 |
| Prss38 | 0 | 0,0413366 | inf | 1 | 1 |
| Prss42 | 0 | 0,031668 | inf | 1 | 1 |
| Psg25 | 0 | 0,0298873 | inf | 1 | 1 |
| Ptx4 | 0 | 0,0626848 | inf | 1 | 1 |
| Raet1c,Raet1e | 0 | 0,51807 | inf | 1 | 1 |
| Rbm46os | 0 | 0,0140161 | inf | 1 | 1 |
| Rbp3 | 0 | 0,00843529 | inf | 1 | 1 |
| Rdh7 | 0 | 0,0307653 | inf | 1 | 1 |
| Ren1 | 0 | 0,0350925 | inf | 1 | 1 |
| Ren2 | 0 | 3,09E-005 | inf | 1 | 1 |

| | | | | | |
|---|---|---|---|---|---|
| Rfpl4 | 0 | 0,0580904 | inf | 1 | 1 |
| Rnu12 | 0 | 2,77208 | inf | 1 | 1 |
| Rptn | 0 | 0,0108533 | inf | 1 | 1 |
| Rs1 | 0 | 0,0295085 | inf | 1 | 1 |
| Scg2 | 0 | 0,0117992 | inf | 1 | 1 |
| Scgb3a1 | 0 | 0,140728 | inf | 1 | 1 |
| Sept14 | 0 | 0,0181239 | inf | 1 | 1 |
| Serpina3b | 0 | 0,0144234 | inf | 1 | 1 |
| Serpinb6d | 0 | 0,067749 | inf | 1 | 1 |
| Sez6 | 0 | 0,0108399 | inf | 1 | 1 |
| Shox2 | 0 | 0,0615148 | inf | 1 | 1 |
| Siglec15 | 0 | 0,0326228 | inf | 1 | 1 |
| Six6 | 0 | 0,0143466 | inf | 1 | 1 |
| Skor2 | 0 | 0,0145672 | inf | 1 | 1 |
| Slc17a2 | 0 | 0,0308908 | inf | 1 | 1 |
| Slc22a12 | 0 | 0,0262704 | inf | 1 | 1 |
| Slc34a1 | 0 | 0,0446129 | inf | 1 | 1 |
| Slc6a19os | 0 | 0,0321439 | inf | 1 | 1 |
| Slco6b1 | 0 | 0,0188391 | inf | 1 | 1 |
| Slitrk3 | 0 | 0,00762872 | inf | 1 | 1 |
| Snora26 | 0 | 35,0204 | inf | 1 | 1 |
| Sox5os3 | 0 | 0,0384482 | inf | 1 | 1 |
| Spata16 | 0 | 0,0266386 | inf | 1 | 1 |
| St8sia5 | 0 | 0,0150249 | inf | 1 | 1 |
| Strc | 0 | 0,00770463 | inf | 1 | 1 |
| Taar4 | 0 | 0,0640805 | inf | 1 | 1 |
| Taar7d | 0 | 0,0308291 | inf | 1 | 1 |
| Tas1r2 | 0 | 0,0189403 | inf | 1 | 1 |
| Tas2r143 | 0 | 0,182189 | inf | 1 | 1 |
| Tbata | 0 | 0,0378497 | inf | 1 | 1 |
| Tcte1 | 0 | 0,0130636 | inf | 1 | 1 |
| Tekt3 | 0 | 0,027872 | inf | 1 | 1 |
| Tepp | 0 | 0,0938207 | inf | 1 | 1 |
| Tescl | 0 | 0,550437 | inf | 1 | 1 |
| Tff2 | 0 | 0,0733332 | inf | 1 | 1 |
| Them5 | 0 | 0,0514115 | inf | 1 | 1 |
| Tktl2 | 0 | 0,00694092 | inf | 1 | 1 |
| Tnip3 | 0 | 0,00754955 | inf | 1 | 1 |
| Tppp2 | 0 | 0,302707 | inf | 1 | 1 |
| Tpsg1 | 0 | 0,053137 | inf | 1 | 1 |
| Treh | 0 | 0,0154585 | inf | 1 | 1 |
| Trpv3 | 0 | 0,0574021 | inf | 1 | 1 |
| Tuba3a | 0 | 0,0412475 | inf | 1 | 1 |
| Ucma | 0 | 0,0736254 | inf | 1 | 1 |
| Ugt2b5 | 0 | 0,0253307 | inf | 1 | 1 |
| Usp9y | 0 | 0,00682236 | inf | 1 | 1 |
| Uts2r | 0 | 0,0285414 | inf | 1 | 1 |
| Vmn1r197 | 0 | 0,123472 | inf | 1 | 1 |

| | | | | | |
|---|---|---|---|---|---|
| Vmn1r221 | 0 | 0,0582084 | inf | 1 | 1 |
| Vmn1r26 | 0 | 0,104861 | inf | 1 | 1 |
| Vmn1r49 | 0 | 0,0586875 | inf | 1 | 1 |
| Vmn1r50 | 0 | 0,0369187 | inf | 1 | 1 |
| Vmn2r109 | 0 | 0,0180294 | inf | 1 | 1 |
| Vmn2r120 | 0 | 0,0361501 | inf | 1 | 1 |
| Vmn2r23 | 0 | 0,0229447 | inf | 1 | 1 |
| Vmn2r28 | 0 | 0,016844 | inf | 1 | 1 |
| Vmn2r3 | 0 | 0,0173494 | inf | 1 | 1 |
| Wfdc8 | 0 | 0,0182443 | inf | 1 | 1 |
| Wt1os | 0 | 0,0139679 | inf | 1 | 1 |
| Xkr7 | 0 | 0,0170657 | inf | 1 | 1 |
| Yipf7 | 0 | 0,0421511 | inf | 1 | 1 |
| Zfp831 | 0 | 0,00445619 | inf | 1 | 1 |
| Zp3 | 0 | 0,115319 | inf | 1 | 1 |
| 1010001N08Rik | 0,088815 | 0 | -inf | 1 | 1 |
| 1110017D15Rik | 0,125825 | 0 | -inf | 1 | 1 |
| 1110028F18Rik | 0,036482 | 0 | -inf | 1 | 1 |
| 1110036E04Rik | 0,020642 | 0 | -inf | 1 | 1 |
| 1600002D24Rik | 0,141905 | 0 | -inf | 1 | 1 |
| 1600014K23Rik | 0,092987 | 0 | -inf | 1 | 1 |
| 1600015I10Rik | 0,077888 | 0 | -inf | 1 | 1 |
| 1600025M17Rik | 0,155412 | 0 | -inf | 1 | 1 |
| 1600029O15Rik | 0,022109 | 0 | -inf | 1 | 1 |
| 1700001D01Rik | 0,065528 | 0 | -inf | 1 | 1 |
| 1700007F19Rik | 0,090592 | 0 | -inf | 1 | 1 |
| 1700007K09Rik | 0,050118 | 0 | -inf | 1 | 1 |
| 1700007P06Rik | 0,06701 | 0 | -inf | 1 | 1 |
| 1700008F21Rik | 0,049146 | 0 | -inf | 1 | 1 |
| 1700009C05Rik | 0,020288 | 0 | -inf | 1 | 1 |
| 1700010B08Rik | 0,034185 | 0 | -inf | 1 | 1 |
| 1700011A15Rik | 0,016922 | 0 | -inf | 1 | 1 |
| 1700012L04Rik | 0,036846 | 0 | -inf | 1 | 1 |
| 1700013F07Rik | 0,030484 | 0 | -inf | 1 | 1 |
| 1700015F17Rik | 0,023375 | 0 | -inf | 1 | 1 |
| 1700016G22Rik | 0,030406 | 0 | -inf | 1 | 1 |
| 1700018A04Rik | 0,127715 | 0 | -inf | 1 | 1 |
| 1700018B08Rik | 0,040507 | 0 | -inf | 1 | 1 |
| 1700018C11Rik | 0,025327 | 0 | -inf | 1 | 1 |
| 1700019A02Rik | 0,111847 | 0 | -inf | 1 | 1 |
| 1700019D03Rik | 0,084615 | 0 | -inf | 1 | 1 |
| 1700019G24Rik | 0,133375 | 0 | -inf | 1 | 1 |
| 1700024G13Rik | 0,060939 | 0 | -inf | 1 | 1 |
| 1700024P16Rik | 0,107447 | 0 | -inf | 1 | 1 |
| 1700025N23Rik | 0,033891 | 0 | -inf | 1 | 1 |
| 1700026F02Rik | 0,042545 | 0 | -inf | 1 | 1 |
| 1700027H10Rik | 0,149627 | 0 | -inf | 1 | 1 |
| 1700027I24Rik | 0,02404 | 0 | -inf | 1 | 1 |

| | | | | | |
|---|---|---|---|---|---|
| 1700028P14Rik | 0,017813 | 0 | -inf | 1 | 1 |
| 1700029H14Rik | 0,036415 | 0 | -inf | 1 | 1 |
| 1700029M20Rik | 0,004495 | 0 | -inf | 1 | 1 |
| 1700029N11Rik | 0,032764 | 0 | -inf | 1 | 1 |
| 1700030A11Rik | 0,060642 | 0 | -inf | 1 | 1 |
| 1700030M09Rik | 0,054578 | 0 | -inf | 1 | 1 |
| 1700030O20Rik | 0,030328 | 0 | -inf | 1 | 1 |
| 1700031A10Rik | 0,091768 | 0 | -inf | 1 | 1 |
| 1700034J05Rik | 0,032351 | 0 | -inf | 1 | 1 |
| 1700049L16Rik | 0,137805 | 0 | -inf | 1 | 1 |
| 1700051A21Rik | 0,046194 | 0 | -inf | 1 | 1 |
| 1700060C20Rik | 0,030642 | 0 | -inf | 1 | 1 |
| 1700063D05Rik | 0,082137 | 0 | -inf | 1 | 1 |
| 1700064M15Rik | 0,080461 | 0 | -inf | 1 | 1 |
| 1700065J18Rik | 0,05912 | 0 | -inf | 1 | 1 |
| 1700080O16Rik | 0,009462 | 0 | -inf | 1 | 1 |
| 1700093K21Rik | 0,029643 | 0 | -inf | 1 | 1 |
| 1700110K17Rik | 0,119052 | 0 | -inf | 1 | 1 |
| 1700125H03Rik | 0,042393 | 0 | -inf | 1 | 1 |
| 1810011H11Rik | 0,256642 | 0 | -inf | 1 | 1 |
| 1810012K16Rik | 0,806293 | 0 | -inf | 1 | 1 |
| 1810046K07Rik | 0,047853 | 0 | -inf | 1 | 1 |
| 1810053B23Rik | 0,070987 | 0 | -inf | 1 | 1 |
| 2010002M12Rik | 0,013122 | 0 | -inf | 1 | 1 |
| 2010005H15Rik | 0,036505 | 0 | -inf | 1 | 1 |
| 2010016I18Rik | 0,008736 | 0 | -inf | 1 | 1 |
| 2200002J24Rik | 0,029133 | 0 | -inf | 1 | 1 |
| 2210019I11Rik | 0,047771 | 0 | -inf | 1 | 1 |
| 2210417A02Rik | 0,068368 | 0 | -inf | 1 | 1 |
| 2310002D06Rik | 0,077814 | 0 | -inf | 1 | 1 |
| 2310002J15Rik | 0,020499 | 0 | -inf | 1 | 1 |
| 2310005E17Rik | 0,02409 | 0 | -inf | 1 | 1 |
| 2310007B03Rik | 0,308629 | 0 | -inf | 1 | 1 |
| 2310007L24Rik | 0,202465 | 0 | -inf | 1 | 1 |
| 2310008N11Rik | 0,148073 | 0 | -inf | 1 | 1 |
| 2310015B20Rik | 0,098975 | 0 | -inf | 1 | 1 |
| 2310061N02Rik | 0,020116 | 0 | -inf | 1 | 1 |
| 2310065F04Rik | 0,01646 | 0 | -inf | 1 | 1 |
| 2410003L11Rik | 0,551204 | 0 | -inf | 1 | 1 |
| 2410012E07Rik | 0,035976 | 0 | -inf | 1 | 1 |
| 2410017I17Rik | 0,00905 | 0 | -inf | 1 | 1 |
| 2410088K16Rik | 0,435438 | 0 | -inf | 1 | 1 |
| 2410137M14Rik | 0,017894 | 0 | -inf | 1 | 1 |
| 2700089I24Rik | 0,007288 | 0 | -inf | 1 | 1 |
| 2810047C21Rik1 | 0,008214 | 0 | -inf | 1 | 1 |
| 2810049E08Rik | 0,013172 | 0 | -inf | 1 | 1 |
| 2810442N19Rik | 0,389347 | 0 | -inf | 1 | 1 |
| 2900055J20Rik | 0,102121 | 0 | -inf | 1 | 1 |

| | | | | | |
|---|---|---|---|---|---|
| 2900057B20Rik | 0,014567 | 0 | -inf | 1 | 1 |
| 3110079O15Rik | 0,550752 | 0 | -inf | 1 | 1 |
| 3300005D01Rik | 0,120904 | 0 | -inf | 1 | 1 |
| 4430402I18Rik | 0,062122 | 0 | -inf | 1 | 1 |
| 4833424O15Rik | 0,010802 | 0 | -inf | 1 | 1 |
| 4833427F10Rik | 0,009306 | 0 | -inf | 1 | 1 |
| 4921508D12Rik | 0,01358 | 0 | -inf | 1 | 1 |
| 4921529L05Rik | 0,009002 | 0 | -inf | 1 | 1 |
| 4921534H16Rik | 0,008981 | 0 | -inf | 1 | 1 |
| 4921539E11Rik | 0,005493 | 0 | -inf | 1 | 1 |
| 4930401O10Rik | 0,071148 | 0 | -inf | 1 | 1 |
| 4930405D11Rik | 0,053159 | 0 | -inf | 1 | 1 |
| 4930429F24Rik | 0,13051 | 0 | -inf | 1 | 1 |
| 4930430J02Rik | 0,169992 | 0 | -inf | 1 | 1 |
| 4930442L01Rik | 0,012808 | 0 | -inf | 1 | 1 |
| 4930444F02Rik | 0,07024 | 0 | -inf | 1 | 1 |
| 4930447J18Rik | 0,019047 | 0 | -inf | 1 | 1 |
| 4930447K03Rik | 0,028536 | 0 | -inf | 1 | 1 |
| 4930452G13Rik | 0,027005 | 0 | -inf | 1 | 1 |
| 4930453H23Rik | 0,027379 | 0 | -inf | 1 | 1 |
| 4930456L15Rik | 0,042283 | 0 | -inf | 1 | 1 |
| 4930467K11Rik | 0,091159 | 0 | -inf | 1 | 1 |
| 4930471C04Rik | 0,017894 | 0 | -inf | 1 | 1 |
| 4930474N09Rik | 0,085349 | 0 | -inf | 1 | 1 |
| 4930483J18Rik | 0,05316 | 0 | -inf | 1 | 1 |
| 4930486L24Rik | 0,192523 | 0 | -inf | 1 | 1 |
| 4930500F04Rik | 0,031795 | 0 | -inf | 1 | 1 |
| 4930505A04Rik | 0,049215 | 0 | -inf | 1 | 1 |
| 4930511E03Rik | 0,017497 | 0 | -inf | 1 | 1 |
| 4930513D17Rik | 0,011852 | 0 | -inf | 1 | 1 |
| 4930515G16Rik | 0,045143 | 0 | -inf | 1 | 1 |
| 4930525D18Rik | 0,016158 | 0 | -inf | 1 | 1 |
| 4930528D03Rik | 0,011428 | 0 | -inf | 1 | 1 |
| 4930533B01Rik | 0,050282 | 0 | -inf | 1 | 1 |
| 4930542D17Rik | 0,148317 | 0 | -inf | 1 | 1 |
| 4930547E08Rik | 0,26347 | 0 | -inf | 1 | 1 |
| 4930548J01Rik | 0,101152 | 0 | -inf | 1 | 1 |
| 4930550L24Rik | 0,008646 | 0 | -inf | 1 | 1 |
| 4930563F08Rik | 0,133371 | 0 | -inf | 1 | 1 |
| 4930565D16Rik | 0,058709 | 0 | -inf | 1 | 1 |
| 4930578C19Rik | 0,19667 | 0 | -inf | 1 | 1 |
| 4930583K01Rik | 0,113348 | 0 | -inf | 1 | 1 |
| 4931423N10Rik | 0,007221 | 0 | -inf | 1 | 1 |
| 4931440F15Rik | 0,029019 | 0 | -inf | 1 | 1 |
| 4932438H23Rik | 0,010598 | 0 | -inf | 1 | 1 |
| 4932443I19Rik | 0,061497 | 0 | -inf | 1 | 1 |
| 4933401B06Rik | 0,011653 | 0 | -inf | 1 | 1 |
| 4933401H06Rik | 0,016136 | 0 | -inf | 1 | 1 |

| | | | | | |
|---|---|---|---|---|---|
| 4933402E13Rik | 0,068482 | 0 | -inf | 1 | 1 |
| 4933402J10Rik | 0,012169 | 0 | -inf | 1 | 1 |
| 4933406J08Rik | 0,006202 | 0 | -inf | 1 | 1 |
| 4933417E11Rik | 0,02661 | 0 | -inf | 1 | 1 |
| 4933434I20Rik | 0,010871 | 0 | -inf | 1 | 1 |
| 4933436H12Rik | 0,0089 | 0 | -inf | 1 | 1 |
| 5031414D18Rik | 0,08469 | 0 | -inf | 1 | 1 |
| 5031434C07Rik | 0,006206 | 0 | -inf | 1 | 1 |
| 5133400J02Rik | 0,056589 | 0 | -inf | 1 | 1 |
| 5430416O09Rik | 0,003047 | 0 | -inf | 1 | 1 |
| 5430419D17Rik | 0,008684 | 0 | -inf | 1 | 1 |
| 5430421F17Rik | 0,005657 | 0 | -inf | 1 | 1 |
| 5430427O19Rik | 0,034272 | 0 | -inf | 1 | 1 |
| 5430435G22Rik | 0,196687 | 0 | -inf | 1 | 1 |
| 5730460C07Rik | 0,127038 | 0 | -inf | 1 | 1 |
| 5730488B01Rik | 0,019687 | 0 | -inf | 1 | 1 |
| 5830432E09Rik | 0,074932 | 0 | -inf | 1 | 1 |
| 5930438M14Rik | 0,037119 | 0 | -inf | 1 | 1 |
| 6030469F06Rik | 0,008736 | 0 | -inf | 1 | 1 |
| 6030498E09Rik | 0,012069 | 0 | -inf | 1 | 1 |
| 6330407A03Rik | 0,014433 | 0 | -inf | 1 | 1 |
| 8430423G03Rik | 0,122544 | 0 | -inf | 1 | 1 |
| 8430426J06Rik | 0,032627 | 0 | -inf | 1 | 1 |
| 9030204H09Rik | 0,052685 | 0 | -inf | 1 | 1 |
| 9030404E10Rik | 0,08835 | 0 | -inf | 1 | 1 |
| 9030619P08Rik | 0,018113 | 0 | -inf | 1 | 1 |
| 9130015A21Rik | 0,032764 | 0 | -inf | 1 | 1 |
| 9230009I02Rik | 0,012503 | 0 | -inf | 1 | 1 |
| 9230104L09Rik | 0,091363 | 0 | -inf | 1 | 1 |
| 9330111N05Rik | 0,03448 | 0 | -inf | 1 | 1 |
| 9330158H04Rik | 0,018388 | 0 | -inf | 1 | 1 |
| 9430041J12Rik | 0,023375 | 0 | -inf | 1 | 1 |
| 9530026P05Rik | 0,25441 | 0 | -inf | 1 | 1 |
| 9530036O11Rik | 0,179196 | 0 | -inf | 1 | 1 |
| 9530080O11Rik | 0,03751 | 0 | -inf | 1 | 1 |
| A130077B15Rik | 0,015145 | 0 | -inf | 1 | 1 |
| A1cf | 0,003264 | 0 | -inf | 1 | 1 |
| A230001M10Rik | 0,024988 | 0 | -inf | 1 | 1 |
| A330048O09Rik | 0,115134 | 0 | -inf | 1 | 1 |
| A430078G23Rik | 0,008562 | 0 | -inf | 1 | 1 |
| A630023P12Rik | 0,02682 | 0 | -inf | 1 | 1 |
| A630033H20Rik | 0,326445 | 0 | -inf | 1 | 1 |
| A730020M07Rik | 0,019124 | 0 | -inf | 1 | 1 |
| A930016O22Rik | 0,044581 | 0 | -inf | 1 | 1 |
| A930018P22Rik | 0,086678 | 0 | -inf | 1 | 1 |
| AA467197 | 0,176472 | 0 | -inf | 1 | 1 |
| AF251705 | 0,057014 | 0 | -inf | 1 | 1 |
| AI115009 | 0,086173 | 0 | -inf | 1 | 1 |

| | | | | | |
|---|---|---|---|---|---|
| AI314278 | 0,010907 | 0 | -inf | 1 | 1 |
| AI607873 | 0,075851 | 0 | -inf | 1 | 1 |
| AI747448 | 0,004235 | 0 | -inf | 1 | 1 |
| AI847159 | 0,029496 | 0 | -inf | 1 | 1 |
| AU015791 | 0,146416 | 0 | -inf | 1 | 1 |
| AU021063 | 0,225922 | 0 | -inf | 1 | 1 |
| AU021092 | 0,354575 | 0 | -inf | 1 | 1 |
| AU022751 | 0,014459 | 0 | -inf | 1 | 1 |
| AU022754 | 0,245626 | 0 | -inf | 1 | 1 |
| AU022793 | 0,032036 | 0 | -inf | 1 | 1 |
| AW112010 | 0,145 | 0 | -inf | 1 | 1 |
| Aadac | 0,07596 | 0 | -inf | 1 | 1 |
| Aanat | 0,020586 | 0 | -inf | 1 | 1 |
| Abca13 | 0,008241 | 0 | -inf | 1 | 1 |
| Abcc12 | 0,008239 | 0 | -inf | 1 | 1 |
| Accsl | 0,011705 | 0 | -inf | 1 | 1 |
| Acot12 | 0,004158 | 0 | -inf | 1 | 1 |
| Acot3 | 0,004951 | 0 | -inf | 1 | 1 |
| Acox2 | 0,005269 | 0 | -inf | 1 | 1 |
| Acsm1 | 0,006456 | 0 | -inf | 1 | 1 |
| Adam3 | 0,004726 | 0 | -inf | 1 | 1 |
| Adam30 | 0,024703 | 0 | -inf | 1 | 1 |
| Adamtsl5 | 0,125132 | 0 | -inf | 1 | 1 |
| Adh6-ps1 | 0,004576 | 0 | -inf | 1 | 1 |
| Adh7 | 0,041048 | 0 | -inf | 1 | 1 |
| Adora3 | 0,045544 | 0 | -inf | 1 | 1 |
| Adtrp | 0,029124 | 0 | -inf | 1 | 1 |
| Agbl4 | 0,008126 | 0 | -inf | 1 | 1 |
| Agmo | 0,087949 | 0 | -inf | 1 | 1 |
| Agr2 | 0,085093 | 0 | -inf | 1 | 1 |
| Agt | 0,028937 | 0 | -inf | 1 | 1 |
| Agtr2 | 0,004433 | 0 | -inf | 1 | 1 |
| Ahsg | 0,026076 | 0 | -inf | 1 | 1 |
| Akr1c14 | 0,005278 | 0 | -inf | 1 | 1 |
| Akr1c21 | 0,011769 | 0 | -inf | 1 | 1 |
| Alb | 0,006432 | 0 | -inf | 1 | 1 |
| Aldh8a1 | 0,011602 | 0 | -inf | 1 | 1 |
| Aldob | 0,118395 | 0 | -inf | 1 | 1 |
| Amer3 | 0,002849 | 0 | -inf | 1 | 1 |
| Ampd1 | 0,011065 | 0 | -inf | 1 | 1 |
| Amtn | 0,01446 | 0 | -inf | 1 | 1 |
| Ang4 | 0,241225 | 0 | -inf | 1 | 1 |
| Ankk1 | 0,009531 | 0 | -inf | 1 | 1 |
| Ankrd2 | 0,026402 | 0 | -inf | 1 | 1 |
| Ankrd22 | 0,29782 | 0 | -inf | 1 | 1 |
| Ankrd36 | 0,022111 | 0 | -inf | 1 | 1 |
| Anks4b | 0,032388 | 0 | -inf | 1 | 1 |
| Aoah | 0,187398 | 0 | -inf | 1 | 1 |

| | | | | | |
|---|---|---|---|---|---|
| Aoc1 | 0,016796 | 0 | -inf | 1 | 1 |
| Aox2 | 0,01166 | 0 | -inf | 1 | 1 |
| Apob | 0,349986 | 0 | -inf | 1 | 1 |
| Apoc3 | 0,046379 | 0 | -inf | 1 | 1 |
| Apol6 | 0,012906 | 0 | -inf | 1 | 1 |
| Apol7c | 0,025713 | 0 | -inf | 1 | 1 |
| Apol7e | 0,026351 | 0 | -inf | 1 | 1 |
| Apol9b | 0,008822 | 0 | -inf | 1 | 1 |
| Aqp6 | 0,07019 | 0 | -inf | 1 | 1 |
| Arhgap15 | 0,075053 | 0 | -inf | 1 | 1 |
| Arhgap27os3 | 0,021095 | 0 | -inf | 1 | 1 |
| Arhgap40 | 0,019454 | 0 | -inf | 1 | 1 |
| Arhgef38 | 0,006933 | 0 | -inf | 1 | 1 |
| Arl11 | 0,148337 | 0 | -inf | 1 | 1 |
| Arrdc5 | 0,027765 | 0 | -inf | 1 | 1 |
| Asb15 | 0,00291 | 0 | -inf | 1 | 1 |
| Ascl1 | 0,04037 | 0 | -inf | 1 | 1 |
| Asmt | 0,011984 | 0 | -inf | 1 | 1 |
| Astl | 0,005776 | 0 | -inf | 1 | 1 |
| Atoh7 | 0,16861 | 0 | -inf | 1 | 1 |
| Atp13a5 | 0,005636 | 0 | -inf | 1 | 1 |
| Atp1a4 | 0,018025 | 0 | -inf | 1 | 1 |
| Atp1b4 | 0,002908 | 0 | -inf | 1 | 1 |
| Atp6v0c-ps2 | 0,000222 | 0 | -inf | 1 | 1 |
| Atp6v0d2 | 0,023188 | 0 | -inf | 1 | 1 |
| Atp8b4 | 0,047833 | 0 | -inf | 1 | 1 |
| B230206H07Rik | 0,041809 | 0 | -inf | 1 | 1 |
| B230312C02Rik | 0,010449 | 0 | -inf | 1 | 1 |
| B430010I23Rik | 0,006043 | 0 | -inf | 1 | 1 |
| B930025P03Rik | 0,004106 | 0 | -inf | 1 | 1 |
| BB019430 | 0,037168 | 0 | -inf | 1 | 1 |
| BB123696 | 0,011243 | 0 | -inf | 1 | 1 |
| BC021614 | 0,088066 | 0 | -inf | 1 | 1 |
| BC021767 | 0,558376 | 0 | -inf | 1 | 1 |
| BC021785 | 0,068968 | 0 | -inf | 1 | 1 |
| BC024386 | 0,029418 | 0 | -inf | 1 | 1 |
| BC055111 | 0,217905 | 0 | -inf | 1 | 1 |
| BC100530 | 0,274823 | 0 | -inf | 1 | 1 |
| BC107364 | 0,110084 | 0 | -inf | 1 | 1 |
| BC147527 | 0,057095 | 0 | -inf | 1 | 1 |
| Bach2os | 0,040279 | 0 | -inf | 1 | 1 |
| Bank1 | 0,132319 | 0 | -inf | 1 | 1 |
| Barhl2 | 0,011212 | 0 | -inf | 1 | 1 |
| Batf2 | 0,010135 | 0 | -inf | 1 | 1 |
| Bcl2a1a | 0,140133 | 0 | -inf | 1 | 1 |
| Bcl2a1b | 0,560441 | 0 | -inf | 1 | 1 |
| Bex6 | 0,053247 | 0 | -inf | 1 | 1 |
| Bfsp2 | 0,039025 | 0 | -inf | 1 | 1 |

| | | | | | |
|---|---|---|---|---|---|
| Bhlhe23 | 0,01791 | 0 | -inf | 1 | 1 |
| Blk | 0,005208 | 0 | -inf | 1 | 1 |
| Bpi | 0,00751 | 0 | -inf | 1 | 1 |
| Bpifb1 | 0,239862 | 0 | -inf | 1 | 1 |
| Bpifb4 | 0,012379 | 0 | -inf | 1 | 1 |
| Bsnd | 0,037764 | 0 | -inf | 1 | 1 |
| Btla | 0,043001 | 0 | -inf | 1 | 1 |
| Btn1a1 | 0,041956 | 0 | -inf | 1 | 1 |
| Btnl10 | 0,028709 | 0 | -inf | 1 | 1 |
| Btnl2 | 0,005014 | 0 | -inf | 1 | 1 |
| C030018K13Rik | 0,22346 | 0 | -inf | 1 | 1 |
| C030023E24Rik | 0,015547 | 0 | -inf | 1 | 1 |
| C130071C03Rik | 0,010162 | 0 | -inf | 1 | 1 |
| C230004F18Rik | 0,035573 | 0 | -inf | 1 | 1 |
| C330022C24Rik | 0,013795 | 0 | -inf | 1 | 1 |
| C4bp-ps1 | 0,021505 | 0 | -inf | 1 | 1 |
| C5ar1 | 0,192406 | 0 | -inf | 1 | 1 |
| C5ar2 | 0,03853 | 0 | -inf | 1 | 1 |
| C86187 | 0,022841 | 0 | -inf | 1 | 1 |
| C8b | 0,020383 | 0 | -inf | 1 | 1 |
| C9 | 0,015135 | 0 | -inf | 1 | 1 |
| C920009B18Rik | 0,026084 | 0 | -inf | 1 | 1 |
| Cabp7 | 0,729281 | 0 | -inf | 1 | 1 |
| Cacna2d4 | 0,00845 | 0 | -inf | 1 | 1 |
| Calcb | 0,054722 | 0 | -inf | 1 | 1 |
| Calhm1 | 0,014031 | 0 | -inf | 1 | 1 |
| Calm4 | 0,016636 | 0 | -inf | 1 | 1 |
| Calml3 | 0,033944 | 0 | -inf | 1 | 1 |
| Car1 | 0,054343 | 0 | -inf | 1 | 1 |
| Card14 | 0,026218 | 0 | -inf | 1 | 1 |
| Casp1 | 0,117592 | 0 | -inf | 1 | 1 |
| Casp12 | 0,192262 | 0 | -inf | 1 | 1 |
| Catsper4 | 0,040878 | 0 | -inf | 1 | 1 |
| Cav3 | 0,055937 | 0 | -inf | 1 | 1 |
| Cblc | 0,347187 | 0 | -inf | 1 | 1 |
| Ccdc108 | 0,00429 | 0 | -inf | 1 | 1 |
| Ccdc113 | 0,038875 | 0 | -inf | 1 | 1 |
| Ccdc129 | 0,013468 | 0 | -inf | 1 | 1 |
| Ccdc135 | 0,081139 | 0 | -inf | 1 | 1 |
| Ccdc185 | 0,00639 | 0 | -inf | 1 | 1 |
| Ccdc27 | 0,078774 | 0 | -inf | 1 | 1 |
| Ccdc83 | 0,009898 | 0 | -inf | 1 | 1 |
| Cckar | 0,015222 | 0 | -inf | 1 | 1 |
| Ccl24 | 0,087211 | 0 | -inf | 1 | 1 |
| Ccl3 | 0,303319 | 0 | -inf | 1 | 1 |
| Ccl4 | 0,051876 | 0 | -inf | 1 | 1 |
| Ccl6 | 0,355755 | 0 | -inf | 1 | 1 |
| Ccl9 | 0,171045 | 0 | -inf | 1 | 1 |

| | | | | | |
|---|---|---|---|---|---|
| Ccnb1ip1 | 0,254352 | 0 | -inf | 1 | 1 |
| Ccnb3 | 0,027229 | 0 | -inf | 1 | 1 |
| Ccr1 | 0,340406 | 0 | -inf | 1 | 1 |
| Ccr2 | 0,045474 | 0 | -inf | 1 | 1 |
| Ccr5 | 0,128398 | 0 | -inf | 1 | 1 |
| Ccr6 | 0,01161 | 0 | -inf | 1 | 1 |
| Ccr8 | 0,012424 | 0 | -inf | 1 | 1 |
| Ccrl2 | 0,378891 | 0 | -inf | 1 | 1 |
| Cd177 | 0,009425 | 0 | -inf | 1 | 1 |
| Cd180 | 0,507852 | 0 | -inf | 1 | 1 |
| Cd200r1 | 0,168465 | 0 | -inf | 1 | 1 |
| Cd200r2 | 0,007651 | 0 | -inf | 1 | 1 |
| Cd200r3 | 0,011001 | 0 | -inf | 1 | 1 |
| Cd200r4 | 0,007906 | 0 | -inf | 1 | 1 |
| Cd207 | 0,01784 | 0 | -inf | 1 | 1 |
| Cd209f | 0,01307 | 0 | -inf | 1 | 1 |
| Cd226 | 0,170719 | 0 | -inf | 1 | 1 |
| Cd244 | 0,061639 | 0 | -inf | 1 | 1 |
| Cd300ld | 0,023873 | 0 | -inf | 1 | 1 |
| Cd300lg | 0,160136 | 0 | -inf | 1 | 1 |
| Cd33 | 0,391519 | 0 | -inf | 1 | 1 |
| Cd3d | 0,036838 | 0 | -inf | 1 | 1 |
| Cd48 | 0,15771 | 0 | -inf | 1 | 1 |
| Cd53 | 0,457899 | 0 | -inf | 1 | 1 |
| Cd5l | 0,041988 | 0 | -inf | 1 | 1 |
| Cd69 | 0,008353 | 0 | -inf | 1 | 1 |
| Cd84 | 0,141944 | 0 | -inf | 1 | 1 |
| Cd8b1 | 0,034224 | 0 | -inf | 1 | 1 |
| Cdh17 | 0,019993 | 0 | -inf | 1 | 1 |
| Cdh7 | 0,016929 | 0 | -inf | 1 | 1 |
| Cdhr2 | 0,081449 | 0 | -inf | 1 | 1 |
| Cdsn | 0,004781 | 0 | -inf | 1 | 1 |
| Ceacam13 | 0,055897 | 0 | -inf | 1 | 1 |
| Ceacam15 | 0,054874 | 0 | -inf | 1 | 1 |
| Ceacam3 | 0,021638 | 0 | -inf | 1 | 1 |
| Ceacam5 | 0,017513 | 0 | -inf | 1 | 1 |
| Ceacam9 | 0,288607 | 0 | -inf | 1 | 1 |
| Cel | 0,006449 | 0 | -inf | 1 | 1 |
| Celf6 | 0,004619 | 0 | -inf | 1 | 1 |
| Ces1d | 0,023541 | 0 | -inf | 1 | 1 |
| Ces1g | 0,019894 | 0 | -inf | 1 | 1 |
| Ces2a | 0,007118 | 0 | -inf | 1 | 1 |
| Ces2e | 0,004741 | 0 | -inf | 1 | 1 |
| Cetn1 | 0,013965 | 0 | -inf | 1 | 1 |
| Cfc1 | 0,241293 | 0 | -inf | 1 | 1 |
| Cfh | 0,192092 | 0 | -inf | 1 | 1 |
| ChkbCpt1b | 0,090381 | 0 | -inf | 1 | 1 |
| Chrm2 | 0,12276 | 0 | -inf | 1 | 1 |

| | | | | | |
|---|---|---|---|---|---|
| Chrna10 | 0,02049 | 0 | -inf | 1 | 1 |
| Chrnb3 | 0,002716 | 0 | -inf | 1 | 1 |
| Chrng | 0,006889 | 0 | -inf | 1 | 1 |
| Chst13 | 0,024523 | 0 | -inf | 1 | 1 |
| Chst4 | 0,006586 | 0 | -inf | 1 | 1 |
| Cib4 | 0,073545 | 0 | -inf | 1 | 1 |
| Cidea | 0,049283 | 0 | -inf | 1 | 1 |
| Cidec | 0,007896 | 0 | -inf | 1 | 1 |
| Cilp | 0,032947 | 0 | -inf | 1 | 1 |
| Ckmt2 | 0,032356 | 0 | -inf | 1 | 1 |
| Clca1 | 0,01472 | 0 | -inf | 1 | 1 |
| Clca3 | 0,054168 | 0 | -inf | 1 | 1 |
| Clca4 | 0,003814 | 0 | -inf | 1 | 1 |
| Clca6 | 0,014846 | 0 | -inf | 1 | 1 |
| Cldn22 | 0,029944 | 0 | -inf | 1 | 1 |
| Cldn24 | 0,025769 | 0 | -inf | 1 | 1 |
| Clec12a | 0,005861 | 0 | -inf | 1 | 1 |
| Clec18a | 0,029296 | 0 | -inf | 1 | 1 |
| Clec1a | 0,395979 | 0 | -inf | 1 | 1 |
| Clec2f | 0,02611 | 0 | -inf | 1 | 1 |
| Clec4a2 | 0,024257 | 0 | -inf | 1 | 1 |
| Clec4a3 | 0,012268 | 0 | -inf | 1 | 1 |
| Clec4e | 0,035836 | 0 | -inf | 1 | 1 |
| Clec4g | 0,011472 | 0 | -inf | 1 | 1 |
| Clec4n | 0,132499 | 0 | -inf | 1 | 1 |
| Clec5a | 0,104363 | 0 | -inf | 1 | 1 |
| Clic3 | 0,487717 | 0 | -inf | 1 | 1 |
| Clrn1 | 0,017886 | 0 | -inf | 1 | 1 |
| Clvs1 | 0,011383 | 0 | -inf | 1 | 1 |
| Cml2 | 0,032712 | 0 | -inf | 1 | 1 |
| Cmtm2a | 0,013441 | 0 | -inf | 1 | 1 |
| Cndp1 | 0,004506 | 0 | -inf | 1 | 1 |
| Cnga4 | 0,015591 | 0 | -inf | 1 | 1 |
| Cngb3 | 0,002626 | 0 | -inf | 1 | 1 |
| Cntnap3 | 0,007608 | 0 | -inf | 1 | 1 |
| Cntnap4 | 0,043315 | 0 | -inf | 1 | 1 |
| Col28a1 | 0,013241 | 0 | -inf | 1 | 1 |
| Col6a6 | 0,016316 | 0 | -inf | 1 | 1 |
| Colq | 0,020262 | 0 | -inf | 1 | 1 |
| Cox7b2 | 0,122356 | 0 | -inf | 1 | 1 |
| Cpa3 | 0,289162 | 0 | -inf | 1 | 1 |
| Cpn2 | 0,009475 | 0 | -inf | 1 | 1 |
| Cpne4 | 0,05495 | 0 | -inf | 1 | 1 |
| Cps1 | 0,158008 | 0 | -inf | 1 | 1 |
| Cryaa | 0,025626 | 0 | -inf | 1 | 1 |
| Cryba4 | 0,043167 | 0 | -inf | 1 | 1 |
| Crybb1 | 0,12376 | 0 | -inf | 1 | 1 |
| Cryge | 0,027004 | 0 | -inf | 1 | 1 |

| | | | | | |
|---|---|---|---|---|---|
| Csn3 | 0,008491 | 0 | -inf | 1 | 1 |
| Csnka2ip | 0,010561 | 0 | -inf | 1 | 1 |
| Cst9 | 0,368764 | 0 | -inf | 1 | 1 |
| Ctcfl | 0,021618 | 0 | -inf | 1 | 1 |
| Ctrb1 | 0,017318 | 0 | -inf | 1 | 1 |
| Cts3 | 0,030507 | 0 | -inf | 1 | 1 |
| Cts7 | 0,03218 | 0 | -inf | 1 | 1 |
| Cts8 | 0,018121 | 0 | -inf | 1 | 1 |
| Ctsg | 0,081623 | 0 | -inf | 1 | 1 |
| Cuzd1 | 0,005764 | 0 | -inf | 1 | 1 |
| Cxcl2 | 0,013456 | 0 | -inf | 1 | 1 |
| Cxcr2 | 0,09594 | 0 | -inf | 1 | 1 |
| Cybb | 0,231746 | 0 | -inf | 1 | 1 |
| Cyp1a1 | 0,026942 | 0 | -inf | 1 | 1 |
| Cyp2ab1 | 0,004673 | 0 | -inf | 1 | 1 |
| Cyp2c37 | 0,007301 | 0 | -inf | 1 | 1 |
| Cyp2c65 | 0,006809 | 0 | -inf | 1 | 1 |
| Cyp2d11 | 0,008534 | 0 | -inf | 1 | 1 |
| Cyp2d9 | 0,028666 | 0 | -inf | 1 | 1 |
| Cyp2j8 | 0,008988 | 0 | -inf | 1 | 1 |
| Cyp4b1-ps2 | 0,575186 | 0 | -inf | 1 | 1 |
| Cyp4f14 | 0,006457 | 0 | -inf | 1 | 1 |
| Cyp4x1 | 0,008961 | 0 | -inf | 1 | 1 |
| Cypt3 | 0,025601 | 0 | -inf | 1 | 1 |
| D030040B21Rik | 0,009155 | 0 | -inf | 1 | 1 |
| D130058E03 | 0,023258 | 0 | -inf | 1 | 1 |
| D3Bwg0562e | 0,127379 | 0 | -inf | 1 | 1 |
| D630024D03Rik | 0,030961 | 0 | -inf | 1 | 1 |
| D6Ertd474e | 0,016222 | 0 | -inf | 1 | 1 |
| D7Ertd143e | 0,005832 | 0 | -inf | 1 | 1 |
| D830013O20Rik | 0,024387 | 0 | -inf | 1 | 1 |
| DXBay18,Gm1468 | 9,05E-005 | 0 | -inf | 1 | 1 |
| Dazl | 0,045865 | 0 | -inf | 1 | 1 |
| Dbpht2 | 0,0191 | 0 | -inf | 1 | 1 |
| Dcstamp | 0,071743 | 0 | -inf | 1 | 1 |
| Ddi1 | 0,017976 | 0 | -inf | 1 | 1 |
| Defa22 | 0,192906 | 0 | -inf | 1 | 1 |
| Defa24,Defa6 | 8,20E-006 | 0 | -inf | 1 | 1 |
| Defa6 | 0,492855 | 0 | -inf | 1 | 1 |
| Dgkeos | 0,135977 | 0 | -inf | 1 | 1 |
| Dhrs7c | 0,012857 | 0 | -inf | 1 | 1 |
| Dhrs9 | 0,015875 | 0 | -inf | 1 | 1 |
| Dkk4 | 0,010656 | 0 | -inf | 1 | 1 |
| Dlx1as | 0,172215 | 0 | -inf | 1 | 1 |
| Dlx6as2 | 0,010285 | 0 | -inf | 1 | 1 |
| Dmbt1 | 0,006869 | 0 | -inf | 1 | 1 |
| Dmkn | 0,090764 | 0 | -inf | 1 | 1 |
| Dmp1 | 0,039366 | 0 | -inf | 1 | 1 |

| | | | | | |
|---|---|---|---|---|---|
| Dmr | 0,031513 | 0 | -inf | 1 | 1 |
| Dmrt3 | 0,010817 | 0 | -inf | 1 | 1 |
| Dnase2b | 0,005646 | 0 | -inf | 1 | 1 |
| Doxl2 | 0,045484 | 0 | -inf | 1 | 1 |
| Dpep2 | 0,015145 | 0 | -inf | 1 | 1 |
| Dppa4 | 0,010073 | 0 | -inf | 1 | 1 |
| Dppa5a | 0,217898 | 0 | -inf | 1 | 1 |
| Dpt | 0,028966 | 0 | -inf | 1 | 1 |
| Dsc1 | 0,016166 | 0 | -inf | 1 | 1 |
| Duoxa1 | 0,073308 | 0 | -inf | 1 | 1 |
| Dydc2 | 0,22884 | 0 | -inf | 1 | 1 |
| Dynlrb2 | 0,038031 | 0 | -inf | 1 | 1 |
| Dyrk4 | 0,066494 | 0 | -inf | 1 | 1 |
| Dytn | 0,005321 | 0 | -inf | 1 | 1 |
| E030019B13Rik | 0,011351 | 0 | -inf | 1 | 1 |
| E030044B06Rik | 0,017497 | 0 | -inf | 1 | 1 |
| E130215H24Rik | 0,01657 | 0 | -inf | 1 | 1 |
| E230016K23Rik | 0,021489 | 0 | -inf | 1 | 1 |
| E230029C05Rik | 0,099949 | 0 | -inf | 1 | 1 |
| E330017L17Rik | 0,018894 | 0 | -inf | 1 | 1 |
| E330021D16Rik | 0,084796 | 0 | -inf | 1 | 1 |
| Eddm3b | 0,080154 | 0 | -inf | 1 | 1 |
| Edn2 | 0,039884 | 0 | -inf | 1 | 1 |
| Efcab10 | 0,591003 | 0 | -inf | 1 | 1 |
| Efcab9 | 0,042017 | 0 | -inf | 1 | 1 |
| Elane | 0,058141 | 0 | -inf | 1 | 1 |
| Emr1 | 0,369883 | 0 | -inf | 1 | 1 |
| Emx1 | 0,29329 | 0 | -inf | 1 | 1 |
| Endou | 0,298805 | 0 | -inf | 1 | 1 |
| Enpep | 0,01477 | 0 | -inf | 1 | 1 |
| Entpd4 | 0,004107 | 0 | -inf | 1 | 1 |
| Epha6 | 0,00295 | 0 | -inf | 1 | 1 |
| Eppin | 0,021466 | 0 | -inf | 1 | 1 |
| Eps8l3 | 0,028896 | 0 | -inf | 1 | 1 |
| Erbb4 | 0,010234 | 0 | -inf | 1 | 1 |
| Erp27 | 0,05538 | 0 | -inf | 1 | 1 |
| Esp4 | 0,119191 | 0 | -inf | 1 | 1 |
| Esp5 | 0,071617 | 0 | -inf | 1 | 1 |
| Esp6-esp5 | 3,49E-005 | 0 | -inf | 1 | 1 |
| Esx1 | 0,008473 | 0 | -inf | 1 | 1 |
| F12 | 0,03034 | 0 | -inf | 1 | 1 |
| F630028O10Rik | 0,386656 | 0 | -inf | 1 | 1 |
| F730035M05Rik | 0,054158 | 0 | -inf | 1 | 1 |
| F9 | 0,004677 | 0 | -inf | 1 | 1 |
| Fabp1 | 0,164273 | 0 | -inf | 1 | 1 |
| Fabp2 | 0,015543 | 0 | -inf | 1 | 1 |
| Fam183b | 0,039686 | 0 | -inf | 1 | 1 |
| Fam187a | 0,008703 | 0 | -inf | 1 | 1 |

| | | | | | |
|---|---|---|---|---|---|
| Fam229a | 0,133647 | 0 | -inf | 1 | 1 |
| Fam24a | 0,141356 | 0 | -inf | 1 | 1 |
| Fam26d | 0,00337 | 0 | -inf | 1 | 1 |
| Fam43b | 0,010937 | 0 | -inf | 1 | 1 |
| Fam71f1 | 0,038435 | 0 | -inf | 1 | 1 |
| Fasl | 0,021065 | 0 | -inf | 1 | 1 |
| Fate1 | 0,032229 | 0 | -inf | 1 | 1 |
| Fbp2 | 0,141218 | 0 | -inf | 1 | 1 |
| Fbxo40 | 0,094669 | 0 | -inf | 1 | 1 |
| Fcamr | 0,042321 | 0 | -inf | 1 | 1 |
| Fcgbp | 0,061302 | 0 | -inf | 1 | 1 |
| Fcgr4 | 0,090619 | 0 | -inf | 1 | 1 |
| Fcna | 0,244865 | 0 | -inf | 1 | 1 |
| Fcrl1 | 0,014344 | 0 | -inf | 1 | 1 |
| Fcrla | 0,028499 | 0 | -inf | 1 | 1 |
| Fezf1 | 0,506575 | 0 | -inf | 1 | 1 |
| Fezf2 | 0,005632 | 0 | -inf | 1 | 1 |
| Ffar1 | 0,059341 | 0 | -inf | 1 | 1 |
| Fga | 0,332655 | 0 | -inf | 1 | 1 |
| Fgb | 0,280857 | 0 | -inf | 1 | 1 |
| Fgf20 | 0,102415 | 0 | -inf | 1 | 1 |
| Fgf22 | 0,125853 | 0 | -inf | 1 | 1 |
| Fgf5 | 0,094442 | 0 | -inf | 1 | 1 |
| Figla | 0,074737 | 0 | -inf | 1 | 1 |
| Fitm1 | 0,058587 | 0 | -inf | 1 | 1 |
| Fmo2 | 0,206056 | 0 | -inf | 1 | 1 |
| Fnd3c2 | 0,056575 | 0 | -inf | 1 | 1 |
| Folr4 | 0,048199 | 0 | -inf | 1 | 1 |
| Foxi1 | 0,011003 | 0 | -inf | 1 | 1 |
| Foxi3 | 0,011804 | 0 | -inf | 1 | 1 |
| Foxr1 | 0,00809 | 0 | -inf | 1 | 1 |
| Foxs1 | 0,03747 | 0 | -inf | 1 | 1 |
| Fpr1 | 0,042281 | 0 | -inf | 1 | 1 |
| Frmpd1os | 0,056048 | 0 | -inf | 1 | 1 |
| Frmpd4 | 0,008182 | 0 | -inf | 1 | 1 |
| Ftcd | 0,01888 | 0 | -inf | 1 | 1 |
| Fut1 | 0,195691 | 0 | -inf | 1 | 1 |
| Fut2 | 0,096824 | 0 | -inf | 1 | 1 |
| Fyb | 0,471662 | 0 | -inf | 1 | 1 |
| G630090E17Rik | 0,041278 | 0 | -inf | 1 | 1 |
| Gabrb1 | 0,025048 | 0 | -inf | 1 | 1 |
| Gabrd | 0,024764 | 0 | -inf | 1 | 1 |
| Gabrg3 | 0,072577 | 0 | -inf | 1 | 1 |
| Gad2 | 0,037123 | 0 | -inf | 1 | 1 |
| Galntl5 | 0,008479 | 0 | -inf | 1 | 1 |
| Gapt | 0,01278 | 0 | -inf | 1 | 1 |
| Gata5 | 0,01909 | 0 | -inf | 1 | 1 |
| Gbgt1 | 0,057591 | 0 | -inf | 1 | 1 |

| | | | | | |
|---|---|---|---|---|---|
| Gbp10 | 0,037437 | 0 | -inf | 1 | 1 |
| Gbp2b | 0,036234 | 0 | -inf | 1 | 1 |
| Gbp6 | 0,036016 | 0 | -inf | 1 | 1 |
| Gc | 0,007356 | 0 | -inf | 1 | 1 |
| Gckr | 0,131088 | 0 | -inf | 1 | 1 |
| Gcnt3 | 0,008661 | 0 | -inf | 1 | 1 |
| Gdf3 | 0,055966 | 0 | -inf | 1 | 1 |
| Gfap | 0,117799 | 0 | -inf | 1 | 1 |
| Gfy | 0,053074 | 0 | -inf | 1 | 1 |
| Ghrh | 0,132519 | 0 | -inf | 1 | 1 |
| Ghrhr | 0,01697 | 0 | -inf | 1 | 1 |
| Ghsr | 0,015402 | 0 | -inf | 1 | 1 |
| Gimap3 | 0,006094 | 0 | -inf | 1 | 1 |
| Gimap5 | 0,469971 | 0 | -inf | 1 | 1 |
| Gimap7 | 0,011106 | 0 | -inf | 1 | 1 |
| Gjb1 | 0,082393 | 0 | -inf | 1 | 1 |
| Gjb4 | 0,019062 | 0 | -inf | 1 | 1 |
| Glipr1 | 0,350587 | 0 | -inf | 1 | 1 |
| Glyat | 0,010694 | 0 | -inf | 1 | 1 |
| Gm10248 | 0,004683 | 0 | -inf | 1 | 1 |
| Gm10494 | 0,009948 | 0 | -inf | 1 | 1 |
| Gm10635 | 0,006726 | 0 | -inf | 1 | 1 |
| Gm10714 | 0,017654 | 0 | -inf | 1 | 1 |
| Gm10768 | 0,070839 | 0 | -inf | 1 | 1 |
| Gm10804 | 0,098095 | 0 | -inf | 1 | 1 |
| Gm10825 | 0,002723 | 0 | -inf | 1 | 1 |
| Gm10863 | 0,024848 | 0 | -inf | 1 | 1 |
| Gm1110 | 0,010847 | 0 | -inf | 1 | 1 |
| Gm11346 | 0,038951 | 0 | -inf | 1 | 1 |
| Gm11747 | 0,017509 | 0 | -inf | 1 | 1 |
| Gm11978 | 0,010733 | 0 | -inf | 1 | 1 |
| Gm12409 | 0,02676 | 0 | -inf | 1 | 1 |
| Gm12429 | 0,009002 | 0 | -inf | 1 | 1 |
| Gm12709 | 0,013851 | 0 | -inf | 1 | 1 |
| Gm12718 | 0,127422 | 0 | -inf | 1 | 1 |
| Gm12830 | 0,030958 | 0 | -inf | 1 | 1 |
| Gm1322 | 0,023541 | 0 | -inf | 1 | 1 |
| Gm13238 | 0,062252 | 0 | -inf | 1 | 1 |
| Gm13272,Gm132[illegible] | 0,019486 | 0 | -inf | 1 | 1 |
| Gm13446 | 0,02263 | 0 | -inf | 1 | 1 |
| Gm13490 | 0,005364 | 0 | -inf | 1 | 1 |
| Gm13629 | 0,03629 | 0 | -inf | 1 | 1 |
| Gm13807 | 0,054207 | 0 | -inf | 1 | 1 |
| Gm14139 | 0,034979 | 0 | -inf | 1 | 1 |
| Gm14207 | 0,331505 | 0 | -inf | 1 | 1 |
| Gm14685 | 0,004511 | 0 | -inf | 1 | 1 |
| Gm15217 | 0,025166 | 0 | -inf | 1 | 1 |
| Gm15284 | 0,89134 | 0 | -inf | 1 | 1 |

| | | | | | |
|---|---|---|---|---|---|
| Gm15350 | 0,06846 | 0 | -inf | 1 | 1 |
| Gm15850 | 0,294777 | 0 | -inf | 1 | 1 |
| Gm1587 | 0,010285 | 0 | -inf | 1 | 1 |
| Gm15915 | 0,033602 | 0 | -inf | 1 | 1 |
| Gm16063 | 0,007912 | 0 | -inf | 1 | 1 |
| Gm16157 | 0,047243 | 0 | -inf | 1 | 1 |
| Gm16548 | 0,22816 | 0 | -inf | 1 | 1 |
| Gm16701 | 0,12949 | 0 | -inf | 1 | 1 |
| Gm17019 | 0,012232 | 0 | -inf | 1 | 1 |
| Gm1720 | 0,018211 | 0 | -inf | 1 | 1 |
| Gm17252 | 0,063882 | 0 | -inf | 1 | 1 |
| Gm17660 | 0,036962 | 0 | -inf | 1 | 1 |
| Gm17745 | 0,136392 | 0 | -inf | 1 | 1 |
| Gm19589 | 0,026938 | 0 | -inf | 1 | 1 |
| Gm19619 | 0,00446 | 0 | -inf | 1 | 1 |
| Gm1966 | 0,027926 | 0 | -inf | 1 | 1 |
| Gm1968 | 0,083722 | 0 | -inf | 1 | 1 |
| Gm20110 | 0,006255 | 0 | -inf | 1 | 1 |
| Gm20125 | 0,0037 | 0 | -inf | 1 | 1 |
| Gm20611 | 0,070265 | 0 | -inf | 1 | 1 |
| Gm20751 | 0,155493 | 0 | -inf | 1 | 1 |
| Gm2087 | 0,144761 | 0 | -inf | 1 | 1 |
| Gm21057 | 0,167632 | 0 | -inf | 1 | 1 |
| Gm2115 | 0,014259 | 0 | -inf | 1 | 1 |
| Gm21949 | 0,026468 | 0 | -inf | 1 | 1 |
| Gm2381 | 0,003056 | 0 | -inf | 1 | 1 |
| Gm2696 | 0,02279 | 0 | -inf | 1 | 1 |
| Gm3336 | 0,043755 | 0 | -inf | 1 | 1 |
| Gm3500 | 0,006652 | 0 | -inf | 1 | 1 |
| Gm4461 | 0,013335 | 0 | -inf | 1 | 1 |
| Gm4489 | 0,037429 | 0 | -inf | 1 | 1 |
| Gm4566 | 0,234639 | 0 | -inf | 1 | 1 |
| Gm4598 | 0,013502 | 0 | -inf | 1 | 1 |
| Gm4827 | 0,03923 | 0 | -inf | 1 | 1 |
| Gm4861 | 0,015708 | 0 | -inf | 1 | 1 |
| Gm4872 | 0,024468 | 0 | -inf | 1 | 1 |
| Gm4952 | 0,01804 | 0 | -inf | 1 | 1 |
| Gm4971 | 0,129548 | 0 | -inf | 1 | 1 |
| Gm4984 | 0,064458 | 0 | -inf | 1 | 1 |
| Gm5039 | 0,028866 | 0 | -inf | 1 | 1 |
| Gm5065 | 0,168211 | 0 | -inf | 1 | 1 |
| Gm5087 | 0,006889 | 0 | -inf | 1 | 1 |
| Gm5136 | 0,02591 | 0 | -inf | 1 | 1 |
| Gm5177 | 0,003303 | 0 | -inf | 1 | 1 |
| Gm5294 | 0,02646 | 0 | -inf | 1 | 1 |
| Gm5420 | 0,020719 | 0 | -inf | 1 | 1 |
| Gm5483 | 0,255014 | 0 | -inf | 1 | 1 |
| Gm5547 | 0,139148 | 0 | -inf | 1 | 1 |

| | | | | | |
|---|---|---|---|---|---|
| Gm572 | 0,215536 | 0 | -inf | 1 | 1 |
| Gm5936 | 0,015553 | 0 | -inf | 1 | 1 |
| Gm6034 | 0,008277 | 0 | -inf | 1 | 1 |
| Gm614 | 0,076389 | 0 | -inf | 1 | 1 |
| Gm6150 | 0,023464 | 0 | -inf | 1 | 1 |
| Gm6213 | 0,122827 | 0 | -inf | 1 | 1 |
| Gm6249 | 0,007607 | 0 | -inf | 1 | 1 |
| Gm6432 | 0,071393 | 0 | -inf | 1 | 1 |
| Gm648 | 0,10621 | 0 | -inf | 1 | 1 |
| Gm6634 | 0,062888 | 0 | -inf | 1 | 1 |
| Gm6792 | 0,021781 | 0 | -inf | 1 | 1 |
| Gm6878 | 0,07512 | 0 | -inf | 1 | 1 |
| Gm6880 | 0,027962 | 0 | -inf | 1 | 1 |
| Gm7168 | 0,008455 | 0 | -inf | 1 | 1 |
| Gm7257 | 0,263355 | 0 | -inf | 1 | 1 |
| Gm7325 | 0,058637 | 0 | -inf | 1 | 1 |
| Gm766 | 0,055837 | 0 | -inf | 1 | 1 |
| Gm7788 | 0,290765 | 0 | -inf | 1 | 1 |
| Gm7854 | 0,033778 | 0 | -inf | 1 | 1 |
| Gm7977 | 0,152497 | 0 | -inf | 1 | 1 |
| Gm805 | 0,008003 | 0 | -inf | 1 | 1 |
| Gm8221 | 0,00648 | 0 | -inf | 1 | 1 |
| Gm826 | 0,07294 | 0 | -inf | 1 | 1 |
| Gm8817 | 0,355925 | 0 | -inf | 1 | 1 |
| Gm8883 | 0,083795 | 0 | -inf | 1 | 1 |
| Gm8884 | 0,06564 | 0 | -inf | 1 | 1 |
| Gm9112 | 0,046749 | 0 | -inf | 1 | 1 |
| Gm9159 | 0,026707 | 0 | -inf | 1 | 1 |
| Gm9513 | 0,533859 | 0 | -inf | 1 | 1 |
| Gm9920 | 0,017731 | 0 | -inf | 1 | 1 |
| Gm9962 | 0,019502 | 0 | -inf | 1 | 1 |
| Gna15 | 0,266238 | 0 | -inf | 1 | 1 |
| Golga7b | 0,009017 | 0 | -inf | 1 | 1 |
| Gp49a | 0,011911 | 0 | -inf | 1 | 1 |
| Gp6 | 0,362171 | 0 | -inf | 1 | 1 |
| Gpa33 | 0,050959 | 0 | -inf | 1 | 1 |
| Gpr110 | 0,003584 | 0 | -inf | 1 | 1 |
| Gpr132 | 0,100625 | 0 | -inf | 1 | 1 |
| Gpr143 | 0,098203 | 0 | -inf | 1 | 1 |
| Gpr149 | 0,003777 | 0 | -inf | 1 | 1 |
| Gpr174 | 0,002487 | 0 | -inf | 1 | 1 |
| Gpr26 | 0,031969 | 0 | -inf | 1 | 1 |
| Gpr52 | 0,07385 | 0 | -inf | 1 | 1 |
| Gprin3 | 0,016337 | 0 | -inf | 1 | 1 |
| Gpx6 | 0,346722 | 0 | -inf | 1 | 1 |
| Gria2 | 0,045643 | 0 | -inf | 1 | 1 |
| Grin1os | 0,013044 | 0 | -inf | 1 | 1 |
| Grin2b | 0,017044 | 0 | -inf | 1 | 1 |

| | | | | | |
|---|---|---|---|---|---|
| Grp | 0,051351 | 0 | -inf | 1 | 1 |
| Grxcr1 | 0,013642 | 0 | -inf | 1 | 1 |
| Gsdma3 | 0,032584 | 0 | -inf | 1 | 1 |
| Gsdmc | 0,190225 | 0 | -inf | 1 | 1 |
| Gsdmc3 | 0,091633 | 0 | -inf | 1 | 1 |
| Gsdmcl-ps | 0,06515 | 0 | -inf | 1 | 1 |
| Gstm3 | 0,243205 | 0 | -inf | 1 | 1 |
| Gstp2 | 0,389103 | 0 | -inf | 1 | 1 |
| Guca1a | 0,55753 | 0 | -inf | 1 | 1 |
| Guca2a | 0,116513 | 0 | -inf | 1 | 1 |
| Guca2b | 0,344102 | 0 | -inf | 1 | 1 |
| Gys2 | 0,009644 | 0 | -inf | 1 | 1 |
| Gzma | 0,061542 | 0 | -inf | 1 | 1 |
| Gzmb | 0,151468 | 0 | -inf | 1 | 1 |
| Gzme | 0,60124 | 0 | -inf | 1 | 1 |
| H2-DMb2 | 0,020323 | 0 | -inf | 1 | 1 |
| H2-Eb2 | 0,014897 | 0 | -inf | 1 | 1 |
| H2-M10.3 | 0,029885 | 0 | -inf | 1 | 1 |
| H2-Q1 | 0,097574 | 0 | -inf | 1 | 1 |
| H2-Q9 | 0,491382 | 0 | -inf | 1 | 1 |
| H2-T3 | 0,005992 | 0 | -inf | 1 | 1 |
| Habp2 | 0,063611 | 0 | -inf | 1 | 1 |
| Hamp | 0,062214 | 0 | -inf | 1 | 1 |
| Havcr2 | 0,077571 | 0 | -inf | 1 | 1 |
| Hbb-bh2 | 0,767223 | 0 | -inf | 1 | 1 |
| Hbb-bs | 0,003709 | 0 | -inf | 1 | 1 |
| Hbq1a | 0,236964 | 0 | -inf | 1 | 1 |
| Hc | 0,010166 | 0 | -inf | 1 | 1 |
| Hcrtr1 | 0,024165 | 0 | -inf | 1 | 1 |
| Heatr9 | 0,00716 | 0 | -inf | 1 | 1 |
| Hephl1 | 0,002339 | 0 | -inf | 1 | 1 |
| Hhatl | 0,007648 | 0 | -inf | 1 | 1 |
| Hic1 | 0,015589 | 0 | -inf | 1 | 1 |
| Higd1c | 0,039164 | 0 | -inf | 1 | 1 |
| Hist1h2ad | 0,035733 | 0 | -inf | 1 | 1 |
| Hist1h3b | 0,483545 | 0 | -inf | 1 | 1 |
| Hist3h2bb-ps | 0,090633 | 0 | -inf | 1 | 1 |
| Hmx1 | 0,009071 | 0 | -inf | 1 | 1 |
| Hnf4g | 0,016382 | 0 | -inf | 1 | 1 |
| Hormad1 | 0,01631 | 0 | -inf | 1 | 1 |
| Hrct1 | 0,167056 | 0 | -inf | 1 | 1 |
| Hrh2 | 0,004576 | 0 | -inf | 1 | 1 |
| Hsd11b1 | 0,113952 | 0 | -inf | 1 | 1 |
| Hsd17b13 | 0,021903 | 0 | -inf | 1 | 1 |
| Hsd17b2 | 0,247712 | 0 | -inf | 1 | 1 |
| Hsd17b3 | 0,010932 | 0 | -inf | 1 | 1 |
| Hsd3b1 | 0,219153 | 0 | -inf | 1 | 1 |
| Hsd3b2 | 0,029644 | 0 | -inf | 1 | 1 |

| | | | | | |
|---|---|---|---|---|---|
| Hsf4 | 0,016555 | 0 | -inf | 1 | 1 |
| Hsh2d | 0,023169 | 0 | -inf | 1 | 1 |
| Hyal4 | 0,022017 | 0 | -inf | 1 | 1 |
| Iapp | 0,024089 | 0 | -inf | 1 | 1 |
| Idi2 | 0,005801 | 0 | -inf | 1 | 1 |
| Ifi202b | 0,014873 | 0 | -inf | 1 | 1 |
| Ifi203 | 0,003461 | 0 | -inf | 1 | 1 |
| Ifi27l2b | 0,02661 | 0 | -inf | 1 | 1 |
| Ifi47 | 0,2283 | 0 | -inf | 1 | 1 |
| Ifitm6 | 0,070194 | 0 | -inf | 1 | 1 |
| Ifna13 | 0,067446 | 0 | -inf | 1 | 1 |
| Ifnk | 0,057603 | 0 | -inf | 1 | 1 |
| Ifnz | 0,009402 | 0 | -inf | 1 | 1 |
| Iigp1 | 0,080124 | 0 | -inf | 1 | 1 |
| Il1b | 0,116003 | 0 | -inf | 1 | 1 |
| Il1f6 | 0,061089 | 0 | -inf | 1 | 1 |
| Il1rl1 | 0,04015 | 0 | -inf | 1 | 1 |
| Il1rn | 0,029531 | 0 | -inf | 1 | 1 |
| Il20ra | 0,04713 | 0 | -inf | 1 | 1 |
| Il21r | 0,137958 | 0 | -inf | 1 | 1 |
| Il27ra | 0,127894 | 0 | -inf | 1 | 1 |
| Il2rb | 0,035631 | 0 | -inf | 1 | 1 |
| Il5 | 0,017787 | 0 | -inf | 1 | 1 |
| Il5ra | 0,014127 | 0 | -inf | 1 | 1 |
| Il7r | 0,003935 | 0 | -inf | 1 | 1 |
| Iltifb | 0,013055 | 0 | -inf | 1 | 1 |
| Impg1 | 0,012261 | 0 | -inf | 1 | 1 |
| Inhbe | 0,024501 | 0 | -inf | 1 | 1 |
| Insc | 0,039188 | 0 | -inf | 1 | 1 |
| Isx | 0,007448 | 0 | -inf | 1 | 1 |
| Itgb6 | 0,047784 | 0 | -inf | 1 | 1 |
| Itgbl1 | 0,005494 | 0 | -inf | 1 | 1 |
| Itih1 | 0,004117 | 0 | -inf | 1 | 1 |
| Itih2 | 0,306459 | 0 | -inf | 1 | 1 |
| Itih3 | 0,031749 | 0 | -inf | 1 | 1 |
| Itk | 0,002948 | 0 | -inf | 1 | 1 |
| Itln1 | 0,044363 | 0 | -inf | 1 | 1 |
| Iyd | 0,031048 | 0 | -inf | 1 | 1 |
| Kcna4 | 0,005098 | 0 | -inf | 1 | 1 |
| Kcne2 | 0,123208 | 0 | -inf | 1 | 1 |
| Kcnj1 | 0,008251 | 0 | -inf | 1 | 1 |
| Kcnj16 | 0,05711 | 0 | -inf | 1 | 1 |
| Kcnmb2 | 0,172295 | 0 | -inf | 1 | 1 |
| Kcnmb4os1 | 0,194665 | 0 | -inf | 1 | 1 |
| Kcns2 | 0,013335 | 0 | -inf | 1 | 1 |
| Khdc3 | 0,063959 | 0 | -inf | 1 | 1 |
| Klhl31 | 0,013706 | 0 | -inf | 1 | 1 |
| Klk4 | 0,01145 | 0 | -inf | 1 | 1 |

| | | | | | |
|---|---|---|---|---|---|
| Klk5 | 0,022856 | 0 | -inf | 1 | 1 |
| Klkb1 | 0,005236 | 0 | -inf | 1 | 1 |
| Klra2 | 0,014941 | 0 | -inf | 1 | 1 |
| Klrb1b | 0,00589 | 0 | -inf | 1 | 1 |
| Klrb1c | 0,009761 | 0 | -inf | 1 | 1 |
| Klrg1 | 0,056152 | 0 | -inf | 1 | 1 |
| Kmo | 0,020624 | 0 | -inf | 1 | 1 |
| Krt12 | 0,007169 | 0 | -inf | 1 | 1 |
| Krt13 | 0,031968 | 0 | -inf | 1 | 1 |
| Krt24 | 0,007965 | 0 | -inf | 1 | 1 |
| Krt75 | 0,085034 | 0 | -inf | 1 | 1 |
| Krt76 | 0,019677 | 0 | -inf | 1 | 1 |
| Krt78 | 0,010986 | 0 | -inf | 1 | 1 |
| Krt85 | 0,115618 | 0 | -inf | 1 | 1 |
| Krtap5-1 | 0,024188 | 0 | -inf | 1 | 1 |
| LOC100861615 | 0,013238 | 0 | -inf | 1 | 1 |
| LOC100862268 | 0,057982 | 0 | -inf | 1 | 1 |
| LOC101055769 | 0,049543 | 0 | -inf | 1 | 1 |
| LOC101056149 | 0,106863 | 0 | -inf | 1 | 1 |
| LOC101243624 | 0,040355 | 0 | -inf | 1 | 1 |
| LOC102631757 | 0,008739 | 0 | -inf | 1 | 1 |
| Lair1 | 0,190723 | 0 | -inf | 1 | 1 |
| Lalba | 0,20123 | 0 | -inf | 1 | 1 |
| Lbx1 | 0,017921 | 0 | -inf | 1 | 1 |
| Lcn2 | 0,191738 | 0 | -inf | 1 | 1 |
| Lcn4 | 0,020751 | 0 | -inf | 1 | 1 |
| Lcn8 | 0,026401 | 0 | -inf | 1 | 1 |
| Ldlrad1 | 0,071746 | 0 | -inf | 1 | 1 |
| Ldlrad2 | 0,040108 | 0 | -inf | 1 | 1 |
| Lect2 | 0,017575 | 0 | -inf | 1 | 1 |
| Lgals7 | 0,647689 | 0 | -inf | 1 | 1 |
| Lgi3 | 0,081532 | 0 | -inf | 1 | 1 |
| Lhcgr | 0,017659 | 0 | -inf | 1 | 1 |
| Lhfpl5 | 0,01134 | 0 | -inf | 1 | 1 |
| Lhx3 | 0,012191 | 0 | -inf | 1 | 1 |
| Lhx8 | 0,046791 | 0 | -inf | 1 | 1 |
| Lilra6 | 0,019969 | 0 | -inf | 1 | 1 |
| Lilrb4 | 0,036577 | 0 | -inf | 1 | 1 |
| Lipk | 0,020503 | 0 | -inf | 1 | 1 |
| Lipm | 0,061645 | 0 | -inf | 1 | 1 |
| Loxhd1 | 0,005552 | 0 | -inf | 1 | 1 |
| Lrrc2 | 0,029157 | 0 | -inf | 1 | 1 |
| Lrrc25 | 0,520798 | 0 | -inf | 1 | 1 |
| Lrrc43 | 0,074824 | 0 | -inf | 1 | 1 |
| Lrrc4c | 0,077738 | 0 | -inf | 1 | 1 |
| Lrriq4 | 0,005855 | 0 | -inf | 1 | 1 |
| Lrtm2 | 0,01996 | 0 | -inf | 1 | 1 |
| Ltb4r1 | 0,134478 | 0 | -inf | 1 | 1 |

| | | | | | |
|---|---|---|---|---|---|
| Ltc4s | 0,123206 | 0 | -inf | 1 | 1 |
| Ltf | 0,032687 | 0 | -inf | 1 | 1 |
| Ly6c2 | 0,109104 | 0 | -inf | 1 | 1 |
| Ly6f | 0,079209 | 0 | -inf | 1 | 1 |
| Ly6g6f | 0,568788 | 0 | -inf | 1 | 1 |
| Ly6k | 0,072899 | 0 | -inf | 1 | 1 |
| Ly9 | 0,021109 | 0 | -inf | 1 | 1 |
| Lyg2 | 0,017576 | 0 | -inf | 1 | 1 |
| Lypd3 | 0,220247 | 0 | -inf | 1 | 1 |
| Lypd4 | 0,013275 | 0 | -inf | 1 | 1 |
| Lyrm7os | 0,057292 | 0 | -inf | 1 | 1 |
| Lyve1 | 0,110852 | 0 | -inf | 1 | 1 |
| Lyzl4 | 0,059737 | 0 | -inf | 1 | 1 |
| Map3k19 | 0,006339 | 0 | -inf | 1 | 1 |
| 03/11/16 | 0,097001 | 0 | -inf | 1 | 1 |
| Marco | 0,006641 | 0 | -inf | 1 | 1 |
| Mat1a | 0,003604 | 0 | -inf | 1 | 1 |
| Matn1 | 0,006578 | 0 | -inf | 1 | 1 |
| Mbl1 | 0,016451 | 0 | -inf | 1 | 1 |
| Mbl2 | 0,013674 | 0 | -inf | 1 | 1 |
| Mc2r | 0,029535 | 0 | -inf | 1 | 1 |
| Mc3r | 0,022037 | 0 | -inf | 1 | 1 |
| Mcemp1 | 0,127633 | 0 | -inf | 1 | 1 |
| Mctp1 | 0,084089 | 0 | -inf | 1 | 1 |
| Me3 | 0,042666 | 0 | -inf | 1 | 1 |
| Meiob | 0,067125 | 0 | -inf | 1 | 1 |
| Mettl11b | 0,013932 | 0 | -inf | 1 | 1 |
| Mettl7b | 0,011562 | 0 | -inf | 1 | 1 |
| Mfsd6l | 0,220084 | 0 | -inf | 1 | 1 |
| Mgat4c | 0,006689 | 0 | -inf | 1 | 1 |
| Mill1 | 0,015529 | 0 | -inf | 1 | 1 |
| Mill2 | 0,013574 | 0 | -inf | 1 | 1 |
| Mir1306 | 40,2487 | 0 | -inf | 1 | 1 |
| Mir147 | 0,036356 | 0 | -inf | 1 | 1 |
| Mir196b | 49,3569 | 0 | -inf | 1 | 1 |
| Mir19b-1 | 25,0412 | 0 | -inf | 1 | 1 |
| Mir3057 | 10,0567 | 0 | -inf | 1 | 1 |
| Mir3068 | 72,619 | 0 | -inf | 1 | 1 |
| Mir3101 | 41,492 | 0 | -inf | 1 | 1 |
| Mir330 | 24,9345 | 0 | -inf | 1 | 1 |
| Mir339 | 7,81552 | 0 | -inf | 1 | 1 |
| Mir431 | 8,36033 | 0 | -inf | 1 | 1 |
| Mir5131 | 3,5507 | 0 | -inf | 1 | 1 |
| Mir6373 | 5,93796 | 0 | -inf | 1 | 1 |
| Mir700 | 72,619 | 0 | -inf | 1 | 1 |
| Mir702 | 15,5767 | 0 | -inf | 1 | 1 |
| Mir7653 | 60,3495 | 0 | -inf | 1 | 1 |
| Mir8112 | 2,07353 | 0 | -inf | 1 | 1 |

| | | | | | |
|---|---|---|---|---|---|
| Mirlet7bhg | 0,009205 | 0 | -inf | 1 | 1 |
| Mlkl | 0,130446 | 0 | -inf | 1 | 1 |
| Mmp13 | 0,057507 | 0 | -inf | 1 | 1 |
| Mmp20 | 0,00385 | 0 | -inf | 1 | 1 |
| Mmp21 | 0,039381 | 0 | -inf | 1 | 1 |
| Mocos | 0,039782 | 0 | -inf | 1 | 1 |
| Mospd4 | 0,061089 | 0 | -inf | 1 | 1 |
| Mrgprd | 0,00406 | 0 | -inf | 1 | 1 |
| Mrgprg | 0,084272 | 0 | -inf | 1 | 1 |
| Mro | 0,026828 | 0 | -inf | 1 | 1 |
| Ms4a15 | 0,015677 | 0 | -inf | 1 | 1 |
| Ms4a2 | 0,029286 | 0 | -inf | 1 | 1 |
| Ms4a3 | 0,029277 | 0 | -inf | 1 | 1 |
| Ms4a4c | 0,062401 | 0 | -inf | 1 | 1 |
| Muc15 | 0,017653 | 0 | -inf | 1 | 1 |
| Muc19 | 0,007319 | 0 | -inf | 1 | 1 |
| Muc4 | 0,005881 | 0 | -inf | 1 | 1 |
| Mug1 | 0,009315 | 0 | -inf | 1 | 1 |
| Musk | 0,013288 | 0 | -inf | 1 | 1 |
| Myadml2 | 0,078614 | 0 | -inf | 1 | 1 |
| Mybpc2 | 0,081347 | 0 | -inf | 1 | 1 |
| Mybpc3 | 0,005995 | 0 | -inf | 1 | 1 |
| Mybph | 0,040084 | 0 | -inf | 1 | 1 |
| Myf6 | 0,033989 | 0 | -inf | 1 | 1 |
| Myh15 | 0,007711 | 0 | -inf | 1 | 1 |
| Myh7 | 0,024291 | 0 | -inf | 1 | 1 |
| Myoc | 0,01261 | 0 | -inf | 1 | 1 |
| Myoz1 | 0,032554 | 0 | -inf | 1 | 1 |
| Myoz3 | 0,016695 | 0 | -inf | 1 | 1 |
| Naip1 | 0,002294 | 0 | -inf | 1 | 1 |
| Naip6 | 0,012735 | 0 | -inf | 1 | 1 |
| Naip7 | 0,035775 | 0 | -inf | 1 | 1 |
| Nat8 | 0,016247 | 0 | -inf | 1 | 1 |
| Nccrp1 | 0,049085 | 0 | -inf | 1 | 1 |
| Ndst4 | 0,025254 | 0 | -inf | 1 | 1 |
| Neu2 | 0,017146 | 0 | -inf | 1 | 1 |
| Neurod6 | 0,021283 | 0 | -inf | 1 | 1 |
| Ngp | 0,043343 | 0 | -inf | 1 | 1 |
| Nipal4 | 0,16853 | 0 | -inf | 1 | 1 |
| Nkain2 | 0,011824 | 0 | -inf | 1 | 1 |
| Nkx2-1 | 0,017217 | 0 | -inf | 1 | 1 |
| Nkx2-2os | 0,003296 | 0 | -inf | 1 | 1 |
| Nlrp1a | 0,015421 | 0 | -inf | 1 | 1 |
| Nlrp1b | 0,035956 | 0 | -inf | 1 | 1 |
| Nlrp1c-ps | 0,010908 | 0 | -inf | 1 | 1 |
| Nlrp4c | 0,003469 | 0 | -inf | 1 | 1 |
| Nnmt | 0,118388 | 0 | -inf | 1 | 1 |
| Nov | 0,208293 | 0 | -inf | 1 | 1 |

| | | | | | |
|---|---|---|---|---|---|
| Nox1 | 0,017236 | 0 | -inf | 1 | 1 |
| Nox3 | 0,007448 | 0 | -inf | 1 | 1 |
| Npffr1 | 0,010724 | 0 | -inf | 1 | 1 |
| Npffr2 | 0,00822 | 0 | -inf | 1 | 1 |
| Npsr1 | 0,014862 | 0 | -inf | 1 | 1 |
| Npy | 0,156324 | 0 | -inf | 1 | 1 |
| Npy4r | 0,006962 | 0 | -inf | 1 | 1 |
| Npy5r | 0,022423 | 0 | -inf | 1 | 1 |
| Nr0b2 | 0,038823 | 0 | -inf | 1 | 1 |
| Nr1h4 | 0,02003 | 0 | -inf | 1 | 1 |
| Nr1i2 | 0,016127 | 0 | -inf | 1 | 1 |
| Nr2e3 | 0,036288 | 0 | -inf | 1 | 1 |
| Nr5a1 | 0,023746 | 0 | -inf | 1 | 1 |
| Nrxn1 | 0,02353 | 0 | -inf | 1 | 1 |
| Nt5c1b | 0,053781 | 0 | -inf | 1 | 1 |
| Ntrk1 | 0,052198 | 0 | -inf | 1 | 1 |
| Nts | 0,242712 | 0 | -inf | 1 | 1 |
| Nupr1 | 0,238431 | 0 | -inf | 1 | 1 |
| Nxf7 | 0,085654 | 0 | -inf | 1 | 1 |
| Nxnl1 | 0,016115 | 0 | -inf | 1 | 1 |
| Nxpe5 | 0,005611 | 0 | -inf | 1 | 1 |
| Oas1d | 0,015462 | 0 | -inf | 1 | 1 |
| Oas1g | 0,043021 | 0 | -inf | 1 | 1 |
| Oas2 | 0,032362 | 0 | -inf | 1 | 1 |
| Ocm | 0,022275 | 0 | -inf | 1 | 1 |
| Odf4 | 0,013055 | 0 | -inf | 1 | 1 |
| Olfm3 | 0,002897 | 0 | -inf | 1 | 1 |
| Olfr1033 | 0,003342 | 0 | -inf | 1 | 1 |
| Olfr1362 | 0,015145 | 0 | -inf | 1 | 1 |
| Olfr1378 | 0,055752 | 0 | -inf | 1 | 1 |
| Olfr1393 | 0,113332 | 0 | -inf | 1 | 1 |
| Olfr1535 | 0,015223 | 0 | -inf | 1 | 1 |
| Olfr31 | 0,031541 | 0 | -inf | 1 | 1 |
| Olfr329-ps | 0,010792 | 0 | -inf | 1 | 1 |
| Olfr330 | 0,013472 | 0 | -inf | 1 | 1 |
| Olfr374 | 0,013502 | 0 | -inf | 1 | 1 |
| Olfr419 | 0,01338 | 0 | -inf | 1 | 1 |
| Olfr420 | 0,042142 | 0 | -inf | 1 | 1 |
| Olfr421-ps1 | 0,013981 | 0 | -inf | 1 | 1 |
| Olfr460 | 0,055978 | 0 | -inf | 1 | 1 |
| Olfr461 | 0,056205 | 0 | -inf | 1 | 1 |
| Olfr463 | 0,056666 | 0 | -inf | 1 | 1 |
| Olfr545 | 0,047502 | 0 | -inf | 1 | 1 |
| Olfr631 | 0,0092 | 0 | -inf | 1 | 1 |
| Olfr632 | 0,031541 | 0 | -inf | 1 | 1 |
| Olfr646 | 0,056435 | 0 | -inf | 1 | 1 |
| Olfr658 | 0,021839 | 0 | -inf | 1 | 1 |
| Olfr666 | 0,015708 | 0 | -inf | 1 | 1 |

| | | | | | |
|---|---|---|---|---|---|
| Olfr70 | 0,019311 | 0 | -inf | 1 | 1 |
| Olfr701 | 0,004137 | 0 | -inf | 1 | 1 |
| Olfr725 | 0,02611 | 0 | -inf | 1 | 1 |
| Olfr727 | 0,028467 | 0 | -inf | 1 | 1 |
| Olfr875 | 0,014785 | 0 | -inf | 1 | 1 |
| Olfr876 | 0,014014 | 0 | -inf | 1 | 1 |
| Olfr90 | 0,070237 | 0 | -inf | 1 | 1 |
| Olfr921 | 0,008988 | 0 | -inf | 1 | 1 |
| Olfr922 | 0,012746 | 0 | -inf | 1 | 1 |
| Olr1 | 0,022872 | 0 | -inf | 1 | 1 |
| Opalin | 0,051575 | 0 | -inf | 1 | 1 |
| Opn4 | 0,012116 | 0 | -inf | 1 | 1 |
| Orm1 | 0,020971 | 0 | -inf | 1 | 1 |
| Orm3 | 0,062361 | 0 | -inf | 1 | 1 |
| Osm | 0,125209 | 0 | -inf | 1 | 1 |
| Otc | 0,011478 | 0 | -inf | 1 | 1 |
| Otogl | 0,006118 | 0 | -inf | 1 | 1 |
| Otor | 0,266255 | 0 | -inf | 1 | 1 |
| Otos | 0,070967 | 0 | -inf | 1 | 1 |
| P2ry10 | 0,009019 | 0 | -inf | 1 | 1 |
| P2ry12 | 0,3883 | 0 | -inf | 1 | 1 |
| P2ry4 | 0,009806 | 0 | -inf | 1 | 1 |
| Padi2 | 0,323777 | 0 | -inf | 1 | 1 |
| Padi6 | 0,005556 | 0 | -inf | 1 | 1 |
| Pah | 0,0061 | 0 | -inf | 1 | 1 |
| Paqr6 | 0,213976 | 0 | -inf | 1 | 1 |
| Pbp2 | 0,123832 | 0 | -inf | 1 | 1 |
| Pcdh20 | 0,011751 | 0 | -inf | 1 | 1 |
| Pcdha10 | 0,034046 | 0 | -inf | 1 | 1 |
| Pcdha2 | 0,019092 | 0 | -inf | 1 | 1 |
| Pcdha7 | 0,00702 | 0 | -inf | 1 | 1 |
| Pcdhb11 | 0,020806 | 0 | -inf | 1 | 1 |
| Pcdhb12 | 0,014657 | 0 | -inf | 1 | 1 |
| Pcdhb14 | 0,029259 | 0 | -inf | 1 | 1 |
| Pcdhb18 | 0,053854 | 0 | -inf | 1 | 1 |
| Pcdhb3 | 0,13553 | 0 | -inf | 1 | 1 |
| Pcdhb9 | 0,020744 | 0 | -inf | 1 | 1 |
| Pdcd1 | 0,092656 | 0 | -inf | 1 | 1 |
| Pde6b | 0,004528 | 0 | -inf | 1 | 1 |
| Pdzd3 | 0,026964 | 0 | -inf | 1 | 1 |
| Peril | 0,012503 | 0 | -inf | 1 | 1 |
| Pga5 | 0,034224 | 0 | -inf | 1 | 1 |
| Pglyrp2 | 0,005462 | 0 | -inf | 1 | 1 |
| Pgr15l | 0,002286 | 0 | -inf | 1 | 1 |
| Phf11a | 0,061711 | 0 | -inf | 1 | 1 |
| Pik3c2g | 0,012404 | 0 | -inf | 1 | 1 |
| Pilrb1 | 0,043485 | 0 | -inf | 1 | 1 |
| Pilrb2 | 0,007379 | 0 | -inf | 1 | 1 |

| | | | | | |
|---|---|---|---|---|---|
| Pinc | 0,065481 | 0 | -inf | 1 | 1 |
| Pirt | 0,018971 | 0 | -inf | 1 | 1 |
| Pitx3 | 0,035332 | 0 | -inf | 1 | 1 |
| Pkd2l1 | 0,017102 | 0 | -inf | 1 | 1 |
| Pkhd1 | 0,001859 | 0 | -inf | 1 | 1 |
| Pkib | 0,017159 | 0 | -inf | 1 | 1 |
| Pla2g12b | 0,068608 | 0 | -inf | 1 | 1 |
| Pla2g2d | 0,020136 | 0 | -inf | 1 | 1 |
| Pla2g2f | 0,087509 | 0 | -inf | 1 | 1 |
| Pla2g4e | 0,022183 | 0 | -inf | 1 | 1 |
| Plekhd1os | 0,136897 | 0 | -inf | 1 | 1 |
| Plet1os | 0,080853 | 0 | -inf | 1 | 1 |
| Plscr5 | 0,045432 | 0 | -inf | 1 | 1 |
| Plxnb3 | 0,011196 | 0 | -inf | 1 | 1 |
| Pmis2 | 0,114897 | 0 | -inf | 1 | 1 |
| Pmp2 | 0,010197 | 0 | -inf | 1 | 1 |
| Pnma5 | 0,036949 | 0 | -inf | 1 | 1 |
| Podnl1 | 0,018915 | 0 | -inf | 1 | 1 |
| Pou2f3 | 0,010205 | 0 | -inf | 1 | 1 |
| Pou4f3 | 0,069146 | 0 | -inf | 1 | 1 |
| Pp2d1 | 0,057338 | 0 | -inf | 1 | 1 |
| Pparg | 0,102077 | 0 | -inf | 1 | 1 |
| Ppp1r1c | 0,01153 | 0 | -inf | 1 | 1 |
| Ppp1r27 | 0,093358 | 0 | -inf | 1 | 1 |
| Ppp2r2b | 0,199015 | 0 | -inf | 1 | 1 |
| Ppp2r2cos | 0,03223 | 0 | -inf | 1 | 1 |
| Pramel5 | 0,006088 | 0 | -inf | 1 | 1 |
| Prf1 | 0,176132 | 0 | -inf | 1 | 1 |
| Prima1 | 0,245007 | 0 | -inf | 1 | 1 |
| Prl2a1 | 0,302404 | 0 | -inf | 1 | 1 |
| Prl2b1 | 0,529538 | 0 | -inf | 1 | 1 |
| Prl2c1 | 0,073807 | 0 | -inf | 1 | 1 |
| Prl2c2 | 0,090968 | 0 | -inf | 1 | 1 |
| Prl2c4 | 0,000794 | 0 | -inf | 1 | 1 |
| Prl6a1 | 0,20522 | 0 | -inf | 1 | 1 |
| Prl7a2 | 0,088339 | 0 | -inf | 1 | 1 |
| Prl7d1 | 0,560256 | 0 | -inf | 1 | 1 |
| Prl8a1 | 0,125695 | 0 | -inf | 1 | 1 |
| Prl8a6 | 0,050823 | 0 | -inf | 1 | 1 |
| Prl8a9 | 0,559775 | 0 | -inf | 1 | 1 |
| Prlh | 0,176295 | 0 | -inf | 1 | 1 |
| Prm1 | 0,120642 | 0 | -inf | 1 | 1 |
| Prm2 | 0,057979 | 0 | -inf | 1 | 1 |
| Prm3 | 0,05825 | 0 | -inf | 1 | 1 |
| Prok1 | 0,253803 | 0 | -inf | 1 | 1 |
| Prok2 | 0,010559 | 0 | -inf | 1 | 1 |
| Prph2 | 0,004872 | 0 | -inf | 1 | 1 |
| Prps1l1 | 0,016071 | 0 | -inf | 1 | 1 |

| | | | | | |
|---|---|---|---|---|---|
| Prr15l | 0,091928 | 0 | -inf | 1 | 1 |
| Prss1 | 0,176807 | 0 | -inf | 1 | 1 |
| Prss27 | 0,05641 | 0 | -inf | 1 | 1 |
| Prss32 | 0,018834 | 0 | -inf | 1 | 1 |
| Prss33 | 0,020065 | 0 | -inf | 1 | 1 |
| Prss43 | 0,192116 | 0 | -inf | 1 | 1 |
| Prss45 | 0,011506 | 0 | -inf | 1 | 1 |
| Prss48 | 0,015876 | 0 | -inf | 1 | 1 |
| Prss57 | 0,00944 | 0 | -inf | 1 | 1 |
| Psapl1 | 0,03029 | 0 | -inf | 1 | 1 |
| Psca | 0,018141 | 0 | -inf | 1 | 1 |
| Psg16 | 0,005689 | 0 | -inf | 1 | 1 |
| Psg17 | 0,050419 | 0 | -inf | 1 | 1 |
| Psg19 | 0,006669 | 0 | -inf | 1 | 1 |
| Psg22 | 0,063957 | 0 | -inf | 1 | 1 |
| Psg29 | 0,179852 | 0 | -inf | 1 | 1 |
| Ptafr | 0,120921 | 0 | -inf | 1 | 1 |
| Ptcra | 0,05481 | 0 | -inf | 1 | 1 |
| Rab11fip4os1 | 0,005965 | 0 | -inf | 1 | 1 |
| Rab44 | 0,098689 | 0 | -inf | 1 | 1 |
| Rag2 | 0,003709 | 0 | -inf | 1 | 1 |
| Rarres2 | 0,042275 | 0 | -inf | 1 | 1 |
| Rasal1 | 0,040774 | 0 | -inf | 1 | 1 |
| Rbakdn | 0,037393 | 0 | -inf | 1 | 1 |
| Rbbp8nl | 0,313565 | 0 | -inf | 1 | 1 |
| Rcvrn | 0,027571 | 0 | -inf | 1 | 1 |
| Rdh9 | 0,040345 | 0 | -inf | 1 | 1 |
| Rem1 | 0,016909 | 0 | -inf | 1 | 1 |
| Rgs1 | 0,010505 | 0 | -inf | 1 | 1 |
| Rgs6 | 0,078998 | 0 | -inf | 1 | 1 |
| Rhbdl2 | 0,099328 | 0 | -inf | 1 | 1 |
| Rhox6 | 0,393915 | 0 | -inf | 1 | 1 |
| Rhox9 | 0,314497 | 0 | -inf | 1 | 1 |
| Rnase10 | 0,049453 | 0 | -inf | 1 | 1 |
| Rnase11 | 0,081491 | 0 | -inf | 1 | 1 |
| Rnase12 | 0,191009 | 0 | -inf | 1 | 1 |
| Rnf133 | 0,010714 | 0 | -inf | 1 | 1 |
| Rnf148 | 0,011395 | 0 | -inf | 1 | 1 |
| Rnf186 | 0,050721 | 0 | -inf | 1 | 1 |
| Rnf222 | 0,030823 | 0 | -inf | 1 | 1 |
| Rnf224 | 0,029338 | 0 | -inf | 1 | 1 |
| Robo3 | 0,030135 | 0 | -inf | 1 | 1 |
| Ros1 | 0,001643 | 0 | -inf | 1 | 1 |
| Rph3a | 0,02415 | 0 | -inf | 1 | 1 |
| Rpl39l | 0,128058 | 0 | -inf | 1 | 1 |
| Rptoros | 0,062302 | 0 | -inf | 1 | 1 |
| Rrh | 0,049698 | 0 | -inf | 1 | 1 |
| Rtdr1 | 0,024021 | 0 | -inf | 1 | 1 |

| | | | | | |
|---|---|---|---|---|---|
| Rtp3 | 0,105237 | 0 | -inf | 1 | 1 |
| S100a2 | 0,31259 | 0 | -inf | 1 | 1 |
| Satl1 | 0,005357 | 0 | -inf | 1 | 1 |
| Scarletltr | 0,620119 | 0 | -inf | 1 | 1 |
| Scarna2 | 120,153 | 0 | -inf | 1 | 1 |
| Scgb1a1 | 0,157114 | 0 | -inf | 1 | 1 |
| Scgb1b30 | 0,041211 | 0 | -inf | 1 | 1 |
| Scgb1c1 | 0,050762 | 0 | -inf | 1 | 1 |
| Scimp | 0,027798 | 0 | -inf | 1 | 1 |
| Scin | 0,077147 | 0 | -inf | 1 | 1 |
| Scn11a | 0,0526 | 0 | -inf | 1 | 1 |
| Scnn1g | 0,192898 | 0 | -inf | 1 | 1 |
| Sctr | 0,020654 | 0 | -inf | 1 | 1 |
| Sdr9c7 | 0,004472 | 0 | -inf | 1 | 1 |
| Sec14l3 | 0,003176 | 0 | -inf | 1 | 1 |
| Sectm1a | 0,008383 | 0 | -inf | 1 | 1 |
| Sel1l2 | 0,00587 | 0 | -inf | 1 | 1 |
| Serpina10 | 0,034636 | 0 | -inf | 1 | 1 |
| Serpina1a | 0,023344 | 0 | -inf | 1 | 1 |
| Serpina1b | 0,397441 | 0 | -inf | 1 | 1 |
| Serpina1c | 0,01939 | 0 | -inf | 1 | 1 |
| Serpina1d | 0,009554 | 0 | -inf | 1 | 1 |
| Serpina5 | 0,006041 | 0 | -inf | 1 | 1 |
| Serpina7 | 0,079736 | 0 | -inf | 1 | 1 |
| Serpinb10 | 0,046859 | 0 | -inf | 1 | 1 |
| Serpinb2 | 0,120576 | 0 | -inf | 1 | 1 |
| Serpinb5 | 0,017569 | 0 | -inf | 1 | 1 |
| Serpinb8 | 0,242056 | 0 | -inf | 1 | 1 |
| Serpinb9c | 0,038729 | 0 | -inf | 1 | 1 |
| Serpinb9d | 0,034336 | 0 | -inf | 1 | 1 |
| Serpinb9e | 0,079364 | 0 | -inf | 1 | 1 |
| Serpinb9f | 0,104312 | 0 | -inf | 1 | 1 |
| Serpinb9g | 0,027523 | 0 | -inf | 1 | 1 |
| Serpinf2 | 0,357148 | 0 | -inf | 1 | 1 |
| Sfta2 | 0,078329 | 0 | -inf | 1 | 1 |
| Sftpc | 0,10874 | 0 | -inf | 1 | 1 |
| Sgk2 | 0,050376 | 0 | -inf | 1 | 1 |
| Sh2d1a | 0,067446 | 0 | -inf | 1 | 1 |
| Sh2d1b1 | 0,077126 | 0 | -inf | 1 | 1 |
| Sh2d2a | 0,010428 | 0 | -inf | 1 | 1 |
| Siah3 | 0,050619 | 0 | -inf | 1 | 1 |
| Siglec1 | 0,005739 | 0 | -inf | 1 | 1 |
| Six3os1 | 0,197496 | 0 | -inf | 1 | 1 |
| Skint4 | 0,003118 | 0 | -inf | 1 | 1 |
| Slamf6 | 0,023809 | 0 | -inf | 1 | 1 |
| Slamf9 | 0,207485 | 0 | -inf | 1 | 1 |
| Slc10a6 | 0,021604 | 0 | -inf | 1 | 1 |
| Slc11a1 | 0,498897 | 0 | -inf | 1 | 1 |

| | | | | | |
|---|---|---|---|---|---|
| Slc13a2os | 0,003455 | 0 | -inf | 1 | 1 |
| Slc13a3 | 0,121468 | 0 | -inf | 1 | 1 |
| Slc15a1 | 0,050735 | 0 | -inf | 1 | 1 |
| Slc16a14 | 0,041395 | 0 | -inf | 1 | 1 |
| Slc17a4 | 0,018201 | 0 | -inf | 1 | 1 |
| Slc18a3 | 0,010704 | 0 | -inf | 1 | 1 |
| Slc22a20 | 0,006986 | 0 | -inf | 1 | 1 |
| Slc22a28 | 0,006571 | 0 | -inf | 1 | 1 |
| Slc22a6 | 0,009284 | 0 | -inf | 1 | 1 |
| Slc22a8 | 0,011433 | 0 | -inf | 1 | 1 |
| Slc23a3 | 0,076645 | 0 | -inf | 1 | 1 |
| Slc24a1 | 0,042537 | 0 | -inf | 1 | 1 |
| Slc25a31 | 0,012411 | 0 | -inf | 1 | 1 |
| Slc26a3 | 0,004371 | 0 | -inf | 1 | 1 |
| Slc26a5 | 0,089995 | 0 | -inf | 1 | 1 |
| Slc28a1 | 0,022385 | 0 | -inf | 1 | 1 |
| Slc2a7 | 0,008608 | 0 | -inf | 1 | 1 |
| Slc32a1 | 0,031833 | 0 | -inf | 1 | 1 |
| Slc34a2 | 0,077587 | 0 | -inf | 1 | 1 |
| Slc34a3 | 0,006394 | 0 | -inf | 1 | 1 |
| Slc35f4 | 0,007053 | 0 | -inf | 1 | 1 |
| Slc38a8 | 0,03183 | 0 | -inf | 1 | 1 |
| Slc45a2 | 0,014926 | 0 | -inf | 1 | 1 |
| Slc47a2 | 0,043546 | 0 | -inf | 1 | 1 |
| Slc51a | 0,066263 | 0 | -inf | 1 | 1 |
| Slc51b | 0,187697 | 0 | -inf | 1 | 1 |
| Slc5a12 | 0,010893 | 0 | -inf | 1 | 1 |
| Slc6a14 | 0,036693 | 0 | -inf | 1 | 1 |
| Slc6a2 | 0,124125 | 0 | -inf | 1 | 1 |
| Slc6a20a | 0,006999 | 0 | -inf | 1 | 1 |
| Slc6a20b | 0,007673 | 0 | -inf | 1 | 1 |
| Slc6a3 | 0,058245 | 0 | -inf | 1 | 1 |
| Slc6a7 | 0,020751 | 0 | -inf | 1 | 1 |
| Slc7a15 | 0,006945 | 0 | -inf | 1 | 1 |
| Slc9a3 | 0,005174 | 0 | -inf | 1 | 1 |
| Slco1a5 | 0,06253 | 0 | -inf | 1 | 1 |
| Slco1b2 | 0,007651 | 0 | -inf | 1 | 1 |
| Slco6d1 | 0,020832 | 0 | -inf | 1 | 1 |
| Slfn1 | 0,025094 | 0 | -inf | 1 | 1 |
| Slfn3 | 0,095489 | 0 | -inf | 1 | 1 |
| Slfn4 | 0,022099 | 0 | -inf | 1 | 1 |
| Slfn5 | 0,068565 | 0 | -inf | 1 | 1 |
| Slfn5os | 0,112105 | 0 | -inf | 1 | 1 |
| Slitrk6 | 0,574851 | 0 | -inf | 1 | 1 |
| Snai3 | 0,04703 | 0 | -inf | 1 | 1 |
| Snord70 | 1280,71 | 0 | -inf | 1 | 1 |
| Sox14 | 0,025423 | 0 | -inf | 1 | 1 |
| Sp110 | 0,200858 | 0 | -inf | 1 | 1 |

| | | | | | |
|---|---|---|---|---|---|
| Sp140 | 0,123134 | 0 | -inf | 1 | 1 |
| Spata19 | 0,072121 | 0 | -inf | 1 | 1 |
| Spata4 | 0,013866 | 0 | -inf | 1 | 1 |
| Spatc1l | 0,068291 | 0 | -inf | 1 | 1 |
| Spem1 | 0,107366 | 0 | -inf | 1 | 1 |
| Spesp1 | 0,012632 | 0 | -inf | 1 | 1 |
| Sphkap | 0,016902 | 0 | -inf | 1 | 1 |
| Spink4 | 0,062541 | 0 | -inf | 1 | 1 |
| Spns3 | 0,033008 | 0 | -inf | 1 | 1 |
| Sppl2c | 0,00586 | 0 | -inf | 1 | 1 |
| Sprr2g | 0,119255 | 0 | -inf | 1 | 1 |
| Sprr2h | 0,185301 | 0 | -inf | 1 | 1 |
| Sptssb | 0,017367 | 0 | -inf | 1 | 1 |
| Sst | 0,059887 | 0 | -inf | 1 | 1 |
| Sstr3 | 0,017491 | 0 | -inf | 1 | 1 |
| Sstr4 | 0,065003 | 0 | -inf | 1 | 1 |
| Sstr5 | 0,004974 | 0 | -inf | 1 | 1 |
| Ssu2 | 0,259798 | 0 | -inf | 1 | 1 |
| St18 | 0,002145 | 0 | -inf | 1 | 1 |
| St6galnac1 | 0,005406 | 0 | -inf | 1 | 1 |
| Stfa1 | 0,913 | 0 | -inf | 1 | 1 |
| Stmnd1 | 0,010205 | 0 | -inf | 1 | 1 |
| Stpg2 | 0,013575 | 0 | -inf | 1 | 1 |
| Stra8 | 0,009455 | 0 | -inf | 1 | 1 |
| Sult1d1 | 0,005401 | 0 | -inf | 1 | 1 |
| Sult1e1 | 0,03928 | 0 | -inf | 1 | 1 |
| Svop | 0,029776 | 0 | -inf | 1 | 1 |
| Syce1 | 0,178902 | 0 | -inf | 1 | 1 |
| Syne1 | 0,0393 | 0 | -inf | 1 | 1 |
| Syt8 | 0,065318 | 0 | -inf | 1 | 1 |
| Tac2 | 0,086957 | 0 | -inf | 1 | 1 |
| Taf7l | 0,027332 | 0 | -inf | 1 | 1 |
| Tarm1 | 0,015708 | 0 | -inf | 1 | 1 |
| Tas2r108 | 0,060115 | 0 | -inf | 1 | 1 |
| Tcp10b | 0,012712 | 0 | -inf | 1 | 1 |
| Tdgf1 | 0,01373 | 0 | -inf | 1 | 1 |
| Tdo2 | 0,115994 | 0 | -inf | 1 | 1 |
| Tectb | 0,004692 | 0 | -inf | 1 | 1 |
| Tenm1 | 0,00145 | 0 | -inf | 1 | 1 |
| Tespa1 | 0,035777 | 0 | -inf | 1 | 1 |
| Tex19.2 | 0,035362 | 0 | -inf | 1 | 1 |
| Tex21 | 0,023343 | 0 | -inf | 1 | 1 |
| Tex43 | 0,032855 | 0 | -inf | 1 | 1 |
| Tgm7 | 0,021615 | 0 | -inf | 1 | 1 |
| Tgtp1 | 0,029545 | 0 | -inf | 1 | 1 |
| Themis | 0,003132 | 0 | -inf | 1 | 1 |
| Timd2 | 0,125402 | 0 | -inf | 1 | 1 |
| Timd4 | 0,105143 | 0 | -inf | 1 | 1 |

| | | | | | |
|---|---|---|---|---|---|
| Timm8a2 | 0,074975 | 0 | -inf | 1 | 1 |
| Tinag | 0,007864 | 0 | -inf | 1 | 1 |
| Tlr1 | 0,119927 | 0 | -inf | 1 | 1 |
| Tlr8 | 0,018046 | 0 | -inf | 1 | 1 |
| Tlr9 | 0,083371 | 0 | -inf | 1 | 1 |
| Tm4sf19 | 0,011233 | 0 | -inf | 1 | 1 |
| Tm4sf20 | 0,068354 | 0 | -inf | 1 | 1 |
| Tmc1 | 0,003057 | 0 | -inf | 1 | 1 |
| Tmco5 | 0,012033 | 0 | -inf | 1 | 1 |
| Tmem151a | 0,003162 | 0 | -inf | 1 | 1 |
| Tmem163 | 0,138212 | 0 | -inf | 1 | 1 |
| Tmem190 | 0,031044 | 0 | -inf | 1 | 1 |
| Tmem213 | 0,036281 | 0 | -inf | 1 | 1 |
| Tmem217 | 0,066577 | 0 | -inf | 1 | 1 |
| Tmem235 | 0,014639 | 0 | -inf | 1 | 1 |
| Tmem255b | 0,012368 | 0 | -inf | 1 | 1 |
| Tmem27 | 0,01118 | 0 | -inf | 1 | 1 |
| Tmem51os1 | 0,022834 | 0 | -inf | 1 | 1 |
| Tmem52 | 0,033596 | 0 | -inf | 1 | 1 |
| Tmem71 | 0,004337 | 0 | -inf | 1 | 1 |
| Tmem72 | 0,054856 | 0 | -inf | 1 | 1 |
| Tmprss3 | 0,009345 | 0 | -inf | 1 | 1 |
| Tmsb15a | 0,119523 | 0 | -inf | 1 | 1 |
| Tnfrsf13b | 0,260252 | 0 | -inf | 1 | 1 |
| Tnfsf10 | 0,002495 | 0 | -inf | 1 | 1 |
| Tnfsf14 | 0,224019 | 0 | -inf | 1 | 1 |
| Tnfsf15 | 0,046356 | 0 | -inf | 1 | 1 |
| Tnfsf18 | 0,006359 | 0 | -inf | 1 | 1 |
| Tnni3k | 0,014616 | 0 | -inf | 1 | 1 |
| Tnp2 | 0,154065 | 0 | -inf | 1 | 1 |
| Tomm20l | 0,060346 | 0 | -inf | 1 | 1 |
| Topaz1 | 0,007447 | 0 | -inf | 1 | 1 |
| Tpbpa | 0,385375 | 0 | -inf | 1 | 1 |
| Tpbpb | 0,081975 | 0 | -inf | 1 | 1 |
| Tph2 | 0,039136 | 0 | -inf | 1 | 1 |
| Tpte | 0,005044 | 0 | -inf | 1 | 1 |
| Trat1 | 0,024641 | 0 | -inf | 1 | 1 |
| Trem1 | 0,035668 | 0 | -inf | 1 | 1 |
| Trem2 | 0,400407 | 0 | -inf | 1 | 1 |
| Trem3 | 0,014014 | 0 | -inf | 1 | 1 |
| Treml4 | 0,152461 | 0 | -inf | 1 | 1 |
| Trhr | 0,025129 | 0 | -inf | 1 | 1 |
| Trhr2 | 0,010412 | 0 | -inf | 1 | 1 |
| Trim30e-ps1 | 0,007015 | 0 | -inf | 1 | 1 |
| Trim34b | 0,008887 | 0 | -inf | 1 | 1 |
| Trim40 | 0,006252 | 0 | -inf | 1 | 1 |
| Trim43a | 0,043486 | 0 | -inf | 1 | 1 |
| Trim50 | 0,009478 | 0 | -inf | 1 | 1 |

| | | | | | |
|---|---|---|---|---|---|
| Trim55 | 0,022416 | 0 | -inf | 1 | 1 |
| Trim58 | 0,41086 | 0 | -inf | 1 | 1 |
| Triml1 | 0,057587 | 0 | -inf | 1 | 1 |
| Triml2 | 0,051572 | 0 | -inf | 1 | 1 |
| Trpc5os | 0,01401 | 0 | -inf | 1 | 1 |
| Trpc7 | 0,007546 | 0 | -inf | 1 | 1 |
| Trpm8 | 0,003249 | 0 | -inf | 1 | 1 |
| Trpv1 | 0,005107 | 0 | -inf | 1 | 1 |
| Trpv5 | 0,017312 | 0 | -inf | 1 | 1 |
| Tsga13 | 0,126407 | 0 | -inf | 1 | 1 |
| Tsix | 0,048688 | 0 | -inf | 1 | 1 |
| Tsks | 0,080143 | 0 | -inf | 1 | 1 |
| Tspo2 | 0,016181 | 0 | -inf | 1 | 1 |
| Tssk5 | 0,009029 | 0 | -inf | 1 | 1 |
| Ttc22 | 0,027943 | 0 | -inf | 1 | 1 |
| Ttc36 | 0,063364 | 0 | -inf | 1 | 1 |
| Ttll10 | 0,202711 | 0 | -inf | 1 | 1 |
| Tubal3 | 0,00818 | 0 | -inf | 1 | 1 |
| Ube2d2b | 0,008377 | 0 | -inf | 1 | 1 |
| Uchl1os | 0,016004 | 0 | -inf | 1 | 1 |
| Ugt1a7c | 0,156103 | 0 | -inf | 1 | 1 |
| Ugt2b34 | 0,008318 | 0 | -inf | 1 | 1 |
| Ugt3a2 | 0,011876 | 0 | -inf | 1 | 1 |
| Umod | 0,017446 | 0 | -inf | 1 | 1 |
| Uox | 0,032525 | 0 | -inf | 1 | 1 |
| Upk2 | 0,079563 | 0 | -inf | 1 | 1 |
| Upk3b | 0,181046 | 0 | -inf | 1 | 1 |
| Uroc1 | 0,086197 | 0 | -inf | 1 | 1 |
| Ush2a | 0,006797 | 0 | -inf | 1 | 1 |
| Usp17lb | 0,009832 | 0 | -inf | 1 | 1 |
| Usp17le | 0,007036 | 0 | -inf | 1 | 1 |
| Usp26 | 0,00301 | 0 | -inf | 1 | 1 |
| Vax2 | 0,011495 | 0 | -inf | 1 | 1 |
| Vax2os | 0,007216 | 0 | -inf | 1 | 1 |
| Vgf | 0,015106 | 0 | -inf | 1 | 1 |
| Vgll1 | 0,012033 | 0 | -inf | 1 | 1 |
| Vip | 0,00894 | 0 | -inf | 1 | 1 |
| Vmn1r21 | 0,017142 | 0 | -inf | 1 | 1 |
| Vmn1r223 | 0,01341 | 0 | -inf | 1 | 1 |
| Vmn1r4 | 0,017142 | 0 | -inf | 1 | 1 |
| Vmn1r51 | 0,005489 | 0 | -inf | 1 | 1 |
| Vmn2r-ps54 | 0,015095 | 0 | -inf | 1 | 1 |
| Vmn2r77 | 0,017569 | 0 | -inf | 1 | 1 |
| Vsx2 | 0,015898 | 0 | -inf | 1 | 1 |
| Vtcn1 | 0,009783 | 0 | -inf | 1 | 1 |
| Wdr27 | 0,009259 | 0 | -inf | 1 | 1 |
| Wdr38 | 0,053201 | 0 | -inf | 1 | 1 |
| Wdr63 | 0,00822 | 0 | -inf | 1 | 1 |

| | | | | | |
|---|---|---|---|---|---|
| Wfdc15a | 0,045143 | 0 | -inf | 1 | 1 |
| Wfdc17 | 0,044632 | 0 | -inf | 1 | 1 |
| Wfdc6a | 0,070035 | 0 | -inf | 1 | 1 |
| Wnt10b | 0,515164 | 0 | -inf | 1 | 1 |
| Wnt2 | 0,117771 | 0 | -inf | 1 | 1 |
| Xcr1 | 0,002784 | 0 | -inf | 1 | 1 |
| Xdh | 0,04431 | 0 | -inf | 1 | 1 |
| Xkr9 | 0,013738 | 0 | -inf | 1 | 1 |
| Xlr3c | 0,008571 | 0 | -inf | 1 | 1 |
| Xlr5a | 0,02454 | 0 | -inf | 1 | 1 |
| Zcchc13 | 0,013518 | 0 | -inf | 1 | 1 |
| Zdhhc11 | 0,010782 | 0 | -inf | 1 | 1 |
| Zfp36l3 | 0,066386 | 0 | -inf | 1 | 1 |
| Zfp534 | 0,026183 | 0 | -inf | 1 | 1 |
| Zfp804a | 0,002973 | 0 | -inf | 1 | 1 |
| Zfp804b | 0,020106 | 0 | -inf | 1 | 1 |
| Zfp936 | 0,004083 | 0 | -inf | 1 | 1 |
| Zic4 | 0,067124 | 0 | -inf | 1 | 1 |
| Zp1 | 0,024102 | 0 | -inf | 1 | 1 |
| Atcayos | 0,000153 | 0,348173 | 11,1555 | 1 | 1 |
| 4933416I08Rik | 0,00015 | 0,0510771 | 8,4069 | 1 | 1 |
| Gm16551 | 0,004726 | 0,218353 | 5,53003 | 1 | 1 |
| Gsx2 | 0,008096 | 0,22106 | 4,77111 | 1 | 1 |
| Gm6537 | 0,014016 | 0,249501 | 4,1539 | 1 | 1 |
| Arhgef33 | 0,004601 | 0,0793179 | 4,10754 | 1 | 1 |
| Pln | 0,011006 | 0,184313 | 4,06581 | 1 | 1 |
| Gpr87 | 0,012336 | 0,206546 | 4,06551 | 1 | 1 |
| Vaultrc5 | 0,083321 | 1,28334 | 3,94507 | 1 | 1 |
| Actrt3 | 0,007007 | 0,105067 | 3,90634 | 1 | 1 |
| Prdm13 | 0,003983 | 0,0576212 | 3,85484 | 1 | 1 |
| Rasgrf1 | 0,002939 | 0,0425286 | 3,85484 | 1 | 1 |
| Shc3 | 0,00888 | 0,128479 | 3,85484 | 1 | 1 |
| Spata31d1d | 0,003265 | 0,0472357 | 3,85484 | 1 | 1 |
| Aqp5 | 0,008781 | 0,123466 | 3,81358 | 1 | 1 |
| Csta1 | 0,004681 | 0,0658131 | 3,81358 | 1 | 1 |
| Cd3e | 0,009601 | 0,126026 | 3,71445 | 1 | 1 |
| 1700011L22Rik | 0,015282 | 0,194365 | 3,6689 | 1 | 1 |
| Gm17746 | 0,012984 | 0,165136 | 3,6689 | 1 | 1 |
| 4930578M01Rik | 0,03365 | 0,419142 | 3,63877 | 1 | 1 |
| Tbc1d22bos | 0,021186 | 0,256469 | 3,59763 | 1 | 1 |
| Pou3f4 | 0,026821 | 0,321557 | 3,58365 | 1 | 1 |
| LOC100504039 | 0,011496 | 0,137126 | 3,57625 | 1 | 1 |
| 4930592I03Rik | 0,030212 | 0,481217 | 3,53371 | 1 | 1 |
| Trim38 | 0,00927 | 0,105462 | 3,50807 | 1 | 1 |
| Hus1b | 0,02409 | 0,272652 | 3,50058 | 1 | 1 |
| Gpr82 | 0,005197 | 0,0586942 | 3,49752 | 1 | 1 |
| Cldn19 | 0,005867 | 0,0642904 | 3,45388 | 1 | 1 |
| Egfem1 | 0,005108 | 0,0554416 | 3,44021 | 1 | 1 |

| | | | | | |
|---|---|---|---|---|---|
| Spata3 | 0,021742 | 0,235935 | 3,43986 | 1 | 1 |
| Actl7a | 0,009098 | 0,0987311 | 3,43981 | 1 | 1 |
| Birc7 | 0,012144 | 0,131776 | 3,43981 | 1 | 1 |
| Cabp4 | 0,009624 | 0,104435 | 3,43981 | 1 | 1 |
| Cyp2c55 | 0,006626 | 0,0719052 | 3,43981 | 1 | 1 |
| Gpr142 | 0,01323 | 0,143568 | 3,43981 | 1 | 1 |
| Nron | 0,010288 | 0,108844 | 3,40319 | 1 | 1 |
| Omg | 0,00585 | 0,0611192 | 3,38519 | 1 | 1 |
| Tex35 | 0,017043 | 0,177989 | 3,38453 | 1 | 1 |
| Vmn2r1 | 0,004666 | 0,0487282 | 3,38453 | 1 | 1 |
| Pcdhb15 | 0,004565 | 0,047675 | 3,38452 | 1 | 1 |
| A630012P03Rik | 0,030152 | 0,297422 | 3,30221 | 1 | 1 |
| 4930447N08Rik | 0,012723 | 0,122075 | 3,26228 | 1 | 1 |
| Crhr2 | 0,005156 | 0,0490496 | 3,24992 | 1 | 1 |
| Efcab8 | 0,009395 | 0,0893415 | 3,2494 | 1 | 1 |
| Gm12216 | 0,012477 | 0,11865 | 3,2494 | 1 | 1 |
| Gm14124 | 0,008407 | 0,0782296 | 3,21812 | 1 | 1 |
| Dnmt3l | 0,012913 | 0,117339 | 3,18382 | 1 | 1 |
| Slc6a18 | 0,006169 | 0,0558067 | 3,17738 | 1 | 1 |
| Pou5f2 | 0,019863 | 0,175628 | 3,14441 | 1 | 1 |
| Agbl1 | 0,011212 | 0,098114 | 3,12948 | 1 | 1 |
| Lrrc23 | 0,028029 | 0,232748 | 3,0538 | 1 | 1 |
| Scn10a | 0,005714 | 0,0474457 | 3,0538 | 1 | 1 |
| AW495222 | 0,021656 | 0,177957 | 3,03869 | 1 | 1 |
| 4930417O22Rik | 0,012057 | 0,0984823 | 3,02998 | 1 | 1 |
| Gm6484 | 0,053682 | 0,438474 | 3,02998 | 1 | 1 |
| Rln3 | 0,043644 | 0,356486 | 3,02998 | 1 | 1 |
| Wdr95 | 0,005192 | 0,0424108 | 3,02998 | 1 | 1 |
| Tfap2e | 0,018549 | 0,148986 | 3,00576 | 1 | 1 |
| Gm17365 | 0,021028 | 0,161936 | 2,94507 | 1 | 1 |
| Morn3 | 0,032719 | 0,251974 | 2,94507 | 1 | 1 |
| Muc6 | 0,010396 | 0,0787647 | 2,92147 | 1 | 1 |
| Nuggc | 0,010015 | 0,0750864 | 2,90634 | 1 | 1 |
| Krt86 | 0,013472 | 0,100999 | 2,90632 | 1 | 1 |
| Tmem132cos | 0,061109 | 0,452054 | 2,88704 | 1 | 1 |
| Pou5f1 | 0,074634 | 0,541734 | 2,85968 | 1 | 1 |
| Sntg1 | 0,001741 | 0,0126008 | 2,85566 | 1 | 1 |
| C130026I21Rik | 0,007025 | 0,0508388 | 2,85542 | 1 | 1 |
| Serpina3a | 0,011828 | 0,0855667 | 2,85488 | 1 | 1 |
| 4930444M15Rik | 0,008135 | 0,0588494 | 2,85484 | 1 | 1 |
| Arhgap33os | 0,00484 | 0,0350112 | 2,85484 | 1 | 1 |
| Arr3 | 0,010241 | 0,0740839 | 2,85484 | 1 | 1 |
| Cyp8b1 | 0,006774 | 0,0490076 | 2,85484 | 1 | 1 |
| I830012O16Rik | 0,006538 | 0,0473008 | 2,85484 | 1 | 1 |
| Mslnl | 0,00488 | 0,0353001 | 2,85484 | 1 | 1 |
| Nobox | 0,007003 | 0,0506615 | 2,85484 | 1 | 1 |
| Nphs1 | 0,002168 | 0,0156803 | 2,85484 | 1 | 1 |
| Onecut3 | 0,002637 | 0,0190786 | 2,85484 | 1 | 1 |

| | | | | | |
|---|---|---|---|---|---|
| Slc7a13 | 0,006129 | 0,0443354 | 2,85484 | 1 | 1 |
| Tekt4 | 0,008907 | 0,064433 | 2,85484 | 1 | 1 |
| Alpk2 | 0,001696 | 0,0121493 | 2,84087 | 1 | 1 |
| Gm101 | 0,010254 | 0,0720891 | 2,81358 | 1 | 1 |
| Fam159b | 0,042778 | 0,296376 | 2,7925 | 1 | 1 |
| Rsph6a | 0,008521 | 0,0586524 | 2,78304 | 1 | 1 |
| Otx2 | 0,006078 | 0,0415127 | 2,77187 | 1 | 1 |
| Sun3 | 0,013183 | 0,0900152 | 2,77152 | 1 | 1 |
| 1700016H13Rik | 0,040574 | 0,276983 | 2,77116 | 1 | 1 |
| 0610043K17Rik | 0,057139 | 0,390049 | 2,77111 | 1 | 1 |
| 1700027J07Rik | 0,027636 | 0,188648 | 2,77111 | 1 | 1 |
| Avpr1a | 0,004795 | 0,0327289 | 2,77111 | 1 | 1 |
| Gm6904 | 0,017043 | 0,116342 | 2,77111 | 1 | 1 |
| Nanp | 0,010101 | 0,0689512 | 2,77111 | 1 | 1 |
| Cygb | 0,069627 | 0,472062 | 2,76127 | 1 | 1 |
| Dusp13 | 0,007676 | 0,0494029 | 2,68625 | 1 | 1 |
| Snord7 | 23,1449 | 148,04 | 2,67723 | 1 | 1 |
| Mfrp | 0,026764 | 0,170844 | 2,67432 | 1 | 1 |
| Gm15412 | 0,02652 | 0,168653 | 2,6689 | 1 | 1 |
| Ube2u | 0,027025 | 0,167659 | 2,63316 | 1 | 1 |
| Mmel1 | 0,025967 | 0,158821 | 2,61263 | 1 | 1 |
| Sftpb | 0,00918 | 0,0541041 | 2,55924 | 1 | 1 |
| 5730420D15Rik | 0,037183 | 0,219149 | 2,55919 | 1 | 1 |
| 1700012B09Rik | 0,02459 | 0,144899 | 2,5589 | 1 | 1 |
| 1700015E13Rik | 0,023237 | 0,136928 | 2,5589 | 1 | 1 |
| AW549542 | 0,007558 | 0,0445352 | 2,5589 | 1 | 1 |
| BC048546 | 0,002632 | 0,0155091 | 2,5589 | 1 | 1 |
| Col6a5 | 0,001301 | 0,00766502 | 2,5589 | 1 | 1 |
| E330020D12Rik | 0,004509 | 0,0265718 | 2,5589 | 1 | 1 |
| Gpr55 | 0,005042 | 0,0297092 | 2,5589 | 1 | 1 |
| Mtnr1b | 0,011373 | 0,0670166 | 2,5589 | 1 | 1 |
| Nlrp4g | 0,005396 | 0,0317964 | 2,5589 | 1 | 1 |
| Olfr1349 | 0,015771 | 0,0929293 | 2,5589 | 1 | 1 |
| Spata31d1a | 0,002904 | 0,0171122 | 2,5589 | 1 | 1 |
| Stx19 | 0,036061 | 0,212492 | 2,5589 | 1 | 1 |
| Wfdc13 | 0,027191 | 0,160223 | 2,5589 | 1 | 1 |
| Eqtn | 0,043686 | 0,25734 | 2,55842 | 1 | 1 |
| Chrm1 | 0,003946 | 0,0230523 | 2,54639 | 1 | 1 |
| Spata21 | 0,017965 | 0,104302 | 2,53754 | 1 | 1 |
| 0610040F04Rik | 0,008016 | 0,0464026 | 2,53319 | 1 | 1 |
| A630019I02Rik | 0,016362 | 0,0938687 | 2,5203 | 1 | 1 |
| Raet1b | 0,122509 | 0,697286 | 2,50887 | 1 | 1 |
| Prdm8 | 0,022345 | 0,12665 | 2,50281 | 1 | 1 |
| Tbx10 | 0,008346 | 0,0472906 | 2,50248 | 1 | 1 |
| Ssmem1 | 0,041652 | 0,23283 | 2,48283 | 1 | 1 |
| Gm20199 | 0,092058 | 0,511967 | 2,47544 | 1 | 1 |
| B130006D01Rik | 0,023783 | 0,131591 | 2,46805 | 1 | 1 |
| Crygs | 0,140808 | 0,766745 | 2,44502 | 1 | 1 |

| | | | | | |
|---|---|---|---|---|---|
| 4931429I11Rik | 0,00826 | 0,0448154 | 2,43981 | 1 | 1 |
| Ifi27l2a | 0,090981 | 0,493636 | 2,43981 | 1 | 1 |
| Barhl1 | 0,022267 | 0,11779 | 2,40323 | 1 | 1 |
| Fam179a | 0,007171 | 0,0374435 | 2,38453 | 1 | 1 |
| Gm5741 | 0,188211 | 0,982784 | 2,38453 | 1 | 1 |
| Serpina6 | 0,018805 | 0,0981919 | 2,38453 | 1 | 1 |
| A230073K19Rik | 0,075299 | 0,391679 | 2,37898 | 1 | 1 |
| Zpbp2 | 0,049458 | 0,2562 | 2,37299 | 1 | 1 |
| Ppp1r42 | 0,040785 | 0,209894 | 2,36354 | 1 | 1 |
| Pcp2 | 0,081477 | 0,418973 | 2,3624 | 1 | 1 |
| Gm9054 | 0,070544 | 0,362178 | 2,36011 | 1 | 1 |
| 5830444B04Rik | 0,064534 | 0,328161 | 2,34626 | 1 | 1 |
| 4931403G20Rik | 0,070829 | 0,357214 | 2,33437 | 1 | 1 |
| Ptprh | 0,01087 | 0,054794 | 2,33366 | 1 | 1 |
| Sumo2 | 0,111828 | 0,560574 | 2,32563 | 1 | 1 |
| Gm9047 | 0,093844 | 0,462134 | 2,29998 | 1 | 1 |
| Kcnip1 | 0,075569 | 0,369368 | 2,2892 | 1 | 1 |
| 4933417D19Rik | 0,080777 | 0,392502 | 2,28069 | 1 | 1 |
| Kcnj8 | 0,017885 | 0,0862573 | 2,26988 | 1 | 1 |
| 1700019B03Rik | 0,016789 | 0,0798304 | 2,2494 | 1 | 1 |
| Ucp1 | 0,016428 | 0,0781119 | 2,2494 | 1 | 1 |
| Gabrr2 | 0,047626 | 0,225469 | 2,2431 | 1 | 1 |
| 1810020O05Rik | 0,070154 | 0,33071 | 2,23697 | 1 | 1 |
| 2010109A12Rik | 0,171914 | 0,810413 | 2,23697 | 1 | 1 |
| Tdrd6 | 0,010361 | 0,0486102 | 2,23017 | 1 | 1 |
| Ccr9 | 0,008609 | 0,0403584 | 2,22903 | 1 | 1 |
| Mdh1b | 0,031395 | 0,146076 | 2,21812 | 1 | 1 |
| Gm16853 | 0,076586 | 0,356314 | 2,218 | 1 | 1 |
| 8430431K14Rik | 0,01339 | 0,0609416 | 2,18624 | 1 | 1 |
| 2610028E06Rik | 0,003542 | 0,0161211 | 2,18614 | 1 | 1 |
| 4932413F04Rik | 0,004619 | 0,0210184 | 2,18614 | 1 | 1 |
| A530065N20Rik | 0,006705 | 0,0305149 | 2,18614 | 1 | 1 |
| Ccdc87 | 0,007995 | 0,0363827 | 2,18614 | 1 | 1 |
| Dbhos | 0,0031 | 0,0141059 | 2,18614 | 1 | 1 |
| Enpp6 | 0,004134 | 0,0188145 | 2,18614 | 1 | 1 |
| Gm13539 | 0,030173 | 0,137313 | 2,18614 | 1 | 1 |
| Hmx3 | 0,01017 | 0,0462836 | 2,18614 | 1 | 1 |
| Mc1r | 0,00342 | 0,0155618 | 2,18614 | 1 | 1 |
| Mmp7 | 0,017549 | 0,0798627 | 2,18614 | 1 | 1 |
| Nps | 0,036061 | 0,164108 | 2,18614 | 1 | 1 |
| Slc5a4b | 0,006135 | 0,0279191 | 2,18614 | 1 | 1 |
| Utf1 | 0,012372 | 0,0563052 | 2,18614 | 1 | 1 |
| Armc4 | 0,019901 | 0,0904538 | 2,18434 | 1 | 1 |
| Tcf21 | 0,063671 | 0,289397 | 2,18434 | 1 | 1 |
| Spag11b | 0,026915 | 0,122067 | 2,18117 | 1 | 1 |
| Rbmxl2 | 0,042041 | 0,18993 | 2,17561 | 1 | 1 |
| 4933408J17Rik | 0,03273 | 0,147012 | 2,16724 | 1 | 1 |
| Ptgir | 0,015116 | 0,0668031 | 2,14386 | 1 | 1 |

| | | | | | |
|---|---|---|---|---|---|
| Tmem232 | 0,014891 | 0,0651675 | 2,12967 | 1 | 1 |
| Emr4 | 0,010795 | 0,0472362 | 2,12948 | 1 | 1 |
| Riiad1 | 0,111583 | 0,481753 | 2,11018 | 1 | 1 |
| Nhlh2 | 0,04137 | 0,177067 | 2,09765 | 1 | 1 |
| Snora24 | 3,42131 | 14,5909 | 2,09245 | 1 | 1 |
| Bcan | 0,029209 | 0,124136 | 2,08742 | 1 | 1 |
| Acer1 | 0,015948 | 0,0676135 | 2,08394 | 1 | 1 |
| B3galt2 | 0,008113 | 0,0343942 | 2,08394 | 1 | 1 |
| Tlx1 | 0,074537 | 0,314048 | 2,07496 | 1 | 1 |
| Scpep1os | 0,123927 | 0,51798 | 2,06341 | 1 | 1 |
| Wdr16 | 0,041461 | 0,17209 | 2,05335 | 1 | 1 |
| 4933416M07Rik | 0,086583 | 0,35828 | 2,04894 | 1 | 1 |
| Slc14a2 | 0,12648 | 0,519187 | 2,03735 | 1 | 1 |
| Best2 | 0,013394 | 0,0547044 | 2,03005 | 1 | 1 |
| 4930415F15Rik | 0,043089 | 0,175975 | 2,02998 | 1 | 1 |
| Gm11413 | 0,01882 | 0,0768587 | 2,02998 | 1 | 1 |
| Gm8580 | 0,058122 | 0,237369 | 2,02998 | 1 | 1 |
| Pcnxl2 | 0,003364 | 0,0137389 | 2,02998 | 1 | 1 |
| Trp53tg5 | 0,065078 | 0,265779 | 2,02998 | 1 | 1 |
| Vwa3a | 0,029773 | 0,121435 | 2,02809 | 1 | 1 |
| Olig1 | 0,030066 | 0,122436 | 2,02581 | 1 | 1 |
| Xlr3b | 0,10045 | 0,405838 | 2,01442 | 1 | 1 |
| Hist1h2bh | 0,204823 | 0,826768 | 2,0131 | 1 | 1 |
| AY074887 | 0,191544 | 0,769677 | 2,00658 | 1 | 1 |
| Otog | 0,006011 | 0,023987 | 1,99662 | 1 | 1 |
| Tmprss11d | 0,025685 | 0,102279 | 1,9935 | 1 | 1 |
| C230024C17Rik | 0,011326 | 0,0444927 | 1,97393 | 1 | 1 |
| Tnfrsf8 | 0,025882 | 0,101674 | 1,97393 | 1 | 1 |
| Pcsk2os1 | 0,014679 | 0,0573797 | 1,96681 | 1 | 1 |
| Kctd4 | 0,029455 | 0,114895 | 1,96375 | 1 | 1 |
| Erich6 | 0,032715 | 0,126508 | 1,95119 | 1 | 1 |
| Card11 | 0,022721 | 0,0875819 | 1,94663 | 1 | 1 |
| Opn1sw | 0,034889 | 0,134204 | 1,94361 | 1 | 1 |
| 4930428G15Rik | 0,127633 | 0,480648 | 1,91297 | 1 | 1 |
| Dpcr1 | 0,083711 | 0,314902 | 1,91142 | 1 | 1 |
| Myt1 | 0,00813 | 0,0304335 | 1,90426 | 1 | 1 |
| Sox15 | 0,126638 | 0,471383 | 1,89619 | 1 | 1 |
| Kctd19 | 0,018877 | 0,0701307 | 1,89343 | 1 | 1 |
| A230028O05Rik | 0,036549 | 0,134604 | 1,88082 | 1 | 1 |
| Plxdc1 | 0,004397 | 0,0159492 | 1,85905 | 1 | 1 |
| Sprn | 0,014052 | 0,0509621 | 1,8587 | 1 | 1 |
| Cd8a | 0,004073 | 0,0147691 | 1,85834 | 1 | 1 |
| Hist1h3f | 0,110105 | 0,398838 | 1,85692 | 1 | 1 |
| Otof | 0,001763 | 0,00638116 | 1,85581 | 1 | 1 |
| Slc17a3 | 0,002994 | 0,0108346 | 1,85564 | 1 | 1 |
| Pla2g10 | 0,019371 | 0,070076 | 1,85502 | 1 | 1 |
| Speer4b | 0,006671 | 0,0241321 | 1,85493 | 1 | 1 |
| Foxg1 | 0,004312 | 0,0155972 | 1,85489 | 1 | 1 |

| | | | | | |
|---|---|---|---|---|---|
| 1700109G14Rik | 0,019912 | 0,0720252 | 1,85484 | 1 | 1 |
| 1700125H20Rik | 0,015341 | 0,0554905 | 1,85484 | 1 | 1 |
| 4930428O21Rik | 0,015401 | 0,0557078 | 1,85484 | 1 | 1 |
| 4930463O16Rik | 0,005566 | 0,0201335 | 1,85484 | 1 | 1 |
| 4930593A02Rik | 0,032229 | 0,116575 | 1,85484 | 1 | 1 |
| 4933406J10Rik | 0,015145 | 0,05478 | 1,85484 | 1 | 1 |
| 4933411E08Rik | 0,007885 | 0,0285224 | 1,85484 | 1 | 1 |
| 5430427M07Rik | 0,01446 | 0,0523023 | 1,85484 | 1 | 1 |
| Adamdec1 | 0,005357 | 0,0193767 | 1,85484 | 1 | 1 |
| Akap14 | 0,014897 | 0,0538836 | 1,85484 | 1 | 1 |
| Alox12b | 0,011039 | 0,0399288 | 1,85484 | 1 | 1 |
| Atp5l | 0,038277 | 0,138454 | 1,85484 | 1 | 1 |
| B930092H01Rik | 0,009562 | 0,0345867 | 1,85484 | 1 | 1 |
| Car6 | 0,010153 | 0,036724 | 1,85484 | 1 | 1 |
| Catsperg2 | 0,003152 | 0,0113999 | 1,85484 | 1 | 1 |
| Cd19 | 0,005299 | 0,0191683 | 1,85484 | 1 | 1 |
| Ces1a | 0,006664 | 0,0241034 | 1,85484 | 1 | 1 |
| Clcnkb | 0,005537 | 0,0200298 | 1,85484 | 1 | 1 |
| EU599041 | 0,019327 | 0,0699076 | 1,85484 | 1 | 1 |
| Entpd8 | 0,022508 | 0,0814151 | 1,85484 | 1 | 1 |
| Gm15880 | 0,004413 | 0,0159641 | 1,85484 | 1 | 1 |
| Gm20750 | 0,003572 | 0,0129217 | 1,85484 | 1 | 1 |
| Gm5150 | 0,016683 | 0,0603429 | 1,85484 | 1 | 1 |
| Gm5544 | 0,05165 | 0,186826 | 1,85484 | 1 | 1 |
| Gm8994 | 0,00864 | 0,0312517 | 1,85484 | 1 | 1 |
| Hrh3 | 0,00489 | 0,0176866 | 1,85484 | 1 | 1 |
| Ifi44 | 0,008749 | 0,0316448 | 1,85484 | 1 | 1 |
| Igll1 | 0,035519 | 0,128478 | 1,85484 | 1 | 1 |
| Ins1 | 0,014531 | 0,0525593 | 1,85484 | 1 | 1 |
| Kcnmb3 | 0,045844 | 0,165825 | 1,85484 | 1 | 1 |
| Kirrel2 | 0,004054 | 0,014662 | 1,85484 | 1 | 1 |
| Klk14 | 0,011096 | 0,0401345 | 1,85484 | 1 | 1 |
| Mirlet7i | 14,0742 | 50,9082 | 1,85484 | 1 | 1 |
| Msmb | 0,037311 | 0,13496 | 1,85484 | 1 | 1 |
| Olfr5 | 0,014081 | 0,0509325 | 1,85484 | 1 | 1 |
| Oxct2a | 0,007388 | 0,026723 | 1,85484 | 1 | 1 |
| Pbld1 | 0,027207 | 0,0984097 | 1,85484 | 1 | 1 |
| Pcdhb10 | 0,008971 | 0,0324488 | 1,85484 | 1 | 1 |
| Pldi | 0,018225 | 0,0659215 | 1,85484 | 1 | 1 |
| Shbg | 0,009832 | 0,035564 | 1,85484 | 1 | 1 |
| Slc22a22 | 0,010932 | 0,0395413 | 1,85484 | 1 | 1 |
| Slc6a1 | 0,002891 | 0,010458 | 1,85484 | 1 | 1 |
| Slurp1 | 0,036846 | 0,133278 | 1,85484 | 1 | 1 |
| Tpo | 0,003843 | 0,0138999 | 1,85484 | 1 | 1 |
| Trcg1 | 0,004737 | 0,017134 | 1,85484 | 1 | 1 |
| Ttc9b | 0,033794 | 0,122238 | 1,85484 | 1 | 1 |
| Ush1g | 0,007941 | 0,0287236 | 1,85484 | 1 | 1 |
| Vmn2r90 | 0,004953 | 0,0179161 | 1,85484 | 1 | 1 |

| | | | | | |
|---|---|---|---|---|---|
| Zfp541 | 0,002715 | 0,00981955 | 1,85484 | 1 | 1 |
| Bcas3os1 | 0,004032 | 0,0145822 | 1,85481 | 1 | 1 |
| Ibsp | 0,06422 | 0,229764 | 1,83907 | 1 | 1 |
| Trim67 | 0,010998 | 0,0392196 | 1,83436 | 1 | 1 |
| Pcdhac1 | 0,019708 | 0,070191 | 1,8325 | 1 | 1 |
| 4930563D23Rik | 0,031097 | 0,110693 | 1,83174 | 1 | 1 |
| Fer1l4 | 0,011038 | 0,0391769 | 1,8275 | 1 | 1 |
| Gm11992 | 0,026931 | 0,0952132 | 1,8219 | 1 | 1 |
| 4930431P03Rik | 0,025654 | 0,0906634 | 1,82136 | 1 | 1 |
| Colec11 | 0,040129 | 0,141059 | 1,81358 | 1 | 1 |
| Pcdha9 | 0,006532 | 0,0228354 | 1,80562 | 1 | 1 |
| 4930453L07Rik | 0,028732 | 0,100021 | 1,79956 | 1 | 1 |
| Pom121l2 | 0,022918 | 0,0790151 | 1,78563 | 1 | 1 |
| Lrrtm2 | 0,016245 | 0,0559287 | 1,78363 | 1 | 1 |
| Fxyd4 | 0,068053 | 0,232558 | 1,77285 | 1 | 1 |
| Aspn | 0,022194 | 0,0757977 | 1,77196 | 1 | 1 |
| Abcb5 | 0,006442 | 0,0219887 | 1,77111 | 1 | 1 |
| Ctsk | 0,051464 | 0,175653 | 1,77111 | 1 | 1 |
| Cyp21a1 | 0,013762 | 0,0469704 | 1,77111 | 1 | 1 |
| Kiss1 | 0,081011 | 0,276504 | 1,77111 | 1 | 1 |
| Toporsl | 0,010768 | 0,0367511 | 1,77111 | 1 | 1 |
| Chrnd | 0,03216 | 0,108975 | 1,76068 | 1 | 1 |
| Myt1l | 0,011996 | 0,0403984 | 1,7517 | 1 | 1 |
| Gm14164 | 0,022293 | 0,0750531 | 1,75132 | 1 | 1 |
| Krt83 | 0,058338 | 0,195283 | 1,74307 | 1 | 1 |
| Dpy19l2 | 0,036591 | 0,122366 | 1,74165 | 1 | 1 |
| Klhl33 | 0,038132 | 0,126862 | 1,73421 | 1 | 1 |
| Gm19299 | 0,07213 | 0,239366 | 1,73055 | 1 | 1 |
| Gm11149 | 0,063455 | 0,209825 | 1,72537 | 1 | 1 |
| Lrrc63 | 0,030656 | 0,0998789 | 1,70401 | 1 | 1 |
| Haao | 0,139313 | 0,451928 | 1,69776 | 1 | 1 |
| Nhlh1 | 0,036338 | 0,117278 | 1,69038 | 1 | 1 |
| LOC381967 | 0,094528 | 0,304565 | 1,68794 | 1 | 1 |
| Prdm14 | 0,032216 | 0,103412 | 1,68255 | 1 | 1 |
| Gm10389 | 0,042085 | 0,134672 | 1,67806 | 1 | 1 |
| Ccdc147 | 0,043332 | 0,138191 | 1,67317 | 1 | 1 |
| Gpr18 | 0,042778 | 0,136021 | 1,6689 | 1 | 1 |
| Cd5 | 0,107904 | 0,342265 | 1,66537 | 1 | 1 |
| Becn2 | 0,022388 | 0,070966 | 1,66444 | 1 | 1 |
| Cryba1 | 0,074078 | 0,234821 | 1,66444 | 1 | 1 |
| Gm1123 | 0,031319 | 0,099277 | 1,66444 | 1 | 1 |
| Hist1h4j | 0,320404 | 1,01565 | 1,66444 | 1 | 1 |
| Pzp | 0,011395 | 0,0360699 | 1,66235 | 1 | 1 |
| E130304I02Rik | 0,173095 | 0,545885 | 1,65703 | 1 | 1 |
| Mroh2a | 0,005967 | 0,0188138 | 1,65662 | 1 | 1 |
| A730085K08Rik | 0,179929 | 0,56426 | 1,64893 | 1 | 1 |
| Tmem8c | 0,135291 | 0,424105 | 1,64835 | 1 | 1 |
| Pirb | 0,021121 | 0,0658913 | 1,64145 | 1 | 1 |

| | | | | | |
|---|---|---|---|---|---|
| 9530091C08Rik | 0,019032 | 0,0593043 | 1,63973 | 1 | 1 |
| Pdxk-ps | 0,066464 | 0,205929 | 1,6315 | 1 | 1 |
| 6530411M01Rik | 0,040867 | 0,126453 | 1,62961 | 1 | 1 |
| Scn7a | 0,005819 | 0,0180041 | 1,62961 | 1 | 1 |
| Grin2a | 0,061833 | 0,190785 | 1,62549 | 1 | 1 |
| C4b | 0,011397 | 0,0351088 | 1,62314 | 1 | 1 |
| Dpys | 0,006205 | 0,0188724 | 1,60477 | 1 | 1 |
| Osr2 | 0,110064 | 0,334497 | 1,60365 | 1 | 1 |
| A230020J21Rik | 0,009759 | 0,0296016 | 1,60086 | 1 | 1 |
| Nlrc5 | 0,026344 | 0,0787227 | 1,5793 | 1 | 1 |
| Rasgrf2 | 0,011551 | 0,0342657 | 1,56874 | 1 | 1 |
| Hist1h3d | 0,295963 | 0,876372 | 1,56613 | 1 | 1 |
| Gjc2 | 0,1196 | 0,353944 | 1,5653 | 1 | 1 |
| 2900060B14Rik | 0,060192 | 0,177343 | 1,5589 | 1 | 1 |
| 5830473C10Rik | 0,011613 | 0,0342159 | 1,5589 | 1 | 1 |
| Insm1 | 0,008395 | 0,0247331 | 1,5589 | 1 | 1 |
| Slc22a30 | 0,013194 | 0,0388721 | 1,5589 | 1 | 1 |
| Igfn1 | 0,039052 | 0,114753 | 1,55507 | 1 | 1 |
| Oca2 | 0,028394 | 0,0830683 | 1,54871 | 1 | 1 |
| Snx31 | 0,066477 | 0,193922 | 1,54456 | 1 | 1 |
| Pnliprp1 | 0,075231 | 0,219455 | 1,54452 | 1 | 1 |
| Gm14379 | 0,095436 | 0,278252 | 1,54378 | 1 | 1 |
| 4933429K18Rik | 0,065731 | 0,191447 | 1,54231 | 1 | 1 |
| Cbln2 | 0,080082 | 0,233247 | 1,54231 | 1 | 1 |
| 4933427G17Rik | 0,049313 | 0,143438 | 1,54039 | 1 | 1 |
| Acaa1b | 0,1899 | 0,551578 | 1,53832 | 1 | 1 |
| Lrrc7 | 0,033911 | 0,0984674 | 1,53789 | 1 | 1 |
| 4931408D14Rik | 0,020713 | 0,0599407 | 1,53298 | 1 | 1 |
| Ppp1r14d | 0,109109 | 0,315737 | 1,53295 | 1 | 1 |
| 9330179D12Rik | 0,065056 | 0,187022 | 1,52345 | 1 | 1 |
| 9630001P10Rik | 0,021616 | 0,0621239 | 1,52304 | 1 | 1 |
| Slain1os | 0,071974 | 0,206373 | 1,51971 | 1 | 1 |
| Fgr | 0,147107 | 0,420026 | 1,51361 | 1 | 1 |
| Ddn | 0,050222 | 0,142465 | 1,50421 | 1 | 1 |
| Kank4os | 0,092143 | 0,260998 | 1,50209 | 1 | 1 |
| 4930519F16Rik | 0,147712 | 0,418206 | 1,50142 | 1 | 1 |
| Btnl9 | 0,081266 | 0,230024 | 1,50106 | 1 | 1 |
| Hist1h3i | 0,19106 | 0,53919 | 1,49676 | 1 | 1 |
| Kbtbd12 | 0,029003 | 0,0818497 | 1,49676 | 1 | 1 |
| A230072E10Rik | 0,028239 | 0,0793792 | 1,49105 | 1 | 1 |
| Mroh8 | 0,148038 | 0,414968 | 1,48703 | 1 | 1 |
| Wscd2 | 0,113246 | 0,317016 | 1,48509 | 1 | 1 |
| 4930478L05Rik | 0,042344 | 0,11819 | 1,48089 | 1 | 1 |
| Iqcj | 0,011156 | 0,0311202 | 1,48004 | 1 | 1 |
| Cyp2t4 | 0,162896 | 0,454238 | 1,4795 | 1 | 1 |
| Zfhx2os | 0,107458 | 0,297326 | 1,46827 | 1 | 1 |
| Ephx1 | 0,093158 | 0,256948 | 1,46372 | 1 | 1 |
| Otud7a | 0,056315 | 0,155096 | 1,46158 | 1 | 1 |

| | | | | | |
|---|---|---|---|---|---|
| Htr2b | 0,167928 | 0,459562 | 1,45242 | 1 | 1 |
| Tbrg3 | 0,074565 | 0,203698 | 1,44987 | 1 | 1 |
| 4933424G05Rik | 0,042269 | 0,11509 | 1,4451 | 1 | 1 |
| 1700019N19Rik | 0,049352 | 0,134368 | 1,44502 | 1 | 1 |
| Il17c | 0,078852 | 0,214688 | 1,44502 | 1 | 1 |
| Iqch | 0,022381 | 0,0609347 | 1,44502 | 1 | 1 |
| Lingo4 | 0,012955 | 0,0352727 | 1,44502 | 1 | 1 |
| Pi15 | 0,005281 | 0,014379 | 1,44502 | 1 | 1 |
| Calr4 | 0,045445 | 0,123534 | 1,44272 | 1 | 1 |
| LOC102632430 | 0,041691 | 0,113325 | 1,44267 | 1 | 1 |
| Tssk1 | 0,066396 | 0,180402 | 1,44204 | 1 | 1 |
| Hist1h2an | 0,31671 | 0,860425 | 1,44189 | 1 | 1 |
| 1700019L03Rik | 0,132982 | 0,360997 | 1,44075 | 1 | 1 |
| A430090L17Rik | 0,013584 | 0,0368508 | 1,43981 | 1 | 1 |
| Cplx4 | 0,026727 | 0,072475 | 1,4392 | 1 | 1 |
| Rad21l | 0,028509 | 0,0773066 | 1,4392 | 1 | 1 |
| Olig2 | 0,071545 | 0,192222 | 1,42584 | 1 | 1 |
| Tmprss9 | 0,038239 | 0,102345 | 1,42032 | 1 | 1 |
| Rp1 | 0,027576 | 0,0735685 | 1,41567 | 1 | 1 |
| Rhcg | 0,034537 | 0,0907394 | 1,3936 | 1 | 1 |
| Foxr2 | 0,019687 | 0,0516862 | 1,39252 | 1 | 1 |
| Ttc16 | 0,088306 | 0,231763 | 1,39207 | 1 | 1 |
| Dear1 | 0,145163 | 0,380117 | 1,38877 | 1 | 1 |
| Asgr1 | 0,056326 | 0,147327 | 1,38715 | 1 | 1 |
| S100a14 | 0,073322 | 0,191735 | 1,3868 | 1 | 1 |
| 4930443O20Rik | 0,043025 | 0,112506 | 1,38676 | 1 | 1 |
| Fscn3 | 0,033612 | 0,087891 | 1,38676 | 1 | 1 |
| Asphd1 | 0,150671 | 0,393381 | 1,38453 | 1 | 1 |
| 1700112J05Rik | 0,197339 | 0,513425 | 1,37948 | 1 | 1 |
| 1700019B21Rik | 0,030326 | 0,0787899 | 1,37746 | 1 | 1 |
| Gm13582 | 0,016924 | 0,0439234 | 1,37595 | 1 | 1 |
| Klk13 | 0,207178 | 0,536907 | 1,3738 | 1 | 1 |
| Foxp3 | 0,085824 | 0,221965 | 1,37089 | 1 | 1 |
| Baiap3 | 0,036384 | 0,0940821 | 1,37061 | 1 | 1 |
| Fscn2 | 0,070312 | 0,180955 | 1,36378 | 1 | 1 |
| Fam196a | 0,088124 | 0,226501 | 1,36191 | 1 | 1 |
| Zmynd10 | 0,107008 | 0,273086 | 1,35163 | 1 | 1 |
| Kcnc4 | 0,067431 | 0,171969 | 1,35067 | 1 | 1 |
| Gm3696 | 0,238564 | 0,607344 | 1,34814 | 1 | 1 |
| Il11 | 0,036036 | 0,0916702 | 1,347 | 1 | 1 |
| 1700016L04Rik | 0,083767 | 0,213085 | 1,34697 | 1 | 1 |
| Slc25a34 | 0,023866 | 0,0607103 | 1,34697 | 1 | 1 |
| H2-M10.2 | 0,18058 | 0,457417 | 1,34087 | 1 | 1 |
| F630111L10Rik | 0,031524 | 0,079767 | 1,33933 | 1 | 1 |
| Tmem239 | 0,151978 | 0,384524 | 1,33922 | 1 | 1 |
| Galnt15 | 0,024945 | 0,0629716 | 1,33594 | 1 | 1 |
| Gm10125 | 0,040849 | 0,103119 | 1,33594 | 1 | 1 |
| Hist1h2bl | 0,339029 | 0,851154 | 1,32801 | 1 | 1 |

| | | | | | |
|---|---|---|---|---|---|
| Psmb11 | 0,019037 | 0,0477923 | 1,32801 | 1 | 1 |
| Fndc9 | 0,201253 | 0,503109 | 1,32186 | 1 | 1 |
| 4930590J08Rik | 0,038946 | 0,0973146 | 1,32118 | 1 | 1 |
| Nmbr | 0,146645 | 0,364614 | 1,31404 | 1 | 1 |
| Gabra5 | 0,110206 | 0,273234 | 1,30993 | 1 | 1 |
| Lrrc4b | 0,232962 | 0,577578 | 1,30992 | 1 | 1 |
| Rad51ap2 | 0,119545 | 0,296382 | 1,3099 | 1 | 1 |
| Sowaha | 0,228902 | 0,566384 | 1,30705 | 1 | 1 |
| 2610206C17Rik | 0,144788 | 0,358038 | 1,30617 | 1 | 1 |
| Ap3b2 | 0,04107 | 0,101542 | 1,30593 | 1 | 1 |
| Inhbc | 0,070544 | 0,173302 | 1,29669 | 1 | 1 |
| Gipr | 0,126828 | 0,311148 | 1,29472 | 1 | 1 |
| Chit1 | 0,043393 | 0,106221 | 1,29154 | 1 | 1 |
| Nxph4 | 0,116758 | 0,285234 | 1,28862 | 1 | 1 |
| Slc35g3 | 0,105307 | 0,257046 | 1,28743 | 1 | 1 |
| Ngb | 0,109014 | 0,265865 | 1,28618 | 1 | 1 |
| Kndc1 | 0,055874 | 0,135896 | 1,28225 | 1 | 1 |
| Xntrpc | 0,102943 | 0,249902 | 1,27951 | 1 | 1 |
| Cr2 | 0,086724 | 0,210026 | 1,27607 | 1 | 1 |
| Grm1 | 0,005292 | 0,0127661 | 1,27043 | 1 | 1 |
| Kif17 | 0,012325 | 0,029721 | 1,2699 | 1 | 1 |
| 4732490B19Rik | 0,015827 | 0,0381656 | 1,26988 | 1 | 1 |
| 4930474M22Rik | 0,173513 | 0,418411 | 1,26988 | 1 | 1 |
| Ankar | 0,008122 | 0,0195852 | 1,26988 | 1 | 1 |
| Pitpnm2os1 | 0,025095 | 0,0605142 | 1,26988 | 1 | 1 |
| Styxl1 | 0,047046 | 0,113284 | 1,26778 | 1 | 1 |
| Pdyn | 0,04764 | 0,113748 | 1,25562 | 1 | 1 |
| Spag17 | 0,079767 | 0,189382 | 1,24744 | 1 | 1 |
| Fosl1 | 0,043423 | 0,102972 | 1,24574 | 1 | 1 |
| 4930570G19Rik | 0,09971 | 0,235998 | 1,24296 | 1 | 1 |
| 1700001K19Rik | 0,244553 | 0,578735 | 1,24275 | 1 | 1 |
| Cysltr2 | 0,088312 | 0,208669 | 1,24053 | 1 | 1 |
| Crym | 0,270306 | 0,638264 | 1,23956 | 1 | 1 |
| Tnf | 0,147908 | 0,348228 | 1,23533 | 1 | 1 |
| Sstr1 | 0,035501 | 0,0834122 | 1,23242 | 1 | 1 |
| Slc25a18 | 0,173035 | 0,405857 | 1,22991 | 1 | 1 |
| Gm5779 | 0,10095 | 0,236568 | 1,22862 | 1 | 1 |
| Gm3558 | 0,099179 | 0,231891 | 1,22534 | 1 | 1 |
| Kcnq3 | 0,181541 | 0,423496 | 1,22205 | 1 | 1 |
| Cplx3 | 0,015623 | 0,0363887 | 1,21979 | 1 | 1 |
| Gm7173 | 0,083358 | 0,194054 | 1,21907 | 1 | 1 |
| Ackr4 | 0,037821 | 0,0879998 | 1,21832 | 1 | 1 |
| Gpbar1 | 0,188208 | 0,437226 | 1,21606 | 1 | 1 |
| Wdr20rt | 0,052595 | 0,122061 | 1,2146 | 1 | 1 |
| 2310005G13Rik | 0,01862 | 0,0431457 | 1,21239 | 1 | 1 |
| Ccr10 | 0,035048 | 0,0812134 | 1,21239 | 1 | 1 |
| Gm6938 | 0,094016 | 0,217855 | 1,21239 | 1 | 1 |
| 1700034P13Rik | 0,106894 | 0,245984 | 1,20238 | 1 | 1 |

| | | | | | |
|---|---|---|---|---|---|
| Galntl6 | 0,020598 | 0,0473251 | 1,20011 | 1 | 1 |
| Kcnj14 | 0,24075 | 0,552915 | 1,19952 | 1 | 1 |
| Spdef | 0,116921 | 0,267935 | 1,19635 | 1 | 1 |
| Il17rb | 0,073293 | 0,167221 | 1,19002 | 1 | 1 |
| 1700001L19Rik | 0,092227 | 0,210376 | 1,18971 | 1 | 1 |
| Pabpc5 | 0,082323 | 0,187528 | 1,18774 | 1 | 1 |
| Gm19522 | 0,005085 | 0,011577 | 1,18689 | 1 | 1 |
| Syce1l | 0,017221 | 0,0392021 | 1,18678 | 1 | 1 |
| 1700007B14Rik | 0,019293 | 0,0439032 | 1,18624 | 1 | 1 |
| Crip3 | 0,047244 | 0,107507 | 1,18622 | 1 | 1 |
| Gm20556 | 0,01145 | 0,0260545 | 1,18615 | 1 | 1 |
| 1700040L02Rik | 0,011439 | 0,0260289 | 1,18614 | 1 | 1 |
| 4833403I15Rik | 0,003923 | 0,00892674 | 1,18614 | 1 | 1 |
| 4930438E09Rik | 0,020678 | 0,0470518 | 1,18614 | 1 | 1 |
| 4930483O08Rik | 0,028639 | 0,0651666 | 1,18614 | 1 | 1 |
| 4930488L21Rik | 0,007134 | 0,0162328 | 1,18614 | 1 | 1 |
| 4930552P12Rik | 0,005161 | 0,0117426 | 1,18614 | 1 | 1 |
| 4930563E18Rik | 0,017536 | 0,0399021 | 1,18614 | 1 | 1 |
| 4933430H16Rik | 0,019139 | 0,0435501 | 1,18614 | 1 | 1 |
| 5430428K19Rik | 0,016897 | 0,0384482 | 1,18614 | 1 | 1 |
| 5730422E09Rik | 0,015656 | 0,0356241 | 1,18614 | 1 | 1 |
| Als2cr11 | 0,006138 | 0,0139668 | 1,18614 | 1 | 1 |
| Aox4 | 0,004961 | 0,0112895 | 1,18614 | 1 | 1 |
| Areg | 0,011887 | 0,0270491 | 1,18614 | 1 | 1 |
| Ccdc121 | 0,007324 | 0,016665 | 1,18614 | 1 | 1 |
| Cckbr | 0,02076 | 0,0472384 | 1,18614 | 1 | 1 |
| Cxcl1 | 0,031753 | 0,0722517 | 1,18614 | 1 | 1 |
| Cyp2b23 | 0,01196 | 0,0272134 | 1,18614 | 1 | 1 |
| Dcpp3 | 0,029133 | 0,0662891 | 1,18614 | 1 | 1 |
| E030018B13Rik | 0,014584 | 0,033186 | 1,18614 | 1 | 1 |
| Fcrlb | 0,010952 | 0,0249203 | 1,18614 | 1 | 1 |
| G630055G22Rik | 0,010486 | 0,0238599 | 1,18614 | 1 | 1 |
| Gabrr3 | 0,009447 | 0,0214968 | 1,18614 | 1 | 1 |
| Glod5 | 0,050546 | 0,115015 | 1,18614 | 1 | 1 |
| Gm11545 | 0,004752 | 0,0108132 | 1,18614 | 1 | 1 |
| Gm136 | 0,010763 | 0,0244895 | 1,18614 | 1 | 1 |
| Gm3317 | 0,015371 | 0,034976 | 1,18614 | 1 | 1 |
| Gm6815 | 0,003693 | 0,00840279 | 1,18614 | 1 | 1 |
| Gtf2a1l | 0,016718 | 0,038041 | 1,18614 | 1 | 1 |
| Hcst | 0,049695 | 0,113079 | 1,18614 | 1 | 1 |
| Hist1h4f | 0,076511 | 0,174095 | 1,18614 | 1 | 1 |
| Irx6 | 0,005436 | 0,0123686 | 1,18614 | 1 | 1 |
| Kif19a | 0,007326 | 0,0166703 | 1,18614 | 1 | 1 |
| Kis2 | 0,012948 | 0,0294626 | 1,18614 | 1 | 1 |
| Klhl40 | 0,005255 | 0,0119565 | 1,18614 | 1 | 1 |
| Mael | 0,008814 | 0,0200551 | 1,18614 | 1 | 1 |
| Mmp12 | 0,003476 | 0,00790894 | 1,18614 | 1 | 1 |
| Mmp27 | 0,008146 | 0,0185358 | 1,18614 | 1 | 1 |

| | | | | | |
|---|---|---|---|---|---|
| Mnda | 0,016348 | 0,0371997 | 1,18614 | 1 | 1 |
| Odf3l2 | 0,01133 | 0,0257796 | 1,18614 | 1 | 1 |
| Olfr91 | 0,016092 | 0,0366174 | 1,18614 | 1 | 1 |
| Pabpc2 | 0,004966 | 0,0112989 | 1,18614 | 1 | 1 |
| Pde6h | 0,133647 | 0,304105 | 1,18614 | 1 | 1 |
| Phxr4 | 0,010058 | 0,022886 | 1,18614 | 1 | 1 |
| Ppef1 | 0,00519 | 0,0118096 | 1,18614 | 1 | 1 |
| Ptchd3 | 0,003901 | 0,00887669 | 1,18614 | 1 | 1 |
| Rbp7 | 0,030328 | 0,0690086 | 1,18614 | 1 | 1 |
| Rnf138rt1 | 0,059737 | 0,135929 | 1,18614 | 1 | 1 |
| Tbx21 | 0,02015 | 0,0458502 | 1,18614 | 1 | 1 |
| Tlx3 | 0,008907 | 0,0202665 | 1,18614 | 1 | 1 |
| Tmem174 | 0,005775 | 0,0131416 | 1,18614 | 1 | 1 |
| Tmem178b | 0,004096 | 0,00931929 | 1,18614 | 1 | 1 |
| Tnmd | 0,010542 | 0,0239875 | 1,18614 | 1 | 1 |
| Ucp3 | 0,005271 | 0,0119938 | 1,18614 | 1 | 1 |
| Vgll2 | 0,009372 | 0,0213264 | 1,18614 | 1 | 1 |
| Zdhhc19 | 0,010694 | 0,0243345 | 1,18614 | 1 | 1 |
| Thy1 | 0,123718 | 0,28127 | 1,1849 | 1 | 1 |
| LOC100505025 | 0,068729 | 0,156057 | 1,18309 | 1 | 1 |
| C330011F03Rik | 0,005526 | 0,0125416 | 1,18237 | 1 | 1 |
| Obscn | 0,004768 | 0,0107824 | 1,17727 | 1 | 1 |
| Csmd2 | 0,030146 | 0,0678657 | 1,1707 | 1 | 1 |
| Galnt9 | 0,008905 | 0,0200328 | 1,16974 | 1 | 1 |
| 4930538K18Rik | 0,113794 | 0,25586 | 1,16893 | 1 | 1 |
| Inpp5j | 0,045253 | 0,10175 | 1,16893 | 1 | 1 |
| Tom1 | 0,036959 | 0,0830196 | 1,16753 | 1 | 1 |
| Ehf | 0,045282 | 0,10146 | 1,16391 | 1 | 1 |
| 1810024B03Rik | 0,105883 | 0,237171 | 1,16346 | 1 | 1 |
| Qrfpr | 0,248269 | 0,554398 | 1,15902 | 1 | 1 |
| Mpz | 0,138934 | 0,308975 | 1,15309 | 1 | 1 |
| E130018N17Rik | 0,123171 | 0,272288 | 1,14447 | 1 | 1 |
| 6820408C15Rik | 0,066585 | 0,147135 | 1,14387 | 1 | 1 |
| Lmo3 | 0,084304 | 0,186249 | 1,14356 | 1 | 1 |
| Tigd4 | 0,029121 | 0,0642377 | 1,14137 | 1 | 1 |
| A930017M01Rik | 0,031984 | 0,069974 | 1,12948 | 1 | 1 |
| Padi4 | 0,131234 | 0,286116 | 1,12446 | 1 | 1 |
| Doc2b | 0,142021 | 0,309264 | 1,12273 | 1 | 1 |
| Kif9 | 0,14963 | 0,325681 | 1,12206 | 1 | 1 |
| Unc80 | 0,013323 | 0,0289931 | 1,12177 | 1 | 1 |
| Abcg8 | 0,035018 | 0,0761393 | 1,12056 | 1 | 1 |
| E130310I04Rik | 0,201857 | 0,438838 | 1,12036 | 1 | 1 |
| Kcnh7 | 0,260176 | 0,565466 | 1,11995 | 1 | 1 |
| Gpr21 | 0,226284 | 0,489618 | 1,11352 | 1 | 1 |
| Fam155a | 0,026196 | 0,0566065 | 1,11161 | 1 | 1 |
| Lamp3 | 0,118774 | 0,256335 | 1,10982 | 1 | 1 |
| Srrm4os | 0,011918 | 0,0256 | 1,103 | 1 | 1 |
| Upp2 | 0,032584 | 0,0697326 | 1,09765 | 1 | 1 |

| | | | | | |
|---|---|---|---|---|---|
| Plk5 | 0,043596 | 0,0931116 | 1,09475 | 1 | 1 |
| Adamts13 | 0,075917 | 0,162127 | 1,09462 | 1 | 1 |
| Kcnq2 | 0,170889 | 0,364909 | 1,09448 | 1 | 1 |
| Napsa | 0,106748 | 0,227321 | 1,09052 | 1 | 1 |
| Sorcs1 | 0,087581 | 0,185156 | 1,08004 | 1 | 1 |
| Tbx4 | 0,089673 | 0,188969 | 1,07541 | 1 | 1 |
| Ecm2 | 0,095966 | 0,202221 | 1,07533 | 1 | 1 |
| 1700084C01Rik | 0,210795 | 0,44218 | 1,06879 | 1 | 1 |
| Chadl | 0,184608 | 0,387167 | 1,06849 | 1 | 1 |
| Gm14858 | 0,038057 | 0,0794888 | 1,0626 | 1 | 1 |
| Nkx2-3 | 0,052736 | 0,109588 | 1,05522 | 1 | 1 |
| Syndig1 | 0,21744 | 0,449995 | 1,04929 | 1 | 1 |
| Ccl7 | 0,069713 | 0,143808 | 1,04465 | 1 | 1 |
| D16Ertd519e | 0,102165 | 0,210752 | 1,04465 | 1 | 1 |
| Il25 | 0,048288 | 0,0996118 | 1,04465 | 1 | 1 |
| Mrgprh | 0,025844 | 0,0533129 | 1,04465 | 1 | 1 |
| Tnxb | 0,004429 | 0,00913697 | 1,04465 | 1 | 1 |
| Rpl3l | 0,049676 | 0,102474 | 1,04464 | 1 | 1 |
| 4930512B01Rik | 0,092668 | 0,190782 | 1,04178 | 1 | 1 |
| Ankfn1 | 0,131334 | 0,270316 | 1,04141 | 1 | 1 |
| Ovol1 | 0,267806 | 0,551092 | 1,04111 | 1 | 1 |
| Psors1c2 | 0,24051 | 0,494914 | 1,04108 | 1 | 1 |
| Cd1d2 | 0,19177 | 0,393468 | 1,03687 | 1 | 1 |
| Alox8 | 0,028932 | 0,0593612 | 1,03685 | 1 | 1 |
| Ccr4 | 0,034376 | 0,0705301 | 1,03685 | 1 | 1 |
| Hist1h4h | 0,384362 | 0,788613 | 1,03685 | 1 | 1 |
| Lrfn5 | 0,025211 | 0,0517262 | 1,03685 | 1 | 1 |
| E030024N20Rik | 0,07787 | 0,159492 | 1,03434 | 1 | 1 |
| Duox1 | 0,086504 | 0,176778 | 1,03111 | 1 | 1 |
| 1700022I11Rik | 0,026867 | 0,054862 | 1,02998 | 1 | 1 |
| 4933433C11Rik | 0,038591 | 0,0788021 | 1,02998 | 1 | 1 |
| Aloxe3 | 0,040542 | 0,0827865 | 1,02998 | 1 | 1 |
| Cacna1f | 0,008059 | 0,0164558 | 1,02998 | 1 | 1 |
| Kcnj12 | 0,012512 | 0,0254914 | 1,02668 | 1 | 1 |
| Avpr2 | 0,081528 | 0,165538 | 1,02179 | 1 | 1 |
| Gdf5 | 0,047665 | 0,0967637 | 1,02154 | 1 | 1 |
| Klhl35 | 0,057563 | 0,116858 | 1,02154 | 1 | 1 |
| Gal | 0,358749 | 0,726315 | 1,01762 | 1 | 1 |
| 1810034E14Rik | 0,211104 | 0,427097 | 1,01661 | 1 | 1 |
| Osr1 | 0,272053 | 0,549547 | 1,01435 | 1 | 1 |
| Hapln4 | 0,06141 | 0,124018 | 1,01401 | 1 | 1 |
| Rsph4a | 0,01547 | 0,031241 | 1,01401 | 1 | 1 |
| C4a | 0,144624 | 0,29118 | 1,00961 | 1 | 1 |
| 4931406H21Rik | 0,043722 | 0,0880064 | 1,00924 | 1 | 1 |
| Tmem150cos | 0,011774 | 0,0236918 | 1,00884 | 1 | 1 |
| Lynx1 | 0,140544 | 0,282259 | 1,006 | 1 | 1 |
| Pou6f2 | 0,086829 | 0,174237 | 1,00481 | 1 | 1 |
| 1700018L02Rik | 0,207183 | 0,414917 | 1,00192 | 1 | 1 |

| | | | | | |
|---|---|---|---|---|---|
| Dntt | 0,060253 | 0,12055 | 1,00053 | 1 | 1 |
| Ces2g | 0,042354 | 0,0847319 | 1,00042 | 1 | 1 |
| Chrna4 | 0,236327 | 0,472607 | 0,999854 | 1 | 1 |
| Shisa9 | 0,055386 | 0,109691 | 0,985861 | 1 | 1 |
| AI197445 | 0,074376 | 0,14709 | 0,983797 | 1 | 1 |
| Cxcr6 | 0,119148 | 0,235449 | 0,982665 | 1 | 1 |
| Trank1 | 0,028696 | 0,0566594 | 0,981456 | 1 | 1 |
| Gareml | 0,182713 | 0,360094 | 0,978792 | 1 | 1 |
| Hcn1 | 0,008452 | 0,0166539 | 0,978565 | 1 | 1 |
| Cxcr3 | 0,025274 | 0,0496421 | 0,973933 | 1 | 1 |
| Dnah5 | 0,004605 | 0,00904465 | 0,973933 | 1 | 1 |
| Tac4 | 0,034021 | 0,0668239 | 0,973933 | 1 | 1 |
| Gsc | 0,323557 | 0,635289 | 0,973395 | 1 | 1 |
| Gm6277 | 0,208988 | 0,409064 | 0,968905 | 1 | 1 |
| Cacna1i | 0,078362 | 0,153272 | 0,967862 | 1 | 1 |
| A730020E08Rik | 0,081293 | 0,15867 | 0,964824 | 1 | 1 |
| Fcgr2b | 0,068938 | 0,134456 | 0,963771 | 1 | 1 |
| Cdh22 | 0,036747 | 0,0716225 | 0,962804 | 1 | 1 |
| C2cd4a | 0,036132 | 0,0703326 | 0,96091 | 1 | 1 |
| U90926 | 0,130436 | 0,253898 | 0,96091 | 1 | 1 |
| Baat | 0,285458 | 0,553802 | 0,956089 | 1 | 1 |
| Clvs2 | 0,164994 | 0,320094 | 0,956086 | 1 | 1 |
| Kl | 0,052988 | 0,102798 | 0,956081 | 1 | 1 |
| Rlbp1 | 0,114268 | 0,221384 | 0,954128 | 1 | 1 |
| Hpd | 0,265621 | 0,513837 | 0,951941 | 1 | 1 |
| Oprd1 | 0,130235 | 0,251846 | 0,951423 | 1 | 1 |
| Espnl | 0,024871 | 0,0480866 | 0,95119 | 1 | 1 |
| Gm17455 | 0,208427 | 0,402631 | 0,949916 | 1 | 1 |
| 4933407L21Rik | 0,094059 | 0,181285 | 0,946626 | 1 | 1 |
| Gm8787 | 0,074867 | 0,144295 | 0,946626 | 1 | 1 |
| Stk32b | 0,171944 | 0,331391 | 0,946597 | 1 | 1 |
| Col4a3 | 0,100727 | 0,193929 | 0,945076 | 1 | 1 |
| Rax | 0,064702 | 0,124354 | 0,942582 | 1 | 1 |
| Ccdc177 | 0,191926 | 0,368485 | 0,941057 | 1 | 1 |
| A230009B12Rik | 0,11474 | 0,219992 | 0,939078 | 1 | 1 |
| Foxo6 | 0,267447 | 0,512514 | 0,938339 | 1 | 1 |
| Lct | 0,1757 | 0,336593 | 0,93789 | 1 | 1 |
| 4732471J01Rik | 0,26402 | 0,505685 | 0,937591 | 1 | 1 |
| Cdnf | 0,109597 | 0,209798 | 0,936795 | 1 | 1 |
| Npb | 0,446155 | 0,854062 | 0,936795 | 1 | 1 |
| Lrrc38 | 0,145087 | 0,277589 | 0,93603 | 1 | 1 |
| 3200001D21Rik | 0,150963 | 0,288776 | 0,935755 | 1 | 1 |
| Gm10406 | 0,307494 | 0,588004 | 0,935268 | 1 | 1 |
| Tbx20 | 0,082876 | 0,157811 | 0,929176 | 1 | 1 |
| Rho | 0,099559 | 0,189133 | 0,925787 | 1 | 1 |
| Gata4 | 0,096486 | 0,183285 | 0,925699 | 1 | 1 |
| Siglecg | 0,051949 | 0,0986679 | 0,92548 | 1 | 1 |
| Gdf9 | 0,155345 | 0,295009 | 0,925278 | 1 | 1 |

| | | | | | |
|---|---|---|---|---|---|
| Ccdc42b | 0,09088 | 0,172412 | 0,923834 | 1 | 1 |
| 4931403E22Rik | 0,280753 | 0,532428 | 0,923284 | 1 | 1 |
| Slc35f3 | 0,075769 | 0,143691 | 0,923284 | 1 | 1 |
| Dbil5 | 0,366848 | 0,695657 | 0,923193 | 1 | 1 |
| Nr0b1 | 0,171303 | 0,323772 | 0,918426 | 1 | 1 |
| Hspb3 | 0,348765 | 0,658963 | 0,917942 | 1 | 1 |
| Nyx | 0,060649 | 0,114508 | 0,916896 | 1 | 1 |
| Trim66 | 0,031817 | 0,0599909 | 0,914962 | 1 | 1 |
| Gm5796 | 0,091826 | 0,173054 | 0,914243 | 1 | 1 |
| Tas1r1 | 0,256824 | 0,482877 | 0,910874 | 1 | 1 |
| 4930465K10Rik | 0,324517 | 0,609206 | 0,908636 | 1 | 1 |
| Crhr1 | 0,295501 | 0,554168 | 0,907163 | 1 | 1 |
| Cym | 0,086206 | 0,161465 | 0,905365 | 1 | 1 |
| Irf4 | 0,019824 | 0,0371311 | 0,905365 | 1 | 1 |
| Psmb9 | 0,294986 | 0,552515 | 0,905365 | 1 | 1 |
| Trim72 | 0,03915 | 0,0733287 | 0,905365 | 1 | 1 |
| 1700023E05Rik | 0,238284 | 0,44547 | 0,902645 | 1 | 1 |
| Ovol3 | 0,052652 | 0,0983198 | 0,901002 | 1 | 1 |
| Prkag2os1 | 0,143063 | 0,266019 | 0,894883 | 1 | 1 |
| Ankef1 | 0,096356 | 0,179166 | 0,894854 | 1 | 1 |
| Lrtm1 | 0,014987 | 0,0278394 | 0,893431 | 1 | 1 |
| Prkag3 | 0,040569 | 0,0753611 | 0,893431 | 1 | 1 |
| Tchh | 0,143287 | 0,265522 | 0,889923 | 1 | 1 |
| Pdcd1lg2 | 0,042387 | 0,078442 | 0,888003 | 1 | 1 |
| Phospho1 | 0,159996 | 0,295762 | 0,886402 | 1 | 1 |
| Glra4 | 0,152099 | 0,279677 | 0,878752 | 1 | 1 |
| Csmd1 | 0,019756 | 0,0361362 | 0,871171 | 1 | 1 |
| Epha10 | 0,028523 | 0,0521158 | 0,869604 | 1 | 1 |
| Zfp781 | 0,113333 | 0,207044 | 0,869375 | 1 | 1 |
| 6030408B16Rik | 0,033545 | 0,0610642 | 0,864216 | 1 | 1 |
| Mboat4 | 0,036171 | 0,0658447 | 0,864216 | 1 | 1 |
| 1700105P06Rik | 0,23588 | 0,428591 | 0,861545 | 1 | 1 |
| Ptf1a | 0,063494 | 0,115368 | 0,861545 | 1 | 1 |
| Dmrtb1 | 0,208367 | 0,378227 | 0,860123 | 1 | 1 |
| Wdr93 | 0,076771 | 0,139277 | 0,859321 | 1 | 1 |
| Gm5424 | 0,403234 | 0,730899 | 0,858053 | 1 | 1 |
| 1600014C23Rik | 0,169765 | 0,30767 | 0,857847 | 1 | 1 |
| CK137956 | 0,026588 | 0,0481854 | 0,857847 | 1 | 1 |
| Pla2g2e | 0,164783 | 0,298641 | 0,857847 | 1 | 1 |
| Pcdhb6 | 0,025205 | 0,0456794 | 0,857846 | 1 | 1 |
| Fam221b | 0,049368 | 0,0894132 | 0,856925 | 1 | 1 |
| Srpk3 | 0,100958 | 0,182816 | 0,856648 | 1 | 1 |
| Erich3 | 0,167236 | 0,30279 | 0,856436 | 1 | 1 |
| Serpina3g | 0,055408 | 0,100319 | 0,856436 | 1 | 1 |
| Htr3a | 0,10812 | 0,195755 | 0,856416 | 1 | 1 |
| Gltpd2 | 0,1244 | 0,225186 | 0,856133 | 1 | 1 |
| Clec2i | 0,01157 | 0,0209276 | 0,855047 | 1 | 1 |
| A930001A20Rik | 0,020357 | 0,0368176 | 0,854876 | 1 | 1 |

| | | | | | |
|---|---|---|---|---|---|
| 2610034M16Rik | 0,005959 | 0,0107768 | 0,854844 | 1 | 1 |
| 4930524C18Rik | 0,027979 | 0,0506015 | 0,854844 | 1 | 1 |
| 4930529C04Rik | 0,014221 | 0,0257191 | 0,854844 | 1 | 1 |
| 4933432I03Rik | 0,027603 | 0,0499221 | 0,854844 | 1 | 1 |
| 5730435O14Rik | 0,057277 | 0,10359 | 0,854844 | 1 | 1 |
| 5830418P13Rik | 0,011984 | 0,0216736 | 0,854844 | 1 | 1 |
| A530016L24Rik | 0,029181 | 0,0527759 | 0,854844 | 1 | 1 |
| Adcy10 | 0,009453 | 0,0170964 | 0,854844 | 1 | 1 |
| Cyp2b19 | 0,009497 | 0,0171753 | 0,854844 | 1 | 1 |
| Cyp2j13 | 0,005673 | 0,0102599 | 0,854844 | 1 | 1 |
| Gabrr1 | 0,011557 | 0,0209007 | 0,854844 | 1 | 1 |
| Gm5105 | 0,016192 | 0,0292837 | 0,854844 | 1 | 1 |
| Gm8298 | 0,023609 | 0,042698 | 0,854844 | 1 | 1 |
| Gtsf1l | 0,034994 | 0,0632886 | 0,854844 | 1 | 1 |
| Hist1h1t | 0,089099 | 0,161141 | 0,854844 | 1 | 1 |
| Khdc1a | 0,013549 | 0,0245038 | 0,854844 | 1 | 1 |
| Mypn | 0,004692 | 0,00848548 | 0,854844 | 1 | 1 |
| Nxph2 | 0,009961 | 0,0180142 | 0,854844 | 1 | 1 |
| R3hdml | 0,02658 | 0,0480711 | 0,854844 | 1 | 1 |
| Scgn | 0,058797 | 0,106338 | 0,854844 | 1 | 1 |
| Sec14l5 | 0,011091 | 0,020058 | 0,854844 | 1 | 1 |
| Sfrp5 | 0,013948 | 0,0252262 | 0,854844 | 1 | 1 |
| Slc46a2 | 0,012746 | 0,0230515 | 0,854844 | 1 | 1 |
| Spata25 | 0,040856 | 0,0738911 | 0,854844 | 1 | 1 |
| Tas2r135 | 0,031045 | 0,0561459 | 0,854844 | 1 | 1 |
| Tex29 | 0,041429 | 0,0749264 | 0,854844 | 1 | 1 |
| Tktl1 | 0,02087 | 0,0377448 | 0,854844 | 1 | 1 |
| Zfp42 | 0,014813 | 0,02679 | 0,854844 | 1 | 1 |
| 9930111J21Rik1 | 0,006467 | 0,0116927 | 0,854502 | 1 | 1 |
| Ccdc11 | 0,102622 | 0,184482 | 0,846136 | 1 | 1 |
| Tex16 | 0,139841 | 0,250909 | 0,843373 | 1 | 1 |
| Rgs20 | 0,285901 | 0,512516 | 0,842081 | 1 | 1 |
| Myo18b | 0,027126 | 0,0486209 | 0,841927 | 1 | 1 |
| Mybphl | 0,022312 | 0,0399099 | 0,838944 | 1 | 1 |
| Ccdc150 | 0,310433 | 0,555244 | 0,838839 | 1 | 1 |
| 1700108F19Rik | 0,356509 | 0,637057 | 0,837485 | 1 | 1 |
| Gm6559 | 0,111246 | 0,197758 | 0,829989 | 1 | 1 |
| Gm4975 | 0,041848 | 0,0743768 | 0,829691 | 1 | 1 |
| A630075F10Rik | 0,200304 | 0,355402 | 0,827259 | 1 | 1 |
| 4933405L10Rik | 0,128743 | 0,227273 | 0,81994 | 1 | 1 |
| Ttc21a | 0,133884 | 0,2361 | 0,818413 | 1 | 1 |
| Ckmt1 | 0,1453 | 0,256106 | 0,81771 | 1 | 1 |
| Abcc8 | 0,271814 | 0,479047 | 0,817548 | 1 | 1 |
| Mageb3 | 0,078338 | 0,177244 | 0,81712 | 1 | 1 |
| Adam21 | 0,107142 | 0,188391 | 0,814204 | 1 | 1 |
| Gm6682 | 0,150648 | 0,26416 | 0,810232 | 1 | 1 |
| Hhipl1 | 0,184839 | 0,323915 | 0,809347 | 1 | 1 |
| Hs3st4 | 0,028793 | 0,0503444 | 0,806102 | 1 | 1 |

| | | | | | |
|---|---|---|---|---|---|
| Mapk10 | 0,069384 | 0,121121 | 0,803776 | 1 | 1 |
| Gja10 | 0,054009 | 0,0940072 | 0,799563 | 1 | 1 |
| Iqub | 0,020589 | 0,0358358 | 0,799563 | 1 | 1 |
| Cnnm1 | 0,094283 | 0,164101 | 0,799508 | 1 | 1 |
| Onecut2 | 0,311516 | 0,539896 | 0,793376 | 1 | 1 |
| Col5a3 | 0,05738 | 0,0993707 | 0,792269 | 1 | 1 |
| Rtn4rl1 | 0,232657 | 0,402621 | 0,791219 | 1 | 1 |
| Wfdc10 | 0,255424 | 0,441084 | 0,788157 | 1 | 1 |
| Adam4 | 0,236566 | 0,406717 | 0,781787 | 1 | 1 |
| Ccdc67 | 0,291665 | 0,500255 | 0,778351 | 1 | 1 |
| Ptgdr2 | 0,060297 | 0,103272 | 0,776289 | 1 | 1 |
| Galnt13 | 0,203101 | 0,347594 | 0,775206 | 1 | 1 |
| Hcn2 | 0,213889 | 0,366023 | 0,775071 | 1 | 1 |
| Sec14l4 | 0,060539 | 0,103585 | 0,774874 | 1 | 1 |
| Syt17 | 0,14242 | 0,243215 | 0,772075 | 1 | 1 |
| 4933400F21Rik | 0,013887 | 0,023699 | 0,771107 | 1 | 1 |
| Dupd1 | 0,052627 | 0,0898126 | 0,771107 | 1 | 1 |
| Slc22a2 | 0,024463 | 0,0417488 | 0,771107 | 1 | 1 |
| Smtnl1 | 0,030387 | 0,0518574 | 0,771107 | 1 | 1 |
| Gm10474 | 0,156287 | 0,266358 | 0,76917 | 1 | 1 |
| Pcdhb2 | 0,080669 | 0,137423 | 0,768537 | 1 | 1 |
| Kcnip2 | 0,236216 | 0,402037 | 0,767222 | 1 | 1 |
| D730005E14Rik | 0,056247 | 0,0954766 | 0,763379 | 1 | 1 |
| Fam19a2 | 0,044979 | 0,0762086 | 0,760716 | 1 | 1 |
| Aox1 | 0,154468 | 0,261627 | 0,760204 | 1 | 1 |
| Stxbp5l | 0,023431 | 0,0396835 | 0,760139 | 1 | 1 |
| Selp | 0,023457 | 0,0396881 | 0,758672 | 1 | 1 |
| Gpr165 | 0,071261 | 0,120245 | 0,754806 | 1 | 1 |
| Elmod1 | 0,244091 | 0,411001 | 0,751724 | 1 | 1 |
| D030018L15Rik | 0,068914 | 0,115931 | 0,750403 | 1 | 1 |
| Hgfac | 0,087286 | 0,146809 | 0,750119 | 1 | 1 |
| Gbx1 | 0,033237 | 0,0558473 | 0,748698 | 1 | 1 |
| Tnn | 0,007401 | 0,012435 | 0,748698 | 1 | 1 |
| Traf1 | 0,019557 | 0,0328608 | 0,748698 | 1 | 1 |
| Tspan10 | 0,022301 | 0,0374719 | 0,748698 | 1 | 1 |
| Mettl7a2Higd1c | 0,033317 | 0,0559782 | 0,748598 | 1 | 1 |
| Nkpd1 | 0,285666 | 0,477833 | 0,742177 | 1 | 1 |
| Dnajb7 | 0,150643 | 0,251578 | 0,739873 | 1 | 1 |
| Chd5 | 0,166132 | 0,277202 | 0,738606 | 1 | 1 |
| 3110035E14Rik | 0,025801 | 0,0430276 | 0,737835 | 1 | 1 |
| A930019D19Rik | 0,102073 | 0,170225 | 0,737835 | 1 | 1 |
| Slc12a3 | 0,024879 | 0,0414907 | 0,737835 | 1 | 1 |
| Steap4 | 0,026214 | 0,0437163 | 0,737835 | 1 | 1 |
| A3galt2 | 0,154147 | 0,25602 | 0,731953 | 1 | 1 |
| Cdhr3 | 0,0646 | 0,107188 | 0,730548 | 1 | 1 |
| Plin1 | 0,039324 | 0,0650048 | 0,725156 | 1 | 1 |
| Lysmd1 | 0,20746 | 0,341741 | 0,720072 | 1 | 1 |
| Trim34a | 0,171849 | 0,2826 | 0,717618 | 1 | 1 |

| | | | | | |
|---|---|---|---|---|---|
| Rpgrip1 | 0,115331 | 0,189462 | 0,716126 | 1 | 1 |
| Abcb9 | 0,041621 | 0,0683446 | 0,715531 | 1 | 1 |
| F730043M19Rik | 0,298797 | 0,490283 | 0,714447 | 1 | 1 |
| Gm4787 | 0,258172 | 0,423623 | 0,714446 | 1 | 1 |
| Dlec1 | 0,142782 | 0,234209 | 0,71399 | 1 | 1 |
| 2310001H17Rik | 0,113931 | 0,186568 | 0,711541 | 1 | 1 |
| Ryr3 | 0,268196 | 0,438228 | 0,708394 | 1 | 1 |
| 1700007K13Rik | 0,098566 | 0,161017 | 0,708055 | 1 | 1 |
| Omd | 0,038229 | 0,0624509 | 0,708055 | 1 | 1 |
| Exd1 | 0,358517 | 0,585355 | 0,70727 | 1 | 1 |
| Ptprv | 0,040315 | 0,0657673 | 0,706065 | 1 | 1 |
| Fam26e | 0,08972 | 0,146133 | 0,70378 | 1 | 1 |
| Gm7104 | 0,07105 | 0,115591 | 0,702121 | 1 | 1 |
| Nrxn3 | 0,016884 | 0,0274344 | 0,700328 | 1 | 1 |
| B230216N24Rik | 0,176168 | 0,286231 | 0,700228 | 1 | 1 |
| Mog | 0,114164 | 0,185489 | 0,700228 | 1 | 1 |
| Efhb | 0,250596 | 0,406129 | 0,696574 | 1 | 1 |
| Gpr37 | 0,03747 | 0,0607062 | 0,696094 | 1 | 1 |
| Mag | 0,122451 | 0,198055 | 0,693691 | 1 | 1 |
| Sowahd | 0,099346 | 0,160684 | 0,693691 | 1 | 1 |
| Gpr75 | 0,141643 | 0,229012 | 0,693169 | 1 | 1 |
| Dpep1 | 0,255516 | 0,413035 | 0,692853 | 1 | 1 |
| Kcnn2 | 0,135025 | 0,217688 | 0,689036 | 1 | 1 |
| 9030624G23Rik | 0,32979 | 0,530941 | 0,687006 | 1 | 1 |
| Slc26a4 | 0,10288 | 0,165627 | 0,686972 | 1 | 1 |
| Paqr9 | 0,222569 | 0,35802 | 0,685787 | 1 | 1 |
| Mrps36 | 0,467827 | 0,751424 | 0,683653 | 1 | 1 |
| Rnase1 | 0,069592 | 0,111705 | 0,682708 | 1 | 1 |
| Six3 | 0,015336 | 0,0246167 | 0,682708 | 1 | 1 |
| Trim69 | 0,034304 | 0,0550625 | 0,682708 | 1 | 1 |
| Wbscr25 | 0,028492 | 0,0457334 | 0,682708 | 1 | 1 |
| Slc26a8 | 0,08363 | 0,134235 | 0,682667 | 1 | 1 |
| 4930509E16Rik | 0,131987 | 0,211846 | 0,682617 | 1 | 1 |
| Fam209 | 0,29274 | 0,469862 | 0,682617 | 1 | 1 |
| Rsph1 | 0,283845 | 0,454064 | 0,677792 | 1 | 1 |
| Kcnk10 | 0,31265 | 0,499857 | 0,676967 | 1 | 1 |
| Adcyap1r1 | 0,292206 | 0,466381 | 0,674521 | 1 | 1 |
| Adamts2 | 0,194287 | 0,309781 | 0,673059 | 1 | 1 |
| Scg3 | 0,062473 | 0,099579 | 0,672618 | 1 | 1 |
| Asb16 | 0,21478 | 0,342051 | 0,671354 | 1 | 1 |
| Vrtn | 0,077543 | 0,123299 | 0,669088 | 1 | 1 |
| Ryr2 | 0,208057 | 0,330147 | 0,666133 | 1 | 1 |
| Tmem169 | 0,258466 | 0,409403 | 0,663546 | 1 | 1 |
| 1700039E22Rik | 0,251056 | 0,397454 | 0,662777 | 1 | 1 |
| Ltbp2 | 0,136498 | 0,215966 | 0,661921 | 1 | 1 |
| Cd163 | 0,108835 | 0,172144 | 0,66147 | 1 | 1 |
| Chrna6 | 0,094208 | 0,148896 | 0,66039 | 1 | 1 |
| Gpnmb | 0,031288 | 0,0493474 | 0,657369 | 1 | 1 |

| | | | | | |
|---|---|---|---|---|---|
| Gm16523 | 0,17839 | 0,28083 | 0,654661 | 1 | 1 |
| Ccdc37 | 0,256721 | 0,403998 | 0,654146 | 1 | 1 |
| Gm14634 | 0,259825 | 0,408463 | 0,652667 | 1 | 1 |
| Sstr2 | 0,207321 | 0,325855 | 0,652364 | 1 | 1 |
| Coro6 | 0,081307 | 0,127646 | 0,650706 | 1 | 1 |
| F7 | 0,053651 | 0,0842267 | 0,650678 | 1 | 1 |
| Nppc | 0,162609 | 0,25527 | 0,650615 | 1 | 1 |
| Lrp2bp | 0,060711 | 0,0950577 | 0,646855 | 1 | 1 |
| Il22ra1 | 0,223315 | 0,349153 | 0,644782 | 1 | 1 |
| Optc | 0,242972 | 0,379251 | 0,642363 | 1 | 1 |
| Scd3 | 0,105076 | 0,16396 | 0,641912 | 1 | 1 |
| Gm14204 | 0,033183 | 0,0517286 | 0,64054 | 1 | 1 |
| Zan | 0,018187 | 0,0283205 | 0,638936 | 1 | 1 |
| Glp1r | 0,278217 | 0,433193 | 0,638802 | 1 | 1 |
| Grm8 | 0,185724 | 0,289027 | 0,638047 | 1 | 1 |
| BC100451 | 0,176734 | 0,275019 | 0,637954 | 1 | 1 |
| Adcyap1 | 0,166149 | 0,258413 | 0,637203 | 1 | 1 |
| 9530051G07Rik | 0,113883 | 0,176862 | 0,635072 | 1 | 1 |
| Gm10190 | 0,130234 | 0,201914 | 0,63264 | 1 | 1 |
| Lrit3 | 0,044949 | 0,0696802 | 0,632451 | 1 | 1 |
| Vwa5b1 | 0,210733 | 0,326438 | 0,631394 | 1 | 1 |
| Scn1a | 0,108408 | 0,16776 | 0,629926 | 1 | 1 |
| Htr2a | 0,293024 | 0,453401 | 0,629771 | 1 | 1 |
| Gpr45 | 0,022692 | 0,035107 | 0,629609 | 1 | 1 |
| Neurog3 | 0,062096 | 0,0960708 | 0,629609 | 1 | 1 |
| E130008D07Rik | 0,342581 | 0,529828 | 0,629076 | 1 | 1 |
| Gm9855 | 3,20114 | 4,93462 | 0,624351 | 1 | 1 |
| Bmp8a | 0,054829 | 0,0844451 | 0,623082 | 1 | 1 |
| Ccdc3 | 0,096532 | 0,148475 | 0,621151 | 1 | 1 |
| Dok7 | 0,264907 | 0,407266 | 0,620488 | 1 | 1 |
| Dscam | 0,066687 | 0,102513 | 0,620326 | 1 | 1 |
| Calb2 | 0,188176 | 0,289084 | 0,619411 | 1 | 1 |
| Rgn | 0,224712 | 0,344535 | 0,616572 | 1 | 1 |
| Gm9899 | 0,108956 | 0,166891 | 0,615166 | 1 | 1 |
| 4632428C04Rik | 0,085168 | 0,130322 | 0,613702 | 1 | 1 |
| 6430531B16Rik | 0,197922 | 0,302856 | 0,613702 | 1 | 1 |
| E130112N10Rik | 0,165142 | 0,252694 | 0,613687 | 1 | 1 |
| Casc1 | 0,162656 | 0,248787 | 0,613084 | 1 | 1 |
| Zfp872 | 0,081603 | 0,124697 | 0,611734 | 1 | 1 |
| Btbd18 | 0,189247 | 0,28802 | 0,605901 | 1 | 1 |
| Sugct | 0,073979 | 0,112588 | 0,605869 | 1 | 1 |
| 4833422C13Rik | 0,24176 | 0,367762 | 0,605198 | 1 | 1 |
| N28178 | 0,051978 | 0,0789832 | 0,603654 | 1 | 1 |
| Nrap | 0,039445 | 0,0599247 | 0,603293 | 1 | 1 |
| Col6a4 | 0,030279 | 0,0459738 | 0,602516 | 1 | 1 |
| 4930471M09Rik | 0,039079 | 0,0592816 | 0,601182 | 1 | 1 |
| 4930479D17Rik | 0,066201 | 0,100424 | 0,601182 | 1 | 1 |
| 4930548H24Rik | 0,029014 | 0,044013 | 0,601182 | 1 | 1 |

| | | | | | |
|---|---|---|---|---|---|
| Adam32 | 0,01582 | 0,0239983 | 0,601182 | 1 | 1 |
| Foxi2 | 0,014866 | 0,0225506 | 0,601182 | 1 | 1 |
| Gm6756 | 0,020885 | 0,0316822 | 0,601182 | 1 | 1 |
| Il23a | 0,030722 | 0,0466042 | 0,601182 | 1 | 1 |
| Nctc1 | 0,028027 | 0,0424512 | 0,598971 | 1 | 1 |
| Selenbp2 | 0,035753 | 0,0541529 | 0,598971 | 1 | 1 |
| Slitrk1 | 0,039517 | 0,0598536 | 0,598971 | 1 | 1 |
| Crtam | 0,194604 | 0,294187 | 0,596195 | 1 | 1 |
| Prdm6 | 0,392739 | 0,59277 | 0,593903 | 1 | 1 |
| Gpr179 | 0,087445 | 0,131978 | 0,59385 | 1 | 1 |
| C1ql2 | 0,108711 | 0,164065 | 0,593768 | 1 | 1 |
| Stk33 | 0,167579 | 0,2524 | 0,590876 | 1 | 1 |
| Snord45c | 224,246 | 337,6 | 0,59023 | 1 | 1 |
| Clec11a | 0,33451 | 0,502597 | 0,587354 | 1 | 1 |
| Fgf1 | 0,311553 | 0,467692 | 0,586082 | 1 | 1 |
| 1700028E10Rik | 0,159067 | 0,238621 | 0,585087 | 1 | 1 |
| AI463170 | 0,299456 | 0,447723 | 0,580266 | 1 | 1 |
| Trpm1 | 0,036939 | 0,0552277 | 0,580266 | 1 | 1 |
| Cpvl | 0,064381 | 0,09605 | 0,577158 | 1 | 1 |
| 4930550C14Rik | 0,292216 | 0,43555 | 0,575802 | 1 | 1 |
| Adipoq | 0,0807 | 0,120161 | 0,574328 | 1 | 1 |
| Pipox | 0,230041 | 0,341966 | 0,571962 | 1 | 1 |
| H2-Q6,H2-Q8 | 1,03599 | 1,53867 | 0,57068 | 1 | 1 |
| Gm10364 | 0,058105 | 0,0861831 | 0,568742 | 1 | 1 |
| 4933433G19Rik | 0,246656 | 0,365677 | 0,568067 | 1 | 1 |
| Nodal | 0,118993 | 0,175989 | 0,564608 | 1 | 1 |
| Tex11 | 0,037 | 0,0546005 | 0,561405 | 1 | 1 |
| Gm11744 | 0,347124 | 0,512061 | 0,560863 | 1 | 1 |
| Clec3b | 0,323028 | 0,476394 | 0,560498 | 1 | 1 |
| Fbxw10 | 0,113957 | 0,168032 | 0,560239 | 1 | 1 |
| AU040972 | 0,056729 | 0,0836381 | 0,560078 | 1 | 1 |
| Asb5 | 0,034409 | 0,0506891 | 0,558896 | 1 | 1 |
| Mrap2 | 0,054789 | 0,0807115 | 0,558879 | 1 | 1 |
| Tcp11 | 0,211067 | 0,310028 | 0,554696 | 1 | 1 |
| Ankrd7 | 0,126105 | 0,18491 | 0,552198 | 1 | 1 |
| Car5a | 0,102425 | 0,150152 | 0,551863 | 1 | 1 |
| Lhx2 | 0,177286 | 0,259831 | 0,551491 | 1 | 1 |
| Dnah11 | 0,264459 | 0,387537 | 0,55129 | 1 | 1 |
| Apc2 | 0,23231 | 0,34025 | 0,550542 | 1 | 1 |
| Btn2a2 | 0,222364 | 0,325489 | 0,549689 | 1 | 1 |
| Atg9b | 0,253267 | 0,37053 | 0,548934 | 1 | 1 |
| Gm5148 | 0,316204 | 0,462103 | 0,547358 | 1 | 1 |
| Stmn3 | 0,177849 | 0,259898 | 0,547293 | 1 | 1 |
| Hrh1 | 0,059523 | 0,0867625 | 0,543624 | 1 | 1 |
| Tmprss7 | 0,223752 | 0,32605 | 0,543193 | 1 | 1 |
| Tspear | 0,308882 | 0,450001 | 0,54287 | 1 | 1 |
| Caln1 | 0,225336 | 0,327686 | 0,540238 | 1 | 1 |
| Pou3f3 | 0,399466 | 0,579328 | 0,536308 | 1 | 1 |

| | | | | | |
|---|---|---|---|---|---|
| Fam83c | 0,316311 | 0,458117 | 0,534373 | 1 | 1 |
| 2010300C02Rik | 0,172203 | 0,249175 | 0,533051 | 1 | 1 |
| D830030K20Rik | 0,048917 | 0,070775 | 0,532916 | 1 | 1 |
| Slc1a7 | 0,021114 | 0,0305489 | 0,532916 | 1 | 1 |
| Slc9a2 | 0,017981 | 0,0260163 | 0,532916 | 1 | 1 |
| 4933411K16Rik | 0,18301 | 0,264618 | 0,531987 | 1 | 1 |
| 4933417O13Rik | 0,067165 | 0,0970881 | 0,531589 | 1 | 1 |
| Cyp4f18 | 0,059274 | 0,0856813 | 0,531589 | 1 | 1 |
| Igsf11 | 0,256582 | 0,369781 | 0,527253 | 1 | 1 |
| Smyd1 | 0,148037 | 0,21334 | 0,527203 | 1 | 1 |
| C1rl | 0,172943 | 0,248885 | 0,525184 | 1 | 1 |
| Gm5468 | 0,064135 | 0,0922525 | 0,524485 | 1 | 1 |
| Slc6a12 | 0,127962 | 0,184 | 0,523997 | 1 | 1 |
| Necab1 | 0,228229 | 0,328136 | 0,523809 | 1 | 1 |
| Ccdc170 | 0,100364 | 0,14319 | 0,512692 | 1 | 1 |
| Adra1d | 0,132522 | 0,188753 | 0,510261 | 1 | 1 |
| Pgr | 0,355085 | 0,505101 | 0,508407 | 1 | 1 |
| Arsj | 0,185084 | 0,263086 | 0,507352 | 1 | 1 |
| Henmt1 | 0,149918 | 0,212885 | 0,505893 | 1 | 1 |
| 4932435O22Rik | 0,08517 | 0,120841 | 0,50469 | 1 | 1 |
| B3gnt9 | 0,261962 | 0,371611 | 0,504438 | 1 | 1 |
| Sp7 | 0,318605 | 0,45154 | 0,503084 | 1 | 1 |
| Slc47a1 | 0,25578 | 0,361793 | 0,500259 | 1 | 1 |
| Rab3c | 0,321815 | 0,455175 | 0,500191 | 1 | 1 |
| Adra1b | 0,049261 | 0,0696293 | 0,499238 | 1 | 1 |
| Rapsn | 0,075178 | 0,106243 | 0,498975 | 1 | 1 |
| BC037032 | 0,29504 | 0,416915 | 0,49884 | 1 | 1 |
| D930015M05Rik | 0,370206 | 0,52269 | 0,497628 | 1 | 1 |
| Vipr1 | 0,167583 | 0,236344 | 0,49601 | 1 | 1 |
| Sdcbp2 | 0,446327 | 0,628812 | 0,494529 | 1 | 1 |
| Ube2ql1 | 0,129075 | 0,181568 | 0,4923 | 1 | 1 |
| Gabra4 | 0,24013 | 0,336677 | 0,487547 | 1 | 1 |
| Rspo4 | 0,371539 | 0,520352 | 0,485972 | 1 | 1 |
| Agap2 | 0,196233 | 0,274557 | 0,484542 | 1 | 1 |
| Ccdc154 | 0,034936 | 0,048868 | 0,484174 | 1 | 1 |
| BC051628 | 0,110681 | 0,154782 | 0,483829 | 1 | 1 |
| Mx1 | 0,036822 | 0,0514817 | 0,483495 | 1 | 1 |
| Atp8b5 | 0,099798 | 0,139471 | 0,48288 | 1 | 1 |
| Chac1 | 0,23391 | 0,326862 | 0,48273 | 1 | 1 |
| Prmt8 | 0,340269 | 0,47546 | 0,482647 | 1 | 1 |
| Ttll7 | 0,339764 | 0,473219 | 0,477976 | 1 | 1 |
| 5830454E08Rik | 0,329161 | 0,458344 | 0,477635 | 1 | 1 |
| 2210011C24Rik | 0,126303 | 0,17571 | 0,476308 | 1 | 1 |
| Calb1 | 0,100029 | 0,139158 | 0,476308 | 1 | 1 |
| Smc2os | 0,210403 | 0,292708 | 0,476308 | 1 | 1 |
| 5730416F02Rik | 0,165491 | 0,229265 | 0,470262 | 1 | 1 |
| Zfp652os | 0,302153 | 0,418591 | 0,470262 | 1 | 1 |
| Pyroxd2 | 0,333009 | 0,460214 | 0,466744 | 1 | 1 |

| | | | | | |
|---|---|---|---|---|---|
| P2ry14 | 0,143259 | 0,19798 | 0,466731 | 1 | 1 |
| Otoa | 0,032641 | 0,0450698 | 0,46547 | 1 | 1 |
| Arhgap36 | 0,407276 | 0,562046 | 0,464681 | 1 | 1 |
| Prkcg | 0,163536 | 0,225493 | 0,46347 | 1 | 1 |
| Egr3 | 0,242952 | 0,334769 | 0,462495 | 1 | 1 |
| Ppp1r32 | 0,174692 | 0,239935 | 0,457832 | 1 | 1 |
| Drc1 | 0,08903 | 0,122152 | 0,456308 | 1 | 1 |
| Tenm2 | 0,256771 | 0,351656 | 0,453683 | 1 | 1 |
| Usp18 | 0,39282 | 0,537709 | 0,452958 | 1 | 1 |
| Zfp114 | 0,241768 | 0,330442 | 0,450769 | 1 | 1 |
| Aph1c | 0,149734 | 0,204488 | 0,449616 | 1 | 1 |
| 4930529M08Rik | 0,043582 | 0,0595003 | 0,449179 | 1 | 1 |
| B3gnt6 | 0,027342 | 0,0373288 | 0,449179 | 1 | 1 |
| Elfn2 | 0,012699 | 0,0173379 | 0,449179 | 1 | 1 |
| Gria1 | 0,034257 | 0,0465685 | 0,442963 | 1 | 1 |
| Kcnb2 | 0,251599 | 0,341994 | 0,442846 | 1 | 1 |
| C1ra | 0,083359 | 0,113306 | 0,44281 | 1 | 1 |
| Vsnl1 | 0,341834 | 0,464568 | 0,442595 | 1 | 1 |
| Rpp25 | 0,394006 | 0,535382 | 0,442353 | 1 | 1 |
| Gpr141 | 0,025642 | 0,0348348 | 0,442043 | 1 | 1 |
| Lrrd1 | 0,030141 | 0,0409471 | 0,442043 | 1 | 1 |
| Nppa | 0,132473 | 0,179968 | 0,442043 | 1 | 1 |
| Piwil1 | 0,022138 | 0,0300753 | 0,442043 | 1 | 1 |
| 0610038B21Rik | 0,276228 | 0,375253 | 0,442007 | 1 | 1 |
| Gja3 | 0,124922 | 0,169699 | 0,441949 | 1 | 1 |
| Gdap1l1 | 0,144435 | 0,196042 | 0,44074 | 1 | 1 |
| Serpina3f | 0,051568 | 0,0698303 | 0,437367 | 1 | 1 |
| Fbxo41 | 0,125105 | 0,169349 | 0,436854 | 1 | 1 |
| Sybu | 0,247651 | 0,335136 | 0,436438 | 1 | 1 |
| Htr7 | 0,399051 | 0,538783 | 0,43313 | 1 | 1 |
| Arl5c | 0,375163 | 0,506503 | 0,433055 | 1 | 1 |
| Dlgap3 | 0,234323 | 0,316103 | 0,431898 | 1 | 1 |
| Kcnd3 | 0,245581 | 0,330446 | 0,428214 | 1 | 1 |
| Iqsec3 | 0,035665 | 0,0479895 | 0,428198 | 1 | 1 |
| Ankrd53 | 0,409248 | 0,549635 | 0,425499 | 1 | 1 |
| Gm9199 | 0,327645 | 0,439886 | 0,424998 | 1 | 1 |
| Slc9a7 | 0,355536 | 0,476819 | 0,423444 | 1 | 1 |
| Cd27 | 0,201148 | 0,269531 | 0,422199 | 1 | 1 |
| Gnb3 | 0,090093 | 0,120614 | 0,420915 | 1 | 1 |
| A730043L09Rik | 0,135968 | 0,181996 | 0,420642 | 1 | 1 |
| Gm5878 | 0,030661 | 0,0410407 | 0,420642 | 1 | 1 |
| 1110046J04Rik | 0,216919 | 0,29024 | 0,420089 | 1 | 1 |
| Gm7904 | 0,107915 | 0,144375 | 0,419924 | 1 | 1 |
| Tnfsf13b | 0,075267 | 0,100696 | 0,419924 | 1 | 1 |
| Fam46b | 0,184655 | 0,247024 | 0,41982 | 1 | 1 |
| Eci3 | 0,088108 | 0,117837 | 0,419439 | 1 | 1 |
| Ifitm10 | 0,033252 | 0,0444714 | 0,419439 | 1 | 1 |
| Mbd3l2 | 0,051574 | 0,068975 | 0,419439 | 1 | 1 |

| | | | | | |
|---|---|---|---|---|---|
| Wnt8b | 0,031722 | 0,0424245 | 0,419439 | 1 | 1 |
| Atp4a | 0,083325 | 0,111281 | 0,417376 | 1 | 1 |
| Dnah2 | 0,145143 | 0,193685 | 0,416236 | 1 | 1 |
| Gsto2 | 0,308671 | 0,411891 | 0,416194 | 1 | 1 |
| Gdnf | 0,17919 | 0,238658 | 0,413459 | 1 | 1 |
| Zfp488 | 0,049119 | 0,065352 | 0,411946 | 1 | 1 |
| Sh2d4b | 0,336076 | 0,446778 | 0,41077 | 1 | 1 |
| Epgn | 0,110395 | 0,146758 | 0,410763 | 1 | 1 |
| 4921511I17Rik | 0,087632 | 0,116472 | 0,410454 | 1 | 1 |
| B230319C09Rik | 0,11162 | 0,14824 | 0,409341 | 1 | 1 |
| Kcnv1 | 0,03862 | 0,0512904 | 0,409341 | 1 | 1 |
| Serpina12 | 0,044156 | 0,0585968 | 0,408205 | 1 | 1 |
| Ubap1l | 0,176071 | 0,232765 | 0,402719 | 1 | 1 |
| Brsk2 | 0,347951 | 0,459571 | 0,401405 | 1 | 1 |
| Ptprn2 | 0,289154 | 0,381514 | 0,399898 | 1 | 1 |
| Muc2 | 0,12793 | 0,168714 | 0,399224 | 1 | 1 |
| Gpat2 | 0,12941 | 0,170529 | 0,398068 | 1 | 1 |
| Ccdc65 | 0,219462 | 0,288738 | 0,395789 | 1 | 1 |
| 1700008I05Rik | 0,041435 | 0,054432 | 0,393603 | 1 | 1 |
| 4932441J04Rik | 0,124779 | 0,163919 | 0,393603 | 1 | 1 |
| Eef1a2 | 0,035362 | 0,0464534 | 0,393603 | 1 | 1 |
| Fam162b | 0,185567 | 0,243774 | 0,393603 | 1 | 1 |
| S100a4 | 0,662376 | 0,870143 | 0,393603 | 1 | 1 |
| Sertm1 | 0,211634 | 0,277615 | 0,391513 | 1 | 1 |
| 0610005C13Rik | 0,490652 | 0,643503 | 0,391248 | 1 | 1 |
| Dcdc2b | 0,32258 | 0,422405 | 0,388971 | 1 | 1 |
| Myh13 | 0,073813 | 0,0965107 | 0,386818 | 1 | 1 |
| Faim3 | 0,04142 | 0,054155 | 0,386759 | 1 | 1 |
| Neurod1 | 0,046557 | 0,0608714 | 0,386759 | 1 | 1 |
| Tacr2 | 0,025953 | 0,0339327 | 0,386759 | 1 | 1 |
| Hrk | 0,049312 | 0,0644363 | 0,385929 | 1 | 1 |
| Ankrd55 | 0,19182 | 0,250549 | 0,385339 | 1 | 1 |
| Gm10767 | 0,437299 | 0,570939 | 0,384718 | 1 | 1 |
| Adarb2 | 0,044749 | 0,0583724 | 0,383417 | 1 | 1 |
| Gm4925 | 0,161312 | 0,210029 | 0,380735 | 1 | 1 |
| Adam33 | 0,0431 | 0,0560093 | 0,377986 | 1 | 1 |
| Trpa1 | 0,296077 | 0,384741 | 0,377912 | 1 | 1 |
| Sez6l2 | 0,201075 | 0,261079 | 0,376753 | 1 | 1 |
| Myo15 | 0,056527 | 0,0733904 | 0,376663 | 1 | 1 |
| Hist2h2aa1,Hist2h | 0,117775 | 0,152851 | 0,376094 | 1 | 1 |
| Ccl17 | 0,133801 | 0,173633 | 0,375947 | 1 | 1 |
| Eras | 0,047514 | 0,0616583 | 0,375947 | 1 | 1 |
| Fam115e | 0,014274 | 0,0185231 | 0,375947 | 1 | 1 |
| Gm10267 | 0,2922 | 0,379186 | 0,375947 | 1 | 1 |
| Hes2 | 0,017902 | 0,0232318 | 0,375947 | 1 | 1 |
| Morn5 | 0,085348 | 0,110755 | 0,375947 | 1 | 1 |
| Olfr726 | 0,054435 | 0,0706398 | 0,375947 | 1 | 1 |
| Spatc1 | 0,029607 | 0,0384211 | 0,375947 | 1 | 1 |

| | | | | | |
|---|---|---|---|---|---|
| Wfikkn2 | 0,215345 | 0,279235 | 0,374832 | 1 | 1 |
| Gm3002 | 0,068518 | 0,088825 | 0,374489 | 1 | 1 |
| Klhl41 | 0,185077 | 0,239493 | 0,371856 | 1 | 1 |
| Gm2897 | 0,393656 | 0,508815 | 0,370207 | 1 | 1 |
| Hyal3 | 0,214844 | 0,277564 | 0,369532 | 1 | 1 |
| Atp8b3 | 0,185716 | 0,239816 | 0,368825 | 1 | 1 |
| Abcc6 | 0,320073 | 0,413122 | 0,368166 | 1 | 1 |
| Cbln3 | 0,244964 | 0,315798 | 0,366436 | 1 | 1 |
| Tcf24 | 0,063541 | 0,0818284 | 0,364923 | 1 | 1 |
| Grk4 | 0,336552 | 0,433034 | 0,363651 | 1 | 1 |
| Hsf3 | 0,087063 | 0,112018 | 0,363596 | 1 | 1 |
| A330076C08Rik | 0,341448 | 0,438808 | 0,361924 | 1 | 1 |
| Dok5 | 0,389951 | 0,50114 | 0,361921 | 1 | 1 |
| Kcna2 | 0,326125 | 0,418912 | 0,361222 | 1 | 1 |
| Chrnb2 | 0,217146 | 0,278833 | 0,36074 | 1 | 1 |
| Has1 | 0,181768 | 0,233197 | 0,359451 | 1 | 1 |
| Dcaf12l2 | 0,393605 | 0,504734 | 0,358775 | 1 | 1 |
| Gm19466 | 0,067602 | 0,0866694 | 0,358448 | 1 | 1 |
| Ogn | 0,060831 | 0,0779877 | 0,358448 | 1 | 1 |
| 5430440P10Rik | 0,211515 | 0,271103 | 0,358078 | 1 | 1 |
| Ckm | 0,027583 | 0,035329 | 0,357096 | 1 | 1 |
| Shank1 | 0,180246 | 0,230775 | 0,356517 | 1 | 1 |
| Aifm3 | 0,186146 | 0,238302 | 0,356356 | 1 | 1 |
| Acot4 | 0,075323 | 0,0963074 | 0,354551 | 1 | 1 |
| Mroh6 | 0,091551 | 0,116975 | 0,353559 | 1 | 1 |
| Rit2 | 0,344439 | 0,440047 | 0,353411 | 1 | 1 |
| Mx2 | 0,045397 | 0,0579335 | 0,351807 | 1 | 1 |
| 9530053A07Rik | 0,013037 | 0,0166302 | 0,351172 | 1 | 1 |
| Cyp4f39 | 0,041682 | 0,0531695 | 0,351172 | 1 | 1 |
| Scml4 | 0,022158 | 0,028265 | 0,351172 | 1 | 1 |
| Vmo1 | 0,214996 | 0,274248 | 0,351172 | 1 | 1 |
| Hrasls | 0,139737 | 0,178185 | 0,350667 | 1 | 1 |
| Ptprcap | 0,543876 | 0,693293 | 0,350188 | 1 | 1 |
| 4933413J09Rik | 0,078088 | 0,0995131 | 0,349789 | 1 | 1 |
| 4921504A21Rik | 0,117518 | 0,14956 | 0,347845 | 1 | 1 |
| Tsx | 0,552112 | 0,702495 | 0,347528 | 1 | 1 |
| Samd15 | 0,266141 | 0,338612 | 0,347445 | 1 | 1 |
| Fam189a1 | 0,312202 | 0,396907 | 0,346324 | 1 | 1 |
| Xrra1 | 0,097538 | 0,123972 | 0,345976 | 1 | 1 |
| Ptchd4 | 0,028354 | 0,0360205 | 0,345254 | 1 | 1 |
| C1qtnf2 | 0,11762 | 0,149393 | 0,344981 | 1 | 1 |
| Bfsp1 | 0,346764 | 0,439858 | 0,343086 | 1 | 1 |
| Fxyd2 | 0,250105 | 0,316832 | 0,341186 | 1 | 1 |
| Ccdc33 | 0,037767 | 0,0478432 | 0,341185 | 1 | 1 |
| Grpr | 0,038791 | 0,04914 | 0,341181 | 1 | 1 |
| Ldb3 | 0,075702 | 0,0957505 | 0,338943 | 1 | 1 |
| Vstm2l | 0,464696 | 0,587665 | 0,338707 | 1 | 1 |
| E330011O21Rik | 0,262643 | 0,332055 | 0,338322 | 1 | 1 |

| | | | | | |
|---|---|---|---|---|---|
| Gm9767 | 0,04962 | 0,0626794 | 0,33706 | 1 | 1 |
| Adrb1 | 0,447832 | 0,565159 | 0,3357 | 1 | 1 |
| Galr1 | 0,062151 | 0,0784002 | 0,335081 | 1 | 1 |
| Npas4 | 0,05394 | 0,0680422 | 0,335081 | 1 | 1 |
| Abcc3 | 0,387346 | 0,487575 | 0,332003 | 1 | 1 |
| Cnn1 | 0,212465 | 0,267436 | 0,331972 | 1 | 1 |
| Qrfp | 0,126869 | 0,159611 | 0,331218 | 1 | 1 |
| Gm1141 | 0,213206 | 0,268013 | 0,330055 | 1 | 1 |
| Rxrg | 0,104575 | 0,131442 | 0,329896 | 1 | 1 |
| Ak8 | 0,15598 | 0,196007 | 0,329548 | 1 | 1 |
| Ccdc63 | 0,041619 | 0,0522449 | 0,328053 | 1 | 1 |
| Gpr22 | 0,01756 | 0,0220425 | 0,328012 | 1 | 1 |
| Vwa5b2 | 0,215066 | 0,269755 | 0,326868 | 1 | 1 |
| Ntsr1 | 0,095682 | 0,119909 | 0,325625 | 1 | 1 |
| Ttll6 | 0,118311 | 0,148268 | 0,325625 | 1 | 1 |
| Fmo4 | 0,186901 | 0,234098 | 0,324835 | 1 | 1 |
| Gabrq | 0,034527 | 0,0432444 | 0,324795 | 1 | 1 |
| Mettl21c | 0,248185 | 0,310616 | 0,323716 | 1 | 1 |
| Atp6v1c2 | 0,329907 | 0,412199 | 0,321282 | 1 | 1 |
| 9330162012Rik | 0,22607 | 0,28197 | 0,318773 | 1 | 1 |
| Kcnc1 | 0,065156 | 0,0812555 | 0,318575 | 1 | 1 |
| Aipl1 | 0,325007 | 0,404652 | 0,316211 | 1 | 1 |
| Slc5a10 | 0,055113 | 0,0685457 | 0,314678 | 1 | 1 |
| Atp6v0a4 | 0,062516 | 0,0777386 | 0,314395 | 1 | 1 |
| Gm996 | 0,088502 | 0,10998 | 0,313468 | 1 | 1 |
| Nrsn2 | 0,187561 | 0,232963 | 0,31274 | 1 | 1 |
| Scn5a | 0,142804 | 0,177321 | 0,312325 | 1 | 1 |
| Htr1d | 0,025144 | 0,0311718 | 0,310057 | 1 | 1 |
| E330013P04Rik | 0,20614 | 0,255541 | 0,309929 | 1 | 1 |
| Gm13283 | 0,18525 | 0,229643 | 0,30992 | 1 | 1 |
| Lta | 0,095376 | 0,118231 | 0,30992 | 1 | 1 |
| 2900092D14Rik | 0,109986 | 0,136342 | 0,309908 | 1 | 1 |
| 1700071K01Rik | 0,0781 | 0,0968123 | 0,309866 | 1 | 1 |
| 2010310C07Rik | 0,101615 | 0,125961 | 0,309866 | 1 | 1 |
| Fa2h | 0,028469 | 0,0352894 | 0,309866 | 1 | 1 |
| Zfyve28 | 0,195972 | 0,242881 | 0,309597 | 1 | 1 |
| Rims1 | 0,134525 | 0,166641 | 0,308873 | 1 | 1 |
| 1700007J10Rik | 0,378543 | 0,467919 | 0,305802 | 1 | 1 |
| Asic4 | 0,200915 | 0,247991 | 0,303703 | 1 | 1 |
| Anks1b | 0,056456 | 0,0696838 | 0,303701 | 1 | 1 |
| Rab26 | 0,346452 | 0,427512 | 0,303312 | 1 | 1 |
| Gpr12 | 0,080737 | 0,099592 | 0,302809 | 1 | 1 |
| Lca5l | 0,373116 | 0,458121 | 0,296104 | 1 | 1 |
| Eml6 | 0,140693 | 0,172735 | 0,296016 | 1 | 1 |
| Shcbp1l | 0,054655 | 0,0670971 | 0,295891 | 1 | 1 |
| 9430018G01Rik | 0,12476 | 0,153056 | 0,294903 | 1 | 1 |
| Cacna2d2 | 0,159419 | 0,195058 | 0,291082 | 1 | 1 |
| Pate2 | 0,453296 | 0,554532 | 0,290816 | 1 | 1 |

| | | | | | |
|---|---|---|---|---|---|
| Dnah8 | 0,04406 | 0,0538489 | 0,289442 | 1 | 1 |
| Gm19897 | 0,147618 | 0,180384 | 0,289201 | 1 | 1 |
| Ascl4 | 0,142816 | 0,17451 | 0,289155 | 1 | 1 |
| Kdm4d | 0,055439 | 0,0676606 | 0,287426 | 1 | 1 |
| AU015228 | 0,11082 | 0,135134 | 0,28618 | 1 | 1 |
| Klhdc8a | 0,117985 | 0,143788 | 0,285335 | 1 | 1 |
| Zpbp | 0,210374 | 0,25635 | 0,285158 | 1 | 1 |
| Gm6498 | 0,199546 | 0,242894 | 0,283604 | 1 | 1 |
| Cyp2u1 | 0,178103 | 0,216723 | 0,283141 | 1 | 1 |
| Gm8773 | 0,50634 | 0,61484 | 0,280104 | 1 | 1 |
| Rplp2-ps1 | 369,442 | 448,318 | 0,279175 | 1 | 1 |
| Smok4a | 0,042002 | 0,0509254 | 0,277927 | 1 | 1 |
| Fbxo47 | 0,336453 | 0,407419 | 0,276108 | 1 | 1 |
| 1700061G19Rik | 0,155192 | 0,187922 | 0,276079 | 1 | 1 |
| Lrrc71 | 0,142257 | 0,171994 | 0,273858 | 1 | 1 |
| Nrg2 | 0,121738 | 0,147162 | 0,273621 | 1 | 1 |
| Fam198a | 0,282671 | 0,341702 | 0,273613 | 1 | 1 |
| Cyp27b1 | 0,176208 | 0,212907 | 0,272943 | 1 | 1 |
| Gpr1 | 0,122593 | 0,14806 | 0,272309 | 1 | 1 |
| Insl3 | 0,092556 | 0,111746 | 0,271836 | 1 | 1 |
| Skor1 | 0,010437 | 0,0125882 | 0,270326 | 1 | 1 |
| 4933405D12Rik | 0,125829 | 0,151713 | 0,269881 | 1 | 1 |
| Actl7b | 0,029157 | 0,0351549 | 0,269881 | 1 | 1 |
| Aicda | 0,016181 | 0,0195092 | 0,269881 | 1 | 1 |
| C030034L19Rik | 0,012107 | 0,014597 | 0,269881 | 1 | 1 |
| Ccdc183 | 0,02333 | 0,0281286 | 0,269881 | 1 | 1 |
| Gbp8 | 0,015736 | 0,0189728 | 0,269881 | 1 | 1 |
| Ifit3 | 0,039559 | 0,0476963 | 0,269881 | 1 | 1 |
| Il12rb1 | 0,0133 | 0,0160359 | 0,269881 | 1 | 1 |
| Kcnk12 | 0,0203 | 0,0244757 | 0,269881 | 1 | 1 |
| Map3k7cl | 0,024506 | 0,0295466 | 0,269881 | 1 | 1 |
| Otop2 | 0,016652 | 0,0200768 | 0,269881 | 1 | 1 |
| Pnpla1 | 0,008293 | 0,00999855 | 0,269881 | 1 | 1 |
| Prlhr | 0,025939 | 0,0312745 | 0,269881 | 1 | 1 |
| Spata45 | 0,158401 | 0,190985 | 0,269881 | 1 | 1 |
| Tbx19 | 0,014436 | 0,0174059 | 0,269881 | 1 | 1 |
| Bglap | 0,073772 | 0,0889455 | 0,269855 | 1 | 1 |
| Cabp5 | 0,138266 | 0,166581 | 0,268775 | 1 | 1 |
| Tmem154 | 0,056592 | 0,0681704 | 0,268554 | 1 | 1 |
| Acvr1c | 0,016807 | 0,0202424 | 0,268296 | 1 | 1 |
| BC006965 | 0,060993 | 0,0734272 | 0,26767 | 1 | 1 |
| Gdf7 | 0,089218 | 0,107406 | 0,26767 | 1 | 1 |
| Ncmap | 0,182523 | 0,219602 | 0,266816 | 1 | 1 |
| Gm1995 | 0,196066 | 0,235864 | 0,26662 | 1 | 1 |
| Pitx2 | 0,150208 | 0,18069 | 0,266551 | 1 | 1 |
| Cdk5r2 | 0,25335 | 0,30474 | 0,26645 | 1 | 1 |
| Gucy1b2 | 0,074306 | 0,0891845 | 0,263318 | 1 | 1 |
| Kcnt1 | 0,056551 | 0,0678726 | 0,263272 | 1 | 1 |

| | | | | | |
|---|---|---|---|---|---|
| Gm10640 | 0,159242 | 0,190702 | 0,2601 | 1 | 1 |
| Il1f5 | 0,068926 | 0,0825407 | 0,260049 | 1 | 1 |
| Trp53cor1 | 0,33478 | 0,400752 | 0,259497 | 1 | 1 |
| Pate4 | 0,247831 | 0,296589 | 0,259107 | 1 | 1 |
| 1700042O10Rik | 0,129102 | 0,1545 | 0,259097 | 1 | 1 |
| Ankrd66 | 0,100669 | 0,120474 | 0,259097 | 1 | 1 |
| A330023F24Rik | 0,096062 | 0,114898 | 0,258304 | 1 | 1 |
| Tctex1d1 | 0,317675 | 0,379715 | 0,257367 | 1 | 1 |
| Pgpep1l | 0,220712 | 0,26371 | 0,256791 | 1 | 1 |
| Akap3 | 0,070274 | 0,0837749 | 0,253536 | 1 | 1 |
| 9230102O04Rik | 0,244409 | 0,291299 | 0,253204 | 1 | 1 |
| Samd12 | 0,178453 | 0,212689 | 0,253204 | 1 | 1 |
| Tmem145 | 0,186636 | 0,222443 | 0,253204 | 1 | 1 |
| Rasd2 | 0,1749 | 0,208406 | 0,252868 | 1 | 1 |
| Mei4 | 0,105657 | 0,125843 | 0,25224 | 1 | 1 |
| Tmem25 | 0,454286 | 0,540824 | 0,25156 | 1 | 1 |
| 5730405O15Rik | 0,251225 | 0,299071 | 0,251508 | 1 | 1 |
| Capn8 | 0,052408 | 0,0623863 | 0,251434 | 1 | 1 |
| Dock3 | 0,226539 | 0,269601 | 0,251064 | 1 | 1 |
| Mitf | 0,444948 | 0,529395 | 0,250708 | 1 | 1 |
| 4732491K20Rik | 0,364704 | 0,433567 | 0,249532 | 1 | 1 |
| Pnmal1 | 0,486962 | 0,578867 | 0,249423 | 1 | 1 |
| Upb1 | 0,129258 | 0,153436 | 0,247381 | 1 | 1 |
| Tnk2os | 0,522492 | 0,620034 | 0,246937 | 1 | 1 |
| Adam1b | 0,035369 | 0,0419707 | 0,246911 | 1 | 1 |
| Rimbp3 | 0,333448 | 0,395663 | 0,246814 | 1 | 1 |
| Rpl34-ps1 | 546,702 | 648,371 | 0,246066 | 1 | 1 |
| Myom3 | 0,039648 | 0,0468697 | 0,2414 | 1 | 1 |
| 6030443J06Rik | 0,382187 | 0,451735 | 0,2412 | 1 | 1 |
| Fam81a | 0,435407 | 0,514403 | 0,240536 | 1 | 1 |
| Ntn5 | 0,085329 | 0,100723 | 0,239283 | 1 | 1 |
| Sspo | 0,100471 | 0,118528 | 0,238441 | 1 | 1 |
| Unc13a | 0,020598 | 0,0242908 | 0,237875 | 1 | 1 |
| 1700001K23Rik | 0,208233 | 0,245405 | 0,236968 | 1 | 1 |
| Pth2r | 0,02676 | 0,0315374 | 0,236968 | 1 | 1 |
| Usp50 | 0,038303 | 0,0451409 | 0,236968 | 1 | 1 |
| Tas1r3 | 0,221934 | 0,261391 | 0,236079 | 1 | 1 |
| Ccdc158 | 0,103488 | 0,121849 | 0,23564 | 1 | 1 |
| Fam46d | 0,042402 | 0,0499038 | 0,235006 | 1 | 1 |
| Gdf15 | 0,134594 | 0,158403 | 0,234977 | 1 | 1 |
| Kcnj3 | 0,080301 | 0,0945051 | 0,234977 | 1 | 1 |
| Xlr4b | 0,109265 | 0,128593 | 0,234977 | 1 | 1 |
| Igfals | 0,189303 | 0,222551 | 0,23344 | 1 | 1 |
| Fhl4 | 0,362431 | 0,425909 | 0,232838 | 1 | 1 |
| Icos | 0,079555 | 0,0934155 | 0,231718 | 1 | 1 |
| Pld5 | 0,341063 | 0,399866 | 0,229479 | 1 | 1 |
| Pik3r5 | 0,242013 | 0,283732 | 0,229441 | 1 | 1 |
| Mip | 0,457928 | 0,536689 | 0,228967 | 1 | 1 |

| | | | | | |
|---|---|---|---|---|---|
| Pstpip1 | 0,219187 | 0,256837 | 0,228692 | 1 | 1 |
| Kcng2 | 0,102021 | 0,119466 | 0,227728 | 1 | 1 |
| Trappc3l | 0,144851 | 0,169529 | 0,226962 | 1 | 1 |
| Celsr3 | 0,118477 | 0,13865 | 0,226847 | 1 | 1 |
| Cpne7 | 0,363002 | 0,424525 | 0,225872 | 1 | 1 |
| Gldn | 0,050596 | 0,0591501 | 0,225368 | 1 | 1 |
| Nrip3 | 0,172384 | 0,201504 | 0,225182 | 1 | 1 |
| Parp14 | 0,153684 | 0,179595 | 0,224787 | 1 | 1 |
| Wbp2nl | 0,074323 | 0,0867247 | 0,222628 | 1 | 1 |
| Ina | 0,100845 | 0,117633 | 0,222149 | 1 | 1 |
| Cxcl5 | 0,057135 | 0,0665369 | 0,219786 | 1 | 1 |
| Pmfbp1 | 0,025974 | 0,030248 | 0,219786 | 1 | 1 |
| Sgcd | 0,227789 | 0,265061 | 0,218627 | 1 | 1 |
| Rnf207 | 0,385913 | 0,448807 | 0,217819 | 1 | 1 |
| Shank2 | 0,352005 | 0,409047 | 0,216672 | 1 | 1 |
| Cacna1b | 0,241943 | 0,280951 | 0,21565 | 1 | 1 |
| Gpr137c | 0,38576 | 0,447626 | 0,214589 | 1 | 1 |
| Kcnab1 | 0,350458 | 0,406598 | 0,214359 | 1 | 1 |
| Gm10432 | 0,0403 | 0,0467497 | 0,214184 | 1 | 1 |
| Slc3a1 | 0,333682 | 0,386954 | 0,213687 | 1 | 1 |
| Pcsk1n | 0,276811 | 0,320927 | 0,213344 | 1 | 1 |
| 4833412C05Rik | 0,121291 | 0,140529 | 0,212389 | 1 | 1 |
| Krt27 | 0,079862 | 0,0925284 | 0,212389 | 1 | 1 |
| Slc6a19 | 0,058287 | 0,067311 | 0,207659 | 1 | 1 |
| Lama2 | 0,031762 | 0,0366485 | 0,206447 | 1 | 1 |
| Dnah17 | 0,189242 | 0,218205 | 0,205446 | 1 | 1 |
| Slc13a5 | 0,050221 | 0,0578644 | 0,204373 | 1 | 1 |
| Jakmip3 | 0,040764 | 0,0469046 | 0,202439 | 1 | 1 |
| Tgoln2 | 13,3383 | 15,3446 | 0,202155 | 1 | 1 |
| Gm17801 | 0,110659 | 0,127296 | 0,202071 | 1 | 1 |
| Avil | 0,15069 | 0,172951 | 0,198784 | 1 | 1 |
| Trim60 | 0,147384 | 0,16905 | 0,197872 | 1 | 1 |
| Oas1b | 0,410302 | 0,46992 | 0,195729 | 1 | 1 |
| Celf5 | 0,10184 | 0,116552 | 0,194667 | 1 | 1 |
| Rell2 | 0,297448 | 0,340288 | 0,19412 | 1 | 1 |
| Irak3 | 0,301968 | 0,345416 | 0,193937 | 1 | 1 |
| Brdt | 0,136607 | 0,155749 | 0,189189 | 1 | 1 |
| Mapt | 0,359309 | 0,409568 | 0,188879 | 1 | 1 |
| Hhipl2 | 0,217383 | 0,247741 | 0,188591 | 1 | 1 |
| Actg2 | 0,122653 | 0,139752 | 0,188288 | 1 | 1 |
| Strip2 | 0,304155 | 0,346465 | 0,187901 | 1 | 1 |
| Fam19a1 | 0,06023 | 0,0686003 | 0,187736 | 1 | 1 |
| 9030612E09Rik | 0,153066 | 0,174284 | 0,187292 | 1 | 1 |
| Efcab5 | 0,07315 | 0,0832761 | 0,187042 | 1 | 1 |
| Mmp19 | 0,209692 | 0,238702 | 0,186936 | 1 | 1 |
| Ppp1r3fos | 0,315797 | 0,359476 | 0,1869 | 1 | 1 |
| Hs3st5 | 0,008672 | 0,00986704 | 0,186189 | 1 | 1 |
| Ms4a7 | 0,022001 | 0,0250313 | 0,186189 | 1 | 1 |

| | | | | | |
|---|---|---|---|---|---|
| 2310034O05Rik | 0,026197 | 0,0298049 | 0,186145 | 1 | 1 |
| Acpt | 0,044052 | 0,050119 | 0,186145 | 1 | 1 |
| Cd4 | 0,008183 | 0,00930962 | 0,186145 | 1 | 1 |
| Dlx4 | 0,014639 | 0,0166547 | 0,186145 | 1 | 1 |
| E230008N13Rik | 0,014552 | 0,0165558 | 0,186145 | 1 | 1 |
| Ermn | 0,007132 | 0,008114 | 0,186145 | 1 | 1 |
| Gm15179 | 0,050118 | 0,0570201 | 0,186145 | 1 | 1 |
| Gsdmc4 | 0,012583 | 0,014316 | 0,186145 | 1 | 1 |
| Il18r1 | 0,005983 | 0,00680682 | 0,186145 | 1 | 1 |
| Khdrbs2 | 0,040963 | 0,0466042 | 0,186145 | 1 | 1 |
| Morc2b | 0,006747 | 0,00767659 | 0,186145 | 1 | 1 |
| Mroh2b | 0,004707 | 0,00535499 | 0,186145 | 1 | 1 |
| Muc5b | 0,001603 | 0,00182358 | 0,186145 | 1 | 1 |
| Nlrc4 | 0,00651 | 0,0074062 | 0,186145 | 1 | 1 |
| Oas3 | 0,005241 | 0,00596237 | 0,186145 | 1 | 1 |
| Patl2 | 0,010608 | 0,0120691 | 0,186145 | 1 | 1 |
| Resp18 | 0,04675 | 0,0531888 | 0,186145 | 1 | 1 |
| Sgcz | 0,010827 | 0,0123177 | 0,186145 | 1 | 1 |
| Slpi | 0,035202 | 0,0400502 | 0,186145 | 1 | 1 |
| Spata18 | 0,04067 | 0,0462705 | 0,186145 | 1 | 1 |
| Tex22 | 0,022254 | 0,0253188 | 0,186145 | 1 | 1 |
| Tnfrsf17 | 0,038908 | 0,0442659 | 0,186145 | 1 | 1 |
| Zmynd12 | 0,018183 | 0,0206871 | 0,186145 | 1 | 1 |
| Kcnn1 | 0,495168 | 0,562959 | 0,185112 | 1 | 1 |
| Syt3 | 0,420385 | 0,477891 | 0,18497 | 1 | 1 |
| Rarb | 0,289016 | 0,328497 | 0,184729 | 1 | 1 |
| Pax5 | 0,117971 | 0,134033 | 0,184154 | 1 | 1 |
| Slc5a1 | 0,106325 | 0,120656 | 0,182418 | 1 | 1 |
| Tdrd5 | 0,089729 | 0,101692 | 0,180571 | 1 | 1 |
| C1qtnf5 | 0,062036 | 0,0702118 | 0,178606 | 1 | 1 |
| Trim54 | 0,01792 | 0,0202665 | 0,17753 | 1 | 1 |
| Htr4 | 0,107705 | 0,121651 | 0,175669 | 1 | 1 |
| Osbp2 | 0,173351 | 0,195698 | 0,174932 | 1 | 1 |
| Oas1a | 0,189739 | 0,213499 | 0,170212 | 1 | 1 |
| Fbxo24 | 0,478464 | 0,537946 | 0,169051 | 1 | 1 |
| Mtl5 | 0,072852 | 0,0819064 | 0,169008 | 1 | 1 |
| Mei1 | 0,039063 | 0,043916 | 0,16893 | 1 | 1 |
| Fbxl21 | 0,423521 | 0,475733 | 0,167717 | 1 | 1 |
| 2900041M22Rik | 0,394554 | 0,44311 | 0,167443 | 1 | 1 |
| Gm7134 | 0,123731 | 0,138957 | 0,167425 | 1 | 1 |
| Neurl1b | 0,479673 | 0,538665 | 0,167338 | 1 | 1 |
| 9230116L04Rik | 0,073808 | 0,0828207 | 0,16622 | 1 | 1 |
| Gnrh1 | 0,781586 | 0,876671 | 0,16563 | 1 | 1 |
| Scd4 | 0,043332 | 0,0486038 | 0,16563 | 1 | 1 |
| Tmem215 | 0,10854 | 0,121727 | 0,165419 | 1 | 1 |
| Hivep3 | 0,057042 | 0,0639451 | 0,164805 | 1 | 1 |
| Atoh1 | 0,065981 | 0,0739106 | 0,163736 | 1 | 1 |
| Duxbl1,Duxbl2 | 0,124527 | 0,139467 | 0,163455 | 1 | 1 |

| | | | | | |
|---|---|---|---|---|---|
| Gm5126 | 0,420062 | 0,47043 | 0,16338 | 1 | 1 |
| Gm960 | 0,146677 | 0,163967 | 0,160764 | 1 | 1 |
| Odf3b | 0,120358 | 0,134546 | 0,160764 | 1 | 1 |
| Cys1 | 0,070795 | 0,0790179 | 0,158529 | 1 | 1 |
| 9030025P20Rik | 0,253362 | 0,282731 | 0,15823 | 1 | 1 |
| A330033J07Rik | 0,009234 | 0,010304 | 0,158128 | 1 | 1 |
| 1700020D05Rik | 0,348509 | 0,388646 | 0,157262 | 1 | 1 |
| Dcdc2a | 0,33328 | 0,371131 | 0,155195 | 1 | 1 |
| Ptger2 | 0,436762 | 0,486195 | 0,154685 | 1 | 1 |
| Poln | 0,387686 | 0,431299 | 0,153799 | 1 | 1 |
| Fcer2a | 0,083968 | 0,0933591 | 0,152952 | 1 | 1 |
| 1810010D01Rik | 0,105285 | 0,117054 | 0,152873 | 1 | 1 |
| Bpifc | 0,036236 | 0,0402859 | 0,152873 | 1 | 1 |
| Gm5607 | 0,015827 | 0,0175956 | 0,152873 | 1 | 1 |
| Ptgds | 0,128058 | 0,142373 | 0,152873 | 1 | 1 |
| Slc27a5 | 0,036201 | 0,040248 | 0,152873 | 1 | 1 |
| Ptgdr | 0,130905 | 0,14551 | 0,152596 | 1 | 1 |
| Smim6 | 0,406856 | 0,451991 | 0,151776 | 1 | 1 |
| Gm14023 | 0,34889 | 0,38758 | 0,151721 | 1 | 1 |
| Chdh | 0,076813 | 0,0852176 | 0,149807 | 1 | 1 |
| Umodl1 | 0,472093 | 0,523663 | 0,149566 | 1 | 1 |
| Shisa7 | 0,240809 | 0,266809 | 0,147919 | 1 | 1 |
| Rab42 | 0,40156 | 0,44445 | 0,146407 | 1 | 1 |
| Sez6l | 0,144131 | 0,159445 | 0,145676 | 1 | 1 |
| Ttc25 | 0,339742 | 0,375665 | 0,145006 | 1 | 1 |
| Gm16325 | 0,042438 | 0,0468853 | 0,143771 | 1 | 1 |
| Epsti1 | 0,021374 | 0,0235257 | 0,138398 | 1 | 1 |
| Gpr39 | 0,487215 | 0,53598 | 0,137621 | 1 | 1 |
| Gm16712 | 0,197623 | 0,21739 | 0,137536 | 1 | 1 |
| 9530052E02Rik | 0,037916 | 0,0416584 | 0,135812 | 1 | 1 |
| Gsg1l | 0,143214 | 0,157123 | 0,133725 | 1 | 1 |
| Cnih3 | 0,139287 | 0,152756 | 0,133172 | 1 | 1 |
| Catip | 0,374539 | 0,410708 | 0,132997 | 1 | 1 |
| Gm12250 | 0,263023 | 0,287857 | 0,130165 | 1 | 1 |
| Drd2 | 0,149621 | 0,163671 | 0,129484 | 1 | 1 |
| Entpd2 | 0,232245 | 0,254054 | 0,129484 | 1 | 1 |
| Tdh | 0,12 | 0,131269 | 0,129484 | 1 | 1 |
| Gm4262 | 0,306036 | 0,334573 | 0,128619 | 1 | 1 |
| Igfbp6 | 0,130967 | 0,143101 | 0,127824 | 1 | 1 |
| Lrguk | 0,518564 | 0,566044 | 0,126391 | 1 | 1 |
| 4833418N02Rik | 0,267726 | 0,291437 | 0,122429 | 1 | 1 |
| 2310043L19Rik | 0,179738 | 0,195567 | 0,121766 | 1 | 1 |
| Ces5a | 0,052571 | 0,0572005 | 0,121766 | 1 | 1 |
| Mrgprf | 0,049274 | 0,0536136 | 0,121766 | 1 | 1 |
| Pin4 | 0,327633 | 0,356486 | 0,121766 | 1 | 1 |
| Sgcg | 0,053635 | 0,0583582 | 0,121766 | 1 | 1 |
| Tmc2 | 0,02948 | 0,0320757 | 0,121766 | 1 | 1 |
| Catsperg1 | 0,158512 | 0,172419 | 0,121324 | 1 | 1 |

| | | | | | |
|---|---|---|---|---|---|
| Tecta | 0,18254 | 0,198493 | 0,120879 | 1 | 1 |
| Slc16a9 | 0,167433 | 0,182022 | 0,12053 | 1 | 1 |
| 4933406I18Rik | 0,056257 | 0,0611171 | 0,119551 | 1 | 1 |
| 6430548M08Rik | 0,467714 | 0,507254 | 0,117081 | 1 | 1 |
| Syt16 | 0,194164 | 0,210253 | 0,11485 | 1 | 1 |
| Nags | 0,065032 | 0,0704188 | 0,114807 | 1 | 1 |
| Prelp | 0,161982 | 0,175262 | 0,113682 | 1 | 1 |
| Gcnt7 | 0,137998 | 0,149246 | 0,11304 | 1 | 1 |
| Tmem30c | 0,130669 | 0,141179 | 0,111612 | 1 | 1 |
| Dnaic1 | 0,489278 | 0,528354 | 0,11085 | 1 | 1 |
| Zfp174 | 0,48273 | 0,520871 | 0,10971 | 1 | 1 |
| Tnfrsf26 | 0,068306 | 0,0736633 | 0,108932 | 1 | 1 |
| Arg2 | 0,225322 | 0,242991 | 0,108915 | 1 | 1 |
| Sh2d5 | 0,348784 | 0,376063 | 0,108642 | 1 | 1 |
| Ttc30a2 | 0,121432 | 0,130824 | 0,107476 | 1 | 1 |
| Slc17a9 | 0,142517 | 0,153389 | 0,10606 | 1 | 1 |
| Zmynd15 | 0,21612 | 0,232599 | 0,106016 | 1 | 1 |
| Rerg | 0,48241 | 0,519089 | 0,105721 | 1 | 1 |
| N4bp2l1 | 0,563294 | 0,604932 | 0,102886 | 1 | 1 |
| Stk32c | 0,553599 | 0,594081 | 0,101817 | 1 | 1 |
| Esrrb | 0,065034 | 0,0697798 | 0,101615 | 1 | 1 |
| Adck3 | 0,417836 | 0,447912 | 0,100279 | 1 | 1 |
| Prss50 | 0,104003 | 0,111427 | 0,099464 | 1 | 1 |
| Krt84 | 0,067593 | 0,0723308 | 0,0977451 | 1 | 1 |
| Nr3c2 | 0,041186 | 0,0440729 | 0,0977451 | 1 | 1 |
| Kcna5 | 0,173369 | 0,185514 | 0,0976844 | 1 | 1 |
| 4833411C07Rik | 0,195799 | 0,209511 | 0,0976546 | 1 | 1 |
| Npw | 0,162026 | 0,173373 | 0,0976546 | 1 | 1 |
| Nwd2 | 0,007818 | 0,0083651 | 0,0976546 | 1 | 1 |
| Ribc2 | 0,044983 | 0,0481337 | 0,0976546 | 1 | 1 |
| Ttc39d | 0,107426 | 0,114949 | 0,0976546 | 1 | 1 |
| 2610100L16Rik | 0,086816 | 0,0928867 | 0,0975107 | 1 | 1 |
| Atp2a1 | 0,214105 | 0,229035 | 0,0972518 | 1 | 1 |
| Cdhr5 | 0,264037 | 0,28182 | 0,0940291 | 1 | 1 |
| Pcdhac2 | 0,485434 | 0,517425 | 0,0920739 | 1 | 1 |
| Capn12 | 0,324305 | 0,345645 | 0,0919412 | 1 | 1 |
| Capn11 | 0,499003 | 0,53175 | 0,0917006 | 1 | 1 |
| Gsap | 0,267503 | 0,284921 | 0,0910047 | 1 | 1 |
| Il33 | 0,194675 | 0,207295 | 0,0906181 | 1 | 1 |
| Pllp | 0,161548 | 0,172009 | 0,0905183 | 1 | 1 |
| Atcay | 0,267651 | 0,284799 | 0,0895918 | 1 | 1 |
| Aoc3 | 0,114372 | 0,121654 | 0,0890543 | 1 | 1 |
| Slfn8 | 0,027964 | 0,029721 | 0,0879195 | 1 | 1 |
| Ccl28 | 0,311455 | 0,330692 | 0,0864618 | 1 | 1 |
| Zfa-ps | 0,379229 | 0,402337 | 0,085333 | 1 | 1 |
| Ky | 0,299162 | 0,317349 | 0,0851439 | 1 | 1 |
| Hepacam2 | 0,203973 | 0,215985 | 0,082553 | 1 | 1 |
| 1700001J11Rik | 0,062765 | 0,0664488 | 0,0822785 | 1 | 1 |

| | | | | | |
|---|---|---|---|---|---|
| Chad | 0,465736 | 0,492919 | 0,0818364 | 1 | 1 |
| Kcnk3 | 0,12469 | 0,131846 | 0,0805094 | 1 | 1 |
| Tunar | 0,349906 | 0,369931 | 0,0802908 | 1 | 1 |
| Qrich2 | 0,235845 | 0,249175 | 0,0793171 | 1 | 1 |
| 2900005J15Rik | 0,547128 | 0,577643 | 0,0783011 | 1 | 1 |
| Cacng8 | 0,458118 | 0,483523 | 0,0778671 | 1 | 1 |
| Insl5 | 0,322162 | 0,339913 | 0,0773796 | 1 | 1 |
| Ccdc162 | 0,041892 | 0,0441971 | 0,0772622 | 1 | 1 |
| Mmp24 | 0,039621 | 0,0417807 | 0,0765891 | 1 | 1 |
| A230057D06Rik | 0,2236 | 0,235781 | 0,0765224 | 1 | 1 |
| Zbtb20 | 0,167569 | 0,176668 | 0,076286 | 1 | 1 |
| Fam71f2 | 0,298386 | 0,314509 | 0,0759235 | 1 | 1 |
| Gpr35 | 0,065633 | 0,0691487 | 0,0752805 | 1 | 1 |
| Trim15 | 0,339963 | 0,357732 | 0,0735044 | 1 | 1 |
| Gadl1 | 0,496099 | 0,521366 | 0,0716681 | 1 | 1 |
| Hist2h2aa2 | 5,62E-005 | 5,90E-005 | 0,069664 | 1 | 1 |
| Gm20755 | 0,138568 | 0,145414 | 0,069568 | 1 | 1 |
| Sell | 0,038305 | 0,0401877 | 0,0692293 | 1 | 1 |
| Tshb | 0,21801 | 0,228719 | 0,0691848 | 1 | 1 |
| 4930469G21Rik | 0,102618 | 0,107655 | 0,0691361 | 1 | 1 |
| Clnk | 0,05569 | 0,058424 | 0,0691361 | 1 | 1 |
| Icam5 | 0,027705 | 0,0290649 | 0,0691361 | 1 | 1 |
| Jakmip1 | 0,106011 | 0,111215 | 0,0691361 | 1 | 1 |
| Slc17a7 | 0,082259 | 0,0862601 | 0,0685209 | 1 | 1 |
| Fam228b | 0,326597 | 0,341952 | 0,0662852 | 1 | 1 |
| Omp | 0,06275 | 0,0655343 | 0,0626338 | 1 | 1 |
| 4930404H11Rik | 0,210377 | 0,219403 | 0,0606069 | 1 | 1 |
| Mpped1 | 0,471593 | 0,491046 | 0,0583185 | 1 | 1 |
| Cass4 | 0,139972 | 0,14566 | 0,0574712 | 1 | 1 |
| Mpp4 | 0,067542 | 0,0702662 | 0,05704 | 1 | 1 |
| Nos1 | 0,161255 | 0,167735 | 0,0568391 | 1 | 1 |
| Myh11 | 0,025014 | 0,0260151 | 0,0566297 | 1 | 1 |
| AI854517 | 0,061488 | 0,063929 | 0,0561586 | 1 | 1 |
| Bco2 | 0,107558 | 0,111755 | 0,0552242 | 1 | 1 |
| BC030500 | 0,143757 | 0,149245 | 0,0540541 | 1 | 1 |
| BC049762 | 0,488231 | 0,506125 | 0,0519321 | 1 | 1 |
| Cd72 | 0,115527 | 0,119761 | 0,0519262 | 1 | 1 |
| Sox30 | 0,133641 | 0,138462 | 0,0511237 | 1 | 1 |
| Jazf1 | 0,169592 | 0,175235 | 0,047229 | 1 | 1 |
| Hba-a2 | 0,01212 | 0,012522 | 0,0470611 | 1 | 1 |
| Ccr7 | 0,143731 | 0,148459 | 0,0466897 | 1 | 1 |
| Hist1h2ah | 0,521229 | 0,538141 | 0,0460671 | 1 | 1 |
| 1700066B17Rik | 0,253691 | 0,261665 | 0,0446466 | 1 | 1 |
| 1810006J02Rik | 0,02797 | 0,0288493 | 0,0446466 | 1 | 1 |
| 9530003J23Rik | 0,040867 | 0,0421511 | 0,0446466 | 1 | 1 |
| Ces2f | 0,025621 | 0,0264259 | 0,0446466 | 1 | 1 |
| Cntn5 | 0,012192 | 0,0125757 | 0,0446466 | 1 | 1 |
| Dpp10 | 0,009372 | 0,00966647 | 0,0446466 | 1 | 1 |

| | | | | | |
|---|---|---|---|---|---|
| Gabra2 | 0,018958 | 0,0195538 | 0,0446466 | 1 | 1 |
| Gjd2 | 0,015507 | 0,0159939 | 0,0446466 | 1 | 1 |
| Gm10421 | 0,065015 | 0,0670581 | 0,0446466 | 1 | 1 |
| Gm19990 | 0,040786 | 0,0420682 | 0,0446466 | 1 | 1 |
| Gm3230 | 0,02197 | 0,0226608 | 0,0446466 | 1 | 1 |
| Lhx4 | 0,05037 | 0,0519534 | 0,0446466 | 1 | 1 |
| Mir1966 | 16,2082 | 16,7176 | 0,0446466 | 1 | 1 |
| Mir6403 | 7,73534 | 7,97847 | 0,0446466 | 1 | 1 |
| Rp1l1 | 0,006328 | 0,0065269 | 0,0446466 | 1 | 1 |
| Tspan2os | 0,107458 | 0,110835 | 0,0446466 | 1 | 1 |
| Wnt1 | 0,153133 | 0,157946 | 0,0446466 | 1 | 1 |
| Etnppl | 0,01693 | 0,0174619 | 0,0446258 | 1 | 1 |
| Tprg | 0,343021 | 0,352872 | 0,0408488 | 1 | 1 |
| Ttn | 0,006952 | 0,00714031 | 0,0386387 | 1 | 1 |
| Gm16796 | 0,135613 | 0,13928 | 0,0384933 | 1 | 1 |
| Dnah7b | 0,322068 | 0,330517 | 0,0373592 | 1 | 1 |
| Caps2 | 0,280256 | 0,287534 | 0,0369865 | 1 | 1 |
| Hsf2bp | 0,506777 | 0,51983 | 0,0366889 | 1 | 1 |
| Mir719 | 302,838 | 310,484 | 0,0359736 | 1 | 1 |
| 4930502E18Rik | 0,314001 | 0,321894 | 0,0358181 | 1 | 1 |
| Gm10941 | 0,619026 | 0,634257 | 0,0350673 | 1 | 1 |
| C2 | 0,055781 | 0,0571293 | 0,034468 | 1 | 1 |
| Cyb5r2 | 0,223232 | 0,228621 | 0,0344155 | 1 | 1 |
| A930011O12Rik | 0,142046 | 0,145393 | 0,0336029 | 1 | 1 |
| Chgb | 0,037416 | 0,0382016 | 0,0299834 | 1 | 1 |
| Gm15772 | 297,055 | 302,984 | 0,0285128 | 1 | 1 |
| Gm16880 | 0,469964 | 0,479233 | 0,0281755 | 1 | 1 |
| C1s2 | 0,038563 | 0,0393051 | 0,027496 | 1 | 1 |
| 9430021M05Rik | 0,319725 | 0,325874 | 0,0274816 | 1 | 1 |
| Kcnu1 | 0,08367 | 0,085247 | 0,0269451 | 1 | 1 |
| Klk7 | 0,157508 | 0,160319 | 0,0255167 | 1 | 1 |
| Cacnb2 | 0,377561 | 0,383868 | 0,0239006 | 1 | 1 |
| Btg3 | 0,000652 | 0,00066239 | 0,0236684 | 1 | 1 |
| 4930412C18Rik | 0,169863 | 0,172476 | 0,0220191 | 1 | 1 |
| Prrt3 | 0,306218 | 0,310666 | 0,020805 | 1 | 1 |
| Opcml | 0,06853 | 0,069509 | 0,0204693 | 1 | 1 |
| Cd28 | 0,04751 | 0,048152 | 0,0193638 | 1 | 1 |
| Zfp575 | 0,254446 | 0,257861 | 0,019237 | 1 | 1 |
| 1700003M02Rik | 0,567746 | 0,574147 | 0,016174 | 1 | 1 |
| Srp54a | 0,400925 | 0,405378 | 0,0159383 | 1 | 1 |
| Pstpip2 | 0,575451 | 0,581778 | 0,0157775 | 1 | 1 |
| B3gat1 | 0,433639 | 0,437995 | 0,0144202 | 1 | 1 |
| Acsm4 | 0,027352 | 0,0276183 | 0,0140083 | 1 | 1 |
| Atp10b | 0,007951 | 0,00802809 | 0,0140083 | 1 | 1 |
| Duox2 | 0,012255 | 0,0123743 | 0,0140083 | 1 | 1 |
| Gm13271 | 0,154511 | 0,156019 | 0,0140083 | 1 | 1 |
| Sp100 | 0,027282 | 0,0275477 | 0,0140083 | 1 | 1 |
| Pfn4 | 0,515788 | 0,52037 | 0,0127621 | 1 | 1 |

| | | | | | |
|---|---|---|---|---|---|
| Wdr17 | 0,432164 | 0,435527 | 0,0111817 | 1 | 1 |
| Kcnq4 | 0,161538 | 0,162653 | 0,00992317 | 1 | 1 |
| Syndig1l | 0,118681 | 0,119375 | 0,00841517 | 1 | 1 |
| Vwa7 | 0,387898 | 0,389951 | 0,00761514 | 1 | 1 |
| Gpr115 | 0,078955 | 0,0793417 | 0,00705605 | 1 | 1 |
| Jph2 | 0,500692 | 0,503123 | 0,00698668 | 1 | 1 |
| Ncam2 | 0,034114 | 0,034264 | 0,00631288 | 1 | 1 |
| 2010009K17Rik | 0,186257 | 0,187005 | 0,005787 | 1 | 1 |
| 4933403O08Rik | 0,161906 | 0,162551 | 0,00574093 | 1 | 1 |
| AA414768 | 0,339432 | 0,340565 | 0,0048061 | 1 | 1 |
| Dcst1 | 0,078632 | 0,0788945 | 0,0048061 | 1 | 1 |
| Hpx | 0,310852 | 0,311513 | 0,00306202 | 1 | 1 |
| Srp54c | 2,98816 | 2,99314 | 0,00240007 | 1 | 1 |
| Rbm46 | 0,264317 | 0,264645 | 0,00178681 | 1 | 1 |
| Marc1 | 0,058023 | 0,0580399 | 0,000423042 | 1 | 1 |
| Rtn4rl2 | 0,643068 | 0,643157 | 0,0002012 | 1 | 1 |
| 0610010B08Rik,Gı | 0 | 0 | 0 | 1 | 1 |
| 0610031O16Rik | 0 | 0 | 0 | 1 | 1 |
| 1110015O18Rik | 0 | 0 | 0 | 1 | 1 |
| 1110025L11Rik | 0 | 0 | 0 | 1 | 1 |
| 1190003K10Rik | 0 | 0 | 0 | 1 | 1 |
| 1500009C09Rik | 0 | 0 | 0 | 1 | 1 |
| 1500012K07Rik | 0 | 0 | 0 | 1 | 1 |
| 1500015L24Rik | 0 | 0 | 0 | 1 | 1 |
| 1500017E21Rik | 0 | 0 | 0 | 1 | 1 |
| 1600010M07Rik | 0 | 0 | 0 | 1 | 1 |
| 1600019K03Rik | 0 | 0 | 0 | 1 | 1 |
| 1600029I14Rik | 0 | 0 | 0 | 1 | 1 |
| 1700001C02Rik | 0 | 0 | 0 | 1 | 1 |
| 1700001F09Rik | 0 | 0 | 0 | 1 | 1 |
| 1700003C15Rik | 0 | 0 | 0 | 1 | 1 |
| 1700003E24Rik | 0 | 0 | 0 | 1 | 1 |
| 1700003G13Rik | 0 | 0 | 0 | 1 | 1 |
| 1700003G18Rik | 0 | 0 | 0 | 1 | 1 |
| 1700003H04Rik | 0 | 0 | 0 | 1 | 1 |
| 1700003L19Rik | 0 | 0 | 0 | 1 | 1 |
| 1700003P14Rik | 0 | 0 | 0 | 1 | 1 |
| 1700006A11Rik | 0 | 0 | 0 | 1 | 1 |
| 1700006E09Rik | 0 | 0 | 0 | 1 | 1 |
| 1700006F04Rik | 0 | 0 | 0 | 1 | 1 |
| 1700006H21Rik | 0 | 0 | 0 | 1 | 1 |
| 1700008K24Rik | 0 | 0 | 0 | 1 | 1 |
| 1700008P02Rik | 0 | 0 | 0 | 1 | 1 |
| 1700009N14Rik | 0 | 0 | 0 | 1 | 1 |
| 1700010D01Rik | 0 | 0 | 0 | 1 | 1 |
| 1700010I02Rik | 0 | 0 | 0 | 1 | 1 |
| 1700010J16Rik | 0 | 0 | 0 | 1 | 1 |
| 1700010K23Rik | 0 | 0 | 0 | 1 | 1 |

| | | | | | |
|---|---|---|---|---|---|
| 1700011B04Rik | 0 | 0 | 0 | 1 | 1 |
| 1700011E24Rik | 0 | 0 | 0 | 1 | 1 |
| 1700011I03Rik | 0 | 0 | 0 | 1 | 1 |
| 1700011M02Rik | 0 | 0 | 0 | 1 | 1 |
| 1700012A03Rik | 0 | 0 | 0 | 1 | 1 |
| 1700012I11Rik | 0 | 0 | 0 | 1 | 1 |
| 1700012P22Rik | 0 | 0 | 0 | 1 | 1 |
| 1700013D24Rik | 0 | 0 | 0 | 1 | 1 |
| 1700013G24Rik | 0 | 0 | 0 | 1 | 1 |
| 1700015G11Rik | 0 | 0 | 0 | 1 | 1 |
| 1700016C15Rik | 0 | 0 | 0 | 1 | 1 |
| 1700016L21Rik | 0 | 0 | 0 | 1 | 1 |
| 1700016P04Rik | 0 | 0 | 0 | 1 | 1 |
| 1700017D01Rik | 0 | 0 | 0 | 1 | 1 |
| 1700017N19Rik | 0 | 0 | 0 | 1 | 1 |
| 1700018B24Rik | 0 | 0 | 0 | 1 | 1 |
| 1700018F24Rik | 0 | 0 | 0 | 1 | 1 |
| 1700018G05Rik | 0 | 0 | 0 | 1 | 1 |
| 1700019E08Rik | 0 | 0 | 0 | 1 | 1 |
| 1700019M22Rik | 0 | 0 | 0 | 1 | 1 |
| 1700019O17Rik | 0 | 0 | 0 | 1 | 1 |
| 1700020G17Rik | 0 | 0 | 0 | 1 | 1 |
| 1700020M21Rik | 0 | 0 | 0 | 1 | 1 |
| 1700020N01Rik | 0 | 0 | 0 | 1 | 1 |
| 1700020N15Rik | 0 | 0 | 0 | 1 | 1 |
| 1700020N18Rik | 0 | 0 | 0 | 1 | 1 |
| 1700021F07Rik | 0 | 0 | 0 | 1 | 1 |
| 1700021N21Rik | 0 | 0 | 0 | 1 | 1 |
| 1700022A21Rik | 0 | 0 | 0 | 1 | 1 |
| 1700022A22Rik | 0 | 0 | 0 | 1 | 1 |
| 1700022E09Rik | 0 | 0 | 0 | 1 | 1 |
| 1700022H16Rik | 0 | 0 | 0 | 1 | 1 |
| 1700023C21Rik | 0 | 0 | 0 | 1 | 1 |
| 1700023F02Rik | 0 | 0 | 0 | 1 | 1 |
| 1700024B18Rik | 0 | 0 | 0 | 1 | 1 |
| 1700024P04Rik | 0 | 0 | 0 | 1 | 1 |
| 1700025B11Rik | 0 | 0 | 0 | 1 | 1 |
| 1700025C18Rik | 0 | 0 | 0 | 1 | 1 |
| 1700025F22Rik | 0 | 0 | 0 | 1 | 1 |
| 1700025K24Rik | 0 | 0 | 0 | 1 | 1 |
| 1700025M24Rik | 0 | 0 | 0 | 1 | 1 |
| 1700026D08Rik | 0 | 0 | 0 | 1 | 1 |
| 1700026D11Rik | 0 | 0 | 0 | 1 | 1 |
| 1700027A15Rik | 0 | 0 | 0 | 1 | 1 |
| 1700027F09Rik | 0 | 0 | 0 | 1 | 1 |
| 1700028B04Rik | 0 | 0 | 0 | 1 | 1 |
| 1700028D13Rik | 0 | 0 | 0 | 1 | 1 |
| 1700028M03Rik | 0 | 0 | 0 | 1 | 1 |

| | | | | | |
|---|---|---|---|---|---|
| 1700028P15Rik | 0 | 0 | 0 | 1 | 1 |
| 1700029B22Rik | 0 | 0 | 0 | 1 | 1 |
| 1700029F12Rik | 0 | 0 | 0 | 1 | 1 |
| 1700029P11Rik | 0 | 0 | 0 | 1 | 1 |
| 1700030F04Rik | 0 | 0 | 0 | 1 | 1 |
| 1700030F18Rik | 0 | 0 | 0 | 1 | 1 |
| 1700030L20Rik | 0 | 0 | 0 | 1 | 1 |
| 1700030N03Rik | 0 | 0 | 0 | 1 | 1 |
| 1700031F05Rik | 0 | 0 | 0 | 1 | 1 |
| 1700031M16Rik | 0 | 0 | 0 | 1 | 1 |
| 1700031P21Rik | 0 | 0 | 0 | 1 | 1 |
| 1700034E13Rik | 0 | 0 | 0 | 1 | 1 |
| 1700034G24Rik | 0 | 0 | 0 | 1 | 1 |
| 1700034I23Rik | 0 | 0 | 0 | 1 | 1 |
| 1700034K08Rik | 0 | 0 | 0 | 1 | 1 |
| 1700034O15Rik | 0 | 0 | 0 | 1 | 1 |
| 1700036G14Rik | 0 | 0 | 0 | 1 | 1 |
| 1700039E15Rik | 0 | 0 | 0 | 1 | 1 |
| 1700041C23Rik | 0 | 0 | 0 | 1 | 1 |
| 1700041M19Rik | 0 | 0 | 0 | 1 | 1 |
| 1700042B14Rik | 0 | 0 | 0 | 1 | 1 |
| 1700042G07Rik | 0 | 0 | 0 | 1 | 1 |
| 1700042G15Rik | 0 | 0 | 0 | 1 | 1 |
| 1700044C05Rik | 0 | 0 | 0 | 1 | 1 |
| 1700044K03Rik | 0 | 0 | 0 | 1 | 1 |
| 1700045H11Rik | 0 | 0 | 0 | 1 | 1 |
| 1700046C09Rik | 0 | 0 | 0 | 1 | 1 |
| 1700047E10Rik | 0 | 0 | 0 | 1 | 1 |
| 1700047G03Rik | 0 | 0 | 0 | 1 | 1 |
| 1700047L14Rik | 0 | 0 | 0 | 1 | 1 |
| 1700047M11Rik | 0 | 0 | 0 | 1 | 1 |
| 1700049E15Rik | 0 | 0 | 0 | 1 | 1 |
| 1700049E22Rik | 0 | 0 | 0 | 1 | 1 |
| 1700052I22Rik | 0 | 0 | 0 | 1 | 1 |
| 1700054K19Rik | 0 | 0 | 0 | 1 | 1 |
| 1700054M17Rik | 0 | 0 | 0 | 1 | 1 |
| 1700054O13Rik | 0 | 0 | 0 | 1 | 1 |
| 1700055C04Rik | 0 | 0 | 0 | 1 | 1 |
| 1700055N04Rik | 0 | 0 | 0 | 1 | 1 |
| 1700057G04Rik | 0 | 0 | 0 | 1 | 1 |
| 1700057H15Rik | 0 | 0 | 0 | 1 | 1 |
| 1700060C16Rik | 0 | 0 | 0 | 1 | 1 |
| 1700061F12Rik | 0 | 0 | 0 | 1 | 1 |
| 1700061I17Rik | 0 | 0 | 0 | 1 | 1 |
| 1700063A18Rik | 0 | 0 | 0 | 1 | 1 |
| 1700063O14Rik | 0 | 0 | 0 | 1 | 1 |
| 1700064J06Rik | 0 | 0 | 0 | 1 | 1 |
| 1700065I16Rik | 0 | 0 | 0 | 1 | 1 |

| | | | | | |
|---|---|---|---|---|---|
| 1700065J11Rik | 0 | 0 | 0 | 1 | 1 |
| 1700065L07Rik | 0 | 0 | 0 | 1 | 1 |
| 1700065O20Rik | 0 | 0 | 0 | 1 | 1 |
| 1700066N21Rik | 0 | 0 | 0 | 1 | 1 |
| 1700066O22Rik | 0 | 0 | 0 | 1 | 1 |
| 1700067G17Rik | 0 | 0 | 0 | 1 | 1 |
| 1700067P10Rik | 0 | 0 | 0 | 1 | 1 |
| 1700069L16Rik | 0 | 0 | 0 | 1 | 1 |
| 1700069P05Rik | 0 | 0 | 0 | 1 | 1 |
| 1700072B07Rik | 0 | 0 | 0 | 1 | 1 |
| 1700072O05Rik | 0 | 0 | 0 | 1 | 1 |
| 1700074H08Rik | 0 | 0 | 0 | 1 | 1 |
| 1700074P13Rik | 0 | 0 | 0 | 1 | 1 |
| 1700080E11Rik | 0 | 0 | 0 | 1 | 1 |
| 1700080N15Rik | 0 | 0 | 0 | 1 | 1 |
| 1700081H04Rik | 0 | 0 | 0 | 1 | 1 |
| 1700084F23Rik | 0 | 0 | 0 | 1 | 1 |
| 1700085C21Rik | 0 | 0 | 0 | 1 | 1 |
| 1700086L19Rik | 0 | 0 | 0 | 1 | 1 |
| 1700091H14Rik | 0 | 0 | 0 | 1 | 1 |
| 1700092C02Rik | 0 | 0 | 0 | 1 | 1 |
| 1700092C10Rik | 0 | 0 | 0 | 1 | 1 |
| 1700092E19Rik | 0 | 0 | 0 | 1 | 1 |
| 1700092K14Rik | 0 | 0 | 0 | 1 | 1 |
| 1700094M24Rik | 0 | 0 | 0 | 1 | 1 |
| 1700095A21Rik | 0 | 0 | 0 | 1 | 1 |
| 1700097N02Rik | 0 | 0 | 0 | 1 | 1 |
| 1700101O22Rik | 0 | 0 | 0 | 1 | 1 |
| 1700104L18Rik | 0 | 0 | 0 | 1 | 1 |
| 1700108J01Rik | 0 | 0 | 0 | 1 | 1 |
| 1700109G15Rik | 0 | 0 | 0 | 1 | 1 |
| 1700109I08Rik | 0 | 0 | 0 | 1 | 1 |
| 1700110I01Rik | 0 | 0 | 0 | 1 | 1 |
| 1700111N16Rik | 0 | 0 | 0 | 1 | 1 |
| 1700112H15Rik | 0 | 0 | 0 | 1 | 1 |
| 1700113H08Rik | 0 | 0 | 0 | 1 | 1 |
| 1700120G07Rik | 0 | 0 | 0 | 1 | 1 |
| 1700121L16Rik | 0 | 0 | 0 | 1 | 1 |
| 1700121N20Rik | 0 | 0 | 0 | 1 | 1 |
| 1700122O11Rik | 0 | 0 | 0 | 1 | 1 |
| 1700123I01Rik | 0 | 0 | 0 | 1 | 1 |
| 1700123K08Rik | 0 | 0 | 0 | 1 | 1 |
| 1700123O12Rik | 0 | 0 | 0 | 1 | 1 |
| 1700123O21Rik | 0 | 0 | 0 | 1 | 1 |
| 1700125G02Rik | 0 | 0 | 0 | 1 | 1 |
| 1700125G22Rik | 0 | 0 | 0 | 1 | 1 |
| 1700126H18Rik | 0 | 0 | 0 | 1 | 1 |
| 1700128A07Rik | 0 | 0 | 0 | 1 | 1 |

| | | | | | |
|---|---|---|---|---|---|
| 1700128F08Rik | 0 | 0 | 0 | 1 | 1 |
| 1700129C05Rik | 0 | 0 | 0 | 1 | 1 |
| 1810007C17Rik | 0 | 0 | 0 | 1 | 1 |
| 1810007D17Rik | 0 | 0 | 0 | 1 | 1 |
| 1810008I18Rik | 0 | 0 | 0 | 1 | 1 |
| 1810009J06Rik | 0 | 0 | 0 | 1 | 1 |
| 1810013A23Rik | 0 | 0 | 0 | 1 | 1 |
| 1810018F18Rik | 0 | 0 | 0 | 1 | 1 |
| 1810064F22Rik | 0 | 0 | 0 | 1 | 1 |
| 1810065E05Rik | 0 | 0 | 0 | 1 | 1 |
| 2010003K11Rik | 0 | 0 | 0 | 1 | 1 |
| 2010106C02Rik | 0 | 0 | 0 | 1 | 1 |
| 2010106E10Rik | 0 | 0 | 0 | 1 | 1 |
| 2010109I03Rik | 0 | 0 | 0 | 1 | 1 |
| 2210010C04Rik | 0 | 0 | 0 | 1 | 1 |
| 2210407C18Rik | 0 | 0 | 0 | 1 | 1 |
| 2210408F21Rik | 0 | 0 | 0 | 1 | 1 |
| 2210409D07Rik | 0 | 0 | 0 | 1 | 1 |
| 2210409E12Rik | 0 | 0 | 0 | 1 | 1 |
| 2210414B05Rik | 0 | 0 | 0 | 1 | 1 |
| 2210420H20Rik | 0 | 0 | 0 | 1 | 1 |
| 2300002M23Rik | 0 | 0 | 0 | 1 | 1 |
| 2300003K06Rik | 0 | 0 | 0 | 1 | 1 |
| 2300005B03Rik | 0 | 0 | 0 | 1 | 1 |
| 2310002L09Rik | 0 | 0 | 0 | 1 | 1 |
| 2310005A03Rik | 0 | 0 | 0 | 1 | 1 |
| 2310015D24Rik | 0 | 0 | 0 | 1 | 1 |
| 2310016D03Rik | 0 | 0 | 0 | 1 | 1 |
| 2310020H05Rik | 0 | 0 | 0 | 1 | 1 |
| 2310030A07Rik | 0 | 0 | 0 | 1 | 1 |
| 2310034C09Rik | 0 | 0 | 0 | 1 | 1 |
| 2310039L15Rik | 0 | 0 | 0 | 1 | 1 |
| 2310042E22Rik | 0 | 0 | 0 | 1 | 1 |
| 2310043O21Rik | 0 | 0 | 0 | 1 | 1 |
| 2310050C09Rik | 0 | 0 | 0 | 1 | 1 |
| 2310057J18Rik | 0 | 0 | 0 | 1 | 1 |
| 2310057N15Rik | 0 | 0 | 0 | 1 | 1 |
| 2310069B03Rik | 0 | 0 | 0 | 1 | 1 |
| 2310079G19Rik | 0 | 0 | 0 | 1 | 1 |
| 2310081J21Rik | 0 | 0 | 0 | 1 | 1 |
| 2410007B07Rik | 0 | 0 | 0 | 1 | 1 |
| 2410114N07Rik | 0 | 0 | 0 | 1 | 1 |
| 2410124H12Rik | 0 | 0 | 0 | 1 | 1 |
| 2410141K09Rik | 0 | 0 | 0 | 1 | 1 |
| 2610020C07Rik | 2,79813 | 0 | 0 | 1 | 1 |
| 2610207O16Rik | 0 | 0 | 0 | 1 | 1 |
| 2700070H01Rik | 0 | 0 | 0 | 1 | 1 |
| 2810007J24Rik | 0 | 0 | 0 | 1 | 1 |

| | | | | | |
|---|---|---|---|---|---|
| 2810404M03Rik | 0 | 0 | 0 | 1 | 1 |
| 2810471M01Rik | 0 | 0 | 0 | 1 | 1 |
| 2900079G21Rik | 0 | 0 | 0 | 1 | 1 |
| 2900092C05Rik | 0 | 0 | 0 | 1 | 1 |
| 3010033K07Rik | 0 | 0 | 0 | 1 | 1 |
| 3100003L05Rik | 0 | 0 | 0 | 1 | 1 |
| 3110009F21Rik | 0 | 0 | 0 | 1 | 1 |
| 3110015C05Rik | 0 | 0 | 0 | 1 | 1 |
| 3110039I08Rik | 0 | 0 | 0 | 1 | 1 |
| 3110039M20Rik | 0 | 0 | 0 | 1 | 1 |
| 3110082J24Rik | 0 | 0 | 0 | 1 | 1 |
| 4833423E24Rik | 0 | 0 | 0 | 1 | 1 |
| 4833428L15Rik | 0 | 0 | 0 | 1 | 1 |
| 4921501E09Rik | 0 | 0 | 0 | 1 | 1 |
| 4921504E06Rik | 0 | 0 | 0 | 1 | 1 |
| 4921509C19Rik | 0 | 0 | 0 | 1 | 1 |
| 4921509O07Rik | 0 | 0 | 0 | 1 | 1 |
| 4921511C20Rik | 0 | 0 | 0 | 1 | 1 |
| 4921511H03Rik | 0 | 0 | 0 | 1 | 1 |
| 4921511M17Rik,G | 0 | 0 | 0 | 1 | 1 |
| 4921515E04Rik | 0 | 0 | 0 | 1 | 1 |
| 4921524L21Rik | 0 | 0 | 0 | 1 | 1 |
| 4921531P14Rik | 0 | 0 | 0 | 1 | 1 |
| 4922502D21Rik | 0 | 0 | 0 | 1 | 1 |
| 4922502H24Rik | 0 | 0 | 0 | 1 | 1 |
| 4922502N22Rik | 0 | 0 | 0 | 1 | 1 |
| 4930401C15Rik | 0 | 0 | 0 | 1 | 1 |
| 4930401O12Rik | 0 | 0 | 0 | 1 | 1 |
| 4930402F06Rik | 0 | 0 | 0 | 1 | 1 |
| 4930402F11Rik | 0 | 0 | 0 | 1 | 1 |
| 4930402K13Rik | 0 | 0 | 0 | 1 | 1 |
| 4930404A05Rik | 0 | 0 | 0 | 1 | 1 |
| 4930404A10Rik | 0 | 0 | 0 | 1 | 1 |
| 4930405A10Rik | 0 | 0 | 0 | 1 | 1 |
| 4930405J17Rik | 0 | 0 | 0 | 1 | 1 |
| 4930405L22Rik | 0 | 0 | 0 | 1 | 1 |
| 4930406D18Rik | 0 | 0 | 0 | 1 | 1 |
| 4930407I10Rik | 0 | 0 | 0 | 1 | 1 |
| 4930412B13Rik | 0 | 0 | 0 | 1 | 1 |
| 4930412D23Rik | 0 | 0 | 0 | 1 | 1 |
| 4930413E15Rik | 0 | 0 | 0 | 1 | 1 |
| 4930413F20Rik | 0 | 0 | 0 | 1 | 1 |
| 4930413M19Rik | 0 | 0 | 0 | 1 | 1 |
| 4930415L06Rik | 0 | 0 | 0 | 1 | 1 |
| 4930417O13Rik | 0 | 0 | 0 | 1 | 1 |
| 4930419G24Rik | 0 | 0 | 0 | 1 | 1 |
| 4930423M02Rik | 0 | 0 | 0 | 1 | 1 |
| 4930425K10Rik | 0 | 0 | 0 | 1 | 1 |

| | | | | | |
|---|---|---|---|---|---|
| 4930425O10Rik | 0 | 0 | 0 | 1 | 1 |
| 4930428D18Rik | 0 | 0 | 0 | 1 | 1 |
| 4930428E07Rik | 0 | 0 | 0 | 1 | 1 |
| 4930429F11Rik | 0 | 0 | 0 | 1 | 1 |
| 4930430D24Rik | 0 | 0 | 0 | 1 | 1 |
| 4930430F21Rik | 0 | 0 | 0 | 1 | 1 |
| 4930432J09Rik | 0 | 0 | 0 | 1 | 1 |
| 4930432M17Rik | 0 | 0 | 0 | 1 | 1 |
| 4930433B08Rik | 0 | 0 | 0 | 1 | 1 |
| 4930433I11Rik | 0 | 0 | 0 | 1 | 1 |
| 4930433N12Rik | 0 | 0 | 0 | 1 | 1 |
| 4930434J06Rik | 0 | 0 | 0 | 1 | 1 |
| 4930435E12Rik | 0 | 0 | 0 | 1 | 1 |
| 4930440C22Rik | 0 | 0 | 0 | 1 | 1 |
| 4930440I19Rik | 0 | 0 | 0 | 1 | 1 |
| 4930441J16Rik | 0 | 0 | 0 | 1 | 1 |
| 4930442J19Rik | 0 | 0 | 0 | 1 | 1 |
| 4930444G20Rik | 0 | 0 | 0 | 1 | 1 |
| 4930447A16Rik | 0 | 0 | 0 | 1 | 1 |
| 4930448C13Rik | 0 | 0 | 0 | 1 | 1 |
| 4930448F12Rik | 0 | 0 | 0 | 1 | 1 |
| 4930448H16Rik | 0 | 0 | 0 | 1 | 1 |
| 4930448I06Rik | 0 | 0 | 0 | 1 | 1 |
| 4930448I18Rik | 0 | 0 | 0 | 1 | 1 |
| 4930448K20Rik | 0 | 0 | 0 | 1 | 1 |
| 4930449E18Rik | 0 | 0 | 0 | 1 | 1 |
| 4930449I24Rik | 0 | 0 | 0 | 1 | 1 |
| 4930451I11Rik | 0 | 0 | 0 | 1 | 1 |
| 4930452A19Rik | 0 | 0 | 0 | 1 | 1 |
| 4930452N14Rik | 0 | 0 | 0 | 1 | 1 |
| 4930455B14Rik | 0 | 0 | 0 | 1 | 1 |
| 4930455F16Rik | 0 | 0 | 0 | 1 | 1 |
| 4930455H04Rik | 0 | 0 | 0 | 1 | 1 |
| 4930455J16Rik | 0 | 0 | 0 | 1 | 1 |
| 4930459C07Rik | 0 | 0 | 0 | 1 | 1 |
| 4930459L07Rik | 0 | 0 | 0 | 1 | 1 |
| 4930465M20Rik | 0 | 0 | 0 | 1 | 1 |
| 4930467D21Rik | 0 | 0 | 0 | 1 | 1 |
| 4930467E23Rik | 0 | 0 | 0 | 1 | 1 |
| 4930468A15Rik | 0 | 0 | 0 | 1 | 1 |
| 4930470H14Rik | 0 | 0 | 0 | 1 | 1 |
| 4930470P17Rik | 0 | 0 | 0 | 1 | 1 |
| 4930471G03Rik | 0 | 0 | 0 | 1 | 1 |
| 4930473O22Rik | 0 | 0 | 0 | 1 | 1 |
| 4930474G06Rik | 0 | 0 | 0 | 1 | 1 |
| 4930474H20Rik | 0 | 0 | 0 | 1 | 1 |
| 4930474N05Rik | 0 | 0 | 0 | 1 | 1 |
| 4930478P22Rik | 0 | 0 | 0 | 1 | 1 |

| | | | | | |
|---|---|---|---|---|---|
| 4930480E11Rik | 0 | 0 | 0 | 1 | 1 |
| 4930480G23Rik | 0 | 0 | 0 | 1 | 1 |
| 4930480K15Rik | 0 | 0 | 0 | 1 | 1 |
| 4930480M12Rik | 0 | 0 | 0 | 1 | 1 |
| 4930482G09Rik | 0 | 0 | 0 | 1 | 1 |
| 4930486F22Rik | 0 | 0 | 0 | 1 | 1 |
| 4930486I03Rik | 0 | 0 | 0 | 1 | 1 |
| 4930487D11Rik | 0 | 0 | 0 | 1 | 1 |
| 4930488B22Rik | 0 | 0 | 0 | 1 | 1 |
| 4930502A04Rik | 0 | 0 | 0 | 1 | 1 |
| 4930503E14Rik | 0 | 0 | 0 | 1 | 1 |
| 4930503H13Rik | 0 | 0 | 0 | 1 | 1 |
| 4930503O07Rik | 0 | 0 | 0 | 1 | 1 |
| 4930504O13Rik | 0 | 0 | 0 | 1 | 1 |
| 4930505G20Rik | 0 | 0 | 0 | 1 | 1 |
| 4930507D10Rik | 0 | 0 | 0 | 1 | 1 |
| 4930509J09Rik | 0 | 0 | 0 | 1 | 1 |
| 4930509K18Rik | 0 | 0 | 0 | 1 | 1 |
| 4930513O06Rik | 0 | 0 | 0 | 1 | 1 |
| 4930515B02Rik | 0 | 0 | 0 | 1 | 1 |
| 4930515L19Rik | 0 | 0 | 0 | 1 | 1 |
| 4930518P08Rik | 0 | 0 | 0 | 1 | 1 |
| 4930519D14Rik | 0 | 0 | 0 | 1 | 1 |
| 4930519F24Rik | 0 | 0 | 0 | 1 | 1 |
| 4930519H02Rik | 0 | 0 | 0 | 1 | 1 |
| 4930520P13Rik | 0 | 0 | 0 | 1 | 1 |
| 4930521E06Rik | 0 | 0 | 0 | 1 | 1 |
| 4930522H14Rik | 0 | 0 | 0 | 1 | 1 |
| 4930522O17Rik | 0 | 0 | 0 | 1 | 1 |
| 4930523O13Rik | 0 | 0 | 0 | 1 | 1 |
| 4930524N10Rik | 0 | 0 | 0 | 1 | 1 |
| 4930525M21Rik | 0 | 0 | 0 | 1 | 1 |
| 4930526L06Rik | 0 | 0 | 0 | 1 | 1 |
| 4930527F14Rik | 0 | 0 | 0 | 1 | 1 |
| 4930527G23Rik | 0 | 0 | 0 | 1 | 1 |
| 4930528P14Rik | 0 | 0 | 0 | 1 | 1 |
| 4930529K09Rik | 0 | 0 | 0 | 1 | 1 |
| 4930529L06Rik | 0 | 0 | 0 | 1 | 1 |
| 4930532M18Rik | 0 | 0 | 0 | 1 | 1 |
| 4930533P14Rik | 0 | 0 | 0 | 1 | 1 |
| 4930539C22Rik | 0 | 0 | 0 | 1 | 1 |
| 4930539M17Rik | 0 | 0 | 0 | 1 | 1 |
| 4930539N22Rik | 0 | 0 | 0 | 1 | 1 |
| 4930540M03Rik | 0 | 0 | 0 | 1 | 1 |
| 4930543E12Rik | 0 | 0 | 0 | 1 | 1 |
| 4930544D05Rik | 0 | 0 | 0 | 1 | 1 |
| 4930544G11Rik | 0 | 0 | 0 | 1 | 1 |
| 4930544M13Rik | 0 | 0 | 0 | 1 | 1 |

| | | | | | |
|---|---|---|---|---|---|
| 4930545H06Rik | 0 | 0 | 0 | 1 | 1 |
| 4930545L23Rik | 0 | 0 | 0 | 1 | 1 |
| 4930546C10Rik | 0 | 0 | 0 | 1 | 1 |
| 4930546K05Rik | 0 | 0 | 0 | 1 | 1 |
| 4930548G14Rik | 0 | 0 | 0 | 1 | 1 |
| 4930548K13Rik | 0 | 0 | 0 | 1 | 1 |
| 4930549C01Rik | 0 | 0 | 0 | 1 | 1 |
| 4930552N02Rik | 0 | 0 | 0 | 1 | 1 |
| 4930553E22Rik | 0 | 0 | 0 | 1 | 1 |
| 4930554C24Rik | 0 | 0 | 0 | 1 | 1 |
| 4930555B11Rik | 0 | 0 | 0 | 1 | 1 |
| 4930555G01Rik | 0 | 0 | 0 | 1 | 1 |
| 4930556C24Rik | 0 | 0 | 0 | 1 | 1 |
| 4930556G01Rik | 0 | 0 | 0 | 1 | 1 |
| 4930556J02Rik | 0 | 0 | 0 | 1 | 1 |
| 4930556N09Rik | 0 | 0 | 0 | 1 | 1 |
| 4930557A04Rik | 0 | 0 | 0 | 1 | 1 |
| 4930557J02Rik | 0 | 0 | 0 | 1 | 1 |
| 4930558C23Rik | 0 | 0 | 0 | 1 | 1 |
| 4930558G05Rik | 0 | 0 | 0 | 1 | 1 |
| 4930558K02Rik | 0 | 0 | 0 | 1 | 1 |
| 4930563M20Rik | 0 | 0 | 0 | 1 | 1 |
| 4930564B18Rik | 0 | 0 | 0 | 1 | 1 |
| 4930564D02Rik | 0 | 0 | 0 | 1 | 1 |
| 4930567H12Rik | 0 | 0 | 0 | 1 | 1 |
| 4930567H17Rik | 0 | 0 | 0 | 1 | 1 |
| 4930567J20Rik | 0 | 0 | 0 | 1 | 1 |
| 4930567K20Rik | 0 | 0 | 0 | 1 | 1 |
| 4930568D16Rik | 0 | 0 | 0 | 1 | 1 |
| 4930568E12Rik | 0 | 0 | 0 | 1 | 1 |
| 4930568G15Rik | 0 | 0 | 0 | 1 | 1 |
| 4930571O06Rik | 0 | 0 | 0 | 1 | 1 |
| 4930572K03Rik | 0 | 0 | 0 | 1 | 1 |
| 4930572O03Rik | 0 | 0 | 0 | 1 | 1 |
| 4930572O13Rik | 0 | 0 | 0 | 1 | 1 |
| 4930573O16Rik | 0 | 0 | 0 | 1 | 1 |
| 4930578E11Rik | 0 | 0 | 0 | 1 | 1 |
| 4930578I06Rik | 0 | 0 | 0 | 1 | 1 |
| 4930578N18Rik | 0 | 0 | 0 | 1 | 1 |
| 4930579F01Rik | 0 | 0 | 0 | 1 | 1 |
| 4930583P06Rik | 0 | 0 | 0 | 1 | 1 |
| 4930584F24Rik | 0 | 0 | 0 | 1 | 1 |
| 4930591A17Rik | 0 | 0 | 0 | 1 | 1 |
| 4930592A05Rik | 0 | 0 | 0 | 1 | 1 |
| 4930593C16Rik | 0 | 0 | 0 | 1 | 1 |
| 4930595M18Rik | 0 | 0 | 0 | 1 | 1 |
| 4930596D02Rik | 0 | 0 | 0 | 1 | 1 |
| 4930597G03Rik | 0 | 0 | 0 | 1 | 1 |

| | | | | | |
|---|---|---|---|---|---|
| 4930598F16Rik | 0 | 0 | 0 | 1 | 1 |
| 4931406B18Rik | 0 | 0 | 0 | 1 | 1 |
| 4931408C20Rik | 0 | 0 | 0 | 1 | 1 |
| 4931412M21 | 0 | 0 | 0 | 1 | 1 |
| 4931417E11Rik | 0 | 0 | 0 | 1 | 1 |
| 4931419H13Rik | 0 | 0 | 0 | 1 | 1 |
| 4931429P17Rik | 0 | 0 | 0 | 1 | 1 |
| 4931431B13Rik | 0 | 0 | 0 | 1 | 1 |
| 4931431F19Rik | 0 | 0 | 0 | 1 | 1 |
| 4931440L10Rik | 0 | 0 | 0 | 1 | 1 |
| 4932411N23Rik | 0 | 0 | 0 | 1 | 1 |
| 4932412D23Rik | 0 | 0 | 0 | 1 | 1 |
| 4932414J04Rik | 0 | 0 | 0 | 1 | 1 |
| 4932414N04Rik | 0 | 0 | 0 | 1 | 1 |
| 4932415M13Rik | 0 | 0 | 0 | 1 | 1 |
| 4932416K20Rik | 0 | 0 | 0 | 1 | 1 |
| 4932418E24Rik | 0 | 0 | 0 | 1 | 1 |
| 4932429P05Rik | 0 | 0 | 0 | 1 | 1 |
| 4933400A11Rik | 0 | 0 | 0 | 1 | 1 |
| 4933400B14Rik | 0 | 0 | 0 | 1 | 1 |
| 4933400C23Rik | 0 | 0 | 0 | 1 | 1 |
| 4933400L20Rik | 0 | 0 | 0 | 1 | 1 |
| 4933401D09Rik | 0 | 0 | 0 | 1 | 1 |
| 4933402C06Rik | 0 | 0 | 0 | 1 | 1 |
| 4933402J07Rik | 0 | 0 | 0 | 1 | 1 |
| 4933402J15Rik | 0 | 0 | 0 | 1 | 1 |
| 4933402N03Rik | 0 | 0 | 0 | 1 | 1 |
| 4933402N22Rik | 0 | 0 | 0 | 1 | 1 |
| 4933402P03Rik | 0 | 0 | 0 | 1 | 1 |
| 4933404G15Rik | 0 | 0 | 0 | 1 | 1 |
| 4933404K08Rik | 0 | 0 | 0 | 1 | 1 |
| 4933405E24Rik | 0 | 0 | 0 | 1 | 1 |
| 4933405O20Rik | 0 | 0 | 0 | 1 | 1 |
| 4933406D12Rik | 0 | 0 | 0 | 1 | 1 |
| 4933406F09Rik | 0 | 0 | 0 | 1 | 1 |
| 4933406K04Rik | 0 | 0 | 0 | 1 | 1 |
| 4933407E24Rik | 0 | 0 | 0 | 1 | 1 |
| 4933407G14Rik | 0 | 0 | 0 | 1 | 1 |
| 4933407I05Rik | 0 | 0 | 0 | 1 | 1 |
| 4933409G03Rik | 0 | 0 | 0 | 1 | 1 |
| 4933409K07Rik | 0 | 0 | 0 | 1 | 1 |
| 4933411G06Rik | 0 | 0 | 0 | 1 | 1 |
| 4933411G11Rik | 0 | 0 | 0 | 1 | 1 |
| 4933412E24Rik | 0 | 0 | 0 | 1 | 1 |
| 4933412O06Rik | 0 | 0 | 0 | 1 | 1 |
| 4933413L06Rik | 0 | 0 | 0 | 1 | 1 |
| 4933415F23Rik | 0 | 0 | 0 | 1 | 1 |
| 4933416C03Rik | 0 | 0 | 0 | 1 | 1 |

| | | | | | |
|---|---|---|---|---|---|
| 4933416E03Rik | 0 | 0 | 0 | 1 | 1 |
| 4933416M06Rik | 0 | 0 | 0 | 1 | 1 |
| 4933417A18Rik | 0 | 0 | 0 | 1 | 1 |
| 4933421I07Rik | 0 | 0 | 0 | 1 | 1 |
| 4933425L06Rik | 0 | 0 | 0 | 1 | 1 |
| 4933428C19Rik | 0 | 0 | 0 | 1 | 1 |
| 4933429O19Rik | 0 | 0 | 0 | 1 | 1 |
| 4933430M04Rik | 0 | 0 | 0 | 1 | 1 |
| 4933430N04Rik | 0 | 0 | 0 | 1 | 1 |
| 4933432G23Rik | 0 | 0 | 0 | 1 | 1 |
| 4933432I09Rik | 0 | 0 | 0 | 1 | 1 |
| 4933432K03Rik | 0 | 0 | 0 | 1 | 1 |
| 4933433F19Rik | 0 | 0 | 0 | 1 | 1 |
| 4933433G08Rik | 0 | 0 | 0 | 1 | 1 |
| 4933433H22Rik | 0 | 0 | 0 | 1 | 1 |
| 4933436E23Rik | 0 | 0 | 0 | 1 | 1 |
| 4933436I01Rik | 0 | 0 | 0 | 1 | 1 |
| 4933438B17Rik | 0 | 0 | 0 | 1 | 1 |
| 4933438K21Rik | 0 | 0 | 0 | 1 | 1 |
| 4933440J02Rik | 0 | 0 | 0 | 1 | 1 |
| 5031410I06Rik | 0 | 0 | 0 | 1 | 1 |
| 5033403H07Rik | 0 | 0 | 0 | 1 | 1 |
| 5330439B14Rik | 0 | 0 | 0 | 1 | 1 |
| 5430401F13Rik | 0 | 0 | 0 | 1 | 1 |
| 5430402E10Rik | 0 | 0 | 0 | 1 | 1 |
| 5430403N17Rik | 0 | 0 | 0 | 1 | 1 |
| 5430425K12Rik | 0 | 0 | 0 | 1 | 1 |
| 5430434I15Rik | 0 | 0 | 0 | 1 | 1 |
| 5430437J10Rik | 0 | 0 | 0 | 1 | 1 |
| 5530400C23Rik | 0 | 0 | 0 | 1 | 1 |
| 5530401A14Rik | 0 | 0 | 0 | 1 | 1 |
| 5730412P04Rik | 0 | 0 | 0 | 1 | 1 |
| 5830411N06Rik | 0 | 0 | 0 | 1 | 1 |
| 5830416I19Rik | 0 | 0 | 0 | 1 | 1 |
| 5830428M24Rik | 0 | 0 | 0 | 1 | 1 |
| 6030440G07Rik | 0 | 0 | 0 | 1 | 1 |
| 6330415B21Rik | 0 | 0 | 0 | 1 | 1 |
| 6430503K07Rik | 0 | 0 | 0 | 1 | 1 |
| 6430710C18Rik | 0 | 0 | 0 | 1 | 1 |
| 7420426K07Rik | 0 | 0 | 0 | 1 | 1 |
| 7420461P10Rik | 0 | 0 | 0 | 1 | 1 |
| 7420700N18Rik | 0 | 0 | 0 | 1 | 1 |
| 7420701I03Rik | 0 | 0 | 0 | 1 | 1 |
| 7530416G11Rik | 0 | 0 | 0 | 1 | 1 |
| 7630403G23Rik | 0 | 0 | 0 | 1 | 1 |
| 8030411F24Rik | 0 | 0 | 0 | 1 | 1 |
| 8030423J24Rik | 0 | 0 | 0 | 1 | 1 |
| 8030442B05Rik | 0 | 0 | 0 | 1 | 1 |

| | | | | | |
|---|---|---|---|---|---|
| 8030443G20Rik | 0 | 0 | 0 | 1 | 1 |
| 8430422H06Rik | 0 | 0 | 0 | 1 | 1 |
| 8430436N08Rik | 0 | 0 | 0 | 1 | 1 |
| 8430437L04Rik | 0 | 0 | 0 | 1 | 1 |
| 9130015L21Rik | 0 | 0 | 0 | 1 | 1 |
| 9130204L05Rik | 0 | 0 | 0 | 1 | 1 |
| 9130209A04Rik | 0 | 0 | 0 | 1 | 1 |
| 9130221F21Rik | 0 | 0 | 0 | 1 | 1 |
| 9130227L01Rik | 0 | 0 | 0 | 1 | 1 |
| 9130230L23Rik | 0 | 0 | 0 | 1 | 1 |
| 9130409I23Rik | 0 | 0 | 0 | 1 | 1 |
| 9230110F15Rik | 0 | 0 | 0 | 1 | 1 |
| 9230112D13Rik | 0 | 0 | 0 | 1 | 1 |
| 9330162B11Rik | 0 | 0 | 0 | 1 | 1 |
| 9330175E14Rik | 0 | 0 | 0 | 1 | 1 |
| 9330175M20Rik | 0 | 0 | 0 | 1 | 1 |
| 9330178D15Rik | 0 | 0 | 0 | 1 | 1 |
| 9330182O14Rik | 0 | 0 | 0 | 1 | 1 |
| 9330188P03Rik | 0 | 0 | 0 | 1 | 1 |
| 9430007A20Rik | 0 | 0 | 0 | 1 | 1 |
| 9430069I07Rik | 0 | 0 | 0 | 1 | 1 |
| 9530002B09Rik | 0 | 0 | 0 | 1 | 1 |
| 9530026F06Rik | 0 | 0 | 0 | 1 | 1 |
| 9630013A20Rik | 0 | 0 | 0 | 1 | 1 |
| 9630028H03Rik | 0 | 0 | 0 | 1 | 1 |
| 9830107B12Rik | 0 | 0 | 0 | 1 | 1 |
| 9830132P13Rik | 0 | 0 | 0 | 1 | 1 |
| 9930111H07Rik | 0 | 0 | 0 | 1 | 1 |
| 9930111J21Rik2 | 0 | 0 | 0 | 1 | 1 |
| A1bg | 0 | 0 | 0 | 1 | 1 |
| A230065H16Rik | 0 | 0 | 0 | 1 | 1 |
| A330049N07Rik | 0 | 0 | 0 | 1 | 1 |
| A330070K13Rik | 0 | 0 | 0 | 1 | 1 |
| A430089I19Rik | 0 | 0 | 0 | 1 | 1 |
| A430107P09Rik | 0 | 0 | 0 | 1 | 1 |
| A4gnt | 0 | 0 | 0 | 1 | 1 |
| A530013C23Rik | 0 | 0 | 0 | 1 | 1 |
| A530032D15Rik | 0 | 0 | 0 | 1 | 1 |
| A530046M15Rik | 0 | 0 | 0 | 1 | 1 |
| A530050N04Rik | 0 | 0 | 0 | 1 | 1 |
| A530053G22Rik | 0 | 0 | 0 | 1 | 1 |
| A530058N18Rik | 0 | 0 | 0 | 1 | 1 |
| A530064D06Rik | 0 | 0 | 0 | 1 | 1 |
| A530099J19Rik | 0 | 0 | 0 | 1 | 1 |
| A630010A05Rik | 0 | 0 | 0 | 1 | 1 |
| A630023A22Rik | 0 | 0 | 0 | 1 | 1 |
| A630073D07Rik | 0 | 0 | 0 | 1 | 1 |
| A630077J23Rik | 0 | 0 | 0 | 1 | 1 |

| | | | | | |
|---|---|---|---|---|---|
| A630095E13Rik | 0 | 0 | 0 | 1 | 1 |
| A630095N17Rik | 0 | 0 | 0 | 1 | 1 |
| A730006G06Rik | 0 | 0 | 0 | 1 | 1 |
| A730018C14Rik | 0 | 0 | 0 | 1 | 1 |
| A730046J19Rik | 0 | 0 | 0 | 1 | 1 |
| A730082K24Rik | 0 | 0 | 0 | 1 | 1 |
| A730090H04Rik | 0 | 0 | 0 | 1 | 1 |
| A830019L24Rik | 0 | 0 | 0 | 1 | 1 |
| A930003A15Rik | 0 | 0 | 0 | 1 | 1 |
| A930006I01Rik | 0 | 0 | 0 | 1 | 1 |
| A930009A15Rik | 0 | 0 | 0 | 1 | 1 |
| AA536875 | 0 | 0 | 0 | 1 | 1 |
| AA543401 | 0 | 0 | 0 | 1 | 1 |
| AA545190 | 0 | 0 | 0 | 1 | 1 |
| AA619741 | 0 | 0 | 0 | 1 | 1 |
| AA792892 | 0 | 0 | 0 | 1 | 1 |
| AB041803 | 0 | 0 | 0 | 1 | 1 |
| AF067061 | 0 | 0 | 0 | 1 | 1 |
| AF067063 | 0 | 0 | 0 | 1 | 1 |
| AF357355 | 0 | 0 | 0 | 1 | 1 |
| AF357359 | 0 | 0 | 0 | 1 | 1 |
| AF357425 | 0 | 0 | 0 | 1 | 1 |
| AF357426 | 0 | 0 | 0 | 1 | 1 |
| AF366264 | 0 | 0 | 0 | 1 | 1 |
| AI427809 | 0 | 0 | 0 | 1 | 1 |
| AI606473 | 0 | 0 | 0 | 1 | 1 |
| AU016765 | 0 | 0 | 0 | 1 | 1 |
| AU018829,Gm163 | 0 | 0 | 0 | 1 | 1 |
| AU019990 | 0 | 0 | 0 | 1 | 1 |
| AU023762 | 0 | 0 | 0 | 1 | 1 |
| AV320801,Gm150 | 0 | 0 | 0 | 1 | 1 |
| AY512915 | 0 | 0 | 0 | 1 | 1 |
| AY761184 | 0 | 0 | 0 | 1 | 1 |
| AY761185 | 0 | 0 | 0 | 1 | 1 |
| Aadacl2 | 0 | 0 | 0 | 1 | 1 |
| Aadacl3 | 0 | 0 | 0 | 1 | 1 |
| Abca15 | 0 | 0 | 0 | 1 | 1 |
| Abca16 | 0 | 0 | 0 | 1 | 1 |
| Abca17 | 0 | 0 | 0 | 1 | 1 |
| Abcb11 | 0 | 0 | 0 | 1 | 1 |
| Abhd12b | 0 | 0 | 0 | 1 | 1 |
| Abo | 0 | 0 | 0 | 1 | 1 |
| Abra | 0 | 0 | 0 | 1 | 1 |
| Ace2 | 0 | 0 | 0 | 1 | 1 |
| Ace3 | 0 | 0 | 0 | 1 | 1 |
| Acnat2 | 0 | 0 | 0 | 1 | 1 |
| Acot5 | 0 | 0 | 0 | 1 | 1 |
| Acrv1 | 0 | 0 | 0 | 1 | 1 |

| | | | | | |
|---|---|---|---|---|---|
| Acsbg2 | 0 | 0 | 0 | 1 | 1 |
| Acsm5 | 0 | 0 | 0 | 1 | 1 |
| Actbl2 | 0 | 0 | 0 | 1 | 1 |
| Actl6b | 0 | 0 | 0 | 1 | 1 |
| Actrt1 | 0 | 0 | 0 | 1 | 1 |
| Actrt2 | 0 | 0 | 0 | 1 | 1 |
| Adam18 | 0 | 0 | 0 | 1 | 1 |
| Adam20 | 0 | 0 | 0 | 1 | 1 |
| Adam24 | 0 | 0 | 0 | 1 | 1 |
| Adam25 | 0 | 0 | 0 | 1 | 1 |
| Adam26b | 0 | 0 | 0 | 1 | 1 |
| Adam29 | 0 | 0 | 0 | 1 | 1 |
| Adam34 | 0 | 0 | 0 | 1 | 1 |
| Adam39 | 0 | 0 | 0 | 1 | 1 |
| Adam6a | 0 | 0 | 0 | 1 | 1 |
| Adam6b | 0 | 0 | 0 | 1 | 1 |
| Adam7 | 0 | 0 | 0 | 1 | 1 |
| Adh4 | 0 | 0 | 0 | 1 | 1 |
| Adig | 0 | 0 | 0 | 1 | 1 |
| Agmat | 0 | 0 | 0 | 1 | 1 |
| Agr3 | 0 | 0 | 0 | 1 | 1 |
| Agxt | 0 | 0 | 0 | 1 | 1 |
| Agxt2 | 0 | 0 | 0 | 1 | 1 |
| Akap4 | 0 | 0 | 0 | 1 | 1 |
| Aknad1 | 0 | 0 | 0 | 1 | 1 |
| Akr1c20 | 0 | 0 | 0 | 1 | 1 |
| Akr1c6 | 0 | 0 | 0 | 1 | 1 |
| Akr1cl | 0 | 0 | 0 | 1 | 1 |
| Allc | 0 | 0 | 0 | 1 | 1 |
| Alox12e | 0 | 0 | 0 | 1 | 1 |
| Alpi | 0 | 0 | 0 | 1 | 1 |
| Alppl2 | 0 | 0 | 0 | 1 | 1 |
| Als2cr12 | 0 | 0 | 0 | 1 | 1 |
| Ambn | 0 | 0 | 0 | 1 | 1 |
| Amelx | 0 | 0 | 0 | 1 | 1 |
| Amy2a4,Amy2a5 | 0 | 0 | 0 | 1 | 1 |
| Amy2a5 | 0 | 0 | 0 | 1 | 1 |
| Amy2b | 0 | 0 | 0 | 1 | 1 |
| Ang2 | 0 | 0 | 0 | 1 | 1 |
| Ang3,Ang5 | 0 | 0 | 0 | 1 | 1 |
| Ang5 | 0 | 0 | 0 | 1 | 1 |
| Ang6 | 0 | 0 | 0 | 1 | 1 |
| Angptl3 | 0 | 0 | 0 | 1 | 1 |
| Ankrd33 | 0 | 0 | 0 | 1 | 1 |
| Ankrd34c | 0 | 0 | 0 | 1 | 1 |
| Ankrd60 | 0 | 0 | 0 | 1 | 1 |
| Ankrd63 | 0 | 0 | 0 | 1 | 1 |
| Ankub1 | 0 | 0 | 0 | 1 | 1 |

| | | | | | |
|---|---|---|---|---|---|
| Antxrl | 0 | 0 | 0 | 1 | 1 |
| Anxa10 | 0 | 0 | 0 | 1 | 1 |
| Anxa13 | 0 | 0 | 0 | 1 | 1 |
| Apcs | 0 | 0 | 0 | 1 | 1 |
| Apobec4 | 0 | 0 | 0 | 1 | 1 |
| Apoc4 | 0 | 0 | 0 | 1 | 1 |
| Apof | 0 | 0 | 0 | 1 | 1 |
| Apoh | 0 | 0 | 0 | 1 | 1 |
| Apol10a | 0 | 0 | 0 | 1 | 1 |
| Apol11a | 0 | 0 | 0 | 1 | 1 |
| Apol11b | 0 | 0 | 0 | 1 | 1 |
| Apol7a | 0 | 0 | 0 | 1 | 1 |
| Apol7b | 0 | 0 | 0 | 1 | 1 |
| Apol9a | 0 | 0 | 0 | 1 | 1 |
| Apoo-ps | 0 | 0 | 0 | 1 | 1 |
| Aqp2 | 0 | 0 | 0 | 1 | 1 |
| Arhgap15os | 0 | 0 | 0 | 1 | 1 |
| Arl14 | 0 | 0 | 0 | 1 | 1 |
| Arl14epl | 0 | 0 | 0 | 1 | 1 |
| Armc12 | 0 | 0 | 0 | 1 | 1 |
| Art2a-ps | 0 | 0 | 0 | 1 | 1 |
| Art2b | 0 | 0 | 0 | 1 | 1 |
| Arx | 0 | 0 | 0 | 1 | 1 |
| Asb11 | 0 | 0 | 0 | 1 | 1 |
| Asb12 | 0 | 0 | 0 | 1 | 1 |
| Asb18 | 0 | 0 | 0 | 1 | 1 |
| Asb9 | 0 | 0 | 0 | 1 | 1 |
| Ascl3 | 0 | 0 | 0 | 1 | 1 |
| Asic5 | 0 | 0 | 0 | 1 | 1 |
| Aspdh | 0 | 0 | 0 | 1 | 1 |
| Asz1 | 0 | 0 | 0 | 1 | 1 |
| Atp13a4 | 0 | 0 | 0 | 1 | 1 |
| Atp6ap1l | 0 | 0 | 0 | 1 | 1 |
| Aurkc | 0 | 0 | 0 | 1 | 1 |
| Avp | 0 | 0 | 0 | 1 | 1 |
| Awat1 | 0 | 0 | 0 | 1 | 1 |
| Aym1 | 0 | 0 | 0 | 1 | 1 |
| Azgp1 | 0 | 0 | 0 | 1 | 1 |
| B020004C17Rik | 0 | 0 | 0 | 1 | 1 |
| B020004J07Rik | 0 | 0 | 0 | 1 | 1 |
| B020031M17Rik | 0 | 0 | 0 | 1 | 1 |
| B230112J18Rik | 0 | 0 | 0 | 1 | 1 |
| B230209E15Rik | 0 | 0 | 0 | 1 | 1 |
| B230214G05Rik | 0 | 0 | 0 | 1 | 1 |
| B430306N03Rik | 0 | 0 | 0 | 1 | 1 |
| B930018H19Rik | 0 | 0 | 0 | 1 | 1 |
| BB014433 | 0 | 0 | 0 | 1 | 1 |
| BB287469 | 0 | 0 | 0 | 1 | 1 |

| | | | | | |
|---|---|---|---|---|---|
| BC018473 | 0 | 0 | 0 | 1 | 1 |
| BC039966 | 0 | 0 | 0 | 1 | 1 |
| BC048562 | 0 | 0 | 0 | 1 | 1 |
| BC048602 | 0 | 0 | 0 | 1 | 1 |
| BC048671 | 0 | 0 | 0 | 1 | 1 |
| BC048679 | 0 | 0 | 0 | 1 | 1 |
| BC049352 | 0 | 0 | 0 | 1 | 1 |
| BC049730 | 0 | 0 | 0 | 1 | 1 |
| BC051537 | 0 | 0 | 0 | 1 | 1 |
| BC051665 | 0 | 0 | 0 | 1 | 1 |
| BC052688 | 0 | 0 | 0 | 1 | 1 |
| BC053393 | 0 | 0 | 0 | 1 | 1 |
| BC055402 | 0 | 0 | 0 | 1 | 1 |
| BC061195 | 0 | 0 | 0 | 1 | 1 |
| BC061212 | 0 | 0 | 0 | 1 | 1 |
| BC061237 | 0 | 0 | 0 | 1 | 1 |
| BC080695 | 0 | 0 | 0 | 1 | 1 |
| BC089597 | 0 | 0 | 0 | 1 | 1 |
| BC094916 | 0 | 0 | 0 | 1 | 1 |
| BC117090 | 0 | 0 | 0 | 1 | 1 |
| Bbox1 | 0 | 0 | 0 | 1 | 1 |
| Bc1 | 0 | 0 | 0 | 1 | 1 |
| Bcas1os2 | 0 | 0 | 0 | 1 | 1 |
| Bcl2a1c | 0 | 0 | 0 | 1 | 1 |
| Bcl2a1d | 0 | 0 | 0 | 1 | 1 |
| Best3 | 0 | 0 | 0 | 1 | 1 |
| Bglap2 | 0 | 0 | 0 | 1 | 1 |
| Bglap3 | 0 | 0 | 0 | 1 | 1 |
| Bmp10 | 0 | 0 | 0 | 1 | 1 |
| Bpifa1 | 0 | 0 | 0 | 1 | 1 |
| Bpifa2 | 0 | 0 | 0 | 1 | 1 |
| Bpifa3 | 0 | 0 | 0 | 1 | 1 |
| Bpifa5 | 0 | 0 | 0 | 1 | 1 |
| Bpifa6 | 0 | 0 | 0 | 1 | 1 |
| Bpifb2 | 0 | 0 | 0 | 1 | 1 |
| Bpifb3 | 0 | 0 | 0 | 1 | 1 |
| Bpifb6 | 0 | 0 | 0 | 1 | 1 |
| Bpifb9a | 0 | 0 | 0 | 1 | 1 |
| Bpifb9b | 0 | 0 | 0 | 1 | 1 |
| Bricd5 | 0 | 0 | 0 | 1 | 1 |
| Brinp3 | 0 | 0 | 0 | 1 | 1 |
| Brs3 | 0 | 0 | 0 | 1 | 1 |
| Bsph1 | 0 | 0 | 0 | 1 | 1 |
| Bsph2 | 0 | 0 | 0 | 1 | 1 |
| Btbd16 | 0 | 0 | 0 | 1 | 1 |
| Btnl1 | 0 | 0 | 0 | 1 | 1 |
| Btnl4 | 0 | 0 | 0 | 1 | 1 |
| Btnl5-ps | 0 | 0 | 0 | 1 | 1 |

| | | | | | |
|---|---|---|---|---|---|
| Btnl6 | 0 | 0 | 0 | 1 | 1 |
| C030007H22Rik | 0 | 0 | 0 | 1 | 1 |
| C030013G03Rik | 0 | 0 | 0 | 1 | 1 |
| C130079G13Rik | 0 | 0 | 0 | 1 | 1 |
| C130080G10Rik | 0 | 0 | 0 | 1 | 1 |
| C1rb | 0 | 0 | 0 | 1 | 1 |
| C230029M16 | 0 | 0 | 0 | 1 | 1 |
| C230079O03Rik | 0 | 0 | 0 | 1 | 1 |
| C330046G13Rik | 0 | 0 | 0 | 1 | 1 |
| C430002N11Rik | 0 | 0 | 0 | 1 | 1 |
| C4bp | 0 | 0 | 0 | 1 | 1 |
| C530044C16Rik | 0 | 0 | 0 | 1 | 1 |
| C630028M04Rik | 0 | 0 | 0 | 1 | 1 |
| C630031E19Rik | 0 | 0 | 0 | 1 | 1 |
| C7 | 0 | 0 | 0 | 1 | 1 |
| C730002L08Rik | 0 | 0 | 0 | 1 | 1 |
| C730027H18Rik | 0 | 0 | 0 | 1 | 1 |
| C87198 | 0 | 0 | 0 | 1 | 1 |
| C87414 | 0 | 0 | 0 | 1 | 1 |
| C87499 | 0 | 0 | 0 | 1 | 1 |
| C87977 | 0 | 0 | 0 | 1 | 1 |
| C8a | 0 | 0 | 0 | 1 | 1 |
| Cabp2 | 0 | 0 | 0 | 1 | 1 |
| Cabs1 | 0 | 0 | 0 | 1 | 1 |
| Cacng3 | 0 | 0 | 0 | 1 | 1 |
| Calm5 | 0 | 0 | 0 | 1 | 1 |
| Capn13 | 0 | 0 | 0 | 1 | 1 |
| Capza3 | 0 | 0 | 0 | 1 | 1 |
| Car10 | 0 | 0 | 0 | 1 | 1 |
| Casp14 | 0 | 0 | 0 | 1 | 1 |
| Catsper1 | 0 | 0 | 0 | 1 | 1 |
| Catsperb | 0 | 0 | 0 | 1 | 1 |
| Cbx5 | 0 | 105,476 | 0 | 1 | 1 |
| Ccdc110 | 0 | 0 | 0 | 1 | 1 |
| Ccdc144b | 0 | 0 | 0 | 1 | 1 |
| Ccdc152 | 0 | 0 | 0 | 1 | 1 |
| Ccdc153 | 0 | 0 | 0 | 1 | 1 |
| Ccdc172 | 0 | 0 | 0 | 1 | 1 |
| Ccdc178 | 0 | 0 | 0 | 1 | 1 |
| Ccdc54 | 0 | 0 | 0 | 1 | 1 |
| Ccdc7 | 0 | 0 | 0 | 1 | 1 |
| Ccdc70 | 0 | 0 | 0 | 1 | 1 |
| Ccer1 | 0 | 0 | 0 | 1 | 1 |
| Ccl1 | 0 | 0 | 0 | 1 | 1 |
| Ccl11 | 0 | 0 | 0 | 1 | 1 |
| Ccl19 | 0 | 0 | 0 | 1 | 1 |
| Ccl20 | 0 | 0 | 0 | 1 | 1 |
| Ccl21a,Gm1987 | 0 | 0 | 0 | 1 | 1 |

| | | | | | |
|---|---|---|---|---|---|
| Ccl21b,Gm13304 | 0 | 0 | 0 | 1 | 1 |
| Ccl21c | 0 | 0 | 0 | 1 | 1 |
| Ccl22 | 0 | 0 | 0 | 1 | 1 |
| Ccl26 | 0 | 0 | 0 | 1 | 1 |
| Ccl27b | 0 | 0 | 0 | 1 | 1 |
| Ccl5 | 0 | 0 | 0 | 1 | 1 |
| Ccl8 | 0 | 0 | 0 | 1 | 1 |
| Ccr1l1 | 0 | 0 | 0 | 1 | 1 |
| Ccr3 | 0 | 0 | 0 | 1 | 1 |
| Cct8l1 | 0 | 0 | 0 | 1 | 1 |
| Cd163l1 | 0 | 0 | 0 | 1 | 1 |
| Cd209a | 0 | 0 | 0 | 1 | 1 |
| Cd209b | 0 | 0 | 0 | 1 | 1 |
| Cd209c | 0 | 0 | 0 | 1 | 1 |
| Cd209d | 0 | 0 | 0 | 1 | 1 |
| Cd209e | 0 | 0 | 0 | 1 | 1 |
| Cd209g | 0 | 0 | 0 | 1 | 1 |
| Cd300c | 0 | 0 | 0 | 1 | 1 |
| Cd300lb | 0 | 0 | 0 | 1 | 1 |
| Cd300lh,Gm11710 | 0 | 0 | 0 | 1 | 1 |
| Cd3g | 0 | 0 | 0 | 1 | 1 |
| Cd40lg | 0 | 0 | 0 | 1 | 1 |
| Cd7 | 0 | 0 | 0 | 1 | 1 |
| Cd96 | 0 | 0 | 0 | 1 | 1 |
| Cdc20b | 0 | 0 | 0 | 1 | 1 |
| Cdh19 | 0 | 0 | 0 | 1 | 1 |
| Cdh9 | 0 | 0 | 0 | 1 | 1 |
| Ceacam-ps1 | 0 | 0 | 0 | 1 | 1 |
| Ceacam11 | 0 | 0 | 0 | 1 | 1 |
| Ceacam14 | 0 | 0 | 0 | 1 | 1 |
| Ceacam16 | 0 | 0 | 0 | 1 | 1 |
| Ceacam20 | 0 | 0 | 0 | 1 | 1 |
| Cebpe | 0 | 0 | 0 | 1 | 1 |
| Cela2a | 0 | 0 | 0 | 1 | 1 |
| Cela3b | 0 | 0 | 0 | 1 | 1 |
| Celrr | 0 | 0 | 0 | 1 | 1 |
| Cerkl | 0 | 0 | 0 | 1 | 1 |
| Ces1b | 0 | 0 | 0 | 1 | 1 |
| Ces1c | 0 | 0 | 0 | 1 | 1 |
| Ces1e | 0 | 0 | 0 | 1 | 1 |
| Ces1f | 0 | 0 | 0 | 1 | 1 |
| Ces2b | 0 | 0 | 0 | 1 | 1 |
| Ces2c | 0 | 0 | 0 | 1 | 1 |
| Ces2d-ps | 0 | 0 | 0 | 1 | 1 |
| Ces2h | 0 | 0 | 0 | 1 | 1 |
| Ces3a | 0 | 0 | 0 | 1 | 1 |
| Ces3b | 0 | 0 | 0 | 1 | 1 |
| Ces4a | 0 | 0 | 0 | 1 | 1 |

| | | | | | |
|---|---|---|---|---|---|
| Cfd | 0 | 0 | 0 | 1 | 1 |
| Cfhr1 | 0 | 0 | 0 | 1 | 1 |
| Cfhr2 | 0 | 0 | 0 | 1 | 1 |
| Cga | 0 | 0 | 0 | 1 | 1 |
| Chat | 0 | 0 | 0 | 1 | 1 |
| Chia1 | 0 | 0 | 0 | 1 | 1 |
| Chil3 | 0 | 0 | 0 | 1 | 1 |
| Chil4 | 0 | 0 | 0 | 1 | 1 |
| Chil6 | 0 | 0 | 0 | 1 | 1 |
| Chrm5 | 0 | 0 | 0 | 1 | 1 |
| Chrna2 | 0 | 0 | 0 | 1 | 1 |
| Chrna9 | 0 | 0 | 0 | 1 | 1 |
| Cib3 | 0 | 0 | 0 | 1 | 1 |
| Clca2 | 0 | 0 | 0 | 1 | 1 |
| Cldn16 | 0 | 0 | 0 | 1 | 1 |
| Cldn17 | 0 | 0 | 0 | 1 | 1 |
| Cldn18 | 0 | 0 | 0 | 1 | 1 |
| Clec10a | 0 | 0 | 0 | 1 | 1 |
| Clec12b | 0 | 0 | 0 | 1 | 1 |
| Clec2e | 0 | 0 | 0 | 1 | 1 |
| Clec2g | 0 | 0 | 0 | 1 | 1 |
| Clec4a4 | 0 | 0 | 0 | 1 | 1 |
| Clec4b1 | 0 | 0 | 0 | 1 | 1 |
| Clec4b2 | 0 | 0 | 0 | 1 | 1 |
| Clec4d | 0 | 0 | 0 | 1 | 1 |
| Clec4f | 0 | 0 | 0 | 1 | 1 |
| Clps | 0 | 0 | 0 | 1 | 1 |
| Clpsl2 | 0 | 0 | 0 | 1 | 1 |
| Clrn2 | 0 | 0 | 0 | 1 | 1 |
| Clrn3 | 0 | 0 | 0 | 1 | 1 |
| Cma2 | 0 | 0 | 0 | 1 | 1 |
| Cml3 | 0 | 0 | 0 | 1 | 1 |
| Cml5 | 0 | 0 | 0 | 1 | 1 |
| Cmtm2b | 0 | 0 | 0 | 1 | 1 |
| Cnga1 | 0 | 0 | 0 | 1 | 1 |
| Cntn6 | 0 | 0 | 0 | 1 | 1 |
| Cntnap5a | 0 | 0 | 0 | 1 | 1 |
| Cntnap5b | 0 | 0 | 0 | 1 | 1 |
| Cntnap5c | 0 | 0 | 0 | 1 | 1 |
| Colec10 | 0 | 0 | 0 | 1 | 1 |
| Cox6a2 | 0 | 0 | 0 | 1 | 1 |
| Cox8b | 0 | 0 | 0 | 1 | 1 |
| Cox8c | 0 | 0 | 0 | 1 | 1 |
| Cpb2 | 0 | 0 | 0 | 1 | 1 |
| Cphx1,Cphx2 | 0 | 0 | 0 | 1 | 1 |
| Cphx2 | 0 | 0 | 0 | 1 | 1 |
| Cpxcr1 | 0 | 0 | 0 | 1 | 1 |
| Crct1 | 0 | 0 | 0 | 1 | 1 |

| | | | | | |
|---|---|---|---|---|---|
| Crh | 0 | 0 | 0 | 1 | 1 |
| Crisp1 | 0 | 0 | 0 | 1 | 1 |
| Crisp3 | 0 | 0 | 0 | 1 | 1 |
| Crisp4 | 0 | 0 | 0 | 1 | 1 |
| Crnn | 0 | 0 | 0 | 1 | 1 |
| Crp | 0 | 0 | 0 | 1 | 1 |
| Crx | 0 | 0 | 0 | 1 | 1 |
| Crxos | 0 | 0 | 0 | 1 | 1 |
| Cryba2 | 0 | 0 | 0 | 1 | 1 |
| Crybb2 | 0 | 0 | 0 | 1 | 1 |
| Cryga | 0 | 0 | 0 | 1 | 1 |
| Crygb | 0 | 0 | 0 | 1 | 1 |
| Crygc | 0 | 0 | 0 | 1 | 1 |
| Crygd | 0 | 0 | 0 | 1 | 1 |
| Crygf | 0 | 0 | 0 | 1 | 1 |
| Csf2 | 0 | 0 | 0 | 1 | 1 |
| Csmd2os | 0 | 0 | 0 | 1 | 1 |
| Csmd3 | 0 | 0 | 0 | 1 | 1 |
| Csn1s1 | 0 | 0 | 0 | 1 | 1 |
| Csn1s2a | 0 | 0 | 0 | 1 | 1 |
| Csn1s2b | 0 | 0 | 0 | 1 | 1 |
| Csn2 | 0 | 0 | 0 | 1 | 1 |
| Csprs | 0 | 0 | 0 | 1 | 1 |
| Csrp3 | 0 | 0 | 0 | 1 | 1 |
| Cst10 | 0 | 0 | 0 | 1 | 1 |
| Cst11 | 0 | 0 | 0 | 1 | 1 |
| Cst12 | 0 | 0 | 0 | 1 | 1 |
| Cst13 | 0 | 0 | 0 | 1 | 1 |
| Cst8 | 0 | 0 | 0 | 1 | 1 |
| Cstl1 | 0 | 0 | 0 | 1 | 1 |
| Ctag2 | 0 | 0 | 0 | 1 | 1 |
| Ctcflos | 0 | 0 | 0 | 1 | 1 |
| Ctla4 | 0 | 0 | 0 | 1 | 1 |
| Ctnna3 | 0 | 0 | 0 | 1 | 1 |
| Ctrc | 0 | 0 | 0 | 1 | 1 |
| Ctrcos | 0 | 0 | 0 | 1 | 1 |
| Cts6 | 0 | 0 | 0 | 1 | 1 |
| Cts8-ps | 0 | 0 | 0 | 1 | 1 |
| Ctsll3 | 0 | 0 | 0 | 1 | 1 |
| Ctsm | 0 | 0 | 0 | 1 | 1 |
| Ctsq | 0 | 0 | 0 | 1 | 1 |
| Ctsr | 0 | 0 | 0 | 1 | 1 |
| Ctxn2 | 0 | 0 | 0 | 1 | 1 |
| Cwh43 | 0 | 0 | 0 | 1 | 1 |
| Cxcl11 | 0 | 0 | 0 | 1 | 1 |
| Cxcl13 | 0 | 0 | 0 | 1 | 1 |
| Cxcl15 | 0 | 0 | 0 | 1 | 1 |
| Cxcl3 | 0 | 0 | 0 | 1 | 1 |

| | | | | | |
|---|---|---|---|---|---|
| Cxcl9 | 0 | 0 | 0 | 1 | 1 |
| Cxcr1 | 0 | 0 | 0 | 1 | 1 |
| Cylc1 | 0 | 0 | 0 | 1 | 1 |
| Cylc2 | 0 | 0 | 0 | 1 | 1 |
| Cyp11b1 | 0 | 0 | 0 | 1 | 1 |
| Cyp11b2 | 0 | 0 | 0 | 1 | 1 |
| Cyp19a1 | 0 | 0 | 0 | 1 | 1 |
| Cyp2a12 | 0 | 0 | 0 | 1 | 1 |
| Cyp2a22 | 0 | 0 | 0 | 1 | 1 |
| Cyp2a4 | 0 | 0 | 0 | 1 | 1 |
| Cyp2a5 | 0 | 0 | 0 | 1 | 1 |
| Cyp2b10 | 0 | 0 | 0 | 1 | 1 |
| Cyp2b13 | 0 | 0 | 0 | 1 | 1 |
| Cyp2c29 | 0 | 0 | 0 | 1 | 1 |
| Cyp2c38 | 0 | 0 | 0 | 1 | 1 |
| Cyp2c39 | 0 | 0 | 0 | 1 | 1 |
| Cyp2c40 | 0 | 0 | 0 | 1 | 1 |
| Cyp2c50 | 0 | 0 | 0 | 1 | 1 |
| Cyp2c53-ps | 0 | 0 | 0 | 1 | 1 |
| Cyp2c54 | 0 | 0 | 0 | 1 | 1 |
| Cyp2c66 | 0 | 0 | 0 | 1 | 1 |
| Cyp2c67 | 0 | 0 | 0 | 1 | 1 |
| Cyp2c68 | 0 | 0 | 0 | 1 | 1 |
| Cyp2c69 | 0 | 0 | 0 | 1 | 1 |
| Cyp2c70 | 0 | 0 | 0 | 1 | 1 |
| Cyp2d10 | 0 | 0 | 0 | 1 | 1 |
| Cyp2d12 | 0 | 0 | 0 | 1 | 1 |
| Cyp2d13 | 0 | 0 | 0 | 1 | 1 |
| Cyp2d34 | 0 | 0 | 0 | 1 | 1 |
| Cyp2d37-ps | 0 | 0 | 0 | 1 | 1 |
| Cyp2d40 | 0 | 0 | 0 | 1 | 1 |
| Cyp2e1 | 0 | 0 | 0 | 1 | 1 |
| Cyp2f2 | 0 | 0 | 0 | 1 | 1 |
| Cyp2g1 | 0 | 0 | 0 | 1 | 1 |
| Cyp2j12 | 0 | 0 | 0 | 1 | 1 |
| Cyp2j5 | 0 | 0 | 0 | 1 | 1 |
| Cyp3a11 | 0 | 0 | 0 | 1 | 1 |
| Cyp3a13 | 0 | 0 | 0 | 1 | 1 |
| Cyp3a16 | 0 | 0 | 0 | 1 | 1 |
| Cyp3a25 | 0 | 0 | 0 | 1 | 1 |
| Cyp3a41a | 0 | 0 | 0 | 1 | 1 |
| Cyp3a44 | 0 | 0 | 0 | 1 | 1 |
| Cyp3a57 | 0 | 0 | 0 | 1 | 1 |
| Cyp3a59 | 0 | 0 | 0 | 1 | 1 |
| Cyp4a10 | 0 | 0 | 0 | 1 | 1 |
| Cyp4a12a | 0 | 0 | 0 | 1 | 1 |
| Cyp4a12b | 0 | 0 | 0 | 1 | 1 |
| Cyp4a14 | 0 | 0 | 0 | 1 | 1 |

| | | | | | |
|---|---|---|---|---|---|
| Cyp4a29 | 0 | 0 | 0 | 1 | 1 |
| Cyp4a30b | 0 | 0 | 0 | 1 | 1 |
| Cyp4a31 | 0 | 0 | 0 | 1 | 1 |
| Cyp4a32 | 0 | 0 | 0 | 1 | 1 |
| Cyp4b1 | 0 | 0 | 0 | 1 | 1 |
| Cyp4f40 | 0 | 0 | 0 | 1 | 1 |
| Cyp7a1 | 0 | 0 | 0 | 1 | 1 |
| Cypt1,Cypt8 | 0 | 0 | 0 | 1 | 1 |
| Cypt10,Cypt4 | 0 | 0 | 0 | 1 | 1 |
| Cypt12 | 0 | 0 | 0 | 1 | 1 |
| Cypt14 | 0 | 0 | 0 | 1 | 1 |
| Cypt15 | 0 | 0 | 0 | 1 | 1 |
| Cypt2 | 0 | 0 | 0 | 1 | 1 |
| Cypt4 | 0 | 0 | 0 | 1 | 1 |
| Cypt7 | 0 | 0 | 0 | 1 | 1 |
| Cypt8 | 0 | 0 | 0 | 1 | 1 |
| D030024E09Rik | 0 | 0 | 0 | 1 | 1 |
| D030025E07Rik | 0 | 0 | 0 | 1 | 1 |
| D130009I18Rik | 0 | 0 | 0 | 1 | 1 |
| D17Ertd648e | 0 | 0 | 0 | 1 | 1 |
| D430036J16Rik | 0 | 0 | 0 | 1 | 1 |
| D5Ertd577e,Gm32 | 0 | 0 | 0 | 1 | 1 |
| D630010B17Rik | 0 | 0 | 0 | 1 | 1 |
| D630013N20Rik | 0 | 0 | 0 | 1 | 1 |
| D730001G18Rik | 0 | 0 | 0 | 1 | 1 |
| D730045A05Rik | 0 | 0 | 0 | 1 | 1 |
| D730048I06Rik | 0 | 0 | 0 | 1 | 1 |
| D730050B12Rik | 0 | 0 | 0 | 1 | 1 |
| D830005E20Rik | 0 | 0 | 0 | 1 | 1 |
| D830015G02Rik | 0 | 0 | 0 | 1 | 1 |
| D830032E09Rik | 0 | 0 | 0 | 1 | 1 |
| D930032P07Rik | 0 | 0 | 0 | 1 | 1 |
| DQ267100 | 0 | 0 | 0 | 1 | 1 |
| DQ267101 | 0 | 0 | 0 | 1 | 1 |
| Daf2 | 0 | 0 | 0 | 1 | 1 |
| Dao | 0 | 0 | 0 | 1 | 1 |
| Dcdc2c | 0 | 0 | 0 | 1 | 1 |
| Dcpp1 | 0 | 0 | 0 | 1 | 1 |
| Dcpp2 | 0 | 0 | 0 | 1 | 1 |
| Defa-ps1 | 0 | 0 | 0 | 1 | 1 |
| Defa-ps12 | 0 | 0 | 0 | 1 | 1 |
| Defa-ps13 | 0 | 0 | 0 | 1 | 1 |
| Defa-rs1 | 0 | 0 | 0 | 1 | 1 |
| Defa-rs7,Defa23 | 0 | 0 | 0 | 1 | 1 |
| Defa17 | 0 | 0 | 0 | 1 | 1 |
| Defa2 | 0 | 0 | 0 | 1 | 1 |
| Defa20 | 0 | 0 | 0 | 1 | 1 |
| Defa21 | 0 | 0 | 0 | 1 | 1 |

| | | | | | |
|---|---|---|---|---|---|
| Defa23 | 0 | 0 | 0 | 1 | 1 |
| Defa25 | 0 | 0 | 0 | 1 | 1 |
| Defa26 | 0 | 0 | 0 | 1 | 1 |
| Defa3 | 0 | 0 | 0 | 1 | 1 |
| Defa4 | 0 | 0 | 0 | 1 | 1 |
| Defa5 | 0 | 0 | 0 | 1 | 1 |
| Defb1 | 0 | 0 | 0 | 1 | 1 |
| Defb10 | 0 | 0 | 0 | 1 | 1 |
| Defb11 | 0 | 0 | 0 | 1 | 1 |
| Defb12 | 0 | 0 | 0 | 1 | 1 |
| Defb13 | 0 | 0 | 0 | 1 | 1 |
| Defb14 | 0 | 0 | 0 | 1 | 1 |
| Defb15 | 0 | 0 | 0 | 1 | 1 |
| Defb18 | 0 | 0 | 0 | 1 | 1 |
| Defb19 | 0 | 0 | 0 | 1 | 1 |
| Defb2 | 0 | 0 | 0 | 1 | 1 |
| Defb20 | 0 | 0 | 0 | 1 | 1 |
| Defb21 | 0 | 0 | 0 | 1 | 1 |
| Defb22 | 0 | 0 | 0 | 1 | 1 |
| Defb23 | 0 | 0 | 0 | 1 | 1 |
| Defb25 | 0 | 0 | 0 | 1 | 1 |
| Defb26 | 0 | 0 | 0 | 1 | 1 |
| Defb28 | 0 | 0 | 0 | 1 | 1 |
| Defb29 | 0 | 0 | 0 | 1 | 1 |
| Defb3 | 0 | 0 | 0 | 1 | 1 |
| Defb30 | 0 | 0 | 0 | 1 | 1 |
| Defb33 | 0 | 0 | 0 | 1 | 1 |
| Defb34 | 0 | 0 | 0 | 1 | 1 |
| Defb35 | 0 | 0 | 0 | 1 | 1 |
| Defb36 | 0 | 0 | 0 | 1 | 1 |
| Defb38 | 0 | 0 | 0 | 1 | 1 |
| Defb39 | 0 | 0 | 0 | 1 | 1 |
| Defb4 | 0 | 0 | 0 | 1 | 1 |
| Defb41 | 0 | 0 | 0 | 1 | 1 |
| Defb42 | 0 | 0 | 0 | 1 | 1 |
| Defb43 | 0 | 0 | 0 | 1 | 1 |
| Defb44-ps | 0 | 0 | 0 | 1 | 1 |
| Defb45 | 0 | 0 | 0 | 1 | 1 |
| Defb46 | 0 | 0 | 0 | 1 | 1 |
| Defb48 | 0 | 0 | 0 | 1 | 1 |
| Defb5 | 0 | 0 | 0 | 1 | 1 |
| Defb50 | 0 | 0 | 0 | 1 | 1 |
| Defb6 | 0 | 0 | 0 | 1 | 1 |
| Defb7 | 0 | 0 | 0 | 1 | 1 |
| Defb8 | 0 | 0 | 0 | 1 | 1 |
| Defb9 | 0 | 0 | 0 | 1 | 1 |
| Dgat2l6 | 0 | 0 | 0 | 1 | 1 |
| Dgkb | 0 | 0 | 0 | 1 | 1 |

| | | | | | |
|---|---|---|---|---|---|
| Dhrsx | 0 | 0 | 0 | 1 | 1 |
| Dio1 | 0 | 0 | 0 | 1 | 1 |
| Dmrtc1b | 0 | 0 | 0 | 1 | 1 |
| Dmrtc1c1 | 0 | 0 | 0 | 1 | 1 |
| Dmrtc1c2 | 0 | 0 | 0 | 1 | 1 |
| Dmrtc2 | 0 | 0 | 0 | 1 | 1 |
| Dnajb8 | 0 | 0 | 0 | 1 | 1 |
| Dnajc5b | 0 | 0 | 0 | 1 | 1 |
| Dnajc5g | 0 | 0 | 0 | 1 | 1 |
| Dpep3 | 0 | 0 | 0 | 1 | 1 |
| Dppa1 | 0 | 0 | 0 | 1 | 1 |
| Dppa2 | 0 | 0 | 0 | 1 | 1 |
| Dppa3 | 0 | 0 | 0 | 1 | 1 |
| Drd3 | 0 | 0 | 0 | 1 | 1 |
| Dreh | 0 | 0 | 0 | 1 | 1 |
| Dsg1a | 0 | 0 | 0 | 1 | 1 |
| Dsg1b | 0 | 0 | 0 | 1 | 1 |
| Dsg1c | 0 | 0 | 0 | 1 | 1 |
| Dsg3 | 0 | 0 | 0 | 1 | 1 |
| Dsg4 | 0 | 0 | 0 | 1 | 1 |
| Dspp | 0 | 0 | 0 | 1 | 1 |
| Dthd1 | 0 | 0 | 0 | 1 | 1 |
| Duoxa2 | 0 | 0 | 0 | 1 | 1 |
| Dusp21 | 0 | 0 | 0 | 1 | 1 |
| Dux | 0 | 0 | 0 | 1 | 1 |
| Duxbl2 | 0 | 0 | 0 | 1 | 1 |
| Duxbl3 | 0 | 0 | 0 | 1 | 1 |
| Dydc1 | 0 | 0 | 0 | 1 | 1 |
| Dynap | 0 | 0 | 0 | 1 | 1 |
| Dynlt1b | 0 | 0 | 0 | 1 | 1 |
| Dynlt1c,Dynlt1f | 0 | 0 | 0 | 1 | 1 |
| Dynlt1f | 0 | 0 | 0 | 1 | 1 |
| E030002O03Rik | 0 | 0 | 0 | 1 | 1 |
| E030003E18Rik | 0 | 0 | 0 | 1 | 1 |
| E030019B06Rik | 0 | 0 | 0 | 1 | 1 |
| E030025P04Rik | 0 | 0 | 0 | 1 | 1 |
| E230019M04Rik | 0 | 0 | 0 | 1 | 1 |
| E330012B07Rik | 0 | 0 | 0 | 1 | 1 |
| E330014E10Rik,Gr | 0 | 0 | 0 | 1 | 1 |
| E330017A01Rik | 0 | 0 | 0 | 1 | 1 |
| E330023G01Rik | 0 | 0 | 0 | 1 | 1 |
| E330034G19Rik | 0 | 0 | 0 | 1 | 1 |
| E530001F21Rik | 0 | 0 | 0 | 1 | 1 |
| Ear1 | 0 | 0 | 0 | 1 | 1 |
| Ear10 | 0 | 0 | 0 | 1 | 1 |
| Ear2,Ear3 | 0 | 0 | 0 | 1 | 1 |
| Ear3 | 0 | 0 | 0 | 1 | 1 |
| Ear6,Ear7 | 0 | 0 | 0 | 1 | 1 |

| | | | | | |
|---|---|---|---|---|---|
| Ear7 | 0 | 0 | 0 | 1 | 1 |
| Efcab3 | 0 | 0 | 0 | 1 | 1 |
| Eftud2 | 0 | 125,794 | 0 | 1 | 1 |
| Egfbp2 | 0 | 0 | 0 | 1 | 1 |
| Enam | 0 | 0 | 0 | 1 | 1 |
| Enthd1 | 0 | 0 | 0 | 1 | 1 |
| Epyc | 0 | 0 | 0 | 1 | 1 |
| Erdr1 | 0 | 0 | 0 | 1 | 1 |
| Ern2 | 0 | 0 | 0 | 1 | 1 |
| Esp1 | 0 | 0 | 0 | 1 | 1 |
| Esp15 | 0 | 0 | 0 | 1 | 1 |
| Esp16 | 0 | 0 | 0 | 1 | 1 |
| Esp18 | 0 | 0 | 0 | 1 | 1 |
| Esp23 | 0 | 0 | 0 | 1 | 1 |
| Esp24 | 0 | 0 | 0 | 1 | 1 |
| Esp3 | 0 | 0 | 0 | 1 | 1 |
| Esp31 | 0 | 0 | 0 | 1 | 1 |
| Esp34 | 0 | 0 | 0 | 1 | 1 |
| Esp36 | 0 | 0 | 0 | 1 | 1 |
| Esp38 | 0 | 0 | 0 | 1 | 1 |
| Esp6 | 0 | 0 | 0 | 1 | 1 |
| Esp8 | 0 | 0 | 0 | 1 | 1 |
| Etd | 0 | 0 | 0 | 1 | 1 |
| F11 | 0 | 0 | 0 | 1 | 1 |
| F13b | 0 | 0 | 0 | 1 | 1 |
| F630206G17Rik | 0 | 0 | 0 | 1 | 1 |
| F830002L21Rik | 0 | 0 | 0 | 1 | 1 |
| F830016B08Rik | 0 | 0 | 0 | 1 | 1 |
| F830045P16Rik | 0 | 0 | 0 | 1 | 1 |
| Fabp12 | 0 | 0 | 0 | 1 | 1 |
| Fabp6 | 0 | 0 | 0 | 1 | 1 |
| Fabp9 | 0 | 0 | 0 | 1 | 1 |
| Fam122c | 0 | 0 | 0 | 1 | 1 |
| Fam151a | 0 | 0 | 0 | 1 | 1 |
| Fam154a | 0 | 0 | 0 | 1 | 1 |
| Fam170a | 0 | 0 | 0 | 1 | 1 |
| Fam170b | 0 | 0 | 0 | 1 | 1 |
| Fam177a | 0 | 0 | 0 | 1 | 1 |
| Fam180a | 0 | 0 | 0 | 1 | 1 |
| Fam216b | 0 | 0 | 0 | 1 | 1 |
| Fam227b | 0 | 0 | 0 | 1 | 1 |
| Fam25c | 0 | 0 | 0 | 1 | 1 |
| Fam3b | 0 | 0 | 0 | 1 | 1 |
| Fam47c | 0 | 0 | 0 | 1 | 1 |
| Fam47e | 0 | 0 | 0 | 1 | 1 |
| Fam50b | 0 | 0 | 0 | 1 | 1 |
| Fam71a | 0 | 0 | 0 | 1 | 1 |
| Fam71b | 0 | 0 | 0 | 1 | 1 |

| | | | | | |
|---|---|---|---|---|---|
| Fam71e2 | 0 | 0 | 0 | 1 | 1 |
| Fbll1 | 0 | 0 | 0 | 1 | 1 |
| Fbxo39 | 0 | 0 | 0 | 1 | 1 |
| Fbxw14 | 0 | 0 | 0 | 1 | 1 |
| Fbxw15 | 0 | 0 | 0 | 1 | 1 |
| Fbxw16 | 0 | 0 | 0 | 1 | 1 |
| Fbxw18 | 0 | 0 | 0 | 1 | 1 |
| Fbxw20 | 0 | 0 | 0 | 1 | 1 |
| Fbxw21 | 0 | 0 | 0 | 1 | 1 |
| Fbxw22 | 0 | 0 | 0 | 1 | 1 |
| Fbxw24 | 0 | 0 | 0 | 1 | 1 |
| Fbxw26 | 0 | 0 | 0 | 1 | 1 |
| Fbxw28 | 0 | 0 | 0 | 1 | 1 |
| Fcer1a | 0 | 0 | 0 | 1 | 1 |
| Fcnb | 0 | 0 | 0 | 1 | 1 |
| Fcrl5 | 0 | 0 | 0 | 1 | 1 |
| Ffar2 | 0 | 0 | 0 | 1 | 1 |
| Ffar3 | 0 | 0 | 0 | 1 | 1 |
| Fgf16 | 0 | 0 | 0 | 1 | 1 |
| Fgf6 | 0 | 0 | 0 | 1 | 1 |
| Fhad1os1 | 0 | 0 | 0 | 1 | 1 |
| Fhl5 | 0 | 0 | 0 | 1 | 1 |
| Flg2 | 0 | 0 | 0 | 1 | 1 |
| Fmo3 | 0 | 0 | 0 | 1 | 1 |
| Fmo9 | 0 | 0 | 0 | 1 | 1 |
| Foxb2 | 0 | 0 | 0 | 1 | 1 |
| Foxe3 | 0 | 0 | 0 | 1 | 1 |
| Fpr-rs3 | 0 | 0 | 0 | 1 | 1 |
| Fpr-rs4 | 0 | 0 | 0 | 1 | 1 |
| Fpr-rs6 | 0 | 0 | 0 | 1 | 1 |
| Fpr2 | 0 | 0 | 0 | 1 | 1 |
| Fpr3 | 0 | 0 | 0 | 1 | 1 |
| Fscb | 0 | 0 | 0 | 1 | 1 |
| Fshb | 0 | 0 | 0 | 1 | 1 |
| Fshr | 0 | 0 | 0 | 1 | 1 |
| Fthl17 | 0 | 0 | 0 | 1 | 1 |
| Ftmt | 0 | 0 | 0 | 1 | 1 |
| Fut4-ps1 | 0 | 0 | 0 | 1 | 1 |
| G630093K05Rik | 0 | 0 | 0 | 1 | 1 |
| G6pc | 0 | 0 | 0 | 1 | 1 |
| G730013B05Rik | 0 | 0 | 0 | 1 | 1 |
| Gabra6 | 0 | 0 | 0 | 1 | 1 |
| Gabrg2 | 0 | 0 | 0 | 1 | 1 |
| Gal3st2 | 0 | 0 | 0 | 1 | 1 |
| Galp | 0 | 0 | 0 | 1 | 1 |
| Gas2l2 | 0 | 0 | 0 | 1 | 1 |
| Gast | 0 | 0 | 0 | 1 | 1 |
| Gata5os | 0 | 0 | 0 | 1 | 1 |

| | | | | | |
|---|---|---|---|---|---|
| Gcg | 0 | 0 | 0 | 1 | 1 |
| Gcm1 | 0 | 0 | 0 | 1 | 1 |
| Gcm2 | 0 | 0 | 0 | 1 | 1 |
| Gdf2 | 0 | 0 | 0 | 1 | 1 |
| Gfral | 0 | 0 | 0 | 1 | 1 |
| Ggt6 | 0 | 0 | 0 | 1 | 1 |
| Gh | 0 | 0 | 0 | 1 | 1 |
| Gif | 0 | 0 | 0 | 1 | 1 |
| Gip | 0 | 0 | 0 | 1 | 1 |
| Gjd3 | 0 | 0 | 0 | 1 | 1 |
| Gjd4 | 0 | 0 | 0 | 1 | 1 |
| Gje1 | 0 | 0 | 0 | 1 | 1 |
| Gkn1 | 0 | 0 | 0 | 1 | 1 |
| Gkn3 | 0 | 0 | 0 | 1 | 1 |
| Glb1l3 | 0 | 0 | 0 | 1 | 1 |
| Gldnos | 0 | 0 | 0 | 1 | 1 |
| Glipr1l1 | 0 | 0 | 0 | 1 | 1 |
| Glipr1l2 | 0 | 0 | 0 | 1 | 1 |
| Glra2 | 0 | 0 | 0 | 1 | 1 |
| Glra3 | 0 | 0 | 0 | 1 | 1 |
| Glrp1 | 0 | 0 | 0 | 1 | 1 |
| Glt6d1 | 0 | 0 | 0 | 1 | 1 |
| Glycam1 | 0 | 0 | 0 | 1 | 1 |
| Gm10007 | 0 | 0 | 0 | 1 | 1 |
| Gm10024 | 0 | 0 | 0 | 1 | 1 |
| Gm10052 | 0 | 0 | 0 | 1 | 1 |
| Gm10057,Samt1 | 0 | 0 | 0 | 1 | 1 |
| Gm10058,Gm100[illegible] | 0 | 0 | 0 | 1 | 1 |
| Gm10081 | 0 | 0 | 0 | 1 | 1 |
| Gm10094,Sap18 | 0 | 0 | 0 | 1 | 1 |
| Gm10096,Gm104[illegible] | 0 | 0 | 0 | 1 | 1 |
| Gm10100 | 0 | 0 | 0 | 1 | 1 |
| Gm10104 | 0 | 0 | 0 | 1 | 1 |
| Gm10142 | 0 | 0 | 0 | 1 | 1 |
| Gm10147,Gm146[illegible] | 0 | 0 | 0 | 1 | 1 |
| Gm10220 | 0 | 0 | 0 | 1 | 1 |
| Gm10228 | 0 | 0 | 0 | 1 | 1 |
| Gm10229 | 0 | 0 | 0 | 1 | 1 |
| Gm10230 | 0 | 0 | 0 | 1 | 1 |
| Gm10272 | 0 | 0 | 0 | 1 | 1 |
| Gm10280 | 0 | 0 | 0 | 1 | 1 |
| Gm10318 | 0 | 0 | 0 | 1 | 1 |
| Gm10319 | 0 | 0 | 0 | 1 | 1 |
| Gm10324 | 0 | 0 | 0 | 1 | 1 |
| Gm10334 | 0 | 0 | 0 | 1 | 1 |
| Gm10352,Rbmy | 0 | 0 | 0 | 1 | 1 |
| Gm10354 | 0 | 0 | 0 | 1 | 1 |
| Gm10375 | 0 | 0 | 0 | 1 | 1 |

| | | | | | |
|---|---|---|---|---|---|
| Gm10377 | 0 | 0 | 0 | 1 | 1 |
| Gm10408 | 0 | 0 | 0 | 1 | 1 |
| Gm10409,Gm302( | 0 | 0 | 0 | 1 | 1 |
| Gm10413 | 0 | 0 | 0 | 1 | 1 |
| Gm10415 | 0 | 0 | 0 | 1 | 1 |
| Gm10436 | 0 | 0 | 0 | 1 | 1 |
| Gm10439 | 0 | 0 | 0 | 1 | 1 |
| Gm10445 | 0 | 0 | 0 | 1 | 1 |
| Gm10466 | 0 | 0 | 0 | 1 | 1 |
| Gm10471 | 0 | 0 | 0 | 1 | 1 |
| Gm10486 | 0 | 0 | 0 | 1 | 1 |
| Gm10487 | 0 | 0 | 0 | 1 | 1 |
| Gm10488 | 0 | 0 | 0 | 1 | 1 |
| Gm10512 | 0 | 0 | 0 | 1 | 1 |
| Gm10538 | 0 | 0 | 0 | 1 | 1 |
| Gm10556 | 0 | 0 | 0 | 1 | 1 |
| Gm10591 | 0 | 0 | 0 | 1 | 1 |
| Gm10619 | 0 | 0 | 0 | 1 | 1 |
| Gm10637 | 0 | 0 | 0 | 1 | 1 |
| Gm10639 | 0 | 0 | 0 | 1 | 1 |
| Gm10649 | 0 | 0 | 0 | 1 | 1 |
| Gm10662,Gm589: | 0 | 0 | 0 | 1 | 1 |
| Gm10665 | 0 | 0 | 0 | 1 | 1 |
| Gm10666 | 0 | 0 | 0 | 1 | 1 |
| Gm10670 | 0 | 0 | 0 | 1 | 1 |
| Gm10681,Hsd3b4 | 0 | 0 | 0 | 1 | 1 |
| Gm10696 | 0 | 0 | 0 | 1 | 1 |
| Gm10731 | 0 | 0 | 0 | 1 | 1 |
| Gm10745 | 0 | 0 | 0 | 1 | 1 |
| Gm10754 | 0 | 0 | 0 | 1 | 1 |
| Gm10782 | 0 | 0 | 0 | 1 | 1 |
| Gm10787 | 0 | 0 | 0 | 1 | 1 |
| Gm10790 | 0 | 0 | 0 | 1 | 1 |
| Gm10823 | 0 | 0 | 0 | 1 | 1 |
| Gm10921 | 0 | 0 | 0 | 1 | 1 |
| Gm10922 | 0 | 0 | 0 | 1 | 1 |
| Gm11127 | 0 | 0 | 0 | 1 | 1 |
| Gm11128 | 0 | 0 | 0 | 1 | 1 |
| Gm11166 | 0 | 0 | 0 | 1 | 1 |
| Gm11190 | 0 | 0 | 0 | 1 | 1 |
| Gm11213 | 0 | 0 | 0 | 1 | 1 |
| Gm11237,LOC101( | 0 | 0 | 0 | 1 | 1 |
| Gm11240 | 0 | 0 | 0 | 1 | 1 |
| Gm11351 | 0 | 0 | 0 | 1 | 1 |
| Gm1140,Gm1469: | 0 | 0 | 0 | 1 | 1 |
| Gm11426 | 0 | 0 | 0 | 1 | 1 |
| Gm11437 | 0 | 0 | 0 | 1 | 1 |
| Gm11487 | 0 | 0 | 0 | 1 | 1 |

| | | | | | |
|---|---|---|---|---|---|
| Gm11541 | 0 | 0 | 0 | 1 | 1 |
| Gm11544 | 0 | 0 | 0 | 1 | 1 |
| Gm11548 | 0 | 0 | 0 | 1 | 1 |
| Gm11554 | 0 | 0 | 0 | 1 | 1 |
| Gm11559 | 0 | 0 | 0 | 1 | 1 |
| Gm11562 | 0 | 0 | 0 | 1 | 1 |
| Gm11563 | 0 | 0 | 0 | 1 | 1 |
| Gm11564 | 0 | 0 | 0 | 1 | 1 |
| Gm11565 | 0 | 0 | 0 | 1 | 1 |
| Gm11567 | 0 | 0 | 0 | 1 | 1 |
| Gm11568 | 0 | 0 | 0 | 1 | 1 |
| Gm11569 | 0 | 0 | 0 | 1 | 1 |
| Gm11595 | 0 | 0 | 0 | 1 | 1 |
| Gm11596 | 0 | 0 | 0 | 1 | 1 |
| Gm11651 | 0 | 0 | 0 | 1 | 1 |
| Gm11710 | 0 | 0 | 0 | 1 | 1 |
| Gm11757,Gm1175 | 0 | 0 | 0 | 1 | 1 |
| Gm11758 | 0 | 0 | 0 | 1 | 1 |
| Gm11780 | 0 | 0 | 0 | 1 | 1 |
| Gm11937 | 0 | 0 | 0 | 1 | 1 |
| Gm11938 | 0 | 0 | 0 | 1 | 1 |
| Gm11981 | 0 | 0 | 0 | 1 | 1 |
| Gm12 | 0 | 0 | 0 | 1 | 1 |
| Gm12130 | 0 | 0 | 0 | 1 | 1 |
| Gm12159 | 0 | 0 | 0 | 1 | 1 |
| Gm12169 | 0 | 0 | 0 | 1 | 1 |
| Gm12171 | 0 | 0 | 0 | 1 | 1 |
| Gm12185 | 0 | 0 | 0 | 1 | 1 |
| Gm12298 | 0 | 0 | 0 | 1 | 1 |
| Gm12505 | 0 | 0 | 0 | 1 | 1 |
| Gm12530 | 0 | 0 | 0 | 1 | 1 |
| Gm12603 | 0 | 0 | 0 | 1 | 1 |
| Gm12669 | 0 | 0 | 0 | 1 | 1 |
| Gm12695 | 0 | 0 | 0 | 1 | 1 |
| Gm12789 | 0 | 0 | 0 | 1 | 1 |
| Gm12794 | 0 | 0 | 0 | 1 | 1 |
| Gm12886 | 0 | 0 | 0 | 1 | 1 |
| Gm12887 | 0 | 0 | 0 | 1 | 1 |
| Gm12888 | 0 | 0 | 0 | 1 | 1 |
| Gm13011 | 0 | 0 | 0 | 1 | 1 |
| Gm13023 | 0 | 0 | 0 | 1 | 1 |
| Gm13031 | 0 | 0 | 0 | 1 | 1 |
| Gm13040 | 0 | 0 | 0 | 1 | 1 |
| Gm13043 | 0 | 0 | 0 | 1 | 1 |
| Gm13051 | 0 | 0 | 0 | 1 | 1 |
| Gm13078 | 0 | 0 | 0 | 1 | 1 |
| Gm13083 | 0 | 0 | 0 | 1 | 1 |
| Gm13084 | 0 | 0 | 0 | 1 | 1 |

| | | | | | |
|---|---|---|---|---|---|
| Gm13088 | 0 | 0 | 0 | 1 | 1 |
| Gm13102 | 0 | 0 | 0 | 1 | 1 |
| Gm13103 | 0 | 0 | 0 | 1 | 1 |
| Gm13119 | 0 | 0 | 0 | 1 | 1 |
| Gm13124 | 0 | 0 | 0 | 1 | 1 |
| Gm13125 | 0 | 0 | 0 | 1 | 1 |
| Gm13128 | 0 | 0 | 0 | 1 | 1 |
| Gm13177 | 0 | 0 | 0 | 1 | 1 |
| Gm13178 | 0 | 0 | 0 | 1 | 1 |
| Gm13242 | 0 | 0 | 0 | 1 | 1 |
| Gm13275,Gm1328 | 0 | 0 | 0 | 1 | 1 |
| Gm13276 | 0 | 0 | 0 | 1 | 1 |
| Gm13278,Gm1327 | 0 | 0 | 0 | 1 | 1 |
| Gm13279 | 0 | 0 | 0 | 1 | 1 |
| Gm13285,Gm1328 | 0 | 0 | 0 | 1 | 1 |
| Gm13287 | 0 | 0 | 0 | 1 | 1 |
| Gm13290 | 0 | 0 | 0 | 1 | 1 |
| Gm13293 | 0 | 0 | 0 | 1 | 1 |
| Gm13298 | 0 | 0 | 0 | 1 | 1 |
| Gm13304 | 0 | 0 | 0 | 1 | 1 |
| Gm13305,Gm200 | 0 | 0 | 0 | 1 | 1 |
| Gm13306 | 0 | 0 | 0 | 1 | 1 |
| Gm13308,Gm215 | 0 | 0 | 0 | 1 | 1 |
| Gm13363 | 0 | 0 | 0 | 1 | 1 |
| Gm13497 | 0 | 0 | 0 | 1 | 1 |
| Gm13544 | 0 | 0 | 0 | 1 | 1 |
| Gm13546 | 0 | 0 | 0 | 1 | 1 |
| Gm13580 | 0 | 0 | 0 | 1 | 1 |
| Gm13749 | 0 | 0 | 0 | 1 | 1 |
| Gm13752 | 0 | 0 | 0 | 1 | 1 |
| Gm13769 | 0 | 0 | 0 | 1 | 1 |
| Gm13871 | 0 | 0 | 0 | 1 | 1 |
| Gm14015 | 0 | 0 | 0 | 1 | 1 |
| Gm14085 | 0 | 0 | 0 | 1 | 1 |
| Gm14151 | 0 | 0 | 0 | 1 | 1 |
| Gm14169 | 0 | 0 | 0 | 1 | 1 |
| Gm14288 | 0 | 0 | 0 | 1 | 1 |
| Gm14306,Gm144 | 0 | 0 | 0 | 1 | 1 |
| Gm14308,OTTMU | 0 | 0 | 0 | 1 | 1 |
| Gm14327 | 0 | 0 | 0 | 1 | 1 |
| Gm14345,Gm143 | 0 | 0 | 0 | 1 | 1 |
| Gm14346,Gm370 | 0 | 0 | 0 | 1 | 1 |
| Gm14347 | 0 | 0 | 0 | 1 | 1 |
| Gm14351,Gm291 | 0 | 0 | 0 | 1 | 1 |
| Gm14374 | 0 | 0 | 0 | 1 | 1 |
| Gm14391 | 0 | 0 | 0 | 1 | 1 |
| Gm14393 | 0 | 0 | 0 | 1 | 1 |
| Gm14405 | 0 | 0 | 0 | 1 | 1 |

| | | | | | |
|---|---|---|---|---|---|
| Gm14430,OTTMU | 0 | 0 | 0 | 1 | 1 |
| Gm14431 | 0 | 0 | 0 | 1 | 1 |
| Gm14436 | 0 | 0 | 0 | 1 | 1 |
| Gm14440 | 0 | 0 | 0 | 1 | 1 |
| Gm14458 | 0 | 0 | 0 | 1 | 1 |
| Gm14459 | 0 | 0 | 0 | 1 | 1 |
| Gm14461 | 0 | 0 | 0 | 1 | 1 |
| Gm14474 | 0 | 0 | 0 | 1 | 1 |
| Gm14475,Gm144 | 0 | 0 | 0 | 1 | 1 |
| Gm14477,Gm144 | 0 | 0 | 0 | 1 | 1 |
| Gm14478 | 0 | 0 | 0 | 1 | 1 |
| Gm14479,Gm144 | 0 | 0 | 0 | 1 | 1 |
| Gm14482 | 0 | 0 | 0 | 1 | 1 |
| Gm14483 | 0 | 0 | 0 | 1 | 1 |
| Gm14484 | 0 | 0 | 0 | 1 | 1 |
| Gm14499 | 0 | 0 | 0 | 1 | 1 |
| Gm14501 | 0 | 0 | 0 | 1 | 1 |
| Gm14511 | 0 | 0 | 0 | 1 | 1 |
| Gm14525 | 0 | 0 | 0 | 1 | 1 |
| Gm14548 | 0 | 0 | 0 | 1 | 1 |
| Gm14625 | 0 | 0 | 0 | 1 | 1 |
| Gm14632 | 0 | 0 | 0 | 1 | 1 |
| Gm14635 | 0 | 0 | 0 | 1 | 1 |
| Gm14692 | 0 | 0 | 0 | 1 | 1 |
| Gm14718 | 0 | 0 | 0 | 1 | 1 |
| Gm14725 | 0 | 0 | 0 | 1 | 1 |
| Gm14743 | 0 | 0 | 0 | 1 | 1 |
| Gm14744 | 0 | 0 | 0 | 1 | 1 |
| Gm14781 | 0 | 0 | 0 | 1 | 1 |
| Gm14812 | 0 | 0 | 0 | 1 | 1 |
| Gm14819 | 0 | 0 | 0 | 1 | 1 |
| Gm14850,Gm214 | 0 | 0 | 0 | 1 | 1 |
| Gm14851 | 0 | 0 | 0 | 1 | 1 |
| Gm14920 | 0 | 0 | 0 | 1 | 1 |
| Gm15008 | 0 | 0 | 0 | 1 | 1 |
| Gm15023 | 0 | 0 | 0 | 1 | 1 |
| Gm15056 | 0 | 0 | 0 | 1 | 1 |
| Gm15080 | 0 | 0 | 0 | 1 | 1 |
| Gm15085 | 0 | 0 | 0 | 1 | 1 |
| Gm15091 | 0 | 0 | 0 | 1 | 1 |
| Gm15093 | 0 | 0 | 0 | 1 | 1 |
| Gm15097 | 0 | 0 | 0 | 1 | 1 |
| Gm15104 | 0 | 0 | 0 | 1 | 1 |
| Gm15107 | 0 | 0 | 0 | 1 | 1 |
| Gm15114 | 0 | 0 | 0 | 1 | 1 |
| Gm15127 | 0 | 0 | 0 | 1 | 1 |
| Gm15140 | 0 | 0 | 0 | 1 | 1 |
| Gm1527 | 0 | 0 | 0 | 1 | 1 |

| | | | | | |
|---|---|---|---|---|---|
| Gm15292 | 0 | 0 | 0 | 1 | 1 |
| Gm15293 | 0 | 0 | 0 | 1 | 1 |
| Gm15299 | 0 | 0 | 0 | 1 | 1 |
| Gm15308 | 0 | 0 | 0 | 1 | 1 |
| Gm15315 | 0 | 0 | 0 | 1 | 1 |
| Gm15319 | 0 | 0 | 0 | 1 | 1 |
| Gm15386 | 0 | 0 | 0 | 1 | 1 |
| Gm1553 | 0 | 0 | 0 | 1 | 1 |
| Gm156 | 0 | 0 | 0 | 1 | 1 |
| Gm15645 | 0 | 0 | 0 | 1 | 1 |
| Gm15679 | 0 | 0 | 0 | 1 | 1 |
| Gm15698 | 0 | 0 | 0 | 1 | 1 |
| Gm15881 | 0 | 0 | 0 | 1 | 1 |
| Gm15941 | 0 | 0 | 0 | 1 | 1 |
| Gm15987 | 0 | 0 | 0 | 1 | 1 |
| Gm15997 | 0 | 0 | 0 | 1 | 1 |
| Gm1604b | 0 | 0 | 0 | 1 | 1 |
| Gm16287 | 0 | 0 | 0 | 1 | 1 |
| Gm16291 | 0 | 0 | 0 | 1 | 1 |
| Gm16294 | 0 | 0 | 0 | 1 | 1 |
| Gm16336 | 0 | 0 | 0 | 1 | 1 |
| Gm16367 | 0 | 0 | 0 | 1 | 1 |
| Gm16381 | 0 | 0 | 0 | 1 | 1 |
| Gm16390 | 0 | 0 | 0 | 1 | 1 |
| Gm16404 | 0 | 0 | 0 | 1 | 1 |
| Gm16405 | 0 | 0 | 0 | 1 | 1 |
| Gm16442 | 0 | 0 | 0 | 1 | 1 |
| Gm16445 | 0 | 0 | 0 | 1 | 1 |
| Gm16451 | 0 | 0 | 0 | 1 | 1 |
| Gm1647 | 0 | 0 | 0 | 1 | 1 |
| Gm16497 | 0 | 0 | 0 | 1 | 1 |
| Gm16501 | 0 | 0 | 0 | 1 | 1 |
| Gm16677 | 0 | 0 | 0 | 1 | 1 |
| Gm16833 | 0 | 0 | 0 | 1 | 1 |
| Gm16894 | 0 | 0 | 0 | 1 | 1 |
| Gm1715 | 0 | 0 | 0 | 1 | 1 |
| Gm17359 | 0 | 0 | 0 | 1 | 1 |
| Gm17644 | 0 | 0 | 0 | 1 | 1 |
| Gm17677 | 0 | 0 | 0 | 1 | 1 |
| Gm17689 | 0 | 0 | 0 | 1 | 1 |
| Gm17727 | 0 | 0 | 0 | 1 | 1 |
| Gm17751 | 0 | 0 | 0 | 1 | 1 |
| Gm17757,Gm188[illegible] | 0 | 0 | 0 | 1 | 1 |
| Gm17821 | 0 | 0 | 0 | 1 | 1 |
| Gm17830 | 0 | 0 | 0 | 1 | 1 |
| Gm18409 | 0 | 0 | 0 | 1 | 1 |
| Gm18853 | 0 | 0 | 0 | 1 | 1 |
| Gm19276 | 0 | 0 | 0 | 1 | 1 |

| | | | | | |
|---|---|---|---|---|---|
| Gm19277 | 0 | 0 | 0 | 1 | 1 |
| Gm19303 | 0 | 0 | 0 | 1 | 1 |
| Gm19345 | 0 | 0 | 0 | 1 | 1 |
| Gm19402 | 0 | 0 | 0 | 1 | 1 |
| Gm19510 | 0 | 0 | 0 | 1 | 1 |
| Gm19668 | 0 | 0 | 0 | 1 | 1 |
| Gm19689 | 0 | 0 | 0 | 1 | 1 |
| Gm19757 | 0 | 0 | 0 | 1 | 1 |
| Gm19782 | 0 | 0 | 0 | 1 | 1 |
| Gm19784 | 0 | 0 | 0 | 1 | 1 |
| Gm1979 | 0 | 0 | 0 | 1 | 1 |
| Gm1987 | 0 | 0 | 0 | 1 | 1 |
| Gm1993 | 0 | 0 | 0 | 1 | 1 |
| Gm2002 | 0 | 0 | 0 | 1 | 1 |
| Gm20063 | 0 | 0 | 0 | 1 | 1 |
| Gm20098 | 0 | 0 | 0 | 1 | 1 |
| Gm2012 | 0 | 0 | 0 | 1 | 1 |
| Gm20139 | 0 | 0 | 0 | 1 | 1 |
| Gm2016 | 0 | 0 | 0 | 1 | 1 |
| Gm20172 | 0 | 0 | 0 | 1 | 1 |
| Gm20187 | 0 | 0 | 0 | 1 | 1 |
| Gm2022 | 0 | 0 | 0 | 1 | 1 |
| Gm20268 | 0 | 0 | 0 | 1 | 1 |
| Gm2030 | 0 | 0 | 0 | 1 | 1 |
| Gm20356 | 0 | 0 | 0 | 1 | 1 |
| Gm20362 | 0 | 0 | 0 | 1 | 1 |
| Gm2042 | 0 | 0 | 0 | 1 | 1 |
| Gm20735 | 0 | 0 | 0 | 1 | 1 |
| Gm20736 | 0 | 0 | 0 | 1 | 1 |
| Gm20738,Gm208[illegible] | 0 | 0 | 0 | 1 | 1 |
| Gm20740 | 0 | 0 | 0 | 1 | 1 |
| Gm20741 | 0 | 0 | 0 | 1 | 1 |
| Gm20743 | 0 | 0 | 0 | 1 | 1 |
| Gm20744 | 0 | 0 | 0 | 1 | 1 |
| Gm20745 | 0 | 0 | 0 | 1 | 1 |
| Gm20747 | 0 | 0 | 0 | 1 | 1 |
| Gm20752 | 0 | 0 | 0 | 1 | 1 |
| Gm20753 | 0 | 0 | 0 | 1 | 1 |
| Gm20754 | 0 | 0 | 0 | 1 | 1 |
| Gm20756 | 0 | 0 | 0 | 1 | 1 |
| Gm20757 | 0 | 0 | 0 | 1 | 1 |
| Gm20758 | 0 | 0 | 0 | 1 | 1 |
| Gm20759 | 0 | 0 | 0 | 1 | 1 |
| Gm20765,Gm213[illegible] | 0 | 0 | 0 | 1 | 1 |
| Gm20767 | 0 | 0 | 0 | 1 | 1 |
| Gm20806 | 0 | 0 | 0 | 1 | 1 |
| Gm20809 | 0 | 0 | 0 | 1 | 1 |
| Gm20815 | 0 | 0 | 0 | 1 | 1 |

| | | | | | |
|---|---|---|---|---|---|
| Gm20816,Gm208[illegible] | 0 | 0 | 0 | 1 | 1 |
| Gm20822 | 0 | 0 | 0 | 1 | 1 |
| Gm20823 | 0 | 0 | 0 | 1 | 1 |
| Gm20826 | 0 | 0 | 0 | 1 | 1 |
| Gm2083,Mup19 | 0 | 0 | 0 | 1 | 1 |
| Gm20831 | 0 | 0 | 0 | 1 | 1 |
| Gm20854 | 0 | 0 | 0 | 1 | 1 |
| Gm20857 | 0 | 0 | 0 | 1 | 1 |
| Gm20865 | 0 | 0 | 0 | 1 | 1 |
| Gm20867 | 0 | 0 | 0 | 1 | 1 |
| Gm20871 | 0 | 0 | 0 | 1 | 1 |
| Gm20877 | 0 | 0 | 0 | 1 | 1 |
| Gm20878 | 0 | 0 | 0 | 1 | 1 |
| Gm20917 | 0 | 0 | 0 | 1 | 1 |
| Gm21002 | 0 | 0 | 0 | 1 | 1 |
| Gm2109 | 0 | 0 | 0 | 1 | 1 |
| Gm21119 | 0 | 0 | 0 | 1 | 1 |
| Gm21221 | 0 | 0 | 0 | 1 | 1 |
| Gm21269 | 0 | 0 | 0 | 1 | 1 |
| Gm21276 | 0 | 0 | 0 | 1 | 1 |
| Gm21283 | 0 | 0 | 0 | 1 | 1 |
| Gm21293,Gm213[illegible] | 0 | 0 | 0 | 1 | 1 |
| Gm21304,Gm213[illegible] | 0 | 0 | 0 | 1 | 1 |
| Gm21312 | 0 | 0 | 0 | 1 | 1 |
| Gm21498 | 0 | 0 | 0 | 1 | 1 |
| Gm21541 | 0 | 0 | 0 | 1 | 1 |
| Gm21586 | 0 | 0 | 0 | 1 | 1 |
| Gm21637 | 0 | 0 | 0 | 1 | 1 |
| Gm21671 | 0 | 0 | 0 | 1 | 1 |
| Gm21677,Rbmy | 0 | 0 | 0 | 1 | 1 |
| Gm21693 | 0 | 0 | 0 | 1 | 1 |
| Gm21704 | 0 | 0 | 0 | 1 | 1 |
| Gm21943 | 0 | 0 | 0 | 1 | 1 |
| Gm21944 | 0 | 0 | 0 | 1 | 1 |
| Gm21950 | 0 | 0 | 0 | 1 | 1 |
| Gm21951 | 0 | 0 | 0 | 1 | 1 |
| Gm2447 | 0 | 0 | 0 | 1 | 1 |
| Gm2506 | 0 | 0 | 0 | 1 | 1 |
| Gm2516 | 0 | 0 | 0 | 1 | 1 |
| Gm2663 | 0 | 0 | 0 | 1 | 1 |
| Gm2799,Gmcl1l | 0 | 0 | 0 | 1 | 1 |
| Gm2825 | 0 | 0 | 0 | 1 | 1 |
| Gm2837 | 0 | 0 | 0 | 1 | 1 |
| Gm2848 | 0 | 0 | 0 | 1 | 1 |
| Gm2863 | 0 | 0 | 0 | 1 | 1 |
| Gm2913 | 0 | 0 | 0 | 1 | 1 |
| Gm2927 | 0 | 0 | 0 | 1 | 1 |
| Gm2933 | 0 | 0 | 0 | 1 | 1 |

| | | | | | |
|---|---|---|---|---|---|
| Gm3020 | 0 | 0 | 0 | 1 | 1 |
| Gm3139 | 0 | 0 | 0 | 1 | 1 |
| Gm3143 | 0 | 0 | 0 | 1 | 1 |
| Gm3238 | 0 | 0 | 0 | 1 | 1 |
| Gm3259 | 0 | 0 | 0 | 1 | 1 |
| Gm3264 | 0 | 0 | 0 | 1 | 1 |
| Gm3279 | 0 | 0 | 0 | 1 | 1 |
| Gm3285 | 0 | 0 | 0 | 1 | 1 |
| Gm3286 | 0 | 0 | 0 | 1 | 1 |
| Gm3376,Rbmy | 0 | 0 | 0 | 1 | 1 |
| Gm3383 | 0 | 0 | 0 | 1 | 1 |
| Gm3402,Gm3415 | 0 | 0 | 0 | 1 | 1 |
| Gm3404 | 0 | 0 | 0 | 1 | 1 |
| Gm3409 | 0 | 0 | 0 | 1 | 1 |
| Gm3415 | 0 | 0 | 0 | 1 | 1 |
| Gm3417,Tcte3 | 0 | 0 | 0 | 1 | 1 |
| Gm3428 | 0 | 0 | 0 | 1 | 1 |
| Gm3434 | 0 | 0 | 0 | 1 | 1 |
| Gm3488 | 0 | 0 | 0 | 1 | 1 |
| Gm362 | 0 | 0 | 0 | 1 | 1 |
| Gm364 | 0 | 0 | 0 | 1 | 1 |
| Gm3646 | 0 | 0 | 0 | 1 | 1 |
| Gm3701 | 0 | 0 | 0 | 1 | 1 |
| Gm3706 | 0 | 0 | 0 | 1 | 1 |
| Gm3750 | 0 | 0 | 0 | 1 | 1 |
| Gm3763 | 0 | 0 | 0 | 1 | 1 |
| Gm3776 | 0 | 0 | 0 | 1 | 1 |
| Gm382 | 0 | 0 | 0 | 1 | 1 |
| Gm3893 | 0 | 0 | 0 | 1 | 1 |
| Gm3985 | 0 | 0 | 0 | 1 | 1 |
| Gm4027 | 0 | 0 | 0 | 1 | 1 |
| Gm4070,Gvin1 | 0 | 0 | 0 | 1 | 1 |
| Gm41 | 0 | 0 | 0 | 1 | 1 |
| Gm4133 | 0 | 0 | 0 | 1 | 1 |
| Gm4141 | 0 | 0 | 0 | 1 | 1 |
| Gm4175 | 0 | 0 | 0 | 1 | 1 |
| Gm4177 | 0 | 0 | 0 | 1 | 1 |
| Gm4201 | 0 | 0 | 0 | 1 | 1 |
| Gm4216 | 0 | 0 | 0 | 1 | 1 |
| Gm4224 | 0 | 0 | 0 | 1 | 1 |
| Gm4251 | 0 | 0 | 0 | 1 | 1 |
| Gm4278 | 0 | 0 | 0 | 1 | 1 |
| Gm428 | 0 | 0 | 0 | 1 | 1 |
| Gm4297 | 0 | 0 | 0 | 1 | 1 |
| Gm4301 | 0 | 0 | 0 | 1 | 1 |
| Gm4302 | 0 | 0 | 0 | 1 | 1 |
| Gm4303 | 0 | 0 | 0 | 1 | 1 |
| Gm4307 | 0 | 0 | 0 | 1 | 1 |

| | | | | | |
|---|---|---|---|---|---|
| Gm4312 | 0 | 0 | 0 | 1 | 1 |
| Gm4340 | 0 | 0 | 0 | 1 | 1 |
| Gm436 | 0 | 0 | 0 | 1 | 1 |
| Gm438 | 0 | 0 | 0 | 1 | 1 |
| Gm44 | 0 | 0 | 0 | 1 | 1 |
| Gm4477 | 0 | 0 | 0 | 1 | 1 |
| Gm4498 | 0 | 0 | 0 | 1 | 1 |
| Gm4541 | 0 | 0 | 0 | 1 | 1 |
| Gm4567,Gm6902 | 0 | 0 | 0 | 1 | 1 |
| Gm4710 | 0 | 0 | 0 | 1 | 1 |
| Gm4719 | 0 | 0 | 0 | 1 | 1 |
| Gm4736 | 0 | 0 | 0 | 1 | 1 |
| Gm4745 | 0 | 0 | 0 | 1 | 1 |
| Gm4759 | 0 | 0 | 0 | 1 | 1 |
| Gm4763 | 0 | 0 | 0 | 1 | 1 |
| Gm4776 | 0 | 0 | 0 | 1 | 1 |
| Gm4788 | 0 | 0 | 0 | 1 | 1 |
| Gm4791 | 0 | 0 | 0 | 1 | 1 |
| Gm4792 | 0 | 0 | 0 | 1 | 1 |
| Gm4794 | 0 | 0 | 0 | 1 | 1 |
| Gm4814 | 0 | 0 | 0 | 1 | 1 |
| Gm4836 | 0 | 0 | 0 | 1 | 1 |
| Gm4841 | 0 | 0 | 0 | 1 | 1 |
| Gm4846 | 0 | 0 | 0 | 1 | 1 |
| Gm4847 | 0 | 0 | 0 | 1 | 1 |
| Gm4850 | 0 | 0 | 0 | 1 | 1 |
| Gm4858 | 0 | 0 | 0 | 1 | 1 |
| Gm4871 | 0 | 0 | 0 | 1 | 1 |
| Gm4884 | 0 | 0 | 0 | 1 | 1 |
| Gm4894 | 0 | 0 | 0 | 1 | 1 |
| Gm4906 | 0 | 0 | 0 | 1 | 1 |
| Gm4922 | 0 | 0 | 0 | 1 | 1 |
| Gm4926 | 0 | 0 | 0 | 1 | 1 |
| Gm4937 | 0 | 0 | 0 | 1 | 1 |
| Gm4951 | 0 | 0 | 0 | 1 | 1 |
| Gm4956 | 0 | 0 | 0 | 1 | 1 |
| Gm4981 | 0 | 0 | 0 | 1 | 1 |
| Gm5 | 0 | 0 | 0 | 1 | 1 |
| Gm5071 | 0 | 0 | 0 | 1 | 1 |
| Gm5072 | 0 | 0 | 0 | 1 | 1 |
| Gm5082 | 0 | 0 | 0 | 1 | 1 |
| Gm5084 | 0 | 0 | 0 | 1 | 1 |
| Gm5091 | 0 | 0 | 0 | 1 | 1 |
| Gm5095 | 0 | 0 | 0 | 1 | 1 |
| Gm5108 | 0 | 0 | 0 | 1 | 1 |
| Gm5111 | 0 | 0 | 0 | 1 | 1 |
| Gm5114 | 0 | 0 | 0 | 1 | 1 |
| Gm5122 | 0 | 0 | 0 | 1 | 1 |

| | | | | | |
|---|---|---|---|---|---|
| Gm5132 | 0 | 0 | 0 | 1 | 1 |
| Gm5166 | 0 | 0 | 0 | 1 | 1 |
| Gm5168 | 0 | 0 | 0 | 1 | 1 |
| Gm5169 | 0 | 0 | 0 | 1 | 1 |
| Gm525 | 0 | 0 | 0 | 1 | 1 |
| Gm5334 | 0 | 0 | 0 | 1 | 1 |
| Gm5346 | 0 | 0 | 0 | 1 | 1 |
| Gm5347 | 0 | 0 | 0 | 1 | 1 |
| Gm5382 | 0 | 0 | 0 | 1 | 1 |
| Gm5409 | 0 | 0 | 0 | 1 | 1 |
| Gm5414 | 0 | 0 | 0 | 1 | 1 |
| Gm5415 | 0 | 0 | 0 | 1 | 1 |
| Gm5416 | 0 | 0 | 0 | 1 | 1 |
| Gm5431 | 0 | 0 | 0 | 1 | 1 |
| Gm5441 | 0 | 0 | 0 | 1 | 1 |
| Gm5458 | 0 | 0 | 0 | 1 | 1 |
| Gm5460 | 0 | 0 | 0 | 1 | 1 |
| Gm5476 | 0 | 0 | 0 | 1 | 1 |
| Gm5477 | 0 | 0 | 0 | 1 | 1 |
| Gm5478 | 0 | 0 | 0 | 1 | 1 |
| Gm5485 | 0 | 0 | 0 | 1 | 1 |
| Gm5531 | 0 | 0 | 0 | 1 | 1 |
| Gm5535 | 0 | 0 | 0 | 1 | 1 |
| Gm5538 | 0 | 0 | 0 | 1 | 1 |
| Gm5549 | 0 | 0 | 0 | 1 | 1 |
| Gm5592 | 0 | 0 | 0 | 1 | 1 |
| Gm5615 | 0 | 0 | 0 | 1 | 1 |
| Gm5627 | 0 | 0 | 0 | 1 | 1 |
| Gm5634 | 0 | 0 | 0 | 1 | 1 |
| Gm5635 | 0 | 0 | 0 | 1 | 1 |
| Gm5640 | 0 | 0 | 0 | 1 | 1 |
| Gm5643 | 0 | 0 | 0 | 1 | 1 |
| Gm5662 | 0 | 0 | 0 | 1 | 1 |
| Gm5712 | 0 | 0 | 0 | 1 | 1 |
| Gm5725 | 0 | 0 | 0 | 1 | 1 |
| Gm5726 | 0 | 0 | 0 | 1 | 1 |
| Gm5728 | 0 | 0 | 0 | 1 | 1 |
| Gm5766 | 0 | 0 | 0 | 1 | 1 |
| Gm5771 | 0 | 0 | 0 | 1 | 1 |
| Gm5795 | 0 | 0 | 0 | 1 | 1 |
| Gm5797 | 0 | 0 | 0 | 1 | 1 |
| Gm5833 | 0 | 0 | 0 | 1 | 1 |
| Gm5860 | 0 | 0 | 0 | 1 | 1 |
| Gm5862 | 0 | 0 | 0 | 1 | 1 |
| Gm5885 | 0 | 0 | 0 | 1 | 1 |
| Gm5886 | 0 | 0 | 0 | 1 | 1 |
| Gm5891 | 0 | 0 | 0 | 1 | 1 |
| Gm5893 | 0 | 0 | 0 | 1 | 1 |

| | | | | | |
|---|---|---|---|---|---|
| Gm590 | 0 | 0 | 0 | 1 | 1 |
| Gm5901 | 0 | 0 | 0 | 1 | 1 |
| Gm5916 | 0 | 0 | 0 | 1 | 1 |
| Gm5925 | 0 | 0 | 0 | 1 | 1 |
| Gm5934 | 0 | 0 | 0 | 1 | 1 |
| Gm5935 | 0 | 0 | 0 | 1 | 1 |
| Gm5938 | 0 | 0 | 0 | 1 | 1 |
| Gm5941 | 0 | 0 | 0 | 1 | 1 |
| Gm595 | 0 | 0 | 0 | 1 | 1 |
| Gm597 | 0 | 0 | 0 | 1 | 1 |
| Gm6026 | 0 | 0 | 0 | 1 | 1 |
| Gm6040 | 0 | 0 | 0 | 1 | 1 |
| Gm6042 | 0 | 0 | 0 | 1 | 1 |
| Gm609 | 0 | 0 | 0 | 1 | 1 |
| Gm6116 | 0 | 0 | 0 | 1 | 1 |
| Gm6121 | 0 | 0 | 0 | 1 | 1 |
| Gm6164 | 0 | 0 | 0 | 1 | 1 |
| Gm6260 | 0 | 0 | 0 | 1 | 1 |
| Gm6268 | 0 | 0 | 0 | 1 | 1 |
| Gm6300 | 0 | 0 | 0 | 1 | 1 |
| Gm6307 | 0 | 0 | 0 | 1 | 1 |
| Gm6313 | 0 | 0 | 0 | 1 | 1 |
| Gm6329 | 0 | 0 | 0 | 1 | 1 |
| Gm6367 | 0 | 0 | 0 | 1 | 1 |
| Gm6370 | 0 | 0 | 0 | 1 | 1 |
| Gm6406 | 0 | 0 | 0 | 1 | 1 |
| Gm6408 | 0 | 0 | 0 | 1 | 1 |
| Gm6416 | 0 | 0 | 0 | 1 | 1 |
| Gm6455 | 0 | 0 | 0 | 1 | 1 |
| Gm6460 | 0 | 0 | 0 | 1 | 1 |
| Gm6567 | 0 | 0 | 0 | 1 | 1 |
| Gm6588 | 0 | 0 | 0 | 1 | 1 |
| Gm6592 | 0 | 0 | 0 | 1 | 1 |
| Gm6602 | 0 | 0 | 0 | 1 | 1 |
| Gm6607 | 0 | 0 | 0 | 1 | 1 |
| Gm6614 | 0 | 0 | 0 | 1 | 1 |
| Gm6639 | 0 | 0 | 0 | 1 | 1 |
| Gm6696 | 0 | 0 | 0 | 1 | 1 |
| Gm6710 | 0 | 0 | 0 | 1 | 1 |
| Gm6760 | 0 | 0 | 0 | 1 | 1 |
| Gm6763 | 0 | 0 | 0 | 1 | 1 |
| Gm6793 | 0 | 0 | 0 | 1 | 1 |
| Gm6812 | 0 | 0 | 0 | 1 | 1 |
| Gm6890 | 0 | 0 | 0 | 1 | 1 |
| Gm6902 | 0 | 0 | 0 | 1 | 1 |
| Gm6927 | 0 | 0 | 0 | 1 | 1 |
| Gm6936 | 0 | 0 | 0 | 1 | 1 |
| Gm694 | 0 | 0 | 0 | 1 | 1 |

| | | | | | |
|---|---|---|---|---|---|
| Gm6994 | 0 | 0 | 0 | 1 | 1 |
| Gm7030 | 0 | 0 | 0 | 1 | 1 |
| Gm7056 | 0 | 0 | 0 | 1 | 1 |
| Gm7073 | 0 | 0 | 0 | 1 | 1 |
| Gm7157 | 0 | 0 | 0 | 1 | 1 |
| Gm7271 | 0 | 0 | 0 | 1 | 1 |
| Gm732 | 0 | 0 | 0 | 1 | 1 |
| Gm7337 | 0 | 0 | 0 | 1 | 1 |
| Gm7361 | 0 | 0 | 0 | 1 | 1 |
| Gm7367 | 0 | 0 | 0 | 1 | 1 |
| Gm7534 | 0 | 0 | 0 | 1 | 1 |
| Gm7538 | 0 | 0 | 0 | 1 | 1 |
| Gm7550 | 0 | 0 | 0 | 1 | 1 |
| Gm7609 | 0 | 0 | 0 | 1 | 1 |
| Gm765 | 0 | 0 | 0 | 1 | 1 |
| Gm7714 | 0 | 0 | 0 | 1 | 1 |
| Gm773 | 0 | 0 | 0 | 1 | 1 |
| Gm7849,Gm7861 | 0 | 0 | 0 | 1 | 1 |
| Gm7861 | 0 | 0 | 0 | 1 | 1 |
| Gm7903 | 0 | 0 | 0 | 1 | 1 |
| Gm7978 | 0 | 0 | 0 | 1 | 1 |
| Gm806 | 0 | 0 | 0 | 1 | 1 |
| Gm8096 | 0 | 0 | 0 | 1 | 1 |
| Gm813 | 0 | 0 | 0 | 1 | 1 |
| Gm815 | 0 | 0 | 0 | 1 | 1 |
| Gm8179 | 0 | 0 | 0 | 1 | 1 |
| Gm8267 | 0 | 0 | 0 | 1 | 1 |
| Gm829 | 0 | 0 | 0 | 1 | 1 |
| Gm8300 | 0 | 0 | 0 | 1 | 1 |
| Gm833 | 0 | 0 | 0 | 1 | 1 |
| Gm8369 | 0 | 0 | 0 | 1 | 1 |
| Gm839 | 0 | 0 | 0 | 1 | 1 |
| Gm8439 | 0 | 0 | 0 | 1 | 1 |
| Gm8453 | 0 | 0 | 0 | 1 | 1 |
| Gm853 | 0 | 0 | 0 | 1 | 1 |
| Gm8579 | 0 | 0 | 0 | 1 | 1 |
| Gm8633 | 0 | 0 | 0 | 1 | 1 |
| Gm8653 | 0 | 0 | 0 | 1 | 1 |
| Gm8693 | 0 | 0 | 0 | 1 | 1 |
| Gm8709 | 0 | 0 | 0 | 1 | 1 |
| Gm8765 | 0 | 0 | 0 | 1 | 1 |
| Gm884 | 0 | 0 | 0 | 1 | 1 |
| Gm8882 | 0 | 0 | 0 | 1 | 1 |
| Gm8979 | 0 | 0 | 0 | 1 | 1 |
| Gm8989 | 0 | 0 | 0 | 1 | 1 |
| Gm9 | 0 | 0 | 0 | 1 | 1 |
| Gm904 | 0 | 0 | 0 | 1 | 1 |
| Gm906 | 0 | 0 | 0 | 1 | 1 |

| | | | | | |
|---|---|---|---|---|---|
| Gm9125 | 0 | 0 | 0 | 1 | 1 |
| Gm9268 | 0 | 0 | 0 | 1 | 1 |
| Gm933 | 0 | 0 | 0 | 1 | 1 |
| Gm9376 | 0 | 0 | 0 | 1 | 1 |
| Gm94 | 0 | 0 | 0 | 1 | 1 |
| Gm9573 | 0 | 0 | 0 | 1 | 1 |
| Gm9696 | 0 | 0 | 0 | 1 | 1 |
| Gm9731 | 0 | 0 | 0 | 1 | 1 |
| Gm9733 | 0 | 0 | 0 | 1 | 1 |
| Gm9758 | 0 | 0 | 0 | 1 | 1 |
| Gm9839 | 0 | 0 | 0 | 1 | 1 |
| Gm9866 | 0 | 0 | 0 | 1 | 1 |
| Gm9961 | 0 | 0 | 0 | 1 | 1 |
| Gm9992 | 0 | 0 | 0 | 1 | 1 |
| Gm9994 | 0 | 0 | 0 | 1 | 1 |
| Gm9999 | 0 | 0 | 0 | 1 | 1 |
| Gmcl1l | 0 | 0 | 0 | 1 | 1 |
| Gml | 0 | 0 | 0 | 1 | 1 |
| Gnat3 | 0 | 0 | 0 | 1 | 1 |
| Gngt1 | 0 | 0 | 0 | 1 | 1 |
| Gnrhr | 0 | 0 | 0 | 1 | 1 |
| Gp2 | 0 | 0 | 0 | 1 | 1 |
| Gphb5 | 0 | 0 | 0 | 1 | 1 |
| Gpr113 | 0 | 0 | 0 | 1 | 1 |
| Gpr119 | 0 | 0 | 0 | 1 | 1 |
| Gpr152 | 0 | 0 | 0 | 1 | 1 |
| Gpr25 | 0 | 0 | 0 | 1 | 1 |
| Gpr31b | 0 | 0 | 0 | 1 | 1 |
| Gpr37l1 | 0 | 0 | 0 | 1 | 1 |
| Gpr6 | 0 | 0 | 0 | 1 | 1 |
| Gprc6a | 0 | 0 | 0 | 1 | 1 |
| Gpx5 | 0 | 0 | 0 | 1 | 1 |
| Grip1os2 | 0 | 0 | 0 | 1 | 1 |
| Grm7 | 0 | 0 | 0 | 1 | 1 |
| Grxcr2 | 0 | 0 | 0 | 1 | 1 |
| Gsc2 | 0 | 0 | 0 | 1 | 1 |
| Gsdma | 0 | 0 | 0 | 1 | 1 |
| Gsdma2 | 0 | 0 | 0 | 1 | 1 |
| Gsdmcl1 | 0 | 0 | 0 | 1 | 1 |
| Gsdmcl2 | 0 | 0 | 0 | 1 | 1 |
| Gspt1 | 68,799 | 0 | 0 | 1 | 1 |
| Gsta1 | 0 | 0 | 0 | 1 | 1 |
| Gsta2 | 0 | 0 | 0 | 1 | 1 |
| Gsx1 | 0 | 0 | 0 | 1 | 1 |
| Gtsf1 | 0 | 0 | 0 | 1 | 1 |
| Gucy2d | 0 | 0 | 0 | 1 | 1 |
| Gucy2f | 0 | 0 | 0 | 1 | 1 |
| Gvin1 | 0 | 0 | 0 | 1 | 1 |

| | | | | | |
|---|---|---|---|---|---|
| Gzmn | 0 | 0 | 0 | 1 | 1 |
| H1fnt | 0 | 0 | 0 | 1 | 1 |
| H1foo | 0 | 0 | 0 | 1 | 1 |
| H2-Ea-ps | 0 | 0 | 0 | 1 | 1 |
| H2-M1 | 0 | 0 | 0 | 1 | 1 |
| H2-M10.1 | 0 | 0 | 0 | 1 | 1 |
| H2-M10.4 | 0 | 0 | 0 | 1 | 1 |
| H2-M10.5 | 0 | 0 | 0 | 1 | 1 |
| H2-M10.6 | 0 | 0 | 0 | 1 | 1 |
| H2-M11 | 0 | 0 | 0 | 1 | 1 |
| H2-M2 | 0 | 0 | 0 | 1 | 1 |
| H2-M9 | 0 | 0 | 0 | 1 | 1 |
| H2-Ob | 0 | 0 | 0 | 1 | 1 |
| H2afb1 | 0 | 0 | 0 | 1 | 1 |
| H2afb2 | 0 | 0 | 0 | 1 | 1 |
| H2afb3 | 0 | 0 | 0 | 1 | 1 |
| H2bfm | 0 | 0 | 0 | 1 | 1 |
| Hamp2 | 0 | 0 | 0 | 1 | 1 |
| Hao1 | 0 | 0 | 0 | 1 | 1 |
| Hao2 | 0 | 0 | 0 | 1 | 1 |
| Havcr1 | 0 | 0 | 0 | 1 | 1 |
| Hcrt | 0 | 0 | 0 | 1 | 1 |
| Hdgfl1 | 0 | 0 | 0 | 1 | 1 |
| Hdhd1a | 0 | 0 | 0 | 1 | 1 |
| Hemt1 | 0 | 0 | 0 | 1 | 1 |
| Hhla1 | 0 | 0 | 0 | 1 | 1 |
| Higd1b | 0 | 0 | 0 | 1 | 1 |
| Hils1 | 0 | 0 | 0 | 1 | 1 |
| Hist1h2aa | 0 | 0 | 0 | 1 | 1 |
| Hist1h2ao,Hist1h2 | 0 | 0 | 0 | 1 | 1 |
| Hist1h2ap | 0 | 0 | 0 | 1 | 1 |
| Hist1h2bn | 0 | 0 | 0 | 1 | 1 |
| Hist1h2bq | 0 | 0 | 0 | 1 | 1 |
| Hist1h2br | 0 | 0 | 0 | 1 | 1 |
| Hist1h4a | 0 | 0 | 0 | 1 | 1 |
| Hist1h4m,Hist1h4 | 0 | 0 | 0 | 1 | 1 |
| Hist1h4n | 0 | 0 | 0 | 1 | 1 |
| Hist2h3c1,Hist2h3 | 0 | 0 | 0 | 1 | 1 |
| Hist2h3c2 | 0 | 0 | 0 | 1 | 1 |
| Hmgb4 | 0 | 0 | 0 | 1 | 1 |
| Hnf4aos | 0 | 0 | 0 | 1 | 1 |
| Hrasls5 | 0 | 0 | 0 | 1 | 1 |
| Hrg | 0 | 0 | 0 | 1 | 1 |
| Hrnr | 0 | 0 | 0 | 1 | 1 |
| Hs3st2 | 0 | 0 | 0 | 1 | 1 |
| Hsd17b6 | 0 | 0 | 0 | 1 | 1 |
| Hsd3b4 | 0 | 0 | 0 | 1 | 1 |
| Hsd3b5 | 0 | 0 | 0 | 1 | 1 |

| | | | | | |
|---|---|---|---|---|---|
| Hsfy2 | 0 | 0 | 0 | 1 | 1 |
| Hspb7 | 0 | 0 | 0 | 1 | 1 |
| Htr1a | 0 | 0 | 0 | 1 | 1 |
| Htr2c | 0 | 0 | 0 | 1 | 1 |
| Htr3b | 0 | 0 | 0 | 1 | 1 |
| Htr5a | 0 | 0 | 0 | 1 | 1 |
| Hyal5 | 0 | 0 | 0 | 1 | 1 |
| Hyal6 | 0 | 0 | 0 | 1 | 1 |
| I730028E13Rik | 0 | 0 | 0 | 1 | 1 |
| I730030J21Rik | 0 | 0 | 0 | 1 | 1 |
| Ifi205 | 0 | 0 | 0 | 1 | 1 |
| Ifi44l | 0 | 0 | 0 | 1 | 1 |
| Ifltd1 | 0 | 0 | 0 | 1 | 1 |
| Ifna1 | 0 | 0 | 0 | 1 | 1 |
| Ifna11 | 0 | 0 | 0 | 1 | 1 |
| Ifna12 | 0 | 0 | 0 | 1 | 1 |
| Ifna14 | 0 | 0 | 0 | 1 | 1 |
| Ifna15 | 0 | 0 | 0 | 1 | 1 |
| Ifna16 | 0 | 0 | 0 | 1 | 1 |
| Ifna2 | 0 | 0 | 0 | 1 | 1 |
| Ifna4 | 0 | 0 | 0 | 1 | 1 |
| Ifna5 | 0 | 0 | 0 | 1 | 1 |
| Ifna6 | 0 | 0 | 0 | 1 | 1 |
| Ifna7 | 0 | 0 | 0 | 1 | 1 |
| Ifna9 | 0 | 0 | 0 | 1 | 1 |
| Ifnab | 0 | 0 | 0 | 1 | 1 |
| Ifnb1 | 0 | 0 | 0 | 1 | 1 |
| Ifne | 0 | 0 | 0 | 1 | 1 |
| Ifng | 0 | 0 | 0 | 1 | 1 |
| Ifnl2 | 0 | 0 | 0 | 1 | 1 |
| Ifnl3 | 0 | 0 | 0 | 1 | 1 |
| Igbp1b | 0 | 0 | 0 | 1 | 1 |
| Igfl3 | 0 | 0 | 0 | 1 | 1 |
| Il11ra2 | 0 | 0 | 0 | 1 | 1 |
| Il12b | 0 | 0 | 0 | 1 | 1 |
| Il13 | 0 | 0 | 0 | 1 | 1 |
| Il13ra2 | 0 | 0 | 0 | 1 | 1 |
| Il17a | 0 | 0 | 0 | 1 | 1 |
| Il1f10 | 0 | 0 | 0 | 1 | 1 |
| Il1f8 | 0 | 0 | 0 | 1 | 1 |
| Il1f9 | 0 | 0 | 0 | 1 | 1 |
| Il1rapl1 | 0 | 0 | 0 | 1 | 1 |
| Il1rapl2 | 0 | 0 | 0 | 1 | 1 |
| Il2 | 0 | 0 | 0 | 1 | 1 |
| Il20 | 0 | 0 | 0 | 1 | 1 |
| Il21 | 0 | 0 | 0 | 1 | 1 |
| Il22ra2 | 0 | 0 | 0 | 1 | 1 |
| Il23r | 0 | 0 | 0 | 1 | 1 |

| | | | | | |
|---|---|---|---|---|---|
| Il24 | 0 | 0 | 0 | 1 | 1 |
| Il27 | 0 | 0 | 0 | 1 | 1 |
| Il2ra | 0 | 0 | 0 | 1 | 1 |
| Il3 | 0 | 0 | 0 | 1 | 1 |
| Il31 | 0 | 0 | 0 | 1 | 1 |
| Il34 | 0 | 0 | 0 | 1 | 1 |
| Il4 | 0 | 0 | 0 | 1 | 1 |
| Il4i1 | 0 | 0 | 0 | 1 | 1 |
| Il6 | 0 | 0 | 0 | 1 | 1 |
| Il9 | 0 | 0 | 0 | 1 | 1 |
| Inhba | 0 | 0 | 0 | 1 | 1 |
| Inmt | 0 | 0 | 0 | 1 | 1 |
| Ins2 | 0 | 0 | 0 | 1 | 1 |
| Insm2 | 0 | 0 | 0 | 1 | 1 |
| Ip6k3 | 0 | 0 | 0 | 1 | 1 |
| Ipw | 0 | 0 | 0 | 1 | 1 |
| Iqcf1 | 0 | 0 | 0 | 1 | 1 |
| Iqcf3 | 0 | 0 | 0 | 1 | 1 |
| Iqcf4 | 0 | 0 | 0 | 1 | 1 |
| Iqcf5 | 0 | 0 | 0 | 1 | 1 |
| Iqcf6 | 0 | 0 | 0 | 1 | 1 |
| Irgc1 | 0 | 0 | 0 | 1 | 1 |
| Itgad | 0 | 0 | 0 | 1 | 1 |
| Ivl | 0 | 0 | 0 | 1 | 1 |
| Izumo2 | 0 | 0 | 0 | 1 | 1 |
| Izumo3 | 0 | 0 | 0 | 1 | 1 |
| Kap | 0 | 0 | 0 | 1 | 1 |
| Kcna10 | 0 | 0 | 0 | 1 | 1 |
| Kcng4 | 0 | 0 | 0 | 1 | 1 |
| Kcnk15 | 0 | 0 | 0 | 1 | 1 |
| Kcnk18 | 0 | 0 | 0 | 1 | 1 |
| Kcnk7 | 0 | 0 | 0 | 1 | 1 |
| Kcnk9 | 0 | 0 | 0 | 1 | 1 |
| Kcnv2 | 0 | 0 | 0 | 1 | 1 |
| Keg1 | 0 | 0 | 0 | 1 | 1 |
| Kera | 0 | 0 | 0 | 1 | 1 |
| Khdc1b | 0 | 0 | 0 | 1 | 1 |
| Khdc1c | 0 | 0 | 0 | 1 | 1 |
| Kir3dl1 | 0 | 0 | 0 | 1 | 1 |
| Kir3dl2 | 0 | 0 | 0 | 1 | 1 |
| Klf17 | 0 | 0 | 0 | 1 | 1 |
| Klhdc7b | 0 | 0 | 0 | 1 | 1 |
| Klhl1 | 0 | 0 | 0 | 1 | 1 |
| Klhl38 | 0 | 0 | 0 | 1 | 1 |
| Klk12 | 0 | 0 | 0 | 1 | 1 |
| Klk1b1 | 0 | 0 | 0 | 1 | 1 |
| Klk1b11 | 0 | 0 | 0 | 1 | 1 |
| Klk1b16 | 0 | 0 | 0 | 1 | 1 |

| | | | | | |
|---|---|---|---|---|---|
| Klk1b21 | 0 | 0 | 0 | 1 | 1 |
| Klk1b22 | 0 | 0 | 0 | 1 | 1 |
| Klk1b24 | 0 | 0 | 0 | 1 | 1 |
| Klk1b26 | 0 | 0 | 0 | 1 | 1 |
| Klk1b27 | 0 | 0 | 0 | 1 | 1 |
| Klk1b3 | 0 | 0 | 0 | 1 | 1 |
| Klk1b4 | 0 | 0 | 0 | 1 | 1 |
| Klk1b5 | 0 | 0 | 0 | 1 | 1 |
| Klk1b7-ps | 0 | 0 | 0 | 1 | 1 |
| Klk1b8 | 0 | 0 | 0 | 1 | 1 |
| Klk1b9 | 0 | 0 | 0 | 1 | 1 |
| Klk6 | 0 | 0 | 0 | 1 | 1 |
| Klra1 | 0 | 0 | 0 | 1 | 1 |
| Klra10 | 0 | 0 | 0 | 1 | 1 |
| Klra12 | 0 | 0 | 0 | 1 | 1 |
| Klra13-ps | 0 | 0 | 0 | 1 | 1 |
| Klra14-ps | 0 | 0 | 0 | 1 | 1 |
| Klra15 | 0 | 0 | 0 | 1 | 1 |
| Klra17 | 0 | 0 | 0 | 1 | 1 |
| Klra18,Klra33,Klra4 | 0 | 0 | 0 | 1 | 1 |
| Klra19,Klra6 | 0 | 0 | 0 | 1 | 1 |
| Klra21 | 0 | 0 | 0 | 1 | 1 |
| Klra22 | 0 | 0 | 0 | 1 | 1 |
| Klra23 | 0 | 0 | 0 | 1 | 1 |
| Klra3 | 0 | 0 | 0 | 1 | 1 |
| Klra33 | 0 | 0 | 0 | 1 | 1 |
| Klra4 | 0 | 0 | 0 | 1 | 1 |
| Klra5 | 0 | 0 | 0 | 1 | 1 |
| Klra6 | 0 | 0 | 0 | 1 | 1 |
| Klra7 | 0 | 0 | 0 | 1 | 1 |
| Klra8 | 0 | 0 | 0 | 1 | 1 |
| Klra9 | 0 | 0 | 0 | 1 | 1 |
| Klrb1 | 0 | 0 | 0 | 1 | 1 |
| Klrb1-ps1 | 0 | 0 | 0 | 1 | 1 |
| Klrb1a | 0 | 0 | 0 | 1 | 1 |
| Klrc1 | 0 | 0 | 0 | 1 | 1 |
| Klrc2 | 0 | 0 | 0 | 1 | 1 |
| Klrc3 | 0 | 0 | 0 | 1 | 1 |
| Klrd1 | 0 | 0 | 0 | 1 | 1 |
| Klre1 | 0 | 0 | 0 | 1 | 1 |
| Klri1 | 0 | 0 | 0 | 1 | 1 |
| Klri2 | 0 | 0 | 0 | 1 | 1 |
| Klrk1 | 0 | 0 | 0 | 1 | 1 |
| Kncn | 0 | 0 | 0 | 1 | 1 |
| Kpna7 | 0 | 0 | 0 | 1 | 1 |
| Kprp | 0 | 0 | 0 | 1 | 1 |
| Krt16 | 0 | 0 | 0 | 1 | 1 |
| Krt26 | 0 | 0 | 0 | 1 | 1 |

| | | | | | |
|---|---|---|---|---|---|
| Krt31 | 0 | 0 | 0 | 1 | 1 |
| Krt32 | 0 | 0 | 0 | 1 | 1 |
| Krt33a | 0 | 0 | 0 | 1 | 1 |
| Krt33b | 0 | 0 | 0 | 1 | 1 |
| Krt34 | 0 | 0 | 0 | 1 | 1 |
| Krt40 | 0 | 0 | 0 | 1 | 1 |
| Krt6a | 0 | 0 | 0 | 1 | 1 |
| Krt6b | 0 | 0 | 0 | 1 | 1 |
| Krt72 | 0 | 0 | 0 | 1 | 1 |
| Krt73 | 0 | 0 | 0 | 1 | 1 |
| Krt74 | 0 | 0 | 0 | 1 | 1 |
| Krt77 | 0 | 0 | 0 | 1 | 1 |
| Krtap1-3 | 0 | 0 | 0 | 1 | 1 |
| Krtap1-4 | 0 | 0 | 0 | 1 | 1 |
| Krtap1-5 | 0 | 0 | 0 | 1 | 1 |
| Krtap10-10 | 0 | 0 | 0 | 1 | 1 |
| Krtap10-4 | 0 | 0 | 0 | 1 | 1 |
| Krtap11-1 | 0 | 0 | 0 | 1 | 1 |
| Krtap12-1 | 0 | 0 | 0 | 1 | 1 |
| Krtap13 | 0 | 0 | 0 | 1 | 1 |
| Krtap13-1 | 0 | 0 | 0 | 1 | 1 |
| Krtap14 | 0 | 0 | 0 | 1 | 1 |
| Krtap15 | 0 | 0 | 0 | 1 | 1 |
| Krtap16-1 | 0 | 0 | 0 | 1 | 1 |
| Krtap16-3 | 0 | 0 | 0 | 1 | 1 |
| Krtap19-1 | 0 | 0 | 0 | 1 | 1 |
| Krtap19-3 | 0 | 0 | 0 | 1 | 1 |
| Krtap19-4 | 0 | 0 | 0 | 1 | 1 |
| Krtap19-5 | 0 | 0 | 0 | 1 | 1 |
| Krtap19-9b | 0 | 0 | 0 | 1 | 1 |
| Krtap2-4 | 0 | 0 | 0 | 1 | 1 |
| Krtap20-2 | 0 | 0 | 0 | 1 | 1 |
| Krtap21-1 | 0 | 0 | 0 | 1 | 1 |
| Krtap22-2 | 0 | 0 | 0 | 1 | 1 |
| Krtap24-1 | 0 | 0 | 0 | 1 | 1 |
| Krtap27-1 | 0 | 0 | 0 | 1 | 1 |
| Krtap3-2 | 0 | 0 | 0 | 1 | 1 |
| Krtap31-1 | 0 | 0 | 0 | 1 | 1 |
| Krtap31-2 | 0 | 0 | 0 | 1 | 1 |
| Krtap4-1 | 0 | 0 | 0 | 1 | 1 |
| Krtap4-13 | 0 | 0 | 0 | 1 | 1 |
| Krtap4-16 | 0 | 0 | 0 | 1 | 1 |
| Krtap4-2 | 0 | 0 | 0 | 1 | 1 |
| Krtap4-6 | 0 | 0 | 0 | 1 | 1 |
| Krtap4-7 | 0 | 0 | 0 | 1 | 1 |
| Krtap4-8 | 0 | 0 | 0 | 1 | 1 |
| Krtap4-9 | 0 | 0 | 0 | 1 | 1 |
| Krtap5-2 | 0 | 0 | 0 | 1 | 1 |

| | | | | | |
|---|---|---|---|---|---|
| Krtap5-3 | 0 | 0 | 0 | 1 | 1 |
| Krtap5-4 | 0 | 0 | 0 | 1 | 1 |
| Krtap5-5 | 0 | 0 | 0 | 1 | 1 |
| Krtap6-1 | 0 | 0 | 0 | 1 | 1 |
| Krtap6-2 | 0 | 0 | 0 | 1 | 1 |
| Krtap6-5 | 0 | 0 | 0 | 1 | 1 |
| Krtap7-1 | 0 | 0 | 0 | 1 | 1 |
| Krtap8-1 | 0 | 0 | 0 | 1 | 1 |
| Krtap9-1 | 0 | 0 | 0 | 1 | 1 |
| Krtap9-5 | 0 | 0 | 0 | 1 | 1 |
| Krtdap | 0 | 0 | 0 | 1 | 1 |
| LOC100038947 | 0 | 0 | 0 | 1 | 1 |
| LOC100040786 | 0 | 0 | 0 | 1 | 1 |
| LOC100043315 | 0 | 0 | 0 | 1 | 1 |
| LOC100048884 | 0 | 0 | 0 | 1 | 1 |
| LOC100502896 | 0 | 0 | 0 | 1 | 1 |
| LOC100503280 | 0 | 0 | 0 | 1 | 1 |
| LOC100861978 | 0 | 0 | 0 | 1 | 1 |
| LOC100862015 | 0 | 0 | 0 | 1 | 1 |
| LOC101055863 | 0 | 0 | 0 | 1 | 1 |
| LOC101056136 | 0 | 0 | 0 | 1 | 1 |
| LOC102633035 | 0 | 0 | 0 | 1 | 1 |
| LOC102634753 | 0 | 0 | 0 | 1 | 1 |
| LOC102636514 | 0 | 0 | 0 | 1 | 1 |
| LOC666331 | 0 | 0 | 0 | 1 | 1 |
| Lao1 | 0 | 0 | 0 | 1 | 1 |
| Lce1a1 | 0 | 0 | 0 | 1 | 1 |
| Lce1a2 | 0 | 0 | 0 | 1 | 1 |
| Lce1c | 0 | 0 | 0 | 1 | 1 |
| Lce1d | 0 | 0 | 0 | 1 | 1 |
| Lce1e | 0 | 0 | 0 | 1 | 1 |
| Lce1f | 0 | 0 | 0 | 1 | 1 |
| Lce1g | 0 | 0 | 0 | 1 | 1 |
| Lce1h | 0 | 0 | 0 | 1 | 1 |
| Lce1j | 0 | 0 | 0 | 1 | 1 |
| Lce1k | 0 | 0 | 0 | 1 | 1 |
| Lce1l | 0 | 0 | 0 | 1 | 1 |
| Lce1m | 0 | 0 | 0 | 1 | 1 |
| Lce3a | 0 | 0 | 0 | 1 | 1 |
| Lce3b | 0 | 0 | 0 | 1 | 1 |
| Lce3c | 0 | 0 | 0 | 1 | 1 |
| Lce3d | 0 | 0 | 0 | 1 | 1 |
| Lce3e | 0 | 0 | 0 | 1 | 1 |
| Lce3f | 0 | 0 | 0 | 1 | 1 |
| Lce6a | 0 | 0 | 0 | 1 | 1 |
| Lcn10 | 0 | 0 | 0 | 1 | 1 |
| Lcn11 | 0 | 0 | 0 | 1 | 1 |
| Lcn12 | 0 | 0 | 0 | 1 | 1 |

| | | | | | |
|---|---|---|---|---|---|
| Lcn3 | 0 | 0 | 0 | 1 | 1 |
| Lcn5 | 0 | 0 | 0 | 1 | 1 |
| Lcn6 | 0 | 0 | 0 | 1 | 1 |
| Lcn9 | 0 | 0 | 0 | 1 | 1 |
| Lelp1 | 0 | 0 | 0 | 1 | 1 |
| Lep | 0 | 0 | 0 | 1 | 1 |
| Lgsn | 0 | 0 | 0 | 1 | 1 |
| Lilra5 | 0 | 0 | 0 | 1 | 1 |
| Lim2 | 0 | 0 | 0 | 1 | 1 |
| Lincrna-cox2 | 0 | 0 | 0 | 1 | 1 |
| Lingo2 | 0 | 0 | 0 | 1 | 1 |
| Lipf | 0 | 0 | 0 | 1 | 1 |
| Lipi | 0 | 0 | 0 | 1 | 1 |
| Lipn | 0 | 0 | 0 | 1 | 1 |
| Lmod2 | 0 | 0 | 0 | 1 | 1 |
| Lmod3 | 0 | 0 | 0 | 1 | 1 |
| Lpcat2b | 0 | 0 | 0 | 1 | 1 |
| Lrcol1 | 0 | 0 | 0 | 1 | 1 |
| Lrg1 | 0 | 0 | 0 | 1 | 1 |
| Lrit1 | 0 | 0 | 0 | 1 | 1 |
| Lrit2 | 0 | 0 | 0 | 1 | 1 |
| Lrrc26 | 0 | 0 | 0 | 1 | 1 |
| Lrrc30 | 0 | 0 | 0 | 1 | 1 |
| Lrrc52 | 0 | 0 | 0 | 1 | 1 |
| Lrrc66 | 0 | 0 | 0 | 1 | 1 |
| Lrrc69 | 0 | 0 | 0 | 1 | 1 |
| Lrrc72 | 0 | 0 | 0 | 1 | 1 |
| Lsmem1 | 0 | 0 | 0 | 1 | 1 |
| Luzp4 | 0 | 0 | 0 | 1 | 1 |
| Ly6d | 0 | 0 | 0 | 1 | 1 |
| Ly6g5c | 0 | 0 | 0 | 1 | 1 |
| Ly6i | 0 | 0 | 0 | 1 | 1 |
| Lyg1 | 0 | 0 | 0 | 1 | 1 |
| Lypd5 | 0 | 0 | 0 | 1 | 1 |
| Lypd8 | 0 | 0 | 0 | 1 | 1 |
| Lyzl1 | 0 | 0 | 0 | 1 | 1 |
| Lyzl4os | 0 | 0 | 0 | 1 | 1 |
| Lyzl6 | 0 | 0 | 0 | 1 | 1 |
| Magea1 | 0 | 0 | 0 | 1 | 1 |
| Magea10 | 0 | 0 | 0 | 1 | 1 |
| Magea2 | 0 | 0 | 0 | 1 | 1 |
| Magea3 | 0 | 0 | 0 | 1 | 1 |
| Magea4 | 0 | 0 | 0 | 1 | 1 |
| Magea6 | 0 | 0 | 0 | 1 | 1 |
| Magea8 | 0 | 0 | 0 | 1 | 1 |
| Mageb1 | 0 | 0 | 0 | 1 | 1 |
| Mageb16 | 0 | 0 | 0 | 1 | 1 |
| Mageb16-ps1 | 0 | 0 | 0 | 1 | 1 |

| | | | | | |
|---|---|---|---|---|---|
| Mageb18 | 0 | 0 | 0 | 1 | 1 |
| Mageb2 | 0 | 0 | 0 | 1 | 1 |
| Mageb4 | 0 | 0 | 0 | 1 | 1 |
| Mageb5 | 0 | 0 | 0 | 1 | 1 |
| Manr | 0 | 0 | 0 | 1 | 1 |
| 03/04/16 | 0 | 0 | 0 | 1 | 1 |
| Mbd3l1 | 0 | 0 | 0 | 1 | 1 |
| Mcidas | 0 | 0 | 0 | 1 | 1 |
| Mcpt-ps1 | 0 | 0 | 0 | 1 | 1 |
| Mcpt1 | 0 | 0 | 0 | 1 | 1 |
| Mcpt2 | 0 | 0 | 0 | 1 | 1 |
| Mcpt4 | 0 | 0 | 0 | 1 | 1 |
| Mcpt8 | 0 | 0 | 0 | 1 | 1 |
| Mcpt9 | 0 | 0 | 0 | 1 | 1 |
| Mefv | 0 | 0 | 0 | 1 | 1 |
| Meig1 | 0 | 0 | 0 | 1 | 1 |
| Mep1b | 0 | 0 | 0 | 1 | 1 |
| Mettl21e | 0 | 0 | 0 | 1 | 1 |
| Mettl7a3 | 0 | 0 | 0 | 1 | 1 |
| Mgl2 | 0 | 0 | 0 | 1 | 1 |
| Mir100 | 0 | 0 | 0 | 1 | 1 |
| Mir101a | 0 | 0 | 0 | 1 | 1 |
| Mir101b | 0 | 0 | 0 | 1 | 1 |
| Mir101c | 0 | 0 | 0 | 1 | 1 |
| Mir103-1 | 0 | 0 | 0 | 1 | 1 |
| Mir103-2 | 0 | 0 | 0 | 1 | 1 |
| Mir105 | 0 | 0 | 0 | 1 | 1 |
| Mir106a | 0 | 0 | 0 | 1 | 1 |
| Mir107 | 0 | 0 | 0 | 1 | 1 |
| Mir10a | 0 | 0 | 0 | 1 | 1 |
| Mir10b | 0 | 0 | 0 | 1 | 1 |
| Mir1187 | 0 | 0 | 0 | 1 | 1 |
| Mir1190 | 0 | 0 | 0 | 1 | 1 |
| Mir1191 | 0 | 0 | 0 | 1 | 1 |
| Mir1191b | 0 | 0 | 0 | 1 | 1 |
| Mir1192 | 0 | 0 | 0 | 1 | 1 |
| Mir1193 | 0 | 0 | 0 | 1 | 1 |
| Mir1195 | 0 | 0 | 0 | 1 | 1 |
| Mir1197 | 0 | 0 | 0 | 1 | 1 |
| Mir1198 | 0 | 0 | 0 | 1 | 1 |
| Mir1199 | 0 | 0 | 0 | 1 | 1 |
| Mir1224 | 0 | 0 | 0 | 1 | 1 |
| Mir122a | 0 | 0 | 0 | 1 | 1 |
| Mir1231 | 0 | 0 | 0 | 1 | 1 |
| Mir1247 | 0 | 0 | 0 | 1 | 1 |
| Mir1249 | 0 | 0 | 0 | 1 | 1 |
| Mir124a-1 | 0 | 0 | 0 | 1 | 1 |
| Mir124a-2 | 0 | 0 | 0 | 1 | 1 |

| | | | | | |
|---|---|---|---|---|---|
| Mir124a-3 | 0 | 0 | 0 | 1 | 1 |
| Mir1251 | 0 | 0 | 0 | 1 | 1 |
| Mir1258 | 0 | 0 | 0 | 1 | 1 |
| Mir125a | 0 | 0 | 0 | 1 | 1 |
| Mir125b-1 | 0 | 0 | 0 | 1 | 1 |
| Mir125b-2 | 0 | 0 | 0 | 1 | 1 |
| Mir126 | 0 | 0 | 0 | 1 | 1 |
| Mir1264 | 0 | 0 | 0 | 1 | 1 |
| Mir126b | 0 | 0 | 0 | 1 | 1 |
| Mir127 | 0 | 0 | 0 | 1 | 1 |
| Mir128-1 | 0 | 0 | 0 | 1 | 1 |
| Mir128-2 | 0 | 0 | 0 | 1 | 1 |
| Mir129-1 | 0 | 0 | 0 | 1 | 1 |
| Mir129-2 | 0 | 0 | 0 | 1 | 1 |
| Mir1298 | 0 | 0 | 0 | 1 | 1 |
| Mir129b | 0 | 0 | 0 | 1 | 1 |
| Mir130a | 0 | 0 | 0 | 1 | 1 |
| Mir130b | 0 | 0 | 0 | 1 | 1 |
| Mir132 | 0 | 0 | 0 | 1 | 1 |
| Mir133a-1 | 0 | 0 | 0 | 1 | 1 |
| Mir133a-2 | 0 | 0 | 0 | 1 | 1 |
| Mir133b | 0 | 0 | 0 | 1 | 1 |
| Mir133c | 0 | 0 | 0 | 1 | 1 |
| Mir134 | 0 | 0 | 0 | 1 | 1 |
| Mir135a-1 | 0 | 0 | 0 | 1 | 1 |
| Mir135a-2 | 0 | 0 | 0 | 1 | 1 |
| Mir135b | 0 | 0 | 0 | 1 | 1 |
| Mir136 | 0 | 0 | 0 | 1 | 1 |
| Mir137 | 0 | 0 | 0 | 1 | 1 |
| Mir138-1 | 0 | 0 | 0 | 1 | 1 |
| Mir138-2 | 0 | 0 | 0 | 1 | 1 |
| Mir139 | 0 | 0 | 0 | 1 | 1 |
| Mir140 | 0 | 0 | 0 | 1 | 1 |
| Mir141 | 0 | 0 | 0 | 1 | 1 |
| Mir142 | 0 | 0 | 0 | 1 | 1 |
| Mir143 | 0 | 0 | 0 | 1 | 1 |
| Mir144 | 0 | 0 | 0 | 1 | 1 |
| Mir145 | 0 | 0 | 0 | 1 | 1 |
| Mir145b | 0 | 0 | 0 | 1 | 1 |
| Mir146 | 0 | 0 | 0 | 1 | 1 |
| Mir146b | 0 | 0 | 0 | 1 | 1 |
| Mir148a | 0 | 0 | 0 | 1 | 1 |
| Mir148b | 0 | 0 | 0 | 1 | 1 |
| Mir149 | 0 | 0 | 0 | 1 | 1 |
| Mir152 | 0 | 0 | 0 | 1 | 1 |
| Mir153 | 0 | 0 | 0 | 1 | 1 |
| Mir154 | 0 | 0 | 0 | 1 | 1 |
| Mir155 | 0 | 0 | 0 | 1 | 1 |

| | | | | | |
|---|---|---|---|---|---|
| Mir15a | 0 | 0 | 0 | 1 | 1 |
| Mir15b | 0 | 0 | 0 | 1 | 1 |
| Mir16-1 | 0 | 0 | 0 | 1 | 1 |
| Mir16-2 | 0 | 0 | 0 | 1 | 1 |
| Mir17 | 0 | 0 | 0 | 1 | 1 |
| Mir18 | 0 | 0 | 0 | 1 | 1 |
| Mir181a-1 | 0 | 0 | 0 | 1 | 1 |
| Mir181a-2 | 0 | 0 | 0 | 1 | 1 |
| Mir181b-1 | 0 | 0 | 0 | 1 | 1 |
| Mir181b-2 | 0 | 0 | 0 | 1 | 1 |
| Mir181c | 0 | 0 | 0 | 1 | 1 |
| Mir181d | 0 | 0 | 0 | 1 | 1 |
| Mir182 | 0 | 0 | 0 | 1 | 1 |
| Mir183 | 0 | 0 | 0 | 1 | 1 |
| Mir184 | 0 | 0 | 0 | 1 | 1 |
| Mir1843 | 0 | 0 | 0 | 1 | 1 |
| Mir1843b | 0 | 0 | 0 | 1 | 1 |
| Mir185 | 0 | 0 | 0 | 1 | 1 |
| Mir186 | 0 | 0 | 0 | 1 | 1 |
| Mir187 | 0 | 0 | 0 | 1 | 1 |
| Mir188 | 0 | 0 | 0 | 1 | 1 |
| Mir1893 | 0 | 0 | 0 | 1 | 1 |
| Mir1896 | 0 | 0 | 0 | 1 | 1 |
| Mir1897 | 0 | 0 | 0 | 1 | 1 |
| Mir1899 | 0 | 0 | 0 | 1 | 1 |
| Mir18b | 0 | 0 | 0 | 1 | 1 |
| Mir190 | 0 | 0 | 0 | 1 | 1 |
| Mir1900 | 0 | 0 | 0 | 1 | 1 |
| Mir1902 | 0 | 0 | 0 | 1 | 1 |
| Mir1904 | 0 | 0 | 0 | 1 | 1 |
| Mir1906-1 | 0 | 0 | 0 | 1 | 1 |
| Mir1906-2 | 0 | 0 | 0 | 1 | 1 |
| Mir190b | 0 | 0 | 0 | 1 | 1 |
| Mir191 | 0 | 0 | 0 | 1 | 1 |
| Mir1912 | 0 | 0 | 0 | 1 | 1 |
| Mir192 | 0 | 0 | 0 | 1 | 1 |
| Mir1928 | 0 | 0 | 0 | 1 | 1 |
| Mir1929 | 0 | 0 | 0 | 1 | 1 |
| Mir193 | 0 | 0 | 0 | 1 | 1 |
| Mir1930 | 0 | 0 | 0 | 1 | 1 |
| Mir1931 | 0 | 0 | 0 | 1 | 1 |
| Mir1933 | 0 | 0 | 0 | 1 | 1 |
| Mir1936 | 0 | 0 | 0 | 1 | 1 |
| Mir193b | 0 | 0 | 0 | 1 | 1 |
| Mir194-2 | 0 | 0 | 0 | 1 | 1 |
| Mir1941 | 0 | 0 | 0 | 1 | 1 |
| Mir1942 | 0 | 0 | 0 | 1 | 1 |
| Mir1945 | 0 | 0 | 0 | 1 | 1 |

| | | | | | |
|---|---|---|---|---|---|
| Mir1946a | 0 | 0 | 0 | 1 | 1 |
| Mir1946b | 0 | 0 | 0 | 1 | 1 |
| Mir1947 | 0 | 0 | 0 | 1 | 1 |
| Mir1948 | 0 | 0 | 0 | 1 | 1 |
| Mir195 | 0 | 0 | 0 | 1 | 1 |
| Mir1950 | 0 | 0 | 0 | 1 | 1 |
| Mir1951 | 0 | 0 | 0 | 1 | 1 |
| Mir1952 | 0 | 0 | 0 | 1 | 1 |
| Mir1953 | 0 | 0 | 0 | 1 | 1 |
| Mir1954 | 0 | 0 | 0 | 1 | 1 |
| Mir1955 | 0 | 0 | 0 | 1 | 1 |
| Mir1956 | 0 | 0 | 0 | 1 | 1 |
| Mir1957 | 0 | 0 | 0 | 1 | 1 |
| Mir1958 | 0 | 0 | 0 | 1 | 1 |
| Mir195b | 0 | 0 | 0 | 1 | 1 |
| Mir1960 | 0 | 0 | 0 | 1 | 1 |
| Mir1961 | 0 | 0 | 0 | 1 | 1 |
| Mir1962 | 0 | 0 | 0 | 1 | 1 |
| Mir1963 | 0 | 0 | 0 | 1 | 1 |
| Mir1964 | 0 | 0 | 0 | 1 | 1 |
| Mir1967 | 0 | 0 | 0 | 1 | 1 |
| Mir1968 | 0 | 0 | 0 | 1 | 1 |
| Mir1969 | 0 | 0 | 0 | 1 | 1 |
| Mir196a-2 | 0 | 0 | 0 | 1 | 1 |
| Mir1970 | 0 | 0 | 0 | 1 | 1 |
| Mir1971 | 0 | 0 | 0 | 1 | 1 |
| Mir1981 | 0 | 0 | 0 | 1 | 1 |
| Mir1983 | 0 | 0 | 0 | 1 | 1 |
| Mir199a-2 | 0 | 0 | 0 | 1 | 1 |
| Mir19a | 0 | 0 | 0 | 1 | 1 |
| Mir19b-2 | 0 | 0 | 0 | 1 | 1 |
| Mir1a-1 | 0 | 0 | 0 | 1 | 1 |
| Mir1a-2 | 0 | 0 | 0 | 1 | 1 |
| Mir200a | 0 | 0 | 0 | 1 | 1 |
| Mir200b | 0 | 0 | 0 | 1 | 1 |
| Mir200c | 0 | 0 | 0 | 1 | 1 |
| Mir201 | 0 | 0 | 0 | 1 | 1 |
| Mir202 | 0 | 0 | 0 | 1 | 1 |
| Mir203 | 0 | 0 | 0 | 1 | 1 |
| Mir204 | 0 | 0 | 0 | 1 | 1 |
| Mir205 | 0 | 0 | 0 | 1 | 1 |
| Mir206 | 0 | 0 | 0 | 1 | 1 |
| Mir207 | 0 | 0 | 0 | 1 | 1 |
| Mir208a | 0 | 0 | 0 | 1 | 1 |
| Mir208b | 0 | 0 | 0 | 1 | 1 |
| Mir20a | 0 | 0 | 0 | 1 | 1 |
| Mir20b | 0 | 0 | 0 | 1 | 1 |
| Mir21 | 0 | 0 | 0 | 1 | 1 |

| | | | | | |
|---|---|---|---|---|---|
| Mir210 | 0 | 0 | 0 | 1 | 1 |
| Mir211 | 0 | 0 | 0 | 1 | 1 |
| Mir212 | 0 | 0 | 0 | 1 | 1 |
| Mir2136 | 0 | 0 | 0 | 1 | 1 |
| Mir2137 | 0 | 0 | 0 | 1 | 1 |
| Mir214 | 0 | 0 | 0 | 1 | 1 |
| Mir216a | 0 | 0 | 0 | 1 | 1 |
| Mir216b | 0 | 0 | 0 | 1 | 1 |
| Mir216c | 0 | 0 | 0 | 1 | 1 |
| Mir217 | 0 | 0 | 0 | 1 | 1 |
| Mir218-1 | 0 | 0 | 0 | 1 | 1 |
| Mir218-2 | 0 | 0 | 0 | 1 | 1 |
| Mir219-2 | 0 | 0 | 0 | 1 | 1 |
| Mir219b | 0 | 0 | 0 | 1 | 1 |
| Mir219c | 0 | 0 | 0 | 1 | 1 |
| Mir21b | 0 | 0 | 0 | 1 | 1 |
| Mir21c | 0 | 0 | 0 | 1 | 1 |
| Mir22 | 0 | 0 | 0 | 1 | 1 |
| Mir221 | 0 | 0 | 0 | 1 | 1 |
| Mir222 | 0 | 0 | 0 | 1 | 1 |
| Mir223 | 0 | 0 | 0 | 1 | 1 |
| Mir23a | 0 | 0 | 0 | 1 | 1 |
| Mir23b | 0 | 0 | 0 | 1 | 1 |
| Mir24-1 | 0 | 0 | 0 | 1 | 1 |
| Mir24-2 | 0 | 0 | 0 | 1 | 1 |
| Mir26a-1 | 0 | 0 | 0 | 1 | 1 |
| Mir26a-2 | 0 | 0 | 0 | 1 | 1 |
| Mir27a | 0 | 0 | 0 | 1 | 1 |
| Mir27b | 0 | 0 | 0 | 1 | 1 |
| Mir28 | 0 | 0 | 0 | 1 | 1 |
| Mir28b | 0 | 0 | 0 | 1 | 1 |
| Mir28c | 0 | 0 | 0 | 1 | 1 |
| Mir290 | 0 | 0 | 0 | 1 | 1 |
| Mir290b | 0 | 0 | 0 | 1 | 1 |
| Mir291a | 0 | 0 | 0 | 1 | 1 |
| Mir291b | 0 | 0 | 0 | 1 | 1 |
| Mir292 | 0 | 0 | 0 | 1 | 1 |
| Mir292b | 0 | 0 | 0 | 1 | 1 |
| Mir293 | 0 | 0 | 0 | 1 | 1 |
| Mir294 | 0 | 0 | 0 | 1 | 1 |
| Mir295 | 0 | 0 | 0 | 1 | 1 |
| Mir296 | 0 | 0 | 0 | 1 | 1 |
| Mir297-1 | 0 | 0 | 0 | 1 | 1 |
| Mir297-2 | 0 | 0 | 0 | 1 | 1 |
| Mir297a-3,Mir297 | 0 | 0 | 0 | 1 | 1 |
| Mir297a-4 | 0 | 0 | 0 | 1 | 1 |
| Mir297b | 0 | 0 | 0 | 1 | 1 |
| Mir297c | 0 | 0 | 0 | 1 | 1 |

| | | | | | |
|---|---|---|---|---|---|
| Mir298 | 0 | 0 | 0 | 1 | 1 |
| Mir299 | 0 | 0 | 0 | 1 | 1 |
| Mir299b | 0 | 0 | 0 | 1 | 1 |
| Mir29a | 0 | 0 | 0 | 1 | 1 |
| Mir29b-1 | 0 | 0 | 0 | 1 | 1 |
| Mir29b-2 | 0 | 0 | 0 | 1 | 1 |
| Mir29c | 0 | 0 | 0 | 1 | 1 |
| Mir300 | 0 | 0 | 0 | 1 | 1 |
| Mir301 | 0 | 0 | 0 | 1 | 1 |
| Mir301b | 0 | 0 | 0 | 1 | 1 |
| Mir302a | 0 | 0 | 0 | 1 | 1 |
| Mir302b | 0 | 0 | 0 | 1 | 1 |
| Mir302c | 0 | 0 | 0 | 1 | 1 |
| Mir302d | 0 | 0 | 0 | 1 | 1 |
| Mir3058 | 0 | 0 | 0 | 1 | 1 |
| Mir3059 | 0 | 0 | 0 | 1 | 1 |
| Mir3060 | 0 | 0 | 0 | 1 | 1 |
| Mir3061 | 0 | 0 | 0 | 1 | 1 |
| Mir3063 | 0 | 0 | 0 | 1 | 1 |
| Mir3065 | 0 | 0 | 0 | 1 | 1 |
| Mir3066 | 0 | 0 | 0 | 1 | 1 |
| Mir3067 | 0 | 0 | 0 | 1 | 1 |
| Mir3069 | 0 | 0 | 0 | 1 | 1 |
| Mir3070a | 0 | 0 | 0 | 1 | 1 |
| Mir3071 | 0 | 0 | 0 | 1 | 1 |
| Mir3073 | 0 | 0 | 0 | 1 | 1 |
| Mir3073b | 0 | 0 | 0 | 1 | 1 |
| Mir3074-1 | 0 | 0 | 0 | 1 | 1 |
| Mir3074-2 | 0 | 0 | 0 | 1 | 1 |
| Mir3075 | 0 | 0 | 0 | 1 | 1 |
| Mir3076 | 0 | 0 | 0 | 1 | 1 |
| Mir3077 | 0 | 0 | 0 | 1 | 1 |
| Mir3078 | 0 | 0 | 0 | 1 | 1 |
| Mir3079 | 0 | 0 | 0 | 1 | 1 |
| Mir3081 | 0 | 0 | 0 | 1 | 1 |
| Mir3082 | 0 | 0 | 0 | 1 | 1 |
| Mir3083 | 0 | 0 | 0 | 1 | 1 |
| Mir3084 | 0 | 0 | 0 | 1 | 1 |
| Mir3084-2 | 0 | 0 | 0 | 1 | 1 |
| Mir3085 | 0 | 0 | 0 | 1 | 1 |
| Mir3086 | 0 | 0 | 0 | 1 | 1 |
| Mir3087 | 0 | 0 | 0 | 1 | 1 |
| Mir3088 | 0 | 0 | 0 | 1 | 1 |
| Mir3089 | 0 | 0 | 0 | 1 | 1 |
| Mir3091 | 0 | 0 | 0 | 1 | 1 |
| Mir3092 | 0 | 0 | 0 | 1 | 1 |
| Mir3093 | 0 | 0 | 0 | 1 | 1 |
| Mir3094 | 0 | 0 | 0 | 1 | 1 |

| | | | | | |
|---|---|---|---|---|---|
| Mir3095 | 0 | 0 | 0 | 1 | 1 |
| Mir3097 | 0 | 0 | 0 | 1 | 1 |
| Mir3098 | 0 | 0 | 0 | 1 | 1 |
| Mir3099 | 0 | 0 | 0 | 1 | 1 |
| Mir30a | 0 | 0 | 0 | 1 | 1 |
| Mir30b | 0 | 0 | 0 | 1 | 1 |
| Mir30c-1 | 0 | 0 | 0 | 1 | 1 |
| Mir30c-2 | 0 | 0 | 0 | 1 | 1 |
| Mir30d | 0 | 0 | 0 | 1 | 1 |
| Mir30f | 0 | 0 | 0 | 1 | 1 |
| Mir31 | 0 | 0 | 0 | 1 | 1 |
| Mir3100 | 0 | 0 | 0 | 1 | 1 |
| Mir3102 | 0 | 0 | 0 | 1 | 1 |
| Mir3103 | 0 | 0 | 0 | 1 | 1 |
| Mir3104 | 0 | 0 | 0 | 1 | 1 |
| Mir3106 | 0 | 0 | 0 | 1 | 1 |
| Mir3107 | 0 | 0 | 0 | 1 | 1 |
| Mir3108 | 0 | 0 | 0 | 1 | 1 |
| Mir3109 | 0 | 0 | 0 | 1 | 1 |
| Mir3110 | 0 | 0 | 0 | 1 | 1 |
| Mir3112 | 0 | 0 | 0 | 1 | 1 |
| Mir32 | 0 | 0 | 0 | 1 | 1 |
| Mir320 | 0 | 0 | 0 | 1 | 1 |
| Mir322 | 0 | 0 | 0 | 1 | 1 |
| Mir323 | 0 | 0 | 0 | 1 | 1 |
| Mir324 | 0 | 0 | 0 | 1 | 1 |
| Mir325 | 0 | 0 | 0 | 1 | 1 |
| Mir326 | 0 | 0 | 0 | 1 | 1 |
| Mir328 | 0 | 0 | 0 | 1 | 1 |
| Mir329 | 0 | 0 | 0 | 1 | 1 |
| Mir33 | 0 | 0 | 0 | 1 | 1 |
| Mir331 | 0 | 0 | 0 | 1 | 1 |
| Mir335 | 0 | 0 | 0 | 1 | 1 |
| Mir337 | 0 | 0 | 0 | 1 | 1 |
| Mir338 | 0 | 0 | 0 | 1 | 1 |
| Mir340 | 0 | 0 | 0 | 1 | 1 |
| Mir341 | 0 | 0 | 0 | 1 | 1 |
| Mir344 | 0 | 0 | 0 | 1 | 1 |
| Mir344-2 | 0 | 0 | 0 | 1 | 1 |
| Mir344b | 0 | 0 | 0 | 1 | 1 |
| Mir344c | 0 | 0 | 0 | 1 | 1 |
| Mir344d-1 | 0 | 0 | 0 | 1 | 1 |
| Mir344d-2 | 0 | 0 | 0 | 1 | 1 |
| Mir344d-3 | 0 | 0 | 0 | 1 | 1 |
| Mir344e | 0 | 0 | 0 | 1 | 1 |
| Mir344f | 0 | 0 | 0 | 1 | 1 |
| Mir344g | 0 | 0 | 0 | 1 | 1 |
| Mir344h-2 | 0 | 0 | 0 | 1 | 1 |

| | | | | | |
|---|---|---|---|---|---|
| Mir344i | 0 | 0 | 0 | 1 | 1 |
| Mir346 | 0 | 0 | 0 | 1 | 1 |
| Mir3470a | 0 | 0 | 0 | 1 | 1 |
| Mir3470b | 0 | 0 | 0 | 1 | 1 |
| Mir3471-1 | 0 | 0 | 0 | 1 | 1 |
| Mir3473 | 0 | 0 | 0 | 1 | 1 |
| Mir3473c | 0 | 0 | 0 | 1 | 1 |
| Mir3473d | 0 | 0 | 0 | 1 | 1 |
| Mir3473e | 0 | 0 | 0 | 1 | 1 |
| Mir3473f | 0 | 0 | 0 | 1 | 1 |
| Mir3473g | 0 | 0 | 0 | 1 | 1 |
| Mir3474 | 0 | 0 | 0 | 1 | 1 |
| Mir3475 | 0 | 0 | 0 | 1 | 1 |
| Mir34a | 0 | 0 | 0 | 1 | 1 |
| Mir34b | 0 | 0 | 0 | 1 | 1 |
| Mir34c | 0 | 0 | 0 | 1 | 1 |
| Mir350 | 0 | 0 | 0 | 1 | 1 |
| Mir351 | 0 | 0 | 0 | 1 | 1 |
| Mir3544 | 0 | 0 | 0 | 1 | 1 |
| Mir3547 | 0 | 0 | 0 | 1 | 1 |
| Mir3569 | 0 | 0 | 0 | 1 | 1 |
| Mir3572 | 0 | 0 | 0 | 1 | 1 |
| Mir362 | 0 | 0 | 0 | 1 | 1 |
| Mir3620 | 0 | 0 | 0 | 1 | 1 |
| Mir363 | 0 | 0 | 0 | 1 | 1 |
| Mir365-1 | 0 | 0 | 0 | 1 | 1 |
| Mir365-2 | 0 | 0 | 0 | 1 | 1 |
| Mir367 | 0 | 0 | 0 | 1 | 1 |
| Mir369 | 0 | 0 | 0 | 1 | 1 |
| Mir370 | 0 | 0 | 0 | 1 | 1 |
| Mir374 | 0 | 0 | 0 | 1 | 1 |
| Mir374c | 0 | 0 | 0 | 1 | 1 |
| Mir375 | 0 | 0 | 0 | 1 | 1 |
| Mir376a | 0 | 0 | 0 | 1 | 1 |
| Mir376b | 0 | 0 | 0 | 1 | 1 |
| Mir376c | 0 | 0 | 0 | 1 | 1 |
| Mir377 | 0 | 0 | 0 | 1 | 1 |
| Mir378 | 0 | 0 | 0 | 1 | 1 |
| Mir378c | 0 | 0 | 0 | 1 | 1 |
| Mir379 | 0 | 0 | 0 | 1 | 1 |
| Mir380 | 0 | 0 | 0 | 1 | 1 |
| Mir381 | 0 | 0 | 0 | 1 | 1 |
| Mir382 | 0 | 0 | 0 | 1 | 1 |
| Mir383 | 0 | 0 | 0 | 1 | 1 |
| Mir384 | 0 | 0 | 0 | 1 | 1 |
| Mir3962 | 0 | 0 | 0 | 1 | 1 |
| Mir3963 | 0 | 0 | 0 | 1 | 1 |
| Mir3964 | 0 | 0 | 0 | 1 | 1 |

| | | | | | |
|---|---|---|---|---|---|
| Mir3965 | 0 | 0 | 0 | 1 | 1 |
| Mir3966 | 0 | 0 | 0 | 1 | 1 |
| Mir3967 | 0 | 0 | 0 | 1 | 1 |
| Mir3968 | 0 | 0 | 0 | 1 | 1 |
| Mir3969 | 0 | 0 | 0 | 1 | 1 |
| Mir3970 | 0 | 0 | 0 | 1 | 1 |
| Mir3971 | 0 | 0 | 0 | 1 | 1 |
| Mir409 | 0 | 0 | 0 | 1 | 1 |
| Mir410 | 0 | 0 | 0 | 1 | 1 |
| Mir411 | 0 | 0 | 0 | 1 | 1 |
| Mir412 | 0 | 0 | 0 | 1 | 1 |
| Mir421 | 0 | 0 | 0 | 1 | 1 |
| Mir423 | 0 | 0 | 0 | 1 | 1 |
| Mir425 | 0 | 0 | 0 | 1 | 1 |
| Mir429 | 0 | 0 | 0 | 1 | 1 |
| Mir432 | 0 | 0 | 0 | 1 | 1 |
| Mir433 | 0 | 0 | 0 | 1 | 1 |
| Mir434 | 0 | 0 | 0 | 1 | 1 |
| Mir448 | 0 | 0 | 0 | 1 | 1 |
| Mir449a | 0 | 0 | 0 | 1 | 1 |
| Mir449b | 0 | 0 | 0 | 1 | 1 |
| Mir449c | 0 | 0 | 0 | 1 | 1 |
| Mir450-1 | 0 | 0 | 0 | 1 | 1 |
| Mir450-2 | 0 | 0 | 0 | 1 | 1 |
| Mir450b | 0 | 0 | 0 | 1 | 1 |
| Mir451 | 0 | 0 | 0 | 1 | 1 |
| Mir452 | 0 | 0 | 0 | 1 | 1 |
| Mir453 | 0 | 0 | 0 | 1 | 1 |
| Mir455 | 0 | 0 | 0 | 1 | 1 |
| Mir463 | 0 | 0 | 0 | 1 | 1 |
| Mir465 | 0 | 0 | 0 | 1 | 1 |
| Mir465b-1 | 0 | 0 | 0 | 1 | 1 |
| Mir465c-1 | 0 | 0 | 0 | 1 | 1 |
| Mir465d | 0 | 0 | 0 | 1 | 1 |
| Mir466 | 0 | 0 | 0 | 1 | 1 |
| Mir4660 | 0 | 0 | 0 | 1 | 1 |
| Mir466b-2,Mir466 | 0 | 0 | 0 | 1 | 1 |
| Mir466b-3 | 0 | 0 | 0 | 1 | 1 |
| Mir466d | 0 | 0 | 0 | 1 | 1 |
| Mir466f-1 | 0 | 0 | 0 | 1 | 1 |
| Mir466f-2 | 0 | 0 | 0 | 1 | 1 |
| Mir466f-3 | 0 | 0 | 0 | 1 | 1 |
| Mir466g | 0 | 0 | 0 | 1 | 1 |
| Mir466h | 0 | 0 | 0 | 1 | 1 |
| Mir466n | 0 | 0 | 0 | 1 | 1 |
| Mir466p | 0 | 0 | 0 | 1 | 1 |
| Mir467a-1 | 0 | 0 | 0 | 1 | 1 |
| Mir467a-10 | 0 | 0 | 0 | 1 | 1 |

| | | | | | |
|---|---|---|---|---|---|
| Mir467a-2 | 0 | 0 | 0 | 1 | 1 |
| Mir467a-3 | 0 | 0 | 0 | 1 | 1 |
| Mir467a-5 | 0 | 0 | 0 | 1 | 1 |
| Mir467a-7 | 0 | 0 | 0 | 1 | 1 |
| Mir467a-8 | 0 | 0 | 0 | 1 | 1 |
| Mir467b | 0 | 0 | 0 | 1 | 1 |
| Mir467c | 0 | 0 | 0 | 1 | 1 |
| Mir467d | 0 | 0 | 0 | 1 | 1 |
| Mir467e | 0 | 0 | 0 | 1 | 1 |
| Mir467f | 0 | 0 | 0 | 1 | 1 |
| Mir468 | 0 | 0 | 0 | 1 | 1 |
| Mir470 | 0 | 0 | 0 | 1 | 1 |
| Mir471 | 0 | 0 | 0 | 1 | 1 |
| Mir483 | 0 | 0 | 0 | 1 | 1 |
| Mir484 | 0 | 0 | 0 | 1 | 1 |
| Mir485 | 0 | 0 | 0 | 1 | 1 |
| Mir486 | 0 | 0 | 0 | 1 | 1 |
| Mir487b | 0 | 0 | 0 | 1 | 1 |
| Mir488 | 0 | 0 | 0 | 1 | 1 |
| Mir489 | 0 | 0 | 0 | 1 | 1 |
| Mir490 | 0 | 0 | 0 | 1 | 1 |
| Mir491 | 0 | 0 | 0 | 1 | 1 |
| Mir493 | 0 | 0 | 0 | 1 | 1 |
| Mir494 | 0 | 0 | 0 | 1 | 1 |
| Mir495 | 0 | 0 | 0 | 1 | 1 |
| Mir496 | 0 | 0 | 0 | 1 | 1 |
| Mir496b | 0 | 0 | 0 | 1 | 1 |
| Mir497 | 0 | 0 | 0 | 1 | 1 |
| Mir497b | 0 | 0 | 0 | 1 | 1 |
| Mir499 | 0 | 0 | 0 | 1 | 1 |
| Mir500 | 0 | 0 | 0 | 1 | 1 |
| Mir501 | 0 | 0 | 0 | 1 | 1 |
| Mir503 | 0 | 0 | 0 | 1 | 1 |
| Mir504 | 0 | 0 | 0 | 1 | 1 |
| Mir5046 | 0 | 0 | 0 | 1 | 1 |
| Mir505 | 0 | 0 | 0 | 1 | 1 |
| Mir509 | 0 | 0 | 0 | 1 | 1 |
| Mir5098 | 0 | 0 | 0 | 1 | 1 |
| Mir5100 | 0 | 0 | 0 | 1 | 1 |
| Mir5101 | 0 | 0 | 0 | 1 | 1 |
| Mir5103 | 0 | 0 | 0 | 1 | 1 |
| Mir5104 | 0 | 0 | 0 | 1 | 1 |
| Mir5106 | 0 | 0 | 0 | 1 | 1 |
| Mir5107 | 0 | 0 | 0 | 1 | 1 |
| Mir5108 | 0 | 0 | 0 | 1 | 1 |
| Mir511 | 0 | 0 | 0 | 1 | 1 |
| Mir5112 | 0 | 0 | 0 | 1 | 1 |
| Mir5113 | 0 | 0 | 0 | 1 | 1 |

| | | | | | |
|---|---|---|---|---|---|
| Mir5114 | 0 | 0 | 0 | 1 | 1 |
| Mir5116 | 0 | 0 | 0 | 1 | 1 |
| Mir5119 | 0 | 0 | 0 | 1 | 1 |
| Mir5120 | 0 | 0 | 0 | 1 | 1 |
| Mir5122 | 0 | 0 | 0 | 1 | 1 |
| Mir5123 | 0 | 0 | 0 | 1 | 1 |
| Mir5124 | 0 | 0 | 0 | 1 | 1 |
| Mir5125 | 0 | 0 | 0 | 1 | 1 |
| Mir5126 | 0 | 0 | 0 | 1 | 1 |
| Mir5127 | 0 | 0 | 0 | 1 | 1 |
| Mir5128 | 0 | 0 | 0 | 1 | 1 |
| Mir5129 | 0 | 0 | 0 | 1 | 1 |
| Mir5130 | 0 | 0 | 0 | 1 | 1 |
| Mir5132 | 0 | 0 | 0 | 1 | 1 |
| Mir5134 | 0 | 0 | 0 | 1 | 1 |
| Mir5135 | 0 | 0 | 0 | 1 | 1 |
| Mir5136 | 0 | 0 | 0 | 1 | 1 |
| Mir532 | 0 | 0 | 0 | 1 | 1 |
| Mir539 | 0 | 0 | 0 | 1 | 1 |
| Mir540 | 0 | 0 | 0 | 1 | 1 |
| Mir541 | 0 | 0 | 0 | 1 | 1 |
| Mir542 | 0 | 0 | 0 | 1 | 1 |
| Mir543 | 0 | 0 | 0 | 1 | 1 |
| Mir544 | 0 | 0 | 0 | 1 | 1 |
| Mir547 | 0 | 0 | 0 | 1 | 1 |
| Mir551b | 0 | 0 | 0 | 1 | 1 |
| Mir5615-1 | 0 | 0 | 0 | 1 | 1 |
| Mir5615-2 | 0 | 0 | 0 | 1 | 1 |
| Mir5616 | 0 | 0 | 0 | 1 | 1 |
| Mir5617 | 0 | 0 | 0 | 1 | 1 |
| Mir5618 | 0 | 0 | 0 | 1 | 1 |
| Mir5619 | 0 | 0 | 0 | 1 | 1 |
| Mir5620 | 0 | 0 | 0 | 1 | 1 |
| Mir5621 | 0 | 0 | 0 | 1 | 1 |
| Mir5622 | 0 | 0 | 0 | 1 | 1 |
| Mir5623 | 0 | 0 | 0 | 1 | 1 |
| Mir5624 | 0 | 0 | 0 | 1 | 1 |
| Mir5625 | 0 | 0 | 0 | 1 | 1 |
| Mir5626 | 0 | 0 | 0 | 1 | 1 |
| Mir5627 | 0 | 0 | 0 | 1 | 1 |
| Mir568 | 0 | 0 | 0 | 1 | 1 |
| Mir5709 | 0 | 0 | 0 | 1 | 1 |
| Mir5710 | 0 | 0 | 0 | 1 | 1 |
| Mir574 | 0 | 0 | 0 | 1 | 1 |
| Mir582 | 0 | 0 | 0 | 1 | 1 |
| Mir592 | 0 | 0 | 0 | 1 | 1 |
| Mir598 | 0 | 0 | 0 | 1 | 1 |
| Mir6237 | 0 | 0 | 0 | 1 | 1 |

| | | | | | |
|---|---|---|---|---|---|
| Mir6238 | 0 | 0 | 0 | 1 | 1 |
| Mir6239 | 0 | 0 | 0 | 1 | 1 |
| Mir6241 | 0 | 0 | 0 | 1 | 1 |
| Mir6335 | 0 | 0 | 0 | 1 | 1 |
| Mir6336 | 0 | 0 | 0 | 1 | 1 |
| Mir6337 | 0 | 0 | 0 | 1 | 1 |
| Mir6338 | 0 | 0 | 0 | 1 | 1 |
| Mir6339 | 0 | 0 | 0 | 1 | 1 |
| Mir6341 | 0 | 0 | 0 | 1 | 1 |
| Mir6342 | 0 | 0 | 0 | 1 | 1 |
| Mir6343 | 0 | 0 | 0 | 1 | 1 |
| Mir6344 | 0 | 0 | 0 | 1 | 1 |
| Mir6345 | 0 | 0 | 0 | 1 | 1 |
| Mir6348 | 0 | 0 | 0 | 1 | 1 |
| Mir6349 | 0 | 0 | 0 | 1 | 1 |
| Mir6350 | 0 | 0 | 0 | 1 | 1 |
| Mir6353 | 0 | 0 | 0 | 1 | 1 |
| Mir6354 | 0 | 0 | 0 | 1 | 1 |
| Mir6355 | 0 | 0 | 0 | 1 | 1 |
| Mir6356 | 0 | 0 | 0 | 1 | 1 |
| Mir6358 | 0 | 0 | 0 | 1 | 1 |
| Mir6359 | 0 | 0 | 0 | 1 | 1 |
| Mir6360 | 0 | 0 | 0 | 1 | 1 |
| Mir6362 | 0 | 0 | 0 | 1 | 1 |
| Mir6364 | 0 | 0 | 0 | 1 | 1 |
| Mir6365 | 0 | 0 | 0 | 1 | 1 |
| Mir6366 | 0 | 0 | 0 | 1 | 1 |
| Mir6367 | 0 | 0 | 0 | 1 | 1 |
| Mir6368 | 0 | 0 | 0 | 1 | 1 |
| Mir6369 | 0 | 0 | 0 | 1 | 1 |
| Mir6370 | 0 | 0 | 0 | 1 | 1 |
| Mir6372 | 0 | 0 | 0 | 1 | 1 |
| Mir6374 | 0 | 0 | 0 | 1 | 1 |
| Mir6375 | 0 | 0 | 0 | 1 | 1 |
| Mir6376 | 0 | 0 | 0 | 1 | 1 |
| Mir6378 | 0 | 0 | 0 | 1 | 1 |
| Mir6380 | 0 | 0 | 0 | 1 | 1 |
| Mir6381 | 0 | 0 | 0 | 1 | 1 |
| Mir6382 | 0 | 0 | 0 | 1 | 1 |
| Mir6383 | 0 | 0 | 0 | 1 | 1 |
| Mir6384 | 0 | 0 | 0 | 1 | 1 |
| Mir6388 | 0 | 0 | 0 | 1 | 1 |
| Mir6389 | 0 | 0 | 0 | 1 | 1 |
| Mir6390 | 0 | 0 | 0 | 1 | 1 |
| Mir6391 | 0 | 0 | 0 | 1 | 1 |
| Mir6392 | 0 | 0 | 0 | 1 | 1 |
| Mir6393 | 0 | 0 | 0 | 1 | 1 |
| Mir6394 | 0 | 0 | 0 | 1 | 1 |

| | | | | | |
|---|---|---|---|---|---|
| Mir6396 | 0 | 0 | 0 | 1 | 1 |
| Mir6397 | 0 | 0 | 0 | 1 | 1 |
| Mir6398 | 0 | 0 | 0 | 1 | 1 |
| Mir6399 | 0 | 0 | 0 | 1 | 1 |
| Mir6400 | 0 | 0 | 0 | 1 | 1 |
| Mir6401 | 0 | 0 | 0 | 1 | 1 |
| Mir6402 | 0 | 0 | 0 | 1 | 1 |
| Mir6404 | 0 | 0 | 0 | 1 | 1 |
| Mir6405 | 0 | 0 | 0 | 1 | 1 |
| Mir6407 | 0 | 0 | 0 | 1 | 1 |
| Mir6408 | 0 | 0 | 0 | 1 | 1 |
| Mir6409 | 0 | 0 | 0 | 1 | 1 |
| Mir6410 | 0 | 0 | 0 | 1 | 1 |
| Mir6411 | 0 | 0 | 0 | 1 | 1 |
| Mir6412 | 0 | 0 | 0 | 1 | 1 |
| Mir6413 | 0 | 0 | 0 | 1 | 1 |
| Mir6414 | 0 | 0 | 0 | 1 | 1 |
| Mir6415 | 0 | 0 | 0 | 1 | 1 |
| Mir6416 | 0 | 0 | 0 | 1 | 1 |
| Mir6417 | 0 | 0 | 0 | 1 | 1 |
| Mir6419 | 0 | 0 | 0 | 1 | 1 |
| Mir6420 | 0 | 0 | 0 | 1 | 1 |
| Mir653 | 0 | 0 | 0 | 1 | 1 |
| Mir6537 | 0 | 0 | 0 | 1 | 1 |
| Mir6538 | 0 | 0 | 0 | 1 | 1 |
| Mir6539 | 0 | 0 | 0 | 1 | 1 |
| Mir654 | 0 | 0 | 0 | 1 | 1 |
| Mir6540 | 0 | 0 | 0 | 1 | 1 |
| Mir6541 | 0 | 0 | 0 | 1 | 1 |
| Mir6546 | 0 | 0 | 0 | 1 | 1 |
| Mir664 | 0 | 0 | 0 | 1 | 1 |
| Mir665 | 0 | 0 | 0 | 1 | 1 |
| Mir666 | 0 | 0 | 0 | 1 | 1 |
| Mir667 | 0 | 0 | 0 | 1 | 1 |
| Mir668 | 0 | 0 | 0 | 1 | 1 |
| Mir669a-1 | 0 | 0 | 0 | 1 | 1 |
| Mir669a-2 | 0 | 0 | 0 | 1 | 1 |
| Mir669a-3 | 0 | 0 | 0 | 1 | 1 |
| Mir669a-4,Mir669 | 0 | 0 | 0 | 1 | 1 |
| Mir669b | 0 | 0 | 0 | 1 | 1 |
| Mir669c | 0 | 0 | 0 | 1 | 1 |
| Mir669e | 0 | 0 | 0 | 1 | 1 |
| Mir669g | 0 | 0 | 0 | 1 | 1 |
| Mir669h | 0 | 0 | 0 | 1 | 1 |
| Mir669i | 0 | 0 | 0 | 1 | 1 |
| Mir669j | 0 | 0 | 0 | 1 | 1 |
| Mir669k | 0 | 0 | 0 | 1 | 1 |
| Mir669m-1 | 0 | 0 | 0 | 1 | 1 |

| | | | | | |
|---|---|---|---|---|---|
| Mir669m-2 | 0 | 0 | 0 | 1 | 1 |
| Mir669p-1 | 0 | 0 | 0 | 1 | 1 |
| Mir670 | 0 | 0 | 0 | 1 | 1 |
| Mir671 | 0 | 0 | 0 | 1 | 1 |
| Mir6715 | 0 | 0 | 0 | 1 | 1 |
| Mir672 | 0 | 0 | 0 | 1 | 1 |
| Mir673 | 0 | 0 | 0 | 1 | 1 |
| Mir674 | 0 | 0 | 0 | 1 | 1 |
| Mir675 | 0 | 0 | 0 | 1 | 1 |
| Mir676 | 0 | 0 | 0 | 1 | 1 |
| Mir6769b | 0 | 0 | 0 | 1 | 1 |
| Mir679 | 0 | 0 | 0 | 1 | 1 |
| Mir680-2 | 0 | 0 | 0 | 1 | 1 |
| Mir680-3 | 0 | 0 | 0 | 1 | 1 |
| Mir683-1 | 0 | 0 | 0 | 1 | 1 |
| Mir684-1 | 0 | 0 | 0 | 1 | 1 |
| Mir687 | 0 | 0 | 0 | 1 | 1 |
| Mir688 | 0 | 0 | 0 | 1 | 1 |
| Mir6896 | 0 | 0 | 0 | 1 | 1 |
| Mir6897 | 0 | 0 | 0 | 1 | 1 |
| Mir6898 | 0 | 0 | 0 | 1 | 1 |
| Mir6899 | 0 | 0 | 0 | 1 | 1 |
| Mir690 | 0 | 0 | 0 | 1 | 1 |
| Mir6900 | 0 | 0 | 0 | 1 | 1 |
| Mir6901 | 0 | 0 | 0 | 1 | 1 |
| Mir6902 | 0 | 0 | 0 | 1 | 1 |
| Mir6903 | 0 | 0 | 0 | 1 | 1 |
| Mir6906 | 0 | 0 | 0 | 1 | 1 |
| Mir6908 | 0 | 0 | 0 | 1 | 1 |
| Mir6909 | 0 | 0 | 0 | 1 | 1 |
| Mir691 | 0 | 0 | 0 | 1 | 1 |
| Mir6911 | 0 | 0 | 0 | 1 | 1 |
| Mir6913 | 0 | 0 | 0 | 1 | 1 |
| Mir6914 | 0 | 0 | 0 | 1 | 1 |
| Mir6916 | 0 | 0 | 0 | 1 | 1 |
| Mir6917 | 0 | 0 | 0 | 1 | 1 |
| Mir6918 | 0 | 0 | 0 | 1 | 1 |
| Mir6919 | 0 | 0 | 0 | 1 | 1 |
| Mir692-1 | 0 | 0 | 0 | 1 | 1 |
| Mir692-2b | 0 | 0 | 0 | 1 | 1 |
| Mir6923 | 0 | 0 | 0 | 1 | 1 |
| Mir6924 | 0 | 0 | 0 | 1 | 1 |
| Mir6925 | 0 | 0 | 0 | 1 | 1 |
| Mir6926 | 0 | 0 | 0 | 1 | 1 |
| Mir6927 | 0 | 0 | 0 | 1 | 1 |
| Mir6928 | 0 | 0 | 0 | 1 | 1 |
| Mir6929 | 0 | 0 | 0 | 1 | 1 |
| Mir693 | 0 | 0 | 0 | 1 | 1 |

| | | | | | |
|---|---|---|---|---|---|
| Mir6930 | 0 | 0 | 0 | 1 | 1 |
| Mir6931 | 0 | 0 | 0 | 1 | 1 |
| Mir6932 | 0 | 0 | 0 | 1 | 1 |
| Mir6933 | 0 | 0 | 0 | 1 | 1 |
| Mir6934 | 0 | 0 | 0 | 1 | 1 |
| Mir6936 | 0 | 0 | 0 | 1 | 1 |
| Mir6937 | 0 | 0 | 0 | 1 | 1 |
| Mir694 | 0 | 0 | 0 | 1 | 1 |
| Mir6940 | 0 | 0 | 0 | 1 | 1 |
| Mir6941 | 0 | 0 | 0 | 1 | 1 |
| Mir6942 | 0 | 0 | 0 | 1 | 1 |
| Mir6946 | 0 | 0 | 0 | 1 | 1 |
| Mir6947 | 0 | 0 | 0 | 1 | 1 |
| Mir6949 | 0 | 0 | 0 | 1 | 1 |
| Mir695 | 0 | 0 | 0 | 1 | 1 |
| Mir6950 | 0 | 0 | 0 | 1 | 1 |
| Mir6951 | 0 | 0 | 0 | 1 | 1 |
| Mir6952 | 0 | 0 | 0 | 1 | 1 |
| Mir6953 | 0 | 0 | 0 | 1 | 1 |
| Mir6954 | 0 | 0 | 0 | 1 | 1 |
| Mir6956 | 0 | 0 | 0 | 1 | 1 |
| Mir6957 | 0 | 0 | 0 | 1 | 1 |
| Mir6959 | 0 | 0 | 0 | 1 | 1 |
| Mir6960 | 0 | 0 | 0 | 1 | 1 |
| Mir6961 | 0 | 0 | 0 | 1 | 1 |
| Mir6962 | 0 | 0 | 0 | 1 | 1 |
| Mir6964 | 0 | 0 | 0 | 1 | 1 |
| Mir6968 | 0 | 0 | 0 | 1 | 1 |
| Mir6969 | 0 | 0 | 0 | 1 | 1 |
| Mir6971 | 0 | 0 | 0 | 1 | 1 |
| Mir6973a | 0 | 0 | 0 | 1 | 1 |
| Mir6974 | 0 | 0 | 0 | 1 | 1 |
| Mir6975 | 0 | 0 | 0 | 1 | 1 |
| Mir6977 | 0 | 0 | 0 | 1 | 1 |
| Mir6978 | 0 | 0 | 0 | 1 | 1 |
| Mir6979 | 0 | 0 | 0 | 1 | 1 |
| Mir698 | 0 | 0 | 0 | 1 | 1 |
| Mir6980 | 0 | 0 | 0 | 1 | 1 |
| Mir6981 | 0 | 0 | 0 | 1 | 1 |
| Mir6982 | 0 | 0 | 0 | 1 | 1 |
| Mir6983 | 0 | 0 | 0 | 1 | 1 |
| Mir6984 | 0 | 0 | 0 | 1 | 1 |
| Mir6985 | 0 | 0 | 0 | 1 | 1 |
| Mir6986 | 0 | 0 | 0 | 1 | 1 |
| Mir6987 | 0 | 0 | 0 | 1 | 1 |
| Mir6988 | 0 | 0 | 0 | 1 | 1 |
| Mir6989 | 0 | 0 | 0 | 1 | 1 |
| Mir6994 | 0 | 0 | 0 | 1 | 1 |

| | | | | | |
|---|---|---|---|---|---|
| Mir6995 | 0 | 0 | 0 | 1 | 1 |
| Mir6996 | 0 | 0 | 0 | 1 | 1 |
| Mir6997 | 0 | 0 | 0 | 1 | 1 |
| Mir6999 | 0 | 0 | 0 | 1 | 1 |
| Mir7-2 | 0 | 0 | 0 | 1 | 1 |
| Mir7000 | 0 | 0 | 0 | 1 | 1 |
| Mir7001 | 0 | 0 | 0 | 1 | 1 |
| Mir7002 | 0 | 0 | 0 | 1 | 1 |
| Mir7003 | 0 | 0 | 0 | 1 | 1 |
| Mir7004 | 0 | 0 | 0 | 1 | 1 |
| Mir7005 | 0 | 0 | 0 | 1 | 1 |
| Mir7006 | 0 | 0 | 0 | 1 | 1 |
| Mir7008 | 0 | 0 | 0 | 1 | 1 |
| Mir7009 | 0 | 0 | 0 | 1 | 1 |
| Mir701 | 0 | 0 | 0 | 1 | 1 |
| Mir7010 | 0 | 0 | 0 | 1 | 1 |
| Mir7011 | 0 | 0 | 0 | 1 | 1 |
| Mir7014 | 0 | 0 | 0 | 1 | 1 |
| Mir7015 | 0 | 0 | 0 | 1 | 1 |
| Mir7016 | 0 | 0 | 0 | 1 | 1 |
| Mir7017 | 0 | 0 | 0 | 1 | 1 |
| Mir7018 | 0 | 0 | 0 | 1 | 1 |
| Mir7020 | 0 | 0 | 0 | 1 | 1 |
| Mir7021 | 0 | 0 | 0 | 1 | 1 |
| Mir7023 | 0 | 0 | 0 | 1 | 1 |
| Mir7025 | 0 | 0 | 0 | 1 | 1 |
| Mir7026 | 0 | 0 | 0 | 1 | 1 |
| Mir7028 | 0 | 0 | 0 | 1 | 1 |
| Mir7029 | 0 | 0 | 0 | 1 | 1 |
| Mir7032 | 0 | 0 | 0 | 1 | 1 |
| Mir7034 | 0 | 0 | 0 | 1 | 1 |
| Mir7035 | 0 | 0 | 0 | 1 | 1 |
| Mir7037 | 0 | 0 | 0 | 1 | 1 |
| Mir7038 | 0 | 0 | 0 | 1 | 1 |
| Mir7039 | 0 | 0 | 0 | 1 | 1 |
| Mir7040 | 0 | 0 | 0 | 1 | 1 |
| Mir7041 | 0 | 0 | 0 | 1 | 1 |
| Mir7042 | 0 | 0 | 0 | 1 | 1 |
| Mir7043 | 0 | 0 | 0 | 1 | 1 |
| Mir7044 | 0 | 0 | 0 | 1 | 1 |
| Mir7045 | 0 | 0 | 0 | 1 | 1 |
| Mir7046 | 0 | 0 | 0 | 1 | 1 |
| Mir7047 | 0 | 0 | 0 | 1 | 1 |
| Mir7048 | 0 | 0 | 0 | 1 | 1 |
| Mir7049 | 0 | 0 | 0 | 1 | 1 |
| Mir705 | 0 | 0 | 0 | 1 | 1 |
| Mir7050 | 0 | 0 | 0 | 1 | 1 |
| Mir7051 | 0 | 0 | 0 | 1 | 1 |

| | | | | | |
|---|---|---|---|---|---|
| Mir7053 | 0 | 0 | 0 | 1 | 1 |
| Mir7054 | 0 | 0 | 0 | 1 | 1 |
| Mir7055 | 0 | 0 | 0 | 1 | 1 |
| Mir7056 | 0 | 0 | 0 | 1 | 1 |
| Mir7057 | 0 | 0 | 0 | 1 | 1 |
| Mir7058 | 0 | 0 | 0 | 1 | 1 |
| Mir7059 | 0 | 0 | 0 | 1 | 1 |
| Mir706 | 0 | 0 | 0 | 1 | 1 |
| Mir7061 | 0 | 0 | 0 | 1 | 1 |
| Mir7064 | 0 | 0 | 0 | 1 | 1 |
| Mir7065 | 0 | 0 | 0 | 1 | 1 |
| Mir7066 | 0 | 0 | 0 | 1 | 1 |
| Mir707 | 0 | 0 | 0 | 1 | 1 |
| Mir7070 | 0 | 0 | 0 | 1 | 1 |
| Mir7071 | 0 | 0 | 0 | 1 | 1 |
| Mir7072 | 0 | 0 | 0 | 1 | 1 |
| Mir7073 | 0 | 0 | 0 | 1 | 1 |
| Mir7074 | 0 | 0 | 0 | 1 | 1 |
| Mir7076 | 0 | 0 | 0 | 1 | 1 |
| Mir7077 | 0 | 0 | 0 | 1 | 1 |
| Mir708 | 0 | 0 | 0 | 1 | 1 |
| Mir7080 | 0 | 0 | 0 | 1 | 1 |
| Mir7081 | 0 | 0 | 0 | 1 | 1 |
| Mir7083 | 0 | 0 | 0 | 1 | 1 |
| Mir7084 | 0 | 0 | 0 | 1 | 1 |
| Mir7085 | 0 | 0 | 0 | 1 | 1 |
| Mir7086 | 0 | 0 | 0 | 1 | 1 |
| Mir7087 | 0 | 0 | 0 | 1 | 1 |
| Mir7088 | 0 | 0 | 0 | 1 | 1 |
| Mir709 | 0 | 0 | 0 | 1 | 1 |
| Mir7090 | 0 | 0 | 0 | 1 | 1 |
| Mir7092 | 0 | 0 | 0 | 1 | 1 |
| Mir7093 | 0 | 0 | 0 | 1 | 1 |
| Mir7094-1 | 0 | 0 | 0 | 1 | 1 |
| Mir7094-2 | 0 | 0 | 0 | 1 | 1 |
| Mir710 | 0 | 0 | 0 | 1 | 1 |
| Mir711 | 0 | 0 | 0 | 1 | 1 |
| Mir7117 | 0 | 0 | 0 | 1 | 1 |
| Mir7118 | 0 | 0 | 0 | 1 | 1 |
| Mir7119 | 0 | 0 | 0 | 1 | 1 |
| Mir713 | 0 | 0 | 0 | 1 | 1 |
| Mir718 | 0 | 0 | 0 | 1 | 1 |
| Mir721 | 0 | 0 | 0 | 1 | 1 |
| Mir7210 | 0 | 0 | 0 | 1 | 1 |
| Mir7211 | 0 | 0 | 0 | 1 | 1 |
| Mir7212 | 0 | 0 | 0 | 1 | 1 |
| Mir7213 | 0 | 0 | 0 | 1 | 1 |
| Mir7214 | 0 | 0 | 0 | 1 | 1 |

| | | | | | |
|---|---|---|---|---|---|
| Mir7215 | 0 | 0 | 0 | 1 | 1 |
| Mir7216 | 0 | 0 | 0 | 1 | 1 |
| Mir7217 | 0 | 0 | 0 | 1 | 1 |
| Mir7218 | 0 | 0 | 0 | 1 | 1 |
| Mir7219 | 0 | 0 | 0 | 1 | 1 |
| Mir7220 | 0 | 0 | 0 | 1 | 1 |
| Mir7221 | 0 | 0 | 0 | 1 | 1 |
| Mir7222 | 0 | 0 | 0 | 1 | 1 |
| Mir7223 | 0 | 0 | 0 | 1 | 1 |
| Mir7224 | 0 | 0 | 0 | 1 | 1 |
| Mir7225 | 0 | 0 | 0 | 1 | 1 |
| Mir7226 | 0 | 0 | 0 | 1 | 1 |
| Mir7227 | 0 | 0 | 0 | 1 | 1 |
| Mir7228 | 0 | 0 | 0 | 1 | 1 |
| Mir7229 | 0 | 0 | 0 | 1 | 1 |
| Mir7230 | 0 | 0 | 0 | 1 | 1 |
| Mir7231 | 0 | 0 | 0 | 1 | 1 |
| Mir7232 | 0 | 0 | 0 | 1 | 1 |
| Mir7233 | 0 | 0 | 0 | 1 | 1 |
| Mir7234 | 0 | 0 | 0 | 1 | 1 |
| Mir7235 | 0 | 0 | 0 | 1 | 1 |
| Mir7236 | 0 | 0 | 0 | 1 | 1 |
| Mir7237 | 0 | 0 | 0 | 1 | 1 |
| Mir7238 | 0 | 0 | 0 | 1 | 1 |
| Mir7239 | 0 | 0 | 0 | 1 | 1 |
| Mir7240 | 0 | 0 | 0 | 1 | 1 |
| Mir7241 | 0 | 0 | 0 | 1 | 1 |
| Mir7242 | 0 | 0 | 0 | 1 | 1 |
| Mir7243 | 0 | 0 | 0 | 1 | 1 |
| Mir741 | 0 | 0 | 0 | 1 | 1 |
| Mir742 | 0 | 0 | 0 | 1 | 1 |
| Mir743 | 0 | 0 | 0 | 1 | 1 |
| Mir743b | 0 | 0 | 0 | 1 | 1 |
| Mir744 | 0 | 0 | 0 | 1 | 1 |
| Mir7578 | 0 | 0 | 0 | 1 | 1 |
| Mir758 | 0 | 0 | 0 | 1 | 1 |
| Mir759 | 0 | 0 | 0 | 1 | 1 |
| Mir760 | 0 | 0 | 0 | 1 | 1 |
| Mir761 | 0 | 0 | 0 | 1 | 1 |
| Mir764 | 0 | 0 | 0 | 1 | 1 |
| Mir7646 | 0 | 0 | 0 | 1 | 1 |
| Mir7647 | 0 | 0 | 0 | 1 | 1 |
| Mir7648 | 0 | 0 | 0 | 1 | 1 |
| Mir7649 | 0 | 0 | 0 | 1 | 1 |
| Mir7652 | 0 | 0 | 0 | 1 | 1 |
| Mir7654 | 0 | 0 | 0 | 1 | 1 |
| Mir7655 | 0 | 0 | 0 | 1 | 1 |
| Mir7656 | 0 | 0 | 0 | 1 | 1 |

| | | | | | |
|---|---|---|---|---|---|
| Mir7657 | 0 | 0 | 0 | 1 | 1 |
| Mir7658 | 0 | 0 | 0 | 1 | 1 |
| Mir7661 | 0 | 0 | 0 | 1 | 1 |
| Mir7662 | 0 | 0 | 0 | 1 | 1 |
| Mir7663 | 0 | 0 | 0 | 1 | 1 |
| Mir7665 | 0 | 0 | 0 | 1 | 1 |
| Mir7666 | 0 | 0 | 0 | 1 | 1 |
| Mir7667 | 0 | 0 | 0 | 1 | 1 |
| Mir7668 | 0 | 0 | 0 | 1 | 1 |
| Mir7669 | 0 | 0 | 0 | 1 | 1 |
| Mir767 | 0 | 0 | 0 | 1 | 1 |
| Mir7670 | 0 | 0 | 0 | 1 | 1 |
| Mir7671 | 0 | 0 | 0 | 1 | 1 |
| Mir7672 | 0 | 0 | 0 | 1 | 1 |
| Mir7673 | 0 | 0 | 0 | 1 | 1 |
| Mir7674 | 0 | 0 | 0 | 1 | 1 |
| Mir7675 | 0 | 0 | 0 | 1 | 1 |
| Mir7676-1 | 0 | 0 | 0 | 1 | 1 |
| Mir7676-2 | 0 | 0 | 0 | 1 | 1 |
| Mir7677 | 0 | 0 | 0 | 1 | 1 |
| Mir7678 | 0 | 0 | 0 | 1 | 1 |
| Mir7679 | 0 | 0 | 0 | 1 | 1 |
| Mir7680 | 0 | 0 | 0 | 1 | 1 |
| Mir7681 | 0 | 0 | 0 | 1 | 1 |
| Mir7682 | 0 | 0 | 0 | 1 | 1 |
| Mir7684 | 0 | 0 | 0 | 1 | 1 |
| Mir7685 | 0 | 0 | 0 | 1 | 1 |
| Mir7686 | 0 | 0 | 0 | 1 | 1 |
| Mir7687 | 0 | 0 | 0 | 1 | 1 |
| Mir770 | 0 | 0 | 0 | 1 | 1 |
| Mir7b | 0 | 0 | 0 | 1 | 1 |
| Mir802 | 0 | 0 | 0 | 1 | 1 |
| Mir804 | 0 | 0 | 0 | 1 | 1 |
| Mir8092 | 0 | 0 | 0 | 1 | 1 |
| Mir8093 | 0 | 0 | 0 | 1 | 1 |
| Mir8095 | 0 | 0 | 0 | 1 | 1 |
| Mir8096 | 0 | 0 | 0 | 1 | 1 |
| Mir8098 | 0 | 0 | 0 | 1 | 1 |
| Mir8099-1 | 0 | 0 | 0 | 1 | 1 |
| Mir8099-2 | 0 | 0 | 0 | 1 | 1 |
| Mir8100 | 0 | 0 | 0 | 1 | 1 |
| Mir8101 | 0 | 0 | 0 | 1 | 1 |
| Mir8103 | 0 | 0 | 0 | 1 | 1 |
| Mir8104 | 0 | 0 | 0 | 1 | 1 |
| Mir8105 | 0 | 0 | 0 | 1 | 1 |
| Mir8106 | 0 | 0 | 0 | 1 | 1 |
| Mir8107 | 0 | 0 | 0 | 1 | 1 |
| Mir8108 | 0 | 0 | 0 | 1 | 1 |

| | | | | | |
|---|---|---|---|---|---|
| Mir8109 | 0 | 0 | 0 | 1 | 1 |
| Mir8110 | 0 | 0 | 0 | 1 | 1 |
| Mir8111 | 0 | 0 | 0 | 1 | 1 |
| Mir8113 | 0 | 0 | 0 | 1 | 1 |
| Mir8114 | 0 | 0 | 0 | 1 | 1 |
| Mir8115 | 0 | 0 | 0 | 1 | 1 |
| Mir8118 | 0 | 0 | 0 | 1 | 1 |
| Mir8119 | 0 | 0 | 0 | 1 | 1 |
| Mir8120 | 0 | 0 | 0 | 1 | 1 |
| Mir871 | 0 | 0 | 0 | 1 | 1 |
| Mir872 | 0 | 0 | 0 | 1 | 1 |
| Mir873 | 0 | 0 | 0 | 1 | 1 |
| Mir874 | 0 | 0 | 0 | 1 | 1 |
| Mir875 | 0 | 0 | 0 | 1 | 1 |
| Mir876 | 0 | 0 | 0 | 1 | 1 |
| Mir877 | 0 | 0 | 0 | 1 | 1 |
| Mir878 | 0 | 0 | 0 | 1 | 1 |
| Mir879 | 0 | 0 | 0 | 1 | 1 |
| Mir880 | 0 | 0 | 0 | 1 | 1 |
| Mir881 | 0 | 0 | 0 | 1 | 1 |
| Mir882 | 0 | 0 | 0 | 1 | 1 |
| Mir883a | 0 | 0 | 0 | 1 | 1 |
| Mir883b | 0 | 0 | 0 | 1 | 1 |
| Mir9-1 | 0 | 0 | 0 | 1 | 1 |
| Mir9-2 | 0 | 0 | 0 | 1 | 1 |
| Mir9-3 | 0 | 0 | 0 | 1 | 1 |
| Mir92-1 | 0 | 0 | 0 | 1 | 1 |
| Mir92-2 | 0 | 0 | 0 | 1 | 1 |
| Mir92b | 0 | 0 | 0 | 1 | 1 |
| Mir93 | 0 | 0 | 0 | 1 | 1 |
| Mir96 | 0 | 0 | 0 | 1 | 1 |
| Mir98 | 0 | 0 | 0 | 1 | 1 |
| Mir99a | 0 | 0 | 0 | 1 | 1 |
| Mir99b | 0 | 0 | 0 | 1 | 1 |
| Mirlet7a-1 | 0 | 0 | 0 | 1 | 1 |
| Mirlet7a-2 | 0 | 0 | 0 | 1 | 1 |
| Mirlet7b | 0 | 0 | 0 | 1 | 1 |
| Mirlet7c-1 | 0 | 0 | 0 | 1 | 1 |
| Mirlet7c-2 | 0 | 0 | 0 | 1 | 1 |
| Mirlet7e | 0 | 0 | 0 | 1 | 1 |
| Mirlet7f-1 | 0 | 0 | 0 | 1 | 1 |
| Mirlet7f-2 | 0 | 0 | 0 | 1 | 1 |
| Mirlet7g | 0 | 0 | 0 | 1 | 1 |
| Mirlet7j | 0 | 0 | 0 | 1 | 1 |
| Mirlet7k | 0 | 0 | 0 | 1 | 1 |
| Mlana | 0 | 0 | 0 | 1 | 1 |
| Mlc1 | 0 | 0 | 0 | 1 | 1 |
| Mlip | 0 | 0 | 0 | 1 | 1 |

| | | | | | |
|---|---|---|---|---|---|
| Mmp10 | 0 | 0 | 0 | 1 | 1 |
| Mmp1a | 0 | 0 | 0 | 1 | 1 |
| Mmp1b | 0 | 0 | 0 | 1 | 1 |
| Mmp3 | 0 | 0 | 0 | 1 | 1 |
| Mmp8 | 0 | 0 | 0 | 1 | 1 |
| Mov10l1 | 0 | 0 | 0 | 1 | 1 |
| Mptx1 | 0 | 0 | 0 | 1 | 1 |
| Mptx2 | 0 | 0 | 0 | 1 | 1 |
| Mrgpra1 | 0 | 0 | 0 | 1 | 1 |
| Mrgpra2a | 0 | 0 | 0 | 1 | 1 |
| Mrgpra2b | 0 | 0 | 0 | 1 | 1 |
| Mrgpra3 | 0 | 0 | 0 | 1 | 1 |
| Mrgpra4 | 0 | 0 | 0 | 1 | 1 |
| Mrgpra6 | 0 | 0 | 0 | 1 | 1 |
| Mrgpra9 | 0 | 0 | 0 | 1 | 1 |
| Mrgprb1 | 0 | 0 | 0 | 1 | 1 |
| Mrgprb2 | 0 | 0 | 0 | 1 | 1 |
| Mrgprb3 | 0 | 0 | 0 | 1 | 1 |
| Mrgprb4 | 0 | 0 | 0 | 1 | 1 |
| Mrgprb8 | 0 | 0 | 0 | 1 | 1 |
| Mrgprx1 | 0 | 0 | 0 | 1 | 1 |
| Mrgprx2 | 0 | 0 | 0 | 1 | 1 |
| Mroh4 | 0 | 0 | 0 | 1 | 1 |
| Mroh5 | 0 | 0 | 0 | 1 | 1 |
| Mroh9 | 0 | 0 | 0 | 1 | 1 |
| Ms4a1 | 0 | 0 | 0 | 1 | 1 |
| Ms4a18 | 0 | 0 | 0 | 1 | 1 |
| Ms4a4b | 0 | 0 | 0 | 1 | 1 |
| Ms4a5 | 0 | 0 | 0 | 1 | 1 |
| Ms4a8a | 0 | 0 | 0 | 1 | 1 |
| Msln | 0 | 0 | 0 | 1 | 1 |
| Mstn | 0 | 0 | 0 | 1 | 1 |
| Mt4 | 0 | 0 | 0 | 1 | 1 |
| Muc20 | 0 | 0 | 0 | 1 | 1 |
| Mug-ps1 | 0 | 0 | 0 | 1 | 1 |
| Mup1 | 0 | 0 | 0 | 1 | 1 |
| Mup10 | 0 | 0 | 0 | 1 | 1 |
| Mup11 | 0 | 0 | 0 | 1 | 1 |
| Mup12 | 0 | 0 | 0 | 1 | 1 |
| Mup13 | 0 | 0 | 0 | 1 | 1 |
| Mup14 | 0 | 0 | 0 | 1 | 1 |
| Mup15 | 0 | 0 | 0 | 1 | 1 |
| Mup16 | 0 | 0 | 0 | 1 | 1 |
| Mup17 | 0 | 0 | 0 | 1 | 1 |
| Mup19 | 0 | 0 | 0 | 1 | 1 |
| Mup2 | 0 | 0 | 0 | 1 | 1 |
| Mup20 | 0 | 0 | 0 | 1 | 1 |
| Mup21 | 0 | 0 | 0 | 1 | 1 |

| | | | | | |
|---|---|---|---|---|---|
| Mup3 | 0 | 0 | 0 | 1 | 1 |
| Mup4 | 0 | 0 | 0 | 1 | 1 |
| Mup5 | 0 | 0 | 0 | 1 | 1 |
| Mup6 | 0 | 0 | 0 | 1 | 1 |
| Mup7 | 0 | 0 | 0 | 1 | 1 |
| Mup8 | 0 | 0 | 0 | 1 | 1 |
| Mup9 | 0 | 0 | 0 | 1 | 1 |
| Myl10 | 0 | 0 | 0 | 1 | 1 |
| Myo3a | 0 | 0 | 0 | 1 | 1 |
| Myot | 0 | 0 | 0 | 1 | 1 |
| Myoz2 | 0 | 0 | 0 | 1 | 1 |
| Naa11 | 0 | 0 | 0 | 1 | 1 |
| Nat3 | 0 | 0 | 0 | 1 | 1 |
| Ncr1 | 0 | 0 | 0 | 1 | 1 |
| Nell1os | 0 | 0 | 0 | 1 | 1 |
| Neu4 | 0 | 0 | 0 | 1 | 1 |
| Nipsnap3a | 0 | 0 | 0 | 1 | 1 |
| Nkx2-5 | 0 | 0 | 0 | 1 | 1 |
| Nkx6-3 | 0 | 0 | 0 | 1 | 1 |
| Nlrp12 | 0 | 0 | 0 | 1 | 1 |
| Nlrp14 | 0 | 0 | 0 | 1 | 1 |
| Nlrp2 | 0 | 0 | 0 | 1 | 1 |
| Nlrp4a | 0 | 0 | 0 | 1 | 1 |
| Nlrp4b | 0 | 0 | 0 | 1 | 1 |
| Nlrp4e | 0 | 0 | 0 | 1 | 1 |
| Nlrp5 | 0 | 0 | 0 | 1 | 1 |
| Nlrp9a | 0 | 0 | 0 | 1 | 1 |
| Nlrp9b | 0 | 0 | 0 | 1 | 1 |
| Nlrp9c | 0 | 0 | 0 | 1 | 1 |
| Nme8 | 0 | 0 | 0 | 1 | 1 |
| Nme9 | 0 | 0 | 0 | 1 | 1 |
| Nmrk2 | 0 | 0 | 0 | 1 | 1 |
| Nms | 0 | 0 | 0 | 1 | 1 |
| Nmu | 0 | 0 | 0 | 1 | 1 |
| Nmur1 | 0 | 0 | 0 | 1 | 1 |
| Nmur2 | 0 | 0 | 0 | 1 | 1 |
| Nphs1os | 0 | 0 | 0 | 1 | 1 |
| Nppb | 0 | 0 | 0 | 1 | 1 |
| Npvf | 0 | 0 | 0 | 1 | 1 |
| Npy2r | 0 | 0 | 0 | 1 | 1 |
| Npy6r | 0 | 0 | 0 | 1 | 1 |
| Nrg3 | 0 | 0 | 0 | 1 | 1 |
| Nrg3os | 0 | 0 | 0 | 1 | 1 |
| Nrl | 0 | 0 | 0 | 1 | 1 |
| Ntsr2 | 0 | 0 | 0 | 1 | 1 |
| Nutf2-ps1 | 0 | 0 | 0 | 1 | 1 |
| Nxph1 | 0 | 0 | 0 | 1 | 1 |
| Nyap2 | 0 | 0 | 0 | 1 | 1 |

| | | | | | |
|---|---|---|---|---|---|
| OTTMUSG000000 | 0 | 0 | 0 | 1 | 1 |
| Oas1e | 0 | 0 | 0 | 1 | 1 |
| Oas1f | 0 | 0 | 0 | 1 | 1 |
| Oaz1-ps | 0 | 0 | 0 | 1 | 1 |
| Obox1 | 0 | 0 | 0 | 1 | 1 |
| Obox2 | 0 | 0 | 0 | 1 | 1 |
| Obox3 | 0 | 0 | 0 | 1 | 1 |
| Obox5 | 0 | 0 | 0 | 1 | 1 |
| Obox6 | 0 | 0 | 0 | 1 | 1 |
| Obp1a | 0 | 0 | 0 | 1 | 1 |
| Obp2a | 0 | 0 | 0 | 1 | 1 |
| Obp2b | 0 | 0 | 0 | 1 | 1 |
| Oc90 | 0 | 0 | 0 | 1 | 1 |
| Odam | 0 | 0 | 0 | 1 | 1 |
| Odf1 | 0 | 0 | 0 | 1 | 1 |
| Odf3 | 0 | 0 | 0 | 1 | 1 |
| Olah | 0 | 0 | 0 | 1 | 1 |
| Olfr1 | 0 | 0 | 0 | 1 | 1 |
| Olfr10 | 0 | 0 | 0 | 1 | 1 |
| Olfr100,Olfr102 | 0 | 0 | 0 | 1 | 1 |
| Olfr1000 | 0 | 0 | 0 | 1 | 1 |
| Olfr1002 | 0 | 0 | 0 | 1 | 1 |
| Olfr1006 | 0 | 0 | 0 | 1 | 1 |
| Olfr1008 | 0 | 0 | 0 | 1 | 1 |
| Olfr1009 | 0 | 0 | 0 | 1 | 1 |
| Olfr101 | 0 | 0 | 0 | 1 | 1 |
| Olfr1010 | 0 | 0 | 0 | 1 | 1 |
| Olfr1012 | 0 | 0 | 0 | 1 | 1 |
| Olfr1013 | 0 | 0 | 0 | 1 | 1 |
| Olfr1014 | 0 | 0 | 0 | 1 | 1 |
| Olfr1015 | 0 | 0 | 0 | 1 | 1 |
| Olfr1016 | 0 | 0 | 0 | 1 | 1 |
| Olfr1018 | 0 | 0 | 0 | 1 | 1 |
| Olfr1019 | 0 | 0 | 0 | 1 | 1 |
| Olfr102 | 0 | 0 | 0 | 1 | 1 |
| Olfr1020 | 0 | 0 | 0 | 1 | 1 |
| Olfr1022 | 0 | 0 | 0 | 1 | 1 |
| Olfr1023 | 0 | 0 | 0 | 1 | 1 |
| Olfr1024 | 0 | 0 | 0 | 1 | 1 |
| Olfr1026 | 0 | 0 | 0 | 1 | 1 |
| Olfr1028 | 0 | 0 | 0 | 1 | 1 |
| Olfr1029 | 0 | 0 | 0 | 1 | 1 |
| Olfr103 | 0 | 0 | 0 | 1 | 1 |
| Olfr1030 | 0 | 0 | 0 | 1 | 1 |
| Olfr1031 | 0 | 0 | 0 | 1 | 1 |
| Olfr1032 | 0 | 0 | 0 | 1 | 1 |
| Olfr1034 | 0 | 0 | 0 | 1 | 1 |
| Olfr1036 | 0 | 0 | 0 | 1 | 1 |

| | | | | | |
|---|---|---|---|---|---|
| Olfr1037 | 0 | 0 | 0 | 1 | 1 |
| Olfr1038-ps | 0 | 0 | 0 | 1 | 1 |
| Olfr1039 | 0 | 0 | 0 | 1 | 1 |
| Olfr1040 | 0 | 0 | 0 | 1 | 1 |
| Olfr1042 | 0 | 0 | 0 | 1 | 1 |
| Olfr1043 | 0 | 0 | 0 | 1 | 1 |
| Olfr1044 | 0 | 0 | 0 | 1 | 1 |
| Olfr1045 | 0 | 0 | 0 | 1 | 1 |
| Olfr1046 | 0 | 0 | 0 | 1 | 1 |
| Olfr1047 | 0 | 0 | 0 | 1 | 1 |
| Olfr1048 | 0 | 0 | 0 | 1 | 1 |
| Olfr1049 | 0 | 0 | 0 | 1 | 1 |
| Olfr1051 | 0 | 0 | 0 | 1 | 1 |
| Olfr1052 | 0 | 0 | 0 | 1 | 1 |
| Olfr1053 | 0 | 0 | 0 | 1 | 1 |
| Olfr1054 | 0 | 0 | 0 | 1 | 1 |
| Olfr1055 | 0 | 0 | 0 | 1 | 1 |
| Olfr1056 | 0 | 0 | 0 | 1 | 1 |
| Olfr1057 | 0 | 0 | 0 | 1 | 1 |
| Olfr1058 | 0 | 0 | 0 | 1 | 1 |
| Olfr1061 | 0 | 0 | 0 | 1 | 1 |
| Olfr1062 | 0 | 0 | 0 | 1 | 1 |
| Olfr1065 | 0 | 0 | 0 | 1 | 1 |
| Olfr1066 | 0 | 0 | 0 | 1 | 1 |
| Olfr107 | 0 | 0 | 0 | 1 | 1 |
| Olfr1076 | 0 | 0 | 0 | 1 | 1 |
| Olfr1077-ps1 | 0 | 0 | 0 | 1 | 1 |
| Olfr1079 | 0 | 0 | 0 | 1 | 1 |
| Olfr108 | 0 | 0 | 0 | 1 | 1 |
| Olfr1080 | 0 | 0 | 0 | 1 | 1 |
| Olfr1082 | 0 | 0 | 0 | 1 | 1 |
| Olfr1084 | 0 | 0 | 0 | 1 | 1 |
| Olfr1085 | 0 | 0 | 0 | 1 | 1 |
| Olfr1086 | 0 | 0 | 0 | 1 | 1 |
| Olfr1087 | 0 | 0 | 0 | 1 | 1 |
| Olfr1089 | 0 | 0 | 0 | 1 | 1 |
| Olfr109 | 0 | 0 | 0 | 1 | 1 |
| Olfr1090 | 0 | 0 | 0 | 1 | 1 |
| Olfr1093 | 0 | 0 | 0 | 1 | 1 |
| Olfr1094 | 0 | 0 | 0 | 1 | 1 |
| Olfr1095 | 0 | 0 | 0 | 1 | 1 |
| Olfr1097 | 0 | 0 | 0 | 1 | 1 |
| Olfr1098 | 0 | 0 | 0 | 1 | 1 |
| Olfr1099 | 0 | 0 | 0 | 1 | 1 |
| Olfr11 | 0 | 0 | 0 | 1 | 1 |
| Olfr110 | 0 | 0 | 0 | 1 | 1 |
| Olfr1100 | 0 | 0 | 0 | 1 | 1 |
| Olfr1101 | 0 | 0 | 0 | 1 | 1 |

| | | | | | |
|---|---|---|---|---|---|
| Olfr1102 | 0 | 0 | 0 | 1 | 1 |
| Olfr1104 | 0 | 0 | 0 | 1 | 1 |
| Olfr1105 | 0 | 0 | 0 | 1 | 1 |
| Olfr1106 | 0 | 0 | 0 | 1 | 1 |
| Olfr1107 | 0 | 0 | 0 | 1 | 1 |
| Olfr1109 | 0 | 0 | 0 | 1 | 1 |
| Olfr111 | 0 | 0 | 0 | 1 | 1 |
| Olfr1110 | 0 | 0 | 0 | 1 | 1 |
| Olfr1111 | 0 | 0 | 0 | 1 | 1 |
| Olfr1112 | 0 | 0 | 0 | 1 | 1 |
| Olfr1113 | 0 | 0 | 0 | 1 | 1 |
| Olfr1115 | 0 | 0 | 0 | 1 | 1 |
| Olfr1116-ps | 0 | 0 | 0 | 1 | 1 |
| Olfr1118 | 0 | 0 | 0 | 1 | 1 |
| Olfr112 | 0 | 0 | 0 | 1 | 1 |
| Olfr1120 | 0 | 0 | 0 | 1 | 1 |
| Olfr1121 | 0 | 0 | 0 | 1 | 1 |
| Olfr1122 | 0 | 0 | 0 | 1 | 1 |
| Olfr1123 | 0 | 0 | 0 | 1 | 1 |
| Olfr1124 | 0 | 0 | 0 | 1 | 1 |
| Olfr1126 | 0 | 0 | 0 | 1 | 1 |
| Olfr1128 | 0 | 0 | 0 | 1 | 1 |
| Olfr1129 | 0 | 0 | 0 | 1 | 1 |
| Olfr113 | 0 | 0 | 0 | 1 | 1 |
| Olfr1130 | 0 | 0 | 0 | 1 | 1 |
| Olfr1131 | 0 | 0 | 0 | 1 | 1 |
| Olfr1132 | 0 | 0 | 0 | 1 | 1 |
| Olfr1133 | 0 | 0 | 0 | 1 | 1 |
| Olfr1134 | 0 | 0 | 0 | 1 | 1 |
| Olfr1135 | 0 | 0 | 0 | 1 | 1 |
| Olfr1136 | 0 | 0 | 0 | 1 | 1 |
| Olfr1137 | 0 | 0 | 0 | 1 | 1 |
| Olfr1138 | 0 | 0 | 0 | 1 | 1 |
| Olfr114 | 0 | 0 | 0 | 1 | 1 |
| Olfr1140 | 0 | 0 | 0 | 1 | 1 |
| Olfr1141 | 0 | 0 | 0 | 1 | 1 |
| Olfr1143 | 0 | 0 | 0 | 1 | 1 |
| Olfr1145 | 0 | 0 | 0 | 1 | 1 |
| Olfr1148 | 0 | 0 | 0 | 1 | 1 |
| Olfr115 | 0 | 0 | 0 | 1 | 1 |
| Olfr1151 | 0 | 0 | 0 | 1 | 1 |
| Olfr1152 | 0 | 0 | 0 | 1 | 1 |
| Olfr1153 | 0 | 0 | 0 | 1 | 1 |
| Olfr1154 | 0 | 0 | 0 | 1 | 1 |
| Olfr1155 | 0 | 0 | 0 | 1 | 1 |
| Olfr1156 | 0 | 0 | 0 | 1 | 1 |
| Olfr1157 | 0 | 0 | 0 | 1 | 1 |
| Olfr1158 | 0 | 0 | 0 | 1 | 1 |

| | | | | | |
|---|---|---|---|---|---|
| Olfr116 | 0 | 0 | 0 | 1 | 1 |
| Olfr1160 | 0 | 0 | 0 | 1 | 1 |
| Olfr1161 | 0 | 0 | 0 | 1 | 1 |
| Olfr1162 | 0 | 0 | 0 | 1 | 1 |
| Olfr1163 | 0 | 0 | 0 | 1 | 1 |
| Olfr1164 | 0 | 0 | 0 | 1 | 1 |
| Olfr1166 | 0 | 0 | 0 | 1 | 1 |
| Olfr1167 | 0 | 0 | 0 | 1 | 1 |
| Olfr117 | 0 | 0 | 0 | 1 | 1 |
| Olfr1170 | 0 | 0 | 0 | 1 | 1 |
| Olfr1173 | 0 | 0 | 0 | 1 | 1 |
| Olfr1176 | 0 | 0 | 0 | 1 | 1 |
| Olfr1178 | 0 | 0 | 0 | 1 | 1 |
| Olfr1179 | 0 | 0 | 0 | 1 | 1 |
| Olfr118 | 0 | 0 | 0 | 1 | 1 |
| Olfr1180 | 0 | 0 | 0 | 1 | 1 |
| Olfr1181 | 0 | 0 | 0 | 1 | 1 |
| Olfr1182 | 0 | 0 | 0 | 1 | 1 |
| Olfr1183 | 0 | 0 | 0 | 1 | 1 |
| Olfr1184 | 0 | 0 | 0 | 1 | 1 |
| Olfr1186 | 0 | 0 | 0 | 1 | 1 |
| Olfr1189 | 0 | 0 | 0 | 1 | 1 |
| Olfr119 | 0 | 0 | 0 | 1 | 1 |
| Olfr1193 | 0 | 0 | 0 | 1 | 1 |
| Olfr1195 | 0 | 0 | 0 | 1 | 1 |
| Olfr1196 | 0 | 0 | 0 | 1 | 1 |
| Olfr1197 | 0 | 0 | 0 | 1 | 1 |
| Olfr1199 | 0 | 0 | 0 | 1 | 1 |
| Olfr12 | 0 | 0 | 0 | 1 | 1 |
| Olfr120 | 0 | 0 | 0 | 1 | 1 |
| Olfr1200 | 0 | 0 | 0 | 1 | 1 |
| Olfr1201 | 0 | 0 | 0 | 1 | 1 |
| Olfr1202 | 0 | 0 | 0 | 1 | 1 |
| Olfr1204 | 0 | 0 | 0 | 1 | 1 |
| Olfr1205 | 0 | 0 | 0 | 1 | 1 |
| Olfr1206 | 0 | 0 | 0 | 1 | 1 |
| Olfr1208 | 0 | 0 | 0 | 1 | 1 |
| Olfr1209 | 0 | 0 | 0 | 1 | 1 |
| Olfr121 | 0 | 0 | 0 | 1 | 1 |
| Olfr1211 | 0 | 0 | 0 | 1 | 1 |
| Olfr1212 | 0 | 0 | 0 | 1 | 1 |
| Olfr1213 | 0 | 0 | 0 | 1 | 1 |
| Olfr1214 | 0 | 0 | 0 | 1 | 1 |
| Olfr1215 | 0 | 0 | 0 | 1 | 1 |
| Olfr1216 | 0 | 0 | 0 | 1 | 1 |
| Olfr1217 | 0 | 0 | 0 | 1 | 1 |
| Olfr1218 | 0 | 0 | 0 | 1 | 1 |
| Olfr1219 | 0 | 0 | 0 | 1 | 1 |

| | | | | | |
|---|---|---|---|---|---|
| Olfr122 | 0 | 0 | 0 | 1 | 1 |
| Olfr1221 | 0 | 0 | 0 | 1 | 1 |
| Olfr1222 | 0 | 0 | 0 | 1 | 1 |
| Olfr1223 | 0 | 0 | 0 | 1 | 1 |
| Olfr1225 | 0 | 0 | 0 | 1 | 1 |
| Olfr1226 | 0 | 0 | 0 | 1 | 1 |
| Olfr1228 | 0 | 0 | 0 | 1 | 1 |
| Olfr1229 | 0 | 0 | 0 | 1 | 1 |
| Olfr123 | 0 | 0 | 0 | 1 | 1 |
| Olfr1230 | 0 | 0 | 0 | 1 | 1 |
| Olfr1231 | 0 | 0 | 0 | 1 | 1 |
| Olfr1232 | 0 | 0 | 0 | 1 | 1 |
| Olfr1233 | 0 | 0 | 0 | 1 | 1 |
| Olfr1234 | 0 | 0 | 0 | 1 | 1 |
| Olfr1238 | 0 | 0 | 0 | 1 | 1 |
| Olfr1239 | 0 | 0 | 0 | 1 | 1 |
| Olfr124 | 0 | 0 | 0 | 1 | 1 |
| Olfr1240 | 0 | 0 | 0 | 1 | 1 |
| Olfr1241 | 0 | 0 | 0 | 1 | 1 |
| Olfr1242 | 0 | 0 | 0 | 1 | 1 |
| Olfr1243 | 0 | 0 | 0 | 1 | 1 |
| Olfr1245 | 0 | 0 | 0 | 1 | 1 |
| Olfr1246 | 0 | 0 | 0 | 1 | 1 |
| Olfr1247 | 0 | 0 | 0 | 1 | 1 |
| Olfr1248 | 0 | 0 | 0 | 1 | 1 |
| Olfr125 | 0 | 0 | 0 | 1 | 1 |
| Olfr1250 | 0 | 0 | 0 | 1 | 1 |
| Olfr1251 | 0 | 0 | 0 | 1 | 1 |
| Olfr1252 | 0 | 0 | 0 | 1 | 1 |
| Olfr1253 | 0 | 0 | 0 | 1 | 1 |
| Olfr1254 | 0 | 0 | 0 | 1 | 1 |
| Olfr1255 | 0 | 0 | 0 | 1 | 1 |
| Olfr1256 | 0 | 0 | 0 | 1 | 1 |
| Olfr1257 | 0 | 0 | 0 | 1 | 1 |
| Olfr1258 | 0 | 0 | 0 | 1 | 1 |
| Olfr1259 | 0 | 0 | 0 | 1 | 1 |
| Olfr126 | 0 | 0 | 0 | 1 | 1 |
| Olfr1260 | 0 | 0 | 0 | 1 | 1 |
| Olfr1261 | 0 | 0 | 0 | 1 | 1 |
| Olfr1262 | 0 | 0 | 0 | 1 | 1 |
| Olfr1263 | 0 | 0 | 0 | 1 | 1 |
| Olfr1264 | 0 | 0 | 0 | 1 | 1 |
| Olfr1265 | 0 | 0 | 0 | 1 | 1 |
| Olfr1269 | 0 | 0 | 0 | 1 | 1 |
| Olfr127 | 0 | 0 | 0 | 1 | 1 |
| Olfr1271 | 0 | 0 | 0 | 1 | 1 |
| Olfr1272 | 0 | 0 | 0 | 1 | 1 |
| Olfr1273-ps | 0 | 0 | 0 | 1 | 1 |

| | | | | | |
|---|---|---|---|---|---|
| Olfr1274-ps | 0 | 0 | 0 | 1 | 1 |
| Olfr1275 | 0 | 0 | 0 | 1 | 1 |
| Olfr1276 | 0 | 0 | 0 | 1 | 1 |
| Olfr1277 | 0 | 0 | 0 | 1 | 1 |
| Olfr1278 | 0 | 0 | 0 | 1 | 1 |
| Olfr1279 | 0 | 0 | 0 | 1 | 1 |
| Olfr128 | 0 | 0 | 0 | 1 | 1 |
| Olfr1280 | 0 | 0 | 0 | 1 | 1 |
| Olfr1281 | 0 | 0 | 0 | 1 | 1 |
| Olfr1282 | 0 | 0 | 0 | 1 | 1 |
| Olfr1283 | 0 | 0 | 0 | 1 | 1 |
| Olfr1284 | 0 | 0 | 0 | 1 | 1 |
| Olfr1286 | 0 | 0 | 0 | 1 | 1 |
| Olfr1287 | 0 | 0 | 0 | 1 | 1 |
| Olfr1288 | 0 | 0 | 0 | 1 | 1 |
| Olfr1289 | 0 | 0 | 0 | 1 | 1 |
| Olfr129 | 0 | 0 | 0 | 1 | 1 |
| Olfr1290 | 0 | 0 | 0 | 1 | 1 |
| Olfr1294 | 0 | 0 | 0 | 1 | 1 |
| Olfr1295 | 0 | 0 | 0 | 1 | 1 |
| Olfr1297 | 0 | 0 | 0 | 1 | 1 |
| Olfr1298 | 0 | 0 | 0 | 1 | 1 |
| Olfr1299 | 0 | 0 | 0 | 1 | 1 |
| Olfr13 | 0 | 0 | 0 | 1 | 1 |
| Olfr130 | 0 | 0 | 0 | 1 | 1 |
| Olfr1300-ps1 | 0 | 0 | 0 | 1 | 1 |
| Olfr1301 | 0 | 0 | 0 | 1 | 1 |
| Olfr1302 | 0 | 0 | 0 | 1 | 1 |
| Olfr1303 | 0 | 0 | 0 | 1 | 1 |
| Olfr1305 | 0 | 0 | 0 | 1 | 1 |
| Olfr1306 | 0 | 0 | 0 | 1 | 1 |
| Olfr1307 | 0 | 0 | 0 | 1 | 1 |
| Olfr1308 | 0 | 0 | 0 | 1 | 1 |
| Olfr1309 | 0 | 0 | 0 | 1 | 1 |
| Olfr131 | 0 | 0 | 0 | 1 | 1 |
| Olfr1310 | 0 | 0 | 0 | 1 | 1 |
| Olfr1311 | 0 | 0 | 0 | 1 | 1 |
| Olfr1312 | 0 | 0 | 0 | 1 | 1 |
| Olfr1313 | 0 | 0 | 0 | 1 | 1 |
| Olfr1314 | 0 | 0 | 0 | 1 | 1 |
| Olfr1316 | 0 | 0 | 0 | 1 | 1 |
| Olfr1317 | 0 | 0 | 0 | 1 | 1 |
| Olfr1318 | 0 | 0 | 0 | 1 | 1 |
| Olfr132 | 0 | 0 | 0 | 1 | 1 |
| Olfr1320 | 0 | 0 | 0 | 1 | 1 |
| Olfr1321 | 0 | 0 | 0 | 1 | 1 |
| Olfr1322 | 0 | 0 | 0 | 1 | 1 |
| Olfr1323 | 0 | 0 | 0 | 1 | 1 |

| | | | | | |
|---|---|---|---|---|---|
| Olfr1324 | 0 | 0 | 0 | 1 | 1 |
| Olfr1325 | 0 | 0 | 0 | 1 | 1 |
| Olfr1328 | 0 | 0 | 0 | 1 | 1 |
| Olfr1329 | 0 | 0 | 0 | 1 | 1 |
| Olfr133 | 0 | 0 | 0 | 1 | 1 |
| Olfr1330 | 0 | 0 | 0 | 1 | 1 |
| Olfr1331 | 0 | 0 | 0 | 1 | 1 |
| Olfr1333 | 0 | 0 | 0 | 1 | 1 |
| Olfr1335 | 0 | 0 | 0 | 1 | 1 |
| Olfr1336 | 0 | 0 | 0 | 1 | 1 |
| Olfr1337 | 0 | 0 | 0 | 1 | 1 |
| Olfr1338 | 0 | 0 | 0 | 1 | 1 |
| Olfr1339 | 0 | 0 | 0 | 1 | 1 |
| Olfr134 | 0 | 0 | 0 | 1 | 1 |
| Olfr1340 | 0 | 0 | 0 | 1 | 1 |
| Olfr1341 | 0 | 0 | 0 | 1 | 1 |
| Olfr1342 | 0 | 0 | 0 | 1 | 1 |
| Olfr1344 | 0 | 0 | 0 | 1 | 1 |
| Olfr1346 | 0 | 0 | 0 | 1 | 1 |
| Olfr1348 | 0 | 0 | 0 | 1 | 1 |
| Olfr135 | 0 | 0 | 0 | 1 | 1 |
| Olfr1350 | 0 | 0 | 0 | 1 | 1 |
| Olfr1351 | 0 | 0 | 0 | 1 | 1 |
| Olfr1352 | 0 | 0 | 0 | 1 | 1 |
| Olfr1353 | 0 | 0 | 0 | 1 | 1 |
| Olfr1354 | 0 | 0 | 0 | 1 | 1 |
| Olfr1355 | 0 | 0 | 0 | 1 | 1 |
| Olfr1356 | 0 | 0 | 0 | 1 | 1 |
| Olfr1357 | 0 | 0 | 0 | 1 | 1 |
| Olfr1359 | 0 | 0 | 0 | 1 | 1 |
| Olfr136 | 0 | 0 | 0 | 1 | 1 |
| Olfr1360 | 0 | 0 | 0 | 1 | 1 |
| Olfr1361 | 0 | 0 | 0 | 1 | 1 |
| Olfr1364 | 0 | 0 | 0 | 1 | 1 |
| Olfr1367 | 0 | 0 | 0 | 1 | 1 |
| Olfr1368 | 0 | 0 | 0 | 1 | 1 |
| Olfr137 | 0 | 0 | 0 | 1 | 1 |
| Olfr1370 | 0 | 0 | 0 | 1 | 1 |
| Olfr1371 | 0 | 0 | 0 | 1 | 1 |
| Olfr1373 | 0 | 0 | 0 | 1 | 1 |
| Olfr1377 | 0 | 0 | 0 | 1 | 1 |
| Olfr138 | 0 | 0 | 0 | 1 | 1 |
| Olfr1380 | 0 | 0 | 0 | 1 | 1 |
| Olfr1381 | 0 | 0 | 0 | 1 | 1 |
| Olfr1382 | 0 | 0 | 0 | 1 | 1 |
| Olfr1383 | 0 | 0 | 0 | 1 | 1 |
| Olfr1384 | 0 | 0 | 0 | 1 | 1 |
| Olfr1385 | 0 | 0 | 0 | 1 | 1 |

| | | | | | |
|---|---|---|---|---|---|
| Olfr1386 | 0 | 0 | 0 | 1 | 1 |
| Olfr1387 | 0 | 0 | 0 | 1 | 1 |
| Olfr1388 | 0 | 0 | 0 | 1 | 1 |
| Olfr1389 | 0 | 0 | 0 | 1 | 1 |
| Olfr139 | 0 | 0 | 0 | 1 | 1 |
| Olfr1390 | 0 | 0 | 0 | 1 | 1 |
| Olfr1391 | 0 | 0 | 0 | 1 | 1 |
| Olfr1394 | 0 | 0 | 0 | 1 | 1 |
| Olfr1395 | 0 | 0 | 0 | 1 | 1 |
| Olfr140 | 0 | 0 | 0 | 1 | 1 |
| Olfr1402 | 0 | 0 | 0 | 1 | 1 |
| Olfr1404 | 0 | 0 | 0 | 1 | 1 |
| Olfr1406 | 0 | 0 | 0 | 1 | 1 |
| Olfr1408 | 0 | 0 | 0 | 1 | 1 |
| Olfr141 | 0 | 0 | 0 | 1 | 1 |
| Olfr1410 | 0 | 0 | 0 | 1 | 1 |
| Olfr1411 | 0 | 0 | 0 | 1 | 1 |
| Olfr1412 | 0 | 0 | 0 | 1 | 1 |
| Olfr1413 | 0 | 0 | 0 | 1 | 1 |
| Olfr1415 | 0 | 0 | 0 | 1 | 1 |
| Olfr1416 | 0 | 0 | 0 | 1 | 1 |
| Olfr1418 | 0 | 0 | 0 | 1 | 1 |
| Olfr1419 | 0 | 0 | 0 | 1 | 1 |
| Olfr142 | 0 | 0 | 0 | 1 | 1 |
| Olfr1423 | 0 | 0 | 0 | 1 | 1 |
| Olfr1424 | 0 | 0 | 0 | 1 | 1 |
| Olfr1425 | 0 | 0 | 0 | 1 | 1 |
| Olfr1426 | 0 | 0 | 0 | 1 | 1 |
| Olfr1427 | 0 | 0 | 0 | 1 | 1 |
| Olfr1428 | 0 | 0 | 0 | 1 | 1 |
| Olfr143 | 0 | 0 | 0 | 1 | 1 |
| Olfr1431 | 0 | 0 | 0 | 1 | 1 |
| Olfr1433,Olfr1434 | 0 | 0 | 0 | 1 | 1 |
| Olfr1434 | 0 | 0 | 0 | 1 | 1 |
| Olfr1436 | 0 | 0 | 0 | 1 | 1 |
| Olfr1437 | 0 | 0 | 0 | 1 | 1 |
| Olfr1440 | 0 | 0 | 0 | 1 | 1 |
| Olfr1441 | 0 | 0 | 0 | 1 | 1 |
| Olfr1442 | 0 | 0 | 0 | 1 | 1 |
| Olfr1443 | 0 | 0 | 0 | 1 | 1 |
| Olfr1444 | 0 | 0 | 0 | 1 | 1 |
| Olfr1445 | 0 | 0 | 0 | 1 | 1 |
| Olfr1446 | 0 | 0 | 0 | 1 | 1 |
| Olfr1447 | 0 | 0 | 0 | 1 | 1 |
| Olfr1448 | 0 | 0 | 0 | 1 | 1 |
| Olfr1449 | 0 | 0 | 0 | 1 | 1 |
| Olfr145 | 0 | 0 | 0 | 1 | 1 |
| Olfr1450 | 0 | 0 | 0 | 1 | 1 |

| | | | | | |
|---|---|---|---|---|---|
| Olfr1451 | 0 | 0 | 0 | 1 | 1 |
| Olfr1453 | 0 | 0 | 0 | 1 | 1 |
| Olfr1454 | 0 | 0 | 0 | 1 | 1 |
| Olfr1457 | 0 | 0 | 0 | 1 | 1 |
| Olfr1459 | 0 | 0 | 0 | 1 | 1 |
| Olfr146 | 0 | 0 | 0 | 1 | 1 |
| Olfr1461 | 0 | 0 | 0 | 1 | 1 |
| Olfr1462 | 0 | 0 | 0 | 1 | 1 |
| Olfr1463 | 0 | 0 | 0 | 1 | 1 |
| Olfr1465 | 0 | 0 | 0 | 1 | 1 |
| Olfr1466 | 0 | 0 | 0 | 1 | 1 |
| Olfr1467 | 0 | 0 | 0 | 1 | 1 |
| Olfr1469 | 0 | 0 | 0 | 1 | 1 |
| Olfr147 | 0 | 0 | 0 | 1 | 1 |
| Olfr1471 | 0 | 0 | 0 | 1 | 1 |
| Olfr1472 | 0 | 0 | 0 | 1 | 1 |
| Olfr1475 | 0 | 0 | 0 | 1 | 1 |
| Olfr1477 | 0 | 0 | 0 | 1 | 1 |
| Olfr148 | 0 | 0 | 0 | 1 | 1 |
| Olfr1480 | 0 | 0 | 0 | 1 | 1 |
| Olfr1484 | 0 | 0 | 0 | 1 | 1 |
| Olfr1487 | 0 | 0 | 0 | 1 | 1 |
| Olfr1489 | 0 | 0 | 0 | 1 | 1 |
| Olfr149 | 0 | 0 | 0 | 1 | 1 |
| Olfr1490 | 0 | 0 | 0 | 1 | 1 |
| Olfr1491 | 0 | 0 | 0 | 1 | 1 |
| Olfr1494 | 0 | 0 | 0 | 1 | 1 |
| Olfr1495 | 0 | 0 | 0 | 1 | 1 |
| Olfr1496 | 0 | 0 | 0 | 1 | 1 |
| Olfr1497 | 0 | 0 | 0 | 1 | 1 |
| Olfr1499 | 0 | 0 | 0 | 1 | 1 |
| Olfr15 | 0 | 0 | 0 | 1 | 1 |
| Olfr150 | 0 | 0 | 0 | 1 | 1 |
| Olfr1500 | 0 | 0 | 0 | 1 | 1 |
| Olfr1501 | 0 | 0 | 0 | 1 | 1 |
| Olfr1502 | 0 | 0 | 0 | 1 | 1 |
| Olfr1504 | 0 | 0 | 0 | 1 | 1 |
| Olfr1505 | 0 | 0 | 0 | 1 | 1 |
| Olfr1507 | 0 | 0 | 0 | 1 | 1 |
| Olfr1508 | 0 | 0 | 0 | 1 | 1 |
| Olfr1509 | 0 | 0 | 0 | 1 | 1 |
| Olfr151 | 0 | 0 | 0 | 1 | 1 |
| Olfr1510 | 0 | 0 | 0 | 1 | 1 |
| Olfr1511 | 0 | 0 | 0 | 1 | 1 |
| Olfr1512 | 0 | 0 | 0 | 1 | 1 |
| Olfr1513 | 0 | 0 | 0 | 1 | 1 |
| Olfr152 | 0 | 0 | 0 | 1 | 1 |
| Olfr153 | 0 | 0 | 0 | 1 | 1 |

| | | | | | |
|---|---|---|---|---|---|
| Olfr1532-ps1 | 0 | 0 | 0 | 1 | 1 |
| Olfr1537 | 0 | 0 | 0 | 1 | 1 |
| Olfr154 | 0 | 0 | 0 | 1 | 1 |
| Olfr155 | 0 | 0 | 0 | 1 | 1 |
| Olfr156 | 0 | 0 | 0 | 1 | 1 |
| Olfr157 | 0 | 0 | 0 | 1 | 1 |
| Olfr159 | 0 | 0 | 0 | 1 | 1 |
| Olfr16 | 0 | 0 | 0 | 1 | 1 |
| Olfr160 | 0 | 0 | 0 | 1 | 1 |
| Olfr161 | 0 | 0 | 0 | 1 | 1 |
| Olfr164 | 0 | 0 | 0 | 1 | 1 |
| Olfr165 | 0 | 0 | 0 | 1 | 1 |
| Olfr166 | 0 | 0 | 0 | 1 | 1 |
| Olfr167 | 0 | 0 | 0 | 1 | 1 |
| Olfr168 | 0 | 0 | 0 | 1 | 1 |
| Olfr169 | 0 | 0 | 0 | 1 | 1 |
| Olfr17 | 0 | 0 | 0 | 1 | 1 |
| Olfr170 | 0 | 0 | 0 | 1 | 1 |
| Olfr171 | 0 | 0 | 0 | 1 | 1 |
| Olfr172 | 0 | 0 | 0 | 1 | 1 |
| Olfr173 | 0 | 0 | 0 | 1 | 1 |
| Olfr175-ps1 | 0 | 0 | 0 | 1 | 1 |
| Olfr176 | 0 | 0 | 0 | 1 | 1 |
| Olfr177 | 0 | 0 | 0 | 1 | 1 |
| Olfr178 | 0 | 0 | 0 | 1 | 1 |
| Olfr18 | 0 | 0 | 0 | 1 | 1 |
| Olfr180 | 0 | 0 | 0 | 1 | 1 |
| Olfr181 | 0 | 0 | 0 | 1 | 1 |
| Olfr183 | 0 | 0 | 0 | 1 | 1 |
| Olfr186 | 0 | 0 | 0 | 1 | 1 |
| Olfr187 | 0 | 0 | 0 | 1 | 1 |
| Olfr19 | 0 | 0 | 0 | 1 | 1 |
| Olfr190 | 0 | 0 | 0 | 1 | 1 |
| Olfr191 | 0 | 0 | 0 | 1 | 1 |
| Olfr192 | 0 | 0 | 0 | 1 | 1 |
| Olfr193 | 0 | 0 | 0 | 1 | 1 |
| Olfr194 | 0 | 0 | 0 | 1 | 1 |
| Olfr195 | 0 | 0 | 0 | 1 | 1 |
| Olfr196 | 0 | 0 | 0 | 1 | 1 |
| Olfr197 | 0 | 0 | 0 | 1 | 1 |
| Olfr198 | 0 | 0 | 0 | 1 | 1 |
| Olfr199 | 0 | 0 | 0 | 1 | 1 |
| Olfr2 | 0 | 0 | 0 | 1 | 1 |
| Olfr20 | 0 | 0 | 0 | 1 | 1 |
| Olfr201 | 0 | 0 | 0 | 1 | 1 |
| Olfr202 | 0 | 0 | 0 | 1 | 1 |
| Olfr203 | 0 | 0 | 0 | 1 | 1 |
| Olfr204 | 0 | 0 | 0 | 1 | 1 |

| | | | | | |
|---|---|---|---|---|---|
| Olfr205 | 0 | 0 | 0 | 1 | 1 |
| Olfr206 | 0 | 0 | 0 | 1 | 1 |
| Olfr207 | 0 | 0 | 0 | 1 | 1 |
| Olfr211 | 0 | 0 | 0 | 1 | 1 |
| Olfr212 | 0 | 0 | 0 | 1 | 1 |
| Olfr213 | 0 | 0 | 0 | 1 | 1 |
| Olfr214 | 0 | 0 | 0 | 1 | 1 |
| Olfr215 | 0 | 0 | 0 | 1 | 1 |
| Olfr220 | 0 | 0 | 0 | 1 | 1 |
| Olfr221 | 0 | 0 | 0 | 1 | 1 |
| Olfr222 | 0 | 0 | 0 | 1 | 1 |
| Olfr223 | 0 | 0 | 0 | 1 | 1 |
| Olfr224 | 0 | 0 | 0 | 1 | 1 |
| Olfr225 | 0 | 0 | 0 | 1 | 1 |
| Olfr228 | 0 | 0 | 0 | 1 | 1 |
| Olfr229 | 0 | 0 | 0 | 1 | 1 |
| Olfr23 | 0 | 0 | 0 | 1 | 1 |
| Olfr231 | 0 | 0 | 0 | 1 | 1 |
| Olfr235 | 0 | 0 | 0 | 1 | 1 |
| Olfr237-ps1 | 0 | 0 | 0 | 1 | 1 |
| Olfr239 | 0 | 0 | 0 | 1 | 1 |
| Olfr24 | 0 | 0 | 0 | 1 | 1 |
| Olfr242 | 0 | 0 | 0 | 1 | 1 |
| Olfr243 | 0 | 0 | 0 | 1 | 1 |
| Olfr247 | 0 | 0 | 0 | 1 | 1 |
| Olfr248 | 0 | 0 | 0 | 1 | 1 |
| Olfr25 | 0 | 0 | 0 | 1 | 1 |
| Olfr259 | 0 | 0 | 0 | 1 | 1 |
| Olfr26 | 0 | 0 | 0 | 1 | 1 |
| Olfr262 | 0 | 0 | 0 | 1 | 1 |
| Olfr263 | 0 | 0 | 0 | 1 | 1 |
| Olfr266 | 0 | 0 | 0 | 1 | 1 |
| Olfr27 | 0 | 0 | 0 | 1 | 1 |
| Olfr270 | 0 | 0 | 0 | 1 | 1 |
| Olfr272 | 0 | 0 | 0 | 1 | 1 |
| Olfr273 | 0 | 0 | 0 | 1 | 1 |
| Olfr275 | 0 | 0 | 0 | 1 | 1 |
| Olfr279 | 0 | 0 | 0 | 1 | 1 |
| Olfr281 | 0 | 0 | 0 | 1 | 1 |
| Olfr282 | 0 | 0 | 0 | 1 | 1 |
| Olfr283 | 0 | 0 | 0 | 1 | 1 |
| Olfr284 | 0 | 0 | 0 | 1 | 1 |
| Olfr285 | 0 | 0 | 0 | 1 | 1 |
| Olfr286 | 0 | 0 | 0 | 1 | 1 |
| Olfr287 | 0 | 0 | 0 | 1 | 1 |
| Olfr288 | 0 | 0 | 0 | 1 | 1 |
| Olfr29-ps1 | 0 | 0 | 0 | 1 | 1 |
| Olfr290 | 0 | 0 | 0 | 1 | 1 |

| | | | | | |
|---|---|---|---|---|---|
| Olfr291 | 0 | 0 | 0 | 1 | 1 |
| Olfr292 | 0 | 0 | 0 | 1 | 1 |
| Olfr293 | 0 | 0 | 0 | 1 | 1 |
| Olfr294 | 0 | 0 | 0 | 1 | 1 |
| Olfr295 | 0 | 0 | 0 | 1 | 1 |
| Olfr297 | 0 | 0 | 0 | 1 | 1 |
| Olfr298 | 0 | 0 | 0 | 1 | 1 |
| Olfr299 | 0 | 0 | 0 | 1 | 1 |
| Olfr3 | 0 | 0 | 0 | 1 | 1 |
| Olfr30 | 0 | 0 | 0 | 1 | 1 |
| Olfr301 | 0 | 0 | 0 | 1 | 1 |
| Olfr303 | 0 | 0 | 0 | 1 | 1 |
| Olfr304 | 0 | 0 | 0 | 1 | 1 |
| Olfr305 | 0 | 0 | 0 | 1 | 1 |
| Olfr307 | 0 | 0 | 0 | 1 | 1 |
| Olfr308 | 0 | 0 | 0 | 1 | 1 |
| Olfr309 | 0 | 0 | 0 | 1 | 1 |
| Olfr310 | 0 | 0 | 0 | 1 | 1 |
| Olfr311 | 0 | 0 | 0 | 1 | 1 |
| Olfr312 | 0 | 0 | 0 | 1 | 1 |
| Olfr313 | 0 | 0 | 0 | 1 | 1 |
| Olfr314 | 0 | 0 | 0 | 1 | 1 |
| Olfr315 | 0 | 0 | 0 | 1 | 1 |
| Olfr316 | 0 | 0 | 0 | 1 | 1 |
| Olfr317 | 0 | 0 | 0 | 1 | 1 |
| Olfr318 | 0 | 0 | 0 | 1 | 1 |
| Olfr319 | 0 | 0 | 0 | 1 | 1 |
| Olfr32 | 0 | 0 | 0 | 1 | 1 |
| Olfr320 | 0 | 0 | 0 | 1 | 1 |
| Olfr322 | 0 | 0 | 0 | 1 | 1 |
| Olfr323 | 0 | 0 | 0 | 1 | 1 |
| Olfr324 | 0 | 0 | 0 | 1 | 1 |
| Olfr325 | 0 | 0 | 0 | 1 | 1 |
| Olfr328 | 0 | 0 | 0 | 1 | 1 |
| Olfr33 | 0 | 0 | 0 | 1 | 1 |
| Olfr331 | 0 | 0 | 0 | 1 | 1 |
| Olfr332 | 0 | 0 | 0 | 1 | 1 |
| Olfr338 | 0 | 0 | 0 | 1 | 1 |
| Olfr339 | 0 | 0 | 0 | 1 | 1 |
| Olfr340 | 0 | 0 | 0 | 1 | 1 |
| Olfr341 | 0 | 0 | 0 | 1 | 1 |
| Olfr342 | 0 | 0 | 0 | 1 | 1 |
| Olfr344 | 0 | 0 | 0 | 1 | 1 |
| Olfr345 | 0 | 0 | 0 | 1 | 1 |
| Olfr346 | 0 | 0 | 0 | 1 | 1 |
| Olfr347 | 0 | 0 | 0 | 1 | 1 |
| Olfr348 | 0 | 0 | 0 | 1 | 1 |
| Olfr350 | 0 | 0 | 0 | 1 | 1 |

| | | | | | |
|---|---|---|---|---|---|
| Olfr351 | 0 | 0 | 0 | 1 | 1 |
| Olfr352 | 0 | 0 | 0 | 1 | 1 |
| Olfr353 | 0 | 0 | 0 | 1 | 1 |
| Olfr354 | 0 | 0 | 0 | 1 | 1 |
| Olfr355 | 0 | 0 | 0 | 1 | 1 |
| Olfr356 | 0 | 0 | 0 | 1 | 1 |
| Olfr357 | 0 | 0 | 0 | 1 | 1 |
| Olfr358 | 0 | 0 | 0 | 1 | 1 |
| Olfr360 | 0 | 0 | 0 | 1 | 1 |
| Olfr361 | 0 | 0 | 0 | 1 | 1 |
| Olfr362 | 0 | 0 | 0 | 1 | 1 |
| Olfr365 | 0 | 0 | 0 | 1 | 1 |
| Olfr366 | 0 | 0 | 0 | 1 | 1 |
| Olfr367-ps | 0 | 0 | 0 | 1 | 1 |
| Olfr368 | 0 | 0 | 0 | 1 | 1 |
| Olfr371 | 0 | 0 | 0 | 1 | 1 |
| Olfr372 | 0 | 0 | 0 | 1 | 1 |
| Olfr373 | 0 | 0 | 0 | 1 | 1 |
| Olfr376 | 0 | 0 | 0 | 1 | 1 |
| Olfr378 | 0 | 0 | 0 | 1 | 1 |
| Olfr380 | 0 | 0 | 0 | 1 | 1 |
| Olfr381 | 0 | 0 | 0 | 1 | 1 |
| Olfr382 | 0 | 0 | 0 | 1 | 1 |
| Olfr384 | 0 | 0 | 0 | 1 | 1 |
| Olfr385 | 0 | 0 | 0 | 1 | 1 |
| Olfr389 | 0 | 0 | 0 | 1 | 1 |
| Olfr39 | 0 | 0 | 0 | 1 | 1 |
| Olfr390 | 0 | 0 | 0 | 1 | 1 |
| Olfr391-ps | 0 | 0 | 0 | 1 | 1 |
| Olfr392 | 0 | 0 | 0 | 1 | 1 |
| Olfr393 | 0 | 0 | 0 | 1 | 1 |
| Olfr394 | 0 | 0 | 0 | 1 | 1 |
| Olfr395 | 0 | 0 | 0 | 1 | 1 |
| Olfr397 | 0 | 0 | 0 | 1 | 1 |
| Olfr398 | 0 | 0 | 0 | 1 | 1 |
| Olfr399 | 0 | 0 | 0 | 1 | 1 |
| Olfr401 | 0 | 0 | 0 | 1 | 1 |
| Olfr402 | 0 | 0 | 0 | 1 | 1 |
| Olfr403 | 0 | 0 | 0 | 1 | 1 |
| Olfr406 | 0 | 0 | 0 | 1 | 1 |
| Olfr410 | 0 | 0 | 0 | 1 | 1 |
| Olfr411 | 0 | 0 | 0 | 1 | 1 |
| Olfr414 | 0 | 0 | 0 | 1 | 1 |
| Olfr417 | 0 | 0 | 0 | 1 | 1 |
| Olfr418-ps1 | 0 | 0 | 0 | 1 | 1 |
| Olfr424 | 0 | 0 | 0 | 1 | 1 |
| Olfr426 | 0 | 0 | 0 | 1 | 1 |
| Olfr427 | 0 | 0 | 0 | 1 | 1 |

| | | | | | |
|---|---|---|---|---|---|
| Olfr429 | 0 | 0 | 0 | 1 | 1 |
| Olfr43 | 0 | 0 | 0 | 1 | 1 |
| Olfr430 | 0 | 0 | 0 | 1 | 1 |
| Olfr432 | 0 | 0 | 0 | 1 | 1 |
| Olfr433 | 0 | 0 | 0 | 1 | 1 |
| Olfr434 | 0 | 0 | 0 | 1 | 1 |
| Olfr435 | 0 | 0 | 0 | 1 | 1 |
| Olfr437 | 0 | 0 | 0 | 1 | 1 |
| Olfr44 | 0 | 0 | 0 | 1 | 1 |
| Olfr441 | 0 | 0 | 0 | 1 | 1 |
| Olfr446 | 0 | 0 | 0 | 1 | 1 |
| Olfr447 | 0 | 0 | 0 | 1 | 1 |
| Olfr448 | 0 | 0 | 0 | 1 | 1 |
| Olfr449 | 0 | 0 | 0 | 1 | 1 |
| Olfr45 | 0 | 0 | 0 | 1 | 1 |
| Olfr450 | 0 | 0 | 0 | 1 | 1 |
| Olfr452 | 0 | 0 | 0 | 1 | 1 |
| Olfr453 | 0 | 0 | 0 | 1 | 1 |
| Olfr455 | 0 | 0 | 0 | 1 | 1 |
| Olfr458 | 0 | 0 | 0 | 1 | 1 |
| Olfr459 | 0 | 0 | 0 | 1 | 1 |
| Olfr46 | 0 | 0 | 0 | 1 | 1 |
| Olfr462 | 0 | 0 | 0 | 1 | 1 |
| Olfr464 | 0 | 0 | 0 | 1 | 1 |
| Olfr466 | 0 | 0 | 0 | 1 | 1 |
| Olfr467 | 0 | 0 | 0 | 1 | 1 |
| Olfr47 | 0 | 0 | 0 | 1 | 1 |
| Olfr470 | 0 | 0 | 0 | 1 | 1 |
| Olfr472 | 0 | 0 | 0 | 1 | 1 |
| Olfr473 | 0 | 0 | 0 | 1 | 1 |
| Olfr474 | 0 | 0 | 0 | 1 | 1 |
| Olfr476 | 0 | 0 | 0 | 1 | 1 |
| Olfr477 | 0 | 0 | 0 | 1 | 1 |
| Olfr478 | 0 | 0 | 0 | 1 | 1 |
| Olfr479 | 0 | 0 | 0 | 1 | 1 |
| Olfr48 | 0 | 0 | 0 | 1 | 1 |
| Olfr480 | 0 | 0 | 0 | 1 | 1 |
| Olfr481 | 0 | 0 | 0 | 1 | 1 |
| Olfr482 | 0 | 0 | 0 | 1 | 1 |
| Olfr483 | 0 | 0 | 0 | 1 | 1 |
| Olfr484 | 0 | 0 | 0 | 1 | 1 |
| Olfr485 | 0 | 0 | 0 | 1 | 1 |
| Olfr486 | 0 | 0 | 0 | 1 | 1 |
| Olfr487 | 0 | 0 | 0 | 1 | 1 |
| Olfr488 | 0 | 0 | 0 | 1 | 1 |
| Olfr49 | 0 | 0 | 0 | 1 | 1 |
| Olfr490 | 0 | 0 | 0 | 1 | 1 |
| Olfr491 | 0 | 0 | 0 | 1 | 1 |

| | | | | | |
|---|---|---|---|---|---|
| Olfr492 | 0 | 0 | 0 | 1 | 1 |
| Olfr493 | 0 | 0 | 0 | 1 | 1 |
| Olfr494 | 0 | 0 | 0 | 1 | 1 |
| Olfr495 | 0 | 0 | 0 | 1 | 1 |
| Olfr497 | 0 | 0 | 0 | 1 | 1 |
| Olfr498 | 0 | 0 | 0 | 1 | 1 |
| Olfr50 | 0 | 0 | 0 | 1 | 1 |
| Olfr502 | 0 | 0 | 0 | 1 | 1 |
| Olfr503 | 0 | 0 | 0 | 1 | 1 |
| Olfr504 | 0 | 0 | 0 | 1 | 1 |
| Olfr506 | 0 | 0 | 0 | 1 | 1 |
| Olfr507 | 0 | 0 | 0 | 1 | 1 |
| Olfr508 | 0 | 0 | 0 | 1 | 1 |
| Olfr509 | 0 | 0 | 0 | 1 | 1 |
| Olfr51 | 0 | 0 | 0 | 1 | 1 |
| Olfr510 | 0 | 0 | 0 | 1 | 1 |
| Olfr512 | 0 | 0 | 0 | 1 | 1 |
| Olfr513 | 0 | 0 | 0 | 1 | 1 |
| Olfr514 | 0 | 0 | 0 | 1 | 1 |
| Olfr516 | 0 | 0 | 0 | 1 | 1 |
| Olfr517 | 0 | 0 | 0 | 1 | 1 |
| Olfr518 | 0 | 0 | 0 | 1 | 1 |
| Olfr519 | 0 | 0 | 0 | 1 | 1 |
| Olfr52 | 0 | 0 | 0 | 1 | 1 |
| Olfr520 | 0 | 0 | 0 | 1 | 1 |
| Olfr522 | 0 | 0 | 0 | 1 | 1 |
| Olfr523 | 0 | 0 | 0 | 1 | 1 |
| Olfr524 | 0 | 0 | 0 | 1 | 1 |
| Olfr525 | 0 | 0 | 0 | 1 | 1 |
| Olfr527 | 0 | 0 | 0 | 1 | 1 |
| Olfr53 | 0 | 0 | 0 | 1 | 1 |
| Olfr530 | 0 | 0 | 0 | 1 | 1 |
| Olfr531 | 0 | 0 | 0 | 1 | 1 |
| Olfr532 | 0 | 0 | 0 | 1 | 1 |
| Olfr533 | 0 | 0 | 0 | 1 | 1 |
| Olfr535 | 0 | 0 | 0 | 1 | 1 |
| Olfr536 | 0 | 0 | 0 | 1 | 1 |
| Olfr538 | 0 | 0 | 0 | 1 | 1 |
| Olfr539 | 0 | 0 | 0 | 1 | 1 |
| Olfr54 | 0 | 0 | 0 | 1 | 1 |
| Olfr541 | 0 | 0 | 0 | 1 | 1 |
| Olfr547 | 0 | 0 | 0 | 1 | 1 |
| Olfr549 | 0 | 0 | 0 | 1 | 1 |
| Olfr55 | 0 | 0 | 0 | 1 | 1 |
| Olfr550 | 0 | 0 | 0 | 1 | 1 |
| Olfr551 | 0 | 0 | 0 | 1 | 1 |
| Olfr552 | 0 | 0 | 0 | 1 | 1 |
| Olfr553 | 0 | 0 | 0 | 1 | 1 |

| | | | | | |
|---|---|---|---|---|---|
| Olfr554 | 0 | 0 | 0 | 1 | 1 |
| Olfr555 | 0 | 0 | 0 | 1 | 1 |
| Olfr556 | 0 | 0 | 0 | 1 | 1 |
| Olfr557 | 0 | 0 | 0 | 1 | 1 |
| Olfr559 | 0 | 0 | 0 | 1 | 1 |
| Olfr56 | 0 | 0 | 0 | 1 | 1 |
| Olfr560 | 0 | 0 | 0 | 1 | 1 |
| Olfr561 | 0 | 0 | 0 | 1 | 1 |
| Olfr564 | 0 | 0 | 0 | 1 | 1 |
| Olfr566 | 0 | 0 | 0 | 1 | 1 |
| Olfr568 | 0 | 0 | 0 | 1 | 1 |
| Olfr569 | 0 | 0 | 0 | 1 | 1 |
| Olfr57 | 0 | 0 | 0 | 1 | 1 |
| Olfr570 | 0 | 0 | 0 | 1 | 1 |
| Olfr571 | 0 | 0 | 0 | 1 | 1 |
| Olfr572 | 0 | 0 | 0 | 1 | 1 |
| Olfr574 | 0 | 0 | 0 | 1 | 1 |
| Olfr575 | 0 | 0 | 0 | 1 | 1 |
| Olfr576 | 0 | 0 | 0 | 1 | 1 |
| Olfr577 | 0 | 0 | 0 | 1 | 1 |
| Olfr578 | 0 | 0 | 0 | 1 | 1 |
| Olfr58 | 0 | 0 | 0 | 1 | 1 |
| Olfr582 | 0 | 0 | 0 | 1 | 1 |
| Olfr583 | 0 | 0 | 0 | 1 | 1 |
| Olfr584 | 0 | 0 | 0 | 1 | 1 |
| Olfr585 | 0 | 0 | 0 | 1 | 1 |
| Olfr586 | 0 | 0 | 0 | 1 | 1 |
| Olfr589 | 0 | 0 | 0 | 1 | 1 |
| Olfr59 | 0 | 0 | 0 | 1 | 1 |
| Olfr591 | 0 | 0 | 0 | 1 | 1 |
| Olfr592 | 0 | 0 | 0 | 1 | 1 |
| Olfr593 | 0 | 0 | 0 | 1 | 1 |
| Olfr594 | 0 | 0 | 0 | 1 | 1 |
| Olfr596 | 0 | 0 | 0 | 1 | 1 |
| Olfr597 | 0 | 0 | 0 | 1 | 1 |
| Olfr598 | 0 | 0 | 0 | 1 | 1 |
| Olfr599 | 0 | 0 | 0 | 1 | 1 |
| Olfr6 | 0 | 0 | 0 | 1 | 1 |
| Olfr60 | 0 | 0 | 0 | 1 | 1 |
| Olfr600 | 0 | 0 | 0 | 1 | 1 |
| Olfr601 | 0 | 0 | 0 | 1 | 1 |
| Olfr603 | 0 | 0 | 0 | 1 | 1 |
| Olfr605 | 0 | 0 | 0 | 1 | 1 |
| Olfr606 | 0 | 0 | 0 | 1 | 1 |
| Olfr608 | 0 | 0 | 0 | 1 | 1 |
| Olfr609 | 0 | 0 | 0 | 1 | 1 |
| Olfr610 | 0 | 0 | 0 | 1 | 1 |
| Olfr611 | 0 | 0 | 0 | 1 | 1 |

| | | | | | |
|---|---|---|---|---|---|
| Olfr612 | 0 | 0 | 0 | 1 | 1 |
| Olfr613 | 0 | 0 | 0 | 1 | 1 |
| Olfr615 | 0 | 0 | 0 | 1 | 1 |
| Olfr616 | 0 | 0 | 0 | 1 | 1 |
| Olfr617 | 0 | 0 | 0 | 1 | 1 |
| Olfr618 | 0 | 0 | 0 | 1 | 1 |
| Olfr619 | 0 | 0 | 0 | 1 | 1 |
| Olfr62 | 0 | 0 | 0 | 1 | 1 |
| Olfr620 | 0 | 0 | 0 | 1 | 1 |
| Olfr622 | 0 | 0 | 0 | 1 | 1 |
| Olfr623 | 0 | 0 | 0 | 1 | 1 |
| Olfr624 | 0 | 0 | 0 | 1 | 1 |
| Olfr628 | 0 | 0 | 0 | 1 | 1 |
| Olfr629 | 0 | 0 | 0 | 1 | 1 |
| Olfr63 | 0 | 0 | 0 | 1 | 1 |
| Olfr630 | 0 | 0 | 0 | 1 | 1 |
| Olfr633 | 0 | 0 | 0 | 1 | 1 |
| Olfr635 | 0 | 0 | 0 | 1 | 1 |
| Olfr638 | 0 | 0 | 0 | 1 | 1 |
| Olfr639 | 0 | 0 | 0 | 1 | 1 |
| Olfr64 | 0 | 0 | 0 | 1 | 1 |
| Olfr640 | 0 | 0 | 0 | 1 | 1 |
| Olfr641 | 0 | 0 | 0 | 1 | 1 |
| Olfr642 | 0 | 0 | 0 | 1 | 1 |
| Olfr643 | 0 | 0 | 0 | 1 | 1 |
| Olfr644 | 0 | 0 | 0 | 1 | 1 |
| Olfr645 | 0 | 0 | 0 | 1 | 1 |
| Olfr648 | 0 | 0 | 0 | 1 | 1 |
| Olfr649 | 0 | 0 | 0 | 1 | 1 |
| Olfr65 | 0 | 0 | 0 | 1 | 1 |
| Olfr651 | 0 | 0 | 0 | 1 | 1 |
| Olfr652 | 0 | 0 | 0 | 1 | 1 |
| Olfr653 | 0 | 0 | 0 | 1 | 1 |
| Olfr654 | 0 | 0 | 0 | 1 | 1 |
| Olfr656 | 0 | 0 | 0 | 1 | 1 |
| Olfr657 | 0 | 0 | 0 | 1 | 1 |
| Olfr659 | 0 | 0 | 0 | 1 | 1 |
| Olfr66 | 0 | 0 | 0 | 1 | 1 |
| Olfr661 | 0 | 0 | 0 | 1 | 1 |
| Olfr663 | 0 | 0 | 0 | 1 | 1 |
| Olfr665 | 0 | 0 | 0 | 1 | 1 |
| Olfr667 | 0 | 0 | 0 | 1 | 1 |
| Olfr668 | 0 | 0 | 0 | 1 | 1 |
| Olfr669 | 0 | 0 | 0 | 1 | 1 |
| Olfr67 | 0 | 0 | 0 | 1 | 1 |
| Olfr670 | 0 | 0 | 0 | 1 | 1 |
| Olfr671 | 0 | 0 | 0 | 1 | 1 |
| Olfr672 | 0 | 0 | 0 | 1 | 1 |

| | | | | | |
|---|---|---|---|---|---|
| Olfr675 | 0 | 0 | 0 | 1 | 1 |
| Olfr676 | 0 | 0 | 0 | 1 | 1 |
| Olfr677 | 0 | 0 | 0 | 1 | 1 |
| Olfr678 | 0 | 0 | 0 | 1 | 1 |
| Olfr679 | 0 | 0 | 0 | 1 | 1 |
| Olfr68 | 0 | 0 | 0 | 1 | 1 |
| Olfr681 | 0 | 0 | 0 | 1 | 1 |
| Olfr683 | 0 | 0 | 0 | 1 | 1 |
| Olfr684 | 0 | 0 | 0 | 1 | 1 |
| Olfr685 | 0 | 0 | 0 | 1 | 1 |
| Olfr686 | 0 | 0 | 0 | 1 | 1 |
| Olfr688 | 0 | 0 | 0 | 1 | 1 |
| Olfr689 | 0 | 0 | 0 | 1 | 1 |
| Olfr69 | 0 | 0 | 0 | 1 | 1 |
| Olfr690 | 0 | 0 | 0 | 1 | 1 |
| Olfr691 | 0 | 0 | 0 | 1 | 1 |
| Olfr692 | 0 | 0 | 0 | 1 | 1 |
| Olfr693 | 0 | 0 | 0 | 1 | 1 |
| Olfr694 | 0 | 0 | 0 | 1 | 1 |
| Olfr695 | 0 | 0 | 0 | 1 | 1 |
| Olfr697 | 0 | 0 | 0 | 1 | 1 |
| Olfr699 | 0 | 0 | 0 | 1 | 1 |
| Olfr700 | 0 | 0 | 0 | 1 | 1 |
| Olfr702 | 0 | 0 | 0 | 1 | 1 |
| Olfr703 | 0 | 0 | 0 | 1 | 1 |
| Olfr704 | 0 | 0 | 0 | 1 | 1 |
| Olfr705 | 0 | 0 | 0 | 1 | 1 |
| Olfr706 | 0 | 0 | 0 | 1 | 1 |
| Olfr707 | 0 | 0 | 0 | 1 | 1 |
| Olfr71 | 0 | 0 | 0 | 1 | 1 |
| Olfr710 | 0 | 0 | 0 | 1 | 1 |
| Olfr711 | 0 | 0 | 0 | 1 | 1 |
| Olfr713 | 0 | 0 | 0 | 1 | 1 |
| Olfr714 | 0 | 0 | 0 | 1 | 1 |
| Olfr715 | 0 | 0 | 0 | 1 | 1 |
| Olfr716 | 0 | 0 | 0 | 1 | 1 |
| Olfr720 | 0 | 0 | 0 | 1 | 1 |
| Olfr722 | 0 | 0 | 0 | 1 | 1 |
| Olfr723 | 0 | 0 | 0 | 1 | 1 |
| Olfr724 | 0 | 0 | 0 | 1 | 1 |
| Olfr728 | 0 | 0 | 0 | 1 | 1 |
| Olfr729 | 0 | 0 | 0 | 1 | 1 |
| Olfr73 | 0 | 0 | 0 | 1 | 1 |
| Olfr730 | 0 | 0 | 0 | 1 | 1 |
| Olfr731 | 0 | 0 | 0 | 1 | 1 |
| Olfr732 | 0 | 0 | 0 | 1 | 1 |
| Olfr733 | 0 | 0 | 0 | 1 | 1 |
| Olfr734 | 0 | 0 | 0 | 1 | 1 |

| | | | | | |
|---|---|---|---|---|---|
| Olfr735 | 0 | 0 | 0 | 1 | 1 |
| Olfr736 | 0 | 0 | 0 | 1 | 1 |
| Olfr738 | 0 | 0 | 0 | 1 | 1 |
| Olfr739 | 0 | 0 | 0 | 1 | 1 |
| Olfr74 | 0 | 0 | 0 | 1 | 1 |
| Olfr740 | 0 | 0 | 0 | 1 | 1 |
| Olfr741 | 0 | 0 | 0 | 1 | 1 |
| Olfr742 | 0 | 0 | 0 | 1 | 1 |
| Olfr743 | 0 | 0 | 0 | 1 | 1 |
| Olfr744 | 0 | 0 | 0 | 1 | 1 |
| Olfr745 | 0 | 0 | 0 | 1 | 1 |
| Olfr746 | 0 | 0 | 0 | 1 | 1 |
| Olfr747 | 0 | 0 | 0 | 1 | 1 |
| Olfr748 | 0 | 0 | 0 | 1 | 1 |
| Olfr749 | 0 | 0 | 0 | 1 | 1 |
| Olfr75-ps1 | 0 | 0 | 0 | 1 | 1 |
| Olfr76 | 0 | 0 | 0 | 1 | 1 |
| Olfr761 | 0 | 0 | 0 | 1 | 1 |
| Olfr763 | 0 | 0 | 0 | 1 | 1 |
| Olfr765 | 0 | 0 | 0 | 1 | 1 |
| Olfr768 | 0 | 0 | 0 | 1 | 1 |
| Olfr769 | 0 | 0 | 0 | 1 | 1 |
| Olfr77 | 0 | 0 | 0 | 1 | 1 |
| Olfr770 | 0 | 0 | 0 | 1 | 1 |
| Olfr771 | 0 | 0 | 0 | 1 | 1 |
| Olfr772 | 0 | 0 | 0 | 1 | 1 |
| Olfr773 | 0 | 0 | 0 | 1 | 1 |
| Olfr774 | 0 | 0 | 0 | 1 | 1 |
| Olfr775 | 0 | 0 | 0 | 1 | 1 |
| Olfr776 | 0 | 0 | 0 | 1 | 1 |
| Olfr777 | 0 | 0 | 0 | 1 | 1 |
| Olfr78 | 0 | 0 | 0 | 1 | 1 |
| Olfr780 | 0 | 0 | 0 | 1 | 1 |
| Olfr781 | 0 | 0 | 0 | 1 | 1 |
| Olfr782 | 0 | 0 | 0 | 1 | 1 |
| Olfr784 | 0 | 0 | 0 | 1 | 1 |
| Olfr786 | 0 | 0 | 0 | 1 | 1 |
| Olfr787 | 0 | 0 | 0 | 1 | 1 |
| Olfr788 | 0 | 0 | 0 | 1 | 1 |
| Olfr790 | 0 | 0 | 0 | 1 | 1 |
| Olfr791 | 0 | 0 | 0 | 1 | 1 |
| Olfr792 | 0 | 0 | 0 | 1 | 1 |
| Olfr794 | 0 | 0 | 0 | 1 | 1 |
| Olfr796 | 0 | 0 | 0 | 1 | 1 |
| Olfr798 | 0 | 0 | 0 | 1 | 1 |
| Olfr799 | 0 | 0 | 0 | 1 | 1 |
| Olfr8 | 0 | 0 | 0 | 1 | 1 |
| Olfr800 | 0 | 0 | 0 | 1 | 1 |

| | | | | | |
|---|---|---|---|---|---|
| Olfr801 | 0 | 0 | 0 | 1 | 1 |
| Olfr802 | 0 | 0 | 0 | 1 | 1 |
| Olfr803 | 0 | 0 | 0 | 1 | 1 |
| Olfr804 | 0 | 0 | 0 | 1 | 1 |
| Olfr805 | 0 | 0 | 0 | 1 | 1 |
| Olfr806 | 0 | 0 | 0 | 1 | 1 |
| Olfr807 | 0 | 0 | 0 | 1 | 1 |
| Olfr808 | 0 | 0 | 0 | 1 | 1 |
| Olfr809 | 0 | 0 | 0 | 1 | 1 |
| Olfr810 | 0 | 0 | 0 | 1 | 1 |
| Olfr811 | 0 | 0 | 0 | 1 | 1 |
| Olfr812 | 0 | 0 | 0 | 1 | 1 |
| Olfr813 | 0 | 0 | 0 | 1 | 1 |
| Olfr814 | 0 | 0 | 0 | 1 | 1 |
| Olfr815 | 0 | 0 | 0 | 1 | 1 |
| Olfr816 | 0 | 0 | 0 | 1 | 1 |
| Olfr818 | 0 | 0 | 0 | 1 | 1 |
| Olfr819 | 0 | 0 | 0 | 1 | 1 |
| Olfr820 | 0 | 0 | 0 | 1 | 1 |
| Olfr821 | 0 | 0 | 0 | 1 | 1 |
| Olfr822 | 0 | 0 | 0 | 1 | 1 |
| Olfr823 | 0 | 0 | 0 | 1 | 1 |
| Olfr824 | 0 | 0 | 0 | 1 | 1 |
| Olfr825 | 0 | 0 | 0 | 1 | 1 |
| Olfr826 | 0 | 0 | 0 | 1 | 1 |
| Olfr827 | 0 | 0 | 0 | 1 | 1 |
| Olfr828 | 0 | 0 | 0 | 1 | 1 |
| Olfr829 | 0 | 0 | 0 | 1 | 1 |
| Olfr830 | 0 | 0 | 0 | 1 | 1 |
| Olfr832 | 0 | 0 | 0 | 1 | 1 |
| Olfr834 | 0 | 0 | 0 | 1 | 1 |
| Olfr835 | 0 | 0 | 0 | 1 | 1 |
| Olfr836 | 0 | 0 | 0 | 1 | 1 |
| Olfr837 | 0 | 0 | 0 | 1 | 1 |
| Olfr843 | 0 | 0 | 0 | 1 | 1 |
| Olfr845 | 0 | 0 | 0 | 1 | 1 |
| Olfr846 | 0 | 0 | 0 | 1 | 1 |
| Olfr847 | 0 | 0 | 0 | 1 | 1 |
| Olfr849 | 0 | 0 | 0 | 1 | 1 |
| Olfr850 | 0 | 0 | 0 | 1 | 1 |
| Olfr851 | 0 | 0 | 0 | 1 | 1 |
| Olfr853 | 0 | 0 | 0 | 1 | 1 |
| Olfr854 | 0 | 0 | 0 | 1 | 1 |
| Olfr855 | 0 | 0 | 0 | 1 | 1 |
| Olfr857 | 0 | 0 | 0 | 1 | 1 |
| Olfr859 | 0 | 0 | 0 | 1 | 1 |
| Olfr860 | 0 | 0 | 0 | 1 | 1 |
| Olfr862 | 0 | 0 | 0 | 1 | 1 |

| | | | | | |
|---|---|---|---|---|---|
| Olfr866 | 0 | 0 | 0 | 1 | 1 |
| Olfr867 | 0 | 0 | 0 | 1 | 1 |
| Olfr868 | 0 | 0 | 0 | 1 | 1 |
| Olfr869 | 0 | 0 | 0 | 1 | 1 |
| Olfr870 | 0 | 0 | 0 | 1 | 1 |
| Olfr871 | 0 | 0 | 0 | 1 | 1 |
| Olfr872 | 0 | 0 | 0 | 1 | 1 |
| Olfr873 | 0 | 0 | 0 | 1 | 1 |
| Olfr874 | 0 | 0 | 0 | 1 | 1 |
| Olfr877 | 0 | 0 | 0 | 1 | 1 |
| Olfr878 | 0 | 0 | 0 | 1 | 1 |
| Olfr881 | 0 | 0 | 0 | 1 | 1 |
| Olfr883 | 0 | 0 | 0 | 1 | 1 |
| Olfr884 | 0 | 0 | 0 | 1 | 1 |
| Olfr885 | 0 | 0 | 0 | 1 | 1 |
| Olfr887 | 0 | 0 | 0 | 1 | 1 |
| Olfr888 | 0 | 0 | 0 | 1 | 1 |
| Olfr889 | 0 | 0 | 0 | 1 | 1 |
| Olfr890 | 0 | 0 | 0 | 1 | 1 |
| Olfr891 | 0 | 0 | 0 | 1 | 1 |
| Olfr893 | 0 | 0 | 0 | 1 | 1 |
| Olfr894 | 0 | 0 | 0 | 1 | 1 |
| Olfr895 | 0 | 0 | 0 | 1 | 1 |
| Olfr898 | 0 | 0 | 0 | 1 | 1 |
| Olfr899 | 0 | 0 | 0 | 1 | 1 |
| Olfr9 | 0 | 0 | 0 | 1 | 1 |
| Olfr900 | 0 | 0 | 0 | 1 | 1 |
| Olfr901 | 0 | 0 | 0 | 1 | 1 |
| Olfr902 | 0 | 0 | 0 | 1 | 1 |
| Olfr904 | 0 | 0 | 0 | 1 | 1 |
| Olfr905 | 0 | 0 | 0 | 1 | 1 |
| Olfr906 | 0 | 0 | 0 | 1 | 1 |
| Olfr907 | 0 | 0 | 0 | 1 | 1 |
| Olfr908 | 0 | 0 | 0 | 1 | 1 |
| Olfr910 | 0 | 0 | 0 | 1 | 1 |
| Olfr911-ps1 | 0 | 0 | 0 | 1 | 1 |
| Olfr912 | 0 | 0 | 0 | 1 | 1 |
| Olfr913 | 0 | 0 | 0 | 1 | 1 |
| Olfr914 | 0 | 0 | 0 | 1 | 1 |
| Olfr915 | 0 | 0 | 0 | 1 | 1 |
| Olfr916 | 0 | 0 | 0 | 1 | 1 |
| Olfr917 | 0 | 0 | 0 | 1 | 1 |
| Olfr918 | 0 | 0 | 0 | 1 | 1 |
| Olfr919 | 0 | 0 | 0 | 1 | 1 |
| Olfr92 | 0 | 0 | 0 | 1 | 1 |
| Olfr923 | 0 | 0 | 0 | 1 | 1 |
| Olfr924 | 0 | 0 | 0 | 1 | 1 |
| Olfr926 | 0 | 0 | 0 | 1 | 1 |

| | | | | | |
|---|---|---|---|---|---|
| Olfr93 | 0 | 0 | 0 | 1 | 1 |
| Olfr930 | 0 | 0 | 0 | 1 | 1 |
| Olfr933 | 0 | 0 | 0 | 1 | 1 |
| Olfr934 | 0 | 0 | 0 | 1 | 1 |
| Olfr935 | 0 | 0 | 0 | 1 | 1 |
| Olfr936 | 0 | 0 | 0 | 1 | 1 |
| Olfr937 | 0 | 0 | 0 | 1 | 1 |
| Olfr938 | 0 | 0 | 0 | 1 | 1 |
| Olfr943 | 0 | 0 | 0 | 1 | 1 |
| Olfr944 | 0 | 0 | 0 | 1 | 1 |
| Olfr945 | 0 | 0 | 0 | 1 | 1 |
| Olfr947-ps1 | 0 | 0 | 0 | 1 | 1 |
| Olfr948 | 0 | 0 | 0 | 1 | 1 |
| Olfr951 | 0 | 0 | 0 | 1 | 1 |
| Olfr952 | 0 | 0 | 0 | 1 | 1 |
| Olfr954 | 0 | 0 | 0 | 1 | 1 |
| Olfr955 | 0 | 0 | 0 | 1 | 1 |
| Olfr957 | 0 | 0 | 0 | 1 | 1 |
| Olfr958 | 0 | 0 | 0 | 1 | 1 |
| Olfr959 | 0 | 0 | 0 | 1 | 1 |
| Olfr96 | 0 | 0 | 0 | 1 | 1 |
| Olfr961 | 0 | 0 | 0 | 1 | 1 |
| Olfr963 | 0 | 0 | 0 | 1 | 1 |
| Olfr965 | 0 | 0 | 0 | 1 | 1 |
| Olfr967 | 0 | 0 | 0 | 1 | 1 |
| Olfr968 | 0 | 0 | 0 | 1 | 1 |
| Olfr969 | 0 | 0 | 0 | 1 | 1 |
| Olfr97 | 0 | 0 | 0 | 1 | 1 |
| Olfr970 | 0 | 0 | 0 | 1 | 1 |
| Olfr971 | 0 | 0 | 0 | 1 | 1 |
| Olfr972 | 0 | 0 | 0 | 1 | 1 |
| Olfr974 | 0 | 0 | 0 | 1 | 1 |
| Olfr975 | 0 | 0 | 0 | 1 | 1 |
| Olfr976 | 0 | 0 | 0 | 1 | 1 |
| Olfr978 | 0 | 0 | 0 | 1 | 1 |
| Olfr979 | 0 | 0 | 0 | 1 | 1 |
| Olfr98 | 0 | 0 | 0 | 1 | 1 |
| Olfr980 | 0 | 0 | 0 | 1 | 1 |
| Olfr981 | 0 | 0 | 0 | 1 | 1 |
| Olfr982 | 0 | 0 | 0 | 1 | 1 |
| Olfr983 | 0 | 0 | 0 | 1 | 1 |
| Olfr984 | 0 | 0 | 0 | 1 | 1 |
| Olfr985 | 0 | 0 | 0 | 1 | 1 |
| Olfr986 | 0 | 0 | 0 | 1 | 1 |
| Olfr987 | 0 | 0 | 0 | 1 | 1 |
| Olfr988 | 0 | 0 | 0 | 1 | 1 |
| Olfr99 | 0 | 0 | 0 | 1 | 1 |
| Olfr992 | 0 | 0 | 0 | 1 | 1 |

| | | | | | |
|---|---|---|---|---|---|
| Olfr993 | 0 | 0 | 0 | 1 | 1 |
| Olfr994 | 0 | 0 | 0 | 1 | 1 |
| Olfr996 | 0 | 0 | 0 | 1 | 1 |
| Olfr998 | 0 | 0 | 0 | 1 | 1 |
| Omt2a | 0 | 0 | 0 | 1 | 1 |
| Omt2b | 0 | 0 | 0 | 1 | 1 |
| Oog1 | 0 | 0 | 0 | 1 | 1 |
| Oog2 | 0 | 0 | 0 | 1 | 1 |
| Oog3 | 0 | 0 | 0 | 1 | 1 |
| Oog4 | 0 | 0 | 0 | 1 | 1 |
| Oosp1 | 0 | 0 | 0 | 1 | 1 |
| Oosp2 | 0 | 0 | 0 | 1 | 1 |
| Oosp3 | 0 | 0 | 0 | 1 | 1 |
| Opn1mw | 0 | 0 | 0 | 1 | 1 |
| Opn5 | 0 | 0 | 0 | 1 | 1 |
| Oprm1 | 0 | 0 | 0 | 1 | 1 |
| Ostn | 0 | 0 | 0 | 1 | 1 |
| Otol1 | 0 | 0 | 0 | 1 | 1 |
| Otop3 | 0 | 0 | 0 | 1 | 1 |
| Otp | 0 | 0 | 0 | 1 | 1 |
| Otud6a | 0 | 0 | 0 | 1 | 1 |
| Otx2os1 | 0 | 0 | 0 | 1 | 1 |
| Oxgr1 | 0 | 0 | 0 | 1 | 1 |
| Pabpc6 | 0 | 0 | 0 | 1 | 1 |
| Pabpn1l | 0 | 0 | 0 | 1 | 1 |
| Panx3 | 0 | 0 | 0 | 1 | 1 |
| Papl | 0 | 0 | 0 | 1 | 1 |
| Pax6os1 | 0 | 0 | 0 | 1 | 1 |
| Pbsn | 0 | 0 | 0 | 1 | 1 |
| Pcdha1 | 0 | 0 | 0 | 1 | 1 |
| Pcdha12 | 0 | 0 | 0 | 1 | 1 |
| Pcdha3 | 0 | 0 | 0 | 1 | 1 |
| Pcdha4 | 0 | 0 | 0 | 1 | 1 |
| Pcdha4-g | 0 | 0 | 0 | 1 | 1 |
| Pcdha6 | 0 | 0 | 0 | 1 | 1 |
| Pcdha8 | 0 | 0 | 0 | 1 | 1 |
| Pcdhb1 | 0 | 0 | 0 | 1 | 1 |
| Pcdhb13 | 0 | 0 | 0 | 1 | 1 |
| Pcsk2os2 | 0 | 0 | 0 | 1 | 1 |
| Pdc | 0 | 0 | 0 | 1 | 1 |
| Pdcl2 | 0 | 0 | 0 | 1 | 1 |
| Pde6c | 0 | 0 | 0 | 1 | 1 |
| Pdha2 | 0 | 0 | 0 | 1 | 1 |
| Pdilt | 0 | 0 | 0 | 1 | 1 |
| Pea15b | 0 | 0 | 0 | 1 | 1 |
| Pebp4 | 0 | 0 | 0 | 1 | 1 |
| Pet2 | 0 | 0 | 0 | 1 | 1 |
| Pfpl | 0 | 0 | 0 | 1 | 1 |

| | | | | | |
|---|---|---|---|---|---|
| Pgc | 0 | 0 | 0 | 1 | 1 |
| Pgk2 | 0 | 0 | 0 | 1 | 1 |
| Pglyrp4 | 0 | 0 | 0 | 1 | 1 |
| Phf11b | 0 | 0 | 0 | 1 | 1 |
| Phgr1 | 0 | 0 | 0 | 1 | 1 |
| Phox2b | 0 | 0 | 0 | 1 | 1 |
| Pigr | 0 | 0 | 0 | 1 | 1 |
| Pih1d3 | 0 | 0 | 0 | 1 | 1 |
| Pinlyp | 0 | 0 | 0 | 1 | 1 |
| Pip | 0 | 0 | 0 | 1 | 1 |
| Pira1 | 0 | 0 | 0 | 1 | 1 |
| Pira11 | 0 | 0 | 0 | 1 | 1 |
| Pira2 | 0 | 0 | 0 | 1 | 1 |
| Pira4,Pira7 | 0 | 0 | 0 | 1 | 1 |
| Pira6 | 0 | 0 | 0 | 1 | 1 |
| Pira7 | 0 | 0 | 0 | 1 | 1 |
| Pisd-ps3 | 0 | 0 | 0 | 1 | 1 |
| Pla2g10os | 0 | 0 | 0 | 1 | 1 |
| Pla2g2a | 0 | 0 | 0 | 1 | 1 |
| Plac8l1 | 0 | 0 | 0 | 1 | 1 |
| Plac9a,Plac9b | 0 | 0 | 0 | 1 | 1 |
| Plac9b | 0 | 0 | 0 | 1 | 1 |
| Plcz1 | 0 | 0 | 0 | 1 | 1 |
| Pnlip | 0 | 0 | 0 | 1 | 1 |
| Pnmt | 0 | 0 | 0 | 1 | 1 |
| Pnpla5 | 0 | 0 | 0 | 1 | 1 |
| Pom121l12 | 0 | 0 | 0 | 1 | 1 |
| Poteg | 0 | 0 | 0 | 1 | 1 |
| Pou1f1 | 0 | 0 | 0 | 1 | 1 |
| Pou2af1 | 0 | 0 | 0 | 1 | 1 |
| Ppp1r2-ps7 | 0 | 0 | 0 | 1 | 1 |
| Ppp1r2-ps9 | 0 | 0 | 0 | 1 | 1 |
| Ppp1r3a | 0 | 0 | 0 | 1 | 1 |
| Ppp3r2 | 0 | 0 | 0 | 1 | 1 |
| Ppy | 0 | 0 | 0 | 1 | 1 |
| Prame | 0 | 0 | 0 | 1 | 1 |
| Pramef17 | 0 | 0 | 0 | 1 | 1 |
| Pramef25 | 0 | 0 | 0 | 1 | 1 |
| Pramef6 | 0 | 0 | 0 | 1 | 1 |
| Pramel3 | 0 | 0 | 0 | 1 | 1 |
| Pramel4 | 0 | 0 | 0 | 1 | 1 |
| Pramel6 | 0 | 0 | 0 | 1 | 1 |
| Pramel7 | 0 | 0 | 0 | 1 | 1 |
| Prb1 | 0 | 0 | 0 | 1 | 1 |
| Prg3 | 0 | 0 | 0 | 1 | 1 |
| Prh1 | 0 | 0 | 0 | 1 | 1 |
| Prl | 0 | 0 | 0 | 1 | 1 |
| Prl3a1 | 0 | 0 | 0 | 1 | 1 |

| | | | | | |
|---|---|---|---|---|---|
| Prl3c1 | 0 | 0 | 0 | 1 | 1 |
| Prl3d2 | 0 | 0 | 0 | 1 | 1 |
| Prl3d3 | 0 | 0 | 0 | 1 | 1 |
| Prl5a1 | 0 | 0 | 0 | 1 | 1 |
| Prl7b1 | 0 | 0 | 0 | 1 | 1 |
| Prl7c1 | 0 | 0 | 0 | 1 | 1 |
| Prl8a8 | 0 | 0 | 0 | 1 | 1 |
| Prn | 0 | 0 | 0 | 1 | 1 |
| Prol1 | 0 | 0 | 0 | 1 | 1 |
| Prop1 | 0 | 0 | 0 | 1 | 1 |
| Prp2 | 0 | 0 | 0 | 1 | 1 |
| Prpmp5 | 0 | 0 | 0 | 1 | 1 |
| Prr27 | 0 | 0 | 0 | 1 | 1 |
| Prr30 | 0 | 0 | 0 | 1 | 1 |
| Prr32 | 0 | 0 | 0 | 1 | 1 |
| Prrxl1 | 0 | 0 | 0 | 1 | 1 |
| Prss2 | 0 | 0 | 0 | 1 | 1 |
| Prss21 | 0 | 0 | 0 | 1 | 1 |
| Prss28 | 0 | 0 | 0 | 1 | 1 |
| Prss29 | 0 | 0 | 0 | 1 | 1 |
| Prss3 | 0 | 0 | 0 | 1 | 1 |
| Prss34 | 0 | 0 | 0 | 1 | 1 |
| Prss37 | 0 | 0 | 0 | 1 | 1 |
| Prss39 | 0 | 0 | 0 | 1 | 1 |
| Prss40 | 0 | 0 | 0 | 1 | 1 |
| Prss46 | 0 | 0 | 0 | 1 | 1 |
| Prss51 | 0 | 0 | 0 | 1 | 1 |
| Prss52 | 0 | 0 | 0 | 1 | 1 |
| Prss54 | 0 | 0 | 0 | 1 | 1 |
| Prss55 | 0 | 0 | 0 | 1 | 1 |
| Prss58 | 0 | 0 | 0 | 1 | 1 |
| Psg-ps1 | 0 | 0 | 0 | 1 | 1 |
| Psg18 | 0 | 0 | 0 | 1 | 1 |
| Psg20 | 0 | 0 | 0 | 1 | 1 |
| Psg21 | 0 | 0 | 0 | 1 | 1 |
| Psg23 | 0 | 0 | 0 | 1 | 1 |
| Psg26 | 0 | 0 | 0 | 1 | 1 |
| Psg27 | 0 | 0 | 0 | 1 | 1 |
| Psg28 | 0 | 0 | 0 | 1 | 1 |
| Ptgs2os | 0 | 0 | 0 | 1 | 1 |
| Pth | 0 | 0 | 0 | 1 | 1 |
| Pth2 | 0 | 0 | 0 | 1 | 1 |
| Ptprtos | 0 | 0 | 0 | 1 | 1 |
| Pvalb | 0 | 0 | 0 | 1 | 1 |
| Pydc4 | 0 | 0 | 0 | 1 | 1 |
| Pyhin1 | 0 | 0 | 0 | 1 | 1 |
| Pyy | 0 | 0 | 0 | 1 | 1 |
| Rab11fip4os2 | 0 | 0 | 0 | 1 | 1 |

| | | | | | |
|---|---|---|---|---|---|
| Rag1 | 0 | 0 | 0 | 1 | 1 |
| Rbm31y | 0 | 0 | 0 | 1 | 1 |
| Rbmy | 0 | 0 | 0 | 1 | 1 |
| Rbpjl | 0 | 0 | 0 | 1 | 1 |
| Rd3l | 0 | 0 | 0 | 1 | 1 |
| Rdh16 | 0 | 0 | 0 | 1 | 1 |
| Rdh18-ps | 0 | 0 | 0 | 1 | 1 |
| Rdh19 | 0 | 0 | 0 | 1 | 1 |
| Rdh8 | 0 | 0 | 0 | 1 | 1 |
| Reg1 | 0 | 0 | 0 | 1 | 1 |
| Reg2 | 0 | 0 | 0 | 1 | 1 |
| Reg3a | 0 | 0 | 0 | 1 | 1 |
| Reg3b | 0 | 0 | 0 | 1 | 1 |
| Reg3d | 0 | 0 | 0 | 1 | 1 |
| Reg3g | 0 | 0 | 0 | 1 | 1 |
| Reg4 | 0 | 0 | 0 | 1 | 1 |
| Rergl | 0 | 0 | 0 | 1 | 1 |
| Retn | 0 | 0 | 0 | 1 | 1 |
| Retnla | 0 | 0 | 0 | 1 | 1 |
| Retnlb | 0 | 0 | 0 | 1 | 1 |
| Retnlg | 0 | 0 | 0 | 1 | 1 |
| Rex2 | 0 | 0 | 0 | 1 | 1 |
| Rfpl4b | 0 | 0 | 0 | 1 | 1 |
| Rfx6 | 0 | 0 | 0 | 1 | 1 |
| Rgr | 0 | 0 | 0 | 1 | 1 |
| Rgs13 | 0 | 0 | 0 | 1 | 1 |
| Rgs21 | 0 | 0 | 0 | 1 | 1 |
| Rhox1 | 0 | 0 | 0 | 1 | 1 |
| Rhox10 | 0 | 0 | 0 | 1 | 1 |
| Rhox11 | 0 | 0 | 0 | 1 | 1 |
| Rhox12 | 0 | 0 | 0 | 1 | 1 |
| Rhox13 | 0 | 0 | 0 | 1 | 1 |
| Rhox2a | 0 | 0 | 0 | 1 | 1 |
| Rhox2b | 0 | 0 | 0 | 1 | 1 |
| Rhox2c | 0 | 0 | 0 | 1 | 1 |
| Rhox2d | 0 | 0 | 0 | 1 | 1 |
| Rhox2e | 0 | 0 | 0 | 1 | 1 |
| Rhox2f | 0 | 0 | 0 | 1 | 1 |
| Rhox2g | 0 | 0 | 0 | 1 | 1 |
| Rhox2h | 0 | 0 | 0 | 1 | 1 |
| Rhox3a,Rhox3e | 0 | 0 | 0 | 1 | 1 |
| Rhox3c | 0 | 0 | 0 | 1 | 1 |
| Rhox3e | 0 | 0 | 0 | 1 | 1 |
| Rhox3f | 0 | 0 | 0 | 1 | 1 |
| Rhox3g | 0 | 0 | 0 | 1 | 1 |
| Rhox3h | 0 | 0 | 0 | 1 | 1 |
| Rhox4a | 0 | 0 | 0 | 1 | 1 |
| Rhox4b | 0 | 0 | 0 | 1 | 1 |

| | | | | | |
|---|---|---|---|---|---|
| Rhox4c | 0 | 0 | 0 | 1 | 1 |
| Rhox4d | 0 | 0 | 0 | 1 | 1 |
| Rhox4e | 0 | 0 | 0 | 1 | 1 |
| Rhox4f | 0 | 0 | 0 | 1 | 1 |
| Rhox4g | 0 | 0 | 0 | 1 | 1 |
| Rhox7 | 0 | 0 | 0 | 1 | 1 |
| Rhox8 | 0 | 0 | 0 | 1 | 1 |
| Rn4.5s | 0 | 0 | 0 | 1 | 1 |
| Rnase13 | 0 | 0 | 0 | 1 | 1 |
| Rnase2a | 0 | 0 | 0 | 1 | 1 |
| Rnase2b | 0 | 0 | 0 | 1 | 1 |
| Rnase9 | 0 | 0 | 0 | 1 | 1 |
| Rnf183 | 0 | 0 | 0 | 1 | 1 |
| Rnu6 | 0 | 0 | 0 | 1 | 1 |
| Rnu7 | 0 | 0 | 0 | 1 | 1 |
| Ropn1 | 0 | 0 | 0 | 1 | 1 |
| Rpe65 | 0 | 0 | 0 | 1 | 1 |
| Rpl10l | 0 | 0 | 0 | 1 | 1 |
| Rpl26 | 0 | 0 | 0 | 1 | 1 |
| Rprl1 | 0 | 0 | 0 | 1 | 1 |
| Rprl2 | 0 | 0 | 0 | 1 | 1 |
| Rprl3 | 0 | 0 | 0 | 1 | 1 |
| Rprml | 0 | 0 | 0 | 1 | 1 |
| Rps27rt | 0 | 0 | 0 | 1 | 1 |
| Rtp1 | 0 | 0 | 0 | 1 | 1 |
| Rtp2 | 0 | 0 | 0 | 1 | 1 |
| Rxfp1 | 0 | 0 | 0 | 1 | 1 |
| Rxfp4 | 0 | 0 | 0 | 1 | 1 |
| S100a5 | 0 | 0 | 0 | 1 | 1 |
| S100a7a | 0 | 0 | 0 | 1 | 1 |
| S100z | 0 | 0 | 0 | 1 | 1 |
| Saa4 | 0 | 0 | 0 | 1 | 1 |
| Samd7 | 0 | 0 | 0 | 1 | 1 |
| Samt1 | 0 | 0 | 0 | 1 | 1 |
| Samt2 | 0 | 0 | 0 | 1 | 1 |
| Samt3 | 0 | 0 | 0 | 1 | 1 |
| Samt4 | 0 | 0 | 0 | 1 | 1 |
| Sbk3 | 0 | 0 | 0 | 1 | 1 |
| Sbp | 0 | 0 | 0 | 1 | 1 |
| Sbpl | 0 | 0 | 0 | 1 | 1 |
| Scarna17 | 0 | 0 | 0 | 1 | 1 |
| Scarna3b | 0 | 0 | 0 | 1 | 1 |
| Scarna9 | 0 | 0 | 0 | 1 | 1 |
| Scgb1b19 | 0 | 0 | 0 | 1 | 1 |
| Scgb1b2 | 0 | 0 | 0 | 1 | 1 |
| Scgb1b20 | 0 | 0 | 0 | 1 | 1 |
| Scgb1b24 | 0 | 0 | 0 | 1 | 1 |
| Scgb1b27 | 0 | 0 | 0 | 1 | 1 |

| | | | | | |
|---|---|---|---|---|---|
| Scgb1b29 | 0 | 0 | 0 | 1 | 1 |
| Scgb1b3 | 0 | 0 | 0 | 1 | 1 |
| Scgb1b7 | 0 | 0 | 0 | 1 | 1 |
| Scgb2b12 | 0 | 0 | 0 | 1 | 1 |
| Scgb2b17 | 0 | 0 | 0 | 1 | 1 |
| Scgb2b19 | 0 | 0 | 0 | 1 | 1 |
| Scgb2b2 | 0 | 0 | 0 | 1 | 1 |
| Scgb2b20 | 0 | 0 | 0 | 1 | 1 |
| Scgb2b23-ps | 0 | 0 | 0 | 1 | 1 |
| Scgb2b24 | 0 | 0 | 0 | 1 | 1 |
| Scgb2b26 | 0 | 0 | 0 | 1 | 1 |
| Scgb2b27 | 0 | 0 | 0 | 1 | 1 |
| Scgb2b3 | 0 | 0 | 0 | 1 | 1 |
| Scgb2b7 | 0 | 0 | 0 | 1 | 1 |
| Scgb3a2 | 0 | 0 | 0 | 1 | 1 |
| Scp2d1 | 0 | 0 | 0 | 1 | 1 |
| Scrg1 | 0 | 0 | 0 | 1 | 1 |
| Sdr16c5 | 0 | 0 | 0 | 1 | 1 |
| Sdr16c6 | 0 | 0 | 0 | 1 | 1 |
| Sebox | 0 | 0 | 0 | 1 | 1 |
| Sectm1b | 0 | 0 | 0 | 1 | 1 |
| Sele | 0 | 0 | 0 | 1 | 1 |
| Sept12 | 0 | 0 | 0 | 1 | 1 |
| Serpina11 | 0 | 0 | 0 | 1 | 1 |
| Serpina1e | 0 | 0 | 0 | 1 | 1 |
| Serpina1f | 0 | 0 | 0 | 1 | 1 |
| Serpina3c | 0 | 0 | 0 | 1 | 1 |
| Serpina3j | 0 | 0 | 0 | 1 | 1 |
| Serpina3k | 0 | 0 | 0 | 1 | 1 |
| Serpina3n | 0 | 0 | 0 | 1 | 1 |
| Serpina4-ps1 | 0 | 0 | 0 | 1 | 1 |
| Serpina9 | 0 | 0 | 0 | 1 | 1 |
| Serpinb11 | 0 | 0 | 0 | 1 | 1 |
| Serpinb12 | 0 | 0 | 0 | 1 | 1 |
| Serpinb13 | 0 | 0 | 0 | 1 | 1 |
| Serpinb1b | 0 | 0 | 0 | 1 | 1 |
| Serpinb1c | 0 | 0 | 0 | 1 | 1 |
| Serpinb3a | 0 | 0 | 0 | 1 | 1 |
| Serpinb3b | 0 | 0 | 0 | 1 | 1 |
| Serpinb3c | 0 | 0 | 0 | 1 | 1 |
| Serpinb3d | 0 | 0 | 0 | 1 | 1 |
| Serpinb6e | 0 | 0 | 0 | 1 | 1 |
| Serpinb7 | 0 | 0 | 0 | 1 | 1 |
| Serpini2 | 0 | 0 | 0 | 1 | 1 |
| Sftpa1 | 0 | 0 | 0 | 1 | 1 |
| Sftpd | 0 | 0 | 0 | 1 | 1 |
| Sgca | 0 | 0 | 0 | 1 | 1 |
| Sh2d1b2 | 0 | 0 | 0 | 1 | 1 |

| | | | | | |
|---|---|---|---|---|---|
| Sirpb1a | 0 | 0 | 0 | 1 | 1 |
| Sirpb1b | 0 | 0 | 0 | 1 | 1 |
| Sis | 0 | 0 | 0 | 1 | 1 |
| Sit1 | 0 | 0 | 0 | 1 | 1 |
| Skint1 | 0 | 0 | 0 | 1 | 1 |
| Skint10 | 0 | 0 | 0 | 1 | 1 |
| Skint11 | 0 | 0 | 0 | 1 | 1 |
| Skint2 | 0 | 0 | 0 | 1 | 1 |
| Skint3 | 0 | 0 | 0 | 1 | 1 |
| Skint5 | 0 | 0 | 0 | 1 | 1 |
| Skint6 | 0 | 0 | 0 | 1 | 1 |
| Skint7 | 0 | 0 | 0 | 1 | 1 |
| Skint8 | 0 | 0 | 0 | 1 | 1 |
| Skint9 | 0 | 0 | 0 | 1 | 1 |
| Slamf7 | 0 | 0 | 0 | 1 | 1 |
| Slc10a2 | 0 | 0 | 0 | 1 | 1 |
| Slc13a1 | 0 | 0 | 0 | 1 | 1 |
| Slc13a2 | 0 | 0 | 0 | 1 | 1 |
| Slc15a5 | 0 | 0 | 0 | 1 | 1 |
| Slc17a1 | 0 | 0 | 0 | 1 | 1 |
| Slc17a6 | 0 | 0 | 0 | 1 | 1 |
| Slc22a13 | 0 | 0 | 0 | 1 | 1 |
| Slc22a14 | 0 | 0 | 0 | 1 | 1 |
| Slc22a16 | 0 | 0 | 0 | 1 | 1 |
| Slc22a19 | 0 | 0 | 0 | 1 | 1 |
| Slc22a26 | 0 | 0 | 0 | 1 | 1 |
| Slc22a27 | 0 | 0 | 0 | 1 | 1 |
| Slc22a29 | 0 | 0 | 0 | 1 | 1 |
| Slc25a54 | 0 | 0 | 0 | 1 | 1 |
| Slc26a7 | 0 | 0 | 0 | 1 | 1 |
| Slc26a9 | 0 | 0 | 0 | 1 | 1 |
| Slc30a8 | 0 | 0 | 0 | 1 | 1 |
| Slc36a1os | 0 | 0 | 0 | 1 | 1 |
| Slc36a3 | 0 | 0 | 0 | 1 | 1 |
| Slc38a11 | 0 | 0 | 0 | 1 | 1 |
| Slc39a12 | 0 | 0 | 0 | 1 | 1 |
| Slc5a4a | 0 | 0 | 0 | 1 | 1 |
| Slc6a5 | 0 | 0 | 0 | 1 | 1 |
| Slc7a12 | 0 | 0 | 0 | 1 | 1 |
| Slc9a4 | 0 | 0 | 0 | 1 | 1 |
| Slc9c1 | 0 | 0 | 0 | 1 | 1 |
| Slco1a1 | 0 | 0 | 0 | 1 | 1 |
| Slco1a4 | 0 | 0 | 0 | 1 | 1 |
| Slco1a6 | 0 | 0 | 0 | 1 | 1 |
| Slco6c1 | 0 | 0 | 0 | 1 | 1 |
| Sln | 0 | 0 | 0 | 1 | 1 |
| Slx | 0 | 0 | 0 | 1 | 1 |
| Slxl1 | 0 | 0 | 0 | 1 | 1 |

| | | | | | |
|---|---|---|---|---|---|
| Sly | 0 | 0 | 0 | 1 | 1 |
| Smco1 | 0 | 0 | 0 | 1 | 1 |
| Smco2 | 0 | 0 | 0 | 1 | 1 |
| Smcp | 0 | 0 | 0 | 1 | 1 |
| Smgc | 0 | 0 | 0 | 1 | 1 |
| Smim23 | 0 | 0 | 0 | 1 | 1 |
| Smim9 | 0 | 0 | 0 | 1 | 1 |
| Smlr1 | 0 | 0 | 0 | 1 | 1 |
| Smok2a | 0 | 0 | 0 | 1 | 1 |
| Smok2b | 0 | 0 | 0 | 1 | 1 |
| Smok3a | 0 | 0 | 0 | 1 | 1 |
| Smok3b | 0 | 0 | 0 | 1 | 1 |
| Smr2 | 0 | 0 | 0 | 1 | 1 |
| Smr3a | 0 | 0 | 0 | 1 | 1 |
| Sncb | 0 | 0 | 0 | 1 | 1 |
| Snora19 | 0 | 0 | 0 | 1 | 1 |
| Snora20 | 0 | 0 | 0 | 1 | 1 |
| Snora2b | 0 | 0 | 0 | 1 | 1 |
| Snora35 | 0 | 0 | 0 | 1 | 1 |
| Snora36b | 0 | 0 | 0 | 1 | 1 |
| Snord110 | 0 | 0 | 0 | 1 | 1 |
| Snord111 | 0 | 0 | 0 | 1 | 1 |
| Snord116 | 0 | 0 | 0 | 1 | 1 |
| Snord116l1 | 0 | 0 | 0 | 1 | 1 |
| Snord116l2 | 0 | 0 | 0 | 1 | 1 |
| Snord118 | 0 | 0 | 0 | 1 | 1 |
| Snord14a | 0 | 0 | 0 | 1 | 1 |
| Snord14c | 0 | 0 | 0 | 1 | 1 |
| Snord14d | 0 | 0 | 0 | 1 | 1 |
| Snord19 | 0 | 0 | 0 | 1 | 1 |
| Snord1a | 0 | 0 | 0 | 1 | 1 |
| Snord1c | 0 | 0 | 0 | 1 | 1 |
| Snord38a | 0 | 0 | 0 | 1 | 1 |
| Snord42a | 0 | 0 | 0 | 1 | 1 |
| Snord43 | 0 | 0 | 0 | 1 | 1 |
| Snord45b | 0 | 0 | 0 | 1 | 1 |
| Snord58b | 0 | 0 | 0 | 1 | 1 |
| Snord61 | 0 | 0 | 0 | 1 | 1 |
| Snord64 | 0 | 0 | 0 | 1 | 1 |
| Snord68 | 0 | 0 | 0 | 1 | 1 |
| Snord73a | 0 | 0 | 0 | 1 | 1 |
| Snord90 | 0 | 0 | 0 | 1 | 1 |
| Snord91a | 0 | 0 | 0 | 1 | 1 |
| Snord93 | 0 | 0 | 0 | 1 | 1 |
| Snord98 | 0 | 0 | 0 | 1 | 1 |
| Sntn | 0 | 0 | 0 | 1 | 1 |
| Sohlh1 | 0 | 0 | 0 | 1 | 1 |
| Sohlh2 | 0 | 0 | 0 | 1 | 1 |

| | | | | | |
|---|---|---|---|---|---|
| Sorbs2os | 0 | 0 | 0 | 1 | 1 |
| Spaca3 | 0 | 0 | 0 | 1 | 1 |
| Spaca5 | 0 | 0 | 0 | 1 | 1 |
| Spaca7 | 0 | 0 | 0 | 1 | 1 |
| Spag11a | 0 | 0 | 0 | 1 | 1 |
| Spam1 | 0 | 0 | 0 | 1 | 1 |
| Spata20 | 0 | 0 | 0 | 1 | 1 |
| Spata22 | 0 | 0 | 0 | 1 | 1 |
| Spata31 | 0 | 0 | 0 | 1 | 1 |
| Spata31d1b | 0 | 0 | 0 | 1 | 1 |
| Spata31d1c | 0 | 0 | 0 | 1 | 1 |
| Spata32 | 0 | 0 | 0 | 1 | 1 |
| Spdyb | 0 | 0 | 0 | 1 | 1 |
| Speer1-ps1 | 0 | 0 | 0 | 1 | 1 |
| Speer2 | 0 | 0 | 0 | 1 | 1 |
| Speer3 | 0 | 0 | 0 | 1 | 1 |
| Speer4a | 0 | 0 | 0 | 1 | 1 |
| Speer4c | 0 | 0 | 0 | 1 | 1 |
| Speer4d | 0 | 0 | 0 | 1 | 1 |
| Speer4e | 0 | 0 | 0 | 1 | 1 |
| Speer4f | 0 | 0 | 0 | 1 | 1 |
| Speer5-ps1 | 0 | 0 | 0 | 1 | 1 |
| Speer6-ps1 | 0 | 0 | 0 | 1 | 1 |
| Speer7-ps1 | 0 | 0 | 0 | 1 | 1 |
| Speer8-ps1 | 0 | 0 | 0 | 1 | 1 |
| Speer9-ps1 | 0 | 0 | 0 | 1 | 1 |
| Spert | 0 | 0 | 0 | 1 | 1 |
| Spic | 0 | 0 | 0 | 1 | 1 |
| Spin2d | 0 | 0 | 0 | 1 | 1 |
| Spink11 | 0 | 0 | 0 | 1 | 1 |
| Spink12 | 0 | 0 | 0 | 1 | 1 |
| Spink13 | 0 | 0 | 0 | 1 | 1 |
| Spink14 | 0 | 0 | 0 | 1 | 1 |
| Spink5 | 0 | 0 | 0 | 1 | 1 |
| Spink6 | 0 | 0 | 0 | 1 | 1 |
| Spink7 | 0 | 0 | 0 | 1 | 1 |
| Spinkl | 0 | 0 | 0 | 1 | 1 |
| Spint3 | 0 | 0 | 0 | 1 | 1 |
| Spint4 | 0 | 0 | 0 | 1 | 1 |
| Spint5 | 0 | 0 | 0 | 1 | 1 |
| Spn-ps | 0 | 0 | 0 | 1 | 1 |
| Sprr1a | 0 | 0 | 0 | 1 | 1 |
| Sprr1b | 0 | 0 | 0 | 1 | 1 |
| Sprr2a2 | 0 | 0 | 0 | 1 | 1 |
| Sprr2b | 0 | 0 | 0 | 1 | 1 |
| Sprr2d | 0 | 0 | 0 | 1 | 1 |
| Sprr2e | 0 | 0 | 0 | 1 | 1 |
| Sprr2f | 0 | 0 | 0 | 1 | 1 |

| | | | | | |
|---|---|---|---|---|---|
| Sprr2i | 0 | 0 | 0 | 1 | 1 |
| Sprr2j-ps | 0 | 0 | 0 | 1 | 1 |
| Sprr2k | 0 | 0 | 0 | 1 | 1 |
| Sprr3 | 0 | 0 | 0 | 1 | 1 |
| Sprr4 | 0 | 0 | 0 | 1 | 1 |
| Spry3 | 0 | 0 | 0 | 1 | 1 |
| Spz1 | 0 | 0 | 0 | 1 | 1 |
| Srp54b,Srp54c | 0 | 0 | 0 | 1 | 1 |
| Sry | 0 | 0 | 0 | 1 | 1 |
| Ssty1 | 0 | 0 | 0 | 1 | 1 |
| Ssty2 | 0 | 0 | 0 | 1 | 1 |
| Ssx9 | 0 | 0 | 0 | 1 | 1 |
| Ssxb1 | 0 | 0 | 0 | 1 | 1 |
| Ssxb10 | 0 | 0 | 0 | 1 | 1 |
| Ssxb2 | 0 | 0 | 0 | 1 | 1 |
| Ssxb3 | 0 | 0 | 0 | 1 | 1 |
| Ssxb5 | 0 | 0 | 0 | 1 | 1 |
| Ssxb6 | 0 | 0 | 0 | 1 | 1 |
| Ssxb8 | 0 | 0 | 0 | 1 | 1 |
| Ssxb9 | 0 | 0 | 0 | 1 | 1 |
| St8sia3os | 0 | 0 | 0 | 1 | 1 |
| Stfa2 | 0 | 0 | 0 | 1 | 1 |
| Stfa2l1 | 0 | 0 | 0 | 1 | 1 |
| Stfa3 | 0 | 0 | 0 | 1 | 1 |
| Stoml3 | 0 | 0 | 0 | 1 | 1 |
| Sucnr1 | 0 | 0 | 0 | 1 | 1 |
| Sult1b1 | 0 | 0 | 0 | 1 | 1 |
| Sult1c1 | 0 | 0 | 0 | 1 | 1 |
| Sult1c2 | 0 | 0 | 0 | 1 | 1 |
| Sult2a1 | 0 | 0 | 0 | 1 | 1 |
| Sult2a2 | 0 | 0 | 0 | 1 | 1 |
| Sult2a3 | 0 | 0 | 0 | 1 | 1 |
| Sult2a4 | 0 | 0 | 0 | 1 | 1 |
| Sult2a5 | 0 | 0 | 0 | 1 | 1 |
| Sult2a6 | 0 | 0 | 0 | 1 | 1 |
| Sult2a7 | 0 | 0 | 0 | 1 | 1 |
| Sult3a1 | 0 | 0 | 0 | 1 | 1 |
| Sun5 | 0 | 0 | 0 | 1 | 1 |
| Sva | 0 | 0 | 0 | 1 | 1 |
| Sval1 | 0 | 0 | 0 | 1 | 1 |
| Sval2 | 0 | 0 | 0 | 1 | 1 |
| Sval3 | 0 | 0 | 0 | 1 | 1 |
| Svs2 | 0 | 0 | 0 | 1 | 1 |
| Svs3a | 0 | 0 | 0 | 1 | 1 |
| Svs3b | 0 | 0 | 0 | 1 | 1 |
| Svs4 | 0 | 0 | 0 | 1 | 1 |
| Svs5 | 0 | 0 | 0 | 1 | 1 |
| Svs6 | 0 | 0 | 0 | 1 | 1 |

| | | | | | |
|---|---|---|---|---|---|
| Syce3 | 0 | 0 | 0 | 1 | 1 |
| Sycn | 0 | 0 | 0 | 1 | 1 |
| Synb | 0 | 0 | 0 | 1 | 1 |
| Syt10 | 0 | 0 | 0 | 1 | 1 |
| Taar1 | 0 | 0 | 0 | 1 | 1 |
| Taar3 | 0 | 0 | 0 | 1 | 1 |
| Taar5 | 0 | 0 | 0 | 1 | 1 |
| Taar6 | 0 | 0 | 0 | 1 | 1 |
| Taar7a | 0 | 0 | 0 | 1 | 1 |
| Taar7b | 0 | 0 | 0 | 1 | 1 |
| Taar7e | 0 | 0 | 0 | 1 | 1 |
| Taar7f | 0 | 0 | 0 | 1 | 1 |
| Taar8a | 0 | 0 | 0 | 1 | 1 |
| Taar8b | 0 | 0 | 0 | 1 | 1 |
| Taar8c | 0 | 0 | 0 | 1 | 1 |
| Taar9 | 0 | 0 | 0 | 1 | 1 |
| Tas2r102 | 0 | 0 | 0 | 1 | 1 |
| Tas2r103 | 0 | 0 | 0 | 1 | 1 |
| Tas2r104 | 0 | 0 | 0 | 1 | 1 |
| Tas2r105 | 0 | 0 | 0 | 1 | 1 |
| Tas2r106 | 0 | 0 | 0 | 1 | 1 |
| Tas2r107 | 0 | 0 | 0 | 1 | 1 |
| Tas2r109 | 0 | 0 | 0 | 1 | 1 |
| Tas2r110 | 0 | 0 | 0 | 1 | 1 |
| Tas2r113 | 0 | 0 | 0 | 1 | 1 |
| Tas2r114 | 0 | 0 | 0 | 1 | 1 |
| Tas2r115 | 0 | 0 | 0 | 1 | 1 |
| Tas2r116 | 0 | 0 | 0 | 1 | 1 |
| Tas2r117 | 0 | 0 | 0 | 1 | 1 |
| Tas2r118 | 0 | 0 | 0 | 1 | 1 |
| Tas2r119 | 0 | 0 | 0 | 1 | 1 |
| Tas2r120 | 0 | 0 | 0 | 1 | 1 |
| Tas2r121 | 0 | 0 | 0 | 1 | 1 |
| Tas2r122 | 0 | 0 | 0 | 1 | 1 |
| Tas2r123 | 0 | 0 | 0 | 1 | 1 |
| Tas2r124 | 0 | 0 | 0 | 1 | 1 |
| Tas2r125 | 0 | 0 | 0 | 1 | 1 |
| Tas2r126 | 0 | 0 | 0 | 1 | 1 |
| Tas2r129 | 0 | 0 | 0 | 1 | 1 |
| Tas2r130 | 0 | 0 | 0 | 1 | 1 |
| Tas2r131 | 0 | 0 | 0 | 1 | 1 |
| Tas2r134 | 0 | 0 | 0 | 1 | 1 |
| Tas2r136 | 0 | 0 | 0 | 1 | 1 |
| Tas2r138 | 0 | 0 | 0 | 1 | 1 |
| Tas2r139 | 0 | 0 | 0 | 1 | 1 |
| Tas2r140 | 0 | 0 | 0 | 1 | 1 |
| Tas2r144 | 0 | 0 | 0 | 1 | 1 |
| Tat | 0 | 0 | 0 | 1 | 1 |

| | | | | | |
|---|---|---|---|---|---|
| Tbc1d21 | 0 | 0 | 0 | 1 | 1 |
| Tbpl2 | 0 | 0 | 0 | 1 | 1 |
| Tcf23 | 0 | 0 | 0 | 1 | 1 |
| Tchhl1 | 0 | 0 | 0 | 1 | 1 |
| Tcl1 | 0 | 0 | 0 | 1 | 1 |
| Tcl1b1 | 0 | 0 | 0 | 1 | 1 |
| Tcl1b2 | 0 | 0 | 0 | 1 | 1 |
| Tcl1b3 | 0 | 0 | 0 | 1 | 1 |
| Tcl1b4 | 0 | 0 | 0 | 1 | 1 |
| Tcl1b5 | 0 | 0 | 0 | 1 | 1 |
| Tcp10a | 0 | 0 | 0 | 1 | 1 |
| Tcp10c | 0 | 0 | 0 | 1 | 1 |
| Tcstv1 | 0 | 0 | 0 | 1 | 1 |
| Tcstv3 | 0 | 0 | 0 | 1 | 1 |
| Tcte3 | 0 | 0 | 0 | 1 | 1 |
| Tctex1d4 | 0 | 0 | 0 | 1 | 1 |
| Tdpoz1 | 0 | 0 | 0 | 1 | 1 |
| Tdpoz2 | 0 | 0 | 0 | 1 | 1 |
| Tdpoz3 | 0 | 0 | 0 | 1 | 1 |
| Tdpoz4 | 0 | 0 | 0 | 1 | 1 |
| Tdpoz5 | 0 | 0 | 0 | 1 | 1 |
| Teddm1 | 0 | 0 | 0 | 1 | 1 |
| Tex101 | 0 | 0 | 0 | 1 | 1 |
| Tex12 | 0 | 0 | 0 | 1 | 1 |
| Tex13 | 0 | 0 | 0 | 1 | 1 |
| Tex13a | 0 | 0 | 0 | 1 | 1 |
| Tex24 | 0 | 0 | 0 | 1 | 1 |
| Tex28 | 0 | 0 | 0 | 1 | 1 |
| Tex33 | 0 | 0 | 0 | 1 | 1 |
| Tex36 | 0 | 0 | 0 | 1 | 1 |
| Tex37 | 0 | 0 | 0 | 1 | 1 |
| Tfap2d | 0 | 0 | 0 | 1 | 1 |
| Tff1 | 0 | 0 | 0 | 1 | 1 |
| Tgif2lx1,Tgif2lx2 | 0 | 0 | 0 | 1 | 1 |
| Tgif2lx2 | 0 | 0 | 0 | 1 | 1 |
| Tgm5 | 0 | 0 | 0 | 1 | 1 |
| Tgm6 | 0 | 0 | 0 | 1 | 1 |
| Tgtp2 | 0 | 0 | 0 | 1 | 1 |
| Theg | 0 | 0 | 0 | 1 | 1 |
| Tigit | 0 | 0 | 0 | 1 | 1 |
| Tlr11 | 0 | 0 | 0 | 1 | 1 |
| Tmbim7 | 0 | 0 | 0 | 1 | 1 |
| Tmco2 | 0 | 0 | 0 | 1 | 1 |
| Tmco5b | 0 | 0 | 0 | 1 | 1 |
| Tmed11 | 0 | 0 | 0 | 1 | 1 |
| Tmed6 | 0 | 0 | 0 | 1 | 1 |
| Tmem179 | 0 | 0 | 0 | 1 | 1 |
| Tmem182 | 0 | 0 | 0 | 1 | 1 |

| | | | | | |
|---|---|---|---|---|---|
| Tmem202 | 0 | 0 | 0 | 1 | 1 |
| Tmem207 | 0 | 0 | 0 | 1 | 1 |
| Tmem210 | 0 | 0 | 0 | 1 | 1 |
| Tmem212 | 0 | 0 | 0 | 1 | 1 |
| Tmem225 | 0 | 0 | 0 | 1 | 1 |
| Tmem233 | 0 | 0 | 0 | 1 | 1 |
| Tmem247 | 0 | 0 | 0 | 1 | 1 |
| Tmem254a,Tmem | 0 | 0 | 0 | 1 | 1 |
| Tmem254b | 0 | 0 | 0 | 1 | 1 |
| Tmem254c | 0 | 0 | 0 | 1 | 1 |
| Tmem45b | 0 | 0 | 0 | 1 | 1 |
| Tmem63c | 0 | 0 | 0 | 1 | 1 |
| Tmem92 | 0 | 0 | 0 | 1 | 1 |
| Tmevpg1 | 0 | 0 | 0 | 1 | 1 |
| Tmlhe | 0 | 0 | 0 | 1 | 1 |
| Tmprss11a | 0 | 0 | 0 | 1 | 1 |
| Tmprss11bnl | 0 | 0 | 0 | 1 | 1 |
| Tmprss11c | 0 | 0 | 0 | 1 | 1 |
| Tmprss11e | 0 | 0 | 0 | 1 | 1 |
| Tmprss11f | 0 | 0 | 0 | 1 | 1 |
| Tmprss11g | 0 | 0 | 0 | 1 | 1 |
| Tmprss12 | 0 | 0 | 0 | 1 | 1 |
| Tmprss15 | 0 | 0 | 0 | 1 | 1 |
| Tmprss5 | 0 | 0 | 0 | 1 | 1 |
| Tnfsf8 | 0 | 0 | 0 | 1 | 1 |
| Tnp1 | 0 | 0 | 0 | 1 | 1 |
| Tnr | 0 | 0 | 0 | 1 | 1 |
| Tpsab1 | 0 | 0 | 0 | 1 | 1 |
| Tpsb2 | 0 | 0 | 0 | 1 | 1 |
| Trdn | 0 | 0 | 0 | 1 | 1 |
| Trex2 | 0 | 0 | 0 | 1 | 1 |
| Trim30b | 0 | 0 | 0 | 1 | 1 |
| Trim31 | 0 | 0 | 0 | 1 | 1 |
| Trim42 | 0 | 0 | 0 | 1 | 1 |
| Trim43b | 0 | 0 | 0 | 1 | 1 |
| Trim52 | 0 | 0 | 0 | 1 | 1 |
| Trpd52l3 | 0 | 0 | 0 | 1 | 1 |
| Try10 | 0 | 0 | 0 | 1 | 1 |
| Try4 | 0 | 0 | 0 | 1 | 1 |
| Try5 | 0 | 0 | 0 | 1 | 1 |
| Tsga8 | 0 | 0 | 0 | 1 | 1 |
| Tspy-ps | 0 | 0 | 0 | 1 | 1 |
| Tssk3 | 0 | 0 | 0 | 1 | 1 |
| Ttc23l | 0 | 0 | 0 | 1 | 1 |
| Ttc24 | 0 | 0 | 0 | 1 | 1 |
| Ttc29 | 0 | 0 | 0 | 1 | 1 |
| Ttll2 | 0 | 0 | 0 | 1 | 1 |
| Ttll8 | 0 | 0 | 0 | 1 | 1 |

| | | | | | |
|---|---|---|---|---|---|
| Tuba3b | 0 | 0 | 0 | 1 | 1 |
| Txndc8 | 0 | 0 | 0 | 1 | 1 |
| Tyr | 0 | 0 | 0 | 1 | 1 |
| Tyrp1 | 0 | 0 | 0 | 1 | 1 |
| Uba1y | 0 | 0 | 0 | 1 | 1 |
| Ubd | 0 | 0 | 0 | 1 | 1 |
| Ube2dnl1 | 0 | 0 | 0 | 1 | 1 |
| Ube2dnl2 | 0 | 0 | 0 | 1 | 1 |
| Ubqln3 | 0 | 0 | 0 | 1 | 1 |
| Ubqlnl | 0 | 0 | 0 | 1 | 1 |
| Ubtfl1 | 0 | 0 | 0 | 1 | 1 |
| Ucn | 0 | 0 | 0 | 1 | 1 |
| Ucn3 | 0 | 0 | 0 | 1 | 1 |
| Ugt1a1 | 0 | 0 | 0 | 1 | 1 |
| Ugt1a10 | 0 | 0 | 0 | 1 | 1 |
| Ugt1a2 | 0 | 0 | 0 | 1 | 1 |
| Ugt1a5 | 0 | 0 | 0 | 1 | 1 |
| Ugt1a6a | 0 | 0 | 0 | 1 | 1 |
| Ugt1a9 | 0 | 0 | 0 | 1 | 1 |
| Ugt2a1 | 0 | 0 | 0 | 1 | 1 |
| Ugt2a2 | 0 | 0 | 0 | 1 | 1 |
| Ugt2a3 | 0 | 0 | 0 | 1 | 1 |
| Ugt2b1 | 0 | 0 | 0 | 1 | 1 |
| Ugt2b35 | 0 | 0 | 0 | 1 | 1 |
| Ugt2b36 | 0 | 0 | 0 | 1 | 1 |
| Ugt2b37 | 0 | 0 | 0 | 1 | 1 |
| Ugt2b38 | 0 | 0 | 0 | 1 | 1 |
| Ugt3a1 | 0 | 0 | 0 | 1 | 1 |
| Unc13c | 0 | 0 | 0 | 1 | 1 |
| Unc93a | 0 | 0 | 0 | 1 | 1 |
| Usp17la | 0 | 0 | 0 | 1 | 1 |
| Usp17lc | 0 | 0 | 0 | 1 | 1 |
| Usp17ld | 0 | 0 | 0 | 1 | 1 |
| Uts2 | 0 | 0 | 0 | 1 | 1 |
| Uts2b | 0 | 0 | 0 | 1 | 1 |
| V1ra8 | 0 | 0 | 0 | 1 | 1 |
| V1rd18,V1rd19 | 0 | 0 | 0 | 1 | 1 |
| V1rd19 | 0 | 0 | 0 | 1 | 1 |
| Vamp7 | 0 | 0 | 0 | 1 | 1 |
| Vmn1r-ps103 | 0 | 0 | 0 | 1 | 1 |
| Vmn1r-ps79 | 0 | 0 | 0 | 1 | 1 |
| Vmn1r1 | 0 | 0 | 0 | 1 | 1 |
| Vmn1r10 | 0 | 0 | 0 | 1 | 1 |
| Vmn1r100,Vmn1r1 | 0 | 0 | 0 | 1 | 1 |
| Vmn1r101 | 0 | 0 | 0 | 1 | 1 |
| Vmn1r103 | 0 | 0 | 0 | 1 | 1 |
| Vmn1r107 | 0 | 0 | 0 | 1 | 1 |
| Vmn1r11 | 0 | 0 | 0 | 1 | 1 |

| | | | | | |
|---|---|---|---|---|---|
| Vmn1r112 | 0 | 0 | 0 | 1 | 1 |
| Vmn1r113 | 0 | 0 | 0 | 1 | 1 |
| Vmn1r114 | 0 | 0 | 0 | 1 | 1 |
| Vmn1r115 | 0 | 0 | 0 | 1 | 1 |
| Vmn1r116 | 0 | 0 | 0 | 1 | 1 |
| Vmn1r117 | 0 | 0 | 0 | 1 | 1 |
| Vmn1r118 | 0 | 0 | 0 | 1 | 1 |
| Vmn1r119 | 0 | 0 | 0 | 1 | 1 |
| Vmn1r12 | 0 | 0 | 0 | 1 | 1 |
| Vmn1r120 | 0 | 0 | 0 | 1 | 1 |
| Vmn1r121 | 0 | 0 | 0 | 1 | 1 |
| Vmn1r122 | 0 | 0 | 0 | 1 | 1 |
| Vmn1r123 | 0 | 0 | 0 | 1 | 1 |
| Vmn1r124 | 0 | 0 | 0 | 1 | 1 |
| Vmn1r125 | 0 | 0 | 0 | 1 | 1 |
| Vmn1r126 | 0 | 0 | 0 | 1 | 1 |
| Vmn1r127 | 0 | 0 | 0 | 1 | 1 |
| Vmn1r128 | 0 | 0 | 0 | 1 | 1 |
| Vmn1r129 | 0 | 0 | 0 | 1 | 1 |
| Vmn1r13 | 0 | 0 | 0 | 1 | 1 |
| Vmn1r130 | 0 | 0 | 0 | 1 | 1 |
| Vmn1r131 | 0 | 0 | 0 | 1 | 1 |
| Vmn1r132,Vmn1r1 | 0 | 0 | 0 | 1 | 1 |
| Vmn1r138 | 0 | 0 | 0 | 1 | 1 |
| Vmn1r139 | 0 | 0 | 0 | 1 | 1 |
| Vmn1r14 | 0 | 0 | 0 | 1 | 1 |
| Vmn1r142 | 0 | 0 | 0 | 1 | 1 |
| Vmn1r143 | 0 | 0 | 0 | 1 | 1 |
| Vmn1r148 | 0 | 0 | 0 | 1 | 1 |
| Vmn1r15 | 0 | 0 | 0 | 1 | 1 |
| Vmn1r151 | 0 | 0 | 0 | 1 | 1 |
| Vmn1r152 | 0 | 0 | 0 | 1 | 1 |
| Vmn1r155 | 0 | 0 | 0 | 1 | 1 |
| Vmn1r157 | 0 | 0 | 0 | 1 | 1 |
| Vmn1r158 | 0 | 0 | 0 | 1 | 1 |
| Vmn1r159 | 0 | 0 | 0 | 1 | 1 |
| Vmn1r16 | 0 | 0 | 0 | 1 | 1 |
| Vmn1r160 | 0 | 0 | 0 | 1 | 1 |
| Vmn1r163 | 0 | 0 | 0 | 1 | 1 |
| Vmn1r165 | 0 | 0 | 0 | 1 | 1 |
| Vmn1r166 | 0 | 0 | 0 | 1 | 1 |
| Vmn1r167 | 0 | 0 | 0 | 1 | 1 |
| Vmn1r168 | 0 | 0 | 0 | 1 | 1 |
| Vmn1r169 | 0 | 0 | 0 | 1 | 1 |
| Vmn1r17 | 0 | 0 | 0 | 1 | 1 |
| Vmn1r170 | 0 | 0 | 0 | 1 | 1 |
| Vmn1r171 | 0 | 0 | 0 | 1 | 1 |
| Vmn1r172 | 0 | 0 | 0 | 1 | 1 |

| | | | | | |
|---|---|---|---|---|---|
| Vmn1r173 | 0 | 0 | 0 | 1 | 1 |
| Vmn1r174 | 0 | 0 | 0 | 1 | 1 |
| Vmn1r175 | 0 | 0 | 0 | 1 | 1 |
| Vmn1r176 | 0 | 0 | 0 | 1 | 1 |
| Vmn1r177 | 0 | 0 | 0 | 1 | 1 |
| Vmn1r178 | 0 | 0 | 0 | 1 | 1 |
| Vmn1r179 | 0 | 0 | 0 | 1 | 1 |
| Vmn1r18 | 0 | 0 | 0 | 1 | 1 |
| Vmn1r180 | 0 | 0 | 0 | 1 | 1 |
| Vmn1r181 | 0 | 0 | 0 | 1 | 1 |
| Vmn1r183 | 0 | 0 | 0 | 1 | 1 |
| Vmn1r184 | 0 | 0 | 0 | 1 | 1 |
| Vmn1r185 | 0 | 0 | 0 | 1 | 1 |
| Vmn1r186,Vmn1r( | 0 | 0 | 0 | 1 | 1 |
| Vmn1r187,Vmn1r( | 0 | 0 | 0 | 1 | 1 |
| Vmn1r188 | 0 | 0 | 0 | 1 | 1 |
| Vmn1r189 | 0 | 0 | 0 | 1 | 1 |
| Vmn1r19 | 0 | 0 | 0 | 1 | 1 |
| Vmn1r191 | 0 | 0 | 0 | 1 | 1 |
| Vmn1r192 | 0 | 0 | 0 | 1 | 1 |
| Vmn1r193 | 0 | 0 | 0 | 1 | 1 |
| Vmn1r194 | 0 | 0 | 0 | 1 | 1 |
| Vmn1r195 | 0 | 0 | 0 | 1 | 1 |
| Vmn1r196 | 0 | 0 | 0 | 1 | 1 |
| Vmn1r198 | 0 | 0 | 0 | 1 | 1 |
| Vmn1r199 | 0 | 0 | 0 | 1 | 1 |
| Vmn1r2 | 0 | 0 | 0 | 1 | 1 |
| Vmn1r20 | 0 | 0 | 0 | 1 | 1 |
| Vmn1r200 | 0 | 0 | 0 | 1 | 1 |
| Vmn1r201 | 0 | 0 | 0 | 1 | 1 |
| Vmn1r202 | 0 | 0 | 0 | 1 | 1 |
| Vmn1r203 | 0 | 0 | 0 | 1 | 1 |
| Vmn1r204 | 0 | 0 | 0 | 1 | 1 |
| Vmn1r205 | 0 | 0 | 0 | 1 | 1 |
| Vmn1r206 | 0 | 0 | 0 | 1 | 1 |
| Vmn1r207-ps | 0 | 0 | 0 | 1 | 1 |
| Vmn1r208 | 0 | 0 | 0 | 1 | 1 |
| Vmn1r209 | 0 | 0 | 0 | 1 | 1 |
| Vmn1r210 | 0 | 0 | 0 | 1 | 1 |
| Vmn1r211 | 0 | 0 | 0 | 1 | 1 |
| Vmn1r212 | 0 | 0 | 0 | 1 | 1 |
| Vmn1r213 | 0 | 0 | 0 | 1 | 1 |
| Vmn1r214 | 0 | 0 | 0 | 1 | 1 |
| Vmn1r215 | 0 | 0 | 0 | 1 | 1 |
| Vmn1r216 | 0 | 0 | 0 | 1 | 1 |
| Vmn1r217 | 0 | 0 | 0 | 1 | 1 |
| Vmn1r218 | 0 | 0 | 0 | 1 | 1 |
| Vmn1r219 | 0 | 0 | 0 | 1 | 1 |

| | | | | | |
|---|---|---|---|---|---|
| Vmn1r22 | 0 | 0 | 0 | 1 | 1 |
| Vmn1r220 | 0 | 0 | 0 | 1 | 1 |
| Vmn1r222 | 0 | 0 | 0 | 1 | 1 |
| Vmn1r224 | 0 | 0 | 0 | 1 | 1 |
| Vmn1r225 | 0 | 0 | 0 | 1 | 1 |
| Vmn1r226 | 0 | 0 | 0 | 1 | 1 |
| Vmn1r227 | 0 | 0 | 0 | 1 | 1 |
| Vmn1r228 | 0 | 0 | 0 | 1 | 1 |
| Vmn1r229 | 0 | 0 | 0 | 1 | 1 |
| Vmn1r23 | 0 | 0 | 0 | 1 | 1 |
| Vmn1r230 | 0 | 0 | 0 | 1 | 1 |
| Vmn1r231 | 0 | 0 | 0 | 1 | 1 |
| Vmn1r232 | 0 | 0 | 0 | 1 | 1 |
| Vmn1r233 | 0 | 0 | 0 | 1 | 1 |
| Vmn1r234 | 0 | 0 | 0 | 1 | 1 |
| Vmn1r235 | 0 | 0 | 0 | 1 | 1 |
| Vmn1r236 | 0 | 0 | 0 | 1 | 1 |
| Vmn1r237 | 0 | 0 | 0 | 1 | 1 |
| Vmn1r238 | 0 | 0 | 0 | 1 | 1 |
| Vmn1r24 | 0 | 0 | 0 | 1 | 1 |
| Vmn1r25 | 0 | 0 | 0 | 1 | 1 |
| Vmn1r27 | 0 | 0 | 0 | 1 | 1 |
| Vmn1r28 | 0 | 0 | 0 | 1 | 1 |
| Vmn1r29 | 0 | 0 | 0 | 1 | 1 |
| Vmn1r3 | 0 | 0 | 0 | 1 | 1 |
| Vmn1r30 | 0 | 0 | 0 | 1 | 1 |
| Vmn1r31 | 0 | 0 | 0 | 1 | 1 |
| Vmn1r33 | 0 | 0 | 0 | 1 | 1 |
| Vmn1r34 | 0 | 0 | 0 | 1 | 1 |
| Vmn1r35 | 0 | 0 | 0 | 1 | 1 |
| Vmn1r36 | 0 | 0 | 0 | 1 | 1 |
| Vmn1r37 | 0 | 0 | 0 | 1 | 1 |
| Vmn1r38 | 0 | 0 | 0 | 1 | 1 |
| Vmn1r39 | 0 | 0 | 0 | 1 | 1 |
| Vmn1r40 | 0 | 0 | 0 | 1 | 1 |
| Vmn1r41 | 0 | 0 | 0 | 1 | 1 |
| Vmn1r42 | 0 | 0 | 0 | 1 | 1 |
| Vmn1r43 | 0 | 0 | 0 | 1 | 1 |
| Vmn1r44 | 0 | 0 | 0 | 1 | 1 |
| Vmn1r45 | 0 | 0 | 0 | 1 | 1 |
| Vmn1r46 | 0 | 0 | 0 | 1 | 1 |
| Vmn1r47 | 0 | 0 | 0 | 1 | 1 |
| Vmn1r48 | 0 | 0 | 0 | 1 | 1 |
| Vmn1r5 | 0 | 0 | 0 | 1 | 1 |
| Vmn1r52 | 0 | 0 | 0 | 1 | 1 |
| Vmn1r54 | 0 | 0 | 0 | 1 | 1 |
| Vmn1r55 | 0 | 0 | 0 | 1 | 1 |
| Vmn1r56 | 0 | 0 | 0 | 1 | 1 |

| | | | | | |
|---|---|---|---|---|---|
| Vmn1r57 | 0 | 0 | 0 | 1 | 1 |
| Vmn1r58 | 0 | 0 | 0 | 1 | 1 |
| Vmn1r59 | 0 | 0 | 0 | 1 | 1 |
| Vmn1r6 | 0 | 0 | 0 | 1 | 1 |
| Vmn1r60 | 0 | 0 | 0 | 1 | 1 |
| Vmn1r61 | 0 | 0 | 0 | 1 | 1 |
| Vmn1r62 | 0 | 0 | 0 | 1 | 1 |
| Vmn1r63 | 0 | 0 | 0 | 1 | 1 |
| Vmn1r64 | 0 | 0 | 0 | 1 | 1 |
| Vmn1r65 | 0 | 0 | 0 | 1 | 1 |
| Vmn1r66 | 0 | 0 | 0 | 1 | 1 |
| Vmn1r67 | 0 | 0 | 0 | 1 | 1 |
| Vmn1r68 | 0 | 0 | 0 | 1 | 1 |
| Vmn1r69 | 0 | 0 | 0 | 1 | 1 |
| Vmn1r7 | 0 | 0 | 0 | 1 | 1 |
| Vmn1r70 | 0 | 0 | 0 | 1 | 1 |
| Vmn1r71 | 0 | 0 | 0 | 1 | 1 |
| Vmn1r72 | 0 | 0 | 0 | 1 | 1 |
| Vmn1r73 | 0 | 0 | 0 | 1 | 1 |
| Vmn1r74 | 0 | 0 | 0 | 1 | 1 |
| Vmn1r75 | 0 | 0 | 0 | 1 | 1 |
| Vmn1r76 | 0 | 0 | 0 | 1 | 1 |
| Vmn1r77 | 0 | 0 | 0 | 1 | 1 |
| Vmn1r78 | 0 | 0 | 0 | 1 | 1 |
| Vmn1r79 | 0 | 0 | 0 | 1 | 1 |
| Vmn1r8 | 0 | 0 | 0 | 1 | 1 |
| Vmn1r80 | 0 | 0 | 0 | 1 | 1 |
| Vmn1r81 | 0 | 0 | 0 | 1 | 1 |
| Vmn1r82 | 0 | 0 | 0 | 1 | 1 |
| Vmn1r83 | 0 | 0 | 0 | 1 | 1 |
| Vmn1r84 | 0 | 0 | 0 | 1 | 1 |
| Vmn1r85 | 0 | 0 | 0 | 1 | 1 |
| Vmn1r86 | 0 | 0 | 0 | 1 | 1 |
| Vmn1r87 | 0 | 0 | 0 | 1 | 1 |
| Vmn1r88 | 0 | 0 | 0 | 1 | 1 |
| Vmn1r89 | 0 | 0 | 0 | 1 | 1 |
| Vmn1r9 | 0 | 0 | 0 | 1 | 1 |
| Vmn1r91 | 0 | 0 | 0 | 1 | 1 |
| Vmn1r93 | 0 | 0 | 0 | 1 | 1 |
| Vmn1r94 | 0 | 0 | 0 | 1 | 1 |
| Vmn2r-ps11 | 0 | 0 | 0 | 1 | 1 |
| Vmn2r-ps129 | 0 | 0 | 0 | 1 | 1 |
| Vmn2r-ps159 | 0 | 0 | 0 | 1 | 1 |
| Vmn2r-ps60 | 0 | 0 | 0 | 1 | 1 |
| Vmn2r10 | 0 | 0 | 0 | 1 | 1 |
| Vmn2r100 | 0 | 0 | 0 | 1 | 1 |
| Vmn2r101 | 0 | 0 | 0 | 1 | 1 |
| Vmn2r102 | 0 | 0 | 0 | 1 | 1 |

| | | | | | |
|---|---|---|---|---|---|
| Vmn2r103 | 0 | 0 | 0 | 1 | 1 |
| Vmn2r104 | 0 | 0 | 0 | 1 | 1 |
| Vmn2r105 | 0 | 0 | 0 | 1 | 1 |
| Vmn2r106 | 0 | 0 | 0 | 1 | 1 |
| Vmn2r107 | 0 | 0 | 0 | 1 | 1 |
| Vmn2r108 | 0 | 0 | 0 | 1 | 1 |
| Vmn2r11 | 0 | 0 | 0 | 1 | 1 |
| Vmn2r110 | 0 | 0 | 0 | 1 | 1 |
| Vmn2r111 | 0 | 0 | 0 | 1 | 1 |
| Vmn2r112 | 0 | 0 | 0 | 1 | 1 |
| Vmn2r113 | 0 | 0 | 0 | 1 | 1 |
| Vmn2r114 | 0 | 0 | 0 | 1 | 1 |
| Vmn2r115 | 0 | 0 | 0 | 1 | 1 |
| Vmn2r116 | 0 | 0 | 0 | 1 | 1 |
| Vmn2r117 | 0 | 0 | 0 | 1 | 1 |
| Vmn2r118 | 0 | 0 | 0 | 1 | 1 |
| Vmn2r12 | 0 | 0 | 0 | 1 | 1 |
| Vmn2r121 | 0 | 0 | 0 | 1 | 1 |
| Vmn2r122 | 0 | 0 | 0 | 1 | 1 |
| Vmn2r123 | 0 | 0 | 0 | 1 | 1 |
| Vmn2r124 | 0 | 0 | 0 | 1 | 1 |
| Vmn2r13 | 0 | 0 | 0 | 1 | 1 |
| Vmn2r14 | 0 | 0 | 0 | 1 | 1 |
| Vmn2r15 | 0 | 0 | 0 | 1 | 1 |
| Vmn2r16 | 0 | 0 | 0 | 1 | 1 |
| Vmn2r17 | 0 | 0 | 0 | 1 | 1 |
| Vmn2r19 | 0 | 0 | 0 | 1 | 1 |
| Vmn2r2 | 0 | 0 | 0 | 1 | 1 |
| Vmn2r20 | 0 | 0 | 0 | 1 | 1 |
| Vmn2r21 | 0 | 0 | 0 | 1 | 1 |
| Vmn2r22 | 0 | 0 | 0 | 1 | 1 |
| Vmn2r24 | 0 | 0 | 0 | 1 | 1 |
| Vmn2r25 | 0 | 0 | 0 | 1 | 1 |
| Vmn2r26 | 0 | 0 | 0 | 1 | 1 |
| Vmn2r27 | 0 | 0 | 0 | 1 | 1 |
| Vmn2r30 | 0 | 0 | 0 | 1 | 1 |
| Vmn2r31 | 0 | 0 | 0 | 1 | 1 |
| Vmn2r32 | 0 | 0 | 0 | 1 | 1 |
| Vmn2r33 | 0 | 0 | 0 | 1 | 1 |
| Vmn2r34 | 0 | 0 | 0 | 1 | 1 |
| Vmn2r35 | 0 | 0 | 0 | 1 | 1 |
| Vmn2r36 | 0 | 0 | 0 | 1 | 1 |
| Vmn2r37 | 0 | 0 | 0 | 1 | 1 |
| Vmn2r38 | 0 | 0 | 0 | 1 | 1 |
| Vmn2r39 | 0 | 0 | 0 | 1 | 1 |
| Vmn2r4 | 0 | 0 | 0 | 1 | 1 |
| Vmn2r40 | 0 | 0 | 0 | 1 | 1 |
| Vmn2r41 | 0 | 0 | 0 | 1 | 1 |

| | | | | | |
|---|---|---|---|---|---|
| Vmn2r42 | 0 | 0 | 0 | 1 | 1 |
| Vmn2r43 | 0 | 0 | 0 | 1 | 1 |
| Vmn2r44 | 0 | 0 | 0 | 1 | 1 |
| Vmn2r45 | 0 | 0 | 0 | 1 | 1 |
| Vmn2r46 | 0 | 0 | 0 | 1 | 1 |
| Vmn2r47 | 0 | 0 | 0 | 1 | 1 |
| Vmn2r48 | 0 | 0 | 0 | 1 | 1 |
| Vmn2r49 | 0 | 0 | 0 | 1 | 1 |
| Vmn2r5 | 0 | 0 | 0 | 1 | 1 |
| Vmn2r50 | 0 | 0 | 0 | 1 | 1 |
| Vmn2r51 | 0 | 0 | 0 | 1 | 1 |
| Vmn2r52 | 0 | 0 | 0 | 1 | 1 |
| Vmn2r53 | 0 | 0 | 0 | 1 | 1 |
| Vmn2r54 | 0 | 0 | 0 | 1 | 1 |
| Vmn2r55 | 0 | 0 | 0 | 1 | 1 |
| Vmn2r56 | 0 | 0 | 0 | 1 | 1 |
| Vmn2r58 | 0 | 0 | 0 | 1 | 1 |
| Vmn2r59 | 0 | 0 | 0 | 1 | 1 |
| Vmn2r6 | 0 | 0 | 0 | 1 | 1 |
| Vmn2r60 | 0 | 0 | 0 | 1 | 1 |
| Vmn2r61 | 0 | 0 | 0 | 1 | 1 |
| Vmn2r62 | 0 | 0 | 0 | 1 | 1 |
| Vmn2r63 | 0 | 0 | 0 | 1 | 1 |
| Vmn2r65 | 0 | 0 | 0 | 1 | 1 |
| Vmn2r66 | 0 | 0 | 0 | 1 | 1 |
| Vmn2r67 | 0 | 0 | 0 | 1 | 1 |
| Vmn2r68 | 0 | 0 | 0 | 1 | 1 |
| Vmn2r69 | 0 | 0 | 0 | 1 | 1 |
| Vmn2r7 | 0 | 0 | 0 | 1 | 1 |
| Vmn2r70 | 0 | 0 | 0 | 1 | 1 |
| Vmn2r71 | 0 | 0 | 0 | 1 | 1 |
| Vmn2r72 | 0 | 0 | 0 | 1 | 1 |
| Vmn2r73 | 0 | 0 | 0 | 1 | 1 |
| Vmn2r74 | 0 | 0 | 0 | 1 | 1 |
| Vmn2r75 | 0 | 0 | 0 | 1 | 1 |
| Vmn2r76 | 0 | 0 | 0 | 1 | 1 |
| Vmn2r78 | 0 | 0 | 0 | 1 | 1 |
| Vmn2r79 | 0 | 0 | 0 | 1 | 1 |
| Vmn2r8 | 0 | 0 | 0 | 1 | 1 |
| Vmn2r80 | 0 | 0 | 0 | 1 | 1 |
| Vmn2r81 | 0 | 0 | 0 | 1 | 1 |
| Vmn2r82 | 0 | 0 | 0 | 1 | 1 |
| Vmn2r83 | 0 | 0 | 0 | 1 | 1 |
| Vmn2r84 | 0 | 0 | 0 | 1 | 1 |
| Vmn2r85 | 0 | 0 | 0 | 1 | 1 |
| Vmn2r86 | 0 | 0 | 0 | 1 | 1 |
| Vmn2r88 | 0 | 0 | 0 | 1 | 1 |
| Vmn2r89 | 0 | 0 | 0 | 1 | 1 |

| | | | | | |
|---|---|---|---|---|---|
| Vmn2r9 | 0 | 0 | 0 | 1 | 1 |
| Vmn2r91 | 0 | 0 | 0 | 1 | 1 |
| Vmn2r92 | 0 | 0 | 0 | 1 | 1 |
| Vmn2r93 | 0 | 0 | 0 | 1 | 1 |
| Vmn2r94 | 0 | 0 | 0 | 1 | 1 |
| Vmn2r95 | 0 | 0 | 0 | 1 | 1 |
| Vmn2r96 | 0 | 0 | 0 | 1 | 1 |
| Vmn2r97 | 0 | 0 | 0 | 1 | 1 |
| Vmn2r98 | 0 | 0 | 0 | 1 | 1 |
| Vmn2r99 | 0 | 0 | 0 | 1 | 1 |
| Vnn3 | 0 | 0 | 0 | 1 | 1 |
| Vpreb2 | 0 | 0 | 0 | 1 | 1 |
| Vsig1 | 0 | 0 | 0 | 1 | 1 |
| Vsig4 | 0 | 0 | 0 | 1 | 1 |
| Vstm2a | 0 | 0 | 0 | 1 | 1 |
| Vsx1 | 0 | 0 | 0 | 1 | 1 |
| Vwc2l | 0 | 0 | 0 | 1 | 1 |
| Wap | 0 | 0 | 0 | 1 | 1 |
| Wbscr28 | 0 | 0 | 0 | 1 | 1 |
| Wdr64 | 0 | 0 | 0 | 1 | 1 |
| Wfdc11 | 0 | 0 | 0 | 1 | 1 |
| Wfdc12 | 0 | 0 | 0 | 1 | 1 |
| Wfdc15b | 0 | 0 | 0 | 1 | 1 |
| Wfdc16 | 0 | 0 | 0 | 1 | 1 |
| Wfdc18 | 0 | 0 | 0 | 1 | 1 |
| Wfdc5 | 0 | 0 | 0 | 1 | 1 |
| Wfdc6b | 0 | 0 | 0 | 1 | 1 |
| Wfdc9 | 0 | 0 | 0 | 1 | 1 |
| Xcl1 | 0 | 0 | 0 | 1 | 1 |
| Xirp1 | 0 | 0 | 0 | 1 | 1 |
| Xirp2 | 0 | 0 | 0 | 1 | 1 |
| Xlr4c | 0 | 0 | 0 | 1 | 1 |
| Xlr5b | 0 | 0 | 0 | 1 | 1 |
| Xlr5c | 0 | 0 | 0 | 1 | 1 |
| Zar1l | 0 | 0 | 0 | 1 | 1 |
| Zbbx | 0 | 0 | 0 | 1 | 1 |
| Zc2hc1b | 0 | 0 | 0 | 1 | 1 |
| Zcchc5 | 0 | 0 | 0 | 1 | 1 |
| Zdhhc22 | 0 | 0 | 0 | 1 | 1 |
| Zdhhc25 | 0 | 0 | 0 | 1 | 1 |
| Zf12 | 0 | 0 | 0 | 1 | 1 |
| Zfp352 | 0 | 0 | 0 | 1 | 1 |
| Zfp474 | 0 | 0 | 0 | 1 | 1 |
| Zfp600 | 0 | 0 | 0 | 1 | 1 |
| Zfp616 | 0 | 0 | 0 | 1 | 1 |
| Zfp648 | 0 | 0 | 0 | 1 | 1 |
| Zfp735 | 0 | 0 | 0 | 1 | 1 |
| Zfp91Cntf | 0 | 0 | 0 | 1 | 1 |

| | | | | | |
|---|---|---|---|---|---|
| Zfy1 | 0 | 0 | 0 | 1 | 1 |
| Zfy2 | 0 | 0 | 0 | 1 | 1 |
| Zg16 | 0 | 0 | 0 | 1 | 1 |
| Zim3 | 0 | 0 | 0 | 1 | 1 |
| Znrf4 | 0 | 0 | 0 | 1 | 1 |
| Zp2 | 0 | 0 | 0 | 1 | 1 |
| Zp3r | 0 | 0 | 0 | 1 | 1 |
| Zp4-ps | 0 | 0 | 0 | 1 | 1 |
| Zscan4a | 0 | 0 | 0 | 1 | 1 |
| Zscan4b | 0 | 0 | 0 | 1 | 1 |
| Zscan4c | 0 | 0 | 0 | 1 | 1 |
| Zscan4d | 0 | 0 | 0 | 1 | 1 |
| Zscan4e | 0 | 0 | 0 | 1 | 1 |
| Zscan4f | 0 | 0 | 0 | 1 | 1 |
| Zscan5b | 0 | 0 | 0 | 1 | 1 |
| Zswim2 | 0 | 0 | 0 | 1 | 1 |
| 9130008F23Rik | 0,587451 | 0,587364 | -0,000213231 | 1 | 1 |
| Zar1 | 0,168326 | 0,168107 | -0,00187371 | 1 | 1 |
| Gsg1 | 0,517437 | 0,516538 | -0,00250884 | 1 | 1 |
| Lrp1b | 0,016282 | 0,0162328 | -0,00438025 | 1 | 1 |
| AF529169 | 0,147952 | 0,147411 | -0,00528567 | 1 | 1 |
| Pde4c | 0,16241 | 0,161613 | -0,00709708 | 1 | 1 |
| Hipk4 | 0,527586 | 0,524589 | -0,00821758 | 1 | 1 |
| Gm13032 | 0,212497 | 0,211262 | -0,00840891 | 1 | 1 |
| Alpk3 | 0,267002 | 0,265399 | -0,00868672 | 1 | 1 |
| Pcdhb8 | 0,058182 | 0,0578305 | -0,00874039 | 1 | 1 |
| Tbx3os2 | 0,108396 | 0,1076 | -0,0106345 | 1 | 1 |
| Gabre | 0,480641 | 0,476949 | -0,0111276 | 1 | 1 |
| Efcab4b | 0,432379 | 0,428785 | -0,0120423 | 1 | 1 |
| AI839979 | 0,26269 | 0,259933 | -0,015223 | 1 | 1 |
| C130026L21Rik | 0,202884 | 0,200608 | -0,0162755 | 1 | 1 |
| Rgag1 | 0,034562 | 0,0341724 | -0,016355 | 1 | 1 |
| 9230116N13Rik | 0,260682 | 0,25773 | -0,0164284 | 1 | 1 |
| 1810010H24Rik | 0,383436 | 0,378587 | -0,018358 | 1 | 1 |
| Akr1b3 | 0,000252 | 0,00024876 | -0,01917 | 1 | 1 |
| Ccdc69 | 0,17043 | 0,168121 | -0,0196761 | 1 | 1 |
| Drd4 | 0,533465 | 0,526093 | -0,0200751 | 1 | 1 |
| Fhad1 | 0,030289 | 0,0298422 | -0,0214347 | 1 | 1 |
| Isg15 | 0,739982 | 0,728822 | -0,0219229 | 1 | 1 |
| 1700001P01Rik | 0,479133 | 0,471533 | -0,0230665 | 1 | 1 |
| Nkx2-2 | 0,075667 | 0,0744056 | -0,0242573 | 1 | 1 |
| Sdr42e1 | 0,560238 | 0,550809 | -0,0244871 | 1 | 1 |
| 9230102K24Rik | 0,376837 | 0,370464 | -0,0246082 | 1 | 1 |
| Ptpn20 | 0,094377 | 0,0927744 | -0,0247133 | 1 | 1 |
| 9430083A17Rik | 0,401585 | 0,394579 | -0,0253941 | 1 | 1 |
| AI317395 | 0,028674 | 0,0281607 | -0,0260668 | 1 | 1 |
| Gm19395 | 0,156315 | 0,153516 | -0,0260668 | 1 | 1 |
| Artn | 0,525844 | 0,516042 | -0,0271459 | 1 | 1 |

| | | | | | |
|---|---|---|---|---|---|
| Gprc5d | 0,255337 | 0,250477 | -0,027725 | 1 | 1 |
| Ar | 0,050841 | 0,0498732 | -0,0277256 | 1 | 1 |
| A930003O13Rik | 0,166283 | 0,163032 | -0,0284879 | 1 | 1 |
| Plscr4 | 0,166644 | 0,163316 | -0,0291082 | 1 | 1 |
| LOC171588 | 0,218793 | 0,214216 | -0,0305027 | 1 | 1 |
| Mir686 | 2247,24 | 2195,14 | -0,0338455 | 1 | 1 |
| Hid1 | 0,474338 | 0,463238 | -0,0341596 | 1 | 1 |
| Lonrf3 | 0,517045 | 0,504805 | -0,0345624 | 1 | 1 |
| Txk | 0,169594 | 0,165409 | -0,0360535 | 1 | 1 |
| Cyp2c44 | 0,047043 | 0,0458762 | -0,0362478 | 1 | 1 |
| Ppp1r1b | 0,157929 | 0,153873 | -0,037535 | 1 | 1 |
| 1700013H16Rik | 0,162505 | 0,158317 | -0,0376697 | 1 | 1 |
| Cilp2 | 0,186093 | 0,181287 | -0,037748 | 1 | 1 |
| Cd46 | 0,469385 | 0,456855 | -0,039036 | 1 | 1 |
| Abcb4 | 0,062703 | 0,0609682 | -0,0404652 | 1 | 1 |
| Psma8 | 0,326127 | 0,317 | -0,0409504 | 1 | 1 |
| Tmc3 | 0,254867 | 0,247703 | -0,0411328 | 1 | 1 |
| D630045M09Rik | 0,365374 | 0,355055 | -0,041334 | 1 | 1 |
| Arhgap20 | 0,503714 | 0,488826 | -0,0432847 | 1 | 1 |
| Sgip1 | 0,231187 | 0,224035 | -0,0453323 | 1 | 1 |
| Gm21284 | 0,075354 | 0,0730118 | -0,0455591 | 1 | 1 |
| Ltb | 0,298961 | 0,289668 | -0,0455591 | 1 | 1 |
| Fam151b | 0,521954 | 0,505121 | -0,0472941 | 1 | 1 |
| Srrm4 | 0,123479 | 0,119394 | -0,0485271 | 1 | 1 |
| a | 0,71517 | 0,69147 | -0,0486197 | 1 | 1 |
| Eva1c | 0,049398 | 0,0476945 | -0,0506347 | 1 | 1 |
| C130060C02Rik | 0,471802 | 0,455459 | -0,0508584 | 1 | 1 |
| Gm10536 | 0,110964 | 0,107042 | -0,0519058 | 1 | 1 |
| Atp2b2 | 0,098942 | 0,0952556 | -0,0547765 | 1 | 1 |
| Atp6v1g3 | 0,24261 | 0,233313 | -0,0563733 | 1 | 1 |
| Lhx1 | 0,200024 | 0,192176 | -0,0577487 | 1 | 1 |
| Zfp92 | 0,262638 | 0,251642 | -0,061705 | 1 | 1 |
| Zyg11a | 0,397654 | 0,380618 | -0,063169 | 1 | 1 |
| Pla2g5 | 0,173829 | 0,166377 | -0,0632068 | 1 | 1 |
| Grid2ip | 0,105481 | 0,100704 | -0,0668734 | 1 | 1 |
| Trim43c | 0,151368 | 0,144511 | -0,0668827 | 1 | 1 |
| Ndp | 0,254135 | 0,242567 | -0,0672092 | 1 | 1 |
| Sypl2 | 0,07199 | 0,068713 | -0,0672092 | 1 | 1 |
| Oasl1 | 0,120256 | 0,114655 | -0,0688153 | 1 | 1 |
| Ccdc169 | 0,043098 | 0,0410403 | -0,0705728 | 1 | 1 |
| Rhbdl3 | 0,202731 | 0,192907 | -0,07166 | 1 | 1 |
| Fgf7 | 0,047926 | 0,045576 | -0,0725297 | 1 | 1 |
| Proca1 | 0,556396 | 0,528762 | -0,0734932 | 1 | 1 |
| Ccdc146 | 0,037188 | 0,0353161 | -0,07452 | 1 | 1 |
| Cpa4 | 0,197717 | 0,187662 | -0,0752987 | 1 | 1 |
| 5330413P13Rik | 0,038014 | 0,036045 | -0,0767158 | 1 | 1 |
| Wfdc3 | 0,211478 | 0,200526 | -0,0767158 | 1 | 1 |
| Fkbp6 | 0,040188 | 0,0380952 | -0,0771714 | 1 | 1 |

| | | | | | |
|---|---|---|---|---|---|
| Baiap2l2 | 0,282077 | 0,267092 | -0,0787536 | 1 | 1 |
| H2-K2 | 0,37527 | 0,354901 | -0,0805123 | 1 | 1 |
| Extl1 | 0,178689 | 0,168904 | -0,0812493 | 1 | 1 |
| A530072M11Rik | 0,168813 | 0,159539 | -0,0815163 | 1 | 1 |
| Cfi | 0,151973 | 0,143256 | -0,0852126 | 1 | 1 |
| Srcin1 | 0,13574 | 0,127893 | -0,0859136 | 1 | 1 |
| Phf11d | 0,054903 | 0,0517128 | -0,086351 | 1 | 1 |
| Xkr4 | 0,136332 | 0,12836 | -0,0869226 | 1 | 1 |
| Efcab6 | 0,02076 | 0,0195444 | -0,0870534 | 1 | 1 |
| Sytl5 | 0,098771 | 0,0929741 | -0,0872552 | 1 | 1 |
| 6430571L13Rik | 0,385822 | 0,362965 | -0,0881033 | 1 | 1 |
| Fam124a | 0,333628 | 0,313656 | -0,0890578 | 1 | 1 |
| Sv2c | 0,122844 | 0,115413 | -0,0900197 | 1 | 1 |
| Sardh | 0,543847 | 0,510674 | -0,0907994 | 1 | 1 |
| Syna | 0,124359 | 0,11671 | -0,0915817 | 1 | 1 |
| Smco3 | 0,387197 | 0,363342 | -0,0917373 | 1 | 1 |
| 8030423F21Rik | 0,125288 | 0,117358 | -0,0943316 | 1 | 1 |
| Zfp957 | 0,065079 | 0,06096 | -0,0943316 | 1 | 1 |
| Gm10778 | 0,03928 | 0,0367861 | -0,0946348 | 1 | 1 |
| Igfbp1 | 0,269884 | 0,252749 | -0,0946348 | 1 | 1 |
| Slc16a5 | 0,067263 | 0,0629809 | -0,0949002 | 1 | 1 |
| Cds1 | 0,422395 | 0,394092 | -0,100063 | 1 | 1 |
| Lrrc19 | 0,352568 | 0,327985 | -0,104273 | 1 | 1 |
| Kcnh6 | 0,093891 | 0,0872716 | -0,105471 | 1 | 1 |
| Noxa1 | 0,147736 | 0,137319 | -0,105486 | 1 | 1 |
| Dach2 | 0,531656 | 0,494049 | -0,10584 | 1 | 1 |
| Slc5a7 | 0,392506 | 0,364458 | -0,106963 | 1 | 1 |
| Pax4 | 0,078323 | 0,0725953 | -0,109568 | 1 | 1 |
| Aqp7 | 0,104621 | 0,0969668 | -0,10961 | 1 | 1 |
| Pou3f1 | 0,086501 | 0,0801722 | -0,10961 | 1 | 1 |
| 0610039K10Rik | 0,342758 | 0,317606 | -0,109954 | 1 | 1 |
| Ccdc62 | 0,452385 | 0,41858 | -0,112047 | 1 | 1 |
| Prune2 | 0,479009 | 0,442938 | -0,112948 | 1 | 1 |
| Hpse2 | 0,165236 | 0,152653 | -0,114269 | 1 | 1 |
| 6820431F20Rik | 0,419897 | 0,38756 | -0,115615 | 1 | 1 |
| Gm16938 | 0,129229 | 0,119259 | -0,115832 | 1 | 1 |
| Lin7a | 0,230934 | 0,213088 | -0,11603 | 1 | 1 |
| Prdm16 | 0,47014 | 0,433181 | -0,118119 | 1 | 1 |
| Grin2d | 0,383925 | 0,353551 | -0,118908 | 1 | 1 |
| Gpr61 | 0,115589 | 0,106427 | -0,119146 | 1 | 1 |
| 1810041L15Rik | 0,07357 | 0,0676437 | -0,121166 | 1 | 1 |
| Hspb2 | 0,165455 | 0,152052 | -0,121868 | 1 | 1 |
| 4930525G20Rik | 0,399864 | 0,366492 | -0,125726 | 1 | 1 |
| Eaf2 | 0,371034 | 0,339142 | -0,129661 | 1 | 1 |
| Myh14 | 0,339321 | 0,309874 | -0,130972 | 1 | 1 |
| 1700028K03Rik | 0,583296 | 0,532279 | -0,132047 | 1 | 1 |
| 4930539E08Rik | 0,252273 | 0,229819 | -0,134491 | 1 | 1 |
| Akap6 | 0,315427 | 0,287177 | -0,135364 | 1 | 1 |

| | | | | | |
|---|---|---|---|---|---|
| 4833427G06Rik | 0,103572 | 0,0942684 | -0,135784 | 1 | 1 |
| Gm6377 | 0,022773 | 0,0207271 | -0,135784 | 1 | 1 |
| Il19 | 0,07221 | 0,0657236 | -0,135784 | 1 | 1 |
| Ctnna2 | 0,255918 | 0,232819 | -0,136472 | 1 | 1 |
| A930011G23Rik | 0,028188 | 0,0256325 | -0,137111 | 1 | 1 |
| Gpha2 | 0,231223 | 0,21026 | -0,137111 | 1 | 1 |
| Myh3 | 0,015341 | 0,01395 | -0,137111 | 1 | 1 |
| Irs4 | 0,336045 | 0,305539 | -0,137298 | 1 | 1 |
| Alpk1 | 0,545525 | 0,495732 | -0,138084 | 1 | 1 |
| Kcnb1 | 0,255736 | 0,232191 | -0,139338 | 1 | 1 |
| Irs3 | 0,119415 | 0,108384 | -0,139837 | 1 | 1 |
| Klk10 | 0,195703 | 0,177624 | -0,139847 | 1 | 1 |
| Pde6g | 0,238495 | 0,216458 | -0,139877 | 1 | 1 |
| Kcnt2 | 0,378502 | 0,343463 | -0,140143 | 1 | 1 |
| Itgb7 | 0,311596 | 0,282742 | -0,140192 | 1 | 1 |
| Gm5868 | 0,18118 | 0,164305 | -0,141048 | 1 | 1 |
| 4930405A21Rik | 0,538737 | 0,488553 | -0,141065 | 1 | 1 |
| Scn4a | 0,028686 | 0,0260104 | -0,141256 | 1 | 1 |
| Chrna5 | 0,331668 | 0,300651 | -0,14165 | 1 | 1 |
| 1700101E01Rik | 0,14665 | 0,13289 | -0,142153 | 1 | 1 |
| Erich5 | 0,049087 | 0,0444806 | -0,142153 | 1 | 1 |
| Adh6a | 0,161459 | 0,14627 | -0,142539 | 1 | 1 |
| Tnfrsf13c | 0,306162 | 0,277329 | -0,142698 | 1 | 1 |
| Ttc34 | 0,086964 | 0,0787743 | -0,142698 | 1 | 1 |
| Fank1 | 0,043483 | 0,0393832 | -0,142854 | 1 | 1 |
| Adssl1 | 0,339476 | 0,307425 | -0,143075 | 1 | 1 |
| Gm19461 | 0,026466 | 0,0239676 | -0,143075 | 1 | 1 |
| Htra4 | 0,03893 | 0,0352543 | -0,143075 | 1 | 1 |
| Plxna4os1 | 0,04514 | 0,0408784 | -0,143075 | 1 | 1 |
| Pon1 | 0,058796 | 0,0532448 | -0,143075 | 1 | 1 |
| Vmn1r53 | 0,083838 | 0,0759228 | -0,143075 | 1 | 1 |
| 1700003M07Rik | 0,043284 | 0,0391974 | -0,143088 | 1 | 1 |
| Apobec2 | 0,548724 | 0,496795 | -0,143429 | 1 | 1 |
| Nek11 | 0,083123 | 0,0752496 | -0,143564 | 1 | 1 |
| Gabra1 | 0,038276 | 0,0346343 | -0,144222 | 1 | 1 |
| Ccno | 0,028297 | 0,0255883 | -0,145156 | 1 | 1 |
| Ephx3 | 0,021389 | 0,0193416 | -0,145156 | 1 | 1 |
| Epx | 0,019024 | 0,0172029 | -0,145156 | 1 | 1 |
| Pdx1 | 0,043686 | 0,0395045 | -0,145156 | 1 | 1 |
| Slfnl1 | 0,030367 | 0,0274606 | -0,145156 | 1 | 1 |
| Smim18 | 0,123527 | 0,111704 | -0,145156 | 1 | 1 |
| Grin1 | 0,081386 | 0,0735886 | -0,145301 | 1 | 1 |
| Ccin | 0,094059 | 0,0850301 | -0,145595 | 1 | 1 |
| Evx2 | 0,14032 | 0,12685 | -0,145595 | 1 | 1 |
| 4930500L23Rik | 0,262658 | 0,237243 | -0,146815 | 1 | 1 |
| Acss2os | 0,120351 | 0,108553 | -0,148845 | 1 | 1 |
| Efhc1 | 0,469676 | 0,423558 | -0,149104 | 1 | 1 |
| 9430037G07Rik | 0,296756 | 0,267402 | -0,150268 | 1 | 1 |

| | | | | | |
|---|---|---|---|---|---|
| Nudt15 | 0,307368 | 0,276555 | -0,1524 | 1 | 1 |
| Gm10548 | 0,312433 | 0,280869 | -0,153649 | 1 | 1 |
| Carns1 | 0,555138 | 0,499047 | -0,153671 | 1 | 1 |
| C030016D13Rik | 0,282316 | 0,253695 | -0,154214 | 1 | 1 |
| Ccdc17 | 0,399701 | 0,359147 | -0,154345 | 1 | 1 |
| Styx | 0,503797 | 0,452451 | -0,15508 | 1 | 1 |
| Chsy3 | 0,469922 | 0,421972 | -0,155275 | 1 | 1 |
| Zfp433 | 0,287863 | 0,257985 | -0,158096 | 1 | 1 |
| Catsperd | 0,406276 | 0,363601 | -0,160104 | 1 | 1 |
| AI854703 | 0,423471 | 0,378941 | -0,160292 | 1 | 1 |
| Mir678 | 70,2953 | 62,835 | -0,161861 | 1 | 1 |
| Kbtbd13 | 0,451809 | 0,403819 | -0,162005 | 1 | 1 |
| Ttc18 | 0,149154 | 0,132978 | -0,165618 | 1 | 1 |
| Galr3 | 0,110137 | 0,0981316 | -0,166503 | 1 | 1 |
| Acoxl | 0,12916 | 0,115057 | -0,166815 | 1 | 1 |
| Lctl | 0,054212 | 0,0482218 | -0,168935 | 1 | 1 |
| Hmga1-rs1 | 29,2096 | 25,9732 | -0,169417 | 1 | 1 |
| Cdh12 | 0,107831 | 0,0958638 | -0,16972 | 1 | 1 |
| 03/10/16 | 0,052781 | 0,0468887 | -0,170783 | 1 | 1 |
| C130050O18Rik | 0,174169 | 0,154712 | -0,170902 | 1 | 1 |
| Plin4 | 0,05654 | 0,0502237 | -0,170902 | 1 | 1 |
| 5430402O13Rik | 0,188951 | 0,167824 | -0,171061 | 1 | 1 |
| 2510049J12Rik | 0,119146 | 0,105804 | -0,171334 | 1 | 1 |
| Ms4a4d | 0,239571 | 0,212496 | -0,173017 | 1 | 1 |
| Vsig10l | 0,442369 | 0,39223 | -0,173553 | 1 | 1 |
| 4930412O13Rik | 0,557453 | 0,493946 | -0,174497 | 1 | 1 |
| Gm15787 | 0,517091 | 0,458126 | -0,174675 | 1 | 1 |
| 1700110C19Rik | 0,479823 | 0,424968 | -0,175147 | 1 | 1 |
| Lgals6 | 0,097138 | 0,0860079 | -0,17557 | 1 | 1 |
| Gpr176 | 0,135717 | 0,120071 | -0,176719 | 1 | 1 |
| Mab21l3 | 0,577154 | 0,510466 | -0,177141 | 1 | 1 |
| 1700028J19Rik | 0,511965 | 0,452795 | -0,177187 | 1 | 1 |
| Paqr5 | 0,217335 | 0,192203 | -0,17729 | 1 | 1 |
| Ksr2 | 0,163669 | 0,144742 | -0,1773 | 1 | 1 |
| Zfp536 | 0,220919 | 0,195365 | -0,177345 | 1 | 1 |
| Prrt4 | 0,223278 | 0,197414 | -0,177618 | 1 | 1 |
| Col4a4 | 0,03111 | 0,0274979 | -0,17807 | 1 | 1 |
| Gm12657 | 0,305843 | 0,270143 | -0,179065 | 1 | 1 |
| Adrb3 | 0,175976 | 0,155275 | -0,180552 | 1 | 1 |
| Cldn15 | 0,409945 | 0,36159 | -0,181074 | 1 | 1 |
| Slc36a2 | 0,090291 | 0,0796381 | -0,181119 | 1 | 1 |
| Ihh | 0,179679 | 0,158352 | -0,18229 | 1 | 1 |
| Cnpy1 | 0,504657 | 0,444552 | -0,182952 | 1 | 1 |
| Thsd7b | 0,105672 | 0,0929756 | -0,184667 | 1 | 1 |
| Cldn20 | 0,454097 | 0,399507 | -0,184782 | 1 | 1 |
| Tbc1d30 | 0,113164 | 0,0995337 | -0,185161 | 1 | 1 |
| Plcxd2 | 0,515417 | 0,452963 | -0,186347 | 1 | 1 |
| Celf3 | 0,13863 | 0,121661 | -0,188381 | 1 | 1 |

| | | | | | |
|---|---|---|---|---|---|
| Cabyr | 0,548079 | 0,480564 | -0,189656 | 1 | 1 |
| Stard6 | 0,386493 | 0,338609 | -0,190824 | 1 | 1 |
| Hsbp1l1 | 0,163728 | 0,143412 | -0,19114 | 1 | 1 |
| Abcg5 | 0,094904 | 0,0831154 | -0,19136 | 1 | 1 |
| Serpinb6c | 0,17385 | 0,152254 | -0,19136 | 1 | 1 |
| Lrfn2 | 0,249677 | 0,218595 | -0,191803 | 1 | 1 |
| Trpm3 | 0,099668 | 0,0872296 | -0,192315 | 1 | 1 |
| Ect2l | 0,057095 | 0,0499149 | -0,193898 | 1 | 1 |
| Krt20 | 0,089182 | 0,0779665 | -0,193898 | 1 | 1 |
| Serpine3 | 0,42509 | 0,371535 | -0,194273 | 1 | 1 |
| Padi1 | 0,049142 | 0,0429123 | -0,19557 | 1 | 1 |
| Cadps2 | 0,299536 | 0,26144 | -0,19625 | 1 | 1 |
| Brinp2 | 0,297699 | 0,259773 | -0,196605 | 1 | 1 |
| Wdr96 | 0,456608 | 0,398216 | -0,197405 | 1 | 1 |
| Lrrc48 | 0,568964 | 0,496071 | -0,197789 | 1 | 1 |
| Myo7b | 0,025039 | 0,021825 | -0,198203 | 1 | 1 |
| Nptxr | 0,349462 | 0,304211 | -0,200061 | 1 | 1 |
| Grid1 | 0,181781 | 0,158181 | -0,200631 | 1 | 1 |
| Rbfox1 | 0,230374 | 0,200392 | -0,201157 | 1 | 1 |
| Rai2 | 0,285052 | 0,247792 | -0,202098 | 1 | 1 |
| Agtr1b | 0,36859 | 0,320357 | -0,202335 | 1 | 1 |
| Gm14827 | 0,250318 | 0,217428 | -0,203222 | 1 | 1 |
| Kcnc2 | 0,073065 | 0,0634562 | -0,203415 | 1 | 1 |
| Gm14057 | 0,514856 | 0,446836 | -0,204424 | 1 | 1 |
| Wbscr17 | 0,48053 | 0,416958 | -0,204727 | 1 | 1 |
| Fam228a | 0,331939 | 0,287837 | -0,20567 | 1 | 1 |
| Gm16982 | 0,353242 | 0,305903 | -0,207586 | 1 | 1 |
| Smc1b | 0,061712 | 0,0532504 | -0,212748 | 1 | 1 |
| Fibcd1 | 0,23697 | 0,204433 | -0,213076 | 1 | 1 |
| Fam178b | 0,489548 | 0,422327 | -0,213091 | 1 | 1 |
| Neurog1 | 0,153939 | 0,132164 | -0,220039 | 1 | 1 |
| C920025E04Rik | 0,42008 | 0,360618 | -0,220194 | 1 | 1 |
| Oprl1 | 0,208878 | 0,179165 | -0,221373 | 1 | 1 |
| Best1 | 0,583325 | 0,500108 | -0,22206 | 1 | 1 |
| BC030499 | 0,107043 | 0,0916776 | -0,22355 | 1 | 1 |
| Gpr98 | 0,465833 | 0,39874 | -0,224364 | 1 | 1 |
| Olfr1372-ps1 | 0,420199 | 0,359594 | -0,224703 | 1 | 1 |
| Naalad2 | 0,552505 | 0,472732 | -0,224965 | 1 | 1 |
| Slc12a8 | 0,357817 | 0,305515 | -0,227978 | 1 | 1 |
| Adgb | 0,018872 | 0,0161034 | -0,228893 | 1 | 1 |
| Rps13 | 0,501077 | 0,427195 | -0,230137 | 1 | 1 |
| 9430014N10Rik | 0,102252 | 0,0870999 | -0,23138 | 1 | 1 |
| Tcerg1l | 0,122636 | 0,104462 | -0,231403 | 1 | 1 |
| Armc2 | 0,105401 | 0,0897206 | -0,232381 | 1 | 1 |
| Lpo | 0,071762 | 0,0610759 | -0,232621 | 1 | 1 |
| Dzank1 | 0,440345 | 0,374771 | -0,232624 | 1 | 1 |
| A330040F15Rik | 0,111552 | 0,0948632 | -0,23379 | 1 | 1 |
| Cxcl10 | 0,224597 | 0,190997 | -0,23379 | 1 | 1 |

| | | | | | |
|---|---|---|---|---|---|
| Cyct | 0,224594 | 0,190994 | -0,23379 | 1 | 1 |
| Krt80 | 0,064089 | 0,0545014 | -0,23379 | 1 | 1 |
| Prg4 | 0,353837 | 0,300785 | -0,23435 | 1 | 1 |
| Plin5 | 0,156973 | 0,133313 | -0,235702 | 1 | 1 |
| Ccdc149 | 0,583892 | 0,494562 | -0,239549 | 1 | 1 |
| Tmem132b | 0,242523 | 0,205336 | -0,240138 | 1 | 1 |
| Ccdc155 | 0,218409 | 0,184735 | -0,24158 | 1 | 1 |
| Gm1653 | 0,117289 | 0,0991611 | -0,242218 | 1 | 1 |
| Dio2 | 0,046435 | 0,0392334 | -0,243123 | 1 | 1 |
| A930012L18Rik | 0,259953 | 0,219422 | -0,244537 | 1 | 1 |
| Myl4 | 0,51448 | 0,433957 | -0,245564 | 1 | 1 |
| 1700034H15Rik | 0,294528 | 0,248075 | -0,247628 | 1 | 1 |
| Ccl12 | 0,249379 | 0,209926 | -0,248459 | 1 | 1 |
| Foxl2 | 0,03575 | 0,030094 | -0,248459 | 1 | 1 |
| Syt2 | 0,02602 | 0,0219038 | -0,248459 | 1 | 1 |
| Snx20 | 0,264647 | 0,222735 | -0,248738 | 1 | 1 |
| Cd160 | 0,283095 | 0,238183 | -0,249214 | 1 | 1 |
| Hpcal4 | 0,037266 | 0,031339 | -0,249881 | 1 | 1 |
| Hsd17b1 | 0,148927 | 0,125242 | -0,249881 | 1 | 1 |
| Bcl2l14 | 0,452253 | 0,380295 | -0,25001 | 1 | 1 |
| Mb21d1 | 0,502226 | 0,4223 | -0,250067 | 1 | 1 |
| E030013I19Rik | 0,096815 | 0,081391 | -0,250355 | 1 | 1 |
| Myog | 0,065015 | 0,0546218 | -0,251302 | 1 | 1 |
| 6720468P15Rik | 0,072963 | 0,0612981 | -0,251318 | 1 | 1 |
| A430088P11Rik | 0,074528 | 0,0625639 | -0,252461 | 1 | 1 |
| Slc4a5 | 0,131172 | 0,110105 | -0,252589 | 1 | 1 |
| Myo16 | 0,220651 | 0,185096 | -0,253491 | 1 | 1 |
| Mccc1os | 0,225099 | 0,18855 | -0,255611 | 1 | 1 |
| Gabbr2 | 0,099849 | 0,0834872 | -0,25819 | 1 | 1 |
| BC016579 | 0,077347 | 0,0645628 | -0,260633 | 1 | 1 |
| Pla2g2c | 0,422836 | 0,352848 | -0,26105 | 1 | 1 |
| Mc4r | 0,176795 | 0,147432 | -0,26202 | 1 | 1 |
| P2rx2 | 0,383514 | 0,319723 | -0,26246 | 1 | 1 |
| Gpr68 | 0,37923 | 0,316073 | -0,262814 | 1 | 1 |
| Gm6644 | 11,967 | 9,97398 | -0,262821 | 1 | 1 |
| Vstm5 | 0,577366 | 0,4812 | -0,262851 | 1 | 1 |
| Ccdc116 | 0,405855 | 0,338213 | -0,263032 | 1 | 1 |
| Adamts8 | 0,169214 | 0,140916 | -0,264017 | 1 | 1 |
| Lrrc75b | 0,413386 | 0,344021 | -0,264992 | 1 | 1 |
| Trpm2 | 0,030907 | 0,0257026 | -0,265998 | 1 | 1 |
| Plcb1 | 0,301011 | 0,249962 | -0,268109 | 1 | 1 |
| Bdnf | 0,407174 | 0,338107 | -0,268167 | 1 | 1 |
| Dmrta1 | 0,302995 | 0,250955 | -0,271866 | 1 | 1 |
| Them7 | 0,188475 | 0,156042 | -0,272437 | 1 | 1 |
| Gucy2e | 0,192315 | 0,159041 | -0,274078 | 1 | 1 |
| Pnoc | 0,034327 | 0,0283705 | -0,274932 | 1 | 1 |
| Eomes | 0,075423 | 0,0623325 | -0,275023 | 1 | 1 |
| Slc24a4 | 0,072089 | 0,059577 | -0,275029 | 1 | 1 |

| | | | | | |
|---|---|---|---|---|---|
| Ccdc184 | 0,094702 | 0,0782633 | -0,275051 | 1 | 1 |
| Gpr15 | 0,059978 | 0,0495652 | -0,275096 | 1 | 1 |
| Krt25 | 0,045014 | 0,0371997 | -0,275096 | 1 | 1 |
| Lrrc10 | 0,053216 | 0,043977 | -0,275096 | 1 | 1 |
| Mafa | 0,074356 | 0,0614471 | -0,275096 | 1 | 1 |
| Tnfsf11 | 0,032071 | 0,0265033 | -0,275096 | 1 | 1 |
| Abcc2 | 0,078781 | 0,0650811 | -0,275606 | 1 | 1 |
| Svs1 | 0,131173 | 0,108353 | -0,27574 | 1 | 1 |
| Hormad2 | 0,358769 | 0,296262 | -0,276183 | 1 | 1 |
| Jsrp1 | 0,56479 | 0,465353 | -0,279389 | 1 | 1 |
| Trim17 | 0,573469 | 0,472206 | -0,280301 | 1 | 1 |
| Fndc1 | 0,373485 | 0,307125 | -0,282224 | 1 | 1 |
| Gnmt | 0,314467 | 0,258353 | -0,283565 | 1 | 1 |
| Dgki | 0,057931 | 0,0475862 | -0,283795 | 1 | 1 |
| E030011O05Rik | 0,455357 | 0,374042 | -0,283795 | 1 | 1 |
| Lrrn4cl | 0,288712 | 0,236991 | -0,284798 | 1 | 1 |
| Trpv6 | 0,325903 | 0,267517 | -0,284811 | 1 | 1 |
| Dpyd | 0,309133 | 0,253339 | -0,287157 | 1 | 1 |
| Fgf12 | 0,235323 | 0,192546 | -0,289439 | 1 | 1 |
| Slc23a1 | 0,432716 | 0,353526 | -0,291605 | 1 | 1 |
| Gm6083 | 0,501525 | 0,409363 | -0,29294 | 1 | 1 |
| Tppp | 0,195026 | 0,159174 | -0,293068 | 1 | 1 |
| Ido2 | 0,17068 | 0,139243 | -0,293685 | 1 | 1 |
| Adad2 | 0,068703 | 0,0560393 | -0,293935 | 1 | 1 |
| Kiss1r | 0,120252 | 0,0980862 | -0,293935 | 1 | 1 |
| Nefh | 0,031268 | 0,0255048 | -0,293935 | 1 | 1 |
| Mfi2 | 0,134104 | 0,109275 | -0,295386 | 1 | 1 |
| Zfp641 | 0,30547 | 0,248858 | -0,295709 | 1 | 1 |
| Arpp21 | 0,146539 | 0,119373 | -0,295813 | 1 | 1 |
| Hlf | 0,179951 | 0,146459 | -0,297112 | 1 | 1 |
| Apobr | 0,406292 | 0,330305 | -0,298719 | 1 | 1 |
| Palm2 | 0,547469 | 0,444396 | -0,30093 | 1 | 1 |
| L1td1 | 0,583734 | 0,473829 | -0,300943 | 1 | 1 |
| Tnnc2 | 0,351944 | 0,285641 | -0,301144 | 1 | 1 |
| Apon | 0,523214 | 0,424585 | -0,301346 | 1 | 1 |
| Frmd3 | 0,112466 | 0,0911564 | -0,303078 | 1 | 1 |
| Slitrk4 | 0,447439 | 0,362522 | -0,303625 | 1 | 1 |
| Adamtsl4 | 0,181552 | 0,147012 | -0,304446 | 1 | 1 |
| Gpc5 | 0,083946 | 0,067952 | -0,304953 | 1 | 1 |
| Tdrd9 | 0,327896 | 0,26524 | -0,305942 | 1 | 1 |
| Gm5820 | 0,298484 | 0,241167 | -0,307622 | 1 | 1 |
| Pak7 | 0,132171 | 0,106782 | -0,307736 | 1 | 1 |
| Synpo2l | 0,162423 | 0,131223 | -0,307736 | 1 | 1 |
| Gm5069 | 0,496167 | 0,400524 | -0,308938 | 1 | 1 |
| Pla1a | 0,214025 | 0,172714 | -0,309392 | 1 | 1 |
| Stox1 | 0,388134 | 0,313135 | -0,309769 | 1 | 1 |
| Slco4c1 | 0,440438 | 0,355123 | -0,310621 | 1 | 1 |
| Gm6568 | 0,679324 | 0,547369 | -0,311587 | 1 | 1 |

| | | | | | |
|---|---|---|---|---|---|
| Rmst | 0,250321 | 0,2015 | -0,313001 | 1 | 1 |
| Btbd17 | 0,331232 | 0,266504 | -0,313686 | 1 | 1 |
| Klb | 0,322107 | 0,25897 | -0,314754 | 1 | 1 |
| Vipr2 | 0,422368 | 0,339076 | -0,316889 | 1 | 1 |
| Gm13247 | 0,019387 | 0,015561 | -0,317161 | 1 | 1 |
| 1700028I16Rik | 0,062568 | 0,0502151 | -0,317292 | 1 | 1 |
| 5830403L16Rik | 0,036764 | 0,0295059 | -0,317292 | 1 | 1 |
| BC033916 | 0,088403 | 0,0709503 | -0,317292 | 1 | 1 |
| E130114P18Rik | 0,051906 | 0,0416586 | -0,317292 | 1 | 1 |
| E230025N22Rik | 0,058839 | 0,0472225 | -0,317292 | 1 | 1 |
| Elovl3 | 0,063651 | 0,0510849 | -0,317292 | 1 | 1 |
| Fev | 0,04181 | 0,0335557 | -0,317292 | 1 | 1 |
| Gm15133 | 0,084884 | 0,0681259 | -0,317292 | 1 | 1 |
| Grid2 | 0,018904 | 0,0151715 | -0,317292 | 1 | 1 |
| Hist1h2af | 0,201916 | 0,162052 | -0,317292 | 1 | 1 |
| Nsun7 | 0,022704 | 0,0182213 | -0,317292 | 1 | 1 |
| Serpina3m | 0,036638 | 0,0294045 | -0,317292 | 1 | 1 |
| Tmem89 | 0,1596 | 0,128091 | -0,317292 | 1 | 1 |
| Atp2b3 | 0,015376 | 0,0123384 | -0,317527 | 1 | 1 |
| Oacyl | 0,143157 | 0,114853 | -0,31781 | 1 | 1 |
| Plcd4 | 0,337981 | 0,270855 | -0,31942 | 1 | 1 |
| Mir546 | 489,792 | 392,475 | -0,319567 | 1 | 1 |
| Zfp354b | 0,305026 | 0,244094 | -0,321494 | 1 | 1 |
| Gm6225 | 0,372043 | 0,297529 | -0,322437 | 1 | 1 |
| Nkg7 | 0,660879 | 0,528029 | -0,323768 | 1 | 1 |
| Gm6402 | 0,344312 | 0,274955 | -0,32452 | 1 | 1 |
| Sarm1 | 0,486756 | 0,388246 | -0,326227 | 1 | 1 |
| Scn3b | 0,115829 | 0,0923604 | -0,326654 | 1 | 1 |
| Gm10390 | 0,089246 | 0,0711062 | -0,327806 | 1 | 1 |
| Tlx2 | 0,51981 | 0,413477 | -0,330179 | 1 | 1 |
| Fndc8 | 0,337526 | 0,268382 | -0,330711 | 1 | 1 |
| Pdzrn4 | 0,222746 | 0,176795 | -0,333324 | 1 | 1 |
| 5930403L14Rik | 0,497697 | 0,394409 | -0,335576 | 1 | 1 |
| A230056J06Rik | 0,04817 | 0,0381294 | -0,337223 | 1 | 1 |
| Sh2d7 | 0,108887 | 0,0861597 | -0,337741 | 1 | 1 |
| Tap2 | 0,533437 | 0,422014 | -0,338025 | 1 | 1 |
| Tssk2 | 0,194872 | 0,153867 | -0,340843 | 1 | 1 |
| Acnat1 | 0,4888 | 0,385214 | -0,343585 | 1 | 1 |
| A830009L08Rik | 0,034112 | 0,026776 | -0,349322 | 1 | 1 |
| Saa2 | 0,218695 | 0,171665 | -0,349322 | 1 | 1 |
| Phkg1 | 0,099148 | 0,0777104 | -0,351476 | 1 | 1 |
| Ankrd29 | 0,483011 | 0,378283 | -0,352592 | 1 | 1 |
| Asprv1 | 0,36594 | 0,286531 | -0,352918 | 1 | 1 |
| Nol4 | 0,208257 | 0,16297 | -0,35376 | 1 | 1 |
| Scn3a | 0,1373 | 0,10732 | -0,355418 | 1 | 1 |
| Luzp2 | 0,220674 | 0,172453 | -0,35572 | 1 | 1 |
| Bnc2 | 0,507689 | 0,396669 | -0,356011 | 1 | 1 |
| Tmem171 | 0,666003 | 0,519558 | -0,358244 | 1 | 1 |

| | | | | | |
|---|---|---|---|---|---|
| Irgm2 | 0,52579 | 0,410089 | -0,358551 | 1 | 1 |
| Pnpla3 | 0,047971 | 0,0374151 | -0,358553 | 1 | 1 |
| Ntrk2 | 0,104088 | 0,0810958 | -0,360104 | 1 | 1 |
| Adra1a | 0,085843 | 0,0668602 | -0,36056 | 1 | 1 |
| Tusc5 | 0,177629 | 0,138339 | -0,360666 | 1 | 1 |
| C2cd4c | 0,071959 | 0,0560241 | -0,361136 | 1 | 1 |
| Glt28d2 | 0,542189 | 0,42201 | -0,36152 | 1 | 1 |
| Ankmy1 | 0,081842 | 0,0635814 | -0,36423 | 1 | 1 |
| Cybrd1 | 0,410516 | 0,318705 | -0,365218 | 1 | 1 |
| Tmem102 | 0,294826 | 0,228832 | -0,365576 | 1 | 1 |
| Bai3 | 0,292862 | 0,227219 | -0,366139 | 1 | 1 |
| Msh4 | 0,036332 | 0,0281627 | -0,367443 | 1 | 1 |
| AB124611 | 0,389317 | 0,301774 | -0,367479 | 1 | 1 |
| Hnf1a | 0,095251 | 0,0738223 | -0,367674 | 1 | 1 |
| Grk1 | 0,079502 | 0,0616059 | -0,367917 | 1 | 1 |
| Slc41a2 | 0,077838 | 0,060229 | -0,370015 | 1 | 1 |
| Zfp300 | 0,518395 | 0,400816 | -0,371114 | 1 | 1 |
| Lgals12 | 0,204269 | 0,15771 | -0,373197 | 1 | 1 |
| Npas2 | 0,321615 | 0,24733 | -0,378898 | 1 | 1 |
| Slc24a2 | 0,105429 | 0,0809428 | -0,381292 | 1 | 1 |
| A330074K22Rik | 0,235777 | 0,180902 | -0,382211 | 1 | 1 |
| Galr2 | 0,286539 | 0,219828 | -0,382356 | 1 | 1 |
| Tbx15 | 0,079208 | 0,06074 | -0,383004 | 1 | 1 |
| Gm10653 | 0,530537 | 0,406815 | -0,383079 | 1 | 1 |
| Dclk3 | 0,291103 | 0,223114 | -0,383748 | 1 | 1 |
| Kng1 | 0,10671 | 0,0816872 | -0,385517 | 1 | 1 |
| Gm17769 | 0,466591 | 0,357031 | -0,386111 | 1 | 1 |
| Mast1 | 0,271053 | 0,207264 | -0,387111 | 1 | 1 |
| Hydin | 0,031535 | 0,0240968 | -0,388088 | 1 | 1 |
| Pbld2 | 0,486817 | 0,371689 | -0,389282 | 1 | 1 |
| Gm973 | 0,151852 | 0,115797 | -0,391071 | 1 | 1 |
| Sh3rf3 | 0,239923 | 0,182885 | -0,391631 | 1 | 1 |
| Zfp947 | 0,573803 | 0,437383 | -0,391658 | 1 | 1 |
| Fbxl13 | 0,048318 | 0,0367688 | -0,394072 | 1 | 1 |
| Itih4 | 0,044721 | 0,0340108 | -0,394952 | 1 | 1 |
| Gsta3 | 0,12632 | 0,0960498 | -0,395224 | 1 | 1 |
| 1700016K19Rik | 0,206319 | 0,156876 | -0,395254 | 1 | 1 |
| Ccdc105 | 0,09771 | 0,074294 | -0,395254 | 1 | 1 |
| Lsamp | 0,115132 | 0,087516 | -0,395674 | 1 | 1 |
| Spib | 0,114651 | 0,0871274 | -0,39605 | 1 | 1 |
| 1700001C19Rik | 0,099136 | 0,075322 | -0,396341 | 1 | 1 |
| Prr19 | 0,369736 | 0,280719 | -0,397367 | 1 | 1 |
| Sag | 0,168129 | 0,127624 | -0,39767 | 1 | 1 |
| 4930502E09Rik | 0,064399 | 0,048845 | -0,398818 | 1 | 1 |
| 4930519G04Rik | 0,019316 | 0,0146511 | -0,398818 | 1 | 1 |
| Afm | 0,019518 | 0,0148042 | -0,398818 | 1 | 1 |
| Atp4b | 0,031598 | 0,0239661 | -0,398818 | 1 | 1 |
| Cdr1 | 0,016825 | 0,0127615 | -0,398818 | 1 | 1 |

| | | | | | |
|---|---|---|---|---|---|
| Clec3a | 0,013017 | 0,00987311 | -0,398818 | 1 | 1 |
| Cngb1 | 0,005927 | 0,00449544 | -0,398818 | 1 | 1 |
| Erich4 | 0,05026 | 0,0381214 | -0,398818 | 1 | 1 |
| Foxe1 | 0,038908 | 0,0295108 | -0,398818 | 1 | 1 |
| Grifin | 0,10592 | 0,0803379 | -0,398818 | 1 | 1 |
| Nek5 | 0,032258 | 0,0244673 | -0,398818 | 1 | 1 |
| Orm2 | 0,062252 | 0,0472169 | -0,398818 | 1 | 1 |
| Stap1 | 0,038073 | 0,0288775 | -0,398818 | 1 | 1 |
| Tbx5 | 0,018991 | 0,0144041 | -0,398818 | 1 | 1 |
| Cmtm5 | 0,051726 | 0,0392328 | -0,398833 | 1 | 1 |
| 4933431G14Rik | 0,13372 | 0,101371 | -0,399577 | 1 | 1 |
| Wisp2 | 0,135626 | 0,102791 | -0,399924 | 1 | 1 |
| Cacna1s | 0,023738 | 0,0179863 | -0,400287 | 1 | 1 |
| Hist2h2ac | 0,538236 | 0,407771 | -0,400477 | 1 | 1 |
| Ppp1r2-ps3 | 0,23683 | 0,179424 | -0,400477 | 1 | 1 |
| Ucn2 | 0,178069 | 0,134906 | -0,400477 | 1 | 1 |
| Rgs8 | 0,28228 | 0,213804 | -0,400836 | 1 | 1 |
| Cdk15 | 0,167503 | 0,126853 | -0,401029 | 1 | 1 |
| Cyp46a1 | 0,39297 | 0,297088 | -0,403529 | 1 | 1 |
| Plekha4 | 0,274711 | 0,207644 | -0,403805 | 1 | 1 |
| AI182371 | 0,429879 | 0,324885 | -0,404 | 1 | 1 |
| Lpar3 | 0,248139 | 0,187437 | -0,404741 | 1 | 1 |
| Gbp3 | 0,227084 | 0,171284 | -0,406834 | 1 | 1 |
| Tmem181c-ps | 0,192699 | 0,145166 | -0,40864 | 1 | 1 |
| Amd2 | 7,7754 | 5,85524 | -0,409188 | 1 | 1 |
| Ptchd2 | 0,135334 | 0,101907 | -0,409264 | 1 | 1 |
| Gm3458 | 0,207376 | 0,156141 | -0,409395 | 1 | 1 |
| Hand1 | 0,20622 | 0,155207 | -0,409987 | 1 | 1 |
| Wt1 | 0,21735 | 0,16344 | -0,411255 | 1 | 1 |
| Map3k8 | 0,57319 | 0,430585 | -0,412714 | 1 | 1 |
| Fam186b | 0,081368 | 0,0610726 | -0,413939 | 1 | 1 |
| Hgd | 0,16729 | 0,125332 | -0,416603 | 1 | 1 |
| Cytl1 | 0,568992 | 0,425702 | -0,418566 | 1 | 1 |
| Gm10584 | 0,455054 | 0,340368 | -0,418944 | 1 | 1 |
| Zxda | 0,550941 | 0,411962 | -0,419387 | 1 | 1 |
| Nxnl2 | 0,210013 | 0,156871 | -0,420904 | 1 | 1 |
| Pcdhb22 | 0,198477 | 0,148238 | -0,42105 | 1 | 1 |
| Mtus2 | 0,118559 | 0,0884823 | -0,422142 | 1 | 1 |
| Col20a1 | 0,126249 | 0,0942042 | -0,422411 | 1 | 1 |
| Fgf10 | 0,451296 | 0,33651 | -0,423423 | 1 | 1 |
| Stpg1 | 0,27216 | 0,202332 | -0,427728 | 1 | 1 |
| Gdap1 | 0,114198 | 0,0848514 | -0,428529 | 1 | 1 |
| Bsn | 0,360488 | 0,26778 | -0,428903 | 1 | 1 |
| Efcab12 | 0,456472 | 0,338723 | -0,430419 | 1 | 1 |
| 2610005L07Rik | 0,068474 | 0,0507816 | -0,431258 | 1 | 1 |
| Ppfia2 | 0,020906 | 0,0155038 | -0,431258 | 1 | 1 |
| Slc10a5 | 0,035029 | 0,025978 | -0,431258 | 1 | 1 |
| Ak5 | 0,537063 | 0,398247 | -0,431428 | 1 | 1 |

| | | | | | |
|---|---|---|---|---|---|
| Scn2a1 | 0,057205 | 0,042368 | -0,433172 | 1 | 1 |
| Actl10 | 0,196259 | 0,145257 | -0,434154 | 1 | 1 |
| Zc3h12a | 0,346051 | 0,255763 | -0,436178 | 1 | 1 |
| 3110070M22Rik | 0,665552 | 0,490966 | -0,438928 | 1 | 1 |
| Gm10789 | 0,319813 | 0,235793 | -0,439707 | 1 | 1 |
| Vdr | 0,198262 | 0,146165 | -0,439808 | 1 | 1 |
| Ttll13 | 0,137201 | 0,101075 | -0,44086 | 1 | 1 |
| 4930562C15Rik | 0,08658 | 0,0637745 | -0,441056 | 1 | 1 |
| Alms1-ps2 | 0,097954 | 0,0721503 | -0,441104 | 1 | 1 |
| Pcdhb16 | 0,037357 | 0,027516 | -0,441104 | 1 | 1 |
| 1700007G11Rik | 0,144272 | 0,106083 | -0,443592 | 1 | 1 |
| Rdh1 | 0,025716 | 0,0189088 | -0,443592 | 1 | 1 |
| Gm1564 | 0,063619 | 0,0467113 | -0,445684 | 1 | 1 |
| Got1l1 | 0,298037 | 0,218642 | -0,446923 | 1 | 1 |
| Serpind1 | 0,125196 | 0,0918241 | -0,447248 | 1 | 1 |
| Foxq1 | 0,377821 | 0,276458 | -0,450642 | 1 | 1 |
| Car5b | 0,178757 | 0,130788 | -0,450772 | 1 | 1 |
| Rorb | 0,046376 | 0,033899 | -0,452128 | 1 | 1 |
| Zfp112 | 0,546575 | 0,399466 | -0,452349 | 1 | 1 |
| Ocstamp | 0,087859 | 0,0642016 | -0,452573 | 1 | 1 |
| Kcnq5 | 0,112777 | 0,0822583 | -0,455245 | 1 | 1 |
| Rgs22 | 0,108957 | 0,0790959 | -0,462089 | 1 | 1 |
| Pde8b | 0,4502 | 0,326784 | -0,462229 | 1 | 1 |
| Oscar | 0,200387 | 0,145383 | -0,462931 | 1 | 1 |
| Ugt1a6b | 0,035525 | 0,025767 | -0,463289 | 1 | 1 |
| Krt222 | 0,257289 | 0,186493 | -0,464264 | 1 | 1 |
| Medag | 0,166153 | 0,12035 | -0,46528 | 1 | 1 |
| Hsd3b3 | 0,045969 | 0,0332672 | -0,466552 | 1 | 1 |
| Gm7008 | 0,300081 | 0,217086 | -0,467084 | 1 | 1 |
| Lipc | 0,034464 | 0,0249322 | -0,467084 | 1 | 1 |
| Vit | 0,024135 | 0,0174586 | -0,467177 | 1 | 1 |
| Rnf112 | 0,020931 | 0,0151407 | -0,467185 | 1 | 1 |
| Hecw1 | 0,047317 | 0,0342212 | -0,467475 | 1 | 1 |
| Creb3l1 | 0,47825 | 0,345545 | -0,468892 | 1 | 1 |
| Cd80 | 0,233404 | 0,168366 | -0,471225 | 1 | 1 |
| Gpr123 | 0,063688 | 0,0459143 | -0,472071 | 1 | 1 |
| Tex14 | 0,114268 | 0,0823115 | -0,47326 | 1 | 1 |
| Tff3 | 0,675516 | 0,486301 | -0,474142 | 1 | 1 |
| Park2 | 0,229123 | 0,164866 | -0,474829 | 1 | 1 |
| Gprin2 | 0,122943 | 0,088422 | -0,475515 | 1 | 1 |
| B430319G15Rik | 0,493862 | 0,355131 | -0,475756 | 1 | 1 |
| Mzf1 | 0,548144 | 0,393965 | -0,476486 | 1 | 1 |
| D7Ertd443e | 0,050626 | 0,036352 | -0,477833 | 1 | 1 |
| Amdhd1 | 0,050721 | 0,0364196 | -0,477868 | 1 | 1 |
| Prss30 | 0,088892 | 0,0638276 | -0,477868 | 1 | 1 |
| Impg2 | 0,07674 | 0,0550365 | -0,479581 | 1 | 1 |
| Zpld1 | 0,153877 | 0,11024 | -0,481137 | 1 | 1 |
| Itga11 | 0,100066 | 0,0715909 | -0,483111 | 1 | 1 |

| | | | | | |
|---|---|---|---|---|---|
| Nol3 | 0,167954 | 0,120147 | -0,483266 | 1 | 1 |
| 2010107G12Rik | 0,430199 | 0,307249 | -0,485596 | 1 | 1 |
| 9030617O03Rik | 0,064598 | 0,0460812 | -0,487308 | 1 | 1 |
| C330013E15Rik | 0,36901 | 0,263133 | -0,487869 | 1 | 1 |
| Runx2 | 0,221331 | 0,157803 | -0,488085 | 1 | 1 |
| Mmp9 | 0,207311 | 0,147772 | -0,488429 | 1 | 1 |
| Mos | 0,352061 | 0,250742 | -0,489621 | 1 | 1 |
| Snap25 | 0,200951 | 0,143063 | -0,490191 | 1 | 1 |
| 1700067K01Rik | 0,376901 | 0,268188 | -0,490944 | 1 | 1 |
| Nfasc | 0,467039 | 0,332265 | -0,491208 | 1 | 1 |
| Mal | 0,310755 | 0,220644 | -0,494053 | 1 | 1 |
| 5730522E02Rik | 0,310392 | 0,220239 | -0,495021 | 1 | 1 |
| Rcan2 | 0,447768 | 0,317401 | -0,496444 | 1 | 1 |
| C430002E04Rik | 0,11154 | 0,0790555 | -0,496622 | 1 | 1 |
| Tmem88b | 0,123246 | 0,0872393 | -0,498493 | 1 | 1 |
| Thsd7a | 0,173812 | 0,122792 | -0,501316 | 1 | 1 |
| Xlr3a | 0,410295 | 0,289803 | -0,50159 | 1 | 1 |
| Tulp2 | 0,365147 | 0,25782 | -0,502114 | 1 | 1 |
| Dfnb59 | 0,607969 | 0,428483 | -0,50476 | 1 | 1 |
| A830082K12Rik | 0,130565 | 0,0919674 | -0,505575 | 1 | 1 |
| Ciita | 0,132527 | 0,0933355 | -0,505787 | 1 | 1 |
| Gm19583 | 0,137487 | 0,0967993 | -0,506227 | 1 | 1 |
| Rspo2 | 0,10194 | 0,0717357 | -0,506964 | 1 | 1 |
| Lrriq3 | 0,196079 | 0,137958 | -0,507209 | 1 | 1 |
| 1700023F06Rik | 0,160727 | 0,113046 | -0,5077 | 1 | 1 |
| Npas3 | 0,322376 | 0,226661 | -0,508208 | 1 | 1 |
| Krt15 | 0,398448 | 0,27995 | -0,50922 | 1 | 1 |
| Gm11201 | 0,101431 | 0,0712413 | -0,509712 | 1 | 1 |
| Kcnj4 | 0,243128 | 0,170749 | -0,509838 | 1 | 1 |
| BB031773 | 0,017259 | 0,0121156 | -0,510468 | 1 | 1 |
| Ppp1r3d | 0,180281 | 0,126538 | -0,510674 | 1 | 1 |
| Tbxas1 | 0,520614 | 0,365261 | -0,511285 | 1 | 1 |
| Lrriq1 | 0,082039 | 0,0575579 | -0,511287 | 1 | 1 |
| H2-T23 | 0,419992 | 0,294512 | -0,512036 | 1 | 1 |
| Gm17762 | 0,212615 | 0,14908 | -0,512157 | 1 | 1 |
| Bcl2l15 | 0,42639 | 0,29895 | -0,512268 | 1 | 1 |
| 6030407O03Rik | 0,048193 | 0,0337057 | -0,515826 | 1 | 1 |
| Chp2 | 0,07959 | 0,0556647 | -0,515826 | 1 | 1 |
| Rgsl1 | 0,022399 | 0,015666 | -0,515826 | 1 | 1 |
| Kcnma1 | 0,28086 | 0,196363 | -0,51633 | 1 | 1 |
| Dnah1 | 0,537485 | 0,375712 | -0,516596 | 1 | 1 |
| Synpr | 0,277459 | 0,193753 | -0,518054 | 1 | 1 |
| 4933402D24Rik | 0,51125 | 0,356157 | -0,521515 | 1 | 1 |
| Car7 | 0,133039 | 0,0926367 | -0,522194 | 1 | 1 |
| Krt1 | 0,079173 | 0,0550717 | -0,523692 | 1 | 1 |
| 9230112J17Rik | 0,065319 | 0,0453743 | -0,525629 | 1 | 1 |
| Kcna1 | 0,078372 | 0,0543967 | -0,526824 | 1 | 1 |
| Mustn1 | 0,594935 | 0,41263 | -0,527885 | 1 | 1 |

| | | | | | |
|---|---|---|---|---|---|
| Hcar1 | 0,032658 | 0,0226218 | -0,529738 | 1 | 1 |
| Hist1h2bg | 0,386984 | 0,268057 | -0,529738 | 1 | 1 |
| Htr5b | 0,1677 | 0,116163 | -0,529738 | 1 | 1 |
| Gcgr | 0,466834 | 0,323235 | -0,530323 | 1 | 1 |
| Rps6kl1 | 0,416828 | 0,288179 | -0,532486 | 1 | 1 |
| Il9r | 0,10277 | 0,0709143 | -0,535273 | 1 | 1 |
| Pthlh | 0,149269 | 0,102882 | -0,536925 | 1 | 1 |
| Cemip | 0,247595 | 0,170571 | -0,537606 | 1 | 1 |
| Rassf10 | 0,194161 | 0,133701 | -0,538244 | 1 | 1 |
| Abhd16b | 0,079106 | 0,0544667 | -0,538422 | 1 | 1 |
| Cers1 | 1,01314 | 0,69689 | -0,539835 | 1 | 1 |
| Cftr | 0,347409 | 0,238924 | -0,540081 | 1 | 1 |
| Cntn3 | 0,148656 | 0,102145 | -0,541352 | 1 | 1 |
| Ica1l | 0,170023 | 0,116749 | -0,542318 | 1 | 1 |
| Fmn1 | 0,519618 | 0,356485 | -0,543608 | 1 | 1 |
| Nod1 | 0,511228 | 0,350657 | -0,543907 | 1 | 1 |
| Cxcl16 | 0,507033 | 0,347703 | -0,544225 | 1 | 1 |
| Flrt1 | 0,240611 | 0,164897 | -0,545137 | 1 | 1 |
| Rasl10a | 0,563337 | 0,385493 | -0,547293 | 1 | 1 |
| Gm7457 | 0,048182 | 0,0329339 | -0,548923 | 1 | 1 |
| Phactr3 | 0,314769 | 0,215131 | -0,549081 | 1 | 1 |
| Myh6 | 0,020374 | 0,0138903 | -0,552635 | 1 | 1 |
| Tnfaip8l3 | 0,225607 | 0,153652 | -0,554151 | 1 | 1 |
| Psd | 0,386094 | 0,262702 | -0,555525 | 1 | 1 |
| Kazald1 | 0,120833 | 0,0821525 | -0,556638 | 1 | 1 |
| Msantd1 | 0,372687 | 0,253221 | -0,55757 | 1 | 1 |
| Slc16a11 | 0,213603 | 0,14495 | -0,559378 | 1 | 1 |
| Thrb | 0,02861 | 0,0194103 | -0,559701 | 1 | 1 |
| H2-DMb1 | 0,175381 | 0,118843 | -0,561438 | 1 | 1 |
| Igj | 0,099796 | 0,0676243 | -0,561438 | 1 | 1 |
| Pla2g4f | 0,064901 | 0,0439408 | -0,562681 | 1 | 1 |
| C1s1 | 0,111535 | 0,075472 | -0,563488 | 1 | 1 |
| Cct6b | 0,158159 | 0,106955 | -0,564373 | 1 | 1 |
| Ptger4 | 0,47503 | 0,320903 | -0,565882 | 1 | 1 |
| LOC102634101 | 0,223405 | 0,150456 | -0,570313 | 1 | 1 |
| Slc5a8 | 0,066968 | 0,0450913 | -0,570612 | 1 | 1 |
| Scrt2 | 0,050117 | 0,0337375 | -0,570954 | 1 | 1 |
| Cpb1 | 0,568986 | 0,383024 | -0,570955 | 1 | 1 |
| Tmem211 | 0,113111 | 0,0761095 | -0,571597 | 1 | 1 |
| Ankrd35 | 0,053828 | 0,0362067 | -0,572093 | 1 | 1 |
| Gm15328 | 0,224201 | 0,150807 | -0,572093 | 1 | 1 |
| Oas1c | 0,39519 | 0,265493 | -0,573873 | 1 | 1 |
| Tbr1 | 0,268389 | 0,1802 | -0,574727 | 1 | 1 |
| Ttbk1 | 0,144616 | 0,0968462 | -0,578461 | 1 | 1 |
| 4931409K22Rik | 0,043233 | 0,028947 | -0,578728 | 1 | 1 |
| Fgf11 | 0,475165 | 0,318086 | -0,579013 | 1 | 1 |
| D5Ertd605e | 0,289686 | 0,193832 | -0,579683 | 1 | 1 |
| Pcdhb4 | 0,070712 | 0,0473144 | -0,579683 | 1 | 1 |

| | | | | | |
|---|---|---|---|---|---|
| Creb3l3 | 0,108175 | 0,0723617 | -0,580076 | 1 | 1 |
| Lin7b | 0,485765 | 0,324817 | -0,580631 | 1 | 1 |
| Gm14295 | 0,465303 | 0,310444 | -0,583836 | 1 | 1 |
| Rps15a-ps4 | 0,636273 | 0,424116 | -0,585189 | 1 | 1 |
| Rgs17 | 0,213092 | 0,141653 | -0,589113 | 1 | 1 |
| Lmod1 | 0,31357 | 0,208058 | -0,591802 | 1 | 1 |
| Cntnap1 | 0,303131 | 0,200626 | -0,595438 | 1 | 1 |
| Dnah9 | 0,104669 | 0,0692121 | -0,596736 | 1 | 1 |
| Tagap | 0,276275 | 0,182608 | -0,597359 | 1 | 1 |
| Scn1b | 0,397597 | 0,262137 | -0,600984 | 1 | 1 |
| Ano4 | 0,027009 | 0,017798 | -0,601728 | 1 | 1 |
| Syt15 | 0,276178 | 0,181951 | -0,602044 | 1 | 1 |
| St6galnac5 | 0,471555 | 0,31063 | -0,602229 | 1 | 1 |
| Fmr1nb | 0,335531 | 0,22054 | -0,605405 | 1 | 1 |
| Calcr | 0,039874 | 0,0261907 | -0,606387 | 1 | 1 |
| 4931440J10Rik | 0,094811 | 0,0622753 | -0,606397 | 1 | 1 |
| A330032B11Rik | 0,03798 | 0,0249467 | -0,606397 | 1 | 1 |
| Csf3 | 0,0562 | 0,0369141 | -0,606397 | 1 | 1 |
| Gm9871 | 0,048035 | 0,0315513 | -0,606397 | 1 | 1 |
| Grm3 | 0,019549 | 0,0128403 | -0,606397 | 1 | 1 |
| Kcnh4 | 0,018195 | 0,0119508 | -0,606397 | 1 | 1 |
| Neto1 | 0,019578 | 0,0128596 | -0,606397 | 1 | 1 |
| Nlrp4f | 0,019602 | 0,012875 | -0,606397 | 1 | 1 |
| Slc12a1 | 0,01467 | 0,00963599 | -0,606397 | 1 | 1 |
| Gm15612 | 0,469172 | 0,307486 | -0,609597 | 1 | 1 |
| Tuba8 | 0,313371 | 0,205111 | -0,611468 | 1 | 1 |
| Kif4-ps | 0,061563 | 0,0402762 | -0,612135 | 1 | 1 |
| Gpr128 | 0,298753 | 0,195388 | -0,612609 | 1 | 1 |
| Kcnj10 | 0,143377 | 0,0937701 | -0,612609 | 1 | 1 |
| Gm9926 | 0,022742 | 0,0148673 | -0,613241 | 1 | 1 |
| Mep1a | 0,039212 | 0,0256336 | -0,613241 | 1 | 1 |
| Spef2 | 0,0764 | 0,0499447 | -0,613241 | 1 | 1 |
| Stmn4 | 0,103711 | 0,0677989 | -0,613241 | 1 | 1 |
| Clcnka | 0,067112 | 0,0438548 | -0,613845 | 1 | 1 |
| Arhgef37 | 0,168019 | 0,109775 | -0,614071 | 1 | 1 |
| Syt4 | 0,068853 | 0,0449851 | -0,614071 | 1 | 1 |
| Slc22a18 | 0,11378 | 0,0743028 | -0,614763 | 1 | 1 |
| Bace2 | 0,251394 | 0,16412 | -0,6152 | 1 | 1 |
| 5330434G04Rik | 0,091226 | 0,0595452 | -0,615465 | 1 | 1 |
| D130043K22Rik | 0,132734 | 0,0866044 | -0,616026 | 1 | 1 |
| En2 | 0,303691 | 0,198144 | -0,616052 | 1 | 1 |
| Lrrc73 | 0,446541 | 0,291239 | -0,61659 | 1 | 1 |
| 1810062G17Rik | 0,267907 | 0,174716 | -0,616719 | 1 | 1 |
| D330050G23Rik | 0,125766 | 0,0820165 | -0,616753 | 1 | 1 |
| Magix | 0,073858 | 0,0481604 | -0,616896 | 1 | 1 |
| Myh8 | 0,045796 | 0,0298376 | -0,618084 | 1 | 1 |
| Cyp2w1 | 0,0633 | 0,0411529 | -0,62121 | 1 | 1 |
| Zbp1 | 0,046411 | 0,0301727 | -0,62121 | 1 | 1 |

| | | | | | |
|---|---|---|---|---|---|
| Sds | 0,129454 | 0,0841059 | -0,622158 | 1 | 1 |
| Plekhb1 | 0,291035 | 0,189042 | -0,622482 | 1 | 1 |
| Smpx | 0,19499 | 0,126645 | -0,622608 | 1 | 1 |
| 4930426L09Rik | 0,129295 | 0,0839749 | -0,622632 | 1 | 1 |
| 1700029J03Rik | 0,070905 | 0,0460062 | -0,624053 | 1 | 1 |
| 5330411J11Rik | 0,028805 | 0,0186903 | -0,624053 | 1 | 1 |
| AY512931 | 0,018932 | 0,0122839 | -0,624053 | 1 | 1 |
| Actl9 | 0,035605 | 0,0231021 | -0,624053 | 1 | 1 |
| Il17b | 0,084998 | 0,0551507 | -0,624053 | 1 | 1 |
| Klk1 | 0,061725 | 0,0400502 | -0,624053 | 1 | 1 |
| Taar2 | 0,101665 | 0,0659652 | -0,624053 | 1 | 1 |
| Daw1 | 0,0361 | 0,023423 | -0,624072 | 1 | 1 |
| AI593442 | 0,007632 | 0,0049491 | -0,624804 | 1 | 1 |
| Ifih1 | 0,450466 | 0,291773 | -0,626568 | 1 | 1 |
| Sidt1 | 0,108175 | 0,0700476 | -0,626959 | 1 | 1 |
| Itgae | 0,338815 | 0,219376 | -0,627093 | 1 | 1 |
| Syn2 | 0,315365 | 0,203258 | -0,633708 | 1 | 1 |
| Rbfox3 | 0,401396 | 0,258599 | -0,634309 | 1 | 1 |
| 1700030C10Rik | 0,38951 | 0,250149 | -0,638872 | 1 | 1 |
| Klhl14 | 0,293964 | 0,188138 | -0,643849 | 1 | 1 |
| Tacr1 | 0,53394 | 0,341608 | -0,644337 | 1 | 1 |
| Cd79a | 0,268943 | 0,172059 | -0,644393 | 1 | 1 |
| Ldhd | 0,129174 | 0,0824539 | -0,647658 | 1 | 1 |
| Fndc7 | 0,014431 | 0,00918003 | -0,652557 | 1 | 1 |
| A330076H08Rik | 0,290104 | 0,184362 | -0,654024 | 1 | 1 |
| Pcdh9 | 0,126259 | 0,0800311 | -0,657751 | 1 | 1 |
| B830017H08Rik | 0,19408 | 0,122929 | -0,658819 | 1 | 1 |
| Tdrd12 | 0,263226 | 0,166554 | -0,660316 | 1 | 1 |
| Gpr64 | 0,088044 | 0,0556502 | -0,661846 | 1 | 1 |
| 2610027K06Rik | 0,143118 | 0,0904526 | -0,661974 | 1 | 1 |
| Larp6 | 0,194221 | 0,122669 | -0,662922 | 1 | 1 |
| Fgf14 | 0,264869 | 0,167181 | -0,663868 | 1 | 1 |
| Piwil2 | 0,061591 | 0,0386876 | -0,670859 | 1 | 1 |
| Pydc3 | 0,048964 | 0,0307465 | -0,671314 | 1 | 1 |
| BC030870 | 0,120728 | 0,0757737 | -0,671988 | 1 | 1 |
| Mchr1 | 0,077546 | 0,0486709 | -0,671988 | 1 | 1 |
| 9430076C15Rik | 0,060797 | 0,0381085 | -0,673895 | 1 | 1 |
| Plekhd1 | 0,340347 | 0,213068 | -0,675695 | 1 | 1 |
| Fam20a | 0,410448 | 0,256896 | -0,676017 | 1 | 1 |
| Bai1 | 0,212337 | 0,132793 | -0,67718 | 1 | 1 |
| Cd274 | 0,270648 | 0,168804 | -0,68107 | 1 | 1 |
| Slc27a2 | 0,539016 | 0,335999 | -0,681869 | 1 | 1 |
| Nptx1 | 0,30537 | 0,190181 | -0,683183 | 1 | 1 |
| Lgr6 | 0,116061 | 0,0722586 | -0,683643 | 1 | 1 |
| Tbc1d10c | 0,616859 | 0,38388 | -0,684285 | 1 | 1 |
| Wdr52 | 0,07704 | 0,0479239 | -0,684866 | 1 | 1 |
| Dram1 | 0,302397 | 0,187432 | -0,69008 | 1 | 1 |
| Pla2g4d | 0,036486 | 0,0226145 | -0,69008 | 1 | 1 |

| | | | | | |
|---|---|---|---|---|---|
| Hepacam | 0,045171 | 0,0279967 | -0,690134 | 1 | 1 |
| Rab9b | 0,073249 | 0,0453992 | -0,690134 | 1 | 1 |
| Fam154b | 0,361997 | 0,224331 | -0,690347 | 1 | 1 |
| Ddo | 0,154613 | 0,0957607 | -0,691154 | 1 | 1 |
| Lum | 0,221525 | 0,1372 | -0,691183 | 1 | 1 |
| Nipal1 | 0,429152 | 0,265617 | -0,692141 | 1 | 1 |
| Themis2 | 0,230432 | 0,142267 | -0,695743 | 1 | 1 |
| Tcam1 | 0,14694 | 0,0905707 | -0,698113 | 1 | 1 |
| Mfsd7a | 0,287843 | 0,177316 | -0,698956 | 1 | 1 |
| Folh1 | 0,114313 | 0,0704114 | -0,699106 | 1 | 1 |
| Tmem196 | 0,325571 | 0,200438 | -0,699815 | 1 | 1 |
| Pkdrej | 0,120944 | 0,0744055 | -0,700856 | 1 | 1 |
| Nanog | 0,210018 | 0,129043 | -0,702658 | 1 | 1 |
| Myo3b | 0,185858 | 0,114138 | -0,70343 | 1 | 1 |
| 6430550D23Rik | 0,418822 | 0,256953 | -0,704833 | 1 | 1 |
| C1qtnf7 | 0,376453 | 0,230886 | -0,705286 | 1 | 1 |
| Pnma3 | 0,320062 | 0,195964 | -0,707763 | 1 | 1 |
| Padi3 | 0,26658 | 0,163125 | -0,708588 | 1 | 1 |
| Mcf2 | 0,222027 | 0,135689 | -0,710428 | 1 | 1 |
| Unc79 | 0,031411 | 0,0191911 | -0,710845 | 1 | 1 |
| Slc25a2 | 0,347327 | 0,212106 | -0,711505 | 1 | 1 |
| Mkln1os | 0,358381 | 0,218826 | -0,71171 | 1 | 1 |
| D630023F18Rik | 0,092732 | 0,0565417 | -0,713753 | 1 | 1 |
| Cdkl4 | 0,358285 | 0,218387 | -0,71422 | 1 | 1 |
| Gm12253 | 0,313509 | 0,191091 | -0,714251 | 1 | 1 |
| Apol8 | 0,340419 | 0,207367 | -0,715127 | 1 | 1 |
| Faah | 0,403146 | 0,244735 | -0,720081 | 1 | 1 |
| Islr2 | 0,157794 | 0,0957227 | -0,72111 | 1 | 1 |
| Maats1 | 0,034671 | 0,0210184 | -0,722073 | 1 | 1 |
| Pde6a | 0,022232 | 0,0134773 | -0,722073 | 1 | 1 |
| Ptk6 | 0,042648 | 0,0258541 | -0,722073 | 1 | 1 |
| Dnaic2 | 0,19988 | 0,121098 | -0,722957 | 1 | 1 |
| Dnah7a | 0,02061 | 0,0124661 | -0,725313 | 1 | 1 |
| Gata3 | 0,513158 | 0,310322 | -0,725635 | 1 | 1 |
| Spn | 0,477412 | 0,288632 | -0,726004 | 1 | 1 |
| Ctxn3 | 0,126123 | 0,076192 | -0,727115 | 1 | 1 |
| Slfn10-ps | 0,09158 | 0,0552685 | -0,728573 | 1 | 1 |
| 4931431C16Rik | 0,221131 | 0,133366 | -0,729515 | 1 | 1 |
| Cst7 | 0,090752 | 0,05471 | -0,730119 | 1 | 1 |
| Tex19.1 | 0,046233 | 0,027872 | -0,730119 | 1 | 1 |
| A230108P19Rik | 0,06189 | 0,037268 | -0,731777 | 1 | 1 |
| Amhr2 | 0,081526 | 0,0490919 | -0,731777 | 1 | 1 |
| Klf14 | 0,154181 | 0,0928063 | -0,73233 | 1 | 1 |
| Dnaaf1 | 0,053406 | 0,0320873 | -0,734992 | 1 | 1 |
| Bhlha15 | 0,116967 | 0,0702683 | -0,735157 | 1 | 1 |
| Lrrc17 | 0,489438 | 0,292173 | -0,744305 | 1 | 1 |
| Masp2 | 0,432195 | 0,257837 | -0,745225 | 1 | 1 |
| Adad1 | 0,117215 | 0,0698675 | -0,746464 | 1 | 1 |

| | | | | | |
|---|---|---|---|---|---|
| C1qtnf9 | 0,121975 | 0,0727049 | -0,746464 | 1 | 1 |
| Fut9 | 0,519333 | 0,309092 | -0,748625 | 1 | 1 |
| A230070E04Rik | 0,106524 | 0,0633362 | -0,750072 | 1 | 1 |
| Gm16532 | 0,095374 | 0,0566371 | -0,751846 | 1 | 1 |
| Mzb1 | 0,269815 | 0,160228 | -0,751846 | 1 | 1 |
| Tmeff2 | 0,285194 | 0,169272 | -0,752597 | 1 | 1 |
| Liph | 0,232141 | 0,137735 | -0,753102 | 1 | 1 |
| Dkkl1 | 0,389422 | 0,231019 | -0,753321 | 1 | 1 |
| Ptprq | 0,019882 | 0,0117804 | -0,75505 | 1 | 1 |
| Ccdc79 | 0,211379 | 0,12517 | -0,755947 | 1 | 1 |
| Nacad | 0,133798 | 0,0790315 | -0,75956 | 1 | 1 |
| Gm5464 | 0,373034 | 0,219893 | -0,762505 | 1 | 1 |
| Mir8097 | 58,8194 | 34,6402 | -0,763845 | 1 | 1 |
| Ak7 | 0,081997 | 0,0482505 | -0,765023 | 1 | 1 |
| Drd5 | 0,319134 | 0,187792 | -0,765023 | 1 | 1 |
| Pcsk2 | 0,026095 | 0,0153554 | -0,765023 | 1 | 1 |
| Thrsp | 0,142018 | 0,0835696 | -0,765023 | 1 | 1 |
| Trpm5 | 0,056686 | 0,0333563 | -0,765023 | 1 | 1 |
| Aatk | 0,315236 | 0,185411 | -0,765707 | 1 | 1 |
| Xkr6 | 0,246065 | 0,14464 | -0,766578 | 1 | 1 |
| Spink10 | 0,324231 | 0,190497 | -0,767252 | 1 | 1 |
| Prss16 | 0,206696 | 0,121154 | -0,770676 | 1 | 1 |
| Scn9a | 0,022775 | 0,0133466 | -0,770953 | 1 | 1 |
| Lingo3 | 0,252537 | 0,147832 | -0,772538 | 1 | 1 |
| Zbtb7c | 0,275132 | 0,160989 | -0,773167 | 1 | 1 |
| Sbspon | 0,176806 | 0,10335 | -0,774632 | 1 | 1 |
| Myo1h | 0,097155 | 0,0567886 | -0,774688 | 1 | 1 |
| Cacnb4 | 0,48373 | 0,282256 | -0,777198 | 1 | 1 |
| Kcnj11 | 0,126074 | 0,0734914 | -0,778626 | 1 | 1 |
| Aldh3b1 | 0,101816 | 0,0593438 | -0,778793 | 1 | 1 |
| Dscaml1 | 0,02481 | 0,0144607 | -0,778793 | 1 | 1 |
| Gm5523 | 0,159227 | 0,0928061 | -0,778793 | 1 | 1 |
| Acr | 0,141123 | 0,0822434 | -0,778986 | 1 | 1 |
| Adam5 | 0,151319 | 0,0881846 | -0,778989 | 1 | 1 |
| Fas | 0,320015 | 0,186072 | -0,782282 | 1 | 1 |
| Acta1 | 0,296773 | 0,17238 | -0,783768 | 1 | 1 |
| Adora1 | 0,062558 | 0,0363179 | -0,784501 | 1 | 1 |
| Frmpd1 | 0,160553 | 0,0931494 | -0,785433 | 1 | 1 |
| Pigz | 0,10761 | 0,0623868 | -0,786506 | 1 | 1 |
| Fam212b | 0,280817 | 0,16276 | -0,786878 | 1 | 1 |
| Ddx60 | 0,03686 | 0,0213529 | -0,787611 | 1 | 1 |
| Ano3 | 0,05594 | 0,0323599 | -0,78967 | 1 | 1 |
| Il31ra | 0,17943 | 0,103792 | -0,789736 | 1 | 1 |
| Cpeb1 | 0,153535 | 0,0887452 | -0,790821 | 1 | 1 |
| Epha8 | 0,162893 | 0,0941244 | -0,791287 | 1 | 1 |
| Mmd2 | 0,134079 | 0,0774665 | -0,791437 | 1 | 1 |
| Tsnaxip1 | 0,129882 | 0,074995 | -0,792341 | 1 | 1 |
| 4933433G15Rik | 0,274716 | 0,158524 | -0,793245 | 1 | 1 |

| | | | | | |
|---|---|---|---|---|---|
| Kcna7 | 0,059345 | 0,0341889 | -0,795599 | 1 | 1 |
| Kcne1 | 0,104323 | 0,0600997 | -0,795627 | 1 | 1 |
| Ppara | 0,133392 | 0,0766633 | -0,799059 | 1 | 1 |
| Tnfaip8l2 | 0,264399 | 0,151846 | -0,800113 | 1 | 1 |
| Agbl2 | 0,241703 | 0,138527 | -0,803066 | 1 | 1 |
| Hr | 0,530041 | 0,3037 | -0,803457 | 1 | 1 |
| Esr2 | 0,088675 | 0,050735 | -0,805546 | 1 | 1 |
| 1700112E06Rik | 0,544294 | 0,311365 | -0,805781 | 1 | 1 |
| Alx1 | 0,336235 | 0,192344 | -0,805781 | 1 | 1 |
| Cp | 0,151903 | 0,0865008 | -0,812367 | 1 | 1 |
| Ltk | 0,023301 | 0,0132567 | -0,813663 | 1 | 1 |
| 6330409D20Rik | 0,051764 | 0,0294501 | -0,813668 | 1 | 1 |
| Cacng1 | 0,047359 | 0,0269408 | -0,813855 | 1 | 1 |
| Ecel1 | 0,017733 | 0,0100878 | -0,813855 | 1 | 1 |
| Gm11961 | 0,042432 | 0,024138 | -0,813855 | 1 | 1 |
| Gm19424 | 0,023261 | 0,0132321 | -0,813855 | 1 | 1 |
| Kcng3 | 0,01501 | 0,00853875 | -0,813855 | 1 | 1 |
| Kcnh3 | 0,014002 | 0,00796512 | -0,813855 | 1 | 1 |
| Krt42 | 0,035229 | 0,0200402 | -0,813855 | 1 | 1 |
| Lyz1 | 0,044592 | 0,0253665 | -0,813855 | 1 | 1 |
| Prodh2 | 0,058664 | 0,0333714 | -0,813855 | 1 | 1 |
| Ranbp3l | 0,011808 | 0,00671678 | -0,813855 | 1 | 1 |
| Tesc | 0,064108 | 0,0364685 | -0,813855 | 1 | 1 |
| Nkx6-2 | 0,214817 | 0,122114 | -0,814883 | 1 | 1 |
| Morc1 | 0,066554 | 0,0378273 | -0,8151 | 1 | 1 |
| Olfr558 | 0,052262 | 0,0296956 | -0,815514 | 1 | 1 |
| Crisp2 | 0,075979 | 0,0431502 | -0,81624 | 1 | 1 |
| Gm14446 | 0,047128 | 0,0267634 | -0,816321 | 1 | 1 |
| Il17f | 0,097318 | 0,0552646 | -0,816343 | 1 | 1 |
| Slc16a8 | 0,190311 | 0,107835 | -0,819533 | 1 | 1 |
| Gm10791 | 0,391949 | 0,221794 | -0,821446 | 1 | 1 |
| D10Bwg1379e | 0,260781 | 0,147284 | -0,824237 | 1 | 1 |
| Trpc5 | 0,151212 | 0,0853427 | -0,82523 | 1 | 1 |
| Gpx2-ps1 | 0,210795 | 0,118687 | -0,828684 | 1 | 1 |
| Pkd1l2 | 0,023892 | 0,013452 | -0,828684 | 1 | 1 |
| Npy1r | 0,122491 | 0,0689347 | -0,82937 | 1 | 1 |
| Serpina3i | 0,291801 | 0,164102 | -0,830389 | 1 | 1 |
| Alk | 0,051779 | 0,0291057 | -0,83107 | 1 | 1 |
| Cideb | 0,256393 | 0,143951 | -0,832784 | 1 | 1 |
| 4930449E01Rik | 0,313031 | 0,175556 | -0,83437 | 1 | 1 |
| Akr1c12 | 0,242017 | 0,13573 | -0,83437 | 1 | 1 |
| Rasgef1a | 0,040988 | 0,0229874 | -0,83437 | 1 | 1 |
| Trpc4 | 0,077056 | 0,0432016 | -0,834819 | 1 | 1 |
| Crb1 | 0,138219 | 0,0774766 | -0,835128 | 1 | 1 |
| Ace | 0,284988 | 0,159675 | -0,835756 | 1 | 1 |
| Krt23 | 0,093213 | 0,0522079 | -0,836264 | 1 | 1 |
| Mc5r | 0,129894 | 0,0727528 | -0,836264 | 1 | 1 |
| Sycp2 | 0,022696 | 0,0127117 | -0,836264 | 1 | 1 |

| | | | | | |
|---|---|---|---|---|---|
| Tldc2 | 0,286062 | 0,160221 | -0,836264 | 1 | 1 |
| Nutm1 | 0,057057 | 0,0318916 | -0,839236 | 1 | 1 |
| Il6ra | 0,563775 | 0,314816 | -0,840608 | 1 | 1 |
| Aldh1a7 | 0,440257 | 0,245768 | -0,841051 | 1 | 1 |
| Apobec1 | 0,283405 | 0,15797 | -0,843216 | 1 | 1 |
| Casz1 | 0,136471 | 0,0759242 | -0,845965 | 1 | 1 |
| Cpa5 | 0,043754 | 0,0243227 | -0,847127 | 1 | 1 |
| Fgf21 | 0,103585 | 0,0575817 | -0,847127 | 1 | 1 |
| Gm1045 | 0,138672 | 0,0770864 | -0,847127 | 1 | 1 |
| Gm13547 | 0,181088 | 0,100665 | -0,847127 | 1 | 1 |
| Gm2762 | 0,036049 | 0,0200392 | -0,847127 | 1 | 1 |
| Gm6583 | 0,037416 | 0,0207991 | -0,847127 | 1 | 1 |
| Gpr158 | 0,011095 | 0,00616748 | -0,847127 | 1 | 1 |
| Leap2 | 0,262666 | 0,146014 | -0,847127 | 1 | 1 |
| Ptgs2 | 0,019529 | 0,0108561 | -0,847127 | 1 | 1 |
| Tnfsf13 | 0,503038 | 0,279464 | -0,848008 | 1 | 1 |
| Bmp8b | 0,318019 | 0,176557 | -0,84898 | 1 | 1 |
| 2310069G16Rik | 0,352756 | 0,195062 | -0,854738 | 1 | 1 |
| 1700066B19Rik | 0,270237 | 0,149419 | -0,854864 | 1 | 1 |
| Mucl1 | 2,84E-006 | 1,57E-006 | -0,855 | 1 | 1 |
| Pla2r1 | 0,36682 | 0,20269 | -0,855802 | 1 | 1 |
| Sync | 0,182456 | 0,100662 | -0,858024 | 1 | 1 |
| Asb14 | 0,085979 | 0,0472486 | -0,863716 | 1 | 1 |
| 4930524B15Rik | 0,128146 | 0,070385 | -0,86445 | 1 | 1 |
| Zfp78 | 0,494016 | 0,270802 | -0,867318 | 1 | 1 |
| Tcap | 0,342351 | 0,187357 | -0,869683 | 1 | 1 |
| Astn2 | 0,105671 | 0,0577913 | -0,870653 | 1 | 1 |
| Gm128 | 0,318043 | 0,173884 | -0,871101 | 1 | 1 |
| Xkrx | 0,106442 | 0,0581518 | -0,872176 | 1 | 1 |
| Apol10b | 0,083107 | 0,0453098 | -0,875137 | 1 | 1 |
| Tlr3 | 0,574396 | 0,312658 | -0,877459 | 1 | 1 |
| Gm4907 | 0,321128 | 0,174704 | -0,878234 | 1 | 1 |
| Mylk4 | 0,136133 | 0,0739964 | -0,879492 | 1 | 1 |
| Nbeal2 | 0,5143 | 0,279523 | -0,879641 | 1 | 1 |
| 4930431F12Rik | 0,100698 | 0,0546557 | -0,881591 | 1 | 1 |
| Pglyrp1 | 0,434449 | 0,235805 | -0,881591 | 1 | 1 |
| Sphk1 | 0,178294 | 0,0967711 | -0,881611 | 1 | 1 |
| 1700034F02Rik | 0,165782 | 0,0899668 | -0,881823 | 1 | 1 |
| Gm12522 | 0,328292 | 0,177849 | -0,884329 | 1 | 1 |
| Prss53 | 0,312595 | 0,169271 | -0,884966 | 1 | 1 |
| Bcas1 | 0,111966 | 0,0605164 | -0,887665 | 1 | 1 |
| Cd300a | 0,410771 | 0,221269 | -0,892533 | 1 | 1 |
| Rgs7bp | 0,176839 | 0,095216 | -0,89316 | 1 | 1 |
| A330102I10Rik | 0,23249 | 0,125176 | -0,893215 | 1 | 1 |
| Ccdc148 | 0,156748 | 0,0843528 | -0,893943 | 1 | 1 |
| Kcnmb1 | 0,190297 | 0,102195 | -0,89693 | 1 | 1 |
| Osmr | 0,359224 | 0,192528 | -0,899813 | 1 | 1 |
| Gbp5 | 0,056713 | 0,0303754 | -0,900781 | 1 | 1 |

| | | | | | |
|---|---|---|---|---|---|
| A830052D11Rik | 0,249166 | 0,133396 | -0,901387 | 1 | 1 |
| Cnksr1 | 0,434035 | 0,232365 | -0,901416 | 1 | 1 |
| Slc5a11 | 0,121844 | 0,0652295 | -0,901441 | 1 | 1 |
| S100b | 0,109762 | 0,0587283 | -0,902255 | 1 | 1 |
| Siglech | 0,072925 | 0,0390184 | -0,902255 | 1 | 1 |
| Micalcl | 0,059404 | 0,0317819 | -0,902345 | 1 | 1 |
| Batf | 0,287483 | 0,153793 | -0,902489 | 1 | 1 |
| Sntg2 | 0,047173 | 0,0252359 | -0,902489 | 1 | 1 |
| Tmem95 | 0,307697 | 0,164607 | -0,902489 | 1 | 1 |
| Amz1 | 0,360372 | 0,192759 | -0,90269 | 1 | 1 |
| Efhc2 | 0,433275 | 0,231109 | -0,906706 | 1 | 1 |
| 1700088E04Rik | 0,390253 | 0,207784 | -0,909327 | 1 | 1 |
| D030025P21Rik | 0,046463 | 0,0247305 | -0,909791 | 1 | 1 |
| 4933427E11Rik | 0,280141 | 0,148949 | -0,911328 | 1 | 1 |
| Bspry | 0,615692 | 0,327325 | -0,911488 | 1 | 1 |
| Cpxm2 | 0,570049 | 0,302198 | -0,91559 | 1 | 1 |
| Nfia | 0,523651 | 0,277253 | -0,917403 | 1 | 1 |
| Grb14 | 0,443924 | 0,234864 | -0,918485 | 1 | 1 |
| B930059L03Rik | 0,230987 | 0,122114 | -0,919581 | 1 | 1 |
| Igsf23 | 0,150531 | 0,0795802 | -0,919581 | 1 | 1 |
| Pcdhb5 | 0,128106 | 0,0675768 | -0,922738 | 1 | 1 |
| Cspg5 | 0,22126 | 0,11661 | -0,924052 | 1 | 1 |
| Esyt3 | 0,323154 | 0,170309 | -0,924069 | 1 | 1 |
| Adhfe1 | 0,063365 | 0,0333697 | -0,92514 | 1 | 1 |
| Fmo5 | 0,073596 | 0,0387483 | -0,925498 | 1 | 1 |
| Asic2 | 0,453774 | 0,238865 | -0,925779 | 1 | 1 |
| 1700084J12Rik | 0,348343 | 0,183335 | -0,926026 | 1 | 1 |
| 4933422H20Rik | 0,106238 | 0,0559095 | -0,926139 | 1 | 1 |
| Chil1 | 0,123623 | 0,0650583 | -0,926139 | 1 | 1 |
| St8sia3 | 0,030214 | 0,0159006 | -0,926139 | 1 | 1 |
| Tnfsf12Tnfsf13 | 0,237296 | 0,12464 | -0,928926 | 1 | 1 |
| Unc5a | 0,168004 | 0,0882281 | -0,929189 | 1 | 1 |
| Abca6 | 0,081109 | 0,0424864 | -0,932868 | 1 | 1 |
| Myrip | 0,131174 | 0,0684347 | -0,938683 | 1 | 1 |
| A430035B10Rik | 0,403296 | 0,210368 | -0,938923 | 1 | 1 |
| Bcmo1 | 0,125881 | 0,0656571 | -0,939034 | 1 | 1 |
| Tmem240 | 0,201778 | 0,105217 | -0,939393 | 1 | 1 |
| Ddr2 | 0,407014 | 0,212233 | -0,939429 | 1 | 1 |
| Foxf1 | 0,343903 | 0,17897 | -0,942288 | 1 | 1 |
| Prr9 | 0,211513 | 0,109965 | -0,943707 | 1 | 1 |
| Prr33 | 0,394556 | 0,204896 | -0,945338 | 1 | 1 |
| Gja6 | 0,159974 | 0,0829784 | -0,947029 | 1 | 1 |
| Gm16793 | 0,078572 | 0,0406325 | -0,951387 | 1 | 1 |
| Rab40b | 0,049847 | 0,0257577 | -0,952511 | 1 | 1 |
| Vmn2r57 | 0,034672 | 0,0179161 | -0,952511 | 1 | 1 |
| Gm13710 | 0,416199 | 0,214965 | -0,953168 | 1 | 1 |
| Gm7444 | 0,069776 | 0,0360393 | -0,953168 | 1 | 1 |
| Fam163a | 0,436665 | 0,225474 | -0,953562 | 1 | 1 |

| | | | | | |
|---|---|---|---|---|---|
| En1 | 0,035302 | 0,0182058 | -0,955353 | 1 | 1 |
| Olfr94 | 0,086687 | 0,0447057 | -0,955353 | 1 | 1 |
| Plcxd3 | 0,544037 | 0,279961 | -0,95848 | 1 | 1 |
| Slco1c1 | 0,073697 | 0,0378408 | -0,961669 | 1 | 1 |
| Ccdc176 | 0,362206 | 0,185918 | -0,962143 | 1 | 1 |
| Plxna4 | 0,380439 | 0,195147 | -0,963103 | 1 | 1 |
| Adcy8 | 0,282035 | 0,144553 | -0,96427 | 1 | 1 |
| Lgi1 | 0,06717 | 0,0343763 | -0,966397 | 1 | 1 |
| Irf7 | 0,343336 | 0,17558 | -0,967491 | 1 | 1 |
| Dcc | 0,103532 | 0,0529093 | -0,968488 | 1 | 1 |
| Trpv4 | 0,368294 | 0,188163 | -0,968878 | 1 | 1 |
| Ccdc175 | 0,034982 | 0,0178626 | -0,969689 | 1 | 1 |
| Ccser1 | 0,153525 | 0,078354 | -0,970396 | 1 | 1 |
| Nwd1 | 0,025188 | 0,0128474 | -0,971261 | 1 | 1 |
| BC048644 | 0,112297 | 0,0572258 | -0,972588 | 1 | 1 |
| Trhde | 0,164085 | 0,0835396 | -0,973914 | 1 | 1 |
| Pdk4 | 0,129415 | 0,0658414 | -0,974937 | 1 | 1 |
| Prr16 | 0,12024 | 0,0611514 | -0,975453 | 1 | 1 |
| Trim14 | 0,282202 | 0,143077 | -0,979937 | 1 | 1 |
| Disc1 | 0,394851 | 0,199759 | -0,983044 | 1 | 1 |
| Angptl7 | 0,057294 | 0,028971 | -0,98378 | 1 | 1 |
| Tal2 | 0,169957 | 0,0858953 | -0,984518 | 1 | 1 |
| Cmya5 | 0,034728 | 0,0175437 | -0,985157 | 1 | 1 |
| Bcl3 | 0,196005 | 0,0990093 | -0,985255 | 1 | 1 |
| Sytl3 | 0,041004 | 0,0207019 | -0,985988 | 1 | 1 |
| Cacng6 | 0,092068 | 0,046483 | -0,985992 | 1 | 1 |
| Ccdc13 | 0,109417 | 0,0552424 | -0,985992 | 1 | 1 |
| Ceacam18 | 0,030761 | 0,0155303 | -0,985992 | 1 | 1 |
| Fam150a | 0,08151 | 0,0411525 | -0,985992 | 1 | 1 |
| Ferd3l | 0,078278 | 0,0395209 | -0,985992 | 1 | 1 |
| Hist2h2ab | 0,227801 | 0,115012 | -0,985992 | 1 | 1 |
| Neurod4 | 0,016972 | 0,00856865 | -0,985992 | 1 | 1 |
| Slc25a41 | 0,05216 | 0,0263345 | -0,985992 | 1 | 1 |
| Tacr3 | 0,014974 | 0,00756015 | -0,985992 | 1 | 1 |
| Vwde | 0,014125 | 0,00713146 | -0,985992 | 1 | 1 |
| Itpka | 0,490836 | 0,247812 | -0,985993 | 1 | 1 |
| Gpr20 | 0,501145 | 0,252596 | -0,988398 | 1 | 1 |
| Ephx4 | 0,241661 | 0,121774 | -0,988772 | 1 | 1 |
| Gm3435 | 0,329414 | 0,165683 | -0,991479 | 1 | 1 |
| Gm14322 | 0,296377 | 0,149011 | -0,992007 | 1 | 1 |
| Mir143hg | 0,426222 | 0,214268 | -0,992193 | 1 | 1 |
| Fbxo15 | 0,411406 | 0,20623 | -0,99631 | 1 | 1 |
| Grip2 | 0,056373 | 0,0282292 | -0,99781 | 1 | 1 |
| Rd3 | 0,134163 | 0,0671382 | -0,998782 | 1 | 1 |
| Pde11a | 0,150019 | 0,0749051 | -1,00201 | 1 | 1 |
| Hkdc1 | 0,144389 | 0,0719755 | -1,00438 | 1 | 1 |
| Ambp | 0,419793 | 0,209243 | -1,0045 | 1 | 1 |
| B430212C06Rik | 0,266723 | 0,132937 | -1,0046 | 1 | 1 |

| | | | | | |
|---|---|---|---|---|---|
| Klk9 | 0,237626 | 0,118363 | -1,00548 | 1 | 1 |
| Gpr160 | 0,231065 | 0,115084 | -1,00561 | 1 | 1 |
| Wdfy4 | 0,196738 | 0,0979386 | -1,00633 | 1 | 1 |
| B3galt5 | 0,141971 | 0,0706186 | -1,00748 | 1 | 1 |
| Camk1d | 0,314416 | 0,156195 | -1,00933 | 1 | 1 |
| Slc2a9 | 0,500499 | 0,247712 | -1,0147 | 1 | 1 |
| Cd79b | 0,589595 | 0,291583 | -1,01582 | 1 | 1 |
| Gm20257 | 0,266948 | 0,131982 | -1,01622 | 1 | 1 |
| Scrt1 | 0,051342 | 0,0253841 | -1,01622 | 1 | 1 |
| 1700025F24Rik | 0,489137 | 0,241025 | -1,02105 | 1 | 1 |
| Panx2 | 0,105209 | 0,0518422 | -1,02105 | 1 | 1 |
| Gm13139,Gm132! | 0,424291 | 0,208827 | -1,02275 | 1 | 1 |
| Cpeb2 | 0,523002 | 0,257155 | -1,02418 | 1 | 1 |
| Dhrs2 | 0,337698 | 0,165905 | -1,02537 | 1 | 1 |
| Abcb1a | 0,059544 | 0,0292051 | -1,02773 | 1 | 1 |
| Asic3 | 0,163521 | 0,0802044 | -1,02773 | 1 | 1 |
| 5830416P10Rik | 0,068659 | 0,0336351 | -1,02949 | 1 | 1 |
| 4921525O09Rik | 0,171748 | 0,0841254 | -1,02968 | 1 | 1 |
| Slc5a5 | 0,345593 | 0,169226 | -1,03013 | 1 | 1 |
| Tmc5 | 0,080858 | 0,0395647 | -1,03117 | 1 | 1 |
| 4933428G20Rik | 0,248785 | 0,121622 | -1,03249 | 1 | 1 |
| Vmn1r32 | 0,166591 | 0,0814027 | -1,03316 | 1 | 1 |
| Tm4sf4 | 0,515944 | 0,251992 | -1,03384 | 1 | 1 |
| Nckap5 | 0,339625 | 0,165036 | -1,04116 | 1 | 1 |
| Zfp345 | 0,101673 | 0,0493709 | -1,0422 | 1 | 1 |
| AI118078 | 0,26066 | 0,126551 | -1,04245 | 1 | 1 |
| Lims2 | 0,171528 | 0,0832382 | -1,04313 | 1 | 1 |
| B020014A21Rik | 0,344709 | 0,167148 | -1,04425 | 1 | 1 |
| Fam217a | 0,156705 | 0,0759058 | -1,04577 | 1 | 1 |
| Adamtsl1 | 0,139599 | 0,067609 | -1,046 | 1 | 1 |
| Tmem132e | 0,210959 | 0,101895 | -1,04988 | 1 | 1 |
| Tekt5 | 0,134682 | 0,0649535 | -1,05208 | 1 | 1 |
| Gab3 | 0,341399 | 0,164545 | -1,05298 | 1 | 1 |
| 4933408N05Rik | 0,064435 | 0,0310475 | -1,05337 | 1 | 1 |
| Aldh3a1 | 0,063829 | 0,0307555 | -1,05337 | 1 | 1 |
| Chrna1 | 0,064718 | 0,0311835 | -1,05337 | 1 | 1 |
| Fam26f | 0,202259 | 0,0974562 | -1,05337 | 1 | 1 |
| Gm13939 | 0,077547 | 0,0373654 | -1,05337 | 1 | 1 |
| Gm8909 | 0,077009 | 0,0371062 | -1,05337 | 1 | 1 |
| Slc28a2 | 0,023998 | 0,0115632 | -1,05337 | 1 | 1 |
| Faim2 | 0,020036 | 0,00965356 | -1,05342 | 1 | 1 |
| Cd101 | 0,058701 | 0,0282826 | -1,05346 | 1 | 1 |
| Cd22 | 0,053217 | 0,0256361 | -1,0537 | 1 | 1 |
| Asb10 | 0,084081 | 0,0404765 | -1,0547 | 1 | 1 |
| Lrrn4 | 0,470536 | 0,226158 | -1,05697 | 1 | 1 |
| Ubl4b | 0,247467 | 0,118938 | -1,05703 | 1 | 1 |
| Nlrp10 | 0,067819 | 0,0325767 | -1,05785 | 1 | 1 |
| Cacna1e | 0,057339 | 0,0275086 | -1,05963 | 1 | 1 |

| | | | | | |
|---|---|---|---|---|---|
| C330024C12Rik | 0,094336 | 0,0451576 | -1,06283 | 1 | 1 |
| Lrrc6 | 0,059813 | 0,028632 | -1,06283 | 1 | 1 |
| Rbp2 | 0,480538 | 0,23003 | -1,06283 | 1 | 1 |
| Stab2 | 0,014182 | 0,0067888 | -1,06283 | 1 | 1 |
| 4933406M09Rik | 0,143716 | 0,0687459 | -1,06388 | 1 | 1 |
| Nxf2 | 0,10916 | 0,0520824 | -1,06758 | 1 | 1 |
| Ceacam2 | 0,015609 | 0,00742869 | -1,07118 | 1 | 1 |
| Nat1 | 0,207432 | 0,0984946 | -1,07452 | 1 | 1 |
| Edaradd | 0,283364 | 0,13454 | -1,07462 | 1 | 1 |
| H2-Aa | 0,301205 | 0,142803 | -1,07672 | 1 | 1 |
| Slc37a1 | 0,478741 | 0,226518 | -1,07962 | 1 | 1 |
| Atp1a3 | 0,057574 | 0,0271963 | -1,082 | 1 | 1 |
| Hrc | 0,449476 | 0,212046 | -1,08387 | 1 | 1 |
| Slc10a1 | 0,105646 | 0,0497742 | -1,08576 | 1 | 1 |
| Ascl2 | 0,210247 | 0,0990022 | -1,08655 | 1 | 1 |
| Asgr2 | 0,126794 | 0,0596724 | -1,08735 | 1 | 1 |
| Pkd1l3 | 0,102221 | 0,0481049 | -1,08744 | 1 | 1 |
| Mybpc1 | 0,33135 | 0,155876 | -1,08796 | 1 | 1 |
| Sptbn4 | 0,136169 | 0,0640468 | -1,0882 | 1 | 1 |
| Ralyl | 0,230578 | 0,108223 | -1,09125 | 1 | 1 |
| Ablim3 | 0,37474 | 0,175789 | -1,09205 | 1 | 1 |
| Bcas3os2 | 0,508565 | 0,237123 | -1,10079 | 1 | 1 |
| AU018091 | 0,056591 | 0,0263772 | -1,10128 | 1 | 1 |
| Cacng2 | 0,068424 | 0,0318924 | -1,10128 | 1 | 1 |
| Clec4a1 | 0,135583 | 0,0631957 | -1,10128 | 1 | 1 |
| Hist1h2ac | 0,816419 | 0,380396 | -1,10181 | 1 | 1 |
| Prkg2 | 0,279405 | 0,13012 | -1,10252 | 1 | 1 |
| Rnf152 | 0,275716 | 0,128389 | -1,10266 | 1 | 1 |
| Adamts5 | 0,346233 | 0,160991 | -1,10476 | 1 | 1 |
| M1ap | 0,230719 | 0,107227 | -1,10546 | 1 | 1 |
| AA986860 | 0,282309 | 0,131131 | -1,10627 | 1 | 1 |
| Dlgap1 | 0,267395 | 0,124167 | -1,10669 | 1 | 1 |
| Slc6a11 | 0,075099 | 0,0348448 | -1,10786 | 1 | 1 |
| Armc3 | 0,044769 | 0,0206935 | -1,11332 | 1 | 1 |
| Phex | 0,103538 | 0,0478199 | -1,11448 | 1 | 1 |
| Ebi3 | 0,300806 | 0,138802 | -1,11581 | 1 | 1 |
| Calca | 0,533042 | 0,245743 | -1,1171 | 1 | 1 |
| Samd9l | 0,213732 | 0,0983129 | -1,12035 | 1 | 1 |
| Rasef | 0,321034 | 0,147608 | -1,12096 | 1 | 1 |
| Dnah10 | 0,054791 | 0,0251909 | -1,12104 | 1 | 1 |
| Wee2 | 0,147893 | 0,0679267 | -1,12251 | 1 | 1 |
| Syne3 | 0,318164 | 0,146105 | -1,12276 | 1 | 1 |
| Phldb3 | 0,264877 | 0,121211 | -1,1278 | 1 | 1 |
| Grin3a | 0,170101 | 0,0778306 | -1,12798 | 1 | 1 |
| Cutal | 0,300539 | 0,137493 | -1,1282 | 1 | 1 |
| Sntb1 | 0,580282 | 0,264134 | -1,13548 | 1 | 1 |
| Fbxo43 | 0,015437 | 0,00702536 | -1,13578 | 1 | 1 |
| BC048609 | 0,710862 | 0,323282 | -1,13678 | 1 | 1 |

| | | | | | |
|---|---|---|---|---|---|
| Crygn | 0,44395 | 0,20185 | -1,13711 | 1 | 1 |
| 1700096J18Rik | 0,473755 | 0,215303 | -1,13777 | 1 | 1 |
| BC049635 | 0,081573 | 0,0370716 | -1,13777 | 1 | 1 |
| Gm10845 | 0,037188 | 0,0169006 | -1,13777 | 1 | 1 |
| Spink8 | 0,249351 | 0,11332 | -1,13777 | 1 | 1 |
| Gm5127 | 0,317296 | 0,144038 | -1,13937 | 1 | 1 |
| Spata2l | 0,223014 | 0,101166 | -1,14041 | 1 | 1 |
| Ano2 | 0,171735 | 0,0778502 | -1,14142 | 1 | 1 |
| Hist1h3h | 0,54198 | 0,245405 | -1,14308 | 1 | 1 |
| Olfr856-ps1 | 0,107043 | 0,0484684 | -1,14308 | 1 | 1 |
| Plekhs1 | 0,070577 | 0,0319568 | -1,14308 | 1 | 1 |
| Slc5a9 | 0,059806 | 0,0270649 | -1,14387 | 1 | 1 |
| Gm1661 | 0,077434 | 0,0350397 | -1,14398 | 1 | 1 |
| Cyp2d26 | 0,172599 | 0,0780576 | -1,14481 | 1 | 1 |
| Cntn1 | 0,08662 | 0,0391235 | -1,14667 | 1 | 1 |
| Vwc2 | 0,084233 | 0,0380413 | -1,14681 | 1 | 1 |
| Cyp24a1 | 0,030661 | 0,0138394 | -1,14762 | 1 | 1 |
| Hcrtr2 | 0,026898 | 0,0121407 | -1,14764 | 1 | 1 |
| Lamp5 | 0,054134 | 0,0244338 | -1,14764 | 1 | 1 |
| Boll | 0,114467 | 0,0516656 | -1,14766 | 1 | 1 |
| Slc4a10 | 0,018225 | 0,00822522 | -1,14781 | 1 | 1 |
| Elovl7 | 0,52269 | 0,235786 | -1,14848 | 1 | 1 |
| Fmo1 | 0,331924 | 0,149269 | -1,15293 | 1 | 1 |
| Sptlc3 | 0,495792 | 0,222555 | -1,15558 | 1 | 1 |
| Slc7a10 | 0,462648 | 0,207576 | -1,15628 | 1 | 1 |
| Nebl | 0,328438 | 0,146665 | -1,16309 | 1 | 1 |
| C330024D21Rik | 0,142537 | 0,0636019 | -1,1642 | 1 | 1 |
| Cers3 | 0,458491 | 0,204285 | -1,16631 | 1 | 1 |
| Ccdc78 | 0,54846 | 0,244169 | -1,16751 | 1 | 1 |
| Phf19 | 0,184053 | 0,0818977 | -1,16822 | 1 | 1 |
| Zfp385c | 0,042998 | 0,0191278 | -1,16861 | 1 | 1 |
| Fam46a | 0,228014 | 0,100953 | -1,17544 | 1 | 1 |
| Ccna1 | 0,339048 | 0,149976 | -1,17676 | 1 | 1 |
| C130030K03Rik | 0,129855 | 0,0574177 | -1,17733 | 1 | 1 |
| Dcx | 0,029026 | 0,0128211 | -1,17884 | 1 | 1 |
| Gm13483 | 0,304353 | 0,134257 | -1,18074 | 1 | 1 |
| 9030625G05Rik | 0,259861 | 0,114604 | -1,18108 | 1 | 1 |
| Lancl3 | 0,157669 | 0,0694556 | -1,18274 | 1 | 1 |
| L3mbtl1 | 0,264605 | 0,116513 | -1,18335 | 1 | 1 |
| Cela1 | 0,459319 | 0,201971 | -1,18535 | 1 | 1 |
| Tmem52b | 0,14189 | 0,062265 | -1,18828 | 1 | 1 |
| Msr1 | 0,15463 | 0,067749 | -1,19055 | 1 | 1 |
| Sdk1 | 0,51314 | 0,224741 | -1,19109 | 1 | 1 |
| 4933406C10Rik | 0,136744 | 0,059879 | -1,19136 | 1 | 1 |
| Fbxl8 | 0,296657 | 0,129721 | -1,19338 | 1 | 1 |
| Ntng1 | 0,111304 | 0,0486202 | -1,19488 | 1 | 1 |
| Slc5a2 | 0,039646 | 0,0173146 | -1,19517 | 1 | 1 |
| 4932411E22Rik | 0,255061 | 0,111323 | -1,19609 | 1 | 1 |

| | | | | | |
|---|---|---|---|---|---|
| D630032N06Rik | 0,366415 | 0,159853 | -1,19673 | 1 | 1 |
| Golt1a | 0,097534 | 0,0425504 | -1,19673 | 1 | 1 |
| Kctd8 | 0,059501 | 0,0259582 | -1,19673 | 1 | 1 |
| Tmem221 | 0,119793 | 0,0522079 | -1,1982 | 1 | 1 |
| Gm16863 | 0,268944 | 0,117062 | -1,20003 | 1 | 1 |
| Lypd1 | 0,435695 | 0,189375 | -1,20207 | 1 | 1 |
| 2310002F09Rik | 0,120936 | 0,0525642 | -1,2021 | 1 | 1 |
| Gpr151 | 0,19888 | 0,0864419 | -1,2021 | 1 | 1 |
| Nalcn | 0,059129 | 0,0256627 | -1,2042 | 1 | 1 |
| B130024G19Rik | 0,438328 | 0,190118 | -1,20511 | 1 | 1 |
| Kcnj9 | 0,040534 | 0,0175565 | -1,20712 | 1 | 1 |
| P2rx7 | 0,26081 | 0,112865 | -1,20841 | 1 | 1 |
| Ppargc1b | 0,298432 | 0,12894 | -1,2107 | 1 | 1 |
| Lif | 0,094363 | 0,0407347 | -1,21196 | 1 | 1 |
| Slc7a4 | 0,357944 | 0,154341 | -1,21361 | 1 | 1 |
| Kcnrg | 0,179772 | 0,0774866 | -1,21415 | 1 | 1 |
| Mtag2 | 0,334149 | 0,143209 | -1,22237 | 1 | 1 |
| C6 | 0,123442 | 0,0528614 | -1,22355 | 1 | 1 |
| 9630028B13Rik | 0,325325 | 0,139007 | -1,22672 | 1 | 1 |
| Tmtc1 | 0,353183 | 0,150787 | -1,2279 | 1 | 1 |
| Fam84a | 0,397066 | 0,169045 | -1,23197 | 1 | 1 |
| Cdkl2 | 0,219656 | 0,0934868 | -1,23241 | 1 | 1 |
| Syt1 | 0,086269 | 0,0367045 | -1,23289 | 1 | 1 |
| 3300002I08Rik | 0,679851 | 0,289072 | -1,23379 | 1 | 1 |
| Kcnk4 | 0,065722 | 0,0279448 | -1,23379 | 1 | 1 |
| Nkx2-6 | 0,105693 | 0,0449405 | -1,23379 | 1 | 1 |
| Spag6 | 0,088966 | 0,0378282 | -1,23379 | 1 | 1 |
| Amy1 | 0,06896 | 0,0293215 | -1,2338 | 1 | 1 |
| Gcsam | 0,065034 | 0,0276518 | -1,23382 | 1 | 1 |
| Creb5 | 0,312891 | 0,132899 | -1,23532 | 1 | 1 |
| Pcdhb20 | 0,142753 | 0,0604959 | -1,23861 | 1 | 1 |
| Srms | 0,156636 | 0,0662677 | -1,24104 | 1 | 1 |
| Sod3 | 0,366657 | 0,154813 | -1,24391 | 1 | 1 |
| 4931429L15Rik | 0,090429 | 0,0380238 | -1,24988 | 1 | 1 |
| Bmx | 0,118414 | 0,0497664 | -1,25059 | 1 | 1 |
| Fabp4 | 0,37967 | 0,159488 | -1,2513 | 1 | 1 |
| Dnajc6 | 0,460529 | 0,193001 | -1,25468 | 1 | 1 |
| P2ry2 | 0,188514 | 0,0787332 | -1,25963 | 1 | 1 |
| Fgf23 | 0,118506 | 0,04945 | -1,26092 | 1 | 1 |
| Fam19a3 | 0,087811 | 0,0365768 | -1,26347 | 1 | 1 |
| Sult6b1 | 0,194593 | 0,0810559 | -1,26347 | 1 | 1 |
| Gm13212 | 0,256352 | 0,106711 | -1,26441 | 1 | 1 |
| Bpifb5 | 0,305742 | 0,127131 | -1,266 | 1 | 1 |
| Rnaset2a,Rnaset2 | 3,22278 | 1,33723 | -1,26906 | 1 | 1 |
| Tnfrsf11b | 0,366829 | 0,152063 | -1,27044 | 1 | 1 |
| Npc1l1 | 0,029794 | 0,0123262 | -1,27329 | 1 | 1 |
| Klk11 | 0,061441 | 0,0253905 | -1,2749 | 1 | 1 |
| 4930461G14Rik | 0,113082 | 0,0467251 | -1,2751 | 1 | 1 |

| | | | | | |
|---|---|---|---|---|---|
| Ankrd1 | 0,166908 | 0,0689661 | -1,2751 | 1 | 1 |
| Camp | 0,164897 | 0,0681351 | -1,2751 | 1 | 1 |
| Cpne9 | 0,025296 | 0,0104521 | -1,2751 | 1 | 1 |
| Cstad | 0,106256 | 0,0439048 | -1,2751 | 1 | 1 |
| Gm1965 | 0,07663 | 0,0316633 | -1,2751 | 1 | 1 |
| Gm4832 | 0,231792 | 0,0957758 | -1,2751 | 1 | 1 |
| Myod1 | 0,039476 | 0,0163115 | -1,2751 | 1 | 1 |
| Plscr2 | 0,038463 | 0,0158927 | -1,2751 | 1 | 1 |
| Tmigd1 | 0,11211 | 0,0463235 | -1,2751 | 1 | 1 |
| Spt1 | 0,150076 | 0,0620103 | -1,27511 | 1 | 1 |
| Il12a | 0,060971 | 0,0251913 | -1,27519 | 1 | 1 |
| Camk2b | 0,248895 | 0,10282 | -1,27542 | 1 | 1 |
| 6720483E21Rik | 0,368456 | 0,152177 | -1,27574 | 1 | 1 |
| Pi16 | 0,234987 | 0,0970351 | -1,276 | 1 | 1 |
| Aldh1a1 | 0,113368 | 0,0468092 | -1,27615 | 1 | 1 |
| Zmat4 | 0,329738 | 0,135891 | -1,27888 | 1 | 1 |
| 9130019P16Rik | 0,091282 | 0,0375796 | -1,28037 | 1 | 1 |
| Prkcb | 0,431067 | 0,177383 | -1,28105 | 1 | 1 |
| Nr5a2 | 0,350264 | 0,144005 | -1,28232 | 1 | 1 |
| Lox | 0,137141 | 0,0563479 | -1,28323 | 1 | 1 |
| Soat2 | 0,19086 | 0,0782726 | -1,28594 | 1 | 1 |
| Mobp | 0,043844 | 0,0179213 | -1,29069 | 1 | 1 |
| Pabpc1l | 0,343905 | 0,139991 | -1,29667 | 1 | 1 |
| 03/01/16 | 0,197913 | 0,0804906 | -1,29798 | 1 | 1 |
| Dennd2d | 0,535171 | 0,21759 | -1,29839 | 1 | 1 |
| Pcdhb19 | 0,114175 | 0,0463623 | -1,30023 | 1 | 1 |
| H2-Oa | 0,406317 | 0,164964 | -1,30046 | 1 | 1 |
| Psd4 | 0,371305 | 0,150679 | -1,30113 | 1 | 1 |
| 2410012M07Rik | 0,106678 | 0,0432904 | -1,30114 | 1 | 1 |
| Col10a1 | 0,058699 | 0,0238202 | -1,30114 | 1 | 1 |
| Aim1l | 0,204023 | 0,0823796 | -1,30837 | 1 | 1 |
| Spaca1 | 0,320895 | 0,129434 | -1,30988 | 1 | 1 |
| Tmem130 | 0,269111 | 0,108287 | -1,31334 | 1 | 1 |
| G6b | 0,488601 | 0,19644 | -1,31457 | 1 | 1 |
| Gm15408 | 0,090598 | 0,0364114 | -1,31508 | 1 | 1 |
| Ddx4 | 0,259673 | 0,104296 | -1,31601 | 1 | 1 |
| Gda | 0,079477 | 0,031903 | -1,31684 | 1 | 1 |
| C730036E19Rik | 0,043767 | 0,0175631 | -1,31729 | 1 | 1 |
| Enkur | 0,107151 | 0,0429984 | -1,31729 | 1 | 1 |
| Prg2 | 0,168162 | 0,0674812 | -1,31729 | 1 | 1 |
| Syt6 | 0,049565 | 0,0198871 | -1,31748 | 1 | 1 |
| F630042J09Rik | 0,067987 | 0,0272778 | -1,31753 | 1 | 1 |
| Rhov | 0,1385 | 0,0555692 | -1,31753 | 1 | 1 |
| Angpt4 | 0,151888 | 0,0608913 | -1,3187 | 1 | 1 |
| Slc4a11 | 0,365474 | 0,14644 | -1,31945 | 1 | 1 |
| Kcns1 | 0,224245 | 0,0896982 | -1,32192 | 1 | 1 |
| Tmem116 | 0,165201 | 0,0660541 | -1,3225 | 1 | 1 |
| Mapk15 | 0,473511 | 0,189107 | -1,32419 | 1 | 1 |

| | | | | | |
|---|---|---|---|---|---|
| Hsf5 | 0,061386 | 0,024434 | -1,32903 | 1 | 1 |
| Isl2 | 0,147371 | 0,0586592 | -1,32903 | 1 | 1 |
| Pramef12 | 0,215178 | 0,0856488 | -1,32903 | 1 | 1 |
| Zfp296 | 0,509061 | 0,20192 | -1,33406 | 1 | 1 |
| Srrm3 | 0,036509 | 0,0144389 | -1,33828 | 1 | 1 |
| Cbfa2t3 | 0,424553 | 0,167866 | -1,33864 | 1 | 1 |
| BC024139 | 0,222208 | 0,0878515 | -1,33877 | 1 | 1 |
| Gucy2c | 0,296624 | 0,117269 | -1,33882 | 1 | 1 |
| B4galnt2 | 0,489597 | 0,193407 | -1,33995 | 1 | 1 |
| Shpk | 0,179195 | 0,0706943 | -1,34186 | 1 | 1 |
| Zdhhc23 | 0,429965 | 0,169626 | -1,34186 | 1 | 1 |
| Lepr | 0,327889 | 0,129015 | -1,34567 | 1 | 1 |
| Zcchc16 | 0,051774 | 0,0203201 | -1,34932 | 1 | 1 |
| S1pr4 | 0,189343 | 0,0737709 | -1,35988 | 1 | 1 |
| Epha3 | 0,571186 | 0,221984 | -1,3635 | 1 | 1 |
| P2rx5 | 0,139171 | 0,0539738 | -1,36653 | 1 | 1 |
| Trpm6 | 0,101407 | 0,0391836 | -1,37184 | 1 | 1 |
| Pcdhb7 | 0,109328 | 0,0422156 | -1,37282 | 1 | 1 |
| Smpd5 | 0,294341 | 0,113551 | -1,37415 | 1 | 1 |
| Fam107a | 0,241556 | 0,0931858 | -1,37418 | 1 | 1 |
| Mas1 | 0,208548 | 0,0802198 | -1,37835 | 1 | 1 |
| Cdh23 | 0,079434 | 0,0305525 | -1,37847 | 1 | 1 |
| Pcdh10 | 0,273436 | 0,104949 | -1,38152 | 1 | 1 |
| Gal3st1 | 0,405642 | 0,155434 | -1,38391 | 1 | 1 |
| Myh1 | 0,110955 | 0,0424854 | -1,38494 | 1 | 1 |
| Slc9a9 | 0,401428 | 0,153634 | -1,38564 | 1 | 1 |
| Pde1a | 0,129321 | 0,0494484 | -1,38696 | 1 | 1 |
| Elfn1 | 0,282147 | 0,107789 | -1,38824 | 1 | 1 |
| Tmem91 | 0,423903 | 0,161629 | -1,39105 | 1 | 1 |
| Il16 | 0,285694 | 0,108886 | -1,39165 | 1 | 1 |
| Csgalnact1 | 0,511454 | 0,194895 | -1,39191 | 1 | 1 |
| 5031425F14Rik | 0,078525 | 0,0298768 | -1,39413 | 1 | 1 |
| Xlr4a | 0,113929 | 0,0433469 | -1,39413 | 1 | 1 |
| Casp4 | 0,183273 | 0,0696799 | -1,39518 | 1 | 1 |
| Hand2 | 0,110812 | 0,0421306 | -1,39518 | 1 | 1 |
| Mlph | 0,052584 | 0,0199924 | -1,39518 | 1 | 1 |
| Slc30a2 | 0,367631 | 0,139613 | -1,39682 | 1 | 1 |
| Cyp2s1 | 0,433532 | 0,164457 | -1,39843 | 1 | 1 |
| Proz | 0,213135 | 0,0808359 | -1,3987 | 1 | 1 |
| 4631405J19Rik | 0,069102 | 0,0262063 | -1,39882 | 1 | 1 |
| 4930451C15Rik | 0,036749 | 0,0139351 | -1,399 | 1 | 1 |
| Oxct2b | 0,137552 | 0,0521253 | -1,39992 | 1 | 1 |
| Gm8234 | 0,197755 | 0,0749277 | -1,40015 | 1 | 1 |
| Gm16998 | 0,138498 | 0,0524637 | -1,40048 | 1 | 1 |
| Lrat | 0,027619 | 0,0104623 | -1,40048 | 1 | 1 |
| Slc17a8 | 0,033451 | 0,0126714 | -1,40048 | 1 | 1 |
| Hs3st6 | 0,588631 | 0,221419 | -1,41059 | 1 | 1 |
| Ccdc88b | 0,221663 | 0,0833015 | -1,41196 | 1 | 1 |

| | | | | | |
|---|---|---|---|---|---|
| Grem2 | 0,128764 | 0,0483296 | -1,41375 | 1 | 1 |
| Cacna2d3 | 0,396837 | 0,148847 | -1,41472 | 1 | 1 |
| Cmah | 0,048996 | 0,0183525 | -1,41667 | 1 | 1 |
| C1ql3 | 0,106782 | 0,0399909 | -1,41692 | 1 | 1 |
| 4933413G19Rik | 0,423166 | 0,157942 | -1,42183 | 1 | 1 |
| Rxfp2 | 0,171126 | 0,0636076 | -1,42779 | 1 | 1 |
| Ly6c1 | 0,229313 | 0,0851055 | -1,42999 | 1 | 1 |
| 4831440E17Rik | 0,136275 | 0,0502937 | -1,43807 | 1 | 1 |
| Ipcef1 | 0,040863 | 0,0150723 | -1,43888 | 1 | 1 |
| Frmpd3 | 0,18301 | 0,0674535 | -1,43996 | 1 | 1 |
| Gm2011 | 0,37129 | 0,136849 | -1,43996 | 1 | 1 |
| Spock1 | 0,531655 | 0,195879 | -1,44053 | 1 | 1 |
| Plbd1 | 0,535155 | 0,196886 | -1,4426 | 1 | 1 |
| Frat1 | 0,393108 | 0,144601 | -1,44285 | 1 | 1 |
| Fam83a | 0,100785 | 0,0370535 | -1,44359 | 1 | 1 |
| Vmn2r29 | 0,274881 | 0,0999247 | -1,45989 | 1 | 1 |
| Atp5g2 | 0,334295 | 0,121505 | -1,4601 | 1 | 1 |
| Wisp1 | 0,571377 | 0,206822 | -1,46606 | 1 | 1 |
| Aldh1a3 | 0,403961 | 0,14618 | -1,46647 | 1 | 1 |
| Slc2a2 | 0,049992 | 0,0180827 | -1,46708 | 1 | 1 |
| A730056A06Rik | 0,254704 | 0,0920875 | -1,46774 | 1 | 1 |
| Il1a | 0,200873 | 0,0726245 | -1,46775 | 1 | 1 |
| 6030466F02Rik | 0,096296 | 0,0347833 | -1,46907 | 1 | 1 |
| B020018J22Rik | 0,039233 | 0,0141715 | -1,46907 | 1 | 1 |
| Htr1b | 0,123765 | 0,0447057 | -1,46907 | 1 | 1 |
| Mndal | 0,069509 | 0,0251077 | -1,46907 | 1 | 1 |
| Sox10 | 0,094416 | 0,0341042 | -1,46907 | 1 | 1 |
| Gja8 | 0,531059 | 0,191004 | -1,47527 | 1 | 1 |
| F10 | 0,355294 | 0,127289 | -1,4809 | 1 | 1 |
| Esrrg | 0,338638 | 0,121107 | -1,48346 | 1 | 1 |
| C130060K24Rik | 0,366204 | 0,130787 | -1,48543 | 1 | 1 |
| Lgals2 | 0,503195 | 0,179628 | -1,4861 | 1 | 1 |
| Chrm3 | 0,065927 | 0,0235148 | -1,48731 | 1 | 1 |
| Agrp | 0,422264 | 0,150522 | -1,48817 | 1 | 1 |
| Cpne8 | 0,347501 | 0,123868 | -1,48821 | 1 | 1 |
| BC037704 | 0,598939 | 0,213316 | -1,48942 | 1 | 1 |
| Ptk2b | 0,436353 | 0,15535 | -1,48997 | 1 | 1 |
| Gm13003 | 0,230202 | 0,0815326 | -1,49745 | 1 | 1 |
| Dnali1 | 0,185706 | 0,0656354 | -1,50048 | 1 | 1 |
| 9330159M07Rik | 0,257276 | 0,0908278 | -1,50211 | 1 | 1 |
| Gm2373 | 0,077513 | 0,0273059 | -1,50523 | 1 | 1 |
| Runx3 | 0,173412 | 0,0610153 | -1,50696 | 1 | 1 |
| Adam28 | 0,031402 | 0,0110479 | -1,50711 | 1 | 1 |
| Mr1 | 0,377625 | 0,132627 | -1,50958 | 1 | 1 |
| F2 | 0,086191 | 0,0301727 | -1,5143 | 1 | 1 |
| Calcoco2 | 0,042172 | 0,0147474 | -1,51583 | 1 | 1 |
| Cst6 | 0,02054 | 0,00718285 | -1,51583 | 1 | 1 |
| Foxn4 | 0,027825 | 0,00973034 | -1,51583 | 1 | 1 |

| | | | | | |
|---|---|---|---|---|---|
| Gm5134 | 0,029242 | 0,0102257 | -1,51583 | 1 | 1 |
| Krt28 | 0,119788 | 0,0418894 | -1,51583 | 1 | 1 |
| Pxt1 | 0,094899 | 0,033186 | -1,51583 | 1 | 1 |
| Tas2r137 | 0,096445 | 0,0337266 | -1,51583 | 1 | 1 |
| 1700023L04Rik | 0,338895 | 0,118486 | -1,51612 | 1 | 1 |
| Adra2c | 0,067215 | 0,0232792 | -1,52974 | 1 | 1 |
| Atxn1 | 0,454895 | 0,157504 | -1,53014 | 1 | 1 |
| Plcb2 | 0,165248 | 0,0572153 | -1,53016 | 1 | 1 |
| Clca5 | 0,141716 | 0,0490125 | -1,53178 | 1 | 1 |
| Ggt5 | 0,21365 | 0,0736934 | -1,53564 | 1 | 1 |
| Cacng5 | 0,339784 | 0,117191 | -1,53575 | 1 | 1 |
| Nphs2 | 0,042795 | 0,0147327 | -1,53842 | 1 | 1 |
| Vmn2r87 | 0,053042 | 0,0182602 | -1,53842 | 1 | 1 |
| Xaf1 | 0,118198 | 0,0405716 | -1,54266 | 1 | 1 |
| A430105I19Rik | 0,149642 | 0,0512373 | -1,54624 | 1 | 1 |
| Cd6 | 0,133119 | 0,0455289 | -1,54787 | 1 | 1 |
| Gpr3 | 0,421411 | 0,143781 | -1,55136 | 1 | 1 |
| Ptpro | 0,094637 | 0,0322722 | -1,55211 | 1 | 1 |
| Gm15663 | 0,205238 | 0,0696979 | -1,55811 | 1 | 1 |
| Bst1 | 0,074935 | 0,0252477 | -1,56948 | 1 | 1 |
| Krtap26-1 | 0,623672 | 0,209991 | -1,57046 | 1 | 1 |
| Slc22a3 | 0,145557 | 0,049009 | -1,57046 | 1 | 1 |
| Misp | 0,258629 | 0,0870565 | -1,57086 | 1 | 1 |
| Slc22a7 | 0,11494 | 0,0386565 | -1,57209 | 1 | 1 |
| Nkx3-2 | 0,542991 | 0,182451 | -1,57342 | 1 | 1 |
| Tecrl | 0,117433 | 0,0394501 | -1,57373 | 1 | 1 |
| Cadm2 | 0,02221 | 0,00745719 | -1,57448 | 1 | 1 |
| Rgs11 | 0,505961 | 0,169647 | -1,57649 | 1 | 1 |
| Dlg2 | 0,062279 | 0,0208218 | -1,58066 | 1 | 1 |
| Prph | 0,215654 | 0,071867 | -1,58532 | 1 | 1 |
| Synpo2 | 0,526787 | 0,175517 | -1,58561 | 1 | 1 |
| Adap2 | 0,174243 | 0,0579626 | -1,5879 | 1 | 1 |
| Diras1 | 0,089367 | 0,0297284 | -1,5879 | 1 | 1 |
| Col6a3 | 0,331508 | 0,110025 | -1,59122 | 1 | 1 |
| Fbxo2 | 0,316034 | 0,10452 | -1,5963 | 1 | 1 |
| Stk32a | 0,224504 | 0,0738338 | -1,60439 | 1 | 1 |
| Zdhhc14 | 0,561232 | 0,184434 | -1,60549 | 1 | 1 |
| Alox15 | 0,058947 | 0,0193591 | -1,6064 | 1 | 1 |
| Gramd2 | 0,111917 | 0,0367555 | -1,6064 | 1 | 1 |
| Lrrc10b | 0,069553 | 0,0228423 | -1,6064 | 1 | 1 |
| Tmprss6 | 0,043599 | 0,0143185 | -1,6064 | 1 | 1 |
| Btc | 0,079803 | 0,0260845 | -1,61324 | 1 | 1 |
| Tfcp2l1 | 0,361736 | 0,117908 | -1,61727 | 1 | 1 |
| Dnajb13 | 0,088175 | 0,0286623 | -1,62121 | 1 | 1 |
| Rilp | 0,193727 | 0,0629321 | -1,62216 | 1 | 1 |
| Oit1 | 0,395709 | 0,128529 | -1,62235 | 1 | 1 |
| 2610037D02Rik | 0,048368 | 0,0157071 | -1,62263 | 1 | 1 |
| Nlrp6 | 0,039168 | 0,0127194 | -1,62263 | 1 | 1 |

| | | | | | |
|---|---|---|---|---|---|
| Gm8801 | 0,366931 | 0,119091 | -1,62344 | 1 | 1 |
| 2010001E11Rik | 0,039865 | 0,0129332 | -1,62405 | 1 | 1 |
| Cyp17a1 | 0,050678 | 0,0164411 | -1,62405 | 1 | 1 |
| Fam92b | 0,084223 | 0,0273237 | -1,62405 | 1 | 1 |
| Fendrr | 0,038125 | 0,0123686 | -1,62405 | 1 | 1 |
| Xpnpep2 | 0,025184 | 0,00816931 | -1,62421 | 1 | 1 |
| Npas1 | 0,342006 | 0,110807 | -1,62597 | 1 | 1 |
| Crybb3 | 0,300445 | 0,0971575 | -1,6287 | 1 | 1 |
| Hdac9 | 0,499592 | 0,161135 | -1,63248 | 1 | 1 |
| Nav3 | 0,512433 | 0,165101 | -1,63402 | 1 | 1 |
| Adamts16 | 0,320291 | 0,102282 | -1,64683 | 1 | 1 |
| 3830417A13Rik | 0,151601 | 0,0482336 | -1,65217 | 1 | 1 |
| Glp2r | 0,167375 | 0,0529134 | -1,66138 | 1 | 1 |
| Neb | 0,069327 | 0,0218931 | -1,66295 | 1 | 1 |
| Aire | 0,077457 | 0,0244416 | -1,66406 | 1 | 1 |
| Dsc3 | 0,526832 | 0,166115 | -1,66516 | 1 | 1 |
| Nanos2 | 0,110206 | 0,0346425 | -1,66958 | 1 | 1 |
| Tlr6 | 0,056805 | 0,0178563 | -1,66958 | 1 | 1 |
| 4833419F23Rik | 0,201279 | 0,0631951 | -1,67131 | 1 | 1 |
| Tac1 | 0,164177 | 0,0515461 | -1,67131 | 1 | 1 |
| Slc9b1 | 0,150836 | 0,0473355 | -1,67199 | 1 | 1 |
| Dbx1 | 0,411001 | 0,128581 | -1,67646 | 1 | 1 |
| Fam83e | 0,491859 | 0,153324 | -1,68166 | 1 | 1 |
| Ovol2 | 0,257343 | 0,0800206 | -1,68525 | 1 | 1 |
| Tceal7 | 0,293213 | 0,0908696 | -1,69008 | 1 | 1 |
| Tnfrsf23 | 0,073042 | 0,0226365 | -1,69008 | 1 | 1 |
| Trim30d | 0,090012 | 0,0277992 | -1,69508 | 1 | 1 |
| Neurl3 | 0,146773 | 0,0452528 | -1,69751 | 1 | 1 |
| Zic1 | 0,557478 | 0,171748 | -1,69862 | 1 | 1 |
| Slco2a1 | 0,105157 | 0,0322607 | -1,70469 | 1 | 1 |
| Olfr543 | 0,396322 | 0,121469 | -1,70608 | 1 | 1 |
| Gm19705 | 0,425501 | 0,129891 | -1,71187 | 1 | 1 |
| Prdm12 | 0,302903 | 0,0922335 | -1,7155 | 1 | 1 |
| Krt35 | 0,487368 | 0,148196 | -1,7175 | 1 | 1 |
| Gkn2 | 0,154958 | 0,0469697 | -1,72207 | 1 | 1 |
| Il1r2 | 0,077888 | 0,0236087 | -1,72207 | 1 | 1 |
| Klrg2 | 0,170919 | 0,0518076 | -1,72207 | 1 | 1 |
| Ldhc | 0,080282 | 0,0243345 | -1,72207 | 1 | 1 |
| Sbk2 | 0,094964 | 0,0287849 | -1,72207 | 1 | 1 |
| Kng2 | 0,063055 | 0,0191114 | -1,72218 | 1 | 1 |
| Zap70 | 0,086931 | 0,0263157 | -1,72395 | 1 | 1 |
| A730036I17Rik | 0,044008 | 0,0132498 | -1,73178 | 1 | 1 |
| Abca8a | 0,026257 | 0,00790536 | -1,73178 | 1 | 1 |
| 4931402G19Rik | 0,098316 | 0,0295223 | -1,73563 | 1 | 1 |
| Sema6b | 0,544466 | 0,163394 | -1,73648 | 1 | 1 |
| 9330117O12Rik | 0,416266 | 0,124858 | -1,73722 | 1 | 1 |
| Gmnc | 0,036304 | 0,0108299 | -1,7451 | 1 | 1 |
| Ptchd1 | 0,091314 | 0,0272292 | -1,74568 | 1 | 1 |

| | | | | | |
|---|---|---|---|---|---|
| Aox3 | 0,099729 | 0,0297224 | -1,74646 | 1 | 1 |
| Nipal2 | 0,114464 | 0,0341138 | -1,74646 | 1 | 1 |
| Elf5 | 0,224899 | 0,0668465 | -1,75035 | 1 | 1 |
| Aqp4 | 0,218532 | 0,0649345 | -1,75079 | 1 | 1 |
| Mogat1 | 0,378229 | 0,112256 | -1,75247 | 1 | 1 |
| Mb | 0,18959 | 0,055996 | -1,75949 | 1 | 1 |
| Paqr7 | 0,519599 | 0,153094 | -1,76298 | 1 | 1 |
| Izumo1 | 0,183776 | 0,054071 | -1,76502 | 1 | 1 |
| Gm4980 | 0,194402 | 0,0570013 | -1,76998 | 1 | 1 |
| Gm5622 | 0,133319 | 0,0390908 | -1,76998 | 1 | 1 |
| Lrrc36 | 0,097639 | 0,0286203 | -1,77042 | 1 | 1 |
| Pitpnm3 | 0,02861 | 0,00838597 | -1,77045 | 1 | 1 |
| Nlgn1 | 0,05385 | 0,0157699 | -1,77177 | 1 | 1 |
| Abcc9 | 0,334989 | 0,097558 | -1,77978 | 1 | 1 |
| Lax1 | 0,180271 | 0,0524921 | -1,78 | 1 | 1 |
| Gfra1 | 0,350857 | 0,101711 | -1,7864 | 1 | 1 |
| Aqp1 | 0,479498 | 0,138775 | -1,78878 | 1 | 1 |
| Kcnh8 | 0,036232 | 0,0104785 | -1,78981 | 1 | 1 |
| Dbh | 0,071269 | 0,0205889 | -1,7914 | 1 | 1 |
| Olfr920 | 0,084337 | 0,0243642 | -1,7914 | 1 | 1 |
| Tnfsf4 | 0,105406 | 0,0304509 | -1,7914 | 1 | 1 |
| Gucy2g | 0,065867 | 0,018914 | -1,80011 | 1 | 1 |
| Hs6st3 | 0,153724 | 0,0441124 | -1,80108 | 1 | 1 |
| A930041C12Rik | 0,284149 | 0,0808204 | -1,81386 | 1 | 1 |
| Kctd16 | 0,081502 | 0,0231817 | -1,81386 | 1 | 1 |
| Rnase6 | 0,109139 | 0,0310425 | -1,81386 | 1 | 1 |
| Slc13a4 | 0,117524 | 0,0334129 | -1,81448 | 1 | 1 |
| Slc22a1 | 0,105876 | 0,0300884 | -1,8151 | 1 | 1 |
| Spats1 | 0,198129 | 0,0563052 | -1,8151 | 1 | 1 |
| Adprhl1 | 0,081643 | 0,0231817 | -1,81634 | 1 | 1 |
| Gnat1 | 0,094599 | 0,0268603 | -1,81634 | 1 | 1 |
| Hal | 0,037114 | 0,010538 | -1,81634 | 1 | 1 |
| Hist2h3b | 0,318068 | 0,0903123 | -1,81634 | 1 | 1 |
| Nr4a3 | 0,056421 | 0,0160203 | -1,81634 | 1 | 1 |
| Olfr1420 | 0,13056 | 0,0370713 | -1,81634 | 1 | 1 |
| Olfr456 | 0,111777 | 0,031738 | -1,81634 | 1 | 1 |
| Apoa5 | 0,142666 | 0,0404191 | -1,81953 | 1 | 1 |
| C3 | 0,069658 | 0,0196008 | -1,82937 | 1 | 1 |
| Prss56 | 0,191276 | 0,0538227 | -1,82937 | 1 | 1 |
| Nfix | 0,251779 | 0,0708399 | -1,82953 | 1 | 1 |
| Slc39a5 | 0,291925 | 0,0819827 | -1,83221 | 1 | 1 |
| Clec7a | 0,259967 | 0,0725711 | -1,84086 | 1 | 1 |
| Slc7a9 | 0,128845 | 0,0359309 | -1,84234 | 1 | 1 |
| Tnfrsf11a | 0,327995 | 0,0913817 | -1,8437 | 1 | 1 |
| Tspan8 | 0,392304 | 0,109134 | -1,84587 | 1 | 1 |
| Ptprr | 0,047189 | 0,0131158 | -1,84714 | 1 | 1 |
| Fap | 0,184845 | 0,0513404 | -1,84815 | 1 | 1 |
| Slc15a3 | 0,520862 | 0,144115 | -1,85368 | 1 | 1 |

| | | | | | |
|---|---|---|---|---|---|
| Matn4 | 0,362996 | 0,100179 | -1,85738 | 1 | 1 |
| Eif3j2 | 6,09421 | 1,67858 | -1,8602 | 1 | 1 |
| Tmem200a | 0,365951 | 0,100781 | -1,86042 | 1 | 1 |
| Vmn2r18 | 0,116787 | 0,0319568 | -1,86968 | 1 | 1 |
| Kcnk13 | 0,103573 | 0,0282403 | -1,87482 | 1 | 1 |
| Slc4a9 | 0,185638 | 0,0506116 | -1,87496 | 1 | 1 |
| Mdga2 | 0,171884 | 0,0467947 | -1,87701 | 1 | 1 |
| Sigirr | 0,55676 | 0,150695 | -1,88543 | 1 | 1 |
| Dnase1l3 | 0,176264 | 0,0475851 | -1,88915 | 1 | 1 |
| Slamf1 | 0,128632 | 0,0344474 | -1,90078 | 1 | 1 |
| A330041J22Rik | 0,097574 | 0,0261035 | -1,90225 | 1 | 1 |
| BC061194 | 0,110521 | 0,0295671 | -1,90225 | 1 | 1 |
| Lman1l | 0,104321 | 0,0279084 | -1,90225 | 1 | 1 |
| Clic5 | 0,045984 | 0,0123011 | -1,90235 | 1 | 1 |
| 1500015O10Rik | 0,147104 | 0,0393476 | -1,90249 | 1 | 1 |
| 1700106J16Rik | 0,099133 | 0,0265162 | -1,90249 | 1 | 1 |
| A630020A06 | 0,030289 | 0,00810179 | -1,90249 | 1 | 1 |
| Atp12a | 0,026854 | 0,00718285 | -1,90249 | 1 | 1 |
| Hist1h2bf | 0,443234 | 0,118557 | -1,90249 | 1 | 1 |
| Nhlrc4 | 0,103893 | 0,0277894 | -1,90249 | 1 | 1 |
| Nrsn1 | 0,052054 | 0,0139235 | -1,90249 | 1 | 1 |
| Pkp1 | 0,023499 | 0,00628542 | -1,90249 | 1 | 1 |
| Tmem236 | 0,029132 | 0,00779216 | -1,90249 | 1 | 1 |
| Trim75 | 0,043748 | 0,0117018 | -1,90249 | 1 | 1 |
| Gk2 | 0,122508 | 0,0327421 | -1,90366 | 1 | 1 |
| Asb2 | 0,316695 | 0,0841084 | -1,91277 | 1 | 1 |
| 4930426D05Rik | 0,403809 | 0,107234 | -1,91292 | 1 | 1 |
| 9430060I03Rik | 0,159875 | 0,0423987 | -1,91485 | 1 | 1 |
| Abhd15 | 0,353859 | 0,0935996 | -1,9186 | 1 | 1 |
| Ifi27 | 0,180805 | 0,0477166 | -1,92187 | 1 | 1 |
| Dcn | 0,370447 | 0,0976451 | -1,92365 | 1 | 1 |
| Gbp4 | 0,21483 | 0,056379 | -1,92997 | 1 | 1 |
| Fbln5 | 0,526535 | 0,1378 | -1,93396 | 1 | 1 |
| Acan | 0,037273 | 0,00974661 | -1,93517 | 1 | 1 |
| Gata6 | 0,16436 | 0,0429268 | -1,93691 | 1 | 1 |
| Slc6a4 | 0,339861 | 0,0884636 | -1,94179 | 1 | 1 |
| Pax7 | 0,203583 | 0,0528895 | -1,94456 | 1 | 1 |
| A430093F15Rik | 0,108444 | 0,0281675 | -1,94484 | 1 | 1 |
| Sycp1-ps1 | 0,071995 | 0,018504 | -1,96006 | 1 | 1 |
| 1110028F11Rik | 0,10884 | 0,027868 | -1,96553 | 1 | 1 |
| Dlx3 | 0,560084 | 0,143193 | -1,96768 | 1 | 1 |
| Pcdh11x | 0,142302 | 0,0363643 | -1,96837 | 1 | 1 |
| Il18rap | 0,270454 | 0,0686394 | -1,97828 | 1 | 1 |
| Cplx1 | 0,169773 | 0,0430419 | -1,97979 | 1 | 1 |
| Wfdc1 | 0,110313 | 0,02789 | -1,98378 | 1 | 1 |
| 4933440M02Rik | 0,072314 | 0,0182801 | -1,98401 | 1 | 1 |
| Lhx1os | 0,507294 | 0,128159 | -1,98489 | 1 | 1 |
| Slc44a4 | 0,149316 | 0,0377124 | -1,98525 | 1 | 1 |

| | | | | | |
|---|---|---|---|---|---|
| Gm13154 | 0,203171 | 0,051304 | -1,98555 | 1 | 1 |
| BB283400 | 0,087822 | 0,0221696 | -1,98599 | 1 | 1 |
| Ch25h | 0,092871 | 0,0234442 | -1,98599 | 1 | 1 |
| Ofcc1 | 0,040818 | 0,010304 | -1,98599 | 1 | 1 |
| Stk31 | 0,035141 | 0,00887084 | -1,98599 | 1 | 1 |
| Rufy4 | 0,050702 | 0,0127944 | -1,98654 | 1 | 1 |
| Unc45b | 0,11499 | 0,0288566 | -1,99453 | 1 | 1 |
| Cd74 | 0,273173 | 0,0683952 | -1,99785 | 1 | 1 |
| Rbm11 | 0,068839 | 0,0171478 | -2,0052 | 1 | 1 |
| Abca12 | 0,101341 | 0,0251594 | -2,01005 | 1 | 1 |
| Siglec5 | 0,098782 | 0,0244203 | -2,01617 | 1 | 1 |
| 1700119H24Rik | 0,573451 | 0,140907 | -2,02492 | 1 | 1 |
| Chst9 | 0,188177 | 0,0462042 | -2,026 | 1 | 1 |
| Oprk1 | 0,201207 | 0,0491954 | -2,03208 | 1 | 1 |
| Anxa8 | 0,523526 | 0,1279 | -2,03324 | 1 | 1 |
| Stx11 | 0,524216 | 0,12721 | -2,04295 | 1 | 1 |
| Sycp1 | 0,232697 | 0,05635 | -2,04597 | 1 | 1 |
| Ptpre | 0,365709 | 0,0885292 | -2,04647 | 1 | 1 |
| Slc46a3 | 0,58205 | 0,140697 | -2,04855 | 1 | 1 |
| Cyp11a1 | 0,297949 | 0,071644 | -2,05615 | 1 | 1 |
| Plch2 | 0,354521 | 0,085246 | -2,05617 | 1 | 1 |
| Ltb4r2 | 0,234843 | 0,0564352 | -2,05703 | 1 | 1 |
| Abhd1 | 0,437234 | 0,104667 | -2,06259 | 1 | 1 |
| BB557941 | 0,12564 | 0,0300713 | -2,06283 | 1 | 1 |
| Mgst2 | 0,584132 | 0,13981 | -2,06283 | 1 | 1 |
| Pcdhb21 | 0,089672 | 0,0214626 | -2,06283 | 1 | 1 |
| Serpinc1 | 0,114919 | 0,0275054 | -2,06283 | 1 | 1 |
| Klrb1f | 0,083288 | 0,0199328 | -2,06296 | 1 | 1 |
| Siglece | 0,136814 | 0,0327222 | -2,06388 | 1 | 1 |
| Abcg3 | 0,389043 | 0,0929052 | -2,0661 | 1 | 1 |
| Galnt6 | 0,086272 | 0,0204803 | -2,07466 | 1 | 1 |
| Veph1 | 0,065598 | 0,015515 | -2,07998 | 1 | 1 |
| Txndc2 | 0,118835 | 0,0280902 | -2,08082 | 1 | 1 |
| Gm10814 | 0,308624 | 0,0728438 | -2,08297 | 1 | 1 |
| 4933427I22Rik | 0,097873 | 0,022925 | -2,09399 | 1 | 1 |
| Pcdhb17 | 0,247114 | 0,0578455 | -2,0949 | 1 | 1 |
| 6330410L21Rik | 0,033266 | 0,00777986 | -2,09621 | 1 | 1 |
| BC021891 | 0,373278 | 0,0871421 | -2,09881 | 1 | 1 |
| A730090N16Rik | 0,451728 | 0,104771 | -2,10821 | 1 | 1 |
| Slc22a13b-ps | 0,241425 | 0,0557134 | -2,11548 | 1 | 1 |
| Arl9 | 0,930994 | 0,214473 | -2,11798 | 1 | 1 |
| A4galt | 0,482727 | 0,110735 | -2,1241 | 1 | 1 |
| Rasal3 | 0,209664 | 0,0478865 | -2,13039 | 1 | 1 |
| 2210039B01Rik | 0,141848 | 0,0322321 | -2,13777 | 1 | 1 |
| Gbp11 | 0,047302 | 0,0107484 | -2,13777 | 1 | 1 |
| Krt79 | 0,06257 | 0,0142177 | -2,13777 | 1 | 1 |
| Vpreb1 | 0,420007 | 0,0954384 | -2,13777 | 1 | 1 |
| Cldn2 | 0,10751 | 0,0243528 | -2,14231 | 1 | 1 |

| | | | | | |
|---|---|---|---|---|---|
| 4921513I03Rik | 0,138055 | 0,0311833 | -2,1464 | 1 | 1 |
| Rfpl3s | 0,322125 | 0,0727602 | -2,1464 | 1 | 1 |
| Gm12295 | 0,126965 | 0,0286658 | -2,14702 | 1 | 1 |
| Unc5cl | 0,143993 | 0,0325104 | -2,14702 | 1 | 1 |
| Rhoh | 0,102104 | 0,022636 | -2,17335 | 1 | 1 |
| Ano7 | 0,117947 | 0,0261422 | -2,17369 | 1 | 1 |
| Slc1a1 | 0,125084 | 0,0275719 | -2,18162 | 1 | 1 |
| Sh3rf2 | 0,141247 | 0,0310817 | -2,18408 | 1 | 1 |
| Lrrc8e | 0,206264 | 0,0452667 | -2,18797 | 1 | 1 |
| Acbd7 | 0,590338 | 0,129252 | -2,19136 | 1 | 1 |
| Rbm44 | 0,297137 | 0,0647129 | -2,19901 | 1 | 1 |
| Gm10549 | 0,177018 | 0,0385215 | -2,20016 | 1 | 1 |
| Tnfrsf9 | 0,316574 | 0,0688143 | -2,20176 | 1 | 1 |
| Xlr | 0,149783 | 0,0325049 | -2,20414 | 1 | 1 |
| Mctp2 | 0,345588 | 0,074877 | -2,20646 | 1 | 1 |
| 5031434O11Rik | 0,241778 | 0,052361 | -2,20712 | 1 | 1 |
| Aldh3b2 | 0,063504 | 0,0137527 | -2,20712 | 1 | 1 |
| Gpr84 | 0,184523 | 0,0399614 | -2,20712 | 1 | 1 |
| Slc7a14 | 0,014391 | 0,00311651 | -2,20712 | 1 | 1 |
| Art3 | 0,27446 | 0,0594128 | -2,20775 | 1 | 1 |
| Gm7334 | 0,491877 | 0,106361 | -2,20933 | 1 | 1 |
| Ccdc85a | 0,109193 | 0,0235465 | -2,2133 | 1 | 1 |
| Gpr65 | 0,371844 | 0,0799099 | -2,21825 | 1 | 1 |
| Nr1i3 | 0,121771 | 0,0261012 | -2,22198 | 1 | 1 |
| Gfi1 | 0,133079 | 0,0284921 | -2,22365 | 1 | 1 |
| Kcnk16 | 0,561457 | 0,119617 | -2,23076 | 1 | 1 |
| Chrdl2 | 0,158074 | 0,0335756 | -2,23512 | 1 | 1 |
| D830026I12Rik | 0,147752 | 0,0313827 | -2,23513 | 1 | 1 |
| Fbp1 | 0,199645 | 0,042218 | -2,24151 | 1 | 1 |
| B230119M05Rik | 0,328657 | 0,0694549 | -2,24243 | 1 | 1 |
| Tmie | 0,114067 | 0,0241058 | -2,24243 | 1 | 1 |
| Pappa2 | 0,072789 | 0,0153146 | -2,24882 | 1 | 1 |
| Arap2 | 0,305906 | 0,0637255 | -2,26315 | 1 | 1 |
| Foxl2os | 0,058694 | 0,0121269 | -2,27501 | 1 | 1 |
| 2810055G20Rik | 0,044205 | 0,00913272 | -2,2751 | 1 | 1 |
| 5033404E19Rik | 0,515919 | 0,106588 | -2,2751 | 1 | 1 |
| Tex38 | 0,559179 | 0,113513 | -2,30046 | 1 | 1 |
| Eppk1 | 0,043788 | 0,00888184 | -2,30161 | 1 | 1 |
| Tmem252 | 0,512995 | 0,103779 | -2,30543 | 1 | 1 |
| Dner | 0,24002 | 0,0485151 | -2,30665 | 1 | 1 |
| C1ql4 | 0,415331 | 0,0833974 | -2,31619 | 1 | 1 |
| Kynu | 0,287351 | 0,0576728 | -2,31685 | 1 | 1 |
| Cnga2 | 0,075241 | 0,0150966 | -2,31729 | 1 | 1 |
| Plcl1 | 0,290869 | 0,0583497 | -2,31757 | 1 | 1 |
| Dlk2 | 0,444976 | 0,08888 | -2,3238 | 1 | 1 |
| St8sia6 | 0,228322 | 0,04554 | -2,32587 | 1 | 1 |
| 2810433D01Rik | 0,503007 | 0,0995405 | -2,33722 | 1 | 1 |
| Gpr171 | 0,067352 | 0,0133217 | -2,33793 | 1 | 1 |

| | | | | | |
|---|---|---|---|---|---|
| Saa1 | 0,338125 | 0,0667826 | -2,34001 | 1 | 1 |
| Il1rl2 | 0,40693 | 0,0798369 | -2,34965 | 1 | 1 |
| Nrn1l | 0,544922 | 0,106424 | -2,35622 | 1 | 1 |
| Dusp8 | 0,25658 | 0,0500938 | -2,35671 | 1 | 1 |
| 4930503B20Rik | 0,070834 | 0,0138159 | -2,35811 | 1 | 1 |
| Gdpd2 | 0,485413 | 0,0940874 | -2,36714 | 1 | 1 |
| Asb17 | 0,524063 | 0,101382 | -2,36994 | 1 | 1 |
| Sec1 | 0,061567 | 0,0118774 | -2,37395 | 1 | 1 |
| Gfod1 | 0,425048 | 0,0815263 | -2,38229 | 1 | 1 |
| 3632454L22Rik | 0,06774 | 0,0129022 | -2,39238 | 1 | 1 |
| Gabrg1 | 0,04928 | 0,00936803 | -2,39518 | 1 | 1 |
| E130218I03Rik | 0,327854 | 0,0621217 | -2,39989 | 1 | 1 |
| Sla2 | 0,297738 | 0,0554782 | -2,42405 | 1 | 1 |
| Slc35d3 | 0,49805 | 0,0927682 | -2,42459 | 1 | 1 |
| Tgm1 | 0,408669 | 0,0757925 | -2,43081 | 1 | 1 |
| Amica1 | 0,102437 | 0,0189812 | -2,43209 | 1 | 1 |
| Serpinb9b | 0,133945 | 0,0248021 | -2,43311 | 1 | 1 |
| D630041G03Rik | 0,095612 | 0,017654 | -2,43719 | 1 | 1 |
| Ldoc1 | 0,38738 | 0,0713897 | -2,43996 | 1 | 1 |
| Plg | 0,115916 | 0,0211008 | -2,45771 | 1 | 1 |
| Zc3h12d | 0,038039 | 0,00692058 | -2,45851 | 1 | 1 |
| Themis3 | 0,185714 | 0,0337692 | -2,4593 | 1 | 1 |
| Dok6 | 0,186988 | 0,0339821 | -2,4601 | 1 | 1 |
| Ly6g6c | 0,195892 | 0,0356002 | -2,4601 | 1 | 1 |
| Tslp | 0,159244 | 0,0289299 | -2,4606 | 1 | 1 |
| Rab17 | 0,174702 | 0,0315734 | -2,46811 | 1 | 1 |
| Dnm3 | 0,198386 | 0,0356685 | -2,47559 | 1 | 1 |
| Adamtsl3 | 0,109584 | 0,0196859 | -2,47681 | 1 | 1 |
| D030045P18Rik | 0,371959 | 0,0666358 | -2,48077 | 1 | 1 |
| Ramp1 | 0,166308 | 0,0297219 | -2,48426 | 1 | 1 |
| Krt9 | 0,127046 | 0,022655 | -2,48745 | 1 | 1 |
| Slc39a4 | 0,424466 | 0,0744015 | -2,51225 | 1 | 1 |
| Ccl2 | 0,255691 | 0,0447071 | -2,51583 | 1 | 1 |
| Cd70 | 0,241264 | 0,0421845 | -2,51583 | 1 | 1 |
| Cxcr5 | 0,063844 | 0,011163 | -2,51583 | 1 | 1 |
| Tspan1 | 0,087113 | 0,0152315 | -2,51583 | 1 | 1 |
| Tfec | 0,492363 | 0,0854413 | -2,52672 | 1 | 1 |
| Sowahb | 0,508012 | 0,0870063 | -2,54567 | 1 | 1 |
| Lhfpl1 | 0,41849 | 0,071644 | -2,54627 | 1 | 1 |
| A630001G21Rik | 0,327987 | 0,0557804 | -2,55581 | 1 | 1 |
| D030047H15Rik | 0,136242 | 0,0230824 | -2,5613 | 1 | 1 |
| Reps2 | 0,183578 | 0,0308957 | -2,57092 | 1 | 1 |
| Cd2 | 0,170131 | 0,0286318 | -2,57095 | 1 | 1 |
| Plekhg6 | 0,147237 | 0,0247594 | -2,57209 | 1 | 1 |
| Gm711 | 0,129541 | 0,0217728 | -2,57281 | 1 | 1 |
| Dmbx1 | 0,152132 | 0,0252882 | -2,58879 | 1 | 1 |
| Gm15816 | 0,346999 | 0,0573122 | -2,59802 | 1 | 1 |
| Grm6 | 0,063077 | 0,0103643 | -2,60549 | 1 | 1 |

| | | | | | |
|---|---|---|---|---|---|
| Slc25a48 | 0,201347 | 0,033063 | -2,6064 | 1 | 1 |
| Filip1 | 0,337257 | 0,0549933 | -2,61652 | 1 | 1 |
| Osgin1 | 0,094341 | 0,0153182 | -2,62263 | 1 | 1 |
| Negr1 | 0,235684 | 0,0379699 | -2,63392 | 1 | 1 |
| Nlrc3 | 0,430064 | 0,0684577 | -2,65126 | 1 | 1 |
| Tnfrsf25 | 0,505359 | 0,0799385 | -2,66035 | 1 | 1 |
| Popdc2 | 0,406084 | 0,0638587 | -2,66882 | 1 | 1 |
| Ceacam10 | 0,644276 | 0,101262 | -2,66958 | 1 | 1 |
| D830046C22Rik | 0,126508 | 0,0198717 | -2,67045 | 1 | 1 |
| Rxfp3 | 0,134612 | 0,0211382 | -2,67088 | 1 | 1 |
| Serpinb6b | 0,37213 | 0,0583756 | -2,67237 | 1 | 1 |
| Rasgef1c | 0,168384 | 0,0263863 | -2,6739 | 1 | 1 |
| Bean1 | 0,057808 | 0,00905782 | -2,67403 | 1 | 1 |
| Acmsd | 0,214567 | 0,0335862 | -2,67549 | 1 | 1 |
| Diras2 | 0,150066 | 0,0233137 | -2,68635 | 1 | 1 |
| Rassf6 | 0,594171 | 0,0920663 | -2,69013 | 1 | 1 |
| Pof1b | 0,216713 | 0,0335123 | -2,69303 | 1 | 1 |
| 4921506M07Rik | 0,273732 | 0,042218 | -2,69684 | 1 | 1 |
| Dock2 | 0,510037 | 0,0785393 | -2,69912 | 1 | 1 |
| Pik3cg | 0,364564 | 0,0559052 | -2,70512 | 1 | 1 |
| Disp2 | 0,313318 | 0,0475807 | -2,71918 | 1 | 1 |
| 1700101I11Rik | 0,294986 | 0,0447071 | -2,72207 | 1 | 1 |
| BC051019 | 0,102177 | 0,0154856 | -2,72207 | 1 | 1 |
| Dnah6 | 0,014267 | 0,0021623 | -2,72207 | 1 | 1 |
| Zfp572 | 0,198639 | 0,030105 | -2,72207 | 1 | 1 |
| Slc44a3 | 0,555345 | 0,0840615 | -2,72387 | 1 | 1 |
| Ms4a10 | 0,353041 | 0,0531469 | -2,73178 | 1 | 1 |
| Myo1g | 0,293774 | 0,0433679 | -2,76001 | 1 | 1 |
| 3830403N18Rik | 0,355429 | 0,0524304 | -2,76108 | 1 | 1 |
| Tmem125 | 0,174929 | 0,0258043 | -2,76108 | 1 | 1 |
| Proc | 0,133712 | 0,0196939 | -2,7633 | 1 | 1 |
| Hnmt | 0,255925 | 0,0375369 | -2,76934 | 1 | 1 |
| Fibin | 0,098434 | 0,0144311 | -2,76998 | 1 | 1 |
| Gpr101 | 0,035263 | 0,00516984 | -2,76998 | 1 | 1 |
| Hesx1 | 0,634219 | 0,0917113 | -2,78981 | 1 | 1 |
| AI467606 | 0,328124 | 0,0471074 | -2,80022 | 1 | 1 |
| Myh7b | 0,524218 | 0,0751566 | -2,8022 | 1 | 1 |
| Gpr34 | 0,450952 | 0,0635604 | -2,82677 | 1 | 1 |
| Bdkrb2 | 0,260718 | 0,0364688 | -2,83775 | 1 | 1 |
| D930020B18Rik | 0,095519 | 0,0132745 | -2,84713 | 1 | 1 |
| Upk3a | 0,341302 | 0,0474316 | -2,84713 | 1 | 1 |
| Naip5 | 0,038406 | 0,00528973 | -2,86006 | 1 | 1 |
| Tm4sf5 | 0,50689 | 0,0698153 | -2,86006 | 1 | 1 |
| Gbp2 | 0,27612 | 0,038013 | -2,86073 | 1 | 1 |
| Hnf4a | 0,075515 | 0,0102994 | -2,8742 | 1 | 1 |
| Gpr126 | 0,298731 | 0,04074 | -2,87433 | 1 | 1 |
| Srd5a2 | 0,610221 | 0,0828114 | -2,88143 | 1 | 1 |
| Ppp1r17 | 0,617699 | 0,0836706 | -2,88411 | 1 | 1 |

| | | | | | |
|---|---|---|---|---|---|
| Aqp9 | 0,213316 | 0,0285377 | -2,90205 | 1 | 1 |
| Popdc3 | 0,122186 | 0,0163412 | -2,90249 | 1 | 1 |
| Col19a1 | 0,038948 | 0,00520685 | -2,90307 | 1 | 1 |
| Slc52a3 | 0,442485 | 0,0581891 | -2,92681 | 1 | 1 |
| Fam110c | 0,429913 | 0,056527 | -2,92703 | 1 | 1 |
| Itgax | 0,085192 | 0,011197 | -2,9276 | 1 | 1 |
| P2ry13 | 0,405359 | 0,0532111 | -2,9294 | 1 | 1 |
| Rgs14 | 0,247638 | 0,0323874 | -2,93473 | 1 | 1 |
| Casq2 | 0,269138 | 0,0349534 | -2,94484 | 1 | 1 |
| Tnni2 | 0,447585 | 0,0581285 | -2,94484 | 1 | 1 |
| Cd300lf | 0,128426 | 0,0161992 | -2,98694 | 1 | 1 |
| Mroh7 | 0,082893 | 0,0103293 | -3,00451 | 1 | 1 |
| Selplg | 0,143059 | 0,0178265 | -3,00451 | 1 | 1 |
| Papln | 0,105262 | 0,0131086 | -3,0054 | 1 | 1 |
| G6bos | 0,226971 | 0,0281526 | -3,01117 | 1 | 1 |
| C2cd4d | 0,196999 | 0,0242032 | -3,02492 | 1 | 1 |
| Tc2n | 0,069734 | 0,00841627 | -3,0506 | 1 | 1 |
| Kcnd2 | 0,051833 | 0,00619856 | -3,06388 | 1 | 1 |
| Sdpr | 0,527522 | 0,0622077 | -3,08406 | 1 | 1 |
| Krt17 | 0,176658 | 0,0205922 | -3,10079 | 1 | 1 |
| Slamf8 | 0,199978 | 0,0233026 | -3,10128 | 1 | 1 |
| Ccdc64b | 0,406488 | 0,0469977 | -3,11255 | 1 | 1 |
| Alox5 | 0,322701 | 0,0368326 | -3,13114 | 1 | 1 |
| Cytip | 0,085629 | 0,00973212 | -3,13728 | 1 | 1 |
| Zfp750 | 0,202604 | 0,0229345 | -3,14308 | 1 | 1 |
| Podn | 0,500707 | 0,0565844 | -3,14549 | 1 | 1 |
| Ifit1 | 0,099721 | 0,0110576 | -3,17286 | 1 | 1 |
| Krt2 | 0,100756 | 0,0111723 | -3,17286 | 1 | 1 |
| P2rx6 | 0,117762 | 0,0130554 | -3,17315 | 1 | 1 |
| Ackr2 | 0,145018 | 0,0158669 | -3,19213 | 1 | 1 |
| AI662270 | 0,206574 | 0,0223538 | -3,20807 | 1 | 1 |
| Samd11 | 0,466449 | 0,0502826 | -3,21359 | 1 | 1 |
| Nos2 | 0,198622 | 0,0213266 | -3,2193 | 1 | 1 |
| Dlx1 | 0,410849 | 0,0436821 | -3,23349 | 1 | 1 |
| Galnt3 | 0,568223 | 0,0600844 | -3,2414 | 1 | 1 |
| Cpa1 | 0,18526 | 0,0195881 | -3,24151 | 1 | 1 |
| Muc13 | 0,184122 | 0,0194677 | -3,24151 | 1 | 1 |
| Lpar5 | 0,214077 | 0,0226315 | -3,24172 | 1 | 1 |
| Ndst3 | 0,049613 | 0,00524234 | -3,24243 | 1 | 1 |
| Gpr111 | 0,132719 | 0,0138953 | -3,2557 | 1 | 1 |
| Cdh26 | 0,095709 | 0,00988274 | -3,27566 | 1 | 1 |
| Fam65b | 0,266696 | 0,0274363 | -3,28104 | 1 | 1 |
| Lamc3 | 0,046549 | 0,00470034 | -3,30792 | 1 | 1 |
| Cubn | 0,278208 | 0,027923 | -3,31664 | 1 | 1 |
| Csf2rb2 | 0,460244 | 0,0459166 | -3,32531 | 1 | 1 |
| Trpv2 | 0,32088 | 0,0317009 | -3,33944 | 1 | 1 |
| Scnn1b | 0,522704 | 0,0515163 | -3,34289 | 1 | 1 |
| Hk3 | 0,162808 | 0,0159028 | -3,35582 | 1 | 1 |

| | | | | | |
|---|---|---|---|---|---|
| 6430411K18Rik | 0,123426 | 0,0119785 | -3,36512 | 1 | 1 |
| Sec16b | 0,3763 | 0,0362963 | -3,37399 | 1 | 1 |
| Mir1892 | 1448,56 | 139,248 | -3,37889 | 1 | 1 |
| Card9 | 0,399834 | 0,0382573 | -3,3856 | 1 | 1 |
| Hdc | 0,129476 | 0,0122615 | -3,40048 | 1 | 1 |
| Hsd17b14 | 0,244788 | 0,0231817 | -3,40048 | 1 | 1 |
| Nlrp3 | 0,148913 | 0,0141022 | -3,40048 | 1 | 1 |
| Ptpn22 | 0,114037 | 0,0105541 | -3,43362 | 1 | 1 |
| Hpca | 0,217681 | 0,0199659 | -3,44661 | 1 | 1 |
| 3110099E03Rik | 0,12139 | 0,0110304 | -3,4601 | 1 | 1 |
| Hmgcs2 | 0,153273 | 0,0138459 | -3,46858 | 1 | 1 |
| Tlr7 | 0,134828 | 0,0119106 | -3,5008 | 1 | 1 |
| Slc28a3 | 0,210453 | 0,0185189 | -3,50643 | 1 | 1 |
| H2-Q2 | 0,344154 | 0,0300713 | -3,51659 | 1 | 1 |
| Mia2 | 0,5613 | 0,0489449 | -3,51954 | 1 | 1 |
| Clec2d | 0,486568 | 0,0423598 | -3,52187 | 1 | 1 |
| Amn | 0,419458 | 0,0363457 | -3,52867 | 1 | 1 |
| Slc16a12 | 0,171179 | 0,0146831 | -3,54328 | 1 | 1 |
| Zfp457 | 0,164149 | 0,0140764 | -3,54365 | 1 | 1 |
| Cldn23 | 0,611128 | 0,0519534 | -3,55619 | 1 | 1 |
| Il1bos | 0,427093 | 0,0358922 | -3,57281 | 1 | 1 |
| Csf3r | 0,150713 | 0,0126609 | -3,57335 | 1 | 1 |
| Vnn1 | 0,382881 | 0,0315089 | -3,60306 | 1 | 1 |
| Mfap5 | 0,43705 | 0,0354755 | -3,6229 | 1 | 1 |
| Gm19434 | 0,14246 | 0,0113657 | -3,64779 | 1 | 1 |
| Ptger3 | 0,372205 | 0,0296737 | -3,64884 | 1 | 1 |
| Scel | 0,191149 | 0,0150171 | -3,67002 | 1 | 1 |
| Gpihbp1 | 0,579256 | 0,0453856 | -3,6739 | 1 | 1 |
| Vmn1r90 | 0,317557 | 0,0248343 | -3,67661 | 1 | 1 |
| Rbm47 | 0,244145 | 0,019069 | -3,67844 | 1 | 1 |
| Tlr13 | 0,144552 | 0,0112293 | -3,68624 | 1 | 1 |
| P2ry6 | 0,608392 | 0,0471349 | -3,69013 | 1 | 1 |
| Oit3 | 0,139067 | 0,0107142 | -3,69819 | 1 | 1 |
| Gdf6 | 0,483753 | 0,0371172 | -3,70411 | 1 | 1 |
| 4933422A05Rik | 0,313651 | 0,0233628 | -3,74688 | 1 | 1 |
| Insrr | 0,073771 | 0,00549493 | -3,74688 | 1 | 1 |
| Cd86 | 0,245994 | 0,0183228 | -3,74692 | 1 | 1 |
| Fam83b | 0,494767 | 0,0367178 | -3,7522 | 1 | 1 |
| Kcnq1 | 0,51839 | 0,0375024 | -3,78898 | 1 | 1 |
| Tmprss4 | 0,180722 | 0,0130461 | -3,79208 | 1 | 1 |
| Traf3ip3 | 0,227719 | 0,016319 | -3,80263 | 1 | 1 |
| Dlx6os1 | 0,185956 | 0,0133139 | -3,80395 | 1 | 1 |
| Pilra | 0,183881 | 0,0131498 | -3,80566 | 1 | 1 |
| Igsf5 | 0,445607 | 0,0310242 | -3,84431 | 1 | 1 |
| Myl3 | 0,563129 | 0,0387806 | -3,86006 | 1 | 1 |
| Kcnj5 | 0,471707 | 0,0324781 | -3,86035 | 1 | 1 |
| Brinp1 | 0,39664 | 0,0271643 | -3,86805 | 1 | 1 |
| Itgam | 0,281712 | 0,0191149 | -3,88145 | 1 | 1 |

| | | | | | |
|---|---|---|---|---|---|
| Dhh | 0,292897 | 0,0197798 | -3,8883 | 1 | 1 |
| Emx2os | 0,174369 | 0,01117 | -3,96444 | 1 | 1 |
| Zbtb32 | 0,271356 | 0,0171318 | -3,98544 | 1 | 1 |
| Gpr150 | 0,22252 | 0,0138589 | -4,00504 | 1 | 1 |
| Casr | 0,162704 | 0,00988078 | -4,04148 | 1 | 1 |
| Cdh15 | 0,341254 | 0,0203972 | -4,0644 | 1 | 1 |
| Wnt9b | 0,436337 | 0,0260676 | -4,06511 | 1 | 1 |
| Arhgap30 | 0,442359 | 0,0262624 | -4,07415 | 1 | 1 |
| Slfn14 | 0,336653 | 0,0199137 | -4,07943 | 1 | 1 |
| Fcgr1 | 0,501444 | 0,0292234 | -4,10089 | 1 | 1 |
| Evi2a-evi2b,Evi2b | 0,209847 | 0,012195 | -4,10497 | 1 | 1 |
| Dusp15 | 0,159477 | 0,00922664 | -4,1114 | 1 | 1 |
| Gabrp | 0,242519 | 0,0139089 | -4,12402 | 1 | 1 |
| Cdh16 | 0,172327 | 0,00985471 | -4,12819 | 1 | 1 |
| Gm16897 | 0,220903 | 0,0124771 | -4,14606 | 1 | 1 |
| Rtp4 | 0,354594 | 0,0196596 | -4,17286 | 1 | 1 |
| Evi2b | 0,16957 | 0,00883008 | -4,26332 | 1 | 1 |
| Il10ra | 0,523777 | 0,0260163 | -4,33147 | 1 | 1 |
| Ulbp1 | 0,557322 | 0,0264914 | -4,39492 | 1 | 1 |
| Trp73 | 0,212421 | 0,0100328 | -4,40413 | 1 | 1 |
| Tifab | 0,536417 | 0,0249934 | -4,42374 | 1 | 1 |
| Rsad2 | 0,325735 | 0,0150316 | -4,43762 | 1 | 1 |
| Fmnl1 | 0,490944 | 0,0226142 | -4,44026 | 1 | 1 |
| Fgg | 0,457586 | 0,0204048 | -4,48706 | 1 | 1 |
| Slfn2 | 0,42172 | 0,0184342 | -4,51583 | 1 | 1 |
| Mab21l2 | 0,414835 | 0,0171203 | -4,59876 | 1 | 1 |
| Clec9a | 0,239893 | 0,00981812 | -4,6108 | 1 | 1 |
| Baalc | 0,180933 | 0,00715231 | -4,6609 | 1 | 1 |
| Itgb2 | 0,52108 | 0,0202514 | -4,68541 | 1 | 1 |
| Corin | 0,153893 | 0,00597656 | -4,68647 | 1 | 1 |
| Pdia2 | 0,476048 | 0,0177416 | -4,7459 | 1 | 1 |
| Trim30a | 0,33542 | 0,0119977 | -4,80514 | 1 | 1 |
| Atp8b1 | 0,331258 | 0,0111767 | -4,88939 | 1 | 1 |
| Ubash3a | 0,297301 | 0,00979051 | -4,9244 | 1 | 1 |
| BC035044 | 0,590428 | 0,0177673 | -5,05446 | 1 | 1 |
| Palmd | 0,427123 | 0,0122895 | -5,11915 | 1 | 1 |
| Pak6 | 0,273969 | 0,00778261 | -5,13762 | 1 | 1 |
| Naip2 | 0,209449 | 0,005875 | -5,15587 | 1 | 1 |
| Flt3 | 0,457881 | 0,0123939 | -5,20727 | 1 | 1 |
| Cnr2 | 0,283376 | 0,00736164 | -5,26655 | 1 | 1 |
| Macc1 | 0,290102 | 0,00745131 | -5,28292 | 1 | 1 |
| Myo1f | 0,476402 | 0,0117894 | -5,33662 | 1 | 1 |
| Fat2 | 0,541062 | 0,0128124 | -5,40018 | 1 | 1 |
| C3ar1 | 0,537886 | 0,00657244 | -6,35473 | 1 | 1 |
| Ass1 | 0,436275 | 0,0001147 | -11,8932 | 1 | 1 |
| G630025P09Rik | 0,187118 | 4,94E-006 | -15,2104 | 1 | 1 |

v

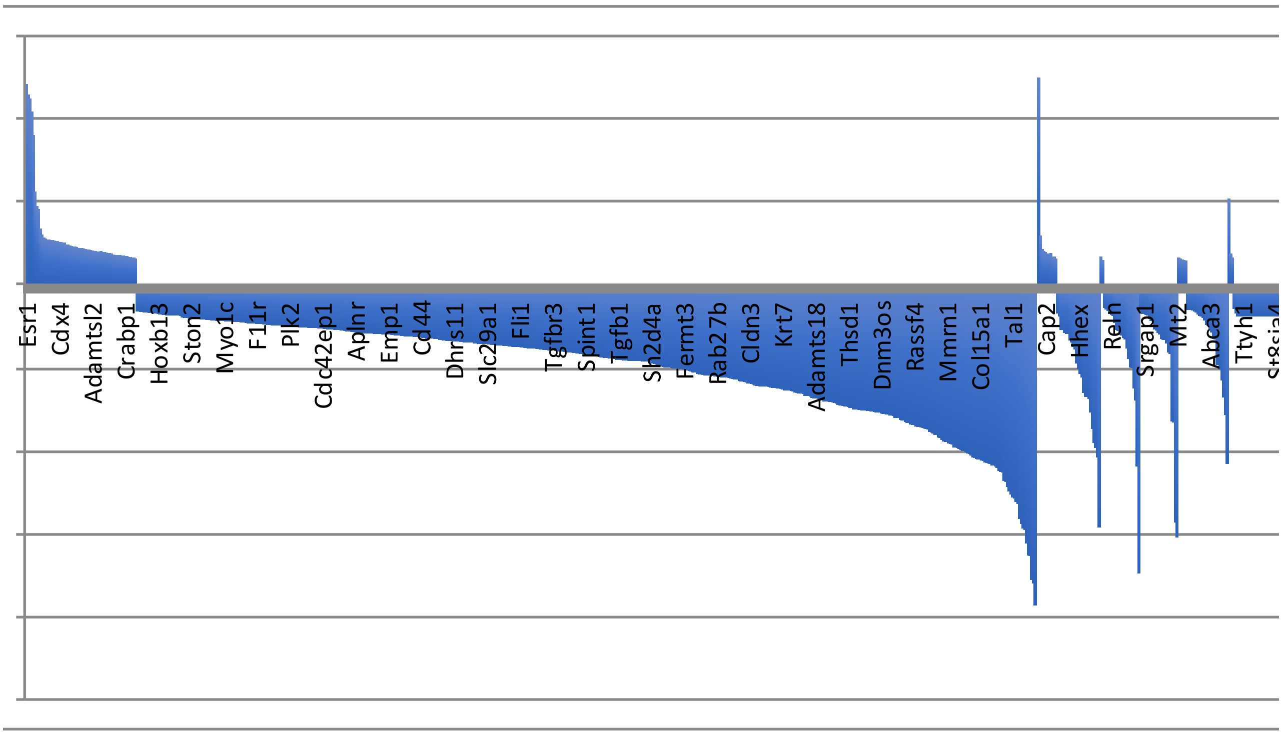
Esr1
Cdx4
Adamtsl2
Crabp1
Hoxb13
Ston2
Myo1c
F11r
Plk2
Cdc42ep1
Aplnr
Emp1
Cd44
Dhrs11
Slc29a1
Fli1
Tgfbr3
Spint1
Tgfb1
Sh2d4a
Fermt3
Rab27b
Cldn3
Krt7
Adamts18
Thsd1
Dnm3os
Rassf4
Mmrn1
Col15a1
Tal1
Cap2
Hhex
Reln
Srgap1
Mt2
Abca3
Ttyh1

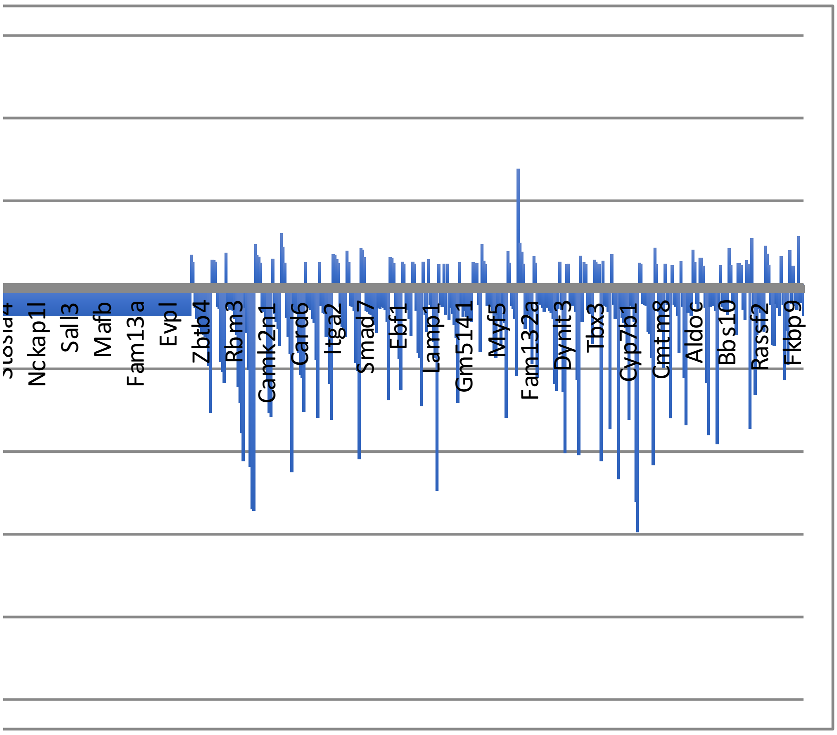
Nckap1l
Sall3
Mafb
Fam13a
Evpl
Zbtb4
Rbm3
Camk2n1
Card6
Itga2
Smad7
Ebf1
Lamp1
Gm5141
Myf5
Fam132a
Dynlt3
Tbx3
Cyp7b1
Cmtm8
Aldoc
Bbs10
Rassf2
Fkbp9
